# Retrotransposon Activity in Chronic Lymphocytic Leukemia: Associations with RNA Splicing and *TP53* Dysfunction

**DOI:** 10.64898/2026.07.16.738931

**Authors:** Anastasiya Volakhava, Sarka Pavlova, Lenka Radova, Kristyna Tausova, Hana Svozilova, Marcela Zenatova, Michael Doubek, Ilgar Mamedov, Sarka Pospisilova, Karla Plevova

**Author notes:** **Correspondence:** Karla Plevova, MSc, PhD, Department of Internal Medicine, Hematology and Oncology, University Hospital Brno, and Faculty of Medicine, Masaryk University. Contributed equally as senior authors (Sarka Pospisilova, Karla Plevova).

## Abstract

Retroelements (RE), particularly autonomous Long Interspersed Element-1 (LINE-1), function as potent drivers of genomic instability in various malignancies. While normally silenced by epigenetic mechanisms, their reactivation in cancer cells can drive tumor evolution. *TP53* is known to repress LINE-1 transcription; however, the consequences of *TP53* dysfunction on retrotransposition and transcriptional activity in chronic lymphocytic leukemia (CLL) remain unknown. To investigate the relationship between *TP53* status, LINE-1 retrotranspositional potential, and transcriptomic alterations, we utilized a multi-omics approach, combining a highly sensitive NGS protocol for detecting novel LINE-1 insertions with transcriptomic profiling of transposable elements and protein-coding genes. We applied these methods to a cohort of CLL patients stratified by the presence or absence of *TP53* clonal evolution and to CLL-derived cell lines MEC1 and HG3, including CRISPR/Cas9-engineered *TP53* mutants. Genomic analysis revealed no evidence of widespread somatic retrotransposition, suggesting that CLL exhibits resistance against *de novo* LINE-1 insertions. Conversely, transcriptomic profiling uncovered distinct transposon expression signatures aligned with patterns of *TP53* mutation status evolution. Notably, differentially expressed genes were significantly enriched in the RNA splicing pathway, indicating that while LINE-1 elements remain largely constrained at the genomic level, their transcriptomic activity may influence cellular rewiring, affecting patterns of TP53 mutation clonal evolution. Based on these results, we argue that the pathogenic contribution of retroelements in CLL lies in transcriptomic dysregulation and splicing alterations, rather than in direct DNA damage caused by LINE-1 insertions.

## Introduction

Transposable Elements (TEs), also known as mobile genetic elements or “jumping genes,” are DNA sequences capable of moving from one location to another within the host genome. In humans, TEs account for nearly half of the genome (∼42%), underscoring their significant contribution to genomic architecture. [1] TEs drive evolutionary processes such as genomic rearrangements, mutagenesis, and gene expression regulation. [2] Although these processes can promote diversification, adaptation, and structural variation, TE mobility must be tightly controlled to prevent genomic instability. [3]

TEs are classified based on their mobilization mechanisms. Retrotransposons (REs) propagate using a “copy-and-paste” mechanism involving an RNA intermediate. These include autonomous Long Interspersed Nuclear Elements (LINEs) and non-autonomous Short Interspersed Nuclear Elements (SINEs). LINE-1 (L1) elements occupy approximately 17% of the human genome and contain two open reading frames (ORF1 and ORF2) essential for retrotransposition. ORF1 encodes an RNA-binding protein with nucleic acid chaperone activity, while ORF2 encodes a protein with both reverse transcriptase and endonuclease activity required for reintegration of the L1 RNA back into the genome. Among LINE-1, the L1HS subfamily is the most active in the human genome. [4] Non-autonomous elements such as Alu and SINE-VNTR-Alu (SVA) rely on the enzymatic machinery of LINE-1 for their propagation. [4–7]

When dysregulated, REs can compromise genomic integrity by generating double-strand breaks, chromosomal rearrangements, and copy-number variations (CNVs). RE insertions can disrupt tumor suppressor genes or activate oncogenes by inserting a new promoter. RE-derived oligonucleotides can activate the immune system, leading to a strong inflammatory response. [3,8] Elevated RE activity has been detected in various solid tumors, including breast, lung, colorectal, and ovarian cancers, often associated with LINE-1 activation. [9] In cancer, global DNA hypomethylation typically leads to TE derepression [10], while hypermethylation of tumor suppressor gene (TSG) promoters facilitates carcinogenesis. [11] In hematological malignancies such as acute myeloid leukemia (AML) and multiple myeloma, enhanced RE activity has been implicated in genomic instability and clonal evolution. [12, 13]

Despite growing evidence of TE dysregulation in hematological malignancies, the role of TE in the pathogenesis and clonal evolution of these diseases, particularly chronic lymphocytic leukemia (CLL), remains unexplored. CLL is the most common leukemia in adults, characterized by abnormal accumulation of B lymphocytes in the blood, bone marrow, and lymphoid tissues. It exhibits an extremely variable clinical course, ranging from indolent to aggressive disease. [14,15] Among genomic changes observed in CLL, *TP53* aberrations associate with poor prognosis and drug resistance. [16–18]

p53, a protein encoded by the *TP53* gene, plays a central role in maintaining genomic stability. It regulates cell cycle checkpoints, DNA repair, and apoptosis. [19, 20] More recently, it has been shown to suppress RE activity by directly binding to their promoters and recruiting chromatin-modifying enzymes such as histone deacetylases (HDACs) and DNA methyltransferases (DNMTs). [21–23] In cancers where *TP53* is mutated or deleted, this regulatory control is lost, leading to RE reactivation and increased genomic instability. [21, 24]

In this study, we aimed to reveal the relationship between *TP53* dysfunction and RE activity in CLL, where *TP53* disruptions are associated with unfavorable disease. We employed a comprehensive sequencing approach to identify TE-related changes in cell-line models and in a unique CLL patient cohort with well-characterized clonal dynamics of *TP53* alterations. This study provides one of the first comprehensive investigations of TEs in CLL, especially in the context of *TP53* dysfunction.

## Materials and methods

### Cell Cultures and Patient Samples

The present research was conducted on CLL-derived cell lines MEC1 and HG3 and CRISPR/Cas9-generated *TP53*-inactivated HG3 clones (Supplementary Table S1 [25–27]), as well as on primary CLL patient samples (Supplementary Table S2; Figure 1). Cell lines were cultured according to the DSMZ recommendations. Long-term cultivation experiments were conducted for MEC1 and all HG3 cell lines, with cells harvested at the initial timepoint (baseline; TP1) and after 12 months of continuous culture (TP2). During cultivation, the cell lines were tested weekly for mycoplasma contamination using the MycoAlert Plus Detection Kit (Lonza). The cell lines were authenticated by comparing their STR profiles at 16 loci against reference sets in the Cellosaurus STR Similarity Search Tool and DSMZCellDive.

**Figure 1.**
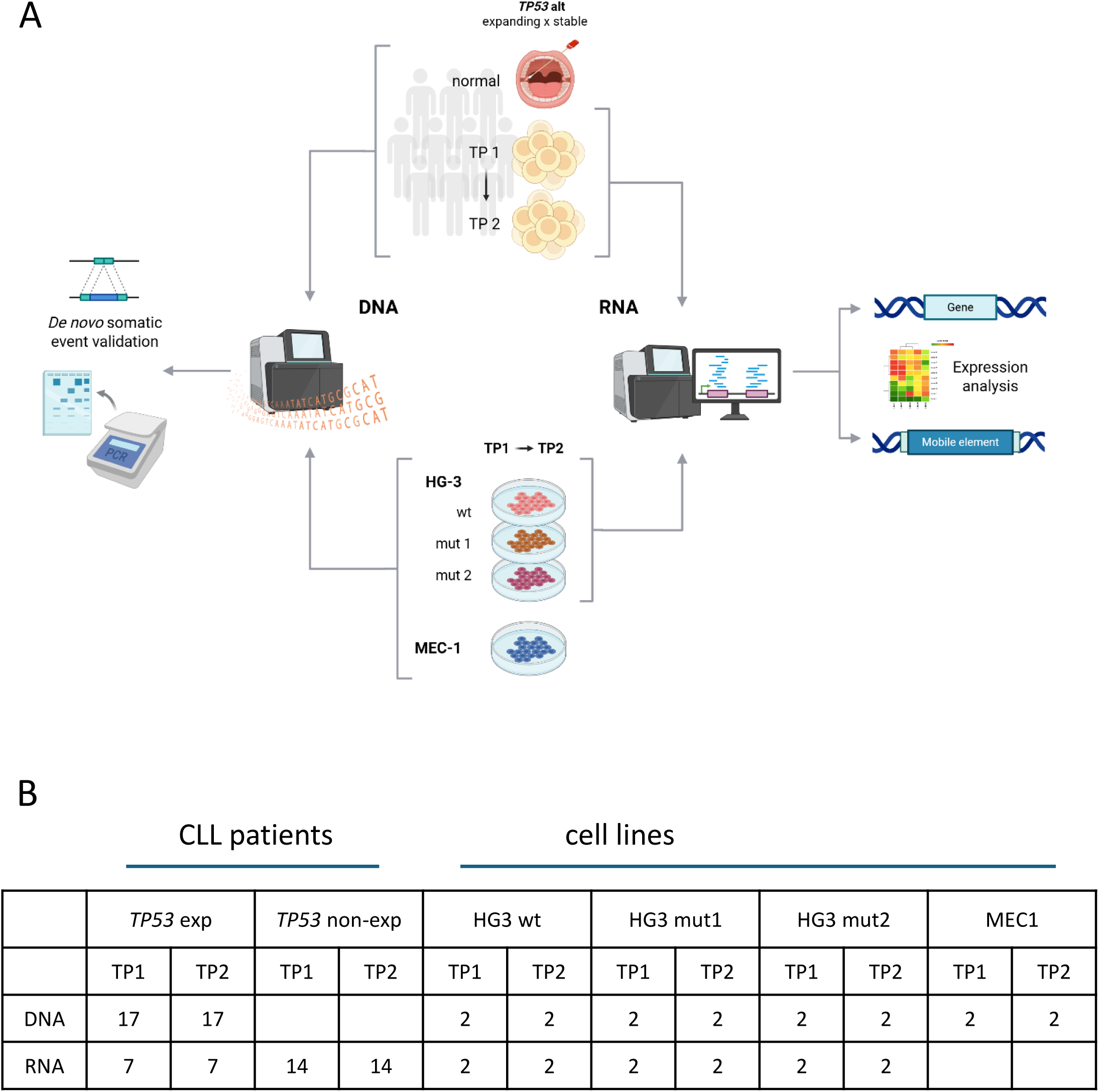
Experimental design and counts of tested samples. (A) The experimental design involved serial samples from CLL patients and CLL-derived cell lines HG3 and MEC1. Somatic retrotransposition was analyzed from gDNA, while RNA samples were used to explore gene and TE expression. (B) The numbers of samples used in specific experiments are summarized and detailed in Supplementary Tables S1 and S2.

The cohort of primary CLL samples included 32 CLL patients monitored and treated at the University Hospital Brno (Supplementary Table S2). The study was approved by the Ethics Committees of both the University Hospital Brno and Masaryk University. Written informed consent was obtained from all patients in line with the Declaration of Helsinki. All patients met the iwCLL/NCI criteria for the diagnosis of CLL. For each patient, paired treatment-naive (TP1) and post-treatment (TP2) samples were collected. The median number of therapies before collecting the post-treatment sample was 1 (range 1 to 10). All patients received chemo(immuno)therapy.

### B cell Separation and genomic DNA and RNA Extraction

Peripheral blood samples were processed by Ficoll-Paque™ PLUS (GE Healthcare) density gradient centrifugation, followed by B cell separation using RosetteSep™ antibody cocktails (StemCell Technologies). Flow cytometry was used to assess purity.

Genomic DNA (gDNA) from patient samples was extracted using the DNeasy® Blood & Tissue Kit (Qiagen). Genomic DNA from cell lines was extracted using the MagCore 101 Genomic DNA Whole Blood Kit (RBC Bioscience) on the MagCore automated nucleic acid extraction system. For total RNA isolation, either the TRI Reagent® (MRC) or RNeasy® Mini Kit (Qiagen) was used. The gDNA/RNA quantity of patient samples was measured on Qubit® 2.0 Fluorometer (Invitrogen); integrity was verified by electrophoresis or the 2100 Bioanalyzer instrument (Agilent), respectively. For cell-line-derived gDNA/RNA, the quantity was assessed using NanoDrop 2000c (Thermo Scientific), and, in case of gDNA, also using Tecan Spark 10M Multimode Plate Reader and the QuantiFluor dsDNA System (Promega).

### *TP53* Status Assessment

The *TP53* mutation status was assessed using amplicon ultra-deep next-generation sequencing (NGS) of exons 2-11 and adjacent splicing sites, enabling the detection of variants with a variant allele frequency (VAF) >0.1%, as described previously [28]. The sequencing libraries were prepared with the Nextera XT Sample Preparation Kit (Illumina) and sequenced on the MiSeq instrument. The variants were annotated using ANNOVAR. The NM_000546.6 sequence was used as a reference for annotation.

### Detection of Somatic Retrotransposition Events

To detect novel insertions of the transpositionally active L1HS subfamily, we applied an NGS protocol described previously [29–31]. Briefly, genomic DNA was digested using TaqI and FspBI endonucleases to generate fragments that consisted of a 3’ part of L1 retroelement and its adjacent genomic region (i.e., flank). Fragments were ligated to a stem-loop adaptor carrying unique molecular identifiers (UMIs), which enabled quantification of the number of cells bearing each insertion after sequencing. Next, a primer specific to transpositionally active L1 subfamily L1HS and a primer corresponding to the ligated adaptor were used to selectively amplify genomic flanks adjacent to the 3’ end of L1. A product of the first PCR was used in the second semi-nested PCR. Finally, an indexing PCR was carried out to introduce sample barcodes and the oligonucleotides necessary for Illumina sequencing on the NextSeq 550 instrument (paired-end, 150+150).

Sequencing data were processed using a custom bioinformatics pipeline [30]. Flanking sequences were mapped to the human reference genome (hg38) to find the coordinates of each transposon insertion. Artificial chimeric sequences were filtered out during a filtration stage in the pipeline. Candidate somatic insertions supported by at least two unique UMIs were identified by comparing insert presence across serial samples: in patient samples, insertions present in the tumor but absent in the matched germline control were considered somatic; in cell lines, insertions that appeared after long-term cultivation but were absent at baseline were retained.

Candidate somatic insertions were validated by locus-specific PCR using initial gDNA as a template, and Encyclo Polymerase Mix (Evrogen) and/or Q5 High-Fidelity 2X Master Mix (New England Biolabs) as described previously. [32, 33] To perform locus-specific PCR, we designed primers that were complementary to the unique genomic 3’ flank and 5’ flank (Supplementary Figure 1A,B). The 3’ flank primers were used in combination with a common universal primer annealing to the L1HS element to detect L1 insertion. A control reaction was performed with primers complementary to the 3’ flank and the 5’ flank in parallel to detect an insertion-free allele.

### RNA Sequencing, Transposable Element and Gene Expression Analyses

Extracted RNA was treated with DNase before library preparation. RNA sequencing (RNAseq) libraries were prepared using the TruSeq Stranded Total RNA kit (Illumina) according to the manufacturer’s instructions. The libraries were sequenced on NextSeq 500/550 and NovaSeq 6000 instruments. Reads were aligned to the human genome reference hg38 and processed with TElocal (https://github.com/mhammell-laboratory/TElocal), a tool designed to accurately quantify transcription of both genes and TEs in RNAseq datasets.

Gene and TE expression matrices were generated by merging the output of TElocal. To avoid bias due to sex chromosome-linked variability, reads mapping to the Y chromosome were excluded from downstream analyses. Expression data were normalized to enable accurate comparison across samples. Differential expression analysis of TE subfamilies was conducted using DESeq2 [34]. TE subfamilies meeting the following criteria were included: a false discovery rate (FDR) < 0.05, a baseMean > 50, and |log2 fold change| > 1. Principal Component Analysis (PCA) and hierarchical clustering were used to visualize data segregation and to confirm the alignment of expression profiles with clinical or biological variables such as *TP53* status and clonal evolution.

### Pathway and Functional Enrichment Analysis

Functional enrichment was performed to identify pathways significantly associated with the differentially expressed genes identified in each pairwise comparison. Pathway analysis was conducted using the top significant differentially expressed genes (*p* < 0.05, |log2 fold change| > 1). Functional annotation was performed using the Gene Ontology (GO) and Kyoto Encyclopedia of Genes and Genomes (KEGG) databases. The enrichment was evaluated across the three main GO domains: Biological Processes, Molecular Function, and Cellular Components. Pathways were considered statistically significant if they met the threshold of *p* < 0.05.

Differential alternative splicing events were identified using the rMATS (Multivariate Analysis of Transcript Splicing) software. [35] Raw RNAseq data were processed to detect five types of alternative splicing events: Skipped Exons (SE), Mutually Exclusive Exons (MXE), Alternative 3’ Splice Sites (A3SS), Alternative 5’ Splice Sites (A5SS), and Retained Introns (RI).

To ensure robust quantification, the analysis was performed using both junction-only (JC) and junction-plus-exon (JCEC) counting modes. Splicing profiles between defined experimental groups were compared. Statistical significance for individual splicing events was defined as *p* < 0.05.

### Statistical Analysis

All statistical analyses were conducted using the R environment for statistical computing (version 4.3.3). The Shapiro-Wilk test did not confirm normality of the data. For comparisons involving more than two groups, the Kruskal-Wallis test was applied, followed by post hoc analysis with the Bonferroni adjustment. Paired longitudinal samples (T1 vs. T2) were analyzed using the Wilcoxon signed-rank test.

## Results

### Targeted DNA Sequencing of CLL Cell Lines Reveals the Absence of *De Novo* Somatic Retrotransposition Events Despite Long-Term Cultivation and *TP53* Inactivation

To investigate *de novo* retrotransposon insertions, we initially performed genomic analysis of CLL-derived cell line models. To simulate the conditions of clonal evolution often observed in patients, we utilized the MEC1 cell line carrying a *TP53* mutation [25] alongside the HG3 *TP53* wild-type line [27] and two CRISPR/Cas9-engineered HG3 isogenic clones bearing *TP53* mutations (mut1 and mut2) [26] [Figure 1, Supplementary Table S1]. These models were subjected to longitudinal cultivation for 12 months, providing conditions to observe the potential accumulation of somatic insertions caused by the loss of p53-mediated repression. The NGS amplicon-based analysis focused on the L1HS subfamily, as these elements represent the only autonomously active retrotransposons in the human genome. We identified five candidate L1HS insertions in MEC1 and no novel insertions in HG3 (Supplementary Figure S1, Supplementary Table S3). To differentiate true novel somatic events from baseline variants or sequencing artifacts, locus-specific PCR validation was performed using gDNA extracted from TP1 and TP2. One of the identified candidates was confirmed to be present at both timepoints. This insertion was most likely a non-reference polymorphic variant, representing an inherent feature of the original donor’s genome or the established cell line. The remaining four candidates in the MEC1 line failed to be validated via PCR, suggesting they were likely chimeric sequences generated during library preparation or misaligned reads resulting from the high sequence similarity.

### Transcriptomic Profiling of HG3 Cell Lines Demonstrates TEs Dysregulation Linked to *TP53* Status and Time in Culture

To gain a deeper understanding of the impact of *TP53* inactivation on the TE regulation during long-term culture, we performed the transcriptomic profiling of the HG3 isogenic cell lines. RNAseq was performed at the same timepoints as the DNA sequencing. Altogether, 4844 differentially expressed TE subfamilies were identified in *TP53*-mutant clones compared to the wt-*TP53* HG3 cell line (Figure 2A, Supplementary Table S4). The analysis comparing TP1 and

**Figure 2.**
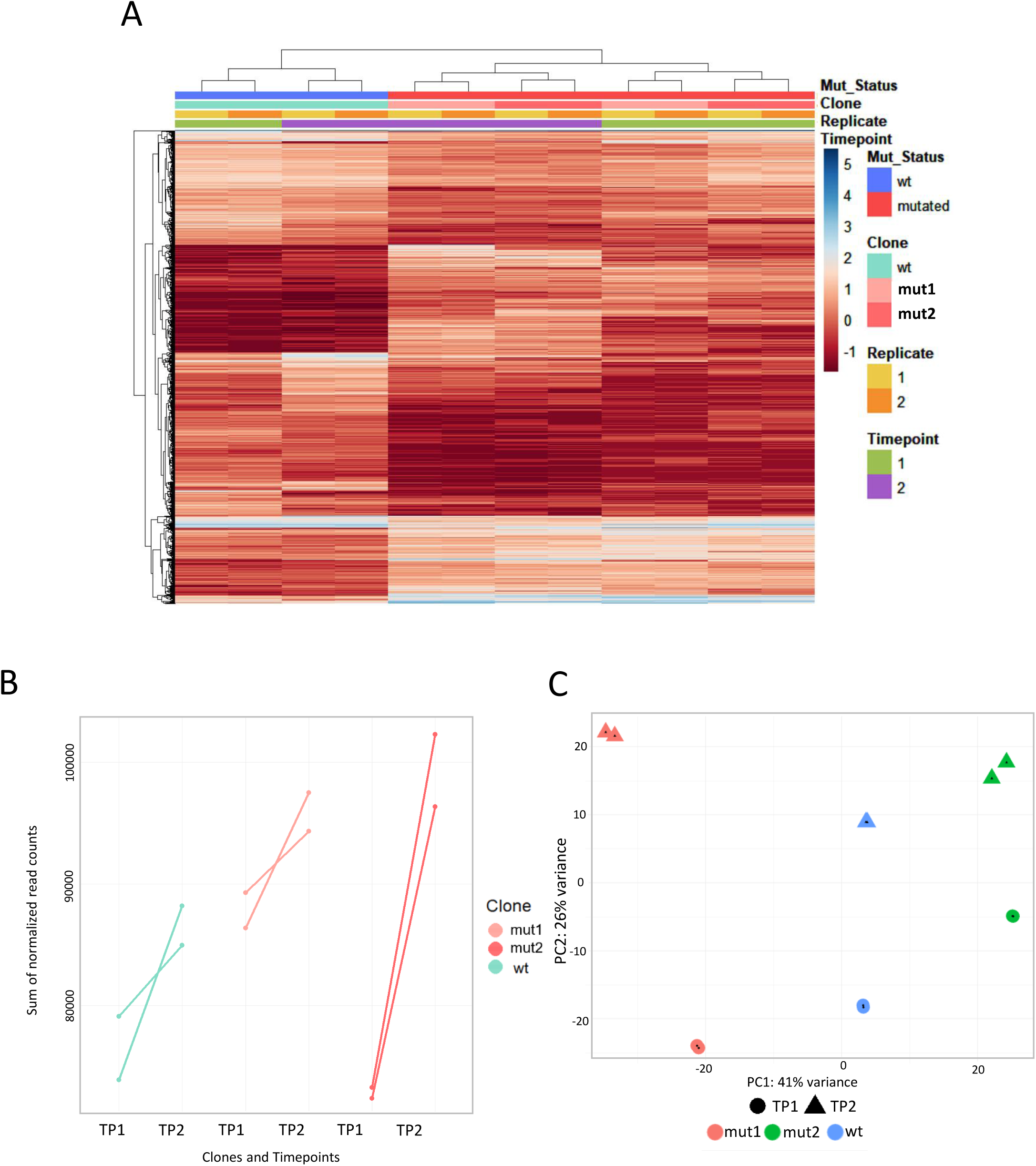
Transcriptomic analysis of the HG3 isogenic cell lines. (A) Overall, 4844 TE subfamilies were differentially expressed between *TP53*-mutant clones and *TP53*-wt HG3 cell lines. (B) Global TE expression assessed as the sum of normalized read counts. Lines indicate changes in the expression levels between TP1 and TP2 for wt, mut1, and mut2 clones. Statistical significance of the global TE expression increase in each clone cannot be evaluated due to the small sample size. (C) PCA analysis for differentially expressed TE subfamilies in the isogenic cell lines.

TP2 showed 7643 differentially expressed TE subfamilies (Supplementary Figure S2, Supplementary Table S5). The global normalized read counts further revealed a consistent increase of TE transcripts from TP1 to TP2 across all HG3 cell lines (Figure 2B). The majority of differentially expressed TEs belonged to evolutionarily older subfamilies, that are generally considered transcriptionally inert within the human genome, as their expression may be influenced by co-regulation with adjacent actively transcribed genes. [36] TE transcriptional data showed high consistency between biological replicates (Figure 2C). We did not observe differences in L1HS transcript abundance between the HG3 isogenic clones.

Next, we applied DESeq2 for differential gene expression analysis in the HG3 cell lines. We identified 2637 differentially expressed genes (DEGs) when comparing isogenic mutant clones against wild-type and 1917 DEGs between the timepoints (Supplementary Figure S3).

Overall, the isogenic HG3 cell lines showed significant transcriptional differences between wild-type and mutated clones, with the latter exhibiting distinct expression profiles and more pronounced changes over time, highlighting the impact of *TP53* mutation status on gene expression dynamics.

### Targeted DNA Sequencing of Serial Patient Samples Shows a Lack of Somatic L1 Insertions During Disease Progression and *TP53* Clonal Evolution

Building on the insights from the cell line models demonstrating TE dysregulation linked to *TP53* status and time, we sought to investigate whether tumor-specific TE changes are present in patient samples. First, we investigated whether tumor-specific L1HS insertions occur during CLL progression, particularly in relation to the expansion of CLL clones bearing dysfunctional *TP53.* For this purpose, we analyzed primary samples from CLL patients, a subset of whom acquired *TP53* mutations after chemoimmunotherapy. The cohort consisted of 34 serial tumor samples and 17 normal samples from 17 CLL patients (Figure 1, Supplementary Table S2): In TP1, the *TP53* mutation was either absent or present in a low frequency (0.1-5.4% VAF), while it expanded in TP2 (16.7-94% VAF).

We identified two candidate tumor-specific L1HS insertions in two different patients. They were subjected to validation using locus-specific PCR (Supplementary Figure S1, Supplementary Table S6). However, as a result, these candidate insertions were present in the germline as well as both tumor samples of the patients, suggesting that they were likely non-reference polymorphic insertions characterized by low UMI counts. To summarize, despite the sensitivity of our approach, no *de novo* somatic retrotransposition events were revealed in the tested CLL cohort.

### Transcriptomic Analysis of Primary CLL Cells Reveals Distinct Patient Clusters Defined by Specific Transposable Element Expression

Following the genomic analyses, we aimed to determine whether primary CLL cells exhibit TE expression changes analogous to those observed in the *TP53*-inactivated HG3 cell lines. We analyzed global TE expression in paired primary CLL samples from 21 CLL patients (N=42). The patient cohort used for the transcriptomic analysis involved 7 cases with *TP53* mutation expanding between TP1 and TP2 (*TP53*exp group; TP1: 0-5.4% VAF; TP2: 33.4-88.6% VAF; six of them involved in the L1HS insertion analysis at the DNA level) and additional 14 cases in which the *TP53* mutation did not expand (*TP53*non-exp group) (Figure 1). The global TE expression did not differ between patient groups categorized by their *TP53* mutation dynamics, nor between TP1 and TP2 (Figure 3A; Supplementary Table S7).

**Figure 3.**
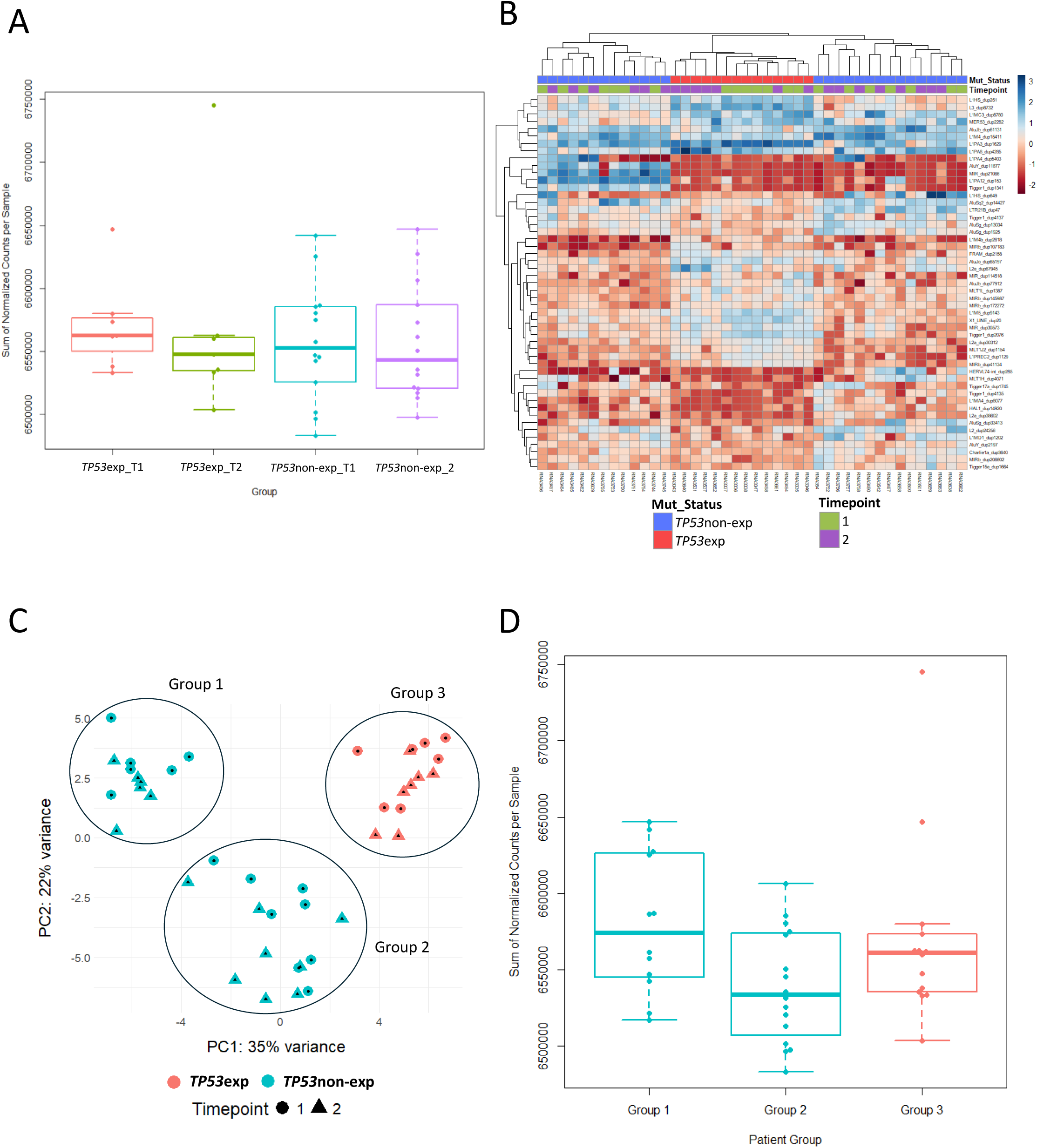
TE expression analysis in CLL samples. (A) Global TE expression was assessed as a sum of normalized TE read counts in RNAseq data. No significant difference was found among the *TP53*exp vs *TP53*non-exp patient groups (Kruskal-Wallis test) or between sample pairs from T1 and T2 (Wilcoxon paired test). (B) Differentially expressed TE subfamilies between *TP53*exp vs. *TP53*non-exp patient groups. (C) PCA analysis for differentially expressed TE subfamilies in the patient cohort. (D) The comparison of global TE expression between groups 1, 2 and 3 did not reach statistical significance (Kruskal-Wallis test, followed by Bonferroni-adjusted pairwise post-hoc testing).

Next, we analyzed the differential expression of TE subfamilies. While we did not observe significant TE expression changes in time, we identified 51 significantly differentially expressed

TE subfamilies when comparing *TP53*exp and *TP53*non-exp patient groups (Figure 3B, Supplementary Table S8). The analysis revealed distinct TE expression profiles across the cohort, with the HERV-K family, L1HS, L1PA2, AluS, and AluY emerging as the most differentially expressed signatures.

Further PCA analysis revealed three distinct patient groups, distinguishing *TP53*non-exp patients (Group 1 and Group 2) from *TP53*exp (Group 3) (Figure 3C). Interestingly, Groups 1 and 2 of *TP53*non-exp cases did not differ in clinical or laboratory parameters (IGHV mutational status, recurrent cytogenetic aberrations, overall survival; data not shown), nor in the global TE expression (Figure 3D), suggesting that their divergence is driven by more subtle regulatory differences.

### Pathway Enrichment Analyses Reveal Differences in RNA Homeostasis and Splicing Machinery among *TP53*non-exp CLL Cases

To better understand TE expression differences among Groups 1, 2, and 3 (Figure 3C), we sought to determine whether these differences are associated with changes in the gene expression landscape. Prior to differential gene expression analysis, TEs and immunoglobulin genes were removed from the dataset to avoid confounding signals. The input data was then filtered to retain only genes with greater than 1 read per million in at least 10 samples, resulting in a baseline of 20,583 evaluable genes. Following the DESeq analysis (Supplementary Table S9), downstream pathway analyses were performed on the top 150 most significant genes in each comparison (Figure 4A). While the comparisons including Group 3 did not show dysregulated KEGG or GO pathways, Spliceosome and Glycosylphosphatidylinositol (GPI)-anchor biosynthesis pathways were significantly dysregulated between the two *TP53*non-exp subgroups (Group 1 vs. Group 2) (adjusted *p* < 0.00579 and *p* < 0.0112, Table 1).

**Figure 4.**
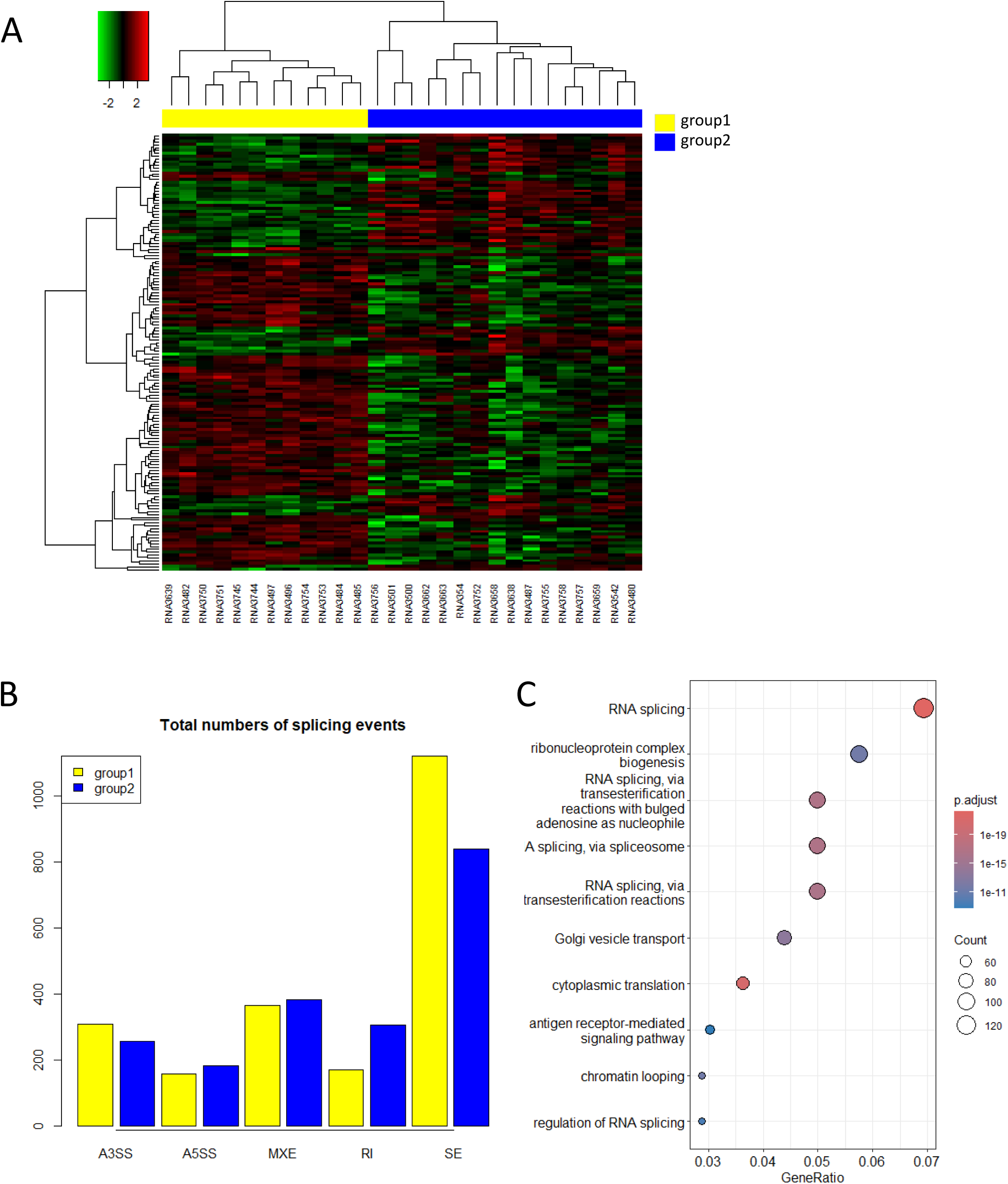
Pathway enrichment analysis between the *TP53*non-exp CLL subgroups. (A) The heatmap shows the expression profiles of 150 differentially expressed genes (DEGs) between\ Groups 1 and 2. (B) Quantitative distribution of differential alternative splicing events between Groups 1 and 2 identified via rMATS analysis (*p* < 0.001). A3SS – alternative 3’ splice sites; A5SS – alternative 5’ splice sites; MXE – mutually exclusive exons; RI – retained introns; SE – skipped exons. (C) Functional enrichment analysis of the most significant alternatively spliced genes between Groups 1 and 2, as represented by the dot plot of the top 10 significantly enriched categories across Gene Ontology (GO) (*p* < 0.05).

**Table 1.** Functional enrichment analysis of 150 differentially expressed genes between Groups 1 and 2 of the TP53non-exp CLL cohort.

|  |  |  |  |  |
| --- | --- | --- | --- | --- |
| ID | Description | p-value | p.adjust | Genes involved |
| hsa03040 | Spliceosome | 7.43E-05 | 0.005794955 | 7 |
| hsa00563 | Glycosylphosphatidylinositol (GPI)-anchor biosynthesis | 0.000286729 | 0.011182438 | 3 |

Given that the spliceosome and RNA metabolism pathways constituted the primary biological signal distinguishing the two *TP53*non-exp subgroups, we performed a deep alternative splicing analysis using rMATS. By comparing Group 1 and Group 2, we identified 15,477 distinct alternative splicing events. These events were categorized into five main types: Skipped Exon (SE), Mutually Exclusive Exons (MXE), Alternative 3’ Splice Site (A3SS), Retained Intron (RI), and Alternative 5’ Splice Site (A5SS). SE represented by far the most frequent class of alternative splicing (Table 2, Figure 4B), consistent with published CLL profiles. [37]

**Table 2.**
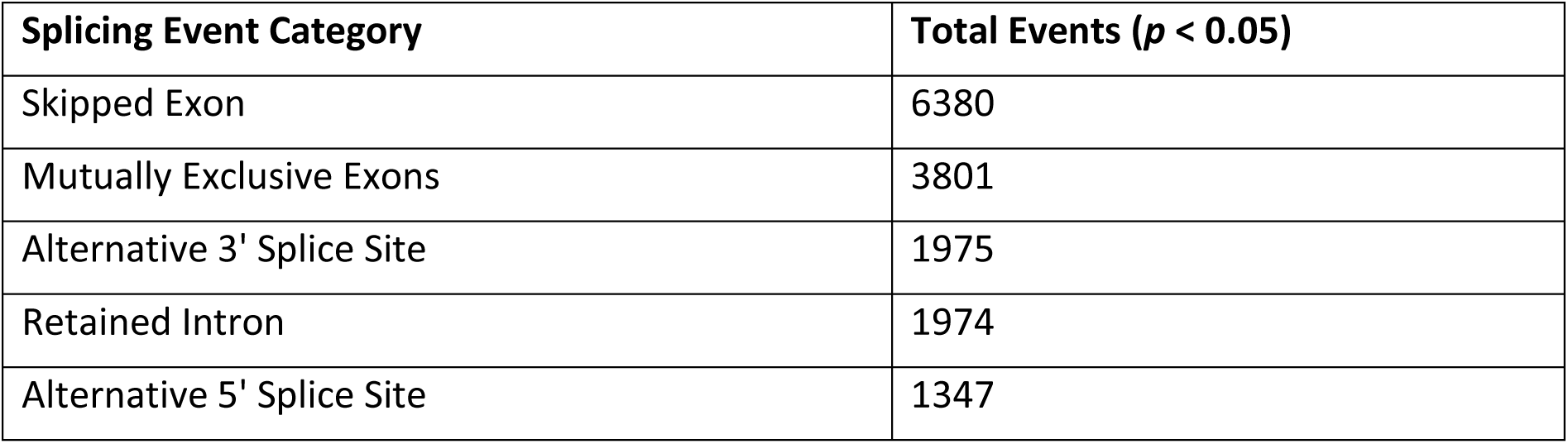
Quantitative profiling of differential alternative splicing events between Groups 1 and 2 of the *TP53*non-exp CLL cohort.

|  |  |
| --- | --- |
| Splicing Event Category | Total Events ($p < 0.05$) |
| Skipped Exon | 6380 |
| Mutually Exclusive Exons | 3801 |
| Alternative 3' Splice Site | 1975 |
| Retained Intron | 1974 |
| Alternative 5' Splice Site | 1347 |

We performed pathway analysis on genes affected by alternative splicing events. As visualized in the functional enrichment dot plot (Figure 4C), the top-ranked biological processes were dominated by RNA processing, spliceosome activity, and translational machinery. Notably, general RNA splicing exhibited the highest gene ratio and statistical significance, encompassing 128 distinct genes (Figure 4C, Supplementary Table S10). To evaluate the biological impact of these transcriptomic alterations, we explored the most significantly different skipped exon events between Group 1 and Group 2, sorted by *p*-value (Supplementary Table S11). Notably, one of the significantly mis-spliced targets was *DDX17*, a DEAD-box RNA helicase essential for global RNA metabolism and alternative splicing. Further differentially spliced genes included, for instance, *CDC42SE1*, *CHEK2*, *PARP15*, or *RAD21*.

## Discussion

In this study, we aimed to determine the extent to which TE dysregulation and *de novo* retrotransposition events contribute to genomic instability and clonal evolution of CLL, particularly in the context of *TP53* dysfunction. By integrating a highly sensitive amplicon-based targeted DNA sequencing approach (limit of detection <1%) with global transcriptomic profiling across both cell line models and serial primary patient samples, we evaluated the insertional and transcriptional landscapes of TEs during disease progression. Our principal findings reveal that while *TP53* dysfunction is a critical driver of genomic instability and poor prognosis in CLL [16, 17], it is insufficient to induce *de novo* somatic LINE-1 (L1HS) insertions, as we detected none in either the *in vitro* models or the primary patient samples. This contrasts with certain solid tumors, such as colorectal [38], breast, and lung cancers [39], where L1 retrotransposition frequently contributes to structural variation.

The absence of detectable somatic retrotransposition events in CLL, even in the setting of *TP53*-mutated clonal expansion, suggests that mature B-cell malignancies may retain secondary defense mechanisms that remain functionally active despite the loss of intact *TP53*. [40] Consistent with recent literature, while LINE-1 is derepressed in approximately 50% of human cancers, its consequences frequently manifest as DNA damage and replication stress rather than as successful genomic insertions. [9] The observed transcriptional activity of TEs in the absence of *de novo* genomic insertions aligns with the characteristically low frequency of LINE-1 mobilization reported in other hematological malignancies, including ALL [32], MDS [33], and AML [41]. These findings distinguish the retrotransposition landscape in hematopoietic tumors from that of solid tumors, like colorectal and lung cancers, which frequently harbor numerous somatic insertions. [42]

The absence of successful insertional mutagenesis does not mean that the transcriptional activity of these elements is benign to the cell. Although TEs may fail to complete their mobilization cycle, the transcriptomic RE derepression is sufficient to induce profound genomic instability and DNA replication stress [43]. When LINE-1 elements are transcribed and translated, the endonuclease activity of the newly synthesized ORF2p generates double- strand breaks to initiate target-primed reverse transcription. These enzymatic nicks stall replication forks and cause widespread DNA damage, triggering a robust cellular response [43]. As noted by Tiwari et al. (2020) [22], p53 normally represses human LINE-1 transposons by recruiting repressive histone marks; when this control is lost, the resulting TE-derived transcripts can induce replication stress or form deleterious R-loops without ever achieving successful integration.

Supporting this view of transcriptional and epigenetic deregulation, Barrow et al. (2020) [44] demonstrated that locus-specific hypomethylation of retrotransposons in CLL is strongly associated with the altered expression of neighboring genes. This suggests that the pathogenic contribution of TEs in CLL is primarily mediated by epigenetic remodeling of the local transcriptome rather than direct insertional mutagenesis. Furthermore, the life cycle of REs is tightly regulated by various post-transcriptional molecular mechanisms.

In our HG3 cell line models, many of the differentially expressed genes encoded non-coding RNAs (ncRNA), which are known to play roles in the regulation of gene expression and genomic stability. [45] We hypothesize that these ncRNAs, particularly miRNAs and piRNAs, may act against the deleterious effects of TE mobilization. Growing evidence suggests that ncRNAs play essential roles in protecting the genome by silencing transposons. [46] In CLL, the upregulation of ncRNAs in *TP53-*mutated samples may be a compensatory mechanism that silences L1 activity post-transcriptionally, explaining why we observed increased RNA transcripts but no new DNA insertions.

Finally, our integrative analysis points to a profound disruption of RNA homeostasis that occurs independently of *TP53* clonal expansion. Specifically, the transcriptomic divergence between the two *TP53*-mutation non-expanding patient subgroups coincided with massive alterations in alternative splicing, particularly affecting genes essential for RNA processing and DNA repair. We detected aberrant splicing of the DEAD-box RNA helicase *DDX17,* suggesting the possibility of an autoregulatory feedback loop that may contribute to the disruption of normal cellular signaling networks and broader transcriptomic instability. [47] Other notable targets included *CDC42SE1*, which regulates B-cell morphology and cytoskeleton organization [48], and *CHEK2*, a vital component of the DNA damage response.[49]. These findings suggest that CLL cell biology and progression may be driven by systemic dysregulation of transcription and splicing, rather than by the direct insertional mutagenesis typically associated with activated transposable elements in solid cancers.

## Conclusion

In this study, our primary aim was to investigate the relationship between *TP53* dysfunction and retrotransposon activity in CLL. Our genomic analysis demonstrates that *TP53* deficiency does not directly promote *de novo* LINE-1 retrotransposition in CLL, distinguishing it from solid tumors where LINE-1 mobilization is a prominent source of genomic instability. While insertional mutagenesis is suppressed, transcriptomic profiling revealed that patients exhibit distinct TE expression signatures. Independent of *TP53* mutation expansion, we identified profound alterations in RNA homeostasis, driven by alternative splicing abnormalities in core spliceosome and DNA repair genes among the patients. Collectively, our findings suggest that the role of REs in CLL is primarily associated with transcriptional activation rather than *de novo* insertional mutagenesis, occurring alongside widespread alterations in alternative splicing and global transcriptomic regulation.

## Data availability

All data supporting the results of this study are presented within the Article or Supplementary Information.

The research was conducted on primary samples of leukemia patients. Due to data protection policies, patient genomic and transcriptomic sequences have not been made publicly available in databases. They will be made available upon request to the corresponding author and the institutional review via a secure environment.

## CRediT authorship contribution statement

**Anastasiya Volakhava:** Investigation, Data curation, Formal analysis, Software, Validation, Visualization, Writing - original draft, Writing - review & editing. **Sarka Pavlova:** Conceptualization, Investigation, Resources, Project administration, Supervision, Writing - review & editing. **Lenka Radova:** Software, Formal analysis, Investigation, Data curation, Visualization, Writing - review & editing. **Kristyna Tausova:** Investigation, Data curation**. Hana Svozilova:** Investigation. **Marcela Zenatova:** Investigation, Resources. **Michael Doubek:** Resources. **Ilgar Mamedov:** Methodology, Investigation, Funding acquisition, Supervision. **Sarka Pospisilova:** Funding acquisition, Resources, Supervision. **Karla Plevova:** Conceptualization, Investigation, Resources, Funding acquisition, Project administration, Supervision, Writing - review & editing.

## Acknowledgments

We gratefully acknowledge Richard Rosenquist and Michal Smida for the generous gift of the HG3 cell line and the CRISPR/Cas9-engineered *TP53*-inactivated HG3 cell lines, respectively. We acknowledge the EATRIS-CZ infrastructure LM2023053 and the Core Facilities Genomics and Bioinformatics, supported by the NCMG research infrastructure LM2023067, funded by the Ministry of Education, Youth and Sports of the Czech Republic, for their support with obtaining scientific data presented in this work. Computational resources were provided by the e-INFRA CZ project ID:90254, supported by the Ministry of Education, Youth and Sports of the Czech Republic.

## Supplementary Materials

**Figure S1.**
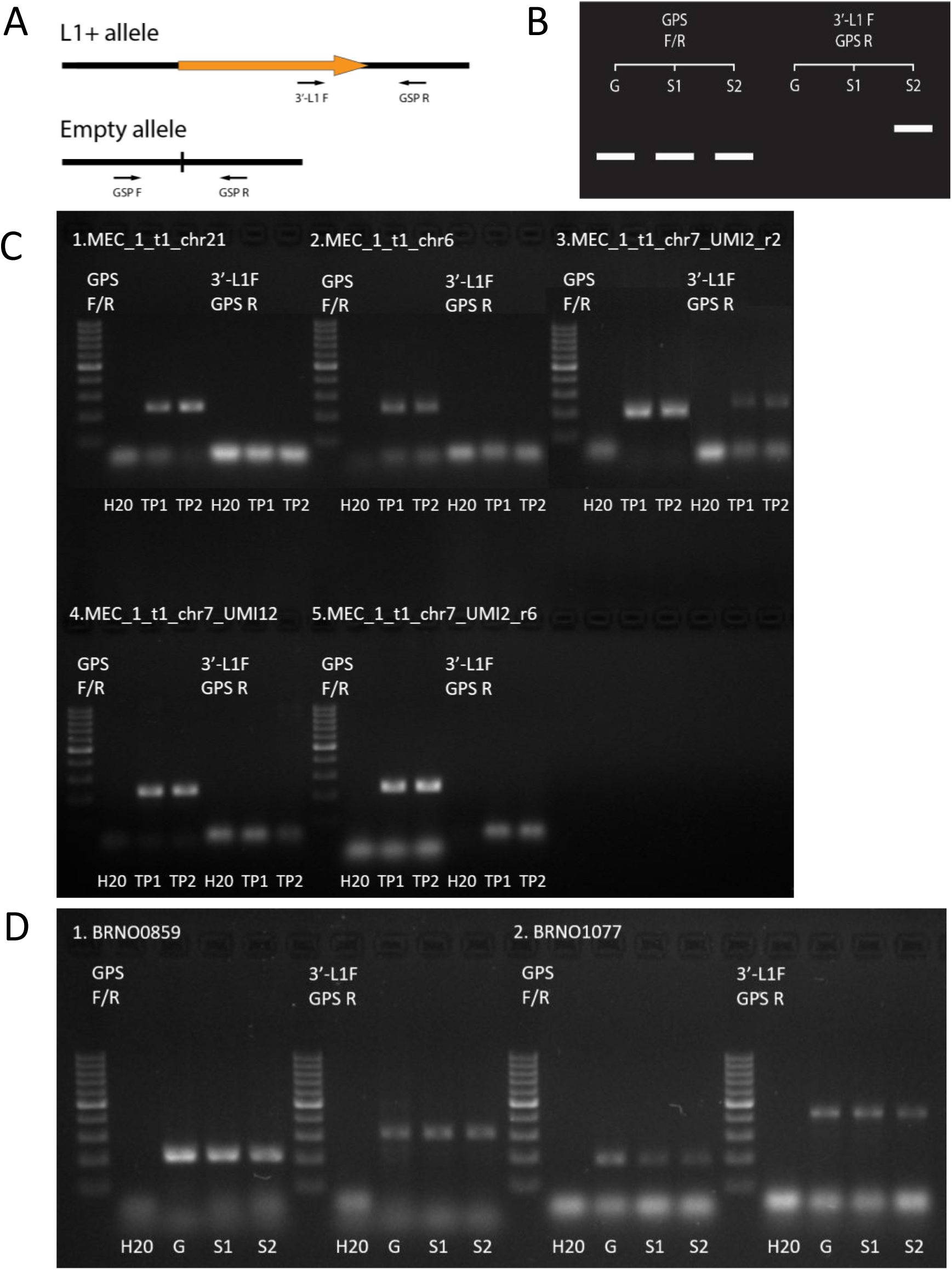
PCR validation of candidate L1 insertions detected by NGS. The approach is based on locus-specific PCR with primers designed for every candidate insertion (A), as described in detail in our previous reports. [32, 33] A hypothetical positive result is illustrated (B), showing a novel insertion in the follow-up sample S2 that was not present in the germinal G or baseline S1 samples. In the cultivated MEC1 cell line (C), one candidate insertion was present in both time points, pointing to non-reference polymorphic insertions. The remaining four candidate insertions were absent in both time points, which may indicate chimeric or incorrectly mapped sequences. The candidate insertions in the two CLL patients (BRNO0859 and BRNO1077) were present in germline as well as both tumor samples (D), suggesting that these were likely non-reference polymorphic insertions. Supplementary Tables S3 and S6 list PCR primers for all validation reactions. 3’-L1 F – a universal L1 forward primer; GSP F – genomic locus-specific forward primer; GSP R – genomic locus-specific reverse primer; H2O – non-template control; TP1 – time point 1; TP2 – time point 2.

**Figure S2.**
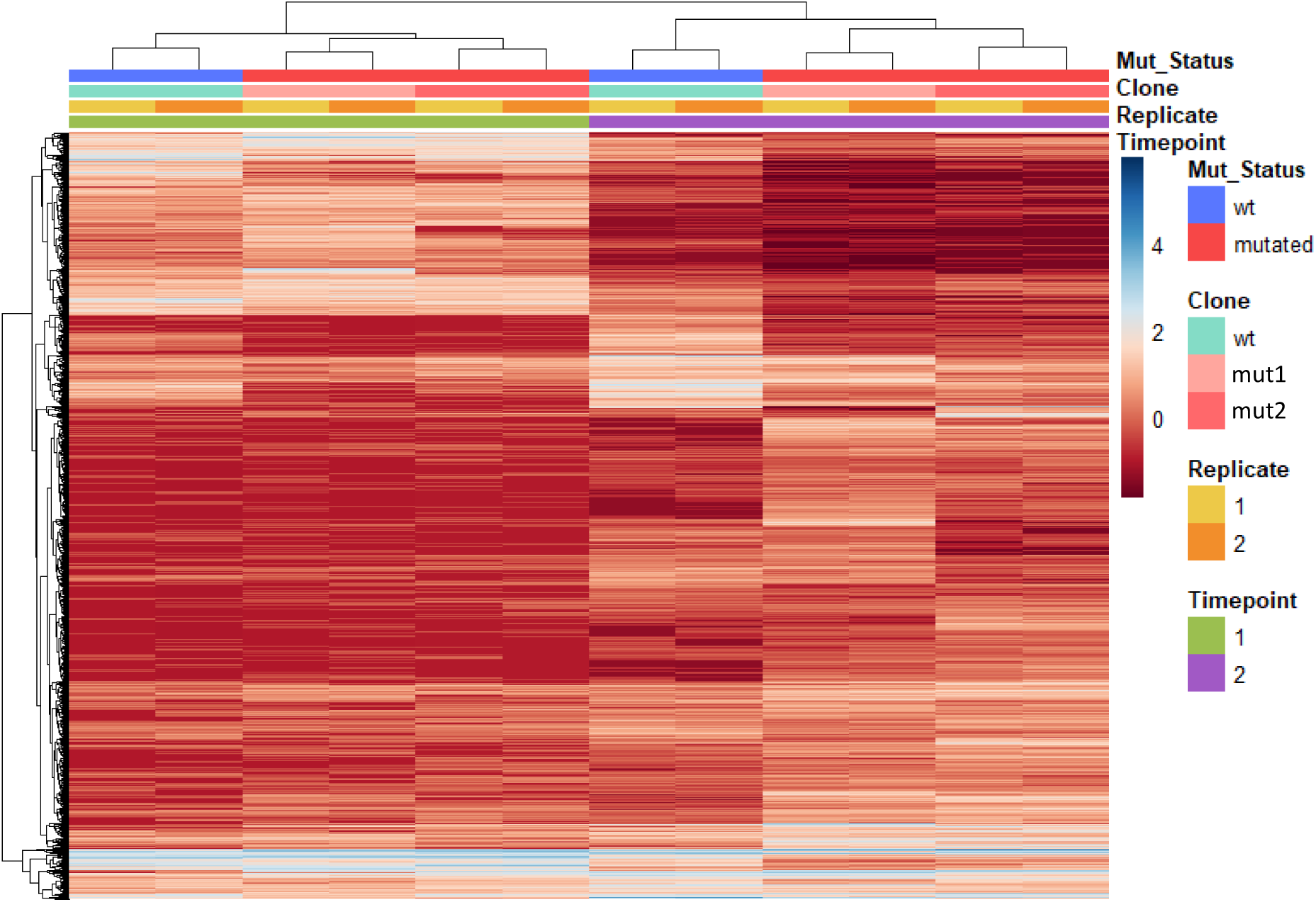
A total of 7643 TE subfamilies differentially expressed in HG3 cell lines between TP1 and TP2.

**Figure S3.**
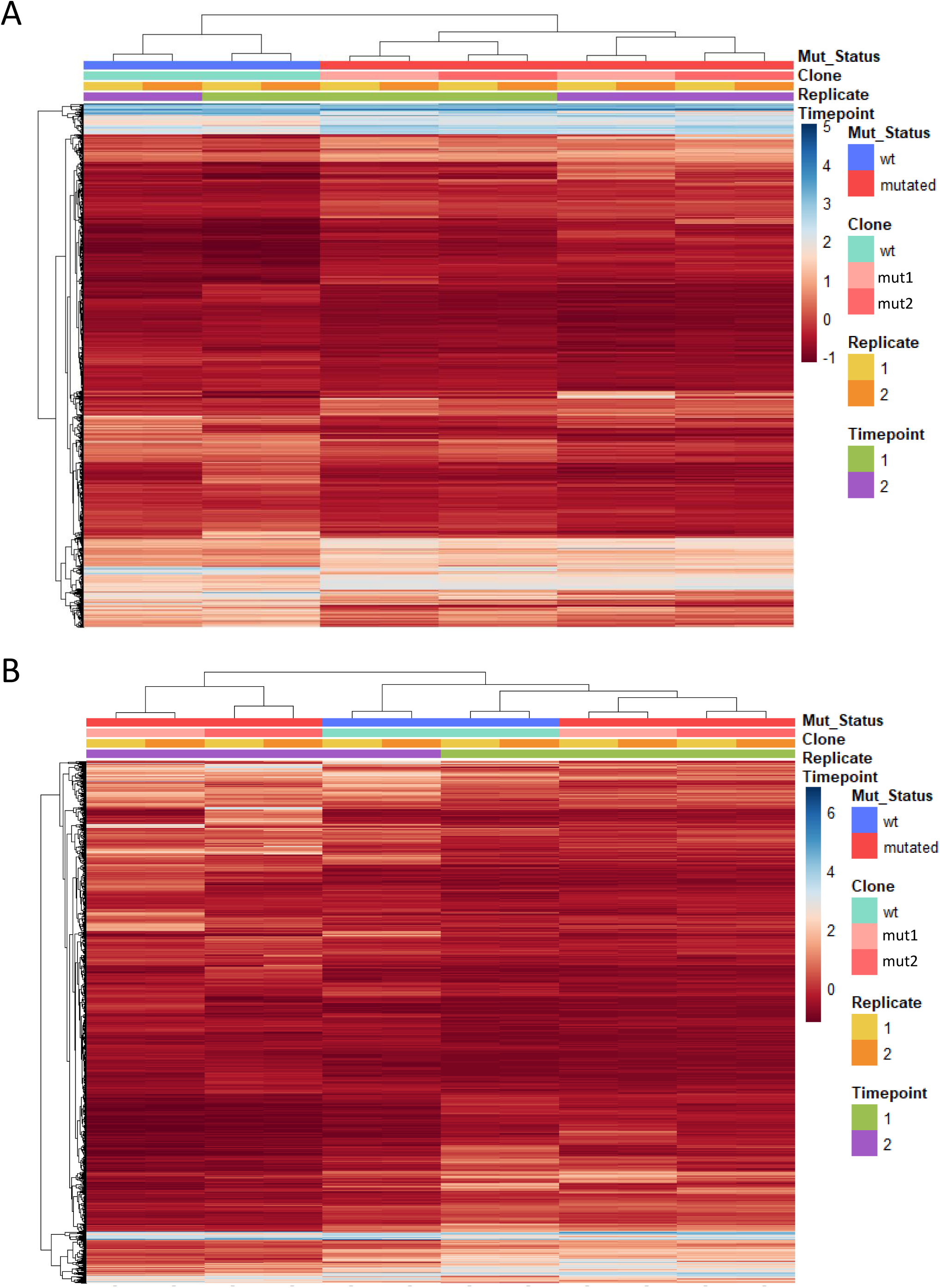
Differentially expressed genes in HG3 cell lines: between wild-type and mutant HG3 cell lines (A) and between T1 and T2 (B)

**Table S1.** CLL-derived cell lines used for L1HS insertion NGS and TE expression analysis.

**Table S2.** A list of CLL samples tested for L1 *de novo* insertions and TE expression activity.

**Table S3.** A list of candidate insertions detected using L1-targeted amplicon-based NGS protocol and validated using site-specific PCR in MEC1 cell lines subjected to long-term cultivation.

**Table S4.** Differentially expressed TE subfamilies based on *TP53* mutation status in HG3 cell lines.

**Table S5.** Differentially expressed TE subfamilies between timepoints in HG3 cell lines.

**Table S6.** A list of candidate insertions detected using L1-targeted amplicon-based NGS protocol and validated using site-specific PCR in paired primary samples from CLL patients.

**Table S7.** Sums of normalized TE read counts per individual patient samples with RNAseq data.

**Table S8.** Differentially expressed TE subfamilies between the groups of patients with different *TP53* mutation evolution.

**Table S9.** Differentially expressed genes between Group 1 and Group 2 of the *TP53*non-exp cases.

**Table S10.** Results of the pathway analysis in Group 1 and Group 2 of the *TP53*non-exp cases.

**Table S11.** Significantly different skipped exon (SE) events between *TP53*non-exp Group 1 and Group 2, sorted by *p*-value.

