## Supplementary Tables 1-11 for "Retrotransposon Activity in Chronic Lymphocytic Leukemia: Associations with RNA Splicing and *TP53* Dysfunction"

**Table S1**. CLL-derived cell lines used for L1HS insertion NGS and TE expression analysis.

| Cell line | Cell type origin | TP53 status | TP53 variant annotation | Cultivation conditions | Span of cultivation [mo] | Cell line source | Research Resource Identifier (RRID) |
|---|---|---|---|---|---|---|---|
| MEC1 | CLL in prolymphocytoid transformation to B-cell prolympho | mut | NM_000546.6: c.949dup, p.Gln317ProfsTer20 | IMDM | 12 | PMID: 10071128 | MEC-1 (CVCL_1870) |
| HG3_wt | chronic lymphocytic leukemia | wt | | RPMI | 12 | PMID: 32217767, 22720208 | HG-3 (CVCL_Y547) |
| HG3_mut1 | chronic lymphocytic leukemia isogenic to HG3 wt | mut | CRISPR/Cas9 generated frameshift mutation in exon 5 | RPMI | 12 | PMID: 32217767 | |
| HG3_mut2 | chronic lymphocytic leukemia isogenic to HG3 wt | mut | CRISPR/Cas9 generated frameshift mutation in exon 2 | RPMI | 12 | PMID: 32217767 | |

**Table S2**. A list of CLL samples tested for L1 *de novo* insertions and TE expression activity.

| Patient ID | Timepoint | Time from previous sampling (mo) | Number of prior therapy lines | TP53 status (mutations >0.1% VAF) | Category based on TP53 status development [stable / expanding mutation] | Material [DNA/RNA] | Experiment / method applied [L1SH insertion NGS / RNAseq] |
|---|---|---|---|---|---|---|---|
| BRNO0029 | N | | 1 | NA | expanding mutation | DNA | L1SH insertion NGS |
| BRNO0029 | T1 | | 0 | wild type | expanding mutation | DNA | L1SH insertion NGS |
| BRNO0029 | T2 | 135 | 2 | p.R248Q 45%; p.H214R 2.2%; p.H193L 0.3% | expanding mutation | DNA | L1SH insertion NGS |
| BRNO0047 | N | | 1 | NA | expanding mutation | DNA | L1SH insertion NGS / RNAseq |
| BRNO0047 | T1 | | 0 | wild type | expanding mutation | DNA/RNA | L1SH insertion NGS / RNAseq |
| BRNO0047 | T2 | 114 | 1 | p.R248Q 33.4%; p.T170M 1.5% | expanding mutation | DNA/RNA | L1SH insertion NGS / RNAseq |
| BRNO0126 | N | | 2 | NA | expanding mutation | DNA | L1SH insertion NGS |
| BRNO0126 | T1 | | 0 | p.V272M 0.7%; p.H193Y 0.6%; c.559+33_54del 0.5%; p.C275F 0.4% | expanding mutation | DNA | L1SH insertion NGS |
| BRNO0126 | T2 | 55 | 2 | c.559+33_54del 94%; p.V272M 0.2% | expanding mutation | DNA | L1SH insertion NGS |
| BRNO0149 | N | | 2 | NA | expanding mutation | DNA | L1SH insertion NGS |
| BRNO0149 | T1 | | 0 | wild type | expanding mutation | DNA | L1SH insertion NGS |
| BRNO0149 | T2 | 52 | 2 | p.V272M 25.4%; p.R248W 16.8%; p.G245S 1.2% | expanding mutation | DNA | L1SH insertion NGS |
| BRNO0178 | T1 | | 0 | wild type | stable | RNA | RNAseq |
| BRNO0178 | T2 | 81 | 3 | p.R175H 5.9%; p.Y163C 4.7%; p.I195T 1.7%; p.H179Y 1.5%; c.376-1G>T 1.3%; p.R24 [illegible] | stable | RNA | RNAseq |
| BRNO0209 | T1 | | 0 | wild type | stable | RNA | RNAseq |
| BRNO0209 | T2 | 141 | 3 | p.Y234C 5.9%; p.P27T 1.2%; p.V216L 0.9%; p.E198Ter 0.7%; c.323del 0.6%; p.R209fs [illegible] | stable | RNA | RNAseq |
| BRNO0311 | T1 | | 0 | p.G266E 0.6% | stable | RNA | RNAseq |
| BRNO0311 | T2 | 51 | 3 | p.G266E 0.2%; p.R248Q 0.2% | stable | RNA | RNAseq |
| BRNO0332 | N | | 1 | NA | expanding mutation | DNA | L1SH insertion NGS / RNAseq |
| BRNO0332 | T1 | | 0 | wild type | expanding mutation | DNA/RNA | L1SH insertion NGS / RNAseq |
| BRNO0332 | T2 | 70 | 1 | p.P278L 42%; p.P75fs 1.1% | expanding mutation | DNA/RNA | L1SH insertion NGS / RNAseq |
| BRNO0462 | N | | 1 | NA | expanding mutation | DNA | L1SH insertion NGS / RNAseq |
| BRNO0462 | T1 | | 0 | p.G105D 2.3% | expanding mutation | DNA/RNA | L1SH insertion NGS / RNAseq |
| BRNO0462 | T2 | 29 | 1 | p.G105D 67.7%; p.R248W 2.1%; p.S241F 1.5%; p.R248Q 1.4%; p.R248G 1.3%; p.R17 [illegible] | expanding mutation | DNA/RNA | L1SH insertion NGS / RNAseq |
| BRNO0467 | T1 | | 0 | wild type | stable | RNA | RNAseq |
| BRNO0467 | T2 | 130 | 3 | p.C277Ter 0.1% | stable | RNA | RNAseq |
| BRNO0475 | T1 | | 3 | wild type | stable | RNA | RNAseq |
| BRNO0475 | T2 | 78 | 10 | p.H168R 4.7% | stable | RNA | RNAseq |
| BRNO0488 | N | | 1 | NA | expanding mutation | DNA | L1SH insertion NGS / RNAseq |
| BRNO0488 | T1 | | 0 | p.R158G 0.8%; p.V173G 0.7%; p.Y220C 0.3% | expanding mutation | DNA/RNA | L1SH insertion NGS / RNAseq |
| BRNO0488 | T2 | 91 | 1 | p.R158G 59.9%; p.V173G 9%; c.R209fs 1.6%; p.G245S 1.3%; p.R213Ter 1%; p.R333fs [illegible] | expanding mutation | DNA/RNA | L1SH insertion NGS / RNAseq |
| BRNO0513 | T1 | | 0 | wild type | stable | RNA | RNAseq |
| BRNO0513 | T2 | 42 | 3 | p.G245S 0.2% | stable | RNA | RNAseq |
| BRNO0823 | T1 | | 2 | p.P278L 0.2%; p.R273H 0.1% | stable | RNA | RNAseq |
| BRNO0823 | T2 | 40 | 5 | p.R273H 3.3%; p.P278L 2% | stable | RNA | RNAseq |
| BRNO0837 | T1 | | 0 | p.C238Y 1.6%; p.T125T 0.9%; p.W146Ter 0.9%; p.M237I 0.4%; p.G262V 0.3%; p.R28 [illegible] | expanding mutation | RNA | RNAseq |
| BRNO0837 | T2 | 35 | 1 | p.W146Ter 71.3%; p.T125T 3.9%; p.C238Y 1.9%; p.R280G 1.5%; p.G262V 0.9%; p.M [illegible] | expanding mutation | RNA | RNAseq |
| BRNO0843 | N | | 1 | NA | expanding mutation | DNA | L1SH insertion NGS |
| BRNO0843 | T1 | | 0 | p.R282W 0.3% | expanding mutation | DNA | L1SH insertion NGS |
| BRNO0843 | T2 | 48 | 1 | p.R282W 16.7%; p.V147D 1.5%; p.Y220C 1%; p.R175H 0.9%; p.R248Q 0.7%; p.G244 [illegible] | expanding mutation | DNA | L1SH insertion NGS |
| BRNO0859 | N | | 2 | NA | expanding mutation | DNA | L1SH insertion NGS |
| BRNO0859 | T1 | | 1 | p.Y163H 0.2% | expanding mutation | DNA | L1SH insertion NGS |
| BRNO0859 | T2 | 42 | 3 | p.Y163H 38% | expanding mutation | DNA | L1SH insertion NGS |
| BRNO0874 | T1 | | 0 | wild type | stable | RNA | RNAseq |

| | | | | | | | |
|---|---|---|---|---|---|---|---|
| BRNO0874 | T2 | 50 | 4 | p.R209fs 0.1% | stable | RNA | RNAseq |
| BRNO1077 | N | | 1 | NA | expanding mutation | DNA | L1SH insertion NGS |
| BRNO1077 | T1 | | 1 | p.Q144Ter 0.8% | expanding mutation | DNA | L1SH insertion NGS |
| BRNO1077 | T2 | 87 | 7 | p.Q144Ter 54.5%; p.Y163C 11.2% | expanding mutation | DNA | L1SH insertion NGS |
| BRNO1162 | N | | 3 | NA | expanding mutation | DNA | L1SH insertion NGS |
| BRNO1162 | T1 | | 0 | wild type | expanding mutation | DNA | L1SH insertion NGS |
| BRNO1162 | T2 | 93 | 3 | p.G262V 76.3%; p.R110_F113delinsL 3.5%; p.V157G 0.8%; p.M246I 0.4%; p.E285K 0 | expanding mutation | DNA | L1SH insertion NGS |
| BRNO1208 | N | | 1 | NA | expanding mutation | DNA | L1SH insertion NGS / RNAseq |
| BRNO1208 | T1 | | 0 | p.I255del 5.4% | expanding mutation | DNA/RNA | L1SH insertion NGS / RNAseq |
| BRNO1208 | T2 | 34 | 1 | p.I255del 64.5%; p.C277F 6.8%; p.I254T 0.3% | expanding mutation | DNA/RNA | L1SH insertion NGS / RNAseq |
| BRNO1266 | T1 | | 0 | wild type | stable | RNA | RNAseq |
| BRNO1266 | T2 | 86 | 3 | p.P177L 1%; p.R248Q 0.7%; p.Q167Ter 0.5%; p.R175H 0.4%; p.C277F 0.1% | stable | RNA | RNAseq |
| BRNO1284 | N | | 2 | NA | expanding mutation | DNA | L1SH insertion NGS |
| BRNO1284 | T1 | | 0 | p.I195fs 0.2% | expanding mutation | DNA | L1SH insertion NGS |
| BRNO1284 | T2 | 38 | 3 | p.I195fs 69% | expanding mutation | DNA | L1SH insertion NGS |
| BRNO1340 | T1 | | 0 | p.P278A 0.1% | stable | RNA | RNAseq |
| BRNO1340 | T2 | 45 | 1 | p.R273H 0.6%; p.P278A 0.2% | stable | RNA | RNAseq |
| BRNO1342 | N | | 0 | NA | expanding mutation | DNA | L1SH insertion NGS |
| BRNO1342 | T1 | | 0 | p.A276D 3.7% | expanding mutation | DNA | L1SH insertion NGS |
| BRNO1342 | T2 | 16 | 1 | p.A276D 33.4%; p.R209fs 0.1% | expanding mutation | DNA | L1SH insertion NGS |
| BRNO1404 | N | | 1 | NA | expanding mutation | DNA | L1SH insertion NGS |
| BRNO1404 | T1 | | 0 | p.R175H 0.2% | expanding mutation | DNA | L1SH insertion NGS |
| BRNO1404 | T2 | 34 | 2 | p.R248Q 41%; p.R175H 5%; p.F134L 0.3% | expanding mutation | DNA | L1SH insertion NGS |
| BRNO1503 | N | | 1 | NA | expanding mutation | DNA | L1SH insertion NGS / RNAseq |
| BRNO1503 | T1 | | 0 | p.K132R 0.9%; p.R249G 0.3%; p.F212fs 0.3% | expanding mutation | DNA/RNA | L1SH insertion NGS / RNAseq |
| BRNO1503 | T2 | 28 | 2 | p.K132R 88.6%; p.I254N 12%; p.F109S 0.2% | expanding mutation | DNA/RNA | L1SH insertion NGS / RNAseq |
| BRNO1530 | T1 | | 0 | p.G244S 0.4% | stable | RNA | RNAseq |
| BRNO1530 | T2 | 41 | 1 | p.G244S 0.4%; p.Y163C 0.2% | stable | RNA | RNAseq |
| BRNO1692 | N | | 1 | NA | expanding mutation | DNA | L1SH insertion NGS |
| BRNO1692 | T1 | | 0 | p.R248Q 0.1% | expanding mutation | DNA | L1SH insertion NGS |
| BRNO1692 | T2 | 55 | 3 | p.I195F 54.2%; p.G187S 1.9%; p.C141fs 0.5%; c.673-2A>C 0.4% | expanding mutation | DNA | L1SH insertion NGS |
| BRNO1794 | T1 | | 0 | c.672+1G>C 0.4% | stable | RNA | RNAseq |
| BRNO1794 | T2 | 55 | 4 | wild type | stable | RNA | RNAseq |
| BRNO1876 | T1 | | 0 | p.R273C 0.1% | stable | RNA | RNAseq |
| BRNO1876 | T2 | 29 | 1 | p.R273C 0.3% | stable | RNA | RNAseq |
| BRNO1921 | T1 | | 0 | wild type | stable | RNA | RNAseq |
| BRNO1921 | T2 | 22 | 1 | p.G245S 1% | stable | RNA | RNAseq |

**Table S3**. A list of candidate insertions detected using L1-targeted amplicon-based NGS protocol and validated using site-specific PCR in MEC1 cell lines subjected to long-term cultivation.

| Cell line | Time point | Chromosome | Coordinate | Sample reads | UMI | Forward Primer | Reverse primer | Product size |
|---|---|---|---|---|---|---|---|---|
| MEC1 | T1 | chr21 | 37686359 | 9 | 2 | GAAGGATTTGCTGTGGGGTA | GAGAGTGAATGCGTGGAGGT | 234 bp |
| MEC1 | T1 | chr6 | 127521564 | 4 | 2 | CAAAGCCAGATTTTGCACCT | GCAATGAAGAAGCCCATGTT | 235 bp |
| MEC1 | T1 | chr7 | 23881024 | 2 | 2 | ACAGAGGCAGTCCAGGCTAA | TTGTCAACGTTATGCCCAAA | 200 bp |
| MEC1 | T1 | chr7 | 34481963 | 144 | 12 | AAGGCCACTTATCCAATCCA | CTTGGCCCAACAGTTCCTT | 231 bp |
| MEC1 | T1 | chr7 | 51089907 | 6 | 2 | TCAGCTTTACTGAGGAACAAACA | TGGCTCATCAATTCTGTGGA | 245 bp |

**Table S4.** Differentially expressed TE subfamilies based on *TP53* mutation status in HG3 cell lines.

| TE families | baseMean | log2FoldChange | lfcSE | stat | pvalue | FDR | Heatmap_order | HG3_mut1_TP1_RNA1 | HG3_mut1_TP1_RNA2 | HG3_mut1_TP2_RNA1 | HG3_mut1_TP2_RNA2 | HG3_mut2_TP1_RNA1 | HG3_mut2_TP1_RNA2 | HG3_mut2_TP2_RNA1 | HG3_mut2_TP2_RNA2 | HG3_wt_TP1_RNA1 | HG3_wt_TP1_RNA2 | HG3_wt_TP2_RNA1 | HG3_wt_TP2_RNA2 |
|---|---|---|---|---|---|---|---|---|---|---|---|---|---|---|---|---|---|---|---|
| THE1B-int_dup2596 | 1167.18054531514 | 1.03777881563599 | 0.293722128250021 | 3.53319929219843 | 0.000410562853576766 | 0.0267795473566823 | 1 | 988 | 911 | 477 | 511 | 1593 | 1711 | 800 | 711 | 1474 | 1326 | 1761 | 2290 |
| L1PB1_dup6906 | 534.493902316509 | 1.76721330064695 | 0.360198344997582 | 4.90622271087563 | 9.28470946345507e-07 | 0.000301811404097169 | 2 | 280 | 236 | 201 | 238 | 666 | 750 | 260 | 148 | 687 | 737 | 957 | 1591 |
| L1M3f_dup660 | 289.287814898852 | 1.20791343355503 | 0.339548463169047 | 3.55741098717228 | 0.00037452795062499 | 0.025202892460241 5 | 3 | 178 | 182 | 129 | 188 | 485 | 467 | 142 | 94 | 334 | 392 | 457 | 590 |
| L1M2_dup5165 | 339.931714161076 | 1.07650806312544 | 0.317237747437653 | 3.39337948217214 | 0.000690359286741005 | 0.0374528570093195 | 4 | 277 | 242 | 163 | 175 | 437 | 491 | 212 | 203 | 346 | 339 | 540 | 876 |
| L1PA7_dup11794 | 377.769985375855 | 1.06519656992876 | 0.255552887101013 | 4.16820401448923 | 3.0700909099366e-05 | 0.0042471851840452 | 5 | 361 | 327 | 174 | 198 | 464 | 517 | 301 | 164 | 405 | 470 | 606 | 788 |
| LTR48B_dup395 | 579.050032186774 | -1.1241577374668 | 0.267286401660799 | -4.20581717020315 | 2.60140476327143e-05 | 0.00376356984672793 | 6 | 966 | 769 | 609 | 638 | 436 | 472 | 1206 | 935 | 225 | 211 | 376 | 553 |
| L1MA2_dup2806 | 324.701109085925 | -1.02904474743249 | 0.164962038625135 | -6.23807001907222 | 4.43002373114257e-10 | 5.19264249620911e-07 | 7 | 665 | 570 | 312 | 439 | 395 | 417 | 347 | 269 | 159 | 226 | 145 | 231 |
| L1PA13_dup7462 | 504.748005542502 | -1.54775132216182 | 0.328121405083889 | -4.71700808963108 | 2.39338177668099e-06 | 0.000647228895125728 | 8 | 902 | 766 | 744 | 1111 | 639 | 622 | 490 | 312 | 336 | 358 | 86 | 152 |
| L1PA15_dup276 | 48.7211887528992 | 3.16761121272008 | 0.602753976665286 | 5.25523071659314 | 1.47839020366614e-07 | 7.09227893954726e-05 | 9 | 75 | 67 | 0 | 9 | 22 | 35 | 5 | 1 | 100 | 92 | 88 | 104 |
| MIRb_dup41760 | 53.9869989065261 | 3.88053008200787 | 1.03746506770824 | 3.74039589649025 | 0.000183730632997595 | 0.0155028259571793 | 10 | 174 | 122 | 0 | 1 | 67 | 83 | 1 | 0 | 74 | 63 | 36 | 57 |
| ERV3-16A3_I-int_dup | 18.8439615872923 | 2.92144466030083 | 0.878793842411795 | 3.32437998459697 | 0.000886153652536653 | 0.0440754007120582 | 11 | 61 | 41 | 0 | 0 | 31 | 31 | 0 | 0 | 26 | 16 | 10 | 21 |
| L1MA2_dup6577 | 38.0198642268624 | 1.5365561702825 | 0.420436708562101 | 3.65466701406151 | 0.000257515984370109 | 0.0195142988588949 | 12 | 36 | 27 | 4 | 9 | 73 | 76 | 1 | 2 | 88 | 100 | 12 | 25 |
| L1PA4_dup10597 | 37.0934731210145 | 1.74483465004926 | 0.389685926906117 | 4.47754083372281 | 7.55077881815563e-06 | 0.00149225715669839 | 13 | 18 | 27 | 3 | 7 | 63 | 59 | 6 | 2 | 107 | 105 | 14 | 21 |
| L1M2_dup7939 | 21.4194352015391 | 1.99449077058646 | 0.52767955793604 | 3.7797385564597 | 0.000156993116554046 | 0.0138252716919204 | 14 | 8 | 15 | 3 | 3 | 45 | 30 | 1 | 0 | 53 | 71 | 6 | 17 |
| L1MCc_dup2887 | 15.1762619448705 | 1.68341902638926 | 0.494871733321166 | 3.40172798937526 | 0.000669612495499507 | 0.0367631783235177 | 15 | 20 | 9 | 3 | 1 | 30 | 27 | 1 | 0 | 28 | 45 | 6 | 14 |
| THE1B_dup21097 | 15.0809618502895 | 1.41759445431014 | 0.393007742021482 | 3.60703951280597 | 0.000309710512371056 | 0.0221304437167097 | 16 | 24 | 14 | 5 | 3 | 22 | 19 | 0 | 0 | 38 | 40 | 7 | 7 |
| L1PA7_dup7006 | 30.2398600220442 | 2.74191523347035 | 0.365031565147209 | 7.51144694121067 | 5.84773723338166e-14 | 1.90608997771813e-10 | 17 | 17 | 10 | 9 | 4 | 13 | 24 | 3 | 2 | 116 | 102 | 17 | 25 |
| L1PA7_dup3319 | 20.5494403989349 | 1.52913099107744 | 0.450952869100246 | 3.39088870668105 | 0.000696663851205123 | 0.0376746310770123 | 18 | 24 | 10 | 3 | 3 | 27 | 18 | 16 | 1 | 54 | 65 | 10 | 13 |
| L1PA8A_dup647 | 20.0530919858095 | 1.70688643508334 | 0.366936101857562 | 4.65172662608685 | 3.29167261032069e-06 | 0.00080996931844402 | 19 | 22 | 13 | 2 | 6 | 16 | 20 | 6 | 4 | 63 | 64 | 8 | 8 |
| SVA_A_dup943 | 51.555955696327 | 1.46687833198659 | 0.421861045336978 | 3.47715995160176 | 0.000506755404958741 | 0.0307681606502456 | 20 | 21 | 15 | 17 | 24 | 40 | 42 | 77 | 42 | 106 | 100 | 63 | 77 |
| L2c_dup114177 | 40.2045336297267 | 1.23891366764297 | 0.312670655935618 | 3.9623598957047 | 7.42125578354335e-05 | 0.00807803352548493 | 21 | 21 | 11 | 38 | 51 | 34 | 34 | 37 | 19 | 63 | 61 | 76 | 55 |
| MIR_dup86401 | 36.3081682694068 | 1.06526321495012 | 0.319143439793761 | 3.33788222511646 | 0.000844195236355861 | 0.0427844259233082 | 22 | 13 | 15 | 42 | 51 | 21 | 28 | 45 | 26 | 50 | 60 | 54 | 48 |
| L2b_dup73142 | 41.6638480182885 | 1.8236480035474 | 0.423446405930372 | 4.30667961283219 | 1.65723371566188e-05 | 0.00269352550346231 | 23 | 9 | 7 | 43 | 50 | 16 | 22 | 52 | 24 | 71 | 98 | 51 | 73 |
| HSMAR2_dup1514 | 46.9903152688242 | 1.70229344414672 | 0.370735285512741 | 4.59166826214663 | 4.39716920197561e-06 | 0.000997415465141361 | 24 | 16 | 10 | 44 | 62 | 36 | 28 | 24 | 20 | 70 | 114 | 72 | 88 |
| THE1D_dup10624 | 42.5625994375959 | 1.67538510027688 | 0.325509683698752 | 5.14695932004096 | 2.64742710330216e-07 | 0.000112289605975461 | 25 | 13 | 13 | 41 | 43 | 31 | 32 | 29 | 15 | 63 | 98 | 78 | 72 |
| L1PA11_dup2632 | 37.1729172989345 | 1.09492017727654 | 0.253071624247391 | 4.32652289853801 | 1.51481578846004e-05 | 0.0025241201512669 | 26 | 33 | 37 | 11 | 15 | 43 | 49 | 26 | 19 | 70 | 66 | 35 | 48 |
| L1PA5_dup1101 | 36.7688750103457 | 1.01777113890091 | 0.304288671542259 | 3.34475527380772 | 0.000823552031214091 | 0.0421330706556156 | 27 | 33 | 31 | 12 | 19 | 60 | 53 | 24 | 16 | 47 | 54 | 45 | 65 |
| L1PA15_dup2846 | 32.8425886812939 | 1.06816507191872 | 0.280674595672165 | 3.80570628189792 | 0.000141400106875756 | 0.0128012209430167 | 28 | 29 | 30 | 12 | 19 | 40 | 53 | 26 | 8 | 45 | 59 | 37 | 53 |
| MLT1D_dup9306 | 35.7731528982347 | 1.02322929802721 | 0.259427010396233 | 3.94418952931844 | 8.0070314591557e-05 | 0.00852838454601729 | 29 | 54 | 33 | 16 | 20 | 37 | 46 | 22 | 13 | 51 | 49 | 56 | 49 |
| MLT1I_dup3698 | 33.4536583706441 | 1.69751274087703 | 0.233395852561391 | 7.27310585105845 | 3.51314815018824e-13 | 9.28821721938544e-10 | 30 | 27 | 15 | 16 | 18 | 28 | 21 | 22 | 17 | 62 | 75 | 50 | 57 |
| AluJr_dup23268 | 34.3022841039851 | 1.41999822907032 | 0.227604564296714 | 6.23888292160589 | 4.4070667466718e-10 | 5.19264249620911e-07 | 31 | 27 | 21 | 19 | 19 | 31 | 37 | 19 | 18 | 51 | 68 | 48 | 66 |
| MER5B_dup7891 | 30.0714270090565 | 1.32036186311373 | 0.243914591261571 | 5.413213929862 | 6.19034578637443e-08 | 3.40964819093669e-05 | 32 | 27 | 15 | 15 | 20 | 32 | 33 | 20 | 13 | 48 | 57 | 41 | 51 |
| LTR101_Mam_dup73 | 36.5882383374282 | 1.17010604271863 | 0.220224507404127 | 5.31324173004691 | 1.07692028184816e-07 | 5.42900155475935e-05 | 33 | 35 | 28 | 23 | 21 | 28 | 41 | 28 | 21 | 48 | 55 | 57 | 72 |
| UCON40_dup47 | 34.812474983624 | 1.50503259644063 | 0.249633827908449 | 6.02896093470388 | 1.65017165965386e-09 | 1.70028388963981e-06 | 34 | 23 | 32 | 14 | 17 | 24 | 27 | 30 | 20 | 51 | 62 | 50 | 83 |
| MIR3_dup27877 | 39.0345296825041 | 1.41208069390508 | 0.251371813584869 | 5.61749813460423 | 1.93742265317313e-08 | 1.36772564567561e-05 | 35 | 32 | 31 | 12 | 25 | 32 | 40 | 33 | 16 | 69 | 58 | 72 | 61 |
| L2c_dup14873 | 39.59115809642 | 1.04581744751123 | 0.236372162960203 | 4.42445266995043 | 9.66870974443876e-06 | 0.00180867201953634 | 36 | 40 | 38 | 15 | 35 | 43 | 36 | 35 | 17 | 46 | 57 | 65 | 73 |
| L3_dup5139 | 40.939935959125 | 1.11757634771842 | 0.217347788351361 | 5.14188046814519 | 2.72002158254019e-07 | 0.000114147840472506 | 37 | 46 | 34 | 23 | 30 | 31 | 38 | 38 | 19 | 49 | 64 | 79 | 63 |
| HAL1M8_dup1417 | 37.340053063941 | 1.14584233604628 | 0.319520989424866 | 3.58612540011465 | 0.000335627611040227 | 0.0233991349360029 | 38 | 41 | 22 | 11 | 27 | 48 | 42 | 29 | 18 | 42 | 44 | 71 | 77 |
| MLT1J_dup1154 | 42.8592773184609 | 1.06880189589966 | 0.289961114113487 | 3.68601803440906 | 0.000227790216094792 | 0.0179229528587944 | 39 | 35 | 29 | 11 | 31 | 50 | 39 | 46 | 31 | 53 | 65 | 69 | 76 |
| MER74B_dup110 | 44.8662786393462 | 1.47230152606438 | 0.26769663166453 | 5.49988812675621 | 3.80032293065263e-08 | 2.34875750664174e-05 | 40 | 26 | 40 | 25 | 17 | 38 | 54 | 27 | 20 | 51 | 104 | 66 | 91 |
| Cheshire_dup761 | 44.9479150608331 | 1.73473084555563 | 0.309715666916067 | 5.60104324985842 | 2.13065562708337e-08 | 1.47807867009323e-05 | 41 | 20 | 23 | 29 | 16 | 28 | 52 | 28 | 22 | 83 | 59 | 82 | 111 |
| LTR2_dup419 | 40.3918829569101 | 1.69611199658149 | 0.343162053461669 | 4.94259775948957 | 7.70884282137831e-07 | 0.00026354717154823 | 42 | 20 | 23 | 15 | 10 | 31 | 47 | 29 | 24 | 80 | 59 | 59 | 96 |
| L2b_dup28428 | 39.4376453251939 | 1.2683025570888 | 0.273299710531279 | 4.64070216036194 | 3.47227291445369e-06 | 0.000837944676371802 | 43 | 48 | 35 | 31 | 37 | 27 | 31 | 16 | 13 | 49 | 55 | 69 | 88 |
| Ricksha_a_dup10 | 40.0344062508976 | 1.07014244822501 | 0.289840625045436 | 3.69217547766899 | 0.000222343962031609 | 0.0176383903523952 | 44 | 39 | 34 | 43 | 45 | 23 | 42 | 17 | 14 | 48 | 68 | 70 | 60 |
| LTR78_dup1511 | 33.9134805518549 | 1.04283472055085 | 0.301672619371152 | 3.45684246294769 | 0.000546544258070263 | 0.0323341670887088 | 45 | 41 | 25 | 29 | 30 | 33 | 45 | 11 | 10 | 40 | 53 | 54 | 56 |
| MIR_dup159331 | 34.0986033196346 | 1.14170166798855 | 0.309744476775304 | 3.68594681614539 | 0.000227853934917388 | 0.0179229528587944 | 46 | 43 | 31 | 21 | 22 | 23 | 53 | 14 | 13 | 34 | 59 | 64 | 52 |
| L1PA4_dup1430 | 34.0445833537428 | 1.05446579934166 | 0.319461366245993 | 3.30076156542101 | 0.000964227881275979 | 0.0466709046863495 | 47 | 34 | 36 | 28 | 31 | 21 | 24 | 25 | 15 | 25 | 27 | 77 | 97 |
| L1ME3G_dup696 | 29.2932323512798 | 1.01296780052438 | 0.293555909196283 | 3.45068100757281 | 0.00055917405885262 | 0.0328445397698705 | 48 | 30 | 26 | 25 | 24 | 20 | 37 | 26 | 7 | 24 | 45 | 48 | 65 |
| L2a_dup24048 | 27.9481063101736 | 1.24243895279414 | 0.289412831241857 | 4.29296430107433 | 1.76303361889296e-05 | 0.00282179735087705 | 49 | 28 | 23 | 24 | 37 | 18 | 15 | 17 | 8 | 30 | 42 | 49 | 67 |
| THE1B_dup2541 | 25.9773437151868 | 1.03453027143177 | 0.296961296048883 | 3.48372089291216 | 0.000494494874735748 | 0.0302456737546355 | 50 | 26 | 17 | 19 | 27 | 23 | 23 | 23 | 10 | 26 | 22 | 63 | 53 |
| L2b_dup74943 | 31.5686990570142 | 1.48999168063731 | 0.332326286317151 | 4.48352038940234 | 7.34215836096911e-06 | 0.00146563524610667 | 51 | 10 | 20 | 29 | 41 | 19 | 20 | 20 | 14 | 46 | 78 | 42 | 53 |
| L2c_dup28344 | 30.0162075787698 | 1.03131025647119 | 0.267680525935906 | 3.85276535476523 | 0.000116791299490787 | 0.0112237570874938 | 52 | 25 | 24 | 25 | 41 | 21 | 24 | 26 | 11 | 29 | 63 | 38 | 56 |
| MLT2D_dup2512 | 32.1542924355104 | 1.28723190963085 | 0.256148230837476 | 5.02533984100632 | 5.02541815745117e-07 | 0.000186840335764512 | 53 | 19 | 17 | 32 | 35 | 29 | 18 | 27 | 14 | 32 | 63 | 53 | 69 |
| L2a_dup137013 | 35.793256074206 | 1.24449019738527 | 0.362092517910268 | 3.4369398311999 | 0.000588326281577377 | 0.0339698824062632 | 54 | 17 | 18 | 36 | 40 | 37 | 38 | 22 | 8 | 53 | 56 | 78 | 42 |
| L2c_dup75991 | 32.3415968560188 | 1.15694049009982 | 0.280235785467264 | 4.12845378819391 | 3.65210817810943e-05 | 0.00480644099860855 | 55 | 18 | 22 | 28 | 30 | 31 | 42 | 18 | 11 | 48 | 53 | 45 | 57 |
| MER57F_dup317 | 33.8839806923607 | 1.28763786209457 | 0.296655026159939 | 4.34052265610479 | 1.42144211075418e-05 | 0.00242978781957984 | 56 | 16 | 18 | 26 | 31 | 35 | 40 | 20 | 13 | 48 | 61 | 64 | 48 |
| MIRb_dup187276 | 37.176211604279 | 1.01368310152196 | 0.250106961919811 | 4.05299834015402 | 5.05653613279981e-05 | 0.00608898049926712 | 57 | 24 | 24 | 38 | 37 | 35 | 40 | 29 | 14 | 52 | 68 | 48 | 55 |
| MSTD_dup6382 | 36.5820807568259 | 1.11785801200966 | 0.266752447451568 | 4.19061951516911 | 2.78193831001861e-05 | 0.00395194682457127 | 58 | 22 | 18 | 23 | 39 | 38 | 42 | 28 | 18 | 53 | 68 | 55 | 50 |
| THE1D-int_dup2166 | 33.900990102684 | 1.18861348551408 | 0.322729206582322 | 3.68300563218744 | 0.000230500064778718 | 0.0180654046158883 | 59 | 12 | 28 | 44 | 24 | 23 | 36 | 20 | 17 | 48 | 54 | 47 | 68 |
| L2c_dup71692 | 33.5109068035992 | 1.16534381021501 | 0.253282812240346 | 4.60095890403011 | 4.20550402977047e-06 | 0.000966843344799773 | 60 | 26 | 21 | 29 | 41 | 21 | 35 | 22 | 13 | 40 | 42 | 61 | 75 |
| L2_dup47005 | 38.9615024346802 | 1.66106559057615 | 0.263725754404028 | 6.29845801116338 | 3.00621133127475e-10 | 3.70630031829794e-07 | 61 | 21 | 19 | 29 | 29 | 30 | 38 | 19 | 12 | 74 | 65 | 57 | 88 |
| MIRc_dup79392 | 34.9771935015089 | 1.46300200079589 | 0.319838029406977 | 4.57419651912091 | 4.78050979005385e-06 | 0.00106209447479379 | 62 | 14 | 17 | 31 | 32 | 33 | 38 | 16 | 11 | 58 | 55 | 58 | 71 |
| MIR_dup137204 | 34.8745529341494 | 1.86877375822993 | 0.319999481105032 | 5.83992746418407 | 5.22235547617085e-09 | 4.55178972942472e-06 | 63 | 12 | 9 | 16 | 27 | 34 | 31 | 19 | 13 | 54 | 74 | 68 | 75 |
| L2b_dup74956 | 39.9203457933733 | 1.66636027774188 | 0.253965016239507 | 6.56137724170005 | 5.33130551862496e-11 | 8.23742092814763e-08 | 64 | 22 | 14 | 29 | 31 | 33 | 35 | 21 | 16 | 62 | 73 | 83 | 75 |
| MLT1C_dup15276 | 36.7270363331262 | 1.5316609526373 | 0.294839822484634 | 5.19489172029033 | 2.04839028760078e-07 | 9.2383040279331e-05 | 65 | 24 | 12 | 25 | 37 | 34 | 30 | 24 | 12 | 43 | 83 | 79 | 57 |
| MER44B_dup2089 | 38.2881922583616 | 1.36922957925625 | 0.368386088115822 | 3.71683302770587 | 0.000201735670804928 | 0.0165410737165229 | 66 | 16 | 10 | 21 | 23 | 39 | 56 | 31 | 21 | 44 | 77 | 67 | 73 |

| | | | | | | | | | | | | | | | | | | | |
|---|---|---|---|---|---|---|---|---|---|---|---|---|---|---|---|---|---|---|---|
| MLT1A0_dup20137 | 34.8186940487488 | 1.51306739210892 | 0.289021055183879 | 5.2351459001708 | 1.64854588735227e-07 | 7.81404182693072e-05 | 67 | 17 | 13 | 16 | 22 | 31 | 41 | 28 | 18 | 53 | 70 | 59 | 62 |
| MLT1D_dup16182 | 39.8699477516521 | 1.26225551223433 | 0.354681113552488 | 3.55884614095059 | 0.000372487649323267 | 0.0251082000583213 | 68 | 15 | 11 | 36 | 35 | 50 | 42 | 27 | 19 | 57 | 68 | 54 | 83 |
| L1MA7_dup7104 | 37.1950069122483 | 1.67183934770244 | 0.278662430456121 | 5.99951469943735 | 1.97908114274302e-09 | 1.92997721963578e-06 | 69 | 14 | 14 | 28 | 32 | 30 | 30 | 19 | 19 | 48 | 83 | 60 | 87 |
| L2b_dup74957 | 34.8793417146289 | 1.43322122652722 | 0.314827142465212 | 4.55240680744541 | 5.30356562595673e-06 | 0.00114723838766718 | 70 | 12 | 14 | 30 | 27 | 37 | 29 | 19 | 22 | 57 | 53 | 63 | 65 |
| MIRb_dup174359 | 33.3409288156409 | 1.04750616701197 | 0.284316669699918 | 3.68429388300572 | 0.000229337518844978 | 0.0179980030537886 | 71 | 16 | 17 | 29 | 27 | 35 | 39 | 25 | 23 | 32 | 48 | 62 | 66 |
| MIR_dup137132 | 33.9980948521151 | 1.1877950401303 | 0.271456383380815 | 4.37563863975892 | 1.21077456038211e-05 | 0.00215642936784941 | 72 | 21 | 14 | 26 | 23 | 39 | 34 | 30 | 19 | 48 | 51 | 60 | 57 |
| MIRb_dup118485 | 33.3417487569992 | 1.11482103096744 | 0.237477732811139 | 4.69442342139096 | 2.67359740589935e-06 | 0.000704044193033047 | 73 | 24 | 19 | 31 | 24 | 36 | 27 | 26 | 20 | 44 | 47 | 58 | 59 |
| L1M1_dup9399 | 37.0015337915979 | 1.12504407476808 | 0.3170816688201 | 3.54812083257449 | 0.000387990182277143 | 0.02578790835534 | 74 | 28 | 27 | 21 | 25 | 58 | 48 | 19 | 13 | 45 | 47 | 58 | 76 |
| L1MEc_dup2479 | 37.4145740684717 | 1.28356181501978 | 0.280736687148441 | 4.5721199749753 | 4.82814344987368e-06 | 0.00106968102544101 | 75 | 23 | 27 | 19 | 25 | 49 | 46 | 23 | 13 | 48 | 61 | 67 | 66 |
| Charlie4z_dup3811 | 31.5339873686863 | 1.20904454321435 | 0.295074536380444 | 4.09742080101249 | 4.17779036153391e-05 | 0.00528489370210285 | 76 | 25 | 22 | 17 | 19 | 43 | 33 | 24 | 12 | 30 | 49 | 55 | 71 |
| AluSx3_dup15832 | 31.8887270451807 | 1.10486264647129 | 0.283908605040763 | 3.89161380407141 | 9.95796616156694e-05 | 0.0100436975964263 | 77 | 24 | 25 | 22 | 22 | 42 | 41 | 19 | 10 | 41 | 46 | 42 | 67 |
| L1M4b_dup4221 | 33.6709852265444 | 1.29331085408722 | 0.230456075696784 | 5.61196249730842 | 2.00044798915256e-08 | 1.40235392691298e-05 | 78 | 26 | 30 | 22 | 22 | 34 | 33 | 19 | 13 | 48 | 59 | 43 | 71 |
| L1P1_dup2348 | 30.561671648318 | 1.55329868688853 | 0.260811708975327 | 5.95563248671274 | 2.59067924849366e-09 | 2.43652871328882e-06 | 79 | 27 | 14 | 19 | 19 | 21 | 29 | 19 | 14 | 44 | 45 | 72 | 57 |
| L2c_dup73170 | 30.90767496551 | 1.44750923232305 | 0.35385245767989 | 4.09071408409583 | 4.30047033078145e-05 | 0.00540274399856453 | 80 | 18 | 17 | 16 | 14 | 23 | 51 | 23 | 12 | 43 | 39 | 67 | 63 |
| MLT1C_dup2135 | 31.2114893156438 | 1.03813531484183 | 0.25211691345543 | 4.11767421952576 | 3.82715189220343e-05 | 0.00497816297128209 | 81 | 32 | 22 | 21 | 24 | 36 | 31 | 15 | 20 | 33 | 52 | 46 | 59 |
| AluSz_dup29103 | 31.1673693956234 | 1.40645548765445 | 0.282830097481624 | 4.97279285400604 | 6.59951678290121e-07 | 0.000232641277099883 | 82 | 24 | 19 | 18 | 22 | 37 | 36 | 11 | 11 | 41 | 62 | 55 | 51 |
| MIR3_dup57106 | 29.1655494774587 | 1.0262066685755 | 0.294966771903782 | 3.47905854599191 | 0.000503178644691087 | 0.0306448287150411 | 83 | 20 | 17 | 28 | 23 | 32 | 36 | 15 | 16 | 34 | 51 | 52 | 38 |
| AluSx1_dup83843 | 31.8236874943729 | 1.36914462527265 | 0.366601648074461 | 3.73469304479111 | 0.00018794427987229 | 0.0157300702140316 | 84 | 15 | 12 | 20 | 25 | 35 | 47 | 13 | 15 | 45 | 43 | 55 | 72 |
| L1MB8_dup16028 | 29.2768452405746 | 1.42520504336271 | 0.387423945869498 | 3.6786705069665 | 0.000234452926492671 | 0.0183028661572261 | 85 | 14 | 11 | 22 | 19 | 41 | 30 | 11 | 14 | 34 | 36 | 59 | 77 |
| AluSp_dup37015 | 26.4673627677635 | 1.09933295401009 | 0.310637483963276 | 3.53895782306832 | 0.000401710030294767 | 0.026397485464932 | 86 | 14 | 16 | 22 | 27 | 32 | 25 | 14 | 15 | 44 | 30 | 40 | 48 |
| AluSx_dup100338 | 26.9374672060784 | 1.14322091706339 | 0.329312802954264 | 3.47153498681972 | 0.000517491893030852 | 0.0311835256753778 | 87 | 18 | 18 | 22 | 23 | 29 | 33 | 15 | 10 | 33 | 24 | 51 | 65 |
| L1ME1_dup21387 | 55.8243203373914 | 1.95119017273247 | 0.494371192039608 | 3.94681203951735 | 7.9198682771481e-05 | 0.00845888137039488 | 88 | 25 | 27 | 35 | 49 | 36 | 21 | 10 | 12 | 26 | 29 | 210 | 255 |
| L1PA5_dup10738 | 52.3136675742783 | 2.68217186578396 | 0.505203997559812 | 5.30908678224862 | 1.10175868218867e-07 | 5.5424750822847e-05 | 89 | 9 | 13 | 19 | 36 | 32 | 28 | 13 | 3 | 48 | 42 | 182 | 258 |
| L1PA4_dup9113 | 39.0489984284459 | 2.03048110583607 | 0.540074474655456 | 3.75963168252199 | 0.000170163692416021 | 0.0146278070648925 | 90 | 18 | 13 | 34 | 30 | 28 | 12 | 11 | 2 | 20 | 24 | 136 | 189 |
| L1PA5_dup10736 | 44.8643474184738 | 2.84797701060507 | 0.534023620874335 | 5.33305438052008 | 9.6574374172701e-08 | 4.9955404428038e-05 | 91 | 12 | 12 | 24 | 16 | 19 | 23 | 7 | 2 | 28 | 34 | 176 | 237 |
| L1PA4_dup3783 | 20.447741931568 | 1.09095723831585 | 0.306407761365146 | 3.56047520942447 | 0.000370184260318616 | 0.0250309360630217 | 92 | 16 | 9 | 25 | 27 | 6 | 11 | 20 | 15 | 21 | 13 | 39 | 61 |
| L1MB4_dup8117 | 26.2608190571334 | 1.34657749536223 | 0.327066127111465 | 4.11714140884824 | 3.83600724070829e-05 | 0.00497816297128209 | 93 | 13 | 8 | 28 | 23 | 11 | 7 | 25 | 21 | 8 | 14 | 87 | 98 |
| L1PA3_dup6649 | 28.6015670092003 | 1.00392836652795 | 0.273615657163721 | 3.66911885428849 | 0.000243387912356935 | 0.0187725057353916 | 94 | 20 | 12 | 35 | 37 | 8 | 8 | 34 | 30 | 14 | 23 | 55 | 96 |
| LTR39_dup1133 | 35.0525874854453 | 1.34520782575643 | 0.394129680439232 | 3.41310967562067 | 0.000642260889685775 | 0.0358656206658464 | 95 | 6 | 7 | 23 | 35 | 15 | 21 | 32 | 34 | 9 | 19 | 97 | 163 |
| L1P2_dup1120 | 35.0572696262656 | 1.15473132185882 | 0.296571370726951 | 3.89360348245471 | 9.87660415362013e-05 | 0.00997918663243013 | 96 | 19 | 16 | 33 | 26 | 31 | 30 | 31 | 22 | 30 | 26 | 85 | 99 |
| LTR41B_dup765 | 36.1224218134288 | 1.02100143224082 | 0.267266983146057 | 3.82015548730482 | 0.000133367572000866 | 0.0123144277405193 | 97 | 12 | 16 | 49 | 55 | 25 | 20 | 41 | 17 | 26 | 29 | 80 | 100 |
| AluSx_dup107084 | 60.2001896369823 | 1.52760363885887 | 0.363483504451396 | 4.20267665561461 | 2.63777248374592e-05 | 0.00380392370556004 | 98 | 73 | 69 | 29 | 38 | 60 | 65 | 5 | 9 | 122 | 102 | 67 | 98 |
| L2a_dup20545 | 71.6941050823992 | 1.36085171616044 | 0.371508893913678 | 3.66303940081886 | 0.000249240183416642 | 0.0191062192922862 | 99 | 67 | 77 | 52 | 67 | 69 | 75 | 16 | 8 | 107 | 116 | 95 | 151 |
| THE1D_dup1163 | 60.3473138516638 | 1.09674280627675 | 0.317891131086452 | 3.45005789412378 | 0.000560466344866057 | 0.0329029492590101 | 100 | 71 | 66 | 39 | 65 | 60 | 66 | 17 | 12 | 97 | 81 | 75 | 106 |
| L2a_dup112240 | 67.7065022700435 | 1.10442038285169 | 0.326472755899441 | 3.3828868194805 | 0.000717281860181256 | 0.0383561600635049 | 101 | 121 | 111 | 22 | 22 | 96 | 104 | 12 | 12 | 91 | 109 | 52 | 92 |
| THE1B-int_dup3441 | 68.2719708347424 | 1.71057054420343 | 0.421526243955246 | 4.05804043931615 | 4.94862001797245e-05 | 0.00598325375404711 | 102 | 110 | 112 | 12 | 30 | 53 | 50 | 21 | 17 | 87 | 83 | 121 | 166 |
| L1PA7_dup730 | 55.1065048438186 | 2.16767443152533 | 0.318631136116218 | 6.80308414911054 | 1.02402791296629e-11 | 1.95568469034522e-08 | 103 | 50 | 42 | 16 | 13 | 63 | 58 | 3 | 6 | 135 | 135 | 58 | 79 |
| L1MA3_dup7784 | 57.1291831539532 | 1.54194334400677 | 0.446215635961382 | 3.45560132756132 | 0.000549066772841674 | 0.0324591906411391 | 104 | 32 | 30 | 30 | 21 | 83 | 109 | 20 | 13 | 61 | 75 | 95 | 155 |
| L1ME2_dup4575 | 45.255085666881 | 1.27768672532817 | 0.292206762805725 | 4.37254330823839 | 1.22807445555214e-05 | 0.00217584068801794 | 105 | 38 | 34 | 13 | 18 | 55 | 72 | 19 | 23 | 72 | 94 | 41 | 76 |
| L1PA15_dup2317 | 50.8436992563856 | 1.40002837023995 | 0.307756379528212 | 4.54914491906289 | 5.38643402124455e-06 | 0.00115821603421951 | 106 | 48 | 51 | 9 | 14 | 70 | 80 | 15 | 21 | 79 | 109 | 57 | 68 |
| MIRb_dup116673 | 39.3796802797282 | 1.42624116494838 | 0.380721869146816 | 3.74614982886204 | 0.000179569366286154 | 0.0152436005816358 | 107 | 38 | 50 | 20 | 15 | 47 | 57 | 14 | 6 | 57 | 42 | 74 | 71 |
| L1PB1_dup10398 | 43.6518346891317 | 1.41389036432159 | 0.403498780161407 | 3.50407593241301 | 0.000458194689013159 | 0.0287742395016957 | 108 | 77 | 55 | 12 | 16 | 42 | 60 | 13 | 13 | 39 | 60 | 82 | 85 |
| Tigger16a_dup991 | 45.9225503225644 | 1.4333929395598 | 0.384780469424772 | 3.72522270088877 | 0.000195142865272658 | 0.0161563897773723 | 109 | 49 | 57 | 17 | 16 | 59 | 70 | 8 | 15 | 61 | 61 | 56 | 105 |
| L1PA3_dup1820 | 74.3143276566154 | 2.66449010582222 | 0.472275118911078 | 5.64181765909184 | 1.68264247701468e-08 | 1.22815351790103e-05 | 110 | 54 | 55 | 13 | 15 | 9 | 9 | 43 | 22 | 212 | 222 | 80 | 145 |
| LOR1a_dup55 | 51.5441780432401 | 1.69801410386437 | 0.508331119463139 | 3.34037016198749 | 0.000836667920810296 | 0.042545655222169 | 111 | 47 | 58 | 27 | 50 | 6 | 4 | 36 | 29 | 88 | 118 | 77 | 95 |
| ERV3-16A3_I-int_dup | 67.7201255315808 | 2.2692912362993 | 0.592736872242184 | 3.82849682982451 | 0.000128928291470011 | 0.0120258609337998 | 112 | 122 | 112 | 16 | 12 | 18 | 13 | 19 | 15 | 75 | 83 | 171 | 206 |
| L1M4a2_dup453 | 39.3369195032358 | 2.19924244603357 | 0.633872444923193 | 3.46953470473077 | 0.000521360706739588 | 0.031306627043134 | 113 | 69 | 75 | 8 | 7 | 5 | 11 | 12 | 9 | 54 | 42 | 78 | 129 |
| LTR12C_dup993 | 39.2148641985077 | -2.06549073606602 | 0.420445300125179 | -4.91262653061186 | 8.98643521327218e-07 | 0.000296161539232308 | 114 | 163 | 117 | 3 | 1 | 76 | 89 | 3 | 3 | 24 | 23 | 0 | 0 |
| L1MA9_dup12019 | 19.6334559174861 | -1.6167333448068 | 0.484004702560278 | -3.34032569571048 | 0.000836801906481937 | 0.042545655222169 | 115 | 67 | 58 | 6 | 1 | 37 | 45 | 2 | 1 | 11 | 22 | 0 | 0 |
| LTR12C_dup1603 | 21.0604579270952 | -2.21291211732611 | 0.574232794269192 | -3.85368467181052 | 0.000116353428657924 | 0.0111997706049508 | 116 | 88 | 84 | 2 | 5 | 29 | 33 | 4 | 3 | 6 | 17 | 0 | 1 |
| L1PA7_dup10206 | 39.7980430240714 | -1.71303436423967 | 0.5223925986236 | -3.27920871917629 | 0.00104098612770519 | 0.0491367754697359 | 117 | 124 | 103 | 14 | 22 | 107 | 69 | 4 | 0 | 27 | 34 | 0 | 4 |
| L1PA8_dup6066 | 24.7804398366232 | -1.83651102761197 | 0.550326772771886 | -3.33712826356209 | 0.000846488737737162 | 0.0428550232318312 | 118 | 81 | 71 | 5 | 11 | 45 | 68 | 0 | 0 | 12 | 24 | 1 | 0 |
| L2a_dup119788 | 18.7336820776164 | -1.59022440106916 | 0.454335932722824 | -3.50010704973077 | 0.000465071352174147 | 0.02906928846143 | 119 | 53 | 48 | 5 | 10 | 45 | 47 | 1 | 0 | 12 | 16 | 1 | 1 |
| THE1D_dup9318 | 28.9247238627343 | -1.57226460009979 | 0.479486646980972 | -3.27905815521528 | 0.00104154169729202 | 0.0491532488504257 | 120 | 80 | 74 | 12 | 11 | 72 | 64 | 0 | 2 | 17 | 33 | 1 | 1 |
| L1MC_dup5888 | 29.0195502495872 | -1.5099921961399 | 0.387911032456545 | -3.8926250346052 | 9.91653613623499e-05 | 0.0100152805919888 | 121 | 75 | 98 | 6 | 5 | 68 | 60 | 3 | 1 | 24 | 24 | 0 | 1 |
| L1M4_dup14479 | 25.1923966365671 | -1.72774471094069 | 0.480073248434893 | -3.5989189495007 | 0.000319542751575233 | 0.0225888055752586 | 122 | 86 | 84 | 8 | 12 | 43 | 47 | 2 | 1 | 16 | 21 | 1 | 1 |
| MER110_dup554 | 21.141832477641 | -1.58210633829669 | 0.457633017212722 | -3.45715077101022 | 0.000545919321191326 | 0.0323179885759807 | 123 | 61 | 52 | 9 | 11 | 43 | 49 | 3 | 1 | 18 | 21 | 0 | 0 |
| HERVL-int_dup831 | 19.7219119204788 | -3.74312823831988 | 0.64365859897231 | -5.81539381948179 | 6.04913008957561e-09 | 5.1572665100993e-06 | 124 | 52 | 47 | 7 | 13 | 44 | 29 | 26 | 24 | 4 | 5 | 0 | 0 |
| L1M3c_dup758 | 20.4742470579324 | -2.73893625403614 | 0.609011573081101 | -4.49734680767948 | 6.88067037234602e-06 | 0.00139220067382504 | 125 | 74 | 65 | 16 | 10 | 19 | 20 | 18 | 19 | 11 | 7 | 1 | 0 |
| L1PB1_dup9381 | 18.7405794222416 | -2.9723649846784 | 0.653407953374676 | -4.54901867864775 | 5.38966595123362e-06 | 0.00115821603421951 | 126 | 54 | 65 | 10 | 23 | 13 | 14 | 31 | 17 | 7 | 7 | 1 | 0 |
| L1ME4c_dup6749 | 14.0977170321448 | -1.91380161126363 | 0.556418270291971 | -3.43950174436112 | 0.000582786028946705 | 0.0337811460277108 | 127 | 25 | 33 | 22 | 29 | 12 | 15 | 10 | 13 | 8 | 10 | 0 | 3 |
| L1PA7_dup9661 | 15.4332398271595 | -2.31195039537832 | 0.679366131579577 | -3.40309928315511 | 0.000666260653189498 | 0.0366552733835441 | 128 | 56 | 48 | 14 | 18 | 11 | 6 | 14 | 11 | 10 | 9 | 0 | 1 |
| AluY_dup35194 | 19.6454381024511 | -2.97681333306801 | 0.483964045722199 | -6.15089769452985 | 7.70456073555086e-10 | 8.72985432753041e-07 | 129 | 36 | 40 | 22 | 28 | 28 | 30 | 21 | 30 | 5 | 5 | 0 | 4 |
| L2a_dup134045 | 12.3853390716428 | -2.69698799836952 | 0.611906511440189 | -4.40751642276508 | 1.04562674764784e-05 | 0.00192404429751195 | 130 | 30 | 12 | 15 | 30 | 25 | 15 | 9 | 13 | 5 | 5 | 0 | 1 |
| L1PA13_dup6200 | 24.0910464880274 | -1.8999360950861 | 0.561549427874306 | -3.38338176619311 | 0.000715990288237267 | 0.0383191689439732 | 131 | 31 | 30 | 10 | 4 | 95 | 73 | 16 | 14 | 15 | 6 | 1 | 7 |
| L1P1_dup2172 | 10.8243410412246 | -3.40586532438196 | 0.985879259253423 | -3.45464750618765 | 0.000551012708867503 | 0.0325257429978137 | 132 | 14 | 10 | 0 | 3 | 29 | 30 | 28 | 16 | 3 | 4 | 0 | 0 |
| L1PA7_dup9299 | 17.3011610318843 | -2.15245712463597 | 0.65125792825994 | -3.30507627045247 | 0.000949505573386435 | 0.0462026693588975 | 133 | 23 | 22 | 1 | 6 | 48 | 48 | 30 | 21 | 5 | 7 | 2 | 6 |
| L1P3_dup2390 | 14.1444762991891 | -2.74846992226823 | 0.712835437364824 | -3.85568642943544 | 0.000115405341175392 | 0.0111310252579326 | 134 | 14 | 27 | 2 | 6 | 35 | 51 | 20 | 14 | 4 | 1 | 4 | 1 |
| L1PA7_dup9366 | 12.5530129957211 | -1.8821838222301 | 0.521614001666314 | -3.60838439194001 | 0.00030810975605985 | 0.022062438764796 | 135 | 20 | 17 | 3 | 10 | 31 | 40 | 16 | 7 | 6 | 6 | 2 | 3 |
| L1PA8_dup5517 | 11.7474498281782 | -2.14365181871971 | 0.604081690149592 | -3.54861247025857 | 0.000387266571876148 | 0.0257723946337853 | 136 | 14 | 11 | 7 | 5 | 34 | 29 | 21 | 13 | 6 | 4 | 2 | 2 |
| L1PA7_dup7904 | 13.7094904839528 | 1.89173185000034 | 0.506133934196929 | 3.73761117796203 | 0.000185776949187006 | 0.0156138688051098 | 137 | 3 | 7 | 6 | 8 | 9 | 3 | 11 | 13 | 28 | 41 | 19 | 14 |
| UCON28a_dup42 | 15.9718946545987 | 1.64981419705938 | 0.416208286575791 | 3.96391482407199 | 7.37305688322941e-05 | 0.008040281361764 | 138 | 10 | 6 | 14 | 6 | 10 | 9 | 16 | 7 | 30 | 49 | 20 | 16 |

| | | | | | | | | | | | | | | | | | | | |
|---|---|---|---|---|---|---|---|---|---|---|---|---|---|---|---|---|---|---|---|
| MIRb_dup71184 | 15.5992796944714 | 1.51120036673828 | 0.362833651632406 | 4.16499506024129 | 3.11359306089343e-05 | 0.00429737247370851 | 139 | 13 | 9 | 14 | 9 | 7 | 9 | 9 | 12 | 22 | 37 | 31 | 18 |
| MER5A1_dup10465 | 14.1379596393689 | 1.29063359592778 | 0.33401019864093 | 3.86405445456246 | 0.000111520343160653 | 0.0108664289818611 | 140 | 12 | 8 | 10 | 14 | 7 | 11 | 14 | 7 | 20 | 33 | 20 | 20 |
| MIR_dup137141 | 13.6414717866802 | 1.58294406902951 | 0.383156060048156 | 4.13132985246419 | 3.60670603953852e-05 | 0.00477910326615774 | 141 | 9 | 7 | 10 | 10 | 3 | 12 | 12 | 8 | 19 | 34 | 27 | 17 |
| L2a_dup12591 | 11.4075992683406 | 1.93763859934414 | 0.54497007461144 | 3.5554954108731 | 0.000377267530410685 | 0.0253156513793291 | 142 | 1 | 3 | 12 | 14 | 6 | 3 | 8 | 8 | 23 | 30 | 12 | 18 |
| MER57A-int_dup941 | 14.5067335118431 | 1.80924274723566 | 0.547587797201479 | 3.30402312922611 | 0.000953079687580933 | 0.0463008369418401 | 143 | 5 | 3 | 16 | 18 | 5 | 8 | 7 | 8 | 34 | 42 | 9 | 19 |
| MIR_dup57152 | 16.8494224946346 | 1.80117039319004 | 0.443112165926849 | 4.06481819207686 | 4.80699181658887e-05 | 0.00587000943116681 | 144 | 2 | 6 | 11 | 8 | 14 | 10 | 17 | 10 | 34 | 44 | 21 | 26 |
| L2a_dup133506 | 16.462931537571 | 2.00029146421176 | 0.378753093037107 | 5.28125446625934 | 1.28302342929769e-07 | 6.26879451555786e-05 | 145 | 4 | 3 | 10 | 14 | 11 | 9 | 12 | 9 | 32 | 36 | 22 | 40 |
| MIRb_dup174455 | 17.0578627742119 | 1.73409095925428 | 0.352969502576504 | 4.91286342473291 | 8.97558015946431e-07 | 0.000296161539232308 | 146 | 3 | 6 | 14 | 13 | 13 | 14 | 11 | 9 | 27 | 33 | 28 | 40 |
| MSTA_dup10681 | 14.4973499944117 | 1.47772520027271 | 0.445013954027908 | 3.32062666102383 | 0.000898155941608577 | 0.0444885652069922 | 147 | 3 | 5 | 15 | 21 | 9 | 12 | 10 | 6 | 23 | 33 | 25 | 17 |
| MIRc_dup32868 | 15.4620827454528 | 1.85890918592205 | 0.419998793912628 | 4.42598696202152 | 9.60023226654841e-06 | 0.0017986904463749 | 148 | 3 | 7 | 9 | 17 | 11 | 9 | 6 | 8 | 38 | 31 | 22 | 24 |
| Tigger3a_dup4402 | 13.1524858469556 | 1.57850538223362 | 0.44413033489773 | 3.55414899231574 | 0.0003792043214684 | 0.0254026327353941 | 149 | 2 | 6 | 9 | 16 | 13 | 6 | 11 | 6 | 27 | 25 | 22 | 17 |
| L2c_dup114237 | 16.463643623012 | 1.57541869126794 | 0.477962899344454 | 3.29611083502234 | 0.000980333310416834 | 0.0471817131635202 | 150 | 5 | 1 | 12 | 20 | 11 | 15 | 15 | 9 | 26 | 44 | 17 | 29 |
| MIR_dup137234 | 13.4698944396593 | 1.47993755265716 | 0.397316755136421 | 3.72483046215609 | 0.00019544652967705 | 0.0161715524870134 | 151 | 5 | 2 | 12 | 12 | 9 | 12 | 17 | 6 | 22 | 24 | 21 | 26 |
| MSTA_dup15814 | 13.3625160302129 | 1.73928756020837 | 0.487490063344164 | 3.56784207718394 | 0.000359933318457692 | 0.0245329972482767 | 152 | 3 | 1 | 11 | 10 | 10 | 9 | 19 | 5 | 20 | 35 | 22 | 21 |
| L1ME3Cz_dup9415 | 16.6489591033736 | 1.89587804971534 | 0.470746925278958 | 4.02738275686425 | 5.64011759459486e-05 | 0.00663064931404875 | 153 | 5 | 2 | 16 | 14 | 8 | 5 | 21 | 12 | 33 | 38 | 25 | 23 |
| L1MEc_dup14689 | 18.0282384583472 | 1.48134847478962 | 0.419397780896764 | 3.53208467536994 | 0.000412297312529658 | 0.0268338337875224 | 154 | 4 | 4 | 16 | 30 | 9 | 11 | 23 | 10 | 30 | 31 | 24 | 34 |
| L1MEf_dup9174 | 19.7982329507831 | 1.80663598785 | 0.513535601877482 | 3.51803454569646 | 0.000434755761780841 | 0.0278087081969634 | 155 | 0 | 4 | 23 | 29 | 12 | 10 | 13 | 12 | 24 | 45 | 28 | 49 |
| L1MB8_dup8943 | 21.6167918719584 | 2.20503453805447 | 0.478261379931841 | 4.61052184135947 | 4.01659435973348e-06 | 0.000936992707373669 | 156 | 3 | 1 | 13 | 8 | 18 | 12 | 15 | 14 | 43 | 46 | 37 | 52 |
| L1MA5A_dup2823 | 22.239813083348 | 2.03419059353456 | 0.442053407683216 | 4.60168513165789 | 4.19086420240843e-06 | 0.000964579257338518 | 157 | 4 | 4 | 17 | 18 | 10 | 23 | 19 | 3 | 41 | 45 | 45 | 47 |
| MIRc_dup72833 | 17.99064520329 | 1.37228992466569 | 0.393997126442296 | 3.48299475444695 | 0.000495838080002915 | 0.0303141541069819 | 158 | 6 | 4 | 16 | 10 | 11 | 21 | 24 | 11 | 22 | 33 | 28 | 40 |
| MIRb_dup174472 | 20.4375040868437 | 1.49234372535821 | 0.395867125142286 | 3.76980969263822 | 0.000163372073231084 | 0.0142013401740939 | 159 | 5 | 4 | 19 | 26 | 11 | 20 | 17 | 12 | 25 | 42 | 43 | 31 |
| MLT2C1_dup8 | 20.0812570495348 | 1.63169923751162 | 0.432106766626322 | 3.77614831225887 | 0.000159272176334597 | 0.0139593168184873 | 160 | 2 | 3 | 15 | 19 | 15 | 20 | 19 | 11 | 32 | 29 | 39 | 46 |
| MARNA_dup2606 | 20.0729887754047 | 1.44845683679481 | 0.418896049581105 | 3.45779540829585 | 0.000544614799354665 | 0.0322898394372453 | 161 | 3 | 8 | 22 | 17 | 20 | 16 | 10 | 13 | 22 | 39 | 48 | 30 |
| MER39_dup2869 | 20.6103869810844 | 1.52744067254653 | 0.458204145428258 | 3.33353743694073 | 0.000857491311098299 | 0.0432372594851866 | 162 | 4 | 6 | 7 | 26 | 23 | 23 | 9 | 11 | 35 | 41 | 29 | 40 |
| L1PA7_dup10222 | 19.6984027175596 | 1.37432484123441 | 0.41279014880266 | 3.32935474652382 | 0.000870474552936817 | 0.0436058891412255 | 163 | 4 | 8 | 13 | 21 | 18 | 27 | 13 | 7 | 35 | 39 | 30 | 27 |
| MIR3_dup60814 | 21.4175079756061 | 1.46440833101775 | 0.440813143004729 | 3.32206140914006 | 0.000893550258533336 | 0.0443403186133173 | 164 | 4 | 6 | 12 | 16 | 22 | 26 | 16 | 10 | 43 | 55 | 21 | 28 |
| AluSx1_dup69530 | 16.564493538919 | 1.39253327077367 | 0.390337556845027 | 3.56751034163628 | 0.00036038916003038 | 0.024535953954961 | 165 | 17 | 1 | 9 | 16 | 12 | 14 | 15 | 10 | 25 | 21 | 36 | 31 |
| MIRb_dup174375 | 19.0986372563769 | 1.80705482483538 | 0.385804683076157 | 4.68385922749071 | 2.81523391320447e-06 | 0.000721069589570882 | 166 | 8 | 3 | 8 | 20 | 11 | 17 | 14 | 10 | 43 | 29 | 32 | 38 |
| L1PA3_dup9429 | 17.0289600445418 | 1.58649457980447 | 0.415221773103371 | 3.82083667710149 | 0.000132999709215659 | 0.0122852285749337 | 167 | 8 | 3 | 9 | 20 | 17 | 15 | 5 | 11 | 21 | 33 | 35 | 35 |
| MER57A1_dup1005 | 19.3622220214282 | 1.52357873971563 | 0.395800990117572 | 3.84935555432302 | 0.000118428996647807 | 0.0113524173231182 | 168 | 11 | 5 | 12 | 19 | 20 | 25 | 7 | 6 | 30 | 29 | 40 | 37 |
| MIR_dup137216 | 18.3984244006708 | 1.35369975485053 | 0.369852888403381 | 3.66010323914009 | 0.000252113650358401 | 0.0192397160513727 | 169 | 7 | 6 | 13 | 24 | 19 | 16 | 13 | 8 | 30 | 27 | 38 | 27 |
| L2c_dup134707 | 16.5350023526718 | 1.59044721901632 | 0.42019755844654 | 3.78499871559503 | 0.000153709372347647 | 0.013581333201869 | 170 | 6 | 3 | 10 | 18 | 16 | 13 | 15 | 6 | 30 | 30 | 37 | 19 |
| MSTA_dup16240 | 17.8588391395001 | 1.59322615377366 | 0.388281018293523 | 4.10328107404224 | 4.07331889059217e-05 | 0.0051858209563448 | 171 | 7 | 3 | 14 | 16 | 15 | 16 | 17 | 6 | 24 | 29 | 49 | 27 |
| AluSc8_dup16285 | 20.510381415337 | 1.90462293580097 | 0.453454003867578 | 4.20025607791783 | 2.66613262242286e-05 | 0.00383552353672932 | 172 | 6 | 1 | 12 | 25 | 19 | 15 | 11 | 6 | 34 | 40 | 44 | 42 |
| MamTip2_dup5625 | 21.0863141681986 | 1.52938846452258 | 0.411735142396877 | 3.71449581791681 | 0.000203609236885255 | 0.0166259449141726 | 173 | 9 | 1 | 15 | 23 | 18 | 18 | 18 | 12 | 32 | 42 | 40 | 33 |
| MIRb_dup174450 | 21.1263970756416 | 1.83353800243276 | 0.465500441276598 | 3.93885341419747 | 8.18719292552789e-05 | 0.00869303796455894 | 174 | 9 | 7 | 16 | 38 | 10 | 8 | 11 | 7 | 23 | 70 | 39 | 27 |
| L1MC5_dup630 | 19.8477167357539 | 1.76646834499966 | 0.488853921690705 | 3.61348915620911 | 0.000302103995639665 | 0.0217645622649209 | 175 | 7 | 6 | 15 | 20 | 9 | 12 | 12 | 10 | 60 | 49 | 17 | 15 |
| L1PA3_dup3966 | 21.6571265153111 | 2.17401656560087 | 0.537755833254286 | 4.04275775577288 | 5.2826190176282e-05 | 0.00629748529443166 | 176 | 10 | 3 | 8 | 22 | 7 | 11 | 9 | 10 | 52 | 91 | 15 | 18 |
| L1PA15_dup2518 | 21.0969715765212 | 1.61647660302671 | 0.348416748460996 | 4.63949167244947 | 3.49267208567747e-06 | 0.000841161287549202 | 177 | 13 | 18 | 19 | 24 | 5 | 4 | 17 | 11 | 31 | 30 | 36 | 58 |
| L2c_dup17585 | 18.1776163112374 | 1.69776600701492 | 0.388858680927382 | 4.36602316030581 | 1.2652906957669e-05 | 0.00222850377420394 | 178 | 17 | 17 | 17 | 12 | 5 | 3 | 14 | 6 | 27 | 25 | 37 | 49 |
| L2c_dup17586 | 19.7691552630518 | 1.95006700047306 | 0.355874662223011 | 5.47964552545478 | 4.26178795601294e-08 | 2.5543460881145e-05 | 179 | 17 | 13 | 14 | 16 | 3 | 5 | 14 | 7 | 34 | 25 | 44 | 56 |
| L1ME2_dup11653 | 20.3121417900412 | 1.86344751267823 | 0.314599993347678 | 5.92322807400333 | 3.15682537143695e-09 | 2.934195194656e-06 | 180 | 12 | 5 | 18 | 15 | 11 | 11 | 11 | 9 | 17 | 28 | 64 | 58 |
| MER117_dup2734 | 17.7633176887207 | 1.43950753358762 | 0.361850511534204 | 3.97818294489699 | 6.94439326629831e-05 | 0.0077106420911928 | 181 | 13 | 6 | 16 | 21 | 7 | 8 | 10 | 18 | 22 | 33 | 38 | 27 |
| MIRb_dup174364 | 17.9919822748722 | 1.4020984426656 | 0.334803364366845 | 4.18782662270177 | 2.81638532396838e-05 | 0.00398423081032687 | 182 | 9 | 6 | 21 | 21 | 9 | 11 | 10 | 15 | 19 | 34 | 33 | 37 |
| AluJr_dup4540 | 15.1829046414589 | 1.32840542022614 | 0.336980721184006 | 3.9420813616835 | 8.07775633674313e-05 | 0.00858708329551212 | 183 | 11 | 10 | 11 | 18 | 11 | 10 | 7 | 9 | 16 | 19 | 40 | 29 |
| Chap1_Mam_dup971 | 15.6813146910755 | 1.36706568198208 | 0.322887915388004 | 4.23387069268083 | 2.29703053409456e-05 | 0.00342461921970968 | 184 | 9 | 14 | 13 | 12 | 8 | 10 | 10 | 12 | 13 | 28 | 34 | 34 |
| MIRb_dup107236 | 20.1662719941933 | 1.03497033697772 | 0.289159628154196 | 3.57923526041407 | 0.00034460114455388 | 0.0238569868903164 | 185 | 12 | 16 | 16 | 25 | 11 | 13 | 23 | 15 | 17 | 36 | 39 | 32 |
| Charlie17_dup431 | 18.4066897498848 | 1.31422401068732 | 0.284335964002145 | 4.62208154110748 | 3.79908616599066e-06 | 0.000895914129685643 | 186 | 11 | 10 | 18 | 20 | 13 | 11 | 14 | 10 | 24 | 23 | 40 | 37 |
| L2a_dup12139 | 17.5517399079303 | 1.22636049798545 | 0.286862104637013 | 4.27508715219548 | 1.91062540485894e-05 | 0.00300678354883972 | 187 | 14 | 11 | 13 | 16 | 11 | 12 | 22 | 8 | 19 | 32 | 29 | 36 |
| L2c_dup114174 | 17.1930610799914 | 1.35450246536036 | 0.321649491728036 | 4.21111333981411 | 2.54115205113577e-05 | 0.00370273708487172 | 188 | 15 | 9 | 18 | 15 | 7 | 14 | 10 | 10 | 25 | 26 | 37 | 27 |
| L2c_dup93587 | 17.0382622160448 | 1.32439404970287 | 0.312633388980871 | 4.23625273685628 | 2.27280945554124e-05 | 0.00340987338403667 | 189 | 17 | 8 | 16 | 15 | 6 | 14 | 14 | 9 | 27 | 27 | 29 | 30 |
| L1MD3_dup2686 | 15.8006131146473 | 1.21318044636618 | 0.345176756303207 | 3.51466436894293 | 0.00044031009144801 | 0.028043368581287 | 190 | 16 | 6 | 17 | 17 | 6 | 8 | 20 | 8 | 18 | 24 | 36 | 24 |
| MER5A_dup28789 | 14.5925146701047 | 1.12185392315957 | 0.330533751663135 | 3.39406767845878 | 0.000688626721544737 | 0.0374260916699391 | 191 | 14 | 10 | 13 | 11 | 10 | 9 | 16 | 8 | 15 | 18 | 37 | 23 |
| ERVL-B4-int_dup2701 | 24.8848719403516 | 2.60125317142633 | 0.344580248730361 | 7.54904896903086 | 4.38448010440362e-14 | 1.49038500417489e-10 | 192 | 9 | 5 | 11 | 12 | 5 | 9 | 27 | 6 | 29 | 55 | 83 | 63 |
| MLT1B_dup16233 | 29.1887455785243 | 2.46875348945354 | 0.286607457182096 | 8.61370989340666 | 7.07335468710754e-18 | 5.80371191164997e-14 | 193 | 12 | 5 | 25 | 18 | 5 | 11 | 16 | 10 | 38 | 40 | 99 | 89 |
| L1ME2_dup11654 | 21.847936013767 | 1.90510986286196 | 0.372511450951896 | 5.11423167796249 | 3.15020528478664e-07 | 0.000128793599088289 | 194 | 2 | 10 | 17 | 14 | 9 | 16 | 19 | 8 | 20 | 20 | 73 | 72 |
| L2a_dup13847 | 19.0268133437674 | 1.74550793679393 | 0.411779278000615 | 4.2389406899473 | 2.24576975339667e-05 | 0.00337143173338627 | 195 | 6 | 5 | 25 | 21 | 12 | 10 | 10 | 3 | 19 | 18 | 46 | 72 |
| L1MB7_dup13708 | 19.0678713464189 | 1.79724382579564 | 0.459373299200479 | 3.91238199722027 | 9.13901980561615e-05 | 0.00942606504840546 | 196 | 3 | 4 | 15 | 19 | 12 | 19 | 14 | 3 | 17 | 20 | 62 | 58 |
| L1PA5_dup7723 | 18.6592024514524 | 2.09035082346974 | 0.453152864478599 | 4.61290435816822 | 3.9708104573342e-06 | 0.000928132087505741 | 197 | 4 | 3 | 16 | 15 | 11 | 13 | 9 | 3 | 20 | 13 | 60 | 74 |
| L2c_dup53042 | 13.1740202817181 | 1.27664793311847 | 0.384385114327608 | 3.32127308142945 | 0.000896078151516513 | 0.0444204608001559 | 198 | 5 | 11 | 12 | 8 | 10 | 11 | 16 | 5 | 12 | 33 | 21 | 22 |
| MER53_dup5640 | 12.2453879232216 | 1.23720898786248 | 0.374878073886493 | 3.30029701400217 | 0.000965825531898948 | 0.0466817635614924 | 199 | 8 | 10 | 9 | 8 | 11 | 17 | 6 | 5 | 13 | 21 | 26 | 19 |
| MIRb_dup143668 | 14.0530751090695 | 1.37691469470551 | 0.36913350560242 | 3.73012656344594 | 0.000191383609753859 | 0.0159338573850566 | 200 | 9 | 8 | 17 | 7 | 10 | 14 | 8 | 6 | 17 | 31 | 27 | 20 |
| L2a_dup142441 | 16.3013687553459 | 1.57769917802771 | 0.450778820487287 | 3.49994078320325 | 0.000465361523729895 | 0.0290784960938639 | 201 | 5 | 12 | 14 | 20 | 9 | 17 | 9 | 1 | 19 | 42 | 33 | 24 |
| MER31A_dup117 | 15.7551771475468 | 1.45405463912827 | 0.421785829152352 | 3.4473766983837 | 0.000566058722654022 | 0.0331181236342842 | 202 | 7 | 14 | 11 | 8 | 10 | 19 | 19 | 1 | 19 | 23 | 31 | 39 |
| L2c_dup114271 | 13.8906839832473 | 1.59455820841152 | 0.357189636067235 | 4.46417826107188 | 8.03766269112479e-06 | 0.00155616736102716 | 203 | 7 | 9 | 11 | 12 | 8 | 14 | 6 | 4 | 26 | 34 | 17 | 22 |
| MLT1F2_dup2843 | 12.4970962212401 | 1.44647564255609 | 0.354740233759467 | 4.07756297397289 | 4.55102058551645e-05 | 0.00561011076983783 | 204 | 13 | 9 | 6 | 11 | 7 | 12 | 9 | 2 | 25 | 28 | 14 | 18 |
| MER102a_dup198 | 14.0870292415571 | 1.31659181698147 | 0.386283937593023 | 3.40835248078212 | 0.000653564123734914 | 0.0362622254961759 | 205 | 21 | 10 | 8 | 14 | 13 | 13 | 5 | 2 | 16 | 26 | 19 | 33 |
| MER4D_dup1051 | 14.9984591005937 | 1.58332550938824 | 0.381225938841037 | 4.15324705921559 | 3.2779050415056e-05 | 0.00445693938860623 | 206 | 11 | 9 | 13 | 10 | 14 | 15 | 7 | 1 | 25 | 29 | 16 | 38 |
| MER52A_dup152 | 14.088525023786 | 1.26262152956214 | 0.343626335840295 | 3.6744026806751 | 0.000238406490274146 | 0.0185449930180485 | 207 | 12 | 12 | 14 | 18 | 6 | 8 | 11 | 4 | 20 | 24 | 21 | 28 |
| THE1C_dup5029 | 14.0587148798614 | 1.13561201571283 | 0.327784312788898 | 3.46450995793747 | 0.000531198472810025 | 0.0317101233846599 | 208 | 12 | 13 | 9 | 16 | 9 | 13 | 13 | 4 | 15 | 29 | 19 | 27 |
| MIR3_dup72712 | 14.7653594313426 | 1.06380011759513 | 0.311811705484397 | 3.41167473473303 | 0.00064565105340056 | 0.0359985079011148 | 209 | 11 | 14 | 14 | 12 | 7 | 15 | 13 | 8 | 19 | 21 | 24 | 27 |
| MIR_dup168993 | 14.8466264048556 | 1.22736466458858 | 0.347992964756344 | 3.52698125793419 | 0.00042032651560437 | 0.0271706093675624 | 210 | 8 | 13 | 12 | 18 | 8 | 13 | 11 | 6 | 26 | 25 | 25 | 18 |

| | | | | | | | | | | | | | | | | | | | |
|---|---|---|---|---|---|---|---|---|---|---|---|---|---|---|---|---|---|---|---|
| MIR1_Amn_dup7070 | 15.0231368014218 | 1.71015140101258 | 0.364918861951011 | 4.68638806958177 | 2.78068781841344e-06 | 0.000716512335013653 | 211 | 15 | 6 | 10 | 15 | 7 | 10 | 10 | 3 | 19 | 30 | 40 | 24 |
| L1PA2_dup565 | 16.5967344956515 | 1.64628892635896 | 0.32383054766075 | 5.08379749301367 | 3.69962381855738e-07 | 0.000147209145339541 | 212 | 18 | 10 | 9 | 15 | 6 | 11 | 12 | 5 | 20 | 28 | 31 | 47 |
| MLT2B2_dup165 | 17.2608268644506 | 1.56904266249138 | 0.330033616874026 | 4.75419042869892 | 1.99243175761754e-06 | 0.000561775339842958 | 213 | 15 | 8 | 12 | 13 | 10 | 20 | 9 | 5 | 21 | 27 | 39 | 39 |
| MIR_dup113595 | 14.2138784924803 | 1.2576871642024 | 0.340211564242385 | 3.69677958185559 | 0.000218351767411536 | 0.017434875720975 | 214 | 9 | 18 | 10 | 9 | 12 | 12 | 9 | 6 | 16 | 24 | 20 | 34 |
| MER113_dup2794 | 14.4756192655054 | 1.27613064333072 | 0.344256653221221 | 3.70691642816464 | 0.000209798135952093 | 0.0169896393973003 | 215 | 16 | 14 | 8 | 10 | 10 | 16 | 6 | 7 | 15 | 22 | 25 | 34 |
| MLT2B3_dup337 | 15.0092435151119 | 1.38308818352073 | 0.358232212718523 | 3.86087050359002 | 0.000112983775324601 | 0.010968599511786 | 216 | 15 | 14 | 12 | 10 | 14 | 13 | 5 | 4 | 16 | 22 | 30 | 35 |
| MIR3_dup27870 | 12.506873485053 | 1.37245726191986 | 0.371492156524432 | 3.69444478925251 | 0.000220367767017055 | 0.0175077224342705 | 217 | 11 | 7 | 9 | 11 | 12 | 11 | 8 | 3 | 24 | 14 | 17 | 29 |
| AluSz6_dup34478 | 14.1677233432583 | 1.13981537515022 | 0.335373393669184 | 3.39864579798642 | 0.000677203572545956 | 0.0370346773783084 | 218 | 17 | 12 | 13 | 13 | 11 | 9 | 10 | 4 | 21 | 17 | 26 | 25 |
| MSTA_dup19190 | 13.4722026285809 | 1.37352208496485 | 0.328890754603226 | 4.17622589185236 | 2.96385504358764e-05 | 0.00414358080024386 | 219 | 13 | 8 | 13 | 10 | 9 | 10 | 10 | 4 | 20 | 24 | 25 | 22 |
| L1MC2_dup1708 | 12.8481338329834 | 1.31572633461983 | 0.359122614110353 | 3.66372454121068 | 0.000248574107132105 | 0.0190674450340605 | 220 | 10 | 15 | 8 | 14 | 11 | 7 | 5 | 5 | 19 | 20 | 22 | 24 |
| LTR78_dup1733 | 13.2503690542728 | 1.30076742476699 | 0.37308746361127 | 3.48649459345622 | 0.000489395275917286 | 0.0300670406205563 | 221 | 10 | 9 | 12 | 12 | 17 | 6 | 9 | 3 | 21 | 22 | 16 | 29 |
| MamTip2_dup5103 | 14.8208036154616 | 1.34417950348028 | 0.38690964303437 | 3.47414319513579 | 0.000512487456114559 | 0.0310180227529039 | 222 | 11 | 10 | 14 | 19 | 16 | 5 | 7 | 4 | 22 | 18 | 26 | 35 |
| AluSz_dup75643 | 15.3554812766166 | 1.38998247688134 | 0.340252814896031 | 4.08514615024144 | 4.40490817778844e-05 | 0.00549046768607673 | 223 | 9 | 8 | 11 | 13 | 13 | 12 | 19 | 3 | 24 | 30 | 21 | 30 |
| L2a_dup137067 | 17.0658274594633 | 1.63824689854013 | 0.43059389054552 | 3.8046217898369 | 0.000142021022329172 | 0.0128302618251898 | 224 | 6 | 5 | 10 | 14 | 15 | 14 | 23 | 2 | 28 | 41 | 22 | 34 |
| L1MA1_dup3692 | 16.4655451818108 | 1.50416817291541 | 0.412217226242663 | 3.64896971100849 | 0.000263294141798323 | 0.019794561726491 | 225 | 12 | 3 | 14 | 23 | 12 | 13 | 11 | 3 | 27 | 36 | 20 | 33 |
| L2a_dup137031 | 18.2074128054269 | 1.52024872020136 | 0.3740486153917 | 4.06430784033077 | 4.81752091742295e-05 | 0.0058785119600878 | 226 | 9 | 8 | 12 | 27 | 12 | 13 | 16 | 4 | 18 | 47 | 33 | 33 |
| AluSx1_dup83840 | 16.3936874813077 | 1.26462622938123 | 0.35818237010216 | 3.53067692589264 | 0.000414497696787624 | 0.0269328424248574 | 227 | 10 | 7 | 15 | 17 | 12 | 12 | 21 | 5 | 21 | 39 | 25 | 22 |
| MLT1H1_dup109 | 15.4687921949511 | 1.43991840283515 | 0.360790356100147 | 3.99101134076727 | 6.57921338166716e-05 | 0.00741047468173125 | 228 | 7 | 9 | 12 | 20 | 9 | 11 | 14 | 5 | 20 | 40 | 23 | 24 |
| MIRc_dup79387 | 14.1788122072398 | 1.48935084345315 | 0.440145599841876 | 3.38376856201266 | 0.000714982442396025 | 0.0382981050831384 | 229 | 4 | 5 | 15 | 15 | 13 | 15 | 9 | 2 | 17 | 34 | 21 | 29 |
| MIRb_dup174424 | 15.6448409056006 | 1.50698222667911 | 0.434248724797975 | 3.47032044223089 | 0.000519837781452825 | 0.0312480190412212 | 230 | 3 | 4 | 16 | 19 | 16 | 12 | 12 | 4 | 25 | 28 | 23 | 34 |
| MIRc_dup20057 | 14.6665174068428 | 1.40260472438946 | 0.414924734243922 | 3.38038349761263 | 0.000723847546703678 | 0.0385575617523961 | 231 | 3 | 7 | 16 | 19 | 11 | 11 | 15 | 3 | 23 | 28 | 22 | 26 |
| AluSg_dup27051 | 13.7392392343647 | 1.69799502939629 | 0.479887597606327 | 3.53831821840337 | 0.000402684442941932 | 0.0264073769498129 | 232 | 3 | 5 | 13 | 17 | 10 | 10 | 11 | 1 | 28 | 29 | 24 | 18 |
| MIRc_dup79328 | 15.262797828298 | 1.90725895663173 | 0.388299818490098 | 4.91182036614928 | 9.02347037623993e-07 | 0.000296533043249678 | 233 | 4 | 8 | 12 | 16 | 8 | 7 | 13 | 3 | 23 | 42 | 27 | 27 |
| MLT1B_dup14514 | 17.1161896854279 | 1.68921581685584 | 0.462529916513416 | 3.65212228776306 | 0.000260081973729978 | 0.0196342318143951 | 234 | 4 | 4 | 19 | 15 | 16 | 18 | 7 | 3 | 23 | 36 | 34 | 35 |
| MamRep38_dup1565 | 18.0392719760499 | 1.70967443351755 | 0.417202075065189 | 4.09795285234478 | 4.16820151763575e-05 | 0.00527837614856496 | 235 | 4 | 5 | 15 | 15 | 17 | 17 | 12 | 4 | 27 | 49 | 30 | 28 |
| MER20_dup12363 | 16.2509467726912 | 1.6754376901064 | 0.376861525531999 | 4.4457647613172 | 8.75797432056373e-06 | 0.00167383530737418 | 236 | 5 | 7 | 11 | 9 | 17 | 15 | 14 | 4 | 26 | 23 | 38 | 33 |
| MLT1C_dup15277 | 15.4079109069503 | 1.6900502304584 | 0.422202532086551 | 4.00293722092585 | 6.25609072298898e-05 | 0.00712521917277737 | 237 | 4 | 4 | 18 | 9 | 17 | 10 | 8 | 6 | 26 | 28 | 30 | 29 |
| MER58B_dup3504 | 15.7245193409334 | 1.38898934506762 | 0.419393653028659 | 3.31189882116005 | 0.000926650532472739 | 0.045415610216221 | 238 | 4 | 5 | 9 | 17 | 16 | 20 | 9 | 8 | 20 | 38 | 25 | 24 |
| MIRb_dup174369 | 15.0573671965682 | 1.36379959037884 | 0.402122818061133 | 3.3915001316128 | 0.000695111297381171 | 0.0376163185732682 | 239 | 3 | 9 | 9 | 19 | 18 | 12 | 9 | 7 | 17 | 30 | 21 | 36 |
| L3_dup37619 | 13.8412119068738 | 1.99130487883764 | 0.37748206894466 | 5.2752303822134 | 1.32589413789769e-07 | 6.4517629148508e-05 | 240 | 4 | 4 | 8 | 10 | 12 | 7 | 9 | 6 | 20 | 40 | 22 | 29 |
| X6B_LINE_dup719 | 14.4793747237785 | 1.79756387920434 | 0.409189691857504 | 4.39298426860256 | 1.118052204427e-05 | 0.00203547092451865 | 241 | 3 | 5 | 9 | 15 | 11 | 8 | 9 | 9 | 17 | 43 | 26 | 24 |
| L2a_dup137047 | 14.4914821169068 | 1.50106610026003 | 0.37336059384516 | 4.02041920064696 | 5.80946725224178e-05 | 0.00676291337965716 | 242 | 7 | 8 | 11 | 13 | 17 | 9 | 4 | 8 | 20 | 33 | 20 | 29 |
| L2c_dup114280 | 14.4430877654422 | 1.72051818929268 | 0.42097958041554 | 4.08693976936932 | 4.37100536041265e-05 | 0.00545964956162073 | 243 | 5 | 4 | 11 | 11 | 19 | 10 | 4 | 6 | 24 | 32 | 23 | 28 |
| MIRb_dup174473 | 14.975083037423 | 1.7181287667223 | 0.366371676943768 | 4.68957857510913 | 2.73768293555875e-06 | 0.000710382446875096 | 244 | 5 | 7 | 14 | 10 | 14 | 12 | 5 | 6 | 22 | 33 | 25 | 32 |
| MER5A_dup27617 | 15.1091945388633 | 1.62635043903647 | 0.431061153093638 | 3.77289956973501 | 0.000161361253178766 | 0.0140768967389611 | 245 | 4 | 6 | 14 | 16 | 11 | 18 | 6 | 3 | 27 | 24 | 30 | 28 |
| MIRc_dup79386 | 15.9668859324154 | 1.50926890972205 | 0.447974003224833 | 3.36909932017767 | 0.000754142488854977 | 0.0397706535135386 | 246 | 2 | 7 | 15 | 24 | 13 | 14 | 6 | 6 | 24 | 27 | 31 | 30 |
| AluSg7_dup6350 | 15.4065126114817 | 1.67087469389412 | 0.404059376152323 | 4.13522069406015 | 3.54613677986745e-05 | 0.00471639796168591 | 247 | 4 | 2 | 13 | 15 | 12 | 16 | 10 | 6 | 21 | 26 | 34 | 34 |
| LTR31_dup52 | 14.383988022192 | 1.44922182561949 | 0.423291008551326 | 3.42370094413136 | 0.000617745615141869 | 0.0349643430400921 | 248 | 4 | 3 | 12 | 20 | 11 | 16 | 7 | 7 | 21 | 24 | 24 | 31 |
| AluJb_dup93866 | 13.2006743153408 | 1.47267322565116 | 0.408953355571348 | 3.60107871860764 | 0.000316899600566577 | 0.0224625569651984 | 249 | 3 | 4 | 9 | 14 | 12 | 14 | 6 | 9 | 16 | 19 | 28 | 31 |
| MER5B_dup19713 | 12.534058712187 | 1.47999421563962 | 0.404948729188245 | 3.65476937933946 | 0.000257413262098355 | 0.0195127289147038 | 250 | 6 | 3 | 9 | 13 | 12 | 14 | 5 | 6 | 19 | 21 | 26 | 21 |
| THE1C_dup8257 | 18.4072064747124 | 1.40612141688053 | 0.408263315340965 | 3.44415323161278 | 0.000572850943433024 | 0.0333841759946398 | 251 | 10 | 3 | 21 | 15 | 19 | 12 | 13 | 9 | 35 | 37 | 27 | 23 |
| AluY_dup78931 | 18.1825834855977 | 1.3278820665321 | 0.35835073584723 | 3.70553743497319 | 0.000210942969854196 | 0.017036181826881 | 252 | 9 | 4 | 15 | 17 | 21 | 15 | 15 | 9 | 25 | 38 | 23 | 36 |
| L2c_dup86869 | 17.9207838854394 | 1.44596992149584 | 0.379564840974682 | 3.80954652644524 | 0.000139221906865316 | 0.0126692279259795 | 253 | 9 | 5 | 19 | 12 | 20 | 14 | 12 | 7 | 36 | 26 | 23 | 37 |
| MLT2B4_dup2514 | 16.946026296588 | 1.5156739939645 | 0.426736068832213 | 3.55178318559344 | 0.000382630007301858 | 0.025567334938907 | 254 | 6 | 5 | 18 | 22 | 16 | 9 | 10 | 6 | 34 | 31 | 29 | 21 |
| MIRb_dup174475 | 14.7456271507188 | 1.35799338428184 | 0.387980311063208 | 3.50016056371634 | 0.000464977994448424 | 0.02906928846143 | 255 | 6 | 5 | 18 | 13 | 14 | 12 | 10 | 6 | 27 | 21 | 27 | 22 |
| MIRb_dup42008 | 15.9682858143671 | 1.37567260887141 | 0.413870947861189 | 3.32391682958332 | 0.000887626634984877 | 0.0441209958822042 | 256 | 9 | 4 | 17 | 15 | 15 | 14 | 11 | 5 | 38 | 21 | 22 | 23 |
| L1ME3Cz_dup6515 | 16.0042825066413 | 1.34716208991791 | 0.368293299389625 | 3.65785120758528 | 0.000254338617406611 | 0.019343153135177 | 257 | 12 | 8 | 14 | 19 | 16 | 12 | 6 | 5 | 21 | 37 | 29 | 20 |
| L1PREC2_dup6491 | 16.2170521117857 | 1.73044586163169 | 0.42773812744259 | 4.04557309861032 | 5.21952750675974e-05 | 0.00624103362876107 | 258 | 7 | 6 | 9 | 24 | 16 | 9 | 5 | 4 | 31 | 35 | 25 | 29 |
| MIRc_dup79388 | 15.7955581623709 | 1.36018865649511 | 0.344809707776918 | 3.9447516291366 | 7.98827319526194e-05 | 0.00851221519802865 | 259 | 9 | 8 | 11 | 25 | 11 | 10 | 11 | 7 | 22 | 28 | 20 | 38 |
| MIR_dup122127 | 16.1797250036055 | 1.28341773454218 | 0.368040763486995 | 3.48716191756171 | 0.00048817570504132 | 0.0300231213005329 | 260 | 11 | 5 | 20 | 23 | 12 | 8 | 9 | 9 | 20 | 32 | 25 | 29 |
| L1ME3Cz_dup6516 | 15.4732819113137 | 1.7279051853981 | 0.376721387629716 | 4.58669255884265 | 4.50323060574845e-06 | 0.00101019849178833 | 261 | 7 | 5 | 17 | 20 | 6 | 7 | 10 | 7 | 25 | 33 | 27 | 28 |
| MER33_dup7538 | 15.0562862510361 | 1.55578725485353 | 0.357782289884756 | 4.34841885369636 | 1.3712253523393e-05 | 0.00236090873869557 | 262 | 6 | 9 | 17 | 15 | 10 | 6 | 10 | 7 | 21 | 35 | 23 | 28 |
| L2c_dup100696 | 18.77131984785 | 1.30104306178835 | 0.394785713568753 | 3.29556773984368 | 0.00098223019918167 | 0.0472443393924564 | 263 | 12 | 9 | 13 | 10 | 14 | 12 | 24 | 10 | 7 | 19 | 42 | 75 |
| MIRc_dup17832 | 15.6758598218046 | 1.71166851892383 | 0.510868600492419 | 3.35050640668457 | 0.000806639434217295 | 0.0415987487677218 | 264 | 12 | 14 | 7 | 4 | 14 | 15 | 9 | 4 | 9 | 13 | 48 | 54 |
| AluJb_dup112350 | 17.1040472810425 | 1.46939984532754 | 0.318839700651105 | 4.6085849482573 | 4.05418730158919e-06 | 0.000941580341451141 | 265 | 12 | 10 | 15 | 8 | 8 | 9 | 20 | 11 | 20 | 22 | 30 | 52 |
| AluSq_dup14387 | 19.016993016525 | 1.59412203328089 | 0.355387243921838 | 4.48559159211549 | 7.27119023376531e-06 | 0.00145513089265225 | 266 | 22 | 16 | 11 | 9 | 6 | 11 | 12 | 11 | 24 | 22 | 46 | 49 |
| L1M5_dup39668 | 18.7917675477038 | 1.50885451217577 | 0.403026216172642 | 3.74381231698691 | 0.00018124906146502 | 0.0153314927761662 | 267 | 24 | 16 | 11 | 5 | 7 | 11 | 12 | 14 | 25 | 25 | 27 | 59 |
| LTR1E_dup166 | 15.34043833635 | 1.18421031111259 | 0.354465666353886 | 3.34083219763891 | 0.000835276893325002 | 0.0424953593455443 | 268 | 13 | 15 | 5 | 7 | 11 | 13 | 22 | 8 | 20 | 30 | 21 | 27 |
| MLT1B_dup17482 | 14.9759115568203 | 1.24617480814397 | 0.376571781224114 | 3.30926232468363 | 0.000935421489185201 | 0.0457144465247499 | 269 | 19 | 18 | 5 | 7 | 9 | 16 | 14 | 5 | 13 | 32 | 20 | 33 |
| AluJr4_dup6370 | 16.4092855190909 | 1.44646283893793 | 0.362931264729038 | 3.98550078075486 | 6.7337955861969e-05 | 0.00752950998380266 | 270 | 19 | 14 | 7 | 4 | 14 | 13 | 16 | 6 | 25 | 24 | 22 | 42 |
| L2b_dup9699 | 19.0362224717999 | 1.17121454851205 | 0.351513852834878 | 3.33191576680827 | 0.000862503578042109 | 0.04336135144323 | 271 | 22 | 24 | 7 | 10 | 14 | 13 | 24 | 6 | 25 | 26 | 30 | 40 |
| MER45B_dup489 | 15.8215121372283 | 1.45375751089354 | 0.403986235277702 | 3.59853228636421 | 0.00032001812698576 | 0.0226057129777587 | 272 | 12 | 5 | 3 | 8 | 13 | 18 | 17 | 9 | 34 | 32 | 17 | 24 |
| MamRTE1_dup3275 | 16.2645625644626 | 1.57501037871842 | 0.440065038866751 | 3.57904000457379 | 0.000344858681077414 | 0.0238678719393969 | 273 | 19 | 9 | 2 | 4 | 7 | 21 | 17 | 8 | 27 | 32 | 21 | 35 |
| AluJo_dup15893 | 16.3474201635756 | 1.43464756524031 | 0.428479118303585 | 3.34823216337893 | 0.000813288498988451 | 0.0418328459101397 | 274 | 11 | 13 | 6 | 3 | 13 | 25 | 12 | 9 | 29 | 23 | 15 | 43 |
| AluJb_dup82347 | 17.75447591921 | 1.46857747297526 | 0.351472837504203 | 4.17835268125862 | 2.93628091832956e-05 | 0.00411712610131318 | 275 | 11 | 10 | 8 | 5 | 14 | 21 | 18 | 10 | 26 | 29 | 27 | 42 |
| L2c_dup47475 | 15.9551893225113 | 1.57146591799634 | 0.335258408006986 | 4.68732738826225 | 2.76795984991537e-06 | 0.000715719428984722 | 276 | 11 | 9 | 7 | 6 | 14 | 17 | 12 | 8 | 26 | 26 | 22 | 40 |
| L1PA11_dup2132 | 16.2036708228409 | 1.38797662353459 | 0.414763818500389 | 3.34642647604347 | 0.000818603854191978 | 0.0419611186319613 | 277 | 10 | 7 | 5 | 6 | 15 | 28 | 14 | 8 | 26 | 23 | 30 | 28 |
| MIRb_dup174368 | 15.1520420207705 | 1.3474325589426 | 0.384033105001056 | 3.50863647273038 | 0.000450410085209541 | 0.028498300031631 | 278 | 11 | 5 | 7 | 6 | 15 | 18 | 17 | 8 | 26 | 21 | 21 | 33 |
| AluSx1_dup65234 | 14.5538496194557 | 1.16864328668903 | 0.341806341595004 | 3.41902166365806 | 0.000628467226531363 | 0.035377634890994 | 279 | 12 | 12 | 8 | 9 | 14 | 20 | 5 | 9 | 25 | 21 | 19 | 24 |
| MIR_dup104597 | 16.7251920827303 | 1.19874556780605 | 0.312991203269137 | 3.82996568365298 | 0.000128161130374232 | 0.0119683784646888 | 280 | 17 | 14 | 8 | 8 | 11 | 22 | 10 | 11 | 28 | 27 | 22 | 27 |
| L2c_dup79236 | 14.5982852017913 | 1.31576634597456 | 0.341875932751234 | 3.84866619707909 | 0.000118762709691198 | 0.0113627308886939 | 281 | 10 | 11 | 9 | 10 | 13 | 11 | 7 | 10 | 26 | 37 | 16 | 16 |
| L3_dup15369 | 14.0786986733748 | 1.30894808348706 | 0.349114038787835 | 3.7493424441821 | 0.000177298845229944 | 0.0150922641332448 | 282 | 10 | 7 | 8 | 6 | 18 | 16 | 10 | 6 | 23 | 32 | 19 | 17 |

| | | | | | | | | | | | | | | | | | | | |
|---|---|---|---|---|---|---|---|---|---|---|---|---|---|---|---|---|---|---|---|
| MIRb_dup116671 | 14.3781179265884 | 1.50611192231287 | 0.326496528477255 | 4.61294926882444 | 3.96995224854615e-06 | 0.000928132087505741 | 283 | 12 | 9 | 9 | 8 | 14 | 15 | 6 | 5 | 19 | 36 | 22 | 23 |
| MER39B_dup566 | 15.2900699808481 | 1.61205372989681 | 0.377687087952273 | 4.2682256855456 | 1.97033930823066e-05 | 0.00306427684337421 | 284 | 7 | 11 | 13 | 2 | 13 | 12 | 11 | 8 | 28 | 29 | 22 | 30 |
| MIR3_dup27874 | 17.0197774324018 | 1.35322972088277 | 0.346298035630864 | 3.90770256151623 | 9.31779146795311e-05 | 0.00956072103766093 | 285 | 15 | 9 | 8 | 6 | 19 | 10 | 14 | 14 | 28 | 31 | 27 | 26 |
| MIRc_dup32866 | 15.6496964593737 | 1.38011809182872 | 0.307502421810786 | 4.48815356868292 | 7.18431346022224e-06 | 0.00144381642787672 | 286 | 16 | 11 | 8 | 9 | 12 | 9 | 12 | 11 | 20 | 32 | 26 | 28 |
| MIRb_dup143671 | 14.7318753995299 | 1.40275395155453 | 0.409656940569064 | 3.42421624690632 | 0.000616575365407112 | 0.034914717253013 | 287 | 14 | 3 | 8 | 12 | 13 | 20 | 8 | 6 | 20 | 16 | 30 | 36 |
| L2_dup42864 | 13.887869079203 | 1.35990282899928 | 0.378581585386719 | 3.59209977846689 | 0.000328024199308865 | 0.0229970672153056 | 288 | 14 | 8 | 6 | 8 | 10 | 18 | 11 | 6 | 17 | 18 | 19 | 42 |
| LTR33_dup4819 | 16.7020546747578 | 1.24685105020299 | 0.367480478121739 | 3.39297221059436 | 0.000691386518722369 | 0.0374744092446271 | 289 | 18 | 8 | 9 | 14 | 12 | 26 | 8 | 7 | 23 | 24 | 20 | 42 |
| L1ME4a_dup27399 | 14.0443557839099 | 1.26680585887535 | 0.341354485127356 | 3.71111531873593 | 0.000206348066839236 | 0.0167919620766515 | 290 | 10 | 4 | 11 | 10 | 14 | 14 | 12 | 8 | 16 | 28 | 22 | 27 |
| MIR_dup137200 | 15.789019876733 | 1.4358607771471 | 0.336472449206581 | 4.26739479126131 | 1.97768989236942e-05 | 0.0030709302228926 | 291 | 8 | 6 | 13 | 12 | 15 | 15 | 9 | 9 | 16 | 33 | 28 | 34 |
| L1HS_dup1188 | 16.6300065288877 | 1.55882299316292 | 0.364034685393515 | 4.28207271369721 | 1.85160450860901e-05 | 0.00293279376840857 | 292 | 8 | 8 | 12 | 8 | 14 | 20 | 9 | 8 | 22 | 21 | 41 | 36 |
| L2c_dup64873 | 14.9656983745697 | 1.35768668539222 | 0.402997296546439 | 3.36897219169252 | 0.000754490404913693 | 0.039780184774561 | 293 | 10 | 7 | 11 | 8 | 17 | 21 | 8 | 5 | 21 | 15 | 28 | 37 |
| L1PA7_dup10264 | 15.5069638993221 | 1.44802628149499 | 0.404291506789519 | 3.58163913210489 | 0.00034144521792036 | 0.0236867416394397 | 294 | 11 | 11 | 5 | 6 | 20 | 24 | 10 | 4 | 16 | 28 | 31 | 29 |
| MIRb_dup110313 | 16.1903903527529 | 1.249333082372 | 0.377649739103931 | 3.30817938690057 | 0.000939046383304171 | 0.0458061358593059 | 295 | 11 | 9 | 6 | 9 | 19 | 20 | 17 | 7 | 14 | 27 | 36 | 30 |
| AluSp_dup23269 | 16.7727540805661 | 1.7357463552488 | 0.344766496680654 | 5.03455635034215 | 4.78956774701844e-07 | 0.000179191586027052 | 296 | 13 | 9 | 4 | 7 | 21 | 13 | 12 | 5 | 25 | 39 | 32 | 27 |
| L1M6_dup614 | 16.6443538715551 | 1.38189967200325 | 0.369852976455061 | 3.73634865737293 | 0.000186711738157767 | 0.0156529341744761 | 297 | 16 | 7 | 8 | 4 | 19 | 21 | 15 | 6 | 24 | 33 | 29 | 24 |
| L3b_dup8255 | 16.8548973628786 | 1.39419458768756 | 0.368140695929687 | 3.78712433344737 | 0.000152400850660272 | 0.0134907636946463 | 298 | 13 | 7 | 5 | 17 | 20 | 17 | 12 | 6 | 18 | 25 | 32 | 43 |
| L1MC1_dup4505 | 14.8967065579018 | 1.17468128931267 | 0.342835468205127 | 3.42637036786936 | 0.000611705668905485 | 0.0348227569164275 | 299 | 13 | 10 | 3 | 13 | 15 | 18 | 12 | 8 | 21 | 25 | 26 | 21 |
| MLT1I_dup5739 | 15.7258436964671 | 1.16567717963581 | 0.33145825835818 | 3.51681441099036 | 0.000436759046952509 | 0.0279068389329113 | 300 | 14 | 13 | 5 | 12 | 16 | 17 | 13 | 8 | 16 | 24 | 33 | 27 |
| AluJb_dup98888 | 13.728082894603 | 1.35753522832716 | 0.381156425604726 | 3.56162230814647 | 0.000368570339774063 | 0.0249359789786406 | 301 | 15 | 14 | 2 | 13 | 8 | 9 | 9 | 9 | 16 | 20 | 26 | 32 |
| LTR84b_dup63 | 12.495456672973 | 1.25048020697351 | 0.346258039069562 | 3.61141133454605 | 0.000304535202474362 | 0.0218649330148206 | 302 | 10 | 9 | 4 | 15 | 8 | 11 | 11 | 7 | 16 | 17 | 22 | 28 |
| MER5B_dup20618 | 13.8977318864313 | 1.43040903183645 | 0.317985406451474 | 4.49834804621681 | 6.84835180060451e-06 | 0.00138738284900527 | 303 | 8 | 10 | 7 | 13 | 9 | 11 | 11 | 8 | 18 | 20 | 30 | 29 |
| L2_dup19474 | 15.9956947364601 | 1.91984531390565 | 0.37025009472808 | 5.18526623285708 | 2.15706499157681e-07 | 9.61170386677407e-05 | 304 | 11 | 11 | 5 | 10 | 18 | 8 | 9 | 2 | 28 | 36 | 37 | 21 |
| MER47B_dup219 | 13.6045595727056 | 1.84133334098582 | 0.347421132210599 | 5.30000385776661 | 1.1580023395928e-07 | 5.7644775041161e-05 | 305 | 10 | 6 | 6 | 7 | 12 | 9 | 11 | 3 | 25 | 29 | 28 | 21 |
| MSTA_dup19087 | 16.2174291333273 | 1.55205755183774 | 0.375849492931372 | 4.12946560000052 | 3.6360739882134e-05 | 0.00479594934146023 | 306 | 6 | 11 | 9 | 17 | 16 | 9 | 9 | 8 | 23 | 36 | 37 | 18 |
| L1MC5a_dup6744 | 14.3844674982361 | 1.21961822738158 | 0.354243023636077 | 3.44288566324606 | 0.00057554258688053 | 0.0334918210755379 | 307 | 10 | 16 | 12 | 11 | 16 | 7 | 10 | 5 | 16 | 24 | 31 | 22 |
| MIRc_dup79394 | 14.0834490477657 | 1.40112350605155 | 0.349349423635123 | 4.01066499973663 | 6.05479693406155e-05 | 0.0069579262646345 | 308 | 8 | 9 | 11 | 13 | 13 | 12 | 11 | 3 | 19 | 25 | 32 | 20 |
| L1MA9_dup608 | 29.6401335035717 | 1.30812890264501 | 0.320365044547427 | 4.08324480123284 | 4.44111956999116e-05 | 0.0055182592020946 | 309 | 49 | 44 | 14 | 14 | 22 | 23 | 12 | 6 | 45 | 57 | 29 | 55 |
| L1MA9_dup610 | 35.8285302788585 | 1.47593833549463 | 0.330520199434044 | 4.46550116459421 | 7.98815377810059e-06 | 0.00155253948901038 | 310 | 45 | 57 | 12 | 13 | 33 | 40 | 11 | 8 | 49 | 63 | 46 | 70 |
| L1MCc_dup1255 | 33.3915556331116 | 1.51534988593507 | 0.340097179703179 | 4.45563790695824 | 8.36441062464206e-06 | 0.00161286713978207 | 311 | 31 | 29 | 5 | 14 | 43 | 48 | 15 | 11 | 51 | 53 | 50 | 64 |
| L3b_dup3380 | 28.4834296460283 | 1.51217655919958 | 0.322453641615539 | 4.68959367809697 | 2.73748088806638e-06 | 0.000710382446875096 | 312 | 25 | 20 | 4 | 16 | 38 | 25 | 16 | 13 | 42 | 59 | 53 | 39 |
| MER21B_dup1166 | 28.5920768478543 | 1.68262656743467 | 0.330216228038391 | 5.0955296092809 | 3.47767658796489e-07 | 0.000139816406724132 | 313 | 32 | 26 | 4 | 7 | 29 | 30 | 11 | 13 | 47 | 60 | 52 | 37 |
| MIR3_dup45882 | 25.158184401657 | 1.44874421397802 | 0.396737626047199 | 3.65164309826682 | 0.000260567841529666 | 0.0196516879938567 | 314 | 29 | 27 | 11 | 7 | 33 | 39 | 9 | 4 | 29 | 36 | 46 | 46 |
| X7A_LINE_dup507 | 29.6580620344896 | 1.48311593046651 | 0.425615875301357 | 3.48463489388498 | 0.000492808985982944 | 0.0301898601598025 | 315 | 38 | 23 | 10 | 7 | 50 | 49 | 6 | 8 | 36 | 43 | 49 | 51 |
| L1ME4b_dup7265 | 29.3576417586832 | 1.28541018555986 | 0.364786802079451 | 3.52372996564687 | 0.000425517666417771 | 0.027387131905178 | 316 | 37 | 20 | 10 | 6 | 54 | 34 | 19 | 7 | 44 | 51 | 31 | 52 |
| L1M5_dup25451 | 31.2146537494163 | 1.73720197133005 | 0.346644865435125 | 5.01147469514495 | 5.40144873122791e-07 | 0.000197731249200116 | 317 | 52 | 29 | 4 | 8 | 35 | 32 | 11 | 6 | 48 | 69 | 46 | 46 |
| L1P2_dup1291 | 28.0523406066101 | 1.83342856313528 | 0.286357572484597 | 6.4025845282436 | 1.52768498754882e-10 | 2.07718018312737e-07 | 318 | 26 | 21 | 6 | 9 | 34 | 27 | 8 | 7 | 59 | 66 | 27 | 50 |
| MIRc_dup83169 | 18.7226186021979 | 1.38693626093912 | 0.390726431270152 | 3.54963511536841 | 0.000385765443465778 | 0.0257227955943545 | 319 | 18 | 25 | 4 | 8 | 17 | 20 | 6 | 12 | 23 | 28 | 28 | 44 |
| AluSq2_dup17516 | 20.1072485321978 | 1.13279733416799 | 0.333126272036943 | 3.40050434101566 | 0.000672616679115761 | 0.0368600756169689 | 320 | 24 | 19 | 8 | 10 | 23 | 25 | 6 | 13 | 25 | 34 | 22 | 41 |
| Tigger6a_dup400 | 19.8586670420456 | 1.26797852862399 | 0.378515240316209 | 3.34987443983691 | 0.000808481997734688 | 0.0416396228209909 | 321 | 23 | 17 | 7 | 8 | 27 | 22 | 8 | 11 | 21 | 26 | 31 | 49 |
| AluSc_dup31642 | 25.3234874960153 | 1.27421802471932 | 0.356560641799866 | 3.57363622156178 | 0.000352057956520268 | 0.0241902346295616 | 322 | 24 | 20 | 14 | 17 | 23 | 37 | 7 | 12 | 35 | 29 | 34 | 66 |
| L1ME3A_dup2031 | 23.6142934197154 | 1.19576161275532 | 0.341808431179265 | 3.49833855364496 | 0.000468166435412181 | 0.0291847866493547 | 323 | 24 | 27 | 17 | 21 | 17 | 20 | 9 | 10 | 23 | 26 | 40 | 68 |
| L1PA14_dup1 | 21.8647342281914 | 1.90121421401979 | 0.481576715892977 | 3.94789480320787 | 7.88414322965259e-05 | 0.00843938667805609 | 324 | 38 | 27 | 7 | 7 | 23 | 16 | 4 | 2 | 24 | 36 | 32 | 63 |
| MER113_dup2322 | 24.9561183448908 | 1.45345518310455 | 0.439376537893709 | 3.30799452804683 | 0.0009396664560868 | 0.0458269880221418 | 325 | 36 | 22 | 13 | 19 | 25 | 23 | 10 | 2 | 19 | 32 | 56 | 66 |
| L2c_dup19449 | 24.8774224075203 | 1.59488790005436 | 0.341953667973062 | 4.66404676840608 | 3.10050867537291e-06 | 0.000775766180094935 | 326 | 22 | 18 | 17 | 18 | 24 | 28 | 5 | 3 | 35 | 52 | 39 | 50 |
| MLT2B1_dup1917 | 23.6086638082079 | 1.12884743023449 | 0.341168391942567 | 3.30876909143601 | 0.000937070867521711 | 0.0457566724078229 | 327 | 26 | 23 | 18 | 24 | 21 | 29 | 10 | 2 | 25 | 48 | 33 | 41 |
| L1MEf_dup3353 | 32.0254364154591 | 1.75655224459036 | 0.458571628899597 | 3.83048608743075 | 0.000127890364661188 | 0.0119528832159892 | 328 | 36 | 33 | 29 | 28 | 9 | 12 | 4 | 7 | 47 | 64 | 57 | 75 |
| L1PA3_dup1014 | 33.8127844872666 | 1.84924874828177 | 0.522846590000959 | 3.53688593106896 | 0.000404874490789771 | 0.0264883875681778 | 329 | 44 | 32 | 17 | 22 | 26 | 40 | 1 | 2 | 48 | 49 | 65 | 80 |
| THE1B-int_dup524 | 26.5810723372806 | 1.35138747997468 | 0.348104820217197 | 3.88212803009017 | 0.000103546332820118 | 0.0103120186895199 | 330 | 41 | 28 | 13 | 18 | 21 | 18 | 4 | 4 | 54 | 81 | 17 | 23 |
| L1MD3_dup2457 | 38.4691163158353 | 1.75335704059107 | 0.291054617905095 | 6.02415125109883 | 1.6999933610698e-09 | 1.73607991542109e-06 | 331 | 35 | 18 | 18 | 19 | 40 | 25 | 11 | 10 | 85 | 126 | 27 | 47 |
| L1PA7_dup4487 | 33.1815520471661 | 2.39863626892641 | 0.264263194524584 | 9.07669444184846 | 1.11922267739871e-19 | 1.47952532886841e-15 | 332 | 28 | 15 | 5 | 8 | 18 | 26 | 12 | 7 | 97 | 88 | 38 | 49 |
| THE1C_dup8258 | 18.5146618056774 | 1.51048147309004 | 0.316534737118728 | 4.7719295734783 | 1.82469312469556e-06 | 0.000523737551566719 | 333 | 7 | 11 | 14 | 20 | 13 | 16 | 8 | 10 | 26 | 37 | 30 | 38 |
| L2_dup47004 | 17.9683033861727 | 1.55777280473897 | 0.349401142326636 | 4.45840787573241 | 8.25706704050555e-06 | 0.00159345991404715 | 334 | 8 | 7 | 14 | 17 | 14 | 11 | 9 | 13 | 36 | 30 | 32 | 26 |
| MER58A_dup10300 | 17.1708597643715 | 1.35627454852576 | 0.348020417334787 | 3.89711201116415 | 9.73466021618511e-05 | 0.00986509139608339 | 335 | 6 | 8 | 15 | 21 | 14 | 10 | 12 | 12 | 28 | 33 | 25 | 27 |
| MIR_dup137210 | 21.0747974799376 | 1.87995016417648 | 0.306987168418651 | 6.1238721274914 | 9.13282008672276e-10 | 1.01077472885744e-06 | 336 | 8 | 8 | 16 | 15 | 17 | 11 | 11 | 10 | 39 | 34 | 35 | 56 |
| MIRc_dup79383 | 21.9732102125199 | 1.80173098610085 | 0.306403893867533 | 5.88024833287461 | 4.09651416231657e-09 | 3.67829871270407e-06 | 337 | 12 | 8 | 16 | 22 | 15 | 11 | 13 | 8 | 33 | 56 | 41 | 37 |
| MIRc_dup79323 | 21.084676777823 | 1.52812702677931 | 0.345733255140296 | 4.41995961932908 | 9.87193461798082e-06 | 0.00183610546772347 | 338 | 12 | 10 | 25 | 15 | 16 | 12 | 16 | 7 | 28 | 48 | 46 | 26 |
| MLT1C_dup10070 | 20.9985082591584 | 1.23478577894399 | 0.311261565862245 | 3.96703581286508 | 7.27720609334086e-05 | 0.00795763824028531 | 339 | 13 | 15 | 25 | 20 | 18 | 17 | 12 | 6 | 28 | 41 | 35 | 32 |
| L3_dup37629 | 19.0786957945381 | 1.24426132853256 | 0.345907923534077 | 3.59708825348716 | 0.000321799323616981 | 0.0226912052413817 | 340 | 9 | 9 | 17 | 14 | 24 | 18 | 9 | 11 | 30 | 39 | 27 | 26 |
| MIRb_dup114607 | 18.1166102959652 | 1.16403000107119 | 0.345660306033686 | 3.36755473727363 | 0.00075837970031098 | 0.0399321567094925 | 341 | 11 | 8 | 14 | 20 | 23 | 16 | 9 | 10 | 18 | 39 | 29 | 30 |
| LTR90B_dup438 | 19.1853541579276 | 1.40143921899107 | 0.291202096263985 | 4.81260003609526 | 1.48979257422945e-06 | 0.000439814123905212 | 342 | 14 | 8 | 16 | 16 | 18 | 10 | 13 | 12 | 27 | 37 | 30 | 37 |
| MIR_dup112778 | 18.3185754757669 | 1.41940972517987 | 0.345725400185381 | 4.10559861791693 | 4.03269191033812e-05 | 0.00515158666763517 | 343 | 15 | 7 | 13 | 17 | 16 | 9 | 12 | 11 | 29 | 48 | 26 | 21 |
| MER2_dup12 | 19.6090904786627 | 1.33891739347581 | 0.344562311319897 | 3.88584981435402 | 0.000101972537228708 | 0.0102250136297607 | 344 | 13 | 10 | 13 | 6 | 16 | 19 | 24 | 11 | 35 | 28 | 27 | 40 |
| AluSz6_dup20707 | 19.0829904422201 | 1.4107435484133 | 0.300542268289529 | 4.69399381472109 | 2.67922137062301e-06 | 0.000704044193033047 | 345 | 11 | 12 | 8 | 11 | 16 | 18 | 21 | 10 | 24 | 32 | 37 | 39 |
| L1MEc_dup2478 | 19.4372362283095 | 1.34126036628577 | 0.336659928830058 | 3.98402141575579 | 6.7758762315646e-05 | 0.00755880284011191 | 346 | 9 | 15 | 10 | 8 | 18 | 22 | 21 | 9 | 21 | 42 | 34 | 34 |
| AluSz_dup387 | 19.2937285378555 | 1.43658079299375 | 0.391143524055159 | 3.6727715138936 | 0.00023993400156925 | 0.0185904708360133 | 347 | 4 | 9 | 14 | 11 | 24 | 15 | 21 | 8 | 30 | 40 | 25 | 38 |
| Charlie17a_dup875 | 16.7229763006962 | 1.02627218851708 | 0.313548134759322 | 3.27309294729132 | 0.00106377489935375 | 0.0498217036967053 | 348 | 9 | 13 | 16 | 10 | 18 | 18 | 14 | 9 | 22 | 29 | 20 | 30 |
| L1PB1_dup5443 | 17.887261777009 | 1.2954958689723 | 0.309033382286279 | 4.19209037997129 | 2.76395834675019e-05 | 0.00393332679863325 | 349 | 10 | 12 | 11 | 13 | 22 | 20 | 11 | 6 | 26 | 37 | 23 | 31 |
| AluJr4_dup6371 | 24.4404495222653 | 1.36117411546694 | 0.339287923649019 | 4.01185547905036 | 6.02433777740923e-05 | 0.00693165897864321 | 350 | 10 | 10 | 19 | 13 | 23 | 22 | 22 | 17 | 40 | 50 | 42 | 29 |
| AluJr_dup62313 | 21.5015118190196 | 1.14031400478926 | 0.31045162467668 | 3.67308113132551 | 0.000239643355132264 | 0.0185904708360133 | 351 | 15 | 10 | 23 | 15 | 18 | 15 | 20 | 15 | 36 | 40 | 28 | 28 |
| MamGypsy2-I_dup81 | 21.4828382718455 | 1.52028706878201 | 0.340207845534473 | 4.46870079199263 | 7.86961192596732e-06 | 0.00153361398799035 | 352 | 9 | 8 | 11 | 13 | 28 | 17 | 18 | 10 | 34 | 44 | 38 | 34 |
| MIRb_dup71176 | 19.0940885496692 | 1.49686324178579 | 0.32306361858829 | 4.63333893282913 | 3.59814725410618e-06 | 0.000857880507540631 | 353 | 9 | 8 | 14 | 15 | 18 | 15 | 14 | 9 | 34 | 37 | 36 | 24 |
| L1MC3_dup6347 | 18.9390568820922 | 1.62180300230906 | 0.295043580289473 | 5.49682525109641 | 3.8668954527306e-08 | 2.37142346751401e-05 | 354 | 11 | 9 | 11 | 13 | 18 | 18 | 9 | 8 | 31 | 37 | 34 | 34 |

| | | | | | | | | | | | | | | | | | | |
|---|---|---|---|---|---|---|---|---|---|---|---|---|---|---|---|---|---|---|
| L2c_dup20171 | 20.1561043724055 | 1.41419832393711 | 0.297464801080162 | 4.75417030452623 | 1.99263020528799e-06 | 0.000561775339842958 | 355 | 10 | 11 | 10 | 17 | 19 | 21 | 14 | 10 | 33 | 36 | 36 | 31 |
| MER21A_dup256 | 21.2575324494577 | 1.00242287496742 | 0.303292186706579 | 3.30513913283635 | 0.000949292626362166 | 0.0462017556294481 | 356 | 11 | 13 | 15 | 28 | 24 | 22 | 16 | 10 | 26 | 37 | 31 | 34 |
| MLT1C_dup10071 | 20.3105416617545 | 1.09783906514682 | 0.326418565622579 | 3.36328622439999 | 0.000770204627211808 | 0.0403938968980694 | 357 | 12 | 10 | 17 | 24 | 22 | 18 | 17 | 8 | 27 | 43 | 32 | 23 |
| AluSx3_dup9560 | 18.7809474700146 | 1.11358880228219 | 0.291686832169677 | 3.81775479543899 | 0.000134671674384929 | 0.01238527951313 | 358 | 14 | 16 | 13 | 14 | 22 | 16 | 11 | 11 | 22 | 33 | 34 | 27 |
| MIRb_dup71185 | 21.5839228267639 | 1.10201450885821 | 0.306682068579338 | 3.5933451015351 | 0.000326459752289755 | 0.0229346891698666 | 359 | 15 | 13 | 13 | 13 | 31 | 24 | 14 | 12 | 29 | 39 | 35 | 28 |
| L2c_dup114253 | 19.5342106669053 | 1.56957769323806 | 0.321924743556298 | 4.87560439094851 | 1.08475812074487e-06 | 0.000342326068698618 | 360 | 12 | 7 | 13 | 19 | 22 | 12 | 14 | 5 | 24 | 38 | 36 | 45 |
| MIRb_dup174463 | 20.3541683281424 | 1.3787837191088 | 0.410753764946965 | 3.35671596165849 | 0.000788740957100599 | 0.0410673426210632 | 361 | 8 | 7 | 17 | 29 | 25 | 11 | 15 | 6 | 18 | 49 | 30 | 45 |
| MER5A_dup27632 | 21.2575779451099 | 1.29651618394925 | 0.383327624139568 | 3.38226650599329 | 0.000718903635378255 | 0.0383973612623376 | 362 | 7 | 9 | 18 | 27 | 22 | 23 | 15 | 5 | 26 | 47 | 31 | 38 |
| MLT1H_dup5226 | 20.5167978032425 | 1.42113101890143 | 0.350094382625787 | 4.05927969549991 | 4.92243242680891e-05 | 0.00596371235351056 | 363 | 8 | 13 | 16 | 21 | 17 | 26 | 11 | 4 | 28 | 38 | 33 | 43 |
| L3_dup37624 | 21.1664479381274 | 1.30529718228048 | 0.363059498246542 | 3.59527071619014 | 0.000324054424200111 | 0.0227993654703488 | 364 | 9 | 10 | 18 | 25 | 27 | 17 | 12 | 6 | 32 | 37 | 39 | 31 |
| MIRb_dup174428 | 21.6342170415166 | 1.46057032545676 | 0.363748022689704 | 4.01533543648345 | 5.9361306694707e-05 | 0.00686669201885209 | 365 | 11 | 12 | 13 | 27 | 26 | 19 | 9 | 4 | 30 | 40 | 37 | 44 |
| MER4D1_dup1228 | 20.2898569924785 | 1.68354319072526 | 0.327300883380922 | 5.14371722231469 | 2.69354878044599e-07 | 0.000113236600373145 | 366 | 6 | 16 | 17 | 15 | 18 | 14 | 12 | 4 | 30 | 38 | 39 | 44 |
| MER91C_dup1286 | 21.9287921954341 | 1.58147890176104 | 0.337017119126798 | 4.69257735588806 | 2.6978447202168e-06 | 0.000706983876428091 | 367 | 6 | 13 | 15 | 21 | 25 | 13 | 12 | 9 | 30 | 48 | 34 | 47 |
| MLT1G_dup1481 | 23.6356345943954 | 1.49065637574942 | 0.283221089697471 | 5.2632251974657 | 1.41550019215288e-07 | 6.84578473012215e-05 | 368 | 14 | 17 | 16 | 22 | 25 | 17 | 12 | 6 | 34 | 51 | 34 | 47 |
| L2c_dup127358 | 24.5209176416706 | 1.11808257294429 | 0.298116842316217 | 3.75048442166956 | 0.000176493270530028 | 0.0150468891972547 | 369 | 16 | 20 | 17 | 36 | 20 | 17 | 25 | 6 | 31 | 44 | 34 | 46 |
| ERVL-B4-int_dup2819 | 22.3177955527307 | 1.24724218550054 | 0.273305044000722 | 4.5635534831082 | 5.02949895998667e-06 | 0.00110056527813885 | 370 | 17 | 15 | 14 | 22 | 20 | 24 | 17 | 6 | 27 | 46 | 36 | 37 |
| L1ME1_dup25045 | 24.9967566131895 | 1.39039533193729 | 0.297186093967738 | 4.6785342926854 | 2.88932852606396e-06 | 0.000734512997289332 | 371 | 15 | 16 | 13 | 29 | 18 | 28 | 19 | 6 | 31 | 40 | 50 | 52 |
| AluSc8_dup16283 | 22.0649459963047 | 1.30690584372699 | 0.35066678868612 | 3.7269165084715 | 0.000193836632668505 | 0.0160762117103312 | 372 | 7 | 14 | 13 | 25 | 22 | 27 | 15 | 7 | 30 | 30 | 40 | 48 |
| L1ME1_dup25046 | 24.4606198819506 | 1.43859316751809 | 0.364623101359567 | 3.9454251860456 | 7.9658501255558e-05 | 0.00849212443537411 | 373 | 7 | 9 | 16 | 26 | 28 | 25 | 16 | 9 | 38 | 34 | 42 | 56 |
| AluSp_dup30491 | 24.3955296415995 | 1.75261447525675 | 0.256329971853559 | 6.83733729061548 | 8.06786380710902e-12 | 1.60832729164361e-08 | 374 | 13 | 14 | 14 | 23 | 16 | 16 | 14 | 9 | 31 | 40 | 53 | 65 |
| AluY_dup102111 | 23.0719765189532 | 1.51058080046723 | 0.326397943031955 | 4.62803406919552 | 3.69153298014177e-06 | 0.000874015429346082 | 375 | 9 | 12 | 9 | 24 | 25 | 21 | 13 | 11 | 33 | 43 | 46 | 40 |
| L2c_dup75995 | 21.8501900090269 | 1.54281846091625 | 0.291536716199935 | 5.29202112525062 | 1.20971914224427e-07 | 5.98436239127764e-05 | 376 | 9 | 16 | 13 | 25 | 17 | 16 | 14 | 7 | 29 | 42 | 43 | 43 |
| MIRc_dup96512 | 21.0295147662564 | 1.64904917218619 | 0.296751348558668 | 5.55700649784975 | 2.74440548088935e-08 | 1.79895401255013e-05 | 377 | 8 | 16 | 10 | 16 | 19 | 20 | 11 | 7 | 35 | 41 | 38 | 38 |
| MIR_dup137135 | 19.4402106239755 | 1.23312378688428 | 0.291653281372069 | 4.22804701899156 | 2.35728537139815e-05 | 0.00349039592397451 | 378 | 13 | 11 | 14 | 16 | 16 | 19 | 17 | 9 | 31 | 44 | 22 | 28 |
| L2a_dup105398 | 20.0968784074471 | 1.10014228104598 | 0.287532361541414 | 3.82615116833561 | 0.000130162376717425 | 0.0121077470251776 | 379 | 17 | 10 | 18 | 18 | 14 | 19 | 20 | 10 | 34 | 34 | 25 | 30 |
| L2a_dup143435 | 19.3469331637216 | 1.00384246498288 | 0.274085341131284 | 3.66251788891566 | 0.000249748305868498 | 0.0191335939991608 | 380 | 16 | 15 | 13 | 18 | 16 | 18 | 21 | 9 | 31 | 29 | 25 | 30 |
| L1PA6_dup5339 | 21.0376936183471 | 1.39742503515934 | 0.336917240691596 | 4.14768039857747 | 3.35860804787815e-05 | 0.0045329968834964 | 381 | 19 | 10 | 12 | 18 | 11 | 23 | 22 | 5 | 32 | 49 | 20 | 44 |
| LOR1-int_dup242 | 20.5273321189165 | 1.26016957168406 | 0.310563627781873 | 4.05768563654579 | 4.95614188781088e-05 | 0.00598930491435779 | 382 | 17 | 10 | 10 | 16 | 17 | 26 | 20 | 8 | 21 | 34 | 33 | 50 |
| MIR3_dup67834 | 19.5815878563294 | 1.38777702352426 | 0.300618560693239 | 4.61640498951227 | 3.90444650484953e-06 | 0.000916222834214355 | 383 | 12 | 9 | 9 | 23 | 16 | 18 | 17 | 8 | 26 | 38 | 29 | 42 |
| MLT1B_dup674 | 19.8780945874764 | 1.18719387588462 | 0.36251301355434 | 3.27490002150409 | 0.0010569937416565 | 0.0495649949257222 | 384 | 10 | 18 | 23 | 29 | 10 | 17 | 11 | 5 | 18 | 30 | 34 | 51 |
| Charlie8_dup2340 | 18.9178857212536 | 1.41080053360048 | 0.321561582253785 | 4.387341683395 | 1.1474445770715e-05 | 0.00207154664139496 | 385 | 8 | 11 | 16 | 19 | 12 | 21 | 15 | 5 | 20 | 29 | 41 | 44 |
| HERVP71A-int_dup23 | 21.650932041519 | 1.40419248138133 | 0.363120049271722 | 3.86701996818296 | 0.000110173409947571 | 0.0107793265638917 | 386 | 14 | 9 | 17 | 23 | 14 | 27 | 17 | 3 | 23 | 29 | 49 | 53 |
| MIRb_dup174365 | 19.672202164694 | 1.7770960591719 | 0.320410737807503 | 5.54630619227108 | 2.91767700464736e-08 | 1.89168820857717e-05 | 387 | 8 | 10 | 19 | 16 | 8 | 15 | 16 | 4 | 34 | 35 | 38 | 42 |
| MER5A_dup27641 | 22.9245168396507 | 1.47397957565891 | 0.302159489945267 | 4.87815085975258 | 1.0708501057585e-06 | 0.000338835770298951 | 388 | 12 | 12 | 20 | 19 | 12 | 26 | 18 | 7 | 39 | 36 | 35 | 50 |
| MLT1C_dup15275 | 23.8162603263797 | 1.58340141992593 | 0.305773953909453 | 5.17833974961389 | 2.23869256137193e-07 | 9.86459148533713e-05 | 389 | 8 | 13 | 25 | 23 | 11 | 21 | 18 | 7 | 34 | 38 | 44 | 58 |
| MIRb_dup130625 | 23.995822775219 | 1.15149747586713 | 0.328356920078168 | 3.5068469870926 | 0.000453449811994817 | 0.0285609855960214 | 390 | 18 | 19 | 10 | 23 | 21 | 16 | 31 | 11 | 38 | 40 | 47 | 23 |
| MIR3_dup27873 | 19.7868020120037 | 1.34548938143754 | 0.326171867060321 | 4.12509329380239 | 3.70584507343063e-05 | 0.00486103093628735 | 391 | 11 | 18 | 11 | 11 | 17 | 19 | 20 | 5 | 39 | 41 | 29 | 20 |
| MIR_dup57154 | 24.9667489631632 | 1.27746175080776 | 0.309692477291746 | 4.12493633032075 | 3.70837330083678e-05 | 0.00486166718149261 | 392 | 22 | 18 | 11 | 13 | 27 | 22 | 29 | 5 | 43 | 57 | 33 | 28 |
| AluSz_dup29100 | 25.0854378692436 | 1.67208731802702 | 0.306033873587531 | 5.46373281632426 | 4.66225104958778e-08 | 2.75096544669864e-05 | 393 | 20 | 17 | 5 | 12 | 23 | 17 | 16 | 15 | 54 | 39 | 42 | 43 |
| L2b_dup31005 | 23.8630282941084 | 1.79648011258298 | 0.31526165453295 | 5.69837811466291 | 1.20952559036267e-08 | 9.25407640271496e-06 | 394 | 10 | 15 | 7 | 14 | 19 | 22 | 10 | 13 | 57 | 48 | 36 | 33 |
| L2c_dup47484 | 26.4841552842569 | 1.7597704849706 | 0.265203299943969 | 6.63555274516719 | 3.23288735514693e-11 | 5.23301098372647e-08 | 395 | 20 | 23 | 7 | 17 | 18 | 20 | 13 | 11 | 41 | 63 | 52 | 40 |
| MIRb_dup71177 | 25.575593600255 | 1.84395773375112 | 0.247030140263379 | 7.4645050672162 | 8.36132968358783e-14 | 2.58382461414414e-10 | 396 | 20 | 21 | 11 | 12 | 18 | 21 | 11 | 7 | 43 | 61 | 44 | 45 |
| MIRc_dup32870 | 26.7162875325485 | 1.63520188506362 | 0.25123456411835 | 6.50866607786228 | 7.5821053274612e-11 | 1.10682922346508e-07 | 397 | 20 | 20 | 10 | 12 | 20 | 31 | 16 | 9 | 46 | 58 | 41 | 45 |
| MLT1C_dup7425 | 21.3250139537263 | 1.06551034896685 | 0.267981449865157 | 3.97606009484235 | 7.00664265118374e-05 | 0.00776526591652802 | 398 | 22 | 23 | 10 | 14 | 18 | 19 | 17 | 12 | 27 | 48 | 28 | 26 |
| MSTA_dup9124 | 21.6862664396159 | 1.55114323533914 | 0.302159480283147 | 5.13352496464979 | 2.84365291282814e-07 | 0.000117880807664774 | 399 | 21 | 24 | 8 | 6 | 12 | 21 | 16 | 8 | 32 | 55 | 32 | 32 |
| L2a_dup57886 | 19.8230575306327 | 1.31956295458123 | 0.328547135394991 | 4.01635811858414 | 5.91044198022434e-05 | 0.00685029725974895 | 400 | 16 | 21 | 7 | 13 | 12 | 27 | 18 | 4 | 32 | 37 | 22 | 39 |
| MSTD_dup3927 | 21.4719671442916 | 1.2208170201141 | 0.287788576148232 | 4.24206212926706 | 2.21475356979446e-05 | 0.00333961820607295 | 401 | 26 | 17 | 9 | 15 | 20 | 29 | 13 | 5 | 32 | 36 | 34 | 32 |
| L1PA10_dup2271 | 21.7083797068965 | 1.52701712992473 | 0.352012036384646 | 4.33796851269071 | 1.43805750693845e-05 | 0.00244993718135918 | 402 | 19 | 14 | 6 | 6 | 21 | 26 | 22 | 5 | 45 | 36 | 31 | 35 |
| L2b_dup31004 | 21.1498775285519 | 1.64131234142561 | 0.309286806569764 | 5.30676481040065 | 1.11588004907589e-07 | 5.58987777173499e-05 | 403 | 14 | 18 | 3 | 14 | 20 | 18 | 17 | 5 | 40 | 47 | 28 | 36 |
| MIRc_dup65542 | 19.587171122388 | 1.63752135595324 | 0.325379882535811 | 5.03264474494062 | 4.83759094476984e-07 | 0.000180704146773031 | 404 | 19 | 17 | 11 | 13 | 12 | 16 | 6 | 7 | 36 | 32 | 44 | 26 |
| MER5A1_dup15477 | 19.3344165551343 | 1.4699689021099 | 0.330371315231145 | 4.4494447136896 | 8.60925873473335e-06 | 0.0016507160990289 | 405 | 16 | 13 | 12 | 10 | 16 | 23 | 12 | 6 | 19 | 32 | 47 | 38 |
| MIR3_dup57101 | 18.5515558604904 | 1.40137056259391 | 0.304647108091752 | 4.59997986316632 | 4.2253178332245e-06 | 0.000970050335570093 | 406 | 19 | 10 | 12 | 10 | 15 | 17 | 14 | 8 | 23 | 35 | 41 | 27 |
| MER5B_dup7892 | 21.877002565652 | 1.71676556299536 | 0.26309172721071 | 6.52534985115821 | 6.78429907270363e-11 | 1.00893551697096e-07 | 407 | 17 | 15 | 10 | 11 | 18 | 16 | 14 | 8 | 31 | 44 | 43 | 45 |
| MIR_dup57149 | 20.9580907608351 | 1.85510435696625 | 0.277096149189829 | 6.69480381589636 | 2.15961161818692e-11 | 3.6444747950433e-08 | 408 | 18 | 13 | 13 | 7 | 14 | 16 | 11 | 6 | 39 | 42 | 37 | 41 |
| L2c_dup67904 | 18.394288236213 | 1.55704382354585 | 0.391235197990456 | 3.97981529152658 | 6.8968834001777e-05 | 0.00767221046067641 | 409 | 16 | 14 | 5 | 11 | 26 | 16 | 9 | 5 | 38 | 17 | 32 | 37 |
| L1MCb_dup780 | 17.0876144605044 | 1.39167880557037 | 0.319788304757541 | 4.35187524017027 | 1.3497806627961e-05 | 0.00233242490624314 | 410 | 11 | 15 | 6 | 8 | 20 | 21 | 13 | 4 | 28 | 37 | 20 | 28 |
| AluYm1_dup1453 | 18.1350804852732 | 1.62728236640216 | 0.345223986700904 | 4.71370017464055 | 2.43258522341084e-06 | 0.000655519732242034 | 411 | 14 | 10 | 8 | 9 | 19 | 20 | 9 | 6 | 33 | 31 | 18 | 48 |
| MIRc_dup34919 | 19.5522971980416 | 1.47100502469147 | 0.302785297501081 | 4.85824456085493 | 1.18431071157651e-06 | 0.000364085266895072 | 412 | 17 | 12 | 9 | 12 | 24 | 23 | 8 | 4 | 37 | 39 | 23 | 32 |
| L2c_dup6035 | 18.8091026411054 | 1.12534161982209 | 0.318878307426272 | 3.52906294851141 | 0.000417033919464464 | 0.0270312048490578 | 413 | 15 | 8 | 12 | 15 | 20 | 23 | 12 | 12 | 19 | 36 | 34 | 29 |
| LTR47B4_dup95 | 19.3938543367215 | 1.18598209296176 | 0.333954247694275 | 3.55133106151561 | 0.000383287965680471 | 0.025596923458267 | 414 | 11 | 8 | 11 | 12 | 22 | 24 | 15 | 14 | 23 | 31 | 35 | 35 |
| LOR1-int_dup250 | 19.6020588248392 | 1.1038338075273 | 0.320764366071454 | 3.44126070188672 | 0.000579010369098612 | 0.0336596290107234 | 415 | 16 | 11 | 10 | 10 | 21 | 20 | 18 | 16 | 17 | 34 | 34 | 39 |
| MER84_dup24 | 18.3885274242748 | 1.29386694242216 | 0.293005881365198 | 4.41583949234623 | 1.00618726827956e-05 | 0.00186028155196619 | 416 | 18 | 10 | 10 | 10 | 17 | 16 | 16 | 11 | 22 | 32 | 27 | 42 |
| L2a_dup112251 | 19.1269783471161 | 1.42908758525606 | 0.33197641429411 | 4.30478649603698 | 1.67146855924138e-05 | 0.00271295537378751 | 417 | 15 | 10 | 8 | 12 | 10 | 22 | 17 | 12 | 24 | 29 | 48 | 31 |
| LOR1-int_dup241 | 17.1541829573547 | 1.21857969328357 | 0.309915049930966 | 3.93197972655735 | 8.42491722065088e-05 | 0.00889385693427238 | 418 | 14 | 11 | 11 | 10 | 10 | 18 | 18 | 11 | 19 | 25 | 38 | 30 |
| L1ME3G_dup2072 | 16.543955722875 | 1.01782321351921 | 0.298271402896136 | 3.4124062972059 | 0.000643920603792224 | 0.0359245795990491 | 419 | 13 | 11 | 12 | 13 | 15 | 20 | 13 | 10 | 19 | 24 | 30 | 27 |
| MIR_dup137146 | 16.8424564407088 | 1.33634584479207 | 0.311243193324971 | 4.29357452131261 | 1.75819263529101e-05 | 0.00281910313205495 | 420 | 10 | 10 | 8 | 14 | 11 | 19 | 12 | 12 | 23 | 33 | 29 | 27 |
| HERVIP10FH-int_dup3 | 18.005338161032 | 1.00455309867529 | 0.306526485804471 | 3.27721467865612 | 0.00104836626736142 | 0.049304134923274 | 421 | 21 | 13 | 12 | 12 | 15 | 21 | 17 | 8 | 21 | 21 | 26 | 42 |
| MSTA_dup7001 | 17.9945360939252 | 1.05540805109593 | 0.316629745749471 | 3.33325616201268 | 0.000858358736426345 | 0.0432547760588506 | 422 | 17 | 11 | 12 | 12 | 16 | 15 | 25 | 8 | 27 | 19 | 32 | 32 |
| AluSx1_dup9092 | 17.373148253007 | 1.04000508240356 | 0.315484836424749 | 3.29652953907224 | 0.00097887320348036 | 0.0471305065308251 | 423 | 21 | 15 | 12 | 10 | 17 | 9 | 19 | 9 | 21 | 30 | 31 | 23 |
| MLT1D_dup8091 | 17.9879990510021 | 1.03877294472736 | 0.284846585777241 | 3.64678039546421 | 0.000265546683311837 | 0.0199034214965659 | 424 | 17 | 19 | 9 | 16 | 15 | 13 | 16 | 11 | 19 | 30 | 31 | 30 |
| L1M1_dup9390 | 17.2219555867977 | 1.27446900254609 | 0.358079215585091 | 3.55918173151614 | 0.000372012054755005 | 0.0250856084628151 | 425 | 11 | 11 | 8 | 15 | 26 | 13 | 9 | 9 | 22 | 22 | 29 | 41 |
| Tigger12_dup824 | 17.4840953135904 | 1.23675561073639 | 0.335141796170785 | 3.69024581495694 | 0.000224037454751532 | 0.0177459441439108 | 426 | 11 | 12 | 9 | 15 | 20 | 10 | 17 | 10 | 23 | 16 | 31 | 47 |

| | | | | | | | | | | | | | | | | | | | |
|---|---|---|---|---|---|---|---|---|---|---|---|---|---|---|---|---|---|---|---|
| MIR_dup137150 | 18.4731941729429 | 1.18785862049886 | 0.34355540664592 | 3.45754599555206 | 0.00054511917924913 | 0.0322898394372453 | 427 | 11 | 11 | 9 | 23 | 20 | 18 | 9 | 11 | 32 | 22 | 30 | 32 |
| MIRc_dup32871 | 20.7591018756782 | 1.20311653998442 | 0.301819197063374 | 3.98621609125745 | 6.71353733590164e-05 | 0.00752099507970081 | 428 | 13 | 18 | 7 | 23 | 23 | 14 | 16 | 12 | 25 | 35 | 36 | 38 |
| MIRb_dup71181 | 19.4322820867325 | 1.36987460709832 | 0.305714810037531 | 4.48089056244986 | 7.43322210896872e-06 | 0.0014788507257029 | 429 | 12 | 21 | 11 | 16 | 19 | 9 | 13 | 8 | 32 | 44 | 23 | 31 |
| AluSg_dup10634 | 18.4112153626092 | 1.5397024301626 | 0.284799386771792 | 5.40627017359539 | 6.43507063656752e-08 | 3.51999843146826e-05 | 430 | 15 | 15 | 9 | 10 | 18 | 11 | 12 | 8 | 31 | 32 | 30 | 36 |
| AluSx3_dup9561 | 18.7916940429611 | 1.79670432589228 | 0.326713836600505 | 5.49932119370024 | 3.81256098155331e-08 | 2.35021667180488e-05 | 431 | 10 | 14 | 7 | 8 | 19 | 11 | 15 | 5 | 44 | 34 | 29 | 31 |
| AluY_dup31630 | 18.6983650489911 | 1.17885543759874 | 0.303810534768971 | 3.88023225888195 | 0.000104356764772532 | 0.0103506772615944 | 432 | 16 | 18 | 5 | 10 | 24 | 18 | 13 | 11 | 26 | 34 | 25 | 31 |
| L2c_dup47482 | 20.619548096567 | 1.43221755217518 | 0.316121390059871 | 4.53059361754651 | 5.88181833506583e-06 | 0.00123854437659785 | 433 | 12 | 13 | 8 | 11 | 24 | 18 | 17 | 10 | 41 | 37 | 21 | 40 |
| L2a_dup57887 | 20.4132824338338 | 1.66910553833584 | 0.33485567174454 | 4.98455208968116 | 6.21055134043837e-07 | 0.000220563559589843 | 434 | 16 | 12 | 8 | 7 | 24 | 9 | 12 | 13 | 39 | 38 | 28 | 42 |
| MER45B_dup491 | 17.5815209251872 | 1.41755689198919 | 0.325001066688425 | 4.36169919820043 | 1.29056221674154e-05 | 0.00225300159372548 | 435 | 11 | 13 | 8 | 6 | 19 | 15 | 10 | 13 | 32 | 37 | 21 | 27 |
| AluJb_dup33722 | 18.45214060531 | 1.22880384648131 | 0.375405397530433 | 3.27327165396362 | 0.00106310250264136 | 0.0498052742849971 | 436 | 22 | 15 | 22 | 19 | 11 | 12 | 7 | 4 | 33 | 24 | 27 | 34 |
| MER83_dup101 | 21.5370621437446 | 1.09865566335239 | 0.297994056284469 | 3.68683750626085 | 0.000227058238814796 | 0.0179017891626989 | 437 | 19 | 21 | 14 | 27 | 19 | 17 | 9 | 10 | 30 | 34 | 38 | 30 |
| L1MB8_dup2561 | 19.0635471587435 | 1.04815329308771 | 0.307822818180719 | 3.40505391797288 | 0.00066150991391548 | 0.0365100077262221 | 438 | 19 | 15 | 11 | 24 | 19 | 21 | 8 | 7 | 25 | 33 | 23 | 35 |
| L2b_dup85687 | 18.8380898545341 | 1.03138016358636 | 0.298593309320479 | 3.45413018775776 | 0.000552070799739038 | 0.0325639659183701 | 439 | 16 | 18 | 13 | 23 | 17 | 18 | 14 | 4 | 29 | 30 | 27 | 27 |
| AluJb_dup37194 | 18.2337482736985 | 1.58087475615415 | 0.384360578764255 | 4.11299920828709 | 3.9055172572035e-05 | 0.00504233428802249 | 440 | 11 | 8 | 4 | 15 | 15 | 20 | 20 | 4 | 38 | 28 | 26 | 37 |
| AluSx3_dup9562 | 16.7700693820232 | 1.51279109399568 | 0.376923115532306 | 4.01352698111722 | 5.98181613923918e-05 | 0.00690276053863922 | 441 | 7 | 15 | 6 | 17 | 10 | 14 | 19 | 3 | 33 | 32 | 22 | 30 |
| L1PA3_dup569 | 19.7298751161957 | 1.43210337498782 | 0.366606683231219 | 3.90637552585095 | 9.36908710733251e-05 | 0.00958442304746922 | 442 | 20 | 9 | 11 | 17 | 14 | 9 | 18 | 8 | 47 | 44 | 16 | 26 |
| Tigger19a_dup1234 | 21.0789067257584 | 1.65675122519041 | 0.321536867181566 | 5.15260113004359 | 2.56897997307269e-07 | 0.000109352148241996 | 443 | 19 | 8 | 10 | 20 | 20 | 11 | 15 | 5 | 33 | 51 | 26 | 46 |
| AluY_dup49917 | 24.8839667627793 | 1.84203285754251 | 0.478032012624358 | 3.8533671572117 | 0.000116504485589402 | 0.0112014561305119 | 444 | 11 | 12 | 5 | 13 | 27 | 41 | 13 | 5 | 37 | 26 | 44 | 81 |
| L1PA3_dup2284 | 23.6846898361064 | 1.47544405608795 | 0.416130602477564 | 3.54562737588495 | 0.000391679638002084 | 0.0259172978709799 | 445 | 11 | 11 | 8 | 8 | 29 | 31 | 18 | 14 | 25 | 38 | 35 | 71 |
| AluSg4_dup3297 | 25.2847974049977 | 1.54927497406817 | 0.314510480971192 | 4.92598837814272 | 8.39349865624429e-07 | 0.000281591563794387 | 446 | 15 | 14 | 8 | 14 | 30 | 26 | 19 | 10 | 41 | 37 | 44 | 56 |
| MIR_dup168516 | 23.0563917510705 | 1.63225612972363 | 0.376991013391493 | 4.32969506365548 | 1.49315957891974e-05 | 0.00250381500469088 | 447 | 20 | 18 | 5 | 7 | 27 | 25 | 14 | 9 | 35 | 27 | 46 | 54 |
| THE1B-int_dup1286 | 21.2985177814027 | 2.2097916907366 | 0.480787979229179 | 4.59618748014341 | 4.30291563796899e-06 | 0.00098259267216139 | 448 | 7 | 6 | 4 | 5 | 25 | 34 | 5 | 6 | 43 | 45 | 30 | 49 |
| L1M5_dup28029 | 19.5974226676715 | 1.82082172152499 | 0.473275995292554 | 3.84727249984325 | 0.000119440099142017 | 0.0114091906184048 | 449 | 6 | 12 | 4 | 7 | 25 | 23 | 18 | 3 | 26 | 30 | 42 | 51 |
| L1PA10_dup4828 | 16.8130171841873 | 1.52500690096353 | 0.457763656921523 | 3.33142851754387 | 0.00086401487008797 | 0.043399668800527 | 450 | 6 | 10 | 8 | 7 | 24 | 24 | 11 | 4 | 24 | 20 | 24 | 50 |
| L1MB1_dup848 | 25.9000738836694 | 1.15454657503392 | 0.309677113338481 | 3.72822699936495 | 0.000192831649238724 | 0.0160151900906657 | 451 | 36 | 18 | 8 | 10 | 22 | 38 | 18 | 13 | 41 | 49 | 22 | 46 |
| L1MC4a_dup3331 | 27.514278780946 | 1.15526080774383 | 0.30454301776686 | 3.79342404963042 | 0.000148584092599605 | 0.0132375650465738 | 452 | 28 | 24 | 9 | 11 | 35 | 46 | 18 | 7 | 39 | 55 | 24 | 47 |
| L1ME1_dup19496 | 23.4093206371064 | 1.17245937499189 | 0.284654720085878 | 4.11888260499659 | 3.80714030051874e-05 | 0.00496108327462887 | 453 | 27 | 11 | 9 | 15 | 24 | 28 | 18 | 12 | 33 | 49 | 33 | 31 |
| MIRb_dup71178 | 20.3003629526662 | 1.13212734872634 | 0.294857138122129 | 3.83957924823043 | 0.000123245343444263 | 0.0116464402268422 | 454 | 23 | 12 | 7 | 16 | 21 | 21 | 20 | 8 | 32 | 34 | 23 | 37 |
| MLT1K_dup10980 | 24.4944841519833 | 1.09799966546955 | 0.285036318706489 | 3.85213951138695 | 0.000117090277320647 | 0.0112346293087562 | 455 | 36 | 24 | 8 | 18 | 33 | 18 | 14 | 7 | 35 | 52 | 32 | 27 |
| L1PA7_dup5556 | 23.5697592026673 | 1.00060640482773 | 0.266862636456961 | 3.74951854674156 | 0.000177174393743532 | 0.0150887395467782 | 456 | 33 | 22 | 12 | 23 | 19 | 24 | 16 | 8 | 34 | 34 | 34 | 37 |
| MLT1A0_dup10525 | 24.5740597916166 | 1.00556931772554 | 0.266236329612743 | 3.77698009579762 | 0.000158741412082863 | 0.0139431096491211 | 457 | 31 | 20 | 17 | 19 | 23 | 31 | 10 | 11 | 35 | 37 | 33 | 39 |
| MIR_dup57155 | 22.7901509439828 | 1.17916403307507 | 0.309148892908002 | 3.81422693118287 | 0.000136609895276196 | 0.012507032759288 | 458 | 16 | 17 | 11 | 8 | 22 | 32 | 17 | 15 | 35 | 41 | 26 | 40 |
| AluJr_dup23267 | 25.3408186111847 | 1.5252872329949 | 0.239874059567995 | 6.35870021019318 | 2.03468063233349e-10 | 2.67482937978578e-07 | 459 | 20 | 19 | 12 | 14 | 17 | 20 | 20 | 13 | 40 | 51 | 39 | 48 |
| LTR54_dup111 | 22.975280455486 | 1.13897966469173 | 0.264148700564657 | 4.31188819879483 | 1.61866280073103e-05 | 0.00265075250366652 | 460 | 21 | 17 | 10 | 15 | 18 | 23 | 25 | 14 | 28 | 38 | 37 | 42 |
| L1M4a1_dup2081 | 26.6899521882483 | 1.03305643168944 | 0.265707709690408 | 3.88794300659591 | 0.000101097356670626 | 0.01016378093529 | 461 | 19 | 22 | 15 | 14 | 27 | 30 | 29 | 15 | 36 | 50 | 30 | 45 |
| THE1D_dup750 | 24.4598996859423 | 1.20432040082338 | 0.291582851913284 | 4.13028541603518 | 3.62313139859366e-05 | 0.0047841821518855 | 462 | 18 | 15 | 16 | 12 | 22 | 25 | 31 | 10 | 29 | 51 | 37 | 41 |
| MER21C_dup656 | 28.3675007361864 | 1.28963753460033 | 0.292260358685379 | 4.41263242268389 | 1.0212129391915e-05 | 0.0018822117275667 | 463 | 20 | 22 | 11 | 14 | 37 | 23 | 25 | 15 | 39 | 46 | 54 | 45 |
| AluSq2_dup59192 | 27.8097745628068 | 1.45452343777624 | 0.322034746386679 | 4.51666614890598 | 6.28208296246434e-06 | 0.00130208755451789 | 464 | 19 | 15 | 12 | 9 | 31 | 30 | 27 | 11 | 37 | 63 | 51 | 39 |
| MIRb_dup71190 | 26.5954350621742 | 1.59030447261857 | 0.259689514488537 | 6.12386863501479 | 9.13302037875607e-10 | 1.01077472885744e-06 | 465 | 21 | 16 | 13 | 12 | 29 | 21 | 18 | 10 | 36 | 56 | 45 | 54 |
| L2_dup47423 | 27.0996067799631 | 1.03809056202982 | 0.231395891061479 | 4.48621000687521 | 7.25012823091702e-06 | 0.00145213721551665 | 466 | 24 | 21 | 16 | 25 | 25 | 30 | 20 | 14 | 35 | 44 | 40 | 45 |
| MER5B_dup15475 | 27.5976981109373 | 1.0227117328879 | 0.269399794179387 | 3.79626026071459 | 0.000146895282897171 | 0.013130407582363 | 467 | 19 | 18 | 20 | 17 | 27 | 35 | 24 | 17 | 42 | 36 | 43 | 43 |
| L2a_dup57879 | 28.9423031296628 | 1.18878932019797 | 0.252787760991341 | 4.70271707592163 | 2.56722043360486e-06 | 0.000681763206801944 | 468 | 25 | 22 | 10 | 20 | 31 | 33 | 27 | 10 | 46 | 54 | 39 | 42 |
| MIR3_dup27875 | 28.4082403163957 | 1.39068329645474 | 0.268233901843111 | 5.18459183160278 | 2.16488456686239e-07 | 9.62851632049791e-05 | 469 | 24 | 16 | 8 | 21 | 27 | 25 | 23 | 16 | 51 | 45 | 41 | 53 |
| L2c_dup77283 | 25.7590731339759 | 1.29637410326946 | 0.381770148030767 | 3.39569269613239 | 0.000684551691852288 | 0.0372908280378856 | 470 | 27 | 21 | 7 | 12 | 35 | 38 | 14 | 9 | 28 | 33 | 58 | 41 |
| MIR3_dup57107 | 27.519774681344 | 1.27425202407815 | 0.325950926548163 | 3.9093370206753 | 9.25497707823747e-05 | 0.00951678814113351 | 471 | 25 | 23 | 10 | 14 | 26 | 41 | 19 | 11 | 36 | 32 | 48 | 61 |
| MIRb_dup213918 | 29.29762632852 | 1.25579513141956 | 0.345776951515432 | 3.63180693772618 | 0.000281443649130851 | 0.0207012026386675 | 472 | 27 | 18 | 13 | 11 | 38 | 38 | 19 | 15 | 35 | 37 | 64 | 50 |
| AluSc_dup26553 | 28.1196691195899 | 1.51109120033775 | 0.301984090528331 | 5.00387685223433 | 5.61886596891056e-07 | 0.000203189465324983 | 473 | 15 | 19 | 15 | 18 | 28 | 33 | 21 | 6 | 32 | 56 | 56 | 55 |
| Charlie2b_dup1971 | 30.0730899342557 | 1.38345012174702 | 0.394054901448018 | 3.51080551634634 | 0.000446751114958235 | 0.0283473708799606 | 474 | 18 | 19 | 13 | 14 | 49 | 41 | 20 | 7 | 36 | 43 | 40 | 81 |
| Tigger2_dup984 | 31.699390522887 | 1.41917453192289 | 0.351069963850148 | 4.04242651908738 | 5.29008932757471e-05 | 0.00630323282493286 | 475 | 22 | 17 | 14 | 15 | 47 | 41 | 19 | 10 | 41 | 49 | 43 | 81 |
| AluSg_dup27054 | 33.050893170901 | 1.46985199621813 | 0.348738180553218 | 4.21477222220532 | 2.50030405523518e-05 | 0.00366340731974747 | 476 | 12 | 8 | 29 | 42 | 27 | 29 | 14 | 19 | 42 | 43 | 68 | 83 |
| L1M5_dup43731 | 30.5826257599625 | 1.21847535998861 | 0.329634783353285 | 3.69644048966372 | 0.000218643481196067 | 0.0174464593483164 | 477 | 14 | 13 | 28 | 38 | 24 | 39 | 18 | 12 | 35 | 57 | 44 | 65 |
| MLT1C_dup15281 | 32.0319601651191 | 1.61472815639309 | 0.266108375639106 | 6.06793436138579 | 1.29565834120986e-09 | 1.36414477724567e-06 | 478 | 11 | 16 | 24 | 29 | 22 | 29 | 20 | 14 | 41 | 59 | 71 | 64 |
| L2a_dup137046 | 29.0286165599906 | 1.60710720837896 | 0.453103547521556 | 3.54688727812819 | 0.000389811339152889 | 0.0258485014139323 | 479 | 7 | 4 | 22 | 27 | 38 | 25 | 13 | 13 | 34 | 63 | 47 | 71 |
| Charlie7_dup3618 | 29.6112601606607 | 1.26986938831103 | 0.379171166318114 | 3.34906633497981 | 0.000810843799555408 | 0.0417251381334367 | 480 | 15 | 8 | 23 | 23 | 47 | 27 | 19 | 13 | 30 | 51 | 60 | 56 |
| LTR16E1_dup2433 | 26.6355266858139 | 1.29077046279813 | 0.35352442994372 | 3.65114926570596 | 0.000261069446815316 | 0.0196832796552336 | 481 | 8 | 13 | 24 | 30 | 30 | 24 | 15 | 12 | 29 | 50 | 57 | 41 |
| AluSx1_dup83846 | 25.8664468647532 | 1.31925741460287 | 0.344279613864642 | 3.83193590754274 | 0.00012713886479007 | 0.0119196943740496 | 482 | 13 | 5 | 27 | 28 | 28 | 18 | 22 | 11 | 37 | 41 | 39 | 56 |
| MIRb_dup174453 | 26.9895172507139 | 1.32479720434501 | 0.379329059736042 | 3.49247485881223 | 0.000478566644376508 | 0.0296235740798159 | 483 | 16 | 5 | 22 | 31 | 28 | 23 | 23 | 9 | 53 | 47 | 40 | 35 |
| AluSg_dup27053 | 27.8714450622492 | 1.21464366216356 | 0.324721079779021 | 3.7405753361936 | 0.000183599503304412 | 0.0154972569752649 | 484 | 13 | 11 | 26 | 39 | 27 | 24 | 19 | 11 | 42 | 49 | 42 | 45 |
| MIR3_dup67862 | 29.3798790511429 | 1.46189564934559 | 0.300948670018646 | 4.85762455522736 | 1.18802404049878e-06 | 0.000364755572052289 | 485 | 11 | 10 | 27 | 34 | 27 | 21 | 18 | 14 | 42 | 45 | 52 | 67 |
| AluJb_dup93865 | 28.8284091048953 | 1.15107144609449 | 0.34226616993267 | 3.36308857612461 | 0.000770756290713804 | 0.040405017922491 | 486 | 17 | 14 | 26 | 40 | 38 | 20 | 16 | 9 | 42 | 40 | 47 | 53 |
| MER102c_dup2586 | 29.3059638624952 | 1.18370610061681 | 0.338938717518262 | 3.49238974314887 | 0.00047871918536214 | 0.0296253095657165 | 487 | 17 | 11 | 32 | 28 | 36 | 25 | 16 | 13 | 49 | 40 | 36 | 61 |
| MamTip2_dup5624 | 30.1803836023041 | 1.55920935653557 | 0.35120726152601 | 4.43957038291505 | 9.01386332008499e-06 | 0.00171037697094174 | 488 | 15 | 13 | 35 | 21 | 25 | 26 | 15 | 8 | 43 | 77 | 33 | 65 |
| L1PA4_dup1883 | 34.5543061163807 | 1.51571982821679 | 0.443656413404525 | 3.41642717747609 | 0.000634486311856751 | 0.0355650129472477 | 489 | 13 | 10 | 27 | 50 | 35 | 33 | 10 | 8 | 66 | 70 | 42 | 63 |
| MER103C_dup7098 | 29.1721301122441 | 1.77546839481575 | 0.379058751209369 | 4.68388709969418 | 2.81485092078338e-06 | 0.000721069589570882 | 490 | 8 | 7 | 21 | 28 | 27 | 28 | 14 | 8 | 44 | 65 | 46 | 68 |
| MIR_dup137218 | 29.8564823546178 | 1.63514843050696 | 0.365537300989974 | 4.4732737974443 | 7.70310403625139e-06 | 0.00151356134848049 | 491 | 9 | 13 | 22 | 43 | 22 | 25 | 15 | 7 | 41 | 63 | 47 | 70 |
| MIR_dup137201 | 24.5757845967605 | 1.85586609745618 | 0.39620727594971 | 4.68407878933486 | 2.81221826417011e-06 | 0.000721069589570882 | 492 | 10 | 6 | 13 | 23 | 23 | 20 | 12 | 5 | 56 | 59 | 43 | 26 |
| L3_dup37635 | 22.7472014150144 | 1.55319169990542 | 0.439932492620964 | 3.53052280965211 | 0.000414739253275703 | 0.0269411810974448 | 493 | 10 | 4 | 19 | 27 | 29 | 18 | 11 | 4 | 36 | 41 | 42 | 43 |
| MIR1_Amn_dup8559 | 25.0487918896564 | 1.72934499618287 | 0.419058263607 | 4.12674118700755 | 3.67940086914458e-05 | 0.00483167063581389 | 494 | 13 | 2 | 14 | 28 | 28 | 17 | 13 | 9 | 43 | 41 | 56 | 46 |
| MIR_dup137199 | 21.2164837710376 | 1.65478686458711 | 0.362024499824229 | 4.57092507659164 | 4.85575881801354e-06 | 0.00107280258840395 | 495 | 10 | 3 | 12 | 24 | 20 | 17 | 12 | 10 | 31 | 40 | 44 | 41 |
| L2c_dup114235 | 24.7989338755458 | 1.270721544629 | 0.343589982634495 | 3.69836610161238 | 0.000216991766213843 | 0.0173554026230316 | 496 | 26 | 10 | 24 | 17 | 25 | 27 | 9 | 10 | 27 | 36 | 42 | 61 |
| MIRb_dup183265 | 26.485527775668 | 1.35354075142355 | 0.40156731572799 | 3.37064471736141 | 0.000749925053533414 | 0.0396096929607241 | 497 | 24 | 10 | 28 | 19 | 34 | 24 | 5 | 9 | 32 | 43 | 48 | 56 |
| AluY_dup78934 | 25.7042227040228 | 1.63103160042928 | 0.410433760279663 | 3.97392163675308 | 7.06988320635865e-05 | 0.00781711542741796 | 498 | 14 | 9 | 22 | 31 | 19 | 27 | 9 | 3 | 40 | 53 | 54 | 39 |

| | | | | | | | | | | | | | | | | | | | |
|---|---|---|---|---|---|---|---|---|---|---|---|---|---|---|---|---|---|---|---|
| L2_dup48801 | 23.8482959179431 | 1.53580067095101 | 0.399866303351855 | 3.84078542772236 | 0.000122641280887197 | 0.0116309295424412 | 499 | 18 | 9 | 23 | 23 | 24 | 21 | 10 | 2 | 25 | 41 | 53 | 55 |
| AluY_dup102112 | 23.7521040590279 | 1.68349184666608 | 0.394182170137539 | 4.27084727368229 | 1.94731735005321e-05 | 0.00304439142034009 | 500 | 13 | 8 | 10 | 16 | 30 | 29 | 7 | 9 | 38 | 33 | 44 | 58 |
| AluSx_dup84765 | 27.221935825076 | 1.44927849091142 | 0.348994360980588 | 4.15272753072373 | 3.28535828222572e-05 | 0.00446197409716028 | 501 | 17 | 10 | 19 | 23 | 32 | 29 | 7 | 13 | 39 | 45 | 47 | 57 |
| MLT1L_dup9421 | 25.3974230521602 | 1.56522776809658 | 0.401422686034299 | 3.89920107296288 | 9.65106054845933e-05 | 0.00979288380922688 | 502 | 11 | 12 | 20 | 19 | 29 | 30 | 6 | 8 | 38 | 34 | 41 | 70 |
| L3_dup37636 | 27.8823797232606 | 1.74295267052622 | 0.372900493542901 | 4.67404227322562 | 2.95328501976337e-06 | 0.000747576975864484 | 503 | 15 | 8 | 19 | 20 | 35 | 23 | 11 | 7 | 36 | 56 | 40 | 82 |
| MIRb_dup174460 | 23.7988015412169 | 1.49502275197913 | 0.420884163661537 | 3.55210027142144 | 0.000382169194706961 | 0.0255440078426185 | 504 | 9 | 8 | 12 | 34 | 29 | 21 | 8 | 9 | 28 | 43 | 37 | 64 |
| L1PA7_dup9484 | 23.6820140676068 | 1.38854764275918 | 0.381140949994728 | 3.64313423361722 | 0.000269338286312476 | 0.0201154952526392 | 505 | 16 | 11 | 8 | 19 | 31 | 36 | 12 | 7 | 25 | 42 | 39 | 54 |
| MIRb_dup71179 | 24.5598557540119 | 1.36219576210038 | 0.318124119802025 | 4.28196316251687 | 1.85251655162015e-05 | 0.00293279376840857 | 506 | 17 | 11 | 8 | 21 | 26 | 32 | 13 | 13 | 33 | 49 | 34 | 49 |
| AluJr_dup26607 | 24.6920365558354 | 1.49135139541546 | 0.331772936773326 | 4.49509658599543 | 6.95383757164548e-06 | 0.00140580954530396 | 507 | 19 | 12 | 10 | 22 | 24 | 36 | 10 | 5 | 37 | 54 | 34 | 45 |
| L2b_dup74944 | 21.7759133335594 | 1.3056571619248 | 0.381256589369695 | 3.42461533342506 | 0.000615670459943731 | 0.0349133277554268 | 508 | 16 | 9 | 13 | 23 | 29 | 27 | 9 | 5 | 24 | 38 | 32 | 52 |
| L1PB1_dup7191 | 27.527643735698 | 1.45687329822058 | 0.352318818295532 | 4.13509929804127 | 3.54801188144275e-05 | 0.00471639796168591 | 509 | 12 | 10 | 19 | 27 | 29 | 34 | 14 | 7 | 44 | 56 | 45 | 44 |
| MER44C_dup529 | 27.2650790804391 | 1.392288219185 | 0.320127523323595 | 4.34916749653429 | 1.36655310479133e-05 | 0.00235456803093902 | 510 | 16 | 11 | 16 | 27 | 35 | 24 | 17 | 9 | 39 | 49 | 55 | 41 |
| L3_dup24808 | 25.4998456858007 | 1.25620525594584 | 0.345611218729857 | 3.63473518181056 | 0.000278266316837578 | 0.0205309634189874 | 511 | 11 | 13 | 23 | 31 | 29 | 20 | 16 | 8 | 39 | 56 | 36 | 34 |
| MIR_dup137214 | 25.0545874860075 | 1.53857680454728 | 0.296270355390264 | 5.1931513786473 | 2.0676392871719e-07 | 9.29980470828435e-05 | 512 | 12 | 12 | 16 | 23 | 25 | 21 | 17 | 7 | 49 | 45 | 38 | 43 |
| HAL1_dup3237 | 23.4643375397846 | 1.04349592920502 | 0.287503447593919 | 3.62950753438928 | 0.000283962443337678 | 0.0208156893229905 | 513 | 16 | 16 | 21 | 17 | 25 | 32 | 14 | 10 | 28 | 41 | 36 | 37 |
| MER20_dup15686 | 24.4702055289822 | 1.09580928500188 | 0.297213604906767 | 3.68694187248131 | 0.000226965174370229 | 0.0179003829568109 | 514 | 17 | 12 | 19 | 21 | 21 | 37 | 17 | 11 | 32 | 40 | 33 | 47 |
| LTR35B_dup16 | 23.7281628846185 | 1.11864431775237 | 0.291746654804626 | 3.8343004086936 | 0.000125922171478161 | 0.0118289289437594 | 515 | 21 | 12 | 19 | 15 | 33 | 25 | 13 | 10 | 34 | 46 | 28 | 38 |
| MIRb_dup174355 | 25.4247033063473 | 1.26306974668913 | 0.275163323748072 | 4.59025472393821 | 4.42705409463137e-06 | 0.00100083295050532 | 516 | 17 | 12 | 21 | 20 | 27 | 25 | 18 | 10 | 42 | 46 | 31 | 46 |
| L1MB7_dup10965 | 28.4972518125949 | 1.35619853787564 | 0.34294039643842 | 3.95461879661987 | 7.66567759452014e-05 | 0.00828722090370599 | 517 | 12 | 9 | 22 | 31 | 25 | 33 | 24 | 9 | 40 | 52 | 57 | 42 |
| L1MA4_dup8531 | 25.6560539844698 | 1.2364709221505 | 0.323776067961207 | 3.81890770969104 | 0.000134043896731112 | 0.0123468775134145 | 518 | 14 | 11 | 27 | 26 | 22 | 30 | 18 | 7 | 40 | 36 | 43 | 47 |
| MSTC_dup1731 | 29.1029988716098 | 1.45503067092375 | 0.350073111481283 | 4.15636226606209 | 3.23354960207033e-05 | 0.00440922747056863 | 519 | 14 | 11 | 28 | 34 | 23 | 31 | 14 | 7 | 45 | 49 | 57 | 50 |
| L2c_dup114183 | 25.9356977785239 | 1.58603025805254 | 0.303055412917612 | 5.23346619281047 | 1.66360445480865e-07 | 7.85412749214084e-05 | 520 | 15 | 11 | 28 | 21 | 18 | 21 | 16 | 6 | 41 | 53 | 47 | 45 |
| L2c_dup114217 | 22.7853814315536 | 1.31634366737998 | 0.360075664138583 | 3.65574183006536 | 0.000256439334416483 | 0.0194574980443446 | 521 | 11 | 7 | 22 | 28 | 16 | 27 | 16 | 7 | 36 | 41 | 38 | 35 |
| MER3_dup8354 | 22.8605023181981 | 1.51521234785039 | 0.31598921295534 | 4.79513947225959 | 1.62561687794947e-06 | 0.000475195373022808 | 522 | 18 | 7 | 16 | 22 | 16 | 22 | 9 | 12 | 38 | 42 | 44 | 35 |
| L1PA4_dup10452 | 25.5666910121252 | 1.7690890449569 | 0.32541220026157 | 5.43645580446858 | 5.435081237983e-08 | 3.10133295024725e-05 | 523 | 8 | 8 | 16 | 22 | 23 | 21 | 12 | 12 | 43 | 56 | 38 | 56 |
| MER50_dup1412 | 25.0232614499155 | 1.60223519724828 | 0.361521871141319 | 4.43191774868294 | 9.33986482575122e-06 | 0.00176239768106915 | 524 | 6 | 11 | 17 | 24 | 20 | 30 | 12 | 9 | 39 | 55 | 43 | 43 |
| L1P1_dup2788 | 25.6325041042343 | 1.73384534232504 | 0.352251318702624 | 4.92218268681281 | 8.55842629521163e-07 | 0.000284817245208451 | 525 | 10 | 8 | 18 | 32 | 20 | 17 | 12 | 9 | 45 | 62 | 38 | 45 |
| MIR1_Amn_dup8556 | 23.1025493927944 | 1.79668714653336 | 0.330168965662738 | 5.44172025049939 | 5.27684975612808e-08 | 3.02555492065458e-05 | 526 | 10 | 7 | 12 | 25 | 17 | 19 | 12 | 8 | 38 | 59 | 39 | 39 |
| MIRb_dup174457 | 24.1119591993666 | 1.51367254690063 | 0.308761913432635 | 4.90239398400049 | 9.46757123752832e-07 | 0.00030523617106533 | 527 | 12 | 9 | 14 | 30 | 19 | 18 | 18 | 11 | 29 | 56 | 46 | 40 |
| THE1D_dup10626 | 25.068008608703 | 1.46981368219893 | 0.339012530453831 | 4.33557332005195 | 1.45380704506973e-05 | 0.00246387158936012 | 528 | 11 | 8 | 17 | 33 | 25 | 21 | 16 | 8 | 34 | 51 | 42 | 49 |
| MER84-int_dup3 | 23.9267689726797 | 1.10036033776131 | 0.314199634981263 | 3.50210571640837 | 0.000461596426311205 | 0.0289191741061206 | 529 | 18 | 17 | 18 | 15 | 10 | 18 | 39 | 17 | 25 | 45 | 41 | 38 |
| AluSg_dup24441 | 22.8066733159568 | 1.20041898357582 | 0.298104904721022 | 4.02683405930277 | 5.65329001603168e-05 | 0.00664285306743048 | 530 | 26 | 19 | 19 | 14 | 14 | 8 | 23 | 15 | 22 | 37 | 44 | 47 |
| MLT1E2_dup228 | 24.8445935107175 | 1.00848830000709 | 0.301063080238481 | 3.34975746347987 | 0.000808823482243223 | 0.0416481956948379 | 531 | 27 | 18 | 12 | 14 | 27 | 16 | 25 | 21 | 22 | 39 | 45 | 46 |
| AluY_dup71334 | 31.4047685789244 | 1.4117466948617 | 0.301468071884111 | 4.68290617324277 | 2.82835997606357e-06 | 0.000722875341422581 | 532 | 38 | 27 | 16 | 13 | 15 | 24 | 27 | 17 | 42 | 39 | 59 | 78 |
| L2c_dup119020 | 29.6916027129233 | 1.64374266128099 | 0.277746646070239 | 5.91813685075176 | 3.25608996822037e-09 | 3.00299838596187e-06 | 533 | 23 | 23 | 18 | 11 | 19 | 22 | 21 | 14 | 44 | 39 | 52 | 86 |
| AluSz6_dup32234 | 26.3432278668332 | 1.29030542076165 | 0.312344502780158 | 4.13103291166236 | 3.61136867988249e-05 | 0.00477925879812747 | 534 | 32 | 28 | 19 | 15 | 14 | 15 | 19 | 13 | 28 | 32 | 49 | 71 |
| L1M5_dup39667 | 30.1119790401902 | 1.23573330490162 | 0.287482465655621 | 4.29846495884002 | 1.71985084325967e-05 | 0.00276882022158501 | 535 | 33 | 24 | 21 | 19 | 14 | 20 | 32 | 16 | 25 | 35 | 70 | 77 |
| AluSc8_dup14791 | 26.3533964318067 | 1.07940070735193 | 0.281789017831289 | 3.83052794484056 | 0.000127868609723228 | 0.0119528832159892 | 536 | 22 | 28 | 12 | 24 | 13 | 18 | 31 | 19 | 26 | 37 | 42 | 63 |
| MIRb_dup22059 | 26.359642572397 | 1.07751809212836 | 0.240502170897239 | 4.48028426566161 | 7.45436919345805e-06 | 0.00147918080419546 | 537 | 22 | 20 | 21 | 21 | 16 | 28 | 23 | 16 | 30 | 35 | 47 | 53 |
| MLT1E1_dup683 | 27.9794459808324 | 1.2322770376915 | 0.25303588276272 | 4.86996952462685 | 1.11615461854163e-06 | 0.000349259924646234 | 538 | 24 | 22 | 19 | 22 | 15 | 27 | 23 | 16 | 37 | 38 | 38 | 72 |
| L1MB2_dup9366 | 24.5129234672352 | 1.05738092464899 | 0.319877099875942 | 3.30558494202642 | 0.000947783711639317 | 0.0461377543064093 | 539 | 19 | 20 | 14 | 17 | 23 | 17 | 24 | 19 | 29 | 16 | 49 | 62 |
| L3_dup38247 | 20.7282104182827 | 1.21878975690356 | 0.342365922071092 | 3.5599038290104 | 0.000370990631848026 | 0.025042762237081 | 540 | 20 | 18 | 13 | 13 | 20 | 12 | 15 | 12 | 19 | 18 | 42 | 63 |
| MER5A_dup28790 | 21.0916167472353 | 1.09840159222026 | 0.300268462967244 | 3.65806512400901 | 0.000254126482934919 | 0.0193431531351772 | 541 | 18 | 23 | 15 | 13 | 15 | 22 | 16 | 11 | 24 | 22 | 40 | 47 |
| L1PA4_dup11161 | 27.2521979097837 | 1.81152626928173 | 0.288304605370053 | 6.28337610825379 | 3.31298246267853e-10 | 4.04262012853592e-07 | 542 | 11 | 14 | 14 | 18 | 16 | 9 | 23 | 19 | 19 | 36 | 77 | 93 |
| L1PA7_dup9226 | 28.7046734094907 | 1.37231886892344 | 0.250430997077977 | 5.4798283157262 | 4.25738746894918e-08 | 2.5543460881145e-05 | 543 | 12 | 14 | 28 | 29 | 14 | 20 | 30 | 15 | 21 | 32 | 63 | 94 |
| AluYj4_dup1763 | 20.6749988430855 | 1.24454773145313 | 0.318524010457455 | 3.90723364830723 | 9.33588660380157e-05 | 0.00956737689934412 | 544 | 9 | 6 | 18 | 26 | 15 | 20 | 14 | 15 | 23 | 26 | 35 | 55 |
| L2a_dup137014 | 20.7837815308914 | 1.4396131319592 | 0.343844162924389 | 4.18681858582479 | 2.828917560781e-05 | 0.00399958180580865 | 545 | 10 | 5 | 12 | 19 | 25 | 16 | 14 | 13 | 30 | 39 | 31 | 44 |
| MIRb_dup174427 | 20.9641323947093 | 1.316350975084 | 0.351665867658547 | 3.74318663294934 | 0.000181701166933718 | 0.0153566509172711 | 546 | 8 | 7 | 13 | 28 | 25 | 14 | 18 | 10 | 31 | 32 | 33 | 44 |
| AluSc8_dup16282 | 25.8736646064005 | 1.3104497960545 | 0.352493198593018 | 3.71765980531024 | 0.00020107679202559 | 0.0164984201225245 | 547 | 12 | 7 | 14 | 26 | 32 | 30 | 18 | 12 | 36 | 44 | 43 | 49 |
| L2b_dup74914 | 24.2526607713389 | 1.41666223565301 | 0.30785972572045 | 4.60164846940519 | 4.19160209496284e-06 | 0.000964579257338518 | 548 | 16 | 6 | 15 | 20 | 27 | 21 | 20 | 11 | 33 | 36 | 43 | 57 |
| MER5A_dup27638 | 25.1630266010239 | 1.3417667175613 | 0.289549048171964 | 4.63398766472347 | 3.58688376966763e-06 | 0.000856835251145652 | 549 | 10 | 11 | 25 | 24 | 19 | 23 | 20 | 13 | 37 | 39 | 35 | 59 |
| MIR_dup137221 | 23.9659513107221 | 1.54818792830318 | 0.302892081535334 | 5.11135160898083 | 3.19862020515848e-07 | 0.000130548693539732 | 550 | 7 | 8 | 21 | 22 | 19 | 19 | 21 | 11 | 31 | 46 | 46 | 49 |
| L1MC5a_dup3492 | 26.8012780701052 | 1.27348914632191 | 0.369499563870121 | 3.44652408512596 | 0.000567847945800621 | 0.0332064751318443 | 551 | 20 | 8 | 16 | 15 | 30 | 28 | 19 | 19 | 54 | 34 | 24 | 61 |
| LTR10E_dup221 | 24.6000123171648 | 1.22999594954586 | 0.345866781803733 | 3.55627083679818 | 0.000376156298751493 | 0.0252767259708339 | 552 | 21 | 6 | 12 | 18 | 14 | 27 | 30 | 19 | 32 | 28 | 55 | 46 |
| L1ME3F_dup401 | 26.9995428277535 | 1.14918571861387 | 0.276206590343065 | 4.16060209565062 | 3.17409682298589e-05 | 0.00436316373565685 | 553 | 29 | 15 | 14 | 19 | 27 | 19 | 34 | 12 | 34 | 42 | 50 | 45 |
| L2a_dup87945 | 25.703819694226 | 1.10761478151649 | 0.319449161827483 | 3.46726463509912 | 0.000525783974190409 | 0.0314728857988515 | 554 | 29 | 11 | 18 | 15 | 28 | 17 | 36 | 10 | 27 | 47 | 39 | 50 |
| AluJb_dup37193 | 28.383981885118 | 1.47782880162606 | 0.308525648658489 | 4.78997064928591 | 1.66805710455637e-06 | 0.00048521701198138 | 555 | 24 | 12 | 14 | 21 | 35 | 14 | 23 | 11 | 50 | 58 | 50 | 36 |
| MIR_dup57150 | 28.6227327941377 | 1.4016219833766 | 0.260202638153255 | 5.386655544019 | 7.17808111839388e-08 | 3.82958674842455e-05 | 556 | 22 | 15 | 14 | 21 | 28 | 21 | 26 | 14 | 34 | 66 | 51 | 45 |
| MIRb_dup174361 | 25.1608270585714 | 1.20936644610096 | 0.272023917216053 | 4.44580924529675 | 8.75616204730224e-06 | 0.00167383530737418 | 557 | 19 | 11 | 16 | 17 | 27 | 22 | 24 | 15 | 37 | 44 | 42 | 37 |
| AluSc_dup16738 | 29.9187555904746 | 1.25484039174696 | 0.292305016179809 | 4.29291432677658 | 1.7634306358393e-05 | 0.00282179735087705 | 558 | 14 | 14 | 30 | 34 | 20 | 19 | 30 | 19 | 49 | 53 | 46 | 41 |
| L2b_dup46681 | 28.7279606514955 | 1.14865611188022 | 0.308661468084163 | 3.72141077086764 | 0.000198112870118912 | 0.016317690929575 | 559 | 13 | 11 | 29 | 36 | 19 | 26 | 24 | 21 | 39 | 52 | 48 | 38 |
| AluSx_dup52338 | 26.1389338842889 | 1.10247249293417 | 0.255633347020622 | 4.31271000353969 | 1.6126558311634e-05 | 0.00264455550931776 | 560 | 14 | 18 | 18 | 25 | 22 | 22 | 29 | 16 | 34 | 49 | 38 | 41 |
| L1ME3G_dup697 | 27.3989514793687 | 1.13818698777047 | 0.249414432167445 | 4.56343675816781 | 5.03229733895533e-06 | 0.00110056527813885 | 561 | 19 | 14 | 19 | 31 | 18 | 22 | 30 | 18 | 30 | 40 | 60 | 44 |
| HAL1_dup15476 | 24.4501306832136 | 1.20123322167376 | 0.366531984080795 | 3.27729440770705 | 0.00104807025557235 | 0.049304134923274 | 562 | 8 | 7 | 21 | 27 | 19 | 17 | 30 | 22 | 37 | 46 | 33 | 34 |
| L1PA13_dup5365 | 24.5464164924593 | 1.20767165308916 | 0.359891389942049 | 3.35565586407505 | 0.000791770262735379 | 0.0411800146310016 | 563 | 12 | 6 | 16 | 28 | 18 | 15 | 33 | 24 | 36 | 43 | 27 | 47 |
| MIR_dup86405 | 23.5600116950659 | 1.15352134866121 | 0.333971123340798 | 3.45395535135567 | 0.000552428826918844 | 0.0325751948712446 | 564 | 15 | 9 | 16 | 30 | 14 | 18 | 27 | 19 | 21 | 55 | 41 | 31 |
| MIRc_dup49395 | 23.5448833206215 | 1.13954511922037 | 0.315872668582999 | 3.60760911772568 | 0.000309031585970257 | 0.0220885640598614 | 565 | 21 | 7 | 25 | 27 | 15 | 16 | 19 | 17 | 32 | 41 | 43 | 29 |
| AluJo_dup53493 | 14.5861698498884 | 2.15433322475289 | 0.475353476326933 | 4.53206578270862 | 5.84096531455404e-06 | 0.00123248081434485 | 566 | 10 | 12 | 11 | 13 | 3 | 1 | 5 | 4 | 35 | 30 | 22 | 31 |
| AluSz_dup71423 | 16.1297756947388 | 1.63350649080448 | 0.458553296073674 | 3.56230454516683 | 0.000367613582658759 | 0.024892481940615 | 567 | 17 | 15 | 15 | 17 | 2 | 3 | 6 | 8 | 25 | 29 | 28 | 36 |
| MIRc_dup10819 | 19.2957110688375 | 2.01886371643972 | 0.580529531777632 | 3.47762448924481 | 0.000505878079267053 | 0.0307619896369226 | 568 | 21 | 15 | 14 | 21 | 1 | 2 | 5 | 5 | 49 | 36 | 24 | 42 |
| MIR_dup129860 | 16.4229265196953 | 1.67705254865366 | 0.492316387772206 | 3.40645282242692 | 0.000658129236261576 | 0.0364093803308099 | 569 | 27 | 21 | 9 | 11 | 2 | 9 | 4 | 4 | 27 | 26 | 30 | 35 |
| AluJb_dup53216 | 18.8748577333059 | 2.16451365069614 | 0.535390761858002 | 4.04286701396282 | 5.2801571342754e-05 | 0.00629748529443166 | 570 | 22 | 18 | 7 | 7 | 2 | 3 | 7 | 9 | 26 | 19 | 48 | 71 |

| | | | | | | | | | | | | | | | | | | | |
|---|---|---|---|---|---|---|---|---|---|---|---|---|---|---|---|---|---|---|---|
| L1MA4_dup4647 | 13.9131336579818 | 2.15285057612912 | 0.558069706883594 | 3.8576732432785 | 0.00011447153949793 | 0.0110678768538701 | 571 | 15 | 15 | 3 | 8 | 1 | 2 | 6 | 6 | 16 | 18 | 39 | 48 |
| MamRep137_dup615 | 13.3684521836565 | 1.9137359780862 | 0.550349326323196 | 3.47731138488276 | 0.000506469252389007 | 0.0307681606502456 | 572 | 21 | 16 | 6 | 6 | 2 | 5 | 5 | 3 | 14 | 16 | 37 | 40 |
| L1MA6_dup1225 | 17.3058331256306 | 2.42580042370055 | 0.721102891428153 | 3.3640142794272 | 0.000768175686178003 | 0.040302291057938 8 | 573 | 34 | 30 | 3 | 6 | 2 | 2 | 2 | 2 | 26 | 24 | 34 | 54 |
| L1ME3E_dup1183 | 14.8085546623699 | 2.86699352108775 | 0.716733427423713 | 4.00008344998379 | 6.33201511334313e-05 | 0.00720209210401312 | 574 | 22 | 18 | 4 | 3 | 1 | 2 | 3 | 0 | 24 | 23 | 36 | 51 |
| L1M4a2_dup454 | 24.0620301307934 | 2.38611482867831 | 0.578184679844786 | 4.12690773702075 | 3.67673818570636e-05 | 0.00483084011229202 | 575 | 35 | 39 | 3 | 3 | 7 | 8 | 7 | 6 | 31 | 38 | 60 | 65 |
| L1M3_dup1353 | 21.996587374596 | 2.19755329141886 | 0.661811650133487 | 3.32051164553482 | 0.000898526103697323 | 0.0444885652069922 | 576 | 43 | 36 | 6 | 8 | 8 | 2 | 5 | 2 | 23 | 29 | 48 | 73 |
| LTR16B_dup175 | 19.5863555863501 | 2.09450760537061 | 0.616187210280358 | 3.39914164141387 | 0.00067597699267579 | 0.0369770325303779 | 577 | 35 | 26 | 7 | 2 | 7 | 2 | 7 | 6 | 25 | 26 | 43 | 62 |
| L1MA3_dup1485 | 18.3309741097954 | 2.31880708129502 | 0.569478659440636 | 4.07180680584702 | 4.66498647340216e-05 | 0.00572467700567381 | 578 | 15 | 9 | 3 | 6 | 5 | 0 | 15 | 13 | 48 | 46 | 25 | 32 |
| L1MA3_dup1483 | 16.9493721387438 | 2.38716505560283 | 0.577254717081253 | 4.13537557158985 | 3.54374588384753e-05 | 0.00471599640983212 | 579 | 12 | 10 | 1 | 3 | 7 | 1 | 15 | 9 | 56 | 32 | 20 | 32 |
| L1ME2_dup2722 | 19.2570373003585 | 2.5248791054738 | 0.678551917843644 | 3.72098145930758 | 0.000198450009576031 | 0.0163335821440948 | 580 | 12 | 15 | 0 | 2 | 4 | 1 | 13 | 12 | 58 | 53 | 21 | 33 |
| Tigger2a_dup676 | 15.0657195103905 | 2.00692380902744 | 0.485295548845313 | 4.13546716800228 | 3.54233259996479e-05 | 0.00471599640983212 | 581 | 11 | 17 | 6 | 7 | 3 | 3 | 12 | 3 | 44 | 37 | 13 | 23 |
| L1ME2_dup2724 | 13.4578280236309 | 2.52690444778479 | 0.51695188672417 | 4.88808438982073 | 1.01821884942605e-06 | 0.000324338825094417 | 582 | 4 | 10 | 1 | 3 | 6 | 2 | 9 | 6 | 39 | 44 | 12 | 21 |
| L2a_dup34097 | 13.4164083411177 | 2.8482267739293 | 0.442619601865859 | 6.43493139915771 | 1.23529108676149e-10 | 1.70891030773575e-07 | 583 | 7 | 7 | 2 | 2 | 6 | 2 | 6 | 3 | 42 | 48 | 15 | 15 |
| L1MC4a_dup1924 | 15.8647083541862 | 2.94331060294178 | 0.466584930842777 | 6.3081990188268 | 2.82301066513246e-10 | 3.5353899775032e-07 | 584 | 12 | 9 | 3 | 3 | 2 | 1 | 7 | 5 | 45 | 38 | 24 | 40 |
| L2a_dup34101 | 14.2744635967173 | 2.81863078121779 | 0.497322439258707 | 5.66761231489805 | 1.44801137269042e-08 | 1.08008938585014e-05 | 585 | 8 | 6 | 2 | 5 | 1 | 2 | 10 | 6 | 34 | 49 | 18 | 30 |
| L2a_dup34107 | 15.9188605951394 | 2.88092449196631 | 0.506722669016507 | 5.68540676808059 | 1.30501664091636e-08 | 9.88928310953771e-06 | 586 | 6 | 8 | 3 | 9 | 2 | 2 | 5 | 5 | 48 | 61 | 15 | 21 |
| Tigger4_dup280 | 19.5211194838109 | 3.38761559672411 | 0.517833777256706 | 6.54189770059122 | 6.07430501993601e-11 | 9.14782647008668e-08 | 587 | 6 | 6 | 2 | 4 | 3 | 1 | 12 | 4 | 60 | 76 | 24 | 29 |
| L1PA15_dup2694 | 15.137744801705 | 2.88206112381073 | 0.568929906431041 | 5.06575782224249 | 4.06778802162584e-07 | 0.00015716498706176 | 588 | 1 | 9 | 0 | 5 | 7 | 7 | 2 | 7 | 45 | 51 | 27 | 12 |
| THE1C-int_dup638 | 18.9622384040269 | 2.84543728182436 | 0.368686034547936 | 7.71777885569572 | 1.18374347030006e-14 | 4.94152673305293e-11 | 589 | 11 | 11 | 0 | 5 | 13 | 5 | 3 | 5 | 56 | 64 | 24 | 23 |
| L1PA16_dup4206 | 16.4015417643601 | 2.72478179737448 | 0.431942845631511 | 6.30819985776308 | 2.8229953667213e-10 | 3.5353899775032e-07 | 590 | 3 | 10 | 9 | 2 | 2 | 7 | 7 | 8 | 30 | 24 | 43 | 57 |
| L1MC3_dup11158 | 12.3562163871034 | 3.34624653854854 | 0.516781950040962 | 6.4751614066306 | 9.47106432751013e-11 | 1.36581931665074e-07 | 591 | 2 | 3 | 0 | 0 | 5 | 5 | 6 | 5 | 23 | 26 | 32 | 45 |
| L1PA4_dup10738 | 16.1778367361239 | 3.69713429681559 | 0.480879063381843 | 7.6882829350378 | 1.49122750737504e-14 | 5.91386034116435e-11 | 592 | 2 | 6 | 0 | 1 | 2 | 4 | 9 | 4 | 31 | 37 | 42 | 62 |
| AluY_dup82072 | 11.9167983425135 | 2.38795972961634 | 0.711128205377124 | 3.35798764774063 | 0.000785121215320213 | 0.0409505595573397 | 593 | 13 | 11 | 0 | 0 | 14 | 24 | 1 | 3 | 24 | 12 | 11 | 34 |
| LTR16C_dup3104 | 14.4395573320463 | 2.44917931887419 | 0.717912100945071 | 3.4115309042013 | 0.000645991780812841 | 0.0359985079011148 | 594 | 18 | 22 | 0 | 0 | 13 | 19 | 5 | 2 | 15 | 15 | 31 | 44 |
| THE1B_dup1362 | 13.2224404275041 | 2.4082046467996 | 0.642517819900665 | 3.74807448480093 | 0.000178197340395426 | 0.0151487475375956 | 595 | 24 | 19 | 0 | 0 | 16 | 14 | 3 | 1 | 18 | 22 | 24 | 24 |
| L1PB2_dup662 | 13.5533386816142 | 1.70170535062446 | 0.49904519193399 | 3.40992234396589 | 0.000649813791093824 | 0.0361516465596472 | 596 | 23 | 21 | 0 | 4 | 11 | 6 | 3 | 5 | 35 | 23 | 11 | 20 |
| LTR41_dup58 | 11.1794445434171 | 2.04010900154797 | 0.555090814204355 | 3.67527069326877 | 0.000237597361400477 | 0.0184973080163121 | 597 | 18 | 14 | 0 | 1 | 9 | 6 | 4 | 4 | 21 | 17 | 15 | 29 |
| Tigger15a_dup1079 | 13.6578769184699 | 1.65801114331922 | 0.494802948448663 | 3.35085138137823 | 0.000805635270337086 | 0.041555970092267 2 | 598 | 25 | 18 | 0 | 0 | 14 | 13 | 7 | 2 | 29 | 32 | 8 | 18 |
| L2a_dup96476 | 12.5486586709693 | 2.06018872810662 | 0.548419584264093 | 3.75659219185458 | 0.000172242836413996 | 0.0147638666978979 | 599 | 8 | 11 | 0 | 0 | 18 | 14 | 3 | 6 | 24 | 27 | 17 | 23 |
| L2c_dup5679 | 13.1373094760684 | 1.82794581287482 | 0.507975424034549 | 3.59849261674221 | 0.000320066935447829 | 0.0226057129777587 | 600 | 12 | 8 | 0 | 1 | 18 | 14 | 9 | 5 | 25 | 23 | 17 | 29 |
| L1MC1_dup12593 | 7.51427164356419 | 1.77409774078768 | 0.493144761425361 | 3.59751918617146 | 0.000321266804453143 | 0.0226702701816155 | 601 | 5 | 6 | 4 | 0 | 5 | 11 | 4 | 2 | 14 | 17 | 13 | 10 |
| MIR3_dup57906 | 10.5681727748034 | 1.86793739306249 | 0.425939140426222 | 4.38545607993037 | 1.15743010812025e-05 | 0.00208374840047616 | 602 | 7 | 7 | 5 | 2 | 6 | 12 | 7 | 3 | 19 | 26 | 23 | 11 |
| MIR_dup6785 | 9.694247839659 | 1.86876432011752 | 0.455098107705518 | 4.10628892644561 | 4.02066519253134e-05 | 0.00514909149570539 | 603 | 12 | 6 | 2 | 1 | 5 | 8 | 9 | 3 | 14 | 27 | 17 | 16 |
| L2c_dup5682 | 11.0946370424715 | 1.85087382036668 | 0.553950028457794 | 3.34122885690527 | 0.000834084402961396 | 0.0424528443523106 | 604 | 10 | 9 | 2 | 0 | 17 | 13 | 8 | 1 | 11 | 27 | 17 | 25 |
| L1ME3C_dup75 | 9.42666945949721 | 1.80757160497203 | 0.497551510252388 | 3.63293361134633 | 0.000280217133586762 | 0.0206365045089556 | 605 | 8 | 8 | 1 | 0 | 12 | 9 | 7 | 3 | 13 | 22 | 13 | 21 |
| L2_dup41533 | 10.2105740140241 | 1.76157518063675 | 0.479529129513514 | 3.67355197467123 | 0.00023920199453074 | 0.0185791657393772 | 606 | 8 | 12 | 1 | 1 | 10 | 10 | 8 | 3 | 12 | 24 | 14 | 25 |
| MLT1F_dup2340 | 10.8973085049735 | 1.82106728744308 | 0.532881854206537 | 3.41739406787393 | 0.000632236930764917 | 0.035497463125953 | 607 | 5 | 8 | 3 | 0 | 15 | 15 | 4 | 5 | 20 | 17 | 14 | 27 |
| HERVL-int_dup1711 | 9.44095254597667 | 2.01442603068624 | 0.530533668498687 | 3.79698056937026 | 0.000146469265264377 | 0.0131021713506006 | 608 | 5 | 2 | 3 | 0 | 12 | 8 | 6 | 5 | 15 | 19 | 17 | 24 |
| MIRb_dup71169 | 8.46851862989082 | 1.71789081461169 | 0.522654787959595 | 3.28685559605836 | 0.00101312771506968 | 0.0482428831879068 | 609 | 6 | 4 | 2 | 0 | 13 | 6 | 7 | 4 | 18 | 15 | 13 | 14 |
| L3b_dup244 | 9.8827668467747 | 1.72037180232369 | 0.524016942894355 | 3.28304614125908 | 0.00102691855809988 | 0.0487065362603376 | 610 | 10 | 6 | 1 | 0 | 14 | 10 | 6 | 5 | 17 | 15 | 17 | 20 |
| Mam_R4_dup311 | 8.96641873250004 | 1.92796186661675 | 0.543471518484288 | 3.54749384474412 | 0.000388914839215385 | 0.0258277226714887 | 611 | 8 | 4 | 1 | 0 | 13 | 8 | 5 | 4 | 18 | 13 | 16 | 19 |
| MIRc_dup22361 | 9.2532116789781 | 2.14936511096991 | 0.647926778137249 | 3.31729631108813 | 0.000908931638021043 | 0.0448679548849407 | 612 | 9 | 11 | 1 | 1 | 6 | 6 | 4 | 3 | 10 | 5 | 34 | 27 |
| LTR5_Hs_dup54 | 10.3279423463107 | 2.48255109604922 | 0.482438815563846 | 5.14583614742475 | 2.6633181872646e-07 | 0.000112762617328623 | 613 | 8 | 8 | 1 | 2 | 6 | 5 | 3 | 4 | 19 | 13 | 29 | 29 |
| THE1A_dup2964 | 10.2504407567305 | 2.19325921930702 | 0.475873567375235 | 4.60891163046591 | 4.04782321015206e-06 | 0.000941580341451141 | 614 | 9 | 7 | 1 | 3 | 3 | 3 | 7 | 7 | 17 | 15 | 29 | 25 |
| THE1D-int_dup762 | 12.9425653238351 | 2.4971302068288 | 0.72721522768776 | 3.43382551926013 | 0.000595127144270024 | 0.0342379408777744 | 615 | 10 | 11 | 0 | 0 | 1 | 4 | 15 | 4 | 17 | 24 | 33 | 45 |
| MSTD_dup190 | 10.0624458925713 | 2.13198241209815 | 0.615775060533949 | 3.4622746985717 | 0.000535630159588725 | 0.0318946581465212 | 616 | 8 | 7 | 0 | 0 | 7 | 8 | 8 | 5 | 7 | 28 | 17 | 33 |
| L1PA10_dup102 | 14.0775563940441 | 2.45933044907428 | 0.580830969225215 | 4.23415860961208 | 2.29408994226364e-05 | 0.00342452650816728 | 617 | 17 | 6 | 0 | 0 | 9 | 9 | 7 | 6 | 22 | 27 | 25 | 48 |
| THE1B_dup403 | 11.9454498630577 | 1.83312969481802 | 0.554256611560936 | 3.30736640137757 | 0.000941776219112789 | 0.0458922556283047 | 618 | 12 | 10 | 0 | 0 | 8 | 9 | 11 | 7 | 24 | 21 | 20 | 23 |
| L1MC1_dup10028 | 8.38061295121581 | 1.80414280414464 | 0.527623824666098 | 3.4193732727789 | 0.000627655609636466 | 0.0353570411199239 | 619 | 6 | 2 | 4 | 8 | 7 | 9 | 0 | 4 | 15 | 12 | 13 | 24 |
| L1PA5_dup9519 | 9.53846620585931 | 2.08192876133934 | 0.623153912463136 | 3.34095432877908 | 0.000834909557890376 | 0.0424857548464036 | 620 | 2 | 5 | 8 | 6 | 5 | 11 | 0 | 2 | 13 | 7 | 26 | 36 |
| L1MEf_dup10018 | 14.7308252058115 | 2.34824842550378 | 0.660583340273343 | 3.55481024473323 | 0.000378251967198173 | 0.0253602543214811 | 621 | 3 | 3 | 1 | 5 | 16 | 24 | 2 | 6 | 22 | 16 | 41 | 44 |
| L1PA4_dup9046 | 13.8830564270709 | 2.45281722597799 | 0.626613657773927 | 3.91440115539729 | 9.06288538960978e-05 | 0.00936784242795868 | 622 | 7 | 2 | 2 | 6 | 19 | 16 | 1 | 3 | 21 | 15 | 37 | 45 |
| FAM_dup1434 | 13.9800443681385 | 1.72593152756039 | 0.501736666258578 | 3.43991508619564 | 0.000581896722443165 | 0.0337547543097392 | 623 | 12 | 9 | 4 | 7 | 11 | 15 | 0 | 11 | 20 | 19 | 30 | 34 |
| L1P3_dup447 | 12.6712585249027 | 1.6651909037912 | 0.442974453179628 | 3.75911272498588 | 0.00017051700262559 | 0.0146475952009923 | 624 | 10 | 3 | 7 | 8 | 9 | 15 | 2 | 9 | 23 | 17 | 23 | 29 |
| MIRb_dup174470 | 14.1552295187783 | 1.44915576928347 | 0.44161259457779 | 3.28150914868938 | 0.00103253174845838 | 0.0488928954067022 | 625 | 6 | 6 | 10 | 15 | 13 | 17 | 2 | 8 | 26 | 19 | 26 | 25 |
| X6A_LINE_dup885 | 10.7918335015449 | 1.79039963316198 | 0.526243078177855 | 3.40222932596346 | 0.000668385268597654 | 0.0367297000281148 | 626 | 2 | 7 | 8 | 16 | 9 | 6 | 5 | 0 | 19 | 21 | 14 | 29 |
| L2a_dup137063 | 7.83224618277894 | 1.79835175521925 | 0.506737770790447 | 3.5488804247097 | 0.000386872717871383 | 0.0257640122380 7 | 627 | 2 | 6 | 6 | 9 | 7 | 5 | 3 | 0 | 10 | 13 | 15 | 24 |
| LTR57_dup42 | 8.79347233763109 | 1.83499874945953 | 0.473402180710863 | 3.8761941204075 | 0.000106103045985972 | 0.0104758487054681 | 628 | 5 | 4 | 5 | 9 | 9 | 4 | 6 | 0 | 13 | 10 | 22 | 25 |
| MLT1I-int_dup67 | 10.9208540048087 | 1.75329278176067 | 0.478060078772905 | 3.66751556888217 | 0.000244918642104647 | 0.0188574298004422 | 629 | 11 | 8 | 10 | 8 | 4 | 12 | 2 | 0 | 18 | 21 | 22 | 20 |
| L1MC5_dup6113 | 11.9455913793826 | 1.78688926323573 | 0.454918898983338 | 3.92792927976636 | 8.56803958824072e-05 | 0.00900897369802708 | 630 | 6 | 14 | 10 | 7 | 9 | 10 | 2 | 0 | 26 | 24 | 18 | 19 |
| MER5B_dup2366 | 10.5204449947709 | 1.73733483593442 | 0.480043689173217 | 3.61911816594578 | 0.000295608640527617 | 0.0213926075361874 | 631 | 10 | 15 | 7 | 8 | 7 | 5 | 2 | 0 | 15 | 24 | 15 | 24 |
| AluSx1_dup41191 | 10.8998043702258 | 1.83640996704352 | 0.473064929488248 | 3.88194062288682 | 0.000103626183029621 | 0.0103126038256655 | 632 | 9 | 9 | 4 | 10 | 5 | 15 | 1 | 1 | 18 | 23 | 22 | 18 |
| AluSx_dup41963 | 9.02362715347307 | 1.71960626998285 | 0.471022389631539 | 3.65079518051792 | 0.000261429662304036 | 0.0196981313515086 | 633 | 11 | 7 | 3 | 7 | 9 | 8 | 1 | 1 | 22 | 12 | 12 | 17 |
| MLT2B3_dup338 | 9.69787317614288 | 1.49594705375971 | 0.445957366700926 | 3.35446203036477 | 0.000795194654948383 | 0.0412888041983632 | 634 | 10 | 7 | 5 | 11 | 10 | 9 | 1 | 1 | 18 | 18 | 14 | 16 |
| L3_dup25439 | 14.051846917197 | 1.65903719060799 | 0.448377185784668 | 3.70009278617656 | 0.000215520653010065 | 0.0172667600340515 | 635 | 11 | 16 | 13 | 6 | 10 | 5 | 9 | 0 | 28 | 37 | 20 | 16 |
| L2b_dup31003 | 12.8298071985646 | 1.60630745111678 | 0.435302306168798 | 3.69009634994651 | 0.000224169130829338 | 0.0177504652260625 | 636 | 11 | 5 | 4 | 12 | 10 | 14 | 12 | 0 | 19 | 36 | 19 | 19 |
| L2a_dup137065 | 13.2036495720833 | 1.71945664006519 | 0.421314546862745 | 4.08117083273973 | 4.48094041453859e-05 | 0.00554457539198024 | 637 | 9 | 3 | 8 | 11 | 14 | 13 | 8 | 1 | 17 | 21 | 28 | 35 |
| L3_dup24811 | 13.3008800015309 | 1.69669127231867 | 0.475333355011719 | 3.56947656719112 | 0.000357695209291372 | 0.0244434647530284 | 638 | 9 | 2 | 10 | 14 | 15 | 10 | 8 | 0 | 21 | 28 | 24 | 26 |
| MER5A_dup27620 | 13.0297073075194 | 2.11845080221082 | 0.544405008231885 | 3.89131394858233 | 9.97028258969587e-05 | 0.0100482374463692 | 639 | 3 | 3 | 8 | 12 | 15 | 11 | 3 | 0 | 24 | 25 | 30 | 27 |
| MLT2B2_dup280 | 13.8199499498126 | 1.97339475653808 | 0.472245699517722 | 4.17874584893711 | 2.93121022204379e-05 | 0.0041140044901846 | 640 | 6 | 8 | 5 | 11 | 10 | 21 | 4 | 0 | 25 | 25 | 25 | 32 |
| MIR3_dup45007 | 11.1573772670738 | 1.77470441271078 | 0.468108465083694 | 3.79122478033691 | 0.000149906203405567 | 0.01331451342872 | 641 | 1 | 3 | 9 | 15 | 8 | 8 | 7 | 4 | 15 | 20 | 32 | 17 |
| L1MA2_dup2673 | 8.46717892184196 | 1.5131282776907 | 0.412581212780844 | 3.66746771500342 | 0.000244964468884398 | 0.0188574298004422 | 642 | 2 | 3 | 8 | 9 | 6 | 5 | 9 | 4 | 8 | 12 | 21 | 21 |

| | | | | | | | | | | | | | | | | | | | |
|---|---|---|---|---|---|---|---|---|---|---|---|---|---|---|---|---|---|---|---|
| MIRb_dup174517 | 8.44421688424053 | 1.68454760979397 | 0.509845313176556 | 3.3040366680994 | 0.000953033660856589 | 0.0463008369418401 | 643 | 1 | 1 | 7 | 7 | 10 | 7 | 5 | 4 | 10 | 18 | 19 | 16 |
| AluY_dup31628 | 9.46277048918064 | 1.35473731242577 | 0.398680135933008 | 3.39805571013806 | 0.00067866598333296 | 0.0370890804019625 | 644 | 4 | 4 | 9 | 7 | 6 | 11 | 8 | 5 | 12 | 17 | 18 | 17 |
| L2a_dup137053 | 8.8397165903874 | 1.46742930982154 | 0.403517733105664 | 3.6365918754735 | 0.000276269132534981 | 0.0204342974852871 | 645 | 5 | 3 | 7 | 6 | 6 | 9 | 7 | 5 | 14 | 15 | 17 | 15 |
| L2b_dup74913 | 10.5793713774368 | 1.66665103908882 | 0.447478744988369 | 3.72453676907523 | 0.000195674192396668 | 0.0161722443153934 | 646 | 4 | 3 | 8 | 9 | 6 | 10 | 7 | 6 | 13 | 29 | 24 | 11 |
| MER58A_dup6785 | 9.51592423673299 | 1.58836995075909 | 0.437714579152831 | 3.62878009188837 | 0.00028476368030754 | 0.020834648649285 | 647 | 3 | 5 | 6 | 10 | 5 | 9 | 6 | 5 | 16 | 22 | 19 | 10 |
| L3_dup37625 | 11.2377044803155 | 1.52638441808797 | 0.419315688561847 | 3.64017960626064 | 0.000272447930660159 | 0.0202777276536948 | 648 | 2 | 5 | 4 | 12 | 9 | 11 | 10 | 7 | 14 | 25 | 20 | 21 |
| AluSg_dup10635 | 8.85862706651825 | 1.56140967897223 | 0.472497742287143 | 3.30458653921639 | 0.000951166059139941 | 0.0462680418532698 | 649 | 2 | 4 | 3 | 9 | 7 | 9 | 10 | 3 | 11 | 25 | 13 | 15 |
| L2a_dup57885 | 9.14340785561987 | 1.67755022335774 | 0.450699494623464 | 3.72210362640688 | 0.000197569903750731 | 0.0162893168114593 | 650 | 2 | 5 | 3 | 11 | 9 | 5 | 7 | 4 | 14 | 19 | 20 | 14 |
| MIR3_dup67878 | 11.5901446547028 | 1.41439027735467 | 0.408763806485891 | 3.46016514895011 | 0.000539844186734678 | 0.0320950622291132 | 651 | 4 | 4 | 10 | 9 | 16 | 8 | 7 | 6 | 18 | 18 | 21 | 22 |
| hAT-N1_Mam | 11.8454545370433 | 1.28249008606332 | 0.380182504280413 | 3.37335377515791 | 0.000742584762170989 | 0.039331664037174 | 652 | 4 | 5 | 9 | 9 | 13 | 7 | 16 | 7 | 14 | 20 | 20 | 25 |
| Charlie7a_dup1230 | 10.3873853618882 | 1.82380853808028 | 0.444036692426304 | 4.10733745473744 | 4.00246258049885e-05 | 0.00513588925876228 | 653 | 4 | 3 | 12 | 8 | 7 | 3 | 8 | 5 | 11 | 30 | 17 | 22 |
| MIR_dup137193 | 10.7606494217071 | 1.65359558731669 | 0.475823177523365 | 3.47523127377605 | 0.000510413097074911 | 0.03095205798549 | 654 | 2 | 3 | 11 | 8 | 9 | 5 | 8 | 8 | 18 | 28 | 17 | 13 |
| MER49_dup1031 | 10.6150357324796 | 1.48679261358228 | 0.417008639420161 | 3.56537604508537 | 0.0003633348548847 | 0.02468172339212 | 655 | 3 | 3 | 11 | 12 | 11 | 6 | 7 | 5 | 15 | 19 | 21 | 19 |
| Tigger19b_dup2880 | 10.3189444752281 | 1.52883602204166 | 0.440280267537199 | 3.47241549250783 | 0.000515797373293227 | 0.03113442967672 | 656 | 3 | 3 | 11 | 10 | 9 | 6 | 10 | 4 | 14 | 23 | 22 | 13 |
| MIRc_dup79399 | 10.7382413700429 | 1.92797771198048 | 0.430437906218281 | 4.47910763463935 | 7.49557329038866e-06 | 0.00148381171560301 | 657 | 3 | 5 | 10 | 6 | 11 | 3 | 7 | 3 | 21 | 19 | 18 | 26 |
| L1PA7_dup11540 | 10.0883007833024 | 1.78657974455577 | 0.397474000121428 | 4.4948342382394 | 6.96241623306914e-06 | 0.0014059862931934 | 658 | 6 | 5 | 8 | 11 | 4 | 4 | 7 | 4 | 15 | 25 | 15 | 22 |
| MamRTE1_dup7772 | 10.1697989904094 | 1.68040519638387 | 0.384406947992115 | 4.37142253843533 | 1.23439644214319e-05 | 0.00218378956001638 | 659 | 4 | 6 | 6 | 11 | 6 | 5 | 7 | 6 | 14 | 22 | 21 | 18 |
| MIRc_dup79351 | 7.58899436368347 | 1.82110151768023 | 0.458959413751775 | 3.96789228658279 | 7.25110917013423e-05 | 0.00795102498892516 | 660 | 6 | 3 | 5 | 3 | 4 | 6 | 2 | 6 | 10 | 16 | 17 | 15 |
| LTR25-int_dup29 | 8.89474889261769 | 1.83394163500144 | 0.457394811609119 | 4.00953746840638 | 6.08377993654278e-05 | 0.00697501986589356 | 661 | 7 | 5 | 5 | 6 | 7 | 7 | 2 | 3 | 10 | 14 | 29 | 16 |
| MLT1C_dup13199 | 8.66821101772847 | 1.51280384461282 | 0.437211706739165 | 3.46011742433818 | 0.000539939877548435 | 0.0320950622291132 | 662 | 6 | 5 | 5 | 7 | 8 | 5 | 4 | 6 | 9 | 19 | 22 | 11 |
| AluSp_dup37009 | 7.84402318692719 | 1.88409306000732 | 0.483136339274664 | 3.89971299371917 | 9.63067826121759e-05 | 0.00977637103047646 | 663 | 5 | 2 | 6 | 7 | 4 | 2 | 5 | 5 | 13 | 10 | 26 | 11 |
| AluSx3_dup21095 | 8.63978469872055 | 1.52161087954995 | 0.450622552136968 | 3.37668603653784 | 0.000733647402400131 | 0.0389922860870006 | 664 | 7 | 5 | 6 | 4 | 7 | 5 | 8 | 4 | 14 | 11 | 25 | 10 |
| L1M1_dup7432 | 8.11342462274704 | 1.53696463675642 | 0.454321214194879 | 3.38299112772036 | 0.00071700948614854 | 0.0383561600635049 | 665 | 5 | 8 | 4 | 3 | 4 | 3 | 7 | 8 | 9 | 15 | 18 | 16 |
| L1PB1_dup4083 | 8.98341707275823 | 1.35838959468215 | 0.400245392363653 | 3.39389189881779 | 0.000689068870004625 | 0.0374260916699391 | 666 | 9 | 8 | 6 | 7 | 4 | 4 | 7 | 6 | 8 | 13 | 20 | 22 |
| MER21C_dup1096 | 8.70764632926045 | 1.61891568687723 | 0.434319481153991 | 3.7274765630493 | 0.000193406539727993 | 0.016054021806127 | 667 | 7 | 6 | 4 | 6 | 3 | 3 | 10 | 5 | 9 | 8 | 26 | 24 |
| AluSp_dup24968 | 7.72362616424666 | 1.61618281485259 | 0.444528180931257 | 3.63572633677982 | 0.000277198486669296 | 0.0204846106275154 | 668 | 6 | 6 | 7 | 4 | 2 | 5 | 4 | 5 | 15 | 12 | 13 | 15 |
| THE1B_dup7229 | 9.22545685925421 | 1.56638647039992 | 0.411388121097869 | 3.80756368516357 | 0.000140342604017069 | 0.0127457867387197 | 669 | 6 | 5 | 9 | 8 | 4 | 7 | 4 | 5 | 18 | 12 | 17 | 18 |
| L1M3a_dup420 | 8.3932649599418 | 1.67938853645124 | 0.469875379118529 | 3.57411477826679 | 0.000351414762961249 | 0.0241627001658766 | 670 | 6 | 5 | 11 | 3 | 5 | 2 | 3 | 6 | 11 | 13 | 17 | 22 |
| L2a_dup152271 | 9.63525470988059 | 1.53767033452316 | 0.427035255874641 | 3.60080418038025 | 0.000317234444588007 | 0.0224790551375634 | 671 | 7 | 7 | 12 | 3 | 5 | 4 | 8 | 5 | 11 | 20 | 14 | 25 |
| L1MEj_dup1977 | 12.8639732097468 | 1.46672101771848 | 0.43948221118244 | 3.33738426811913 | 0.000845709341544974 | 0.0428337920356025 | 672 | 1 | 8 | 9 | 13 | 6 | 15 | 13 | 6 | 17 | 27 | 26 | 19 |
| MLT1C_dup3781 | 14.9033982194794 | 2.05776365086695 | 0.418760439688241 | 4.91393994236637 | 8.92641040037592e-07 | 0.000295000506823312 | 673 | 2 | 5 | 5 | 8 | 5 | 15 | 11 | 11 | 19 | 30 | 35 | 39 |
| MLT2B1_dup961 | 14.8263714185371 | 1.7472069561883 | 0.382694239458771 | 4.5655428695747 | 4.98203366880351e-06 | 0.00109359315807852 | 674 | 4 | 6 | 6 | 10 | 7 | 17 | 11 | 10 | 15 | 25 | 34 | 42 |
| MIRb_dup71166 | 10.7450757932557 | 1.86297107054999 | 0.477719601076145 | 3.89971662530348 | 9.63053381383346e-05 | 0.00977637103047646 | 675 | 6 | 2 | 8 | 3 | 2 | 11 | 10 | 7 | 19 | 27 | 16 | 20 |
| MSTA_dup17784 | 12.4481039681938 | 1.68080242963774 | 0.406108212129958 | 4.13880433695802 | 3.49120531521516e-05 | 0.0046768976735448 | 676 | 9 | 4 | 7 | 4 | 4 | 14 | 9 | 10 | 21 | 24 | 21 | 25 |
| L2b_dup74916 | 13.566579869343 | 1.42196869986998 | 0.406601245469307 | 3.49720694590769 | 0.000470156957219519 | 0.0292781908774027 | 677 | 10 | 3 | 10 | 12 | 10 | 14 | 14 | 4 | 13 | 37 | 16 | 30 |
| THE1D_dup1245 | 12.7390677052917 | 1.45822226961206 | 0.38025294258614 | 3.83487438570373 | 0.000125628481943576 | 0.0118153338990301 | 678 | 9 | 5 | 11 | 10 | 5 | 11 | 16 | 4 | 22 | 26 | 16 | 24 |
| MER5A_dup27615 | 14.082915260099 | 1.4221063223293 | 0.395052816768844 | 3.59978783080392 | 0.000318476932982399 | 0.0225268466989982 | 679 | 9 | 2 | 8 | 12 | 12 | 12 | 15 | 8 | 28 | 25 | 22 | 19 |
| MIRb_dup174423 | 12.278446647832 | 1.85228333294145 | 0.420827348596782 | 4.40152794041962 | 1.07491222668708e-05 | 0.00197050126880804 | 680 | 7 | 2 | 6 | 12 | 8 | 7 | 12 | 4 | 26 | 22 | 28 | 16 |
| MIRc_dup79382 | 13.4382710580504 | 1.97923620439864 | 0.433925373484053 | 4.56123639073479 | 5.0853290499242e-06 | 0.00110910513850895 | 681 | 6 | 2 | 7 | 12 | 7 | 7 | 10 | 8 | 23 | 40 | 26 | 15 |
| AluJo_dup34550 | 10.2387088909732 | 1.62676761303052 | 0.437053924892736 | 3.72212104817447 | 0.000197556268965893 | 0.0162893168114593 | 682 | 4 | 6 | 4 | 8 | 6 | 10 | 9 | 5 | 18 | 24 | 7 | 26 |
| L1PA7_dup7230 | 11.2186005717435 | 1.35721882444386 | 0.412947147085315 | 3.28666473185117 | 0.00101381457471574 | 0.0482511438789805 | 683 | 7 | 6 | 4 | 7 | 8 | 10 | 13 | 8 | 16 | 29 | 9 | 22 |
| MIRb_dup145993 | 11.2026277119083 | 1.68961685257548 | 0.418312065088131 | 4.03913009829039 | 5.36498032248782e-05 | 0.00636061588348125 | 684 | 2 | 10 | 4 | 5 | 10 | 6 | 11 | 7 | 17 | 22 | 19 | 25 |
| AluY_dup66115 | 10.7134347263506 | 1.5032878248115 | 0.447351367278805 | 3.36041853175917 | 0.00077824479135561 | 0.0407078995655973 | 685 | 5 | 5 | 2 | 5 | 10 | 7 | 12 | 10 | 19 | 22 | 14 | 19 |
| SVA_A_dup942 | 11.03843197936 | 1.76136689863772 | 0.491507588163008 | 3.58360062195737 | 0.000338890127671993 | 0.0235626197100543 | 686 | 2 | 4 | 3 | 4 | 8 | 10 | 12 | 9 | 23 | 19 | 13 | 27 |
| MSTD_dup5438 | 12.552255906028 | 1.50358614984586 | 0.43974071658135 | 3.41925615061325 | 0.000627925853547278 | 0.0353638923427599 | 687 | 8 | 7 | 1 | 7 | 11 | 9 | 12 | 9 | 24 | 36 | 13 | 14 |
| AluJb_dup81411 | 11.3724446499996 | 1.52236808723603 | 0.407309574928023 | 3.73761919911912 | 0.00018577102429201 | 0.0156138688051098 | 688 | 7 | 10 | 2 | 7 | 5 | 13 | 8 | 8 | 23 | 17 | 18 | 20 |
| L2_dup42863 | 10.4477544886889 | 1.34368541408326 | 0.394787577621741 | 3.40356558881062 | 0.000665124424921163 | 0.0366258787677276 | 689 | 10 | 5 | 2 | 7 | 8 | 12 | 9 | 7 | 14 | 23 | 13 | 20 |
| Tigger19b_dup2290 | 10.6360962527154 | 1.40509354367118 | 0.400091183373496 | 3.51193328436706 | 0.000444859657884758 | 0.028265040361828 | 690 | 5 | 8 | 5 | 6 | 8 | 13 | 9 | 4 | 25 | 20 | 12 | 13 |
| MIR_dup57146 | 12.3981074883951 | 1.63005026095446 | 0.362185635496787 | 4.50059334550535 | 6.77640378864109e-06 | 0.00137813519307008 | 691 | 12 | 8 | 3 | 6 | 9 | 8 | 12 | 5 | 25 | 27 | 19 | 17 |
| L2a_dup39618 | 12.5203132273149 | 1.62149736894625 | 0.361804276586678 | 4.48169763012125 | 7.40516130921654e-06 | 0.00147681113018408 | 692 | 9 | 6 | 6 | 7 | 7 | 10 | 11 | 7 | 23 | 30 | 21 | 15 |
| L2a_dup57882 | 12.5393341953165 | 1.62200981693156 | 0.386583280634095 | 4.19575780481517 | 2.7196074246029e-05 | 0.00389354078069124 | 693 | 10 | 4 | 4 | 7 | 10 | 11 | 11 | 6 | 28 | 26 | 20 | 14 |
| MER44B_dup1417 | 12.3543437482106 | 1.58867914211506 | 0.379339782462129 | 4.18801089567679 | 2.81410009709379e-05 | 0.00398336622072028 | 694 | 10 | 11 | 6 | 4 | 5 | 13 | 7 | 8 | 14 | 31 | 20 | 24 |
| L1MEj_dup1975 | 12.463680828232 | 1.29823618940795 | 0.358988595940691 | 3.61637167332868 | 0.000298761321709129 | 0.0215826648244835 | 695 | 13 | 9 | 9 | 6 | 7 | 14 | 7 | 7 | 17 | 30 | 20 | 14 |
| UCON84_dup35 | 12.2357794113295 | 1.61061329153775 | 0.378126892364336 | 4.25945185085086 | 2.04928835439574e-05 | 0.00314593526951645 | 696 | 11 | 8 | 11 | 4 | 6 | 9 | 7 | 6 | 17 | 34 | 20 | 17 |
| MamGypsy2-I_dup81 | 11.1676053333125 | 1.2620988383212 | 0.370494840688265 | 3.40652203408991 | 0.000657962392966713 | 0.036409190594618 | 697 | 10 | 16 | 5 | 7 | 6 | 11 | 7 | 5 | 15 | 20 | 17 | 20 |
| L1MC2_dup5623 | 9.76917853704338 | 1.32173066034736 | 0.395189193051207 | 3.34455163144123 | 0.000824156878456794 | 0.0421369844884236 | 698 | 11 | 11 | 4 | 6 | 5 | 6 | 7 | 6 | 14 | 23 | 15 | 12 |
| MIRb_dup29939 | 11.1686259643994 | 1.44242360358192 | 0.378767132932772 | 3.80820688535806 | 0.000139978141028349 | 0.0127175405670606 | 699 | 12 | 11 | 5 | 8 | 5 | 8 | 8 | 5 | 13 | 29 | 20 | 15 |
| MIR3_dup57904 | 10.9652984521257 | 1.50370590838839 | 0.45296266261273 | 3.31971271034763 | 0.000901101275630185 | 0.0445821616253597 | 700 | 7 | 9 | 2 | 7 | 4 | 8 | 14 | 8 | 10 | 30 | 21 | 17 |
| MIRb_dup154838 | 11.0211914596571 | 1.61124011560224 | 0.375717166562174 | 4.28843890830207 | 1.79933272848969e-05 | 0.00286767599071138 | 701 | 7 | 4 | 6 | 8 | 5 | 6 | 13 | 8 | 19 | 18 | 20 | 22 |
| AluSq2_dup46066 | 10.4856213737225 | 1.44230101664102 | 0.386748228514936 | 3.72930219274508 | 0.000192010768648665 | 0.0159693094571392 | 702 | 8 | 9 | 5 | 5 | 6 | 7 | 9 | 8 | 17 | 12 | 22 | 21 |
| Tigger18a_dup165 | 10.8636367601911 | 1.41357630588852 | 0.378993269207725 | 3.7298190251335 | 0.000191617351095633 | 0.0159477377487938 | 703 | 6 | 7 | 4 | 6 | 9 | 8 | 12 | 8 | 18 | 16 | 17 | 23 |
| MIR3_dup27872 | 14.0489875545347 | 1.55271423796251 | 0.402845492045344 | 3.85436667065332 | 0.000116029593304354 | 0.0111731192263852 | 704 | 4 | 11 | 9 | 10 | 7 | 7 | 10 | 14 | 24 | 32 | 16 | 26 |
| MIR_dup72716 | 11.6479985073488 | 1.3195765149144 | 0.38317276539831 | 3.44381603828939 | 0.000573565816730513 | 0.0334122554045634 | 705 | 10 | 9 | 8 | 9 | 4 | 7 | 7 | 12 | 18 | 20 | 14 | 25 |
| AluSx3_dup5118 | 11.8739744296301 | 1.1506355818935 | 0.337913671220765 | 3.40511698664528 | 0.000661357151384567 | 0.0365100077262221 | 706 | 13 | 9 | 10 | 10 | 11 | 8 | 8 | 5 | 14 | 18 | 19 | 25 |
| L1ME3F_dup399 | 12.4643468563899 | 1.19975027024874 | 0.330296305035768 | 3.63234541821115 | 0.000280856824131227 | 0.0206772146889632 | 707 | 10 | 7 | 10 | 10 | 11 | 11 | 11 | 6 | 13 | 23 | 23 | 22 |
| X7A_LINE_dup308 | 10.6691108656985 | 1.18030051786691 | 0.359090892777644 | 3.28691298388837 | 0.00101292127874289 | 0.0482427074843386 | 708 | 10 | 6 | 9 | 9 | 8 | 10 | 7 | 6 | 16 | 18 | 12 | 22 |
| MIR_dup104594 | 11.7500790657406 | 1.28191606626177 | 0.355952850912094 | 3.60136479586267 | 0.00031655103484674 | 0.0224506863003405 | 709 | 11 | 8 | 5 | 10 | 10 | 9 | 9 | 7 | 13 | 30 | 14 | 21 |
| MLT1C_dup1283 | 11.2458633745348 | 1.19404972422277 | 0.355763166214362 | 3.35630508612942 | 0.000789913784706952 | 0.0411194105044586 | 710 | 12 | 6 | 6 | 7 | 9 | 9 | 10 | 9 | 12 | 18 | 20 | 23 |
| MER115_dup338 | 13.9641151269321 | 1.34720237834128 | 0.328539334021145 | 4.10058169246356 | 4.12112918988187e-05 | 0.00523828101610914 | 711 | 15 | 7 | 8 | 9 | 10 | 12 | 8 | 10 | 22 | 27 | 23 | 20 |
| MIRb_dup71174 | 14.0566220690751 | 1.52051977777189 | 0.350522491748802 | 4.33786650946655 | 1.43872489680042e-05 | 0.00244993718135918 | 712 | 9 | 5 | 7 | 12 | 10 | 10 | 10 | 11 | 21 | 33 | 24 | 20 |
| AluSp_dup14784 | 12.0304315107417 | 1.71278728203417 | 0.355830003582149 | 4.81349876286851 | 1.48310519885342e-06 | 0.00043855939092937 | 713 | 11 | 6 | 4 | 10 | 10 | 7 | 5 | 6 | 22 | 27 | 19 | 20 |
| MER20_dup9706 | 11.6870331286906 | 1.33689007940727 | 0.378980837092134 | 3.52759281884814 | 0.000419356704084522 | 0.0271226556972263 | 714 | 10 | 6 | 4 | 13 | 8 | 9 | 7 | 9 | 21 | 24 | 13 | 19 |

| | | | | | | | | | | | | | | | | | | | |
|---|---|---|---|---|---|---|---|---|---|---|---|---|---|---|---|---|---|---|---|
| L1MC5a_dup13425 | 9.12591994393085 | 1.66144998902535 | 0.435356818771467 | 3.81629485835042 | 0.000135470605310266 | 0.0124218453376326 | 715 | 4 | 7 | 11 | 10 | 3 | 5 | 5 | 2 | 10 | 19 | 17 | 23 |
| MER53_dup4541 | 8.32573399202528 | 1.54370523655546 | 0.459337560636996 | 3.36072067438747 | 0.00077739401514481 | 0.0406812835556734 | 716 | 3 | 7 | 9 | 8 | 6 | 6 | 2 | 3 | 9 | 20 | 15 | 16 |
| L2b_dup12526 | 8.5288467315348 | 1.55458037095616 | 0.47460851652825 | 3.27550036886797 | 0.00105474976617431 | 0.049500908001123 | 717 | 8 | 7 | 5 | 12 | 6 | 5 | 1 | 2 | 11 | 18 | 12 | 21 |
| L1ME3A_dup2032 | 10.0492246465028 | 1.40023417841756 | 0.389848242927045 | 3.59174166825627 | 0.000328475375322909 | 0.0230151359412794 | 718 | 8 | 11 | 8 | 8 | 9 | 7 | 3 | 3 | 15 | 16 | 17 | 20 |
| L2b_dup30999 | 10.3953846898333 | 1.48040939318745 | 0.399214067509127 | 3.70830968563903 | 0.000208647387936109 | 0.0169385231558667 | 719 | 7 | 8 | 7 | 11 | 11 | 5 | 4 | 3 | 18 | 25 | 12 | 17 |
| MamRTE1_dup3278 | 9.25321452081361 | 1.98313925921163 | 0.47139222239407 | 4.2069834099931 | 2.58802132387559e-05 | 0.00375493488982256 | 720 | 5 | 7 | 8 | 5 | 2 | 5 | 7 | 1 | 17 | 29 | 15 | 12 |
| L1ME3Cz_dup5108 | 8.68842494055549 | 1.50679863985283 | 0.449788406229935 | 3.35001662777974 | 0.000808067094170073 | 0.0416272640808383 | 721 | 9 | 6 | 4 | 6 | 6 | 7 | 9 | 1 | 8 | 26 | 16 | 12 |
| MIRb_dup145990 | 8.81853080064576 | 1.83028642352344 | 0.447336885488596 | 4.09151689229548 | 4.28560722355059e-05 | 0.00539029296425262 | 722 | 7 | 3 | 6 | 2 | 5 | 7 | 10 | 2 | 11 | 22 | 16 | 20 |
| MIRb_dup185039 | 8.67466363153621 | 1.46415197806095 | 0.427398766124073 | 3.42572813519989 | 0.0006131537694762 | 0.0348608368369612 | 723 | 7 | 6 | 6 | 4 | 3 | 10 | 8 | 3 | 14 | 23 | 10 | 13 |
| L1P1_dup134 | 8.93106494212343 | 1.3977421268755 | 0.421526341703186 | 3.31590695193067 | 0.000913462372860661 | 0.0450009767645348 | 724 | 6 | 7 | 4 | 9 | 10 | 10 | 3 | 2 | 16 | 15 | 15 | 13 |
| Tigger2a_dup1668 | 9.8321967176077 | 1.55094682486823 | 0.383795295399252 | 4.04107826088597 | 5.32059972794169e-05 | 0.00632691365749532 | 725 | 5 | 7 | 4 | 8 | 8 | 11 | 5 | 4 | 17 | 19 | 15 | 18 |
| AluSz_dup33141 | 9.45868105250435 | 1.42510621594974 | 0.410332641872014 | 3.47305105791277 | 0.000514577452860763 | 0.0310844495045461 | 726 | 5 | 10 | 4 | 10 | 6 | 8 | 5 | 4 | 15 | 23 | 15 | 11 |
| AluSz6_dup13998 | 8.94242719125387 | 1.38880989667673 | 0.413720932000275 | 3.35687607093518 | 0.000788284369420807 | 0.0410593057198402 | 727 | 6 | 5 | 4 | 10 | 10 | 5 | 6 | 4 | 15 | 20 | 13 | 12 |
| AluJb_dup8277 | 9.11457265969981 | 1.48392207824917 | 0.404296831907649 | 3.67037770552734 | 0.00024219232434987 | 0.0187227728426752 | 728 | 8 | 6 | 4 | 7 | 9 | 7 | 6 | 3 | 12 | 21 | 19 | 11 |
| AluSx_dup32473 | 9.31938058498951 | 1.51525182359907 | 0.392621429820718 | 3.85932022174891 | 0.000113702867278004 | 0.011015937482627 | 729 | 5 | 8 | 4 | 7 | 9 | 6 | 8 | 3 | 15 | 20 | 16 | 14 |
| MIRb_dup71188 | 10.84885500134 | 1.60655435354069 | 0.449408956331262 | 3.57481605764103 | 0.000350474209020885 | 0.0241326771274171 | 730 | 4 | 6 | 10 | 9 | 9 | 7 | 10 | 1 | 15 | 34 | 15 | 15 |
| MLT1A0_dup17008 | 10.2209245184925 | 1.5990885725564 | 0.41699945069309 | 3.83474982976254 | 0.000125692159166213 | 0.0118166521157502 | 731 | 3 | 6 | 9 | 8 | 11 | 6 | 8 | 2 | 15 | 25 | 16 | 18 |
| LTR33_dup6359 | 11.1285583531118 | 1.57300731247063 | 0.440912572245099 | 3.56761728172317 | 0.000360242153516372 | 0.024535953954961 | 732 | 7 | 6 | 8 | 9 | 11 | 11 | 4 | 1 | 21 | 32 | 17 | 8 |
| Tigger3b_dup4981 | 10.2065074336935 | 1.67633823064969 | 0.423320582462541 | 3.95997336320878 | 7.49581238137778e-05 | 0.0081368548033728 | 733 | 6 | 6 | 6 | 3 | 10 | 15 | 5 | 1 | 17 | 29 | 13 | 14 |
| L2a_dup89293 | 8.23706013210886 | 1.73478387074423 | 0.506220189283093 | 3.42693536818638 | 0.000610434343295503 | 0.0347905174250998 | 734 | 9 | 2 | 4 | 4 | 10 | 11 | 2 | 1 | 9 | 23 | 15 | 13 |
| L1MC2_dup2043 | 8.43784541727086 | 1.53337407000392 | 0.454684270658093 | 3.37239304052584 | 0.000745180237750112 | 0.0394378685168346 | 735 | 6 | 7 | 2 | 3 | 11 | 11 | 3 | 3 | 11 | 25 | 13 | 8 |
| AluSg_dup16292 | 8.24953992604897 | 1.60703129461312 | 0.457482592847811 | 3.51277036489938 | 0.00044346056849426 | 0.0281912018249894 | 736 | 7 | 4 | 2 | 6 | 13 | 8 | 3 | 1 | 16 | 18 | 11 | 12 |
| MER39_dup1579 | 8.53444610630736 | 1.66657987546071 | 0.455273824003035 | 3.66060991780103 | 0.000251615582706003 | 0.0192185382035177 | 737 | 6 | 7 | 3 | 4 | 14 | 6 | 4 | 1 | 14 | 17 | 16 | 13 |
| AluJo_dup34548 | 9.69481035130867 | 2.14025774295352 | 0.43683893400714 | 4.89942076206636 | 9.61196043711602e-07 | 0.000307409615345431 | 738 | 3 | 4 | 3 | 6 | 10 | 8 | 4 | 2 | 17 | 24 | 13 | 26 |
| L3b_dup2442 | 8.33169643977313 | 1.8884086020416 | 0.464222888045869 | 4.0678920636395 | 4.74403455693629e-05 | 0.00580670805907799 | 739 | 5 | 5 | 3 | 3 | 9 | 7 | 5 | 2 | 13 | 21 | 7 | 24 |
| AluSz_dup75651 | 8.41841567546386 | 2.1634176515666 | 0.494966822992151 | 4.37083366211983 | 1.237730582331e-05 | 0.00218806122691926 | 740 | 1 | 3 | 5 | 4 | 7 | 9 | 2 | 2 | 19 | 23 | 11 | 15 |
| AluSq2_dup45244 | 8.62349560234148 | 1.64546944683964 | 0.500825293587662 | 3.28551586333095 | 0.00101795810150156 | 0.0484147628262821 | 741 | 3 | 4 | 8 | 8 | 11 | 5 | 3 | 1 | 19 | 19 | 9 | 15 |
| THE1B-int_dup113 | 9.17701417960316 | 1.73944939636508 | 0.422018875467826 | 4.12173364150319 | 3.76031832650785e-05 | 0.0049162236512046 | 742 | 5 | 4 | 5 | 10 | 8 | 7 | 4 | 2 | 16 | 22 | 11 | 20 |
| Tigger19a_dup2803 | 9.14301344952578 | 1.5431837760247 | 0.461281393704527 | 3.34542818567094 | 0.000821556313265763 | 0.0420671483788111 | 743 | 4 | 5 | 6 | 10 | 10 | 9 | 3 | 2 | 17 | 14 | 10 | 24 |
| AluSx1_dup73503 | 9.1644136857858 | 1.85565437599076 | 0.472137921903891 | 3.93032266611387 | 8.48319413981188e-05 | 0.00892765198050277 | 744 | 8 | 4 | 2 | 3 | 7 | 12 | 3 | 5 | 9 | 21 | 17 | 24 |
| ORSL-2b_dup75 | 8.36424742864235 | 1.57304569970365 | 0.459193275953529 | 3.4256723303214 | 0.000613279748344244 | 0.0348608368369612 | 745 | 11 | 4 | 3 | 2 | 7 | 11 | 5 | 3 | 9 | 18 | 16 | 16 |
| L1PA16_dup11952 | 10.7646955243562 | 2.01272751195692 | 0.38091475803075 | 5.28393156086233 | 1.2644050363086e-07 | 6.20330145916464e-05 | 746 | 7 | 8 | 4 | 6 | 4 | 7 | 5 | 5 | 18 | 31 | 14 | 23 |
| MIR_dup57147 | 10.5794559728141 | 2.11019728711381 | 0.408992429696839 | 5.15950206872526 | 2.4760748806138e-07 | 0.000106541069356697 | 747 | 5 | 3 | 4 | 5 | 5 | 9 | 6 | 6 | 16 | 30 | 23 | 17 |
| MIR_dup114524 | 8.4408001308877 | 1.79688412688559 | 0.454600972981715 | 3.95266229876254 | 7.72864742455045e-05 | 0.00833258151373847 | 748 | 7 | 5 | 2 | 7 | 3 | 6 | 5 | 5 | 14 | 17 | 22 | 10 |
| L1M5_dup45705 | 8.96645436480568 | 1.47755498059636 | 0.421456727696533 | 3.50582843622384 | 0.000455188521615293 | 0.0286005513504813 | 749 | 9 | 4 | 3 | 6 | 10 | 5 | 4 | 7 | 16 | 16 | 14 | 15 |
| L2c_dup51273 | 8.77462174059876 | 1.46253858125442 | 0.425248260729921 | 3.43925823175392 | 0.000583310539379259 | 0.0337856622072427 | 750 | 7 | 3 | 4 | 5 | 6 | 11 | 5 | 6 | 18 | 16 | 12 | 13 |
| MLT1K_dup12283 | 9.03496990152613 | 1.74534159231749 | 0.501164541497663 | 3.48257198544369 | 0.000496621680990627 | 0.0303379081982717 | 751 | 4 | 4 | 6 | 7 | 12 | 8 | 3 | 1 | 14 | 12 | 26 | 15 |
| MIR_dup70189 | 8.02894648415975 | 1.58923743190963 | 0.438642794194019 | 3.62307885355729 | 0.000291117025712668 | 0.0211762179304455 | 752 | 3 | 4 | 3 | 8 | 9 | 7 | 5 | 3 | 11 | 17 | 15 | 15 |
| MIRc_dup245 | 8.44375286256871 | 1.81242580349008 | 0.469014855905894 | 3.86432493697755 | 0.000111396849471642 | 0.0108588425827035 | 753 | 3 | 3 | 3 | 5 | 12 | 7 | 3 | 4 | 13 | 16 | 16 | 19 |
| L2a_dup41599 | 7.55699214272217 | 1.63347059295013 | 0.488480520896301 | 3.34398307214566 | 0.000825847763919131 | 0.0421870270574284 | 754 | 7 | 2 | 3 | 5 | 9 | 8 | 5 | 1 | 15 | 9 | 13 | 17 |
| MER103C_dup2941 | 8.27010719464392 | 1.6195366644485 | 0.488382417300725 | 3.31612401895963 | 0.000912753135118197 | 0.0449846639372068 | 755 | 7 | 1 | 3 | 5 | 10 | 7 | 7 | 3 | 14 | 12 | 21 | 12 |
| L1PB1_dup2754 | 8.29496710633252 | 1.95294587905811 | 0.474304622303084 | 4.11749282470657 | 3.83016450518263e-05 | 0.00497816297128209 | 756 | 7 | 3 | 2 | 7 | 10 | 3 | 5 | 1 | 13 | 15 | 19 | 19 |
| L2b_dup74937 | 7.84556363668249 | 1.75513819006387 | 0.497063901085322 | 3.5310111762926 | 0.000413974255675236 | 0.0269061781592187 | 757 | 4 | 2 | 4 | 8 | 10 | 3 | 4 | 3 | 15 | 8 | 18 | 18 |
| AluSq_dup8572 | 8.86203585855382 | 1.78618890969698 | 0.469861669702902 | 3.80152079829452 | 0.000143810664328695 | 0.0129421983110271 | 758 | 7 | 5 | 7 | 3 | 10 | 4 | 6 | 1 | 10 | 12 | 21 | 27 |
| L1ME4a_dup23221 | 9.42919312850334 | 1.43931701225183 | 0.418503201098839 | 3.43920191882093 | 0.000583431896352787 | 0.0337856622072427 | 759 | 9 | 2 | 9 | 8 | 8 | 7 | 7 | 3 | 9 | 22 | 15 | 21 |
| AluSx4_dup4545 | 8.79688029872817 | 1.52577774166424 | 0.438178812786078 | 3.48208926844926 | 0.000497517807221522 | 0.0303688468697268 | 760 | 8 | 4 | 6 | 12 | 6 | 7 | 3 | 2 | 9 | 15 | 17 | 24 |
| HAL1_dup22254 | 9.91132545738938 | 1.71115590293534 | 0.435211675072552 | 3.93177849066224 | 8.431974220586e-05 | 0.00889735936980735 | 761 | 5 | 5 | 6 | 9 | 11 | 9 | 3 | 2 | 10 | 20 | 19 | 27 |
| AluSx_dup106642 | 11.5870009827105 | 1.33946465175652 | 0.393642259185913 | 3.40274607336787 | 0.000667122506241436 | 0.0366857249526519 | 762 | 11 | 7 | 5 | 4 | 14 | 18 | 6 | 4 | 20 | 18 | 14 | 22 |
| Charlie2b_dup1873 | 11.9216973228183 | 1.31365000332574 | 0.383833329900149 | 3.42244901886836 | 0.000620597345743284 | 0.0350673607290979 | 763 | 16 | 8 | 6 | 7 | 11 | 15 | 5 | 4 | 17 | 20 | 12 | 29 |
| MIRb_dup29938 | 11.2276943164474 | 1.34306397415275 | 0.36776102443887 | 3.65200193849252 | 0.00026020392032434 | 0.0196366895101476 | 764 | 13 | 10 | 6 | 6 | 14 | 9 | 6 | 2 | 20 | 20 | 18 | 14 |
| Tigger19a_dup1863 | 11.2921877074873 | 1.39747591773969 | 0.421555703768513 | 3.31504450122938 | 0.000916285364816121 | 0.0450840441307975 | 765 | 14 | 5 | 7 | 7 | 16 | 11 | 3 | 3 | 18 | 16 | 21 | 19 |
| MER45A_dup1108 | 12.7253042036564 | 1.4364455371114 | 0.347420425696857 | 4.13460300795549 | 3.55568745158188e-05 | 0.0047239620678621 | 766 | 8 | 8 | 4 | 9 | 11 | 14 | 12 | 5 | 21 | 22 | 20 | 24 |
| MIR_dup121267 | 13.1750591651475 | 1.35275975482563 | 0.385584634522758 | 3.50833418582655 | 0.000450922229240323 | 0.028498300031631 | 767 | 7 | 8 | 4 | 8 | 20 | 13 | 11 | 5 | 23 | 21 | 21 | 21 |
| L1MC2_dup2042 | 12.6419951147636 | 1.47400560091076 | 0.378732699679169 | 3.89194173663752 | 9.94451292403263e-05 | 0.0100350172698128 | 768 | 11 | 11 | 3 | 4 | 18 | 9 | 9 | 5 | 22 | 25 | 21 | 16 |
| L2a_dup61521 | 12.5659257049103 | 1.41414859785767 | 0.366612090552016 | 3.85734304541991 | 0.000114626236981946 | 0.0110738345858328 | 769 | 15 | 13 | 3 | 6 | 16 | 9 | 7 | 4 | 16 | 28 | 21 | 18 |
| AluSx_dup32474 | 13.6990761754679 | 1.59913425966784 | 0.345790477976491 | 4.62457575184172 | 3.75365887987299e-06 | 0.000886407198837999 | 770 | 9 | 11 | 4 | 11 | 14 | 10 | 9 | 3 | 28 | 29 | 21 | 18 |
| MER2_dup3118 | 11.5763044558007 | 1.33745024167804 | 0.370070018831616 | 3.61404646045264 | 0.000301455006160895 | 0.0217363675442304 | 771 | 9 | 10 | 2 | 10 | 12 | 11 | 8 | 4 | 23 | 25 | 14 | 13 |
| MamTip2_dup2259 | 11.4019635072652 | 1.85031525361572 | 0.491405924469971 | 3.76534991028337 | 0.000166316005065357 | 0.0143488862006097 | 772 | 5 | 5 | 3 | 3 | 19 | 8 | 10 | 2 | 21 | 19 | 24 | 21 |
| AluSc_dup26564 | 11.2104042820088 | 1.4219568662395 | 0.39479377775246 | 3.60177121922901 | 0.000316056452658111 | 0.0224289796254658 | 773 | 7 | 6 | 6 | 6 | 18 | 9 | 7 | 4 | 17 | 21 | 15 | 23 |
| MIR_dup56567 | 9.88362445276689 | 1.5060324048116 | 0.425011556024941 | 3.54350930806977 | 0.000394839372446924 | 0.026068382163223 | 774 | 6 | 8 | 3 | 6 | 15 | 9 | 5 | 3 | 14 | 15 | 19 | 20 |
| MIRb_dup174366 | 11.3165089844596 | 1.83484036969202 | 0.442886977817459 | 4.14290882683905 | 3.42928301987583e-05 | 0.00461110957597297 | 775 | 4 | 3 | 4 | 4 | 15 | 7 | 8 | 7 | 18 | 24 | 23 | 21 |
| MIRb_dup71170 | 9.89969115143712 | 1.62011091164535 | 0.426729696541106 | 3.79657409544565 | 0.000146709526002764 | 0.0131187316325643 | 776 | 7 | 5 | 3 | 4 | 12 | 5 | 9 | 5 | 22 | 18 | 11 | 19 |
| MIRc_dup79408 | 11.0255824401119 | 1.7987503876254 | 0.406084938851526 | 4.42949298418346 | 9.44548856542587e-06 | 0.00177669266576192 | 777 | 5 | 6 | 5 | 6 | 9 | 11 | 4 | 6 | 17 | 14 | 28 | 25 |
| LTR2C_dup30 | 10.8580926671933 | 1.60903399896309 | 0.472088006702155 | 3.40833483613203 | 0.000653606390171339 | 0.0362622254961759 | 778 | 8 | 8 | 4 | 6 | 14 | 9 | 2 | 6 | 18 | 9 | 23 | 27 |
| MER20_dup4166 | 9.8789997483569 | 1.38015329454998 | 0.401638743435542 | 3.4363051799844 | 0.000589706300604554 | 0.0340330476361026 | 779 | 7 | 7 | 5 | 8 | 11 | 8 | 5 | 5 | 16 | 10 | 17 | 24 |
| AluSz6_dup10229 | 11.1241919649071 | 2.12086925410454 | 0.421210793937437 | 5.03517308822707 | 4.7741724488929e-07 | 0.000178896887799098 | 780 | 7 | 4 | 5 | 5 | 10 | 7 | 4 | 4 | 22 | 16 | 16 | 38 |
| Charlie29a_dup1656 | 11.8369811992339 | 1.432928021969 | 0.412751301709785 | 3.47164991614376 | 0.000517270419247321 | 0.0311835256753778 | 781 | 6 | 6 | 4 | 12 | 12 | 13 | 7 | 6 | 16 | 15 | 16 | 37 |
| HERVL-int_dup852 | 14.7074955888006 | 2.49916832526834 | 0.461307623152222 | 5.41757430365218 | 6.04130456861072e-08 | 3.35865480580058e-05 | 782 | 4 | 4 | 7 | 15 | 7 | 5 | 4 | 3 | 37 | 47 | 19 | 24 |
| MER5A_dup27622 | 14.4578847219349 | 2.05416902097046 | 0.37600093091985 | 5.46320195523223 | 4.67622083626989e-08 | 2.75096544669864e-05 | 783 | 9 | 3 | 6 | 17 | 8 | 8 | 6 | 5 | 24 | 35 | 30 | 28 |
| MLT1C_dup15274 | 13.4451568035024 | 1.60680365661183 | 0.434844586664065 | 3.69512167309824 | 0.00021978151351622 | 0.0174779121709676 | 784 | 7 | 5 | 6 | 21 | 9 | 14 | 7 | 2 | 21 | 32 | 19 | 26 |
| MER44C_dup530 | 12.2739840630317 | 1.68800457901052 | 0.478366272191687 | 3.52868644203687 | 0.000417627647243732 | 0.027047585234365 | 785 | 6 | 1 | 12 | 10 | 16 | 8 | 7 | 2 | 17 | 22 | 21 | 33 |
| MIR3_dup67865 | 12.0457061425704 | 1.75377890317906 | 0.46902691460134 | 3.73918606498249 | 0.000184617045843074 | 0.0155610653879476 | 786 | 6 | 1 | 13 | 9 | 13 | 8 | 5 | 3 | 22 | 28 | 17 | 23 |

| | | | | | | | | | | | | | | | | | | | |
|---|---|---|---|---|---|---|---|---|---|---|---|---|---|---|---|---|---|---|---|
| MIR_dup137205 | 10.596248690023 | 1.88626128695916 | 0.43260936480571 | 4.36019522556176 | 1.29946456909366e-05 | 0.00226323878879064 | 787 | 2 | 2 | 8 | 9 | 6 | 10 | 8 | 4 | 15 | 27 | 18 | 23 |
| Charlie17a_dup966 | 10.7462880390375 | 1.70080285973324 | 0.477514667680466 | 3.56178139615042 | 0.000368347029664794 | 0.0249279585667289 | 788 | 4 | 1 | 10 | 8 | 8 | 10 | 9 | 3 | 25 | 21 | 17 | 14 |
| L2a_dup137066 | 11.0550356770588 | 1.73276285129683 | 0.467443145558379 | 3.70689541126327 | 0.000209815540214093 | 0.0169896393973003 | 789 | 5 | 1 | 7 | 17 | 6 | 8 | 8 | 4 | 20 | 26 | 17 | 18 |
| Charlie16a_dup1459 | 11.4249749655473 | 1.51721507337582 | 0.460422477581333 | 3.29526716711568 | 0.000983281481144137 | 0.0472676432766945 | 790 | 6 | 3 | 9 | 17 | 11 | 7 | 5 | 4 | 15 | 28 | 24 | 13 |
| MIRb_dup174439 | 10.0153808716589 | 1.59148442064361 | 0.460578736435007 | 3.45540142161597 | 0.000549474079373881 | 0.0324681398242244 | 791 | 5 | 4 | 10 | 15 | 6 | 5 | 3 | 5 | 11 | 22 | 25 | 14 |
| MIR3_dup67872 | 10.1621945761031 | 2.16043211201038 | 0.47684908887678 | 4.53064116594893 | 5.88049458966498e-06 | 0.00123854437659785 | 792 | 4 | 3 | 12 | 7 | 5 | 6 | 2 | 2 | 19 | 20 | 27 | 17 |
| HERVP71A-int_dup24 | 12.7116182087813 | 1.84433203753438 | 0.398636299781909 | 4.62660334380837 | 3.71711432373458e-06 | 0.000879197301068934 | 793 | 9 | 4 | 14 | 8 | 6 | 10 | 3 | 5 | 17 | 25 | 30 | 27 |
| HSMAR1_dup32 | 11.7112949478655 | 2.12207179389757 | 0.411705192531823 | 5.15434789842624 | 2.545150451708e-07 | 0.000108531965839088 | 794 | 7 | 4 | 10 | 7 | 6 | 6 | 2 | 5 | 25 | 28 | 15 | 27 |
| L2c_dup114228 | 12.5781569982002 | 1.40465570009123 | 0.350241976650088 | 4.01052927329313 | 6.05827882515699e-05 | 0.0069579262646345 | 795 | 7 | 6 | 9 | 16 | 11 | 9 | 8 | 5 | 19 | 23 | 21 | 23 |
| MIR_dup142238 | 12.9090574619906 | 1.31924775774892 | 0.375761143844399 | 3.51086795258218 | 0.000446646202900952 | 0.0283473708799606 | 796 | 9 | 6 | 14 | 19 | 9 | 6 | 7 | 6 | 16 | 21 | 29 | 20 |
| L2b_dup74967 | 12.9008010599068 | 1.61895736712814 | 0.436221915508733 | 3.71131598292137 | 0.000206184530383566 | 0.0167919620766515 | 797 | 7 | 2 | 13 | 21 | 6 | 8 | 9 | 4 | 18 | 26 | 29 | 19 |
| AluSq2_dup45249 | 12.4031258584121 | 1.55106332213682 | 0.428907000731404 | 3.6163161699199 | 0.000298825357149541 | 0.0215826648244835 | 798 | 8 | 2 | 14 | 16 | 8 | 10 | 6 | 3 | 22 | 19 | 21 | 26 |
| MLT1K_dup8222 | 12.9812081507445 | 1.61154034470983 | 0.367294060297705 | 4.38760252045357 | 1.14606976220681e-05 | 0.00207154664139496 | 799 | 9 | 3 | 12 | 13 | 8 | 10 | 10 | 3 | 22 | 25 | 22 | 25 |
| MIR1_Amn_dup8821 | 13.4749347724662 | 1.39265630721459 | 0.378520317442268 | 3.67921150606928 | 0.000233956180787823 | 0.018270081192563 | 800 | 8 | 5 | 13 | 9 | 7 | 14 | 10 | 9 | 23 | 16 | 31 | 20 |
| MIRb_dup49013 | 13.9280099448717 | 1.10706212070619 | 0.330757806889998 | 3.34704759085058 | 0.000816771868666597 | 0.0419213975543016 | 801 | 9 | 7 | 14 | 16 | 9 | 12 | 10 | 10 | 19 | 19 | 27 | 21 |
| MER5B_dup19716 | 14.449239446713 | 1.79333742681683 | 0.337328655164186 | 5.31629139523876 | 1.05903505849986e-07 | 5.37298840148845e-05 | 802 | 6 | 6 | 8 | 16 | 8 | 11 | 7 | 7 | 22 | 23 | 33 | 33 |
| MIR_dup112776 | 14.2094010472526 | 1.67523180164825 | 0.371454470616347 | 4.50992499529907 | 6.48505505391682e-06 | 0.00132946272810474 | 803 | 8 | 6 | 6 | 12 | 14 | 14 | 7 | 5 | 20 | 21 | 37 | 27 |
| L1PA7_dup551 | 12.4411614541287 | 1.35281267776993 | 0.388492327771346 | 3.48221208261839 | 0.000497289669460241 | 0.0303638921450825 | 804 | 7 | 5 | 14 | 12 | 11 | 12 | 6 | 4 | 14 | 14 | 28 | 31 |
| MIR_dup112774 | 13.2066986977731 | 1.43483390311767 | 0.431794891120247 | 3.32295247726332 | 0.000890700872149175 | 0.0442368419378851 | 805 | 6 | 7 | 13 | 13 | 7 | 16 | 6 | 5 | 22 | 13 | 36 | 19 |
| AluJb_dup8278 | 11.4108048896378 | 1.44254928847178 | 0.430602175473117 | 3.35007431601293 | 0.000807898816600204 | 0.0416272640808383 | 806 | 6 | 3 | 9 | 7 | 11 | 17 | 8 | 3 | 12 | 23 | 22 | 23 |
| L2c_dup134704 | 12.0436076038749 | 1.58380186801153 | 0.370888824157941 | 4.27028738762179 | 1.9522124744824e-05 | 0.00304803903840675 | 807 | 7 | 4 | 8 | 7 | 11 | 14 | 7 | 5 | 18 | 20 | 21 | 28 |
| MER5A1_dup12437 | 11.8879261760978 | 1.63706899262215 | 0.373323428407807 | 4.38512257214638 | 1.15920486578497e-05 | 0.00208374840047616 | 808 | 5 | 5 | 8 | 7 | 12 | 12 | 9 | 3 | 18 | 25 | 19 | 25 |
| L1M5_dup44657 | 10.706255112652 | 1.63655516347975 | 0.360008290938114 | 4.54588187181799 | 5.47057135519643e-06 | 0.00116744445890903 | 809 | 8 | 6 | 7 | 7 | 6 | 8 | 10 | 3 | 18 | 21 | 19 | 20 |
| LTR12D_dup430 | 10.0549894419988 | 1.48924633675734 | 0.398590209539206 | 3.73628428675907 | 0.000186759517375109 | 0.0156529341744761 | 810 | 5 | 5 | 6 | 12 | 8 | 9 | 8 | 2 | 18 | 20 | 15 | 17 |
| AluSz6_dup35678 | 9.68818321934126 | 1.26088421299223 | 0.382959668811203 | 3.29247259092925 | 0.000993105774221881 | 0.0475655286942431 | 811 | 7 | 5 | 7 | 13 | 9 | 7 | 6 | 4 | 11 | 14 | 19 | 21 |
| HERVE_a-int_dup184 | 10.3307424732371 | 1.27418203318497 | 0.388968676593552 | 3.27579599556396 | 0.00105364639447585 | 0.0494694050868097 | 812 | 8 | 8 | 9 | 10 | 10 | 9 | 6 | 2 | 10 | 18 | 23 | 18 |
| MER103C_dup7097 | 11.1368228712809 | 1.68179167884759 | 0.420363265651076 | 4.000805532431 | 6.31272215028824e-05 | 0.00718358194534905 | 813 | 3 | 6 | 9 | 13 | 5 | 12 | 4 | 4 | 17 | 22 | 18 | 26 |
| L2c_dup114223 | 11.479994144647 | 1.52665242717867 | 0.367113551353912 | 4.15852921132544 | 3.20303277537685e-05 | 0.00439263584193471 | 814 | 4 | 8 | 10 | 8 | 10 | 10 | 7 | 4 | 19 | 21 | 17 | 24 |
| MIRb_dup27846 | 11.0351016654355 | 1.50313353536579 | 0.380530162118121 | 3.95010352661401 | 7.81173973397474e-05 | 0.00838796129395466 | 815 | 5 | 8 | 9 | 10 | 9 | 10 | 3 | 5 | 17 | 21 | 19 | 20 |
| Tigger3b_dup1414 | 11.7925075628789 | 1.60768989798708 | 0.371949606224782 | 4.32233257162126 | 1.5438823434002e-05 | 0.00255644139236398 | 816 | 8 | 4 | 7 | 8 | 6 | 12 | 6 | 9 | 18 | 17 | 22 | 29 |
| AluJo_dup75930 | 11.6931247606115 | 1.42421049510353 | 0.405528915152723 | 3.51198260318176 | 0.000444777112746573 | 0.0282650403618282 | 817 | 5 | 5 | 8 | 10 | 9 | 15 | 4 | 8 | 19 | 20 | 15 | 26 |
| MIRb_dup132438 | 11.3564654304395 | 1.35897569157545 | 0.370202118590145 | 3.67090198389703 | 0.000241696020461378 | 0.0186904781555746 | 818 | 8 | 5 | 7 | 9 | 7 | 14 | 6 | 8 | 16 | 24 | 17 | 19 |
| HAL1_dup15475 | 12.2921691335388 | 1.34857705446688 | 0.368682634901145 | 3.65782634386477 | 0.000254363284778772 | 0.0193431531351772 | 819 | 6 | 5 | 11 | 11 | 8 | 10 | 9 | 10 | 13 | 29 | 18 | 23 |
| MER102a_dup1285 | 10.4692120882904 | 1.4350908928271 | 0.373612445123347 | 3.84112176015258 | 0.000122473341725295 | 0.0116213079607753 | 820 | 4 | 5 | 8 | 9 | 7 | 7 | 7 | 10 | 12 | 16 | 22 | 23 |
| MER5A1_dup6630 | 12.605315878889 | 1.444445704859 | 0.416091875748863 | 3.47145856250942 | 0.000517639214909546 | 0.0311835256753778 | 821 | 16 | 12 | 7 | 6 | 8 | 13 | 7 | 3 | 26 | 13 | 28 | 15 |
| L1ME4a_dup1647 | 11.3303260919379 | 1.59186210385984 | 0.458289179359541 | 3.47348830291927 | 0.000513739755767195 | 0.0310471542025556 | 822 | 10 | 13 | 11 | 8 | 5 | 7 | 5 | 1 | 15 | 15 | 33 | 19 |
| LTR16_dup129 | 11.8523233906785 | 1.65401404466972 | 0.423371586113218 | 3.9067667716071 | 9.3539361192595e-05 | 0.009577158708405 | 823 | 7 | 11 | 7 | 6 | 9 | 11 | 8 | 2 | 15 | 11 | 27 | 37 |
| L1MC1_dup1705 | 11.1448719864862 | 1.24037274136285 | 0.36969768479148 | 3.35510010581336 | 0.000793362693269772 | 0.0412267917477111 | 824 | 8 | 12 | 11 | 10 | 5 | 7 | 6 | 7 | 13 | 17 | 23 | 20 |
| L2_dup8008 | 10.9002970554726 | 1.24653482066675 | 0.371956602243131 | 3.3512910193 | 0.000804357238690881 | 0.0415354127014618 | 825 | 8 | 9 | 7 | 15 | 6 | 6 | 8 | 6 | 13 | 11 | 26 | 23 |
| THE1D_dup9835 | 11.1947195939177 | 1.64565638122816 | 0.442887824378322 | 3.71574085049222 | 0.000202609162553299 | 0.0165898278709247 | 826 | 7 | 11 | 7 | 8 | 3 | 13 | 6 | 3 | 8 | 28 | 26 | 22 |
| AluSg_dup22404 | 11.0722829759528 | 1.40688630758019 | 0.387949369123815 | 3.62646886308296 | 0.000287323447442445 | 0.0209844889579926 | 827 | 5 | 9 | 7 | 8 | 5 | 13 | 10 | 5 | 18 | 20 | 12 | 26 |
| AluSx1_dup86778 | 10.1382445620863 | 1.41874079393122 | 0.391290607007337 | 3.62579823927288 | 0.000288070212206171 | 0.021025022039365 | 828 | 9 | 8 | 6 | 10 | 3 | 10 | 6 | 5 | 14 | 16 | 14 | 27 |
| LTR16A1_dup1755 | 11.7860554836352 | 1.72647148213747 | 0.418857871046677 | 4.12185517207356 | 3.75833465362753e-05 | 0.0049162236512046 | 829 | 5 | 8 | 11 | 8 | 2 | 12 | 4 | 7 | 20 | 20 | 20 | 28 |
| MER3_dup6979 | 11.6745968723718 | 1.33732941277757 | 0.380815009623322 | 3.51175604685428 | 0.00044515641920395 | 0.0282763452546458 | 830 | 6 | 7 | 12 | 10 | 4 | 13 | 9 | 6 | 20 | 17 | 21 | 19 |
| L1MA9_dup5789 | 31.6382769825878 | 3.37332391697565 | 0.687130834339655 | 4.90928910244215 | 9.14071414215051e-07 | 0.000299586276483216 | 831 | 11 | 10 | 2 | 2 | 47 | 52 | 1 | 3 | 47 | 55 | 73 | 90 |
| L1PA7_dup5384 | 34.3479090914763 | 2.91842548982831 | 0.627848978229315 | 4.6482921706092 | 3.34694613466728e-06 | 0.000820177595220949 | 832 | 9 | 6 | 3 | 5 | 48 | 38 | 7 | 6 | 42 | 43 | 96 | 132 |
| L1PB1_dup11487 | 19.5156918221876 | 2.51679959195054 | 0.756798720760664 | 3.32558647749944 | 0.000882327248316978 | 0.043940192429475 | 833 | 3 | 1 | 3 | 1 | 26 | 40 | 6 | 4 | 33 | 23 | 50 | 51 |
| THE1B-int_dup3854 | 18.2812328885706 | 2.94294085511279 | 0.829918168426991 | 3.54606148783413 | 0.000391034954115724 | 0.0258818367710765 | 834 | 3 | 0 | 0 | 1 | 29 | 22 | 5 | 4 | 25 | 43 | 46 | 48 |
| HERVK-int_dup18 | 32.3306649216301 | 2.19668811684221 | 0.486680462344891 | 4.51361475712066 | 6.37319390209552e-06 | 0.00131410398286657 | 835 | 42 | 28 | 5 | 4 | 24 | 33 | 11 | 9 | 39 | 39 | 62 | 116 |
| L1M1_dup95 | 26.7182928853444 | 2.12927280808854 | 0.466507290505844 | 4.5642862425145 | 5.01196570207628e-06 | 0.00109813737656192 | 836 | 23 | 25 | 3 | 1 | 20 | 12 | 18 | 12 | 43 | 57 | 44 | 72 |
| L1MC1_dup244 | 28.0243969404682 | 1.72391520203666 | 0.502015597982085 | 3.43398732821481 | 0.000594772000170299 | 0.0342340634621485 | 837 | 45 | 30 | 2 | 3 | 22 | 15 | 23 | 11 | 43 | 46 | 38 | 73 |
| L2c_dup5680 | 26.712881973138 | 1.75358113945069 | 0.519386203220344 | 3.37625668255718 | 0.000734793326075028 | 0.0390183291154315 | 838 | 30 | 21 | 1 | 2 | 17 | 24 | 23 | 16 | 39 | 52 | 39 | 68 |
| AluJb_dup53405 | 21.6571696641122 | 1.59014247520074 | 0.353142655782559 | 4.50283320115211 | 6.70535097495147e-06 | 0.00136602007113511 | 839 | 30 | 21 | 4 | 4 | 21 | 32 | 9 | 5 | 33 | 46 | 33 | 29 |
| L1ME1_dup25854 | 18.7614586741743 | 2.0429765145562 | 0.432317255316212 | 4.72564185082526 | 2.29389638336279e-06 | 0.000626662995218877 | 840 | 20 | 18 | 1 | 2 | 22 | 25 | 5 | 4 | 35 | 37 | 27 | 33 |
| L1PA13_dup7133 | 17.3041830158396 | 1.68785134498138 | 0.427130833720165 | 3.95160267471391 | 7.7629552465228e-05 | 0.00835442853500278 | 841 | 25 | 21 | 6 | 2 | 17 | 15 | 9 | 3 | 23 | 27 | 32 | 37 |
| LTR62_dup11 | 20.7668420473777 | 1.97335015255892 | 0.551939443800927 | 3.57530191893781 | 0.00034982395450904 | 0.0241119967850035 | 842 | 33 | 21 | 4 | 1 | 21 | 29 | 6 | 5 | 20 | 27 | 37 | 61 |
| L1MDa_dup82 | 26.1410339332407 | 2.22720144452571 | 0.543536813966941 | 4.09760919094097 | 4.17439273133574e-05 | 0.00528340453643838 | 843 | 21 | 26 | 1 | 1 | 20 | 29 | 10 | 11 | 35 | 46 | 44 | 83 |
| L2a_dup5965 | 24.2895392957584 | 2.02128051456008 | 0.434016651788387 | 4.65714968822346 | 3.20617333732362e-06 | 0.000796342506182469 | 844 | 28 | 20 | 2 | 2 | 21 | 20 | 19 | 4 | 48 | 49 | 34 | 52 |
| MLT2C1_dup39 | 20.6256969101794 | 2.10622690977123 | 0.45853954633707 | 4.59333753565272 | 4.36212645686367e-06 | 0.000991240230967941 | 845 | 14 | 21 | 1 | 1 | 25 | 14 | 12 | 6 | 36 | 48 | 32 | 42 |
| Tigger1_dup2716 | 29.3575451360945 | 1.22552945803115 | 0.363909223486142 | 3.36767902250716 | 0.000758037935982142 | 0.0399229957315641 | 846 | 50 | 45 | 2 | 3 | 38 | 52 | 6 | 8 | 50 | 57 | 21 | 26 |
| L1ME2_dup4576 | 24.2469479334246 | 1.72317090144784 | 0.432035309340429 | 3.98849553310476 | 6.64936547822856e-05 | 0.00746315999095554 | 847 | 25 | 35 | 6 | 1 | 27 | 36 | 5 | 7 | 38 | 45 | 33 | 39 |
| L1PA4_dup9862 | 22.483948495793 | 1.51078131060743 | 0.402306523302271 | 3.7552990645203 | 0.000173134616812506 | 0.0148136244272091 | 848 | 28 | 17 | 3 | 3 | 25 | 33 | 4 | 13 | 42 | 54 | 17 | 32 |
| L1ME1_dup18928 | 27.5222752686724 | 2.43674610063842 | 0.526341950873702 | 4.62958746988254 | 3.66394933007821e-06 | 0.000870081923447894 | 849 | 26 | 15 | 3 | 3 | 44 | 42 | 4 | 3 | 61 | 33 | 39 | 63 |
| MSTA-int_dup2197 | 30.731740345369 | 2.4052927229381 | 0.572267294165913 | 4.20309311305978 | 2.63292217402842e-05 | 0.00380153701226558 | 850 | 50 | 37 | 1 | 0 | 38 | 45 | 7 | 3 | 48 | 56 | 46 | 49 |
| L1MC1_dup10106 | 22.7464814164961 | 2.19175151958699 | 0.608363334579458 | 3.60270153542714 | 0.000314927059854329 | 0.0223755252863835 | 851 | 39 | 35 | 5 | 3 | 31 | 34 | 2 | 0 | 28 | 27 | 41 | 41 |
| L1PA3_dup1673 | 18.1570984452117 | 2.26818299498638 | 0.690791650695051 | 3.28345455927877 | 0.00102543174719776 | 0.0486544224854446 | 852 | 49 | 41 | 1 | 2 | 15 | 25 | 1 | 0 | 27 | 19 | 25 | 22 |
| L1PA2_dup187 | 27.8615179535395 | 2.24425622607009 | 0.437298501086372 | 5.13209219902362 | 2.86539067799409e-07 | 0.000118575347872345 | 853 | 38 | 38 | 6 | 11 | 17 | 12 | 4 | 0 | 62 | 70 | 36 | 44 |
| THE1B-int_dup3889 | 19.3710316115239 | 2.14713244693347 | 0.479197470643575 | 4.48068401540146 | 7.44041983264315e-06 | 0.00147904606307277 | 854 | 20 | 21 | 6 | 6 | 12 | 28 | 3 | 0 | 24 | 52 | 44 | 24 |
| L1PA3_dup7081 | 18.5007966426459 | 1.09327143631819 | 0.318781925752116 | 3.42952767393819 | 0.000604632770876888 | 0.0346090808994641 | 855 | 17 | 22 | 5 | 8 | 27 | 16 | 14 | 6 | 32 | 45 | 18 | 15 |
| L1M4_dup15347 | 20.0211054882569 | 1.48788934035158 | 0.349541645815248 | 4.25668688742747 | 2.07478608875541e-05 | 0.00317483633874595 | 856 | 14 | 14 | 3 | 8 | 30 | 29 | 8 | 5 | 46 | 42 | 19 | 22 |
| L1MCb_dup782 | 17.5720125628098 | 1.27844906157188 | 0.308519357140821 | 4.14382122865745 | 3.41566051764898e-05 | 0.00459696129826077 | 857 | 21 | 20 | 5 | 6 | 20 | 22 | 7 | 5 | 38 | 31 | 19 | 18 |
| AluJr_dup23266 | 17.9741926313824 | 1.28960047905028 | 0.376779463079999 | 3.4226931279863 | 0.000620040335902848 | 0.0350525345133616 | 858 | 14 | 15 | 1 | 17 | 16 | 24 | 11 | 9 | 24 | 38 | 29 | 25 |

| | | | | | | | | | | | | | | | | | | |
|---|---|---|---|---|---|---|---|---|---|---|---|---|---|---|---|---|---|---|
| L1MA8_dup8321 | 21.2579972813442 | 1.54455259945328 | 0.304338441889063 | 5.07511502610737 | 3.87261661339942e-07 | 0.000152058355229693 | 859 | 21 | 20 | 4 | 13 | 18 | 15 | 7 | 12 | 38 | 60 | 22 | 27 |
| L1M1_dup96 | 13.9938517203938 | 1.80789627518511 | 0.544615986417748 | 3.31957988798067 | 0.000901530058562984 | 0.0445821616253597 | 860 | 17 | 11 | 1 | 1 | 18 | 7 | 8 | 8 | 17 | 20 | 29 | 38 |
| L2a_dup5963 | 13.0569044879026 | 1.59992466169354 | 0.486248489901221 | 3.29034371298213 | 0.0010006507133731 | 0.0478662029861271 | 861 | 11 | 19 | 1 | 3 | 7 | 17 | 7 | 7 | 19 | 18 | 27 | 25 |
| MLT1N2_dup133 | 16.9721404822252 | 1.60843707816547 | 0.453852746573394 | 3.54396242021059 | 0.000394161423670459 | 0.0260404258365742 | 862 | 20 | 17 | 1 | 5 | 15 | 20 | 8 | 9 | 16 | 29 | 32 | 42 |
| Tigger3a_dup119 | 17.8416204819404 | 1.76255745747754 | 0.536695993732896 | 3.28408908965087 | 0.00102312574524726 | 0.048592550614492 | 863 | 19 | 21 | 0 | 2 | 12 | 13 | 20 | 5 | 27 | 39 | 23 | 42 |
| MLT1H1_dup80 | 20.8067289969558 | 1.81118122036529 | 0.479954824015801 | 3.77364937227021 | 0.000160876822663779 | 0.0140580229326315 | 864 | 31 | 18 | 1 | 4 | 14 | 12 | 11 | 13 | 26 | 38 | 38 | 54 |
| L1ME2_dup372 | 21.5835492108194 | 1.5919678557998 | 0.477010442726581 | 3.33738575344419 | 0.00084570482147163 | 0.0428337920356025 | 865 | 21 | 25 | 1 | 3 | 18 | 15 | 18 | 14 | 31 | 37 | 40 | 43 |
| MLT1E1A_dup60 | 20.2337552347503 | 1.99809225983517 | 0.474114284389855 | 4.21436840361506 | 2.50478145840654e-05 | 0.00366732586027206 | 866 | 18 | 19 | 1 | 2 | 19 | 11 | 14 | 9 | 24 | 50 | 47 | 36 |
| L3_dup35386 | 11.5804655731718 | 1.62528803648359 | 0.479372115466828 | 3.39045176814431 | 0.00069777531617564 | 0.0377175932264264 | 867 | 16 | 14 | 10 | 6 | 5 | 2 | 3 | 4 | 18 | 23 | 16 | 27 |
| Tigger1_dup6307 | 12.1396197367392 | 1.48467460374542 | 0.441832699959921 | 3.36026419927745 | 0.000778679695712429 | 0.0407145608885742 | 868 | 15 | 11 | 9 | 3 | 14 | 5 | 3 | 6 | 24 | 19 | 23 | 14 |
| L1PA7_dup11056 | 15.3437513728557 | 1.53487282962713 | 0.383409817864404 | 4.003217335895 | 6.24868491054723e-05 | 0.00712092710596299 | 869 | 26 | 18 | 4 | 7 | 13 | 13 | 2 | 1 | 35 | 42 | 15 | 8 |
| L1MC3_dup1593 | 16.2125347172108 | 1.61736948997629 | 0.374515583675696 | 4.3185639275743 | 1.57047734519305e-05 | 0.00258966599015457 | 870 | 25 | 21 | 4 | 8 | 15 | 10 | 3 | 1 | 28 | 53 | 14 | 16 |
| L2a_dup112235 | 15.9643148691321 | 1.65012687687842 | 0.346264416663546 | 4.76551097215917 | 1.88375645337941e-06 | 0.000538740760884396 | 871 | 25 | 17 | 3 | 4 | 18 | 14 | 4 | 1 | 33 | 46 | 13 | 15 |
| AluJb_dup53403 | 17.1960148850731 | 2.09078510706351 | 0.399262313181482 | 5.23662023195552 | 1.63543719157555e-07 | 7.77593831715409e-05 | 872 | 25 | 12 | 2 | 5 | 17 | 10 | 5 | 3 | 38 | 40 | 15 | 38 |
| HERVH-int_dup5629 | 16.3923367020501 | 2.03666011681741 | 0.381940185632824 | 5.33240594582352 | 9.69199719097048e-08 | 5.00254222039623e-05 | 873 | 18 | 19 | 4 | 9 | 10 | 6 | 4 | 3 | 39 | 32 | 26 | 28 |
| MER101-int_dup157 | 15.2800135939507 | 1.58552750535076 | 0.361051197644035 | 4.39142015231299 | 1.12612691350858e-05 | 0.00204703891949361 | 874 | 15 | 19 | 5 | 10 | 9 | 6 | 9 | 5 | 36 | 33 | 17 | 20 |
| L1MCc_dup1858 | 16.7016241017581 | 1.89880135495406 | 0.364577293061603 | 5.20822714713945 | 1.90653445842749e-07 | 8.74544048961656e-05 | 875 | 13 | 9 | 7 | 13 | 13 | 10 | 5 | 3 | 44 | 45 | 12 | 26 |
| MER21C_dup2763 | 16.8652516893241 | 1.87420204106771 | 0.358375639393515 | 5.22971383947707 | 1.69772623578762e-07 | 7.93773995421099e-05 | 876 | 17 | 13 | 5 | 11 | 11 | 10 | 2 | 5 | 41 | 53 | 17 | 15 |
| MER110_dup98 | 20.0560881769418 | 2.76446411936905 | 0.326587013166708 | 8.46470927476198 | 2.5679243538803e-17 | 1.79713920090707e-13 | 877 | 14 | 10 | 5 | 3 | 10 | 6 | 5 | 7 | 40 | 57 | 37 | 49 |
| L1PREC2_dup2546 | 18.0403176842046 | 2.92920134301219 | 0.387304183384999 | 7.56305113312041 | 3.93722736255903e-14 | 1.37771691472275e-10 | 878 | 17 | 14 | 1 | 2 | 9 | 11 | 1 | 2 | 48 | 52 | 25 | 32 |
| Tigger1_dup4321 | 21.4523834197813 | 2.61626630321398 | 0.375251785488444 | 6.97202892668061 | 3.1240229550111e-12 | 6.99283363762677e-09 | 879 | 25 | 15 | 3 | 6 | 7 | 14 | 1 | 3 | 59 | 64 | 30 | 26 |
| L1PA15_dup277 | 13.3576032782384 | 2.23622598006191 | 0.474809035175865 | 4.70973763006348 | 2.48035876470503e-06 | 0.000663136456883711 | 880 | 18 | 19 | 3 | 3 | 8 | 8 | 2 | 1 | 26 | 26 | 22 | 28 |
| LTR8_dup3217 | 14.789796878176 | 2.83387511383054 | 0.527671155406139 | 5.37053254625859 | 7.85044629672335e-08 | 4.1424527977874e-05 | 881 | 16 | 13 | 1 | 1 | 12 | 9 | 2 | 2 | 23 | 33 | 34 | 37 |
| L2a_dup160290 | 12.7582492130679 | 1.83076361871271 | 0.505667365632722 | 3.62048995671679 | 0.000294045665346181 | 0.0212989314722869 | 882 | 22 | 15 | 1 | 2 | 8 | 16 | 2 | 5 | 16 | 27 | 24 | 20 |
| MIR1_Amn_dup4850 | 11.278490726966 | 1.64041792899803 | 0.47953799330653 | 3.42082994860731 | 0.000624303552029557 | 0.0352349461554139 | 883 | 19 | 9 | 1 | 3 | 14 | 10 | 3 | 5 | 16 | 19 | 24 | 16 |
| L1PA12_dup79 | 14.7486159588683 | 1.84881041901968 | 0.457422697283987 | 4.04179860334273 | 5.30427805916311e-05 | 0.0063138156431497 | 884 | 24 | 14 | 2 | 11 | 9 | 10 | 1 | 4 | 22 | 34 | 27 | 25 |
| L1MB8_dup12110 | 12.1715377631409 | 1.57082756782927 | 0.3995352324723 | 3.93163716278307 | 8.43693368722781e-05 | 0.00889864637917158 | 885 | 23 | 10 | 4 | 5 | 8 | 9 | 6 | 3 | 19 | 23 | 19 | 23 |
| THE1B_dup17349 | 16.4015112755794 | 1.84416310664787 | 0.394344287376681 | 4.67653054876465 | 2.91769152143839e-06 | 0.000740141819573751 | 886 | 26 | 16 | 3 | 5 | 10 | 12 | 5 | 6 | 27 | 31 | 32 | 29 |
| L1PA7_dup5797 | 14.7491878834188 | 1.61147559516998 | 0.491125530864442 | 3.28118880794811 | 0.001033705222219 | 0.0489192567235723 | 887 | 13 | 9 | 4 | 4 | 24 | 23 | 4 | 5 | 15 | 21 | 32 | 31 |
| MLT2A2_dup1396 | 15.0328755501238 | 1.46448163819924 | 0.417602929372002 | 3.50687587465336 | 0.000453400590202545 | 0.0285609855960214 | 888 | 23 | 13 | 3 | 3 | 23 | 16 | 6 | 5 | 21 | 24 | 22 | 28 |
| AluJo_dup32000 | 16.4064880413075 | 1.59041750611422 | 0.346485678214179 | 4.59013923551295 | 4.42950432020663e-06 | 0.00100083295050532 | 889 | 18 | 15 | 6 | 3 | 18 | 19 | 5 | 6 | 24 | 37 | 27 | 23 |
| LTR78_dup824 | 16.6429891386519 | 1.290526122183 | 0.387937580487253 | 3.32663342531055 | 0.000879019261585767 | 0.0438488715340224 | 890 | 15 | 11 | 7 | 4 | 26 | 26 | 8 | 6 | 25 | 26 | 26 | 25 |
| AluJr4_dup6368 | 14.4485947475749 | 1.94081661526288 | 0.390585166954446 | 4.96899723662379 | 6.73000289960474e-07 | 0.000236540217126935 | 891 | 8 | 9 | 7 | 4 | 20 | 10 | 6 | 2 | 30 | 34 | 15 | 31 |
| L1PB1_dup10475 | 17.9135212771373 | 1.76721562383615 | 0.358922396495749 | 4.92367052346113 | 8.49357956993002e-07 | 0.000283452073540893 | 892 | 12 | 14 | 8 | 4 | 25 | 19 | 5 | 2 | 40 | 44 | 24 | 17 |
| MER61E_dup9 | 14.9000019363846 | 1.42486478545363 | 0.334355572130121 | 4.26152546636523 | 2.03036211874458e-05 | 0.0031229253051506 | 893 | 14 | 14 | 8 | 5 | 14 | 17 | 3 | 7 | 25 | 40 | 17 | 16 |
| L1MC2_dup2041 | 12.2227960542448 | 1.37174179700582 | 0.345942633568704 | 3.96522909840599 | 7.33254868294737e-05 | 0.00800206116441897 | 894 | 16 | 10 | 4 | 5 | 14 | 12 | 5 | 5 | 20 | 26 | 15 | 18 |
| LTR10E_dup109 | 13.1397410531489 | 1.71682679506797 | 0.37746657842328 | 4.54828822789914 | 5.40840298634518e-06 | 0.00115833290458046 | 895 | 16 | 10 | 5 | 3 | 10 | 14 | 3 | 6 | 18 | 35 | 22 | 19 |
| MER39_dup1580 | 15.4799523998112 | 1.77422320982224 | 0.36250427776333 | 4.89435109778369 | 9.86306202765407e-07 | 0.000315016933856668 | 896 | 14 | 17 | 8 | 9 | 12 | 14 | 2 | 2 | 22 | 41 | 22 | 29 |
| MIR3_dup84576 | 14.355053046103 | 1.43296352203971 | 0.381582105928493 | 3.75532159337743 | 0.000173119043119885 | 0.0148136244272091 | 897 | 18 | 21 | 6 | 4 | 16 | 17 | 3 | 2 | 23 | 28 | 22 | 16 |
| L1MEc_dup1808 | 18.0740997778299 | 2.04161336413353 | 0.614020001404435 | 3.32499488528678 | 0.000884201570897412 | 0.0440151102486937 | 898 | 22 | 12 | 10 | 22 | 0 | 1 | 8 | 5 | 32 | 42 | 34 | 37 |
| MLT1H_dup765 | 17.6885247397446 | 2.11263071585386 | 0.608913682789896 | 3.46950770784833 | 0.000521413106217125 | 0.031306627043134 | 899 | 17 | 9 | 8 | 22 | 0 | 1 | 17 | 5 | 43 | 32 | 26 | 38 |
| L2_dup5580 | 18.0322706483824 | 2.15424386177867 | 0.629092082741891 | 3.42436969225462 | 0.000616227290665105 | 0.034914717253013 | 900 | 13 | 15 | 12 | 17 | 0 | 0 | 9 | 9 | 39 | 37 | 23 | 47 |
| MER21A_dup176 | 20.6353827892047 | 2.51481779301764 | 0.647722890941652 | 3.8825519804644 | 0.000103365910833019 | 0.0103082585997794 | 901 | 16 | 13 | 9 | 21 | 0 | 0 | 12 | 3 | 43 | 50 | 34 | 55 |
| L1M4_dup1447 | 19.3716403173366 | 2.13101979595865 | 0.635046366881621 | 3.35569165826894 | 0.000791667802330548 | 0.0411800146310016 | 902 | 11 | 11 | 17 | 30 | 2 | 0 | 12 | 6 | 28 | 71 | 28 | 24 |
| MLT1J2-int_dup73 | 18.6804597242545 | 2.51054189818576 | 0.629014458735997 | 3.99123082676142 | 6.57312645359284e-05 | 0.00740904380448414 | 903 | 6 | 6 | 20 | 31 | 1 | 0 | 17 | 5 | 31 | 49 | 42 | 24 |
| MIRc_dup10816 | 16.0064321242726 | 1.99222801405372 | 0.465989580166621 | 4.2752630076865 | 1.90911786293787e-05 | 0.00300639946402789 | 904 | 17 | 7 | 10 | 13 | 1 | 2 | 13 | 9 | 23 | 42 | 30 | 32 |
| MamTip2_dup586 | 13.6855765776588 | 2.18705187928209 | 0.520372063739709 | 4.20286182076073 | 2.63561489272551e-05 | 0.00380311716958437 | 905 | 9 | 4 | 12 | 12 | 1 | 1 | 16 | 6 | 24 | 38 | 22 | 24 |
| MLT1J_dup1242 | 12.3637434406391 | 1.87032301196478 | 0.52565446763148 | 3.55808449682198 | 0.000373569151133441 | 0.0251525991045834 | 906 | 7 | 11 | 9 | 13 | 1 | 1 | 14 | 4 | 24 | 27 | 24 | 17 |
| LTR39_dup82 | 15.0224495947372 | 2.03982383497351 | 0.560132644895724 | 3.64167997270227 | 0.000270864665007683 | 0.0201977949169283 | 907 | 12 | 16 | 9 | 14 | 1 | 0 | 8 | 5 | 26 | 35 | 27 | 33 |
| L1PA10_dup464 | 13.8079062390038 | 2.26231020838656 | 0.563376942736773 | 4.0156244190554 | 5.92886103695368e-05 | 0.00686384130784084 | 908 | 9 | 7 | 15 | 11 | 1 | 0 | 7 | 6 | 26 | 40 | 23 | 23 |
| L2c_dup15950 | 11.8422839800282 | 1.96754589820779 | 0.518675250015139 | 3.79340617881876 | 0.000148594791420737 | 0.0132375650465738 | 909 | 8 | 10 | 11 | 11 | 2 | 0 | 9 | 3 | 21 | 28 | 24 | 19 |
| L2c_dup15969 | 13.334457607179 | 2.00173058257072 | 0.579975150232454 | 3.45140749869788 | 0.000557670880248222 | 0.0327886225034701 | 910 | 10 | 11 | 10 | 15 | 1 | 0 | 9 | 4 | 22 | 41 | 19 | 23 |
| L2c_dup15956 | 8.39438373792545 | 1.9495653953541 | 0.577242666247841 | 3.37737577166038 | 0.000731810009557464 | 0.0389120144210414 | 911 | 11 | 3 | 4 | 8 | 0 | 1 | 9 | 3 | 18 | 15 | 15 | 17 |
| MIRb_dup23822 | 11.1516575420387 | 2.15448167622091 | 0.516774381779061 | 4.16909535802421 | 3.05811030380807e-05 | 0.00423539108090265 | 912 | 12 | 4 | 6 | 7 | 0 | 4 | 7 | 4 | 26 | 34 | 16 | 14 |
| L1MA7_dup595 | 9.67301602687535 | 2.4344138179414 | 0.59964624157788 | 4.05974998114822 | 4.91252892030079e-05 | 0.00595948609287004 | 913 | 5 | 4 | 9 | 10 | 0 | 1 | 3 | 4 | 23 | 27 | 15 | 15 |
| MIR_dup16874 | 10.8468676011802 | 1.99886800051199 | 0.520796798972504 | 3.83809578794574 | 0.000123992116333762 | 0.0116984251043431 | 914 | 8 | 6 | 8 | 15 | 0 | 2 | 7 | 4 | 18 | 29 | 17 | 21 |
| L2a_dup12590 | 13.0183978021273 | 2.51729978208282 | 0.5663712169814 | 4.44461107239758 | 8.80510089251669e-06 | 0.00168014317319228 | 915 | 6 | 2 | 9 | 14 | 1 | 4 | 5 | 5 | 39 | 35 | 13 | 21 |
| MLT1C_dup3239 | 11.0431178236226 | 2.37558083389425 | 0.507650270290374 | 4.67956184192599 | 2.87488637186077e-06 | 0.000731623222073565 | 916 | 6 | 4 | 5 | 8 | 2 | 3 | 5 | 5 | 38 | 25 | 12 | 15 |
| L1MC1_dup11016 | 8.72189477417301 | 2.46829350174437 | 0.538135071858225 | 4.58675457301295 | 4.50189376349367e-06 | 0.00101019849178833 | 917 | 1 | 4 | 6 | 9 | 1 | 3 | 3 | 4 | 21 | 22 | 12 | 19 |
| MER5A_dup18635 | 8.51184208881433 | 2.00090297853942 | 0.545289185646984 | 3.66943455180641 | 0.000243087561068905 | 0.0187554191978281 | 918 | 4 | 2 | 11 | 10 | 2 | 3 | 4 | 3 | 15 | 24 | 16 | 10 |
| L4_A_Mam_dup489 | 11.8134802519234 | 2.24447780679467 | 0.647213295281379 | 3.46791053762094 | 0.000524521875955391 | 0.0314298368914836 | 919 | 4 | 12 | 10 | 7 | 0 | 1 | 5 | 6 | 28 | 31 | 26 | 9 |
| AluSc_dup3021 | 14.0305377543059 | 2.02941054104482 | 0.612632516354255 | 3.31260663916722 | 0.000924308809925422 | 0.0453568950477448 | 920 | 14 | 7 | 5 | 8 | 0 | 2 | 10 | 12 | 36 | 38 | 18 | 15 |
| LTR62_dup14 | 13.2583397275695 | 2.36016112013277 | 0.619313323253903 | 3.81093225595788 | 0.000138443705820888 | 0.0126215042242364 | 921 | 6 | 7 | 9 | 10 | 0 | 1 | 7 | 10 | 34 | 40 | 14 | 18 |
| L1M_dup64 | 11.3235575372582 | 2.06192271917771 | 0.599332445690619 | 3.44036558341461 | 0.00058092891498541 | 0.0337310179612295 | 922 | 12 | 5 | 8 | 8 | 0 | 0 | 7 | 9 | 25 | 19 | 21 | 23 |
| L3_dup4581 | 12.7863091982539 | 2.21677858733644 | 0.582670284640175 | 3.80451628609377 | 0.000142081564343906 | 0.0128302618251898 | 923 | 8 | 6 | 12 | 14 | 0 | 0 | 6 | 9 | 21 | 32 | 23 | 26 |
| MLT1A0_dup1689 | 11.9982580146931 | 2.62410001789855 | 0.698716101839205 | 3.75560261312317 | 0.000172924891169601 | 0.0148063282311054 | 924 | 9 | 1 | 14 | 9 | 0 | 0 | 7 | 5 | 24 | 38 | 17 | 22 |
| L2c_dup15962 | 10.7656595874053 | 2.35572162343358 | 0.689527585561167 | 3.41642839643085 | 0.000634483471390381 | 0.0355650129472477 | 925 | 8 | 2 | 9 | 12 | 0 | 0 | 12 | 4 | 27 | 26 | 18 | 12 |
| L3_dup4580 | 11.5554543597953 | 2.28362382483439 | 0.618820309580997 | 3.69028583819531 | 0.000224002207299765 | 0.0177459441439108 | 926 | 8 | 5 | 10 | 12 | 0 | 0 | 10 | 4 | 19 | 33 | 28 | 13 |
| MLT2B4_dup367 | 12.5447735372528 | 2.35367131457212 | 0.675532509719379 | 3.48417179145065 | 0.000493662514512958 | 0.0302277459285384 | 927 | 8 | 7 | 4 | 13 | 0 | 0 | 10 | 6 | 24 | 49 | 15 | 16 |
| MIRb_dup23835 | 10.6111242485264 | 2.16280936775584 | 0.593700203971803 | 3.64293182533345 | 0.00026955024809444 | 0.0201214572693337 | 928 | 6 | 9 | 6 | 7 | 0 | 0 | 10 | 6 | 22 | 23 | 25 | 14 |
| Charlie18a_dup250 | 11.137000547274 | 2.75749521917938 | 0.542020783462887 | 5.08743447356791 | 3.62939723964225e-07 | 0.000144899422077838 | 929 | 5 | 6 | 7 | 6 | 0 | 0 | 6 | 4 | 27 | 30 | 18 | 25 |
| L2c_dup15417 | 9.92569031535442 | 2.33121755767263 | 0.577744306792074 | 4.03503337076001 | 5.45945377188445e-05 | 0.00644015967881324 | 930 | 5 | 7 | 8 | 7 | 0 | 0 | 9 | 3 | 17 | 29 | 20 | 17 |

| | | | | | | | | | | | | | | | | | | | |
|---|---|---|---|---|---|---|---|---|---|---|---|---|---|---|---|---|---|---|---|
| MIRb_dup23818 | 13.056593951178 | 2.08223950272207 | 0.60880122597499 | 3.42022882655564 | 0.000625684812073237 | 0.0352710728011321 | 931 | 5 | 9 | 12 | 11 | 1 | 0 | 11 | 10 | 21 | 43 | 14 | 22 |
| MER3_dup871 | 8.73984299609049 | 2.36364418431897 | 0.602772225061527 | 3.92128914711972 | 8.80764986885143e-05 | 0.00917576644349265 | 932 | 3 | 4 | 5 | 6 | 1 | 0 | 11 | 5 | 15 | 30 | 11 | 16 |
| MIR_dup16857 | 10.8510484992419 | 2.40073901824045 | 0.604950883387149 | 3.96848584598902 | 7.23307525569426e-05 | 0.00794289530722213 | 933 | 4 | 3 | 6 | 12 | 1 | 1 | 9 | 8 | 19 | 40 | 16 | 12 |
| MLT1A1_dup510 | 15.1528630216772 | 2.45418780574977 | 0.634005388531197 | 3.87092578414104 | 0.000108422800538765 | 0.0106562460541086 | 934 | 10 | 3 | 18 | 19 | 0 | 0 | 12 | 3 | 28 | 29 | 36 | 31 |
| L1M3_dup504 | 10.0019361453692 | 2.04903394309418 | 0.536019685558374 | 3.82268412578859 | 0.000132006838503422 | 0.0122219841223872 | 935 | 8 | 5 | 8 | 13 | 0 | 0 | 9 | 3 | 13 | 18 | 27 | 23 |
| L1MA1_dup4158 | 10.5746088816515 | 2.55399312007411 | 0.6160351474721 | 4.1458561748537 | 3.38546292065239e-05 | 0.00456147995536553 | 936 | 5 | 2 | 14 | 13 | 0 | 0 | 8 | 1 | 16 | 17 | 31 | 27 |
| MIR1_Amn_dup1198 | 11.7235735518562 | 2.64612858176919 | 0.576465907109779 | 4.59026032438944 | 4.426935307354e-06 | 0.00100083295050532 | 937 | 7 | 4 | 7 | 15 | 1 | 0 | 3 | 3 | 25 | 31 | 20 | 28 |
| AluY_dup9214 | 11.6552459450282 | 2.30417175904615 | 0.596352235424319 | 3.86377651021409 | 0.000111647378263875 | 0.0108743541008498 | 938 | 8 | 10 | 12 | 5 | 0 | 0 | 5 | 4 | 22 | 19 | 25 | 34 |
| L2c_dup15955 | 8.6568096746903 | 2.19803583714099 | 0.609816831926363 | 3.60441975699091 | 0.000312851085697647 | 0.02226845624372 | 939 | 8 | 6 | 8 | 5 | 0 | 0 | 3 | 4 | 20 | 20 | 15 | 15 |
| MIRb_dup23834 | 10.573202715619 | 2.20640852404079 | 0.644468605926791 | 3.4236090070948 | 0.000617954620772321 | 0.0349674579690303 | 940 | 14 | 5 | 8 | 7 | 0 | 0 | 7 | 2 | 26 | 18 | 22 | 20 |
| L1PREC2_dup6492 | 9.6351622107009 | 1.89420346087568 | 0.4990865656797 | 3.79534051031005 | 0.000147440954016896 | 0.0131594093190189 | 941 | 6 | 6 | 6 | 11 | 8 | 0 | 7 | 1 | 16 | 22 | 20 | 17 |
| THE1D_dup883 | 11.2493014007268 | 2.2898149955613 | 0.589020315590803 | 3.88749748514965 | 0.00010128303672847 | 0.0101730238317402 | 942 | 11 | 8 | 5 | 6 | 2 | 0 | 10 | 1 | 19 | 39 | 24 | 13 |
| MER5B_dup1705 | 7.82093858223146 | 2.63912717444757 | 0.740740928458469 | 3.56282078261798 | 0.000366891164220515 | 0.0248716943004123 | 943 | 5 | 1 | 5 | 10 | 0 | 0 | 3 | 4 | 24 | 20 | 10 | 10 |
| L2a_dup17015 | 7.61839776291408 | 2.5018237082567 | 0.71323174046684 | 3.50772906800131 | 0.000451949072257633 | 0.0285174950801948 | 944 | 5 | 2 | 7 | 7 | 0 | 0 | 5 | 2 | 16 | 30 | 9 | 9 |
| MLT1J1_dup405 | 8.69733485144256 | 2.42019630618394 | 0.696325361622536 | 3.47566875999539 | 0.000509581263150007 | 0.0309160691589729 | 945 | 4 | 3 | 8 | 13 | 0 | 0 | 6 | 3 | 17 | 28 | 15 | 9 |
| L1ME5_dup282 | 6.82411568460717 | 2.24311166330975 | 0.681189167293531 | 3.29293501865565 | 0.000991473866291906 | 0.0475064922651417 | 946 | 4 | 4 | 5 | 9 | 0 | 0 | 6 | 1 | 16 | 15 | 16 | 7 |
| MIRb_dup23786 | 7.34110489008167 | 2.11999384366651 | 0.641862744979485 | 3.30287722764509 | 0.00095698276241757 | 0.0464525133386803 | 947 | 6 | 5 | 3 | 5 | 0 | 0 | 7 | 4 | 21 | 16 | 10 | 10 |
| MIRb_dup23861 | 7.68222126612305 | 2.43224190637107 | 0.593255525121962 | 4.09982175196943 | 4.13468468847723e-05 | 0.00524710230871681 | 948 | 5 | 7 | 3 | 6 | 0 | 0 | 4 | 2 | 19 | 19 | 11 | 17 |
| AluSx_dup10546 | 7.21322074151355 | 2.39107487722812 | 0.668245088472993 | 3.57814059313472 | 0.000346047303935167 | 0.0239223044108534 | 949 | 7 | 2 | 5 | 3 | 0 | 0 | 8 | 2 | 16 | 22 | 9 | 14 |
| AluJb_dup12392 | 7.11796189742581 | 2.04511007754484 | 0.578771843373939 | 3.53353415678087 | 0.000410043102502455 | 0.0267529794538111 | 950 | 7 | 4 | 5 | 6 | 0 | 0 | 5 | 4 | 12 | 19 | 10 | 16 |
| L2c_dup15948 | 7.55612750160173 | 2.27308166450692 | 0.600478659414564 | 3.78544953907781 | 0.00015343096674178 | 0.0135668096664213 | 951 | 5 | 4 | 3 | 6 | 0 | 0 | 10 | 3 | 12 | 22 | 14 | 15 |
| MER3_dup874 | 7.94482892834215 | 2.2199786199711 | 0.596025784219678 | 3.72463520664213 | 0.000195597858579781 | 0.0161715524870134 | 952 | 3 | 7 | 9 | 6 | 0 | 0 | 3 | 4 | 9 | 19 | 20 | 19 |
| MIRb_dup23840 | 9.74145860157506 | 2.66821840615078 | 0.611507176653406 | 4.36334765644638 | 1.28087147198033e-05 | 0.00224762716277162 | 953 | 6 | 4 | 4 | 8 | 1 | 0 | 1 | 6 | 24 | 28 | 19 | 14 |
| MLT1E2_dup248 | 7.74130589299532 | 2.26295469906642 | 0.600534584207253 | 3.768233768008 | 0.000164406704455025 | 0.0142669284092835 | 954 | 4 | 5 | 5 | 6 | 1 | 0 | 3 | 5 | 20 | 21 | 12 | 9 |
| AluSx1_dup26020 | 9.59150746397239 | 2.74137022504273 | 0.763608226088702 | 3.5900218611897 | 0.000330650229423314 | 0.023122798592195 | 955 | 8 | 11 | 3 | 6 | 0 | 0 | 0 | 1 | 26 | 26 | 17 | 16 |
| L1MA3_dup4146 | 10.6084634039445 | 2.69014170033065 | 0.767302390117521 | 3.50597330984284 | 0.000454940837243877 | 0.0285925389484499 | 956 | 5 | 11 | 3 | 8 | 0 | 0 | 1 | 1 | 41 | 30 | 10 | 11 |
| MLT1H_dup1936 | 10.0452449516046 | 2.85593696959155 | 0.529992605281882 | 5.38863550383423 | 7.09946251367353e-08 | 3.79615439839879e-05 | 957 | 5 | 7 | 4 | 3 | 4 | 1 | 0 | 2 | 29 | 39 | 13 | 9 |
| Tigger1_dup5835 | 15.7988832238336 | 2.49579930106118 | 0.749434007110649 | 3.3302455951838 | 0.000867694141401757 | 0.0435119810684895 | 958 | 10 | 14 | 9 | 8 | 2 | 0 | 0 | 2 | 57 | 54 | 11 | 13 |
| L1MA4_dup2162 | 24.2266685566555 | 4.13410362200808 | 0.698856224807483 | 5.91552808039581 | 3.30812462107493e-09 | 3.03920857562276e-06 | 959 | 2 | 4 | 14 | 17 | 0 | 2 | 2 | 0 | 65 | 70 | 41 | 76 |
| MER31-int_dup259 | 20.4212274582055 | 3.85488850542314 | 0.760615951665293 | 5.06811420005489 | 4.01776339829038e-07 | 0.000155702073219805 | 960 | 5 | 3 | 15 | 14 | 0 | 1 | 1 | 0 | 49 | 77 | 33 | 48 |
| L1MA4_dup2161 | 12.6896663789032 | 3.80760458663026 | 0.780298314059002 | 4.87967808981111 | 1.06259135058547e-06 | 0.000336670255001879 | 961 | 1 | 2 | 11 | 9 | 2 | 0 | 0 | 0 | 26 | 43 | 26 | 35 |
| MER31-int_dup260 | 19.5049766507157 | 3.99011336216621 | 0.805610993845838 | 4.95290331518212 | 7.31143077302657e-07 | 0.000251769277383297 | 962 | 3 | 4 | 17 | 11 | 0 | 0 | 0 | 1 | 42 | 66 | 39 | 54 |
| ERVL-E-int_dup4593 | 38.4513944168659 | 4.65580944167479 | 0.530260385732091 | 8.78023244230637 | 1.63137154432645e-18 | 1.61740972285959e-14 | 963 | 2 | 3 | 0 | 2 | 7 | 12 | 1 | 2 | 181 | 180 | 11 | 14 |
| MER49_dup651 | 24.0704243860995 | 4.89927279016974 | 0.634660827545539 | 7.71951344329376 | 1.16774604497909e-14 | 4.94152673305293e-11 | 964 | 2 | 0 | 0 | 0 | 6 | 10 | 1 | 0 | 110 | 115 | 4 | 14 |
| MLT1D_dup16594 | 6.50617852113221 | 2.44185645061664 | 0.744472448960386 | 3.2799822935376 | 0.00103813601839074 | 0.0490568324622138 | 965 | 2 | 3 | 1 | 0 | 14 | 9 | 0 | 0 | 10 | 28 | 6 | 5 |
| ERV3-16A3_I-int_dup | 10.7966195197041 | 1.90390237110785 | 0.578111356687121 | 3.29331425353447 | 0.000990137399354521 | 0.0474902708378978 | 966 | 6 | 4 | 2 | 1 | 21 | 17 | 1 | 0 | 31 | 32 | 3 | 8 |
| MER2_dup8619 | 9.35404505458004 | 2.38118846529227 | 0.576403807146844 | 4.13111161961086 | 3.61013223039828e-05 | 0.00477925879812747 | 967 | 4 | 3 | 2 | 2 | 9 | 10 | 0 | 0 | 29 | 43 | 2 | 3 |
| HERVE-int_dup188 | 5.84859108322128 | 2.67600552837652 | 0.643394784366524 | 4.15919680015938 | 3.19368647249252e-05 | 0.00438249666311249 | 968 | 1 | 0 | 4 | 3 | 5 | 4 | 1 | 0 | 14 | 16 | 13 | 9 |
| AluY_dup31631 | 7.80244849712004 | 2.38956853496464 | 0.495108602476407 | 4.82635228516053 | 1.39056444838779e-06 | 0.000417249997775637 | 969 | 5 | 1 | 3 | 3 | 5 | 7 | 5 | 0 | 12 | 23 | 17 | 16 |
| MER47B_dup220 | 7.80925304268937 | 1.95266589482152 | 0.590962635857758 | 3.30421210469137 | 0.000952437433159061 | 0.0463008369418401 | 970 | 6 | 0 | 4 | 4 | 8 | 5 | 6 | 0 | 18 | 27 | 9 | 7 |
| L1MEf_dup1996 | 7.63637161574849 | 1.97852080785523 | 0.591949540899295 | 3.34238084693746 | 0.000830630083160418 | 0.0423586287315242 | 971 | 4 | 4 | 2 | 0 | 10 | 8 | 7 | 0 | 14 | 23 | 6 | 16 |
| L2c_dup10773 | 6.33973560843839 | 2.06065633434644 | 0.58736737073244 | 3.50829214734355 | 0.000450993495197216 | 0.028498300031631 | 972 | 4 | 1 | 2 | 0 | 6 | 10 | 4 | 1 | 13 | 16 | 9 | 11 |
| MLT1K_dup12416 | 5.83792436813675 | 2.08804537472179 | 0.627254656535291 | 3.32886388800258 | 0.000872010090106629 | 0.0436358851976175 | 973 | 3 | 1 | 2 | 0 | 10 | 5 | 2 | 1 | 14 | 20 | 4 | 7 |
| MLT1K-int_dup145 | 7.37336376468653 | 2.52367521759639 | 0.56915382184087 | 4.43408288014972 | 9.24650528788914e-06 | 0.0017475527777856 | 974 | 3 | 5 | 0 | 3 | 6 | 6 | 3 | 0 | 24 | 14 | 15 | 7 |
| Tigger1_dup4324 | 8.83418722953734 | 2.66338189329487 | 0.483516585906418 | 5.50835684013195 | 3.62198622473511e-08 | 2.25022750451911e-05 | 975 | 5 | 6 | 0 | 3 | 5 | 5 | 3 | 0 | 30 | 29 | 8 | 8 |
| L1MCc_dup1857 | 7.9114810093607 | 2.08848972788071 | 0.586632264035576 | 3.5601344418282 | 0.000370664977668026 | 0.0250423065551251 | 976 | 5 | 4 | 3 | 6 | 4 | 6 | 2 | 0 | 21 | 30 | 10 | 2 |
| MIRb_dup105954 | 6.91490799407679 | 1.71118195221202 | 0.482426000597479 | 3.54703508951165 | 0.000389592697441289 | 0.0258485014139323 | 977 | 8 | 6 | 3 | 1 | 6 | 6 | 4 | 0 | 16 | 21 | 6 | 6 |
| L1MB7_dup16627 | 6.35749578661697 | 2.17592950042197 | 0.517791163364585 | 4.20233031070457 | 2.64181269217502e-05 | 0.0038074425369611 | 978 | 5 | 5 | 3 | 1 | 5 | 6 | 1 | 0 | 17 | 18 | 8 | 6 |
| L2a_dup57862 | 6.47727203792076 | 2.29057050000639 | 0.529276092672652 | 4.32774223456767 | 1.50645624714756e-05 | 0.00251901080944324 | 979 | 8 | 3 | 2 | 1 | 7 | 4 | 1 | 0 | 17 | 19 | 7 | 8 |
| MLT1H2_dup1576 | 13.5313307493828 | 2.59434730160254 | 0.406661785787762 | 6.37961911414154 | 1.77528959338644e-10 | 2.37316324487601e-07 | 980 | 13 | 10 | 2 | 1 | 10 | 13 | 0 | 1 | 38 | 39 | 15 | 17 |
| L1MB3_dup1405 | 9.69345657546477 | 1.69508110303446 | 0.495952651658934 | 3.41782849101524 | 0.000631228702757658 | 0.0354743374837916 | 981 | 13 | 10 | 5 | 2 | 12 | 13 | 0 | 0 | 17 | 23 | 9 | 15 |
| MLT1B_dup7391 | 8.33040641031804 | 2.10223842015933 | 0.589148635690415 | 3.56826493826256 | 0.000359353041906367 | 0.0245145123020219 | 982 | 8 | 14 | 2 | 1 | 9 | 11 | 0 | 0 | 18 | 13 | 11 | 14 |
| L2_dup50371 | 7.47923306935682 | 2.48719281694744 | 0.662724175137038 | 3.75298338321993 | 0.000174742442288346 | 0.0149276401006521 | 983 | 9 | 5 | 2 | 0 | 4 | 15 | 0 | 0 | 11 | 23 | 12 | 11 |
| L3b_dup8807 | 7.44073308024571 | 2.26787996100996 | 0.595034490813876 | 3.81134202474213 | 0.000138214373088363 | 0.012605426300837 | 984 | 8 | 6 | 1 | 1 | 9 | 12 | 0 | 0 | 14 | 18 | 14 | 7 |
| MIRb_dup38130 | 8.40853076898789 | 2.09385947284456 | 0.619114347832531 | 3.38202382189169 | 0.000719539046103474 | 0.0384140538173968 | 985 | 15 | 15 | 3 | 1 | 3 | 3 | 2 | 0 | 17 | 16 | 14 | 14 |
| L1MEf_dup6663 | 8.64955260608563 | 1.54306680176288 | 0.426403008217882 | 3.61879905165775 | 0.000295973343256332 | 0.0214124880311557 | 986 | 8 | 12 | 3 | 5 | 7 | 9 | 4 | 0 | 15 | 20 | 10 | 14 |
| LTR54B_dup63 | 10.1641688750195 | 1.63573961210971 | 0.468064326348211 | 3.49468976811231 | 0.000474613074326265 | 0.0294554727656853 | 987 | 10 | 18 | 2 | 5 | 4 | 9 | 5 | 0 | 22 | 28 | 10 | 10 |
| L1PB1_dup10474 | 8.57748206881173 | 2.12411929290167 | 0.497040346487469 | 4.27353495126218 | 1.92398109127941e-05 | 0.00302380188075014 | 988 | 7 | 8 | 1 | 1 | 10 | 4 | 3 | 2 | 28 | 16 | 7 | 13 |
| L2b_dup4679 | 9.34278839042286 | 1.95590900903467 | 0.530440659301593 | 3.68732859130732 | 0.000226620645194519 | 0.0178850666804162 | 989 | 13 | 14 | 1 | 0 | 6 | 7 | 5 | 1 | 20 | 21 | 8 | 18 |
| L1MC1_dup243 | 8.69731473233662 | 2.03064719569758 | 0.470447790266804 | 4.3164135058344 | 1.58584779900422e-05 | 0.00261139197496095 | 990 | 9 | 6 | 2 | 1 | 6 | 10 | 4 | 2 | 18 | 16 | 10 | 23 |
| MIRb_dup116669 | 8.48129149288455 | 1.62025284217557 | 0.464771175535859 | 3.48613022377624 | 0.000490062379839632 | 0.0300924859440829 | 991 | 8 | 11 | 2 | 4 | 7 | 11 | 3 | 1 | 18 | 11 | 12 | 16 |
| LTR78_dup2505 | 11.6981314259361 | 1.89796924834619 | 0.473759035812035 | 4.00619113278342 | 6.17057232417144e-05 | 0.00705895674157355 | 992 | 15 | 10 | 3 | 1 | 16 | 16 | 2 | 1 | 24 | 20 | 13 | 22 |
| MLT1A0_dup18776 | 10.035501709431 | 1.5541519885785 | 0.39668683227906 | 3.91783104987259 | 8.9349291957012e-05 | 0.00927992432300445 | 993 | 15 | 11 | 2 | 3 | 10 | 9 | 3 | 2 | 18 | 29 | 10 | 10 |
| L2a_dup57863 | 12.5985016871653 | 1.71910452829377 | 0.405038073873455 | 4.24430353387189 | 2.19273381213204e-05 | 0.00332114729256251 | 994 | 14 | 10 | 3 | 4 | 7 | 13 | 6 | 1 | 33 | 44 | 7 | 7 |
| MLT1D_dup9676 | 12.739057364943 | 2.16356361775152 | 0.501179767167696 | 4.3169412643659 | 1.58206234409565e-05 | 0.00260696264908715 | 995 | 10 | 5 | 4 | 5 | 6 | 5 | 6 | 2 | 40 | 50 | 9 | 5 |
| AluSc_dup16626 | 8.89471097584654 | 2.13153496081328 | 0.461513536887099 | 4.61857516724308 | 3.86384013492824e-06 | 0.000910282479946172 | 996 | 9 | 4 | 5 | 3 | 6 | 4 | 3 | 1 | 27 | 23 | 7 | 13 |
| L1MD2_dup8878 | 9.23929164209228 | 1.87828867603488 | 0.499347210146334 | 3.76148827483074 | 0.000168905340511474 | 0.0145459102979889 | 997 | 5 | 4 | 5 | 4 | 8 | 11 | 1 | 1 | 30 | 25 | 6 | 7 |
| L1PB2_dup904 | 10.3549608225427 | 2.45012338073961 | 0.454480009913243 | 5.3910476309119 | 7.00481031622249e-08 | 3.76245281152117e-05 | 998 | 4 | 9 | 2 | 6 | 9 | 3 | 2 | 1 | 25 | 33 | 11 | 19 |
| Tigger1_dup4322 | 13.1714131501147 | 2.79934359625559 | 0.426901082309585 | 6.55735886428493 | 5.47691037810713e-11 | 8.40779946341341e-08 | 999 | 11 | 12 | 1 | 2 | 8 | 4 | 2 | 1 | 41 | 39 | 12 | 21 |
| L2_dup19468 | 10.3715168813112 | 3.05788790701426 | 0.492981571948026 | 6.20284424614688 | 5.545171381992e-10 | 6.40510363912364e-07 | 1000 | 3 | 3 | 2 | 3 | 9 | 4 | 2 | 0 | 37 | 28 | 11 | 18 |
| L2a_dup57861 | 12.4206538852223 | 2.77891701782982 | 0.476544409565095 | 5.83139149689305 | 5.49670494588581e-09 | 4.77342684326184e-06 | 1001 | 17 | 4 | 4 | 2 | 6 | 5 | 2 | 0 | 30 | 39 | 16 | 25 |
| Tigger2_dup535 | 14.1130944249972 | 4.39614098584154 | 0.50531256128662 | 8.69984505164119 | 3.32337640411701e-18 | 3.04147739174625e-14 | 1002 | 4 | 4 | 0 | 1 | 4 | 2 | 0 | 1 | 54 | 64 | 10 | 14 |

| | | | | | | | | | | | | | | | | | | | |
|---|---|---|---|---|---|---|---|---|---|---|---|---|---|---|---|---|---|---|---|
| L1PA5_dup242 | 7.53863800407546 | 2.84181979073751 | 0.605983257001578 | 4.68960116950906 | 2.73738067340322e-06 | 0.000710382446875096 | 1003 | 7 | 8 | 1 | 0 | 5 | 2 | 0 | 0 | 26 | 29 | 2 | 6 |
| L1PA7_dup267 | 9.16699935405664 | 3.48576966445113 | 0.607901116815624 | 5.73410636702014 | 9.80278013205479e-09 | 7.67280368849312e-06 | 1004 | 8 | 6 | 1 | 0 | 1 | 2 | 0 | 1 | 34 | 35 | 5 | 11 |
| L1M3_dup5631 | 14.4332645168682 | 2.53824772910928 | 0.594220661216341 | 4.27155751184015 | 1.9411245049924e-05 | 0.0030418245330925 | 1005 | 3 | 1 | 2 | 3 | 18 | 17 | 7 | 2 | 21 | 36 | 33 | 36 |
| L1ME1_dup26354 | 10.4758670481618 | 2.40967392245766 | 0.678241763794075 | 3.55282445153181 | 0.00038111870691602 | 0.0255037468213251 | 1006 | 1 | 1 | 2 | 2 | 14 | 13 | 4 | 3 | 7 | 30 | 27 | 28 |
| L1MEc_dup16025 | 10.4644385925396 | 2.48593616462314 | 0.689409573257922 | 3.6058915643939 | 0.000311083026384769 | 0.0221951909433734 | 1007 | 1 | 0 | 2 | 3 | 11 | 13 | 7 | 1 | 13 | 22 | 22 | 38 |
| L1ME3_dup7501 | 9.25534761739218 | 2.81594469959945 | 0.735640652647108 | 3.82788075871914 | 0.000129251342987595 | 0.012046549180778 | 1008 | 0 | 0 | 2 | 2 | 7 | 11 | 2 | 3 | 8 | 21 | 23 | 39 |
| MIRb_dup183944 | 8.38957924173885 | 2.55549044217968 | 0.716581510059263 | 3.56622436709026 | 0.000362161339636233 | 0.0246284201546394 | 1009 | 1 | 0 | 1 | 2 | 7 | 11 | 2 | 4 | 7 | 18 | 29 | 23 |
| L1PA14_dup831 | 9.96666841635427 | 2.45293068855899 | 0.583221696248771 | 4.20582894006862 | 2.60126936740677e-05 | 0.00376356984672793 | 1010 | 3 | 0 | 8 | 5 | 9 | 8 | 1 | 1 | 19 | 22 | 18 | 29 |
| L1PB1_dup11488 | 8.54341507292008 | 2.95085243488603 | 0.76072496295869 | 3.87900039905259 | 0.00010488658287648 | 0.0103834216322931 | 1011 | 2 | 0 | 2 | 2 | 13 | 5 | 2 | 0 | 10 | 13 | 25 | 35 |
| Tigger7_dup1770 | 8.04893463994645 | 3.04289153667449 | 0.712118718271959 | 4.27301159005963 | 1.92850428658004e-05 | 0.00302691214363176 | 1012 | 2 | 0 | 0 | 0 | 8 | 7 | 3 | 2 | 19 | 22 | 22 | 10 |
| LTR16E2_dup524 | 6.62472957237662 | 2.09914978134512 | 0.625519702202381 | 3.3558491826145 | 0.000791217037212702 | 0.0411782434682007 | 1013 | 2 | 0 | 0 | 3 | 5 | 7 | 5 | 5 | 11 | 12 | 11 | 21 |
| HERVL-int_dup1710 | 8.21807069756183 | 2.43196099420479 | 0.72839046189644 | 3.33881499199335 | 0.000841365796943234 | 0.0427138096691817 | 1014 | 0 | 1 | 0 | 1 | 6 | 9 | 9 | 3 | 15 | 18 | 18 | 21 |
| L1ME3Cz_dup10384 | 7.10070688853068 | 3.14126072267874 | 0.654307756789237 | 4.8008917670383 | 1.57960629245754e-06 | 0.000463453759389768 | 1015 | 0 | 1 | 0 | 1 | 5 | 5 | 2 | 3 | 19 | 12 | 14 | 23 |
| L1P4a_dup542 | 6.75656167861862 | 2.79580724339583 | 0.679070156534678 | 4.11711104735179 | 3.8365124360478e-05 | 0.00497816297128209 | 1016 | 0 | 0 | 0 | 2 | 5 | 5 | 4 | 3 | 16 | 20 | 9 | 17 |
| MIR_dup97813 | 9.02374706299304 | 2.92314872600151 | 0.716999128495417 | 4.07692089129254 | 4.56360105152399e-05 | 0.00561764415833382 | 1017 | 0 | 0 | 1 | 2 | 9 | 9 | 3 | 1 | 30 | 20 | 9 | 22 |
| L1MA9_dup14215 | 8.4855730593679 | 2.77542464535115 | 0.641474863342477 | 4.32663040121243 | 1.51407699596289e-05 | 0.002524120151266 9 | 1018 | 0 | 2 | 2 | 3 | 8 | 10 | 1 | 1 | 14 | 19 | 25 | 19 |
| L2a_dup142925 | 7.38545098885317 | 2.69408542561872 | 0.816974669284823 | 3.29763642240843 | 0.000975022966883637 | 0.0469892704244484 | 1019 | 0 | 1 | 2 | 2 | 7 | 14 | 0 | 1 | 11 | 13 | 18 | 23 |
| L1MA7_dup7429 | 7.85042296967704 | 2.89463999376087 | 0.708673967774249 | 4.0845863195062 | 4.41554101900342e-05 | 0.00549508537294869 | 1020 | 0 | 1 | 1 | 0 | 7 | 9 | 3 | 2 | 11 | 25 | 20 | 17 |
| AluJr_dup65984 | 6.83650930088981 | 2.83996702442204 | 0.658985468731387 | 4.30960492936098 | 1.63546447163128e-05 | 0.00266908250457323 | 1021 | 1 | 2 | 1 | 0 | 8 | 7 | 2 | 1 | 13 | 12 | 16 | 21 |
| MLT1C_dup16104 | 6.4219468358444 | 2.73241536476772 | 0.561813198134728 | 4.86356563683373 | 1.15289769821187e-06 | 0.000357428388883535 | 1022 | 1 | 2 | 2 | 1 | 4 | 7 | 2 | 1 | 12 | 14 | 14 | 19 |
| LTR67B_dup114 | 10.0643058144511 | 2.03032227351726 | 0.535200666714825 | 3.79357201847264 | 0.000148495535105023 | 0.0132375650465738 | 1023 | 8 | 4 | 0 | 2 | 8 | 9 | 11 | 2 | 14 | 35 | 19 | 12 |
| MSTA-int_dup2758 | 9.78875497593667 | 2.51997297973413 | 0.473216448021265 | 5.32520158644378 | 1.00841141503453e-07 | 5.18245059528741e-05 | 1024 | 3 | 3 | 0 | 3 | 9 | 7 | 6 | 2 | 21 | 32 | 14 | 18 |
| L1MD2_dup1954 | 10.0726279933262 | 2.28410319325412 | 0.540520605765492 | 4.22574675024524 | 2.38149705128346e-05 | 0.00351967513891115 | 1025 | 12 | 7 | 0 | 4 | 2 | 2 | 4 | 6 | 20 | 34 | 14 | 16 |
| AluSz_dup42776 | 8.02143592238378 | 2.16067056106521 | 0.523312696976122 | 4.12883267222504 | 3.64609615305099e-05 | 0.00480384272001036 | 1026 | 7 | 10 | 0 | 1 | 7 | 2 | 4 | 2 | 19 | 24 | 11 | 8 |
| L1ME1_dup23546 | 8.00582295556208 | 1.53906379259029 | 0.449757095783772 | 3.42198890694149 | 0.00062164849826926 | 0.0351100815497691 | 1027 | 10 | 7 | 0 | 3 | 9 | 6 | 5 | 3 | 15 | 21 | 10 | 8 |
| Arthur1B_dup850 | 8.13910184330928 | 2.92428830577776 | 0.523812626261543 | 5.58269915455921 | 2.36814187586583e-08 | 1.6008235420306e-05 | 1028 | 2 | 6 | 0 | 3 | 6 | 2 | 1 | 2 | 23 | 25 | 13 | 12 |
| SVA_F_dup205 | 6.54264861019653 | 2.19735151368082 | 0.647261986157358 | 3.39484097733899 | 0.000686684729174611 | 0.0373728006789072 | 1029 | 6 | 12 | 0 | 1 | 4 | 2 | 1 | 2 | 11 | 24 | 11 | 4 |
| L1MEd_dup476 | 8.20442401705652 | 1.70212352507972 | 0.515337578778283 | 3.30292917724914 | 0.000956805496096443 | 0.0464525133386803 | 1030 | 9 | 7 | 0 | 1 | 8 | 8 | 6 | 4 | 13 | 15 | 9 | 22 |
| AluY_dup3552 | 6.65757281021453 | 2.03812278445739 | 0.594349873835098 | 3.42916331639176 | 0.000605445090899617 | 0.0346472432898509 | 1031 | 7 | 7 | 0 | 1 | 9 | 5 | 2 | 2 | 7 | 14 | 10 | 20 |
| L1MB3_dup12460 | 6.62166422354348 | 1.93314308377984 | 0.583418059448201 | 3.31347830680492 | 0.000921432546727012 | 0.0452624254259921 | 1032 | 6 | 2 | 0 | 1 | 10 | 9 | 3 | 2 | 10 | 15 | 9 | 15 |
| L2b_dup47838 | 6.33406008172101 | 2.23588236124214 | 0.544899941555889 | 4.10328977987753 | 4.07316555146461e-05 | 0.005185820956344 8 | 1033 | 3 | 3 | 0 | 2 | 3 | 5 | 3 | 5 | 9 | 10 | 18 | 17 |
| L1ME4a_dup18487 | 5.13960049814303 | 2.18988132000601 | 0.634030828101107 | 3.45390353741096 | 0.00055253497222203 | 0.0325751948712446 | 1034 | 5 | 5 | 0 | 0 | 3 | 2 | 3 | 3 | 6 | 10 | 13 | 14 |
| MER33_dup4360 | 6.0249519839811 | 1.975588662545 | 0.596557359193827 | 3.31164913498806 | 0.000927477896956972 | 0.0454374419743306 | 1035 | 7 | 4 | 0 | 0 | 3 | 3 | 4 | 5 | 12 | 14 | 11 | 9 |
| MIRc_dup65156 | 6.23490994027194 | 2.17051155835203 | 0.552969148128827 | 3.925194679842 | 8.66596237318007e-05 | 0.00908383736937756 | 1036 | 5 | 3 | 0 | 0 | 5 | 4 | 4 | 4 | 13 | 14 | 10 | 13 |
| MIRb_dup71171 | 7.60707716117692 | 1.97923192803092 | 0.566089509314875 | 3.49632327655451 | 0.00047171683714724 | 0.0293292747666154 | 1037 | 2 | 3 | 8 | 6 | 8 | 4 | 2 | 0 | 13 | 23 | 14 | 10 |
| MIRb_dup114615 | 6.76124699333593 | 1.7353582464751 | 0.521196918987437 | 3.32956351669633 | 0.0008698222240359 | 0.0435911370936071 | 1038 | 2 | 3 | 5 | 8 | 5 | 5 | 6 | 0 | 9 | 17 | 15 | 10 |
| MIRb_dup174521 | 6.51545062989294 | 2.03433828261122 | 0.558392335588485 | 3.64320595566064 | 0.00026926321654608 | 0.0201154952526392 | 1039 | 1 | 2 | 3 | 7 | 6 | 6 | 3 | 0 | 13 | 17 | 11 | 11 |
| MER5A_dup27635 | 8.14416972986096 | 1.65782166258995 | 0.498832312007579 | 3.32340472476201 | 0.000889257935582789 | 0.0441799381726591 | 1040 | 3 | 5 | 5 | 12 | 7 | 4 | 5 | 1 | 9 | 25 | 12 | 15 |
| HERVK22-int_dup191 | 7.12265781435293 | 1.74964528207403 | 0.483796546390782 | 3.61648981400701 | 0.000298625063133067 | 0.0215826648244835 | 1041 | 5 | 2 | 6 | 8 | 6 | 4 | 3 | 1 | 12 | 17 | 9 | 16 |
| L2c_dup134696 | 6.57203318932183 | 2.15012778688053 | 0.489188674759641 | 4.39529346818379 | 1.1062319924491e-05 | 0.0020158671111376 | 1042 | 3 | 2 | 3 | 5 | 6 | 4 | 3 | 1 | 9 | 16 | 16 | 14 |
| MIR3_dup67868 | 6.36968098099713 | 1.61959408265012 | 0.493017223121515 | 3.28506593014284 | 0.0010195851045351 | 0.0484727666900516 | 1043 | 3 | 2 | 4 | 8 | 5 | 4 | 5 | 2 | 10 | 16 | 11 | 9 |
| MIRb_dup174490 | 6.98138449526502 | 1.84895150325924 | 0.508058171362341 | 3.63925158078127 | 0.00027343156945731 | 0.0203141009821374 | 1044 | 2 | 2 | 4 | 8 | 7 | 4 | 5 | 1 | 10 | 18 | 15 | 11 |
| AluJo_dup53492 | 7.10907314552412 | 1.99494543843126 | 0.586860582545464 | 3.39935156281639 | 0.000675458327572599 | 0.0369647210700528 | 1045 | 5 | 8 | 6 | 7 | 2 | 2 | 1 | 0 | 11 | 23 | 11 | 12 |
| MIRb_dup63769 | 8.93951679163827 | 1.96404696758456 | 0.508130054725039 | 3.86524463436306 | 0.000110977909189573 | 0.010826875596565 | 1046 | 5 | 14 | 3 | 7 | 5 | 3 | 3 | 1 | 15 | 14 | 21 | 20 |
| SVA_E_dup604 | 8.220143253074 | 2.75312049336589 | 0.474733420325324 | 5.79929782798784 | 6.65931694611772e-09 | 5.61899939737917e-06 | 1047 | 5 | 6 | 4 | 2 | 3 | 2 | 2 | 1 | 13 | 21 | 19 | 24 |
| MER74A_dup234 | 7.49701883562681 | 2.08308722572817 | 0.630996347824162 | 3.30126669181397 | 0.000962493466077889 | 0.0466154020515712 | 1048 | 6 | 7 | 3 | 5 | 10 | 0 | 1 | 0 | 22 | 14 | 10 | 11 |
| AluJr_dup76110 | 7.67351162616284 | 1.90762383040907 | 0.554657348046857 | 3.43928343711029 | 0.000583256228285191 | 0.0337856622072427 | 1049 | 2 | 3 | 8 | 6 | 8 | 0 | 5 | 3 | 16 | 12 | 18 | 12 |
| MIR_dup137191 | 5.93070350903372 | 2.39166561973608 | 0.620226786964395 | 3.85611468256913 | 0.000115203457041542 | 0.0111160591197108 | 1050 | 1 | 2 | 5 | 6 | 3 | 0 | 3 | 2 | 14 | 16 | 11 | 8 |
| L2_dup15147 | 7.01775729966697 | 1.80121290546223 | 0.493386801205781 | 3.65071157367866 | 0.000261514784612541 | 0.0196981313515086 | 1051 | 2 | 5 | 3 | 3 | 7 | 4 | 5 | 4 | 7 | 13 | 11 | 25 |
| L2c_dup91349 | 8.43411625136727 | 2.27903088712321 | 0.530140387506452 | 4.29891957080043 | 1.716327375686e-05 | 0.00276548443276155 | 1052 | 1 | 2 | 1 | 4 | 7 | 6 | 5 | 5 | 10 | 12 | 19 | 35 |
| LTR1D1_dup23 | 9.38071292629927 | 1.58606371549731 | 0.444758817979458 | 3.56612089829452 | 0.000362304281840886 | 0.024631101327689 | 1053 | 5 | 2 | 7 | 5 | 7 | 9 | 8 | 5 | 11 | 9 | 17 | 35 |
| AluSq2_dup28362 | 7.85251356143401 | 1.52106608903874 | 0.436369963442763 | 3.48572591256788 | 0.000490803602934893 | 0.0301146864631114 | 1054 | 3 | 5 | 6 | 4 | 6 | 5 | 6 | 6 | 12 | 8 | 17 | 19 |
| MSTD_dup6383 | 9.20898453494298 | 1.82986931584213 | 0.454718483380291 | 4.02418063642196 | 5.717402193924e-05 | 0.0066884610739205 5 | 1055 | 3 | 4 | 8 | 5 | 9 | 4 | 3 | 6 | 15 | 11 | 17 | 29 |
| MER5A1_dup12438 | 6.66833647008116 | 1.6898734001467 | 0.500577027851964 | 3.37585088032935 | 0.000735877919364651 | 0.0390578768691985 | 1056 | 2 | 4 | 5 | 4 | 3 | 9 | 4 | 2 | 14 | 11 | 13 | 10 |
| L2c_dup114270 | 6.38094826465997 | 1.82029497822689 | 0.513445021194715 | 3.54525782330368 | 0.000392229229577803 | 0.0259320300808891 | 1057 | 2 | 1 | 5 | 6 | 4 | 6 | 4 | 2 | 12 | 15 | 12 | 9 |
| MIR_dup137237 | 7.21504821479845 | 1.9667211932457 | 0.506714094134292 | 3.8813232471968 | 0.000103889644898322 | 0.0103178404285482 | 1058 | 3 | 2 | 6 | 5 | 2 | 8 | 5 | 1 | 12 | 20 | 14 | 11 |
| LTR1A2_dup53 | 7.5854892788178 | 1.47360727063915 | 0.434523143724146 | 3.39132055892207 | 0.000695566941454446 | 0.0376324173377261 | 1059 | 5 | 3 | 4 | 7 | 3 | 7 | 9 | 4 | 8 | 15 | 16 | 15 |
| AluSq2_dup45253 | 7.80019434343953 | 1.60925336879618 | 0.455502832203736 | 3.53291627411088 | 0.000411002612257811 | 0.0267935418017252 | 1060 | 2 | 2 | 6 | 9 | 5 | 7 | 7 | 3 | 12 | 15 | 11 | 19 |
| MER102c_dup2587 | 7.12341068010093 | 1.68310278249833 | 0.497606729732532 | 3.38239553834615 | 0.00071856600709654 | 0.0383879450212378 | 1061 | 2 | 1 | 6 | 5 | 3 | 9 | 7 | 3 | 10 | 15 | 13 | 15 |
| MLT2B4_dup227 | 6.7328985196073 | 1.75899274537836 | 0.532969454028437 | 3.30036314855017 | 0.000965597937321186 | 0.04668024518363 | 1062 | 1 | 1 | 6 | 7 | 5 | 2 | 7 | 5 | 9 | 16 | 14 | 10 |
| MER58A_dup6786 | 7.49538104437085 | 2.0134965355341 | 0.470881667752275 | 4.27601385533951 | 1.90269386912762e-05 | 0.00300025444256753 | 1063 | 2 | 2 | 6 | 6 | 4 | 3 | 7 | 3 | 13 | 17 | 17 | 12 |
| MIR3_dup67838 | 6.9479984280931 | 1.8703071618893 | 0.52457941182441 | 3.56534610343293 | 0.000363376339170527 | 0.02468172339212 | 1064 | 2 | 2 | 5 | 9 | 3 | 4 | 5 | 3 | 9 | 20 | 16 | 8 |
| AluSp_dup37019 | 6.86767905610838 | 1.95101771378858 | 0.49318986848886 | 3.95591604459865 | 7.62419350220377e-05 | 0.00824986970020636 | 1065 | 2 | 3 | 3 | 7 | 2 | 5 | 7 | 2 | 13 | 17 | 10 | 14 |
| MIR_dup114522 | 7.02697834994737 | 1.7010716127295 | 0.500523793055683 | 3.39858291719662 | 0.000677359270020817 | 0.037034680345674 | 1066 | 1 | 5 | 3 | 9 | 3 | 4 | 6 | 4 | 13 | 16 | 11 | 11 |
| MIRb_dup174515 | 7.71661883344006 | 1.71213875877291 | 0.518848694529513 | 3.299880633458 | 0.000967259599966974 | 0.0467415826104268 | 1067 | 1 | 3 | 5 | 11 | 5 | 6 | 4 | 3 | 17 | 17 | 9 | 13 |
| L2a_dup64481 | 6.71641235372112 | 1.84284305010414 | 0.530367231876121 | 3.4746548039653 | 0.000511511126576174 | 0.0310014292298582 | 1068 | 1 | 2 | 6 | 9 | 4 | 3 | 4 | 3 | 14 | 15 | 9 | 12 |
| MIRc_dup79342 | 6.33162172003642 | 1.71487978337348 | 0.52316553313694 | 3.27789136469088 | 0.00104585637634097 | 0.049278681450461 | 1069 | 2 | 3 | 5 | 10 | 3 | 3 | 4 | 2 | 12 | 14 | 11 | 9 |
| L3b_dup8256 | 8.47172695989194 | 1.98885734911129 | 0.422326261935907 | 4.70929120058633 | 2.48579715181657e-06 | 0.000663844544428895 | 1070 | 4 | 3 | 5 | 7 | 3 | 7 | 5 | 3 | 17 | 17 | 15 | 18 |
| MLT2A1_dup1855 | 8.39853760348749 | 1.73657037791621 | 0.430143355469065 | 4.03718982482597 | 5.40952966965919e-05 | 0.00639114174169179 | 1071 | 5 | 6 | 4 | 5 | 4 | 9 | 6 | 2 | 18 | 16 | 16 | 11 |
| L1PA3_dup3659 | 9.18324850779551 | 2.40197019393558 | 0.431573137839383 | 5.56561561259521 | 2.61228516609041e-08 | 1.71707957494626e-05 | 1072 | 6 | 2 | 2 | 3 | 5 | 8 | 5 | 2 | 20 | 26 | 16 | 16 |
| MIRb_dup71167 | 7.32550721106195 | 1.86375251526204 | 0.475479067073597 | 3.91973620780568 | 8.86459425990358e-05 | 0.00922701113633866 | 1073 | 6 | 2 | 3 | 3 | 4 | 6 | 8 | 2 | 13 | 18 | 15 | 10 |
| AluSc8_dup6513 | 8.41582906669943 | 2.13924037038679 | 0.483920754876593 | 4.42064191053828 | 9.84081335074703e-06 | 0.00183365871069448 | 1074 | 4 | 1 | 7 | 4 | 4 | 4 | 7 | 3 | 20 | 20 | 15 | 12 |

| | | | | | | | | | | | | | | | | | | |
|---|---|---|---|---|---|---|---|---|---|---|---|---|---|---|---|---|---|---|
| L3_dup37644 | 6.95763591087056 | 1.58953502371243 | 0.473953405282799 | 3.35377909725954 | 0.0007971597554315 | 0.0413428455025945 | 1075 | 7 | 1 | 5 | 5 | 6 | 4 | 5 | 3 | 15 | 11 | 11 | 12 |
| MER2_dup3497 | 8.6456868531084 | 1.69093763967327 | 0.495515804686169 | 3.41247973057935 | 0.000643747141753462 | 0.0359245795990491 | 1076 | 4 | 1 | 3 | 7 | 10 | 9 | 6 | 2 | 18 | 24 | 10 | 11 |
| MIRb_dup174500 | 9.71249705867567 | 1.74616645794712 | 0.427852711987667 | 4.08123265091621 | 4.47974860538943e-05 | 0.00554457539198024 | 1077 | 10 | 2 | 4 | 9 | 5 | 8 | 6 | 3 | 21 | 23 | 11 | 17 |
| L1MA4_dup3514 | 8.3713724747849 | 1.62568983724382 | 0.495008681059561 | 3.28416429740999 | 0.00102285274457314 | 0.0485923564930782 | 1078 | 4 | 1 | 9 | 9 | 6 | 9 | 2 | 3 | 13 | 16 | 15 | 17 |
| LTR88a_dup718 | 7.72659289425953 | 1.72538718044141 | 0.506810874147547 | 3.40440047452317 | 0.000663094591849045 | 0.0365571607395998 | 1079 | 2 | 1 | 5 | 13 | 4 | 6 | 3 | 5 | 11 | 14 | 13 | 20 |
| MIR_dup168501 | 7.06311766412554 | 2.02016060207018 | 0.546698336251131 | 3.69520166445541 | 0.000219712329307143 | 0.0174779121709676 | 1080 | 3 | 2 | 6 | 4 | 3 | 9 | 1 | 2 | 16 | 15 | 7 | 18 |
| AluSx_dup560 | 7.24131504002594 | 2.01327378116065 | 0.484973388687921 | 4.15130773795133 | 3.30580906551473e-05 | 0.00448207432423344 | 1081 | 4 | 1 | 5 | 3 | 3 | 8 | 4 | 3 | 13 | 18 | 10 | 17 |
| MIRc_dup79409 | 6.25391792820206 | 2.44167176470028 | 0.506275469119662 | 4.82281270500026 | 1.4154787208285e-06 | 0.000424190805675387 | 1082 | 4 | 1 | 3 | 3 | 2 | 5 | 2 | 2 | 12 | 13 | 13 | 17 |
| L1MC4_dup7306 | 7.47628040185484 | 2.256289008431 | 0.672397217831893 | 3.35558944712217 | 0.000791960412988075 | 0.0411809030657475 | 1083 | 0 | 6 | 3 | 2 | 3 | 1 | 2 | 10 | 14 | 15 | 17 | 15 |
| MIRb_dup102160 | 9.09284945788888 | 1.73106366642337 | 0.459495531919312 | 3.7673133821187 | 0.000165013807515121 | 0.0143094198017401 | 1084 | 4 | 10 | 3 | 3 | 9 | 5 | 2 | 7 | 18 | 20 | 14 | 13 |
| MSTA-int_dup2754 | 8.95396124414819 | 2.06194860855813 | 0.581573235930824 | 3.54546681512591 | 0.000391918332448581 | 0.025925226644737 | 1085 | 2 | 4 | 2 | 2 | 11 | 5 | 2 | 8 | 21 | 12 | 22 | 14 |
| L2_dup3431 | 9.96649195952662 | 2.14707312510451 | 0.556485209974378 | 3.85827527240728 | 0.000114189995909305 | 0.0110451434010713 | 1086 | 2 | 4 | 2 | 7 | 2 | 7 | 14 | 3 | 21 | 31 | 16 | 12 |
| L1MA8_dup1679 | 9.06232531722984 | 3.03246228231323 | 0.525742279176986 | 5.76796351067741 | 8.02351720819969e-09 | 6.57809325703013e-06 | 1087 | 2 | 6 | 2 | 4 | 1 | 2 | 3 | 2 | 25 | 31 | 17 | 11 |
| MIRb_dup46310 | 6.98005470592855 | 2.47700450136952 | 0.554713534273028 | 4.46537599738886 | 7.99282557402819e-06 | 0.00155253948901038 | 1088 | 2 | 5 | 1 | 3 | 3 | 2 | 6 | 1 | 21 | 19 | 11 | 8 |
| MIR_dup39861 | 6.2267656311849 | 1.86536942020772 | 0.535042672140719 | 3.4863937351844 | 0.00048957984683319 | 0.0300706159614275 | 1089 | 1 | 4 | 5 | 2 | 5 | 4 | 7 | 1 | 7 | 16 | 14 | 12 |
| MER58A_dup10296 | 5.94949000799574 | 1.76568678317778 | 0.535731914528028 | 3.29584020532605 | 0.00098127812442723 | 0.0472080882723336 | 1090 | 3 | 1 | 5 | 4 | 3 | 7 | 4 | 2 | 5 | 16 | 14 | 11 |
| MIRc_dup72153 | 5.55525816049638 | 2.07077569862806 | 0.586619237960707 | 3.53001668650827 | 0.000415533458429948 | 0.026973724727073 | 1091 | 3 | 1 | 4 | 2 | 1 | 5 | 7 | 1 | 6 | 14 | 16 | 10 |
| AluSx3_dup24021 | 5.41400081032023 | 1.82855008655262 | 0.555274190288123 | 3.29305794962991 | 0.000991040460686219 | 0.0475034237087012 | 1092 | 2 | 2 | 5 | 6 | 1 | 3 | 7 | 1 | 7 | 10 | 7 | 19 |
| MER5A1_dup12432 | 4.98325354803008 | 2.02445929737905 | 0.564555281340831 | 3.58593633659916 | 0.000335870897177187 | 0.0234023825767856 | 1093 | 2 | 1 | 3 | 5 | 4 | 1 | 5 | 1 | 10 | 8 | 10 | 12 |
| MIRc_dup79336 | 6.0000644519018 | 2.07441002018113 | 0.624789699116746 | 3.32017320886322 | 0.000899616137565763 | 0.0445216434004208 | 1094 | 2 | 1 | 4 | 13 | 2 | 1 | 5 | 1 | 9 | 15 | 12 | 11 |
| MER45A_dup2222 | 5.73068022632778 | 2.10217929617777 | 0.610314511447588 | 3.44441965043837 | 0.000572286703814187 | 0.0333599035686686 | 1095 | 4 | 2 | 4 | 1 | 5 | 2 | 0 | 4 | 15 | 15 | 5 | 10 |
| MLT1J-int_dup526 | 6.52057395443316 | 2.67973825087963 | 0.608748699087271 | 4.40204349495531 | 1.07236051757597e-05 | 0.00196733921137341 | 1096 | 2 | 3 | 1 | 0 | 9 | 5 | 0 | 2 | 15 | 20 | 9 | 11 |
| MSTB_dup2678 | 6.52805960821326 | 2.34283963765358 | 0.595043135666544 | 3.93726017027189 | 8.24172353250844e-05 | 0.0087392386259637 | 1097 | 10 | 2 | 2 | 2 | 5 | 3 | 0 | 1 | 23 | 12 | 7 | 9 |
| LTR2_dup331 | 5.61956723063387 | 1.96635864329302 | 0.535380470312743 | 3.67282475235671 | 0.000239884001620002 | 0.0185904708360133 | 1098 | 7 | 3 | 3 | 3 | 3 | 4 | 0 | 2 | 11 | 14 | 10 | 8 |
| MamTip2_dup2944 | 4.57360535562976 | 2.08863935047983 | 0.634579238175811 | 3.29137674986645 | 0.000996982932811366 | 0.0477320122597049 | 1099 | 6 | 5 | 2 | 2 | 2 | 2 | 0 | 1 | 12 | 6 | 8 | 9 |
| L1MD_dup2751 | 5.38155785693594 | 2.56707966563231 | 0.650921861378258 | 3.94376010078505 | 8.02139029666543e-05 | 0.0085360363843039 | 1100 | 6 | 3 | 0 | 1 | 1 | 6 | 0 | 2 | 14 | 16 | 4 | 11 |
| AluY_dup34659 | 5.39004569565589 | 2.42794662216573 | 0.563475852525436 | 4.30887430452244 | 1.64087586328601e-05 | 0.00267607846583586 | 1101 | 5 | 3 | 1 | 2 | 1 | 4 | 1 | 2 | 13 | 15 | 10 | 7 |
| MIR3_dup20850 | 6.05881856057555 | 1.70285212682722 | 0.50263428754511 | 3.38785508474567 | 0.00070441471494961 | 0.0379557662507699 | 1102 | 8 | 6 | 1 | 2 | 3 | 7 | 2 | 2 | 10 | 18 | 7 | 8 |
| L3_dup9409 | 7.0610154759115 | 2.23812725021153 | 0.65320503231983 | 3.42637784381867 | 0.000611688830955281 | 0.0348227569164275 | 1103 | 8 | 3 | 0 | 3 | 2 | 1 | 2 | 6 | 18 | 23 | 5 | 12 |
| Tigger6a_dup138 | 5.5513427285271 | 2.25286218469613 | 0.668549879367142 | 3.36977427447705 | 0.000752297816526024 | 0.0396997685187627 | 1104 | 9 | 1 | 0 | 1 | 3 | 2 | 2 | 3 | 15 | 16 | 3 | 11 |
| MLT1H1_dup626 | 5.82936085359987 | 2.60333993360879 | 0.592188455527729 | 4.3961342192813 | 1.10195810026321e-05 | 0.00201232941001711 | 1105 | 3 | 1 | 0 | 3 | 2 | 4 | 4 | 1 | 16 | 20 | 8 | 7 |
| L1PA8_dup1390 | 7.19071657391967 | 3.53552900048325 | 0.650879056112141 | 5.43192927669515 | 5.57480182397582e-08 | 3.15083561712055e-05 | 1106 | 2 | 1 | 0 | 4 | 1 | 1 | 1 | 3 | 24 | 22 | 11 | 13 |
| L2c_dup29818 | 7.21416370227788 | 3.3520391480726 | 0.598753321625971 | 5.59836417937495 | 2.1638382895222e-08 | 1.48808284866662e-05 | 1107 | 5 | 1 | 0 | 1 | 1 | 3 | 1 | 3 | 23 | 22 | 8 | 16 |
| AluJo_dup5295 | 4.57024724226273 | 2.23371923874805 | 0.577175574598533 | 3.87008622168698 | 0.000108796868256973 | 0.010683725028802 | 1108 | 2 | 3 | 2 | 2 | 2 | 2 | 2 | 2 | 12 | 12 | 6 | 7 |
| MER5B_dup9792 | 4.23542025845308 | 2.10827823892233 | 0.632656536158761 | 3.3324214932212 | 0.000860937568481347 | 0.0433519044356457 | 1109 | 2 | 3 | 2 | 1 | 3 | 1 | 4 | 1 | 8 | 14 | 8 | 4 |
| L1ME1_dup22281 | 5.28831647229308 | 2.00856110591285 | 0.575763856683293 | 3.48851544361126 | 0.000485710762435284 | 0.029926204166386 | 1110 | 5 | 2 | 3 | 2 | 3 | 3 | 3 | 1 | 12 | 17 | 9 | 3 |
| MLT1B_dup14507 | 5.23191544318521 | 1.83367329439426 | 0.551291676425331 | 3.32613999595299 | 0.00088057688962737 | 0.0438713878934829 | 1111 | 3 | 5 | 4 | 3 | 2 | 3 | 3 | 1 | 8 | 18 | 8 | 6 |
| AluSz_dup51264 | 4.83484359314541 | 2.19744777346411 | 0.612543394472433 | 3.58741567257731 | 0.000333971691088267 | 0.0233041724362724 | 1112 | 1 | 3 | 1 | 1 | 7 | 3 | 1 | 2 | 12 | 12 | 6 | 8 |
| MER5A1_dup7841 | 6.39386354130089 | 2.1444594064548 | 0.64391776662236 | 3.33033116589322 | 0.000867427502150585 | 0.0435077791793261 | 1113 | 1 | 3 | 4 | 1 | 7 | 2 | 2 | 2 | 21 | 21 | 3 | 6 |
| L2c_dup29820 | 7.96534038835666 | 2.45239936674418 | 0.614683944583813 | 3.98969159411612 | 6.61592607511169e-05 | 0.00743263997105065 | 1114 | 4 | 9 | 2 | 2 | 1 | 1 | 2 | 4 | 18 | 33 | 5 | 13 |
| MER1B_dup1010 | 5.38473263320894 | 2.28938399601928 | 0.580050681388751 | 3.94686890210708 | 7.91798833381367e-05 | 0.00845888137039488 | 1115 | 5 | 5 | 1 | 2 | 2 | 1 | 1 | 3 | 10 | 18 | 6 | 11 |
| SVA_F_dup861 | 6.23176473196616 | 1.93155346876754 | 0.561198423421044 | 3.44183694778196 | 0.000577778399594558 | 0.0336001669821296 | 1116 | 4 | 3 | 2 | 4 | 2 | 4 | 3 | 5 | 12 | 17 | 3 | 17 |
| L1PB1_dup1482 | 5.86602433009109 | 1.71606695580943 | 0.489030486071006 | 3.5091206063588 | 0.000449590981508806 | 0.0284744146090217 | 1117 | 4 | 5 | 2 | 4 | 4 | 5 | 4 | 1 | 12 | 14 | 6 | 11 |
| L2a_dup129398 | 7.56375051984699 | 1.84519805995854 | 0.494847284062091 | 3.72882325393751 | 0.000192376018809603 | 0.0159885100145546 | 1118 | 3 | 8 | 2 | 6 | 4 | 7 | 2 | 2 | 17 | 23 | 5 | 12 |
| L1ME4a_dup18000 | 5.18685906544669 | 1.85579825769307 | 0.566471395751175 | 3.27606702052832 | 0.00105263578257433 | 0.0494512288095621 | 1119 | 5 | 3 | 1 | 1 | 8 | 6 | 2 | 0 | 10 | 14 | 7 | 6 |
| AluSx_dup34165 | 4.57199593949385 | 2.11831359857577 | 0.595738168649587 | 3.55577955224448 | 0.000376859983144644 | 0.0253045658679988 | 1120 | 4 | 4 | 1 | 1 | 5 | 5 | 1 | 0 | 9 | 9 | 8 | 9 |
| L1MB3_dup8961 | 4.74355708896261 | 1.90526725805808 | 0.571193542970591 | 3.3355896289538 | 0.000851187103892304 | 0.0430013941874221 | 1121 | 6 | 3 | 1 | 2 | 4 | 4 | 3 | 0 | 8 | 9 | 11 | 8 |
| MIRc_dup71475 | 4.76141245108647 | 2.13117195407873 | 0.588198433648596 | 3.62321936299467 | 0.000290958861492344 | 0.0211720175096811 | 1122 | 4 | 5 | 3 | 2 | 4 | 1 | 1 | 0 | 10 | 11 | 8 | 9 |
| THE1B_dup17350 | 5.46316567634939 | 2.46415607006958 | 0.552025034206587 | 4.46384840791009 | 8.05005286932423e-06 | 0.00155729908946685 | 1123 | 6 | 3 | 1 | 3 | 2 | 3 | 2 | 0 | 10 | 14 | 10 | 14 |
| Tigger19a_dup3162 | 5.95619611613873 | 1.78365878209841 | 0.519193992185299 | 3.43543802306138 | 0.000591596764243565 | 0.0341007954614097 | 1124 | 9 | 6 | 2 | 4 | 4 | 3 | 2 | 0 | 10 | 16 | 8 | 10 |
| AluSp_dup22688 | 5.69402979680457 | 2.26787764750884 | 0.570034908092832 | 3.97848906323385 | 6.93546009054114e-05 | 0.00770431833195099 | 1125 | 1 | 5 | 1 | 4 | 2 | 4 | 5 | 0 | 12 | 15 | 9 | 12 |
| MIRb_dup199601 | 6.01877756597803 | 1.86871993281273 | 0.534639816725623 | 3.49528762047245 | 0.000473551153188516 | 0.0294125822752792 | 1126 | 3 | 6 | 1 | 4 | 3 | 7 | 4 | 0 | 15 | 14 | 6 | 10 |
| L3_dup16365 | 4.39088312639991 | 2.17937811445272 | 0.615053220923621 | 3.54339761229112 | 0.000395006659569608 | 0.0260721926818169 | 1127 | 1 | 2 | 1 | 3 | 4 | 5 | 2 | 0 | 9 | 11 | 6 | 10 |
| MER112_dup2269 | 4.40843003259411 | 2.07941464912024 | 0.62331899863331 | 3.33603604844319 | 0.000849821435254591 | 0.0429688931647023 | 1128 | 1 | 3 | 1 | 3 | 3 | 5 | 3 | 0 | 6 | 6 | 12 | 13 |
| AluSc_dup4467 | 6.31782907669738 | 1.68451091027648 | 0.482540883634082 | 3.49091852609502 | 0.000481363020650862 | 0.0297734352253158 | 1129 | 5 | 5 | 5 | 4 | 4 | 2 | 2 | 4 | 10 | 13 | 13 | 10 |
| L2b_dup8619 | 5.68426016078799 | 1.76214333488444 | 0.535690533734534 | 3.28948007089049 | 0.00100372664327139 | 0.0479583814979629 | 1130 | 4 | 6 | 4 | 2 | 3 | 3 | 1 | 4 | 6 | 12 | 14 | 11 |
| MIRb_dup71168 | 5.86567045380169 | 1.74867819829046 | 0.532926970580169 | 3.28127172168968 | 0.00103340137387363 | 0.0489192567235723 | 1131 | 7 | 4 | 4 | 2 | 2 | 5 | 3 | 2 | 5 | 19 | 8 | 13 |
| MIRb_dup40635 | 6.35358595356421 | 1.78053649357152 | 0.510238228673478 | 3.48961797355047 | 0.000483711499205628 | 0.0298566060674403 | 1132 | 6 | 4 | 2 | 1 | 4 | 4 | 4 | 5 | 7 | 19 | 11 | 11 |
| MIRb_dup174370 | 6.1180641693571 | 1.75901677626495 | 0.507331902293286 | 3.46719133631 | 0.000525927379229213 | 0.0314728857988515 | 1133 | 5 | 2 | 2 | 2 | 6 | 6 | 4 | 3 | 6 | 12 | 14 | 15 |
| AluSx_dup73908 | 5.17452323017674 | 1.78680452119405 | 0.53109158799168 | 3.36439996715227 | 0.000767102864814314 | 0.0402844975217627 | 1134 | 4 | 4 | 2 | 1 | 5 | 4 | 2 | 3 | 7 | 10 | 9 | 13 |
| MIR3_dup69142 | 5.14102796932351 | 1.84159671381927 | 0.534862776871247 | 3.44311998040309 | 0.00057504413587279 | 0.033478173957623 | 1135 | 3 | 3 | 2 | 1 | 4 | 5 | 3 | 3 | 6 | 10 | 11 | 13 |
| L1PA7_dup11132 | 6.88223519791525 | 1.86786574498953 | 0.538427525247912 | 3.46911266122491 | 0.000522180431808968 | 0.0313289826089805 | 1136 | 4 | 6 | 2 | 2 | 6 | 10 | 2 | 2 | 7 | 11 | 15 | 20 |
| AluSx_dup51591 | 7.56231282312126 | 1.93932894039573 | 0.454038079060941 | 4.27129139566162 | 1.9434426757938e-05 | 0.00304233165088442 | 1137 | 4 | 3 | 3 | 4 | 5 | 9 | 3 | 3 | 14 | 13 | 14 | 18 |
| MADE2_dup2503 | 6.48887255837006 | 1.74633772085441 | 0.518674930960132 | 3.36692139259887 | 0.000760123519267937 | 0.0400037420957086 | 1138 | 2 | 3 | 4 | 3 | 6 | 10 | 2 | 2 | 11 | 12 | 10 | 15 |
| AluJr_dup62323 | 5.67795117790432 | 1.91814806625888 | 0.538371985859437 | 3.56286752773144 | 0.000366825815071643 | 0.0248716943004123 | 1139 | 1 | 2 | 2 | 6 | 4 | 6 | 4 | 1 | 10 | 9 | 12 | 14 |
| AluSz_dup5801 | 6.96571192182081 | 2.09559485426171 | 0.554831822472574 | 3.77699109781197 | 0.000158734402822341 | 0.0139431096491211 | 1140 | 3 | 2 | 1 | 4 | 4 | 10 | 5 | 1 | 9 | 11 | 15 | 24 |
| L2b_dup74972 | 5.70013129106005 | 1.94309755176633 | 0.514302048276321 | 3.77812524425793 | 0.000158013411294412 | 0.0138996891548466 | 1141 | 2 | 2 | 5 | 2 | 5 | 3 | 3 | 3 | 10 | 10 | 10 | 15 |
| MIRc_dup79403 | 6.0368166332452 | 1.9237562797215 | 0.495300048229268 | 3.88402199151618 | 0.000102742608626258 | 0.0102652751583559 | 1142 | 2 | 3 | 3 | 3 | 6 | 2 | 5 | 3 | 9 | 12 | 11 | 16 |
| L2a_dup136997 | 7.00287493532312 | 1.86880429581746 | 0.482098329346062 | 3.87639654000125 | 0.000106014857717238 | 0.0104714866477318 | 1143 | 2 | 3 | 5 | 5 | 7 | 3 | 5 | 2 | 16 | 12 | 12 | 13 |
| L2c_dup70063 | 6.7542426175399 | 1.7435593261956 | 0.498604570077247 | 3.4968779486427 | 0.000470737150027342 | 0.0292836658563152 | 1144 | 2 | 2 | 3 | 4 | 6 | 5 | 8 | 3 | 13 | 9 | 10 | 19 |
| MIR3_dup27868 | 6.99362796976474 | 1.73436192745506 | 0.465950269232059 | 3.7222039388742 | 0.000197491408389658 | 0.0162893168114593 | 1145 | 6 | 4 | 1 | 2 | 7 | 6 | 5 | 3 | 16 | 15 | 9 | 10 |
| LTR1B0_dup43 | 5.7951305267069 | 1.85174438877759 | 0.495337824302646 | 3.73834643333475 | 0.00018523458412136 | 0.0155910252399509 | 1146 | 6 | 3 | 2 | 4 | 3 | 4 | 3 | 2 | 10 | 15 | 11 | 8 |

| | | | | | | | | | | | | | | | | | | | |
|---|---|---|---|---|---|---|---|---|---|---|---|---|---|---|---|---|---|---|---|
| Tigger3b_dup2232 | 7.06441517614746 | 1.97696001542778 | 0.47140636208524 | 4.19374911845229 | 2.74381436918164e-05 | 0.00391180140137384 | 1147 | 6 | 2 | 1 | 6 | 5 | 5 | 3 | 3 | 15 | 15 | 12 | 13 |
| MIRb_dup184035 | 4.20673568660813 | 3.18678743434301 | 0.724312182885882 | 4.39974296945535 | 1.08379170223563e-05 | 0.00198219754327562 | 1148 | 0 | 1 | 1 | 0 | 4 | 2 | 1 | 1 | 7 | 14 | 10 | 10 |
| L1MB5_dup7498 | 3.55959658973571 | 2.53309986919486 | 0.715619389031466 | 3.53973062778974 | 0.000400535631264046 | 0.0263395581133461 | 1149 | 0 | 1 | 1 | 1 | 4 | 3 | 1 | 1 | 8 | 6 | 8 | 9 |
| MLT1E2_dup3350 | 3.24725751778226 | 2.85555238432104 | 0.745901848283057 | 3.8283219044088 | 0.000129019940325813 | 0.0120296938560995 | 1150 | 0 | 1 | 1 | 1 | 3 | 1 | 1 | 1 | 5 | 6 | 11 | 9 |
| LTR53B_dup153 | 4.88964433948315 | 2.33914363167414 | 0.663257680341885 | 3.52674940826677 | 0.000420694729346075 | 0.0271796437873964 | 1151 | 0 | 1 | 1 | 4 | 4 | 4 | 2 | 2 | 4 | 18 | 11 | 10 |
| MLT1D_dup16964 | 4.14750145100568 | 2.68194532733042 | 0.703164811974705 | 3.81410628299031 | 0.000136676642272182 | 0.0125083308931141 | 1152 | 0 | 2 | 1 | 1 | 3 | 4 | 1 | 1 | 4 | 9 | 14 | 12 |
| THE1D-int_dup2689 | 4.43617539762862 | 2.47504804485499 | 0.631580307162984 | 3.91881763377445 | 8.89844077520849e-05 | 0.00925010217867086 | 1153 | 0 | 1 | 3 | 2 | 3 | 2 | 2 | 2 | 5 | 8 | 17 | 10 |
| L2a_dup61522 | 4.72998466405111 | 2.35748957721959 | 0.68787640184087 | 3.42719937900262 | 0.000609841127261899 | 0.0347765478027451 | 1154 | 1 | 5 | 0 | 1 | 5 | 4 | 2 | 1 | 5 | 12 | 6 | 18 |
| MER11D_dup202 | 4.68517386822721 | 2.21503992743167 | 0.671862570937399 | 3.29686460185033 | 0.000977706221610053 | 0.0470898042609459 | 1155 | 2 | 5 | 0 | 1 | 1 | 6 | 1 | 3 | 6 | 11 | 8 | 14 |
| AluSx1_dup74123 | 5.10071090634894 | 2.20083703311032 | 0.61122512785815 | 3.60069789804368 | 0.000317364162088718 | 0.0224815519238947 | 1156 | 2 | 2 | 0 | 1 | 5 | 6 | 4 | 1 | 10 | 11 | 12 | 8 |
| L1ME1_dup23557 | 4.5828549943143 | 2.44574803111699 | 0.71080052873359 | 3.4408359761275 | 0.00057991996538555 | 0.03368887599698 | 1157 | 1 | 1 | 0 | 1 | 8 | 4 | 1 | 1 | 11 | 13 | 8 | 5 |
| LTR48B_dup300 | 5.21815987414675 | 2.27599590550766 | 0.614687923474562 | 3.70268524659221 | 0.000213329485415917 | 0.0171534565551927 | 1158 | 1 | 3 | 0 | 5 | 3 | 1 | 5 | 2 | 9 | 16 | 10 | 9 |
| L1PREC2_dup6715 | 3.55517187136676 | 2.20843589151643 | 0.665683984601481 | 3.3175439737198 | 0.000908126194393964 | 0.0448495216793828 | 1159 | 2 | 2 | 0 | 1 | 2 | 2 | 3 | 2 | 4 | 10 | 9 | 7 |
| THE1B_dup6821 | 4.57903340645186 | 1.97861609795178 | 0.588353939252566 | 3.36296906665633 | 0.000771090036006859 | 0.0404136100677727 | 1160 | 3 | 1 | 0 | 3 | 3 | 3 | 4 | 3 | 6 | 8 | 13 | 10 |
| L1MC4_dup4707 | 5.446120028703 | 3.70536217539087 | 0.684580939090427 | 5.41259910086603 | 6.21164564126081e-08 | 3.4134785998971e-05 | 1161 | 1 | 2 | 1 | 0 | 1 | 0 | 2 | 2 | 17 | 15 | 12 | 10 |
| MIR_dup34587 | 6.0218801209702 | 2.77513317287573 | 0.614761959434103 | 4.51415890376543 | 6.3568542338752e-06 | 0.00131293568783474 | 1162 | 2 | 4 | 1 | 0 | 1 | 2 | 4 | 3 | 18 | 13 | 13 | 9 |
| MLT1G1_dup722 | 5.9001745882354 | 2.94755864437162 | 0.616315533188141 | 4.78254803854153 | 1.73087004385489e-06 | 0.000499822334290165 | 1163 | 1 | 2 | 1 | 1 | 2 | 1 | 3 | 4 | 15 | 18 | 11 | 10 |
| MIRb_dup66793 | 6.55610691838716 | 2.27054866798816 | 0.638303943024229 | 3.55715908197354 | 0.000374887150277781 | 0.0252199315408529 | 1164 | 2 | 3 | 2 | 0 | 1 | 2 | 8 | 7 | 11 | 15 | 15 | 13 |
| MIRb_dup75922 | 6.37986845641342 | 2.25484248815251 | 0.665997582149122 | 3.38566167293928 | 0.000710068685784108 | 0.0381825996627311 | 1165 | 1 | 3 | 2 | 0 | 3 | 3 | 6 | 4 | 20 | 18 | 8 | 5 |
| AluSx_dup21152 | 4.73221878047707 | 2.39530125824372 | 0.631791134691518 | 3.79128659254337 | 0.000149868893645777 | 0.01331451342872 | 1166 | 1 | 4 | 4 | 2 | 1 | 0 | 3 | 2 | 7 | 15 | 9 | 10 |
| AluSz6_dup33777 | 5.22907930571026 | 1.97257117128966 | 0.576990363104713 | 3.41872463982847 | 0.000629153605235098 | 0.0354078958730536 | 1167 | 2 | 6 | 5 | 4 | 2 | 1 | 2 | 1 | 7 | 15 | 9 | 11 |
| MIRb_dup143679 | 5.48202198269744 | 1.95142795127972 | 0.530726493319282 | 3.67689944979956 | 0.000236086044421683 | 0.018400173575487 | 1168 | 2 | 4 | 3 | 6 | 2 | 1 | 5 | 2 | 7 | 14 | 10 | 13 |
| MIRb_dup46312 | 5.97609841289063 | 2.73429729303636 | 0.598435551545768 | 4.56907562723109 | 4.8988000432527e-06 | 0.00107830700748548 | 1169 | 2 | 2 | 3 | 2 | 0 | 2 | 4 | 3 | 10 | 21 | 15 | 8 |
| MER57A-int_dup946 | 4.88033707925813 | 2.03216166545966 | 0.605912021874809 | 3.353889000538 | 0.000796843211128389 | 0.0413428455025945 | 1170 | 1 | 1 | 3 | 4 | 1 | 3 | 4 | 4 | 10 | 13 | 5 | 10 |
| MLT1I_dup2169 | 4.5390362957173 | 2.79089847076263 | 0.740916976324565 | 3.7668167418802 | 0.00016534227582661 | 0.0143119933168183 | 1171 | 1 | 1 | 2 | 3 | 0 | 1 | 2 | 3 | 12 | 17 | 5 | 6 |
| L1PA12_dup1706 | 38.5657936822046 | -3.95003535338244 | 1.15286287342331 | -3.42628377098589 | 0.000611900740991795 | 0.0348240453757555 | 1172 | 0 | 0 | 209 | 259 | 1 | 3 | 34 | 20 | 1 | 0 | 2 | 2 |
| L1PA8_dup6180 | 58.5109754896935 | -5.33299728176616 | 1.2500236668505 | -4.26631704918269 | 1.98726314865935e-05 | 0.00307252317849837 | 1173 | 3 | 1 | 316 | 452 | 4 | 6 | 18 | 13 | 1 | 0 | 0 | 2 |
| L1MA3_dup2025 | 30.4422397414132 | -4.98286667672112 | 1.34737533180428 | -3.6982023932771 | 0.000217131731880469 | 0.0173607617856284 | 1174 | 6 | 7 | 158 | 239 | 2 | 0 | 6 | 4 | 0 | 0 | 2 | 2 |
| L1PA6_dup1343 | 18.6020769012835 | -4.91549562920702 | 1.32987804094374 | -3.69620031151034 | 0.000218850322666923 | 0.0174512596773806 | 1175 | 9 | 7 | 92 | 136 | 3 | 0 | 8 | 2 | 0 | 0 | 1 | 2 |
| L1PA16_dup2993 | 45.8578454870208 | -5.31105697890058 | 1.35355175390702 | -3.92379306042066 | 8.71656166196133e-05 | 0.00910878779629798 | 1176 | 17 | 15 | 245 | 342 | 2 | 1 | 10 | 2 | 0 | 0 | 4 | 3 |
| L1PREC2_dup1646 | 47.3082658427876 | -4.77836674072867 | 1.43148751977362 | -3.33804289225263 | 0.000843707243062223 | 0.0427729368088106 | 1177 | 14 | 8 | 246 | 371 | 1 | 0 | 6 | 6 | 0 | 0 | 7 | 3 |
| L1MA1_dup1056 | 13.3357075466083 | -3.7150608378032 | 1.1199200901373 | -3.31725528501568 | 0.00090906512612231 | 0.0448679548849407 | 1178 | 1 | 0 | 59 | 106 | 3 | 6 | 3 | 3 | 0 | 0 | 1 | 5 |
| L1PA16_dup10755 | 25.846221282442 | -4.91367102263 | 1.48326645599345 | -3.31273656380166 | 0.00092387956747533 | 0.0453535543596254 | 1179 | 1 | 0 | 125 | 222 | 1 | 3 | 9 | 3 | 0 | 0 | 1 | 0 |
| MLT2B3_dup2261 | 8.26558217821925 | -4.19261922544825 | 1.16308391048376 | -3.60474355087968 | 0.000312461311848726 | 0.0222601554817835 | 1180 | 1 | 0 | 49 | 48 | 0 | 3 | 7 | 4 | 0 | 0 | 0 | 1 |
| L1PA10_dup4823 | 9.57670102377904 | -4.19867460448914 | 1.14731385072705 | -3.65956935134048 | 0.000252639465086817 | 0.0192551409864022 | 1181 | 0 | 1 | 51 | 65 | 0 | 2 | 10 | 3 | 0 | 0 | 0 | 1 |
| L1PREC2_dup7594 | 12.0523976016935 | -4.21102246151172 | 1.14342068021584 | -3.6828286687248 | 0.00023066019187628 | 0.0180720019809659 | 1182 | 0 | 0 | 70 | 77 | 0 | 2 | 11 | 5 | 0 | 0 | 0 | 1 |
| L1PB2_dup2770 | 8.78619035193561 | -4.16170236024887 | 1.23620839084962 | -3.36650551076476 | 0.000761270611396487 | 0.0400489270173223 | 1183 | 0 | 0 | 46 | 59 | 1 | 0 | 4 | 9 | 0 | 0 | 0 | 0 |
| L1MA2_dup5853 | 8.90970651926925 | -4.93289675794029 | 1.35052872983399 | -3.65256706426873 | 0.000259631758780876 | 0.0196237807108081 | 1184 | 0 | 0 | 41 | 72 | 3 | 2 | 5 | 2 | 0 | 0 | 0 | 0 |
| REP522_dup82 | 5.53200516343422 | -4.31805374406107 | 1.26700439713054 | -3.40808110361765 | 0.00065421446855306 | 0.0362837448787836 | 1185 | 0 | 0 | 31 | 32 | 2 | 1 | 5 | 4 | 0 | 0 | 0 | 0 |
| MLT1D_dup10293 | 10.1052154109878 | -3.83435641463447 | 1.11708748619973 | -3.43245847975501 | 0.000598135459591803 | 0.0343362943469315 | 1186 | 0 | 3 | 57 | 64 | 0 | 0 | 9 | 4 | 0 | 0 | 1 | 1 |
| MLT1D_dup13027 | 14.9360689906808 | -3.85398818170303 | 1.03290426211486 | -3.73121529560935 | 0.000190558281725431 | 0.0158887770714079 | 1187 | 0 | 3 | 80 | 103 | 0 | 1 | 11 | 5 | 0 | 0 | 1 | 3 |
| ALR/Alpha_dup1273 | 12.1036512462181 | -5.20394979464536 | 1.19251954668931 | -4.36382766982114 | 1.27806270854688e-05 | 0.00224435357378521 | 1188 | 0 | 2 | 68 | 79 | 1 | 2 | 8 | 6 | 0 | 0 | 0 | 0 |
| L1PREC2_dup5949 | 13.1104413640582 | -4.98574826147136 | 1.1673629547083 | -4.27094952890398 | 1.94642458861002e-05 | 0.00304439142034009 | 1189 | 1 | 1 | 72 | 89 | 1 | 0 | 14 | 4 | 0 | 0 | 0 | 0 |
| HERV9NC-int_dup77 | 9.7651404513396 | -4.56126002499258 | 1.12623987303602 | -4.04998982383454 | 5.12198602318456e-05 | 0.00615222658391052 | 1190 | 1 | 11 | 45 | 54 | 1 | 4 | 5 | 8 | 0 | 0 | 2 | 0 |
| LTR12C_dup699 | 6.61173608884047 | -3.67197188141271 | 1.04910628303896 | -3.5000952151159 | 0.000465092000591159 | 0.02906928846143 | 1191 | 3 | 3 | 30 | 39 | 0 | 1 | 7 | 5 | 0 | 1 | 1 | 0 |
| MIR_dup167127 | 8.18828302991744 | -3.06308738432563 | 0.919636935726934 | -3.3307572426986 | 0.000866100972827913 | 0.0434770257891153 | 1192 | 3 | 2 | 38 | 46 | 1 | 1 | 12 | 5 | 0 | 2 | 1 | 1 |
| MSTA-int_dup1462 | 10.5498369068127 | -4.6302692024037 | 0.970040471521756 | -4.77327424817641 | 1.81254702057217e-06 | 0.000520879605503703 | 1193 | 2 | 2 | 40 | 66 | 0 | 1 | 17 | 14 | 0 | 0 | 1 | 0 |
| L1PA2_dup4853 | 8.78441663883705 | -3.91309804400988 | 1.13410131977845 | -3.45039545917671 | 0.000559765917858999 | 0.0328711898052511 | 1194 | 1 | 1 | 51 | 55 | 0 | 0 | 6 | 5 | 0 | 0 | 1 | 0 |
| L1PA4_dup6269 | 8.35313516524596 | -4.68843773913813 | 1.13420089324958 | -4.13369251165492 | 3.56981010841423e-05 | 0.00474007832620944 | 1195 | 1 | 1 | 42 | 52 | 0 | 1 | 10 | 7 | 0 | 0 | 0 | 0 |
| AluSx_dup96641 | 10.4684402671384 | -4.77539465617376 | 1.30550347999741 | -3.65789500322377 | 0.000254295173077225 | 0.0193431531351772 | 1196 | 22 | 11 | 36 | 60 | 0 | 0 | 5 | 7 | 1 | 0 | 0 | 1 |
| L1P2_dup462 | 11.5936292366053 | -5.57537260484729 | 1.35793821940274 | -4.10576307904455 | 4.02982353589699e-05 | 0.00515158666763517 | 1197 | 20 | 13 | 45 | 68 | 1 | 0 | 6 | 5 | 1 | 0 | 0 | 0 |
| ERV3-16A3_I-int_dup | 14.277474553291 | -4.46777855148138 | 1.08166864747825 | -4.13045026487303 | 3.62053418669607e-05 | 0.0047841821518855 | 1198 | 12 | 6 | 74 | 79 | 0 | 1 | 10 | 9 | 1 | 0 | 0 | 2 |
| L1PA5_dup3252 | 11.5876180129923 | -5.38512214756541 | 1.274536339324 | -4.22516171678684 | 2.38769250885258e-05 | 0.00352309507275537 | 1199 | 13 | 11 | 52 | 65 | 0 | 1 | 9 | 6 | 1 | 0 | 0 | 0 |
| MIRb_dup194034 | 12.9865578459895 | -5.75928570739647 | 1.71966267200764 | -3.3490787473294 | 0.000810807474330714 | 0.0417251381334367 | 1200 | 13 | 24 | 64 | 61 | 0 | 0 | 9 | 4 | 0 | 0 | 0 | 1 |
| L1M5_dup48057 | 9.32490559441349 | -4.19748025811515 | 1.19339681150737 | -3.51725446024391 | 0.000436035558150841 | 0.0278755821923052 | 1201 | 8 | 11 | 47 | 47 | 0 | 0 | 5 | 5 | 0 | 0 | 2 | 1 |
| MER9a3_dup497 | 9.21490903517206 | -4.40147870340611 | 1.29376988419566 | -3.40205685506625 | 0.000668807225695793 | 0.0367334858133218 | 1202 | 22 | 12 | 36 | 45 | 0 | 0 | 7 | 2 | 0 | 0 | 1 | 2 |
| SVA_D_dup1221 | 15.6905142573759 | -4.02408672880464 | 1.10463932004057 | -3.64289651454453 | 0.000269587241536466 | 0.0201214572693337 | 1203 | 21 | 19 | 76 | 72 | 0 | 1 | 13 | 4 | 0 | 0 | 4 | 4 |
| MSTA_dup17450 | 17.2561352650646 | -5.00102564253158 | 1.16606213783107 | -4.28881573312527 | 1.79628316882554e-05 | 0.0028647345501968 | 1204 | 20 | 27 | 74 | 89 | 1 | 0 | 11 | 9 | 0 | 1 | 1 | 1 |
| Tigger17c_dup672 | 16.8492103661604 | -3.98874320340116 | 1.10662751946495 | -3.60441352961267 | 0.000312858586497592 | 0.02226845624372 | 1205 | 28 | 25 | 75 | 81 | 1 | 0 | 7 | 5 | 1 | 0 | 3 | 3 |
| L1PA16_dup2992 | 10.2636136957239 | -5.14481583479151 | 1.3130340772876 | -3.91826527870428 | 8.91885203147076e-05 | 0.00926727146497966 | 1206 | 2 | 3 | 54 | 76 | 0 | 1 | 2 | 4 | 0 | 0 | 0 | 0 |
| L1MA3_dup2026 | 10.4852069611417 | -4.70972721355117 | 1.3019402445202 | -3.61746803155841 | 0.000297499061527451 | 0.0215097878134946 | 1207 | 3 | 4 | 55 | 79 | 0 | 1 | 1 | 2 | 0 | 0 | 0 | 1 |
| Tigger17a_dup481 | 8.60630277579723 | -4.39268718118366 | 1.28859574532223 | -3.40889468022045 | 0.000652266564067034 | 0.0362288094896112 | 1208 | 1 | 3 | 43 | 67 | 0 | 2 | 1 | 2 | 0 | 0 | 0 | 1 |
| L1MA3_dup2580 | 10.6546919597818 | -4.40276590727656 | 1.22559268415025 | -3.59235655060161 | 0.000327701054217294 | 0.0229879584453974 | 1209 | 5 | 3 | 60 | 70 | 1 | 2 | 5 | 0 | 0 | 0 | 2 | 0 |
| MLT1K_dup14180 | 6.96776770326144 | -5.13550728340001 | 1.4782385721649 | -3.47407203417713 | 0.000512623393775375 | 0.0310180227529039 | 1210 | 4 | 3 | 49 | 36 | 1 | 1 | 1 | 0 | 0 | 0 | 0 | 0 |
| L1MC4_dup24440 | 6.88842409718332 | -4.27065560290654 | 1.12016303753065 | -3.81253037265094 | 0.000137551319880194 | 0.0125690423810341 | 1211 | 15 | 9 | 19 | 33 | 3 | 2 | 8 | 3 | 1 | 0 | 1 | 0 |
| L1PA7_dup3770 | 12.4130513287232 | -3.83760769929252 | 0.981992699329421 | -3.90797986778632 | 9.30710594358036e-05 | 0.00955387675086787 | 1212 | 11 | 7 | 50 | 80 | 2 | 1 | 13 | 4 | 1 | 1 | 1 | 1 |
| LTR1E_dup103 | 9.39326949062 | -4.2102214866247 | 1.04422534575798 | -4.03190892054875 | 5.53256295018042e-05 | 0.00651063908874199 | 1213 | 5 | 3 | 44 | 59 | 2 | 1 | 6 | 6 | 1 | 0 | 1 | 0 |
| AluSp_dup41981 | 6.63463580012965 | -5.60656103052706 | 1.52769219609355 | -3.66995461838683 | 0.000242593533296257 | 0.0187416106732828 | 1214 | 11 | 13 | 33 | 28 | 0 | 0 | 4 | 1 | 0 | 0 | 0 | 0 |
| AluSq_dup15690 | 6.68792261242614 | -5.38905130257416 | 1.39643799290482 | -3.85914113620186 | 0.000113786212962844 | 0.0110195255309959 | 1215 | 7 | 8 | 33 | 33 | 0 | 0 | 7 | 3 | 0 | 0 | 0 | 0 |
| L1MEc_dup5356 | 6.81231908175982 | -5.18635820989327 | 1.30856378896977 | -3.96339731666921 | 7.38906535560132e-05 | 0.00805404738939034 | 1216 | 3 | 6 | 30 | 44 | 0 | 1 | 7 | 3 | 0 | 0 | 0 | 0 |
| L2b_dup27299 | 6.43007795475272 | -5.15399555302808 | 1.49044245299063 | -3.45803056178813 | 0.000544139653744335 | 0.0322824939682598 | 1217 | 2 | 8 | 31 | 42 | 0 | 0 | 5 | 1 | 0 | 0 | 0 | 0 |
| L1M3_dup1656 | 6.40670128451164 | -5.45927182512014 | 1.35724321898781 | -4.02232389062257 | 5.76267390197597e-05 | 0.00672489065365166 | 1218 | 11 | 6 | 31 | 31 | 0 | 2 | 3 | 3 | 0 | 0 | 0 | 0 |

| | | | | | | | | | | | | | | | | | | | |
|---|---|---|---|---|---|---|---|---|---|---|---|---|---|---|---|---|---|---|---|
| LTR12C_dup700 | 6.16270853514593 | -5.19311766663195 | 1.29534938589907 | -4.00904784698494 | 6.0964064350223e-05 | 0.00698418644962839 | 1219 | 6 | 3 | 34 | 31 | 1 | 2 | 2 | 4 | 0 | 0 | 0 | 0 |
| L1MCb_dup935 | 4.56240119322185 | -5.03697599206297 | 1.23532028604466 | -4.07746561678407 | 4.55292600845553e-05 | 0.00561011076983783 | 1220 | 4 | 8 | 14 | 24 | 2 | 1 | 6 | 3 | 0 | 0 | 0 | 0 |
| AluSc_dup19053 | 4.93214730690802 | -4.95149809070678 | 1.34756753178169 | -3.67439699601558 | 0.000238411797812152 | 0.0185449930180485 | 1221 | 6 | 4 | 21 | 30 | 1 | 0 | 4 | 2 | 0 | 0 | 0 | 0 |
| L1PA3_dup9588 | 4.92350106338671 | -5.18061400565394 | 1.28301304054914 | -4.03784984401762 | 5.39433621353115e-05 | 0.0063858742520641 | 1222 | 9 | 7 | 19 | 20 | 2 | 0 | 5 | 4 | 0 | 0 | 0 | 0 |
| HERVH-int_dup5474 | 7.87552780289223 | -4.10551493945223 | 1.08050964499925 | -3.79960971052238 | 0.000144924144544411 | 0.0130030620278147 | 1223 | 2 | 5 | 48 | 33 | 2 | 0 | 13 | 2 | 0 | 0 | 1 | 1 |
| L1MA1_dup3734 | 8.36930337075385 | -3.92478783120074 | 0.959370864313858 | -4.09100169412352 | 4.29513993630965e-05 | 0.00539889787261033 | 1224 | 4 | 6 | 35 | 44 | 2 | 0 | 14 | 6 | 0 | 0 | 2 | 1 |
| MER11A_dup295 | 8.74813032800759 | -5.62844615835162 | 1.32585957240413 | -4.24512993344066 | 2.18466789325441e-05 | 0.00331103813075358 | 1225 | 6 | 7 | 44 | 52 | 2 | 0 | 6 | 3 | 0 | 0 | 0 | 0 |
| MSTA_dup17064 | 9.33954402500058 | -5.26982758381181 | 1.3898438674966 | -3.7916687673011 | 0.000149638407560279 | 0.0133056279990053 | 1226 | 3 | 4 | 50 | 67 | 1 | 0 | 1 | 3 | 0 | 0 | 0 | 0 |
| BLACKJACK_dup591 | 5.56623654572333 | -4.13207130141502 | 1.16327650548801 | -3.55209727173297 | 0.000382173551645003 | 0.0255440078426185 | 1227 | 3 | 1 | 25 | 33 | 3 | 0 | 3 | 6 | 0 | 0 | 0 | 1 |
| L1MA9_dup4094 | 5.76375102221597 | -3.9980694934892 | 1.17817417936857 | -3.39344518280983 | 0.000690193707656837 | 0.03745241321827 | 1228 | 3 | 2 | 23 | 41 | 1 | 0 | 6 | 3 | 0 | 1 | 0 | 0 |
| L1PA15-16_dup68 | 26.0231173934498 | -3.01640412071669 | 0.713923407917578 | -4.22510886639108 | 2.38825294493474e-05 | 0.00352309507275537 | 1229 | 6 | 6 | 96 | 161 | 1 | 0 | 33 | 29 | 0 | 0 | 12 | 10 |
| L1M2_dup780 | 26.4370541929101 | -2.44097989601107 | 0.537569940772122 | -4.54076709070645 | 5.60499233956456e-06 | 0.00119075421008329 | 1230 | 8 | 6 | 78 | 145 | 2 | 2 | 56 | 35 | 2 | 1 | 10 | 13 |
| HERVL-int_dup220 | 36.4075632269072 | -2.82606029911501 | 0.61957490734371 | -4.56128914457029 | 5.08405137572725e-06 | 0.00110910513850895 | 1231 | 4 | 3 | 144 | 226 | 1 | 3 | 60 | 32 | 1 | 1 | 9 | 16 |
| L1M2_dup781 | 36.6990560355874 | -1.89846070521149 | 0.554766757230885 | -3.42208807659573 | 0.000621421799467073 | 0.0351056105166173 | 1232 | 4 | 6 | 147 | 175 | 1 | 2 | 71 | 43 | 2 | 2 | 12 | 30 |
| PRIMA41-int_dup441 | 23.2101957674641 | -1.6087845703525 | 0.482780044654927 | -3.33233444125139 | 0.00086120694167786 | 0.0433519044356457 | 1233 | 1 | 0 | 67 | 59 | 8 | 8 | 80 | 37 | 1 | 0 | 23 | 23 |
| L1PA16_dup13857 | 26.6485002276535 | -1.55678502335667 | 0.375787395810092 | -4.14272815084892 | 3.43198669406005e-05 | 0.00461110957597297 | 1234 | 3 | 3 | 58 | 84 | 6 | 9 | 77 | 55 | 0 | 1 | 18 | 37 |
| Tigger18a_dup816 | 25.1192889568887 | -1.23533018688388 | 0.367910603310485 | -3.35769117760752 | 0.000785963710284567 | 0.0409765383451734 | 1235 | 9 | 9 | 72 | 77 | 6 | 4 | 50 | 40 | 0 | 2 | 33 | 28 |
| Tigger17c_dup119 | 18.8934466352546 | -3.07858462419335 | 0.74989657188678 | -4.10534564312978 | 4.03710783830311e-05 | 0.00515350676873858 | 1236 | 2 | 0 | 80 | 100 | 0 | 2 | 44 | 19 | 0 | 1 | 8 | 1 |
| L1MDa_dup2999 | 17.8183631025881 | -5.1919958509028 | 0.797449973362129 | -6.51074803979593 | 7.47774810617213e-11 | 1.09833348819212e-07 | 1237 | 2 | 2 | 52 | 58 | 3 | 6 | 65 | 42 | 1 | 0 | 1 | 0 |
| L1MDa_dup2998 | 10.1607769999559 | -4.44250837117425 | 0.850414130292008 | -5.2239352721583 | 1.75160030286592e-07 | 8.15628739071888e-05 | 1238 | 3 | 1 | 20 | 39 | 1 | 4 | 40 | 24 | 0 | 0 | 2 | 0 |
| L2a_dup80477 | 11.6587364808589 | -4.2427783928271 | 0.796172527217852 | -5.32896859384622 | 9.87720729998153e-08 | 5.08710382727577e-05 | 1239 | 3 | 1 | 45 | 47 | 3 | 4 | 29 | 19 | 1 | 0 | 2 | 0 |
| HERV-Fc1-int_dup2 | 26.1327434603562 | -3.74526157507979 | 0.625907064768034 | -5.98373430481699 | 2.18078948469077e-09 | 2.10085074787137e-06 | 1240 | 4 | 7 | 88 | 91 | 5 | 4 | 87 | 49 | 2 | 2 | 2 | 3 |
| L1MCa_dup3376 | 23.6089370240019 | -4.53179991156291 | 0.630768178185241 | -7.18457282452197 | 6.74176397391274e-13 | 1.67101642764234e-09 | 1241 | 6 | 2 | 76 | 77 | 9 | 2 | 77 | 53 | 0 | 1 | 1 | 4 |
| Tigger3_dup566 | 12.1356115754835 | -2.76530654446349 | 0.726357915315075 | -3.80708530348151 | 0.000140614253634424 | 0.0127558517709862 | 1242 | 6 | 1 | 22 | 39 | 3 | 1 | 41 | 35 | 0 | 3 | 3 | 2 |
| MER1B_dup521 | 9.42066077924811 | -2.33970710018514 | 0.602326031046411 | -3.88445290355525 | 0.000102560570209754 | 0.0102580401173309 | 1243 | 1 | 3 | 22 | 34 | 2 | 0 | 30 | 21 | 1 | 1 | 4 | 4 |
| L1MC5a_dup1465 | 10.5217499681133 | -2.22097776448241 | 0.626213843670467 | -3.54667624635133 | 0.000390123694738628 | 0.0258485014139323 | 1244 | 1 | 3 | 37 | 33 | 1 | 1 | 24 | 24 | 0 | 2 | 2 | 8 |
| L1MA3_dup1348 | 11.8986445434875 | -2.56635138870402 | 0.561334376862162 | -4.57187639754015 | 4.83376054792043e-06 | 0.00106992929054463 | 1245 | 2 | 1 | 31 | 47 | 1 | 1 | 38 | 25 | 1 | 1 | 3 | 6 |
| L1ME3Cz_dup1167 | 11.0782417008375 | -2.59724303289475 | 0.583707850667813 | -4.44955987815355 | 8.60464383646472e-06 | 0.0016507160990289 | 1246 | 1 | 2 | 37 | 36 | 1 | 3 | 34 | 21 | 1 | 1 | 5 | 3 |
| LTR62_dup116 | 12.8369412220545 | -3.34642454675442 | 0.781025728251704 | -4.28465340603473 | 1.83024286229724e-05 | 0.00290526329627872 | 1247 | 4 | 4 | 38 | 30 | 0 | 1 | 42 | 38 | 1 | 2 | 2 | 0 |
| L1PA5_dup3579 | 10.1857645414155 | -2.63904484971061 | 0.783472053569422 | -3.36839691688731 | 0.000756066639493818 | 0.0398368096990691 | 1248 | 2 | 3 | 22 | 30 | 2 | 2 | 29 | 32 | 2 | 2 | 0 | 2 |
| L1PB1_dup9598 | 10.6187282725164 | -3.07854652304709 | 0.766418767652445 | -4.01679428137797 | 5.89951808816879e-05 | 0.00684096847469498 | 1249 | 1 | 2 | 25 | 48 | 0 | 2 | 39 | 20 | 1 | 2 | 0 | 2 |
| MER41-int_dup811 | 17.4320725262858 | -3.76190010336094 | 0.761481776141349 | -4.94023655093047 | 7.80278519772313e-07 | 0.000266376115732773 | 1250 | 8 | 14 | 53 | 55 | 1 | 2 | 37 | 45 | 2 | 0 | 1 | 4 |
| L1PA13_dup1660 | 11.4196086680074 | -2.92965623712007 | 0.716877391747302 | -4.08669079377631 | 4.37569663434386e-05 | 0.00546074960142377 | 1251 | 5 | 6 | 21 | 24 | 1 | 1 | 45 | 34 | 2 | 0 | 2 | 4 |
| L1PA2_dup526 | 12.0271537838699 | -4.02944510828681 | 0.729465704649893 | -5.52383077449918 | 3.31686948119073e-08 | 2.11025621810537e-05 | 1252 | 3 | 4 | 29 | 36 | 2 | 3 | 54 | 24 | 1 | 0 | 2 | 1 |
| MLT1C_dup19335 | 18.4541212967639 | -3.73274537934218 | 0.627066622006192 | -5.95270940653787 | 2.63739347870501e-09 | 2.47069775072418e-06 | 1253 | 5 | 3 | 81 | 67 | 2 | 7 | 42 | 28 | 0 | 0 | 4 | 5 |
| L1MB3_dup6407 | 13.00676666318 | -3.67869795562983 | 0.739839621637339 | -4.9722910858552 | 6.61662571391978e-07 | 0.00023289935238526 | 1254 | 6 | 13 | 33 | 62 | 3 | 4 | 35 | 14 | 0 | 0 | 2 | 5 |
| L1ME3A_dup6486 | 11.8749282516558 | -3.94504176558601 | 0.725956326634285 | -5.43426873056704 | 5.5021607138177e-08 | 3.13121932406234e-05 | 1255 | 8 | 8 | 35 | 41 | 5 | 4 | 40 | 14 | 0 | 0 | 2 | 3 |
| L1MA7_dup3347 | 12.7344913752978 | -3.96011087871868 | 0.714579594934092 | -5.54187512041109 | 2.99249695951675e-08 | 1.91412011163756e-05 | 1256 | 2 | 8 | 40 | 51 | 3 | 3 | 36 | 21 | 0 | 0 | 3 | 2 |
| MLT1C_dup9855 | 9.78257887131069 | -4.09903735640664 | 0.792036183781843 | -5.17531577513846 | 2.27525835873928e-07 | 9.98872002635749e-05 | 1257 | 2 | 3 | 29 | 45 | 2 | 3 | 25 | 18 | 0 | 0 | 1 | 2 |
| L1MB5_dup9402 | 14.2630756432611 | -3.29705970209689 | 0.766545654741526 | -4.3011915620455 | 1.69882116727292e-05 | 0.00274424780358399 | 1258 | 4 | 2 | 70 | 64 | 0 | 0 | 28 | 16 | 1 | 0 | 4 | 2 |
| MLT1F1-int_dup174 | 17.932979102599 | -2.98752153138862 | 0.748394154906742 | -3.9919092256418 | 6.55434633259336e-05 | 0.00739614215118511 | 1259 | 3 | 1 | 88 | 90 | 0 | 0 | 30 | 19 | 1 | 0 | 2 | 8 |
| AluJr_dup40173 | 15.8072973026736 | -3.79071158919786 | 0.803127965828689 | -4.71993474325913 | 2.3592031600399e-06 | 0.000640825291231568 | 1260 | 0 | 2 | 75 | 76 | 0 | 0 | 35 | 20 | 1 | 0 | 1 | 2 |
| L1PA8_dup1259 | 12.6075172956437 | -3.18982442246473 | 0.645883244502161 | -4.93870130494469 | 7.86445661502038e-07 | 0.000267058535441124 | 1261 | 1 | 1 | 45 | 54 | 1 | 0 | 32 | 25 | 1 | 0 | 2 | 4 |
| HERVL-int_dup1870 | 11.4047286086596 | -2.74809396959335 | 0.581717402325754 | -4.72410479488193 | 2.31131215493043e-06 | 0.000630696653689307 | 1262 | 2 | 2 | 32 | 34 | 0 | 1 | 36 | 30 | 0 | 0 | 5 | 5 |
| MLT2A2_dup2221 | 14.8338748162907 | -3.47280777815588 | 0.636154713168369 | -5.45906161861091 | 4.78657632559574e-08 | 2.79153600580932e-05 | 1263 | 1 | 2 | 46 | 45 | 0 | 0 | 41 | 46 | 0 | 0 | 2 | 5 |
| L1MC3_dup3059 | 10.1611917954216 | -3.74764245442921 | 0.77478325159301 | -4.83702047859681 | 1.31799853799937e-06 | 0.000397985380866496 | 1264 | 1 | 1 | 22 | 45 | 0 | 0 | 45 | 20 | 0 | 0 | 1 | 2 |
| THE1D-int_dup632 | 11.7734523593608 | -3.90969651529962 | 0.82504936479543 | -4.73874253120481 | 2.15048534682277e-06 | 0.000594308230354346 | 1265 | 5 | 2 | 22 | 61 | 0 | 0 | 53 | 16 | 0 | 0 | 1 | 3 |
| ERVL-E-int_dup2276 | 8.97408366432143 | -4.22828855478562 | 0.867296465647298 | -4.87525168412842 | 1.08669814784789e-06 | 0.000342439823200773 | 1266 | 4 | 3 | 30 | 35 | 0 | 0 | 31 | 15 | 0 | 0 | 1 | 1 |
| L2a_dup46290 | 9.36033877291793 | -3.6777927388385 | 0.790382319576319 | -4.65318194466946 | 3.26851599677068e-06 | 0.000806774177767216 | 1267 | 3 | 4 | 24 | 46 | 1 | 0 | 34 | 12 | 0 | 0 | 2 | 2 |
| L1ME2z_dup596 | 14.1308489873048 | -1.96736700393091 | 0.590411737942416 | -3.3321949370234 | 0.000861638785873185 | 0.0433545152343795 | 1268 | 3 | 0 | 54 | 62 | 1 | 0 | 35 | 16 | 1 | 0 | 8 | 11 |
| L1MA7_dup688 | 11.9259405292238 | -2.63115045859777 | 0.6554309194075 | -4.01438256983099 | 5.96016078764707e-05 | 0.0068877922233 | 1269 | 1 | 2 | 38 | 64 | 1 | 0 | 30 | 15 | 0 | 0 | 7 | 4 |
| L1MC3_dup1030 | 12.0305085579011 | -1.93293297320835 | 0.562987290313345 | -3.43335099471344 | 0.000596169786333916 | 0.034268915854522 | 1270 | 2 | 3 | 38 | 51 | 0 | 0 | 30 | 18 | 0 | 0 | 10 | 9 |
| L1MC3_dup3058 | 12.9000604045492 | -3.4013275176533 | 0.698181562921674 | -4.87169484026446 | 1.10644973747986e-06 | 0.000347328877617918 | 1271 | 4 | 7 | 39 | 47 | 0 | 0 | 38 | 27 | 1 | 0 | 2 | 4 |
| L2b_dup26240 | 13.0362143022265 | -3.34494480595163 | 0.681449791183798 | -4.908571180484 | 9.17423315475371e-07 | 0.000299858802505635 | 1272 | 3 | 5 | 38 | 46 | 0 | 0 | 30 | 37 | 1 | 0 | 3 | 3 |
| L1MEi_dup519 | 11.9611319296872 | -2.2942738284518 | 0.618025020901551 | -3.71226689997924 | 0.000205411210024812 | 0.0167386218426589 | 1273 | 5 | 9 | 31 | 40 | 1 | 0 | 36 | 21 | 1 | 0 | 6 | 8 |
| L1MC4_dup2389 | 14.2555671187349 | -3.06530344409763 | 0.765210762503749 | -4.00582897457955 | 6.18003544081856e-05 | 0.00706638497357526 | 1274 | 6 | 8 | 38 | 71 | 0 | 0 | 37 | 21 | 0 | 0 | 3 | 9 |
| L1PA4_dup1024 | 13.6717467115195 | -2.26013631276269 | 0.584187508063588 | -3.86885423184482 | 0.000109347987275905 | 0.0107161928255159 | 1275 | 5 | 3 | 50 | 49 | 0 | 0 | 37 | 21 | 1 | 0 | 5 | 11 |
| HAL1_dup6239 | 16.9861700715923 | -3.36682034113299 | 0.603072303992441 | -5.58278057016393 | 2.36703315287527e-08 | 1.6008235420306e-05 | 1276 | 8 | 6 | 54 | 55 | 0 | 1 | 52 | 36 | 1 | 0 | 4 | 5 |
| L1MA7_dup685 | 19.0205067580436 | -2.62350017762146 | 0.536847274204289 | -4.88686504278148 | 1.02454272076151e-06 | 0.00032580489270755 | 1277 | 6 | 7 | 54 | 84 | 0 | 2 | 45 | 35 | 1 | 0 | 9 | 9 |
| L1PA16_dup1121 | 10.9868407231347 | -2.01023392999571 | 0.601472592659904 | -3.34218708304864 | 0.000831210167346164 | 0.0423609197856822 | 1278 | 4 | 4 | 30 | 45 | 0 | 1 | 19 | 24 | 1 | 0 | 7 | 8 |
| L1ME3Cz_dup1168 | 13.8027486811892 | -3.19981094077495 | 0.701612492027256 | -4.56065274939638 | 5.09948511513977e-06 | 0.00111117407070243 | 1279 | 11 | 3 | 34 | 55 | 0 | 1 | 38 | 30 | 0 | 0 | 5 | 5 |
| L1P4_dup284 | 14.5077916756004 | -2.61868342042199 | 0.580126479388513 | -4.51398705879145 | 6.36201006982029e-06 | 0.00131293568783474 | 1280 | 6 | 2 | 44 | 64 | 0 | 1 | 35 | 26 | 0 | 0 | 8 | 7 |
| L1MB3_dup16146 | 12.9836733783324 | -2.67542224509977 | 0.787030108284347 | -3.39938995590899 | 0.000675363507539133 | 0.0369647210700528 | 1281 | 6 | 6 | 66 | 61 | 2 | 5 | 11 | 7 | 2 | 0 | 3 | 6 |
| L1PA16_dup3769 | 18.5329412687817 | -2.93022661442491 | 0.849027002449407 | -3.4512761148601 | 0.00055794244670856 | 0.0327964855297715 | 1282 | 8 | 6 | 82 | 122 | 1 | 3 | 12 | 9 | 2 | 0 | 4 | 6 |
| L1PREC2_dup6610 | 13.4226364570491 | -3.40851946565546 | 0.89410771452795 | -3.81220227750189 | 0.000137734084509003 | 0.0125809115057885 | 1283 | 11 | 8 | 65 | 61 | 5 | 1 | 13 | 8 | 0 | 0 | 1 | 9 |
| L1MB5_dup9401 | 11.7858258203644 | -3.15503680653593 | 0.807954060772321 | -3.90497054191428 | 9.423686361934e-05 | 0.00962785948938063 | 1284 | 7 | 3 | 53 | 62 | 2 | 1 | 13 | 10 | 0 | 0 | 3 | 6 |
| L1MC4_dup6651 | 11.0265924584336 | -3.10598623878215 | 0.902718692563875 | -3.4407022523934 | 0.000580206624533311 | 0.0336973018016117 | 1285 | 8 | 5 | 45 | 66 | 1 | 1 | 6 | 9 | 0 | 0 | 5 | 4 |
| L1MA10_dup432 | 18.4734376249467 | -2.00884385469857 | 0.600274489760966 | -3.34654210526005 | 0.00081826251590101 | 0.0419526594717909 | 1286 | 9 | 10 | 57 | 81 | 0 | 2 | 42 | 21 | 1 | 1 | 8 | 18 |
| MER4B-int_dup92 | 14.9750310781189 | -2.07150404874267 | 0.613534826545882 | -3.37634305195836 | 0.000734562677093845 | 0.0390147881169134 | 1287 | 4 | 7 | 51 | 61 | 0 | 1 | 43 | 16 | 1 | 1 | 5 | 13 |
| LTR12D_dup37 | 15.0026524297838 | -2.49576618747718 | 0.541707596631645 | -4.60722021067443 | 4.08087759209288e-06 | 0.000944580252459273 | 1288 | 9 | 10 | 43 | 46 | 3 | 1 | 44 | 26 | 0 | 2 | 5 | 9 |
| L1ME2_dup6728 | 17.4883708443897 | -1.83900018544101 | 0.488280656437364 | -3.76627695813077 | 0.000165699975987164 | 0.0143217023197391 | 1289 | 5 | 9 | 53 | 68 | 4 | 2 | 44 | 22 | 0 | 3 | 8 | 17 |
| THE1A-int_dup137 | 19.0257245933881 | -2.24118186829448 | 0.456691137381367 | -4.90743455444581 | 9.22754314383962e-07 | 0.000300774928890967 | 1290 | 5 | 7 | 50 | 76 | 3 | 3 | 43 | 39 | 0 | 2 | 8 | 14 |

| | | | | | | | | | | | | | | | | | | | |
|---|---|---|---|---|---|---|---|---|---|---|---|---|---|---|---|---|---|---|---|
| L1MA7_dup4440 | 15.3943309798501 | -1.96957324801275 | 0.515067984706297 | -3.82390928284126 | 0.000131352260829905 | 0.0121898381651453 | 1291 | 6 | 2 | 33 | 68 | 5 | 2 | 39 | 29 | 1 | 3 | 8 | 8 |
| L1MB3_dup8238 | 15.7826239655389 | -2.29736997592025 | 0.500947501045871 | -4.58604937867508 | 4.51711811503306e-06 | 0.00101208115536691 | 1292 | 8 | 6 | 32 | 62 | 2 | 2 | 45 | 33 | 1 | 2 | 6 | 9 |
| L1PA6_dup3339 | 12.669306746751 | -2.0426176963928 | 0.577280431961101 | -3.53834563464026 | 0.000402642630124048 | 0.0264073769498129 | 1293 | 4 | 7 | 28 | 50 | 2 | 1 | 30 | 28 | 1 | 3 | 5 | 6 |
| L1PA2_dup3773 | 25.4332080127126 | -3.17732786311898 | 0.589207206420013 | -5.39254752572397 | 6.94657182832656e-08 | 3.73961307751808e-05 | 1294 | 2 | 3 | 42 | 55 | 0 | 1 | 119 | 85 | 0 | 0 | 7 | 10 |
| L1MA9_dup7820 | 35.5578309829605 | -3.74876873163781 | 0.88901632390851 | -4.21676028979599 | 2.47837182140939e-05 | 0.00363350992863265 | 1295 | 1 | 1 | 13 | 30 | 1 | 2 | 221 | 160 | 1 | 0 | 2 | 7 |
| L1P3_dup353 | 22.0539164067497 | -3.80954280837546 | 0.77133095498093 | -4.93892120337586 | 7.85559449046582e-07 | 0.000267058535441124 | 1296 | 1 | 0 | 21 | 41 | 1 | 3 | 101 | 96 | 0 | 0 | 3 | 5 |
| L1PB1_dup9010 | 20.1553849409051 | -4.26204720091434 | 1.00733810119657 | -4.23099969697529 | 2.32654985477404e-05 | 0.0034599576984004 | 1297 | 2 | 0 | 16 | 14 | 6 | 3 | 112 | 90 | 1 | 1 | 1 | 0 |
| L1M4_dup14436 | 13.2988645223534 | -4.73860677261426 | 1.10217530514552 | -4.29932221352811 | 1.71321244043559e-05 | 0.00276374269391109 | 1298 | 1 | 5 | 4 | 5 | 3 | 3 | 78 | 61 | 0 | 0 | 1 | 1 |
| L1PB1_dup9688 | 10.6887359214425 | -5.24597657277458 | 1.1243843417421 | -4.66564356868095 | 3.07652596939312e-06 | 0.000770575840329701 | 1299 | 4 | 7 | 6 | 11 | 4 | 6 | 51 | 42 | 0 | 1 | 0 | 0 |
| L1PB1_dup9690 | 16.7049259179689 | -6.17165618065684 | 1.19444383865162 | -5.16697058576138 | 2.37918687760693e-07 | 0.0001036846155273 | 1300 | 4 | 8 | 10 | 12 | 2 | 2 | 108 | 64 | 0 | 0 | 0 | 0 |
| L1PA5_dup8564 | 7.31523166253564 | -2.72356832753733 | 0.751662877129403 | -3.62339076520398 | 0.000290766031864207 | 0.0211644583107863 | 1301 | 1 | 2 | 12 | 21 | 1 | 0 | 35 | 18 | 0 | 0 | 4 | 2 |
| MER90a_dup520 | 5.86708729736552 | -3.47647812951149 | 1.0394630181272 | -3.34449429069162 | 0.000824327263109274 | 0.0421369844884236 | 1302 | 1 | 2 | 5 | 7 | 1 | 0 | 40 | 18 | 0 | 0 | 1 | 1 |
| L1PA3_dup8337 | 11.1819360652802 | -3.1946933762116 | 0.954590957548695 | -3.34666209746559 | 0.000817908437660595 | 0.041947056743146 | 1303 | 0 | 0 | 11 | 11 | 0 | 1 | 74 | 41 | 0 | 0 | 2 | 2 |
| L1PB1_dup10190 | 9.09974420012603 | -3.40845972387108 | 0.893829525676797 | -3.81332192096724 | 0.000137111329747619 | 0.012533650583222 | 1304 | 1 | 0 | 9 | 16 | 4 | 2 | 50 | 30 | 1 | 0 | 1 | 2 |
| MLT1H2_dup2076 | 9.21695775619112 | -4.2530672870642 | 0.947980617407894 | -4.48644962667439 | 7.24198294082987e-06 | 0.00145172777829714 | 1305 | 1 | 1 | 13 | 15 | 1 | 5 | 45 | 33 | 0 | 0 | 1 | 1 |
| L2a_dup78432 | 15.2655758769994 | -3.12178064980986 | 0.845798707491959 | -3.69092624776746 | 0.000223438922372337 | 0.0177103257237869 | 1306 | 9 | 6 | 15 | 17 | 2 | 0 | 69 | 59 | 0 | 0 | 4 | 9 |
| L2a_dup78444 | 14.2761153836258 | -3.54357478676266 | 1.04393086340026 | -3.39445351315761 | 0.000687657134230394 | 0.0373915138166328 | 1307 | 8 | 4 | 7 | 4 | 0 | 1 | 88 | 59 | 1 | 0 | 2 | 3 |
| L1MEf_dup7433 | 10.7094571525802 | -4.86583005800009 | 1.08870443695498 | -4.46937652941826 | 7.84479277457239e-06 | 0.00153003037831017 | 1308 | 3 | 4 | 6 | 16 | 0 | 2 | 56 | 45 | 0 | 0 | 0 | 1 |
| L2_dup26571 | 9.96956529366746 | -3.79257002477316 | 0.993572100809867 | -3.817105997322 | 0.000135026170841125 | 0.0123906429799315 | 1309 | 3 | 5 | 5 | 11 | 0 | 2 | 59 | 37 | 0 | 0 | 1 | 3 |
| L1MA5_dup3295 | 14.0211618360763 | -2.24868817451725 | 0.558599292281317 | -4.02558364392769 | 5.68341732517715e-05 | 0.00665852495744265 | 1310 | 3 | 5 | 43 | 63 | 6 | 5 | 25 | 18 | 0 | 0 | 8 | 12 |
| L1MA8_dup10812 | 15.5647497811356 | -1.65307411507388 | 0.497966443645338 | -3.31964961930493 | 0.000901304925300557 | 0.0445821616253597 | 1311 | 4 | 1 | 42 | 52 | 8 | 9 | 39 | 20 | 0 | 0 | 16 | 17 |
| L1PA16_dup13858 | 16.3596196633182 | -1.92783316845684 | 0.462704853483596 | -4.16644250420681 | 3.09389878022678e-05 | 0.00427515005319304 | 1312 | 4 | 2 | 48 | 54 | 7 | 5 | 48 | 23 | 0 | 0 | 10 | 18 |
| L2a_dup168107 | 15.3785440266792 | -1.79904367976094 | 0.496656065545849 | -3.62231291343175 | 0.000291980622079944 | 0.0212009829421527 | 1313 | 1 | 3 | 31 | 47 | 5 | 10 | 49 | 29 | 1 | 0 | 15 | 11 |
| MSTA-int_dup3086 | 17.2364650279455 | -1.79919764952958 | 0.497944078153091 | -3.61325242827051 | 0.000302380064756244 | 0.0217645622649209 | 1314 | 3 | 2 | 40 | 67 | 10 | 3 | 35 | 35 | 1 | 0 | 11 | 19 |
| L2b_dup60232 | 11.1957859967299 | -1.98278430454632 | 0.556353143027338 | -3.56389521546909 | 0.000365391865744111 | 0.0247843594316842 | 1315 | 6 | 2 | 27 | 41 | 2 | 2 | 38 | 15 | 0 | 0 | 11 | 7 |
| L1ME4c_dup4308 | 12.1782497875272 | -2.46495729214027 | 0.618600985613316 | -3.98472900862965 | 6.75571775003769e-05 | 0.00755047447510789 | 1316 | 5 | 1 | 36 | 55 | 4 | 4 | 21 | 21 | 0 | 0 | 4 | 11 |
| MER41-int_dup513 | 10.075268275498 | -2.18831569818124 | 0.596478233251553 | -3.66872683057046 | 0.000243761364153466 | 0.0187891291075027 | 1317 | 2 | 1 | 39 | 32 | 4 | 2 | 24 | 15 | 0 | 0 | 7 | 7 |
| HAL1M8_dup1424 | 11.1028144879233 | -2.49801480874284 | 0.638667813697159 | -3.91128964881162 | 9.18045932302225e-05 | 0.00946060447845756 | 1318 | 0 | 3 | 31 | 50 | 3 | 6 | 30 | 14 | 0 | 0 | 6 | 7 |
| HERVL18-int_dup496 | 13.2733612891477 | -2.35620109043975 | 0.604476042595364 | -3.89792303483728 | 9.70212395991922e-05 | 0.00984050122662804 | 1319 | 1 | 0 | 37 | 62 | 3 | 6 | 31 | 21 | 0 | 0 | 7 | 10 |
| MSTA_dup19401 | 13.7163155657443 | -2.08542931291573 | 0.593338879266895 | -3.51473565240221 | 0.000440191927841609 | 0.0280433489858633 | 1320 | 1 | 0 | 28 | 40 | 3 | 7 | 59 | 24 | 0 | 0 | 12 | 9 |
| Tigger3a_dup2811 | 13.9350274788782 | -2.48216523328363 | 0.713403406000883 | -3.47932910384866 | 0.000502670861235525 | 0.0306295827778613 | 1321 | 2 | 0 | 40 | 68 | 4 | 11 | 20 | 24 | 0 | 1 | 5 | 10 |
| L1PA5_dup10951 | 10.2848152161973 | -2.23326069012995 | 0.6520275991161 | -3.4251014729398 | 0.000614569836416498 | 0.0348903560487367 | 1322 | 5 | 0 | 30 | 37 | 6 | 11 | 18 | 15 | 0 | 1 | 6 | 7 |
| L2a_dup168108 | 9.67959829906943 | -2.30292501968831 | 0.657330347324673 | -3.50345154314142 | 0.000459270207299256 | 0.0288341711730947 | 1323 | 2 | 0 | 26 | 37 | 7 | 6 | 28 | 12 | 0 | 1 | 6 | 5 |
| L2a_dup19314 | 11.0568630437885 | -2.44712232783428 | 0.688066308490216 | -3.55652107600482 | 0.000375798344516039 | 0.0252651332902664 | 1324 | 5 | 1 | 42 | 47 | 5 | 4 | 24 | 10 | 0 | 2 | 7 | 2 |
| L1MEf_dup6278 | 10.8295994308164 | -2.48800465068554 | 0.583566317043226 | -4.26344800586091 | 2.01296356412781e-05 | 0.00310017235100295 | 1325 | 3 | 2 | 36 | 40 | 3 | 3 | 33 | 14 | 1 | 1 | 4 | 5 |
| MER31A_dup730 | 9.43407362790743 | -1.93049161223656 | 0.548775107708911 | -3.51781920338223 | 0.000435108698269594 | 0.0278238039017621 | 1326 | 2 | 3 | 28 | 31 | 5 | 3 | 26 | 13 | 1 | 1 | 6 | 6 |
| AluSc_dup16512 | 10.4285725109926 | -1.99517180330082 | 0.602608866647909 | -3.3108902203833 | 0.00092999685340587 | 0.0455327224857023 | 1327 | 1 | 2 | 30 | 41 | 8 | 5 | 18 | 17 | 1 | 1 | 5 | 8 |
| L1MB1_dup6018 | 10.0420760505937 | -2.09819179036944 | 0.621402824242948 | -3.37654047986932 | 0.000734035699540779 | 0.0390042113807348 | 1328 | 1 | 3 | 28 | 41 | 6 | 3 | 16 | 19 | 0 | 1 | 8 | 5 |
| L4_C_Mam_dup2665 | 7.70619414122304 | -3.03486994180251 | 0.887285139927415 | -3.42039983003748 | 0.000625291590675941 | 0.0352592052839397 | 1329 | 1 | 0 | 21 | 46 | 4 | 6 | 10 | 10 | 0 | 0 | 4 | 2 |
| AluSq_dup8355 | 7.6967918416805 | -2.93303314684211 | 0.840347322070523 | -3.49026297794993 | 0.00048254544716736 | 0.0298233140186194 | 1330 | 0 | 0 | 30 | 38 | 4 | 3 | 15 | 8 | 0 | 0 | 3 | 3 |
| MER44B_dup2130 | 7.7288722937473 | -2.60903899942202 | 0.712999449087696 | -3.65924406079186 | 0.000252960340857007 | 0.0192734233959531 | 1331 | 0 | 0 | 28 | 28 | 2 | 2 | 19 | 15 | 0 | 0 | 5 | 2 |
| MLT2B1_dup3987 | 8.68454508709376 | -3.07035664492609 | 0.904177752399457 | -3.39574451680341 | 0.000684422111570655 | 0.0372908280378856 | 1332 | 0 | 0 | 17 | 24 | 10 | 3 | 30 | 22 | 0 | 1 | 1 | 4 |
| THE1D_dup11443 | 7.26192723693284 | -2.54496849920664 | 0.758685929916465 | -3.35444272636881 | 0.000795250139279914 | 0.0412888041983632 | 1333 | 0 | 1 | 17 | 20 | 6 | 2 | 19 | 20 | 0 | 1 | 4 | 2 |
| L2a_dup63141 | 7.76715895547772 | -2.70659046347001 | 0.777629965643655 | -3.48056348526859 | 0.000500360228276518 | 0.0305122283130406 | 1334 | 2 | 1 | 18 | 17 | 8 | 9 | 27 | 12 | 0 | 0 | 6 | 2 |
| MLT2A2_dup1776 | 7.9936112210745 | -2.92395759222676 | 0.836849935434371 | -3.49400468162678 | 0.000475832674025048 | 0.0295080743949868 | 1335 | 1 | 0 | 23 | 10 | 7 | 4 | 36 | 16 | 0 | 0 | 3 | 4 |
| THE1D-int_dup2261 | 8.21823251794345 | -4.37752432532686 | 0.954724157219721 | -4.58511947374912 | 4.53726913153248e-06 | 0.00101563973732044 | 1336 | 2 | 0 | 21 | 26 | 9 | 4 | 31 | 14 | 0 | 0 | 1 | 1 |
| MER11D_dup37 | 16.2244967408518 | -4.42619419352002 | 0.883882141339554 | -5.00767465084425 | 5.50915548796584e-07 | 0.000200134581944965 | 1337 | 8 | 5 | 44 | 65 | 17 | 20 | 50 | 9 | 0 | 3 | 0 | 1 |
| L1MD1_dup1200 | 13.4568385617159 | -4.27909225531961 | 0.932632866745482 | -4.58818513468429 | 4.47116041826649e-06 | 0.00100747607659549 | 1338 | 2 | 4 | 34 | 46 | 30 | 16 | 28 | 14 | 0 | 2 | 1 | 1 |
| L1PA13_dup1288 | 11.2593993588105 | -4.46406385823413 | 1.01029553846149 | -4.41857227740719 | 9.93550543798704e-06 | 0.00184264518857932 | 1339 | 2 | 2 | 27 | 41 | 21 | 17 | 23 | 13 | 0 | 1 | 2 | 0 |
| MER74A_dup306 | 11.8190486192995 | -5.0875763264609 | 1.10507979480213 | -4.60380901939473 | 4.14832904674928e-06 | 0.000958328449861946 | 1340 | 2 | 1 | 26 | 46 | 23 | 15 | 33 | 11 | 0 | 0 | 2 | 0 |
| L2b_dup9247 | 11.5238445894605 | -4.0914751303083 | 0.816572107771002 | -5.01054970084247 | 5.42747784918145e-07 | 0.000198378900814336 | 1341 | 8 | 3 | 20 | 31 | 8 | 8 | 59 | 15 | 0 | 1 | 3 | 0 |
| MER52D_dup54 | 14.1062553392678 | -4.0607182294709 | 0.782984092087541 | -5.18620783041003 | 2.14619295922744e-07 | 9.58120131100062e-05 | 1342 | 1 | 5 | 31 | 42 | 10 | 12 | 49 | 29 | 0 | 1 | 4 | 0 |
| L1MA5_dup1879 | 8.84077367684709 | -3.56146123788381 | 0.821637044263779 | -4.33459185262883 | 1.46030805705319e-05 | 0.00247136885450625 | 1343 | 13 | 8 | 11 | 28 | 4 | 2 | 23 | 21 | 0 | 0 | 2 | 3 |
| MIR_dup72105 | 7.82452785330315 | -3.99498265677929 | 0.892805674721202 | -4.47463851305248 | 7.65406949845767e-06 | 0.001506414574756 | 1344 | 7 | 9 | 14 | 33 | 2 | 2 | 20 | 14 | 0 | 0 | 1 | 2 |
| HERV16-int_dup41 | 9.33529161449393 | -3.43185776723508 | 0.827474584019919 | -4.1473875252614 | 3.36290583409036e-05 | 0.00453622444216817 | 1345 | 10 | 14 | 17 | 16 | 1 | 4 | 33 | 20 | 0 | 0 | 2 | 4 |
| L2a_dup14886 | 8.14961591659992 | -3.401156394411 | 0.812488635739843 | -4.18609718930277 | 2.83791871951105e-05 | 0.00400754544589185 | 1346 | 8 | 4 | 10 | 21 | 1 | 5 | 37 | 17 | 0 | 0 | 3 | 2 |
| L2_dup24554 | 11.1971556912261 | -3.44838699019032 | 0.739539852848532 | -4.66288189461048 | 3.11811720361596e-06 | 0.000778534644419312 | 1347 | 14 | 16 | 18 | 42 | 2 | 8 | 25 | 17 | 0 | 0 | 3 | 4 |
| L1M5_dup5184 | 11.987495854113 | -2.86428397512174 | 0.62046004643827 | -4.61638745566948 | 3.90477624174527e-06 | 0.000916222834214355 | 1348 | 9 | 5 | 30 | 46 | 1 | 6 | 24 | 25 | 0 | 0 | 5 | 6 |
| L1MC3_dup5226 | 8.94014275596513 | -3.31208124057392 | 0.764101697332663 | -4.33460788287185 | 1.46020165411063e-05 | 0.00247136885450625 | 1349 | 12 | 4 | 27 | 29 | 1 | 5 | 22 | 13 | 0 | 0 | 3 | 3 |
| L1MC1_dup5770 | 10.2404285321421 | -3.3023902693424 | 0.728971729083142 | -4.53020348744657 | 5.89269031467178e-06 | 0.0012397365955923 | 1350 | 9 | 11 | 25 | 33 | 2 | 3 | 23 | 21 | 0 | 2 | 2 | 2 |
| LTR33C_dup402 | 9.56278018416067 | -3.87527401050805 | 0.829586580341598 | -4.67133160340218 | 2.99253377553649e-06 | 0.000755903865980687 | 1351 | 8 | 14 | 15 | 29 | 5 | 3 | 31 | 17 | 0 | 1 | 1 | 2 |
| L2c_dup14109 | 9.62545000060403 | -3.82138192885205 | 0.763659690346588 | -5.00403776336251 | 5.61417515133792e-07 | 0.000203189465324983 | 1352 | 6 | 4 | 22 | 24 | 3 | 5 | 29 | 26 | 0 | 1 | 1 | 2 |
| THE1D_dup5549 | 11.6133249370281 | -3.38319149207067 | 0.671113588484306 | -5.04116076640846 | 4.62716526581887e-07 | 0.000174487395162567 | 1353 | 9 | 5 | 20 | 39 | 6 | 3 | 39 | 25 | 0 | 1 | 1 | 5 |
| LTR10B_dup13 | 8.48445840310541 | -2.11528152595354 | 0.573572761765262 | -3.68790442461637 | 0.000226108542812316 | 0.0178683571331848 | 1354 | 5 | 7 | 23 | 27 | 4 | 5 | 20 | 10 | 0 | 1 | 5 | 6 |
| MLT1C_dup16505 | 9.01632581230745 | -1.89235574822671 | 0.563053040306417 | -3.36088363397679 | 0.000776935510404809 | 0.0406662333829262 | 1355 | 6 | 5 | 24 | 33 | 2 | 5 | 20 | 11 | 0 | 1 | 5 | 9 |
| MER2_dup4954 | 8.1800767121198 | -2.30338878623621 | 0.634685188336417 | -3.62918314239167 | 0.000284319481559768 | 0.0208226172235213 | 1356 | 5 | 5 | 16 | 26 | 4 | 3 | 17 | 20 | 0 | 3 | 2 | 4 |
| MER5A1_dup6553 | 8.07857205727963 | -3.02737583145017 | 0.731954691563071 | -4.13601533857961 | 3.53388579981702e-05 | 0.00471341922939048 | 1357 | 5 | 5 | 18 | 31 | 2 | 7 | 14 | 17 | 0 | 1 | 3 | 2 |
| MIRb_dup190728 | 6.90368625667816 | -2.38359921897118 | 0.681198030868869 | -3.49912816972018 | 0.000466782144242327 | 0.0291431426506715 | 1358 | 4 | 4 | 14 | 27 | 4 | 4 | 11 | 14 | 0 | 1 | 3 | 4 |
| ALR/Alpha_dup1618 | 10.6760929984433 | -2.43595570072695 | 0.660577178092382 | -3.68761710442603 | 0.000226363927626063 | 0.0178752355164257 | 1359 | 5 | 8 | 8 | 23 | 9 | 9 | 25 | 33 | 0 | 0 | 5 | 9 |
| MLT1B_dup8113 | 8.79204616948922 | -2.94219240699993 | 0.78436121395387 | -3.75106820003081 | 0.000176082791830986 | 0.0150226590121964 | 1360 | 10 | 11 | 4 | 18 | 4 | 7 | 40 | 15 | 0 | 0 | 4 | 4 |
| L2_dup40333 | 12.2560379774071 | -2.46720737411474 | 0.59927936331763 | -4.11695700725652 | 3.83907653513933e-05 | 0.00497817297128209 | 1361 | 9 | 6 | 24 | 16 | 5 | 5 | 36 | 38 | 1 | 0 | 5 | 8 |
| L1MA10_dup2099 | 10.8377309085577 | -3.67406593724181 | 0.8641051394573 | -4.25187372401153 | 2.1198938670926e-05 | 0.00323346324422574 | 1362 | 13 | 16 | 7 | 9 | 10 | 5 | 41 | 30 | 0 | 0 | 2 | 4 |

| | | | | | | | | | | | | | | | | | | | |
|---|---|---|---|---|---|---|---|---|---|---|---|---|---|---|---|---|---|---|---|
| CR1_Mam_dup784 | 12.0941546712895 | -4.35611899418832 | 0.940065655522215 | -4.63384548579051 | 3.58934942810395e-06 | 0.000856835251145652 | 1363 | 26 | 13 | 12 | 12 | 4 | 7 | 40 | 35 | 0 | 0 | 2 | 2 |
| L1ME5_dup1462 | 12.1061457685157 | -4.00572136421135 | 0.831320130941773 | -4.81850639136263 | 1.44636893321895e-06 | 0.000429660052663816 | 1364 | 13 | 10 | 12 | 17 | 8 | 6 | 46 | 36 | 0 | 0 | 4 | 1 |
| L2a_dup73360 | 11.3471982904757 | -3.52841624912196 | 0.820069217429454 | -4.30258345932399 | 1.68818047853095e-05 | 0.00273347199332482 | 1365 | 26 | 13 | 12 | 26 | 5 | 2 | 38 | 21 | 0 | 0 | 3 | 4 |
| Charlie20a_dup400 | 10.5111504104835 | -4.45663723114549 | 0.958425511743524 | -4.64995680575967 | 3.32004554242837e-06 | 0.000816106979998616 | 1366 | 15 | 11 | 13 | 17 | 6 | 1 | 36 | 31 | 0 | 0 | 2 | 1 |
| L1M5_dup24521 | 11.402647365009 | -4.25713385973927 | 0.92569864427979 | -4.59883341738217 | 4.24863328067996e-06 | 0.000972110993582927 | 1367 | 12 | 14 | 14 | 14 | 11 | 1 | 39 | 34 | 0 | 0 | 2 | 2 |
| L1PA2_dup4611 | 12.3616191598231 | -2.02230834253465 | 0.54405314925417 | -3.71711540555731 | 0.000201510409433645 | 0.0165282991668728 | 1368 | 5 | 6 | 22 | 43 | 13 | 19 | 27 | 13 | 2 | 5 | 4 | 5 |
| L2_dup51125 | 15.246984207456 | -2.3510451223101 | 0.57336626642527 | -4.10042456276336 | 4.12392854053318e-05 | 0.0052389545171399 | 1369 | 4 | 6 | 27 | 40 | 20 | 8 | 53 | 26 | 4 | 2 | 3 | 6 |
| L1PREC2_dup6949 | 16.0866345727394 | -2.38041689202504 | 0.507354003586616 | -4.69182636817144 | 2.70776890374271e-06 | 0.000708681084918037 | 1370 | 8 | 5 | 37 | 40 | 17 | 14 | 36 | 33 | 2 | 5 | 4 | 5 |
| L1ME2_dup11350 | 14.3236358145496 | -2.25186499619093 | 0.616641937859886 | -3.65181940755803 | 0.00026038897620263 | 0.0196444246453744 | 1371 | 5 | 2 | 32 | 36 | 12 | 8 | 41 | 34 | 2 | 5 | 2 | 4 |
| MER61-int_dup1050 | 12.6652025161964 | -2.64461550039399 | 0.558931104565664 | -4.7315590039475 | 2.22802034254299e-06 | 0.000611702896822642 | 1372 | 8 | 3 | 28 | 30 | 10 | 7 | 38 | 28 | 1 | 3 | 4 | 3 |
| Tigger5b_dup1422 | 14.5519797969754 | -2.67816529210452 | 0.639440686798617 | -4.18829353119966 | 2.8105984706126e-05 | 0.0039816402410238 | 1373 | 3 | 3 | 34 | 53 | 2 | 5 | 51 | 30 | 2 | 2 | 5 | 1 |
| L1PA16_dup7469 | 13.7365031884452 | -2.97570690224443 | 0.689738089974278 | -4.31425630322286 | 1.60141073446371e-05 | 0.0026322210337009 | 1374 | 7 | 7 | 35 | 52 | 1 | 5 | 41 | 25 | 3 | 1 | 1 | 3 |
| THE1B-int_dup2450 | 14.0932849931955 | -3.0342663016073 | 0.610762749282169 | -4.96799502781315 | 6.7648696880784e-07 | 0.000237414997463053 | 1375 | 12 | 6 | 30 | 40 | 3 | 5 | 52 | 28 | 2 | 2 | 2 | 3 |
| L1MA4A_dup5681 | 14.7848871369694 | -2.0613145281365 | 0.60218532769065 | -3.42305671252658 | 0.000619211568559483 | 0.035022323720574 | 1376 | 1 | 6 | 18 | 56 | 5 | 11 | 42 | 36 | 1 | 4 | 5 | 7 |
| L1PA13_dup7681 | 11.5679518283906 | -2.29448975449626 | 0.690191989949369 | -3.32442246202332 | 0.000886018674109012 | 0.0440754007120582 | 1377 | 3 | 1 | 19 | 27 | 2 | 6 | 47 | 33 | 1 | 3 | 3 | 3 |
| MLT1A1_dup1241 | 9.87212057543256 | -1.93708764342016 | 0.553216870228672 | -3.50149778082412 | 0.000462650824331163 | 0.0289699771174481 | 1378 | 4 | 3 | 22 | 28 | 2 | 7 | 21 | 25 | 1 | 2 | 4 | 7 |
| L1MC5a_dup13074 | 11.2981595503965 | -2.25058538295738 | 0.590754299295236 | -3.80968092088759 | 0.000139146253293782 | 0.0126692279259795 | 1379 | 6 | 9 | 34 | 28 | 4 | 5 | 24 | 22 | 3 | 1 | 7 | 1 |
| MIR_dup63985 | 10.8176187463057 | -1.71499342256746 | 0.493655823608072 | -3.47406703324753 | 0.000512632948236488 | 0.0310180227529039 | 1380 | 8 | 6 | 24 | 21 | 5 | 6 | 26 | 24 | 4 | 1 | 6 | 6 |
| MLT1A0_dup3769 | 9.10462070584196 | -2.14535758421431 | 0.625412802270532 | -3.43030647346152 | 0.00060289986964837 | 0.0345348128029252 | 1381 | 9 | 6 | 20 | 35 | 3 | 1 | 17 | 17 | 1 | 2 | 5 | 3 |
| MER34A_dup1200 | 9.6698687298949 | -2.37143842762801 | 0.603279131461368 | -3.93091407270047 | 8.46235146818764e-05 | 0.00891628753785281 | 1382 | 6 | 8 | 23 | 38 | 3 | 6 | 20 | 14 | 1 | 3 | 2 | 4 |
| MER4A1_dup816 | 8.53857829210152 | -2.76340309141123 | 0.660613482066394 | -4.18308612589517 | 2.87578382494848e-05 | 0.0040562018850693 | 1383 | 10 | 7 | 16 | 24 | 3 | 4 | 23 | 17 | 1 | 2 | 2 | 2 |
| L1PA5_dup4886 | 9.76194740060888 | -3.00597313569504 | 0.676582371342616 | -4.4428782998439 | 8.8763377888244e-06 | 0.00168967125719969 | 1384 | 7 | 7 | 26 | 28 | 1 | 5 | 21 | 23 | 1 | 1 | 2 | 3 |
| L3_dup19409 | 9.20257677980582 | -2.58735887037027 | 0.686322242523444 | -3.76988928824041 | 0.000163319979731026 | 0.0142013401740939 | 1385 | 5 | 10 | 25 | 19 | 1 | 3 | 25 | 21 | 2 | 1 | 2 | 3 |
| MSTA_dup17270 | 9.60632496328409 | -1.96069607563424 | 0.537343369397506 | -3.64886995410897 | 0.00026339638887256 | 0.0197959940450632 | 1386 | 10 | 11 | 20 | 25 | 2 | 6 | 24 | 14 | 2 | 1 | 5 | 6 |
| L1ME4b_dup21261 | 10.2274271224599 | -2.0830603607746 | 0.63045830602599 | -3.30404095479257 | 0.000953019088252917 | 0.0463008369418401 | 1387 | 14 | 11 | 28 | 33 | 2 | 2 | 16 | 15 | 2 | 2 | 3 | 6 |
| MIRb_dup213181 | 11.1422892419827 | -2.26221352878685 | 0.609073314541942 | -3.71419530123966 | 0.000203851321123361 | 0.0166400022147579 | 1388 | 10 | 12 | 27 | 42 | 2 | 2 | 25 | 15 | 2 | 1 | 4 | 6 |
| L1PA15_dup3421 | 11.988884900842 | -2.49966834977826 | 0.684208146727115 | -3.65337414021645 | 0.000258816679874693 | 0.0195879114851984 | 1389 | 5 | 5 | 43 | 46 | 10 | 8 | 7 | 19 | 1 | 1 | 2 | 9 |
| LTR37-int_dup1170 | 8.3168286761092 | -2.06474917872646 | 0.618740192630243 | -3.33702126242887 | 0.000846814696007959 | 0.0428624057979812 | 1390 | 4 | 6 | 21 | 25 | 9 | 2 | 13 | 16 | 1 | 2 | 2 | 6 |
| THE1B_dup3842 | 11.0679654398349 | -1.74810327732772 | 0.524492598255499 | -3.33294174816201 | 0.000859329322180653 | 0.0432840759728192 | 1391 | 7 | 6 | 34 | 31 | 10 | 3 | 19 | 16 | 3 | 1 | 4 | 10 |
| MIRc_dup8968 | 12.2687864392932 | -3.46429046600728 | 0.753186404190419 | -4.59951274576041 | 4.23480283421183e-06 | 0.000970765313284554 | 1392 | 20 | 10 | 44 | 28 | 5 | 3 | 16 | 24 | 1 | 0 | 3 | 3 |
| L1PA3_dup9589 | 9.58060269673861 | -2.3113555547386 | 0.687851870018309 | -3.36025190231303 | 0.000778714357895915 | 0.0407145608885742 | 1393 | 8 | 8 | 38 | 33 | 3 | 2 | 14 | 9 | 1 | 0 | 7 | 4 |
| MER4B_dup250 | 11.6267473745737 | -3.13587191952296 | 0.67388552062644 | -4.65341934726223 | 3.26475336772411e-06 | 0.000806774177767216 | 1394 | 10 | 6 | 41 | 36 | 6 | 2 | 19 | 22 | 0 | 0 | 3 | 6 |
| MER89_dup1151 | 10.3597727428468 | -2.81613646591716 | 0.671467297916272 | -4.19400389960363 | 2.74073266195322e-05 | 0.0039097504435319 | 1395 | 6 | 5 | 30 | 44 | 7 | 4 | 12 | 18 | 0 | 0 | 4 | 6 |
| MER5B_dup1409 | 10.1440668965233 | -4.86454669003177 | 0.948586867622379 | -5.12820370602926 | 2.9251982885313e-07 | 0.000120840144437998 | 1396 | 7 | 8 | 17 | 21 | 10 | 10 | 28 | 26 | 0 | 0 | 0 | 2 |
| Tigger1_dup5448 | 11.3433701685632 | -5.01187304758297 | 0.953020969600564 | -5.25893260216885 | 1.44893978794758e-07 | 6.97913819398736e-05 | 1397 | 15 | 10 | 25 | 26 | 6 | 5 | 22 | 32 | 0 | 0 | 0 | 2 |
| MIR3_dup7894 | 9.73622411132327 | -4.21932667686183 | 0.85619339263269 | -4.92800658492343 | 8.3072818142127e-07 | 0.000279192722961109 | 1398 | 3 | 5 | 22 | 17 | 2 | 6 | 41 | 26 | 0 | 0 | 0 | 3 |
| MamRTE1_dup8792 | 13.0835025223701 | -3.48145830660393 | 0.690058128151825 | -5.0451667251979 | 4.53126072768371e-07 | 0.000171414207489575 | 1399 | 3 | 9 | 27 | 33 | 4 | 8 | 36 | 38 | 0 | 0 | 1 | 7 |
| MIRc_dup8969 | 11.7228323082854 | -5.55237767809816 | 1.07358056299598 | -5.17183141117035 | 2.31810687599106e-07 | 0.000101394165113491 | 1400 | 6 | 13 | 43 | 42 | 4 | 8 | 12 | 22 | 0 | 1 | 0 | 0 |
| AluJr_dup5793 | 8.62344252364388 | -4.03198902377981 | 0.847937685790283 | -4.75505345657798 | 1.98393914498514e-06 | 0.000561320313665438 | 1401 | 9 | 5 | 25 | 30 | 2 | 4 | 22 | 14 | 0 | 1 | 2 | 0 |
| L1PA12_dup778 | 10.2306280150255 | -5.30071257139515 | 1.03096336584403 | -5.14151399264859 | 2.72533351458005e-07 | 0.000114169402897934 | 1402 | 11 | 6 | 20 | 45 | 2 | 5 | 18 | 25 | 0 | 0 | 1 | 0 |
| MER113_dup1166 | 10.2828264681855 | -2.10124984172243 | 0.60517456610179 | -3.47213838687497 | 0.000516330100330774 | 0.0311549391865464 | 1403 | 36 | 28 | 0 | 3 | 20 | 32 | 1 | 1 | 6 | 4 | 0 | 1 |
| MSTA_dup14325 | 10.9087882579649 | -2.04445085973291 | 0.582991129682319 | -3.50683013109816 | 0.000453478535389411 | 0.0285609855960214 | 1404 | 34 | 28 | 6 | 6 | 21 | 27 | 2 | 1 | 9 | 5 | 0 | 0 |
| L1PA15_dup447 | 7.55883841960496 | -2.90579437468226 | 0.88581530620963 | -3.280361441389 | 0.00103674174807953 | 0.0490337014487243 | 1405 | 33 | 34 | 0 | 0 | 13 | 14 | 0 | 1 | 1 | 1 | 0 | 1 |
| L2a_dup116819 | 6.34608060603111 | -4.5185156912205 | 1.25231121299103 | -3.60814120671206 | 0.00030839863483859 | 0.0220656474689521 | 1406 | 39 | 19 | 5 | 7 | 4 | 9 | 0 | 1 | 0 | 0 | 1 | 0 |
| MSTA-int_dup2279 | 8.42365990221301 | -5.68515797512161 | 1.27644195805625 | -4.45391029277893 | 8.43203420729707e-06 | 0.00162327573745106 | 1407 | 45 | 35 | 7 | 8 | 8 | 5 | 0 | 3 | 0 | 0 | 0 | 0 |
| L1PA7_dup6251 | 15.0075779981633 | -5.8192839019906 | 1.18749131469159 | -4.90048544355201 | 9.56001411802826e-07 | 0.000306986115968739 | 1408 | 95 | 70 | 3 | 2 | 16 | 7 | 2 | 3 | 0 | 0 | 0 | 0 |
| L1PA8A_dup2385 | 9.96497818949336 | -3.65315257001704 | 1.00754579141583 | -3.62579309163064 | 0.000288075951324605 | 0.021025022039365 | 1409 | 53 | 44 | 4 | 6 | 6 | 10 | 4 | 2 | 0 | 0 | 2 | 1 |
| L1MEg_dup11514 | 13.0542066776467 | -2.71789530951129 | 0.785643933682001 | -3.45944924028573 | 0.000541281288704471 | 0.0321346600603977 | 1410 | 32 | 26 | 29 | 46 | 10 | 10 | 13 | 0 | 2 | 3 | 5 | 1 |
| L1PA16_dup8711 | 17.9818300177667 | -3.11576002004326 | 0.773836346113417 | -4.02638107616956 | 5.66418660282968e-05 | 0.00664846903522944 | 1411 | 44 | 32 | 46 | 69 | 20 | 12 | 4 | 3 | 2 | 6 | 2 | 2 |
| L1PA6_dup929 | 22.3168456672619 | -2.44837018425241 | 0.668853122839554 | -3.66054982818124 | 0.000251674602818124 | 0.0192185382035177 | 1412 | 70 | 49 | 39 | 48 | 36 | 27 | 5 | 1 | 6 | 4 | 4 | 7 |
| ERVL-B4-int_dup3001 | 17.0491856236167 | -5.2982558180298 | 1.60927395774839 | -3.29232682385716 | 0.000993620701647434 | 0.0475806141022742 | 1413 | 77 | 49 | 43 | 51 | 2 | 6 | 1 | 0 | 0 | 0 | 1 | 2 |
| HERV9NC-int_dup20 | 15.1952436905044 | -3.04951096932189 | 0.900511993215753 | -3.38641905082464 | 0.000708111639089803 | 0.0381204371210095 | 1414 | 56 | 48 | 31 | 30 | 4 | 13 | 5 | 4 | 0 | 0 | 5 | 7 |
| L1PA13_dup1436 | 9.75865163895834 | -3.6643964390363 | 0.946765375940507 | -3.87043773690618 | 0.000108640102434963 | 0.0106731948034639 | 1415 | 38 | 33 | 15 | 13 | 7 | 13 | 4 | 1 | 1 | 4 | 0 | 0 |
| MER57-int_dup106 | 10.8126373667334 | -4.81859541547606 | 1.13093643674223 | -4.26103471328376 | 2.03482621200173e-05 | 0.00312776975349459 | 1416 | 50 | 47 | 7 | 19 | 9 | 6 | 3 | 1 | 1 | 1 | 0 | 0 |
| L1PA12_dup1370 | 6.47541479394767 | -4.92322911425638 | 1.43143898287932 | -3.43935660069377 | 0.000583098606147316 | 0.0337856622072427 | 1417 | 32 | 19 | 13 | 18 | 3 | 1 | 1 | 0 | 0 | 1 | 0 | 0 |
| L1PA3_dup3700 | 10.9510349034295 | -3.94246238523753 | 1.06942751197288 | -3.68651670272112 | 0.0002273445269742 | 0.017906561011388 | 1418 | 43 | 40 | 21 | 28 | 4 | 2 | 4 | 1 | 1 | 1 | 1 | 1 |
| L1M5_dup10591 | 9.69765963730614 | -2.64823381920529 | 0.783156574681367 | -3.38161474323674 | 0.000720611302637467 | 0.0384540428386128 | 1419 | 21 | 21 | 20 | 26 | 12 | 14 | 6 | 0 | 3 | 4 | 0 | 2 |
| L1PA13_dup1434 | 10.089350355853 | -2.57285294110173 | 0.755136261101213 | -3.40713732558644 | 0.000656480884630969 | 0.03636002461915 | 1420 | 21 | 19 | 27 | 23 | 11 | 15 | 2 | 4 | 2 | 6 | 0 | 2 |
| MER50-int_dup315 | 13.919627743593 | -4.77655599958706 | 1.24615844103559 | -3.83302463177765 | 0.000126577273717176 | 0.0118810871684052 | 1421 | 43 | 26 | 45 | 47 | 13 | 11 | 0 | 0 | 1 | 0 | 1 | 1 |
| L1ME3G_dup9452 | 14.0852458284913 | -4.39628029222881 | 0.986231847255539 | -4.44369541673177 | 0.000573821744485843 | 0.0334161989264387 | 1422 | 40 | 33 | 33 | 55 | 10 | 10 | 0 | 1 | 2 | 4 | 1 | 1 |
| THE1B-int_dup2744 | 10.5300957050063 | -3.60600384763625 | 1.02590921186297 | -3.51493466082445 | 0.000439862196567241 | 0.0280433489858633 | 1423 | 27 | 26 | 30 | 36 | 4 | 11 | 0 | 2 | 1 | 1 | 2 | 1 |
| MSTA-int_dup2464 | 8.79458852210479 | -1.86527556703722 | 0.542241666611387 | -3.43993404028469 | 0.000581855972961668 | 0.0337547543097392 | 1424 | 18 | 12 | 15 | 13 | 17 | 13 | 11 | 4 | 0 | 5 | 3 | 5 |
| AluJb_dup4169 | 9.76149558971945 | -2.02348846480427 | 0.538236310492519 | -3.75947966600888 | 0.000170267115031743 | 0.0146314116841254 | 1425 | 20 | 11 | 18 | 15 | 15 | 13 | 10 | 11 | 0 | 6 | 5 | 2 |
| L2a_dup54686 | 8.78984430092665 | -2.30218048217241 | 0.565969655328253 | -4.06767476047313 | 4.74845942905718e-05 | 0.00580913587304082 | 1426 | 15 | 14 | 15 | 12 | 15 | 11 | 13 | 9 | 0 | 2 | 5 | 3 |
| AluSx1_dup111538 | 11.7375982433592 | -1.91308624172623 | 0.491963821372644 | -3.88867261903217 | 0.000100793969767233 | 0.0101453138452767 | 1427 | 14 | 14 | 15 | 15 | 20 | 14 | 19 | 21 | 0 | 7 | 4 | 6 |
| L2a_dup113971 | 10.6213995277778 | -2.14994685835577 | 0.564767831015599 | -3.80678000035804 | 0.000140787879190034 | 0.0127638717197426 | 1428 | 25 | 20 | 11 | 10 | 18 | 10 | 11 | 17 | 0 | 5 | 3 | 5 |
| AluJo_dup14737 | 7.96310497363673 | -2.35401912143145 | 0.614034616649622 | -3.83369122456933 | 0.000126234583449544 | 0.0118535967614385 | 1429 | 17 | 10 | 10 | 15 | 9 | 7 | 18 | 10 | 0 | 3 | 1 | 5 |
| AluSz6_dup10184 | 7.62468950842611 | -2.09529228890163 | 0.618158916090798 | -3.38956898357487 | 0.000700025934544253 | 0.0377963174542925 | 1430 | 14 | 12 | 7 | 19 | 7 | 5 | 15 | 11 | 0 | 4 | 2 | 4 |
| AluJb_dup4174 | 6.77363306378575 | -2.743089594721 | 0.715961395034852 | -3.83133729520077 | 0.000127448643691368 | 0.0119346300557994 | 1431 | 11 | 14 | 12 | 13 | 7 | 5 | 11 | 9 | 0 | 1 | 2 | 3 |
| LTR56_dup199 | 9.33366180948933 | -2.63172532545245 | 0.606365596648115 | -4.34016266753948 | 1.42377279988137e-05 | 0.00243202471385909 | 1432 | 17 | 14 | 19 | 20 | 7 | 7 | 16 | 13 | 0 | 2 | 2 | 5 |
| AluSx_dup3635 | 7.46038832945888 | -2.59126085654195 | 0.678179902326187 | -3.82090481840262 | 0.000132962963530782 | 0.0122852285749337 | 1433 | 12 | 12 | 13 | 10 | 14 | 5 | 16 | 8 | 0 | 4 | 1 | 2 |
| L2c_dup17404 | 8.22508196069587 | -1.90253177508314 | 0.575839418812932 | -3.30392764532364 | 0.000953404352919078 | 0.0463071549621725 | 1434 | 15 | 19 | 7 | 10 | 11 | 10 | 17 | 7 | 0 | 4 | 2 | 6 |

| | | | | | | | | | | | | | | | | | | | |
|---|---|---|---|---|---|---|---|---|---|---|---|---|---|---|---|---|---|---|---|
| L1PA7_dup12373 | 9.68667895363444 | -1.98716327909869 | 0.537751870125818 | -3.69531635219372 | 0.000219613172006944 | 0.0174779121709676 | 1435 | 14 | 8 | 16 | 30 | 9 | 7 | 18 | 12 | 0 | 3 | 8 | 3 |
| MIR3_dup1074 | 9.96907630478843 | -2.9116296080404 | 0.61189294864953 | -4.75839706024799 | 1.95136363248621e-06 | 0.000556087625861988 | 1436 | 14 | 9 | 14 | 20 | 11 | 12 | 26 | 16 | 0 | 1 | 5 | 2 |
| ERV3-16A3_I-int_dup | 6.96520425727799 | -2.35732928756473 | 0.686575276591106 | -3.43346078418237 | 0.000595928401967604 | 0.03426746726307 | 1437 | 6 | 5 | 7 | 15 | 16 | 13 | 13 | 8 | 0 | 1 | 4 | 3 |
| AluY_dup73757 | 8.16552047570649 | -1.96021708715066 | 0.560222263144609 | -3.49899890830413 | 0.000467008493216993 | 0.0291431426506715 | 1438 | 8 | 10 | 12 | 12 | 9 | 13 | 13 | 15 | 0 | 2 | 5 | 5 |
| MLT1A1_dup4766 | 7.23735978792169 | -2.80799377704491 | 0.714805877100069 | -3.92833056778547 | 8.5537582166398e-05 | 0.00899793347752685 | 1439 | 4 | 6 | 13 | 15 | 6 | 12 | 17 | 14 | 0 | 2 | 2 | 2 |
| L2_dup11234 | 7.96475844652361 | -2.56396647219014 | 0.736240944263464 | -3.48251002904373 | 0.000496736613939577 | 0.0303379081982717 | 1440 | 18 | 19 | 4 | 5 | 17 | 15 | 13 | 6 | 0 | 0 | 2 | 5 |
| AluSz_dup63183 | 10.8953802866533 | -2.32811758458493 | 0.672187966729166 | -3.4634919097309 | 0.000533212629189978 | 0.0317904816500222 | 1441 | 23 | 28 | 11 | 7 | 15 | 10 | 18 | 15 | 1 | 0 | 2 | 10 |
| L1HS_dup651 | 9.01051680143185 | -2.49428875935402 | 0.643990247100906 | -3.87317784792973 | 0.000107425375931187 | 0.0105800656048519 | 1442 | 23 | 19 | 13 | 7 | 13 | 10 | 17 | 7 | 1 | 0 | 3 | 5 |
| L1MC4_dup487 | 10.1872915218714 | -2.79657251547601 | 0.670026446431964 | -4.17382407868878 | 2.99529077486763e-05 | 0.0041716169089471 | 1443 | 25 | 21 | 12 | 10 | 8 | 9 | 22 | 16 | 0 | 1 | 2 | 6 |
| L2c_dup10852 | 10.6882038401585 | -2.62856692212822 | 0.691897555643185 | -3.7990695308711 | 0.000145240344595003 | 0.0130216122965345 | 1444 | 24 | 27 | 16 | 7 | 9 | 10 | 20 | 14 | 0 | 0 | 5 | 6 |
| AluSq2_dup34686 | 6.76057879974943 | -2.81918645676803 | 0.842277821962023 | -3.34709805156801 | 0.000816623201377076 | 0.0419213975543016 | 1445 | 15 | 13 | 4 | 10 | 9 | 3 | 9 | 16 | 0 | 0 | 4 | 2 |
| MIR_dup39524 | 7.35363737060245 | -2.67275457638638 | 0.731837360483872 | -3.65211551186622 | 0.000260088838135431 | 0.0196342318143951 | 1446 | 14 | 15 | 5 | 14 | 7 | 9 | 9 | 14 | 0 | 0 | 4 | 3 |
| HERVL40-int_dup580 | 9.72731776721471 | -2.71059237025204 | 0.802949620048873 | -3.37579382637612 | 0.000736030527436838 | 0.0390578768691985 | 1447 | 31 | 23 | 4 | 10 | 9 | 3 | 26 | 13 | 0 | 2 | 1 | 6 |
| L1ME2_dup5836 | 8.38367765036415 | -3.67456646267573 | 0.946265374104319 | -3.88323039523016 | 0.00010307781272996 | 0.0102838378313808 | 1448 | 22 | 12 | 6 | 5 | 7 | 4 | 33 | 16 | 0 | 1 | 2 | 1 |
| MamGypLTR2c_dup5 | 7.94007098808565 | -2.84306331750638 | 0.778338970783214 | -3.65273155299613 | 0.000259465443961296 | 0.0196237807108081 | 1449 | 14 | 9 | 7 | 9 | 6 | 3 | 28 | 19 | 0 | 1 | 2 | 4 |
| AluJo_dup75785 | 8.53215351370129 | -2.56361552933887 | 0.707836854147579 | -3.62176045838435 | 0.000292605003399854 | 0.0212261380010619 | 1450 | 5 | 4 | 21 | 17 | 12 | 17 | 16 | 10 | 0 | 0 | 2 | 8 |
| L1PA6_dup2791 | 10.9405069663371 | -2.26158727400828 | 0.690586743005757 | -3.27487791637123 | 0.00105707645036384 | 0.0495649949257222 | 1451 | 4 | 7 | 16 | 17 | 17 | 22 | 19 | 21 | 0 | 0 | 4 | 13 |
| MLT1J-int_dup689 | 10.6676405562323 | -2.27804463070709 | 0.684857450566423 | -3.32630480813635 | 0.000880056336086684 | 0.0438702505527443 | 1452 | 11 | 5 | 10 | 19 | 24 | 23 | 15 | 16 | 0 | 0 | 5 | 10 |
| AluJr_dup82689 | 10.0952753410681 | -1.98927906671483 | 0.591308552824254 | -3.36419802692432 | 0.000767664404149488 | 0.0402961999359705 | 1453 | 11 | 19 | 21 | 21 | 7 | 6 | 21 | 10 | 0 | 0 | 10 | 7 |
| AluSg4_dup6967 | 9.49334821276111 | -2.10403800313353 | 0.591119300484134 | -3.55941347442098 | 0.000371683963158315 | 0.0250753366310373 | 1454 | 11 | 11 | 22 | 29 | 5 | 10 | 18 | 7 | 0 | 0 | 5 | 10 |
| L1MC4_dup24225 | 11.004482670997 | -2.06039799663221 | 0.544856135548018 | -3.78154500281042 | 0.00015585803877759 | 0.0137440172797455 | 1455 | 9 | 7 | 28 | 26 | 18 | 8 | 23 | 10 | 0 | 1 | 6 | 10 |
| L1MA2_dup7367 | 10.1757050278462 | -1.80431274635829 | 0.537171788153941 | -3.35891196475348 | 0.000782499912213655 | 0.0408765585316334 | 1456 | 6 | 7 | 23 | 26 | 12 | 5 | 26 | 11 | 0 | 0 | 11 | 8 |
| MIRb_dup215003 | 11.1725668651155 | -2.06368479501219 | 0.509007591793099 | -4.05433008914931 | 5.02781791802642e-05 | 0.00605908178693109 | 1457 | 6 | 8 | 24 | 33 | 8 | 6 | 28 | 17 | 0 | 0 | 11 | 6 |
| L1MD3_dup1542 | 7.73601793879511 | -2.3243857639131 | 0.655049884673064 | -3.54841030935102 | 0.000387563966916336 | 0.0257739786673769 | 1458 | 9 | 11 | 12 | 18 | 14 | 15 | 6 | 7 | 1 | 0 | 2 | 6 |
| L1PA2_dup4638 | 9.3058787102339 | -1.8968935680216 | 0.562306626995921 | -3.37341492515498 | 0.000742419846747896 | 0.039331664037174 | 1459 | 8 | 10 | 14 | 32 | 13 | 9 | 15 | 8 | 2 | 0 | 4 | 9 |
| L3_dup41746 | 6.71722519902612 | -2.75266722943355 | 0.733728726262486 | -3.75161436496463 | 0.000175699573689653 | 0.0149953410190668 | 1460 | 7 | 7 | 14 | 22 | 7 | 6 | 7 | 11 | 1 | 0 | 2 | 3 |
| Tigger3c_dup1861 | 7.63486823055694 | -2.25994455008624 | 0.62170440230219 | -3.63507889234434 | 0.000277895581202794 | 0.0205163332190133 | 1461 | 7 | 9 | 14 | 18 | 13 | 10 | 8 | 10 | 2 | 0 | 4 | 3 |
| AluSc8_dup18153 | 7.3912106788647 | -2.66088404202833 | 0.727319954244202 | -3.65847798689011 | 0.000253717528330439 | 0.0193187427187567 | 1462 | 5 | 18 | 9 | 19 | 7 | 6 | 18 | 8 | 1 | 0 | 4 | 2 |
| LTR12C_dup2601 | 8.28625669615343 | -2.58025518639394 | 0.668182664064795 | -3.86160151282184 | 0.00011264618990642 | 0.0109492280651442 | 1463 | 7 | 13 | 19 | 28 | 4 | 7 | 14 | 9 | 2 | 0 | 4 | 2 |
| THE1D_dup11563 | 8.81711550512614 | -1.81638785239927 | 0.554534622615533 | -3.27551748497154 | 0.00105468585435494 | 0.049500908001123 | 1464 | 14 | 4 | 19 | 18 | 5 | 5 | 16 | 18 | 1 | 2 | 6 | 5 |
| L1MC_dup7361 | 8.97838036491675 | -2.87162058965916 | 0.68045243967562 | -4.22016355915794 | 2.44125136289046e-05 | 0.00358571602959465 | 1465 | 19 | 6 | 16 | 29 | 6 | 6 | 18 | 12 | 2 | 1 | 1 | 3 |
| L1MA3_dup1312 | 8.72350881767305 | -1.88228695993001 | 0.525162176088914 | -3.58420131081817 | 0.00033811123899241 | 0.0235378048195693 | 1466 | 9 | 10 | 17 | 24 | 6 | 7 | 20 | 9 | 3 | 1 | 4 | 5 |
| MIR_dup126688 | 8.06257590845017 | -2.09096034495679 | 0.589767082401299 | -3.54540022214062 | 0.000392017371453034 | 0.025925226644737 | 1467 | 11 | 9 | 17 | 23 | 7 | 4 | 17 | 8 | 3 | 2 | 3 | 2 |
| L1MEg_dup974 | 8.3157905090262 | -1.81608486502755 | 0.549076009268212 | -3.30752907497809 | 0.000941229407463386 | 0.0458844033165914 | 1468 | 14 | 8 | 9 | 15 | 9 | 11 | 17 | 12 | 3 | 1 | 1 | 8 |
| L1ME4b_dup2274 | 8.33382257939612 | -1.6701235040889 | 0.508164253387941 | -3.28658203121166 | 0.00101411232186623 | 0.0482511438789805 | 1469 | 10 | 9 | 10 | 15 | 10 | 8 | 17 | 14 | 2 | 3 | 3 | 6 |
| MER2B_dup1098 | 8.13944761767815 | -2.05106802694663 | 0.603881388409626 | -3.3964749805393 | 0.000682597973188608 | 0.0372317419350031 | 1470 | 10 | 12 | 7 | 17 | 7 | 5 | 15 | 19 | 1 | 2 | 4 | 4 |
| AluSz_dup49527 | 7.78856863479017 | -2.05146635174752 | 0.607039518979904 | -3.37946095370347 | 0.000726281232420266 | 0.0386612335860118 | 1471 | 13 | 7 | 14 | 11 | 8 | 5 | 13 | 17 | 2 | 3 | 1 | 4 |
| MIRc_dup22349 | 7.2501690448605 | -2.31864756803919 | 0.651251386610287 | -3.56029578701947 | 0.000370437296964683 | 0.0250409298476018 | 1472 | 13 | 8 | 12 | 12 | 9 | 3 | 15 | 13 | 1 | 3 | 1 | 3 |
| AluJb_dup106786 | 7.95040595502639 | -2.90067815205304 | 0.691795885539483 | -4.19296820447407 | 2.75328045808738e-05 | 0.00392059887420742 | 1473 | 8 | 10 | 13 | 16 | 9 | 5 | 17 | 17 | 1 | 2 | 1 | 2 |
| LTR22B_dup13 | 7.11706031245768 | -2.50304457175113 | 0.665647988829302 | -3.76031267840728 | 0.000169701109857827 | 0.0145985901251737 | 1474 | 9 | 6 | 10 | 12 | 10 | 5 | 18 | 14 | 1 | 2 | 2 | 2 |
| L1MA3_dup1241 | 8.49053971402471 | -2.07414805353743 | 0.543914423425103 | -3.81337203833691 | 0.000137083516178983 | 0.012533650583222 | 1475 | 12 | 9 | 17 | 17 | 14 | 9 | 15 | 7 | 2 | 3 | 3 | 3 |
| L1PA5_dup4893 | 10.2508332689633 | -2.89357468633065 | 0.60908651433567 | -4.75067928484127 | 2.02734456128058e-06 | 0.000569537814614484 | 1476 | 11 | 14 | 26 | 27 | 11 | 11 | 18 | 9 | 1 | 2 | 3 | 2 |
| HAL1ME_dup3858 | 10.582116984326 | -1.91760580713622 | 0.485339508813336 | -3.95106059225387 | 7.78056206610971e-05 | 0.00836167592889316 | 1477 | 10 | 9 | 24 | 18 | 13 | 12 | 18 | 16 | 2 | 1 | 5 | 8 |
| L1MA3_dup8202 | 9.81950209668786 | -1.88026520439847 | 0.549504212668676 | -3.42174847990142 | 0.000622198426593653 | 0.0351328018068945 | 1478 | 6 | 7 | 23 | 26 | 17 | 11 | 13 | 10 | 4 | 2 | 4 | 4 |
| AluSz_dup35223 | 9.94005544468859 | -2.02355696932545 | 0.562281600149249 | -3.59883191765182 | 0.000319649693493561 | 0.0225896542821559 | 1479 | 5 | 13 | 13 | 18 | 9 | 9 | 25 | 21 | 1 | 4 | 6 | 2 |
| Tigger1_dup9307 | 7.87372171827832 | -1.92446454241998 | 0.58122521065027 | -3.31104795035801 | 0.000929472801705272 | 0.045516430391966 | 1480 | 4 | 7 | 11 | 14 | 9 | 10 | 21 | 14 | 1 | 4 | 3 | 3 |
| MER20_dup7054 | 9.37321836054625 | -1.92067817546198 | 0.540545592602384 | -3.55322141508015 | 0.000380544021677844 | 0.0254780325779843 | 1481 | 10 | 18 | 8 | 18 | 13 | 8 | 19 | 13 | 3 | 5 | 3 | 2 |
| MER58A_dup874 | 8.57048724520091 | -1.97135779489848 | 0.564768767435021 | -3.49055739015401 | 0.000482014074721338 | 0.0298059566069759 | 1482 | 6 | 13 | 10 | 18 | 12 | 11 | 16 | 12 | 2 | 1 | 7 | 2 |
| L1PB1_dup11002 | 13.8279023841368 | -3.14973632719554 | 0.812216057209471 | -3.87795377749251 | 0.000105338724292856 | 0.0104201523291801 | 1483 | 7 | 12 | 34 | 75 | 23 | 18 | 5 | 4 | 0 | 2 | 4 | 4 |
| AluY_dup5085 | 10.3854696029534 | -2.54173118271719 | 0.650236151675862 | -3.90893550930128 | 9.270370520967e-05 | 0.00952849928285967 | 1484 | 25 | 14 | 18 | 27 | 12 | 15 | 5 | 10 | 0 | 1 | 3 | 7 |
| L1MC3_dup11858 | 9.07276486987837 | -2.40987828209381 | 0.702201662695823 | -3.43188917104246 | 0.000599392453456717 | 0.0344001535769927 | 1485 | 12 | 9 | 17 | 34 | 18 | 4 | 6 | 9 | 0 | 1 | 5 | 5 |
| HERV9NC-int_dup194 | 7.66465843756408 | -3.10014335762197 | 0.888786783601001 | -3.48806194558998 | 0.000486535343881699 | 0.0299687131595327 | 1486 | 17 | 19 | 19 | 22 | 3 | 5 | 3 | 7 | 1 | 2 | 2 | 0 |
| MLT1C_dup15027 | 10.9310651234307 | -3.3980090961422 | 0.870573838399923 | -3.90318310321338 | 9.49358259739923e-05 | 0.00967849187969476 | 1487 | 27 | 29 | 22 | 32 | 5 | 12 | 3 | 8 | 0 | 2 | 4 | 0 |
| L1HS_dup895 | 10.030559049722 | -3.09701515773778 | 0.836380539257957 | -3.7028780708904 | 0.00021316734749413 | 0.0171475042822307 | 1488 | 8 | 11 | 34 | 46 | 4 | 10 | 8 | 6 | 0 | 0 | 1 | 8 |
| L1M6_dup4563 | 5.78621411793436 | -3.64227322442995 | 1.0203330133829 | -3.56969065653777 | 0.000357403022351258 | 0.0244410279552528 | 1489 | 6 | 11 | 13 | 28 | 3 | 2 | 4 | 7 | 0 | 0 | 1 | 2 |
| L1PREC2_dup6361 | 6.50854113174505 | -2.98559305272507 | 0.889145984518372 | -3.35782099307606 | 0.000785594703533846 | 0.0409662759866458 | 1490 | 13 | 19 | 16 | 17 | 4 | 4 | 4 | 4 | 0 | 1 | 1 | 3 |
| L1MC3_dup8881 | 10.3487117496787 | -3.91355840335914 | 0.851823915530233 | -4.5943279262394 | 4.34146201238597e-06 | 0.000988655508332668 | 1491 | 30 | 23 | 20 | 22 | 10 | 14 | 8 | 6 | 1 | 0 | 0 | 3 |
| HERVE_a-int_dup146 | 5.98017776664187 | -4.85241749043795 | 1.211216895895 | -4.00623332359675 | 6.16947077984362e-05 | 0.00705895674157355 | 1492 | 18 | 15 | 10 | 17 | 2 | 12 | 5 | 1 | 0 | 0 | 0 | 1 |
| Harlequin-int_dup391 | 8.67532529953238 | -3.58452274745075 | 1.03048776345549 | -3.47847191841554 | 0.000504281271346678 | 0.030686298026668 | 1493 | 29 | 27 | 15 | 20 | 6 | 13 | 1 | 1 | 0 | 0 | 2 | 2 |
| AluSp_dup28604 | 14.7816359131805 | -1.68806610232645 | 0.499279783385438 | -3.3810023127319 | 0.000722219349967115 | 0.0385030227999302 | 1494 | 24 | 20 | 9 | 9 | 22 | 17 | 39 | 24 | 5 | 4 | 8 | 7 |
| L2a_dup34772 | 12.0718275019583 | -2.18448796504824 | 0.582991288154277 | -3.74703363400888 | 0.000178938101019038 | 0.0152008587593988 | 1495 | 15 | 16 | 5 | 11 | 22 | 24 | 29 | 18 | 2 | 1 | 5 | 7 |
| THE1C-int_dup305 | 13.684891553266 | -2.17783718272397 | 0.591457183750476 | -3.68215526424777 | 0.000231270483416118 | 0.0181078928749364 | 1496 | 23 | 15 | 4 | 12 | 34 | 23 | 35 | 15 | 4 | 1 | 6 | 5 |
| L1M4a1_dup1255 | 14.000872774106 | -3.42168041026968 | 0.755611219286388 | -4.52836104458741 | 5.94429514993585e-06 | 0.00124618612664902 | 1497 | 36 | 27 | 5 | 9 | 17 | 13 | 45 | 22 | 1 | 0 | 3 | 4 |
| L2_dup11237 | 9.4108009026546 | -2.47221518882024 | 0.733522693823765 | -3.37033224689052 | 0.000750776024180718 | 0.0396370427889295 | 1498 | 17 | 18 | 5 | 2 | 23 | 12 | 22 | 12 | 2 | 0 | 4 | 3 |
| MLT1L_dup2375 | 11.4625954443949 | -2.32004256288583 | 0.673219022778729 | -3.44619282044318 | 0.000568544529798018 | 0.0332390409551153 | 1499 | 22 | 19 | 6 | 4 | 26 | 13 | 23 | 19 | 5 | 0 | 2 | 5 |
| AluY_dup4980 | 13.2891326110196 | -2.33342554709117 | 0.61980910973194 | -3.76474871126104 | 0.000166716656576586 | 0.0143678238195481 | 1500 | 29 | 21 | 10 | 15 | 23 | 23 | 6 | 25 | 3 | 0 | 7 | 4 |
| L1MEj_dup443 | 11.9548139754044 | -2.32323627907994 | 0.564711359570782 | -4.1140243412949 | 3.88820408489868e-05 | 0.00502277394500201 | 1501 | 30 | 16 | 11 | 16 | 26 | 14 | 13 | 15 | 3 | 0 | 2 | 8 |
| MER5A1_dup3266 | 11.9348487607383 | -2.2292360677487 | 0.643387334497258 | -3.46484294641986 | 0.000530541212322599 | 0.0316867869747272 | 1502 | 32 | 23 | 1 | 6 | 29 | 23 | 9 | 16 | 2 | 3 | 3 | 4 |
| AluJb_dup72424 | 11.5171082672362 | -1.86190692053355 | 0.551271646226626 | -3.37747630098162 | 0.000731542565596183 | 0.0389064880003016 | 1503 | 34 | 22 | 4 | 11 | 13 | 14 | 15 | 17 | 6 | 3 | 3 | 4 |
| L1M5_dup24520 | 11.118996807322 | -2.1320330940991 | 0.620796903251794 | -3.43434879093518 | 0.000593979361264508 | 0.0341967125805576 | 1504 | 34 | 31 | 5 | 12 | 9 | 11 | 14 | 14 | 2 | 2 | 6 | 3 |
| MER5A1_dup3269 | 9.12236611612137 | -2.02497726153791 | 0.59132759987066 | -3.42445923711464 | 0.000616024251996647 | 0.034914717253013 | 1505 | 22 | 21 | 4 | 6 | 18 | 14 | 10 | 10 | 4 | 1 | 1 | 5 |
| L2_dup3485 | 10.1224604820482 | -3.19799361311585 | 0.706099689112676 | -4.52909647522239 | 5.92364487773065e-06 | 0.0012436890883113 | 1506 | 29 | 26 | 7 | 9 | 16 | 10 | 14 | 13 | 1 | 1 | 2 | 2 |

| | | | | | | | | | | | | | | | | | | | |
|---|---|---|---|---|---|---|---|---|---|---|---|---|---|---|---|---|---|---|---|
| MER11B_dup163 | 10.8017038960645 | -3.51303183690212 | 0.750105514262781 | -4.68338356418403 | 2.82177772408343e-06 | 0.000721968518639523 | 1507 | 30 | 30 | 5 | 5 | 18 | 20 | 15 | 11 | 1 | 1 | 1 | 2 |
| L1ME1_dup23528 | 8.29262887580165 | -2.29688870748691 | 0.654786599104759 | -3.50784318223262 | 0.00045175526150938 | 0.0285163983699895 | 1508 | 16 | 17 | 16 | 5 | 15 | 18 | 7 | 3 | 5 | 2 | 0 | 2 |
| MER5B_dup1329 | 5.62054572303784 | -2.66927602518543 | 0.801246073727602 | -3.33140605952335 | 0.000864084586876914 | 0.043399668800527 | 1509 | 10 | 10 | 14 | 6 | 7 | 11 | 9 | 2 | 1 | 1 | 0 | 3 |
| Charlie3_dup284 | 6.91383576474029 | -2.66667796346867 | 0.80260868691566 | -3.32251320841845 | 0.000892104476667996 | 0.0442973063032644 | 1510 | 28 | 11 | 9 | 8 | 5 | 9 | 3 | 10 | 1 | 2 | 0 | 3 |
| L1PA3_dup3548 | 8.17874200957474 | -2.48931553938958 | 0.656378560411097 | -3.79249976999628 | 0.000149138388709528 | 0.0132710856544044 | 1511 | 28 | 17 | 5 | 5 | 16 | 12 | 7 | 7 | 2 | 5 | 0 | 1 |
| THE1B_dup15896 | 7.72647344617338 | -2.09229031296339 | 0.617342696795595 | -3.38918776203837 | 0.000700999925267881 | 0.0378403194686459 | 1512 | 11 | 12 | 7 | 17 | 11 | 15 | 7 | 10 | 1 | 3 | 0 | 6 |
| AluJb_dup72687 | 9.98732139671529 | -1.95203592604539 | 0.502587104634286 | -3.88397535083161 | 0.000102762330222839 | 0.0102652751583559 | 1513 | 14 | 15 | 14 | 19 | 12 | 13 | 18 | 11 | 2 | 6 | 0 | 6 |
| MIR1_Amn_dup7648 | 8.13095694758988 | -1.97885706564721 | 0.602548358369227 | -3.2841464724971 | 0.00102291744208278 | 0.0485923564930782 | 1514 | 7 | 16 | 15 | 16 | 10 | 6 | 12 | 11 | 2 | 5 | 0 | 4 |
| L1MA3_dup1242 | 8.50013357395767 | -2.36325704357069 | 0.630962537793233 | -3.74547917192689 | 0.000180049785347316 | 0.0152721731603586 | 1515 | 19 | 16 | 6 | 10 | 11 | 15 | 20 | 5 | 5 | 2 | 0 | 2 |
| MSTA_dup15120 | 8.01567710037456 | -1.94280098746526 | 0.569782000148263 | -3.40972685511253 | 0.0006502797117739 | 0.0361678479030935 | 1516 | 16 | 9 | 12 | 8 | 13 | 12 | 15 | 7 | 5 | 3 | 0 | 3 |
| FLAM_A_dup10762 | 8.43609754706642 | -2.05928737027077 | 0.605781504846738 | -3.39938963767435 | 0.000675364293437453 | 0.0369647210700528 | 1517 | 24 | 12 | 10 | 19 | 6 | 10 | 11 | 8 | 3 | 3 | 0 | 5 |
| MER21B_dup500 | 7.61120359850752 | -2.40383019501327 | 0.675571937407155 | -3.55821499075165 | 0.000373383647244816 | 0.0251512195776959 | 1518 | 17 | 12 | 13 | 17 | 5 | 7 | 19 | 4 | 2 | 3 | 0 | 3 |
| L1M4_dup3029 | 10.4382729844404 | -2.00246925255275 | 0.565052288519051 | -3.54386539660079 | 0.000394306499328322 | 0.0260404258365742 | 1519 | 20 | 29 | 14 | 14 | 12 | 7 | 14 | 10 | 4 | 5 | 0 | 5 |
| THE1D_dup8867 | 9.93191163843579 | -2.24678078528585 | 0.630362353466255 | -3.56426866695192 | 0.000364872082415005 | 0.024768838791851 | 1520 | 28 | 24 | 11 | 12 | 10 | 7 | 10 | 12 | 5 | 6 | 0 | 1 |
| AluSc_dup2013 | 9.50781168315228 | -2.24442406040721 | 0.63244572051769 | -3.54880108694551 | 0.000386989293800046 | 0.0257645647740755 | 1521 | 28 | 15 | 7 | 9 | 10 | 8 | 23 | 12 | 4 | 4 | 0 | 3 |
| AluSz_dup61448 | 8.83500534751052 | -2.30694476795648 | 0.646279422646733 | -3.56957793659709 | 0.000357556833534647 | 0.0244410279552528 | 1522 | 22 | 18 | 9 | 11 | 10 | 5 | 21 | 9 | 2 | 6 | 1 | 1 |
| L1PREC2_dup2784 | 7.78234772120604 | -2.25827006119612 | 0.669448146575595 | -3.37333081396636 | 0.000742646694992618 | 0.039331664037174 | 1523 | 8 | 15 | 7 | 17 | 12 | 11 | 12 | 8 | 6 | 3 | 0 | 0 |
| LTR16C_dup2210 | 9.18391041021492 | -2.05771669318648 | 0.599603618408041 | -3.43179498924599 | 0.000599600637147411 | 0.0344038035222276 | 1524 | 14 | 14 | 7 | 11 | 9 | 20 | 14 | 14 | 6 | 5 | 1 | 0 |
| L2a_dup134049 | 12.2985324024608 | -2.03489034161576 | 0.573273636556118 | -3.54959693217388 | 0.000385821394250171 | 0.0257227955943545 | 1525 | 21 | 25 | 24 | 18 | 18 | 12 | 15 | 7 | 9 | 7 | 1 | 0 |
| AluJr4_dup10368 | 12.3234030043954 | -3.81602979570912 | 0.732517029427806 | -5.20947587892933 | 1.89374824618325e-07 | 8.73274845322327e-05 | 1526 | 24 | 27 | 20 | 12 | 16 | 16 | 24 | 14 | 4 | 1 | 0 | 0 |
| AluSx_dup55466 | 12.7640853862687 | -2.84088546339815 | 0.596164884864292 | -4.76526802487677 | 1.8860277251327e-06 | 0.000538742800821639 | 1527 | 28 | 29 | 13 | 12 | 11 | 24 | 19 | 16 | 6 | 3 | 1 | 0 |
| MSTD_dup4846 | 11.4985055042827 | -1.97695786974739 | 0.58051407906744 | -3.40552958323294 | 0.000660358584641162 | 0.0364823969772524 | 1528 | 14 | 10 | 14 | 12 | 41 | 29 | 8 | 5 | 2 | 7 | 2 | 4 |
| AluJb_dup89991 | 10.708110117529 | -1.56215712322722 | 0.457137997520754 | -3.41725503392725 | 0.000632559923075228 | 0.0355072194989522 | 1529 | 19 | 27 | 9 | 10 | 17 | 17 | 14 | 7 | 3 | 6 | 4 | 5 |
| L2_dup11231 | 9.33160051366688 | -2.30737935256856 | 0.607303166906506 | -3.7993863333892 | 0.00014505482209828 | 0.0130098811530333 | 1530 | 12 | 20 | 6 | 6 | 15 | 20 | 22 | 9 | 3 | 2 | 2 | 3 |
| AluSz_dup74298 | 10.1997419621702 | -2.48750231344262 | 0.55682757952436 | -4.46727569702535 | 7.9222005481825e-06 | 0.00154240490874888 | 1531 | 20 | 18 | 12 | 11 | 10 | 17 | 19 | 14 | 2 | 3 | 2 | 3 |
| MIR1_Amn_dup1862 | 10.9754355562699 | -2.05742120793106 | 0.498393382513466 | -4.1281069936266 | 3.65761926576485e-05 | 0.00481103302272903 | 1532 | 22 | 20 | 8 | 14 | 13 | 16 | 23 | 12 | 4 | 3 | 3 | 4 |
| L1PA5_dup1894 | 10.0867810690289 | -1.88994432898285 | 0.537761392682295 | -3.51446636872911 | 0.000440638463473456 | 0.028056772766833 | 1533 | 25 | 17 | 8 | 19 | 13 | 18 | 9 | 7 | 8 | 2 | 1 | 3 |
| MLT1D_dup2517 | 9.64288829552282 | -1.80946876619502 | 0.526762869067455 | -3.43507272902204 | 0.000592394818713822 | 0.0341385268911792 | 1534 | 21 | 15 | 11 | 6 | 12 | 17 | 10 | 15 | 4 | 3 | 3 | 4 |
| L1MC5_dup8105 | 8.67825514855158 | -1.77176170335622 | 0.537225228663694 | -3.29798678249593 | 0.000973807182487207 | 0.0469530950038705 | 1535 | 21 | 17 | 15 | 10 | 10 | 11 | 9 | 6 | 3 | 3 | 5 | 2 |
| AluY_dup73756 | 8.98568301870107 | -1.83149989734885 | 0.528186529367565 | -3.4675248146556 | 0.00052527524142345 | 0.0314651412518991 | 1536 | 20 | 16 | 11 | 12 | 9 | 11 | 11 | 11 | 5 | 4 | 3 | 1 |
| Tigger4b_dup407 | 9.98120955902792 | -2.11092986204031 | 0.554428217464481 | -3.80739975987161 | 0.000140435633475007 | 0.0127486837713916 | 1537 | 26 | 24 | 12 | 9 | 13 | 9 | 11 | 11 | 4 | 3 | 3 | 2 |
| Harlequin-int_dup19 | 9.68228198455324 | -2.24888819197223 | 0.643399193907637 | -3.49532329736656 | 0.000473487853079158 | 0.0294125822752792 | 1538 | 32 | 29 | 5 | 8 | 19 | 7 | 14 | 3 | 2 | 3 | 3 | 2 |
| L2a_dup24834 | 7.21419800266861 | -2.07787675166778 | 0.63176483033926 | -3.2890035213767 | 0.00100542765732045 | 0.048014784126543 | 1539 | 10 | 20 | 5 | 8 | 16 | 9 | 12 | 4 | 2 | 4 | 2 | 1 |
| AluY_dup52994 | 7.86031829000525 | -1.794566538124 | 0.540205097810457 | -3.32200963189293 | 0.00089371608713519 | 0.0443403186133173 | 1540 | 15 | 12 | 10 | 12 | 8 | 11 | 17 | 6 | 2 | 3 | 5 | 2 |
| L2_dup2037 | 8.7445584289496 | -1.71332789218988 | 0.522435188248234 | -3.2795032393105 | 0.00103990016353172 | 0.0491147448018496 | 1541 | 22 | 12 | 8 | 13 | 8 | 17 | 14 | 7 | 1 | 5 | 4 | 4 |
| Charlie30a_dup138 | 10.9594746676265 | -1.96090366456218 | 0.541549292058657 | -3.62091446395943 | 0.000293563565695682 | 0.0212769650329043 | 1542 | 26 | 10 | 13 | 21 | 15 | 14 | 27 | 5 | 3 | 6 | 5 | 1 |
| L1ME3D_dup1951 | 9.9978067266145 | -1.9255525651571 | 0.539023048190695 | -3.57230098345605 | 0.000353858380381136 | 0.0242928984934131 | 1543 | 19 | 18 | 21 | 9 | 11 | 9 | 22 | 7 | 3 | 5 | 3 | 3 |
| L1MC4_dup17167 | 10.2843863198087 | -2.12394475093358 | 0.558910904127129 | -3.80014906714089 | 0.000144609073110823 | 0.0129894867914035 | 1544 | 19 | 18 | 20 | 18 | 11 | 10 | 24 | 4 | 1 | 6 | 3 | 3 |
| L2_dup56441 | 9.80052232580836 | -1.73401426376092 | 0.516960878560025 | -3.3542465893956 | 0.000795814087906007 | 0.0413000608420682 | 1545 | 20 | 16 | 17 | 12 | 11 | 10 | 23 | 5 | 1 | 5 | 4 | 6 |
| THE1D-int_dup1267 | 7.29215498742681 | -3.69449540194493 | 0.979220554383332 | -3.77289404864622 | 0.000161364825315157 | 0.0140768967389611 | 1546 | 23 | 28 | 4 | 12 | 6 | 9 | 7 | 4 | 1 | 0 | 0 | 2 |
| AluJr_dup35443 | 6.25621065810962 | -2.8063606818766 | 0.837322052384196 | -3.35159055453723 | 0.000803487564755351 | 0.0415354127014618 | 1547 | 13 | 17 | 9 | 13 | 2 | 12 | 8 | 3 | 3 | 1 | 0 | 1 |
| L1M4a1_dup1753 | 6.8252352574045 | -3.34307339225895 | 0.987722289022435 | -3.38462888750607 | 0.000712745479124516 | 0.0382401208062598 | 1548 | 27 | 25 | 3 | 11 | 3 | 8 | 7 | 2 | 3 | 1 | 0 | 0 |
| L1PA10_dup147 | 7.9670003139803 | -3.02609363810781 | 0.902905768928034 | -3.35150548622644 | 0.000803734464275546 | 0.0415354127014618 | 1549 | 29 | 27 | 5 | 7 | 3 | 8 | 13 | 7 | 1 | 0 | 4 | 0 |
| Harlequin-int_dup389 | 6.05414140296588 | -3.0535562385587 | 0.8360924009966 | -3.65217556686191 | 0.000260028004603735 | 0.0196342318143951 | 1550 | 16 | 13 | 8 | 12 | 10 | 8 | 4 | 4 | 1 | 1 | 2 | 0 |
| L1PA3_dup9223 | 7.7686288588204 | -2.61025320718649 | 0.738424914822011 | -3.53489319603424 | 0.000407940008333429 | 0.0266449885322279 | 1551 | 26 | 20 | 8 | 16 | 9 | 7 | 8 | 2 | 2 | 3 | 2 | 0 |
| GSAT_dup20 | 12.3566798276255 | -2.94915392260394 | 0.680218756076443 | -4.33559630083548 | 1.45365515687999e-05 | 0.00246387158936012 | 1552 | 46 | 43 | 8 | 9 | 14 | 19 | 3 | 9 | 2 | 3 | 2 | 1 |
| LTR49-int_dup399 | 10.4263439667704 | -1.87023284331449 | 0.56651923427279 | -3.30126980722059 | 0.000962482777907315 | 0.0466154020515712 | 1553 | 33 | 23 | 13 | 14 | 14 | 16 | 6 | 3 | 1 | 9 | 2 | 3 |
| L1MA6_dup2050 | 14.1109553299952 | -2.55285234467019 | 0.688233328010258 | -3.70928323400249 | 0.000207846816039186 | 0.0168852651672033 | 1554 | 64 | 55 | 5 | 4 | 13 | 15 | 8 | 6 | 7 | 3 | 0 | 2 |
| AluJr_dup54887 | 15.7496418546116 | -2.38322683367412 | 0.586843162934815 | -4.06109670215045 | 4.8842733906966e-05 | 0.00593561448530487 | 1555 | 56 | 60 | 7 | 13 | 19 | 12 | 17 | 6 | 5 | 5 | 3 | 2 |
| Tigger1_dup4214 | 11.5339347806856 | -2.95199080288432 | 0.716688595138063 | -4.11893090375695 | 3.80634251197203e-05 | 0.00496108327462887 | 1556 | 39 | 35 | 3 | 4 | 15 | 17 | 19 | 9 | 3 | 3 | 2 | 0 |
| AluSz_dup58153 | 4.96897519151999 | -3.17105376446061 | 0.956093716210952 | -3.31667671347916 | 0.000910949582271812 | 0.044930231711484 | 1557 | 6 | 6 | 8 | 5 | 9 | 4 | 11 | 10 | 2 | 1 | 0 | 0 |
| MER44A_dup1149 | 4.77122083901765 | -3.15161948306392 | 0.947215389239488 | -3.327246916453 | 0.000877086192033203 | 0.0438075463999858 | 1558 | 6 | 4 | 9 | 10 | 10 | 5 | 7 | 7 | 1 | 2 | 0 | 0 |
| L1MEb_dup301 | 6.58937124238431 | -2.82277675850832 | 0.818742978564508 | -3.44769583668059 | 0.000565390357036135 | 0.0331044572401402 | 1559 | 4 | 16 | 15 | 16 | 7 | 5 | 8 | 8 | 3 | 1 | 0 | 1 |
| MLT1F_dup3211 | 7.54338504961514 | -2.47903302810201 | 0.756128174246353 | -3.27858835649512 | 0.00104327698307016 | 0.0491961127652817 | 1560 | 11 | 9 | 18 | 21 | 6 | 7 | 6 | 11 | 4 | 3 | 0 | 0 |
| L1MC5_dup12054 | 6.70652855055855 | -2.63195961109271 | 0.775514198862276 | -3.39382517425722 | 0.000689236775140885 | 0.0374260916699391 | 1561 | 6 | 13 | 8 | 13 | 5 | 4 | 27 | 7 | 2 | 1 | 0 | 3 |
| L1M5_dup43755 | 6.29622031247737 | -2.50421366285694 | 0.750505092244393 | -3.33670442577287 | 0.000847780560463205 | 0.0428879957031394 | 1562 | 9 | 13 | 9 | 11 | 4 | 3 | 16 | 10 | 2 | 2 | 0 | 2 |
| MER2_dup1531 | 6.39399881084044 | -2.35928223752186 | 0.692770597814514 | -3.40557501280322 | 0.000660248721664685 | 0.0364823969772524 | 1563 | 8 | 9 | 9 | 12 | 5 | 4 | 18 | 11 | 1 | 2 | 0 | 4 |
| AluJb_dup56718 | 6.7018031820004 | -2.18457075756329 | 0.665108455522216 | -3.28453313053742 | 0.00102151487326089 | 0.0485451124491581 | 1564 | 9 | 14 | 4 | 12 | 10 | 5 | 14 | 10 | 2 | 2 | 1 | 3 |
| L1MC3_dup5746 | 6.97807890617905 | -2.44891766197912 | 0.673209260352613 | -3.63767673174374 | 0.000275108414205328 | 0.0203791964242417 | 1565 | 7 | 7 | 11 | 11 | 12 | 6 | 22 | 8 | 2 | 2 | 1 | 2 |
| MER58A_dup11906 | 5.31551223768078 | -2.59427728620281 | 0.770169493746336 | -3.36844981172061 | 0.000755921581791264 | 0.0398368096990691 | 1566 | 8 | 6 | 5 | 11 | 6 | 5 | 14 | 9 | 1 | 1 | 1 | 2 |
| MIR_dup160645 | 5.60785673310535 | -2.62643563723003 | 0.767805267144428 | -3.42070541792205 | 0.000624589465814831 | 0.0352396741735448 | 1567 | 5 | 10 | 9 | 9 | 5 | 5 | 15 | 9 | 1 | 2 | 1 | 1 |
| L1M3_dup5967 | 5.95056872647426 | -2.53587836316392 | 0.730192209158451 | -3.47289156383431 | 0.000514883336838051 | 0.0310950331135195 | 1568 | 6 | 7 | 15 | 13 | 5 | 7 | 15 | 5 | 1 | 1 | 0 | 4 |
| MIR_dup171454 | 5.58510672704741 | -2.64364234363333 | 0.777032326284919 | -3.40222955236996 | 0.00066838471484762 | 0.0367297000281148 | 1569 | 5 | 4 | 9 | 19 | 7 | 8 | 11 | 6 | 1 | 2 | 0 | 2 |
| AluYe5_dup1065 | 5.64047470604062 | -2.68759755120862 | 0.783924804210441 | -3.42838692789614 | 0.000607179405915628 | 0.0346803590223921 | 1570 | 8 | 10 | 9 | 6 | 5 | 6 | 13 | 10 | 1 | 1 | 0 | 3 |
| L1MC5_dup208 | 5.03996317376237 | -3.23570797799552 | 0.978221408734712 | -3.30774602672085 | 0.000940500602754442 | 0.0458582697588132 | 1571 | 5 | 10 | 9 | 5 | 2 | 8 | 10 | 11 | 1 | 1 | 0 | 1 |
| AluJb_dup10338 | 7.24739288365211 | -2.47376500204068 | 0.670010135559505 | -3.69213071676134 | 0.000222383108172381 | 0.0176383903523952 | 1572 | 12 | 10 | 8 | 13 | 7 | 8 | 17 | 11 | 4 | 1 | 0 | 2 |
| FLAM_A_dup1854 | 6.03158276547083 | -2.46039679685851 | 0.721197190415333 | -3.41154517732048 | 0.000645957961026855 | 0.0359985079011148 | 1573 | 13 | 7 | 6 | 12 | 7 | 8 | 8 | 10 | 2 | 3 | 0 | 1 |
| L1P1_dup1461 | 8.22111663446853 | -2.4443120784073 | 0.740469416949299 | -3.30103043077424 | 0.000963304338175173 | 0.0466423730071497 | 1574 | 17 | 8 | 10 | 5 | 7 | 13 | 24 | 13 | 5 | 0 | 0 | 3 |
| AluSx3_dup13883 | 5.57926082382405 | -3.47807883831171 | 0.924895688402226 | -3.76050930058951 | 0.000169567769815696 | 0.0145923951379984 | 1575 | 9 | 7 | 13 | 10 | 6 | 4 | 14 | 7 | 1 | 0 | 0 | 2 |
| MER63C_dup792 | 7.13948107095251 | -3.45292266165601 | 0.858131148117224 | -4.02377034003703 | 5.7273770682479e-05 | 0.00669397449229353 | 1576 | 11 | 11 | 18 | 20 | 3 | 5 | 13 | 9 | 1 | 0 | 0 | 3 |
| L1PA5_dup252 | 7.62742500859013 | -2.39942188919352 | 0.66630758099713 | -3.60107247407088 | 0.0003169072131163 | 0.0224625569651984 | 1577 | 12 | 12 | 18 | 16 | 4 | 4 | 18 | 8 | 1 | 0 | 2 | 6 |
| L1PA5_dup7769 | 7.71815033640535 | -2.56240183611158 | 0.657048569026049 | -3.89986670225895 | 9.62456624546953e-05 | 0.00977637103047646 | 1578 | 14 | 5 | 12 | 17 | 8 | 5 | 19 | 13 | 1 | 0 | 2 | 5 |

| | | | | | | | | | | | | | | | | | | | |
|---|---|---|---|---|---|---|---|---|---|---|---|---|---|---|---|---|---|---|---|
| AluJo_dup34815 | 9.1167559742261 | -2.25528697370389 | 0.613462679779684 | -3.67632302345408 | 0.000236619873963011 | 0.0184357408415202 | 1579 | 12 | 11 | 19 | 14 | 8 | 10 | 21 | 12 | 3 | 6 | 1 | 0 |
| AluSq_dup10486 | 9.7765362446402 | -2.75304381432022 | 0.627895562254788 | -4.38455689101222 | 1.16222107221501e-05 | 0.00208713853018319 | 1580 | 10 | 8 | 16 | 18 | 9 | 16 | 31 | 12 | 2 | 2 | 4 | 0 |
| L1P1_dup2044 | 9.25687069177898 | -2.01561473825077 | 0.583933950267281 | -3.45178549273967 | 0.000556890264335558 | 0.032757320946675 | 1581 | 12 | 19 | 12 | 16 | 6 | 7 | 16 | 17 | 3 | 5 | 1 | 3 |
| MIRc_dup3051 | 8.20619538160095 | -1.98405542620927 | 0.595175661857422 | -3.33356276702821 | 0.00085741323541125 | 0.0432372594851866 | 1582 | 17 | 12 | 8 | 16 | 6 | 8 | 10 | 16 | 3 | 3 | 1 | 4 |
| AluJr_dup76137 | 6.97140854255423 | -2.2427561299974 | 0.645403373727732 | -3.47496809172789 | 0.000510914119593221 | 0.0309732410447716 | 1583 | 16 | 13 | 13 | 11 | 6 | 5 | 10 | 8 | 2 | 2 | 2 | 2 |
| HAL1_dup22094 | 7.89903439166547 | -2.12468539349867 | 0.617729481284159 | -3.43950783938918 | 0.000582772906284359 | 0.0337811460277108 | 1584 | 22 | 11 | 14 | 11 | 8 | 5 | 10 | 11 | 1 | 3 | 2 | 4 |
| HAL1M8_dup159 | 8.92555728066418 | -2.13322265366895 | 0.56483239068053 | -3.77673569870659 | 0.00015889718986154 | 0.0139516423390383 | 1585 | 16 | 15 | 9 | 18 | 6 | 9 | 17 | 14 | 2 | 3 | 4 | 2 |
| AluSz6_dup10183 | 8.16263879841206 | -2.18156492290589 | 0.586685952315071 | -3.71845433540279 | 0.000200445517434647 | 0.0164636551920968 | 1586 | 15 | 13 | 11 | 13 | 6 | 8 | 16 | 13 | 1 | 2 | 5 | 2 |
| L1MEd_dup421 | 8.22855772515097 | -2.01173412259884 | 0.607364750122766 | -3.31223391247551 | 0.000925541242703464 | 0.0453986469874909 | 1587 | 18 | 15 | 8 | 11 | 6 | 8 | 17 | 12 | 1 | 3 | 6 | 1 |
| L1PA3_dup6399 | 8.7574918043811 | -1.87636224507649 | 0.559899841136269 | -3.35124625373812 | 0.000804487286523488 | 0.0415354127014618 | 1588 | 20 | 10 | 9 | 18 | 5 | 7 | 21 | 12 | 2 | 3 | 3 | 5 |
| L1MA3_dup1722 | 9.07433542649786 | -2.50573681158091 | 0.596528104706346 | -4.2005343785343 | 2.66285728552275e-05 | 0.00383544939262104 | 1589 | 14 | 18 | 12 | 15 | 8 | 8 | 22 | 12 | 1 | 3 | 2 | 3 |
| MLT1J_dup7116 | 7.29579750735031 | -2.36614959103383 | 0.634979053325612 | -3.72634274885362 | 0.000194278182301588 | 0.0161072182459699 | 1590 | 13 | 10 | 9 | 10 | 8 | 8 | 22 | 8 | 1 | 2 | 2 | 3 |
| L1MA4_dup1054 | 9.32624511877883 | -2.52759274284877 | 0.669656624518499 | -3.77446089578547 | 0.000160354057902073 | 0.0140226411839642 | 1591 | 10 | 9 | 11 | 26 | 7 | 4 | 28 | 18 | 2 | 4 | 1 | 1 |
| L1M5_dup25046 | 6.65205936860595 | -2.63840309308985 | 0.748134597445117 | -3.52664226744761 | 0.000420864987561953 | 0.0271832628475615 | 1592 | 9 | 8 | 13 | 7 | 3 | 5 | 18 | 15 | 1 | 1 | 3 | 1 |
| MIRc_dup41282 | 7.38730389363801 | -3.05246256009868 | 0.824259761806562 | -3.70327741513001 | 0.000212831923486388 | 0.0171263120953304 | 1593 | 13 | 8 | 8 | 23 | 1 | 4 | 18 | 16 | 1 | 1 | 2 | 1 |
| L1ME2_dup3613 | 6.70341434678843 | -2.67604001359654 | 0.777674324339519 | -3.44108057813186 | 0.000579395961322315 | 0.0336666548016604 | 1594 | 14 | 4 | 14 | 8 | 17 | 14 | 7 | 4 | 0 | 1 | 1 | 4 |
| MamRep137_dup487 | 5.58834110659637 | -2.69085597250254 | 0.776967938543365 | -3.46327800545704 | 0.000533636731558616 | 0.0317939826815871 | 1595 | 10 | 7 | 5 | 15 | 10 | 5 | 11 | 6 | 0 | 2 | 1 | 2 |
| AluSp_dup28449 | 6.25009551185025 | -2.5922172417434 | 0.753154435757896 | -3.44181368212333 | 0.000577828092470033 | 0.0336001669821296 | 1596 | 12 | 5 | 10 | 16 | 14 | 3 | 12 | 5 | 1 | 1 | 1 | 3 |
| AluY_dup32090 | 6.38332505853176 | -2.35531823053743 | 0.665599621755548 | -3.53864117940028 | 0.000402192148867612 | 0.0264073769498129 | 1597 | 12 | 6 | 9 | 10 | 13 | 9 | 13 | 5 | 1 | 1 | 2 | 3 |
| AluSx_dup92168 | 4.97768952349145 | -2.80446722636404 | 0.849348045431515 | -3.30190578697243 | 0.000960303192781684 | 0.0465567040565669 | 1598 | 12 | 9 | 3 | 12 | 5 | 8 | 9 | 4 | 1 | 1 | 1 | 1 |
| MIR_dup35171 | 5.81481561604532 | -2.68963177434719 | 0.780354754718226 | -3.44667826790954 | 0.000567524001073121 | 0.0331956897638507 | 1599 | 11 | 7 | 3 | 8 | 12 | 12 | 12 | 6 | 0 | 1 | 3 | 1 |
| MLT1I_dup4630 | 5.50185749707545 | -3.00722977543968 | 0.838330956707374 | -3.58716298304296 | 0.000334295386567671 | 0.0233130861817793 | 1600 | 12 | 9 | 4 | 10 | 8 | 8 | 10 | 7 | 0 | 0 | 2 | 2 |
| L1MA5_dup2508 | 7.08574122841918 | -3.12902015961344 | 0.799962418372147 | -3.91145894825999 | 9.17402514929657e-05 | 0.00945807013940434 | 1601 | 13 | 15 | 9 | 8 | 9 | 8 | 21 | 6 | 0 | 0 | 1 | 4 |
| L1PA7_dup5351 | 6.5742668227715 | -3.03313432587539 | 0.79773734343245 | -3.80217166821429 | 0.000143433282841055 | 0.0129228988712222 | 1602 | 7 | 14 | 9 | 20 | 10 | 6 | 10 | 6 | 0 | 0 | 2 | 3 |
| MLT1D_dup7166 | 5.77291666525164 | -3.16109920833782 | 0.900183700519523 | -3.51161569190094 | 0.000445391556529611 | 0.0282837308006391 | 1603 | 17 | 7 | 13 | 9 | 5 | 9 | 12 | 2 | 0 | 0 | 1 | 3 |
| AluSx1_dup78264 | 6.03278771824822 | -2.7922972188051 | 0.785110736984476 | -3.55656480960891 | 0.000375735818554555 | 0.0252651332902664 | 1604 | 12 | 10 | 10 | 6 | 10 | 8 | 16 | 3 | 0 | 1 | 3 | 1 |
| Kanga2_a_dup519 | 5.35123608667132 | -2.60579946583764 | 0.789074363584522 | -3.30234967208958 | 0.000958784650664131 | 0.0465099767532787 | 1605 | 15 | 10 | 10 | 6 | 6 | 5 | 8 | 5 | 0 | 1 | 2 | 2 |
| LTR12C_dup939 | 6.1564443738355 | -3.1204915438958 | 0.857397649060091 | -3.63949160266137 | 0.000273176845293106 | 0.0203084963943936 | 1606 | 14 | 19 | 11 | 9 | 6 | 8 | 7 | 3 | 0 | 1 | 1 | 2 |
| AluSp_dup21844 | 5.7165589944737 | -4.78241966252362 | 1.11767914157568 | -4.27888423844204 | 1.87832514817272e-05 | 0.00296772878955582 | 1607 | 13 | 12 | 10 | 9 | 9 | 4 | 6 | 9 | 0 | 1 | 0 | 0 |
| MIRb_dup215212 | 6.32815136334892 | -4.2055715729237 | 1.06685989534908 | -3.94200924719139 | 8.08018605004986e-05 | 0.00858708329551212 | 1608 | 15 | 9 | 14 | 13 | 6 | 7 | 10 | 7 | 0 | 2 | 0 | 0 |
| ERV3-16A3_I-int_dup | 4.13624765948148 | -4.33010881831951 | 1.13772377191814 | -3.80594035669939 | 0.000141266425507585 | 0.0127955009074576 | 1609 | 6 | 11 | 6 | 10 | 6 | 5 | 8 | 2 | 0 | 1 | 0 | 0 |
| L1M6_dup1105 | 5.01056106585488 | -3.27378388299175 | 0.989280081710622 | -3.30925886765137 | 0.000935433040180082 | 0.0457144465247499 | 1610 | 5 | 9 | 10 | 11 | 11 | 3 | 8 | 5 | 0 | 3 | 0 | 0 |
| Tigger7_dup622 | 5.41287311258609 | -3.94199158093036 | 1.13702237126393 | -3.46694285051612 | 0.000526413799262164 | 0.0314851555133322 | 1611 | 13 | 3 | 3 | 5 | 18 | 8 | 8 | 9 | 0 | 1 | 0 | 1 |
| LTR12D_dup450 | 4.56100451566078 | -4.44944862922941 | 1.13115947551222 | -3.93352902535213 | 8.37077255237396e-05 | 0.00885240820332078 | 1612 | 6 | 5 | 3 | 11 | 10 | 10 | 7 | 6 | 1 | 0 | 0 | 0 |
| MLT1B-int_dup475 | 4.3719960937307 | -4.29104541327388 | 1.12178787891926 | -3.82518432754675 | 0.000130674277814246 | 0.0121411252279533 | 1613 | 9 | 10 | 2 | 6 | 8 | 11 | 7 | 3 | 1 | 0 | 0 | 0 |
| MER5A_dup2591 | 7.42433801338173 | -2.30127655080366 | 0.631898842019731 | -3.64184327897819 | 0.000270692857020215 | 0.0201913111462483 | 1614 | 13 | 13 | 10 | 9 | 7 | 11 | 12 | 11 | 4 | 0 | 2 | 2 |
| FLAM_A_dup3905 | 5.70935949888396 | -2.67297005103069 | 0.77431766767922 | -3.45203288340571 | 0.000556379915662957 | 0.0327369869961271 | 1615 | 11 | 8 | 9 | 7 | 9 | 7 | 6 | 10 | 1 | 0 | 3 | 1 |
| L1PB1_dup5159 | 6.52304327208003 | -2.63615068327153 | 0.699307777733711 | -3.76965732000673 | 0.000163471841344834 | 0.0142013401740939 | 1616 | 13 | 11 | 10 | 13 | 9 | 8 | 8 | 7 | 1 | 0 | 2 | 3 |
| MER6_dup233 | 7.6212533308576 | -2.50794090317327 | 0.690935462158494 | -3.62977592051568 | 0.000283667364920559 | 0.0208132947312326 | 1617 | 20 | 12 | 12 | 10 | 8 | 17 | 9 | 4 | 4 | 0 | 3 | 0 |
| THE1D-int_dup1882 | 7.93434007437476 | -2.90366145919651 | 0.697745518756288 | -4.16149065976404 | 3.16176932105816e-05 | 0.00435124559206769 | 1618 | 16 | 11 | 10 | 21 | 11 | 11 | 10 | 8 | 1 | 1 | 4 | 0 |
| AluJr_dup72980 | 6.89912719549206 | -2.41199447692799 | 0.70398545219834 | -3.42619947884353 | 0.00061209067691017 | 0.0348265251573571 | 1619 | 8 | 14 | 17 | 13 | 6 | 4 | 12 | 8 | 1 | 4 | 1 | 1 |
| MIR_dup154062 | 6.29298132397054 | -2.30805202437974 | 0.685946644703471 | -3.36476902715588 | 0.000766077597340187 | 0.0402395364186993 | 1620 | 6 | 12 | 13 | 12 | 9 | 9 | 10 | 4 | 1 | 4 | 1 | 1 |
| AluSx1_dup28797 | 5.8317531075278 | -2.65843044225768 | 0.776131545139659 | -3.42523179080334 | 0.00061427510712587 | 0.0348840822530244 | 1621 | 5 | 10 | 13 | 9 | 4 | 8 | 8 | 11 | 2 | 1 | 1 | 1 |
| AluYa5_dup1467 | 6.13705891114826 | -2.22740563689263 | 0.679294378353576 | -3.27899907294693 | 0.00104175978132333 | 0.0491537919743726 | 1622 | 5 | 9 | 11 | 12 | 5 | 12 | 8 | 9 | 2 | 2 | 2 | 1 |
| Charlie4a_dup2446 | 6.45503340206896 | -2.28265416166948 | 0.678824485852629 | -3.36265737203393 | 0.0007719611140095 | 0.0404414485323876 | 1623 | 6 | 9 | 9 | 13 | 8 | 9 | 8 | 12 | 2 | 3 | 2 | 0 |
| THE1D_dup11273 | 7.46737870486725 | -2.1593877586517 | 0.638202065176777 | -3.3835486854051 | 0.000715555197149559 | 0.0383104548611121 | 1624 | 7 | 16 | 8 | 13 | 6 | 11 | 13 | 12 | 1 | 5 | 2 | 1 |
| L1M4b_dup3066 | 7.89490122971522 | -2.47614540942761 | 0.679240948100038 | -3.64545956240395 | 0.000266914385385693 | 0.0199846476856622 | 1625 | 14 | 16 | 21 | 16 | 10 | 8 | 5 | 5 | 1 | 2 | 2 | 3 |
| L1MA9_dup13335 | 7.91501204707748 | -2.33074805905266 | 0.656246131269449 | -3.55163702762565 | 0.000382842589686135 | 0.0255743579015882 | 1626 | 9 | 9 | 16 | 28 | 11 | 9 | 5 | 8 | 1 | 2 | 2 | 4 |
| L2c_dup34366 | 7.48311993637812 | -2.19305663855123 | 0.661143485727362 | -3.31706609214869 | 0.000909680945300985 | 0.044879730916564 | 1627 | 12 | 9 | 17 | 22 | 11 | 11 | 5 | 3 | 2 | 2 | 2 | 3 |
| MER4A1_dup2015 | 7.44984555734143 | -2.11314919115127 | 0.633427503327384 | -3.33605531819127 | 0.000849762531844286 | 0.0429688931647023 | 1628 | 14 | 7 | 17 | 22 | 8 | 9 | 7 | 5 | 1 | 1 | 4 | 4 |
| AluJb_dup53141 | 7.57049865420469 | -2.02054946186924 | 0.600852953794732 | -3.36280191202907 | 0.000771557061491382 | 0.0404291822403938 | 1629 | 13 | 12 | 14 | 15 | 12 | 8 | 14 | 2 | 3 | 3 | 2 | 2 |
| L1MB7_dup19280 | 8.33862399636841 | -2.17702116579384 | 0.577972871887806 | -3.76664939079776 | 0.000165453097109638 | 0.0143159645981273 | 1630 | 11 | 14 | 16 | 11 | 13 | 16 | 9 | 7 | 2 | 5 | 1 | 2 |
| L1PA4_dup2159 | 6.76983512772905 | -2.60995301703253 | 0.779008503902467 | -3.35035240816742 | 0.000807088073103966 | 0.0416114172932681 | 1631 | 8 | 22 | 3 | 7 | 14 | 8 | 12 | 7 | 1 | 1 | 0 | 4 |
| L1MB7_dup1121 | 7.95745988696936 | -4.54660475467648 | 1.10394613825536 | -4.1185023409401 | 3.81342696715043e-05 | 0.00496383858383795 | 1632 | 10 | 11 | 6 | 5 | 17 | 15 | 23 | 13 | 0 | 1 | 0 | 1 |
| L2a_dup34793 | 8.62923097722518 | -3.29292977510078 | 0.805675474376487 | -4.08716645824323 | 4.36673816940444e-05 | 0.00545964956162073 | 1633 | 13 | 9 | 4 | 6 | 25 | 19 | 15 | 13 | 2 | 2 | 0 | 1 |
| AluSx_dup3872 | 6.66066086901917 | -2.87111502037623 | 0.812117460776167 | -3.53534452717247 | 0.000407243811350115 | 0.0266068193123324 | 1634 | 21 | 17 | 8 | 3 | 6 | 8 | 10 | 8 | 0 | 1 | 2 | 2 |
| MER9a1_dup18 | 6.72793561411761 | -2.35202974358641 | 0.717216693213992 | -3.2793851089083 | 0.00104033561180084 | 0.0491158129931673 | 1635 | 23 | 13 | 4 | 8 | 7 | 9 | 10 | 7 | 0 | 2 | 4 | 1 |
| L2c_dup126065 | 5.5215742348682 | -2.99337363048718 | 0.871243035625074 | -3.43575042564281 | 0.000590915055288792 | 0.0340697537547242 | 1636 | 9 | 6 | 7 | 5 | 5 | 12 | 14 | 9 | 0 | 1 | 3 | 0 |
| MER58A_dup5452 | 5.46512743433339 | -2.6422222490868 | 0.786783928137047 | -3.35825651058109 | 0.000784357898350327 | 0.0409505595573397 | 1637 | 11 | 11 | 7 | 8 | 3 | 9 | 10 | 7 | 0 | 1 | 3 | 1 |
| L1PA6_dup2024 | 6.66522400523944 | -2.90179396677522 | 0.804091967734344 | -3.60878367551846 | 0.000307635998113612 | 0.0220351460587422 | 1638 | 13 | 13 | 4 | 3 | 12 | 11 | 14 | 10 | 0 | 3 | 1 | 1 |
| L2a_dup105772 | 6.40458414182771 | -2.60314956375624 | 0.793433608033109 | -3.28086627211235 | 0.00103488798386159 | 0.0489654913915154 | 1639 | 12 | 10 | 9 | 3 | 8 | 9 | 13 | 11 | 0 | 5 | 1 | 0 |
| AluJb_dup19335 | 7.18860086481472 | -2.67256450344114 | 0.777247062744865 | -3.43850061523928 | 0.000584945208484658 | 0.0338403531675396 | 1640 | 13 | 21 | 8 | 3 | 17 | 4 | 8 | 10 | 1 | 2 | 2 | 1 |
| MER5C_dup909 | 7.81131803580938 | -2.81091418287301 | 0.766019329045248 | -3.6695081655139 | 0.000243017575773484 | 0.0187554191978292 | 1641 | 22 | 21 | 5 | 8 | 10 | 5 | 9 | 13 | 1 | 1 | 3 | 1 |
| Harlequin-int_dup474 | 6.43552027125875 | -2.45489369544442 | 0.741890528388785 | -3.30897026111909 | 0.000936397828424303 | 0.0457331932845339 | 1642 | 21 | 15 | 7 | 2 | 9 | 10 | 7 | 6 | 1 | 1 | 1 | 3 |
| L2b_dup28626 | 6.32522358198635 | -3.10151674924701 | 0.815999534230668 | -3.80088053870171 | 0.00014418280599413 | 0.0129658813133331 | 1643 | 18 | 15 | 7 | 5 | 9 | 11 | 8 | 5 | 1 | 1 | 1 | 1 |
| Tigger1_dup9404 | 7.43813481663891 | -2.75828977556067 | 0.693426931339221 | -3.97776557399286 | 6.95659056636613e-05 | 0.0077205825508608 | 1644 | 23 | 11 | 9 | 7 | 10 | 13 | 11 | 7 | 1 | 1 | 2 | 2 |
| AluJr_dup14207 | 6.88922704385824 | -2.40979496306034 | 0.67507944496869 | -3.56964647793728 | 0.00035746329856782 | 0.0244410279552528 | 1645 | 15 | 11 | 11 | 6 | 8 | 11 | 7 | 11 | 1 | 2 | 3 | 1 |
| AluSz_dup7080 | 6.24380810533499 | -2.48856498105858 | 0.69257872908363 | -3.59318713751326 | 0.000326657808431189 | 0.0229418296590814 | 1646 | 14 | 9 | 7 | 10 | 9 | 10 | 7 | 8 | 2 | 1 | 2 | 1 |
| AluSz6_dup9130 | 7.15754453701545 | -2.51304629459375 | 0.747797660071908 | -3.36059662763868 | 0.000777743202687525 | 0.0406906077631235 | 1647 | 12 | 8 | 4 | 3 | 11 | 14 | 19 | 13 | 1 | 1 | 3 | 2 |
| Tigger2a_dup694 | 7.28322395296392 | -2.73946238230811 | 0.737340072785264 | -3.71533093537143 | 0.000202937916847619 | 0.0165996100248276 | 1648 | 12 | 12 | 5 | 5 | 17 | 9 | 20 | 8 | 2 | 1 | 1 | 2 |
| L1MC1_dup1269 | 6.50597656593907 | -2.32755695007452 | 0.701794698859361 | -3.31657813012487 | 0.000911271037292586 | 0.04493023171484 | 1649 | 8 | 3 | 8 | 10 | 13 | 9 | 11 | 13 | 1 | 3 | 2 | 1 |
| Tigger15a_dup1039 | 8.30139402575918 | -2.29758569699742 | 0.651525750784594 | -3.52646951287893 | 0.000421139648566477 | 0.0271936213888193 | 1650 | 9 | 13 | 7 | 6 | 10 | 19 | 13 | 17 | 2 | 3 | 3 | 1 |

| | | | | | | | | | | | | | | | | | | | |
|---|---|---|---|---|---|---|---|---|---|---|---|---|---|---|---|---|---|---|---|
| L1PA15_dup143 | 5.26302305138191 | -4.42369984499303 | 1.20169029948838 | -3.6812312181237 | 0.000232110392623995 | 0.0181586635340565 | 1651 | 18 | 26 | 6 | 2 | 4 | 4 | 4 | 3 | 0 | 1 | 0 | 0 |
| L1ME3C | 5.2856947983882 | -4.48224402850964 | 1.26605217916415 | -3.54033119825191 | 0.000399625185217132 | 0.0262895256626142 | 1652 | 25 | 22 | 2 | 5 | 1 | 4 | 4 | 5 | 0 | 0 | 1 | 0 |
| MSTB_dup7397 | 3.39041731815508 | -4.59782423824963 | 1.35041427873505 | -3.40475090544545 | 0.000662244316897325 | 0.0365256185677352 | 1653 | 14 | 16 | 3 | 2 | 0 | 3 | 3 | 3 | 0 | 0 | 0 | 0 |
| LTR12C_dup1462 | 3.16663032289023 | -4.60840655849268 | 1.34940225561631 | -3.4151466245978 | 0.000637476859411763 | 0.0356738167426132 | 1654 | 8 | 10 | 6 | 1 | 6 | 2 | 0 | 6 | 0 | 0 | 0 | 0 |
| AluSc5_dup853 | 3.33845236557028 | -4.69997827857701 | 1.29200508474194 | -3.63773976904724 | 0.000275041109514055 | 0.0203791964242417 | 1655 | 4 | 16 | 3 | 1 | 3 | 3 | 4 | 7 | 0 | 0 | 0 | 0 |
| MER67A_dup190 | 2.81365944037356 | -4.47332889657289 | 1.30375654275196 | -3.43110753418013 | 0.00060112225850264 | 0.0344495753664906 | 1656 | 4 | 11 | 1 | 1 | 4 | 2 | 6 | 6 | 0 | 0 | 0 | 0 |
| MIRc_dup94807 | 3.72437672986154 | -4.15003997351619 | 1.25879743683974 | -3.29682906245428 | 0.000977829939790262 | 0.0470898042609459 | 1657 | 9 | 11 | 8 | 0 | 3 | 4 | 5 | 6 | 0 | 0 | 0 | 1 |
| AluJb_dup113619 | 3.16793520143681 | -4.52146988327119 | 1.20577970014578 | -3.74983082127237 | 0.0001769539116311 | 0.015075358201566 | 1658 | 8 | 7 | 4 | 0 | 8 | 7 | 5 | 2 | 0 | 0 | 0 | 0 |
| L1MC5_dup11567 | 3.13507277224636 | -4.52841600952651 | 1.27599895048363 | -3.54891828697049 | 0.000386817096057147 | 0.02576401223807 | 1659 | 6 | 8 | 1 | 0 | 6 | 8 | 4 | 6 | 0 | 0 | 0 | 0 |
| L1PB1_dup6637 | 5.31953781554903 | -4.68522575922979 | 1.38890799238979 | -3.37331614829884 | 0.000742686254952464 | 0.039331664037174 | 1660 | 31 | 29 | 0 | 3 | 2 | 2 | 1 | 2 | 0 | 0 | 0 | 0 |
| L1P1_dup1713 | 5.69543993302456 | -5.15104453910704 | 1.39092294717577 | -3.70332846227469 | 0.00021278908284526 | 0.0171263120953304 | 1661 | 20 | 32 | 0 | 1 | 8 | 1 | 9 | 3 | 0 | 0 | 0 | 0 |
| L1PA15_dup2718 | 5.65789346979442 | -4.66535940400113 | 1.32969028223567 | -3.50860607641432 | 0.00045046155904974 | 0.028498300031631 | 1662 | 26 | 30 | 0 | 0 | 9 | 3 | 2 | 3 | 0 | 0 | 0 | 0 |
| L1PA3_dup2919 | 6.47724243314523 | -3.42501651890322 | 1.00203991257445 | -3.41804400795137 | 0.000630729076996442 | 0.035463010622636 | 1663 | 31 | 20 | 2 | 0 | 7 | 12 | 7 | 3 | 0 | 2 | 1 | 0 |
| LTR12C_dup1391 | 4.66606673873167 | -3.57283979627866 | 1.02126413023886 | -3.49844833524411 | 0.000467973746214576 | 0.0291804195536618 | 1664 | 12 | 13 | 5 | 3 | 9 | 10 | 3 | 3 | 1 | 0 | 0 | 1 |
| MER5A_dup12151 | 5.94221226734572 | -2.8620756598525 | 0.860979004515935 | -3.3242107471153 | 0.000886691619442053 | 0.0440929412914856 | 1665 | 19 | 19 | 2 | 1 | 14 | 11 | 2 | 3 | 4 | 0 | 0 | 0 |
| L2b_dup86267 | 2.49904732082638 | -4.24467287770893 | 1.28681240372373 | -3.29859493538129 | 0.000971700164206321 | 0.0468800014745007 | 1666 | 0 | 3 | 5 | 10 | 4 | 3 | 2 | 5 | 0 | 0 | 0 | 0 |
| L1MC4a_dup8006 | 3.07347568183366 | -4.63181037634838 | 1.27257587671767 | -3.63971253980951 | 0.000272942571474487 | 0.0203081904665629 | 1667 | 5 | 3 | 4 | 9 | 7 | 5 | 0 | 6 | 0 | 0 | 0 | 0 |
| L1PA15_dup7161 | 3.11975695959253 | -4.64852467607304 | 1.35473945752607 | -3.43130529656371 | 0.000600684161024467 | 0.034441058645573 | 1668 | 1 | 4 | 12 | 8 | 7 | 5 | 0 | 3 | 0 | 0 | 0 | 0 |
| AluSp_dup45070 | 2.49390554973363 | -4.08730915907772 | 1.20752372056313 | -3.38486862781594 | 0.000712123281248733 | 0.0382401208062598 | 1669 | 5 | 7 | 2 | 2 | 4 | 9 | 1 | 2 | 0 | 0 | 0 | 0 |
| L1PB3_dup2669 | 2.79849860578755 | -4.47105371711233 | 1.18680299189865 | -3.76730910490841 | 0.000165016633757133 | 0.0143094198017401 | 1670 | 6 | 8 | 5 | 5 | 3 | 6 | 3 | 1 | 0 | 0 | 0 | 0 |
| MSTA_dup14106 | 3.02634468672958 | -4.32794236788775 | 1.23627645834883 | -3.50078846738547 | 0.000463883891637343 | 0.0290242746462107 | 1671 | 12 | 11 | 2 | 4 | 2 | 7 | 1 | 1 | 0 | 0 | 0 | 0 |
| L1MB8_dup400 | 2.90944656089644 | -4.42916205725393 | 1.22951728316769 | -3.6023585173547 | 0.00031534303928963 | 0.022398392485615 | 1672 | 13 | 8 | 2 | 2 | 3 | 4 | 6 | 1 | 0 | 0 | 0 | 0 |
| MER5B_dup16280 | 3.49059609583946 | -4.61994050423325 | 1.19180140461634 | -3.87643485427883 | 0.000105998173099387 | 0.0104714866477318 | 1673 | 17 | 5 | 1 | 4 | 8 | 5 | 3 | 3 | 0 | 0 | 0 | 0 |
| L1MCc_dup2421 | 3.99575194410174 | -4.26166790334529 | 1.2153304150182 | -3.5065919939817 | 0.00045388451354019 | 0.0285741803351622 | 1674 | 12 | 10 | 8 | 9 | 5 | 2 | 1 | 4 | 0 | 0 | 1 | 0 |
| L1ME3Cz_dup2329 | 3.23508852702098 | -4.65838632074735 | 1.25978196631084 | -3.69777187269083 | 0.000217500218628309 | 0.0173785450039394 | 1675 | 12 | 11 | 3 | 6 | 3 | 1 | 2 | 4 | 0 | 0 | 0 | 0 |
| L1ME3E_dup2329 | 3.23670041260692 | -4.6869070830719 | 1.1808807364828 | -3.96899275114914 | 7.21770772162958e-05 | 0.00793446039108713 | 1676 | 5 | 10 | 4 | 8 | 6 | 4 | 2 | 3 | 0 | 0 | 0 | 0 |
| AluSg_dup36554 | 3.20797321923131 | -4.70720951803064 | 1.20979833257981 | -3.89090428649613 | 9.98713251948227e-05 | 0.0100609578089785 | 1677 | 3 | 11 | 7 | 7 | 5 | 2 | 5 | 2 | 0 | 0 | 0 | 0 |
| L1MC3_dup1230 | 2.75718314993801 | -4.46452050201225 | 1.22438333624039 | -3.64634209717447 | 0.000265999804343827 | 0.0199287120416864 | 1678 | 5 | 9 | 4 | 6 | 6 | 1 | 3 | 2 | 0 | 0 | 0 | 0 |
| L1MC3_dup8882 | 3.00704306070215 | -4.56860175575354 | 1.19451343649766 | -3.82465497344992 | 0.000130955351397512 | 0.0121624910396692 | 1679 | 5 | 8 | 5 | 5 | 9 | 2 | 3 | 2 | 0 | 0 | 0 | 0 |
| AluSq2_dup36062 | 3.06846691169158 | -3.88467088109929 | 1.1484374276876 | -3.38257077612068 | 0.000718107712799124 | 0.0383806958287737 | 1680 | 4 | 6 | 7 | 6 | 5 | 4 | 4 | 3 | 0 | 0 | 0 | 1 |
| MLT1B_dup3336 | 3.20160190602549 | -3.90675772593587 | 1.19112343815312 | -3.27989325102479 | 0.00103846371295312 | 0.0490568324622138 | 1681 | 8 | 7 | 5 | 7 | 8 | 3 | 1 | 2 | 0 | 0 | 0 | 1 |
| AluSz_dup85003 | 3.32404476730924 | -4.01170200082382 | 1.15946065629012 | -3.45997251313374 | 0.000540230530373757 | 0.0321043191259526 | 1682 | 6 | 4 | 6 | 8 | 2 | 7 | 9 | 2 | 0 | 0 | 0 | 1 |
| MER57A-int_dup1376 | 4.22433040709642 | -4.32539039094697 | 1.18894977883582 | -3.63799251065278 | 0.000274771413196015 | 0.0203791964242417 | 1683 | 9 | 2 | 15 | 12 | 3 | 7 | 6 | 2 | 0 | 0 | 0 | 1 |
| AluSz_dup98539 | 4.90168341693017 | -3.34085192809757 | 0.974601916493941 | -3.42791438387074 | 0.000608237248150271 | 0.0347151883541292 | 1684 | 13 | 5 | 7 | 12 | 5 | 7 | 7 | 6 | 0 | 0 | 0 | 3 |
| L1MA3_dup6010 | 4.14241683105858 | -4.30302216845737 | 1.14090129473142 | -3.77159898786015 | 0.000162204786654107 | 0.0141429022224984 | 1685 | 10 | 6 | 9 | 6 | 3 | 4 | 6 | 8 | 0 | 0 | 0 | 1 |
| MIR_dup172248 | 4.07985741845504 | -4.3038612257047 | 1.17516895136186 | -3.66233401649791 | 0.000249927689041566 | 0.0191398102915848 | 1686 | 12 | 7 | 10 | 8 | 5 | 3 | 2 | 5 | 0 | 0 | 0 | 1 |
| AluSc_dup4085 | 3.24993510126796 | -3.96295296249226 | 1.17527102545712 | -3.37194815208763 | 0.000746384979245958 | 0.0394840640888525 | 1687 | 6 | 4 | 2 | 3 | 5 | 5 | 10 | 6 | 0 | 0 | 0 | 1 |
| AluSx3_dup6088 | 3.88030684809615 | -4.16934348492949 | 1.122429370015 | -3.71457090870117 | 0.000203548788932153 | 0.0166259449141726 | 1688 | 11 | 8 | 4 | 4 | 6 | 8 | 6 | 3 | 0 | 0 | 0 | 1 |
| ERVL-B4-int_dup519 | 2.90499570213988 | -4.57387466045656 | 1.22391683528713 | -3.73707961896246 | 0.000186169985875598 | 0.0156365702291405 | 1689 | 8 | 3 | 2 | 11 | 2 | 4 | 8 | 2 | 0 | 0 | 0 | 0 |
| MER103C_dup7113 | 2.74959034901548 | -4.48753217105307 | 1.21237567827295 | -3.7014369815185 | 0.000214381904234174 | 0.0172161041461036 | 1690 | 8 | 2 | 6 | 4 | 4 | 2 | 9 | 2 | 0 | 0 | 0 | 0 |
| AluJb_dup91119 | 2.28480875798535 | -4.19892723359523 | 1.25299490629533 | -3.35111277188668 | 0.000804875178695834 | 0.0415354127014618 | 1691 | 7 | 4 | 4 | 2 | 1 | 2 | 4 | 5 | 0 | 0 | 0 | 0 |
| MER5A_dup24420 | 2.4862027755585 | -4.31711283053348 | 1.21387727480939 | -3.55646564947134 | 0.000375877601840515 | 0.0252651332902664 | 1692 | 7 | 3 | 6 | 5 | 1 | 2 | 6 | 3 | 0 | 0 | 0 | 0 |
| MLT1C_dup17513 | 3.15469503743932 | -4.60307928136434 | 1.20472472532838 | -3.82085565655642 | 0.000132989473447319 | 0.0122852285749337 | 1693 | 4 | 4 | 11 | 3 | 1 | 5 | 6 | 6 | 0 | 0 | 0 | 0 |
| AluJr_dup80270 | 2.42208667805838 | -4.21659389176462 | 1.1816156956564 | -3.5684985459018 | 0.000359032846076468 | 0.0245137531111941 | 1694 | 2 | 3 | 6 | 6 | 2 | 3 | 7 | 3 | 0 | 0 | 0 | 0 |
| AluSx_dup58014 | 2.69602719821103 | -4.44779458765137 | 1.17497107954937 | -3.78545026772677 | 0.000153430517149772 | 0.0135668096664213 | 1695 | 3 | 4 | 7 | 4 | 3 | 6 | 5 | 3 | 0 | 0 | 0 | 0 |
| L1PB3_dup2670 | 3.21844444620101 | -3.93970861382114 | 1.18656843444022 | -3.3202540194656 | 0.000899355752155231 | 0.044518016185215 | 1696 | 9 | 6 | 5 | 2 | 2 | 4 | 6 | 6 | 0 | 0 | 1 | 0 |
| MLT1D_dup1362 | 2.72591600927455 | -4.40021245160588 | 1.25071273957569 | -3.51816393354935 | 0.00043454382939068 | 0.0278026259823057 | 1697 | 9 | 4 | 4 | 2 | 1 | 9 | 2 | 4 | 0 | 0 | 0 | 0 |
| AluSx3_dup27241 | 3.3891121991174 | -4.76169945726843 | 1.30901932731175 | -3.63760821396521 | 0.000275181587885859 | 0.0203791964242417 | 1698 | 13 | 6 | 8 | 8 | 2 | 6 | 0 | 2 | 0 | 0 | 0 | 0 |
| L2b_dup73439 | 3.86934634460254 | -4.83190489935797 | 1.39749513904679 | -3.45754683816197 | 0.000545117474532061 | 0.0322898394372453 | 1699 | 7 | 4 | 10 | 26 | 2 | 1 | 0 | 3 | 0 | 0 | 0 | 0 |
| Arthur1B_dup886 | 6.80391747433031 | -5.79302363846171 | 1.28738255495146 | -4.49984630922713 | 6.80026105585405e-06 | 0.00138180607787895 | 1700 | 11 | 7 | 23 | 24 | 16 | 8 | 1 | 1 | 0 | 0 | 0 | 0 |
| L1PA16_dup11701 | 4.99296262747666 | -4.59640403960001 | 1.26550698148535 | -3.632065335748 | 0.0002811619090277 | 0.0206868743368909 | 1701 | 11 | 12 | 20 | 12 | 4 | 2 | 3 | 1 | 1 | 0 | 0 | 0 |
| AluSq2_dup55381 | 4.6185537902711 | -5.21794965158222 | 1.13887010106282 | -4.58168991065153 | 4.6123347467503e-06 | 0.00103050385319272 | 1702 | 8 | 9 | 14 | 8 | 6 | 3 | 7 | 5 | 0 | 0 | 0 | 0 |
| L1PB4_dup6132 | 4.9323953588839 | -5.25598102824824 | 1.22195843685962 | -4.30127643437356 | 1.69817051505734e-05 | 0.00274424780358399 | 1703 | 10 | 6 | 22 | 14 | 4 | 2 | 3 | 4 | 0 | 0 | 0 | 0 |
| AluSp_dup50605 | 2.34392823454217 | -4.12115503940819 | 1.25739140763067 | -3.27754350347739 | 0.00104714593043624 | 0.0492906400719254 | 1704 | 3 | 6 | 3 | 2 | 4 | 9 | 4 | 0 | 0 | 0 | 0 | 0 |
| L1M5_dup10592 | 2.72528579865788 | -4.41605066096285 | 1.24496853207194 | -3.54711829833437 | 0.000389469665947114 | 0.0258485014139323 | 1705 | 8 | 6 | 3 | 6 | 2 | 8 | 4 | 0 | 0 | 0 | 0 | 0 |
| L1MC1_dup2514 | 3.81058154596478 | -4.15444695587468 | 1.24199635338411 | -3.34497516402114 | 0.000822899387952939 | 0.0421177925932136 | 1706 | 11 | 14 | 4 | 11 | 2 | 5 | 2 | 1 | 0 | 0 | 1 | 0 |
| L1MCa_dup603 | 3.9871393870123 | -4.11640672850396 | 1.24918403084643 | -3.29527645795692 | 0.000983248969936874 | 0.0472676432766945 | 1707 | 14 | 16 | 6 | 6 | 3 | 5 | 2 | 0 | 0 | 0 | 1 | 0 |
| L1M1_dup823 | 3.84054472244023 | -4.45186927075992 | 1.30878949281789 | -3.40151666497171 | 0.000670130426032976 | 0.0367831267249925 | 1708 | 16 | 15 | 1 | 6 | 7 | 6 | 0 | 0 | 0 | 0 | 0 | 0 |
| LTR49_dup310 | 5.18186125928678 | -5.19140347518295 | 1.20856357524685 | -4.29551542137333 | 1.74287920241291e-05 | 0.00280210226146854 | 1709 | 23 | 12 | 6 | 9 | 12 | 5 | 1 | 1 | 0 | 0 | 0 | 0 |
| FLAM_C_dup21363 | 8.97123625019197 | -5.99908890195321 | 1.41958460626305 | -4.22594671390905 | 2.37938296177147e-05 | 0.00351967513891115 | 1710 | 44 | 43 | 7 | 4 | 0 | 1 | 10 | 8 | 0 | 0 | 0 | 0 |
| MER2_dup4902 | 5.60402400902161 | -5.51014501650726 | 1.41988149592402 | -3.88070767336918 | 0.000104152967139216 | 0.0103390829866115 | 1711 | 21 | 15 | 7 | 14 | 0 | 0 | 9 | 8 | 0 | 0 | 0 | 0 |
| MSTB_dup1026 | 4.9633656006181 | -4.58630681909874 | 1.34893536747031 | -3.39994556425601 | 0.000673992697303511 | 0.0369135431765362 | 1712 | 10 | 24 | 7 | 6 | 0 | 1 | 12 | 4 | 0 | 0 | 1 | 0 |
| L1PA13_dup4188 | 10.8666555034313 | -3.65508057309216 | 1.11521877837563 | -3.27745608661284 | 0.00104747022417251 | 0.04929615307772 | 1713 | 50 | 43 | 8 | 4 | 1 | 1 | 21 | 10 | 0 | 0 | 2 | 3 |
| LTR8B_dup31 | 6.0305329181389 | -3.99356429046396 | 1.19015454287665 | -3.3555006065106 | 0.000792214827939013 | 0.041185132062219 | 1714 | 28 | 20 | 9 | 4 | 7 | 1 | 7 | 2 | 0 | 0 | 1 | 1 |
| MLT1J1_dup608 | 5.19518359629384 | -4.56279818161051 | 1.299579334408 | -3.51098086958193 | 0.000446456526046997 | 0.0283437952366005 | 1715 | 24 | 16 | 2 | 4 | 7 | 0 | 11 | 4 | 0 | 0 | 0 | 1 |
| ERVL-E-int_dup6260 | 6.89448016619233 | -4.35936846763848 | 1.30993535593138 | -3.3279264109479 | 0.000874949750797523 | 0.0437467521334876 | 1716 | 22 | 19 | 19 | 21 | 2 | 0 | 3 | 4 | 0 | 1 | 0 | 1 |
| MER39_dup2366 | 4.71953704743458 | -5.24009803531877 | 1.33771840017221 | -3.91719066930991 | 8.95868901857841e-05 | 0.00930054195992434 | 1717 | 16 | 17 | 8 | 13 | 4 | 0 | 3 | 2 | 0 | 0 | 0 | 0 |
| L1MEg_dup11516 | 6.92651832449367 | -4.3068507435893 | 1.20571761622635 | -3.5720227403402 | 0.000354234643630404 | 0.0243047048769551 | 1718 | 18 | 8 | 27 | 26 | 0 | 4 | 6 | 3 | 0 | 1 | 0 | 1 |
| L1MC4_dup19131 | 8.48275252061684 | -5.37937303589401 | 1.4264177113432 | -3.77124666436486 | 0.000162434010427389 | 0.0141577007491412 | 1719 | 37 | 20 | 17 | 34 | 0 | 1 | 3 | 3 | 0 | 0 | 1 | 0 |
| L1PA10_dup2089 | 4.74554501765246 | -5.2594284082833 | 1.49606677189259 | -3.51550379107071 | 0.000438920493237772 | 0.0279998326230442 | 1720 | 18 | 17 | 11 | 15 | 0 | 1 | 1 | 1 | 0 | 0 | 0 | 0 |
| L1PA4_dup8278 | 6.78549323127662 | -5.0524073068534 | 1.44477955383658 | -3.49700914124708 | 0.000470505709402148 | 0.0292836658563152 | 1721 | 23 | 18 | 17 | 28 | 0 | 1 | 1 | 3 | 0 | 0 | 0 | 1 |
| THE1D_dup5996 | 6.6209962955456 | -4.14626724940125 | 1.09637158632523 | -3.78180837693772 | 0.000155693194669145 | 0.013736215387002 | 1722 | 7 | 8 | 5 | 9 | 1 | 9 | 31 | 14 | 1 | 1 | 0 | 0 |

| | | | | | | | | | | | | | | | | | | | |
|---|---|---|---|---|---|---|---|---|---|---|---|---|---|---|---|---|---|---|---|
| AluSx3_dup10115 | 3.93032280912436 | -4.23176224996619 | 1.19495579070899 | -3.54135465334278 | 0.000398078112826867 | 0.026216743602187 | 1723 | 8 | 7 | 5 | 3 | 1 | 6 | 14 | 6 | 1 | 0 | 0 | 0 |
| L1MEb_dup745 | 4.78459438170955 | -5.23450335974421 | 1.2237242189349 | -4.27751880591215 | 1.88987996711577e-05 | 0.00298400383978321 | 1724 | 5 | 11 | 5 | 3 | 2 | 5 | 22 | 9 | 0 | 0 | 0 | 0 |
| LTR82A_dup507 | 5.42996423663829 | -4.63868606414534 | 1.18649887802139 | -3.90955790188426 | 9.2465190660744e-05 | 0.00951391693553835 | 1725 | 8 | 6 | 4 | 4 | 4 | 5 | 26 | 12 | 1 | 0 | 0 | 0 |
| MIR3_dup68043 | 4.84230904269293 | -3.7065027336098 | 1.11116402032691 | -3.33569362020861 | 0.000850868796291226 | 0.0429944418352755 | 1726 | 11 | 6 | 9 | 6 | 2 | 1 | 12 | 12 | 1 | 1 | 0 | 0 |
| MLT1A0_dup19138 | 6.73994656777182 | -3.67136630734491 | 1.12015264330575 | -3.27755893742313 | 0.00104708868417827 | 0.0492906400719254 | 1727 | 19 | 14 | 10 | 5 | 1 | 1 | 22 | 12 | 1 | 2 | 0 | 0 |
| MER67D_dup884 | 6.26210299952828 | -4.10895185458655 | 1.17311736650174 | -3.50259229972831 | 0.000460754116493326 | 0.0289044553132405 | 1728 | 14 | 11 | 5 | 7 | 2 | 2 | 20 | 16 | 2 | 0 | 0 | 0 |
| MIRb_dup175058 | 7.62640043095133 | -5.20347282404533 | 1.18546195946495 | -4.38940514497309 | 1.13661141826281e-05 | 0.00206137302233203 | 1729 | 18 | 22 | 7 | 5 | 7 | 3 | 17 | 16 | 1 | 0 | 0 | 0 |
| AluJb_dup51488 | 9.36636124453507 | -4.7964428338503 | 1.13972903134441 | -4.20840629828694 | 2.57178133996024e-05 | 0.00374212319188565 | 1730 | 12 | 24 | 6 | 8 | 3 | 12 | 42 | 14 | 0 | 1 | 0 | 1 |
| L1MEc_dup8482 | 6.67819109085707 | -4.96847429138115 | 1.17333577413922 | -4.23448632598465 | 2.29074722494405e-05 | 0.00342452650816728 | 1731 | 13 | 9 | 4 | 8 | 3 | 6 | 26 | 16 | 0 | 1 | 0 | 0 |
| MIRc_dup79635 | 6.1521071247751 | -4.11895194255107 | 1.12632920922916 | -3.65696983510711 | 0.000255214400800317 | 0.0193892866579924 | 1732 | 8 | 14 | 4 | 9 | 3 | 5 | 21 | 13 | 0 | 2 | 0 | 0 |
| MER57-int_dup1031 | 6.13164750311421 | -3.57758226318383 | 1.03497734635567 | -3.45667687875595 | 0.000546880170333562 | 0.032337959495574 | 1733 | 12 | 14 | 5 | 3 | 4 | 5 | 19 | 13 | 0 | 2 | 0 | 1 |
| UCON26_dup159 | 6.82523435336726 | -3.74631937203485 | 1.03254007593432 | -3.62825565743287 | 0.00028534262881309 | 0.0208654385849906 | 1734 | 20 | 13 | 3 | 8 | 3 | 6 | 18 | 14 | 0 | 2 | 0 | 1 |
| L3_dup27248 | 6.41501363409251 | -3.61914406278236 | 1.01673239087951 | -3.55958371666677 | 0.000371443113909891 | 0.0250661948900746 | 1735 | 16 | 19 | 5 | 9 | 1 | 8 | 10 | 11 | 1 | 1 | 0 | 1 |
| MIRb_dup175060 | 5.47923507063862 | -4.00749583705711 | 1.15795536138154 | -3.46083793098538 | 0.000538496897220185 | 0.0320413060029893 | 1736 | 10 | 18 | 6 | 11 | 1 | 4 | 14 | 6 | 1 | 0 | 0 | 1 |
| AluSz_dup75770 | 4.32389767522205 | -3.64326830647638 | 1.06905793582108 | -3.40792410252136 | 0.00065459099031795 | 0.0362837448787836 | 1737 | 7 | 8 | 5 | 9 | 2 | 5 | 10 | 8 | 0 | 1 | 0 | 1 |
| L2a_dup23213 | 5.40125500374725 | -4.70714232052771 | 1.21247343532878 | -3.88226428998118 | 0.00010348831185283 | 0.0103120186895199 | 1738 | 12 | 16 | 7 | 13 | 1 | 2 | 9 | 9 | 0 | 1 | 0 | 0 |
| L1MA7_dup3679 | 6.49067375652944 | -4.27400959970743 | 1.13273626855518 | -3.77317273080608 | 0.000161184611247392 | 0.0140745811038062 | 1739 | 9 | 12 | 4 | 11 | 3 | 1 | 15 | 23 | 0 | 0 | 0 | 2 |
| AluY_dup81169 | 5.24659375558073 | -3.95487059216819 | 1.1655369567754 | -3.39317476737057 | 0.000690875446805854 | 0.0374723157204618 | 1740 | 12 | 15 | 2 | 5 | 6 | 4 | 11 | 10 | 0 | 0 | 0 | 2 |
| L1MA10_dup2100 | 5.3943237427226 | -3.41683049447968 | 1.0305528676357 | -3.31553149943545 | 0.000914690319857665 | 0.0450428192981895 | 1741 | 12 | 15 | 4 | 5 | 6 | 2 | 15 | 8 | 0 | 1 | 0 | 2 |
| L1ME1_dup25940 | 7.96994594872624 | -5.56764371174329 | 1.28469010322832 | -4.33384183294652 | 1.46529468978794e-05 | 0.00247628558420654 | 1742 | 7 | 4 | 5 | 3 | 1 | 4 | 41 | 33 | 0 | 0 | 0 | 0 |
| Tigger1_dup6128 | 6.526774583408 | -5.50635358567315 | 1.32652091577135 | -4.15097381443948 | 3.31063645574768e-05 | 0.00448606322380033 | 1743 | 6 | 10 | 2 | 3 | 1 | 3 | 35 | 22 | 0 | 0 | 0 | 0 |
| L1MC4_dup11020 | 5.08956758528161 | -5.30485595490709 | 1.3307597331771 | -3.98633639315347 | 6.7101359461915e-05 | 0.00752072542558871 | 1744 | 11 | 8 | 3 | 1 | 1 | 4 | 15 | 19 | 0 | 0 | 0 | 0 |
| LTR12C_dup118 | 6.35983525693194 | -4.08677911309284 | 1.20274235151477 | -3.39788410040257 | 0.000679091833310888 | 0.0370953134455906 | 1745 | 10 | 5 | 2 | 1 | 3 | 6 | 23 | 25 | 1 | 0 | 1 | 0 |
| L1M3_dup2724 | 5.49862754281892 | -4.54203447981694 | 1.11953729524464 | -4.05706402020704 | 4.96934637772842e-05 | 0.00600221367104044 | 1746 | 5 | 5 | 6 | 12 | 7 | 1 | 21 | 13 | 0 | 0 | 1 | 0 |
| L1MA3_dup1979 | 6.55698834543354 | -4.65621191886466 | 1.11807633923186 | -4.16448479901073 | 3.12056415705774e-05 | 0.00430449715313195 | 1747 | 10 | 2 | 7 | 13 | 3 | 1 | 30 | 18 | 0 | 0 | 1 | 0 |
| MLT1C_dup8341 | 5.90001779009805 | -5.41577901375959 | 1.27621938439696 | -4.24361131007164 | 2.19951194168897e-05 | 0.00332929432873488 | 1748 | 11 | 7 | 3 | 6 | 1 | 2 | 21 | 22 | 0 | 0 | 0 | 0 |
| THE1B_dup10247 | 4.46234122422059 | -4.99478478489071 | 1.31223842061418 | -3.80630890425609 | 0.000141056187258275 | 0.0127813235085139 | 1749 | 8 | 5 | 1 | 7 | 1 | 1 | 20 | 14 | 0 | 0 | 0 | 0 |
| LTR40b_dup998 | 3.93069668797329 | -4.81727377540162 | 1.23976330650326 | -3.88563990411096 | 0.000102060695824916 | 0.0102295426827108 | 1750 | 5 | 6 | 3 | 8 | 1 | 1 | 20 | 8 | 0 | 0 | 0 | 0 |
| MLT2B1_dup3715 | 4.36852628660309 | -5.05757505811081 | 1.25995857153265 | -4.01408044072326 | 5.96779932422108e-05 | 0.00689327173786946 | 1751 | 8 | 6 | 3 | 5 | 3 | 1 | 27 | 6 | 0 | 0 | 0 | 0 |
| L1MC_dup3524 | 5.13385358810575 | -5.06477879163571 | 1.21366318259417 | -4.17313375265276 | 3.00438445818486e-05 | 0.0041757083194349 | 1752 | 6 | 3 | 5 | 6 | 1 | 3 | 27 | 15 | 0 | 0 | 0 | 0 |
| MER4CL34_dup685 | 5.31758353199384 | -5.28391352325904 | 1.18905427475999 | -4.44379506925839 | 8.83857968510118e-06 | 0.0016851800334544 | 1753 | 8 | 6 | 4 | 15 | 1 | 4 | 26 | 8 | 0 | 0 | 0 | 0 |
| L1ME1_dup14052 | 7.17776974257687 | -4.39627908765791 | 1.09002325913492 | -4.03319750364499 | 5.50229965180144e-05 | 0.00647822955441635 | 1754 | 11 | 5 | 3 | 7 | 4 | 9 | 23 | 25 | 0 | 0 | 1 | 1 |
| THE1B_dup10248 | 5.85802947900001 | -4.80709524911975 | 1.13304125079463 | -4.24264804635173 | 2.20897723699951e-05 | 0.00333513513727846 | 1755 | 9 | 8 | 3 | 11 | 5 | 9 | 18 | 12 | 0 | 0 | 0 | 1 |
| L1M5_dup24526 | 6.47838229352827 | -3.83217196408006 | 1.03261248344522 | -3.7111423941868 | 0.000206325993893638 | 0.0167919620766515 | 1756 | 11 | 7 | 3 | 7 | 5 | 5 | 22 | 19 | 0 | 0 | 0 | 3 |
| MLT1C_dup8340 | 5.67831449979745 | -4.12407206078833 | 1.16271729507697 | -3.54692587635013 | 0.000389754233848722 | 0.0258485014139323 | 1757 | 13 | 10 | 2 | 6 | 2 | 4 | 20 | 14 | 0 | 0 | 0 | 2 |
| MIRc_dup44128 | 5.26829721636393 | -3.3557145878606 | 1.02295828468259 | -3.2804021807222 | 0.00103659203726095 | 0.0490337014487243 | 1758 | 7 | 10 | 8 | 4 | 3 | 1 | 16 | 14 | 0 | 1 | 1 | 1 |
| Ricksha_dup91 | 4.81951943098663 | -3.73191711779063 | 1.06895542058903 | -3.49118124658014 | 0.000480889904946411 | 0.0297519057000464 | 1759 | 10 | 5 | 7 | 7 | 3 | 1 | 16 | 11 | 0 | 1 | 1 | 0 |
| ALR/Alpha_dup1290 | 4.7643718646717 | -3.79008083748358 | 1.12063221277892 | -3.38209163922304 | 0.000719361430005395 | 0.038413190041307 | 1760 | 6 | 5 | 2 | 3 | 5 | 3 | 17 | 16 | 0 | 0 | 1 | 1 |
| MIRb_dup96341 | 5.46999931365842 | -3.52220977181482 | 0.998020632166559 | -3.52919534756372 | 0.00041682532163263 | 0.0270255736209899 | 1761 | 7 | 5 | 5 | 7 | 6 | 1 | 16 | 18 | 0 | 0 | 1 | 2 |
| L1PA3_dup8679 | 5.93041948581031 | -3.1211346065506 | 0.908668085362713 | -3.43484563486648 | 0.000592891454221653 | 0.034153947339677 | 1762 | 8 | 5 | 7 | 8 | 2 | 2 | 19 | 19 | 1 | 0 | 1 | 2 |
| SVA_D_dup572 | 5.23536422003788 | -3.04500612889495 | 0.874614662208712 | -3.48154022618971 | 0.000498538893698383 | 0.0304167527179373 | 1763 | 7 | 3 | 6 | 13 | 2 | 2 | 18 | 13 | 0 | 0 | 2 | 2 |
| L1MC4_dup11022 | 3.47174538895585 | -4.81563790827104 | 1.36078186125536 | -3.53887573415219 | 0.000401834966212719 | 0.026398404989081 | 1764 | 12 | 8 | 0 | 3 | 1 | 4 | 13 | 5 | 0 | 0 | 0 | 0 |
| L1PA14_dup1509 | 4.79869110293962 | -5.27468683897403 | 1.26494302541637 | -4.16990072516333 | 3.04732353186206e-05 | 0.00422695046279277 | 1765 | 13 | 5 | 0 | 11 | 5 | 4 | 18 | 8 | 0 | 0 | 0 | 0 |
| L1MC2_dup2649 | 6.70604853951022 | -5.01350355058229 | 1.15032490001456 | -4.35833697985573 | 1.31054487288946e-05 | 0.00227619642572668 | 1766 | 16 | 15 | 5 | 5 | 6 | 8 | 12 | 16 | 0 | 0 | 1 | 0 |
| MER52A_dup880 | 5.19202917548646 | -4.63860982984117 | 1.22414110668446 | -3.7892770731347 | 0.00015108632547556 | 0.0133938633530045 | 1767 | 10 | 12 | 2 | 1 | 6 | 8 | 13 | 12 | 0 | 0 | 1 | 0 |
| L1M5_dup19415 | 4.38825779682676 | -3.66970803278958 | 1.10019201131195 | -3.33551597817325 | 0.000851412608911548 | 0.0430036562598317 | 1768 | 11 | 5 | 3 | 8 | 8 | 4 | 9 | 7 | 0 | 0 | 2 | 0 |
| MLT1H1_dup501 | 5.20626764767757 | -4.64093493177382 | 1.18229574262732 | -3.92535874438713 | 8.66005772342605e-05 | 0.00908164872216102 | 1769 | 12 | 5 | 2 | 6 | 9 | 4 | 14 | 13 | 0 | 0 | 1 | 0 |
| LTR16_dup1398 | 4.54126367696594 | -3.70621705835936 | 1.09537771427096 | -3.38350599074042 | 0.000715666461540195 | 0.0383104548611121 | 1770 | 7 | 7 | 3 | 6 | 2 | 4 | 13 | 13 | 0 | 0 | 2 | 0 |
| L2c_dup63712 | 5.21879283814995 | -4.63459306658735 | 1.15412988262742 | -4.01565988052968 | 5.92796955067017e-05 | 0.00686384130784084 | 1771 | 11 | 6 | 3 | 7 | 4 | 7 | 16 | 12 | 0 | 0 | 1 | 0 |
| MIR3_dup25613 | 4.60393895740563 | -4.45799436758743 | 1.11603398889586 | -3.99449695255062 | 6.4831755287487e-05 | 0.00734243543247805 | 1772 | 8 | 6 | 6 | 7 | 4 | 8 | 10 | 9 | 0 | 0 | 1 | 0 |
| AluJb_dup109107 | 3.4547630899064 | -4.77116295492598 | 1.30536278490625 | -3.65504747806077 | 0.000257134387470484 | 0.0194978008161414 | 1773 | 1 | 10 | 1 | 5 | 2 | 4 | 13 | 8 | 0 | 0 | 0 | 0 |
| L1ME4c_dup1028 | 3.76236451033802 | -4.80085479983629 | 1.21315695814924 | -3.95732371445187 | 7.5794184713177e-05 | 0.00820888624295021 | 1774 | 2 | 6 | 2 | 8 | 4 | 2 | 14 | 10 | 0 | 0 | 0 | 0 |
| L1M2_dup3503 | 2.70473822794093 | -4.37193302706514 | 1.228095656586 | -3.55992874302539 | 0.00037095543725579 | 0.025042762237081 | 1775 | 1 | 5 | 6 | 11 | 1 | 4 | 5 | 3 | 0 | 0 | 0 | 0 |
| L2b_dup39268 | 3.15605578589989 | -4.54022660424468 | 1.24491821736515 | -3.64700792462818 | 0.000265311744790398 | 0.0198959560119433 | 1776 | 1 | 6 | 7 | 8 | 1 | 3 | 15 | 2 | 0 | 0 | 0 | 0 |
| L1PA13_dup186 | 2.80474316677885 | -4.33249580847984 | 1.26712836823242 | -3.41914514511536 | 0.000628182083969724 | 0.0353699512901704 | 1777 | 0 | 5 | 3 | 8 | 2 | 2 | 12 | 5 | 0 | 0 | 0 | 0 |
| MamGypLTR3a_dup6 | 2.77755460977058 | -4.32807473541909 | 1.21009063867204 | -3.57665334901585 | 0.000348021192113247 | 0.0240378085859445 | 1778 | 2 | 2 | 3 | 10 | 3 | 2 | 13 | 3 | 0 | 0 | 0 | 0 |
| AluJo_dup5002 | 2.51448754881272 | -4.29766027882255 | 1.26043245379065 | -3.40967123299443 | 0.000650412336173773 | 0.0361678479030935 | 1779 | 1 | 5 | 3 | 8 | 4 | 1 | 10 | 2 | 0 | 0 | 0 | 0 |
| LTR102_Mam_dup19 | 2.85892488536908 | -4.46156132457976 | 1.2170570361638 | -3.66586050777269 | 0.000246508274344508 | 0.0189456259196312 | 1780 | 2 | 6 | 3 | 9 | 3 | 1 | 10 | 4 | 0 | 0 | 0 | 0 |
| AluJb_dup23763 | 2.4329488589743 | -4.17647666438573 | 1.21801206972134 | -3.42892879981168 | 0.000605968473008012 | 0.0346605226630683 | 1781 | 3 | 1 | 3 | 8 | 1 | 4 | 7 | 5 | 0 | 0 | 0 | 0 |
| L1MB7_dup871 | 3.03097841207046 | -4.51206572113484 | 1.19344031535234 | -3.78072171946257 | 0.000156374386274539 | 0.013775882901326 | 1782 | 2 | 1 | 4 | 10 | 4 | 5 | 6 | 7 | 0 | 0 | 0 | 0 |
| L2a_dup137524 | 3.08785348226769 | -4.62582413977271 | 1.19241166745242 | -3.87938517043845 | 0.000104720822200467 | 0.0103801369157049 | 1783 | 8 | 2 | 3 | 5 | 4 | 4 | 5 | 8 | 0 | 0 | 0 | 0 |
| MER5A_dup20554 | 2.4211026804715 | -4.26993094787778 | 1.25455073529809 | -3.40355382029505 | 0.000665153078615678 | 0.0366258787677276 | 1784 | 4 | 3 | 1 | 7 | 1 | 5 | 4 | 6 | 0 | 0 | 0 | 0 |
| L1MA7_dup3678 | 3.5748219338439 | -4.71110876714043 | 1.2219717423574 | -3.85533364139164 | 0.00011557190067753 | 0.0111425735326643 | 1785 | 6 | 2 | 2 | 6 | 2 | 3 | 15 | 10 | 0 | 0 | 0 | 0 |
| L2c_dup63708 | 3.75536579648525 | -4.72921626854411 | 1.28427787914184 | -3.68239330860718 | 0.000231054576140469 | 0.0180969427828572 | 1786 | 6 | 1 | 1 | 6 | 3 | 2 | 20 | 10 | 0 | 0 | 0 | 0 |
| L2a_dup161215 | 2.38905588804221 | -4.28750049871318 | 1.28628309551883 | -3.33324795579608 | 0.000858384055835747 | 0.0432547760588506 | 1787 | 7 | 3 | 1 | 2 | 2 | 3 | 11 | 3 | 0 | 0 | 0 | 0 |
| MIRb_dup96475 | 2.63016449165685 | -4.35590972613829 | 1.26081199348276 | -3.45484477357 | 0.000550609728204626 | 0.0325100204435181 | 1788 | 3 | 2 | 1 | 3 | 4 | 3 | 12 | 6 | 0 | 0 | 0 | 0 |
| ERVL-B4-int_dup1947 | 2.55578969504734 | -4.05500964970651 | 1.23290805037171 | -3.2889797811637 | 0.00100551246628463 | 0.048014784126543 | 1789 | 3 | 2 | 8 | 10 | 0 | 1 | 6 | 4 | 0 | 0 | 0 | 0 |
| L2c_dup2070 | 2.54241435172651 | -4.22366840203281 | 1.23666001886089 | -3.41538364434497 | 0.000636922346860948 | 0.0356679512229172 | 1790 | 5 | 2 | 5 | 7 | 0 | 2 | 9 | 4 | 0 | 0 | 0 | 0 |
| AluYj4_dup228 | 2.98362417761897 | -4.4974684259193 | 1.23078610105697 | -3.6541430083237 | 0.0002580424198133 | 0.0195479661333638 | 1791 | 5 | 4 | 11 | 5 | 0 | 3 | 7 | 4 | 0 | 0 | 0 | 0 |
| MLT1A_dup7128 | 3.41988489381541 | -4.45723358183162 | 1.20040576491774 | -3.71310577814248 | 0.000204731266424274 | 0.016694649047495 | 1792 | 2 | 2 | 13 | 8 | 0 | 4 | 8 | 7 | 0 | 0 | 0 | 0 |
| L2b_dup9246 | 3.52316697001724 | -4.34546957339224 | 1.23859486677855 | -3.50838655152377 | 0.000450833470719483 | 0.028498300031631 | 1793 | 2 | 3 | 5 | 7 | 0 | 1 | 19 | 9 | 0 | 0 | 0 | 0 |
| MSTA_dup8132 | 3.76776616163662 | -4.68706421195264 | 1.28399437734436 | -3.65037752084765 | 0.000261855151935129 | 0.0197175272096064 | 1794 | 6 | 2 | 3 | 5 | 0 | 3 | 17 | 12 | 0 | 0 | 0 | 0 |

| | | | | | | | | | | | | | | | | | | | |
|---|---|---|---|---|---|---|---|---|---|---|---|---|---|---|---|---|---|---|---|
| AluSx1_dup83992 | 3.8535567697787 | -4.15773434827594 | 1.17512793561313 | -3.53811208318065 | 0.000402998952068345 | 0.0264092505229563 | 1795 | 7 | 7 | 7 | 6 | 1 | 2 | 12 | 7 | 0 | 1 | 0 | 0 |
| L1ME3_dup1134 | 3.43063677298109 | -3.77815150528599 | 1.13488076320706 | -3.32911758466087 | 0.000871216145585075 | 0.0436058891412255 | 1796 | 2 | 3 | 7 | 6 | 1 | 3 | 10 | 10 | 0 | 1 | 0 | 0 |
| MIR_dup137714 | 3.03370842625403 | -4.61951920261501 | 1.29312030600506 | -3.5723816114886 | 0.00035374941838695 | 0.0242924268708517 | 1797 | 10 | 6 | 3 | 5 | 0 | 2 | 9 | 5 | 0 | 0 | 0 | 0 |
| L1PA4_dup335 | 2.62380329811677 | -4.39201619932525 | 1.28157829645344 | -3.42703696799441 | 0.000610205991254287 | 0.034785834881407 | 1798 | 3 | 7 | 2 | 7 | 0 | 4 | 6 | 5 | 0 | 0 | 0 | 0 |
| LTR33_dup1194 | 2.71553634907707 | -4.45523972994937 | 1.32709547940751 | -3.35713578945987 | 0.000787544245232071 | 0.0410409555356965 | 1799 | 4 | 10 | 2 | 7 | 0 | 2 | 5 | 5 | 0 | 0 | 0 | 0 |
| AluSx1_dup1605 | 4.34641626566574 | -4.34811977778863 | 1.32130910804871 | -3.29076652185488 | 0.000999148029933371 | 0.0478164274196553 | 1800 | 12 | 5 | 1 | 8 | 0 | 3 | 20 | 8 | 0 | 1 | 0 | 0 |
| MLT2B1_dup3714 | 4.69378873362532 | -4.41948684548407 | 1.25951277692373 | -3.50888607599389 | 0.000449987608708316 | 0.0284919509158353 | 1801 | 7 | 10 | 3 | 13 | 0 | 2 | 18 | 8 | 1 | 0 | 0 | 0 |
| MER20_dup6442 | 4.7652897456588 | -3.77982347413319 | 1.06711509935254 | -3.54209539010979 | 0.000396961894845727 | 0.0261794609287587 | 1802 | 10 | 5 | 7 | 9 | 6 | 8 | 9 | 6 | 1 | 0 | 1 | 0 |
| MER44D_dup12 | 5.40852554658292 | -2.9596642348947 | 0.901388740245101 | -3.28344930744301 | 0.00102545085340583 | 0.0486544224854446 | 1803 | 15 | 9 | 10 | 11 | 2 | 4 | 10 | 6 | 1 | 0 | 3 | 0 |
| MIR_dup157631 | 4.58473541949169 | -3.22479918490912 | 0.974674134463473 | -3.3085921446805 | 0.000937663236132298 | 0.0457661655404614 | 1804 | 8 | 4 | 8 | 14 | 5 | 3 | 5 | 9 | 0 | 0 | 3 | 0 |
| AluSq2_dup59019 | 3.67297036490504 | -4.11421143328176 | 1.14670072235014 | -3.58786852845941 | 0.000333392316027688 | 0.0232705685038205 | 1805 | 7 | 6 | 6 | 9 | 3 | 2 | 11 | 4 | 0 | 0 | 1 | 0 |
| MER63A_dup247 | 3.84747355974964 | -4.18311636795968 | 1.13399224014577 | -3.68884038167842 | 0.000225278490690826 | 0.0178145947975803 | 1806 | 4 | 5 | 4 | 10 | 4 | 6 | 12 | 5 | 0 | 0 | 1 | 0 |
| AluSz6_dup17863 | 6.70822967575613 | -4.24577134061384 | 0.969204766849767 | -4.380675256493 | 1.18312088185035e-05 | 0.00211827600716903 | 1807 | 9 | 8 | 16 | 17 | 3 | 3 | 19 | 11 | 0 | 0 | 2 | 0 |
| MER9a1_dup287 | 6.69238974000711 | -4.30543618114567 | 1.10748471622897 | -3.88758067542986 | 0.000101248341075769 | 0.0101730238317402 | 1808 | 15 | 13 | 14 | 21 | 1 | 2 | 13 | 8 | 0 | 0 | 2 | 0 |
| L1MC_dup3523 | 5.48598296097115 | -3.55194189569452 | 1.02747296242612 | -3.45696872383629 | 0.000546288248173342 | 0.0323270588161786 | 1809 | 15 | 9 | 4 | 9 | 2 | 4 | 19 | 8 | 0 | 0 | 1 | 2 |
| L2a_dup165166 | 5.47833085527566 | -4.04446033683991 | 1.07635426329445 | -3.7575549935212 | 0.000171581666489009 | 0.0147272393349269 | 1810 | 12 | 8 | 6 | 22 | 5 | 2 | 14 | 4 | 0 | 0 | 1 | 1 |
| AluJb_dup35006 | 2.4941967193448 | -4.24664869680783 | 1.29559526450102 | -3.27775873620792 | 0.00104634786912656 | 0.0492906400719254 | 1811 | 6 | 2 | 10 | 9 | 1 | 1 | 4 | 1 | 0 | 0 | 0 | 0 |
| L1MB7_dup17176 | 2.63159474697098 | -4.41392173262921 | 1.30422768615126 | -3.38431838205689 | 0.000713552084484036 | 0.0382595763784293 | 1812 | 8 | 4 | 9 | 8 | 1 | 2 | 1 | 2 | 0 | 0 | 0 | 0 |
| MER49_dup337 | 3.34495037318006 | -4.69609019280587 | 1.27613901274598 | -3.679920561868 | 0.000233306620720744 | 0.0182373118180086 | 1813 | 8 | 1 | 6 | 15 | 5 | 1 | 2 | 6 | 0 | 0 | 0 | 0 |
| AluSq_dup16653 | 2.37487817310236 | -4.05918762576463 | 1.22202220150438 | -3.32169711873282 | 0.00089471758860656 | 0.0443807528327239 | 1814 | 3 | 2 | 6 | 11 | 1 | 1 | 5 | 3 | 0 | 0 | 0 | 0 |
| L1PA17_dup1931 | 2.74696502544886 | -4.11803233134075 | 1.2382707762822 | -3.32563152600983 | 0.000882184673594773 | 0.043940192429475 | 1815 | 3 | 1 | 10 | 12 | 1 | 1 | 3 | 5 | 0 | 0 | 0 | 0 |
| L1PB4_dup2966 | 3.69528698160961 | -3.88622799967675 | 1.12201672878456 | -3.4636096770915 | 0.00053297926902289 | 0.0317845326182758 | 1816 | 4 | 3 | 12 | 16 | 2 | 1 | 5 | 5 | 0 | 0 | 1 | 0 |
| L2_dup24562 | 3.40366961886684 | -3.86918129654192 | 1.17692133120068 | -3.28754454012201 | 0.00101065200616697 | 0.0481731975679899 | 1817 | 6 | 2 | 6 | 16 | 3 | 1 | 4 | 6 | 0 | 0 | 1 | 0 |
| AluSx4_dup4241 | 4.19611005882105 | -4.96304301365056 | 1.14153862047066 | -4.34767858454432 | 1.37586032012748e-05 | 0.00236717613689842 | 1818 | 6 | 5 | 9 | 11 | 4 | 1 | 9 | 9 | 0 | 0 | 0 | 0 |
| Chap1_Mam_dup116 | 3.3716586299655 | -4.74061919108927 | 1.20304700885587 | -3.94051035096103 | 8.13084419679789e-05 | 0.00863706184487174 | 1819 | 8 | 5 | 5 | 4 | 3 | 1 | 12 | 6 | 0 | 0 | 0 | 0 |
| MER57A1_dup404 | 3.17007511161525 | -3.69895004373845 | 1.12792193468209 | -3.27943799123031 | 0.00104014065792905 | 0.0491158129931673 | 1820 | 3 | 2 | 4 | 8 | 3 | 1 | 11 | 8 | 0 | 0 | 0 | 1 |
| L1M5_dup22422 | 3.22493003983761 | -4.46568962091965 | 1.16560975521127 | -3.83120474151337 | 0.000127517335700671 | 0.0119363650513894 | 1821 | 4 | 2 | 8 | 12 | 2 | 1 | 9 | 5 | 0 | 0 | 0 | 0 |
| MLT1O_dup969 | 3.11223882468424 | -4.47745197707834 | 1.16141842264339 | -3.85515839062346 | 0.000115654724930712 | 0.0111460425995801 | 1822 | 2 | 2 | 7 | 8 | 4 | 2 | 11 | 5 | 0 | 0 | 0 | 0 |
| L1PB4_dup1241 | 3.28725045050843 | -3.83715726662464 | 1.15111415939113 | -3.33342895256738 | 0.000857825770320218 | 0.0432449599035201 | 1823 | 4 | 3 | 8 | 10 | 3 | 1 | 11 | 3 | 1 | 0 | 0 | 0 |
| MER47B_dup598 | 3.43841411345401 | -4.51725975489222 | 1.24298105649081 | -3.63421448082674 | 0.000278828842280051 | 0.020566090423177 | 1824 | 6 | 2 | 10 | 7 | 1 | 0 | 15 | 5 | 0 | 0 | 0 | 0 |
| L1MC1_dup5773 | 3.47394188288894 | -4.34076881514566 | 1.20271185176774 | -3.6091511102727 | 0.000307200632166182 | 0.0220155277341164 | 1825 | 3 | 2 | 12 | 13 | 1 | 0 | 12 | 4 | 0 | 0 | 0 | 0 |
| L2c_dup59402 | 2.73755453482514 | -4.1238003912451 | 1.23487456744608 | -3.33944879905801 | 0.000839448238940204 | 0.0426693520431741 | 1826 | 3 | 2 | 9 | 8 | 1 | 0 | 11 | 3 | 0 | 0 | 0 | 0 |
| FLAM_A_dup2026 | 4.41829953087349 | -4.4045030246886 | 1.26763339781545 | -3.47458739433579 | 0.000511639668604024 | 0.0310014292298582 | 1827 | 25 | 5 | 4 | 7 | 1 | 3 | 11 | 3 | 1 | 0 | 0 | 0 |
| THE1C-int_dup200 | 4.95746035900616 | -4.57289382954742 | 1.27241025269943 | -3.59388319910656 | 0.00032578592580099 | 0.0229048854870021 | 1828 | 19 | 6 | 7 | 10 | 0 | 2 | 16 | 6 | 0 | 0 | 0 | 1 |
| L2_dup24563 | 3.36047957790569 | -3.91292077233478 | 1.17599205410691 | -3.32733606376822 | 0.000876805622992774 | 0.0438027274341042 | 1829 | 7 | 4 | 9 | 11 | 1 | 1 | 7 | 4 | 0 | 0 | 0 | 1 |
| L2c_dup59385 | 4.64912364250228 | -4.43746686291232 | 1.17099507454113 | -3.78948379834236 | 0.000150960656368413 | 0.0133931708949434 | 1830 | 12 | 5 | 11 | 12 | 3 | 1 | 14 | 4 | 0 | 0 | 0 | 1 |
| L2c_dup114611 | 3.95177875698435 | -4.14298338738737 | 1.2366492766877 | -3.35016844750366 | 0.000807624303238514 | 0.0416224761659934 | 1831 | 8 | 6 | 6 | 8 | 1 | 0 | 12 | 9 | 0 | 1 | 0 | 0 |
| HAL1_dup3208 | 3.02283137880614 | -4.62124623628681 | 1.35305476364232 | -3.41541699601779 | 0.000636844355958564 | 0.0356679512229172 | 1832 | 8 | 10 | 2 | 8 | 1 | 0 | 7 | 4 | 0 | 0 | 0 | 0 |
| AluSz_dup36955 | 3.34065358180838 | -4.58392176487486 | 1.27275160679866 | -3.60158395431515 | 0.000316284247974436 | 0.0224384530913909 | 1833 | 4 | 6 | 3 | 11 | 1 | 0 | 12 | 7 | 0 | 0 | 0 | 0 |
| L2a_dup137525 | 2.73125789026728 | -4.33276696778379 | 1.32265602054352 | -3.2758078445848 | 0.0010536021924752 | 0.0494694050868097 | 1834 | 4 | 5 | 3 | 3 | 1 | 0 | 12 | 7 | 0 | 0 | 0 | 0 |
| ALR/Alpha_dup1292 | 4.17610052770248 | -4.1395508496075 | 1.1802047022102 | -3.50748547421922 | 0.000452363049739425 | 0.0285326097886149 | 1835 | 4 | 5 | 5 | 5 | 3 | 1 | 21 | 9 | 1 | 0 | 0 | 0 |
| AluSc_dup14924 | 4.44883955369207 | -4.40698640576591 | 1.20417370000797 | -3.65975972215366 | 0.000252451855056294 | 0.019247007082097 | 1836 | 7 | 10 | 4 | 4 | 6 | 1 | 16 | 8 | 1 | 0 | 0 | 0 |
| AluSx3_dup24069 | 4.27902320589987 | -4.33738697573442 | 1.27718016086979 | -3.39606510390872 | 0.000683620973544261 | 0.0372571864798357 | 1837 | 13 | 6 | 4 | 7 | 2 | 0 | 18 | 6 | 1 | 0 | 0 | 0 |
| MIR_dup137713 | 3.69014741320135 | -4.10726642314846 | 1.23480356327222 | -3.32625086719399 | 0.000880226675260415 | 0.0438702505527443 | 1838 | 10 | 6 | 3 | 8 | 1 | 1 | 8 | 9 | 1 | 0 | 0 | 0 |
| MIRb_dup175062 | 3.68295915658716 | -4.06251540917199 | 1.17765935057448 | -3.44965240346476 | 0.000561308792550569 | 0.0329292854911829 | 1839 | 7 | 5 | 4 | 9 | 2 | 1 | 12 | 7 | 1 | 0 | 0 | 0 |
| MER4-int_dup1137 | 39.7407884658694 | -4.71533884707934 | 1.35197188222867 | -3.48774919734743 | 0.000487104764772938 | 0.0299727515796901 | 1840 | 30 | 21 | 5 | 2 | 1 | 11 | 219 | 180 | 0 | 0 | 7 | 8 |
| AluSz_dup144 | 10.4116396301892 | -5.61442991567764 | 1.37498000706191 | -4.08328112906507 | 4.44042506450095e-05 | 0.0055182592020946 | 1841 | 8 | 3 | 0 | 3 | 17 | 21 | 36 | 38 | 0 | 0 | 0 | 1 |
| L1P3_dup2884 | 14.4656334699843 | -4.02853257587781 | 1.08859926170845 | -3.70065708987844 | 0.000215041907415338 | 0.0172400140504885 | 1842 | 10 | 2 | 2 | 2 | 12 | 8 | 73 | 61 | 0 | 0 | 2 | 5 |
| AluY_dup13399 | 10.752609490338 | -3.34713486973328 | 0.892363572515298 | -3.75086452744674 | 0.000176225900367992 | 0.0150294796017785 | 1843 | 38 | 33 | 7 | 5 | 4 | 3 | 21 | 20 | 1 | 1 | 2 | 2 |
| MER1A_dup368 | 14.0889756464943 | -3.0980094702495 | 0.766967072093008 | -4.03929918633302 | 5.36111451366057e-05 | 0.00635920116683688 | 1844 | 35 | 32 | 14 | 11 | 5 | 4 | 59 | 17 | 1 | 2 | 2 | 5 |
| AluSg_dup15059 | 12.033047065921 | -2.43850381590507 | 0.690323121400678 | -3.53240930270061 | 0.000411791453804772 | 0.0268205162266047 | 1845 | 35 | 23 | 6 | 7 | 6 | 11 | 35 | 20 | 1 | 2 | 5 | 5 |
| L1ME4c_dup1140 | 15.039441144105 | -2.95523618153632 | 0.713596247594212 | -4.14132808503334 | 3.45300631251221e-05 | 0.0046341175411 | 1846 | 50 | 41 | 11 | 9 | 8 | 8 | 39 | 19 | 2 | 1 | 4 | 4 |
| AluSx3_dup17387 | 10.4376193611102 | -1.95469477733865 | 0.593455897374741 | -3.29374901485619 | 0.000988607303157957 | 0.0474264422091176 | 1847 | 25 | 26 | 12 | 7 | 7 | 10 | 23 | 10 | 5 | 4 | 4 | 1 |
| L2c_dup3881 | 9.4929288451132 | -2.37220513920657 | 0.647427953655633 | -3.66404497336293 | 0.000248263164391121 | 0.019049734574076 | 1848 | 24 | 19 | 13 | 8 | 9 | 6 | 26 | 9 | 2 | 2 | 5 | 1 |
| AluJo_dup74476 | 9.55444061886517 | -2.14954972398583 | 0.615181839128353 | -3.49416967027426 | 0.000475538692180895 | 0.0294975311912605 | 1849 | 20 | 14 | 18 | 14 | 8 | 5 | 19 | 13 | 5 | 2 | 4 | 0 |
| L2b_dup20935 | 11.2738209158093 | -1.73916043299808 | 0.516377183257689 | -3.36800402764927 | 0.000757144900148873 | 0.0398847908812981 | 1850 | 16 | 17 | 19 | 25 | 10 | 5 | 21 | 15 | 4 | 5 | 7 | 1 |
| THE1B_dup10240 | 11.821513302262 | -2.56824232952729 | 0.758491297954045 | -3.38598786361144 | 0.000709225201401934 | 0.0381466941258146 | 1851 | 30 | 20 | 5 | 12 | 13 | 3 | 34 | 23 | 2 | 6 | 3 | 0 |
| L2_dup26672 | 12.2976367974018 | -2.23694024713947 | 0.625757890144864 | -3.57476954325197 | 0.000350536521010152 | 0.0241326771274171 | 1852 | 25 | 17 | 14 | 16 | 9 | 4 | 34 | 24 | 3 | 2 | 8 | 1 |
| L1M5_dup24524 | 12.5550510169365 | -2.37485203707532 | 0.684643224535449 | -3.4687439413232 | 0.000522897570669753 | 0.0313482956287692 | 1853 | 20 | 13 | 10 | 11 | 8 | 4 | 47 | 33 | 4 | 2 | 4 | 2 |
| MLT1C_dup8339 | 12.1459806196442 | -3.07597756839142 | 0.761165335225958 | -4.04114247724943 | 5.31914277039506e-05 | 0.00632691365749532 | 1854 | 18 | 18 | 7 | 11 | 11 | 4 | 47 | 30 | 2 | 1 | 4 | 1 |
| L1MA8_dup569 | 10.9612086084246 | -3.18381454432615 | 0.750897133613493 | -4.24001424669833 | 2.23505600806437e-05 | 0.00336171072626349 | 1855 | 17 | 20 | 17 | 15 | 2 | 17 | 26 | 20 | 0 | 4 | 0 | 3 |
| MLT1B_dup14964 | 11.1378762108164 | -3.55693449511523 | 0.934966918372941 | -3.80434261920744 | 0.00014218127387111 | 0.0128343950654534 | 1856 | 19 | 17 | 11 | 11 | 1 | 3 | 37 | 35 | 1 | 2 | 0 | 2 |
| L1PA5_dup5266 | 11.3783292435308 | -2.72363752634952 | 0.762786627687461 | -3.57064141856651 | 0.000356108126691233 | 0.0244171151047913 | 1857 | 22 | 18 | 14 | 9 | 1 | 6 | 44 | 23 | 2 | 1 | 2 | 5 |
| THE1B-int_dup1873 | 13.3120522265846 | -3.13825689256526 | 0.74264483846385 | -4.2257842915285 | 2.38110001478629e-05 | 0.00351967513891115 | 1858 | 15 | 18 | 20 | 9 | 3 | 9 | 57 | 31 | 2 | 2 | 1 | 3 |
| L1MCb_dup1862 | 12.5316222858827 | -3.00527032854205 | 0.834804118771561 | -3.59997065295317 | 0.000318253097241524 | 0.0225251502695944 | 1859 | 36 | 20 | 8 | 17 | 4 | 7 | 37 | 23 | 6 | 3 | 0 | 0 |
| L2a_dup22925 | 11.0766083478841 | -3.0256824865738 | 0.852144703354576 | -3.55066748013902 | 0.000384255564658322 | 0.0256471457470376 | 1860 | 30 | 15 | 14 | 10 | 3 | 3 | 35 | 24 | 4 | 2 | 1 | 0 |
| L1MCa_dup3788 | 9.21758652370092 | -4.62959715304421 | 0.953180463790885 | -4.85699962274918 | 1.19177821168471e-06 | 0.000365436673141146 | 1861 | 12 | 8 | 19 | 27 | 4 | 4 | 34 | 13 | 1 | 1 | 0 | 0 |
| L2_dup47144 | 7.77963054969883 | -5.20899328897532 | 1.15135128665843 | -4.5242432516781 | 6.06119817241593e-06 | 0.00126512093011726 | 1862 | 14 | 16 | 14 | 11 | 2 | 6 | 30 | 9 | 1 | 0 | 0 | 0 |
| L2a_dup32944 | 14.5872951460974 | -2.86403902631267 | 0.806845238532788 | -3.54967581084174 | 0.000385705819724437 | 0.0257227955943545 | 1863 | 13 | 18 | 3 | 8 | 9 | 11 | 77 | 35 | 0 | 1 | 4 | 10 |
| L1MA10_dup2101 | 14.8571041641805 | -4.0137831035975 | 0.873681006328876 | -4.59410594315542 | 4.34608550560199e-06 | 0.000988655508332668 | 1864 | 27 | 29 | 5 | 13 | 12 | 5 | 57 | 36 | 0 | 1 | 2 | 3 |
| Tigger1_dup5862 | 18.2598727008434 | -4.11121028176149 | 0.839095304416472 | -4.89957488752786 | 9.60442379658172e-07 | 0.000307409615345431 | 1865 | 25 | 33 | 11 | 17 | 7 | 6 | 73 | 52 | 0 | 1 | 3 | 3 |
| L1MA5_dup2029 | 12.9760655735547 | -4.05962503569705 | 0.946926771662386 | -4.28715837083172 | 1.80973273676581e-05 | 0.00288232038676357 | 1866 | 18 | 15 | 5 | 8 | 4 | 10 | 56 | 42 | 0 | 1 | 1 | 3 |

| | | | | | | | | | | | | | | | | | | |
|---|---|---|---|---|---|---|---|---|---|---|---|---|---|---|---|---|---|---|
| MLT1A0_dup9731 | 13.4867925139584 | -3.36150883164174 | 0.716768799494062 | -4.68980909048286 | 2.73460066226331e-06 | 0.000710382446875096 | 1867 | 17 | 16 | 14 | 14 | 8 | 8 | 38 | 44 | 0 | 2 | 2 | 4 |
| L1M2_dup4146 | 15.119415854909 | -3.92682885432053 | 0.808981404939967 | -4.8540409338728 | 1.20970749070969e-06 | 0.000369505851841346 | 1868 | 29 | 20 | 12 | 12 | 6 | 17 | 41 | 45 | 0 | 0 | 3 | 4 |
| ALR/Alpha_dup1295 | 14.152023526822 | -3.83768376091031 | 0.827599480920269 | -4.63712683415764 | 3.53285642155348e-06 | 0.000848953661159664 | 1869 | 26 | 24 | 8 | 10 | 12 | 11 | 33 | 44 | 0 | 0 | 2 | 5 |
| L1PA14_dup1511 | 14.15356607549 | -4.27566565345799 | 0.871627632306062 | -4.90538103082607 | 9.32461454431279e-07 | 0.000302282661084612 | 1870 | 27 | 20 | 9 | 13 | 12 | 8 | 52 | 35 | 0 | 0 | 2 | 3 |
| Ricksha_0_dup198 | 9.72420534923268 | -3.40220058913162 | 0.768993935063509 | -4.42422291516596 | 9.67900408312051e-06 | 0.00180917541678098 | 1871 | 21 | 6 | 12 | 11 | 11 | 9 | 25 | 23 | 0 | 0 | 3 | 3 |
| THE1B-int_dup636 | 10.7181008437336 | -2.94586621945915 | 0.685399353987725 | -4.2980288824607 | 1.72323712702715e-05 | 0.00277239608808385 | 1872 | 19 | 15 | 10 | 8 | 14 | 17 | 27 | 19 | 0 | 0 | 2 | 7 |
| L2c_dup63710 | 13.9175128681014 | -5.38121816921312 | 1.06847374069296 | -5.03635977588287 | 4.74468396259991e-07 | 0.00017807232967899 | 1873 | 32 | 18 | 8 | 16 | 9 | 13 | 54 | 29 | 0 | 0 | 2 | 0 |
| L1ME1_dup14051 | 14.0941922745234 | -3.89260875161239 | 0.835750726244444 | -4.6576193467451 | 3.19886982972591e-06 | 0.000796190669983223 | 1874 | 22 | 12 | 11 | 14 | 7 | 10 | 46 | 47 | 0 | 0 | 1 | 6 |
| L2a_dup78627 | 15.9421290387821 | -4.23343531491028 | 0.81018188254808 | -5.22529003190719 | 1.73882336573714e-07 | 8.11266793301354e-05 | 1875 | 19 | 10 | 16 | 24 | 6 | 8 | 71 | 45 | 0 | 0 | 1 | 5 |
| THE1B_dup10243 | 17.0120325240564 | -4.18937074033986 | 0.848720553904681 | -4.93610143063692 | 7.9699668381383e-07 | 0.000269761270166096 | 1876 | 29 | 9 | 12 | 21 | 6 | 15 | 70 | 49 | 0 | 0 | 1 | 6 |
| L1M3f_dup330 | 11.5630440028414 | -4.27818371061908 | 1.11154943081169 | -3.84884701663247 | 0.000118675090519667 | 0.0113627308886939 | 1877 | 13 | 10 | 2 | 3 | 2 | 5 | 59 | 45 | 0 | 0 | 3 | 1 |
| MSTA_dup8758 | 7.6620623580627 | -5.15148823500376 | 1.23567394044241 | -4.16897052401977 | 3.05978552615789e-05 | 0.00423539108090265 | 1878 | 13 | 11 | 4 | 2 | 5 | 6 | 31 | 23 | 0 | 0 | 1 | 0 |
| MSTB_dup3730 | 7.71651735712587 | -4.46515877251028 | 1.18214225258125 | -3.77717551568811 | 0.000158616955602401 | 0.0139431096491211 | 1879 | 15 | 9 | 3 | 5 | 2 | 2 | 45 | 18 | 0 | 0 | 2 | 0 |
| AluJr_dup34428 | 6.55873834259309 | -4.88598385877198 | 1.13398616072888 | -4.30868032430965 | 1.64231544708918e-05 | 0.00267615631641256 | 1880 | 12 | 9 | 9 | 10 | 2 | 4 | 21 | 16 | 0 | 0 | 1 | 0 |
| MIR_dup77361 | 8.35925869471157 | -4.6305832198381 | 1.10295503259568 | -4.19834270934921 | 2.68875510663069e-05 | 0.00386106531443782 | 1881 | 23 | 10 | 7 | 9 | 4 | 4 | 27 | 21 | 0 | 0 | 2 | 0 |
| MLT2B4_dup2131 | 8.68823720125802 | -4.67580010099798 | 1.09632803119022 | -4.26496447046214 | 1.99934014858297e-05 | 0.00308518152396059 | 1882 | 18 | 16 | 10 | 13 | 2 | 3 | 23 | 23 | 0 | 0 | 2 | 0 |
| HUERS-P3-int_dup101 | 7.94054518065965 | -3.05341788439714 | 0.875448611232843 | -3.48783223277628 | 0.000486953521553788 | 0.0299711957164091 | 1883 | 8 | 6 | 12 | 7 | 2 | 2 | 31 | 26 | 0 | 2 | 2 | 1 |
| L2a_dup78625 | 7.56086578752971 | -3.99037333804254 | 0.998100037814946 | -3.99796932858386 | 6.38882271709669e-05 | 0.00725629981022572 | 1884 | 9 | 9 | 10 | 7 | 3 | 1 | 38 | 18 | 0 | 0 | 1 | 2 |
| L1MC4_dup11023 | 8.01920578909172 | -5.28760492563999 | 1.17180126074695 | -4.51237347386834 | 6.41061775318529e-06 | 0.00131725462167481 | 1885 | 24 | 14 | 5 | 6 | 8 | 5 | 23 | 17 | 0 | 0 | 0 | 1 |
| MER57B1_dup812 | 9.30083924344731 | -6.21663526641932 | 1.14612438745715 | -5.4240493740927 | 5.82637766909342e-08 | 3.27745451737613e-05 | 1886 | 13 | 22 | 13 | 8 | 12 | 4 | 24 | 21 | 0 | 0 | 0 | 0 |
| ALR/Alpha_dup1294 | 10.0075473978964 | -4.89502754258198 | 1.04597590947905 | -4.67986642734433 | 2.87061876428297e-06 | 0.000731319328143551 | 1887 | 6 | 15 | 9 | 11 | 15 | 10 | 27 | 29 | 0 | 0 | 1 | 1 |
| LTR7B_dup366 | 10.6342127996162 | -5.61133936133108 | 1.1668815877611 | -4.80883357847608 | 1.51813526370358e-06 | 0.000447072046357935 | 1888 | 14 | 16 | 4 | 14 | 11 | 5 | 38 | 31 | 0 | 0 | 1 | 0 |
| ALR/Alpha_dup1285 | 8.96628940575894 | -4.67754504371135 | 1.11130016708503 | -4.2090743637524 | 2.56418992705128e-05 | 0.00373401919450517 | 1889 | 14 | 9 | 6 | 4 | 14 | 6 | 32 | 25 | 1 | 0 | 1 | 0 |
| ALR/Alpha_dup1299 | 7.6287208584826 | -3.188322826493 | 0.836049750252579 | -3.81355634103087 | 0.000136981279667997 | 0.0125313908388624 | 1890 | 7 | 8 | 8 | 4 | 13 | 10 | 21 | 19 | 1 | 0 | 2 | 2 |
| L1ME1_dup14053 | 11.7816932245128 | -3.77873919280417 | 0.816056831405524 | -4.63048533800754 | 3.648096163487e-06 | 0.000868049889717078 | 1891 | 11 | 21 | 5 | 14 | 14 | 12 | 36 | 30 | 1 | 1 | 2 | 1 |
| THE1B_dup10242 | 11.5020517904374 | -3.7662352216617 | 0.747687095927465 | -5.03718098409856 | 4.72438035746156e-07 | 0.00017759042788887 | 1892 | 11 | 15 | 12 | 20 | 12 | 10 | 36 | 26 | 1 | 1 | 1 | 2 |
| LTR1B1_dup165 | 10.196851176498 | -2.77518970409966 | 0.821019679771063 | -3.38017440078111 | 0.000724398482627388 | 0.0385782724590995 | 1893 | 6 | 6 | 5 | 7 | 8 | 7 | 42 | 37 | 2 | 2 | 1 | 2 |
| ALR/Alpha_dup1286 | 9.82065758480646 | -2.87516263804972 | 0.765298709201387 | -3.75691557228685 | 0.000172020499567754 | 0.0147501224469005 | 1894 | 12 | 13 | 7 | 5 | 10 | 8 | 22 | 34 | 1 | 1 | 2 | 4 |
| L1MEb_dup744 | 11.4008660353732 | -3.28250493717863 | 0.80607963241225 | -4.07218444082937 | 4.65742751939632e-05 | 0.00571835009561546 | 1895 | 17 | 15 | 8 | 9 | 6 | 5 | 36 | 38 | 1 | 1 | 1 | 4 |
| L1MEc_dup8480 | 7.94796365749649 | -3.45706705773249 | 0.937977528308162 | -3.68566085369661 | 0.000228109953369049 | 0.0179253140569193 | 1896 | 14 | 20 | 4 | 7 | 7 | 3 | 29 | 14 | 2 | 1 | 1 | 0 |
| MLT1C_dup12029 | 8.79596440558302 | -4.08302317610137 | 1.04427383535395 | -3.90991619043815 | 9.23281495723873e-05 | 0.00950983647660379 | 1897 | 10 | 15 | 2 | 15 | 3 | 5 | 35 | 24 | 1 | 0 | 2 | 0 |
| MLT1H_dup4355 | 8.93465241814851 | -3.24022395236834 | 0.84212339154942 | -3.84768311257411 | 0.000119240148249641 | 0.0113994489363673 | 1898 | 21 | 7 | 7 | 10 | 6 | 4 | 35 | 20 | 1 | 0 | 1 | 4 |
| AluSx4_dup4240 | 7.16354491250734 | -3.13268645804129 | 0.807089838876976 | -3.88145942018086 | 0.000103831479501961 | 0.0103178404285482 | 1899 | 12 | 9 | 7 | 17 | 5 | 3 | 23 | 13 | 1 | 0 | 1 | 3 |
| L1MC2_dup2647 | 7.60064717107066 | -3.11506746798207 | 0.792185404362132 | -3.93224547035214 | 8.4156065902654e-05 | 0.00889191796504125 | 1900 | 11 | 8 | 6 | 13 | 5 | 7 | 24 | 18 | 2 | 0 | 1 | 2 |
| LTR33C_dup432 | 10.3120259714448 | -4.0021323813534 | 0.945232945814766 | -4.23401702096161 | 2.29553558649889e-05 | 0.00342453611702235 | 1901 | 22 | 14 | 5 | 14 | 5 | 7 | 29 | 30 | 1 | 0 | 0 | 3 |
| L1P4_dup1640 | 9.27945070829945 | -3.27624166678324 | 0.857875949557918 | -3.8190156379504 | 0.000133985269660774 | 0.0123468775134145 | 1902 | 19 | 15 | 8 | 10 | 1 | 8 | 25 | 25 | 1 | 0 | 2 | 3 |
| MSTD_dup3674 | 11.9098956988625 | -3.64255428737084 | 0.883096612698793 | -4.12475173722951 | 3.71134865655322e-05 | 0.00486288858718179 | 1903 | 20 | 19 | 6 | 13 | 5 | 3 | 43 | 35 | 1 | 0 | 3 | 2 |
| L1MA2_dup6123 | 3.27294700307748 | -4.71018173290065 | 1.43309299590015 | -3.28672441102967 | 0.00101359976200933 | 0.0482511438789805 | 1904 | 0 | 1 | 1 | 3 | 10 | 7 | 8 | 10 | 0 | 0 | 0 | 0 |
| L1MC4_dup1142 | 3.0744078402254 | -4.63254808973695 | 1.30808074127639 | -3.54148482089618 | 0.000397881750894479 | 0.0262110661955531 | 1905 | 3 | 1 | 1 | 2 | 6 | 9 | 10 | 7 | 0 | 0 | 0 | 0 |
| MER5A_dup34245 | 2.34783988301961 | -4.23036281721842 | 1.25312005688501 | -3.3758639437423 | 0.000735842981446773 | 0.0390578768691985 | 1906 | 2 | 2 | 1 | 5 | 6 | 7 | 3 | 4 | 0 | 0 | 0 | 0 |
| AluSx1_dup116766 | 2.6319983560171 | -4.41852799428202 | 1.27551058435135 | -3.46412491475248 | 0.00053195942599729 | 0.0317475840427267 | 1907 | 2 | 1 | 9 | 6 | 4 | 8 | 4 | 1 | 0 | 0 | 0 | 0 |
| L1ME1_dup24775 | 2.5759565869386 | -4.38316600239313 | 1.26190872820739 | -3.47344138638272 | 0.00051382958002935 | 0.0310471542025556 | 1908 | 5 | 0 | 4 | 8 | 3 | 6 | 7 | 2 | 0 | 0 | 0 | 0 |
| HAL1ME_dup1613 | 4.56690113782188 | -5.16001977853784 | 1.31763870507749 | -3.9161112668092 | 8.99887286375745e-05 | 0.00932598346010292 | 1909 | 1 | 0 | 16 | 12 | 8 | 12 | 7 | 4 | 0 | 0 | 0 | 0 |
| MER5A_dup6428 | 3.51899342694656 | -4.67094536603252 | 1.24858379193867 | -3.74099471432347 | 0.000183293376108095 | 0.0154769076193814 | 1910 | 1 | 0 | 11 | 9 | 5 | 6 | 6 | 7 | 0 | 0 | 0 | 0 |
| MIR1_Amn_dup2207 | 3.50890606573755 | -4.776638371494 | 1.38862015631614 | -3.43984519435892 | 0.000582047006181486 | 0.0337547543097392 | 1911 | 0 | 0 | 5 | 11 | 9 | 7 | 6 | 7 | 0 | 0 | 0 | 0 |
| AluSz_dup99902 | 3.41918565905417 | -4.78103947667379 | 1.44352703868555 | -3.31205398204893 | 0.000926136733202007 | 0.0454029130489933 | 1912 | 0 | 0 | 3 | 11 | 7 | 11 | 11 | 3 | 0 | 0 | 0 | 0 |
| L2_dup10653 | 4.10017729215415 | -5.01044507282278 | 1.40095963686599 | -3.57643785086583 | 0.000348308075857049 | 0.0240436650472531 | 1913 | 0 | 0 | 9 | 11 | 7 | 12 | 13 | 3 | 0 | 0 | 0 | 0 |
| L1MCa_dup203 | 2.8763708583768 | -4.39727313494878 | 1.32958574103882 | -3.30725052113835 | 0.000942165918850871 | 0.0459018451529257 | 1914 | 0 | 1 | 8 | 2 | 6 | 3 | 12 | 5 | 0 | 0 | 0 | 0 |
| L1MB1_dup586 | 3.50886768867544 | -4.34559205165111 | 1.28705969419132 | -3.37637179632254 | 0.000734485930353194 | 0.0390147881169134 | 1915 | 0 | 1 | 2 | 9 | 3 | 2 | 17 | 11 | 0 | 0 | 0 | 0 |
| L1PA17_dup510 | 3.83297941616703 | -4.21964684873759 | 1.18201412951985 | -3.56987851782421 | 0.000357146815073247 | 0.0244406914428247 | 1916 | 0 | 1 | 7 | 11 | 2 | 1 | 21 | 8 | 0 | 0 | 0 | 0 |
| L1M5_dup9950 | 4.44969238502083 | -4.31984333771839 | 1.21342437643184 | -3.56004331347059 | 0.000370793630372154 | 0.025042762237081 | 1917 | 1 | 1 | 5 | 16 | 9 | 7 | 7 | 10 | 0 | 0 | 1 | 0 |
| MIRc_dup20441 | 5.13952585291326 | -4.33197870515614 | 1.14324566123496 | -3.78919321720988 | 0.00015113732991022 | 0.0133938633530045 | 1918 | 0 | 1 | 14 | 18 | 7 | 5 | 9 | 11 | 0 | 0 | 1 | 0 |
| HAL1_dup1602 | 4.04475262012634 | -4.85359764852274 | 1.19037944151095 | -4.07735338772489 | 4.55512343444592e-05 | 0.00561011076983783 | 1919 | 0 | 3 | 11 | 14 | 6 | 4 | 9 | 6 | 0 | 0 | 0 | 0 |
| L1ME3G_dup2996 | 3.63197401655912 | -4.67838350858759 | 1.21798445005991 | -3.84108640168393 | 0.000122490986886203 | 0.0116213079607753 | 1920 | 1 | 1 | 11 | 16 | 6 | 3 | 8 | 3 | 0 | 0 | 0 | 0 |
| MIR_dup32488 | 4.16747600609693 | -4.0215154688721 | 1.21521187122874 | -3.3093121982143 | 0.000935254861273556 | 0.0457144465247499 | 1921 | 0 | 0 | 12 | 13 | 6 | 4 | 13 | 6 | 0 | 1 | 0 | 0 |
| MIR_dup32478 | 3.17971884656524 | -4.40008671135082 | 1.31230882777258 | -3.35293539007828 | 0.000799593697132039 | 0.0413968933546084 | 1922 | 0 | 0 | 12 | 12 | 4 | 4 | 6 | 4 | 0 | 0 | 0 | 0 |
| MIRb_dup43698 | 3.86888367505423 | -4.6931508886237 | 1.30826200391761 | -3.58731727633302 | 0.000334097701867859 | 0.02330613068562 | 1923 | 0 | 0 | 12 | 17 | 5 | 5 | 9 | 4 | 0 | 0 | 0 | 0 |
| MIRc_dup20437 | 3.56037926640157 | -4.56579154354612 | 1.33288500678538 | -3.42549546307658 | 0.000613679183263152 | 0.034866883223671 | 1924 | 0 | 0 | 8 | 16 | 7 | 2 | 8 | 6 | 0 | 0 | 0 | 0 |
| MIRc_dup20438 | 3.32711771740187 | -4.30023701469344 | 1.25513593121428 | -3.42611258888364 | 0.000612286523928305 | 0.0348293400006322 | 1925 | 0 | 0 | 11 | 11 | 4 | 2 | 8 | 7 | 0 | 0 | 0 | 0 |
| L2_dup4739 | 4.07330536129953 | -4.38221059676968 | 1.15769003413861 | -3.78530562373746 | 0.000153519789902364 | 0.0135696210728484 | 1926 | 2 | 0 | 8 | 14 | 2 | 1 | 18 | 9 | 0 | 0 | 0 | 0 |
| Tigger1_dup1606 | 4.35021407078017 | -3.99343920065157 | 1.15895331995031 | -3.4457291177377 | 0.000569520941330264 | 0.0332656588926594 | 1927 | 0 | 0 | 9 | 8 | 1 | 1 | 24 | 13 | 0 | 0 | 0 | 0 |
| HAL1_dup10566 | 4.87091113236467 | -4.82469630008849 | 1.28078637450131 | -3.76697972131928 | 0.000165234416607554 | 0.0143119933168183 | 1928 | 4 | 0 | 5 | 5 | 1 | 4 | 24 | 18 | 0 | 0 | 0 | 0 |
| LTR1D_dup141 | 3.48672603742985 | -4.12794683402182 | 1.19822530285238 | -3.44505062962302 | 0.000570952437438823 | 0.0333130639552934 | 1929 | 2 | 0 | 8 | 10 | 0 | 2 | 16 | 8 | 0 | 0 | 0 | 0 |
| LTR62_dup315 | 4.31951793569661 | -4.59016227300062 | 1.21601791585096 | -3.77474888582415 | 0.000160168925950443 | 0.0140116011963986 | 1930 | 4 | 0 | 11 | 9 | 0 | 3 | 18 | 11 | 0 | 0 | 0 | 0 |
| L2_dup4738 | 4.09635639816185 | -3.83396447430249 | 1.14521392641891 | -3.34781509887093 | 0.00081451334913767 | 0.0418777427342074 | 1931 | 4 | 0 | 7 | 10 | 3 | 0 | 11 | 15 | 0 | 0 | 0 | 1 |
| L2a_dup81493 | 3.12052228338809 | -4.13915793280416 | 1.23953525866506 | -3.33928212519095 | 0.000839952110858041 | 0.0426693520431741 | 1932 | 3 | 0 | 7 | 10 | 2 | 0 | 9 | 9 | 0 | 0 | 0 | 0 |
| L2a_dup14878 | 4.55018572057903 | -4.14950491412971 | 1.08187190214034 | -3.83548635094455 | 0.000125316066017533 | 0.0117999432705215 | 1933 | 3 | 1 | 11 | 15 | 4 | 3 | 9 | 11 | 1 | 0 | 0 | 0 |
| L1ME3D_dup3749 | 4.41199799939545 | -3.92395716276218 | 1.06347407819963 | -3.68975346291946 | 0.000224471482891242 | 0.0177684935023418 | 1934 | 2 | 0 | 10 | 19 | 2 | 3 | 15 | 7 | 1 | 0 | 0 | 0 |
| MIR_dup153551 | 3.64834367451208 | -3.88809921608557 | 1.16322593138047 | -3.34251421946151 | 0.0008302310149629 | 0.0423563784490485 | 1935 | 2 | 0 | 6 | 9 | 5 | 3 | 13 | 8 | 1 | 0 | 0 | 0 |
| MER61C_dup160 | 4.07233479559877 | -3.91482016446826 | 1.07878246266919 | -3.6289245514633 | 0.000284604397854444 | 0.020834648649285 | 1936 | 2 | 0 | 11 | 9 | 3 | 3 | 18 | 7 | 0 | 0 | 0 | 1 |
| MamGypLTR1b_dup7 | 3.58014286570384 | -3.72723323360628 | 1.10322662933558 | -3.37848374440618 | 0.000728867414913536 | 0.0387902227486057 | 1937 | 2 | 0 | 14 | 9 | 2 | 3 | 13 | 4 | 0 | 0 | 0 | 1 |
| L1PREC2_dup7589 | 3.54879657668429 | -3.68058066642865 | 1.11875864701125 | -3.28987907826345 | 0.00100230446063969 | 0.0479184559634305 | 1938 | 4 | 0 | 14 | 7 | 1 | 2 | 13 | 5 | 0 | 0 | 1 | 0 |

| | | | | | | | | | | | | | | | | | | | |
|---|---|---|---|---|---|---|---|---|---|---|---|---|---|---|---|---|---|---|---|
| MamRep434_dup451 | 4.14854078803128 | -3.50738940420945 | 1.04564449557409 | -3.35428476796388 | 0.000795704284766277 | 0.0413000608420682 | 1939 | 7 | 0 | 9 | 10 | 4 | 1 | 12 | 9 | 0 | 0 | 1 | 1 |
| AluY_dup103478 | 9.33149451932899 | -6.23094107553893 | 1.27106571750278 | -4.90213919684707 | 9.47986225518728e-07 | 0.00030523617106533 | 1940 | 0 | 5 | 11 | 16 | 30 | 16 | 27 | 15 | 0 | 0 | 0 | 0 |
| HERV17-int_dup775 | 7.2768812354149 | -5.12891309933585 | 1.30687564130435 | -3.92456094308778 | 8.68880617467069e-05 | 0.00909300946410297 | 1941 | 1 | 2 | 6 | 3 | 19 | 18 | 18 | 21 | 0 | 0 | 1 | 0 |
| L1PA3_dup3976 | 8.31302678416323 | -3.99558711220352 | 1.02262859612048 | -3.90717326638572 | 9.33821912362928e-05 | 0.00956737689934412 | 1942 | 4 | 3 | 7 | 12 | 20 | 24 | 19 | 14 | 1 | 0 | 2 | 0 |
| LTR1A2_dup289 | 10.2820531626301 | -5.65221692717644 | 1.29054170916898 | -4.37972433360259 | 1.18829535420189e-05 | 0.00212594079963101 | 1943 | 3 | 0 | 16 | 16 | 25 | 35 | 15 | 19 | 0 | 1 | 0 | 0 |
| L2b_dup19081 | 10.0917147648016 | -4.93842498610527 | 1.15504605129837 | -4.27552215823262 | 1.90689832915867e-05 | 0.00300489291278139 | 1944 | 2 | 0 | 22 | 31 | 16 | 21 | 19 | 18 | 0 | 0 | 0 | 2 |
| L2b_dup19086 | 11.0506441355157 | -6.38273631070325 | 1.21990916817863 | -5.23214061931588 | 1.67558202414353e-07 | 7.86386667291629e-05 | 1945 | 2 | 1 | 29 | 32 | 23 | 18 | 20 | 18 | 0 | 0 | 0 | 0 |
| L1MEf_dup4044 | 9.35683587815356 | -4.18690232164671 | 1.06162147105706 | -3.94387494582019 | 8.01754783118819e-05 | 0.00853576481539107 | 1946 | 5 | 2 | 13 | 13 | 34 | 30 | 16 | 6 | 0 | 0 | 2 | 1 |
| Tigger1_dup11380 | 6.62159109776581 | -5.00334784206533 | 1.25524517256819 | -3.98595266598688 | 6.7209910635612e-05 | 0.00752473348877803 | 1947 | 3 | 0 | 11 | 17 | 14 | 21 | 10 | 9 | 0 | 0 | 1 | 0 |
| L1ME4a_dup6658 | 7.44625114679623 | -5.86236944247737 | 1.26415839430237 | -4.63736939049678 | 3.5287144870143e-06 | 0.000848953661159664 | 1948 | 3 | 0 | 23 | 22 | 18 | 13 | 12 | 7 | 0 | 0 | 0 | 0 |
| Tigger18a_dup220 | 6.52926285333037 | -4.89561016861055 | 1.23196821985207 | -3.97381205920913 | 7.07313823944545e-05 | 0.00781711542741796 | 1949 | 2 | 0 | 19 | 23 | 11 | 13 | 12 | 6 | 0 | 0 | 0 | 1 |
| L2b_dup19082 | 10.0382005116237 | -4.1910825673422 | 1.0951898119275 | -3.82680930894164 | 0.000129815000893581 | 0.0120943462030635 | 1950 | 2 | 0 | 22 | 37 | 25 | 16 | 22 | 7 | 0 | 0 | 1 | 3 |
| L2b_dup19087 | 9.79543475763966 | -4.43693844522015 | 1.09618167858351 | -4.04763054510599 | 5.17387268068496e-05 | 0.00620202674497865 | 1951 | 2 | 1 | 28 | 27 | 10 | 32 | 19 | 8 | 0 | 0 | 2 | 1 |
| L2c_dup28751 | 10.7941477276135 | -5.0509361479389 | 1.19109912206298 | -4.24056743421212 | 2.22955438653293e-05 | 0.00335768068391117 | 1952 | 1 | 1 | 30 | 32 | 19 | 32 | 15 | 10 | 0 | 0 | 1 | 1 |
| L2a_dup32315 | 7.01990667147371 | -3.7385806348441 | 0.995607795663129 | -3.75507368577202 | 0.000173290488618145 | 0.0148216314179486 | 1953 | 1 | 1 | 17 | 32 | 6 | 10 | 21 | 5 | 1 | 0 | 0 | 2 |
| L2c_dup28754 | 6.12558450419194 | -4.06964700615308 | 1.13046941685763 | -3.59996205599749 | 0.000318263619503485 | 0.0225251502695944 | 1954 | 3 | 0 | 14 | 23 | 5 | 13 | 21 | 3 | 1 | 0 | 0 | 1 |
| HERVIP10B3-int_dup4 | 5.67097333298646 | -4.02020524065879 | 1.18171405005736 | -3.40201188304706 | 0.000668917292281121 | 0.0367334858133218 | 1955 | 0 | 0 | 14 | 18 | 8 | 10 | 21 | 4 | 0 | 0 | 1 | 1 |
| L1M4_dup3476 | 7.65519257388867 | -5.1522953942912 | 1.3647669050882 | -3.77522005778579 | 0.000159866470712095 | 0.0139954331273216 | 1956 | 0 | 0 | 20 | 27 | 10 | 23 | 14 | 7 | 0 | 0 | 0 | 1 |
| L2b_dup19075 | 5.16628282711407 | -5.23737973125199 | 1.36695166712292 | -3.83143007702337 | 0.000127400583019997 | 0.0119346300557994 | 1957 | 0 | 0 | 13 | 22 | 5 | 13 | 10 | 6 | 0 | 0 | 0 | 0 |
| L1PA2_dup732 | 8.26075450785746 | -5.08532419889947 | 1.20724225378 | -4.21234775619956 | 2.52730051184522e-05 | 0.00369120158319118 | 1958 | 3 | 0 | 30 | 33 | 6 | 14 | 5 | 15 | 0 | 0 | 0 | 1 |
| MIRb_dup43702 | 6.85642244218681 | -4.33361125978682 | 1.04464161033288 | -4.14841914865512 | 3.34779038323041e-05 | 0.00452499294019949 | 1959 | 2 | 3 | 19 | 28 | 12 | 7 | 13 | 6 | 0 | 0 | 0 | 2 |
| MIRb_dup43703 | 8.05906732920388 | -4.58156221322201 | 1.12324505711136 | -4.07886256361936 | 4.52565802536893e-05 | 0.00558538498186948 | 1960 | 0 | 2 | 22 | 28 | 12 | 14 | 15 | 11 | 0 | 0 | 0 | 2 |
| L1ME4a_dup6656 | 4.95374448149094 | -3.90342561384666 | 1.17109858137976 | -3.33313153641408 | 0.000858743330085499 | 0.0432637180649003 | 1961 | 1 | 0 | 8 | 17 | 10 | 9 | 10 | 8 | 0 | 0 | 0 | 2 |
| L1PB1_dup319 | 6.00637224951925 | -4.00405502803223 | 1.0022332034429 | -3.99513308307627 | 6.4657922397287e-05 | 0.00732623523940231 | 1962 | 2 | 1 | 13 | 20 | 9 | 5 | 14 | 12 | 0 | 1 | 0 | 1 |
| L1ME3G_dup2994 | 5.28476869379173 | -3.94658241562046 | 1.12655198635797 | -3.50324038607341 | 0.000459634460680613 | 0.0288494279559771 | 1963 | 0 | 3 | 15 | 20 | 6 | 10 | 14 | 2 | 0 | 0 | 1 | 1 |
| MLT1K_dup3033 | 6.36127186955739 | -5.63869086908091 | 1.32762854010099 | -4.24719015806333 | 2.16468224209444e-05 | 0.0032849329131212 | 1964 | 0 | 2 | 19 | 21 | 10 | 17 | 15 | 2 | 0 | 0 | 0 | 0 |
| L2_dup10656 | 7.42507758091105 | -4.93425460906806 | 1.11233011629718 | -4.43596243307125 | 9.16618333962452e-06 | 0.00173512860853643 | 1965 | 1 | 4 | 18 | 35 | 9 | 7 | 15 | 9 | 1 | 0 | 0 | 0 |
| MER103C_dup1638 | 6.21312333020546 | -5.51302958371528 | 1.24862366981814 | -4.41528517917511 | 1.00876913718214e-05 | 0.00186360699624178 | 1966 | 0 | 3 | 17 | 22 | 15 | 4 | 11 | 9 | 0 | 0 | 0 | 0 |
| L1PA6_dup5436 | 5.55119615240898 | -3.31708768516977 | 1.00108391108254 | -3.31349614997085 | 0.000921373755908553 | 0.0452624254259921 | 1967 | 0 | 3 | 10 | 22 | 9 | 6 | 15 | 6 | 1 | 1 | 0 | 1 |
| L2a_dup32314 | 6.01128508692127 | -3.98399614674889 | 1.12494372196829 | -3.54150707181883 | 0.000397848193721764 | 0.0262110661955531 | 1968 | 0 | 1 | 11 | 22 | 12 | 6 | 19 | 7 | 1 | 0 | 1 | 0 |
| LTR102_Mam_dup42 | 6.37739871886771 | -4.64965946715435 | 1.16625377332522 | -3.98683337494997 | 6.6961016667778e-05 | 0.00750853255043879 | 1969 | 0 | 1 | 17 | 21 | 10 | 5 | 17 | 11 | 1 | 0 | 0 | 0 |
| MER5B_dup4428 | 6.3057837041043 | -4.72695871046887 | 1.21143806507572 | -3.90194005516362 | 9.54247920680995e-05 | 0.00972001180369692 | 1970 | 1 | 0 | 16 | 20 | 6 | 12 | 20 | 8 | 0 | 1 | 0 | 0 |
| L1MA7_dup6578 | 4.98789767957709 | -4.35205558726057 | 1.16219391761188 | -3.74468969533357 | 0.000180616868384396 | 0.0152997726467047 | 1971 | 5 | 0 | 12 | 23 | 2 | 7 | 13 | 5 | 0 | 1 | 0 | 0 |
| MLT1A_dup1561 | 4.63328986131869 | -4.28560336771412 | 1.18255829682936 | -3.62401023205752 | 0.00029007012164065 | 0.0211202647380373 | 1972 | 3 | 0 | 11 | 23 | 5 | 6 | 7 | 6 | 0 | 1 | 0 | 0 |
| LTR16C_dup1074 | 6.03650896373986 | -4.12583713813302 | 1.07515402918747 | -3.8374381959495 | 0.000124324510364232 | 0.0117251367194323 | 1973 | 5 | 2 | 11 | 21 | 13 | 10 | 10 | 6 | 0 | 1 | 1 | 0 |
| AluSz6_dup15111 | 4.92258854377016 | -3.79250697253094 | 1.08384521599188 | -3.49912230692481 | 0.000466792408335715 | 0.0291431426506715 | 1974 | 2 | 2 | 10 | 7 | 6 | 12 | 13 | 9 | 0 | 1 | 1 | 0 |
| L1M3_dup6572 | 4.13898918063353 | -4.23806366043073 | 1.12828230174924 | -3.75620857817252 | 0.000172506936485574 | 0.0147811795135025 | 1975 | 3 | 4 | 12 | 8 | 4 | 7 | 7 | 7 | 0 | 0 | 1 | 0 |
| AluJo_dup13737 | 8.696938593639 | -2.83708100562251 | 0.844821702793221 | -3.35820090350699 | 0.000784515713487232 | 0.0409505595573397 | 1976 | 0 | 2 | 20 | 35 | 11 | 12 | 22 | 8 | 1 | 1 | 1 | 4 |
| L1M4_dup3477 | 10.6241968471828 | -3.93206729658688 | 0.948631838154048 | -4.14498769537224 | 3.3983195784287e-05 | 0.00457621137752573 | 1977 | 1 | 3 | 23 | 35 | 14 | 25 | 22 | 13 | 1 | 1 | 0 | 2 |
| AluJr_dup25105 | 8.89839986857303 | -2.65998750281627 | 0.663296237044619 | -4.01025568103341 | 6.06530325369338e-05 | 0.00695860486018961 | 1978 | 5 | 3 | 20 | 16 | 10 | 14 | 21 | 17 | 1 | 2 | 1 | 4 |
| L1ME2_dup11349 | 8.88909323283677 | -2.55750553596257 | 0.725052534431282 | -3.52733824724664 | 0.000419760148834782 | 0.027141371840935 | 1979 | 3 | 1 | 13 | 16 | 9 | 11 | 33 | 20 | 2 | 1 | 2 | 3 |
| ERVL-E-int_dup7594 | 11.1648858364669 | -3.14207060103522 | 0.650193613309924 | -4.83251532576574 | 1.34818729615707e-06 | 0.000405045169660846 | 1980 | 3 | 5 | 28 | 30 | 9 | 6 | 36 | 21 | 2 | 1 | 2 | 2 |
| L1M1_dup8761 | 10.4234104395547 | -3.42698397014215 | 0.796327321318523 | -4.30348661712108 | 1.68131008460679e-05 | 0.00272521123563929 | 1981 | 2 | 2 | 24 | 23 | 16 | 7 | 35 | 20 | 1 | 0 | 2 | 3 |
| LTR16E1_dup413 | 9.08670698530128 | -3.73012263513948 | 0.940462162903529 | -3.96626550463584 | 7.30075339432042e-05 | 0.00797605632307147 | 1982 | 1 | 3 | 30 | 30 | 9 | 14 | 11 | 15 | 1 | 0 | 3 | 0 |
| MSTA-int_dup1618 | 9.43052034404987 | -3.14591213530288 | 0.857940972029155 | -3.66681652685538 | 0.000245588872571126 | 0.0188932718631778 | 1983 | 0 | 4 | 42 | 40 | 3 | 5 | 13 | 12 | 1 | 1 | 3 | 0 |
| L1MA10_dup4471 | 5.36439432441093 | -3.35555580431273 | 0.997747976542301 | -3.36312965117847 | 0.000770641614405744 | 0.040405017922491 | 1984 | 6 | 1 | 16 | 22 | 8 | 2 | 4 | 8 | 1 | 0 | 1 | 1 |
| MLT1C_dup2758 | 5.33175848418245 | -3.35535780855327 | 0.938765385964846 | -3.57422403799506 | 0.000351268069028037 | 0.0241627001658766 | 1985 | 3 | 2 | 20 | 12 | 5 | 7 | 7 | 9 | 1 | 0 | 1 | 1 |
| MSTD_dup1335 | 7.332961033598 | -3.84293556927661 | 1.03684584251396 | -3.70637119975225 | 0.000210250082860889 | 0.0170048151653355 | 1986 | 1 | 1 | 25 | 19 | 13 | 12 | 9 | 11 | 1 | 0 | 1 | 1 |
| MLT1H1_dup1564 | 9.64082385706514 | -2.87350604202092 | 0.755177572643953 | -3.80507333124379 | 0.0001417621852982 | 0.0128160125163242 | 1987 | 0 | 0 | 21 | 25 | 0 | 4 | 34 | 31 | 0 | 0 | 5 | 2 |
| MLT2D_dup4369 | 8.48155849555134 | -2.26053866279656 | 0.654813252099908 | -3.45218832323153 | 0.000556059477661966 | 0.03272622519707 | 1988 | 1 | 1 | 24 | 25 | 0 | 5 | 28 | 17 | 0 | 1 | 6 | 3 |
| L1MCa_dup1079 | 6.12519795600859 | -3.2660271391484 | 0.885689547319581 | -3.68755299081104 | 0.000226420952008048 | 0.0178752355164257 | 1989 | 1 | 0 | 17 | 14 | 0 | 0 | 25 | 19 | 0 | 0 | 1 | 1 |
| L1PA11_dup3215 | 6.22326535202345 | -2.85808942157838 | 0.770906310310689 | -3.70744068812528 | 0.000209364430634971 | 0.01696778910486 | 1990 | 1 | 0 | 16 | 24 | 1 | 0 | 19 | 16 | 0 | 0 | 2 | 2 |
| L1PA16_dup1122 | 7.61818867272257 | -2.38025172110485 | 0.697330975055587 | -3.41337443229897 | 0.000641637192789596 | 0.035843779007008 | 1991 | 1 | 0 | 20 | 25 | 0 | 0 | 24 | 21 | 0 | 0 | 5 | 2 |
| MLT1L_dup10543 | 7.38085515194266 | -3.08311591333648 | 0.925233439094066 | -3.3322573342737 | 0.000861445606408236 | 0.0433539628304598 | 1992 | 0 | 0 | 30 | 46 | 0 | 1 | 16 | 6 | 0 | 0 | 1 | 2 |
| L1PA3_dup2692 | 6.59936088633577 | -3.0069330031317 | 0.826705811414757 | -3.63724672261089 | 0.000275567945276892 | 0.020401459336296 | 1993 | 1 | 1 | 31 | 21 | 0 | 1 | 23 | 7 | 0 | 0 | 1 | 3 |
| L1MA9_dup13759 | 6.76249768706634 | -3.69972801813706 | 0.926549338077837 | -3.99301782008963 | 6.52376635901851e-05 | 0.00736923588814351 | 1994 | 1 | 1 | 31 | 34 | 1 | 1 | 15 | 6 | 0 | 0 | 1 | 1 |
| MamRTE1_dup1581 | 7.52249284003416 | -3.7394859968604 | 0.900042867246195 | -4.15478654733621 | 3.25591350030323e-05 | 0.0044346399183924 | 1995 | 0 | 1 | 33 | 37 | 1 | 1 | 19 | 8 | 0 | 0 | 1 | 1 |
| AluSg4_dup2976 | 8.30072909567192 | -3.72951244781984 | 0.825285177419609 | -4.51905904754134 | 6.21150710967983e-06 | 0.00129196090097891 | 1996 | 3 | 1 | 31 | 31 | 3 | 0 | 21 | 16 | 0 | 1 | 0 | 2 |
| L1MCa_dup3375 | 9.61257735546296 | -3.54628885764733 | 0.820043880178855 | -4.32451109430128 | 1.52870511132073e-05 | 0.00253837589963938 | 1997 | 1 | 0 | 36 | 34 | 1 | 1 | 33 | 18 | 0 | 2 | 0 | 1 |
| L1PA5_dup6227 | 9.9725986900855 | -3.59768908383577 | 0.866391503699259 | -4.15249811254451 | 3.2886546747436e-05 | 0.00446390316735049 | 1998 | 4 | 0 | 34 | 50 | 3 | 6 | 25 | 10 | 0 | 2 | 0 | 2 |
| L1ME1_dup3212 | 8.16879760270501 | -4.12804651217831 | 0.903040248332414 | -4.57127632993248 | 4.84762532122136e-06 | 0.00107200098018898 | 1999 | 2 | 0 | 34 | 36 | 3 | 2 | 18 | 12 | 0 | 0 | 0 | 2 |
| MER41-int_dup233 | 10.6216771272403 | -4.9244701536109 | 1.06124971880426 | -4.64025579121892 | 3.4797818057087e-06 | 0.000838905938744846 | 2000 | 3 | 0 | 57 | 46 | 1 | 6 | 11 | 15 | 0 | 0 | 0 | 1 |
| Arthur1B_dup397 | 11.9786532549018 | -3.91949892524934 | 0.782686846446904 | -5.0077485561976 | 5.50704113986451e-07 | 0.000200134581944965 | 2001 | 0 | 1 | 46 | 58 | 0 | 1 | 25 | 24 | 0 | 0 | 1 | 2 |
| L1PA12_dup1703 | 12.4010881095064 | -3.77348800944123 | 0.905521149273865 | -4.16720030500357 | 3.0836352232258e-05 | 0.00426344373518702 | 2002 | 0 | 0 | 62 | 68 | 0 | 2 | 15 | 16 | 0 | 0 | 2 | 1 |
| MLT1K_dup16703 | 8.20144814499443 | -3.56783047724985 | 0.912783908745497 | -3.90873507197708 | 9.27806408233369e-05 | 0.00953228944790576 | 2003 | 1 | 0 | 37 | 41 | 0 | 0 | 17 | 12 | 0 | 0 | 0 | 2 |
| MLT1H1_dup508 | 10.1790939684755 | -3.10003647929903 | 0.914456307786308 | -3.39003236448061 | 0.000698843726583965 | 0.0377496184712255 | 2004 | 0 | 0 | 43 | 59 | 0 | 0 | 20 | 13 | 0 | 1 | 0 | 2 |
| THE1B_dup20516 | 9.02511136496458 | -2.9722028152803 | 0.906461374493348 | -3.27890729700598 | 0.00104209862779447 | 0.049156072268067 | 2005 | 0 | 0 | 38 | 53 | 0 | 0 | 19 | 10 | 0 | 1 | 1 | 1 |
| L2b_dup85159 | 11.5996239316699 | -3.73583460035446 | 0.920362764598743 | -4.0590892461661 | 4.92644838417977e-05 | 0.00596554039298748 | 2006 | 0 | 0 | 51 | 65 | 1 | 0 | 24 | 14 | 0 | 1 | 1 | 0 |
| MER47A_dup384 | 8.56667878411533 | -2.89836604479453 | 0.798209327948851 | -3.63108515938097 | 0.000282232030607215 | 0.0207335544164447 | 2007 | 0 | 0 | 33 | 37 | 2 | 1 | 23 | 13 | 0 | 1 | 4 | 0 |
| AluJb_dup60101 | 13.3182203481428 | -3.73362284166243 | 0.907536107514314 | -4.11402126124611 | 3.88825599366819e-05 | 0.00502277394500201 | 2008 | 0 | 0 | 59 | 63 | 0 | 1 | 28 | 23 | 1 | 1 | 0 | 0 |
| L1PA16_dup7523 | 13.1413135241136 | -3.87278143357673 | 0.912725260548871 | -4.24309658225939 | 2.20456498605244e-05 | 0.003330586794738 | 2009 | 0 | 0 | 50 | 62 | 1 | 1 | 36 | 23 | 0 | 2 | 0 | 0 |
| MIRb_dup38502 | 12.6319100349812 | -2.44190407473447 | 0.740509513834215 | -3.29759986754359 | 0.000975149896850236 | 0.0469892704244484 | 2010 | 0 | 0 | 59 | 40 | 1 | 1 | 31 | 24 | 1 | 2 | 2 | 3 |

| | | | | | | | | | | | | | | | | | | | |
|---|---|---|---|---|---|---|---|---|---|---|---|---|---|---|---|---|---|---|---|
| LTR8_dup475 | 12.8612463200713 | -3.28568193252232 | 0.769658421325062 | -4.26901316413275 | 1.96339682475051e-05 | 0.00305947885305884 | 2011 | 0 | 0 | 55 | 64 | 1 | 0 | 28 | 19 | 1 | 0 | 1 | 3 |
| MER74A_dup270 | 14.1127895138423 | -3.15055506985414 | 0.832209093956735 | -3.7857734224879 | 0.00015323124593034 | 0.0135592272384309 | 2012 | 0 | 0 | 64 | 77 | 2 | 0 | 25 | 17 | 1 | 1 | 1 | 2 |
| AluSx3_dup12248 | 3.4955428458535 | -3.62090683266464 | 1.09641703256669 | -3.30249049870026 | 0.000958303344229037 | 0.0465071277882771 | 2013 | 2 | 3 | 8 | 8 | 0 | 1 | 10 | 11 | 0 | 0 | 0 | 1 |
| Charlie7_dup1348 | 3.4255176098843 | -4.20989444656439 | 1.26069239247281 | -3.33935103574855 | 0.00083974375328329 | 0.0426693520431741 | 2014 | 3 | 2 | 9 | 8 | 0 | 0 | 7 | 13 | 0 | 0 | 0 | 0 |
| L1MA4_dup4710 | 5.16765116039266 | -3.97758219842922 | 1.06987696050946 | -3.71779405038794 | 0.000200969999894116 | 0.0164953458416024 | 2015 | 4 | 1 | 7 | 12 | 1 | 0 | 26 | 15 | 0 | 0 | 1 | 0 |
| MER47A_dup2176 | 4.51487865050949 | -3.9770663565138 | 1.107742918711 | -3.59024308739578 | 0.000330369715294427 | 0.0231206330221905 | 2016 | 5 | 3 | 9 | 15 | 0 | 0 | 18 | 9 | 0 | 0 | 1 | 0 |
| L1MEg_dup4932 | 4.39642806951556 | -4.29055363005638 | 1.22605857973384 | -3.49946870482144 | 0.000466186324955058 | 0.0291223867885944 | 2017 | 4 | 0 | 9 | 15 | 0 | 0 | 19 | 11 | 0 | 0 | 0 | 0 |
| LTR33A__dup451 | 3.68031974538592 | -4.17717494407036 | 1.19096850168458 | -3.50737650757506 | 0.000452548348542243 | 0.0285326097886149 | 2018 | 2 | 2 | 7 | 14 | 0 | 0 | 16 | 8 | 0 | 0 | 0 | 0 |
| MIRb_dup71042 | 3.80242716119317 | -3.898729410859 | 1.14147800853651 | -3.41550987553197 | 0.000636627209555098 | 0.0356679512229172 | 2019 | 1 | 1 | 7 | 15 | 0 | 0 | 16 | 10 | 0 | 0 | 0 | 0 |
| L1MA2_dup2220 | 4.09725949749614 | -4.13490623959525 | 1.1406956712659 | -3.62489868573489 | 0.000289074754381556 | 0.0210626434775512 | 2020 | 2 | 1 | 10 | 20 | 0 | 0 | 14 | 8 | 0 | 0 | 0 | 0 |
| MER52A_dup550 | 3.86570386979305 | -4.08439528728536 | 1.15576590197482 | -3.53392956160626 | 0.000409430176822119 | 0.0267276468735572 | 2021 | 1 | 2 | 9 | 18 | 0 | 0 | 12 | 9 | 0 | 0 | 0 | 0 |
| AluJb_dup31103 | 3.76095267115035 | -4.06169397828313 | 1.15337699700823 | -3.52156665931334 | 0.000429004793967493 | 0.0275593884193815 | 2022 | 1 | 2 | 11 | 11 | 0 | 0 | 15 | 9 | 0 | 0 | 0 | 0 |
| L1ME3A_dup4081 | 3.63790406783956 | -3.86695857334843 | 1.13909930508404 | -3.39475105997289 | 0.000686910277630964 | 0.0373765270800771 | 2023 | 1 | 1 | 14 | 14 | 0 | 0 | 11 | 7 | 0 | 0 | 0 | 0 |
| MER11B_dup165 | 3.2501184711127 | -3.93144667663976 | 1.18803537771554 | -3.30920000395904 | 0.000935629741939482 | 0.0457144465247499 | 2024 | 2 | 1 | 12 | 11 | 0 | 0 | 11 | 6 | 0 | 0 | 0 | 0 |
| MLT1H1_dup1187 | 4.66085860044762 | -3.83117655767943 | 1.15158363604019 | -3.32687651836842 | 0.000878252819641593 | 0.0438290153990014 | 2025 | 4 | 2 | 6 | 10 | 0 | 0 | 20 | 16 | 0 | 1 | 0 | 0 |
| THE1B_dup20032 | 4.06989670634787 | -3.86668459793075 | 1.17434810385431 | -3.29262216649386 | 0.000992577649997748 | 0.0475498050143677 | 2026 | 6 | 2 | 12 | 11 | 0 | 0 | 15 | 7 | 0 | 1 | 0 | 0 |
| THE1B-int_dup2551 | 5.52486119099442 | -3.95732261245867 | 1.05123966578586 | -3.76443425914712 | 0.000166926575475178 | 0.014380706346132 | 2027 | 3 | 2 | 11 | 15 | 0 | 0 | 29 | 12 | 1 | 0 | 0 | 0 |
| LTR1D1_dup158 | 4.02585529626589 | -3.76285024904613 | 1.12833440110825 | -3.33487151091933 | 0.000853388212919297 | 0.0430760101211911 | 2028 | 3 | 3 | 6 | 11 | 0 | 0 | 22 | 8 | 0 | 0 | 0 | 1 |
| THE1B-int_dup2550 | 4.60665289564272 | -4.56946534219313 | 1.20081037806727 | -3.80531799662474 | 0.000141622121443377 | 0.0128086531580676 | 2029 | 3 | 3 | 12 | 9 | 0 | 0 | 22 | 11 | 0 | 0 | 0 | 0 |
| L1PB1_dup7302 | 4.77231703553499 | -4.60107191534068 | 1.1760019905696 | -3.91246949600156 | 9.13570809091578e-05 | 0.00942606504840546 | 2030 | 3 | 3 | 9 | 17 | 0 | 0 | 17 | 13 | 0 | 0 | 0 | 0 |
| MER110A_dup207 | 4.87059261587436 | -4.52030095912002 | 1.19720577117567 | -3.77570929572202 | 0.000159552987363507 | 0.0139731303390493 | 2031 | 1 | 4 | 8 | 17 | 0 | 0 | 19 | 14 | 0 | 0 | 0 | 0 |
| MER21C_dup1742 | 4.62119693012967 | -4.23286419865501 | 1.13425371539994 | -3.73184953347276 | 0.000190079032327557 | 0.015864098711404 | 2032 | 1 | 2 | 9 | 13 | 0 | 0 | 22 | 13 | 0 | 0 | 0 | 0 |
| ERVL-E-int_dup4370 | 6.05987797185954 | -3.543541407702 | 1.00792168799576 | -3.51569119893459 | 0.000438610813579883 | 0.0279950881566735 | 2033 | 0 | 1 | 14 | 27 | 0 | 0 | 31 | 9 | 0 | 1 | 0 | 0 |
| MLT2B1_dup422 | 6.42902502791714 | -3.28180101263455 | 0.899223851442906 | -3.64959293213645 | 0.000262656205495165 | 0.0197590874083948 | 2034 | 0 | 1 | 20 | 31 | 0 | 0 | 23 | 10 | 0 | 1 | 0 | 1 |
| L1MB7_dup22773 | 6.53715952845919 | -3.52018270082764 | 0.915574631368364 | -3.84477963917215 | 0.000120660818287706 | 0.0115073182638423 | 2035 | 2 | 1 | 21 | 15 | 0 | 0 | 34 | 12 | 0 | 1 | 0 | 1 |
| L1MB7_dup22777 | 8.30264191299679 | -3.76755757107809 | 0.844308127035377 | -4.46230167688558 | 8.1083962179678e-06 | 0.00156706758989073 | 2036 | 1 | 1 | 24 | 24 | 0 | 0 | 32 | 23 | 0 | 1 | 0 | 1 |
| MLT1H_dup3382 | 6.75190435271859 | -5.03426729708175 | 1.1838074863474 | -4.25260640360945 | 2.11296764209777e-05 | 0.00322703593431704 | 2037 | 4 | 4 | 16 | 18 | 0 | 0 | 15 | 26 | 0 | 0 | 0 | 0 |
| L1MA7_dup3348 | 5.95883231112245 | -4.04184812350078 | 1.03540632465743 | -3.90363476371272 | 9.47587472881852e-05 | 0.00966872422051222 | 2038 | 2 | 3 | 22 | 26 | 0 | 0 | 12 | 12 | 0 | 1 | 0 | 0 |
| L1P2_dup654 | 5.25621040892716 | -4.68766830266973 | 1.13000926366656 | -4.14834502104841 | 3.34887435287473e-05 | 0.00452499294019949 | 2039 | 2 | 3 | 13 | 23 | 0 | 1 | 12 | 14 | 0 | 0 | 0 | 0 |
| THE1C_dup2506 | 4.7910104579997 | -4.12740649110607 | 1.08076179413637 | -3.81897890312108 | 0.00013400522145716 | 0.0123468775134145 | 2040 | 1 | 1 | 14 | 20 | 0 | 0 | 13 | 13 | 0 | 0 | 0 | 0 |
| SVA_C_dup50 | 7.98377246892943 | -3.09824863628825 | 0.785083106909881 | -3.9463957497227 | 7.93364415977558e-05 | 0.00846537620287874 | 2041 | 3 | 1 | 26 | 31 | 0 | 0 | 27 | 14 | 0 | 0 | 0 | 5 |
| L1PA10_dup536 | 7.42502373506036 | -3.5024326769939 | 0.872217850220681 | -4.015548037807 | 5.93078166009127e-05 | 0.00686384130784084 | 2042 | 2 | 2 | 16 | 40 | 0 | 0 | 28 | 11 | 0 | 0 | 0 | 3 |
| L2a_dup46293 | 6.54100249371228 | -3.41388432442287 | 0.905411532594652 | -3.77053328958562 | 0.00016289907000293 | 0.0141774623668315 | 2043 | 3 | 2 | 17 | 24 | 0 | 0 | 33 | 8 | 0 | 0 | 0 | 3 |
| THE1D_dup3189 | 4.86754740825031 | -2.9792004325783 | 0.908322685938143 | -3.27989213381948 | 0.00103846782510389 | 0.0490568324622138 | 2044 | 2 | 2 | 22 | 17 | 0 | 0 | 10 | 8 | 0 | 0 | 1 | 2 |
| Tigger1_dup4283 | 4.9051023390276 | -2.92118079210918 | 0.860934018779283 | -3.39303678143781 | 0.000691223561595263 | 0.0374744092446271 | 2045 | 1 | 2 | 15 | 13 | 0 | 0 | 16 | 13 | 0 | 0 | 1 | 2 |
| LTR16C_dup1633 | 5.73694295808498 | -2.97216327961071 | 0.895348076096089 | -3.319561809492 | 0.000901588434944193 | 0.0445821616253597 | 2046 | 1 | 4 | 14 | 21 | 0 | 0 | 20 | 12 | 0 | 0 | 0 | 4 |
| L2c_dup39573 | 4.32411320222925 | -3.68996292892336 | 1.02965245939044 | -3.58369748478805 | 0.000338764416181544 | 0.0235626197100543 | 2047 | 2 | 2 | 11 | 20 | 0 | 0 | 14 | 8 | 0 | 0 | 0 | 1 |
| THE1B_dup6490 | 4.48864401743883 | -3.33442916195315 | 0.99994602459292 | -3.33460914883942 | 0.000854193696610491 | 0.0430983828103647 | 2048 | 3 | 2 | 14 | 22 | 0 | 0 | 12 | 6 | 0 | 0 | 0 | 2 |
| L1ME1_dup5863 | 4.68045044479398 | -4.80181174976417 | 1.34970492887031 | -3.55767519778063 | 0.000374151549996742 | 0.0251846859166972 | 2049 | 2 | 0 | 17 | 27 | 0 | 7 | 7 | 4 | 0 | 0 | 0 | 0 |
| MIRc_dup89853 | 5.7253624041955 | -3.44511242538738 | 0.994169152143581 | -3.46531816840141 | 0.000529604520416142 | 0.0316387841363142 | 2050 | 2 | 0 | 18 | 34 | 0 | 3 | 12 | 7 | 0 | 1 | 1 | 0 |
| MER5B_dup24170 | 6.49411949474687 | -3.0034798361667 | 0.896727676267878 | -3.34937787207259 | 0.000809932530652202 | 0.0416962799520919 | 2051 | 3 | 0 | 30 | 30 | 0 | 1 | 8 | 10 | 0 | 0 | 2 | 2 |
| MLT1G1_dup1571 | 4.32849707463246 | -3.75881885145931 | 1.08576500420105 | -3.46190827381217 | 0.000536359921596369 | 0.0319301220675931 | 2052 | 3 | 0 | 21 | 18 | 1 | 1 | 8 | 5 | 0 | 0 | 0 | 1 |
| THE1B-int_dup380 | 5.30713505966296 | -3.8248041602636 | 1.11742777927998 | -3.42286475348603 | 0.000619648998200249 | 0.035038735866387 | 2053 | 4 | 0 | 21 | 27 | 0 | 0 | 15 | 5 | 0 | 0 | 0 | 1 |
| LTR67B_dup2052 | 5.90333462988251 | -4.22987753231835 | 1.1553296242518 | -3.66118676741944 | 0.000251049660079428 | 0.0192074485017261 | 2054 | 1 | 1 | 27 | 36 | 0 | 0 | 11 | 5 | 0 | 0 | 0 | 0 |
| L2a_dup165195 | 5.07053023712732 | -4.22747087541582 | 1.26772811334467 | -3.33468259551521 | 0.000853968135296187 | 0.0430961403862538 | 2055 | 3 | 0 | 19 | 35 | 0 | 0 | 8 | 5 | 0 | 0 | 0 | 0 |
| L1PA3_dup5643 | 4.66513319364221 | -3.84833663576535 | 1.11874194930764 | -3.43987872998504 | 0.000581974892260286 | 0.0337547543097392 | 2056 | 1 | 0 | 18 | 25 | 0 | 0 | 12 | 7 | 0 | 0 | 0 | 0 |
| THE1B_dup21736 | 5.11975690963224 | -4.08513881911949 | 1.19156455297369 | -3.42838229697631 | 0.000607189764430944 | 0.0346803590223921 | 2057 | 2 | 0 | 26 | 24 | 0 | 0 | 14 | 4 | 0 | 0 | 0 | 0 |
| L1ME3A_dup13965 | 6.47585274942659 | -3.77621442048015 | 1.11697729583958 | -3.38074411587906 | 0.00072289829108112 | 0.0385242456370858 | 2058 | 0 | 0 | 30 | 35 | 0 | 0 | 10 | 11 | 0 | 0 | 0 | 0 |
| L1MEd_dup12001 | 6.28475849405913 | -3.55911846287619 | 1.01807272752699 | -3.49593733988108 | 0.000472399617849248 | 0.02936405419769 | 2059 | 1 | 0 | 35 | 25 | 0 | 0 | 12 | 9 | 0 | 0 | 1 | 0 |
| MSTD_dup6960 | 5.98709048113218 | -3.4731153920271 | 1.05274804050608 | -3.29909461560954 | 0.000969972122727405 | 0.046817703382766 | 2060 | 0 | 1 | 24 | 38 | 0 | 0 | 9 | 8 | 0 | 0 | 1 | 0 |
| ERV3-16A3_I-int_dup | 6.38242652242305 | -4.05213011241145 | 1.1508510233129 | -3.52098580122629 | 0.000429945638031034 | 0.0275974763385305 | 2061 | 0 | 1 | 22 | 46 | 0 | 0 | 9 | 9 | 0 | 0 | 0 | 0 |
| L1M6_dup3143 | 5.26862281447327 | -4.14713870258099 | 1.16025755623895 | -3.57432595916394 | 0.000351131279601739 | 0.0241614469219535 | 2062 | 0 | 2 | 13 | 35 | 0 | 0 | 11 | 10 | 0 | 0 | 0 | 0 |
| MIR_dup28627 | 5.14027634243306 | -3.67217495482688 | 1.06410691264672 | -3.45094549352488 | 0.000558626376181648 | 0.0328285778480905 | 2063 | 0 | 1 | 21 | 29 | 1 | 1 | 11 | 5 | 0 | 1 | 0 | 0 |
| L1PA6_dup5780 | 5.6323542676294 | -4.34142015854305 | 1.25245065573268 | -3.46634028148864 | 0.000527595091749993 | 0.0315424979149608 | 2064 | 0 | 2 | 28 | 36 | 1 | 0 | 6 | 4 | 0 | 0 | 0 | 0 |
| LTR16A_dup3757 | 6.87598003682379 | -4.10218623274926 | 1.15388989110957 | -3.55509331033713 | 0.000377844971920927 | 0.0253401070148526 | 2065 | 0 | 0 | 36 | 39 | 1 | 0 | 10 | 7 | 0 | 0 | 0 | 0 |
| MLT1H_dup4915 | 6.27997401528895 | -4.32896309322961 | 1.10570956937368 | -3.9150996004147 | 9.03668955734584e-05 | 0.00934482456937077 | 2066 | 0 | 1 | 29 | 33 | 1 | 0 | 14 | 7 | 0 | 0 | 0 | 0 |
| LTR46_dup95 | 6.96061170877219 | -4.18566898572541 | 1.01554420485519 | -4.12160196051956 | 3.76246880058902e-05 | 0.00491633388920897 | 2067 | 3 | 0 | 25 | 38 | 1 | 1 | 13 | 11 | 1 | 0 | 0 | 0 |
| L2a_dup152019 | 7.03582225211887 | -3.51331358889858 | 1.07150000247207 | -3.27887408380119 | 0.00104222127958612 | 0.049156072268067 | 2068 | 0 | 0 | 24 | 49 | 0 | 1 | 15 | 7 | 1 | 0 | 0 | 0 |
| MLT1D_dup10306 | 7.24523197403236 | -3.54795527773343 | 0.941458065850445 | -3.76857494393918 | 0.00016418219303676 | 0.0142526435988051 | 2069 | 0 | 0 | 32 | 33 | 1 | 2 | 21 | 7 | 1 | 0 | 1 | 0 |
| L2b_dup26246 | 4.99259344671035 | -3.68928617880556 | 1.06079683196889 | -3.47784426538876 | 0.000505463503707056 | 0.0307446367211347 | 2070 | 0 | 0 | 13 | 22 | 0 | 0 | 19 | 12 | 0 | 0 | 0 | 0 |
| MSTA-int_dup1348 | 4.59313312069126 | -3.88170828286489 | 1.08322331073243 | -3.58347927376142 | 0.000339047678593969 | 0.0235626197100543 | 2071 | 0 | 0 | 18 | 15 | 1 | 0 | 16 | 10 | 0 | 0 | 0 | 0 |
| L1ME1_dup30805 | 4.81954240050733 | -3.6936753297704 | 1.01343172724266 | -3.64472043895855 | 0.000267682614950275 | 0.0200295621059617 | 2072 | 1 | 0 | 15 | 22 | 0 | 2 | 19 | 6 | 0 | 0 | 0 | 1 |
| MLT1K_dup1452 | 4.43385604426241 | -3.48476744670698 | 0.995079918956559 | -3.50199755850878 | 0.000461783850284753 | 0.0289233008791408 | 2073 | 0 | 0 | 18 | 14 | 1 | 1 | 14 | 9 | 0 | 0 | 0 | 1 |
| L1MEf_dup6281 | 6.75559159348771 | -3.67923199161443 | 0.962424599566157 | -3.82287816964878 | 0.000131902960173155 | 0.0122171201873731 | 2074 | 0 | 0 | 13 | 38 | 3 | 2 | 26 | 9 | 0 | 1 | 0 | 1 |
| AluJb_dup47100 | 4.62262121767709 | -3.28771220061979 | 0.945808662640666 | -3.4760859468559 | 0.000508789203624987 | 0.0308758877443895 | 2075 | 1 | 0 | 15 | 18 | 2 | 2 | 10 | 10 | 0 | 1 | 1 | 0 |
| ERVL-E-int_dup4369 | 4.78822336591231 | -3.20161282296917 | 0.909845256330073 | -3.51885422350072 | 0.000433414789593857 | 0.0277527759754305 | 2076 | 0 | 1 | 13 | 18 | 1 | 1 | 17 | 10 | 0 | 1 | 1 | 0 |
| AluSx_dup20521 | 6.51900773981039 | -4.57145392741244 | 1.08679479214895 | -4.20636348318635 | 2.59512725478729e-05 | 0.00376295033374537 | 2077 | 1 | 0 | 19 | 26 | 5 | 7 | 13 | 13 | 0 | 0 | 1 | 0 |
| Charlie19a_dup1397 | 5.61562640893742 | -3.77267818247047 | 0.953515614529493 | -3.95659821924581 | 7.60246381115491e-05 | 0.00823009942679284 | 2078 | 3 | 0 | 9 | 26 | 4 | 2 | 15 | 13 | 0 | 0 | 1 | 1 |
| MIR_dup170843 | 6.25413274426963 | -3.46598237314086 | 0.849240783541069 | -4.08127169621882 | 4.47899599548479e-05 | 0.00554457539198024 | 2079 | 2 | 0 | 16 | 21 | 3 | 3 | 15 | 17 | 0 | 0 | 2 | 1 |
| L1ME1_dup25579 | 5.5452376016681 | -3.19332343234324 | 0.888128194768855 | -3.59556587793538 | 0.0003236872005368 | 0.022787004325127 | 2080 | 0 | 1 | 15 | 27 | 1 | 3 | 20 | 6 | 0 | 0 | 2 | 1 |
| L2b_dup94897 | 5.36552206927397 | -3.09713150171551 | 0.920676483917193 | -3.36397372564375 | 0.000768288570679105 | 0.0403022910579388 | 2081 | 0 | 0 | 19 | 22 | 2 | 2 | 22 | 4 | 0 | 0 | 1 | 2 |
| L1MC2_dup6215 | 5.48020286393523 | -3.3570233589989 | 0.95631939455491 | -3.51035791819461 | 0.000447503891082002 | 0.0283875662136492 | 2082 | 0 | 0 | 18 | 16 | 5 | 4 | 11 | 13 | 0 | 0 | 1 | 2 |

| | | | | | | | | | | | | | | | | | | | |
|---|---|---|---|---|---|---|---|---|---|---|---|---|---|---|---|---|---|---|---|
| L2_dup4735 | 4.2569594572749 | -3.37075781353424 | 0.975568786977449 | -3.45517185310702 | 0.000549942170305838 | 0.0324867278191641 | 2083 | 0 | 1 | 13 | 10 | 4 | 2 | 14 | 9 | 0 | 0 | 1 | 1 |
| MLT2A2_dup1686 | 5.2757208085083 | -3.5882017204539 | 1.0000697063011 | -3.58795161761812 | 0.000333286115527755 | 0.0232699818208237 | 2084 | 0 | 4 | 15 | 27 | 3 | 1 | 10 | 8 | 1 | 0 | 0 | 1 |
| MLT2A2_dup3203 | 5.64163053056447 | -4.16583851108993 | 1.02566076063721 | -4.06161439626669 | 4.87345271071318e-05 | 0.00592549094891855 | 2085 | 0 | 2 | 17 | 17 | 4 | 1 | 19 | 12 | 1 | 0 | 0 | 0 |
| L1MA4_dup3244 | 3.37214009208548 | -4.11589774593993 | 1.23353157735612 | -3.33667805631836 | 0.000847860992778057 | 0.0428879957031394 | 2086 | 0 | 2 | 9 | 16 | 2 | 0 | 13 | 4 | 0 | 0 | 0 | 0 |
| L1MEc_dup6193 | 3.94726112446478 | -4.23557865905538 | 1.18124878808715 | -3.58567873404063 | 0.000336202644646639 | 0.023418640071162 | 2087 | 1 | 2 | 15 | 17 | 1 | 0 | 6 | 9 | 0 | 0 | 0 | 0 |
| BLACKJACK_dup1902 | 4.8933101787357 | -4.42388875865975 | 1.12536103462822 | -3.9310840010746 | 8.45637171582389e-05 | 0.0089151963858814 | 2088 | 0 | 2 | 16 | 13 | 2 | 0 | 15 | 14 | 0 | 0 | 0 | 0 |
| L1MEi_dup1846 | 5.49328275985616 | -4.73068908754855 | 1.12823031704512 | -4.19301716686574 | 2.75268603414683e-05 | 0.00392059887420742 | 2089 | 1 | 4 | 12 | 20 | 1 | 0 | 21 | 13 | 0 | 0 | 0 | 0 |
| THE1D_dup12207 | 5.60291552984435 | -3.91764716063526 | 1.01778790782881 | -3.84917833126213 | 0.000118514704557476 | 0.0113526972184514 | 2090 | 1 | 2 | 20 | 32 | 0 | 1 | 12 | 7 | 0 | 0 | 1 | 0 |
| Tigger3b_dup5881 | 4.34359165567375 | -3.25377177530096 | 0.977351769328442 | -3.3291716221445 | 0.000871047121709701 | 0.0436058891412255 | 2091 | 2 | 2 | 13 | 22 | 0 | 1 | 9 | 7 | 0 | 0 | 2 | 0 |
| L1PB2_dup674 | 4.48856920210342 | -3.35266838812399 | 0.967407632155032 | -3.465621188718 | 0.000529008053500027 | 0.031611087468136 | 2092 | 2 | 3 | 11 | 18 | 1 | 0 | 10 | 11 | 0 | 0 | 2 | 0 |
| MIRb_dup90020 | 4.79313022835538 | -3.30428513373991 | 0.937271003062559 | -3.52543194331529 | 0.000422792796304976 | 0.0272683996487349 | 2093 | 1 | 2 | 17 | 20 | 1 | 0 | 14 | 7 | 0 | 0 | 2 | 0 |
| L1MC5a_dup14617 | 4.63443392666883 | -3.37651671884164 | 1.01974722350623 | -3.3111310734753 | 0.000929196738484359 | 0.0455122781258541 | 2094 | 2 | 2 | 20 | 22 | 2 | 0 | 7 | 5 | 0 | 0 | 2 | 0 |
| MSTB_dup4365 | 5.45165777245996 | -4.23527641350998 | 1.04784991806602 | -4.04187311607265 | 5.30259244453678e-05 | 0.0063138156431497 | 2095 | 3 | 4 | 21 | 20 | 2 | 0 | 10 | 10 | 0 | 0 | 1 | 0 |
| L1M4_dup9611 | 4.62960955495382 | -3.22100475701217 | 0.938173546272216 | -3.43327177557997 | 0.000596344014937288 | 0.034268915854522 | 2096 | 2 | 1 | 13 | 21 | 1 | 0 | 15 | 7 | 1 | 0 | 1 | 0 |
| L1PB3_dup1395 | 6.12226360888644 | -3.00874163633257 | 0.828556217276416 | -3.63130657111323 | 0.000281989968035043 | 0.0207285711875398 | 2097 | 4 | 2 | 21 | 24 | 1 | 1 | 15 | 9 | 1 | 0 | 3 | 0 |
| L3_dup42627 | 6.14062129340152 | -2.53174918979289 | 0.747201278412847 | -3.38830949964467 | 0.000703248610593301 | 0.0379186933823326 | 2098 | 3 | 3 | 21 | 21 | 3 | 2 | 14 | 8 | 0 | 0 | 1 | 6 |
| LTR73_dup82 | 6.84266050593633 | -3.76860334258065 | 0.891129178790861 | -4.22902025012145 | 2.34711221273894e-05 | 0.00347966331820797 | 2099 | 5 | 5 | 25 | 29 | 3 | 2 | 13 | 7 | 0 | 0 | 1 | 2 |
| L3_dup5400 | 6.22850238016432 | -3.13648238396119 | 0.805204893889521 | -3.89525996148694 | 9.80934650149554e-05 | 0.00992809341831075 | 2100 | 2 | 1 | 17 | 31 | 2 | 2 | 13 | 11 | 0 | 0 | 2 | 2 |
| MIRc_dup48417 | 4.83598083055934 | -3.41137807423272 | 0.938217674638986 | -3.63601983467789 | 0.000276883021619051 | 0.0204733397955769 | 2101 | 1 | 1 | 21 | 19 | 2 | 1 | 10 | 7 | 0 | 0 | 1 | 1 |
| L1PA13_dup4383 | 6.01918152032525 | -3.37700830511442 | 0.92589919952069 | -3.64727424633545 | 0.000265036997998212 | 0.0198894470064334 | 2102 | 3 | 3 | 15 | 38 | 1 | 0 | 11 | 8 | 0 | 0 | 2 | 1 |
| MLT1B-int_dup47 | 6.91017560291446 | -2.70355426083093 | 0.76437399425254 | -3.53695217414436 | 0.000404772956489691 | 0.0264883875681778 | 2103 | 1 | 3 | 23 | 32 | 1 | 0 | 18 | 9 | 0 | 0 | 4 | 2 |
| LTR1A1_dup336 | 5.23747425512583 | -3.60496127849537 | 0.9387005951566 | -3.84037391378661 | 0.000122847055365777 | 0.0116365308264591 | 2104 | 3 | 1 | 18 | 16 | 3 | 0 | 11 | 13 | 0 | 0 | 1 | 1 |
| AluSx1_dup40890 | 4.26154258034639 | -3.46308804100384 | 0.981269884977197 | -3.52919017899373 | 0.000416833463008551 | 0.0270255736209899 | 2105 | 3 | 2 | 11 | 13 | 4 | 0 | 16 | 6 | 0 | 0 | 1 | 1 |
| MIRb_dup89994 | 5.44757284913165 | -3.73962100143041 | 0.944921489504981 | -3.95759969792781 | 7.57066921857351e-05 | 0.00820811751848647 | 2106 | 2 | 4 | 10 | 21 | 3 | 0 | 16 | 13 | 0 | 0 | 1 | 1 |
| L2_dup51444 | 8.29098884039194 | -3.75170500626523 | 1.13392260597687 | -3.30860764790304 | 0.000937611321784775 | 0.0457661655404614 | 2107 | 0 | 2 | 34 | 58 | 0 | 0 | 15 | 5 | 1 | 0 | 0 | 0 |
| MSTA-int_dup1588 | 8.69015761803722 | -3.75115156554725 | 1.03095120464633 | -3.63853453843539 | 0.000274193859973307 | 0.0203440387294071 | 2108 | 1 | 0 | 48 | 37 | 0 | 0 | 22 | 8 | 1 | 0 | 0 | 0 |
| MSTA-int_dup1584 | 13.0522679507192 | -3.20092357700858 | 0.908030948153841 | -3.52512608024707 | 0.000423281279674892 | 0.0272874796460368 | 2109 | 3 | 2 | 55 | 75 | 1 | 0 | 26 | 12 | 1 | 2 | 0 | 1 |
| L1P4_dup1851 | 11.7477740101322 | -4.69360686566333 | 1.03048180623819 | -4.55476927127663 | 5.24431082240187e-06 | 0.00113648705186451 | 2110 | 4 | 1 | 44 | 81 | 0 | 0 | 22 | 10 | 0 | 0 | 0 | 1 |
| LTR16_dup2938 | 9.92254341222918 | -4.54234185900009 | 1.03844782314325 | -4.37416474642992 | 1.21898298291178e-05 | 0.00216780362370648 | 2111 | 3 | 2 | 49 | 56 | 0 | 0 | 15 | 9 | 0 | 0 | 0 | 1 |
| L1MC_dup7881 | 9.96834488043521 | -3.7197646552952 | 0.957403768951862 | -3.88526218083253 | 0.000102219514154586 | 0.0102398985115826 | 2112 | 3 | 3 | 51 | 50 | 0 | 0 | 17 | 8 | 1 | 0 | 0 | 2 |
| L3_dup5399 | 7.76088287464777 | -3.48993744167193 | 0.931758807236524 | -3.74553738002502 | 0.000180008040655807 | 0.0152721731603586 | 2113 | 5 | 1 | 25 | 43 | 0 | 0 | 17 | 11 | 1 | 0 | 0 | 2 |
| MLT1D_dup17476 | 10.0977936525359 | -4.80289259052075 | 1.10099274318447 | -4.36232901647385 | 1.28685149306792e-05 | 0.00225282475182657 | 2114 | 18 | 10 | 26 | 41 | 2 | 0 | 30 | 9 | 0 | 0 | 2 | 0 |
| L1M1_dup3085 | 10.6245812795923 | -4.9029803096234 | 0.956850027625541 | -5.1240844103755 | 2.98987026052381e-07 | 0.000123084371801142 | 2115 | 6 | 3 | 26 | 36 | 0 | 0 | 35 | 30 | 0 | 0 | 1 | 0 |
| MIR_dup149071 | 10.2499862209599 | -3.87541229854774 | 0.891652327933293 | -4.34632667592573 | 1.38436347220229e-05 | 0.00237836643145592 | 2116 | 6 | 7 | 37 | 25 | 1 | 0 | 24 | 26 | 0 | 0 | 4 | 0 |
| L1MEc_dup8722 | 11.9999219468983 | -5.01297635731953 | 0.909398952815614 | -5.51240612472527 | 3.53960993630517e-08 | 2.22225864354636e-05 | 2117 | 2 | 3 | 53 | 38 | 3 | 0 | 35 | 22 | 0 | 0 | 1 | 0 |
| MER11A_dup252 | 8.3925403661785 | -5.30407739742709 | 1.09106531343366 | -4.86137477942064 | 1.16573311711658e-06 | 0.000359969672019978 | 2118 | 3 | 5 | 18 | 40 | 1 | 0 | 28 | 17 | 0 | 0 | 0 | 0 |
| L1MEi_dup515 | 8.99335205641679 | -3.41036185722441 | 0.753549310457224 | -4.52573150807494 | 6.01869533485152e-06 | 0.00125845736392494 | 2119 | 3 | 2 | 26 | 44 | 0 | 4 | 20 | 16 | 0 | 0 | 2 | 3 |
| MLT1F2_dup544 | 8.99978885715428 | -2.93574097611164 | 0.702644852771222 | -4.17812919931473 | 2.93916688760106e-05 | 0.00411874560799248 | 2120 | 1 | 4 | 30 | 38 | 0 | 1 | 24 | 15 | 0 | 0 | 2 | 5 |
| L1ME3Cz_dup5052 | 8.57320753332686 | -3.07292397343595 | 0.721510587683405 | -4.25901438716563 | 2.05330255925862e-05 | 0.00315006531763605 | 2121 | 3 | 6 | 18 | 37 | 2 | 2 | 19 | 19 | 0 | 1 | 3 | 2 |
| MER84_dup28 | 8.6204730623638 | -2.4880482224564 | 0.666885230895893 | -3.73084918841875 | 0.000190835440175351 | 0.0158993451148334 | 2122 | 2 | 6 | 26 | 32 | 1 | 1 | 23 | 14 | 0 | 1 | 5 | 3 |
| L1MEi_dup523 | 7.59638983661382 | -2.63487167909295 | 0.714882612997792 | -3.68574033161035 | 0.000228038770750382 | 0.0179253140569193 | 2123 | 5 | 5 | 19 | 31 | 1 | 1 | 15 | 15 | 0 | 0 | 3 | 5 |
| THE1B_dup1942 | 8.15499346067151 | -2.3743608188123 | 0.664008505581152 | -3.57579880205633 | 0.000349160116151924 | 0.0240815226080828 | 2124 | 1 | 5 | 20 | 36 | 1 | 2 | 21 | 13 | 0 | 0 | 5 | 5 |
| L1P4_dup283 | 10.0369552507645 | -3.07222225332574 | 0.830863300781951 | -3.69762661370935 | 0.000217624679616548 | 0.0173826525733599 | 2125 | 2 | 7 | 40 | 50 | 1 | 1 | 23 | 6 | 0 | 0 | 2 | 6 |
| L1ME2z_dup597 | 11.7036428903729 | -3.031126143641 | 0.770832684975065 | -3.93227506140201 | 8.41457043668044e-05 | 0.00889191796504125 | 2126 | 1 | 2 | 47 | 65 | 1 | 0 | 24 | 12 | 0 | 0 | 1 | 7 |
| L1P4_dup285 | 10.3192895030764 | -2.56606183795244 | 0.725003424626951 | -3.53937891986207 | 0.00040106970882174 | 0.0263626886561596 | 2127 | 1 | 4 | 37 | 49 | 1 | 0 | 19 | 16 | 0 | 0 | 2 | 9 |
| LTR8B_dup147 | 6.55054997410921 | -2.59683961929296 | 0.781387512000526 | -3.32336974857005 | 0.000889369452930652 | 0.0441799381726591 | 2128 | 4 | 3 | 24 | 25 | 0 | 0 | 15 | 10 | 0 | 1 | 2 | 3 |
| Ricksha_c_dup103 | 7.89608527298435 | -2.45654034951918 | 0.721514583716024 | -3.40469950983837 | 0.000662368958166043 | 0.0365256185677352 | 2129 | 1 | 2 | 28 | 35 | 0 | 0 | 26 | 8 | 0 | 0 | 3 | 5 |
| L1MC1_dup2193 | 7.09178429722196 | -2.4994719936569 | 0.762863058144982 | -3.27643600901942 | 0.00105126132087555 | 0.0493964111882019 | 2130 | 1 | 3 | 24 | 32 | 0 | 0 | 12 | 14 | 0 | 0 | 4 | 3 |
| L1MB8_dup1418 | 6.38312835577209 | -3.14239914074498 | 0.851056296058712 | -3.69235167555613 | 0.000222189929087048 | 0.0176348248387414 | 2131 | 1 | 1 | 22 | 34 | 0 | 0 | 15 | 10 | 0 | 0 | 1 | 2 |
| Tigger20a_dup110 | 6.55885471845977 | -2.76622746797082 | 0.76132924050558 | -3.63341813343966 | 0.000279691216475922 | 0.0206105314944502 | 2132 | 2 | 1 | 23 | 28 | 0 | 0 | 16 | 12 | 0 | 0 | 2 | 3 |
| MER50_dup226 | 8.92241371399076 | -2.6480689876089 | 0.786646304359601 | -3.36627652470148 | 0.000761902890457848 | 0.0400733300558981 | 2133 | 2 | 2 | 34 | 47 | 0 | 0 | 18 | 11 | 1 | 1 | 3 | 1 |
| L1MC1_dup1302 | 7.54823366239941 | -2.62525043197696 | 0.745136798813842 | -3.52317914798465 | 0.000426403033106279 | 0.0274218638150018 | 2134 | 4 | 3 | 21 | 38 | 1 | 0 | 18 | 10 | 1 | 0 | 2 | 4 |
| L1PA13_dup648 | 8.32152901888677 | -2.77241359057151 | 0.794869511702557 | -3.48788518084331 | 0.000486857103225204 | 0.0299711957164091 | 2135 | 1 | 3 | 24 | 47 | 0 | 0 | 15 | 15 | 1 | 0 | 1 | 4 |
| L1MEd_dup3282 | 5.74999486274556 | -3.39258179566017 | 1.02490168865141 | -3.31015338663772 | 0.000932448583291021 | 0.0456339799670435 | 2136 | 3 | 0 | 24 | 26 | 6 | 0 | 9 | 6 | 0 | 0 | 2 | 1 |
| L1PA3_dup940 | 6.71477889185969 | -3.18049573676997 | 0.857833128721681 | -3.70759257282295 | 0.000209238938156914 | 0.0169633963811533 | 2137 | 5 | 0 | 16 | 30 | 1 | 0 | 27 | 9 | 0 | 0 | 2 | 2 |
| L1MA8_dup8018 | 7.81172020635532 | -3.07044677945226 | 0.753565710923122 | -4.07455744727418 | 4.6101932327823e-05 | 0.0056662037136757 | 2138 | 2 | 0 | 22 | 30 | 3 | 0 | 28 | 14 | 0 | 0 | 4 | 1 |
| MSTB1_dup4474 | 7.6693318910277 | -2.73689098982894 | 0.759308250055067 | -3.60445311851999 | 0.000312810905016087 | 0.02226845624372 | 2139 | 3 | 0 | 20 | 30 | 5 | 0 | 21 | 15 | 0 | 0 | 5 | 2 |
| SVA_D_dup758 | 14.7588466758036 | -3.95969704267274 | 0.869170737743084 | -4.55571830795249 | 5.22068620872796e-06 | 0.00113343193487407 | 2140 | 3 | 0 | 66 | 92 | 1 | 0 | 22 | 14 | 0 | 0 | 1 | 3 |
| Charlie1_dup581 | 10.2668640971469 | -4.00676116215976 | 0.938442904003541 | -4.26958437755382 | 1.95837553924759e-05 | 0.00305565656433971 | 2141 | 3 | 2 | 43 | 70 | 1 | 2 | 9 | 8 | 0 | 0 | 2 | 1 |
| THE1B-int_dup3036 | 10.4609023004639 | -3.59294883599691 | 0.939699098115157 | -3.82350993334315 | 0.000131565288785054 | 0.012202838846748 | 2142 | 2 | 0 | 51 | 68 | 2 | 1 | 10 | 6 | 0 | 0 | 2 | 2 |
| ERV3-16A3_I-int_dup | 7.33544889423214 | -4.32468707134194 | 1.00698509150403 | -4.29468828071984 | 1.74938962346991e-05 | 0.00281067024541642 | 2143 | 9 | 6 | 8 | 17 | 6 | 3 | 15 | 25 | 0 | 1 | 0 | 1 |
| L1M5_dup22412 | 6.25128280515886 | -3.52907277905171 | 0.934556611262547 | -3.77620011085694 | 0.000159239074749332 | 0.0139593168184873 | 2144 | 10 | 6 | 10 | 17 | 4 | 1 | 14 | 15 | 1 | 1 | 0 | 1 |
| MER5A_dup14421 | 5.54589164795631 | -3.05839849263196 | 0.846460529389658 | -3.61316137781076 | 0.000302486309541213 | 0.0217645622649209 | 2145 | 8 | 6 | 16 | 12 | 4 | 6 | 8 | 8 | 0 | 1 | 1 | 2 |
| MIRc_dup60274 | 6.22323733390127 | -3.68216274265691 | 0.93147158411909 | -3.95305965896876 | 7.7158189514656e-05 | 0.00832252156040541 | 2146 | 12 | 5 | 11 | 10 | 6 | 4 | 13 | 15 | 0 | 0 | 2 | 1 |
| LTR12_dup565 | 6.19509928347225 | -3.57103522559609 | 0.869720196301898 | -4.10595872187439 | 4.02641384141777e-05 | 0.00515090896725802 | 2147 | 3 | 1 | 14 | 19 | 2 | 5 | 17 | 16 | 0 | 0 | 0 | 3 |
| MLT1A0_dup9050 | 5.01920488488755 | -4.20739412937581 | 1.06049312467874 | -3.96739406552057 | 7.26627927457698e-05 | 0.00795763824028531 | 2148 | 5 | 1 | 12 | 12 | 3 | 1 | 12 | 16 | 0 | 0 | 0 | 1 |
| Tigger16a_dup533 | 5.0317629284138 | -3.68775704852114 | 1.03346804737925 | -3.56833194589116 | 0.000359261170162569 | 0.0245145123020219 | 2149 | 4 | 2 | 5 | 7 | 2 | 2 | 18 | 20 | 0 | 0 | 0 | 2 |
| AluSz_dup7850 | 4.82926921131967 | -4.47122404182385 | 1.15226892511424 | -3.88036502970048 | 0.000104299811586931 | 0.0103493423552393 | 2150 | 7 | 3 | 7 | 8 | 5 | 5 | 26 | 4 | 0 | 0 | 0 | 1 |
| L1M6_dup3229 | 4.77657662258155 | -2.9562650562463 | 0.89314683400467 | -3.30994293848758 | 0.000933149924020252 | 0.0456495254566042 | 2151 | 4 | 4 | 4 | 16 | 4 | 6 | 15 | 7 | 0 | 0 | 1 | 3 |
| MIR_dup157632 | 4.16252358677775 | -4.22097751116849 | 1.14082754783823 | -3.69992600473829 | 0.000215662339015536 | 0.0172722958328478 | 2152 | 4 | 2 | 3 | 13 | 5 | 5 | 14 | 8 | 0 | 0 | 0 | 1 |
| THE1A_dup1917 | 4.81675254665237 | -3.60001923991103 | 1.00564745245051 | -3.57980247564758 | 0.000343854024973129 | 0.0238121914511805 | 2153 | 2 | 3 | 13 | 6 | 1 | 5 | 20 | 10 | 1 | 0 | 0 | 1 |
| L1MA8_dup595 | 4.79065606203173 | -3.66048093067398 | 1.07010815322338 | -3.42066446241709 | 0.000624683523421331 | 0.0352396741735448 | 2154 | 1 | 3 | 18 | 13 | 1 | 9 | 8 | 7 | 1 | 0 | 0 | 1 |

| | | | | | | | | | | | | | | | | | | | |
|---|---|---|---|---|---|---|---|---|---|---|---|---|---|---|---|---|---|---|---|
| THE1C-int_dup899 | 5.44952718467377 | -3.78965238206701 | 1.00790230917535 | -3.7599401723443 | 0.000169953996655615 | 0.0146150609642996 | 2155 | 0 | 4 | 14 | 12 | 3 | 6 | 12 | 15 | 1 | 0 | 0 | 1 |
| L1ME3A_dup6482 | 6.11279616638827 | -3.29733588314227 | 0.935840110082119 | -3.52339662258431 | 0.000426053266095947 | 0.0274067776302964 | 2156 | 7 | 3 | 11 | 27 | 1 | 4 | 15 | 10 | 2 | 1 | 0 | 0 |
| MSTA_dup8134 | 6.51657213093197 | -4.72723949085419 | 1.09299251393658 | -4.32504288051188 | 1.52502194573711e-05 | 0.00253579924458674 | 2157 | 9 | 2 | 17 | 29 | 5 | 2 | 13 | 9 | 1 | 0 | 0 | 0 |
| L1PA3_dup1147 | 6.62085186662749 | -3.51193548593725 | 0.864139047195817 | -4.0640860950951 | 4.82210256884112e-05 | 0.00588108671371332 | 2158 | 3 | 5 | 12 | 18 | 1 | 3 | 23 | 17 | 1 | 1 | 0 | 1 |
| L1PA8A_dup269 | 7.87868756826222 | -2.35812934015701 | 0.718904337669068 | -3.28017125032638 | 0.00103744093593207 | 0.0490472721917925 | 2159 | 4 | 6 | 15 | 19 | 4 | 4 | 31 | 13 | 3 | 2 | 0 | 2 |
| L1PA16_dup12736 | 7.72478002449578 | -2.62243573978981 | 0.774791010647447 | -3.38470078221273 | 0.000712558837832981 | 0.0382401208062598 | 2160 | 4 | 3 | 14 | 20 | 6 | 9 | 16 | 19 | 4 | 0 | 0 | 2 |
| L1PA3_dup9975 | 5.63984343774257 | -2.89933810213677 | 0.844922998559712 | -3.43148204875367 | 0.000600292858348895 | 0.0344352180503101 | 2161 | 3 | 1 | 14 | 17 | 5 | 2 | 13 | 13 | 2 | 0 | 0 | 2 |
| LTR40c_dup269 | 5.03051234531841 | -3.76989255328086 | 1.10473756790703 | -3.41247791583939 | 0.000643751427952141 | 0.0359245795990491 | 2162 | 7 | 1 | 15 | 26 | 0 | 4 | 12 | 3 | 0 | 0 | 0 | 2 |
| THE1B_dup12172 | 4.14891914583227 | -3.73101858114011 | 1.06699169106455 | -3.49676441942824 | 0.000470937515875373 | 0.0292884731187876 | 2163 | 2 | 2 | 19 | 14 | 0 | 1 | 13 | 4 | 0 | 0 | 0 | 1 |
| L1MC1_dup2192 | 4.57333825646713 | -3.75635965265541 | 1.00585017702594 | -3.73451209578953 | 0.000188079452271825 | 0.0157358485760449 | 2164 | 1 | 1 | 12 | 22 | 0 | 2 | 13 | 9 | 0 | 0 | 0 | 1 |
| L2_dup4736 | 4.21869593706964 | -3.39200961196423 | 0.973687614495903 | -3.48367336860944 | 0.000494582680951651 | 0.0302456737546355 | 2165 | 2 | 2 | 11 | 15 | 0 | 3 | 16 | 6 | 0 | 0 | 0 | 2 |
| L1ME4a_dup14289 | 4.04996670025386 | -3.76956766160769 | 1.08522068244064 | -3.4735494103651 | 0.000513622784184058 | 0.0310471542025556 | 2166 | 2 | 2 | 16 | 13 | 1 | 1 | 4 | 11 | 0 | 0 | 0 | 1 |
| Charlie15b_dup783 | 3.94538955907421 | -3.196533091055 | 0.972165639304311 | -3.28805397127846 | 0.00100882497652029 | 0.04813882739989 | 2167 | 1 | 2 | 8 | 19 | 1 | 1 | 9 | 9 | 0 | 0 | 0 | 2 |
| L1MB7_dup4203 | 3.94692811727554 | -3.7550619256895 | 1.06428725596504 | -3.52824099381383 | 0.000418331111407026 | 0.0270784044164462 | 2168 | 1 | 1 | 10 | 18 | 2 | 2 | 6 | 10 | 0 | 0 | 0 | 1 |
| L2a_dup165187 | 4.58157158499093 | -3.60156242154258 | 1.07961835256485 | -3.33595887193503 | 0.000850057385159211 | 0.042971692069066 | 2169 | 5 | 2 | 22 | 20 | 2 | 1 | 5 | 3 | 0 | 0 | 0 | 2 |
| L2c_dup129297 | 4.87671300342516 | -3.78727578014332 | 1.01981351620248 | -3.71369443527884 | 0.0002042553996936 | 0.0166672686335711 | 2170 | 7 | 1 | 19 | 12 | 4 | 3 | 9 | 7 | 0 | 0 | 0 | 2 |
| L2c_dup59401 | 7.57314352737526 | -3.54157537747673 | 0.947988990137833 | -3.73588239348835 | 0.000187058083433361 | 0.0156669210561895 | 2171 | 14 | 4 | 26 | 30 | 3 | 2 | 17 | 4 | 0 | 1 | 0 | 3 |
| L2a_dup73368 | 5.55214604303179 | -3.05185244054266 | 0.875937479641878 | -3.4840984790266 | 0.000493797760493553 | 0.0302282484997167 | 2172 | 10 | 2 | 12 | 17 | 1 | 3 | 16 | 9 | 0 | 1 | 1 | 2 |
| L1PA8_dup4439 | 6.8925120196216 | -3.0245715452687 | 0.74915034624293 | -4.03733584378256 | 5.40616486569786e-05 | 0.00639033932008616 | 2173 | 3 | 3 | 21 | 27 | 3 | 2 | 18 | 10 | 0 | 1 | 1 | 3 |
| LTR16E2_dup1018 | 5.59762639589519 | -2.95347648573021 | 0.823806803699657 | -3.58515670478365 | 0.000336875868290831 | 0.0234586670636026 | 2174 | 3 | 2 | 17 | 16 | 1 | 2 | 25 | 6 | 0 | 1 | 1 | 2 |
| MER11C_dup368 | 8.37907854026951 | -3.50805223624481 | 0.810752094726539 | -4.3269110978098 | 1.51214957909992e-05 | 0.00252321138673569 | 2175 | 10 | 6 | 23 | 17 | 5 | 3 | 32 | 11 | 0 | 0 | 0 | 5 |
| L1ME3F_dup1936 | 9.49226149342623 | -4.29152098913436 | 0.909801786222436 | -4.71698457193964 | 2.3936583414331e-06 | 0.000647228895125728 | 2176 | 8 | 11 | 21 | 42 | 2 | 2 | 23 | 15 | 0 | 0 | 0 | 3 |
| L2a_dup73362 | 6.97092580944177 | -4.33797268676562 | 1.0211751949904 | -4.24802003421795 | 2.15668112415503e-05 | 0.00327487968582127 | 2177 | 8 | 12 | 14 | 14 | 2 | 2 | 21 | 15 | 0 | 0 | 0 | 2 |
| L1MA7_dup3346 | 10.3221552027802 | -4.70971992665567 | 0.941129229394645 | -5.00432860818175 | 5.6057061357086e-07 | 0.000203189465324983 | 2178 | 10 | 5 | 32 | 48 | 1 | 2 | 27 | 13 | 0 | 0 | 0 | 2 |
| L1PA14_dup1397 | 8.87692807250339 | -4.34400842323663 | 0.894889524881723 | -4.85423988375635 | 1.20849378607854e-06 | 0.000369505851841346 | 2179 | 6 | 2 | 30 | 39 | 1 | 1 | 21 | 16 | 0 | 0 | 0 | 2 |
| L1ME2z_dup2690 | 9.19888689876266 | -3.45885286924326 | 0.841516414401254 | -4.11026191533562 | 3.95210584423767e-05 | 0.0050941916425405 | 2180 | 10 | 5 | 24 | 48 | 2 | 1 | 13 | 15 | 0 | 1 | 1 | 3 |
| L1ME3_dup3690 | 8.92652619318237 | -3.87230650056762 | 0.87230047638779 | -4.43918879490116 | 9.02985806141842e-06 | 0.00171204669823288 | 2181 | 9 | 8 | 26 | 44 | 2 | 4 | 12 | 11 | 0 | 0 | 1 | 3 |
| LTR81B_dup569 | 7.00080966208047 | -3.29671809793292 | 0.950262472792528 | -3.46927106175617 | 0.000521872633511056 | 0.0313184125229311 | 2182 | 3 | 8 | 24 | 12 | 0 | 5 | 7 | 22 | 1 | 0 | 2 | 1 |
| L1M3_dup2446 | 4.32326897998333 | -3.0928011893982 | 0.93091098782841 | -3.32233825772425 | 0.000892664070796284 | 0.0443158449801153 | 2183 | 4 | 3 | 14 | 15 | 0 | 3 | 8 | 7 | 0 | 0 | 1 | 2 |
| MIR_dup72084 | 5.18293470833531 | -3.88572966137045 | 1.02440053953197 | -3.7931741652008 | 0.000148733758193526 | 0.0132449860879928 | 2184 | 9 | 5 | 10 | 13 | 0 | 4 | 15 | 10 | 0 | 0 | 1 | 1 |
| AluJr_dup29924 | 5.74259649905507 | -3.4903899894148 | 0.942393606032666 | -3.70374965096465 | 0.0002124359147351 | 0.0171118064209743 | 2185 | 3 | 2 | 8 | 20 | 0 | 7 | 15 | 16 | 0 | 0 | 1 | 2 |
| LTR12C_dup1843 | 5.91241167748612 | -4.37063815645675 | 1.01836310318465 | -4.29182689630916 | 1.77209076624936e-05 | 0.00283374939157238 | 2186 | 2 | 3 | 16 | 20 | 0 | 5 | 17 | 13 | 0 | 0 | 1 | 0 |
| L1ME1_dup27656 | 5.86946779606021 | -2.553179611792 | 0.730794598767883 | -3.49370345114298 | 0.00047636985311984 | 0.0295183075704306 | 2187 | 3 | 3 | 13 | 17 | 4 | 4 | 20 | 8 | 1 | 0 | 2 | 3 |
| THE1D_dup8578 | 7.57013582926312 | -2.45549366459952 | 0.735371388666762 | -3.33912048040292 | 0.00084044104731119 | 0.0426850769356475 | 2188 | 2 | 2 | 11 | 13 | 2 | 5 | 28 | 24 | 1 | 0 | 4 | 3 |
| L1ME1_dup19838 | 6.16451409815238 | -2.95274369089832 | 0.79530697780948 | -3.71270939811327 | 0.000205052284056877 | 0.0167150979041444 | 2189 | 2 | 1 | 17 | 15 | 0 | 2 | 19 | 18 | 1 | 0 | 1 | 2 |
| L1PB1_dup9599 | 6.3870685775065 | -3.01985889279402 | 0.857793064407526 | -3.52049814587837 | 0.000430737004310368 | 0.027633364040883 | 2190 | 1 | 3 | 7 | 24 | 1 | 1 | 27 | 16 | 1 | 0 | 0 | 3 |
| L1MEg_dup13058 | 6.56244896570777 | -2.85889901572384 | 0.831099576640897 | -3.43989949709614 | 0.000581930239509729 | 0.0337547543097392 | 2191 | 2 | 2 | 18 | 24 | 4 | 0 | 20 | 12 | 1 | 2 | 1 | 0 |
| L1MDa_dup2656 | 7.53469294904553 | -3.40991127283789 | 0.863431312510831 | -3.94925597836147 | 7.83944831001194e-05 | 0.00840633333742271 | 2192 | 6 | 7 | 24 | 24 | 3 | 0 | 17 | 13 | 1 | 0 | 3 | 0 |
| MLT1D_dup8027 | 6.47665115794259 | -3.81141177291028 | 1.0001819302883 | -3.81071848779718 | 0.000138563486334066 | 0.0126274180013465 | 2193 | 1 | 7 | 18 | 27 | 1 | 0 | 20 | 10 | 1 | 0 | 1 | 0 |
| UCON17_dup16 | 6.6084533274524 | -3.27367457758222 | 0.823576233757894 | -3.97495027587765 | 7.03939619092194e-05 | 0.00779430510025646 | 2194 | 6 | 5 | 13 | 27 | 2 | 1 | 19 | 11 | 1 | 0 | 1 | 2 |
| OldhAT1_dup420 | 7.61364417132623 | -3.6419684306335 | 0.797845426819661 | -4.56475441007536 | 5.00079418444303e-06 | 0.0010966995142963 | 2195 | 3 | 2 | 23 | 26 | 1 | 1 | 23 | 17 | 1 | 0 | 1 | 1 |
| THE1D_dup5554 | 6.51651757808552 | -3.51426891522641 | 0.860771263405459 | -4.08269776725927 | 4.45159007946593e-05 | 0.00552549845095775 | 2196 | 4 | 4 | 23 | 25 | 1 | 1 | 15 | 10 | 1 | 0 | 1 | 1 |
| L1ME3_dup3693 | 9.42931146520005 | -3.81002198262731 | 0.770160891558681 | -4.9470468111104 | 7.53478616150559e-07 | 0.000258338937750088 | 2197 | 5 | 6 | 23 | 39 | 5 | 4 | 24 | 15 | 1 | 0 | 2 | 1 |
| L1PBa_dup994 | 8.17971392960817 | -3.14210938863236 | 0.806638744611778 | -3.89531176083614 | 9.80725029254768e-05 | 0.00992809341831075 | 2198 | 11 | 3 | 25 | 26 | 4 | 2 | 21 | 11 | 2 | 0 | 3 | 0 |
| L1PA6_dup5433 | 8.33491435316859 | -2.81084669531386 | 0.767967627863484 | -3.66011091266144 | 0.00025210610035782 | 0.0192397160513727 | 2199 | 1 | 1 | 17 | 33 | 6 | 1 | 32 | 14 | 2 | 0 | 2 | 2 |
| MER6A_dup60 | 8.11812223079383 | -3.50870682245345 | 0.866064125694773 | -4.05132451322667 | 5.09285155033108e-05 | 0.00612960877589823 | 2200 | 1 | 9 | 16 | 26 | 4 | 2 | 34 | 12 | 1 | 0 | 3 | 0 |
| AluJo_dup25315 | 4.28371965401295 | -3.63658671421621 | 1.06194398675998 | -3.42446189211122 | 0.000616018232869576 | 0.034914717253013 | 2201 | 2 | 9 | 9 | 5 | 4 | 4 | 10 | 9 | 0 | 0 | 1 | 1 |
| Charlie5_dup1080 | 5.08865878788065 | -3.90252655308522 | 1.06815620748909 | -3.65351671012507 | 0.000258672945832066 | 0.019583262096391 | 2202 | 3 | 13 | 11 | 10 | 3 | 2 | 17 | 6 | 0 | 0 | 1 | 1 |
| AluSz_dup4247 | 4.86750025767868 | -3.23402207055741 | 0.943980350733446 | -3.42594214810156 | 0.000612670861051235 | 0.034842873017136 | 2203 | 5 | 3 | 16 | 5 | 2 | 1 | 13 | 13 | 0 | 0 | 2 | 1 |
| MLT1C_dup16504 | 6.68820712817825 | -3.15625363107527 | 0.916185350761511 | -3.44499464922885 | 0.000571070696181563 | 0.0333130639552934 | 2204 | 7 | 12 | 18 | 9 | 2 | 1 | 17 | 14 | 0 | 0 | 4 | 1 |
| MSTB1_dup270 | 5.13091471765301 | -4.82747845911584 | 1.13583639356608 | -4.25015300307421 | 2.13624541003558e-05 | 0.00325007065432433 | 2205 | 2 | 3 | 15 | 11 | 1 | 2 | 25 | 9 | 0 | 0 | 0 | 0 |
| AluY_dup40190 | 4.77879991245842 | -4.92975940904987 | 1.13686725412497 | -4.33626651762809 | 1.44923210554663e-05 | 0.00246137746314346 | 2206 | 5 | 3 | 10 | 17 | 2 | 1 | 18 | 8 | 0 | 0 | 0 | 0 |
| L1MB7_dup22775 | 4.8055384705871 | -4.76444638235631 | 1.1285554052754 | -4.22172129085116 | 2.42443774653882e-05 | 0.00356983455469013 | 2207 | 3 | 3 | 13 | 15 | 1 | 1 | 14 | 12 | 0 | 0 | 0 | 0 |
| L1ME3_dup3691 | 7.25370998988303 | -4.68753698675304 | 0.988261499610775 | -4.74321522046464 | 2.10352579609467e-06 | 0.00058458332848473 | 2208 | 2 | 4 | 19 | 30 | 2 | 4 | 26 | 10 | 0 | 0 | 0 | 1 |
| MLT1C_dup7745 | 5.55870446719234 | -4.33461402793237 | 1.04264000856047 | -4.15734480965965 | 3.21967848822153e-05 | 0.00440039987109913 | 2209 | 3 | 2 | 17 | 18 | 4 | 1 | 25 | 5 | 0 | 0 | 0 | 1 |
| L2c_dup14107 | 4.59444827677208 | -4.20103430685635 | 1.08685619285594 | -3.86530834021127 | 0.000110948945031584 | 0.010826875596565 | 2210 | 1 | 3 | 10 | 15 | 5 | 3 | 19 | 5 | 0 | 0 | 1 | 0 |
| AluJb_dup57790 | 3.89819361048935 | -3.35817346925219 | 0.97332016931487 | -3.45077968704742 | 0.000558969660472504 | 0.0328406406989606 | 2211 | 2 | 2 | 9 | 13 | 1 | 4 | 13 | 6 | 0 | 0 | 1 | 1 |
| MLT1G1_dup1958 | 3.91667493540642 | -3.7235801741524 | 1.05425711320352 | -3.53194692975582 | 0.00041251213344491 | 0.0268404736408758 | 2212 | 1 | 1 | 8 | 8 | 2 | 2 | 16 | 11 | 0 | 0 | 1 | 0 |
| AluSc_dup13636 | 6.45624485216855 | -3.49428625574729 | 0.850632592808692 | -4.10786782129938 | 3.99328514601813e-05 | 0.00512782637536119 | 2213 | 1 | 5 | 18 | 19 | 2 | 2 | 21 | 13 | 0 | 1 | 2 | 0 |
| L1ME1_dup12944 | 4.02502226117758 | -3.33632908605712 | 0.98599498763259 | -3.38371809989396 | 0.000715113853077521 | 0.0382981050831384 | 2214 | 3 | 3 | 7 | 15 | 2 | 1 | 15 | 6 | 0 | 1 | 1 | 0 |
| L1MB7_dup8332 | 5.12373429561689 | -3.61142569791613 | 0.948281630172456 | -3.80838938877193 | 0.000139874889764373 | 0.0127130139495315 | 2215 | 3 | 3 | 14 | 10 | 1 | 2 | 22 | 10 | 0 | 1 | 1 | 0 |
| MIR_dup72087 | 5.58298009764548 | -3.89420924662546 | 0.962320445663369 | -4.04668659402847 | 5.19477181004868e-05 | 0.00622081113796599 | 2216 | 6 | 5 | 13 | 13 | 3 | 2 | 16 | 12 | 0 | 1 | 1 | 0 |
| L1MC4_dup3155 | 5.93268820375272 | -3.67892698662367 | 1.06527435340024 | -3.45350188416809 | 0.000553358437948165 | 0.032607584664889 | 2217 | 8 | 14 | 5 | 17 | 4 | 0 | 20 | 8 | 0 | 0 | 2 | 1 |
| AluSz_dup36962 | 6.5645211515619 | -3.39421419394282 | 0.918877005231603 | -3.69387216637042 | 0.000220864865431326 | 0.0175413589018432 | 2218 | 8 | 7 | 9 | 26 | 3 | 0 | 15 | 14 | 0 | 0 | 3 | 1 |
| L1MC3_dup11464 | 4.68872907910643 | -3.24282870314281 | 0.986683013384981 | -3.28659626156707 | 0.00101406108254626 | 0.0482511438789805 | 2219 | 7 | 5 | 7 | 15 | 1 | 0 | 15 | 9 | 0 | 0 | 2 | 1 |
| ERVL-B4-int_dup3636 | 4.81912286149982 | -3.83519707114441 | 1.08531885005517 | -3.53370539077014 | 0.000409777563809296 | 0.0267429874377857 | 2220 | 9 | 7 | 14 | 8 | 6 | 0 | 11 | 6 | 0 | 0 | 1 | 1 |
| MIR_dup167813 | 5.06235156882941 | -3.8411952588229 | 1.12958109407201 | -3.40054846790666 | 0.000672508125589243 | 0.0368600756169689 | 2221 | 11 | 6 | 12 | 18 | 1 | 0 | 12 | 6 | 0 | 0 | 2 | 0 |
| MER39_dup1572 | 5.48260665042099 | -3.84414403580005 | 1.03341622860221 | -3.7198409792728 | 0.000199348251412662 | 0.0163905041570965 | 2222 | 5 | 6 | 16 | 21 | 4 | 0 | 7 | 10 | 1 | 0 | 0 | 1 |
| MER58A_dup5343 | 4.22626796251527 | -4.01570235231251 | 1.1669471989298 | -3.44120312898073 | 0.000579133589847704 | 0.0336596290107234 | 2223 | 7 | 2 | 11 | 18 | 1 | 0 | 9 | 7 | 1 | 0 | 0 | 0 |
| L1M5_dup22419 | 6.23635500877897 | -3.31312935216033 | 0.935042410880731 | -3.54329313152721 | 0.000395163200685425 | 0.0260752919995269 | 2224 | 8 | 3 | 17 | 21 | 1 | 0 | 15 | 13 | 0 | 0 | 0 | 4 |
| L1M5_dup22410 | 5.02148125189685 | -3.56429297014805 | 0.953156709188264 | -3.73946166017496 | 0.000184414771362873 | 0.0155529272708604 | 2225 | 4 | 3 | 14 | 17 | 1 | 0 | 15 | 10 | 0 | 1 | 0 | 1 |
| L3_dup19415 | 5.4720118422102 | -4.22036553006324 | 1.0093442071366 | -4.18129464678452 | 2.89853961682983e-05 | 0.00407862748472023 | 2226 | 4 | 2 | 14 | 20 | 2 | 0 | 20 | 10 | 0 | 0 | 0 | 1 |

| | | | | | | | | | | | | | | | | | | |
|---|---|---|---|---|---|---|---|---|---|---|---|---|---|---|---|---|---|---|
| HAL1_dup21086 | 4.59836308090206 | -4.35348098356718 | 1.10596762769032 | -3.93635480331274 | 8.27286352968483e-05 | 0.00876444695206761 | 2227 | 6 | 6 | 9 | 16 | 3 | 2 | 11 | 7 | 0 | 0 | 0 | 1 |
| L1MC4_dup10111 | 4.91185383455579 | -4.23849114873117 | 1.08068718276154 | -3.92203332873838 | 8.78048428564632e-05 | 0.00915147224630923 | 2228 | 5 | 3 | 10 | 25 | 2 | 1 | 8 | 10 | 0 | 0 | 0 | 1 |
| AluSp_dup18904 | 4.80069868615464 | -3.19004330676919 | 0.894616375267622 | -3.56582261957244 | 0.000362716649379371 | 0.0246520925030631 | 2229 | 4 | 4 | 9 | 14 | 2 | 2 | 16 | 9 | 1 | 0 | 0 | 2 |
| MER5C_dup1168 | 4.83905586606546 | -3.65012293116451 | 0.976911447682945 | -3.73639078528758 | 0.000186680474849288 | 0.0156529341744761 | 2230 | 3 | 5 | 10 | 13 | 1 | 1 | 14 | 13 | 0 | 0 | 0 | 2 |
| L1MC2_dup6137 | 11.8764754272336 | -3.6090292196407 | 1.00254766080518 | -3.59985800250353 | 0.000318391001823018 | 0.0225268466989982 | 2231 | 19 | 10 | 33 | 45 | 0 | 2 | 34 | 13 | 2 | 2 | 0 | 0 |
| LTR16E2_dup1847 | 9.1578968827753 | -4.16901111015213 | 1.00802628822596 | -4.13581586001019 | 3.53695736341394e-05 | 0.00471397839482619 | 2232 | 22 | 13 | 18 | 19 | 1 | 3 | 23 | 17 | 1 | 0 | 2 | 0 |
| MSTB_dup1027 | 7.68041655211267 | -3.92486659234042 | 1.09141125269706 | -3.59613901967878 | 0.000322975241961013 | 0.0227503454480922 | 2233 | 18 | 21 | 11 | 16 | 0 | 2 | 17 | 12 | 1 | 0 | 1 | 1 |
| L2b_dup39269 | 9.13375586905718 | -4.2570867568913 | 1.01471462090591 | -4.19535371737394 | 2.72446075994171e-05 | 0.00389354078069124 | 2234 | 18 | 8 | 24 | 25 | 1 | 1 | 12 | 24 | 0 | 0 | 2 | 1 |
| L2c_dup122477 | 8.19123928037085 | -3.53278125605502 | 0.962572170332885 | -3.67014688865696 | 0.000242411128254595 | 0.0187336012743319 | 2235 | 16 | 8 | 21 | 16 | 1 | 0 | 27 | 14 | 0 | 0 | 2 | 3 |
| MLT1C-int_dup406 | 9.29149657407917 | -3.317514808572 | 0.926440657752578 | -3.58092531972836 | 0.000342379513763294 | 0.0237307998199594 | 2236 | 17 | 9 | 16 | 28 | 0 | 1 | 27 | 18 | 0 | 0 | 2 | 5 |
| L2c_dup122476 | 7.46297326044515 | -3.65610231424574 | 1.00942090047699 | -3.62198000112549 | 0.000292356727927806 | 0.0212153443072613 | 2237 | 12 | 7 | 10 | 24 | 2 | 0 | 24 | 16 | 0 | 2 | 1 | 0 |
| LTR12C_dup2636 | 7.56244084075876 | -4.81430752980597 | 1.20218211278206 | -4.00464079328616 | 6.21117894134334e-05 | 0.00708305811651604 | 2238 | 7 | 9 | 7 | 14 | 1 | 0 | 33 | 24 | 0 | 1 | 0 | 0 |
| L1PA10_dup1447 | 9.22083391853567 | -3.20881575543303 | 0.783339524435641 | -4.09632816337824 | 4.19754798187798e-05 | 0.00530424722302675 | 2239 | 3 | 5 | 17 | 23 | 0 | 2 | 36 | 27 | 0 | 2 | 1 | 2 |
| MIRb_dup183496 | 8.29298791261681 | -3.61890854683257 | 1.01002312554086 | -3.58299573081029 | 0.000339676161410695 | 0.0235915306196816 | 2240 | 12 | 13 | 9 | 19 | 0 | 1 | 34 | 17 | 0 | 1 | 2 | 1 |
| L1PA11_dup2003 | 5.57241604755677 | -4.47719500103491 | 1.35887599306519 | -3.2947782018989 | 0.000984993910186987 | 0.047304144699605 | 2241 | 8 | 7 | 30 | 16 | 0 | 0 | 3 | 7 | 1 | 0 | 0 | 0 |
| MIR3_dup75191 | 6.14652165908603 | -5.35832701190616 | 1.47296084646182 | -3.63779324126457 | 0.000274984029526673 | 0.0203791964242417 | 2242 | 13 | 4 | 34 | 24 | 0 | 0 | 2 | 5 | 0 | 0 | 0 | 0 |
| HERVIP10B3-int_dup2 | 3.52580705390173 | -4.48665687320954 | 1.32339682265536 | -3.39025815719218 | 0.000698268341874637 | 0.0377356708779701 | 2243 | 4 | 4 | 18 | 9 | 0 | 0 | 9 | 3 | 0 | 0 | 0 | 0 |
| HERVIP10B3-int_dup2 | 4.61235561118041 | -5.08574864769491 | 1.36840067410767 | -3.71656397422583 | 0.000201950522986015 | 0.0165472862060711 | 2244 | 11 | 6 | 15 | 16 | 0 | 0 | 12 | 3 | 0 | 0 | 0 | 0 |
| L2c_dup134280 | 4.67159425586395 | -4.93528580850636 | 1.36257583864403 | -3.62202650930441 | 0.000292304158311045 | 0.0212153443072613 | 2245 | 9 | 3 | 19 | 21 | 0 | 0 | 9 | 3 | 0 | 0 | 0 | 0 |
| MIRc_dup41293 | 3.10257643335209 | -4.40588146778116 | 1.33965740780049 | -3.28881208145218 | 0.00100611174132759 | 0.0480337609153162 | 2246 | 6 | 3 | 9 | 8 | 0 | 0 | 4 | 9 | 0 | 0 | 0 | 0 |
| MLT1H2-int_dup173 | 3.0227029115777 | -4.53348833145585 | 1.38554446238272 | -3.27199050953559 | 0.0010679316081454 | 0.0499724787476433 | 2247 | 9 | 4 | 7 | 11 | 0 | 0 | 4 | 5 | 0 | 0 | 0 | 0 |
| L1MC2_dup6136 | 5.18923628207918 | -4.99634894375377 | 1.26706809833732 | -3.94323631879779 | 8.03893698928215e-05 | 0.00855088467077215 | 2248 | 5 | 6 | 10 | 25 | 0 | 0 | 10 | 12 | 0 | 0 | 0 | 0 |
| MLT1F1_dup3044 | 4.37121680722121 | -4.43187581135705 | 1.22999086976547 | -3.60317781237036 | 0.000314350328601763 | 0.022347894618905 | 2249 | 2 | 3 | 12 | 27 | 0 | 0 | 8 | 7 | 0 | 0 | 0 | 0 |
| L1MC2_dup6128 | 4.35308425999722 | -4.27545969539151 | 1.25703346703332 | -3.40122980614182 | 0.000670834078255243 | 0.0368058632062167 | 2250 | 9 | 5 | 20 | 14 | 1 | 1 | 4 | 3 | 0 | 1 | 0 | 0 |
| THE1D_dup5556 | 5.0254487785987 | -4.32059938819867 | 1.11922342021864 | -3.86035469786241 | 0.00011322255217889 | 0.0109828183451929 | 2251 | 7 | 3 | 15 | 21 | 2 | 1 | 7 | 9 | 0 | 1 | 0 | 0 |
| L1P4_dup972 | 4.85437359845217 | -4.60508977781257 | 1.38448741615821 | -3.32620558631812 | 0.000880369690557328 | 0.0438702505527443 | 2252 | 3 | 3 | 23 | 31 | 0 | 0 | 2 | 4 | 0 | 0 | 0 | 0 |
| MSTB2_dup430 | 4.10740696425636 | -4.10247029123695 | 1.23606392554632 | -3.3189790644717 | 0.000903472026591391 | 0.0446474672563479 | 2253 | 1 | 1 | 15 | 29 | 0 | 1 | 4 | 5 | 0 | 0 | 0 | 0 |
| L2a_dup147214 | 5.69158064306064 | -3.95937389662961 | 1.12188126303883 | -3.52922722490691 | 0.00041677511268311 | 0.0270255736209899 | 2254 | 6 | 8 | 18 | 11 | 0 | 0 | 18 | 11 | 0 | 0 | 0 | 2 |
| Tigger2a_dup2829 | 5.6498496500659 | -3.63117400048117 | 1.08800422428373 | -3.33746314530323 | 0.000845469337100685 | 0.0428337920356025 | 2255 | 6 | 11 | 15 | 21 | 0 | 1 | 13 | 6 | 0 | 0 | 0 | 3 |
| L1M1_dup9219 | 5.78556837896401 | -3.82047187389708 | 1.01432915918576 | -3.7665010803435 | 0.000165551367933547 | 0.0143192605577302 | 2256 | 6 | 5 | 17 | 17 | 0 | 3 | 19 | 8 | 0 | 2 | 0 | 0 |
| MLT1G1-int_dup137 | 6.66440283612612 | -4.07973853180536 | 0.993920290565057 | -4.10469387790239 | 4.04850620317915e-05 | 0.00516528609663091 | 2257 | 6 | 6 | 19 | 31 | 0 | 3 | 15 | 8 | 0 | 1 | 0 | 1 |
| MSTB-int_dup386 | 5.63624695766201 | -5.152741729819 | 1.23291256401891 | -4.17932453622069 | 2.92376201210472e-05 | 0.00410683279653052 | 2258 | 8 | 4 | 11 | 11 | 0 | 1 | 32 | 9 | 0 | 0 | 0 | 0 |
| L1MA9_dup8574 | 4.79489737732595 | -3.56739018903426 | 1.00253673391057 | -3.55836356750645 | 0.000373172542520925 | 0.0251472426515673 | 2259 | 5 | 4 | 12 | 14 | 0 | 1 | 19 | 7 | 1 | 0 | 0 | 1 |
| MLT1C_dup7742 | 6.03944117685312 | -4.42788494623166 | 1.05281051737355 | -4.20577575277072 | 2.60188126628203e-05 | 0.00376356984672793 | 2260 | 5 | 4 | 18 | 23 | 0 | 2 | 20 | 8 | 1 | 0 | 0 | 0 |
| MIR_dup149072 | 7.33586917584176 | -3.34689597240944 | 0.937801584783766 | -3.56887429784111 | 0.000358518378879399 | 0.0244856527499533 | 2261 | 8 | 7 | 21 | 22 | 0 | 0 | 15 | 17 | 1 | 0 | 2 | 1 |
| L1MC5_dup13213 | 6.83862925247435 | -4.02446652041238 | 1.08199364433212 | -3.71949183019143 | 0.000199624003963476 | 0.0164075071665262 | 2262 | 8 | 7 | 25 | 14 | 0 | 0 | 18 | 14 | 0 | 1 | 1 | 0 |
| L1MEb_dup1540 | 6.31556462129486 | -3.17350660963062 | 0.949264947085016 | -3.34311997864849 | 0.000828420740899749 | 0.0422912254053061 | 2263 | 8 | 6 | 19 | 13 | 0 | 0 | 22 | 11 | 0 | 1 | 1 | 2 |
| L2c_dup18343 | 6.47384021739821 | -3.6354026842378 | 0.952377651101417 | -3.81718605012779 | 0.000134982383334667 | 0.0123906429799315 | 2264 | 2 | 2 | 24 | 37 | 0 | 0 | 16 | 6 | 0 | 0 | 1 | 1 |
| L2c_dup59406 | 5.61242722430476 | -3.91226087316285 | 1.02378224913483 | -3.82137986517055 | 0.00013270705683355 | 0.0122772602431244 | 2265 | 2 | 2 | 24 | 25 | 0 | 0 | 16 | 6 | 0 | 0 | 1 | 0 |
| L2a_dup73380 | 5.86690167321706 | -3.44544938388781 | 0.959266565197001 | -3.59175385538454 | 0.000328460011458748 | 0.0230151359412794 | 2266 | 6 | 2 | 21 | 26 | 0 | 1 | 18 | 4 | 0 | 0 | 1 | 2 |
| L1MA6_dup455 | 6.12897124074969 | -2.8282759421508 | 0.856633750349802 | -3.30161628700234 | 0.000961294779277927 | 0.0465763077886511 | 2267 | 5 | 1 | 18 | 29 | 0 | 0 | 19 | 7 | 0 | 0 | 2 | 3 |
| MER67D_dup986 | 6.23202379465853 | -3.35433859802859 | 0.940037976198345 | -3.56830115693202 | 0.00035930338106662 | 0.0245145123020219 | 2268 | 5 | 1 | 27 | 25 | 0 | 0 | 16 | 7 | 0 | 0 | 2 | 1 |
| MER45C_dup529 | 5.37801771677475 | -3.24418425170303 | 0.988708863356325 | -3.28123310302908 | 0.00103354288676602 | 0.0489192567235723 | 2269 | 9 | 2 | 14 | 18 | 0 | 1 | 17 | 8 | 0 | 1 | 2 | 0 |
| Tigger3b_dup2755 | 7.00170945274909 | -3.34513622521319 | 0.837600027871517 | -3.99371551325487 | 6.50459015983839e-05 | 0.0073561844589967 | 2270 | 7 | 1 | 18 | 28 | 1 | 1 | 17 | 15 | 0 | 0 | 3 | 1 |
| L1P2_dup413 | 7.6102783366065 | -3.51582546096638 | 0.880392091692955 | -3.99347687711006 | 6.51114309829882e-05 | 0.00736009717656918 | 2271 | 9 | 2 | 23 | 32 | 0 | 1 | 19 | 12 | 0 | 0 | 2 | 2 |
| LTR9B_dup291 | 6.52056931120328 | -3.31755853071858 | 0.857251126523096 | -3.86999611674375 | 0.00010883708685927 | 0.010683725028802 | 2272 | 6 | 4 | 15 | 28 | 0 | 1 | 17 | 12 | 0 | 0 | 1 | 3 |
| L1M4_dup12830 | 37.6422835775506 | 1.75375011325303 | 0.373803459586175 | 4.69163692383839 | 2.71027790874994e-06 | 0.000708681084918037 | 2273 | 0 | 0 | 58 | 55 | 2 | 3 | 35 | 24 | 2 | 2 | 133 | 193 |
| L1P2_dup1119 | 46.8142063784341 | 2.6207019139618 | 0.376763497218205 | 6.95582755046994 | 3.50498398887809e-12 | 7.58179018379623e-09 | 2274 | 0 | 0 | 50 | 59 | 0 | 1 | 36 | 17 | 2 | 5 | 202 | 262 |
| MER21B_dup1655 | 13.5234047061733 | 1.63279418097711 | 0.479790967808545 | 3.40313655431015 | 0.000666169769745819 | 0.0366552733835441 | 2275 | 0 | 0 | 14 | 30 | 0 | 0 | 18 | 13 | 0 | 3 | 50 | 53 |
| L1MC3_dup8112 | 15.6245717759221 | 1.8227220837067 | 0.527123975228375 | 3.45786222855261 | 0.00054447974461987 | 0.0322898394372453 | 2276 | 0 | 0 | 32 | 35 | 0 | 0 | 20 | 8 | 4 | 4 | 47 | 60 |
| MLT1F2_dup4804 | 21.0909501651882 | 1.68103989752211 | 0.479227770187125 | 3.5078098601542 | 0.000451811847456399 | 0.0285163983699895 | 2277 | 0 | 0 | 44 | 46 | 0 | 1 | 20 | 17 | 3 | 6 | 53 | 93 |
| L1PB_dup1655 | 15.4666822447547 | 2.52077387490129 | 0.444905180059831 | 5.66586766771808 | 1.46282392008622e-08 | 1.08772843902761e-05 | 2278 | 4 | 2 | 11 | 14 | 0 | 0 | 9 | 16 | 11 | 13 | 49 | 70 |
| L2_dup55459 | 20.696496999132 | 3.02819819184863 | 0.510581199867846 | 5.93088463232179 | 3.01306872429401e-09 | 2.81155941439554e-06 | 2279 | 2 | 2 | 5 | 22 | 0 | 0 | 13 | 20 | 18 | 11 | 68 | 107 |
| L1PA10_dup1402 | 21.0208861065525 | 3.09351459530505 | 0.687636794623012 | 4.49876245642299 | 6.83501775334008e-06 | 0.00138724214323772 | 2280 | 0 | 1 | 32 | 37 | 0 | 0 | 13 | 23 | 19 | 13 | 56 | 75 |
| L1M4a2_dup2015 | 16.1723563716104 | 2.532867769012 | 0.506250358446489 | 5.00319205063778 | 5.63887132699444e-07 | 0.000203603471437484 | 2281 | 0 | 3 | 15 | 22 | 0 | 0 | 14 | 16 | 8 | 17 | 59 | 54 |
| L1ME4a_dup26458 | 15.6052386420729 | 3.01267843424118 | 0.445051311986721 | 6.76928334576187 | 1.29421859415674e-11 | 2.3507949435513e-08 | 2282 | 0 | 1 | 16 | 16 | 0 | 0 | 10 | 6 | 7 | 13 | 61 | 76 |
| AluSz6_dup34553 | 18.4773311225177 | 2.82191130989222 | 0.384256859551419 | 7.34381505430123 | 2.0758992799524e-13 | 5.88038011985184e-10 | 2283 | 0 | 2 | 6 | 16 | 2 | 2 | 14 | 11 | 7 | 10 | 72 | 103 |
| HAL1_dup18027 | 13.6348005996859 | 2.69380289467309 | 0.395807380092435 | 6.80584301900583 | 1.00459209129774e-11 | 1.95568469034522e-08 | 2284 | 1 | 1 | 6 | 10 | 2 | 2 | 12 | 8 | 8 | 8 | 57 | 63 |
| L3b_dup8003 | 16.8436960930612 | 3.11662477905888 | 0.420169327964491 | 7.41754471740562 | 1.19311411880801e-13 | 3.59362952042901e-10 | 2285 | 1 | 1 | 9 | 16 | 1 | 0 | 9 | 6 | 9 | 5 | 81 | 84 |
| L1MEg_dup11427 | 14.5070334261982 | 2.49292038255124 | 0.453911459349506 | 5.4920851439261 | 3.97215798155327e-08 | 2.41728670864111e-05 | 2286 | 0 | 2 | 12 | 14 | 1 | 0 | 20 | 11 | 14 | 8 | 54 | 50 |
| AluSz6_dup34552 | 14.6057452638758 | 2.78267968763446 | 0.438158206213015 | 6.3508560336803 | 2.14120028145474e-10 | 2.77402747421019e-07 | 2287 | 0 | 3 | 13 | 16 | 1 | 0 | 10 | 6 | 12 | 12 | 48 | 70 |
| MIRc_dup77483 | 17.8482738158156 | 2.83925746035689 | 0.390532877388183 | 7.27021366125526 | 3.58919343874815e-13 | 9.38499145028975e-10 | 2288 | 1 | 2 | 16 | 14 | 1 | 0 | 17 | 6 | 14 | 10 | 68 | 85 |
| AluJb_dup90755 | 20.8816423258661 | 3.10933814008087 | 0.409611574488773 | 7.59094306346583 | 3.17585490799869e-14 | 1.16258764913639e-10 | 2289 | 0 | 1 | 17 | 17 | 4 | 0 | 13 | 5 | 12 | 17 | 80 | 110 |
| L2c_dup110919 | 19.9008651791695 | 3.06906897622182 | 0.383831492269365 | 7.99587589354968 | 1.2865592667737e-15 | 6.24758431207622e-12 | 2290 | 0 | 0 | 12 | 19 | 1 | 1 | 19 | 7 | 13 | 13 | 82 | 95 |
| MIR_dup133535 | 19.0362080765701 | 3.10293878072027 | 0.409067460327627 | 7.5853962528212 | 3.31472462495828e-14 | 1.19503858425806e-10 | 2291 | 0 | 0 | 13 | 17 | 3 | 1 | 9 | 7 | 11 | 10 | 95 | 82 |
| L1M3_dup5205 | 22.5712532119373 | 3.23049421273548 | 0.421014959080539 | 7.6731102851799 | 1.67875030048268e-14 | 6.44277288707504e-11 | 2292 | 0 | 1 | 12 | 22 | 2 | 2 | 6 | 14 | 19 | 20 | 82 | 112 |
| MIRb_dup170296 | 14.6360950471523 | 2.73189859738269 | 0.49567653912782 | 5.51145430887181 | 3.55880819244428e-08 | 2.22263910659415e-05 | 2293 | 0 | 1 | 12 | 20 | 1 | 1 | 6 | 12 | 6 | 17 | 59 | 54 |
| L1PA5_dup9445 | 17.6084197101488 | 2.71263015431109 | 0.356651624975652 | 7.60582586577665 | 2.8309024491089e-14 | 1.05250299086823e-10 | 2294 | 0 | 2 | 14 | 15 | 2 | 2 | 13 | 8 | 8 | 14 | 67 | 87 |
| THE1A_dup3517 | 17.6083230842194 | 2.81545249947442 | 0.400955074498762 | 7.02186523763054 | 2.18925648078265e-12 | 5.05752254928456e-09 | 2295 | 0 | 1 | 14 | 23 | 1 | 1 | 11 | 8 | 10 | 14 | 70 | 78 |
| L1PA13_dup7407 | 29.3735051488104 | 3.28692865728831 | 0.384687999110395 | 8.54440134573849 | 1.29204282707939e-17 | 9.60738820413226e-14 | 2296 | 0 | 0 | 19 | 24 | 1 | 4 | 18 | 7 | 18 | 14 | 119 | 166 |
| HAL1_dup18029 | 28.7912559565787 | 3.26780657377989 | 0.375160176943561 | 8.71043030313823 | 3.02723410276764e-18 | 2.88127298326859e-14 | 2297 | 0 | 2 | 16 | 30 | 2 | 1 | 17 | 6 | 17 | 18 | 114 | 160 |
| MLT1J_dup11706 | 29.9131770584209 | 3.37785195955785 | 0.446453391977006 | 7.5659677365198 | 3.84987604662304e-14 | 1.36725762058174e-10 | 2298 | 0 | 2 | 27 | 24 | 1 | 3 | 7 | 6 | 18 | 18 | 135 | 152 |

| | | | | | | | | | | | | | | | | | | | |
|---|---|---|---|---|---|---|---|---|---|---|---|---|---|---|---|---|---|---|---|
| L1ME1_dup24293 | 26.0446008726889 | 2.69336218452752 | 0.332079739545602 | 8.11058870442669 | 5.03750973879269e-16 | 2.72421657342446e-12 | 2299 | 4 | 4 | 19 | 24 | 2 | 4 | 19 | 7 | 10 | 30 | 97 | 125 |
| THE1C-int_dup1566 | 32.2352114524276 | 2.71080573071197 | 0.370091527225699 | 7.32469005986944 | 2.394514786037e-13 | 6.7031201797454e-10 | 2300 | 3 | 3 | 18 | 33 | 7 | 5 | 15 | 7 | 11 | 21 | 124 | 185 |
| L1ME3A_dup12026 | 24.0285029374954 | 2.74728456506836 | 0.341867436404877 | 8.0361107040763 | 9.27349666226656e-16 | 4.59706549333266e-12 | 2301 | 1 | 2 | 17 | 23 | 1 | 4 | 20 | 10 | 19 | 14 | 93 | 110 |
| L3_dup36531 | 30.8139947681344 | 3.31938323690272 | 0.305686417824025 | 10.8587854852406 | 1.81142254783013e-27 | 1.07755187391497e-22 | 2302 | 1 | 2 | 17 | 20 | 2 | 1 | 16 | 12 | 14 | 25 | 126 | 171 |
| L1MA9_dup15923 | 28.2433795694613 | 2.73502525510197 | 0.316966625942676 | 8.62874836417825 | 6.20269867331526e-18 | 5.27109763757383e-14 | 2303 | 1 | 2 | 19 | 21 | 3 | 1 | 26 | 17 | 15 | 14 | 100 | 154 |
| MER65A_dup1087 | 30.988875526773 | 3.04045837920866 | 0.319982922321313 | 9.50193953212156 | 2.06016803328332e-21 | 4.0850728570636e-17 | 2304 | 1 | 1 | 18 | 30 | 2 | 1 | 24 | 10 | 14 | 16 | 121 | 176 |
| L2a_dup155582 | 21.6759807420504 | 1.64943703081598 | 0.433546218264894 | 3.80452408838262 | 0.000142077086265035 | 0.0128302618251898 | 2305 | 1 | 3 | 22 | 14 | 7 | 10 | 23 | 15 | 7 | 7 | 67 | 111 |
| L1MC1_dup10505 | 18.0564601705964 | 2.32854914536155 | 0.423476977379163 | 5.49864401076205 | 3.82722902623616e-08 | 2.35316237177465e-05 | 2306 | 3 | 4 | 14 | 21 | 1 | 1 | 14 | 6 | 9 | 3 | 70 | 95 |
| L1MD2_dup320 | 16.8532395223643 | 1.55029709668105 | 0.404180581742288 | 3.83565457300851 | 0.000125230314955264 | 0.0117965370238896 | 2307 | 5 | 3 | 15 | 29 | 2 | 2 | 20 | 9 | 4 | 4 | 50 | 85 |
| L1PA14_dup684 | 40.4499901647835 | 2.85569099572946 | 0.515017700680285 | 5.54484048209875 | 2.94222278892998e-08 | 1.90241886884438e-05 | 2308 | 0 | 0 | 32 | 61 | 4 | 2 | 44 | 42 | 26 | 32 | 108 | 176 |
| L1PA7_dup2717 | 32.999299069905 | 3.01917114450308 | 0.552730466816672 | 5.46228465004167 | 4.70045572212289e-08 | 2.75481437747846e-05 | 2309 | 0 | 0 | 39 | 42 | 3 | 2 | 42 | 30 | 33 | 36 | 78 | 118 |
| L2c_dup110923 | 35.3678379784329 | 3.0975741985378 | 0.444919667070149 | 6.96209771740527 | 3.35242826038992e-12 | 7.31832013620862e-09 | 2310 | 0 | 0 | 23 | 47 | 4 | 3 | 24 | 14 | 19 | 34 | 103 | 201 |
| MLT1G_dup2162 | 28.4579094435088 | 2.86291630223399 | 0.367518290015038 | 7.78986074983326 | 6.70830850881716e-15 | 3.06964457007501e-11 | 2311 | 0 | 0 | 24 | 27 | 6 | 1 | 17 | 11 | 12 | 17 | 108 | 155 |
| AluJr_dup16778 | 22.484143683763 | 3.02401515031159 | 0.554763101682278 | 5.45100267328792 | 5.00866173800757e-08 | 2.90680738027305e-05 | 2312 | 0 | 0 | 20 | 24 | 4 | 2 | 17 | 18 | 36 | 23 | 50 | 89 |
| HERVL-int_dup506 | 26.8361738095356 | 2.90869728274012 | 0.529218478493594 | 5.4962126247361 | 3.88034613156047e-08 | 2.37355486020639e-05 | 2313 | 0 | 0 | 34 | 38 | 1 | 3 | 30 | 20 | 29 | 24 | 80 | 84 |
| L1PA17_dup4652 | 45.3908234830969 | 2.83503071820084 | 0.372513065663265 | 7.61055377521598 | 2.72923980883884e-14 | 1.03081221516503e-10 | 2314 | 3 | 3 | 33 | 55 | 2 | 0 | 41 | 31 | 25 | 31 | 150 | 224 |
| L1MC1_dup12648 | 23.6305361374569 | 2.18019283681416 | 0.368646363772613 | 5.91404948228091 | 3.33797533735088e-09 | 3.05483799854344e-06 | 2315 | 2 | 3 | 20 | 35 | 3 | 0 | 30 | 16 | 14 | 15 | 69 | 105 |
| L1MD2_dup8423 | 27.9193953076239 | 2.72226751285812 | 0.353896388705607 | 7.69227265306362 | 1.44544176258569e-14 | 5.82944212949515e-11 | 2316 | 1 | 4 | 23 | 26 | 4 | 0 | 19 | 17 | 14 | 28 | 109 | 118 |
| L2a_dup133447 | 35.6920944301383 | 2.5888114970582 | 0.327900706044651 | 7.89510802915344 | 2.90063883990578e-15 | 1.3533243321573e-11 | 2317 | 3 | 2 | 26 | 41 | 11 | 2 | 21 | 14 | 20 | 28 | 126 | 177 |
| L1MD2_dup8424 | 32.6413718741051 | 2.62660375361555 | 0.294795696758331 | 8.90991212727505 | 5.10730266813659e-19 | 5.52391927578376e-15 | 2318 | 5 | 2 | 18 | 35 | 4 | 3 | 24 | 13 | 13 | 18 | 121 | 180 |
| L1MEd_dup17354 | 32.0226256016591 | 2.40633963403437 | 0.266548768297222 | 9.02776497301654 | 1.75213653225971e-19 | 2.19428357528983e-15 | 2319 | 4 | 2 | 22 | 35 | 5 | 4 | 26 | 18 | 15 | 19 | 116 | 157 |
| L1MB4_dup5480 | 29.2722190273536 | 1.87547876357189 | 0.367109093339788 | 5.10877773827299 | 3.24249507098989e-07 | 0.000131886962762694 | 2320 | 1 | 0 | 36 | 48 | 1 | 5 | 24 | 28 | 9 | 7 | 104 | 122 |
| L1P3_dup2991 | 36.5005716354005 | 1.33744697747458 | 0.399701246232481 | 3.34611660604299 | 0.000819519244197639 | 0.0419989933404806 | 2321 | 6 | 3 | 61 | 106 | 2 | 3 | 46 | 19 | 11 | 12 | 94 | 130 |
| L1P1_dup2812 | 26.5731680020379 | 1.39888819949919 | 0.396170922931105 | 3.5310218860824 | 0.000413957494196649 | 0.0269061781592187 | 2322 | 3 | 1 | 43 | 58 | 10 | 4 | 31 | 13 | 10 | 15 | 68 | 99 |
| PRIMA41-int_dup315 | 23.5446270572566 | 1.77939193141848 | 0.527279796440386 | 3.37466359119196 | 0.000739059749373036 | 0.0392098798493459 | 2323 | 1 | 1 | 46 | 67 | 1 | 1 | 37 | 15 | 9 | 14 | 53 | 70 |
| L1PA5_dup10740 | 41.3027756191438 | 3.37074210252005 | 0.906908144975957 | 3.71674035699548 | 0.000201809648581697 | 0.0165414394217776 | 2324 | 6 | 3 | 14 | 21 | 15 | 13 | 1 | 0 | 24 | 21 | 189 | 240 |
| L1MEc_dup17223 | 22.530293239922 | 3.00918384901382 | 0.446779798899495 | 6.73527284900978 | 1.63622597202019e-11 | 2.90547332192772e-08 | 2325 | 6 | 2 | 9 | 11 | 6 | 11 | 4 | 4 | 22 | 19 | 98 | 98 |
| L1PA10_dup6569 | 29.0400312157961 | 2.88675650560572 | 0.603219778975904 | 4.7855799929283 | 1.70494263258574e-06 | 0.000493754345008578 | 2326 | 9 | 3 | 14 | 15 | 9 | 17 | 3 | 1 | 20 | 17 | 101 | 177 |
| L1PA7_dup9931 | 23.9113717146002 | 1.77564996826335 | 0.520495097531114 | 3.41146338685199 | 0.000646151783743711 | 0.0359985079011148 | 2327 | 8 | 5 | 25 | 36 | 9 | 16 | 5 | 3 | 14 | 17 | 78 | 100 |
| L1M4a1_dup1806 | 27.0556696928771 | 2.05213698525329 | 0.514079265471998 | 3.99186880912059 | 6.55546375936111e-05 | 0.00739614215118511 | 2328 | 13 | 5 | 18 | 17 | 18 | 12 | 8 | 7 | 18 | 8 | 111 | 118 |
| L1PA5_dup10739 | 29.7878463931015 | 2.67261508669213 | 0.569935296016281 | 4.68933071064927 | 2.74100090711837e-06 | 0.000710468629460987 | 2329 | 5 | 7 | 17 | 15 | 19 | 11 | 3 | 3 | 18 | 19 | 104 | 173 |
| L1PA5_dup8380 | 29.8989081716459 | 2.31526381431925 | 0.476287040835543 | 4.86106825467604 | 1.16753985903996e-06 | 0.000359969672019978 | 2330 | 9 | 10 | 16 | 24 | 16 | 14 | 10 | 2 | 20 | 20 | 108 | 145 |
| L1MD1_dup4827 | 29.1181972485046 | 1.85066932099692 | 0.52630404333589 | 3.51635018660842 | 0.000437523496870235 | 0.0279406779351275 | 2331 | 10 | 13 | 11 | 31 | 20 | 9 | 12 | 7 | 6 | 17 | 118 | 133 |
| L1M4a1_dup1805 | 28.894715316417 | 1.86366727060036 | 0.563333707827694 | 3.30828289645042 | 0.000938699346386598 | 0.045798596406665 | 2332 | 8 | 11 | 9 | 36 | 20 | 15 | 5 | 9 | 13 | 11 | 113 | 131 |
| L1MC2_dup4324 | 28.8840660588026 | 2.01328207633708 | 0.601731528591017 | 3.34581450476972 | 0.00082041259873396 | 0.0420246410372884 | 2333 | 7 | 14 | 13 | 35 | 19 | 18 | 1 | 6 | 17 | 19 | 91 | 141 |
| L1MA7_dup6197 | 35.9959050973067 | 2.19628267665852 | 0.496913767043591 | 4.41984670645251 | 9.87709395697362e-06 | 0.00183610546772347 | 2334 | 14 | 26 | 15 | 28 | 20 | 20 | 6 | 6 | 26 | 24 | 123 | 162 |
| L1MD1_dup4829 | 32.5887026722209 | 2.23722458267399 | 0.523759222362665 | 4.27147530230002 | 1.94184036116073e-05 | 0.0030418245330925 | 2335 | 8 | 16 | 11 | 21 | 19 | 21 | 8 | 8 | 20 | 14 | 102 | 183 |
| L1ME2_dup11813 | 30.9421649546203 | 2.49838590522001 | 0.558666801194989 | 4.47205006611448 | 7.74732825928355e-06 | 0.00151723931685883 | 2336 | 10 | 12 | 12 | 13 | 19 | 27 | 4 | 5 | 19 | 20 | 113 | 151 |
| ERV3-16A3_I-int_dup | 33.3050437178351 | 3.56472967797857 | 0.57821262340874 | 6.16508449255812 | 7.04455149424099e-10 | 8.02020502319936e-07 | 2337 | 10 | 6 | 7 | 2 | 10 | 4 | 5 | 6 | 17 | 17 | 168 | 185 |
| L1MA4A_dup2485 | 23.3604699664659 | 3.79318477844514 | 0.702344225979847 | 5.40074886093529 | 6.636327755504e-08 | 3.60522293176063e-05 | 2338 | 10 | 3 | 0 | 4 | 4 | 11 | 4 | 2 | 11 | 15 | 112 | 133 |
| L1ME3A_dup10622 | 13.3441017220695 | 2.09924267127151 | 0.57339789777787 | 3.66105749499058 | 0.000251176379877471 | 0.0192074485017261 | 2339 | 2 | 3 | 8 | 9 | 9 | 7 | 5 | 3 | 6 | 5 | 43 | 78 |
| L1PA10_dup6571 | 14.093466664967 | 2.44471009055165 | 0.576367914195576 | 4.24157908575408 | 2.21952652047032e-05 | 0.00334469574059424 | 2340 | 4 | 2 | 9 | 14 | 2 | 7 | 2 | 2 | 6 | 8 | 44 | 90 |
| L1PA16_dup10248 | 11.1256139475417 | 2.05707816604497 | 0.490320577378234 | 4.1953739266752 | 2.72421783846016e-05 | 0.00389354078069124 | 2341 | 4 | 2 | 6 | 13 | 3 | 5 | 6 | 3 | 4 | 6 | 44 | 52 |
| L1PA5_dup10742 | 18.7191729153274 | 3.12860794242012 | 0.617820304780769 | 5.06394483672773 | 4.10668525481355e-07 | 0.000158118014505156 | 2342 | 3 | 2 | 4 | 12 | 5 | 4 | 5 | 0 | 6 | 8 | 80 | 124 |
| L1M6_dup5342 | 12.8589790009991 | 2.35398880869209 | 0.688978907418115 | 3.41663407013931 | 0.000634004370077274 | 0.035563131504575 | 2343 | 2 | 2 | 8 | 11 | 8 | 9 | 2 | 0 | 10 | 5 | 50 | 62 |
| L4_A_Mam_dup6118 | 14.1124144332642 | 3.0474026814421 | 0.55993051547637 | 5.44246580104582 | 5.25480511678894e-08 | 3.02019289449145e-05 | 2344 | 2 | 2 | 7 | 10 | 5 | 4 | 2 | 0 | 11 | 10 | 61 | 71 |
| L1MA7_dup6196 | 19.177067666464 | 1.87628263498799 | 0.535758405373523 | 3.50210583010801 | 0.000461596229321445 | 0.0289191741061206 | 2345 | 7 | 11 | 12 | 17 | 8 | 12 | 10 | 1 | 8 | 10 | 65 | 95 |
| L2b_dup87314 | 18.1158622459843 | 2.18563775185532 | 0.562397040117115 | 3.88628957115453 | 0.000101788080102261 | 0.0102151271649146 | 2346 | 9 | 4 | 8 | 19 | 7 | 13 | 5 | 1 | 13 | 9 | 63 | 89 |
| LTR80A_dup389 | 15.9781703388378 | 2.20702123043047 | 0.616071624033167 | 3.58241013598714 | 0.000340438744950835 | 0.0236267334177332 | 2347 | 5 | 2 | 7 | 12 | 21 | 5 | 5 | 2 | 14 | 9 | 60 | 65 |
| L1ME4a_dup23657 | 17.7768490187802 | 2.41855129414724 | 0.609472900519016 | 3.96826715689516 | 7.23971467092883e-05 | 0.00794289530722213 | 2348 | 8 | 4 | 5 | 10 | 10 | 14 | 3 | 3 | 7 | 10 | 83 | 76 |
| MLT1E2_dup3616 | 14.7691110356377 | 3.04122076054391 | 0.546123147257279 | 5.56874539344729 | 2.56580083717562e-08 | 1.69589457222942e-05 | 2349 | 3 | 6 | 4 | 6 | 4 | 7 | 3 | 1 | 9 | 14 | 65 | 71 |
| THE1A_dup3971 | 14.8754679860783 | 2.99580565022952 | 0.638236481605693 | 4.69388030388452 | 2.68070923187589e-06 | 0.000704044193033047 | 2350 | 5 | 4 | 4 | 3 | 10 | 8 | 3 | 1 | 12 | 7 | 51 | 88 |
| L1PA10_dup6570 | 14.6198082144836 | 2.86851741766756 | 0.645046806766966 | 4.44699111378418 | 8.70814386824791e-06 | 0.00166833172373117 | 2351 | 5 | 5 | 5 | 12 | 6 | 6 | 0 | 0 | 13 | 13 | 44 | 84 |
| L2_dup52921 | 13.2657224626892 | 2.73719425695576 | 0.731062827658018 | 3.74412998910701 | 0.000181019923620031 | 0.0153284579166163 | 2352 | 4 | 1 | 11 | 6 | 6 | 7 | 0 | 0 | 9 | 9 | 54 | 67 |
| L1MB7_dup13706 | 18.6858634797576 | 2.2502557501709 | 0.566184147836985 | 3.97442379615826 | 7.05498453247111e-05 | 0.00780793185843428 | 2353 | 4 | 1 | 15 | 23 | 6 | 14 | 1 | 6 | 15 | 25 | 62 | 69 |
| Charlie17b_dup471 | 17.6455404373531 | 2.45195537222529 | 0.572210526529277 | 4.28505813602825 | 1.8269140864443e-05 | 0.00290385370215815 | 2354 | 8 | 3 | 16 | 12 | 7 | 6 | 0 | 3 | 13 | 14 | 66 | 82 |
| MLT1E2_dup3617 | 19.8895463442642 | 2.78081941180379 | 0.605743488561981 | 4.59075411343732 | 4.41647388673113e-06 | 0.00100083295050532 | 2355 | 7 | 3 | 13 | 17 | 5 | 7 | 0 | 1 | 17 | 15 | 79 | 96 |
| MER31-int_dup258 | 49.5629997112256 | 3.82872410397663 | 0.937926767178568 | 4.08211412442576 | 4.46278711559163e-05 | 0.00553597972569969 | 2356 | 10 | 7 | 35 | 39 | 4 | 3 | 1 | 0 | 141 | 108 | 91 | 161 |
| PABL_B-int_dup108 | 43.7720935953442 | 2.85534457724928 | 0.660440178752005 | 4.32339622741436 | 1.53645425552527e-05 | 0.00254591326103911 | 2357 | 17 | 18 | 44 | 50 | 1 | 4 | 5 | 3 | 91 | 112 | 80 | 115 |
| L1PA10_dup6914 | 33.9023085025502 | 2.89155697659014 | 0.641588994832034 | 4.50686810385073 | 6.57915011306804e-06 | 0.00134607261634057 | 2358 | 7 | 6 | 47 | 66 | 0 | 1 | 15 | 14 | 62 | 53 | 71 | 82 |
| MLT1D-int_dup579 | 33.2770242056959 | 2.56255543138384 | 0.619519925078801 | 4.13635676214593 | 3.52863444960695e-05 | 0.00471169726569123 | 2359 | 10 | 10 | 50 | 44 | 0 | 0 | 21 | 17 | 53 | 53 | 69 | 90 |
| L1MA9_dup14595 | 24.2005890130569 | 2.33201543834003 | 0.614867298828355 | 3.79271339162733 | 0.000149010105809151 | 0.0132646309902223 | 2360 | 9 | 9 | 14 | 11 | 0 | 0 | 39 | 26 | 38 | 30 | 56 | 68 |
| LTR16D1_dup28 | 27.0033857152314 | 2.18412686476497 | 0.649645623411671 | 3.36202813665524 | 0.000773722392245993 | 0.0405247960258343 | 2361 | 19 | 14 | 26 | 25 | 1 | 0 | 13 | 18 | 52 | 81 | 43 | 33 |
| L1MEg_dup1227 | 19.9942299082719 | 2.13443934196335 | 0.513283116455858 | 4.15840551448746 | 3.20476739327354e-05 | 0.00439263584193471 | 2362 | 18 | 9 | 13 | 21 | 4 | 0 | 15 | 6 | 38 | 53 | 26 | 45 |
| MER66B_dup116 | 22.2019887194175 | 1.95662294435762 | 0.587226513941688 | 3.33197309369433 | 0.000862325929553641 | 0.04336135144323 | 2363 | 19 | 20 | 13 | 17 | 1 | 1 | 21 | 7 | 45 | 67 | 32 | 27 |
| L1PA7_dup3990 | 34.0679673484998 | 2.03940363230946 | 0.513930746126953 | 3.96824600917276 | 7.24035702275508e-05 | 0.00794289530722213 | 2364 | 16 | 12 | 12 | 3 | 8 | 9 | 70 | 21 | 41 | 44 | 82 | 117 |
| L1ME4a_dup10302 | 20.7295650742327 | 1.8008226095541 | 0.447699008423958 | 4.02239579643824 | 5.76091437215136e-05 | 0.00672489065365166 | 2365 | 10 | 11 | 5 | 7 | 6 | 6 | 37 | 18 | 25 | 30 | 48 | 58 |
| LTR1A1_dup118 | 26.2737622328925 | 2.02494269966074 | 0.484326097415847 | 4.18094897315043 | 2.90295009978159e-05 | 0.00408241941159947 | 2366 | 10 | 10 | 6 | 6 | 6 | 7 | 31 | 35 | 28 | 40 | 61 | 87 |
| L1PA4_dup2047 | 34.8266503027518 | 2.57385449201632 | 0.420452269478322 | 6.12163301011514 | 9.26211545325077e-10 | 1.02031635353667e-06 | 2367 | 28 | 25 | 6 | 10 | 5 | 6 | 18 | 11 | 103 | 94 | 51 | 52 |
| MER52A_dup224 | 22.3440098283979 | 2.76471445622138 | 0.488131829321814 | 5.66386842681932 | 1.47997904796447e-08 | 1.09364936194706e-05 | 2368 | 10 | 7 | 4 | 9 | 3 | 4 | 17 | 10 | 57 | 85 | 25 | 33 |
| L1PA5_dup1955 | 23.8890546724659 | 2.49554410894877 | 0.478188263418552 | 5.2187481372048 | 1.80136523962967e-07 | 8.37163385368987e-05 | 2369 | 15 | 18 | 4 | 5 | 4 | 3 | 14 | 15 | 50 | 70 | 36 | 53 |
| L2c_dup29819 | 23.6337675304853 | 2.3030223870383 | 0.522538970726529 | 4.40736962419515 | 1.04633543786425e-05 | 0.00192404429751195 | 2370 | 20 | 19 | 3 | 5 | 1 | 8 | 18 | 10 | 55 | 79 | 29 | 35 |

| | | | | | | | | | | | | | | | | | | | |
|---|---|---|---|---|---|---|---|---|---|---|---|---|---|---|---|---|---|---|---|
| L1PA8_dup805 | 30.2862679776957 | 7.66929390513159 | 0.723166363708599 | 10.605158494653 | 2.81985468868789e-26 | 1.11828857292421e-21 | 2371 | 1 | 0 | 0 | 0 | 0 | 1 | 0 | 1 | 27 | 33 | 144 | 188 |
| THE1A-int_dup39 | 35.2637319831957 | 5.45684676008243 | 0.513971850535666 | 10.6170148314451 | 2.48377207640297e-26 | 1.11828857292421e-21 | 2372 | 1 | 5 | 0 | 3 | 0 | 1 | 7 | 3 | 109 | 107 | 79 | 100 |
| L1PA13_dup1610 | 12.3751882342154 | 3.16315931549536 | 0.609142117677347 | 5.1928100581133 | 2.07143490346159e-07 | 9.29980470828435e-05 | 2373 | 0 | 0 | 9 | 15 | 1 | 1 | 12 | 8 | 16 | 26 | 25 | 44 |
| L1PA8_dup1656 | 12.8461243621459 | 3.05303901794729 | 0.680288941307336 | 4.48785631011457 | 7.19434237290226e-06 | 0.00144461180612878 | 2374 | 0 | 0 | 14 | 18 | 0 | 2 | 19 | 8 | 18 | 26 | 21 | 37 |
| LTR33B_dup361 | 13.3295994461853 | 3.07049236347234 | 0.751837816869807 | 4.08398233578618 | 4.42703975459911e-05 | 0.00550651542837345 | 2375 | 0 | 0 | 16 | 19 | 0 | 1 | 19 | 19 | 23 | 13 | 28 | 25 |
| THE1C-int_dup384 | 15.6268241764652 | 2.9599938621977 | 0.637474704606834 | 4.6433118691718 | 3.42868198355529e-06 | 0.000829950318676547 | 2376 | 0 | 0 | 21 | 20 | 0 | 2 | 17 | 13 | 18 | 19 | 47 | 40 |
| L1PBa_dup498 | 16.7253686972978 | 3.26205637457069 | 0.620205036090859 | 5.25964186800445 | 1.44336235364489e-07 | 6.96637522515998e-05 | 2377 | 0 | 0 | 17 | 24 | 1 | 0 | 17 | 10 | 12 | 26 | 38 | 74 |
| MER51B_dup123 | 16.6640997432255 | 2.91119820685356 | 0.667929991694429 | 4.35853793519337 | 1.30934228836564e-05 | 0.00227619642572668 | 2378 | 0 | 0 | 12 | 26 | 2 | 1 | 28 | 18 | 28 | 20 | 37 | 35 |
| MLT1A0_dup4491 | 18.7666751501383 | 3.37994613574797 | 0.694133827296808 | 4.86930041849512 | 1.11994033312212e-06 | 0.000349259924646234 | 2379 | 0 | 0 | 13 | 26 | 2 | 0 | 26 | 17 | 31 | 22 | 49 | 48 |
| MLT1D-int_dup580 | 15.3154453046886 | 2.56175978807749 | 0.746577324274742 | 3.43133886441849 | 0.000600609828602439 | 0.034441058645573 | 2380 | 5 | 0 | 25 | 40 | 0 | 1 | 9 | 8 | 28 | 21 | 29 | 26 |
| L1PA10_dup5521 | 13.8639377394893 | 3.11407293737264 | 0.808127630215135 | 3.85344198235572 | 0.000116468871030265 | 0.0112014561305119 | 2381 | 0 | 0 | 29 | 36 | 0 | 1 | 15 | 3 | 11 | 23 | 24 | 41 |
| L1MCc_dup3254 | 12.7926444671103 | 2.42632124892837 | 0.497440589888376 | 4.87761010711416 | 1.07378909916134e-06 | 0.000339314503305503 | 2382 | 0 | 0 | 18 | 18 | 0 | 1 | 13 | 10 | 7 | 12 | 32 | 57 |
| MLT1E1A_dup2913 | 13.6138262432825 | 2.5959955955006 | 0.618870662886477 | 4.19473041974975 | 2.73196309456845e-05 | 0.00389957822735564 | 2383 | 0 | 1 | 18 | 29 | 0 | 1 | 17 | 5 | 15 | 9 | 32 | 52 |
| L1PA2_dup4911 | 15.9612434281279 | 2.92549263485017 | 0.52968998922979 | 5.52302798681179 | 3.33206634567869e-08 | 2.1142716231703e-05 | 2384 | 3 | 0 | 19 | 11 | 1 | 6 | 4 | 3 | 14 | 28 | 44 | 74 |
| THE1C_dup7987 | 20.5231626896492 | 2.75609946407745 | 0.373899615216269 | 7.37122840440284 | 1.69062901359274e-13 | 4.90583428376023e-10 | 2385 | 2 | 0 | 16 | 14 | 3 | 1 | 20 | 13 | 13 | 22 | 70 | 93 |
| THE1C_dup7988 | 21.5662806913652 | 2.62728691849957 | 0.420462568935224 | 6.24856316021874 | 4.14245709572579e-10 | 4.92840548049784e-07 | 2386 | 1 | 0 | 26 | 23 | 4 | 2 | 17 | 10 | 16 | 22 | 68 | 92 |
| L1PREC2_dup6316 | 22.9867049093828 | 2.98112090959538 | 0.367304879110627 | 8.116202858001 | 4.80996416293169e-16 | 2.66165519235568e-12 | 2387 | 1 | 0 | 19 | 17 | 4 | 3 | 15 | 6 | 19 | 16 | 97 | 102 |
| LTR105_Mam_dup49 | 21.0200460142407 | 2.77010236953281 | 0.379166431700222 | 7.30576901840014 | 2.75685407956539e-13 | 7.4543454638212e-10 | 2388 | 2 | 0 | 17 | 26 | 2 | 1 | 11 | 11 | 13 | 15 | 71 | 108 |
| L2a_dup133434 | 22.9694311688531 | 2.92651886612694 | 0.369776577344179 | 7.91428945323113 | 2.48669155309731e-15 | 1.18339661658659e-11 | 2389 | 1 | 0 | 15 | 26 | 2 | 2 | 20 | 6 | 14 | 17 | 85 | 117 |
| Tigger1_dup11835 | 19.1268826449923 | 2.8759099197862 | 0.357855299239142 | 8.03651622848913 | 9.24287127531953e-16 | 4.59706549333266e-12 | 2390 | 2 | 0 | 12 | 18 | 1 | 1 | 18 | 8 | 13 | 15 | 74 | 89 |
| MSTD_dup6173 | 16.1590849299157 | 2.47869476354467 | 0.412047184933508 | 6.015560484765 | 1.79265350666768e-09 | 1.79224677015776e-06 | 2391 | 2 | 0 | 17 | 17 | 1 | 2 | 17 | 6 | 7 | 13 | 76 | 53 |
| MSTD_dup6174 | 19.5050115818357 | 3.04420290520287 | 0.365497045383147 | 8.3289398468643 | 8.15694971943411e-17 | 5.39142654983463e-13 | 2392 | 2 | 1 | 14 | 10 | 3 | 2 | 14 | 5 | 10 | 20 | 71 | 106 |
| L1MC4_dup18972 | 18.1263341918065 | 2.78614062507302 | 0.374079455363719 | 7.44799155667089 | 9.47718849271112e-14 | 2.89110140139313e-10 | 2393 | 2 | 1 | 12 | 19 | 0 | 3 | 11 | 10 | 12 | 18 | 73 | 74 |
| L1PA11_dup3442 | 20.820997160009 | 3.41237941122892 | 0.386766381051589 | 8.82284391407266 | 1.11588837998292e-18 | 1.15443989766703e-14 | 2394 | 1 | 0 | 11 | 10 | 0 | 2 | 11 | 10 | 12 | 10 | 82 | 126 |
| HUERS-P2-int_dup24 | 14.8074007174008 | 2.56651533520519 | 0.427170418127474 | 6.00817665805524 | 1.87621346891944e-09 | 1.86750702110292e-06 | 2395 | 2 | 0 | 9 | 15 | 0 | 2 | 13 | 12 | 12 | 6 | 49 | 73 |
| MIRb_dup170299 | 15.132551608552 | 2.54416596170729 | 0.414046803012792 | 6.14463375443255 | 8.01482283604151e-10 | 8.99573129502232e-07 | 2396 | 1 | 0 | 14 | 13 | 0 | 3 | 14 | 11 | 12 | 9 | 53 | 66 |
| AluSc8_dup15772 | 15.2264561004933 | 3.33646482756104 | 0.524862247442801 | 6.35683904456215 | 2.0594758531198e-10 | 2.69254967772771e-07 | 2397 | 0 | 1 | 9 | 7 | 0 | 2 | 14 | 4 | 16 | 4 | 61 | 81 |
| L1ME3Cz_dup9410 | 14.6349974808952 | 3.15442528828809 | 0.509716680331336 | 6.18858556137027 | 6.07064546086422e-10 | 6.97819229386859e-07 | 2398 | 0 | 0 | 9 | 13 | 0 | 3 | 10 | 4 | 13 | 4 | 62 | 74 |
| L2c_dup110921 | 19.510666265592 | 3.12584626011954 | 0.441508616405395 | 7.07992130610964 | 1.44236344514808e-12 | 3.50208788080822e-09 | 2399 | 1 | 2 | 17 | 11 | 0 | 2 | 18 | 3 | 16 | 13 | 62 | 114 |
| AluJr_dup60454 | 17.3968983256183 | 3.09003368523687 | 0.419013047976239 | 7.37455241587632 | 1.64898093437291e-13 | 4.84405453592959e-10 | 2400 | 0 | 2 | 6 | 19 | 0 | 1 | 14 | 8 | 12 | 14 | 64 | 89 |
| AluJr_dup60455 | 20.6769137665835 | 3.66709507905649 | 0.385617880604588 | 9.50966037494699 | 1.9128601710778e-21 | 4.0850728570636e-17 | 2401 | 0 | 1 | 7 | 13 | 0 | 1 | 12 | 7 | 16 | 15 | 86 | 113 |
| AluSq2_dup43750 | 9.51604082243269 | 2.72009872665538 | 0.504230536168162 | 5.39455374386175 | 6.86940641803249e-08 | 3.71129473126804e-05 | 2402 | 0 | 0 | 4 | 9 | 0 | 3 | 10 | 6 | 7 | 9 | 33 | 43 |
| L1MC3_dup12191 | 11.7556462278224 | 2.60173068328681 | 0.541581801152183 | 4.80394776514237 | 1.55567429817047e-06 | 0.000456995650064779 | 2403 | 0 | 0 | 7 | 18 | 0 | 3 | 10 | 9 | 13 | 7 | 34 | 51 |
| L1MC4_dup18971 | 15.2334413895296 | 3.24785668133895 | 0.502836733042784 | 6.45906805910021 | 1.05349728368285e-10 | 1.49211586109047e-07 | 2404 | 0 | 0 | 7 | 18 | 0 | 2 | 5 | 9 | 12 | 10 | 56 | 80 |
| MIR_dup133534 | 11.75468704932 | 3.29395550157741 | 0.476861459571126 | 6.90757333280801 | 4.93014121420717e-12 | 1.02904156259275e-08 | 2405 | 0 | 0 | 4 | 10 | 1 | 3 | 4 | 5 | 10 | 11 | 44 | 61 |
| L1P1_dup674 | 14.9714552547641 | 2.39338883980272 | 0.484925020666342 | 4.93558537465016 | 7.99107140575834e-07 | 0.000270091402942411 | 2406 | 0 | 0 | 13 | 18 | 2 | 1 | 18 | 18 | 11 | 16 | 38 | 57 |
| L3b_dup8000 | 17.4016960165173 | 2.81890070034217 | 0.407644764302477 | 6.91509114600209 | 4.67562653164225e-12 | 9.93345205980488e-09 | 2407 | 0 | 0 | 11 | 11 | 2 | 3 | 18 | 11 | 13 | 16 | 71 | 68 |
| THE1B_dup4577 | 12.4834578857266 | 3.21193695604163 | 0.534993580021832 | 6.00369252264777 | 1.92879812764915e-09 | 1.89648677389092e-06 | 2408 | 0 | 0 | 8 | 15 | 1 | 0 | 12 | 4 | 16 | 10 | 39 | 57 |
| L2c_dup110930 | 11.5290597634198 | 2.67058372629207 | 0.525222027386806 | 5.08467578859805 | 3.68254563383818e-07 | 0.000146775042443762 | 2409 | 0 | 0 | 9 | 10 | 1 | 1 | 12 | 12 | 13 | 9 | 46 | 31 |
| L1ME3Cz_dup9408 | 11.9869510843251 | 2.36586015395738 | 0.437310806781564 | 5.41001986977907 | 6.30177566932437e-08 | 3.45502837191948e-05 | 2410 | 0 | 0 | 12 | 10 | 3 | 2 | 14 | 5 | 8 | 6 | 46 | 51 |
| L2b_dup73109 | 11.0150249395678 | 2.93709645163691 | 0.477940966900189 | 6.14531219344141 | 7.98063899110433e-10 | 8.99573129502232e-07 | 2411 | 0 | 0 | 10 | 9 | 1 | 1 | 10 | 4 | 8 | 13 | 48 | 38 |
| AluY_dup80270 | 15.9084628539134 | 1.63593155375621 | 0.474924106984906 | 3.44461679181135 | 0.000571869517817832 | 0.033351486344775 | 2412 | 1 | 0 | 24 | 25 | 1 | 4 | 20 | 5 | 3 | 2 | 63 | 66 |
| Charlie6_dup225 | 12.1271412951844 | 1.79555984157172 | 0.543945822075641 | 3.30099022494566 | 0.000963442391677695 | 0.0466423730071497 | 2413 | 0 | 0 | 16 | 32 | 2 | 3 | 15 | 5 | 6 | 11 | 36 | 35 |
| MER66C_dup450 | 12.8628635045072 | 1.717908809289 | 0.488707236447722 | 3.51521050061792 | 0.00043940554686676 | 0.0280232624644219 | 2414 | 0 | 1 | 17 | 33 | 1 | 1 | 10 | 10 | 5 | 5 | 37 | 51 |
| L3_dup36526 | 15.7379026410953 | 2.94270723515235 | 0.440852139279605 | 6.67504356440465 | 2.47158734547716e-11 | 4.05589187935799e-08 | 2415 | 1 | 0 | 15 | 15 | 2 | 0 | 7 | 5 | 6 | 8 | 72 | 77 |
| Tigger3a_dup4267 | 14.7954505106374 | 3.17701382378541 | 0.566246108744179 | 5.61065899566426 | 2.01557590272994e-08 | 1.40644640396181e-05 | 2416 | 0 | 0 | 15 | 19 | 2 | 0 | 5 | 3 | 6 | 14 | 61 | 71 |
| AluSg_dup26192 | 11.1982051315981 | 2.77230409009996 | 0.535832293484309 | 5.17382793797803 | 2.2934603904751e-07 | 0.000100500870363165 | 2417 | 0 | 0 | 11 | 17 | 1 | 1 | 5 | 3 | 4 | 8 | 46 | 53 |
| L1ME4b_dup16165 | 13.934363818057 | 3.01384795826534 | 0.456102898404159 | 6.60782461328437 | 3.9000833578634e-11 | 6.18672823113443e-08 | 2418 | 1 | 0 | 10 | 12 | 1 | 1 | 6 | 5 | 2 | 7 | 65 | 76 |
| MLT1G_dup2161 | 15.14096474612 | 2.96852417690975 | 0.484703475693159 | 6.12441281272959 | 9.10186363169149e-10 | 1.01077472885744e-06 | 2419 | 1 | 0 | 13 | 15 | 1 | 1 | 12 | 1 | 7 | 6 | 70 | 75 |
| L1PA13_dup7409 | 16.2313688726456 | 3.14624241503874 | 0.445426418274355 | 7.06343918088138 | 1.6243131266053e-12 | 3.86498811223225e-09 | 2420 | 0 | 0 | 15 | 13 | 1 | 1 | 9 | 5 | 10 | 8 | 56 | 98 |
| MIR_dup133527 | 13.4415774641464 | 3.05189569712329 | 0.487325843424408 | 6.26253612096131 | 3.78766101732014e-10 | 4.57491770775258e-07 | 2421 | 0 | 0 | 15 | 10 | 0 | 1 | 11 | 3 | 6 | 8 | 53 | 72 |
| AluY_dup76592 | 10.2258007311832 | 2.32101478958944 | 0.506802070137078 | 4.57972633963721 | 4.65584704921038e-06 | 0.00103924970166174 | 2422 | 0 | 1 | 14 | 6 | 3 | 2 | 8 | 4 | 8 | 4 | 32 | 52 |
| THE1B_dup18272 | 12.8357340493957 | 2.40393140417083 | 0.424991384041833 | 5.65642385807568 | 1.54559498599584e-08 | 1.1350868658573e-05 | 2423 | 0 | 2 | 16 | 12 | 1 | 2 | 9 | 6 | 8 | 7 | 52 | 52 |
| AluJr_dup60452 | 13.0506129590695 | 2.66272970609718 | 0.485545614695979 | 5.48399496464287 | 4.15826611604159e-08 | 2.51127611484171e-05 | 2424 | 0 | 0 | 4 | 11 | 2 | 4 | 17 | 4 | 7 | 8 | 54 | 62 |
| L1MD2_dup8421 | 13.2059169081985 | 2.26278249043376 | 0.467064268209231 | 4.8446919288206 | 1.26808284222784e-06 | 0.000384376101881206 | 2425 | 1 | 0 | 11 | 12 | 7 | 2 | 8 | 6 | 5 | 8 | 46 | 69 |
| L1MEg1_dup716 | 12.9284176633078 | 2.69501518422864 | 0.536752068138249 | 5.02096842137271 | 5.14116121286867e-07 | 0.000190251748982465 | 2426 | 1 | 0 | 7 | 7 | 6 | 2 | 7 | 4 | 7 | 4 | 48 | 79 |
| L1ME3Cz_dup10794 | 9.024889881715 | 2.42070855013802 | 0.66775360413371 | 3.62515235433054 | 0.000288791147907649 | 0.021059362083982 | 2427 | 2 | 3 | 5 | 3 | 0 | 1 | 10 | 12 | 10 | 18 | 7 | 43 |
| THE1B_dup6813 | 7.49028533101998 | 3.01769776338585 | 0.850706149586913 | 3.54728570476561 | 0.000389222250846425 | 0.0258409256975177 | 2428 | 0 | 1 | 2 | 3 | 0 | 0 | 12 | 14 | 9 | 16 | 15 | 19 |
| THE1B-int_dup1420 | 8.62153468975249 | 2.83543412876235 | 0.76329834807209 | 3.71471278029617 | 0.000203434628551381 | 0.0166259449141726 | 2429 | 1 | 3 | 0 | 3 | 0 | 1 | 9 | 11 | 18 | 15 | 26 | 15 |
| THE1B_dup6819 | 10.0306809739249 | 2.38878149961178 | 0.644422537224744 | 3.70685592390865 | 0.000209848243671529 | 0.0169896393973003 | 2430 | 3 | 3 | 0 | 4 | 1 | 2 | 12 | 13 | 19 | 12 | 23 | 30 |
| THE1C_dup6766 | 8.74909164075662 | 2.8876857291984 | 0.568236112951808 | 5.08184126876023 | 3.73793715489984e-07 | 0.000148237865709966 | 2431 | 1 | 4 | 0 | 5 | 1 | 1 | 8 | 5 | 16 | 11 | 30 | 26 |
| MARNA_dup1023 | 13.1173735132658 | 3.07142095903149 | 0.671431806551629 | 4.57443470663959 | 4.77507489306356e-06 | 0.00106187847710739 | 2432 | 1 | 1 | 2 | 5 | 0 | 2 | 20 | 14 | 17 | 23 | 39 | 39 |
| MLT1D_dup5763 | 7.77924038555136 | 2.23712176401432 | 0.667083882081257 | 3.35358389567838 | 0.000797722264315021 | 0.0413539917003708 | 2433 | 2 | 5 | 2 | 1 | 0 | 1 | 13 | 7 | 8 | 12 | 28 | 18 |
| MSTA_dup5774 | 9.65813711596093 | 2.3498378190535 | 0.676453250034455 | 3.47376233159323 | 0.00051321540530119 | 0.0310336855984236 | 2434 | 3 | 4 | 3 | 3 | 0 | 0 | 18 | 9 | 14 | 10 | 38 | 17 |
| L1PB1_dup8279 | 10.1868358406907 | 2.81707635062452 | 0.61612425780222 | 4.57225359162668 | 4.82506479479178e-06 | 0.00106968102544101 | 2435 | 8 | 4 | 3 | 0 | 2 | 1 | 8 | 2 | 10 | 12 | 29 | 54 |
| ERVL-B4-int_dup2699 | 11.5930380221137 | 2.88916544610237 | 0.488770966277437 | 5.91108237894478 | 3.39866996382595e-09 | 3.09846713874532e-06 | 2436 | 3 | 2 | 4 | 2 | 1 | 3 | 13 | 6 | 17 | 28 | 26 | 40 |
| ERVL-B4-int_dup2700 | 11.3695320506048 | 2.81818614055896 | 0.462600043435187 | 6.09205766526003 | 1.11468513843365e-09 | 1.20780925714484e-06 | 2437 | 6 | 1 | 7 | 1 | 2 | 4 | 9 | 3 | 14 | 24 | 32 | 41 |
| HAL1_dup7060 | 8.58773421343524 | 2.12199329305184 | 0.563592025480316 | 3.76512299165943 | 0.000166467121806327 | 0.0143567182911664 | 2438 | 1 | 4 | 2 | 3 | 2 | 2 | 17 | 6 | 10 | 11 | 19 | 33 |
| MIRb_dup66796 | 10.3298688020057 | 2.30451492254461 | 0.561614902960701 | 4.10337209784811 | 4.07171592504826e-05 | 0.0051858209563448 | 2439 | 2 | 5 | 2 | 4 | 2 | 1 | 18 | 8 | 15 | 19 | 29 | 23 |
| AluJb_dup75643 | 7.96328000499234 | 2.92483987923637 | 0.514016973163823 | 5.69016206066834 | 1.26918858094527e-08 | 9.64850945947613e-06 | 2440 | 4 | 3 | 3 | 1 | 2 | 1 | 4 | 3 | 11 | 12 | 32 | 23 |
| L3_dup14176 | 6.47126442692107 | 2.65684629711507 | 0.546293618113371 | 4.86340350504278 | 1.15384288982594e-06 | 0.000357428388883535 | 2441 | 4 | 4 | 1 | 1 | 2 | 1 | 3 | 4 | 8 | 17 | 17 | 18 |
| AluSx_dup30073 | 6.91665961024443 | 2.10825585108893 | 0.51577868559955 | 4.08752030657927 | 4.36008522619597e-05 | 0.0054545995753545 | 2442 | 2 | 1 | 2 | 3 | 3 | 3 | 9 | 6 | 10 | 13 | 14 | 20 |

| | | | | | | | | | | | | | | | | | | |
|---|---|---|---|---|---|---|---|---|---|---|---|---|---|---|---|---|---|---|
| Ricksha_c_dup789 | 6.7179594192599 | 2.30270052601434 | 0.512413963080898 | 4.4938286071857 | 6.99539380870533e-06 | 0.00140942080881134 | 2443 | 3 | 1 | 2 | 4 | 1 | 3 | 8 | 4 | 12 | 12 | 17 | 16 |
| AluSx_dup30070 | 8.31846933357876 | 2.36890978712879 | 0.470897708549382 | 5.03062500436944 | 4.88883516054547e-07 | 0.000182332095785447 | 2444 | 2 | 4 | 4 | 3 | 2 | 3 | 7 | 5 | 16 | 9 | 19 | 29 |
| MSTA1_dup133 | 6.85422128566483 | 2.09224600870604 | 0.492757471204646 | 4.24599550685881 | 2.1762499110369e-05 | 0.00330038216272521 | 2445 | 1 | 4 | 2 | 4 | 2 | 5 | 6 | 4 | 7 | 10 | 19 | 23 |
| L2_dup17667 | 5.94648053982455 | 2.49760484269076 | 0.63640898944909 | 3.92452791223585 | 8.68999836995375e-05 | 0.00909300946410297 | 2446 | 3 | 1 | 1 | 1 | 2 | 0 | 8 | 4 | 5 | 8 | 16 | 28 |
| L2c_dup44109 | 6.92972746374856 | 2.43056792871955 | 0.63299528460213 | 3.83978836469104 | 0.000123140415968012 | 0.0116418460383675 | 2447 | 1 | 1 | 4 | 1 | 3 | 0 | 13 | 4 | 7 | 12 | 21 | 21 |
| L1M1_dup6231 | 7.13984393114743 | 2.31615683232604 | 0.606508405021359 | 3.81883715567713 | 0.000134082235022497 | 0.0123468775134145 | 2448 | 4 | 0 | 4 | 2 | 2 | 0 | 8 | 5 | 2 | 7 | 25 | 35 |
| MLT1E1A_dup1123 | 5.83528529756732 | 2.41721148786531 | 0.619156239788886 | 3.90404122986713 | 9.4599654256459e-05 | 0.00965663205993454 | 2449 | 1 | 0 | 2 | 3 | 4 | 0 | 6 | 5 | 4 | 10 | 20 | 19 |
| L2_dup17673 | 9.59354101015174 | 2.95806935929992 | 0.674509917999483 | 4.38550906423023 | 1.15714839167268e-05 | 0.00208374840047616 | 2450 | 0 | 2 | 3 | 0 | 5 | 0 | 10 | 5 | 8 | 15 | 37 | 37 |
| MLT1F2_dup5010 | 7.63627639676121 | 3.31303356345798 | 0.652992613618677 | 5.07361568011957 | 3.90327001851529e-07 | 0.000153009470811473 | 2451 | 0 | 1 | 1 | 2 | 2 | 0 | 9 | 4 | 15 | 13 | 17 | 31 |
| L1MB7_dup637 | 11.7985852471931 | 2.04045985774693 | 0.522207114723909 | 3.90737659487025 | 9.3303668510708e-05 | 0.00956737689934412 | 2452 | 2 | 0 | 6 | 5 | 2 | 6 | 22 | 10 | 8 | 14 | 42 | 34 |
| MLT1N2_dup1708 | 9.79768452250882 | 1.97018244469403 | 0.577928269258617 | 3.40904321434464 | 0.000651911519252821 | 0.036224555790675 | 2453 | 9 | 0 | 3 | 4 | 1 | 3 | 16 | 8 | 11 | 11 | 27 | 32 |
| MSTA-int_dup2760 | 7.94772723820736 | 2.20167784202157 | 0.548627468906704 | 4.01306527070006 | 5.99353317654164e-05 | 0.00690957967647954 | 2454 | 3 | 0 | 2 | 5 | 5 | 3 | 9 | 5 | 11 | 26 | 14 | 15 |
| MER3_dup8141 | 9.33418774275512 | 1.69677108191321 | 0.499573369128653 | 3.39644021632435 | 0.000682684685022731 | 0.0372317419350031 | 2455 | 2 | 1 | 8 | 8 | 7 | 4 | 7 | 10 | 13 | 23 | 13 | 18 |
| MLT1I_dup6464 | 9.40237373287901 | 2.29885242855396 | 0.513964167975111 | 4.47278734938052 | 7.72065472158539e-06 | 0.00151575817523297 | 2456 | 1 | 1 | 6 | 4 | 6 | 4 | 5 | 8 | 15 | 25 | 23 | 15 |
| MIR3_dup48853 | 10.6331359934606 | 2.37589379167765 | 0.504089126708777 | 4.71324149995058 | 2.43806959033533e-06 | 0.000656254419391324 | 2457 | 2 | 0 | 4 | 3 | 6 | 9 | 8 | 6 | 22 | 18 | 26 | 25 |
| MIR_dup137142 | 11.5199338782132 | 2.0801244434332 | 0.444692792410689 | 4.67766619772901 | 2.90158377997135e-06 | 0.000736841253051294 | 2458 | 8 | 0 | 4 | 12 | 6 | 7 | 8 | 4 | 22 | 28 | 21 | 22 |
| MER5A1_dup14498 | 13.326290895847 | 1.515202692873 | 0.382341734383542 | 3.96295396660266 | 7.40280599809477e-05 | 0.00806533001841877 | 2459 | 6 | 3 | 15 | 21 | 2 | 6 | 14 | 6 | 5 | 14 | 34 | 51 |
| L1ME3G_dup15138 | 13.7813365780402 | 1.97061879465812 | 0.351862850704286 | 5.60053097595765 | 2.13696215798048e-08 | 1.47814417919426e-05 | 2460 | 10 | 4 | 8 | 7 | 3 | 7 | 11 | 10 | 17 | 20 | 36 | 40 |
| MIRb_dup201083 | 14.9269623623624 | 1.90345028410519 | 0.349697143676733 | 5.44313935221838 | 5.23496613692637e-08 | 3.01607567171206e-05 | 2461 | 6 | 2 | 12 | 13 | 4 | 5 | 15 | 13 | 14 | 24 | 43 | 37 |
| MIR_dup137140 | 10.0033349700134 | 1.64163291970225 | 0.456224619040373 | 3.59829972164868 | 0.000320304367843684 | 0.0226120505216138 | 2462 | 4 | 4 | 11 | 13 | 4 | 1 | 12 | 6 | 13 | 20 | 22 | 15 |
| MamRTE1_dup7778 | 9.37319820278694 | 1.81246875083823 | 0.499745774939135 | 3.62678153919154 | 0.000286975891144978 | 0.0209655404956656 | 2463 | 2 | 1 | 10 | 12 | 8 | 2 | 8 | 4 | 17 | 16 | 18 | 18 |
| LTR49-int_dup877 | 13.150130466299 | 1.87879838594192 | 0.557566535846918 | 3.36963979211578 | 0.000752665026652714 | 0.0397103397853452 | 2464 | 1 | 1 | 23 | 17 | 5 | 5 | 6 | 12 | 22 | 20 | 23 | 26 |
| MLT1J1_dup3980 | 11.0535502556974 | 1.60543353665366 | 0.48540560490594 | 3.30740626071831 | 0.000941642208562463 | 0.0458922556283047 | 2465 | 0 | 3 | 15 | 9 | 6 | 4 | 16 | 9 | 16 | 16 | 22 | 21 |
| MLT1H_dup8426 | 16.8628990948168 | 2.80765714980267 | 0.775325033383165 | 3.62126466825318 | 0.00029316640703656 | 0.021254653835686 | 2466 | 0 | 1 | 4 | 6 | 4 | 3 | 31 | 23 | 37 | 43 | 20 | 28 |
| MLT1D_dup5758 | 16.1647110946182 | 2.35219440463928 | 0.599563792739352 | 3.92317620430734 | 8.73891887554391e-05 | 0.00912413686160672 | 2467 | 2 | 6 | 6 | 7 | 2 | 0 | 31 | 19 | 22 | 24 | 37 | 45 |
| THE1B_dup6816 | 11.4673130358258 | 2.38779349942513 | 0.699167980188474 | 3.41519286793062 | 0.000637368636940461 | 0.0356738167426132 | 2468 | 3 | 2 | 4 | 2 | 2 | 0 | 24 | 15 | 13 | 25 | 30 | 21 |
| L1PA7_dup3988 | 14.7789758810712 | 2.03370547188411 | 0.471591684336644 | 4.31242861023893 | 1.6147102750892e-05 | 0.00264610090300534 | 2469 | 9 | 8 | 4 | 8 | 1 | 2 | 20 | 11 | 14 | 19 | 53 | 38 |
| L1PA7_dup3989 | 12.5386427997606 | 2.12536880749399 | 0.619424607212496 | 3.43119853933228 | 0.00060092062040974 | 0.0344463160549304 | 2470 | 5 | 4 | 4 | 1 | 1 | 3 | 28 | 10 | 19 | 14 | 29 | 40 |
| L1PA8_dup2434 | 12.3250938644695 | 2.19813500706816 | 0.572633514212727 | 3.8386419105948 | 0.000123716703835362 | 0.0116817042900036 | 2471 | 5 | 4 | 5 | 3 | 0 | 2 | 24 | 11 | 17 | 16 | 30 | 38 |
| MSTB_dup5677 | 12.3162522509529 | 1.9877389146572 | 0.51660730404225 | 3.84767868960411 | 0.000119242300364865 | 0.0113994489363673 | 2472 | 7 | 1 | 1 | 6 | 7 | 5 | 19 | 8 | 16 | 15 | 39 | 31 |
| MER20B_dup1244 | 17.9560307641739 | 2.01370690944374 | 0.542883493805583 | 3.7092800433621 | 0.000207849435059797 | 0.0168852651672033 | 2473 | 11 | 3 | 2 | 9 | 4 | 4 | 33 | 18 | 23 | 34 | 37 | 47 |
| LTR16B2_dup455 | 13.2781486484186 | 1.82477546577455 | 0.50025552406351 | 3.64768678804811 | 0.00026461193207335 | 0.0198685234424504 | 2474 | 8 | 4 | 4 | 4 | 5 | 2 | 22 | 14 | 12 | 21 | 26 | 47 |
| MIR_dup52893 | 13.3201054961024 | 1.89958627956724 | 0.547222234654938 | 3.47132510206766 | 0.000517896578137506 | 0.0311835256753778 | 2475 | 5 | 5 | 3 | 5 | 3 | 3 | 28 | 13 | 9 | 29 | 34 | 32 |
| L1ME3G_dup15139 | 16.6845903244744 | 1.93124215610264 | 0.397724254869806 | 4.85573140801994 | 1.19943190177137e-06 | 0.000367310197295869 | 2476 | 7 | 6 | 6 | 14 | 16 | 2 | 17 | 7 | 22 | 22 | 39 | 54 |
| L2_dup29432 | 11.6468532173274 | 1.44330019274495 | 0.401604179020603 | 3.59383758472022 | 0.000325842995358199 | 0.0229048854870021 | 2477 | 7 | 5 | 7 | 11 | 12 | 2 | 14 | 7 | 11 | 20 | 19 | 34 |
| L1MC3_dup1885 | 14.6206710626074 | 1.71849400414244 | 0.381109200125845 | 4.50919055109397 | 6.50754397374075e-06 | 0.00133256803646791 | 2478 | 3 | 4 | 8 | 12 | 11 | 4 | 15 | 15 | 15 | 22 | 35 | 39 |
| MLT1A0-int_dup142 | 14.1213673008169 | 1.36149881452589 | 0.354353718974305 | 3.84220269640971 | 0.000121935070210401 | 0.011587045613532 | 2479 | 6 | 5 | 8 | 12 | 9 | 7 | 17 | 15 | 14 | 14 | 39 | 31 |
| AluSz_dup29098 | 13.3322752715689 | 1.77965858769133 | 0.381318390770524 | 4.66711973711838 | 3.05451343440041e-06 | 0.000766945003482527 | 2480 | 6 | 4 | 6 | 9 | 8 | 9 | 17 | 6 | 16 | 34 | 27 | 25 |
| MIRc_dup246 | 14.0293076947912 | 1.8093457164147 | 0.415648901236743 | 4.35306267147846 | 1.34248751995153e-05 | 0.00232150825161619 | 2481 | 5 | 4 | 5 | 5 | 10 | 9 | 18 | 10 | 22 | 30 | 29 | 25 |
| L1M2_dup1898 | 15.0946510916258 | 1.57733985934344 | 0.361769702493115 | 4.36006622023153 | 1.30023090292621e-05 | 0.00226323878879064 | 2482 | 9 | 9 | 5 | 8 | 12 | 9 | 15 | 11 | 19 | 29 | 38 | 22 |
| L2a_dup136979 | 13.0212101146636 | 1.51794419837573 | 0.361797386056742 | 4.19556430442996 | 2.72193045495549e-05 | 0.00389354078069124 | 2483 | 5 | 10 | 7 | 10 | 7 | 6 | 16 | 9 | 16 | 27 | 26 | 23 |
| MLT1D_dup13855 | 8.29996599186707 | 1.97998905410821 | 0.579526819894567 | 3.41656155701031 | 0.000634173245492333 | 0.0355642203799007 | 2484 | 2 | 2 | 7 | 19 | 3 | 1 | 4 | 1 | 9 | 5 | 25 | 31 |
| L1MD2_dup2979 | 7.40601512729287 | 1.74052508842945 | 0.471834836635938 | 3.68884396251653 | 0.000225275320529357 | 0.0178145947975803 | 2485 | 3 | 3 | 6 | 12 | 2 | 3 | 3 | 5 | 10 | 10 | 18 | 18 |
| L1PA3_dup3361 | 9.93580702249196 | 2.05738437028527 | 0.446433545162981 | 4.60848964549512 | 4.05604569939154e-06 | 0.000941580341451141 | 2486 | 3 | 4 | 9 | 13 | 3 | 2 | 3 | 6 | 15 | 11 | 23 | 33 |
| MLT1H_dup6675 | 6.92837406429207 | 2.16913043387976 | 0.616766453297677 | 3.51693971402308 | 0.000436552921111146 | 0.0279011607216526 | 2487 | 1 | 4 | 5 | 2 | 3 | 4 | 2 | 3 | 3 | 5 | 32 | 25 |
| L1PA10_dup5994 | 8.69612088086339 | 2.54022507673069 | 0.597524595527937 | 4.2512477239306 | 2.12582873824322e-05 | 0.00324034983410281 | 2488 | 5 | 3 | 1 | 3 | 5 | 7 | 4 | 1 | 7 | 8 | 32 | 37 |
| L2c_dup91350 | 7.72171646947007 | 2.27181721706719 | 0.572660796316252 | 3.96712544612985 | 7.27447079370057e-05 | 0.00795763824028531 | 2489 | 6 | 1 | 3 | 2 | 4 | 5 | 4 | 3 | 10 | 4 | 28 | 28 |
| THE1A-int_dup107 | 8.39350275133906 | 2.41152860941129 | 0.529115093413718 | 4.55766361502316 | 5.17257929497535e-06 | 0.00112401329033991 | 2490 | 3 | 2 | 6 | 4 | 5 | 6 | 2 | 1 | 6 | 13 | 30 | 30 |
| MLT1K_dup10937 | 9.72040044925449 | 2.07243158035018 | 0.618456988508061 | 3.35097123786995 | 0.0008052866596622 | 0.0415469946921036 | 2491 | 3 | 2 | 7 | 8 | 5 | 8 | 2 | 1 | 5 | 5 | 44 | 37 |
| MLT1C_dup17823 | 10.3662481323783 | 2.38847555482606 | 0.668158616500496 | 3.57471339266084 | 0.000350611755732312 | 0.0241326771274171 | 2492 | 5 | 1 | 6 | 4 | 6 | 8 | 2 | 1 | 5 | 5 | 41 | 53 |
| MamRep1527_dup85 | 10.9393918378635 | 2.58486122187215 | 0.617462239510007 | 4.18626606855084 | 2.83580910657328e-05 | 0.00400694438047913 | 2493 | 5 | 1 | 7 | 4 | 5 | 6 | 2 | 1 | 5 | 8 | 52 | 47 |
| Kanga1a_dup562 | 11.3168157537753 | 2.23371166052927 | 0.633805334076176 | 3.52428662309238 | 0.000424624657206299 | 0.0273444488994885 | 2494 | 7 | 1 | 3 | 13 | 4 | 7 | 1 | 3 | 7 | 6 | 50 | 46 |
| L2a_dup115545 | 9.02776757147869 | 2.11910839367222 | 0.578733975211382 | 3.66162776757356 | 0.000250617818994093 | 0.0191809287740008 | 2495 | 7 | 1 | 3 | 9 | 2 | 5 | 3 | 3 | 4 | 6 | 37 | 39 |
| L1M5_dup37747 | 8.661921497145 | 2.02080743142155 | 0.579367905222042 | 3.48795197871217 | 0.000486735489833139 | 0.0299711957164091 | 2496 | 3 | 0 | 8 | 9 | 3 | 7 | 3 | 2 | 3 | 10 | 26 | 41 |
| THE1C_dup7985 | 6.69567853588021 | 2.10239040251563 | 0.626863735509487 | 3.35382361974872 | 0.000797031507341995 | 0.0413428455025945 | 2497 | 3 | 1 | 2 | 7 | 2 | 5 | 4 | 1 | 2 | 5 | 23 | 35 |
| L2b_dup73094 | 11.1874801251131 | 2.78720142680864 | 0.591962268838974 | 4.70841060913431 | 2.49655806021298e-06 | 0.000665970856721343 | 2498 | 3 | 4 | 8 | 4 | 2 | 1 | 6 | 0 | 3 | 9 | 45 | 65 |
| L1PA4_dup10148 | 12.5060703339714 | 2.51609671579398 | 0.493485488530983 | 5.09862351430829 | 3.42132256789479e-07 | 0.000137981359278016 | 2499 | 1 | 4 | 7 | 8 | 5 | 1 | 8 | 3 | 4 | 8 | 52 | 65 |
| L1MB2_dup5729 | 10.8710624695741 | 2.74227163239998 | 0.47619938251235 | 5.75866272218204 | 8.4782912755721e-09 | 6.81545775627459e-06 | 2500 | 5 | 4 | 4 | 6 | 1 | 3 | 4 | 3 | 8 | 10 | 46 | 47 |
| MLT1C_dup13031 | 10.5472751670188 | 2.37188432338828 | 0.525635014385524 | 4.51241690236533 | 6.40930487134469e-06 | 0.00131725462167481 | 2501 | 3 | 6 | 9 | 8 | 4 | 1 | 2 | 2 | 7 | 9 | 42 | 44 |
| L1PA16_dup11606 | 8.64717698326586 | 3.16352578422176 | 0.568136621854417 | 5.56824830952791 | 2.57312965843464e-08 | 1.69602745070883e-05 | 2502 | 1 | 0 | 3 | 7 | 1 | 1 | 4 | 3 | 6 | 2 | 33 | 54 |
| L3_dup36525 | 8.65600163117459 | 2.45638823183067 | 0.496287164367252 | 4.94953004670688 | 7.43928928591945e-07 | 0.000255801897171588 | 2503 | 2 | 0 | 6 | 10 | 1 | 2 | 6 | 3 | 4 | 5 | 28 | 49 |
| AluY_dup76596 | 11.2960622364877 | 3.01552647407392 | 0.549706951353591 | 5.48569827368661 | 4.11839330976357e-08 | 2.49352471879135e-05 | 2504 | 1 | 0 | 4 | 15 | 2 | 1 | 5 | 2 | 8 | 4 | 48 | 60 |
| L2b_dup73108 | 10.6956239767801 | 2.87244437575716 | 0.503793175514483 | 5.70163415338798 | 1.1866429243953e-08 | 9.137764960782e-06 | 2505 | 1 | 0 | 9 | 10 | 2 | 0 | 8 | 2 | 8 | 4 | 45 | 52 |
| THE1B-int_dup2767 | 10.2018646035019 | 2.46260780692095 | 0.664501428875714 | 3.70594809869332 | 0.000210601426723367 | 0.0170274437931085 | 2506 | 3 | 0 | 7 | 8 | 2 | 1 | 3 | 2 | 1 | 1 | 47 | 63 |
| L2c_dup110912 | 6.82513193772205 | 2.33225950115898 | 0.610943135159666 | 3.81747394632638 | 0.000134825019615411 | 0.0123865151032465 | 2507 | 3 | 0 | 8 | 6 | 2 | 0 | 5 | 1 | 4 | 4 | 17 | 42 |
| MER103C_dup6943 | 9.74427531372281 | 3.13292408339244 | 0.695468450420149 | 4.50476809048602 | 6.64454657111193e-06 | 0.00135595478422796 | 2508 | 1 | 0 | 8 | 12 | 1 | 0 | 2 | 0 | 3 | 3 | 42 | 60 |
| L1PB4_dup7311 | 10.9532940784743 | 2.13784995072803 | 0.523724151103126 | 4.08201521015415 | 4.46468740538513e-05 | 0.00553597972569969 | 2509 | 4 | 3 | 6 | 20 | 0 | 0 | 9 | 4 | 3 | 9 | 39 | 50 |
| MIR3_dup66178 | 11.6611230087294 | 2.32711118837047 | 0.447347967735337 | 5.20201578237022 | 1.97138479844996e-07 | 8.93491670956141e-05 | 2510 | 2 | 1 | 12 | 16 | 0 | 0 | 15 | 6 | 9 | 11 | 39 | 41 |
| MLT1D-int_dup506 | 11.3111897355812 | 2.5336483480214 | 0.420645700909431 | 6.02323604530769 | 1.70963823718093e-09 | 1.738468290531e-06 | 2511 | 2 | 1 | 10 | 12 | 0 | 1 | 10 | 5 | 7 | 10 | 43 | 47 |
| L1MB1_dup5919 | 11.3903693976293 | 2.67481874202342 | 0.547370491811924 | 4.88666959954152 | 1.02555985239489e-06 | 0.00032580489270755 | 2512 | 1 | 2 | 11 | 10 | 0 | 0 | 4 | 12 | 11 | 7 | 30 | 58 |
| L1MC5a_dup16544 | 11.753721936 | 2.65994455807455 | 0.548965153920597 | 4.84537960028568 | 1.26369827997964e-06 | 0.000383535651693924 | 2513 | 2 | 1 | 4 | 16 | 0 | 0 | 11 | 11 | 14 | 6 | 40 | 45 |
| LTR8A_dup1384 | 9.54123302947042 | 2.40146836817397 | 0.47547813732507 | 5.05063888254479 | 4.40334822987489e-07 | 0.00016737365781243 | 2514 | 1 | 1 | 7 | 10 | 0 | 0 | 11 | 7 | 6 | 4 | 40 | 37 |

| | | | | | | | | | | | | | | | | | | | |
|---|---|---|---|---|---|---|---|---|---|---|---|---|---|---|---|---|---|---|---|
| AluSp_dup49092 | 9.09237692403508 | 2.20793016279042 | 0.544594078635677 | 4.05426766358119 | 5.02916062013604e-05 | 0.00605908178693109 | 2515 | 1 | 0 | 11 | 17 | 0 | 0 | 10 | 3 | 3 | 4 | 29 | 45 |
| Charlie4a_dup2050 | 8.9127069721716 | 2.27406511476667 | 0.626685831610753 | 3.62871633609731 | 0.000284834004589798 | 0.020834648649285 | 2516 | 0 | 0 | 10 | 20 | 0 | 0 | 6 | 4 | 3 | 1 | 33 | 43 |
| L1ME3Cz_dup9407 | 8.87564807633591 | 2.26817364471523 | 0.529362759447192 | 4.28472461331406 | 1.82965678645068e-05 | 0.00290526329627872 | 2517 | 2 | 0 | 8 | 10 | 0 | 0 | 5 | 10 | 3 | 4 | 36 | 37 |
| AluSc_dup25765 | 7.419856282204 | 2.2445329336997 | 0.555728650205917 | 4.03890087881563 | 5.37022511539995e-05 | 0.00636366327345098 | 2518 | 0 | 1 | 7 | 9 | 0 | 0 | 5 | 9 | 3 | 4 | 26 | 32 |
| MIRb_dup170318 | 9.07890428496578 | 2.57941413316038 | 0.545167895786881 | 4.73141238340406 | 2.22963050233383e-06 | 0.000611702896822642 | 2519 | 0 | 1 | 11 | 12 | 0 | 0 | 4 | 7 | 6 | 6 | 30 | 41 |
| AluSg4_dup5063 | 8.44676900211947 | 2.94143926799353 | 0.599472469168439 | 4.90671285050632 | 9.26154715041505e-07 | 0.000301470328078339 | 2520 | 0 | 0 | 2 | 10 | 1 | 0 | 7 | 5 | 3 | 4 | 26 | 56 |
| L1MD2_dup8422 | 11.2071541668101 | 2.83210638552486 | 0.526965863695493 | 5.37436403501345 | 7.68534494250441e-08 | 4.08191314216329e-05 | 2521 | 0 | 0 | 4 | 11 | 1 | 0 | 15 | 6 | 6 | 4 | 54 | 46 |
| L1M4_dup20180 | 8.51329303589271 | 2.68022791854706 | 0.545145670262096 | 4.91653527626562 | 8.80893304453801e-07 | 0.000292336176041233 | 2522 | 1 | 0 | 5 | 7 | 2 | 0 | 5 | 5 | 1 | 4 | 44 | 38 |
| MER1B_dup3417 | 10.433398564857 | 2.23512606460236 | 0.491001203785725 | 4.55218041701131 | 5.30927743900971e-06 | 0.00114743081698692 | 2523 | 0 | 0 | 9 | 14 | 1 | 0 | 8 | 8 | 1 | 1 | 45 | 52 |
| MIRb_dup170322 | 14.8253718162838 | 4.03490631939486 | 0.518482810248097 | 7.78214096907886 | 7.13073454031797e-15 | 3.20137690741604e-11 | 2524 | 0 | 1 | 2 | 8 | 0 | 0 | 7 | 4 | 6 | 5 | 60 | 107 |
| L2_dup45754 | 13.9847820218942 | 3.31612606990175 | 0.493415410872747 | 6.72075901325464 | 1.80780369624434e-11 | 3.11709897323592e-08 | 2525 | 0 | 0 | 10 | 9 | 0 | 0 | 8 | 9 | 9 | 6 | 66 | 64 |
| MIRb_dup170316 | 12.5172002204665 | 2.74678745125613 | 0.484292963257052 | 5.67174759836061 | 1.41348125030328e-08 | 1.05764845781341e-05 | 2526 | 0 | 0 | 11 | 16 | 0 | 0 | 11 | 8 | 6 | 7 | 58 | 46 |
| L3b_dup8001 | 10.4713590180129 | 2.96547384561987 | 0.575034585367732 | 5.157035630689 | 2.50890075459966e-07 | 0.000107371024991722 | 2527 | 0 | 0 | 15 | 11 | 0 | 0 | 5 | 7 | 7 | 10 | 33 | 48 |
| MER66B_dup965 | 12.844137842089 | 2.97292289975494 | 0.496630642593401 | 5.98618499299674 | 2.14820028052309e-09 | 2.07786855263962e-06 | 2528 | 1 | 0 | 11 | 17 | 0 | 0 | 6 | 8 | 8 | 13 | 43 | 61 |
| AluSx_dup82053 | 12.7405634338578 | 2.64972679572751 | 0.499612864291363 | 5.30355998636227 | 1.13565861980167e-07 | 5.67700474679262e-05 | 2529 | 1 | 0 | 7 | 20 | 0 | 0 | 16 | 8 | 6 | 11 | 37 | 64 |
| AluJb_dup90754 | 11.2027770189623 | 3.28878152860934 | 0.516673671361867 | 6.36529730640358 | 1.9491181716057e-10 | 2.59097694112229e-07 | 2530 | 1 | 0 | 8 | 8 | 0 | 0 | 7 | 4 | 7 | 6 | 34 | 75 |
| LTR8A_dup1385 | 10.5037645444482 | 2.8902296200201 | 0.477708309537559 | 6.05019749984663 | 1.44668367965438e-09 | 1.50319910410061e-06 | 2531 | 1 | 0 | 8 | 10 | 1 | 0 | 6 | 7 | 10 | 9 | 32 | 52 |
| L1M6_dup5550 | 5.59928062677694 | 2.9158873112328 | 0.672801500269734 | 4.33394888397809 | 1.46458195079608e-05 | 0.00247628558420654 | 2532 | 0 | 2 | 4 | 1 | 0 | 0 | 5 | 3 | 2 | 4 | 26 | 26 |
| AluJb_dup90750 | 6.10460244728476 | 2.75120695821511 | 0.595964064465784 | 4.61639740087561 | 3.90458921150469e-06 | 0.000916222834214355 | 2533 | 0 | 1 | 7 | 4 | 0 | 0 | 5 | 3 | 3 | 5 | 23 | 29 |
| MIR3_dup66176 | 6.57137469104739 | 2.90311181830309 | 0.583020225985802 | 4.97943585643938 | 6.3769883267977e-07 | 0.000225800426251221 | 2534 | 0 | 1 | 5 | 5 | 0 | 0 | 4 | 5 | 4 | 7 | 29 | 24 |
| MLT1A0_dup16556 | 5.22435021829403 | 2.55365961161121 | 0.73009742310552 | 3.49769706178257 | 0.000469293865796582 | 0.0292397481515668 | 2535 | 0 | 1 | 4 | 2 | 0 | 1 | 3 | 5 | 1 | 3 | 33 | 13 |
| Tigger5b_dup2135 | 6.67988563561698 | 2.93051206172789 | 0.616967607771019 | 4.74986372836532 | 2.03553768722783e-06 | 0.000571165153921125 | 2536 | 0 | 2 | 6 | 3 | 0 | 1 | 3 | 2 | 3 | 3 | 35 | 29 |
| LTR78_dup3867 | 7.39564548962586 | 2.70751754732142 | 0.580550458488342 | 4.66370753434827 | 3.10562675496712e-06 | 0.000776230529240972 | 2537 | 0 | 0 | 5 | 7 | 0 | 0 | 8 | 6 | 5 | 2 | 26 | 38 |
| MLT1C_dup14894 | 5.61093305457504 | 2.27277564703116 | 0.669407090887369 | 3.39520700926302 | 0.000685767286056606 | 0.037339950262706 | 2538 | 0 | 0 | 2 | 10 | 0 | 0 | 9 | 4 | 2 | 2 | 21 | 25 |
| MLT1A1_dup5436 | 5.49963719184825 | 2.93545290983511 | 0.645050803669942 | 4.55073134260772 | 5.34597721413348e-06 | 0.00115212843387852 | 2539 | 0 | 0 | 3 | 6 | 0 | 1 | 4 | 2 | 4 | 2 | 20 | 31 |
| THE1D_dup10313 | 6.65207029894254 | 2.9552588365189 | 0.686016603309891 | 4.30785322433944 | 1.64846711718476e-05 | 0.00268294224805503 | 2540 | 0 | 0 | 3 | 8 | 0 | 1 | 7 | 2 | 7 | 1 | 23 | 36 |
| MIRb_dup170301 | 5.75539051231403 | 3.4531293420463 | 0.676433922929713 | 5.10490267414503 | 3.30964808159058e-07 | 0.000134388655703439 | 2541 | 0 | 0 | 2 | 6 | 0 | 0 | 5 | 1 | 5 | 4 | 23 | 30 |
| MSTB_dup6759 | 6.23901543760564 | 2.99419417883355 | 0.616985983184555 | 4.85293711759724 | 1.21646271386143e-06 | 0.000371092867836501 | 2542 | 0 | 0 | 5 | 5 | 0 | 0 | 7 | 2 | 4 | 3 | 28 | 28 |
| ERV3-16A3_I-int_dup | 5.1184891597612 | 2.87695894446876 | 0.658430892269943 | 4.36941671213275 | 1.24578841979586e-05 | 0.00220067090821637 | 2543 | 1 | 0 | 4 | 6 | 0 | 0 | 4 | 1 | 4 | 3 | 18 | 27 |
| MIRb_dup170300 | 5.83021513153477 | 2.93878205053772 | 0.727411057914929 | 4.04005688195273 | 5.34382393723337e-05 | 0.00634501761760944 | 2544 | 1 | 0 | 3 | 10 | 0 | 0 | 5 | 1 | 7 | 3 | 29 | 16 |
| L1PA16_dup2866 | 4.15242984141799 | 2.34066525867444 | 0.704910693970933 | 3.32051319222994 | 0.000898521124926495 | 0.0444885652069922 | 2545 | 0 | 0 | 5 | 7 | 0 | 0 | 5 | 3 | 5 | 4 | 12 | 12 |
| L1ME3A_dup3515 | 5.56213856302342 | 2.88245258348253 | 0.717530803232617 | 4.01718305401875 | 5.88979722272218e-05 | 0.0068349722875298 | 2546 | 0 | 0 | 4 | 11 | 0 | 0 | 5 | 2 | 7 | 4 | 14 | 26 |
| AluSx1_dup113008 | 5.59943097157504 | 2.86509839508746 | 0.72398868194541 | 3.95738008968419 | 7.57763048197596e-05 | 0.00820888624295021 | 2547 | 0 | 0 | 2 | 12 | 0 | 0 | 5 | 3 | 6 | 4 | 19 | 22 |
| L2c_dup110918 | 5.69727341196766 | 2.92866850685773 | 0.653726463418267 | 4.47996015266696 | 7.46569754658283e-06 | 0.00147918080419546 | 2548 | 0 | 0 | 2 | 8 | 0 | 0 | 5 | 4 | 5 | 4 | 21 | 25 |
| MLT1D_dup4082 | 6.26004639547645 | 2.41048494105082 | 0.617967325915091 | 3.90066730062362 | 9.59279069072136e-05 | 0.00975872669386227 | 2549 | 0 | 0 | 7 | 9 | 0 | 0 | 8 | 6 | 5 | 8 | 19 | 18 |
| L1M8_dup355 | 7.44471492028145 | 2.68936533275879 | 0.672705596162476 | 3.99783404226243 | 6.39247430944971e-05 | 0.00725698326353206 | 2550 | 0 | 0 | 6 | 15 | 0 | 0 | 4 | 8 | 5 | 8 | 17 | 34 |
| L1PB_dup323 | 6.94287427200797 | 2.26509932709061 | 0.67579043464736 | 3.35177772717747 | 0.000802944568701794 | 0.0415252006833986 | 2551 | 0 | 0 | 13 | 13 | 0 | 0 | 4 | 7 | 6 | 4 | 19 | 23 |
| L1MD3_dup4076 | 5.44233707666809 | 2.76747144375398 | 0.842979966950254 | 3.28296229122286 | 0.00102722405374009 | 0.048709471036915 | 2552 | 0 | 0 | 11 | 11 | 0 | 0 | 1 | 3 | 4 | 7 | 12 | 22 |
| MER58B_dup1495 | 5.810931074556 | 3.13554016086129 | 0.720204728984729 | 4.35367893971129 | 1.3387172645551e-05 | 0.00231667213259512 | 2553 | 0 | 0 | 5 | 10 | 0 | 0 | 3 | 3 | 8 | 10 | 14 | 21 |
| THE1C_dup2045 | 4.93095162392492 | 2.5821867270395 | 0.740405075436648 | 3.48753245041801 | 0.000487499760575769 | 0.0299893014555227 | 2554 | 0 | 0 | 6 | 8 | 0 | 0 | 3 | 6 | 5 | 8 | 11 | 15 |
| AluSx1_dup81388 | 6.57255471843141 | 3.37943062451787 | 0.698415817461602 | 4.83870860313623 | 1.30685484842177e-06 | 0.000395624534049066 | 2555 | 0 | 0 | 7 | 6 | 0 | 0 | 4 | 1 | 2 | 9 | 22 | 37 |
| MIR_dup133521 | 6.37193455578424 | 2.8312324928527 | 0.624276449528996 | 4.53522232816697 | 5.75428374076756e-06 | 0.00121773511727169 | 2556 | 0 | 0 | 7 | 8 | 0 | 0 | 4 | 3 | 2 | 6 | 26 | 28 |
| LTR78_dup3866 | 6.72091535923511 | 2.98979134926448 | 0.702196850990703 | 4.25776809600654 | 2.06477972307555e-05 | 0.00316155776053368 | 2557 | 0 | 0 | 3 | 11 | 0 | 0 | 3 | 4 | 1 | 5 | 21 | 43 |
| L2c_dup110932 | 6.15548077194233 | 3.46544725510358 | 0.671098664942693 | 5.1638416765432 | 2.4193234912272e-07 | 0.000104857622485163 | 2558 | 0 | 0 | 4 | 3 | 0 | 0 | 2 | 5 | 3 | 7 | 22 | 34 |
| L1MB1_dup5918 | 6.90768878783878 | 3.48790079326426 | 0.619834860181324 | 5.62714525647028 | 1.83216437903495e-08 | 1.31708817321405e-05 | 2559 | 0 | 0 | 4 | 6 | 0 | 0 | 4 | 2 | 3 | 7 | 26 | 40 |
| L2b_dup73100 | 5.52930900620751 | 3.54031543414421 | 0.661514635641268 | 5.3518323607644 | 8.70680417371298e-08 | 4.56332428616368e-05 | 2560 | 0 | 0 | 3 | 4 | 0 | 0 | 3 | 2 | 4 | 5 | 19 | 33 |
| MLT1B_dup17325 | 5.99652078694613 | 2.74452879070739 | 0.65415126054668 | 4.19555683254934 | 2.72202019495494e-05 | 0.00389354078069124 | 2561 | 1 | 0 | 3 | 6 | 0 | 0 | 3 | 7 | 4 | 5 | 15 | 34 |
| L2a_dup38937 | 4.93763523735604 | 2.69943181045894 | 0.660548901566362 | 4.08664945783389 | 4.37647596089664e-05 | 0.00546074960142377 | 2562 | 0 | 0 | 4 | 5 | 0 | 0 | 3 | 5 | 3 | 3 | 17 | 24 |
| MLT1A0-int_dup930 | 4.3201902130227 | 2.40607406474939 | 0.673049272736885 | 3.57488546858589 | 0.000350381243423756 | 0.0241326771274171 | 2563 | 0 | 0 | 5 | 5 | 0 | 0 | 4 | 3 | 2 | 2 | 15 | 21 |
| L1PA13_dup8700 | 4.85238494820155 | 2.56722070817831 | 0.739733686376465 | 3.47046613593288 | 0.000519555852349682 | 0.031242415173919 | 2564 | 0 | 0 | 4 | 3 | 0 | 0 | 6 | 6 | 1 | 5 | 11 | 28 |
| MLT1F2_dup4610 | 4.20493205084582 | 2.47889280658649 | 0.690065185841164 | 3.59225890169319 | 0.000327823909243334 | 0.0229897989710623 | 2565 | 0 | 0 | 3 | 4 | 0 | 0 | 6 | 4 | 2 | 4 | 13 | 19 |
| MLT1B_dup14132 | 7.8898640148932 | 3.00331303652219 | 0.611447182194917 | 4.91181106721463 | 9.02389842508815e-07 | 0.000296533043249678 | 2566 | 0 | 1 | 9 | 7 | 0 | 0 | 7 | 1 | 8 | 4 | 31 | 35 |
| MIR_dup133531 | 8.80433620901545 | 2.60240397386067 | 0.519570029038607 | 5.0087646099912 | 5.4780521926331e-07 | 0.000199920338501269 | 2567 | 0 | 1 | 9 | 12 | 0 | 0 | 8 | 3 | 4 | 6 | 33 | 41 |
| L2a_dup165791 | 8.08440031948257 | 2.78103258561177 | 0.553558943023122 | 5.02391411188096 | 5.06288856822574e-07 | 0.00018793979457957 | 2568 | 0 | 2 | 8 | 5 | 0 | 0 | 6 | 4 | 5 | 4 | 25 | 48 |
| MamRTE1_dup7519 | 8.27411810667567 | 2.9709226732468 | 0.548098918073572 | 5.42041331460539 | 5.94613999931485e-08 | 3.3134899959648e-05 | 2569 | 0 | 1 | 7 | 7 | 0 | 0 | 6 | 4 | 7 | 5 | 24 | 48 |
| MER30_dup3206 | 7.81081677251838 | 2.8659035546971 | 0.531202868279601 | 5.39512063249745 | 6.84775300214076e-08 | 3.71129473126804e-05 | 2570 | 0 | 2 | 3 | 6 | 1 | 0 | 6 | 4 | 4 | 5 | 30 | 42 |
| AluJb_dup90749 | 6.89762428322074 | 2.69391949769812 | 0.554235041149536 | 4.86060840200699 | 1.17025541213827e-06 | 0.000360229746286199 | 2571 | 1 | 1 | 5 | 6 | 1 | 0 | 5 | 2 | 2 | 4 | 29 | 36 |
| MIR_dup133520 | 7.59213038580901 | 2.16139663558858 | 0.55036750488036 | 3.92718795427144 | 8.59448177219224e-05 | 0.00902879717336007 | 2572 | 0 | 2 | 10 | 9 | 2 | 0 | 6 | 2 | 2 | 5 | 29 | 34 |
| MIRc_dup77479 | 5.91393049804205 | 2.60479384260882 | 0.665146355743707 | 3.91612134700246 | 8.99849681339303e-05 | 0.00932598346010292 | 2573 | 0 | 1 | 4 | 7 | 3 | 0 | 2 | 2 | 7 | 2 | 18 | 31 |
| L1PA16_dup11605 | 6.31051783928249 | 2.48700512540422 | 0.604714725452031 | 4.11269152333797 | 3.91072791842471e-05 | 0.00504632356441153 | 2574 | 0 | 0 | 4 | 7 | 1 | 0 | 5 | 2 | 5 | 4 | 21 | 31 |
| L2a_dup38936 | 5.47837509722663 | 2.9461168304958 | 0.683053574307306 | 4.31315630473568 | 1.60940250849101e-05 | 0.00264104061576139 | 2575 | 0 | 0 | 4 | 9 | 1 | 0 | 3 | 1 | 6 | 5 | 16 | 27 |
| L2a_dup133435 | 8.57562697115658 | 2.72460616670142 | 0.607985967732493 | 4.48136356972669 | 7.41676389000014e-06 | 0.00147681113018408 | 2576 | 0 | 0 | 11 | 8 | 2 | 0 | 5 | 3 | 8 | 2 | 31 | 42 |
| MamGypsy2-I_dup15 | 7.48527568100772 | 2.26525458282484 | 0.591378833111277 | 3.8304627355484 | 0.000127902503066438 | 0.0119528832159892 | 2577 | 0 | 0 | 9 | 13 | 1 | 0 | 4 | 5 | 4 | 2 | 30 | 30 |
| MER3_dup8137 | 9.24439421844207 | 2.82477554349925 | 0.545605940342662 | 5.17731816065855 | 2.25098160730304e-07 | 9.90040793958096e-05 | 2578 | 0 | 0 | 8 | 8 | 3 | 0 | 4 | 3 | 3 | 4 | 39 | 51 |
| AluSg_dup26193 | 8.7536879394621 | 3.24361262008947 | 0.609851125379306 | 5.318695801491 | 1.04513706083705e-07 | 5.31380733072507e-05 | 2579 | 0 | 0 | 5 | 12 | 1 | 0 | 2 | 3 | 4 | 7 | 36 | 46 |
| L1MB8_dup12601 | 6.64074703004715 | 3.01113464167251 | 0.58647572987908 | 5.13428687371828 | 2.83215826115807e-07 | 0.000117609202375134 | 2580 | 0 | 0 | 6 | 6 | 1 | 0 | 3 | 3 | 5 | 5 | 28 | 29 |
| L1PA2_dup4204 | 7.65402592475383 | 3.49289550549335 | 0.608158065701261 | 5.74340077437882 | 9.27937488773018e-09 | 7.31122561932399e-06 | 2581 | 0 | 1 | 1 | 7 | 1 | 1 | 2 | 2 | 7 | 5 | 32 | 41 |
| L2b_dup73103 | 8.10236112010659 | 3.38411280845982 | 0.586753831074381 | 5.76751719245413 | 8.04478805189315e-09 | 6.57809325703013e-06 | 2582 | 0 | 0 | 7 | 4 | 0 | 1 | 3 | 3 | 5 | 4 | 31 | 49 |
| MER2_dup7270 | 6.85258431977557 | 2.83900403474843 | 0.613905197971999 | 4.62449909876463 | 3.75504718057334e-06 | 0.000886407198837999 | 2583 | 0 | 1 | 4 | 7 | 0 | 1 | 2 | 5 | 5 | 3 | 21 | 41 |
| MLT1A0_dup20088 | 6.75945911951715 | 2.45003018017323 | 0.584764220190658 | 4.18977443485584 | 2.79231887856481e-05 | 0.00396196247989852 | 2584 | 0 | 0 | 3 | 6 | 1 | 1 | 6 | 6 | 2 | 2 | 33 | 28 |
| MIRb_dup170315 | 6.92467747865912 | 3.04681901617676 | 0.587126581068105 | 5.18937332156545 | 2.1100299115827e-07 | 9.45523874466023e-05 | 2585 | 0 | 0 | 6 | 2 | 1 | 1 | 5 | 3 | 5 | 4 | 29 | 34 |
| MER103C_dup6944 | 7.56526022377275 | 2.63094373766885 | 0.55335672733519 | 4.7545165852392 | 1.98921812215705e-06 | 0.000561775339842958 | 2586 | 0 | 0 | 9 | 6 | 1 | 1 | 5 | 3 | 2 | 5 | 32 | 36 |

| | | | | | | | | | | | | | | | | | | | |
|---|---|---|---|---|---|---|---|---|---|---|---|---|---|---|---|---|---|---|---|
| MIR_dup133517 | 7.59666541014869 | 2.92756476681294 | 0.549801144620799 | 5.32477022912001 | 1.01080729019711e-07 | 5.18356791968192e-05 | 2587 | 0 | 0 | 5 | 7 | 1 | 1 | 4 | 3 | 3 | 4 | 36 | 36 |
| MLT1-int_dup231 | 7.57190919091268 | 3.10352047418132 | 0.612284747731328 | 5.06875352633013 | 4.00429355368557e-07 | 0.000155433219237401 | 2588 | 1 | 0 | 8 | 3 | 0 | 0 | 4 | 5 | 8 | 4 | 33 | 30 |
| MER2_dup8992 | 8.12923312306599 | 2.89960502194371 | 0.556809353257322 | 5.20753648440186 | 1.91364220077138e-07 | 8.75660590586052e-05 | 2589 | 2 | 0 | 5 | 8 | 0 | 0 | 5 | 4 | 7 | 3 | 28 | 45 |
| AluY_dup76595 | 7.28778476091662 | 2.93196513399738 | 0.560116105938908 | 5.23456673162898 | 1.65372318168771e-07 | 7.8229983337945e-05 | 2590 | 1 | 0 | 6 | 6 | 0 | 0 | 8 | 2 | 6 | 6 | 25 | 36 |
| MLT-int_dup355 | 6.57232068874664 | 2.46625406214538 | 0.558315039788154 | 4.41731618600347 | 9.99339920017509e-06 | 0.00185049756115553 | 2591 | 1 | 0 | 7 | 7 | 0 | 0 | 6 | 4 | 3 | 4 | 22 | 33 |
| MIR3_dup66180 | 7.45977805418792 | 3.11360891785397 | 0.640241328871511 | 4.86318014387171 | 1.15514625608936e-06 | 0.000357428388883535 | 2592 | 0 | 0 | 4 | 11 | 0 | 0 | 4 | 5 | 4 | 9 | 35 | 24 |
| MLT1A0-int_dup927 | 8.16960555012092 | 2.91058414974354 | 0.566200118233419 | 5.14055729770023 | 2.73924772734833e-07 | 0.000114550622096947 | 2593 | 1 | 0 | 3 | 8 | 0 | 0 | 6 | 8 | 6 | 8 | 35 | 29 |
| AluSp_dup35805 | 7.84816834558075 | 3.86998822357009 | 0.683802708302676 | 5.6595099384384 | 1.51805858555696e-08 | 1.11831569101838e-05 | 2594 | 0 | 0 | 1 | 7 | 0 | 0 | 5 | 2 | 6 | 5 | 48 | 27 |
| AluSc_dup25764 | 11.0834941082489 | 3.84880456348091 | 0.586262963850147 | 6.56497988241449 | 5.2039887637925e-11 | 8.09325693064948e-08 | 2595 | 0 | 0 | 3 | 10 | 0 | 0 | 3 | 5 | 8 | 6 | 50 | 60 |
| L1PA11_dup3443 | 7.6657482146073 | 3.35133119729352 | 0.709824619232563 | 4.72135103022612 | 2.34283193886091e-06 | 0.000637523445339182 | 2596 | 0 | 0 | 2 | 13 | 0 | 0 | 2 | 4 | 5 | 4 | 32 | 39 |
| MIRc_dup23820 | 6.77348818540409 | 3.1276509554646 | 0.649777446740093 | 4.81341876538791 | 1.48369928344306e-06 | 0.00043855939092937 | 2597 | 0 | 0 | 6 | 2 | 0 | 1 | 8 | 4 | 11 | 10 | 22 | 20 |
| LTR33_dup7068 | 6.93226821981619 | 2.62302165766008 | 0.581618254129725 | 4.50986818077933 | 6.48679207605758e-06 | 0.00132946272810474 | 2598 | 0 | 0 | 9 | 7 | 0 | 1 | 7 | 3 | 8 | 5 | 24 | 25 |
| MIRb_dup170321 | 7.75479141307835 | 2.92820371546772 | 0.559627538116697 | 5.23241534060661 | 1.67309287792391e-07 | 7.86386667291629e-05 | 2599 | 0 | 0 | 6 | 5 | 0 | 2 | 6 | 4 | 9 | 4 | 32 | 31 |
| Kanga2_a_dup1175 | 10.147081723583 | 3.05232217628395 | 0.508096037294123 | 6.00737252850694 | 1.88553943680572e-09 | 1.86750702110292e-06 | 2600 | 0 | 1 | 5 | 8 | 0 | 0 | 11 | 6 | 7 | 12 | 31 | 52 |
| L1PA6_dup5016 | 10.0707468076195 | 3.77789204179895 | 0.55260033493533 | 6.83657211724468 | 8.11105355825412e-12 | 1.60832729164361e-08 | 2601 | 0 | 0 | 7 | 5 | 0 | 0 | 4 | 4 | 7 | 13 | 44 | 46 |
| MIRc_dup77477 | 7.33561426568654 | 3.6289383691344 | 0.595712005563487 | 6.09176638248505 | 1.11671571100949e-09 | 1.20780925714484e-06 | 2602 | 0 | 0 | 4 | 3 | 0 | 0 | 6 | 3 | 8 | 10 | 27 | 33 |
| L1MA8_dup10203 | 8.21731411603679 | 2.47663273056576 | 0.561871812264639 | 4.40782519518754 | 1.04413760219318e-05 | 0.00192297496820014 | 2603 | 0 | 0 | 9 | 8 | 1 | 1 | 7 | 8 | 10 | 7 | 16 | 38 |
| L1PA10_dup1403 | 8.29989063560232 | 2.139941109888 | 0.60315535924884 | 3.54791029719615 | 0.000388300442207302 | 0.0258013228208486 | 2604 | 0 | 0 | 8 | 7 | 2 | 1 | 8 | 13 | 10 | 5 | 16 | 34 |
| L1MA6_dup1085 | 7.50675079719635 | 2.64657326738586 | 0.681527657486893 | 3.88329547350287 | 0.00010305021622422 | 0.0102838378313808 | 2605 | 0 | 0 | 5 | 10 | 0 | 0 | 13 | 8 | 9 | 5 | 14 | 33 |
| L1PBa_dup500 | 8.8160960746577 | 2.98937594188094 | 0.706969872279968 | 4.22843470293896 | 2.35322790398549e-05 | 0.00348655770138064 | 2606 | 0 | 0 | 6 | 15 | 0 | 0 | 13 | 8 | 11 | 11 | 18 | 31 |
| L1P1_dup2815 | 7.23615593607627 | 2.18625242752429 | 0.613851533533478 | 3.56153289206544 | 0.000368695907714828 | 0.0249373838706971 | 2607 | 1 | 0 | 7 | 18 | 1 | 1 | 4 | 3 | 8 | 5 | 23 | 23 |
| L1MA6_dup1086 | 7.63956516448611 | 1.97153747562514 | 0.55996529030146 | 3.5208207718799 | 0.000430213295290013 | 0.0276072094824912 | 2608 | 2 | 0 | 12 | 12 | 0 | 1 | 8 | 6 | 10 | 6 | 19 | 21 |
| L1ME4a_dup26456 | 8.06538258241429 | 2.3956905164681 | 0.50458887411526 | 4.74780685695593 | 2.05634263393825e-06 | 0.000574294019219568 | 2609 | 2 | 0 | 6 | 10 | 0 | 3 | 6 | 4 | 9 | 7 | 31 | 25 |
| L3_dup36520 | 7.21396250420683 | 1.99602101753152 | 0.492794165158543 | 4.05041528218857 | 5.11268173250501e-05 | 0.00614780985184786 | 2610 | 1 | 1 | 6 | 8 | 0 | 2 | 8 | 6 | 3 | 5 | 27 | 27 |
| L1MDa_dup6638 | 9.85881040063329 | 2.67869768530095 | 0.480373694605696 | 5.57627887492818 | 2.45717905096738e-08 | 1.64234810803788e-05 | 2611 | 1 | 0 | 8 | 12 | 0 | 1 | 8 | 4 | 6 | 6 | 33 | 52 |
| THE1D_dup10315 | 9.63156477864619 | 2.88166288253666 | 0.499794440679335 | 5.76569615024093 | 8.13214579451757e-09 | 6.62675192884341e-06 | 2612 | 0 | 0 | 6 | 9 | 0 | 1 | 9 | 5 | 7 | 4 | 40 | 45 |
| AluSx1_dup81389 | 8.8293680349261 | 2.43812253970931 | 0.506692933887959 | 4.81183449905467 | 1.49551174916869e-06 | 0.000440955438249929 | 2613 | 0 | 0 | 9 | 13 | 0 | 1 | 9 | 6 | 6 | 9 | 28 | 34 |
| L1PA7_dup2715 | 7.68083509458806 | 2.14655513044729 | 0.568452313329072 | 3.77613931743237 | 0.000159277925084654 | 0.0139593168184873 | 2614 | 0 | 0 | 12 | 10 | 0 | 1 | 5 | 8 | 4 | 5 | 20 | 35 |
| L1MA8_dup10204 | 9.52875168198304 | 2.08919043891245 | 0.564458749214199 | 3.70122784316989 | 0.000214558705997583 | 0.0172181197299823 | 2615 | 0 | 0 | 4 | 7 | 6 | 3 | 7 | 9 | 7 | 4 | 36 | 39 |
| MADE1_dup6316 | 7.5333569895614 | 3.08031796784399 | 0.598336030932938 | 5.14814052404816 | 2.63081378063153e-07 | 0.000111784217115384 | 2616 | 0 | 0 | 4 | 5 | 2 | 2 | 1 | 4 | 8 | 4 | 33 | 33 |
| MIR_dup133522 | 8.34901573665292 | 2.49538946832345 | 0.542638429133657 | 4.59862282939939 | 4.25292942230517e-06 | 0.000972110993582927 | 2617 | 0 | 0 | 7 | 8 | 2 | 1 | 3 | 8 | 6 | 4 | 32 | 36 |
| L1MA6_dup1087 | 7.58554736279654 | 2.05094379568134 | 0.500335795221699 | 4.09913465170439 | 4.14697730265114e-05 | 0.00525709462576787 | 2618 | 0 | 3 | 8 | 7 | 3 | 3 | 9 | 1 | 8 | 12 | 18 | 26 |
| AluSx1_dup22521 | 8.05169560647736 | 2.42526364604878 | 0.523050131747762 | 4.63677093043607 | 3.53894233491249e-06 | 0.000848953661159664 | 2619 | 0 | 1 | 7 | 9 | 1 | 1 | 11 | 4 | 10 | 10 | 17 | 33 |
| L2b_dup1841 | 6.88756234564659 | 2.15546843495909 | 0.500453014055948 | 4.30703457551385 | 1.65457755273543e-05 | 0.00269104655067113 | 2620 | 0 | 1 | 4 | 6 | 2 | 3 | 7 | 5 | 4 | 8 | 23 | 26 |
| AluSx1_dup81390 | 10.4301253278615 | 2.42363645982902 | 0.45009553186511 | 5.3847156619972 | 7.25592585083065e-08 | 3.86245756711801e-05 | 2621 | 0 | 2 | 10 | 5 | 2 | 2 | 9 | 7 | 5 | 10 | 34 | 50 |
| L1P1_dup672 | 10.1375905901275 | 2.42321099016767 | 0.476287390103288 | 5.08770763307877 | 3.62417505724226e-07 | 0.000144899422077838 | 2622 | 0 | 3 | 7 | 11 | 1 | 1 | 10 | 6 | 6 | 12 | 23 | 54 |
| AluSx_dup82052 | 7.92186435336834 | 2.65596622529039 | 0.535372616280948 | 4.96096764108049 | 7.01428773688099e-07 | 0.000243652804356187 | 2623 | 0 | 1 | 6 | 5 | 2 | 0 | 11 | 3 | 5 | 11 | 29 | 30 |
| L2c_dup110936 | 9.93273156429786 | 2.72919525203998 | 0.504112462513848 | 5.41386189802391 | 6.16797446602526e-08 | 3.40783262835684e-05 | 2624 | 0 | 1 | 11 | 6 | 3 | 0 | 6 | 4 | 4 | 10 | 39 | 46 |
| HERVKC4-int_dup34 | 11.4433884429209 | 3.23684531669386 | 0.947109305404143 | 3.4176048088902 | 0.00063174764695525 | 0.0354867388107682 | 2625 | 0 | 0 | 18 | 10 | 0 | 1 | 0 | 0 | 2 | 4 | 41 | 80 |
| AluSz_dup73300 | 8.52306921330146 | 3.50801803411012 | 0.664069723558482 | 5.28260498809081 | 1.27359804430336e-07 | 6.23554650719769e-05 | 2626 | 0 | 0 | 6 | 9 | 1 | 1 | 1 | 1 | 4 | 13 | 41 | 34 |
| MER51B_dup124 | 8.60695315920767 | 2.5200318613343 | 0.763503530581973 | 3.30061585813685 | 0.000964728720776952 | 0.0466769083293685 | 2627 | 0 | 0 | 13 | 18 | 0 | 3 | 2 | 2 | 3 | 13 | 27 | 33 |
| AluSx_dup82055 | 9.78334423172859 | 2.63389491183688 | 0.617684516121717 | 4.26414268626068 | 2.00671185000632e-05 | 0.00309254571153888 | 2628 | 1 | 1 | 10 | 11 | 1 | 2 | 0 | 3 | 4 | 5 | 43 | 48 |
| AluSx1_dup81391 | 10.6677041068802 | 3.35024702568725 | 0.515797039136638 | 6.49528161560413 | 8.28782302306332e-11 | 1.20247215673526e-07 | 2629 | 0 | 1 | 7 | 6 | 1 | 1 | 5 | 2 | 6 | 6 | 36 | 72 |
| LTR56_dup554 | 9.54196994249717 | 2.98658765176024 | 0.68841026581534 | 4.33838206672086 | 1.43535471656889e-05 | 0.0024482932859405 | 2630 | 0 | 1 | 13 | 6 | 1 | 1 | 1 | 1 | 2 | 6 | 43 | 52 |
| FRAM_dup5323 | 8.1124569703023 | 3.77195656445027 | 0.828826333567407 | 4.5509612951306 | 5.34013718534641e-06 | 0.00115200750925153 | 2631 | 0 | 0 | 5 | 10 | 0 | 0 | 2 | 2 | 0 | 12 | 38 | 39 |
| MIR_dup133525 | 8.9047626266988 | 3.27605035179043 | 0.867640676967284 | 3.77581461860617 | 0.00015948557654798 | 0.0139723692920786 | 2632 | 0 | 0 | 10 | 15 | 0 | 0 | 7 | 1 | 0 | 11 | 48 | 26 |
| L1MEg_dup8069 | 4.71459212385237 | 2.85518957523486 | 0.780732807448601 | 3.65706365608676 | 0.000255121040454993 | 0.0193888795471302 | 2633 | 0 | 0 | 4 | 4 | 1 | 0 | 1 | 1 | 0 | 1 | 21 | 31 |
| L1MC2_dup4896 | 6.52153758103253 | 2.74254560147924 | 0.730240448843585 | 3.75567473127839 | 0.000172875098945478 | 0.0148063282311054 | 2634 | 0 | 0 | 5 | 11 | 1 | 0 | 4 | 1 | 0 | 5 | 25 | 37 |
| L3_dup36524 | 6.4097757261143 | 2.93936343342719 | 0.627229704126037 | 4.68626312512225 | 2.78238506868203e-06 | 0.000716512335013653 | 2635 | 0 | 0 | 6 | 6 | 1 | 0 | 3 | 2 | 1 | 4 | 26 | 37 |
| MLT1A_dup7133 | 4.54570310349778 | 3.0773330442574 | 0.84775625418444 | 3.62996799083754 | 0.000283456368682874 | 0.0208042285942675 | 2636 | 0 | 1 | 2 | 7 | 0 | 0 | 1 | 1 | 0 | 5 | 21 | 23 |
| MamGypLTR3a_dup9 | 4.64154939608569 | 2.88185982832885 | 0.833480098873632 | 3.45762284213313 | 0.000544963728352728 | 0.0322898394372453 | 2637 | 0 | 1 | 2 | 4 | 0 | 3 | 1 | 1 | 0 | 8 | 17 | 25 |
| L1PA13_dup9196 | 7.27608716624045 | 3.96592097562854 | 0.868638977346474 | 4.56567236683722 | 4.97895887691454e-06 | 0.00109359315807852 | 2638 | 0 | 1 | 6 | 6 | 0 | 0 | 0 | 0 | 10 | 3 | 36 | 31 |
| LTR78_dup3865 | 7.0682297950004 | 2.91768822733986 | 0.69333898525119 | 4.20816986986937 | 2.57447305906248e-05 | 0.00374212319188565 | 2639 | 0 | 1 | 7 | 9 | 0 | 0 | 2 | 1 | 3 | 1 | 35 | 35 |
| L2a_dup165790 | 7.13492661365219 | 3.63713472632385 | 0.717135955329582 | 5.07175062035803 | 3.94172713672531e-07 | 0.000154161275032637 | 2640 | 0 | 0 | 6 | 8 | 0 | 0 | 1 | 2 | 8 | 5 | 35 | 26 |
| MER5A_dup26804 | 5.83537309149672 | 2.97536718357445 | 0.713641968190727 | 4.16927159023144 | 3.05574682189454e-05 | 0.00423472762540779 | 2641 | 0 | 0 | 8 | 7 | 0 | 0 | 2 | 2 | 5 | 2 | 24 | 26 |
| MIR_dup133529 | 4.8415605175486 | 3.22739062847175 | 0.717474552155072 | 4.49826494720637 | 6.85102858455782e-06 | 0.00138738284900527 | 2642 | 0 | 0 | 5 | 5 | 0 | 0 | 2 | 1 | 3 | 4 | 24 | 19 |
| MIRb_dup149762 | 6.13443243772941 | 2.5218483016672 | 0.72728748532933 | 3.46747105173308 | 0.000525380327209484 | 0.0314651412518991 | 2643 | 1 | 2 | 6 | 12 | 0 | 0 | 2 | 0 | 4 | 5 | 26 | 23 |
| Kanga2_a_dup1177 | 4.91692639911376 | 2.31479359765281 | 0.626328957517349 | 3.69581123444814 | 0.000219185785810977 | 0.0174663030778897 | 2644 | 1 | 3 | 5 | 4 | 0 | 0 | 5 | 1 | 3 | 6 | 19 | 17 |
| Tigger13a_dup2856 | 5.01806455379895 | 2.59606343477731 | 0.661047246971316 | 3.9271980129583 | 8.59412247554126e-05 | 0.00902879717336007 | 2645 | 1 | 1 | 3 | 6 | 0 | 0 | 6 | 1 | 4 | 4 | 25 | 14 |
| L2b_dup73107 | 7.1606330547685 | 3.54293608393514 | 0.680741903820581 | 5.2045218078259 | 1.94496784293168e-07 | 8.84889710046311e-05 | 2646 | 0 | 0 | 4 | 6 | 0 | 0 | 7 | 0 | 3 | 6 | 37 | 32 |
| L1MC4a_dup7789 | 5.70717252368939 | 3.09908317859375 | 0.686297886278196 | 4.51565310130871 | 6.31219227063748e-06 | 0.00130605295828618 | 2647 | 1 | 0 | 4 | 5 | 0 | 1 | 4 | 0 | 6 | 2 | 24 | 28 |
| L1PA3_dup9152 | 4.5472752597129 | 3.13664470316549 | 0.751320521412433 | 4.17484231266411 | 2.98192528853322e-05 | 0.00416395067315333 | 2648 | 1 | 0 | 4 | 4 | 0 | 1 | 1 | 0 | 2 | 4 | 17 | 27 |
| L2a_dup155965 | 5.95134603911111 | 3.52426997844987 | 0.706479709424596 | 4.98849424476949 | 6.08517178133113e-07 | 0.00021746742009862 | 2649 | 1 | 0 | 4 | 4 | 0 | 1 | 1 | 0 | 3 | 3 | 24 | 39 |
| THE1D_dup12312 | 4.42675839017028 | 3.33735079718176 | 0.906595620489703 | 3.68119007168717 | 0.000232147859054575 | 0.0181586635340565 | 2650 | 0 | 0 | 2 | 3 | 0 | 0 | 6 | 2 | 8 | 0 | 18 | 17 |
| L1PA8_dup4527 | 5.16465035815322 | 3.00906382789776 | 0.885843248785902 | 3.39683553723737 | 0.000681699245817874 | 0.0372016212954467 | 2651 | 0 | 0 | 7 | 5 | 0 | 0 | 1 | 1 | 3 | 0 | 15 | 38 |
| MIRb_dup170320 | 6.80622335274706 | 3.1011207669567 | 0.70556134209281 | 4.39525322880966 | 1.10643694247674e-05 | 0.00201586711111376 | 2652 | 0 | 0 | 5 | 4 | 0 | 2 | 3 | 2 | 5 | 0 | 34 | 34 |
| L1PA10_dup6573 | 5.13418399349685 | 3.36212739974421 | 0.983947677683381 | 3.41697782920739 | 0.000633204360612532 | 0.0355350105637522 | 2653 | 0 | 0 | 2 | 2 | 3 | 0 | 0 | 1 | 3 | 0 | 24 | 33 |
| LTR88a_dup489 | 3.55072862494715 | 3.11906930097277 | 0.898527423800407 | 3.47131230316863 | 0.000517921265619482 | 0.0311835256753778 | 2654 | 0 | 0 | 3 | 1 | 1 | 1 | 1 | 0 | 2 | 0 | 18 | 20 |
| L1PA15_dup4374 | 3.82536034473367 | 3.71598921319558 | 1.07607198517343 | 3.45329054598208 | 0.000553792180372325 | 0.0326187018391823 | 2655 | 0 | 0 | 1 | 3 | 0 | 0 | 0 | 0 | 1 | 0 | 14 | 34 |
| L1MB3_dup8398 | 4.86292538050753 | 2.93401865291387 | 0.8069984433464 | 3.63571783678878 | 0.000277207627867666 | 0.0204846106275154 | 2656 | 0 | 1 | 3 | 5 | 0 | 0 | 2 | 0 | 1 | 0 | 24 | 30 |
| L1MB4_dup4305 | 5.52694249006911 | 3.38633999093304 | 0.750023049022465 | 4.51498123337223 | 6.3322373063616e-06 | 0.00130906215299698 | 2657 | 0 | 0 | 2 | 5 | 0 | 1 | 2 | 0 | 1 | 1 | 28 | 35 |
| AluSx1_dup81392 | 4.1175210776351 | 2.82190192529648 | 0.780725576857475 | 3.61446071314202 | 0.000300973448649527 | 0.0217082231622796 | 2658 | 0 | 0 | 5 | 5 | 1 | 0 | 2 | 0 | 3 | 2 | 14 | 23 |

| | | | | | | | | | | | | | | | | | | | |
|---|---|---|---|---|---|---|---|---|---|---|---|---|---|---|---|---|---|---|---|
| Tigger19a_dup2728 | 3.46478270155727 | 3.33365699197446 | 0.861524465449393 | 3.86948615584067 | 0.000109064973863121 | 0.0107016801116842 | 2659 | 0 | 0 | 4 | 1 | 1 | 0 | 1 | 0 | 2 | 2 | 14 | 21 |
| L1M1_dup8973 | 5.04293100085171 | 2.4488179327631 | 0.715705926761617 | 3.42154206245485 | 0.000622670925920573 | 0.0351511402465235 | 2660 | 0 | 0 | 7 | 5 | 1 | 1 | 4 | 0 | 1 | 3 | 20 | 26 |
| MIR_dup133514 | 4.46415669352424 | 2.65239172448828 | 0.786221011244371 | 3.37359557497742 | 0.000741932850908825 | 0.039331664037174 | 2661 | 0 | 0 | 4 | 2 | 2 | 1 | 4 | 0 | 2 | 2 | 11 | 33 |
| LTR16E2_dup1587 | 4.13187961099927 | 2.45870726949243 | 0.716145119315367 | 3.43325284663385 | 0.000596385652852224 | 0.034268915854522 | 2662 | 0 | 0 | 5 | 4 | 1 | 2 | 4 | 0 | 7 | 5 | 14 | 10 |
| AluSx_dup82054 | 4.26296730967369 | 2.93463736372633 | 0.780233559766217 | 3.76122934856281 | 0.000169080307487776 | 0.0145557101467027 | 2663 | 0 | 0 | 5 | 6 | 0 | 1 | 2 | 0 | 5 | 4 | 12 | 21 |
| MER33_dup7091 | 4.85406657088522 | 3.07887795135547 | 0.679818398644639 | 4.52897120391837 | 5.92715753215021e-06 | 0.0012436890883113 | 2664 | 0 | 0 | 4 | 5 | 0 | 1 | 3 | 1 | 5 | 6 | 22 | 15 |
| L1MC5a_dup13064 | 4.72519364740837 | 3.32423569971854 | 0.855626644932116 | 3.88514747572206 | 0.000102267789463798 | 0.0102398985115826 | 2665 | 0 | 0 | 1 | 5 | 0 | 0 | 7 | 0 | 4 | 2 | 26 | 17 |
| MER5A_dup26802 | 3.02743569382584 | 2.93415753321835 | 0.881606044855826 | 3.32819579713543 | 0.000874104093161516 | 0.0437136554332514 | 2666 | 0 | 0 | 1 | 3 | 1 | 0 | 4 | 0 | 3 | 2 | 17 | 8 |
| MIR_dup133516 | 5.13857036231768 | 2.93095158621791 | 0.667661475310103 | 4.38987674832758 | 1.13414924730479e-05 | 0.0020600479144976 | 2667 | 1 | 0 | 3 | 7 | 1 | 0 | 4 | 0 | 6 | 6 | 18 | 21 |
| MLT1A1_dup5437 | 4.11153339568941 | 2.92013120581239 | 0.791993835139624 | 3.68706305055718 | 0.000226857163537671 | 0.0178977966296865 | 2668 | 1 | 0 | 4 | 2 | 0 | 0 | 7 | 0 | 5 | 5 | 16 | 13 |
| AluY_dup76586 | 7.01104474167045 | 2.71615901411821 | 0.574758588759845 | 4.72573888800662 | 2.29280113251807e-06 | 0.000626662995218877 | 2669 | 1 | 0 | 5 | 4 | 1 | 2 | 7 | 1 | 3 | 5 | 36 | 27 |
| L3_dup36521 | 5.38628402517941 | 2.09942278463009 | 0.581244370949322 | 3.61194514658471 | 0.000303908859563456 | 0.0218405006033483 | 2670 | 1 | 0 | 6 | 4 | 1 | 3 | 5 | 2 | 2 | 5 | 19 | 23 |
| LTR45C_dup202 | 7.35012327955573 | 2.6421008786894 | 0.63071631484823 | 4.18904793881093 | 2.80127224133384e-05 | 0.00397229752524685 | 2671 | 1 | 0 | 8 | 9 | 1 | 1 | 2 | 1 | 3 | 3 | 37 | 31 |
| MIR_dup133518 | 5.87676543612853 | 2.6873715127815 | 0.694054522320351 | 3.87198905324776 | 0.000107950801562354 | 0.0106144386082852 | 2672 | 1 | 0 | 6 | 5 | 1 | 3 | 1 | 0 | 3 | 5 | 28 | 24 |
| L1MC_dup1698 | 6.15165909970295 | 2.86926925494189 | 0.696986490496489 | 4.11667843504112 | 3.84371768493964e-05 | 0.00498146649379437 | 2673 | 0 | 0 | 4 | 8 | 1 | 1 | 6 | 2 | 2 | 17 | 15 | 25 |
| MIRb_dup170293 | 4.2448131458222 | 2.62405090467999 | 0.703133552461132 | 3.73193811544791 | 0.000190012187323427 | 0.0158640841841614 | 2674 | 0 | 0 | 2 | 4 | 1 | 1 | 5 | 1 | 1 | 4 | 16 | 22 |
| MIR_dup133532 | 4.99089891711361 | 2.47659472939778 | 0.634888688282538 | 3.90083297293784 | 9.58622757884729e-05 | 0.00975622116114797 | 2675 | 0 | 0 | 5 | 6 | 1 | 1 | 2 | 3 | 4 | 3 | 16 | 24 |
| L2_dup55460 | 4.74469394819255 | 2.51249096763868 | 0.669662211720311 | 3.75187807175842 | 0.000175514823982065 | 0.0149849480829697 | 2676 | 0 | 0 | 3 | 8 | 0 | 1 | 5 | 2 | 4 | 4 | 12 | 24 |
| THE1D_dup9105 | 4.41201546590215 | 2.69446951748808 | 0.708708852554555 | 3.80194138647459 | 0.000143566695970732 | 0.0129251309267695 | 2677 | 0 | 0 | 4 | 6 | 0 | 1 | 2 | 2 | 2 | 4 | 12 | 26 |
| L1M6_dup5341 | 6.07716196706494 | 3.40462967527129 | 0.846832129890544 | 4.02043044317573 | 5.80918999935551e-05 | 0.00676291337965716 | 2678 | 3 | 0 | 2 | 4 | 0 | 2 | 0 | 0 | 6 | 2 | 21 | 41 |
| L1MC5a_dup6752 | 6.75182209791214 | 3.74994747872248 | 0.858650575531315 | 4.36725669973737 | 1.25816825306797e-05 | 0.00221760076403341 | 2679 | 2 | 0 | 3 | 1 | 0 | 3 | 0 | 0 | 3 | 4 | 31 | 43 |
| L3_dup42913 | 6.3086606181158 | 3.45572077319565 | 0.733604027759534 | 4.71060768811427 | 2.46979255347717e-06 | 0.0006617987150109 | 2680 | 0 | 1 | 3 | 4 | 1 | 2 | 0 | 0 | 4 | 3 | 26 | 40 |
| MIRb_dup149751 | 5.89548862592442 | 2.38186336283518 | 0.648087050604732 | 3.67522134659635 | 0.000237643291359662 | 0.0184973080163121 | 2681 | 1 | 3 | 5 | 6 | 1 | 3 | 1 | 0 | 4 | 5 | 19 | 30 |
| MIRb_dup149763 | 6.3512156955908 | 2.2529598690218 | 0.639260490973796 | 3.52432208909051 | 0.000424567820760681 | 0.0273444488994885 | 2682 | 1 | 2 | 7 | 7 | 3 | 2 | 1 | 0 | 4 | 5 | 21 | 31 |
| CER_dup50 | 7.57284574743106 | 3.44404632309564 | 0.607934838284567 | 5.6651570303387 | 1.46889952745697e-08 | 1.08884351077968e-05 | 2683 | 0 | 0 | 3 | 4 | 2 | 4 | 2 | 0 | 9 | 10 | 31 | 32 |
| L1PA4_dup11271 | 8.73095407530126 | 3.53837926311511 | 0.614898387901789 | 5.75441297738491 | 8.69433776074237e-09 | 6.94926420988751e-06 | 2684 | 1 | 1 | 2 | 6 | 3 | 2 | 1 | 0 | 7 | 13 | 23 | 57 |
| MER52D_dup585 | 7.05661075498692 | 3.43490501979015 | 0.618938557745135 | 5.54967044273976 | 2.86208599758549e-08 | 1.86581346515473e-05 | 2685 | 1 | 1 | 5 | 2 | 1 | 3 | 1 | 0 | 10 | 9 | 19 | 39 |
| MER31A_dup1227 | 5.67384086338447 | 2.79961472024989 | 0.762501089939311 | 3.67162061430327 | 0.000241017284886416 | 0.018650113453523 | 2686 | 1 | 3 | 3 | 3 | 3 | 0 | 1 | 0 | 3 | 3 | 19 | 37 |
| MIRb_dup203248 | 5.45129385798591 | 3.25831496557498 | 0.822493492337558 | 3.961508505453 | 7.44777285087585e-05 | 0.00809948701450871 | 2687 | 2 | 1 | 2 | 2 | 4 | 0 | 0 | 0 | 4 | 4 | 20 | 33 |
| MIRb_dup149764 | 5.37952206199737 | 2.16360368009125 | 0.635159610789433 | 3.40639367387062 | 0.000658271852366268 | 0.0364093803308099 | 2688 | 3 | 3 | 4 | 8 | 2 | 0 | 2 | 0 | 4 | 8 | 14 | 23 |
| THE1D_dup11828 | 4.83703684154093 | 3.0454245412299 | 0.659725466733365 | 4.61619976004465 | 3.90830766950304e-06 | 0.000916222834214355 | 2689 | 1 | 2 | 2 | 4 | 1 | 0 | 1 | 1 | 3 | 7 | 19 | 22 |
| MIR3_dup79382 | 8.92965952119588 | 3.23647737123181 | 0.778611105060077 | 4.156731582941 | 3.22832910415197e-05 | 0.00440716005173003 | 2690 | 0 | 1 | 7 | 1 | 5 | 2 | 1 | 0 | 7 | 5 | 42 | 45 |
| MER58D_dup443 | 6.0394258583478 | 3.73587271474118 | 0.92530278180756 | 4.03745972474353 | 5.40331175600063e-05 | 0.00639033932008616 | 2691 | 0 | 3 | 1 | 0 | 3 | 1 | 1 | 0 | 2 | 4 | 29 | 36 |
| UCON1_dup219 | 5.21965467173419 | 2.88827985026368 | 0.766768211099025 | 3.76682263095369 | 0.000165338377305285 | 0.0143119933168183 | 2692 | 0 | 1 | 2 | 2 | 5 | 1 | 2 | 0 | 3 | 5 | 24 | 23 |
| L1MA4A_dup3282 | 21.1822235811987 | 2.74751016362769 | 0.749568092329225 | 3.66545773725508 | 0.000246896583794667 | 0.018966350703973 | 2693 | 1 | 0 | 16 | 28 | 3 | 1 | 3 | 0 | 3 | 3 | 90 | 142 |
| L1PA7_dup7467 | 15.0202951356662 | 2.55773578082005 | 0.618004162786664 | 4.13870315903194 | 3.49274506227846e-05 | 0.0046768976735448 | 2694 | 0 | 1 | 18 | 19 | 3 | 0 | 5 | 2 | 5 | 2 | 68 | 78 |
| L1MB4_dup4075 | 19.1290203250204 | 2.63905762314527 | 0.594997371082465 | 4.43541056046028 | 9.18969799963664e-06 | 0.00173819704310138 | 2695 | 0 | 0 | 18 | 21 | 3 | 1 | 4 | 5 | 2 | 2 | 93 | 108 |
| L1PA10_dup3708 | 16.5795150311953 | 2.51714946548595 | 0.577541508992318 | 4.35838710515859 | 1.31024480756624e-05 | 0.00227619642572668 | 2696 | 1 | 1 | 15 | 22 | 3 | 1 | 2 | 5 | 5 | 3 | 65 | 100 |
| L1M5_dup38550 | 10.8396411959007 | 2.43734885606353 | 0.576183898121576 | 4.23015787843007 | 2.3352735715191e-05 | 0.00346859553838129 | 2697 | 1 | 0 | 9 | 5 | 0 | 1 | 8 | 10 | 2 | 0 | 36 | 73 |
| L1PA13_dup8699 | 7.59240812338471 | 2.23804354359383 | 0.640629347385477 | 3.4935076776106 | 0.000476719275891312 | 0.0295243270435662 | 2698 | 0 | 1 | 5 | 5 | 1 | 0 | 5 | 10 | 3 | 0 | 31 | 37 |
| L1MC4_dup17258 | 11.3941379430131 | 2.60001221743653 | 0.603738707392509 | 4.3065190049943 | 1.65843686871199e-05 | 0.00269364108643374 | 2699 | 2 | 1 | 6 | 7 | 0 | 0 | 4 | 8 | 0 | 0 | 56 | 68 |
| L1MEd_dup15777 | 12.2784542478632 | 2.25994647061086 | 0.52420826266477 | 4.31116148212278 | 1.62399248859806e-05 | 0.00265544184995128 | 2700 | 2 | 0 | 5 | 10 | 1 | 0 | 15 | 9 | 0 | 1 | 54 | 68 |
| L1PA3_dup6147 | 10.0182210088425 | 3.19989120030822 | 0.614388878592729 | 5.20825052634032 | 1.90629430567631e-07 | 8.74544048961656e-05 | 2701 | 0 | 0 | 4 | 4 | 0 | 1 | 4 | 5 | 0 | 0 | 49 | 68 |
| L1PB1_dup7168 | 8.7818546065612 | 2.30935423116844 | 0.699674694064913 | 3.30061134232501 | 0.000964744249383618 | 0.0466769083293685 | 2702 | 0 | 0 | 10 | 12 | 0 | 1 | 3 | 2 | 0 | 0 | 42 | 49 |
| L1MEd_dup15775 | 12.2069574614109 | 2.56930509509507 | 0.593315083887838 | 4.33042267905796 | 1.48823397482721e-05 | 0.00250090202376348 | 2703 | 0 | 0 | 11 | 4 | 0 | 0 | 14 | 6 | 0 | 0 | 59 | 70 |
| MIRb_dup170307 | 7.84928364287441 | 2.50615037939376 | 0.58082452704038 | 4.31481499612968 | 1.59736618530931e-05 | 0.00262854007143575 | 2704 | 0 | 0 | 9 | 7 | 0 | 0 | 8 | 3 | 0 | 2 | 34 | 43 |
| AluY_dup92078 | 4.40826548921708 | 2.40389273790348 | 0.663748759002651 | 3.62169074563027 | 0.000292683881138931 | 0.0212261380010619 | 2705 | 1 | 3 | 2 | 2 | 0 | 4 | 4 | 0 | 8 | 6 | 14 | 11 |
| AluJr_dup21555 | 4.74739276198072 | 2.63840502336072 | 0.631399889401315 | 4.17865930553523 | 2.93232566017209e-05 | 0.0041140044901846 | 2706 | 1 | 4 | 1 | 2 | 0 | 2 | 4 | 1 | 6 | 8 | 16 | 15 |
| L1ME4a_dup23653 | 5.62315525943135 | 2.84500871354986 | 0.642768586432576 | 4.42617883574541 | 9.59170135253505e-06 | 0.0017985082506149 | 2707 | 2 | 3 | 2 | 1 | 0 | 3 | 3 | 1 | 6 | 5 | 18 | 29 |
| MLT1L_dup10858 | 4.75546508826083 | 2.65490475124979 | 0.692888666638332 | 3.83164695726734 | 0.000127288306407708 | 0.0119290048666752 | 2708 | 2 | 1 | 4 | 3 | 0 | 1 | 1 | 2 | 4 | 3 | 27 | 12 |
| MER5A1_dup4768 | 5.23959339047063 | 2.42816789268209 | 0.667176748155514 | 3.63946720174981 | 0.000273202730697108 | 0.0203084963943936 | 2709 | 1 | 2 | 4 | 1 | 0 | 1 | 2 | 7 | 7 | 5 | 17 | 17 |
| AluSz_dup61166 | 3.6848900464206 | 2.47382112237717 | 0.737377562952192 | 3.35489069191756 | 0.000793963503746438 | 0.0412490043367794 | 2710 | 1 | 2 | 1 | 1 | 0 | 2 | 1 | 4 | 7 | 4 | 13 | 8 |
| Charlie4a_dup2149 | 4.31191521334874 | 2.92535052811828 | 0.690749599456951 | 4.23503760323294 | 2.28513463401069e-05 | 0.00342189204294719 | 2711 | 1 | 3 | 1 | 1 | 0 | 1 | 1 | 3 | 5 | 6 | 15 | 17 |
| LTR82B_dup259 | 3.59094990405248 | 2.53363609786215 | 0.689614096763882 | 3.67399116368361 | 0.000238790993744266 | 0.0185623527597103 | 2712 | 2 | 3 | 2 | 2 | 1 | 1 | 0 | 1 | 6 | 6 | 7 | 14 |
| MLT1A0_dup13462 | 3.00720704482261 | 2.75562505982491 | 0.752736100768533 | 3.66081161380656 | 0.00025141757143909 | 0.019211241314594 | 2713 | 2 | 2 | 1 | 1 | 1 | 1 | 0 | 1 | 8 | 5 | 6 | 8 |
| AluSg_dup2167 | 3.87586613898683 | 2.66506237300445 | 0.701295164072524 | 3.80020069941455 | 0.000144578945372345 | 0.0129894867914035 | 2714 | 3 | 1 | 1 | 2 | 1 | 2 | 1 | 1 | 6 | 4 | 7 | 21 |
| L1MDa_dup4965 | 3.74248216010883 | 3.29106065744284 | 0.737193452532709 | 4.46431075335251 | 8.03269105585347e-06 | 0.00155616736102716 | 2715 | 1 | 1 | 1 | 2 | 1 | 1 | 0 | 1 | 4 | 6 | 12 | 18 |
| BSR/Beta_dup1535 | 3.46815837168202 | 2.1752090429575 | 0.662629547590203 | 3.28269249517792 | 0.00102820758792305 | 0.0487269234646362 | 2716 | 3 | 1 | 1 | 2 | 3 | 2 | 1 | 1 | 4 | 5 | 10 | 11 |
| Tigger15a_dup1216 | 4.23735125195572 | 2.14813115792424 | 0.628433839601172 | 3.41822960916225 | 0.00063029909872962 | 0.0354555908620137 | 2717 | 2 | 2 | 1 | 2 | 4 | 1 | 4 | 1 | 4 | 6 | 15 | 12 |
| AluSz_dup61165 | 3.89227091783876 | 2.24877445276801 | 0.632359346695188 | 3.55616543745335 | 0.000376307162429203 | 0.0252797244696158 | 2718 | 1 | 2 | 2 | 2 | 2 | 0 | 4 | 2 | 5 | 5 | 12 | 12 |
| L1MA9_dup4656 | 4.19801723252119 | 2.33878386461533 | 0.67150134738455 | 3.48291760504238 | 0.000495980990124896 | 0.0303151021516205 | 2719 | 4 | 2 | 1 | 2 | 1 | 0 | 2 | 3 | 5 | 6 | 17 | 9 |
| FLAM_A_dup13972 | 3.05992112780931 | 2.68777955448138 | 0.814537585390068 | 3.2997612420724 | 0.000967671162870557 | 0.0467519761462736 | 2720 | 0 | 0 | 2 | 2 | 0 | 0 | 5 | 2 | 3 | 2 | 14 | 9 |
| L1MEc_dup17904 | 3.15071025511421 | 2.70886607525204 | 0.788370538923004 | 3.43603158858858 | 0.000590302141164661 | 0.0340509171591676 | 2721 | 0 | 0 | 3 | 1 | 0 | 1 | 3 | 2 | 3 | 2 | 14 | 11 |
| L4_A_Mam_dup1954 | 3.32714560450139 | 3.4345360433008 | 0.814440389689414 | 4.21705024306389 | 2.47518840919804e-05 | 0.00363108003215189 | 2722 | 0 | 0 | 2 | 1 | 0 | 0 | 3 | 2 | 6 | 6 | 10 | 11 |
| MIR_dup15452 | 2.32469769247943 | 3.24276058330302 | 0.970248498705083 | 3.34219592983744 | 0.000831183673925115 | 0.0423609197856822 | 2723 | 0 | 0 | 2 | 0 | 0 | 0 | 2 | 2 | 4 | 4 | 9 | 5 |
| MIRc_dup92367 | 2.82163356059973 | 3.2628035296961 | 0.940470316121336 | 3.46933175217319 | 0.000521754746857887 | 0.0313184125229311 | 2724 | 0 | 0 | 4 | 2 | 0 | 0 | 1 | 1 | 1 | 8 | 11 | 8 |
| MER1B_dup4268 | 2.85848685889938 | 3.28151640394089 | 0.882152221620867 | 3.71989813494029 | 0.000199303144851285 | 0.0163905041570965 | 2725 | 0 | 0 | 2 | 2 | 0 | 0 | 2 | 0 | 1 | 2 | 15 | 14 |
| L1MB4_dup7375 | 2.56647737481595 | 3.00298790396554 | 0.876345261245093 | 3.42671779807305 | 0.00061092361349438 | 0.0348017309400368 | 2726 | 0 | 0 | 2 | 3 | 0 | 0 | 1 | 1 | 2 | 2 | 13 | 9 |
| MER3_dup8136 | 3.24167923906521 | 2.72994833818434 | 0.800721205678836 | 3.4093618588132 | 0.000651150461712705 | 0.0361921625234037 | 2727 | 0 | 0 | 3 | 5 | 0 | 0 | 2 | 1 | 1 | 3 | 14 | 14 |
| Tigger3a_dup4746 | 3.14765527824522 | 3.56232300481253 | 0.923582224607565 | 3.8570718555418 | 0.000114753436713898 | 0.0110816238848722 | 2728 | 2 | 0 | 2 | 1 | 0 | 0 | 0 | 1 | 8 | 4 | 5 | 16 |
| MLT2A2_dup3761 | 4.49988704500971 | 3.66881637037074 | 0.780549896964867 | 4.70029704012103 | 2.59783336090752e-06 | 0.000687590719571191 | 2729 | 0 | 0 | 2 | 5 | 0 | 0 | 1 | 2 | 8 | 6 | 11 | 22 |
| THE1B-int_dup2384 | 3.8119841861857 | 2.62453745098257 | 0.772471600185283 | 3.39758439061456 | 0.000679836159672991 | 0.0371272880838366 | 2730 | 1 | 0 | 4 | 6 | 0 | 0 | 1 | 2 | 5 | 2 | 12 | 16 |

| | | | | | | | | | | | | | | | | | | |
|---|---|---|---|---|---|---|---|---|---|---|---|---|---|---|---|---|---|---|
| L1P3_dup2963 | 3.70994812168584 | 2.53823702236482 | 0.755856404695255 | 3.35809421815799 | 0.000784818573365228 | 0.0409505595573397 | 2731 | 0 | 2 | 4 | 1 | 1 | 0 | 3 | 2 | 5 | 8 | 4 | 17 |
| MLT2A2_dup3767 | 5.07261030048706 | 2.03806034629017 | 0.619503715336083 | 3.28982747937922 | 0.00100248826954837 | 0.0479184559634305 | 2732 | 3 | 1 | 7 | 6 | 0 | 0 | 3 | 3 | 5 | 5 | 10 | 23 |
| Tigger1_dup11834 | 3.42141840052705 | 2.76060770188699 | 0.831246599881558 | 3.32104540611696 | 0.000896809459824458 | 0.0444381973609726 | 2733 | 1 | 0 | 5 | 1 | 0 | 0 | 2 | 2 | 3 | 3 | 6 | 22 |
| AluJb_dup90752 | 5.21802052956634 | 3.97216663651889 | 0.827796179278297 | 4.79848389730667 | 1.59871158348063e-06 | 0.000467905321577961 | 2734 | 0 | 0 | 0 | 4 | 0 | 0 | 3 | 1 | 5 | 1 | 23 | 32 |
| AluSc_dup25760 | 3.69681959113949 | 3.09951563413613 | 0.843837670704272 | 3.67311835171954 | 0.000239608437570354 | 0.0185904708360133 | 2735 | 0 | 0 | 0 | 3 | 0 | 0 | 4 | 2 | 1 | 2 | 13 | 25 |
| MIR_dup133526 | 4.78457515511275 | 3.89435813953643 | 0.80768854320288 | 4.82160874053431 | 1.4240505594836e-06 | 0.000425687354807643 | 2736 | 0 | 0 | 0 | 5 | 0 | 0 | 2 | 1 | 2 | 5 | 20 | 29 |
| MSTA_dup3859 | 3.70812214959331 | 2.79998471936779 | 0.763719381288869 | 3.66624808531436 | 0.000246135153298207 | 0.0189291775005479 | 2737 | 0 | 0 | 1 | 3 | 0 | 1 | 6 | 2 | 4 | 7 | 8 | 16 |
| AluSz6_dup34554 | 4.84568676850602 | 3.25264495205507 | 0.747506603372418 | 4.3513260449882 | 1.35316655986331e-05 | 0.00233657888427601 | 2738 | 0 | 0 | 4 | 3 | 0 | 0 | 6 | 1 | 4 | 5 | 10 | 32 |
| AluJb_dup90744 | 3.63373229077895 | 3.49574733940685 | 0.786136524186937 | 4.44674332237945 | 8.71819049798823e-06 | 0.00166891275642342 | 2739 | 0 | 0 | 1 | 3 | 0 | 0 | 3 | 1 | 2 | 5 | 14 | 19 |
| MLT1A1_dup6518 | 3.48096206486262 | 2.85352585511082 | 0.798135336471496 | 3.575240594817 | 0.00034990596563345 | 0.0241119967850035 | 2740 | 0 | 0 | 2 | 4 | 0 | 0 | 6 | 1 | 2 | 6 | 11 | 14 |
| MIR_dup133519 | 3.88287933963815 | 3.17143366932298 | 0.790681726651269 | 4.01101171612348 | 6.04591096201281e-05 | 0.00695311904189028 | 2741 | 0 | 0 | 2 | 5 | 0 | 0 | 1 | 3 | 3 | 5 | 19 | 11 |
| MIR_dup133512 | 3.275150541639 | 2.70415221985449 | 0.779877615981802 | 3.46740586527821 | 0.000525507767876103 | 0.0314651412518991 | 2742 | 0 | 0 | 2 | 3 | 0 | 0 | 2 | 4 | 2 | 3 | 11 | 15 |
| MLT1A0_dup20087 | 4.1102525484637 | 2.71944755430256 | 0.756325257748131 | 3.59560589368573 | 0.000323637445180039 | 0.022787004325127 | 2743 | 0 | 0 | 3 | 4 | 0 | 0 | 2 | 6 | 4 | 4 | 15 | 13 |
| L1M3_dup5203 | 4.34174599887318 | 2.57430818098733 | 0.740279402646565 | 3.47748184237458 | 0.000506147331352792 | 0.0307681606502456 | 2744 | 0 | 0 | 3 | 7 | 0 | 0 | 3 | 3 | 2 | 2 | 24 | 12 |
| MER123_dup25 | 4.40140154301321 | 2.61801413181575 | 0.687626736874479 | 3.80731869693666 | 0.000140481659067977 | 0.0127486837713916 | 2745 | 0 | 0 | 2 | 5 | 0 | 1 | 4 | 3 | 2 | 3 | 22 | 15 |
| L1ME2z_dup2143 | 2.91088727719621 | 2.83551358640585 | 0.81672246009414 | 3.47182026324954 | 0.000516942315511107 | 0.0311798114592157 | 2746 | 1 | 1 | 1 | 2 | 0 | 0 | 2 | 1 | 3 | 1 | 11 | 15 |
| AluY_dup101976 | 3.36427135607245 | 2.64804216809023 | 0.777337207051864 | 3.40655528137296 | 0.000657882260254359 | 0.036409190594618 | 2747 | 0 | 0 | 3 | 2 | 0 | 0 | 4 | 2 | 1 | 2 | 11 | 20 |
| THE1B_dup13745 | 3.7230591428259 | 2.92314196096822 | 0.78593266950809 | 3.71932873435302 | 0.000199752937301375 | 0.0164124352275942 | 2748 | 0 | 0 | 1 | 4 | 0 | 0 | 2 | 3 | 2 | 1 | 15 | 21 |
| L3_dup36518 | 2.69025855070956 | 3.02334157551132 | 0.838620444443341 | 3.6051369788846 | 0.000311988326251917 | 0.0222464411981836 | 2749 | 0 | 0 | 2 | 1 | 1 | 0 | 1 | 2 | 3 | 3 | 9 | 12 |
| MLT1C_dup13030 | 3.04578416673501 | 3.14806569553027 | 0.807140240162342 | 3.90027102960086 | 9.60850619015444e-05 | 0.00977014097876609 | 2750 | 0 | 0 | 1 | 2 | 1 | 1 | 1 | 1 | 2 | 3 | 11 | 17 |
| L1PA11_dup3441 | 3.47421043878475 | 3.75648357374203 | 0.848938699222219 | 4.42491733983408 | 9.64792181250055e-06 | 0.00180620330731491 | 2751 | 0 | 0 | 1 | 2 | 0 | 0 | 1 | 2 | 2 | 5 | 9 | 24 |
| MSTA1_dup132 | 3.4149467792974 | 3.70737733736238 | 0.81257515936931 | 4.56250390454946 | 5.05471533021254e-06 | 0.0011044529788455 | 2752 | 0 | 1 | 1 | 1 | 0 | 0 | 1 | 2 | 4 | 6 | 12 | 15 |
| L2c_dup96715 | 4.08660330155065 | 3.26442576790009 | 0.813056763320095 | 4.01500352148834 | 5.94449072297893e-05 | 0.00687302132930002 | 2753 | 0 | 1 | 2 | 3 | 0 | 0 | 0 | 2 | 2 | 1 | 23 | 19 |
| L1ME3A_dup15341 | 4.26841242374451 | 3.11433665122838 | 0.866925089973043 | 3.59239418405254 | 0.000327653717945632 | 0.0229879584453974 | 2754 | 0 | 0 | 3 | 7 | 0 | 0 | 0 | 2 | 1 | 4 | 16 | 24 |
| L1M3_dup5202 | 4.91550027959087 | 3.76012463096837 | 0.826299669013406 | 4.55055807472115 | 5.35038168086779e-06 | 0.00115212843387852 | 2755 | 0 | 0 | 3 | 3 | 0 | 0 | 0 | 2 | 4 | 1 | 18 | 34 |
| AluSx3_dup23211 | 3.18268030603488 | 3.76219866781955 | 0.901231272282956 | 4.17450967750967 | 2.98628525922002e-05 | 0.00416515024789195 | 2756 | 0 | 0 | 2 | 2 | 0 | 0 | 0 | 1 | 3 | 2 | 10 | 22 |
| L1MC4a_dup7786 | 4.27213377704361 | 3.80891846837962 | 0.804852256700557 | 4.73244429231527 | 2.21832196840162e-06 | 0.000610220623229239 | 2757 | 0 | 0 | 3 | 3 | 0 | 0 | 0 | 1 | 3 | 3 | 19 | 24 |
| Charlie5_dup1946 | 2.5369738763253 | 3.68182919237137 | 1.04454409266955 | 3.5248193141964 | 0.000423771734473087 | 0.0273116980311303 | 2758 | 0 | 0 | 2 | 1 | 0 | 0 | 0 | 0 | 1 | 1 | 12 | 17 |
| MSTA-int_dup2915 | 2.22478387369706 | 3.60960178367491 | 1.04033060154955 | 3.46966798659817 | 0.00052110208581576 | 0.0313037507981613 | 2759 | 0 | 0 | 2 | 2 | 0 | 0 | 0 | 0 | 3 | 2 | 7 | 13 |
| MER5A1_dup12054 | 4.15755421907759 | 3.71727576202446 | 0.930774428708466 | 3.99374504430952 | 6.50377967247278e-05 | 0.0073561844589967 | 2760 | 0 | 0 | 3 | 5 | 0 | 0 | 0 | 0 | 4 | 2 | 22 | 18 |
| THE1C-int_dup1743 | 3.78055477061187 | 3.5535687136394 | 0.905009264480727 | 3.92655506756426 | 8.61711707075022e-05 | 0.00904458993611262 | 2761 | 0 | 1 | 2 | 4 | 0 | 0 | 0 | 0 | 3 | 2 | 11 | 28 |
| L3_dup42915 | 3.53397445756499 | 3.3335007070082 | 0.83660145625535 | 3.98457435387339 | 6.76011882839158e-05 | 0.00755184617249043 | 2762 | 1 | 0 | 0 | 2 | 2 | 1 | 0 | 1 | 3 | 4 | 18 | 13 |
| MER74A_dup1402 | 3.69711240994311 | 2.80536208016315 | 0.84322602103746 | 3.32693964627845 | 0.000878053886614652 | 0.0438282798624732 | 2763 | 0 | 2 | 0 | 4 | 2 | 1 | 0 | 1 | 5 | 2 | 15 | 15 |
| AluSc_dup31475 | 3.48180692594977 | 3.1159001386467 | 0.807083361991888 | 3.86069182612888 | 0.000113066434896563 | 0.0109721476011001 | 2764 | 0 | 0 | 2 | 2 | 2 | 1 | 0 | 1 | 4 | 3 | 16 | 13 |
| THE1B_dup21074 | 3.68343141632474 | 2.32435107792678 | 0.701273928140479 | 3.31446954557417 | 0.00091817181082311 | 0.0451577899575352 | 2765 | 1 | 0 | 3 | 2 | 3 | 2 | 1 | 1 | 3 | 4 | 15 | 12 |
| MIR_dup116621 | 3.29469424515803 | 2.96402625718219 | 0.873814576603765 | 3.39205403130536 | 0.000693707590504533 | 0.0375659868753281 | 2766 | 1 | 0 | 1 | 3 | 1 | 2 | 0 | 0 | 1 | 4 | 13 | 18 |
| L3_dup42914 | 4.63676845428927 | 3.16758513369272 | 0.82594779235349 | 3.83509122854711 | 0.000125517697068905 | 0.0118095824226009 | 2767 | 0 | 0 | 3 | 6 | 1 | 1 | 0 | 0 | 2 | 4 | 19 | 26 |
| THE1B-int_dup4010 | 3.68704627201467 | 3.1101004299802 | 0.875396476212264 | 3.5527906662787 | 0.000381167655359015 | 0.0255037468213251 | 2768 | 0 | 0 | 2 | 4 | 1 | 1 | 0 | 0 | 2 | 1 | 18 | 20 |
| MIRc_dup30831 | 6.17692596092318 | 3.06808810275741 | 0.7892647487963 | 3.88727370307177 | 0.00010137642383489 | 0.0101781074032982 | 2769 | 1 | 0 | 1 | 0 | 0 | 2 | 7 | 6 | 11 | 8 | 12 | 29 |
| L1MB8_dup6873 | 7.4285089736841 | 3.24059325632731 | 0.687621737762006 | 4.71275568872275 | 2.44389137226383e-06 | 0.000657078165496823 | 2770 | 1 | 1 | 1 | 0 | 1 | 5 | 4 | 2 | 6 | 7 | 27 | 42 |
| AluSp_dup43963 | 5.43268664171317 | 3.18520936073096 | 0.821854853065364 | 3.87563491150625 | 0.000106347036793412 | 0.0104912321794549 | 2771 | 0 | 0 | 0 | 0 | 1 | 2 | 6 | 4 | 9 | 5 | 17 | 24 |
| L1MCa_dup7417 | 7.66964389241182 | 4.06604836837388 | 0.686772130545759 | 5.92052034077548 | 3.20924597257171e-09 | 2.97131222641848e-06 | 2772 | 0 | 0 | 0 | 0 | 2 | 2 | 3 | 3 | 10 | 10 | 25 | 43 |
| AluY_dup64148 | 4.14106466811544 | 2.96082493917161 | 0.80313923902974 | 3.68656491338729 | 0.000227301481691436 | 0.017906561011388 | 2773 | 0 | 0 | 1 | 0 | 3 | 0 | 4 | 3 | 5 | 7 | 12 | 17 |
| L1M2_dup5813 | 5.96412458689195 | 2.6054836083768 | 0.721073455186434 | 3.61334006908235 | 0.000302277831623745 | 0.0217645622649209 | 2774 | 0 | 0 | 2 | 1 | 3 | 0 | 8 | 5 | 4 | 4 | 17 | 34 |
| L1PA13_dup8790 | 5.720158089585 | 3.98413459719845 | 0.707652842524886 | 5.63006937551917 | 1.8013715013672e-08 | 1.30282414366054e-05 | 2775 | 0 | 0 | 2 | 0 | 2 | 1 | 1 | 2 | 8 | 8 | 21 | 27 |
| MSTA_dup19316 | 3.74357705908448 | 3.15986626868216 | 0.905546516743351 | 3.48945770344974 | 0.000484001646091176 | 0.0298667675520775 | 2776 | 0 | 0 | 1 | 0 | 3 | 1 | 1 | 2 | 6 | 2 | 7 | 25 |
| AluSx_dup68892 | 4.11425109394856 | 3.25254138360557 | 0.932018033613007 | 3.48978374484553 | 0.000483411563842971 | 0.0298483738175653 | 2777 | 0 | 0 | 1 | 1 | 0 | 0 | 8 | 4 | 2 | 10 | 16 | 10 |
| MSTB_dup2433 | 5.2374569167337 | 2.95001550135212 | 0.770162408907567 | 3.83038105629767 | 0.00012794496882118 | 0.0119528832159892 | 2778 | 1 | 0 | 0 | 2 | 0 | 1 | 7 | 5 | 3 | 11 | 17 | 20 |
| L1MA4_dup3037 | 5.08218876710548 | 2.95114364894623 | 0.737810894019181 | 3.99986456267953 | 6.33787447017876e-05 | 0.00720531236828072 | 2779 | 2 | 0 | 2 | 1 | 0 | 0 | 8 | 3 | 8 | 8 | 19 | 12 |
| L2b_dup29048 | 3.91726415230996 | 3.08204260683035 | 0.832174106341944 | 3.70360310822255 | 0.000212558728821124 | 0.0171159050044234 | 2780 | 4 | 0 | 1 | 0 | 0 | 0 | 4 | 1 | 7 | 7 | 12 | 13 |
| AluSq2_dup16093 | 3.62309936213719 | 2.51065902707042 | 0.709069223708465 | 3.54078127088856 | 0.000398944156970709 | 0.0262592438103879 | 2781 | 2 | 0 | 2 | 1 | 2 | 0 | 2 | 3 | 6 | 4 | 9 | 14 |
| MER20_dup4684 | 4.63725138344152 | 2.63709736177414 | 0.747322194497865 | 3.52872881494713 | 0.000417560788344804 | 0.027047585234365 | 2782 | 2 | 0 | 1 | 1 | 2 | 0 | 9 | 1 | 6 | 8 | 16 | 13 |
| THE1B_dup17353 | 5.12915079838118 | 3.09019682510351 | 0.628143274694882 | 4.91957320820566 | 8.67331242152707e-07 | 0.000288237429811827 | 2783 | 3 | 0 | 1 | 3 | 1 | 0 | 3 | 2 | 6 | 10 | 18 | 18 |
| MIR3_dup72035 | 4.07922596687627 | 3.94903222213813 | 0.804997353146164 | 4.90564621946167 | 9.31202384327164e-07 | 0.000302282661084612 | 2784 | 1 | 0 | 1 | 0 | 0 | 1 | 2 | 1 | 13 | 10 | 8 | 11 |
| MSTD_dup1402 | 5.03320707502935 | 3.04647339006175 | 0.659743362267556 | 4.61766432873373 | 3.88083336850127e-06 | 0.000913381579328786 | 2785 | 3 | 1 | 1 | 1 | 0 | 1 | 5 | 1 | 13 | 12 | 10 | 13 |
| AluSx4_dup8316 | 2.64314637706478 | 2.77000270610581 | 0.806133124882957 | 3.43616038170865 | 0.000590021579505135 | 0.0340429861195269 | 2786 | 1 | 0 | 2 | 2 | 0 | 1 | 1 | 1 | 6 | 7 | 5 | 6 |
| L1PA4_dup9311 | 4.06514784231764 | 2.61949818306085 | 0.692259402373476 | 3.78398353865567 | 0.000154338035857795 | 0.0136267600297659 | 2787 | 1 | 0 | 2 | 3 | 0 | 4 | 1 | 2 | 8 | 6 | 11 | 12 |
| MLT1F-int_dup426 | 3.84893108816592 | 2.70511376567613 | 0.743058255786625 | 3.64051370751867 | 0.000272094620537053 | 0.0202641084752143 | 2788 | 1 | 0 | 1 | 1 | 0 | 4 | 4 | 1 | 6 | 12 | 7 | 11 |
| L1MA6_dup1721 | 3.35524801003802 | 2.8332481621894 | 0.854366924936139 | 3.31619598031744 | 0.000912518123574553 | 0.0449823984736007 | 2789 | 0 | 2 | 0 | 5 | 0 | 0 | 3 | 1 | 4 | 9 | 7 | 12 |
| L1PA4_dup607 | 3.42853397655899 | 3.41763144740353 | 0.814332959713983 | 4.19684774714743 | 2.70655751470656e-05 | 0.00388428549091899 | 2790 | 1 | 3 | 0 | 1 | 0 | 0 | 1 | 1 | 5 | 8 | 9 | 14 |
| L1ME4b_dup14913 | 3.32598919287448 | 3.63777821371081 | 1.00030158682454 | 3.63668143850392 | 0.000276173132988153 | 0.0204335486013679 | 2791 | 0 | 0 | 0 | 2 | 0 | 0 | 6 | 1 | 7 | 9 | 8 | 8 |
| MER20_dup13030 | 2.9362777528169 | 4.18782107887686 | 0.936511900115439 | 4.47172222623189 | 7.75921717466654e-06 | 0.00151831800151579 | 2792 | 0 | 0 | 0 | 1 | 0 | 0 | 2 | 1 | 5 | 8 | 11 | 8 |
| MIR3_dup25959 | 3.06731813781923 | 3.04473001211517 | 0.910589525370635 | 3.34369101256231 | 0.000826717594722322 | 0.0422223963927447 | 2793 | 1 | 0 | 0 | 1 | 0 | 0 | 4 | 3 | 3 | 8 | 11 | 7 |
| MIRc_dup64074 | 3.05493407798871 | 3.30247641453718 | 0.872681228929642 | 3.78428721170928 | 0.000154149728530126 | 0.0136151860819708 | 2794 | 0 | 0 | 0 | 3 | 0 | 0 | 2 | 3 | 4 | 5 | 11 | 10 |
| Tigger1_dup3862 | 3.1437052719634 | 3.18823574402152 | 0.853757438152386 | 3.73435779478674 | 0.000188194790279806 | 0.0157399639957535 | 2795 | 0 | 1 | 0 | 1 | 0 | 0 | 4 | 2 | 3 | 5 | 12 | 12 |
| MSTA-int_dup3070 | 7.27791362021408 | 4.58018668973836 | 0.736165101670838 | 6.22168407513877 | 4.91846655884001e-10 | 5.7089241161451e-07 | 2796 | 0 | 0 | 0 | 3 | 0 | 1 | 3 | 1 | 4 | 16 | 30 | 37 |
| L1MB3_dup10702 | 7.27575856067949 | 3.28204910626468 | 0.559672417183255 | 5.8642323714696 | 4.5121521905872e-09 | 4.02115567468712e-06 | 2797 | 1 | 0 | 1 | 6 | 1 | 3 | 4 | 1 | 12 | 14 | 21 | 28 |
| L3_dup14175 | 5.5761041580218 | 2.9505279961283 | 0.622629108576723 | 4.73882116252637 | 2.14965115562676e-06 | 0.000594308230354346 | 2798 | 3 | 0 | 0 | 3 | 1 | 3 | 3 | 2 | 5 | 14 | 18 | 19 |
| L1MB7_dup6269 | 5.37045515436497 | 2.54678982378396 | 0.687412847374131 | 3.70489122150178 | 0.000211481469823195 | 0.0170580236673051 | 2799 | 1 | 5 | 1 | 2 | 0 | 2 | 1 | 5 | 6 | 13 | 9 | 22 |
| AluJr_dup21558 | 6.05305953866317 | 2.66052364742297 | 0.674097328770365 | 3.94679452635732 | 7.92044736757071e-05 | 0.00845888137039488 | 2800 | 1 | 6 | 1 | 1 | 0 | 1 | 5 | 3 | 6 | 9 | 25 | 18 |
| MIR3_dup25957 | 5.78357490197167 | 2.58223665734516 | 0.697078159958713 | 3.7043717701586 | 0.000211915272580579 | 0.017083559495959 | 2801 | 1 | 4 | 1 | 1 | 0 | 1 | 3 | 7 | 7 | 11 | 21 | 13 |
| MIRb_dup66787 | 5.97170700821066 | 2.07903771759437 | 0.608153828108612 | 3.41860500008736 | 0.000629430272182546 | 0.0354150899845703 | 2802 | 2 | 1 | 2 | 5 | 1 | 2 | 5 | 7 | 4 | 9 | 27 | 9 |

| | | | | | | | | | | | | | | | | | | | |
|---|---|---|---|---|---|---|---|---|---|---|---|---|---|---|---|---|---|---|---|
| L4_A_Mam_dup1951 | 5.9393437047652 | 2.34412773664725 | 0.710772493057538 | 3.29800007673832 | 0.000973761077799339 | 0.0469530950038705 | 2803 | 3 | 1 | 1 | 2 | 0 | 0 | 9 | 8 | 6 | 10 | 17 | 17 |
| AluJb_dup20491 | 4.20827987695539 | 2.31612191554952 | 0.691317345213628 | 3.35030204519721 | 0.00080723484414836 | 0.0416114172932681 | 2804 | 1 | 1 | 3 | 3 | 0 | 0 | 6 | 4 | 5 | 8 | 14 | 7 |
| L2a_dup148884 | 4.69100135704249 | 2.6808058432193 | 0.633107493350771 | 4.23436125993537 | 2.29202235529209e-05 | 0.00342452650816728 | 2805 | 2 | 2 | 2 | 2 | 0 | 0 | 5 | 2 | 4 | 9 | 16 | 16 |
| MER41C_dup264 | 5.36460653016356 | 4.84466764473934 | 1.32483625460971 | 3.65680485258653 | 0.000255378651012626 | 0.0193955724525536 | 2806 | 2 | 2 | 0 | 0 | 0 | 2 | 0 | 0 | 0 | 1 | 32 | 33 |
| HERVIP10B3-int_dup1 | 3.90910426544495 | 4.58690481037792 | 1.38914017757951 | 3.30197404438356 | 0.000960069537624527 | 0.046554861666926 | 2807 | 2 | 3 | 0 | 0 | 0 | 0 | 0 | 0 | 0 | 3 | 17 | 28 |
| MER66B_dup445 | 2.43946359318646 | 4.18257202187527 | 1.24945950834627 | 3.34750505633522 | 0.000815425002169067 | 0.0418974557473809 | 2808 | 1 | 1 | 0 | 0 | 0 | 0 | 0 | 0 | 0 | 1 | 13 | 17 |
| HERVH-int_dup3388 | 8.71037840299146 | 5.46961369306256 | 0.868024948073006 | 6.30121715419007 | 2.95317298327526e-10 | 3.65987343061674e-07 | 2809 | 0 | 0 | 0 | 2 | 0 | 0 | 0 | 0 | 1 | 2 | 49 | 64 |
| AluSx1_dup14119 | 5.19163889468822 | 3.62446713570383 | 1.03897264119826 | 3.4885106613815 | 0.000485719451012343 | 0.029926204166386 | 2810 | 0 | 0 | 0 | 0 | 0 | 0 | 5 | 1 | 0 | 1 | 25 | 39 |
| MER50_dup1041 | 4.2738150933523 | 3.67450538811975 | 0.898881519587206 | 4.08786398212658 | 4.35363275355179e-05 | 0.00544939242070823 | 2811 | 0 | 0 | 0 | 2 | 0 | 0 | 2 | 1 | 1 | 0 | 26 | 25 |
| L2_dup23725 | 3.7042133247935 | 4.89346940051158 | 1.15569536989697 | 4.23422082321558 | 2.29345500479513e-05 | 0.00342452650816728 | 2812 | 1 | 0 | 0 | 0 | 1 | 0 | 0 | 0 | 3 | 0 | 19 | 25 |
| MER61B_dup175 | 3.25968878876166 | 4.15768572035417 | 1.22400535907261 | 3.39678718686684 | 0.000681819700559973 | 0.0372016212954467 | 2813 | 0 | 0 | 0 | 1 | 2 | 0 | 0 | 0 | 2 | 0 | 24 | 13 |
| Tigger5b_dup1775 | 2.26241333824066 | 4.09494255917593 | 1.20544886858585 | 3.39702717045132 | 0.000681222026661793 | 0.0371860647754226 | 2814 | 0 | 0 | 0 | 1 | 0 | 0 | 1 | 0 | 3 | 0 | 14 | 10 |
| L1MCb_dup911 | 2.83244225816684 | 4.72247363115337 | 1.25229861985919 | 3.771044346982 | 0.000162565777057349 | 0.0141588127334144 | 2815 | 0 | 0 | 1 | 0 | 0 | 0 | 0 | 0 | 3 | 0 | 15 | 18 |
| L1M5_dup21532 | 2.43775870139255 | 4.49073605611951 | 1.33313488462622 | 3.36855340589081 | 0.000755637562136487 | 0.0398318421267453 | 2816 | 1 | 0 | 0 | 0 | 0 | 0 | 0 | 0 | 1 | 0 | 12 | 19 |
| MIR_dup70220 | 2.35006988674416 | 4.89285664874234 | 1.4357119365643 | 3.4079654310398 | 0.000654491856271279 | 0.0362837448787836 | 2817 | 0 | 0 | 0 | 0 | 0 | 0 | 0 | 0 | 2 | 0 | 13 | 16 |
| AluJr4_dup2610 | 2.8470324053581 | 4.6621029170326 | 1.33937605209341 | 3.48080205685762 | 0.000499914790705248 | 0.0304928799767114 | 2818 | 0 | 2 | 0 | 0 | 0 | 0 | 0 | 0 | 3 | 0 | 15 | 17 |
| Charlie10b_dup125 | 3.62034510645842 | 4.55716506570223 | 1.36206836956286 | 3.34576822099232 | 0.000820549545944214 | 0.0420246410372884 | 2819 | 0 | 3 | 0 | 0 | 1 | 0 | 0 | 0 | 3 | 0 | 15 | 26 |
| L1PA17_dup2481 | 3.56536280008535 | 3.11215960511707 | 0.911755752145403 | 3.41336986116514 | 0.00064164795639418 | 0.035843779007008 | 2820 | 0 | 2 | 0 | 2 | 1 | 0 | 2 | 0 | 1 | 1 | 17 | 22 |
| MSTA_dup7809 | 4.56390119910653 | 4.16649003659943 | 1.04602068548868 | 3.98318130262685 | 6.79988396335056e-05 | 0.00758202994162798 | 2821 | 0 | 3 | 0 | 0 | 1 | 0 | 1 | 0 | 2 | 3 | 30 | 19 |
| L1MD_dup5428 | 3.8852250306844 | 3.88193911481526 | 0.93804225465566 | 4.13834141857529 | 3.49825538139798e-05 | 0.00467900997741497 | 2822 | 1 | 0 | 1 | 0 | 1 | 2 | 0 | 0 | 2 | 2 | 21 | 21 |
| L1M3_dup2375 | 4.76741622503489 | 3.81787276692746 | 0.993219812501631 | 3.84393536946404 | 0.000121076905144148 | 0.0115331326146635 | 2823 | 1 | 0 | 0 | 0 | 3 | 3 | 1 | 0 | 2 | 4 | 22 | 27 |
| L1PA8_dup3308 | 3.56096573236864 | 3.89278214153126 | 0.8846774221831 | 4.40022774846579 | 1.08137322869669e-05 | 0.00197929564827278 | 2824 | 1 | 1 | 0 | 0 | 1 | 1 | 1 | 0 | 2 | 4 | 15 | 21 |
| MER4-int_dup2801 | 4.41853207413462 | 4.20835394854553 | 0.827995582914598 | 5.08258019171051 | 3.72342030140716e-07 | 0.000147908675632492 | 2825 | 0 | 0 | 1 | 1 | 0 | 1 | 2 | 0 | 4 | 2 | 27 | 19 |
| MSTA_dup19311 | 3.55964991531929 | 4.32243091723155 | 0.966431842805491 | 4.47256674064443 | 7.72862671596947e-06 | 0.00151607239287557 | 2826 | 0 | 0 | 1 | 0 | 0 | 1 | 1 | 0 | 1 | 3 | 26 | 13 |
| ERV3-16A3_I-int_dup | 4.6540338486121 | 4.731491151819 | 0.932288069952665 | 5.07513858035235 | 3.87213691655862e-07 | 0.000152058355229693 | 2827 | 0 | 0 | 0 | 0 | 0 | 1 | 2 | 0 | 3 | 2 | 17 | 38 |
| LOR1-int_dup479 | 5.07125778289462 | 5.11494643876345 | 0.930776873524094 | 5.49535187675787 | 3.89932119171357e-08 | 2.37904584688071e-05 | 2828 | 0 | 0 | 0 | 0 | 0 | 1 | 0 | 1 | 2 | 3 | 22 | 39 |
| THE1D_dup5316 | 3.87823052752921 | 4.64476422818956 | 0.999854933992529 | 4.64543812335107 | 3.39355479766285e-06 | 0.000827223543608194 | 2829 | 0 | 0 | 0 | 0 | 0 | 1 | 0 | 1 | 1 | 2 | 18 | 29 |
| L1PA13_dup1920 | 3.36258064461333 | 4.43305468845481 | 1.08667200806064 | 4.0794781273205 | 4.51369163207807e-05 | 0.0055792980212283 | 2830 | 0 | 0 | 1 | 1 | 0 | 0 | 0 | 0 | 0 | 3 | 25 | 14 |
| L1M2_dup2632 | 2.48423986742474 | 3.65200335047248 | 1.0588021896277 | 3.44918379112592 | 0.000562283850864243 | 0.0329702299600156 | 2831 | 1 | 0 | 1 | 1 | 1 | 0 | 0 | 0 | 0 | 5 | 11 | 13 |
| THE1C-int_dup1244 | 2.60120705621663 | 3.81437824691693 | 1.11110018763392 | 3.43297417223881 | 0.000596998964886344 | 0.0342958753497938 | 2832 | 1 | 0 | 1 | 0 | 1 | 0 | 0 | 0 | 0 | 2 | 13 | 17 |
| MER101_dup150 | 2.86949411258793 | 4.36715303754954 | 1.13954658044743 | 3.83236026721683 | 0.000126919690790971 | 0.0119038363204369 | 2833 | 0 | 0 | 0 | 0 | 1 | 0 | 1 | 0 | 0 | 3 | 18 | 15 |
| MamTip3_dup1269 | 2.6034389583925 | 4.7582683371561 | 1.25409078158747 | 3.79419768250981 | 0.000148121632047022 | 0.013210251071612 | 2834 | 0 | 0 | 0 | 0 | 1 | 0 | 0 | 0 | 0 | 3 | 13 | 18 |
| L1ME4b_dup8595 | 2.75660197070427 | 3.72203496519924 | 1.12297761410005 | 3.31443380390273 | 0.000918289199006515 | 0.0451577899575352 | 2835 | 0 | 1 | 1 | 0 | 1 | 0 | 0 | 0 | 0 | 1 | 15 | 18 |
| THE1B_dup9109 | 3.81216095294601 | 4.91027377949582 | 1.22184131484591 | 4.01874917784643 | 5.85079130103824e-05 | 0.00680103755210964 | 2836 | 0 | 1 | 0 | 0 | 0 | 0 | 0 | 0 | 0 | 1 | 25 | 24 |
| L1M5_dup21539 | 3.13616141867894 | 4.43610731328742 | 1.20049604147849 | 3.69522860552213 | 0.000219689032688062 | 0.0174779121709676 | 2837 | 0 | 0 | 0 | 0 | 0 | 0 | 1 | 0 | 0 | 1 | 21 | 19 |
| L1PA6_dup3882 | 3.12164238727076 | 3.94274385089403 | 1.1044494584605 | 3.56987259189738 | 0.000357154894292895 | 0.0244406914428247 | 2838 | 0 | 0 | 0 | 0 | 0 | 1 | 2 | 0 | 0 | 2 | 18 | 19 |
| L1M5_dup21538 | 2.32172761710149 | 4.64030650107988 | 1.28420711886944 | 3.61336301045038 | 0.000302251075831126 | 0.0217645622649209 | 2839 | 1 | 0 | 0 | 0 | 0 | 0 | 0 | 0 | 0 | 3 | 13 | 14 |
| MER47A_dup986 | 2.69058010106211 | 4.50620753535048 | 1.21912713731412 | 3.69625726261676 | 0.000218801259753482 | 0.0174512596773806 | 2840 | 1 | 0 | 0 | 0 | 0 | 1 | 0 | 0 | 0 | 3 | 20 | 10 |
| L1MB8_dup5902 | 4.27451733832316 | 4.42382096279312 | 0.854112193652603 | 5.17943777839617 | 2.22555628309737e-07 | 9.82490195432073e-05 | 2841 | 0 | 0 | 0 | 2 | 1 | 0 | 1 | 0 | 3 | 3 | 18 | 29 |
| MER52A_dup24 | 2.84555367151783 | 4.59938397484595 | 1.0153962845479 | 4.52964428257072 | 5.90830754618604e-06 | 0.00124192415846712 | 2842 | 0 | 0 | 0 | 1 | 0 | 0 | 1 | 0 | 3 | 3 | 8 | 22 |
| L1MA3_dup5547 | 3.15910748889814 | 5.43344634015104 | 1.11670287288539 | 4.86561508175568 | 1.14101394160557e-06 | 0.000354901573005595 | 2843 | 0 | 0 | 0 | 0 | 0 | 0 | 0 | 0 | 1 | 4 | 16 | 21 |
| L1PREC2_dup4835 | 3.5981424689958 | 5.68893752224936 | 1.10245566548278 | 5.16024154110371 | 2.46631433783581e-07 | 0.000106463686417179 | 2844 | 0 | 0 | 0 | 0 | 0 | 0 | 0 | 0 | 1 | 6 | 16 | 25 |
| L1MA3_dup4936 | 3.11779411228582 | 5.68721353508737 | 1.04447324302638 | 5.44505431140446 | 5.17895811553273e-08 | 2.99104943630716e-05 | 2845 | 0 | 0 | 0 | 0 | 0 | 0 | 0 | 0 | 5 | 5 | 10 | 20 |
| L2a_dup100066 | 3.36848961838793 | 5.76231971293961 | 1.03936594726909 | 5.54407206439656 | 2.95517116822573e-08 | 1.90323406797792e-05 | 2846 | 0 | 0 | 0 | 0 | 0 | 0 | 0 | 0 | 3 | 7 | 12 | 22 |
| AluJr_dup9790 | 3.51386503969096 | 5.68366414348134 | 1.06281555715168 | 5.347742705906 | 8.90579443658969e-08 | 4.65735860001927e-05 | 2847 | 0 | 0 | 0 | 0 | 0 | 0 | 0 | 0 | 3 | 4 | 11 | 29 |
| L2a_dup100071 | 3.03658439118576 | 5.46950369400826 | 1.07666931594072 | 5.08002189068553 | 3.77391378585545e-07 | 0.000149167390313814 | 2848 | 0 | 0 | 0 | 0 | 0 | 0 | 0 | 0 | 2 | 4 | 14 | 20 |
| THE1D-int_dup584 | 2.87818890408005 | 5.47004070073649 | 1.09719833151982 | 4.98546210251658 | 6.18138951014373e-07 | 0.000220185165925249 | 2849 | 0 | 0 | 0 | 0 | 0 | 0 | 0 | 0 | 2 | 5 | 11 | 20 |
| L2a_dup155959 | 3.6571512410965 | 4.68364306937956 | 0.94354890801761 | 4.9638582903136 | 6.91063746854443e-07 | 0.0002411082907757 | 2850 | 0 | 1 | 0 | 2 | 0 | 0 | 0 | 0 | 4 | 5 | 12 | 24 |
| L2c_dup135341 | 2.91883360331419 | 4.50655346281227 | 0.986067769880172 | 4.57022691590449 | 4.87196400132631e-06 | 0.00107538436573246 | 2851 | 0 | 0 | 0 | 1 | 0 | 0 | 0 | 1 | 2 | 3 | 14 | 17 |
| L1PA5_dup1796 | 2.29819063711016 | 4.25725650621827 | 1.06069353694435 | 4.01365366897831 | 5.97860491749881e-05 | 0.00690240235661898 | 2852 | 0 | 1 | 0 | 1 | 0 | 0 | 0 | 0 | 2 | 2 | 11 | 13 |
| MSTA-int_dup688 | 2.42604045642283 | 4.54070572878007 | 1.13787136762235 | 3.9905263969056 | 6.59268107906032e-05 | 0.00741703116802878 | 2853 | 0 | 0 | 0 | 1 | 0 | 0 | 0 | 0 | 1 | 2 | 9 | 20 |
| L1MC1_dup5602 | 3.50611385694917 | 3.9097994207503 | 0.954879097122995 | 4.09454917646678 | 4.22990630573408e-05 | 0.005342298185998 | 2854 | 0 | 1 | 0 | 1 | 2 | 1 | 0 | 0 | 1 | 6 | 16 | 18 |
| L1MC5_dup14135 | 3.31751700358909 | 3.15857697254985 | 0.853547675077784 | 3.70052788470428 | 0.00021515143464542 | 0.0172429852705083 | 2855 | 0 | 1 | 2 | 2 | 1 | 1 | 0 | 0 | 1 | 3 | 15 | 18 |
| L2_dup23724 | 4.08360919430705 | 3.69539473859956 | 0.880938647835161 | 4.19483779906887 | 2.73066922216725e-05 | 0.00389957822735564 | 2856 | 0 | 2 | 1 | 1 | 1 | 1 | 0 | 0 | 2 | 3 | 13 | 31 |
| L1PA10_dup6572 | 4.92019863174569 | 4.30930457845252 | 0.918836243216874 | 4.68995929390585 | 2.73259405050645e-06 | 0.000710382446875096 | 2857 | 0 | 0 | 1 | 1 | 2 | 0 | 0 | 0 | 1 | 2 | 26 | 33 |
| L2c_dup56369 | 4.34858485871849 | 4.71932208405171 | 0.940292238458587 | 5.01899504327298 | 5.19424899309678e-07 | 0.000191323648747896 | 2858 | 0 | 1 | 1 | 0 | 1 | 0 | 0 | 0 | 1 | 5 | 20 | 29 |
| MLT1E2_dup3618 | 5.16631900096732 | 4.10421939227187 | 0.810409244390081 | 5.06437879464311 | 4.09734223682429e-07 | 0.000158013970159383 | 2859 | 0 | 1 | 2 | 2 | 1 | 0 | 0 | 0 | 1 | 5 | 25 | 32 |
| LTR78_dup623 | 8.27865009982484 | 3.55797044575862 | 0.976258760940357 | 3.64449527943953 | 0.000267917052643733 | 0.0200408019517025 | 2860 | 0 | 0 | 0 | 0 | 0 | 0 | 9 | 7 | 1 | 4 | 36 | 53 |
| L1PA5_dup8844 | 4.1735761931 | 3.57202092721896 | 0.863612370419841 | 4.13613914015896 | 3.53198078174339e-05 | 0.00471341922939048 | 2861 | 1 | 0 | 1 | 0 | 2 | 0 | 1 | 1 | 1 | 2 | 23 | 23 |
| MER61-int_dup923 | 5.11781093616265 | 2.76103376298178 | 0.79802411001468 | 3.45983752662685 | 0.000540501408062462 | 0.0321123965150638 | 2862 | 3 | 0 | 3 | 1 | 2 | 0 | 4 | 0 | 1 | 4 | 21 | 30 |
| LTR16A_dup1806 | 7.6138983817008 | 4.11313660785459 | 0.803600994843 | 5.11838167728772 | 3.0816847466633e-07 | 0.000126426648056818 | 2863 | 1 | 0 | 0 | 1 | 2 | 2 | 2 | 0 | 3 | 4 | 47 | 38 |
| MER20B_dup1677 | 7.59750561970215 | 3.17501982460885 | 0.694867106248483 | 4.56924755260104 | 4.89478356436426e-06 | 0.00107830700748548 | 2864 | 2 | 0 | 0 | 3 | 4 | 1 | 3 | 2 | 4 | 5 | 40 | 35 |
| MIRb_dup87640 | 6.67258862759582 | 2.62701969105108 | 0.750406166029344 | 3.50079704828592 | 0.000463868956333206 | 0.0290242746462107 | 2865 | 2 | 0 | 3 | 1 | 3 | 3 | 2 | 2 | 1 | 3 | 29 | 40 |
| L2c_dup129978 | 5.86478896185881 | 2.85391846801562 | 0.739382626154249 | 3.85986682275685 | 0.000113448837336925 | 0.0109958032786037 | 2866 | 1 | 0 | 1 | 3 | 1 | 5 | 1 | 2 | 3 | 4 | 25 | 31 |
| THE1B_dup9103 | 6.37713675484398 | 3.43751835372537 | 0.850806377050182 | 4.04030628642388 | 5.33814410771925e-05 | 0.00634143803222849 | 2867 | 4 | 0 | 0 | 1 | 1 | 4 | 1 | 1 | 3 | 5 | 30 | 34 |
| L1PA16_dup13778 | 6.08869635111439 | 4.77742192396638 | 0.850458806910019 | 5.61746422654407 | 1.93780277611618e-08 | 1.36772564567561e-05 | 2868 | 0 | 0 | 0 | 1 | 1 | 0 | 0 | 2 | 1 | 6 | 29 | 41 |
| MLT1A0_dup16558 | 5.99353306494102 | 2.96186809231842 | 0.863749524411986 | 3.42908216862379 | 0.000605626144718025 | 0.0346492711303379 | 2869 | 0 | 0 | 1 | 5 | 2 | 1 | 0 | 4 | 0 | 5 | 29 | 32 |
| AluSx3_dup23212 | 6.1113975114588 | 3.52392940494786 | 0.795630671216995 | 4.42910200980269 | 9.46262604618247e-06 | 0.0017785102821366 | 2870 | 1 | 0 | 1 | 1 | 2 | 0 | 1 | 3 | 2 | 2 | 17 | 53 |
| L1PA16_dup8977 | 6.0603888871577 | 3.195178166426 | 0.641094080382329 | 4.98394582667257 | 6.23005290010808e-07 | 0.00022092640348273 | 2871 | 1 | 0 | 2 | 3 | 2 | 1 | 3 | 1 | 4 | 3 | 21 | 40 |
| MLT2A2_dup1589 | 7.5194971318696 | 3.49001150974449 | 0.716694380803041 | 4.86959519039901 | 1.11827103225128e-06 | 0.000349259924646234 | 2872 | 1 | 0 | 4 | 1 | 3 | 0 | 2 | 1 | 3 | 3 | 35 | 47 |
| L1MA3_dup7285 | 6.88226372883044 | 4.11272071190594 | 0.712216855774672 | 5.7745343690758 | 7.71661783483771e-09 | 6.37548037266769e-06 | 2873 | 0 | 0 | 3 | 1 | 1 | 1 | 0 | 2 | 2 | 5 | 33 | 43 |
| MSTA_dup19305 | 8.0592296174809 | 3.94721741699249 | 0.657561927068987 | 6.00280711899912 | 1.93934954065785e-09 | 1.89901426255708e-06 | 2874 | 1 | 0 | 3 | 3 | 1 | 0 | 0 | 3 | 5 | 3 | 42 | 44 |

| | | | | | | | | | | | | | | | | | | | |
|---|---|---|---|---|---|---|---|---|---|---|---|---|---|---|---|---|---|---|---|
| MSTA_dup18514 | 6.22658522679349 | 3.63441837409815 | 0.732104164841962 | 4.9643459887743 | 6.89329615823556e-07 | 0.000240856424033409 | 2875 | 1 | 5 | 0 | 0 | 5 | 2 | 1 | 0 | 10 | 23 | 19 | 9 |
| AluJb_dup99979 | 5.47209426508661 | 3.35168020612974 | 0.814685441064492 | 4.11407892812021 | 3.88728422971681e-05 | 0.00502277394500201 | 2876 | 1 | 3 | 0 | 0 | 4 | 4 | 2 | 0 | 4 | 13 | 13 | 27 |
| L2c_dup119697 | 6.81525451062423 | 3.22256416964625 | 0.673126787961489 | 4.78745494501197 | 1.68909641588181e-06 | 0.000490737162116498 | 2877 | 2 | 1 | 0 | 0 | 7 | 5 | 3 | 0 | 16 | 18 | 10 | 21 |
| L1MB7_dup7456 | 5.20906635221542 | 2.5842157757307 | 0.710098571297075 | 3.63923528392732 | 0.000273448872625664 | 0.0203141009821374 | 2878 | 2 | 1 | 1 | 0 | 8 | 4 | 3 | 0 | 6 | 15 | 13 | 12 |
| PABL_A_dup269 | 3.8053247880656 | 2.72224678104796 | 0.741342062355921 | 3.67205224049597 | 0.00024061048045095 | 0.0186307521579505 | 2879 | 1 | 1 | 1 | 0 | 4 | 5 | 1 | 0 | 6 | 11 | 7 | 10 |
| MLT1E1A_dup2986 | 6.39717509491517 | 2.6280820215103 | 0.788662133428532 | 3.33232940966152 | 0.000861222513794295 | 0.0433519044356457 | 2880 | 0 | 1 | 2 | 0 | 9 | 8 | 3 | 0 | 11 | 16 | 11 | 18 |
| MIRb_dup122110 | 5.26764130574676 | 3.01518621202997 | 0.815783663084681 | 3.69606103734512 | 0.000218970349743743 | 0.0174549811859714 | 2881 | 0 | 1 | 0 | 0 | 7 | 6 | 2 | 0 | 11 | 19 | 9 | 8 |
| MSTB1_dup3868 | 4.90656385624879 | 2.95651598518223 | 0.894810242609686 | 3.30407034295858 | 0.00095291918880714 | 0.0463008369418401 | 2882 | 0 | 1 | 0 | 0 | 7 | 7 | 2 | 0 | 9 | 10 | 13 | 11 |
| L1MD2_dup9070 | 6.13393445339001 | 3.11340526373452 | 0.787816870502307 | 3.95194033068806 | 7.75200723934064e-05 | 0.00834642133290565 | 2883 | 0 | 2 | 0 | 2 | 9 | 4 | 1 | 0 | 11 | 11 | 16 | 20 |
| AluSz6_dup35675 | 5.25026667757214 | 2.28256162770738 | 0.610548281096555 | 3.73854402408252 | 0.000185089084200227 | 0.0155842913054166 | 2884 | 0 | 2 | 4 | 3 | 4 | 5 | 2 | 0 | 12 | 10 | 10 | 12 |
| L1MD2_dup9069 | 6.10058787542124 | 3.02185623870667 | 0.658691210451265 | 4.58766746961208 | 4.48225834963362e-06 | 0.0010090212348741 | 2885 | 0 | 1 | 3 | 1 | 3 | 6 | 2 | 0 | 16 | 10 | 13 | 19 |
| L1PA13_dup7902 | 4.99236607358718 | 3.23817848061662 | 0.716787501489968 | 4.51762687530894 | 6.2536557169394e-06 | 0.00129845755953129 | 2886 | 0 | 1 | 1 | 1 | 6 | 2 | 1 | 0 | 13 | 11 | 10 | 14 |
| L2a_dup142895 | 5.2602200395416 | 2.65702271113004 | 0.738656906351742 | 3.59709993676657 | 0.000321784875274934 | 0.0226912052413817 | 2887 | 0 | 2 | 1 | 1 | 7 | 7 | 1 | 0 | 14 | 9 | 9 | 12 |
| L1MB8_dup6875 | 4.94337674327099 | 2.8829196851028 | 0.871181235424542 | 3.30920773758161 | 0.000935603896706126 | 0.0457144465247499 | 2888 | 0 | 0 | 0 | 2 | 2 | 5 | 4 | 0 | 3 | 6 | 20 | 23 |
| L1ME1_dup26352 | 6.5981741729996 | 3.34110890833774 | 0.858560012431412 | 3.89152634639463 | 9.96155696148121e-05 | 0.0100436975964263 | 2889 | 0 | 0 | 0 | 1 | 4 | 8 | 3 | 0 | 10 | 14 | 24 | 18 |
| L1M5_dup1223 | 5.06188693034332 | 4.10823162206602 | 0.829041689483029 | 4.95539811107426 | 7.21823403896106e-07 | 0.000249643883231777 | 2890 | 1 | 0 | 0 | 1 | 2 | 1 | 2 | 0 | 6 | 15 | 6 | 32 |
| MIR_dup143638 | 4.98779969823622 | 2.86693444877981 | 0.864775288768341 | 3.31523632325694 | 0.000915656790686758 | 0.0450628962230779 | 2891 | 1 | 0 | 0 | 1 | 3 | 9 | 1 | 1 | 4 | 9 | 15 | 20 |
| ERV3-16A3_I-int_dup | 3.78197939135473 | 2.44404121018467 | 0.742190687637799 | 3.29300980313215 | 0.000991210185141592 | 0.0475034237087012 | 2892 | 2 | 0 | 0 | 2 | 4 | 4 | 1 | 1 | 5 | 9 | 5 | 15 |
| L2b_dup79137 | 4.51398385533599 | 2.80237081484086 | 0.6622023629035 | 4.2318949188788 | 2.31730674982736e-05 | 0.00344836692867055 | 2893 | 1 | 0 | 0 | 3 | 3 | 3 | 1 | 2 | 7 | 9 | 13 | 14 |
| L1ME1_dup26361 | 7.10011515031925 | 3.90292867566353 | 1.14629207377939 | 3.40482915736769 | 0.000662054587326918 | 0.0365252123431697 | 2894 | 0 | 0 | 1 | 0 | 8 | 13 | 0 | 0 | 9 | 10 | 20 | 29 |
| L1MEd_dup19749 | 5.65980042599765 | 3.46779658097744 | 0.942028150122716 | 3.68120271196322 | 0.000232136348680561 | 0.0181586635340565 | 2895 | 0 | 0 | 2 | 0 | 4 | 8 | 0 | 0 | 12 | 12 | 14 | 17 |
| L2c_dup119715 | 4.96382886044811 | 3.45803858455426 | 0.986180678640231 | 3.50649597933949 | 0.000454048296030395 | 0.0285741803351622 | 2896 | 0 | 0 | 0 | 1 | 4 | 9 | 0 | 0 | 8 | 18 | 8 | 13 |
| L1PA6_dup2144 | 5.39197076215854 | 4.40646743557019 | 0.898591646176888 | 4.90374849835047 | 9.4024861639505e-07 | 0.00030397880064774 | 2897 | 0 | 0 | 0 | 0 | 4 | 4 | 0 | 0 | 13 | 15 | 12 | 17 |
| Tigger3b_dup1084 | 4.64973827984659 | 4.57340055309259 | 1.02085915023722 | 4.4799525498007 | 7.46596347843104e-06 | 0.00147918080419546 | 2898 | 0 | 0 | 0 | 0 | 2 | 4 | 0 | 0 | 7 | 10 | 12 | 24 |
| L1M4_dup12908 | 3.39483932802767 | 2.80284000960042 | 0.831582859301222 | 3.37048795348625 | 0.000750351866508376 | 0.0396234410182428 | 2899 | 1 | 0 | 1 | 1 | 5 | 3 | 0 | 0 | 6 | 6 | 9 | 10 |
| MLT2B5_dup319 | 5.15511080661233 | 3.01842762816281 | 0.767655640492842 | 3.93200735973874 | 8.42394860801434e-05 | 0.00889385693427238 | 2900 | 1 | 0 | 1 | 2 | 6 | 5 | 0 | 0 | 12 | 9 | 11 | 16 |
| L1ME4a_dup33049 | 3.75333104494801 | 3.91375914225609 | 0.856501290180053 | 4.56947255903531 | 4.88953178310259e-06 | 0.00107826184398714 | 2901 | 1 | 0 | 0 | 1 | 2 | 2 | 0 | 0 | 4 | 9 | 12 | 17 |
| MER66D_dup192 | 4.86485209254506 | 3.60593612811408 | 0.795115857860273 | 4.5351077990294 | 5.7574071719241e-06 | 0.00121773511727169 | 2902 | 0 | 0 | 1 | 2 | 3 | 3 | 0 | 0 | 5 | 11 | 17 | 20 |
| HERVL-int_dup825 | 4.59097739705512 | 3.86476838359525 | 0.856316130142283 | 4.51324954366221 | 6.38418308202795e-06 | 0.0013152301537976 | 2903 | 0 | 0 | 0 | 1 | 5 | 2 | 0 | 0 | 12 | 17 | 9 | 8 |
| Charlie4a_dup2783 | 5.03972507815467 | 3.86954686444953 | 0.830874974639659 | 4.65719510462775 | 3.20546638359947e-06 | 0.000796342506182469 | 2904 | 0 | 0 | 0 | 0 | 4 | 4 | 1 | 0 | 14 | 17 | 11 | 8 |
| L2a_dup142885 | 4.54720788476153 | 3.77768755129836 | 0.811655659911557 | 4.65429829160558 | 3.25085897700584e-06 | 0.00080492080139712 | 2905 | 0 | 0 | 0 | 1 | 3 | 3 | 1 | 0 | 8 | 16 | 15 | 8 |
| L1P4_dup3225 | 4.35337834490597 | 3.68943235510865 | 0.876882004231144 | 4.20744448774903 | 2.58274820991595e-05 | 0.00374957809375751 | 2906 | 0 | 0 | 2 | 0 | 4 | 2 | 0 | 0 | 7 | 13 | 15 | 10 |
| LTR33_dup7691 | 3.29003769530399 | 3.93662582600643 | 1.11665191461246 | 3.52538313371599 | 0.000422870713119076 | 0.0272683996487349 | 2907 | 0 | 0 | 0 | 0 | 4 | 3 | 0 | 0 | 5 | 8 | 14 | 6 |
| L2b_dup79071 | 4.03249935609458 | 4.44631306716317 | 0.934481253873097 | 4.75805485528443 | 1.95467385948105e-06 | 0.000556087625861988 | 2908 | 0 | 0 | 0 | 0 | 2 | 3 | 0 | 0 | 6 | 18 | 11 | 9 |
| L2c_dup119724 | 3.50045773001268 | 4.15347929216952 | 1.05891498882362 | 3.92239163295226 | 8.7674330094366e-05 | 0.00914187386004997 | 2909 | 0 | 0 | 0 | 0 | 2 | 4 | 0 | 0 | 4 | 10 | 11 | 13 |
| L3_dup39458 | 3.99036711790107 | 4.27296002728968 | 0.953727586425966 | 4.48027307598633 | 7.45476001937067e-06 | 0.00147918080419546 | 2910 | 0 | 0 | 0 | 0 | 2 | 4 | 0 | 0 | 7 | 13 | 10 | 13 |
| L2a_dup142908 | 2.9266622375224 | 3.91106867103591 | 1.15156316656794 | 3.39631275520237 | 0.00068300269689281 | 0.0372405498888305 | 2911 | 0 | 0 | 0 | 0 | 4 | 2 | 0 | 0 | 4 | 4 | 10 | 13 |
| L2c_dup8971 | 4.03354045252871 | 4.21766463437667 | 1.23038730690473 | 3.42791624288371 | 0.000608233083185099 | 0.0347151883541292 | 2912 | 0 | 0 | 0 | 0 | 7 | 2 | 0 | 0 | 4 | 8 | 9 | 22 |
| L1MB7_dup18720 | 4.9576966983004 | 3.59053976153106 | 0.869674370300906 | 4.12860247944152 | 3.6497476660623e-05 | 0.00480599257414976 | 2913 | 1 | 0 | 0 | 0 | 4 | 6 | 1 | 0 | 9 | 10 | 16 | 14 |
| L1MB7_dup7457 | 4.72218339731775 | 3.49102596936874 | 0.950345961369693 | 3.6734264270848 | 0.000239319606092101 | 0.0185791657393772 | 2914 | 1 | 0 | 0 | 0 | 5 | 6 | 1 | 0 | 9 | 6 | 12 | 19 |
| L2a_dup9637 | 5.15563273089965 | 3.35478824958907 | 0.973978800382263 | 3.44441608818631 | 0.000572294244775307 | 0.0333599035686686 | 2915 | 0 | 0 | 0 | 1 | 8 | 4 | 1 | 0 | 5 | 11 | 19 | 16 |
| L1MD2_dup9072 | 4.38302226127913 | 3.69293308958428 | 1.03428607092201 | 3.57051418694273 | 0.000356281155870425 | 0.0244171151047913 | 2916 | 0 | 0 | 0 | 0 | 6 | 3 | 1 | 0 | 6 | 7 | 16 | 16 |
| PABL_A_dup430 | 4.44905784926561 | 3.76573144763046 | 0.927107946773924 | 4.06180473453404 | 4.86948003677683e-05 | 0.00592368761160991 | 2917 | 0 | 0 | 0 | 0 | 6 | 2 | 1 | 0 | 9 | 12 | 12 | 12 |
| L2b_dup2386 | 2.81012186293032 | 3.24086605414972 | 0.878860354640933 | 3.68757793776442 | 0.000226398761911103 | 0.0178752355164257 | 2918 | 2 | 1 | 0 | 1 | 2 | 1 | 0 | 0 | 2 | 7 | 10 | 10 |
| MER57D_dup173 | 2.83444849206594 | 3.25377672966217 | 0.986517023779102 | 3.29824691437939 | 0.000972905407230703 | 0.0469286336973275 | 2919 | 2 | 3 | 0 | 0 | 2 | 2 | 0 | 0 | 1 | 6 | 8 | 13 |
| AluSx1_dup88557 | 3.01259641542389 | 3.5969083827872 | 1.07110372121736 | 3.35813265469672 | 0.000784709446671221 | 0.0409505595573397 | 2920 | 0 | 4 | 0 | 0 | 2 | 2 | 0 | 0 | 5 | 3 | 13 | 8 |
| L2b_dup52622 | 4.00253442988419 | 3.63735829163877 | 0.953593222511766 | 3.81437095584421 | 0.000136530255770926 | 0.0125045528251227 | 2921 | 0 | 3 | 0 | 0 | 5 | 3 | 0 | 0 | 4 | 11 | 12 | 12 |
| MER39_dup1221 | 3.13189721966307 | 3.45604256781879 | 0.878220221947979 | 3.9352801056584 | 8.30997197804901e-05 | 0.00879734300557002 | 2922 | 0 | 1 | 1 | 2 | 1 | 1 | 0 | 0 | 7 | 14 | 7 | 3 |
| AluSx1_dup89212 | 2.82647848948605 | 3.63912674573716 | 0.88357703010517 | 4.11862986671802 | 3.81131756621088e-05 | 0.0049638082627784 | 2923 | 0 | 1 | 1 | 0 | 1 | 1 | 0 | 1 | 6 | 9 | 5 | 9 |
| L2_dup48977 | 2.18105909619415 | 3.28637769484363 | 0.953929419188552 | 3.44509523318732 | 0.000570858228605526 | 0.0333130639552934 | 2924 | 0 | 1 | 1 | 0 | 1 | 2 | 0 | 0 | 6 | 5 | 4 | 6 |
| L1PB1_dup6110 | 3.03931756625846 | 3.6335171834609 | 1.07379305908555 | 3.38381511476266 | 0.000714861232174909 | 0.0382981050831384 | 2925 | 0 | 1 | 0 | 0 | 1 | 6 | 0 | 0 | 5 | 7 | 6 | 12 |
| L2a_dup142921 | 3.164912497967 | 3.74831573039506 | 1.09194709278508 | 3.43268987587552 | 0.000597625254769921 | 0.0343152844767094 | 2926 | 0 | 1 | 0 | 0 | 0 | 6 | 0 | 0 | 10 | 7 | 7 | 6 |
| L1ME3A_dup8679 | 3.05336087252926 | 3.89039382076382 | 0.904730840480538 | 4.30005659881947 | 1.70754493794679e-05 | 0.00275646870966545 | 2927 | 0 | 2 | 0 | 0 | 2 | 1 | 0 | 0 | 10 | 9 | 7 | 4 |
| MIR_dup162939 | 2.76022577836377 | 3.22764003536911 | 0.976786707533427 | 3.3043447566148 | 0.000951986840137685 | 0.0462950052449216 | 2928 | 0 | 5 | 0 | 0 | 1 | 2 | 0 | 0 | 5 | 10 | 6 | 4 |
| Charlie1a_dup2378 | 3.39166521497375 | 3.0730333997498 | 0.877418503268444 | 3.50235764153883 | 0.000461160147056158 | 0.0289070633170244 | 2929 | 1 | 2 | 0 | 0 | 4 | 5 | 0 | 0 | 6 | 8 | 10 | 5 |
| AluSc8_dup6511 | 2.97629399296406 | 3.6519351401471 | 0.843893803692574 | 4.32748187528757 | 1.50823752029159e-05 | 0.00252021829356252 | 2930 | 2 | 1 | 0 | 0 | 1 | 2 | 0 | 0 | 9 | 8 | 6 | 6 |
| MIR_dup57134 | 3.57652239496628 | 3.52152238905889 | 0.787704245123696 | 4.47061496857351 | 7.79950028338751e-06 | 0.00152494650323001 | 2931 | 1 | 2 | 0 | 0 | 3 | 2 | 0 | 0 | 9 | 13 | 7 | 5 |
| UCON27_dup35 | 3.38738734825875 | 3.2508639408756 | 0.767690417906341 | 4.23460273184263 | 2.28956099754357e-05 | 0.00342452650816728 | 2932 | 2 | 2 | 0 | 0 | 3 | 2 | 0 | 0 | 9 | 11 | 5 | 6 |
| MLT1D_dup16988 | 3.11044549087745 | 3.47646446702561 | 0.990597889816938 | 3.50946080418973 | 0.000449016234098461 | 0.0284455848878574 | 2933 | 0 | 0 | 1 | 0 | 2 | 4 | 0 | 0 | 9 | 4 | 9 | 8 |
| L2c_dup119704 | 2.9892116243664 | 3.18624965787992 | 0.941330383547421 | 3.38483673061999 | 0.000712206034905279 | 0.0382401208062598 | 2934 | 1 | 0 | 1 | 0 | 3 | 3 | 0 | 0 | 4 | 5 | 13 | 7 |
| LTR16A2_dup973 | 2.90885680380757 | 3.32108773706402 | 0.936408958919419 | 3.54662106276346 | 0.000390205412483693 | 0.0258485014139323 | 2935 | 1 | 0 | 1 | 0 | 3 | 2 | 0 | 0 | 8 | 6 | 10 | 3 |
| MIR_dup143648 | 2.43871679637105 | 3.67906187303241 | 0.998053245642873 | 3.68623807306255 | 0.000227593453644302 | 0.0179143076152322 | 2936 | 1 | 0 | 0 | 0 | 2 | 2 | 0 | 0 | 3 | 8 | 9 | 5 |
| L1ME3A_dup13060 | 2.08564091164093 | 3.68406185539727 | 1.01902411819391 | 3.61528426032428 | 0.000300018236403966 | 0.0216524535272606 | 2937 | 1 | 0 | 0 | 0 | 1 | 2 | 0 | 0 | 3 | 6 | 6 | 7 |
| L2b_dup79108 | 2.45445454040243 | 3.51022631480279 | 1.01929058876162 | 3.44379351041347 | 0.000573613606984953 | 0.0334122554045634 | 2938 | 1 | 0 | 0 | 0 | 2 | 3 | 0 | 0 | 2 | 9 | 6 | 8 |
| AluSz_dup80372 | 2.19257244471197 | 3.79834910805961 | 1.11190109734122 | 3.41608540286743 | 0.000635283198101262 | 0.0356012943606695 | 2939 | 0 | 0 | 0 | 0 | 2 | 2 | 0 | 0 | 2 | 6 | 6 | 10 |
| L2a_dup143001 | 2.84706904659538 | 4.50194578615282 | 1.02952298975874 | 4.372846289919 | 1.22637072320404e-05 | 0.00217444119302167 | 2940 | 0 | 0 | 0 | 0 | 1 | 2 | 0 | 0 | 3 | 9 | 9 | 12 |
| MLT2C1_dup82 | 2.83349139303282 | 3.9837984776297 | 0.981307733416853 | 4.05968315745191 | 4.91393497441059e-05 | 0.00595948609287004 | 2941 | 0 | 0 | 1 | 0 | 1 | 2 | 0 | 0 | 3 | 5 | 12 | 12 |
| MER20B_dup3403 | 3.19311652656671 | 3.73844708716156 | 1.00829770876375 | 3.70768182320395 | 0.000209165229310548 | 0.0169632002909093 | 2942 | 0 | 0 | 1 | 0 | 1 | 4 | 0 | 0 | 3 | 12 | 9 | 10 |
| L1MB5_dup7500 | 2.31806642246847 | 3.41599213133149 | 0.99757253360219 | 3.42430451547869 | 0.000616375115028594 | 0.034914717253013 | 2943 | 0 | 0 | 1 | 0 | 1 | 3 | 0 | 0 | 5 | 7 | 5 | 6 |
| LTR16C_dup4766 | 2.79763212633946 | 3.73528889405257 | 0.96218215796076 | 3.88210159910791 | 0.00010355759097903 | 0.0103120186895199 | 2944 | 0 | 0 | 1 | 0 | 1 | 3 | 0 | 0 | 5 | 10 | 7 | 7 |
| AluSx1_dup89240 | 2.20478886579679 | 3.51545934079742 | 1.04829240041225 | 3.3535102795889 | 0.000797934498116277 | 0.0413559839008442 | 2945 | 1 | 0 | 0 | 0 | 1 | 3 | 0 | 0 | 6 | 4 | 8 | 3 |
| L2c_dup119746 | 2.16341523645251 | 4.06896652857925 | 1.13704341144158 | 3.57854984922739 | 0.000345505974597271 | 0.0238918234907068 | 2946 | 0 | 0 | 0 | 0 | 1 | 2 | 0 | 0 | 7 | 2 | 5 | 9 |

| | | | | | | | | | | | | | | | | | | | |
|---|---|---|---|---|---|---|---|---|---|---|---|---|---|---|---|---|---|---|---|
| MLT2A1_dup3372 | 2.3613035438292 | 3.91127633110754 | 1.11151884711138 | 3.51885740963563 | 0.000433409584699785 | 0.0277527759754305 | 2947 | 0 | 0 | 0 | 0 | 1 | 3 | 0 | 0 | 7 | 3 | 8 | 6 |
| AluJb_dup64988 | 2.10109128729676 | 3.68375568724486 | 1.05435850109878 | 3.49383599924113 | 0.000476133412742304 | 0.0295183075704306 | 2948 | 0 | 0 | 0 | 0 | 2 | 1 | 0 | 0 | 7 | 9 | 2 | 3 |
| L2c_dup119786 | 1.81530604464029 | 3.72788930017438 | 1.11360066360567 | 3.34759974738524 | 0.000815146471026541 | 0.041892190538513 | 2949 | 0 | 0 | 0 | 0 | 2 | 1 | 0 | 0 | 4 | 6 | 3 | 6 |
| L2c_dup119791 | 2.43156413749398 | 3.18107325735192 | 0.899046032370043 | 3.53827628710629 | 0.000402748400691211 | 0.0264073769498129 | 2950 | 0 | 1 | 0 | 1 | 2 | 2 | 0 | 0 | 6 | 8 | 4 | 5 |
| L2b_dup79112 | 1.90670881433498 | 3.79727572565196 | 1.05570496060472 | 3.59690999602468 | 0.00032201984430508 | 0.0226964851519599 | 2951 | 0 | 0 | 0 | 1 | 1 | 1 | 0 | 0 | 4 | 6 | 6 | 4 |
| MER103C_dup7503 | 2.20950186914634 | 3.62982688699942 | 1.0301032298184 | 3.52375061248895 | 0.000425484512745235 | 0.027387131905178 | 2952 | 0 | 0 | 0 | 1 | 1 | 2 | 0 | 0 | 6 | 6 | 7 | 3 |
| MER41C_dup660 | 2.93080156868798 | 3.4413789191715 | 0.906637734065779 | 3.79575964011423 | 0.000147192055685645 | 0.0131421241584152 | 2953 | 0 | 0 | 0 | 2 | 2 | 2 | 0 | 0 | 8 | 9 | 5 | 7 |
| MLT1A1_dup5864 | 3.05623846857362 | 3.13173971628127 | 0.93784206873141 | 3.33930394113955 | 0.000839886143091594 | 0.0426693520431741 | 2954 | 0 | 0 | 0 | 2 | 4 | 2 | 0 | 0 | 5 | 9 | 6 | 10 |
| L2_dup48978 | 2.52537524645463 | 3.59646796009622 | 1.03190489214217 | 3.48527077202839 | 0.00049163926217691 | 0.03013487786655 | 2955 | 0 | 0 | 0 | 1 | 2 | 2 | 0 | 0 | 3 | 10 | 3 | 11 |
| MIR_dup143663 | 2.98041456039211 | 3.06212909875542 | 0.893194004513552 | 3.42829114759129 | 0.00060739368230682 | 0.0346836806167935 | 2956 | 0 | 0 | 0 | 1 | 3 | 3 | 1 | 0 | 4 | 12 | 5 | 8 |
| L2b_dup79076 | 2.70557741016755 | 3.66743004281071 | 1.01902123249787 | 3.59897313799924 | 0.00031947618342112 | 0.0225888055752586 | 2957 | 1 | 0 | 0 | 0 | 4 | 1 | 0 | 0 | 5 | 6 | 9 | 7 |
| AluSx_dup90134 | 3.05342692130457 | 3.10536057541815 | 0.927896724463437 | 3.34666616827842 | 0.000817896427821833 | 0.041947056743146 | 2958 | 1 | 0 | 0 | 0 | 5 | 2 | 1 | 0 | 5 | 6 | 8 | 10 |
| MER113_dup2542 | 2.47693684006089 | 3.27290424551803 | 0.937859135504235 | 3.4897610116666 | 0.000483452685453408 | 0.0298483738175653 | 2959 | 1 | 0 | 0 | 0 | 3 | 1 | 1 | 0 | 4 | 5 | 7 | 9 |
| AluJb_dup99998 | 3.82793509723009 | 3.45531212488869 | 0.969848514961639 | 3.56273384099099 | 0.000367012736597715 | 0.0248716943004123 | 2960 | 1 | 0 | 0 | 0 | 5 | 6 | 0 | 0 | 7 | 10 | 7 | 11 |
| L1PA7_dup7761 | 3.32758201973522 | 3.64556746588861 | 0.847365295835374 | 4.30223834254934 | 1.69081286799973e-05 | 0.00273502487212144 | 2961 | 2 | 0 | 0 | 0 | 2 | 3 | 0 | 0 | 7 | 14 | 5 | 7 |
| MER113_dup1928 | 3.95997306433563 | 3.19334667518643 | 0.86424328477368 | 3.69496267017299 | 0.00021991909525639 | 0.0174779121709676 | 2962 | 0 | 1 | 0 | 1 | 1 | 1 | 6 | 0 | 6 | 10 | 17 | 6 |
| AluSc_dup28157 | 3.9734818995484 | 3.12963254463731 | 0.813971676762609 | 3.84489120934124 | 0.00012060593340633 | 0.0115066958421421 | 2963 | 0 | 1 | 0 | 0 | 3 | 3 | 3 | 0 | 11 | 9 | 11 | 6 |
| MIRb_dup124345 | 4.47790352696375 | 3.26241161166467 | 0.753242409654398 | 4.33115763245662 | 1.48327442666438e-05 | 0.00249764515492702 | 2964 | 0 | 1 | 0 | 1 | 4 | 1 | 3 | 0 | 12 | 14 | 10 | 7 |
| MER21C_dup4430 | 3.49517917173121 | 3.54975183812701 | 1.00675797534075 | 3.52592373248949 | 0.000422008480265959 | 0.0272343337277868 | 2965 | 0 | 0 | 0 | 0 | 2 | 5 | 1 | 0 | 7 | 8 | 14 | 5 |
| MLT1C_dup15989 | 3.67243669911643 | 3.81875745228286 | 0.929973129077944 | 4.10630945441306 | 4.02030807111914e-05 | 0.00514909149570539 | 2966 | 0 | 0 | 0 | 0 | 1 | 3 | 2 | 0 | 10 | 12 | 11 | 4 |
| Charlie7_dup2289 | 3.60022821536058 | 3.09934955957322 | 0.92627486498184 | 3.34603655647504 | 0.000819755873721459 | 0.0420020738864182 | 2967 | 0 | 0 | 0 | 1 | 4 | 2 | 2 | 0 | 10 | 10 | 11 | 2 |
| MIRb_dup183945 | 2.49103697803431 | 3.13936018958501 | 0.94368282627841 | 3.32671115989857 | 0.000878774106757214 | 0.0438458342643011 | 2968 | 0 | 0 | 0 | 1 | 2 | 1 | 2 | 0 | 5 | 9 | 7 | 3 |
| MIR_dup143689 | 2.91420874902581 | 3.71847612850796 | 0.912253629499799 | 4.07614287108602 | 4.57888916507402e-05 | 0.00563063752595713 | 2969 | 0 | 0 | 0 | 1 | 2 | 1 | 1 | 0 | 6 | 11 | 8 | 5 |
| UCON78_dup159 | 3.15381395723159 | 3.80908307896328 | 0.888097697358269 | 4.2890360939948 | 1.79450211960382e-05 | 0.0028638135570171 | 2970 | 0 | 0 | 0 | 1 | 2 | 1 | 1 | 0 | 9 | 11 | 7 | 5 |
| AluSz6_dup1822 | 2.27947528434872 | 3.11404901176568 | 0.920025036877418 | 3.38474376994654 | 0.000712447261796644 | 0.0382401208062598 | 2971 | 1 | 1 | 0 | 0 | 2 | 1 | 1 | 0 | 5 | 7 | 2 | 8 |
| LTR9D_dup389 | 2.0461107618653 | 3.3735148198463 | 0.973064627348702 | 3.46689698200014 | 0.000526503634385738 | 0.0314851555133322 | 2972 | 0 | 1 | 0 | 0 | 1 | 1 | 1 | 0 | 6 | 8 | 2 | 4 |
| MLT1E2_dup3355 | 2.39877301934742 | 3.39818258585809 | 1.01673359741537 | 3.34225464221561 | 0.000831007868196121 | 0.0423609197856822 | 2973 | 0 | 0 | 0 | 0 | 1 | 3 | 1 | 0 | 5 | 11 | 2 | 6 |
| AluSx_dup90123 | 2.21858602841349 | 4.05301866938573 | 1.01011038129096 | 4.01245125726342 | 6.00914894870609e-05 | 0.00692422738861414 | 2974 | 0 | 0 | 0 | 0 | 1 | 1 | 1 | 0 | 5 | 6 | 6 | 7 |
| L2a_dup12154 | 2.35248882230927 | 3.45514156569586 | 0.979943531444404 | 3.5258578222391 | 0.000422113516462044 | 0.0272343337277868 | 2975 | 0 | 0 | 0 | 0 | 2 | 2 | 1 | 0 | 4 | 7 | 6 | 7 |
| L2_dup48952 | 2.13316653301884 | 3.59830726946222 | 1.05564870112946 | 3.40862188871384 | 0.000652919092836703 | 0.0362565896065629 | 2976 | 0 | 0 | 0 | 0 | 2 | 1 | 1 | 0 | 6 | 4 | 3 | 9 |
| L2a_dup142922 | 1.86397675117406 | 3.7016357613605 | 1.05949701118505 | 3.49376706331639 | 0.000476256367541889 | 0.0295183075704306 | 2977 | 0 | 0 | 0 | 0 | 1 | 1 | 1 | 0 | 6 | 5 | 3 | 5 |
| MER20_dup13031 | 2.05117415907979 | 3.45765341074749 | 1.02619531433883 | 3.36939115043148 | 0.000753344392332249 | 0.0397373719303678 | 2978 | 0 | 0 | 0 | 0 | 2 | 1 | 1 | 0 | 7 | 5 | 3 | 5 |
| LTR37-int_dup926 | 3.2992728152327 | 3.49853932453431 | 0.934468731232946 | 3.74388056828644 | 0.000181199808611256 | 0.0153314927761662 | 2979 | 0 | 0 | 0 | 0 | 4 | 2 | 1 | 0 | 12 | 7 | 6 | 6 |
| AluSx1_dup89259 | 3.92362741494536 | 3.70066562680812 | 0.919077087659841 | 4.02650188596342 | 5.66127855880241e-05 | 0.0066484690352294 | 2980 | 0 | 0 | 0 | 0 | 2 | 5 | 1 | 0 | 9 | 10 | 7 | 14 |
| L1MA1_dup397 | 3.0578358210385 | 4.19079683911909 | 1.01158056520926 | 4.14282063460972 | 3.43060249444861e-05 | 0.00461110957597297 | 2981 | 0 | 0 | 0 | 0 | 1 | 2 | 1 | 0 | 10 | 4 | 5 | 14 |
| MER41E_dup296 | 3.02587091110107 | 2.99907214146679 | 0.820971372606448 | 3.65307761212822 | 0.000259115869109937 | 0.0196043194248754 | 2982 | 0 | 1 | 1 | 0 | 3 | 1 | 2 | 0 | 7 | 7 | 5 | 10 |
| AluSz_dup29097 | 3.51538055214209 | 2.91568580516593 | 0.787761329674977 | 3.70122992248034 | 0.000214556947512608 | 0.0172181197299823 | 2983 | 0 | 2 | 0 | 0 | 3 | 2 | 3 | 0 | 7 | 10 | 7 | 9 |
| AluSz_dup80332 | 2.96655364778126 | 3.00810340542125 | 0.855456790999984 | 3.51637094598891 | 0.000437489285216904 | 0.0279406779351275 | 2984 | 0 | 1 | 0 | 0 | 3 | 2 | 2 | 0 | 6 | 11 | 4 | 7 |
| AluSp_dup39460 | 2.84208463815076 | 2.98875103491582 | 0.895184167198065 | 3.33869961560048 | 0.000841715300427323 | 0.042716510562465 | 2985 | 0 | 1 | 0 | 0 | 1 | 4 | 2 | 0 | 6 | 7 | 5 | 9 |
| L1PA7_dup9585 | 3.50001804555189 | 4.07911069610331 | 0.870710058175073 | 4.684809435477 | 2.80220525324368e-06 | 0.00072083624993332 | 2986 | 0 | 1 | 0 | 0 | 1 | 2 | 1 | 0 | 10 | 9 | 6 | 12 |
| MER20_dup13040 | 2.31866175183249 | 3.03405372940524 | 0.920828224294543 | 3.29491825875523 | 0.000984503128224591 | 0.047304144699605 | 2987 | 0 | 0 | 1 | 0 | 1 | 2 | 2 | 0 | 4 | 5 | 6 | 8 |
| MIR_dup143656 | 2.83177067876508 | 3.37614930030251 | 0.946951284220798 | 3.56528298399277 | 0.000363463805930709 | 0.02468172339212 | 2988 | 0 | 0 | 1 | 0 | 0 | 3 | 2 | 0 | 4 | 6 | 8 | 12 |
| MIRb_dup184112 | 2.21442920619838 | 3.63073673755783 | 1.03038844021788 | 3.52365825919989 | 0.00042563282780243 | 0.0273871359773599 | 2989 | 0 | 0 | 1 | 0 | 0 | 2 | 1 | 0 | 4 | 7 | 7 | 5 |
| MLT1C_dup16096 | 2.76207545467328 | 3.59339703056078 | 0.961710061609701 | 3.73646608682266 | 0.000186624605510907 | 0.0156529341744761 | 2990 | 0 | 0 | 0 | 1 | 0 | 2 | 2 | 0 | 7 | 11 | 5 | 5 |
| AluSx_dup90162 | 2.08174191629039 | 3.9547148392614 | 1.09479100669012 | 3.61230117446587 | 0.000303491788688116 | 0.0218171169641034 | 2991 | 0 | 0 | 0 | 0 | 0 | 1 | 2 | 0 | 5 | 6 | 7 | 4 |
| MLT1D_dup16968 | 2.52168933481565 | 4.66745645234103 | 1.00719348102394 | 4.63412099093015 | 3.58457311014366e-06 | 0.000856835251145652 | 2992 | 0 | 0 | 0 | 0 | 0 | 1 | 1 | 0 | 5 | 8 | 7 | 9 |
| MIR_dup143771 | 3.42190764332137 | 3.49889911983859 | 0.857882502946916 | 4.07852952801754 | 4.53214468423259e-05 | 0.0055904909293628 | 2993 | 0 | 1 | 0 | 0 | 1 | 3 | 2 | 0 | 4 | 10 | 11 | 11 |
| MIR_dup143783 | 3.36320916877011 | 3.97729363961436 | 0.97280550404349 | 4.088477730731 | 4.34213221333619e-05 | 0.00544072138828063 | 2994 | 0 | 0 | 0 | 0 | 1 | 3 | 1 | 0 | 3 | 7 | 14 | 14 |
| MLT1O_dup4109 | 2.48911400200918 | 3.82754851738296 | 1.02431300711141 | 3.73669814871995 | 0.000186452527981308 | 0.0156529341744761 | 2995 | 0 | 0 | 0 | 0 | 1 | 2 | 1 | 0 | 2 | 6 | 9 | 11 |
| MLT1C_dup16099 | 3.47880340472996 | 3.38687659504643 | 0.968540387282833 | 3.49688731571437 | 0.000470720621788208 | 0.0292836658563152 | 2996 | 1 | 0 | 0 | 0 | 1 | 4 | 2 | 0 | 3 | 13 | 15 | 4 |
| THE1B_dup19557 | 2.57606709102357 | 3.91653730323361 | 0.957003827523738 | 4.09249910041396 | 4.26748900531712e-05 | 0.00537549994102269 | 2997 | 1 | 0 | 0 | 0 | 1 | 1 | 1 | 0 | 4 | 7 | 9 | 8 |
| L1MB7_dup18715 | 3.19802298811478 | 3.20757304237475 | 0.871485327616722 | 3.68058180755217 | 0.000232702384497246 | 0.0181960570429122 | 2998 | 2 | 0 | 0 | 0 | 2 | 2 | 2 | 0 | 5 | 10 | 12 | 4 |
| MIRb_dup122104 | 2.6920052440908 | 3.00816323201375 | 0.901950185724942 | 3.33517668672128 | 0.000852452172081036 | 0.0430470255810684 | 2999 | 1 | 0 | 0 | 0 | 3 | 3 | 1 | 0 | 4 | 9 | 7 | 5 |
| MER52A_dup1359 | 3.26272573822461 | 3.66478973546439 | 0.948553307973681 | 3.86355696053941 | 0.000111747820434835 | 0.0108796836666069 | 3000 | 0 | 0 | 0 | 3 | 0 | 2 | 1 | 0 | 2 | 8 | 10 | 17 |
| THE1D-int_dup2394 | 2.95667420274537 | 4.11640966188746 | 1.00104433448984 | 4.1121152381181 | 3.92050509728306e-05 | 0.00505619786383802 | 3001 | 0 | 0 | 0 | 3 | 0 | 1 | 0 | 0 | 5 | 4 | 9 | 16 |
| L1ME2_dup8358 | 3.07915552802745 | 3.25658854797213 | 0.892377721499256 | 3.64933869314984 | 0.00026291627224222 | 0.0197724005420188 | 3002 | 0 | 1 | 0 | 1 | 1 | 1 | 3 | 0 | 2 | 7 | 6 | 19 |
| L2c_dup119765 | 3.33209242988445 | 3.1979275329849 | 0.8023151221157 | 3.985874682945 | 6.72319912801312e-05 | 0.00752473348877803 | 3003 | 1 | 2 | 0 | 2 | 0 | 2 | 1 | 0 | 4 | 8 | 7 | 16 |
| MIRb_dup184038 | 2.44898304165448 | 3.18298658974221 | 0.951488931582839 | 3.3452691713893 | 0.000822027511585515 | 0.0420822199379791 | 3004 | 0 | 2 | 0 | 1 | 0 | 2 | 1 | 0 | 4 | 7 | 3 | 11 |
| THE1B_dup15421 | 3.60052419094221 | 3.63222691456653 | 0.997449995415645 | 3.64151278887214 | 0.000271040658290162 | 0.0201982588404356 | 3005 | 0 | 1 | 0 | 0 | 1 | 0 | 5 | 0 | 12 | 5 | 9 | 10 |
| MLT1K_dup15305 | 3.28661849435585 | 4.29403230486999 | 0.943457476856236 | 4.55137874276904 | 5.329551010201e-06 | 0.00115076710042948 | 3006 | 0 | 2 | 0 | 0 | 1 | 0 | 1 | 0 | 5 | 10 | 9 | 13 |
| MSTA_dup5773 | 3.22185364445281 | 3.57422387995382 | 0.868471224687072 | 4.11553518222976 | 3.86282067543308e-05 | 0.00500077654209249 | 3007 | 0 | 2 | 1 | 0 | 1 | 0 | 2 | 0 | 5 | 6 | 8 | 16 |
| THE1D_dup2602 | 3.94235768225031 | 3.08840997211836 | 0.763559196085415 | 4.04475512566923 | 5.23778420740487e-05 | 0.006256575306301 | 3008 | 1 | 0 | 0 | 2 | 4 | 1 | 2 | 0 | 8 | 6 | 12 | 13 |
| UCON56_dup12 | 3.2296492062747 | 3.1292414220154 | 0.868083212113799 | 3.60477126886908 | 0.000312427966857475 | 0.0222601554817835 | 3009 | 0 | 0 | 0 | 2 | 2 | 3 | 1 | 0 | 5 | 6 | 14 | 7 |
| L1MC4_dup20736 | 3.01129463214985 | 3.02292040494445 | 0.881262819098231 | 3.43021439170407 | 0.000603104518882947 | 0.0345382209025564 | 3010 | 0 | 0 | 0 | 1 | 2 | 3 | 2 | 0 | 8 | 4 | 6 | 11 |
| L1PA7_dup11883 | 2.77463954435751 | 2.84565831048839 | 0.869649477775549 | 3.27218998367851 | 0.00106717838519265 | 0.0499470550832123 | 3011 | 0 | 0 | 0 | 1 | 2 | 2 | 3 | 0 | 5 | 4 | 8 | 10 |
| MIRb_dup76263 | 3.38760042561608 | 2.74203609284808 | 0.827697313942173 | 3.31284884783334 | 0.000923508754497013 | 0.0453456900737817 | 3012 | 0 | 0 | 1 | 3 | 3 | 0 | 4 | 0 | 7 | 7 | 9 | 8 |
| L2a_dup142606 | 2.92489793838035 | 3.72022799279908 | 0.9413607782909 | 3.95196834050532 | 7.75109971749109e-05 | 0.00834642133290565 | 3013 | 0 | 1 | 0 | 1 | 2 | 0 | 1 | 0 | 10 | 5 | 9 | 5 |
| L1MB3_dup14047 | 2.62684271612201 | 3.97273814162301 | 0.959690461409876 | 4.13960365489794 | 3.47906376840592e-05 | 0.00466382708415276 | 3014 | 0 | 0 | 0 | 2 | 1 | 0 | 1 | 0 | 5 | 6 | 7 | 11 |
| MER57F_dup294 | 3.49141082252214 | 3.48941398376671 | 0.909644281309656 | 3.83602036033563 | 0.000125044046012443 | 0.0117836556722681 | 3015 | 0 | 0 | 0 | 1 | 4 | 0 | 2 | 0 | 7 | 8 | 10 | 11 |
| L1MD_dup7450 | 2.86346064362405 | 4.07900480260703 | 0.939689995727056 | 4.34079837090425 | 1.41965949993991e-05 | 0.00242848525789145 | 3016 | 0 | 0 | 0 | 1 | 1 | 0 | 1 | 1 | 4 | 9 | 6 | 13 |
| MLT1L_dup9901 | 3.51308587837713 | 3.44129908944695 | 0.929547853179651 | 3.70212149667765 | 0.000213804183728442 | 0.0171755065163565 | 3017 | 0 | 0 | 0 | 1 | 4 | 0 | 1 | 1 | 4 | 9 | 6 | 19 |
| L1MA6_dup4633 | 3.11855521168873 | 2.84484703094166 | 0.83968560976065 | 3.38799069303161 | 0.000704066533530495 | 0.0379533299781096 | 3018 | 0 | 0 | 2 | 2 | 3 | 0 | 1 | 1 | 3 | 12 | 6 | 9 |

| | | | | | | | | | | | | | | | | | | | |
|---|---|---|---|---|---|---|---|---|---|---|---|---|---|---|---|---|---|---|---|
| L1ME1_dup26732 | 2.46521469473275 | 3.39593446026389 | 1.01196791553897 | 3.35577285417713 | 0.000791435425738884 | 0.0411800146310016 | 3019 | 0 | 0 | 1 | 0 | 3 | 0 | 0 | 1 | 4 | 8 | 5 | 8 |
| L1MA9_dup15261 | 2.95325140032093 | 3.17271477689567 | 0.884100847601868 | 3.58863447026625 | 0.000332414524972741 | 0.0232227558893611 | 3020 | 0 | 0 | 3 | 1 | 1 | 1 | 1 | 0 | 6 | 12 | 3 | 8 |
| Charlie18a_dup2059 | 3.02029226976522 | 3.11782394000083 | 0.910334114080785 | 3.42492266495919 | 0.000614974445995783 | 0.0348988098084695 | 3021 | 0 | 0 | 1 | 0 | 2 | 2 | 1 | 1 | 3 | 17 | 3 | 7 |
| L2a_dup142879 | 3.29651628652079 | 3.3537962416937 | 0.849400626307843 | 3.94842685279372 | 7.86664449216047e-05 | 0.00842790000149309 | 3022 | 0 | 0 | 1 | 0 | 1 | 1 | 3 | 1 | 6 | 14 | 5 | 8 |
| MIR1_Amn_dup1051 | 3.12539561322431 | 2.65605572698822 | 0.754285654328927 | 3.52128628158951 | 0.000429458695133355 | 0.027573658573179 | 3023 | 2 | 0 | 1 | 1 | 3 | 2 | 0 | 1 | 5 | 10 | 7 | 6 |
| L1MC4a_dup1496 | 2.22069304754094 | 2.94256801271501 | 0.887589983935878 | 3.31523345910986 | 0.00091566617315938 | 0.0450628962230779 | 3024 | 1 | 0 | 0 | 1 | 1 | 1 | 1 | 1 | 3 | 6 | 4 | 9 |
| L1MC4a_dup1925 | 2.69168828747659 | 2.95612792797302 | 0.856997531124328 | 3.44940075159232 | 0.000561832217587649 | 0.0329518680912277 | 3025 | 2 | 0 | 1 | 0 | 1 | 1 | 1 | 1 | 8 | 7 | 2 | 8 |
| AluSc_dup32567 | 3.48045728249135 | 2.94043689992025 | 0.832445334965729 | 3.53228827937549 | 0.000411979973642088 | 0.026820516226047 | 3026 | 0 | 0 | 0 | 1 | 1 | 1 | 2 | 4 | 4 | 3 | 10 | 18 |
| MIR_dup143726 | 3.8238439636557 | 2.76238757868829 | 0.810581458028665 | 3.40790867016181 | 0.000654628011248846 | 0.0362837448787836 | 3027 | 0 | 0 | 0 | 1 | 4 | 2 | 2 | 2 | 3 | 9 | 8 | 18 |
| AluSz_dup80386 | 4.05218962316321 | 2.7235307903948 | 0.753779284473737 | 3.61316747023139 | 0.000302479199337982 | 0.0217645622649209 | 3028 | 0 | 0 | 0 | 3 | 2 | 4 | 1 | 2 | 11 | 8 | 9 | 8 |
| MSTA-int_dup2778 | 3.22727283474067 | 2.49870743237088 | 0.750906152606774 | 3.32758950462265 | 0.000876008435305589 | 0.0437720922190726 | 3029 | 0 | 0 | 1 | 2 | 3 | 2 | 2 | 1 | 7 | 5 | 5 | 12 |
| L1PA6_dup111 | 3.11712668772142 | 2.96964951899373 | 0.888763463216316 | 3.34132718310333 | 0.000833789045891762 | 0.0424521638085235 | 3030 | 0 | 1 | 0 | 0 | 1 | 3 | 1 | 2 | 10 | 3 | 9 | 6 |
| MIR_dup143755 | 3.21571195663649 | 3.54974074970163 | 0.893795885754925 | 3.9715340004093 | 7.1411306088696e-05 | 0.00788127766167944 | 3031 | 0 | 0 | 0 | 0 | 2 | 2 | 1 | 1 | 6 | 5 | 14 | 8 |
| MIRb_dup184077 | 3.85767401666538 | 3.45648941138343 | 0.865402852843656 | 3.9940813691862 | 6.49455589055313e-05 | 0.00735182490929379 | 3032 | 0 | 0 | 0 | 0 | 2 | 4 | 1 | 1 | 8 | 6 | 13 | 12 |
| AluJo_dup60238 | 3.20623378212514 | 3.70082186143244 | 1.01533543792619 | 3.64492533520875 | 0.000267469442649368 | 0.0200199068891621 | 3033 | 0 | 0 | 0 | 0 | 4 | 1 | 0 | 1 | 7 | 6 | 13 | 6 |
| HAL1_dup19465 | 2.90097548694247 | 4.08248348346626 | 0.960749983808018 | 4.24926729354183 | 2.14470880402503e-05 | 0.00326086185995232 | 3034 | 0 | 0 | 0 | 0 | 1 | 2 | 0 | 1 | 7 | 7 | 6 | 11 |
| L1M5_dup46214 | 2.59264945878395 | 3.4519567914422 | 0.996359135127033 | 3.46457082566126 | 0.000531078274025241 | 0.0317101233846599 | 3035 | 0 | 0 | 0 | 0 | 2 | 1 | 0 | 2 | 8 | 4 | 6 | 7 |
| L1MB1_dup5250 | 2.73288132680464 | 3.0233848186669 | 0.904047819592087 | 3.34427532830185 | 0.000824978197626078 | 0.0421607092380444 | 3036 | 0 | 0 | 0 | 2 | 1 | 3 | 0 | 1 | 5 | 8 | 8 | 5 |
| L2a_dup142945 | 2.60989890817626 | 3.4646825569039 | 0.987656012862561 | 3.50798507960486 | 0.000451514372273306 | 0.0285127491573631 | 3037 | 0 | 0 | 0 | 1 | 1 | 2 | 0 | 1 | 3 | 5 | 14 | 5 |
| L1PA6_dup5305 | 2.06735701720783 | 3.47510326252148 | 1.01088079927135 | 3.43769835674626 | 0.000586680847694459 | 0.0339077874630813 | 3038 | 0 | 0 | 1 | 0 | 1 | 1 | 0 | 1 | 3 | 7 | 6 | 5 |
| AluSz_dup80330 | 1.86506052785928 | 3.7536938166048 | 1.06388343003615 | 3.52829427607238 | 0.000418246908361449 | 0.0270784044164462 | 3039 | 0 | 0 | 0 | 0 | 1 | 1 | 0 | 1 | 3 | 5 | 4 | 8 |
| MIRb_dup184056 | 2.32101228022436 | 3.40128447722734 | 0.987866162804926 | 3.44306203136851 | 0.000575167370490627 | 0.033478173957623 | 3040 | 0 | 0 | 0 | 0 | 2 | 2 | 0 | 1 | 4 | 7 | 6 | 6 |
| MLT2B1_dup3832 | 4.83769871941739 | 2.6889357589708 | 0.814694921341444 | 3.30054317086365 | 0.000964978662458682 | 0.046677252330751 | 3041 | 0 | 0 | 0 | 1 | 8 | 2 | 3 | 1 | 10 | 15 | 10 | 8 |
| L2b_dup79097 | 5.00665598434667 | 2.84751838700358 | 0.822838038534329 | 3.46060616263644 | 0.000538960674197883 | 0.0320608841456723 | 3042 | 0 | 0 | 0 | 0 | 4 | 4 | 4 | 2 | 7 | 18 | 7 | 16 |
| MSTA_dup16904 | 5.01485625629632 | 3.38356685671513 | 0.890988807032686 | 3.79754137202197 | 0.000146138391235158 | 0.0130774898987743 | 3043 | 0 | 0 | 0 | 0 | 7 | 3 | 1 | 1 | 7 | 18 | 8 | 17 |
| L1MB5_dup7505 | 4.40292317118644 | 3.39286278498682 | 0.749458140155578 | 4.52708777608648 | 5.98021045651566e-06 | 0.0012526119342307 | 3044 | 0 | 0 | 0 | 1 | 3 | 1 | 2 | 2 | 6 | 11 | 17 | 11 |
| THE1A-int_dup1015 | 4.15655272268698 | 2.70202124183475 | 0.823383593366579 | 3.2816068520226 | 0.00103217408654983 | 0.0488856877384926 | 3045 | 0 | 0 | 0 | 1 | 3 | 3 | 1 | 4 | 5 | 11 | 15 | 7 |
| AluSx_dup90112 | 3.26414377225945 | 2.80653539469769 | 0.835712019750186 | 3.35825658644545 | 0.000784357683064317 | 0.0409505595573397 | 3046 | 0 | 0 | 0 | 2 | 4 | 1 | 1 | 1 | 7 | 13 | 4 | 6 |
| AluJb_dup99996 | 3.20210093364408 | 2.83251960448318 | 0.86288904900422 | 3.28260001416396 | 0.00102854492548013 | 0.0487327202425138 | 3047 | 0 | 0 | 0 | 0 | 4 | 2 | 2 | 1 | 7 | 11 | 6 | 5 |
| MER5A_dup29079 | 2.80191172620151 | 2.82520001225919 | 0.830032154230356 | 3.40372357608103 | 0.000664739872475868 | 0.0366258787677276 | 3048 | 0 | 0 | 0 | 1 | 2 | 2 | 2 | 1 | 5 | 7 | 9 | 5 |
| MIRc_dup55665 | 3.81425642930246 | 3.04350637011022 | 0.857479394976907 | 3.54936385403428 | 0.000386163093916457 | 0.0257311575315164 | 3049 | 0 | 0 | 1 | 0 | 3 | 5 | 0 | 1 | 12 | 10 | 6 | 6 |
| L1M3c_dup364 | 4.92096236152699 | 3.27457970916433 | 0.959347751795647 | 3.41333963939059 | 0.000641719123540023 | 0.035843779007008 | 3050 | 0 | 0 | 1 | 0 | 4 | 7 | 0 | 1 | 8 | 8 | 9 | 24 |
| L2b_dup79080 | 4.90817448811937 | 3.53794088145306 | 0.760453722920588 | 4.65240786495894 | 3.28081341182593e-06 | 0.000808970391803454 | 3051 | 1 | 0 | 1 | 0 | 5 | 2 | 0 | 1 | 10 | 12 | 10 | 18 |
| AluSx4_dup8702 | 3.97923794175109 | 3.4605859483405 | 1.03207969717365 | 3.35302201740553 | 0.000799343475723843 | 0.0413968933546084 | 3052 | 0 | 0 | 0 | 0 | 5 | 4 | 0 | 1 | 4 | 11 | 15 | 9 |
| L1MC_dup275 | 4.26692311593287 | 3.4304973052454 | 0.928364199604517 | 3.69520637127842 | 0.000219708259031519 | 0.0174779121709676 | 3053 | 0 | 0 | 0 | 0 | 4 | 3 | 0 | 2 | 5 | 11 | 12 | 16 |
| MIR_dup143757 | 2.97834571390573 | 3.40472525323516 | 0.972428131684746 | 3.5012615763557 | 0.000463061101087422 | 0.0289880391368976 | 3054 | 0 | 0 | 0 | 0 | 1 | 3 | 0 | 2 | 4 | 9 | 10 | 7 |
| MLT1F2_dup5008 | 3.56041083061456 | 3.32435912545238 | 0.895302769670018 | 3.71311162890476 | 0.000204726531585097 | 0.016694649047495 | 3055 | 0 | 0 | 1 | 0 | 2 | 4 | 0 | 1 | 5 | 12 | 12 | 6 |
| L1MB5_dup7499 | 4.79637272700491 | 4.50825713548316 | 0.821108795375189 | 5.49045042614993 | 4.00910008330343e-08 | 2.43354420515744e-05 | 3056 | 0 | 0 | 0 | 1 | 2 | 1 | 0 | 1 | 11 | 17 | 13 | 11 |
| L2a_dup142864 | 3.07066534048061 | 3.2784881003354 | 0.884131426265367 | 3.70814564773923 | 0.000208782564962246 | 0.0169437163037199 | 3057 | 0 | 0 | 0 | 1 | 3 | 2 | 0 | 1 | 4 | 10 | 7 | 10 |
| MER63B_dup1318 | 3.71033457033433 | 3.84532587574883 | 0.887857780659609 | 4.33101557424215 | 1.48423182111353e-05 | 0.00249764515492702 | 3058 | 0 | 0 | 0 | 0 | 3 | 2 | 0 | 1 | 7 | 19 | 5 | 7 |
| Charlie1_dup614 | 7.32575365597125 | 3.4663086138094 | 0.812362980382842 | 4.26694556191598 | 1.98167491052724e-05 | 0.0030709302228926 | 3059 | 0 | 2 | 6 | 0 | 3 | 0 | 0 | 3 | 21 | 19 | 11 | 21 |
| L1PA6_dup349 | 5.83824387018085 | 2.79648547067001 | 0.566761497571878 | 4.93414863686175 | 8.05011199989525e-07 | 0.000271700985805259 | 3060 | 2 | 1 | 4 | 4 | 2 | 1 | 1 | 2 | 12 | 17 | 10 | 15 |
| MIRc_dup21225 | 5.67391375153434 | 2.8780424038362 | 0.709734925005859 | 4.05509480009377 | 5.0113974169903e-05 | 0.00604380116463849 | 3061 | 2 | 1 | 2 | 2 | 3 | 0 | 1 | 4 | 13 | 19 | 3 | 18 |
| LTR33_dup3035 | 5.0389346843059 | 3.08843512696103 | 0.701810999999225 | 4.40066503227285 | 1.07919612113166e-05 | 0.00197683141061428 | 3062 | 0 | 2 | 2 | 1 | 3 | 2 | 0 | 1 | 18 | 19 | 3 | 6 |
| MER113_dup2554 | 2.83697976920421 | 2.76920977682802 | 0.813622934661167 | 3.40355422500627 | 0.000665152093215622 | 0.0366258787677276 | 3063 | 0 | 1 | 1 | 2 | 1 | 2 | 0 | 1 | 6 | 11 | 3 | 6 |
| L1PA3_dup8392 | 3.53451073777825 | 3.18515463391934 | 0.797230952902504 | 3.99527216338382 | 6.46199752831732e-05 | 0.00732542383931869 | 3064 | 0 | 1 | 1 | 1 | 2 | 2 | 0 | 1 | 9 | 12 | 9 | 3 |
| MIRb_dup174545 | 3.90408576413292 | 2.43603980434278 | 0.710524668251226 | 3.42850841525104 | 0.000606907719804407 | 0.0346803590223921 | 3065 | 0 | 1 | 2 | 3 | 3 | 2 | 0 | 2 | 9 | 12 | 7 | 5 |
| AluJr_dup27806 | 3.72743715303149 | 2.88613461832738 | 0.756064018238971 | 3.81731513298276 | 0.000134911805412302 | 0.0123897045351739 | 3066 | 1 | 2 | 1 | 0 | 4 | 2 | 0 | 1 | 8 | 5 | 11 | 10 |
| LTR22B_dup11 | 4.11382163109596 | 2.51822871893291 | 0.757606618693932 | 3.32392650327446 | 0.000887595846330715 | 0.0441209958822042 | 3067 | 0 | 5 | 2 | 1 | 3 | 2 | 0 | 1 | 6 | 5 | 13 | 13 |
| MER58A_dup7345 | 4.94713376019862 | 2.84905689025069 | 0.763001794519295 | 3.73401073328491 | 0.000188454457252402 | 0.0157506091624095 | 3068 | 0 | 3 | 1 | 0 | 3 | 6 | 0 | 2 | 7 | 10 | 13 | 16 |
| L1ME3C_dup1855 | 4.52964334792836 | 3.43338084903874 | 0.827733459572804 | 4.14793048333544 | 3.35494229484665e-05 | 0.00453061917871499 | 3069 | 0 | 2 | 0 | 0 | 5 | 3 | 0 | 1 | 10 | 9 | 9 | 16 |
| MamRTE1_dup4066 | 3.42774556397149 | 3.21915279456799 | 0.864787809629295 | 3.72247707324636 | 0.000197277826904235 | 0.0162893168114593 | 3070 | 0 | 1 | 0 | 0 | 3 | 4 | 0 | 1 | 8 | 7 | 8 | 9 |
| MER31-int_dup22 | 5.54553005928965 | 5.15398109937018 | 0.899293661564821 | 5.73114358484632 | 9.97557762215611e-09 | 7.78245505862806e-06 | 3071 | 0 | 0 | 0 | 0 | 0 | 0 | 3 | 1 | 12 | 22 | 12 | 17 |
| MLT2B1_dup83 | 5.30646715779784 | 4.89397592491111 | 0.905397397272855 | 5.4053346515599 | 6.46875015799266e-08 | 3.52222705971104e-05 | 3072 | 0 | 0 | 0 | 0 | 1 | 0 | 2 | 1 | 12 | 29 | 5 | 13 |
| LTR33_dup7661 | 4.88183691978449 | 3.75504337312086 | 0.831161263996216 | 4.51782768973935 | 6.24772932664017e-06 | 0.00129836000205827 | 3073 | 1 | 0 | 0 | 0 | 3 | 0 | 3 | 1 | 18 | 9 | 13 | 8 |
| MIR3_dup31965 | 4.96159688668696 | 4.86842907695473 | 0.865296500561403 | 5.62631314676079 | 1.84102005439598e-08 | 1.31946794537141e-05 | 3074 | 1 | 0 | 0 | 1 | 1 | 0 | 1 | 0 | 19 | 12 | 12 | 10 |
| THE1A_dup111 | 6.36698989222534 | 5.98462801718978 | 0.942499482066811 | 6.34974143865423 | 2.15677121165679e-10 | 2.77402747421019e-07 | 3075 | 0 | 0 | 0 | 0 | 1 | 0 | 0 | 1 | 30 | 9 | 15 | 15 |
| L1PA8A_dup1780 | 6.5872021897527 | 6.45823913241757 | 0.894240561084215 | 7.22203779773457 | 5.12142321571544e-13 | 1.29640656221981e-09 | 3076 | 0 | 0 | 0 | 0 | 0 | 0 | 0 | 1 | 18 | 23 | 10 | 27 |
| L2b_dup4668 | 4.36157130251357 | 5.05066686171263 | 0.949203762002774 | 5.32095116337926 | 1.03226113202057e-07 | 5.25958045652603e-05 | 3077 | 1 | 0 | 0 | 0 | 0 | 0 | 0 | 2 | 14 | 13 | 6 | 15 |
| L3_dup1452 | 4.20984068695907 | 5.74131606969549 | 0.942594858634622 | 6.09096900656977 | 1.12229279577981e-09 | 1.20834878545078e-06 | 3078 | 0 | 0 | 0 | 0 | 0 | 0 | 0 | 1 | 13 | 17 | 6 | 12 |
| MER31-int_dup20 | 4.34863587119236 | 5.81204505021572 | 0.929140368724894 | 6.25529279089648 | 3.96770820998856e-10 | 4.75503876091944e-07 | 3079 | 0 | 0 | 0 | 0 | 0 | 0 | 0 | 1 | 14 | 15 | 11 | 9 |
| MER61-int_dup946 | 5.81473826538681 | 5.84899052768511 | 0.874622560636271 | 6.68744529460809 | 2.27100061118151e-11 | 3.7788497302671e-08 | 3080 | 1 | 0 | 0 | 1 | 0 | 0 | 0 | 0 | 15 | 26 | 11 | 15 |
| Cheshire_dup378 | 6.00063849749969 | 6.63737724746513 | 0.908486765341703 | 7.30597021407204 | 2.752731528479e-13 | 7.4543454638212e-10 | 3081 | 0 | 0 | 0 | 0 | 0 | 0 | 0 | 0 | 19 | 24 | 16 | 10 |
| Tigger1_dup4829 | 6.04679452100545 | 6.67190459813396 | 0.889139095668833 | 7.50378048905294 | 6.20031021060701e-14 | 1.96711868449746e-10 | 3082 | 0 | 0 | 0 | 0 | 0 | 0 | 0 | 0 | 21 | 20 | 13 | 16 |
| L1M1_dup7940 | 6.47470316424671 | 6.74861703174192 | 0.879455030127553 | 7.67363514967118 | 1.67189199071496e-14 | 6.44277288707504e-11 | 3083 | 0 | 0 | 0 | 0 | 0 | 0 | 0 | 0 | 22 | 24 | 12 | 17 |
| MLT1L_dup4597 | 6.35726576629388 | 6.71498282496045 | 0.891906198062452 | 7.52879937323887 | 5.12089944498071e-14 | 1.69235769352136e-10 | 3084 | 0 | 0 | 0 | 0 | 0 | 0 | 0 | 0 | 24 | 21 | 11 | 17 |
| Tigger1_dup4830 | 5.3262172952669 | 6.52408783811596 | 0.959003693876564 | 6.80298509773591 | 1.02473252214751e-11 | 1.95568469034522e-08 | 3085 | 0 | 0 | 0 | 0 | 0 | 0 | 0 | 0 | 21 | 11 | 10 | 20 |
| L1MEj_dup29 | 4.21458276396267 | 6.19626730916565 | 0.990620196033313 | 6.25493739576179 | 3.97675402579984e-10 | 4.75503876091944e-07 | 3086 | 0 | 0 | 0 | 0 | 0 | 0 | 0 | 0 | 16 | 8 | 11 | 14 |
| LTR16D_dup31 | 3.87720051058146 | 6.03942170351643 | 1.00394928063815 | 6.01566415753346 | 1.7915064916817e-09 | 1.79224677015776e-06 | 3087 | 0 | 0 | 0 | 0 | 0 | 0 | 0 | 0 | 16 | 9 | 10 | 9 |
| MLT1J2_dup2569 | 5.10746761380287 | 6.08078846636415 | 0.903136913111446 | 6.73296415868431 | 1.66241024368972e-11 | 2.92535001668019e-08 | 3088 | 0 | 0 | 0 | 0 | 1 | 0 | 0 | 0 | 14 | 18 | 10 | 18 |
| MLT1J1-int_dup8 | 4.53766285540255 | 6.29577959351944 | 0.960638484909183 | 6.55374492321597 | 5.6111781471641e-11 | 8.55870125259684e-08 | 3089 | 0 | 0 | 0 | 0 | 0 | 0 | 0 | 0 | 10 | 20 | 8 | 17 |
| LTR78_dup1972 | 4.63471729764398 | 6.20249382848551 | 0.947243529577241 | 6.54794003317574 | 5.83361329818973e-11 | 8.84130541306404e-08 | 3090 | 0 | 0 | 0 | 0 | 0 | 0 | 0 | 0 | 13 | 24 | 7 | 10 |

| | | | | | | | | | | | | | | | | | | | |
|---|---|---|---|---|---|---|---|---|---|---|---|---|---|---|---|---|---|---|---|
| L2a_dup5942 | 4.62667069700749 | 6.29740134643679 | 0.929378008261804 | 6.77593109634118 | 1.23607582398651e-11 | 2.29780701573668e-08 | 3091 | 0 | 0 | 0 | 0 | 0 | 0 | 0 | 0 | 15 | 16 | 10 | 13 |
| L1MC1_dup241 | 4.08976258286937 | 6.11765522378011 | 0.951694333234148 | 6.42817237651343 | 1.29147243772785e-10 | 1.77630462813637e-07 | 3092 | 0 | 0 | 0 | 0 | 0 | 0 | 0 | 0 | 12 | 16 | 9 | 11 |
| MamRTE1_dup162 | 3.70620029503747 | 5.95795876994152 | 0.975080109661427 | 6.11022490450582 | 9.9490833731032e-10 | 1.09094220843153e-06 | 3093 | 0 | 0 | 0 | 0 | 0 | 0 | 0 | 0 | 12 | 14 | 8 | 9 |
| LTR8_dup76 | 4.18766107308204 | 4.70410192848914 | 0.90774442011577 | 5.18218765573816 | 2.19298410331165e-07 | 9.71716565077456e-05 | 3094 | 0 | 0 | 2 | 1 | 0 | 0 | 1 | 0 | 12 | 14 | 7 | 13 |
| LTR88b_dup257 | 3.7100579887915 | 5.41885710794222 | 0.992474329609817 | 5.45994686842187 | 4.76277094482597e-08 | 2.78447738387607e-05 | 3095 | 0 | 0 | 1 | 0 | 0 | 0 | 0 | 0 | 16 | 14 | 6 | 4 |
| L1HS_dup1346 | 4.23181299473976 | 5.33727170593032 | 0.930007242730942 | 5.73895714000846 | 9.52613223751833e-09 | 7.48087479006118e-06 | 3096 | 0 | 0 | 1 | 0 | 1 | 0 | 0 | 0 | 17 | 12 | 8 | 9 |
| MER21B_dup2143 | 3.83807930738878 | 5.0288771996252 | 0.944805838460097 | 5.32265677763124 | 1.02262582495897e-07 | 5.22807148722699e-05 | 3097 | 0 | 0 | 1 | 0 | 1 | 0 | 0 | 0 | 15 | 16 | 5 | 5 |
| MIR1_Amn_dup2286 | 2.71413027749176 | 3.50032470194926 | 0.945440154964998 | 3.70232286366009 | 0.000213634511511735 | 0.0171676722310609 | 3098 | 0 | 1 | 2 | 0 | 0 | 0 | 1 | 1 | 9 | 9 | 4 | 4 |
| MLT1E2_dup684 | 3.65914636700901 | 3.36122103681761 | 0.842051964164867 | 3.99170262627581 | 6.56006023984861e-05 | 0.00739782034991004 | 3099 | 0 | 2 | 3 | 0 | 0 | 0 | 2 | 1 | 6 | 13 | 6 | 12 |
| L1M5_dup10214 | 4.21119868807947 | 2.99322301802242 | 0.852747130814543 | 3.51009450499505 | 0.000447947455850734 | 0.0284081304184059 | 3100 | 0 | 1 | 1 | 0 | 0 | 3 | 2 | 3 | 8 | 20 | 3 | 9 |
| MER53_dup1144 | 4.24744836493648 | 2.85372557620933 | 0.76559921259574 | 3.72744058413261 | 0.00019343414269694 | 0.016054021806127 | 3101 | 2 | 2 | 1 | 0 | 0 | 1 | 4 | 1 | 12 | 17 | 3 | 7 |
| MIR_dup34578 | 3.44123292515823 | 2.81834094873822 | 0.854262324176702 | 3.29915164110088 | 0.000969775092805701 | 0.046817703382766 | 3102 | 1 | 3 | 1 | 0 | 0 | 0 | 2 | 2 | 6 | 17 | 3 | 6 |
| MER53_dup1145 | 4.31203379746956 | 3.44447676371964 | 0.954699577909276 | 3.60791692320929 | 0.000308665285115016 | 0.0220756447045319 | 3103 | 1 | 0 | 1 | 1 | 0 | 0 | 4 | 3 | 14 | 16 | 6 | 3 |
| HERVK11-int_dup20 | 3.88502332203327 | 3.4868981257463 | 0.878027458668831 | 3.97128596756274 | 7.14857076195004e-05 | 0.00788217710158927 | 3104 | 0 | 0 | 1 | 1 | 1 | 0 | 3 | 2 | 10 | 16 | 8 | 3 |
| LTR9B_dup16 | 3.5843095591428 | 4.18868975216314 | 0.96295414251797 | 4.3498330473042 | 1.36241218564329e-05 | 0.0023491342748194 | 3105 | 0 | 0 | 1 | 1 | 0 | 0 | 2 | 1 | 8 | 17 | 9 | 3 |
| L1MB5_dup1567 | 6.73399165274347 | 3.6020885216088 | 0.761339826142626 | 4.73124930277062 | 2.23142273642782e-06 | 0.000611702896822642 | 3106 | 0 | 2 | 1 | 1 | 0 | 1 | 5 | 2 | 20 | 28 | 9 | 9 |
| L1P3_dup2569 | 4.62330595166432 | 3.1558076949586 | 0.749225225070444 | 4.21209482724221 | 2.530132795754e-05 | 0.00369120158319118 | 3107 | 0 | 3 | 1 | 3 | 0 | 0 | 2 | 2 | 15 | 11 | 7 | 10 |
| Tigger7_dup594 | 6.0580701433139 | 4.42914346564885 | 0.768794905784862 | 5.76115090295394 | 8.35423052500429e-09 | 6.74270898513513e-06 | 3108 | 0 | 2 | 1 | 1 | 0 | 0 | 1 | 1 | 25 | 20 | 7 | 10 |
| MSTD_dup5439 | 5.22308819907792 | 2.80121840222555 | 0.703558498305786 | 3.98150034285857 | 6.84816223361551e-05 | 0.00762514183827738 | 3109 | 0 | 1 | 1 | 1 | 2 | 3 | 6 | 1 | 17 | 11 | 6 | 13 |
| L1PB3_dup3060 | 3.19440085489291 | 2.57308588719747 | 0.761049230024408 | 3.38097167132663 | 0.000722299892033702 | 0.0385030227999302 | 3110 | 0 | 1 | 1 | 1 | 2 | 2 | 2 | 1 | 9 | 9 | 2 | 8 |
| Tigger1_dup6876 | 4.21212486619881 | 2.80801728621317 | 0.701193570186672 | 4.00462497889936 | 6.21159445351965e-05 | 0.00708305811651604 | 3111 | 0 | 3 | 1 | 1 | 2 | 1 | 3 | 1 | 11 | 10 | 4 | 14 |
| MIR_dup28909 | 4.16740610636574 | 3.18784872405486 | 0.829915739289111 | 3.84117154686752 | 0.000122448500407646 | 0.0116213079607753 | 3112 | 0 | 1 | 0 | 0 | 2 | 4 | 1 | 1 | 16 | 15 | 1 | 6 |
| Tigger1_dup8169 | 4.219289257641 | 3.33892064012603 | 0.919894761697156 | 3.62967676211777 | 0.000283776351617162 | 0.02081487292903 | 3113 | 0 | 0 | 0 | 0 | 2 | 1 | 4 | 1 | 16 | 15 | 3 | 6 |
| MIR_dup64591 | 5.4405121503059 | 3.89473413905305 | 0.758136151316444 | 5.13724893911225 | 2.78789547094034e-07 | 0.000116176633227385 | 3114 | 2 | 0 | 2 | 0 | 0 | 2 | 2 | 0 | 17 | 21 | 5 | 13 |
| MLT1A0_dup422 | 4.90985902065114 | 5.14570765336837 | 0.884047885995631 | 5.82062095830158 | 5.8629394765094e-09 | 5.01821221826441e-06 | 3115 | 0 | 0 | 0 | 0 | 0 | 2 | 1 | 0 | 16 | 20 | 7 | 11 |
| THE1D_dup11065 | 3.95471305574506 | 2.9991437012127 | 0.817264184654537 | 3.66973587920099 | 0.000242801205722409 | 0.0187454820560754 | 3116 | 0 | 1 | 4 | 0 | 0 | 2 | 4 | 0 | 7 | 11 | 9 | 11 |
| L1ME3_dup4996 | 3.75279661793049 | 3.81130236912919 | 0.845955318305939 | 4.50532349245286 | 6.6271905844594e-06 | 0.00135357381185388 | 3117 | 0 | 0 | 1 | 0 | 1 | 2 | 2 | 0 | 11 | 11 | 10 | 6 |
| MLT1A0_dup17822 | 4.39703466259355 | 3.06395669882585 | 0.707754320273548 | 4.32912468502011 | 1.49703163075578e-05 | 0.00250854005923813 | 3118 | 0 | 0 | 3 | 1 | 2 | 3 | 2 | 0 | 10 | 13 | 10 | 9 |
| L3_dup15368 | 6.50090023426936 | 3.00954432108668 | 0.568322125201123 | 5.2954903348548 | 1.1869747237684e-07 | 5.88408099212076e-05 | 3119 | 3 | 5 | 1 | 2 | 5 | 2 | 0 | 0 | 18 | 20 | 8 | 13 |
| LTR16E1_dup836 | 4.92369030146068 | 2.66391891310557 | 0.662514439207157 | 4.0209220440441 | 5.79707883862137e-05 | 0.00675841117753356 | 3120 | 2 | 4 | 1 | 3 | 2 | 3 | 0 | 0 | 18 | 13 | 3 | 8 |
| MSTA_dup6485 | 5.13169334743252 | 2.758588061799 | 0.619865746945262 | 4.45029923881662 | 8.57507225954541e-06 | 0.00164681528964471 | 3121 | 4 | 2 | 3 | 1 | 2 | 3 | 0 | 0 | 15 | 18 | 6 | 6 |
| AluSx1_dup20428 | 4.38368934583011 | 2.69987161495761 | 0.693619920027489 | 3.89243667461368 | 9.92424087533884e-05 | 0.0100188095855892 | 3122 | 2 | 4 | 3 | 1 | 1 | 1 | 1 | 0 | 13 | 13 | 3 | 10 |
| MER20_dup7480 | 5.06894628270697 | 2.49193676941748 | 0.592786395217974 | 4.20376848983042 | 2.62507436323753e-05 | 0.00379423495328771 | 3123 | 3 | 3 | 3 | 2 | 3 | 2 | 1 | 0 | 12 | 18 | 4 | 10 |
| AluSx1_dup73994 | 5.02697962198341 | 2.32517475852707 | 0.689145883880528 | 3.37399498845468 | 0.000740857164928736 | 0.0392877198498179 | 3124 | 3 | 7 | 3 | 3 | 3 | 0 | 0 | 0 | 10 | 15 | 8 | 9 |
| MIRb_dup165984 | 4.70230485930236 | 2.22849576742724 | 0.676486788664764 | 3.29421919949951 | 0.000986955002649167 | 0.0473662757281885 | 3125 | 6 | 4 | 2 | 5 | 1 | 1 | 0 | 0 | 10 | 11 | 8 | 10 |
| L1PA15_dup4718 | 3.72121955754302 | 3.15073739929739 | 0.87861472428565 | 3.58602845161634 | 0.00033575234335561 | 0.0234009745436713 | 3126 | 0 | 3 | 2 | 3 | 1 | 0 | 0 | 0 | 10 | 5 | 5 | 17 |
| L1PA11_dup1797 | 4.04031573823875 | 3.12894960663161 | 0.754154985702231 | 4.14894771758101 | 3.34007076336404e-05 | 0.00452079907769863 | 3127 | 2 | 2 | 2 | 3 | 1 | 0 | 0 | 0 | 5 | 15 | 7 | 14 |
| MLT1D_dup19353 | 3.26046801331291 | 3.13096455045304 | 0.875336083980421 | 3.57687133862412 | 0.000347731216475764 | 0.0240247537850006 | 3128 | 1 | 4 | 1 | 2 | 0 | 0 | 0 | 0 | 6 | 12 | 5 | 9 |
| LTR62_dup47 | 4.36488151618092 | 2.67057050892808 | 0.699418112834592 | 3.81827473427137 | 0.000134388218392512 | 0.0123702665429882 | 3129 | 1 | 1 | 5 | 4 | 1 | 2 | 0 | 0 | 8 | 10 | 11 | 11 |
| AluSz6_dup43057 | 2.67466526433356 | 2.76903306421283 | 0.837551280058738 | 3.30610570378287 | 0.000946023921142719 | 0.0460628393317399 | 3130 | 0 | 1 | 1 | 2 | 2 | 2 | 0 | 0 | 5 | 4 | 9 | 7 |
| L1P4d_dup18 | 3.68140679274457 | 3.20818053590708 | 0.740310830355611 | 4.33355883010117 | 1.46718049813972e-05 | 0.00247771278076901 | 3131 | 1 | 2 | 1 | 1 | 2 | 2 | 0 | 0 | 9 | 6 | 8 | 13 |
| L3_dup23318 | 3.44892235875468 | 3.11820007300459 | 0.812079138834674 | 3.83977364260234 | 0.000123147800251195 | 0.0116418460383675 | 3132 | 1 | 2 | 2 | 1 | 0 | 2 | 0 | 0 | 7 | 17 | 5 | 4 |
| L1PA4_dup587 | 3.28583279504021 | 3.12897432932429 | 0.787848444987836 | 3.97154344751242 | 7.14084737234058e-05 | 0.00788127766167944 | 3133 | 2 | 1 | 1 | 2 | 0 | 2 | 0 | 0 | 9 | 10 | 7 | 5 |
| L2b_dup46217 | 3.55004081962529 | 2.48756281103792 | 0.747102991116593 | 3.32961163402665 | 0.000869671939616766 | 0.0435911370936071 | 3134 | 3 | 2 | 3 | 1 | 0 | 3 | 0 | 0 | 10 | 10 | 5 | 5 |
| MIRc_dup81770 | 3.88060085885207 | 2.93992371382785 | 0.807401911932356 | 3.64121470407684 | 0.000271354716075483 | 0.020215331644113 | 3135 | 4 | 3 | 1 | 1 | 0 | 3 | 0 | 0 | 10 | 4 | 12 | 9 |
| L2a_dup80970 | 2.77270234816298 | 3.43300864729676 | 0.862453977823685 | 3.98051227725752 | 6.87669128486991e-05 | 0.00765332640069997 | 3136 | 2 | 1 | 1 | 0 | 1 | 1 | 0 | 0 | 6 | 5 | 8 | 9 |
| MIR_dup43656 | 3.31698696731741 | 3.526456092615 | 0.8395750222016 | 4.20028704923552 | 2.66576792973442e-05 | 0.00383552353672932 | 3137 | 2 | 3 | 1 | 0 | 0 | 1 | 0 | 0 | 8 | 8 | 8 | 9 |
| HERV9NC-int_dup97 | 3.50993219733645 | 3.57995957584289 | 0.955138325866131 | 3.74810588047186 | 0.000178175041348622 | 0.0151487475375956 | 3138 | 4 | 4 | 0 | 0 | 0 | 1 | 0 | 0 | 4 | 16 | 9 | 5 |
| L1MC3_dup277 | 3.23536574762033 | 3.02953807392579 | 0.924432440406922 | 3.27718710584435 | 0.00104846865554271 | 0.049304134923274 | 3139 | 3 | 5 | 0 | 0 | 0 | 1 | 0 | 0 | 7 | 17 | 3 | 2 |
| AluSx3_dup13302 | 3.62466442517267 | 4.43695270475191 | 0.917723000705613 | 4.83474065849985 | 1.33319329070747e-06 | 0.000401046789823868 | 3140 | 1 | 1 | 0 | 1 | 1 | 0 | 0 | 0 | 8 | 13 | 13 | 5 |
| LOR1a_dup282 | 4.69737314570342 | 3.41644502287321 | 0.733977412780216 | 4.65470048994033 | 3.2445199384269e-06 | 0.000804188063821799 | 3141 | 1 | 4 | 0 | 3 | 1 | 1 | 0 | 0 | 9 | 18 | 10 | 10 |
| AluJo_dup27559 | 2.9493306934308 | 2.62485801330515 | 0.792192717463166 | 3.31340841116378 | 0.000921662876889936 | 0.0452643952331172 | 3142 | 1 | 3 | 0 | 0 | 2 | 3 | 0 | 1 | 9 | 8 | 3 | 4 |
| MIR_dup95917 | 2.65357668735793 | 2.88804569486949 | 0.851376120491098 | 3.39220894897026 | 0.000693315465885398 | 0.0375532988494348 | 3143 | 3 | 1 | 0 | 0 | 2 | 1 | 0 | 1 | 7 | 7 | 6 | 3 |
| AluSx1_dup45451 | 3.96959303797918 | 2.95462258387182 | 0.725963018994256 | 4.06993539142687 | 4.702617735856e-05 | 0.00576194170842427 | 3144 | 3 | 2 | 1 | 0 | 2 | 2 | 0 | 1 | 14 | 9 | 3 | 9 |
| L2c_dup47466 | 3.4949930494571 | 2.68700110998208 | 0.771258285478128 | 3.48391863085959 | 0.000494129688837523 | 0.0302407877932441 | 3145 | 2 | 2 | 1 | 0 | 2 | 1 | 0 | 2 | 6 | 18 | 4 | 3 |
| THE1B_dup2554 | 3.63492738644257 | 2.37676916431648 | 0.700311703559395 | 3.39387325991604 | 0.000689115768956805 | 0.0374260916699391 | 3146 | 3 | 2 | 3 | 0 | 2 | 2 | 0 | 1 | 8 | 11 | 7 | 4 |
| L1ME3A_dup209 | 3.22960601246092 | 2.82176270037118 | 0.841801247242856 | 3.35205336130502 | 0.000802145561936971 | 0.0415109456025782 | 3147 | 1 | 1 | 0 | 1 | 0 | 4 | 0 | 2 | 9 | 9 | 3 | 8 |
| AluJo_dup71260 | 3.36254148973314 | 3.39715305087362 | 0.823249473290746 | 4.12651712644807 | 3.68298587869643e-05 | 0.00483371074402814 | 3148 | 0 | 1 | 0 | 1 | 1 | 3 | 0 | 1 | 11 | 8 | 6 | 7 |
| L1MEj_dup770 | 3.96760947511078 | 2.43116078682843 | 0.686689797667358 | 3.54040615586095 | 0.00039951168764051 | 0.0262893274411805 | 3149 | 1 | 3 | 0 | 3 | 1 | 4 | 0 | 2 | 8 | 9 | 8 | 9 |
| THE1B-int_dup2524 | 3.63721726840166 | 3.18098372400909 | 0.891953349238388 | 3.56631176588466 | 0.000362040639234365 | 0.0246272504125958 | 3150 | 3 | 2 | 0 | 0 | 3 | 5 | 0 | 0 | 6 | 8 | 3 | 16 |
| L2a_dup129401 | 3.30690892392014 | 2.55399860555618 | 0.725854480709876 | 3.51860968476547 | 0.000433814444082295 | 0.027770893653916 | 3151 | 1 | 3 | 0 | 1 | 2 | 2 | 1 | 1 | 6 | 13 | 3 | 7 |
| LTR40a_dup1537 | 3.47071643812386 | 2.53097959228661 | 0.739969251798545 | 3.42038481482155 | 0.000625326108857578 | 0.0352592052839397 | 3152 | 5 | 2 | 0 | 1 | 1 | 3 | 1 | 0 | 6 | 13 | 3 | 8 |
| MIRb_dup71161 | 4.29455690707889 | 3.75488653578017 | 0.808965385673931 | 4.6415911017652 | 3.45736525858968e-06 | 0.000835194146010539 | 3153 | 1 | 1 | 0 | 0 | 2 | 5 | 0 | 0 | 14 | 10 | 9 | 8 |
| MIRc_dup32859 | 5.00334796704729 | 3.44238962920756 | 0.725787499790505 | 4.74297177920699 | 2.10605618163107e-06 | 0.00058458322848473 | 3154 | 1 | 1 | 0 | 1 | 3 | 5 | 0 | 0 | 18 | 16 | 7 | 5 |
| THE1C_dup9354 | 5.64780303101 | 3.14608743434314 | 0.792039541809946 | 3.97213430424671 | 7.12315381474e-05 | 0.00786873703622156 | 3155 | 9 | 1 | 0 | 1 | 3 | 5 | 0 | 0 | 11 | 18 | 16 | 4 |
| MIR1_Amn_dup3443 | 5.14108296622225 | 2.36167362511985 | 0.648757783827569 | 3.64030102449999 | 0.000272319481874957 | 0.0202745092097054 | 3156 | 4 | 1 | 1 | 1 | 4 | 8 | 0 | 1 | 11 | 18 | 8 | 4 |
| THE1C_dup3401 | 4.83900781298439 | 2.86486714347086 | 0.701284460788737 | 4.08517128733857 | 4.40443132016547e-05 | 0.00549046768607673 | 3157 | 6 | 1 | 1 | 0 | 3 | 6 | 0 | 0 | 9 | 16 | 8 | 9 |
| Tigger1_dup3611 | 4.65028440661053 | 3.31696068405261 | 0.995926566565759 | 3.33052736557721 | 0.000866816428598395 | 0.0434862959981602 | 3158 | 1 | 0 | 0 | 0 | 10 | 1 | 0 | 1 | 6 | 21 | 9 | 7 |
| L1MEd_dup7862 | 4.62048081534002 | 3.34907414255648 | 0.921834868379927 | 3.63305214136919 | 0.000280088391432537 | 0.0206334094079896 | 3159 | 2 | 1 | 0 | 0 | 10 | 4 | 0 | 0 | 10 | 7 | 11 | 11 |
| MLT1I_dup9170 | 4.31655243535886 | 3.5203845954975 | 0.925530629515234 | 3.80363921326016 | 0.000142585803480461 | 0.0128514096950613 | 3160 | 1 | 1 | 0 | 0 | 7 | 4 | 0 | 0 | 11 | 7 | 14 | 6 |
| L1MA1_dup1664 | 4.24359669211296 | 3.0257110041291 | 0.86203490364864 | 3.50996344964978 | 0.000448168294411544 | 0.0284091139766903 | 3161 | 1 | 1 | 0 | 0 | 7 | 4 | 0 | 1 | 7 | 10 | 14 | 6 |
| L2a_dup73069 | 4.62685859313322 | 2.66894667302849 | 0.708681071048665 | 3.7660758584664 | 0.000165833425364128 | 0.0143280320376517 | 3162 | 3 | 2 | 0 | 0 | 5 | 5 | 1 | 1 | 6 | 10 | 15 | 9 |

| | | | | | | | | | | | | | | | | | | | |
|---|---|---|---|---|---|---|---|---|---|---|---|---|---|---|---|---|---|---|---|
| L2b_dup45721 | 4.80523888765117 | 1.9518645021433 | 0.583113598533319 | 3.34731432614972 | 0.000815986297896078 | 0.0419172443089767 | 3163 | 4 | 3 | 1 | 2 | 3 | 6 | 1 | 2 | 11 | 6 | 10 | 9 |
| MSTD_dup192 | 5.51019519913104 | 2.13921259671393 | 0.606402849237187 | 3.52770868310549 | 0.000419173202494407 | 0.0271185852620177 | 3164 | 4 | 3 | 1 | 1 | 3 | 5 | 1 | 4 | 17 | 6 | 10 | 9 |
| AmnSINE1_dup2448 | 3.54173971232941 | 2.16640883666599 | 0.634645147256427 | 3.41357504430843 | 0.000641164979016974 | 0.0358409616513724 | 3165 | 2 | 2 | 2 | 1 | 2 | 2 | 1 | 2 | 7 | 7 | 6 | 9 |
| HERVK-int_dup17 | 4.17937711812729 | 2.02927621652306 | 0.593727483187701 | 3.41785798027734 | 0.000631160317044187 | 0.0354743374837916 | 3166 | 3 | 3 | 2 | 1 | 2 | 4 | 1 | 2 | 6 | 11 | 9 | 7 |
| MER1A_dup78 | 5.94896503537425 | 2.25310655285991 | 0.646456352151216 | 3.48531891652428 | 0.000491550803928472 | 0.03013487786655 | 3167 | 7 | 1 | 1 | 0 | 4 | 6 | 1 | 4 | 7 | 15 | 10 | 18 |
| L4_C_Mam_dup4800 | 4.82185957352723 | 2.10713184625994 | 0.633090487808529 | 3.32832649808698 | 0.000873694069568817 | 0.0437023353116716 | 3168 | 3 | 1 | 2 | 0 | 3 | 2 | 3 | 5 | 11 | 7 | 11 | 9 |
| MIRb_dup52755 | 5.0474164918946 | 2.18346004964496 | 0.617301493169906 | 3.53710475967371 | 0.000404539170987726 | 0.0264883875681778 | 3169 | 2 | 1 | 2 | 0 | 5 | 4 | 4 | 2 | 9 | 11 | 14 | 7 |
| L2b_dup79116 | 4.54053387857067 | 2.64420673362902 | 0.741066091414813 | 3.56811189212667 | 0.000359562960345956 | 0.0245147759777876 | 3170 | 2 | 0 | 1 | 0 | 6 | 3 | 2 | 1 | 7 | 7 | 16 | 11 |
| L3_dup26266 | 5.32386390526548 | 2.40648949083477 | 0.719598694136047 | 3.34421047515103 | 0.000825171085514492 | 0.0421615115125255 | 3171 | 7 | 0 | 1 | 0 | 6 | 7 | 1 | 1 | 8 | 12 | 11 | 12 |
| LTR16C_dup4758 | 6.04352481952362 | 2.83107348533993 | 0.646008838340065 | 4.3824067370571 | 1.17375416643392e-05 | 0.00210467301346107 | 3172 | 2 | 0 | 3 | 0 | 5 | 6 | 1 | 1 | 11 | 16 | 12 | 17 |
| MER91C_dup1287 | 4.80021289200289 | 2.49896376164316 | 0.645086871915673 | 3.87384067237665 | 0.000107133468318402 | 0.0105644344187694 | 3173 | 2 | 0 | 2 | 1 | 2 | 3 | 5 | 1 | 11 | 16 | 6 | 9 |
| L1ME2_dup5292 | 3.88661125260275 | 2.26409675888626 | 0.677745075622805 | 3.34063181028014 | 0.000835879925811128 | 0.0425169484452874 | 3174 | 1 | 0 | 1 | 1 | 3 | 4 | 4 | 1 | 8 | 7 | 7 | 11 |
| MLT1C_dup16103 | 5.20645154921438 | 2.45609251265133 | 0.602151304797599 | 4.0788627261663 | 4.52565486153229e-05 | 0.00558538498186948 | 3175 | 1 | 0 | 1 | 2 | 4 | 4 | 4 | 2 | 11 | 11 | 13 | 10 |
| AluSp_dup9434 | 4.47511112701707 | 2.30048835681216 | 0.702469049692593 | 3.2748608039299 | 0.0010571404824962 | 0.0495649949257222 | 3176 | 2 | 0 | 2 | 0 | 3 | 2 | 3 | 4 | 7 | 16 | 5 | 10 |
| L3_dup26265 | 3.86555981283875 | 2.70559103817326 | 0.762107705377199 | 3.55014260987448 | 0.00038502251893365 | 0.0256911296383024 | 3177 | 1 | 0 | 1 | 0 | 6 | 2 | 1 | 1 | 9 | 12 | 5 | 8 |
| AluSx1_dup89209 | 4.49683235303831 | 2.71837759057716 | 0.701798030985513 | 3.87344716080185 | 0.000107306680253459 | 0.0105727516934119 | 3178 | 2 | 0 | 0 | 0 | 3 | 5 | 2 | 2 | 10 | 10 | 10 | 10 |
| L2_dup10919 | 5.43004321788859 | 2.85687393378399 | 0.675539554218004 | 4.22902540043132 | 2.34705848797941e-05 | 0.00347966331820797 | 3179 | 5 | 0 | 0 | 0 | 2 | 3 | 2 | 3 | 11 | 21 | 7 | 11 |
| MIRb_dup122102 | 3.23868933441385 | 2.90323883630895 | 0.754228470037685 | 3.84928301123915 | 0.000118464072576561 | 0.0113524173231182 | 3180 | 1 | 1 | 0 | 0 | 1 | 3 | 2 | 1 | 7 | 9 | 7 | 7 |
| MIR_dup34577 | 3.45247664006707 | 2.92948656747087 | 0.748594143395559 | 3.9133175076457 | 9.10367063066975e-05 | 0.00940183164880792 | 3181 | 2 | 1 | 0 | 0 | 2 | 1 | 2 | 1 | 11 | 11 | 4 | 5 |
| Tigger13a_dup607 | 3.77867938041668 | 2.9620331393078 | 0.763597203384107 | 3.87905184327636 | 0.000104864406218559 | 0.0103834216322931 | 3182 | 4 | 1 | 0 | 0 | 1 | 1 | 1 | 2 | 10 | 11 | 9 | 4 |
| MER135_dup135 | 4.36455049226057 | 3.01291949407464 | 0.731073799871769 | 4.12122482655391 | 3.76863421416409e-05 | 0.00492168735852628 | 3183 | 4 | 1 | 2 | 0 | 1 | 0 | 2 | 1 | 9 | 17 | 10 | 5 |
| LTR79_dup1390 | 4.22467094613067 | 2.46450864363082 | 0.69312307774465 | 3.55565803933419 | 0.000377034220132345 | 0.0253071324523585 | 3184 | 3 | 2 | 2 | 0 | 1 | 0 | 2 | 4 | 9 | 10 | 9 | 8 |
| MLT1D_dup3506 | 3.41535196259563 | 2.58160466280757 | 0.740613306834506 | 3.48576597123503 | 0.000490730116810873 | 0.0301146864631114 | 3185 | 3 | 1 | 1 | 0 | 3 | 0 | 1 | 2 | 6 | 10 | 8 | 6 |
| MLT1C_dup6087 | 5.71874641414259 | 3.23767538206386 | 0.635695280130591 | 5.09312477732845 | 3.52209728728103e-07 | 0.000141326974893654 | 3186 | 5 | 0 | 2 | 0 | 2 | 2 | 1 | 1 | 17 | 18 | 9 | 10 |
| HAL1_dup12058 | 5.76241575457435 | 3.35297247086241 | 0.691790449741872 | 4.84680364135339 | 1.25466497989322e-06 | 0.000381280349049389 | 3187 | 2 | 1 | 2 | 0 | 1 | 1 | 2 | 2 | 11 | 32 | 7 | 7 |
| MER39_dup1220 | 4.51051845346464 | 2.7334724170898 | 0.640702039896196 | 4.26637071037367 | 1.9867854504825e-05 | 0.00307252317849837 | 3188 | 2 | 2 | 3 | 0 | 2 | 1 | 1 | 2 | 11 | 13 | 9 | 7 |
| LTR8_dup2572 | 2.91093028100068 | 4.24863144655827 | 1.13435645695046 | 3.74541125986098 | 0.000180098500869803 | 0.0152721731603586 | 3189 | 0 | 0 | 3 | 1 | 0 | 0 | 0 | 0 | 10 | 2 | 12 | 6 |
| CER_dup79 | 3.28178926052899 | 3.90952728792633 | 0.936871180766302 | 4.1729614147471 | 3.0066587595276e-05 | 0.00417642980265356 | 3190 | 0 | 0 | 3 | 3 | 0 | 0 | 0 | 0 | 5 | 6 | 9 | 16 |
| L1PA11_dup1794 | 2.45139302704507 | 3.95968719313896 | 1.0310304691768 | 3.84051423456038 | 0.000122776852464626 | 0.0116365308264591 | 3191 | 0 | 0 | 2 | 2 | 0 | 0 | 0 | 0 | 4 | 7 | 5 | 11 |
| L1PA12_dup1681 | 3.03144735178734 | 4.83732535309786 | 1.0111804227948 | 4.78383999932277 | 1.71977599011786e-06 | 0.000497222135777138 | 3192 | 0 | 0 | 1 | 1 | 0 | 0 | 0 | 0 | 3 | 7 | 17 | 9 |
| LTR16B_dup674 | 2.85491614670336 | 3.7832073661776 | 1.03243866632271 | 3.66434103020613 | 0.000247976199387912 | 0.0190338531417923 | 3193 | 0 | 0 | 4 | 1 | 0 | 0 | 0 | 0 | 3 | 4 | 15 | 9 |
| THE1A_dup110 | 3.54907905715453 | 5.19517771668137 | 1.00531595646391 | 5.16770641436432 | 2.36984176746324e-07 | 0.000103466856734093 | 3194 | 0 | 0 | 2 | 0 | 0 | 0 | 0 | 0 | 5 | 14 | 10 | 13 |
| L1ME1_dup2689 | 2.24489959231822 | 3.66520856932789 | 1.09312271790309 | 3.35297081407179 | 0.000799491366808393 | 0.0413968933546084 | 3195 | 0 | 1 | 3 | 0 | 0 | 0 | 0 | 0 | 3 | 6 | 7 | 8 |
| LTR103b_Mam_dup5 | 1.79106339560459 | 4.55462399923273 | 1.14532394972881 | 3.97671244045075 | 6.98745763212855e-05 | 0.00775122421321426 | 3196 | 0 | 0 | 1 | 0 | 0 | 0 | 0 | 0 | 3 | 6 | 6 | 6 |
| MIR_dup128893 | 2.02032491979296 | 4.71712986267601 | 1.11195576304374 | 4.24219201829024 | 2.21347180457148e-05 | 0.00333961820607295 | 3197 | 0 | 0 | 1 | 0 | 0 | 0 | 0 | 0 | 3 | 6 | 8 | 7 |
| MER39_dup60 | 2.72692714239151 | 5.18392489085601 | 1.01609123335037 | 5.10183015137627 | 3.3638457967032e-07 | 0.000136124770738493 | 3198 | 0 | 0 | 1 | 0 | 0 | 0 | 0 | 0 | 6 | 9 | 8 | 9 |
| THE1D-int_dup2677 | 2.66902112120657 | 5.11414535847514 | 1.03824011666655 | 4.92578284770483 | 8.40232701091907e-07 | 0.000281591563794387 | 3199 | 0 | 0 | 1 | 0 | 0 | 0 | 0 | 0 | 4 | 7 | 12 | 9 |
| MLT1H2_dup365 | 3.02601132924563 | 3.77704269878014 | 0.956725567529908 | 3.94788518982699 | 7.88445974520214e-05 | 0.00843938667805609 | 3200 | 0 | 3 | 1 | 1 | 0 | 0 | 0 | 0 | 6 | 6 | 11 | 9 |
| AluJr_dup79740 | 2.63253586985897 | 4.29280718667923 | 1.0494379202074 | 4.09057754062369 | 4.30300314318064e-05 | 0.00540297504057472 | 3201 | 0 | 2 | 1 | 0 | 0 | 0 | 0 | 0 | 9 | 5 | 5 | 9 |
| MER45R_dup130 | 3.39873334148083 | 4.30396052069388 | 0.964546205208582 | 4.46216106336052 | 8.11372021696258e-06 | 0.00156706758989073 | 3202 | 0 | 1 | 2 | 0 | 0 | 1 | 0 | 0 | 10 | 6 | 7 | 14 |
| L2b_dup79121 | 1.87638001662612 | 3.70648223044805 | 1.10581716699059 | 3.35180384342831 | 0.000802868831388413 | 0.0415252006833986 | 3203 | 0 | 0 | 1 | 0 | 0 | 1 | 0 | 1 | 5 | 2 | 8 | 4 |
| MIR3_dup72181 | 1.92515331825289 | 4.2313598308032 | 1.10057958607113 | 3.84466501501121 | 0.000120717230048316 | 0.0115080857456237 | 3204 | 0 | 0 | 1 | 0 | 0 | 1 | 0 | 0 | 6 | 3 | 5 | 7 |
| MIRb_dup124344 | 2.50158526195774 | 3.10196980199662 | 0.936575279510338 | 3.31203467554514 | 0.000926200650314833 | 0.0454029130489933 | 3205 | 0 | 0 | 1 | 1 | 0 | 3 | 0 | 1 | 6 | 7 | 4 | 7 |
| L1ME3_dup5133 | 2.75791124590073 | 3.60171441516462 | 0.972423929328342 | 3.70385210250054 | 0.000212350092053067 | 0.017110685744551 | 3206 | 0 | 0 | 1 | 0 | 0 | 3 | 0 | 1 | 6 | 7 | 8 | 7 |
| THE1B_dup19537 | 3.1176062228997 | 3.77269209849334 | 0.96624393606333 | 3.90449239336398 | 9.4423361822364e-05 | 0.00964277306960696 | 3207 | 0 | 0 | 1 | 0 | 0 | 3 | 0 | 1 | 6 | 5 | 12 | 10 |
| LTR33_dup7686 | 3.82064027199255 | 3.86663946998725 | 0.881772685416209 | 4.38507512643368 | 1.15945755853447e-05 | 0.00208374840047616 | 3208 | 1 | 0 | 3 | 0 | 0 | 2 | 0 | 0 | 7 | 13 | 8 | 13 |
| AluSq4_dup1391 | 2.98560997536402 | 3.51785921586565 | 0.869804416393129 | 4.04442556230443 | 5.24515696660574e-05 | 0.00625909789155451 | 3209 | 0 | 0 | 1 | 2 | 0 | 2 | 1 | 0 | 5 | 9 | 9 | 8 |
| LTR89_dup59 | 2.22137204895799 | 3.28241868145664 | 0.970796978819404 | 3.38115873150782 | 0.000721808327425012 | 0.0385029393461205 | 3210 | 1 | 0 | 1 | 1 | 0 | 2 | 0 | 0 | 5 | 4 | 8 | 5 |
| MLT1A1_dup5691 | 2.66298111700931 | 4.09670448148145 | 1.0985504760008 | 3.72919093931417 | 0.000192095555011826 | 0.0159707788025311 | 3211 | 0 | 0 | 0 | 3 | 0 | 0 | 1 | 0 | 10 | 8 | 4 | 5 |
| L1PA3_dup1823 | 1.99801766231155 | 3.91728907090757 | 1.1284095244037 | 3.47151365367785 | 0.000517533012643181 | 0.0311835256753778 | 3212 | 0 | 0 | 0 | 1 | 0 | 0 | 2 | 0 | 8 | 5 | 4 | 3 |
| MSTD_dup189 | 2.09631581949509 | 4.39496888517889 | 1.07005001013521 | 4.1072555895061 | 4.00388095408371e-05 | 0.00513588925876228 | 3213 | 0 | 0 | 0 | 1 | 0 | 0 | 1 | 0 | 7 | 5 | 4 | 7 |
| AluJo_dup275 | 2.17358096841999 | 3.44399969374473 | 0.97564094114895 | 3.52998685119646 | 0.000415580320112335 | 0.026973724727073 | 3214 | 0 | 0 | 1 | 2 | 0 | 0 | 2 | 0 | 4 | 3 | 5 | 11 |
| THE1B_dup20769 | 2.89316160296101 | 3.75712693810182 | 0.959319284488305 | 3.91645096565097 | 8.98620827924177e-05 | 0.0093209778344048 | 3215 | 0 | 0 | 2 | 3 | 0 | 0 | 1 | 0 | 9 | 4 | 7 | 9 |
| L1MEc_dup16192 | 2.08750788985638 | 4.7879063228297 | 1.09106177351055 | 4.388299946963 | 1.14240149891005e-05 | 0.00206787341108031 | 3216 | 0 | 0 | 0 | 0 | 0 | 0 | 1 | 0 | 6 | 6 | 5 | 7 |
| MLT1H_dup273 | 2.34755610367854 | 4.96018663208901 | 1.05825088122823 | 4.68715568309481 | 2.77028230814945e-06 | 0.000715719428984722 | 3217 | 0 | 0 | 0 | 0 | 0 | 0 | 1 | 0 | 6 | 8 | 7 | 6 |
| L2c_dup5661 | 2.13185424526575 | 4.76649539377165 | 1.09994397728917 | 4.33339832953927 | 1.46825103257396e-05 | 0.00247776212905563 | 3218 | 0 | 0 | 0 | 0 | 0 | 0 | 1 | 0 | 7 | 8 | 3 | 6 |
| MSTA_dup17700 | 2.1117665271088 | 4.81252762646876 | 1.11092205438995 | 4.33201196020141 | 1.47752913403769e-05 | 0.00249165235526388 | 3219 | 0 | 0 | 0 | 0 | 0 | 0 | 1 | 0 | 4 | 9 | 4 | 8 |
| Tigger2a_dup60 | 1.75616627357202 | 4.50819688314572 | 1.15589534040669 | 3.90017739976317 | 9.61222295449027e-05 | 0.00977014097876609 | 3220 | 0 | 0 | 0 | 0 | 0 | 0 | 1 | 0 | 4 | 8 | 4 | 4 |
| L1MEd_dup18673 | 2.09359565698521 | 4.38449059297196 | 1.05049029604598 | 4.17375639687018 | 2.99618119127474e-05 | 0.0041716169089471 | 3221 | 1 | 0 | 0 | 0 | 0 | 0 | 1 | 0 | 4 | 6 | 6 | 8 |
| MIR_dup6762 | 2.19601994546871 | 4.80653531112719 | 1.09826442342337 | 4.37648275644301 | 1.20609728156359e-05 | 0.00215258700619384 | 3222 | 0 | 0 | 0 | 0 | 0 | 0 | 1 | 0 | 4 | 4 | 6 | 13 |
| MER41C_dup36 | 1.60026399550832 | 3.9442708755404 | 1.17337322408855 | 3.361480213258 | 0.000775259109827222 | 0.0405963477436065 | 3223 | 0 | 0 | 0 | 0 | 0 | 1 | 1 | 0 | 5 | 2 | 6 | 4 |
| MLT1A_dup192 | 1.91643147867911 | 4.65130223361089 | 1.17869858014606 | 3.94613373762994 | 7.94232628916276e-05 | 0.00847084164590373 | 3224 | 0 | 0 | 0 | 0 | 0 | 0 | 1 | 0 | 7 | 4 | 7 | 3 |
| LTR16A2_dup1407 | 2.13815911030164 | 4.07854097355357 | 1.12332967379941 | 3.63076046923859 | 0.000282587356386793 | 0.0207468469925368 | 3225 | 0 | 0 | 0 | 1 | 0 | 0 | 2 | 0 | 4 | 6 | 10 | 3 |
| AluJb_dup111094 | 2.15821044462684 | 3.97472130806358 | 1.19442835042414 | 3.32771849115284 | 0.000875602971605672 | 0.0437702152692612 | 3226 | 0 | 0 | 0 | 0 | 0 | 0 | 3 | 0 | 4 | 2 | 11 | 7 |
| MSTA_dup20263 | 2.63311781411783 | 4.30302369940323 | 1.10731141823835 | 3.88601041091857 | 0.000101905138043341 | 0.0102225632280189 | 3227 | 0 | 0 | 0 | 0 | 0 | 0 | 3 | 0 | 5 | 3 | 12 | 10 |
| L2b_dup53510 | 2.96969787267314 | 4.07125028410681 | 0.952516131222027 | 4.27420612696986 | 1.91819518257926e-05 | 0.00301670106354266 | 3228 | 0 | 0 | 0 | 0 | 0 | 2 | 1 | 1 | 8 | 9 | 7 | 7 |
| MIR_dup143647 | 2.72441856638653 | 3.94747517686089 | 0.970650220403987 | 4.06683591460778 | 4.76557730468624e-05 | 0.00582708148684929 | 3229 | 0 | 0 | 0 | 0 | 0 | 1 | 1 | 2 | 6 | 6 | 6 | 11 |
| L1PREC2_dup103 | 2.94541336262989 | 4.4629499633234 | 0.963466923412648 | 4.63217766471464 | 3.61839428657015e-06 | 0.000861844291203423 | 3230 | 1 | 0 | 0 | 0 | 0 | 0 | 1 | 1 | 8 | 10 | 5 | 9 |
| AluJo_dup76133 | 2.8622678315291 | 4.2946034243226 | 0.988444410636822 | 4.34481026763632 | 1.3939609249598e-05 | 0.00239312717352442 | 3231 | 0 | 0 | 0 | 0 | 1 | 0 | 1 | 1 | 12 | 8 | 4 | 5 |
| AluSz_dup80365 | 2.42313676256641 | 4.16935590362816 | 0.997031302067759 | 4.18177031651992 | 2.89248091340179e-05 | 0.00407492163067083 | 3232 | 0 | 0 | 0 | 0 | 1 | 0 | 1 | 1 | 6 | 7 | 5 | 8 |
| L1MC3_dup4611 | 3.29167903307346 | 5.84468125814611 | 1.07559748195492 | 5.43389265612942 | 5.51377580115708e-08 | 3.13121932406234e-05 | 3233 | 0 | 0 | 0 | 0 | 0 | 0 | 0 | 0 | 11 | 4 | 12 | 12 |
| L1PA15_dup2745 | 2.67554315043 | 5.5158562384302 | 1.11998981671134 | 4.92491641988907 | 8.43964196050125e-07 | 0.000282444310257866 | 3234 | 0 | 0 | 0 | 0 | 0 | 0 | 0 | 0 | 7 | 3 | 9 | 14 |

| | | | | | | | | | | | | | | | | | | | |
|---|---|---|---|---|---|---|---|---|---|---|---|---|---|---|---|---|---|---|---|
| MSTA_dup20255 | 2.91797301397445 | 5.67639992412918 | 1.1110743180232 | 5.1089291076663 | 3.23989878863737e-07 | 0.000131886962762694 | 3235 | 0 | 0 | 0 | 0 | 0 | 0 | 0 | 0 | 10 | 4 | 7 | 14 |
| AluSz_dup33311 | 2.71342150336297 | 5.55462471727799 | 1.0553315576313 | 5.26339298499266 | 1.41420839264422e-07 | 6.84578473012215e-05 | 3236 | 0 | 0 | 0 | 0 | 0 | 0 | 0 | 0 | 8 | 9 | 7 | 8 |
| Tigger3b_dup133 | 2.44729661348391 | 5.40238008675403 | 1.09713813609695 | 4.92406553834043 | 8.47644266345995e-07 | 0.000283277475561747 | 3237 | 0 | 0 | 0 | 0 | 0 | 0 | 0 | 0 | 6 | 10 | 7 | 6 |
| L1PA8A_dup180 | 2.85846564815148 | 5.49787992908067 | 1.06446510485652 | 5.1649226489409 | 2.40538373398047e-07 | 0.000104443693059437 | 3238 | 0 | 0 | 0 | 0 | 0 | 0 | 0 | 0 | 12 | 10 | 5 | 5 |
| LTR49-int_dup56 | 3.2545966254616 | 5.73034740566847 | 1.01910582998816 | 5.62291691112694 | 1.87759723828168e-08 | 1.3416418992798e-05 | 3239 | 0 | 0 | 0 | 0 | 0 | 0 | 0 | 0 | 12 | 12 | 7 | 6 |
| L2c_dup5656 | 2.97728534882117 | 5.64905380069209 | 1.04034031303182 | 5.43000567211442 | 5.63522608774936e-08 | 3.1774395892787e-05 | 3240 | 0 | 0 | 0 | 0 | 0 | 0 | 0 | 0 | 9 | 12 | 5 | 9 |
| L2a_dup120007 | 2.6403304608195 | 5.43815796739355 | 1.09540344907139 | 4.96452514551023 | 6.88693633970972e-07 | 0.000240856424033409 | 3241 | 0 | 0 | 0 | 0 | 0 | 0 | 0 | 0 | 11 | 8 | 4 | 7 |
| MER57A-int_dup31 | 2.99737306697812 | 5.57730228410997 | 1.05198748772392 | 5.30168119791714 | 1.14741084780166e-07 | 5.72372791595419e-05 | 3242 | 0 | 0 | 0 | 0 | 0 | 0 | 0 | 0 | 12 | 11 | 4 | 7 |
| L3_dup1454 | 3.35798928459081 | 5.46901941923964 | 0.986698760086026 | 5.54274479757411 | 2.977666883621e-08 | 1.90976799000022e-05 | 3243 | 0 | 0 | 0 | 0 | 0 | 0 | 1 | 0 | 11 | 10 | 10 | 7 |
| LTR78_dup1971 | 3.65803830799207 | 5.99890775832297 | 1.00421362424061 | 5.9737366766552 | 2.31879974461245e-09 | 2.21585993586969e-06 | 3244 | 0 | 0 | 0 | 0 | 0 | 0 | 0 | 0 | 10 | 11 | 14 | 8 |
| L1PB1_dup9121 | 2.78516628214964 | 5.58684652819446 | 1.08290336568105 | 5.15913672932472 | 2.48091087908802e-07 | 0.000106556465710375 | 3245 | 0 | 0 | 0 | 0 | 0 | 0 | 0 | 0 | 10 | 7 | 9 | 6 |
| LTR82A_dup13 | 3.50112121246824 | 5.83876280846444 | 1.04288624884508 | 5.598657394256 | 2.16018225744046e-08 | 1.48808284866662e-05 | 3246 | 0 | 0 | 0 | 0 | 0 | 0 | 0 | 0 | 15 | 10 | 9 | 5 |
| L1M5_dup1224 | 3.78824088439585 | 5.24585710267767 | 0.967684806029833 | 5.42103903046706 | 5.92536199734356e-08 | 3.3134899959648e-05 | 3247 | 0 | 0 | 0 | 0 | 0 | 1 | 0 | 1 | 14 | 7 | 8 | 13 |
| L2a_dup164349 | 3.47329525392924 | 5.541034291717 | 0.97206138542611 | 5.70029256875383 | 1.19601984682157e-08 | 9.18026253134853e-06 | 3248 | 0 | 0 | 0 | 0 | 0 | 1 | 0 | 0 | 9 | 7 | 12 | 13 |
| MLT1E1A_dup1270 | 3.04976585324182 | 5.34774224965429 | 1.00628251465412 | 5.31435473813477 | 1.07035926100732e-07 | 5.40738226580994e-05 | 3249 | 0 | 0 | 0 | 0 | 0 | 1 | 0 | 0 | 8 | 9 | 6 | 13 |
| L1PA16_dup174 | 3.41228817268763 | 5.07286373008486 | 0.985990899779201 | 5.14493970605699 | 2.6760674609437e-07 | 0.000113101163066023 | 3250 | 0 | 0 | 0 | 1 | 0 | 1 | 0 | 0 | 13 | 8 | 10 | 6 |
| MER117_dup3414 | 2.95529662548384 | 4.09284644973176 | 1.00061347734863 | 4.09033712056001 | 4.30746625367531e-05 | 0.00540297504057472 | 3251 | 0 | 0 | 0 | 1 | 0 | 3 | 0 | 0 | 9 | 9 | 5 | 8 |
| MIR3_dup72168 | 3.01052941136391 | 4.53422891375037 | 0.975850938189977 | 4.64643598351253 | 3.37718869351338e-06 | 0.000825274058964267 | 3252 | 1 | 0 | 0 | 0 | 0 | 2 | 0 | 0 | 7 | 11 | 8 | 7 |
| MIRb_dup184098 | 2.96419001228789 | 4.92656537059705 | 1.07128102056756 | 4.59876099362518 | 4.25011030643817e-06 | 0.000972110993582927 | 3253 | 0 | 0 | 0 | 0 | 0 | 2 | 0 | 0 | 8 | 6 | 13 | 6 |
| MIRc_dup84136 | 2.47850563883295 | 4.22953953569124 | 1.10722422201581 | 3.81994852676808 | 0.000133479526598822 | 0.0123199842653543 | 3254 | 0 | 0 | 0 | 1 | 0 | 1 | 0 | 1 | 1 | 13 | 7 | 7 |
| AluSp_dup39463 | 2.74430541937597 | 3.56617689108939 | 1.03369394289483 | 3.44993497891883 | 0.000560721589122264 | 0.0329029492590101 | 3255 | 0 | 1 | 0 | 0 | 0 | 1 | 1 | 2 | 1 | 13 | 8 | 7 |
| AluSz6_dup37859 | 2.48523047034493 | 3.83242848107694 | 0.998059501249413 | 3.83987976295936 | 0.000123094581900609 | 0.0116418460383675 | 3256 | 0 | 1 | 0 | 0 | 0 | 2 | 0 | 1 | 4 | 8 | 8 | 6 |
| MSTA-int_dup3305 | 3.46528548779334 | 5.04931422982973 | 0.998033865737794 | 5.05926141704323 | 4.20883524620093e-07 | 0.000161268198307975 | 3257 | 0 | 1 | 0 | 0 | 0 | 0 | 1 | 0 | 2 | 9 | 14 | 18 |
| THE1A_dup3872 | 3.33949770631804 | 4.72389581947334 | 1.09302452089059 | 4.32185713054666 | 1.54721366799215e-05 | 0.00255839682726935 | 3258 | 0 | 0 | 0 | 0 | 0 | 0 | 2 | 1 | 1 | 14 | 10 | 15 |
| ERVL-B4-int_dup2703 | 1.89975305170237 | 3.77578913979254 | 1.10867495772679 | 3.40567730287184 | 0.000660001414349377 | 0.0364823969772524 | 3259 | 0 | 0 | 0 | 0 | 1 | 1 | 1 | 0 | 1 | 5 | 5 | 11 |
| L1PA10_dup212 | 2.1878674624071 | 4.05170020986241 | 1.13572127364785 | 3.56751282543882 | 0.000360385745004246 | 0.024535953954961 | 3260 | 0 | 0 | 0 | 0 | 0 | 2 | 1 | 0 | 2 | 9 | 4 | 10 |
| L2b_dup79110 | 2.23339289533018 | 4.47375256685536 | 1.11553183527023 | 4.01042124070948 | 6.0610516201362e-05 | 0.0069579262646345 | 3261 | 0 | 0 | 1 | 0 | 0 | 0 | 1 | 0 | 2 | 8 | 4 | 13 |
| MER113_dup100 | 2.01888569107102 | 4.34690819085738 | 1.15945670964113 | 3.74909054793674 | 0.000177477002922557 | 0.0150982992268183 | 3262 | 0 | 0 | 0 | 1 | 0 | 0 | 1 | 0 | 1 | 9 | 6 | 8 |
| L2c_dup119705 | 2.13728178422632 | 3.49656487928638 | 1.0680308470574 | 3.27384259445312 | 0.00106095693180902 | 0.0497145447215891 | 3263 | 0 | 0 | 0 | 0 | 1 | 0 | 2 | 1 | 1 | 5 | 11 | 6 |
| LTR10E_dup79 | 1.71772373608966 | 3.60276703883737 | 1.08288590629401 | 3.3270051977749 | 0.000877847360427475 | 0.0438282798624732 | 3264 | 1 | 0 | 0 | 0 | 0 | 0 | 1 | 1 | 4 | 3 | 4 | 7 |
| AluSz_dup89724 | 2.1212439564899 | 3.45271385091067 | 1.01531591503429 | 3.40063008939838 | 0.000672307377102569 | 0.0368600756169689 | 3265 | 1 | 0 | 0 | 0 | 0 | 0 | 2 | 1 | 3 | 2 | 8 | 10 |
| Charlie4a_dup259 | 2.43446131924724 | 4.14901846222928 | 0.993377511753539 | 4.17667846628148 | 2.95796681734229e-05 | 0.00413777996660393 | 3266 | 1 | 0 | 0 | 0 | 0 | 0 | 1 | 1 | 5 | 4 | 8 | 10 |
| L2b_dup38128 | 2.27119917430316 | 3.56008030153265 | 1.04360154211827 | 3.41134059107129 | 0.000646442880242705 | 0.0360062026175634 | 3267 | 1 | 0 | 0 | 0 | 0 | 0 | 1 | 1 | 1 | 1 | 11 | 14 |
| MSTB_dup3277 | 2.24734478334323 | 4.13835162291183 | 1.11065218658911 | 3.72605544101164 | 0.000194499641503865 | 0.0161143494767683 | 3268 | 0 | 0 | 0 | 0 | 0 | 0 | 1 | 1 | 3 | 1 | 14 | 8 |
| L1M4b_dup4263 | 2.43169961972802 | 3.70106746730638 | 1.03658666937578 | 3.57043706681577 | 0.000356386073978718 | 0.0244171151047913 | 3269 | 0 | 2 | 0 | 1 | 0 | 0 | 0 | 1 | 4 | 3 | 5 | 15 |
| LTR78_dup1559 | 2.07406051077465 | 4.3347673623499 | 1.14439436426272 | 3.78782655500281 | 0.000151970875241889 | 0.0134577081802406 | 3270 | 0 | 1 | 0 | 0 | 0 | 0 | 0 | 1 | 4 | 5 | 2 | 13 |
| MER34A1_dup233 | 2.13902687446715 | 3.91105000292771 | 1.1166430542504 | 3.50250690051818 | 0.000460901844668501 | 0.0289061018269613 | 3271 | 0 | 0 | 0 | 0 | 0 | 1 | 0 | 2 | 5 | 3 | 4 | 11 |
| AluSx1_dup100500 | 2.01868290903272 | 4.27505103073957 | 1.11779026090958 | 3.82455562572253 | 0.000131008165971603 | 0.012162648872524 | 3272 | 0 | 0 | 0 | 0 | 0 | 0 | 1 | 1 | 6 | 2 | 7 | 7 |
| MIRb_dup11009 | 2.09046138454297 | 4.74954266762276 | 1.12513207150847 | 4.22132013467097 | 2.42875712367435e-05 | 0.00357398294712318 | 3273 | 0 | 0 | 0 | 0 | 0 | 0 | 0 | 1 | 6 | 3 | 6 | 9 |
| MER103C_dup7499 | 1.59935082690048 | 4.39587668121034 | 1.25987423120098 | 3.48913929053058 | 0.000484578570356173 | 0.0298946156344231 | 3274 | 0 | 0 | 0 | 0 | 1 | 0 | 0 | 0 | 4 | 3 | 2 | 10 |
| MIR_dup143695 | 1.97340124521399 | 4.30677365712357 | 1.11094930558286 | 3.87666083004933 | 0.000105899818416989 | 0.0104688152027623 | 3275 | 0 | 0 | 0 | 0 | 1 | 1 | 0 | 0 | 4 | 3 | 4 | 12 |
| THE1B_dup19559 | 1.93939857382203 | 4.29191522047764 | 1.21856907425847 | 3.52209432451695 | 0.000428151775625429 | 0.0275120179322086 | 3276 | 0 | 0 | 0 | 0 | 1 | 1 | 0 | 0 | 6 | 1 | 3 | 12 |
| AluJr_dup65995 | 2.08664407187525 | 3.887315269071 | 1.05318760543588 | 3.69099982662837 | 0.000223374289856398 | 0.0177103257237869 | 3277 | 0 | 1 | 1 | 0 | 1 | 0 | 0 | 0 | 2 | 4 | 6 | 12 |
| L2c_dup56368 | 2.32703193292001 | 3.81129222231529 | 1.05196845693682 | 3.62300998398109 | 0.000291194578048875 | 0.0211762179304455 | 3278 | 0 | 1 | 1 | 0 | 1 | 0 | 0 | 0 | 1 | 2 | 9 | 16 |
| L2c_dup12701 | 1.90505801526574 | 3.7028508572662 | 1.06490461893691 | 3.47716667898646 | 0.000506742689505664 | 0.0307681606502456 | 3279 | 0 | 1 | 0 | 0 | 1 | 0 | 0 | 1 | 2 | 3 | 7 | 9 |
| Tigger3a_dup2847 | 1.54597435284813 | 3.88245063757709 | 1.17819292307134 | 3.29525883371989 | 0.000983310642833408 | 0.0472676432766945 | 3280 | 0 | 1 | 0 | 0 | 1 | 0 | 0 | 0 | 1 | 3 | 6 | 8 |
| MIRc_dup6106 | 2.24690496222427 | 4.49884034342516 | 1.10507056158718 | 4.07108876103226 | 4.67939136185026e-05 | 0.00573643718179712 | 3281 | 0 | 0 | 0 | 0 | 2 | 0 | 0 | 0 | 4 | 4 | 8 | 10 |
| AluJb_dup99368 | 2.17377788072075 | 4.46399487697712 | 1.12473249358755 | 3.96893919436643 | 7.21932990960286e-05 | 0.00793446039108713 | 3282 | 0 | 0 | 0 | 0 | 2 | 0 | 0 | 0 | 3 | 7 | 7 | 8 |
| L2b_dup45740 | 2.58202617118462 | 4.72008139951289 | 1.09009750237412 | 4.32996258521191 | 1.49134678327334e-05 | 0.00250381500469088 | 3283 | 0 | 0 | 0 | 0 | 2 | 0 | 0 | 0 | 3 | 7 | 7 | 14 |
| L2_dup47802 | 1.86774785227234 | 4.14693969483008 | 1.15033320285358 | 3.60499000162991 | 0.00031216494622816 | 0.0222523667750766 | 3284 | 1 | 0 | 0 | 0 | 1 | 0 | 0 | 0 | 3 | 1 | 7 | 11 |
| MLT1G1_dup1519 | 2.53994086019611 | 4.53140392144054 | 1.18038003121984 | 3.83893644554257 | 0.000123568407807641 | 0.011672333610241 | 3285 | 0 | 0 | 0 | 0 | 2 | 0 | 0 | 0 | 3 | 1 | 11 | 16 |
| L1ME1_dup1399 | 2.11368570927411 | 4.77407953893484 | 1.23759641190123 | 3.85754151597834 | 0.000114533230029902 | 0.0110693436038566 | 3286 | 0 | 0 | 0 | 0 | 1 | 0 | 0 | 0 | 5 | 2 | 14 | 3 |
| L1ME4a_dup11900 | 1.97900775387504 | 3.9262400749614 | 1.12597083929463 | 3.48698202292789 | 0.000488504192871422 | 0.0300259608191217 | 3287 | 1 | 1 | 0 | 0 | 0 | 1 | 0 | 0 | 5 | 1 | 9 | 6 |
| MIRb_dup29976 | 1.93442923384452 | 4.25683371329708 | 1.10470956001496 | 3.85335102308648 | 0.00011651216630251 | 0.0112014561305119 | 3288 | 1 | 1 | 0 | 0 | 0 | 0 | 0 | 0 | 4 | 2 | 8 | 8 |
| L1PB1_dup8282 | 2.84585742775842 | 4.65093322446892 | 1.07509077742598 | 4.32608419877282 | 1.51783423724114e-05 | 0.00252561248541665 | 3289 | 0 | 1 | 0 | 0 | 0 | 0 | 1 | 0 | 5 | 1 | 14 | 14 |
| L1M1_dup3222 | 2.15786553007614 | 4.38818199640542 | 1.21142278125737 | 3.62233735760756 | 0.000291953024275723 | 0.0212009829421527 | 3290 | 0 | 2 | 0 | 0 | 0 | 0 | 0 | 0 | 4 | 1 | 10 | 10 |
| L3_dup14171 | 2.90690089283177 | 5.1397643849874 | 1.03033457174128 | 4.98844212933779 | 6.08681388994482e-07 | 0.00021746742009862 | 3291 | 0 | 1 | 0 | 0 | 0 | 0 | 0 | 0 | 4 | 3 | 14 | 15 |
| AluSq2_dup18975 | 1.96127912950853 | 3.84982462271844 | 1.13817533653571 | 3.38245303613434 | 0.000718415604845395 | 0.0383879450212378 | 3292 | 2 | 0 | 1 | 0 | 0 | 0 | 0 | 0 | 5 | 2 | 7 | 7 |
| L1ME4a_dup503 | 2.41841296519575 | 3.67439777048638 | 1.05977868315717 | 3.46713689271427 | 0.000526033918712785 | 0.0314728857988515 | 3293 | 1 | 0 | 3 | 0 | 0 | 0 | 0 | 0 | 3 | 3 | 9 | 12 |
| LTR8_dup2490 | 1.66144863459303 | 4.32954332531989 | 1.19112961278029 | 3.6348213316719 | 0.000278173349725624 | 0.0205304701841853 | 3294 | 0 | 0 | 1 | 0 | 0 | 0 | 0 | 0 | 3 | 2 | 6 | 9 |
| THE1D_dup7183 | 1.70336166108321 | 4.27731781050355 | 1.18476082294512 | 3.61027958357944 | 0.0003058671312056 | 0.0219414713300716 | 3295 | 0 | 0 | 1 | 0 | 0 | 0 | 0 | 0 | 2 | 2 | 7 | 10 |
| L1ME4c_dup7939 | 1.83385516379879 | 4.53982736658831 | 1.18993091098788 | 3.81520248332683 | 0.000136071311031795 | 0.0124673177415369 | 3296 | 0 | 0 | 1 | 0 | 0 | 0 | 0 | 0 | 4 | 3 | 10 | 4 |
| L2b_dup3516 | 2.5655347154488 | 4.98465095542493 | 1.06023951075555 | 4.70143859463674 | 2.5833495407314e-06 | 0.000684518585544402 | 3297 | 0 | 0 | 1 | 0 | 0 | 0 | 0 | 0 | 5 | 3 | 11 | 12 |
| MLT1B_dup427 | 2.42906062554135 | 4.99490655508652 | 1.07501095965666 | 4.64637733245233 | 3.37814854391835e-06 | 0.000825274058964267 | 3298 | 0 | 0 | 1 | 0 | 0 | 0 | 0 | 0 | 7 | 4 | 9 | 8 |
| CR1-3_Croc_dup279 | 1.72638628242979 | 4.1120599751301 | 1.19272388036088 | 3.44762106539355 | 0.000565546883247333 | 0.0331044572401402 | 3299 | 2 | 0 | 0 | 0 | 0 | 0 | 0 | 0 | 4 | 4 | 6 | 5 |
| MLT1A0_dup17830 | 1.80397347848887 | 4.1696057776757 | 1.09265262290259 | 3.81603969118684 | 0.00013561069971326 | 0.0124299012149351 | 3300 | 1 | 0 | 0 | 0 | 1 | 0 | 0 | 0 | 4 | 4 | 6 | 6 |
| L1M3d_dup99 | 1.87914020062269 | 4.20681699318636 | 1.14566641698976 | 3.67193882163342 | 0.000240717314432669 | 0.0186329609954443 | 3301 | 1 | 0 | 0 | 1 | 0 | 0 | 0 | 0 | 6 | 2 | 4 | 9 |
| AluJr_dup74984 | 1.41868147137613 | 4.63696728775642 | 1.38369922608401 | 3.35113816669497 | 0.000804801369175408 | 0.0415354127014618 | 3302 | 0 | 0 | 0 | 0 | 0 | 0 | 0 | 0 | 5 | 2 | 3 | 7 |
| HAL1_dup13726 | 1.38280908198279 | 4.60478622463976 | 1.39581691794647 | 3.29899012215323 | 0.000970333256375227 | 0.0468235483718231 | 3303 | 0 | 0 | 0 | 0 | 0 | 0 | 0 | 0 | 4 | 3 | 2 | 8 |
| L1PA16_dup14388 | 2.01893659391817 | 5.14008589598749 | 1.18791015323387 | 4.32699887444731 | 1.51154733685195e-05 | 0.00252321138673569 | 3304 | 0 | 0 | 0 | 0 | 0 | 0 | 0 | 0 | 5 | 4 | 5 | 11 |
| L2b_dup79123 | 1.86463583027465 | 4.9994635576671 | 1.25437041083886 | 3.98563575357593 | 6.72996860967596e-05 | 0.0075287687390595 | 3305 | 0 | 0 | 0 | 0 | 0 | 0 | 0 | 0 | 5 | 2 | 6 | 10 |
| MER53_dup237 | 1.81337009606441 | 4.93125960543967 | 1.27036667978438 | 3.88176082064484 | 0.000103702847549901 | 0.0103140051945626 | 3306 | 0 | 0 | 0 | 0 | 0 | 0 | 0 | 0 | 4 | 2 | 5 | 12 |

| | | | | | | | | | | | | | | | | | | | |
|---|---|---|---|---|---|---|---|---|---|---|---|---|---|---|---|---|---|---|---|
| L1PA16_dup14377 | 2.21193947921096 | 4.07099812294122 | 1.10037696889632 | 3.69963952173992 | 0.000215905919376854 | 0.0172859858317782 | 3307 | 0 | 0 | 1 | 0 | 0 | 0 | 0 | 1 | 1 | 3 | 14 | 8 |
| L2_dup52917 | 2.34128573772827 | 3.39839331709047 | 0.968655247430879 | 3.50836205771235 | 0.000450874985075126 | 0.028498300031631 | 3308 | 0 | 0 | 1 | 1 | 1 | 0 | 0 | 1 | 2 | 1 | 12 | 11 |
| MLT1K_dup1101 | 1.85097521054703 | 4.07234930991033 | 1.10968282129337 | 3.66983180397791 | 0.000242710113582592 | 0.0187445312192546 | 3309 | 0 | 0 | 1 | 0 | 0 | 1 | 0 | 0 | 2 | 3 | 6 | 11 |
| MSTA_dup19307 | 2.8925263576076 | 4.64403653411762 | 1.01470241751813 | 4.57674728466352 | 4.72261388983656e-06 | 0.00105231283297733 | 3310 | 0 | 0 | 1 | 0 | 0 | 1 | 0 | 0 | 3 | 3 | 10 | 20 |
| AluJo_dup68997 | 2.52881095767188 | 3.2593931985307 | 0.97902003608375 | 3.32924054503403 | 0.000870831581885517 | 0.0436058891412255 | 3311 | 0 | 0 | 2 | 2 | 0 | 1 | 0 | 0 | 1 | 2 | 15 | 10 |
| MLT1A0_dup19217 | 2.61715171708095 | 3.6096478610652 | 0.969552320800987 | 3.72300471426145 | 0.000196865844619053 | 0.0162650834249046 | 3312 | 0 | 0 | 3 | 1 | 0 | 1 | 0 | 0 | 3 | 6 | 10 | 9 |
| L1MEi_dup3292 | 2.10877787412172 | 3.91179168796949 | 1.17409631317639 | 3.33174684569664 | 0.000863027239719804 | 0.0433785127972895 | 3313 | 0 | 2 | 0 | 0 | 0 | 1 | 0 | 0 | 1 | 2 | 8 | 14 |
| MSTA_dup19317 | 1.95801071663616 | 4.52670614240808 | 1.1972365341652 | 3.78096225201182 | 0.000156223362644011 | 0.0137676756472933 | 3314 | 0 | 0 | 0 | 0 | 0 | 1 | 0 | 0 | 1 | 3 | 13 | 7 |
| AluJb_dup121788 | 2.15263315851893 | 4.66102399132652 | 1.16631738212913 | 3.99635987831867 | 6.4323925918617e-05 | 0.00729881777617131 | 3315 | 1 | 0 | 0 | 0 | 0 | 0 | 0 | 0 | 3 | 1 | 9 | 14 |
| HERVIP10B3-int_dup1 | 2.99090286309153 | 4.98823739388897 | 1.06749588747643 | 4.67283991667755 | 2.97063303402115e-06 | 0.000751169232638892 | 3316 | 1 | 0 | 0 | 0 | 0 | 0 | 0 | 0 | 2 | 2 | 18 | 16 |
| L2b_dup13130 | 2.7891519968738 | 4.84258050194979 | 1.11804440652529 | 4.33129531679318 | 1.48234707172019e-05 | 0.00249764515492702 | 3317 | 1 | 0 | 0 | 0 | 0 | 0 | 0 | 0 | 2 | 1 | 14 | 19 |
| MSTA-int_dup3312 | 3.01195893863657 | 4.64702349547644 | 0.995402977607329 | 4.66848462383203 | 3.03429482065915e-06 | 0.000764019381372023 | 3318 | 0 | 0 | 1 | 0 | 0 | 0 | 1 | 0 | 4 | 2 | 13 | 18 |
| MSTA_dup19306 | 2.95824138754237 | 4.93977159363063 | 1.05515341978208 | 4.68156715508807 | 2.84690102812139e-06 | 0.000726833382014349 | 3319 | 0 | 0 | 1 | 0 | 0 | 0 | 0 | 0 | 2 | 3 | 19 | 13 |
| MIRb_dup82624 | 2.58094604805268 | 5.49563940989566 | 1.08399091062208 | 5.06982056403206 | 3.98190933465584e-07 | 0.000154816895186931 | 3320 | 0 | 0 | 0 | 0 | 0 | 0 | 0 | 0 | 4 | 8 | 10 | 10 |
| L2a_dup21560 | 2.34741751590381 | 5.29453051869645 | 1.11853881128266 | 4.73343478589274 | 2.20751911997307e-06 | 0.000607951787640175 | 3321 | 0 | 0 | 0 | 0 | 0 | 0 | 0 | 0 | 3 | 5 | 9 | 13 |
| MER4-int_dup1591 | 2.796196530141 | 5.52512851409129 | 1.07076611826264 | 5.15997697336187 | 2.46980217790098e-07 | 0.000106463686417179 | 3322 | 0 | 0 | 0 | 0 | 0 | 0 | 0 | 0 | 3 | 6 | 11 | 16 |
| L1M4_dup11537 | 2.83107794323698 | 5.50083190580167 | 1.07039213920184 | 5.13908100063541 | 2.76085377226508e-07 | 0.00011525159854305 | 3323 | 0 | 0 | 0 | 0 | 0 | 0 | 0 | 0 | 3 | 5 | 15 | 13 |
| L1ME1_dup589 | 2.71177082728277 | 5.40540262828063 | 1.10227183781356 | 4.90387438274997 | 9.39645928232361e-07 | 0.00030397880064774 | 3324 | 0 | 0 | 0 | 0 | 0 | 0 | 0 | 0 | 2 | 5 | 14 | 14 |
| L1PA13_dup5430 | 2.86042228024895 | 5.46139203867423 | 1.08909055027987 | 5.01463541049986 | 5.31341216604451e-07 | 0.000195410381956975 | 3325 | 0 | 0 | 0 | 0 | 0 | 0 | 0 | 0 | 2 | 5 | 15 | 15 |
| L1PBb_dup183 | 3.5184711893826 | 5.84556967261987 | 1.00965422031534 | 5.78967487581457 | 7.05227985448494e-09 | 5.90866824737772e-06 | 3326 | 0 | 0 | 0 | 0 | 0 | 0 | 0 | 0 | 5 | 6 | 19 | 14 |
| MSTA_dup20256 | 3.04257284359425 | 5.57742869993587 | 1.10463334930876 | 5.04912213941939 | 4.43844908225999e-07 | 0.000168438788728459 | 3327 | 0 | 0 | 0 | 0 | 0 | 0 | 0 | 0 | 3 | 5 | 21 | 9 |
| MSTA-int_dup3066 | 2.88733597544989 | 5.06581897634685 | 1.06971666917535 | 4.73566423925327 | 2.18338804271126e-06 | 0.000602701219502288 | 3328 | 0 | 0 | 0 | 1 | 0 | 0 | 0 | 0 | 4 | 3 | 19 | 9 |
| MIRb_dup183961 | 2.66566967557354 | 5.11426213939073 | 1.08381162853677 | 4.71877400530884 | 2.37270226671939e-06 | 0.000643757142026011 | 3329 | 0 | 0 | 0 | 1 | 0 | 0 | 0 | 0 | 2 | 9 | 11 | 11 |
| MSTA-int_dup3012 | 2.3874034615958 | 4.84964858410921 | 1.08139690031206 | 4.48461483726258 | 7.30457584365067e-06 | 0.00145935734986843 | 3330 | 0 | 0 | 0 | 1 | 0 | 0 | 0 | 0 | 2 | 5 | 10 | 13 |
| L1M4_dup11538 | 2.11953829443229 | 5.09496906202318 | 1.2046086440223 | 4.22956375691496 | 2.34144914458665e-05 | 0.00347559861608119 | 3331 | 0 | 0 | 0 | 0 | 0 | 0 | 0 | 0 | 2 | 4 | 6 | 16 |
| Tigger6a_dup512 | 2.42940988984285 | 5.24079471667288 | 1.14179478196794 | 4.58996204873184 | 4.43326607647367e-06 | 0.00100083295050532 | 3332 | 0 | 0 | 0 | 0 | 0 | 0 | 0 | 0 | 2 | 4 | 9 | 17 |
| HERVFH19-int_dup59 | 1.74854469607861 | 4.76473620015623 | 1.27282496673394 | 3.74343395571705 | 0.000181522330391785 | 0.0153491515378122 | 3333 | 0 | 0 | 0 | 0 | 0 | 0 | 0 | 0 | 2 | 2 | 6 | 13 |
| THE1B_dup9107 | 2.00294359922341 | 4.9067525985402 | 1.20850677249506 | 4.06017799007434 | 4.90353211794794e-05 | 0.00595597679089965 | 3334 | 0 | 0 | 0 | 0 | 0 | 0 | 0 | 0 | 2 | 2 | 10 | 12 |
| MSTA-int_dup691 | 1.75780400204179 | 4.45504278670732 | 1.1563614157864 | 3.85263873896864 | 0.000116851728222477 | 0.0112250308129292 | 3335 | 0 | 0 | 0 | 1 | 0 | 0 | 0 | 0 | 3 | 3 | 8 | 7 |
| MER31B_dup919 | 1.65299265636399 | 4.762227324117 | 1.26223362267807 | 3.77285728929721 | 0.000161388610457356 | 0.0140768967389611 | 3336 | 0 | 0 | 0 | 0 | 0 | 0 | 0 | 0 | 2 | 3 | 8 | 8 |
| MLT2B4_dup180 | 1.58777592995869 | 4.7593222476032 | 1.27774087991596 | 3.72479453574049 | 0.000195474365469377 | 0.0161715524870134 | 3337 | 0 | 0 | 0 | 0 | 0 | 0 | 0 | 0 | 2 | 4 | 7 | 7 |
| L1MEd_dup19752 | 1.44703483209073 | 4.61414055001604 | 1.32726316063374 | 3.47643232093692 | 0.000508132459456749 | 0.0308438995402795 | 3338 | 0 | 0 | 0 | 0 | 0 | 0 | 0 | 0 | 3 | 2 | 6 | 7 |
| L1PA13_dup5428 | 1.81520377510573 | 4.92436600572546 | 1.21580997609235 | 4.05027603207577 | 5.1157252093579e-05 | 0.00614780985184786 | 3339 | 0 | 0 | 0 | 0 | 0 | 0 | 0 | 0 | 3 | 3 | 7 | 10 |
| AluSq_dup9861 | 2.51372335810774 | 5.29330533536523 | 1.139018810637 | 4.64725014717266 | 3.36389153109311e-06 | 0.000823482031129095 | 3340 | 0 | 0 | 0 | 0 | 0 | 0 | 0 | 0 | 4 | 2 | 12 | 14 |
| L1MCa_dup7713 | 1.82113217760924 | 4.72117250115722 | 1.32168575259735 | 3.57208397826737 | 0.000354151800564163 | 0.0243047048769551 | 3341 | 0 | 0 | 0 | 0 | 0 | 0 | 0 | 0 | 2 | 1 | 13 | 7 |
| L1P3_dup593 | 2.9050734952908 | 2.79236383423514 | 0.834833897696509 | 3.34481367124632 | 0.000823378658382176 | 0.0421330706556156 | 3342 | 3 | 2 | 0 | 0 | 3 | 1 | 1 | 0 | 3 | 6 | 8 | 10 |
| THE1B_dup8009 | 2.48374061684962 | 3.07158533058645 | 0.93714045216274 | 3.27761470919094 | 0.00104688184449349 | 0.0492906400719254 | 3343 | 3 | 1 | 0 | 0 | 2 | 0 | 1 | 0 | 4 | 4 | 9 | 7 |
| AluSx_dup8133 | 3.03235300958072 | 4.580380914212 | 0.971183915362386 | 4.71628580514833 | 2.40188975421478e-06 | 0.000647981926821305 | 3344 | 1 | 1 | 0 | 0 | 1 | 0 | 0 | 0 | 8 | 5 | 8 | 13 |
| L1MB7_dup7044 | 4.13757296890857 | 4.24908937610263 | 0.982499085651448 | 4.32477692667293 | 1.52686289032779e-05 | 0.00253708741132637 | 3345 | 1 | 3 | 0 | 0 | 2 | 0 | 0 | 0 | 5 | 5 | 21 | 15 |
| AmnSINE1_dup1837 | 3.97676691495082 | 3.00702432349679 | 0.90923842057604 | 3.30719012246723 | 0.000942369095928573 | 0.0459023453223788 | 3346 | 8 | 5 | 0 | 0 | 3 | 0 | 0 | 0 | 10 | 9 | 5 | 8 |
| L1M4_dup18419 | 2.63040620515988 | 2.93244470470937 | 0.869262963974608 | 3.3734840045423 | 0.000742233587470676 | 0.039331664037174 | 3347 | 2 | 2 | 0 | 0 | 4 | 1 | 0 | 0 | 5 | 7 | 5 | 6 |
| MER5A_dup19623 | 2.80078083549988 | 2.99023252725283 | 0.881134327725116 | 3.39361710600121 | 0.00068976060060055 | 0.0374374488755699 | 3348 | 1 | 2 | 0 | 0 | 5 | 1 | 0 | 0 | 7 | 8 | 5 | 4 |
| MSTA_dup1145 | 2.74850391661058 | 3.18064290340595 | 0.96313338125866 | 3.30239088925499 | 0.000958643758613719 | 0.0465099767532787 | 3349 | 2 | 1 | 0 | 0 | 5 | 0 | 0 | 0 | 9 | 6 | 4 | 5 |
| MamRTE1_dup7779 | 2.75075122461644 | 3.28485236281712 | 0.955422271324942 | 3.43811575405482 | 0.000585777235671069 | 0.0338718235226471 | 3350 | 2 | 2 | 0 | 0 | 4 | 0 | 0 | 0 | 6 | 6 | 8 | 5 |
| L1MB7_dup7042 | 4.41756736526596 | 4.23896991650681 | 0.907552751827128 | 4.67076972437438 | 3.00073180340574e-06 | 0.000757170869239853 | 3351 | 2 | 4 | 0 | 0 | 0 | 1 | 0 | 0 | 6 | 14 | 13 | 15 |
| L2a_dup61549 | 4.44635297931068 | 3.97965448550413 | 0.812075584553137 | 4.90059615287414 | 9.55462809634312e-07 | 0.000306986115968739 | 3352 | 2 | 3 | 0 | 0 | 0 | 1 | 0 | 1 | 11 | 9 | 15 | 11 |
| L1MB3_dup15227 | 3.27974972240748 | 2.9046324959405 | 0.871828120297531 | 3.33165727087265 | 0.000863305044579968 | 0.0433833119614837 | 3353 | 4 | 3 | 0 | 0 | 0 | 3 | 0 | 1 | 5 | 8 | 5 | 12 |
| L1PA5_dup10093 | 3.95756126955888 | 4.19651831174949 | 0.883615188781352 | 4.74926004558292 | 2.04162279270264e-06 | 0.000572197852806151 | 3354 | 1 | 1 | 0 | 0 | 0 | 1 | 0 | 2 | 5 | 18 | 10 | 10 |
| L2a_dup14287 | 3.15032913394933 | 3.5489106318365 | 0.886074133440704 | 4.00520734992654 | 6.19631039967925e-05 | 0.00707479498254356 | 3355 | 2 | 1 | 2 | 0 | 1 | 0 | 0 | 0 | 3 | 8 | 11 | 12 |
| L2a_dup92224 | 4.19896967553525 | 3.46646818431568 | 0.848165769937677 | 4.08701731097967 | 4.36954527236094e-05 | 0.00545964956162073 | 3356 | 5 | 2 | 1 | 0 | 0 | 0 | 0 | 1 | 7 | 10 | 12 | 14 |
| MIRc_dup14175 | 2.91103105736917 | 3.46287073105597 | 1.05091303325554 | 3.29510684659475 | 0.000983842643517408 | 0.0472836642404349 | 3357 | 5 | 1 | 0 | 1 | 0 | 0 | 0 | 0 | 5 | 6 | 12 | 6 |
| MLT1K_dup6055 | 2.91288559826171 | 3.55793490667559 | 1.05271003594465 | 3.37978625185509 | 0.000725422224385605 | 0.0386241478191222 | 3358 | 4 | 3 | 0 | 0 | 0 | 1 | 0 | 0 | 5 | 4 | 12 | 7 |
| L2b_dup33110 | 3.04713447988136 | 3.98083576867683 | 1.13273991464794 | 3.51434227504387 | 0.000440844382264163 | 0.0280623749016128 | 3359 | 3 | 3 | 0 | 0 | 0 | 0 | 0 | 0 | 5 | 3 | 14 | 10 |
| L2c_dup50046 | 2.83705135476801 | 4.21721415351826 | 1.09066492314043 | 3.86664507498357 | 0.000110342834602931 | 0.0107914657305504 | 3360 | 2 | 2 | 0 | 0 | 0 | 0 | 0 | 0 | 7 | 3 | 14 | 6 |
| AluSq2_dup16092 | 3.12241791948238 | 4.5920090636334 | 0.996353466541729 | 4.6088152626918 | 4.04969955311856e-06 | 0.000941580341451141 | 3361 | 2 | 1 | 0 | 0 | 0 | 0 | 0 | 0 | 4 | 5 | 12 | 16 |
| L1MB7_dup7046 | 4.06294496532656 | 4.40102432035225 | 1.07647976037768 | 4.08834841335814 | 4.34455298838775e-05 | 0.00544088950197322 | 3362 | 2 | 4 | 0 | 0 | 0 | 0 | 0 | 0 | 4 | 9 | 14 | 19 |
| L1M7_dup1878 | 3.00024798822498 | 3.90598640457335 | 1.05419631474066 | 3.70517933894906 | 0.000211241217682862 | 0.0170444227815416 | 3363 | 3 | 3 | 0 | 0 | 0 | 0 | 0 | 0 | 4 | 9 | 6 | 13 |
| LTR16_dup928 | 2.53068389298834 | 4.04215123289087 | 1.07322617641985 | 3.76635542600629 | 0.000165647932252706 | 0.0143217023197391 | 3364 | 3 | 1 | 0 | 0 | 0 | 0 | 0 | 0 | 4 | 5 | 8 | 11 |
| LTR48_dup336 | 2.91206830539141 | 4.05398205442954 | 1.05977527997368 | 3.8253223405345 | 0.000130601089907624 | 0.0121390652106092 | 3365 | 4 | 1 | 0 | 0 | 0 | 0 | 0 | 0 | 7 | 6 | 6 | 12 |
| L2a_dup61553 | 4.00958701829312 | 4.2697605171153 | 0.930913499415336 | 4.58663508456687 | 4.50446992085353e-06 | 0.00101019849178833 | 3366 | 4 | 2 | 0 | 0 | 0 | 0 | 0 | 0 | 9 | 14 | 6 | 14 |
| MER34C__dup237 | 3.19993716341274 | 5.01804907511972 | 0.972722187088533 | 5.15876901105679 | 2.48578757963564e-07 | 0.000106573551607925 | 3367 | 1 | 1 | 0 | 0 | 0 | 0 | 0 | 0 | 7 | 12 | 6 | 12 |
| AluSx3_dup23986 | 3.27263431719298 | 2.9417935184932 | 0.840465348201756 | 3.50019608159623 | 0.000464916041449827 | 0.02906928846143 | 3368 | 0 | 2 | 0 | 0 | 1 | 1 | 2 | 2 | 11 | 9 | 6 | 3 |
| MIR3_dup17929 | 2.99658347793088 | 3.54556419442986 | 0.953604276806327 | 3.71806658240266 | 0.000200753364194394 | 0.0164832505164249 | 3369 | 0 | 1 | 0 | 0 | 1 | 0 | 1 | 2 | 8 | 14 | 5 | 2 |
| MER58A_dup3816 | 2.72715824233713 | 2.98089497240431 | 0.881623610810402 | 3.38114240119344 | 0.000721851228526823 | 0.0385029393461205 | 3370 | 1 | 2 | 0 | 0 | 0 | 0 | 2 | 2 | 6 | 5 | 6 | 9 |
| OldhAT1_dup864 | 2.62372303219771 | 3.79488935345887 | 0.954956155345799 | 3.97388857301485 | 7.07086522508883e-05 | 0.00781711542741796 | 3371 | 1 | 1 | 0 | 0 | 0 | 0 | 1 | 1 | 10 | 7 | 3 | 6 |
| MER5B_dup8295 | 2.52553033844674 | 3.84333482073388 | 1.06218693227987 | 3.6183224477113 | 0.000296518819326318 | 0.0214454306940486 | 3372 | 1 | 2 | 0 | 0 | 0 | 0 | 0 | 1 | 7 | 6 | 10 | 2 |
| AluSx1_dup88605 | 3.37464759697838 | 3.8217594119592 | 1.03129143122549 | 3.70579963746794 | 0.000210724839814222 | 0.0170316347603379 | 3373 | 0 | 2 | 0 | 0 | 0 | 0 | 0 | 3 | 8 | 10 | 10 | 6 |
| MIRb_dup75957 | 1.98995739699349 | 3.85286732702644 | 1.10733656131258 | 3.47940044755629 | 0.000502537042862228 | 0.030629272336295 | 3374 | 0 | 2 | 0 | 0 | 0 | 0 | 0 | 1 | 3 | 6 | 7 | 5 |
| MIRb_dup46316 | 3.78062235731364 | 3.52207681995356 | 0.843737913956814 | 4.17437306264495 | 2.98807767486179e-05 | 0.00416520873123997 | 3375 | 1 | 2 | 0 | 2 | 0 | 0 | 1 | 1 | 10 | 14 | 10 | 3 |
| MIRc_dup21227 | 4.36623311889163 | 3.33354946630702 | 0.752077575723928 | 4.43245427587473 | 9.31664642800558e-06 | 0.0017594117071097 | 3376 | 2 | 2 | 0 | 1 | 0 | 0 | 3 | 1 | 13 | 15 | 8 | 6 |
| AluSx_dup21158 | 3.04061743314183 | 3.1964466764927 | 0.892145354783967 | 3.58287655632833 | 0.000339831225247587 | 0.0235954131084804 | 3377 | 3 | 0 | 0 | 1 | 0 | 0 | 2 | 1 | 6 | 13 | 4 | 7 |
| Charlie4a_dup683 | 3.681043412976 | 3.24825460275208 | 0.816852761206152 | 3.97654847607512 | 6.99227502619427e-05 | 0.00775295374362917 | 3378 | 2 | 1 | 0 | 1 | 0 | 0 | 2 | 2 | 9 | 14 | 3 | 10 |

| | | | | | | | | | | | | | | | | | | | |
|---|---|---|---|---|---|---|---|---|---|---|---|---|---|---|---|---|---|---|---|
| MLT1K_dup12182 | 2.49420905139594 | 4.65167639545434 | 1.0471378946037 | 4.44227681896166 | 8.90119408964532e-06 | 0.0016930483843763 | 3379 | 0 | 1 | 0 | 0 | 0 | 0 | 1 | 0 | 3 | 11 | 7 | 8 |
| AluSq2_dup53631 | 1.88325495737704 | 4.1475667551076 | 1.08387745710628 | 3.82660117886453 | 0.000129924760293206 | 0.0120972195307501 | 3380 | 0 | 1 | 0 | 0 | 0 | 0 | 1 | 0 | 6 | 6 | 4 | 4 |
| MER5A_dup6848 | 2.46434460071707 | 4.05195773888953 | 1.03732095939903 | 3.90617552086965 | 9.37684124735853e-05 | 0.00958517281496258 | 3381 | 0 | 2 | 0 | 0 | 0 | 0 | 1 | 0 | 6 | 12 | 4 | 4 |
| MER41E_dup293 | 3.39077501585535 | 3.90173755126266 | 0.907616759387102 | 4.29888222193851 | 1.71661658829684e-05 | 0.00276548443276155 | 3382 | 0 | 2 | 0 | 0 | 0 | 1 | 2 | 0 | 10 | 13 | 4 | 8 |
| MLT2D_dup2368 | 2.70203679649754 | 3.98407240656375 | 0.929486419752477 | 4.28631588573905 | 1.81660626614114e-05 | 0.0028913324053727 | 3383 | 1 | 1 | 0 | 0 | 0 | 1 | 1 | 0 | 6 | 9 | 5 | 9 |
| AluSg_dup28592 | 2.74778695993805 | 3.38020684887013 | 0.896671476504576 | 3.76972719378443 | 0.000163426083342213 | 0.0142013401740939 | 3384 | 2 | 1 | 0 | 0 | 1 | 0 | 2 | 0 | 6 | 8 | 4 | 10 |
| PRIMA41-int_dup13 | 2.82608906983489 | 3.11988698908074 | 0.865035733817562 | 3.60665677394864 | 0.000310167492613562 | 0.0221484413675961 | 3385 | 1 | 3 | 0 | 0 | 2 | 0 | 1 | 0 | 8 | 11 | 3 | 4 |
| MER2_dup7912 | 4.16520217777093 | 4.30422671793483 | 0.836742523246905 | 5.14402770069897 | 2.68909856969606e-07 | 0.000113236600373145 | 3386 | 2 | 2 | 0 | 0 | 0 | 0 | 1 | 0 | 12 | 17 | 6 | 9 |
| MLT1G3_dup1294 | 3.94024766683918 | 3.51162492018856 | 0.862455783356375 | 4.07165791911389 | 4.6679698700568e-05 | 0.00572538535412647 | 3387 | 2 | 4 | 0 | 0 | 0 | 0 | 2 | 0 | 11 | 13 | 5 | 10 |
| L1MA7_dup3805 | 3.7787366967306 | 4.5194028427147 | 0.903772296738833 | 5.00059899935248 | 5.71524720327111e-07 | 0.000206048516822659 | 3388 | 2 | 2 | 0 | 0 | 0 | 0 | 0 | 0 | 11 | 14 | 9 | 6 |
| L2c_dup29830 | 3.46264754860259 | 4.34930983192906 | 0.907078566434194 | 4.79485459459877 | 1.62792867589513e-06 | 0.000475287259772446 | 3389 | 1 | 2 | 0 | 0 | 0 | 0 | 1 | 0 | 9 | 12 | 9 | 7 |
| MLT1A0_dup3785 | 3.28884953107912 | 3.35621823170528 | 0.870311524435802 | 3.8563412496244 | 0.000115096785133521 | 0.011102716573553 | 3390 | 1 | 2 | 1 | 0 | 0 | 0 | 3 | 0 | 9 | 9 | 9 | 5 |
| AluSz_dup18885 | 3.34054888850407 | 2.96244055692619 | 0.759014627775937 | 3.90300851724917 | 9.50043581079687e-05 | 0.00968133061856904 | 3391 | 2 | 2 | 1 | 2 | 0 | 0 | 2 | 0 | 7 | 11 | 6 | 8 |
| L2_dup26433 | 3.12768148140307 | 3.09195324634214 | 0.855009920008113 | 3.61627762905113 | 0.000298869830115544 | 0.0215826648244835 | 3392 | 1 | 2 | 2 | 1 | 0 | 0 | 1 | 0 | 11 | 11 | 3 | 4 |
| AluY_dup48501 | 2.85495832135189 | 2.81571507657529 | 0.850539283557864 | 3.31050561803208 | 0.00093127582754036 | 0.0455859613371567 | 3393 | 0 | 2 | 2 | 1 | 2 | 0 | 1 | 0 | 6 | 10 | 7 | 3 |
| MIR_dup34597 | 2.07000701211624 | 3.41615076559297 | 1.01486464155015 | 3.36611467749529 | 0.000762350078974701 | 0.0400879893683346 | 3394 | 0 | 1 | 1 | 0 | 1 | 0 | 1 | 0 | 6 | 7 | 5 | 2 |
| MIR_dup57117 | 3.56631751016757 | 3.30519256306866 | 0.912417284472186 | 3.62245720167461 | 0.000291817754061559 | 0.0212009829421527 | 3395 | 0 | 4 | 1 | 1 | 0 | 0 | 1 | 0 | 11 | 15 | 5 | 3 |
| MLT1A1-int_dup242 | 3.02257630556629 | 3.4724466134067 | 0.974862309663943 | 3.56198673287895 | 0.000368058987719728 | 0.0249155516051091 | 3396 | 0 | 3 | 1 | 1 | 0 | 0 | 0 | 0 | 11 | 13 | 3 | 2 |
| L1MB8_dup6681 | 2.89066012678283 | 3.61278237464932 | 0.961587273386758 | 3.75710294285086 | 0.000171891798431084 | 0.0147444000971459 | 3397 | 0 | 2 | 0 | 2 | 0 | 0 | 1 | 0 | 8 | 12 | 5 | 4 |
| L1PA8_dup4394 | 3.31815861076813 | 4.01808704003566 | 1.0120975862191 | 3.9700589100761 | 7.18548636270973e-05 | 0.00791188217520281 | 3398 | 0 | 2 | 0 | 1 | 0 | 0 | 1 | 0 | 13 | 13 | 6 | 1 |
| L2a_dup61550 | 3.02980877796439 | 3.94799318703964 | 1.14511204627899 | 3.44769160351472 | 0.000565399217668014 | 0.0331044572401402 | 3399 | 0 | 5 | 0 | 0 | 1 | 0 | 0 | 0 | 6 | 6 | 9 | 10 |
| L2b_dup33106 | 2.9322155209638 | 4.31643809518732 | 1.22629671229014 | 3.51989698082634 | 0.000431714445553963 | 0.0276811440209602 | 3400 | 0 | 4 | 0 | 0 | 0 | 0 | 0 | 0 | 3 | 8 | 10 | 12 |
| AluJb_dup86234 | 2.67457026150988 | 4.06651984328955 | 1.02922513240069 | 3.95104988721399 | 7.78091014529819e-05 | 0.00836167592889316 | 3401 | 0 | 3 | 0 | 0 | 0 | 1 | 0 | 0 | 7 | 6 | 7 | 8 |
| L2b_dup42806 | 2.63124029227154 | 5.13155000475281 | 1.04871874618009 | 4.89316132036759 | 9.92290039079024e-07 | 0.000316503278336055 | 3402 | 0 | 1 | 0 | 0 | 0 | 0 | 0 | 0 | 7 | 7 | 5 | 12 |
| MER57A1_dup303 | 2.33741590726945 | 4.57432277771583 | 1.13297466407223 | 4.03744489861266 | 5.40365314232471e-05 | 0.00639033932008616 | 3403 | 0 | 2 | 0 | 0 | 0 | 0 | 0 | 0 | 3 | 6 | 5 | 14 |
| L1MC4a_dup2967 | 1.53686882378557 | 4.32097548110508 | 1.24679356179534 | 3.46567035113897 | 0.000528911340952008 | 0.031611087468136 | 3404 | 0 | 1 | 0 | 0 | 0 | 0 | 0 | 0 | 5 | 2 | 6 | 4 |
| MIR_dup60766 | 1.80791458721671 | 4.16720932590018 | 1.21660899678871 | 3.42526591279508 | 0.000614197957928605 | 0.0348840822530244 | 3405 | 0 | 2 | 0 | 0 | 0 | 0 | 0 | 0 | 4 | 2 | 8 | 6 |
| AluSz_dup3829 | 2.10269390955611 | 4.78340229974869 | 1.09582540195918 | 4.36511354016493 | 1.27056745976343e-05 | 0.00223614234305376 | 3406 | 0 | 1 | 0 | 0 | 0 | 0 | 0 | 0 | 5 | 7 | 7 | 5 |
| MLT1I_dup9171 | 1.86550404248253 | 4.17198687082985 | 1.0928426844043 | 3.81755483233525 | 0.000134780838531164 | 0.0123865151032465 | 3407 | 0 | 1 | 1 | 0 | 0 | 0 | 0 | 0 | 5 | 5 | 6 | 4 |
| L1M5_dup24335 | 1.80355732458244 | 4.48402370764136 | 1.14486507624874 | 3.91663943696639 | 8.97918899312516e-05 | 0.00931775884935961 | 3408 | 0 | 1 | 0 | 0 | 0 | 0 | 0 | 0 | 6 | 6 | 3 | 5 |
| L1MC4_dup20988 | 1.90521005129889 | 4.44409993620681 | 1.14477993098254 | 3.88205611919886 | 0.000103576965647469 | 0.0103120186895199 | 3409 | 0 | 1 | 0 | 0 | 0 | 0 | 0 | 0 | 6 | 9 | 2 | 4 |
| L1MA4_dup7544 | 1.87763934342327 | 4.61808620050902 | 1.15179417835468 | 4.00947173314062 | 6.08547369638281e-05 | 0.00697501986589356 | 3410 | 0 | 1 | 0 | 0 | 0 | 0 | 0 | 0 | 4 | 7 | 3 | 8 |
| MIRc_dup84138 | 1.58033643749092 | 4.28142106085879 | 1.2720101760293 | 3.36587013338498 | 0.000763026223750332 | 0.0401146791507946 | 3411 | 0 | 1 | 0 | 0 | 0 | 0 | 0 | 0 | 4 | 7 | 1 | 6 |
| AluJr_dup65589 | 2.05918744065164 | 4.3625119135303 | 1.05962283010387 | 4.11704220557673 | 3.8376581518756e-05 | 0.00497816297128209 | 3412 | 0 | 1 | 0 | 0 | 1 | 0 | 0 | 0 | 5 | 5 | 5 | 8 |
| MSTB_dup2136 | 2.06109965756272 | 3.91955652825086 | 1.03187485525127 | 3.79848051176362 | 0.000145585873711395 | 0.013047674691575 | 3413 | 0 | 1 | 0 | 0 | 1 | 0 | 1 | 0 | 5 | 6 | 4 | 7 |
| MER31B_dup296 | 2.17673739843659 | 4.02754250785016 | 1.10184749656605 | 3.65526311073189 | 0.000256918348199645 | 0.0194876293531122 | 3414 | 0 | 1 | 0 | 2 | 0 | 0 | 0 | 0 | 4 | 9 | 3 | 8 |
| MIRb_dup207174 | 2.56593016410937 | 4.67741565150235 | 1.00322302298079 | 4.66238866568746 | 3.12560184923958e-06 | 0.000779585385345032 | 3415 | 0 | 1 | 0 | 1 | 0 | 0 | 0 | 0 | 6 | 9 | 6 | 8 |
| L3_dup12646 | 2.00104202788915 | 3.86569372117961 | 1.03853234698164 | 3.72226607328579 | 0.000197442802380159 | 0.0162893168114593 | 3416 | 0 | 1 | 1 | 1 | 0 | 0 | 0 | 0 | 4 | 4 | 5 | 9 |
| MIRb_dup5888 | 1.99556500171011 | 3.83275830994495 | 1.06730189712754 | 3.5910723294507 | 0.000329320219665552 | 0.0230675387072533 | 3417 | 0 | 1 | 1 | 1 | 0 | 0 | 0 | 0 | 5 | 8 | 2 | 6 |
| MLT1B_dup1767 | 2.25541330056557 | 3.69662884709074 | 1.03891198583916 | 3.5581732595999 | 0.000373442960873473 | 0.0251512195776959 | 3418 | 1 | 2 | 0 | 1 | 0 | 0 | 0 | 0 | 7 | 3 | 6 | 7 |
| UCON69_dup10 | 2.72147421045751 | 3.63277301022677 | 0.972926993257464 | 3.73385982237357 | 0.000188567471882551 | 0.0157545209496367 | 3419 | 1 | 2 | 0 | 2 | 0 | 0 | 0 | 0 | 8 | 6 | 4 | 10 |
| MIRb_dup122094 | 1.861255225136 | 3.80148130953023 | 1.08026329554845 | 3.51903218890744 | 0.000433124153344153 | 0.0277527759754305 | 3420 | 1 | 1 | 0 | 0 | 0 | 1 | 0 | 0 | 4 | 6 | 2 | 8 |
| MIRc_dup71469 | 1.9705853571158 | 3.54784903209402 | 1.01402612733867 | 3.49877477161799 | 0.000467401220510818 | 0.0291600028357806 | 3421 | 1 | 2 | 0 | 0 | 1 | 0 | 0 | 0 | 4 | 6 | 4 | 6 |
| MLT1G_dup91 | 1.92882342041807 | 3.81565185491062 | 1.08020958472357 | 3.53232549393371 | 0.000411921995389156 | 0.0268205162266047 | 3422 | 1 | 2 | 0 | 0 | 0 | 0 | 0 | 0 | 5 | 6 | 4 | 5 |
| L2a_dup61551 | 1.7914270915557 | 4.16081073654951 | 1.09459489593444 | 3.80123345358513 | 0.000143977567280886 | 0.0129523199335416 | 3423 | 1 | 1 | 0 | 0 | 0 | 0 | 0 | 0 | 4 | 4 | 5 | 7 |
| MIR1_Amn_dup3703 | 1.86229312863835 | 4.21993454571435 | 1.09094709222878 | 3.8681385887313 | 0.00010966933172052 | 0.0107344215572073 | 3424 | 1 | 1 | 0 | 0 | 0 | 0 | 0 | 0 | 4 | 5 | 4 | 8 |
| LTR79_dup1389 | 3.00272353489924 | 2.48369005398648 | 0.758493664172158 | 3.27450336278979 | 0.00105847879304109 | 0.0496081928085395 | 3425 | 2 | 2 | 2 | 1 | 0 | 0 | 1 | 2 | 6 | 9 | 5 | 6 |
| UCON35_dup27 | 3.01095503493835 | 2.68685781876328 | 0.814351875123739 | 3.29938187758825 | 0.000968979974941793 | 0.0468057062763905 | 3426 | 3 | 2 | 1 | 1 | 0 | 0 | 1 | 1 | 11 | 5 | 4 | 6 |
| L1ME4a_dup2850 | 2.83606063663096 | 3.06217271310812 | 0.90373217200572 | 3.38836306592031 | 0.000703111268859241 | 0.0379186933823326 | 3427 | 1 | 2 | 2 | 1 | 0 | 0 | 0 | 1 | 5 | 10 | 2 | 11 |
| MIRc_dup21222 | 3.92140165747316 | 3.11025771491161 | 0.844261641497434 | 3.68399742690556 | 0.000229604558860023 | 0.0180070818597584 | 3428 | 3 | 1 | 1 | 2 | 0 | 0 | 1 | 1 | 10 | 17 | 1 | 10 |
| AluSx1_dup20429 | 3.25044009156464 | 2.89070643687392 | 0.883225617826195 | 3.272896956939 | 0.00106451277949322 | 0.0498320987269906 | 3429 | 2 | 0 | 1 | 3 | 0 | 0 | 1 | 2 | 6 | 16 | 3 | 5 |
| L1MC_dup1457 | 4.29191416043155 | 3.21535287133737 | 0.753550697970825 | 4.2669363587556 | 1.98175662968611e-05 | 0.0030709302228926 | 3430 | 4 | 0 | 1 | 3 | 0 | 0 | 1 | 1 | 10 | 14 | 8 | 10 |
| AluSc8_dup16286 | 3.5712468977934 | 2.8601561228078 | 0.758670707587137 | 3.76995723467984 | 0.000163275522707027 | 0.0142013401740939 | 3431 | 1 | 1 | 1 | 3 | 0 | 0 | 3 | 2 | 6 | 9 | 4 | 15 |
| MIRc_dup10829 | 3.51244710108933 | 2.71990131641439 | 0.825167922178786 | 3.29617916948677 | 0.000980094876490556 | 0.0471797805950681 | 3432 | 1 | 0 | 4 | 4 | 0 | 0 | 1 | 3 | 7 | 8 | 4 | 11 |
| L1PA6_dup2403 | 3.57654585578254 | 2.54763711401762 | 0.725209564575398 | 3.51296678706828 | 0.00044313286559784 | 0.0281778976049021 | 3433 | 2 | 1 | 1 | 3 | 2 | 1 | 1 | 0 | 11 | 13 | 3 | 4 |
| AluSq_dup3423 | 3.6574968012196 | 2.70615581056001 | 0.713733258781659 | 3.79155066302979 | 0.000149709599507232 | 0.0133069855675562 | 3434 | 1 | 2 | 1 | 3 | 2 | 0 | 2 | 0 | 9 | 12 | 6 | 6 |
| MIR_dup34586 | 3.24721229458786 | 2.62916445507568 | 0.792125488269527 | 3.31912619150703 | 0.000902996127822515 | 0.0446332190757906 | 3435 | 3 | 3 | 1 | 1 | 1 | 0 | 1 | 0 | 9 | 12 | 5 | 2 |
| MIR_dup34582 | 3.49321140532782 | 2.52881462447392 | 0.745572400119767 | 3.39177606905472 | 0.000694411679964385 | 0.0375870067335773 | 3436 | 2 | 2 | 3 | 1 | 1 | 0 | 1 | 1 | 6 | 16 | 4 | 5 |
| L2_dup10918 | 4.14623664486391 | 3.55065691162584 | 0.741907465470626 | 4.78584874378301 | 1.70266253613675e-06 | 0.000493754345008578 | 3437 | 2 | 2 | 1 | 0 | 2 | 0 | 1 | 0 | 11 | 16 | 6 | 8 |
| L2b_dup30991 | 3.17037952260162 | 3.21741203135519 | 0.828180442155273 | 3.88491670122291 | 0.000102364979464522 | 0.0102427827601619 | 3438 | 1 | 2 | 2 | 0 | 1 | 0 | 1 | 0 | 9 | 11 | 6 | 4 |
| L1PA5_dup8661 | 3.37513660399533 | 2.28514738930677 | 0.684446588627303 | 3.33867890829841 | 0.000841778042191922 | 0.0427165105624651 | 3439 | 2 | 2 | 3 | 3 | 1 | 1 | 1 | 0 | 7 | 7 | 5 | 10 |
| L2c_dup99089 | 3.4630264823754 | 2.20189380129747 | 0.650245443146609 | 3.38625026058201 | 0.00070854735459297 | 0.0381266415278107 | 3440 | 2 | 2 | 2 | 2 | 2 | 1 | 3 | 0 | 4 | 8 | 9 | 9 |
| L1P3_dup1512 | 3.49141500369065 | 3.05157770362471 | 0.836678119797573 | 3.64725410097136 | 0.000265057771332152 | 0.0198894470064334 | 3441 | 3 | 0 | 3 | 1 | 1 | 0 | 1 | 0 | 4 | 15 | 5 | 11 |
| MLT1G3_dup1295 | 4.11619924591671 | 3.34160100003873 | 0.737334015093427 | 4.53200439913968 | 5.84266328738869e-06 | 0.00123248081434485 | 3442 | 3 | 0 | 1 | 1 | 1 | 1 | 2 | 0 | 6 | 18 | 8 | 10 |
| L2c_dup5667 | 2.33652256721002 | 4.94286651513807 | 1.13953816963604 | 4.33760504636435 | 1.44043695490636e-05 | 0.00244993718135918 | 3443 | 0 | 0 | 0 | 0 | 0 | 0 | 0 | 1 | 3 | 10 | 10 | 4 |
| MIRb_dup165043 | 2.14169107979315 | 4.79551540973205 | 1.11164664580636 | 4.31388465734408 | 1.60410659737385e-05 | 0.00263416665575375 | 3444 | 0 | 0 | 0 | 0 | 0 | 0 | 0 | 1 | 3 | 7 | 9 | 6 |
| L1PA2_dup68 | 1.73951946538619 | 4.48265016037634 | 1.19402794454941 | 3.75422550271046 | 0.000173878275254173 | 0.0148665612948723 | 3445 | 0 | 0 | 0 | 0 | 0 | 0 | 0 | 1 | 5 | 4 | 7 | 3 |
| L2a_dup142891 | 1.6832151614484 | 4.00269509874286 | 1.14093418700557 | 3.50826116381707 | 0.000451046026909981 | 0.028498300031631 | 3446 | 1 | 0 | 0 | 0 | 0 | 0 | 0 | 1 | 3 | 6 | 6 | 3 |
| LTR82B_dup10 | 1.53982226637166 | 4.7520861105917 | 1.30883501633779 | 3.63077549979399 | 0.000282570898397945 | 0.0207468469925368 | 3447 | 0 | 0 | 0 | 0 | 0 | 0 | 0 | 0 | 5 | 3 | 6 | 4 |
| MIR_dup6759 | 1.43308383989567 | 4.64914513994994 | 1.33010637460204 | 3.49531829087049 | 0.000473496735431997 | 0.0294125822752792 | 3448 | 0 | 0 | 0 | 0 | 0 | 0 | 0 | 0 | 4 | 3 | 6 | 4 |
| MER30_dup1384 | 1.52400155617522 | 4.72457769715489 | 1.35868859785052 | 3.47730723922265 | 0.000506477084138587 | 0.0307681606502456 | 3449 | 0 | 0 | 0 | 0 | 0 | 0 | 0 | 0 | 3 | 7 | 6 | 2 |
| MER61E_dup253 | 1.54448060075165 | 4.75407848059373 | 1.35099885954935 | 3.51893596873911 | 0.000433281268410408 | 0.0277527759754305 | 3450 | 0 | 0 | 0 | 0 | 0 | 0 | 0 | 0 | 4 | 5 | 7 | 2 |

| | | | | | | | | | | | | | | | | | | |
|---|---|---|---|---|---|---|---|---|---|---|---|---|---|---|---|---|---|---|
| MIRb_dup183581 | 1.89410618006005 | 4.19657349604747 | 1.08481186420089 | 3.86848045687516 | 0.000109515712056932 | 0.0107237965518925 | 3451 | 1 | 0 | 0 | 1 | 0 | 0 | 0 | 0 | 4 | 8 | 4 | 5 |
| MLT1H1_dup79 | 1.82884232362276 | 4.52897373697589 | 1.18051295955406 | 3.8364455894552 | 0.000124827835951524 | 0.011767941463281 | 3452 | 0 | 0 | 0 | 1 | 0 | 0 | 0 | 0 | 4 | 10 | 2 | 5 |
| L2c_dup5657 | 1.76772032067965 | 4.52006915538647 | 1.16928343956776 | 3.86567448270487 | 0.000110782614882587 | 0.0108166926880805 | 3453 | 0 | 0 | 0 | 1 | 0 | 0 | 0 | 0 | 4 | 8 | 5 | 3 |
| MER57A-int_dup30 | 1.79384867593344 | 4.5060650023728 | 1.15329642215931 | 3.90711781966349 | 9.34036148419237e-05 | 0.00956737689934412 | 3454 | 0 | 0 | 0 | 1 | 0 | 0 | 0 | 0 | 5 | 8 | 4 | 3 |
| MIR_dup107109 | 1.45550879810677 | 4.68224861507138 | 1.32198100552037 | 3.5418425798246 | 0.000397342526133666 | 0.026196842555987 | 3455 | 0 | 0 | 0 | 0 | 0 | 0 | 0 | 0 | 3 | 5 | 3 | 7 |
| MLT1L_dup8752 | 1.63028860632673 | 4.84355284862623 | 1.27593835699309 | 3.79607119895719 | 0.000147007294284931 | 0.0131305546718927 | 3456 | 0 | 0 | 0 | 0 | 0 | 0 | 0 | 0 | 4 | 5 | 3 | 8 |
| L1MCa_dup5844 | 1.51242384299435 | 4.67522853707575 | 1.31651776282491 | 3.55120809539546 | 0.000383467096253064 | 0.0256017008094926 | 3457 | 0 | 0 | 0 | 0 | 0 | 0 | 0 | 0 | 4 | 7 | 2 | 5 |
| MIRb_dup11013 | 1.52216787713389 | 4.71604872264969 | 1.27925839068362 | 3.68654898572094 | 0.000227315701990772 | 0.017906561011388 | 3458 | 0 | 0 | 0 | 0 | 0 | 0 | 0 | 0 | 4 | 6 | 4 | 4 |
| MSTA_dup367 | 1.506774532317 | 4.71943195466424 | 1.27943952531216 | 3.68867137625187 | 0.000225428160979343 | 0.0178205080373391 | 3459 | 0 | 0 | 0 | 0 | 0 | 0 | 0 | 0 | 4 | 5 | 4 | 5 |
| L1P5_dup489 | 1.83486412788314 | 5.00649137847522 | 1.20447608821253 | 4.15657183025106 | 3.23058631311325e-05 | 0.00440771267694981 | 3460 | 0 | 0 | 0 | 0 | 0 | 0 | 0 | 0 | 4 | 7 | 6 | 5 |
| MIRb_dup11007 | 2.00681928076443 | 5.13893262259597 | 1.16237549973782 | 4.42106068456801 | 9.82175829556548e-06 | 0.00183297890148755 | 3461 | 0 | 0 | 0 | 0 | 0 | 0 | 0 | 0 | 5 | 6 | 7 | 6 |
| L2a_dup5943 | 2.1812077625649 | 5.19761116766814 | 1.13983812697967 | 4.55995552758067 | 5.11644551171678e-06 | 0.00111384971979045 | 3462 | 0 | 0 | 0 | 0 | 0 | 0 | 0 | 0 | 8 | 7 | 5 | 5 |
| L2c_dup5658 | 1.81907945664675 | 4.92861286603871 | 1.20493118700131 | 4.09036874404792 | 4.30687895078117e-05 | 0.00540297504057472 | 3463 | 0 | 0 | 0 | 0 | 0 | 0 | 0 | 0 | 6 | 7 | 4 | 4 |
| MIRb_dup11017 | 1.7328200747472 | 4.89713431904787 | 1.23453796422485 | 3.96677498866768 | 7.28517108784182e-05 | 0.00796268865258433 | 3464 | 0 | 0 | 0 | 0 | 0 | 0 | 0 | 0 | 6 | 5 | 5 | 4 |
| L1PB3_dup2139 | 2.56332769777497 | 5.49349165918961 | 1.08538721419496 | 5.06131967222791 | 4.16364428606289e-07 | 0.000160052100693299 | 3465 | 0 | 0 | 0 | 0 | 0 | 0 | 0 | 0 | 7 | 7 | 6 | 11 |
| L1PA10_dup4455 | 2.16298976240933 | 5.22580073030546 | 1.15725430175961 | 4.51568918115892 | 6.31111755560236e-06 | 0.00130605295828618 | 3466 | 0 | 0 | 0 | 0 | 0 | 0 | 0 | 0 | 8 | 5 | 6 | 6 |
| MER52A_dup908 | 2.06698634637024 | 5.16572008369832 | 1.19088115552604 | 4.33772930214562 | 1.43962308701299e-05 | 0.00244993718135918 | 3467 | 0 | 0 | 0 | 0 | 0 | 0 | 0 | 0 | 8 | 4 | 5 | 7 |
| L1MB3_dup14038 | 1.87413773645079 | 4.23663060385819 | 1.15992730578744 | 3.65249665450549 | 0.000259702980753341 | 0.0196237807108081 | 3468 | 0 | 0 | 0 | 0 | 0 | 2 | 0 | 0 | 4 | 4 | 6 | 7 |
| Tigger1_dup3855 | 1.848009381197 | 4.21418508830042 | 1.16849172221396 | 3.60651685261042 | 0.00031033471258002 | 0.0221484413675961 | 3469 | 0 | 0 | 0 | 0 | 0 | 2 | 0 | 0 | 3 | 4 | 7 | 7 |
| L1MA9_dup481 | 1.9507000114158 | 4.69292054467396 | 1.12696741201863 | 4.16420252673319 | 3.12442687839468e-05 | 0.00430600629491679 | 3470 | 0 | 0 | 0 | 0 | 0 | 1 | 0 | 0 | 3 | 6 | 8 | 6 |
| L1PA6_dup2143 | 2.06164539594732 | 4.7752915205045 | 1.10768860748762 | 4.31104146799476 | 1.62487427127544e-05 | 0.00265544184995128 | 3471 | 0 | 0 | 0 | 0 | 0 | 1 | 0 | 0 | 3 | 6 | 6 | 10 |
| L1MC_dup276 | 1.89017765835055 | 4.60821219450866 | 1.13812722377072 | 4.0489429461508 | 5.14494841900274e-05 | 0.00617526689724256 | 3472 | 0 | 0 | 0 | 0 | 0 | 1 | 0 | 0 | 6 | 5 | 6 | 4 |
| MER2_dup7919 | 1.68926949487679 | 4.46392429711709 | 1.19395505521749 | 3.73877080013195 | 0.000184922225508698 | 0.0155767044856915 | 3473 | 0 | 0 | 0 | 0 | 0 | 1 | 0 | 0 | 4 | 5 | 7 | 3 |
| L2c_dup8973 | 1.87195965819498 | 4.62072965112407 | 1.16359894127021 | 3.97106725284571 | 7.15513755272577e-05 | 0.00788576359481651 | 3474 | 0 | 0 | 0 | 0 | 0 | 1 | 0 | 0 | 6 | 3 | 7 | 5 |
| MER5B_dup787 | 2.29421899075889 | 4.9207481517116 | 1.08226423510826 | 4.54671603485017 | 5.44894362545802e-06 | 0.00116491854438745 | 3475 | 0 | 0 | 0 | 0 | 0 | 1 | 0 | 0 | 7 | 5 | 8 | 6 |
| AluSz6_dup37855 | 2.13360749510095 | 3.99799673007126 | 1.1565784118045 | 3.45674507605028 | 0.000546741798649217 | 0.0323378135772773 | 3476 | 0 | 0 | 0 | 0 | 1 | 2 | 0 | 0 | 2 | 11 | 7 | 3 |
| L1MD2_dup8132 | 2.19171153328336 | 4.80858251515114 | 1.27697622253521 | 3.7656006668664 | 0.000166149163514243 | 0.0143396912809431 | 3477 | 0 | 0 | 0 | 0 | 0 | 1 | 0 | 0 | 3 | 12 | 9 | 1 |
| AluJb_dup100005 | 1.76490610093279 | 4.45903015458013 | 1.31440112544889 | 3.39244243499662 | 0.000692724859736231 | 0.0375383988783414 | 3478 | 0 | 0 | 0 | 0 | 0 | 1 | 0 | 0 | 2 | 11 | 1 | 7 |
| MamTip3_dup1153 | 1.7113031266684 | 4.3289908745389 | 1.17732040941148 | 3.67698617974597 | 0.000236005821528479 | 0.0183999479716303 | 3479 | 0 | 0 | 0 | 0 | 0 | 1 | 0 | 0 | 5 | 8 | 2 | 4 |
| MIRc_dup4364 | 1.91360448110396 | 5.07990838510794 | 1.33611728943143 | 3.80199285294004 | 0.000143536868870693 | 0.0129251309267695 | 3480 | 0 | 0 | 0 | 0 | 0 | 0 | 0 | 0 | 1 | 10 | 7 | 6 |
| L1M5_dup35159 | 1.729508004124 | 4.91999018254119 | 1.26544354597352 | 3.88795707101747 | 0.000101091500274085 | 0.01016378093529 | 3481 | 0 | 0 | 0 | 0 | 0 | 0 | 0 | 0 | 2 | 6 | 5 | 9 |
| L1ME3A_dup9024 | 1.53512789303161 | 4.74344258935448 | 1.30845374872835 | 3.62522755883767 | 0.000288707117898824 | 0.021059362083982 | 3482 | 0 | 0 | 0 | 0 | 0 | 0 | 0 | 0 | 2 | 5 | 7 | 5 |
| MER5A1_dup4766 | 1.49321486654142 | 4.71476353852284 | 1.30254298746248 | 3.61966060537305 | 0.000294989674284568 | 0.0213543088035643 | 3483 | 0 | 0 | 0 | 0 | 0 | 0 | 0 | 0 | 3 | 5 | 6 | 4 |
| MLT1D_dup15019 | 2.59713516840854 | 5.12181089429743 | 1.08484954214332 | 4.72121773142692 | 2.34436811155382e-06 | 0.000637523445339182 | 3484 | 0 | 0 | 0 | 0 | 0 | 1 | 0 | 0 | 3 | 11 | 10 | 7 |
| AluY_dup3542 | 2.20077202647727 | 5.2575846065376 | 1.1898805312872 | 4.41858192338855 | 9.93506209274298e-06 | 0.00184264518857932 | 3485 | 0 | 0 | 0 | 0 | 0 | 0 | 0 | 0 | 3 | 12 | 5 | 7 |
| Helitron1Nb_Mam_du | 2.69107274078516 | 5.55024798376934 | 1.11303803726904 | 4.98657529924805 | 6.14589599165452e-07 | 0.000219249080604232 | 3486 | 0 | 0 | 0 | 0 | 0 | 0 | 0 | 0 | 4 | 14 | 6 | 9 |
| MIR1_Amn_dup9187 | 1.90230926949511 | 3.33497210616594 | 1.00317955786004 | 3.32440197772771 | 0.000886083763667691 | 0.0440754007120582 | 3487 | 1 | 0 | 1 | 0 | 1 | 1 | 0 | 0 | 4 | 8 | 2 | 5 |
| L1MA1_dup1812 | 1.65726287124337 | 3.77194227819636 | 1.14909021778543 | 3.28254667894205 | 0.00102873951979923 | 0.0487327202425138 | 3488 | 0 | 0 | 1 | 0 | 1 | 0 | 0 | 0 | 5 | 8 | 2 | 2 |
| MIRb_dup183948 | 1.88508204280991 | 4.2029109401998 | 1.08351559852555 | 3.87895748424768 | 0.000104905086044869 | 0.0103834216322931 | 3489 | 0 | 0 | 1 | 0 | 1 | 0 | 0 | 0 | 4 | 6 | 4 | 7 |
| L1PA7_dup11426 | 2.272462649386 | 3.67542922501024 | 1.06919429580236 | 3.43756905497899 | 0.000586961032705775 | 0.0339097792839445 | 3490 | 0 | 0 | 1 | 1 | 1 | 0 | 0 | 0 | 10 | 9 | 2 | 1 |
| MIR_dup28910 | 2.60661086983598 | 4.36059580007101 | 1.05988849380489 | 4.11420241427182 | 3.88520409818248e-05 | 0.00502277394500201 | 3491 | 0 | 0 | 0 | 1 | 1 | 0 | 0 | 0 | 7 | 16 | 3 | 2 |
| MIR_dup143799 | 2.11174443769125 | 3.8311871728301 | 1.08798442199148 | 3.52136215867629 | 0.000429335814115916 | 0.0275732090757424 | 3492 | 0 | 1 | 0 | 0 | 2 | 0 | 0 | 0 | 7 | 7 | 5 | 2 |
| MLT1B_dup8338 | 2.17491207591512 | 3.81583211809249 | 1.09338585189536 | 3.48992271253358 | 0.000483160258777234 | 0.0298483738175653 | 3493 | 1 | 0 | 0 | 0 | 2 | 0 | 0 | 0 | 10 | 5 | 4 | 2 |
| MER57-int_dup485 | 1.64884293197437 | 4.12896690762353 | 1.21854907212488 | 3.38842891277536 | 0.000702942474374946 | 0.0379186933823326 | 3494 | 0 | 0 | 0 | 0 | 1 | 0 | 0 | 0 | 8 | 5 | 2 | 2 |
| MIRb_dup183974 | 1.67719629268898 | 4.36498973035651 | 1.21030137552532 | 3.60653124802236 | 0.000310317504731237 | 0.0221484413675961 | 3495 | 0 | 0 | 0 | 0 | 1 | 0 | 0 | 0 | 6 | 5 | 5 | 2 |
| L1MA6_dup788 | 2.4658263144716 | 3.99255793417314 | 1.05050168189974 | 3.80062022076249 | 0.000144334371204968 | 0.0129746075900027 | 3496 | 0 | 1 | 0 | 0 | 0 | 1 | 0 | 0 | 11 | 11 | 1 | 2 |
| L2b_dup39689 | 2.52795018195702 | 4.66807483643851 | 1.09409075927222 | 4.26662486350189 | 1.9845244347555e-05 | 0.00307228139981998 | 3497 | 0 | 1 | 0 | 0 | 0 | 0 | 0 | 0 | 7 | 16 | 2 | 3 |
| AluY_dup16811 | 1.8508571225779 | 4.78846434985284 | 1.28364147801168 | 3.73037521136355 | 0.0001911948233895 | 0.0159237113917529 | 3498 | 0 | 0 | 0 | 0 | 0 | 0 | 0 | 0 | 7 | 9 | 1 | 4 |
| L2a_dup132176 | 1.93429184913877 | 4.83507696239397 | 1.26920693541036 | 3.80952611232831 | 0.000139233401807286 | 0.0126692279259795 | 3499 | 0 | 0 | 0 | 0 | 0 | 0 | 0 | 0 | 7 | 10 | 1 | 4 |
| L1M1_dup7939 | 1.63133963252907 | 4.48635959022945 | 1.33091209330785 | 3.37089099482075 | 0.000749254981803292 | 0.0395830874556319 | 3500 | 0 | 0 | 0 | 0 | 0 | 0 | 0 | 0 | 7 | 8 | 1 | 2 |
| L3b_dup4309 | 1.90494751212681 | 4.64517375089201 | 1.27840450620239 | 3.63357116496006 | 0.000279525303041784 | 0.0206046864180856 | 3501 | 0 | 0 | 0 | 0 | 0 | 0 | 0 | 0 | 8 | 10 | 1 | 2 |
| L1M5_dup1225 | 1.74920439586145 | 4.83802005373663 | 1.25940079185946 | 3.84152533888237 | 0.000122272110967249 | 0.0116144350164523 | 3502 | 0 | 0 | 0 | 0 | 0 | 0 | 0 | 0 | 7 | 6 | 2 | 5 |
| L1MB8_dup8233 | 2.09085365720316 | 5.07754468100816 | 1.19750797418185 | 4.24009258433304 | 2.2342761314125e-05 | 0.00336171072626349 | 3503 | 0 | 0 | 0 | 0 | 0 | 0 | 0 | 0 | 8 | 8 | 2 | 6 |
| AluSx3_dup11069 | 1.66859432463925 | 4.73533343388941 | 1.34058002695634 | 3.53230194294371 | 0.000411958685662244 | 0.0268205162266047 | 3504 | 0 | 0 | 0 | 0 | 0 | 0 | 0 | 0 | 7 | 6 | 1 | 5 |
| L2c_dup53779 | 2.11698201245696 | 5.04396156184019 | 1.26384653138959 | 3.99096048180343 | 6.58062459654633e-05 | 0.00741047468173125 | 3505 | 0 | 0 | 0 | 0 | 0 | 0 | 0 | 0 | 9 | 8 | 1 | 6 |
| HERV17-int_dup543 | 1.72864369406959 | 3.83450521620105 | 1.15191508870652 | 3.3288089146456 | 0.000872182217813574 | 0.0436358851976175 | 3506 | 1 | 1 | 0 | 0 | 0 | 0 | 0 | 0 | 6 | 7 | 1 | 4 |
| L1MB7_dup9040 | 1.69476980992503 | 4.2401807791833 | 1.24324155889323 | 3.41058481262326 | 0.000648237196257096 | 0.036080806526454 | 3507 | 1 | 0 | 0 | 0 | 0 | 0 | 0 | 0 | 8 | 5 | 1 | 4 |
| L1PA17_dup779 | 1.83941705093108 | 4.10121113433143 | 1.16085160871065 | 3.53293315317589 | 0.000410976372925792 | 0.0267935418017252 | 3508 | 0 | 0 | 0 | 0 | 0 | 0 | 1 | 1 | 7 | 6 | 1 | 5 |
| L3_dup1450 | 1.57489445193073 | 4.36015098112949 | 1.28424133054434 | 3.39511809613026 | 0.000685990038137215 | 0.0373435336569658 | 3509 | 0 | 0 | 0 | 0 | 0 | 0 | 1 | 0 | 5 | 5 | 1 | 7 |
| L1PA7_dup4205 | 1.63177808898054 | 4.33740684468272 | 1.17882988303032 | 3.67941711278384 | 0.000233767651711499 | 0.0182653743020935 | 3510 | 1 | 0 | 0 | 0 | 0 | 0 | 0 | 0 | 5 | 6 | 4 | 3 |
| MamTip1_dup413 | 1.58195470739214 | 4.10783117853025 | 1.25412764386121 | 3.27544903314869 | 0.00105494147610674 | 0.049500908001123 | 3511 | 1 | 0 | 0 | 0 | 0 | 0 | 0 | 0 | 5 | 8 | 3 | 1 |
| L1MC4_dup18000 | 1.90904223558899 | 4.91060813583139 | 1.3138631892627 | 3.73753384367751 | 0.000185834081918044 | 0.0156138688051098 | 3512 | 0 | 0 | 0 | 0 | 0 | 0 | 0 | 0 | 8 | 7 | 5 | 1 |
| L2c_dup110079 | 1.75287175394409 | 4.86598446299176 | 1.35881013063128 | 3.58106283821353 | 0.000342199332568108 | 0.0237252221408074 | 3513 | 0 | 0 | 0 | 0 | 0 | 0 | 0 | 0 | 5 | 8 | 6 | 1 |
| L1MC1_dup10368 | 1.56368957720459 | 4.67565664086216 | 1.27757560445944 | 3.65978860628017 | 0.000252423401229847 | 0.019247007082097 | 3514 | 0 | 0 | 0 | 0 | 0 | 0 | 0 | 0 | 5 | 7 | 3 | 3 |
| L2a_dup5944 | 1.5762582666828 | 4.69093794112738 | 1.30234063086528 | 3.60192858147313 | 0.00031586515071737 | 0.0224220910359772 | 3515 | 0 | 0 | 0 | 0 | 0 | 0 | 0 | 0 | 5 | 7 | 4 | 2 |
| ERVL-B4-int_dup3213 | 2.04794965441562 | 5.02220632003345 | 1.1774290881428 | 4.26540024414988 | 1.99544157108454e-05 | 0.00308116379022247 | 3516 | 0 | 0 | 0 | 0 | 0 | 0 | 0 | 0 | 8 | 8 | 4 | 3 |
| MER39_dup59 | 2.03721464397873 | 5.02535730079142 | 1.21478126510034 | 4.13684129412083 | 3.52119470399738e-05 | 0.00470440311643662 | 3517 | 0 | 0 | 0 | 0 | 0 | 0 | 0 | 0 | 7 | 9 | 5 | 2 |
| L1MC3_dup9211 | 1.91790752802452 | 4.95331465596384 | 1.19914102453548 | 4.13071903522161 | 3.61630347005122e-05 | 0.00478312921336747 | 3518 | 0 | 0 | 0 | 0 | 0 | 0 | 0 | 0 | 6 | 9 | 4 | 3 |
| MER96B_dup43 | 1.74312771980455 | 4.80507211571798 | 1.26772554760485 | 3.7903094441824 | 0.000150459723537293 | 0.0133586900659719 | 3519 | 0 | 0 | 0 | 0 | 0 | 0 | 0 | 0 | 5 | 9 | 4 | 2 |
| MLT1K_dup588 | 1.84986614628054 | 4.86853718978499 | 1.24048899030622 | 3.92469197859078 | 8.68407818206176e-05 | 0.00909300946410297 | 3520 | 0 | 0 | 0 | 0 | 0 | 0 | 0 | 0 | 6 | 9 | 4 | 2 |
| AluJb_dup50990 | 2.18078039718536 | 5.10222736575016 | 1.20346830335441 | 4.23960261481654 | 2.23915819671123e-05 | 0.003365753229783 | 3521 | 0 | 0 | 0 | 0 | 0 | 0 | 0 | 0 | 6 | 12 | 5 | 2 |
| L2a_dup5947 | 2.05864862589247 | 5.07851752363235 | 1.19388548951752 | 4.25377271792182 | 2.10198657104313e-05 | 0.00321439136653874 | 3522 | 0 | 0 | 0 | 0 | 0 | 0 | 0 | 0 | 5 | 11 | 5 | 3 |

| | | | | | | | | | | | | | | | | | | | |
|---|---|---|---|---|---|---|---|---|---|---|---|---|---|---|---|---|---|---|---|
| THE1B_dup10145 | 1.57998049656197 | 4.13479833534623 | 1.25767048109597 | 3.28766429481835 | 0.00101022224140979 | 0.0481676941991385 | 3523 | 0 | 0 | 0 | 1 | 0 | 0 | 0 | 0 | 5 | 9 | 2 | 1 |
| L1PA7_dup172 | 1.62286566651304 | 4.72011977896954 | 1.30541637733116 | 3.61579635504459 | 0.000299425702668698 | 0.0216162464342252 | 3524 | 0 | 0 | 0 | 0 | 0 | 0 | 0 | 0 | 7 | 5 | 4 | 2 |
| MER84_dup273 | 1.55482428476906 | 4.62426174751104 | 1.38659939457435 | 3.33496593580338 | 0.00085309848938882 | 0.0430705226301957 | 3525 | 0 | 0 | 0 | 0 | 0 | 0 | 0 | 0 | 7 | 5 | 4 | 1 |
| L2b_dup39690 | 1.58377729536153 | 4.46826761914402 | 1.36288989621819 | 3.27852428251379 | 0.00104351385900725 | 0.0491975329295304 | 3526 | 0 | 0 | 0 | 0 | 0 | 0 | 0 | 0 | 8 | 6 | 2 | 1 |
| MER50_dup2006 | 2.11277505440143 | 4.93760916324143 | 1.20464843634094 | 4.09879680601187 | 4.1530342633872e-05 | 0.00526196959976534 | 3527 | 0 | 0 | 0 | 0 | 0 | 0 | 0 | 0 | 10 | 8 | 3 | 2 |
| AluSc5_dup2894 | 1.5334956407336 | 3.80420869500041 | 1.15493471788302 | 3.29387335586678 | 0.00098817010097615 | 0.0474150277973121 | 3528 | 1 | 0 | 1 | 0 | 0 | 0 | 0 | 0 | 5 | 5 | 2 | 4 |
| L2c_dup101110 | 1.55858588965806 | 4.28362648686725 | 1.19542356331184 | 3.58335456848428 | 0.000339209659536438 | 0.0235660092403087 | 3529 | 0 | 0 | 1 | 0 | 0 | 0 | 0 | 0 | 5 | 6 | 3 | 3 |
| L2c_dup5660 | 1.99949058775708 | 4.67742945813892 | 1.13915245113716 | 4.10606100480203 | 4.02463232821851e-05 | 0.00515090896725802 | 3530 | 0 | 0 | 1 | 0 | 0 | 0 | 0 | 0 | 8 | 5 | 3 | 6 |
| L2a_dup45374 | 1.70593357777714 | 3.86946116733777 | 1.17869454406581 | 3.2828362418565 | 0.0010276834548758 | 0.0487194744485838 | 3531 | 0 | 0 | 1 | 0 | 0 | 0 | 1 | 0 | 7 | 7 | 2 | 1 |
| MER67D_dup883 | 1.90171354258165 | 4.21282345580892 | 1.26651187148612 | 3.32631975321763 | 0.000880009146678411 | 0.0438702505527443 | 3532 | 0 | 0 | 1 | 0 | 0 | 0 | 0 | 0 | 11 | 6 | 1 | 1 |
| L1MB8_dup13570 | 2.79362872032835 | 4.83357107934132 | 1.03117120475217 | 4.68745738541352 | 2.76620277089292e-06 | 0.000715719428984722 | 3533 | 0 | 0 | 1 | 0 | 0 | 0 | 1 | 0 | 4 | 14 | 8 | 6 |
| MLT1A0_dup9661 | 2.3873623824592 | 4.43171166009136 | 1.06826280472096 | 4.14852191848894 | 3.34628812896857e-05 | 0.00452499294019949 | 3534 | 0 | 0 | 1 | 0 | 0 | 0 | 1 | 0 | 6 | 14 | 2 | 4 |
| L1PA7_dup11662 | 3.19857827285399 | 3.31435985988803 | 0.97125438749711 | 3.41245290889138 | 0.000643810494074716 | 0.0359245795990491 | 3535 | 0 | 1 | 2 | 2 | 0 | 0 | 2 | 0 | 7 | 18 | 3 | 3 |
| MLT1D_dup16487 | 3.06264234778651 | 3.6926757174659 | 1.07796602956003 | 3.4255956275107 | 0.000613452943357403 | 0.0348623534894007 | 3536 | 0 | 0 | 3 | 2 | 0 | 0 | 2 | 0 | 12 | 10 | 3 | 3 |
| L1MA2_dup1530 | 2.70867663543353 | 4.15821535072616 | 1.12099682951126 | 3.70939082186261 | 0.000207758521244435 | 0.0168852651672033 | 3537 | 0 | 0 | 2 | 2 | 0 | 0 | 0 | 0 | 6 | 11 | 9 | 2 |
| L1MA5_dup3752 | 2.39697136098165 | 3.87599222460712 | 1.10438495860789 | 3.50963873094843 | 0.000448715908818624 | 0.028434129331493 | 3538 | 0 | 0 | 2 | 2 | 0 | 0 | 0 | 0 | 7 | 10 | 4 | 3 |
| L1ME4b_dup1127 | 1.93285636472437 | 3.82098394521483 | 1.16354795861828 | 3.28390756643352 | 0.00102378494393989 | 0.0486141553124567 | 3539 | 0 | 0 | 2 | 1 | 0 | 0 | 0 | 0 | 4 | 11 | 2 | 3 |
| MER57A1_dup362 | 3.07278520384743 | 5.63734621472686 | 1.22889476492006 | 4.58733031960924 | 4.48950050533459e-06 | 0.00100969630174135 | 3540 | 0 | 0 | 0 | 0 | 0 | 0 | 0 | 0 | 12 | 11 | 10 | 1 |
| MIRb_dup10999 | 2.81273699002527 | 5.49844355273525 | 1.16540838816786 | 4.71804013816938 | 2.38127519839108e-06 | 0.000645347276032305 | 3541 | 0 | 0 | 0 | 0 | 0 | 0 | 0 | 0 | 12 | 9 | 8 | 2 |
| ERVL-B4-int_dup66 | 2.52434187543815 | 4.8538660490177 | 1.10352527442634 | 4.39850917917666 | 1.08997017506586e-05 | 0.00199196653821982 | 3542 | 0 | 0 | 0 | 0 | 0 | 0 | 1 | 0 | 7 | 15 | 4 | 2 |
| MIRb_dup82623 | 2.55925698525771 | 5.31826336728447 | 1.16202490265904 | 4.5767206495444 | 4.72321495473672e-06 | 0.00105231283297733 | 3543 | 0 | 0 | 0 | 0 | 0 | 0 | 0 | 0 | 5 | 17 | 5 | 3 |
| L1PA3_dup345 | 3.2298660378611 | 4.5911787087131 | 1.0985101882802 | 4.17945937843409 | 2.9220290613665e-05 | 0.00410682295945607 | 3544 | 0 | 0 | 0 | 0 | 0 | 1 | 1 | 0 | 19 | 8 | 4 | 1 |
| MLT1E2_dup3054 | 2.55820343620484 | 4.06726213346093 | 1.08638748128897 | 3.74384112806163 | 0.000181228268711978 | 0.0153314927761662 | 3545 | 0 | 0 | 1 | 0 | 0 | 1 | 1 | 0 | 12 | 7 | 5 | 1 |
| AluSc5_dup195 | 2.56540764470579 | 3.80650517470138 | 1.0428364666237 | 3.65014582490137 | 0.000262091471446103 | 0.0197290785399286 | 3546 | 1 | 0 | 1 | 0 | 0 | 0 | 2 | 0 | 11 | 6 | 5 | 3 |
| MIR_dup34589 | 2.31828440616249 | 3.40976842590445 | 0.997740301458755 | 3.41749092516276 | 0.000632012010498477 | 0.0354932097828819 | 3547 | 1 | 1 | 1 | 0 | 0 | 0 | 1 | 0 | 9 | 9 | 3 | 1 |
| L1M3a_dup247 | 2.79086560212807 | 4.12736505151475 | 1.06843365887468 | 3.86300545404183 | 0.000112000506250013 | 0.0108909164120007 | 3548 | 0 | 2 | 0 | 0 | 0 | 0 | 0 | 1 | 4 | 19 | 3 | 4 |
| L2a_dup34140 | 2.02338474066363 | 3.6222625288451 | 1.06833415498389 | 3.39057074225968 | 0.000697472511913198 | 0.0377097919381272 | 3549 | 0 | 1 | 0 | 0 | 0 | 0 | 1 | 1 | 6 | 10 | 2 | 2 |
| MIRb_dup11012 | 3.17427012709037 | 5.34719362435045 | 1.15117547514109 | 4.64498570358709 | 3.40100005828286e-06 | 0.000827249705078879 | 3550 | 0 | 0 | 0 | 0 | 0 | 0 | 0 | 1 | 8 | 15 | 1 | 13 |
| LTR82A_dup12 | 2.39746062095131 | 4.5535922407551 | 1.0947252437205 | 4.15957544313074 | 3.18839695934374e-05 | 0.00437907830751988 | 3551 | 0 | 0 | 0 | 0 | 0 | 0 | 0 | 2 | 6 | 9 | 4 | 7 |
| L1ME1_dup21141 | 2.4022788935882 | 4.97033559795903 | 1.08298763039069 | 4.5894666369975 | 4.44380014325209e-06 | 0.00100226016008177 | 3552 | 0 | 0 | 0 | 0 | 0 | 0 | 0 | 1 | 4 | 7 | 5 | 13 |
| MIRb_dup82625 | 2.80267687885882 | 5.21161612469823 | 1.03340863214086 | 5.04313198342615 | 4.57973126222217e-07 | 0.00017297281506678 | 3553 | 0 | 0 | 0 | 0 | 0 | 0 | 0 | 1 | 5 | 12 | 6 | 10 |
| MER5A_dup8252 | 3.35109367867749 | 4.55362820755496 | 1.041070429227 | 4.37398669649667 | 1.21997816822334e-05 | 0.00216795313828283 | 3554 | 0 | 0 | 0 | 1 | 0 | 0 | 0 | 2 | 14 | 12 | 3 | 5 |
| L1PA12_dup1541 | 2.76225333502245 | 4.67491742917938 | 1.0100857726865 | 4.62823807204574 | 3.68789919013333e-06 | 0.000874015429346082 | 3555 | 1 | 0 | 0 | 0 | 0 | 0 | 0 | 1 | 7 | 13 | 6 | 4 |
| AluY_dup19481 | 2.29485207433719 | 4.04046666000021 | 1.06436222702034 | 3.7961387180297 | 0.000146967282756853 | 0.0131305546718927 | 3556 | 2 | 0 | 0 | 0 | 0 | 0 | 0 | 1 | 7 | 7 | 3 | 7 |
| L1MA8_dup7327 | 1.87116330902445 | 3.97159624889805 | 1.10544741299706 | 3.59275005052511 | 0.000327206416669384 | 0.0229668017760511 | 3557 | 1 | 0 | 0 | 0 | 0 | 0 | 0 | 1 | 7 | 7 | 2 | 3 |
| AluSg_dup1376 | 3.01088148503484 | 4.94548571442022 | 1.08463159998117 | 4.55959951241146 | 5.1251266430232e-06 | 0.00111471972961682 | 3558 | 0 | 0 | 0 | 0 | 0 | 0 | 2 | 0 | 10 | 9 | 3 | 12 |
| L1MA8_dup1678 | 3.6878848031756 | 3.96559262023673 | 1.04868727031942 | 3.78148255678629 | 0.000155897147411737 | 0.0137440172797455 | 3559 | 2 | 0 | 0 | 0 | 0 | 0 | 3 | 0 | 13 | 15 | 2 | 8 |
| L1PA11_dup1796 | 2.60527732480101 | 4.2838030717867 | 1.00488735302346 | 4.26296844009211 | 2.01729018945438e-05 | 0.0031048262058209 | 3560 | 1 | 1 | 1 | 0 | 0 | 0 | 0 | 0 | 4 | 12 | 5 | 8 |
| MIR_dup76880 | 2.23952372291821 | 3.47945238109802 | 1.01977849883686 | 3.41196876092857 | 0.000644955039933726 | 0.0359738565241609 | 3561 | 1 | 1 | 2 | 0 | 0 | 0 | 0 | 0 | 7 | 9 | 2 | 4 |
| L1PA7_dup11878 | 2.67946680705248 | 4.34072649765399 | 1.00687455606151 | 4.31108967003117 | 1.62452006047652e-05 | 0.00265544184995128 | 3562 | 1 | 0 | 1 | 1 | 0 | 0 | 0 | 0 | 9 | 6 | 4 | 10 |
| MLT1A0_dup17340 | 2.08669467422911 | 3.81410526447714 | 1.05359774534588 | 3.620077284073 | 0.000294515035601908 | 0.0213264378153779 | 3563 | 1 | 0 | 1 | 1 | 0 | 0 | 0 | 0 | 8 | 7 | 2 | 4 |
| MER72B_dup150 | 3.28043899217419 | 4.86005592275166 | 1.00826965458398 | 4.82019457855942 | 1.43418269153193e-06 | 0.000427399328953033 | 3564 | 1 | 0 | 1 | 0 | 0 | 0 | 0 | 0 | 11 | 15 | 2 | 8 |
| MamGypsy2-I_dup21 | 2.29597986980438 | 4.87021240621852 | 1.14140209237614 | 4.26686830061773 | 1.98236104929817e-05 | 0.0030709302228926 | 3565 | 1 | 0 | 0 | 0 | 0 | 0 | 0 | 0 | 5 | 11 | 2 | 9 |
| MLT1D_dup16966 | 2.17595805313991 | 4.46015935463517 | 1.08516223497218 | 4.11013138026271 | 3.95434066840218e-05 | 0.00509431264040945 | 3566 | 1 | 0 | 0 | 0 | 1 | 0 | 0 | 0 | 5 | 6 | 3 | 11 |
| L1MB3_dup5884 | 2.31535288725106 | 4.47282535121435 | 1.07603273175459 | 4.1567744355889 | 3.22772387705086e-05 | 0.00440716005173003 | 3567 | 1 | 0 | 0 | 0 | 1 | 0 | 0 | 0 | 9 | 5 | 3 | 8 |
| L2a_dup77725 | 2.41457228504832 | 4.59566681760988 | 1.02813330245356 | 4.46991339220575 | 7.82512771055103e-06 | 0.0015274544259124 | 3568 | 1 | 0 | 0 | 0 | 1 | 0 | 0 | 0 | 7 | 6 | 5 | 9 |
| L1MEj_dup28 | 2.34029204638306 | 4.44918408276706 | 1.12707874781823 | 3.9475361339034 | 7.89596036950795e-05 | 0.00844507275200178 | 3569 | 0 | 0 | 0 | 0 | 2 | 0 | 0 | 0 | 9 | 6 | 3 | 7 |
| MIR3_dup31964 | 2.32462242597908 | 4.8197205384291 | 1.06479884407127 | 4.52641413471189 | 5.99929588377054e-06 | 0.00125550436091439 | 3570 | 0 | 0 | 0 | 0 | 1 | 0 | 0 | 0 | 7 | 10 | 4 | 5 |
| L2a_dup120002 | 1.82496900395114 | 4.53693805465094 | 1.15212014485195 | 3.93790359011033 | 8.21966055451218e-05 | 0.00872361886843869 | 3571 | 0 | 0 | 0 | 0 | 1 | 0 | 0 | 0 | 4 | 8 | 3 | 6 |
| LTR78_dup1789 | 1.89018573035644 | 4.62781473671711 | 1.1289370290672 | 4.09926737945775 | 4.14460002871804e-05 | 0.00525688165476195 | 3572 | 0 | 0 | 0 | 0 | 1 | 0 | 0 | 0 | 4 | 7 | 4 | 7 |
| L2c_dup5664 | 3.9447156218418 | 5.69821427062085 | 1.08485833623634 | 5.25249618340905 | 1.50051574705312e-07 | 7.18393802712881e-05 | 3573 | 0 | 0 | 0 | 0 | 1 | 0 | 0 | 0 | 11 | 14 | 2 | 20 |
| Tigger1_dup8185 | 3.26944250508899 | 5.85158811880239 | 1.07622625785796 | 5.43713561723439 | 5.41439273204505e-08 | 3.0969545505269e-05 | 3574 | 0 | 0 | 0 | 0 | 0 | 0 | 0 | 0 | 6 | 12 | 5 | 18 |
| L1MA9_dup13840 | 3.62639190559861 | 4.62709321722657 | 1.2748879641383 | 3.62941164038208 | 0.000284067944183764 | 0.0208170098696489 | 3575 | 0 | 0 | 0 | 4 | 0 | 0 | 0 | 0 | 16 | 13 | 2 | 6 |
| MLT1A0_dup423 | 3.3714892453331 | 5.46260217309107 | 0.971582896505758 | 5.6223737498231 | 1.88351211449005e-08 | 1.34183884309715e-05 | 3576 | 0 | 0 | 0 | 1 | 0 | 0 | 0 | 0 | 9 | 14 | 7 | 9 |
| MLT2D_dup720 | 3.2800291128416 | 4.95688464060943 | 0.977502264932319 | 5.07096997975001 | 3.95793208873983e-07 | 0.00015438919816185 | 3577 | 0 | 0 | 0 | 1 | 1 | 0 | 0 | 0 | 12 | 11 | 4 | 9 |
| AluSx_dup37593 | 4.71680576044806 | 6.11010583032528 | 0.947806756423036 | 6.44657340636022 | 1.14407216972939e-10 | 1.61081299703212e-07 | 3578 | 0 | 0 | 0 | 0 | 0 | 0 | 0 | 0 | 18 | 22 | 6 | 7 |
| AluY_dup76063 | 4.33419840995346 | 5.91451383224901 | 0.982325413704982 | 6.02093130212479 | 1.73416354743043e-09 | 1.75590331683779e-06 | 3579 | 0 | 0 | 0 | 0 | 0 | 0 | 0 | 0 | 15 | 24 | 4 | 6 |
| HAL1_dup8728 | 3.87971003513393 | 5.93505262352708 | 1.03189090244993 | 5.75162801555475 | 8.83881133423562e-09 | 7.01053267245343e-06 | 3580 | 0 | 0 | 0 | 0 | 0 | 0 | 0 | 0 | 13 | 18 | 3 | 11 |
| Ricksha_0_dup316 | 3.84471638744461 | 5.92875629772141 | 0.98524058684677 | 6.01757213098194 | 1.77052429012197e-09 | 1.78512361329391e-06 | 3581 | 0 | 0 | 0 | 0 | 0 | 0 | 0 | 0 | 14 | 16 | 5 | 9 |
| MER34D_dup14 | 3.25883962247718 | 5.74377614395638 | 1.1069108543792 | 5.1890142022121 | 2.11410274123143e-07 | 9.45568215911753e-05 | 3582 | 0 | 0 | 0 | 0 | 0 | 0 | 0 | 0 | 15 | 8 | 3 | 11 |
| MIRb_dup82622 | 3.15775050667854 | 5.65721603786005 | 1.20512037199657 | 4.69431616070645 | 2.67500048930042e-06 | 0.000704044193033047 | 3583 | 0 | 0 | 0 | 0 | 0 | 0 | 0 | 0 | 14 | 10 | 1 | 11 |
| Cheshire_dup377 | 2.64411005349549 | 5.40847549443582 | 1.14250962182798 | 4.73385553268463 | 2.20294553641721e-06 | 0.000607395224343371 | 3584 | 0 | 0 | 0 | 0 | 0 | 0 | 0 | 0 | 8 | 13 | 2 | 8 |
| MamGypsy2-I_dup20 | 2.85758690644747 | 5.49005744281154 | 1.11842007599909 | 4.90876152943449 | 9.16533445357326e-07 | 0.000299858802505635 | 3585 | 0 | 0 | 0 | 0 | 0 | 0 | 0 | 0 | 10 | 13 | 2 | 8 |
| Zaphod3_dup508 | 2.6828070982275 | 5.3868835073271 | 1.12812697165481 | 4.77506844768126 | 1.79646141090387e-06 | 0.000516881265875373 | 3586 | 0 | 0 | 0 | 0 | 0 | 0 | 0 | 0 | 9 | 13 | 2 | 7 |
| L1M1_dup7938 | 2.9242459854746 | 5.4346774721596 | 1.06778782998381 | 5.08966043585829 | 3.58705262921365e-07 | 0.000143691047964793 | 3587 | 0 | 0 | 0 | 0 | 0 | 0 | 0 | 0 | 11 | 14 | 3 | 5 |
| LTR79_dup139 | 2.22837877331295 | 5.04800167973921 | 1.16627312627569 | 4.32831861251849 | 1.50252003445541e-05 | 0.00251419572516544 | 3588 | 0 | 0 | 0 | 0 | 0 | 0 | 0 | 0 | 9 | 10 | 2 | 4 |
| Charlie8_dup2260 | 2.24981275522669 | 5.11513071162188 | 1.16954199396323 | 4.37361867981178 | 1.22203760437946e-05 | 0.00216999223740056 | 3589 | 0 | 0 | 0 | 0 | 0 | 0 | 0 | 0 | 7 | 12 | 2 | 5 |
| L1MC1_dup10369 | 2.43716125292488 | 5.24851950819626 | 1.11482119276412 | 4.70794737511485 | 2.50223675702914e-06 | 0.000666546133848742 | 3590 | 0 | 0 | 0 | 0 | 0 | 0 | 0 | 0 | 8 | 12 | 3 | 5 |
| LTR77_dup88 | 3.46867312829144 | 5.59897845190773 | 1.07781517928256 | 5.19474819016248 | 2.04997122319714e-07 | 9.2383040279331e-05 | 3591 | 0 | 0 | 0 | 0 | 0 | 0 | 0 | 0 | 17 | 13 | 2 | 6 |
| MER61C_dup132 | 2.61154973472535 | 5.13733260933247 | 1.17913791259333 | 4.356854744865 | 1.31944767424951e-05 | 0.00228635571176538 | 3592 | 0 | 0 | 0 | 0 | 0 | 0 | 0 | 0 | 11 | 13 | 1 | 4 |
| L1PBb_dup182 | 4.54693353329804 | 5.86954830374225 | 0.924287496011618 | 6.35034913819549 | 2.14826796623347e-10 | 2.77402747421019e-07 | 3593 | 0 | 1 | 0 | 0 | 0 | 0 | 0 | 0 | 8 | 7 | 19 | 22 |
| L1PB3_dup2140 | 4.99389542114799 | 6.32311265029189 | 0.988224403579231 | 6.39845831310209 | 1.56953580198029e-10 | 2.10997042902826e-07 | 3594 | 0 | 0 | 0 | 0 | 0 | 0 | 0 | 0 | 10 | 4 | 26 | 22 |

| | | | | | | | | | | | | | | | | | | | |
|---|---|---|---|---|---|---|---|---|---|---|---|---|---|---|---|---|---|---|---|
| L1PB3_dup2181 | 4.69006446283427 | 6.22205862488798 | 0.96212418724247 | 6.46700156527707 | 9.99666145088668e-11 | 1.42615767093198e-07 | 3595 | 0 | 0 | 0 | 0 | 0 | 0 | 0 | 0 | 8 | 5 | 23 | 23 |
| L1MA9_dup9915 | 4.333134091269 | 6.07455651186733 | 0.973701333256391 | 6.23862400552737 | 4.41436610177847e-10 | 5.19264249620911e-07 | 3596 | 0 | 0 | 0 | 0 | 0 | 0 | 0 | 0 | 6 | 5 | 17 | 28 |
| L1PA6_dup3791 | 6.06673681885518 | 6.56137413885062 | 0.902737206160605 | 7.26831030567194 | 3.6401185534553e-13 | 9.41469184043995e-10 | 3597 | 0 | 0 | 0 | 0 | 0 | 0 | 0 | 0 | 8 | 8 | 26 | 36 |
| L1PA13_dup5429 | 9.14706791557245 | 7.07552470310118 | 0.85818139957913 | 8.24478916295688 | 1.65451686101372e-16 | 1.03428304407981e-12 | 3598 | 0 | 0 | 0 | 0 | 0 | 0 | 0 | 0 | 8 | 13 | 43 | 55 |
| MER58A_dup7646 | 5.99334922920493 | 6.48463036793286 | 0.942571009069396 | 6.87972609547493 | 5.99677504917127e-12 | 1.23009365159492e-08 | 3599 | 0 | 0 | 0 | 0 | 0 | 0 | 0 | 0 | 5 | 9 | 39 | 23 |
| L1PA5_dup1106 | 13.9157418564205 | 5.7562935974989 | 0.890128778644257 | 6.46680989942403 | 1.0009343760586e-10 | 1.42615767093198e-07 | 3600 | 2 | 3 | 0 | 0 | 0 | 2 | 0 | 0 | 16 | 10 | 61 | 87 |
| MSTA_dup19525 | 15.0833904766994 | 6.48210972895552 | 0.750785296126306 | 8.63377288074249 | 5.93608067647763e-18 | 5.2313579727598e-14 | 3601 | 0 | 0 | 0 | 3 | 0 | 0 | 0 | 0 | 6 | 12 | 63 | 120 |
| L1ME1_dup18307 | 17.2844691573433 | 7.87642883193837 | 0.817159192406276 | 9.63879364649227 | 5.48285010898999e-22 | 1.30462225203373e-17 | 3602 | 0 | 0 | 0 | 0 | 0 | 0 | 0 | 0 | 14 | 18 | 92 | 101 |
| L1ME1_dup18308 | 13.059212742401 | 7.50395065167324 | 0.837280951317322 | 8.96228516827837 | 3.18018145597693e-19 | 3.78355728361943e-15 | 3603 | 0 | 0 | 0 | 0 | 0 | 0 | 0 | 0 | 13 | 12 | 64 | 81 |
| L1PA13_dup9197 | 9.52498151340925 | 6.21298280815186 | 0.807591218385865 | 7.69322729953623 | 1.43469304055375e-14 | 5.82944212949515e-11 | 3604 | 0 | 0 | 0 | 0 | 1 | 1 | 1 | 0 | 14 | 20 | 39 | 44 |
| MSTA_dup20229 | 12.5716755315099 | 6.11918166360726 | 0.717099713061835 | 8.5332368039584 | 1.42308981249785e-17 | 1.02611675310489e-13 | 3605 | 0 | 0 | 0 | 0 | 1 | 0 | 1 | 2 | 11 | 15 | 59 | 74 |
| L1PA3_dup9836 | 16.3311558019084 | 5.07279405275858 | 0.748323147498941 | 6.77888164987674 | 1.21109683107433e-11 | 2.26909957926624e-08 | 3606 | 1 | 0 | 1 | 2 | 0 | 1 | 1 | 7 | 48 | 55 | 29 | 46 |
| L2b_dup4666 | 7.78974049835209 | 5.24884902596147 | 0.776784221043405 | 6.75715196545963 | 1.40730565708978e-11 | 2.53683902940822e-08 | 3607 | 0 | 0 | 0 | 1 | 0 | 2 | 1 | 1 | 16 | 34 | 16 | 23 |
| MLT1B_dup15459 | 9.51270376377967 | 4.51320745926233 | 0.611962199515557 | 7.37497751141342 | 1.64372795832847e-13 | 4.84405453592959e-10 | 3608 | 2 | 0 | 0 | 1 | 0 | 4 | 1 | 2 | 21 | 31 | 23 | 30 |
| L1PA7_dup170 | 7.73253436792457 | 5.23489123324402 | 0.779723948144113 | 6.71377510682341 | 1.89652538516585e-11 | 3.24655128991852e-08 | 3609 | 0 | 0 | 1 | 1 | 0 | 0 | 1 | 2 | 24 | 26 | 16 | 19 |
| MER89_dup1018 | 12.9287823963839 | 6.02890076664554 | 0.803048116868423 | 7.50752120577272 | 6.02574260013193e-14 | 1.93756939017702e-10 | 3610 | 0 | 0 | 0 | 2 | 0 | 0 | 1 | 2 | 44 | 42 | 19 | 41 |
| THE1D-int_dup2538 | 9.82452335524453 | 3.85261806276332 | 0.634305713193501 | 6.0737558918818 | 1.2495262865274e-09 | 1.33721899853131e-06 | 3611 | 1 | 0 | 8 | 1 | 0 | 2 | 1 | 3 | 14 | 10 | 44 | 39 |
| L1PA12_dup1725 | 9.72858976497914 | 4.15495365582595 | 0.591372001698348 | 7.02595598691422 | 2.1260547635039e-12 | 4.95966888977157e-09 | 3612 | 0 | 0 | 5 | 2 | 0 | 2 | 3 | 1 | 6 | 10 | 48 | 50 |
| L1PA13_dup8785 | 12.131522720844 | 3.80947724534288 | 0.504068127178972 | 7.557465032877 | 4.11000461996473e-14 | 1.41733211493062e-10 | 3613 | 1 | 0 | 5 | 4 | 1 | 4 | 3 | 1 | 11 | 13 | 50 | 65 |
| MIRc_dup27325 | 11.9874083520586 | 6.16031385125697 | 0.734024542554753 | 8.39251754419023 | 4.75840971308012e-17 | 3.23498445025303e-13 | 3614 | 0 | 0 | 1 | 2 | 0 | 0 | 1 | 0 | 17 | 12 | 58 | 62 |
| MSTA-int_dup3282 | 9.96270367683817 | 4.89449872934342 | 0.702799214384171 | 6.96429169123679 | 3.30060114494064e-12 | 7.27189666698191e-09 | 3615 | 0 | 1 | 5 | 3 | 0 | 0 | 0 | 0 | 11 | 17 | 35 | 57 |
| L1PREC2_dup2667 | 9.46751685995031 | 4.92419533187559 | 0.99291677501183 | 4.95932333484539 | 7.07391446191248e-07 | 0.000245008682758985 | 3616 | 6 | 8 | 0 | 0 | 0 | 2 | 0 | 0 | 14 | 19 | 32 | 38 |
| L2a_dup61546 | 10.6615876667129 | 4.6221615682216 | 1.03055485021118 | 4.48511941627795 | 7.28731100465371e-06 | 0.00145713151622969 | 3617 | 8 | 11 | 0 | 0 | 0 | 0 | 1 | 0 | 16 | 22 | 34 | 42 |
| L1MB3_dup5881 | 7.71889801258391 | 4.35468581938322 | 0.929139888997101 | 4.68679245283893 | 2.77520146936636e-06 | 0.00071621267768964 | 3618 | 4 | 7 | 0 | 0 | 0 | 0 | 0 | 1 | 16 | 20 | 10 | 38 |
| MER90a_dup388 | 7.62588300400372 | 4.15456585686119 | 1.06359640684288 | 3.9061488268782 | 9.37787662462613e-05 | 0.00958517281496258 | 3619 | 12 | 4 | 0 | 0 | 0 | 0 | 0 | 1 | 13 | 11 | 24 | 31 |
| MER4D1_dup1149 | 6.291899562496 | 4.42915670774231 | 0.789275856668526 | 5.61167134446181 | 2.0038174003783e-08 | 1.40235392691298e-05 | 3620 | 3 | 1 | 3 | 0 | 0 | 0 | 0 | 0 | 15 | 19 | 21 | 13 |
| MLT1B_dup13514 | 8.36764241717924 | 5.14879076236886 | 0.782610499342973 | 6.57899525586666 | 4.73637882393454e-11 | 7.41448944499953e-08 | 3621 | 3 | 1 | 0 | 2 | 0 | 0 | 0 | 0 | 28 | 17 | 16 | 33 |
| MLT1C_dup14136 | 12.081374376727 | 5.70756606572907 | 0.73403689407424 | 7.77558473123806 | 7.50996911411401e-15 | 3.2490311105945e-11 | 3622 | 4 | 1 | 0 | 0 | 0 | 0 | 0 | 1 | 30 | 30 | 37 | 43 |
| HAL1_dup19466 | 11.6999060641842 | 3.96376426737074 | 0.582682542039877 | 6.80261374142811 | 1.02737842322692e-11 | 1.95568469034522e-08 | 3623 | 1 | 3 | 0 | 1 | 6 | 2 | 5 | 0 | 30 | 25 | 40 | 27 |
| AluSx1_dup76488 | 7.31297857426982 | 4.12585201953418 | 0.686856394094206 | 6.00686264990684 | 1.89147618403252e-09 | 1.86750702110292e-06 | 3624 | 0 | 2 | 0 | 0 | 1 | 4 | 2 | 1 | 22 | 12 | 20 | 23 |
| Tigger2a_dup3003 | 6.78387632126183 | 4.00021516709907 | 0.767240511333191 | 5.21376948689547 | 1.85041386702474e-07 | 8.54948695151592e-05 | 3625 | 0 | 1 | 0 | 0 | 4 | 4 | 2 | 0 | 21 | 9 | 18 | 22 |
| L1MC3_dup11155 | 7.25603404894378 | 3.37062974587813 | 0.616406651922797 | 5.46819171299322 | 4.54649986518851e-08 | 2.70455364230536e-05 | 3626 | 2 | 2 | 0 | 0 | 3 | 1 | 4 | 3 | 22 | 12 | 15 | 22 |
| THE1B_dup19511 | 8.95455009457105 | 3.6214366857636 | 0.619865465512849 | 5.84229463851061 | 5.14866378633007e-09 | 4.52067879447267e-06 | 3627 | 1 | 6 | 0 | 0 | 3 | 2 | 4 | 1 | 23 | 20 | 14 | 35 |
| MER77B_dup23 | 9.07367035111542 | 5.50043962424866 | 0.747493143033212 | 7.3585151589869 | 1.85967141748374e-13 | 5.33134186872996e-10 | 3628 | 1 | 0 | 1 | 1 | 1 | 0 | 1 | 0 | 25 | 27 | 20 | 32 |
| THE1B_dup17354 | 8.1229062896963 | 5.65289495656791 | 0.814715544289718 | 6.93848914039831 | 3.96315450978545e-12 | 8.49564651338206e-09 | 3629 | 1 | 0 | 0 | 0 | 1 | 0 | 2 | 0 | 20 | 29 | 23 | 21 |
| CER_dup80 | 8.02007322516518 | 7.12769472591182 | 0.877624927974009 | 8.12157277980475 | 4.60180733629108e-16 | 2.60709916295504e-12 | 3630 | 0 | 0 | 0 | 0 | 0 | 0 | 0 | 0 | 20 | 17 | 36 | 23 |
| MLT1H1-int_dup13 | 7.59813445065299 | 7.05421939516229 | 0.860148036559077 | 8.20116897944903 | 2.38060619404503e-16 | 1.4161393036206e-12 | 3631 | 0 | 0 | 0 | 0 | 0 | 0 | 0 | 0 | 23 | 20 | 22 | 25 |
| L1MEc_dup469 | 11.8268229403138 | 7.32594846213092 | 0.820204229814783 | 8.93185891492568 | 4.18909507345112e-19 | 4.74656388736857e-15 | 3632 | 0 | 0 | 0 | 0 | 0 | 0 | 1 | 0 | 34 | 34 | 26 | 47 |
| Tigger1_dup4828 | 11.5970153562029 | 7.65585650214005 | 0.834068220508748 | 9.17893322619376 | 4.35383051337504e-20 | 6.90651036890358e-16 | 3633 | 0 | 0 | 0 | 0 | 0 | 0 | 0 | 0 | 40 | 28 | 29 | 39 |
| Cheshire_dup380 | 11.1153488643942 | 7.52455785928994 | 0.822487491624466 | 9.14853774180621 | 5.77093712315532e-20 | 8.58232127941447e-16 | 3634 | 0 | 0 | 0 | 0 | 0 | 0 | 0 | 0 | 34 | 46 | 25 | 24 |
| MER61E_dup2 | 12.4787650207222 | 7.34992314008928 | 0.809350349243709 | 9.08126270280523 | 1.07322445015446e-19 | 1.47952532886841e-15 | 3635 | 0 | 0 | 0 | 0 | 0 | 0 | 0 | 1 | 40 | 44 | 28 | 32 |
| L1MC4_dup20648 | 7.08344577990573 | 4.17965689700848 | 0.723001858838921 | 5.78097669585616 | 7.42681791225728e-09 | 6.17895669562927e-06 | 3636 | 0 | 3 | 0 | 0 | 1 | 2 | 0 | 3 | 11 | 28 | 16 | 22 |
| L1MEi_dup4739 | 6.24481025199139 | 5.31783912736125 | 0.919381551304892 | 5.78414818071296 | 7.28806790708582e-09 | 6.0847951095419e-06 | 3637 | 0 | 1 | 0 | 0 | 3 | 0 | 0 | 0 | 12 | 12 | 19 | 31 |
| L1P3_dup1513 | 5.06165038848856 | 4.93402034901232 | 0.885365217581662 | 5.5728644530326 | 2.50584538885787e-08 | 1.66554144823475e-05 | 3638 | 0 | 1 | 2 | 0 | 1 | 0 | 0 | 0 | 11 | 16 | 10 | 21 |
| L1ME3E_dup1182 | 8.14270604718405 | 2.56690811595777 | 0.773954266522178 | 3.31661472388073 | 0.000911151702178732 | 0.044930231711484 | 3639 | 14 | 13 | 0 | 1 | 2 | 4 | 1 | 2 | 10 | 8 | 18 | 31 |
| L1PB4_dup1007 | 7.61918436591387 | 3.18032263664684 | 0.883848841552235 | 3.59826532222579 | 0.00032034672702361 | 0.0226120505216138 | 3640 | 16 | 10 | 0 | 0 | 4 | 5 | 0 | 1 | 9 | 7 | 21 | 24 |
| MIR_dup89593 | 6.23850793381833 | 2.04631228736224 | 0.575684589210744 | 3.55457194045738 | 0.000378594921737188 | 0.025376097816247 | 3641 | 7 | 7 | 1 | 0 | 4 | 5 | 3 | 2 | 8 | 10 | 15 | 16 |
| MLT1E1A_dup2975 | 7.9559374832876 | 2.52856316980645 | 0.706745338206954 | 3.57775712567349 | 0.000346555242115923 | 0.0239504599594875 | 3642 | 14 | 9 | 1 | 0 | 1 | 8 | 2 | 1 | 9 | 15 | 18 | 23 |
| MER58B_dup5548 | 6.34905073535418 | 2.46816584200341 | 0.748832868598797 | 3.29601696920942 | 0.000980660916377489 | 0.0471879357751988 | 3643 | 7 | 12 | 0 | 1 | 9 | 6 | 0 | 0 | 12 | 7 | 11 | 13 |
| THE1A_dup315 | 6.83373016263219 | 2.31580920953958 | 0.672025371415074 | 3.4460145524911 | 0.000568919721236725 | 0.0332528056962402 | 3644 | 10 | 7 | 1 | 1 | 6 | 8 | 0 | 1 | 13 | 6 | 12 | 20 |
| L1MEd_dup10521 | 4.75770256879617 | 3.26028262770408 | 0.746361841557709 | 4.36823326993733 | 1.25255669695823e-05 | 0.00220934659610395 | 3645 | 3 | 4 | 0 | 0 | 5 | 3 | 0 | 0 | 7 | 13 | 10 | 14 |
| L1PA7_dup2503 | 6.35864131959452 | 3.60076784726439 | 0.871436401694973 | 4.13199154896534 | 3.59633646444359e-05 | 0.00477263734728664 | 3646 | 6 | 8 | 0 | 0 | 5 | 5 | 0 | 0 | 6 | 10 | 19 | 22 |
| L1PA4_dup9045 | 7.67643635270969 | 3.51255763060667 | 0.795875358717906 | 4.41345192074439 | 1.01735319551146e-05 | 0.00187655134464473 | 3647 | 5 | 4 | 0 | 1 | 4 | 11 | 0 | 0 | 10 | 13 | 17 | 33 |
| HUERS-P1-int_dup107 | 5.1334761858797 | 2.53797184185013 | 0.752864013490354 | 3.371089328714 | 0.000748715759162683 | 0.039563384416994 | 3648 | 3 | 3 | 0 | 1 | 6 | 11 | 0 | 0 | 12 | 8 | 9 | 9 |
| L1MA4_dup8568 | 5.3996196911379 | 2.95465031022398 | 0.845506484608436 | 3.49453299768873 | 0.000474891901473002 | 0.0294650921480821 | 3649 | 3 | 3 | 0 | 0 | 7 | 9 | 1 | 0 | 8 | 5 | 14 | 18 |
| THE1B-int_dup1412 | 5.9761949949602 | 4.12627190859884 | 1.14373462245813 | 3.60771793349284 | 0.000308902044363408 | 0.0220859392572402 | 3650 | 13 | 2 | 0 | 0 | 6 | 5 | 0 | 0 | 2 | 4 | 17 | 31 |
| L1ME4a_dup33050 | 4.72836671639467 | 3.97309322393741 | 1.02169874167919 | 3.88871304412838 | 0.000100777185386368 | 0.0101453138452767 | 3651 | 2 | 3 | 0 | 0 | 5 | 2 | 0 | 0 | 3 | 7 | 13 | 27 |
| L1MEg_dup6070 | 5.06086570500858 | 4.33921711862407 | 1.05779429510235 | 4.10213700217037 | 4.09351758494127e-05 | 0.00520874938645153 | 3652 | 3 | 1 | 0 | 0 | 3 | 1 | 0 | 0 | 1 | 5 | 20 | 34 |
| L2b_dup13132 | 7.24379464109194 | 3.72206892057819 | 0.895250050943001 | 4.15757465376025 | 3.21644182019297e-05 | 0.00440039987109913 | 3653 | 8 | 1 | 0 | 3 | 0 | 3 | 0 | 0 | 5 | 18 | 31 | 24 |
| L1MB7_dup7043 | 6.24801749636646 | 4.66857842075304 | 1.08797883441811 | 4.29105628994149 | 1.77825227383854e-05 | 0.00284169251545188 | 3654 | 7 | 3 | 0 | 0 | 0 | 1 | 0 | 0 | 7 | 8 | 22 | 33 |
| L2c_dup20202 | 7.33211312976775 | 4.31417475061153 | 0.886613511738055 | 4.86590232778466 | 1.13935778567889e-06 | 0.000354850297998887 | 3655 | 3 | 5 | 0 | 1 | 0 | 2 | 0 | 0 | 10 | 10 | 30 | 32 |
| L1MA4_dup4648 | 5.95021936315292 | 2.94981484171754 | 0.808210267865426 | 3.64981114321689 | 0.00026243318485542 | 0.0197485536380796 | 3656 | 8 | 3 | 0 | 4 | 3 | 1 | 0 | 0 | 3 | 20 | 15 | 20 |
| L1PA7_dup570 | 6.7562522839625 | 2.81407509425925 | 0.810988088208005 | 3.4699339425285 | 0.000520586380653284 | 0.0312806684169006 | 3657 | 13 | 6 | 0 | 1 | 1 | 5 | 2 | 0 | 5 | 12 | 18 | 25 |
| ERV3-16A3_I-int_dup | 11.9132894815603 | 3.75577572115064 | 0.749238915940722 | 5.0127878320829 | 5.36470399390339e-07 | 0.0001969922618107 | 3658 | 1 | 1 | 2 | 3 | 1 | 5 | 0 | 2 | 2 | 10 | 67 | 63 |
| MLT2A2_dup1590 | 13.6135558322335 | 4.39787456658177 | 0.792088263966911 | 5.55225315996538 | 2.8201077633986e-08 | 1.84349824689462e-05 | 3659 | 1 | 1 | 2 | 2 | 1 | 2 | 0 | 0 | 1 | 2 | 82 | 89 |
| LOR1-int_dup476 | 12.9161543591902 | 3.41375944848907 | 0.708237060179169 | 4.82008022515153 | 1.43500502883343e-06 | 0.000427399328953033 | 3660 | 3 | 5 | 1 | 1 | 0 | 3 | 4 | 3 | 5 | 6 | 68 | 71 |
| LTR16D1_dup99 | 11.598428310768 | 3.11582833483359 | 0.73169027863056 | 4.25839788477881 | 2.05897235063412e-05 | 0.00315672831793805 | 3661 | 5 | 6 | 2 | 2 | 2 | 6 | 2 | 1 | 5 | 4 | 57 | 61 |
| MER77_dup454 | 10.9308119661086 | 3.84626562465671 | 0.754364924012589 | 5.09868036307653 | 3.42029538728196e-07 | 0.000137981359278016 | 3662 | 3 | 2 | 0 | 1 | 2 | 4 | 2 | 1 | 6 | 5 | 61 | 56 |
| LOR1-int_dup478 | 15.2308474334351 | 4.52241265031826 | 0.959970540108958 | 4.71099107875223 | 2.46515028541871e-06 | 0.000661299492462503 | 3663 | 7 | 4 | 0 | 0 | 2 | 0 | 2 | 1 | 2 | 11 | 83 | 91 |
| MER50_dup1040 | 12.0251890697461 | 4.07659707618447 | 0.867534907372518 | 4.69905826444628 | 2.61363887810378e-06 | 0.00069100546276587 | 3664 | 1 | 5 | 1 | 0 | 6 | 1 | 0 | 2 | 7 | 7 | 53 | 75 |
| L1MB8_dup5901 | 12.3623502847813 | 3.33574664997698 | 0.784733934802975 | 4.25079954113938 | 2.13008750310357e-05 | 0.00324277543834602 | 3665 | 2 | 2 | 1 | 1 | 7 | 4 | 3 | 2 | 1 | 9 | 58 | 75 |
| Tigger2b_Pri_dup922 | 9.83720696778853 | 3.39152296323599 | 0.740193754984068 | 4.58193944544829 | 4.60683308888064e-06 | 0.00103024201707405 | 3666 | 2 | 2 | 1 | 2 | 5 | 1 | 1 | 2 | 2 | 7 | 50 | 55 |

| | | | | | | | | | | | | | | | | | | | |
|---|---|---|---|---|---|---|---|---|---|---|---|---|---|---|---|---|---|---|---|
| MLT1D_dup2061 | 4.83926059724478 | 4.12203058160117 | 0.764142826087779 | 5.39431954456072 | 6.87837146448861e-08 | 3.71129473126804e-05 | 3667 | 1 | 1 | 0 | 1 | 0 | 0 | 2 | 1 | 4 | 4 | 23 | 26 |
| LOR1a_dup332 | 5.70847074757578 | 3.05415670706948 | 0.70699071254947 | 4.31993893675904 | 1.56072381347433e-05 | 0.00257715467398309 | 3668 | 3 | 1 | 1 | 1 | 0 | 2 | 2 | 2 | 3 | 3 | 26 | 31 |
| L1MA7_dup3214 | 6.87504717546122 | 3.49925180032668 | 0.703390809305779 | 4.97483298620337 | 6.53039184638689e-07 | 0.000230546085798275 | 3669 | 1 | 2 | 0 | 2 | 0 | 2 | 2 | 2 | 3 | 4 | 26 | 48 |
| MLT1L_dup4970 | 4.92804898706983 | 2.94579690155734 | 0.739615154081039 | 3.98287796741732 | 6.80857206835843e-05 | 0.0075881615427523 | 3670 | 1 | 4 | 0 | 1 | 0 | 1 | 3 | 2 | 4 | 5 | 17 | 26 |
| L2b_dup29049 | 5.00177317547911 | 2.93180772589435 | 0.686215367172724 | 4.27243088124607 | 1.93353496649404e-05 | 0.00303280758824912 | 3671 | 1 | 2 | 1 | 1 | 2 | 1 | 3 | 2 | 11 | 2 | 15 | 21 |
| L1ME4a_dup23655 | 4.08512897672601 | 3.32417789463222 | 0.811425275853493 | 4.09671474817654 | 4.19054744980023e-05 | 0.00529821468379472 | 3672 | 2 | 2 | 0 | 0 | 1 | 0 | 2 | 1 | 4 | 3 | 16 | 22 |
| MLT1B_dup10390 | 3.90559899872434 | 2.67930180526025 | 0.814925068615061 | 3.28778915810461 | 0.00100977432355866 | 0.0481658507396003 | 3673 | 3 | 1 | 0 | 0 | 2 | 1 | 3 | 1 | 3 | 2 | 16 | 19 |
| L1PA3_dup3873 | 5.38376808881193 | 2.88438133843179 | 0.805505488998113 | 3.58083387118737 | 0.000342499381748318 | 0.0237321950755636 | 3674 | 2 | 3 | 1 | 0 | 2 | 0 | 0 | 5 | 6 | 6 | 14 | 29 |
| MLT1C_dup17820 | 4.82114491339987 | 2.87918521074214 | 0.725287424154502 | 3.9697161633521 | 7.19582995410337e-05 | 0.00791929759195084 | 3675 | 2 | 1 | 4 | 0 | 2 | 1 | 0 | 2 | 5 | 4 | 20 | 20 |
| L1MC1_dup5601 | 4.88259941914851 | 4.91173994433395 | 0.859120219139438 | 5.71717419158629 | 1.08310074023649e-08 | 8.36751586806211e-06 | 3676 | 0 | 1 | 1 | 0 | 0 | 0 | 0 | 1 | 3 | 5 | 22 | 31 |
| Tigger2a_dup399 | 5.36705623797928 | 4.02417478981273 | 0.825469058458939 | 4.8750159058965 | 1.08799688307341e-06 | 0.000342439823200773 | 3677 | 2 | 2 | 2 | 0 | 0 | 1 | 0 | 0 | 4 | 7 | 17 | 36 |
| L1PA13_dup5757 | 5.46830409824366 | 3.16568330383031 | 0.896346174434124 | 3.53176417116842 | 0.000412797315628037 | 0.0268516867316645 | 3678 | 2 | 3 | 0 | 1 | 1 | 2 | 2 | 0 | 2 | 1 | 27 | 32 |
| LTR24B_dup155 | 6.45415986991004 | 3.21017777973853 | 0.948829813987304 | 3.38330197092803 | 0.00071619836914782 | 0.0383216858882948 | 3679 | 2 | 2 | 0 | 2 | 4 | 1 | 2 | 0 | 4 | 0 | 29 | 40 |
| L1M5_dup21530 | 8.06199975667539 | 3.8359826014418 | 0.939088404043397 | 4.08479391815015 | 4.41159527000156e-05 | 0.00549508537294869 | 3680 | 5 | 3 | 0 | 0 | 5 | 1 | 2 | 0 | 3 | 6 | 36 | 46 |
| L1MD_dup3367 | 7.75008967779211 | 4.1141870254013 | 0.950067070957015 | 4.33041745279839 | 1.48826929876908e-05 | 0.00250090202376348 | 3681 | 2 | 3 | 0 | 0 | 2 | 0 | 2 | 0 | 4 | 2 | 39 | 49 |
| LTR48B_dup437 | 6.44190092660125 | 3.48067232537457 | 0.886815622276541 | 3.92491092617352 | 8.67618358985112e-05 | 0.00909054680964647 | 3682 | 3 | 5 | 1 | 1 | 2 | 2 | 0 | 0 | 3 | 4 | 34 | 29 |
| THE1C_dup4023 | 7.83655457934151 | 4.42134572908305 | 0.958591570440882 | 4.61233529004386 | 3.98170033314032e-06 | 0.00092976414864515 | 3683 | 4 | 1 | 0 | 1 | 1 | 1 | 0 | 0 | 2 | 4 | 46 | 44 |
| L1MCb_dup912 | 7.4176373181208 | 3.90074484409804 | 1.0749361543474 | 3.62881537505472 | 0.000284724769057694 | 0.020834648649285 | 3684 | 0 | 4 | 0 | 1 | 5 | 0 | 0 | 1 | 2 | 4 | 37 | 44 |
| THE1B_dup9102 | 7.83225320179215 | 4.18821945883833 | 0.804718809106384 | 5.20457507820555 | 1.94441003064538e-07 | 8.84889710046311e-05 | 3685 | 0 | 3 | 0 | 2 | 1 | 0 | 1 | 1 | 2 | 7 | 39 | 48 |
| L1MC2_dup2336 | 5.53663178034989 | 3.5037872788797 | 1.06656389646069 | 3.28511708534928 | 0.00101940000104118 | 0.0484727666900516 | 3686 | 3 | 2 | 0 | 0 | 2 | 0 | 1 | 1 | 0 | 3 | 18 | 46 |
| MER67D_dup372 | 9.12114758709514 | 3.7549931203767 | 0.791452290762805 | 4.7444339528762 | 2.09090184369737e-06 | 0.000581896760351363 | 3687 | 3 | 3 | 0 | 1 | 2 | 0 | 1 | 2 | 3 | 6 | 43 | 57 |
| THE1B-int_dup1959 | 9.61778872452862 | 2.97498681194333 | 0.902268971289698 | 3.29722832836742 | 0.000976440865915911 | 0.0470419514641076 | 3688 | 6 | 2 | 0 | 1 | 4 | 1 | 1 | 4 | 1 | 3 | 41 | 65 |
| L1PA2_dup4041 | 10.9889485414469 | 6.75206006471693 | 1.54285041412559 | 4.37635431335296 | 1.20680787500634e-05 | 0.00215258700619384 | 3689 | 0 | 0 | 0 | 0 | 0 | 0 | 0 | 0 | 59 | 52 | 1 | 4 |
| L1PA3_dup205 | 20.3011091855389 | 6.94307974552941 | 0.810253920059939 | 8.56901716071401 | 1.04371982950624e-17 | 8.27829861838974e-14 | 3690 | 0 | 0 | 0 | 0 | 1 | 0 | 0 | 0 | 100 | 105 | 3 | 8 |
| ERVL-E-int_dup4596 | 19.0029888037232 | 5.39025095762411 | 0.654007245494744 | 8.2418826316618 | 1.69521818896358e-16 | 1.03428304407981e-12 | 3691 | 4 | 0 | 2 | 0 | 0 | 2 | 1 | 0 | 90 | 88 | 8 | 11 |
| L1MC4_dup18842 | 17.488251076344 | 6.43252647676735 | 1.32641793137587 | 4.8495472841618 | 1.2374357020897e-06 | 0.000377007523136281 | 3692 | 0 | 0 | 4 | 0 | 0 | 1 | 0 | 0 | 58 | 96 | 14 | 26 |
| MLT1A0_dup16374 | 8.74577292158397 | 6.54188118375974 | 0.867035873263903 | 7.54511016843309 | 4.5190531812201e-14 | 1.51449384261774e-10 | 3693 | 0 | 0 | 1 | 0 | 0 | 0 | 0 | 0 | 33 | 44 | 9 | 11 |
| L1PA5_dup8867 | 9.52973318938316 | 4.69878426549422 | 0.732438973324714 | 6.41525702020651 | 1.40585474477437e-10 | 1.92251444310392e-07 | 3694 | 0 | 1 | 0 | 0 | 2 | 4 | 1 | 0 | 32 | 55 | 2 | 11 |
| L2a_dup28904 | 13.3036608321324 | 4.11926798108781 | 0.817701510423397 | 5.03761816332576 | 4.71360571748331e-07 | 0.000177465763615868 | 3695 | 0 | 0 | 0 | 1 | 6 | 12 | 0 | 0 | 55 | 59 | 7 | 7 |
| LTR33C_dup456 | 12.1438939301305 | 4.55411915132123 | 0.65893068780833 | 6.91137813973833 | 4.79967742242995e-12 | 1.01067614509515e-08 | 3696 | 1 | 1 | 0 | 0 | 3 | 5 | 0 | 0 | 67 | 46 | 3 | 2 |
| MLT1G_dup1343 | 11.015221124189 | 4.42827602612509 | 0.643072710154137 | 6.88612027256402 | 5.73345222921214e-12 | 1.18630610794097e-08 | 3697 | 1 | 0 | 1 | 1 | 4 | 3 | 0 | 0 | 46 | 57 | 3 | 5 |
| ERVL-E-int_dup4595 | 6.83169946216604 | 4.19147177609695 | 1.05646582717266 | 3.96744662088531 | 7.26467762525341e-05 | 0.00795763824028531 | 3698 | 0 | 0 | 0 | 0 | 3 | 0 | 1 | 0 | 33 | 35 | 1 | 0 |
| L2_dup9697 | 5.67087504787338 | 3.14341541575782 | 0.888372278122798 | 3.53839881451514 | 0.000402561536449611 | 0.0264073769498129 | 3699 | 0 | 0 | 0 | 0 | 4 | 7 | 0 | 1 | 21 | 26 | 2 | 2 |
| MIR3_dup15374 | 6.89032306299382 | 3.85376711238979 | 0.85401398627895 | 4.51253395647669 | 6.40576750314236e-06 | 0.00131725462167481 | 3700 | 0 | 0 | 0 | 0 | 3 | 3 | 0 | 2 | 33 | 28 | 1 | 4 |
| AluSz_dup16191 | 7.12207985224739 | 3.27249577352598 | 0.956895845348205 | 3.41990801761204 | 0.000626423130881612 | 0.0353043292990895 | 3701 | 0 | 0 | 0 | 0 | 7 | 7 | 0 | 0 | 23 | 40 | 2 | 1 |
| LTR16A_dup618 | 7.31297361145935 | 3.10070128588797 | 0.792294335665923 | 3.91357245193711 | 9.09405972824431e-05 | 0.00939598409073739 | 3702 | 0 | 1 | 1 | 1 | 7 | 4 | 0 | 0 | 35 | 27 | 2 | 1 |
| MER20_dup2539 | 5.08240046811163 | 3.12982208944476 | 0.863429067012484 | 3.6248745948224 | 0.000289101702057115 | 0.0210626434775512 | 3703 | 0 | 0 | 1 | 1 | 5 | 3 | 0 | 0 | 15 | 29 | 1 | 3 |
| L1MA2_dup6546 | 4.11854087433684 | 4.17903278241063 | 0.96380418051602 | 4.3359770240602 | 1.45114102236134e-05 | 0.0024628616384221 9 | 3704 | 0 | 0 | 2 | 0 | 1 | 0 | 0 | 1 | 17 | 19 | 3 | 2 |
| MIRb_dup38848 | 3.75901872340998 | 4.32424663756732 | 1.01309218746509 | 4.26836441053528 | 1.96911460197935e-05 | 0.00306427684337421 | 3705 | 0 | 0 | 0 | 0 | 3 | 0 | 0 | 0 | 13 | 21 | 3 | 2 |
| L1MA7_dup6863 | 2.70690579722303 | 4.50897472600398 | 1.34306580999626 | 3.35722545570314 | 0.000787288871667452 | 0.0410366347990763 | 3706 | 0 | 0 | 0 | 0 | 0 | 0 | 0 | 1 | 8 | 20 | 1 | 0 |
| L1MB2_dup5377 | 2.36940754676872 | 3.80842692886227 | 1.14076157807703 | 3.33849509139509 | 0.000842335185523354 | 0.0427356690947846 | 3707 | 0 | 0 | 0 | 0 | 1 | 0 | 0 | 2 | 8 | 11 | 3 | 1 |
| L1MC4_dup11961 | 2.59072439959806 | 3.89876666652889 | 1.16572793951437 | 3.34449105522261 | 0.000824336878088356 | 0.0421369844884236 | 3708 | 0 | 0 | 0 | 0 | 1 | 1 | 0 | 0 | 8 | 17 | 2 | 0 |
| MIRc_dup47229 | 3.61803738891499 | 4.01357946500024 | 1.20478126453959 | 3.33137606230457 | 0.000864177715855264 | 0.043399668800527 | 3709 | 0 | 0 | 0 | 0 | 2 | 0 | 0 | 0 | 16 | 20 | 1 | 0 |
| FLAM_A_dup6660 | 3.18859724621731 | 5.16607812696944 | 1.15967296396414 | 4.45477154982567 | 8.39825717710515e-06 | 0.0016180823500109 | 3710 | 0 | 0 | 0 | 0 | 0 | 0 | 0 | 0 | 16 | 15 | 2 | 1 |
| THE1D-int_dup2178 | 3.02506203260502 | 4.62940770456947 | 1.30070342268541 | 3.55915700983678 | 0.000372047070618016 | 0.0250856084628151 | 3711 | 0 | 0 | 1 | 0 | 0 | 0 | 0 | 0 | 12 | 19 | 1 | 0 |
| L1MA7_dup6861 | 4.08894331743877 | 4.7189908845456 | 0.98610400742842 | 4.78549001829114 | 1.70570663762808e-06 | 0.000493754345008578 | 3712 | 1 | 0 | 0 | 0 | 0 | 0 | 0 | 1 | 12 | 28 | 2 | 2 |
| L1PREC2_dup6281 | 4.27950279662157 | 4.28688572538952 | 1.01275362615078 | 4.23290088990634 | 2.30696186594586e-05 | 0.00343624067165728 | 3713 | 3 | 0 | 0 | 0 | 0 | 0 | 0 | 0 | 22 | 18 | 1 | 2 |
| L2a_dup83773 | 4.02694974897736 | 3.41361764526694 | 1.01983327837761 | 3.34723107947356 | 0.000816231395093949 | 0.0419207847047323 | 3714 | 3 | 0 | 0 | 0 | 0 | 1 | 0 | 1 | 20 | 17 | 1 | 0 |
| MIR_dup82093 | 5.99015256069901 | 3.58066712182884 | 0.815971903782235 | 4.38822354695247 | 1.14280279373077e-05 | 0.00206787341108031 | 3715 | 3 | 0 | 0 | 0 | 1 | 3 | 0 | 1 | 34 | 19 | 1 | 1 |
| Charlie2b_dup1874 | 4.80036697203194 | 3.58708527274417 | 1.05971591729004 | 3.38494988535914 | 0.000711912508138504 | 0.0382401208062598 | 3716 | 1 | 0 | 0 | 0 | 0 | 4 | 0 | 0 | 15 | 33 | 0 | 1 |
| ERVL-E-int_dup4594 | 5.30823422271848 | 4.23486853015162 | 0.97947197786866 | 4.32362397887761 | 1.53486817796273e-05 | 0.0025450574458085 | 3717 | 0 | 1 | 1 | 0 | 0 | 1 | 0 | 0 | 25 | 28 | 0 | 1 |
| MIRb_dup102161 | 3.17208434068263 | 3.61992566005184 | 1.0121494751483 | 3.57647338553571 | 0.000348260754800399 | 0.0240436650472531 | 3718 | 1 | 1 | 0 | 0 | 0 | 1 | 0 | 0 | 16 | 14 | 0 | 1 |
| MLT1E3_dup1531 | 3.06729728942344 | 4.27304207359099 | 1.25411080053921 | 3.40722850943775 | 0.000656261594856619 | 0.0363598180026449 | 3719 | 0 | 0 | 0 | 0 | 0 | 1 | 0 | 0 | 18 | 12 | 0 | 1 |
| L1PB1_dup10170 | 4.82488687209163 | 3.60230383251117 | 0.965865057792876 | 3.72961399053289 | 0.000191773334572315 | 0.0159551391147357 | 3720 | 0 | 0 | 1 | 0 | 2 | 1 | 0 | 1 | 25 | 20 | 0 | 1 |
| MIRb_dup111528 | 4.49942847684711 | 3.33062964525148 | 0.810585862834374 | 4.10891652317408 | 3.97519721703091e-05 | 0.00511272984696482 | 3721 | 1 | 1 | 0 | 0 | 2 | 1 | 1 | 1 | 20 | 20 | 0 | 2 |
| MLT1H_dup6411 | 4.29114397110056 | 3.31851673352492 | 0.833466012759259 | 3.98158615075219 | 6.84568993946038e-05 | 0.00762514183827738 | 3722 | 1 | 0 | 0 | 0 | 2 | 3 | 1 | 0 | 19 | 19 | 1 | 1 |
| L2a_dup78647 | 2.84329867441572 | 3.80562842821346 | 1.0762990320217 | 3.53584674425011 | 0.000406470425371584 | 0.0265708823723811 | 3723 | 0 | 1 | 0 | 1 | 1 | 0 | 0 | 0 | 14 | 11 | 0 | 3 |
| PRIMA41-int_dup16 | 3.10457265638805 | 3.54322745445998 | 1.04215049619566 | 3.39991917424061 | 0.000674057748825574 | 0.0369135431765362 | 3724 | 3 | 1 | 0 | 0 | 1 | 0 | 0 | 0 | 16 | 9 | 0 | 4 |
| L3_dup9412 | 2.41320785658123 | 3.63581550284306 | 1.08412507564518 | 3.35368638224637 | 0.000797426884701616 | 0.0413476874062346 | 3725 | 1 | 1 | 0 | 0 | 0 | 1 | 1 | 0 | 6 | 12 | 0 | 7 |
| L2b_dup52618 | 3.18054550124 | 3.21537137802986 | 0.968017430151191 | 3.32160483673075 | 0.00089501352082894 | 0.0443861790802757 | 3726 | 1 | 0 | 1 | 0 | 2 | 2 | 1 | 0 | 8 | 15 | 0 | 8 |
| MLT1H_dup274 | 3.21924443730838 | 4.0196383155922 | 0.954065580987202 | 4.21316772735157 | 2.51813924412764e-05 | 0.00368047395935623 | 3727 | 1 | 0 | 1 | 0 | 1 | 0 | 1 | 0 | 12 | 13 | 1 | 7 |
| MER47A_dup2035 | 1.79947935660131 | 3.79796245458894 | 1.11765199197353 | 3.39816193400468 | 0.000678402512949659 | 0.0370831987930897 | 3728 | 1 | 0 | 0 | 0 | 0 | 1 | 0 | 0 | 8 | 6 | 2 | 2 |
| PRIMA41-int_dup17 | 2.2978867544955 | 3.48546652669854 | 1.02224819175198 | 3.40960889422066 | 0.000650561005544283 | 0.0361678479030935 | 3729 | 2 | 0 | 0 | 0 | 0 | 3 | 0 | 0 | 7 | 7 | 4 | 4 |
| L2c_dup65643 | 2.11011973930495 | 3.3698173561546 | 0.970012251047619 | 3.47399463513495 | 0.000512771286108403 | 0.0310180227529039 | 3730 | 1 | 1 | 0 | 1 | 0 | 1 | 0 | 0 | 8 | 7 | 3 | 2 |
| MIRc_dup32850 | 2.82720552424024 | 3.74927196659235 | 0.980633875100727 | 3.82331475771958 | 0.000131669521157059 | 0.0122050003432948 | 3731 | 2 | 0 | 0 | 1 | 0 | 1 | 0 | 0 | 10 | 13 | 4 | 1 |
| L1ME4c_dup2852 | 3.39184201917938 | 2.99037815064107 | 0.827742188652779 | 3.61269268576024 | 0.000303033769383859 | 0.0217973619376698 | 3732 | 3 | 3 | 0 | 0 | 0 | 2 | 0 | 1 | 13 | 10 | 4 | 2 |
| MIRc_dup60503 | 3.22461576682159 | 3.22017555096153 | 0.878414155669532 | 3.66589669596922 | 0.000246473413510364 | 0.0189456259196312 | 3733 | 1 | 2 | 0 | 0 | 0 | 2 | 0 | 1 | 15 | 11 | 1 | 2 |
| L1ME1_dup5754 | 4.18410412452609 | 3.16284360904907 | 0.758462922421701 | 4.17007017159179 | 3.04505864149719e-05 | 0.00422695046279277 | 3734 | 4 | 2 | 0 | 0 | 0 | 3 | 1 | 0 | 14 | 18 | 3 | 3 |
| L1MEj_dup781 | 3.44836284157047 | 3.17708699865749 | 0.779404713817733 | 4.07629944024237 | 4.5758086796175e-05 | 0.00562975890424129 | 3735 | 2 | 2 | 0 | 1 | 1 | 2 | 0 | 0 | 14 | 11 | 2 | 4 |
| L1PA16_dup6311 | 3.06616000273064 | 3.04180209747574 | 0.857635289589735 | 3.54673150043865 | 0.000390041888618312 | 0.0258485014139323 | 3736 | 4 | 2 | 0 | 1 | 0 | 1 | 0 | 0 | 9 | 13 | 2 | 4 |
| AluSx1_dup17655 | 2.43002534952886 | 3.94602338832841 | 1.12841314622103 | 3.49696687028448 | 0.000470580269222264 | 0.0292836658563152 | 3737 | 0 | 0 | 0 | 1 | 0 | 1 | 0 | 0 | 13 | 9 | 1 | 1 |
| DNA1_Mam_dup129 | 2.43520445104527 | 4.60647275107184 | 1.3681734176861 | 3.36687783253563 | 0.000760243592062041 | 0.0400037420957086 | 3738 | 0 | 0 | 0 | 0 | 0 | 1 | 0 | 0 | 15 | 5 | 0 | 5 |

| | | | | | | | | | | | | | | | | | | | |
|---|---|---|---|---|---|---|---|---|---|---|---|---|---|---|---|---|---|---|---|
| AluSz6_dup35625 | 3.50430644342937 | 3.03116800030417 | 0.92406442074357 | 3.28025615125953 | 0.00103712876588803 | 0.0490422578155789 | 3739 | 0 | 3 | 0 | 1 | 0 | 4 | 1 | 0 | 15 | 7 | 2 | 7 |
| LTR16A_dup2128 | 4.60303528019936 | 3.74137891235018 | 0.967692248082315 | 3.86629005219842 | 0.000110503505838219 | 0.0107981242841968 | 3740 | 0 | 1 | 0 | 0 | 0 | 6 | 0 | 0 | 19 | 20 | 2 | 3 |
| HUERS-P3-int_dup46 | 10.7609508149171 | 2.73319768508019 | 0.792755362943597 | 3.44771894690374 | 0.000565341986257845 | 0.0331044572401402 | 3741 | 2 | 0 | 0 | 2 | 2 | 9 | 5 | 4 | 42 | 44 | 4 | 6 |
| MER61E_dup221 | 6.03080174798073 | 3.01054104714843 | 0.849875801266452 | 3.54233058837802 | 0.000396608086144365 | 0.0261633788926274 | 3742 | 1 | 0 | 1 | 0 | 2 | 2 | 3 | 2 | 26 | 25 | 2 | 2 |
| MLT1C_dup8688 | 5.6930551747851 | 3.83012131892747 | 0.980472068735396 | 3.90640533377716 | 9.36793198058381e-05 | 0.00958442304746922 | 3743 | 1 | 0 | 0 | 0 | 0 | 1 | 3 | 3 | 21 | 25 | 2 | 8 |
| AmnSINE1_dup2106 | 6.53517833800025 | 2.37816937951082 | 0.599093165389067 | 3.96961527338803 | 7.19887734670457e-05 | 0.00791929759195084 | 3744 | 1 | 0 | 3 | 6 | 4 | 5 | 1 | 3 | 16 | 17 | 10 | 12 |
| L1MCa_dup1794 | 6.45859753359388 | 2.98018516113665 | 0.704394666223689 | 4.23084572334092 | 2.32814315382694e-05 | 0.00346016459013433 | 3745 | 1 | 0 | 1 | 6 | 1 | 4 | 0 | 4 | 14 | 19 | 10 | 18 |
| AluJr_dup77939 | 7.45101149945425 | 3.44261726575905 | 0.766467747069765 | 4.4915357220448 | 7.07114376012132e-06 | 0.00142227419538954 | 3746 | 0 | 0 | 5 | 4 | 1 | 1 | 5 | 3 | 27 | 17 | 10 | 13 |
| L2c_dup114190 | 6.08513221104011 | 2.18125128828177 | 0.583795884942593 | 3.73632522006602 | 0.000186729133215581 | 0.0156529341744761 | 3747 | 1 | 0 | 3 | 7 | 4 | 2 | 4 | 4 | 12 | 16 | 9 | 12 |
| MER92D_dup196 | 5.65220897612568 | 2.0928640570474 | 0.637327010510452 | 3.28381509418716 | 0.00102412090654907 | 0.0486204056723314 | 3748 | 0 | 0 | 4 | 8 | 3 | 2 | 5 | 4 | 11 | 13 | 9 | 10 |
| MER4E_dup363 | 7.34532569152511 | 2.35955977852371 | 0.697502387649551 | 3.38286982281878 | 0.000717326251681461 | 0.0383561600635049 | 3749 | 2 | 1 | 5 | 12 | 1 | 2 | 6 | 2 | 20 | 21 | 4 | 13 |
| THE1C_dup3400 | 4.77021082676636 | 2.44049213252657 | 0.730403575111087 | 3.3412926985679 | 0.000833892621187324 | 0.0424521638085235 | 3750 | 1 | 0 | 2 | 4 | 2 | 3 | 2 | 1 | 14 | 19 | 2 | 6 |
| MLT1C_dup8794 | 4.60162085503892 | 4.48991532480685 | 1.36691361317246 | 3.28471037345678 | 0.00102087253494948 | 0.0485242781064899 | 3751 | 0 | 0 | 2 | 1 | 0 | 0 | 7 | 3 | 15 | 22 | 2 | 0 |
| MSTA-int_dup2764 | 5.43051958257132 | 2.63661694638582 | 0.774269608245323 | 3.40529567260301 | 0.000660924523181502 | 0.0365004750531395 | 3752 | 0 | 3 | 0 | 2 | 4 | 0 | 5 | 2 | 11 | 24 | 5 | 9 |
| MER5B_dup4653 | 4.91023984443749 | 2.89111217066367 | 0.828707717087206 | 3.48869946671371 | 0.000485376530803645 | 0.0299244120663339 | 3753 | 1 | 1 | 0 | 1 | 1 | 0 | 4 | 6 | 12 | 15 | 8 | 8 |
| MADE1_dup1403 | 3.20894882253804 | 2.74751067759063 | 0.78646927773688 | 3.49347489516283 | 0.000476777810400793 | 0.0295243270435662 | 3754 | 1 | 2 | 0 | 1 | 1 | 0 | 2 | 2 | 11 | 7 | 4 | 6 |
| MIRb_dup46323 | 5.46143234920406 | 2.78653543766521 | 0.729231876170701 | 3.82119258458324 | 0.000132807889028515 | 0.0122818134328718 | 3755 | 2 | 3 | 0 | 3 | 1 | 0 | 3 | 2 | 21 | 17 | 4 | 6 |
| AluJb_dup90191 | 6.27933016265309 | 3.79865591607339 | 1.00898287479519 | 3.76483685795407 | 0.000166657857040337 | 0.0143678238195481 | 3756 | 0 | 1 | 0 | 4 | 1 | 0 | 0 | 3 | 19 | 35 | 4 | 4 |
| MER81_dup2714 | 4.8731267472897 | 2.85442343848714 | 0.803503332389174 | 3.5524724334368 | 0.000381629002987185 | 0.0255219490569951 | 3757 | 2 | 0 | 1 | 5 | 3 | 0 | 0 | 2 | 11 | 21 | 5 | 8 |
| MIRb_dup71162 | 3.3849282465087 | 2.84809604408387 | 0.832840151180861 | 3.41973911805961 | 0.000626812166730294 | 0.0353178896104207 | 3758 | 1 | 1 | 0 | 2 | 4 | 0 | 0 | 1 | 10 | 12 | 5 | 3 |
| L1PA3_dup9744 | 6.75313803161097 | 3.8608615923813 | 0.640743032433831 | 6.0256005870497 | 1.68482793645635e-09 | 1.72800891451743e-06 | 3759 | 1 | 0 | 2 | 1 | 1 | 2 | 4 | 0 | 8 | 21 | 23 | 22 |
| MIR3_dup3593 | 6.07200037414503 | 2.55438244842775 | 0.731193919048026 | 3.49344049763627 | 0.000476839235912454 | 0.0295243270435662 | 3760 | 0 | 0 | 1 | 2 | 4 | 3 | 9 | 1 | 3 | 11 | 24 | 21 |
| THE1D_dup530 | 6.95756380744309 | 2.3632198749397 | 0.618745162756323 | 3.81937511141464 | 0.000133790176399733 | 0.0123438686753047 | 3761 | 0 | 0 | 6 | 4 | 5 | 3 | 8 | 1 | 7 | 18 | 20 | 16 |
| MER5B_dup19178 | 4.53541591839832 | 2.49997464940459 | 0.653322025920799 | 3.82655803756363 | 0.000129947522213432 | 0.0120972195307501 | 3762 | 0 | 0 | 3 | 3 | 3 | 2 | 4 | 1 | 7 | 10 | 15 | 8 |
| MLT1F2_dup5018 | 6.41351000296106 | 2.88491076951708 | 0.663045423430484 | 4.35100019933936 | 1.35517929075856e-05 | 0.00233835744393644 | 3763 | 0 | 0 | 1 | 2 | 4 | 5 | 5 | 1 | 9 | 20 | 20 | 12 |
| L2c_dup114262 | 8.30781084336948 | 2.17741613248723 | 0.582801615574837 | 3.73611890272399 | 0.000186882326708502 | 0.0156577120109089 | 3764 | 1 | 0 | 4 | 7 | 8 | 5 | 8 | 1 | 16 | 20 | 19 | 13 |
| L2c_dup114263 | 7.27776717191253 | 1.93612719344355 | 0.540998102423127 | 3.57880588632686 | 0.000345167713297961 | 0.0238823136692052 | 3765 | 2 | 0 | 9 | 8 | 4 | 3 | 7 | 2 | 14 | 13 | 15 | 13 |
| LTR16E1_dup107 | 8.09434509489346 | 2.14902575999484 | 0.481075529796905 | 4.46712756498359 | 7.92768612794447e-06 | 0.00154240490874888 | 3766 | 4 | 0 | 6 | 6 | 4 | 3 | 8 | 3 | 14 | 19 | 12 | 22 |
| MIRc_dup31474 | 6.58051050444133 | 1.83919190236844 | 0.519980869838089 | 3.53703762782874 | 0.000404642012196358 | 0.0264883875681778 | 3767 | 1 | 0 | 8 | 7 | 5 | 2 | 6 | 3 | 6 | 10 | 18 | 18 |
| MSTD_dup1163 | 6.67305119549434 | 1.83661889347777 | 0.517821585147738 | 3.5468179507306 | 0.000389913927296854 | 0.0258485014139323 | 3768 | 1 | 0 | 6 | 5 | 5 | 4 | 5 | 5 | 9 | 7 | 14 | 23 |
| L2_dup45793 | 7.43823985203989 | 2.38797403857449 | 0.571592448906354 | 4.17775644717399 | 2.94398648071448e-05 | 0.00412064592435346 | 3769 | 1 | 0 | 9 | 7 | 1 | 3 | 7 | 3 | 9 | 18 | 21 | 14 |
| MIR_dup137227 | 6.32303370111657 | 2.07126817179769 | 0.60648868385417 | 3.41518024480689 | 0.000637398176922468 | 0.0356738167426132 | 3770 | 1 | 0 | 6 | 6 | 2 | 4 | 5 | 4 | 10 | 20 | 11 | 8 |
| AluJb_dup27565 | 5.99846381437839 | 1.88272263089656 | 0.574332723674519 | 3.27810440410064 | 0.00104506734499352 | 0.0492524138742215 | 3771 | 0 | 1 | 6 | 3 | 3 | 7 | 4 | 3 | 7 | 8 | 13 | 21 |
| Charlie29a_dup1197 | 6.38303079418527 | 2.31059425082349 | 0.59875138729409 | 3.85902112271614 | 0.000113842098994625 | 0.0110204524358727 | 3772 | 0 | 1 | 3 | 2 | 5 | 5 | 2 | 5 | 11 | 14 | 9 | 21 |
| L1ME1_dup26353 | 5.69485933782546 | 2.54694370916826 | 0.576138533329585 | 4.4207140502287 | 9.83752833511216e-06 | 0.00183365871069448 | 3773 | 0 | 1 | 2 | 3 | 5 | 3 | 4 | 1 | 12 | 11 | 12 | 16 |
| L2c_dup34378 | 6.81108978969913 | 2.46204941052589 | 0.596090665830416 | 4.13032706542385 | 3.62247504084883e-05 | 0.0047841821518855 | 3774 | 0 | 1 | 2 | 6 | 6 | 2 | 5 | 2 | 18 | 10 | 16 | 14 |
| L2_dup15153 | 6.76674804359938 | 2.59571450978354 | 0.64159042670792 | 4.04575006379455 | 5.21558568476127e-05 | 0.00623945576342989 | 3775 | 0 | 1 | 6 | 4 | 7 | 1 | 2 | 1 | 8 | 16 | 13 | 27 |
| MER58D_dup258 | 6.1167883237531 | 3.2198091280311 | 0.694853052694306 | 4.63379863633933 | 3.59016224434206e-06 | 0.000856835251145652 | 3776 | 0 | 0 | 3 | 8 | 2 | 1 | 1 | 1 | 14 | 11 | 12 | 23 |
| MLT1C_dup3701 | 5.18438317710956 | 3.17998352364495 | 0.914118165856992 | 3.47874448011182 | 0.000503768683832638 | 0.0306729128053329 | 3777 | 0 | 0 | 7 | 1 | 0 | 0 | 6 | 7 | 12 | 4 | 10 | 15 |
| L2c_dup5210 | 5.05089049909371 | 2.19617744231796 | 0.668139867757016 | 3.28700253988832 | 0.00101259920440149 | 0.0482370230811846 | 3778 | 0 | 0 | 2 | 2 | 1 | 3 | 7 | 6 | 7 | 6 | 11 | 18 |
| MIRb_dup10297 | 4.11209526876102 | 2.29820283836115 | 0.6986982114185 | 3.28926394944583 | 0.00100449774501522 | 0.0479855893265189 | 3779 | 0 | 0 | 1 | 4 | 1 | 1 | 5 | 5 | 6 | 4 | 12 | 12 |
| THE1C_dup9572 | 4.10850623957249 | 3.29638603878932 | 1.00500073055884 | 3.27998372394849 | 0.00103813075496869 | 0.0490568324622138 | 3780 | 0 | 0 | 7 | 7 | 0 | 0 | 1 | 2 | 10 | 7 | 12 | 3 |
| L1M5_dup53126 | 5.68598742299355 | 3.04889616003591 | 0.825143742081446 | 3.69498792094695 | 0.000219897240916659 | 0.0174779121709676 | 3781 | 0 | 1 | 8 | 7 | 0 | 0 | 1 | 4 | 15 | 6 | 13 | 13 |
| MLT1H1_dup3508 | 5.34860474945692 | 2.64883795899195 | 0.748326210105703 | 3.53968352734537 | 0.000400607115951516 | 0.0263395581133461 | 3782 | 0 | 1 | 7 | 8 | 0 | 0 | 4 | 3 | 11 | 3 | 17 | 12 |
| L2a_dup133438 | 5.03016244735573 | 2.4870216365889 | 0.653765569982818 | 3.80414899587671 | 0.000142292518934142 | 0.0128346989045881 | 3783 | 0 | 0 | 5 | 3 | 2 | 0 | 5 | 3 | 4 | 3 | 24 | 15 |
| MER49_dup1008 | 4.79406723894177 | 2.33673237826753 | 0.647053710663118 | 3.61134221125598 | 0.000304616395922311 | 0.0218649330148206 | 3784 | 0 | 0 | 5 | 7 | 1 | 0 | 5 | 2 | 2 | 4 | 20 | 17 |
| MIRb_dup170297 | 5.59010816476587 | 2.04625078116318 | 0.580830863608575 | 3.52297184837987 | 0.000426736685120673 | 0.027435905776202 | 3785 | 0 | 0 | 6 | 8 | 1 | 0 | 8 | 4 | 3 | 4 | 20 | 19 |
| L1PA15_dup1633 | 5.12624404863965 | 2.60197043241755 | 0.78088788722556 | 3.33206658085346 | 0.000862036297909384 | 0.043359797441665 | 3786 | 0 | 0 | 7 | 4 | 1 | 0 | 11 | 1 | 9 | 5 | 14 | 13 |
| L1PB_dup322 | 6.68419999781966 | 2.77473812314109 | 0.598423047947844 | 4.63675009285893 | 3.53929896644779e-06 | 0.000848953661159664 | 3787 | 0 | 0 | 7 | 6 | 1 | 0 | 9 | 2 | 8 | 8 | 21 | 24 |
| L1MB7_dup22217 | 7.21379150683232 | 2.56499025965038 | 0.642378523229407 | 3.99295768288687 | 6.52542174120111e-05 | 0.00736923588814351 | 3788 | 1 | 0 | 11 | 10 | 1 | 0 | 6 | 3 | 8 | 13 | 24 | 14 |
| HSMAR2_dup261 | 5.27647357492933 | 2.31182482669078 | 0.657597690955721 | 3.51556104664979 | 0.000438825860363607 | 0.0279998326230442 | 3789 | 0 | 1 | 6 | 9 | 2 | 0 | 2 | 3 | 8 | 9 | 12 | 14 |
| HUERS-P2-int_dup61 | 6.2625324521937 | 2.90814830402479 | 0.666685473000732 | 4.36209940338927 | 1.28820313232141e-05 | 0.00225282475182657 | 3790 | 0 | 0 | 4 | 8 | 1 | 0 | 5 | 5 | 11 | 12 | 16 | 15 |
| L1M5_dup53128 | 5.26286389820274 | 2.44969866555566 | 0.70281846491559 | 3.48553543744738 | 0.000491153162832173 | 0.0301206006400165 | 3791 | 1 | 2 | 5 | 7 | 0 | 0 | 5 | 2 | 13 | 12 | 7 | 10 |
| MER58C_dup210 | 4.88597979896052 | 2.16644980715395 | 0.65311128810198 | 3.31712197694502 | 0.000909499001132407 | 0.044879730916564 | 3792 | 2 | 3 | 5 | 3 | 0 | 0 | 6 | 2 | 9 | 13 | 8 | 9 |
| MER53_dup495 | 5.47780963616915 | 2.26763225782402 | 0.610294049230311 | 3.71563881490227 | 0.000202690948885289 | 0.0165908154535463 | 3793 | 4 | 3 | 4 | 4 | 0 | 0 | 2 | 4 | 11 | 13 | 11 | 10 |
| L2a_dup17012 | 3.80663235285474 | 2.22308103338621 | 0.67581363001027 | 3.28948830663926 | 0.00100369726970709 | 0.0479583814979629 | 3794 | 3 | 2 | 3 | 3 | 0 | 0 | 2 | 2 | 8 | 9 | 8 | 6 |
| MIRb_dup23888 | 4.41773640919399 | 2.47871335403022 | 0.647823599826399 | 3.82621651124543 | 0.000130127848686677 | 0.0121077470251776 | 3795 | 3 | 1 | 3 | 4 | 0 | 0 | 3 | 2 | 9 | 11 | 9 | 9 |
| MIRb_dup23859 | 5.41807465971092 | 2.61260034061246 | 0.761974489107551 | 3.42872416066373 | 0.00060642552017859 | 0.0346783289652523 | 3796 | 2 | 4 | 7 | 1 | 0 | 0 | 2 | 1 | 14 | 17 | 8 | 8 |
| MLT2D_dup375 | 6.413784524938 | 2.40873594084861 | 0.655935702708828 | 3.67221349730045 | 0.000240458662488217 | 0.0186250575860747 | 3797 | 5 | 6 | 5 | 2 | 0 | 0 | 4 | 1 | 8 | 23 | 11 | 15 |
| MIRc_dup10805 | 5.82653962612742 | 2.74274344174074 | 0.827212657669757 | 3.31564491465472 | 0.000914319225284854 | 0.0450338651168764 | 3798 | 0 | 2 | 5 | 8 | 0 | 0 | 7 | 3 | 15 | 16 | 6 | 8 |
| HAL1_dup22243 | 6.09901997401042 | 2.23984264983508 | 0.670133953373992 | 3.34238048759941 | 0.000830631158587628 | 0.0423586287315242 | 3799 | 0 | 4 | 7 | 9 | 0 | 0 | 5 | 3 | 5 | 14 | 14 | 17 |
| L1MB3_dup14662 | 6.18667487273241 | 2.77988710269418 | 0.595641453589397 | 4.66704774481745 | 3.05558346558226e-06 | 0.000766945003482527 | 3800 | 1 | 2 | 4 | 2 | 0 | 0 | 7 | 4 | 9 | 10 | 14 | 25 |
| L1ME3B_dup517 | 5.96876892288022 | 3.00840798209575 | 0.647630248501889 | 4.64525551277888 | 3.3965580514285e-06 | 0.000827223543608194 | 3801 | 0 | 1 | 5 | 4 | 0 | 0 | 4 | 5 | 11 | 11 | 14 | 18 |
| LOR1-int_dup1315 | 5.60126563934207 | 2.89864042837011 | 0.81980518294127 | 3.53576738557624 | 0.000406592541917323 | 0.0265715652235813 | 3802 | 1 | 1 | 7 | 9 | 0 | 0 | 4 | 0 | 7 | 18 | 7 | 18 |
| AluSz_dup100066 | 4.32857702946762 | 3.70017363828693 | 0.924385149953488 | 4.00284842143247 | 6.2584401801689e-05 | 0.00712521917277737 | 3803 | 0 | 0 | 4 | 5 | 0 | 0 | 3 | 0 | 13 | 7 | 12 | 8 |
| MLT1H2_dup363 | 5.05418031787271 | 2.97287342919044 | 0.873350677310337 | 3.40398594336243 | 0.000664101708220637 | 0.036604203165223 | 3804 | 1 | 1 | 5 | 7 | 0 | 0 | 4 | 0 | 17 | 9 | 10 | 6 |
| AluJr_dup17203 | 7.0748749551149 | 3.93562830505887 | 0.779603767982434 | 5.04824176933371 | 4.45894646984055e-07 | 0.000168947209667624 | 3805 | 2 | 1 | 4 | 4 | 0 | 0 | 0 | 0 | 21 | 24 | 10 | 18 |
| MER31-int_dup261 | 7.7171240680424 | 4.01185638423483 | 0.903556227495871 | 4.44007385722231 | 8.99280098080158e-06 | 0.00170774223637495 | 3806 | 0 | 2 | 7 | 5 | 0 | 0 | 0 | 0 | 22 | 19 | 18 | 19 |
| L1MA2_dup1529 | 5.32349766764631 | 4.17107949874306 | 0.965664478467562 | 4.31938793623459 | 1.56462534320709e-05 | 0.00258180542243242 | 3807 | 0 | 0 | 6 | 5 | 0 | 0 | 0 | 0 | 7 | 14 | 18 | 17 |
| LTR88a_dup209 | 6.84252826328194 | 3.89789465654593 | 1.08625062558678 | 3.5883934745173 | 0.000332721887211912 | 0.0232374059801954 | 3808 | 1 | 0 | 6 | 14 | 0 | 0 | 0 | 0 | 15 | 20 | 16 | 12 |
| L1M4_dup16501 | 13.4206069684879 | 3.209959438958 | 0.800027427867372 | 4.01231173725477 | 6.01270263063095e-05 | 0.00692496873256588 | 3809 | 0 | 2 | 14 | 16 | 2 | 2 | 5 | 1 | 40 | 51 | 10 | 15 |
| L1M1_dup9379 | 11.3872860161728 | 3.64495296703631 | 0.830768848216655 | 4.38744540657806 | 1.14689768651091e-05 | 0.00207154664139496 | 3810 | 1 | 0 | 14 | 25 | 0 | 0 | 3 | 3 | 15 | 28 | 19 | 38 |

| | | | | | | | | | | | | | | | | | | | |
|---|---|---|---|---|---|---|---|---|---|---|---|---|---|---|---|---|---|---|---|
| L1MC5a_dup16549 | 11.5700930164459 | 2.94431574535979 | 0.678876582506354 | 4.33704125496512 | 1.4441352700013e-05 | 0.00245447293539806 | 3811 | 3 | 1 | 14 | 19 | 0 | 0 | 3 | 6 | 20 | 24 | 23 | 31 |
| L1PA7_dup12827 | 10.9567722838525 | 2.84530337587762 | 0.777193143060883 | 3.66099907247216 | 0.00025123366845655 | 0.0192074485017261 | 3812 | 1 | 2 | 18 | 27 | 0 | 0 | 3 | 3 | 16 | 20 | 19 | 31 |
| MLT1D_dup20000 | 12.4654610319126 | 3.64261605926803 | 0.724140762848059 | 5.03025964860969 | 4.89816059651843e-07 | 0.000182394009592985 | 3813 | 0 | 2 | 14 | 13 | 0 | 0 | 5 | 4 | 33 | 21 | 30 | 28 |
| MLT1D_dup20002 | 9.35201417144006 | 2.54427473706006 | 0.655453001657792 | 3.88170430316896 | 0.000103726956664908 | 0.0103140051945626 | 3814 | 2 | 2 | 14 | 13 | 0 | 0 | 6 | 6 | 17 | 19 | 12 | 25 |
| L1M5_dup53131 | 11.5379356349203 | 2.49452681060707 | 0.708248282953902 | 3.52210781253589 | 0.000428129991799524 | 0.0275120179322086 | 3815 | 2 | 4 | 21 | 19 | 0 | 0 | 4 | 5 | 24 | 15 | 19 | 30 |
| MLT1D-int_dup581 | 10.6795298474335 | 2.33873627155066 | 0.600494017423119 | 3.89468704715295 | 9.83255934973082e-05 | 0.00994312863166616 | 3816 | 2 | 5 | 13 | 18 | 0 | 0 | 6 | 5 | 18 | 17 | 21 | 29 |
| LOR1a_dup791 | 8.61154040870153 | 3.28422721244118 | 0.829281097421947 | 3.96033048703404 | 7.4846107452837e-05 | 0.00813211501551268 | 3817 | 0 | 0 | 9 | 22 | 0 | 1 | 5 | 2 | 16 | 17 | 15 | 22 |
| AluYm1_dup986 | 8.28578500413203 | 3.3544515051881 | 0.712779930041254 | 4.70615313901157 | 2.52434919698427e-06 | 0.000671127144162704 | 3818 | 0 | 0 | 8 | 7 | 0 | 0 | 7 | 8 | 13 | 15 | 21 | 23 |
| LOR1-int_dup1312 | 7.5032180817586 | 2.66272244592933 | 0.738561619377248 | 3.60528136863452 | 0.000311814906739996 | 0.0222407403474685 | 3819 | 0 | 1 | 15 | 11 | 0 | 0 | 6 | 6 | 10 | 14 | 12 | 19 |
| L1MEd_dup1698 | 10.2766294747595 | 3.14579011732887 | 0.579095594843269 | 5.43224667108765 | 5.56489236755858e-08 | 3.15083561712055e-05 | 3820 | 1 | 0 | 8 | 11 | 2 | 0 | 3 | 6 | 13 | 24 | 26 | 35 |
| L1MEj_dup499 | 9.51236998803563 | 2.79501380260939 | 0.664899201820051 | 4.20366545027953 | 2.62627023276914e-05 | 0.00379423495328771 | 3821 | 0 | 0 | 10 | 13 | 2 | 0 | 9 | 8 | 16 | 14 | 16 | 31 |
| MER57C1_dup175 | 9.15597771821002 | 3.09384308916614 | 0.754539773009014 | 4.10030484785217 | 4.12606253734888e-05 | 0.0052389545171399 | 3822 | 1 | 0 | 8 | 19 | 0 | 0 | 5 | 9 | 17 | 15 | 17 | 22 |
| MER90a_dup1188 | 7.71968851046052 | 3.03658268465944 | 0.757579130195696 | 4.00827129949452 | 6.11648316085153e-05 | 0.00700381473624628 | 3823 | 1 | 0 | 12 | 12 | 0 | 0 | 3 | 5 | 16 | 11 | 15 | 20 |
| L1MEg1_dup849 | 7.77496670129747 | 2.74562989742329 | 0.710488854177309 | 3.86442360253845 | 0.000111351834027138 | 0.0108588425827035 | 3824 | 1 | 1 | 10 | 16 | 0 | 0 | 6 | 2 | 15 | 12 | 15 | 20 |
| MLT1D_dup20003 | 8.29812348915978 | 2.34529165699651 | 0.654674573414982 | 3.58237779842702 | 0.000340480902765364 | 0.0236267334177332 | 3825 | 2 | 1 | 14 | 12 | 1 | 0 | 4 | 6 | 18 | 10 | 17 | 16 |
| L2b_dup20087 | 9.35958868197302 | 2.74255196637228 | 0.608010805782349 | 4.51069609337509 | 6.46152380276143e-06 | 0.00132656923448824 | 3826 | 14 | 8 | 1 | 0 | 1 | 3 | 1 | 3 | 23 | 29 | 14 | 14 |
| L2c_dup29842 | 11.8281605320458 | 2.50101302871705 | 0.493713818264055 | 5.06571405578813 | 4.06872282072588e-07 | 0.00015716498706176 | 3827 | 11 | 7 | 3 | 0 | 4 | 3 | 4 | 5 | 36 | 39 | 9 | 16 |
| MIRc_dup21221 | 10.6720262461313 | 2.80836354622681 | 0.482023921172226 | 5.82619123838749 | 5.67066323843816e-09 | 4.88881027149785e-06 | 3828 | 8 | 7 | 2 | 0 | 3 | 3 | 2 | 5 | 29 | 34 | 12 | 20 |
| MLT1A0_dup3784 | 7.59478303258474 | 2.55039319266854 | 0.512646414458498 | 4.97495568239268 | 6.52625690352681e-07 | 0.000230546085798275 | 3829 | 5 | 5 | 3 | 1 | 1 | 4 | 5 | 1 | 18 | 23 | 8 | 18 |
| MLT1J_dup2949 | 10.813284055707 | 2.79845285784591 | 0.533209986049345 | 5.24831291810602 | 1.53498410736115e-07 | 7.33420338173004e-05 | 3830 | 10 | 5 | 2 | 2 | 1 | 2 | 7 | 1 | 34 | 35 | 12 | 16 |
| L3_dup9413 | 10.0819620949188 | 2.43378921046711 | 0.55862831581736 | 4.35672367754237 | 1.32023767912284e-05 | 0.00228635571176538 | 3831 | 7 | 4 | 5 | 3 | 1 | 2 | 5 | 5 | 29 | 36 | 10 | 10 |
| L2a_dup34098 | 11.5681012956293 | 2.53165571771666 | 0.453889073129171 | 5.57769699160875 | 2.43723712356695e-08 | 1.63360795663172e-05 | 3832 | 9 | 7 | 2 | 3 | 2 | 4 | 5 | 4 | 30 | 44 | 12 | 14 |
| L2c_dup29838 | 10.5574657478966 | 2.49468244353579 | 0.533580901471792 | 4.67535932537058 | 2.9343936924069e-06 | 0.000743585986723591 | 3833 | 8 | 4 | 2 | 4 | 1 | 4 | 5 | 5 | 31 | 35 | 6 | 19 |
| L1MA10_dup798 | 9.85698983294902 | 2.10176730767225 | 0.568384306446548 | 3.69779264457912 | 0.000217482426294875 | 0.0173785450039394 | 3834 | 7 | 9 | 1 | 1 | 2 | 4 | 5 | 7 | 26 | 34 | 10 | 8 |
| MER58C_dup530 | 7.92172375793035 | 2.50123814228788 | 0.578805149544028 | 4.32138197847464 | 1.55054981355359e-05 | 0.00256213281899876 | 3835 | 9 | 3 | 1 | 1 | 1 | 3 | 2 | 5 | 23 | 25 | 8 | 11 |
| MLT1F1_dup628 | 9.25308069094942 | 2.91088005230571 | 0.726716925243601 | 4.00552120253696 | 6.1880882566474e-05 | 0.00706879907977063 | 3836 | 6 | 7 | 1 | 0 | 1 | 0 | 8 | 1 | 28 | 31 | 15 | 10 |
| L1PA5_dup1953 | 8.88310199711787 | 2.74455604064948 | 0.703552112170676 | 3.90099893550412 | 9.5796572198871e-05 | 0.00975370610544825 | 3837 | 7 | 5 | 0 | 1 | 1 | 0 | 6 | 4 | 31 | 26 | 9 | 12 |
| Tigger7_dup592 | 9.12553475936206 | 2.29551839368414 | 0.588931877011379 | 3.89776557066852 | 9.70843298825211e-05 | 0.00984270471164311 | 3838 | 7 | 7 | 1 | 3 | 3 | 0 | 8 | 4 | 30 | 17 | 11 | 16 |
| Charlie7_dup713 | 10.2896033296002 | 2.26582976620403 | 0.649778514046896 | 3.48708016227281 | 0.000488324964758911 | 0.0300245405851483 | 3839 | 4 | 7 | 3 | 0 | 7 | 0 | 7 | 8 | 23 | 33 | 13 | 16 |
| MIRb_dup46307 | 9.46272533821609 | 2.72889186004216 | 0.828242753242883 | 3.2947971465582 | 0.000984927511779143 | 0.047304144699605 | 3840 | 2 | 5 | 2 | 1 | 1 | 0 | 6 | 8 | 34 | 34 | 8 | 4 |
| L1MD1_dup393 | 12.0273177682697 | 2.92617017178493 | 0.86085736867985 | 3.39913472108893 | 0.000675994097430274 | 0.0369770325303779 | 3841 | 8 | 3 | 2 | 0 | 0 | 1 | 3 | 4 | 54 | 50 | 2 | 4 |
| L1MA9_dup7192 | 15.2638432300241 | 3.13330679186684 | 0.700159180525914 | 4.47513491076886 | 7.63630794803347e-06 | 0.00150540922204041 | 3842 | 9 | 8 | 1 | 6 | 0 | 0 | 3 | 3 | 59 | 60 | 10 | 13 |
| MER45B_dup686 | 10.036794485674 | 2.51842657924855 | 0.708869990895549 | 3.55273408607254 | 0.000381249642538448 | 0.0255037468213251 | 3843 | 11 | 4 | 3 | 6 | 0 | 1 | 2 | 1 | 36 | 39 | 6 | 6 |
| L1MD2_dup1953 | 7.99036966398641 | 2.26948041982568 | 0.677191361761414 | 3.35131330370581 | 0.000804292507835927 | 0.0415354127014618 | 3844 | 9 | 9 | 0 | 0 | 0 | 4 | 5 | 3 | 19 | 27 | 8 | 11 |
| AluSx1_dup20425 | 7.89089205475397 | 2.28488104414628 | 0.621853668181853 | 3.67430661111433 | 0.000238496201456 | 0.0185454958011926 | 3845 | 8 | 6 | 0 | 2 | 1 | 2 | 7 | 2 | 21 | 28 | 5 | 12 |
| Tigger2a_dup677 | 8.00809558484846 | 2.56183460443422 | 0.662114370004259 | 3.86917233712619 | 0.0001092054341863 | 0.0107066321561159 | 3846 | 5 | 8 | 0 | 2 | 0 | 2 | 8 | 1 | 21 | 24 | 11 | 14 |
| AluJb_dup24729 | 5.99123347113764 | 3.00381969678074 | 0.638819106498113 | 4.70214441964193 | 2.57443298257768e-06 | 0.000682916421931359 | 3847 | 4 | 5 | 1 | 1 | 1 | 0 | 1 | 2 | 21 | 15 | 7 | 11 |
| Cheshire_dup186 | 5.50968502791288 | 2.7347515968037 | 0.704953120941773 | 3.87933823620818 | 0.000104741028361929 | 0.0103801369157049 | 3848 | 5 | 3 | 1 | 1 | 0 | 0 | 2 | 3 | 16 | 20 | 5 | 8 |
| THE1C_dup1698 | 4.89168951997245 | 2.84313351875383 | 0.736317654057753 | 3.86128663775163 | 0.00011279148470442 | 0.0109588740790028 | 3849 | 5 | 2 | 1 | 1 | 0 | 0 | 1 | 3 | 17 | 11 | 8 | 7 |
| MLT1H1_dup666 | 6.14774338182526 | 2.52281305656165 | 0.742544852610393 | 3.397522786257 | 0.000679989247431602 | 0.0371272880838366 | 3850 | 3 | 6 | 1 | 0 | 0 | 1 | 4 | 4 | 21 | 13 | 5 | 13 |
| MER6A_dup207 | 6.45495572161663 | 2.72902230691293 | 0.749174364866953 | 3.64270647113985 | 0.000269786422759426 | 0.0201300094543475 | 3851 | 3 | 4 | 4 | 0 | 0 | 0 | 3 | 4 | 17 | 23 | 8 | 9 |
| OldhAT1_dup863 | 5.81014597656171 | 2.61898958050919 | 0.746939592785912 | 3.50629368934771 | 0.000454393544323805 | 0.0285813618321663 | 3852 | 6 | 5 | 1 | 0 | 0 | 0 | 2 | 4 | 13 | 15 | 12 | 11 |
| MIR_dup34576 | 5.93674143181475 | 2.86456475920864 | 0.74929472858488 | 3.82301469625799 | 0.000131829918966455 | 0.0122151097735171 | 3853 | 2 | 5 | 0 | 0 | 0 | 1 | 5 | 3 | 14 | 18 | 14 | 8 |
| L1MB5_dup1568 | 5.99088513869896 | 3.23175730249756 | 0.625518396597405 | 5.16652638847579 | 2.38484545811237e-07 | 0.000103741213414261 | 3854 | 3 | 2 | 0 | 2 | 0 | 2 | 2 | 2 | 16 | 23 | 11 | 7 |
| MER131_dup239 | 6.36182787978015 | 2.81478958486432 | 0.643616952881204 | 4.3733925470168 | 1.22330469777406e-05 | 0.0021706223685052 | 3855 | 5 | 3 | 0 | 1 | 0 | 1 | 5 | 3 | 17 | 16 | 10 | 15 |
| Eulor4_dup10 | 5.47194766646375 | 2.88787002160376 | 0.873903133160349 | 3.30456535973293 | 0.000951237931069217 | 0.0462680418532698 | 3856 | 2 | 2 | 1 | 1 | 0 | 0 | 9 | 1 | 17 | 19 | 8 | 4 |
| L1MA4A_dup1103 | 5.32958861654133 | 2.52545277387917 | 0.674025239440109 | 3.74682226436651 | 0.000179088883440881 | 0.0152082382081456 | 3857 | 2 | 3 | 1 | 5 | 0 | 1 | 5 | 1 | 14 | 17 | 7 | 8 |
| MLT1B_dup3166 | 5.28467761156434 | 2.31999672117375 | 0.605941448694294 | 3.82874735863171 | 0.00012879713850622 | 0.0120183387917651 | 3858 | 5 | 5 | 2 | 2 | 0 | 1 | 2 | 2 | 12 | 17 | 8 | 7 |
| Eulor2C_dup4 | 5.96373317702539 | 2.97918840689152 | 0.639103567368008 | 4.66151115250472 | 3.13896055237123e-06 | 0.000782097494863376 | 3859 | 4 | 4 | 1 | 1 | 0 | 1 | 3 | 1 | 22 | 15 | 6 | 11 |
| Tigger4_dup281 | 5.54398042210442 | 3.21871259985863 | 0.636430095961083 | 5.05744876033558 | 4.24902544384385e-07 | 0.000162285490892595 | 3860 | 2 | 2 | 2 | 1 | 0 | 1 | 3 | 1 | 16 | 21 | 7 | 9 |
| L2a_dup17024 | 6.22585774079864 | 2.93332777471552 | 0.777017555052417 | 3.77511132874937 | 0.000159936218437524 | 0.0139963896404322 | 3861 | 1 | 5 | 4 | 5 | 0 | 0 | 2 | 0 | 12 | 26 | 13 | 7 |
| L1PA2_dup2312 | 5.88109873098536 | 3.54178536406255 | 0.739174249800944 | 4.79154321868807 | 1.65503359293158e-06 | 0.000482017898780537 | 3862 | 1 | 3 | 2 | 2 | 0 | 0 | 2 | 0 | 21 | 23 | 8 | 5 |
| THE1B-int_dup2100 | 6.68619375764659 | 3.28308806066006 | 0.769429545161218 | 4.26691187166742 | 1.98197407760661e-05 | 0.0030709302228926 | 3863 | 1 | 5 | 3 | 3 | 0 | 0 | 1 | 0 | 22 | 28 | 6 | 8 |
| L1M4_dup16499 | 8.84987914021874 | 3.57830728206316 | 0.841734726218806 | 4.25111043967194 | 2.12713239174796e-05 | 0.00324034983410281 | 3864 | 0 | 3 | 5 | 8 | 1 | 0 | 1 | 0 | 31 | 30 | 13 | 10 |
| L1MDa_dup5690 | 7.54351624554103 | 3.91129588894932 | 0.889765835647585 | 4.3958710620785 | 1.10329414139332e-05 | 0.00201322413932496 | 3865 | 0 | 1 | 4 | 6 | 0 | 1 | 1 | 0 | 22 | 36 | 8 | 9 |
| MLT1B_dup5947 | 7.19788848666024 | 4.31818160227536 | 0.689031293864611 | 6.26703263077605 | 3.67993063352984e-10 | 4.46747333941781e-07 | 3866 | 0 | 2 | 2 | 1 | 1 | 2 | 0 | 0 | 21 | 35 | 10 | 9 |
| THE1B_dup18029 | 8.81581919568095 | 4.15322872917773 | 0.815080855749905 | 5.09548065063632 | 3.47857550791718e-07 | 0.000139816406724132 | 3867 | 0 | 1 | 1 | 3 | 3 | 0 | 1 | 0 | 27 | 52 | 9 | 3 |
| L1PA8A_dup939 | 7.3953502151749 | 2.42454556201612 | 0.664851138211238 | 3.64674950928006 | 0.000265578590317117 | 0.0199034214965659 | 3868 | 10 | 8 | 1 | 2 | 0 | 6 | 0 | 0 | 20 | 31 | 3 | 6 |
| L1MEd_dup10520 | 5.27603551906858 | 2.65622455209875 | 0.683868216442335 | 3.88411756568705 | 0.000102702207101995 | 0.0102652751583559 | 3869 | 5 | 6 | 2 | 0 | 0 | 4 | 0 | 1 | 13 | 15 | 6 | 11 |
| L4_C_Mam_dup3286 | 5.01457143459309 | 2.44203304413626 | 0.704143657152026 | 3.46808924476191 | 0.000524173179879396 | 0.0314168542719352 | 3870 | 2 | 6 | 3 | 1 | 0 | 3 | 0 | 1 | 14 | 19 | 5 | 4 |
| L1PA10_dup3032 | 6.09840212965496 | 3.1196186810627 | 0.936646172399991 | 3.33062662613485 | 0.000866507428790264 | 0.0434799613350755 | 3871 | 4 | 5 | 2 | 1 | 0 | 0 | 1 | 0 | 23 | 23 | 0 | 11 |
| L1ME1_dup13950 | 5.87504162977579 | 3.37304421345034 | 0.811657794012649 | 4.15574671780676 | 3.24226854461757e-05 | 0.00441858437066192 | 3872 | 3 | 4 | 2 | 0 | 0 | 0 | 0 | 1 | 27 | 21 | 3 | 3 |
| Tigger1_dup5836 | 5.49752595688329 | 3.2527833261809 | 0.715103522849301 | 4.54868871743257 | 5.39812218717457e-06 | 0.00115821603421951 | 3873 | 2 | 4 | 2 | 0 | 2 | 0 | 0 | 1 | 19 | 24 | 3 | 5 |
| L1M3_dup2697 | 3.26037764178826 | 3.08179620176597 | 0.909704538589508 | 3.38768915734363 | 0.000704840959959651 | 0.0379701351728683 | 3874 | 3 | 1 | 2 | 1 | 0 | 0 | 0 | 0 | 9 | 17 | 2 | 3 |
| L1PA16_dup6308 | 2.91950769723576 | 3.03551904267229 | 0.899676894530597 | 3.37401022647809 | 0.000740816155132388 | 0.0392877198498179 | 3875 | 3 | 1 | 1 | 2 | 0 | 0 | 0 | 0 | 10 | 10 | 3 | 4 |
| L1ME3F_dup2082 | 4.28262258627799 | 3.98846984540537 | 0.83955172240815 | 4.75071367129703 | 2.02699980992394e-06 | 0.000569537814614484 | 3876 | 2 | 3 | 1 | 0 | 0 | 0 | 0 | 0 | 15 | 18 | 3 | 7 |
| AluSx_dup46230 | 3.6197285392506 | 3.06477987568522 | 0.863686026446593 | 3.54848843426869 | 0.000387449013552827 | 0.0257735373158627 | 3877 | 2 | 3 | 1 | 2 | 0 | 0 | 0 | 0 | 11 | 17 | 2 | 4 |
| MLT1E1A_dup1617 | 4.49253248122011 | 3.13430660631498 | 0.859235648907403 | 3.64778464476019 | 0.000264511198384552 | 0.0198672290438165 | 3878 | 2 | 5 | 1 | 1 | 0 | 0 | 0 | 0 | 18 | 19 | 2 | 2 |
| L1PA15_dup3630 | 3.18537265579391 | 4.08229067911426 | 0.910918353827337 | 4.48151106184435 | 7.41163905427547e-06 | 0.00147681113018408 | 3879 | 1 | 2 | 0 | 1 | 0 | 0 | 0 | 0 | 9 | 15 | 5 | 4 |
| L2c_dup5659 | 3.25536541275468 | 4.86088309137152 | 0.968065681900061 | 5.02123273477774 | 5.13409051543791e-07 | 0.000190251748982465 | 3880 | 0 | 1 | 0 | 1 | 0 | 0 | 0 | 0 | 12 | 13 | 5 | 5 |
| L1MA9_dup7189 | 2.72131799291529 | 4.38016809476538 | 1.07523719442951 | 4.0736761316087 | 4.6276823951624e-05 | 0.00568476259782814 | 3881 | 0 | 1 | 0 | 1 | 0 | 0 | 0 | 0 | 13 | 10 | 1 | 4 |
| MLT1C_dup8279 | 2.22732497263664 | 3.68489281739982 | 1.03914583194844 | 3.54607861967796 | 0.000391009532585596 | 0.0258818367710765 | 3882 | 1 | 1 | 0 | 1 | 0 | 0 | 0 | 0 | 6 | 13 | 1 | 3 |

| | | | | | | | | | | | | | | | | | | | |
|---|---|---|---|---|---|---|---|---|---|---|---|---|---|---|---|---|---|---|---|
| L2a_dup57856 | 4.24528155219849 | 3.92399248729109 | 0.804146112426679 | 4.87970087357584 | 1.0624686088181e-06 | 0.000336670255001879 | 3883 | 2 | 1 | 0 | 2 | 1 | 0 | 0 | 0 | 14 | 19 | 4 | 6 |
| Tigger1_dup5834 | 4.54893823675363 | 3.60903027521858 | 0.913455924265182 | 3.95096268943882 | 7.78374596862244e-05 | 0.00836167592889316 | 3884 | 3 | 1 | 0 | 3 | 0 | 0 | 0 | 0 | 16 | 22 | 3 | 4 |
| L1PA13_dup3929 | 3.38878557263744 | 3.76615433562149 | 0.991183083145319 | 3.79965558297298 | 0.000144897322488401 | 0.0130030620278147 | 3885 | 1 | 1 | 0 | 2 | 0 | 0 | 0 | 0 | 16 | 14 | 2 | 1 |
| MLT1A0_dup9658 | 3.38525809287414 | 3.94111942669037 | 0.990491206823669 | 3.97895448191695 | 6.92189902458623e-05 | 0.0076964401182439 | 3886 | 2 | 0 | 0 | 2 | 0 | 0 | 0 | 0 | 14 | 15 | 2 | 3 |
| LTR84a_dup106 | 8.22023277854913 | 4.57757002999818 | 0.679985846577361 | 6.73186074833602 | 1.67506918637639e-11 | 2.92535001668019e-08 | 3887 | 4 | 1 | 0 | 1 | 0 | 2 | 0 | 0 | 36 | 33 | 7 | 7 |
| MLT1J2_dup2311 | 5.48473433658803 | 3.95001137737391 | 0.725051966636692 | 5.44790105969491 | 5.09676949363401e-08 | 2.95074431613683e-05 | 3888 | 3 | 2 | 0 | 0 | 2 | 0 | 0 | 1 | 18 | 26 | 6 | 4 |
| L2a_dup77724 | 5.33544855362922 | 5.0149014377701 | 0.828434044421306 | 6.05347096916232 | 1.4175762934209e-09 | 1.47941495050145e-06 | 3889 | 2 | 1 | 0 | 0 | 1 | 0 | 0 | 0 | 12 | 27 | 11 | 9 |
| MLT2D_dup833 | 4.47424106568733 | 4.02856222182631 | 0.774332632171538 | 5.20262488554643 | 1.96493229191033e-07 | 8.92266754066593e-05 | 3890 | 2 | 3 | 0 | 0 | 1 | 1 | 0 | 0 | 10 | 23 | 5 | 8 |
| L1PA7_dup5784 | 5.30771336208756 | 3.42645102984484 | 0.72167711555321 | 4.74790035044724 | 2.05539254641101e-06 | 0.000574294019219568 | 3891 | 1 | 3 | 0 | 1 | 1 | 2 | 0 | 1 | 16 | 31 | 2 | 2 |
| MLT1O_dup2415 | 3.5150120012662 | 2.68974351414585 | 0.800633454183542 | 3.35951926576533 | 0.000780782070647228 | 0.0408137017975011 | 3892 | 2 | 3 | 0 | 2 | 3 | 1 | 0 | 0 | 6 | 18 | 5 | 2 |
| THE1C_dup3397 | 3.70079796776038 | 2.96188256698785 | 0.803603237099499 | 3.68575240896039 | 0.000228027955766441 | 0.0179253140569193 | 3893 | 1 | 2 | 0 | 1 | 3 | 1 | 0 | 0 | 15 | 16 | 1 | 1 |
| L1MC4_dup4499 | 4.78042472223181 | 2.56418150206912 | 0.724715021713226 | 3.53819284165988 | 0.00040287570814991 | 0.0264084471767048 | 3894 | 4 | 4 | 1 | 2 | 2 | 0 | 1 | 0 | 20 | 14 | 2 | 4 |
| LTR7_dup163 | 4.43572691319056 | 2.91562556733765 | 0.792918911884169 | 3.67707911066141 | 0.00023591989129175 | 0.0183992771069507 | 3895 | 2 | 5 | 0 | 1 | 2 | 0 | 1 | 0 | 20 | 13 | 3 | 2 |
| L1PA7_dup7258 | 5.41195279695065 | 3.00168125737437 | 0.745650214810509 | 4.02558893936239 | 5.68328941776135e-05 | 0.00665852495744265 | 3896 | 9 | 3 | 0 | 0 | 2 | 1 | 1 | 0 | 18 | 20 | 2 | 7 |
| L1MC1_dup7233 | 3.98201486833396 | 2.71886524003696 | 0.769343770975202 | 3.53400565860252 | 0.000409312315442195 | 0.0267272854583448 | 3897 | 7 | 2 | 1 | 0 | 1 | 2 | 0 | 0 | 14 | 13 | 2 | 4 |
| L1PB1_dup5380 | 4.14057211037262 | 2.36460936851346 | 0.687734837147078 | 3.43825736430997 | 0.0005854709619763 | 0.0338625361007323 | 3898 | 4 | 3 | 1 | 0 | 4 | 2 | 1 | 0 | 9 | 20 | 2 | 3 |
| LTR39_dup79 | 6.46638790853702 | 2.83150411287818 | 0.654152533517354 | 4.32850744711374 | 1.5012325752656e-05 | 0.00251380919320302 | 3899 | 7 | 2 | 1 | 5 | 0 | 0 | 3 | 1 | 14 | 18 | 10 | 19 |
| MLT1E1A_dup861 | 8.0817893652892 | 2.59227224826176 | 0.746854265468268 | 3.47092112627414 | 0.000518676326397296 | 0.0312131909865784 | 3900 | 11 | 10 | 2 | 2 | 0 | 0 | 3 | 1 | 15 | 12 | 19 | 26 |
| L2a_dup120267 | 6.91263463984729 | 2.6896746612405 | 0.694638739183921 | 3.87204817341514 | 0.000107924614330313 | 0.0106144386082852 | 3901 | 6 | 7 | 2 | 5 | 0 | 3 | 0 | 0 | 17 | 10 | 14 | 21 |
| AluJb_dup88604 | 6.31166483922726 | 2.40707686474948 | 0.608124754325833 | 3.95819582680525 | 7.55180327097195e-05 | 0.00819389594671542 | 3902 | 6 | 7 | 3 | 4 | 0 | 1 | 1 | 1 | 12 | 15 | 11 | 17 |
| AluSx1_dup28489 | 7.97584075296403 | 2.23559211595382 | 0.499530817136991 | 4.47538377865631 | 7.62741807242268e-06 | 0.00150490349971865 | 3903 | 9 | 5 | 3 | 5 | 0 | 5 | 4 | 1 | 13 | 20 | 14 | 21 |
| AluY_dup19474 | 7.54578280289274 | 3.16281951841958 | 0.869633422475043 | 3.63695717836827 | 0.000275877773114139 | 0.0204180443550286 | 3904 | 6 | 3 | 0 | 2 | 0 | 0 | 0 | 5 | 26 | 22 | 15 | 6 |
| L1MA2_dup3011 | 6.52343830083626 | 2.76589848380791 | 0.781097417360836 | 3.54104164516802 | 0.000398550666396279 | 0.0262406023426478 | 3905 | 5 | 3 | 2 | 5 | 0 | 0 | 0 | 2 | 18 | 29 | 6 | 6 |
| MER57F_dup63 | 7.61508327061096 | 2.79256234256716 | 0.720541827560923 | 3.87564223997952 | 0.000106343835855224 | 0.0104912321794549 | 3906 | 6 | 4 | 1 | 1 | 0 | 2 | 0 | 5 | 27 | 25 | 5 | 10 |
| L1P4_dup1620 | 9.33917752988109 | 3.03350373389752 | 0.825707741795639 | 3.67382256499214 | 0.000238948692941201 | 0.0185685452941172 | 3907 | 5 | 7 | 0 | 4 | 0 | 0 | 3 | 0 | 35 | 41 | 4 | 7 |
| AluJr4_dup4065 | 6.14021298838835 | 3.272931598372 | 0.679517718418367 | 4.81655078250206 | 1.4606100522122e-06 | 0.000432809862370714 | 3908 | 6 | 1 | 1 | 2 | 1 | 0 | 2 | 0 | 22 | 21 | 7 | 8 |
| L1PA5_dup5214 | 4.01418689091168 | 2.86777553477924 | 0.808358695368191 | 3.54765223806126 | 0.000388681052790341 | 0.0258194030673508 | 3909 | 3 | 2 | 1 | 3 | 0 | 0 | 1 | 0 | 12 | 18 | 3 | 4 |
| MLT1C_dup8281 | 5.02473947135601 | 2.89316995006076 | 0.842306349722885 | 3.43481911422442 | 0.00059294947786559 | 0.034153947339677 | 3910 | 4 | 1 | 2 | 5 | 0 | 0 | 1 | 0 | 19 | 16 | 5 | 5 |
| Tigger1_dup5832 | 5.77667672180655 | 2.99593687273124 | 0.753632856719508 | 3.97532677353302 | 7.02826860108548e-05 | 0.0077856070789287 | 3911 | 7 | 3 | 1 | 2 | 0 | 0 | 1 | 0 | 19 | 24 | 4 | 6 |
| Tigger3b_dup2961 | 5.74977989628413 | 3.9977145310954 | 0.766582404183686 | 5.2149834241923 | 1.83833673151644e-07 | 8.52680062217174e-05 | 3912 | 3 | 1 | 1 | 1 | 0 | 0 | 1 | 0 | 20 | 30 | 5 | 3 |
| L2c_dup63218 | 4.93951445189748 | 2.88339409732434 | 0.716360013570258 | 4.02506287718912 | 5.69600939897005e-05 | 0.00666999336834315 | 3913 | 2 | 4 | 2 | 1 | 0 | 1 | 2 | 0 | 16 | 21 | 3 | 5 |
| MIRb_dup71189 | 5.09620970658441 | 2.88539851837493 | 0.789660088699855 | 3.65397537455088 | 0.000258211043804124 | 0.0195545146495914 | 3914 | 2 | 1 | 2 | 4 | 0 | 1 | 5 | 0 | 12 | 20 | 11 | 3 |
| LTR1F2_dup30 | 4.48308913877805 | 3.08550596214973 | 0.759085920574904 | 4.06476510565876 | 4.80808602848718e-05 | 0.00587000943116681 | 3915 | 1 | 1 | 3 | 3 | 0 | 2 | 1 | 0 | 15 | 11 | 11 | 4 |
| L1MB4_dup1421 | 4.6522800693159 | 2.27234027005273 | 0.654105502153076 | 3.47396599260061 | 0.000512826025694587 | 0.0310180227529039 | 3916 | 1 | 3 | 4 | 4 | 0 | 3 | 3 | 0 | 9 | 14 | 9 | 7 |
| L1PREC2_dup1317 | 3.73137848650897 | 2.4441972258188 | 0.714139581170471 | 3.42257632858379 | 0.000620306791508197 | 0.0350592683639452 | 3917 | 2 | 2 | 2 | 4 | 0 | 1 | 2 | 0 | 10 | 11 | 5 | 6 |
| L1M4_dup8100 | 4.43141506389849 | 2.82547184070368 | 0.814145525435527 | 3.47047518217606 | 0.000519538351836905 | 0.031242415173919 | 3918 | 5 | 2 | 0 | 3 | 0 | 0 | 2 | 0 | 15 | 15 | 4 | 6 |
| L2c_dup29846 | 4.48038036280712 | 3.13405346062003 | 0.791418589882323 | 3.96004529169075 | 7.4935549802252e-05 | 0.0081368548033728 | 3919 | 3 | 4 | 0 | 2 | 0 | 0 | 2 | 0 | 13 | 13 | 4 | 13 |
| L2c_dup101329 | 4.89485624439817 | 2.39204102889176 | 0.697963002492252 | 3.42717453554183 | 0.000609896926112881 | 0.0347765478027451 | 3920 | 7 | 4 | 0 | 4 | 0 | 2 | 2 | 0 | 10 | 10 | 10 | 12 |
| MER57F_dup64 | 5.44909372637161 | 2.69887700710106 | 0.690280425719361 | 3.90982694357657 | 9.23622674784672e-05 | 0.00950983647660379 | 3921 | 5 | 5 | 0 | 2 | 0 | 2 | 3 | 0 | 17 | 15 | 4 | 12 |
| L1MC4a_dup1922 | 4.45803615400055 | 2.59013954120644 | 0.785090861661296 | 3.29915894795357 | 0.00096974984943109 | 0.046817703382766 | 3922 | 6 | 1 | 0 | 2 | 0 | 1 | 5 | 0 | 11 | 12 | 6 | 11 |
| MamRep1527_dup43 | 5.18181444423317 | 3.19316777804067 | 0.747717531949841 | 4.27055357350492 | 1.94988373245981e-05 | 0.00304640206568537 | 3923 | 6 | 1 | 0 | 3 | 0 | 1 | 1 | 0 | 16 | 19 | 7 | 7 |
| FRAM_dup3384 | 4.99644573346354 | 2.49328567162742 | 0.681231996433658 | 3.65996559862149 | 0.000252249111295427 | 0.0192438817044892 | 3924 | 0 | 1 | 1 | 5 | 3 | 2 | 6 | 0 | 10 | 14 | 9 | 11 |
| LTR88c_dup156 | 5.44233756686984 | 2.80682603985621 | 0.777897188627777 | 3.608222372943 | 0.000308302189606805 | 0.0220656474689521 | 3925 | 0 | 2 | 1 | 5 | 2 | 1 | 4 | 0 | 19 | 18 | 7 | 4 |
| Tigger4_dup279 | 8.84127564989002 | 3.35202259175488 | 0.611442062604189 | 5.48215897591066 | 4.20166397458097e-08 | 2.5310611040396e-05 | 3926 | 5 | 2 | 2 | 3 | 1 | 0 | 6 | 0 | 27 | 25 | 11 | 24 |
| L2a_dup34125 | 6.10934997206363 | 3.19172542555152 | 0.660162080163454 | 4.83476031334799 | 1.33306157615398e-06 | 0.000401046789823868 | 3927 | 2 | 3 | 2 | 1 | 1 | 0 | 5 | 0 | 15 | 23 | 11 | 10 |
| L2b_dup43481 | 5.29179909349776 | 3.08401237708798 | 0.622497414506177 | 4.95425732737301 | 7.26070676778243e-07 | 0.000250748350154827 | 3928 | 2 | 1 | 3 | 1 | 2 | 1 | 3 | 0 | 11 | 20 | 10 | 10 |
| L1PA10_dup3602 | 5.11622652101726 | 2.65880879871167 | 0.787309055123216 | 3.37708398170977 | 0.000732586789689107 | 0.0389446148926194 | 3929 | 10 | 7 | 0 | 0 | 1 | 2 | 1 | 0 | 16 | 11 | 8 | 4 |
| MLT1D_dup15018 | 3.80772651883098 | 2.42893699373017 | 0.713713910360082 | 3.40323616854368 | 0.000665926923068486 | 0.0366538625113241 | 3930 | 5 | 4 | 0 | 2 | 2 | 1 | 1 | 0 | 7 | 7 | 11 | 7 |
| L2a_dup160291 | 4.82253997558393 | 2.69480329042624 | 0.617512551177304 | 4.36396521056702 | 1.27725898266906e-05 | 0.00224435357378521 | 3931 | 4 | 3 | 0 | 2 | 2 | 2 | 2 | 0 | 15 | 14 | 5 | 8 |
| MLT1M_dup807 | 5.77855096072379 | 2.63113981227992 | 0.586418533883742 | 4.48679511347359 | 7.23025436420648e-06 | 0.0014505987394144 | 3932 | 6 | 4 | 0 | 1 | 4 | 5 | 1 | 0 | 18 | 12 | 9 | 8 |
| L1MA1_dup1527 | 5.30244125967858 | 3.87612839135967 | 0.841988469582113 | 4.60354094074878 | 4.15367496175121e-06 | 0.000958632727884436 | 3933 | 6 | 1 | 0 | 0 | 1 | 1 | 0 | 1 | 15 | 6 | 15 | 18 |
| L1PA4_dup608 | 4.00036934385772 | 3.22187457150684 | 0.738759021626613 | 4.36119827601274 | 1.29352080511461e-05 | 0.00225485788640147 | 3934 | 2 | 3 | 0 | 0 | 1 | 2 | 2 | 0 | 10 | 7 | 8 | 14 |
| Tigger2a_dup2815 | 5.32923803026192 | 3.80126052930123 | 0.729051097980093 | 5.2139836834935 | 1.84827732462297e-07 | 8.54948695151592e-05 | 3935 | 1 | 4 | 0 | 0 | 1 | 3 | 1 | 0 | 13 | 13 | 9 | 20 |
| AluSg_dup14950 | 5.41894500564453 | 3.70425357987832 | 0.718434822049583 | 5.15600506293759 | 2.52274077262391e-07 | 0.000107769492977158 | 3936 | 3 | 5 | 0 | 0 | 1 | 1 | 1 | 0 | 10 | 17 | 15 | 13 |
| L1ME3A_dup5499 | 5.76243884047622 | 3.64074685009928 | 0.884535019368855 | 4.11600080310791 | 3.85502960992587e-05 | 0.00499340705260436 | 3937 | 6 | 7 | 0 | 0 | 0 | 1 | 1 | 0 | 12 | 10 | 16 | 18 |
| L1M5_dup49103 | 3.68415017338467 | 3.88082781592676 | 0.842763690996088 | 4.60488255176239 | 4.12698709698943e-06 | 0.000954324656734934 | 3938 | 3 | 0 | 0 | 1 | 0 | 1 | 1 | 0 | 6 | 11 | 11 | 12 |
| THE1A-int_dup1324 | 3.25024487588564 | 3.95085360336681 | 0.982811566234171 | 4.0199502520155 | 5.8210432013153e-05 | 0.00677307552850939 | 3939 | 1 | 0 | 0 | 0 | 0 | 1 | 3 | 0 | 11 | 4 | 8 | 11 |
| MIR_dup34594 | 4.01959402477275 | 3.29795009212321 | 0.832483043300145 | 3.96158230328562 | 7.4454708304022e-05 | 0.00809948701450871 | 3940 | 2 | 1 | 0 | 1 | 0 | 1 | 4 | 0 | 14 | 7 | 12 | 5 |
| MIR_dup34603 | 3.16686936876988 | 3.35110666395809 | 0.871671143240023 | 3.84446208865186 | 0.000120817160318975 | 0.0115129996112371 | 3941 | 3 | 1 | 0 | 0 | 0 | 1 | 2 | 0 | 11 | 7 | 7 | 5 |
| THE1C_dup2999 | 2.84891513376278 | 3.22308273001494 | 0.937610176998303 | 3.43755092370416 | 0.000587000331464145 | 0.0339097792839445 | 3942 | 3 | 1 | 0 | 0 | 0 | 1 | 2 | 0 | 9 | 3 | 9 | 6 |
| L2a_dup34096 | 4.09276985872822 | 2.74532673028138 | 0.709023171595379 | 3.87198449960965 | 0.000107952818841461 | 0.0106144386082852 | 3943 | 1 | 3 | 0 | 1 | 0 | 2 | 3 | 2 | 6 | 16 | 7 | 9 |
| MER103C_dup2692 | 3.76634458290553 | 2.96210246671338 | 0.786575464471488 | 3.76582108202887 | 0.000166002639750051 | 0.0143374461408224 | 3944 | 1 | 1 | 0 | 0 | 0 | 3 | 4 | 1 | 5 | 7 | 10 | 16 |
| L2b_dup29046 | 5.01405809038744 | 2.760402861985 | 0.691720144236478 | 3.99063535301829 | 6.58965291611742e-05 | 0.00741703116802878 | 3945 | 4 | 2 | 0 | 2 | 0 | 1 | 4 | 2 | 11 | 6 | 8 | 23 |
| AluSg_dup31955 | 4.61884077779769 | 2.72213637286644 | 0.634558400592386 | 4.28981220692251 | 1.78824263697994e-05 | 0.00285573947985791 | 3946 | 3 | 1 | 0 | 2 | 3 | 1 | 2 | 2 | 10 | 7 | 9 | 17 |
| AluY_dup19473 | 5.46859518684465 | 3.2080447565091 | 0.626844445804813 | 5.11776849580321 | 3.09171759025016e-07 | 0.000126619936958634 | 3947 | 4 | 3 | 0 | 1 | 1 | 1 | 2 | 1 | 14 | 16 | 7 | 16 |
| L1PA10_dup6908 | 263.368410949215 | 3.29635935499962 | 0.349650095560187 | 9.42759460631179 | 4.19600797524336e-21 | 7.1315950976947e-17 | 3948 | 17 | 16 | 206 | 268 | 2 | 6 | 176 | 137 | 129 | 158 | 914 | 1466 |
| L1PA4_dup10537 | 152.248254985836 | 1.54672018119632 | 0.297143606590627 | 5.20529517341166 | 1.93688482464977e-07 | 8.84595002852427e-05 | 3949 | 18 | 17 | 259 | 340 | 15 | 15 | 136 | 91 | 54 | 38 | 435 | 616 |
| L1PA4_dup8359 | 74.1954449305104 | 1.85168530556886 | 0.35742134703546 | 5.18067910864079 | 2.21079530918399e-07 | 9.7778791940352e-05 | 3950 | 0 | 2 | 91 | 111 | 8 | 8 | 60 | 43 | 5 | 15 | 287 | 366 |
| L1M4_dup12829 | 110.165051307909 | 2.26412711564988 | 0.301926782088015 | 7.49892772013136 | 6.43419927838375e-14 | 2.01446313354513e-10 | 3951 | 0 | 0 | 124 | 157 | 8 | 5 | 88 | 65 | 14 | 17 | 406 | 599 |
| L1PA4_dup8363 | 94.5787751168963 | 2.26488152920255 | 0.331370473930771 | 6.83489238596352 | 8.20666105742617e-12 | 1.6138365057606e-08 | 3952 | 1 | 0 | 97 | 148 | 4 | 8 | 67 | 70 | 20 | 14 | 365 | 466 |
| L1MA4A_dup6036 | 89.3860885884918 | 3.27532591247907 | 0.320113304758901 | 10.2317706380431 | 1.42882721641505e-24 | 4.24979651046369e-20 | 3953 | 5 | 4 | 47 | 73 | 1 | 4 | 74 | 46 | 54 | 51 | 323 | 500 |
| HSMAR2_dup1463 | 57.9134514914616 | 2.73767113144057 | 0.283029909161589 | 9.67272730839753 | 3.93742531622623e-22 | 1.04099400477196e-17 | 3954 | 3 | 4 | 49 | 47 | 10 | 4 | 41 | 36 | 33 | 57 | 216 | 252 |

| | | | | | | | | | | | | | | | | | | | |
|---|---|---|---|---|---|---|---|---|---|---|---|---|---|---|---|---|---|---|---|
| L1MC2_dup4897 | 63.9671531112043 | 3.02093350541085 | 0.268424866740295 | 11.2542982403104 | 2.20574797032155e-29 | 2.62424453273065e-24 | 3955 | 4 | 2 | 35 | 53 | 5 | 7 | 46 | 34 | 41 | 55 | 260 | 291 |
| L1PA6_dup26 | 68.6641148121592 | 1.17088660849277 | 0.252084019092064 | 4.64482680302376 | 3.40361872556052e-06 | 0.000827249705078879 | 3956 | 16 | 14 | 78 | 83 | 39 | 30 | 90 | 54 | 28 | 46 | 190 | 229 |
| L1M3f_dup115 | 130.289748908571 | 1.64147579819864 | 0.345970765293267 | 4.74455058885458 | 2.08969751986664e-06 | 0.000581896760351363 | 3957 | 39 | 19 | 75 | 106 | 56 | 69 | 155 | 159 | 161 | 128 | 236 | 434 |
| THE1B-int_dup3980 | 92.8448612556543 | 2.16684640294082 | 0.196795694602629 | 11.0106392688932 | 3.39585557090298e-28 | 2.69343416558027e-23 | 3958 | 29 | 19 | 67 | 75 | 32 | 46 | 66 | 46 | 132 | 128 | 232 | 298 |
| L1MA4_dup1393 | 85.1175439817247 | 1.40895462686882 | 0.316999013911258 | 4.44466564575261 | 8.80286619782007e-06 | 0.00168014317319228 | 3959 | 22 | 21 | 62 | 57 | 41 | 49 | 114 | 110 | 81 | 92 | 177 | 243 |
| L1PA8_dup1199 | 81.701073309614 | 1.42853330108471 | 0.235216052689124 | 6.07328149908522 | 1.25322483535122e-09 | 1.33721899853131e-06 | 3960 | 25 | 30 | 63 | 77 | 34 | 40 | 123 | 77 | 91 | 88 | 158 | 230 |
| L1MC4_dup7046 | 45.3965894489185 | 1.04746066713124 | 0.261008930518963 | 4.01312194586056 | 5.99209373573524e-05 | 0.00690957967647954 | 3961 | 60 | 48 | 32 | 41 | 35 | 29 | 26 | 23 | 58 | 49 | 83 | 87 |
| L1PA2_dup563 | 42.1172259138474 | 1.40497250201825 | 0.217983525206246 | 6.44531507915074 | 1.15360504986299e-10 | 1.61468063055706e-07 | 3962 | 35 | 25 | 37 | 38 | 22 | 26 | 27 | 24 | 46 | 49 | 90 | 116 |
| AluSq2_dup40955 | 40.370316032347 | 1.08560244811345 | 0.213716680142551 | 5.07963368787752 | 3.78163330360816e-07 | 0.000149224629860754 | 3963 | 34 | 29 | 37 | 34 | 24 | 35 | 42 | 21 | 56 | 52 | 76 | 65 |
| MER57-int_dup391 | 45.3617119086833 | 1.3138586594448 | 0.243742397036091 | 5.39035750620872 | 7.03176534531857e-08 | 3.76842440733597e-05 | 3964 | 31 | 37 | 41 | 39 | 22 | 35 | 35 | 23 | 73 | 44 | 81 | 106 |
| AluSp_dup33457 | 51.094369028039 | 1.21263644373283 | 0.181864458681795 | 6.66780333289064 | 2.59660406265961e-11 | 4.23185993351783e-08 | 3965 | 42 | 40 | 37 | 43 | 31 | 46 | 38 | 31 | 59 | 86 | 93 | 94 |
| THE1A-int_dup80 | 49.0898780783104 | 1.006783664777 | 0.201964072372791 | 4.98496417183868 | 6.19732956509175e-07 | 0.00022042298665102 | 3966 | 27 | 33 | 44 | 51 | 45 | 39 | 51 | 30 | 55 | 76 | 67 | 102 |
| MER30_dup2981 | 50.5733274960922 | 1.29081196387643 | 0.218708607449675 | 5.90197148127078 | 3.59183264496918e-09 | 3.26206950587724e-06 | 3967 | 49 | 37 | 22 | 33 | 36 | 36 | 56 | 29 | 67 | 80 | 77 | 114 |
| MLT1J_dup1153 | 44.7334606353953 | 1.20814069543343 | 0.25875812520496 | 4.66899616959457 | 3.0267502155266e-06 | 0.000762927019897978 | 3968 | 33 | 33 | 24 | 24 | 31 | 34 | 46 | 40 | 61 | 54 | 91 | 82 |
| MSTB2_dup119 | 42.0320314905833 | 1.09742241084314 | 0.250799526221819 | 4.37569570953867 | 1.21045778997988e-05 | 0.00215642936784941 | 3969 | 38 | 23 | 22 | 25 | 40 | 32 | 47 | 35 | 47 | 72 | 66 | 78 |
| AluJr_dup76560 | 51.9956483130824 | 1.14634256451005 | 0.257464745734778 | 4.45242536502816 | 8.49057566591742e-06 | 0.00163190510290984 | 3970 | 46 | 34 | 52 | 31 | 45 | 59 | 29 | 22 | 71 | 114 | 81 | 57 |
| AluY_dup65414 | 41.4345513343402 | 1.00266766510271 | 0.255523890579785 | 3.92396837269363 | 8.71021754631346e-05 | 0.00910878779629798 | 3971 | 32 | 31 | 31 | 49 | 32 | 51 | 35 | 13 | 55 | 61 | 75 | 56 |
| THE1B_dup18811 | 47.0653055135281 | 1.30377709469565 | 0.244119966219873 | 5.34072290310482 | 9.25766696388075e-08 | 4.80966118643574e-05 | 3972 | 25 | 31 | 30 | 45 | 44 | 53 | 35 | 14 | 71 | 84 | 74 | 81 |
| L1PA3_dup4529 | 51.451557537688 | 1.07460203019556 | 0.31029229482485 | 3.46319276410695 | 0.000533805824905401 | 0.0317939826815871 | 3973 | 45 | 46 | 40 | 51 | 65 | 58 | 16 | 15 | 62 | 73 | 68 | 111 |
| L1PA8_dup2340 | 48.8703448594748 | 1.12807031390897 | 0.277921295525747 | 4.05895601405782 | 4.92925966242408e-05 | 0.00596590854341383 | 3974 | 44 | 38 | 48 | 56 | 43 | 46 | 23 | 12 | 51 | 86 | 79 | 95 |
| L1PA3_dup3880 | 44.4984623563507 | 1.52217969963452 | 0.337998354978194 | 4.50351215387639 | 6.68395433892831e-06 | 0.00136282793413079 | 3975 | 20 | 29 | 30 | 61 | 34 | 40 | 17 | 11 | 68 | 77 | 60 | 114 |
| Tigger1_dup10137 | 41.1521926576586 | 1.15222493976999 | 0.253761319338166 | 4.54058539250624 | 5.60982521035598e-06 | 0.00119075421008329 | 3976 | 26 | 20 | 39 | 45 | 44 | 36 | 27 | 18 | 50 | 63 | 76 | 73 |
| AluSx_dup70127 | 43.2172724479073 | 1.28591035721982 | 0.260158405009382 | 4.94279766657334 | 7.70093959250136e-07 | 0.00026354717154823 | 3977 | 29 | 23 | 30 | 36 | 45 | 53 | 24 | 16 | 68 | 64 | 65 | 85 |
| L2_dup47003 | 45.4049752895306 | 1.26094370352352 | 0.301549529603219 | 4.18154757257517 | 2.89531654864868e-05 | 0.00407650290819384 | 3978 | 22 | 22 | 34 | 45 | 55 | 45 | 25 | 21 | 67 | 55 | 73 | 104 |
| L2a_dup137041 | 48.5930197257992 | 1.42027259096124 | 0.320868715592419 | 4.4263355133859 | 9.58474066601848e-06 | 0.0017985082506149 | 3979 | 27 | 21 | 42 | 53 | 56 | 39 | 19 | 15 | 76 | 85 | 87 | 83 |
| L1MA3_dup7539 | 48.5620697889684 | 1.27901020184232 | 0.299032703083442 | 4.27715827952579 | 1.89294216086414e-05 | 0.00298685686610728 | 3980 | 33 | 19 | 35 | 67 | 53 | 37 | 28 | 18 | 70 | 71 | 71 | 111 |
| L2a_dup137051 | 48.1567037285725 | 1.3999007175968 | 0.301537498355727 | 4.64254271933145 | 3.44147453783321e-06 | 0.000832200305263477 | 3981 | 28 | 18 | 36 | 55 | 48 | 49 | 28 | 13 | 64 | 78 | 86 | 105 |
| L2c_dup114239 | 45.8559941131607 | 1.27896653148262 | 0.336592140492143 | 3.79975162109431 | 0.000144841183227271 | 0.0130030620278147 | 3982 | 30 | 9 | 44 | 52 | 31 | 54 | 35 | 19 | 67 | 80 | 79 | 73 |
| MLT1A_dup7308 | 48.8529530061693 | 1.36068537530074 | 0.317859366141413 | 4.28077797995538 | 1.86241091229005e-05 | 0.00294649751951974 | 3983 | 37 | 10 | 35 | 45 | 43 | 56 | 40 | 17 | 68 | 77 | 98 | 87 |
| ERV3-16A3_I-int_dup | 44.0021971172428 | 1.15835592775245 | 0.351114610903473 | 3.29908209963642 | 0.000970015371878645 | 0.046817703382766 | 3984 | 26 | 16 | 28 | 33 | 65 | 58 | 22 | 24 | 44 | 76 | 79 | 80 |
| L1PA7_dup2773 | 42.7398690880679 | 1.46112932806043 | 0.294389371644499 | 4.96325434542137 | 6.93217046923068e-07 | 0.000241505451606378 | 3985 | 22 | 18 | 23 | 29 | 43 | 43 | 32 | 23 | 57 | 52 | 71 | 126 |
| L2b_dup14136 | 45.4187103483984 | 1.19262509373598 | 0.263827087113248 | 4.52048008711115 | 6.16995488490747e-06 | 0.00128556574872521 | 3986 | 35 | 29 | 30 | 30 | 47 | 65 | 30 | 16 | 58 | 72 | 59 | 101 |
| MLT1I_dup8718 | 47.4364068231582 | 1.19862201632753 | 0.252857828002655 | 4.74030021453379 | 2.13401783227038e-06 | 0.000591130392453328 | 3987 | 33 | 24 | 31 | 39 | 51 | 59 | 34 | 20 | 54 | 80 | 71 | 103 |
| MIRb_dup178808 | 53.6696284790144 | 1.03878592988414 | 0.316886805181139 | 3.27809777150661 | 0.00104509190173798 | 0.0492524138742215 | 3988 | 28 | 22 | 40 | 49 | 59 | 74 | 38 | 32 | 54 | 60 | 109 | 113 |
| L1PB1_dup2756 | 50.635966725568 | 1.30874413168616 | 0.330060787950557 | 3.96516090206452 | 7.33464542358256e-05 | 0.00800206116441897 | 3989 | 24 | 26 | 18 | 35 | 63 | 59 | 34 | 34 | 55 | 78 | 94 | 114 |
| LTR33_dup4818 | 59.9865818772263 | 1.24633177304777 | 0.305334833092441 | 4.08185256960329 | 4.46781363995104e-05 | 0.00553697075193641 | 3990 | 33 | 32 | 30 | 36 | 71 | 84 | 41 | 33 | 70 | 87 | 97 | 139 |
| L1PA3_dup9430 | 51.3286001314316 | 1.41349339169796 | 0.312758201159894 | 4.5194446906776 | 6.2002042276189e-06 | 0.00129073822847332 | 3991 | 18 | 30 | 52 | 55 | 41 | 51 | 24 | 17 | 67 | 97 | 93 | 97 |
| L2c_dup114233 | 55.8101578041027 | 1.59636960111294 | 0.295304650521826 | 5.40583969230433 | 6.45054714401351e-08 | 3.52037130901247e-05 | 3992 | 19 | 20 | 47 | 53 | 43 | 50 | 39 | 20 | 92 | 117 | 93 | 97 |
| LTR32_dup978 | 63.0668472695832 | 1.41619661027036 | 0.249649748339613 | 5.67273397906186 | 1.40536376852389e-08 | 1.05489175793434e-05 | 3993 | 31 | 35 | 54 | 62 | 56 | 59 | 37 | 20 | 88 | 124 | 97 | 126 |
| L1PA5_dup9767 | 57.9867645244415 | 1.45593423411695 | 0.320394856726736 | 4.54418728499973 | 5.51476101603101e-06 | 0.00117476752436931 | 3994 | 25 | 29 | 47 | 59 | 62 | 55 | 24 | 19 | 95 | 86 | 81 | 142 |
| L2c_dup114255 | 63.2635547665235 | 1.34383412302043 | 0.300084449078475 | 4.47818648099623 | 7.52798279429735e-06 | 0.00148898868991843 | 3995 | 33 | 34 | 43 | 59 | 76 | 73 | 27 | 22 | 92 | 105 | 86 | 141 |
| L1ME3Cz_dup9745 | 52.8049637991442 | 1.28451845654365 | 0.311779344891434 | 4.11996008584513 | 3.7893803750115e-05 | 0.00494335472978337 | 3996 | 25 | 25 | 44 | 38 | 68 | 58 | 38 | 17 | 62 | 85 | 82 | 125 |
| L2a_dup137035 | 54.6688386763295 | 1.46616566361683 | 0.308178167876192 | 4.7575260561801 | 1.95979966538836e-06 | 0.000556475526468375 | 3997 | 21 | 27 | 29 | 53 | 57 | 66 | 30 | 19 | 83 | 94 | 86 | 117 |
| MER33_dup9248 | 45.1216718796219 | 1.1619763949892 | 0.295135076387991 | 3.93710029051796 | 8.24721451258689e-05 | 0.00874116572121158 | 3998 | 40 | 37 | 19 | 30 | 55 | 61 | 27 | 19 | 45 | 60 | 87 | 90 |
| MLT1A0_dup2303 | 48.5441643942906 | 1.2610146027596 | 0.328375393400651 | 3.84016168111912 | 0.000122953307908083 | 0.0116373300729899 | 3999 | 44 | 37 | 14 | 19 | 66 | 72 | 33 | 20 | 64 | 66 | 90 | 79 |
| L1PA15_dup2489 | 48.2005803630935 | 1.07826349838448 | 0.288450546528701 | 3.73812257026593 | 0.000185399560107246 | 0.0155993931150207 | 4000 | 62 | 60 | 16 | 21 | 52 | 43 | 38 | 25 | 52 | 69 | 77 | 91 |
| LTR33C_dup68 | 54.463890893853 | 1.00990795101213 | 0.297813038565682 | 3.39108037672233 | 0.000696176808787054 | 0.0376568508623879 | 4001 | 48 | 48 | 20 | 24 | 74 | 60 | 59 | 28 | 61 | 74 | 91 | 97 |
| L1MEf_dup7410 | 58.6442782319639 | 1.01736414182081 | 0.234968851632489 | 4.32978301060964 | 1.49256339578533e-05 | 0.00250381500469088 | 4002 | 46 | 38 | 34 | 32 | 72 | 66 | 52 | 36 | 92 | 90 | 83 | 80 |
| L1PA2_dup1805 | 55.7334855646383 | 1.06891932880548 | 0.176078352449937 | 6.07070269532084 | 1.27351763332119e-09 | 1.35280547668859e-06 | 4003 | 51 | 57 | 26 | 38 | 53 | 62 | 39 | 26 | 86 | 116 | 64 | 68 |
| L1MB2_dup2513 | 63.0379937259137 | 1.41587748356869 | 0.246058046779362 | 5.75424174133307 | 8.70315422216166e-09 | 6.94926420988751e-06 | 4004 | 59 | 57 | 23 | 27 | 84 | 59 | 34 | 24 | 93 | 108 | 84 | 131 |
| L2c_dup27084 | 54.7299540988406 | 1.48690163421458 | 0.248142611086476 | 5.99212536574948 | 2.07116159097645e-09 | 2.01152904459788e-06 | 4005 | 39 | 42 | 17 | 32 | 43 | 70 | 35 | 23 | 104 | 88 | 70 | 110 |
| MER61A_dup782 | 69.9326491733404 | 1.17998845231208 | 0.332966060338714 | 3.54387006024494 | 0.000394299524819977 | 0.0260404258365742 | 4006 | 44 | 65 | 21 | 59 | 86 | 108 | 44 | 23 | 75 | 95 | 116 | 150 |
| MLT1J2_dup6567 | 66.1943799656623 | 1.31629568344126 | 0.304267706814887 | 4.32611037569649 | 1.51765396764441e-05 | 0.00252561248541665 | 4007 | 53 | 52 | 23 | 30 | 77 | 96 | 35 | 34 | 78 | 96 | 123 | 129 |
| MSTB_dup8303 | 67.0792809505891 | 1.14964609307835 | 0.320813728121951 | 3.58353147731051 | 0.000338979892367686 | 0.0235626197100543 | 4008 | 49 | 39 | 29 | 41 | 102 | 88 | 43 | 34 | 65 | 94 | 125 | 136 |
| L1MD1_dup6545 | 77.1305668048208 | 1.13427024349752 | 0.314837808887237 | 3.60271292544718 | 0.000314913255945937 | 0.0223755252863835 | 4009 | 65 | 81 | 31 | 29 | 76 | 89 | 78 | 40 | 70 | 99 | 145 | 175 |
| L1MEc_dup2477 | 71.5956025340729 | 1.22016848778818 | 0.211796368269358 | 5.761045374661 | 8.35945613969079e-09 | 6.74270898513513e-06 | 4010 | 73 | 59 | 27 | 38 | 69 | 87 | 54 | 34 | 92 | 123 | 105 | 134 |
| L1PA6_dup2629 | 79.6010024810452 | 1.17467864070901 | 0.20093610891719 | 5.84603059668644 | 5.03441484765394e-09 | 4.4532300198508e-06 | 4011 | 88 | 80 | 42 | 50 | 81 | 93 | 51 | 22 | 103 | 131 | 113 | 146 |
| MIR3_dup5787 | 59.8781557916587 | 1.00357046405945 | 0.216008676203556 | 4.6459729382061 | 3.38477375764297e-06 | 0.000826044488754988 | 4012 | 51 | 35 | 46 | 54 | 57 | 69 | 50 | 30 | 59 | 85 | 119 | 102 |
| MIR_dup11121 | 54.7923192780003 | 1.02150923219503 | 0.288814155466588 | 3.53690846816268 | 0.000404839944312389 | 0.0264883875681778 | 4013 | 47 | 48 | 34 | 43 | 61 | 72 | 33 | 24 | 39 | 81 | 117 | 97 |
| MIR3_dup62797 | 64.2295279287866 | 1.54930738281774 | 0.181318403481236 | 8.54467805292621 | 1.28895094279667e-17 | 9.60738820413226e-14 | 4014 | 44 | 32 | 35 | 47 | 53 | 48 | 55 | 28 | 81 | 118 | 118 | 149 |
| L2a_dup123000 | 63.7907072592859 | 1.16927614983065 | 0.195460324641278 | 5.98216621187232 | 2.20189409149232e-09 | 2.1126285947348e-06 | 4015 | 59 | 51 | 46 | 44 | 49 | 55 | 51 | 36 | 61 | 88 | 115 | 155 |
| L2c_dup103558 | 63.3274067576901 | 1.13549245878816 | 0.169268777933099 | 6.7082215199601 | 1.97010706515275e-11 | 3.34842211232026e-08 | 4016 | 65 | 54 | 42 | 44 | 51 | 46 | 57 | 35 | 84 | 105 | 85 | 125 |
| AluJr_dup41600 | 61.9298553983684 | 1.09696417740833 | 0.269942826350472 | 4.06369079052362 | 4.83028051330816e-05 | 0.00588502778811891 | 4017 | 62 | 49 | 30 | 36 | 59 | 62 | 46 | 44 | 58 | 69 | 121 | 147 |
| THE1B-int_dup516 | 65.1299855379758 | 1.12607622597394 | 0.182465394278874 | 6.17145092319744 | 6.76661355241438e-10 | 7.74081071318649e-07 | 4018 | 54 | 42 | 40 | 70 | 43 | 52 | 62 | 42 | 69 | 73 | 134 | 146 |
| L1MEf_dup3351 | 62.236423406755 | 1.63699416484082 | 0.369551120664768 | 4.42968258869329 | 9.43718835928356e-06 | 0.00177653577637507 | 4019 | 58 | 59 | 46 | 79 | 16 | 29 | 23 | 15 | 86 | 110 | 134 | 129 |
| L1PA3_dup3252 | 87.809604817363 | 1.88560514905309 | 0.377821613878606 | 4.99072863962445 | 6.01519526878388e-07 | 0.000215555971901514 | 4020 | 82 | 69 | 57 | 104 | 24 | 28 | 30 | 19 | 139 | 170 | 134 | 253 |
| L1ME2_dup10166 | 93.4039571832844 | 1.25553865668663 | 0.318881469658261 | 3.93732084223072 | 8.23964069281839e-05 | 0.0087392386259637 | 4021 | 44 | 38 | 84 | 91 | 121 | 85 | 58 | 35 | 130 | 159 | 105 | 224 |
| HERVE-int_dup87 | 102.630411700222 | 1.1487510608719 | 0.328378300499604 | 3.49825508909741 | 0.000468312982319815 | 0.0291862757703171 | 4022 | 47 | 31 | 109 | 146 | 85 | 118 | 62 | 45 | 117 | 152 | 179 | 210 |
| THE1C-int_dup1628 | 101.563162411089 | 1.46740918368651 | 0.352684689018257 | 4.16068298221631 | 3.17297275591753e-05 | 0.00436316373565685 | 4023 | 37 | 39 | 90 | 140 | 87 | 96 | 42 | 32 | 165 | 180 | 170 | 186 |
| THE1B_dup13144 | 72.1150776145668 | 1.25568011623274 | 0.342847079642527 | 3.66250783743582 | 0.000249758108820461 | 0.0191335939991608 | 4024 | 41 | 32 | 29 | 41 | 80 | 101 | 49 | 52 | 90 | 89 | 155 | 136 |
| Tigger3b_dup2231 | 72.8165391821916 | 1.4093617343631 | 0.199111114412306 | 7.07826752174512 | 1.45967739827436e-12 | 3.50832725464434e-09 | 4025 | 44 | 52 | 37 | 49 | 63 | 80 | 40 | 39 | 109 | 155 | 97 | 133 |
| MIRc_dup47222 | 75.9406083808134 | 1.21072134013734 | 0.260743901030745 | 4.64333522414616 | 3.42829425619124e-06 | 0.000829950318676547 | 4026 | 39 | 38 | 46 | 50 | 92 | 83 | 63 | 43 | 110 | 136 | 122 | 115 |

| | | | | | | | | | | | | | | | | | | | |
|---|---|---|---|---|---|---|---|---|---|---|---|---|---|---|---|---|---|---|---|
| L1PA5_dup5345 | 83.0751217343296 | 1.10952598445595 | 0.206596131378956 | 5.37050707121307 | 7.85155544452153e-08 | 4.1424527977874e-05 | 4027 | 56 | 60 | 46 | 58 | 95 | 104 | 55 | 45 | 120 | 145 | 107 | 137 |
| L2c_dup114176 | 74.4831715247977 | 1.17610983053488 | 0.210354711026676 | 5.59107911011191 | 2.25662729915358e-08 | 1.53855426740515e-05 | 4028 | 52 | 43 | 48 | 67 | 84 | 78 | 48 | 37 | 98 | 120 | 110 | 147 |
| L1PA10_dup5876 | 67.845923070243 | 1.21125947246974 | 0.357318068622024 | 3.38986348252941 | 0.000699274374318222 | 0.0377643078237684 | 4029 | 33 | 20 | 59 | 84 | 79 | 76 | 32 | 27 | 105 | 120 | 91 | 118 |
| L1PA3_dup9425 | 67.3301385928243 | 1.56062506581879 | 0.305838388006647 | 5.10277691427302 | 3.34705471368837e-07 | 0.000135676027411123 | 4030 | 38 | 19 | 48 | 59 | 73 | 62 | 42 | 18 | 109 | 112 | 116 | 144 |
| MLT1I_dup8723 | 62.5765923673053 | 1.39573959852607 | 0.278054439600519 | 5.01966305782183 | 5.17621916751581e-07 | 0.000190955138919956 | 4031 | 32 | 22 | 47 | 67 | 66 | 50 | 40 | 29 | 104 | 86 | 105 | 130 |
| MIRb_dup119281 | 75.401402398172 | 1.10744257658682 | 0.275592687102694 | 4.01840334817793 | 5.85938343977031e-05 | 0.0068065457987524 | 4032 | 45 | 51 | 50 | 80 | 87 | 98 | 48 | 23 | 94 | 116 | 136 | 119 |
| MLT1C_dup15280 | 80.3181907863025 | 1.54880858525684 | 0.280595988023057 | 5.51971037137414 | 3.39558860537135e-08 | 2.14884767631301e-05 | 4033 | 39 | 27 | 49 | 78 | 83 | 74 | 53 | 26 | 123 | 150 | 142 | 157 |
| L1PA5_dup10091 | 145.770326092511 | 3.63518840001292 | 0.439404760205702 | 8.27298365705266 | 1.30647613183053e-16 | 8.40191269363641e-13 | 4034 | 45 | 37 | 6 | 6 | 43 | 50 | 54 | 31 | 276 | 352 | 366 | 533 |
| HERVI-int_dup36 | 161.441877609244 | 1.86453082007766 | 0.333317238579209 | 5.5938625557603 | 2.22072778513803e-08 | 1.52280488058344e-05 | 4035 | 208 | 175 | 27 | 34 | 172 | 181 | 58 | 43 | 245 | 275 | 265 | 319 |
| L1PA4_dup191 | 149.088717114295 | 1.86369371738585 | 0.511786829785659 | 3.6415429411624 | 0.000271008909258893 | 0.0201982588404356 | 4036 | 124 | 113 | 10 | 14 | 150 | 151 | 104 | 76 | 203 | 212 | 301 | 402 |
| L1MA4_dup917 | 93.1599724148366 | 1.51560969030408 | 0.460629061727078 | 3.29030409983571 | 0.00100079160757475 | 0.047866202986127l | 4037 | 119 | 85 | 57 | 62 | 105 | 123 | 15 | 8 | 101 | 111 | 164 | 240 |
| L1MB7_dup2384 | 94.9448074418486 | 1.44200104884319 | 0.397007400923866 | 3.63217674402932 | 0.000281040518363829 | 0.0206843387511907 | 4038 | 106 | 89 | 61 | 101 | 85 | 87 | 17 | 12 | 141 | 134 | 152 | 212 |
| L1MA3_dup846 | 142.160009249824 | 1.38376398708199 | 0.414651974410911 | 3.33716965666902 | 0.000846362672856071 | 0.0428550232318312 | 4039 | 156 | 129 | 100 | 167 | 123 | 129 | 34 | 12 | 214 | 202 | 225 | 304 |
| L1MA4_dup915 | 179.150550947347 | 1.61108917934278 | 0.444467530601764 | 3.62476236939407 | 0.000289227266487952 | 0.0210653416442431 | 4040 | 226 | 164 | 91 | 111 | 214 | 238 | 29 | 17 | 207 | 225 | 277 | 486 |
| L1PA8_dup738 | 152.028600430663 | 1.36913409385934 | 0.413818373373252 | 3.30853867772667 | 0.000937842296887906 | 0.0457661655404614 | 4041 | 197 | 179 | 92 | 130 | 161 | 155 | 25 | 19 | 185 | 202 | 240 | 344 |
| L1PA4_dup8634 | 212.237992473133 | 2.3953452374532 | 0.474173299003831 | 5.05162404227626 | 4.38069303279352e-07 | 0.000166778941500974 | 4042 | 81 | 51 | 97 | 124 | 89 | 110 | 65 | 28 | 109 | 86 | 865 | 1105 |
| MER61-int_dup385 | 180.808443962734 | 2.07739177931092 | 0.360018764827852 | 5.77023194972699 | 7.91624935072244e-09 | 6.51778501040485e-06 | 4043 | 131 | 109 | 141 | 194 | 57 | 51 | 50 | 41 | 363 | 395 | 295 | 401 |
| L1P2_dup1538 | 110.274829943405 | 1.25717521933931 | 0.337946937150737 | 3.72003732283735 | 0.000199193339552937 | 0.0163890934900633 | 4044 | 110 | 93 | 38 | 46 | 187 | 152 | 57 | 38 | 131 | 126 | 178 | 230 |
| LTR85a_dup89 | 106.375807270778 | 1.16407593589559 | 0.265589328666291 | 4.38299212449999 | 1.17060346260194e-05 | 0.00210060642166125 | 4045 | 92 | 84 | 74 | 61 | 106 | 127 | 79 | 42 | 101 | 120 | 225 | 241 |
| MIRc_dup6929 | 89.1311639145533 | 1.13374658520196 | 0.309384434024201 | 3.66452368160602 | 0.000247799312387028 | 0.0190264134189234 | 4046 | 82 | 88 | 56 | 50 | 100 | 125 | 59 | 24 | 91 | 95 | 152 | 214 |
| L1PA8_dup4353 | 85.5266048422003 | 1.25777314980312 | 0.331202678437661 | 3.79759353316901 | 0.000146107651916983 | 0.0130774898987743 | 4047 | 54 | 52 | 33 | 39 | 110 | 125 | 74 | 37 | 96 | 122 | 128 | 211 |
| AluY_dup49916 | 98.6242490969446 | 1.64704561298471 | 0.419005840993954 | 3.93084165384816 | 8.4649010901673e-05 | 0.00891628753785281 | 4048 | 61 | 42 | 28 | 29 | 136 | 162 | 49 | 39 | 135 | 114 | 189 | 249 |
| Charlie2b_dup1969 | 93.7079071316373 | 1.58170324068398 | 0.380923868921669 | 4.15228178050777 | 3.2917659172481e-05 | 0.00446557886513978 | 4049 | 33 | 42 | 39 | 37 | 120 | 137 | 52 | 42 | 134 | 134 | 168 | 227 |
| L1PA4_dup6356 | 115.864120186827 | 1.81382815867522 | 0.457914097523982 | 3.96106642814207 | 7.46157693547839e-05 | 0.00811079207624185 | 4050 | 79 | 49 | 19 | 41 | 162 | 207 | 57 | 28 | 159 | 128 | 203 | 330 |
| L1PA5_dup9838 | 104.17358736893 | 1.71247602434232 | 0.400073492610884 | 4.28040361576241 | 1.86554669636593e-05 | 0.00294949750307966 | 4051 | 68 | 57 | 50 | 50 | 136 | 160 | 28 | 23 | 125 | 131 | 216 | 270 |
| AluSq2_dup40954 | 99.6917954750379 | 1.57501304618964 | 0.238265336967552 | 6.61033227172334 | 3.83458360639068e-11 | 6.13330620216037e-08 | 4052 | 55 | 48 | 63 | 39 | 93 | 82 | 85 | 52 | 145 | 189 | 167 | 218 |
| L2c_dup103557 | 93.4502260421895 | 1.28254503738281 | 0.17544116398119 | 7.31039972762789 | 2.66348846574902e-13 | 7.36937705198973e-10 | 4053 | 73 | 62 | 70 | 58 | 57 | 81 | 101 | 49 | 124 | 159 | 138 | 201 |
| AluSz_dup12569 | 112.868021517954 | 1.21468528834728 | 0.23196878975618 | 5.23641688877209 | 1.63723916220243e-07 | 7.77593831715409e-05 | 4054 | 71 | 58 | 120 | 81 | 70 | 99 | 83 | 79 | 125 | 99 | 239 | 302 |
| L1PA3_dup7525 | 98.83517442492 | 1.16146492866266 | 0.216129413253877 | 5.37393273398817 | 7.70376067230612e-08 | 4.08258137401459e-05 | 4055 | 59 | 59 | 96 | 103 | 66 | 72 | 83 | 54 | 92 | 74 | 223 | 288 |
| L1PB1_dup1477 | 81.8936042145512 | 1.01033887265349 | 0.239401222819715 | 4.22027448629345 | 2.44005039465898e-05 | 0.00358571602959465 | 4056 | 89 | 81 | 65 | 84 | 68 | 66 | 56 | 30 | 80 | 88 | 148 | 194 |
| HERVI-int_dup60 | 81.6610541041286 | 1.06530640759468 | 0.198426993912269 | 5.36875747896321 | 7.92809443704184e-08 | 4.17358044008044e-05 | 4057 | 59 | 57 | 69 | 61 | 70 | 78 | 68 | 52 | 70 | 105 | 151 | 198 |
| L1PA3_dup561 | 87.2747151707567 | 1.10733773833633 | 0.24801105243676 | 4.46487254280205 | 8.01164313006271e-06 | 0.00155366479458223 | 4058 | 91 | 63 | 44 | 55 | 71 | 74 | 88 | 57 | 78 | 95 | 180 | 216 |
| MIRb_dup158864 | 85.8511019095241 | 1.18297949155919 | 0.202473377841027 | 5.84264214966579 | 5.13793105934933e-09 | 4.52067879447267e-06 | 4059 | 68 | 68 | 47 | 55 | 75 | 69 | 78 | 59 | 100 | 104 | 163 | 192 |
| HAL1_dup22256 | 134.540187423102 | 1.24059830558283 | 0.366086024831335 | 3.38881634761777 | 0.000701950070642336 | 0.0378830146312228 | 4060 | 157 | 117 | 46 | 45 | 144 | 138 | 118 | 67 | 112 | 130 | 271 | 378 |
| L1PA5_dup8571 | 138.562804736108 | 1.14399366045476 | 0.254530314517919 | 4.49452813752847 | 6.97243839345153e-06 | 0.0014059862931934 | 4061 | 130 | 136 | 69 | 59 | 108 | 131 | 153 | 78 | 140 | 204 | 218 | 330 |
| L1PA3_dup3758 | 125.372339509226 | 1.09861286123508 | 0.207640114918453 | 5.29094708730316 | 1.21684570403513e-07 | 5.99469084663238e-05 | 4062 | 140 | 115 | 85 | 88 | 142 | 155 | 52 | 39 | 162 | 215 | 168 | 207 |
| AluJb_dup84451 | 117.460091335288 | 1.40891933791951 | 0.134346301406281 | 10.4872208849185 | 9.88944687335534e-26 | 3.36164903675344e-21 | 4063 | 87 | 77 | 94 | 80 | 74 | 78 | 97 | 68 | 157 | 187 | 223 | 244 |
| L1PA7_dup11683 | 127.664033709676 | 1.05430642737887 | 0.157702073237646 | 6.68543162264015 | 2.30245008743204e-11 | 3.8045749201674e-08 | 4064 | 142 | 114 | 102 | 101 | 111 | 104 | 90 | 60 | 138 | 197 | 195 | 260 |
| HERVL-int_dup1998 | 159.656172109825 | 1.6481348640928 | 0.428847186147605 | 3.84317518531072 | 0.000121452708637885 | 0.0115557182680308 | 4065 | 28 | 20 | 124 | 149 | 135 | 121 | 143 | 108 | 249 | 348 | 265 | 273 |
| L1M3f_dup661 | 191.628613115634 | 1.29251714099714 | 0.34640025000661 | 3.73128235609667 | 0.000190507555145284 | 0.0158887770714079 | 4066 | 129 | 87 | 95 | 115 | 273 | 310 | 120 | 57 | 215 | 257 | 345 | 419 |
| MIRb_dup110316 | 137.804995457233 | 1.72674010706677 | 0.38360410085311 | 4.5013598739602 | 6.75200725647498e-06 | 0.00137434826231753 | 4067 | 50 | 60 | 39 | 55 | 187 | 182 | 67 | 62 | 205 | 215 | 258 | 326 |
| Charlie2a_dup1232 | 160.030403917729 | 1.69656075145767 | 0.389296024906532 | 4.35802228359513 | 1.31243023891105e-05 | 0.00227780835614828 | 4068 | 73 | 47 | 52 | 63 | 213 | 225 | 84 | 69 | 243 | 244 | 320 | 346 |
| MER112_dup2337 | 171.653705631788 | 1.57763380060897 | 0.397256819715344 | 3.97131961570713 | 7.14756100080529e-05 | 0.00788217710158927 | 4069 | 92 | 69 | 54 | 65 | 262 | 252 | 85 | 74 | 248 | 209 | 292 | 438 |
| L1PA2_dup2463 | 41.2069727082669 | -3.2637983772054 | 0.949701247949607 | -3.43665798507888 | 0.000588938771106218 | 0.0339969977752645 | 4070 | 198 | 196 | 14 | 21 | 5 | 10 | 61 | 20 | 2 | 6 | 3 | 9 |
| L1PA7_dup7400 | 93.9941209607165 | -3.47317726944131 | 0.788634653313596 | -4.40403836535474 | 1.06254139733047e-05 | 0.0019508292849475 | 4071 | 522 | 420 | 17 | 16 | 51 | 43 | 59 | 62 | 9 | 13 | 5 | 8 |
| L1MA3_dup4191 | 57.920113788413 | -3.63798522942988 | 0.609501241890195 | -5.96879051164473 | 2.39018760885765e-09 | 2.27494232310897e-06 | 4072 | 105 | 89 | 36 | 68 | 30 | 19 | 204 | 154 | 0 | 5 | 11 | 17 |
| Ricksha_dup92 | 58.2840074029327 | -3.13642626287592 | 0.447024274097974 | -7.01623255069256 | 2.2793057055206e-12 | 5.16525405148385e-09 | 4073 | 67 | 78 | 46 | 77 | 36 | 52 | 185 | 155 | 4 | 6 | 17 | 13 |
| L1M1_dup4055 | 61.7716968926329 | -4.04181997023236 | 0.589418218844517 | -6.85730410260452 | 7.01721517983994e-12 | 1.42710964374547e-08 | 4074 | 97 | 96 | 38 | 54 | 36 | 35 | 207 | 186 | 2 | 4 | 12 | 5 |
| L1MA3_dup4193 | 56.1114297934412 | -3.57818421339681 | 0.555531020534738 | -6.44101603894693 | 1.18676329324329e-10 | 1.65137765247993e-07 | 4075 | 83 | 83 | 49 | 52 | 24 | 28 | 188 | 166 | 2 | 4 | 14 | 10 |
| MER57A-int_dup1411 | 35.7513517168984 | -4.19215510910866 | 0.577012722045384 | -7.26527327551528 | 3.72284864650526e-13 | 9.52512843055206e-10 | 4076 | 60 | 61 | 55 | 46 | 31 | 18 | 105 | 71 | 3 | 4 | 2 | 2 |
| MER57A-int_dup1412 | 28.580438734753 | -4.56260224266686 | 0.677842939386737 | -6.73106110214081 | 1.6843021201667e-11 | 2.92535001668019e-08 | 4077 | 60 | 57 | 30 | 42 | 22 | 24 | 68 | 56 | 2 | 4 | 1 | 0 |
| L1MA3_dup4188 | 39.3725925246846 | -3.41983437647225 | 0.626835029706601 | -5.45571675863896 | 4.87756902362286e-08 | 2.8376480168581e-05 | 4078 | 74 | 78 | 14 | 23 | 30 | 32 | 120 | 101 | 2 | 3 | 6 | 13 |
| L1PA4_dup5557 | 38.9033044385458 | -3.59888494655969 | 0.642481765749076 | -5.60153632121172 | 2.124602562505e-08 | 1.47807867009323e-05 | 4079 | 75 | 75 | 28 | 23 | 17 | 19 | 148 | 91 | 2 | 2 | 8 | 9 |
| L1MEc_dup8479 | 34.8337526882261 | -3.62364044632742 | 0.566214939382389 | -6.39976128196121 | 1.55620063471006e-10 | 2.10393020583364e-07 | 4080 | 69 | 59 | 25 | 43 | 25 | 22 | 123 | 68 | 2 | 2 | 4 | 10 |
| THE1B-int_dup2122 | 36.5896633905368 | -3.46784669948123 | 0.572700458078063 | -6.0552539299847 | 1.40196331938728e-09 | 1.46956636121112e-06 | 4081 | 61 | 48 | 21 | 41 | 30 | 21 | 142 | 85 | 3 | 1 | 9 | 8 |
| L1ME3A_dup7082 | 31.9780066821587 | -4.02229937525118 | 0.633567261868984 | -6.34865406931798 | 2.17206836539048e-10 | 2.77868268425378e-07 | 4082 | 64 | 31 | 25 | 33 | 28 | 18 | 99 | 92 | 2 | 1 | 6 | 3 |
| L1MC2_dup2648 | 31.6500050353015 | -3.78914791870417 | 0.650194254046011 | -5.82771670946823 | 5.61908578438162e-09 | 4.86195994927443e-06 | 4083 | 34 | 53 | 21 | 27 | 24 | 21 | 109 | 91 | 0 | 1 | 6 | 10 |
| L1MA3_dup4192 | 25.925319114145 | -3.34945773765578 | 0.633845182186107 | -5.28434676446326 | 1.26154093428255e-07 | 6.20203758571894e-05 | 4084 | 54 | 46 | 14 | 20 | 25 | 17 | 75 | 61 | 1 | 1 | 7 | 8 |
| LTR1B0_dup41 | 29.0971742017961 | -3.46070023867535 | 0.601892880810115 | -5.74969458687969 | 8.94048026129915e-09 | 7.06761301081425e-06 | 4085 | 50 | 45 | 19 | 27 | 21 | 20 | 85 | 81 | 1 | 2 | 8 | 6 |
| HSATI_dup17 | 41.9223485188593 | -1.44065820727851 | 0.427339179262242 | -3.37122893755181 | 0.00074833641211442 | 0.0395615328898235 | 4086 | 19 | 22 | 78 | 100 | 46 | 44 | 58 | 66 | 3 | 3 | 44 | 61 |
| MER52A_dup584 | 42.4578589691748 | -1.52159189939861 | 0.35669255484591 | -4.26583588226542 | 1.99155144283076e-05 | 0.00307715389360914 | 4087 | 29 | 30 | 52 | 74 | 66 | 73 | 74 | 63 | 7 | 11 | 36 | 30 |
| MER54A_dup401 | 39.1165439795907 | -1.4666375069459 | 0.427601005973356 | -3.42992061865562 | 0.00060375785383763 | 0.0345673162389915 | 4088 | 31 | 26 | 54 | 78 | 75 | 64 | 68 | 31 | 5 | 6 | 35 | 42 |
| L1MB8_dup16409 | 29.6303921660244 | -1.24040320115954 | 0.375826564182795 | -3.30046707543596 | 0.000965240385035278 | 0.04667725233075l | 4089 | 30 | 16 | 72 | 89 | 27 | 23 | 28 | 34 | 9 | 5 | 29 | 24 |
| L1MEd_dup10967 | 40.9680388517886 | -1.17671173977047 | 0.349639676606405 | -3.36549830726195 | 0.000764055360048699 | 0.0401510416745026 | 4090 | 47 | 28 | 64 | 138 | 39 | 23 | 48 | 49 | 6 | 10 | 35 | 55 |
| MIRb_dup101982 | 45.8942306863637 | -1.20261370079274 | 0.365052178193569 | -3.29436111501588 | 0.000986456792467318 | 0.0473519160662555 | 4091 | 62 | 43 | 97 | 127 | 34 | 35 | 44 | 43 | 7 | 9 | 45 | 60 |
| LTR16D2_dup157 | 33.464121306684 | -1.21726915450696 | 0.360075412296306 | -3.3805950446438 | 0.000723290551052095 | 0.0385365189119216 | 4092 | 40 | 37 | 66 | 101 | 38 | 17 | 38 | 27 | 7 | 12 | 26 | 33 |
| L1PA8A_dup2278 | 40.785820172198 | -1.7365167696748 | 0.323910176281312 | -5.36110593872375 | 8.27139900593645e-08 | 4.34469383635001e-05 | 4093 | 59 | 46 | 93 | 101 | 48 | 26 | 62 | 34 | 8 | 12 | 27 | 20 |
| L2a_dup64558 | 33.5611114706709 | -2.01167697252067 | 0.342789021001999 | -5.86855718610937 | 4.39603778541331e-09 | 3.93240453717277e-06 | 4094 | 45 | 45 | 75 | 91 | 34 | 33 | 45 | 27 | 5 | 7 | 16 | 20 |
| AluJo_dup9148 | 50.038046076839 | -1.88603291836909 | 0.372895954807059 | -5.05779935141678 | 4.24122337345134e-07 | 0.00016224793196451 | 4095 | 149 | 126 | 53 | 54 | 42 | 59 | 49 | 46 | 8 | 24 | 16 | 22 |
| HERVE_a-int_dup61 | 44.6047715419901 | -1.35994693052398 | 0.346680573508234 | -3.922766472785 | 8.75379903869981e-05 | 0.00913165921114628 | 4096 | 112 | 130 | 32 | 42 | 49 | 55 | 36 | 35 | 19 | 18 | 20 | 23 |
| L3_dup29059 | 47.367880943591 | -1.22380975887447 | 0.291378561906536 | -4.20006794894892 | 2.66834889357136e-05 | 0.00383639242193191 | 4097 | 55 | 70 | 33 | 44 | 86 | 82 | 79 | 54 | 19 | 29 | 19 | 33 |
| MER20_dup3595 | 52.1649768483944 | -1.46876555416641 | 0.301633993576992 | -4.86936348502612 | 1.11958298422846e-06 | 0.000349259924646234 | 4098 | 102 | 92 | 49 | 48 | 69 | 57 | 80 | 70 | 27 | 15 | 23 | 29 |

| | | | | | | | | | | | | | | | | | | | |
|---|---|---|---|---|---|---|---|---|---|---|---|---|---|---|---|---|---|---|---|
| MER47A_dup1607 | 44.4836196781259 | -1.33432814608276 | 0.280436349838271 | -4.75804276746674 | 1.95479088629678e-06 | 0.000556087625861988 | 4099 | 70 | 74 | 54 | 45 | 51 | 48 | 82 | 54 | 27 | 18 | 18 | 24 |
| L1PA4_dup8533 | 54.6056211296408 | -1.19558790578875 | 0.307848130240098 | -3.88369390080845 | 0.000102881414703966 | 0.0102728582052664 | 4100 | 119 | 102 | 42 | 60 | 65 | 61 | 78 | 58 | 15 | 23 | 35 | 47 |
| L1PA11_dup580 | 53.6020607307268 | -1.0477798211467 | 0.224474375548174 | -4.66770346765855 | 3.04585063297143e-06 | 0.00076611836650425 | 4101 | 97 | 72 | 68 | 67 | 74 | 92 | 59 | 37 | 32 | 29 | 19 | 44 |
| Harlequin-int_dup372 | 53.2227414990327 | -1.38663218448477 | 0.304531688379415 | -4.55332642676308 | 5.28042413378447e-06 | 0.00114327188438351 | 4102 | 143 | 109 | 58 | 62 | 58 | 63 | 58 | 39 | 24 | 17 | 24 | 34 |
| MIR_dup27239 | 53.387282447716 | -1.06924799858075 | 0.28889709572443 | -3.70113792905927 | 0.00021463475983227 | 0.0172181197299823 | 4103 | 124 | 95 | 61 | 41 | 80 | 78 | 51 | 38 | 26 | 21 | 25 | 46 |
| L1PA4_dup9828 | 42.3374120907064 | -1.00243053404021 | 0.245412109523208 | -4.08468243880775 | 4.41371369979729e-05 | 0.00549508537294869 | 4104 | 62 | 59 | 62 | 73 | 65 | 45 | 44 | 33 | 18 | 30 | 28 | 26 |
| AluJb_dup27133 | 47.0168187794419 | -1.00259800030853 | 0.249126645165483 | -4.0244510965198 | 5.71083593430709e-05 | 0.00668406575122792 | 4105 | 80 | 50 | 61 | 74 | 45 | 55 | 57 | 64 | 23 | 29 | 20 | 42 |
| L1MB1_dup998 | 48.3496289615529 | -1.2077649525606 | 0.244972037707745 | -4.93021556199601 | 8.21389309180733e-07 | 0.000276444555250805 | 4106 | 102 | 84 | 48 | 72 | 52 | 57 | 60 | 45 | 23 | 29 | 19 | 32 |
| L1PA7_dup7021 | 46.1515120427365 | -1.13345453455741 | 0.233065148048918 | -4.86325194498625 | 1.15472712486246e-06 | 0.000357428388883535 | 4107 | 83 | 71 | 47 | 80 | 57 | 52 | 66 | 38 | 25 | 28 | 18 | 32 |
| L1MD2_dup8533 | 48.3615859358762 | -1.09387752013935 | 0.226199909306531 | -4.83588841168368 | 1.32552270186008e-06 | 0.00039975009482484 | 4108 | 66 | 51 | 58 | 64 | 56 | 65 | 73 | 70 | 21 | 22 | 30 | 39 |
| L1PA7_dup12782 | 48.0805777126276 | -1.21079886863731 | 0.223286104515419 | -5.42263420854163 | 5.87270908178801e-08 | 3.29572555465832e-05 | 4109 | 52 | 51 | 61 | 82 | 53 | 53 | 99 | 63 | 24 | 18 | 29 | 32 |
| HERVL-int_dup2209 | 45.8347959368344 | -1.12142494474651 | 0.261786381498531 | -4.2837405762943 | 1.8377718220549e-05 | 0.00291526969313784 | 4110 | 67 | 76 | 49 | 84 | 46 | 40 | 78 | 48 | 19 | 33 | 27 | 24 |
| L2a_dup155929 | 45.4322182710822 | -1.09551238402767 | 0.260325911978737 | -4.20823411584609 | 2.57374135861057e-05 | 0.00374212319188565 | 4111 | 83 | 62 | 54 | 87 | 44 | 35 | 66 | 51 | 22 | 26 | 24 | 32 |
| L1PA8A_dup2328 | 52.7212606878175 | -1.19658409966944 | 0.257067245279482 | -4.65475132146268 | 3.24371962834119e-06 | 0.000804188063821799 | 4112 | 100 | 86 | 57 | 88 | 45 | 53 | 84 | 54 | 25 | 27 | 34 | 27 |
| Tigger1_dup10604 | 51.7057776429886 | -1.45003056933853 | 0.295227627945482 | -4.91156799730918 | 9.03509436410011e-07 | 0.000296533043249678 | 4113 | 93 | 86 | 64 | 69 | 39 | 44 | 111 | 64 | 20 | 23 | 28 | 25 |
| L1PA7_dup5800 | 47.5696905478255 | -1.08248092391853 | 0.318262384679713 | -3.40122168382513 | 0.000670854011941205 | 0.0368058632062167 | 4114 | 56 | 59 | 50 | 84 | 40 | 49 | 134 | 42 | 29 | 31 | 19 | 27 |
| L2a_dup118113 | 45.7259266733035 | -1.23381486085582 | 0.311113636660811 | -3.96580128758861 | 7.31497867531132e-05 | 0.00798792985716213 | 4115 | 65 | 65 | 71 | 63 | 32 | 48 | 124 | 35 | 15 | 31 | 23 | 28 |
| L2a_dup134924 | 40.4755790513889 | -1.15052734549442 | 0.288360123385494 | -3.989897535022 | 6.6101844758477e-05 | 0.00742969747420905 | 4116 | 65 | 52 | 67 | 64 | 25 | 35 | 92 | 38 | 15 | 24 | 25 | 26 |
| MIR_dup38960 | 41.4826508767012 | -1.32346463904009 | 0.255349652172048 | -5.18295062390911 | 2.18402872288399e-07 | 9.69553915103272e-05 | 4117 | 74 | 53 | 46 | 64 | 62 | 44 | 56 | 50 | 14 | 17 | 19 | 35 |
| L1MEd_dup723 | 37.5207723863574 | -1.12323111035131 | 0.292643835339837 | -3.83821893615816 | 0.000123929961625216 | 0.0116971989880498 | 4118 | 82 | 69 | 30 | 46 | 48 | 40 | 47 | 37 | 17 | 15 | 24 | 27 |
| L3_dup42328 | 46.2938626109489 | -1.23581433936252 | 0.256512757403709 | -4.81775000928141 | 1.45186114706818e-06 | 0.00043075380611008 | 4119 | 97 | 74 | 40 | 62 | 70 | 64 | 54 | 40 | 22 | 15 | 29 | 29 |
| AluSz_dup32431 | 35.6480461732535 | -1.46310359053558 | 0.307685739056704 | -4.75518818331043 | 1.9826165106457e-06 | 0.000561320313665438 | 4120 | 61 | 58 | 38 | 37 | 41 | 43 | 61 | 49 | 10 | 10 | 23 | 25 |
| MER44A_dup1047 | 39.1569568503893 | -1.08865220239902 | 0.294282450444682 | -3.69934462878772 | 0.000216156920040069 | 0.0172944433409059 | 4121 | 68 | 56 | 37 | 45 | 36 | 53 | 60 | 53 | 17 | 12 | 29 | 34 |
| Tigger3b_dup4476 | 38.3893380961546 | -1.24254146215831 | 0.329630656691818 | -3.76949606152681 | 0.000163577489821293 | 0.0142053318952618 | 4122 | 77 | 59 | 28 | 40 | 40 | 47 | 56 | 59 | 13 | 15 | 26 | 28 |
| L1MC1_dup639 | 45.2435294907341 | -1.41090599028437 | 0.251184283670157 | -5.61701540267185 | 1.9428410994022e-08 | 1.36772564567561e-05 | 4123 | 66 | 73 | 50 | 60 | 56 | 52 | 72 | 61 | 16 | 16 | 29 | 26 |
| MLT1B_dup6553 | 44.0532678331184 | -1.19160091543419 | 0.301434014274402 | -3.9531070118366 | 7.71429154329794e-05 | 0.00832252156040541 | 4124 | 75 | 66 | 49 | 47 | 39 | 40 | 84 | 64 | 14 | 16 | 32 | 38 |
| L2_dup23972 | 41.2417904726306 | -1.2309455573127 | 0.329642846860744 | -3.73417948860484 | 0.000188328154648282 | 0.0157455836563388 | 4125 | 95 | 69 | 51 | 43 | 24 | 56 | 66 | 41 | 15 | 22 | 29 | 21 |
| AluYj4_dup279 | 42.086372671875 | -1.51656542026916 | 0.326594711672375 | -4.6435700458938 | 3.42439820123154e-06 | 0.000829950318676547 | 4126 | 100 | 84 | 42 | 54 | 39 | 64 | 62 | 30 | 13 | 11 | 22 | 29 |
| L1MEd_dup15712 | 39.277124005785 | -1.10136541145251 | 0.256508743420949 | -4.29367590657561 | 1.75738955626209e-05 | 0.00281910313205495 | 4127 | 73 | 59 | 46 | 48 | 40 | 44 | 62 | 43 | 12 | 22 | 29 | 28 |
| L1PA5_dup10250 | 37.0249025427358 | -1.05322440950497 | 0.295510804620817 | -3.56408088312173 | 0.000365133360288885 | 0.0247738872390359 | 4128 | 76 | 57 | 53 | 46 | 38 | 43 | 58 | 24 | 11 | 18 | 23 | 38 |
| L1M5_dup47153 | 31.0247767762668 | -1.01716348904134 | 0.259757033193753 | -3.91582655736076 | 9.00950032997693e-05 | 0.00932605117680356 | 4129 | 52 | 47 | 26 | 47 | 46 | 35 | 38 | 32 | 18 | 18 | 17 | 20 |
| L1MA2_dup2805 | 34.9702061970128 | -1.14457928925185 | 0.289500469562133 | -3.95363534637098 | 7.69726902906441e-05 | 0.00831381923009424 | 4130 | 53 | 54 | 24 | 61 | 52 | 39 | 46 | 40 | 21 | 19 | 21 | 15 |
| MIR_dup27347 | 32.9998178315789 | -1.30278529452215 | 0.326036504108682 | -3.99582647373706 | 6.44689448027288e-05 | 0.00731178624405629 | 4131 | 47 | 52 | 25 | 48 | 35 | 32 | 64 | 50 | 15 | 9 | 20 | 24 |
| Tigger3b_dup1229 | 33.4945131899353 | -1.49320689463245 | 0.337334266605931 | -4.42649040566278 | 9.5778640409566e-06 | 0.0017985082506149 | 4132 | 71 | 55 | 31 | 39 | 39 | 35 | 51 | 45 | 20 | 8 | 16 | 15 |
| Tigger2_dup601 | 41.7899579834475 | -1.72872491136177 | 0.289698706396393 | -5.96732009219386 | 2.41181981567223e-09 | 2.2863859675695e-06 | 4133 | 90 | 61 | 51 | 63 | 36 | 48 | 84 | 45 | 11 | 19 | 14 | 23 |
| L1MC3_dup5134 | 38.3633085424641 | -1.6853160436396 | 0.31810618456161 | -5.29796692246703 | 1.17099172181444e-07 | 5.81696860623923e-05 | 4134 | 68 | 61 | 56 | 56 | 35 | 36 | 84 | 39 | 20 | 15 | 10 | 16 |
| L1MC5a_dup8465 | 36.6658751353596 | -1.81076257135063 | 0.298399859552772 | -6.0682420362547 | 1.29317919121533e-09 | 1.36414477724567e-06 | 4135 | 80 | 49 | 60 | 61 | 31 | 43 | 62 | 36 | 13 | 15 | 12 | 15 |
| L1PA7_dup5427 | 36.4593128628588 | -1.23425027353345 | 0.282968324912683 | -4.36179658594053 | 1.28998776728599e-05 | 0.00225300159372548 | 4136 | 69 | 60 | 39 | 56 | 45 | 42 | 51 | 31 | 25 | 16 | 9 | 26 |
| AluSx1_dup93749 | 38.0903456683604 | -1.83427658003891 | 0.31142050866203 | -5.89003141738994 | 3.86122205294099e-09 | 3.48016038867082e-06 | 4137 | 86 | 70 | 39 | 46 | 47 | 39 | 53 | 51 | 12 | 14 | 12 | 18 |
| L1MA10_dup4338 | 38.8099888884225 | -1.52601164955563 | 0.285469167195806 | -5.34562686592675 | 9.01046598365519e-08 | 4.70176390119916e-05 | 4138 | 73 | 65 | 30 | 54 | 52 | 50 | 52 | 50 | 20 | 16 | 11 | 21 |
| L1ME1_dup12897 | 38.4058586156695 | -1.03265658038618 | 0.295863979153729 | -3.49030856456378 | 0.000482463134034984 | 0.0298233140186194 | 4139 | 82 | 52 | 38 | 54 | 33 | 37 | 54 | 49 | 23 | 28 | 17 | 22 |
| MER44A_dup506 | 38.5379583456571 | -1.10637350700357 | 0.330115763338075 | -3.35147130150983 | 0.000803833700723053 | 0.0415354127014618 | 4140 | 82 | 81 | 52 | 56 | 28 | 32 | 49 | 31 | 14 | 25 | 24 | 25 |
| MIRc_dup75680 | 33.8273262110901 | -1.30940720743994 | 0.287886329687342 | -4.54834798464386 | 5.40686780702043e-06 | 0.00115833290458046 | 4141 | 64 | 58 | 48 | 45 | 33 | 31 | 52 | 35 | 15 | 21 | 16 | 16 |
| MLT1D_dup19387 | 36.5290306728296 | -1.30074840669605 | 0.2862409651402 | -4.5442426665203 | 5.51331144882937e-06 | 0.00117476752436931 | 4142 | 66 | 66 | 39 | 59 | 36 | 34 | 52 | 43 | 14 | 19 | 15 | 27 |
| L2_dup40987 | 38.387369435202 | -1.62233294489587 | 0.303794588784555 | -5.34022989476747 | 9.28287887112436e-08 | 4.81225249644566e-05 | 4143 | 89 | 58 | 51 | 65 | 53 | 36 | 56 | 29 | 8 | 22 | 20 | 15 |
| L1PA5_dup7974 | 34.7183175820991 | -1.14294334521398 | 0.248835794428876 | -4.59316292431819 | 4.36577947120104e-06 | 0.000991240230967941 | 4144 | 54 | 49 | 53 | 56 | 37 | 51 | 45 | 27 | 12 | 24 | 19 | 23 |
| THE1D-int_dup2243 | 34.1398605247299 | -1.0529809317457 | 0.265412855481342 | -3.96733206398785 | 7.26816922972576e-05 | 0.00795763824028531 | 4145 | 47 | 43 | 40 | 63 | 41 | 43 | 65 | 23 | 15 | 26 | 22 | 17 |
| MLT1A-int_dup364 | 20.746315339036 | -2.08626570196392 | 0.539286012517749 | -3.86857002321241 | 0.000109475498679443 | 0.0107237965518925 | 4146 | 31 | 29 | 26 | 51 | 27 | 44 | 12 | 17 | 16 | 9 | 2 | 1 |
| L1MA4_dup8337 | 24.047397831005 | -1.80506659513428 | 0.500280022288882 | -3.60811248643458 | 0.000308432768282445 | 0.0220656474689521 | 4147 | 34 | 32 | 37 | 58 | 44 | 25 | 28 | 15 | 9 | 24 | 5 | 1 |
| AluSp_dup36086 | 20.5813154642696 | -1.8228179853785 | 0.461237095184942 | -3.95201948066992 | 7.74944302839729e-05 | 0.00834642133290565 | 4148 | 32 | 20 | 35 | 52 | 25 | 24 | 23 | 23 | 12 | 12 | 4 | 2 |
| LTR80A_dup353 | 20.9383777073342 | -1.61018758359363 | 0.489417116963924 | -3.29001076542306 | 0.00100183549746538 | 0.0479065010610725 | 4149 | 32 | 31 | 40 | 49 | 17 | 23 | 20 | 20 | 13 | 17 | 3 | 3 |
| ERVL-E-int_dup6960 | 18.8787564039808 | -2.02239787337145 | 0.599538435668652 | -3.37325808163727 | 0.000742842906302522 | 0.039331664037174 | 4150 | 27 | 27 | 33 | 64 | 26 | 10 | 20 | 14 | 11 | 11 | 2 | 1 |
| L1PA14_dup69 | 17.7886420832128 | -1.9756559009441 | 0.589992234806039 | -3.34861339589257 | 0.00081217037685161 | 0.0417843659438558 | 4151 | 27 | 18 | 43 | 59 | 20 | 20 | 15 | 8 | 9 | 11 | 2 | 2 |
| L2a_dup134055 | 19.0567972415245 | -1.92418703954988 | 0.532249450560093 | -3.61519779405133 | 0.000300118392959537 | 0.0216531143514706 | 4152 | 31 | 25 | 40 | 54 | 27 | 17 | 19 | 9 | 11 | 11 | 2 | 3 |
| ERVL-E-int_dup6961 | 24.3096317338402 | -1.355822626154 | 0.410568967791962 | -3.30230176295497 | 0.000958948441880029 | 0.0465099767532787 | 4153 | 45 | 28 | 34 | 62 | 26 | 23 | 37 | 15 | 14 | 21 | 6 | 7 |
| MER94_dup3664 | 22.9062000581109 | -1.41847878464589 | 0.427972694182699 | -3.31441422297926 | 0.000918353515522484 | 0.0451577899575352 | 4154 | 22 | 32 | 46 | 57 | 23 | 28 | 28 | 17 | 15 | 15 | 7 | 5 |
| AluSx3_dup23378 | 21.7772785390228 | -1.69417412397384 | 0.444816657145836 | -3.8087020725449 | 0.000139698155714427 | 0.012705330511537 | 4155 | 39 | 51 | 33 | 48 | 16 | 22 | 21 | 16 | 13 | 11 | 4 | 7 |
| AluY_dup77113 | 24.2306693609913 | -1.52908166010435 | 0.458049499871466 | -3.33824545280245 | 0.000843092380025374 | 0.0427558523993004 | 4156 | 48 | 39 | 40 | 51 | 32 | 20 | 18 | 18 | 21 | 12 | 6 | 4 |
| L1ME3B_dup6300 | 27.6291273458902 | -1.77806067560496 | 0.397784237796991 | -4.46991234608043 | 7.82516598382498e-06 | 0.0015274544259124 | 4157 | 42 | 45 | 41 | 64 | 41 | 35 | 24 | 21 | 19 | 11 | 4 | 8 |
| AluY_dup77110 | 24.2107266095228 | -1.42144207642478 | 0.386565066934268 | -3.67710949077166 | 0.000235891806146299 | 0.0183992771069507 | 4158 | 38 | 43 | 32 | 54 | 34 | 31 | 26 | 10 | 17 | 15 | 5 | 9 |
| L1MA9_dup13333 | 28.2238451540226 | -1.66540108085293 | 0.390170235477516 | -4.26839602158453 | 1.96883563149143e-05 | 0.00306427684337421 | 4159 | 37 | 42 | 38 | 70 | 50 | 48 | 25 | 12 | 15 | 13 | 6 | 12 |
| L2_dup46013 | 22.6527530040986 | -1.53109045563209 | 0.425855448609849 | -3.59532902685674 | 0.000323981846451489 | 0.0227993654703488 | 4160 | 37 | 29 | 37 | 49 | 32 | 31 | 26 | 12 | 15 | 15 | 1 | 10 |
| MERX_dup348 | 27.4592441426311 | -1.58967352610638 | 0.465256616839371 | -3.4167671529435 | 0.000633694543125322 | 0.0355541338737321 | 4161 | 40 | 36 | 43 | 64 | 49 | 36 | 28 | 10 | 15 | 27 | 3 | 6 |
| L1MDa_dup5495 | 21.8373608907659 | -2.13672937898645 | 0.525787507692253 | -4.06386486503802 | 4.82667769217231e-05 | 0.00588365087162722 | 4162 | 41 | 27 | 47 | 72 | 17 | 28 | 19 | 12 | 11 | 3 | 3 | 9 |
| L1M3_dup5255 | 21.4568604376549 | -1.7611223414359 | 0.457198814505735 | -3.85198361316795 | 0.000117164865209974 | 0.0112369653435117 | 4163 | 25 | 15 | 41 | 55 | 26 | 35 | 25 | 23 | 7 | 16 | 2 | 8 |
| L2c_dup68262 | 20.015008502361 | -1.39798759042014 | 0.419907978464982 | -3.32927132161344 | 0.000870735351478984 | 0.0436058891412255 | 4164 | 22 | 18 | 40 | 44 | 27 | 17 | 30 | 22 | 12 | 12 | 2 | 11 |
| L1PA12_dup698 | 20.7778506729766 | -1.49939927838094 | 0.440351411002497 | -3.4050061857811 | 0.000661625550820227 | 0.0365100077262221 | 4165 | 80 | 57 | 10 | 15 | 25 | 30 | 11 | 9 | 8 | 11 | 8 | 5 |
| MLT1E2_dup1342 | 21.2753005593187 | -1.29100719413874 | 0.360755321071027 | -3.57862273605817 | 0.000345409649203649 | 0.0238918234907068 | 4166 | 64 | 46 | 12 | 20 | 27 | 40 | 18 | 8 | 8 | 19 | 8 | 6 |
| L1M5_dup40758 | 13.9319982404314 | -1.64445778994857 | 0.46063876725858 | -3.56995091779899 | 0.000357048121055539 | 0.0244406914428247 | 4167 | 30 | 34 | 11 | 13 | 11 | 21 | 21 | 15 | 3 | 10 | 4 | 6 |
| AluSz_dup85219 | 15.8942832593417 | -1.50102349275055 | 0.387491538323795 | -3.87369360178459 | 0.000107198173462961 | 0.0105664360326503 | 4168 | 34 | 32 | 16 | 15 | 16 | 24 | 21 | 17 | 6 | 8 | 8 | 6 |
| L2a_dup67723 | 16.1943812206067 | -1.72642137297263 | 0.405461274883768 | -4.25791926360301 | 2.06338436428386e-05 | 0.00316145560813837 | 4169 | 43 | 32 | 14 | 16 | 25 | 18 | 21 | 15 | 4 | 9 | 5 | 7 |
| L1PREC2_dup2786 | 16.99696085713 | -1.93354890312096 | 0.393444809570394 | -4.91440948282484 | 8.90504554825552e-07 | 0.000294703750768457 | 4170 | 34 | 28 | 15 | 20 | 23 | 28 | 27 | 19 | 9 | 5 | 5 | 4 |

| | | | | | | | | | | | | | | | | | | | |
|---|---|---|---|---|---|---|---|---|---|---|---|---|---|---|---|---|---|---|---|
| LTR42_dup26 | 15.2701793110697 | -1.69358969207705 | 0.387416258310174 | -4.37149875811646 | 1.23396552331922e-05 | 0.00218378956001638 | 4171 | 34 | 28 | 16 | 21 | 20 | 23 | 19 | 12 | 8 | 5 | 4 | 7 |
| L1ME3F_dup91 | 19.2334324997926 | -1.84386480340329 | 0.40323948225048 | -4.57262962721972 | 4.81641066439865e-06 | 0.00106907243652146 | 4172 | 47 | 36 | 23 | 20 | 15 | 29 | 32 | 19 | 7 | 5 | 6 | 10 |
| L2_dup15846 | 16.0730633176284 | -1.59468412628365 | 0.423310005583026 | -3.76717796709598 | 0.000165103307481978 | 0.0143117200736272 | 4173 | 46 | 25 | 21 | 15 | 16 | 26 | 17 | 15 | 5 | 5 | 6 | 11 |
| L1PA5_dup7729 | 18.4070273099182 | -1.57125914786783 | 0.461420175407273 | -3.40526754488132 | 0.000660992607583142 | 0.0365004750531395 | 4174 | 38 | 24 | 12 | 12 | 21 | 23 | 51 | 24 | 7 | 11 | 5 | 9 |
| LTR33_dup3438 | 15.2262071652189 | -1.40470271294025 | 0.401458068673541 | -3.49900231817865 | 0.00046700252088915 | 0.0291431426506715 | 4175 | 36 | 16 | 11 | 15 | 22 | 26 | 27 | 15 | 5 | 11 | 5 | 8 |
| L1MC4_dup4778 | 19.5820224842645 | -1.53861232733381 | 0.458189684494932 | -3.35802480806577 | 0.000785015673985181 | 0.0409505595573397 | 4176 | 30 | 37 | 11 | 11 | 33 | 30 | 34 | 28 | 4 | 7 | 8 | 16 |
| L2a_dup11725 | 19.6259216007244 | -1.20892879545658 | 0.363582058629876 | -3.32505074648709 | 0.000884024429970552 | 0.0440151102486937 | 4177 | 49 | 45 | 9 | 13 | 29 | 31 | 22 | 14 | 11 | 11 | 6 | 11 |
| MER11B_dup29 | 21.8198599241874 | -1.49736694384656 | 0.429593322947529 | -3.48554519789277 | 0.000491135244822992 | 0.0301206006400165 | 4178 | 56 | 53 | 11 | 13 | 31 | 23 | 34 | 21 | 8 | 12 | 5 | 13 |
| L1PA5_dup9013 | 20.5318520763499 | -1.76721813031595 | 0.449597621499805 | -3.93066610188176 | 8.47108469205655e-05 | 0.00891885273511543 | 4179 | 44 | 48 | 37 | 24 | 27 | 37 | 11 | 8 | 5 | 8 | 10 | 7 |
| L1MA5A_dup3127 | 24.785426982449 | -1.19538476103118 | 0.360305376273878 | -3.31769892915098 | 0.000907622587785953 | 0.0448339556307487 | 4180 | 54 | 48 | 30 | 47 | 28 | 34 | 18 | 12 | 9 | 14 | 12 | 18 |
| L1PA11_dup2078 | 19.1620107680544 | -1.22521255909134 | 0.371541961273747 | -3.29764249209154 | 0.00097500189251956 | 0.0469892704244484 | 4181 | 36 | 43 | 22 | 26 | 27 | 26 | 13 | 14 | 6 | 12 | 6 | 16 |
| L1PA13_dup7154 | 21.8775995825536 | -1.05975202359776 | 0.318536259124616 | -3.3269431445893 | 0.000878042863717834 | 0.0438282798624732 | 4182 | 32 | 30 | 33 | 40 | 29 | 32 | 26 | 11 | 13 | 14 | 8 | 16 |
| L2c_dup15840 | 21.0771993494784 | -1.17806035176336 | 0.341585090427342 | -3.44880495307786 | 0.000563073265498909 | 0.0330083841420062 | 4183 | 26 | 26 | 36 | 50 | 25 | 24 | 31 | 12 | 10 | 9 | 9 | 19 |
| L1PA4_dup6418 | 19.0381266301168 | -1.08908828325952 | 0.324043478889835 | -3.36093257297048 | 0.000776797864183026 | 0.0406662333829262 | 4184 | 39 | 28 | 23 | 25 | 21 | 36 | 20 | 11 | 10 | 14 | 8 | 11 |
| HAL1_dup3095 | 16.9433954763414 | -1.19684555933568 | 0.360441711730986 | -3.32049682482071 | 0.000898573812481842 | 0.0444885652069922 | 4185 | 34 | 30 | 20 | 18 | 21 | 23 | 30 | 8 | 10 | 9 | 7 | 10 |
| MIR3_dup14565 | 16.9082571391299 | -1.25837092865457 | 0.35146055596164 | -3.58040442180349 | 0.000343062816292081 | 0.0237643158327323 | 4186 | 28 | 33 | 23 | 25 | 17 | 21 | 23 | 13 | 9 | 8 | 7 | 11 |
| MER112_dup1640 | 18.5656755747317 | -1.18136360700482 | 0.344999423473298 | -3.42424806137755 | 0.000616503182573167 | 0.034914717253013 | 4187 | 33 | 28 | 25 | 20 | 24 | 32 | 24 | 14 | 5 | 7 | 12 | 17 |
| AluJr_dup13518 | 18.6912245885478 | -1.05079290171729 | 0.317219724706746 | -3.31250808154725 | 0.000924634545896623 | 0.0453635240531789 | 4188 | 34 | 27 | 26 | 25 | 23 | 23 | 26 | 14 | 6 | 14 | 13 | 11 |
| L1MEg2_dup540 | 18.9594627330054 | -1.10926350453468 | 0.312978720253067 | -3.54421381631873 | 0.000393785752776886 | 0.0260277068695136 | 4189 | 32 | 29 | 20 | 25 | 24 | 23 | 33 | 16 | 9 | 10 | 11 | 13 |
| MIRb_dup70715 | 20.0969037394019 | -1.17579129154556 | 0.323948987639491 | -3.62955692534546 | 0.000283908118629317 | 0.0208156893229905 | 4190 | 40 | 30 | 24 | 30 | 24 | 19 | 31 | 18 | 8 | 10 | 15 | 11 |
| L1ME3Cz_dup10639 | 17.1571796122918 | -1.21552426564714 | 0.351545785235045 | -3.45765563604877 | 0.000544897402989777 | 0.0322898394372453 | 4191 | 33 | 29 | 17 | 27 | 15 | 22 | 24 | 18 | 5 | 11 | 9 | 12 |
| L1P1_dup712 | 18.1224782940277 | -1.45622295807368 | 0.366017018812554 | -3.97856625027441 | 6.93320932757728e-05 | 0.00770431833195099 | 4192 | 38 | 32 | 25 | 27 | 17 | 19 | 29 | 15 | 5 | 8 | 12 | 9 |
| L2a_dup39515 | 16.5069734001903 | -1.13553667108614 | 0.34465460864727 | -3.29470908728885 | 0.000985236182469593 | 0.047304144699605 | 4193 | 31 | 28 | 23 | 23 | 17 | 20 | 23 | 12 | 6 | 10 | 11 | 10 |
| L1MA4_dup3690 | 21.6012275547767 | -1.42426201795613 | 0.387782734015982 | -3.67283505174686 | 0.000239874329873867 | 0.0185904708360133 | 4194 | 32 | 41 | 15 | 17 | 43 | 25 | 36 | 23 | 13 | 14 | 9 | 4 |
| L1MB7_dup7558 | 21.8005189834889 | -1.2578772469375 | 0.364223750554387 | -3.45358380672012 | 0.000553190388352137 | 0.0326057567864349 | 4195 | 29 | 20 | 26 | 17 | 40 | 42 | 36 | 21 | 14 | 16 | 7 | 8 |
| MIRb_dup132550 | 17.9906726390141 | -1.65714336466769 | 0.417352232267338 | -3.97061100084447 | 7.16885464363485e-05 | 0.0078972235510849 | 4196 | 25 | 27 | 18 | 14 | 20 | 27 | 39 | 28 | 8 | 8 | 8 | 5 |
| L1MEd_dup17076 | 18.0887854621579 | -1.5911568723154 | 0.405511838815329 | -3.92382347446092 | 8.71546074049395e-05 | 0.00910878779629798 | 4197 | 24 | 27 | 18 | 24 | 20 | 19 | 34 | 31 | 11 | 8 | 5 | 6 |
| L4_A_Mam_dup5132 | 17.1772264173973 | -1.37815606986087 | 0.379976342946544 | -3.62695229701379 | 0.000286786250976053 | 0.0209581207846277 | 4198 | 22 | 23 | 18 | 21 | 22 | 27 | 30 | 21 | 11 | 13 | 5 | 4 |
| AluSz_dup88002 | 19.2759581848161 | -1.38422945210403 | 0.40310836701367 | -3.43388916077022 | 0.000594987438056322 | 0.0342379408777744 | 4199 | 27 | 44 | 21 | 19 | 22 | 22 | 37 | 16 | 15 | 11 | 4 | 7 |
| L1ME3D_dup1950 | 18.2619689433285 | -1.37515669620736 | 0.35119274700165 | -3.91567510419257 | 9.01515851412555e-05 | 0.0093266126426179 | 4200 | 33 | 27 | 24 | 18 | 32 | 19 | 27 | 18 | 10 | 8 | 6 | 11 |
| L1PA5_dup8113 | 17.7945544274514 | -1.3498214846131 | 0.348861545050847 | -3.86921832962805 | 0.000109184837997608 | 0.0107066321561159 | 4201 | 32 | 34 | 19 | 13 | 31 | 26 | 23 | 14 | 11 | 10 | 5 | 8 |
| AluJr_dup48555 | 23.2575731687273 | -1.45658525550274 | 0.393078818900768 | -3.70558062521922 | 0.000210907024683258 | 0.017036181826881 | 4202 | 40 | 33 | 10 | 31 | 29 | 37 | 52 | 25 | 14 | 9 | 11 | 8 |
| L1PB1_dup4899 | 21.186701149423 | -1.56771523605722 | 0.369795303186355 | -4.23941359597849 | 2.24104429642449e-05 | 0.00336646165503171 | 4203 | 24 | 31 | 14 | 31 | 34 | 39 | 38 | 23 | 13 | 9 | 7 | 7 |
| L1PA8_dup5492 | 22.5172460782799 | -1.71994764401239 | 0.416530890052564 | -4.1292199092253 | 3.63996129593306e-05 | 0.00479841678959605 | 4204 | 35 | 34 | 20 | 21 | 18 | 29 | 51 | 41 | 10 | 9 | 7 | 9 |
| L2a_dup105462 | 22.3834722629379 | -1.50476012348116 | 0.419056338129529 | -3.5908301260821 | 0.00032962643125588 | 0.0230821927050064 | 4205 | 38 | 28 | 12 | 20 | 23 | 40 | 43 | 39 | 7 | 8 | 11 | 15 |
| MER57B1_dup635 | 20.1739552940447 | -1.32145288097674 | 0.3479074614166 | -3.79828841725926 | 0.000145698727055727 | 0.0130528724804224 | 4206 | 32 | 27 | 16 | 21 | 24 | 33 | 41 | 24 | 10 | 9 | 11 | 10 |
| L1MB7_dup19278 | 22.2375639744914 | -1.23529526720874 | 0.321058334099132 | -3.84757265583806 | 0.000119293904915807 | 0.0113998022084726 | 4207 | 42 | 32 | 26 | 38 | 19 | 31 | 33 | 20 | 12 | 13 | 6 | 16 |
| AluSp_dup10295 | 19.8793772872866 | -1.13668498714974 | 0.33327543730767 | -3.41064735022876 | 0.000648088548276478 | 0.036080806526454 | 4208 | 34 | 31 | 21 | 33 | 20 | 18 | 33 | 22 | 8 | 13 | 7 | 17 |
| L2_dup4528 | 19.8418972962405 | -1.23642564104696 | 0.337037144476828 | -3.66851446883171 | 0.00024396388976383 | 0.018798650166368 | 4209 | 42 | 27 | 23 | 37 | 21 | 20 | 32 | 15 | 10 | 11 | 8 | 13 |
| Looper_dup57 | 19.3770220080086 | -1.18716014855606 | 0.332312606533333 | -3.57241983968184 | 0.000353697767170801 | 0.0242924268708517 | 4210 | 23 | 28 | 35 | 30 | 16 | 20 | 37 | 19 | 9 | 12 | 8 | 13 |
| HERVL-int_dup2210 | 21.2203338579506 | -1.29967388979263 | 0.334528875786304 | -3.88508730894389 | 0.00010229312012823 | 0.0102398985115826 | 4211 | 35 | 27 | 30 | 34 | 14 | 21 | 41 | 28 | 8 | 13 | 11 | 11 |
| AluSg_dup3986 | 21.0984361577351 | -1.11092257832737 | 0.327111540414384 | -3.39615831627357 | 0.000683388202124866 | 0.0372530330224063 | 4212 | 30 | 23 | 23 | 32 | 15 | 26 | 47 | 28 | 8 | 14 | 12 | 14 |
| MIR3_dup27530 | 20.9092556515355 | -1.09025043634613 | 0.316228948782183 | -3.4476617037901 | 0.000565461805823008 | 0.0331044572401402 | 4213 | 33 | 28 | 25 | 26 | 19 | 23 | 37 | 28 | 8 | 15 | 12 | 13 |
| L1PA13_dup8064 | 21.5104929861837 | -1.0347960972209 | 0.305811975685619 | -3.3837657760162 | 0.000714989696949985 | 0.0382981050831384 | 4214 | 24 | 28 | 25 | 28 | 31 | 26 | 34 | 26 | 15 | 11 | 10 | 14 |
| AluJb_dup36815 | 22.3941336579215 | -1.01756419692674 | 0.3106248576541 | -3.27586209491276 | 0.00105339983706848 | 0.0494694050868097 | 4215 | 36 | 29 | 24 | 24 | 21 | 29 | 44 | 26 | 13 | 14 | 12 | 14 |
| MER20_dup15383 | 23.8507547677319 | -1.09901579311275 | 0.304513025110244 | -3.60909288761907 | 0.000307269580514069 | 0.0220155277341164 | 4216 | 41 | 26 | 27 | 31 | 25 | 26 | 42 | 32 | 14 | 13 | 10 | 17 |
| L2a_dup125413 | 24.8567288524708 | -1.04381968939929 | 0.301715494731082 | -3.4596157891382 | 0.00054094664379316 | 0.032130826286572 | 4217 | 42 | 34 | 30 | 29 | 28 | 31 | 28 | 33 | 18 | 14 | 12 | 13 |
| FLAM_A_dup3224 | 23.6626404379677 | -1.11454374649122 | 0.322551940353065 | -3.45539309195053 | 0.000549491057097626 | 0.0324681398242244 | 4218 | 50 | 38 | 26 | 23 | 20 | 33 | 32 | 27 | 12 | 14 | 10 | 17 |
| L2a_dup109372 | 23.5040811555523 | -1.18006185542626 | 0.320290857140992 | -3.68434449225255 | 0.00022929196047809 | 0.0179980030537886 | 4219 | 50 | 36 | 27 | 36 | 21 | 31 | 25 | 25 | 12 | 10 | 11 | 18 |
| Charlie29a_dup1062 | 22.8810550963711 | -1.6921901506165 | 0.329470646640682 | -5.13608774520659 | 2.80516724480782e-07 | 0.000116692014900881 | 4220 | 39 | 38 | 35 | 29 | 22 | 28 | 36 | 29 | 7 | 10 | 9 | 11 |
| Charlie15b_dup1834 | 22.0793324786259 | -1.45769402020549 | 0.362454644568864 | -4.02172807563109 | 5.77727304390868e-05 | 0.00673862260640144 | 4221 | 44 | 39 | 24 | 22 | 20 | 27 | 33 | 31 | 8 | 13 | 6 | 14 |
| L1PA3_dup2394 | 22.16671382219 | -1.42428556138695 | 0.326340357756206 | -4.36441747867105 | 1.27461953126145e-05 | 0.00224161580920574 | 4222 | 39 | 39 | 30 | 25 | 24 | 30 | 27 | 27 | 7 | 12 | 8 | 15 |
| AluJb_dup101957 | 20.4202430110326 | -1.77675083887242 | 0.378553212606911 | -4.69352994427602 | 2.68530662861892e-06 | 0.000704474058493228 | 4223 | 40 | 30 | 19 | 32 | 24 | 21 | 28 | 34 | 8 | 8 | 6 | 9 |
| AluSg_dup34713 | 20.9836218101212 | -1.35225920081311 | 0.367525562931397 | -3.67936094030424 | 0.000233819144361229 | 0.0182653743020935 | 4224 | 35 | 36 | 19 | 19 | 27 | 21 | 33 | 33 | 9 | 13 | 8 | 11 |
| MER2_dup1443 | 21.9976853668637 | -1.62270736664339 | 0.355410777519314 | -4.56572357757273 | 4.97774342721687e-06 | 0.00109359315807852 | 4225 | 42 | 41 | 21 | 31 | 28 | 19 | 35 | 28 | 7 | 12 | 7 | 11 |
| MER8_dup307 | 21.2470577033367 | -1.43004863013311 | 0.372958482104695 | -3.83433732908552 | 0.0001259032607884 | 0.0118289289437594 | 4226 | 41 | 42 | 18 | 32 | 18 | 20 | 33 | 28 | 8 | 13 | 8 | 11 |
| AluJb_dup4989 | 20.1837794554289 | -1.16822750180416 | 0.348193616575411 | -3.35510889973812 | 0.000793337472649372 | 0.0412267917477111 | 4227 | 37 | 35 | 22 | 22 | 19 | 24 | 29 | 25 | 8 | 9 | 17 | 10 |
| L1M1_dup8265 | 19.4193043028721 | -1.18269542133873 | 0.360843130461527 | -3.27758885093928 | 0.00104697773979387 | 0.0492906400719254 | 4228 | 32 | 29 | 19 | 20 | 14 | 29 | 32 | 29 | 9 | 10 | 11 | 12 |
| MER58A_dup12520 | 19.2498392115503 | -1.26401436198429 | 0.386200606750788 | -3.27294763366322 | 0.00106432194233899 | 0.0498320987269906 | 4229 | 39 | 27 | 24 | 22 | 15 | 21 | 23 | 32 | 7 | 14 | 7 | 12 |
| MIRb_dup82898 | 20.3353045238377 | -1.31040916352609 | 0.384280513195831 | -3.41003282375212 | 0.000649550615110989 | 0.0361454561887744 | 4230 | 46 | 30 | 28 | 24 | 20 | 19 | 18 | 31 | 8 | 9 | 12 | 12 |
| HAL1_dup20635 | 22.4166649537065 | -1.43941357160572 | 0.341589507137211 | -4.21386940034879 | 2.51032477280399e-05 | 0.00367130755002838 | 4231 | 56 | 40 | 20 | 31 | 29 | 25 | 30 | 18 | 12 | 8 | 8 | 13 |
| AluSz_dup5301 | 19.667359989709 | -1.21169277026753 | 0.364128664981584 | -3.32765005009649 | 0.000875818091497559 | 0.0437717730727742 | 4232 | 53 | 33 | 18 | 27 | 18 | 24 | 24 | 15 | 14 | 11 | 8 | 8 |
| MIR_dup38499 | 20.6540487388967 | -1.37281584502614 | 0.351147412472242 | -3.90951434145812 | 9.24818650950224e-05 | 0.00951391693553835 | 4233 | 49 | 39 | 19 | 32 | 24 | 25 | 25 | 15 | 11 | 8 | 12 | 8 |
| MIRc_dup23435 | 20.8972949121136 | -1.2964519185929 | 0.36885141576644 | -3.51483514276063 | 0.00044002705632643 | 0.0280433489858633 | 4234 | 34 | 45 | 25 | 20 | 29 | 16 | 36 | 20 | 11 | 8 | 10 | 13 |
| AluJr4_dup1970 | 19.500431693449 | -1.49446831106111 | 0.391028709808385 | -3.8218889651183 | 0.000132433320972674 | 0.0122567012804994 | 4235 | 32 | 42 | 18 | 28 | 19 | 16 | 31 | 27 | 8 | 8 | 11 | 8 |
| L1M5_dup10658 | 19.7654959811977 | -1.33561849322143 | 0.33231152356687 | -4.01917597947117 | 5.84020386339587e-05 | 0.00679204862404493 | 4236 | 31 | 33 | 17 | 32 | 25 | 25 | 25 | 25 | 9 | 8 | 11 | 11 |
| L1MC4a_dup9366 | 17.2952015252444 | -1.40063815478828 | 0.367468951727469 | -3.81158230703271 | 0.000138080062293262 | 0.0126028379372583 | 4237 | 28 | 29 | 19 | 29 | 19 | 17 | 22 | 24 | 9 | 5 | 8 | 11 |
| L1ME3E_dup3205 | 17.8089843303889 | -1.31032518375847 | 0.346980870803746 | -3.77636144817793 | 0.000159136014577262 | 0.0139593168184873 | 4238 | 23 | 29 | 16 | 32 | 21 | 22 | 27 | 22 | 9 | 6 | 9 | 12 |
| L1ME3F_dup4184 | 20.5928040105799 | -1.19925051099943 | 0.353342070365977 | -3.39402129431487 | 0.000688743368583695 | 0.0374260916699391 | 4239 | 38 | 28 | 24 | 44 | 15 | 20 | 33 | 21 | 9 | 14 | 13 | 8 |
| MER1B_dup3872 | 20.656066494667 | -1.38619738562455 | 0.415547199100643 | -3.33583619050895 | 0.000850432581223296 | 0.0429815273941713 | 4240 | 49 | 34 | 17 | 28 | 13 | 18 | 45 | 24 | 7 | 11 | 13 | 9 |
| Charlie1a_dup2684 | 22.7852362181013 | -1.09610166945134 | 0.315484296198309 | -3.47434621202934 | 0.0005120998207909 | 0.0310135158966433 | 4241 | 37 | 26 | 21 | 36 | 25 | 29 | 41 | 26 | 10 | 11 | 21 | 10 |
| L2a_dup87550 | 24.4420690042106 | -1.54638031748874 | 0.349846138069426 | -4.42017261080032 | 9.86220938491978e-06 | 0.00183610546772347 | 4242 | 47 | 35 | 21 | 48 | 29 | 24 | 39 | 29 | 14 | 8 | 12 | 8 |

| | | | | | | | | | | | | | | | | | | | |
|---|---|---|---|---|---|---|---|---|---|---|---|---|---|---|---|---|---|---|---|
| Tigger10_dup143 | 24.3246686548242 | -1.46137135733504 | 0.303933448382402 | -4.80819523192584 | 1.52298992933059e-06 | 0.000447947295085904 | 4243 | 37 | 36 | 28 | 40 | 29 | 23 | 43 | 31 | 10 | 10 | 13 | 12 |
| LTR80A_dup354 | 41.9876968385906 | -1.66763264431671 | 0.408818857154697 | -4.07914805085832 | 4.52010449366117e-05 | 0.00558432390367964 | 4244 | 77 | 66 | 63 | 105 | 51 | 48 | 26 | 34 | 27 | 25 | 7 | 11 |
| L1MC4a_dup7864 | 37.8437363653214 | -1.43265993975616 | 0.361546155830729 | -3.96259209689097 | 7.41403924772759e-05 | 0.00807387177501048 | 4245 | 75 | 52 | 55 | 88 | 50 | 33 | 37 | 28 | 21 | 29 | 9 | 13 |
| LTR16C_dup4473 | 36.4636207237936 | -1.23142463598203 | 0.348012150797074 | -3.53845299125798 | 0.000402478938346943 | 0.0264073769498129 | 4246 | 76 | 40 | 55 | 84 | 39 | 39 | 44 | 22 | 23 | 25 | 9 | 20 |
| L1PA4_dup5951 | 34.7641464786627 | -2.43828437630184 | 0.360029708566587 | -6.77245326784161 | 1.26616710727097e-11 | 2.33549921323021e-08 | 4247 | 75 | 57 | 66 | 66 | 50 | 46 | 41 | 19 | 11 | 10 | 7 | 7 |
| L1ME4a_dup26617 | 35.2029421797488 | -1.6602859262162 | 0.396064176503185 | -4.19196187061076 | 2.76552483929691e-05 | 0.00393332679863325 | 4248 | 52 | 38 | 67 | 88 | 56 | 36 | 34 | 27 | 15 | 26 | 9 | 8 |
| MER74C_dup183 | 34.0483583931098 | -1.87631843805399 | 0.463430302334453 | -4.0487607922968 | 5.14895376435378e-05 | 0.00617526689724256 | 4249 | 43 | 67 | 58 | 83 | 55 | 33 | 34 | 16 | 22 | 19 | 5 | 5 |
| L1MC5_dup11733 | 33.1349339342799 | -1.7542720782022 | 0.434146392304793 | -4.04073858333622 | 5.32831271279851e-05 | 0.00633292056323454 | 4250 | 58 | 55 | 61 | 67 | 46 | 39 | 24 | 21 | 18 | 27 | 4 | 6 |
| L1PA7_dup11257 | 29.0833767185361 | -1.43530956371919 | 0.432650334213491 | -3.31748169414551 | 0.000908328676544642 | 0.0448502127555699 | 4251 | 45 | 38 | 52 | 75 | 39 | 32 | 24 | 16 | 21 | 20 | 6 | 8 |
| Tigger4b_dup1990 | 32.5676636997339 | -1.46938002306662 | 0.413297966515263 | -3.55525587376042 | 0.000377611421270486 | 0.0253315836610169 | 4252 | 62 | 44 | 58 | 68 | 48 | 41 | 24 | 14 | 21 | 26 | 7 | 8 |
| L1MC3_dup10157 | 32.3649702042817 | -3.55632645959619 | 0.62569417666068 | -5.68380942679737 | 1.31727059489685e-08 | 9.95045298328019e-06 | 4253 | 100 | 85 | 70 | 82 | 24 | 15 | 19 | 18 | 3 | 4 | 3 | 6 |
| LTR49-int_dup397 | 33.1240490592683 | -2.39132429554406 | 0.470657931450974 | -5.08081163781078 | 3.75825636133931e-07 | 0.000148795685217179 | 4254 | 85 | 87 | 51 | 46 | 51 | 53 | 9 | 15 | 7 | 10 | 10 | 5 |
| L1MB7_dup15534 | 26.1756817581949 | -1.84463966702589 | 0.386994348172977 | -4.76658037962193 | 1.87378995536751e-06 | 0.000536669815358667 | 4255 | 64 | 53 | 37 | 39 | 27 | 26 | 20 | 31 | 9 | 8 | 9 | 12 |
| L1MC1_dup1101 | 28.0512062874656 | -2.75809545952484 | 0.518201918475167 | -5.32243390306362 | 1.02387991941356e-07 | 5.22807148722699e-05 | 4256 | 86 | 80 | 42 | 41 | 25 | 19 | 19 | 27 | 7 | 7 | 2 | 7 |
| L2a_dup121085 | 35.64876441265 | -1.82081559261779 | 0.45260458438586 | -4.0229720498489 | 5.74683186240819e-05 | 0.00671298799377801 | 4257 | 81 | 76 | 45 | 88 | 39 | 40 | 25 | 21 | 13 | 3 | 18 | 19 |
| LTR37-int_dup377 | 38.9300449648314 | -2.15229586555077 | 0.544695316608508 | -3.95137575067072 | 7.77032114264552e-05 | 0.00835857520166334 | 4258 | 91 | 76 | 73 | 70 | 47 | 52 | 18 | 27 | 3 | 2 | 22 | 29 |
| AluSg_dup21506 | 26.8240866273023 | -1.59759197232368 | 0.455644601780806 | -3.5062238553464 | 0.00045451278687394 | 0.0285813618321663 | 4259 | 58 | 40 | 37 | 27 | 36 | 40 | 41 | 21 | 4 | 2 | 15 | 31 |
| L1PB_dup1645 | 27.1314044982156 | -1.95344216517229 | 0.573514075678096 | -3.40609280227801 | 0.000658997744217865 | 0.036441059085676 | 4260 | 58 | 53 | 50 | 34 | 45 | 35 | 16 | 19 | 2 | 1 | 12 | 29 |
| AluY_dup73723 | 26.91649551792 | -1.61755202747074 | 0.473231019698827 | -3.41810228015099 | 0.000630594049687288 | 0.035463010622636 | 4261 | 44 | 27 | 44 | 73 | 27 | 30 | 29 | 24 | 3 | 1 | 18 | 38 |
| AluJb_dup73823 | 21.1103728502562 | -1.50168795297749 | 0.372970915095093 | -4.02628701649461 | 5.66645171738348e-05 | 0.00664846903522944 | 4262 | 35 | 30 | 37 | 39 | 26 | 26 | 27 | 16 | 2 | 6 | 12 | 22 |
| L4_A_Mam_dup5264 | 25.9255067192378 | -1.41618067475091 | 0.384347820867939 | -3.68463302732632 | 0.000229032383860235 | 0.0179918585704878 | 4263 | 44 | 38 | 32 | 48 | 35 | 29 | 28 | 28 | 3 | 5 | 19 | 29 |
| LTR12C_dup1137 | 21.0070799213787 | -2.41609781323736 | 0.603331924150678 | -4.00459136426196 | 6.21247774249997e-05 | 0.00708305811651604 | 4264 | 79 | 65 | 22 | 12 | 18 | 20 | 19 | 18 | 2 | 2 | 5 | 11 |
| L2a_dup36339 | 21.8808772514916 | -2.47774178456743 | 0.5185668913324 | -4.77805626618612 | 1.76997857373757e-06 | 0.000510496147523102 | 4265 | 50 | 50 | 31 | 33 | 33 | 25 | 31 | 13 | 2 | 2 | 2 | 17 |
| MER67A_dup408 | 16.5035869450702 | -2.16729411164024 | 0.427031606714634 | -5.07525456561473 | 3.86977564081908e-07 | 0.000152058355229693 | 4266 | 33 | 22 | 23 | 31 | 21 | 19 | 22 | 22 | 3 | 1 | 6 | 11 |
| L1PA2_dup3939 | 21.8260028388169 | -2.37232473385759 | 0.406194682607225 | -5.84036383398828 | 5.20869428672596e-09 | 4.55178972942472e-06 | 4267 | 42 | 39 | 26 | 34 | 24 | 33 | 34 | 27 | 4 | 1 | 6 | 13 |
| MSTA-int_dup1343 | 19.0230125212176 | -2.07277005273796 | 0.427468747684465 | -4.84893939958383 | 1.24123330024967e-06 | 0.000377680944835306 | 4268 | 41 | 31 | 20 | 24 | 28 | 20 | 33 | 24 | 5 | 2 | 4 | 14 |
| AluJb_dup71242 | 24.2914749857904 | -1.56532661167589 | 0.366647340920893 | -4.2692975973706 | 1.96089496684003e-05 | 0.00305758265910693 | 4269 | 41 | 37 | 35 | 37 | 21 | 18 | 42 | 36 | 3 | 6 | 15 | 23 |
| L1PA3_dup9703 | 26.9453414889685 | -1.21799086716534 | 0.370462587088977 | -3.28779673877374 | 0.00100989060265718 | 0.0481658507396003 | 4270 | 58 | 27 | 27 | 32 | 24 | 26 | 62 | 33 | 6 | 8 | 22 | 27 |
| L1PA4_dup10556 | 28.8630982193605 | -1.26308745825804 | 0.333078829020415 | -3.79215773627156 | 0.000149344001480678 | 0.0132844141219893 | 4271 | 42 | 42 | 30 | 39 | 30 | 26 | 52 | 44 | 8 | 6 | 23 | 28 |
| AluSg_dup1573 | 29.5414955044694 | -1.82446835968631 | 0.323604774848208 | -5.63795253188742 | 1.72083925692767e-08 | 1.25219210345233e-05 | 4272 | 45 | 39 | 46 | 41 | 35 | 54 | 37 | 37 | 5 | 6 | 16 | 19 |
| L1ME4b_dup7704 | 24.5797814264134 | -1.86733308886793 | 0.398072341670389 | -4.69093904146226 | 2.71953994069934e-06 | 0.000710322338890938 | 4273 | 50 | 35 | 34 | 23 | 27 | 33 | 33 | 40 | 4 | 4 | 13 | 17 |
| AluSx4_dup3766 | 27.6151602621701 | -1.39469297405697 | 0.391473730728801 | -3.56267321299053 | 0.000367097536344582 | 0.0248716943004123 | 4274 | 45 | 44 | 31 | 21 | 27 | 37 | 49 | 41 | 4 | 8 | 23 | 23 |
| Charlie20a_dup532 | 25.6452227555011 | -1.31808457726941 | 0.340925626638238 | -3.86619389767393 | 0.000110547060073728 | 0.0107981242841968 | 4275 | 36 | 33 | 35 | 27 | 33 | 23 | 39 | 44 | 6 | 9 | 21 | 18 |
| L2a_dup113970 | 30.0788825355253 | -1.87258004370293 | 0.449691859483372 | -4.16414040906643 | 3.12527752991069e-05 | 0.00430600629491679 | 4276 | 81 | 61 | 20 | 36 | 28 | 40 | 37 | 40 | 7 | 3 | 17 | 17 |
| L1PB3_dup2432 | 25.595409682566 | -1.21943144269936 | 0.336786879105129 | -3.62078073213388 | 0.000293715360822831 | 0.0212814845451734 | 4277 | 35 | 34 | 22 | 24 | 37 | 39 | 52 | 30 | 9 | 8 | 17 | 22 |
| L2a_dup160556 | 26.9148715828497 | -1.49606043207138 | 0.365498054007013 | -4.09321038968561 | 4.25441366049476e-05 | 0.00536470966009584 | 4278 | 58 | 52 | 36 | 25 | 21 | 36 | 49 | 23 | 10 | 7 | 15 | 17 |
| MLT1B_dup12024 | 24.3167035580499 | -1.52868312942116 | 0.348248342661997 | -4.38963504531267 | 1.13541050604491e-05 | 0.00206076573814922 | 4279 | 34 | 55 | 23 | 33 | 30 | 37 | 34 | 23 | 12 | 5 | 14 | 11 |
| AluSc_dup7856 | 25.7135956621682 | -1.87183058556777 | 0.335035062090694 | -5.59161119996641 | 2.24972139636534e-08 | 1.53825346948146e-05 | 4280 | 56 | 31 | 40 | 35 | 34 | 31 | 50 | 21 | 11 | 7 | 8 | 11 |
| L1M5_dup48435 | 26.9882946324018 | -2.06172868191992 | 0.365853389799362 | -5.63539587005221 | 1.74656654407253e-08 | 1.26703817956062e-05 | 4281 | 63 | 53 | 37 | 35 | 36 | 31 | 40 | 22 | 6 | 4 | 10 | 15 |
| MER44C_dup452 | 24.1197142802652 | -2.45285467657712 | 0.393931094375854 | -6.22660843887796 | 4.7664036253887e-10 | 5.55954253454284e-07 | 4282 | 52 | 50 | 21 | 28 | 34 | 33 | 46 | 25 | 5 | 4 | 6 | 9 |
| L1M5_dup49829 | 24.288799008161 | -1.58459390059591 | 0.371353565697827 | -4.26707603471703 | 1.98051672676827e-05 | 0.0030709302228076 | 4283 | 49 | 36 | 24 | 30 | 34 | 43 | 25 | 29 | 8 | 5 | 7 | 22 |
| L1ME4c_dup5879 | 22.2140894935821 | -1.49667312207945 | 0.357160197247572 | -4.19048128434647 | 2.78363375873583e-05 | 0.00395199593291263 | 4284 | 42 | 31 | 19 | 21 | 38 | 39 | 23 | 29 | 11 | 7 | 6 | 15 |
| AluSz_dup20575 | 22.1579374416439 | -1.33855094446664 | 0.326355817040988 | -4.10150784687416 | 4.1046658469627e-05 | 0.0052201433437808 | 4285 | 47 | 33 | 31 | 33 | 36 | 22 | 24 | 17 | 7 | 12 | 12 | 13 |
| AluSx1_dup11829 | 22.1946985754352 | -1.20153681000115 | 0.327486999126227 | -3.6689603349354 | 0.000243538857254489 | 0.0187780612210877 | 4286 | 41 | 32 | 20 | 28 | 31 | 23 | 35 | 27 | 5 | 11 | 15 | 18 |
| THE1B-int_dup3050 | 21.1060299717883 | -1.16113071790063 | 0.324024269144601 | -3.58346836477998 | 0.000339061845491103 | 0.0235626197100543 | 4287 | 37 | 39 | 20 | 35 | 22 | 24 | 28 | 21 | 6 | 13 | 13 | 15 |
| AluJb_dup41363 | 26.4204622488948 | -1.6750460208617 | 0.414670855311219 | -4.03945924679114 | 5.35745753161617e-05 | 0.0063580328669224 | 4288 | 58 | 45 | 33 | 18 | 31 | 24 | 45 | 39 | 5 | 10 | 8 | 22 |
| AluJb_dup41684 | 24.8792998235101 | -1.9294061611092 | 0.447222186483971 | -4.31420045655223 | 1.60181556184971e-05 | 0.0026322210337009 | 4289 | 68 | 57 | 18 | 18 | 30 | 30 | 32 | 32 | 5 | 6 | 7 | 16 |
| L1ME1_dup9610 | 24.4080497308542 | -1.65161692489091 | 0.45915903709718 | -3.59704762718489 | 0.000321849569606815 | 0.0226912052413817 | 4290 | 66 | 46 | 17 | 22 | 26 | 22 | 35 | 37 | 3 | 8 | 14 | 17 |
| L1MA8_dup298 | 20.5681152735793 | -1.28800916516742 | 0.379353228903699 | -3.39527666309751 | 0.000685592830831719 | 0.0373389955882546 | 4291 | 43 | 32 | 19 | 24 | 23 | 20 | 39 | 23 | 7 | 5 | 11 | 21 |
| L1ME3Cz_dup11135 | 20.4852551344974 | -1.43688425324581 | 0.38568581640131 | -3.72553045028422 | 0.000194904921484208 | 0.0161423064557882 | 4292 | 52 | 29 | 19 | 23 | 21 | 20 | 34 | 27 | 7 | 8 | 8 | 16 |
| L2a_dup139455 | 20.5157290542238 | -1.53002413973438 | 0.422369638057641 | -3.62247662206624 | 0.000291795839435036 | 0.0212009829421527 | 4293 | 48 | 35 | 15 | 21 | 16 | 25 | 45 | 24 | 7 | 5 | 12 | 13 |
| L2a_dup36341 | 19.1952947071916 | -1.34556300845552 | 0.40398755136822 | -3.33070413654674 | 0.000866266208577558 | 0.0434770257891153 | 4294 | 49 | 35 | 16 | 22 | 16 | 25 | 23 | 23 | 6 | 6 | 12 | 14 |
| L2a_dup139038 | 22.5484863874672 | -1.32313265825576 | 0.376592111524397 | -3.5134369992507 | 0.000442349300374404 | 0.0281506409807136 | 4295 | 48 | 41 | 19 | 24 | 33 | 27 | 37 | 19 | 6 | 9 | 7 | 24 |
| AluSq2_dup52756 | 21.1057064620348 | -1.71737988513635 | 0.451311479287867 | -3.80530955667355 | 0.000141626950900103 | 0.0128086531580676 | 4296 | 49 | 46 | 19 | 28 | 23 | 15 | 31 | 27 | 4 | 7 | 5 | 19 |
| L1M6_dup1814 | 22.570779411606 | -1.66741778774147 | 0.449987734335372 | -3.70547386186923 | 0.000210995889167367 | 0.017036181826881 | 4297 | 67 | 55 | 26 | 27 | 18 | 22 | 28 | 16 | 4 | 8 | 8 | 17 |
| L1PA5_dup5368 | 31.0514387646069 | -1.45405677101051 | 0.407242537035703 | -3.57049335168795 | 0.000356309498353186 | 0.0244171151047913 | 4298 | 13 | 23 | 38 | 50 | 21 | 20 | 105 | 71 | 7 | 10 | 10 | 29 |
| SAR_dup12 | 29.3289793717251 | -1.67285880676294 | 0.399166461870346 | -4.1908801629389 | 2.77874404593254e-05 | 0.00394976720880206 | 4299 | 12 | 23 | 65 | 59 | 13 | 11 | 75 | 69 | 2 | 10 | 10 | 25 |
| MER21B_dup2392 | 24.7897801539404 | -1.5228359872302 | 0.346748853078103 | -4.39175493649633 | 1.12439393010489e-05 | 0.00204545136156528 | 4300 | 39 | 24 | 39 | 42 | 19 | 18 | 46 | 45 | 9 | 10 | 8 | 17 |
| L1MC5a_dup8468 | 26.8559988283266 | -1.56763906773155 | 0.325252270249053 | -4.81976364540415 | 1.43728398032339e-06 | 0.000427494967477538 | 4301 | 45 | 34 | 38 | 49 | 21 | 22 | 45 | 43 | 7 | 12 | 16 | 12 |
| L1MEc_dup16663 | 27.5952853416048 | -1.77064545285437 | 0.317837347441804 | -5.57091690799043 | 2.53402168011149e-08 | 1.67955521642287e-05 | 4302 | 49 | 39 | 40 | 44 | 24 | 27 | 52 | 38 | 8 | 11 | 8 | 16 |
| L1M1_dup4132 | 26.8722796503826 | -1.37934257968275 | 0.413222274229114 | -3.33801604053408 | 0.000843788781406526 | 0.0427729368088106 | 4303 | 37 | 24 | 27 | 34 | 20 | 16 | 79 | 55 | 11 | 13 | 12 | 13 |
| AluJr_dup73289 | 23.7463452618109 | -1.56756151427373 | 0.344826116669356 | -4.54594776467243 | 5.46885993565426e-06 | 0.00116744445890903 | 4304 | 23 | 27 | 28 | 40 | 25 | 21 | 54 | 43 | 10 | 7 | 9 | 15 |
| MER44C_dup439 | 25.2178000799141 | -1.72206336642327 | 0.423891959931882 | -4.06250537684177 | 4.85488290874865e-05 | 0.00590895124606193 | 4305 | 25 | 29 | 30 | 32 | 23 | 13 | 69 | 58 | 7 | 9 | 7 | 16 |
| HERVL-int_dup1128 | 23.6564953721734 | -2.37850559033146 | 0.506426137759236 | -4.69664855936453 | 2.64464905167836e-06 | 0.000698428039123927 | 4306 | 21 | 25 | 20 | 25 | 28 | 22 | 90 | 48 | 3 | 9 | 7 | 5 |
| L1MEc_dup8655 | 20.3987257194238 | -1.76290508935238 | 0.482117807316661 | -3.65658571950337 | 0.000255596964702861 | 0.019405958954431 | 4307 | 29 | 27 | 18 | 21 | 22 | 11 | 63 | 38 | 8 | 8 | 9 | 5 |
| MIRb_dup165624 | 19.4459084535639 | -1.37999491073351 | 0.400555680588141 | -3.44520119826348 | 0.000570634473900078 | 0.0333121174010373 | 4308 | 24 | 24 | 26 | 20 | 16 | 23 | 34 | 38 | 5 | 6 | 19 | 9 |
| AluSz6_dup29385 | 19.858404097161 | -1.26912507287305 | 0.38202129445709 | -3.32213175361513 | 0.00089332500978811 | 0.0443394060865752 | 4309 | 25 | 29 | 26 | 22 | 19 | 14 | 42 | 33 | 5 | 9 | 18 | 10 |
| L1MC5a_dup10811 | 19.9639375323349 | -1.40884728954932 | 0.405974841292439 | -3.47028225952177 | 0.000519911691661952 | 0.0312480190412212 | 4310 | 30 | 33 | 20 | 20 | 17 | 16 | 51 | 30 | 5 | 9 | 14 | 11 |
| AluJb_dup102063 | 18.5746783691258 | -1.40440962562196 | 0.375635594302402 | -3.73875545055869 | 0.000184933515049963 | 0.0155767044856915 | 4311 | 10 | 20 | 30 | 32 | 15 | 19 | 42 | 33 | 8 | 6 | 8 | 13 |
| L1MEf_dup4906 | 16.6563838106329 | -1.36831514332674 | 0.388210312459961 | -3.5246748976249 | 0.000424002810051647 | 0.0273191910751555 | 4312 | 16 | 15 | 28 | 24 | 18 | 12 | 47 | 23 | 5 | 10 | 8 | 9 |
| THE1A-int_dup185 | 24.0468508218641 | -2.23085933054052 | 0.671055686216642 | -3.32440269319216 | 0.000886081490179924 | 0.0440754007120582 | 4313 | 66 | 46 | 31 | 49 | 2 | 4 | 57 | 33 | 6 | 8 | 7 | 6 |
| L1MA3_dup4700 | 29.3036631641987 | -2.23717114620282 | 0.555480155476596 | -4.02745466988527 | 5.63839335620895e-05 | 0.00663064931404875 | 4314 | 55 | 49 | 67 | 101 | 5 | 14 | 42 | 23 | 5 | 5 | 16 | 9 |

| | | | | | | | | | | | | | | | | | | |
|---|---|---|---|---|---|---|---|---|---|---|---|---|---|---|---|---|---|---|
| HERVK9-int_dup583 | 32.6631657732151 | -2.00857724873463 | 0.530259729023414 | -3.78791210947481 | 0.000151918567720984 | 0.0134577081802406 | 4315 | 74 | 45 | 47 | 71 | 7 | 10 | 62 | 60 | 3 | 9 | 20 | 15 |
| L1PA4_dup10951 | 34.1601822215423 | -1.7868155026284 | 0.514534211654609 | -3.47268551275233 | 0.000515278760573886 | 0.0311110174989885 | 4316 | 50 | 60 | 59 | 79 | 8 | 6 | 89 | 45 | 6 | 11 | 15 | 21 |
| L2_dup26674 | 28.0264437055253 | -2.01493756099173 | 0.565617404200622 | -3.56236838899858 | 0.000367524168115208 | 0.024892481940615 | 4317 | 59 | 59 | 13 | 28 | 9 | 25 | 78 | 51 | 5 | 14 | 5 | 14 |
| L2a_dup22924 | 29.8495545251205 | -3.10462462899921 | 0.613639855593629 | -5.05935949351892 | 4.20667118763102e-07 | 0.000161268198307975 | 4318 | 75 | 67 | 14 | 40 | 11 | 19 | 113 | 37 | 3 | 6 | 5 | 6 |
| L1PA13_dup888 | 34.9967103379793 | -2.16526793036997 | 0.593700489083458 | -3.64707115824119 | 0.000265246486621568 | 0.0198959560119433 | 4319 | 91 | 77 | 32 | 31 | 7 | 14 | 94 | 61 | 11 | 9 | 13 | 8 |
| L1PA13_dup889 | 35.2842529819119 | -1.87975821470498 | 0.555102554451467 | -3.386326003422 | 0.000708351801055551 | 0.0381247404781643 | 4320 | 79 | 81 | 40 | 29 | 10 | 13 | 103 | 50 | 10 | 16 | 14 | 10 |
| Harlequin-int_dup68 | 31.036126096331 | -2.11684561585061 | 0.606778101249456 | -3.48866515039299 | 0.000485438841485249 | 0.0299244120663339 | 4321 | 53 | 49 | 21 | 38 | 5 | 11 | 95 | 80 | 7 | 3 | 12 | 19 |
| Harlequin-int_dup69 | 30.4955066677907 | -2.55634566332665 | 0.546201484547847 | -4.68022467101646 | 2.8656071189539e-06 | 0.000730825028431516 | 4322 | 59 | 40 | 32 | 26 | 17 | 11 | 97 | 74 | 2 | 5 | 11 | 14 |
| L1M3d_dup360 | 35.2875052537872 | -1.97503181773601 | 0.505362958883255 | -3.9081451915281 | 9.3007409825804e-05 | 0.00955146359016433 | 4323 | 106 | 111 | 15 | 27 | 22 | 28 | 58 | 38 | 13 | 19 | 5 | 11 |
| L1MA6 | 25.8784253224921 | -1.58996653119024 | 0.460866122179005 | -3.44995315271336 | 0.000560683842817132 | 0.0329029492590101 | 4324 | 74 | 59 | 12 | 14 | 25 | 29 | 48 | 29 | 7 | 14 | 11 | 11 |
| L1MB1_dup4415 | 25.8229251199051 | -1.8184299483627 | 0.494675632124341 | -3.67600469938981 | 0.000236915159716994 | 0.0184527052680916 | 4325 | 97 | 67 | 19 | 8 | 29 | 25 | 36 | 19 | 6 | 11 | 9 | 9 |
| L1PA10_dup5892 | 30.3374597530539 | -1.85444309811567 | 0.5453406207348 | -3.40052258644691 | 0.000672571792811356 | 0.0368600756169689 | 4326 | 43 | 35 | 12 | 16 | 45 | 39 | 86 | 59 | 18 | 14 | 5 | 7 |
| L1PA3_dup9449 | 24.1961192349108 | -1.78711340685667 | 0.545306209561026 | -3.27726582885478 | 0.00104817635200825 | 0.049304134923274 | 4327 | 37 | 20 | 16 | 11 | 40 | 35 | 65 | 42 | 16 | 12 | 3 | 6 |
| L1MA4_dup3692 | 34.5960928701454 | -1.6844367361345 | 0.371321121235161 | -4.5363342934315 | 5.72404249550534e-06 | 0.00121283438614026 | 4328 | 68 | 57 | 24 | 26 | 56 | 75 | 37 | 35 | 27 | 18 | 1 | 11 |
| L1PB1_dup4898 | 29.3402876738623 | -1.57413221881281 | 0.325407988947086 | -4.83741110323134 | 1.31541183821237e-06 | 0.000397709002865667 | 4329 | 46 | 55 | 17 | 31 | 63 | 44 | 45 | 24 | 12 | 21 | 10 | 7 |
| L2c_dup87479 | 28.8715266653415 | -1.36793289682503 | 0.343336030488208 | -3.98423927392618 | 6.7696636543705e-05 | 0.00755544663785905 | 4330 | 60 | 51 | 13 | 21 | 37 | 61 | 44 | 28 | 13 | 17 | 12 | 12 |
| MIRc_dup75650 | 29.417172049602 | -1.22338513340986 | 0.28720878383871 | -4.2595672634333 | 2.04823056485637e-05 | 0.00314593526951645 | 4331 | 46 | 49 | 32 | 31 | 43 | 57 | 24 | 27 | 22 | 20 | 8 | 11 |
| L1MA4_dup3691 | 25.9162013228167 | -1.47852117768706 | 0.311364736580386 | -4.74851839012073 | 2.04912258118085e-06 | 0.000573624143178422 | 4332 | 32 | 40 | 22 | 36 | 49 | 47 | 36 | 22 | 14 | 17 | 6 | 10 |
| THE1A_dup3380 | 25.8329502909394 | -1.40015618729776 | 0.345816991716501 | -4.04883571610502 | 5.14730592179515e-05 | 0.00617526689724256 | 4333 | 23 | 41 | 35 | 32 | 39 | 40 | 40 | 27 | 18 | 16 | 6 | 9 |
| MIRb_dup192741 | 23.5939121361219 | -1.22803771723452 | 0.350011310402532 | -3.5085658112654 | 0.000450529753457801 | 0.028498300031631 | 4334 | 28 | 27 | 40 | 39 | 23 | 20 | 43 | 32 | 11 | 14 | 18 | 5 |
| THE1B-int_dup204 | 27.4678163638699 | -1.0416917484928 | 0.295355456358212 | -3.52690876727674 | 0.000420441609819937 | 0.0271706679223831 | 4335 | 30 | 31 | 50 | 59 | 30 | 24 | 38 | 27 | 13 | 11 | 26 | 15 |
| L1ME4b_dup16385 | 39.214343670589 | -1.17347637672641 | 0.245737382991203 | -4.77532706844369 | 1.7941541295607e-06 | 0.000516842371080448 | 4336 | 55 | 43 | 57 | 55 | 34 | 56 | 63 | 52 | 18 | 21 | 18 | 29 |
| MLT1E2_dup1701 | 38.471180604581 | -1.05122958077565 | 0.237949613986955 | -4.41786630018775 | 9.96800466935706e-06 | 0.00184723274069691 | 4337 | 39 | 36 | 62 | 71 | 38 | 48 | 65 | 45 | 21 | 15 | 20 | 35 |
| Charlie1a_dup477 | 32.8794856086181 | -1.09915961845211 | 0.264838719161278 | -4.15029804528982 | 3.3204262589469e-05 | 0.00449676805128844 | 4338 | 22 | 33 | 49 | 76 | 33 | 34 | 60 | 42 | 9 | 16 | 20 | 33 |
| MER57F_dup406 | 34.8854642468511 | -1.05751473706658 | 0.289670975985176 | -3.65074455067497 | 0.00026148120676763 | 0.0196981313515086 | 4339 | 39 | 45 | 71 | 58 | 36 | 30 | 40 | 42 | 9 | 13 | 26 | 40 |
| MER5B_dup15415 | 29.0917241833648 | -1.25535460559641 | 0.341732442204255 | -3.67350140214688 | 0.000239249363776486 | 0.0185791657393772 | 4340 | 33 | 30 | 49 | 38 | 33 | 33 | 60 | 35 | 22 | 15 | 9 | 12 |
| L4_C_Mam_dup1038 | 32.1449154212893 | -1.23283901910762 | 0.299429869367078 | -4.11728803714052 | 3.83356833106858e-05 | 0.00497816297128209 | 4341 | 51 | 32 | 43 | 57 | 33 | 30 | 44 | 51 | 17 | 17 | 11 | 22 |
| SVA_B_dup16 | 26.0192545291399 | -1.27261170738259 | 0.302182055357426 | -4.21140727856036 | 2.53784719283807e-05 | 0.00370018742737161 | 4342 | 26 | 26 | 40 | 51 | 34 | 23 | 42 | 37 | 12 | 11 | 9 | 22 |
| AluSp_dup18970 | 32.7522317969129 | -1.01264630733785 | 0.297218115996939 | -3.40708137504067 | 0.000656615474891309 | 0.03636002461915 | 4343 | 51 | 29 | 41 | 57 | 39 | 23 | 64 | 39 | 19 | 8 | 22 | 31 |
| ERVL-E-int_dup3472 | 31.6584722987479 | -1.63545711710996 | 0.357409077497363 | -4.57586899740125 | 4.7424725498476e-06 | 0.00105561494232557 | 4344 | 36 | 38 | 44 | 43 | 25 | 17 | 87 | 60 | 10 | 6 | 18 | 20 |
| L1MA7_dup1469 | 33.4008698179942 | -1.30577910930987 | 0.303492068887047 | -4.30251477113838 | 1.68870408840136e-05 | 0.00273347199332482 | 4345 | 45 | 36 | 47 | 60 | 22 | 22 | 74 | 55 | 15 | 11 | 18 | 23 |
| L1MA3_dup374 | 37.177230994718 | -1.9015290007557 | 0.538257227518641 | -3.53275144956571 | 0.000411258922331582 | 0.0268029075686417 | 4346 | 86 | 95 | 20 | 15 | 35 | 18 | 95 | 57 | 13 | 12 | 13 | 14 |
| L1MCa_dup3304 | 42.723261410535 | -1.78302234638619 | 0.496179526692944 | -3.59350245317476 | 0.000326262575692341 | 0.0229276062716154 | 4347 | 109 | 83 | 19 | 38 | 28 | 32 | 94 | 76 | 10 | 18 | 26 | 11 |
| AluYe5_dup184 | 34.8535415278564 | -1.31569738071527 | 0.381863398530882 | -3.44546606398275 | 0.000570075543464473 | 0.033287655279803 | 4348 | 66 | 61 | 27 | 18 | 51 | 48 | 51 | 49 | 14 | 10 | 22 | 23 |
| L1PB4_dup2804 | 34.6186320034113 | -1.61851767850494 | 0.409813066463247 | -3.94940476757612 | 7.83457728550043e-05 | 0.00840489777626549 | 4349 | 61 | 50 | 19 | 17 | 67 | 63 | 62 | 44 | 16 | 9 | 11 | 20 |
| MER33_dup2078 | 31.9112743324987 | -1.21031445598947 | 0.292648265236857 | -4.13573084060448 | 3.53826725944525e-05 | 0.00471397839482619 | 4350 | 56 | 60 | 34 | 45 | 33 | 60 | 28 | 27 | 15 | 15 | 13 | 24 |
| MIRb_dup51041 | 31.1888603549576 | -1.32057226993786 | 0.364222200787927 | -3.62573249813178 | 0.000288143515210885 | 0.021025022039365 | 4351 | 74 | 67 | 30 | 41 | 26 | 46 | 35 | 24 | 13 | 15 | 6 | 28 |
| MIRb_dup166259 | 30.0883722345155 | -1.11019613664691 | 0.337688573350328 | -3.28763311601653 | 0.00101033411676557 | 0.0481676941991385 | 4352 | 71 | 49 | 21 | 55 | 41 | 31 | 42 | 17 | 14 | 17 | 11 | 26 |
| AluSz_dup1521 | 30.4775114935735 | -1.0016899981146 | 0.305888288206903 | -3.27469222174684 | 0.00105777148302305 | 0.0495848095546496 | 4353 | 68 | 40 | 31 | 50 | 25 | 42 | 41 | 25 | 20 | 15 | 12 | 26 |
| L1M4_dup2083 | 28.5689824507766 | -1.11989648362074 | 0.337523934277312 | -3.31797650444732 | 0.000906721112493159 | 0.0447987254637245 | 4354 | 68 | 38 | 27 | 36 | 28 | 30 | 49 | 28 | 22 | 11 | 11 | 19 |
| L1MA10_dup4339 | 31.6929423589978 | -1.32974745980696 | 0.283879760992121 | -4.6841925439125 | 2.81065708132816e-06 | 0.000721069589570882 | 4355 | 67 | 37 | 31 | 45 | 41 | 40 | 44 | 39 | 11 | 13 | 17 | 23 |
| L1MC1_dup2534 | 31.0917752556637 | -1.16360141903533 | 0.289182943676727 | -4.0237553579097 | 5.72774161629514e-05 | 0.00669397449229353 | 4356 | 69 | 44 | 26 | 38 | 48 | 43 | 32 | 31 | 19 | 11 | 15 | 21 |
| L1M5_dup38901 | 30.4255863203929 | -1.1913741988895 | 0.359478565979289 | -3.31417311528427 | 0.000919145812575752 | 0.0451780767442987 | 4357 | 73 | 60 | 31 | 20 | 25 | 37 | 44 | 34 | 11 | 18 | 17 | 19 |
| L2a_dup39245 | 27.0437161370609 | -1.25264106391371 | 0.356883530502304 | -3.50994360022904 | 0.000448201751105273 | 0.0284091139766903 | 4358 | 57 | 47 | 20 | 27 | 21 | 40 | 43 | 35 | 8 | 14 | 15 | 20 |
| L1PB1_dup4086 | 26.8100392158832 | -1.38109228756352 | 0.342225691867033 | -4.03561836643205 | 5.4458675515638e-05 | 0.0064279730691898 | 4359 | 64 | 47 | 26 | 26 | 25 | 33 | 44 | 29 | 7 | 15 | 15 | 15 |
| L2a_dup3816 | 26.9786111654636 | -1.2908497576664 | 0.326923066997721 | -3.94848173156101 | 7.8648416584555e-05 | 0.00842790000149309 | 4360 | 55 | 47 | 20 | 27 | 29 | 35 | 46 | 32 | 9 | 18 | 12 | 16 |
| AluY_dup40398 | 33.0034407049831 | -1.12313145499933 | 0.309098152523782 | -3.63357543818679 | 0.000279520671426533 | 0.0206046864180856 | 4361 | 59 | 53 | 34 | 50 | 27 | 34 | 47 | 44 | 8 | 21 | 17 | 31 |
| L1MA3_dup8242 | 32.5753699830433 | -1.67100588608832 | 0.330533601764868 | -5.05547961588796 | 4.29310483632757e-07 | 0.000163706269773205 | 4362 | 78 | 67 | 37 | 39 | 39 | 32 | 38 | 36 | 8 | 15 | 12 | 18 |
| AluJr4_dup4524 | 27.9599812543214 | -1.27241060317128 | 0.314379434563215 | -4.04737226192647 | 5.17958315249177e-05 | 0.00620574568380064 | 4363 | 56 | 54 | 31 | 33 | 36 | 29 | 38 | 26 | 6 | 16 | 19 | 17 |
| MER4B_dup1644 | 32.2406241063596 | -1.32554497178877 | 0.335267752577326 | -3.95369062964994 | 7.69548990024555e-05 | 0.00831381923009424 | 4364 | 68 | 58 | 36 | 38 | 38 | 26 | 50 | 36 | 6 | 16 | 19 | 26 |
| L1M5_dup48426 | 31.295043597757 | -1.84637858905842 | 0.345581587527635 | -5.34281528789717 | 9.15140043690039e-08 | 4.76485586074114e-05 | 4365 | 72 | 56 | 37 | 37 | 33 | 40 | 61 | 27 | 6 | 13 | 7 | 21 |
| L1MB7_dup20116 | 31.5425348931394 | -1.80951084921523 | 0.315875081948361 | -5.72856471632366 | 1.01283913564896e-08 | 7.87585035853357e-06 | 4366 | 65 | 55 | 35 | 58 | 33 | 33 | 49 | 33 | 12 | 15 | 11 | 9 |
| L1PA6_dup3033 | 28.7961817833273 | -1.55498704779426 | 0.330063320303881 | -4.71117798355363 | 2.46289019542856e-06 | 0.000661299492462503 | 4367 | 59 | 50 | 34 | 45 | 30 | 22 | 43 | 36 | 13 | 12 | 11 | 14 |
| L1MA3_dup1720 | 31.0914532048544 | -1.88382664542638 | 0.376900368641364 | -4.9982085510214 | 5.78653833157986e-07 | 0.00020830312403118 | 4368 | 57 | 62 | 30 | 39 | 28 | 26 | 70 | 43 | 7 | 8 | 16 | 15 |
| L1MB7_dup20111 | 28.8624615422819 | -1.49509869029997 | 0.321601157445074 | -4.64892198205263 | 3.33674385828265e-06 | 0.000818520468147345 | 4369 | 47 | 57 | 44 | 44 | 25 | 27 | 47 | 29 | 7 | 11 | 18 | 18 |
| MER101-int_dup267 | 28.5083388206938 | -1.31067674016605 | 0.34104132587526 | -3.8431610503574 | 0.000121459706775256 | 0.0115557182680308 | 4370 | 47 | 52 | 27 | 41 | 32 | 22 | 41 | 42 | 12 | 13 | 20 | 12 |
| Tigger1_dup6669 | 28.0879924285093 | -1.20147038738191 | 0.335239101718597 | -3.58392079331616 | 0.000338474765883143 | 0.0235562201353701 | 4371 | 36 | 43 | 18 | 39 | 39 | 28 | 50 | 43 | 9 | 16 | 19 | 17 |
| AluSx_dup76022 | 25.3547611107396 | -1.04863791085788 | 0.286439173487208 | -3.66094447938601 | 0.000251287213005781 | 0.0192074485017261 | 4372 | 41 | 31 | 37 | 38 | 27 | 29 | 27 | 33 | 11 | 13 | 16 | 20 |
| FLAM_C_dup19619 | 27.3606293772431 | -1.08491300480048 | 0.267238902279802 | -4.0597120985947 | 4.91332596975873e-05 | 0.00595948609287004 | 4373 | 40 | 31 | 30 | 44 | 30 | 31 | 45 | 36 | 12 | 16 | 18 | 17 |
| L3_dup36404 | 26.5797060258009 | -1.13052672102506 | 0.278490766137802 | -4.05947650151409 | 4.91828568827832e-05 | 0.00596172392451897 | 4374 | 43 | 26 | 34 | 35 | 38 | 33 | 37 | 33 | 13 | 16 | 16 | 14 |
| AluYg6_dup171 | 26.2764817256966 | -1.27475951167041 | 0.301938334858745 | -4.22192005618224 | 2.42230028516823e-05 | 0.00356889575018353 | 4375 | 60 | 40 | 24 | 41 | 34 | 28 | 35 | 24 | 10 | 18 | 12 | 14 |
| L1M5_dup37266 | 25.5657466565278 | -1.27538135810216 | 0.346837830686889 | -3.6771691126552 | 0.000235836697320223 | 0.0183992771069507 | 4376 | 65 | 30 | 26 | 30 | 25 | 24 | 49 | 29 | 10 | 14 | 13 | 16 |
| L1ME4a_dup7136 | 24.2508409418959 | -1.10053465092339 | 0.325437919619796 | -3.38170380455088 | 0.000720377733234256 | 0.0384502019094119 | 4377 | 49 | 33 | 25 | 24 | 34 | 20 | 39 | 30 | 12 | 15 | 11 | 17 |
| L1MB7_dup19279 | 31.3250971248083 | -1.28484491793112 | 0.276701184883392 | -4.64343843873519 | 3.42658124369906e-06 | 0.000829950318676547 | 4378 | 47 | 55 | 38 | 45 | 47 | 38 | 46 | 25 | 11 | 17 | 10 | 27 |
| L3_dup29058 | 27.8955509083368 | -1.24728683572388 | 0.302181572778985 | -4.12760720070824 | 3.66557561887056e-05 | 0.00481883456468384 | 4379 | 39 | 50 | 27 | 29 | 47 | 30 | 44 | 31 | 13 | 16 | 10 | 19 |
| AluJr_dup16687 | 34.3803333048037 | -1.02257408573181 | 0.272878369490879 | -3.74736219525085 | 0.000178703955890869 | 0.0151863898172888 | 4380 | 55 | 47 | 29 | 46 | 34 | 43 | 53 | 48 | 20 | 18 | 20 | 23 |
| MER20_dup9716 | 31.4429566302281 | -1.21456181228642 | 0.310157821672155 | -3.91594771248506 | 9.00497647473975e-05 | 0.00932605117680356 | 4381 | 46 | 37 | 25 | 31 | 38 | 52 | 57 | 46 | 14 | 17 | 15 | 21 |
| L1MC4a_dup6256 | 31.3642647734965 | -1.15957890330755 | 0.336358037040355 | -3.44745412807967 | 0.000565896495459312 | 0.0331167750881853 | 4382 | 54 | 60 | 21 | 30 | 37 | 42 | 57 | 33 | 15 | 14 | 11 | 29 |
| AluSz_dup20808 | 29.6701194441282 | -1.23622901903729 | 0.352911778632373 | -3.50294066077367 | 0.000460151960278713 | 0.0288742928113077 | 4383 | 58 | 52 | 24 | 33 | 28 | 37 | 48 | 37 | 10 | 8 | 18 | 29 |
| L2b_dup86452 | 27.0999316164218 | -1.10681823048967 | 0.309565322457782 | -3.57539475578895 | 0.00034969983439882 | 0.0241117579819941 | 4384 | 47 | 47 | 26 | 27 | 33 | 33 | 38 | 33 | 11 | 11 | 15 | 25 |
| HERVIP10F-int_dup24 | 101.296237095206 | -5.19642690617042 | 0.920036836183679 | -5.64806397070496 | 1.62264801166445e-08 | 1.18800801164157e-05 | 4385 | 155 | 114 | 380 | 384 | 14 | 13 | 194 | 83 | 0 | 0 | 9 | 22 |
| L1M4_dup7343 | 70.6262205234589 | -3.6707307502733 | 0.387748637858601 | -9.46677922724745 | 2.88605213066618e-21 | 5.28249661756534e-17 | 4386 | 74 | 59 | 177 | 258 | 25 | 31 | 157 | 117 | 2 | 4 | 10 | 19 |

| | | | | | | | | | | | | | | | | | | | |
|---|---|---|---|---|---|---|---|---|---|---|---|---|---|---|---|---|---|---|---|
| L1PB1_dup5583 | 78.1416066963819 | -3.65006922620582 | 0.451286123907059 | -8.08814858875993 | 6.05784765286569e-16 | 3.20320137246395e-12 | 4387 | 85 | 65 | 208 | 341 | 29 | 24 | 194 | 78 | 3 | 6 | 8 | 19 |
| AluJb_dup122348 | 43.7133137485681 | -2.20860010592264 | 0.524926794055879 | -4.20744403016229 | 2.58275343803575e-05 | 0.00374957809375751 | 4388 | 33 | 28 | 101 | 101 | 32 | 53 | 90 | 51 | 0 | 0 | 40 | 49 |
| L1MA4_dup9912 | 76.7059856513465 | -2.11061886770468 | 0.389864509965937 | -5.41372403425253 | 6.17272768956737e-08 | 3.40783262835684e-05 | 4389 | 14 | 12 | 197 | 243 | 33 | 32 | 224 | 129 | 3 | 0 | 50 | 79 |
| L1PA8_dup688 | 63.0887050215275 | -2.35865348962713 | 0.42856468576604 | -5.50361139861803 | 3.72089231460707e-08 | 2.30565479867576e-05 | 4390 | 15 | 20 | 206 | 304 | 10 | 6 | 111 | 95 | 1 | 3 | 23 | 50 |
| L1PA8_dup535 | 72.453512219252 | -2.41606389361171 | 0.483463781647925 | -4.99740411862159 | 5.81072139479441e-07 | 0.000208857690786367 | 4391 | 38 | 29 | 225 | 296 | 4 | 7 | 175 | 114 | 5 | 2 | 27 | 43 |
| L1PB1_dup1068 | 93.8373004268277 | -1.98678177822911 | 0.398036693752655 | -4.99145382677639 | 5.99265094186634e-07 | 0.000215071994119657 | 4392 | 41 | 28 | 252 | 429 | 7 | 11 | 219 | 135 | 4 | 5 | 51 | 81 |
| L1MC3_dup11469 | 49.0418840000239 | -1.35524632845735 | 0.345174283453855 | -3.92626679744677 | 8.62744576040247e-05 | 0.00905143831086739 | 4393 | 42 | 35 | 125 | 178 | 25 | 16 | 69 | 62 | 11 | 15 | 32 | 35 |
| L1MC3_dup11472 | 58.6798186264731 | -1.32147581106575 | 0.388029984135156 | -3.40560231192201 | 0.000660182711992413 | 0.0364823969772524 | 4394 | 37 | 42 | 158 | 230 | 17 | 15 | 105 | 72 | 13 | 18 | 27 | 44 |
| L1P2_dup868 | 78.2757026612886 | -1.44402096929307 | 0.391728058824784 | -3.68628423918686 | 0.000227552191331477 | 0.0179143076152322 | 4395 | 22 | 18 | 220 | 322 | 20 | 20 | 188 | 124 | 7 | 21 | 36 | 40 |
| L1PA15_dup4275 | 62.8374356633359 | -1.63386360787629 | 0.406602823168449 | -4.01832824264332 | 5.86125101562693e-05 | 0.0068065457987524 | 4396 | 47 | 28 | 133 | 208 | 12 | 17 | 188 | 113 | 12 | 16 | 20 | 33 |
| L1PA15_dup4276 | 58.1737110904116 | -1.50496566735691 | 0.421560205445278 | -3.56998988025273 | 0.000356995018871404 | 0.0244406914428247 | 4397 | 39 | 32 | 130 | 196 | 25 | 14 | 158 | 92 | 10 | 25 | 14 | 32 |
| L1MA2_dup1202 | 34.8356285286904 | -4.34663840713616 | 0.720965392457726 | -6.02891408187949 | 1.65065005718439e-09 | 1.70028388963981e-06 | 4398 | 6 | 9 | 74 | 91 | 52 | 50 | 102 | 58 | 0 | 0 | 3 | 11 |
| L1PB3_dup541 | 43.2241519116699 | -4.65254050751023 | 0.679403362883278 | -6.84797980358178 | 7.49001362122676e-12 | 1.51035489925121e-08 | 4399 | 11 | 11 | 95 | 119 | 78 | 66 | 114 | 61 | 1 | 0 | 6 | 5 |
| L1M2c_dup375 | 48.7417681237779 | -3.29533777625986 | 0.949994792352827 | -3.46879562160376 | 0.00052279700023578 | 0.0313482956287692 | 4400 | 8 | 5 | 70 | 93 | 108 | 130 | 99 | 72 | 0 | 2 | 16 | 28 |
| L1PA5_dup11042 | 29.7468713522079 | -3.81486284339274 | 1.05607044845243 | -3.61231852380972 | 0.000303471478403466 | 0.0218171169641034 | 4401 | 3 | 2 | 37 | 70 | 51 | 68 | 73 | 58 | 0 | 0 | 9 | 13 |
| MER45C_dup222 | 21.459615170846 | -4.91129948686782 | 0.958269491130227 | -5.12517567586881 | 2.97260447136852e-07 | 0.000122585674791032 | 4402 | 4 | 2 | 57 | 65 | 44 | 37 | 39 | 28 | 1 | 1 | 2 | 0 |
| L1MD1_dup1198 | 26.2934263422833 | -4.30333870552687 | 0.744560666364314 | -5.77970191004052 | 7.48330960674628e-09 | 6.20426337173118e-06 | 4403 | 10 | 4 | 63 | 86 | 47 | 41 | 66 | 25 | 0 | 1 | 5 | 3 |
| LTR1F1_dup33 | 25.9424315095883 | -3.99271260405 | 0.827953128850517 | -4.82238965579278 | 1.4184850251426e-06 | 0.000424557028669913 | 4404 | 6 | 1 | 63 | 85 | 50 | 69 | 37 | 22 | 2 | 1 | 2 | 5 |
| THE1B_dup3667 | 23.2775915957463 | -4.45416641675905 | 0.839314853001567 | -5.3069076531054 | 1.11500630021994e-07 | 5.58987777173499e-05 | 4405 | 6 | 2 | 46 | 50 | 40 | 53 | 55 | 41 | 1 | 0 | 3 | 3 |
| L1PA12_dup1702 | 14.3209567820246 | -2.59260288740256 | 0.752414481706952 | -3.44571104150055 | 0.000569559035784145 | 0.0332656588926594 | 4406 | 4 | 0 | 32 | 31 | 25 | 17 | 40 | 22 | 2 | 0 | 8 | 5 |
| MER5B_dup24217 | 12.2577571797915 | -2.7387134002217 | 0.662918674530211 | -4.13129619883245 | 3.60723419011847e-05 | 0.00477910326615774 | 4407 | 5 | 4 | 13 | 39 | 16 | 19 | 26 | 25 | 2 | 0 | 6 | 3 |
| L1MA7_dup1312 | 17.1417649599828 | -3.85067040861674 | 0.779766706698696 | -4.93823393014477 | 7.88332427855478e-07 | 0.00026720875766168 | 4408 | 7 | 4 | 32 | 50 | 28 | 33 | 42 | 22 | 0 | 0 | 1 | 8 |
| L2a_dup32318 | 15.7004404583022 | -4.04507432202008 | 0.878845637310157 | -4.60271309350827 | 4.17022525764022e-06 | 0.000961519786002385 | 4409 | 5 | 1 | 27 | 55 | 33 | 24 | 36 | 20 | 0 | 0 | 4 | 3 |
| L1M2_dup9092 | 13.2532772419058 | -3.03062846030111 | 0.868672039621821 | -3.48880627218128 | 0.000485182643950187 | 0.0299241237421906 | 4410 | 2 | 1 | 17 | 21 | 34 | 35 | 37 | 15 | 1 | 0 | 5 | 5 |
| L1M2c_dup376 | 13.9525916303881 | -4.10188979664967 | 0.98058332520574 | -4.18311192043678 | 2.87545742015851e-05 | 0.0040562018850693 | 4411 | 2 | 2 | 21 | 20 | 33 | 41 | 34 | 21 | 0 | 0 | 1 | 5 |
| Tigger1_dup11781 | 16.0795269747475 | -3.99437273982193 | 0.972580940613951 | -4.1069823322884 | 4.00861878464862e-05 | 0.00513919614941811 | 4412 | 1 | 1 | 27 | 24 | 29 | 38 | 58 | 24 | 0 | 0 | 4 | 4 |
| BSR/Beta_dup2021 | 14.3923585569718 | -2.89414767747085 | 0.88161171867244 | -3.28279175080494 | 0.0010278456529292 | 0.0487194744485838 | 4413 | 1 | 1 | 43 | 32 | 28 | 21 | 19 | 23 | 0 | 0 | 6 | 11 |
| L1M2_dup9094 | 17.5854675024899 | -2.76271972834427 | 0.825207342417129 | -3.34790977532015 | 0.000814235149837962 | 0.0418724869166509 | 4414 | 2 | 2 | 24 | 33 | 38 | 27 | 55 | 26 | 0 | 0 | 11 | 12 |
| L1PA2_dup2773 | 15.3870731302446 | -2.66728490916911 | 0.693965000656505 | -3.84354384824279 | 0.000121270319854693 | 0.0115469337847718 | 4415 | 3 | 2 | 28 | 36 | 19 | 21 | 43 | 29 | 0 | 0 | 9 | 10 |
| MLT2B4_dup302 | 19.7101992236926 | -3.656474968514 | 0.620205467048738 | -5.89558648348171 | 3.73352898154811e-09 | 3.37786420929067e-06 | 4416 | 14 | 13 | 27 | 31 | 10 | 16 | 72 | 57 | 2 | 1 | 2 | 4 |
| L2_dup26675 | 16.9149745245142 | -3.65568144063167 | 0.699025216420203 | -5.22968464478703 | 1.69799435026998e-07 | 7.93773995421099e-05 | 4417 | 26 | 29 | 13 | 23 | 15 | 7 | 61 | 36 | 1 | 2 | 1 | 4 |
| L1PA6_dup2932 | 15.2961737734749 | -2.94039525348735 | 0.662800585935864 | -4.43631963501595 | 9.15099403279903e-06 | 0.0017336325048793 | 4418 | 31 | 25 | 12 | 15 | 7 | 9 | 42 | 40 | 3 | 2 | 3 | 3 |
| MER30_dup1868 | 14.5669570063128 | -3.33862268310605 | 0.677051001349397 | -4.93112435614453 | 8.17576637335143e-07 | 0.00027555111975545 | 4419 | 23 | 16 | 10 | 16 | 13 | 14 | 50 | 35 | 3 | 1 | 2 | 2 |
| MER50B_dup290 | 21.6145064650643 | -2.28864240455663 | 0.544148763641242 | -4.20591308384473 | 2.60030160598804e-05 | 0.00376356984672793 | 4420 | 38 | 21 | 22 | 34 | 8 | 9 | 72 | 50 | 6 | 3 | 5 | 9 |
| L1M4_dup8159 | 21.6412567481681 | -1.85438759294733 | 0.509633666529912 | -3.6386677622258 | 0.000274052078759917 | 0.0203440387294071 | 4421 | 43 | 47 | 19 | 19 | 11 | 14 | 48 | 41 | 5 | 6 | 8 | 14 |
| L1MA3_dup1721 | 18.7169742510815 | -2.59613109323121 | 0.524149544742422 | -4.95303510090237 | 7.30647885827344e-07 | 0.000251769277383297 | 4422 | 48 | 33 | 20 | 27 | 10 | 16 | 36 | 33 | 3 | 2 | 5 | 8 |
| MLT1C_dup8338 | 20.0967887823864 | -2.45904978267189 | 0.665461413779873 | -3.69525524959335 | 0.000219665995169416 | 0.0174779121709676 | 4423 | 35 | 41 | 9 | 12 | 23 | 4 | 66 | 44 | 5 | 4 | 4 | 7 |
| L1M5_dup24516 | 19.9470275812932 | -1.8640843178626 | 0.563832205656625 | -3.3060976282682 | 0.000946051187303897 | 0.0460628393317399 | 4424 | 30 | 31 | 10 | 23 | 17 | 8 | 65 | 42 | 2 | 11 | 8 | 8 |
| LTR40a_dup1034 | 18.8564905549712 | -2.11372180322426 | 0.56713160087868 | -3.72703936784582 | 0.000193742206139451 | 0.0160739829086673 | 4425 | 38 | 23 | 11 | 19 | 12 | 11 | 55 | 45 | 4 | 2 | 11 | 8 |
| MLT1C_dup8337 | 18.392431225846 | -2.2116882307443 | 0.596909610988112 | -3.70523139522435 | 0.000211197836929492 | 0.0170444227815416 | 4426 | 33 | 24 | 12 | 14 | 11 | 10 | 61 | 46 | 3 | 6 | 8 | 4 |
| L1MC3_dup5722 | 17.8730456681077 | -2.0906215829863 | 0.629494637386918 | -3.32111102910207 | 0.000896598617688703 | 0.0444370036835151 | 4427 | 30 | 33 | 10 | 17 | 8 | 12 | 64 | 33 | 10 | 3 | 3 | 5 |
| L1PA3_dup4979 | 21.681498831941 | -2.48658796404606 | 0.555572271014845 | -4.47572367048466 | 7.61529269138325e-06 | 0.00150375803713185 | 4428 | 44 | 42 | 12 | 17 | 15 | 22 | 65 | 39 | 7 | 5 | 3 | 6 |
| L1M5_dup24519 | 15.7288663198766 | -2.05698785425469 | 0.537630968157976 | -3.82602189249311 | 0.00013023071316762 | 0.0121093697832679 | 4429 | 39 | 28 | 11 | 16 | 12 | 10 | 38 | 27 | 5 | 5 | 6 | 4 |
| L2b_dup16136 | 14.6795847503799 | -2.28837300276589 | 0.564987963233807 | -4.05030399173107 | 5.11511398009686e-05 | 0.00614780985184786 | 4430 | 29 | 24 | 10 | 20 | 12 | 10 | 49 | 21 | 4 | 6 | 4 | 2 |
| MER68-int_dup78 | 14.1872750566009 | -2.27601295365908 | 0.596309427884864 | -3.81683207949974 | 0.000135176100432238 | 0.0123996192727252 | 4431 | 30 | 28 | 10 | 12 | 7 | 13 | 44 | 23 | 4 | 4 | 2 | 6 |
| MIRc_dup92882 | 15.3120022295159 | -1.89819578695611 | 0.539203916511068 | -3.52036720956782 | 0.000430949719012498 | 0.0276395584474792 | 4432 | 29 | 31 | 14 | 9 | 10 | 15 | 42 | 24 | 5 | 5 | 3 | 9 |
| L1MB7_dup9095 | 17.2811614183926 | -2.14533564617271 | 0.540609623343882 | -3.96836377588504 | 7.23678060155807e-05 | 0.00794289530722213 | 4433 | 36 | 24 | 11 | 28 | 8 | 14 | 51 | 31 | 3 | 5 | 3 | 11 |
| MLT1A_dup4307 | 15.913746509381 | -1.80240289574331 | 0.538888181847111 | -3.34466955568655 | 0.000823806576207788 | 0.0421369844884236 | 4434 | 22 | 23 | 9 | 26 | 10 | 13 | 50 | 29 | 3 | 9 | 2 | 10 |
| L1MA9_dup7257 | 15.0998000043808 | -2.67957208315981 | 0.634151877497139 | -4.2254421665288 | 2.38472065730472e-05 | 0.00352225165439497 | 4435 | 30 | 25 | 11 | 19 | 12 | 6 | 51 | 28 | 0 | 4 | 6 | 4 |
| L2a_dup78626 | 17.2798618474887 | -2.81374980535913 | 0.571587974018514 | -4.92268895298344 | 8.53630756817792e-07 | 0.000284479025296592 | 4436 | 33 | 21 | 18 | 25 | 12 | 11 | 63 | 29 | 1 | 4 | 7 | 2 |
| L1M5_dup45847 | 17.2494497502003 | -1.9964653831422 | 0.463802099392241 | -4.30456305773159 | 1.67315631845469e-05 | 0.00271384358112488 | 4437 | 31 | 33 | 30 | 24 | 13 | 14 | 37 | 19 | 0 | 7 | 6 | 12 |
| L1M5_dup48428 | 15.223651506566 | -1.85561098426199 | 0.503630285768026 | -3.68447060611579 | 0.000229178469923286 | 0.0179973928067215 | 4438 | 40 | 35 | 22 | 20 | 14 | 15 | 17 | 13 | 0 | 6 | 8 | 9 |
| AluSx_dup10435 | 10.9779625601689 | -2.18928473908169 | 0.52652091550907 | -4.15802045957655 | 3.21017278682578e-05 | 0.004394981438055 | 4439 | 12 | 17 | 28 | 15 | 11 | 13 | 16 | 15 | 0 | 3 | 5 | 6 |
| L1PA2_dup4707 | 13.5420517610742 | -1.9017784958152 | 0.453206951253439 | -4.19626947591037 | 2.71347373902262e-05 | 0.00389186390780877 | 4440 | 18 | 13 | 23 | 24 | 9 | 12 | 26 | 27 | 0 | 2 | 9 | 11 |
| MIRb_dup119936 | 12.8430769482878 | -2.1472677778597 | 0.546850966530871 | -3.92660507026549 | 8.6153266684478e-05 | 0.00904458993611262 | 4441 | 18 | 22 | 23 | 20 | 8 | 11 | 18 | 25 | 0 | 1 | 6 | 12 |
| L1PA15_dup3669 | 13.5003439822226 | -2.12807688901856 | 0.563096848069272 | -3.77923779242459 | 0.000157309145445193 | 0.013847977033704 | 4442 | 17 | 34 | 14 | 22 | 8 | 12 | 28 | 21 | 1 | 5 | 9 | 2 |
| AluJo_dup14742 | 10.797420171258 | -2.133200034901 | 0.57764133348846 | -3.69294908662145 | 0.000221668415084903 | 0.0175993035354662 | 4443 | 24 | 15 | 13 | 14 | 5 | 13 | 29 | 14 | 2 | 1 | 8 | 3 |
| MIRb_dup3053 | 10.1206468828902 | -2.01693056176658 | 0.532272958440568 | -3.78927865822021 | 0.000151085361520968 | 0.0133938633530045 | 4444 | 16 | 17 | 14 | 15 | 5 | 12 | 26 | 13 | 1 | 3 | 6 | 4 |
| L2c_dup116248 | 9.1589653433159 | -2.23383448972362 | 0.645016136579299 | -3.46322264365395 | 0.000533746547137536 | 0.0317939826815871 | 4445 | 15 | 23 | 11 | 6 | 5 | 14 | 15 | 16 | 1 | 2 | 2 | 6 |
| MER102a_dup2062 | 10.5460396930679 | -2.02046038889459 | 0.511969232118037 | -3.9464488530607 | 7.93188560181365e-05 | 0.00846537620287874 | 4446 | 16 | 20 | 11 | 15 | 12 | 16 | 13 | 17 | 2 | 5 | 1 | 6 |
| L1P4_dup3347 | 10.5637991406214 | -1.8573819967836 | 0.55093510746729 | -3.37132626258325 | 0.000748072064801292 | 0.0395615328898235 | 4447 | 23 | 16 | 7 | 15 | 7 | 17 | 17 | 18 | 1 | 5 | 2 | 8 |
| MER46C_dup1683 | 10.5150124969458 | -2.14266619893055 | 0.547599329134032 | -3.91283569013669 | 9.12186041557806e-05 | 0.00941653014509821 | 4448 | 14 | 15 | 9 | 13 | 10 | 19 | 22 | 19 | 1 | 6 | 2 | 4 |
| L1MA4A_dup110 | 12.1397615668815 | -1.6930465559553 | 0.50432274322047 | -3.35706961209791 | 0.000787732770698178 | 0.0410417915166518 | 4449 | 21 | 22 | 14 | 13 | 8 | 9 | 24 | 23 | 2 | 6 | 5 | 7 |
| MIR_dup153011 | 10.6970892746019 | -1.64652672568316 | 0.479984612440552 | -3.43037398076401 | 0.000602749877541992 | 0.0345345346404062 | 4450 | 21 | 16 | 11 | 13 | 10 | 10 | 21 | 17 | 2 | 5 | 6 | 5 |
| L1M4b_dup1817 | 11.9605785496162 | -2.28422232176333 | 0.582619155754617 | -3.92060971425626 | 8.83252118846006e-05 | 0.00919765027006266 | 4451 | 23 | 21 | 10 | 15 | 7 | 8 | 34 | 22 | 2 | 2 | 5 | 5 |
| L1MB7_dup18000 | 10.9338473003866 | -2.4596770412297 | 0.591538585602402 | -4.1581007580847 | 3.20904484354737e-05 | 0.004394981438055 | 4452 | 29 | 16 | 15 | 11 | 7 | 10 | 22 | 19 | 2 | 3 | 3 | 3 |
| AluJo_dup53567 | 10.2941979904036 | -1.81061356072961 | 0.490764317085111 | -3.68937491520111 | 0.000224805724525153 | 0.0177890332317466 | 4453 | 22 | 16 | 15 | 17 | 11 | 9 | 22 | 8 | 2 | 4 | 3 | 7 |
| L1MB8_dup3119 | 9.78012405846528 | -1.9679370469533 | 0.521015418691558 | -3.77711863479097 | 0.000158653171691557 | 0.0139431096491211 | 4454 | 22 | 13 | 14 | 16 | 9 | 8 | 23 | 10 | 1 | 3 | 5 | 5 |
| AluY_dup35580 | 12.6655064300488 | -1.957365916802 | 0.536904557132769 | -3.64564966118917 | 0.000266717135064356 | 0.0199761647529189 | 4455 | 44 | 19 | 16 | 16 | 10 | 12 | 23 | 10 | 2 | 3 | 4 | 9 |
| L1ME1_dup9614 | 11.9882548198146 | -1.83194656321467 | 0.480605308835498 | -3.81174849619006 | 0.000137987239333848 | 0.012599198637963 | 4456 | 31 | 22 | 11 | 14 | 15 | 14 | 21 | 11 | 1 | 4 | 6 | 7 |
| MLT1G_dup995 | 9.74671921122201 | -1.71260182686427 | 0.52260586877053 | -3.27704285237611 | 0.00104900447356352 | 0.0493098416567651 | 4457 | 13 | 18 | 17 | 20 | 8 | 9 | 15 | 10 | 4 | 0 | 8 | 4 |
| AluJb_dup95927 | 9.51417163559173 | -2.32357088499248 | 0.60147918880336 | -3.86309439835353 | 0.000111959717878561 | 0.0108909164120007 | 4458 | 21 | 12 | 8 | 13 | 8 | 11 | 20 | 18 | 2 | 0 | 4 | 5 |

| | | | | | | | | | | | | | | | | | | | |
|---|---|---|---|---|---|---|---|---|---|---|---|---|---|---|---|---|---|---|---|
| AluSz6_dup640 | 10.0662498176517 | -2.18668244428691 | 0.563883734954897 | -3.87789593622845 | 0.000105363765324937 | 0.0104201523291801 | 4459 | 20 | 16 | 16 | 11 | 7 | 12 | 21 | 14 | 4 | 0 | 5 | 3 |
| L2_dup4527 | 11.3985676525873 | -1.75807655292711 | 0.53267974398593 | -3.300438157779 | 0.000965339861647708 | 0.046677252330751 | 4460 | 29 | 24 | 16 | 16 | 6 | 12 | 24 | 6 | 3 | 3 | 6 | 6 |
| MIR3_dup11563 | 12.4704640624221 | -1.67284503210065 | 0.475761799063268 | -3.51613987376526 | 0.000437870235248563 | 0.0279553182174549 | 4461 | 31 | 21 | 14 | 11 | 12 | 18 | 28 | 8 | 6 | 3 | 5 | 6 |
| FAM_dup3727 | 9.09440391331498 | -1.81277172679337 | 0.542298008536487 | -3.34275932837287 | 0.000829498080492618 | 0.0423280613898556 | 4462 | 13 | 9 | 15 | 11 | 10 | 8 | 32 | 8 | 3 | 2 | 4 | 5 |
| FRAM_dup3087 | 11.3183103353383 | -2.19762134959507 | 0.528988885576711 | -4.15438095112186 | 3.26169379113824e-05 | 0.00443743276630177 | 4463 | 13 | 14 | 18 | 22 | 10 | 13 | 36 | 10 | 5 | 3 | 2 | 3 |
| AluJr_dup54295 | 9.84236210982285 | -1.89451083237614 | 0.519913545962016 | -3.64389588824936 | 0.000268542081513648 | 0.0200749337505016 | 4464 | 15 | 13 | 21 | 21 | 7 | 12 | 19 | 7 | 4 | 2 | 6 | 2 |
| MER2B_dup73 | 9.90704336531874 | -1.75255075905656 | 0.493056337017168 | -3.55446351153081 | 0.000378751062933666 | 0.025379414368013 | 4465 | 16 | 15 | 13 | 14 | 11 | 11 | 18 | 13 | 5 | 3 | 6 | 1 |
| L1MEd_dup1892 | 10.519394943733 | -1.87977314865812 | 0.507138491480133 | -3.70662684895367 | 0.000210038058187872 | 0.0169992230590379 | 4466 | 22 | 19 | 16 | 17 | 10 | 10 | 20 | 8 | 6 | 2 | 2 | 5 |
| AluJo_dup3423 | 11.038032771731 | -1.67138954149781 | 0.462979694727416 | -3.61007093946497 | 0.000306113273853575 | 0.0219525102653294 | 4467 | 19 | 14 | 13 | 14 | 9 | 14 | 23 | 17 | 5 | 3 | 4 | 6 |
| MIRb_dup25111 | 9.94603679409916 | -1.67069959037585 | 0.483884844766065 | -3.45268013339735 | 0.000555046746798614 | 0.032679057648076 | 4468 | 17 | 14 | 15 | 14 | 10 | 13 | 13 | 14 | 6 | 1 | 4 | 5 |
| L1PA6_dup4532 | 11.155416401699 | -2.21815434219234 | 0.578339668688796 | -3.83538336082205 | 0.000125368592462222 | 0.0118002195814936 | 4469 | 19 | 27 | 14 | 18 | 8 | 5 | 25 | 15 | 3 | 4 | 2 | 4 |
| THE1A_dup3638 | 11.870654180122 | -3.31465094085139 | 0.68020169936706 | -4.87304125222817 | 1.098932648318e-06 | 0.000345424871778962 | 4470 | 29 | 23 | 17 | 23 | 9 | 6 | 27 | 15 | 1 | 2 | 1 | 3 |
| HERVK3-int_dup224 | 11.7026514446821 | -2.7088015338922 | 0.710699684723143 | -3.81145734565428 | 0.00013814989670959 | 0.0126043770408206 | 4471 | 24 | 30 | 25 | 25 | 2 | 5 | 23 | 10 | 3 | 1 | 3 | 3 |
| HERVK3-int_dup227 | 13.4853987964743 | -2.18233532160567 | 0.620902860649591 | -3.51477736682112 | 0.000440122793325709 | 0.0280433489858633 | 4472 | 26 | 26 | 26 | 32 | 3 | 7 | 34 | 10 | 2 | 7 | 2 | 5 |
| L1ME4a_dup30963 | 9.21680409436081 | -1.91742445179821 | 0.572055775562237 | -3.35181381555618 | 0.000802839913895569 | 0.0415252006833986 | 4473 | 14 | 14 | 15 | 22 | 4 | 7 | 10 | 18 | 2 | 3 | 5 | 3 |
| MER20_dup14094 | 11.401878022512 | -2.05570919077637 | 0.497563797290797 | -4.131548963107 | 3.6032691739292e-05 | 0.00477910326615774 | 4474 | 19 | 15 | 24 | 23 | 9 | 11 | 13 | 17 | 3 | 2 | 6 | 4 |
| L1MC5a_dup1092 | 12.0263168023656 | -1.65238421634673 | 0.477915733464887 | -3.45748026407701 | 0.000545252178472644 | 0.0322898394372453 | 4475 | 21 | 8 | 18 | 31 | 5 | 11 | 24 | 19 | 2 | 6 | 6 | 6 |
| MER44B_dup745 | 12.5434525027716 | -1.53971009385742 | 0.467919974671904 | -3.29054149683828 | 0.000999947519612779 | 0.0478450658559788 | 4476 | 21 | 20 | 20 | 22 | 5 | 13 | 18 | 19 | 2 | 5 | 10 | 6 |
| MLT1K_dup13641 | 12.1456711887507 | -1.78278545926018 | 0.513108317742477 | -3.47448169833593 | 0.000511841278742005 | 0.0310057497228985 | 4477 | 16 | 16 | 33 | 21 | 7 | 4 | 24 | 16 | 2 | 1 | 7 | 11 |
| MLT2A2_dup3549 | 11.5575212803917 | -2.25708601844999 | 0.510200819231198 | -4.42391688404403 | 9.69273230304786e-06 | 0.00181031937251258 | 4478 | 14 | 14 | 27 | 23 | 6 | 7 | 24 | 20 | 1 | 2 | 6 | 5 |
| L1MC4_dup22132 | 13.5406037658706 | -2.08864202338174 | 0.495613803708778 | -4.21425312965863 | 2.50606097551645e-05 | 0.00366732586027206 | 4479 | 27 | 15 | 22 | 35 | 8 | 8 | 25 | 19 | 1 | 2 | 9 | 7 |
| L1PA2_dup2011 | 13.5768722786194 | -2.63471928001711 | 0.602219489198573 | -4.37501496924878 | 1.21424145407814e-05 | 0.00216098651482481 | 4480 | 32 | 28 | 21 | 28 | 5 | 8 | 23 | 19 | 2 | 1 | 4 | 6 |
| L1PA15_dup849 | 13.1497728167027 | -2.10738765173115 | 0.483116183363158 | -4.36207215635131 | 1.28836361447661e-05 | 0.00225282475182657 | 4481 | 10 | 10 | 19 | 29 | 14 | 13 | 38 | 21 | 4 | 3 | 7 | 2 |
| MER21A_dup874 | 13.228106316025 | -1.41555693274254 | 0.416592889198853 | -3.39793829766222 | 0.000678957315849298 | 0.0370953134455906 | 4482 | 11 | 13 | 23 | 20 | 14 | 11 | 30 | 21 | 7 | 2 | 9 | 7 |
| AluJb_dup100465 | 14.2513578162583 | -2.04515661961828 | 0.569683289301589 | -3.58998878504153 | 0.000330692189022886 | 0.023122798592195 | 4483 | 22 | 10 | 15 | 14 | 11 | 13 | 43 | 34 | 6 | 5 | 2 | 4 |
| MIRc_dup1175 | 11.1766373484292 | -1.79096814941597 | 0.533529500894544 | -3.35683059027314 | 0.000788414042824633 | 0.0410593057198402 | 4484 | 13 | 6 | 21 | 12 | 13 | 9 | 29 | 22 | 5 | 4 | 2 | 5 |
| SVA_D_dup260 | 11.5889374146893 | -1.5719342702535 | 0.469562823873972 | -3.34765486178138 | 0.000814984394177843 | 0.041892190538513 | 4485 | 15 | 11 | 17 | 32 | 12 | 7 | 27 | 12 | 4 | 6 | 2 | 8 |
| L1PA16_dup4101 | 12.1734904236911 | -1.60767144371041 | 0.477731551179842 | -3.36521931561771 | 0.000764828396209551 | 0.0401827903653076 | 4486 | 28 | 13 | 22 | 19 | 16 | 8 | 24 | 9 | 4 | 7 | 2 | 8 |
| L2_dup21205 | 12.1243250960005 | -1.63364884394853 | 0.46812320574097 | -3.48978393703577 | 0.000483411216207645 | 0.0298483738175653 | 4487 | 21 | 17 | 23 | 28 | 15 | 8 | 18 | 9 | 2 | 7 | 3 | 9 |
| AluJb_dup109944 | 10.046715745702 | -2.39388529576438 | 0.558960240076028 | -4.28274700797818 | 1.84600023927697e-05 | 0.00292637157185209 | 4488 | 18 | 17 | 17 | 17 | 16 | 8 | 16 | 11 | 1 | 2 | 1 | 7 |
| L1M5_dup22736 | 10.8999567194696 | -1.60552798665527 | 0.454903671828731 | -3.52938014371478 | 0.000416534333851146 | 0.0270255736209899 | 4489 | 15 | 18 | 17 | 23 | 15 | 9 | 13 | 12 | 3 | 4 | 4 | 8 |
| L1ME2z_dup2431 | 9.53540188742062 | -1.56479358486933 | 0.471891345028301 | -3.31600399404546 | 0.000913145237515796 | 0.0449946690175882 | 4490 | 13 | 16 | 17 | 14 | 10 | 13 | 13 | 10 | 3 | 3 | 3 | 8 |
| L2b_dup21177 | 9.44675895710632 | -1.5450284804741 | 0.469803063043351 | -3.28867264182041 | 0.00100661028068309 | 0.0480479217346878 | 4491 | 16 | 17 | 13 | 13 | 13 | 10 | 13 | 10 | 2 | 3 | 5 | 7 |
| MIR_dup52031 | 13.0784416352389 | -1.56538510505112 | 0.476387542684205 | -3.28594886472254 | 0.00101639459496017 | 0.0483500656322254 | 4492 | 18 | 28 | 19 | 15 | 16 | 6 | 28 | 15 | 5 | 5 | 4 | 9 |
| L1ME3_dup8032 | 12.7962515764301 | -1.9729986883963 | 0.482072533765424 | -4.09274237838317 | 4.26301264042197e-05 | 0.00537270553886571 | 4493 | 14 | 21 | 18 | 22 | 20 | 6 | 31 | 16 | 2 | 4 | 6 | 6 |
| L2a_dup39246 | 12.5702190059217 | -1.77512523970815 | 0.499225427103025 | -3.55575886831145 | 0.000376889636456119 | 0.0253045658679988 | 4494 | 17 | 23 | 12 | 16 | 26 | 6 | 25 | 16 | 5 | 5 | 5 | 4 |
| AluSz6_dup31632 | 13.7591705532678 | -1.67039368134829 | 0.488751608893025 | -3.41767403105142 | 0.000631587008633966 | 0.0354860926461435 | 4495 | 21 | 20 | 23 | 39 | 15 | 5 | 17 | 16 | 7 | 3 | 6 | 6 |
| Charlie5_dup932 | 12.7585681898637 | -2.28489959160247 | 0.5261981296548 | -4.34227995660384 | 1.41011688526846e-05 | 0.00241389692361214 | 4496 | 15 | 13 | 31 | 39 | 17 | 7 | 18 | 13 | 3 | 4 | 4 | 3 |
| L1MA8_dup7141 | 17.2869365959478 | -1.47423557093456 | 0.387524828910519 | -3.80423513785994 | 0.000142243016482705 | 0.0128346989045881 | 4497 | 18 | 18 | 19 | 28 | 21 | 17 | 33 | 32 | 3 | 3 | 12 | 17 |
| MIR_dup168043 | 15.7155810654699 | -1.43055009354298 | 0.400735929688828 | -3.56980741570591 | 0.000357243764711928 | 0.0244406914428247 | 4498 | 18 | 12 | 32 | 25 | 17 | 12 | 35 | 20 | 1 | 2 | 12 | 20 |
| AluJo_dup68437 | 12.5862034161846 | -1.66775203367801 | 0.499502470608983 | -3.33882639588294 | 0.00084133125906634 | 0.0427138096691817 | 4499 | 23 | 10 | 15 | 20 | 11 | 18 | 30 | 16 | 2 | 3 | 2 | 16 |
| L2a_dup95091 | 14.6895834236028 | -1.59856191190013 | 0.44817314947593 | -3.56683998978832 | 0.000361311946196684 | 0.0245917426629623 | 4500 | 22 | 21 | 24 | 27 | 11 | 13 | 22 | 22 | 1 | 3 | 6 | 19 |
| L2c_dup15837 | 13.2630735666218 | -1.53036486659739 | 0.433325979434006 | -3.53167116496523 | 0.000412942516089267 | 0.0268537906349759 | 4501 | 18 | 21 | 16 | 27 | 13 | 16 | 23 | 14 | 3 | 1 | 7 | 15 |
| L1MA8_dup8901 | 13.1290166800121 | -1.68521882412939 | 0.482167824363887 | -3.49508768311668 | 0.00047390604027787 | 0.0294269432828701 | 4502 | 23 | 24 | 13 | 14 | 9 | 19 | 19 | 23 | 6 | 2 | 7 | 6 |
| MER11C_dup693 | 13.7394961987318 | -1.681212382987 | 0.447973488051565 | -3.75292830452832 | 0.000174780854985656 | 0.0149276401006521 | 4503 | 35 | 16 | 10 | 19 | 13 | 19 | 25 | 18 | 6 | 5 | 6 | 5 |
| L2c_dup48289 | 15.3675776487504 | -1.49248599940229 | 0.428948559629393 | -3.47940555084689 | 0.000502527471965926 | 0.030629272336295 | 4504 | 28 | 24 | 12 | 17 | 13 | 22 | 32 | 21 | 5 | 5 | 12 | 6 |
| Tigger1_dup370 | 13.5293679197856 | -1.43253798469963 | 0.436270030225374 | -3.2836039275024 | 0.00102488848267813 | 0.0486471404147875 | 4505 | 25 | 21 | 11 | 19 | 11 | 15 | 24 | 21 | 4 | 4 | 11 | 7 |
| L1M5_dup40760 | 11.6150462009847 | -1.5523050727016 | 0.456209062231499 | -3.40261779349288 | 0.000667435773335985 | 0.0366944714700102 | 4506 | 14 | 15 | 9 | 17 | 11 | 18 | 25 | 19 | 2 | 5 | 7 | 7 |
| AluSx1_dup64394 | 14.1603051885143 | -1.4583129146074 | 0.4254601073861 | -3.42761375106786 | 0.000608911140454248 | 0.0347453166010855 | 4507 | 22 | 17 | 20 | 13 | 10 | 20 | 30 | 22 | 3 | 6 | 10 | 8 |
| L1ME4b_dup14363 | 14.5038368547923 | -1.46044710395541 | 0.419423923206412 | -3.48203100288268 | 0.000497626074592827 | 0.0303688468697268 | 4508 | 23 | 16 | 19 | 21 | 12 | 17 | 25 | 24 | 3 | 4 | 15 | 6 |
| L2a_dup135481 | 16.7157694200854 | -2.17354473271388 | 0.447138723544746 | -4.86100759845366 | 1.16789770283772e-06 | 0.000359969672019978 | 4509 | 34 | 25 | 22 | 31 | 10 | 23 | 40 | 15 | 5 | 6 | 5 | 4 |
| L2a_dup168232 | 15.2548374129653 | -1.53329607317091 | 0.377118023927501 | -4.06582548668022 | 4.78627424796052e-05 | 0.00584938270264619 | 4510 | 30 | 25 | 17 | 19 | 18 | 20 | 29 | 13 | 6 | 6 | 7 | 8 |
| MSTB1_dup759 | 15.20363389208 | -1.41883011000568 | 0.383196394033802 | -3.70261863654314 | 0.000213385521939308 | 0.0171534565551927 | 4511 | 28 | 22 | 20 | 24 | 14 | 14 | 33 | 14 | 6 | 7 | 9 | 7 |
| AluSz_dup49526 | 12.8534620516443 | -1.56837482786836 | 0.422927255526389 | -3.70837965010393 | 0.00020858975804881 | 0.0169385231558667 | 4512 | 22 | 18 | 23 | 18 | 13 | 11 | 21 | 16 | 6 | 5 | 7 | 4 |
| MIR3_dup14567 | 13.8051834942132 | -1.75798357419629 | 0.43573838088934 | -4.03449329069486 | 5.47202535362073e-05 | 0.00644896753240534 | 4513 | 24 | 15 | 18 | 32 | 11 | 13 | 24 | 20 | 5 | 6 | 7 | 3 |
| AluSx1_dup110051 | 13.8386679120732 | -1.35138501639647 | 0.407116567955364 | -3.3194056021435 | 0.000902092982541912 | 0.044597842681055 | 4514 | 20 | 18 | 24 | 28 | 8 | 11 | 26 | 17 | 5 | 9 | 6 | 7 |
| AluSz_dup26460 | 12.7321843468516 | -1.57476354556948 | 0.450123303877117 | -3.4985159222047 | 0.000467855154052204 | 0.0291804195536618 | 4515 | 28 | 15 | 15 | 24 | 8 | 11 | 22 | 19 | 5 | 5 | 6 | 6 |
| L2a_dup155915 | 13.1552139276544 | -1.55277709457566 | 0.424385784336228 | -3.65888102732829 | 0.000253318898580092 | 0.0192945642258446 | 4516 | 23 | 23 | 15 | 26 | 10 | 14 | 18 | 17 | 5 | 7 | 5 | 6 |
| AluJr4_dup13382 | 15.4683166391387 | -1.67090397930998 | 0.411251893270392 | -4.06296969485653 | 4.84523220804841e-05 | 0.00590022324962276 | 4517 | 32 | 16 | 26 | 31 | 14 | 10 | 26 | 20 | 5 | 4 | 7 | 10 |
| AluSx4_dup9316 | 14.6594953465709 | -1.52056681141724 | 0.415789736808518 | -3.65705710556656 | 0.000255127557773884 | 0.0193888795471302 | 4518 | 27 | 24 | 20 | 26 | 18 | 9 | 27 | 13 | 8 | 4 | 7 | 7 |
| AluY_dup3754 | 15.1764840854777 | -1.66818634809684 | 0.413436199346922 | -4.03493054244395 | 5.46184522196008e-05 | 0.00644015967881324 | 4519 | 22 | 18 | 25 | 23 | 16 | 12 | 31 | 22 | 9 | 4 | 6 | 5 |
| L1PA15_dup7709 | 15.9572373768245 | -1.80357326210594 | 0.403165030256796 | -4.47353596356596 | 7.69366110238046e-06 | 0.00151295527658431 | 4520 | 30 | 24 | 20 | 23 | 15 | 15 | 29 | 24 | 6 | 5 | 6 | 7 |
| AluSz6_dup46096 | 14.4435496297955 | -1.53561560421498 | 0.448380112435333 | -3.42480757202729 | 0.000615235011242697 | 0.0349052718133416 | 4521 | 15 | 28 | 25 | 20 | 8 | 17 | 25 | 20 | 7 | 8 | 4 | 6 |
| L1MEd_dup15713 | 12.901155400084 | -1.54464805286602 | 0.455149080720717 | -3.39371893362963 | 0.000689504196662784 | 0.0374320706317871 | 4522 | 20 | 25 | 23 | 12 | 10 | 17 | 26 | 11 | 4 | 8 | 3 | 8 |
| L2a_dup87554 | 13.3253585459728 | -1.5350465863039 | 0.450380687945573 | -3.40833127926971 | 0.000653614910676205 | 0.0362622254961759 | 4523 | 33 | 27 | 17 | 19 | 13 | 14 | 15 | 11 | 6 | 3 | 9 | 5 |
| L1M5_dup25856 | 14.3351354405185 | -1.47413717715822 | 0.442729421666704 | -3.32965713371536 | 0.000869529853000362 | 0.0435911370936071 | 4524 | 31 | 36 | 17 | 15 | 16 | 12 | 17 | 14 | 5 | 5 | 6 | 10 |
| L1MB8_dup9888 | 13.531168733222 | -1.42023934376813 | 0.417647342218488 | -3.40057076916617 | 0.000672453269992211 | 0.0368600756169689 | 4525 | 26 | 25 | 14 | 15 | 14 | 16 | 22 | 16 | 5 | 3 | 7 | 11 |
| L1ME3D_dup1953 | 13.7954626015268 | -1.50378697782988 | 0.453588036981835 | -3.31531445986991 | 0.000915400862416491 | 0.0450628962230779 | 4526 | 37 | 20 | 13 | 29 | 8 | 20 | 17 | 12 | 5 | 6 | 6 | 8 |
| L1PA16_dup12781 | 13.4060577121092 | -1.49527022877858 | 0.44987925827031 | -3.32371453293396 | 0.000888270715526751 | 0.0441437894061672 | 4527 | 28 | 22 | 13 | 22 | 7 | 20 | 24 | 14 | 5 | 3 | 6 | 11 |
| AluSx1_dup72927 | 14.9223183106513 | -1.35444803987989 | 0.395627066063055 | -3.42354746695721 | 0.000618094560184838 | 0.0349674579690303 | 4528 | 24 | 21 | 22 | 36 | 9 | 17 | 19 | 16 | 6 | 5 | 9 | 10 |
| L1ME3B_dup471 | 13.7255817064348 | -1.43688571348677 | 0.395282994338678 | -3.63508102819026 | 0.000277893278868418 | 0.0205163332190133 | 4529 | 27 | 18 | 21 | 27 | 12 | 15 | 18 | 14 | 5 | 6 | 8 | 7 |
| AluJb_dup98947 | 15.1582551904863 | -1.59864663101614 | 0.426263870945263 | -3.75036858617891 | 0.00017657482616732 | 0.0150484504252182 | 4530 | 34 | 22 | 25 | 28 | 9 | 15 | 22 | 16 | 3 | 5 | 10 | 9 |

| | | | | | | | | | | | | | | | | | | |
|---|---|---|---|---|---|---|---|---|---|---|---|---|---|---|---|---|---|---|
| MIRb_dup207502 | 14.1618559409701 | -1.74187107603302 | 0.466851274705719 | -3.73110489444634 | 0.000190641820156127 | 0.0158887770714079 | 4531 | 36 | 27 | 19 | 24 | 11 | 12 | 19 | 14 | 3 | 3 | 8 | 9 |
| AluSz_dup32432 | 18.0445305133578 | -1.48181367537774 | 0.448516159596169 | -3.30381334913756 | 0.00095379311867915 | 0.0463165831463733 | 4532 | 44 | 23 | 15 | 27 | 11 | 21 | 37 | 23 | 2 | 7 | 9 | 18 |
| AluJr_dup47568 | 17.1887384360982 | -1.6915253257945 | 0.405315763943014 | -4.17335193020601 | 3.00150756730661e-05 | 0.00417414798135791 | 4533 | 26 | 25 | 24 | 22 | 10 | 28 | 32 | 25 | 3 | 5 | 11 | 10 |
| L1PA3_dup1074 | 17.7905658547164 | -1.47677112360422 | 0.435891245373504 | -3.38793481006671 | 0.000704209996609961 | 0.0379533299781096 | 4534 | 35 | 33 | 16 | 23 | 10 | 17 | 35 | 26 | 4 | 6 | 13 | 11 |
| L1M5_dup10657 | 16.5084825803844 | -2.10033825607023 | 0.509243581266571 | -4.12442754967348 | 3.71657954566029e-05 | 0.00486706239169887 | 4535 | 43 | 34 | 15 | 25 | 16 | 16 | 23 | 21 | 2 | 3 | 3 | 14 |
| MER20_dup10166 | 16.6216747666521 | -1.80905326071146 | 0.450894449111782 | -4.01214356103767 | 6.0169888532551e-05 | 0.0069265526351071 | 4536 | 37 | 33 | 15 | 24 | 12 | 18 | 22 | 26 | 6 | 5 | 4 | 10 |
| AluSc5_dup1372 | 16.8761200494993 | -1.5604181718042 | 0.406314705057406 | -3.84041766734417 | 0.000122825161271372 | 0.0116365308264591 | 4537 | 34 | 32 | 18 | 29 | 19 | 14 | 19 | 22 | 5 | 6 | 6 | 13 |
| L1MC3_dup5974 | 17.8108102151111 | -1.86230735265377 | 0.463197958138569 | -4.02054309595348 | 5.80641255215112e-05 | 0.00676291337965716 | 4538 | 55 | 35 | 15 | 28 | 15 | 15 | 23 | 20 | 4 | 6 | 6 | 10 |
| AluSx1_dup91542 | 17.3956069439166 | -1.67894114671199 | 0.453236216930239 | -3.70434021818341 | 0.000211941649080341 | 0.017083559495959 | 4539 | 37 | 34 | 18 | 15 | 19 | 16 | 37 | 20 | 3 | 4 | 8 | 15 |
| L1PA6_dup1325 | 16.6610817840357 | -2.08693687341556 | 0.504458352374778 | -4.13698546885216 | 3.51898385045262e-05 | 0.00470409062516741 | 4540 | 47 | 36 | 10 | 16 | 17 | 24 | 25 | 20 | 2 | 3 | 6 | 10 |
| MER34A1_dup294 | 17.4414640264243 | -2.23128792269729 | 0.479903740433102 | -4.6494489096576 | 3.32823111136973e-06 | 0.000817274798788423 | 4541 | 49 | 38 | 21 | 14 | 19 | 23 | 27 | 16 | 2 | 3 | 7 | 8 |
| AluSx3_dup3805 | 17.2494607676977 | -1.30881931409169 | 0.343645391742998 | -3.80863339226884 | 0.000139736956909099 | 0.012705330511537 | 4542 | 30 | 25 | 20 | 23 | 18 | 23 | 29 | 19 | 8 | 9 | 5 | 13 |
| L1PREC2_dup5401 | 17.2628950347603 | -1.39491520860632 | 0.358231776280902 | -3.89389021568124 | 9.86493091553434e-05 | 0.00997162638754348 | 4543 | 27 | 19 | 21 | 27 | 20 | 25 | 38 | 15 | 7 | 7 | 5 | 15 |
| AluJb_dup78043 | 17.1776499101097 | -1.35357437203321 | 0.34751987544016 | -3.89495527505817 | 9.82168507428533e-05 | 0.00993635491788222 | 4544 | 29 | 28 | 18 | 28 | 28 | 19 | 22 | 16 | 6 | 9 | 6 | 13 |
| L1MEg_dup12756 | 16.8543453566534 | -1.44566888744374 | 0.374672738740722 | -3.85848431968293 | 0.000114092386191901 | 0.0110401898840252 | 4545 | 33 | 22 | 15 | 30 | 23 | 19 | 29 | 17 | 3 | 6 | 9 | 15 |
| MIR_dup78811 | 13.6724054213488 | -2.03563523393869 | 0.441818057696488 | -4.60740614485497 | 4.0772313782003e-06 | 0.000944580252459273 | 4546 | 21 | 16 | 18 | 18 | 27 | 18 | 30 | 12 | 3 | 6 | 4 | 5 |
| L2c_dup109609 | 12.7995512258409 | -1.41002041026044 | 0.406178036126082 | -3.47143440770085 | 0.000517685785863906 | 0.0311835256753778 | 4547 | 15 | 16 | 12 | 23 | 19 | 15 | 29 | 13 | 3 | 7 | 7 | 8 |
| L1PA5_dup4810 | 14.4899195845165 | -1.44187565457877 | 0.388522430897603 | -3.71117737333108 | 0.000206297480837522 | 0.0167919620766515 | 4548 | 18 | 23 | 13 | 18 | 19 | 23 | 29 | 16 | 5 | 8 | 8 | 6 |
| MER58A_dup6364 | 13.6499124800961 | -1.57061408641508 | 0.397874778735851 | -3.94750853875512 | 7.89687024501524e-05 | 0.00844507275200178 | 4549 | 19 | 25 | 18 | 19 | 19 | 16 | 22 | 14 | 3 | 5 | 9 | 7 |
| AluSx1_dup93744 | 14.3536829386367 | -1.70461028679174 | 0.425549785867603 | -4.00566594885346 | 6.18429975132042e-05 | 0.00706786449869206 | 4550 | 17 | 23 | 17 | 16 | 18 | 19 | 21 | 26 | 5 | 4 | 6 | 8 |
| L2b_dup8352 | 14.710956466331 | -1.41718971808206 | 0.404109018848398 | -3.50694899638882 | 0.000453276019658408 | 0.0285609855960214 | 4551 | 15 | 19 | 19 | 15 | 19 | 22 | 23 | 25 | 5 | 8 | 5 | 10 |
| L1MA3_dup4334 | 14.5564892264495 | -1.80489647689986 | 0.414238433591423 | -4.35714393097596 | 1.31770620745391e-05 | 0.00228529825975823 | 4552 | 26 | 22 | 23 | 18 | 15 | 19 | 21 | 20 | 3 | 9 | 4 | 6 |
| AluJr_dup12163 | 14.4946416619317 | -1.6542090005977 | 0.396709612466682 | -4.16982333831558 | 3.0483584486196e-05 | 0.00422695046279277 | 4553 | 30 | 21 | 17 | 21 | 17 | 20 | 17 | 19 | 4 | 6 | 5 | 9 |
| Tigger1_dup9454 | 15.3562045947826 | -1.48073847619362 | 0.395386251144169 | -3.74504290907602 | 0.000180362946783948 | 0.0152837043217426 | 4554 | 32 | 18 | 21 | 18 | 14 | 19 | 26 | 21 | 4 | 6 | 6 | 13 |
| L2_dup4529 | 14.6014261151252 | -1.58637095062426 | 0.448329577817445 | -3.53840350740858 | 0.000402554381017039 | 0.0264073769498129 | 4555 | 27 | 22 | 22 | 26 | 9 | 12 | 39 | 11 | 2 | 8 | 8 | 8 |
| MER20_dup6276 | 13.025580134923 | -1.49727864687194 | 0.439479147482339 | -3.40693899915266 | 0.000656958078934292 | 0.0363705321196135 | 4556 | 30 | 18 | 17 | 24 | 14 | 10 | 20 | 13 | 2 | 8 | 9 | 5 |
| Tigger19a_dup1651 | 13.3741807983959 | -1.52961742451573 | 0.418329712225428 | -3.65648764554275 | 0.000255694728620643 | 0.0194071891216484 | 4557 | 27 | 22 | 17 | 20 | 13 | 14 | 21 | 15 | 2 | 9 | 8 | 5 |
| AluSx1_dup14776 | 15.6992769647303 | -1.27630815205701 | 0.364410977884236 | -3.50238667195824 | 0.000461109897445958 | 0.0289070633170244 | 4558 | 18 | 24 | 26 | 20 | 16 | 18 | 29 | 18 | 3 | 7 | 9 | 15 |
| AluJo_dup65108 | 14.7753268874338 | -1.47086267055666 | 0.381794681189908 | -3.85249649359322 | 0.000116919651505759 | 0.0112270231627076 | 4559 | 23 | 22 | 24 | 24 | 15 | 23 | 25 | 10 | 3 | 8 | 6 | 11 |
| AluY_dup31327 | 15.4039117438302 | -1.22639823787651 | 0.359150555039598 | -3.41471903820752 | 0.000638478342467962 | 0.0357214484833515 | 4560 | 24 | 21 | 20 | 26 | 16 | 21 | 27 | 13 | 4 | 8 | 6 | 16 |
| AluSx1_dup106299 | 13.9629403634202 | -1.39436053003252 | 0.410087664961206 | -3.40015233124465 | 0.000673483217596792 | 0.0368990646314268 | 4561 | 24 | 16 | 22 | 14 | 20 | 24 | 22 | 12 | 1 | 7 | 8 | 12 |
| MIR_dup152473 | 13.2309706450916 | -1.45325761679142 | 0.429335726171111 | -3.38489794397457 | 0.000712047231679805 | 0.0382401208062598 | 4562 | 17 | 19 | 14 | 23 | 19 | 17 | 28 | 11 | 1 | 4 | 7 | 15 |
| L1ME4c_dup6178 | 17.3039921623814 | -1.71939917591919 | 0.418409366720295 | -4.10937066107462 | 3.96738843238118e-05 | 0.00510559333656773 | 4563 | 34 | 35 | 21 | 26 | 20 | 23 | 16 | 20 | 1 | 7 | 11 | 9 |
| AluSz6_dup42178 | 19.8683114378591 | -1.67772492470001 | 0.404648949314132 | -4.14612450506471 | 3.38150000795276e-05 | 0.00455872181808684 | 4564 | 36 | 26 | 21 | 22 | 28 | 29 | 31 | 28 | 1 | 5 | 13 | 16 |
| AluSz_dup99900 | 20.696623014517 | -1.86087208599403 | 0.409879556209911 | -4.54004611306113 | 5.62419264962083e-06 | 0.00119273987897208 | 4565 | 27 | 29 | 22 | 49 | 23 | 33 | 30 | 24 | 1 | 3 | 16 | 14 |
| HAL1_dup17353 | 21.1116907346405 | -1.97507800703719 | 0.40561385621779 | -4.8693553653569 | 1.11962898591305e-06 | 0.000349259924646234 | 4566 | 44 | 42 | 22 | 25 | 25 | 18 | 46 | 23 | 5 | 5 | 10 | 9 |
| L1PA16_dup5143 | 21.070981466571 | -2.13611260596312 | 0.410927194022086 | -5.19827511305643 | 2.01146280419787e-07 | 9.09923057809253e-05 | 4567 | 40 | 39 | 33 | 24 | 20 | 21 | 49 | 22 | 2 | 6 | 10 | 9 |
| L1ME4a_dup26821 | 23.2744631698524 | -1.83824721511878 | 0.336199236935198 | -5.4677316696977 | 4.55831229609842e-08 | 2.70481839802353e-05 | 4568 | 29 | 33 | 24 | 37 | 37 | 35 | 43 | 28 | 3 | 12 | 8 | 12 |
| MIR_dup27346 | 21.776400737248 | -1.52364326870267 | 0.377755202265164 | -4.03341438997086 | 5.49722136578979e-05 | 0.00647545462922879 | 4569 | 41 | 33 | 23 | 28 | 20 | 27 | 36 | 32 | 3 | 15 | 6 | 16 |
| L1MA4A_dup394 | 19.0412035712681 | -1.43274838115808 | 0.412645754774882 | -3.47210255910597 | 0.000516399015540245 | 0.0311549391865464 | 4570 | 33 | 21 | 20 | 48 | 14 | 14 | 46 | 20 | 2 | 13 | 9 | 13 |
| L1MA2_dup6916 | 23.2887469716184 | -2.1391078625108 | 0.365097269384351 | -5.85900811068209 | 4.65639957976872e-09 | 4.13422259107331e-06 | 4571 | 36 | 37 | 34 | 40 | 23 | 20 | 49 | 33 | 3 | 9 | 10 | 7 |
| MIRb_dup210761 | 22.3604864724185 | -1.61279959973223 | 0.374324085978802 | -4.30856485100338 | 1.64317297970075e-05 | 0.00267615631641256 | 4572 | 27 | 29 | 33 | 40 | 16 | 16 | 39 | 45 | 4 | 10 | 15 | 9 |
| AluJr_dup58157 | 22.7287624836675 | -1.57520294233512 | 0.341683844402473 | -4.61011829543709 | 4.02439908058978e-06 | 0.000937893891900114 | 4573 | 25 | 32 | 47 | 45 | 18 | 17 | 37 | 31 | 9 | 5 | 10 | 16 |
| MIRb_dup171876 | 20.2435155837979 | -1.27610226141932 | 0.318198839171995 | -4.01039257320972 | 6.06178761004065e-05 | 0.0069579262646345 | 4574 | 21 | 25 | 35 | 39 | 21 | 21 | 29 | 26 | 10 | 7 | 10 | 15 |
| AluJr_dup71399 | 21.9851191134073 | -1.26687603628595 | 0.355850841253271 | -3.56013219421975 | 0.000370668150282629 | 0.0250423065551251 | 4575 | 31 | 33 | 45 | 38 | 20 | 14 | 35 | 22 | 11 | 8 | 8 | 19 |
| AluSx_dup78990 | 21.682472723408 | -1.66964851520183 | 0.32457190235917 | -5.14415605006437 | 2.68726096231576e-07 | 0.000113236600373145 | 4576 | 35 | 24 | 36 | 44 | 24 | 20 | 39 | 24 | 8 | 7 | 8 | 13 |
| AluJo_dup14740 | 19.9033203683884 | -1.6099570629285 | 0.390735410915986 | -4.1203254630911 | 3.78337580377655e-05 | 0.00493822895779164 | 4577 | 41 | 26 | 37 | 34 | 12 | 18 | 32 | 24 | 5 | 8 | 6 | 16 |
| MER39B_dup425 | 17.7763669125907 | -1.62512719064115 | 0.365774367782782 | -4.44297723892517 | 8.87225547201781e-06 | 0.00168967125719969 | 4578 | 32 | 25 | 29 | 28 | 17 | 17 | 34 | 19 | 4 | 7 | 6 | 14 |
| MLT1A0_dup11366 | 18.3177118938807 | -1.61639697695471 | 0.35036805539432 | -4.61342565929859 | 3.96085972333481e-06 | 0.000927628669024238 | 4579 | 29 | 21 | 30 | 33 | 20 | 17 | 34 | 22 | 6 | 5 | 6 | 15 |
| L1PA14_dup2417 | 21.8621016723732 | -1.72133264568392 | 0.383082370670778 | -4.49337473470749 | 7.01032649283114e-06 | 0.00141123447348832 | 4580 | 46 | 23 | 27 | 44 | 14 | 21 | 49 | 27 | 7 | 5 | 8 | 16 |
| Tigger4a_dup3312 | 20.0452543865867 | -1.24625723763278 | 0.336885040693854 | -3.69935463761161 | 0.000216148396453895 | 0.0172944433409059 | 4581 | 35 | 20 | 25 | 34 | 19 | 16 | 34 | 31 | 8 | 7 | 12 | 16 |
| L1MB7_dup20360 | 18.9442262497311 | -1.75276297704542 | 0.389001644139065 | -4.50579837759971 | 6.61238510395229e-06 | 0.00135171012538233 | 4582 | 22 | 17 | 38 | 49 | 26 | 9 | 33 | 23 | 4 | 7 | 9 | 10 |
| L1MC3_dup11862 | 24.3080527403723 | -1.94305808927918 | 0.352136901204676 | -5.51790534485846 | 3.43064116310506e-08 | 2.1652714647114e-05 | 4583 | 38 | 26 | 48 | 63 | 20 | 22 | 36 | 30 | 5 | 7 | 7 | 16 |
| L1ME1_dup11259 | 22.440716248743 | -2.24843606071918 | 0.414781048449833 | -5.42077818917304 | 5.93401512875445e-08 | 3.3134899959648e-05 | 4584 | 36 | 26 | 56 | 58 | 23 | 17 | 27 | 24 | 5 | 6 | 5 | 10 |
| MIR_dup127186 | 21.1570293434924 | -1.65296669330306 | 0.395884392585027 | -4.17537726736232 | 2.9749261280073e-05 | 0.00415661639726849 | 4585 | 24 | 13 | 43 | 41 | 31 | 35 | 31 | 20 | 3 | 4 | 13 | 19 |
| AluJr_dup15976 | 19.5215230507167 | -1.11630071141348 | 0.334067144263071 | -3.3415459454295 | 0.0008331322648705 | 0.0424316117073793 | 4586 | 24 | 17 | 31 | 42 | 16 | 27 | 26 | 24 | 5 | 8 | 11 | 23 |
| L1PA15_dup7588 | 21.4887738710216 | -1.41468897200952 | 0.348863597284894 | -4.05513496684561 | 5.01053633016732e-05 | 0.00604380116463849 | 4587 | 30 | 24 | 39 | 55 | 23 | 25 | 21 | 21 | 6 | 10 | 11 | 15 |
| MER72_dup415 | 19.3864097509577 | -1.55689128061081 | 0.417832337789927 | -3.72611485469457 | 0.000194453825511154 | 0.0161143494767683 | 4588 | 35 | 24 | 43 | 37 | 9 | 22 | 26 | 20 | 3 | 4 | 13 | 18 |
| MIR_dup168039 | 19.537985702307 | -1.526385135699 | 0.415587051282619 | -3.67284093907195 | 0.000239868801486227 | 0.0185904708360133 | 4589 | 21 | 14 | 49 | 55 | 15 | 20 | 28 | 16 | 2 | 2 | 16 | 22 |
| LTR39_dup1120 | 15.3235048274193 | -1.94822755260808 | 0.423539971929378 | -4.59986702962933 | 4.22760709566955e-06 | 0.000970050335570093 | 4590 | 17 | 14 | 29 | 44 | 11 | 15 | 21 | 25 | 3 | 3 | 9 | 7 |
| Tigger17a_dup416 | 20.0822717767569 | -1.71960664004262 | 0.395601671261225 | -4.34681338569758 | 1.38129643661615e-05 | 0.00237481186349036 | 4591 | 31 | 11 | 48 | 49 | 11 | 18 | 33 | 28 | 5 | 6 | 12 | 9 |
| L1MA3_dup5689 | 12.981455738111 | -2.12383383690551 | 0.56739472946083 | -3.74313282557929 | 0.000181740096421186 | 0.0153566509172711 | 4592 | 25 | 15 | 16 | 31 | 36 | 19 | 7 | 6 | 1 | 4 | 5 | 6 |
| AluSp_dup38994 | 11.7540086086295 | -2.19907265073094 | 0.515709297574252 | -4.26417103797575 | 2.00645709437805e-05 | 0.00309254571153888 | 4593 | 19 | 24 | 16 | 27 | 19 | 15 | 9 | 10 | 1 | 4 | 5 | 4 |
| L1MEg2_dup545 | 9.97786080114284 | -2.03866040127261 | 0.531911522026775 | -3.83270584834219 | 0.000126741467608984 | 0.0118918080645454 | 4594 | 20 | 15 | 13 | 24 | 17 | 15 | 8 | 6 | 3 | 2 | 3 | 5 |
| AluJr_dup4695 | 11.20879753463 | -2.25158044362263 | 0.53325402055026 | -4.22234124235808 | 2.41777690152777e-05 | 0.00356443830613958 | 4595 | 25 | 17 | 7 | 25 | 17 | 14 | 20 | 10 | 1 | 3 | 6 | 3 |
| L1PA7_dup11551 | 12.2917034349528 | -1.67700342813464 | 0.512364502730044 | -3.27306716058397 | 0.00106387195617056 | 0.0498217036967053 | 4596 | 18 | 32 | 9 | 24 | 23 | 8 | 14 | 11 | 2 | 5 | 7 | 6 |
| L1MA9_dup14853 | 13.744727403602 | -1.94977008864861 | 0.450901270687617 | -4.32416188509569 | 1.53112835515695e-05 | 0.00254062669174459 | 4597 | 21 | 13 | 28 | 34 | 16 | 14 | 26 | 10 | 1 | 2 | 10 | 8 |
| L2b_dup20934 | 11.7510324522581 | -1.76100049455593 | 0.466569394829714 | -3.77435921445009 | 0.000160419470970396 | 0.0140232077294349 | 4598 | 20 | 13 | 19 | 22 | 16 | 9 | 28 | 9 | 3 | 2 | 9 | 5 |
| MIR_dup15778 | 12.3901200821402 | -1.4891393034457 | 0.44489827244875 | -3.34714561881613 | 0.000816483081811598 | 0.0419213975543016 | 4599 | 17 | 12 | 20 | 24 | 19 | 7 | 21 | 16 | 3 | 2 | 12 | 7 |
| MIR3_dup14533 | 12.254483700906 | -2.07458217564236 | 0.473505811510499 | -4.38132357663019 | 1.1796053675824e-05 | 0.00211357212947863 | 4600 | 19 | 18 | 12 | 32 | 13 | 14 | 20 | 16 | 2 | 5 | 3 | 6 |
| Charlie1a_dup696 | 11.4119195587325 | -1.48635667189841 | 0.420219514625444 | -3.53709578010258 | 0.000404552925636918 | 0.0264883875681778 | 4601 | 15 | 18 | 14 | 17 | 15 | 19 | 16 | 12 | 4 | 4 | 4 | 9 |
| MIR_dup44721 | 11.6842187236358 | -1.5109091063987 | 0.429454761486087 | -3.51820317737389 | 0.000434479568561473 | 0.0278026259823057 | 4602 | 20 | 12 | 18 | 20 | 10 | 18 | 16 | 15 | 5 | 6 | 3 | 7 |

| | | | | | | | | | | | | | | | | | | | |
|---|---|---|---|---|---|---|---|---|---|---|---|---|---|---|---|---|---|---|---|
| L1PA8A_dup2277 | 12.4103179172109 | -1.53659320563952 | 0.443583575059326 | -3.46404441470587 | 0.000532118644965057 | 0.031748158018036 | 4603 | 15 | 11 | 18 | 30 | 23 | 12 | 18 | 12 | 6 | 5 | 3 | 8 |
| MER1B_dup168 | 13.5940155557693 | -1.42678625896849 | 0.39841795983482 | -3.58112937368591 | 0.00034211218747266 | 0.0237252221408074 | 4604 | 15 | 16 | 15 | 31 | 16 | 14 | 22 | 20 | 3 | 5 | 6 | 13 |
| AluY_dup72309 | 13.4793788676156 | -2.22227938526873 | 0.449979678404914 | -4.93862165764076 | 7.86766885529607e-07 | 0.000267058535441124 | 4605 | 20 | 17 | 21 | 28 | 15 | 13 | 21 | 22 | 3 | 3 | 4 | 6 |
| L1MCb_dup2016 | 14.2825264568017 | -1.8199172207897 | 0.429649859108889 | -4.23581477383533 | 2.27724443267689e-05 | 0.00341222420514946 | 4606 | 18 | 19 | 24 | 38 | 18 | 10 | 20 | 17 | 4 | 4 | 4 | 10 |
| MLT1A_dup699 | 11.3553914722813 | -1.65590562625558 | 0.488710894814687 | -3.38831330306945 | 0.000703238857945981 | 0.0379186933823326 | 4607 | 14 | 7 | 28 | 25 | 14 | 12 | 14 | 14 | 1 | 5 | 3 | 11 |
| L1ME1_dup23408 | 12.4028140325104 | -1.55562604411626 | 0.436027476520659 | -3.56772480608238 | 0.000360094400360968 | 0.024535953954961 | 4608 | 17 | 8 | 23 | 20 | 19 | 21 | 20 | 11 | 3 | 6 | 8 | 5 |
| L1PA6_dup2642 | 14.199405210509 | -1.76769711277035 | 0.422614862917842 | -4.18276134579297 | 2.87989661548438e-05 | 0.00405959644590075 | 4609 | 21 | 14 | 17 | 22 | 16 | 27 | 27 | 18 | 1 | 6 | 5 | 11 |
| MER92B_dup458 | 14.2193361391659 | -1.96352068300739 | 0.455493591293734 | -4.31075369783014 | 1.62699047128261e-05 | 0.00265707532381477 | 4610 | 20 | 11 | 20 | 19 | 22 | 19 | 24 | 26 | 1 | 7 | 4 | 8 |
| AluSx1_dup65040 | 16.1335715663034 | -1.72601412692411 | 0.436130471457418 | -3.95756371059396 | 7.57180955051894e-05 | 0.00820811751848647 | 4611 | 23 | 19 | 21 | 18 | 28 | 22 | 22 | 24 | 11 | 6 | 1 | 7 |
| L1MC1_dup10581 | 15.1212590911241 | -1.76769439636569 | 0.404660242957923 | -4.36834215154044 | 1.25193252466497e-05 | 0.00220934659610395 | 4612 | 20 | 19 | 23 | 28 | 19 | 26 | 17 | 18 | 8 | 5 | 3 | 7 |
| AluJb_dup54542 | 16.103118392908 | -1.17202152960879 | 0.346990307926468 | -3.37767800090013 | 0.000731006245445063 | 0.0388866559532016 | 4613 | 20 | 21 | 20 | 26 | 21 | 16 | 25 | 21 | 8 | 11 | 6 | 10 |
| MER21B_dup2087 | 16.8648321994131 | -1.18718462313328 | 0.361062849260747 | -3.28802762611541 | 0.00100891938609286 | 0.04813882739989 | 4614 | 19 | 27 | 24 | 32 | 15 | 16 | 22 | 23 | 9 | 10 | 8 | 9 |
| L3_dup33361 | 15.1983916893101 | -1.54691884275032 | 0.416903960182374 | -3.71049208089466 | 0.000206856767778057 | 0.0168276035780231 | 4615 | 22 | 25 | 29 | 24 | 15 | 14 | 21 | 17 | 10 | 7 | 4 | 5 |
| MIR_dup42426 | 17.9240826379226 | -1.47605334204612 | 0.387747223929676 | -3.80674122457115 | 0.000140809945383928 | 0.0127638717197426 | 4616 | 28 | 26 | 31 | 30 | 18 | 13 | 30 | 21 | 12 | 6 | 5 | 9 |
| LTR79_dup717 | 16.8966838274727 | -1.42272009778471 | 0.359884777047658 | -3.95326556865257 | 7.70917923555441e-05 | 0.00832252156040541 | 4617 | 29 | 23 | 27 | 18 | 21 | 19 | 24 | 22 | 7 | 7 | 7 | 11 |
| Tigger19b_dup1456 | 17.3217807334213 | -1.62951062971797 | 0.362222239886156 | -4.49864875837033 | 6.83867362350215e-06 | 0.00138724214323772 | 4618 | 24 | 28 | 21 | 21 | 24 | 20 | 31 | 23 | 6 | 7 | 7 | 9 |
| AluSg_dup27348 | 16.6527498984224 | -1.54026062282466 | 0.409903875309359 | -3.7576142007982 | 0.000171541086000093 | 0.0147272393349269 | 4619 | 32 | 22 | 20 | 20 | 28 | 14 | 24 | 23 | 6 | 5 | 4 | 15 |
| AluSx_dup27928 | 16.8050079087827 | -1.4464800704631 | 0.429772811083258 | -3.36568538809421 | 0.000763537398773921 | 0.0401326860809939 | 4620 | 28 | 30 | 19 | 16 | 22 | 13 | 22 | 29 | 9 | 4 | 7 | 11 |
| LTR78_dup4183 | 15.1285752827176 | -1.3513640003007 | 0.372580671438129 | -3.62703732076212 | 0.0002866918691185 | 0.0209576600581476 | 4621 | 22 | 20 | 16 | 24 | 15 | 20 | 27 | 20 | 8 | 5 | 5 | 12 |
| AluSx_dup85793 | 17.0126580025773 | -1.40721109460525 | 0.371320640723468 | -3.78974648935079 | 0.000150801107387714 | 0.0133840060792528 | 4622 | 23 | 19 | 17 | 35 | 16 | 20 | 29 | 26 | 10 | 5 | 8 | 9 |
| Tigger1_dup10663 | 17.1318846371134 | -1.22346891465605 | 0.353779019633144 | -3.45828567201285 | 0.000543624620793487 | 0.0322657281165695 | 4623 | 21 | 21 | 20 | 25 | 18 | 18 | 35 | 24 | 11 | 5 | 9 | 11 |
| L1ME4c_dup5390 | 15.8645500376726 | -1.87731476954255 | 0.452062229212163 | -4.15277952509827 | 3.28461163806542e-05 | 0.00446197409716028 | 4624 | 25 | 19 | 14 | 21 | 20 | 12 | 38 | 30 | 5 | 5 | 4 | 9 |
| L1MEc_dup4259 | 16.5192441928782 | -1.71658377809808 | 0.410875050528703 | -4.17787299542581 | 2.94247873401259e-05 | 0.00412064592435346 | 4625 | 30 | 16 | 15 | 22 | 25 | 21 | 29 | 26 | 8 | 6 | 3 | 9 |
| AluSx_dup10429 | 12.5449628478155 | -2.07822766306721 | 0.502188480096618 | -4.13834196807417 | 3.49824700473531e-05 | 0.00467900997741497 | 4626 | 33 | 24 | 13 | 19 | 16 | 10 | 24 | 9 | 5 | 6 | 2 | 3 |
| AluJo_dup2397 | 11.0905194517709 | -1.95878873017596 | 0.49090091289846 | -3.99019166334516 | 6.60199240201676e-05 | 0.00742399661668374 | 4627 | 18 | 18 | 17 | 12 | 14 | 12 | 21 | 14 | 5 | 6 | 2 | 2 |
| AluSx1_dup72928 | 11.746983065554 | -1.8517142007626 | 0.478942310888772 | -3.86625728958125 | 0.000110518344202718 | 0.0107981242841968 | 4628 | 27 | 19 | 17 | 18 | 13 | 13 | 13 | 13 | 7 | 5 | 1 | 4 |
| L1MA4_dup3694 | 12.9448430007341 | -1.72270953881967 | 0.422222043043656 | -4.08010327078436 | 4.50156972667651e-05 | 0.00556720639388653 | 4629 | 19 | 26 | 12 | 18 | 25 | 21 | 13 | 11 | 6 | 7 | 3 | 4 |
| L1PREC2_dup2785 | 13.9573591675143 | -1.87742710871397 | 0.437285210082155 | -4.29336978573149 | 1.75981542481305e-05 | 0.00281980499038765 | 4630 | 29 | 22 | 14 | 22 | 25 | 18 | 15 | 13 | 8 | 8 | 3 | 1 |
| L1ME3G_dup12352 | 12.1164841246317 | -1.6136682611687 | 0.432795522611496 | -3.72847725279549 | 0.000192640293672908 | 0.0160048838401863 | 4631 | 21 | 17 | 12 | 13 | 16 | 24 | 17 | 14 | 5 | 7 | 6 | 2 |
| MIRb_dup133289 | 13.8508277746633 | -1.91959677030249 | 0.451441351484695 | -4.25215094715924 | 2.11727066116475e-05 | 0.00323153357755938 | 4632 | 36 | 23 | 18 | 10 | 20 | 17 | 20 | 14 | 6 | 6 | 4 | 3 |
| L1MA3_dup6014 | 14.7972575823203 | -1.91815441131771 | 0.552591403626824 | -3.47119842749685 | 0.00051814096590554 | 0.0311888616932354 | 4633 | 22 | 29 | 22 | 16 | 13 | 11 | 26 | 25 | 7 | 11 | 2 | 1 |
| L1PA15_dup6863 | 16.1792151401303 | -1.55888076609087 | 0.451420794374511 | -3.4532763787518 | 0.000553821267978013 | 0.0326187018391823 | 4634 | 33 | 34 | 21 | 21 | 13 | 11 | 23 | 21 | 9 | 10 | 4 | 5 |
| AluSx4_dup10970 | 13.0125737150995 | -2.19472632221828 | 0.477719859426896 | -4.59417015832504 | 4.34474753875309e-06 | 0.000988655508332668 | 4635 | 17 | 14 | 27 | 25 | 13 | 17 | 19 | 19 | 5 | 5 | 2 | 3 |
| L1P3_dup1255 | 14.5599729746731 | -1.47925534796648 | 0.440953078371909 | -3.35467744873945 | 0.00079457573437909 | 0.041271800413134 | 4636 | 18 | 21 | 17 | 24 | 16 | 21 | 21 | 19 | 11 | 9 | 5 | 1 |
| HAL1_dup20832 | 16.3702567656106 | -2.00344530191999 | 0.44798660470804 | -4.47210983735925 | 7.74516256916858e-06 | 0.00151723931685883 | 4637 | 20 | 17 | 44 | 42 | 20 | 14 | 27 | 10 | 5 | 4 | 6 | 7 |
| L1MC3_dup9646 | 17.1121185088103 | -1.52270982345992 | 0.405247109911444 | -3.75748472035412 | 0.000171629843223893 | 0.0147272393349269 | 4638 | 18 | 19 | 24 | 48 | 27 | 22 | 27 | 10 | 6 | 9 | 5 | 11 |
| LTR12C_dup2253 | 16.9049860195419 | -2.19093790902352 | 0.482321450462061 | -4.54248490695286 | 5.55949780852383e-06 | 0.00118323816238552 | 4639 | 18 | 25 | 38 | 45 | 24 | 22 | 12 | 15 | 5 | 6 | 3 | 6 |
| L1PA12_dup1461 | 13.4710101929762 | -1.90710788409478 | 0.529197296867552 | -3.60377480267457 | 0.000313628820052601 | 0.0223166038326065 | 4640 | 23 | 17 | 28 | 38 | 20 | 9 | 12 | 10 | 4 | 9 | 3 | 3 |
| THE1B-int_dup2231 | 14.5884438032914 | -1.84907380564636 | 0.523977490916623 | -3.5289183938258 | 0.000417261780110567 | 0.0270386088045177 | 4641 | 26 | 24 | 32 | 43 | 11 | 13 | 11 | 10 | 5 | 7 | 6 | 3 |
| L1PA4_dup9457 | 17.1977694006049 | -1.70195077147227 | 0.45730143622424 | -3.72172627648978 | 0.00019786544590146 | 0.0163080330413816 | 4642 | 38 | 39 | 25 | 34 | 19 | 15 | 24 | 6 | 3 | 8 | 8 | 9 |
| Tigger4_dup774 | 14.5450227270986 | -1.42378713452987 | 0.434469195881598 | -3.27707268553484 | 0.00104889363989448 | 0.0493098416567651 | 4643 | 37 | 25 | 21 | 24 | 16 | 19 | 8 | 11 | 5 | 8 | 8 | 6 |
| LTR62_dup211 | 18.2678403832365 | -1.27935239687404 | 0.350697471606157 | -3.64802287001028 | 0.000264266119607242 | 0.0198550887578354 | 4644 | 38 | 25 | 24 | 33 | 23 | 20 | 20 | 16 | 7 | 6 | 13 | 12 |
| MER63A_dup786 | 16.1587686222724 | -1.49409714402672 | 0.386733282412746 | -3.86337874698905 | 0.00011182941434008 | 0.0108831745703741 | 4645 | 29 | 25 | 22 | 33 | 20 | 15 | 21 | 15 | 5 | 4 | 13 | 8 |
| L1MEb_dup1538 | 17.6458137703437 | -1.57079195752482 | 0.377354115860225 | -4.16264694488361 | 3.14579569410151e-05 | 0.00433176795271226 | 4646 | 28 | 28 | 16 | 35 | 30 | 25 | 22 | 15 | 2 | 13 | 7 | 9 |
| L1MEc_dup16624 | 17.6121161072388 | -1.28950041342262 | 0.355928300756372 | -3.62292183757893 | 0.000291293865781813 | 0.0211769661433912 | 4647 | 20 | 20 | 19 | 32 | 26 | 26 | 24 | 22 | 6 | 15 | 7 | 8 |
| L2a_dup134056 | 17.301565159659 | -1.63729937254136 | 0.405713179815428 | -4.03560804528515 | 5.44610697699757e-05 | 0.0064279730691898 | 4648 | 34 | 29 | 18 | 36 | 33 | 21 | 15 | 10 | 6 | 13 | 4 | 6 |
| MLT1J2-int_dup503 | 15.6931344155776 | -1.67785106508195 | 0.430572136684295 | -3.89679433974192 | 9.74743246112232e-05 | 0.00987382956319374 | 4649 | 23 | 22 | 26 | 33 | 23 | 22 | 16 | 12 | 5 | 14 | 3 | 4 |
| L1PA4_dup10231 | 19.5277863301985 | -2.20384779486762 | 0.416520901375786 | -5.29108572364128 | 1.21592353250227e-07 | 5.99469084663238e-05 | 4650 | 41 | 26 | 28 | 42 | 34 | 27 | 19 | 14 | 8 | 8 | 2 | 5 |
| L2c_dup111474 | 20.741556134934 | -1.99539563923185 | 0.39809893067662 | -5.01231097466256 | 5.37801965707481e-07 | 0.000197176928400974 | 4651 | 35 | 39 | 34 | 40 | 29 | 32 | 19 | 13 | 6 | 13 | 5 | 4 |
| L1P1_dup620 | 16.6963065050661 | -2.45921749059349 | 0.522373300234485 | -4.70777792335403 | 2.50431712935193e-06 | 0.000666546133848742 | 4652 | 47 | 24 | 27 | 24 | 12 | 12 | 31 | 23 | 3 | 6 | 1 | 7 |
| Tigger2_dup1512 | 16.4047758392113 | -1.76949125580133 | 0.430659063819948 | -4.10879836152973 | 3.97723136624776e-05 | 0.00511272984696482 | 4653 | 19 | 26 | 24 | 22 | 12 | 14 | 38 | 29 | 2 | 8 | 5 | 11 |
| AluSx1_dup66943 | 16.731944253385 | -1.65505523427987 | 0.437980076468659 | -3.77883680834123 | 0.000157562635423926 | 0.0138651622960731 | 4654 | 30 | 20 | 34 | 29 | 13 | 11 | 31 | 21 | 5 | 12 | 4 | 6 |
| L1MC5a_dup8466 | 18.2054088791396 | -1.70044052515492 | 0.469092868498302 | -3.62495497021413 | 0.000289011804713238 | 0.0210626434775512 | 4655 | 33 | 32 | 36 | 38 | 10 | 14 | 29 | 17 | 2 | 13 | 4 | 11 |
| L2c_dup86835 | 15.0194832505651 | -1.70283135873536 | 0.489992396521176 | -3.47521996427912 | 0.000510434617665616 | 0.03095205798549 | 4656 | 28 | 19 | 19 | 17 | 20 | 23 | 42 | 7 | 2 | 11 | 2 | 10 |
| L1PA7_dup10835 | 11.632751883313 | -1.59799860072399 | 0.466084464667666 | -3.42856010415068 | 0.000606792160553442 | 0.0346803590223921 | 4657 | 34 | 14 | 14 | 15 | 12 | 19 | 16 | 8 | 4 | 9 | 2 | 5 |
| Tigger4b_dup1151 | 13.1454565615808 | -1.65199049446382 | 0.478597565466129 | -3.45173192190158 | 0.00055700083443406 | 0.032757320946675 | 4658 | 38 | 15 | 18 | 22 | 16 | 16 | 20 | 7 | 5 | 6 | 1 | 10 |
| REP522_dup208 | 14.2618079463292 | -2.83835231821706 | 0.549900522168419 | -5.1615741462194 | 2.44881860401125e-07 | 0.000105943016645466 | 4659 | 9 | 21 | 25 | 36 | 17 | 23 | 29 | 15 | 0 | 0 | 6 | 7 |
| L1PA4_dup11629 | 17.8151391544725 | -2.36773346694191 | 0.523742269058423 | -4.52079888682384 | 6.16066954902609e-06 | 0.00128475607056316 | 4660 | 19 | 7 | 24 | 50 | 16 | 33 | 41 | 23 | 0 | 1 | 9 | 14 |
| AluJo_dup76472 | 18.1895490882472 | -2.08846503481938 | 0.53289456254455 | -3.91909616200069 | 8.88816502571453e-05 | 0.00924750028512754 | 4661 | 12 | 7 | 30 | 52 | 26 | 21 | 36 | 24 | 0 | 0 | 16 | 16 |
| AluSg_dup35426 | 13.8379972080319 | -2.40331987192652 | 0.629446833551131 | -3.81814593993235 | 0.000134458381013701 | 0.0123719388742019 | 4662 | 8 | 5 | 21 | 26 | 23 | 24 | 39 | 17 | 0 | 0 | 11 | 8 |
| MIR3_dup82835 | 17.391235796332 | -1.47281453342162 | 0.444676995386481 | -3.31209967842288 | 0.000925985464700629 | 0.0454029130489933 | 4663 | 7 | 9 | 52 | 30 | 13 | 13 | 39 | 23 | 1 | 0 | 19 | 21 |
| MLT1A1_dup6595 | 18.8364058443141 | -1.93863082480977 | 0.437638664899977 | -4.42975216838495 | 9.43414415775954e-06 | 0.00177653577637507 | 4664 | 14 | 13 | 44 | 35 | 11 | 19 | 37 | 36 | 0 | 0 | 13 | 21 |
| Cheshire_dup1051 | 19.9095333599643 | -2.29376895006834 | 0.506126419512024 | -4.53200793643583 | 5.84256542699703e-06 | 0.00123248081434485 | 4665 | 13 | 18 | 42 | 59 | 26 | 6 | 38 | 30 | 0 | 1 | 17 | 10 |
| MIRb_dup213465 | 15.737043362142 | -1.58843358708662 | 0.476495456748628 | -3.33357551386809 | 0.000857373947938534 | 0.0432372594851866 | 4666 | 11 | 12 | 45 | 51 | 6 | 5 | 30 | 16 | 0 | 1 | 18 | 15 |
| CER_dup66 | 24.3982810634173 | -3.58186766384956 | 0.490990926835228 | -7.29518096584217 | 2.98257916465373e-13 | 7.97407620126625e-10 | 4667 | 24 | 20 | 44 | 54 | 35 | 37 | 43 | 44 | 2 | 3 | 2 | 5 |
| AluSx3_dup20954 | 20.5801798441557 | -2.4217154484165 | 0.570726924093876 | -4.24321220216021 | 2.20342899284762e-05 | 0.003330586794738 | 4668 | 15 | 22 | 31 | 28 | 48 | 54 | 24 | 20 | 1 | 1 | 7 | 15 |
| L1MA2_dup7369 | 23.2029922394232 | -2.83506828101744 | 0.647369838464463 | -4.37936417881642 | 1.19026077994729e-05 | 0.00212785718666671 | 4669 | 5 | 11 | 32 | 55 | 53 | 45 | 64 | 20 | 1 | 1 | 9 | 12 |
| L2_dup43585 | 26.0470811548413 | -1.82374005993725 | 0.506312528012606 | -3.60200460987181 | 0.000315772763448316 | 0.0224220910359772 | 4670 | 35 | 35 | 49 | 65 | 13 | 33 | 42 | 21 | 0 | 2 | 21 | 32 |
| L2a_dup165376 | 20.2245964235239 | -2.23705462188911 | 0.536956744860442 | -4.16617286830158 | 3.09755851824244e-05 | 0.0042777229203814 | 4671 | 26 | 19 | 61 | 64 | 14 | 23 | 25 | 10 | 0 | 2 | 13 | 15 |
| AluJr_dup83492 | 27.314506471586 | -2.30752261482255 | 0.387094201350104 | -5.96113970907958 | 2.50484689394067e-09 | 2.36515198026035e-06 | 4672 | 20 | 12 | 71 | 88 | 11 | 25 | 63 | 37 | 0 | 2 | 16 | 18 |
| AluJb_dup123575 | 29.2400216938139 | -1.72338223273578 | 0.396764211401517 | -4.34359295322568 | 1.40171127903423e-05 | 0.00240469785148579 | 4673 | 21 | 15 | 71 | 85 | 25 | 24 | 44 | 39 | 1 | 2 | 34 | 22 |
| L1PA16_dup13067 | 26.8317219999142 | -2.07231222096569 | 0.365190961917845 | -5.67459887310106 | 1.39014007817801e-08 | 1.04676668051312e-05 | 4674 | 17 | 17 | 63 | 75 | 26 | 15 | 69 | 33 | 1 | 2 | 22 | 15 |

| | | | | | | | | | | | | | | | | | | |
|---|---|---|---|---|---|---|---|---|---|---|---|---|---|---|---|---|---|---|
| L1MC3_dup11859 | 26.6558226600293 | -2.29893006166316 | 0.364369919432437 | -6.30932999421055 | 2.80246012390459e-10 | 3.5353899775032e-07 | 4675 | 36 | 34 | 50 | 75 | 25 | 20 | 47 | 32 | 1 | 6 | 13 | 12 |
| AluSg7_dup8350 | 20.4430399293767 | -2.19096620865819 | 0.441694600804691 | -4.96036447958982 | 7.03610344942443e-07 | 0.000244054325273578 | 4676 | 12 | 25 | 45 | 55 | 20 | 21 | 35 | 26 | 0 | 3 | 16 | 9 |
| L1MA5A_dup3135 | 18.9511528802459 | -2.17153505809385 | 0.397200618817014 | -5.46709887955701 | 4.57460892935237e-08 | 2.70773606045691e-05 | 4677 | 15 | 19 | 40 | 49 | 20 | 20 | 35 | 25 | 1 | 5 | 12 | 6 |
| L1PA16_dup12584 | 19.724641357186 | -1.47626563716549 | 0.417423652756649 | -3.53661232998247 | 0.000405294105910914 | 0.0265085517661018 | 4678 | 13 | 5 | 46 | 74 | 15 | 9 | 33 | 28 | 2 | 6 | 16 | 13 |
| LTR43_dup80 | 16.1666767922484 | -1.54078825540358 | 0.446066762060082 | -3.4541651305462 | 0.000551999270378009 | 0.0325639659183701 | 4679 | 12 | 9 | 44 | 46 | 10 | 9 | 36 | 20 | 1 | 10 | 7 | 9 |
| L1MC1_dup11598 | 27.8027961756159 | -1.54799533343004 | 0.403095900329425 | -3.84026563446803 | 0.000122901253697118 | 0.0116370321178728 | 4680 | 28 | 16 | 75 | 98 | 12 | 10 | 46 | 34 | 5 | 7 | 10 | 28 |
| L1PB1_dup11673 | 24.7137083343036 | -1.70084699954126 | 0.450412676688092 | -3.77619700237498 | 0.000159241061023583 | 0.0139593168184873 | 4681 | 19 | 13 | 64 | 97 | 19 | 4 | 46 | 29 | 3 | 8 | 11 | 16 |
| Arthur1_dup2110 | 27.2543230599907 | -1.32690267939135 | 0.357996624157738 | -3.70646701631103 | 0.00021017059310581 | 0.0170041659119875 | 4682 | 8 | 15 | 72 | 90 | 10 | 12 | 66 | 35 | 6 | 7 | 18 | 21 |
| L1M4_dup17743 | 24.9235266077092 | -1.75532350469896 | 0.422512065092261 | -4.15449320794109 | 3.26009300572339e-05 | 0.00443743276630177 | 4683 | 14 | 12 | 68 | 97 | 17 | 11 | 34 | 36 | 7 | 3 | 13 | 14 |
| L1MEc_dup16318 | 20.5985847026903 | -1.47018079575922 | 0.434267675614503 | -3.38542534550577 | 0.00071068037834073 | 0.0382068579540586 | 4684 | 9 | 14 | 56 | 86 | 8 | 11 | 34 | 19 | 4 | 4 | 12 | 19 |
| L1MC5_dup15126 | 27.7056995288594 | -1.98059454086524 | 0.42228289297793 | -4.69020785307719 | 2.72927657883371e-06 | 0.000710382446875096 | 4685 | 7 | 6 | 66 | 82 | 13 | 18 | 74 | 53 | 1 | 2 | 9 | 33 |
| L1PA7_dup6921 | 28.1802990982546 | -1.90246374493283 | 0.421187423144042 | -4.51690539743919 | 6.27499222030663e-06 | 0.00130175178627121 | 4686 | 4 | 5 | 66 | 111 | 14 | 14 | 64 | 51 | 2 | 3 | 12 | 26 |
| L1PA5_dup5827 | 31.7551823346335 | -1.56984793474739 | 0.409906646392155 | -3.82976940863146 | 0.000128263392367792 | 0.0119732291723604 | 4687 | 5 | 5 | 94 | 122 | 12 | 18 | 69 | 42 | 4 | 4 | 22 | 25 |
| L1PA6_dup3196 | 27.1983740366468 | -1.6886537431715 | 0.464210077082398 | -3.63769298974474 | 0.000275091054121984 | 0.0203791964242417 | 4688 | 6 | 5 | 83 | 119 | 12 | 10 | 38 | 42 | 3 | 3 | 11 | 28 |
| L1MA4A_dup5679 | 31.3692941412707 | -2.26131176019385 | 0.543649009286087 | -4.15950681702405 | 3.18935502464086e-05 | 0.00437907830751988 | 4689 | 8 | 5 | 69 | 99 | 23 | 22 | 88 | 66 | 6 | 7 | 4 | 9 |
| L1P2_dup1460 | 23.0782391488212 | -1.59016169590393 | 0.461218244757721 | -3.44774239522819 | 0.000565292911756624 | 0.0331044572401402 | 4690 | 8 | 6 | 47 | 66 | 27 | 23 | 62 | 28 | 8 | 7 | 9 | 12 |
| L1PA3_dup1626 | 28.3349747431205 | -2.40623881192864 | 0.457757492320414 | -5.25657985351851 | 1.4675906888548e-07 | 7.05469361717664e-05 | 4691 | 12 | 18 | 67 | 62 | 3 | 6 | 97 | 69 | 1 | 3 | 10 | 17 |
| L1PA8_dup4438 | 23.9904267170679 | -1.86185246962638 | 0.524548204708287 | -3.54944017139815 | 0.000386051179203331 | 0.0257309058506207 | 4692 | 14 | 8 | 61 | 64 | 4 | 4 | 71 | 56 | 5 | 4 | 4 | 14 |
| L1MA9_dup8577 | 26.6370855843567 | -1.60114267630074 | 0.464654343624998 | -3.44587906745783 | 0.000569205023846227 | 0.0332613110520909 | 4693 | 13 | 11 | 58 | 82 | 7 | 7 | 68 | 62 | 5 | 9 | 10 | 10 |
| MLT2C2_dup712 | 21.9475048356635 | -1.90378665380991 | 0.426392250046936 | -4.46487161434183 | 8.01167787209029e-06 | 0.00155366479458223 | 4694 | 16 | 13 | 55 | 63 | 8 | 4 | 61 | 38 | 3 | 6 | 10 | 10 |
| L1MEj_dup2046 | 20.5344711220069 | -2.78259336347036 | 0.698400340942901 | -3.98423826614062 | 6.76969238058866e-05 | 0.00755544663785905 | 4695 | 20 | 24 | 75 | 69 | 1 | 3 | 41 | 24 | 1 | 4 | 6 | 4 |
| L1MC5_dup13211 | 24.6812236895945 | -2.26302903935901 | 0.604264608147423 | -3.74509611988875 | 0.000180324723199603 | 0.0152837043217426 | 4696 | 30 | 25 | 60 | 70 | 4 | 2 | 57 | 46 | 6 | 1 | 10 | 9 |
| Tigger1_dup10677 | 19.9380728182471 | -2.80050434376243 | 0.679297050733827 | -4.12265052635973 | 3.74537705170686e-05 | 0.00490207639133906 | 4697 | 29 | 20 | 31 | 74 | 1 | 3 | 50 | 37 | 1 | 1 | 8 | 9 |
| L1PA3_dup5641 | 19.5504370493723 | -2.43366425882778 | 0.645341205841859 | -3.77112795029573 | 0.000162511315138693 | 0.0141588127334144 | 4698 | 6 | 6 | 77 | 93 | 5 | 3 | 32 | 24 | 3 | 2 | 2 | 8 |
| L1PA13_dup5751 | 21.883378524121 | -2.0428183066941 | 0.45147659805414 | -4.52474904679141 | 6.04672114460004e-06 | 0.00126320729541089 | 4699 | 10 | 4 | 68 | 83 | 14 | 10 | 49 | 24 | 3 | 1 | 10 | 17 |
| L1PA8_dup5175 | 16.5936631285241 | -2.08403567666709 | 0.580066301455875 | -3.59275426177403 | 0.000327201126823267 | 0.0229668017760511 | 4700 | 2 | 4 | 40 | 84 | 10 | 12 | 33 | 18 | 3 | 1 | 6 | 11 |
| L1PB_dup871 | 17.5626615718498 | -1.83404889998205 | 0.508156269032617 | -3.60922222503237 | 0.000307116436400303 | 0.0220155277341164 | 4701 | 16 | 13 | 32 | 46 | 1 | 9 | 43 | 41 | 4 | 4 | 6 | 10 |
| L1ME1_dup12948 | 14.1563089482404 | -2.69017127112023 | 0.565649624148659 | -4.75589685959594 | 1.97567325425328e-06 | 0.000560314121759894 | 4702 | 9 | 17 | 37 | 53 | 6 | 8 | 25 | 19 | 2 | 1 | 2 | 8 |
| L1PA8_dup4886 | 14.6811501221215 | -2.14627688556444 | 0.634813406843156 | -3.38095708507103 | 0.000722338235477328 | 0.0385030227999302 | 4703 | 21 | 20 | 41 | 56 | 2 | 6 | 17 | 14 | 2 | 3 | 3 | 11 |
| L1PREC2_dup3229 | 23.6582898849347 | -2.78890138008036 | 0.506019548796877 | -5.51144987720595 | 3.55889781552273e-08 | 2.22263910659415e-05 | 4704 | 10 | 20 | 70 | 95 | 7 | 7 | 57 | 30 | 0 | 3 | 5 | 13 |
| THE1B-int_dup2005 | 21.1195387960532 | -3.43706645762124 | 0.615907905610983 | -5.58048764483978 | 2.39845190067767e-08 | 1.61671964860807e-05 | 4705 | 7 | 17 | 68 | 94 | 7 | 7 | 33 | 34 | 0 | 2 | 2 | 8 |
| L1MB7_dup10650 | 17.9948127345959 | -2.12504609391986 | 0.509032376078808 | -4.17467767038626 | 2.9840825583874e-05 | 0.0041645191110736 | 4706 | 8 | 12 | 35 | 62 | 7 | 5 | 43 | 41 | 3 | 5 | 4 | 7 |
| MER41-int_dup657 | 18.8225047271898 | -2.99238195200242 | 0.519323357773443 | -5.76207849543302 | 8.30843387767483e-09 | 6.74270898513513e-06 | 4707 | 13 | 7 | 42 | 64 | 8 | 6 | 61 | 34 | 3 | 1 | 6 | 3 |
| LTR1A2_dup338 | 20.9334342845745 | -2.68889137789927 | 0.46312384920393 | -5.80598771261994 | 6.39876546235294e-09 | 5.41836529076524e-06 | 4708 | 20 | 26 | 43 | 61 | 11 | 9 | 52 | 34 | 2 | 5 | 4 | 7 |
| L1PA7_dup5842 | 23.4817309203332 | -3.5698927808352 | 0.501516609892428 | -7.11819451324039 | 1.09349980130454e-12 | 2.68241137856917e-09 | 4709 | 15 | 17 | 63 | 86 | 15 | 11 | 52 | 38 | 2 | 1 | 3 | 6 |
| L1MA4_dup4178 | 18.023111726733 | -2.95106368815131 | 0.51292722805059 | -5.75337694465353 | 8.74781312391343e-09 | 6.96156234643045e-06 | 4710 | 15 | 12 | 50 | 61 | 11 | 7 | 40 | 28 | 2 | 3 | 2 | 6 |
| L1P3_dup1465 | 19.7037559774048 | -3.0598453072928 | 0.560106514535517 | -5.46297039560459 | 4.68232710746535e-08 | 2.75096544669864e-05 | 4711 | 17 | 10 | 44 | 87 | 11 | 5 | 33 | 38 | 2 | 1 | 2 | 9 |
| MLT1D_dup15897 | 32.4778678118851 | -4.3510619902685 | 1.25963849886012 | -3.45421483561028 | 0.000551897536890243 | 0.0325639659183701 | 4712 | 90 | 72 | 114 | 113 | 0 | 4 | 17 | 14 | 0 | 1 | 8 | 4 |
| ALR/Alpha_dup313 | 16.5638338750883 | -2.89035782016874 | 0.767322530978486 | -3.76680952725703 | 0.000165347051969495 | 0.0143119933168183 | 4713 | 33 | 36 | 42 | 68 | 2 | 5 | 14 | 10 | 3 | 4 | 2 | 3 |
| L1MB3_dup12893 | 14.6480386694483 | -3.51311723130871 | 0.982437432384799 | -3.57591956037431 | 0.000348998960501999 | 0.024077386678924 | 4714 | 44 | 35 | 34 | 60 | 2 | 1 | 5 | 8 | 0 | 1 | 4 | 4 |
| L1PA4_dup9288 | 20.9570442300782 | -3.05857125645558 | 0.821709934804196 | -3.72220308761924 | 0.000197492074380628 | 0.0162893168114593 | 4715 | 42 | 59 | 53 | 86 | 7 | 5 | 12 | 4 | 0 | 4 | 6 | 6 |
| L1P2_dup606 | 15.4918770125424 | -2.52479281607848 | 0.762945766177889 | -3.30926905686479 | 0.000935398995302179 | 0.0457144465247499 | 4716 | 50 | 35 | 25 | 35 | 5 | 0 | 20 | 18 | 3 | 3 | 5 | 4 |
| L1MC2_dup4933 | 9.49216544316566 | -3.23019355998969 | 0.809795314314874 | -3.98890127281434 | 6.63800394575187e-05 | 0.00745392395882905 | 4717 | 30 | 18 | 18 | 31 | 4 | 7 | 9 | 5 | 1 | 1 | 1 | 3 |
| L1PB4_dup2567 | 8.75393199104613 | -2.33792819909147 | 0.680696221727284 | -3.4346131570393 | 0.000593400264794843 | 0.0341716407083431 | 4718 | 21 | 13 | 25 | 21 | 5 | 5 | 13 | 4 | 2 | 1 | 2 | 5 |
| L1MB7_dup17177 | 12.0566992989147 | -3.03344965320764 | 0.718689308226833 | -4.22080809952752 | 2.43428100713805e-05 | 0.00357989758049734 | 4719 | 22 | 22 | 34 | 40 | 11 | 10 | 5 | 7 | 1 | 1 | 3 | 4 |
| L1MCc_dup2748 | 14.4956071227519 | -2.49576097590855 | 0.692592909599912 | -3.60350350301776 | 0.000313956514085281 | 0.0223332426608479 | 4720 | 36 | 25 | 41 | 49 | 9 | 5 | 10 | 5 | 2 | 3 | 3 | 7 |
| L1M3_dup5262 | 10.0157675699472 | -3.26418586369403 | 0.897879844114662 | -3.63543728605761 | 0.000277509500877663 | 0.0205005512871271 | 4721 | 19 | 11 | 34 | 44 | 9 | 3 | 2 | 6 | 1 | 1 | 3 | 1 |
| MSTB1_dup3288 | 11.581247689436 | -3.20876587287824 | 0.914701347213311 | -3.50799294507866 | 0.000451501023116177 | 0.0285127491573631 | 4722 | 30 | 23 | 34 | 42 | 4 | 1 | 8 | 6 | 3 | 0 | 1 | 3 |
| AluJb_dup104236 | 13.6425606810759 | -3.43942776409784 | 0.951748852057946 | -3.61379764909707 | 0.000301744589361429 | 0.0217506567889108 | 4723 | 22 | 14 | 23 | 42 | 0 | 0 | 39 | 31 | 0 | 2 | 3 | 2 |
| L1PREC2_dup6836 | 12.9177509070136 | -3.38556919895119 | 0.780181474179302 | -4.33946371581378 | 1.42830846440446e-05 | 0.00243802213680907 | 4724 | 13 | 12 | 23 | 45 | 0 | 3 | 37 | 29 | 0 | 2 | 2 | 3 |
| L1MA4_dup4714 | 14.6951973552988 | -2.41531351439071 | 0.687476158971653 | -3.51330512756051 | 0.000442568921789996 | 0.0281558906032628 | 4725 | 9 | 12 | 20 | 30 | 0 | 2 | 52 | 45 | 1 | 1 | 7 | 7 |
| THE1B_dup20031 | 13.3090873775604 | -2.7623802463715 | 0.794147557316206 | -3.47842188888281 | 0.000504375411153364 | 0.030686298026668 | 4726 | 17 | 10 | 21 | 38 | 0 | 1 | 41 | 33 | 1 | 1 | 7 | 2 |
| MER57C2_dup385 | 12.666748371329 | -2.02106816175733 | 0.58345173993397 | -3.46398514808792 | 0.000532235895261721 | 0.031748158018036 | 4727 | 8 | 10 | 32 | 42 | 1 | 4 | 20 | 29 | 1 | 3 | 8 | 4 |
| MER58B_dup6660 | 10.5928583567798 | -2.54421724910884 | 0.756296084753779 | -3.36404921352613 | 0.000768078456836805 | 0.0403022910579388 | 4728 | 13 | 17 | 25 | 37 | 0 | 2 | 23 | 13 | 1 | 1 | 4 | 5 |
| L1M4b_dup1637 | 15.1384046656024 | -3.53554694849233 | 0.641767379312487 | -5.50907862016904 | 3.60716782448355e-08 | 2.24688784074493e-05 | 4729 | 19 | 22 | 34 | 35 | 5 | 9 | 50 | 19 | 1 | 1 | 2 | 4 |
| L2_dup24556 | 15.9736005783124 | -3.99498409047396 | 0.730068282543001 | -5.47206910093188 | 4.44811313988808e-08 | 2.65265846913236e-05 | 4730 | 26 | 22 | 37 | 48 | 7 | 4 | 41 | 21 | 1 | 1 | 2 | 2 |
| MSTC_dup1472 | 17.4345676885385 | -4.00100698611601 | 0.721151170407398 | -5.54808360618167 | 2.88817768695891e-08 | 1.87767849153312e-05 | 4731 | 29 | 24 | 21 | 30 | 9 | 7 | 53 | 43 | 1 | 0 | 2 | 4 |
| THE1A-int_dup677 | 15.2612799484268 | -3.51103123495033 | 0.754776717469844 | -4.65174819742715 | 3.29132822734916e-06 | 0.00080996931844402 | 4732 | 28 | 23 | 29 | 35 | 4 | 2 | 34 | 33 | 2 | 0 | 3 | 3 |
| L1M3_dup5821 | 14.4597836668923 | -3.78741632311088 | 0.914451620006323 | -4.14173504672089 | 3.44688390051269e-05 | 0.00462851149317941 | 4733 | 22 | 26 | 23 | 38 | 1 | 1 | 39 | 30 | 0 | 0 | 5 | 3 |
| L1MA5A_dup3053 | 17.4869970453854 | -3.09038071904946 | 0.690618905598998 | -4.47479889993609 | 7.64832639856994e-06 | 0.001506414574756 | 4734 | 23 | 21 | 58 | 48 | 2 | 5 | 32 | 25 | 0 | 0 | 6 | 10 |
| L1M4b_dup1638 | 13.2715333756146 | -3.19621458361524 | 0.690384248178794 | -4.62961690109056 | 3.66342863395398e-06 | 0.000870081923447894 | 4735 | 18 | 19 | 21 | 47 | 7 | 3 | 33 | 20 | 0 | 4 | 2 | 2 |
| L1MD3_dup1979 | 14.8799776413106 | -4.1340215962637 | 0.740259366453295 | -5.58455830970498 | 2.34294900656962e-08 | 1.59284384090633e-05 | 4736 | 14 | 12 | 37 | 57 | 5 | 4 | 46 | 20 | 0 | 2 | 1 | 2 |
| L1PA3_dup4639 | 13.7493443639576 | -3.11174359862123 | 0.656099511254587 | -4.74279213022272 | 2.10792536978936e-06 | 0.00058458322848473 | 4737 | 16 | 13 | 34 | 39 | 3 | 5 | 36 | 24 | 0 | 1 | 8 | 1 |
| L1M3_dup5824 | 10.8686057907145 | -3.18954418970733 | 0.785970414590885 | -4.05809701039136 | 4.94742170373913e-05 | 0.00598325375404711 | 4738 | 11 | 21 | 23 | 22 | 2 | 2 | 36 | 18 | 0 | 2 | 3 | 2 |
| L1ME3E_dup1977 | 13.2565275930814 | -3.2938062283052 | 0.770773218753351 | -4.27337918361072 | 1.92532626972893e-05 | 0.00302391871007867 | 4739 | 15 | 22 | 21 | 33 | 2 | 2 | 41 | 28 | 0 | 2 | 4 | 2 |
| LTR8_dup1550 | 15.8774972632524 | -4.15489829225963 | 0.874293723367104 | -4.7522911136296 | 2.01124511032239e-06 | 0.000566352342036417 | 4740 | 19 | 15 | 56 | 67 | 4 | 2 | 19 | 23 | 0 | 2 | 0 | 3 |
| MER57-int_dup491 | 16.0169326260735 | -3.8568025867294 | 0.673516245730745 | -5.72636905371879 | 1.02602890119422e-08 | 7.952425828129e-06 | 4741 | 9 | 11 | 48 | 62 | 10 | 5 | 24 | 32 | 0 | 1 | 2 | 4 |
| L1MB5_dup2904 | 13.4311205471823 | -3.34987169259719 | 0.643200404413652 | -5.20813057580548 | 1.90752675569709e-07 | 8.74544048961656e-05 | 4742 | 14 | 14 | 26 | 41 | 9 | 3 | 37 | 24 | 0 | 0 | 2 | 7 |
| L2a_dup73367 | 18.0370169627912 | -4.11368971409412 | 0.673929481792422 | -6.10403584534263 | 1.03423075528803e-09 | 1.12885812521911e-06 | 4743 | 24 | 16 | 46 | 53 | 6 | 7 | 46 | 32 | 0 | 0 | 3 | 4 |
| L1PA3_dup2913 | 38.2372271641099 | -2.49537941586537 | 0.74019969350405 | -3.37122460028648 | 0.000748348194704243 | 0.0395615328898235 | 4744 | 22 | 19 | 172 | 233 | 5 | 6 | 31 | 12 | 6 | 2 | 4 | 14 |
| L1PA4_dup9346 | 35.4489142356542 | -2.47592867622486 | 0.748007656611267 | -3.31003119331916 | 0.000932855745973749 | 0.0456445184724387 | 4745 | 78 | 70 | 101 | 140 | 8 | 6 | 12 | 16 | 2 | 3 | 18 | 24 |
| L1PA15_dup6367 | 42.3999402003134 | -2.22349134914377 | 0.657082771029987 | -3.38388319885242 | 0.000714683994801707 | 0.0382981050831384 | 4746 | 92 | 90 | 108 | 163 | 7 | 11 | 31 | 17 | 9 | 7 | 12 | 24 |

| | | | | | | | | | | | | | | | | | | | |
|---|---|---|---|---|---|---|---|---|---|---|---|---|---|---|---|---|---|---|---|
| ERVL47-int_dup117 | 47.6959306072115 | -2.55769166521799 | 0.644717003959211 | -3.96715403737018 | 7.27359849175129e-05 | 0.00795763824028531 | 4747 | 89 | 75 | 151 | 213 | 13 | 11 | 32 | 16 | 5 | 8 | 18 | 16 |
| L1PA13_dup2478 | 44.0622076961081 | -2.30106762625419 | 0.577009988649252 | -3.98791645122273 | 6.66561282160624e-05 | 0.00747786849811371 | 4748 | 66 | 66 | 134 | 191 | 13 | 13 | 45 | 20 | 9 | 8 | 11 | 19 |
| L1PA15_dup462 | 26.5649098645704 | -4.89473659089263 | 0.726055772390629 | -6.74154352464702 | 1.56712774422334e-11 | 2.80369758065388e-08 | 4749 | 32 | 30 | 84 | 88 | 9 | 20 | 39 | 40 | 0 | 0 | 4 | 2 |
| MER41-int_dup656 | 23.4972762050275 | -5.07480647874001 | 0.778470829157778 | -6.51894237865073 | 7.08048448272877e-11 | 1.04644283274993e-07 | 4750 | 20 | 21 | 50 | 81 | 10 | 5 | 62 | 52 | 2 | 0 | 2 | 0 |
| L1M4a2_dup804 | 24.4430761937064 | -4.12071807769158 | 0.625252157046426 | -6.59048998272486 | 4.38377378042388e-11 | 6.90795652951483e-08 | 4751 | 25 | 16 | 65 | 105 | 6 | 9 | 54 | 37 | 1 | 0 | 5 | 3 |
| L1MA3_dup3847 | 21.7746381156597 | -3.3794489442077 | 0.611354148504626 | -5.52780896714913 | 3.24254845868152e-08 | 2.06850250817542e-05 | 4752 | 20 | 28 | 40 | 81 | 7 | 4 | 47 | 44 | 1 | 1 | 7 | 4 |
| L1ME2_dup5464 | 26.1238247663036 | -3.86874642722521 | 0.612906069188647 | -6.31213594009043 | 2.75210311984158e-10 | 3.50188197301511e-07 | 4753 | 32 | 17 | 63 | 83 | 7 | 4 | 66 | 57 | 1 | 1 | 4 | 5 |
| HERVIP10B3-int_dup2 | 48.5597657202074 | -5.02763048839808 | 0.798672057108425 | -6.29498734011116 | 3.07424949030924e-10 | 3.77064623309857e-07 | 4754 | 97 | 60 | 176 | 151 | 8 | 8 | 59 | 67 | 0 | 0 | 8 | 4 |
| REP522_dup224 | 30.5723027114488 | -2.59521702567526 | 0.465102258772075 | -5.57988480324077 | 2.40677928912426e-08 | 1.61775001336147e-05 | 4755 | 27 | 25 | 92 | 96 | 10 | 23 | 62 | 36 | 2 | 0 | 8 | 26 |
| L1PREC2_dup3226 | 34.2249201238605 | -3.44725747205186 | 0.489540426319778 | -7.04182389586768 | 1.89739544882588e-12 | 4.47007581649826e-09 | 4756 | 45 | 19 | 74 | 112 | 13 | 11 | 99 | 60 | 1 | 1 | 5 | 14 |
| L1MA3_dup3848 | 30.6615519808897 | -3.23519851937865 | 0.477863682785004 | -6.77012846116243 | 1.28668098606528e-11 | 2.3507949435513e-08 | 4757 | 20 | 22 | 77 | 144 | 9 | 13 | 71 | 37 | 1 | 1 | 9 | 10 |
| L1PREC2_dup3228 | 34.9274690331052 | -3.32845040025915 | 0.509508269726048 | -6.53267198596951 | 6.46065389014967e-11 | 9.66847012921732e-08 | 4758 | 19 | 30 | 86 | 151 | 12 | 11 | 97 | 44 | 2 | 2 | 2 | 15 |
| L1PA7_dup10205 | 191.972643557466 | -1.97524578717939 | 0.483764022443812 | -4.083077069686 | 4.44432755175417e-05 | 0.00551936306696085 | 4759 | 597 | 514 | 59 | 88 | 422 | 432 | 15 | 5 | 147 | 152 | 6 | 5 |
| L1PA13_dup6826 | 61.2571452597168 | -1.87248794982251 | 0.426871748381026 | -4.38653519921193 | 1.15170534397964e-05 | 0.00207766246989067 | 4760 | 196 | 138 | 23 | 22 | 153 | 132 | 4 | 7 | 53 | 45 | 1 | 1 |
| L1PA13_dup6827 | 65.2183400413548 | -1.72171073575698 | 0.420514713440588 | -4.09429368516076 | 4.23457287342403e-05 | 0.00534535637633822 | 4761 | 204 | 168 | 14 | 31 | 147 | 125 | 6 | 8 | 54 | 66 | 2 | 0 |
| MER57A-int_dup1410 | 87.9685992925309 | -4.81519041995943 | 0.592460102181708 | -8.12745094940184 | 4.38411938384832e-16 | 2.54435041685164e-12 | 4762 | 158 | 137 | 133 | 132 | 47 | 61 | 214 | 214 | 8 | 7 | 0 | 2 |
| CER_dup61 | 100.153688245821 | -3.80150669993151 | 0.30438619177611 | -12.4890905127776 | 8.56268694011012e-36 | 2.03745710665144e-30 | 4763 | 106 | 98 | 177 | 191 | 150 | 147 | 181 | 186 | 6 | 6 | 12 | 20 |
| L1MA7_dup1311 | 100.856035447002 | -4.14504207006572 | 0.575287770976787 | -7.20516284055162 | 5.79745239714765e-13 | 1.45208485062284e-09 | 4764 | 23 | 23 | 242 | 316 | 169 | 182 | 204 | 132 | 3 | 10 | 5 | 15 |
| L1MCb_dup1043 | 83.5539784320716 | -2.2284750464492 | 0.454799609309278 | -4.89990536674751 | 9.58828273798426e-07 | 0.000307409615345431 | 4765 | 173 | 168 | 52 | 61 | 58 | 56 | 224 | 170 | 20 | 23 | 24 | 30 |
| L1PA12_dup268 | 92.9883334555886 | -2.54490852932181 | 0.545437782017479 | -4.66580903124942 | 3.07405104958744e-06 | 0.000770575840329701 | 4766 | 229 | 213 | 64 | 72 | 40 | 32 | 287 | 169 | 19 | 14 | 18 | 40 |
| AluSx1_dup35690 | 44.4282247791636 | -1.47271455985808 | 0.347070058298684 | -4.24327747278817 | 2.20278793937536e-05 | 0.003330586794738 | 4767 | 55 | 51 | 50 | 54 | 29 | 35 | 108 | 96 | 13 | 14 | 33 | 22 |
| Tigger1_dup10931 | 48.6574957678871 | -1.50282507650417 | 0.302569596309254 | -4.9668740509147 | 6.80407464231359e-07 | 0.000238439226044175 | 4768 | 75 | 60 | 68 | 72 | 31 | 48 | 100 | 80 | 23 | 17 | 24 | 21 |
| L1MA4A_dup2821 | 47.0184362681621 | -1.50602688731391 | 0.390672205828709 | -3.85496297111607 | 0.000115747147027199 | 0.0111504334601351 | 4769 | 77 | 57 | 43 | 60 | 46 | 28 | 137 | 74 | 23 | 20 | 11 | 28 |
| L1MA8_dup4592 | 57.4396947786575 | -1.70799217565277 | 0.490912822051209 | -3.47921687707436 | 0.000502881430628333 | 0.0306345685848155 | 4770 | 111 | 94 | 40 | 59 | 31 | 18 | 178 | 113 | 17 | 22 | 19 | 33 |
| L1PA4_dup5556 | 53.5782357409116 | -2.01553192477887 | 0.434791603410996 | -4.63562752584633 | 3.55856252169657e-06 | 0.000852714720833447 | 4771 | 122 | 103 | 38 | 57 | 46 | 27 | 129 | 92 | 18 | 16 | 17 | 19 |
| L1M4_dup8158 | 43.2151982090751 | -2.40088132400699 | 0.501052440367649 | -4.79167673995427 | 1.65393232563082e-06 | 0.000482017898780537 | 4772 | 103 | 92 | 37 | 38 | 22 | 26 | 106 | 82 | 8 | 12 | 7 | 19 |
| L1MB7_dup9096 | 58.736225949027 | -2.56892932384245 | 0.469147182374289 | -5.47574283797562 | 4.35680026366213e-08 | 2.60473164707876e-05 | 4773 | 121 | 107 | 44 | 63 | 26 | 42 | 164 | 126 | 10 | 11 | 14 | 21 |
| HERVK9-int_dup471 | 122.503761769489 | -2.05556855766219 | 0.29486971606286 | -6.97110773228398 | 3.14455044096587e-12 | 6.99283363762677e-09 | 4774 | 132 | 135 | 160 | 177 | 173 | 198 | 221 | 199 | 47 | 51 | 19 | 39 |
| L1PA7_dup9221 | 96.5236466467754 | -1.39484426753416 | 0.296137907778893 | -4.71011724907438 | 2.47574324738313e-06 | 0.000662647022206778 | 4775 | 143 | 119 | 125 | 136 | 104 | 100 | 183 | 133 | 63 | 56 | 23 | 39 |
| L1PA5_dup4270 | 111.270276147156 | -1.24429541973578 | 0.237804650345768 | -5.23242677519793 | 1.67298935093698e-07 | 7.86386667291629e-05 | 4776 | 185 | 199 | 79 | 79 | 191 | 216 | 141 | 91 | 77 | 72 | 22 | 57 |
| L1MA4_dup3689 | 81.9159091460581 | -1.45362059360907 | 0.262205730017553 | -5.54381703829185 | 2.95948074416814e-08 | 1.90323406797792e-05 | 4777 | 153 | 120 | 53 | 81 | 149 | 159 | 81 | 91 | 39 | 57 | 14 | 41 |
| HERV3-int_dup89 | 82.3073723905069 | -1.39007043858139 | 0.22047795848236 | -6.30480456255044 | 2.88557775178383e-10 | 3.59482556924584e-07 | 4778 | 141 | 103 | 64 | 103 | 147 | 130 | 113 | 89 | 47 | 49 | 31 | 29 |
| L2_dup9153 | 78.2850166031951 | -1.09931716249657 | 0.231050531676764 | -4.75790795424136 | 1.95609652125475e-06 | 0.000556087625861988 | 4779 | 179 | 123 | 64 | 96 | 116 | 83 | 102 | 68 | 38 | 57 | 27 | 55 |
| L1M3d_dup361 | 131.180828583216 | -2.03802066114256 | 0.405976870004049 | -5.02004131694063 | 5.16603666678541e-07 | 0.000190875428682441 | 4780 | 461 | 415 | 70 | 91 | 121 | 103 | 156 | 116 | 47 | 42 | 24 | 44 |
| L1MA2_dup6801 | 99.2809358315157 | -1.17341362753291 | 0.208784005262273 | -5.6202275938661 | 1.90706063410247e-08 | 1.35455955117059e-05 | 4781 | 164 | 175 | 89 | 138 | 123 | 106 | 159 | 105 | 38 | 53 | 55 | 73 |
| L2b_dup16138 | 95.891721792761 | -1.47821192931666 | 0.262625426649197 | -5.6285941090205 | 1.81684367084688e-08 | 1.3100323760707e-05 | 4782 | 224 | 188 | 89 | 135 | 93 | 108 | 146 | 87 | 37 | 37 | 38 | 63 |
| Harlequin-int_dup373 | 92.7939349702142 | -1.22756727499146 | 0.301719022258299 | -4.06857766475376 | 4.73009950099623e-05 | 0.00579263127053036 | 4783 | 247 | 230 | 86 | 98 | 85 | 120 | 98 | 53 | 47 | 39 | 39 | 59 |
| L1PB4_dup2933 | 102.937469546951 | -1.17543967559419 | 0.206424203573081 | -5.6942919253073 | 1.23884963389307e-08 | 9.44805496751028e-06 | 4784 | 244 | 182 | 104 | 125 | 116 | 150 | 118 | 73 | 50 | 54 | 51 | 63 |
| L2a_dup134925 | 65.7946667998922 | -1.09588990476859 | 0.232341701192487 | -4.71671636707473 | 2.39681456400523e-06 | 0.000647346694945277 | 4785 | 97 | 91 | 115 | 101 | 67 | 58 | 119 | 54 | 31 | 31 | 31 | 60 |
| L1M5_dup45846 | 57.2552931015139 | -1.17953567748138 | 0.270349854439941 | -4.3629972722749 | 1.28292543710895e-05 | 0.00224957241015716 | 4786 | 97 | 91 | 95 | 79 | 55 | 63 | 82 | 51 | 19 | 25 | 24 | 63 |
| L1PA4_dup10376 | 59.006884907156 | -1.24636261617457 | 0.214022859089511 | -5.82350231875607 | 5.76270145576355e-09 | 4.95022296243001e-06 | 4787 | 102 | 79 | 65 | 80 | 79 | 73 | 103 | 58 | 17 | 29 | 32 | 49 |
| L1PA5_dup266 | 55.8241797360953 | -1.05213183489762 | 0.257113130146103 | -4.09209686918653 | 4.27489992582859e-05 | 0.00538198591402757 | 4788 | 99 | 85 | 62 | 95 | 59 | 50 | 104 | 43 | 18 | 36 | 29 | 52 |
| Harlequin-int_dup371 | 74.9429028106344 | -1.1943227766318 | 0.286168306432345 | -4.17349772768831 | 2.9995865401936e-05 | 0.00417391589995852 | 4789 | 177 | 152 | 71 | 85 | 81 | 106 | 82 | 58 | 31 | 27 | 35 | 65 |
| L1P2_dup1489 | 74.0102492952069 | -1.10869210142968 | 0.243412900859247 | -4.55477954338491 | 5.24405456906755e-06 | 0.00113648705186451 | 4790 | 146 | 119 | 84 | 98 | 66 | 77 | 127 | 70 | 43 | 39 | 31 | 54 |
| L2_dup41470 | 76.3927008452725 | -1.16075552259778 | 0.346842179762864 | -3.34663887590428 | 0.00081797694976254 | 0.041947056743146 | 4791 | 209 | 174 | 59 | 66 | 67 | 66 | 121 | 61 | 43 | 38 | 27 | 55 |
| AluSg4_dup373 | 72.0047124002275 | -1.5608994807389 | 0.36849772539325 | -4.23584563262407 | 2.2769316755447e-05 | 0.00341222420514946 | 4792 | 184 | 134 | 52 | 68 | 64 | 74 | 108 | 108 | 25 | 18 | 35 | 49 |
| L1MC3_dup12898 | 72.7673270805569 | -1.20806924500065 | 0.281355045944582 | -4.29375361278789 | 1.7567742770893e-05 | 0.00281910313205495 | 4793 | 136 | 104 | 83 | 61 | 74 | 87 | 100 | 115 | 37 | 37 | 32 | 49 |
| MER70-int_dup184 | 75.906697694281 | -1.20058769262259 | 0.255670278070191 | -4.69584380978763 | 2.65508369674116e-06 | 0.000700406369517485 | 4794 | 157 | 118 | 83 | 86 | 88 | 84 | 87 | 99 | 34 | 29 | 48 | 52 |
| L2_dup19568 | 82.1402822435663 | -1.00631423722524 | 0.261235576050513 | -3.85213320650727 | 0.00011709329295603 | 0.0112346293087562 | 4795 | 259 | 190 | 47 | 69 | 117 | 115 | 48 | 34 | 49 | 64 | 26 | 38 |
| L2b_dup71189 | 53.0247097130348 | -1.24058699062617 | 0.222612180517338 | -5.57286213064854 | 2.50587880640162e-08 | 1.66554144823475e-05 | 4796 | 121 | 93 | 43 | 45 | 87 | 112 | 48 | 24 | 34 | 39 | 9 | 25 |
| Charlie1a_dup4571 | 91.231380562795 | -1.82159942232157 | 0.336443782419674 | -5.41427577950998 | 6.15372608502474e-08 | 3.40783262835684e-05 | 4797 | 148 | 122 | 149 | 210 | 138 | 108 | 86 | 76 | 42 | 61 | 18 | 19 |
| L1MB2_dup6624 | 69.1655063561275 | -1.48443929163321 | 0.405334239696809 | -3.66225980006914 | 0.000250000127702275 | 0.0191398102915848 | 4798 | 101 | 108 | 96 | 164 | 95 | 75 | 68 | 46 | 58 | 50 | 12 | 13 |
| ERV3-16A3_I-int_dup | 75.9059228027096 | -1.52667521473402 | 0.405418684176822 | -3.76567552093421 | 0.000166099389688529 | 0.0143396912809431 | 4799 | 103 | 106 | 151 | 183 | 83 | 79 | 83 | 54 | 46 | 60 | 11 | 21 |
| MLT1D_dup15888 | 77.3595700426806 | -1.47125442427455 | 0.348841593389556 | -4.21754301136786 | 2.46978720097196e-05 | 0.00362539164295172 | 4800 | 149 | 104 | 124 | 158 | 103 | 79 | 71 | 59 | 57 | 50 | 15 | 23 |
| MER41A_dup1904 | 70.081130719045 | -1.42050022637443 | 0.346176125156703 | -4.10340321918912 | 4.07116800314479e-05 | 0.0051858209563448 | 4801 | 106 | 108 | 105 | 167 | 88 | 80 | 72 | 44 | 47 | 51 | 16 | 21 |
| MER76_dup608 | 72.8195535391339 | -1.38752049426752 | 0.378543850956506 | -3.66541548822289 | 0.000246937349150262 | 0.018966350703973 | 4802 | 111 | 114 | 121 | 164 | 97 | 62 | 70 | 51 | 49 | 61 | 16 | 19 |
| HAL1b_dup4214 | 44.502100107739 | -1.43230140582475 | 0.388879425554353 | -3.68315038468025 | 0.000230369161664039 | 0.0180610940828045 | 4803 | 75 | 80 | 74 | 102 | 53 | 39 | 36 | 30 | 20 | 41 | 13 | 12 |
| L1ME5_dup2540 | 49.5685240159692 | -1.51101449455487 | 0.402568719054583 | -3.75343245273362 | 0.000174429550660651 | 0.0149083383123202 | 4804 | 64 | 79 | 80 | 131 | 59 | 43 | 55 | 39 | 28 | 40 | 11 | 11 |
| HERVL74-int_dup252 | 49.9264982964393 | -1.71384392859651 | 0.386878714072027 | -4.42992562335036 | 9.42655936243423e-06 | 0.00177653577637507 | 4805 | 91 | 62 | 97 | 105 | 83 | 58 | 39 | 29 | 26 | 33 | 13 | 9 |
| L1MC4a_dup7863 | 52.8713113345977 | -1.39865971803839 | 0.411122607154027 | -3.40205012738299 | 0.000668823690265657 | 0.0367334858133218 | 4806 | 87 | 83 | 78 | 132 | 66 | 52 | 42 | 36 | 39 | 39 | 17 | 8 |
| ERVL-E-int_dup6962 | 60.3656541752655 | -1.46764344750375 | 0.429807231712617 | -3.41465508073411 | 0.000638628267960702 | 0.0357214484833515 | 4807 | 69 | 76 | 114 | 175 | 58 | 51 | 77 | 54 | 37 | 43 | 16 | 10 |
| L1M1_dup8058 | 66.4212579934895 | -1.49420468835646 | 0.336425808300292 | -4.4414092245347 | 8.93716476936076e-06 | 0.00169853243467278 | 4808 | 126 | 101 | 122 | 127 | 83 | 72 | 54 | 47 | 40 | 44 | 14 | 24 |
| L1MB2_dup6601 | 61.9698776204548 | -1.55070115012654 | 0.374481150049007 | -4.14093246061545 | 3.45896806388222e-05 | 0.00463950177524531 | 4809 | 135 | 64 | 113 | 130 | 80 | 67 | 56 | 45 | 38 | 42 | 10 | 21 |
| L1MA9_dup13336 | 57.7880265051001 | -1.80678711846345 | 0.343100486616871 | -5.26605816354039 | 1.39384140021667e-07 | 6.76855073093789e-05 | 4810 | 79 | 90 | 93 | 131 | 83 | 85 | 64 | 36 | 28 | 35 | 7 | 19 |
| L1P4_dup3003 | 56.5214535272673 | -1.47820687068884 | 0.374246180947868 | -3.94982486379667 | 7.82083971947228e-05 | 0.00839394464541971 | 4811 | 112 | 84 | 78 | 108 | 77 | 63 | 58 | 35 | 48 | 40 | 10 | 11 |
| L1M5_dup42772 | 56.6827666741459 | -1.45725849343653 | 0.320364403630361 | -4.54875284807836 | 5.39647765509592e-06 | 0.00115821603421951 | 4812 | 87 | 89 | 70 | 122 | 88 | 69 | 56 | 40 | 35 | 44 | 11 | 18 |
| L1MA9_dup13338 | 53.8585867778555 | -1.47087588771324 | 0.33517782013882 | -4.38834493017481 | 1.14216528529344e-05 | 0.00206787341108031 | 4813 | 91 | 84 | 76 | 116 | 90 | 60 | 38 | 37 | 29 | 41 | 13 | 18 |
| L1PA5_dup9511 | 137.912259597435 | -1.2845623507938 | 0.293379883620402 | -4.37849499066496 | 1.19501687454186e-05 | 0.00213475589511815 | 4814 | 222 | 226 | 203 | 276 | 184 | 159 | 129 | 87 | 101 | 92 | 34 | 58 |
| HSMAR2_dup1480 | 142.810984118364 | -1.43371651290369 | 0.348975159397509 | -4.10836265646801 | 3.98474055629462e-05 | 0.00511961704324018 | 4815 | 217 | 171 | 240 | 330 | 198 | 137 | 132 | 132 | 90 | 108 | 37 | 36 |
| L1MEc_dup14790 | 142.925801758685 | -1.69510373964543 | 0.372754606502689 | -4.5475058123345 | 5.42854215861442e-06 | 0.00116160062272812 | 4816 | 240 | 214 | 227 | 365 | 199 | 150 | 115 | 100 | 67 | 114 | 22 | 40 |
| Tigger1_dup9878 | 223.67017456967 | -1.46296296493513 | 0.373605650660433 | -3.91579453455537 | 9.01069639980138e-05 | 0.00932605117680356 | 4817 | 373 | 298 | 355 | 543 | 241 | 224 | 256 | 168 | 154 | 177 | 48 | 48 |
| L1MB2_dup6625 | 186.86646598994 | -1.47667601465626 | 0.307693299882485 | -4.79918157210518 | 1.59315313146663e-06 | 0.000466852727857091 | 4818 | 312 | 300 | 295 | 370 | 249 | 244 | 164 | 115 | 122 | 133 | 38 | 61 |

| | | | | | | | | | | | | | | | | | | |
|---|---|---|---|---|---|---|---|---|---|---|---|---|---|---|---|---|---|---|
| THE1B-int_dup3625 | 175.20693039588 | -1.6020798713084 | 0.37670507506346 | -4.25287573053938 | 2.11042703875906e-05 | 0.00322522589701068 | 4819 | 308 | 260 | 288 | 429 | 214 | 176 | 144 | 128 | 105 | 135 | 33 | 40 |
| L1MD1_dup3272 | 173.100157130905 | -1.18466629459884 | 0.342212848825428 | -3.46178204198047 | 0.000536611536091637 | 0.0319371106970637 | 4820 | 237 | 213 | 321 | 426 | 129 | 126 | 204 | 148 | 30 | 31 | 179 | 243 |
| L3_dup9633 | 136.889783217343 | -1.63779635557342 | 0.434329034453143 | -3.77086546294459 | 0.000162682365629085 | 0.014163782719348 | 4821 | 340 | 292 | 104 | 79 | 166 | 157 | 188 | 184 | 29 | 21 | 71 | 121 |
| LTR49-int_dup1623 | 127.5291536403 | -1.13081519217906 | 0.345287493443139 | -3.27499609355323 | 0.00105663434741123 | 0.0495649949257222 | 4822 | 210 | 158 | 138 | 164 | 233 | 225 | 101 | 115 | 35 | 31 | 93 | 143 |
| L1MA4A_dup392 | 140.117208339901 | -1.22936192906601 | 0.242462568895708 | -5.07031635713968 | 3.97154976297093e-07 | 0.000154666837954154 | 4823 | 262 | 245 | 158 | 205 | 138 | 122 | 258 | 130 | 51 | 65 | 66 | 124 |
| L1PA8A_dup2327 | 160.280622899522 | -1.07710789953317 | 0.185930911962085 | -5.79305446397644 | 6.91176805650695e-09 | 5.81140481262757e-06 | 4824 | 300 | 265 | 152 | 200 | 186 | 211 | 216 | 160 | 73 | 80 | 85 | 132 |
| LTR12C_dup2128 | 120.976063253531 | -1.49376644151484 | 0.22008090690662 | -6.78735135414831 | 1.14210831505964e-11 | 2.15682623123159e-08 | 4825 | 154 | 164 | 198 | 206 | 115 | 99 | 226 | 172 | 42 | 63 | 46 | 65 |
| AluJr_dup72799 | 127.981888986963 | -1.23098451712301 | 0.267148321883971 | -4.60786917335626 | 4.06816484178742e-06 | 0.000943473247021394 | 4826 | 222 | 190 | 125 | 139 | 117 | 119 | 241 | 200 | 52 | 57 | 88 | 75 |
| Tigger1_dup1914 | 119.087063025579 | -1.22780608100192 | 0.23273806678892 | -5.27548457346028 | 1.32405749800619e-07 | 6.4517629148508e-05 | 4827 | 213 | 180 | 126 | 175 | 114 | 102 | 201 | 160 | 54 | 69 | 59 | 69 |
| HERVIP10F-int_dup19 | 222.438422163457 | -1.59261583340102 | 0.320979185614707 | -4.96174177260436 | 6.98638370662324e-07 | 0.000243038312493592 | 4828 | 141 | 92 | 575 | 627 | 100 | 103 | 566 | 398 | 54 | 61 | 72 | 98 |
| L1PA6_dup3338 | 192.756273831246 | -1.46769732966358 | 0.332149191369182 | -4.41878941090735 | 9.92553019687668e-06 | 0.00184264518857932 | 4829 | 108 | 109 | 453 | 625 | 52 | 57 | 507 | 339 | 45 | 43 | 85 | 96 |
| L2_dup37746 | 81.508188486637 | -1.18010366504634 | 0.238919576011374 | -4.93933433478956 | 7.83897089065035e-07 | 0.000267058535441124 | 4830 | 138 | 128 | 101 | 155 | 64 | 75 | 133 | 84 | 31 | 53 | 47 | 47 |
| ERVL-E-int_dup3473 | 81.1588795892337 | -1.60699255445577 | 0.25262054985112 | -6.36128990853216 | 2.00066420989359e-10 | 2.644722478263e-07 | 4831 | 132 | 112 | 101 | 118 | 76 | 75 | 141 | 137 | 26 | 22 | 47 | 46 |
| L1PA4_dup9454 | 88.450044878621 | -1.74154479653068 | 0.26036007332076 | -6.68898565866172 | 2.24722751212195e-11 | 3.76562533520683e-08 | 4832 | 168 | 145 | 107 | 168 | 81 | 71 | 152 | 112 | 25 | 24 | 42 | 52 |
| HERVL-int_dup409 | 103.991099042224 | -1.47586418750557 | 0.210325597423921 | -7.01704502724362 | 2.26609598558883e-12 | 5.16525405148385e-09 | 4833 | 141 | 109 | 190 | 225 | 81 | 84 | 151 | 157 | 25 | 29 | 68 | 81 |
| L1PA8_dup7440 | 94.8104038922963 | -2.00369113306249 | 0.303125736221509 | -6.61009902372094 | 3.8406303284018e-11 | 6.13330620216037e-08 | 4834 | 131 | 133 | 177 | 264 | 122 | 91 | 115 | 76 | 12 | 19 | 54 | 57 |
| L1PA11_dup2177 | 105.755736817602 | -1.64224183122609 | 0.378150886047485 | -4.34282158741227 | 1.4066436377104e-05 | 0.00241142094393832 | 4835 | 66 | 53 | 218 | 298 | 23 | 26 | 349 | 213 | 13 | 30 | 36 | 57 |
| Harlequin-int_dup70 | 131.049362997178 | -2.25643479848188 | 0.473391517888915 | -4.76652984519967 | 1.87425977559216e-06 | 0.000536669815358667 | 4836 | 199 | 182 | 132 | 164 | 34 | 50 | 458 | 311 | 22 | 29 | 30 | 64 |
| L1M1_dup4131 | 102.546289032537 | -1.51577994148834 | 0.353727003832442 | -4.28516885922103 | 1.82600442853461e-05 | 0.00290385370215815 | 4837 | 138 | 118 | 95 | 130 | 72 | 69 | 300 | 208 | 37 | 46 | 36 | 55 |
| L1MA4A_dup2822 | 120.143758901565 | -1.56314930079826 | 0.318798716643573 | -4.90324841095867 | 9.42646528699904e-07 | 0.000304340530417948 | 4838 | 150 | 141 | 122 | 188 | 69 | 85 | 337 | 241 | 30 | 51 | 47 | 77 |
| L1PA13_dup2638 | 144.18477684894 | -1.89751133064298 | 0.461746933185773 | -4.10941837242167 | 3.96656889360013e-05 | 0.00510559333656773 | 4839 | 285 | 249 | 335 | 496 | 59 | 72 | 146 | 79 | 44 | 47 | 37 | 74 |
| L1PA7_dup11190 | 194.026812486509 | -3.02371760781485 | 0.87932443745554 | -3.43868256017591 | 0.000584552246850924 | 0.0338258436111844 | 4840 | 608 | 512 | 520 | 611 | 27 | 36 | 75 | 65 | 23 | 13 | 40 | 75 |
| L1M1_dup4054 | 164.322565552873 | -3.76819746445578 | 0.484457990742258 | -7.77817176404165 | 7.35801242001969e-15 | 3.24224004313705e-11 | 4841 | 333 | 272 | 113 | 156 | 125 | 110 | 567 | 354 | 11 | 4 | 36 | 29 |
| REP522_dup205 | 129.367359238224 | -1.75733717856992 | 0.379712767023362 | -4.62806977059531 | 3.6908968031473e-06 | 0.000874015429346082 | 4842 | 80 | 50 | 288 | 348 | 96 | 125 | 208 | 197 | 11 | 5 | 134 | 147 |
| Cheshire_dup1041 | 92.5300891868491 | -2.02981371499423 | 0.441741974851641 | -4.59502114481185 | 4.32705388232061e-06 | 0.000987157395093634 | 4843 | 87 | 66 | 199 | 249 | 84 | 81 | 158 | 103 | 1 | 6 | 101 | 88 |
| Tigger4b_dup1877 | 70.6936338827394 | -1.80054926679654 | 0.530661150039415 | -3.39303012225937 | 0.000691240365680788 | 0.0374744092446271 | 4844 | 100 | 100 | 106 | 152 | 80 | 64 | 83 | 74 | 2 | 3 | 73 | 92 |

**Table S5**. Differentially expressed TE subfamilies between timepoints in HG3 cell lines.

| TE families | baseMean | log2FoldChange | lfcSE | stat | pvalue | FDR | Heatmap_order | HG3_mut1_TP1_RNA1 | HG3_mut1_TP1_RNA2 | HG3_mut1_TP2_RNA1 | HG3_mut1_TP2_RNA2 | HG3_mut2_TP1_RNA1 | HG3_mut2_TP1_RNA2 | HG3_mut2_TP2_RNA1 | HG3_mut2_TP2_RNA2 | HG3_wt_TP1_RNA1 | HG3_wt_TP1_RNA2 | HG3_wt_TP2_RNA1 | HG3_wt_TP2_RNA2 |
|---|---|---|---|---|---|---|---|---|---|---|---|---|---|---|---|---|---|---|---|
| L1PA6_dup4183 | 16.2444107440251 | 1.97494091505406 | 0.530406440574931 | 3.72344821626476 | 0.000196520183560731 | 0.0109742294291344 | 1 | 25 | 14 | 3 | 5 | 60 | 58 | 15 | 8 | 4 | 8 | 2 | 6 |
| L1PA7_dup9366 | 12.5530129957211 | 1.62021940750059 | 0.437183012857918 | 3.7060438302692 | 0.000210521882822823 | 0.0115341560972046 | 2 | 20 | 17 | 3 | 10 | 31 | 40 | 16 | 7 | 6 | 6 | 2 | 3 |
| L1PA13_dup6200 | 24.0910464880274 | 2.21242983361569 | 0.50235688509172 | 4.40409975313018 | 1.06224060121012e-05 | 0.00127719000553584 | 3 | 31 | 30 | 10 | 4 | 95 | 73 | 16 | 14 | 15 | 6 | 1 | 7 |
| HAL1b_dup4408 | 23.1335556643535 | 1.98266078183426 | 0.441461269975778 | 4.49113187651329 | 7.08456658846826e-06 | 0.000938610401703601 | 4 | 42 | 39 | 3 | 4 | 66 | 60 | 17 | 11 | 13 | 16 | 8 | 14 |
| L1P4a_dup492 | 23.0447764445462 | 1.60049088598272 | 0.474388967119371 | 3.37379449547776 | 0.000741396944198924 | 0.0285134050887922 | 5 | 37 | 27 | 5 | 5 | 70 | 67 | 26 | 21 | 9 | 12 | 4 | 8 |
| HERVL-int_dup831 | 19.7219119204788 | 1.57885732343077 | 0.448918090450315 | 3.51702762044408 | 0.000436408367605094 | 0.0196744269492538 | 6 | 52 | 47 | 7 | 13 | 44 | 29 | 26 | 24 | 4 | 5 | 0 | 0 |
| L1M3c_dup758 | 20.4742470579324 | 1.9078183145903 | 0.51382477882601 | 3.71297452596447 | 0.00020483751175342 | 0.0113086465363525 | 7 | 74 | 65 | 16 | 10 | 19 | 20 | 18 | 19 | 11 | 7 | 1 | 0 |
| L1PA3_dup3548 | 8.17874200957474 | 1.75864388055762 | 0.488059778509612 | 3.60333704598234 | 0.000314157730764481 | 0.0154383674936979 | 8 | 28 | 17 | 5 | 5 | 16 | 12 | 7 | 7 | 2 | 5 | 0 | 1 |
| GSAT_dup20 | 12.3566798276255 | 1.86587437941419 | 0.513774772607084 | 3.63169715388331 | 0.000281563430222317 | 0.014279130225429 | 9 | 46 | 43 | 8 | 9 | 14 | 19 | 3 | 9 | 2 | 3 | 2 | 1 |
| L1MA6_dup2050 | 14.1109553299952 | 2.66100527612817 | 0.579749941838012 | 4.58991900489345 | 4.434180378792e-06 | 0.000654525734746924 | 10 | 64 | 55 | 5 | 4 | 13 | 15 | 8 | 6 | 7 | 3 | 0 | 2 |
| L1PA15-16_dup971 | 12.7587072558042 | 2.04408193388007 | 0.531340763199679 | 3.84702638203556 | 0.00011956009951669 | 0.00773276636031481 | 11 | 42 | 33 | 5 | 12 | 11 | 11 | 14 | 5 | 12 | 16 | 0 | 2 |
| L2a_dup111512 | 12.3820699067571 | 2.06864892677684 | 0.626329690289858 | 3.30281153655593 | 0.000957206962013662 | 0.0340810366277574 | 12 | 39 | 28 | 2 | 17 | 16 | 9 | 12 | 9 | 12 | 13 | 0 | 0 |
| AluJr_dup54887 | 15.7496418546116 | 1.71253253747656 | 0.489590737973912 | 3.49788589662375 | 0.000468961723293213 | 0.0206359044478966 | 13 | 56 | 60 | 7 | 13 | 19 | 12 | 17 | 6 | 5 | 5 | 3 | 2 |
| MLT2B3_dup2317 | 12.5654461974101 | 2.16812583235567 | 0.533188839506389 | 4.06633761194787 | 4.77577358201286e-05 | 0.00393209764963887 | 14 | 47 | 47 | 6 | 4 | 14 | 8 | 8 | 6 | 8 | 7 | 3 | 2 |
| MIRc_dup32976 | 11.2397955210328 | 1.55989149209033 | 0.432404975445301 | 3.60747812969524 | 0.000309187590397213 | 0.0152726087858589 | 15 | 29 | 35 | 7 | 10 | 16 | 11 | 10 | 4 | 8 | 7 | 2 | 5 |
| X7D_LINE_dup14 | 10.4205803947993 | 1.84719486649078 | 0.44191428664325 | 4.17998449545939 | 2.91528971844649e-05 | 0.00269600282683819 | 16 | 28 | 28 | 7 | 9 | 18 | 10 | 7 | 5 | 8 | 9 | 2 | 1 |
| L1MC1_dup9607 | 21.0201089723079 | 4.57568377431839 | 0.58457278413423 | 7.82739788526952 | 4.98070886719563e-15 | 1.66921091847004e-11 | 17 | 77 | 47 | 0 | 10 | 46 | 49 | 1 | 0 | 18 | 19 | 1 | 0 |
| L2a_dup119788 | 18.7336820776164 | 3.72319802554832 | 0.459064616085933 | 8.11040079127198 | 5.04530668649718e-16 | 2.03476363529705e-12 | 18 | 53 | 48 | 5 | 10 | 45 | 47 | 1 | 0 | 12 | 16 | 1 | 1 |
| L1PA8_dup6066 | 24.7804398366232 | 4.30800995057281 | 0.539964838497601 | 7.97831570396217 | 1.48343572769357e-15 | 5.51527496346521e-12 | 19 | 81 | 71 | 5 | 11 | 45 | 68 | 0 | 0 | 12 | 24 | 1 | 0 |
| MLT1J2_dup4801 | 16.9854881150525 | 3.78371586632976 | 0.587623119534442 | 6.439018038173 | 1.2024891085422e-10 | 1.19219780592159e-07 | 20 | 58 | 33 | 5 | 11 | 30 | 38 | 0 | 0 | 15 | 25 | 1 | 0 |
| L4_A_Mam_dup4849 | 17.9884984320972 | 3.59392672023165 | 0.516211425282705 | 6.96212161182488 | 3.35185953982447e-12 | 5.08001000040174e-09 | 21 | 46 | 22 | 9 | 6 | 38 | 51 | 1 | 1 | 27 | 20 | 2 | 0 |
| L2a_dup119784 | 15.7763353780105 | 3.18469969348415 | 0.433938028953321 | 7.33906567526656 | 2.15090055667032e-13 | 4.61079444916646e-10 | 22 | 46 | 35 | 8 | 6 | 29 | 25 | 2 | 1 | 20 | 21 | 2 | 2 |
| LTR16D_dup830 | 16.019916033508 | 3.49073043171472 | 0.463302932028264 | 7.53444493958386 | 4.90416381447781e-14 | 1.23859929820496e-10 | 23 | 53 | 46 | 6 | 4 | 24 | 39 | 2 | 1 | 12 | 13 | 2 | 2 |
| HAL1_dup16428 | 12.5447247035239 | 3.98906081338768 | 0.554633017054172 | 7.19225269814412 | 6.3730934112047e-13 | 1.26371007068543e-09 | 24 | 32 | 33 | 3 | 4 | 30 | 24 | 1 | 1 | 16 | 11 | 0 | 1 |
| L2_dup41259 | 12.3028370449679 | 3.33877852541874 | 0.544194542070433 | 6.13526646687062 | 8.50165704917701e-10 | 6.34148993173502e-07 | 25 | 20 | 36 | 7 | 6 | 29 | 20 | 2 | 1 | 15 | 16 | 0 | 0 |
| MER20B_dup2837 | 13.4177607592099 | 3.4210051224728 | 0.552348590406236 | 6.19356178669118 | 5.88196156425829e-10 | 4.66529742123001e-07 | 26 | 31 | 35 | 5 | 10 | 23 | 27 | 1 | 1 | 18 | 16 | 0 | 0 |
| L1PREC2_dup5815 | 15.7328724384445 | 4.66438174856699 | 0.582949179186932 | 8.00135228781459 | 1.23060151591698e-15 | 4.64788425883146e-12 | 27 | 36 | 42 | 3 | 3 | 32 | 46 | 1 | 0 | 16 | 17 | 0 | 1 |
| L2a_dup119789 | 15.1782215640878 | 3.96062990434152 | 0.573333889646122 | 6.90806871156027 | 4.91296025151204e-12 | 7.26098906836201e-09 | 28 | 47 | 44 | 3 | 8 | 21 | 35 | 1 | 0 | 19 | 12 | 0 | 1 |
| L1PA16_dup10543 | 14.2055360107362 | 3.3598017429137 | 0.527835013047436 | 6.36524986001971 | 1.94972080316623e-10 | 1.7912288271436e-07 | 29 | 32 | 35 | 9 | 7 | 34 | 27 | 1 | 0 | 14 | 18 | 1 | 0 |
| L2_dup41260 | 14.1881968820553 | 4.36958259017633 | 0.576532082428595 | 7.57907967891364 | 3.48014741041339e-14 | 9.51824316917499e-11 | 30 | 45 | 28 | 4 | 4 | 37 | 37 | 1 | 0 | 12 | 12 | 0 | 0 |
| L2_dup41262 | 13.0138561058955 | 3.91376784038734 | 0.547330796627801 | 7.15064429865949 | 8.63715344371152e-13 | 1.67087488887592e-09 | 31 | 40 | 22 | 5 | 3 | 38 | 27 | 2 | 0 | 13 | 13 | 1 | 0 |
| L1P2_dup1236 | 11.1851471362497 | 4.40929229321385 | 0.668565679221654 | 6.59515202507427 | 4.24820320497578e-11 | 4.85982192216907e-08 | 32 | 33 | 18 | 3 | 4 | 26 | 33 | 0 | 0 | 8 | 17 | 0 | 0 |
| MLT1H2_dup3256 | 12.7071873138518 | 4.02277134006762 | 0.615526595241155 | 6.53549557593292 | 6.33992563381155e-11 | 6.87825930901967e-08 | 33 | 26 | 32 | 6 | 3 | 38 | 29 | 0 | 0 | 15 | 8 | 0 | 1 |
| L1PA7_dup10206 | 39.7980430240714 | 3.62612067921244 | 0.481341762544157 | 7.53335979003024 | 4.94511079528429e-14 | 1.23859929820496e-10 | 34 | 124 | 103 | 14 | 22 | 107 | 69 | 4 | 0 | 27 | 34 | 0 | 4 |
| THE1B-int_dup3278 | 29.3644622627598 | 3.61773806104757 | 0.463936528318485 | 7.79791596527199 | 6.29378965478164e-15 | 2.02375955837388e-11 | 35 | 91 | 70 | 12 | 15 | 64 | 55 | 2 | 0 | 26 | 34 | 2 | 1 |
| THE1D_dup9318 | 28.9247238627343 | 3.8144577254915 | 0.458185376385901 | 8.32514070086519 | 8.42286367208717e-17 | 4.00837343863691e-13 | 36 | 80 | 74 | 12 | 11 | 72 | 64 | 0 | 2 | 17 | 33 | 1 | 1 |
| L1MC_dup5888 | 29.0195502495872 | 4.56276637344559 | 0.445851275689304 | 10.2338304771953 | 1.39874923209361e-24 | 3.69807538644164e-20 | 37 | 75 | 98 | 6 | 5 | 68 | 60 | 3 | 1 | 24 | 24 | 0 | 1 |
| L1M4_dup14479 | 25.1923966365671 | 3.71412866251887 | 0.457308687880493 | 8.1217102603777 | 4.59659636098812e-16 | 1.88576158226152e-12 | 38 | 86 | 84 | 8 | 12 | 43 | 47 | 2 | 1 | 16 | 21 | 1 | 1 |
| L1PA2_dup3869 | 30.0784888893889 | 4.09585411883106 | 0.422505052639457 | 9.6942133431153 | 3.19087062542563e-22 | 3.99607843072382e-18 | 39 | 94 | 84 | 10 | 8 | 49 | 64 | 2 | 1 | 29 | 36 | 1 | 1 |
| L2a_dup119787 | 27.568341777455 | 3.95335347404154 | 0.512408521146949 | 7.71523757097668 | 1.2075708647888e-14 | 3.73164489601344e-11 | 40 | 93 | 77 | 11 | 11 | 50 | 43 | 1 | 1 | 23 | 38 | 0 | 1 |
| L2_dup41267 | 25.126973426885 | 3.64161509608027 | 0.431635878776927 | 8.43677570641037 | 3.26211610820004e-17 | 1.76410790791311e-13 | 41 | 56 | 77 | 5 | 13 | 52 | 49 | 1 | 5 | 23 | 31 | 0 | 2 |
| L2a_dup119781 | 24.9420874984647 | 3.43454095227988 | 0.48925050521527 | 7.02000491704895 | 2.218604377325e-12 | 3.70903703555628e-09 | 42 | 65 | 68 | 8 | 17 | 49 | 31 | 4 | 1 | 26 | 42 | 0 | 2 |
| LTR12C_dup993 | 39.2148641985077 | 5.68988955698646 | 0.537580878637959 | 10.5842484044496 | 3.52599089521458e-26 | 1.19856489936104e-21 | 43 | 163 | 117 | 3 | 1 | 76 | 89 | 3 | 3 | 24 | 23 | 0 | 0 |
| L1MEc_dup13334 | 18.704503233564 | 3.90908893375781 | 0.559502587687218 | 6.98672181288128 | 2.8138338713874e-12 | 4.43404314146455e-09 | 44 | 39 | 41 | 1 | 13 | 39 | 65 | 3 | 1 | 15 | 20 | 0 | 0 |
| MER110_dup554 | 21.141832477641 | 3.57250912254984 | 0.443418107118202 | 8.05675064955685 | 7.83492452413744e-16 | 3.05621139478755e-12 | 45 | 61 | 52 | 9 | 11 | 43 | 49 | 3 | 1 | 18 | 21 | 0 | 0 |
| L1MA9_dup12019 | 19.6334559174861 | 4.69353775492915 | 0.560884270961497 | 8.36810372108178 | 5.85528441075147e-17 | 2.90258646750139e-13 | 46 | 67 | 58 | 6 | 1 | 37 | 45 | 2 | 1 | 11 | 22 | 0 | 0 |
| LTR12C_dup1603 | 21.0604579270952 | 4.17355547709961 | 0.55144545520767 | 7.56839219126012 | 3.77871724119399e-14 | 1.02173937803766e-10 | 47 | 88 | 84 | 2 | 5 | 29 | 33 | 4 | 3 | 6 | 17 | 0 | 1 |
| L1PA7_dup9651 | 45.5639086290989 | 3.97977038221394 | 0.880235424080096 | 4.52125678351681 | 6.14735627811709e-06 | 0.000843563343110063 | 48 | 287 | 236 | 7 | 8 | 9 | 15 | 2 | 3 | 10 | 15 | 2 | 5 |
| L1PA15_dup7571 | 20.9909326428902 | 2.99254216845982 | 0.682355543529276 | 4.38560541764167 | 1.15663625027017e-05 | 0.0013560464468823 | 49 | 86 | 87 | 2 | 11 | 7 | 9 | 2 | 1 | 18 | 28 | 8 | 9 |
| L1PA15_dup3244 | 22.7616997515476 | 2.51266413761673 | 0.751499372820874 | 3.3435345769951 | 0.000827183850745903 | 0.0307779653713189 | 50 | 100 | 91 | 12 | 21 | 5 | 5 | 2 | 1 | 20 | 25 | 2 | 9 |
| L1PA15_dup7573 | 20.6580804640505 | 3.32879037320431 | 0.759358872683059 | 4.38368536004935 | 1.16688273302163e-05 | 0.00136372828483087 | 51 | 93 | 82 | 6 | 10 | 3 | 5 | 3 | 0 | 25 | 28 | 2 | 5 |
| L1M3f_dup451 | 24.1599620974286 | 2.59415507651798 | 0.464389095584936 | 5.58616707666323 | 2.32135917460331e-08 | 1.04415525550125e-05 | 52 | 77 | 52 | 11 | 7 | 16 | 14 | 7 | 2 | 50 | 41 | 8 | 11 |
| L1PA8A_dup490 | 33.7589794986997 | 2.24422889328457 | 0.645657520767399 | 3.47588128551059 | 0.000509177624520803 | 0.0218457949953529 | 53 | 129 | 100 | 26 | 27 | 13 | 8 | 4 | 2 | 50 | 44 | 8 | 14 |
| THE1B-int_dup3020 | 29.3045795513933 | 1.41534027060603 | 0.383269168712888 | 3.69281013486969 | 0.000221789611293387 | 0.0119886303609306 | 54 | 42 | 57 | 19 | 12 | 64 | 74 | 12 | 11 | 17 | 17 | 20 | 27 |
| L1PA10_dup5794 | 25.1685511176764 | 1.34431952425238 | 0.359859159061767 | 3.73568239240409 | 0.000187206831149714 | 0.0106236862973408 | 55 | 49 | 45 | 11 | 19 | 44 | 54 | 12 | 6 | 14 | 22 | 25 | 22 |
| L1P3_dup2923 | 23.8635799016123 | 1.1305409630671 | 0.325683722714514 | 3.47128482088158 | 0.00051797427923426 | 0.0220957167168654 | 56 | 55 | 44 | 15 | 12 | 36 | 42 | 14 | 13 | 12 | 20 | 19 | 23 |
| L1PB1_dup10964 | 23.9158410461923 | 1.27056515083832 | 0.310455209621677 | 4.09258763087482 | 4.26585951069015e-05 | 0.00361869592559957 | 57 | 37 | 43 | 11 | 14 | 56 | 37 | 17 | 12 | 13 | 26 | 19 | 18 |
| L1PA16_dup8724 | 28.1714124273037 | 1.51921152030721 | 0.438238585705221 | 3.46663112254815 | 0.000527024611052291 | 0.0223455805597734 | 58 | 87 | 68 | 11 | 15 | 38 | 36 | 8 | 10 | 20 | 15 | 23 | 30 |
| L1PA5_dup1372 | 29.6269416598997 | 1.63121091822086 | 0.38887474145562 | 4.19469496042601 | 2.73239049145852e-05 | 0.00256879252422201 | 59 | 69 | 59 | 10 | 12 | 48 | 53 | 10 | 9 | 19 | 29 | 22 | 39 |
| L1PB4_dup998 | 23.9212780093702 | 1.07578350869078 | 0.315837742200261 | 3.40612714996127 | 0.000658914838344075 | 0.0261979648476901 | 60 | 50 | 32 | 12 | 17 | 36 | 25 | 14 | 7 | 30 | 22 | 27 | 30 |
| L2_dup11044 | 22.2861176075327 | 1.15118386234203 | 0.35036606082023 | 3.28566031666148 | 0.00101743625375628 | 0.035490037228246 | 61 | 48 | 37 | 6 | 12 | 39 | 26 | 18 | 10 | 20 | 19 | 22 | 26 |
| L2c_dup23167 | 21.2339217224663 | 1.10334946559012 | 0.336838085575532 | 3.27560781526503 | 0.00105434861886403 | 0.0364330578658465 | 62 | 51 | 35 | 9 | 13 | 33 | 23 | 16 | 8 | 16 | 22 | 19 | 27 |
| L1ME3F_dup4188 | 22.4892847391683 | 1.54570151621518 | 0.36918768820339 | 4.18676344202365 | 2.82960465330167e-05 | 0.00264347510339427 | 63 | 39 | 42 | 8 | 10 | 35 | 33 | 7 | 5 | 18 | 33 | 24 | 31 |
| MIRc_dup31569 | 23.4007548757943 | 1.19534525062345 | 0.378146066038619 | 3.16106752913139 | 0.00157192051055219 | 0.0479344095609191 | 64 | 42 | 35 | 11 | 9 | 47 | 37 | 13 | 8 | 14 | 24 | 25 | 34 |
| LTR1A2_dup757 | 27.6422112251756 | 1.6631853208712 | 0.418463019698615 | 3.97450967607379 | 7.05243951908682e-05 | 0.00527372650474115 | 65 | 59 | 45 | 2 | 8 | 42 | 55 | 13 | 20 | 25 | 33 | 15 | 29 |
| Tigger1_dup10259 | 25.4792775367861 | 1.29345717776451 | 0.395000731412358 | 3.27456907013728 | 0.00105823265784444 | 0.0365460418002109 | 66 | 34 | 23 | 7 | 7 | 53 | 54 | 20 | 20 | 32 | 23 | 15 | 27 |
| L1MA7_dup4426 | 23.9318477803221 | 1.44301212523605 | 0.452537973524181 | 3.18870947778915 | 0.00142909424515648 | 0.0449263124928002 | 67 | 47 | 41 | 3 | 4 | 39 | 26 | 16 | 10 | 27 | 24 | 36 | 26 |
| L1ME1_dup20216 | 27.9126877181115 | 1.74172921295555 | 0.415200162004252 | 4.19491457938717 | 2.72974441122476e-05 | 0.00256731922400509 | 68 | 66 | 72 | 8 | 3 | 31 | 31 | 27 | 12 | 26 | 40 | 15 | 20 |
| THE1D_dup8676 | 27.4934717662513 | 1.52653498143695 | 0.425977524988717 | 3.5836045140583 | 0.000338885075544994 | 0.0163204674240049 | 69 | 67 | 76 | 13 | 4 | 25 | 36 | 13 | 18 | 23 | 27 | 21 | 21 |
| MIRb_dup58213 | 23.8343567695349 | 1.05999923826821 | 0.290824387247577 | 3.64480863623668 | 0.000267590835743584 | 0.0137758911730513 | 70 | 37 | 21 | 14 | 12 | 33 | 36 | 13 | 14 | 37 | 27 | 28 | 21 |
| AluJo_dup1387 | 24.8653442508924 | 1.08423262118199 | 0.253303629063987 | 4.28036749883319 | 1.86584948820283e-05 | 0.00194233018546363 | 71 | 44 | 27 | 17 | 19 | 42 | 41 | 16 | 11 | 23 | 31 | 22 | 21 |
| L1ME4a_dup13891 | 21.7962297194289 | 1.10028981408477 | 0.264652044590376 | 4.15749598982988 | 3.2175492187184e-05 | 0.00291103789504627 | 72 | 28 | 25 | 17 | 15 | 30 | 33 | 11 | 7 | 26 | 37 | 22 | 21 |
| L1ME3_dup6674 | 22.1449263544701 | 1.00985140796176 | 0.319983388687661 | 3.15594947632574 | 0.00159976568822052 | 0.0485471045082667 | 73 | 39 | 23 | 7 | 14 | 25 | 42 | 15 | 13 | 19 | 32 | 29 | 21 |
| Tigger7_dup538 | 23.9173686361725 | 1.07409669578268 | 0.305580428779086 | 3.51493942224676 | 0.000439854310329733 | 0.0197847965455045 | 74 | 51 | 35 | 13 | 15 | 27 | 21 | 17 | 15 | 20 | 45 | 25 | 16 |

| | | | | | | | | | | | | | | | | | | | |
|---|---|---|---|---|---|---|---|---|---|---|---|---|---|---|---|---|---|---|---|
| MIR_dup77130 | 19.0597570127423 | 1.12874354074629 | 0.345955295142424 | 3.26268612330852 | 0.0011036167681503 | 0.0375681252524023 | 75 | 31 | 29 | 7 | 16 | 16 | 18 | 11 | 15 | 31 | 31 | 18 | 9 |
| L1MA5A_dup2325 | 21.2285795153203 | 1.53709009233213 | 0.356507393949108 | 4.31152373953724 | 1.62133363135051e-05 | 0.0017504076780641 | 76 | 54 | 47 | 4 | 12 | 22 | 23 | 17 | 11 | 23 | 26 | 13 | 15 |
| L1PB1_dup9144 | 19.4429974733123 | 1.21161572340209 | 0.320200761649188 | 3.78392517607295 | 0.000154374251227881 | 0.00929942672978969 | 77 | 36 | 26 | 5 | 12 | 19 | 32 | 19 | 10 | 26 | 26 | 10 | 23 |
| THE1B_dup17466 | 19.9555192006303 | 1.237100498645 | 0.281538747410465 | 4.39406834769137 | 1.11248813829933e-05 | 0.00132157814556053 | 78 | 34 | 21 | 11 | 12 | 32 | 37 | 18 | 13 | 23 | 24 | 11 | 13 |
| L1ME3G_dup9704 | 19.7236437777847 | 1.08283827415599 | 0.29410237146043 | 3.68184135605205 | 0.000231555489000035 | 0.0123898588679115 | 79 | 30 | 28 | 10 | 9 | 25 | 31 | 11 | 15 | 24 | 23 | 17 | 21 |
| L2c_dup62353 | 20.3252191801337 | 1.09721663718532 | 0.316416782252295 | 3.46763098143908 | 0.000525067783824034 | 0.0222983721024083 | 80 | 36 | 43 | 9 | 10 | 21 | 32 | 20 | 12 | 18 | 21 | 18 | 16 |
| LTR2B_dup112 | 21.7815305579098 | 1.05349831303224 | 0.26820748309328 | 3.92792289343338 | 8.5682670523982e-05 | 0.00608410883930153 | 81 | 37 | 36 | 13 | 11 | 28 | 32 | 22 | 11 | 23 | 24 | 17 | 20 |
| HSMAR2_dup1405 | 18.5948339992953 | 1.08005438786502 | 0.32022572902411 | 3.37279078466459 | 0.000744104690564017 | 0.0285852009526874 | 82 | 31 | 24 | 19 | 19 | 36 | 31 | 15 | 9 | 7 | 29 | 9 | 9 |
| L2c_dup107630 | 16.4995116573317 | 1.15110750444084 | 0.308418650570971 | 3.73228889468846 | 0.000189747702078078 | 0.010698982634756 | 83 | 31 | 20 | 13 | 17 | 31 | 20 | 14 | 6 | 18 | 20 | 6 | 13 |
| L2a_dup91049 | 16.7222259813424 | 1.1174807566643 | 0.290206849133069 | 3.85063536578317 | 0.000117811792156258 | 0.00765136822309103 | 84 | 30 | 19 | 11 | 14 | 26 | 21 | 15 | 8 | 18 | 26 | 12 | 10 |
| L2c_dup107618 | 17.5041877999832 | 1.18182026111526 | 0.295713028372352 | 3.996510629309 | 6.42829966272749e-05 | 0.00492145492775855 | 85 | 23 | 25 | 13 | 17 | 34 | 27 | 13 | 10 | 14 | 26 | 9 | 9 |
| L2a_dup43487 | 17.272525935581 | 1.04644769808171 | 0.304627079794432 | 3.43517621213411 | 0.000592168638452462 | 0.0243526026348444 | 86 | 36 | 27 | 10 | 19 | 27 | 21 | 11 | 10 | 16 | 17 | 15 | 9 |
| MLT1L_dup3968 | 17.837511925045 | 1.20526323873032 | 0.348349739298249 | 3.4599228957608 | 0.000540330082949291 | 0.0227493479883854 | 87 | 43 | 37 | 11 | 17 | 19 | 21 | 11 | 12 | 17 | 17 | 14 | 5 |
| MamRTE1_dup4113 | 17.2669479732855 | 1.19121545798242 | 0.311203725012738 | 3.82776735057931 | 0.00012931089431615 | 0.00818107153920514 | 88 | 28 | 39 | 14 | 13 | 21 | 20 | 14 | 6 | 19 | 20 | 11 | 12 |
| THE1D_dup4449 | 16.6421443822302 | 1.55891608036484 | 0.330551104838809 | 4.71611214588145 | 2.40393965769202e-06 | 0.000406833446507243 | 89 | 38 | 37 | 11 | 14 | 24 | 19 | 8 | 5 | 17 | 19 | 9 | 9 |
| L1MDa_dup125 | 19.3569898065902 | 1.04381686908101 | 0.313679292986483 | 3.32765627958102 | 0.000875798509322366 | 0.0320802276048004 | 90 | 40 | 31 | 11 | 17 | 40 | 22 | 11 | 12 | 15 | 15 | 15 | 16 |
| MLT1E1A_dup2812 | 18.3801161390981 | 1.07795530399797 | 0.312883989418425 | 3.44522359869427 | 0.000570587183851615 | 0.0236985404169587 | 91 | 29 | 32 | 10 | 13 | 40 | 27 | 14 | 16 | 13 | 14 | 13 | 9 |
| L1PB1_dup10966 | 17.6659090601122 | 1.43211054878642 | 0.336674985284167 | 4.25368860587506 | 2.10277668062368e-05 | 0.00211742403744259 | 92 | 46 | 29 | 9 | 9 | 31 | 28 | 13 | 5 | 13 | 16 | 14 | 13 |
| MER112_dup3333 | 19.5415018250645 | 1.34086617833369 | 0.313181264183123 | 4.28143804142021 | 1.85689427103108e-05 | 0.00193619879147573 | 93 | 44 | 32 | 15 | 14 | 41 | 34 | 11 | 6 | 10 | 18 | 11 | 15 |
| L1MC5_dup14322 | 20.2572372347623 | 1.53834123598373 | 0.336406296198641 | 4.57286695691144 | 4.81095638353158e-06 | 0.000701438619874881 | 94 | 44 | 38 | 10 | 9 | 36 | 43 | 8 | 11 | 12 | 18 | 11 | 17 |
| L1PB1_dup1568 | 21.6698427221845 | 1.16307709287947 | 0.348644157744788 | 3.33600052386611 | 0.000849930036154425 | 0.031383838048231 | 95 | 36 | 52 | 11 | 18 | 36 | 37 | 11 | 9 | 15 | 13 | 16 | 22 |
| L1ME3B_dup2530 | 22.3822583289749 | 1.67501743110263 | 0.339410245882326 | 4.93508210616414 | 8.01170485478991e-07 | 0.000173620503039876 | 96 | 64 | 37 | 8 | 16 | 38 | 27 | 20 | 7 | 22 | 25 | 15 | 5 |
| MLT1B_dup5969 | 23.1051005179002 | 1.4033107193918 | 0.347376058042663 | 4.03974507425452 | 5.35093296565457e-05 | 0.00430437151942408 | 97 | 60 | 50 | 11 | 21 | 34 | 23 | 30 | 9 | 14 | 29 | 10 | 6 |
| L1MA5_dup800 | 27.6477745659467 | 1.43798158967698 | 0.291101840692193 | 4.93978872224817 | 7.820726358248e-07 | 0.000170431939185663 | 98 | 75 | 49 | 18 | 20 | 30 | 37 | 30 | 10 | 28 | 32 | 10 | 13 |
| LTR16E1_dup1794 | 24.5445433494987 | 1.14154756211395 | 0.268793515429782 | 4.24693118168677 | 2.16718490058107e-05 | 0.00216396549875645 | 99 | 44 | 45 | 26 | 17 | 44 | 36 | 25 | 7 | 19 | 23 | 12 | 15 |
| Tigger4_dup539 | 24.2429052635755 | 1.41484817459889 | 0.372353882165645 | 3.79974062945174 | 0.000144847607374118 | 0.00888067734713783 | 100 | 77 | 46 | 16 | 13 | 19 | 31 | 19 | 22 | 24 | 21 | 8 | 8 |
| L1MA9_dup5162 | 23.4586149621837 | 1.42543024919314 | 0.369578648643814 | 3.85690638359067 | 0.000114831115570833 | 0.00750648478725751 | 101 | 80 | 56 | 11 | 18 | 26 | 32 | 11 | 15 | 13 | 15 | 9 | 15 |
| LTR104_Mam_dup707 | 25.0495417280598 | 1.43517998498297 | 0.335591575509892 | 4.27656738046056 | 1.89797128815395e-05 | 0.0019651271227963 | 102 | 83 | 50 | 13 | 26 | 34 | 31 | 15 | 18 | 12 | 23 | 10 | 6 |
| MLT1C_dup6115 | 23.2386006833017 | 1.18227477082312 | 0.287712706861249 | 4.10921986630671 | 3.9699796938889e-05 | 0.00341765842345184 | 103 | 61 | 42 | 14 | 22 | 28 | 33 | 19 | 14 | 21 | 17 | 14 | 10 |
| THE1D-int_dup387 | 23.1473265702668 | 1.10026243135683 | 0.321242562915283 | 3.42502071136502 | 0.000614752554324908 | 0.0249748866811328 | 104 | 43 | 41 | 14 | 25 | 19 | 21 | 20 | 14 | 31 | 36 | 8 | 17 |
| MIR_dup129954 | 25.6193654894195 | 1.10425418052314 | 0.25777242197366 | 4.28383367029065 | 1.83700263994456e-05 | 0.00192050716240882 | 105 | 51 | 51 | 21 | 21 | 36 | 35 | 17 | 19 | 16 | 29 | 11 | 17 |
| MSTA-int_dup341 | 23.3316316987127 | 1.12483044107102 | 0.278538586872808 | 4.03832895721793 | 5.3833324993146e-05 | 0.00431729839865828 | 106 | 50 | 38 | 13 | 15 | 40 | 27 | 18 | 19 | 16 | 28 | 13 | 17 |
| LTR1B_dup215 | 19.5815194904746 | 1.4064642730348 | 0.398107533968533 | 3.53287529882812 | 0.000411066316848902 | 0.0188606991571402 | 107 | 57 | 33 | 6 | 9 | 22 | 13 | 16 | 6 | 26 | 22 | 18 | 19 |
| HAL1_dup10653 | 16.7135507039514 | 1.0978606756955 | 0.309412350937972 | 3.54821219116619 | 0.000387855621837507 | 0.0180797385743576 | 108 | 29 | 27 | 6 | 17 | 20 | 18 | 11 | 6 | 21 | 23 | 16 | 16 |
| L2c_dup63484 | 18.5899145690594 | 1.39817501693359 | 0.338425873781557 | 4.13140697934953 | 3.60549590895928e-05 | 0.00317981219256199 | 109 | 35 | 35 | 7 | 17 | 27 | 21 | 8 | 4 | 22 | 24 | 12 | 23 |
| HAL1b_dup966 | 19.8077675224228 | 1.43730636253015 | 0.346921766994555 | 4.14302733144072 | 3.42751079033255e-05 | 0.00304983549603661 | 110 | 46 | 34 | 7 | 14 | 31 | 30 | 9 | 5 | 17 | 21 | 21 | 17 |
| L2_dup11041 | 18.8556167007388 | 1.24938207893831 | 0.36036779237299 | 3.46696376696498 | 0.000526372838393838 | 0.0223338643731206 | 111 | 54 | 27 | 6 | 14 | 24 | 20 | 13 | 6 | 16 | 24 | 16 | 22 |
| L1PA8A_dup310 | 19.7679630235613 | 2.05525678364239 | 0.455927881379707 | 4.50785500860985 | 6.54863005373723e-06 | 0.000884347518028694 | 112 | 53 | 57 | 6 | 9 | 23 | 17 | 7 | 2 | 26 | 18 | 11 | 20 |
| MIRc_dup46857 | 24.7335625501049 | 1.5969140646549 | 0.507353711940427 | 3.14753598342139 | 0.00164652838803347 | 0.0495804661881818 | 113 | 63 | 73 | 3 | 19 | 21 | 18 | 16 | 4 | 33 | 17 | 16 | 31 |
| L1M2c_dup3 | 20.8649841710474 | 1.8057359599494 | 0.485859989737605 | 3.71657678773798 | 0.00020194028590544 | 0.0112016186696749 | 114 | 82 | 55 | 4 | 4 | 25 | 22 | 17 | 13 | 15 | 11 | 9 | 9 |
| Charlie2b_dup1179 | 19.66513599139 | 1.84554464553547 | 0.356697300828064 | 5.17397984579946 | 2.29159553752697e-07 | 6.40011649340632e-05 | 115 | 66 | 39 | 13 | 12 | 28 | 26 | 6 | 9 | 17 | 18 | 9 | 6 |
| MIR3_dup27963 | 17.2241312473917 | 1.9926692804641 | 0.442945822542573 | 4.49867495989891 | 6.83783096843904e-06 | 0.000912526375555915 | 116 | 62 | 44 | 9 | 7 | 25 | 23 | 4 | 8 | 7 | 17 | 10 | 4 |
| L1MEd_dup15112 | 18.3666437237096 | 1.48290476266182 | 0.455094983116293 | 3.25845113147046 | 0.00112022168151952 | 0.0379541888410713 | 117 | 51 | 44 | 5 | 6 | 19 | 18 | 7 | 11 | 10 | 26 | 17 | 19 |
| L1MEd_dup15397 | 16.97790600413 | 2.14650255228697 | 0.438762815272316 | 4.89217061604175 | 9.97299312388759e-07 | 0.000205021688222428 | 118 | 55 | 41 | 4 | 7 | 24 | 24 | 3 | 6 | 16 | 14 | 8 | 13 |
| L1PB4_dup3480 | 18.2314726381998 | 1.52017003512289 | 0.445499575433197 | 3.41228166972932 | 0.000644215094456635 | 0.0257820043373982 | 119 | 54 | 43 | 6 | 6 | 20 | 27 | 9 | 10 | 17 | 9 | 17 | 12 |
| MLT1I_dup8584 | 16.0061135462489 | 1.46956758597664 | 0.361314254324155 | 4.06728372431774 | 4.75643182655983e-05 | 0.00392295988701077 | 120 | 35 | 32 | 6 | 7 | 22 | 26 | 7 | 12 | 16 | 14 | 13 | 9 |
| L1ME4b_dup21322 | 23.0304591292808 | 1.2361519556477 | 0.375091836714403 | 3.29559812998253 | 0.000982123964708492 | 0.0346825666932116 | 121 | 50 | 59 | 16 | 13 | 23 | 33 | 15 | 5 | 15 | 23 | 16 | 28 |
| L1P1_dup2737 | 23.1638207849869 | 1.45795025562773 | 0.327504889455488 | 4.45169004362569 | 8.51970842069575e-06 | 0.00107967250302574 | 122 | 45 | 59 | 11 | 13 | 38 | 32 | 22 | 5 | 19 | 20 | 15 | 16 |
| L2_dup19566 | 20.3724926891793 | 1.49022914757334 | 0.372061497919456 | 4.00533018306545 | 6.19309125165102e-05 | 0.00478642346165923 | 123 | 58 | 44 | 11 | 16 | 27 | 31 | 19 | 3 | 12 | 21 | 5 | 18 |
| MER50_dup890 | 21.9955905975221 | 1.33944745610074 | 0.351923608928121 | 3.80607444945337 | 0.000141189898056727 | 0.00871254447173391 | 124 | 60 | 54 | 11 | 26 | 32 | 26 | 14 | 6 | 11 | 20 | 11 | 14 |
| L2_dup19567 | 25.3657798450467 | 2.29463840794716 | 0.373101414865293 | 6.15017342878645 | 7.73982674622271e-10 | 5.80965556768678e-07 | 125 | 75 | 83 | 10 | 14 | 40 | 30 | 8 | 5 | 18 | 22 | 8 | 10 |
| L1MD1_dup2146 | 23.2484901617521 | 1.9744058498793 | 0.335355796265835 | 5.88749582343343 | 3.9209107652762e-09 | 2.27552447062051e-06 | 126 | 63 | 55 | 12 | 17 | 28 | 26 | 15 | 5 | 28 | 29 | 7 | 8 |
| L2c_dup47670 | 22.8541249558386 | 1.74294822047508 | 0.355183606661845 | 4.90717529690064 | 9.23974453796405e-07 | 0.000193876565593509 | 127 | 84 | 40 | 13 | 19 | 33 | 21 | 16 | 8 | 20 | 24 | 6 | 9 |
| MLT1M_dup813 | 23.1340439791482 | 1.69038501995387 | 0.361434921210055 | 4.67687243472377 | 2.91283329794314e-06 | 0.00047149457953223 | 128 | 59 | 68 | 13 | 14 | 25 | 20 | 16 | 9 | 24 | 23 | 11 | 9 |
| L2_dup19562 | 26.5511565220863 | 1.6513046978389 | 0.353514088420796 | 4.67111425520703 | 2.99570241654878e-06 | 0.000480897165161573 | 129 | 78 | 71 | 11 | 22 | 29 | 28 | 12 | 10 | 23 | 25 | 12 | 17 |
| L2a_dup58142 | 23.1868566220572 | 1.7396314922476 | 0.335353081825971 | 5.18746236884254 | 2.13178984363581e-07 | 6.04589828526538e-05 | 130 | 71 | 54 | 11 | 17 | 24 | 28 | 11 | 11 | 20 | 27 | 9 | 11 |
| HAL1ME_dup2900 | 18.5098806878976 | 1.8383846777598 | 0.461893675649013 | 3.98010359240503 | 6.88852434873759e-05 | 0.00518373439179227 | 131 | 67 | 55 | 4 | 15 | 18 | 27 | 9 | 5 | 6 | 17 | 8 | 10 |
| MER77B_dup397 | 16.440491816328 | 1.84561882192854 | 0.403675270140303 | 4.57203836461686 | 4.83002475576732e-06 | 0.000702496054992057 | 132 | 38 | 49 | 6 | 13 | 20 | 26 | 10 | 3 | 11 | 19 | 3 | 13 |
| L1PA12_dup698 | 20.7778506729766 | 1.76614787947799 | 0.390646489860126 | 4.52108984803722 | 6.1522067076343e-06 | 0.000843742350002739 | 133 | 80 | 57 | 10 | 15 | 25 | 30 | 11 | 9 | 8 | 11 | 8 | 5 |
| MLT1E2_dup1342 | 21.2753005593187 | 1.53742369794848 | 0.3178746232009 | 4.8365726161689 | 1.32097028441227e-06 | 0.000255960582487591 | 134 | 64 | 46 | 12 | 20 | 27 | 40 | 18 | 8 | 8 | 19 | 8 | 6 |
| L2a_dup11725 | 19.6259216007244 | 1.23485784840191 | 0.319623326058014 | 3.86347850024491 | 0.000111783736044032 | 0.00737413164866463 | 135 | 49 | 45 | 9 | 13 | 29 | 31 | 22 | 14 | 11 | 11 | 6 | 11 |
| L2c_dup47666 | 19.4505534404824 | 1.41980576142502 | 0.301366115807536 | 4.71123224195436 | 2.46223446469779e-06 | 0.000414903913254598 | 136 | 48 | 44 | 15 | 14 | 26 | 33 | 14 | 8 | 15 | 13 | 8 | 10 |
| MER112_dup1514 | 16.8091051807716 | 1.74418414307941 | 0.397235347337072 | 4.39080800530924 | 1.12930223901278e-05 | 0.00133688035106534 | 137 | 55 | 44 | 7 | 11 | 15 | 30 | 12 | 5 | 11 | 12 | 5 | 10 |
| MLT2B4_dup1629 | 15.8235431310583 | 1.69314213253117 | 0.374437222376916 | 4.52183178206257 | 6.13067727958201e-06 | 0.000841760032295107 | 138 | 45 | 43 | 10 | 9 | 18 | 19 | 11 | 6 | 14 | 13 | 5 | 8 |
| Tigger3b_dup2237 | 15.9404573580934 | 1.80529523921657 | 0.393387471997515 | 4.58910201194199 | 4.4515685750211e-06 | 0.000655871787091003 | 139 | 47 | 46 | 7 | 8 | 19 | 22 | 12 | 4 | 12 | 12 | 7 | 8 |
| L1MC2_dup1939 | 17.8593069358517 | 1.47428201421777 | 0.366575336309895 | 4.02177088360207 | 5.77622296142266e-05 | 0.00453307766747585 | 140 | 55 | 42 | 7 | 13 | 24 | 19 | 17 | 8 | 14 | 13 | 6 | 11 |
| MLT1O_dup1617 | 16.8052171991829 | 1.55193145769412 | 0.395242603110589 | 3.92652878379079 | 8.61805833144018e-05 | 0.00611034716249364 | 141 | 59 | 41 | 8 | 9 | 20 | 23 | 11 | 10 | 11 | 8 | 7 | 8 |
| L2c_dup47679 | 20.0313630432068 | 1.99147204623654 | 0.377500628343005 | 5.2754138581911 | 1.32456819865828e-07 | 4.13074317952742e-05 | 142 | 64 | 50 | 9 | 18 | 31 | 25 | 14 | 5 | 20 | 11 | 3 | 6 |
| HERVK9-int_dup210 | 16.6428678796305 | 1.13881001802017 | 0.322789827327186 | 3.52802325726903 | 0.000418675369492034 | 0.0190846991320214 | 143 | 38 | 35 | 15 | 16 | 22 | 26 | 15 | 4 | 13 | 11 | 9 | 10 |
| LTR16E2_dup793 | 16.9675179306581 | 1.43898406982839 | 0.35773004546451 | 4.0225418246876 | 5.75734263786413e-05 | 0.00452538390995712 | 144 | 45 | 41 | 17 | 13 | 20 | 26 | 15 | 4 | 11 | 14 | 8 | 4 |
| MamRTE1_dup308 | 15.4938041211302 | 1.82374935307761 | 0.443954473471637 | 4.10796480732865 | 3.99160906631451e-05 | 0.00343326645742806 | 145 | 23 | 17 | 6 | 7 | 31 | 32 | 2 | 4 | 27 | 12 | 11 | 19 |
| L1PA8_dup740 | 16.5023683930523 | 1.42938931837742 | 0.438010271429913 | 3.26336940389796 | 0.0011009591259568 | 0.037504483920532 | 146 | 35 | 18 | 7 | 10 | 32 | 17 | 3 | 3 | 20 | 19 | 15 | 30 |
| L2c_dup127339 | 13.407276179644 | 1.60074913458957 | 0.43610479101707 | 3.67056076328891 | 0.000242018925664399 | 0.0128193214538876 | 147 | 24 | 19 | 7 | 5 | 25 | 15 | 2 | 3 | 20 | 15 | 10 | 22 |
| MIR_dup71406 | 16.4007718891984 | 1.83130126009349 | 0.380110224344346 | 4.81781636695594 | 1.45137851181108e-06 | 0.000275179052885578 | 148 | 42 | 18 | 7 | 7 | 23 | 25 | 5 | 2 | 25 | 21 | 15 | 15 |
| MIRb_dup199606 | 15.82211694185 | 1.42422437727798 | 0.368148936184981 | 3.86860924287002 | 0.000109457894274644 | 0.00727922529655516 | 149 | 24 | 19 | 11 | 6 | 16 | 29 | 6 | 2 | 24 | 24 | 21 | 14 |
| AluSq2_dup21230 | 15.466907029892 | 1.48343056303134 | 0.376274561154026 | 3.94241523658068 | 8.06651627386781e-05 | 0.00581811240164217 | 150 | 31 | 22 | 10 | 14 | 27 | 16 | 5 | 1 | 22 | 20 | 13 | 13 |
| L1PA8A_dup1941 | 13.8821923497373 | 1.59936759737786 | 0.409532801178152 | 3.90534675800515 | 9.40903679806844e-05 | 0.00652343435301047 | 151 | 17 | 21 | 6 | 11 | 21 | 24 | 2 | 1 | 18 | 23 | 11 | 19 |
| L2a_dup88091 | 18.3857780942107 | 2.01380682880934 | 0.539761482130401 | 3.73091985159997 | 0.000190781915704916 | 0.0107318661263173 | 152 | 35 | 27 | 7 | 7 | 28 | 32 | 0 | 1 | 20 | 26 | 16 | 34 |
| AluSx_dup101416 | 19.2835980718695 | 1.74661041497055 | 0.399255463238099 | 4.37466879176791 | 1.21616989097781e-05 | 0.00141024737270276 | 153 | 31 | 30 | 7 | 9 | 28 | 21 | 2 | 2 | 22 | 40 | 24 | 25 |

| | | | | | | | | | | | | | | | | | | | |
|---|---|---|---|---|---|---|---|---|---|---|---|---|---|---|---|---|---|---|---|
| MLT1B_dup11970 | 18.4420751869247 | 1.72798169938092 | 0.4740102597652 | 3.64545210527062 | 0.000266922125844117 | 0.0137533677254448 | 154 | 26 | 34 | 9 | 10 | 27 | 47 | 2 | 2 | 16 | 20 | 22 | 18 |
| MLT1D_dup5601 | 20.5112772372424 | 1.43865093757898 | 0.403984007878941 | 3.56115813874022 | 0.000369222612949344 | 0.017417732724196 | 155 | 36 | 39 | 17 | 13 | 31 | 29 | 2 | 4 | 23 | 23 | 21 | 19 |
| L1PREC2_dup5704 | 26.6122950872001 | 2.41204226849701 | 0.462549642825315 | 5.21466680584583 | 1.84147931658924e-07 | 5.39621474710767e-05 | 156 | 87 | 86 | 9 | 6 | 34 | 34 | 5 | 5 | 16 | 27 | 12 | 19 |
| L2a_dup116313 | 23.1193977383432 | 2.09738035557791 | 0.493300595768525 | 4.25172881113259 | 2.1212663268397e-05 | 0.00213009056667282 | 157 | 97 | 50 | 9 | 10 | 29 | 23 | 4 | 3 | 18 | 20 | 16 | 19 |
| L1MA1_dup2676 | 19.6320071519423 | 1.51171580211493 | 0.458542818413866 | 3.29678220093832 | 0.000977993094195524 | 0.0346087960724938 | 158 | 50 | 49 | 8 | 13 | 22 | 30 | 8 | 2 | 13 | 17 | 14 | 28 |
| L2_dup15059 | 21.3092034068872 | 1.49811563875334 | 0.455405341595529 | 3.28963124039046 | 0.00100318760991565 | 0.0351668451897249 | 159 | 51 | 49 | 10 | 7 | 21 | 35 | 6 | 6 | 15 | 20 | 18 | 35 |
| L1PA2_dup2681 | 20.1636447184304 | 2.97412574283301 | 0.628983255346523 | 4.72846569054448 | 2.26222843162964e-06 | 0.000385975727989599 | 160 | 45 | 48 | 2 | 1 | 39 | 44 | 2 | 2 | 19 | 13 | 21 | 18 |
| L1M5_dup8492 | 18.0045231783839 | 1.99408930709287 | 0.593954444132859 | 3.35731018900638 | 0.000787047617979041 | 0.0297261811985617 | 161 | 49 | 38 | 4 | 2 | 23 | 21 | 3 | 3 | 13 | 16 | 25 | 34 |
| THE1D_dup4767 | 17.3576046756897 | 1.84805737543756 | 0.43280632520528 | 4.26994077445848 | 1.95524880931846e-05 | 0.00201142945604017 | 162 | 40 | 32 | 4 | 4 | 21 | 22 | 7 | 2 | 21 | 23 | 18 | 25 |
| L1MEg_dup7718 | 13.7757413291908 | 2.15420792598651 | 0.515402139146082 | 4.17966430941788 | 2.9193972205895e-05 | 0.00269875249048325 | 163 | 34 | 32 | 2 | 1 | 18 | 24 | 4 | 4 | 11 | 17 | 13 | 14 |
| THE1A_dup3351 | 17.5164339224429 | 1.96257462180926 | 0.50381012590144 | 3.89546482079479 | 9.80105875399532e-05 | 0.00671694333605464 | 164 | 42 | 36 | 1 | 3 | 25 | 33 | 4 | 10 | 15 | 19 | 12 | 20 |
| L1PA17_dup533 | 13.7620761675456 | 1.2531042354847 | 0.385580961045543 | 3.24991211206792 | 0.00115440679339873 | 0.0388006947890204 | 165 | 24 | 18 | 12 | 8 | 9 | 14 | 5 | 6 | 19 | 33 | 14 | 7 |
| AluSq2_dup58960 | 15.4722553360786 | 1.797629882963 | 0.345880712189526 | 5.19725390752052 | 2.02254025191074e-07 | 5.82881131502447e-05 | 166 | 29 | 28 | 9 | 7 | 13 | 20 | 6 | 4 | 27 | 27 | 8 | 12 |
| L2c_dup22380 | 11.8398781844294 | 1.61464820760444 | 0.403636344240139 | 4.00025476061646 | 6.32743290574617e-05 | 0.00486615174592979 | 167 | 30 | 18 | 8 | 7 | 11 | 12 | 3 | 4 | 15 | 23 | 6 | 11 |
| AluSc8_dup2558 | 12.2516231567717 | 1.41892128044996 | 0.350515977753655 | 4.04809301288737 | 5.16366272417952e-05 | 0.00418233988027831 | 168 | 23 | 18 | 9 | 8 | 15 | 15 | 7 | 2 | 14 | 24 | 9 | 10 |
| L2a_dup13951 | 13.4852291800651 | 1.38003177658673 | 0.409960595708331 | 3.36625468650788 | 0.000761963215800977 | 0.0290461549738833 | 169 | 33 | 18 | 12 | 8 | 8 | 15 | 6 | 3 | 14 | 32 | 10 | 11 |
| L1MB8_dup8613 | 13.7979526640129 | 1.49267128265994 | 0.378395955935057 | 3.94473370882464 | 7.98887058609817e-05 | 0.00577260795772765 | 170 | 12 | 17 | 10 | 13 | 19 | 24 | 6 | 2 | 19 | 32 | 8 | 9 |
| AluSz_dup66415 | 14.4385964921779 | 1.12462400413598 | 0.342240585966342 | 3.28606264204613 | 0.00101598413012476 | 0.0354607182207161 | 171 | 24 | 19 | 9 | 9 | 20 | 19 | 8 | 4 | 19 | 18 | 9 | 24 |
| L1ME4a_dup10569 | 15.005478152249 | 1.07709070910282 | 0.328281851015722 | 3.28099377340002 | 0.0010344202780504 | 0.0359115830516252 | 172 | 25 | 26 | 15 | 9 | 22 | 17 | 10 | 4 | 14 | 21 | 10 | 17 |
| L1MA2_dup2265 | 13.0480436597672 | 1.17185217022287 | 0.353995441931893 | 3.3103594888895 | 0.000931762199100404 | 0.0334838602028075 | 173 | 14 | 15 | 7 | 9 | 26 | 18 | 13 | 3 | 17 | 19 | 12 | 10 |
| L1ME3C_dup953 | 14.8449664159961 | 1.09816143213352 | 0.343282918686967 | 3.19899818008396 | 0.00137906039417922 | 0.0438809714567221 | 174 | 15 | 19 | 7 | 9 | 21 | 28 | 9 | 4 | 15 | 23 | 17 | 20 |
| L2c_dup111816 | 13.9005033056761 | 1.12927959327404 | 0.339958940294312 | 3.32181172319337 | 0.000894350198427488 | 0.0325144464958025 | 175 | 17 | 17 | 9 | 11 | 16 | 26 | 9 | 2 | 14 | 26 | 13 | 16 |
| L2a_dup13953 | 17.9340182673443 | 1.43940026125192 | 0.415066956932697 | 3.46787485057578 | 0.000524591534947791 | 0.0222861020129775 | 176 | 27 | 35 | 6 | 14 | 16 | 9 | 12 | 2 | 19 | 53 | 15 | 17 |
| L2b_dup43152 | 19.5166477956392 | 1.3964603390066 | 0.442302666560765 | 3.15725055393658 | 0.00159264435054641 | 0.0484112611950838 | 177 | 33 | 22 | 17 | 25 | 26 | 19 | 4 | 1 | 28 | 44 | 11 | 14 |
| L2b_dup71186 | 15.9311775652405 | 1.4834106575386 | 0.468984726083252 | 3.16302552095326 | 0.00156138643158116 | 0.0477290937039548 | 178 | 12 | 7 | 7 | 12 | 41 | 43 | 9 | 5 | 20 | 19 | 8 | 16 |
| L2b_dup25516 | 14.7390953574589 | 1.49378468560543 | 0.459140689695388 | 3.2534356443043 | 0.0011401854863505 | 0.0384458929751536 | 179 | 11 | 11 | 6 | 9 | 33 | 35 | 8 | 1 | 18 | 21 | 14 | 18 |
| L2b_dup94578 | 14.3901744039513 | 1.48190373740046 | 0.359303837829993 | 4.12437491998523 | 3.71742940573434e-05 | 0.00325920212740186 | 180 | 19 | 12 | 6 | 9 | 19 | 29 | 8 | 3 | 26 | 21 | 9 | 17 |
| MER68-int_dup235 | 15.0026630846284 | 1.25324712534671 | 0.397996699600823 | 3.14888823601722 | 0.00163892862814776 | 0.0494204171021731 | 181 | 14 | 8 | 7 | 8 | 22 | 32 | 7 | 3 | 23 | 24 | 19 | 19 |
| L1PA10_dup3322 | 20.0411383810971 | 1.67738949415606 | 0.362555841211666 | 4.6265686647061 | 3.71773649004955e-06 | 0.000567760348935386 | 182 | 20 | 20 | 8 | 10 | 30 | 41 | 5 | 9 | 40 | 28 | 9 | 23 |
| L1ME4a_dup17255 | 18.3899834472397 | 1.37695505074978 | 0.394418909519441 | 3.49109796086466 | 0.000481039841632907 | 0.0209790150581349 | 183 | 24 | 27 | 8 | 6 | 41 | 32 | 7 | 11 | 17 | 20 | 10 | 27 |
| L1ME4a_dup18521 | 15.2299510706798 | 1.32040397871108 | 0.387598967500556 | 3.40662408681259 | 0.000657716453978038 | 0.026166360033148 | 184 | 14 | 17 | 8 | 8 | 30 | 39 | 8 | 10 | 14 | 19 | 7 | 16 |
| L2b_dup90432 | 18.3711995585783 | 1.3089692661549 | 0.40704845756811 | 3.21575783378537 | 0.00130100566711484 | 0.0421182441452119 | 185 | 33 | 31 | 7 | 5 | 27 | 31 | 14 | 5 | 21 | 14 | 14 | 31 |
| L2a_dup23936 | 17.569206200126 | 1.50022746569214 | 0.348747183982679 | 4.3017622352089 | 1.69445081346577e-05 | 0.00180919355428757 | 186 | 22 | 24 | 6 | 4 | 29 | 27 | 9 | 5 | 24 | 26 | 18 | 24 |
| MLT2C2_dup1233 | 17.6268579714194 | 1.49232901929073 | 0.408502125931929 | 3.65317320169689 | 0.000259019386285104 | 0.0134745576932653 | 187 | 22 | 16 | 4 | 6 | 26 | 39 | 10 | 4 | 29 | 18 | 19 | 26 |
| L1MEf_dup6335 | 22.3612823849764 | 1.37245327155441 | 0.401056979234112 | 3.42209048244304 | 0.000621416300729342 | 0.025142581719664 | 188 | 40 | 40 | 4 | 12 | 28 | 22 | 9 | 5 | 29 | 25 | 26 | 42 |
| L2a_dup87204 | 21.5011042284809 | 1.58759248967289 | 0.443101840012919 | 3.58290656077304 | 0.000339792178737799 | 0.0163469853946511 | 189 | 46 | 20 | 6 | 9 | 24 | 43 | 6 | 3 | 28 | 23 | 23 | 42 |
| L1M5_dup14759 | 17.8982714149584 | 1.22660069789783 | 0.380693362451097 | 3.2220175576489 | 0.00127291334557585 | 0.0415066741916022 | 190 | 31 | 20 | 7 | 14 | 32 | 28 | 6 | 4 | 14 | 26 | 17 | 30 |
| L2b_dup44535 | 19.7832278676359 | 1.07588546249204 | 0.332656091621211 | 3.23422744868034 | 0.00121972276305988 | 0.040275902383992 | 191 | 26 | 20 | 7 | 17 | 33 | 25 | 9 | 4 | 27 | 26 | 27 | 27 |
| L1PA7_dup1258 | 19.3579007082721 | 1.00444859572452 | 0.300971185390871 | 3.33735800794367 | 0.000845789259010433 | 0.0312989379509326 | 192 | 32 | 25 | 9 | 19 | 29 | 29 | 11 | 6 | 21 | 21 | 22 | 21 |
| MIRb_dup64442 | 19.5994074225883 | 1.08440630308355 | 0.300113287717178 | 3.61332319315857 | 0.0003022975148963 | 0.0150387799455394 | 193 | 39 | 24 | 12 | 18 | 25 | 22 | 9 | 5 | 21 | 31 | 21 | 21 |
| L1MEc_dup6581 | 17.4920891990816 | 1.09831058730906 | 0.312752049635123 | 3.51176143718459 | 0.000445147391068522 | 0.0199437094925985 | 194 | 25 | 14 | 13 | 7 | 18 | 26 | 7 | 6 | 24 | 34 | 20 | 23 |
| MIRb_dup88064 | 16.0075986544475 | 1.07037611172894 | 0.305711668457857 | 3.50126024671675 | 0.000463063411572246 | 0.020460932242512 | 195 | 21 | 19 | 11 | 8 | 22 | 19 | 9 | 5 | 25 | 22 | 18 | 19 |
| Charlie15a_dup2576 | 15.93659200725 | 1.07171086907203 | 0.333002724487308 | 3.21832462698928 | 0.00128941793811805 | 0.0418455865662082 | 196 | 29 | 17 | 10 | 14 | 18 | 27 | 10 | 3 | 20 | 20 | 20 | 13 |
| MER57D_dup332 | 16.5165323505199 | 1.04488402747486 | 0.310300139453849 | 3.36733341246295 | 0.000758988662088375 | 0.0289689233821965 | 197 | 20 | 15 | 11 | 15 | 19 | 19 | 8 | 2 | 28 | 31 | 17 | 20 |
| MIR_dup49845 | 15.8433705417732 | 1.01535821006846 | 0.32260985017026 | 3.14732550643634 | 0.00164771419654391 | 0.0495973437331863 | 198 | 24 | 16 | 10 | 14 | 15 | 22 | 9 | 2 | 26 | 23 | 18 | 19 |
| Charlie1a_dup4014 | 16.7732735523176 | 1.18415246445652 | 0.341629469484765 | 3.46618945444731 | 0.000527891163442068 | 0.0223703634508257 | 199 | 23 | 29 | 10 | 19 | 20 | 16 | 6 | 4 | 27 | 24 | 15 | 15 |
| L1ME3Cz_dup2026 | 17.3829188797993 | 1.36627890259695 | 0.349874152154523 | 3.90505813071188 | 9.42027379947655e-05 | 0.00652570515807461 | 200 | 20 | 28 | 14 | 9 | 20 | 15 | 6 | 4 | 37 | 26 | 17 | 14 |
| L1M4_dup13435 | 19.3660101279428 | 1.01821126520995 | 0.317352376480266 | 3.20845640578735 | 0.0013344953860414 | 0.0428931296943141 | 201 | 33 | 24 | 17 | 15 | 21 | 22 | 4 | 9 | 27 | 28 | 20 | 20 |
| AluY_dup28112 | 16.3594666125528 | 1.0194100109978 | 0.314954247919635 | 3.2366923695467 | 0.00120923692473707 | 0.040067614860817 | 202 | 23 | 21 | 14 | 16 | 19 | 28 | 7 | 5 | 20 | 22 | 11 | 20 |
| L1MA9_dup15293 | 18.3978229932146 | 1.36517002584372 | 0.314197056120269 | 4.34494849411052 | 1.39308345811867e-05 | 0.00156097022104272 | 203 | 26 | 19 | 10 | 14 | 26 | 30 | 5 | 6 | 29 | 28 | 12 | 23 |
| L1MB3_dup6155 | 20.2098520175323 | 1.10423554026386 | 0.313693468597533 | 3.52011007816228 | 0.000431367731439887 | 0.0195280259354976 | 204 | 35 | 33 | 21 | 9 | 17 | 29 | 11 | 8 | 21 | 33 | 18 | 18 |
| MLT2E_dup50 | 20.098950323212 | 1.10477328599806 | 0.327251159397665 | 3.37591863091177 | 0.000735696738090802 | 0.0283491653508913 | 205 | 29 | 34 | 17 | 12 | 14 | 21 | 14 | 4 | 35 | 30 | 21 | 17 |
| THE1D_dup9678 | 21.3223111023471 | 1.05381105621292 | 0.320904646262125 | 3.28387596903826 | 0.00102389972954031 | 0.0356640033214319 | 206 | 33 | 30 | 16 | 21 | 26 | 19 | 8 | 3 | 29 | 35 | 25 | 22 |
| L1ME3Cz_dup1667 | 19.7011309828152 | 1.08049926944192 | 0.290454905213991 | 3.72002417602783 | 0.000199203708636503 | 0.0110824703425815 | 207 | 30 | 29 | 15 | 14 | 23 | 30 | 14 | 3 | 21 | 30 | 16 | 25 |
| L2b_dup27999 | 19.1683326666199 | 1.27152826599795 | 0.297330262838172 | 4.27648451879925 | 1.89867753831067e-05 | 0.0019651271227963 | 208 | 32 | 26 | 10 | 13 | 25 | 31 | 11 | 3 | 23 | 27 | 18 | 23 |
| L1P4_dup1462 | 18.5380432152192 | 1.01343134045652 | 0.32012796259773 | 3.16570702613064 | 0.00154706527478143 | 0.0474562322899501 | 209 | 27 | 26 | 14 | 15 | 21 | 22 | 8 | 3 | 24 | 28 | 15 | 31 |
| MIRb_dup80101 | 17.7392676424722 | 1.17202014759025 | 0.297369159188414 | 3.94129690781973 | 8.10422364575556e-05 | 0.0058399987874408 | 210 | 26 | 24 | 12 | 12 | 19 | 22 | 10 | 2 | 25 | 31 | 18 | 21 |
| L1PA3_dup7081 | 18.5007966426459 | 1.42599643833386 | 0.318222597903232 | 4.48112876876044 | 7.42492939487586e-06 | 0.000973942805839653 | 211 | 17 | 22 | 5 | 8 | 27 | 16 | 14 | 6 | 32 | 45 | 18 | 15 |
| LTR12C_dup1716 | 16.0274811787567 | 1.47945510678496 | 0.368329984372832 | 4.01665671966424 | 5.9029613168989e-05 | 0.00461124764777028 | 212 | 17 | 19 | 5 | 7 | 33 | 13 | 12 | 5 | 32 | 24 | 11 | 17 |
| MIRb_dup75921 | 14.6221801076248 | 1.41898948482789 | 0.437729437414243 | 3.24170449492761 | 0.00118817155024071 | 0.0395584078303696 | 213 | 17 | 18 | 3 | 1 | 27 | 13 | 12 | 5 | 17 | 32 | 19 | 17 |
| THE1B-int_dup2842 | 17.9145545472876 | 1.2901024176611 | 0.36469888375723 | 3.53744547932285 | 0.00040401758907037 | 0.0186062714994523 | 214 | 19 | 15 | 5 | 3 | 30 | 20 | 15 | 11 | 28 | 38 | 16 | 18 |
| MIRc_dup34919 | 19.5522971980416 | 1.05625613003659 | 0.302365383903466 | 3.49331036642016 | 0.000477071685012318 | 0.0208709871597612 | 215 | 17 | 12 | 9 | 12 | 24 | 23 | 8 | 4 | 37 | 39 | 23 | 32 |
| L1M4_dup15347 | 20.0211054882569 | 1.7071572660559 | 0.350963528978247 | 4.86420133461135 | 1.1491989061948e-06 | 0.000229209792903124 | 216 | 14 | 14 | 3 | 8 | 30 | 29 | 8 | 5 | 46 | 42 | 19 | 22 |
| L1MCb_dup782 | 17.5720125628098 | 1.59421101678172 | 0.314049509463391 | 5.07630475050164 | 3.84845870002269e-07 | 9.76229390903269e-05 | 217 | 21 | 20 | 5 | 6 | 20 | 22 | 7 | 5 | 38 | 31 | 19 | 18 |
| L1MA8_dup8321 | 21.2579972813442 | 1.15896729822847 | 0.304345447722572 | 3.80806516706941 | 0.000140058367629489 | 0.0086584381252186 | 218 | 21 | 20 | 4 | 13 | 18 | 15 | 7 | 12 | 38 | 60 | 22 | 27 |
| MER47A_dup1771 | 16.0474493537166 | 1.17762397693686 | 0.35628832565263 | 3.30525558136028 | 0.000948898272565308 | 0.0338917064490881 | 219 | 20 | 17 | 4 | 11 | 25 | 17 | 4 | 8 | 17 | 35 | 17 | 25 |
| Tigger6a_dup401 | 18.0004915903137 | 1.13412191576383 | 0.305444651146088 | 3.7130194014149 | 0.000204801180382328 | 0.0113086465363525 | 220 | 18 | 22 | 4 | 12 | 20 | 28 | 10 | 10 | 26 | 32 | 17 | 23 |
| L1PA16_dup13188 | 38.3182706778114 | 3.32767540505907 | 0.956550223202406 | 3.47882978263121 | 0.000503608361243627 | 0.0216921930596596 | 221 | 4 | 1 | 9 | 18 | 85 | 74 | 3 | 2 | 115 | 123 | 4 | 4 |
| MLT1A0_dup19224 | 28.671533744248 | 3.88162333837589 | 1.22218405120301 | 3.17597282876926 | 0.00149334937729054 | 0.0463764697114037 | 222 | 2 | 2 | 12 | 7 | 79 | 51 | 0 | 0 | 77 | 99 | 1 | 2 |
| ERVL-E-int_dup4593 | 38.4513944168659 | 3.50174303083681 | 0.566760263175133 | 6.17852601595456 | 6.47028278928534e-10 | 5.00103397123045e-07 | 223 | 2 | 3 | 0 | 2 | 7 | 12 | 1 | 2 | 181 | 180 | 11 | 14 |
| MER49_dup651 | 24.0704243860995 | 3.99440740582474 | 0.680320876028647 | 5.87135798204807 | 4.32239780865545e-09 | 2.46642030930055e-06 | 224 | 2 | 0 | 0 | 0 | 6 | 10 | 1 | 0 | 110 | 115 | 4 | 14 |
| L1MD3_dup2524 | 7.6386528217044 | 3.01737419888185 | 0.885595356594124 | 3.40717030234469 | 0.000656401570335071 | 0.0261315255236655 | 225 | 1 | 2 | 0 | 0 | 15 | 6 | 3 | 3 | 21 | 34 | 1 | 1 |
| L1M1_dup4591 | 10.8667654198683 | 2.86312900824524 | 0.906781035331392 | 3.15746458812836 | 0.00159147565170839 | 0.0483880993382834 | 226 | 0 | 4 | 0 | 0 | 17 | 20 | 7 | 3 | 28 | 43 | 1 | 3 |
| MER49_dup677 | 10.4988443474117 | 3.48003303246591 | 0.880183821786101 | 3.95375709747096 | 7.69335134479604e-05 | 0.00561023848805617 | 227 | 0 | 4 | 0 | 0 | 19 | 23 | 4 | 2 | 30 | 35 | 1 | 3 |
| HAL1b_dup2772 | 11.5687491868759 | 4.27069399765074 | 0.915367139489871 | 4.66555310258436 | 3.07787993148356e-06 | 0.000489879075703537 | 228 | 1 | 1 | 0 | 0 | 23 | 22 | 2 | 1 | 43 | 33 | 2 | 2 |
| Tigger2a_dup1756 | 12.3586766910245 | 3.82964752892169 | 0.827623678747289 | 4.62728124782309 | 3.70497229941759e-06 | 0.000567001141486893 | 229 | 2 | 3 | 0 | 0 | 24 | 26 | 3 | 3 | 42 | 34 | 1 | 2 |
| MER34B-int_dup365 | 16.9164347902653 | 2.8243319493624 | 0.743870804957937 | 3.79680440546676 | 0.00014657334762291 | 0.00895878288555892 | 230 | 4 | 1 | 1 | 1 | 35 | 28 | 10 | 2 | 48 | 56 | 7 | 3 |
| L1M1_dup4592 | 21.9743509547603 | 3.56095446294132 | 0.691564300756372 | 5.14912996383223 | 2.6169751150148e-07 | 7.14923950306903e-05 | 231 | 5 | 5 | 0 | 0 | 33 | 47 | 3 | 5 | 72 | 67 | 6 | 7 |
| L1PA3_dup5793 | 18.9028938184537 | 2.82670564964298 | 0.739451868381565 | 3.8227040467553 | 0.000131996170577432 | 0.0082942519616654 | 232 | 2 | 4 | 0 | 1 | 26 | 26 | 6 | 6 | 59 | 75 | 2 | 8 |

| | | | | | | | | | | | | | | | | | | | |
|---|---|---|---|---|---|---|---|---|---|---|---|---|---|---|---|---|---|---|---|
| L1PA15_dup276 | 48.7211887528992 | 2.57825128100465 | 0.584270500878267 | 4.41276990217556 | 1.02056445144602e-05 | 0.00124085522951809 | 233 | 75 | 67 | 0 | 9 | 22 | 35 | 5 | 1 | 100 | 92 | 88 | 104 |
| MIRb_dup41760 | 53.9869989065261 | 5.6170735888566 | 1.01291034473517 | 5.54547953632089 | 2.93149621550736e-08 | 1.25456438578258e-05 | 234 | 174 | 122 | 0 | 1 | 67 | 83 | 1 | 0 | 74 | 63 | 36 | 57 |
| L1PA2_dup187 | 27.8615179535395 | 1.84947365008101 | 0.431719968629228 | 4.28396596051219 | 1.83591012952104e-05 | 0.00192020866672093 | 235 | 38 | 38 | 6 | 11 | 17 | 12 | 4 | 0 | 62 | 70 | 36 | 44 |
| THE1B-int_dup3889 | 19.3710316115239 | 1.63274692914926 | 0.472859203208524 | 3.45292407987509 | 0.000554545052178882 | 0.0232186832633743 | 236 | 20 | 21 | 6 | 6 | 12 | 28 | 3 | 0 | 24 | 52 | 44 | 24 |
| L1MC1_dup10106 | 22.7464814164961 | 2.35554367877495 | 0.591866063219799 | 3.97985933837902 | 6.89560567639065e-05 | 0.00518742266289741 | 237 | 39 | 35 | 5 | 3 | 31 | 34 | 2 | 0 | 28 | 27 | 41 | 41 |
| L1PA3_dup1673 | 18.1570984452117 | 3.16262825387857 | 0.681089490715607 | 4.64348414854509 | 3.42582287798044e-06 | 0.000533657813334721 | 238 | 49 | 41 | 1 | 2 | 15 | 25 | 1 | 0 | 27 | 19 | 25 | 22 |
| L2_dup39050 | 23.1375015644119 | 2.20573898097294 | 0.441885786063991 | 4.99164954052972 | 5.98658064141421e-07 | 0.000138108076988327 | 239 | 45 | 32 | 4 | 9 | 31 | 41 | 1 | 4 | 39 | 31 | 15 | 35 |
| MER101_dup342 | 23.8087278773093 | 2.98745562103311 | 0.391423833441851 | 7.63227827688452 | 2.30640910829693e-14 | 6.53334311527169e-11 | 240 | 46 | 46 | 1 | 7 | 41 | 35 | 1 | 3 | 34 | 51 | 12 | 16 |
| L1MA4A_dup5838 | 22.1740468591736 | 2.9230509399773 | 0.411544527423783 | 7.10263591226746 | 1.22399558479594e-12 | 2.27535041734261e-09 | 241 | 50 | 51 | 8 | 6 | 42 | 38 | 2 | 0 | 30 | 29 | 7 | 13 |
| MLT2D_dup4128 | 20.8254652197988 | 2.5387801235476 | 0.367919296167778 | 6.90037230988252 | 5.18664240720478e-12 | 7.57141603818863e-09 | 242 | 30 | 41 | 6 | 12 | 30 | 40 | 2 | 1 | 36 | 36 | 11 | 10 |
| L1PA5_dup4671 | 36.6027231598451 | 2.08986298471998 | 0.619430260187064 | 3.37384709634439 | 0.000741255293644339 | 0.0285125658101351 | 243 | 123 | 120 | 16 | 16 | 33 | 35 | 1 | 3 | 21 | 31 | 37 | 35 |
| L1PA7_dup11139 | 45.6645847634044 | 2.57959988972968 | 0.808009383745571 | 3.19253704427517 | 0.00141028853882235 | 0.0445292617646792 | 244 | 194 | 139 | 19 | 19 | 23 | 29 | 3 | 0 | 41 | 60 | 24 | 35 |
| L1PA7_dup9678 | 32.4883035975904 | 3.50683334819008 | 0.562261376139312 | 6.23701626504963 | 4.45995633015083e-10 | 3.70502118484417e-07 | 245 | 91 | 78 | 5 | 5 | 54 | 79 | 1 | 0 | 29 | 35 | 13 | 22 |
| MLT2B5_dup346 | 40.5535089012444 | 3.1348512599204 | 0.481732034822019 | 6.50745857306044 | 7.64328198168996e-11 | 8.01184305909779e-08 | 246 | 115 | 96 | 11 | 13 | 69 | 61 | 0 | 1 | 45 | 62 | 15 | 21 |
| L2_dup52958 | 29.3673080585274 | 3.2252668955035 | 0.427250432423797 | 7.54889088632754 | 4.38980444876242e-14 | 1.1605960104058e-10 | 247 | 74 | 59 | 8 | 9 | 64 | 48 | 0 | 1 | 37 | 44 | 13 | 8 |
| L2b_dup87330 | 23.8205126662156 | 3.07476369172727 | 0.402534912260223 | 7.63850189903417 | 2.1976371460798e-14 | 6.30022853447115e-11 | 248 | 53 | 49 | 7 | 6 | 43 | 37 | 0 | 1 | 28 | 50 | 9 | 13 |
| L1MA4_dup9425 | 30.7322204601276 | 3.22278522363392 | 0.398088809159479 | 8.09564386006851 | 5.69622504869784e-16 | 2.25898994239576e-12 | 249 | 73 | 56 | 6 | 11 | 41 | 47 | 0 | 2 | 43 | 78 | 6 | 17 |
| THE1B_dup21099 | 29.6702856183993 | 3.3705446269735 | 0.431254544235061 | 7.81567330021302 | 5.46700030048401e-15 | 1.80673729652634e-11 | 250 | 71 | 59 | 6 | 9 | 28 | 46 | 0 | 1 | 70 | 47 | 9 | 12 |
| L1MB5_dup1608 | 38.70178501556 | 1.81475827508188 | 0.572151252876108 | 3.17181560987484 | 0.00151489159271828 | 0.0468315440978231 | 251 | 74 | 71 | 0 | 2 | 52 | 54 | 29 | 17 | 37 | 71 | 31 | 47 |
| LTR7_dup482 | 41.0764256368113 | 3.47748537208408 | 0.491028722539316 | 7.08204064744021 | 1.42046975420677e-12 | 2.58011523766781e-09 | 252 | 101 | 84 | 0 | 2 | 58 | 65 | 6 | 4 | 68 | 64 | 22 | 32 |
| MSTA-int_dup2197 | 30.731740345369 | 2.62001670911895 | 0.558152195576198 | 4.69409012431496 | 2.6779595988941e-06 | 0.000442947302143162 | 253 | 50 | 37 | 1 | 0 | 38 | 45 | 7 | 3 | 48 | 56 | 46 | 49 |
| L1M5_dup10731 | 22.7742488351862 | 2.89881816495636 | 0.524131014178665 | 5.53071290676994 | 3.18931938032086e-08 | 1.3290469164095e-05 | 254 | 40 | 41 | 0 | 1 | 22 | 32 | 12 | 0 | 44 | 55 | 17 | 13 |
| L1PA7_dup6922 | 21.3624591201147 | 3.16383877130577 | 0.519103984498028 | 6.09480733299551 | 1.09569336066638e-09 | 7.75940036896195e-07 | 255 | 60 | 62 | 0 | 1 | 37 | 40 | 8 | 3 | 20 | 21 | 9 | 8 |
| L1MC5a_dup3390 | 19.7403690046365 | 2.60541278442768 | 0.532870371335161 | 4.88939322691099 | 1.01147267024593e-06 | 0.000207479203443395 | 256 | 57 | 50 | 1 | 1 | 31 | 35 | 8 | 4 | 16 | 21 | 8 | 19 |
| L1PA5_dup6022 | 24.8640392784219 | 3.02291283040431 | 0.580238032488328 | 5.20978057477664 | 1.89064095903296e-07 | 5.51311830438795e-05 | 257 | 73 | 56 | 1 | 1 | 51 | 53 | 6 | 3 | 20 | 18 | 16 | 18 |
| L1M2_dup146 | 20.807767201082 | 2.57282566903263 | 0.658086394016575 | 3.90955608932986 | 9.24658844292721e-05 | 0.00644838433071735 | 258 | 61 | 47 | 0 | 0 | 18 | 19 | 6 | 4 | 24 | 31 | 23 | 29 |
| L2_dup38870 | 17.0937250885972 | 1.81147807488581 | 0.550952265899707 | 3.28790384758226 | 0.00100936306385367 | 0.0352989276295891 | 259 | 39 | 23 | 0 | 1 | 27 | 23 | 13 | 6 | 22 | 23 | 10 | 28 |
| THE1A-int_dup194 | 25.4913818885765 | 1.48239616710651 | 0.354605623672598 | 4.18040794659021 | 2.90986591752258e-05 | 0.00269390483312078 | 260 | 51 | 38 | 9 | 21 | 14 | 18 | 15 | 6 | 50 | 54 | 18 | 19 |
| L1ME1_dup4350 | 25.6100416465658 | 1.11075495593185 | 0.333102669466822 | 3.33457236385939 | 0.000854306687340613 | 0.0314805513293488 | 261 | 40 | 44 | 10 | 27 | 19 | 20 | 9 | 9 | 31 | 56 | 27 | 28 |
| MLT1K_dup10980 | 24.4944841519833 | 1.11281338055768 | 0.280954804554627 | 3.96082701743339 | 7.46906275066659e-05 | 0.00549546569347592 | 262 | 36 | 24 | 8 | 18 | 33 | 18 | 14 | 7 | 35 | 52 | 32 | 27 |
| L1ME2_dup1836 | 31.2243204588562 | 1.1838635031634 | 0.321067436800563 | 3.68727366113672 | 0.000226669552749511 | 0.012180468247185 | 263 | 49 | 38 | 18 | 33 | 20 | 33 | 9 | 7 | 51 | 69 | 22 | 39 |
| THE1B-int_dup524 | 26.5810723372806 | 1.72358955614192 | 0.344065127448208 | 5.00948634034809 | 5.45754999398683e-07 | 0.000128320374591817 | 264 | 41 | 28 | 13 | 18 | 21 | 18 | 4 | 4 | 54 | 81 | 17 | 23 |
| L1MD3_dup2457 | 38.4691163158353 | 1.36570433728197 | 0.28807584623357 | 4.74078044076859 | 2.1289654485842e-06 | 0.000369355273216689 | 265 | 35 | 18 | 18 | 19 | 40 | 25 | 11 | 10 | 85 | 126 | 27 | 47 |
| L1P2_dup1291 | 28.0523406066101 | 1.49763994579878 | 0.288665175382305 | 5.18815594508524 | 2.12386716954234e-07 | 6.04237541856086e-05 | 266 | 26 | 21 | 6 | 9 | 34 | 27 | 8 | 7 | 59 | 66 | 27 | 50 |
| L1PA7_dup4487 | 33.1815520471661 | 1.45457550067405 | 0.269079125001646 | 5.40575379329466 | 6.45363966684725e-08 | 2.3588598220701e-05 | 267 | 28 | 15 | 5 | 8 | 18 | 26 | 12 | 7 | 97 | 88 | 38 | 49 |
| L1PA7_dup7006 | 30.2398600220442 | 2.28167281942689 | 0.377244661079633 | 6.04825741707515 | 1.46420878216238e-09 | 9.93609408656041e-07 | 268 | 17 | 10 | 9 | 4 | 13 | 24 | 3 | 2 | 116 | 102 | 17 | 25 |
| L1PA2_dup1230 | 23.3511116402892 | 1.55545085327249 | 0.419849550835004 | 3.7047815108507 | 0.000211573021550474 | 0.0115757540091628 | 269 | 19 | 15 | 6 | 15 | 13 | 32 | 17 | 8 | 55 | 77 | 11 | 10 |
| THE1C-int_dup1784 | 19.7642410901778 | 1.22902455952479 | 0.371270194469921 | 3.31032379606857 | 0.000931881033699902 | 0.0334838602028075 | 270 | 18 | 13 | 6 | 17 | 19 | 17 | 13 | 8 | 50 | 48 | 16 | 10 |
| L1MA2_dup6577 | 38.0198642268624 | 3.32835402120267 | 0.422519541310212 | 7.87739665455853 | 3.34272455661081e-15 | 1.16968814315782e-11 | 271 | 36 | 27 | 4 | 9 | 73 | 76 | 1 | 2 | 88 | 100 | 12 | 25 |
| L1PA4_dup10597 | 37.0934731210145 | 3.14359569733449 | 0.396752291359028 | 7.9233208372067 | 2.31249872270572e-15 | 8.21268389661099e-12 | 272 | 18 | 27 | 3 | 7 | 63 | 59 | 6 | 2 | 107 | 105 | 14 | 21 |
| L1PA3_dup9593 | 18.4902446516392 | 3.04225395347835 | 0.536814391547536 | 5.66723620189857 | 1.45119230731967e-08 | 6.99777351904828e-06 | 273 | 8 | 12 | 0 | 6 | 45 | 32 | 3 | 2 | 49 | 43 | 8 | 9 |
| L1ME1_dup25709 | 22.2866033024594 | 2.73692268347141 | 0.467245366099217 | 5.85757052300062 | 4.69687387071921e-09 | 2.64834680104776e-06 | 274 | 28 | 15 | 2 | 5 | 58 | 44 | 3 | 3 | 44 | 35 | 15 | 18 |
| L1MA2_dup6576 | 24.0245444542518 | 2.99693863021947 | 0.4846167671165 | 6.18414143623516 | 6.24413661415417e-10 | 4.87136829767714e-07 | 275 | 12 | 15 | 4 | 6 | 51 | 47 | 2 | 1 | 61 | 58 | 15 | 11 |
| L1PA14_dup2727 | 21.1741059524389 | 3.28003729364862 | 0.51350772356465 | 6.38751306578092 | 1.68605206959052e-10 | 1.59836392729397e-07 | 276 | 11 | 11 | 2 | 4 | 46 | 40 | 1 | 2 | 50 | 63 | 7 | 13 |
| L1MA9_dup5789 | 31.6382769825878 | 2.34206002524472 | 0.668207658869671 | 3.50498829840788 | 0.000456627352665897 | 0.0202899443618001 | 277 | 11 | 10 | 2 | 2 | 47 | 52 | 1 | 3 | 47 | 55 | 73 | 90 |
| L1ME1_dup18928 | 27.5222752686724 | 2.12151161563806 | 0.515120385133733 | 4.11847730523669 | 3.81384121225707e-05 | 0.00332048394106008 | 278 | 26 | 15 | 3 | 3 | 44 | 42 | 4 | 3 | 61 | 33 | 39 | 63 |
| HAL1_dup1668 | 25.819994103506 | 2.15655084384968 | 0.413488141597356 | 5.21550832272638 | 1.83313828655349e-07 | 5.38503608311428e-05 | 279 | 31 | 28 | 1 | 3 | 53 | 46 | 11 | 10 | 43 | 45 | 20 | 23 |
| Tigger1_dup2716 | 29.3575451360945 | 2.52912432128842 | 0.363213717712689 | 6.96318502840529 | 3.32664430707162e-12 | 5.08001000040174e-09 | 280 | 50 | 45 | 2 | 3 | 38 | 52 | 6 | 8 | 50 | 57 | 21 | 26 |
| L1PREC2_dup4993 | 21.1511778403831 | 2.00688659856836 | 0.487968878267392 | 4.11273482377426 | 3.90999422445958e-05 | 0.00338192470277448 | 281 | 8 | 19 | 3 | 3 | 49 | 44 | 11 | 6 | 41 | 33 | 18 | 20 |
| L2c_dup50028 | 22.3962197928431 | 1.46485945723157 | 0.385530007966537 | 3.79959906352792 | 0.000144930370611324 | 0.00888267567018937 | 282 | 20 | 13 | 7 | 5 | 33 | 48 | 11 | 9 | 40 | 35 | 25 | 27 |
| AluJb_dup53405 | 21.6571696641122 | 1.56857511069904 | 0.351411312837091 | 4.46364432048381 | 8.05772808672907e-06 | 0.00103536301943529 | 283 | 30 | 21 | 4 | 4 | 21 | 32 | 9 | 5 | 33 | 46 | 33 | 29 |
| L1ME1_dup25854 | 18.7614586741743 | 1.83811888241005 | 0.431826645261092 | 4.25661293155888 | 2.07547221966177e-05 | 0.00209756508641218 | 284 | 20 | 18 | 1 | 2 | 22 | 25 | 5 | 4 | 35 | 37 | 27 | 33 |
| L1ME2_dup4576 | 24.2469479334246 | 1.80445453878403 | 0.423775390562792 | 4.25804465990258 | 2.06222756983936e-05 | 0.00208896892862068 | 285 | 25 | 35 | 6 | 1 | 27 | 36 | 5 | 7 | 38 | 45 | 33 | 39 |
| L1PA4_dup9862 | 22.483948495793 | 1.76794989374489 | 0.396858275076369 | 4.45486463248038 | 8.39461438780532e-06 | 0.00106759215132054 | 286 | 28 | 17 | 3 | 3 | 25 | 33 | 4 | 13 | 42 | 54 | 17 | 32 |
| L2a_dup94038 | 2.52612973936977 | 3.24120391574012 | 1.0233309442176 | 3.16730763791984 | 0.00153857461701757 | 0.0473055531490971 | 287 | 4 | 0 | 0 | 0 | 3 | 2 | 0 | 0 | 5 | 13 | 2 | 1 |
| Tigger1_dup5332 | 2.62420890159482 | 3.22084200595275 | 1.02438430438699 | 3.14417352175282 | 0.00166556645311497 | 0.0499580282577669 | 288 | 2 | 0 | 0 | 0 | 2 | 4 | 0 | 0 | 5 | 15 | 2 | 1 |
| AluSx_dup103295 | 2.70187415170001 | 4.57001785303884 | 1.42854060958057 | 3.19908151185189 | 0.00137866183108891 | 0.0438800251552009 | 289 | 0 | 0 | 0 | 0 | 7 | 2 | 0 | 0 | 10 | 11 | 0 | 0 |
| THE1B_dup21101 | 2.99136279311571 | 4.63723873285606 | 1.35564220133437 | 3.42069517184668 | 0.000624612995513927 | 0.0252333045552728 | 290 | 0 | 0 | 0 | 0 | 5 | 4 | 0 | 0 | 12 | 12 | 0 | 0 |
| AluSq2_dup9716 | 3.60819934114139 | 3.04811287930469 | 0.949510516272799 | 3.21019391261692 | 0.00132645455584164 | 0.0426928901250588 | 291 | 0 | 1 | 0 | 1 | 4 | 5 | 0 | 0 | 16 | 9 | 2 | 2 |
| SATR2_dup129 | 2.71789625239512 | 3.4970185343771 | 1.04512797446443 | 3.34601945390385 | 0.000819806437780826 | 0.0306039633894252 | 292 | 1 | 1 | 0 | 0 | 1 | 1 | 0 | 0 | 14 | 10 | 1 | 0 |
| L1PA5_dup9310 | 3.52728843049091 | 3.46639642558993 | 0.905016613216759 | 3.83020198189412 | 0.000128038117672526 | 0.00812647584628086 | 293 | 2 | 2 | 0 | 1 | 3 | 2 | 0 | 0 | 15 | 12 | 2 | 0 |
| THE1C_dup3397 | 3.70079796776038 | 3.41636338705295 | 0.889271062310104 | 3.84175706581309 | 0.00012215670915604 | 0.00784950049064087 | 294 | 1 | 2 | 0 | 1 | 3 | 1 | 0 | 0 | 15 | 16 | 1 | 1 |
| L1ME1_dup25707 | 2.68858426690013 | 4.42685235812543 | 1.15617135762364 | 3.82888948851339 | 0.000128722788781025 | 0.00815687688396529 | 295 | 3 | 1 | 0 | 1 | 9 | 4 | 0 | 0 | 2 | 13 | 0 | 0 |
| LOR1a_dup670 | 3.34177064398679 | 4.591018619806 | 1.07011507897211 | 4.29021019329609 | 1.78504088335611e-05 | 0.00187773358988088 | 296 | 3 | 1 | 0 | 1 | 8 | 4 | 0 | 0 | 12 | 9 | 0 | 0 |
| MSTB1_dup3790 | 3.94868571672615 | 4.94058505606382 | 1.10674702021466 | 4.46405995753714 | 8.04210439419072e-06 | 0.00103493054201195 | 297 | 5 | 1 | 0 | 0 | 15 | 3 | 0 | 0 | 11 | 10 | 0 | 1 |
| L2_dup15105 | 2.97897762170037 | 3.35160465097321 | 0.949980864860379 | 3.52807595915714 | 0.00041859201952688 | 0.0190845557919799 | 298 | 3 | 3 | 1 | 1 | 3 | 3 | 1 | 0 | 11 | 8 | 0 | 0 |
| AluSz_dup91665 | 3.39141100024656 | 4.28973288238618 | 0.985476722402785 | 4.35295201283595 | 1.34316558812332e-05 | 0.00152263401158452 | 299 | 3 | 3 | 0 | 0 | 2 | 2 | 1 | 0 | 13 | 14 | 0 | 0 |
| L1PA10_dup5290 | 2.76886509189612 | 3.80815701277187 | 1.05588009822298 | 3.60661879997636 | 0.000310212866958184 | 0.015307737627796 | 300 | 4 | 1 | 0 | 0 | 4 | 4 | 2 | 0 | 6 | 12 | 0 | 0 |
| AluSp_dup38142 | 2.35245210900598 | 4.15619576716977 | 1.19371841870155 | 3.48172207285753 | 0.000498200486446139 | 0.0215170420094504 | 301 | 1 | 1 | 1 | 0 | 2 | 9 | 0 | 0 | 8 | 5 | 0 | 0 |
| L2c_dup116499 | 2.29667535541891 | 3.40365527422229 | 1.03257944626881 | 3.2962647925264 | 0.000979796194731131 | 0.0346270985938937 | 302 | 1 | 2 | 1 | 0 | 2 | 4 | 0 | 0 | 8 | 7 | 1 | 0 |
| AluSz_dup91719 | 3.24390030012394 | 4.24349214533205 | 1.12358777095658 | 3.7767340078108 | 0.000158898268133038 | 0.00949741454639135 | 303 | 2 | 0 | 1 | 0 | 2 | 6 | 0 | 0 | 10 | 16 | 0 | 0 |
| THE1C_dup8453 | 2.59100226840458 | 3.74900948916211 | 1.05779931096863 | 3.54415951143808 | 0.000393866874313295 | 0.0182688201316474 | 304 | 3 | 1 | 1 | 1 | 3 | 6 | 0 | 0 | 5 | 11 | 0 | 0 |
| AluSz6_dup38068 | 4.95650413224838 | 3.93987508522257 | 1.04341332940739 | 3.77594858545675 | 0.000159399871186419 | 0.00951783230848776 | 305 | 2 | 1 | 0 | 0 | 12 | 16 | 2 | 0 | 14 | 9 | 2 | 0 |
| MER106B_dup414 | 4.35437558647507 | 3.42636901269531 | 1.07358946195584 | 3.19150767971702 | 0.00141532348396199 | 0.044634931970155 | 306 | 1 | 0 | 0 | 0 | 6 | 14 | 2 | 0 | 16 | 8 | 1 | 2 |
| L1MD3_dup2523 | 5.81909018742934 | 3.28337559789192 | 0.955537303952354 | 3.43615637433621 | 0.000590030307254228 | 0.0242994321478529 | 307 | 4 | 0 | 0 | 0 | 9 | 8 | 4 | 0 | 19 | 21 | 1 | 1 |
| AluJr4_dup1696 | 3.32268338469321 | 3.24022295404824 | 0.943015003487806 | 3.43602481621613 | 0.000590316897470174 | 0.0243058564605361 | 308 | 4 | 0 | 1 | 0 | 6 | 5 | 1 | 0 | 9 | 11 | 1 | 1 |
| MER44B_dup1823 | 4.66163632251274 | 3.35589727801915 | 0.90862692336284 | 3.69337204493009 | 0.000221299885810213 | 0.0119703165785399 | 309 | 2 | 1 | 1 | 0 | 9 | 7 | 2 | 0 | 14 | 16 | 0 | 2 |
| MLT1F2_dup5473 | 8.2021971494765 | 3.60587592516505 | 1.08377145068058 | 3.32715529911834 | 0.000877374621571317 | 0.0321280058024636 | 310 | 0 | 0 | 3 | 0 | 25 | 7 | 1 | 0 | 27 | 27 | 1 | 2 |
| AluJo_dup35590 | 4.9226774566871 | 4.41998468613785 | 1.16736717983615 | 3.78628486605066 | 0.000152916365502181 | 0.00923264082866835 | 311 | 0 | 0 | 0 | 0 | 8 | 8 | 1 | 0 | 19 | 18 | 0 | 1 |

| | | | | | | | | | | | | | | | | | | | |
|---|---|---|---|---|---|---|---|---|---|---|---|---|---|---|---|---|---|---|---|
| MER2B_dup1320 | 5.2576128362446 | 5.30077366992315 | 1.2114179082288 | 4.37567715807781 | 1.21056075267787e-05 | 0.00140648480887055 | 312 | 1 | 0 | 0 | 0 | 12 | 14 | 1 | 0 | 14 | 19 | 0 | 0 |
| AluSq_dup16728 | 5.23426802689481 | 4.47192064540326 | 1.17009931419125 | 3.82182998585396 | 0.000132465006021804 | 0.0083077275495l613 | 313 | 0 | 0 | 0 | 1 | 7 | 10 | 0 | 0 | 18 | 22 | 1 | 0 |
| L1MCc_dup3166 | 5.4084909352353 | 4.31348673982388 | 1.09997086555584 | 3.92145544477142 | 8.8015724639866e-05 | 0.00621453697779156 | 314 | 0 | 1 | 1 | 1 | 12 | 13 | 0 | 0 | 13 | 21 | 1 | 0 |
| L1M1_dup4593 | 14.2234404861109 | 4.47604760418312 | 0.872684942916258 | 5.12905332046348 | 2.91202860867775e-07 | 7.81178759098575e-05 | 315 | 1 | 4 | 0 | 0 | 23 | 31 | 3 | 0 | 47 | 48 | 3 | 2 |
| THE1B-int_dup4015 | 15.6643757302686 | 3.53273464023101 | 0.78627375891451 | 4.49300844671216 | 7.02239981929329e-06 | 0.000932971494919912 | 316 | 3 | 4 | 5 | 3 | 26 | 35 | 1 | 0 | 55 | 41 | 5 | 0 |
| MER68_dup1286 | 11.5894792174746 | 3.70780196407653 | 0.797053484092567 | 4.65188602531209 | 3.28912864297866e-06 | 0.000516250002692744 | 317 | 2 | 1 | 2 | 2 | 19 | 19 | 0 | 1 | 40 | 41 | 2 | 2 |
| THE1C_dup9232 | 14.2886290196295 | 3.36699726099345 | 0.712396656016252 | 4.72629571259054 | 2.2865259993876e-06 | 0.00038862122532163 | 318 | 2 | 4 | 4 | 3 | 20 | 30 | 1 | 1 | 48 | 45 | 1 | 4 |
| LTR50_dup2414 | 8.32879409778047 | 3.73947110585016 | 0.928986712496674 | 4.02532248905933 | 5.68972871732573e-05 | 0.00449186526002916 | 319 | 3 | 1 | 2 | 4 | 18 | 20 | 0 | 0 | 27 | 20 | 1 | 0 |
| L1MB3_dup15600 | 7.20141536757149 | 3.44823729862957 | 1.07505771749015 | 3.20749039100886 | 0.00133898533565126 | 0.0429909870026818 | 320 | 0 | 0 | 2 | 3 | 13 | 19 | 0 | 0 | 26 | 17 | 1 | 1 |
| MER58A_dup12197 | 8.56897274171188 | 3.10694976425043 | 0.960622675922393 | 3.23430816503173 | 0.00121937806771108 | 0.0402706929491742 | 321 | 1 | 0 | 1 | 5 | 14 | 17 | 0 | 0 | 29 | 27 | 3 | 1 |
| L1MEf_dup9738 | 8.45948302331888 | 3.99231412112417 | 0.758156029885107 | 5.26582123435617 | 1.39564043433704e-07 | 4.32968785904513e-05 | 322 | 9 | 3 | 2 | 1 | 13 | 16 | 0 | 1 | 17 | 36 | 2 | 0 |
| Tigger2a_dup3086 | 8.58089192426285 | 5.35925716937848 | 0.930237884145889 | 5.76116847176048 | 8.35336085051878e-09 | 4.33116788005429e-06 | 323 | 4 | 2 | 0 | 1 | 13 | 15 | 0 | 0 | 23 | 40 | 1 | 0 |
| MIR3_dup14280 | 4.98465400617641 | 3.00586456368609 | 0.737763207957694 | 4.07429447723077 | 4.61540513834381e-05 | 0.00384260738645331 | 324 | 4 | 7 | 0 | 0 | 4 | 4 | 1 | 2 | 12 | 20 | 1 | 3 |
| AluSx4_dup2142 | 4.48630160786004 | 2.91475878474932 | 0.756917908082587 | 3.850825503829 | 0.000117720355004366 | 0.00764912277221976 | 325 | 10 | 11 | 0 | 0 | 3 | 4 | 1 | 2 | 8 | 11 | 2 | 2 |
| MLT1K_dup3543 | 4.70075245500998 | 3.1431814783671 | 0.749069518647496 | 4.1961145128991 | 2.7153299766803e-05 | 0.00255578285850938 | 326 | 10 | 7 | 0 | 0 | 5 | 7 | 1 | 3 | 9 | 12 | 1 | 1 |
| L2c_dup116500 | 4.65883968344802 | 2.98435598169701 | 0.733881140146496 | 4.06653859656522 | 4.77165854509285e-05 | 0.00393007285137394 | 327 | 3 | 8 | 1 | 0 | 5 | 9 | 1 | 1 | 11 | 12 | 1 | 3 |
| L2_dup4853 | 5.105354212971 | 2.18775682658677 | 0.656271515994547 | 3.33361539129324 | 0.000857251051137833 | 0.0315659948334947 | 328 | 4 | 8 | 0 | 0 | 6 | 9 | 1 | 4 | 11 | 10 | 3 | 4 |
| L2c_dup14359 | 4.57647173328085 | 2.52646812669391 | 0.694661351321293 | 3.63697810722936 | 0.000275855367171303 | 0.0140885772047527 | 329 | 4 | 5 | 0 | 0 | 10 | 7 | 2 | 2 | 11 | 8 | 2 | 3 |
| MER89_dup1101 | 6.31125564078628 | 2.07101886110254 | 0.635897581666498 | 3.25684343015587 | 0.00112658555647988 | 0.0381338169859754 | 330 | 11 | 12 | 1 | 0 | 3 | 7 | 4 | 3 | 6 | 23 | 2 | 6 |
| MER97b_dup34 | 5.69160904287824 | 2.49682758195266 | 0.697188117636351 | 3.5812824670873 | 0.00034191175162826 | 0.0164223927438308 | 331 | 12 | 12 | 0 | 0 | 4 | 11 | 1 | 2 | 4 | 15 | 6 | 4 |
| MIR_dup9659 | 5.53265465344512 | 2.08576541331006 | 0.611741110257582 | 3.40955573908024 | 0.000650687798179185 | 0.0259648765429389 | 332 | 5 | 11 | 2 | 1 | 8 | 12 | 1 | 1 | 11 | 5 | 5 | 5 |
| MamTip2_dup2460 | 4.99145394733023 | 2.25196753620031 | 0.64410842204276 | 3.49625538051233 | 0.000471836888805776 | 0.0207219823447359 | 333 | 4 | 8 | 1 | 1 | 10 | 13 | 2 | 1 | 9 | 5 | 2 | 5 |
| AluSq2_dup12027 | 4.79813637719065 | 2.16411687659592 | 0.608901096509405 | 3.55413529225349 | 0.000379224076359655 | 0.0177662634521509 | 334 | 5 | 5 | 1 | 2 | 9 | 6 | 2 | 0 | 10 | 11 | 4 | 3 |
| HAL1M8_dup81 | 5.7643642195005 | 2.12171056384573 | 0.571227979947354 | 3.71429733543737 | 0.000203769096147047 | 0.0112679622012097 | 335 | 11 | 7 | 0 | 3 | 8 | 10 | 4 | 0 | 12 | 8 | 4 | 4 |
| Tigger16a_dup123 | 5.91749787692887 | 2.09379550758342 | 0.655665422983904 | 3.19339015630053 | 0.00140612822079721 | 0.0444619624779882 | 336 | 7 | 4 | 0 | 0 | 8 | 10 | 3 | 0 | 8 | 17 | 10 | 6 |
| MLT1G3_dup622 | 6.88206197315206 | 2.75874295402984 | 0.669722674559168 | 4.11923182359882 | 3.80137555597772e-05 | 0.00331205458821924 | 337 | 10 | 15 | 0 | 0 | 13 | 9 | 2 | 0 | 11 | 11 | 4 | 10 |
| AluY_dup20323 | 4.72120224314469 | 2.67168641399895 | 0.69874140615748 | 3.82356962168751 | 0.00013153342811047 | 0.00827110282377745 | 338 | 6 | 8 | 0 | 0 | 5 | 9 | 2 | 0 | 8 | 12 | 2 | 6 |
| MER2_dup6635 | 4.45657040669442 | 2.50365861801156 | 0.740582190604803 | 3.38066274044063 | 0.000723112394589672 | 0.0279899139833741 | 339 | 5 | 7 | 0 | 0 | 11 | 5 | 2 | 0 | 7 | 9 | 2 | 7 |
| HAL1_dup1667 | 7.24829840659604 | 2.37400483295003 | 0.636040583447589 | 3.73247383065086 | 0.000189608400913281 | 0.0106987338306169 | 340 | 10 | 7 | 0 | 0 | 21 | 16 | 5 | 1 | 9 | 10 | 6 | 5 |
| L1MD1_dup5544 | 6.17752655632688 | 1.98621430002639 | 0.599284412298537 | 3.31430996579458 | 0.000918696034361499 | 0.0331941792996004 | 341 | 11 | 8 | 1 | 0 | 11 | 16 | 6 | 1 | 5 | 10 | 6 | 3 |
| L1MC4_dup1731 | 5.86881723949848 | 2.54170947890414 | 0.650695719458568 | 3.90614138513013 | 9.37816528623392e-05 | 0.00651002269111798 | 342 | 10 | 6 | 0 | 0 | 14 | 11 | 3 | 2 | 6 | 14 | 1 | 6 |
| Tigger16a_dup909 | 5.55502752409094 | 2.19998954910006 | 0.656488852286942 | 3.35114532628571 | 0.000804780561098094 | 0.0301708390406565 | 343 | 8 | 5 | 0 | 1 | 15 | 14 | 2 | 1 | 5 | 8 | 4 | 7 |
| THE1A_dup856 | 7.87589969799441 | 2.6388739477069 | 0.675594890124178 | 3.90600045423946 | 9.38363350793642e-05 | 0.00651151373193187 | 344 | 17 | 8 | 0 | 2 | 8 | 9 | 7 | 0 | 15 | 25 | 5 | 0 |
| LTR7_dup481 | 7.7867396386466 | 2.81912407769903 | 0.622623019309249 | 4.52781858407134 | 5.95957111654441e-06 | 0.000823972172514397 | 345 | 11 | 17 | 0 | 0 | 4 | 11 | 5 | 1 | 18 | 19 | 4 | 3 |
| MLT2D_dup929 | 6.55909081279361 | 3.53103802591907 | 0.710723763264945 | 4.96822845728146 | 6.75673317354369e-07 | 0.000151007122028644 | 346 | 13 | 14 | 0 | 0 | 6 | 11 | 4 | 0 | 15 | 13 | 2 | 1 |
| Tigger1_dup4324 | 8.83418722953734 | 2.07527228568919 | 0.509904673795093 | 4.06992207042045 | 4.70288662962593e-05 | 0.00389771181460457 | 347 | 5 | 6 | 0 | 3 | 5 | 5 | 3 | 0 | 30 | 29 | 8 | 8 |
| L3b_dup9091 | 7.01747611684068 | 1.95986726643958 | 0.558237314397341 | 3.51081379888659 | 0.000446737196423093 | 0.0199888784947755 | 348 | 10 | 11 | 0 | 2 | 8 | 5 | 2 | 1 | 18 | 10 | 9 | 8 |
| AluSz_dup42776 | 8.02143592238378 | 1.70713426419543 | 0.537022926844769 | 3.17888525584102 | 0.00147842603982447 | 0.046045230690062 | 349 | 7 | 10 | 0 | 1 | 7 | 2 | 4 | 2 | 19 | 24 | 11 | 8 |
| L1ME1_dup23546 | 8.00582295556208 | 1.44681211521137 | 0.458645284736644 | 3.15453393583265 | 0.00160754681372585 | 0.0486900883577916 | 350 | 10 | 7 | 0 | 3 | 9 | 6 | 5 | 3 | 15 | 21 | 10 | 8 |
| L1MB7_dup16627 | 6.35749578661697 | 1.85462894619291 | 0.543479053144891 | 3.41251228628028 | 0.000643670253591798 | 0.0257756247326075 | 351 | 5 | 5 | 3 | 1 | 5 | 6 | 1 | 0 | 17 | 18 | 8 | 6 |
| L2a_dup57862 | 6.47727203792076 | 1.98255053918734 | 0.557740993075445 | 3.55460789829224 | 0.000378543154566508 | 0.0177448442585662 | 352 | 8 | 3 | 2 | 1 | 7 | 4 | 1 | 0 | 17 | 19 | 7 | 8 |
| L2a_dup112237 | 6.8303932656104 | 2.1078513124202 | 0.549048614295652 | 3.83909777301644 | 0.000123487251740398 | 0.00791028362999303 | 353 | 12 | 4 | 2 | 1 | 6 | 10 | 2 | 0 | 11 | 22 | 7 | 7 |
| MIRb_dup105954 | 6.91490799407679 | 1.84110895232768 | 0.505723520637835 | 3.64054444216005 | 0.000272062140412292 | 0.013945734179781 | 354 | 8 | 6 | 3 | 1 | 6 | 6 | 4 | 0 | 16 | 21 | 6 | 6 |
| L2a_dup46423 | 5.69511034436287 | 2.57490186164447 | 0.60892717089176 | 4.22858756306371 | 2.35162991023773e-05 | 0.00231032589026188 | 355 | 7 | 7 | 1 | 0 | 6 | 8 | 2 | 0 | 11 | 18 | 3 | 6 |
| THE1B_dup11594 | 6.92640017390965 | 2.19477075608029 | 0.576869276629996 | 3.80462410635194 | 0.00014201969329837 | 0.00874942656525124 | 356 | 9 | 5 | 2 | 0 | 8 | 4 | 5 | 0 | 19 | 21 | 4 | 5 |
| L1ME1_dup14496 | 5.89213862241754 | 1.82466900519252 | 0.571405558568388 | 3.19329936125244 | 0.00140657045663215 | 0.0444619624779882 | 357 | 2 | 8 | 2 | 1 | 9 | 6 | 2 | 1 | 10 | 18 | 6 | 6 |
| MER34_dup182 | 6.69372345225683 | 1.81390105206298 | 0.566468965220268 | 3.20211902757577 | 0.0013642061654709 | 0.0435538777972854 | 358 | 4 | 7 | 4 | 0 | 10 | 9 | 6 | 1 | 12 | 20 | 3 | 5 |
| L1MCb_dup781 | 6.77303574996973 | 2.67069579869123 | 0.570107175363954 | 4.68455040402934 | 2.80575118330254e-06 | 0.000458100744702134 | 359 | 9 | 10 | 2 | 0 | 15 | 10 | 1 | 1 | 12 | 13 | 4 | 5 |
| MER1B_dup654 | 5.84257868076232 | 2.62802129724274 | 0.624517461978366 | 4.20808297163959 | 2.57546306188323e-05 | 0.00246335178259389 | 360 | 6 | 10 | 1 | 0 | 6 | 5 | 2 | 1 | 10 | 22 | 4 | 3 |
| AluJr4_dup6367 | 6.53596213556993 | 1.71024459817649 | 0.518537489062273 | 3.29820819950609 | 0.000973039567519163 | 0.0344745194956692 | 361 | 7 | 6 | 2 | 0 | 9 | 5 | 3 | 2 | 11 | 20 | 7 | 7 |
| THE1B_dup14929 | 5.88628893966476 | 2.05580514517901 | 0.588770414206904 | 3.49169233978622 | 0.000479970753796787 | 0.0209523670744882 | 362 | 13 | 7 | 1 | 0 | 7 | 5 | 1 | 2 | 9 | 14 | 7 | 6 |
| AluSx1_dup65979 | 7.50223093064582 | 2.45925294022227 | 0.759173953217823 | 3.23938002588013 | 0.00119789850274392 | 0.0397871520287414 | 363 | 3 | 2 | 0 | 0 | 14 | 22 | 3 | 2 | 17 | 14 | 3 | 10 |
| L1MA3_dup4708 | 6.48967335283623 | 3.80249436179396 | 0.90021478275824 | 4.2239856916626 | 2.40019294837326e-05 | 0.0023406406200558 | 364 | 1 | 2 | 0 | 0 | 11 | 19 | 1 | 1 | 19 | 17 | 2 | 2 |
| L1MA9_dup13956 | 8.92872106946453 | 2.90593964521874 | 0.642626710770556 | 4.521971459503 | 6.12663219453499e-06 | 0.000841760032295107 | 365 | 6 | 6 | 0 | 4 | 16 | 23 | 1 | 0 | 22 | 18 | 3 | 8 |
| L1ME1_dup25710 | 6.59206062654668 | 2.59588384208872 | 0.621847412003425 | 4.17447076562638 | 2.98679568717208e-05 | 0.00274824472768696 | 366 | 7 | 4 | 0 | 2 | 16 | 12 | 2 | 0 | 13 | 14 | 5 | 5 |
| L1PA5_dup506 | 6.62825242518163 | 2.26385446331313 | 0.605329275378974 | 3.73987275255408 | 0.000184113435168419 | 0.0104963343132694 | 367 | 7 | 3 | 0 | 3 | 17 | 14 | 3 | 1 | 12 | 12 | 4 | 5 |
| L1MC1_dup9908 | 6.67963256467731 | 2.75492788450656 | 0.8322816465745 | 3.31009087590154 | 0.000932656855930261 | 0.0335027125967968 | 368 | 5 | 3 | 0 | 1 | 25 | 15 | 0 | 1 | 8 | 9 | 10 | 5 |
| L1PB1_dup732 | 8.25697521113088 | 2.9744011246045 | 0.857219198146963 | 3.46982560707252 | 0.00052079639242672 | 0.0221803147292587 | 369 | 5 | 3 | 0 | 0 | 23 | 17 | 0 | 1 | 12 | 15 | 8 | 18 |
| L1MC1_dup11112 | 5.88458259446834 | 4.0445478788211 | 0.903736188980495 | 4.47536341704293 | 7.62814504315353e-06 | 0.000995113267784106 | 370 | 6 | 1 | 0 | 0 | 16 | 14 | 0 | 1 | 12 | 16 | 2 | 2 |
| L1ME1_dup2356 | 6.10460560034778 | 3.68277209222205 | 0.799755815328077 | 4.60487066381793 | 4.12722285354671e-06 | 0.000617137837130519 | 371 | 9 | 6 | 0 | 0 | 21 | 6 | 0 | 2 | 14 | 10 | 2 | 2 |
| MLT1K_dup12417 | 5.99561833119929 | 2.60487008624977 | 0.65261802168914 | 3.99141611123105 | 6.56799217340769e-05 | 0.00500585350958894 | 372 | 6 | 6 | 1 | 0 | 5 | 10 | 0 | 2 | 20 | 10 | 4 | 5 |
| AluJr_dup74015 | 6.01432852698335 | 2.09401115147567 | 0.61260446566269 | 3.41821071971856 | 0.000630342847023325 | 0.0254060090880041 | 373 | 9 | 3 | 0 | 2 | 6 | 11 | 0 | 2 | 14 | 12 | 7 | 6 |
| L1PA3_dup9591 | 4.75641606917768 | 2.58318628979166 | 0.764113798724152 | 3.3806303381837 | 0.000723197663208391 | 0.0279899139833741 | 374 | 3 | 3 | 0 | 2 | 6 | 16 | 0 | 1 | 11 | 8 | 3 | 4 |
| MSTB-int_dup47 | 7.62608331946345 | 3.6878415656309 | 0.706810180802211 | 5.21758410645033 | 1.81271918975417e-07 | 5.33823366739165e-05 | 375 | 10 | 8 | 0 | 0 | 16 | 15 | 0 | 0 | 12 | 22 | 3 | 7 |
| L1MCc_dup2885 | 5.86656990994596 | 3.53659362913049 | 0.737355032708117 | 4.79632398539613 | 1.61603828297465e-06 | 0.000299077259378741 | 376 | 8 | 5 | 1 | 0 | 13 | 6 | 0 | 0 | 12 | 19 | 3 | 3 |
| MLT1D_dup16594 | 6.50617852113221 | 3.14247824111529 | 0.797744157932629 | 3.93920558347816 | 8.17518563291372e-05 | 0.00588120626801949 | 377 | 2 | 3 | 1 | 0 | 14 | 9 | 0 | 0 | 10 | 28 | 6 | 5 |
| L2_dup50371 | 7.47923306935682 | 2.31133620805189 | 0.68023486164844 | 3.39785026961238 | 0.000679175813805388 | 0.0267783211585314 | 378 | 9 | 5 | 2 | 0 | 4 | 15 | 0 | 0 | 11 | 23 | 12 | 11 |
| L1MA2_dup6983 | 9.41304924277363 | 2.40070096878315 | 0.484022043344851 | 4.95990007437063 | 7.05294502586327e-07 | 0.000155968406795917 | 379 | 15 | 10 | 3 | 1 | 14 | 14 | 1 | 0 | 17 | 22 | 8 | 10 |
| L1MB3_dup1405 | 9.69345657546477 | 1.98285879866283 | 0.50559561940587 | 3.92182748931429 | 8.7879903116286e-05 | 0.0062086316588206 | 380 | 13 | 10 | 5 | 2 | 12 | 13 | 0 | 0 | 17 | 23 | 9 | 15 |
| L3b_dup8807 | 7.44073308024571 | 2.31316027993602 | 0.617847535471642 | 3.7439014435337 | 0.000181184746715267 | 0.0103934874016178 | 381 | 8 | 6 | 1 | 1 | 9 | 12 | 0 | 0 | 14 | 18 | 14 | 7 |
| MLT1B_dup7391 | 8.33040641031804 | 2.21798629320425 | 0.602319356696427 | 3.68240912158187 | 0.000231040240405656 | 0.0123706347982818 | 382 | 8 | 14 | 2 | 1 | 9 | 11 | 0 | 0 | 18 | 13 | 11 | 14 |
| MER110A_dup543 | 9.10631943534643 | 3.0929796432299 | 0.578957562933726 | 5.3423253123372 | 9.17617856157237e-08 | 3.10147014774418e-05 | 383 | 12 | 8 | 3 | 2 | 23 | 13 | 0 | 1 | 20 | 20 | 1 | 6 |
| L1MCa_dup6311 | 6.88522252276077 | 3.66217237872815 | 0.759140446437065 | 4.8241038873849 | 1.4063410624269e-06 | 0.00026899777366578 | 384 | 4 | 6 | 1 | 2 | 21 | 13 | 0 | 0 | 15 | 16 | 1 | 3 |
| L1MD1_dup5562 | 7.83749378890534 | 2.73441221712947 | 0.613912908845578 | 4.454071868714 | 8.42568759679269e-06 | 0.00107080208931867 | 385 | 10 | 7 | 3 | 2 | 20 | 21 | 0 | 0 | 11 | 13 | 3 | 7 |
| L1ME3G_dup13318 | 8.06286302347716 | 2.71123399956326 | 0.643734650255613 | 4.21172605589383 | 2.53426768665646e-05 | 0.00243742465226013 | 386 | 7 | 4 | 2 | 3 | 24 | 13 | 0 | 0 | 17 | 16 | 5 | 6 |
| MER2_dup8619 | 9.35404505458004 | 3.48716956262742 | 0.670112323113996 | 5.20385828217965 | 1.95192882100025e-07 | 5.6640689663625e-05 | 387 | 4 | 3 | 2 | 2 | 9 | 10 | 0 | 0 | 29 | 43 | 2 | 3 |
| AluJb_dup96545 | 5.22193466997173 | 2.78652048480129 | 0.795197406464435 | 3.50418708882688 | 0.0004580034673197 | 0.020323182520728 | 388 | 1 | 3 | 2 | 1 | 5 | 11 | 0 | 0 | 15 | 17 | 1 | 5 |
| Charlie25_dup896 | 6.40431148018406 | 3.04807555186515 | 0.712809492960553 | 4.27614332015333 | 1.90158829592328e-05 | 0.00196642906849961 | 389 | 5 | 2 | 3 | 1 | 10 | 12 | 0 | 0 | 14 | 24 | 2 | 3 |
| Charlie25_dup897 | 9.40981438634527 | 3.07379396599542 | 0.648511921858753 | 4.73976477901189 | 2.13966464136114e-06 | 0.000370272467456958 | 390 | 5 | 3 | 2 | 1 | 17 | 14 | 2 | 0 | 31 | 25 | 6 | 2 |

| | | | | | | | | | | | | | | | | | | |
|---|---|---|---|---|---|---|---|---|---|---|---|---|---|---|---|---|---|---|
| L1MC1_dup11111 | 7.51395988059127 | 3.31673153159964 | 0.634954455105511 | 5.22357391924826 | 1.75502357957765e-07 | 5.20050860107326e-05 | 391 | 9 | 4 | 1 | 1 | 13 | 9 | 2 | 0 | 21 | 23 | 3 | 2 |
| AluSg_dup35379 | 8.86258680917595 | 2.10486234205629 | 0.592967763079267 | 3.54970788146355 | 0.000385658838491235 | 0.018001454778783 2 | 392 | 5 | 6 | 7 | 2 | 7 | 17 | 3 | 0 | 29 | 19 | 4 | 4 |
| L1MC1_dup12206 | 8.73966240726681 | 2.89776739433235 | 0.747925970161302 | 3.87440403186883 | 0.000106885953595684 | 0.00715216116824484 | 393 | 2 | 3 | 5 | 2 | 13 | 20 | 1 | 0 | 23 | 29 | 3 | 1 |
| ERVL-E-int_dup4596 | 19.0029888037232 | 2.85149751518837 | 0.638475471747537 | 4.46610346264938 | 7.96570981909139e-06 | 0.00102843667314895 | 394 | 4 | 0 | 2 | 0 | 0 | 2 | 1 | 0 | 90 | 88 | 8 | 11 |
| L1PA3_dup205 | 20.3011091855389 | 3.61051855385655 | 0.732501510797012 | 4.92902540218389 | 8.26408301226442e-07 | 0.00017779434868 3207 | 395 | 0 | 0 | 0 | 0 | 1 | 0 | 0 | 0 | 100 | 105 | 3 | 8 |
| L2_dup9697 | 5.67087504787338 | 3.54517206150908 | 0.948269829137338 | 3.73856886782342 | 0.00018507079761721 | 0.0105351330167045 | 396 | 0 | 0 | 0 | 0 | 4 | 7 | 0 | 1 | 21 | 26 | 2 | 2 |
| MIR3_dup15374 | 6.89032306299382 | 2.94562224256797 | 0.889837942219068 | 3.31029067520116 | 0.00093199131787546 | 0.0334838602028075 | 397 | 0 | 0 | 0 | 0 | 3 | 3 | 0 | 2 | 33 | 28 | 1 | 4 |
| AluSz_dup16191 | 7.12207985224739 | 4.45815075255843 | 1.02110645598792 | 4.36599996642386 | 1.2654249847626e-05 | 0.00145319890648804 | 398 | 0 | 0 | 0 | 0 | 7 | 7 | 0 | 0 | 23 | 40 | 2 | 1 |
| LTR16A_dup618 | 7.31297361145935 | 3.69884378162559 | 0.868800537110318 | 4.2574142436976 | 2.06804948526594e-05 | 0.00209397490562166 | 399 | 0 | 1 | 1 | 1 | 7 | 4 | 0 | 0 | 35 | 27 | 2 | 1 |
| MER20_dup2539 | 5.08240046811163 | 2.95434471476741 | 0.913838506981645 | 3.23289584778545 | 0.00122542230640794 | 0.04040429327663 8 | 400 | 0 | 0 | 1 | 1 | 5 | 3 | 0 | 0 | 15 | 29 | 1 | 3 |
| L1PA5_dup8867 | 9.52973318938316 | 2.91734342627706 | 0.74218170666475 | 3.93076708854379 | 8.46752703992904e-05 | 0.00603092803958492 | 401 | 0 | 1 | 0 | 0 | 2 | 4 | 1 | 0 | 32 | 55 | 2 | 11 |
| L2a_dup28904 | 13.3036608321324 | 3.600651127114 | 0.864697623444297 | 4.16405808168148 | 3.12640527638733e-05 | 0.00284697906580659 | 402 | 0 | 0 | 0 | 1 | 6 | 12 | 0 | 0 | 55 | 59 | 7 | 7 |
| LTR33C_dup456 | 12.1438939301305 | 4.45658624613643 | 0.73439000535846 | 6.06841897849786 | 1.29175553971776e-09 | 8.9351181294675e-07 | 403 | 1 | 1 | 0 | 0 | 3 | 5 | 0 | 0 | 67 | 46 | 3 | 2 |
| MLT1G_dup1343 | 11.015221124189 | 3.38831751855649 | 0.674518115078959 | 5.02331582030211 | 5.07869274001094e-07 | 0.000120604253763936 | 404 | 1 | 0 | 1 | 1 | 4 | 3 | 0 | 0 | 46 | 57 | 3 | 5 |
| Tigger2_dup535 | 14.1130944249972 | 2.68461842306022 | 0.500049011139967 | 5.36871059286782 | 7.93015546053047e-08 | 2.75869703393477e-05 | 405 | 4 | 4 | 0 | 1 | 4 | 2 | 0 | 1 | 54 | 64 | 10 | 14 |
| L1PA5_dup242 | 7.53863800407546 | 3.52117758548074 | 0.706462472817102 | 4.98423868353493 | 6.22062526514305e-07 | 0.000141914947204193 | 406 | 7 | 8 | 1 | 0 | 5 | 2 | 0 | 0 | 26 | 29 | 2 | 6 |
| L1PA7_dup267 | 9.16699935405664 | 2.67907161825327 | 0.646220805063655 | 4.14575265491393 | 3.38699296384611e-05 | 0.00302750348525667 | 407 | 8 | 6 | 1 | 0 | 1 | 2 | 0 | 1 | 34 | 35 | 5 | 11 |
| L1PA10_dup3602 | 5.11622652101726 | 2.63777825999304 | 0.828379348218351 | 3.18426366575444 | 0.00145122759837132 | 0.0454420058062985 | 408 | 10 | 7 | 0 | 0 | 1 | 2 | 1 | 0 | 16 | 11 | 8 | 4 |
| MLT1M_dup807 | 5.77855096072379 | 1.97059349275515 | 0.613264601421811 | 3.21328426292088 | 0.00131226341194713 | 0.0423846653752101 | 409 | 6 | 4 | 0 | 1 | 4 | 5 | 1 | 0 | 18 | 12 | 9 | 8 |
| L1ME1_dup5754 | 4.18410412452609 | 2.79884657136228 | 0.812628173928032 | 3.4441909118575 | 0.000572771110250316 | 0.0237581897226402 | 410 | 4 | 2 | 0 | 0 | 0 | 3 | 1 | 0 | 14 | 18 | 3 | 3 |
| L1MC1_dup7233 | 3.98201486833396 | 2.86161320776838 | 0.837831791724332 | 3.41549847598756 | 0.000636653857257406 | 0.0255764373997925 | 411 | 7 | 2 | 1 | 0 | 1 | 2 | 0 | 0 | 14 | 13 | 2 | 4 |
| L1PA7_dup7258 | 5.41195279695065 | 2.92534844699538 | 0.806845117361069 | 3.62566294825363 | 0.000288221084074541 | 0.0145355274542157 | 412 | 9 | 3 | 0 | 0 | 2 | 1 | 1 | 0 | 18 | 20 | 2 | 7 |
| LTR84a_dup106 | 8.22023277854913 | 2.6182597863675 | 0.669060892120894 | 3.9133355681091 | 9.10298947080871e-05 | 0.00636316078913352 | 413 | 4 | 1 | 0 | 1 | 0 | 2 | 0 | 0 | 36 | 33 | 7 | 7 |
| L1PA7_dup5784 | 5.30771336208756 | 3.03147900419744 | 0.777715638910104 | 3.89792727898049 | 9.70195396580548e-05 | 0.00666432509233256 | 414 | 1 | 3 | 0 | 1 | 1 | 2 | 0 | 1 | 16 | 31 | 2 | 2 |
| MLT1J2_dup2311 | 5.48473433658803 | 2.47328236540764 | 0.736284375990546 | 3.35914009051226 | 0.00078185421345835 3 | 0.0295816636469329 | 415 | 3 | 2 | 0 | 0 | 2 | 0 | 0 | 1 | 18 | 26 | 6 | 4 |
| L2a_dup151134 | 14.0000491837285 | 3.47438536114408 | 0.817246523112706 | 4.25133085658283 | 2.12503967903972e-05 | 0.00213061060222488 | 416 | 38 | 33 | 1 | 1 | 0 | 3 | 5 | 2 | 44 | 32 | 2 | 3 |
| MLT1B_dup16467 | 10.5485015268151 | 3.55412544228406 | 0.733226024895261 | 4.84724398972575 | 1.2518842586255e-06 | 0.000244967805758966 | 417 | 20 | 18 | 0 | 1 | 4 | 1 | 1 | 1 | 31 | 37 | 3 | 6 |
| THE1B-int_dup3927 | 9.58681700377032 | 2.34245951662304 | 0.697750078683079 | 3.35716123607489 | 0.000787471764295209 | 0.0297327445933018 | 418 | 23 | 15 | 0 | 2 | 4 | 1 | 5 | 4 | 24 | 28 | 2 | 6 |
| L1MA3_dup8367 | 12.7962012317966 | 2.93172158890663 | 0.795744254116888 | 3.68425103133196 | 0.000229376100498388 | 0.0122953650842959 | 419 | 15 | 21 | 4 | 5 | 2 | 1 | 0 | 0 | 41 | 52 | 3 | 3 |
| MER11C_dup837 | 10.4984477312837 | 3.01679692855091 | 0.833998164247032 | 3.61727046638598 | 0.00029772615286877 | 0.0148879301019452 | 420 | 16 | 5 | 3 | 2 | 0 | 2 | 0 | 0 | 28 | 60 | 3 | 3 |
| L1PA8A_dup939 | 7.3953502151749 | 2.92564599713432 | 0.717338802840836 | 4.07847168666752 | 4.53327218012774e-05 | 0.00379844169994105 | 421 | 10 | 8 | 1 | 2 | 0 | 6 | 0 | 0 | 20 | 31 | 3 | 6 |
| Tigger1_dup3878 | 7.37085310470212 | 3.4022293428546 | 0.826664755248276 | 4.11560952762863 | 3.86157567893273e-05 | 0.00334856591289842 | 422 | 20 | 15 | 1 | 1 | 0 | 4 | 0 | 1 | 18 | 22 | 2 | 4 |
| L2c_dup129951 | 5.71078359044994 | 2.82110715242395 | 0.755947354486493 | 3.7318830943463 | 0.000190053704296103 | 0.0107136978731202 | 423 | 6 | 7 | 1 | 2 | 1 | 2 | 1 | 0 | 17 | 25 | 3 | 1 |
| L1MA4_dup6448 | 7.59338598336277 | 2.16578783489097 | 0.657101190415566 | 3.29597308067769 | 0.000980814128805178 | 0.0346416503922632 | 424 | 11 | 14 | 5 | 4 | 2 | 3 | 1 | 1 | 18 | 26 | 3 | 2 |
| MamRep1879_dup128 | 6.04035428573529 | 2.51339589494575 | 0.701795598436548 | 3.58137882389839 | 0.000341785653875936 | 0.0164201487401409 | 425 | 12 | 13 | 1 | 2 | 1 | 5 | 1 | 1 | 14 | 15 | 6 | 1 |
| ERVL-E-int_dup4595 | 6.83169946216604 | 3.50552050122423 | 1.08449170079966 | 3.23240878527646 | 0.00122751317737752 | 0.0404489475840289 | 426 | 0 | 0 | 0 | 0 | 3 | 0 | 1 | 0 | 33 | 35 | 1 | 0 |
| MIRb_dup111528 | 4.49942847684711 | 3.09122028482842 | 0.867319120171373 | 3.56410946436604 | 0.000365093581754399 | 0.0172708861638434 | 427 | 1 | 1 | 0 | 0 | 2 | 1 | 1 | 1 | 20 | 20 | 0 | 2 |
| MIR_dup82093 | 5.99015256069901 | 3.81502723816639 | 0.88906285156145 | 4.29106584699395 | 1.77817573403404e-05 | 0.00187382552351844 | 428 | 3 | 0 | 0 | 0 | 1 | 3 | 0 | 1 | 34 | 19 | 1 | 1 |
| MLT1H_dup6411 | 4.29114397110056 | 3.45530145924769 | 0.902684287222794 | 3.82780725017192 | 0.000129289939841044 | 0.00818107153920514 | 429 | 1 | 0 | 0 | 0 | 2 | 3 | 1 | 0 | 19 | 19 | 1 | 1 |
| LTR16A_dup2128 | 4.60303528019936 | 3.28975390704929 | 1.00554808439718 | 3.2716027787189 | 0.00106939710362309 | 0.0367980857872308 | 430 | 0 | 1 | 0 | 0 | 0 | 6 | 0 | 0 | 19 | 20 | 2 | 3 |
| Charlie2b_dup1874 | 4.80036697203194 | 4.03182443395262 | 1.10594635071224 | 3.64558771893056 | 0.00026678139258373 | 0.0137520721923155 | 431 | 1 | 0 | 0 | 0 | 0 | 4 | 0 | 0 | 15 | 33 | 0 | 1 |
| ERVL-E-int_dup4594 | 5.30823422271848 | 3.28230829099345 | 1.00631451249341 | 3.26171216875395 | 0.00110741525214464 | 0.0376381987697197 | 432 | 0 | 1 | 1 | 0 | 0 | 1 | 0 | 0 | 25 | 28 | 0 | 1 |
| MIRb_dup102161 | 3.17208434068263 | 3.46886029442336 | 1.05657902682215 | 3.28310538668988 | 0.00102670275652482 | 0.0357215695429235 | 433 | 1 | 1 | 0 | 0 | 0 | 1 | 0 | 0 | 16 | 14 | 0 | 1 |
| L1PA10_dup4290 | 9.31437229743356 | 5.78345107714621 | 1.82130714252408 | 3.17543973891803 | 0.00149609592169325 | 0.0464314647428227 | 434 | 10 | 17 | 0 | 0 | 0 | 0 | 0 | 1 | 34 | 42 | 0 | 0 |
| L1MA5_dup3774 | 5.04856565957593 | 4.63702520942352 | 1.11907403997362 | 4.14362682341629 | 3.41855874505026e-05 | 0.00304314395491855 | 435 | 9 | 2 | 1 | 0 | 1 | 0 | 0 | 0 | 18 | 26 | 0 | 0 |
| L1PA7_dup8518 | 5.09312627566562 | 5.51046664513717 | 1.1912266376273 | 4.62587594256033 | 3.73018532334965e-06 | 0.000568234748367321 | 436 | 9 | 9 | 0 | 0 | 1 | 0 | 0 | 0 | 17 | 22 | 0 | 0 |
| MSTA_dup18182 | 3.76268111852181 | 4.83918054017232 | 1.15380629457917 | 4.19410135211414 | 2.73955479479103e-05 | 0.0025724787103447 | 437 | 4 | 5 | 0 | 0 | 1 | 0 | 0 | 0 | 14 | 18 | 0 | 0 |
| L1MA9_dup15194 | 4.97287916353436 | 3.27854932248305 | 0.927701659260701 | 3.53405568455679 | 0.000409234850963645 | 0.0187911609122724 | 438 | 8 | 4 | 1 | 2 | 4 | 0 | 0 | 0 | 13 | 24 | 2 | 0 |
| L1MC3_dup278 | 2.54429723274215 | 3.489860106109 | 1.05875993411979 | 3.29617696480961 | 0.000980102568254094 | 0.0346270985938937 | 439 | 4 | 3 | 0 | 1 | 2 | 0 | 0 | 0 | 9 | 9 | 1 | 0 |
| THE1D-int_dup1754 | 3.32396935876297 | 3.62027215917582 | 1.04245352906294 | 3.47283793305404 | 0.00051498623011545 | 0.02202748759860 7 | 440 | 9 | 4 | 0 | 1 | 1 | 0 | 0 | 0 | 10 | 12 | 1 | 1 |
| L1PA4_dup5807 | 3.7233120414511 | 3.46437917157685 | 1.04138723977725 | 3.32669639040121 | 0.000878820681061896 | 0.0321710562732237 | 441 | 10 | 11 | 0 | 0 | 3 | 0 | 2 | 1 | 7 | 10 | 0 | 1 |
| MIR3_dup14279 | 4.26135167558313 | 3.14229395013113 | 0.947466879437444 | 3.3165211558602 | 0.000911456863678586 | 0.0329801573730025 | 442 | 4 | 8 | 0 | 0 | 5 | 0 | 2 | 1 | 12 | 15 | 2 | 0 |
| LTR7_dup163 | 4.43572691319056 | 2.69504158926312 | 0.847861437857908 | 3.1786344665846 | 0.00147970566155878 | 0.0460723121802446 | 443 | 2 | 5 | 0 | 1 | 2 | 0 | 1 | 0 | 20 | 13 | 3 | 2 |
| MIR3_dup79372 | 3.48319812483575 | 3.17262344575509 | 0.944367901662142 | 3.35952062768238 | 0.000780778222198673 | 0.0295503507013338 | 444 | 2 | 2 | 1 | 0 | 2 | 0 | 1 | 0 | 13 | 17 | 0 | 1 |
| THE1D-int_dup2210 | 3.72959668069458 | 4.34864847033272 | 1.12333328763737 | 3.87120057616998 | 0.000108300630546865 | 0.00721436221615462 | 445 | 2 | 7 | 0 | 0 | 1 | 0 | 0 | 0 | 16 | 14 | 0 | 1 |
| L1MA4A_dup5324 | 2.879380846955 | 3.49539818313057 | 1.08258847300742 | 3.22874136413109 | 0.00124336287286496 | 0.0408355034018946 | 446 | 2 | 1 | 0 | 0 | 0 | 4 | 0 | 0 | 8 | 16 | 2 | 0 |
| L2b_dup84648 | 3.57586092106082 | 4.14613167610104 | 1.09272758836747 | 3.79429577896477 | 0.000148063089075897 | 0.00902155378692947 | 447 | 6 | 7 | 0 | 0 | 0 | 2 | 0 | 0 | 8 | 17 | 2 | 0 |
| AluSz_dup94881 | 2.56322242089782 | 3.62738456349089 | 1.12327572120397 | 3.22929134407259 | 0.00124097401047254 | 0.0407739301154237 | 448 | 3 | 2 | 0 | 0 | 0 | 6 | 1 | 1 | 6 | 11 | 0 | 0 |
| L1M3b_dup529 | 4.17222535865517 | 3.84000411968032 | 0.982686396462951 | 3.90765979207803 | 9.31944054626264e-05 | 0.00648588359233989 | 449 | 3 | 4 | 0 | 0 | 0 | 1 | 0 | 1 | 16 | 20 | 1 | 0 |
| L1MA9_dup937 | 3.17565503626868 | 3.5681766193645 | 1.11828320978685 | 3.19076293745357 | 0.00141897657907399 | 0.0446792014336328 | 450 | 2 | 1 | 0 | 0 | 0 | 1 | 0 | 1 | 8 | 23 | 0 | 0 |
| L1PA7_dup8519 | 3.11852930849499 | 4.72374192768461 | 1.37103534846413 | 3.44538303332316 | 0.00057025070326471 | 0.0236887000417292 | 451 | 4 | 6 | 0 | 0 | 0 | 0 | 0 | 0 | 15 | 9 | 0 | 0 |
| L1MC3_dup7507 | 4.04851942098165 | 5.01741103032749 | 1.34131001125951 | 3.7406796252986 | 0.00018352333207264 | 0.01047460848485 4 | 452 | 8 | 4 | 0 | 0 | 0 | 0 | 0 | 0 | 21 | 11 | 0 | 0 |
| L1PA7_dup8520 | 4.06496692787709 | 5.26340190063984 | 1.36578940434221 | 3.85374339843762 | 0.000116325509831782 | 0.00758274803259302 | 453 | 10 | 7 | 0 | 0 | 0 | 0 | 0 | 0 | 16 | 13 | 0 | 0 |
| L1MB3_dup9540 | 2.43739823498099 | 4.38841648927834 | 1.38425364427071 | 3.17024015608813 | 0.00152312993624038 | 0.0469763675707911 | 454 | 4 | 4 | 0 | 0 | 0 | 0 | 0 | 0 | 7 | 13 | 0 | 0 |
| LOR1a_dup484 | 2.2434913103505 | 4.3211648169939 | 1.24093179073226 | 3.48219366226732 | 0.000497323880609542 | 0.0215039484091438 | 455 | 4 | 3 | 0 | 0 | 0 | 1 | 0 | 0 | 6 | 12 | 0 | 0 |
| L1MD2_dup8397 | 3.27380858919882 | 3.96647220278664 | 1.1531770499101 | 3.43960383455071 | 0.000582566263930023 | 0.0240658528189398 | 456 | 13 | 8 | 0 | 0 | 0 | 1 | 0 | 0 | 7 | 8 | 1 | 2 |
| L1ME3A_dup4657 | 2.55418529954 | 4.33543858261937 | 1.25433748041849 | 3.45635736020016 | 0.000547528905146194 | 0.0229758146641934 | 457 | 7 | 8 | 0 | 1 | 0 | 1 | 0 | 0 | 8 | 5 | 0 | 0 |
| THE1D_dup10241 | 2.60674113614137 | 3.85925056766014 | 1.20380809775007 | 3.20586858891638 | 0.00134655467810644 | 0.0431350699295524 | 458 | 11 | 5 | 1 | 0 | 0 | 1 | 0 | 0 | 5 | 8 | 0 | 1 |
| L1PA12_dup1442 | 2.99127247593572 | 4.10893707832666 | 1.24217432686393 | 3.30785863905298 | 0.000940122510953349 | 0.0336686160117661 | 459 | 16 | 7 | 1 | 0 | 0 | 1 | 0 | 0 | 6 | 5 | 1 | 0 |
| hAT-N1_Mam_dup77 | 2.42800581796603 | 4.29086534400023 | 1.27704891835595 | 3.35998510497484 | 0.000779466748127833 | 0.029510102601436 | 460 | 10 | 6 | 1 | 0 | 0 | 1 | 0 | 0 | 7 | 4 | 0 | 0 |
| AluSc5_dup6066 | 3.38183452793212 | 2.74300023196145 | 0.825444810836508 | 3.32305709109939 | 0.000890366899416166 | 0.0324340542327739 | 461 | 6 | 5 | 0 | 1 | 4 | 1 | 0 | 1 | 8 | 10 | 2 | 2 |
| L2a_dup120978 | 3.58391076203072 | 2.91685086966358 | 0.841406723311064 | 3.46663603802133 | 0.000527014974361618 | 0.0223455805597734 | 462 | 5 | 3 | 0 | 2 | 2 | 3 | 0 | 1 | 9 | 15 | 1 | 1 |
| L3b_dup3379 | 3.39575208642675 | 2.69402824775992 | 0.815259976773936 | 3.3045020294268 | 0.00095145287053696 | 0.033952370235871 | 463 | 4 | 5 | 0 | 1 | 4 | 2 | 0 | 0 | 10 | 8 | 2 | 4 |
| SVA_C_dup322 | 3.13156945445471 | 3.21898554473056 | 0.90645460008211 | 3.55118231452404 | 0.000383504662390097 | 0.0179279765023721 | 464 | 5 | 3 | 0 | 1 | 4 | 1 | 0 | 0 | 11 | 8 | 2 | 1 |
| THE1B_dup18142 | 3.75132405507869 | 3.89453375827287 | 0.972609501465028 | 4.00421109644373 | 6.22247829264916e-05 | 0.0048009527231605 | 465 | 6 | 10 | 0 | 1 | 2 | 1 | 0 | 0 | 10 | 12 | 1 | 1 |
| MLT1J_dup10545 | 3.49421184368505 | 3.12041757012731 | 0.875245605098337 | 3.56519078982033 | 0.000363591597915557 | 0.0172203754692705 | 466 | 4 | 9 | 0 | 2 | 5 | 2 | 0 | 0 | 7 | 10 | 1 | 2 |
| L1M3c_dup638 | 3.55896118149831 | 2.56266640975684 | 0.804116563008026 | 3.18693399396036 | 0.00143789585274487 | 0.0451136031879259 | 467 | 4 | 7 | 0 | 3 | 2 | 4 | 0 | 0 | 7 | 12 | 2 | 2 |
| L1PA7_dup8473 | 3.56658722794773 | 3.11882076418005 | 0.830564238300822 | 3.75506266747118 | 0.000173298112251115 | 0.0100883407835225 | 468 | 6 | 5 | 0 | 2 | 4 | 4 | 0 | 0 | 7 | 12 | 2 | 1 |
| L1MA9_dup15193 | 2.90375284075359 | 3.68235021612468 | 1.08483732849069 | 3.3943800783919 | 0.000687841575060439 | 0.0269820092735891 | 469 | 7 | 1 | 0 | 1 | 1 | 1 | 0 | 0 | 5 | 18 | 0 | 1 |

| | | | | | | | | | | | | | | | | | | | |
|---|---|---|---|---|---|---|---|---|---|---|---|---|---|---|---|---|---|---|---|
| MSTA-int_dup1403 | 4.08816907387592 | 3.92994776182513 | 0.967088100554314 | 4.06369157016054 | 4.83026437147632e-05 | 0.00397146539784141 | 470 | 8 | 4 | 0 | 2 | 6 | 1 | 0 | 0 | 8 | 19 | 0 | 1 |
| L1MEc_dup14517 | 4.1043836226299 | 3.91441185867799 | 0.955549342840701 | 4.09650416067583 | 4.19435953218894e-05 | 0.00357333001520311 | 471 | 10 | 1 | 0 | 1 | 3 | 3 | 0 | 0 | 12 | 16 | 2 | 0 |
| HAL1_dup17923 | 3.39025720555484 | 4.2282477175137 | 1.0772712597846 | 3.9249610338246 | 8.67437781221231e-05 | 0.00614112913687792 | 472 | 13 | 6 | 0 | 1 | 1 | 2 | 0 | 0 | 8 | 9 | 1 | 0 |
| L1MEf_dup3220 | 2.79980441068863 | 4.38384418447883 | 1.09622417328928 | 3.99903987824395 | 6.35999643586233e-05 | 0.00488218438317827 | 473 | 9 | 4 | 0 | 0 | 2 | 1 | 0 | 0 | 8 | 8 | 1 | 0 |
| L1MD2_dup3221 | 4.79813845993274 | 2.80734115936399 | 0.849860264901517 | 3.30329734816971 | 0.0009555500670566 | 0.0340373227927919 | 474 | 17 | 8 | 1 | 0 | 1 | 3 | 0 | 2 | 9 | 12 | 4 | 1 |
| MLT1H_dup457 | 3.88726239292677 | 3.12703514499465 | 0.987396830203065 | 3.16694873767374 | 0.00154047470976748 | 0.0473333929868714 | 475 | 8 | 7 | 1 | 0 | 0 | 1 | 0 | 1 | 15 | 8 | 2 | 1 |
| L2b_dup71059 | 3.70401222165766 | 3.29338257113719 | 0.98461724787754 | 3.34483534412633 | 0.000823314323617907 | 0.0306820887888357 | 476 | 12 | 8 | 0 | 0 | 1 | 2 | 0 | 0 | 9 | 6 | 3 | 4 |
| AluJb_dup111327 | 3.45387361770204 | 2.83285425073827 | 0.857253748805735 | 3.30456910183805 | 0.000951225231984098 | 0.0339493383905337 | 477 | 8 | 4 | 0 | 0 | 2 | 5 | 0 | 0 | 8 | 7 | 3 | 5 |
| HERV4_I-int_dup118 | 3.94200905942964 | 3.43514051564782 | 0.918284006529866 | 3.7408258133875 | 0.000183416608446783 | 0.0104710288659977 | 478 | 12 | 5 | 0 | 0 | 2 | 5 | 0 | 0 | 12 | 5 | 3 | 3 |
| LTR79_dup804 | 3.71125519018433 | 2.74907091534713 | 0.862703777652562 | 3.1865757245522 | 0.00143967794958302 | 0.0451510461579603 | 479 | 4 | 10 | 0 | 0 | 1 | 4 | 1 | 0 | 6 | 12 | 4 | 3 |
| L1M1_dup7919 | 2.87444576969479 | 3.06068744764322 | 0.906046474377784 | 3.37806893376535 | 0.000729967794654555 | 0.0281786042937821 | 480 | 4 | 5 | 0 | 0 | 3 | 2 | 0 | 0 | 4 | 12 | 2 | 3 |
| L2b_dup10963 | 3.01239825118412 | 2.89377868395967 | 0.867743985951936 | 3.3348300083983 | 0.000853515583819669 | 0.0314625281344005 | 481 | 4 | 5 | 0 | 0 | 3 | 2 | 0 | 0 | 7 | 9 | 2 | 4 |
| L1PA16_dup11427 | 2.61156067589556 | 3.22462248629081 | 0.97300792839282 | 3.31407627029014 | 0.000919464230387005 | 0.0331941792996004 | 482 | 6 | 7 | 0 | 0 | 2 | 2 | 0 | 0 | 4 | 7 | 2 | 2 |
| L2c_dup34883 | 2.7077995667114 | 3.55553918811279 | 0.953925970328211 | 3.72726951430986 | 0.000193565437855028 | 0.0108602031775177 | 483 | 7 | 4 | 0 | 0 | 3 | 3 | 0 | 0 | 5 | 8 | 1 | 2 |
| L1MEj_dup1271 | 2.64627738318526 | 3.56105228218232 | 1.05375994414068 | 3.37937715509416 | 0.000726502670670083 | 0.0280807298024787 | 484 | 4 | 9 | 0 | 0 | 1 | 2 | 0 | 0 | 8 | 4 | 1 | 2 |
| MER53_dup1193 | 3.02688848972723 | 3.19645260440582 | 0.935829058372491 | 3.41563726388749 | 0.000636329495682408 | 0.0255677234345907 | 485 | 2 | 8 | 0 | 0 | 2 | 5 | 0 | 0 | 7 | 7 | 2 | 3 |
| L1ME4a_dup16041 | 5.96810897586685 | 2.05715548113331 | 0.610896139063632 | 3.3674389959093 | 0.000758698099033822 | 0.0289689233821965 | 486 | 10 | 9 | 5 | 5 | 4 | 9 | 1 | 0 | 15 | 10 | 2 | 2 |
| MIRb_dup85866 | 5.99152261539514 | 2.09903044608795 | 0.618783065123541 | 3.39219116423115 | 0.000693360471806024 | 0.0271486507856436 | 487 | 12 | 12 | 5 | 4 | 2 | 9 | 2 | 0 | 10 | 14 | 2 | 2 |
| L1PA10_dup5289 | 5.19729605798434 | 2.3980456305961 | 0.689392183921904 | 3.47849262948383 | 0.000504242304440802 | 0.0217044933741807 | 488 | 11 | 7 | 3 | 2 | 2 | 8 | 1 | 0 | 16 | 7 | 1 | 4 |
| MamTip2_dup2255 | 4.92354742362024 | 2.84209761819602 | 0.70193540097472 | 4.04894469526602 | 5.14490997236864e-05 | 0.00417107580335682 | 489 | 6 | 8 | 1 | 1 | 3 | 8 | 2 | 0 | 15 | 10 | 1 | 3 |
| L1MEd_dup15968 | 4.9740729274438 | 2.70850931734637 | 0.665682022055539 | 4.06877341975205 | 4.72612784794909e-05 | 0.00390985380164496 | 490 | 10 | 9 | 2 | 1 | 7 | 3 | 1 | 0 | 11 | 11 | 2 | 3 |
| LTR8_dup1511 | 5.16283760451979 | 2.95586425979296 | 0.72740255184906 | 4.06358797103357 | 4.83240976640309e-05 | 0.00397167527681051 | 491 | 15 | 13 | 1 | 2 | 6 | 2 | 2 | 0 | 8 | 12 | 1 | 2 |
| L1MA4_dup6449 | 5.21622270881836 | 2.68365323211792 | 0.754036511467429 | 3.55904945092813 | 0.000372199453256276 | 0.0175234212711749 | 492 | 10 | 9 | 1 | 3 | 4 | 1 | 1 | 0 | 18 | 10 | 1 | 3 |
| MSTB_dup2749 | 5.5971637564875 | 2.62637297506281 | 0.766145877771004 | 3.42803250825271 | 0.000607972652949744 | 0.024809579982641 | 493 | 16 | 13 | 1 | 4 | 3 | 2 | 0 | 0 | 14 | 9 | 2 | 4 |
| L1MA3_dup8134 | 5.36334356575921 | 3.81399411137553 | 0.937877984245098 | 4.06662079230427 | 4.76997660642599e-05 | 0.00393007285137394 | 494 | 20 | 15 | 1 | 0 | 1 | 5 | 1 | 0 | 14 | 5 | 1 | 2 |
| LTR81B_dup665 | 5.39739773708971 | 3.74897365048417 | 0.904903984048107 | 4.14295186734957 | 3.42863925087958e-05 | 0.00304983549603661 | 495 | 16 | 13 | 1 | 0 | 5 | 1 | 2 | 0 | 13 | 12 | 0 | 2 |
| L2a_dup83685 | 5.0462193666753 | 3.48834050468069 | 0.934026672715334 | 3.73473328608447 | 0.000187914231263014 | 0.0106422634544002 | 496 | 15 | 4 | 1 | 3 | 1 | 5 | 1 | 0 | 15 | 15 | 0 | 0 |
| L1MA8_dup2990 | 4.25925015214829 | 2.45671644804417 | 0.703539880687666 | 3.49193630024625 | 0.000479532592765374 | 0.020942686771975 5 | 497 | 10 | 6 | 1 | 2 | 3 | 6 | 2 | 0 | 7 | 12 | 0 | 4 |
| LTR45_dup84 | 5.03412190279849 | 2.94637840270705 | 0.801147900486188 | 3.67769596714789 | 0.000235350247654672 | 0.0125474317163188 | 498 | 7 | 9 | 1 | 2 | 1 | 6 | 3 | 0 | 16 | 13 | 0 | 1 |
| L1MB8_dup14578 | 7.64014725675635 | 3.38566578049543 | 0.815651949624565 | 4.15087070171758 | 3.31212846517657e-05 | 0.00297736199386061 | 499 | 19 | 12 | 0 | 1 | 2 | 2 | 0 | 3 | 22 | 24 | 1 | 3 |
| L1PA7_dup6714 | 5.34989430457884 | 2.98992371610802 | 0.786284174101723 | 3.80259938402525 | 0.000143185796556069 | 0.00880708577964302 | 500 | 7 | 8 | 0 | 3 | 2 | 2 | 0 | 1 | 16 | 20 | 1 | 2 |
| AluJr4_dup3261 | 4.52663830566419 | 2.67954530880836 | 0.832485358595935 | 3.21872965228802 | 0.00128759818695487 | 0.0418195153023083 | 501 | 7 | 9 | 0 | 1 | 2 | 4 | 1 | 4 | 13 | 10 | 0 | 1 |
| HAL1_dup16568 | 5.34722172872987 | 2.7615589593963 | 0.75680299740194 | 3.64897994441958 | 0.000263283655041099 | 0.0136367637314779 | 502 | 3 | 9 | 0 | 2 | 6 | 2 | 1 | 2 | 14 | 20 | 0 | 3 |
| L1PA4_dup5808 | 5.31193548667835 | 3.64805250123551 | 0.86910730586208 | 4.19747075721214 | 2.69912496560735e-05 | 0.00254767794676055 | 503 | 13 | 14 | 1 | 0 | 2 | 6 | 2 | 2 | 13 | 10 | 0 | 0 |
| LTR12__dup504 | 5.45143034461059 | 3.88332251421639 | 0.922826838080761 | 4.20807279759298 | 2.57557899511059e-05 | 0.00246335178259389 | 504 | 15 | 8 | 1 | 0 | 3 | 2 | 1 | 2 | 14 | 18 | 0 | 0 |
| L1PA3_dup6929 | 6.53232436437385 | 2.69209936557671 | 0.80638491695837 | 3.3384793154752 | 0.00084238301778025 | 0.031197147011477 | 505 | 7 | 7 | 1 | 2 | 1 | 1 | 1 | 2 | 26 | 23 | 0 | 2 |
| L2c_dup129954 | 5.80499829893522 | 2.95158686243431 | 0.870574380067889 | 3.39039021824205 | 0.000697932016455786 | 0.0272721203650101 | 506 | 9 | 6 | 1 | 0 | 0 | 3 | 1 | 2 | 24 | 16 | 0 | 3 |
| L1ME1_dup25395 | 4.26134995148754 | 3.34031923545652 | 0.85920614453171 | 3.88768080479345 | 0.000101206595622887 | 0.00687852173724179 | 507 | 3 | 3 | 0 | 0 | 6 | 5 | 2 | 0 | 15 | 12 | 0 | 3 |
| LTR16A_dup3479 | 3.83011179851774 | 4.22571595256713 | 0.950598592230172 | 4.44532107148749 | 8.77606983173664e-06 | 0.00110429968914987 | 508 | 6 | 2 | 0 | 0 | 6 | 2 | 1 | 0 | 13 | 13 | 0 | 1 |
| L1PB1_dup5380 | 4.14057211037262 | 2.78450897474639 | 0.771559615983907 | 3.60893561179396 | 0.000307455901911034 | 0.0152158698078459 | 509 | 4 | 3 | 1 | 0 | 4 | 2 | 1 | 0 | 9 | 20 | 2 | 3 |
| L1MDa_dup5404 | 4.09449075897786 | 3.43428442089802 | 0.815967944140289 | 4.2088472293068 | 2.56676852414638e-05 | 0.00245973541379997 | 510 | 6 | 4 | 0 | 0 | 6 | 5 | 1 | 0 | 9 | 14 | 1 | 3 |
| L1PA7_dup10149 | 4.29599609189268 | 3.37665819857848 | 0.839163295951096 | 4.02383923947891 | 5.72570087647087e-05 | 0.00450870574053039 | 511 | 8 | 2 | 1 | 0 | 7 | 3 | 1 | 0 | 14 | 11 | 1 | 2 |
| L2b_dup76711 | 4.47690023800077 | 2.73858802824799 | 0.807494505773749 | 3.391463358161 | 0.000695204582992844 | 0.0271950564365606 | 512 | 2 | 3 | 0 | 0 | 9 | 5 | 1 | 0 | 15 | 8 | 3 | 6 |
| L1MCc_dup2889 | 4.24190977022994 | 3.2633190270709 | 0.981389501189511 | 3.3252027081149 | 0.000883542712346552 | 0.0322782225708344 | 513 | 1 | 3 | 0 | 1 | 16 | 3 | 0 | 0 | 11 | 9 | 1 | 5 |
| L1PA5_dup9922 | 6.0317709043547 | 3.06398232427394 | 0.793844167564586 | 3.85967731384089 | 0.000113536849921507 | 0.00744719955235963 | 514 | 2 | 3 | 0 | 2 | 14 | 5 | 0 | 0 | 15 | 22 | 2 | 6 |
| HAL1_dup18988 | 4.93970893057774 | 3.60063759284816 | 0.873707342210144 | 4.1211025922478 | 3.77063456669047e-05 | 0.00329129645123158 | 515 | 6 | 1 | 0 | 2 | 13 | 10 | 1 | 0 | 8 | 17 | 1 | 1 |
| L1M3de_dup401 | 6.40817070570302 | 2.35812081962606 | 0.634710487765168 | 3.71526997754372 | 0.000202986848156916 | 0.0112429954775478 | 516 | 6 | 4 | 1 | 2 | 16 | 8 | 3 | 0 | 13 | 16 | 1 | 8 |
| L1M5_dup3602 | 6.03541565569249 | 3.71440673768071 | 0.790081927351237 | 4.70129313061662 | 2.58519084144141e-06 | 0.000431069250145492 | 517 | 5 | 5 | 1 | 0 | 20 | 14 | 1 | 0 | 10 | 13 | 2 | 2 |
| L1PA16_dup12141 | 3.8475099106002 | 4.40433667958231 | 1.09975939138928 | 4.00481843034637 | 6.2065134567735e-05 | 0.00479252837891568 | 518 | 3 | 1 | 0 | 1 | 12 | 8 | 0 | 0 | 14 | 4 | 1 | 0 |
| L1MA6_dup4707 | 3.27976672447857 | 3.14198442045419 | 0.911844538681282 | 3.44574572437331 | 0.000569485946053943 | 0.0236796912027431 | 519 | 2 | 1 | 1 | 1 | 5 | 4 | 0 | 0 | 13 | 8 | 1 | 1 |
| MLT1D_dup16595 | 3.12926468538184 | 2.92095585818712 | 0.920219321101868 | 3.17419531540543 | 0.0015025254663955 1 | 0.0465578616528964 | 520 | 2 | 1 | 1 | 1 | 9 | 5 | 0 | 0 | 8 | 7 | 1 | 2 |
| AluJo_dup49443 | 4.22102220606238 | 3.05279749870957 | 0.833737314527246 | 3.66158194615603 | 0.000250662656298339 | 0.013151968338603 | 521 | 6 | 3 | 1 | 3 | 7 | 11 | 0 | 0 | 13 | 4 | 1 | 1 |
| THE1B_dup16114 | 4.18884173740556 | 2.81515976346185 | 0.831205965365684 | 3.38683777639076 | 0.000707031814674179 | 0.0275479600744166 | 522 | 3 | 2 | 1 | 3 | 14 | 6 | 0 | 0 | 10 | 8 | 1 | 2 |
| L1ME2z_dup2540 | 3.63233947513003 | 3.73847017439217 | 1.00153227513483 | 3.73275057350387 | 0.000189400126649168 | 0.0106895167304704 | 523 | 1 | 1 | 0 | 0 | 8 | 8 | 0 | 0 | 8 | 13 | 1 | 3 |
| L1PBb_dup271 | 3.58861448975684 | 4.3641405612717 | 1.020409380944 | 4.27685264637057 | 1.89554181023674e-05 | 0.00196358986320675 | 524 | 2 | 3 | 0 | 0 | 8 | 9 | 0 | 0 | 10 | 8 | 0 | 2 |
| LTR85a_dup1517 | 2.68545466191337 | 3.95974034862691 | 1.09924865791298 | 3.60222441039393 | 0.000315505811173662 | 0.0154822325728043 | 525 | 1 | 2 | 0 | 0 | 5 | 9 | 0 | 0 | 6 | 7 | 0 | 2 |
| Tigger15a_dup4062 | 4.38896596951817 | 3.41077490281036 | 0.961183764724592 | 3.54851489172588 | 0.000387410091151498 | 0.0180679501272313 | 526 | 3 | 2 | 2 | 0 | 12 | 3 | 0 | 0 | 16 | 9 | 0 | 3 |
| MIR_dup140066 | 4.81272500502778 | 3.88354339499315 | 0.933748859258837 | 4.15908769952952 | 3.19521211765854e-05 | 0.00289302870071682 | 527 | 8 | 1 | 2 | 0 | 7 | 14 | 0 | 0 | 13 | 10 | 1 | 1 |
| L1M5_dup44897 | 3.69708842990974 | 3.59320887354091 | 0.955761513337363 | 3.75952455021338 | 0.000170236572450481 | 0.00997466423745434 | 528 | 6 | 1 | 1 | 0 | 11 | 7 | 0 | 0 | 9 | 6 | 2 | 1 |
| MER75_dup414 | 3.7746062567374 | 3.93528522237348 | 0.911060783851041 | 4.31945408267831 | 1.56415648222304e-05 | 0.00170258361536617 | 529 | 4 | 4 | 1 | 0 | 7 | 7 | 0 | 0 | 11 | 8 | 1 | 1 |
| FLAM_A_dup1205 | 3.24520853930489 | 3.0746095773479 | 0.958679023545957 | 3.20713137748185 | 0.0013406575475003 | 0.0430330636446117 | 530 | 1 | 6 | 0 | 0 | 6 | 11 | 0 | 1 | 5 | 5 | 3 | 1 |
| LTR81A_dup545 | 4.15166437843134 | 2.70393746888461 | 0.79920290070641 | 3.38329286154319 | 0.000716222127126231 | 0.0277968015431705 | 531 | 4 | 6 | 0 | 0 | 11 | 10 | 0 | 1 | 5 | 6 | 4 | 4 |
| L1M7_dup2365 | 3.99653156864986 | 3.57913412854908 | 0.899884850560293 | 3.97732457249459 | 6.96950047104709e-05 | 0.00522813606268528 | 532 | 12 | 3 | 1 | 0 | 3 | 5 | 0 | 1 | 12 | 8 | 1 | 1 |
| LTR16A1_dup577 | 3.5537045237557 | 2.73986273014641 | 0.777389053697635 | 3.52444212729046 | 0.000424375504860017 | 0.019304049155399 | 533 | 7 | 5 | 1 | 0 | 3 | 4 | 1 | 1 | 9 | 8 | 2 | 1 |
| L1MC4a_dup7209 | 2.86355579017552 | 3.06680434895696 | 0.890356867923149 | 3.44446643749782 | 0.0005721876684854 | 0.0237438109496211 | 534 | 3 | 2 | 0 | 0 | 4 | 3 | 0 | 1 | 8 | 9 | 2 | 1 |
| L1ME2z_dup5500 | 2.36960070019328 | 3.09677647054045 | 0.971757184794643 | 3.18678011235376 | 0.00143866103841859 | 0.0451257104465528 | 535 | 4 | 2 | 0 | 0 | 3 | 1 | 0 | 1 | 8 | 6 | 1 | 1 |
| L1ME1_dup5561 | 3.37133350339902 | 2.72102854525024 | 0.824420788274276 | 3.30053364004327 | 0.000965011439156842 | 0.0343024517331362 | 536 | 8 | 5 | 0 | 0 | 5 | 4 | 0 | 2 | 8 | 4 | 2 | 2 |
| MamRep564_dup182 | 3.6265240776294 | 3.00218572017382 | 0.814698879536263 | 3.68502497742811 | 0.000228680213104742 | 0.0122694694361575 | 537 | 8 | 5 | 0 | 0 | 6 | 3 | 0 | 1 | 6 | 10 | 2 | 3 |
| L2a_dup104995 | 3.0896912164749 | 3.16487101896669 | 0.911205134470552 | 3.47328049331681 | 0.000514137729754818 | 0.0220027799207569 | 538 | 5 | 3 | 0 | 0 | 4 | 4 | 0 | 2 | 8 | 8 | 0 | 2 |
| LTR33_dup7243 | 4.1525265801076 | 4.08991325627397 | 0.909392110031915 | 4.49741449387595 | 6.87848096730179e-06 | 0.000916922707140387 | 539 | 10 | 6 | 0 | 0 | 4 | 4 | 0 | 1 | 11 | 11 | 0 | 2 |
| L2c_dup130011 | 2.44668841907297 | 3.4501498727955 | 1.01591009458616 | 3.39611732493017 | 0.000683490557291083 | 0.0268905165583968 | 540 | 5 | 5 | 0 | 0 | 2 | 3 | 0 | 0 | 4 | 8 | 0 | 3 |
| LTR39_dup589 | 3.03383818275976 | 4.59168961887269 | 1.01955437131738 | 4.5036240813128 | 6.68043331507878e-06 | 0.000896044185789028 | 541 | 8 | 3 | 0 | 0 | 4 | 5 | 0 | 0 | 5 | 11 | 0 | 1 |
| L1PB2_dup145 | 15.2092948687503 | 4.38411581340114 | 0.82764202157954 | 5.29711602249742 | 1.17645949581432e-07 | 3.74243089827587e-05 | 542 | 74 | 70 | 0 | 0 | 23 | 12 | 1 | 3 | 6 | 4 | 0 | 3 |
| L1MB1_dup283 | 13.9467708003201 | 4.9285941108883 | 0.94775417418008 | 5.20028742173794 | 1.98980580907598e-07 | 5.76694680933487e-05 | 543 | 62 | 70 | 0 | 0 | 22 | 19 | 0 | 1 | 2 | 2 | 0 | 3 |
| L1PA15_dup447 | 7.55883841960496 | 4.88518431828959 | 0.914898634390888 | 5.33959078596942 | 9.31566114903933e-08 | 3.12200606728072e-05 | 544 | 33 | 34 | 0 | 0 | 13 | 14 | 0 | 1 | 1 | 1 | 0 | 1 |
| L1PA3_dup2919 | 6.47724243314523 | 2.34351925959542 | 0.740157874514803 | 3.16624241974277 | 0.00154422041727142 | 0.0474137103232176 | 545 | 31 | 20 | 2 | 0 | 7 | 12 | 7 | 3 | 0 | 2 | 1 | 0 |
| L1MB1_dup284 | 11.0261459266517 | 4.35600071563413 | 0.956857583613198 | 4.5524023535304 | 5.30367794112918e-06 | 0.000753876315041771 | 546 | 59 | 50 | 0 | 0 | 18 | 14 | 2 | 0 | 0 | 0 | 2 | 0 |
| L1PA7_dup6251 | 15.0075779981633 | 3.68443263666205 | 0.799805584195857 | 4.60666030528714 | 4.09187635428392e-06 | 0.000613126959065769 | 547 | 95 | 70 | 3 | 2 | 16 | 7 | 2 | 3 | 0 | 0 | 0 | 0 |
| MER34_dup1190 | 5.81934907697467 | 3.20623711729313 | 0.741531153467128 | 4.32380635972196 | 1.53359919028641e-05 | 0.00168085579425099 | 548 | 22 | 8 | 2 | 1 | 10 | 7 | 0 | 0 | 12 | 5 | 1 | 4 |

| | | | | | | | | | | | | | | | | | | | |
|---|---|---|---|---|---|---|---|---|---|---|---|---|---|---|---|---|---|---|---|
| L1MEc_dup17245 | 6.75007939215673 | 2.91592702721725 | 0.627905137190067 | 4.64389738912839 | 3.41897415612872e-06 | 0.000533113515435258 | 549 | 18 | 15 | 1 | 3 | 13 | 12 | 0 | 0 | 8 | 8 | 2 | 5 |
| MLT1G3_dup2476 | 6.40847629606326 | 2.69432256573047 | 0.607006382583617 | 4.43870549476359 | 9.05015512236405e-06 | 0.00113339379512949 | 550 | 14 | 14 | 2 | 3 | 7 | 12 | 0 | 0 | 12 | 8 | 2 | 5 |
| MLT1A0_dup14671 | 6.84371198101908 | 2.41509231686165 | 0.719867030791122 | 3.35491446831161 | 0.000793895267799509 | 0.0299100861793581 | 551 | 32 | 15 | 3 | 3 | 8 | 6 | 0 | 0 | 3 | 10 | 5 | 4 |
| L1MC3_dup8993 | 5.86095578622159 | 2.88789833470538 | 0.786195611421467 | 3.67325674775005 | 0.000239478645993353 | 0.0127137407183254 | 552 | 26 | 10 | 1 | 1 | 10 | 8 | 0 | 0 | 6 | 4 | 5 | 4 |
| MIRb_dup150793 | 5.44277139991936 | 2.70233150703859 | 0.778178616321187 | 3.47263655202166 | 0.000515372760610011 | 0.0220390753561513 | 553 | 21 | 8 | 2 | 0 | 11 | 5 | 0 | 0 | 6 | 6 | 5 | 5 |
| L2a_dup164346 | 7.25904032668603 | 2.93672095813332 | 0.716572500910222 | 4.09828866500315 | 4.16216011945944e-05 | 0.00355353194038355 | 554 | 24 | 21 | 2 | 1 | 8 | 6 | 0 | 0 | 8 | 11 | 7 | 3 |
| THE1C_dup2054 | 7.14655199163009 | 3.35080616561783 | 0.80893486678384 | 4.14224470128226 | 3.43923109652037e-05 | 0.00305697154461201 | 555 | 22 | 22 | 1 | 0 | 11 | 8 | 0 | 0 | 7 | 8 | 6 | 5 |
| MIRc_dup24599 | 6.44576699413884 | 2.14721708439828 | 0.664180331426999 | 3.2328826717662 | 0.00122547882535322 | 0.040404293276638 | 556 | 12 | 11 | 1 | 2 | 7 | 13 | 0 | 0 | 9 | 7 | 8 | 11 |
| MER5A_dup30606 | 6.52918346832005 | 2.26390562003644 | 0.70261549833163 | 3.22211170321764 | 0.00127249514721122 | 0.0415001549202743 | 557 | 10 | 16 | 3 | 1 | 4 | 15 | 0 | 0 | 11 | 6 | 5 | 10 |
| MLT1H1_dup3305 | 6.17361285442187 | 2.31317116476001 | 0.708034394296745 | 3.26703219983765 | 0.00108681318163521 | 0.0371929885398206 | 558 | 9 | 16 | 2 | 1 | 3 | 9 | 0 | 0 | 13 | 7 | 8 | 7 |
| MLT1F1_dup1789 | 7.46262623007694 | 2.37440699322562 | 0.749578140161422 | 3.16765773440817 | 0.0015367232131339 | 0.0472975218823386 | 559 | 19 | 10 | 2 | 1 | 14 | 16 | 0 | 0 | 6 | 6 | 13 | 8 |
| AluSq2_dup38787 | 7.48632218169675 | 3.01870418318927 | 0.792177162663637 | 3.81064277722814 | 0.000138605932570758 | 0.00859544624224174 | 560 | 21 | 27 | 3 | 0 | 12 | 11 | 0 | 0 | 8 | 3 | 5 | 4 |
| THE1A-int_dup257 | 6.77442899134419 | 3.0551932351185 | 0.813708803500206 | 3.7546518139861 | 0.000173582609685713 | 0.0100961348433822 | 561 | 17 | 16 | 1 | 0 | 8 | 16 | 0 | 0 | 7 | 6 | 5 | 10 |
| HERVL-int_dup92 | 8.54371987342951 | 2.29764065756992 | 0.493432736048706 | 4.65644147562825 | 3.21721677236004e-06 | 0.000507978674264088 | 562 | 20 | 15 | 5 | 5 | 17 | 12 | 0 | 3 | 11 | 12 | 1 | 5 |
| AluYc_dup3332 | 7.29120508379714 | 2.08758323680316 | 0.57287504446812 | 3.64404638840805 | 0.000268385015647333 | 0.0137988636415774 | 563 | 5 | 15 | 5 | 3 | 16 | 17 | 0 | 3 | 10 | 8 | 2 | 5 |
| FLAM_A_dup4771 | 7.28326864486061 | 2.1020928578917 | 0.531282324975247 | 3.95663992396066 | 7.60113726885307e-05 | 0.00557535440855308 | 564 | 10 | 11 | 2 | 3 | 13 | 14 | 0 | 3 | 9 | 14 | 2 | 9 |
| Chap1_Mam_dup1315 | 7.37618124820669 | 2.05288038573172 | 0.526493858170452 | 3.89915353023377 | 9.65295554286603e-05 | 0.00664222718219433 | 565 | 12 | 8 | 3 | 2 | 13 | 8 | 0 | 1 | 9 | 20 | 8 | 7 |
| L1MCc_dup248 | 7.02304000662631 | 1.8623967527757 | 0.559070151669131 | 3.33123982243629 | 0.000864600801530422 | 0.031753094971594 | 566 | 13 | 7 | 3 | 6 | 15 | 5 | 0 | 1 | 11 | 15 | 3 | 8 |
| MIRb_dup203282 | 9.24679675283717 | 2.16847584433646 | 0.501262642053544 | 4.32602724083481 | 1.51822655343639e-05 | 0.00166657317094875 | 567 | 14 | 15 | 5 | 8 | 18 | 13 | 0 | 0 | 12 | 20 | 4 | 6 |
| AluSx1_dup23380 | 7.0309355106721 | 1.97754268674767 | 0.560477544974922 | 3.52831742230842 | 0.000418210334821705 | 0.0190797204147751 | 568 | 15 | 13 | 4 | 5 | 6 | 15 | 0 | 0 | 7 | 13 | 5 | 6 |
| L1MB2_dup6109 | 7.75177588793776 | 1.9193174531445 | 0.582305873180696 | 3.29606404733773 | 0.000980496593881285 | 0.03463557638475 | 569 | 10 | 18 | 5 | 6 | 9 | 12 | 0 | 0 | 10 | 15 | 2 | 10 |
| L1M1_dup7918 | 10.0543771727539 | 2.97987855716663 | 0.568823072765179 | 5.23867385104609 | 1.61734574386925e-07 | 4.84686335479488e-05 | 570 | 19 | 16 | 3 | 2 | 5 | 10 | 0 | 1 | 21 | 34 | 7 | 2 |
| L1ME4a_dup1852 | 8.25967566617543 | 1.84790046775413 | 0.514219999702623 | 3.59359898258097 | 0.000326141670314372 | 0.0158667155764922 | 571 | 11 | 12 | 7 | 6 | 6 | 14 | 0 | 2 | 16 | 18 | 4 | 4 |
| MER34A_dup328 | 6.60911777640825 | 2.02677557683572 | 0.6134635469938 | 3.3038239790574 | 0.000953756955997454 | 0.0339956542184604 | 572 | 14 | 21 | 7 | 1 | 5 | 9 | 1 | 1 | 8 | 8 | 4 | 3 |
| ERVL-B4-int_dup1607 | 9.53646977447586 | 1.96333999728313 | 0.496576081447606 | 3.95375466244698 | 7.69342968006492e-05 | 0.00561023848805617 | 573 | 23 | 21 | 7 | 3 | 10 | 6 | 2 | 2 | 17 | 14 | 5 | 7 |
| L1PB4_dup2291 | 8.6149026096 | 2.31433889880677 | 0.553579609658522 | 4.18067945138799 | 2.90639337901461e-05 | 0.00269390483312078 | 574 | 31 | 19 | 6 | 5 | 12 | 6 | 1 | 2 | 11 | 10 | 3 | 2 |
| AluYa5_dup3523 | 7.50312775096764 | 1.70279777535865 | 0.538107841220788 | 3.16441732477929 | 0.00155393803596883 | 0.0476056830058759 | 575 | 12 | 16 | 2 | 4 | 7 | 9 | 2 | 0 | 14 | 8 | 8 | 11 |
| L1MCc_dup1878 | 9.95244240039528 | 2.13358218894889 | 0.528325491931394 | 4.03838584647728 | 5.38202734648778e-05 | 0.00431729839865828 | 576 | 20 | 26 | 3 | 5 | 10 | 17 | 2 | 0 | 16 | 8 | 8 | 9 |
| SVA_F_dup230 | 8.23855726992883 | 1.96745515412465 | 0.512343582729661 | 3.8401089043459 | 0.000122979743613847 | 0.00789240605506488 | 577 | 17 | 11 | 2 | 5 | 7 | 16 | 1 | 0 | 12 | 15 | 9 | 8 |
| L2c_dup34882 | 8.95264619338871 | 1.98351282193598 | 0.50351465355708 | 3.93933484939009 | 8.17078245937416e-05 | 0.00588083787984949 | 578 | 20 | 20 | 5 | 4 | 11 | 11 | 1 | 0 | 9 | 16 | 7 | 9 |
| MIR_dup9661 | 8.69511264790711 | 1.87529701160166 | 0.510329938626325 | 3.67467567481827 | 0.000238151737077005 | 0.0126659373906513 | 579 | 22 | 15 | 4 | 5 | 15 | 10 | 1 | 0 | 10 | 11 | 8 | 9 |
| L1MB3_dup7403 | 6.89006741793023 | 1.74560801417044 | 0.545748478465997 | 3.19855772951863 | 0.00138116876754821 | 0.0439186935137013 | 580 | 7 | 6 | 6 | 4 | 11 | 6 | 2 | 0 | 18 | 14 | 5 | 3 |
| THE1A_dup567 | 6.81854698144624 | 1.86570794569344 | 0.526853415278196 | 3.54122777150127 | 0.000398269605295595 | 0.0184084420166406 | 581 | 13 | 10 | 4 | 4 | 6 | 5 | 1 | 0 | 15 | 14 | 5 | 6 |
| MER50_dup1912 | 6.60519341539363 | 1.99373821911825 | 0.560747880840579 | 3.55549844634201 | 0.000377263174426233 | 0.0176987900832067 | 582 | 14 | 19 | 3 | 6 | 6 | 7 | 3 | 0 | 8 | 11 | 4 | 2 |
| L1ME2_dup5543 | 8.38890095547543 | 2.0638470183802 | 0.4811519161849 | 4.28938750726516 | 1.79166533018362e-05 | 0.00188303709653654 | 583 | 15 | 18 | 5 | 5 | 7 | 14 | 2 | 0 | 13 | 15 | 5 | 5 |
| L1PA7_dup3813 | 7.95331904865008 | 2.25688449316941 | 0.533121614369148 | 4.23333894619902 | 2.30247087098284e-05 | 0.00227518161904852 | 584 | 16 | 16 | 2 | 4 | 6 | 13 | 3 | 0 | 18 | 9 | 6 | 4 |
| AluSc8_dup7843 | 6.23959427058379 | 2.23300904689871 | 0.702629162228264 | 3.17807624126661 | 0.00148255760326911 | 0.046128510414423 | 585 | 16 | 18 | 2 | 8 | 5 | 7 | 1 | 0 | 6 | 12 | 4 | 0 |
| LTR7_dup485 | 8.26127702249574 | 2.41205108145644 | 0.761826994325288 | 3.16614021217858 | 0.00154476313312562 | 0.0474222949909314 | 586 | 29 | 25 | 3 | 12 | 5 | 3 | 1 | 0 | 10 | 14 | 2 | 1 |
| L2_dup19134 | 6.59032662997805 | 2.4058153447454 | 0.683265582111665 | 3.52105448851984 | 0.000429834281535682 | 0.0194888239242167 | 587 | 29 | 21 | 3 | 5 | 4 | 5 | 2 | 0 | 6 | 6 | 2 | 2 |
| LTR9B_dup371 | 5.55209540659578 | 2.37647034399811 | 0.677994232848303 | 3.50514831669642 | 0.00045635297647336 | 0.0202828835590077 | 588 | 19 | 19 | 3 | 5 | 5 | 6 | 1 | 0 | 5 | 5 | 2 | 1 |
| MIRb_dup79028 | 6.88912817481815 | 2.33338139695625 | 0.647024487143661 | 3.60632625707435 | 0.000310562629072178 | 0.0153154684636701 | 589 | 30 | 20 | 3 | 4 | 8 | 5 | 2 | 0 | 6 | 5 | 2 | 4 |
| MLT1J_dup10178 | 6.50755594432205 | 2.59985882271854 | 0.608297920111157 | 4.27398933444233 | 1.92006224064136e-05 | 0.0019820873315039 | 590 | 24 | 17 | 4 | 2 | 12 | 7 | 2 | 0 | 6 | 5 | 2 | 2 |
| MER34_dup1082 | 5.98628922994071 | 2.84867850261081 | 0.612365410257844 | 4.6519258842712 | 3.28849279983638e-06 | 0.000516250002692744 | 591 | 15 | 12 | 1 | 3 | 8 | 16 | 4 | 0 | 6 | 9 | 1 | 1 |
| MER31A_dup346 | 7.06128086525408 | 2.53835586602603 | 0.568043762839489 | 4.46859209110493 | 7.87361140210325e-06 | 0.00102015500907472 | 592 | 17 | 21 | 4 | 6 | 14 | 11 | 1 | 0 | 5 | 8 | 1 | 2 |
| Tigger1_dup5556 | 8.54208405407447 | 3.09967571971588 | 0.582096636571265 | 5.32501912049168 | 1.00942420869274e-07 | 3.32670987204437e-05 | 593 | 20 | 22 | 3 | 5 | 10 | 16 | 1 | 0 | 16 | 9 | 2 | 1 |
| Tigger13a_dup2867 | 5.91226967539383 | 2.70176207321337 | 0.660250854588282 | 4.0920235913941 | 4.27625134863406e-05 | 0.00362234568672866 | 594 | 6 | 15 | 2 | 3 | 11 | 11 | 0 | 0 | 4 | 16 | 2 | 4 |
| L1MA2_dup7115 | 4.97753172250639 | 2.31025859361085 | 0.622729248115823 | 3.70989254254709 | 0.000207347236034849 | 0.0114180618897358 | 595 | 7 | 7 | 2 | 2 | 10 | 10 | 0 | 0 | 6 | 10 | 3 | 5 |
| L1MC4_dup22742 | 4.35558977096441 | 2.35189365402579 | 0.689504820778734 | 3.41098942770194 | 0.000647276012311259 | 0.025876468082227 | 596 | 8 | 7 | 1 | 4 | 9 | 6 | 0 | 0 | 4 | 11 | 1 | 4 |
| L1M5_dup50449 | 5.93379282757678 | 3.03040326170275 | 0.677826338515514 | 4.47076646260094 | 7.79397700050516e-06 | 0.00101120264523566 | 597 | 16 | 10 | 0 | 4 | 10 | 14 | 0 | 0 | 7 | 9 | 1 | 4 |
| L1PA3_dup10257 | 7.37752683153126 | 3.15017339310938 | 0.629001139383598 | 5.00821571833154 | 5.49369425420095e-07 | 0.000128788430838433 | 598 | 18 | 15 | 0 | 3 | 16 | 8 | 0 | 0 | 9 | 15 | 3 | 5 |
| THE1B_dup21096 | 6.47693342275555 | 3.20446607001428 | 0.653526434390515 | 4.9033457583132 | 9.42179288430484e-07 | 0.000196722507389869 | 599 | 17 | 8 | 0 | 3 | 9 | 11 | 0 | 0 | 10 | 16 | 4 | 2 |
| LTR16A_dup3477 | 5.12956461880099 | 2.64792109653274 | 0.706875185775448 | 3.74595282139936 | 0.000179710365637756 | 0.0103380447382833 | 600 | 5 | 3 | 1 | 3 | 10 | 15 | 0 | 0 | 8 | 12 | 3 | 3 |
| MER51E_dup214 | 5.56139650895046 | 1.92579126737135 | 0.604268461851765 | 3.18697961080049 | 0.00143766909232374 | 0.0451136031879259 | 601 | 8 | 3 | 2 | 5 | 9 | 9 | 0 | 0 | 11 | 12 | 5 | 4 |
| ERV3-16A3_I-int_dup53 | 5.13624419407587 | 3.44761703036525 | 0.797645402274268 | 4.32224271654461 | 1.544511416202e-05 | 0.00168846459493511 | 602 | 4 | 4 | 0 | 2 | 12 | 8 | 0 | 0 | 11 | 16 | 3 | 1 |
| L2a_dup93877 | 5.26188721510347 | 2.54438893905882 | 0.742938961532789 | 3.42476175136836 | 0.000615338775751757 | 0.0249922467841018 | 603 | 3 | 8 | 0 | 2 | 11 | 8 | 0 | 0 | 11 | 9 | 8 | 3 |
| L1MD2_dup6921 | 4.52012270679664 | 2.59923901515677 | 0.710280597189934 | 3.65945377846456 | 0.000252753425687175 | 0.0132412299930781 | 604 | 8 | 7 | 0 | 1 | 7 | 6 | 0 | 0 | 7 | 10 | 3 | 7 |
| L2a_dup148587 | 5.99329913059523 | 2.43951768184077 | 0.737992081532342 | 3.30561498271829 | 0.000947682113905563 | 0.0338736920948435 | 605 | 12 | 6 | 0 | 1 | 10 | 8 | 0 | 0 | 8 | 10 | 10 | 10 |
| MIRb_dup203281 | 6.55361931823331 | 2.75584267753973 | 0.666933091552286 | 4.13211266984146 | 3.59444142330199e-05 | 0.00317241225296341 | 606 | 17 | 9 | 1 | 2 | 5 | 10 | 0 | 0 | 9 | 18 | 8 | 2 |
| AluJb_dup118870 | 4.84391065114223 | 2.15395493981371 | 0.681504629466216 | 3.16058739248885 | 0.00157451363720777 | 0.0479827384629918 | 607 | 7 | 6 | 1 | 2 | 4 | 13 | 0 | 0 | 7 | 9 | 7 | 4 |
| L1P4a_dup527 | 5.51817227625537 | 2.41501056743333 | 0.678087400682325 | 3.56150337700306 | 0.000368737364805707 | 0.0174017415720069 | 608 | 9 | 12 | 0 | 4 | 6 | 12 | 0 | 0 | 6 | 11 | 6 | 3 |
| MLT1L_dup10863 | 4.44851308803715 | 2.71375082002364 | 0.814600325629597 | 3.33138931404944 | 0.000864136573511601 | 0.0317409449090447 | 609 | 14 | 4 | 2 | 0 | 8 | 4 | 0 | 0 | 6 | 10 | 6 | 1 |
| L2_dup10156 | 6.56592160266665 | 2.95077059251124 | 0.702137947995159 | 4.202551081218 | 2.63923666404847e-05 | 0.00251198322905471 | 610 | 19 | 9 | 1 | 0 | 9 | 6 | 0 | 0 | 11 | 13 | 7 | 6 |
| MIRb_dup199602 | 5.66484305004291 | 2.81634979452889 | 0.694226437258654 | 4.05681726217461 | 4.97459729836285e-05 | 0.00406067076760291 | 611 | 16 | 6 | 2 | 0 | 8 | 7 | 0 | 0 | 8 | 14 | 6 | 3 |
| L1MEc_dup14514 | 4.97394255053335 | 2.88473343507179 | 0.705078007142201 | 4.09136777186413 | 4.28836433322908e-05 | 0.00362873093753387 | 612 | 5 | 11 | 1 | 1 | 5 | 6 | 0 | 1 | 14 | 9 | 3 | 2 |
| MIRb_dup203280 | 4.18901141078096 | 2.24945844653762 | 0.665457374877688 | 3.38031935847292 | 0.000724016501435684 | 0.0280094018773596 | 613 | 7 | 8 | 1 | 2 | 8 | 3 | 0 | 1 | 7 | 8 | 3 | 3 |
| ERV3-16A3_I-int_dup58 | 6.30028756746881 | 2.25956109296367 | 0.579438065422996 | 3.89957309986904 | 9.6362441396424e-05 | 0.00663648552257988 | 614 | 18 | 12 | 2 | 3 | 8 | 13 | 0 | 1 | 6 | 8 | 3 | 6 |
| L1MA5A_dup2455 | 5.52310087467638 | 2.05369467336295 | 0.629767932461885 | 3.26103405318631 | 0.00111006708030682 | 0.0377014018684966 | 615 | 14 | 15 | 2 | 3 | 6 | 6 | 0 | 1 | 8 | 5 | 3 | 6 |
| L1ME4b_dup10762 | 5.4993757315701 | 2.44921996000046 | 0.688962868066918 | 3.5549375351279 | 0.000378068896794943 | 0.0177261047717772 | 616 | 9 | 14 | 1 | 1 | 13 | 12 | 0 | 1 | 4 | 5 | 5 | 4 |
| MLT1E3_dup286 | 5.22003414706243 | 3.02903662790114 | 0.705101347248426 | 4.2958882999183 | 1.73995183230291e-05 | 0.00184416293402739 | 617 | 14 | 14 | 1 | 0 | 7 | 8 | 0 | 1 | 6 | 8 | 2 | 4 |
| HAL1_dup716 | 11.0711286650288 | 2.54701843607803 | 0.792706179574001 | 3.21306746649407 | 0.00131325437094973 | 0.0423994063161473 | 618 | 40 | 48 | 0 | 1 | 12 | 6 | 4 | 2 | 6 | 6 | 3 | 15 |
| L2c_dup70163 | 8.11313967993282 | 2.95405608392141 | 0.822170572845311 | 3.59299661346212 | 0.000326896838121338 | 0.0158839687652889 | 619 | 34 | 34 | 0 | 0 | 12 | 7 | 2 | 2 | 1 | 6 | 3 | 4 |
| L1PA16_dup7044 | 8.80966400727522 | 2.94096886707268 | 0.677910445552497 | 4.33828522095686 | 1.43598721919201e-05 | 0.00159816377389084 | 620 | 29 | 32 | 1 | 0 | 11 | 19 | 3 | 2 | 7 | 2 | 2 | 4 |
| L2a_dup86245 | 8.37545986906696 | 3.23855406773913 | 0.74067854797317 | 4.37241510045251 | 1.22879607592656e-05 | 0.00142211629903901 | 621 | 26 | 31 | 1 | 0 | 11 | 16 | 1 | 1 | 5 | 7 | 1 | 7 |
| AluSx1_dup85349 | 7.68612832889728 | 2.45751517938726 | 0.580918011563147 | 4.23039935149287 | 2.33276802164777e-05 | 0.0022947716355269 | 622 | 22 | 22 | 2 | 0 | 7 | 14 | 2 | 4 | 7 | 9 | 3 | 4 |
| MLT1F2_dup2919 | 8.14701837579689 | 2.82987726489617 | 0.588475079841746 | 4.80883109894336 | 1.51815409189136e-06 | 0.000285564184623858 | 623 | 23 | 25 | 2 | 0 | 8 | 13 | 4 | 3 | 11 | 8 | 1 | 3 |
| AluJr_dup23334 | 5.70755209908987 | 2.25489989955302 | 0.669574813496673 | 3.36765937741508 | 0.000758091947296335 | 0.0289542450226924 | 624 | 20 | 13 | 2 | 0 | 3 | 12 | 3 | 2 | 4 | 8 | 1 | 5 |
| L1ME3G_dup10618 | 8.24016740846184 | 2.2426147712869 | 0.59878583370885 | 3.74527025363351 | 0.000180199688781583 | 0.0103544542735625 | 625 | 27 | 17 | 2 | 0 | 6 | 14 | 3 | 2 | 7 | 13 | 4 | 10 |
| L1M5_dup27947 | 7.65287054316375 | 2.22613471492645 | 0.63362739248295 | 3.51331830242226 | 0.000442546975569796 | 0.0198646071776892 | 626 | 13 | 17 | 1 | 0 | 14 | 19 | 2 | 2 | 6 | 7 | 9 | 6 |
| MIR3_dup57838 | 8.5584195636715 | 2.27868735875125 | 0.599259470078998 | 3.80250537959936 | 0.000143240155191592 | 0.00880708577964302 | 627 | 20 | 25 | 3 | 0 | 19 | 10 | 2 | 2 | 6 | 8 | 5 | 8 |

| | | | | | | | | | | | | | | | | | | | |
|---|---|---|---|---|---|---|---|---|---|---|---|---|---|---|---|---|---|---|---|
| L2c_dup70162 | 9.23715669809904 | 2.92292534991878 | 0.719033118490344 | 4.06507749748112 | 4.80165043192063e-05 | 0.00395203567510822 | 628 | 37 | 19 | 0 | 0 | 21 | 11 | 2 | 2 | 8 | 6 | 3 | 9 |
| MER45C_dup644 | 6.93957594199054 | 2.19035540291511 | 0.675968542592658 | 3.24032150152145 | 0.00119394998210044 | 0.0397003385188472 | 629 | 11 | 16 | 0 | 0 | 20 | 8 | 3 | 2 | 6 | 7 | 5 | 9 |
| L1ME1_dup15527 | 9.52272386506987 | 3.52370889427398 | 0.613428295611729 | 5.74428815801533 | 9.2308477574867e-09 | 4.76451909002805e-06 | 630 | 35 | 24 | 1 | 0 | 16 | 14 | 2 | 1 | 10 | 11 | 3 | 3 |
| L1PA6_dup3201 | 10.6333048007233 | 3.17209176603318 | 0.577566757807095 | 5.49216471196676 | 3.97036831642892e-08 | 1.58597375599916e-05 | 631 | 39 | 27 | 0 | 1 | 17 | 18 | 5 | 1 | 8 | 13 | 2 | 5 |
| L1PA8_dup6861 | 8.90992999240161 | 1.84574798821775 | 0.526449786029393 | 3.50602856568488 | 0.00045484640177680 5 | 0.0202562009951682 | 632 | 23 | 14 | 2 | 1 | 18 | 17 | 10 | 2 | 4 | 13 | 3 | 8 |
| LTR12C_dup2332 | 8.82479028871538 | 1.79642501788961 | 0.50684023148375 | 3.54436152913644 | 0.000393565175884788 | 0.0182673636422175 | 633 | 22 | 15 | 1 | 1 | 16 | 19 | 7 | 6 | 4 | 11 | 4 | 6 |
| MLT1B_dup2293 | 7.61435793181915 | 1.97038813672447 | 0.501798811584381 | 3.92664966762907 | 8.61373010682341e-05 | 0.006109098730248 | 634 | 18 | 17 | 2 | 1 | 17 | 12 | 8 | 4 | 6 | 6 | 2 | 3 |
| L1MC3_dup8408 | 7.73293104362454 | 1.83176029863648 | 0.532040118379895 | 3.44289882540124 | 0.000575514577034203 | 0.0238284593385645 | 635 | 25 | 17 | 1 | 3 | 12 | 9 | 5 | 4 | 8 | 5 | 2 | 7 |
| AluSc8_dup13661 | 6.57514170347841 | 1.7369807933478 | 0.510248729524119 | 3.40418445523184 | 0.000663619240115361 | 0.0263185572256973 | 636 | 16 | 14 | 2 | 2 | 7 | 13 | 5 | 3 | 7 | 6 | 2 | 6 |
| L1PA3_dup3688 | 7.13768043923224 | 1.98297818443982 | 0.50745464155696 | 3.90769543137037 | 9.31806636869136e-05 | 0.00648588359233989 | 637 | 25 | 13 | 2 | 3 | 10 | 13 | 6 | 2 | 5 | 7 | 2 | 4 |
| L1PA11_dup3571 | 8.8412060022618 | 2.3829859428739 | 0.563946901639793 | 4.22555019975262 | 2.38357679713935e-05 | 0.00232825354914663 | 638 | 23 | 27 | 2 | 7 | 17 | 26 | 2 | 1 | 2 | 3 | 1 | 4 |
| Charlie2b_dup1178 | 7.89134503796862 | 3.05627994485002 | 0.743505653892275 | 4.11063443680664 | 3.94573468461335e-05 | 0.00340294231701707 | 639 | 28 | 25 | 1 | 4 | 22 | 9 | 1 | 0 | 5 | 2 | 4 | 0 |
| MIR_dup84478 | 5.52867532917365 | 2.71363202035858 | 0.800945570162192 | 3.38803549385892 | 0.000703951540195221 | 0.0274692154898946 | 640 | 23 | 16 | 0 | 1 | 9 | 9 | 1 | 1 | 3 | 2 | 5 | 1 |
| L1MB5_dup3727 | 4.91096709095755 | 2.14694180397179 | 0.680360726956573 | 3.15559337702458 | 0.0016017198701858 | 0.0485799583339667 | 641 | 21 | 12 | 1 | 3 | 9 | 5 | 1 | 2 | 3 | 2 | 3 | 1 |
| MLT1A_dup6362 | 5.54032960762474 | 2.72530087430656 | 0.727370541654074 | 3.7467847791981 | 0.000179115636289481 | 0.010312085456699 | 642 | 16 | 27 | 2 | 2 | 6 | 5 | 1 | 1 | 4 | 3 | 2 | 1 |
| MLT1G_dup392 | 8.53449443349656 | 2.74965741281388 | 0.765184919156483 | 3.59345478978469 | 0.000326322290691163 | 0.015866717029654 | 643 | 33 | 33 | 1 | 1 | 7 | 7 | 3 | 0 | 3 | 11 | 3 | 9 |
| Tigger1_dup746 | 9.1653881099864 | 2.72007724773793 | 0.829694881178973 | 3.27840668833918 | 0.00104394872393025 | 0.0361471803062146 | 644 | 52 | 36 | 2 | 1 | 4 | 8 | 3 | 0 | 3 | 3 | 3 | 6 |
| L1MEf_dup3595 | 9.60953836127474 | 2.07293063951655 | 0.617998054904773 | 3.3542672554786 | 0.000795754649651784 | 0.0299514972021163 | 645 | 48 | 23 | 3 | 7 | 11 | 9 | 4 | 0 | 4 | 8 | 3 | 7 |
| L2_dup40187 | 9.2242472127415 | 2.42925894478751 | 0.661915029923116 | 3.67004650894492 | 0.000242506341508879 | 0.0128315352316371 | 646 | 37 | 33 | 4 | 4 | 15 | 11 | 1 | 0 | 3 | 4 | 4 | 4 |
| THE1D_dup9157 | 8.91731065370664 | 2.19157226790348 | 0.573713657282367 | 3.8199757668046 | 0.000133464786135084 | 0.00835062108906092 | 647 | 21 | 32 | 4 | 6 | 13 | 14 | 1 | 0 | 6 | 7 | 4 | 6 |
| Tigger3a_dup3740 | 10.6141941777304 | 2.77148331768186 | 0.587617714504516 | 4.71647339634543 | 2.39967728142363e-06 | 0.000406497324436657 | 648 | 37 | 35 | 5 | 3 | 19 | 14 | 1 | 0 | 7 | 7 | 3 | 5 |
| L1PA5_dup10626 | 5.80123375520207 | 2.52313897381018 | 0.699785123251286 | 3.60559104498731 | 0.000311443273823877 | 0.0153429981849474 | 649 | 17 | 17 | 1 | 1 | 8 | 9 | 1 | 0 | 5 | 5 | 4 | 6 |
| L2_dup4734 | 6.77352258094397 | 2.56858163785574 | 0.780384336348743 | 3.29143156546889 | 0.000996788659083619 | 0.0350285370994135 | 650 | 28 | 13 | 1 | 1 | 9 | 16 | 1 | 0 | 2 | 5 | 5 | 8 |
| L1MD2_dup9409 | 6.15459520156226 | 2.19463115149242 | 0.615507998633492 | 3.56556073416559 | 0.000363079065270253 | 0.0172063754759601 | 651 | 18 | 13 | 2 | 3 | 5 | 16 | 3 | 0 | 2 | 11 | 3 | 4 |
| L1M4_dup16311 | 6.52050868741876 | 1.99913416314389 | 0.632038280446466 | 3.16299538333615 | 0.0015615480794783 8 | 0.0477290937039548 | 652 | 24 | 17 | 4 | 3 | 8 | 14 | 2 | 0 | 2 | 4 | 5 | 2 |
| L1MEg_dup7498 | 7.10288162854241 | 3.0062172511272 | 0.633379903600827 | 4.74630981190997 | 2.07161328869254e-06 | 0.00036291185593887 | 653 | 21 | 24 | 2 | 1 | 11 | 14 | 2 | 0 | 3 | 8 | 3 | 2 |
| AluSx3_dup28791 | 5.28097588586279 | 1.86436603361733 | 0.579065170123981 | 3.21961349051293 | 0.00128363538954919 | 0.0417262167215399 | 654 | 8 | 9 | 4 | 1 | 9 | 10 | 1 | 0 | 6 | 8 | 5 | 5 |
| L1MB3_dup3747 | 8.34933770139182 | 2.42776725054897 | 0.496067795640644 | 4.89402309902751 | 9.87952354313207e-07 | 0.000204209296418747 | 655 | 16 | 17 | 2 | 2 | 13 | 12 | 2 | 0 | 11 | 16 | 5 | 8 |
| MER5A_dup24135 | 7.45117047615108 | 2.3250417590092 | 0.59200947951033 | 3.9273725159474 | 8.58789148105643e-05 | 0.00609440628199061 | 656 | 24 | 17 | 2 | 1 | 12 | 8 | 3 | 0 | 7 | 9 | 5 | 7 |
| L1PA3_dup8485 | 7.99161095985045 | 2.07569792809008 | 0.583065175226058 | 3.55997582480435 | 0.000370888936064401 | 0.0174761064389853 | 657 | 18 | 21 | 5 | 2 | 15 | 6 | 4 | 0 | 3 | 18 | 5 | 5 |
| L1PA4_dup6837 | 7.67626129183103 | 2.09362775191244 | 0.646415921419649 | 3.23882454397851 | 0.00120023383223649 | 0.0398480311769699 | 658 | 14 | 17 | 1 | 5 | 17 | 11 | 1 | 0 | 3 | 14 | 4 | 12 |
| L3_dup32207 | 7.90807108640548 | 2.66960858044033 | 0.675068155612366 | 3.95457637609744 | 7.66703773782304e-05 | 0.0055985622377827 1 | 659 | 24 | 36 | 1 | 2 | 14 | 4 | 3 | 2 | 6 | 3 | 2 | 3 |
| MLT1C_dup13086 | 9.52643643091453 | 2.86337517696646 | 0.607793335825908 | 4.7110999877541 | 2.46383309418417e-06 | 0.000414903913254598 | 660 | 27 | 36 | 2 | 2 | 19 | 4 | 4 | 1 | 8 | 11 | 3 | 3 |
| L1PA2_dup1749 | 9.50136870219408 | 2.15279050906301 | 0.651800631532816 | 3.30283587482934 | 0.000957123891329652 | 0.0340810366277574 | 661 | 35 | 29 | 1 | 3 | 7 | 3 | 3 | 2 | 9 | 12 | 7 | 10 |
| L1PA5_dup10157 | 7.2381734132111 | 2.12671579281039 | 0.663754768141269 | 3.20406857304526 | 0.00135500199495025 | 0.0433355596657024 | 662 | 18 | 23 | 2 | 1 | 7 | 6 | 1 | 1 | 9 | 6 | 6 | 11 |
| THE1B-int_dup3198 | 8.11024320127696 | 2.02881095630611 | 0.57329741140269 | 3.53884548570036 | 0.000401881012250742 | 0.0185447917256933 | 663 | 32 | 13 | 3 | 2 | 13 | 5 | 3 | 2 | 10 | 8 | 3 | 9 |
| L1PB1_dup9547 | 8.32139979222439 | 2.13995536087961 | 0.567570907026339 | 3.77037535643155 | 0.000163002198206297 | 0.00966020449673615 | 664 | 24 | 24 | 2 | 3 | 9 | 5 | 3 | 1 | 10 | 11 | 4 | 9 |
| THE1D_dup9391 | 8.02154905974042 | 1.83240234214955 | 0.555917875961103 | 3.29617452754435 | 0.000980111071545115 | 0.0346270985938937 | 665 | 22 | 18 | 2 | 2 | 9 | 8 | 4 | 1 | 12 | 6 | 6 | 11 |
| L2c_dup97438 | 10.8778174330857 | 2.81625642732016 | 0.789928873452172 | 3.56520254160665 | 0.000363575306219222 | 0.0172203754692705 | 666 | 51 | 42 | 0 | 5 | 6 | 9 | 2 | 0 | 6 | 8 | 4 | 9 |
| MSTB2_dup557 | 9.2953038854259 | 2.90969161033026 | 0.812887822012826 | 3.57945036392038 | 0.000344317637840684 | 0.0165056884562976 | 667 | 44 | 32 | 0 | 4 | 4 | 2 | 3 | 0 | 11 | 10 | 3 | 6 |
| FordPrefect_dup387 | 6.1077714711126 | 2.59443141706621 | 0.711182181407824 | 3.64805458417193 | 0.000264233509118715 | 0.0136592019467221 | 668 | 30 | 8 | 0 | 1 | 8 | 7 | 5 | 0 | 6 | 8 | 3 | 3 |
| L2_dup19132 | 7.11390818394199 | 2.60120243384426 | 0.644614199660478 | 4.03528565646603 | 5.45359063407253e-05 | 0.00435656947443041 | 669 | 23 | 19 | 0 | 6 | 9 | 6 | 3 | 0 | 10 | 9 | 1 | 4 |
| MLT1D_dup13913 | 5.98572500968854 | 3.15659744828076 | 0.794335527347062 | 3.9738842587379 | 7.0709933719851e-05 | 0.00528593964464457 | 670 | 28 | 20 | 0 | 3 | 7 | 3 | 1 | 0 | 5 | 6 | 2 | 2 |
| L1PREC2_dup3851 | 7.85461103037065 | 2.78908923042513 | 0.721182116951857 | 3.86738545627486 | 0.000110008472005641 | 0.00730155533608211 | 671 | 27 | 24 | 0 | 1 | 9 | 11 | 0 | 3 | 4 | 11 | 3 | 7 |
| L2_dup29520 | 7.43781829647174 | 2.80969986526282 | 0.709662028537333 | 3.95920840101001 | 7.51985975134301e-05 | 0.00552429931581681 | 672 | 26 | 17 | 0 | 0 | 13 | 14 | 0 | 5 | 6 | 6 | 3 | 3 |
| L2c_dup31081 | 7.55920992653241 | 3.95719094802775 | 0.720168142142144 | 5.49481533056582 | 3.91119476518724e-08 | 1.57204923918453e-05 | 673 | 19 | 23 | 0 | 0 | 13 | 12 | 0 | 2 | 10 | 10 | 1 | 3 |
| L4_A_Mam_dup3450 | 6.03171018486289 | 3.82910315869959 | 0.81240255373782 | 4.71330763435208 | 2.43727809048173e-06 | 0.000411013871380416 | 674 | 17 | 15 | 0 | 0 | 8 | 6 | 0 | 3 | 13 | 8 | 1 | 1 |
| L1MEd_dup15967 | 6.75374187938929 | 2.98104052203772 | 0.606796114213495 | 4.91275479886951 | 8.98055608742627e-07 | 0.000190114537257894 | 675 | 16 | 11 | 1 | 2 | 13 | 7 | 0 | 2 | 11 | 14 | 4 | 1 |
| MamRep605_dup4006 | 7.8023502629874 | 3.78692802534429 | 0.672991705257991 | 5.62700549762731 | 1.83364885804385e-08 | 8.48850994506036e-06 | 676 | 12 | 19 | 1 | 1 | 23 | 10 | 0 | 1 | 11 | 13 | 2 | 2 |
| HAL1_dup11971 | 8.52859457814018 | 3.82451895118718 | 0.635969722871747 | 6.01368086190866 | 1.81357380477029e-09 | 1.18228118506814e-06 | 677 | 18 | 16 | 0 | 0 | 15 | 18 | 0 | 1 | 14 | 15 | 4 | 3 |
| MIR_dup158603 | 8.36168732498075 | 3.71450158090686 | 0.617856548361569 | 6.01191585774557 | 1.83343484506026e-09 | 1.19172786770006e-06 | 678 | 17 | 17 | 0 | 1 | 13 | 16 | 1 | 1 | 17 | 13 | 3 | 2 |
| L1MC3_dup10306 | 6.38791469809131 | 2.60295783195896 | 0.805845715646896 | 3.23009452233598 | 0.00123749298347141 | 0.0407045210734155 | 679 | 11 | 8 | 0 | 0 | 13 | 19 | 2 | 0 | 7 | 4 | 7 | 10 |
| L2a_dup165065 | 7.10382775012091 | 2.20749372757143 | 0.694067761960791 | 3.18051615210466 | 0.00147012943502101 | 0.045858864518289 | 680 | 14 | 8 | 0 | 2 | 15 | 8 | 1 | 0 | 12 | 6 | 8 | 15 |
| ERV3-16A3_I-int_dup5 | 9.34795019979327 | 2.57785761966271 | 0.577518388465975 | 4.46368058774735 | 8.05636365479209e-06 | 0.00103536301943529 | 681 | 19 | 18 | 0 | 3 | 19 | 19 | 1 | 1 | 12 | 9 | 6 | 10 |
| L1M2_dup1431 | 8.67149612620137 | 3.08081023553541 | 0.716869718120887 | 4.29758735466057 | 1.72667221538713e-05 | 0.00183186988031498 | 682 | 20 | 17 | 0 | 1 | 18 | 16 | 1 | 0 | 14 | 5 | 6 | 10 |
| L1MB3_dup8103 | 8.64220794054745 | 2.94167372620077 | 0.783825107621601 | 3.75297205664519 | 0.00017475034096122 | 0.0101318578534012 | 683 | 12 | 16 | 0 | 1 | 24 | 18 | 1 | 0 | 7 | 9 | 5 | 17 |
| L1PA5_dup6021 | 9.86123534108172 | 2.55237043469948 | 0.75138048459296 | 3.39690807391965 | 0.000681518573084245 | 0.0268218399901984 | 684 | 24 | 26 | 0 | 1 | 23 | 14 | 2 | 1 | 4 | 7 | 10 | 15 |
| THE1B-int_dup3748 | 7.09579408633273 | 2.25280328923814 | 0.704642908451051 | 3.19708502309384 | 0.00138824000714194 | 0.0440846332229276 | 685 | 11 | 20 | 0 | 1 | 11 | 8 | 1 | 1 | 4 | 12 | 9 | 12 |
| L3_dup23104 | 6.94364694153823 | 3.23471128212403 | 0.77746377396053 | 4.16059421733038 | 3.17420632690248e-05 | 0.0028761983955108 | 686 | 23 | 19 | 0 | 0 | 9 | 9 | 2 | 0 | 3 | 15 | 4 | 5 |
| HAL1_dup11965 | 6.15486717665711 | 3.44028698544572 | 0.817111104654251 | 4.21030501953761 | 2.55026140472607e-05 | 0.00244786002504618 | 687 | 20 | 18 | 0 | 0 | 13 | 8 | 1 | 0 | 5 | 6 | 4 | 3 |
| MIR3_dup16561 | 8.07368194358131 | 3.45072191054128 | 0.739294590985639 | 4.66758711969031 | 3.047575401873e-06 | 0.000486357060076507 | 688 | 24 | 28 | 0 | 1 | 12 | 13 | 1 | 0 | 7 | 8 | 5 | 3 |
| L1M4b_dup692 | 5.63498152423367 | 3.65839417531788 | 0.93136684889432 | 3.92798410171135 | 8.56608721140034e-05 | 0.00608410883930153 | 689 | 14 | 17 | 0 | 0 | 10 | 11 | 0 | 0 | 3 | 8 | 2 | 7 |
| AluJb_dup22195 | 6.35322980647926 | 3.93533990118154 | 0.94015774723594 | 4.18582935975524 | 2.84126746359879e-05 | 0.0026522880654903 | 690 | 16 | 16 | 0 | 0 | 16 | 18 | 0 | 0 | 4 | 4 | 2 | 5 |
| L2_dup10157 | 7.29627239510742 | 4.16244417022191 | 0.97967115255308 | 4.24881773784432 | 2.14901673433828e-05 | 0.00215194824733185 | 691 | 21 | 28 | 0 | 0 | 12 | 15 | 0 | 0 | 5 | 4 | 2 | 6 |
| L1MC2_dup1058 | 9.08706783890958 | 4.24736432035475 | 0.892706949891938 | 4.7578483855972 | 1.95667369665081e-06 | 0.000347449760763638 | 692 | 28 | 30 | 0 | 0 | 10 | 12 | 0 | 0 | 11 | 12 | 3 | 8 |
| LTR8A_dup240 | 8.8331015181609 | 4.07727358074395 | 0.875657135721311 | 4.65624433858504 | 3.22029729251194e-06 | 0.00050812789095759 | 693 | 27 | 24 | 0 | 0 | 16 | 16 | 0 | 0 | 8 | 9 | 4 | 8 |
| MLT1I_dup1931 | 9.83637008498029 | 4.28730128192576 | 0.82735882421076 | 5.1819128006709 | 2.19621891510338e-07 | 6.19172400442167e-05 | 694 | 27 | 27 | 0 | 0 | 15 | 19 | 0 | 0 | 12 | 13 | 3 | 7 |
| AluJo_dup12879 | 9.61145455081341 | 4.46752195837766 | 0.960108328511207 | 4.65314363568247 | 3.26912355118778e-06 | 0.000513787894657152 | 695 | 40 | 25 | 0 | 0 | 23 | 16 | 0 | 0 | 5 | 6 | 3 | 6 |
| L2a_dup30879 | 7.7274117679706 | 4.18063864106482 | 0.865596645844618 | 4.82977684945336 | 1.36686147205263e-06 | 0.00026313852736977 | 696 | 20 | 24 | 0 | 0 | 17 | 13 | 0 | 0 | 7 | 9 | 3 | 4 |
| MER70A_dup96 | 8.73201214696688 | 4.32668219609689 | 0.935634895309853 | 4.62432752111499 | 3.75815649000083e-06 | 0.000571398277424752 | 697 | 31 | 29 | 0 | 0 | 12 | 14 | 0 | 0 | 10 | 5 | 5 | 4 |
| AluJr_dup13620 | 8.38948385917794 | 3.90797678106888 | 0.949540730785947 | 4.11564944437318 | 3.86090738574637e-05 | 0.00334856591289842 | 698 | 31 | 16 | 0 | 0 | 14 | 11 | 0 | 0 | 9 | 7 | 13 | 5 |
| THE1C-int_dup1742 | 8.6410455184683 | 4.10085098275221 | 0.960101336798593 | 4.27126890212056 | 1.94363874060251e-05 | 0.00200381743401822 | 699 | 38 | 22 | 0 | 0 | 5 | 5 | 0 | 0 | 15 | 8 | 8 | 7 |
| L1MB7_dup4916 | 9.42052791285512 | 3.24482695949277 | 0.7013778400452 | 4.62636082041551 | 3.72146744847976e-06 | 0.000567760348935386 | 700 | 30 | 17 | 0 | 0 | 7 | 18 | 0 | 2 | 17 | 11 | 7 | 7 |
| L2c_dup93037 | 7.60668853080768 | 2.40007367019114 | 0.729409376142461 | 3.2904343550999 | 0.00100032839077125 | 0.0351223460632221 | 701 | 15 | 16 | 0 | 0 | 15 | 9 | 0 | 3 | 9 | 8 | 10 | 9 |
| L1PA5_dup6024 | 9.16548928436608 | 2.85581846996307 | 0.640385027962271 | 4.45953347636873 | 8.21382439169069e-06 | 0.00105021314277552 | 702 | 19 | 16 | 0 | 0 | 16 | 15 | 1 | 1 | 13 | 13 | 7 | 13 |
| MSTD_dup1280 | 9.42135408199349 | 3.21649418469365 | 0.757333524544943 | 4.24713033352951 | 2.16526012131134e-05 | 0.0021629512377227 | 703 | 22 | 25 | 0 | 0 | 11 | 10 | 0 | 1 | 14 | 12 | 8 | 14 |
| L1MD_dup6403 | 9.78033780197107 | 4.64652841836123 | 0.75906590608808 | 6.12137678835763 | 9.27702402886777e-10 | 6.70951598654398e-07 | 704 | 20 | 29 | 3 | 0 | 23 | 22 | 1 | 0 | 5 | 19 | 0 | 1 |
| L2c_dup100945 | 8.4333766187418 | 3.26579510029103 | 0.755575947893251 | 4.32225921086681 | 1.54439592161902e-05 | 0.00168846459493511 | 705 | 25 | 19 | 9 | 0 | 20 | 14 | 1 | 0 | 4 | 14 | 1 | 0 |
| L1PA5_dup8624 | 9.55362173631834 | 2.78833168820009 | 0.666626559707702 | 4.18274916832402 | 2.88005093122687e-05 | 0.00267903283378307 | 706 | 16 | 23 | 4 | 14 | 25 | 19 | 2 | 0 | 6 | 13 | 0 | 0 |

| | | | | | | | | | | | | | | | | | | |
|---|---|---|---|---|---|---|---|---|---|---|---|---|---|---|---|---|---|---|
| AluSc8_dup14405 | 10.8765106707016 | 3.08963406258314 | 0.663019480593961 | 4.65994462155971 | 3.16294469027493e-06 | 0.000500405609888403 | 707 | 20 | 26 | 10 | 7 | 35 | 16 | 1 | 0 | 11 | 10 | 0 | 0 |
| MIRb_dup155672 | 8.78564057717145 | 3.6811825076247 | 0.655847907569338 | 5.61286003224081 | 1.99009570615314e-08 | 9.14160835707172e-06 | 708 | 26 | 13 | 4 | 4 | 24 | 14 | 1 | 0 | 14 | 9 | 0 | 0 |
| L1PA11_dup3122 | 10.4295427374435 | 2.8267582156322 | 0.519800744462167 | 5.43815730498235 | 5.38344353576214e-08 | 2.03651646353014e-05 | 709 | 38 | 15 | 7 | 4 | 23 | 16 | 5 | 0 | 10 | 13 | 1 | 1 |
| L1MEd_dup15859 | 9.19559776237243 | 4.10752524542514 | 0.679143791966843 | 6.0480936349716 | 1.46569768955255e-09 | 9.93609408656041e-07 | 710 | 36 | 27 | 3 | 2 | 20 | 14 | 2 | 0 | 5 | 9 | 0 | 0 |
| MIRb_dup155680 | 9.46751792140847 | 3.94315992742154 | 0.645467596972445 | 6.10899748634458 | 1.00258907190115e-09 | 7.18560419586118e-07 | 711 | 33 | 25 | 5 | 2 | 17 | 23 | 1 | 0 | 6 | 9 | 0 | 0 |
| HAL1b_dup3806 | 8.37931108668849 | 3.51096234621129 | 0.667703883825098 | 5.25826257906112 | 1.45422777370308e-07 | 4.46487331408456e-05 | 712 | 22 | 31 | 6 | 1 | 18 | 14 | 1 | 0 | 5 | 7 | 1 | 0 |
| THE1B_dup16444 | 8.58990233084474 | 3.4408146457101 | 0.669650935782987 | 5.13822121623251 | 2.77351271445928e-07 | 7.51647216804928e-05 | 713 | 28 | 32 | 5 | 3 | 12 | 10 | 1 | 0 | 6 | 11 | 1 | 0 |
| AluJo_dup49447 | 10.4771422388356 | 3.02859029858127 | 0.644531626035747 | 4.69890099452357 | 2.61565207257763e-06 | 0.000435387066031795 | 714 | 32 | 28 | 6 | 10 | 20 | 17 | 1 | 1 | 3 | 17 | 0 | 0 |
| L1ME3F_dup3232 | 9.00908632689009 | 3.06914541588313 | 0.574640835080657 | 5.34098036289458 | 9.24452711309644e-08 | 3.11131293982015e-05 | 715 | 26 | 22 | 1 | 8 | 20 | 15 | 3 | 2 | 5 | 13 | 0 | 0 |
| L2a_dup119783 | 11.5883998607662 | 3.67005073940294 | 0.553191637446699 | 6.63432071450385 | 3.26000320549739e-11 | 3.95767715681267e-08 | 716 | 36 | 24 | 2 | 7 | 27 | 21 | 2 | 1 | 9 | 18 | 0 | 0 |
| L2c_dup100941 | 7.95301204221551 | 2.65792474455012 | 0.520194383530438 | 5.10948374050369 | 3.23040288471351e-07 | 8.49349662769105e-05 | 717 | 20 | 16 | 3 | 6 | 22 | 18 | 4 | 1 | 4 | 7 | 1 | 0 |
| Charlie19a_dup1117 | 8.44576092443796 | 3.57622969377784 | 0.633751246679386 | 5.64295488571568 | 1.67156158547718e-08 | 7.90738355900502e-06 | 718 | 19 | 30 | 4 | 3 | 20 | 16 | 1 | 1 | 7 | 5 | 0 | 0 |
| MSTA_dup14325 | 10.9087882579649 | 3.21066003404146 | 0.530570911127795 | 6.05133068305009 | 1.43654219897463e-09 | 9.82239856543726e-07 | 719 | 34 | 28 | 6 | 6 | 21 | 27 | 2 | 1 | 9 | 5 | 0 | 0 |
| L2_dup41266 | 11.8080917566643 | 4.01508664502801 | 0.582050040172067 | 6.89818120077969 | 5.26725356443343e-12 | 7.64220680879681e-09 | 720 | 35 | 38 | 2 | 4 | 23 | 26 | 0 | 1 | 7 | 12 | 1 | 1 |
| Tigger4b_dup1775 | 10.9997875123887 | 3.36043014293072 | 0.5368312553999 | 6.25975128893612 | 3.85591933054578e-10 | 3.24205152306024e-07 | 721 | 33 | 33 | 2 | 7 | 26 | 20 | 1 | 1 | 8 | 7 | 1 | 1 |
| FLAM_A_dup9702 | 9.97792296573525 | 3.31018741829968 | 0.615561696849922 | 5.37750713736616 | 7.55242201269432e-08 | 2.65445880093436e-05 | 722 | 23 | 25 | 2 | 9 | 27 | 14 | 0 | 1 | 10 | 13 | 1 | 0 |
| MIRb_dup155674 | 8.81741170222776 | 3.29817964781453 | 0.600641434375541 | 5.49109578369913 | 3.99447636902998e-08 | 1.5894141707445e-05 | 723 | 19 | 20 | 4 | 4 | 25 | 17 | 0 | 2 | 4 | 15 | 1 | 0 |
| LTR17_dup487 | 8.58673600188312 | 4.7822418981472 | 0.863481344574293 | 5.53832683033228 | 3.05374987076335e-08 | 1.28934561829224e-05 | 724 | 14 | 9 | 0 | 0 | 39 | 20 | 2 | 2 | 8 | 12 | 0 | 0 |
| L1PA16_dup7719 | 10.8638682223713 | 3.97675328166397 | 0.675448336353159 | 5.88757580355445 | 3.91901436337755e-09 | 2.27552447062051e-06 | 725 | 9 | 17 | 1 | 5 | 25 | 34 | 1 | 2 | 20 | 17 | 0 | 0 |
| L1PA3_dup6036 | 10.748017966811 | 5.20099715249002 | 0.894082026449623 | 5.81713645798591 | 5.98642711408088e-09 | 3.28213453015458e-06 | 726 | 11 | 10 | 0 | 3 | 32 | 48 | 1 | 0 | 13 | 15 | 0 | 0 |
| HAL1_dup16425 | 8.99331393710914 | 5.55097943739707 | 0.899788778976964 | 6.16920278079972 | 6.86351601462685e-10 | 5.26821348908516e-07 | 727 | 21 | 23 | 0 | 2 | 32 | 24 | 0 | 0 | 8 | 3 | 0 | 0 |
| Tigger19a_dup2488 | 8.89843817942822 | 3.94371775201391 | 0.651449330059684 | 6.05375977845061 | 1.41503580500975e-09 | 9.70323082590349e-07 | 728 | 24 | 23 | 1 | 4 | 24 | 26 | 0 | 0 | 4 | 6 | 1 | 1 |
| L1MEc_dup13436 | 7.94275243871867 | 3.57256807386507 | 0.710945645877163 | 5.02509312010378 | 5.03188317994233e-07 | 0.00012021249750347 | 729 | 24 | 15 | 2 | 7 | 17 | 27 | 0 | 0 | 6 | 4 | 0 | 0 |
| HAL1_dup16426 | 7.86812386985881 | 4.91964130608023 | 0.827068633850007 | 5.94828664119359 | 2.70963784191901e-09 | 1.6815482144846e-06 | 730 | 31 | 21 | 1 | 2 | 23 | 14 | 0 | 0 | 4 | 5 | 0 | 0 |
| L2c_dup100955 | 7.62683057465858 | 4.58556238906687 | 0.800720489830707 | 5.72679536405567 | 1.02345498119877e-08 | 5.17042503091981e-06 | 731 | 29 | 18 | 1 | 3 | 21 | 14 | 0 | 0 | 7 | 4 | 0 | 0 |
| LTR48_dup681 | 6.17888721694406 | 4.216821165932 | 0.855198438759658 | 4.93081017786689 | 8.18892780369941e-07 | 0.000176496613693756 | 732 | 17 | 24 | 0 | 2 | 11 | 13 | 0 | 0 | 6 | 3 | 0 | 2 |
| AluSz_dup66186 | 6.42278792614642 | 3.86480318587986 | 0.791142118997222 | 4.88509345296712 | 1.03379808160929e-06 | 0.000210786733784578 | 733 | 15 | 21 | 0 | 5 | 12 | 15 | 1 | 0 | 3 | 10 | 0 | 0 |
| L2c_dup100956 | 7.30295112255444 | 4.2070625489999 | 0.756293764447994 | 5.56273599858458 | 2.65577511300724e-08 | 1.16377728368254e-05 | 734 | 24 | 24 | 0 | 4 | 12 | 21 | 1 | 0 | 4 | 4 | 0 | 0 |
| L2c_dup100944 | 6.86507218811884 | 4.13411785464594 | 0.778003456763467 | 5.31375255303372 | 1.07390423644666e-07 | 3.49563908954224e-05 | 735 | 20 | 17 | 0 | 4 | 17 | 13 | 0 | 0 | 5 | 10 | 1 | 0 |
| MLT1D_dup14180 | 5.91079273440318 | 4.21507704436468 | 0.82642203618016 | 5.10039284993823 | 3.38949223066389e-07 | 8.85308582126839e-05 | 736 | 15 | 19 | 1 | 2 | 10 | 13 | 0 | 0 | 6 | 7 | 1 | 0 |
| MER113_dup1166 | 10.2828264681855 | 4.39406680637784 | 0.670003307109883 | 6.55827629468283 | 5.44332865788317e-11 | 6.08083699919562e-08 | 737 | 36 | 28 | 0 | 3 | 20 | 32 | 1 | 1 | 6 | 4 | 0 | 1 |
| CR1_Mam_dup1239 | 6.12992236326623 | 4.25685919912882 | 0.80267331039424 | 5.30335211599103 | 1.13695314074011e-07 | 3.6707388334674e-05 | 738 | 24 | 12 | 0 | 2 | 16 | 12 | 1 | 1 | 5 | 5 | 0 | 0 |
| THE1B_dup16439 | 7.36331595424768 | 3.99598704872132 | 0.736919193699783 | 5.42255797227784 | 5.87521510518196e-08 | 2.18434989596504e-05 | 739 | 26 | 16 | 0 | 3 | 14 | 19 | 2 | 1 | 9 | 3 | 0 | 0 |
| L1MC4_dup17369 | 10.3337821822476 | 4.27414123765893 | 0.64851583372532 | 6.59065055221034 | 4.37903486644117e-11 | 4.96178014442958e-08 | 740 | 24 | 19 | 2 | 2 | 33 | 26 | 3 | 0 | 9 | 12 | 0 | 0 |
| MLT1A0_dup15047 | 10.6358963092383 | 4.50302345693459 | 0.673525284077771 | 6.68575265604228 | 2.29740778088129e-11 | 2.89237561814592e-08 | 741 | 29 | 20 | 1 | 3 | 30 | 32 | 1 | 0 | 8 | 10 | 0 | 1 |
| AluSz_dup66183 | 7.74516115759973 | 4.01647833107308 | 0.69905267133542 | 5.74560186344798 | 9.15945960554254e-09 | 4.73794951152266e-06 | 742 | 18 | 18 | 2 | 1 | 19 | 22 | 2 | 0 | 9 | 5 | 0 | 1 |
| L1MB5_dup6175 | 8.70746755281573 | 3.57811916300432 | 0.610449988284687 | 5.86144521528869 | 4.58855868620439e-09 | 2.59341849203703e-06 | 743 | 24 | 21 | 4 | 3 | 28 | 18 | 1 | 0 | 5 | 6 | 0 | 1 |
| L1ME4a_dup24281 | 8.27008446543422 | 3.58076609453314 | 0.634902805326037 | 5.63986497538679 | 1.70183562980474e-08 | 8.01871249048553e-06 | 744 | 20 | 14 | 1 | 6 | 20 | 24 | 1 | 1 | 9 | 8 | 0 | 0 |
| MLT1I_dup7673 | 8.35867501821166 | 3.55274769432188 | 0.597702813257301 | 5.94400363444915 | 2.78143429088918e-09 | 1.71458850720186e-06 | 745 | 19 | 19 | 3 | 4 | 19 | 18 | 1 | 1 | 10 | 10 | 0 | 0 |
| AluSz_dup66185 | 8.32134124432899 | 3.72884393818615 | 0.645976123970624 | 5.77241758606503 | 7.81421667184926e-09 | 4.10455099381864e-06 | 746 | 25 | 22 | 4 | 2 | 11 | 19 | 1 | 1 | 10 | 9 | 0 | 0 |
| MER20B_dup2839 | 8.3767550203126 | 4.19670695846623 | 0.726373155291211 | 5.77761847047297 | 7.57653719712246e-09 | 4.00623715534778e-06 | 747 | 36 | 17 | 1 | 3 | 11 | 19 | 1 | 1 | 7 | 11 | 0 | 0 |
| AluJb_dup82015 | 9.81055874115621 | 3.12031918876758 | 0.526087280841821 | 5.93118157841526 | 3.00762374964111e-09 | 1.82564296105128e-06 | 748 | 25 | 22 | 4 | 5 | 13 | 30 | 3 | 1 | 10 | 10 | 0 | 1 |
| L2c_dup100954 | 10.2658575811467 | 3.89023812216482 | 0.603355448761869 | 6.44767214773296 | 1.13581129166782e-10 | 1.14034495192908e-07 | 749 | 25 | 29 | 1 | 2 | 19 | 24 | 5 | 1 | 13 | 9 | 0 | 0 |
| MSTB-int_dup297 | 5.02251681682377 | 2.40550581437666 | 0.739173661104416 | 3.25431754532831 | 0.00113665147939187 | 0.0383742441707402 | 750 | 4 | 7 | 0 | 1 | 12 | 3 | 5 | 2 | 13 | 10 | 1 | 1 |
| L2a_dup120974 | 6.80290443287258 | 2.1257338682966 | 0.614919075427077 | 3.4569327139839 | 0.000546361251307132 | 0.0229367928456737 | 751 | 5 | 12 | 2 | 1 | 9 | 4 | 2 | 5 | 13 | 22 | 1 | 4 |
| L1PA8_dup6108 | 7.04498708391687 | 2.23543764395464 | 0.546553595824894 | 4.09006117795415 | 4.31259421017803e-05 | 0.00364664016323746 | 752 | 10 | 6 | 2 | 3 | 10 | 6 | 3 | 3 | 18 | 18 | 2 | 2 |
| L1PB1_dup9995 | 6.85716053982691 | 2.37689065405327 | 0.606873642795129 | 3.91661539806841 | 8.98008399178106e-05 | 0.0062992343273615 | 753 | 7 | 10 | 1 | 2 | 4 | 8 | 4 | 3 | 19 | 19 | 1 | 2 |
| MIRc_dup71789 | 4.64022570832787 | 2.46911366310743 | 0.701513214868944 | 3.51969657986943 | 0.00043204073974539 | 0.0195293248213253 | 754 | 9 | 10 | 0 | 2 | 7 | 3 | 2 | 3 | 7 | 11 | 2 | 0 |
| L2a_dup159668 | 5.11629236380427 | 2.14467735569253 | 0.673978234308865 | 3.18211664192955 | 0.00146202924424654 | 0.0456523962667711 | 755 | 10 | 12 | 1 | 7 | 6 | 4 | 2 | 2 | 8 | 10 | 1 | 0 |
| LTR16E1_dup1715 | 4.10503648068221 | 2.11134914876051 | 0.655971503279269 | 3.21865986282278 | 0.00128791157714488 | 0.0418195153023083 | 756 | 9 | 6 | 1 | 3 | 6 | 4 | 2 | 2 | 7 | 8 | 2 | 0 |
| L1M5_dup50450 | 5.57931187213729 | 2.69145428716008 | 0.597995734046768 | 4.50079178482833 | 6.77007992377592e-06 | 0.000905516266184814 | 757 | 11 | 10 | 1 | 3 | 8 | 9 | 1 | 1 | 7 | 14 | 2 | 2 |
| L1M7_dup2361 | 5.10286508035564 | 2.19012848210967 | 0.607158089269739 | 3.60717994343755 | 0.000309542999987147 | 0.0152841914660597 | 758 | 11 | 8 | 3 | 2 | 3 | 9 | 2 | 1 | 7 | 13 | 3 | 1 |
| MER5C1_dup276 | 4.73796577913605 | 1.97669956597065 | 0.597487027917104 | 3.30835561880164 | 0.000938455600909253 | 0.0336247186288139 | 759 | 11 | 6 | 2 | 1 | 7 | 6 | 2 | 1 | 5 | 11 | 5 | 2 |
| L1PA13_dup6878 | 4.28525769543539 | 2.12742420601096 | 0.662946071738172 | 3.2090456474583 | 0.00133176346889783 | 0.0428226743743735 | 760 | 7 | 7 | 1 | 2 | 4 | 3 | 2 | 2 | 6 | 15 | 2 | 1 |
| MER44B_dup1579 | 5.82845128651731 | 2.11091863049162 | 0.573789242758482 | 3.67890938551482 | 0.000234233466164565 | 0.0124994205741183 | 761 | 8 | 11 | 1 | 3 | 5 | 4 | 2 | 2 | 15 | 12 | 3 | 3 |
| MIR_dup24127 | 5.55493362817881 | 2.63699057801582 | 0.668934669905107 | 3.94207490903394 | 8.07797371412537e-05 | 0.00582461070721599 | 762 | 9 | 16 | 2 | 1 | 4 | 3 | 2 | 1 | 10 | 15 | 2 | 2 |
| MER47A_dup1584 | 4.49673422329667 | 2.73451694646675 | 0.818700603624817 | 3.34006953745926 | 0.000837574149520689 | 0.0310627157876414 | 763 | 10 | 2 | 1 | 0 | 3 | 5 | 3 | 1 | 17 | 8 | 0 | 2 |
| L1ME1_dup2357 | 4.48736034349071 | 3.48750196428503 | 0.851625597426243 | 4.09511171907568 | 4.2196486151656e-05 | 0.00358973367674006 | 764 | 5 | 2 | 0 | 0 | 9 | 8 | 1 | 1 | 11 | 13 | 0 | 3 |
| L1PA7_dup4383 | 4.41460929652312 | 3.12841034979652 | 0.760082705686847 | 4.11588150393388 | 3.85702437733911e-05 | 0.00334827990693298 | 765 | 8 | 6 | 0 | 0 | 8 | 8 | 1 | 2 | 9 | 8 | 0 | 3 |
| Ricksha_b_dup86 | 3.83642793570106 | 2.47440049769331 | 0.752638238547803 | 3.28763590655134 | 0.00101032410333895 | 0.0353169647558528 | 766 | 5 | 6 | 0 | 0 | 7 | 5 | 3 | 1 | 6 | 10 | 0 | 4 |
| MER4B-int_dup600 | 5.8876318782941 | 2.9735973885311 | 0.729788700185512 | 4.0746004806257 | 4.60934086962348e-05 | 0.00383890172405821 | 767 | 10 | 4 | 5 | 2 | 12 | 13 | 1 | 1 | 12 | 11 | 0 | 0 |
| GSAT_dup14 | 5.59251075086154 | 2.06967774999648 | 0.58744955272501 | 3.5231582701797 | 0.000426436625231827 | 0.0193582707786507 | 768 | 10 | 4 | 2 | 3 | 11 | 9 | 3 | 4 | 9 | 11 | 0 | 2 |
| L1MB8_dup5161 | 5.62261234511292 | 2.5148219701589 | 0.630611050948377 | 3.98791293996015 | 6.665711451854e-05 | 0.00506734625278867 | 769 | 9 | 5 | 2 | 1 | 7 | 13 | 4 | 1 | 13 | 10 | 0 | 3 |
| MLT1H1_dup862 | 4.63662335733857 | 2.48477646731677 | 0.688459519597486 | 3.60918310602999 | 0.000307162748364322 | 0.0152108527205613 | 770 | 15 | 5 | 2 | 5 | 8 | 12 | 1 | 2 | 4 | 5 | 0 | 0 |
| L1HS_dup1250 | 3.16949250399689 | 3.27102653031325 | 0.899580872509739 | 3.63616727553069 | 0.000276724672396381 | 0.0141133601948738 | 771 | 12 | 3 | 1 | 1 | 9 | 5 | 1 | 1 | 3 | 4 | 0 | 0 |
| MER20B_dup2245 | 3.29720527484344 | 2.99670306937252 | 0.824768728834092 | 3.63338589910982 | 0.000279726175996937 | 0.014213052460766 | 772 | 8 | 8 | 1 | 2 | 8 | 5 | 1 | 1 | 4 | 3 | 0 | 0 |
| L1MA9_dup323 | 4.11760402345039 | 2.22658372575621 | 0.701470963782729 | 3.1741637797083 | 0.00150268873164461 | 0.0465578616528964 | 773 | 10 | 8 | 1 | 5 | 3 | 11 | 1 | 2 | 4 | 6 | 0 | 1 |
| THE1B_dup712 | 3.82338511158126 | 2.91036144850903 | 0.780172253033412 | 3.7304088131732 | 0.000191169324585002 | 0.0107485765849959 | 774 | 7 | 7 | 1 | 2 | 5 | 15 | 1 | 1 | 4 | 4 | 0 | 1 |
| L1HS_dup382 | 4.78548177634606 | 2.46538385115834 | 0.668193279514664 | 3.68962682915495 | 0.00022458324319404 | 0.0120902000871152 | 775 | 8 | 10 | 5 | 2 | 12 | 8 | 1 | 1 | 3 | 9 | 0 | 1 |
| MER50_dup892 | 4.71358414710235 | 2.23421518212214 | 0.647295786357819 | 3.45161397495501 | 0.000557244348761213 | 0.0232907190954392 | 776 | 13 | 11 | 5 | 1 | 7 | 7 | 2 | 1 | 6 | 4 | 0 | 2 |
| MSTC_dup1665 | 3.00721975693223 | 2.79356070803436 | 0.870647845000878 | 3.20860003740265 | 0.00133382898673197 | 0.042877502307069 | 777 | 7 | 7 | 0 | 1 | 1 | 8 | 1 | 1 | 5 | 4 | 1 | 1 |
| AluSq2_dup45958 | 3.36241629552831 | 2.53942924193049 | 0.805432456830598 | 3.15287671907738 | 0.00161670067553214 | 0.0488759731481423 | 778 | 10 | 7 | 0 | 0 | 4 | 6 | 1 | 1 | 5 | 3 | 1 | 4 |
| Charlie26a_dup286 | 3.25971296218134 | 2.65552362296518 | 0.830800526426139 | 3.19634321175562 | 0.00139181446399053 | 0.0441561342197164 | 779 | 9 | 7 | 0 | 0 | 3 | 8 | 2 | 1 | 4 | 4 | 0 | 3 |
| L1M5_dup27952 | 3.12013249628029 | 2.93361733714591 | 0.90884301785594 | 3.22785924467642 | 0.00124720326702856 | 0.0409333832519142 | 780 | 8 | 4 | 0 | 0 | 2 | 10 | 1 | 1 | 4 | 6 | 0 | 3 |
| L1PA2_dup115 | 4.16473613107694 | 2.95062995407946 | 0.862296743931535 | 3.42182662157162 | 0.000622019643666612 | 0.025155349593716 | 781 | 21 | 8 | 0 | 1 | 2 | 9 | 1 | 2 | 1 | 7 | 1 | 1 |
| MIR_dup84500 | 3.26653233699991 | 2.81037194268393 | 0.865851626113733 | 3.24578929914117 | 0.00117125491656097 | 0.0391701226108246 | 782 | 11 | 4 | 0 | 0 | 7 | 11 | 1 | 1 | 1 | 3 | 2 | 1 |
| LTR12C_dup926 | 4.37282191266522 | 3.78678842418678 | 0.871724350478273 | 4.34402047173416 | 1.39898469876909e-05 | 0.0015650343823851 | 783 | 11 | 11 | 0 | 0 | 9 | 7 | 3 | 1 | 6 | 6 | 0 | 0 |
| THE1D_dup6698 | 4.31875338152624 | 3.12791446638579 | 0.820091790286638 | 3.8141028887663 | 0.000136678520525385 | 0.00850473515819386 | 784 | 15 | 8 | 0 | 0 | 6 | 9 | 4 | 1 | 7 | 3 | 0 | 1 |
| L1M4b_dup2451 | 5.27837300818515 | 2.29850311657692 | 0.726025890832161 | 3.1658693520453 | 0.00154620223340031 | 0.0474432678799888 | 785 | 9 | 13 | 0 | 0 | 13 | 10 | 3 | 6 | 3 | 6 | 0 | 2 |

| | | | | | | | | | | | | | | | | | | | |
|---|---|---|---|---|---|---|---|---|---|---|---|---|---|---|---|---|---|---|---|
| LTR16A1_dup1262 | 5.38374628348155 | 3.39820710863782 | 0.754220444332616 | 4.50558869647823 | 6.61891842073533e-06 | 0.00089056609194507 | 786 | 16 | 13 | 0 | 0 | 13 | 7 | 1 | 2 | 6 | 6 | 0 | 3 |
| MER50_dup301 | 2.92098525414107 | 2.73387637894595 | 0.836000932471591 | 3.27018340860385 | 0.00107477774250085 | 0.0369351624374795 | 787 | 4 | 6 | 1 | 0 | 4 | 8 | 1 | 1 | 3 | 6 | 1 | 1 |
| MIR_dup24727 | 3.78797791096281 | 2.3887934902388 | 0.691155887662918 | 3.45622967680459 | 0.000547788347677423 | 0.0229802620198258 | 788 | 6 | 11 | 2 | 1 | 7 | 6 | 2 | 1 | 4 | 5 | 1 | 1 |
| MLT1A0_dup10795 | 3.98687163070252 | 2.72539413996644 | 0.766593335239762 | 3.55520197565251 | 0.000377688840326176 | 0.0177117754829035 | 789 | 9 | 11 | 1 | 0 | 3 | 7 | 2 | 1 | 7 | 5 | 0 | 3 |
| AluJr_dup58388 | 3.2489434216039 | 3.24657480004592 | 0.903197279510601 | 3.59453562770371 | 0.000324970675496986 | 0.0158324062964385 | 790 | 8 | 10 | 1 | 0 | 4 | 7 | 1 | 1 | 5 | 2 | 0 | 1 |
| MLT1J1_dup2597 | 3.94161189254184 | 3.55404584549183 | 0.846731086539817 | 4.1973725802551 | 2.70029493319077e-05 | 0.0025476794676055 | 791 | 8 | 10 | 1 | 0 | 9 | 10 | 1 | 1 | 4 | 4 | 0 | 1 |
| HAL1_dup18819 | 5.47502280729364 | 2.28422628171763 | 0.616389214611211 | 3.70581805711577 | 0.000210709524209396 | 0.0115410061057596 | 792 | 15 | 10 | 1 | 0 | 6 | 12 | 4 | 4 | 6 | 7 | 1 | 2 |
| AluJb_dup82824 | 5.97588832992504 | 2.61922195189414 | 0.593411229355374 | 4.4138395472216 | 1.0155323757741e-05 | 0.00123717139960142 | 793 | 18 | 14 | 1 | 1 | 8 | 11 | 4 | 2 | 7 | 6 | 1 | 2 |
| L1MA5_dup3339 | 5.16990274948589 | 2.23578786473911 | 0.607621588373029 | 3.67957279254291 | 0.000233624997520543 | 0.01248073869833 8 | 794 | 16 | 8 | 1 | 1 | 6 | 10 | 5 | 2 | 6 | 7 | 2 | 1 |
| L1ME1_dup4381 | 5.09442927322369 | 2.36949546970637 | 0.635182772379952 | 3.73041520132569 | 0.000191164477283512 | 0.0107485765849959 | 795 | 14 | 10 | 1 | 0 | 11 | 9 | 3 | 1 | 3 | 7 | 2 | 4 |
| L1ME3_dup7229 | 4.6896185104289 | 2.55295391219516 | 0.665820594802892 | 3.83429700451205 | 0.000125923915240747 | 0.00802224683691426 | 796 | 10 | 7 | 0 | 0 | 11 | 10 | 2 | 2 | 6 | 5 | 2 | 3 |
| MER44C_dup785 | 4.3765978513301 | 2.66606114342807 | 0.696155257596414 | 3.82969332535543 | 0.000128303053482752 | 0.00813677994776304 | 797 | 12 | 9 | 0 | 0 | 7 | 8 | 3 | 1 | 5 | 6 | 1 | 3 |
| L3_dup17562 | 5.83674556512516 | 2.74529240816642 | 0.867502482806542 | 3.16459314247131 | 0.00155299945899696 | 0.0475892907099108 | 798 | 28 | 23 | 0 | 4 | 3 | 3 | 1 | 4 | 2 | 6 | 0 | 1 |
| HUERS-P3-int_dup161 | 5.11087389477598 | 2.63832932664359 | 0.7020926397911 | 3.75780798304422 | 0.000171408331347521 | 0.0100186506536029 | 799 | 12 | 17 | 0 | 5 | 7 | 6 | 3 | 1 | 7 | 5 | 0 | 1 |
| LTR12F_dup500 | 5.43041122374916 | 2.2100759318383 | 0.611676661230698 | 3.6131441199532 | 0.000302506451297425 | 0.0150428840251655 | 800 | 16 | 14 | 0 | 4 | 9 | 7 | 4 | 3 | 4 | 6 | 0 | 2 |
| L1ME4a_dup23747 | 5.3491227326924 | 2.44724480686437 | 0.75061370034849 | 3.26032525882245 | 0.00111284515571027 | 0.0377634132088756 | 801 | 26 | 9 | 0 | 4 | 3 | 12 | 2 | 1 | 4 | 5 | 0 | 4 |
| MamGypLTR2c_dup57 | 3.74440182533411 | 3.13690896422253 | 0.866089064305373 | 3.62192422639398 | 0.00029241978349126 | 0.0146948506446909 | 802 | 14 | 12 | 0 | 1 | 4 | 4 | 1 | 1 | 5 | 3 | 0 | 2 |
| MER33_dup3203 | 7.4988109253024 | 3.317051604676 | 0.779944800169897 | 4.252931238151 | 2.10990378829476e-05 | 0.0021228040879897 9 | 803 | 28 | 18 | 0 | 0 | 12 | 16 | 2 | 7 | 4 | 7 | 0 | 0 |
| L2_dup29517 | 7.9265420425738 | 3.17845742725551 | 0.657725672580322 | 4.83249713332021 | 1.348310540314e-06 | 0.00026040998362464 | 804 | 29 | 26 | 0 | 1 | 11 | 12 | 3 | 2 | 6 | 7 | 0 | 4 |
| L1PBa_dup1863 | 6.79571279312519 | 2.45010307837394 | 0.586802709386997 | 4.17534384074921 | 2.97536301240211e-05 | 0.00274090486778564 | 805 | 12 | 16 | 1 | 1 | 17 | 16 | 6 | 5 | 4 | 6 | 0 | 1 |
| MER34_dup1083 | 7.42056188418309 | 2.32921597078538 | 0.591656853293782 | 3.93676834438496 | 8.25862605910983e-05 | 0.00592435042586961 | 806 | 15 | 15 | 0 | 2 | 20 | 21 | 4 | 5 | 3 | 5 | 0 | 4 |
| AluSx3_dup20581 | 3.7453209073484 | 2.89296643888439 | 0.886960191012299 | 3.26166435449894 | 0.00110760204169541 | 0.0376391710101764 | 807 | 13 | 14 | 0 | 1 | 6 | 8 | 0 | 1 | 2 | 0 | 1 | 2 |
| MLT1H1_dup584 | 3.82993239054864 | 3.01373090628873 | 0.8767042252353 | 3.43756858874481 | 0.000586962043217622 | 0.0242096152427561 | 808 | 16 | 14 | 0 | 1 | 5 | 6 | 1 | 1 | 3 | 0 | 1 | 1 |
| L1MCc_dup3045 | 5.17935377580091 | 2.25860212976341 | 0.681503427859516 | 3.31414639667666 | 0.000919233650796266 | 0.0331941792996004 | 809 | 11 | 19 | 1 | 2 | 15 | 9 | 1 | 2 | 1 | 1 | 2 | 2 |
| MLT1J_dup10176 | 3.84380097043114 | 2.70355552344531 | 0.763011995225 | 3.54326739338885 | 0.000395201772441164 | 0.0183057584086598 | 810 | 11 | 10 | 0 | 1 | 12 | 8 | 1 | 2 | 1 | 1 | 1 | 1 |
| MER5A_dup12151 | 5.94221226734572 | 3.08601677412224 | 0.699820932713808 | 4.4097234447605 | 1.03502713892757e-05 | 0.00125461318186072 | 811 | 19 | 19 | 2 | 1 | 14 | 11 | 2 | 3 | 4 | 0 | 0 | 0 |
| Zaphod3_dup901 | 5.17983063754629 | 3.69513661105958 | 0.914172756502277 | 4.04205505444897 | 5.29847879722275e-05 | 0.00427178019361617 | 812 | 13 | 8 | 3 | 0 | 14 | 16 | 1 | 1 | 8 | 0 | 0 | 0 |
| THE1B-int_dup1707 | 5.1052182302133 | 2.49257706871982 | 0.76801342396552 | 3.24548632997817 | 0.00117250193730315 | 0.0392039778235067 | 813 | 18 | 17 | 3 | 1 | 3 | 5 | 2 | 2 | 9 | 1 | 1 | 1 |
| L1MEc_dup11923 | 3.87987728032755 | 2.60420729501477 | 0.825228854572165 | 3.15573950254673 | 0.00160091770497288 | 0.0485682941115323 | 814 | 16 | 12 | 1 | 2 | 4 | 7 | 2 | 0 | 4 | 0 | 1 | 1 |
| L2_dup22986 | 3.99606003508126 | 2.76487755544817 | 0.845847262027009 | 3.26876692704821 | 0.00108017238763097 | 0.0370342518601747 | 815 | 15 | 11 | 1 | 2 | 4 | 4 | 2 | 0 | 8 | 1 | 0 | 2 |
| L1MB5_dup3176 | 4.8934405320781 | 2.43967457776809 | 0.761448988928747 | 3.20398951635667 | 0.00135537411928865 | 0.0433359110707143 | 816 | 11 | 12 | 0 | 3 | 7 | 10 | 6 | 1 | 9 | 1 | 0 | 1 |
| MER57A-int_dup1221 | 4.87626264721429 | 3.17656375013282 | 0.826058513302955 | 3.8454464169027 | 0.000120333159156025 | 0.00777019161082808 | 817 | 14 | 12 | 1 | 1 | 6 | 10 | 4 | 1 | 10 | 1 | 0 | 0 |
| ERV3-16A3_I-int_dup4 | 5.28010633648741 | 3.92101393276371 | 0.796232237910655 | 4.92446015882578 | 8.45935611736648e-07 | 0.000180885133352533 | 818 | 14 | 20 | 1 | 1 | 9 | 15 | 1 | 1 | 3 | 2 | 0 | 0 |
| L1MB7_dup7655 | 5.06234336827148 | 2.52300989481928 | 0.67588674060924 | 3.73288857915019 | 0.000189296345168466 | 0.0106861941037855 | 819 | 21 | 11 | 1 | 2 | 6 | 11 | 4 | 2 | 4 | 2 | 1 | 0 |
| Tigger15a_dup285 | 4.33848951989218 | 2.50460285011562 | 0.701733490227188 | 3.56916533840326 | 0.000358120369574968 | 0.0170154371922694 | 820 | 12 | 11 | 1 | 3 | 13 | 5 | 3 | 2 | 3 | 2 | 0 | 0 |
| L1MB2_dup6329 | 4.10578996954798 | 2.57794884930704 | 0.750320041268913 | 3.43579900244609 | 0.000590809119186615 | 0.0243135016731197 | 821 | 13 | 4 | 0 | 4 | 6 | 11 | 1 | 0 | 6 | 4 | 2 | 1 |
| Tigger2_dup707 | 3.62646648613442 | 3.1521312647125 | 0.905345116931594 | 3.48169024801917 | 0.000498259695359752 | 0.0215170420094504 | 822 | 7 | 6 | 0 | 2 | 3 | 18 | 2 | 0 | 4 | 3 | 1 | 0 |
| L1PA13_dup4377 | 3.47392988399058 | 2.82292746082102 | 0.861528834726108 | 3.27664884451426 | 0.00105046927576329 | 0.0363306631236588 | 823 | 11 | 7 | 0 | 1 | 2 | 12 | 1 | 0 | 2 | 5 | 1 | 3 |
| L4_B_Mam_dup2204 | 3.95274532827812 | 2.94593928860078 | 0.79346980737999 | 3.71273016465261 | 0.000205035454059627 | 0.0113143242466772 | 824 | 12 | 12 | 0 | 2 | 4 | 10 | 1 | 0 | 2 | 5 | 1 | 2 |
| L2c_dup46775 | 4.1749944215306 | 2.7388898934699 | 0.748051151388249 | 3.66136712494461 | 0.000250872964098004 | 0.0131573514487513 | 825 | 18 | 10 | 1 | 2 | 5 | 8 | 1 | 0 | 3 | 3 | 2 | 1 |
| LTR16C_dup2766 | 4.49114191376486 | 2.93148894546263 | 0.756481981451359 | 3.87516030433187 | 0.000106554529918862 | 0.00714205686121825 | 826 | 15 | 14 | 1 | 1 | 3 | 8 | 3 | 0 | 5 | 5 | 1 | 1 |
| L1M5_dup38754 | 3.84232510863681 | 3.21797021641918 | 0.820863021923424 | 3.92022801670237 | 8.84652270196748e-05 | 0.00623886986023223 | 827 | 11 | 11 | 0 | 2 | 8 | 6 | 1 | 0 | 3 | 5 | 0 | 2 |
| L2c_dup13127 | 4.360808051671 | 3.76921352425173 | 0.854214564886842 | 4.41249034983504 | 1.02188350716858e-05 | 0.00124121027563417 | 828 | 10 | 13 | 1 | 1 | 9 | 8 | 1 | 0 | 3 | 8 | 0 | 1 |
| AluSc5_dup2626 | 4.6364476879044 | 2.25661413747537 | 0.662545666584448 | 3.40597524259521 | 0.000659281574424316 | 0.0261979648476901 | 829 | 15 | 10 | 1 | 2 | 6 | 9 | 5 | 0 | 4 | 5 | 0 | 3 |
| AluSg_dup5039 | 3.4428725966429 | 2.52570221183298 | 0.771366524486358 | 3.27432178044648 | 0.00105915926344407 | 0.0365621224574875 | 830 | 5 | 7 | 1 | 2 | 5 | 11 | 2 | 0 | 2 | 7 | 0 | 2 |
| MER34A_dup773 | 5.15132780893704 | 2.82983232836017 | 0.710451092132263 | 3.98314867792947 | 6.80081789385858e-05 | 0.00512746329078604 | 831 | 18 | 6 | 1 | 1 | 15 | 14 | 2 | 0 | 2 | 4 | 1 | 3 |
| LTR81B_dup464 | 5.15770377764478 | 2.24091283811637 | 0.625422218589352 | 3.58304001922218 | 0.000339618552431678 | 0.0163419365170694 | 832 | 7 | 10 | 3 | 3 | 12 | 12 | 1 | 0 | 8 | 3 | 2 | 3 |
| MLT1K_dup10179 | 5.46517925432428 | 2.1557417035112 | 0.590628417518055 | 3.64991192359162 | 0.000262330243139153 | 0.0135992226653571 | 833 | 11 | 9 | 2 | 3 | 16 | 14 | 3 | 0 | 4 | 3 | 2 | 3 |
| MIRb_dup155669 | 4.11361503691954 | 4.53845092794365 | 0.969236030167382 | 4.68250331878385 | 2.83392598100718e-06 | 0.000460655673537113 | 834 | 12 | 11 | 0 | 0 | 11 | 7 | 2 | 0 | 3 | 6 | 0 | 0 |
| MLT2B3_dup1890 | 3.38584442593659 | 3.44804630144055 | 0.912857558985891 | 3.77720079929124 | 0.000158600860020375 | 0.00948678739024838 | 835 | 7 | 6 | 0 | 2 | 12 | 6 | 2 | 0 | 2 | 6 | 0 | 0 |
| L1ME1_dup13186 | 2.91810422559687 | 4.06754432188029 | 1.0342800303487 | 3.93273021089749 | 8.39864820433908e-05 | 0.00599587382427143 | 836 | 9 | 7 | 0 | 2 | 7 | 4 | 0 | 0 | 2 | 6 | 0 | 0 |
| THE1B_dup16440 | 3.1359894127602 | 4.13670678036156 | 1.02450640586207 | 4.03775589561173 | 5.39649640873406e-05 | 0.00432303096906091 | 837 | 4 | 9 | 0 | 2 | 10 | 7 | 0 | 0 | 3 | 4 | 0 | 0 |
| AluY_dup69488 | 4.87046315006157 | 4.69872294946203 | 0.966692538802289 | 4.86061778782698 | 1.17019992565672e-06 | 0.000232618539273445 | 838 | 19 | 13 | 0 | 2 | 15 | 6 | 0 | 0 | 4 | 3 | 0 | 0 |
| MIRc_dup71283 | 4.64682987137286 | 4.27091499110677 | 0.94754142205193 | 4.50736494649276 | 6.56376826988737e-06 | 0.00088538685076339 | 839 | 11 | 17 | 0 | 3 | 12 | 7 | 0 | 0 | 2 | 7 | 0 | 0 |
| AluJr_dup53302 | 4.84075213721186 | 3.09553987767319 | 0.762537067744758 | 4.05952708217636 | 4.91722047954823e-05 | 0.00402211393683941 | 840 | 15 | 18 | 1 | 5 | 10 | 9 | 1 | 0 | 1 | 3 | 0 | 0 |
| L1MA6_dup2049 | 4.94260281838578 | 3.54983083522263 | 0.850310266824154 | 4.17474770530642 | 2.98316472573349e-05 | 0.00274685290231121 | 841 | 19 | 23 | 2 | 2 | 9 | 4 | 1 | 0 | 1 | 3 | 0 | 0 |
| AluSz_dup66184 | 4.71305985809415 | 3.13390821315865 | 0.775505648463892 | 4.04111590852529 | 5.31974552266057e-05 | 0.00428507842970545 | 842 | 12 | 14 | 2 | 4 | 4 | 13 | 1 | 0 | 4 | 6 | 0 | 0 |
| L2a_dup119790 | 4.81662296232951 | 3.9725194550726 | 0.877542451844275 | 4.52686869646455 | 5.98641097691333e-06 | 0.00082720240784705 | 843 | 15 | 13 | 1 | 2 | 8 | 11 | 1 | 0 | 1 | 10 | 0 | 0 |
| MIR3_dup59601 | 3.51610725062556 | 3.04384694974712 | 0.872143632815544 | 3.49007529863017 | 0.000482884466927859 | 0.0210153400367565 | 844 | 12 | 12 | 2 | 2 | 3 | 8 | 0 | 0 | 3 | 2 | 0 | 1 |
| AluJb_dup82013 | 4.8238683251794 | 3.78441846554705 | 0.847075284688171 | 4.46762942321023 | 7.90911614451075e-06 | 0.00102336027135371 | 845 | 16 | 15 | 1 | 1 | 7 | 15 | 0 | 0 | 3 | 2 | 0 | 2 |
| HAL1b_dup1706 | 4.47386629671569 | 3.09195444537767 | 0.792329827689681 | 3.90235775219187 | 9.52602218472721e-05 | 0.00657580178348448 | 846 | 18 | 9 | 1 | 2 | 8 | 11 | 0 | 0 | 3 | 3 | 0 | 3 |
| L1PA15_dup6474 | 2.87566364105927 | 2.82331218708568 | 0.884703009131988 | 3.19125419258575 | 0.00141656590955173 | 0.0446681939987006 | 847 | 9 | 4 | 2 | 3 | 6 | 8 | 0 | 0 | 2 | 3 | 0 | 0 |
| L2a_dup119797 | 3.81924601943113 | 3.93692157140262 | 0.90618288675144 | 4.34451105727236 | 1.39586212948989e-05 | 0.00156301087182872 | 848 | 11 | 11 | 1 | 1 | 10 | 6 | 1 | 0 | 4 | 3 | 0 | 0 |
| LTR85a_dup1310 | 3.79644292175235 | 3.93131133188607 | 0.908097799641135 | 4.32917174057646 | 1.49671182911105e-05 | 0.00164801755154863 | 849 | 13 | 11 | 1 | 1 | 7 | 7 | 1 | 0 | 4 | 3 | 0 | 0 |
| HAL1_dup16427 | 3.73712286436096 | 4.3838530168685 | 0.983776445506296 | 4.45614756979913 | 8.34456023783925e-06 | 0.00106293079783346 | 850 | 8 | 11 | 1 | 1 | 12 | 6 | 0 | 0 | 2 | 6 | 0 | 0 |
| AluYf1_dup1253 | 3.71184462378737 | 4.3483325504917 | 0.961869789730207 | 4.52070810094926 | 6.16331240876162e-06 | 0.000844778533649304 | 851 | 12 | 11 | 1 | 1 | 7 | 8 | 0 | 0 | 3 | 4 | 0 | 0 |
| L1M5_dup38756 | 4.01626728996001 | 4.04033473879541 | 0.926882279059681 | 4.35905921396439 | 1.30622768598309e-05 | 0.00148998208170002 | 852 | 15 | 10 | 2 | 1 | 8 | 6 | 0 | 0 | 3 | 6 | 0 | 0 |
| MIR_dup120678 | 3.61296935791629 | 3.46771625427319 | 1.03382112835897 | 3.3542710234387 | 0.000795743812968679 | 0.0299514972021163 | 853 | 13 | 6 | 0 | 3 | 2 | 15 | 0 | 0 | 5 | 1 | 1 | 0 |
| THE1B_dup16442 | 4.06142470035174 | 3.98483055950951 | 0.952345619652287 | 4.18422731966197 | 2.86137676876683e-05 | 0.00266478730575731 | 854 | 15 | 10 | 0 | 3 | 7 | 12 | 0 | 0 | 4 | 1 | 0 | 0 |
| L4_C_Mam_dup247 | 4.7600973243112 | 3.21994768771919 | 0.935287145235575 | 3.44273702907374 | 0.000575858977320787 | 0.0238384377553187 | 855 | 20 | 15 | 0 | 0 | 9 | 11 | 2 | 0 | 1 | 1 | 3 | 0 |
| MLT1D_dup10461 | 5.44397379114831 | 5.34888567561137 | 0.969211639131187 | 5.51880049687205 | 3.41321419029442e-08 | 1.40027700642034e-05 | 856 | 17 | 16 | 0 | 0 | 10 | 19 | 0 | 0 | 1 | 6 | 1 | 0 |
| THE1B_dup12474 | 4.79577389186067 | 4.70515166796188 | 0.985464401903828 | 4.77455264631777 | 1.80107164218419e-06 | 0.000324419222536836 | 857 | 14 | 7 | 0 | 0 | 14 | 18 | 0 | 0 | 3 | 3 | 2 | 0 |
| L2a_dup86234 | 7.01538896714709 | 4.13510511875467 | 0.770627306479711 | 5.36589488068386 | 8.05488459589041e-08 | 2.79238268338246e-05 | 858 | 17 | 22 | 1 | 0 | 10 | 8 | 1 | 1 | 10 | 14 | 0 | 2 |
| AluSz_dup23986 | 7.40758195318119 | 2.73640954768836 | 0.669212598745367 | 4.08899885151378 | 4.33238997940072e-05 | 0.0036620776768685 | 859 | 22 | 24 | 2 | 2 | 4 | 8 | 1 | 2 | 11 | 10 | 0 | 6 |
| MER77_dup572 | 6.92574519703638 | 2.8965554652681 | 0.691416621120671 | 4.18930551680008 | 2.79809472267757e-05 | 0.0026212419168592 | 860 | 18 | 21 | 2 | 3 | 4 | 4 | 1 | 1 | 12 | 15 | 0 | 4 |
| L1M5_dup46985 | 4.82068970945473 | 2.75742736827146 | 0.843386173353292 | 3.26947186875019 | 0.00107748450667021 | 0.0369752755268131 | 861 | 14 | 22 | 2 | 3 | 4 | 1 | 3 | 0 | 5 | 6 | 0 | 1 |
| L2b_dup31125 | 5.66290627462064 | 2.00142218029492 | 0.62184518048247 | 3.21852165637446 | 0.00128853240216089 | 0.0418282579760677 | 862 | 11 | 16 | 5 | 5 | 7 | 5 | 4 | 0 | 8 | 8 | 0 | 2 |
| L2_dup7890 | 6.55950541862571 | 2.69151019064068 | 0.720010351968865 | 3.73815485191395 | 0.000185375761613016 | 0.0105449249277482 | 863 | 23 | 15 | 3 | 2 | 2 | 4 | 4 | 2 | 11 | 14 | 0 | 1 |
| LTR12_dup719 | 4.80732216476077 | 2.78019372145982 | 0.754605408982552 | 3.68430134261614 | 0.000229330803181257 | 0.0122953650842959 | 864 | 10 | 13 | 4 | 1 | 3 | 5 | 1 | 1 | 6 | 14 | 0 | 1 |

| | | | | | | | | | | | | | | | | | | | |
|---|---|---|---|---|---|---|---|---|---|---|---|---|---|---|---|---|---|---|---|
| MLT1E1_dup360 | 4.90403984971095 | 2.51154510078036 | 0.798786863714214 | 3.14419930380695 | 0.00166541970927215 | 0.0499580282577669 | 865 | 18 | 16 | 4 | 2 | 1 | 4 | 1 | 2 | 7 | 5 | 0 | 1 |
| L1M4c_dup2506 | 5.44445450055613 | 2.31082135299784 | 0.731343190345776 | 3.15969490589676 | 0.00157934425513569 | 0.0480991486154507 | 866 | 20 | 18 | 3 | 5 | 3 | 4 | 4 | 2 | 3 | 8 | 0 | 0 |
| AluSz_dup29170 | 6.27597809868817 | 2.16563732166543 | 0.620055954315574 | 3.49264821439525 | 0.000478256103427021 | 0.0209035868453427 | 867 | 23 | 17 | 5 | 6 | 6 | 7 | 2 | 1 | 4 | 8 | 0 | 2 |
| L1P3_dup1273 | 5.8887678602857 | 2.05647588938383 | 0.651344575198101 | 3.1572779872441 | 0.0015924945113092 | 0.0484112611950838 | 868 | 25 | 16 | 3 | 6 | 3 | 8 | 3 | 3 | 4 | 4 | 0 | 1 |
| L1MEd_dup15858 | 4.21168130927432 | 3.75299947913292 | 0.905549229571253 | 4.14444555478209 | 3.4063687279765e-05 | 0.00303631442785359 | 869 | 16 | 8 | 0 | 3 | 6 | 10 | 0 | 1 | 2 | 8 | 0 | 0 |
| AluY_dup69490 | 4.71757285228213 | 3.15831978725542 | 0.843226718041005 | 3.74551673907211 | 0.000180022842541105 | 0.0103492909618956 | 870 | 15 | 12 | 0 | 6 | 6 | 12 | 0 | 1 | 5 | 3 | 0 | 0 |
| MIRb_dup155668 | 3.65597748583254 | 3.21093477179464 | 0.887912314250661 | 3.61627462561374 | 0.000298873296093536 | 0.0149138761266151 | 871 | 7 | 5 | 0 | 4 | 9 | 9 | 0 | 1 | 6 | 4 | 0 | 0 |
| L2_dup28618 | 4.92176218514804 | 4.29680411795797 | 0.935031077744881 | 4.59535968400221 | 4.32003421088933e-06 | 0.00064045785691232 | 872 | 10 | 11 | 0 | 2 | 5 | 5 | 0 | 1 | 13 | 11 | 0 | 0 |
| L1MEc_dup17243 | 4.03067386231428 | 3.73421870011882 | 0.956383818393056 | 3.90451890580203 | 9.44130117270768e-05 | 0.00653438001408116 | 873 | 12 | 5 | 0 | 4 | 10 | 6 | 0 | 0 | 7 | 6 | 0 | 0 |
| L2c_dup100947 | 4.7869846144116 | 4.8165421822035 | 0.987094090688784 | 4.87951678329121 | 1.06346073906197e-06 | 0.00021572568543635 | 874 | 20 | 11 | 0 | 2 | 6 | 7 | 0 | 0 | 7 | 7 | 0 | 0 |
| AluSq2_dup39871 | 5.00435221927135 | 2.84481787675596 | 0.749846956024548 | 3.79386467318382 | 0.000148320531253154 | 0.00902615783364784 | 875 | 13 | 8 | 1 | 6 | 9 | 7 | 2 | 0 | 10 | 6 | 0 | 0 |
| L2_dup41258 | 4.21121834326656 | 2.58471985979411 | 0.736241889641973 | 3.5106938305983 | 0.000446938838696323 | 0.0199900956602322 | 876 | 9 | 9 | 1 | 6 | 10 | 8 | 2 | 0 | 3 | 6 | 0 | 0 |
| MIR1_Amn_dup7697 | 6.82678276617809 | 3.90698945497293 | 0.743772420355699 | 5.25293671564813 | 1.49692981719541e-07 | 4.56651874720998e-05 | 877 | 23 | 12 | 1 | 4 | 17 | 14 | 1 | 0 | 9 | 5 | 0 | 0 |
| THE1B_dup16441 | 6.537523614173 | 3.81265483810582 | 0.752327686670863 | 5.06781141470049 | 4.02415797190682e-07 | 0.000100703974779425 | 878 | 20 | 17 | 1 | 4 | 21 | 11 | 1 | 0 | 6 | 2 | 0 | 0 |
| AluY_dup69489 | 6.03772759676332 | 3.28364029722213 | 0.693943491511135 | 4.73185545709444 | 2.22476815931755e-06 | 0.000380571304411915 | 879 | 18 | 17 | 3 | 4 | 10 | 9 | 1 | 0 | 7 | 7 | 0 | 0 |
| L3b_dup7254 | 6.63033590877832 | 3.39792674933812 | 0.659446091060479 | 5.15269829543427 | 2.5676487953138e-07 | 7.0387299567942e-05 | 880 | 17 | 13 | 4 | 3 | 16 | 14 | 1 | 0 | 8 | 7 | 0 | 0 |
| AluSx3_dup21147 | 6.84461458034123 | 3.47799882205221 | 0.663446231086608 | 5.24232207387155 | 1.58568271627674e-07 | 4.7760361975593e-05 | 881 | 18 | 15 | 2 | 5 | 16 | 15 | 1 | 0 | 5 | 10 | 0 | 0 |
| MLT1L_dup8182 | 7.26871554803833 | 3.33937808907365 | 0.630357806597385 | 5.29759139035545 | 1.17340180476864e-07 | 3.73770101522728e-05 | 882 | 23 | 16 | 2 | 5 | 19 | 13 | 1 | 0 | 5 | 8 | 0 | 1 |
| L1MA7_dup6243 | 4.42126532007209 | 3.82629091259082 | 0.914930920267282 | 4.18205443474687 | 2.88886778578289e-05 | 0.00268461998497634 | 883 | 10 | 12 | 2 | 2 | 5 | 11 | 0 | 0 | 9 | 3 | 0 | 0 |
| L1PB1_dup5668 | 7.33222585227522 | 4.03098171980283 | 0.816879494366132 | 4.93460999768481 | 8.03110741925721e-07 | 0.000173882610189497 | 884 | 27 | 19 | 3 | 3 | 5 | 13 | 0 | 0 | 13 | 8 | 0 | 0 |
| MIRc_dup71286 | 6.61687174402565 | 3.62161709980879 | 0.755322102301561 | 4.79479825729086 | 1.62838623037488e-06 | 0.000300829184761477 | 885 | 19 | 20 | 4 | 2 | 11 | 5 | 1 | 0 | 9 | 11 | 0 | 0 |
| L1MEc_dup13433 | 4.63066499406554 | 3.35764373694482 | 0.889006078779266 | 3.77685126917845 | 0.000158823507760054 | 0.00949533125062153 | 886 | 20 | 8 | 3 | 3 | 8 | 3 | 0 | 0 | 4 | 10 | 0 | 0 |
| L1MC5_dup6227 | 4.56004653733112 | 3.24634756873956 | 0.828754898979775 | 3.91713831524366 | 8.96063413068513e-05 | 0.00628896961191906 | 887 | 13 | 7 | 4 | 2 | 9 | 5 | 0 | 0 | 8 | 8 | 0 | 0 |
| MIRb_dup155675 | 4.34818220744545 | 3.15778913363022 | 0.861254858188414 | 3.66649790547759 | 0.000245894932186034 | 0.0129532246034842 | 888 | 11 | 8 | 4 | 2 | 8 | 3 | 0 | 0 | 8 | 9 | 0 | 0 |
| L1PA7_dup7528 | 6.5466439706517 | 4.0325589274864 | 0.893692253140846 | 4.51224558936718 | 6.41448530927757e-06 | 0.000872670738365558 | 889 | 12 | 3 | 4 | 0 | 21 | 19 | 0 | 0 | 11 | 9 | 0 | 1 |
| AluSq2_dup39872 | 3.05748968555281 | 4.65746427385417 | 1.09195129586331 | 4.26526740844421 | 1.99662919617417e-05 | 0.00203901257816678 | 890 | 6 | 1 | 1 | 0 | 11 | 8 | 0 | 0 | 6 | 4 | 0 | 0 |
| L1M1_dup4808 | 4.34355185096631 | 4.66725598194776 | 0.978466712632326 | 4.76996909725385 | 1.84254183627941e-06 | 0.000330637601640529 | 891 | 10 | 6 | 2 | 0 | 14 | 7 | 0 | 0 | 7 | 7 | 0 | 0 |
| L2_dup25063 | 4.63416228035725 | 5.27593383860304 | 0.985016070089103 | 5.35619062349489 | 8.49948380343877e-08 | 2.90995420588927e-05 | 892 | 19 | 6 | 1 | 0 | 8 | 7 | 0 | 0 | 9 | 7 | 0 | 0 |
| Tigger2a_dup3088 | 5.09070171030885 | 2.9452732868263 | 0.888438974223472 | 3.31511040406639 | 0.000916069365301533 | 0.0331269059568448 | 893 | 3 | 5 | 4 | 4 | 11 | 10 | 0 | 0 | 14 | 9 | 0 | 0 |
| MIRb_dup155677 | 6.11607586113119 | 3.27977526190471 | 0.779130304437202 | 4.20953368547746 | 2.55898289679389e-05 | 0.00245524090467951 | 894 | 8 | 9 | 4 | 4 | 15 | 10 | 0 | 0 | 8 | 17 | 0 | 0 |
| Tigger2_dup1591 | 5.20769994354221 | 4.09088212185486 | 0.909244744208095 | 4.49920898406491 | 6.82067809316645e-06 | 0.00091125944388354 | 895 | 4 | 11 | 1 | 3 | 13 | 14 | 0 | 0 | 6 | 12 | 0 | 0 |
| HERVL40-int_dup703 | 6.45795903469584 | 4.79106551911813 | 0.959030601844048 | 4.99573789397935 | 5.86112216889435e-07 | 0.000135796550691308 | 896 | 11 | 4 | 0 | 2 | 22 | 13 | 1 | 0 | 13 | 12 | 0 | 0 |
| L1PB_dup907 | 5.53569128368414 | 5.02690340425172 | 0.966856875855052 | 5.19922185980848 | 2.00124543928809e-07 | 5.79304558755283e-05 | 897 | 9 | 7 | 0 | 1 | 18 | 10 | 0 | 0 | 13 | 7 | 1 | 0 |
| L4_B_Mam_dup2265 | 6.07943755011358 | 2.93915018755221 | 0.789735595117609 | 3.72168888641079 | 0.000197894752587068 | 0.0110354499177601 | 898 | 7 | 3 | 4 | 4 | 21 | 22 | 1 | 0 | 3 | 11 | 1 | 0 |
| MIR_dup120671 | 5.65378552579754 | 2.75562979897094 | 0.705391391899422 | 3.90652598063438 | 9.36325800683571e-05 | 0.00650870669965695 | 899 | 9 | 9 | 3 | 6 | 13 | 14 | 2 | 0 | 3 | 13 | 0 | 0 |
| L1ME1_dup22062 | 5.41216828890106 | 4.49165398342659 | 0.946478179450082 | 4.74564980043841 | 2.07838029894783e-06 | 0.000362977083394936 | 900 | 13 | 9 | 1 | 2 | 22 | 11 | 0 | 0 | 1 | 10 | 0 | 0 |
| X9_LINE_dup365 | 4.66807589282553 | 4.24003972411132 | 0.915764447158189 | 4.63005496366349 | 3.65568682426588e-06 | 0.000560474263586835 | 901 | 12 | 5 | 1 | 1 | 15 | 15 | 0 | 0 | 4 | 5 | 0 | 1 |
| AluJr_dup54738 | 5.18509885283355 | 3.70964520988808 | 0.776224032157311 | 4.77909090185999 | 1.76089573491796e-06 | 0.000320019705297638 | 902 | 15 | 10 | 2 | 3 | 14 | 14 | 0 | 0 | 4 | 4 | 0 | 0 |
| L2a_dup119775 | 4.69607274730773 | 4.59574455752242 | 0.92286988735403 | 4.97984019253129 | 6.36367988977438e-07 | 0.000144623894465354 | 903 | 13 | 14 | 1 | 1 | 10 | 15 | 0 | 0 | 2 | 4 | 0 | 0 |
| HAL1_dup16424 | 7.97536777653545 | 3.8660751843145 | 0.726160761285613 | 5.32399351552665 | 1.01513526717563e-07 | 3.33628972767088e-05 | 904 | 29 | 20 | 6 | 1 | 16 | 20 | 0 | 0 | 5 | 5 | 0 | 0 |
| Tigger19b_dup2530 | 6.26241943218139 | 4.65235308480232 | 0.861433455640025 | 5.40071093633777 | 6.63773100737428e-08 | 2.40702843694485e-05 | 905 | 19 | 15 | 3 | 0 | 16 | 15 | 0 | 0 | 5 | 6 | 0 | 0 |
| AluSx_dup74467 | 7.42448587798141 | 3.53377188376781 | 0.688683100269607 | 5.13120168388683 | 2.87898224858673e-07 | 7.73185451605212e-05 | 906 | 14 | 24 | 2 | 5 | 22 | 15 | 0 | 0 | 4 | 7 | 1 | 0 |
| Charlie8_dup1994 | 6.69017043136843 | 3.60147097026358 | 0.696932414936525 | 5.16760433734682 | 2.37113603585958e-07 | 6.57381529552576e-05 | 907 | 19 | 18 | 2 | 5 | 18 | 17 | 0 | 0 | 3 | 4 | 0 | 0 |
| L1PA2_dup3871 | 6.77172226061012 | 3.01294107615785 | 0.714449960053316 | 4.21714779847284 | 2.47411821804734e-05 | 0.00239403773781171 | 908 | 13 | 21 | 4 | 7 | 18 | 17 | 0 | 0 | 2 | 5 | 0 | 0 |
| L1M5_dup38755 | 5.65828588541941 | 3.04291579164051 | 0.746360939883848 | 4.07700300087255 | 4.56199042350054e-05 | 0.0038101346904537 | 909 | 12 | 20 | 4 | 5 | 12 | 10 | 0 | 0 | 3 | 6 | 0 | 0 |
| MIR_dup120676 | 5.22070565596462 | 3.25222315941887 | 0.752901049746063 | 4.31958908878633 | 1.56319994396796e-05 | 0.00170232116186454 | 910 | 9 | 14 | 4 | 3 | 12 | 12 | 0 | 0 | 6 | 5 | 0 | 0 |
| Tigger19b_dup2529 | 5.31333351944835 | 3.77270510617069 | 0.79573974520914 | 4.74112940680012 | 2.12530124200056e-06 | 0.000369355273216689 | 911 | 14 | 10 | 3 | 2 | 14 | 11 | 0 | 0 | 4 | 9 | 0 | 0 |
| MIR_dup120672 | 6.18569416937391 | 3.98745833306525 | 0.768167436303391 | 5.19087134473425 | 2.09312222816904e-07 | 5.96467139765163e-05 | 912 | 14 | 13 | 3 | 2 | 15 | 14 | 0 | 0 | 7 | 9 | 0 | 0 |
| MIR_dup120677 | 5.77711896737822 | 3.40048338905865 | 0.728891148237624 | 4.66528287149683 | 3.08192775351037e-06 | 0.000490195442003194 | 913 | 13 | 9 | 4 | 3 | 15 | 13 | 0 | 0 | 7 | 8 | 0 | 0 |
| MER20_dup10906 | 7.37925653158552 | 3.58087086991971 | 0.667993433100439 | 5.36063783336818 | 8.29286251125004e-08 | 2.85564900593618e-05 | 914 | 17 | 18 | 4 | 4 | 17 | 17 | 0 | 0 | 6 | 10 | 0 | 0 |
| MIR_dup120670 | 6.24404478295883 | 3.53027079286913 | 0.719929846534187 | 4.90363166614664 | 9.40808298950161e-07 | 0.000196714913446393 | 915 | 17 | 15 | 3 | 4 | 14 | 10 | 0 | 0 | 8 | 7 | 0 | 0 |
| THE1A-int_dup1098 | 6.53436499428132 | 2.84736560341533 | 0.720430219918543 | 3.95231283293096 | 7.73994632845973e-05 | 0.00564070220236349 | 916 | 15 | 15 | 7 | 5 | 12 | 12 | 0 | 0 | 7 | 9 | 0 | 0 |
| L1MEg_dup9402 | 4.09746110011094 | 2.73407171790959 | 0.84407192779368 | 3.23914541863296 | 0.00119888431436586 | 0.0398143373434891 | 917 | 5 | 3 | 0 | 2 | 19 | 14 | 3 | 1 | 2 | 2 | 1 | 0 |
| L1PREC2_dup4181 | 3.68439937437165 | 3.86903317369639 | 1.0166883334041 | 3.80552529873338 | 0.000141503548991318 | 0.00872511103091169 | 918 | 4 | 7 | 0 | 1 | 18 | 6 | 0 | 1 | 1 | 7 | 0 | 1 |
| L1ME1_dup25253 | 3.07286803108682 | 2.82159686482761 | 0.882053560440641 | 3.19889515940284 | 0.00137955327321882 | 0.0438907852853759 | 919 | 2 | 7 | 1 | 2 | 9 | 9 | 0 | 1 | 3 | 3 | 0 | 1 |
| AluSx_dup116243 | 4.10413280866147 | 2.68235453031541 | 0.852517868500897 | 3.1463909783289 | 0.00165298874078822 | 0.0497029240600507 | 920 | 2 | 8 | 2 | 4 | 10 | 12 | 0 | 2 | 2 | 9 | 0 | 0 |
| L1MA3_dup4870 | 3.82433060954375 | 3.17568558518349 | 0.8566829423121 | 3.70695554718603 | 0.000209765744792028 | 0.0115139376955672 | 921 | 3 | 5 | 2 | 2 | 10 | 7 | 0 | 1 | 6 | 10 | 0 | 0 |
| MER20B_dup2838 | 8.30078979299667 | 3.88727152151274 | 0.670943134272574 | 5.79374215629654 | 6.88351139074938e-09 | 3.68068314917585e-06 | 922 | 13 | 18 | 2 | 2 | 22 | 23 | 0 | 3 | 9 | 10 | 0 | 0 |
| L1PA16_dup7721 | 6.15280916850266 | 3.45671201071653 | 0.687748896633598 | 5.02612512740695 | 5.00489395834761e-07 | 0.000119846261266864 | 923 | 12 | 13 | 3 | 3 | 13 | 15 | 0 | 1 | 8 | 8 | 0 | 0 |
| MER4B-int_dup601 | 5.56156619563576 | 3.09191107126132 | 0.749983244384073 | 4.12264019818278 | 3.74554504221983e-05 | 0.00327781338954042 | 924 | 6 | 15 | 5 | 1 | 10 | 15 | 0 | 2 | 6 | 8 | 0 | 0 |
| L2c_dup100942 | 7.71922306684562 | 4.21719677266115 | 0.739868991101004 | 5.69992366673661 | 1.19861086025619e-08 | 5.96662468105689e-06 | 925 | 22 | 23 | 1 | 2 | 22 | 18 | 0 | 1 | 1 | 8 | 0 | 1 |
| HAL1b_dup3805 | 6.33950740318729 | 3.97283162683496 | 0.803950406989805 | 4.94163768348629 | 7.74690794924298e-07 | 0.000169425161662736 | 926 | 16 | 8 | 0 | 2 | 15 | 19 | 0 | 2 | 3 | 14 | 0 | 1 |
| MIR_dup120665 | 6.75532224517538 | 3.01317405715941 | 0.605317611000299 | 4.97783973636597 | 6.42978590908032e-07 | 0.000145310429698672 | 927 | 15 | 13 | 1 | 4 | 15 | 17 | 1 | 3 | 4 | 11 | 0 | 1 |
| MIR_dup120666 | 6.57314350300083 | 4.03903922955504 | 0.76820850983894 | 5.25773820235583 | 1.45837929813398e-07 | 4.4718494906416e-05 | 928 | 16 | 21 | 1 | 1 | 13 | 13 | 1 | 1 | 3 | 12 | 0 | 1 |
| MIRb_dup155670 | 5.05048626986665 | 3.0073846744417 | 0.707263530565144 | 4.25214159146398 | 2.11735913841339e-05 | 0.00212760615518966 | 929 | 20 | 9 | 2 | 2 | 12 | 13 | 0 | 2 | 2 | 2 | 0 | 1 |
| AluSz6_dup31319 | 5.63319038436823 | 3.55144329150063 | 0.804710493036684 | 4.41331798483051 | 1.01798306471617e-05 | 0.00123900254893583 | 930 | 16 | 26 | 1 | 3 | 9 | 6 | 0 | 2 | 3 | 5 | 0 | 0 |
| Zaphod3_dup880 | 6.73882362009723 | 2.82658585381928 | 0.684968402146836 | 4.12659305883331 | 3.68177057543038e-05 | 0.00323389657195038 | 931 | 24 | 26 | 2 | 2 | 12 | 6 | 0 | 2 | 3 | 5 | 1 | 3 |
| LTR33A__dup1154 | 8.72623665960308 | 3.37806484181386 | 0.625368682195859 | 5.40171731969763 | 6.60059093055153e-08 | 2.40150490758565e-05 | 932 | 26 | 22 | 6 | 2 | 20 | 25 | 0 | 1 | 2 | 7 | 1 | 0 |
| MIRb_dup155683 | 6.90841645636047 | 3.0309518447721 | 0.623209319237963 | 4.86345719039991 | 1.15352983381612e-06 | 0.000229688765160212 | 933 | 18 | 18 | 5 | 3 | 15 | 16 | 0 | 1 | 6 | 4 | 1 | 0 |
| Charlie1b_dup2209 | 5.63556586828145 | 3.25487750779306 | 0.730400188894669 | 4.45629335435789 | 8.33889049678812e-06 | 0.00106277752445032 | 934 | 14 | 21 | 2 | 3 | 11 | 12 | 0 | 1 | 1 | 6 | 1 | 0 |
| MIR_dup120669 | 6.04872832167965 | 3.21873126799043 | 0.768569537232224 | 4.18795061743111 | 2.81484743241639e-05 | 0.00263330083315559 | 935 | 25 | 14 | 3 | 4 | 8 | 15 | 0 | 0 | 1 | 8 | 1 | 0 |
| L1PA2_dup1941 | 12.1970361067473 | 5.0271736670921 | 1.46666113121382 | 3.42763134585259 | 0.000608871681212077 | 0.0248334900681674 | 936 | 77 | 69 | 0 | 1 | 0 | 2 | 1 | 0 | 4 | 4 | 0 | 2 |
| L1M1_dup3446 | 6.67744456016981 | 3.84685635058067 | 1.04804199348978 | 3.67051737857503 | 0.000242060010642402 | 0.0128193214538876 | 937 | 30 | 40 | 2 | 2 | 0 | 5 | 2 | 0 | 1 | 5 | 0 | 0 |
| L1PA4_dup6083 | 9.72990664540026 | 3.94002453361503 | 0.955387504372178 | 4.1240067674992 | 3.72337946441913e-05 | 0.00326201491178452 | 938 | 59 | 46 | 2 | 4 | 1 | 2 | 2 | 1 | 5 | 5 | 0 | 0 |
| L1M1_dup3443 | 8.24520847404973 | 4.50731307180199 | 1.04596964981474 | 4.3092197489672 | 1.63831519668661e-05 | 0.00176393913027508 | 939 | 57 | 39 | 2 | 1 | 1 | 3 | 1 | 0 | 2 | 3 | 0 | 0 |
| L1MA5_dup1396 | 8.8207338046159 | 3.85207286945158 | 1.01626341784825 | 3.79042756218428 | 0.00015038818749721 | 0.00910309530964414 | 940 | 47 | 51 | 4 | 3 | 1 | 2 | 1 | 0 | 3 | 3 | 0 | 0 |
| L3_dup15760 | 7.8142530638486 | 3.38143932539944 | 0.8995047352031 | 3.75922348495023 | 0.000170441538537488 | 0.00998421524589887 | 941 | 37 | 24 | 2 | 1 | 0 | 3 | 1 | 0 | 13 | 11 | 2 | 4 |
| L1PA7_dup11137 | 7.58140282332428 | 3.83079058472201 | 0.948530473009251 | 4.03865842345441 | 5.37577804670611e-05 | 0.00431560351923593 | 942 | 23 | 29 | 2 | 1 | 0 | 3 | 0 | 0 | 12 | 19 | 2 | 2 |
| L2c_dup110461 | 7.66387025653744 | 4.31956089141929 | 1.07724663969608 | 4.00981607391038 | 6.0766062348637e-05 | 0.00472054896232739 | 943 | 37 | 36 | 1 | 1 | 0 | 2 | 0 | 0 | 9 | 8 | 1 | 2 |

| | | | | | | | | | | | | | | | | | | | |
|---|---|---|---|---|---|---|---|---|---|---|---|---|---|---|---|---|---|---|---|
| L1PA16_dup5634 | 5.29862233133685 | 3.13991169928241 | 0.99179831599097 | 3.1658772238841 | 0.00154616039231453 | 0.0474432678799888 | 944 | 30 | 18 | 2 | 2 | 0 | 3 | 0 | 0 | 6 | 4 | 2 | 1 |
| L2a_dup132831 | 7.07202860722522 | 3.5317206442919 | 1.06753827097998 | 3.30828480842177 | 0.000938692937229704 | 0.0336281586332519 | 945 | 30 | 29 | 4 | 3 | 0 | 2 | 0 | 0 | 12 | 7 | 1 | 0 |
| THE1D_dup10240 | 8.35544970768629 | 3.61436237140473 | 1.02279654881663 | 3.53380383966539 | 0.000409624968302464 | 0.0188018176519479 | 946 | 44 | 32 | 4 | 0 | 1 | 2 | 0 | 1 | 13 | 5 | 0 | 3 |
| THE1A-int_dup499 | 10.1559228348173 | 5.23707759524962 | 1.1673121327922 | 4.48644149934654 | 7.24225906606209e-06 | 0.000956307755678807 | 947 | 56 | 54 | 2 | 0 | 0 | 6 | 0 | 0 | 6 | 6 | 1 | 0 |
| L1M4b_dup3732 | 9.95384834747618 | 4.86782710499349 | 1.16801708211138 | 4.16759924109508 | 3.07824509194984e-05 | 0.00281173169539001 | 948 | 60 | 52 | 1 | 0 | 1 | 4 | 0 | 0 | 4 | 5 | 0 | 3 |
| ERV3-16A3_I-int_dup29 | 12.1271370344138 | 4.92527957649308 | 1.02008847586723 | 4.82828665651366 | 1.37712763191616e-06 | 0.000264472971351027 | 949 | 66 | 63 | 1 | 0 | 4 | 5 | 0 | 0 | 6 | 8 | 1 | 3 |
| MLT1B_dup7374 | 8.604375837619 | 4.94865465418877 | 1.13164396248223 | 4.3729784439746 | 1.22562829871225e-05 | 0.00141914039496538 | 950 | 48 | 39 | 0 | 0 | 2 | 5 | 0 | 0 | 7 | 5 | 1 | 4 |
| L1MA10_dup4323 | 4.16268550339334 | 3.53758033764401 | 1.11134763826965 | 3.18314469372696 | 0.00145684791882301 | 0.0455700190472277 | 951 | 16 | 15 | 2 | 3 | 3 | 0 | 0 | 0 | 6 | 7 | 0 | 0 |
| L1MA1_dup3916 | 5.2963823935163 | 4.13324020090282 | 1.13241387334767 | 3.64993779940547 | 0.000262303818499803 | 0.0135992226653571 | 952 | 18 | 18 | 3 | 1 | 4 | 0 | 0 | 0 | 12 | 8 | 0 | 0 |
| THE1B_dup18145 | 6.47578879353727 | 3.68074135711233 | 0.975395269983274 | 3.77358950815441 | 0.000160915449193759 | 0.00957708541117015 | 953 | 16 | 22 | 1 | 2 | 2 | 0 | 0 | 1 | 16 | 15 | 0 | 2 |
| L2a_dup132835 | 5.57744335340422 | 3.7096665676139 | 1.0505212648285 | 3.53126270910804 | 0.000413580758707297 | 0.0189577898692673 | 954 | 16 | 23 | 1 | 1 | 1 | 0 | 0 | 0 | 7 | 16 | 1 | 3 |
| MER5B_dup19078 | 3.78844733397888 | 3.55286768549611 | 1.09269928530522 | 3.25145969552244 | 0.00114814053073075 | 0.0386524401139303 | 955 | 12 | 10 | 1 | 1 | 1 | 0 | 0 | 0 | 8 | 11 | 0 | 2 |
| L2b_dup12591 | 4.78641023626556 | 3.98027877025838 | 1.17015805725261 | 3.40148815417603 | 0.000670200331049722 | 0.0265010156440533 | 956 | 16 | 17 | 0 | 2 | 0 | 1 | 0 | 0 | 8 | 13 | 0 | 2 |
| MLT1A0_dup16447 | 4.17879413138807 | 4.23466422843015 | 1.3053833628337 | 3.24400045917365 | 0.00117863556246337 | 0.0393174845851546 | 957 | 15 | 18 | 0 | 3 | 0 | 1 | 0 | 0 | 6 | 9 | 0 | 0 |
| L1ME3B_dup6216 | 4.67435965382358 | 3.28747269463797 | 0.985422525875625 | 3.33610467420236 | 0.000849611678371025 | 0.0313769517960068 | 958 | 11 | 18 | 1 | 2 | 0 | 2 | 0 | 0 | 12 | 7 | 1 | 2 |
| AluJb_dup108883 | 3.60758425683463 | 3.45749628134803 | 1.04174983247955 | 3.31893144932751 | 0.000903626092855244 | 0.0327431694672748 | 959 | 7 | 13 | 1 | 1 | 0 | 2 | 0 | 0 | 5 | 13 | 1 | 1 |
| AluJb_dup90415 | 4.48080426702057 | 3.75245522360465 | 1.0654639429668 | 3.52189790032301 | 0.000428469127829866 | 0.0194417458220071 | 960 | 12 | 16 | 2 | 1 | 0 | 3 | 0 | 0 | 6 | 14 | 1 | 0 |
| L1PREC2_dup6298 | 5.12597473371187 | 3.15510706330769 | 0.972268753059963 | 3.24509766808592 | 0.00117410346570097 | 0.039226793491952 | 961 | 16 | 11 | 1 | 2 | 1 | 1 | 0 | 0 | 7 | 20 | 4 | 0 |
| L1PA4_dup8730 | 7.17442182102943 | 4.28101279526337 | 1.06195322789962 | 4.03126303757327 | 5.54779123710202e-05 | 0.00441348958108819 | 962 | 31 | 21 | 0 | 1 | 1 | 1 | 0 | 0 | 18 | 9 | 4 | 1 |
| L3_dup36357 | 5.4483666996536 | 4.91262462762924 | 1.2881628741833 | 3.8136672978903 | 0.000136919764249507 | 0.00851751901284004 | 963 | 21 | 14 | 0 | 1 | 0 | 0 | 0 | 0 | 15 | 13 | 1 | 0 |
| MIR_dup132994 | 5.55723774470528 | 4.19613465588 | 1.22328354387633 | 3.43022243443522 | 0.000603086641552292 | 0.02467791126583 | 964 | 24 | 19 | 0 | 2 | 0 | 0 | 0 | 0 | 9 | 13 | 1 | 1 |
| L1ME3_dup8971 | 3.50136695853066 | 4.79214501265119 | 1.47163857567576 | 3.25633283325064 | 0.00112861367001034 | 0.0381841473516679 | 965 | 19 | 20 | 0 | 0 | 0 | 0 | 0 | 0 | 2 | 3 | 1 | 0 |
| L1PA14_dup1022 | 3.64097050789384 | 4.14953250764899 | 1.31911480680843 | 3.14569473879888 | 0.00165692846114977 | 0.0497865260281276 | 966 | 20 | 21 | 1 | 0 | 0 | 1 | 0 | 0 | 2 | 1 | 1 | 0 |
| MER21-int_dup247 | 4.6407060790136 | 4.53089275932569 | 1.30165532835127 | 3.48086982831677 | 0.000499788321903827 | 0.0215634872246107 | 967 | 24 | 29 | 0 | 0 | 0 | 1 | 1 | 0 | 2 | 2 | 1 | 0 |
| AluJb_dup94550 | 3.48989297469364 | 4.31231007087453 | 1.28156368261919 | 3.36488161248552 | 0.000765765082912989 | 0.0291439346716127 | 968 | 14 | 18 | 0 | 0 | 0 | 2 | 2 | 0 | 4 | 4 | 0 | 0 |
| L1PA7_dup4633 | 2.2313443204931 | 4.72584600725721 | 1.32303996322715 | 3.57196013620779 | 0.000354319353642358 | 0.0168685219931542 | 969 | 11 | 8 | 0 | 0 | 0 | 2 | 0 | 0 | 3 | 4 | 0 | 0 |
| MIRb_dup4946 | 2.58167014996582 | 4.86225560444977 | 1.31684035288565 | 3.69236528467167 | 0.000222178036108569 | 0.012000062632745 | 970 | 13 | 12 | 0 | 0 | 0 | 3 | 0 | 0 | 2 | 3 | 0 | 0 |
| MLT1D-int_dup209 | 2.64453288083592 | 4.93346321804061 | 1.34964609539414 | 3.65537546092768 | 0.000256805853609439 | 0.0134033616238104 | 971 | 12 | 13 | 0 | 0 | 0 | 2 | 0 | 0 | 4 | 2 | 0 | 0 |
| L1MDa_dup5985 | 2.20586750699944 | 4.66649135012835 | 1.40657686968548 | 3.31762269855313 | 0.000907870305592026 | 0.0328803816947337 | 972 | 11 | 11 | 0 | 0 | 0 | 1 | 0 | 0 | 2 | 3 | 0 | 0 |
| LTR2B_dup125 | 2.56704801684669 | 4.80963987612027 | 1.41571122525179 | 3.39733117201556 | 0.000680465616589251 | 0.0268101135786938 | 973 | 15 | 13 | 0 | 0 | 0 | 1 | 0 | 0 | 2 | 2 | 0 | 0 |
| L1M1_dup3449 | 5.6981683775729 | 4.60909625689807 | 1.35154937546536 | 3.41023150213145 | 0.000649077588998769 | 0.0259093132011242 | 974 | 31 | 24 | 0 | 2 | 0 | 0 | 0 | 0 | 5 | 10 | 0 | 1 |
| Tigger2_dup2216 | 4.49515429402832 | 4.82066877121524 | 1.46512869969663 | 3.29026983923897 | 0.00100091347888654 | 0.0351242695261776 | 975 | 22 | 24 | 0 | 0 | 0 | 0 | 0 | 0 | 5 | 4 | 0 | 2 |
| L1PA4_dup8305 | 4.65627515153532 | 4.75585323675449 | 1.3583434691945 | 3.50121552067734 | 0.000463141137309177 | 0.020460932242512 | 976 | 23 | 21 | 0 | 0 | 0 | 0 | 1 | 1 | 7 | 5 | 0 | 0 |
| MLT1C_dup14754 | 2.67418550885331 | 4.46709866407122 | 1.41590454354434 | 3.15494337837848 | 0.0016052925637723 | 0.0486403851240753 | 977 | 11 | 10 | 1 | 0 | 0 | 0 | 0 | 0 | 4 | 7 | 0 | 0 |
| Tigger1_dup9768 | 3.12958288573031 | 4.69889051633572 | 1.4242159675042 | 3.29928228832462 | 0.000969323830677861 | 0.0343992137530909 | 978 | 16 | 12 | 1 | 0 | 0 | 0 | 0 | 0 | 5 | 5 | 0 | 0 |
| MIRb_dup169605 | 3.08003785451184 | 4.69440346232967 | 1.41783414917695 | 3.31096797538327 | 0.000929738481242141 | 0.0334483750616333 | 979 | 12 | 14 | 0 | 0 | 0 | 0 | 0 | 0 | 5 | 6 | 0 | 1 |
| AluJb_dup105776 | 3.45661170917599 | 5.35602694496389 | 1.50799674324938 | 3.5517496764767 | 0.000382678735558472 | 0.0178963982726407 | 980 | 16 | 16 | 0 | 0 | 0 | 0 | 0 | 0 | 5 | 6 | 0 | 0 |
| MLT1B_dup14011 | 2.89690395304046 | 5.10080782507277 | 1.53210603554589 | 3.329278592167 | 0.00087071261977755 | 0.031925409490887 | 981 | 15 | 12 | 0 | 0 | 0 | 0 | 0 | 0 | 5 | 4 | 0 | 0 |
| L2c_dup110458 | 3.80741030845579 | 4.99725183719706 | 1.41084997528626 | 3.54201504393344 | 0.000397082827141842 | 0.0183714311466251 | 982 | 19 | 12 | 0 | 1 | 0 | 0 | 0 | 0 | 4 | 12 | 0 | 0 |
| LTR85a_dup1431 | 2.87127815754192 | 4.56567427462075 | 1.40249247074981 | 3.25540020345337 | 0.00113232683250239 | 0.0382825576138981 | 983 | 9 | 12 | 0 | 1 | 0 | 0 | 0 | 0 | 5 | 8 | 0 | 0 |
| L1P4_dup3273 | 3.20605259585446 | 5.25316316527655 | 1.35972184552822 | 3.8634101397671 | 0.000111815037285807 | 0.00737415212361657 | 984 | 12 | 15 | 0 | 0 | 0 | 1 | 0 | 0 | 3 | 9 | 0 | 0 |
| MLT1L_dup9059 | 2.85915688096848 | 5.0843415303506 | 1.52874078191099 | 3.32583626374837 | 0.000881536965055516 | 0.0322412843617357 | 985 | 15 | 9 | 0 | 0 | 0 | 0 | 0 | 0 | 3 | 9 | 0 | 0 |
| AluSx1_dup28342 | 2.93362689362136 | 4.08154775066847 | 1.17858260106666 | 3.46309859569836 | 0.000533992684908582 | 0.0225646285567852 | 986 | 13 | 11 | 0 | 0 | 1 | 1 | 0 | 1 | 2 | 7 | 0 | 1 |
| L1PA16_dup10923 | 3.49940162611729 | 4.29901287519501 | 1.15019130663918 | 3.73765029380772 | 0.000185748057618252 | 0.0105540194781703 | 987 | 17 | 15 | 0 | 0 | 2 | 1 | 0 | 1 | 4 | 3 | 0 | 1 |
| L1PA13_dup6605 | 2.32007666869831 | 3.78241438803483 | 1.14507454928296 | 3.3032036127284 | 0.000955869551448747 | 0.0340436066889722 | 988 | 5 | 9 | 0 | 0 | 2 | 1 | 0 | 0 | 5 | 4 | 0 | 2 |
| MLT2B2_dup989 | 2.44552303727283 | 4.30951112080422 | 1.08955106039167 | 3.95530900521085 | 7.64357923771879e-05 | 0.00558930272064608 | 989 | 6 | 7 | 0 | 0 | 2 | 3 | 0 | 0 | 7 | 3 | 0 | 1 |
| L3_dup15759 | 3.32330841526643 | 4.26885168393018 | 1.18775559880998 | 3.59404888363161 | 0.000325578711502663 | 0.0158479505340356 | 990 | 17 | 15 | 0 | 0 | 1 | 3 | 0 | 0 | 2 | 3 | 0 | 2 |
| L1PB_dup1490 | 3.51898707376433 | 4.80818201130491 | 1.12123470936986 | 4.2882921578544 | 1.8005216812491e-05 | 0.00188960146141805 | 991 | 9 | 22 | 0 | 0 | 3 | 3 | 0 | 0 | 3 | 3 | 0 | 1 |
| MIRb_dup4947 | 3.12082065999086 | 4.69657259487648 | 1.09690579915137 | 4.28165536047857 | 1.85508137718238e-05 | 0.00193515648125839 | 992 | 12 | 13 | 0 | 0 | 3 | 2 | 0 | 0 | 4 | 4 | 0 | 1 |
| L1MA4A_dup4318 | 4.46222998876106 | 4.28727337494536 | 1.05321074245432 | 4.07067000185988 | 4.68781166191758e-05 | 0.00388903453551815 | 993 | 13 | 17 | 0 | 0 | 1 | 4 | 1 | 0 | 7 | 10 | 0 | 2 |
| MER50_dup2337 | 3.17925257516584 | 4.22905319335369 | 1.07028484902999 | 3.95133426132912 | 7.77166859034214e-05 | 0.00565406288034185 | 994 | 7 | 10 | 0 | 0 | 1 | 4 | 0 | 0 | 8 | 6 | 0 | 2 |
| MLT1J2_dup3261 | 3.31659866888364 | 4.79019671093941 | 1.09161499197805 | 4.38817416959379 | 1.14306222232474e-05 | 0.00134647071065981 | 995 | 14 | 8 | 0 | 0 | 1 | 4 | 0 | 0 | 6 | 7 | 0 | 1 |
| L1PA6_dup994 | 4.30010660179127 | 5.14827425093291 | 1.07739263872662 | 4.77845686510138 | 1.76645647556351e-06 | 0.000320122812288222 | 996 | 8 | 25 | 0 | 0 | 3 | 6 | 0 | 0 | 5 | 5 | 0 | 1 |
| MER117_dup65 | 3.41697647433737 | 5.34417815076511 | 1.02268349729066 | 5.22564230763783 | 1.73551579169042e-07 | 5.15554357764756e-05 | 997 | 10 | 10 | 0 | 0 | 3 | 6 | 0 | 0 | 5 | 8 | 0 | 0 |
| MIRb_dup92010 | 2.81220130153226 | 5.04966881303749 | 1.06165174436709 | 4.75642680363877 | 1.97049639828294e-06 | 0.000349122662684909 | 998 | 6 | 9 | 0 | 0 | 2 | 5 | 0 | 0 | 5 | 7 | 0 | 0 |
| MER33_dup7057 | 2.64243805534754 | 3.95721288771461 | 1.22643098683369 | 3.22660869645103 | 0.00125266642510557 | 0.0410447487177319 | 999 | 9 | 8 | 0 | 0 | 1 | 0 | 0 | 0 | 2 | 11 | 1 | 1 |
| L2b_dup72748 | 2.77224525814163 | 3.68940768864018 | 1.12683709249211 | 3.27412694631901 | 0.00105988984430599 | 0.0365714253035432 | 1000 | 6 | 10 | 0 | 0 | 2 | 0 | 0 | 0 | 7 | 5 | 1 | 2 |
| AluJo_dup19325 | 2.42314001115731 | 4.33104583569606 | 1.23588694855621 | 3.50440292354871 | 0.000457632380837506 | 0.020323182520728 | 1001 | 11 | 8 | 0 | 0 | 3 | 0 | 0 | 0 | 4 | 3 | 1 | 0 |
| L1PA2_dup3607 | 2.37509847704722 | 4.26020086136094 | 1.29770256168282 | 3.28287928771323 | 0.00102752654801938 | 0.0357450046776343 | 1002 | 5 | 11 | 0 | 0 | 1 | 0 | 0 | 0 | 6 | 4 | 1 | 0 |
| AluSx_dup50342 | 2.45237236572132 | 4.30912292977566 | 1.29158870563826 | 3.33629654003999 | 0.000849025488902851 | 0.0313677686235493 | 1003 | 9 | 7 | 0 | 1 | 1 | 0 | 0 | 0 | 8 | 3 | 0 | 0 |
| MSTA_dup5815 | 2.864086222848 | 4.51970149337558 | 1.37034290808369 | 3.29822664583714 | 0.000972975642531086 | 0.0344745194956692 | 1004 | 16 | 12 | 0 | 1 | 1 | 0 | 0 | 0 | 5 | 1 | 0 | 0 |
| L2b_dup84646 | 3.07807342858469 | 4.62152696362817 | 1.16396389783185 | 3.97050713706569 | 7.1719807535753e-05 | 0.00533127328184734 | 1005 | 10 | 7 | 0 | 1 | 3 | 0 | 0 | 0 | 7 | 9 | 0 | 0 |
| MLT1C_dup9087 | 3.61838251134885 | 4.40554893978579 | 1.04929405537139 | 4.19858372134442 | 2.68589551481406e-05 | 0.00253811792759312 | 1006 | 10 | 11 | 0 | 1 | 6 | 1 | 1 | 0 | 7 | 7 | 0 | 0 |
| LTR10F_dup102 | 3.03846960766233 | 5.01174946321784 | 1.59034024213826 | 3.15136932992363 | 0.00162506858516908 | 0.0490771125227367 | 1007 | 25 | 11 | 0 | 0 | 0 | 0 | 0 | 0 | 2 | 2 | 0 | 0 |
| AluY_dup48017 | 2.24883434999612 | 4.65092658429685 | 1.42422299929867 | 3.26558873616498 | 0.00109236773647327 | 0.0373111042710915 | 1008 | 15 | 9 | 0 | 0 | 1 | 0 | 0 | 0 | 2 | 2 | 0 | 0 |
| L1PA7_dup5238 | 3.42785549354432 | 5.25022029427327 | 1.3867255083816 | 3.7860559011427 | 0.000153057256935896 | 0.00923880316054507 | 1009 | 21 | 16 | 0 | 0 | 1 | 0 | 0 | 0 | 3 | 3 | 0 | 0 |
| L1MB4_dup7456 | 3.07786057198711 | 4.07606974322133 | 1.24753752492477 | 3.26729229524949 | 0.00108581509716718 | 0.0371846020672193 | 1010 | 15 | 15 | 1 | 0 | 2 | 0 | 0 | 0 | 3 | 2 | 1 | 0 |
| L1MCb_dup1999 | 3.41470306719237 | 4.64118820754476 | 1.33597426778067 | 3.47401018079092 | 0.000512741578630294 | 0.0219591086516854 | 1011 | 25 | 14 | 0 | 0 | 2 | 0 | 0 | 0 | 1 | 2 | 1 | 0 |
| MER4B-int_dup550 | 3.96060053960835 | 4.58547724628617 | 1.31432784060873 | 3.48883825222968 | 0.000485124603756846 | 0.02109915170271 | 1012 | 25 | 21 | 0 | 1 | 3 | 0 | 0 | 0 | 1 | 1 | 0 | 0 |
| L1MB7_dup10096 | 2.82134687738837 | 4.415692623434 | 1.39050878402966 | 3.17559491471707 | 0.00149529595711416 | 0.0464187464855167 | 1013 | 10 | 21 | 0 | 0 | 1 | 0 | 0 | 0 | 1 | 2 | 0 | 1 |
| MER89_dup420 | 3.63672449181222 | 4.73244785230621 | 1.35829905572823 | 3.48409861020554 | 0.000493797518465201 | 0.0213747761194689 | 1014 | 19 | 21 | 1 | 0 | 1 | 0 | 0 | 0 | 1 | 4 | 0 | 0 |
| L1M3_dup3052 | 5.93867729950817 | 5.30326331005756 | 1.22842167019123 | 4.31713591411325 | 1.5806683549208e-05 | 0.00171601401853916 | 1015 | 24 | 43 | 1 | 0 | 1 | 2 | 0 | 0 | 2 | 3 | 0 | 0 |
| L1M1_dup3442 | 3.64356404613129 | 4.6740430174774 | 1.36146252901453 | 3.43310441372236 | 0.00059671225385949 | 0.0244802230960087 | 1016 | 26 | 16 | 1 | 0 | 0 | 1 | 0 | 0 | 1 | 3 | 0 | 0 |
| L1M6_dup5058 | 4.80509296641279 | 5.52269653402188 | 1.42054664613657 | 3.88772628413283 | 0.000101187639961404 | 0.00687852173724179 | 1017 | 31 | 27 | 0 | 0 | 0 | 1 | 0 | 0 | 1 | 3 | 0 | 0 |
| L1PA13_dup3350 | 4.34747949855777 | 5.52301898812289 | 1.39218749966261 | 3.96715168715518 | 7.27367019169823e-05 | 0.00538454815896774 | 1018 | 23 | 26 | 0 | 0 | 0 | 1 | 0 | 0 | 2 | 4 | 0 | 0 |
| L1MA3_dup8113 | 2.94969538957079 | 4.25095281202877 | 1.14239047194302 | 3.72110317481781 | 0.000198354371367607 | 0.0110558981610299 | 1019 | 13 | 16 | 1 | 0 | 3 | 3 | 0 | 0 | 1 | 1 | 0 | 0 |
| L1PA16_dup6971 | 3.49684400328247 | 3.93510092772032 | 1.16450696014708 | 3.37919914812988 | 0.000726973263108868 | 0.0280812305298219 | 1020 | 22 | 17 | 1 | 1 | 1 | 2 | 0 | 0 | 1 | 1 | 0 | 0 |
| L2c_dup110468 | 3.00778084927814 | 4.60326827329419 | 1.23586382510142 | 3.7247374506786 | 0.000195518602646427 | 0.0109414086136657 | 1021 | 19 | 9 | 1 | 0 | 1 | 1 | 0 | 0 | 5 | 2 | 0 | 0 |
| AluSx1_dup2436 | 2.5090661735438 | 4.6144803359405 | 1.25783909817577 | 3.66857759679502 | 0.00024390366926773 | 0.0128854135185567 | 1022 | 18 | 9 | 0 | 0 | 3 | 1 | 0 | 0 | 1 | 1 | 0 | 0 |

| | | | | | | | | | | | | | | | | | | | |
|---|---|---|---|---|---|---|---|---|---|---|---|---|---|---|---|---|---|---|---|
| MIRb_dup104907 | 2.1348594974341 | 4.52658500962633 | 1.26516630855397 | 3.57785769271711 | 0.00034642196432435 | 0.0165754515831735 | 1023 | 9 | 12 | 0 | 0 | 1 | 2 | 0 | 0 | 2 | 1 | 0 | 0 |
| AluYe5_dup105 | 1.75293023080025 | 4.04473006261747 | 1.26336717949924 | 3.20154752177485 | 0.00136691526214131 | 0.043611158348817 | 1024 | 9 | 7 | 0 | 0 | 2 | 4 | 0 | 0 | 0 | 1 | 0 | 0 |
| L2a_dup119796 | 1.97658039621471 | 4.32471161005546 | 1.28340270832803 | 3.36972298873325 | 0.000752437834923174 | 0.0288215668172295 | 1025 | 10 | 6 | 0 | 0 | 1 | 7 | 0 | 0 | 0 | 2 | 0 | 0 |
| L1M1_dup4368 | 2.69855569737247 | 4.14918575633466 | 1.31479459034158 | 3.15576728624714 | 0.0016007652264553 | 0.0485682941115323 | 1026 | 19 | 12 | 0 | 0 | 1 | 2 | 0 | 0 | 0 | 1 | 0 | 1 |
| L1PA6_dup749 | 2.01140974277201 | 4.19295840485126 | 1.3236369588618 | 3.16775561212554 | 0.00153620597671308 | 0.0472938371503392 | 1027 | 8 | 13 | 0 | 0 | 2 | 2 | 0 | 0 | 0 | 1 | 0 | 0 |
| ERVL-B4-int_dup2616 | 2.05657200189228 | 4.36870490171076 | 1.30002218582369 | 3.36048488198896 | 0.00077805788769222 | 0.0294849119517141 | 1028 | 12 | 9 | 0 | 0 | 2 | 2 | 0 | 0 | 0 | 2 | 0 | 0 |
| MSTA-int_dup908 | 2.0728496354772 | 4.46848749398165 | 1.4167822617454 | 3.15396911341671 | 0.00161066131426767 | 0.0487571602230815 | 1029 | 11 | 11 | 0 | 0 | 1 | 1 | 0 | 0 | 0 | 3 | 0 | 0 |
| L1PB_dup118 | 4.49685276915802 | 3.54253944014298 | 1.05667537390555 | 3.35253335851813 | 0.000800755909517365 | 0.0300697256780142 | 1030 | 21 | 26 | 1 | 3 | 3 | 3 | 0 | 0 | 0 | 2 | 0 | 0 |
| L1MA5A_dup1734 | 2.97169329888967 | 4.04821898780628 | 1.26028015161128 | 3.2121580131454 | 0.00131741893817952 | 0.0424992633763646 | 1031 | 11 | 21 | 0 | 0 | 2 | 2 | 0 | 1 | 0 | 1 | 0 | 0 |
| L1PA8_dup1589 | 3.13913062492654 | 4.45151694291116 | 1.4007034937219 | 3.17805799932915 | 0.00148265088570309 | 0.046128510414423 | 1032 | 16 | 21 | 0 | 0 | 2 | 2 | 0 | 0 | 0 | 0 | 0 | 0 |
| MLT1E3-int_dup74 | 5.44955453942106 | 3.53803246611562 | 1.02670467805734 | 3.4460079336641 | 0.000568933655981514 | 0.023671181623741 5 | 1033 | 33 | 26 | 1 | 3 | 4 | 1 | 0 | 0 | 1 | 2 | 0 | 1 |
| L1MA3_dup3705 | 5.0566876503226 | 2.98659427543893 | 0.914181850889177 | 3.26695861718763 | 0.00108709570003869 | 0.0371973070810189 | 1034 | 22 | 21 | 3 | 4 | 3 | 2 | 0 | 0 | 4 | 5 | 0 | 1 |
| MLT1E3_dup1660 | 4.07347022149829 | 3.73009332243083 | 1.06571533509422 | 3.50008412152673 | 0.000465111356881956 | 0.0205289161425772 | 1035 | 16 | 21 | 1 | 3 | 3 | 1 | 0 | 0 | 3 | 4 | 0 | 0 |
| L1PA7_dup4004 | 6.36974744196434 | 4.14790395682381 | 1.13403221727589 | 3.65765971516018 | 0.000254528655527802 | 0.01331371190772 | 1036 | 36 | 32 | 1 | 1 | 0 | 4 | 0 | 0 | 3 | 4 | 0 | 2 |
| L1MD2_dup3904 | 4.84931251283416 | 5.22643805114259 | 1.25732858540372 | 4.15677978836728 | 3.22764828494317e-05 | 0.00291795592252693 | 1037 | 33 | 21 | 0 | 0 | 1 | 1 | 0 | 0 | 4 | 2 | 0 | 1 |
| ERV3-16A3_I-int_dup2 | 4.9421797891436 | 4.65039094591963 | 1.22805210979311 | 3.78680261923338 | 0.000152598220912075 | 0.00921654526569828 | 1038 | 31 | 25 | 0 | 1 | 1 | 1 | 0 | 0 | 1 | 4 | 0 | 1 |
| L1MA5_dup1394 | 6.53431642227667 | 4.09945507075251 | 1.21068713047904 | 3.38605653562241 | 0.000709047743667102 | 0.0275903637636324 | 1039 | 43 | 33 | 0 | 2 | 1 | 1 | 0 | 0 | 3 | 1 | 0 | 2 |
| L1PA3_dup4243 | 8.0671477231908 | 5.94890531640343 | 1.22555533157197 | 4.85404874276301 | 1.20965983004521e-06 | 0.000238470354531845 | 1040 | 57 | 34 | 0 | 0 | 1 | 2 | 0 | 0 | 6 | 4 | 1 | 0 |
| THE1C_dup4277 | 7.09721082492505 | 5.34090114969719 | 1.16174790150597 | 4.59729786709647 | 4.28005521417956e-06 | 0.000636116188627838 | 1041 | 46 | 30 | 0 | 0 | 2 | 3 | 0 | 0 | 2 | 8 | 1 | 1 |
| L1PB1_dup4817 | 4.09613232277454 | 4.50637117144844 | 1.1092333313004 | 4.06259985549247 | 4.85291772480671e-05 | 0.00398046315390161 | 1042 | 25 | 14 | 0 | 0 | 3 | 2 | 1 | 0 | 3 | 4 | 1 | 0 |
| L2a_dup39588 | 4.53728684658687 | 4.63146026160039 | 1.22368045534673 | 3.78486086082668 | 0.000153794599316005 | 0.00927106273045412 | 1043 | 19 | 29 | 0 | 0 | 2 | 2 | 0 | 0 | 2 | 2 | 2 | 0 |
| MER4CL34_dup202 | 4.99993036115929 | 3.94808241161684 | 1.05142759249024 | 3.75497318105002 | 0.000173360040164413 | 0.0100883407835225 | 1044 | 29 | 17 | 0 | 0 | 1 | 4 | 0 | 1 | 5 | 4 | 1 | 2 |
| THE1D_dup1514 | 5.18032737502579 | 3.14143424464093 | 0.890777569894101 | 3.52662028188967 | 0.000420899932919668 | 0.0191714118373858 | 1045 | 22 | 20 | 0 | 1 | 2 | 8 | 0 | 1 | 5 | 2 | 2 | 3 |
| L1M2_dup8279 | 5.84263226350804 | 4.31978553262605 | 1.11474628804085 | 3.87512887817552 | 0.000106568282570836 | 0.00714205686121825 | 1046 | 44 | 22 | 0 | 1 | 2 | 1 | 0 | 1 | 2 | 3 | 1 | 0 |
| L1MB4_dup3002 | 3.16628158080836 | 3.60663913226311 | 1.09405628948201 | 3.29657547507972 | 0.000978713137837747 | 0.0346180581465968 | 1047 | 18 | 10 | 0 | 1 | 1 | 3 | 0 | 1 | 1 | 5 | 1 | 0 |
| L1MA1_dup1517 | 8.91016201773381 | 4.44415410389348 | 1.08542374022234 | 4.09439552426145 | 4.23271218955297e-05 | 0.0035995601667 4543 | 1048 | 56 | 47 | 0 | 0 | 2 | 2 | 1 | 3 | 2 | 3 | 0 | 0 |
| L1PA5_dup5763 | 4.64828813062582 | 3.72410934564626 | 1.05323257351483 | 3.53588508302419 | 0.0004064114422216 | 0.0186939835744946 | 1049 | 29 | 18 | 0 | 0 | 2 | 2 | 2 | 2 | 3 | 2 | 0 | 0 |
| L2a_dup28141 | 5.04156220006654 | 3.83214699031686 | 1.02709942137374 | 3.73103801888173 | 0.000190692440344138 | 0.0107298041547547 | 1050 | 25 | 23 | 0 | 0 | 2 | 2 | 1 | 2 | 4 | 4 | 0 | 1 |
| L1M1_dup7022 | 4.49958746035101 | 4.25810651361541 | 1.29379574054375 | 3.29117369935523 | 0.000997702876078342 | 0.035050850221665 | 1051 | 34 | 19 | 1 | 0 | 1 | 0 | 0 | 1 | 2 | 1 | 0 | 0 |
| MER4E_dup302 | 4.59232244632756 | 4.57383496429452 | 1.23320428462563 | 3.70890291358582 | 0.00020815921852884 | 0.011449526909862 1 | 1052 | 22 | 25 | 1 | 0 | 2 | 0 | 0 | 1 | 4 | 3 | 0 | 0 |
| L1M2b_dup81 | 4.49487883908138 | 3.2604871300249 | 1.03682949166459 | 3.14467051355794 | 0.00166273981696937 | 0.0499107213937925 | 1053 | 26 | 21 | 2 | 1 | 1 | 2 | 0 | 2 | 2 | 1 | 0 | 0 |
| MER4B-int_dup543 | 4.16840867713147 | 4.75662768701847 | 1.23207664057888 | 3.86065893172329 | 0.000113081658686574 | 0.0074329636347 6118 | 1054 | 26 | 21 | 0 | 0 | 1 | 2 | 0 | 1 | 2 | 1 | 0 | 0 |
| LTR12C_dup224 | 4.61216612662857 | 4.11814293776889 | 1.06705508312255 | 3.85935365746807 | 0.000113687312810687 | 0.00745218769533108 | 1055 | 31 | 18 | 1 | 0 | 6 | 3 | 1 | 0 | 1 | 0 | 0 | 0 |
| L1PA2_dup2091 | 5.79079614877466 | 4.39563181403724 | 1.17494432917609 | 3.74114049907336 | 0.000183187072233221 | 0.0104629455327907 | 1056 | 33 | 34 | 0 | 2 | 4 | 2 | 0 | 0 | 1 | 0 | 0 | 0 |
| L1PA7_dup1919 | 5.96286204550678 | 5.06479233052712 | 1.34650001404025 | 3.76144989061674 | 0.000168931267523962 | 0.00992016766590737 | 1057 | 45 | 29 | 0 | 0 | 1 | 1 | 1 | 0 | 2 | 0 | 0 | 0 |
| L1MDa_dup5873 | 5.46163494272818 | 3.89840392005385 | 1.196739992623 | 3.25751954817635 | 0.00112390517038064 | 0.0380573131736717 | 1058 | 33 | 28 | 0 | 0 | 1 | 1 | 2 | 0 | 1 | 4 | 0 | 2 |
| L1MB7_dup5776 | 4.68882140337624 | 4.59617631672318 | 1.25759856039276 | 3.65472453728616 | 0.000257458255806073 | 0.0134226910902787 | 1059 | 38 | 19 | 0 | 0 | 2 | 2 | 1 | 0 | 0 | 1 | 0 | 0 |
| L2a_dup85080 | 3.81142162498082 | 3.68971695122363 | 1.1366619792537 | 3.24609868067039 | 0.00116998276739074 | 0.0391495879017801 | 1060 | 22 | 18 | 0 | 0 | 2 | 2 | 2 | 1 | 0 | 3 | 0 | 0 |
| MER45A_dup2285 | 4.75551761215993 | 2.68485444055871 | 0.701821429727711 | 3.82555209464089 | 0.000130479337609464 | 0.00823137565103151 | 1061 | 14 | 15 | 2 | 2 | 13 | 6 | 1 | 0 | 2 | 3 | 2 | 1 |
| L1MA6_dup4933 | 5.59633290714535 | 3.21222767190837 | 0.797690667635561 | 4.02690892878282 | 5.6514909311192e-05 | 0.00446315851674773 | 1062 | 19 | 21 | 3 | 2 | 11 | 5 | 0 | 0 | 1 | 8 | 1 | 1 |
| L2_dup41261 | 5.23829672548186 | 3.6203721015291 | 0.809606364475944 | 4.47176833135761 | 7.75754414669752e-06 | 0.00100812484955221 | 1063 | 21 | 16 | 1 | 3 | 14 | 7 | 0 | 0 | 1 | 4 | 1 | 0 |
| LTR78B_dup2315 | 5.90246007208926 | 2.86315397670131 | 0.882321060473665 | 3.24502508776596 | 0.00117440276559636 | 0.0392312846360511 | 1064 | 15 | 26 | 1 | 2 | 13 | 5 | 0 | 0 | 5 | 1 | 1 | 6 |
| L1MA4_dup3752 | 5.71328951448744 | 3.19019424349839 | 0.841260493972719 | 3.79215982012088 | 0.000149342747971894 | 0.00906339296725752 | 1065 | 17 | 26 | 1 | 4 | 12 | 6 | 0 | 0 | 4 | 1 | 0 | 2 |
| L1MC1_dup1296 | 5.04547826947527 | 3.11931844164434 | 0.898862046394428 | 3.47029719872671 | 0.0005198827727005 | 0.0221493332556837 | 1066 | 26 | 21 | 2 | 2 | 6 | 4 | 0 | 0 | 2 | 1 | 0 | 2 |
| L1M2_dup8278 | 6.91895808995454 | 4.55475604903605 | 0.984990351585762 | 4.62416311155051 | 3.7611382157265e-06 | 0.000571486458415873 | 1067 | 32 | 39 | 0 | 1 | 5 | 5 | 0 | 1 | 3 | 2 | 0 | 1 |
| THE1B-int_dup2281 | 6.29389346055272 | 4.18168690152334 | 1.03760717975193 | 4.03012525657648 | 5.5747138659089e-05 | 0.00442750622676088 | 1068 | 32 | 30 | 0 | 0 | 5 | 4 | 1 | 0 | 5 | 1 | 1 | 2 |
| AluJr4_dup16648 | 4.49525859475762 | 4.15386761064446 | 1.15182247018262 | 3.60634361472898 | 0.000310541866090177 | 0.0153154684636701 | 1069 | 26 | 16 | 0 | 0 | 5 | 6 | 0 | 0 | 2 | 1 | 0 | 3 |
| THE1A-int_dup91 | 7.18753047702307 | 5.10798953044801 | 1.06310730096804 | 4.80477325835012 | 1.54926976770487e-06 | 0.000290727558475002 | 1070 | 41 | 29 | 0 | 0 | 13 | 5 | 0 | 0 | 2 | 2 | 0 | 2 |
| L1P2_dup116 | 7.11606965181874 | 3.55610471584263 | 1.10280071274723 | 3.22461227557957 | 0.00126143382247808 | 0.0412468231858415 | 1071 | 43 | 29 | 1 | 1 | 8 | 0 | 0 | 0 | 1 | 5 | 1 | 5 |
| L1P2_dup117 | 11.3141161184228 | 4.09856623978959 | 1.03460992608486 | 3.96146038855364 | 7.44927415294299e-05 | 0.00548429761013667 | 1072 | 66 | 62 | 1 | 1 | 6 | 4 | 0 | 0 | 1 | 3 | 1 | 4 |
| L1PA4_dup4987 | 3.09109474981384 | 3.22688524785979 | 1.01409464563892 | 3.18203558389437 | 0.00146243849379645 | 0.0456548661565062 | 1073 | 16 | 12 | 0 | 0 | 5 | 3 | 1 | 1 | 1 | 0 | 1 | 0 |
| Tigger1_dup738 | 4.68200073333051 | 3.078684832773 | 0.928169915093362 | 3.3169409853834 | 0.000910088377766391 | 0.0329456700343833 | 1074 | 21 | 20 | 0 | 0 | 9 | 4 | 2 | 3 | 1 | 0 | 0 | 0 |
| L1ME4c_dup6081 | 3.30751355459866 | 3.35519842757163 | 1.05201487621635 | 3.18930701782359 | 0.00142614324760479 | 0.0448513192168346 | 1075 | 17 | 10 | 0 | 0 | 8 | 4 | 0 | 1 | 1 | 0 | 0 | 2 |
| L2c_dup117725 | 2.7234284175153 | 3.68259617186447 | 1.15516597515472 | 3.18793684290367 | 0.00143291830708784 | 0.0450049075367376 | 1076 | 11 | 12 | 0 | 0 | 8 | 3 | 0 | 1 | 0 | 0 | 0 | 0 |
| LTR31_dup163 | 3.10276002810174 | 3.3363547365333 | 1.05877277456209 | 3.15115274654962 | 0.00162627416564287 | 0.0490867014268497 | 1077 | 9 | 16 | 0 | 0 | 9 | 2 | 1 | 0 | 0 | 1 | 1 | 1 |
| THE1A-int_dup1319 | 3.29681924406989 | 4.37025437918014 | 1.07227271943924 | 4.07569296500019 | 4.58775197968669e-05 | 0.00382493774547488 | 1078 | 13 | 14 | 0 | 0 | 6 | 8 | 0 | 0 | 0 | 1 | 0 | 1 |
| HERV16-int_dup91 | 4.21361943977713 | 4.08620802403863 | 1.24804641180269 | 3.27408338776158 | 0.00106005324211695 | 0.03657176000417 | 1079 | 17 | 23 | 0 | 0 | 5 | 5 | 0 | 0 | 0 | 2 | 0 | 3 |
| L2-3_Crp_dup222 | 3.92022778706281 | 4.16121976387038 | 1.11530119098143 | 3.73102781340045 | 0.00019070016630016 | 0.0107298041547547 | 1080 | 15 | 19 | 0 | 0 | 9 | 5 | 0 | 0 | 0 | 1 | 0 | 2 |
| L1MC1_dup1300 | 5.64542830913529 | 2.56256663810931 | 0.753380798723198 | 3.40142281626006 | 0.000670360557275839 | 0.0265010156440533 | 1081 | 27 | 18 | 1 | 4 | 10 | 10 | 1 | 0 | 0 | 1 | 1 | 2 |
| L1MEj_dup134 | 4.55376459329301 | 3.66646652261954 | 1.00933367207549 | 3.63256138584994 | 0.000280621789312692 | 0.0142463905013437 | 1082 | 22 | 17 | 0 | 0 | 11 | 7 | 1 | 0 | 0 | 0 | 0 | 2 |
| L1PA10_dup2295 | 4.2838211194128 | 3.12435091597467 | 0.961271666810433 | 3.25022678171873 | 0.00115313012184445 | 0.0387765262821367 | 1083 | 28 | 16 | 0 | 1 | 4 | 5 | 1 | 1 | 0 | 0 | 0 | 1 |
| L1MB5_dup1406 | 5.14911135481794 | 2.70500454524064 | 0.792082586717907 | 3.41505367066478 | 0.000637694449605314 | 0.025609593840639 | 1084 | 10 | 7 | 0 | 0 | 4 | 16 | 0 | 1 | 7 | 7 | 6 | 6 |
| L1ME1_dup15524 | 5.0059963651169 | 3.07092333897535 | 0.854436197068162 | 3.59409321551761 | 0.000325523288497391 | 0.0158479505340356 | 1085 | 16 | 12 | 0 | 0 | 6 | 15 | 0 | 1 | 3 | 4 | 3 | 4 |
| AluY_dup22676 | 5.69469598843236 | 3.58520070719218 | 1.00877503414204 | 3.55401411201795 | 0.000379398854854167 | 0.017770952739592 5 | 1086 | 18 | 5 | 0 | 0 | 14 | 16 | 0 | 0 | 2 | 6 | 5 | 8 |
| MLT1L_dup2092 | 4.91737351289752 | 3.21199631325264 | 0.905206789595271 | 3.54835640891378 | 0.000387643294847334 | 0.0180752834481175 | 1087 | 13 | 10 | 0 | 0 | 7 | 10 | 0 | 0 | 4 | 7 | 8 | 3 |
| Penelope1_Vert_dup1 | 5.16928820839607 | 3.52662633558413 | 0.905824691708694 | 3.89327688664758 | 9.88991612858537e-05 | 0.0067661471625 4276 | 1088 | 16 | 12 | 0 | 0 | 9 | 9 | 0 | 0 | 5 | 6 | 6 | 2 |
| AluSp_dup32646 | 5.2509182535818 | 3.05741580723767 | 0.835701698731484 | 3.65850136702909 | 0.000253694387989192 | 0.013278830806088 | 1089 | 20 | 8 | 0 | 1 | 9 | 11 | 0 | 0 | 3 | 7 | 3 | 6 |
| L1MC5a_dup3897 | 5.23477881719078 | 2.91610838901619 | 0.7779528603571 | 3.74843841782088 | 0.000177939014841724 | 0.0102667014610884 | 1090 | 15 | 12 | 0 | 1 | 7 | 9 | 0 | 0 | 3 | 10 | 4 | 6 |
| L1M2_dup1432 | 3.83077592287229 | 3.23384250392642 | 0.934510600337724 | 3.46046636898258 | 0.000539240586331693 | 0.0227137795282848 | 1091 | 8 | 14 | 0 | 0 | 3 | 7 | 0 | 0 | 4 | 5 | 3 | 4 |
| L1PB1_dup9548 | 4.82164347304659 | 3.55299309753578 | 1.05442022614209 | 3.36961773820999 | 0.000752725261716552 | 0.0288279357998401 | 1092 | 15 | 20 | 0 | 0 | 3 | 4 | 0 | 0 | 4 | 5 | 6 | 4 |
| HERVIP10F-int_dup109 | 5.12145877985239 | 4.55966111964114 | 0.98271368007925 | 4.63986735106144 | 3.48632887482647e-06 | 0.000539725445965817 | 1093 | 19 | 15 | 0 | 0 | 3 | 5 | 0 | 0 | 6 | 13 | 1 | 2 |
| L1M4b_dup696 | 5.22506545744683 | 4.50539975567979 | 0.935127608271506 | 4.81795181302324 | 1.45039385890075e-06 | 0.000275179052885578 | 1094 | 13 | 17 | 0 | 0 | 6 | 11 | 0 | 0 | 6 | 9 | 1 | 2 |
| MER57C2_dup232 | 4.51827161946531 | 3.45332833247447 | 0.894212799430249 | 3.86186412750385 | 0.000112525144994498 | 0.00741679721293155 | 1095 | 18 | 14 | 0 | 1 | 4 | 7 | 1 | 0 | 2 | 8 | 3 | 0 |
| MIRb_dup53569 | 4.63188976951534 | 3.43615887178995 | 0.914624667310771 | 3.75690596875451 | 0.000172027098485387 | 0.0100418153174827 | 1096 | 15 | 11 | 0 | 1 | 4 | 9 | 0 | 0 | 5 | 8 | 5 | 0 |
| HAL1b_dup2237 | 4.00462771214103 | 3.53872567169176 | 0.926293164214981 | 3.82030852477549 | 0.000133284843808152 | 0.00834284266306204 | 1097 | 17 | 14 | 1 | 1 | 3 | 8 | 0 | 0 | 1 | 5 | 1 | 1 |
| L1P3_dup1731 | 3.17407304648781 | 4.13241064383831 | 1.06558387071248 | 3.87807169141483 | 0.000105287693591624 | 0.00709107997151219 | 1098 | 12 | 12 | 0 | 0 | 2 | 9 | 0 | 0 | 1 | 3 | 1 | 1 |
| L1ME3_dup8304 | 3.75829254790168 | 3.24216330143446 | 0.832441171715703 | 3.89476567425444 | 9.82937053730469e-05 | 0.00673053065286187 | 1099 | 6 | 7 | 0 | 2 | 4 | 10 | 0 | 0 | 7 | 7 | 1 | 2 |
| LTR10E_dup41 | 4.33120502539853 | 2.83442671676069 | 0.800813711810161 | 3.53943329760644 | 0.000400987091357821 | 0.0185124707877819 | 1100 | 7 | 12 | 0 | 2 | 10 | 7 | 0 | 0 | 6 | 4 | 1 | 5 |
| L1MA4_dup270 | 4.01888189075053 | 3.23526225161004 | 0.974794203561601 | 3.31891822888296 | 0.000903668873994914 | 0.0327431694672748 | 1101 | 9 | 9 | 0 | 0 | 6 | 13 | 0 | 0 | 4 | 2 | 3 | 5 |

| | | | | | | | | | | | | | | | | | | | |
|---|---|---|---|---|---|---|---|---|---|---|---|---|---|---|---|---|---|---|---|
| L2b_dup18162 | 3.67655564238641 | 3.04235142642957 | 0.895601893623846 | 3.39699083721161 | 0.000681312482509209 | 0.0268218399901984 | 1102 | 12 | 8 | 0 | 1 | 3 | 11 | 0 | 0 | 4 | 3 | 1 | 4 |
| L1ME4b_dup3583 | 4.36394422144792 | 3.39029531635192 | 1.00258621928898 | 3.38154988680802 | 0.000720781437339708 | 0.0279372959585004 | 1103 | 10 | 15 | 0 | 0 | 4 | 9 | 0 | 0 | 2 | 7 | 2 | 7 |
| L2b_dup18160 | 4.1538067932645 | 3.30391117223095 | 1.03464482718803 | 3.19328052043749 | 0.00140666224073653 | 0.0444619624779882 | 1104 | 12 | 12 | 0 | 0 | 5 | 11 | 0 | 0 | 1 | 4 | 3 | 6 |
| L1M6_dup3091 | 4.10207466828673 | 4.07446857491941 | 0.974855358711518 | 4.17956216633481 | 2.92070871864404e-05 | 0.00269891633695718 | 1105 | 18 | 10 | 0 | 0 | 6 | 9 | 0 | 0 | 2 | 5 | 1 | 2 |
| MER112_dup883 | 3.69735128061517 | 3.31523990756072 | 1.03540115423342 | 3.20188932956639 | 0.00136529440222693 | 0.0435711295375923 | 1106 | 16 | 10 | 0 | 0 | 5 | 4 | 0 | 0 | 2 | 4 | 2 | 5 |
| L1MEd_dup3797 | 5.12276258087488 | 3.70879821113128 | 1.15062229090336 | 3.22329772372087 | 0.0012672376493096 | 0.0413627064063952 | 1107 | 16 | 14 | 0 | 0 | 12 | 6 | 0 | 0 | 0 | 6 | 4 | 9 |
| L1MEf_dup10405 | 4.23145294017901 | 3.18739911550063 | 0.969982656679525 | 3.28603722298688 | 0.00101607581900449 | 0.0354607182207161 | 1108 | 13 | 8 | 0 | 0 | 10 | 2 | 0 | 0 | 2 | 9 | 5 | 5 |
| MIR_dup31094 | 6.20235867971752 | 3.78361731525717 | 1.08862054090278 | 3.47560713131453 | 0.000509698367067471 | 0.0218584340791056 | 1109 | 22 | 20 | 0 | 0 | 10 | 4 | 0 | 0 | 2 | 6 | 4 | 13 |
| MIRb_dup41763 | 6.74983427470396 | 3.5297302591755 | 0.938729020014922 | 3.76011626775892 | 0.000169834404911238 | 0.00996091134114112 | 1110 | 20 | 16 | 0 | 0 | 11 | 7 | 0 | 0 | 3 | 12 | 6 | 12 |
| L1MC5_dup14139 | 5.12969181123802 | 2.86236473614853 | 0.859325959520551 | 3.33094177411508 | 0.000865527044545582 | 0.0317822065033091 | 1111 | 13 | 14 | 0 | 1 | 8 | 7 | 1 | 0 | 3 | 10 | 0 | 9 |
| L1ME1_dup15529 | 7.44274439737319 | 3.93802723642974 | 0.769210623461011 | 5.11956948632724 | 3.06233922907839e-07 | 8.11438051450207e-05 | 1112 | 26 | 24 | 0 | 0 | 14 | 10 | 3 | 0 | 5 | 10 | 0 | 3 |
| MamGypLTR2b_dup47 | 6.32929905411907 | 3.41453203221652 | 0.862112016965164 | 3.96065936331159 | 7.47430913215585e-05 | 0.00549613312487247 | 1113 | 16 | 22 | 0 | 0 | 14 | 10 | 2 | 0 | 3 | 8 | 0 | 6 |
| L1MCc_dup3044 | 6.04308126270251 | 2.68976588478498 | 0.841124348877994 | 3.19782192534666 | 0.00138469758841128 | 0.0439887504630519 | 1114 | 11 | 24 | 0 | 1 | 15 | 13 | 2 | 0 | 1 | 3 | 2 | 6 |
| MLT1A0_dup11128 | 5.54513417930279 | 2.59173552846913 | 0.685995977301711 | 3.77806228348952 | 0.000158053355335144 | 0.00946117325498771 | 1115 | 8 | 17 | 0 | 1 | 14 | 9 | 3 | 0 | 4 | 7 | 3 | 4 |
| L1MEf_dup7693 | 5.03069582867552 | 2.08358305722993 | 0.623045876180082 | 3.3441888260371 | 0.000825235484172964 | 0.0307294964815368 | 1116 | 12 | 15 | 0 | 2 | 7 | 6 | 4 | 1 | 4 | 7 | 2 | 4 |
| L1ME2_dup8821 | 3.77674155406165 | 2.90797824266912 | 0.770243005940468 | 3.77540363267366 | 0.000159748775672002 | 0.009533880656145 | 1117 | 9 | 10 | 0 | 1 | 6 | 8 | 3 | 0 | 3 | 6 | 1 | 1 |
| HAL1_dup7780 | 4.37328822681696 | 2.42288214440228 | 0.725309269938565 | 3.34048142609221 | 0.000836332747284276 | 0.0310358752160487 | 1118 | 18 | 9 | 0 | 2 | 8 | 7 | 2 | 0 | 2 | 4 | 3 | 2 |
| L1PA15_dup4052 | 5.29411272723299 | 2.55729462163366 | 0.709584787114731 | 3.60393101440628 | 0.00031344028227719 | 0.0154190327489618 | 1119 | 18 | 16 | 0 | 1 | 8 | 8 | 5 | 0 | 4 | 4 | 3 | 1 |
| L1ME3E_dup1528 | 4.51999720485208 | 3.46875789407457 | 0.941890848981755 | 3.68275994806035 | 0.000230722402570415 | 0.0123563972095476 | 1120 | 12 | 19 | 0 | 0 | 6 | 2 | 3 | 0 | 6 | 6 | 2 | 0 |
| LTR84b_dup649 | 4.19664047760466 | 4.58036438384882 | 0.985461661927766 | 4.64793767307871 | 3.352701768952e-06 | 0.000524843404680956 | 1121 | 10 | 17 | 0 | 0 | 8 | 6 | 1 | 0 | 5 | 4 | 1 | 0 |
| ERV3-16A3_LTR_dup39 | 4.02944739154495 | 2.74504352028056 | 0.812330124648664 | 3.37922162060382 | 0.00072691383759294 | 0.0280812305298219 | 1122 | 11 | 11 | 0 | 0 | 5 | 4 | 2 | 1 | 4 | 8 | 4 | 0 |
| L1PA15_dup4053 | 5.27501083407707 | 3.67083860214702 | 0.849025575242637 | 4.32359013578355 | 1.53510376491813e-05 | 0.00168119844781165 | 1123 | 19 | 21 | 1 | 0 | 6 | 5 | 2 | 1 | 6 | 4 | 1 | 0 |
| L2a_dup66760 | 5.08863715296263 | 2.7755314192054 | 0.882156315037854 | 3.14630340665453 | 0.00165348379694051 | 0.0497029240600507 | 1124 | 16 | 27 | 0 | 0 | 3 | 2 | 3 | 2 | 4 | 4 | 1 | 2 |
| HAL1_dup11964 | 4.95309634796727 | 2.69062276738687 | 0.793763989791981 | 3.38970122352362 | 0.000699688365882338 | 0.0273334506498504 | 1125 | 15 | 20 | 0 | 0 | 7 | 6 | 2 | 2 | 5 | 1 | 2 | 2 |
| L1MA6_dup107 | 6.23438620763636 | 2.96257813843148 | 0.733379663681665 | 4.03962406532919 | 5.35369430943131e-05 | 0.00430513736448781 | 1126 | 21 | 24 | 0 | 0 | 6 | 6 | 3 | 2 | 7 | 5 | 2 | 2 |
| LTR8B_dup412 | 4.98902238650444 | 2.82709480158664 | 0.84356624101242 | 3.35136076355267 | 0.000804154664472189 | 0.0301568771934593 | 1127 | 23 | 18 | 0 | 1 | 2 | 3 | 2 | 1 | 3 | 7 | 3 | 1 |
| L1PA3_dup275 | 5.61548508615954 | 2.78668430064507 | 0.742742118535738 | 3.75188673309495 | 0.000175508759056988 | 0.0101634478419504 | 1128 | 24 | 17 | 0 | 1 | 7 | 3 | 2 | 2 | 5 | 6 | 3 | 1 |
| Tigger8_dup306 | 5.66973648004144 | 3.0386212240333 | 0.742545052065289 | 4.0921708596425 | 4.27353577086003e-05 | 0.00362234568672866 | 1129 | 21 | 19 | 0 | 1 | 4 | 5 | 1 | 2 | 6 | 8 | 2 | 2 |
| L1MC5a_dup3389 | 4.08022301235046 | 2.48813002177685 | 0.761936701923245 | 3.26553375824583 | 0.00109257981368881 | 0.0373111042710915 | 1130 | 8 | 13 | 0 | 0 | 5 | 4 | 1 | 3 | 4 | 8 | 1 | 3 |
| MER5A_dup11613 | 5.37036834150824 | 2.66580683743947 | 0.744586226369028 | 3.58025268670261 | 0.000343262098889038 | 0.016470627824612 | 1131 | 18 | 15 | 1 | 0 | 5 | 2 | 1 | 2 | 9 | 7 | 2 | 4 |
| Tigger1_dup8799 | 5.39216613144385 | 2.69121700823922 | 0.729258857854533 | 3.69034531326329 | 0.000223949838729711 | 0.0120670218130389 | 1132 | 23 | 11 | 1 | 2 | 9 | 6 | 0 | 2 | 5 | 5 | 4 | 0 |
| Charlie2b_dup97 | 6.00822121689923 | 2.84251248346473 | 0.751031564469005 | 3.78481094263254 | 0.000153825471576882 | 0.00927106273045412 | 1133 | 24 | 16 | 0 | 1 | 6 | 6 | 0 | 4 | 5 | 9 | 3 | 1 |
| HAL1_dup11966 | 5.22226163459883 | 3.63528156717605 | 0.816746580995025 | 4.45092964178346 | 8.54993530554162e-06 | 0.00108213984373 | 1134 | 19 | 11 | 0 | 1 | 6 | 7 | 0 | 2 | 5 | 12 | 2 | 0 |
| MLT1B_dup9103 | 4.46610516359328 | 4.18597591619418 | 0.952616030255672 | 4.39419008629396 | 1.11186496510531e-05 | 0.00132157043027922 | 1135 | 16 | 8 | 0 | 0 | 10 | 7 | 0 | 1 | 7 | 4 | 2 | 0 |
| L1M1_dup4411 | 3.98699970762254 | 3.01515287981706 | 0.888130597466325 | 3.39494314059074 | 0.000686428546714326 | 0.0269585101851577 | 1136 | 20 | 9 | 0 | 0 | 8 | 5 | 0 | 2 | 2 | 2 | 2 | 1 |
| L1MEf_dup5982 | 4.10357752751385 | 2.99413571239069 | 0.815291917928831 | 3.67247073906607 | 0.000240216663871708 | 0.0127472333411279 | 1137 | 12 | 9 | 0 | 0 | 11 | 4 | 0 | 2 | 4 | 5 | 1 | 3 |
| L1PA7_dup1617 | 5.64383092774559 | 3.18992323280522 | 0.986616349358677 | 3.23319518765196 | 0.00122413892717105 | 0.0403769006327479 | 1138 | 22 | 23 | 0 | 1 | 7 | 5 | 0 | 0 | 2 | 3 | 6 | 4 |
| L1PREC2_dup2102 | 5.70815672380635 | 3.56069921436614 | 1.02126732724698 | 3.48654962257988 | 0.000489294599831766 | 0.021222328263137 | 1139 | 28 | 22 | 0 | 0 | 3 | 2 | 1 | 0 | 5 | 5 | 5 | 2 |
| L2c_dup48718 | 4.3526514882622 | 3.60732221343549 | 1.05190996376982 | 3.42930701075176 | 0.000605124609071605 | 0.0247268959871324 | 1140 | 17 | 20 | 0 | 1 | 1 | 2 | 0 | 0 | 4 | 6 | 3 | 1 |
| AluJb_dup79853 | 5.81737994906868 | 3.75027254724542 | 0.93632488986851 | 4.00531117758931 | 6.19358923388028e-05 | 0.00478642346165923 | 1141 | 27 | 12 | 0 | 0 | 6 | 4 | 0 | 0 | 10 | 5 | 4 | 5 |
| L2a_dup30881 | 6.08293828745711 | 4.0408963110293 | 0.957552298988041 | 4.22002674454418 | 2.44273338008092e-05 | 0.00237046752388554 | 1142 | 21 | 17 | 0 | 0 | 9 | 3 | 0 | 0 | 14 | 3 | 4 | 3 |
| L1MC5a_dup2908 | 4.41495568707554 | 3.93003927939648 | 0.935020167865493 | 4.2031599044202 | 2.63214507201875e-05 | 0.0025065245920303 | 1143 | 13 | 10 | 0 | 0 | 14 | 3 | 0 | 0 | 5 | 6 | 2 | 2 |
| AluSx1_dup18484 | 5.39391471390469 | 3.93056146187935 | 0.915819849068292 | 4.29185004657641 | 1.77190597963842e-05 | 0.00186887384854186 | 1144 | 19 | 18 | 0 | 0 | 10 | 7 | 0 | 0 | 6 | 3 | 2 | 3 |
| L1MC3_dup2107 | 4.77854397820348 | 3.82093748000064 | 0.938507432892237 | 4.07129165533138 | 4.67531677581992e-05 | 0.0038816222105347 | 1145 | 16 | 18 | 0 | 0 | 6 | 5 | 0 | 0 | 5 | 5 | 3 | 2 |
| Charlie4a_dup1630 | 3.74155923378763 | 3.63782736464855 | 0.971686712688048 | 3.74382742621329 | 0.000181238156961326 | 0.0103940454317473 | 1146 | 13 | 10 | 0 | 0 | 7 | 7 | 0 | 0 | 5 | 1 | 2 | 2 |
| LTR85b_dup206 | 4.15147858581311 | 3.66417603304987 | 1.0545087853772 | 3.47477051292578 | 0.000511290553736254 | 0.0219087956238658 | 1147 | 16 | 14 | 0 | 0 | 4 | 9 | 0 | 0 | 3 | 2 | 4 | 1 |
| LTR12D_dup445 | 4.59759411424975 | 4.1247764731909 | 1.06744096404178 | 3.86417292584759 | 0.000111466237044041 | 0.00735930778015576 | 1148 | 23 | 15 | 0 | 0 | 4 | 2 | 0 | 0 | 6 | 4 | 3 | 1 |
| AluJr_dup13621 | 3.8177138407665 | 3.5708773808739 | 0.957979937415046 | 3.72750747840224 | 0.000193382824474126 | 0.0108525164038492 | 1149 | 17 | 8 | 0 | 0 | 4 | 4 | 0 | 0 | 6 | 4 | 2 | 3 |
| Charlie2b_dup547 | 3.62821453203481 | 3.77467673159014 | 1.06852102701061 | 3.53261811061456 | 0.000411466380028429 | 0.0188681401546048 | 1150 | 17 | 12 | 0 | 0 | 3 | 3 | 0 | 0 | 5 | 2 | 1 | 3 |
| L1MD_dup4331 | 3.7290833271654 | 3.55461204296001 | 0.947979839165565 | 3.74967050574501 | 0.000177067070510142 | 0.0102287936779816 | 1151 | 9 | 11 | 0 | 0 | 9 | 2 | 2 | 0 | 6 | 5 | 0 | 2 |
| LTR101_Mam_dup748 | 3.46848618405863 | 3.48668300010314 | 1.02234068027421 | 3.41049032614837 | 0.000648461844850812 | 0.0258927309568128 | 1152 | 19 | 8 | 0 | 0 | 4 | 3 | 1 | 0 | 3 | 4 | 0 | 3 |
| L2c_dup70170 | 4.35030708728309 | 4.71079964077675 | 1.07090186502721 | 4.39890880258862 | 1.08796530110625e-05 | 0.00130023601977413 | 1153 | 21 | 10 | 0 | 0 | 5 | 4 | 0 | 0 | 9 | 3 | 0 | 2 |
| L2a_dup86241 | 2.55392508743928 | 4.36763079134878 | 1.08141522683183 | 4.03881014709256 | 5.37230249847679e-05 | 0.00431560351923593 | 1154 | 9 | 9 | 0 | 0 | 6 | 2 | 0 | 0 | 3 | 2 | 0 | 1 |
| MamRTE1_dup6565 | 3.17029978184614 | 4.19328049465488 | 1.11130780533129 | 3.77328448026588 | 0.000161112399966562 | 0.00958125193957281 | 1155 | 13 | 9 | 0 | 0 | 9 | 2 | 0 | 0 | 4 | 1 | 0 | 2 |
| L1MB4_dup2429 | 2.77450255289451 | 2.76246425920804 | 0.871288240096161 | 3.17055152598313 | 0.00152149845984657 | 0.0469443040101988 | 1156 | 4 | 5 | 1 | 1 | 4 | 5 | 0 | 0 | 5 | 6 | 0 | 3 |
| L1M5_dup49482 | 2.81764009670447 | 2.71485223604719 | 0.86000072928155 | 3.15680465793867 | 0.00159508163089301 | 0.048448212119539 | 1157 | 6 | 5 | 2 | 1 | 7 | 4 | 0 | 0 | 4 | 4 | 0 | 2 |
| MSTB_dup5951 | 2.94577545296887 | 2.78956795646725 | 0.858765720799689 | 3.24834572328945 | 0.00116078137444323 | 0.0389398022710481 | 1158 | 7 | 8 | 2 | 1 | 5 | 4 | 0 | 0 | 3 | 5 | 0 | 2 |
| LTR80B_dup461 | 3.54437096038133 | 3.13168245695816 | 0.856814404278741 | 3.65503012241535 | 0.000257151783247013 | 0.013412590577925 | 1159 | 5 | 9 | 1 | 0 | 10 | 5 | 1 | 0 | 4 | 6 | 0 | 3 |
| THE1D-int_dup1264 | 3.07570350868812 | 3.24783008894069 | 0.982580079265723 | 3.30541007036065 | 0.000948375328614239 | 0.033880845984224 | 1160 | 8 | 10 | 1 | 0 | 8 | 3 | 0 | 0 | 2 | 4 | 0 | 3 |
| ERV3-16A3_I-int_dup10 | 4.37775569685318 | 2.78683686022807 | 0.75837657539955 | 3.67474016290631 | 0.00023809159526486 | 0.0126659373906513 | 1161 | 8 | 11 | 1 | 0 | 7 | 7 | 0 | 0 | 7 | 5 | 3 | 5 |
| Tigger1_dup5557 | 4.04532839711527 | 3.31941829591427 | 0.812888805315675 | 4.08348383469891 | 4.43655168469572e-05 | 0.00373552628155205 | 1162 | 7 | 9 | 2 | 0 | 6 | 8 | 0 | 0 | 8 | 6 | 1 | 2 |
| MER1B_dup1759 | 3.43059523923786 | 2.62723342945224 | 0.81502762491085 | 3.22349003782493 | 0.00126638703408655 | 0.0413519595461451 | 1163 | 10 | 9 | 1 | 0 | 4 | 2 | 1 | 0 | 4 | 7 | 2 | 3 |
| L1PB1_dup10963 | 3.24692134370715 | 2.65900198372337 | 0.785093757358446 | 3.38685916019756 | 0.000706976710437898 | 0.0275479600744166 | 1164 | 4 | 9 | 1 | 1 | 6 | 6 | 1 | 0 | 5 | 4 | 1 | 2 |
| AluSg_dup23158 | 3.83904444159079 | 2.8990856722164 | 0.827307920808112 | 3.50424019799614 | 0.000457912130234947 | 0.020323182520728 | 1165 | 13 | 13 | 1 | 0 | 7 | 4 | 1 | 0 | 3 | 3 | 1 | 3 |
| THE1B_dup11445 | 3.46217508491612 | 3.03154228101763 | 0.848686856654011 | 3.57203868217027 | 0.000354213075674235 | 0.0168668370030781 | 1166 | 8 | 7 | 1 | 0 | 11 | 5 | 1 | 0 | 5 | 2 | 1 | 2 |
| FLAM_A_dup9701 | 3.2716464514024 | 2.99505994542347 | 0.910781614857001 | 3.28845015815753 | 0.00100740620140039 | 0.0352460338183233 | 1167 | 11 | 8 | 1 | 2 | 3 | 4 | 1 | 0 | 1 | 10 | 1 | 0 |
| L3_dup32208 | 3.91544740153498 | 2.95752306865992 | 0.834652287580274 | 3.54341935278704 | 0.00039497409357016 | 0.0183036480359665 | 1168 | 16 | 12 | 3 | 1 | 4 | 4 | 1 | 0 | 3 | 5 | 1 | 0 |
| Tigger1_dup10646 | 3.26242811063186 | 2.99156313599479 | 0.94059041494711 | 3.18051628897687 | 0.00147012874053644 | 0.045858864518289 | 1169 | 17 | 7 | 1 | 2 | 2 | 2 | 1 | 0 | 3 | 6 | 1 | 0 |
| L1M4_dup14599 | 4.5966667922613 | 2.53520554759088 | 0.703891923021084 | 3.60169717065348 | 0.000316146509462169 | 0.0155072763018935 | 1170 | 11 | 5 | 2 | 3 | 11 | 4 | 1 | 0 | 6 | 11 | 3 | 0 |
| MER58A_dup9101 | 4.01671594080208 | 3.63304689097646 | 0.943060244797429 | 3.85240170075968 | 0.000116964936499753 | 0.0076062691392102 | 1171 | 9 | 11 | 1 | 2 | 8 | 2 | 0 | 0 | 3 | 13 | 1 | 0 |
| AluSz_dup37559 | 4.56004621350637 | 2.80429698061553 | 0.839251194621549 | 3.34142745174179 | 0.000833487953938521 | 0.0309592764108422 | 1172 | 19 | 13 | 3 | 4 | 4 | 3 | 0 | 0 | 5 | 6 | 1 | 0 |
| MLT2D_dup2092 | 3.75330833837574 | 3.14955882986683 | 0.921233072866747 | 3.41885123605675 | 0.000628860974604306 | 0.0253661558676379 | 1173 | 8 | 16 | 3 | 1 | 5 | 4 | 0 | 0 | 6 | 2 | 1 | 0 |
| L2b_dup40046 | 4.46407937625548 | 3.80917986289868 | 0.933557137986622 | 4.08028572425019 | 4.49803766959094e-05 | 0.00378060781112146 | 1174 | 19 | 12 | 2 | 1 | 4 | 5 | 0 | 0 | 3 | 10 | 0 | 1 |
| L2b_dup44485 | 4.20307490721664 | 3.05516667558987 | 0.90838918291583 | 3.36327945449892 | 0.000770223516871603 | 0.0292532489937 | 1175 | 11 | 14 | 3 | 1 | 2 | 2 | 0 | 0 | 9 | 7 | 0 | 2 |
| L3_dup22439 | 3.36424989614046 | 3.76111728885451 | 1.03601762096729 | 3.63036034593978 | 0.000283025809253139 | 0.0143378452647535 | 1176 | 9 | 10 | 1 | 1 | 1 | 2 | 0 | 0 | 6 | 10 | 0 | 1 |
| LOR1a_dup749 | 4.17639928830152 | 2.60596951734593 | 0.780520791487044 | 3.33875733454973 | 0.000841540438698245 | 0.0311772319921954 | 1177 | 12 | 9 | 2 | 2 | 1 | 8 | 0 | 0 | 6 | 8 | 3 | 1 |
| AluJr4_dup19948 | 4.65173718040559 | 2.57742280592222 | 0.66532980338545 | 3.87390252594626 | 0.000107106266246146 | 0.00716488828456719 | 1178 | 8 | 6 | 3 | 1 | 6 | 5 | 0 | 0 | 10 | 12 | 2 | 3 |
| L1MC5a_dup15542 | 4.19060067103949 | 2.82231851430042 | 0.766167604482728 | 3.68368291453133 | 0.000229888182514063 | 0.0123172649125177 | 1179 | 10 | 5 | 3 | 1 | 5 | 12 | 0 | 0 | 3 | 11 | 1 | 2 |
| AluSc8_dup14004 | 4.42968877498034 | 2.87299494787482 | 0.830086285235669 | 3.46107988889269 | 0.000538013127291736 | 0.0226838114398232 | 1180 | 15 | 16 | 1 | 2 | 8 | 2 | 0 | 0 | 4 | 4 | 3 | 1 |

| | | | | | | | | | | | | | | | | | | | |
|---|---|---|---|---|---|---|---|---|---|---|---|---|---|---|---|---|---|---|---|
| L1MA3_dup6404 | 4.16813419161602 | 2.58242916329581 | 0.709848965881515 | 3.63799806355829 | 0.00027476549056737 | 0.0140493119277348 | 1181 | 13 | 10 | 2 | 2 | 7 | 6 | 0 | 0 | 4 | 5 | 1 | 3 |
| L1ME4b_dup7112 | 4.83012320041827 | 2.61015205950739 | 0.691410588292908 | 3.77511149482372 | 0.000159936111881977 | 0.00953788423004232 | 1182 | 14 | 10 | 3 | 2 | 13 | 5 | 0 | 0 | 5 | 5 | 1 | 3 |
| L1MEd_dup15399 | 4.70726043241069 | 2.29080879461965 | 0.71344296032246 | 3.21092073511281 | 0.00132310424886879 | 0.04260953189588 | 1183 | 11 | 17 | 2 | 3 | 6 | 3 | 1 | 0 | 3 | 9 | 1 | 4 |
| Arthur1_dup866 | 4.69425062837455 | 2.40649534732303 | 0.662970082302618 | 3.62987020313922 | 0.000283563773645826 | 0.014358983971894 | 1184 | 12 | 13 | 4 | 2 | 6 | 5 | 1 | 0 | 5 | 8 | 1 | 2 |
| L2a_dup59538 | 4.32686570416327 | 2.64562409284835 | 0.724271929227921 | 3.65280495637682 | 0.000259391257923627 | 0.0134909534989935 | 1185 | 11 | 15 | 2 | 2 | 5 | 4 | 1 | 0 | 6 | 5 | 1 | 2 |
| L1PA15_dup2749 | 4.28938311433524 | 2.46049686530556 | 0.711537310925711 | 3.45800118633898 | 0.000544198987969754 | 0.02287808699456 | 1186 | 7 | 13 | 2 | 3 | 4 | 9 | 0 | 0 | 4 | 8 | 1 | 3 |
| L1PB4_dup6664 | 3.79093459958923 | 2.68780196728749 | 0.774263844412323 | 3.47142900535097 | 0.000517696202236974 | 0.0220878143334192 | 1187 | 13 | 10 | 1 | 2 | 4 | 4 | 0 | 0 | 4 | 6 | 2 | 2 |
| MSTB_dup7895 | 3.52677214519456 | 2.7982295803468 | 0.798922596325944 | 3.50250398876587 | 0.000460906882354251 | 0.0204076942740351 | 1188 | 6 | 10 | 2 | 1 | 5 | 3 | 0 | 0 | 6 | 7 | 1 | 2 |
| AluSx1_dup51193 | 1.61151691631773 | 4.23357804307734 | 1.23524445181794 | 3.42732002305024 | 0.000609570226148632 | 0.0248534607661347 | 1189 | 6 | 3 | 0 | 0 | 1 | 6 | 0 | 0 | 3 | 1 | 0 | 0 |
| L1MEi_dup4708 | 2.10495564353411 | 4.52378126595838 | 1.33656877084102 | 3.38462289756467 | 0.000712761031281462 | 0.0276850532728205 | 1190 | 13 | 4 | 0 | 0 | 1 | 6 | 0 | 0 | 3 | 0 | 0 | 0 |
| L1MEf_dup3588 | 2.53406817238545 | 4.08591973368821 | 1.2334518090956 | 3.31258968008171 | 0.000924364852650683 | 0.0333058452868393 | 1191 | 13 | 13 | 0 | 0 | 2 | 3 | 0 | 0 | 1 | 0 | 0 | 1 |
| L1M3e_dup41 | 2.84776773863486 | 4.33013610415077 | 1.17013115827045 | 3.70055619282121 | 0.000215127433373698 | 0.0117136641330751 | 1192 | 13 | 14 | 0 | 0 | 1 | 6 | 0 | 0 | 1 | 1 | 0 | 1 |
| L4_A_Mam_dup4771 | 1.93144381061384 | 3.86094299174908 | 1.16135822255243 | 3.32450652759276 | 0.000885751599267702 | 0.0323352332064057 | 1193 | 9 | 8 | 0 | 0 | 2 | 3 | 0 | 0 | 1 | 1 | 0 | 1 |
| MLT1A0_dup1168 | 2.21068723326693 | 4.47929341743733 | 1.17427845619907 | 3.81450702241103 | 0.000136455056934049 | 0.00849527341110128 | 1194 | 8 | 9 | 0 | 0 | 4 | 5 | 0 | 0 | 2 | 0 | 0 | 0 |
| L1PA15_dup225 | 2.42996138157613 | 4.02873256238095 | 1.18376424977835 | 3.40332339241981 | 0.000665714350081011 | 0.0263786955444423 | 1195 | 16 | 6 | 0 | 0 | 3 | 5 | 0 | 0 | 1 | 0 | 1 | 0 |
| LTR24C_dup321 | 1.87128136411326 | 4.28817998245154 | 1.29189140094771 | 3.31930375827705 | 0.000902422077752772 | 0.0327129377990495 | 1196 | 11 | 6 | 0 | 0 | 3 | 2 | 0 | 0 | 2 | 0 | 0 | 0 |
| L1PB2_dup143 | 2.21125874771829 | 4.33213560349258 | 1.23049560417358 | 3.52064289283025 | 0.000430501967391402 | 0.0195079453690563 | 1197 | 14 | 7 | 0 | 0 | 3 | 4 | 0 | 0 | 1 | 0 | 0 | 0 |
| LTR86A1_dup470 | 2.34143869395956 | 4.39467997716764 | 1.32914312019144 | 3.30640087617853 | 0.000945027800745154 | 0.0337957598933435 | 1198 | 18 | 8 | 0 | 0 | 2 | 2 | 0 | 0 | 1 | 0 | 0 | 0 |
| MLT1H2_dup2287 | 2.09908682143652 | 4.09492791642268 | 1.07721625479675 | 3.80139818554384 | 0.000143881861124936 | 0.00883512601993136 | 1199 | 3 | 6 | 0 | 0 | 4 | 7 | 0 | 0 | 2 | 3 | 1 | 0 |
| L3_dup32213 | 2.82920252131927 | 3.62016532309454 | 1.02725986389773 | 3.52409886760159 | 0.000424925665321561 | 0.0193214909918984 | 1200 | 9 | 6 | 0 | 0 | 4 | 5 | 0 | 0 | 4 | 4 | 3 | 0 |
| MER57B2_dup154 | 3.27970982775764 | 4.25869301660696 | 1.02728192860032 | 4.145593237885 | 3.38935046058756e-05 | 0.00302847309310915 | 1201 | 6 | 12 | 0 | 0 | 6 | 6 | 0 | 0 | 5 | 3 | 2 | 0 |
| AluJo_dup49169 | 2.25864858434364 | 3.68132768379893 | 1.07327305840322 | 3.43000101882356 | 0.00060357898288958 | 0.0246880493255903 | 1202 | 8 | 6 | 0 | 1 | 2 | 4 | 0 | 0 | 4 | 2 | 1 | 0 |
| L2a_dup86246 | 2.46717601440264 | 3.81017968448548 | 1.03113339614961 | 3.69513750472365 | 0.000219767819182712 | 0.0119009725769799 | 1203 | 9 | 6 | 0 | 1 | 4 | 4 | 0 | 0 | 3 | 3 | 1 | 0 |
| L1M5_dup29310 | 2.33315571132817 | 3.76189793069595 | 1.05933198749501 | 3.55119827882443 | 0.000383481399889019 | 0.0179279765023721 | 1204 | 6 | 8 | 0 | 0 | 2 | 5 | 0 | 0 | 3 | 3 | 1 | 1 |
| L1MEh_dup501 | 2.50489130243685 | 4.36041511448779 | 1.09523141120996 | 3.98127287973835 | 6.85471998198321e-05 | 0.00516156076212967 | 1205 | 9 | 9 | 0 | 0 | 2 | 4 | 0 | 0 | 4 | 2 | 1 | 0 |
| MamRep564_dup119 | 1.78168671007765 | 3.85398173151562 | 1.21026694026981 | 3.18440635142561 | 0.00145051236182803 | 0.0454255875819339 | 1206 | 7 | 7 | 0 | 0 | 2 | 1 | 0 | 0 | 3 | 1 | 1 | 0 |
| L1PA11_dup2072 | 2.10935860875376 | 4.1032763518889 | 1.10299071295926 | 3.72013680956573 | 0.000199114889332136 | 0.0110814798446162 | 1207 | 3 | 7 | 0 | 0 | 4 | 3 | 0 | 0 | 5 | 2 | 1 | 0 |
| L2c_dup30509 | 1.94466064151993 | 3.96764476867888 | 1.10769528626758 | 3.58189189560244 | 0.000341114952670528 | 0.0163977030702758 | 1208 | 6 | 4 | 0 | 0 | 6 | 3 | 0 | 0 | 3 | 1 | 1 | 0 |
| AluJr_dup38524 | 2.40285615176666 | 4.29586043134621 | 1.12656938685815 | 3.81322311919623 | 0.000137166177203668 | 0.00852745751931099 | 1209 | 7 | 8 | 0 | 0 | 3 | 3 | 1 | 0 | 6 | 1 | 0 | 0 |
| ERV3-16A3_LTR_dup18 | 1.93664316811382 | 4.51273966544997 | 1.2220935957677 | 3.69263015621576 | 0.000221946684371275 | 0.0119916725157601 | 1210 | 5 | 8 | 0 | 0 | 2 | 1 | 0 | 0 | 5 | 2 | 0 | 0 |
| AluSq2_dup1252 | 2.12525403544498 | 4.65631293719295 | 1.10418470372875 | 4.21696924569676 | 2.47607729180344e-05 | 0.00239403773781171 | 1211 | 8 | 4 | 0 | 0 | 4 | 2 | 0 | 0 | 4 | 4 | 0 | 0 |
| MamGypLTR1a_dup55 | 2.26359839003629 | 4.6995066986216 | 1.11927245521941 | 4.19871558234707 | 2.68433221671838e-05 | 0.00253811792759312 | 1212 | 7 | 9 | 0 | 0 | 5 | 2 | 0 | 0 | 3 | 2 | 0 | 0 |
| MIRb_dup34508 | 3.13969384962982 | 2.7958002518351 | 0.847010112565482 | 3.30078733460099 | 0.000964139329524352 | 0.0342816940978783 | 1213 | 7 | 6 | 1 | 1 | 8 | 5 | 0 | 2 | 5 | 2 | 1 | 0 |
| MIRc_dup68708 | 3.0338097813069 | 3.07419608592159 | 0.892909605988839 | 3.44289731603584 | 0.000575517788988804 | 0.0238284593385645 | 1214 | 14 | 7 | 1 | 2 | 6 | 5 | 0 | 1 | 2 | 1 | 0 | 0 |
| MamRTE1_dup6750 | 2.91681602097005 | 3.42586195724642 | 0.974892048051145 | 3.51409365179958 | 0.000441257213419214 | 0.0198348119869389 | 1215 | 12 | 11 | 1 | 1 | 4 | 3 | 0 | 1 | 2 | 2 | 0 | 0 |
| MER4B-int_dup599 | 2.74859390544421 | 4.43717714096556 | 1.03611247675808 | 4.28252457189704 | 1.8478471862648e-05 | 0.00192930165240441 | 1216 | 6 | 4 | 0 | 0 | 6 | 11 | 0 | 1 | 3 | 3 | 0 | 0 |
| MIR3_dup41766 | 2.96710409124319 | 3.57533183268581 | 0.97536915422348 | 3.66561913220665 | 0.000246740914724657 | 0.0129748096563698 | 1217 | 7 | 7 | 0 | 1 | 3 | 9 | 0 | 1 | 3 | 5 | 0 | 1 |
| L1PA4_dup11237 | 2.93348253212261 | 3.40026658502754 | 0.976240465556741 | 3.48302155564551 | 0.000495788442947913 | 0.0214531509084714 | 1218 | 6 | 12 | 0 | 0 | 5 | 8 | 0 | 1 | 1 | 2 | 1 | 1 |
| L1PA16_dup10555 | 2.81098110556578 | 3.40521905092486 | 1.05760608417696 | 3.2197423046926 | 0.00128305877619285 | 0.0417177726826414 | 1219 | 13 | 9 | 0 | 0 | 2 | 5 | 0 | 1 | 2 | 2 | 0 | 2 |
| Tigger15a_dup3450 | 2.79573373559542 | 3.93865532068284 | 1.04501918211137 | 3.76897897005598 | 0.000163916696004752 | 0.00970231894217579 | 1220 | 11 | 9 | 0 | 0 | 5 | 3 | 0 | 1 | 3 | 2 | 0 | 1 |
| L1MA3_dup8115 | 2.69308274578944 | 4.41545407818326 | 1.10431392031648 | 3.99836857704181 | 6.37805787599591e-05 | 0.00489287380606462 | 1221 | 11 | 8 | 0 | 0 | 3 | 4 | 0 | 1 | 5 | 1 | 0 | 0 |
| MIR3_dup1829 | 2.35813707589834 | 4.25009232045516 | 1.07179754613357 | 3.96538724667456 | 7.32768849359417e-05 | 0.00541320759483626 | 1222 | 7 | 4 | 0 | 0 | 4 | 4 | 0 | 1 | 6 | 2 | 0 | 0 |
| LTR82B_dup852 | 1.79032704967404 | 3.85039641799395 | 1.16453300864815 | 3.30638667122341 | 0.000945075716046109 | 0.0337957598933435 | 1223 | 7 | 4 | 0 | 0 | 2 | 1 | 0 | 1 | 2 | 5 | 0 | 0 |
| THE1D-int_dup1263 | 2.93045454004274 | 4.07007604764603 | 1.02056368226285 | 3.98806671095884 | 6.66139336736653e-05 | 0.00506568202681814 | 1224 | 10 | 7 | 1 | 0 | 3 | 4 | 0 | 1 | 4 | 6 | 0 | 0 |
| HERVL40-int_dup701 | 2.510630896188 | 4.34594900593977 | 1.07090705436687 | 4.05819439531954 | 4.94535949621271e-05 | 0.00404235146233539 | 1225 | 3 | 4 | 0 | 0 | 10 | 4 | 0 | 1 | 4 | 4 | 0 | 0 |
| L2a_dup86263 | 2.50259606860043 | 3.77684018639875 | 1.06210673870022 | 3.55598928881738 | 0.000376559419577315 | 0.0176797173738642 | 1226 | 3 | 6 | 1 | 0 | 9 | 4 | 0 | 1 | 4 | 2 | 0 | 0 |
| MIR_dup84486 | 2.65527456121945 | 3.39123318483464 | 0.98409071256027 | 3.44605750420284 | 0.0005688293018884 | 0.0236709788504962 | 1227 | 9 | 4 | 0 | 0 | 6 | 4 | 1 | 1 | 3 | 4 | 1 | 0 |
| MLT1A1_dup3465 | 2.7855442856848 | 3.83067864915779 | 1.01011357023548 | 3.79232470687902 | 0.000149243594172595 | 0.00906339296725752 | 1228 | 6 | 9 | 0 | 0 | 9 | 5 | 1 | 1 | 1 | 3 | 0 | 0 |
| L2a_dup74861 | 3.8505940595803 | 3.22551391030469 | 0.971314478157489 | 3.32077198769161 | 0.000897688429137949 | 0.0325860215041432 | 1229 | 10 | 10 | 1 | 2 | 3 | 13 | 0 | 0 | 0 | 9 | 0 | 2 |
| Tigger3b_dup5748 | 3.47271296434262 | 3.30389931881714 | 0.978849453121473 | 3.37528851682071 | 0.000737383414507531 | 0.028386577244525 | 1230 | 12 | 10 | 3 | 0 | 6 | 8 | 0 | 0 | 0 | 5 | 0 | 1 |
| MLT1H_dup6702 | 1.87738451221187 | 4.12595047304765 | 1.19665584505174 | 3.44790065590602 | 0.000564961795071993 | 0.0235306142639945 | 1231 | 4 | 6 | 0 | 0 | 8 | 5 | 0 | 0 | 0 | 1 | 0 | 0 |
| MER117_dup1967 | 2.48934750143303 | 4.70414285581314 | 1.06075821531251 | 4.43469849010526 | 9.22012367144457e-06 | 0.00114878335870386 | 1232 | 10 | 7 | 0 | 0 | 6 | 6 | 0 | 0 | 1 | 2 | 0 | 0 |
| AluSx3_dup21146 | 3.00892471216222 | 5.04176265092214 | 1.04876377390087 | 4.80733867472303 | 1.5295275557442e-06 | 0.000287476274707037 | 1233 | 6 | 12 | 0 | 0 | 7 | 8 | 0 | 0 | 1 | 4 | 0 | 0 |
| Tigger2_dup1593 | 2.38868016959529 | 4.71527608281856 | 1.08656015682724 | 4.33963646944978 | 1.42718614452672e-05 | 0.00159060062925319 | 1234 | 4 | 7 | 0 | 0 | 9 | 6 | 0 | 0 | 1 | 3 | 0 | 0 |
| MER21C_dup2029 | 2.17864083068625 | 3.57465114120679 | 1.1332802055054 | 3.15425181154791 | 0.00160910178827147 | 0.0487185817676604 | 1235 | 6 | 9 | 0 | 1 | 3 | 4 | 0 | 0 | 0 | 4 | 1 | 0 |
| THE1B_dup11443 | 2.70919339126865 | 4.90832343259669 | 1.21971271692726 | 4.02416353004986 | 5.71781774463979e-05 | 0.00450507238763595 | 1236 | 12 | 10 | 0 | 0 | 6 | 2 | 0 | 0 | 0 | 5 | 0 | 0 |
| L3_dup19763 | 2.21542301925847 | 3.66754439025402 | 1.10891960608549 | 3.30731314526987 | 0.00094195529832532 | 0.0337196472717491 | 1237 | 4 | 7 | 0 | 2 | 4 | 4 | 0 | 0 | 1 | 6 | 0 | 0 |
| MIR3_dup18247 | 2.31041636607356 | 4.25199596444936 | 1.12936936488301 | 3.76492943465827 | 0.000166596123405726 | 0.00981939092888254 | 1238 | 6 | 5 | 0 | 1 | 4 | 2 | 0 | 0 | 1 | 10 | 0 | 0 |
| AluJo_dup34587 | 2.36481341978769 | 4.80253023705947 | 1.24649019565597 | 3.85284236795148 | 0.000116754558546182 | 0.0075967405490374 | 1239 | 9 | 11 | 0 | 0 | 1 | 2 | 0 | 0 | 1 | 6 | 0 | 0 |
| L1MA4_dup8955 | 2.08769323753549 | 4.06488431437899 | 1.24426086009601 | 3.26690683982884 | 0.00108729453838524 | 0.0371987614997288 | 1240 | 11 | 8 | 0 | 1 | 1 | 1 | 0 | 0 | 1 | 4 | 0 | 0 |
| L1M2_dup5013 | 3.27042888832999 | 5.08884202881816 | 1.11802797799939 | 4.55162315161755 | 5.32336230449232e-06 | 0.000755320671976583 | 1241 | 6 | 3 | 0 | 0 | 20 | 8 | 0 | 0 | 2 | 2 | 0 | 0 |
| L1M2_dup5014 | 3.7918131079747 | 5.45516340598223 | 0.989812450179049 | 5.511310152742 | 3.56172463344569e-08 | 1.45368461343031e-05 | 1242 | 14 | 6 | 0 | 0 | 10 | 9 | 0 | 0 | 3 | 6 | 0 | 0 |
| L2c_dup100959 | 3.93342745475754 | 5.36504827329143 | 0.994221526316776 | 5.39623024776676 | 6.80556029215489e-08 | 2.45356946860165e-05 | 1243 | 14 | 7 | 0 | 0 | 13 | 11 | 0 | 0 | 3 | 2 | 0 | 0 |
| MLT1D_dup3235 | 2.90883203645911 | 4.96251449860301 | 1.06213296696974 | 4.6722158646116 | 2.97967556963985e-06 | 0.000479055326414544 | 1244 | 13 | 5 | 0 | 0 | 8 | 7 | 0 | 0 | 3 | 1 | 0 | 0 |
| HAL1b_dup3807 | 5.41758858965154 | 4.7131470440233 | 0.914656657415273 | 5.1529139440609 | 2.56469676265621e-07 | 7.0387299567942e-05 | 1245 | 21 | 14 | 1 | 0 | 18 | 10 | 1 | 0 | 1 | 4 | 0 | 0 |
| L1MC1_dup10938 | 3.6997910865387 | 4.35399612121476 | 1.01471754332472 | 4.29084541787747 | 1.77994188913568e-05 | 0.00187402678208973 | 1246 | 14 | 11 | 1 | 0 | 13 | 8 | 0 | 0 | 0 | 1 | 0 | 0 |
| MLT1I_dup7505 | 3.91507236085977 | 4.37536929958428 | 1.00565054286468 | 4.35078500243302 | 1.35651011583501e-05 | 0.00153318123621987 | 1247 | 20 | 11 | 0 | 0 | 4 | 9 | 1 | 0 | 2 | 3 | 0 | 1 |
| L2a_dup116309 | 2.62744333426392 | 3.68748091601226 | 1.03083919386608 | 3.5771640600729 | 0.000347342186022023 | 0.0166020943228561 | 1248 | 9 | 10 | 0 | 0 | 4 | 7 | 1 | 0 | 1 | 1 | 0 | 1 |
| L2b_dup14926 | 2.73923577699889 | 3.88326402216825 | 1.01746059183368 | 3.81662351675928 | 0.000135290362919082 | 0.00843600647147374 | 1249 | 8 | 8 | 0 | 0 | 4 | 9 | 1 | 0 | 2 | 2 | 0 | 1 |
| L1MB5_dup3650 | 3.39960920768172 | 3.31316173439212 | 0.862112747716315 | 3.84307243242661 | 0.000121503589687569 | 0.00782416132779684 | 1250 | 11 | 7 | 0 | 1 | 7 | 11 | 2 | 0 | 2 | 2 | 0 | 1 |
| L1PREC2_dup5830 | 3.37296128044633 | 2.6536515183668 | 0.804368159380476 | 3.29905092266537 | 0.000970123112180542 | 0.0344121815818294 | 1251 | 11 | 10 | 1 | 1 | 5 | 8 | 2 | 0 | 1 | 3 | 0 | 2 |
| L1MD3_dup156 | 2.66913679898149 | 3.98074262638677 | 1.10740397381384 | 3.59466167768687 | 0.00032481338840965 | 0.0158279841324027 | 1252 | 12 | 11 | 0 | 0 | 4 | 6 | 1 | 0 | 0 | 1 | 0 | 0 |
| L1PB4_dup1832 | 2.67279859674252 | 3.47611745078354 | 1.00643803944493 | 3.45388122720469 | 0.000552580682441152 | 0.023152740137846 | 1253 | 11 | 9 | 1 | 0 | 5 | 7 | 1 | 0 | 0 | 1 | 0 | 0 |
| L2c_dup100958 | 2.42284831142738 | 3.49873789757315 | 1.01330618482401 | 3.4527943774278 | 0.000554811742587567 | 0.0232257626498489 | 1254 | 7 | 9 | 1 | 0 | 6 | 5 | 1 | 0 | 1 | 1 | 0 | 0 |
| MamGypLTR1d_dup26 | 2.9964939124947 | 3.16721286875986 | 0.940705882285806 | 3.36684709684593 | 0.00076032832522072 | 0.0290024180302933 | 1255 | 11 | 9 | 1 | 0 | 3 | 8 | 3 | 0 | 1 | 3 | 0 | 0 |
| L1PA16_dup7046 | 3.65723907219036 | 2.9826038457648 | 0.884870748614615 | 3.37066611189766 | 0.000749866821212738 | 0.028753614264793 | 1256 | 13 | 11 | 0 | 2 | 3 | 10 | 4 | 0 | 1 | 4 | 0 | 0 |
| MSTA_dup6514 | 3.75036017037235 | 3.31012082171974 | 0.975142965803103 | 3.39449797393925 | 0.000687545487618765 | 0.0269820092735891 | 1257 | 9 | 18 | 1 | 0 | 5 | 4 | 4 | 0 | 1 | 6 | 0 | 0 |
| L1ME3Cz_dup1726 | 2.39777393926242 | 3.7173353375962 | 1.12197359655533 | 3.31321106754126 | 0.000922313479947792 | 0.0332566757538502 | 1258 | 12 | 8 | 0 | 0 | 1 | 4 | 1 | 0 | 2 | 2 | 0 | 1 |
| L1MCc_dup2830 | 1.80418003998929 | 3.83422756774659 | 1.13825056039206 | 3.36852684388392 | 0.000755710376593617 | 0.0289096883069043 | 1259 | 6 | 7 | 0 | 0 | 2 | 3 | 1 | 0 | 1 | 3 | 0 | 0 |

| | | | | | | | | | | | | | | | | | | | |
|---|---|---|---|---|---|---|---|---|---|---|---|---|---|---|---|---|---|---|---|
| L2_dup52641 | 2.12729689695212 | 3.98025375362622 | 1.12094900936853 | 3.55078930474139 | 0.000384077755612201 | 0.0179512405493814 | 1260 | 8 | 9 | 0 | 0 | 3 | 3 | 1 | 0 | 2 | 1 | 0 | 0 |
| MLT2F_dup1024 | 3.16156095632145 | 3.2329020037068 | 0.988803246852867 | 3.26950989895753 | 0.00107733967685266 | 0.0369752755268131 | 1261 | 6 | 10 | 0 | 0 | 2 | 11 | 2 | 0 | 2 | 5 | 2 | 0 |
| L1PA5_dup5830 | 3.14403536831026 | 3.20504857475968 | 0.982530222900048 | 3.26203560975419 | 0.00110615247323978 | 0.0376113970273667 | 1262 | 7 | 15 | 0 | 0 | 2 | 4 | 2 | 1 | 3 | 4 | 1 | 0 |
| THE1C_dup8360 | 3.42031392318566 | 3.62669371869654 | 1.00224568241418 | 3.61856756515095 | 0.000296238163453895 | 0.0148303568359353 | 1263 | 15 | 14 | 0 | 0 | 2 | 6 | 1 | 1 | 1 | 3 | 1 | 0 |
| L1ME3_dup8485 | 2.20631151858898 | 4.07790260250694 | 1.20025615744667 | 3.39752691723903 | 0.000679978980876574 | 0.0268011062752455 | 1264 | 12 | 2 | 1 | 0 | 4 | 6 | 0 | 0 | 0 | 4 | 0 | 0 |
| L1ME4a_dup31258 | 2.95907310497066 | 5.12224866897805 | 1.4583330819496 | 3.51239969275763 | 0.000444079599372685 | 0.0199108657155329 | 1265 | 10 | 0 | 0 | 0 | 11 | 8 | 0 | 0 | 0 | 9 | 0 | 0 |
| L1M3_dup663 | 3.00237054791876 | 3.52414720334861 | 1.00528281366805 | 3.50562762581188 | 0.00045553204688366 | 0.0202601922294915 | 1266 | 3 | 4 | 2 | 1 | 7 | 3 | 0 | 0 | 5 | 11 | 0 | 0 |
| MIR_dup120680 | 2.76340122021171 | 3.11497982628604 | 0.96490756529277 | 3.22826759612036 | 0.00124542410910524 | 0.0408806297510215 | 1267 | 3 | 7 | 1 | 3 | 6 | 5 | 0 | 0 | 4 | 5 | 0 | 0 |
| L1M5_dup42312 | 2.32942063768536 | 3.68557092509882 | 1.11378489671581 | 3.3090509091714 | 0.000936128135752768 | 0.0335817797964463 | 1268 | 3 | 8 | 1 | 1 | 1 | 2 | 0 | 0 | 4 | 8 | 0 | 0 |
| MER96B_dup777 | 1.90780381649485 | 3.43818486835587 | 1.09354987730452 | 3.14405857447545 | 0.00166622084385493 | 0.0499676762030735 | 1269 | 7 | 3 | 1 | 1 | 3 | 2 | 0 | 0 | 2 | 5 | 0 | 0 |
| L1MC1_dup456 | 1.93968475926253 | 3.98883292315786 | 1.19030860022112 | 3.35109140807425 | 0.000804937277069569 | 0.0301719605119086 | 1270 | 8 | 6 | 1 | 0 | 1 | 1 | 0 | 0 | 3 | 4 | 0 | 0 |
| L1MEc_dup2975 | 2.61189111954431 | 4.34514231822338 | 1.116606897253 | 3.89138051082526 | 9.96754733461039e-05 | 0.00679807388604637 | 1271 | 10 | 12 | 1 | 0 | 3 | 2 | 0 | 0 | 2 | 3 | 0 | 0 |
| L1PA15_dup2751 | 2.68528105724437 | 4.47010867453053 | 1.10021312774493 | 4.06294795235971 | 4.84568371268216e-05 | 0.00397727166849903 | 1272 | 6 | 12 | 1 | 0 | 3 | 2 | 0 | 0 | 3 | 6 | 0 | 0 |
| AluJo_dup49445 | 3.65203090491961 | 3.88871139805709 | 0.968332901595772 | 4.01588275235578 | 5.92236955364223e-05 | 0.00462337318179446 | 1273 | 10 | 11 | 2 | 0 | 5 | 5 | 0 | 0 | 5 | 6 | 1 | 0 |
| L1MEg_dup1278 | 2.41592645368938 | 3.76855902764336 | 1.04164265336723 | 3.6179000691466 | 0.000297003020980927 | 0.0148592684672682 | 1274 | 5 | 7 | 1 | 0 | 3 | 3 | 0 | 0 | 5 | 4 | 1 | 0 |
| L1MC4_dup11975 | 3.00077504474883 | 3.59221458257516 | 1.02298962465718 | 3.51148681862629 | 0.000445607560627558 | 0.0199568109582317 | 1275 | 10 | 7 | 3 | 0 | 5 | 4 | 0 | 0 | 5 | 3 | 0 | 0 |
| L1ME4a_dup24282 | 2.968482709925 | 3.19570488154827 | 0.935110819638097 | 3.41746113341417 | 0.00063208118443719 | 0.0254505833552071 | 1276 | 7 | 8 | 3 | 1 | 6 | 4 | 0 | 0 | 3 | 5 | 0 | 0 |
| AluSg_dup9460 | 2.55618797727107 | 3.88746506049644 | 1.03351954540127 | 3.76138514050755 | 0.000168975012136026 | 0.00992028824024642 | 1277 | 9 | 5 | 1 | 0 | 3 | 4 | 0 | 0 | 3 | 6 | 0 | 1 |
| AluY_dup94574 | 2.21525376499237 | 4.18078171803306 | 1.04400172537156 | 4.00457357150929 | 6.21294532928248e-05 | 0.00479515241427652 | 1278 | 6 | 4 | 1 | 0 | 4 | 4 | 0 | 0 | 4 | 4 | 0 | 0 |
| L1MA5A_dup2504 | 2.85636455031048 | 4.53024458368089 | 1.023499036798 | 4.42623238596656 | 9.58932174445846e-06 | 0.00118593594168758 | 1279 | 10 | 4 | 1 | 0 | 5 | 9 | 0 | 0 | 3 | 4 | 0 | 0 |
| MIRc_dup81311 | 2.31036887463935 | 3.90190750744381 | 1.0718462414681 | 3.64036123511467 | 0.000272255805907071 | 0.0139496511611464 | 1280 | 2 | 1 | 0 | 0 | 4 | 2 | 0 | 0 | 9 | 7 | 0 | 1 |
| L1ME3A_dup4660 | 1.79630446090625 | 4.21306613373164 | 1.15416307655114 | 3.65032136214327 | 0.000261912412954092 | 0.0135834810402734 | 1281 | 4 | 2 | 0 | 0 | 1 | 2 | 0 | 0 | 5 | 7 | 0 | 0 |
| LTR40a_dup1867 | 1.80494305030276 | 3.96936520458638 | 1.19907197710399 | 3.31036441546504 | 0.000931745797811407 | 0.0334838602028075 | 1282 | 3 | 1 | 0 | 0 | 1 | 1 | 0 | 0 | 8 | 6 | 0 | 0 |
| MIRb_dup17027 | 1.77010356574681 | 3.76100232252162 | 1.12156049449793 | 3.35336554824467 | 0.000798351908653603 | 0.030024441798086 | 1283 | 3 | 4 | 0 | 0 | 1 | 3 | 0 | 0 | 4 | 5 | 1 | 0 |
| HAL1_dup10086 | 1.6944526683329 | 4.29142264675865 | 1.19553694604443 | 3.58953578219168 | 0.00033126735911997 | 0.0160537154853687 | 1284 | 1 | 3 | 0 | 0 | 2 | 6 | 0 | 0 | 4 | 4 | 0 | 0 |
| LTR42_dup154 | 2.10112139685235 | 4.60209740223739 | 1.1224562609851 | 4.10002381580416 | 4.13107623563721e-05 | 0.00353460289811194 | 1285 | 4 | 5 | 0 | 0 | 1 | 5 | 0 | 0 | 5 | 5 | 0 | 0 |
| L1ME3D_dup3963 | 1.96553917225512 | 4.5428948427426 | 1.15024111903747 | 3.94951525167535 | 7.8309621357872e-05 | 0.00568786970806478 | 1286 | 6 | 1 | 0 | 0 | 4 | 6 | 0 | 0 | 4 | 3 | 0 | 0 |
| L1PB1_dup8936 | 1.95580343482303 | 4.00004098190459 | 1.0967964499941 | 3.64702218166927 | 0.000265297029966356 | 0.0137052468719875 | 1287 | 7 | 3 | 0 | 1 | 2 | 3 | 0 | 0 | 4 | 4 | 0 | 0 |
| MIR_dup84484 | 2.15128802834504 | 4.12176688364835 | 1.07454460528426 | 3.83582669661067 | 0.000125142632338218 | 0.00798744334612383 | 1288 | 4 | 2 | 0 | 1 | 6 | 3 | 0 | 0 | 4 | 6 | 0 | 0 |
| HERVL40-int_dup700 | 1.72850569046254 | 4.35648477106691 | 1.35003628760899 | 3.22693901715971 | 0.00125122123921731 | 0.0410249165514042 | 1289 | 3 | 3 | 0 | 0 | 8 | 0 | 0 | 0 | 2 | 5 | 0 | 0 |
| AluSp_dup22715 | 2.16579608214866 | 3.60436380779978 | 1.10753184468299 | 3.25441099062163 | 0.00113627761366931 | 0.0383738463920174 | 1290 | 3 | 6 | 2 | 0 | 6 | 2 | 0 | 0 | 4 | 3 | 0 | 0 |
| L1PA7_dup5956 | 2.15970937885343 | 4.13774537108931 | 1.11672436526211 | 3.70525216409882 | 0.000211180531660002 | 0.0115618534821766 | 1291 | 7 | 5 | 1 | 0 | 4 | 1 | 0 | 0 | 5 | 3 | 0 | 0 |
| LTR16_dup2558 | 2.00627597005764 | 3.98970056293268 | 1.12598459351571 | 3.54329942515062 | 0.000395153769454387 | 0.0183057584086598 | 1292 | 4 | 1 | 1 | 0 | 6 | 2 | 0 | 0 | 4 | 6 | 0 | 0 |
| MIR_dup140068 | 2.34860322833705 | 4.11437101224784 | 1.07213644774179 | 3.83754420523043 | 0.000124270868960817 | 0.00794224679894756 | 1293 | 2 | 3 | 1 | 0 | 5 | 2 | 0 | 0 | 7 | 7 | 0 | 0 |
| MLT2B4_dup2342 | 2.8716593784651 | 3.08051647730911 | 0.927724298670703 | 3.32050856242858 | 0.000898536028211337 | 0.0326068405930723 | 1294 | 4 | 7 | 0 | 1 | 3 | 3 | 2 | 0 | 5 | 9 | 0 | 1 |
| MER39B_dup800 | 2.5699112218643 | 3.41183614579038 | 1.00557093936005 | 3.39293431447182 | 0.000691482173611107 | 0.0270974007381536 | 1295 | 5 | 7 | 0 | 0 | 2 | 3 | 1 | 0 | 6 | 5 | 0 | 2 |
| MIR_dup84495 | 2.24254361380061 | 3.69048503442884 | 1.08328614664319 | 3.40674995786168 | 0.000657413232447518 | 0.0261586704026684 | 1296 | 5 | 8 | 0 | 0 | 3 | 2 | 1 | 0 | 2 | 6 | 0 | 1 |
| L1ME1_dup25252 | 2.0900643765089 | 3.55364577046411 | 1.07607583973473 | 3.30241200410199 | 0.000958571589460666 | 0.0341143098154065 | 1297 | 7 | 3 | 1 | 0 | 6 | 2 | 1 | 0 | 3 | 3 | 0 | 0 |
| MER41A_dup1153 | 2.98753342527145 | 4.08832797663095 | 1.00351660478384 | 4.07400132408532 | 4.62122184063291e-05 | 0.00384610441444994 | 1298 | 8 | 9 | 1 | 0 | 6 | 4 | 1 | 0 | 4 | 4 | 0 | 0 |
| L1MEc_dup764 | 3.06548740829657 | 3.63140913543392 | 0.990053791219143 | 3.66789074254464 | 0.000244559639444322 | 0.0129114683752426 | 1299 | 9 | 3 | 1 | 0 | 10 | 5 | 1 | 0 | 5 | 3 | 0 | 1 |
| L2c_dup70173 | 2.86432436185221 | 4.05577468700119 | 1.01331656967375 | 4.00247544388523 | 6.26831754609555e-05 | 0.00483161997675171 | 1300 | 11 | 4 | 0 | 0 | 6 | 5 | 1 | 0 | 4 | 4 | 0 | 1 |
| L1M5_dup17279 | 1.67969087282894 | 3.75572068378309 | 1.16926424520657 | 3.21203756907795 | 0.00131797138838128 | 0.0424997994280759 | 1301 | 2 | 1 | 0 | 0 | 6 | 3 | 1 | 0 | 4 | 3 | 0 | 0 |
| HERVL40-int_dup704 | 2.0892444078296 | 3.5285423205009 | 1.1058855890609 | 3.19069382529643 | 0.00141931602707678 | 0.0446792014336328 | 1302 | 2 | 6 | 0 | 0 | 6 | 5 | 2 | 0 | 2 | 3 | 0 | 0 |
| AluSz_dup49505 | 2.93451160255039 | 4.06658736021061 | 1.02423117281106 | 3.97038038692929 | 7.17579744805386e-05 | 0.00533127328184734 | 1303 | 6 | 7 | 0 | 0 | 8 | 5 | 2 | 0 | 5 | 3 | 0 | 0 |
| L2a_dup86233 | 2.44908399815707 | 3.77754709257769 | 1.10113928458867 | 3.43058062267643 | 0.000602290962963018 | 0.0246580738942186 | 1304 | 4 | 10 | 0 | 0 | 5 | 3 | 2 | 0 | 4 | 2 | 0 | 0 |
| L2c_dup70195 | 2.69087103756985 | 3.43861388076328 | 1.00855416128498 | 3.40944890493756 | 0.000650942702548613 | 0.0259706928740161 | 1305 | 4 | 7 | 0 | 0 | 8 | 6 | 1 | 0 | 1 | 5 | 1 | 1 |
| MIRb_dup115598 | 2.81887442109862 | 4.04485302269508 | 1.05826309061373 | 3.82216204889969 | 0.000132286706996524 | 0.00830092109256193 | 1306 | 2 | 6 | 0 | 0 | 9 | 9 | 0 | 0 | 2 | 5 | 1 | 1 |
| AluSc_dup481 | 3.25499163364489 | 4.2404205633875 | 1.07502634715084 | 3.94448059308125 | 7.99731295506418e-05 | 0.00577695394172951 | 1307 | 1 | 8 | 0 | 0 | 7 | 13 | 1 | 0 | 4 | 5 | 0 | 1 |
| MIRb_dup4005 | 2.38340212788462 | 3.23567735907153 | 1.01493216355031 | 3.18807253851615 | 0.0014322460161344 | 0.0449956707888983 | 1308 | 2 | 3 | 1 | 0 | 4 | 7 | 1 | 0 | 4 | 6 | 0 | 1 |
| AluJr_dup1542 | 3.14789721562999 | 5.20033778753442 | 1.07513312289944 | 4.83692454150242 | 1.31863457732382e-06 | 0.000255716237274567 | 1309 | 13 | 7 | 0 | 0 | 3 | 8 | 0 | 0 | 6 | 2 | 0 | 0 |
| L1MC1_dup7554 | 2.7459570345384 | 4.99810790281947 | 1.09470802015735 | 4.56569953886064 | 4.97831393323829e-06 | 0.000719228832518712 | 1310 | 5 | 7 | 0 | 0 | 8 | 6 | 0 | 0 | 6 | 1 | 0 | 0 |
| L1MEd_dup12204 | 3.70233530408211 | 5.43573626359006 | 1.00908355342162 | 5.38680493320741 | 7.17212000240517e-08 | 2.56242832746592e-05 | 1311 | 7 | 7 | 0 | 0 | 9 | 12 | 0 | 0 | 7 | 3 | 0 | 0 |
| AluJb_dup40167 | 2.0378915863795 | 4.04527532136613 | 1.14777874794495 | 3.52443824962696 | 0.000424381716080345 | 0.019304049155399 | 1312 | 7 | 4 | 0 | 0 | 1 | 9 | 0 | 0 | 2 | 2 | 0 | 1 |
| AluSx_dup20025 | 2.15341617301748 | 4.06687621614145 | 1.11303073234609 | 3.65387594246309 | 0.000258311112053689 | 0.0134527908002825 | 1313 | 9 | 3 | 0 | 0 | 3 | 9 | 0 | 0 | 1 | 2 | 0 | 1 |
| L1M1_dup7258 | 2.70507349290847 | 4.46794892113749 | 1.0361871278675 | 4.31191316797443 | 1.61847997514631e-05 | 0.00174825352846227 | 1314 | 8 | 8 | 0 | 0 | 3 | 8 | 0 | 0 | 3 | 3 | 0 | 1 |
| MLT1J_dup7895 | 2.57849789962392 | 3.56513930254834 | 1.09917938483424 | 3.24345539202955 | 0.00118089300448815 | 0.0393596815864879 | 1315 | 11 | 5 | 0 | 0 | 2 | 6 | 0 | 0 | 3 | 3 | 0 | 3 |
| L1MC4_dup16967 | 2.93302709760709 | 3.21313192963734 | 0.936105554331313 | 3.43244617529545 | 0.00059816260101788 | 0.0245228115544108 | 1316 | 6 | 7 | 0 | 1 | 6 | 7 | 0 | 0 | 3 | 4 | 0 | 3 |
| L1MEd_dup12205 | 2.27290152970551 | 4.23570484152763 | 1.04732825947799 | 4.04429538036029 | 5.24807201399113e-05 | 0.00424314557744184 | 1317 | 7 | 4 | 0 | 0 | 5 | 4 | 0 | 0 | 4 | 3 | 0 | 1 |
| L1MA4_dup1000 | 2.50299119585734 | 3.38323667358557 | 0.998504711027956 | 3.38830316594354 | 0.000703264851591297 | 0.0274506329358174 | 1318 | 6 | 3 | 0 | 1 | 6 | 4 | 0 | 0 | 4 | 5 | 0 | 2 |
| L1ME1_dup6211 | 2.40778544669357 | 3.76371790572151 | 1.03483857042953 | 3.63700968756829 | 0.000275821561236833 | 0.0140885772047527 | 1319 | 6 | 3 | 0 | 1 | 5 | 3 | 0 | 0 | 6 | 4 | 0 | 1 |
| L1PB1_dup4338 | 2.28824963780091 | 3.20344243007214 | 1.01216476517343 | 3.16494165801478 | 0.00155114050274673 | 0.0475632726773244 | 1320 | 7 | 3 | 0 | 1 | 3 | 7 | 1 | 0 | 3 | 3 | 0 | 1 |
| L2b_dup74472 | 2.727468287243 | 3.46501626125587 | 1.01066780973284 | 3.42844229121318 | 0.000607055581100255 | 0.0247849085965136 | 1321 | 6 | 4 | 0 | 2 | 4 | 6 | 0 | 0 | 7 | 3 | 0 | 1 |
| MamTip2_dup6053 | 2.0646069471916 | 3.53636571823721 | 1.09512119757895 | 3.22920031687385 | 0.00124136909825012 | 0.0407812800569134 | 1322 | 4 | 1 | 0 | 1 | 4 | 6 | 0 | 0 | 5 | 3 | 0 | 1 |
| AluSz_dup16986 | 3.32183573894553 | 3.63198393442891 | 1.10178427372093 | 3.29645650337975 | 0.00097912774884468 | 0.0346180581465968 | 1323 | 12 | 9 | 0 | 0 | 7 | 3 | 0 | 0 | 2 | 5 | 4 | 0 |
| MER44B_dup1150 | 2.64924301692153 | 4.46436271730251 | 1.06255574668453 | 4.20153270191475 | 2.65113941145678e-05 | 0.0025188649054405 | 1324 | 10 | 6 | 0 | 0 | 5 | 2 | 0 | 0 | 3 | 6 | 1 | 0 |
| MIR3_dup61688 | 2.92928151816244 | 4.60512636418647 | 1.03289880888649 | 4.45844871207761 | 8.25549441738868e-06 | 0.00105384220742487 | 1325 | 8 | 5 | 0 | 0 | 10 | 3 | 0 | 0 | 4 | 5 | 1 | 0 |
| THE1D_dup6702 | 2.89712760409997 | 3.1622774679016 | 0.976346613281726 | 3.23888814165336 | 0.00119996624499918 | 0.0398447066888886 | 1326 | 10 | 5 | 0 | 0 | 6 | 2 | 0 | 0 | 5 | 3 | 2 | 3 |
| L2_dup12783 | 2.60525929362966 | 3.91620336817944 | 1.04332682799707 | 3.75357295824317 | 0.000174331760950436 | 0.0101215782210759 | 1327 | 6 | 8 | 0 | 0 | 7 | 3 | 0 | 0 | 4 | 2 | 1 | 1 |
| MER5A1_dup13794 | 2.4510285449499 | 3.83337135800928 | 1.05618585196677 | 3.62944774432547 | 0.00028402821897071 | 0.0143794422534478 | 1328 | 6 | 8 | 0 | 0 | 5 | 2 | 0 | 0 | 4 | 3 | 1 | 1 |
| L1PREC2_dup1170 | 3.009379191755 | 3.16293639086336 | 0.948212939063546 | 3.33568153371443 | 0.000850905786260155 | 0.0314006179973985 | 1329 | 10 | 9 | 0 | 1 | 7 | 3 | 0 | 0 | 3 | 2 | 2 | 1 |
| Tigger13a_dup1580 | 2.66809635319639 | 3.43169967428932 | 1.02188958107335 | 3.35819029555501 | 0.000784545822633554 | 0.0296599206088916 | 1330 | 10 | 5 | 0 | 0 | 7 | 5 | 0 | 0 | 2 | 2 | 2 | 1 |
| THE1A-int_dup1072 | 3.46127154233936 | 4.85448779377161 | 1.06353538449128 | 4.56448169431961 | 5.0072988849573e-06 | 0.000722977391067992 | 1331 | 5 | 14 | 0 | 0 | 9 | 2 | 0 | 0 | 5 | 6 | 0 | 1 |
| Tigger19a_dup2316 | 2.63613667308164 | 4.96237502343407 | 1.14068858285901 | 4.35033285859356 | 1.35931033542815e-05 | 0.0015343570069914 | 1332 | 9 | 9 | 0 | 0 | 6 | 1 | 0 | 0 | 2 | 6 | 0 | 0 |
| L1ME1_dup21485 | 3.18772302076732 | 2.91750180782091 | 0.904960300469873 | 3.22390032613153 | 0.00126457406931697 | 0.041321112537448 | 1333 | 11 | 7 | 0 | 1 | 8 | 1 | 0 | 1 | 3 | 5 | 1 | 2 |
| MIR_dup82019 | 2.91940597014591 | 3.52895149246733 | 0.997554797963652 | 3.53760164320909 | 0.000403778739611712 | 0.0186016526574344 | 1334 | 5 | 9 | 0 | 1 | 5 | 2 | 0 | 1 | 5 | 6 | 1 | 0 |
| L1M5_dup13726 | 3.78264421848463 | 3.25107338986368 | 0.85569578130955 | 3.7993331986378 | 0.000145085922719015 | 0.00888384327516693 | 1335 | 6 | 5 | 0 | 0 | 10 | 5 | 0 | 0 | 6 | 8 | 4 | 2 |
| L1MCa_dup7053 | 3.33825719689213 | 3.28341766035052 | 0.899875278999734 | 3.64874748420719 | 0.000263521966824858 | 0.0136431675191705 | 1336 | 5 | 5 | 0 | 0 | 10 | 6 | 0 | 0 | 5 | 5 | 2 | 3 |
| L3_dup42416 | 3.45194537671991 | 2.87682916904255 | 0.888942013662757 | 3.23623940012576 | 0.00121115760156076 | 0.0400932257458232 | 1337 | 5 | 6 | 0 | 0 | 10 | 5 | 0 | 0 | 4 | 5 | 3 | 5 |
| L1PB4_dup3736 | 3.27083884155509 | 3.39080458171852 | 1.00696646725479 | 3.36734607554769 | 0.000758953808137552 | 0.0289689233821965 | 1338 | 6 | 3 | 0 | 0 | 12 | 2 | 0 | 0 | 4 | 8 | 4 | 1 |

| | | | | | | | | | | | | | | | | | | |
|---|---|---|---|---|---|---|---|---|---|---|---|---|---|---|---|---|---|---|
| L2a_dup15181 | 2.74594570850087 | 3.35519365578496 | 1.01665073321573 | 3.30024220331036 | 0.000966014194353504 | 0.034322713676219 | 1339 | 4 | 3 | 0 | 0 | 11 | 3 | 0 | 0 | 3 | 6 | 1 | 3 |
| L1MA7_dup7247 | 2.45439942078028 | 3.46866200950828 | 1.06741204193948 | 3.24959984825142 | 0.00115567499535791 | 0.0388182160425513 | 1340 | 7 | 7 | 0 | 0 | 2 | 3 | 0 | 0 | 1 | 8 | 2 | 1 |
| FLAM_C_dup10182 | 3.39126732734725 | 2.60044973773575 | 0.798218527893275 | 3.25781680938812 | 0.00112272858211016 | 0.0380282954019621 | 1341 | 6 | 3 | 0 | 2 | 5 | 5 | 0 | 0 | 4 | 12 | 3 | 2 |
| LTR79_dup805 | 3.52031758564569 | 3.26997400367231 | 0.957971715533111 | 3.41343481300235 | 0.000641495030691487 | 0.0257014946241651 | 1342 | 12 | 3 | 0 | 1 | 8 | 5 | 0 | 0 | 1 | 11 | 3 | 1 |
| AluSc_dup27279 | 2.90912810491682 | 3.9472874640831 | 1.01144495101901 | 3.90262214479029 | 9.51561912373039e-05 | 0.00657052672087972 | 1343 | 2 | 4 | 0 | 0 | 5 | 5 | 1 | 0 | 4 | 13 | 1 | 0 |
| L2a_dup15180 | 2.51929997032064 | 3.42016726944478 | 1.01493005049354 | 3.36985516172433 | 0.000752077030397654 | 0.0288170243276973 | 1344 | 3 | 2 | 0 | 0 | 5 | 5 | 0 | 0 | 5 | 7 | 3 | 0 |
| L2a_dup30877 | 3.02675563505381 | 3.70254047212627 | 0.969901769434716 | 3.81743861987612 | 0.000134844319703745 | 0.00841260264714927 | 1345 | 6 | 4 | 0 | 0 | 6 | 4 | 0 | 0 | 7 | 6 | 3 | 0 |
| Tigger17a_dup1051 | 3.2396254010671 | 4.18319957466495 | 1.00037059575926 | 4.18164987295534 | 2.89401388869721e-05 | 0.00268676952305871 | 1346 | 4 | 6 | 0 | 0 | 8 | 3 | 0 | 0 | 8 | 7 | 2 | 0 |
| LTR81B_dup666 | 2.55039568353593 | 3.2761955889807 | 0.936649859405912 | 3.4977804737607 | 0.000469147125019716 | 0.0206359044478966 | 1347 | 4 | 3 | 0 | 0 | 3 | 2 | 1 | 0 | 7 | 8 | 1 | 1 |
| MLT1J-int_dup525 | 2.70397236551367 | 3.03227110856368 | 0.896988704606759 | 3.38050088366836 | 0.000723538424291365 | 0.0279985483666341 | 1348 | 3 | 2 | 0 | 0 | 3 | 4 | 1 | 0 | 7 | 9 | 1 | 2 |
| L2a_dup61517 | 2.540726030337 | 2.91068836537824 | 0.908739186139495 | 3.20299642600802 | 0.00136005668910469 | 0.0434564319787437 | 1349 | 6 | 3 | 1 | 0 | 4 | 4 | 1 | 0 | 5 | 5 | 1 | 1 |
| MLT2B4_dup2343 | 2.55680566189861 | 3.40091729598469 | 0.996784938604736 | 3.41188672126725 | 0.000645149172504295 | 0.025813126786734 | 1350 | 7 | 5 | 0 | 0 | 2 | 6 | 1 | 0 | 3 | 6 | 1 | 1 |
| L1MB7_dup5155 | 3.18777956521024 | 2.94718383871357 | 0.864861468803426 | 3.40769469449382 | 0.0006551415215558 | 0.0260985469773396 | 1351 | 7 | 3 | 2 | 0 | 3 | 5 | 0 | 0 | 5 | 11 | 1 | 2 |
| MIR_dup90346 | 2.59808285384871 | 3.02744629702385 | 0.938515641946397 | 3.22578139533743 | 0.00125629271627968 | 0.0411352451724074 | 1352 | 7 | 2 | 1 | 0 | 4 | 5 | 0 | 0 | 4 | 6 | 2 | 1 |
| L2c_dup13125 | 2.23547840669517 | 3.12264825068259 | 0.990962362399277 | 3.15112699449267 | 0.00162641756553707 | 0.0490867014268497 | 1353 | 5 | 2 | 0 | 1 | 1 | 4 | 0 | 0 | 5 | 7 | 1 | 1 |
| L1PB1_dup6112 | 2.45639200075492 | 3.03442833660334 | 0.959993787273272 | 3.16088330657036 | 0.00157291499627337 | 0.0479524448050304 | 1354 | 3 | 1 | 0 | 0 | 4 | 4 | 0 | 0 | 6 | 7 | 3 | 1 |
| LTR44_dup149 | 2.15050922649091 | 3.40045164081769 | 0.995011182568856 | 3.41750092902335 | 0.000631988783953666 | 0.0254505833552071 | 1355 | 2 | 2 | 0 | 0 | 2 | 3 | 0 | 0 | 6 | 8 | 1 | 1 |
| L1PA3_dup4242 | 23.7254007376521 | 4.24946060752654 | 1.27226761966113 | 3.3400681915165 | 0.000837578208894064 | 0.0310627157876414 | 1356 | 150 | 128 | 4 | 1 | 2 | 1 | 3 | 0 | 9 | 8 | 0 | 5 |
| L1PA7_dup2766 | 31.210643378997 | 4.35048322438917 | 1.1257299420591 | 3.86458870982112 | 0.000111276543412519 | 0.00735484190475015 | 1357 | 202 | 174 | 3 | 1 | 4 | 5 | 2 | 1 | 7 | 4 | 1 | 7 |
| LTR16B1_dup547 | 17.3738140815204 | 6.35863267658485 | 1.50401899624026 | 4.22776088099959 | 2.36028433621954e-05 | 0.0023130980917055 | 1358 | 119 | 82 | 0 | 0 | 1 | 7 | 0 | 0 | 5 | 9 | 2 | 4 |
| MER1A_dup2548 | 14.9394321180421 | 4.97780076380886 | 1.01018378500208 | 4.92761895183124 | 8.32377486418803e-07 | 0.000178861074740485 | 1359 | 94 | 62 | 0 | 1 | 3 | 7 | 0 | 0 | 11 | 9 | 3 | 4 |
| ERV3-16A3_I-int_dup2 | 16.4309638376452 | 6.15348475770626 | 1.40349708679177 | 4.38439439284652 | 1.16308889499104e-05 | 0.00136099802516487 | 1360 | 89 | 89 | 0 | 0 | 8 | 3 | 0 | 0 | 7 | 10 | 2 | 5 |
| L2_dup24185 | 24.4832709373524 | 6.74369342230914 | 1.3388884368846 | 5.03678516934596 | 4.73415602336025e-07 | 0.000114642320552505 | 1361 | 149 | 119 | 0 | 0 | 9 | 10 | 0 | 0 | 14 | 12 | 1 | 4 |
| L1ME4b_dup14764 | 12.3056433986596 | 2.74310496330353 | 0.781846380509385 | 3.50849608271686 | 0.000450647870858528 | 0.0201257241515209 | 1362 | 60 | 62 | 2 | 0 | 7 | 9 | 5 | 4 | 2 | 4 | 1 | 4 |
| L1PA4_dup4703 | 10.9861741080847 | 3.34177036795634 | 0.949654368220536 | 3.51893328750559 | 0.000433285647279197 | 0.0195707263909445 | 1363 | 57 | 62 | 2 | 0 | 2 | 3 | 4 | 3 | 1 | 7 | 0 | 2 |
| L1PREC2_dup2103 | 14.3957240670584 | 3.24607920531029 | 0.822754528313482 | 3.94537993241343 | 7.96735477672981e-05 | 0.00576055970739518 | 1364 | 76 | 73 | 0 | 2 | 3 | 5 | 7 | 2 | 7 | 7 | 2 | 3 |
| Harlequin-int_dup237 | 18.8193339280632 | 4.38140781934482 | 0.863300999357431 | 5.07517983021678 | 3.87129697556236e-07 | 9.79957053348044e-05 | 1365 | 119 | 93 | 1 | 1 | 4 | 4 | 4 | 2 | 7 | 9 | 1 | 1 |
| L1MA6_dup2086 | 13.5523136608382 | 4.0116885135335 | 0.913876156802971 | 4.3897507158603 | 1.13480674507569e-05 | 0.00134006315515524 | 1366 | 94 | 53 | 1 | 0 | 3 | 2 | 2 | 2 | 8 | 8 | 1 | 3 |
| HERVL-int_dup951 | 15.0952256330576 | 5.10096663168449 | 1.02584320874601 | 4.97246225173132 | 6.61078462462688e-07 | 0.000148537276514775 | 1367 | 101 | 75 | 2 | 1 | 5 | 1 | 0 | 2 | 3 | 8 | 0 | 0 |
| L1P1_dup951 | 12.0160115124758 | 3.62600553554576 | 0.960920655079165 | 3.77347028225451 | 0.00016099240413034 | 0.00957926946566586 | 1368 | 73 | 61 | 4 | 2 | 2 | 1 | 0 | 2 | 6 | 3 | 1 | 1 |
| L1PREC2_dup2899 | 8.78858996615667 | 5.05930676085011 | 1.21768553554923 | 4.1548549384453 | 3.2549397944641e-05 | 0.00293816352175097 | 1369 | 45 | 55 | 1 | 0 | 2 | 0 | 2 | 0 | 2 | 7 | 0 | 0 |
| L1MA1_dup1519 | 11.2179463914217 | 4.4446263809533 | 1.05406196757399 | 4.21666516550539 | 2.47941704422616e-05 | 0.00239629312756067 | 1370 | 62 | 66 | 2 | 2 | 3 | 0 | 1 | 0 | 4 | 5 | 0 | 1 |
| L1PA10_dup2580 | 11.0616760395808 | 4.949584269819 | 1.08540298575907 | 4.56013511549128 | 5.11207175576092e-06 | 0.000734095972236746 | 1371 | 51 | 74 | 1 | 0 | 3 | 1 | 2 | 0 | 6 | 3 | 0 | 1 |
| L1M1_dup3441 | 14.6249232895513 | 4.42944880698006 | 0.960090309883404 | 4.61357516202615 | 3.95801038853686e-06 | 0.00059531778755423 | 1372 | 88 | 67 | 2 | 2 | 4 | 0 | 3 | 1 | 11 | 10 | 0 | 1 |
| Tigger3b_dup4869 | 9.97785702840521 | 4.11832197493473 | 1.03725373760116 | 3.9704093855175 | 7.17492407439358e-05 | 0.00533127328184734 | 1373 | 56 | 48 | 1 | 1 | 2 | 0 | 1 | 0 | 9 | 6 | 3 | 1 |
| HAL1M8_dup1560 | 21.4523836885297 | 3.23388503649524 | 1.02862654010906 | 3.14388644507692 | 0.00166720121270374 | 0.0499815874710852 | 1374 | 87 | 80 | 4 | 10 | 3 | 1 | 0 | 1 | 34 | 33 | 9 | 5 |
| Charlie1_dup1454 | 14.9857092089455 | 3.67120474680929 | 0.897890841880763 | 4.08869828666412 | 4.33800643685296e-05 | 0.0036642217949003 | 1375 | 60 | 48 | 1 | 4 | 3 | 1 | 0 | 0 | 16 | 41 | 10 | 4 |
| L2b_dup72752 | 14.8863217297045 | 3.24647088772924 | 0.884561932249931 | 3.67014537859624 | 0.00024241256033393 | 0.0128294259522281 | 1376 | 60 | 49 | 4 | 3 | 2 | 0 | 1 | 0 | 28 | 23 | 7 | 7 |
| L2c_dup115296 | 13.2445210621032 | 4.34848571255333 | 0.951445433320159 | 4.57039947880025 | 4.8679537882618e-06 | 0.000705426389830538 | 1377 | 70 | 64 | 0 | 2 | 3 | 3 | 0 | 1 | 12 | 9 | 4 | 1 |
| L1PA4_dup10898 | 11.0311576669409 | 3.36187517195585 | 0.914267456363745 | 3.67712440003805 | 0.000235878024298677 | 0.0125674501499716 | 1378 | 56 | 50 | 2 | 4 | 1 | 4 | 0 | 0 | 8 | 10 | 5 | 2 |
| L1M4_dup17598 | 11.5151069735657 | 3.22801850558098 | 0.908193930446058 | 3.55432732741953 | 0.000378947257777374 | 0.0177567909017517 | 1379 | 62 | 50 | 3 | 5 | 5 | 1 | 0 | 0 | 11 | 5 | 4 | 2 |
| L1M4b_dup3735 | 10.8098349535692 | 2.95930264327963 | 0.804263279679492 | 3.6795197767315 | 0.000233673568377261 | 0.012480738698338 | 1380 | 50 | 39 | 3 | 10 | 7 | 3 | 0 | 0 | 13 | 9 | 2 | 2 |
| L1PA13_dup7382 | 9.49738861474708 | 4.19287377004526 | 1.09455394221707 | 3.8306689221295 | 0.000127795363948421 | 0.00811539783028319 | 1381 | 43 | 41 | 0 | 4 | 1 | 1 | 0 | 0 | 8 | 20 | 0 | 3 |
| L1PA7_dup4632 | 8.52725917922353 | 3.92263518878308 | 0.951378803093005 | 4.1231055138398 | 3.73798365482004e-05 | 0.00327239977457619 | 1382 | 26 | 31 | 0 | 3 | 2 | 1 | 0 | 0 | 17 | 19 | 1 | 4 |
| MSTB_dup6723 | 9.68826893968525 | 3.48108923865347 | 1.06625902990094 | 3.26476882355395 | 0.00109553450111333 | 0.0373731974769768 | 1383 | 33 | 38 | 0 | 5 | 1 | 0 | 0 | 0 | 18 | 17 | 5 | 2 |
| L1PA4_dup10100 | 7.66677248439842 | 3.28964021208153 | 0.950575022585175 | 3.4606844635314 | 0.000538803949622082 | 0.0226994059130269 | 1384 | 35 | 21 | 1 | 3 | 2 | 0 | 0 | 0 | 13 | 14 | 2 | 5 |
| L1ME3G_dup12596 | 8.84114947613698 | 3.29199154103856 | 0.957951714401183 | 3.43649005638702 | 0.00058930398488417 | 0.0242822821841326 | 1385 | 37 | 26 | 1 | 2 | 2 | 0 | 0 | 0 | 13 | 17 | 5 | 8 |
| L3_dup36359 | 11.1846687257347 | 3.57213340384271 | 0.939072025100856 | 3.80389715417096 | 0.000142437335651209 | 0.00876905414459572 | 1386 | 49 | 36 | 1 | 3 | 3 | 0 | 0 | 0 | 17 | 20 | 5 | 6 |
| HAL1ME_dup5439 | 10.2269341978519 | 2.3029783860386 | 0.714547568829689 | 3.22298820470483 | 0.00126860777561916 | 0.0414017481521709 | 1387 | 35 | 19 | 3 | 7 | 1 | 5 | 1 | 0 | 17 | 26 | 4 | 10 |
| ERVL-E-int_dup1830 | 8.84785207613393 | 2.42458243121146 | 0.768991538592029 | 3.1529377236721 | 0.0016163628596097 | 0.0488759731481423 | 1388 | 32 | 23 | 3 | 7 | 3 | 3 | 0 | 0 | 15 | 15 | 2 | 8 |
| MLT1H1_dup2788 | 12.1676252417276 | 3.15301475065506 | 0.791315271122832 | 3.9845240774655 | 6.76155015047068e-05 | 0.00511568779683274 | 1389 | 43 | 32 | 4 | 5 | 3 | 2 | 0 | 0 | 22 | 31 | 2 | 6 |
| L1MA6_dup3465 | 7.09575830337762 | 2.6530785772237 | 0.828558319144442 | 3.20204204812431 | 0.00136457078067613 | 0.0435538777972854 | 1390 | 18 | 15 | 3 | 2 | 1 | 1 | 0 | 0 | 12 | 26 | 2 | 7 |
| L1PA3_dup3766 | 6.52925318903596 | 2.93946255124902 | 0.75923462804277 | 3.87161286205643 | 0.000108117575659125 | 0.00721094694314257 | 1391 | 19 | 16 | 2 | 5 | 3 | 4 | 0 | 0 | 9 | 20 | 2 | 1 |
| Tigger1_dup10645 | 7.91471946791754 | 2.96688136542989 | 0.843254783471047 | 3.51836885314479 | 0.000434208376200599 | 0.0196049613441039 | 1392 | 28 | 32 | 5 | 3 | 2 | 2 | 0 | 0 | 10 | 13 | 2 | 2 |
| HSMAR1_dup633 | 7.38771271167415 | 3.16295274965089 | 0.902244437976622 | 3.50564948534806 | 0.00045549464016432 | 0.0202601922294915 | 1393 | 31 | 36 | 1 | 5 | 1 | 4 | 1 | 0 | 6 | 7 | 0 | 3 |
| L1M1_dup8668 | 10.916360230157 | 2.89618523894941 | 0.849172624867078 | 3.41059656674961 | 0.000648209255013609 | 0.0258894554666196 | 1394 | 51 | 51 | 3 | 7 | 2 | 3 | 1 | 0 | 5 | 13 | 1 | 5 |
| L1MA4_dup8952 | 8.96941444776124 | 3.09154892150513 | 0.82478263638049 | 3.74831959978232 | 0.000178023315069949 | 0.0102690753279113 | 1395 | 39 | 29 | 3 | 6 | 3 | 3 | 0 | 1 | 16 | 9 | 0 | 3 |
| L1MB3_dup5418 | 10.6373534802222 | 2.60947967523968 | 0.821926582521414 | 3.17483304559224 | 0.00149922734197563 | 0.0464862065563896 | 1396 | 50 | 47 | 3 | 8 | 2 | 6 | 0 | 1 | 7 | 7 | 1 | 6 |
| L1MD2_dup2863 | 10.7196141565095 | 2.82723503804376 | 0.635507797645333 | 4.44878103544781 | 8.63589993880015e-06 | 0.00108896547262201 | 1397 | 41 | 32 | 3 | 5 | 4 | 6 | 2 | 1 | 17 | 16 | 2 | 5 |
| MSTA_dup6348 | 8.03530813547285 | 2.33602735850044 | 0.702964841127657 | 3.32310696329153 | 0.000890207726613191 | 0.0324332212091108 | 1398 | 30 | 29 | 6 | 6 | 4 | 3 | 1 | 2 | 10 | 7 | 2 | 1 |
| L1PA15_dup2750 | 10.1018539155022 | 4.42491019553129 | 0.75352813354577 | 5.87225612228919 | 4.29903868766994e-09 | 2.45898812398152e-06 | 1399 | 42 | 29 | 1 | 2 | 13 | 9 | 0 | 0 | 16 | 11 | 1 | 2 |
| L1PA8_dup3546 | 11.7898243162619 | 4.59212865930988 | 0.715368859050444 | 6.41924596131472 | 1.36950985253264e-10 | 1.32467232264525e-07 | 1400 | 47 | 38 | 2 | 1 | 13 | 22 | 0 | 0 | 9 | 15 | 2 | 1 |
| L1PB1_dup6490 | 9.4912167274838 | 4.59230928817618 | 0.847154508034469 | 5.42086389746193 | 5.93117049800471e-08 | 2.2017134092328e-05 | 1401 | 29 | 29 | 2 | 1 | 4 | 6 | 0 | 0 | 19 | 23 | 1 | 1 |
| L1PREC2_dup2140 | 14.1129160055364 | 5.59724429582546 | 0.880010875823122 | 6.36042627381174 | 2.01194608650798e-10 | 1.84128662884703e-07 | 1402 | 48 | 37 | 1 | 0 | 8 | 9 | 0 | 0 | 25 | 41 | 2 | 1 |
| MER34C__dup520 | 7.39052417619353 | 3.58690360301381 | 0.960744043628112 | 3.73346431529087 | 0.000188863962769877 | 0.0106719127250632 | 1403 | 23 | 26 | 0 | 0 | 0 | 3 | 1 | 3 | 14 | 16 | 2 | 1 |
| L1PA4_dup10102 | 9.68999464950095 | 3.35317861010956 | 0.942805851265269 | 3.55659503556275 | 0.000375692610172399 | 0.0176494676841227 | 1404 | 43 | 32 | 2 | 1 | 0 | 1 | 0 | 3 | 18 | 14 | 2 | 3 |
| L2a_dup132839 | 7.16327815525165 | 3.50890448913701 | 1.03626495404873 | 3.38610745777666 | 0.000708916180596277 | 0.0275897562165786 | 1405 | 22 | 37 | 0 | 3 | 0 | 1 | 0 | 1 | 9 | 12 | 1 | 3 |
| L1MEd_dup20872 | 12.4890371632337 | 5.41374987758682 | 1.50067642215339 | 3.60753977184393 | 0.000309114166505571 | 0.0152725248055097 | 1406 | 68 | 25 | 0 | 0 | 0 | 4 | 0 | 0 | 26 | 18 | 4 | 11 |
| L3_dup37943 | 18.3195477397757 | 4.84266706236941 | 1.38159386974267 | 3.50513068161731 | 0.000456383207008847 | 0.0202828835590077 | 1407 | 99 | 94 | 1 | 0 | 0 | 4 | 0 | 1 | 16 | 9 | 3 | 8 |
| L1MB5_dup2339 | 7.97131647086269 | 1.95717283396352 | 0.59313129541154 | 3.29972950188971 | 0.000967780604094517 | 0.0343649490556445 | 1408 | 28 | 23 | 6 | 5 | 5 | 13 | 5 | 1 | 3 | 10 | 0 | 5 |
| L1MEd_dup8662 | 8.44447236745739 | 1.82327442992681 | 0.569122439175879 | 3.20365936118599 | 0.0013569291976388 | 0.043379803152138 | 1409 | 25 | 32 | 8 | 5 | 6 | 9 | 5 | 4 | 4 | 7 | 0 | 3 |
| L1PA8_dup3061 | 9.09684184187327 | 1.91284991048407 | 0.585327334992792 | 3.26800030705489 | 0.00108310248127653 | 0.0371025376875275 | 1410 | 41 | 24 | 4 | 6 | 6 | 8 | 2 | 6 | 4 | 10 | 2 | 4 |
| L1PA3_dup4163 | 8.56107265477709 | 1.94125509134635 | 0.602811339432311 | 3.22033605601132 | 0.0012804040455419 | 0.041683817351281 | 1411 | 36 | 30 | 4 | 3 | 10 | 6 | 4 | 5 | 4 | 3 | 1 | 4 |
| L2a_dup58141 | 8.71215732049662 | 1.92728659050252 | 0.596562269026761 | 3.23065451934585 | 0.00123507125763294 | 0.0406360986544148 | 1412 | 41 | 25 | 9 | 3 | 8 | 7 | 4 | 3 | 4 | 5 | 1 | 3 |
| L2c_dup47661 | 8.30987497964934 | 1.95698291082238 | 0.55923961733724 | 3.49936386864068 | 0.000466369676570671 | 0.0205627678101205 | 1413 | 34 | 22 | 5 | 3 | 12 | 7 | 10 | 2 | 4 | 6 | 2 | 1 |
| L1MA5_dup3338 | 9.99396804166379 | 2.51598088297747 | 0.521903161508524 | 4.82078107307341 | 1.42997221378694e-06 | 0.000272422873003802 | 1414 | 34 | 28 | 6 | 4 | 12 | 9 | 4 | 2 | 7 | 17 | 2 | 2 |
| L2c_dup47663 | 7.26713694203399 | 1.99160670787103 | 0.590991468627544 | 3.36994155346459 | 0.000751841285911724 | 0.0288126311189485 | 1415 | 25 | 21 | 4 | 6 | 7 | 4 | 5 | 1 | 6 | 10 | 3 | 1 |
| Tigger17c_dup259 | 6.58202160282974 | 2.17849011161899 | 0.575784775031027 | 3.78351461533799 | 0.000154629240064651 | 0.00931007316711119 | 1416 | 22 | 18 | 3 | 4 | 7 | 6 | 4 | 1 | 5 | 10 | 3 | 1 |
| MIR_dup44719 | 6.5551857740621 | 1.91078990545958 | 0.557604558255758 | 3.42678315155227 | 0.000610776608700385 | 0.0248855909133257 | 1417 | 12 | 9 | 7 | 3 | 14 | 14 | 2 | 1 | 6 | 9 | 5 | 0 |

| | | | | | | | | | | | | | | | | | | | |
|---|---|---|---|---|---|---|---|---|---|---|---|---|---|---|---|---|---|---|---|
| Arthur1B_dup855 | 6.75377193478453 | 1.77164704260567 | 0.557277142110546 | 3.17911306373703 | 0.0014772645615707 | 0.046021104132954 | 1418 | 10 | 14 | 4 | 7 | 12 | 8 | 1 | 5 | 10 | 8 | 3 | 0 |
| BLACKJACK_dup1919 | 6.64097505181869 | 1.75744765880172 | 0.530264699680167 | 3.31428371502335 | 0.000918782295356161 | 0.0331941792996004 | 1419 | 9 | 15 | 5 | 3 | 10 | 17 | 5 | 5 | 7 | 4 | 2 | 0 |
| MamGyp-int_dup234 | 5.74453857817325 | 1.87071989543165 | 0.552664017525084 | 3.38491350279873 | 0.000712006873322405 | 0.0276693103837291 | 1420 | 11 | 16 | 4 | 4 | 7 | 9 | 5 | 3 | 5 | 7 | 1 | 0 |
| L1MEd_dup20406 | 5.68118803086114 | 2.06772249103287 | 0.61195123173907 | 3.37890077475084 | 0.000727762700316407 | 0.0281026003715494 | 1421 | 14 | 14 | 4 | 0 | 11 | 6 | 3 | 5 | 5 | 6 | 1 | 1 |
| L1PB1_dup4337 | 5.76846100613944 | 1.67908254361792 | 0.510340149183893 | 3.2901243343347 | 0.00100143121997728 | 0.0351299650698384 | 1422 | 13 | 12 | 4 | 3 | 10 | 11 | 5 | 2 | 5 | 4 | 3 | 1 |
| MIR_dup139242 | 4.8647615553138 | 2.4017672456918 | 0.616942665994435 | 3.89301531256628 | 9.90059004593065e-05 | 0.00677150272799372 | 1423 | 12 | 10 | 2 | 1 | 8 | 9 | 2 | 2 | 7 | 4 | 2 | 1 |
| THE1B-int_dup1637 | 5.56583814918063 | 1.94444264739454 | 0.57562479172219 | 3.37796890501716 | 0.000730233374530742 | 0.0281842839474602 | 1424 | 10 | 15 | 4 | 2 | 6 | 8 | 3 | 2 | 10 | 4 | 2 | 2 |
| AluJo_dup21446 | 5.98805744452415 | 1.93085926008565 | 0.612123407016801 | 3.15436272809064 | 0.00160849028836271 | 0.048712464064497 | 1425 | 19 | 15 | 2 | 5 | 8 | 4 | 5 | 3 | 8 | 4 | 2 | 0 |
| L1MCc_dup1879 | 6.96668291839721 | 1.85531771107959 | 0.547532248413758 | 3.3885085608283 | 0.000702738351665499 | 0.0274345824159801 | 1426 | 21 | 20 | 3 | 4 | 8 | 9 | 7 | 2 | 5 | 6 | 4 | 0 |
| MIRc_dup32973 | 7.89015413533989 | 2.95360742399804 | 0.624786965343687 | 4.72738323273646 | 2.27431771748732e-06 | 0.000387376380533457 | 1427 | 30 | 22 | 1 | 2 | 12 | 13 | 1 | 3 | 5 | 7 | 4 | 0 |
| L1M5_dup37955 | 6.04960693186279 | 2.29484895490132 | 0.559900291733769 | 4.09867433323739 | 4.15523205043006e-05 | 0.00354889032832603 | 1428 | 18 | 12 | 2 | 3 | 13 | 10 | 1 | 3 | 5 | 5 | 3 | 1 |
| L1PA7_dup9993 | 6.93649270011716 | 2.50671575836237 | 0.577646711001363 | 4.33953091157903 | 1.42787181706041e-05 | 0.00159061979111543 | 1429 | 20 | 22 | 4 | 2 | 13 | 10 | 1 | 2 | 3 | 7 | 3 | 1 |
| MER20_dup5113 | 6.49550546370473 | 1.98341147561086 | 0.56771515338316 | 3.49367365621862 | 0.000476423016622837 | 0.0208502760922085 | 1430 | 15 | 17 | 1 | 3 | 12 | 9 | 4 | 7 | 3 | 8 | 1 | 1 |
| MamGypLTR1c_dup38 | 6.77592508857165 | 2.26388292483946 | 0.573601014523651 | 3.94679030810206 | 7.92058685444645e-05 | 0.00574244960288883 | 1431 | 21 | 16 | 1 | 5 | 14 | 5 | 5 | 3 | 5 | 9 | 1 | 1 |
| L1PA5_dup756 | 7.04645372142635 | 2.99316245458249 | 0.667937710403597 | 4.48119998012073 | 7.42245202065986e-06 | 0.000973942805839653 | 1432 | 30 | 24 | 1 | 3 | 13 | 6 | 2 | 2 | 2 | 6 | 1 | 1 |
| L1PB1_dup4339 | 6.82629296006611 | 1.9072437933637 | 0.538209595474396 | 3.54368225576244 | 0.000394580478734122 | 0.0182912227922987 | 1433 | 18 | 23 | 3 | 5 | 12 | 9 | 2 | 4 | 2 | 5 | 2 | 2 |
| L2c_dup23051 | 6.22410238942505 | 1.73609849083868 | 0.545993331915852 | 3.17970639814745 | 0.00147424339260174 | 0.0459450318658827 | 1434 | 11 | 16 | 3 | 4 | 8 | 15 | 3 | 2 | 2 | 9 | 1 | 6 |
| AluSx3_dup6133 | 4.70531248332276 | 2.3653199856435 | 0.630080529976459 | 3.75399631176014 | 0.000174037424927854 | 0.0101102317167683 | 1435 | 12 | 8 | 1 | 3 | 9 | 12 | 1 | 2 | 2 | 7 | 1 | 2 |
| L2_dup13911 | 5.35420080667361 | 2.15343719030262 | 0.593202043235872 | 3.63019179528745 | 0.000283210696854124 | 0.0143441576146555 | 1436 | 11 | 12 | 2 | 4 | 7 | 12 | 1 | 2 | 2 | 11 | 2 | 2 |
| AluJb_dup74432 | 6.04549289977388 | 2.12848056821631 | 0.555010982587871 | 3.83502423373996 | 0.000125551914840263 | 0.00801141751906176 | 1437 | 17 | 9 | 1 | 1 | 11 | 9 | 2 | 3 | 5 | 10 | 3 | 5 |
| MLT1K_dup2005 | 6.8500856109547 | 1.87556497184813 | 0.513734349279242 | 3.65084595663012 | 0.000261377978496738 | 0.0135646334724939 | 1438 | 17 | 14 | 2 | 1 | 11 | 8 | 4 | 2 | 8 | 8 | 4 | 7 |
| Charlie10a_dup47 | 6.6803137887248 | 1.62016845314847 | 0.477231169475725 | 3.3949342724792 | 0.00068645078068623 | 0.0269585101851577 | 1439 | 13 | 9 | 1 | 5 | 11 | 12 | 7 | 3 | 8 | 9 | 2 | 4 |
| L2a_dup139020 | 5.46215585797432 | 1.93858122877074 | 0.556651163009111 | 3.48257824216387 | 0.000496610075766763 | 0.0214808909449915 | 1440 | 7 | 8 | 1 | 2 | 10 | 10 | 5 | 2 | 5 | 13 | 3 | 2 |
| L2b_dup25163 | 6.9347613885708 | 1.93650481673823 | 0.540638248211494 | 3.58188645206744 | 0.000341122062139583 | 0.0163977030702758 | 1441 | 18 | 19 | 3 | 4 | 6 | 5 | 4 | 3 | 10 | 9 | 2 | 3 |
| MLT1J1_dup3295 | 8.17287879333571 | 1.68280733570418 | 0.518385471737461 | 3.24624710268972 | 0.00116937292264369 | 0.0391401898229534 | 1442 | 24 | 23 | 2 | 5 | 8 | 7 | 8 | 2 | 9 | 7 | 4 | 5 |
| L2_dup19570 | 6.32631328223335 | 1.90820213466187 | 0.505704363217462 | 3.77335509332219 | 0.000161066786276903 | 0.00958125193957281 | 1443 | 14 | 13 | 4 | 6 | 12 | 9 | 3 | 1 | 7 | 7 | 1 | 3 |
| MER131_dup180 | 6.83679597535241 | 1.63463816761716 | 0.478875150427166 | 3.41349549284198 | 0.000641352193813693 | 0.0257014946241651 | 1444 | 16 | 13 | 3 | 6 | 14 | 8 | 4 | 4 | 7 | 6 | 1 | 4 |
| AluJb_dup17750 | 6.65633662283634 | 1.69119269609592 | 0.516829750611947 | 3.27224331434767 | 0.00106697708956728 | 0.036757337563946 | 1445 | 21 | 13 | 2 | 5 | 10 | 6 | 3 | 2 | 5 | 9 | 5 | 4 |
| MLT1G1_dup1279 | 8.1044010961008 | 1.45793955470966 | 0.44344040995026 | 3.28779137398233 | 0.00100976637627573 | 0.0353026550351607 | 1446 | 16 | 19 | 5 | 9 | 12 | 10 | 3 | 2 | 7 | 10 | 4 | 6 |
| MLT1C_dup6116 | 7.78061246870958 | 1.48848665445228 | 0.463675521145692 | 3.21019028732504 | 0.00132647128631941 | 0.0426928901250588 | 1447 | 16 | 18 | 3 | 7 | 15 | 9 | 7 | 3 | 9 | 4 | 4 | 3 |
| L1MC2_dup1941 | 9.49341193217303 | 1.81180942480395 | 0.451392115587931 | 4.01382603336811 | 5.97423853074704e-05 | 0.00465927945407124 | 1448 | 22 | 17 | 2 | 6 | 25 | 15 | 4 | 4 | 8 | 6 | 5 | 6 |
| L3_dup37030 | 8.51812605998729 | 1.4405982272659 | 0.454654157718224 | 3.16855834882461 | 0.00153196994748702 | 0.0471971330372196 | 1449 | 19 | 13 | 1 | 7 | 20 | 12 | 6 | 4 | 7 | 7 | 7 | 5 |
| LTR9_dup392 | 7.24626890122857 | 1.95852158466028 | 0.594696707888249 | 3.29331163041902 | 0.000990146637780967 | 0.0348832442809342 | 1450 | 30 | 15 | 1 | 3 | 14 | 11 | 4 | 2 | 4 | 2 | 4 | 4 |
| MIR3_dup27960 | 7.24031952401729 | 1.83288266127671 | 0.500045349347062 | 3.66543287257847 | 0.000246920574523842 | 0.0129813882071697 | 1451 | 16 | 18 | 5 | 4 | 19 | 11 | 3 | 2 | 4 | 4 | 4 | 2 |
| MLT1L_dup4354 | 7.21884835511339 | 1.85594026551773 | 0.516092779614697 | 3.59613685528275 | 0.000322977927821754 | 0.0157643704643026 | 1452 | 20 | 18 | 3 | 7 | 12 | 13 | 2 | 2 | 6 | 3 | 4 | 2 |
| L1MA2_dup5241 | 10.9929080190115 | 2.08261819434224 | 0.59169246273234 | 3.51976461678259 | 0.000431929935777546 | 0.0195280259354976 | 1453 | 14 | 11 | 4 | 3 | 52 | 28 | 11 | 4 | 6 | 2 | 3 | 1 |
| Tigger1_dup8115 | 7.89958001381824 | 2.0095308469475 | 0.599341568214894 | 3.35289750205842 | 0.000799703158813065 | 0.0300562577518454 | 1454 | 10 | 6 | 4 | 5 | 34 | 20 | 8 | 4 | 4 | 4 | 1 | 0 |
| L2c_dup59217 | 6.73426208220238 | 1.90373264507277 | 0.605552938954409 | 3.14379226424018 | 0.00166773784746421 | 0.0499913769025849 | 1455 | 12 | 9 | 3 | 5 | 10 | 18 | 3 | 6 | 12 | 2 | 0 | 2 |
| AluSq2_dup48571 | 11.1539376948903 | 2.27252198869183 | 0.446688945795796 | 5.08748203885645 | 3.62848737867089e-07 | 9.35410680179007e-05 | 1456 | 20 | 21 | 8 | 4 | 24 | 36 | 5 | 4 | 8 | 7 | 0 | 4 |
| L1ME5_dup2204 | 7.92715328058627 | 1.62171446780834 | 0.47897774285588 | 3.38578251703086 | 0.000709756090886355 | 0.0276088969759759 | 1457 | 14 | 11 | 4 | 8 | 12 | 17 | 3 | 5 | 6 | 14 | 0 | 6 |
| MIR_dup129970 | 10.8904584820172 | 1.67023258114954 | 0.505509049949794 | 3.30406069152555 | 0.000952951995936827 | 0.0339854789606092 | 1458 | 19 | 24 | 13 | 5 | 18 | 18 | 5 | 9 | 15 | 4 | 2 | 1 |
| L2c_dup107694 | 11.3664675197441 | 1.51340260121013 | 0.473377628141974 | 3.19703025922517 | 0.00138850360002219 | 0.0440871200441526 | 1459 | 7 | 14 | 9 | 15 | 23 | 25 | 7 | 4 | 16 | 15 | 2 | 4 |
| L1MA4A_dup1439 | 8.65041503039373 | 1.58814117915488 | 0.448196919425516 | 3.54340047939309 | 0.000395002364673867 | 0.0183036480359665 | 1460 | 18 | 11 | 10 | 8 | 15 | 19 | 7 | 2 | 8 | 9 | 2 | 1 |
| LTR78B_dup2623 | 9.96118458973361 | 1.36997579904602 | 0.376727184701366 | 3.63651962130636 | 0.000276346602094237 | 0.0141076096507006 | 1461 | 16 | 18 | 9 | 7 | 15 | 18 | 9 | 5 | 10 | 11 | 3 | 4 |
| GSAT_dup12 | 8.66599542384205 | 1.27953338008619 | 0.397707914420101 | 3.21726909043758 | 0.00129417155257065 | 0.0419312287919356 | 1462 | 16 | 14 | 6 | 7 | 14 | 14 | 6 | 6 | 10 | 7 | 4 | 4 |
| Zaphod3_dup201 | 9.82781397291217 | 1.63104774008617 | 0.400891757047001 | 4.06854895720629 | 4.73068221148321e-05 | 0.00390985380164496 | 1463 | 18 | 13 | 5 | 8 | 19 | 20 | 8 | 4 | 13 | 8 | 3 | 4 |
| AluJr_dup59100 | 8.40209778594234 | 1.41570836591706 | 0.440394591732781 | 3.21463612971902 | 0.00130609968871546 | 0.0422494408805948 | 1464 | 12 | 11 | 8 | 8 | 21 | 13 | 6 | 3 | 8 | 10 | 1 | 5 |
| Charlie14a_dup130 | 7.8299274244976 | 1.42366719552467 | 0.443366708757181 | 3.21103765213994 | 0.00132256604643839 | 0.0426017734514456 | 1465 | 12 | 12 | 9 | 6 | 13 | 13 | 3 | 4 | 10 | 9 | 2 | 4 |
| L1PA16_dup12632 | 15.3943968798558 | 1.98727476931366 | 0.540992750862057 | 3.67338520922322 | 0.000239358230348014 | 0.01271018377112 | 1466 | 29 | 24 | 2 | 5 | 30 | 23 | 5 | 0 | 12 | 22 | 18 | 27 |
| LTR8_dup2488 | 10.9498687242225 | 1.48863227716054 | 0.461310956886626 | 3.22696058902932 | 0.00125112691371949 | 0.0410249165514042 | 1467 | 26 | 13 | 5 | 6 | 10 | 23 | 6 | 0 | 13 | 13 | 10 | 15 |
| MIRb_dup192929 | 13.8069606187638 | 1.83820378756811 | 0.493361446587193 | 3.72587643457715 | 0.000194637741027477 | 0.0109049380566339 | 1468 | 25 | 21 | 4 | 7 | 21 | 25 | 4 | 0 | 24 | 9 | 16 | 16 |
| L1MC4a_dup8502 | 13.1025266121979 | 2.22066936782273 | 0.650381603226184 | 3.41440987384516 | 0.00063920337158038 | 0.0256572006501459 | 1469 | 32 | 20 | 5 | 0 | 26 | 26 | 2 | 1 | 9 | 12 | 13 | 22 |
| L1MA4_dup5145 | 10.6967059820714 | 2.32026543114052 | 0.692812961289508 | 3.34905026433382 | 0.000810890833170302 | 0.0303330027023331 | 1470 | 22 | 28 | 1 | 2 | 20 | 20 | 3 | 0 | 8 | 5 | 12 | 16 |
| MER31-int_dup425 | 13.7558542884538 | 2.49404912064947 | 0.701901111043107 | 3.55327706625656 | 0.00038046352007772 | 0.0178138080968936 | 1471 | 42 | 43 | 1 | 1 | 15 | 20 | 4 | 1 | 7 | 10 | 13 | 21 |
| L1MEf_dup3599 | 13.2252285799074 | 2.41626232993078 | 0.504145992765639 | 4.79278297279658 | 1.64483530247199e-06 | 0.00030316187519907 | 1472 | 47 | 36 | 10 | 8 | 15 | 18 | 3 | 0 | 9 | 17 | 3 | 4 |
| L2c_dup47668 | 12.1761630895926 | 1.88277640064135 | 0.474336947512519 | 3.96928050938232 | 7.20899767844551e-05 | 0.00534710773564649 | 1473 | 42 | 25 | 7 | 8 | 11 | 22 | 5 | 1 | 8 | 15 | 3 | 11 |
| L2c_dup97441 | 12.8319942032326 | 2.45084380644365 | 0.515755308235767 | 4.75195071637208 | 2.01463484202738e-06 | 0.000355092075645219 | 1474 | 46 | 32 | 5 | 5 | 15 | 19 | 2 | 1 | 9 | 17 | 3 | 11 |
| AluSq2_dup29420 | 8.04802426408721 | 2.03270159560616 | 0.50373518892877 | 4.03525828705525 | 5.45422641252974e-05 | 0.00435656947443041 | 1475 | 21 | 14 | 5 | 4 | 17 | 15 | 1 | 1 | 5 | 10 | 3 | 7 |
| L2a_dup40044 | 9.45757640405334 | 1.63470855570178 | 0.515379024809617 | 3.17185697711631 | 0.00151467582973849 | 0.0468315440978231 | 1476 | 27 | 12 | 6 | 7 | 10 | 22 | 1 | 1 | 7 | 12 | 6 | 11 |
| L1MA3_dup6579 | 12.3092629865857 | 2.29157208349225 | 0.621804633817005 | 3.68535703798992 | 0.000228382251400238 | 0.01225866076961 | 1477 | 53 | 33 | 3 | 4 | 16 | 10 | 1 | 2 | 8 | 9 | 8 | 12 |
| L1MC3_dup8994 | 10.0148453166352 | 2.18465379875581 | 0.61240996155746 | 3.56730611174233 | 0.000360670063057131 | 0.017115222141225 | 1478 | 35 | 31 | 5 | 3 | 16 | 9 | 1 | 1 | 8 | 5 | 6 | 8 |
| L1PA14_dup2391 | 11.9818590035747 | 2.56833774241591 | 0.52432503534596 | 4.89836946412691 | 9.66352033649255e-07 | 0.000200202324766804 | 1479 | 52 | 27 | 4 | 3 | 17 | 17 | 2 | 2 | 8 | 11 | 6 | 5 |
| AluSx_dup72475 | 8.82677773840439 | 2.00521015780242 | 0.52079712449429 | 3.85027117757138 | 0.000117987116842249 | 0.00765600286450665 | 1480 | 32 | 17 | 5 | 4 | 13 | 11 | 1 | 2 | 9 | 7 | 6 | 5 |
| L1M5_dup19664 | 8.99399620637639 | 1.97109862112165 | 0.581986069039642 | 3.38684845906061 | 0.000707004285851244 | 0.0275479600744166 | 1481 | 34 | 20 | 2 | 5 | 14 | 13 | 1 | 2 | 5 | 6 | 8 | 6 |
| L2a_dup40049 | 16.178579976028 | 2.7665457351474 | 0.594271882635521 | 4.65535357802579 | 3.23425187989105e-06 | 0.000509991582380753 | 1482 | 52 | 33 | 5 | 1 | 22 | 26 | 2 | 2 | 22 | 16 | 18 | 2 |
| L2a_dup164345 | 13.7431061470106 | 3.08632756924877 | 0.654579398466143 | 4.71497816228387 | 2.41736674401396e-06 | 0.000408234739014297 | 1483 | 65 | 40 | 3 | 3 | 13 | 15 | 1 | 0 | 8 | 16 | 6 | 8 |
| LTR16B2_dup1052 | 16.8242836223189 | 3.29956995933237 | 0.695202322911567 | 4.74620099874449 | 2.07272747691041e-06 | 0.00036291185593887 | 1484 | 61 | 63 | 3 | 2 | 22 | 20 | 1 | 0 | 14 | 9 | 12 | 7 |
| Charlie13a_dup418 | 11.9772425984387 | 3.01939197883133 | 0.628185517965991 | 4.80652911039378 | 1.53573130634121e-06 | 0.000288414460472507 | 1485 | 37 | 44 | 1 | 3 | 11 | 18 | 1 | 2 | 16 | 6 | 3 | 8 |
| L1M4_dup14209 | 12.0099720536554 | 2.53594564790733 | 0.664136718978632 | 3.81840903452429 | 0.000134315093082115 | 0.00839058522932971 | 1486 | 48 | 36 | 0 | 6 | 12 | 19 | 2 | 1 | 9 | 6 | 8 | 8 |
| L1PBa_dup1864 | 12.9395524845444 | 1.99708789442392 | 0.48373322575461 | 4.12849022580271 | 3.65152958758994e-05 | 0.00320970394993969 | 1487 | 28 | 29 | 2 | 1 | 17 | 29 | 10 | 9 | 11 | 15 | 2 | 9 |
| L2_dup47548 | 15.3682132395786 | 1.818725371707 | 0.508104224912977 | 3.57943367233069 | 0.000344339629567471 | 0.0165056884562976 | 1488 | 33 | 23 | 2 | 2 | 22 | 47 | 18 | 5 | 11 | 16 | 4 | 14 |
| L3b_dup5090 | 12.47348814912 | 2.70027622975019 | 0.546947261309998 | 4.93699561322006 | 7.93352548776205e-07 | 0.00017255490454397 | 1489 | 37 | 38 | 1 | 0 | 22 | 23 | 6 | 6 | 10 | 7 | 2 | 5 |
| MLT1I_dup8883 | 10.7568678281492 | 2.01647795157479 | 0.497971141658222 | 4.04938716902349 | 5.13519286728253e-05 | 0.00416461691206002 | 1490 | 30 | 19 | 3 | 0 | 14 | 15 | 7 | 8 | 16 | 11 | 4 | 5 |
| MER67D_dup700 | 12.4630334101008 | 1.97362572340666 | 0.463993495889051 | 4.25356333847961 | 2.10395390860337e-05 | 0.00211771326876708 | 1491 | 29 | 19 | 4 | 3 | 21 | 24 | 12 | 11 | 12 | 16 | 0 | 4 |
| L1PA5_dup9867 | 10.7705932330119 | 1.8012045929056 | 0.520098289106102 | 3.46320038083829 | 0.000533790713557232 | 0.0225606934097291 | 1492 | 22 | 26 | 2 | 6 | 15 | 13 | 17 | 2 | 8 | 20 | 1 | 6 |
| L2c_dup101876 | 7.80174378445448 | 2.15958508824117 | 0.554672777530059 | 3.89343983647032 | 9.88327221290403e-05 | 0.0067635464192455 | 1493 | 20 | 17 | 2 | 4 | 7 | 10 | 9 | 3 | 9 | 15 | 0 | 2 |
| L1PB1_dup6261 | 10.1762921107093 | 1.74978892235452 | 0.452365319731224 | 3.86808812707872 | 0.000109692024054244 | 0.00728895891770291 | 1494 | 21 | 26 | 3 | 4 | 10 | 9 | 8 | 2 | 11 | 19 | 6 | 9 |
| L1MA1_dup3205 | 10.0393822101391 | 1.9151773998181 | 0.512286922913923 | 3.73848582533484 | 0.000185131928852835 | 0.010536092308734 | 1495 | 26 | 26 | 4 | 2 | 7 | 6 | 6 | 4 | 12 | 20 | 4 | 8 |
| L1MA5A_dup2323 | 10.7308885534369 | 2.20677054005273 | 0.536873822893121 | 4.11040815542248 | 3.94960356628598e-05 | 0.00340504481950538 | 1496 | 30 | 37 | 3 | 3 | 8 | 7 | 7 | 5 | 11 | 16 | 3 | 4 |

| | | | | | | | | | | | | | | | | | | | |
|---|---|---|---|---|---|---|---|---|---|---|---|---|---|---|---|---|---|---|---|
| L1M1_dup6856 | 10.1723219785516 | 1.53521588237595 | 0.427962730965568 | 3.58726536516907 | 0.000334164199629648 | 0.0161523898134645 | 1497 | 26 | 13 | 4 | 3 | 14 | 12 | 5 | 6 | 9 | 19 | 5 | 12 |
| L2a_dup96123 | 9.32881168302067 | 1.49026660909009 | 0.426463687736883 | 3.49447479807366 | 0.000474995452439624 | 0.0208184321101858 | 1498 | 17 | 10 | 2 | 3 | 12 | 11 | 6 | 7 | 11 | 22 | 5 | 9 |
| L2a_dup134699 | 10.156511346858 | 1.22694081442021 | 0.38770974647673 | 3.16458594494954 | 0.00155303787166129 | 0.0475892907099108 | 1499 | 26 | 13 | 5 | 5 | 14 | 14 | 9 | 6 | 11 | 10 | 7 | 8 |
| MLT1A_dup1639 | 9.95815186807829 | 1.35358425341909 | 0.391575556043764 | 3.45676391829681 | 0.000546703573654867 | 0.0229468916099631 | 1500 | 21 | 19 | 7 | 3 | 14 | 15 | 8 | 4 | 8 | 12 | 7 | 8 |
| L1MD_dup7225 | 10.6662952914581 | 1.7031292796845 | 0.436626983640705 | 3.90065054038433 | 9.59345488419155e-05 | 0.0066108433706164 | 1501 | 25 | 19 | 4 | 3 | 23 | 12 | 10 | 6 | 7 | 16 | 2 | 8 |
| L1PREC2_dup3600 | 11.1760738619042 | 1.39321818612873 | 0.409664070884256 | 3.40087961124215 | 0.000671694022254739 | 0.026536095935485 | 1502 | 23 | 15 | 6 | 2 | 14 | 16 | 11 | 9 | 11 | 20 | 4 | 8 |
| THE1B_dup625 | 9.73943973053766 | 1.72631085721325 | 0.505158109808239 | 3.41736740179935 | 0.000632298867318117 | 0.0254505833552071 | 1503 | 13 | 19 | 9 | 4 | 9 | 14 | 2 | 5 | 21 | 12 | 1 | 8 |
| AluJr_dup61992 | 8.17846329683872 | 1.37771549330947 | 0.434969898712172 | 3.16738123118062 | 0.00153818526572535 | 0.0473055531490971 | 1504 | 12 | 10 | 5 | 3 | 10 | 12 | 5 | 7 | 14 | 12 | 3 | 6 |
| L1PB1_dup6980 | 9.49391970464801 | 1.41057555153627 | 0.411249988212473 | 3.429971044297 | 0.000603645663202089 | 0.0246880493255903 | 1505 | 19 | 14 | 5 | 8 | 13 | 9 | 4 | 6 | 15 | 13 | 3 | 8 |
| MER58A_dup5945 | 7.11346983151719 | 1.57708053805343 | 0.499981439870019 | 3.15427816373229 | 0.00160895648526078 | 0.0487185817676604 | 1506 | 10 | 9 | 3 | 2 | 7 | 15 | 2 | 3 | 7 | 16 | 4 | 11 |
| MIR1_Amn_dup9647 | 8.42168466037849 | 1.54583683712933 | 0.457951722075783 | 3.37554541802449 | 0.000736695313463953 | 0.0283738593488983 | 1507 | 11 | 8 | 4 | 1 | 8 | 18 | 4 | 4 | 14 | 15 | 6 | 10 |
| AluSx1_dup29074 | 7.8504829124278 | 1.62691502099142 | 0.468560804358317 | 3.47215346622824 | 0.000516301097549435 | 0.0220559750372528 | 1508 | 16 | 15 | 4 | 4 | 6 | 22 | 5 | 4 | 7 | 8 | 4 | 4 |
| L1MA4_dup4482 | 8.50855554710462 | 1.6359797831377 | 0.436594996689755 | 3.74713360332031 | 0.000178866828604312 | 0.0103002532427594 | 1509 | 19 | 11 | 5 | 4 | 9 | 16 | 7 | 2 | 8 | 17 | 3 | 7 |
| L3_dup17154 | 8.83573684084 | 1.38225562246663 | 0.431945700703351 | 3.20006801830845 | 0.00137395158783195 | 0.0437769529349572 | 1510 | 18 | 12 | 8 | 6 | 7 | 16 | 4 | 2 | 8 | 18 | 5 | 8 |
| L1M3c_dup295 | 13.8411577399098 | 1.83500318771946 | 0.478672350730111 | 3.83352659688942 | 0.000126319135873208 | 0.00804096658761006 | 1511 | 44 | 44 | 7 | 6 | 13 | 17 | 6 | 9 | 8 | 12 | 1 | 9 |
| L1PA2_dup1195 | 10.3124868804506 | 1.51560581495454 | 0.461202582867997 | 3.28620409176749 | 0.00101547404786326 | 0.0354553173577214 | 1512 | 30 | 25 | 4 | 11 | 11 | 19 | 3 | 7 | 6 | 7 | 2 | 7 |
| AluJr_dup23333 | 13.5185582340042 | 1.44337315650779 | 0.40527831649271 | 3.56143691327673 | 0.000368830736240514 | 0.0174026964833403 | 1513 | 33 | 36 | 6 | 12 | 19 | 24 | 7 | 8 | 8 | 7 | 2 | 11 |
| L3_dup15434 | 13.0471841473689 | 1.35288203355997 | 0.361932010401647 | 3.73794523468272 | 0.000185530345676119 | 0.0105511958968092 | 1514 | 32 | 29 | 9 | 13 | 15 | 21 | 6 | 8 | 9 | 12 | 4 | 8 |
| L1PA6_dup4677 | 9.56838635753278 | 1.86863739102026 | 0.47312251797428 | 3.94958455797246 | 7.82869516977217e-05 | 0.00568786970806478 | 1515 | 30 | 18 | 4 | 3 | 9 | 16 | 8 | 5 | 13 | 7 | 2 | 5 |
| L3b_dup3404 | 11.9275358503661 | 1.93370262737985 | 0.4793731016618 | 4.03381545747239 | 5.48784226395349e-05 | 0.00437457325071583 | 1516 | 43 | 32 | 7 | 6 | 9 | 14 | 5 | 5 | 13 | 8 | 3 | 6 |
| AluSg4_dup4919 | 10.8468356413331 | 1.9011247697067 | 0.433077861114862 | 4.38979901861683 | 1.13455471181852e-05 | 0.00134006315515524 | 1517 | 28 | 26 | 6 | 8 | 20 | 17 | 4 | 3 | 10 | 7 | 1 | 8 |
| L1PA4_dup4587 | 9.44792335459919 | 2.30824278374757 | 0.525295680756751 | 4.39417811397625 | 1.11192623596068e-05 | 0.00132157043027922 | 1518 | 31 | 31 | 3 | 4 | 13 | 10 | 2 | 4 | 6 | 9 | 2 | 5 |
| MIR_dup57368 | 9.5767383584205 | 2.06996771745728 | 0.517573729864895 | 3.99936781566872 | 6.35119085237191e-05 | 0.00487969150325633 | 1519 | 27 | 32 | 3 | 4 | 9 | 11 | 5 | 4 | 9 | 9 | 1 | 7 |
| MER117_dup3101 | 8.65214903631864 | 1.4134582633562 | 0.440442027350617 | 3.20918117614377 | 0.00133113584392734 | 0.0428140645470579 | 1520 | 23 | 13 | 6 | 5 | 15 | 14 | 8 | 4 | 3 | 12 | 6 | 2 |
| L1MC2_dup1938 | 8.46109374278142 | 1.5405852900558 | 0.47420500819744 | 3.24877481980191 | 0.0011590318943292 | 0.0389085543089673 | 1521 | 23 | 21 | 9 | 6 | 14 | 10 | 3 | 6 | 4 | 7 | 3 | 1 |
| LTR16E2_dup895 | 9.61147070641377 | 1.57461529267687 | 0.432721024198471 | 3.63886939765298 | 0.000273837622283911 | 0.0140095822128505 | 1522 | 19 | 23 | 8 | 7 | 13 | 17 | 5 | 9 | 6 | 10 | 2 | 1 |
| MER5A_dup11340 | 10.9131939579272 | 1.37976461873565 | 0.422889501410583 | 3.26270719451142 | 0.00110353472244293 | 0.0375681252524023 | 1523 | 32 | 22 | 8 | 9 | 13 | 10 | 9 | 2 | 11 | 11 | 4 | 9 |
| MER5B_dup4221 | 10.0963675750175 | 1.46202291630324 | 0.438220493349837 | 3.33627235259417 | 0.000849099365840491 | 0.0313677686235493 | 1524 | 29 | 16 | 9 | 8 | 16 | 7 | 5 | 2 | 11 | 13 | 6 | 6 |
| L1MD1_dup2143 | 12.7106594451683 | 1.31880342388761 | 0.409912520364955 | 3.21728017166553 | 0.00129412156429949 | 0.0419312287919356 | 1525 | 37 | 25 | 14 | 13 | 21 | 13 | 9 | 2 | 6 | 14 | 5 | 6 |
| MIRb_dup166118 | 10.1867884899368 | 1.41648461431521 | 0.379637288630851 | 3.73115248879718 | 0.000190605802209011 | 0.0107298041547547 | 1526 | 21 | 16 | 8 | 8 | 19 | 21 | 8 | 2 | 6 | 11 | 5 | 6 |
| MamRep38_dup671 | 11.2290974727624 | 1.23550014892033 | 0.382325670387401 | 3.23153856676281 | 0.00123125707183855 | 0.0405403638233296 | 1527 | 28 | 20 | 13 | 8 | 20 | 17 | 5 | 7 | 5 | 10 | 5 | 5 |
| HERV9N-int_dup85 | 13.5810377646689 | 1.22884323500859 | 0.365248398623113 | 3.36440416889164 | 0.000767091185002514 | 0.0291761955093683 | 1528 | 38 | 30 | 13 | 14 | 22 | 17 | 10 | 6 | 6 | 9 | 7 | 3 |
| MIR_dup57364 | 12.3874373797806 | 1.27451730342568 | 0.360097148845115 | 3.53937071568947 | 0.000401082175006429 | 0.0185132685187352 | 1529 | 34 | 25 | 7 | 9 | 17 | 19 | 11 | 7 | 6 | 11 | 7 | 6 |
| L1P4_dup1384 | 11.0117418954059 | 1.47097288837778 | 0.4202215200547 | 3.50047015247174 | 0.000464438245031288 | 0.0205068143722796 | 1530 | 31 | 24 | 8 | 6 | 10 | 15 | 6 | 7 | 6 | 16 | 4 | 7 |
| MLT1A_dup3090 | 12.8496804675406 | 1.36257529573409 | 0.406316989906204 | 3.35347851451801 | 0.000798026092401151 | 0.030016932750946 | 1531 | 45 | 21 | 10 | 10 | 16 | 13 | 6 | 6 | 8 | 15 | 7 | 8 |
| L2c_dup47667 | 12.6143868570441 | 1.60169197542067 | 0.472996929748752 | 3.38626294312621 | 0.000708514607191483 | 0.0275831506418169 | 1532 | 38 | 37 | 6 | 13 | 7 | 16 | 9 | 3 | 7 | 16 | 6 | 5 |
| AluSx_dup32535 | 9.72004992531737 | 1.64583076116742 | 0.410824130560467 | 4.00616867106149 | 6.17115884490705e-05 | 0.00478151274018318 | 1533 | 24 | 19 | 4 | 9 | 11 | 17 | 9 | 4 | 8 | 13 | 2 | 4 |
| MIR3_dup27961 | 12.2981206698144 | 1.39859992447173 | 0.367096714202247 | 3.80989496871714 | 0.000139025841034062 | 0.00861026620788416 | 1534 | 25 | 24 | 8 | 8 | 11 | 26 | 12 | 7 | 9 | 16 | 4 | 6 |
| L1ME3E_dup1531 | 17.6154632874706 | 2.47235586342512 | 0.579796181125052 | 4.26418100689745 | 2.00636752549595e-05 | 0.00204632287707526 | 1535 | 74 | 60 | 8 | 13 | 8 | 8 | 5 | 4 | 16 | 22 | 2 | 5 |
| L1MD3_dup1530 | 12.2345394003106 | 2.16310452214469 | 0.515086287017846 | 4.19949933955386 | 2.6750581051891e-05 | 0.00253365714629041 | 1536 | 35 | 29 | 6 | 14 | 16 | 8 | 3 | 2 | 19 | 15 | 2 | 4 |
| L2_dup19564 | 13.1538109816838 | 1.88968660922061 | 0.492917826568155 | 3.83367471689386 | 0.000126243059349345 | 0.00803827428416893 | 1537 | 45 | 32 | 10 | 7 | 21 | 5 | 6 | 5 | 11 | 16 | 2 | 7 |
| L1ME3C_dup1040 | 12.5593046802298 | 1.62621661664934 | 0.448147286597913 | 3.62875479843844 | 0.000284791577688368 | 0.0144027665769684 | 1538 | 27 | 38 | 9 | 13 | 20 | 10 | 6 | 2 | 12 | 11 | 7 | 4 |
| MLT1A_dup3091 | 14.5105480456928 | 1.48567438729687 | 0.397373631505714 | 3.73873420253731 | 0.000184949143942187 | 0.0105351330167045 | 1539 | 34 | 34 | 7 | 19 | 22 | 22 | 8 | 2 | 13 | 10 | 9 | 6 |
| L1PA15_dup2701 | 17.2414733572649 | 1.74970193293795 | 0.455366497416137 | 3.84240374042929 | 0.000121835203058148 | 0.00783095603103028 | 1540 | 48 | 42 | 11 | 21 | 18 | 13 | 9 | 4 | 13 | 31 | 7 | 3 |
| THE1C-int_dup624 | 18.1482163872114 | 1.99438013232331 | 0.463749295471066 | 4.30055668396748 | 1.70369584011832e-05 | 0.00181543936575367 | 1541 | 58 | 43 | 10 | 20 | 18 | 19 | 3 | 5 | 21 | 21 | 8 | 3 |
| AluJo_dup21449 | 15.2020117463937 | 1.62276126614334 | 0.400355904527439 | 4.05329669874302 | 5.050088761062e-05 | 0.00411101067512712 | 1542 | 35 | 40 | 7 | 8 | 23 | 13 | 10 | 6 | 20 | 10 | 7 | 11 |
| MSTB1_dup1495 | 15.5892775252021 | 1.56311568230545 | 0.44023936464358 | 3.55060407551459 | 0.00038434813728726 | 0.0179568234586598 | 1543 | 51 | 40 | 12 | 10 | 20 | 9 | 10 | 4 | 15 | 12 | 6 | 10 |
| L2b_dup31121 | 15.4386689638374 | 1.35213199772378 | 0.418496344765098 | 3.2309290502471 | 0.00123388563895892 | 0.0406083198129627 | 1544 | 47 | 38 | 8 | 13 | 13 | 13 | 7 | 11 | 14 | 14 | 7 | 10 |
| L3_dup15435 | 15.7451328894841 | 1.53304540444974 | 0.394229250781821 | 3.88871551618622 | 0.00010077615908084 | 0.00685906291437348 | 1545 | 54 | 34 | 11 | 12 | 17 | 14 | 9 | 8 | 11 | 18 | 7 | 6 |
| L4_A_Mam_dup2186 | 15.4222672250154 | 1.41237995890304 | 0.418959390818145 | 3.37116195473013 | 0.000748518396475261 | 0.0287130353647755 | 1546 | 45 | 36 | 12 | 14 | 11 | 12 | 9 | 9 | 17 | 17 | 6 | 6 |
| L1ME1_dup20213 | 14.3021802894041 | 1.70973554038746 | 0.445634511598629 | 3.83663180451197 | 0.000124733264884444 | 0.00796345088440942 | 1547 | 45 | 36 | 5 | 6 | 14 | 10 | 12 | 6 | 17 | 14 | 8 | 7 |
| MSTA_dup6516 | 13.086544714085 | 1.70007517843135 | 0.421441441145619 | 4.0339535044536 | 5.48461749626399e-05 | 0.00437391430415276 | 1548 | 35 | 38 | 4 | 9 | 12 | 15 | 7 | 7 | 12 | 13 | 6 | 7 |
| L1PA5_dup3882 | 17.6797414178757 | 2.01146225155454 | 0.399141482386609 | 5.03947181717945 | 4.66818361944223e-07 | 0.000113808977408996 | 1549 | 51 | 56 | 10 | 9 | 25 | 18 | 12 | 10 | 11 | 16 | 2 | 4 |
| MLT1L_dup3966 | 17.3643017148179 | 1.35742387498954 | 0.412657918046088 | 3.28946523410206 | 0.00100377956206605 | 0.0351729810384137 | 1550 | 35 | 55 | 10 | 13 | 14 | 18 | 24 | 9 | 12 | 20 | 5 | 6 |
| L1ME3C_dup1039 | 17.8758480962073 | 1.40282386320954 | 0.356720349680357 | 3.93255911659247 | 8.40463015505358e-05 | 0.00599654610756935 | 1551 | 51 | 29 | 10 | 18 | 28 | 14 | 18 | 8 | 16 | 23 | 6 | 7 |
| L1PA6_dup4676 | 17.7018631124621 | 2.00805123645479 | 0.374747328104622 | 5.35841375203664 | 8.39557982514441e-08 | 2.88684196108932e-05 | 1552 | 53 | 34 | 8 | 10 | 28 | 14 | 11 | 8 | 19 | 27 | 5 | 5 |
| L2c_dup47676 | 14.4244056674967 | 1.36714129293682 | 0.414950476336131 | 3.29470953981835 | 0.000985234596006802 | 0.0347668146494787 | 1553 | 32 | 28 | 13 | 12 | 17 | 12 | 14 | 9 | 16 | 20 | 6 | 1 |
| L1MC5_dup4604 | 16.6729222818087 | 1.58634173219773 | 0.362837408142391 | 4.37204570586943 | 1.2308774847653e-05 | 0.00142244960655641 | 1554 | 46 | 37 | 12 | 17 | 21 | 19 | 9 | 9 | 14 | 18 | 6 | 3 |
| MER44A_dup817 | 15.9901192813103 | 1.56590913009694 | 0.422847710223434 | 3.70324609129255 | 0.000212858215612536 | 0.0116220194979671 | 1555 | 43 | 40 | 8 | 13 | 15 | 14 | 14 | 13 | 13 | 21 | 4 | 3 |
| L2a_dup73107 | 13.1954456677199 | 1.59149840282184 | 0.4806078890001 | 3.31142796289287 | 0.000928211348311596 | 0.0334085883354033 | 1556 | 40 | 25 | 4 | 2 | 19 | 27 | 9 | 6 | 9 | 7 | 8 | 13 |
| L1PREC2_dup919 | 10.9762300688544 | 1.42573732745806 | 0.450697365314073 | 3.16340284453298 | 0.00155936390029521 | 0.0476912808759073 | 1557 | 26 | 26 | 3 | 6 | 16 | 18 | 9 | 3 | 9 | 6 | 11 | 7 |
| MLT1D_dup15946 | 12.684872126648 | 1.18953777388766 | 0.36934674586535 | 3.22065318621034 | 0.00127898819803534 | 0.0416434216981004 | 1558 | 30 | 26 | 7 | 11 | 19 | 17 | 11 | 4 | 12 | 7 | 9 | 9 |
| L1M2a_dup382 | 14.1780883118945 | 1.57105612537005 | 0.419099918787622 | 3.74864335434572 | 0.000177793702594223 | 0.0102608053256088 | 1559 | 28 | 24 | 7 | 11 | 30 | 29 | 5 | 3 | 10 | 13 | 4 | 18 |
| MLT1D-int_dup510 | 12.4823993818992 | 1.43282955210343 | 0.43430375908006 | 3.29914149289992 | 0.000969810153347617 | 0.0344113401056445 | 1560 | 25 | 26 | 1 | 10 | 28 | 20 | 7 | 8 | 6 | 11 | 6 | 11 |
| L1PBa_dup1862 | 15.2674307672979 | 1.4270571882644 | 0.410604690105003 | 3.47550143155807 | 0.000509899271179303 | 0.0218609895459514 | 1561 | 32 | 23 | 5 | 4 | 23 | 28 | 21 | 5 | 9 | 25 | 7 | 14 |
| MER31A_dup1080 | 15.6437443509332 | 1.51836932835883 | 0.36899412906303 | 4.11488749757171 | 3.87368300570081e-05 | 0.00335539634683104 | 1562 | 35 | 33 | 8 | 4 | 25 | 19 | 15 | 5 | 11 | 22 | 9 | 13 |
| AluSx1_dup51184 | 16.2611539570725 | 1.2298103054841 | 0.339753480006715 | 3.61971363901791 | 0.000294929223895259 | 0.0147928391882338 | 1563 | 24 | 25 | 6 | 15 | 28 | 29 | 14 | 3 | 15 | 20 | 9 | 20 |
| L2_dup43906 | 17.2039224626589 | 1.39195483510332 | 0.344654066132538 | 4.03870132948915 | 5.37479497798522e-05 | 0.00431560351923593 | 1564 | 32 | 30 | 5 | 19 | 30 | 33 | 14 | 3 | 16 | 15 | 12 | 11 |
| AluJo_dup38610 | 14.0649537856343 | 1.40245090107503 | 0.320947534244978 | 4.36972013003391 | 1.24405875803312e-05 | 0.00143514303733388 | 1565 | 25 | 23 | 6 | 8 | 26 | 22 | 13 | 7 | 13 | 17 | 8 | 9 |
| L1ME4a_dup32374 | 15.5668419226235 | 1.0309415822811 | 0.321023453087959 | 3.21142138483766 | 0.00132080103458741 | 0.0425679700631091 | 1566 | 27 | 31 | 9 | 10 | 23 | 15 | 15 | 8 | 12 | 21 | 12 | 14 |
| LTR84b_dup201 | 17.0521905584743 | 1.37167943963046 | 0.37395804776676 | 3.66800353093614 | 0.000244451809083884 | 0.0129086396281123 | 1567 | 21 | 43 | 7 | 8 | 32 | 22 | 11 | 7 | 16 | 16 | 17 | 13 |
| L2c_dup131837 | 15.2633847823024 | 1.03887725693942 | 0.326367385514449 | 3.18315279972553 | 0.00145680713235719 | 0.0455700190472277 | 1568 | 27 | 26 | 9 | 7 | 20 | 27 | 13 | 8 | 13 | 14 | 14 | 15 |
| L2c_dup64705 | 15.7093717517582 | 1.1885581134671 | 0.327788652479389 | 3.62598919906732 | 0.000287857386357586 | 0.014526938208747 | 1569 | 23 | 29 | 9 | 8 | 24 | 24 | 13 | 4 | 13 | 21 | 16 | 15 |
| MIR3_dup67409 | 12.9854642991782 | 1.37641616110093 | 0.342692745073808 | 4.01647300938479 | 5.90756263196984e-05 | 0.00461332752880439 | 1570 | 27 | 16 | 9 | 11 | 19 | 18 | 11 | 6 | 18 | 16 | 6 | 5 |
| MSTC_dup2903 | 11.4751976019949 | 1.32207410836758 | 0.389752715787124 | 3.3920844033058 | 0.000693630697240024 | 0.0271547630611179 | 1571 | 24 | 9 | 5 | 8 | 20 | 17 | 9 | 6 | 14 | 16 | 11 | 4 |
| MLT1G1-int_dup237 | 15.602565467813 | 1.097832758156 | 0.34309946510606 | 3.19975071315437 | 0.00137546499370154 | 0.043805586956091 | 1572 | 35 | 32 | 9 | 10 | 28 | 18 | 20 | 7 | 9 | 13 | 11 | 8 |
| THE1D_dup4448 | 15.9202897061163 | 1.55175844900084 | 0.371104898582917 | 4.18145504121965 | 2.89649529306166e-05 | 0.00268697648734055 | 1573 | 47 | 30 | 6 | 13 | 30 | 19 | 14 | 7 | 9 | 16 | 10 | 3 |
| L1ME3A_dup11561 | 13.0853098790419 | 1.05889162577322 | 0.333662737207573 | 3.17353874944232 | 0.00150592798528175 | 0.0466271360293886 | 1574 | 30 | 20 | 10 | 11 | 17 | 22 | 9 | 10 | 12 | 9 | 7 | 8 |
| MamGypsy2-I_dup818 | 14.2070453174894 | 1.15245538003012 | 0.345223956146093 | 3.33828333611477 | 0.000842977433152063 | 0.0312143025690633 | 1575 | 38 | 22 | 10 | 13 | 18 | 17 | 12 | 12 | 13 | 13 | 6 | 5 |

| | | | | | | | | | | | | | | | | | | | |
|---|---|---|---|---|---|---|---|---|---|---|---|---|---|---|---|---|---|---|---|
| L2c_dup46657 | 13.5169437161942 | 1.25724068378426 | 0.344209069495689 | 3.65254955549626 | 0.000259649467814124 | 0.0134995606015218 | 1576 | 25 | 17 | 5 | 14 | 28 | 26 | 9 | 10 | 9 | 14 | 7 | 7 |
| MER103C_dup6759 | 12.1999909265388 | 1.52248325093237 | 0.360153504815651 | 4.22731760367478 | 2.36493742167463e-05 | 0.00231575061620491 | 1577 | 29 | 21 | 5 | 12 | 27 | 18 | 11 | 5 | 9 | 10 | 4 | 5 |
| AluSz_dup42778 | 12.391887995537 | 1.59189354714276 | 0.469927827046447 | 3.38752773409483 | 0.000705255864403141 | 0.0275040126812403 | 1578 | 26 | 16 | 4 | 1 | 18 | 11 | 3 | 5 | 12 | 25 | 15 | 19 |
| L1MC2_dup2041 | 12.2227960542448 | 1.16481915944446 | 0.349038090831744 | 3.33722648055042 | 0.000846189641100169 | 0.0313088851409144 | 1579 | 16 | 10 | 4 | 5 | 14 | 12 | 5 | 5 | 20 | 26 | 15 | 18 |
| MLT2A2_dup1396 | 15.0328755501238 | 1.32607582596071 | 0.412373972129645 | 3.21571174609393 | 0.00130121460452308 | 0.0421192776884571 | 1580 | 23 | 13 | 3 | 3 | 23 | 16 | 6 | 5 | 21 | 24 | 22 | 28 |
| L2c_dup127336 | 16.1282593124598 | 1.59720653430737 | 0.400920229605273 | 3.98385119124547 | 6.78073421213369e-05 | 0.00512574933197374 | 1581 | 28 | 27 | 7 | 11 | 24 | 8 | 2 | 5 | 23 | 34 | 13 | 17 |
| AluSz_dup12320 | 13.0177845703346 | 1.36335789080586 | 0.38927038027476 | 3.50234171385851 | 0.000461187718919001 | 0.0204149852232464 | 1582 | 22 | 15 | 8 | 10 | 19 | 10 | 3 | 3 | 26 | 18 | 14 | 11 |
| MLT1D_dup653 | 11.9423456081914 | 1.43445062946462 | 0.404236886139884 | 3.54853967722343 | 0.000387373631737818 | 0.0180679501272313 | 1583 | 18 | 13 | 5 | 13 | 24 | 16 | 2 | 3 | 16 | 18 | 10 | 11 |
| AluJr_dup74018 | 12.0144947720527 | 1.66660325199297 | 0.384519747779788 | 4.33424619051664 | 1.4626042362853e-05 | 0.0016208237693096 | 1584 | 20 | 21 | 4 | 6 | 12 | 16 | 3 | 4 | 17 | 23 | 7 | 16 |
| AluSz_dup89930 | 12.7542048069009 | 1.57762231463072 | 0.44696438448945 | 3.52963763865162 | 0.000416129187845991 | 0.0190351623914442 | 1585 | 21 | 28 | 4 | 5 | 10 | 14 | 2 | 4 | 16 | 22 | 12 | 21 |
| MER68_dup933 | 12.2869600727708 | 1.50376317641914 | 0.39872386324795 | 3.77144012442518 | 0.000162308106449991 | 0.00963827419449703 | 1586 | 20 | 20 | 4 | 3 | 18 | 18 | 4 | 6 | 14 | 18 | 10 | 18 |
| MIRc_dup90365 | 10.5765648184912 | 1.51113067894082 | 0.425669514941216 | 3.55000916415051 | 0.000385217741322882 | 0.0179868564907407 | 1587 | 24 | 15 | 4 | 4 | 11 | 13 | 2 | 4 | 13 | 17 | 12 | 13 |
| L1ME3Cz_dup6836 | 10.4324814260143 | 1.30707534685244 | 0.404166822407349 | 3.2339996120094 | 0.00122069621436185 | 0.0402968620175563 | 1588 | 16 | 18 | 5 | 4 | 15 | 13 | 3 | 5 | 13 | 13 | 13 | 11 |
| MIRc_dup29456 | 11.2452793724462 | 1.19617407689794 | 0.368889347918338 | 3.24263653490679 | 0.00118429188109756 | 0.0394619123287551 | 1589 | 17 | 16 | 5 | 7 | 14 | 14 | 3 | 6 | 14 | 18 | 14 | 11 |
| HERVL-int_dup331 | 12.3411240813595 | 1.49237317740212 | 0.416642753903561 | 3.58190119333638 | 0.0003411028097879 | 0.0163977030702758 | 1590 | 18 | 21 | 4 | 5 | 10 | 6 | 3 | 6 | 26 | 24 | 13 | 12 |
| THE1C-int_dup79 | 13.5662519051483 | 1.71016079359153 | 0.38424565673115 | 4.45069648448387 | 8.55922410057629e-06 | 0.0010822421789763 | 1591 | 23 | 16 | 4 | 3 | 15 | 10 | 5 | 6 | 31 | 25 | 14 | 10 |
| AluJr_dup74020 | 11.9127409106258 | 2.02935889026039 | 0.481769224005708 | 4.21230495669093 | 2.52777956620114e-05 | 0.00243216755624463 | 1592 | 23 | 28 | 3 | 4 | 11 | 7 | 2 | 2 | 21 | 21 | 11 | 13 |
| ERVL-B4-int_dup748 | 13.4701648534516 | 2.15893305479513 | 0.456128163717863 | 4.73317200410922 | 2.21038022361933e-06 | 0.000378927329026892 | 1593 | 28 | 26 | 5 | 2 | 18 | 8 | 2 | 3 | 19 | 31 | 9 | 15 |
| Tigger15a_dup4456 | 11.9999171163241 | 1.94270167561699 | 0.47635116902792 | 4.07829727715672 | 4.53667353861321e-05 | 0.00379947246968682 | 1594 | 24 | 22 | 4 | 2 | 13 | 9 | 1 | 4 | 21 | 21 | 8 | 18 |
| L2a_dup58146 | 12.4284491795324 | 1.49349091168913 | 0.385315253219657 | 3.8760233320889 | 0.000106177507276584 | 0.00713283826833257 | 1595 | 26 | 25 | 9 | 9 | 18 | 20 | 2 | 7 | 9 | 16 | 9 | 6 |
| MamRTE1_dup7617 | 12.4652780883353 | 1.5345425189575 | 0.414407068640274 | 3.70298345535598 | 0.000213078782998232 | 0.0116274926429384 | 1596 | 25 | 26 | 6 | 6 | 21 | 26 | 3 | 8 | 10 | 8 | 10 | 7 |
| MIRb_dup71437 | 14.1266047008059 | 1.77333286296631 | 0.442340775139609 | 4.00897444375691 | 6.09830151599654e-05 | 0.00473431142748878 | 1597 | 33 | 47 | 9 | 6 | 19 | 13 | 4 | 9 | 12 | 12 | 6 | 6 |
| L1MC2_dup1940 | 12.7439975178412 | 2.0609858240582 | 0.389593057227973 | 5.29009895279575 | 1.22250201640929e-07 | 3.86307390168027e-05 | 1598 | 35 | 28 | 5 | 4 | 21 | 17 | 6 | 5 | 12 | 15 | 7 | 5 |
| L3_dup6994 | 12.0278116718356 | 1.67627068730995 | 0.409775328824267 | 4.09070671023443 | 4.30060707604973e-05 | 0.00363779684080245 | 1599 | 30 | 21 | 6 | 5 | 26 | 14 | 3 | 7 | 10 | 13 | 6 | 10 |
| L1ME4a_dup1296 | 13.8797897283145 | 1.25333503378302 | 0.333617733544072 | 3.75679979738683 | 0.000172100068354561 | 0.0100418153174827 | 1600 | 17 | 18 | 9 | 10 | 23 | 22 | 6 | 8 | 12 | 27 | 10 | 11 |
| MLT1A0_dup7772 | 13.0082505265004 | 1.34524504052455 | 0.353961967133691 | 3.80053555306538 | 0.000144383699701745 | 0.00885908298329846 | 1601 | 21 | 18 | 10 | 9 | 21 | 15 | 6 | 8 | 11 | 28 | 7 | 8 |
| L1MA9_dup6268 | 12.7918835183869 | 1.20168683054798 | 0.357213009754418 | 3.36406233181186 | 0.000768041948690582 | 0.0291983558912173 | 1602 | 15 | 18 | 13 | 12 | 20 | 22 | 4 | 6 | 15 | 18 | 7 | 9 |
| THE1D-int_dup533 | 13.0361285334531 | 1.5905242678409 | 0.367498878782031 | 4.32797039575258 | 1.50489690963436e-05 | 0.00165473290230988 | 1603 | 23 | 23 | 9 | 13 | 17 | 26 | 3 | 6 | 15 | 16 | 4 | 8 |
| Tigger2a_dup715 | 12.2556220524781 | 1.42088970242761 | 0.374191677526211 | 3.79722422428297 | 0.000146325422097723 | 0.00894592725757062 | 1604 | 17 | 25 | 6 | 6 | 20 | 17 | 11 | 4 | 14 | 16 | 4 | 14 |
| AluSz_dup46182 | 11.1589517058962 | 1.25116778229563 | 0.38036036320597 | 3.28942735186661 | 0.00100391468942728 | 0.0351729810384137 | 1605 | 25 | 15 | 5 | 6 | 19 | 12 | 8 | 4 | 9 | 17 | 9 | 13 |
| MIR3_dup53985 | 13.1484037746341 | 1.11142990020974 | 0.343638041608123 | 3.23430402236196 | 0.00121939575659519 | 0.0402706929491742 | 1606 | 25 | 16 | 8 | 7 | 21 | 22 | 8 | 6 | 10 | 17 | 11 | 16 |
| L2c_dup98863 | 11.8908684830979 | 1.2913858540697 | 0.409683127290518 | 3.15215777278993 | 0.00162068677428859 | 0.0489696425644282 | 1607 | 17 | 22 | 7 | 6 | 19 | 24 | 4 | 4 | 11 | 10 | 11 | 15 |
| THE1A-int_dup460 | 12.0937894397993 | 1.19942529812511 | 0.356096776136394 | 3.36825654851112 | 0.000756451708159637 | 0.028924097404749 | 1608 | 17 | 25 | 7 | 10 | 17 | 21 | 7 | 4 | 10 | 14 | 11 | 10 |
| L3b_dup6488 | 13.9700816456054 | 1.41639428871553 | 0.368532677311923 | 3.84333432532146 | 0.000121373945525588 | 0.00782032083456042 | 1609 | 26 | 28 | 4 | 7 | 16 | 22 | 8 | 6 | 10 | 23 | 11 | 16 |
| MER1A_dup324 | 12.6228319793536 | 1.33615290225563 | 0.409082756654353 | 3.26621662859428 | 0.00108994834149091 | 0.0372664459746642 | 1610 | 14 | 29 | 2 | 9 | 15 | 20 | 5 | 8 | 12 | 19 | 11 | 13 |
| AluJo_dup64074 | 11.0408418839273 | 1.40204060066658 | 0.431026487553574 | 3.25279452922789 | 0.00114276097400037 | 0.0385258434003247 | 1611 | 13 | 23 | 6 | 7 | 18 | 19 | 4 | 8 | 17 | 6 | 5 | 9 |
| MLT1N2_dup4954 | 13.0923997987607 | 1.61427545047184 | 0.449773053590611 | 3.58908884733049 | 0.000331835742092191 | 0.0160700778452441 | 1612 | 13 | 20 | 5 | 5 | 26 | 26 | 5 | 6 | 24 | 7 | 12 | 10 |
| MER74B_dup579 | 10.8902517337218 | 1.539896391918 | 0.484320137292113 | 3.17950106416746 | 0.00147528827730128 | 0.0459715747028196 | 1613 | 27 | 18 | 4 | 4 | 8 | 17 | 4 | 7 | 20 | 7 | 5 | 13 |
| MER20_dup12261 | 10.6627044612213 | 1.71625449902341 | 0.527958532684304 | 3.25073730752572 | 0.00115106160225188 | 0.0387343380016158 | 1614 | 19 | 25 | 1 | 3 | 10 | 22 | 3 | 5 | 13 | 8 | 10 | 14 |
| THE1D_dup6472 | 13.1768674398705 | 1.71276879681753 | 0.474852251125889 | 3.60695098897921 | 0.00030981615144329 | 0.0152945049733039 | 1615 | 38 | 24 | 3 | 4 | 13 | 18 | 4 | 5 | 18 | 10 | 14 | 14 |
| HERV17-int_dup729 | 11.3166292033852 | 2.20258331308384 | 0.408441592024768 | 5.39265186526418 | 6.94253798831779e-08 | 2.49539145644753e-05 | 1616 | 21 | 23 | 3 | 3 | 16 | 21 | 10 | 3 | 14 | 19 | 3 | 5 |
| L1PA4_dup9446 | 12.7166690258169 | 2.57014792541454 | 0.433105551742129 | 5.93422992403663 | 2.9522786576097e-09 | 1.80123819862461e-06 | 1617 | 28 | 33 | 4 | 2 | 13 | 20 | 6 | 4 | 20 | 18 | 4 | 4 |
| HAL1_dup1662 | 12.8264314006317 | 2.63911609092355 | 0.417297704210966 | 6.32430052763803 | 2.54382219297565e-10 | 2.28412194539541e-07 | 1618 | 20 | 17 | 1 | 3 | 26 | 28 | 4 | 3 | 20 | 22 | 7 | 6 |
| L1ME4a_dup11898 | 9.56813937084417 | 1.92486770889464 | 0.4452852793693 | 4.3227741811295 | 1.54079419663906e-05 | 0.00168641129675013 | 1619 | 16 | 13 | 3 | 2 | 13 | 23 | 4 | 3 | 14 | 12 | 9 | 6 |
| MLT1L_dup10816 | 9.44858591912997 | 1.74661656756427 | 0.415015652059345 | 4.20855589156071 | 2.57007961505148e-05 | 0.00246092621361385 | 1620 | 15 | 14 | 2 | 3 | 13 | 14 | 7 | 3 | 15 | 16 | 9 | 5 |
| CR1-16_AMi_dup52 | 9.66122040510856 | 1.39833243780677 | 0.407987922720253 | 3.4273868414619 | 0.000609420236419419 | 0.0248516036975244 | 1621 | 14 | 14 | 7 | 7 | 16 | 12 | 6 | 1 | 17 | 11 | 5 | 10 |
| MLT1H1_dup1158 | 8.92192935625168 | 1.3578146900074 | 0.41378976835564 | 3.28141194839887 | 0.0010328876826877 | 0.0358696951469707 | 1622 | 21 | 11 | 5 | 5 | 10 | 13 | 4 | 3 | 12 | 11 | 7 | 10 |
| AluJb_dup18078 | 10.1841152052844 | 1.31560052503929 | 0.3933096901343 | 3.34494816181636 | 0.000822979505763588 | 0.0306820887888357 | 1623 | 15 | 11 | 5 | 7 | 18 | 17 | 6 | 4 | 17 | 9 | 7 | 10 |
| AluSx3_dup21016 | 10.7893522065012 | 1.6451541325714 | 0.409397480494121 | 4.01847644637623 | 5.85756631954864e-05 | 0.00458330968586426 | 1624 | 17 | 15 | 5 | 6 | 20 | 19 | 3 | 2 | 16 | 11 | 10 | 10 |
| HAL1_dup6363 | 12.2871251259025 | 1.42573855970634 | 0.405764499036491 | 3.51370946224186 | 0.000441895857937782 | 0.019842866920714 | 1625 | 29 | 20 | 8 | 10 | 18 | 20 | 2 | 3 | 12 | 12 | 10 | 12 |
| L1PA3_dup5095 | 10.351369038256 | 1.43671498193148 | 0.408180799995503 | 3.51980049514163 | 0.000431871515467275 | 0.0195280259354976 | 1626 | 23 | 18 | 6 | 8 | 12 | 17 | 3 | 2 | 11 | 12 | 10 | 9 |
| L1MCb_dup742 | 10.301520879889 | 1.60252865615758 | 0.418874532561434 | 3.82579634612316 | 0.000130350020099276 | 0.00822712622879108 | 1627 | 16 | 15 | 5 | 9 | 14 | 20 | 1 | 2 | 12 | 17 | 9 | 9 |
| MIRc_dup53623 | 10.4793060871056 | 1.33135777217406 | 0.409669022321968 | 3.24983755087958 | 0.00115470949297816 | 0.0388006947890204 | 1628 | 16 | 14 | 6 | 6 | 14 | 22 | 3 | 3 | 9 | 16 | 13 | 10 |
| AluSz_dup91824 | 10.0745463051866 | 1.79568516224229 | 0.471048304837578 | 3.81210407467968 | 0.000137788832625549 | 0.00855815754839963 | 1629 | 11 | 17 | 6 | 4 | 25 | 24 | 3 | 2 | 6 | 14 | 8 | 7 |
| L2b_dup75915 | 10.3757418260605 | 1.41206345103961 | 0.414261278584764 | 3.4086300700457 | 0.000652899513919197 | 0.026035667460871 | 1630 | 14 | 10 | 8 | 4 | 18 | 23 | 4 | 5 | 13 | 13 | 10 | 6 |
| THE1B-int_dup1179 | 12.935325223741 | 4.20464483212366 | 1.33637883045805 | 3.14629709502544 | 0.00165351948282115 | 0.0497029240600507 | 1631 | 17 | 13 | 0 | 0 | 34 | 35 | 1 | 0 | 4 | 2 | 25 | 38 |
| L1MA5_dup746 | 13.8054016750579 | 4.71951026527751 | 0.889145406247312 | 5.3079172789032 | 1.1088494108656e-07 | 3.59463599343087e-05 | 1632 | 41 | 48 | 0 | 0 | 24 | 19 | 0 | 0 | 11 | 17 | 8 | 6 |
| L1MD2_dup1658 | 14.6208479835467 | 4.22367555503415 | 0.772726296311224 | 5.46593997796733 | 4.6046013986326e-08 | 1.78469992384381e-05 | 1633 | 53 | 35 | 0 | 1 | 14 | 25 | 0 | 0 | 20 | 19 | 10 | 6 |
| L1M4b_dup695 | 16.9984655915564 | 4.1714788198202 | 0.838534780025446 | 4.97472367180001 | 6.53407794084305e-07 | 0.000146952524547433 | 1634 | 51 | 47 | 0 | 1 | 28 | 16 | 0 | 0 | 15 | 23 | 12 | 23 |
| L1MC4_dup4260 | 14.4925230664671 | 3.61315820721146 | 0.775877831633874 | 4.65686485667818 | 3.21061044865979e-06 | 0.000507272187129351 | 1635 | 41 | 47 | 0 | 1 | 17 | 21 | 0 | 1 | 10 | 19 | 13 | 14 |
| ERV3-16A3_I-int_dup1( | 18.8439615872923 | 4.86951076882252 | 0.890279755446873 | 5.4696411313748 | 4.50947717899096e-08 | 1.7561572124913e-05 | 1636 | 61 | 41 | 0 | 0 | 31 | 31 | 0 | 0 | 26 | 16 | 10 | 21 |
| L2a_dup30882 | 15.3802606896523 | 4.79914748198308 | 0.952223522826965 | 5.03993796302716 | 4.65682774577663e-07 | 0.000113765249979114 | 1637 | 66 | 26 | 0 | 0 | 23 | 31 | 0 | 0 | 14 | 12 | 9 | 17 |
| L2b_dup18163 | 25.1657871323847 | 6.15711260169776 | 1.21202043078142 | 5.08004027434433 | 3.7735485992172e-07 | 9.64447685273187e-05 | 1638 | 76 | 87 | 0 | 0 | 43 | 40 | 0 | 0 | 23 | 19 | 13 | 17 |
| L1M4b_dup697 | 20.5230220610529 | 4.34909603938135 | 0.844700595194419 | 5.14868352659365 | 2.62321042143494e-07 | 7.15805535480229e-05 | 1639 | 65 | 57 | 0 | 0 | 25 | 29 | 0 | 1 | 18 | 25 | 15 | 26 |
| MLT1H_dup1639 | 20.4830723941702 | 4.38436665310346 | 0.816104449840611 | 5.3723106815062 | 7.77340256654909e-08 | 2.71209684325526e-05 | 1640 | 61 | 60 | 0 | 0 | 30 | 27 | 0 | 1 | 22 | 23 | 18 | 15 |
| L1M2_dup6814 | 12.8135424306503 | 3.282308430248 | 0.870941001010019 | 3.76869205427411 | 0.000164105194899313 | 0.00970557867204541 | 1641 | 48 | 40 | 1 | 3 | 28 | 21 | 0 | 0 | 1 | 8 | 4 | 14 |
| HERV1_I-int_dup2 | 17.9970758230422 | 2.95979958704494 | 0.61078889538944 | 4.84586345525776 | 1.26062198677321e-06 | 0.000245667452305273 | 1642 | 41 | 44 | 3 | 2 | 37 | 36 | 0 | 2 | 14 | 19 | 9 | 21 |
| L1PA4_dup2559 | 12.6669310028354 | 3.05422466726368 | 0.628654974453171 | 4.85834804682864 | 1.18369200334233e-06 | 0.000234908071248785 | 1643 | 24 | 25 | 1 | 2 | 19 | 36 | 0 | 1 | 11 | 20 | 6 | 15 |
| MER21A_dup423 | 11.5533702919211 | 3.31318332091446 | 0.787411775939032 | 4.20768830509717 | 2.57996392589485e-05 | 0.00246642063604249 | 1644 | 28 | 33 | 1 | 1 | 17 | 22 | 0 | 0 | 11 | 8 | 8 | 18 |
| MER31B_dup534 | 10.4630703348238 | 2.64360076154109 | 0.687656447657451 | 3.84436264728804 | 0.000120866158288875 | 0.007796047411278 | 1645 | 31 | 35 | 1 | 2 | 8 | 17 | 0 | 2 | 8 | 10 | 8 | 11 |
| L2a_dup57861 | 12.4206538852223 | 1.53763508850023 | 0.484776830446064 | 3.17184112756664 | 0.00151475849436754 | 0.0468315440978231 | 1646 | 17 | 4 | 4 | 2 | 6 | 5 | 2 | 0 | 30 | 39 | 16 | 25 |
| MLT1H2_dup1576 | 13.5313307493828 | 2.33107667388858 | 0.432451865267824 | 5.39037257347685 | 7.03117577353785e-08 | 2.51584985054171e-05 | 1647 | 13 | 10 | 2 | 1 | 10 | 13 | 0 | 1 | 38 | 39 | 15 | 17 |
| L1PA15_dup277 | 13.3576032782384 | 1.57462083391202 | 0.476583215669772 | 3.30397878510914 | 0.000953230454161172 | 0.0339903152473901 | 1648 | 18 | 19 | 3 | 3 | 8 | 8 | 2 | 1 | 26 | 26 | 22 | 28 |
| Tigger1_dup4322 | 13.1714131501147 | 2.02840107531151 | 0.445076395102691 | 4.55742227094184 | 5.17852452234145e-06 | 0.000742294696381361 | 1649 | 11 | 12 | 1 | 2 | 8 | 4 | 2 | 1 | 41 | 39 | 12 | 21 |
| L1PB2_dup662 | 13.5533386816142 | 1.88863228733924 | 0.497535034386715 | 3.79597848755968 | 0.000147062251393746 | 0.00897943917632444 | 1650 | 23 | 21 | 0 | 4 | 11 | 6 | 3 | 5 | 35 | 23 | 11 | 20 |
| THE1C-int_dup638 | 18.9622384040269 | 1.66254986268787 | 0.378000976993044 | 4.39826869208982 | 1.09117837628541e-05 | 0.0013026741686044 | 1651 | 11 | 11 | 0 | 5 | 13 | 5 | 3 | 5 | 56 | 64 | 24 | 23 |
| L1PREC2_dup2546 | 18.0403176842046 | 1.94732658980224 | 0.400039014345828 | 4.86784168535824 | 1.12823653262706e-06 | 0.000226245953966858 | 1652 | 17 | 14 | 1 | 2 | 9 | 11 | 1 | 2 | 48 | 52 | 25 | 32 |
| Tigger1_dup4321 | 21.4523834197813 | 1.84323758753505 | 0.383795946032086 | 4.80264996697217 | 1.56579466457537e-06 | 0.000292674451890849 | 1653 | 25 | 15 | 3 | 6 | 7 | 14 | 1 | 3 | 59 | 64 | 30 | 26 |
| AluJo_dup32000 | 16.4064880413075 | 1.25701051620523 | 0.347250586224544 | 3.61989458353975 | 0.000294723061637871 | 0.0147887333666142 | 1654 | 18 | 15 | 6 | 3 | 18 | 19 | 5 | 6 | 24 | 37 | 27 | 23 |

| | | | | | | | | | | | | | | | | | | | |
|---|---|---|---|---|---|---|---|---|---|---|---|---|---|---|---|---|---|---|---|
| MER61E_dup9 | 14.9000019363846 | 1.32373595466496 | 0.337837963306538 | 3.91825697061718 | 8.91915937877181e-05 | 0.00627336180177723 | 1655 | 14 | 14 | 8 | 5 | 14 | 17 | 3 | 7 | 25 | 40 | 17 | 16 |
| L1PB1_dup10475 | 17.9135212771373 | 1.66174878346811 | 0.363466195388173 | 4.57194865589469 | 4.83209355758074e-06 | 0.000702496054992057 | 1656 | 12 | 14 | 8 | 4 | 25 | 19 | 5 | 2 | 40 | 44 | 24 | 17 |
| MIR3_dup84576 | 14.355053046103 | 1.59619508958257 | 0.383768728593905 | 4.15926304217358 | 3.19276049283712e-05 | 0.0028919093575509 | 1657 | 18 | 21 | 6 | 4 | 16 | 17 | 3 | 2 | 23 | 28 | 22 | 16 |
| AluJb_dup53403 | 17.1960148850731 | 1.56958051674579 | 0.401921869475694 | 3.9051881371703 | 9.41521074664802e-05 | 0.00652570515807461 | 1658 | 25 | 12 | 2 | 5 | 17 | 10 | 5 | 3 | 38 | 40 | 15 | 38 |
| MER21C_dup2763 | 16.8652516893241 | 1.48282488267901 | 0.363431856024369 | 4.0800630382263 | 4.50234892952601e-05 | 0.00378289519203741 | 1659 | 17 | 13 | 5 | 11 | 11 | 10 | 2 | 5 | 41 | 53 | 17 | 15 |
| L1ME3E_dup4223 | 18.3775532043498 | 3.25935005856294 | 0.551146213025535 | 5.91376658594937 | 3.3437164079675e-09 | 1.99905513670913e-06 | 1660 | 38 | 28 | 4 | 2 | 8 | 9 | 6 | 2 | 50 | 63 | 3 | 3 |
| L1PA7_dup6715 | 19.0538293047692 | 3.88336039729988 | 0.475612282113976 | 8.16497080361198 | 3.215129704576e-16 | 1.39095864124553e-12 | 1661 | 47 | 49 | 3 | 4 | 21 | 22 | 3 | 1 | 34 | 43 | 3 | 2 |
| L1MA2_dup4509 | 17.3581079928136 | 2.37146834906604 | 0.387052011321346 | 6.1270017457606 | 8.95504900093746e-10 | 6.57659904190452e-07 | 1662 | 37 | 30 | 2 | 7 | 14 | 20 | 3 | 5 | 35 | 36 | 12 | 9 |
| MSTB1_dup919 | 18.280343857737 | 2.4468991372794 | 0.397708309229983 | 6.15249689406019 | 7.62724659160886e-10 | 5.76150100789512e-07 | 1663 | 42 | 23 | 0 | 5 | 19 | 20 | 7 | 5 | 27 | 55 | 8 | 14 |
| HERVH-int_dup1188 | 21.6706470589871 | 2.71980490937247 | 0.505742671448489 | 5.37784344275859 | 7.53833211402225e-08 | 2.65342599586263e-05 | 1664 | 32 | 38 | 1 | 0 | 12 | 13 | 11 | 3 | 45 | 81 | 10 | 13 |
| L1PA7_dup3319 | 20.5494403989349 | 2.18617594589392 | 0.448538660427105 | 4.87399668918664 | 1.09362823064409e-06 | 0.00022071625357832 | 1665 | 24 | 10 | 3 | 3 | 27 | 18 | 16 | 1 | 54 | 65 | 10 | 13 |
| L1PA8A_dup647 | 20.0530919858095 | 2.57267300858559 | 0.38798498506509 | 6.63085714039678 | 3.33743126601051e-11 | 3.99059507548812e-08 | 1666 | 22 | 13 | 2 | 6 | 16 | 20 | 6 | 4 | 63 | 64 | 8 | 8 |
| L1PA10_dup1468 | 14.1449582304155 | 2.14560930507365 | 0.563284644905424 | 3.80910313192347 | 0.000139471778206954 | 0.00862890060770462 | 1667 | 16 | 6 | 9 | 3 | 28 | 9 | 10 | 1 | 35 | 43 | 3 | 5 |
| MLT1D_dup9676 | 12.739057364943 | 1.61072736052753 | 0.505721552650846 | 3.18500833528759 | 0.00144749838928126 | 0.0453550766046771 | 1668 | 10 | 5 | 4 | 5 | 6 | 5 | 6 | 2 | 40 | 50 | 9 | 5 |
| L1MA6_dup3469 | 15.1908590946441 | 2.18019452612393 | 0.643895060100789 | 3.38594696748049 | 0.000709330902123396 | 0.0275968690053391 | 1669 | 39 | 30 | 5 | 12 | 4 | 3 | 4 | 0 | 31 | 43 | 7 | 8 |
| LTR54B_dup63 | 10.1641688750195 | 1.75709581245333 | 0.475445624464523 | 3.69568194981769 | 0.000219297362138663 | 0.0118917343052521 | 1670 | 10 | 18 | 2 | 5 | 4 | 9 | 5 | 0 | 22 | 28 | 10 | 10 |
| LTR8A_dup184 | 14.1003929651102 | 2.08914815339457 | 0.422924422360124 | 4.93976711426622 | 7.82159303332455e-07 | 0.000170431939185663 | 1671 | 25 | 21 | 5 | 3 | 8 | 15 | 7 | 0 | 30 | 35 | 13 | 9 |
| L2a_dup95585 | 13.6801445151078 | 2.09084656372728 | 0.552710317800163 | 3.7828976525154 | 0.000155013165585849 | 0.00932375194602892 | 1672 | 25 | 25 | 8 | 3 | 2 | 9 | 2 | 4 | 32 | 38 | 5 | 10 |
| HERVL-int_dup1838 | 14.8015630213181 | 1.83900009578825 | 0.426923077016762 | 4.30756779099118 | 1.65059515965619e-05 | 0.00177394993613167 | 1673 | 16 | 15 | 8 | 10 | 9 | 15 | 5 | 1 | 40 | 42 | 5 | 10 |
| L2a_dup57863 | 12.5985016871653 | 2.17988063959548 | 0.426572172167392 | 5.11022701860653 | 3.21771929677684e-07 | 8.46950703308477e-05 | 1674 | 14 | 10 | 3 | 4 | 7 | 13 | 6 | 1 | 33 | 44 | 7 | 7 |
| L1MC3_dup1593 | 16.2125347172108 | 1.98420335396283 | 0.383590257837816 | 5.17271571271698 | 2.307159045915e-07 | 6.42832864565914e-05 | 1675 | 25 | 21 | 4 | 8 | 15 | 10 | 3 | 1 | 28 | 53 | 14 | 16 |
| L2a_dup112235 | 15.9643148691321 | 2.23828897057611 | 0.366230028547385 | 6.1117024714059 | 9.85738349645107e-10 | 7.08617816751223e-07 | 1676 | 25 | 17 | 3 | 4 | 18 | 14 | 4 | 1 | 33 | 46 | 13 | 15 |
| L1PA7_dup11056 | 15.3437513728557 | 2.21649532038326 | 0.397807270721226 | 5.57178182380815 | 2.5214706800088e-08 | 1.12144647182313e-05 | 1677 | 26 | 18 | 4 | 7 | 13 | 13 | 2 | 1 | 35 | 42 | 15 | 8 |
| L1PB1_dup9996 | 13.8622815092077 | 2.28082472675056 | 0.402115929282 | 5.67205763477985 | 1.41092489032484e-08 | 6.87958881051711e-06 | 1678 | 27 | 17 | 5 | 7 | 14 | 14 | 2 | 2 | 34 | 29 | 9 | 6 |
| ERV3-16A3_I-int_dup5 | 10.7966195197041 | 3.25842231292824 | 0.623118649349739 | 5.22921648441849 | 1.70229937792239e-07 | 5.06952850787384e-05 | 1679 | 6 | 4 | 2 | 1 | 21 | 17 | 1 | 0 | 31 | 32 | 3 | 8 |
| L1MCc_dup2888 | 14.4672137727477 | 3.83341132636356 | 0.587532299294129 | 6.52459674296899 | 6.81847293204931e-11 | 7.24298375129198e-08 | 1680 | 11 | 11 | 1 | 2 | 39 | 18 | 1 | 0 | 37 | 40 | 3 | 7 |
| L1M2_dup7939 | 21.4194352015391 | 3.37528882384618 | 0.543369938353176 | 6.21176952496832 | 5.23912326732299e-10 | 4.25470452206975e-07 | 1681 | 8 | 15 | 3 | 3 | 45 | 30 | 1 | 0 | 53 | 71 | 6 | 17 |
| L1MA6_dup4708 | 18.5251257404838 | 3.75377845455419 | 0.637373995772237 | 5.88944399905451 | 3.87497085622847e-09 | 2.25988680234348e-06 | 1682 | 9 | 8 | 2 | 1 | 47 | 43 | 1 | 0 | 38 | 53 | 8 | 11 |
| L1MCc_dup2887 | 15.1762619448705 | 3.16722185446084 | 0.520107577114609 | 6.0895514578572 | 1.13227467907273e-09 | 7.94172994201091e-07 | 1683 | 20 | 9 | 3 | 1 | 30 | 27 | 1 | 0 | 28 | 45 | 6 | 14 |
| L1MEf_dup9739 | 15.0313477556763 | 3.20528055170651 | 0.5083542105997 | 6.30521098256524 | 2.87801579339258e-10 | 2.53634202212811e-07 | 1684 | 9 | 12 | 2 | 1 | 26 | 30 | 4 | 0 | 36 | 44 | 6 | 8 |
| L2a_dup156037 | 11.2391034288953 | 2.39054451610832 | 0.567776858027078 | 4.21035919712372 | 2.54964988135986e-05 | 0.00244786002504618 | 1685 | 28 | 12 | 5 | 5 | 25 | 25 | 0 | 0 | 16 | 9 | 7 | 9 |
| L1MC1_dup12203 | 13.3980309364553 | 2.76796746835682 | 0.497167180858617 | 5.56747825465166 | 2.58452319093326e-08 | 1.14304142603315e-05 | 1686 | 34 | 26 | 7 | 2 | 28 | 26 | 0 | 1 | 13 | 18 | 7 | 6 |
| MER50_dup2338 | 12.7222122070685 | 3.54133185584427 | 0.541899507938743 | 6.53503427104898 | 6.35949773471429e-11 | 6.87825930901967e-08 | 1687 | 34 | 24 | 2 | 2 | 27 | 17 | 0 | 1 | 26 | 13 | 3 | 6 |
| MLT1O_dup4551 | 11.3697152369157 | 2.53995625634226 | 0.463136299631085 | 5.48425217018291 | 4.15222128173875e-08 | 1.64453021177503e-05 | 1688 | 26 | 21 | 3 | 6 | 23 | 15 | 0 | 1 | 16 | 17 | 7 | 6 |
| L1MC_dup7540 | 17.9529136773438 | 2.89807728348039 | 0.496754503391281 | 5.83402317179931 | 5.41066023295443e-09 | 2.99405804602459e-06 | 1689 | 50 | 36 | 9 | 7 | 37 | 24 | 0 | 0 | 22 | 27 | 3 | 10 |
| MLT1H1-int_dup362 | 20.1843715032507 | 3.07064048206197 | 0.504986188466782 | 6.08064250506517 | 1.19701911045108e-09 | 8.32824296068398e-07 | 1690 | 45 | 40 | 5 | 6 | 51 | 32 | 0 | 0 | 27 | 24 | 9 | 12 |
| MLT2B2_dup2047 | 18.4524047435419 | 3.18494632169725 | 0.500312494263617 | 6.36591402016654 | 1.94130163010618e-10 | 1.7912288271436e-07 | 1691 | 42 | 38 | 4 | 4 | 20 | 35 | 0 | 0 | 25 | 40 | 14 | 6 |
| L2a_dup156040 | 15.4627741289019 | 2.47461400715054 | 0.403853340332197 | 6.12750659711024 | 8.92668993190029e-10 | 6.57606861466237e-07 | 1692 | 25 | 26 | 7 | 5 | 23 | 18 | 0 | 1 | 28 | 35 | 9 | 11 |
| THE1B_dup21097 | 15.0809618502895 | 3.07845215153483 | 0.438442856262217 | 7.02133039132862 | 2.19765484159462e-12 | 3.7086750279296e-09 | 1693 | 24 | 14 | 5 | 3 | 22 | 19 | 0 | 0 | 38 | 40 | 7 | 7 |
| MLT1G3_dup2478 | 12.1485761575842 | 3.23174216268726 | 0.555613606226441 | 5.81652811679015 | 6.00824405302034e-09 | 3.28652330905742e-06 | 1694 | 32 | 20 | 1 | 7 | 10 | 14 | 0 | 0 | 25 | 31 | 4 | 4 |
| L1MB1_dup5646 | 11.1618614465049 | 3.32793406567181 | 0.518060187244594 | 6.42383674254547 | 1.32881716312154e-10 | 1.29584724875458e-07 | 1695 | 26 | 15 | 3 | 3 | 21 | 15 | 0 | 0 | 22 | 23 | 2 | 6 |
| L1MC1_dup12204 | 8.89732703352742 | 3.61839003811137 | 0.587033661537678 | 6.16385443491152 | 7.0995211357745e-10 | 5.43184133817685e-07 | 1696 | 23 | 14 | 2 | 3 | 14 | 15 | 0 | 0 | 16 | 18 | 2 | 2 |
| L1PA6_dup4968 | 10.9647527585019 | 2.7522245917839 | 0.473283045333202 | 5.81517681421752 | 6.05698288225659e-09 | 3.29802024920235e-06 | 1697 | 22 | 22 | 4 | 4 | 11 | 11 | 2 | 0 | 22 | 26 | 5 | 4 |
| MLT1G3_dup2477 | 10.8244063229715 | 2.82627993166061 | 0.500974793876179 | 5.64156114480914 | 1.68515173952384e-08 | 7.95585547247497e-06 | 1698 | 18 | 20 | 2 | 9 | 19 | 15 | 1 | 0 | 21 | 21 | 3 | 3 |
| L1MB5_dup8584 | 8.85641142292066 | 2.97400355922862 | 0.621913684941867 | 4.78201980634437 | 1.73542574923696e-06 | 0.00031666995040486 | 1699 | 18 | 29 | 2 | 4 | 10 | 7 | 0 | 0 | 16 | 14 | 5 | 3 |
| MLT1A0_dup19223 | 8.69028935703503 | 2.91723950473001 | 0.573563395342237 | 5.08616750723663 | 3.65371383688985e-07 | 9.37849614490391e-05 | 1700 | 15 | 13 | 4 | 5 | 12 | 9 | 0 | 0 | 20 | 22 | 3 | 1 |
| MamRep605_dup4007 | 10.6305469981246 | 2.78992668366157 | 0.534986252132608 | 5.21495023945032 | 1.83866586222502e-07 | 5.39461389954371e-05 | 1701 | 21 | 17 | 3 | 4 | 14 | 17 | 0 | 0 | 17 | 25 | 10 | 2 |
| L1PA5_dup10748 | 15.2147016872776 | 2.76015615163962 | 0.493903347765039 | 5.5884540247188 | 2.29100012695458e-08 | 1.03245135645518e-05 | 1702 | 26 | 31 | 5 | 7 | 25 | 28 | 1 | 0 | 14 | 39 | 2 | 13 |
| L2a_dup40051 | 10.9609652900276 | 2.33857826650747 | 0.540534891799305 | 4.32641500481667 | 1.51555761393531e-05 | 0.00166491630658103 | 1703 | 29 | 18 | 3 | 5 | 7 | 20 | 1 | 1 | 11 | 27 | 3 | 14 |
| LTR16B1_dup1119 | 10.6544548843133 | 2.23847640477309 | 0.508225897994726 | 4.40449102181785 | 1.06032531987073e-05 | 0.00127578556111878 | 1704 | 30 | 19 | 4 | 9 | 10 | 15 | 1 | 0 | 12 | 23 | 4 | 8 |
| L1PA3_dup10255 | 13.2908374580809 | 2.79471934438572 | 0.49374922720191 | 5.66019993635933 | 1.5119673901281e-08 | 7.22422876729761e-06 | 1705 | 35 | 24 | 2 | 4 | 22 | 17 | 1 | 0 | 22 | 19 | 7 | 12 |
| MLT1A_dup8346 | 11.3303957287597 | 2.75971680878897 | 0.599744000400465 | 4.60149131453793 | 4.19476652496924e-06 | 0.000626178115150772 | 1706 | 22 | 27 | 4 | 2 | 10 | 15 | 0 | 0 | 22 | 19 | 3 | 15 |
| L1M4_dup19199 | 9.34728318461637 | 1.35903105425261 | 0.428202733718403 | 3.17380284439367 | 0.00150455851472143 | 0.0465968606460894 | 1707 | 12 | 11 | 5 | 7 | 20 | 21 | 5 | 4 | 9 | 10 | 3 | 11 |
| LTR5B_dup79 | 8.65319066451149 | 1.72385392927222 | 0.494351294213652 | 3.48710309743255 | 0.000488283087999626 | 0.0211989812677116 | 1708 | 12 | 11 | 3 | 5 | 22 | 21 | 2 | 3 | 8 | 8 | 4 | 10 |
| AluJb_dup24784 | 9.24096667147349 | 1.83404893632501 | 0.524544938214561 | 3.49645721979077 | 0.000471480086456135 | 0.0207139587614276 | 1709 | 20 | 15 | 2 | 2 | 20 | 22 | 3 | 4 | 4 | 10 | 5 | 11 |
| L1M5_dup44511 | 10.6803122801401 | 1.88012266493711 | 0.55891426823525 | 3.36388382224258 | 0.000768538878465291 | 0.02912580187748 | 1710 | 13 | 11 | 2 | 2 | 34 | 31 | 5 | 6 | 8 | 8 | 3 | 11 |
| AluSp_dup7560 | 12.9004414037022 | 1.96171653981977 | 0.581968307496311 | 3.37083053243789 | 0.000749419436344167 | 0.0287429653772323 | 1711 | 18 | 16 | 3 | 5 | 32 | 40 | 1 | 3 | 10 | 8 | 14 | 13 |
| L1PB3_dup1412 | 8.65254269377057 | 1.98205507089722 | 0.601829437456923 | 3.29338338661623 | 0.000989893946314194 | 0.0348795061379649 | 1712 | 14 | 6 | 2 | 2 | 18 | 25 | 1 | 2 | 10 | 8 | 12 | 8 |
| L2a_dup68134 | 10.5337207996138 | 1.96896232366383 | 0.529015714089267 | 3.72193541935425 | 0.000197701592997215 | 0.0110272628334072 | 1713 | 23 | 8 | 6 | 5 | 31 | 23 | 2 | 1 | 12 | 7 | 6 | 9 |
| MER31B_dup1397 | 10.8111288930438 | 1.98475657685224 | 0.546982284927052 | 3.62855732542953 | 0.000285009468364084 | 0.0144099802399453 | 1714 | 11 | 3 | 2 | 3 | 26 | 20 | 3 | 2 | 22 | 16 | 12 | 11 |
| MER117_dup1471 | 9.32651044984168 | 1.49577105478295 | 0.432614341325232 | 3.45751611054073 | 0.000545179643995873 | 0.0229071720943391 | 1715 | 9 | 6 | 3 | 5 | 16 | 12 | 5 | 1 | 16 | 22 | 7 | 13 |
| MIRc_dup97686 | 8.00989860469773 | 1.6905558760471 | 0.452296854044867 | 3.73771309910416 | 0.000185701677489652 | 0.0105540194781703 | 1716 | 13 | 6 | 2 | 6 | 13 | 13 | 4 | 1 | 11 | 18 | 6 | 7 |
| L1PA7_dup7719 | 10.0146048019155 | 1.4591039460949 | 0.426987663368386 | 3.41720398801323 | 0.000632678547293245 | 0.025459720888591 | 1717 | 10 | 8 | 4 | 5 | 13 | 21 | 3 | 3 | 12 | 23 | 12 | 10 |
| MIRb_dup21854 | 9.4863268042135 | 1.64388273581199 | 0.507894767261325 | 3.2366601150001 | 0.00120937359707828 | 0.040067614860817 | 1718 | 7 | 7 | 4 | 5 | 14 | 27 | 3 | 1 | 16 | 13 | 10 | 10 |
| L1ME3Cz_dup2025 | 10.4674548355705 | 1.7841165254638 | 0.469691925350157 | 3.79848242895327 | 0.000145584747799167 | 0.00890751051679626 | 1719 | 7 | 17 | 7 | 2 | 13 | 8 | 3 | 2 | 27 | 21 | 5 | 12 |
| MIRb_dup199604 | 11.3042793845639 | 1.93632782888103 | 0.402247258487394 | 4.81377507994058 | 1.48105494665444e-06 | 0.000279913503047368 | 1720 | 18 | 14 | 2 | 3 | 14 | 11 | 3 | 4 | 19 | 29 | 12 | 8 |
| MLT1A0_dup18776 | 10.035501709431 | 1.87208522076106 | 0.415386605992782 | 4.5068502300086 | 6.57970411476004e-06 | 0.000887033583734103 | 1721 | 15 | 11 | 2 | 3 | 10 | 9 | 3 | 2 | 18 | 29 | 10 | 10 |
| L1ME2_dup1837 | 11.1561039803949 | 1.33840158894927 | 0.3868193281767 | 3.46001735553887 | 0.000540140573021929 | 0.0227476617324382 | 1722 | 11 | 17 | 6 | 7 | 13 | 9 | 2 | 3 | 20 | 23 | 11 | 14 |
| AluSq2_dup21767 | 10.0207125529968 | 1.43385540566052 | 0.390495873632002 | 3.67188362920261 | 0.000240769318593176 | 0.0127708640842559 | 1723 | 13 | 18 | 4 | 6 | 14 | 10 | 6 | 2 | 16 | 16 | 8 | 11 |
| L1ME5_dup1527 | 11.3288236730942 | 1.67353795755664 | 0.372517693931759 | 4.4925059529205 | 7.0389950240719e-06 | 0.00093465441406128 | 1724 | 18 | 14 | 4 | 5 | 18 | 12 | 5 | 3 | 17 | 24 | 7 | 13 |
| L2c_dup8615 | 12.9930484762617 | 2.07894788173196 | 0.461761074470763 | 4.50221553238263 | 6.7248732034256e-06 | 0.000901496720711159 | 1725 | 24 | 15 | 4 | 3 | 19 | 23 | 3 | 1 | 24 | 17 | 7 | 21 |
| LTR78_dup2505 | 11.6981314259361 | 1.92071261381158 | 0.4802960955317 | 3.99901775525638 | 6.360590885265e-05 | 0.00488218438317827 | 1726 | 15 | 10 | 3 | 1 | 16 | 16 | 2 | 1 | 24 | 20 | 13 | 22 |
| LTR78_dup2361 | 10.7266084423928 | 1.45669491121311 | 0.44294666455959 | 3.28864630386473 | 0.00100670447260685 | 0.0352370259545322 | 1727 | 13 | 15 | 7 | 3 | 23 | 10 | 2 | 5 | 17 | 14 | 8 | 14 |
| L1M2_dup1094 | 11.4821487677116 | 1.45425431700861 | 0.420048087320724 | 3.46211388863728 | 0.000535950310477117 | 0.0226272591513109 | 1728 | 15 | 13 | 8 | 3 | 25 | 18 | 5 | 2 | 11 | 19 | 12 | 13 |
| L1MB3_dup11291 | 11.4647103144343 | 1.38948415043013 | 0.416804299742927 | 3.33366078825753 | 0.000857111163752665 | 0.0315657286751728 | 1729 | 23 | 9 | 8 | 3 | 16 | 10 | 4 | 2 | 16 | 24 | 11 | 17 |
| L1PA5_dup7396 | 12.5303129271452 | 1.41933927289696 | 0.396679636918726 | 3.57804923873056 | 0.000346168248078095 | 0.0165754515831735 | 1730 | 26 | 18 | 3 | 8 | 8 | 15 | 12 | 4 | 18 | 26 | 6 | 13 |
| MER50_dup78 | 12.9208055718994 | 1.49555119728147 | 0.430025592628755 | 3.47781904825509 | 0.000505511056050942 | 0.0217473031536969 | 1731 | 10 | 14 | 2 | 13 | 13 | 22 | 4 | 6 | 25 | 29 | 5 | 14 |
| MIR_dup21438 | 11.1790280521261 | 1.40040199696549 | 0.40229672659176 | 3.481017628032 | 0.000499512615264658 | 0.0215594102578931 | 1732 | 9 | 12 | 3 | 7 | 21 | 16 | 8 | 7 | 19 | 19 | 5 | 10 |
| L1PA16_dup12633 | 14.5003948275308 | 1.34656259531323 | 0.398743958011445 | 3.37701065623313 | 0.000732782111350785 | 0.0282597361859763 | 1733 | 22 | 21 | 3 | 7 | 22 | 15 | 7 | 3 | 12 | 30 | 22 | 19 |

| | | | | | | | | | | | | | | | | | | |
|---|---|---|---|---|---|---|---|---|---|---|---|---|---|---|---|---|---|---|
| L1PB4_dup5500 | 12.7497879152419 | 1.48937264436587 | 0.437513610973688 | 3.40417442339969 | 0.000663643613904768 | 0.0263185572256973 | 1734 | 20 | 11 | 1 | 8 | 19 | 23 | 6 | 2 | 17 | 20 | 11 | 23 |
| L2c_dup11066 | 11.882181898393 | 1.75101107421182 | 0.43560494916021 | 4.01972263535468 | 5.82666976698607e-05 | 0.00456664283391063 | 1735 | 19 | 18 | 1 | 6 | 14 | 15 | 3 | 2 | 15 | 26 | 9 | 21 |
| L2a_dup114014 | 10.4753962394787 | 1.37440380111384 | 0.424624000017644 | 3.23675487267967 | 0.00120897212053014 | 0.0400654707787832 | 1736 | 22 | 18 | 2 | 8 | 16 | 19 | 7 | 3 | 10 | 9 | 6 | 14 |
| MIRb_dup216517 | 9.64158927096417 | 1.60944643805005 | 0.456345115433348 | 3.52681859325198 | 0.00042058482118729 | 0.0191619052937071 | 1737 | 15 | 13 | 2 | 4 | 12 | 25 | 5 | 2 | 9 | 14 | 10 | 11 |
| L1MA9_dup14260 | 11.6831660387202 | 1.40505798041036 | 0.431523194299495 | 3.25604277816684 | 0.0011297672844649 | 0.038217743285355 | 1738 | 18 | 15 | 3 | 2 | 18 | 19 | 8 | 3 | 16 | 13 | 14 | 17 |
| L1MEg2_dup199 | 10.8614530700405 | 1.73002136541346 | 0.548996140447223 | 3.15124504154828 | 0.00162576031808114 | 0.0490855430333884 | 1739 | 20 | 19 | 1 | 2 | 17 | 15 | 4 | 2 | 8 | 16 | 11 | 24 |
| L1M1_dup6275 | 8.13333850836593 | 1.91405819108658 | 0.525912855685402 | 3.63949686795935 | 0.000273171259968669 | 0.0139835390025421 | 1740 | 6 | 9 | 1 | 3 | 17 | 10 | 3 | 3 | 22 | 9 | 6 | 7 |
| L1PB1_dup10474 | 8.57748206881173 | 1.8027293270026 | 0.513016219333399 | 3.51398115510855 | 0.000441444129229685 | 0.0198356726501429 | 1741 | 7 | 8 | 1 | 1 | 10 | 4 | 3 | 2 | 28 | 16 | 7 | 13 |
| L1MD2_dup8878 | 9.23929164209228 | 1.8546837321916 | 0.513239072125798 | 3.61368382284232 | 0.000301877154758196 | 0.0150304376367637 | 1742 | 5 | 4 | 5 | 4 | 8 | 11 | 1 | 1 | 30 | 25 | 6 | 7 |
| L1PB4_dup6392 | 8.21827961861659 | 2.89787547482059 | 0.577140545813855 | 5.02109147561994 | 5.1378681982085e-07 | 0.000121524372394724 | 1743 | 8 | 5 | 1 | 3 | 18 | 11 | 1 | 1 | 18 | 25 | 2 | 5 |
| MLT1D_dup16593 | 9.43511770790872 | 2.29629993235192 | 0.483390065389078 | 4.75040779024624 | 2.0300684874193e-06 | 0.000357018977315205 | 1744 | 7 | 10 | 3 | 2 | 19 | 16 | 2 | 2 | 17 | 23 | 4 | 9 |
| L1MA3_dup3082 | 7.46666470459768 | 1.64671644601185 | 0.502127979328198 | 3.2794755795425 | 0.00104000210690265 | 0.0360682613801281 | 1745 | 5 | 11 | 5 | 8 | 10 | 17 | 3 | 2 | 11 | 14 | 3 | 3 |
| L1PA16_dup10635 | 6.60345123123149 | 1.83115150251671 | 0.52962794397919 | 3.45742992478634 | 0.000545354054021721 | 0.0229104547560474 | 1746 | 7 | 12 | 4 | 6 | 8 | 7 | 3 | 1 | 11 | 17 | 1 | 4 |
| L1M4_dup15345 | 6.7630860358771 | 1.83988989886386 | 0.506729112776142 | 3.63091413632015 | 0.00028241913807126 | 0.0143162556939719 | 1747 | 11 | 4 | 1 | 4 | 10 | 8 | 2 | 2 | 11 | 19 | 4 | 7 |
| MADE1_dup3210 | 5.84882672325359 | 1.81580962212965 | 0.561165281615347 | 3.23578396885627 | 0.00121309155772941 | 0.0401366384017768 | 1748 | 10 | 3 | 1 | 4 | 7 | 12 | 2 | 1 | 12 | 10 | 3 | 7 |
| L2b_dup30359 | 6.06402857288688 | 1.78724338903061 | 0.515606322999877 | 3.46629455324006 | 0.000527684839409234 | 0.0223695878849224 | 1749 | 7 | 9 | 2 | 3 | 6 | 10 | 1 | 1 | 11 | 12 | 6 | 6 |
| HERV4_I-int_dup120 | 6.23158177056803 | 1.79831644877904 | 0.498784855055023 | 3.60539505270396 | 0.000311678429834937 | 0.01535140460881 | 1750 | 8 | 8 | 2 | 4 | 12 | 9 | 2 | 1 | 9 | 12 | 5 | 5 |
| MIR1_Amn_dup10080 | 6.33097750331569 | 1.85676141725851 | 0.506396123760463 | 3.66661854255584 | 0.000245779009204992 | 0.0129532246034842 | 1751 | 10 | 9 | 2 | 4 | 11 | 12 | 1 | 1 | 8 | 10 | 5 | 6 |
| MIRc_dup24602 | 6.9773818602391 | 2.05306496077204 | 0.515002523232116 | 3.98651437256494 | 6.705106785729e-05 | 0.0050875425358325 | 1752 | 12 | 9 | 2 | 2 | 11 | 14 | 2 | 1 | 7 | 15 | 8 | 4 |
| Tigger15a_dup4310 | 7.34637709983985 | 2.03301782170255 | 0.505161587447458 | 4.02449012795139 | 5.70988891496399e-05 | 0.00450096777238542 | 1753 | 11 | 8 | 2 | 5 | 14 | 17 | 1 | 1 | 8 | 14 | 5 | 6 |
| L1MA10_dup286 | 7.50959817525746 | 1.87884927736926 | 0.534404882642006 | 3.51577865097443 | 0.000438466374413402 | 0.019741025530023 | 1754 | 22 | 12 | 2 | 2 | 4 | 9 | 4 | 1 | 12 | 12 | 6 | 8 |
| L1MEc_dup7923 | 6.79675445267578 | 1.6654131443902 | 0.528733124743965 | 3.14981805839507 | 0.0016337217007959 | 0.0492821429789023 | 1755 | 13 | 13 | 2 | 2 | 4 | 11 | 4 | 1 | 12 | 8 | 8 | 6 |
| Helitron3Na_Mam_dup | 8.12642869497223 | 1.62707824822239 | 0.493747742939102 | 3.29536341480163 | 0.000982944732284979 | 0.0346963013304082 | 1756 | 20 | 10 | 2 | 5 | 11 | 5 | 2 | 3 | 13 | 14 | 10 | 5 |
| L1PA6_dup3797 | 7.26524364866892 | 1.93588108317479 | 0.508064151103805 | 3.81030836158968 | 0.000138793565322213 | 0.0086025979927479 | 1757 | 17 | 7 | 4 | 5 | 9 | 9 | 1 | 1 | 14 | 13 | 6 | 3 |
| THE1B-int_dup3307 | 7.88096923609828 | 2.15120468468151 | 0.533766277889737 | 4.03023715395878 | 5.5720606278859e-05 | 0.00442750622676088 | 1758 | 16 | 9 | 3 | 4 | 5 | 6 | 1 | 3 | 19 | 21 | 2 | 5 |
| L2a_dup11645 | 9.68565689631769 | 1.85980909193395 | 0.433238296591462 | 4.29280861494968 | 1.76427073597536e-05 | 0.00186412595267493 | 1759 | 17 | 13 | 4 | 7 | 7 | 11 | 3 | 3 | 18 | 25 | 4 | 6 |
| L2_dup8042 | 8.07441186953853 | 1.66107447068056 | 0.454012678364097 | 3.65865216950717 | 0.000253545179759994 | 0.0132739408895867 | 1760 | 9 | 10 | 3 | 7 | 10 | 8 | 3 | 2 | 19 | 16 | 5 | 5 |
| THE1B-int_dup3308 | 8.59612595837831 | 1.8405900640041 | 0.481304516873231 | 3.82416952153576 | 0.000131213614742052 | 0.00826190917528771 | 1761 | 9 | 10 | 4 | 5 | 5 | 8 | 2 | 2 | 25 | 21 | 4 | 6 |
| L1PA16_dup10636 | 9.13240122744549 | 2.4837174519134 | 0.510847528954172 | 4.8619545189896 | 1.16232332390963e-06 | 0.00023124597460786 | 1762 | 22 | 11 | 1 | 4 | 9 | 10 | 6 | 2 | 16 | 26 | 2 | 3 |
| L1M2c_dup99 | 8.52180112800614 | 2.2252810497849 | 0.521127739835188 | 4.27012588216598 | 1.95362670478013e-05 | 0.00201142945604017 | 1763 | 21 | 21 | 3 | 4 | 5 | 7 | 3 | 2 | 14 | 17 | 3 | 5 |
| MER11B_dup28 | 8.30904777249806 | 2.10001240545225 | 0.515658300437315 | 4.07248831963199 | 4.65135334521509e-05 | 0.00386577339532151 | 1764 | 19 | 16 | 2 | 6 | 6 | 6 | 4 | 1 | 13 | 22 | 3 | 5 |
| L1MEh_dup242 | 7.85823804372628 | 2.0325612754973 | 0.599124879130406 | 3.39255027841181 | 0.000692452226214504 | 0.0271264796540725 | 1765 | 10 | 6 | 0 | 0 | 17 | 16 | 3 | 5 | 12 | 13 | 8 | 5 |
| L1MB4_dup488 | 9.96850260967176 | 2.23659812788495 | 0.529732119696984 | 4.22213047825818 | 2.42003942446046e-05 | 0.00235324356719522 | 1766 | 12 | 13 | 0 | 1 | 20 | 20 | 3 | 7 | 14 | 19 | 3 | 9 |
| L1MEg_dup984 | 10.3757402858673 | 2.39628204936272 | 0.552882349983844 | 4.33416268295188 | 1.46315948008246e-05 | 0.0016208237693096 | 1767 | 9 | 10 | 0 | 1 | 25 | 24 | 3 | 4 | 17 | 17 | 6 | 10 |
| HERVL-int_dup835 | 9.43347127938505 | 2.66979760386167 | 0.552526579711854 | 4.83198040038903 | 1.35181565973661e-06 | 0.00026087520597866 | 1768 | 13 | 13 | 1 | 0 | 13 | 15 | 6 | 5 | 17 | 26 | 1 | 3 |
| MER104_dup722 | 11.0273394553867 | 1.84396176860123 | 0.521842289981463 | 3.5335613920189 | 0.000410000857095336 | 0.018815441454659 | 1769 | 6 | 14 | 1 | 1 | 16 | 19 | 8 | 7 | 25 | 20 | 8 | 5 |
| L1PA10_dup5291 | 9.24925819906844 | 1.90367892846503 | 0.517292193098757 | 3.68008439690794 | 0.000233156773704217 | 0.0124587293231133 | 1770 | 10 | 17 | 0 | 6 | 14 | 6 | 5 | 5 | 19 | 20 | 2 | 7 |
| Tigger3b_dup4005 | 8.216368975136 | 1.71062076534299 | 0.476861535126122 | 3.58724837156462 | 0.000334185970983693 | 0.0161523898134645 | 1771 | 7 | 10 | 0 | 4 | 8 | 14 | 3 | 4 | 17 | 17 | 5 | 10 |
| L1M3_dup3306 | 7.90288857432388 | 1.61303302367538 | 0.443577726039883 | 3.6364157372734 | 0.000276458020278689 | 0.0141102702902688 | 1772 | 9 | 10 | 1 | 5 | 10 | 11 | 3 | 2 | 14 | 16 | 6 | 10 |
| L2a_dup112236 | 7.98016002919858 | 1.77434423572754 | 0.452975300867616 | 3.91708826580393 | 8.96249399192149e-05 | 0.00628896961191906 | 1773 | 10 | 11 | 1 | 3 | 7 | 11 | 4 | 3 | 17 | 16 | 7 | 6 |
| MIR_dup67407 | 9.5848806484879 | 1.7802252540402 | 0.502126290180989 | 3.54537352226374 | 0.000392057086845978 | 0.0182326782024546 | 1774 | 20 | 24 | 1 | 3 | 16 | 10 | 3 | 7 | 12 | 8 | 6 | 8 |
| MIRc_dup32974 | 7.30148798984568 | 1.89182812332622 | 0.50863851991805 | 3.71939609220125 | 0.000199699678931488 | 0.0110970900987931 | 1775 | 20 | 15 | 1 | 6 | 9 | 11 | 2 | 5 | 8 | 8 | 4 | 2 |
| AluY_dup14391 | 7.57268597653738 | 1.65200688582632 | 0.457810372854187 | 3.60849597078152 | 0.000307977296803209 | 0.0152353359386978 | 1776 | 11 | 18 | 2 | 6 | 10 | 13 | 5 | 5 | 8 | 10 | 3 | 3 |
| MLT1N2_dup1907 | 9.29587605412625 | 1.97394017444399 | 0.425529286439524 | 4.63878806312083 | 3.50458208426093e-06 | 0.000541341739817781 | 1777 | 22 | 18 | 2 | 7 | 15 | 13 | 5 | 4 | 12 | 11 | 3 | 4 |
| MLT1J1_dup2519 | 11.1737682098382 | 2.44944929709784 | 0.49746188500684 | 4.92389340957078 | 8.48390601794093e-07 | 0.000181212881628813 | 1778 | 22 | 22 | 2 | 4 | 14 | 7 | 2 | 5 | 29 | 16 | 5 | 4 |
| L1ME3Cz_dup8254 | 9.1452540843779 | 2.4926210337037 | 0.551086514453007 | 4.52310293997632 | 6.09395832014943e-06 | 0.000839139471323077 | 1779 | 24 | 23 | 0 | 2 | 8 | 10 | 4 | 4 | 14 | 15 | 6 | 2 |
| HERV17-int_dup730 | 8.88532935470688 | 2.62373314565578 | 0.540264398944668 | 4.85638726294178 | 1.19546792480812e-06 | 0.000236456201859014 | 1780 | 14 | 30 | 2 | 1 | 15 | 9 | 3 | 4 | 9 | 16 | 3 | 3 |
| L1PA8A_dup1337 | 8.03777621834224 | 2.71194085162029 | 0.553106583522582 | 4.90310716308725 | 9.43324875945975e-07 | 0.000196722507389869 | 1781 | 16 | 17 | 1 | 2 | 17 | 9 | 3 | 2 | 17 | 7 | 4 | 2 |
| L1MA10_dup2492 | 8.3912480265583 | 2.19399509294224 | 0.681751413417123 | 3.2181746157963 | 0.00129009253043496 | 0.0418617697050152 | 1782 | 6 | 9 | 3 | 0 | 25 | 14 | 2 | 0 | 13 | 9 | 12 | 10 |
| L1P1_dup2443 | 10.5333549909802 | 1.85469542114524 | 0.483327678057944 | 3.83734577046691 | 0.00012437129608484 | 0.00794460467602773 | 1783 | 10 | 10 | 4 | 4 | 14 | 23 | 2 | 0 | 21 | 16 | 11 | 14 |
| L2c_dup133602 | 10.6204374207416 | 1.9509379844142 | 0.604869755403321 | 3.22538524531341 | 0.00125803258179241 | 0.041169553116102 | 1784 | 10 | 12 | 1 | 3 | 24 | 21 | 3 | 0 | 11 | 14 | 16 | 19 |
| L1MB3_dup1641 | 8.39428969801076 | 2.30429477072695 | 0.645248578655414 | 3.57117372583556 | 0.000355385065020811 | 0.0169057286448304 | 1785 | 16 | 11 | 0 | 2 | 12 | 11 | 0 | 1 | 8 | 19 | 15 | 10 |
| L3_dup41818 | 10.6707403548515 | 1.64108881831423 | 0.449402613666721 | 3.65171178005492 | 0.000260498150337651 | 0.0135307777516356 | 1786 | 14 | 14 | 3 | 6 | 13 | 15 | 0 | 2 | 19 | 17 | 13 | 15 |
| MIRc_dup34917 | 9.38247455903274 | 1.67630705929301 | 0.518054801644425 | 3.23577168664787 | 0.00121314375272364 | 0.0401366384017768 | 1787 | 12 | 8 | 2 | 5 | 14 | 19 | 0 | 2 | 11 | 18 | 13 | 13 |
| HUERS-P2-int_dup237 | 10.1841196920485 | 2.86136359288236 | 0.5416348191488 | 5.28282800832322 | 1.27204802905193e-07 | 3.99839815483212e-05 | 1788 | 14 | 6 | 1 | 2 | 16 | 21 | 0 | 1 | 23 | 23 | 8 | 7 |
| THE1C-int_dup179 | 9.01780617714775 | 2.5228565151048 | 0.529477134736384 | 4.76480729684555 | 1.89034221924588e-06 | 0.000337705615241662 | 1789 | 9 | 11 | 2 | 2 | 18 | 10 | 0 | 1 | 23 | 16 | 6 | 9 |
| L2c_dup31080 | 9.15275494967519 | 2.89934064072143 | 0.558329692967421 | 5.19288276665344 | 2.07062579237481e-07 | 5.9289666040002e-05 | 1790 | 20 | 21 | 0 | 2 | 15 | 10 | 2 | 1 | 11 | 21 | 2 | 9 |
| L1PA5_dup689 | 7.31103060775302 | 2.04574924012089 | 0.535303040523861 | 3.8216656459094 | 0.00013255333099349 | 0.00830981871543592 | 1791 | 7 | 16 | 1 | 2 | 10 | 10 | 3 | 1 | 12 | 14 | 3 | 11 |
| L1MA2_dup7118 | 8.54374187224362 | 2.29914996632342 | 0.480855794962364 | 4.78137102726062 | 1.74103687317076e-06 | 0.000317450390669341 | 1792 | 15 | 18 | 2 | 5 | 9 | 15 | 1 | 1 | 12 | 17 | 4 | 7 |
| L1PB1_dup10721 | 9.13719832337701 | 2.21311835553806 | 0.496018100675647 | 4.46176934374668 | 8.12856938176518e-06 | 0.00104211237613874 | 1793 | 15 | 17 | 2 | 2 | 9 | 9 | 1 | 2 | 15 | 23 | 4 | 13 |
| L1PA3_dup7687 | 9.55407591915169 | 1.34238742379394 | 0.413861452527702 | 3.24356717832785 | 0.0011804297066155 8 | 0.0393508583211298 | 1794 | 16 | 11 | 1 | 4 | 13 | 12 | 6 | 5 | 10 | 20 | 11 | 10 |
| HERVIP10FH-int_dup15 | 8.94250497580521 | 1.85989541844848 | 0.512957817812787 | 3.62582527034861 | 0.000288040076954357 | 0.01452999452003 | 1795 | 16 | 18 | 1 | 2 | 14 | 11 | 2 | 3 | 7 | 17 | 11 | 10 |
| L1PA17_dup2460 | 8.10659694886209 | 1.77591292226162 | 0.472797659327328 | 3.75617959866446 | 0.000172526902985663 | 0.0100543929605257 | 1796 | 14 | 14 | 1 | 2 | 10 | 10 | 3 | 2 | 10 | 16 | 8 | 11 |
| L3_dup41817 | 10.2227441437148 | 2.0563940487254 | 0.464824773777567 | 4.42401989896831 | 9.68810909131534e-06 | 0.00119442839680939 | 1797 | 21 | 15 | 2 | 2 | 12 | 18 | 2 | 2 | 11 | 21 | 10 | 12 |
| MIRb_dup165033 | 8.79882876006149 | 1.70345354576892 | 0.485709230903694 | 3.50714674003526 | 0.000452939302580388 | 0.020193947028629 | 1798 | 15 | 10 | 3 | 2 | 10 | 12 | 1 | 1 | 11 | 19 | 11 | 15 |
| L1M5_dup49217 | 9.19199498054106 | 1.56203153048327 | 0.469724568890736 | 3.32542011624396 | 0.000882853952733029 | 0.0322740154612096 | 1799 | 14 | 13 | 1 | 5 | 12 | 9 | 4 | 1 | 12 | 20 | 16 | 7 |
| MLT1L_dup8751 | 8.98946743591198 | 1.43439547075104 | 0.438409951673435 | 3.27181320879207 | 0.00106860151619107 | 0.0367813476597136 | 1800 | 11 | 13 | 2 | 5 | 11 | 9 | 2 | 2 | 12 | 20 | 14 | 10 |
| MIR_dup155486 | 7.99911597961372 | 1.91372533661814 | 0.486309161162671 | 3.93520313712123 | 8.31263566772443e-05 | 0.00595232743482503 | 1801 | 17 | 12 | 3 | 2 | 10 | 8 | 2 | 1 | 15 | 12 | 9 | 7 |
| MIRb_dup53568 | 8.60788065953344 | 2.10271127418959 | 0.472553656561377 | 4.4496772905967 | 8.59994129050773e-06 | 0.00108626664002925 | 1802 | 12 | 16 | 3 | 1 | 12 | 12 | 3 | 1 | 11 | 20 | 9 | 6 |
| MLT2B1_dup944 | 7.62232577170456 | 2.3462327308195 | 0.662354083302481 | 3.54226355655761 | 0.000396708892065159 | 0.018364843196758 | 1803 | 14 | 13 | 0 | 0 | 12 | 10 | 2 | 2 | 11 | 13 | 2 | 16 |
| MSTC_dup2193 | 6.5707499113968 | 2.14852207381891 | 0.591612214714842 | 3.63163913857745 | 0.000281626747866904 | 0.014279130225429 | 1804 | 15 | 6 | 0 | 0 | 8 | 10 | 2 | 2 | 11 | 12 | 6 | 9 |
| L2b_dup4679 | 9.34278839042286 | 1.83197682091862 | 0.537642774819446 | 3.40742386342649 | 0.000655792013619564 | 0.0261159977360202 | 1805 | 13 | 14 | 1 | 0 | 6 | 7 | 5 | 1 | 20 | 21 | 8 | 18 |
| L1ME1_dup14553 | 7.8170534131102 | 1.89135917651339 | 0.525286242612733 | 3.60062575997787 | 0.000317452234822116 | 0.01556169951936 2 | 1806 | 7 | 11 | 1 | 0 | 8 | 11 | 4 | 2 | 20 | 12 | 8 | 9 |
| Tigger1_dup873 | 7.90540764407737 | 1.83314093871229 | 0.4861950215868 | 3.77038196057509 | 0.000162997884559923 | 0.00966020449673615 | 1807 | 9 | 11 | 1 | 0 | 8 | 12 | 5 | 3 | 14 | 18 | 6 | 9 |
| L4_A_Mam_dup1279 | 11.0239773656108 | 2.679399897173 | 0.618255788167676 | 4.33380479156359 | 1.46554138598491e-05 | 0.00162195214246309 | 1808 | 29 | 23 | 0 | 0 | 12 | 11 | 3 | 5 | 11 | 30 | 2 | 11 |
| Tigger15a_dup1079 | 13.6578769184699 | 2.3623516244999 | 0.50101900232001 | 4.71509386582312 | 2.41599344916053e-06 | 0.000408234739014297 | 1809 | 25 | 18 | 0 | 0 | 14 | 13 | 7 | 2 | 29 | 32 | 8 | 18 |
| Tigger1_dup872 | 10.0291037587355 | 2.45823755662036 | 0.505019873284209 | 4.8676055867547 | 1.12958483568805e-06 | 0.000226245953966858 | 1810 | 16 | 12 | 0 | 0 | 13 | 15 | 7 | 2 | 20 | 24 | 8 | 4 |
| L1PA7_dup800 | 14.5450244476598 | 2.54527083179581 | 0.548675145972358 | 4.63893954460995 | 3.50201467150001e-06 | 0.000541341739817781 | 1811 | 13 | 20 | 0 | 1 | 32 | 31 | 5 | 2 | 19 | 28 | 9 | 20 |
| AluSz_dup81446 | 12.3249535490603 | 2.25016750753574 | 0.59938454080038 | 3.75413003567128 | 0.000173944550709232 | 0.0101073040447031 | 1812 | 16 | 17 | 1 | 1 | 35 | 31 | 7 | 1 | 11 | 12 | 8 | 16 |

| | | | | | | | | | | | | | | | | | | |
|---|---|---|---|---|---|---|---|---|---|---|---|---|---|---|---|---|---|---|
| L2c_dup67919 | 13.9184823727432 | 1.79044844130728 | 0.55150432860853 | 3.24648121225932 | 0.00116841159712709 | 0.0391190186984667 | 1813 | 17 | 23 | 1 | 5 | 33 | 32 | 6 | 2 | 10 | 12 | 17 | 19 |
| AluY_dup82072 | 11.9167983425135 | 2.27940486649643 | 0.702344395909833 | 3.24542329912612 | 0.00117276152634514 | 0.0392039778235067 | 1814 | 13 | 11 | 0 | 0 | 14 | 24 | 1 | 3 | 24 | 12 | 11 | 34 |
| THE1B_dup1362 | 13.2224404275041 | 2.39976378471946 | 0.638499871466611 | 3.7584405133979 | 0.000170975675145444 | 0.010003191049461 | 1815 | 24 | 19 | 0 | 0 | 16 | 14 | 3 | 1 | 18 | 22 | 24 | 24 |
| L1MCb_dup1309 | 9.46559920781217 | 2.45997580414194 | 0.632031091836538 | 3.89217529946796 | 9.93494161258412e-05 | 0.00678652633207143 | 1816 | 23 | 23 | 0 | 0 | 10 | 15 | 3 | 2 | 11 | 12 | 12 | 7 |
| L1PB1_dup9546 | 11.2900405859438 | 2.51813236575567 | 0.646902093835924 | 3.89260197138016 | 9.91747921851297e-05 | 0.00678042504119536 | 1817 | 28 | 34 | 0 | 1 | 14 | 18 | 3 | 2 | 11 | 9 | 13 | 9 |
| L1P1_dup2291 | 12.1900030717505 | 1.96078175236461 | 0.608306375721716 | 3.22334571956159 | 0.00126702531184524 | 0.0413614494241086 | 1818 | 43 | 23 | 0 | 2 | 17 | 14 | 4 | 6 | 14 | 7 | 9 | 15 |
| L1PA11_dup2123 | 11.9887962729388 | 2.40443617862316 | 0.569763455808316 | 4.2200603673537 | 2.44236908835277e-05 | 0.00237046752388554 | 1819 | 26 | 20 | 0 | 1 | 29 | 20 | 5 | 2 | 15 | 10 | 10 | 12 |
| L1PA8_dup805 | 30.2862679776957 | -1.80033971284391 | 0.480249929918179 | -3.74875580544205 | 0.000177714015464329 | 0.0102586945957485 | 1820 | 1 | 0 | 0 | 0 | 0 | 1 | 0 | 1 | 27 | 33 | 144 | 188 |
| MSTA_dup20229 | 12.5716755315099 | -1.88685450425227 | 0.57105772988894 | -3.30413967887139 | 0.000952683533078945 | 0.0339809977457656 | 1821 | 0 | 0 | 0 | 0 | 1 | 0 | 1 | 2 | 11 | 15 | 59 | 74 |
| L1ME1_dup18307 | 17.2844691573433 | -1.8977760549171 | 0.592980204898649 | -3.20040372214695 | 0.0013723521004103 | 0.0437552851245114 | 1822 | 0 | 0 | 0 | 0 | 0 | 0 | 0 | 0 | 14 | 18 | 92 | 101 |
| MSTA_dup19525 | 15.0833904766994 | -2.7684731629466 | 0.653032074156393 | -4.23941376313407 | 2.24104262781662e-05 | 0.00222464384279705 | 1823 | 0 | 0 | 0 | 3 | 0 | 0 | 0 | 0 | 6 | 12 | 63 | 120 |
| L1M4a2_dup1031 | 7.84750592833776 | -3.24487892960498 | 0.832477172694641 | -3.8978593480247 | 9.70467519728523e-05 | 0.0066643250923325 6 | 1824 | 0 | 2 | 10 | 7 | 2 | 1 | 2 | 1 | 1 | 0 | 33 | 47 |
| LTR78_dup3865 | 7.0682297950004 | -3.82735191648305 | 0.81390996759991 | -4.70242664280093 | 2.57087598258893e-06 | 0.000429283969510952 | 1825 | 0 | 1 | 7 | 9 | 0 | 0 | 2 | 1 | 3 | 1 | 35 | 35 |
| SST1_dup440 | 7.35416847722182 | -4.96784757035559 | 1.02377386271221 | -4.85248525215778 | 1.21923853228592e-06 | 0.000239648601203287 | 1826 | 0 | 1 | 13 | 15 | 0 | 0 | 1 | 1 | 1 | 0 | 34 | 33 |
| L1MB5_dup7703 | 8.12639899337505 | -4.70924587435136 | 0.957556979771433 | -4.91797979006477 | 8.7441937544001e-07 | 0.000185440813465641 | 1827 | 0 | 1 | 12 | 14 | 0 | 0 | 0 | 3 | 1 | 1 | 35 | 42 |
| LTR57-int_dup256 | 6.16575381855247 | -5.12868283318912 | 1.11726631991079 | -4.59038524816412 | 4.42428642963765e-06 | 0.000653470675845165 | 1828 | 0 | 0 | 5 | 17 | 0 | 0 | 0 | 2 | 1 | 0 | 31 | 27 |
| L1MD2_dup5522 | 7.85788495928712 | -3.75871347094842 | 0.905111435954362 | -4.1527632086376 | 3.28484592663411e-05 | 0.0029606664729503 | 1829 | 2 | 1 | 6 | 18 | 0 | 1 | 1 | 2 | 0 | 1 | 24 | 53 |
| L1MA4A_dup3283 | 8.29172156036966 | -4.42257386378787 | 0.953135085886967 | -4.64002839605083 | 3.48361306665805e-06 | 0.000539656116379568 | 1830 | 1 | 0 | 6 | 14 | 0 | 2 | 1 | 2 | 0 | 0 | 38 | 49 |
| L1PB1_dup7168 | 8.7818546065612 | -5.52501449690186 | 0.936222843523199 | -5.9013882593487 | 3.6045551796862e-09 | 2.12299377917231e-06 | 1831 | 0 | 0 | 10 | 12 | 0 | 1 | 3 | 2 | 0 | 0 | 42 | 49 |
| L1MB3_dup8398 | 4.86292538050753 | -3.95894112599904 | 0.929326805369987 | -4.26000961461871 | 2.04418110175716e-05 | 0.00207776470072067 | 1832 | 0 | 1 | 3 | 5 | 0 | 0 | 2 | 0 | 1 | 0 | 24 | 30 |
| L1PA15_dup7074 | 3.74561790019014 | -2.96705347407711 | 0.92945079716547 | -3.19226524214696 | 0.0014116164012842 | 0.0445534522111648 | 1833 | 1 | 1 | 3 | 5 | 0 | 0 | 2 | 0 | 2 | 0 | 13 | 24 |
| ERVL-E-int_dup5058 | 3.88443584459471 | -4.39345870762928 | 1.16847531051046 | -3.75999275988978 | 0.000169918274563348 | 0.00996337450942592 | 1834 | 0 | 0 | 2 | 10 | 0 | 1 | 1 | 0 | 0 | 0 | 16 | 24 |
| MSTB-int_dup467 | 3.80699885317331 | -4.14176543971433 | 1.02572113271238 | -4.03790592552381 | 5.39304709841127e-05 | 0.00432303096906091 | 1835 | 0 | 0 | 3 | 4 | 0 | 1 | 2 | 0 | 0 | 0 | 19 | 23 |
| HAL1b_dup1101 | 3.12145206052081 | -4.86621121251878 | 1.17435878007928 | -4.14371765687337 | 3.41720429336818e-05 | 0.00304307669457255 | 1836 | 0 | 0 | 5 | 9 | 0 | 0 | 2 | 0 | 0 | 0 | 13 | 14 |
| MamGypsy2-I_dup157 | 3.72634752965587 | -5.04758440132568 | 1.14780582147104 | -4.39759435516423 | 1.09457305191616e-05 | 0.00130498206640708 | 1837 | 0 | 0 | 6 | 9 | 0 | 0 | 2 | 0 | 0 | 0 | 17 | 17 |
| L1MB3_dup8399 | 3.28328233952304 | -4.51957659003269 | 1.1462441226049 | -3.94294417821024 | 8.04873945229332e-05 | 0.00580705081175072 | 1838 | 0 | 0 | 3 | 5 | 0 | 0 | 1 | 0 | 0 | 0 | 16 | 20 |
| L3_dup28927 | 2.74064700973739 | -4.51559022454051 | 1.24305907487104 | -3.63264330378584 | 0.00028053268729349 | 0.0142463905013437 | 1839 | 0 | 0 | 2 | 8 | 0 | 0 | 1 | 0 | 0 | 0 | 12 | 15 |
| L1MEg_dup8069 | 4.71459212385237 | -3.99050890482363 | 0.916993645654426 | -4.35173016054625 | 1.3506743210891e-05 | 0.0015282337232804 | 1840 | 0 | 0 | 4 | 4 | 1 | 0 | 1 | 1 | 0 | 1 | 21 | 31 |
| L1MB3_dup8400 | 4.46392553974084 | -3.83060156489367 | 1.09182404768653 | -3.50844220093004 | 0.000450739164266405 | 0.020126023865741 | 1841 | 0 | 0 | 3 | 6 | 2 | 0 | 0 | 1 | 0 | 0 | 24 | 24 |
| L1PA15_dup4372 | 5.52684780612126 | -4.82551236501568 | 1.1132666702747 | -4.33455208339702 | 1.46057206272418e-05 | 0.00161946542421 7 | 1842 | 0 | 0 | 8 | 7 | 1 | 0 | 0 | 1 | 0 | 0 | 19 | 40 |
| L2a_dup92237 | 3.39311883487761 | -3.93689320446375 | 1.1824563133682 | -3.32941958189525 | 0.000870271918045769 | 0.0319170348044572 | 1843 | 0 | 0 | 2 | 6 | 0 | 0 | 0 | 0 | 0 | 1 | 19 | 18 |
| AluSq2_dup49094 | 4.24760940276751 | -4.27715447524011 | 1.15860883489953 | -3.69162943212929 | 0.000222821954317145 | 0.0120280382808411 | 1844 | 0 | 0 | 5 | 5 | 0 | 0 | 0 | 0 | 0 | 1 | 14 | 34 |
| L1MB4_dup4307 | 4.0399746501042 | -3.78833674875508 | 1.10670862920264 | -3.42306606164669 | 0.000619190271446358 | 0.025094098408732 | 1845 | 0 | 0 | 4 | 6 | 0 | 1 | 0 | 0 | 0 | 1 | 14 | 30 |
| L1ME3_dup4712 | 3.53474756098671 | -3.80065605068745 | 1.20849819556035 | -3.14494143611459 | 0.00166120080693317 | 0.0498708159483372 | 1846 | 0 | 1 | 6 | 6 | 1 | 0 | 0 | 0 | 0 | 0 | 10 | 25 |
| L1PA8A_dup209 | 2.80438851221383 | -4.13222000702669 | 1.30489308350527 | -3.16671155611195 | 0.00154173158417509 | 0.0473597809873645 | 1847 | 0 | 0 | 3 | 3 | 0 | 0 | 0 | 0 | 0 | 0 | 10 | 23 |
| L1PB_dup1450 | 2.64598106047907 | -4.31168478183628 | 1.35062657947436 | -3.19235890020342 | 0.00141115871398306 | 0.0445449152769185 | 1848 | 0 | 0 | 5 | 3 | 0 | 0 | 0 | 0 | 0 | 0 | 12 | 16 |
| HERVKC4-int_dup34 | 11.4433884429209 | -4.24085574302406 | 1.00350145241784 | -4.22605840061927 | 2.37820294605598e-05 | 0.00232395843204204 | 1849 | 0 | 0 | 18 | 10 | 0 | 1 | 0 | 0 | 2 | 4 | 41 | 80 |
| SST1_dup444 | 12.8215950545675 | -5.93815060738449 | 1.80273850890556 | -3.29396114747087 | 0.000987861519976139 | 0.0348204360034377 | 1850 | 0 | 0 | 18 | 20 | 0 | 0 | 0 | 0 | 1 | 0 | 58 | 78 |
| ALR/Alpha_dup1202 | 6.5192002770356 | -3.7847175235894 | 1.10306256964376 | -3.43109958378127 | 0.000601139877015372 | 0.0246151831313543 | 1851 | 0 | 2 | 12 | 16 | 0 | 0 | 0 | 0 | 1 | 2 | 22 | 34 |
| L1PB3_dup3147 | 5.81700120360736 | -3.84821737446243 | 1.1246285127081 | -3.42176757122753 | 0.000622154742500477 | 0.025155349593716 | 1852 | 0 | 1 | 18 | 6 | 0 | 0 | 0 | 0 | 1 | 3 | 19 | 30 |
| LTR19-int_dup171 | 5.38146905484379 | -4.45762153261679 | 1.25341029229852 | -3.55639455013755 | 0.00037597929340638 | 0.0176559638738651 | 1853 | 0 | 0 | 10 | 15 | 0 | 0 | 0 | 0 | 0 | 3 | 18 | 28 |
| L1MA8_dup9552 | 7.5079981293443 | -5.40889199791634 | 1.12132987131584 | -4.82364033660248 | 1.40961505100813e-06 | 0.000269407440102153 | 1854 | 0 | 0 | 9 | 18 | 0 | 0 | 1 | 0 | 0 | 1 | 34 | 40 |
| L1MD2_dup9189 | 6.48527765935571 | -4.89288683322562 | 1.10783158103765 | -4.41663418607608 | 1.00249676733863e-05 | 0.00122516741551699 | 1855 | 0 | 0 | 13 | 12 | 0 | 0 | 1 | 0 | 0 | 2 | 23 | 38 |
| HERVH-int_dup3388 | 8.71037840299146 | -3.53445245233345 | 0.869801092506406 | -4.06351806497348 | 4.83385793484372e-05 | 0.00397167527681051 | 1856 | 0 | 0 | 0 | 2 | 0 | 0 | 0 | 0 | 1 | 2 | 49 | 64 |
| L1ME3A_dup15341 | 4.26841242374451 | -3.31957083667414 | 0.939024457095595 | -3.53512713283483 | 0.000407579012463662 | 0.0187368229713444 | 1857 | 0 | 0 | 3 | 7 | 0 | 0 | 0 | 2 | 1 | 4 | 16 | 24 |
| L2c_dup96715 | 4.08660330155065 | -3.02791603300567 | 0.884895291939377 | -3.42177889360169 | 0.000622128836394778 | 0.025155349593716 | 1858 | 0 | 1 | 2 | 3 | 0 | 0 | 0 | 2 | 2 | 1 | 23 | 19 |
| L1M3_dup5202 | 4.91550027959087 | -3.17010137850209 | 0.883194791996718 | -3.58935696544944 | 0.000331494657317736 | 0.0160614595255805 | 1859 | 0 | 0 | 3 | 3 | 0 | 0 | 0 | 2 | 4 | 1 | 18 | 34 |
| L1MC4a_dup7786 | 4.27213377704361 | -2.79053302591795 | 0.847604230134949 | -3.29225943748967 | 0.000993858829954538 | 0.0349570928532687 | 1860 | 0 | 0 | 3 | 3 | 0 | 0 | 0 | 1 | 3 | 3 | 19 | 24 |
| L1MA3_dup7285 | 6.88226372883044 | -2.60059767925444 | 0.736681191794696 | -3.53015348867383 | 0.000415318649398498 | 0.0190154726476381 | 1861 | 0 | 0 | 3 | 1 | 1 | 1 | 0 | 2 | 2 | 5 | 33 | 43 |
| MSTA_dup19305 | 8.0592296174809 | -2.76954086521451 | 0.698127707484942 | -3.96709776094118 | 7.27531554683889e-05 | 0.00538454815896774 | 1862 | 1 | 0 | 3 | 3 | 1 | 0 | 0 | 3 | 5 | 3 | 42 | 44 |
| MLT1D_dup9420 | 2.05528613323758 | -3.90055003474258 | 1.17869309109728 | -3.30921599880715 | 0.000935576288990692 | 0.0335776495423195 | 1863 | 0 | 0 | 0 | 4 | 1 | 0 | 5 | 4 | 0 | 0 | 6 | 7 |
| L1MD_dup4940 | 2.37677861786164 | -3.64789190571477 | 1.1235512168479 | -3.24675177331822 | 0.00116730150040371 | 0.0391001328500764 | 1864 | 0 | 0 | 0 | 2 | 0 | 0 | 2 | 3 | 0 | 1 | 12 | 11 |
| L1MC4_dup17256 | 3.16118652111884 | -3.94934311379014 | 1.09331211819694 | -3.61227416037726 | 0.000303523415693634 | 0.0150799956713082 | 1865 | 1 | 0 | 0 | 4 | 0 | 0 | 2 | 2 | 0 | 0 | 19 | 14 |
| L1MEi_dup3802 | 2.49792375332362 | -4.34896940297307 | 1.2460456670331 | -3.4902167055628 | 0.000482629011956074 | 0.0210136583492955 | 1866 | 0 | 0 | 0 | 6 | 0 | 0 | 1 | 2 | 0 | 0 | 10 | 15 |
| AluSx_dup82049 | 3.53185938152142 | -2.74769013973262 | 0.865365026423939 | -3.17518047971879 | 0.00149743333831391 | 0.0464487385110731 | 1867 | 1 | 0 | 2 | 6 | 1 | 0 | 2 | 1 | 1 | 2 | 11 | 21 |
| MLT1F2_dup4822 | 3.28287097579089 | -2.94779939498004 | 0.928307243299942 | -3.17545663491886 | 0.00149600880005446 | 0.0464314647428227 | 1868 | 1 | 0 | 3 | 6 | 2 | 0 | 3 | 1 | 0 | 1 | 10 | 18 |
| AluSx1_dup81385 | 3.11720528107287 | -2.97171313479365 | 0.936540689967258 | -3.17307423652628 | 0.00150833951442092 | 0.0466714374640312 | 1869 | 0 | 0 | 5 | 5 | 1 | 1 | 1 | 2 | 0 | 2 | 13 | 11 |
| MSTB_dup4465 | 2.15593454967724 | -3.38087555343423 | 1.04022547301173 | -3.25013724538557 | 0.00115349325401308 | 0.0387832564390839 | 1870 | 0 | 0 | 2 | 5 | 1 | 0 | 3 | 2 | 0 | 1 | 6 | 9 |
| AluJo_dup54822 | 3.20175729405535 | -3.51074463274094 | 0.960275852089816 | -3.65597512954285 | 0.00025620619508091 | 0.0133808690286923 | 1871 | 0 | 0 | 6 | 6 | 1 | 0 | 2 | 2 | 0 | 2 | 9 | 15 |
| L1MA9_dup15237 | 2.9478220648381 | -3.12376351664761 | 0.986247488041142 | -3.16732215242641 | 0.00153849781962367 | 0.0473055531490971 | 1872 | 0 | 0 | 5 | 9 | 1 | 0 | 3 | 1 | 0 | 3 | 8 | 10 |
| AluSq2_dup43748 | 2.91445443833289 | -2.90061087688222 | 0.906823634500362 | -3.19864940273683 | 0.00138072969624548 | 0.0439105998800891 | 1873 | 0 | 0 | 3 | 2 | 0 | 1 | 4 | 2 | 1 | 2 | 7 | 17 |
| LTR85b_dup948 | 3.06922112884689 | -3.34591740267655 | 0.924469072298562 | -3.61928538545632 | 0.000295417700272726 | 0.0148048568047797 | 1874 | 0 | 0 | 2 | 4 | 0 | 1 | 4 | 3 | 1 | 1 | 8 | 17 |
| MER3_dup8136 | 3.24167923906521 | -3.13912340187584 | 0.898623033852817 | -3.49325944653005 | 0.000477162670300431 | 0.0208711302844313 | 1875 | 0 | 0 | 3 | 5 | 0 | 0 | 2 | 1 | 1 | 3 | 14 | 14 |
| MIR_dup104346 | 3.13976190099499 | -3.82914649196902 | 0.955069295044929 | -4.00928656364027 | 6.09024724465433e-05 | 0.00472960173262572 | 1876 | 0 | 0 | 4 | 4 | 0 | 0 | 5 | 1 | 1 | 1 | 13 | 13 |
| AluY_dup101976 | 3.36427135607245 | -3.44564319361014 | 0.900872184879661 | -3.82478585912874 | 0.000130885801456787 | 0.00825218678151476 | 1877 | 0 | 0 | 3 | 2 | 0 | 0 | 4 | 2 | 1 | 2 | 11 | 20 |
| L2a_dup165789 | 2.95218376789932 | -3.64523040810907 | 0.943172243573819 | -3.86486183509467 | 0.00011115210090512 | 0.00735080539243182 | 1878 | 0 | 0 | 3 | 3 | 0 | 0 | 3 | 2 | 1 | 1 | 12 | 14 |
| L1MC4_dup19250 | 2.42141322477256 | -3.52948684866339 | 0.99581080800317 | -3.54433474742137 | 0.000393605159967594 | 0.0182673636422175 | 1879 | 0 | 0 | 4 | 4 | 0 | 0 | 3 | 2 | 1 | 1 | 7 | 10 |
| L1ME3F_dup1038 | 2.51751004256852 | -3.14282323220959 | 0.961100973514257 | -3.27002398168205 | 0.00107538367009349 | 0.0369476013824392 | 1880 | 0 | 0 | 4 | 3 | 0 | 0 | 3 | 2 | 2 | 1 | 6 | 12 |
| AluSz_dup20912 | 2.87548309457465 | -3.13757159103189 | 0.943394640350691 | -3.3258314779757 | 0.000881552100297768 | 0.0322412843617357 | 1881 | 0 | 0 | 3 | 9 | 0 | 0 | 4 | 2 | 2 | 2 | 8 | 8 |
| L1MA10_dup922 | 2.8134483202299 | -3.05303613923766 | 0.927064613151879 | -3.29322907586538 | 0.000990437429843725 | 0.0348883234169645 | 1882 | 0 | 0 | 2 | 6 | 0 | 0 | 4 | 3 | 2 | 2 | 6 | 12 |
| MER68_dup910 | 2.71015816127704 | -4.74660095011888 | 1.14160873089298 | -4.15781766700928 | 3.21302306619367e-05 | 0.00290804863639604 | 1883 | 0 | 0 | 1 | 9 | 0 | 0 | 5 | 1 | 0 | 0 | 8 | 14 |
| MSTA1_dup508 | 3.40708216984729 | -4.46321304708491 | 1.08845270197593 | -4.10051170710735 | 4.12237579411515e-05 | 0.00353096771312643 | 1884 | 0 | 0 | 1 | 7 | 0 | 0 | 8 | 1 | 0 | 1 | 8 | 22 |
| L2c_dup75808 | 2.46096783613565 | -3.94928687684288 | 1.06233280603814 | -3.7175608758345 | 0.000201155524667988 | 0.0111649527577908 | 1885 | 0 | 0 | 4 | 3 | 0 | 0 | 2 | 2 | 0 | 1 | 13 | 7 |
| L2a_dup79862 | 2.15516090343901 | -3.37273391867042 | 1.04469755746985 | -3.22843094114131 | 0.00124471308303694 | 0.0408629276015878 | 1886 | 0 | 0 | 3 | 2 | 0 | 0 | 3 | 3 | 0 | 2 | 7 | 8 |
| MER91A_dup3340 | 2.76222111598001 | -4.14848933809689 | 1.02132175705062 | -4.06188285861738 | 4.86785034696359e-05 | 0.00398858649675826 | 1887 | 0 | 0 | 2 | 4 | 0 | 0 | 4 | 3 | 0 | 1 | 9 | 14 |
| L2b_dup22466 | 1.83083021781714 | -3.74220120017216 | 1.15713544321288 | -3.23402175788655 | 0.00122060156276798 | 0.0402968620175563 | 1888 | 0 | 0 | 2 | 8 | 0 | 0 | 2 | 1 | 0 | 1 | 6 | 5 |
| L1MB4_dup5479 | 2.82694355606765 | -4.36784129006376 | 1.10037657790922 | -3.96940590862351 | 7.20520512826133e-05 | 0.00534710773564649 | 1889 | 0 | 0 | 6 | 10 | 0 | 0 | 1 | 2 | 0 | 1 | 9 | 9 |
| L4_A_Mam_dup4412 | 2.91492682846933 | -4.31355508963347 | 1.03379152336006 | -4.17255799855412 | 3.01198894441624e-05 | 0.00276501049910519 | 1890 | 0 | 0 | 4 | 7 | 0 | 0 | 2 | 3 | 0 | 1 | 9 | 13 |
| L1MD3_dup4802 | 2.50509854384959 | -3.65801517569848 | 1.08798076992571 | -3.36220572717315 | 0.000773224925233936 | 0.0293330517194483 | 1891 | 0 | 0 | 4 | 7 | 0 | 0 | 1 | 2 | 0 | 2 | 5 | 13 |

| | | | | | | | | | | | | | | | | | | | |
|---|---|---|---|---|---|---|---|---|---|---|---|---|---|---|---|---|---|---|---|
| L1PA5_dup9444 | 2.75532611964115 | -3.33050939810628 | 0.994021275782657 | -3.3505413608818 | 0.000806537635517396 | 0.0302271860483261 | 1892 | 0 | 0 | 2 | 6 | 0 | 0 | 2 | 2 | 0 | 3 | 9 | 13 |
| LTR16_dup1697 | 3.05226358552211 | -3.36028681471476 | 1.01904902541533 | -3.297473164596 | 0.00097558996770474 | 0.0345484261373344 | 1893 | 1 | 0 | 1 | 4 | 0 | 1 | 1 | 1 | 0 | 0 | 17 | 15 |
| L1MD2_dup7172 | 2.83186590993295 | -4.08491773929874 | 1.04684949019496 | -3.90210605971446 | 9.53593551121225e-05 | 0.00658073582120334 | 1894 | 0 | 0 | 2 | 5 | 0 | 1 | 3 | 1 | 0 | 0 | 10 | 17 |
| Tigger3b_dup3593 | 3.9010716559369 | -3.8308497037875 | 0.99833246789782 | -3.83724843874315 | 0.000124420583282944 | 0.00794561999727413 | 1895 | 0 | 0 | 2 | 5 | 0 | 2 | 3 | 1 | 0 | 0 | 24 | 15 |
| AluY_dup21861 | 2.78644269282189 | -3.80608588039239 | 1.07931381922494 | -3.52639409650621 | 0.000421259604887942 | 0.019184122094673 | 1896 | 1 | 0 | 8 | 5 | 0 | 1 | 2 | 1 | 0 | 0 | 12 | 7 |
| L1ME4b_dup19871 | 2.45301623211867 | -4.03762395749396 | 1.07298377656466 | -3.76298695812633 | 0.000167895960658162 | 0.00987155232388609 | 1897 | 0 | 0 | 5 | 2 | 0 | 1 | 3 | 2 | 0 | 0 | 12 | 7 |
| LTR43-int_dup44 | 2.64185532970827 | -4.15131850900186 | 1.03658653826148 | -4.00479685561445 | 6.20707992149012e-05 | 0.00479252837891568 | 1898 | 0 | 0 | 4 | 4 | 0 | 1 | 2 | 3 | 0 | 0 | 9 | 12 |
| MLT1F2_dup4606 | 2.90517729634319 | -4.2810116603503 | 1.06357215122614 | -4.02512575702076 | 5.69448757107395e-05 | 0.004494131142908 | 1899 | 0 | 0 | 6 | 4 | 0 | 1 | 1 | 3 | 0 | 0 | 12 | 11 |
| L1MA5A_dup2751 | 1.99647987276295 | -4.1093890452562 | 1.25591530565897 | -3.27202720337899 | 0.00106779301371034 | 0.0367747975742252 | 1900 | 0 | 0 | 3 | 4 | 0 | 0 | 0 | 1 | 0 | 0 | 8 | 11 |
| L1MC5_dup7884 | 2.64826462448023 | -3.63230799982675 | 1.08438837282474 | -3.34963753840785 | 0.000809173713757954 | 0.0302830526099167 | 1901 | 0 | 0 | 2 | 2 | 0 | 0 | 0 | 2 | 0 | 1 | 13 | 15 |
| AluSx3_dup24351 | 3.32462866195592 | -3.96130392985175 | 1.1013297645359 | -3.59683725747761 | 0.000322109869128728 | 0.0157316820442743 | 1902 | 2 | 0 | 6 | 8 | 0 | 0 | 1 | 1 | 0 | 0 | 14 | 13 |
| L1PA8_dup4318 | 2.32602778940777 | -4.09131734428518 | 1.14305076949073 | -3.57929626004975 | 0.000344520724776868 | 0.016511042976386 | 1903 | 1 | 0 | 6 | 7 | 0 | 0 | 1 | 1 | 0 | 0 | 5 | 11 |
| THE1B_dup5135 | 2.66985007025628 | -4.29168634704219 | 1.08684453636221 | -3.948758266207 | 7.85576310338169e-05 | 0.00570414222580793 | 1904 | 1 | 0 | 5 | 8 | 0 | 0 | 1 | 3 | 0 | 0 | 6 | 12 |
| L1P4_dup3032 | 3.17931694693633 | -4.98552520462214 | 1.13647905355488 | -4.38681662370066 | 1.15021682703814e-05 | 0.00135155305247614 | 1905 | 0 | 0 | 2 | 13 | 0 | 0 | 2 | 2 | 0 | 0 | 15 | 9 |
| L1MD2_dup7170 | 3.16019216700647 | -4.77088466878299 | 1.05179284916019 | -4.53595465361103 | 5.73435011794195e-06 | 0.000797933142201062 | 1906 | 0 | 0 | 3 | 6 | 0 | 0 | 2 | 2 | 0 | 0 | 17 | 12 |
| MIRc_dup70777 | 2.97804075861168 | -4.61909258058977 | 1.05347972674173 | -4.38460509807434 | 1.16196374090656e-05 | 0.00136099802516487 | 1907 | 0 | 0 | 2 | 5 | 0 | 0 | 2 | 2 | 0 | 0 | 14 | 15 |
| L1MD2_dup7173 | 3.3135773998531 | -4.97590885122175 | 1.06101605954602 | -4.68975828070952 | 2.73527976500104e-06 | 0.000448325344332882 | 1908 | 0 | 0 | 3 | 9 | 0 | 0 | 2 | 3 | 0 | 0 | 16 | 11 |
| MIR3_dup52641 | 3.19685921060852 | -4.83497452544543 | 1.03003494225374 | -4.69399078332859 | 2.67926109477801e-06 | 0.000442947302143162 | 1909 | 0 | 0 | 3 | 5 | 0 | 0 | 3 | 3 | 0 | 0 | 17 | 11 |
| L1MD_dup5727 | 3.40706347476144 | -4.98916300241781 | 1.06529538691023 | -4.68336112567646 | 2.82208677607374e-06 | 0.000459799907440628 | 1910 | 0 | 0 | 4 | 6 | 0 | 0 | 1 | 5 | 0 | 0 | 11 | 18 |
| MER21C_dup1161 | 3.94739193897733 | -5.32219999522276 | 1.08197267839056 | -4.91897817894956 | 8.69971662608741e-07 | 0.000184992204853529 | 1911 | 0 | 0 | 8 | 12 | 0 | 0 | 1 | 3 | 0 | 0 | 14 | 15 |
| THE1B_dup4553 | 1.9970045387751 | -3.74668981773274 | 1.17855128603924 | -3.17906387453384 | 0.0014775152814587 | 0.0460228892737231 | 1912 | 0 | 0 | 5 | 2 | 0 | 0 | 4 | 0 | 0 | 1 | 8 | 7 |
| UCON48_dup19 | 2.8303321237676 | -3.93590811211283 | 1.20308727995218 | -3.27150671252153 | 0.00106976048983489 | 0.036805267425427 | 1913 | 0 | 0 | 8 | 9 | 0 | 0 | 2 | 0 | 0 | 2 | 12 | 5 |
| AluJo_dup15711 | 2.66233939812666 | -4.1483802523916 | 1.20107788837919 | -3.45388112838352 | 0.000552580884918542 | 0.023152740137846 | 1914 | 0 | 0 | 8 | 5 | 0 | 0 | 1 | 0 | 0 | 1 | 10 | 11 |
| AluJr_dup48404 | 2.25256562781417 | -3.73354100377794 | 1.17838965725399 | -3.16834162689297 | 0.00153311252434477 | 0.047222045830686 | 1915 | 0 | 0 | 4 | 4 | 0 | 0 | 1 | 0 | 0 | 1 | 13 | 7 |
| AluSc_dup26972 | 2.73397700153058 | -3.75780563937945 | 1.11329005156291 | -3.37540574812825 | 0.000737069340462423 | 0.0283836707049153 | 1916 | 0 | 1 | 7 | 6 | 0 | 0 | 3 | 0 | 0 | 1 | 5 | 15 |
| LTR78_dup3864 | 2.21250102402254 | -3.62838935512201 | 1.14657174220047 | -3.16455501350358 | 0.00155320296055898 | 0.0475892907099108 | 1917 | 0 | 0 | 1 | 4 | 0 | 0 | 3 | 0 | 0 | 1 | 7 | 15 |
| L1PA4_dup8364 | 2.88731530615987 | -4.21298401704357 | 1.09779559383891 | -3.83767619462845 | 0.000124204111937209 | 0.00794224679894756 | 1918 | 0 | 0 | 5 | 6 | 0 | 0 | 3 | 0 | 0 | 1 | 9 | 16 |
| L2c_dup33178 | 2.75602549066718 | -3.70536799279601 | 1.13832789753549 | -3.25509723588277 | 0.00113353549254441 | 0.0383126177962095 | 1919 | 0 | 0 | 2 | 9 | 0 | 0 | 2 | 0 | 0 | 2 | 11 | 12 |
| UCON84_dup63 | 1.67677277543467 | -3.95928220348762 | 1.18192757278861 | -3.34985179687973 | 0.000808548087991349 | 0.0302786879674519 | 1920 | 0 | 0 | 1 | 4 | 0 | 0 | 2 | 1 | 0 | 0 | 6 | 9 |
| AluY_dup80264 | 1.78125583577605 | -4.08070177814348 | 1.13511117995491 | -3.59497981361224 | 0.000324416730345705 | 0.0158170015902206 | 1921 | 0 | 0 | 2 | 3 | 0 | 0 | 3 | 1 | 0 | 0 | 8 | 7 |
| L2c_dup87254 | 1.91961763147185 | -4.0688657823964 | 1.14358077622799 | -3.55800470502595 | 0.000373682621751413 | 0.0175758618531848 | 1922 | 0 | 0 | 1 | 3 | 0 | 0 | 3 | 1 | 0 | 0 | 9 | 9 |
| Arthur1_dup1389 | 1.82712594419437 | -3.93401325964284 | 1.21559645721232 | -3.23628226810119 | 0.00121097571251045 | 0.0400932257458232 | 1923 | 0 | 0 | 2 | 3 | 0 | 0 | 2 | 0 | 0 | 0 | 9 | 9 |
| MLT1F-int_dup355 | 2.10258514574873 | -3.93175906813031 | 1.20890069368893 | -3.25234247002758 | 0.00114458021785802 | 0.0385660757328165 | 1924 | 0 | 0 | 1 | 2 | 0 | 0 | 3 | 0 | 0 | 0 | 10 | 13 |
| AluJb_dup90748 | 3.25165605307204 | -4.17771774642288 | 1.06420845035957 | -3.92565737004939 | 8.64932000148724e-05 | 0.00612794935546866 | 1925 | 0 | 1 | 4 | 3 | 0 | 0 | 4 | 0 | 0 | 0 | 17 | 15 |
| MLT1I_dup5979 | 2.76743978492191 | -3.75498054778811 | 1.13475997166573 | -3.30905269973185 | 0.000936122148811309 | 0.0335817797964463 | 1926 | 0 | 0 | 4 | 2 | 1 | 0 | 1 | 0 | 0 | 0 | 16 | 13 |
| MLT1H1_dup2120 | 2.31583321608798 | -3.62502003495515 | 1.12550241793288 | -3.22080164129095 | 0.0012783259078389 | 0.041633251637919 | 1927 | 0 | 0 | 2 | 3 | 1 | 0 | 2 | 0 | 0 | 0 | 9 | 15 |
| Tigger15a_dup3515 | 2.40648319711387 | -3.82850533573633 | 1.16551783783076 | -3.28481058931015 | 0.00102050951252281 | 0.0355632918082531 | 1928 | 0 | 0 | 4 | 1 | 1 | 0 | 4 | 0 | 0 | 0 | 9 | 14 |
| FRAM_dup5550 | 1.97141103255651 | -3.8406607226912 | 1.10840546450112 | -3.46503228799926 | 0.000530167823506362 | 0.0224388674727935 | 1929 | 0 | 0 | 2 | 7 | 0 | 0 | 3 | 2 | 1 | 0 | 7 | 4 |
| L2c_dup87256 | 2.95110067468294 | -3.90415792436686 | 1.08629467993201 | -3.59401366543671 | 0.000325622747013607 | 0.0158479505340356 | 1930 | 0 | 0 | 5 | 10 | 0 | 0 | 2 | 3 | 2 | 0 | 12 | 4 |
| LTR84b_dup250 | 2.68208540648407 | -3.75398463417841 | 1.07967558707517 | -3.47695611451948 | 0.000507140819205476 | 0.0217865777291502 | 1931 | 0 | 0 | 7 | 6 | 0 | 0 | 2 | 3 | 2 | 0 | 10 | 4 |
| MIR_dup133508 | 2.32234749078704 | -3.88933025190501 | 1.11486500088225 | -3.48861095184365 | 0.000485537268995937 | 0.0211093826070724 | 1932 | 0 | 0 | 6 | 1 | 0 | 0 | 2 | 2 | 1 | 0 | 9 | 9 |
| MIRb_dup4574 | 2.67475771311238 | -4.20579739875432 | 1.05476489917933 | -3.98742639428626 | 6.67939172765814e-05 | 0.00507288395796152 | 1933 | 0 | 0 | 5 | 3 | 0 | 0 | 2 | 5 | 1 | 0 | 8 | 10 |
| L1M4a2_dup2013 | 2.53613482740522 | -3.91553882037272 | 1.11943141376482 | -3.49779251522356 | 0.000469125944864289 | 0.0206359044478966 | 1934 | 0 | 0 | 1 | 3 | 0 | 0 | 1 | 5 | 1 | 0 | 11 | 10 |
| LTR83_dup700 | 3.2286903849367 | -3.40318855168739 | 0.974741837687334 | -3.49137424916711 | 0.000480542615913852 | 0.0209689669946588 | 1935 | 0 | 0 | 4 | 5 | 1 | 0 | 1 | 4 | 2 | 0 | 14 | 10 |
| AluSq2_dup35047 | 2.94789824604858 | -3.88264486768047 | 1.03975428801134 | -3.73419461929463 | 0.000188316834178557 | 0.0106485830383676 | 1936 | 0 | 0 | 8 | 5 | 0 | 0 | 5 | 1 | 2 | 0 | 9 | 9 |
| L1MD3_dup4031 | 2.80276654905428 | -3.40542380540368 | 1.03860642554923 | -3.27883953115624 | 0.00104234889192249 | 0.0361180645751257 | 1937 | 0 | 0 | 6 | 7 | 0 | 0 | 3 | 1 | 3 | 0 | 7 | 10 |
| L4_A_Mam_dup4413 | 2.99314615897561 | -4.37884729063452 | 1.05881609380368 | -4.13560703908836 | 3.54017549621845e-05 | 0.00313148921421263 | 1938 | 0 | 0 | 4 | 8 | 0 | 0 | 5 | 1 | 1 | 0 | 14 | 7 |
| MLT1L_dup11504 | 2.67141821458765 | -4.02965920373981 | 1.05320820122876 | -3.82608035053134 | 0.000130199807501468 | 0.00821982578820492 | 1939 | 0 | 0 | 2 | 3 | 0 | 0 | 6 | 1 | 1 | 0 | 11 | 12 |
| L1M2_dup4822 | 2.49576585470569 | -3.61566125335223 | 1.03983557531501 | -3.47714709823895 | 0.000506779699971241 | 0.0217800690544299 | 1940 | 0 | 0 | 3 | 2 | 0 | 1 | 7 | 3 | 1 | 0 | 6 | 10 |
| L1PA2_dup1435 | 2.88202268151246 | -3.22086987188896 | 0.960003708018096 | -3.35505982423585 | 0.000793478228715437 | 0.0299025927478498 | 1941 | 1 | 0 | 2 | 6 | 0 | 1 | 3 | 2 | 1 | 0 | 8 | 15 |
| L1PA5_dup1724 | 3.09859235516094 | -3.33224860863977 | 0.940882972053623 | -3.54161857278235 | 0.000397680076152373 | 0.018391911253674 | 1942 | 1 | 0 | 3 | 5 | 0 | 1 | 5 | 1 | 1 | 0 | 11 | 14 |
| AluSx1_dup85186 | 2.79469249991205 | -3.33176831370823 | 1.05656439482936 | -3.15339825003881 | 0.00161381476807683 | 0.0488177941526582 | 1943 | 0 | 1 | 10 | 6 | 0 | 1 | 1 | 3 | 1 | 0 | 7 | 6 |
| L1MD3_dup4054 | 2.13891485762883 | -3.42010541709444 | 1.06554394767893 | -3.20972722386949 | 0.0013286098944729 | 0.042745404667471 | 1944 | 0 | 0 | 6 | 4 | 0 | 1 | 2 | 2 | 1 | 0 | 5 | 7 |
| LTR47B2_dup122 | 3.35917116863408 | -4.12532787691173 | 1.03541368439202 | -3.98423155797295 | 6.76988359512698e-05 | 0.00512036465965062 | 1945 | 0 | 0 | 11 | 7 | 0 | 1 | 2 | 3 | 1 | 0 | 9 | 10 |
| MIRb_dup57572 | 2.24988144729916 | -4.58989617573586 | 1.14821960455995 | -3.99740272462508 | 6.40412945051612e-05 | 0.00490610748948006 | 1946 | 0 | 0 | 5 | 4 | 0 | 0 | 1 | 6 | 0 | 0 | 7 | 5 |
| MLT1D_dup2310 | 2.98197234143876 | -5.00486898039752 | 1.07060685410545 | -4.67479631874703 | 2.94245502819611e-06 | 0.000473725551966953 | 1947 | 0 | 0 | 10 | 4 | 0 | 0 | 3 | 5 | 0 | 0 | 10 | 6 |
| L1MC5_dup11825 | 2.73089892817015 | -4.86053670662871 | 1.10935926146488 | -4.38139102044406 | 1.17924022676098e-05 | 0.00137614269248096 | 1948 | 0 | 0 | 9 | 6 | 0 | 0 | 2 | 2 | 0 | 0 | 5 | 13 |
| THE1C-int_dup1599 | 1.87253981534163 | -4.35779248994794 | 1.15223382245631 | -3.78203833719973 | 0.00015554939827976 | 0.00934891566634905 | 1949 | 0 | 0 | 7 | 4 | 0 | 0 | 2 | 2 | 0 | 0 | 3 | 7 |
| Kanga1a_dup698 | 2.24279411884333 | -4.49134923268155 | 1.08161452924232 | -4.15244905763958 | 3.28935992721629e-05 | 0.0029636124090928 | 1950 | 0 | 0 | 4 | 5 | 0 | 0 | 2 | 2 | 0 | 0 | 8 | 9 |
| MLT1J1_dup690 | 1.80655210080085 | -4.295289364023 | 1.18365133528943 | -3.6288468030771 | 0.000284690113562046 | 0.0144027665769684 | 1951 | 0 | 0 | 7 | 4 | 0 | 0 | 2 | 1 | 0 | 0 | 6 | 4 |
| Kanga1_dup693 | 3.47278739837836 | -4.5937270307595 | 1.11249374237605 | -4.12921606277828 | 3.64002218559443e-05 | 0.00320371357378448 | 1952 | 0 | 1 | 10 | 7 | 0 | 0 | 1 | 1 | 0 | 0 | 11 | 16 |
| L1MB1_dup1213 | 2.83423635805112 | -3.90423241416199 | 1.11941188847669 | -3.48775321608823 | 0.000487097443909295 | 0.0211617470126787 | 1953 | 0 | 2 | 10 | 7 | 0 | 0 | 2 | 1 | 0 | 0 | 8 | 8 |
| Kanga1a_dup703 | 2.23581140557597 | -4.05152936750796 | 1.1987415955967 | -3.37981878862828 | 0.000725336357150885 | 0.0280453176533351 | 1954 | 0 | 1 | 10 | 3 | 0 | 0 | 1 | 1 | 0 | 0 | 4 | 10 |
| L1PA10_dup6055 | 2.234406520505 | -3.98108443688292 | 1.1553295118926 | -3.44584328185412 | 0.000569280402941906 | 0.0236796912027431 | 1955 | 0 | 1 | 4 | 7 | 0 | 0 | 1 | 1 | 0 | 0 | 4 | 13 |
| L1MB4_dup4300 | 2.46080502692588 | -3.43958923466589 | 1.05968628025607 | -3.24585615455429 | 0.00117097990486833 | 0.039169394739703 | 1956 | 0 | 1 | 4 | 1 | 1 | 0 | 3 | 2 | 0 | 0 | 12 | 8 |
| MIRc_dup4146 | 2.0562061247308 | -3.77679914109814 | 1.1036121346862 | -3.42221603260283 | 0.000621129409591623 | 0.0251395234724763 | 1957 | 0 | 0 | 3 | 4 | 1 | 0 | 1 | 2 | 0 | 0 | 9 | 7 |
| AluSz6_dup31037 | 3.57816199416854 | -3.55174892910466 | 1.05867413661975 | -3.35490289811469 | 0.00079392847246361 | 0.0299100861793581 | 1958 | 0 | 0 | 0 | 1 | 1 | 1 | 5 | 2 | 0 | 0 | 18 | 20 |
| AluSx1_dup14119 | 5.19163889468822 | -4.06396938597438 | 1.09710917075742 | -3.70425249765135 | 0.000212014997001721 | 0.0115919394477416 | 1959 | 0 | 0 | 0 | 0 | 0 | 0 | 5 | 1 | 0 | 1 | 25 | 39 |
| MIRb_dup154782 | 4.16156691768388 | -4.36708160469719 | 0.981633623925709 | -4.44878975032714 | 8.63554959838298e-06 | 0.00108896547262201 | 1960 | 0 | 0 | 2 | 1 | 0 | 0 | 6 | 2 | 0 | 1 | 20 | 24 |
| L1MEd_dup15778 | 4.50239087353282 | -5.19156375029415 | 1.12215030879951 | -4.62644238439693 | 3.72000288709867e-06 | 0.000567760348935386 | 1961 | 0 | 0 | 5 | 0 | 0 | 0 | 4 | 6 | 0 | 0 | 18 | 26 |
| MIRc_dup70776 | 2.55140719581411 | -4.55085183769425 | 1.13490624858306 | -4.00989230905728 | 6.07464468262123e-05 | 0.00472054896232739 | 1962 | 0 | 0 | 4 | 1 | 0 | 0 | 6 | 1 | 0 | 0 | 8 | 15 |
| MamRTE1_dup5846 | 2.81206245382481 | -4.38111429233749 | 1.17243251847234 | -3.73677309637062 | 0.000186396984994848 | 0.0105827766622725 | 1963 | 0 | 0 | 2 | 0 | 0 | 0 | 4 | 2 | 0 | 0 | 11 | 19 |
| MamRTE1_dup6692 | 2.28990267319864 | -4.37819345525899 | 1.22365397665833 | -3.57796692428964 | 0.000346277258072858 | 0.0165754515831735 | 1964 | 0 | 0 | 2 | 0 | 0 | 0 | 5 | 3 | 0 | 0 | 10 | 10 |
| MLT1E2_dup1316 | 2.78084134204703 | -3.54183437628359 | 1.02182290817726 | -3.46619198683023 | 0.000527886191126728 | 0.0223703634508257 | 1965 | 1 | 1 | 1 | 2 | 0 | 0 | 4 | 3 | 0 | 0 | 11 | 14 |
| L1MEi_dup3808 | 3.40326825944579 | -4.71145247594877 | 1.07085726039131 | -4.3997016691348 | 1.08399798051571e-05 | 0.00129744961504925 | 1966 | 0 | 0 | 1 | 2 | 0 | 0 | 6 | 2 | 0 | 0 | 16 | 19 |
| X7B_LINE_dup661 | 3.1116115718215 | -4.80902434256208 | 1.05192200939937 | -4.57165483713754 | 4.83887535042396e-06 | 0.000702496054992057 | 1967 | 0 | 0 | 1 | 5 | 0 | 0 | 5 | 3 | 0 | 0 | 12 | 16 |
| L3_dup36519 | 2.52581232287307 | -3.63097001080397 | 1.08825747127608 | -3.33649904240613 | 0.000848407209536702 | 0.0313568036471606 | 1968 | 1 | 0 | 2 | 1 | 0 | 0 | 1 | 2 | 0 | 0 | 15 | 11 |
| L1MEi_dup3807 | 2.88453286752535 | -4.46805978491555 | 1.06451263704125 | -4.19728205137542 | 2.70137418653405e-05 | 0.00254768601739608 | 1969 | 0 | 0 | 2 | 2 | 0 | 0 | 3 | 2 | 0 | 0 | 17 | 12 |
| Charlie15b_dup1463 | 3.01483695490276 | -4.4509566252913 | 1.11411896432329 | -3.99504610173724 | 6.46816653820205e-05 | 0.00494243530860317 | 1970 | 0 | 0 | 3 | 1 | 0 | 0 | 1 | 3 | 0 | 0 | 18 | 13 |

| | | | | | | | | | | | | | | | | | | | |
|---|---|---|---|---|---|---|---|---|---|---|---|---|---|---|---|---|---|---|---|
| ORSL_dup699 | 2.35751428205197 | -4.25450336789626 | 1.1113833643492 | -3.82811503606373 | 0.000129128404312674 | 0.00817530521991442 | 1971 | 0 | 0 | 3 | 2 | 0 | 0 | 1 | 2 | 0 | 0 | 12 | 11 |
| Charlie25_dup604 | 3.04654629482808 | -4.32046217703565 | 1.07221134416502 | -4.02948747049507 | 5.58985950072297e-05 | 0.00443066192124926 | 1972 | 0 | 0 | 2 | 2 | 0 | 0 | 2 | 1 | 0 | 0 | 11 | 24 |
| MIRc_dup70779 | 2.72205066037239 | -3.65128331671807 | 1.06677954902914 | -3.42271589293314 | 0.000619988414412477 | 0.0251062493330676 | 1973 | 0 | 0 | 1 | 1 | 1 | 0 | 3 | 1 | 0 | 0 | 12 | 18 |
| L1PA8_dup4527 | 5.16465035815322 | -3.95784916291204 | 0.974043476955481 | -4.06331879074114 | 4.83798834257806e-05 | 0.00397232565273664 | 1974 | 0 | 0 | 7 | 5 | 0 | 0 | 1 | 1 | 3 | 0 | 15 | 38 |
| HERV16-int_dup1042 | 3.72374968836827 | -3.4632405405357 | 1.00315972062353 | -3.45233213548794 | 0.000555763162903024 | 0.023261498955167 | 1975 | 1 | 0 | 7 | 3 | 0 | 0 | 1 | 1 | 2 | 0 | 22 | 11 |
| MLT2D_dup3014 | 3.744416615153 | -4.51041991246923 | 1.00416228376885 | -4.4917240822276 | 7.06489145408386e-06 | 0.000937047191713176 | 1976 | 0 | 0 | 7 | 4 | 0 | 0 | 3 | 2 | 1 | 0 | 19 | 13 |
| LTR78B_dup1887 | 2.88159773214283 | -3.70417628235892 | 1.07258305400284 | -3.45350998091484 | 0.000553341826789826 | 0.0231723819636276 | 1977 | 0 | 0 | 1 | 4 | 0 | 0 | 2 | 0 | 1 | 0 | 15 | 16 |
| L1MC1_dup7430 | 2.63272445252843 | -3.67852125028457 | 1.04172470504844 | -3.53118365385556 | 0.000413704394780528 | 0.0189598046842156 | 1978 | 0 | 0 | 3 | 2 | 0 | 0 | 1 | 1 | 1 | 0 | 12 | 15 |
| LTR78_dup3863 | 3.25054056255137 | -3.93091737129129 | 1.00449169977391 | -3.91333982368998 | 9.10282897634941e-05 | 0.00636316078913352 | 1979 | 0 | 0 | 3 | 3 | 0 | 0 | 1 | 1 | 1 | 0 | 18 | 16 |
| AluSq2_dup59146 | 2.91461231307993 | -3.54093478578891 | 1.09716806108341 | -3.22734037873138 | 0.00124946731429112 | 0.040996380249078 | 1980 | 0 | 0 | 0 | 6 | 0 | 0 | 2 | 2 | 2 | 0 | 10 | 17 |
| MER50_dup1041 | 4.2738150933523 | -3.83239731055767 | 0.982186405493954 | -3.90190425068071 | 9.5438911313513e-05 | 0.00658431637906789 | 1981 | 0 | 0 | 0 | 2 | 0 | 0 | 2 | 1 | 1 | 0 | 26 | 25 |
| L1PB4_dup4513 | 4.88651233568444 | -3.34454261053979 | 0.892301335656713 | -3.74822100661576 | 0.00017809329445877 | 0.0102699739702481 | 1982 | 0 | 0 | 3 | 2 | 1 | 2 | 7 | 1 | 0 | 1 | 18 | 32 |
| L1PA13_dup8831 | 3.7950037158273 | -4.14766267576033 | 0.970105540270495 | -4.27547571226511 | 1.90729594176848e-05 | 0.00197061849830674 | 1983 | 0 | 0 | 4 | 5 | 1 | 1 | 6 | 3 | 0 | 0 | 19 | 11 |
| MSTA_dup12899 | 3.78482039022221 | -4.04572436671669 | 0.952231039422819 | -4.24867936374868 | 2.15034438050978e-05 | 0.00215194824733185 | 1984 | 0 | 0 | 5 | 4 | 1 | 1 | 3 | 3 | 0 | 0 | 17 | 16 |
| L2a_dup92269 | 2.60164306146752 | -3.620872271981 | 1.04690342077723 | -3.45864976665454 | 0.000542890349666996 | 0.0228392127195656 | 1985 | 0 | 0 | 2 | 4 | 1 | 1 | 6 | 1 | 0 | 0 | 7 | 14 |
| L1M5_dup28422 | 2.97356882690325 | -3.73800730464553 | 1.01828449038871 | -3.6708870064579 | 0.000241710185484131 | 0.0128064956123819 | 1986 | 0 | 0 | 1 | 5 | 1 | 1 | 5 | 2 | 0 | 0 | 13 | 12 |
| L1MC3_dup12190 | 3.4669112210031 | -4.06617469314586 | 0.987168044345504 | -4.11902990219031 | 3.80470777309198e-05 | 0.00331374449405616 | 1987 | 0 | 0 | 2 | 6 | 1 | 1 | 7 | 3 | 0 | 0 | 11 | 16 |
| L1PA5_dup6330 | 3.98202696970653 | -3.11515733704837 | 0.89960139052984 | -3.46281961082078 | 0.000534546637254417 | 0.0225825927050459 | 1988 | 0 | 0 | 3 | 6 | 2 | 1 | 3 | 1 | 1 | 0 | 20 | 16 |
| THE1C_dup7984 | 3.95828180924805 | -2.79753359812641 | 0.852012286006151 | -3.28344279076067 | 0.00102547456159091 | 0.0356954922472825 | 1989 | 0 | 0 | 2 | 6 | 1 | 2 | 3 | 1 | 1 | 1 | 18 | 18 |
| L1PA7_dup12470 | 4.12604258194754 | -2.57083512725452 | 0.731917272639299 | -3.51246680923935 | 0.000443967453562634 | 0.0199095890888456 | 1990 | 0 | 0 | 5 | 7 | 2 | 1 | 6 | 2 | 2 | 2 | 11 | 17 |
| MIRb_dup170293 | 4.2448131458222 | -2.57736594066097 | 0.787298617333173 | -3.27368279826441 | 0.00106155703523879 | 0.0366053288558306 | 1991 | 0 | 0 | 2 | 4 | 1 | 1 | 5 | 1 | 1 | 4 | 16 | 22 |
| L1MA3_dup6021 | 5.35897411764046 | -5.16250665352517 | 0.956560812445055 | -5.39694558501653 | 6.77849348221693e-08 | 2.44751959046979e-05 | 1992 | 0 | 0 | 9 | 7 | 1 | 0 | 4 | 6 | 0 | 0 | 18 | 26 |
| MER20_dup10847 | 4.2810603503324 | -4.5810482767801 | 0.966308802752075 | -4.74077051117939 | 2.12906979979707e-06 | 0.000369355273216689 | 1993 | 0 | 0 | 3 | 3 | 1 | 0 | 4 | 4 | 0 | 0 | 18 | 24 |
| L1MCb_dup420 | 3.67475717092757 | -5.22974480340566 | 1.03326095165272 | -5.06139789279812 | 4.16193614561402e-07 | 0.000103589545826807 | 1994 | 0 | 0 | 9 | 7 | 0 | 0 | 6 | 1 | 0 | 0 | 13 | 14 |
| L1MA6_dup4584 | 3.26014766000192 | -4.55921771793744 | 1.00125771654151 | -4.55349071734065 | 5.2763000832866e-06 | 0.000751331238550397 | 1995 | 0 | 0 | 7 | 5 | 1 | 0 | 6 | 2 | 0 | 0 | 10 | 13 |
| L1MEc_dup10170 | 3.19116566788926 | -4.28726167177863 | 1.02160350358508 | -4.19660039999224 | 2.70951377553835e-05 | 0.0025523276517587 | 1996 | 0 | 0 | 4 | 5 | 1 | 0 | 3 | 1 | 0 | 0 | 15 | 14 |
| L1M4a2_dup2014 | 4.19258789135819 | -4.11327320617197 | 0.937238194059004 | -4.38871701158289 | 1.14021321681855e-05 | 0.00134404309541751 | 1997 | 1 | 0 | 3 | 3 | 0 | 0 | 3 | 5 | 0 | 1 | 13 | 27 |
| L1M7_dup754 | 3.06160652991708 | -3.77580364161969 | 1.00170614301779 | -3.76937255295701 | 0.000163658449965049 | 0.00969185503618306 | 1998 | 1 | 0 | 1 | 5 | 0 | 0 | 3 | 4 | 0 | 1 | 9 | 17 |
| Charlie7a_dup970 | 4.53046213413544 | -4.81345708168291 | 0.991976363194903 | -4.85239090393242 | 1.21981888382721e-06 | 0.000239648601203287 | 1999 | 0 | 0 | 2 | 9 | 0 | 0 | 4 | 5 | 1 | 0 | 13 | 27 |
| AluSx1_dup85191 | 4.33222592746798 | -4.87596668199184 | 1.01627187931575 | -4.79789589895453 | 1.60341070339886e-06 | 0.000298066533774175 | 2000 | 0 | 0 | 4 | 12 | 0 | 0 | 2 | 5 | 0 | 1 | 14 | 20 |
| MIRb_dup132046 | 5.17157929370602 | -5.03260558109752 | 0.943666698491926 | -5.3330329332805 | 9.65857858576805e-08 | 3.2142939023345e-05 | 2001 | 0 | 0 | 4 | 8 | 0 | 0 | 5 | 6 | 0 | 1 | 18 | 27 |
| AluSg_dup32524 | 3.46074978762621 | -3.60308512542112 | 0.991359175992591 | -3.63449011486029 | 0.000278530935781794 | 0.0141795725386253 | 2002 | 1 | 1 | 1 | 8 | 0 | 1 | 5 | 5 | 0 | 0 | 11 | 13 |
| SVA_D_dup696 | 3.4792222117002 | -4.17150604379692 | 0.977065091170591 | -4.2694249149759 | 1.95977607264594e-05 | 0.002014345042686 | 2003 | 0 | 1 | 4 | 5 | 0 | 1 | 8 | 5 | 0 | 0 | 10 | 12 |
| MLT1A0_dup16557 | 3.71189361945617 | -3.56164345851273 | 0.909147459996446 | -3.91756410838639 | 8.94482619090961e-05 | 0.00628212990797573 | 2004 | 0 | 1 | 2 | 4 | 1 | 0 | 5 | 4 | 1 | 0 | 13 | 18 |
| L1MB8_dup7858 | 3.23595674370748 | -3.97583035340032 | 0.991274198458183 | -4.0108280429212 | 6.0506167715563e-05 | 0.00470635401913803 | 2005 | 0 | 1 | 2 | 6 | 1 | 0 | 5 | 4 | 0 | 0 | 11 | 13 |
| MER47A_dup1590 | 2.80952012876175 | -4.19610068747523 | 1.0300023725979 | -4.07387477845485 | 4.62373488818107e-05 | 0.00384685042553543 | 2006 | 0 | 0 | 2 | 4 | 1 | 0 | 4 | 3 | 0 | 0 | 13 | 10 |
| MIRb_dup10534 | 3.22509197086519 | -4.4402697394863 | 1.00157907707507 | -4.43326926562136 | 9.28148294127089e-06 | 0.00115446510190468 | 2007 | 0 | 0 | 3 | 4 | 1 | 0 | 5 | 4 | 0 | 0 | 10 | 16 |
| L2c_dup110912 | 6.82513193772205 | -2.2253566767478 | 0.648524547226683 | -3.43141471863201 | 0.000600441889094761 | 0.024599301952917 | 2008 | 3 | 0 | 8 | 6 | 2 | 0 | 5 | 1 | 4 | 4 | 17 | 42 |
| MER103C_dup6943 | 9.74427531372281 | -3.57943502420971 | 0.785654679386134 | -4.55599020552704 | 5.21393659407984e-06 | 0.000745574132701276 | 2009 | 1 | 0 | 8 | 12 | 1 | 0 | 2 | 0 | 3 | 3 | 42 | 60 |
| L1PA16_dup11606 | 8.64717698326586 | -2.82530675211115 | 0.636916392853331 | -4.43591464093743 | 9.16821743184812e-06 | 0.00114396469063374 | 2010 | 1 | 0 | 3 | 7 | 1 | 1 | 4 | 3 | 6 | 2 | 33 | 54 |
| L3_dup36525 | 8.65600163117459 | -2.53448465049088 | 0.554056049975098 | -4.57441923178133 | 4.77542781426848e-06 | 0.000697969254726 | 2011 | 2 | 0 | 6 | 10 | 1 | 2 | 6 | 3 | 4 | 5 | 28 | 49 |
| AluY_dup76596 | 11.2960622364877 | -2.71567895131711 | 0.596819939038987 | -4.55024836417155 | 5.35826317444108e-06 | 0.000759207675028269 | 2012 | 1 | 0 | 4 | 15 | 2 | 1 | 5 | 2 | 8 | 4 | 48 | 60 |
| L2b_dup73108 | 10.6956239767801 | -2.85788948025648 | 0.565392827140163 | -5.05469709389857 | 4.31074383935729e-07 | 0.000106513421972971 | 2013 | 1 | 0 | 9 | 10 | 2 | 0 | 8 | 2 | 8 | 4 | 45 | 52 |
| AluSc_dup25763 | 6.37549976361781 | -3.16777866149251 | 0.772985347146567 | -4.09810958666447 | 4.16538077258595e-05 | 0.00355373142098866 | 2014 | 0 | 0 | 8 | 12 | 2 | 1 | 2 | 2 | 2 | 2 | 31 | 22 |
| LTR45C_dup202 | 7.35012327955573 | -2.95600623977771 | 0.709946017429774 | -4.16370564409865 | 3.13123744990658e-05 | 0.00284702111675763 | 2015 | 1 | 0 | 8 | 9 | 1 | 1 | 2 | 1 | 3 | 3 | 37 | 31 |
| L2a_dup133435 | 8.57562697115658 | -2.95229730438588 | 0.671496932903705 | -4.39659089970743 | 1.09964321302936e-05 | 0.00130958810794536 | 2016 | 0 | 0 | 11 | 8 | 2 | 0 | 5 | 3 | 8 | 2 | 31 | 42 |
| MamGypsy2-I_dup1572 | 7.48527568100772 | -3.62617063409459 | 0.729486109817941 | -4.97085631280845 | 6.6657837576397e-07 | 0.000149490724033491 | 2017 | 0 | 0 | 9 | 13 | 1 | 0 | 4 | 5 | 4 | 2 | 30 | 30 |
| MER3_dup8137 | 9.24439421844207 | -3.16709089717499 | 0.6380251461084 | -4.96389666848162 | 6.90927131663527e-07 | 0.000153218403793858 | 2018 | 0 | 0 | 8 | 8 | 3 | 0 | 4 | 3 | 3 | 4 | 39 | 51 |
| AluSg_dup26193 | 8.7536879394621 | -3.10086549201598 | 0.685581638627074 | -4.52297044918777 | 6.09777563246611e-06 | 0.000839179479839665 | 2019 | 0 | 0 | 5 | 12 | 1 | 0 | 2 | 3 | 4 | 7 | 36 | 46 |
| L1MB8_dup12601 | 6.64074703004715 | -2.72131893642622 | 0.659142919791284 | -4.12857189953268 | 3.65023301167546e-05 | 0.00320970394993969 | 2020 | 0 | 0 | 6 | 6 | 1 | 0 | 3 | 3 | 5 | 5 | 28 | 29 |
| L1PA12_dup1725 | 9.72858976497914 | -2.33149839783601 | 0.601448361368762 | -3.87647310656901 | 0.000105981517946879 | 0.00712168265162043 | 2021 | 0 | 0 | 5 | 2 | 0 | 2 | 3 | 1 | 6 | 10 | 48 | 50 |
| L1PA13_dup8785 | 12.131522720844 | -1.60711767314569 | 0.506121056646315 | -3.17536220246367 | 0.00149649578648558 | 0.0464351751305307 | 2022 | 1 | 0 | 5 | 4 | 1 | 4 | 3 | 1 | 11 | 13 | 50 | 65 |
| AluSx1_dup81391 | 10.6677041068802 | -2.7614184091006 | 0.570653125790414 | -4.83904894987783 | 1.30461914691111e-06 | 0.000253204655408573 | 2023 | 0 | 1 | 7 | 6 | 1 | 1 | 5 | 2 | 6 | 6 | 36 | 72 |
| L1PA2_dup4204 | 7.65402592475383 | -2.13109453976241 | 0.636139134738043 | -3.3500447046698 | 0.000807985189335889 | 0.0302719010961608 | 2024 | 0 | 1 | 1 | 7 | 1 | 1 | 2 | 2 | 7 | 5 | 32 | 41 |
| MER51B_dup124 | 8.60695315920767 | -2.59835106430487 | 0.777659436230435 | -3.34124546459554 | 0.000834034509153636 | 0.0309650765041459 | 2025 | 0 | 0 | 13 | 18 | 0 | 3 | 2 | 2 | 3 | 13 | 27 | 33 |
| MLT1F1-int_dup14 | 7.39774667875656 | -1.93929756423443 | 0.590111516485327 | -3.28632387279065 | 0.00101504229052198 | 0.0354454436249697 | 2026 | 0 | 2 | 11 | 17 | 1 | 4 | 4 | 3 | 4 | 8 | 21 | 22 |
| AluSx_dup82055 | 9.78334423172859 | -2.61640716048847 | 0.656770273677501 | -3.98374784205479 | 6.78368526792244e-05 | 0.00512574933197374 | 2027 | 1 | 1 | 10 | 11 | 1 | 2 | 0 | 3 | 4 | 5 | 43 | 48 |
| AluSz_dup73300 | 8.52306921330146 | -2.31192845972195 | 0.696633043264269 | -3.31871776981574 | 0.000904317786283552 | 0.0327616930534449 | 2028 | 0 | 0 | 6 | 9 | 1 | 1 | 1 | 1 | 4 | 13 | 41 | 34 |
| LTR56_dup554 | 9.54196994249717 | -3.04319180072562 | 0.748058279841916 | -4.06812127173932 | 4.73937150541403e-05 | 0.00391295798829718 | 2029 | 0 | 1 | 13 | 6 | 1 | 1 | 1 | 1 | 2 | 6 | 43 | 52 |
| FRAM_dup5323 | 8.1124569703023 | -3.43717607943967 | 0.884815015966293 | -3.88462674956526 | 0.000102487215330727 | 0.00693387061105634 | 2030 | 0 | 0 | 5 | 10 | 0 | 0 | 2 | 2 | 0 | 12 | 38 | 39 |
| MIR_dup133525 | 8.9047626266988 | -3.93989910684487 | 0.931062784778971 | -4.23161485052823 | 2.32019467372379e-05 | 0.00228415822024775 | 2031 | 0 | 0 | 10 | 15 | 0 | 0 | 7 | 1 | 0 | 11 | 48 | 26 |
| MLT1A_dup7133 | 4.54570310349778 | -2.99444691636157 | 0.913196734136401 | -3.27908193757764 | 0.0010414539239379 | 0.0361028256680255 | 2032 | 0 | 1 | 2 | 7 | 0 | 0 | 1 | 1 | 0 | 5 | 21 | 23 |
| L1MC2_dup4896 | 6.52153758103253 | -3.65756465176178 | 0.838340733095914 | -4.36286167111877 | 1.28372117928992e-05 | 0.00147066114457063 | 2033 | 0 | 0 | 5 | 11 | 1 | 0 | 4 | 1 | 0 | 5 | 25 | 37 |
| L3_dup36524 | 6.4097757261143 | -3.51182614138548 | 0.751956012513872 | -4.67025475286122 | 3.00826435209335e-06 | 0.000482023211800138 | 2034 | 0 | 0 | 6 | 6 | 1 | 0 | 3 | 2 | 1 | 4 | 26 | 37 |
| L1ME3Cz_dup9407 | 8.87564807633591 | -3.44717063998018 | 0.645450785481556 | -5.34071801835104 | 9.25791643939018e-08 | 3.11141834051855e-05 | 2035 | 2 | 0 | 8 | 10 | 0 | 0 | 5 | 10 | 3 | 4 | 36 | 37 |
| L1M8_dup355 | 7.44471492028145 | -3.16297351491272 | 0.736322496998335 | -4.29563612113819 | 1.74193110742104e-05 | 0.0018446174423071 | 2036 | 0 | 0 | 6 | 15 | 0 | 0 | 4 | 8 | 5 | 8 | 17 | 34 |
| AluSc_dup25765 | 7.419856282204 | -3.42435845289974 | 0.68367178184858 | -5.00877547357683 | 5.47774304094345e-07 | 0.000128668020298157 | 2037 | 0 | 1 | 7 | 9 | 0 | 0 | 5 | 9 | 3 | 4 | 26 | 32 |
| MIRb_dup170318 | 9.07890428496578 | -3.09970096302468 | 0.620111308222908 | -4.99862028303869 | 5.7741983004543e-07 | 0.000134437122191771 | 2038 | 0 | 1 | 11 | 12 | 0 | 0 | 4 | 7 | 6 | 6 | 30 | 41 |
| MLT1-int_dup231 | 7.57190919091268 | -2.72413635665629 | 0.671913406876646 | -4.05429677213809 | 5.02853448629972e-05 | 0.00409767009204477 | 2039 | 1 | 0 | 8 | 3 | 0 | 0 | 4 | 5 | 8 | 4 | 33 | 30 |
| MER2_dup8992 | 8.12923312306599 | -2.80990985388877 | 0.626686364702623 | -4.48375776489431 | 7.33399139460533e-06 | 0.0009652068121575 | 2040 | 2 | 0 | 5 | 8 | 0 | 0 | 5 | 4 | 7 | 3 | 28 | 45 |
| AluY_dup76595 | 7.28778476091662 | -2.67374120976059 | 0.62478310104386 | -4.27947107611172 | 1.87337980376765e-05 | 0.00194571466952116 | 2041 | 1 | 0 | 6 | 6 | 0 | 0 | 8 | 2 | 6 | 6 | 25 | 36 |
| MLT-int_dup355 | 6.57232068874664 | -3.21519468028675 | 0.683040131463482 | -4.70718268544173 | 2.51163807867718e-06 | 0.00042086917906262 | 2042 | 1 | 0 | 7 | 7 | 0 | 0 | 6 | 4 | 3 | 4 | 22 | 33 |
| MIR3_dup66180 | 7.45977805418792 | -3.01149020557571 | 0.71061978854188 | -4.23783611733495 | 2.25684403368598e-05 | 0.00223752921016435 | 2043 | 0 | 0 | 4 | 11 | 0 | 0 | 4 | 5 | 4 | 9 | 35 | 24 |
| MLT1A0-int_dup927 | 8.16960555012092 | -2.70833717192168 | 0.623700730289354 | -4.3423665235492 | 1.40956121970611e-05 | 0.00157612525368511 | 2044 | 1 | 0 | 3 | 8 | 0 | 0 | 6 | 8 | 6 | 8 | 35 | 29 |
| L2c_dup110932 | 6.15548077194233 | -2.91436990190251 | 0.737633227518378 | -3.95097426902437 | 7.78336932409966e-05 | 0.00566021270535519 | 2045 | 0 | 0 | 4 | 3 | 0 | 0 | 2 | 5 | 3 | 7 | 22 | 34 |
| MLT1B_dup17325 | 5.99652078694613 | -2.80235670779929 | 0.7292826583901 | -3.84262079395323 | 0.000121727469872316 | 0.00783068194619217 | 2046 | 1 | 0 | 3 | 6 | 0 | 0 | 3 | 7 | 4 | 5 | 15 | 34 |
| AluSg4_dup5063 | 8.44676900211947 | -3.58269505787384 | 0.718017898141223 | -4.98970160374635 | 6.04726327152542e-07 | 0.000138973836176245 | 2047 | 0 | 0 | 2 | 10 | 1 | 0 | 7 | 5 | 3 | 4 | 26 | 56 |
| MER30_dup3206 | 7.81081677251838 | -2.64513629099161 | 0.598530322062976 | -4.41938560752365 | 9.89818974116485e-06 | 0.00121341301192747 | 2048 | 0 | 2 | 3 | 6 | 1 | 0 | 6 | 4 | 4 | 5 | 30 | 42 |
| AluSp_dup35805 | 7.84816834558075 | -3.10989356643819 | 0.741421325181961 | -4.19450245199645 | 2.73471193602203e-05 | 0.0025693180362804 | 2049 | 0 | 0 | 1 | 7 | 0 | 0 | 5 | 2 | 6 | 5 | 48 | 27 |

| | | | | | | | | | | | | | | | | | | | |
|---|---|---|---|---|---|---|---|---|---|---|---|---|---|---|---|---|---|---|---|
| AluSc_dup25764 | 11.0834941082489 | -3.2816621348161 | 0.653358342089074 | -5.02275998240595 | 5.09341809960816e-07 | 0.000120833346274114 | 2050 | 0 | 0 | 3 | 10 | 0 | 0 | 3 | 5 | 8 | 6 | 50 | 60 |
| L1PA11_dup3443 | 7.6657482146073 | -3.49278554431933 | 0.792431208283458 | -4.40768297337166 | 1.04482325680294e-05 | 0.00126391212335145 | 2051 | 0 | 0 | 2 | 13 | 0 | 0 | 2 | 4 | 5 | 4 | 32 | 39 |
| MLT1B_dup14132 | 7.8898640148932 | -2.84044084757797 | 0.673762070217867 | -4.21579215146304 | 2.4890293002817e-05 | 0.00240460643883406 | 2052 | 0 | 1 | 9 | 7 | 0 | 0 | 7 | 1 | 8 | 4 | 31 | 35 |
| MIR_dup133531 | 8.80433620901545 | -3.2425333458633 | 0.618251853785768 | -5.24468034508617 | 1.56553508382562e-07 | 4.73934874117012e-05 | 2053 | 0 | 1 | 9 | 12 | 0 | 0 | 8 | 3 | 4 | 6 | 33 | 41 |
| L2a_dup165791 | 8.08440031948257 | -2.94529925941978 | 0.634530290034014 | -4.6417000191778 | 3.4555429264168e-06 | 0.000536705363687448 | 2054 | 0 | 2 | 8 | 5 | 0 | 0 | 6 | 4 | 5 | 4 | 25 | 48 |
| MamRTE1_dup7519 | 8.27411810667567 | -2.85265318408076 | 0.618161218692937 | -4.61473980867405 | 3.93588084700629e-06 | 0.000593489926503016 | 2055 | 0 | 1 | 7 | 7 | 0 | 0 | 6 | 4 | 7 | 5 | 24 | 48 |
| Kanga2_a_dup1175 | 10.147081723583 | -2.63467964707847 | 0.555554957754652 | -4.74242846779169 | 2.11171403538197e-06 | 0.000367305488203946 | 2056 | 0 | 1 | 5 | 8 | 0 | 0 | 11 | 6 | 7 | 12 | 31 | 52 |
| L1PA6_dup5016 | 10.0707468076195 | -2.63961340330085 | 0.590923051166612 | -4.4669325356147 | 7.93491393283032e-06 | 0.00102613099492459 | 2057 | 0 | 0 | 7 | 5 | 0 | 0 | 4 | 4 | 7 | 13 | 44 | 46 |
| MIRc_dup77477 | 7.33561426568654 | -2.27084610194971 | 0.624902110593689 | -3.63392291921095 | 0.00027914428833029 | 0.0142016820250244 | 2058 | 0 | 0 | 4 | 3 | 0 | 0 | 6 | 3 | 8 | 10 | 27 | 33 |
| AluSx3_dup23209 | 5.41064125010217 | -3.52041440732329 | 0.790811445112724 | -4.4516482773229 | 8.52136602709161e-06 | 0.00107967250302574 | 2059 | 0 | 0 | 4 | 8 | 0 | 2 | 6 | 5 | 1 | 2 | 18 | 26 |
| L1MEd_dup4656 | 5.17317787235856 | -2.84407320194342 | 0.734618533647403 | -3.87149666347581 | 0.000108169138249819 | 0.00721094694314257 | 2060 | 0 | 0 | 4 | 13 | 0 | 2 | 4 | 5 | 3 | 3 | 14 | 20 |
| MIR_dup133532 | 4.99089891711361 | -2.47667823655688 | 0.706951660242906 | -3.50332048970067 | 0.000459496247379336 | 0.020365026952049 | 2061 | 0 | 0 | 5 | 6 | 1 | 1 | 2 | 3 | 4 | 3 | 16 | 24 |
| THE1D_dup9105 | 4.41201546590215 | -2.79784095974728 | 0.798808276786383 | -3.50251874079602 | 0.000460881360076532 | 0.0204076942740351 | 2062 | 0 | 0 | 4 | 6 | 0 | 1 | 2 | 2 | 2 | 4 | 12 | 26 |
| L2_dup55460 | 4.74469394819255 | -2.59081610171328 | 0.744299133350032 | -3.48088018059651 | 0.000499769006067632 | 0.0215634872246107 | 2063 | 0 | 0 | 3 | 8 | 0 | 1 | 5 | 2 | 4 | 4 | 12 | 24 |
| MLT1A1_dup5436 | 5.49963719184825 | -3.03973299131875 | 0.74456536405773 | -4.08256029363605 | 4.45422507375111e-05 | 0.00374908043650082 | 2064 | 0 | 0 | 3 | 6 | 0 | 1 | 4 | 2 | 4 | 2 | 20 | 31 |
| L2b_dup73103 | 8.10236112010659 | -3.10859382395852 | 0.669529467126433 | -4.64295296411726 | 3.43464563289439e-06 | 0.000534506337321575 | 2065 | 0 | 0 | 7 | 4 | 0 | 1 | 3 | 3 | 5 | 4 | 31 | 49 |
| MER2_dup7270 | 6.85258431977557 | -2.7877120143081 | 0.688339109565431 | -4.04991082966078 | 5.12371530141334e-05 | 0.00415956179157318 | 2066 | 0 | 1 | 4 | 7 | 0 | 1 | 2 | 5 | 5 | 3 | 21 | 41 |
| THE1D_dup10313 | 6.65207029894254 | -3.04908061136317 | 0.767593168436504 | -3.97226126643855 | 7.11935727308943e-05 | 0.00530345863762657 | 2067 | 0 | 0 | 3 | 8 | 0 | 1 | 7 | 2 | 7 | 1 | 23 | 36 |
| LTR33_dup7068 | 6.93226821981619 | -2.55837308221025 | 0.636114843601529 | -4.02187294942742 | 5.77372000894123e-05 | 0.00453302647607606 | 2068 | 0 | 0 | 9 | 7 | 0 | 1 | 7 | 3 | 8 | 5 | 24 | 25 |
| MIRb_dup170321 | 7.75479141307835 | -2.41863100993082 | 0.607207519499986 | -3.98320332383643 | 6.79925364206186e-05 | 0.00512746329078604 | 2069 | 0 | 0 | 6 | 5 | 0 | 2 | 6 | 4 | 9 | 4 | 32 | 31 |
| AluSx1_dup81388 | 6.57255471843141 | -3.05069615956297 | 0.76872794970783 | -3.96849907788893 | 7.23267371917882e-05 | 0.00536132641988699 | 2070 | 0 | 0 | 7 | 6 | 0 | 0 | 4 | 1 | 2 | 9 | 22 | 37 |
| MIR_dup133521 | 6.37193455578424 | -3.34895689118384 | 0.738784969837114 | -4.53306040040611 | 5.81351825390195e-06 | 0.000805994778747697 | 2071 | 0 | 0 | 7 | 8 | 0 | 0 | 4 | 3 | 2 | 6 | 26 | 28 |
| L1MB1_dup5918 | 6.90768878783878 | -3.05772903718408 | 0.69494034627183 | -4.39998778828713 | 1.08256970123696e-05 | 0.00129639219995234 | 2072 | 0 | 0 | 4 | 6 | 0 | 0 | 4 | 2 | 3 | 7 | 26 | 40 |
| L2b_dup73100 | 5.52930900620751 | -2.79648725552777 | 0.721237215086724 | -3.87734742055914 | 0.000105601511831786 | 0.00710618137282982 | 2073 | 0 | 0 | 3 | 4 | 0 | 0 | 3 | 2 | 4 | 5 | 19 | 33 |
| MIRb_dup170301 | 5.75539051231403 | -2.91294037745778 | 0.743765638169621 | -3.91647614243972 | 8.98527031369637e-05 | 0.0062992343273615 | 2074 | 0 | 0 | 2 | 6 | 0 | 0 | 5 | 1 | 5 | 4 | 23 | 30 |
| MSTB_dup6759 | 6.23901543760564 | -3.38807327037415 | 0.732990019764102 | -4.62226384946487 | 3.79574799427431e-06 | 0.000575642480717397 | 2075 | 0 | 0 | 5 | 5 | 0 | 0 | 7 | 2 | 4 | 3 | 28 | 28 |
| AluSz6_dup34554 | 4.84568676850602 | -2.78790622778004 | 0.811390353909473 | -3.43596175915481 | 0.000590454308828616 | 0.0243073081260439 | 2076 | 0 | 0 | 4 | 3 | 0 | 0 | 6 | 1 | 4 | 5 | 10 | 32 |
| ERV3-16A3_I-int_dup6( | 5.1184891597612 | -2.7866962937708 | 0.745304842147607 | -3.73900199781461 | 0.000184752259015858 | 0.0105270739999491 | 2077 | 1 | 0 | 4 | 6 | 0 | 0 | 4 | 1 | 4 | 3 | 18 | 27 |
| L1M5_dup34383 | 5.78459782681027 | -4.41427068630148 | 0.878238706124002 | -5.0262766324469 | 5.00094354471221e-07 | 0.000119846261266864 | 2078 | 0 | 0 | 7 | 13 | 0 | 0 | 3 | 4 | 1 | 2 | 23 | 24 |
| LTR78_dup3866 | 6.72091535923511 | -3.82044360861611 | 0.815901842371447 | -4.68247944815501 | 2.83425611717925e-06 | 0.000460655673537113 | 2079 | 0 | 0 | 3 | 11 | 0 | 0 | 3 | 4 | 1 | 5 | 21 | 43 |
| L1ME3A_dup3515 | 5.56213856302342 | -2.82922236715353 | 0.783267672803897 | -3.61207600592738 | 0.000303755501302402 | 0.0150860794224382 | 2080 | 0 | 0 | 4 | 11 | 0 | 0 | 5 | 2 | 7 | 4 | 14 | 26 |
| AluSx1_dup113008 | 5.59943097157504 | -2.97228720448474 | 0.799059701053899 | -3.71973107962336 | 0.000199435009839376 | 0.0110927449395138 | 2081 | 0 | 0 | 2 | 12 | 0 | 0 | 5 | 3 | 6 | 4 | 19 | 22 |
| L2c_dup110918 | 5.69727341196766 | -2.98253969038278 | 0.743634826854845 | -4.01075848343096 | 6.05239982960939e-05 | 0.00470635401913803 | 2082 | 0 | 0 | 2 | 8 | 0 | 0 | 5 | 4 | 5 | 4 | 21 | 25 |
| LTR78_dup3867 | 7.39564548962586 | -3.66160177687842 | 0.722429184690702 | -5.06845771803376 | 4.01052045750387e-07 | 0.000100557144444807 | 2083 | 0 | 0 | 5 | 7 | 0 | 0 | 8 | 6 | 5 | 2 | 26 | 38 |
| MLT1C_dup14894 | 5.61093305457504 | -4.03828579362823 | 0.839710477182931 | -4.80914065425967 | 1.51580523627415e-06 | 0.000285347937302602 | 2084 | 0 | 0 | 2 | 10 | 0 | 0 | 9 | 4 | 2 | 2 | 21 | 25 |
| L1PB_dup322 | 6.68419999781966 | -2.26344848312894 | 0.637583465414377 | -3.55004263113674 | 0.000385168772616082 | 0.0179868564907407 | 2085 | 0 | 0 | 7 | 6 | 1 | 0 | 9 | 2 | 8 | 8 | 21 | 24 |
| L1PA16_dup11605 | 6.31051783928249 | -2.13631537219581 | 0.64458975646959 | -3.31422482401268 | 0.000918975840677721 | 0.0331941792996004 | 2086 | 0 | 0 | 4 | 7 | 4 | 0 | 5 | 2 | 5 | 4 | 21 | 31 |
| L2a_dup38936 | 5.47837509722663 | -2.39943095108633 | 0.733938385381231 | -3.26925393040997 | 0.00107831482506941 | 0.0369899544029713 | 2087 | 0 | 0 | 4 | 9 | 1 | 0 | 3 | 1 | 6 | 5 | 16 | 27 |
| Kanga1_dup688 | 4.58077438454988 | -2.99491888054794 | 0.818697664501822 | -3.65815002339154 | 0.000254042336778852 | 0.0132941187304114 | 2088 | 0 | 0 | 11 | 7 | 1 | 0 | 6 | 1 | 3 | 3 | 10 | 19 |
| L1MC4a_dup7788 | 3.81918794905628 | -2.81755981473681 | 0.822496213912155 | -3.42562040660984 | 0.0006133969871539 | 0.0249474846462427 | 2089 | 0 | 0 | 5 | 4 | 1 | 0 | 5 | 1 | 2 | 3 | 11 | 19 |
| ERVL47-int_dup141 | 5.82070475382904 | -3.16157428189633 | 0.758737193365213 | -4.16688981315632 | 3.08783656641469e-05 | 0.00281871810251086 | 2090 | 0 | 0 | 14 | 11 | 1 | 0 | 6 | 3 | 5 | 2 | 15 | 19 |
| L2a_dup156080 | 5.66287248294098 | -2.54109850884265 | 0.657602822317707 | -3.86418431095931 | 0.000111461038740962 | 0.00735930778015576 | 2091 | 0 | 0 | 9 | 9 | 2 | 0 | 9 | 4 | 5 | 4 | 11 | 21 |
| MIRb_dup170297 | 5.59010816476587 | -3.00574266655579 | 0.691786196091197 | -4.34490118990399 | 1.39338368768323e-05 | 0.00156097022104272 | 2092 | 0 | 0 | 6 | 8 | 1 | 0 | 8 | 4 | 3 | 4 | 20 | 19 |
| L2a_dup165790 | 7.13492661365219 | -2.87675335345487 | 0.772425136787333 | -3.72431348547232 | 0.000195847442357869 | 0.0109520835532986 | 2093 | 0 | 0 | 6 | 8 | 0 | 0 | 1 | 2 | 8 | 5 | 35 | 26 |
| MER5A_dup26804 | 5.83537309149672 | -3.37519972652006 | 0.809261289629001 | -4.17071688683811 | 3.03642907640898e-05 | 0.0027799390266072 | 2094 | 0 | 0 | 8 | 7 | 0 | 0 | 2 | 2 | 5 | 2 | 24 | 26 |
| MIR_dup133529 | 4.8415605175486 | -3.02025385898928 | 0.796444735231237 | -3.79217003438711 | 0.000149336603874167 | 0.00906339296725752 | 2095 | 0 | 0 | 5 | 5 | 0 | 0 | 2 | 1 | 3 | 4 | 24 | 19 |
| L1MD3_dup4076 | 5.44233707666809 | -3.12618811321481 | 0.900503824058826 | -3.47159893127849 | 0.000517368658385044 | 0.0220817583476391 | 2096 | 0 | 0 | 11 | 11 | 0 | 0 | 1 | 3 | 4 | 7 | 12 | 22 |
| L1PB_dup323 | 6.94287427200797 | -3.34487487536283 | 0.752373808578348 | -4.44576198323962 | 8.758087511081e-06 | 0.00110320375379125 | 2097 | 0 | 0 | 13 | 13 | 0 | 0 | 4 | 7 | 6 | 4 | 19 | 23 |
| MIRb_dup129977 | 5.54567250166119 | -3.60141149484083 | 0.826889839866944 | -4.35537035431508 | 1.32842122795381e-05 | 0.0015096525313337 | 2098 | 0 | 0 | 12 | 10 | 0 | 0 | 5 | 4 | 4 | 2 | 12 | 24 |
| FLAM_C_dup17836 | 6.4413857491286 | -3.14510868255743 | 0.67800242533287 | -4.63878677279556 | 3.50460396133367e-06 | 0.000541341739817781 | 2099 | 0 | 2 | 11 | 14 | 1 | 0 | 9 | 3 | 2 | 3 | 14 | 28 |
| L1MA9_dup15128 | 5.85775004758945 | -2.63997510630959 | 0.653066006450835 | -4.04243228132 | 5.28995928766622e-05 | 0.00426686322936619 | 2100 | 1 | 3 | 10 | 5 | 1 | 0 | 7 | 3 | 2 | 2 | 21 | 22 |
| AluSx_dup82052 | 7.92186435336834 | -2.151999485585 | 0.570450056094635 | -3.77245906559783 | 0.00016164649270029 | 0.00960617791010568 | 2101 | 0 | 1 | 6 | 5 | 2 | 0 | 11 | 3 | 5 | 11 | 29 | 30 |
| L2c_dup110936 | 9.93273156429786 | -2.47440899654946 | 0.548604354247856 | -4.51037068406413 | 6.47144419136801e-06 | 0.000877134904411177 | 2102 | 0 | 1 | 11 | 6 | 3 | 0 | 6 | 4 | 4 | 10 | 39 | 46 |
| AluJb_dup90749 | 6.89762428322074 | -2.88262332799592 | 0.648787936794168 | -4.44309020639275 | 8.86759652488398e-06 | 0.00111404916721755 | 2103 | 1 | 1 | 5 | 6 | 1 | 0 | 5 | 2 | 2 | 4 | 29 | 36 |
| MIR_dup133520 | 7.59213038580901 | -2.78157235290151 | 0.620747145054337 | -4.48100708164841 | 7.4291646016748e-06 | 0.000973961432677747 | 2104 | 0 | 2 | 10 | 9 | 2 | 0 | 6 | 2 | 2 | 5 | 29 | 34 |
| L1M1_dup6231 | 7.13984393114743 | -2.04419566056993 | 0.634825397325034 | -3.22009117654015 | 0.00128149831495576 | 0.0417098372636057 | 2105 | 4 | 0 | 4 | 2 | 2 | 0 | 8 | 5 | 2 | 7 | 25 | 35 |
| L1M4_dup20180 | 8.51329303589271 | -3.37333111421378 | 0.667869146521959 | -5.05088628780199 | 4.39764817703213e-07 | 0.000108323270510568 | 2106 | 1 | 0 | 5 | 7 | 2 | 0 | 5 | 5 | 1 | 4 | 44 | 38 |
| AluY_dup76586 | 7.01104474167045 | -2.3947914659683 | 0.628471397954316 | -3.81050191586027 | 0.000138684937427911 | 0.00859810477363774 | 2107 | 1 | 0 | 5 | 4 | 1 | 2 | 7 | 1 | 3 | 5 | 36 | 27 |
| L3_dup36521 | 5.38628402517941 | -2.00624489861632 | 0.624327315600735 | -3.21345045857218 | 0.00131150421232001 | 0.0423716471561029 | 2108 | 1 | 0 | 6 | 4 | 1 | 3 | 5 | 2 | 2 | 5 | 19 | 23 |
| MLT1A0_dup20088 | 6.75945911951715 | -3.51581592007667 | 0.736513185950148 | -4.77359535055853 | 1.80965809647613e-06 | 0.000325227269957786 | 2109 | 0 | 0 | 3 | 6 | 1 | 1 | 6 | 6 | 2 | 2 | 33 | 28 |
| MIRb_dup170315 | 6.92467747865912 | -2.63436286431884 | 0.653033995180479 | -4.03403633464868 | 5.48268345061139e-05 | 0.00437391430415276 | 2110 | 0 | 0 | 6 | 2 | 1 | 1 | 5 | 3 | 5 | 4 | 29 | 34 |
| MER103C_dup6944 | 7.56526022377275 | -3.15843702986135 | 0.660231427526491 | -4.78383321087001 | 1.71983410349718e-06 | 0.000315032829554072 | 2111 | 0 | 0 | 9 | 6 | 1 | 1 | 5 | 3 | 2 | 5 | 32 | 36 |
| MIR_dup133517 | 7.59666541014869 | -3.09568481416084 | 0.653390065910683 | -4.73788166620827 | 2.15963852819722e-06 | 0.000372644923299795 | 2112 | 0 | 0 | 5 | 7 | 1 | 1 | 4 | 3 | 3 | 4 | 36 | 36 |
| AluJb_dup90752 | 5.21802052956634 | -3.02838155016429 | 0.872916117954446 | -3.46926982773656 | 0.000521875030762476 | 0.0222024094528532 | 2113 | 0 | 0 | 0 | 4 | 0 | 0 | 3 | 1 | 5 | 1 | 23 | 32 |
| AluSc_dup25760 | 3.69681959113949 | -3.44380204754325 | 0.933904230308435 | -3.68753233552212 | 0.000226439326251817 | 0.0121735951026468 | 2114 | 0 | 0 | 0 | 3 | 0 | 0 | 4 | 2 | 1 | 2 | 13 | 25 |
| MIR_dup133526 | 4.78457515511275 | -2.85129231275198 | 0.850051726480518 | -3.3542574221421 | 0.000795782931050655 | 0.0299514972021163 | 2115 | 0 | 0 | 0 | 5 | 0 | 0 | 2 | 1 | 2 | 5 | 20 | 29 |
| AluSx1_dup81392 | 4.1175210776351 | -2.83275618628619 | 0.858652339133875 | -3.29907234532616 | 0.000970049079296642 | 0.0344121815818294 | 2116 | 0 | 0 | 5 | 5 | 1 | 0 | 2 | 0 | 3 | 2 | 14 | 23 |
| L1M1_dup8973 | 5.04293100085171 | -3.08533982853158 | 0.821540338938241 | -3.75555487940991 | 0.000172957855167501 | 0.0100745727798497 | 2117 | 0 | 0 | 7 | 5 | 1 | 1 | 4 | 0 | 1 | 3 | 20 | 26 |
| L1MB4_dup4305 | 5.52694249006911 | -3.72890634843237 | 0.856207882872188 | -4.3551413424548 | 1.32981084181934e-05 | 0.0015096525313337 | 2118 | 0 | 0 | 2 | 5 | 0 | 1 | 2 | 0 | 1 | 1 | 28 | 35 |
| L1MC4a_dup7789 | 5.70717252368939 | -2.43158908228869 | 0.741011893422889 | -3.2814440684031 | 0.00103277005104609 | 0.0358696951469707 | 2119 | 1 | 0 | 4 | 5 | 0 | 1 | 4 | 0 | 6 | 2 | 24 | 28 |
| L2a_dup155965 | 5.95134603911111 | -2.68234543106901 | 0.759908364198988 | -3.52982748636599 | 0.000415830714627174 | 0.0190279336966687 | 2120 | 1 | 0 | 4 | 4 | 0 | 1 | 1 | 0 | 3 | 3 | 24 | 39 |
| MIRb_dup170300 | 5.83021513153477 | -2.66018391198766 | 0.783990773666262 | -3.39313165580706 | 0.000690984192364008 | 0.0270868080125612 | 2121 | 1 | 0 | 3 | 10 | 0 | 0 | 5 | 1 | 7 | 3 | 29 | 16 |
| Tigger13a_dup2856 | 5.01806455379895 | -2.36104054738584 | 0.719725178912464 | -3.28047512656633 | 0.00103632402242074 | 0.0359564240068424 | 2122 | 1 | 1 | 3 | 6 | 0 | 0 | 6 | 1 | 4 | 4 | 25 | 14 |
| L1MC5a_dup13064 | 4.72519364740837 | -3.20502467176691 | 0.919047822728383 | -3.48733177154169 | 0.000487865740536914 | 0.0211873884828977 | 2123 | 0 | 0 | 1 | 5 | 0 | 0 | 7 | 0 | 4 | 2 | 26 | 17 |
| L2b_dup73107 | 7.1606330547685 | -3.31694305925879 | 0.758291875051718 | -4.37422998767139 | 1.21861852006409e-05 | 0.0014117108197428 | 2124 | 0 | 0 | 4 | 6 | 0 | 0 | 7 | 0 | 3 | 6 | 37 | 32 |
| L1M6_dup5550 | 5.59928062677694 | -2.76200493942077 | 0.753849213171927 | -3.66386923427199 | 0.000248433653867094 | 0.0130551665642799 | 2125 | 0 | 2 | 4 | 1 | 0 | 0 | 5 | 3 | 2 | 4 | 26 | 26 |
| AluJb_dup90750 | 6.10460244728476 | -2.91953225151083 | 0.691963530940989 | -4.21919959790455 | 2.45171152830108e-05 | 0.0023772410403958 | 2126 | 0 | 1 | 7 | 4 | 0 | 0 | 5 | 3 | 3 | 5 | 23 | 29 |
| MIR3_dup66176 | 6.57137469104739 | -2.61946273851406 | 0.649357585344956 | -4.03392952917079 | 5.48517742832064e-05 | 0.00437391430415276 | 2127 | 0 | 1 | 5 | 5 | 0 | 0 | 4 | 5 | 4 | 7 | 29 | 24 |
| MLT1A0_dup16556 | 5.22435021829403 | -3.08118973461179 | 0.829443396493647 | -3.71476793671163 | 0.000203390261858448 | 0.011255148151518 | 2128 | 0 | 1 | 4 | 2 | 0 | 1 | 3 | 5 | 1 | 3 | 33 | 13 |

| | | | | | | | | | | | | | | | | | | |
|---|---|---|---|---|---|---|---|---|---|---|---|---|---|---|---|---|---|---|
| Tigger5b_dup2135 | 6.67988563561698 | -2.72221249603527 | 0.69254436472382 | -3.93074095277761 | 8.46844763958281e-05 | 0.00603092803958492 | 2129 | 0 | 2 | 6 | 3 | 0 | 1 | 3 | 2 | 3 | 3 | 35 | 29 |
| L1ME3_dup8925 | 5.34538967069741 | -2.57590995874549 | 0.662579781996854 | -3.88769779690881 | 0.000101199512964537 | 0.00687852173724179 | 2130 | 1 | 2 | 4 | 7 | 0 | 0 | 6 | 6 | 4 | 2 | 16 | 21 |
| MLT1J_dup11703 | 5.30852872091152 | -2.29796874431353 | 0.674248338377535 | -3.40819341111496 | 0.000653945255046261 | 0.0260642642641939 | 2131 | 1 | 3 | 3 | 12 | 0 | 1 | 4 | 3 | 3 | 2 | 19 | 19 |
| L1MEc_dup15461 | 6.54261919338926 | -3.12578915101675 | 0.651292074422829 | -4.79936617344347 | 1.5916855031961e-06 | 0.00029634992076956 | 2132 | 2 | 0 | 11 | 10 | 0 | 1 | 7 | 5 | 3 | 2 | 21 | 24 |
| L3_dup36520 | 7.21396250420683 | -2.59696336691903 | 0.565894880713807 | -4.58912680680779 | 4.4510399019212e-06 | 0.000655871787091003 | 2133 | 1 | 1 | 6 | 8 | 0 | 2 | 8 | 6 | 3 | 5 | 27 | 27 |
| AluJb_dup90747 | 5.53273046316079 | -2.33821805272895 | 0.651245946997664 | -3.59037636012703 | 0.000330200833400459 | 0.0160152871556368 | 2134 | 0 | 2 | 5 | 6 | 0 | 3 | 8 | 2 | 3 | 2 | 21 | 21 |
| FLAM_C_dup21324 | 4.65283024440465 | -2.37934538069096 | 0.66075039302758 | -3.60097459766724 | 0.000317026554024647 | 0.0155440347051203 | 2135 | 1 | 2 | 8 | 7 | 0 | 2 | 11 | 2 | 2 | 2 | 12 | 13 |
| L1MD3_dup4030 | 4.48091343837632 | -2.74221037057058 | 0.736993409975819 | -3.72080717880579 | 0.000198587026267819 | 0.0110636826392701 | 2136 | 1 | 2 | 8 | 13 | 0 | 1 | 3 | 4 | 1 | 2 | 13 | 11 |
| L1ME3F_dup3794 | 4.11379331366834 | -2.766277032765 | 0.765033004741737 | -3.61589240675812 | 0.000299314685444018 | 0.0149247133576409 | 2137 | 1 | 1 | 7 | 5 | 0 | 1 | 3 | 4 | 2 | 1 | 15 | 13 |
| AluJb_dup95348 | 5.85967031088536 | -3.46480594058113 | 0.772816611134588 | -4.48334817168898 | 7.34808899975146e-06 | 0.000966527576083395 | 2138 | 0 | 1 | 9 | 18 | 0 | 1 | 4 | 4 | 2 | 2 | 18 | 19 |
| MER58A_dup12160 | 6.78627918640383 | -3.54335120437249 | 0.738767100798877 | -4.79630346362316 | 1.61620377002604e-06 | 0.000299077259378741 | 2139 | 1 | 1 | 9 | 16 | 0 | 2 | 4 | 5 | 1 | 1 | 27 | 23 |
| L1MA2_dup6049 | 3.7157708676803 | -2.96960061573178 | 0.938301843774826 | -3.16486707921723 | 0.00155153812814386 | 0.0475689075427547 | 2140 | 0 | 0 | 10 | 4 | 0 | 1 | 3 | 4 | 1 | 4 | 3 | 19 |
| L1ME3Cz_dup11907 | 4.07104424290363 | -3.95841952801633 | 0.988284787313901 | -4.00534297282373 | 6.19275615524253e-05 | 0.00478642346165923 | 2141 | 0 | 0 | 6 | 1 | 0 | 0 | 6 | 8 | 2 | 1 | 8 | 20 |
| L1PA7_dup2718 | 5.43585522327949 | -2.90450697992821 | 0.804510985042796 | -3.61027634666008 | 0.000305870948464101 | 0.0151658196922774 | 2142 | 0 | 0 | 6 | 2 | 0 | 1 | 5 | 10 | 5 | 3 | 12 | 24 |
| MLT2A1_dup833 | 4.24399535522791 | -2.40230668853578 | 0.739069609527256 | -3.25044712645188 | 0.00115223692045376 | 0.038751966964564 | 2143 | 0 | 0 | 6 | 4 | 0 | 2 | 9 | 6 | 2 | 5 | 11 | 9 |
| AluSx_dup41844 | 4.07597206592823 | -2.91792327254928 | 0.838931272379526 | -3.47814340532681 | 0.000504899728900576 | 0.0217249314453845 | 2144 | 1 | 0 | 3 | 2 | 0 | 1 | 10 | 8 | 1 | 3 | 11 | 12 |
| Tigger17a_dup1843 | 3.71984318744737 | -2.98366351639817 | 0.80482249017781 | -3.70723178441371 | 0.000209537149704791 | 0.0115040439833079 | 2145 | 2 | 0 | 6 | 5 | 0 | 1 | 7 | 6 | 1 | 1 | 10 | 9 |
| MLT1D_dup15800 | 5.89108346303413 | -3.35779163449002 | 0.732021638420103 | -4.58701144646082 | 4.49636037900383e-06 | 0.000661242871905096 | 2146 | 1 | 0 | 9 | 9 | 0 | 0 | 10 | 4 | 1 | 5 | 15 | 25 |
| AluSx_dup86063 | 5.34979271773951 | -4.3259156466517 | 0.883320850722559 | -4.89733220167178 | 9.71465276610149e-07 | 0.000201005458007199 | 2147 | 1 | 0 | 10 | 8 | 0 | 0 | 9 | 7 | 1 | 1 | 16 | 17 |
| L1MC3_dup12772 | 5.18034054415987 | -4.67410503529785 | 0.924300170292433 | -5.05691244633119 | 4.26098740014057e-07 | 0.000105503112165853 | 2148 | 0 | 0 | 4 | 7 | 0 | 0 | 8 | 9 | 1 | 1 | 18 | 19 |
| AluJb_dup113220 | 3.6304062231088 | -2.93782607716074 | 0.854549180160221 | -3.4378665913762 | 0.000586316485147003 | 0.0241913754768144 | 2149 | 1 | 0 | 4 | 4 | 1 | 0 | 7 | 3 | 1 | 2 | 17 | 7 |
| L2a_dup133438 | 5.03016244735573 | -2.48734218668776 | 0.722982683220212 | -3.44038971391261 | 0.000580877117517745 | 0.0240169220859908 | 2150 | 0 | 0 | 5 | 3 | 2 | 0 | 5 | 3 | 4 | 3 | 24 | 15 |
| L2b_dup73105 | 4.36047922427489 | -3.56096826829137 | 0.831359072662385 | -4.28330956548962 | 1.84133701655213e-05 | 0.0019233484536458 | 2151 | 0 | 0 | 4 | 6 | 1 | 0 | 6 | 3 | 1 | 2 | 16 | 19 |
| MER49_dup1008 | 4.79406723894177 | -2.95792144191882 | 0.760873009873811 | -3.88753629519515 | 0.00010126684906789 | 0.00687915340718343 | 2152 | 0 | 0 | 5 | 7 | 1 | 0 | 5 | 2 | 2 | 4 | 20 | 17 |
| L1M3_dup5203 | 4.34174599887318 | -3.59211876378807 | 0.86817957932287 | -4.13752966476097 | 3.51065072780428e-05 | 0.00310999738673908 | 2153 | 0 | 0 | 3 | 7 | 0 | 0 | 3 | 3 | 2 | 2 | 24 | 12 |
| MER123_dup25 | 4.40140154301321 | -2.96640343702098 | 0.794374380275066 | -3.73426373090458 | 0.000188265134431128 | 0.0106481900825646 | 2154 | 0 | 0 | 2 | 5 | 0 | 1 | 4 | 3 | 2 | 3 | 22 | 15 |
| THE1B_dup13745 | 3.7230591428259 | -3.47782012965011 | 0.896643745120593 | -3.87870896169846 | 0.000105012299736341 | 0.00707653828180779 | 2155 | 0 | 0 | 1 | 4 | 0 | 0 | 2 | 3 | 2 | 1 | 15 | 21 |
| L2a_dup38937 | 4.93763523735604 | -3.27094789719163 | 0.781126595305 | -4.18747475358261 | 2.82075386486142e-05 | 0.00263727740325468 | 2156 | 0 | 0 | 4 | 5 | 0 | 0 | 3 | 5 | 3 | 3 | 17 | 24 |
| MLT1A0-int_dup930 | 4.3201902130227 | -3.5811276880632 | 0.829518820640938 | -4.31711445111781 | 1.58082200526079e-05 | 0.00171601401853916 | 2157 | 0 | 0 | 5 | 5 | 0 | 0 | 4 | 3 | 2 | 2 | 15 | 21 |
| MLT1B_dup3705 | 4.54234638474602 | -3.02777414506088 | 0.845749177019676 | -3.5799906489184 | 0.000343606503475555 | 0.016483829249l924 | 2158 | 0 | 0 | 3 | 8 | 0 | 0 | 4 | 9 | 3 | 5 | 11 | 14 |
| MLT1A0_dup20087 | 4.1102525484637 | -2.71706337778768 | 0.833753254965384 | -3.25883390752158 | 0.00111871141933726 | 0.0379138167477033 | 2159 | 0 | 0 | 3 | 4 | 0 | 0 | 2 | 6 | 4 | 4 | 15 | 13 |
| MIR_dup133512 | 3.275150541639 | -2.88239324039307 | 0.869949533548995 | -3.31328787387727 | 0.00092206021401913 | 0.0332525825530458 | 2160 | 0 | 0 | 2 | 3 | 0 | 0 | 2 | 4 | 2 | 3 | 11 | 15 |
| Tigger17a_dup1748 | 3.70986801918676 | -3.37437113758723 | 0.868197272897177 | -3.88664102379284 | 0.000101640888860562 | 0.00689425397400665 | 2161 | 0 | 0 | 4 | 5 | 0 | 0 | 2 | 4 | 2 | 2 | 13 | 16 |
| MIRc_dup77475 | 3.65787841056305 | -2.99200224069457 | 0.861688118010748 | -3.47225658350932 | 0.000516102808253371 | 0.0220514632452247 | 2162 | 0 | 0 | 4 | 3 | 1 | 0 | 2 | 5 | 1 | 3 | 12 | 16 |
| L1PA13_dup8700 | 4.85238494820155 | -3.41197571660606 | 0.848892325794999 | -4.01932684856198 | 5.83646567766246e-05 | 0.0045713089602932 | 2163 | 0 | 0 | 4 | 3 | 0 | 0 | 6 | 6 | 1 | 5 | 11 | 28 |
| MLT1F2_dup4610 | 4.20493205084582 | -3.09872866364102 | 0.807242991998161 | -3.83865663047847 | 0.000123709288508722 | 0.00792145596434242 | 2164 | 0 | 0 | 3 | 4 | 0 | 0 | 6 | 4 | 2 | 4 | 13 | 19 |
| MLT1B_dup3704 | 4.38314513964623 | -3.07257814989553 | 0.935663755215156 | -3.2838486398236 | 0.00102399901949191 | 0.0356640033214319 | 2165 | 0 | 0 | 5 | 16 | 0 | 0 | 8 | 1 | 4 | 5 | 7 | 13 |
| L1PA16_dup2866 | 4.15242984141799 | -2.57608354082371 | 0.777017358156604 | -3.31534876767131 | 0.000915288511759459 | 0.0331036996837082 | 2166 | 0 | 0 | 5 | 7 | 0 | 0 | 5 | 3 | 5 | 4 | 12 | 12 |
| MIRb_dup51834 | 3.9594237569416 | -2.75537049568898 | 0.855701197121663 | -3.22001477263006 | 0.00128183991043341 | 0.0417098372636057 | 2167 | 0 | 0 | 4 | 8 | 0 | 0 | 6 | 1 | 5 | 3 | 10 | 15 |
| L1P3_dup2993 | 4.78463008276399 | -3.31025019173336 | 0.854378425901729 | -3.87445433004661 | 0.000106863881105655 | 0.00715216116824484 | 2168 | 0 | 0 | 7 | 18 | 0 | 1 | 3 | 4 | 2 | 3 | 15 | 10 |
| MIRb_dup203318 | 3.48443579445834 | -3.00553503895612 | 0.890334256832981 | -3.37573783766015 | 0.000736180314786798 | 0.0283611662517124 | 2169 | 0 | 0 | 7 | 6 | 0 | 1 | 2 | 3 | 1 | 3 | 13 | 9 |
| AluSq2_dup29101 | 4.24409001752271 | -2.7462883335406 | 0.808200374591883 | -3.39802902829313 | 0.000678732178094166 | 0.026765549806863 | 2170 | 0 | 1 | 12 | 8 | 0 | 1 | 4 | 1 | 3 | 2 | 10 | 14 |
| L1MEd_dup19960 | 4.34752548419057 | -2.57367253855326 | 0.740936207787813 | -3.47354132717765 | 0.000513638255558293 | 0.0219856392079643 | 2171 | 1 | 0 | 10 | 7 | 0 | 1 | 4 | 2 | 3 | 3 | 11 | 15 |
| MIRb_dup129976 | 4.37145964045232 | -2.59283669827396 | 0.770885990952878 | -3.36345027501279 | 0.0007697470178304 | 0.0292444872911819 | 2172 | 0 | 0 | 10 | 8 | 0 | 1 | 4 | 3 | 3 | 5 | 8 | 15 |
| L1MB1_dup3906 | 9.5596336231668 | -6.51468974251491 | 0.92835597627162 | -7.01744794995409 | 2.25957291360668e-12 | 3.70903703555628e-09 | 2173 | 0 | 0 | 9 | 24 | 0 | 0 | 10 | 8 | 0 | 0 | 42 | 35 |
| L1MD2_dup7171 | 11.9909859996243 | -6.81200710712641 | 0.909332037203083 | -7.49122083950626 | 6.82358903398301e-14 | 1.62364571628012e-10 | 2174 | 0 | 0 | 20 | 21 | 0 | 0 | 9 | 10 | 0 | 0 | 47 | 53 |
| MER1B_dup3417 | 10.433398564857 | -5.12296050065198 | 0.785307024798477 | -6.5235128922559 | 6.86795059172795e-11 | 7.26311720666355e-08 | 2175 | 0 | 0 | 9 | 14 | 1 | 0 | 8 | 8 | 1 | 1 | 45 | 52 |
| MLT1H_dup6075 | 7.75316239558538 | -5.23773824399792 | 0.887982570430842 | -5.8984696529084 | 3.66888426690518e-09 | 2.15554650808153e-06 | 2176 | 0 | 0 | 11 | 15 | 1 | 0 | 7 | 7 | 0 | 1 | 31 | 30 |
| LTR39_dup973 | 10.6389441997659 | -4.78702969653118 | 0.853280033692108 | -5.61015083854464 | 2.02150340821109e-08 | 9.23241170768132e-06 | 2177 | 1 | 0 | 26 | 30 | 0 | 0 | 7 | 6 | 0 | 4 | 34 | 35 |
| MER21B_dup1655 | 13.5234047061733 | -5.70020820168076 | 0.819166712603474 | -6.95854471864997 | 3.43805614261478e-12 | 5.17766903107986e-09 | 2178 | 0 | 0 | 14 | 30 | 0 | 0 | 18 | 13 | 0 | 3 | 50 | 53 |
| L2a_dup105205 | 7.53514029062064 | -5.15093526736489 | 0.9230912154124 | -5.58009347436338 | 2.4038936274919e-08 | 1.07720691730167e-05 | 2179 | 0 | 0 | 5 | 17 | 2 | 0 | 10 | 5 | 0 | 0 | 25 | 39 |
| Charlie7_dup2487 | 5.95061640416822 | -4.76818575912348 | 0.895474644255333 | -5.32475798138166 | 1.01087539782424e-07 | 3.32688461148943e-05 | 2180 | 0 | 0 | 9 | 6 | 1 | 0 | 7 | 5 | 1 | 0 | 27 | 22 |
| MIR_dup104344 | 6.42830739031726 | -5.00660431478311 | 0.945886294842237 | -5.293029766985 | 1.2030632850192e-07 | 3.8168546188957e-05 | 2181 | 0 | 0 | 14 | 14 | 1 | 0 | 8 | 2 | 1 | 0 | 22 | 25 |
| L4_A_Mam_dup4414 | 6.15355823006049 | -5.29242603322137 | 0.940417022873861 | -5.62774376100511 | 1.8258204212972e-08 | 8.48528644464812e-06 | 2182 | 0 | 0 | 10 | 8 | 0 | 0 | 6 | 4 | 0 | 1 | 32 | 20 |
| MIRb_dup170307 | 7.84928364287441 | -5.06320029929928 | 0.841821933936546 | -6.01457397958572 | 1.80360386635308e-09 | 1.17901188347047e-06 | 2183 | 0 | 0 | 9 | 7 | 0 | 0 | 8 | 3 | 0 | 2 | 34 | 43 |
| AluJb_dup95343 | 6.35724653732064 | -5.42089178190063 | 0.97244911383517 | -5.57447346578534 | 2.48279643013115e-08 | 1.11047270557141e-05 | 2184 | 1 | 0 | 10 | 14 | 0 | 0 | 3 | 5 | 0 | 0 | 20 | 33 |
| L1PA4_dup8362 | 8.48194300267466 | -6.24209870583997 | 0.96321534766794 | -6.48048094432137 | 9.1430701767092e-11 | 9.45894337507499e-08 | 2185 | 0 | 0 | 13 | 12 | 0 | 0 | 3 | 9 | 0 | 0 | 29 | 47 |
| L1MEd_dup15777 | 12.2784542478632 | -4.88628951521464 | 0.762652782381379 | -6.40696477885681 | 1.4844509396506e-10 | 1.41855085657069e-07 | 2186 | 2 | 0 | 5 | 10 | 1 | 0 | 15 | 9 | 0 | 1 | 54 | 68 |
| L1MEd_dup15775 | 12.2069574614109 | -6.43460051444129 | 0.916922219143696 | -7.01760779714826 | 2.25699019096467e-12 | 3.70903703555628e-09 | 2187 | 0 | 0 | 11 | 4 | 0 | 0 | 14 | 6 | 0 | 0 | 59 | 70 |
| L1MC4_dup17258 | 11.3941379430131 | -4.88299232447057 | 0.808516956180976 | -6.03944331302012 | 1.54646829502392e-09 | 1.03655195754299e-06 | 2188 | 2 | 1 | 6 | 7 | 0 | 0 | 4 | 8 | 0 | 0 | 56 | 68 |
| L1PA3_dup6147 | 10.0182210088425 | -5.34668889576276 | 0.850182409241263 | -6.28887264385341 | 3.19779669131037e-10 | 2.74694199101277e-07 | 2189 | 0 | 0 | 4 | 4 | 0 | 1 | 4 | 5 | 0 | 0 | 49 | 68 |
| L1M1_dup4752 | 10.0126040768394 | -2.90714926099114 | 0.848878844744888 | -3.42469279213198 | 0.000615494970097571 | 0.0249922467841018 | 2190 | 4 | 0 | 4 | 6 | 2 | 1 | 1 | 7 | 1 | 0 | 37 | 72 |
| THE1B-int_dup2767 | 10.2018646035019 | -3.12643414626147 | 0.727862248221668 | -4.29536516545549 | 1.74406014763306e-05 | 0.00184605042655113 | 2191 | 3 | 0 | 7 | 8 | 2 | 1 | 3 | 2 | 1 | 1 | 47 | 63 |
| L1PB3_dup3023 | 5.32785956726661 | -3.39681506576882 | 0.816250458798011 | -4.16148625603333 | 3.1618303039334e-05 | 0.00286826105032305 | 2192 | 0 | 0 | 4 | 5 | 0 | 2 | 5 | 5 | 3 | 0 | 18 | 28 |
| MIRb_dup170320 | 6.80622335274706 | -3.11152998280418 | 0.788031250347527 | -3.94848552190281 | 7.86471715557683e-05 | 0.0057071606840527l | 2193 | 0 | 0 | 5 | 4 | 0 | 2 | 3 | 2 | 5 | 0 | 34 | 34 |
| L1M5_dup38550 | 10.8396411959007 | -4.69917599201751 | 0.780241352702861 | -6.02272101541264 | 1.71508929319651e-09 | 1.13360732488594e-06 | 2194 | 1 | 0 | 9 | 5 | 0 | 1 | 8 | 10 | 2 | 0 | 36 | 73 |
| L1PA13_dup8699 | 7.59240812338471 | -3.91752846671194 | 0.798018037344364 | -4.90907258155298 | 9.15081083392677e-07 | 0.000193031811585952 | 2195 | 0 | 1 | 5 | 5 | 1 | 0 | 5 | 10 | 3 | 0 | 31 | 37 |
| THE1B_dup19120 | 9.78810578363745 | -2.97859656198817 | 0.729857671229978 | -4.08106495197694 | 4.48298241673754e-05 | 0.00376928527962201 | 2196 | 0 | 0 | 10 | 10 | 5 | 6 | 12 | 13 | 1 | 0 | 31 | 40 |
| L1M5_dup52602 | 8.07964272122816 | -2.43321155090903 | 0.694508579937656 | -3.50350106708179 | 0.000459184815760869 | 0.0203579635123972 | 2197 | 6 | 0 | 13 | 11 | 6 | 4 | 12 | 6 | 0 | 0 | 23 | 27 |
| L1PA15-16_dup664 | 7.29820588923722 | -3.46775238302653 | 0.724211908508758 | -4.78831173898681 | 1.68190240659935e-06 | 0.000309275077311197 | 2198 | 2 | 0 | 8 | 11 | 4 | 1 | 13 | 9 | 0 | 0 | 21 | 28 |
| L1MDa_dup3423 | 5.34811704889735 | -3.0225204503991 | 0.817392191420354 | -3.69776036781924 | 0.00021751007380972 | 0.0118217569718428 | 2199 | 3 | 0 | 5 | 4 | 3 | 1 | 13 | 7 | 0 | 0 | 9 | 27 |
| MLT1H-int_dup441 | 5.10669030563345 | -2.63013464896838 | 0.771631234369317 | -3.40853834295353 | 0.000653119059213368 | 0.0260356957050735 | 2200 | 2 | 0 | 6 | 2 | 3 | 2 | 6 | 10 | 1 | 0 | 14 | 19 |
| MLT2B3_dup413 | 4.88431999390999 | -2.65372137994515 | 0.702326500903218 | -3.77847251461018 | 0.000157793264103642 | 0.00944798088082667 | 2201 | 2 | 0 | 7 | 8 | 3 | 2 | 5 | 9 | 1 | 0 | 10 | 16 |
| L1MEf_dup8081 | 7.49619224126923 | -2.81455988965709 | 0.800966121216807 | -3.51395622748848 | 0.000441485557038811 | 0.0198356726501429 | 2202 | 0 | 0 | 6 | 12 | 6 | 1 | 8 | 2 | 0 | 2 | 27 | 38 |
| L1MA4_dup8609 | 4.09341763718795 | -2.88421404125523 | 0.78053543989035 | -3.69517371518763 | 0.000219736500194259 | 0.0119009725769799 | 2203 | 0 | 0 | 6 | 10 | 1 | 3 | 10 | 5 | 0 | 2 | 11 | 6 |
| L1PA4_dup7765 | 4.86449058533516 | -2.55033816351557 | 0.703015512400551 | -3.62771250211401 | 0.000285943406614718 | 0.0144487343024731 | 2204 | 0 | 0 | 8 | 12 | 3 | 5 | 9 | 7 | 0 | 1 | 9 | 10 |
| L1MA3_dup4821 | 4.86923946772013 | -3.39862153317309 | 0.787917466652209 | -4.31342326705045 | 1.6074594719002e-05 | 0.00174016629436199 | 2205 | 0 | 0 | 8 | 7 | 2 | 1 | 7 | 7 | 0 | 2 | 11 | 19 |
| L1MC3_dup10666 | 5.51537089752227 | -3.30877992729643 | 0.74096767622035 | -4.46548484297507 | 7.98876282536776e-06 | 0.00103040517355623 | 2206 | 0 | 0 | 5 | 14 | 2 | 2 | 10 | 5 | 0 | 2 | 17 | 17 |
| L1MCa_dup6162 | 3.51700034335806 | -2.59877997739536 | 0.776585973042009 | -3.34641632428092 | 0.000818633828576658 | 0.0305697810694447 | 2207 | 1 | 0 | 7 | 5 | 2 | 1 | 6 | 3 | 0 | 2 | 7 | 13 |

| | | | | | | | | | | | | | | | | | | |
|---|---|---|---|---|---|---|---|---|---|---|---|---|---|---|---|---|---|---|
| AluSq2_dup41167 | 3.95796666557509 | -2.87492625461353 | 0.8288548465903 | -3.4685521432857 | 0.000523270969557695 | 0.0222498631383801 | 2208 | 0 | 0 | 9 | 14 | 2 | 3 | 6 | 3 | 0 | 1 | 4 | 12 |
| L1MA8_dup5167 | 3.63026830395764 | -3.35491914689833 | 0.892685293906307 | -3.75823279469243 | 0.00017111764312474 | 0.010004801484638 | 2209 | 0 | 0 | 6 | 14 | 2 | 1 | 6 | 4 | 0 | 1 | 6 | 9 |
| L1MEd_dup19961 | 3.69298516494373 | -3.37315563863731 | 0.883236731728897 | -3.81908441696544 | 0.000133947921223997 | 0.0083764446947609 | 2210 | 0 | 0 | 6 | 13 | 2 | 1 | 6 | 5 | 0 | 1 | 8 | 7 |
| L1PA2_dup4014 | 5.49472596956389 | -2.52975210182659 | 0.682579083922055 | -3.70616703824383 | 0.000210419550153757 | 0.01153386092658 | 2211 | 2 | 1 | 8 | 8 | 1 | 2 | 9 | 2 | 3 | 0 | 24 | 12 |
| L1PREC2_dup2051 | 5.70449268109968 | -2.15805181998968 | 0.623732048523333 | -3.45990209273166 | 0.00054037182734887 | 0.0227493479883854 | 2212 | 2 | 3 | 6 | 6 | 2 | 2 | 6 | 4 | 2 | 0 | 20 | 22 |
| L1ME3E_dup3758 | 4.46191778272434 | -2.86372060135374 | 0.781103648322151 | -3.66624916873088 | 0.000246134111035614 | 0.0129600856792388 | 2213 | 1 | 2 | 5 | 11 | 1 | 1 | 6 | 1 | 1 | 0 | 13 | 19 |
| L1PA13_dup7544 | 4.62838956689064 | -4.06506234815254 | 0.916922851531697 | -4.43337445605369 | 9.27695365250072e-06 | 0.00115446510190468 | 2214 | 0 | 1 | 7 | 11 | 1 | 0 | 8 | 2 | 1 | 0 | 13 | 19 |
| L1PA16_dup14391 | 5.43173825157827 | -2.62359839534033 | 0.69282681153084 | -3.7868026347643 | 0.000152598211378118 | 0.00921654526569828 | 2215 | 0 | 0 | 4 | 10 | 2 | 2 | 5 | 11 | 3 | 2 | 11 | 19 |
| MLT2B1_dup864 | 5.19795459315638 | -2.14004564986282 | 0.658088347317473 | -3.25191238925011 | 0.00114631349749452 | 0.0386073193878035 | 2216 | 1 | 0 | 8 | 11 | 4 | 2 | 3 | 5 | 3 | 1 | 16 | 13 |
| LTR24B_dup58 | 3.94741669193976 | -2.50965384881403 | 0.750092883282124 | -3.3457907743808 | 0.00082048281097957 | 0.0306229041398287 | 2217 | 1 | 1 | 6 | 10 | 1 | 1 | 4 | 6 | 2 | 1 | 13 | 4 |
| MER103C_dup6547 | 3.79226394062244 | -3.00861665091568 | 0.817243229666815 | -3.68142132195121 | 0.000231937364737436 | 0.0124075018412352 | 2218 | 0 | 1 | 5 | 9 | 1 | 1 | 7 | 8 | 1 | 1 | 10 | 4 |
| MIR_dup119058 | 4.34057185474069 | -2.30681508839169 | 0.675845046411168 | -3.41323074074627 | 0.000641975622245625 | 0.0257164194294373 | 2219 | 1 | 1 | 5 | 5 | 1 | 1 | 12 | 7 | 1 | 4 | 9 | 9 |
| MSTA_dup19337 | 4.41466837572027 | -1.95093869480802 | 0.616227109052343 | -3.16594104048464 | 0.00154582122722981 | 0.0474432678799888 | 2220 | 1 | 1 | 7 | 6 | 2 | 4 | 9 | 6 | 1 | 2 | 10 | 8 |
| HERVK14-int_dup307 | 4.6355485262853 | -2.14410858905407 | 0.644281646994471 | -3.32790573665443 | 0.000875014682909717 | 0.0320712020547806 | 2221 | 1 | 2 | 6 | 4 | 1 | 1 | 7 | 4 | 1 | 4 | 17 | 12 |
| MIR3_dup1694 | 4.10560014980462 | -2.41131922661896 | 0.688957868865989 | -3.49995164521154 | 0.000465342561987371 | 0.0205353118053871 | 2222 | 1 | 1 | 5 | 12 | 1 | 1 | 7 | 3 | 1 | 3 | 9 | 11 |
| ERVL-E-int_dup6620 | 18.5785124386782 | -5.47607633961838 | 1.51155223435451 | -3.62281647643944 | 0.000291412585729656 | 0.0146535205249427 | 2223 | 1 | 1 | 70 | 107 | 0 | 0 | 0 | 0 | 1 | 3 | 38 | 36 |
| L1PREC2_dup6176 | 30.5019225705984 | -5.59485529579124 | 1.74221815381551 | -3.21134025812918 | 0.00132117400264269 | 0.0425742238939354 | 2224 | 0 | 3 | 121 | 183 | 0 | 0 | 0 | 0 | 3 | 3 | 50 | 60 |
| SST1_dup442 | 50.0355794366797 | -4.85994144101209 | 1.28274826806644 | -3.78869460360898 | 0.000151440940988774 | 0.00915750092617911 | 2225 | 1 | 2 | 92 | 85 | 0 | 2 | 0 | 0 | 9 | 7 | 203 | 278 |
| SST1_dup443 | 52.4159747857841 | -5.08681408410508 | 1.31772420022167 | -3.86030254528934 | 0.000113246721132834 | 0.00744175761023845 | 2226 | 0 | 1 | 101 | 94 | 2 | 1 | 0 | 0 | 9 | 9 | 189 | 309 |
| ALR/Alpha_dup1152 | 49.2335168673066 | -4.91403900558821 | 1.05033876589996 | -4.67852769518387 | 2.88942147814448e-06 | 0.00046802333767091 | 2227 | 2 | 0 | 87 | 103 | 2 | 1 | 1 | 1 | 7 | 9 | 207 | 248 |
| SST1_dup441 | 60.6907516756389 | -5.4602922730905 | 1.0228957879082 | -5.3380728883992 | 9.39396960085708e-08 | 3.13945906587526e-05 | 2228 | 1 | 2 | 103 | 118 | 1 | 0 | 2 | 2 | 8 | 5 | 246 | 339 |
| HERVK22-int_dup385 | 46.7999119151625 | -4.00970484332328 | 0.967477348825621 | -4.14449480206487 | 3.40563680756678e-05 | 0.00303631442785359 | 2229 | 0 | 2 | 105 | 96 | 4 | 5 | 2 | 2 | 13 | 8 | 177 | 216 |
| HERVK22-int_dup386 | 41.4779013081349 | -4.75729573196161 | 1.09463294617286 | -4.346019136912 | 1.38630479848561e-05 | 0.00155597019613423 | 2230 | 0 | 1 | 89 | 78 | 3 | 1 | 0 | 3 | 6 | 8 | 177 | 192 |
| LTR22B_dup128 | 21.4884070523874 | -4.97869461496076 | 1.49417276770739 | -3.33207425711546 | 0.000862012520165829 | 0.0317009810791307 | 2231 | 0 | 2 | 43 | 44 | 0 | 0 | 0 | 0 | 3 | 4 | 106 | 86 |
| HERVK22-int_dup387 | 28.0089892242697 | -5.82155943574622 | 1.26763834835948 | -4.59244503235502 | 4.38082918921971e-06 | 0.000648662590079697 | 2232 | 0 | 0 | 38 | 40 | 0 | 0 | 3 | 0 | 6 | 3 | 119 | 172 |
| MER4E_dup633 | 21.007287557076 | -4.64124649577108 | 1.24268831897007 | -3.73484358460674 | 0.000187831893311857 | 0.0106422634544002 | 2233 | 0 | 0 | 34 | 29 | 2 | 0 | 1 | 0 | 5 | 3 | 95 | 114 |
| L1MA4A_dup3282 | 21.1822235811987 | -3.87147652128509 | 0.786010513207579 | -4.92547676682623 | 8.41549098664621e-07 | 0.000180399316964731 | 2234 | 1 | 0 | 16 | 28 | 3 | 1 | 3 | 0 | 3 | 3 | 90 | 142 |
| L1PA7_dup7467 | 15.0202951356662 | -3.68507754815462 | 0.682360561205474 | -5.40048437389827 | 6.64612005695731e-08 | 2.40702843694485e-05 | 2235 | 0 | 1 | 18 | 19 | 3 | 0 | 5 | 2 | 5 | 2 | 68 | 78 |
| L1MB4_dup4075 | 19.1290203250204 | -4.39189568550381 | 0.70462816055134 | -6.23292671423656 | 4.57800374568489e-10 | 3.75626785955426e-07 | 2236 | 0 | 0 | 18 | 21 | 3 | 1 | 4 | 5 | 2 | 2 | 93 | 108 |
| L1PA10_dup3708 | 16.5795150311953 | -3.35984797610019 | 0.623280643316929 | -5.39058610615597 | 7.02282553343009e-08 | 2.51584985054171e-05 | 2237 | 1 | 1 | 15 | 22 | 3 | 1 | 2 | 5 | 5 | 3 | 65 | 100 |
| L1PA15_dup130 | 18.6681569968835 | -2.060867799235 | 0.474408948591684 | -4.34407446434733 | 1.39864071063726e-05 | 0.0015650343823851 | 2238 | 8 | 11 | 31 | 55 | 15 | 9 | 20 | 11 | 2 | 2 | 32 | 56 |
| MLT1A1_dup6547 | 17.5042599527546 | -1.65014387092591 | 0.517113302140172 | -3.19106830958801 | 0.00141747762351643 | 0.04467326233182 | 2239 | 8 | 11 | 27 | 61 | 15 | 11 | 6 | 11 | 5 | 3 | 39 | 35 |
| MER121_dup889 | 17.8113134756354 | -1.80321475442718 | 0.551566507520907 | -3.26926078693933 | 0.00107828869351729 | 0.0369899544029713 | 2240 | 8 | 11 | 43 | 49 | 16 | 6 | 9 | 4 | 4 | 4 | 35 | 50 |
| MIRb_dup214048 | 18.7804369935284 | -1.71413693316404 | 0.508729959027899 | -3.36944365619725 | 0.000753200882995667 | 0.0288415090610375 | 2241 | 8 | 11 | 46 | 61 | 17 | 7 | 10 | 7 | 7 | 4 | 38 | 33 |
| MER6_dup414 | 17.7680015240523 | -1.92953374043061 | 0.485651220194289 | -3.97308533407718 | 7.0947617290832e-05 | 0.00529538950560989 | 2242 | 5 | 15 | 51 | 69 | 14 | 10 | 22 | 9 | 1 | 4 | 20 | 19 |
| L1MB4_dup3284 | 17.7619705278782 | -2.16808899119037 | 0.499504898993302 | -4.34047593038611 | 1.42174442341757e-05 | 0.00158676547173788 | 2243 | 8 | 9 | 51 | 72 | 9 | 5 | 14 | 7 | 5 | 4 | 21 | 35 |
| Tigger3_dup599 | 14.1567769850665 | -2.10575676142632 | 0.528090476323394 | -3.98749240108769 | 6.67753424827041e-05 | 0.00507288395796152 | 2244 | 7 | 11 | 37 | 59 | 7 | 3 | 14 | 5 | 2 | 4 | 20 | 23 |
| L1PB2_dup1329 | 19.1297906923574 | -1.7488920101669 | 0.515968843871267 | -3.38953026125593 | 0.000700124809576889 | 0.027346010823963 | 2245 | 19 | 19 | 47 | 87 | 10 | 5 | 16 | 9 | 3 | 3 | 21 | 20 |
| LTR33B_dup722 | 15.4078156407898 | -1.52280454131539 | 0.479308251737099 | -3.17708809684888 | 0.00148761839862429 | 0.0462430492641567 | 2246 | 12 | 15 | 34 | 52 | 10 | 6 | 12 | 8 | 4 | 4 | 31 | 16 |
| FAM_dup4072 | 16.1995878240992 | -2.26343037782114 | 0.602515494427764 | -3.75663430858458 | 0.000172213864194557 | 0.0100435294435387 | 2247 | 11 | 10 | 31 | 72 | 6 | 4 | 8 | 4 | 3 | 2 | 40 | 29 |
| L1PB2_dup681 | 22.0617054079543 | -2.33952965500871 | 0.636650837059635 | -3.67474527452724 | 0.000238086828759233 | 0.0126659373906513 | 2248 | 15 | 16 | 64 | 80 | 4 | 5 | 7 | 4 | 4 | 3 | 43 | 54 |
| THE1B_dup21984 | 19.3966091502961 | -2.34170855519524 | 0.536603950785647 | -4.36394206894437 | 1.27739417808675e-05 | 0.00146482330168208 | 2249 | 11 | 6 | 64 | 61 | 4 | 6 | 7 | 10 | 7 | 5 | 26 | 52 |
| L1M4_dup20220 | 17.8249754827768 | -2.14452442275623 | 0.574576124577199 | -3.73235909225131 | 0.000189694815143445 | 0.010698982634756 | 2250 | 10 | 10 | 42 | 64 | 10 | 5 | 3 | 11 | 2 | 5 | 31 | 46 |
| MLT2A2_dup3783 | 22.4461299087252 | -2.03829521450723 | 0.54881439085043 | -3.71399738871412 | 0.000204010899191872 | 0.0112760923157048 | 2251 | 14 | 8 | 51 | 74 | 13 | 8 | 5 | 8 | 5 | 6 | 46 | 64 |
| ERVL-B4-int_dup3878 | 20.2133537021427 | -2.3202152781275 | 0.624231316675246 | -3.71691585498358 | 0.000201669572468991 | 0.0111908740883177 | 2252 | 3 | 7 | 74 | 69 | 12 | 7 | 5 | 8 | 9 | 2 | 36 | 35 |
| MSTD_dup7398 | 18.4319256765414 | -2.3178065974299 | 0.605628009638622 | -3.82711261788062 | 0.000129655204453209 | 0.0081941400474962 | 2253 | 2 | 7 | 56 | 63 | 13 | 9 | 8 | 6 | 5 | 2 | 27 | 50 |
| L1PA2_dup2173 | 21.6590707732538 | -1.80259946117847 | 0.569045813052814 | -3.16775806065227 | 0.00153619303948896 | 0.0472938371503392 | 2254 | 7 | 6 | 54 | 80 | 13 | 16 | 7 | 6 | 11 | 5 | 28 | 58 |
| LTR22C2_dup68 | 21.3600616927231 | -2.22382341936839 | 0.604464565532879 | -3.67899715909391 | 0.000234152876059645 | 0.0124979228907331 | 2255 | 8 | 2 | 54 | 101 | 10 | 11 | 9 | 7 | 11 | 3 | 39 | 32 |
| L1PA8_dup7931 | 24.2701549762739 | -1.90492943438112 | 0.60194376928244 | -3.16463020566179 | 0.00155280166922482 | 0.0475892907099108 | 2256 | 6 | 14 | 62 | 89 | 18 | 10 | 8 | 3 | 4 | 11 | 50 | 52 |
| L1MA9_dup7556 | 26.8962275277262 | -2.37652568374371 | 0.595978616538613 | -3.98760226926655 | 6.67444356175719e-05 | 0.00507236393403346 | 2257 | 8 | 9 | 86 | 123 | 16 | 14 | 7 | 7 | 4 | 5 | 33 | 54 |
| L1MB2_dup8675 | 24.7094659002023 | -2.21366100404033 | 0.54188473507777 | -4.08511415942107 | 4.40551512323932e-05 | 0.00371070690801523 | 2258 | 12 | 8 | 65 | 96 | 11 | 11 | 8 | 8 | 3 | 10 | 52 | 49 |
| L1ME3Cz_dup11923 | 24.5954292334982 | -2.36731927400309 | 0.610498315586696 | -3.87768354729704 | 0.000105455762372796 | 0.00709838099959188 | 2259 | 10 | 7 | 68 | 99 | 14 | 11 | 7 | 4 | 3 | 6 | 61 | 42 |
| AluSp_dup49156 | 20.482936848283 | -1.9353384953155 | 0.503156140899816 | -3.84639744603823 | 0.000119867268299426 | 0.00774421314764467 | 2260 | 11 | 5 | 60 | 62 | 10 | 14 | 8 | 10 | 4 | 9 | 34 | 46 |
| L1MA4_dup9827 | 21.3411349591634 | -2.20934684422687 | 0.565722345354855 | -3.90535544930797 | 9.40869861974484e-05 | 0.00652343435301047 | 2261 | 9 | 8 | 60 | 81 | 12 | 14 | 6 | 10 | 2 | 5 | 25 | 57 |
| L1ME4b_dup21347 | 19.7384502169785 | -1.96327386197718 | 0.605686855047 | -3.24140080904485 | 0.00118943819410103 | 0.0395835049697292 | 2262 | 12 | 7 | 58 | 66 | 12 | 12 | 8 | 2 | 2 | 7 | 38 | 43 |
| L1PA13_dup8729 | 18.3030268509797 | -1.99202001500607 | 0.573639304194933 | -3.47260029157477 | 0.000515442387587863 | 0.0220390753561513 | 2263 | 8 | 9 | 43 | 66 | 14 | 13 | 8 | 5 | 2 | 3 | 37 | 39 |
| L1PA3_dup10182 | 18.8767536079925 | -1.34017662893312 | 0.41623472591037 | -3.21976170056919 | 0.00128297197477607 | 0.0417177726826414 | 2264 | 14 | 12 | 29 | 41 | 10 | 5 | 15 | 5 | 11 | 10 | 38 | 61 |
| MIRb_dup202618 | 22.7569048372984 | -1.47244944047318 | 0.387260070650037 | -3.80222375625145 | 0.000143403121967229 | 0.00881255146167725 | 2265 | 6 | 7 | 49 | 36 | 14 | 8 | 16 | 10 | 19 | 18 | 49 | 62 |
| L1MA3_dup8736 | 20.894511276188 | -1.87556274937021 | 0.472694488663 | -3.96781175654316 | 7.25355916096692e-05 | 0.00537346011243909 | 2266 | 11 | 10 | 61 | 67 | 13 | 10 | 10 | 10 | 5 | 7 | 34 | 40 |
| MLT1B_dup7630 | 21.8156061668464 | -1.51654039927075 | 0.42578978164375 | -3.56171158785489 | 0.000368445003234167 | 0.0173948441943566 | 2267 | 15 | 11 | 46 | 71 | 17 | 10 | 12 | 12 | 7 | 10 | 34 | 45 |
| L1MA4_dup686 | 18.0386035518894 | -1.59715533921233 | 0.413557216101285 | -3.86199364206275 | 0.000112465494056152 | 0.00741499430553759 | 2268 | 13 | 11 | 49 | 52 | 9 | 10 | 14 | 10 | 5 | 8 | 27 | 31 |
| MIRb_dup91532 | 18.2567496249325 | -1.37878401942641 | 0.400167442965103 | -3.44551772930374 | 0.000569966576636517 | 0.0236880521540317 | 2269 | 9 | 10 | 47 | 49 | 12 | 14 | 12 | 13 | 8 | 9 | 28 | 27 |
| L2_dup24967 | 21.6789114141283 | -1.66956183581771 | 0.47320750906455 | -3.52818119712037 | 0.000418425628200469 | 0.0190842830223862 | 2270 | 15 | 11 | 49 | 62 | 5 | 14 | 9 | 9 | 8 | 10 | 39 | 58 |
| MER2_dup4009 | 20.3048560508705 | -1.38453268498518 | 0.401442618978675 | -3.44889311580227 | 0.000562889461797747 | 0.0234648381003726 | 2271 | 17 | 15 | 42 | 47 | 10 | 13 | 13 | 12 | 4 | 11 | 41 | 43 |
| AluY_dup102235 | 15.4774995700512 | -1.78259074876716 | 0.519033317851145 | -3.4344437774193 | 0.000593771231943727 | 0.0244016389561454 | 2272 | 6 | 4 | 39 | 54 | 9 | 12 | 9 | 5 | 4 | 8 | 29 | 28 |
| MIRc_dup41971 | 15.0251506647882 | -1.80089274406073 | 0.47451978580233 | -3.79518999616788 | 0.000147530432824924 | 0.00900340507026401 | 2273 | 8 | 4 | 38 | 50 | 6 | 9 | 11 | 8 | 6 | 8 | 32 | 18 |
| LTR16_dup2962 | 17.5614929048567 | -2.03007740156664 | 0.514495638887796 | -3.94576211754706 | 7.95465583451759e-05 | 0.00575757698463372 | 2274 | 10 | 7 | 44 | 61 | 8 | 6 | 10 | 6 | 2 | 10 | 37 | 35 |
| MIRb_dup91531 | 14.180325797595 | -1.80474508265824 | 0.423316823425609 | -4.26334363008228 | 2.01390448572062e-05 | 0.00205197453834503 | 2275 | 7 | 6 | 30 | 45 | 6 | 7 | 15 | 8 | 3 | 10 | 26 | 26 |
| MLT1J2_dup2902 | 15.9950112424262 | -1.37779818560956 | 0.414802414380704 | -3.32157706378493 | 0.000895102601562849 | 0.0325368291523791 | 2276 | 11 | 9 | 41 | 42 | 8 | 8 | 15 | 7 | 10 | 8 | 19 | 33 |
| AluJr4_dup8160 | 15.4044906024572 | -1.49815319594987 | 0.44759471847559 | -3.34711991475739 | 0.000816558795877513 | 0.0305018680131665 | 2277 | 9 | 13 | 34 | 41 | 7 | 8 | 13 | 5 | 7 | 5 | 26 | 37 |
| MIR_dup44937 | 14.3125314500669 | -1.53749878933466 | 0.410104538278584 | -3.74904114884541 | 0.00017751196096621 | 0.0102513694335678 | 2278 | 13 | 10 | 33 | 37 | 8 | 6 | 15 | 8 | 5 | 4 | 22 | 29 |
| HAL1_dup22284 | 19.578495914733 | -2.21183232237271 | 0.496760917076223 | -4.45250873476693 | 8.48727864959212e-06 | 0.00107650000296154 | 2279 | 13 | 6 | 50 | 67 | 7 | 10 | 13 | 7 | 4 | 4 | 39 | 44 |
| L2a_dup74584 | 18.3659265886343 | -1.83336649614097 | 0.431534803666965 | -4.24847887253112 | 2.15226940183062e-05 | 0.00215268556153001 | 2280 | 18 | 7 | 42 | 57 | 9 | 7 | 19 | 9 | 5 | 5 | 28 | 41 |
| L1PA2_dup2172 | 18.4250622768336 | -1.75031963987593 | 0.512338508004282 | -3.41633434249158 | 0.000634702675113339 | 0.0255239078473075 | 2281 | 10 | 10 | 60 | 60 | 8 | 8 | 11 | 5 | 5 | 11 | 21 | 38 |
| MER20_dup6550 | 17.2209846385219 | -1.62191451349866 | 0.489290527295339 | -3.31482917207523 | 0.000916991444880787 | 0.0331552114182653 | 2282 | 10 | 5 | 39 | 63 | 7 | 11 | 14 | 4 | 7 | 12 | 29 | 30 |
| L1PA4_dup5221 | 29.9459278340894 | -1.96510166880627 | 0.509966845843945 | -3.85339102888977 | 0.000116493122260046 | 0.00758595305672931 | 2283 | 8 | 8 | 68 | 124 | 12 | 13 | 13 | 9 | 16 | 18 | 54 | 57 |
| LTR49-int_dup825 | 30.3102810088501 | -1.79424860653937 | 0.496773088626784 | -3.61180717638945 | 0.000304070629781048 | 0.0150985788968867 | 2284 | 5 | 13 | 88 | 109 | 18 | 17 | 12 | 18 | 10 | 19 | 43 | 48 |
| L1MA8_dup10712 | 28.2296749926816 | -1.72364276880869 | 0.52953476586754 | -3.25501341915641 | 0.00113387008188224 | 0.0383129580379935 | 2285 | 12 | 13 | 61 | 112 | 23 | 10 | 11 | 6 | 9 | 13 | 51 | 59 |
| MLT1D_dup20059 | 31.0722811390851 | -1.69335282479503 | 0.475161567638186 | -3.56374113590861 | 0.000365606520948153 | 0.0172836648461867 | 2286 | 13 | 14 | 73 | 103 | 26 | 16 | 13 | 13 | 6 | 16 | 54 | 68 |

| | | | | | | | | | | | | | | | | | | | |
|---|---|---|---|---|---|---|---|---|---|---|---|---|---|---|---|---|---|---|---|
| L2c_dup60365 | 31.9687198871549 | -1.57223463616441 | 0.489279068371019 | -3.21336991054805 | 0.00131187211373247 | 0.0423777792525367 | 2287 | 21 | 16 | 89 | 118 | 19 | 16 | 21 | 7 | 7 | 21 | 34 | 63 |
| SATR2_dup364 | 26.2023565750575 | -1.42435756908895 | 0.445885444428426 | -3.1944473336976 | 0.00140098845042651 | 0.0443592279208497 | 2288 | 12 | 16 | 75 | 93 | 17 | 21 | 20 | 9 | 10 | 11 | 30 | 35 |
| L1ME2_dup12445 | 29.0227648986764 | -2.00363464936661 | 0.48953117284579 | -4.09296641461847 | 4.25889426770262e-05 | 0.00361407580393284 | 2289 | 19 | 10 | 56 | 108 | 13 | 18 | 13 | 13 | 7 | 6 | 75 | 50 |
| THE1D_dup12374 | 29.8224605188141 | -2.41708851152203 | 0.479852347612552 | -5.03715053921892 | 4.7251315805536e-07 | 0.00011464232052505 | 2290 | 11 | 7 | 90 | 112 | 11 | 15 | 18 | 12 | 8 | 6 | 50 | 61 |
| L2b_dup95983 | 25.3740059873902 | -1.98506252858063 | 0.513617300692035 | -3.86486694647163 | 0.000111149773289563 | 0.00735080539243182 | 2291 | 12 | 10 | 73 | 96 | 27 | 14 | 16 | 12 | 2 | 5 | 39 | 34 |
| L1PA15_dup8146 | 25.7067558039985 | -1.99562135120386 | 0.543133640634053 | -3.67427314734947 | 0.000238527457853529 | 0.0126773854113058 | 2292 | 10 | 10 | 52 | 86 | 19 | 19 | 10 | 10 | 5 | 3 | 52 | 70 |
| AluSp_dup49157 | 27.5639541822719 | -1.80403664233563 | 0.480265240538415 | -3.75633397976743 | 0.000172420561482212 | 0.0100531200496071 | 2293 | 18 | 14 | 72 | 75 | 21 | 13 | 16 | 10 | 7 | 5 | 40 | 79 |
| L1PREC2_dup7640 | 22.6590212805348 | -2.00960289448795 | 0.521103713771223 | -3.85643556432265 | 0.000115052407517302 | 0.00751475711202635 | 2294 | 17 | 7 | 59 | 75 | 13 | 14 | 11 | 8 | 4 | 4 | 37 | 57 |
| L2_dup55584 | 24.9674420201451 | -2.13906993698649 | 0.533716682961565 | -4.00787534899023 | 6.12674407834224e-05 | 0.00475174135092967 | 2295 | 14 | 8 | 72 | 91 | 15 | 14 | 17 | 5 | 5 | 4 | 32 | 63 |
| MLT2B1_dup4328 | 28.227863234627 | -1.88178906311632 | 0.539485302182967 | -3.48811924162137 | 0.000486431092272734 | 0.021140517384827 | 2296 | 7 | 15 | 73 | 99 | 14 | 20 | 17 | 5 | 12 | 5 | 44 | 69 |
| MSTA_dup19128 | 24.6960965927007 | -2.10799467999446 | 0.528507599351708 | -3.98857969607292 | 6.64700723321842e-05 | 0.00505779429530543 | 2297 | 12 | 3 | 65 | 98 | 12 | 16 | 16 | 7 | 5 | 10 | 49 | 40 |
| L1M5_dup22821 | 21.6627129568609 | -1.74641183014231 | 0.51656940677808 | -3.38078834562607 | 0.000722781944751205 | 0.0279874813060651 | 2298 | 9 | 8 | 64 | 80 | 17 | 13 | 15 | 5 | 4 | 11 | 28 | 38 |
| MLT1A0_dup20177 | 20.8866314646886 | -2.00805875892877 | 0.533702344500644 | -3.76250690974124 | 0.000168218659241615 | 0.00988564018076201 | 2299 | 6 | 8 | 58 | 69 | 13 | 10 | 12 | 4 | 4 | 10 | 45 | 41 |
| L2_dup55587 | 22.8913865270636 | -1.86892576287965 | 0.522858513316167 | -3.57443881142188 | 0.000350979877575268 | 0.016753110922673 | 2300 | 17 | 6 | 55 | 76 | 15 | 8 | 12 | 5 | 5 | 10 | 44 | 56 |
| MLT1H1_dup3517 | 23.6606357596144 | -2.03405205371213 | 0.499657396645288 | -4.07089351097133 | 4.68331561300395e-05 | 0.00388690693007268 | 2301 | 9 | 8 | 62 | 71 | 19 | 11 | 16 | 7 | 4 | 7 | 56 | 46 |
| LTR78B_dup3038 | 26.1117550753891 | -2.12439870181671 | 0.527034890983653 | -4.03085021155194 | 5.55754542221008e-05 | 0.00441977173473663 | 2302 | 4 | 11 | 76 | 88 | 22 | 10 | 16 | 9 | 6 | 7 | 53 | 46 |
| THE1D-int_dup1089 | 27.7538368780589 | -2.19016165221191 | 0.482067369048465 | -4.5432688309416 | 5.5388540841569e-06 | 0.000777550545078937 | 2303 | 8 | 11 | 82 | 107 | 14 | 11 | 18 | 9 | 7 | 10 | 46 | 50 |
| LTR40A1_dup546 | 20.2820515148937 | -2.33001435579848 | 0.528879919065243 | -4.40556404545784 | 1.05508971854433e-05 | 0.0012737411373351 | 2304 | 4 | 3 | 59 | 79 | 13 | 9 | 14 | 8 | 5 | 7 | 31 | 41 |
| LTR49_dup587 | 22.5303122948102 | -2.15502602176766 | 0.444129018546064 | -4.85225223252135 | 1.22067235699858e-06 | 0.000239648601203287 | 2305 | 8 | 6 | 68 | 75 | 15 | 10 | 20 | 12 | 6 | 6 | 31 | 44 |
| AluSc8_dup4202 | 12.8128325464452 | -1.3562121569564 | 0.427299210709746 | -3.17391683149549 | 0.00150396778647259 | 0.0465846288622765 | 2306 | 0 | 7 | 18 | 21 | 13 | 6 | 20 | 17 | 7 | 10 | 21 | 23 |
| MIR3_dup70136 | 19.790358209596 | -1.64678972316138 | 0.419228973559183 | -3.92813910064568 | 8.56056950030272e-05 | 0.00608268982234133 | 2307 | 0 | 13 | 33 | 36 | 17 | 10 | 39 | 25 | 10 | 7 | 34 | 30 |
| HERV9-int_dup175 | 13.3073115519393 | -1.9602099266767 | 0.555725115381937 | -3.52730130854261 | 0.000419818719349627 | 0.0191331516939985 | 2308 | 1 | 1 | 33 | 47 | 9 | 6 | 25 | 14 | 9 | 5 | 11 | 13 |
| Harlequin-int | 14.8262148278004 | -2.15063038076604 | 0.594042932246105 | -3.62032820192709 | 0.000294229560270942 | 0.0147670843594663 | 2309 | 1 | 2 | 56 | 53 | 11 | 8 | 13 | 19 | 6 | 2 | 10 | 11 |
| L2c_dup66915 | 12.1028109878466 | -1.84068058132353 | 0.530520825273521 | -3.46957271729066 | 0.000521286934752662 | 0.0221958935541952 | 2310 | 0 | 3 | 36 | 32 | 12 | 10 | 16 | 14 | 3 | 4 | 14 | 13 |
| MLT1N2_dup4325 | 13.5394690640004 | -2.23531407182376 | 0.621980665278379 | -3.59386424146047 | 0.000325809643157109 | 0.0158538039571905 | 2311 | 0 | 2 | 37 | 64 | 7 | 10 | 25 | 7 | 4 | 4 | 15 | 8 |
| THE1D_dup2279 | 12.8951412487464 | -2.14685612328771 | 0.589470618772229 | -3.64200700581017 | 0.000270520709127443 | 0.0138826329534458 | 2312 | 0 | 1 | 18 | 36 | 4 | 7 | 24 | 17 | 7 | 12 | 31 | 8 |
| Tigger19b_dup3295 | 16.3093732820615 | -1.99660077283416 | 0.484441811156513 | -4.1214460165353 | 3.76501702223717e-05 | 0.00328759904724127 | 2313 | 1 | 3 | 24 | 46 | 3 | 12 | 14 | 19 | 13 | 8 | 31 | 36 |
| THE1C_dup4430 | 14.4524157486109 | -2.05519891426939 | 0.629777841745267 | -3.26337126846816 | 0.0011009518817541 | 0.037504483920532 | 2314 | 6 | 0 | 40 | 60 | 4 | 7 | 9 | 5 | 6 | 13 | 18 | 26 |
| L1ME2_dup1614 | 16.6759040266883 | -2.63921033522457 | 0.488167125181616 | -5.40636638373116 | 6.43161664626654e-08 | 2.35442685309621e-05 | 2315 | 1 | 1 | 48 | 34 | 4 | 5 | 17 | 12 | 8 | 11 | 33 | 44 |
| MER45A_dup3067 | 18.7680971767836 | -2.04340120217108 | 0.473552367250436 | -4.3150480147224 | 1.5956821714052e-05 | 0.00172898993605274 | 2316 | 4 | 0 | 45 | 39 | 9 | 6 | 17 | 11 | 12 | 16 | 41 | 44 |
| L1ME1_dup6287 | 20.1748956682817 | -1.60473581304785 | 0.414365017020945 | -3.87275891334894 | 0.000107610261440694 | 0.00718648085006103 | 2317 | 4 | 0 | 31 | 25 | 11 | 13 | 34 | 21 | 15 | 19 | 45 | 39 |
| L1MEd_dup7619 | 18.2490049485141 | -1.56370012008471 | 0.413713809356917 | -3.77966624443924 | 0.000157038715253013 | 0.00941939353153349 | 2318 | 2 | 1 | 25 | 28 | 12 | 10 | 29 | 18 | 11 | 22 | 39 | 37 |
| L1PA3_dup9473 | 17.2793696718816 | -2.81624631261918 | 0.570504131543341 | -4.93641703347634 | 7.95708644685818e-07 | 0.000172909305176632 | 2319 | 1 | 0 | 40 | 30 | 2 | 4 | 36 | 25 | 14 | 12 | 30 | 25 |
| L1PA7_dup1289 | 19.5861217728959 | -2.28198996223446 | 0.505010367208677 | -4.5186992394782 | 6.22207056470944e-06 | 0.00085087172562664 | 2320 | 7 | 0 | 48 | 53 | 4 | 5 | 19 | 23 | 14 | 15 | 34 | 29 |
| L1P1_dup368 | 17.8247965602603 | -2.13774804669401 | 0.422712182392129 | -5.05721892044011 | 4.25414782421232e-07 | 0.000105443485227086 | 2321 | 8 | 8 | 29 | 38 | 2 | 11 | 13 | 20 | 6 | 4 | 35 | 60 |
| X9_LINE_dup591 | 16.6431382673278 | -1.76812583182559 | 0.370083198703755 | -4.77764415682362 | 1.77360891449375e-06 | 0.000320930149633559 | 2322 | 9 | 6 | 30 | 19 | 6 | 8 | 14 | 15 | 4 | 11 | 41 | 54 |
| L2a_dup155582 | 21.6759807420504 | -2.37494819286066 | 0.438503243029183 | -5.41603336033391 | 6.09357365382747e-08 | 2.25146192023856e-05 | 2323 | 1 | 3 | 22 | 14 | 7 | 10 | 23 | 15 | 7 | 7 | 67 | 111 |
| AluSz_dup68846 | 16.1542629019755 | -1.81857773407169 | 0.428346414683379 | -4.24557711173077 | 2.18031507685503e-05 | 0.00217616296676739 | 2324 | 4 | 8 | 20 | 14 | 9 | 15 | 40 | 26 | 5 | 3 | 27 | 37 |
| SST1_dup93 | 17.9438820046564 | -1.98886652045925 | 0.55507465643585 | -3.58306130067228 | 0.000339590873439606 | 0.0163419365170694 | 2325 | 3 | 5 | 16 | 18 | 15 | 16 | 23 | 24 | 2 | 2 | 44 | 66 |
| L1PA7_dup1287 | 17.3366309076952 | -1.63588597644521 | 0.42324298969722 | -3.86512243856772 | 0.000111033486036247 | 0.00734704501345405 | 2326 | 5 | 1 | 22 | 37 | 8 | 11 | 31 | 19 | 10 | 18 | 16 | 49 |
| L2_dup11530 | 17.7247334514395 | -1.38406518126279 | 0.397094639749737 | -3.48547938631224 | 0.000491256072065538 | 0.0212880016979979 | 2327 | 5 | 3 | 26 | 35 | 8 | 14 | 33 | 13 | 7 | 25 | 30 | 33 |
| LTR28C_dup80 | 18.5641084211529 | -1.07095288747222 | 0.329914738203051 | -3.24615048513865 | 0.00116976987689502 | 0.0391479694975615 | 2328 | 9 | 4 | 19 | 37 | 14 | 15 | 25 | 16 | 9 | 22 | 34 | 37 |
| MLT1A0_dup4059 | 19.1512730909907 | -1.28620553107893 | 0.333103250212471 | -3.861281840566 | 0.000112793699666664 | 0.00742343515143953 | 2329 | 7 | 6 | 18 | 30 | 14 | 14 | 40 | 22 | 7 | 20 | 29 | 42 |
| MamRep137_dup275 | 17.2049886715396 | -1.52815453325816 | 0.403796832580504 | -3.78446389361782 | 0.000154040267809325 | 0.00928165752447651 | 2330 | 8 | 8 | 29 | 28 | 1 | 17 | 37 | 27 | 6 | 14 | 21 | 25 |
| L1PB1_dup995 | 16.6773874411159 | -1.81711838389046 | 0.416834746418863 | -4.35932560685453 | 1.30463873839826e-05 | 0.00148889002036889 | 2331 | 5 | 10 | 16 | 31 | 4 | 6 | 49 | 32 | 8 | 11 | 21 | 20 |
| L2c_dup30784 | 15.6249356317178 | -1.42918048331965 | 0.364071416980642 | -3.92554981429824 | 8.6531859473892e-05 | 0.00612794935546866 | 2332 | 4 | 11 | 20 | 32 | 7 | 8 | 33 | 21 | 8 | 13 | 27 | 16 |
| MIR_dup35792 | 14.4780214729229 | -1.48503720870438 | 0.418841540245645 | -3.54558243633958 | 0.000391746432786664 | 0.0182295197674648 | 2333 | 2 | 4 | 18 | 19 | 5 | 13 | 34 | 14 | 7 | 16 | 31 | 24 |
| AluSz_dup91334 | 12.8826108814753 | -1.41056527528503 | 0.421218583816676 | -3.34877265505203 | 0.00081170370597101 | 0.0303538661041926 | 2334 | 3 | 4 | 17 | 26 | 5 | 8 | 31 | 17 | 10 | 13 | 14 | 17 |
| MER103C_dup7245 | 14.3042358533222 | -1.4428447930395 | 0.428441102474343 | -3.36766193697745 | 0.000758084909950612 | 0.0289542450226924 | 2335 | 2 | 4 | 19 | 22 | 5 | 6 | 24 | 20 | 14 | 19 | 22 | 22 |
| MSTB_dup1550 | 15.2306163525623 | -2.06084449700466 | 0.422660638142059 | -4.8758846010922 | 1.08321923033996e-06 | 0.000219274038843957 | 2336 | 2 | 1 | 17 | 39 | 9 | 6 | 26 | 19 | 8 | 10 | 34 | 26 |
| L1MDb_dup219 | 13.9834942775709 | -1.24530170998973 | 0.32929525047381 | -3.78171779944567 | 0.000155749867994819 | 0.00935860052775132 | 2337 | 7 | 4 | 18 | 27 | 9 | 10 | 22 | 14 | 10 | 10 | 25 | 24 |
| L2a_dup140627 | 13.8463677089036 | -1.69137562137565 | 0.361941785058578 | -4.67305984331682 | 2.96745256807273e-06 | 0.000477412757784066 | 2338 | 4 | 4 | 18 | 21 | 7 | 10 | 25 | 18 | 8 | 6 | 32 | 24 |
| AluSx1_dup20906 | 13.3454806017682 | -1.26338994719191 | 0.392363247862456 | -3.21994976357927 | 0.00128213062663166 | 0.041711492218279 5 | 2339 | 4 | 3 | 17 | 25 | 13 | 12 | 24 | 18 | 9 | 6 | 22 | 17 |
| L2a_dup35484 | 14.7439589335798 | -1.65796704100999 | 0.358914264848601 | -4.61939578163427 | 3.84859128009056e-06 | 0.000581802351164186 | 2340 | 5 | 3 | 23 | 36 | 10 | 11 | 28 | 16 | 8 | 6 | 18 | 29 |
| MER5A_dup12477 | 11.6857211370234 | -1.46622442091345 | 0.425750557251167 | -3.44385790210127 | 0.000573477017395267 | 0.0237770626208632 | 2341 | 5 | 2 | 29 | 16 | 5 | 6 | 13 | 9 | 10 | 10 | 22 | 22 |
| L1MB7_dup2685 | 14.1388062302767 | -1.63546206034455 | 0.379807893764125 | -4.30602440653916 | 1.66214725963758e-05 | 0.00178168720078139 | 2342 | 7 | 5 | 34 | 26 | 6 | 4 | 23 | 10 | 10 | 10 | 20 | 29 |
| Tigger3a_dup2843 | 14.7174159627668 | -1.11932945263969 | 0.324433847192845 | -3.45010072877617 | 0.000560377420364748 | 0.023392906257212 3 | 2343 | 10 | 6 | 23 | 25 | 8 | 6 | 18 | 13 | 13 | 13 | 21 | 33 |
| MIR1_Amn_dup8891 | 15.4215562655711 | -1.27674649087356 | 0.36901909483904 | -3.45983855233956 | 0.000540499349285709 | 0.0227506913435587 | 2344 | 4 | 4 | 18 | 21 | 11 | 11 | 16 | 21 | 8 | 16 | 23 | 44 |
| L2_dup11525 | 13.2285988494124 | -1.21846622957919 | 0.358957184698061 | -3.39446118233882 | 0.000687637874749726 | 0.0269820092735891 | 2345 | 5 | 5 | 15 | 17 | 9 | 11 | 30 | 17 | 8 | 10 | 17 | 26 |
| L2a_dup35494 | 14.5898535704841 | -1.22815264346588 | 0.336975600127516 | -3.64463374499855 | 0.000267772858542434 | 0.0137763201294569 | 2346 | 9 | 4 | 17 | 17 | 8 | 9 | 26 | 13 | 12 | 10 | 22 | 42 |
| L2a_dup63435 | 12.86325718328 | -1.42854818861582 | 0.393856299696322 | -3.62707969814697 | 0.000286644838349105 | 0.0144780285938901 | 2347 | 3 | 3 | 25 | 17 | 11 | 7 | 20 | 17 | 7 | 11 | 22 | 20 |
| MIRc_dup21817 | 12.0504687654697 | -1.31798261754341 | 0.387899275571682 | -3.39774446755792 | 0.000679438515826399 | 0.0267842407367177 | 2348 | 4 | 5 | 20 | 12 | 9 | 8 | 26 | 14 | 4 | 12 | 20 | 21 |
| LOR1-int_dup299 | 15.9482982695548 | -1.7317883929898 | 0.368831260458819 | -4.69534060327611 | 2.6616285024201e-06 | 0.000441033325652405 | 2349 | 5 | 3 | 30 | 25 | 11 | 11 | 35 | 11 | 5 | 10 | 27 | 38 |
| L1ME3Cz_dup59 | 14.5882073851304 | -1.56507101617176 | 0.359577449946128 | -4.35252827007433 | 1.34576512206282e-05 | 0.00152412864223874 | 2350 | 6 | 3 | 27 | 26 | 7 | 14 | 30 | 14 | 6 | 9 | 26 | 22 |
| Tigger1_dup10126 | 14.0372567806189 | -1.25342574972145 | 0.385487895476217 | -3.25153075992962 | 0.00114785354286662 | 0.0386524401139303 | 2351 | 4 | 4 | 20 | 25 | 9 | 13 | 30 | 12 | 5 | 16 | 24 | 21 |
| AluJb_dup98409 | 14.6533695958475 | -1.09620575381857 | 0.332741035441444 | -3.29447118647163 | 0.000986070534754398 | 0.0347841935178236 | 2352 | 7 | 9 | 23 | 23 | 15 | 15 | 31 | 15 | 5 | 7 | 22 | 18 |
| AluSg_dup13890 | 12.6300003440374 | -1.12012472088305 | 0.351384450292474 | -3.18774698183348 | 0.00143385944388772 | 0.0450285230612786 | 2353 | 7 | 11 | 24 | 15 | 11 | 10 | 23 | 15 | 6 | 4 | 16 | 20 |
| L1MB1_dup11 | 11.9424820600767 | -1.21822107866114 | 0.352917102610878 | -3.45186183851322 | 0.000556732722145848 | 0.0232795407758408 | 2354 | 9 | 7 | 15 | 22 | 10 | 10 | 25 | 10 | 3 | 6 | 17 | 24 |
| L1ME1_dup70 | 14.6495134581402 | -1.10188754911027 | 0.337808681838261 | -3.26186865036774 | 0.00110680415000232 | 0.0376228028966361 | 2355 | 11 | 10 | 30 | 27 | 10 | 12 | 28 | 8 | 5 | 10 | 16 | 27 |
| AluJb_dup83243 | 11.5537726729124 | -1.25261314426581 | 0.378310514132135 | -3.31107145446214 | 0.000929394733661721 | 0.0334410644633104 | 2356 | 7 | 6 | 20 | 21 | 5 | 12 | 24 | 10 | 6 | 6 | 10 | 25 |
| MLT1E_dup145 | 14.3208833897668 | -1.33482088758602 | 0.351433121090847 | -3.79822164582193 | 0.000145737973807788 | 0.00891459329451617 | 2357 | 11 | 4 | 20 | 33 | 5 | 14 | 26 | 15 | 7 | 9 | 16 | 28 |
| L1MB1_dup4350 | 11.1883182679453 | -1.38694461698045 | 0.357780688835229 | -3.87652173597103 | 0.000105960348136525 | 0.00712168265162043 | 2358 | 8 | 6 | 16 | 25 | 6 | 8 | 21 | 14 | 5 | 5 | 17 | 14 |
| MIR_dup867 | 13.0114268769217 | -1.08526636979629 | 0.328797875102608 | -3.30070980372854 | 0.000964405774987539 | 0.0342860446040916 | 2359 | 12 | 7 | 20 | 26 | 6 | 10 | 20 | 14 | 7 | 9 | 20 | 17 |
| MLT1B_dup3354 | 11.9703767357641 | -1.36437184221075 | 0.390806855183217 | -3.491166605998 | 0.000480916258773532 | 0.0209790150581349 | 2360 | 6 | 4 | 17 | 24 | 5 | 11 | 32 | 13 | 4 | 11 | 14 | 16 |
| MLT1C_dup17666 | 12.0435078542473 | -1.3694515689295 | 0.372970025760779 | -3.67174698861153 | 0.000240898111296463 | 0.0127748477803763 | 2361 | 8 | 5 | 16 | 18 | 5 | 7 | 33 | 11 | 7 | 9 | 20 | 18 |
| MLT1H1_dup3288 | 12.6075353272188 | -1.58518320406762 | 0.385373366361187 | -4.11336989640827 | 3.89924838536114e-05 | 0.00337385656837506 | 2362 | 6 | 8 | 18 | 15 | 6 | 6 | 36 | 17 | 7 | 5 | 21 | 17 |
| LTR37A_dup402 | 15.2423094014831 | -1.28962702455824 | 0.325764150236011 | -3.95877515565762 | 7.53351160346295e-05 | 0.00553120122526093 | 2363 | 9 | 9 | 19 | 23 | 8 | 9 | 31 | 23 | 5 | 14 | 19 | 27 |
| MER103C_dup1255 | 13.5293806242253 | -1.17677132423352 | 0.358106622638482 | -3.28609204589187 | 0.00101587807730765 | 0.0354607182207161 | 2364 | 11 | 5 | 22 | 25 | 8 | 5 | 30 | 16 | 10 | 11 | 14 | 17 |
| AluJb_dup13448 | 14.1262494021597 | -1.06606399647956 | 0.33387105260748 | -3.19304111019441 | 0.00140782902240946 | 0.0444870231827679 | 2365 | 9 | 6 | 26 | 21 | 9 | 11 | 29 | 14 | 11 | 9 | 19 | 17 |

| | | | | | | | | | | | | | | | | | | | |
|---|---|---|---|---|---|---|---|---|---|---|---|---|---|---|---|---|---|---|---|
| Charlie18a_dup577 | 14.0614824982033 | -1.27046400378626 | 0.339138248611559 | -3.74615369687014 | 0.00017956659897241 | 0.0103330481158619 | 2366 | 9 | 7 | 16 | 24 | 11 | 8 | 32 | 17 | 8 | 7 | 25 | 17 |
| L1P2_dup138 | 11.1232535836788 | -1.29586916755061 | 0.408503658378713 | -3.1722339346818 | 0.00151271100089969 | 0.0467884481762741 | 2367 | 15 | 4 | 20 | 24 | 6 | 3 | 19 | 9 | 5 | 7 | 13 | 22 |
| L2a_dup106656 | 9.85857078429614 | -1.35188989984526 | 0.413679329529345 | -3.26796579704223 | 0.00108323455454285 | 0.0371025376875275 | 2368 | 7 | 4 | 19 | 18 | 5 | 4 | 24 | 6 | 6 | 8 | 11 | 18 |
| MLT1H_dup5001 | 10.9493960837911 | -1.36921085864205 | 0.399985752734825 | -3.42314907288657 | 0.000619001203351643 | 0.025094098408732 | 2369 | 8 | 6 | 17 | 26 | 6 | 4 | 26 | 11 | 7 | 6 | 7 | 20 |
| MLT1J2_dup3508 | 12.7484350753641 | -1.16639483565747 | 0.36737448302263 | -3.17494787896203 | 0.00149863417180297 | 0.046473870278096 | 2370 | 10 | 9 | 17 | 31 | 8 | 6 | 12 | 13 | 5 | 10 | 25 | 19 |
| L1M5_dup36154 | 13.1342210962593 | -1.44030047950105 | 0.396377105122225 | -3.63366213862661 | 0.000279426715093034 | 0.0142047047096345 | 2371 | 7 | 11 | 20 | 33 | 3 | 7 | 13 | 12 | 5 | 7 | 16 | 25 |
| MER52D_dup257 | 11.0572172745267 | -1.29608755582704 | 0.381532136136482 | -3.39705999329871 | 0.000681140319944477 | 0.0268202241551396 | 2372 | 6 | 10 | 14 | 28 | 6 | 6 | 14 | 9 | 5 | 6 | 19 | 22 |
| FRAM_dup5608 | 13.6898500432062 | -1.21037735298871 | 0.343692196246822 | -3.52168994875716 | 0.000428805343530286 | 0.0194532919488384 | 2373 | 13 | 9 | 21 | 27 | 5 | 7 | 18 | 12 | 10 | 6 | 20 | 30 |
| MIRc_dup81496 | 14.2158052207769 | -1.28039571293565 | 0.39349930203247 | -3.25387035332022 | 0.00113844222677007 | 0.0384174933393487 | 2374 | 16 | 15 | 22 | 25 | 3 | 4 | 30 | 15 | 8 | 5 | 20 | 22 |
| MLT1B_dup3449 | 12.1840326238695 | -2.03389577421766 | 0.403294408818718 | -5.04320350032897 | 4.57801917638834e-07 | 0.000111954917877174 | 2375 | 5 | 4 | 10 | 16 | 4 | 4 | 19 | 9 | 5 | 5 | 40 | 39 |
| AluSx_dup87770 | 10.0714779360757 | -1.27389747332658 | 0.396555447360938 | -3.21240694537 | 0.00131627781847517 | 0.0424682090569346 | 2376 | 6 | 7 | 7 | 16 | 4 | 7 | 17 | 8 | 4 | 7 | 24 | 25 |
| L1ME1_dup26615 | 9.04669527332931 | -1.3872014658004 | 0.416851398286579 | -3.32780811459993 | 0.000875321346548499 | 0.0320725613067184 | 2377 | 7 | 5 | 8 | 16 | 3 | 7 | 13 | 8 | 3 | 5 | 20 | 24 |
| L2a_dup140626 | 14.811345226055 | -1.37611912200253 | 0.394920721736449 | -3.48454524227493 | 0.000492974112362584 | 0.021350785973831 | 2378 | 13 | 6 | 10 | 14 | 4 | 8 | 26 | 10 | 5 | 11 | 43 | 45 |
| THE1B_dup3158 | 11.5889522884011 | -1.42488691883742 | 0.422360326504138 | -3.37362869905695 | 0.000741843587201052 | 0.028516755121186 | 2379 | 6 | 5 | 9 | 10 | 6 | 5 | 25 | 14 | 3 | 13 | 20 | 35 |
| L1ME3G_dup15238 | 14.6076310787785 | -1.90741052550745 | 0.387210206849472 | -4.92603369375786 | 8.39155337155482e-07 | 0.00018004838219549 | 2380 | 4 | 11 | 22 | 20 | 8 | 4 | 20 | 16 | 5 | 4 | 32 | 44 |
| L1ME4a_dup31834 | 15.5731557028373 | -2.31237991463821 | 0.38304718291006 | -6.03680177744882 | 1.57198627400358e-09 | 1.05069619650016e-06 | 2381 | 7 | 2 | 27 | 30 | 9 | 3 | 22 | 13 | 5 | 5 | 35 | 48 |
| L2c_dup117348 | 9.61195300013719 | -1.99855985172983 | 0.446079094982409 | -4.48028135415905 | 7.45447088269527e-06 | 0.000975666407400335 | 2382 | 5 | 4 | 19 | 11 | 4 | 3 | 11 | 15 | 3 | 4 | 18 | 26 |
| L1MB4_dup5460 | 12.3249531697952 | -1.75964496334437 | 0.430016953681896 | -4.09203625177547 | 4.27601783108923e-05 | 0.00362234568672866 | 2383 | 4 | 3 | 27 | 18 | 12 | 7 | 16 | 12 | 4 | 4 | 19 | 35 |
| L1ME4a_dup7111 | 11.582627098208 | -1.23666217992084 | 0.376869930178291 | -3.28140316033119 | 0.00103291986899146 | 0.0358696951469707 | 2384 | 6 | 4 | 21 | 10 | 8 | 7 | 14 | 12 | 7 | 9 | 21 | 29 |
| MLT1K_dup15133 | 9.71712689962198 | -1.22944028288244 | 0.388477553342733 | -3.16476530575183 | 0.00155208089647353 | 0.0475794178036964 | 2385 | 5 | 4 | 15 | 15 | 10 | 5 | 13 | 11 | 5 | 6 | 15 | 21 |
| L1PA2_dup378 | 15.072322970969 | -2.53274306680043 | 0.690872487771862 | -3.66600655204667 | 0.000246367615215773 | 0.012966642019494 | 2386 | 5 | 3 | 16 | 25 | 1 | 10 | 5 | 5 | 1 | 1 | 50 | 83 |
| L1PB1_dup7169 | 30.5142107764057 | -3.1816817999113 | 0.916302659914156 | -3.47230444601064 | 0.000516010795215287 | 0.0220514632452247 | 2387 | 4 | 1 | 11 | 31 | 8 | 4 | 8 | 8 | 0 | 1 | 141 | 198 |
| BSR/Beta_dup2028 | 17.0427415414914 | -3.86582481144951 | 1.17771528800593 | -3.28247824480143 | 0.00102898925264486 | 0.0357854248333579 | 2388 | 1 | 0 | 63 | 65 | 7 | 11 | 14 | 1 | 0 | 0 | 31 | 41 |
| L1PBa_dup1842 | 20.5414528287503 | -3.47475388578889 | 0.690418427386184 | -5.03282320975088 | 4.83308798526352e-07 | 0.000116398173455619 | 2389 | 0 | 4 | 54 | 74 | 8 | 8 | 12 | 8 | 0 | 2 | 44 | 66 |
| LTR53-int_dup337 | 14.978807795251 | -2.53041285137792 | 0.658542993994837 | -3.84244138112835 | 0.0001218165138875 | 0.00783095603103028 | 2390 | 1 | 4 | 31 | 30 | 12 | 11 | 18 | 7 | 0 | 1 | 34 | 53 |
| MER1A_dup3147 | 23.0294484795589 | -2.27090788277038 | 0.660654129662541 | -3.43736273007231 | 0.000587408377749861 | 0.0242196281150699 | 2391 | 7 | 13 | 59 | 96 | 17 | 16 | 8 | 5 | 0 | 4 | 38 | 51 |
| L1MA8_dup10713 | 15.0595938243623 | -2.25165327637324 | 0.60074280468192 | -3.7481152646771 | 0.000178168376645054 | 0.0102699739702481 | 2392 | 10 | 7 | 31 | 51 | 12 | 5 | 8 | 7 | 0 | 2 | 30 | 41 |
| AluSx1_dup113133 | 14.9055757253537 | -2.17926786437745 | 0.597440877149175 | -3.64767117170877 | 0.000264628010858598 | 0.0136766239512945 | 2393 | 8 | 3 | 45 | 47 | 10 | 8 | 8 | 5 | 0 | 6 | 27 | 34 |
| MLT1L_dup11513 | 16.1287157805527 | -2.60671929020432 | 0.574748163305897 | -4.53541125770758 | 5.74913481093366e-06 | 0.000799055859650946 | 2394 | 7 | 6 | 56 | 49 | 5 | 5 | 10 | 6 | 0 | 6 | 36 | 30 |
| L1ME4a_dup33294 | 12.5828348808473 | -2.12827302675384 | 0.66165139668749 | -3.21660777474194 | 0.00129715802740625 | 0.0420165483241502 | 2395 | 2 | 5 | 39 | 40 | 3 | 7 | 6 | 1 | 2 | 10 | 21 | 34 |
| MER76_dup49 | 11.7512030307078 | -2.50820723708385 | 0.73216292518229 | -3.42575013131042 | 0.000613104120127851 | 0.0249419854621203 | 2396 | 6 | 4 | 34 | 49 | 1 | 2 | 3 | 1 | 2 | 7 | 18 | 34 |
| L2b_dup39900 | 10.4146404988715 | -2.208997711456 | 0.688856461132619 | -3.20676053153941 | 0.00134238689513887 | 0.0430711424151447 | 2397 | 1 | 4 | 39 | 41 | 6 | 3 | 8 | 1 | 4 | 4 | 8 | 22 |
| L2_dup24968 | 10.8084922443484 | -1.98218303337446 | 0.577985738966083 | -3.42946702615923 | 0.000604767912177875 | 0.0247212000740554 | 2398 | 2 | 6 | 31 | 36 | 6 | 3 | 5 | 5 | 6 | 3 | 21 | 18 |
| MIRb_dup214042 | 11.8670556751089 | -2.38877968097197 | 0.641269881902973 | -3.72507698924397 | 0.000195255620899966 | 0.0109318338754502 | 2399 | 1 | 3 | 36 | 46 | 7 | 2 | 5 | 4 | 5 | 5 | 17 | 28 |
| AluSx1_dup113131 | 12.6234907708389 | -2.37568079676342 | 0.579480167047973 | -4.09967576434198 | 4.13729359397008e-05 | 0.0035371493298707 | 2400 | 6 | 2 | 46 | 41 | 5 | 5 | 8 | 4 | 4 | 3 | 25 | 19 |
| L1MA9_dup7557 | 14.9554062372745 | -2.88117462766552 | 0.633693336321688 | -4.54663866972229 | 5.45094605776123e-06 | 0.000768838654807383 | 2401 | 3 | 3 | 50 | 71 | 6 | 5 | 6 | 5 | 3 | 2 | 17 | 33 |
| LTR53-int_dup340 | 12.1455060780175 | -2.53425346998157 | 0.721368085796683 | -3.5131211372939 | 0.000442875512462234 | 0.0198756048073064 | 2402 | 1 | 1 | 26 | 34 | 10 | 6 | 6 | 2 | 1 | 2 | 31 | 45 |
| Zaphod_dup2078 | 17.1842732351634 | -2.60585131053061 | 0.655560292061143 | -3.97499870277007 | 7.03796397473288e-05 | 0.00526757127197664 | 2403 | 2 | 2 | 44 | 67 | 10 | 10 | 5 | 5 | 2 | 4 | 36 | 46 |
| L3_dup44305 | 17.0672371080038 | -2.04577718341936 | 0.620256158078385 | -3.29827790788467 | 0.000972798016471698 | 0.0344745194956692 | 2404 | 6 | 4 | 40 | 54 | 11 | 13 | 7 | 2 | 3 | 4 | 33 | 55 |
| MLT1C_dup18999 | 17.5158593038705 | -2.35786167082097 | 0.611797155559758 | -3.85391898180802 | 0.000116242074613792 | 0.00757997716800584 | 2405 | 8 | 2 | 50 | 61 | 8 | 9 | 7 | 3 | 5 | 3 | 34 | 47 |
| LTR78_dup4517 | 14.6452894697443 | -2.34445895656495 | 0.613358316154539 | -3.82233173467603 | 0.000132195682683767 | 0.00829797593784371 | 2406 | 5 | 4 | 43 | 43 | 5 | 7 | 3 | 4 | 3 | 5 | 45 | 27 |
| THE1D_dup12372 | 17.3746289812229 | -2.26616730213117 | 0.584819676930774 | -3.87498470301884 | 0.000106631397686986 | 0.00714316288120147 | 2407 | 6 | 5 | 48 | 60 | 8 | 10 | 5 | 5 | 3 | 5 | 46 | 31 |
| MERX_dup419 | 18.4274363367766 | -2.11673130990038 | 0.637555747130425 | -3.32007251040803 | 0.000899940702781179 | 0.0326478564512838 | 2408 | 4 | 3 | 48 | 63 | 12 | 6 | 4 | 3 | 4 | 13 | 43 | 44 |
| MLT2B1_dup4329 | 15.0251432981436 | -2.04369111328205 | 0.624726098129841 | -3.27133942282862 | 0.00107039356331102 | 0.0368164016790406 | 2409 | 5 | 5 | 39 | 63 | 10 | 4 | 4 | 3 | 3 | 9 | 27 | 31 |
| Cheshire_dup942 | 21.1226757675443 | -2.04210332648504 | 0.546719533306158 | -3.73519364515093 | 0.000187570797329393 | 0.0106378285158242 | 2410 | 18 | 8 | 74 | 102 | 8 | 10 | 9 | 11 | 3 | 5 | 20 | 17 |
| L1PB1_dup6479 | 14.4010587246808 | -2.05766794410892 | 0.631657523447025 | -3.25756896376377 | 0.00112370949979754 | 0.0380561031367528 | 2411 | 13 | 10 | 53 | 71 | 3 | 2 | 9 | 6 | 2 | 4 | 13 | 9 |
| L1PA16_dup6812 | 17.0025848484354 | -1.59026007201554 | 0.49837629448651 | -3.1908822502363 | 0.00141839074418515 | 0.0446792014336328 | 2412 | 12 | 19 | 48 | 74 | 7 | 13 | 18 | 7 | 3 | 2 | 12 | 15 |
| L2b_dup44443 | 15.8811435254377 | -1.64415851509051 | 0.452831051667979 | -3.63084313461796 | 0.000282496851490435 | 0.0143171450105949 | 2413 | 15 | 10 | 46 | 41 | 4 | 14 | 22 | 8 | 5 | 2 | 24 | 20 |
| L1MB4_dup7651 | 13.5087659041955 | -2.48175073337999 | 0.563732577159647 | -4.40235465171134 | 1.07082326406353e-05 | 0.00128600010060026 | 2414 | 9 | 5 | 44 | 65 | 4 | 6 | 12 | 6 | 2 | 1 | 14 | 16 |
| L1PREC2_dup851 | 12.5902828519062 | -2.43700088973744 | 0.503452570602875 | -4.84057691237762 | 1.29462737514475e-06 | 0.000251470535025463 | 2415 | 4 | 6 | 35 | 48 | 4 | 7 | 16 | 6 | 2 | 2 | 20 | 20 |
| AluSq2_dup26994 | 12.0487946212349 | -1.70982850947381 | 0.520141373020634 | -3.28723804365776 | 0.00101175270375767 | 0.0353565147376006 | 2416 | 9 | 7 | 37 | 50 | 4 | 11 | 9 | 8 | 2 | 3 | 12 | 9 |
| FRAM_dup3286 | 11.7958955489036 | -1.85673083893646 | 0.487227584785659 | -3.81080812522813 | 0.000138513247996557 | 0.00859225011390934 | 2417 | 10 | 5 | 23 | 53 | 2 | 9 | 15 | 11 | 3 | 3 | 14 | 10 |
| L1MA4_dup9832 | 14.5196548751418 | -1.84956128115004 | 0.560840737457859 | -3.29141796923898 | 0.000996836842647679 | 0.0350285370994135 | 2418 | 10 | 5 | 36 | 55 | 9 | 5 | 9 | 2 | 5 | 5 | 27 | 28 |
| MLT1J2_dup6574 | 16.3734379472566 | -2.14621132182095 | 0.585984382524422 | -3.66257426959925 | 0.000249693325914446 | 0.0131068890642045 | 2419 | 8 | 4 | 39 | 61 | 15 | 7 | 8 | 5 | 3 | 2 | 33 | 36 |
| L1MDa_dup6680 | 14.7373853440899 | -2.15091749173748 | 0.590432916129345 | -3.64294983050417 | 0.000269531386761174 | 0.0138428481230896 | 2420 | 6 | 4 | 42 | 47 | 9 | 7 | 5 | 4 | 5 | 2 | 31 | 35 |
| LTR16A2_dup1777 | 16.5517571205383 | -2.01789035154762 | 0.587302318186456 | -3.43586307947619 | 0.000590669940729123 | 0.0243119568910775 | 2421 | 10 | 6 | 52 | 56 | 12 | 5 | 8 | 4 | 5 | 3 | 33 | 29 |
| L1MC_dup7975 | 12.2535007373896 | -2.24184686475126 | 0.627671138268795 | -3.57169021812058 | 0.000354684798334329 | 0.0168757906467627 | 2422 | 4 | 5 | 28 | 38 | 7 | 6 | 4 | 3 | 2 | 2 | 20 | 48 |
| L1PA7_dup12858 | 13.3315991468328 | -1.99434789029483 | 0.57526261984039 | -3.46684769966137 | 0.000526600171152436 | 0.0223355444587407 | 2423 | 7 | 5 | 36 | 44 | 7 | 6 | 3 | 5 | 3 | 5 | 22 | 36 |
| L2a_dup166102 | 15.9378305445909 | -1.77378709542268 | 0.554437228253979 | -3.19925684104701 | 0.00137782360525963 | 0.0438592127862353 | 2424 | 8 | 10 | 28 | 48 | 7 | 10 | 7 | 5 | 6 | 2 | 23 | 62 |
| L1ME3A_dup12783 | 13.4600950899065 | -1.6607330388112 | 0.478325703071821 | -3.47197114465296 | 0.000516651866462854 | 0.0220559807059759 | 2425 | 10 | 7 | 21 | 27 | 8 | 7 | 8 | 8 | 4 | 2 | 30 | 47 |
| L1M8_dup783 | 11.6122532049268 | -1.55253346851295 | 0.43981502541689 | -3.52996914337191 | 0.000415608135745383 | 0.019021406706688 | 2426 | 6 | 8 | 24 | 30 | 7 | 6 | 9 | 6 | 4 | 5 | 20 | 29 |
| MLT1J2-int_dup573 | 13.9543704447232 | -1.6548688894308 | 0.500355932117469 | -3.3073833709286 | 0.000941719163855197 | 0.0337162666510215 | 2427 | 9 | 10 | 38 | 34 | 9 | 9 | 8 | 6 | 3 | 3 | 21 | 36 |
| MLT2A2_dup1707 | 15.4511048076751 | -2.00002548552574 | 0.507536227171219 | -3.94065561915252 | 8.1259214663502e-05 | 0.00585031924124709 | 2428 | 3 | 5 | 41 | 59 | 13 | 4 | 18 | 7 | 6 | 6 | 24 | 20 |
| MSTB_dup3433 | 13.9033041392042 | -1.47113028915008 | 0.459979804848038 | -3.19824973541198 | 0.00138264485668398 | 0.0439421414543243 | 2429 | 11 | 6 | 38 | 44 | 10 | 5 | 9 | 10 | 6 | 7 | 20 | 16 |
| MER5A1_dup14572 | 13.7231266328287 | -1.58685041453639 | 0.463640712809365 | -3.42258643534795 | 0.000620283730639472 | 0.0251138391306346 | 2430 | 12 | 7 | 24 | 33 | 14 | 6 | 14 | 8 | 2 | 3 | 22 | 39 |
| MSTB_dup3434 | 12.1900957043256 | -2.07181722142217 | 0.486712202413798 | -4.25676038354331 | 2.07410443719363e-05 | 0.00209742819554813 | 2431 | 7 | 5 | 26 | 44 | 9 | 4 | 13 | 7 | 1 | 4 | 17 | 28 |
| L1ME4a_dup14590 | 12.6967579277243 | -1.86403708938 | 0.457113063829335 | -4.07784689801809 | 4.54546809333192e-05 | 0.00380026978172835 | 2432 | 5 | 5 | 29 | 33 | 5 | 8 | 15 | 4 | 6 | 4 | 21 | 35 |
| L2a_dup155350 | 12.0040212316398 | -1.91856005080245 | 0.413441575457496 | -4.64046231605869 | 3.47630567926091e-06 | 0.000539367852639763 | 2433 | 6 | 5 | 22 | 26 | 4 | 4 | 15 | 5 | 5 | 6 | 26 | 36 |
| L1MD1_dup696 | 13.3795092159511 | -2.28753218955935 | 0.481437504049765 | -4.7514623815491 | 2.01950734547654e-06 | 0.000355687412899157 | 2434 | 6 | 3 | 30 | 45 | 6 | 3 | 14 | 7 | 3 | 7 | 18 | 39 |
| L1ME4a_dup14593 | 11.7893537963775 | -1.87559072214902 | 0.441455708784466 | -4.24864983015714 | 2.15062784503843e-05 | 0.00215194824733185 | 2435 | 7 | 5 | 32 | 30 | 8 | 2 | 15 | 8 | 4 | 5 | 16 | 24 |
| MSTA-int_dup1643 | 15.4347700290567 | -3.00046628885678 | 0.527322832041578 | -5.68999881389585 | 1.27040254302147e-08 | 6.25853423399146e-06 | 2436 | 4 | 1 | 46 | 58 | 2 | 7 | 26 | 7 | 3 | 4 | 27 | 24 |
| L1ME1_dup29323 | 10.8427969535625 | -2.94965040502001 | 0.548951782276555 | -5.37324133058165 | 7.73337147720967e-08 | 2.70606589634725e-05 | 2437 | 4 | 2 | 42 | 30 | 1 | 3 | 15 | 11 | 3 | 2 | 11 | 19 |
| L1PA3_dup9502 | 11.2963326527674 | -2.95322740857303 | 0.541104360468702 | -5.45777787858697 | 4.82130257971601e-08 | 1.85332740490001e-05 | 2438 | 3 | 1 | 30 | 40 | 1 | 6 | 13 | 12 | 1 | 4 | 15 | 25 |
| L2_dup14394 | 10.7446148141625 | -1.90021505214174 | 0.514870673091314 | -3.69066476583864 | 0.000223668753175779 | 0.0120600691464234 | 2439 | 3 | 3 | 22 | 30 | 3 | 7 | 9 | 7 | 8 | 3 | 30 | 14 |
| THE1B_dup9621 | 12.7219749677038 | -1.86744313140481 | 0.492250455411293 | -3.79368492912369 | 0.000148427992928363 | 0.00902807955146528 | 2440 | 5 | 1 | 26 | 41 | 7 | 9 | 12 | 7 | 7 | 4 | 27 | 22 |
| Charlie19a_dup1201 | 9.74617045056954 | -1.93978470849863 | 0.551705451234719 | -3.51597886908201 | 0.000438135853634656 | 0.0197298777117623 | 2441 | 2 | 2 | 25 | 36 | 7 | 8 | 9 | 8 | 1 | 5 | 16 | 10 |
| L1MA9_dup15859 | 10.7978530004648 | -2.23164531028704 | 0.54878255007857 | -4.06653839479322 | 4.77166267456453e-05 | 0.00393007285137394 | 2442 | 3 | 3 | 30 | 29 | 9 | 8 | 9 | 9 | 1 | 1 | 23 | 17 |
| LTR33_dup8833 | 12.8413187966311 | -3.05779150456427 | 0.687901380335655 | -4.44510156829784 | 8.78503528700468e-06 | 0.00110484355518056 | 2443 | 3 | 3 | 45 | 57 | 4 | 2 | 6 | 3 | 1 | 4 | 32 | 13 |
| MamRep605_dup4109 | 12.8147876031504 | -2.06210504316959 | 0.552082804553564 | -3.73513724057587 | 0.000187612844149359 | 0.0106378285158242 | 2444 | 8 | 6 | 30 | 49 | 7 | 6 | 6 | 6 | 1 | 4 | 28 | 21 |

| | | | | | | | | | | | | | | | | | | |
|---|---|---|---|---|---|---|---|---|---|---|---|---|---|---|---|---|---|---|
| THE1C_dup4432 | 11.6151607315072 | -2.24219665501395 | 0.554478676898042 | -4.04379239172483 | 5.25934941819047e-05 | 0.00424937574417911 | 2445 | 3 | 6 | 36 | 41 | 6 | 3 | 9 | 7 | 1 | 6 | 23 | 13 |
| AluSq2_dup22859 | 12.5891659143201 | -2.42353081175343 | 0.640590473883301 | -3.78327638414888 | 0.000154777381104079 | 0.00931663514398968 | 2446 | 6 | 5 | 24 | 56 | 1 | 11 | 11 | 3 | 2 | 1 | 24 | 29 |
| MLT2A2_dup3780 | 10.8182340598958 | -1.83890640913794 | 0.559692525070555 | -3.28556542524153 | 0.00101777902772659 | 0.035490037228246 | 2447 | 10 | 5 | 32 | 36 | 3 | 6 | 5 | 5 | 3 | 3 | 14 | 23 |
| LTR85a_dup800 | 10.0758184425417 | -1.7564188614472 | 0.550031117605938 | -3.19330816971261 | 0.00140652754761045 | 0.0444619624779882 | 2448 | 7 | 6 | 24 | 34 | 5 | 6 | 6 | 3 | 3 | 2 | 20 | 19 |
| MADE1_dup1861 | 10.5244752280145 | -1.81447170870498 | 0.506392929693178 | -3.58313002080097 | 0.000339501509358686 | 0.0163419365170694 | 2449 | 7 | 8 | 27 | 30 | 3 | 5 | 7 | 6 | 3 | 3 | 22 | 18 |
| Tigger20a_dup543 | 12.0646626617305 | -1.79898892481635 | 0.541283738531055 | -3.32355989429588 | 0.00088876335262815 | 0.0323905168792247 | 2450 | 5 | 7 | 25 | 31 | 3 | 12 | 10 | 3 | 2 | 4 | 29 | 31 |
| Tigger6a_dup773 | 15.2230743665994 | -1.94147936791217 | 0.518308431371029 | -3.74579931639655 | 0.000179820302182729 | 0.0103401453898433 | 2451 | 8 | 9 | 37 | 42 | 4 | 11 | 9 | 5 | 3 | 4 | 33 | 39 |
| L1PA7_dup1610 | 17.8785741688702 | -4.41256280698406 | 0.796085763155736 | -5.54282341326187 | 2.97632981774327e-08 | 1.26465316930846e-05 | 2452 | 3 | 7 | 51 | 73 | 1 | 0 | 12 | 7 | 0 | 0 | 38 | 53 |
| L1PA7_dup1611 | 25.5140848529236 | -4.4010707623217 | 0.693892723143147 | -6.34258094303977 | 2.25947425471215e-10 | 2.05203382065549e-07 | 2453 | 7 | 8 | 83 | 106 | 2 | 0 | 21 | 16 | 0 | 0 | 39 | 66 |
| MIR_dup168630 | 14.2879408271808 | -2.33826675794842 | 0.605617082795816 | -3.86096565696904 | 0.00011293977883601 | 0.00742697260416478 | 2454 | 10 | 7 | 45 | 50 | 0 | 9 | 8 | 10 | 1 | 4 | 17 | 31 |
| L1ME3B_dup7552 | 18.5157216656975 | -3.12196903921701 | 0.758914204726214 | -4.11373119619403 | 3.89314747550609e-05 | 0.00337102936392567 | 2455 | 9 | 11 | 57 | 94 | 0 | 3 | 4 | 6 | 1 | 2 | 34 | 32 |
| L2c_dup129942 | 18.818440527476 | -2.72681550126693 | 0.730482259760174 | -3.73289763691479 | 0.000189289535517915 | 0.0106861941037855 | 2456 | 18 | 11 | 59 | 81 | 1 | 3 | 5 | 9 | 0 | 3 | 34 | 31 |
| L2c_dup110861 | 16.68205463645 | -2.88182032226277 | 0.51914350874554 | -5.5511053759035 | 2.83868901069091e-08 | 1.22364981039467e-05 | 2457 | 4 | 3 | 43 | 83 | 8 | 5 | 23 | 9 | 1 | 4 | 18 | 28 |
| ALR/Alpha_dup1732 | 20.2928373658265 | -3.21710092614901 | 0.616355289304495 | -5.21955596386499 | 1.79352614634237e-07 | 5.30139593065319e-05 | 2458 | 6 | 4 | 85 | 97 | 8 | 5 | 11 | 7 | 1 | 2 | 19 | 32 |
| L1MC1_dup10761 | 19.3146059797237 | -2.97957260915298 | 0.596924261812113 | -4.99154214323228 | 5.98991096517502e-07 | 0.000138108076988327 | 2459 | 5 | 3 | 47 | 89 | 13 | 7 | 15 | 8 | 1 | 1 | 35 | 40 |
| L1ME5_dup2943 | 19.5082589523295 | -2.81042416601912 | 0.602586330221052 | -4.66393614503028 | 3.10217677862528e-06 | 0.00049275738035165 | 2460 | 16 | 10 | 62 | 88 | 3 | 2 | 12 | 14 | 1 | 2 | 18 | 37 |
| L1P4a_dup197 | 22.7607285186062 | -4.18696363694087 | 0.62459001046097 | -6.70353922863854 | 2.03431049455248e-11 | 2.63073937465644e-08 | 2461 | 6 | 3 | 93 | 104 | 2 | 1 | 18 | 15 | 0 | 3 | 26 | 38 |
| MIRb_dup202623 | 19.4160293406764 | -1.82337499690822 | 0.472891901068671 | -3.85579662664477 | 0.000115353360909705 | 0.007528214705162 | 2462 | 12 | 9 | 33 | 38 | 9 | 0 | 16 | 18 | 4 | 18 | 60 | 34 |
| L1MA9_dup8681 | 12.1217038542145 | -2.44693022404976 | 0.661960742466452 | -3.69648842759549 | 0.00021860221901918 | 0.0118648548373033 | 2463 | 2 | 5 | 32 | 38 | 1 | 0 | 9 | 14 | 13 | 4 | 20 | 16 |
| L1MB1_dup508 | 20.9182000441757 | -3.03130538356363 | 0.624953861028298 | -4.85044668509756 | 1.23183742871213e-06 | 0.000241243447582171 | 2464 | 6 | 7 | 62 | 95 | 1 | 0 | 15 | 13 | 9 | 5 | 28 | 40 |
| L1PA4_dup6547 | 18.8262100687321 | -2.5306490303031 | 0.585124140148289 | -4.32497799468974 | 1.52547089308503e-05 | 0.00167348869122181 | 2465 | 4 | 7 | 46 | 68 | 1 | 1 | 15 | 15 | 17 | 7 | 27 | 38 |
| MER34_dup109 | 25.9988340267226 | -2.53649172774045 | 0.620829430173189 | -4.08564994580373 | 4.39536037599512e-05 | 0.00370477654986374 | 2466 | 18 | 7 | 69 | 96 | 0 | 3 | 15 | 8 | 7 | 13 | 49 | 67 |
| L1P4_dup319 | 28.1639702338329 | -2.76981795510246 | 0.591819345869146 | -4.68017474324822 | 2.86630507901633e-06 | 0.000464595250907097 | 2467 | 11 | 9 | 70 | 108 | 3 | 1 | 9 | 10 | 7 | 14 | 56 | 83 |
| L1M8_dup167 | 24.2952130210547 | -2.92279582274191 | 0.549252456980996 | -5.3214069151502 | 1.02967794382412e-07 | 3.37476236943767e-05 | 2468 | 6 | 7 | 58 | 97 | 3 | 1 | 16 | 8 | 7 | 12 | 55 | 58 |
| L1MC3_dup1151 | 26.7075719461143 | -3.18885852287138 | 0.598726821984363 | -5.32606592152083 | 1.00362719531026e-07 | 3.31219246345764e-05 | 2469 | 7 | 6 | 76 | 127 | 2 | 2 | 11 | 9 | 8 | 8 | 47 | 60 |
| L1ME1_dup3530 | 20.6879441250187 | -2.86468257694712 | 0.652978071373753 | -4.38710379801931 | 1.14869979114055e-05 | 0.00135110489620727 | 2470 | 7 | 7 | 65 | 83 | 1 | 1 | 8 | 6 | 9 | 6 | 28 | 59 |
| L1PA10_dup579 | 23.6761553620674 | -2.99058890268973 | 0.667804551584917 | -4.47823977178966 | 7.52610418623379e-06 | 0.000983419212903671 | 2471 | 7 | 8 | 68 | 106 | 0 | 2 | 11 | 5 | 9 | 7 | 46 | 52 |
| L1PA13_dup716 | 21.2249639466183 | -3.23909489158278 | 0.694227566267553 | -4.66575378012927 | 3.07487726023261e-06 | 0.000489879075703537 | 2472 | 5 | 5 | 62 | 95 | 0 | 1 | 10 | 4 | 8 | 8 | 47 | 42 |
| MER52A_dup1195 | 17.0777190097813 | -2.49610910157 | 0.505146048485699 | -4.94136123414746 | 7.75790215323712e-07 | 0.000169509805854376 | 2473 | 8 | 8 | 48 | 48 | 1 | 2 | 15 | 11 | 8 | 4 | 37 | 35 |
| L1MC3_dup1149 | 19.4106244104725 | -3.01142048291561 | 0.561232082491096 | -5.36573117764234 | 8.06219437806792e-08 | 2.79238268338246e-05 | 2474 | 4 | 6 | 44 | 100 | 2 | 3 | 11 | 11 | 6 | 5 | 34 | 37 |
| L1MEi_dup598 | 17.3733339271587 | -2.70214636239268 | 0.559612676991789 | -4.82860105478334 | 1.37495554255918e-06 | 0.000264269120783349 | 2475 | 4 | 7 | 36 | 71 | 2 | 3 | 9 | 6 | 6 | 6 | 32 | 54 |
| L1MB2_dup852 | 22.7132819928773 | -2.89223197964563 | 0.559549752306264 | -5.16885579472582 | 2.35531542232619e-07 | 6.53953189592564e-05 | 2476 | 11 | 5 | 62 | 87 | 3 | 2 | 18 | 5 | 6 | 6 | 41 | 64 |
| L1PBa_dup191 | 19.5286364934663 | -3.82304810281635 | 0.658163990401173 | -5.80865583437051 | 6.29763960420418e-09 | 3.41343542884275e-06 | 2477 | 4 | 2 | 58 | 92 | 1 | 1 | 12 | 5 | 5 | 3 | 38 | 46 |
| L1MCc_dup336 | 15.9552918980588 | -2.54352588200848 | 0.699434947661287 | -3.63654388519377 | 0.000276320584540322 | 0.0141076096507006 | 2478 | 10 | 6 | 41 | 74 | 2 | 0 | 5 | 4 | 4 | 7 | 25 | 40 |
| L1MA5_dup403 | 12.8290551421756 | -2.96582856343049 | 0.652097482721351 | -4.54813680778741 | 5.41229491005371e-06 | 0.000765201381264196 | 2479 | 3 | 3 | 29 | 57 | 2 | 0 | 7 | 5 | 3 | 8 | 20 | 38 |
| MSTB2_dup168 | 12.3080613163777 | -3.02661583467896 | 0.652255712263369 | -4.64022894360926 | 3.48023393644741e-06 | 0.000539483872470304 | 2480 | 5 | 2 | 36 | 54 | 1 | 0 | 10 | 5 | 4 | 5 | 18 | 27 |
| L1MA5A_dup404 | 17.9325780858767 | -2.93403907157017 | 0.625646022936395 | -4.68961515618626 | 2.73719357922952e-06 | 0.000448325344332882 | 2481 | 5 | 5 | 42 | 55 | 0 | 0 | 7 | 10 | 7 | 11 | 38 | 59 |
| L1MCc_dup337 | 20.6387811854549 | -3.87898012792038 | 0.804967130294256 | -4.81880561570559 | 1.44420173590582e-06 | 0.000274474461862497 | 2482 | 3 | 4 | 58 | 93 | 0 | 0 | 7 | 2 | 2 | 9 | 39 | 68 |
| L1M6_dup577 | 14.8154448477326 | -2.591303928815 | 0.666828440100613 | -3.88601291274263 | 0.000101904088410217 | 0.00690247428502043 | 2483 | 7 | 2 | 36 | 52 | 2 | 2 | 1 | 9 | 4 | 10 | 23 | 51 |
| Tigger2b_Pri_dup261 | 11.5766138907475 | -2.34402177454929 | 0.678730899352491 | -3.45353626420352 | 0.000553287907609036 | 0.0231723819636276 | 2484 | 6 | 6 | 35 | 41 | 1 | 1 | 2 | 6 | 6 | 3 | 17 | 30 |
| L1PA4_dup1114 | 24.4850489092993 | -2.48718586033296 | 0.6181893909467 | -4.02333960556014 | 5.7378665548393e-05 | 0.00451572686121649 | 2485 | 11 | 9 | 66 | 90 | 4 | 3 | 5 | 5 | 8 | 10 | 37 | 85 |
| L1PA7_dup1186 | 23.595740388303 | -2.83228954703036 | 0.674546202871306 | -4.19880733888101 | 2.68324489380049e-05 | 0.00253811792759312 | 2486 | 3 | 15 | 65 | 86 | 3 | 2 | 6 | 4 | 6 | 5 | 55 | 69 |
| L2_dup6898 | 16.2466773161214 | -3.02081354684901 | 0.642142621422024 | -4.70427198892269 | 2.54773415569047e-06 | 0.000426017674919131 | 2487 | 3 | 5 | 51 | 76 | 1 | 2 | 9 | 5 | 6 | 5 | 20 | 38 |
| MLT1D_dup1857 | 15.0166734503345 | -2.52739318374247 | 0.628008648728943 | -4.02445601483003 | 5.71071659318988e-05 | 0.00450096777238542 | 2488 | 6 | 8 | 52 | 47 | 1 | 1 | 11 | 3 | 6 | 5 | 28 | 33 |
| L1M1_dup885 | 17.3150208748959 | -3.23577985528345 | 0.666573378604121 | -4.85434906215357 | 1.20782823524591e-06 | 0.000238306714149108 | 2489 | 2 | 6 | 55 | 66 | 2 | 1 | 4 | 7 | 6 | 3 | 36 | 44 |
| L1P1_dup282 | 15.8401070386328 | -2.69092630093327 | 0.694785251706471 | -3.87303313408575 | 0.000107489207582477 | 0.00718242824695875 | 2490 | 7 | 4 | 53 | 64 | 3 | 2 | 2 | 3 | 5 | 5 | 23 | 44 |
| L1M3e_dup71 | 19.4674197373814 | -2.96743929450917 | 0.659772217639885 | -4.49767240142998 | 6.87014469082671e-06 | 0.000916324803028841 | 2491 | 8 | 6 | 59 | 74 | 2 | 1 | 5 | 5 | 4 | 6 | 28 | 68 |
| L1MA4A_dup607 | 17.665869931335 | -2.94361657085686 | 0.646233874520336 | -4.55503291752038 | 5.23773756037641e-06 | 0.000747629694985798 | 2492 | 7 | 6 | 52 | 66 | 1 | 3 | 4 | 6 | 2 | 6 | 33 | 54 |
| L1P3_dup2991 | 36.5005716354005 | -3.53674925443357 | 0.411164676938229 | -8.6017828203781 | 7.84872110458346e-18 | 4.66892947987804e-14 | 2493 | 6 | 3 | 61 | 106 | 2 | 3 | 46 | 19 | 11 | 12 | 94 | 130 |
| L1PA11_dup3580 | 30.9167967700199 | -4.41977741080243 | 0.484473049097984 | -9.12285506702879 | 7.31707891672953e-20 | 6.69642176892356e-16 | 2494 | 3 | 2 | 53 | 96 | 0 | 2 | 44 | 18 | 6 | 4 | 93 | 97 |
| L1P1_dup2812 | 26.5731680020379 | -2.71809708327053 | 0.401304559067132 | -6.77315276354943 | 1.26005776194542e-11 | 1.69393053233822e-08 | 2495 | 3 | 1 | 43 | 58 | 10 | 4 | 31 | 13 | 10 | 15 | 68 | 99 |
| L1MC3_dup10663 | 25.723254656391 | -3.64947984430134 | 0.514217625821785 | -7.09715043016856 | 1.27355415328449e-12 | 2.34912493455373e-09 | 2496 | 2 | 1 | 50 | 75 | 1 | 4 | 17 | 13 | 8 | 9 | 73 | 92 |
| PRIMA41-int_dup315 | 23.5446270572566 | -3.72871413543475 | 0.54299470082796 | -6.8669438757859 | 6.55918596759527e-12 | 9.34570098350554e-09 | 2497 | 1 | 1 | 46 | 67 | 1 | 1 | 37 | 15 | 9 | 14 | 53 | 70 |
| L1PA14_dup373 | 22.6917138896294 | -2.71612650029414 | 0.504779870690905 | -5.38081381212073 | 7.41498558167937e-08 | 2.62945776336554e-05 | 2498 | 4 | 8 | 51 | 100 | 10 | 3 | 25 | 10 | 7 | 4 | 27 | 61 |
| L1MD1_dup695 | 28.9520020240716 | -2.18880277298084 | 0.394072893513965 | -5.5543093904866 | 2.78711452479516e-08 | 1.2079831561328e-05 | 2499 | 10 | 8 | 68 | 89 | 8 | 9 | 40 | 14 | 19 | 9 | 52 | 58 |
| L1PREC2_dup792 | 25.0694574111452 | -2.55529437425691 | 0.460695365849261 | -5.54660316486231 | 2.9127279611744e-08 | 1.24877831972902e-05 | 2500 | 6 | 3 | 69 | 77 | 3 | 8 | 37 | 12 | 13 | 14 | 35 | 58 |
| L1PA10_dup6914 | 33.9023085025502 | -2.13333471283619 | 0.620678751458445 | -3.43709963942437 | 0.000587979260296369 | 0.0242389662284268 | 2501 | 7 | 6 | 47 | 66 | 0 | 1 | 15 | 14 | 62 | 53 | 71 | 82 |
| L1PA17_dup4652 | 45.3908234830969 | -3.28124068005011 | 0.389758139652254 | -8.41865851211641 | 3.80819657896628e-17 | 2.01365587373047e-13 | 2502 | 3 | 3 | 33 | 55 | 2 | 0 | 41 | 31 | 25 | 31 | 150 | 224 |
| L1MC1_dup12648 | 23.6305361374569 | -2.87552672061031 | 0.396499858205964 | -7.25227679429989 | 4.09823274090988e-13 | 8.33468451084224e-10 | 2503 | 2 | 3 | 20 | 35 | 3 | 0 | 30 | 16 | 14 | 15 | 69 | 105 |
| L1MD2_dup8423 | 27.9193953076239 | -2.61475145757341 | 0.375868698516394 | -6.95655548837718 | 3.48692955939415e-12 | 5.21824491156982e-09 | 2504 | 1 | 4 | 23 | 26 | 4 | 0 | 19 | 17 | 14 | 28 | 109 | 118 |
| MER51B_dup123 | 16.6640997432255 | -2.73857241205453 | 0.665366245050787 | -4.11588720110906 | 3.85692909422772e-05 | 0.00334827990693298 | 2505 | 0 | 0 | 12 | 26 | 2 | 1 | 28 | 18 | 28 | 20 | 37 | 35 |
| MLT1A0_dup4491 | 18.7666751501383 | -3.01673811347804 | 0.696200356997069 | -4.33314646158784 | 1.469932486752e-05 | 0.00162454507892565 | 2506 | 0 | 0 | 13 | 26 | 2 | 0 | 26 | 17 | 31 | 22 | 49 | 48 |
| AluJr_dup16778 | 22.484143683763 | -2.39108940264271 | 0.555216959772855 | -4.30658566989907 | 1.65793736439177e-05 | 0.00178023269001608 | 2507 | 0 | 0 | 20 | 24 | 4 | 2 | 17 | 18 | 36 | 23 | 50 | 89 |
| HERVL-int_dup506 | 26.8361738095356 | -3.11283864869411 | 0.535808322132498 | -5.80961235597298 | 6.26176590697105e-09 | 3.40174006963501e-06 | 2508 | 0 | 0 | 34 | 38 | 1 | 3 | 30 | 20 | 29 | 24 | 80 | 84 |
| MSTB1_dup4217 | 29.1127699685048 | -2.57125149782137 | 0.515167386978724 | -4.99109913168389 | 6.00366736558564e-07 | 0.000138191116512148 | 2509 | 0 | 0 | 53 | 68 | 11 | 10 | 37 | 30 | 13 | 22 | 59 | 77 |
| L1PA14_dup684 | 40.4499901647835 | -3.51334546275232 | 0.518328474080597 | -6.77822199327219 | 1.21663812043846e-11 | 1.64485326253324e-08 | 2510 | 0 | 0 | 32 | 61 | 4 | 2 | 44 | 42 | 26 | 32 | 108 | 176 |
| L1PA7_dup2717 | 32.999299069905 | -3.2101415957638 | 0.552008693666308 | -5.8153823166132 | 6.04954609643126e-09 | 3.29802024920235e-06 | 2511 | 0 | 0 | 39 | 42 | 3 | 2 | 42 | 30 | 33 | 36 | 78 | 118 |
| L1M4_dup12830 | 37.6422835775506 | -5.50507064102425 | 0.55480441776874 | -9.92254290829914 | 3.32183312079558e-23 | 5.64583502686304e-19 | 2512 | 0 | 0 | 58 | 55 | 2 | 3 | 35 | 24 | 2 | 2 | 133 | 193 |
| L1P2_dup1119 | 46.8142063784341 | -6.11944911850115 | 0.607512546065304 | -10.0729592469081 | 7.27554324094934e-24 | 1.44265534334244e-19 | 2513 | 0 | 0 | 50 | 59 | 0 | 1 | 36 | 17 | 2 | 5 | 202 | 262 |
| L1MB4_dup5480 | 29.2722190273536 | -3.8515913245337 | 0.424553836849851 | -9.07209166477481 | 1.16753817684003e-19 | 1.02892977417177e-15 | 2514 | 1 | 0 | 36 | 48 | 1 | 5 | 24 | 28 | 9 | 7 | 104 | 122 |
| MLT1F2_dup4804 | 21.0909501651882 | -4.79673187939139 | 0.604153154165345 | -7.93959585631597 | 2.02841079425304e-15 | 7.31291264923232e-12 | 2515 | 0 | 0 | 44 | 46 | 0 | 1 | 20 | 17 | 3 | 6 | 53 | 93 |
| L2a_dup133447 | 35.6920944301383 | -2.38695428369782 | 0.340927425497 | -7.00135602237966 | 2.53496853181201e-12 | 4.07557852885501e-09 | 2516 | 3 | 2 | 26 | 41 | 11 | 2 | 21 | 14 | 20 | 28 | 126 | 177 |
| L1MD2_dup8424 | 32.6413718741051 | -2.8039683991485 | 0.327258023724183 | -8.56806616149377 | 1.05237332049343e-17 | 6.10751273458855e-14 | 2517 | 5 | 2 | 18 | 35 | 4 | 3 | 24 | 13 | 13 | 18 | 121 | 180 |
| L1MEd_dup17354 | 32.0226256016591 | -2.68905552459347 | 0.299367112380569 | -8.98246805806452 | 2.64759854474122e-19 | 2.09995161108998e-15 | 2518 | 4 | 2 | 22 | 35 | 5 | 4 | 26 | 18 | 15 | 19 | 116 | 157 |
| L2c_dup110923 | 35.3678379784329 | -3.00689711154999 | 0.45886364082262 | -6.55292083321185 | 5.64224322919559e-11 | 6.21550558988043e-08 | 2519 | 0 | 0 | 23 | 47 | 4 | 3 | 24 | 14 | 19 | 34 | 103 | 201 |
| MLT1G_dup2162 | 28.4579094435088 | -3.09503276933491 | 0.404361902121101 | -7.65411566495202 | 1.94647026842817e-14 | 5.64822944501718e-11 | 2520 | 0 | 0 | 24 | 27 | 6 | 1 | 17 | 11 | 12 | 17 | 108 | 155 |
| L1PA13_dup7407 | 29.3735051488104 | -3.12577777503043 | 0.421901563256871 | -7.40878452997654 | 1.27462200711027e-13 | 2.91359388978609e-10 | 2521 | 0 | 0 | 19 | 24 | 1 | 4 | 18 | 7 | 18 | 14 | 119 | 166 |
| HAL1_dup18029 | 28.7912559565787 | -3.01364569120966 | 0.409823173090901 | -7.35352681128457 | 1.93044262810218e-13 | 4.21413854666424e-10 | 2522 | 0 | 2 | 16 | 30 | 2 | 1 | 17 | 6 | 17 | 18 | 114 | 160 |
| MLT1J_dup11706 | 29.9131770584209 | -2.96424141877857 | 0.472158457206938 | -6.27806486049956 | 3.42812912677039e-10 | 2.90288118575982e-07 | 2523 | 0 | 2 | 27 | 24 | 1 | 3 | 7 | 6 | 18 | 18 | 135 | 152 |

| | | | | | | | | | | | | | | | | | | | |
|---|---|---|---|---|---|---|---|---|---|---|---|---|---|---|---|---|---|---|---|
| L1ME1_dup24293 | 26.0446008726889 | -2.21102935321688 | 0.350325181153351 | -6.31136290556581 | 2.76588753458898e-10 | 2.45571595263175e-07 | 2524 | 4 | 4 | 19 | 24 | 2 | 4 | 19 | 7 | 10 | 30 | 97 | 125 |
| THE1C-int_dup1566 | 32.2352114524276 | -2.41606629854893 | 0.384491997972509 | -6.28378824862222 | 3.30420694233935e-10 | 2.80793866107814e-07 | 2525 | 3 | 3 | 18 | 33 | 7 | 5 | 15 | 7 | 11 | 21 | 124 | 185 |
| L1ME3A_dup12026 | 24.0285029374954 | -2.59994199665676 | 0.371516256985646 | -6.9981917285445 | 2.59287114680029e-12 | 4.14069340870162e-09 | 2526 | 1 | 2 | 17 | 23 | 1 | 4 | 20 | 10 | 19 | 14 | 93 | 110 |
| L3_dup36531 | 30.8139947681344 | -2.84930955989066 | 0.340190357144963 | -8.37563293622842 | 5.49278619150146e-17 | 2.78082234707023e-13 | 2527 | 1 | 2 | 17 | 20 | 2 | 1 | 16 | 12 | 14 | 25 | 126 | 171 |
| L1MA9_dup15923 | 28.2433795694613 | -3.06215484267409 | 0.360964319762693 | -8.48326184894736 | 2.18967130701258e-17 | 1.2116826251591e-13 | 2528 | 1 | 2 | 19 | 21 | 3 | 1 | 26 | 17 | 15 | 14 | 100 | 154 |
| MER65A_dup1087 | 30.988875526773 | -3.29062160775622 | 0.369644252207308 | -8.90213113853788 | 5.47844874661315e-19 | 3.95022716806549e-15 | 2529 | 1 | 1 | 18 | 30 | 2 | 1 | 24 | 10 | 14 | 16 | 121 | 176 |
| L1PB_dup1655 | 15.4666822447547 | -2.48170284082989 | 0.471896476083023 | -5.25899845963943 | 1.44842102971274e-07 | 4.45855097459285e-05 | 2530 | 4 | 2 | 11 | 14 | 0 | 0 | 9 | 16 | 11 | 13 | 49 | 70 |
| L2_dup55459 | 20.696496999132 | -2.97426392639598 | 0.536133182044987 | -5.54762142319035 | 2.89582046773431e-08 | 1.24377057223016e-05 | 2531 | 2 | 2 | 5 | 22 | 0 | 0 | 13 | 20 | 18 | 11 | 68 | 107 |
| L1PA10_dup1402 | 21.0208861065525 | -3.89154360326197 | 0.704308227899566 | -5.52534167443647 | 3.2884501047452e-08 | 1.36082356282383e-05 | 2532 | 0 | 1 | 32 | 37 | 0 | 0 | 13 | 23 | 19 | 13 | 56 | 75 |
| L1PBa_dup498 | 16.7253686972978 | -3.15761858059124 | 0.640815105193544 | -4.92750335471188 | 8.32869940290893e-07 | 0.000178861074740485 | 2533 | 0 | 0 | 17 | 24 | 1 | 0 | 17 | 10 | 12 | 26 | 38 | 74 |
| L1M4a2_dup2015 | 16.1723563716104 | -3.00720592470017 | 0.535621158794764 | -5.61442705412699 | 1.97214602182717e-08 | 9.07667809109646e-06 | 2534 | 0 | 3 | 15 | 22 | 0 | 0 | 14 | 16 | 8 | 17 | 59 | 54 |
| L1ME4a_dup26458 | 15.6052386420729 | -3.23864557177539 | 0.508030576163881 | -6.37490285767891 | 1.83078958467614e-10 | 1.72185398622668e-07 | 2535 | 0 | 1 | 16 | 16 | 0 | 0 | 10 | 6 | 7 | 13 | 61 | 76 |
| MSTD_dup6173 | 16.1590849299157 | -2.84453836786119 | 0.456267898836025 | -6.23436006591266 | 4.53628613358993e-10 | 3.74331167513952e-07 | 2536 | 2 | 0 | 17 | 17 | 1 | 2 | 17 | 6 | 7 | 13 | 76 | 53 |
| L2c_dup110919 | 19.9008651791695 | -3.07541413625843 | 0.43457531861582 | -7.07682651203946 | 1.47492987922258e-12 | 2.658739886678e-09 | 2537 | 0 | 0 | 12 | 19 | 1 | 1 | 19 | 7 | 13 | 13 | 82 | 95 |
| L2a_dup133434 | 22.9694311688531 | -2.84929686092893 | 0.40603266850255 | -7.017407913105 | 2.26022026071319e-12 | 3.70903703555628e-09 | 2538 | 1 | 0 | 15 | 26 | 2 | 2 | 20 | 6 | 14 | 17 | 85 | 117 |
| Tigger1_dup11835 | 19.1268826449923 | -2.71977693687051 | 0.398941037956339 | -6.8174910026884 | 9.26442044262494e-12 | 1.29672458037696e-08 | 2539 | 2 | 0 | 12 | 18 | 1 | 1 | 18 | 8 | 13 | 15 | 74 | 89 |
| MSTD_dup6174 | 19.5050115818357 | -2.29535260934549 | 0.39147999032274 | -5.86326930133307 | 4.53841253016575e-09 | 2.57118359024481e-06 | 2540 | 2 | 1 | 14 | 10 | 3 | 2 | 14 | 5 | 10 | 20 | 71 | 106 |
| L1PREC2_dup6316 | 22.9867049093828 | -2.41982533470146 | 0.391655723346707 | -6.17845007861496 | 6.47339506921309e-10 | 5.00103397123045e-07 | 2541 | 1 | 0 | 19 | 17 | 4 | 3 | 15 | 6 | 19 | 16 | 97 | 102 |
| LTR105_Mam_dup499 | 21.0200460142407 | -2.84369681580588 | 0.416864287871999 | -6.8216369176701 | 9.00089245990649e-12 | 1.26729370252362e-08 | 2542 | 2 | 0 | 17 | 26 | 2 | 1 | 11 | 11 | 13 | 15 | 71 | 108 |
| THE1C_dup7987 | 20.5231626896492 | -2.49489046896888 | 0.400836148563795 | -6.22421525081541 | 4.83972254317825e-10 | 3.95736295621681e-07 | 2543 | 2 | 0 | 16 | 14 | 3 | 1 | 20 | 13 | 13 | 22 | 70 | 93 |
| THE1C_dup7988 | 21.5662806913652 | -2.52361264515991 | 0.438001988422125 | -5.7616465492567 | 8.32972926690214e-09 | 4.33116788005429e-06 | 2544 | 1 | 0 | 26 | 23 | 4 | 2 | 17 | 10 | 16 | 22 | 68 | 92 |
| AluJb_dup90755 | 20.8816423258661 | -2.74056216994346 | 0.442865180364612 | -6.18825387827317 | 6.08342972616278e-10 | 4.76160450533398e-07 | 2545 | 0 | 1 | 17 | 17 | 4 | 0 | 13 | 5 | 12 | 17 | 80 | 110 |
| L1PA13_dup7409 | 16.2313688726456 | -3.20646023208615 | 0.507576211702236 | -6.31719958138461 | 2.66345520661878e-10 | 2.38255079922598e-07 | 2546 | 0 | 0 | 15 | 13 | 1 | 1 | 9 | 5 | 10 | 8 | 56 | 98 |
| MIR_dup133535 | 19.0362080765701 | -3.00300526802559 | 0.458153196242079 | -6.5545876197246 | 5.57958469922363e-11 | 6.17506912019287e-08 | 2547 | 0 | 0 | 13 | 17 | 3 | 1 | 9 | 7 | 11 | 10 | 95 | 82 |
| L3_dup36526 | 15.7379026410953 | -3.32021086828194 | 0.514575907787316 | -6.4523247552706 | 1.10147221355123e-10 | 1.11055469205789e-07 | 2548 | 1 | 0 | 15 | 15 | 2 | 0 | 7 | 5 | 6 | 8 | 72 | 77 |
| Tigger3a_dup4267 | 14.7954505106374 | -3.13923212604261 | 0.610802202753852 | -5.13952325628349 | 2.75436402137172e-07 | 7.48161987933007e-05 | 2549 | 0 | 0 | 15 | 19 | 2 | 0 | 5 | 3 | 6 | 14 | 61 | 71 |
| AluSz6_dup34553 | 18.4773311225177 | -2.99032685415355 | 0.439008501647039 | -6.81154657127291 | 9.65550288893416e-12 | 1.33574900605252e-08 | 2550 | 0 | 2 | 6 | 16 | 2 | 2 | 14 | 11 | 7 | 10 | 72 | 103 |
| HAL1_dup18027 | 13.6348005996859 | -2.56414735374506 | 0.443707652554394 | -5.7789117203263 | 7.51853613623791e-09 | 3.98542967606232e-06 | 2551 | 1 | 1 | 6 | 10 | 2 | 2 | 12 | 8 | 8 | 8 | 57 | 63 |
| L1MEg_dup11427 | 14.5070334261982 | -2.80381629546273 | 0.492147325533231 | -5.69710765455213 | 1.21856988059128e-08 | 6.04070476681608e-06 | 2552 | 0 | 2 | 12 | 14 | 1 | 0 | 20 | 11 | 14 | 8 | 54 | 50 |
| AluSz6_dup34552 | 14.6057452638758 | -2.52966914958254 | 0.472265078019886 | -5.35646031713576 | 8.48681288380062e-08 | 2.90980285079081e-05 | 2553 | 0 | 3 | 13 | 16 | 1 | 0 | 10 | 6 | 12 | 12 | 48 | 70 |
| MIRc_dup77483 | 17.8482738158156 | -2.7997793900539 | 0.433648316397662 | -6.45633635410326 | 1.07268056803563e-10 | 1.09076944633251e-07 | 2554 | 1 | 2 | 16 | 14 | 1 | 0 | 17 | 6 | 14 | 10 | 68 | 85 |
| L1P1_dup674 | 14.9714552547641 | -2.77669397692445 | 0.510697767607706 | -5.43705916305742 | 5.41671562105041e-08 | 2.04585050026422e-05 | 2555 | 0 | 0 | 13 | 18 | 2 | 1 | 18 | 18 | 11 | 16 | 38 | 57 |
| L3b_dup8000 | 17.4016960165173 | -2.49337097104352 | 0.437399403443876 | -5.70044437969486 | 1.19495517169275e-08 | 5.96089734347176e-06 | 2556 | 0 | 0 | 11 | 11 | 2 | 3 | 18 | 11 | 13 | 16 | 71 | 68 |
| L1M3_dup5205 | 22.5712532119373 | -2.55204609637806 | 0.444682835718774 | -5.73902541629024 | 9.5222930098928e-09 | 4.87266996243431e-06 | 2557 | 0 | 1 | 12 | 22 | 2 | 2 | 6 | 14 | 19 | 20 | 82 | 112 |
| MIRb_dup170296 | 14.6360950471523 | -2.84548195471665 | 0.531463255482238 | -5.35405209177579 | 8.60060823647834e-08 | 2.94034529804177e-05 | 2558 | 0 | 1 | 12 | 20 | 1 | 1 | 6 | 12 | 6 | 17 | 59 | 54 |
| L1PA5_dup9445 | 17.6084197101488 | -2.68435275606154 | 0.401862518777881 | -6.67977885627411 | 2.3930308131022e-11 | 2.96568807215841e-08 | 2559 | 0 | 2 | 14 | 15 | 2 | 2 | 13 | 8 | 8 | 14 | 67 | 87 |
| THE1A_dup3517 | 17.6083230842194 | -2.93410331619901 | 0.447961904784138 | -6.5498947228847 | 5.75776590092347e-11 | 6.31353624452136e-08 | 2560 | 0 | 1 | 14 | 23 | 1 | 1 | 11 | 8 | 10 | 14 | 70 | 78 |
| L1MB7_dup637 | 11.7985852471931 | -1.83036756847479 | 0.527367819486806 | -3.4707608254443 | 0.000518986039884328 | 0.0221309412627807 | 2561 | 2 | 0 | 6 | 5 | 2 | 6 | 22 | 10 | 8 | 14 | 42 | 34 |
| MER5A1_dup14498 | 13.326290895847 | -1.75224185819632 | 0.394170432413004 | -4.44539141982206 | 8.77319835349324e-06 | 0.00110429968914987 | 2562 | 6 | 3 | 15 | 21 | 2 | 6 | 14 | 6 | 5 | 14 | 34 | 51 |
| MIRb_dup201083 | 14.9269623623624 | -1.22429322923978 | 0.356001430357006 | -3.4390121073728 | 0.000583841121963743 | 0.0241017799456601 | 2563 | 6 | 2 | 12 | 13 | 4 | 5 | 15 | 13 | 14 | 24 | 43 | 37 |
| L1PA2_dup423 | 9.7736064418304 | -2.51144973744282 | 0.697794573212195 | -3.59912477662548 | 0.000319289971111838 | 0.0156260327984734 | 2564 | 2 | 2 | 3 | 1 | 3 | 3 | 32 | 18 | 4 | 3 | 21 | 34 |
| SVA_D_dup997 | 12.4348831173277 | -3.31138122236492 | 0.577211425484826 | -5.73686014545458 | 9.6447833895256e-09 | 4.92475885923618e-06 | 2565 | 2 | 3 | 12 | 13 | 0 | 2 | 49 | 19 | 3 | 4 | 19 | 39 |
| L1MC3_dup10662 | 11.7381551592179 | -3.00009984093509 | 0.529388009250232 | -5.66710954633089 | 1.45226500993987e-08 | 6.99777351904828e-06 | 2566 | 3 | 4 | 10 | 31 | 1 | 1 | 19 | 6 | 4 | 2 | 33 | 45 |
| L1MC1_dup10505 | 18.0564601705964 | -2.96401057726171 | 0.467437672023212 | -6.34097496770547 | 2.28315639598206e-10 | 2.06565753535493e-07 | 2567 | 3 | 4 | 14 | 21 | 1 | 1 | 14 | 6 | 9 | 3 | 70 | 95 |
| L1MD2_dup320 | 16.8532395223643 | -3.03041119372074 | 0.448887260977337 | -6.75094050814185 | 1.46889825207357e-11 | 1.94176924159943e-08 | 2568 | 5 | 3 | 15 | 29 | 2 | 2 | 20 | 9 | 4 | 4 | 50 | 85 |
| AluY_dup80270 | 15.9084628539134 | -3.90400626243742 | 0.566274503552484 | -6.89419396060728 | 5.4171070921351e-12 | 7.811993722092e-09 | 2569 | 1 | 0 | 24 | 25 | 1 | 4 | 20 | 5 | 3 | 2 | 63 | 66 |
| L1PA3_dup9508 | 17.0530653636482 | -2.99809267354408 | 0.477728396889595 | -6.27572631868678 | 3.48005861402664e-10 | 2.93640435096874e-07 | 2570 | 3 | 2 | 39 | 37 | 4 | 2 | 18 | 6 | 4 | 8 | 45 | 61 |
| L1PA7_dup13359 | 15.7345743663433 | -2.25980690557788 | 0.475268303164133 | -4.75480247795415 | 1.98640530370902e-06 | 0.000350635902371177 | 2571 | 1 | 2 | 23 | 25 | 7 | 7 | 11 | 6 | 7 | 5 | 53 | 61 |
| L1MD2_dup8421 | 13.2059169081985 | -2.29978472575033 | 0.492920269492693 | -4.66563229001972 | 3.07669474045196e-06 | 0.000489879075703537 | 2572 | 1 | 0 | 11 | 12 | 7 | 2 | 8 | 6 | 5 | 8 | 46 | 69 |
| L1MEg1_dup716 | 12.9284176633078 | -2.2232798661482 | 0.559871722414184 | -3.97105225561554 | 7.15558804541766e-05 | 0.0053224243609095 | 2573 | 1 | 0 | 7 | 7 | 6 | 2 | 7 | 4 | 7 | 4 | 48 | 79 |
| L1PA16_dup10248 | 11.1256139475417 | -1.77571754983564 | 0.503186673098841 | -3.52894391836733 | 0.000417221537484792 | 0.019068334365099 | 2574 | 4 | 2 | 6 | 13 | 3 | 5 | 6 | 3 | 4 | 6 | 44 | 52 |
| L1PA4_dup10148 | 12.5060703339714 | -2.00343868986799 | 0.513509072234526 | -3.90146698119677 | 9.56115046877494e-05 | 0.00659306749425309 | 2575 | 1 | 4 | 7 | 8 | 5 | 1 | 8 | 3 | 4 | 8 | 52 | 65 |
| HERVK9-int_dup720 | 9.39030638779841 | -1.88632370082836 | 0.508979466162054 | -3.7060899824744 | 0.000210483544779038 | 0.0115341560972046 | 2576 | 3 | 3 | 18 | 10 | 6 | 3 | 7 | 4 | 3 | 4 | 26 | 37 |
| THE1B_dup20982 | 9.88144237518565 | -1.86435879077031 | 0.53682744215621 | -3.47292005654921 | 0.000514828679964512 | 0.0220247078538 | 2577 | 2 | 4 | 9 | 11 | 6 | 6 | 9 | 7 | 3 | 1 | 38 | 33 |
| MLT1D_dup4319 | 8.91009482802471 | -1.86502014070362 | 0.52030721689974 | -3.58445948879275 | 0.000337776985079383 | 0.0162892310003661 | 2578 | 1 | 0 | 10 | 7 | 4 | 6 | 11 | 5 | 5 | 5 | 36 | 25 |
| L1MEc_dup12928 | 12.4053449337719 | -1.82868995152336 | 0.523168882236061 | -3.49541039923363 | 0.000473333344820743 | 0.0207685369844581 | 2579 | 1 | 1 | 12 | 19 | 9 | 8 | 11 | 6 | 3 | 7 | 48 | 38 |
| MIRc_dup67692 | 11.4863084034105 | -1.75000832102209 | 0.512975546864013 | -3.41148487821782 | 0.000646100849405716 | 0.0258425134833909 | 2580 | 0 | 1 | 13 | 11 | 11 | 5 | 15 | 8 | 5 | 7 | 32 | 42 |
| L1MEj_dup499 | 9.51236998803563 | -2.20568668324199 | 0.674588648075384 | -3.26967654960525 | 0.00107670523622913 | 0.0369640317616184 | 2581 | 0 | 0 | 10 | 13 | 2 | 0 | 9 | 8 | 16 | 14 | 16 | 31 |
| THE1B_dup4577 | 12.4834578857266 | -2.65821693131645 | 0.569217179595611 | -4.6699520439719 | 3.01270057231385e-06 | 0.000482174281751146 | 2582 | 0 | 0 | 8 | 15 | 1 | 0 | 12 | 4 | 16 | 10 | 39 | 57 |
| L1MA6_dup1085 | 7.50675079719635 | -3.0851157927768 | 0.734366770889524 | -4.20105581443977 | 2.65673077303137e-05 | 0.00251992299657343 | 2583 | 0 | 0 | 5 | 10 | 0 | 0 | 13 | 8 | 9 | 5 | 14 | 33 |
| L1PBa_dup500 | 8.8160960746577 | -2.97902699734387 | 0.738797603081204 | -4.03226402592489 | 5.52420734302342e-05 | 0.00439766825173319 | 2584 | 0 | 0 | 6 | 15 | 0 | 0 | 13 | 8 | 11 | 11 | 18 | 31 |
| MER52-int_dup106 | 6.9513437480026 | -2.6301049732526 | 0.59268865961591 | -4.43758275205912 | 9.09747512488176e-06 | 0.00113692637398378 | 2585 | 0 | 0 | 9 | 14 | 1 | 1 | 9 | 7 | 5 | 6 | 17 | 21 |
| L1MA8_dup10203 | 8.21731411603679 | -2.36432019881196 | 0.596519618541648 | -3.96352462739142 | 7.3851241048017e-05 | 0.00544887051237565 | 2586 | 0 | 0 | 9 | 8 | 1 | 1 | 7 | 8 | 10 | 7 | 16 | 38 |
| L1PA10_dup1403 | 8.29989063560232 | -2.42415142145673 | 0.628694380278426 | -3.85585031058042 | 0.00011532804607884 | 0.007528214705162 | 2587 | 0 | 0 | 8 | 7 | 2 | 1 | 8 | 13 | 10 | 5 | 16 | 34 |
| Charlie6_dup225 | 12.1271412951844 | -2.75708425289249 | 0.563376674224184 | -4.89385588547701 | 9.88792578794834e-07 | 0.000204209296418747 | 2588 | 0 | 0 | 16 | 32 | 2 | 3 | 15 | 5 | 6 | 11 | 36 | 35 |
| ERVL47-int_dup137 | 7.88618293971377 | -2.43478900276425 | 0.604410036679718 | -4.02837288430812 | 5.61642142884275e-05 | 0.00444579061744705 | 2589 | 0 | 0 | 12 | 22 | 3 | 2 | 6 | 5 | 4 | 7 | 19 | 24 |
| MER4A_dup741 | 9.31343755482472 | -2.82193111745754 | 0.596799666037101 | -4.72843950499481 | 2.26252015177627e-06 | 0.000385975727989599 | 2590 | 0 | 0 | 15 | 28 | 2 | 3 | 10 | 5 | 3 | 7 | 21 | 31 |
| L1MA8_dup10204 | 9.52875168198304 | -1.94788683053432 | 0.578348480314619 | -3.36801581889639 | 0.000757112519014157 | 0.0289434757589744 | 2591 | 0 | 0 | 4 | 7 | 6 | 3 | 7 | 9 | 7 | 4 | 36 | 39 |
| MIR_dup133522 | 8.34901573665292 | -2.70659119368388 | 0.603977993866559 | -4.48127451855782 | 7.41985974735478e-06 | 0.000973942805839653 | 2592 | 0 | 0 | 7 | 8 | 2 | 1 | 3 | 8 | 6 | 4 | 32 | 36 |
| AluSx1_dup81390 | 10.4301253278615 | -2.2483517822834 | 0.487727963660346 | -4.60984800914379 | 4.02963463843098e-06 | 0.000605325406361174 | 2593 | 0 | 2 | 10 | 5 | 2 | 2 | 9 | 7 | 5 | 10 | 34 | 50 |
| L1P1_dup672 | 10.1375905901275 | -2.18267398590654 | 0.507183793828568 | -4.30351681671498 | 1.68108081489843e-05 | 0.00179939926037707 | 2594 | 0 | 3 | 7 | 11 | 1 | 1 | 10 | 6 | 6 | 12 | 23 | 54 |
| AluSx1_dup22521 | 8.05169560647736 | -1.94868877674809 | 0.548062683694212 | -3.55559470608903 | 0.00037712506332553 | 0.0176987900832067 | 2595 | 0 | 1 | 7 | 9 | 1 | 1 | 11 | 4 | 10 | 10 | 17 | 33 |
| L2b_dup1841 | 6.88756234564659 | -1.77980689378479 | 0.530333533415811 | -3.35601424696131 | 0.000790744951539857 | 0.0298279325046136 | 2596 | 0 | 1 | 4 | 6 | 2 | 3 | 7 | 5 | 4 | 8 | 23 | 26 |
| AluY_dup76592 | 10.2258007311832 | -2.40092117277407 | 0.542908797048995 | -4.4223287333423 | 9.76427423398802e-06 | 0.00120070800872378 | 2597 | 0 | 1 | 14 | 6 | 3 | 2 | 8 | 4 | 8 | 4 | 32 | 52 |
| THE1B_dup18272 | 12.8357340493957 | -2.71514173146356 | 0.476267535060109 | -5.70087509979216 | 1.19193947139463e-08 | 5.9583451567325e-06 | 2598 | 0 | 2 | 16 | 12 | 1 | 2 | 9 | 6 | 8 | 7 | 52 | 52 |
| L2c_dup110930 | 11.5290597634198 | -2.62279642426084 | 0.558267798459872 | -4.69809727785932 | 2.62596361845728e-06 | 0.000436339063657428 | 2599 | 0 | 0 | 9 | 10 | 1 | 1 | 12 | 12 | 13 | 9 | 46 | 31 |
| L1ME3Cz_dup9408 | 11.9869510843251 | -2.67746123016825 | 0.488435298302075 | -5.48171116927008 | 4.21231535838541e-08 | 1.66219500873362e-05 | 2600 | 0 | 0 | 12 | 10 | 3 | 2 | 14 | 5 | 8 | 6 | 46 | 51 |
| L2b_dup73109 | 11.0150249395678 | -2.47487716701563 | 0.51782990177535 | -4.77932455914705 | 1.75885071778601e-06 | 0.000320019705297638 | 2601 | 0 | 0 | 10 | 9 | 1 | 1 | 10 | 4 | 8 | 13 | 48 | 38 |
| AluJr_dup60452 | 13.0506129590695 | -2.56546112107982 | 0.522432323866668 | -4.91060947778293 | 9.079374826228e-07 | 0.000191865090799436 | 2602 | 0 | 0 | 4 | 11 | 2 | 4 | 17 | 4 | 7 | 8 | 54 | 62 |

| | | | | | | | | | | | | | | | | | | | |
|---|---|---|---|---|---|---|---|---|---|---|---|---|---|---|---|---|---|---|---|
| MIR_dup133534 | 11.75468704932 | -2.15972727100502 | 0.503195582082831 | -4.29202351512202 | 1.77052192895239e-05 | 0.00186824217696898 | 2603 | 0 | 0 | 4 | 10 | 1 | 3 | 4 | 5 | 10 | 11 | 44 | 61 |
| L1MC4_dup18972 | 18.1263341918065 | -2.41356634226434 | 0.403952454839219 | -5.97487727417076 | 2.30263461049082e-09 | 1.46845764841041e-06 | 2604 | 2 | 1 | 12 | 19 | 0 | 3 | 11 | 10 | 12 | 18 | 73 | 74 |
| L1PA11_dup3442 | 20.820997160009 | -3.1344332871755 | 0.44041028420547 | -7.11707559879131 | 1.10241027078884e-12 | 2.08185804994541e-09 | 2605 | 1 | 0 | 11 | 10 | 0 | 2 | 11 | 10 | 12 | 10 | 82 | 126 |
| HUERS-P2-int_dup240 | 14.8074007174008 | -2.86225987571004 | 0.476130284001491 | -6.0115056149234 | 1.83808144472242e-09 | 1.19172786770006e-06 | 2606 | 2 | 0 | 9 | 15 | 0 | 2 | 13 | 12 | 12 | 6 | 49 | 73 |
| MIRb_dup170299 | 15.132551608552 | -2.74534553698526 | 0.457407153494559 | -6.00197333166087 | 1.94933724120936e-09 | 1.25701083793171e-06 | 2607 | 1 | 0 | 14 | 13 | 0 | 3 | 14 | 11 | 12 | 9 | 53 | 66 |
| AluSc8_dup15772 | 15.2264561004933 | -2.80561214130021 | 0.563488361947877 | -4.97900636599079 | 6.3911540794673e-07 | 0.000145052991673874 | 2608 | 0 | 1 | 9 | 7 | 0 | 2 | 14 | 4 | 16 | 4 | 61 | 81 |
| L1ME3Cz_dup9410 | 14.6349974808952 | -2.95080002082281 | 0.5569746720433 | -5.29790701253541 | 1.17137588927059e-07 | 3.73770101522728e-05 | 2609 | 0 | 0 | 9 | 13 | 0 | 3 | 10 | 4 | 13 | 4 | 62 | 74 |
| L2c_dup110921 | 19.510666265592 | -2.63419534397918 | 0.470329460905704 | -5.60074493081205 | 2.13432598968452e-08 | 9.69187656376858e-06 | 2610 | 1 | 2 | 17 | 11 | 0 | 2 | 18 | 3 | 16 | 13 | 62 | 114 |
| AluJr_dup60454 | 17.3968983256183 | -2.78078381298292 | 0.459095322362242 | -6.05709463271068 | 1.38602058066599e-09 | 9.55936385759856e-07 | 2611 | 0 | 2 | 6 | 19 | 0 | 1 | 14 | 8 | 12 | 14 | 64 | 89 |
| AluJr_dup60455 | 20.6769137665835 | -2.77799464071758 | 0.419772149352843 | -6.61786315504822 | 3.64428132880803e-11 | 4.22996178080271e-08 | 2612 | 0 | 1 | 7 | 13 | 0 | 1 | 12 | 7 | 16 | 15 | 86 | 113 |
| L2_dup45754 | 13.9847820218942 | -3.54796376061722 | 0.579317904541539 | -6.12438133329405 | 9.10366315002843e-10 | 6.64472463771983e-07 | 2613 | 0 | 0 | 10 | 9 | 0 | 0 | 8 | 9 | 9 | 6 | 66 | 64 |
| MIRb_dup170316 | 12.5172002204665 | -3.64829074742794 | 0.585223888723393 | -6.23400858667326 | 4.54648144585461e-10 | 3.74331167513952e-07 | 2614 | 0 | 0 | 11 | 16 | 0 | 0 | 11 | 8 | 6 | 7 | 58 | 46 |
| L3b_dup8001 | 10.4713590180129 | -3.19781396625754 | 0.636287359032299 | -5.02573863972553 | 5.01498505501082e-07 | 0.000119929209437146 | 2615 | 0 | 0 | 15 | 11 | 0 | 0 | 5 | 7 | 7 | 10 | 33 | 48 |
| MER66B_dup965 | 12.844137842089 | -2.95651406311573 | 0.546416306089373 | -5.41073542309727 | 6.27664478120434e-08 | 2.30478783813032e-05 | 2616 | 1 | 0 | 11 | 17 | 0 | 0 | 6 | 8 | 8 | 13 | 43 | 61 |
| AluSx_dup82053 | 12.7405634338578 | -3.25799371923397 | 0.562027192964925 | -5.7968613618973 | 6.75675021515116e-09 | 3.62921373971638e-06 | 2617 | 1 | 0 | 7 | 20 | 0 | 0 | 16 | 8 | 6 | 11 | 37 | 64 |
| AluJb_dup90754 | 11.2027770189623 | -3.19111955959758 | 0.593205186471442 | -5.37945323536244 | 7.471239847612e-08 | 2.64153289268928e-05 | 2618 | 1 | 0 | 8 | 8 | 0 | 0 | 7 | 4 | 7 | 6 | 34 | 75 |
| LTR8A_dup1385 | 10.5037645444482 | -2.480623091606 | 0.520614718954304 | -4.76479630961746 | 1.89044522497497e-06 | 0.000337705615241662 | 2619 | 1 | 0 | 8 | 10 | 1 | 0 | 6 | 7 | 10 | 9 | 32 | 52 |
| L1MC3_dup12191 | 11.7556462278224 | -2.63146379122772 | 0.571779982805296 | -4.60223140082153 | 4.17988430532344e-06 | 0.000624348242884175 | 2620 | 0 | 0 | 7 | 18 | 0 | 3 | 10 | 9 | 13 | 7 | 34 | 51 |
| L1MC4_dup18971 | 15.2334413895296 | -2.93706513843252 | 0.546325613060453 | -5.37603412364181 | 7.61443669464875e-08 | 2.67230789637889e-05 | 2621 | 0 | 0 | 7 | 18 | 0 | 2 | 5 | 9 | 12 | 10 | 56 | 80 |
| L1MDa_dup6638 | 9.85881040063329 | -2.95747322532139 | 0.556136877436069 | -5.3178872779595 | 1.04979068867428e-07 | 3.43252716328284e-05 | 2622 | 1 | 0 | 8 | 12 | 0 | 1 | 8 | 4 | 6 | 6 | 33 | 52 |
| THE1D_dup10315 | 9.63156477864619 | -3.17320238837218 | 0.591124859671674 | -5.36807467399468 | 7.95816053836526e-08 | 2.76439776271804e-05 | 2623 | 0 | 0 | 6 | 9 | 0 | 1 | 9 | 5 | 7 | 4 | 40 | 45 |
| AluSq2_dup43750 | 9.51604082243269 | -2.42230938831204 | 0.546611829695644 | -4.43149828949145 | 9.35805555935948e-06 | 0.00116216695622513 | 2624 | 0 | 0 | 4 | 9 | 0 | 3 | 10 | 6 | 7 | 9 | 33 | 43 |
| AluSx1_dup81389 | 8.8293680349261 | -2.77752294569372 | 0.56702476477033 | -4.89841558651981 | 9.66125273438339e-07 | 0.000200202324766804 | 2625 | 0 | 0 | 9 | 13 | 0 | 1 | 9 | 6 | 6 | 9 | 28 | 34 |
| AluSg_dup26192 | 11.1982051315981 | -3.22636727868396 | 0.608011386172673 | -5.30642575461191 | 1.11795665971802e-07 | 3.61431134990849e-05 | 2626 | 0 | 0 | 11 | 17 | 1 | 1 | 5 | 3 | 4 | 8 | 46 | 53 |
| L1ME4b_dup16165 | 13.934363818057 | -3.58009796864564 | 0.561825592002302 | -6.37225861478907 | 1.86264445237484e-10 | 1.73511255455854e-07 | 2627 | 1 | 0 | 10 | 12 | 1 | 1 | 6 | 5 | 2 | 7 | 65 | 76 |
| MIR_dup133527 | 13.4415774641464 | -3.44363105916155 | 0.570087252040709 | -6.04053335140293 | 1.53605628839818e-09 | 1.03248149604292e-06 | 2628 | 0 | 0 | 15 | 10 | 0 | 1 | 11 | 3 | 6 | 8 | 53 | 72 |
| MLT1G_dup2161 | 15.14096474612 | -3.34134059042792 | 0.554504643982801 | -6.02581173428649 | 1.68262958531449e-09 | 1.11836586398671e-06 | 2629 | 1 | 0 | 13 | 15 | 1 | 1 | 12 | 1 | 7 | 6 | 70 | 75 |
| MIRb_dup170322 | 14.8253718162838 | -3.69223898549907 | 0.599087612701368 | -6.16310353814571 | 7.13328316376129e-10 | 5.4401801143729e-07 | 2630 | 0 | 1 | 2 | 8 | 0 | 0 | 7 | 4 | 6 | 5 | 60 | 107 |
| L1MD2_dup8422 | 11.2071541668101 | -3.58481867371175 | 0.630260940625143 | -5.68783251926741 | 1.28661910325587e-08 | 6.325327874862e-06 | 2631 | 0 | 0 | 4 | 11 | 1 | 0 | 15 | 6 | 6 | 4 | 54 | 46 |
| L3b_dup8003 | 16.8436960930612 | -3.35892800371576 | 0.496927853301681 | -6.75938766844809 | 1.38576150898918e-11 | 1.85245173043786e-08 | 2632 | 1 | 1 | 9 | 16 | 1 | 0 | 9 | 6 | 9 | 5 | 81 | 84 |
| L1PA10_dup5521 | 13.8639377394893 | -3.46798739656157 | 0.81345559943296 | -4.2632780436683 | 2.01449594511948e-05 | 0.00205197453834503 | 2633 | 0 | 0 | 29 | 36 | 0 | 1 | 15 | 3 | 11 | 23 | 24 | 41 |
| L1MCc_dup3254 | 12.7926444671103 | -3.17104631637005 | 0.549146669254675 | -5.77449795092798 | 7.71828690717074e-09 | 4.06313162923373e-06 | 2634 | 0 | 0 | 18 | 18 | 0 | 1 | 13 | 10 | 7 | 12 | 32 | 57 |
| MLT1E1A_dup2913 | 13.6138262432825 | -3.06863211157075 | 0.638744554056459 | -4.80416168260514 | 1.5540121957388e-06 | 0.000291158256635642 | 2635 | 0 | 1 | 18 | 29 | 0 | 1 | 17 | 5 | 15 | 9 | 32 | 52 |
| L1PA13_dup1610 | 12.3751882342154 | -2.2201933212765 | 0.620586594231158 | -3.57757215820475 | 0.000346800497919724 | 0.0165902274382804 | 2636 | 0 | 0 | 9 | 15 | 1 | 1 | 12 | 8 | 16 | 26 | 25 | 44 |
| L1PA8_dup1656 | 12.8461243621459 | -2.50468189277998 | 0.683840949819593 | -3.66266731093064 | 0.000249602620938059 | 0.0131050188088542 | 2637 | 0 | 0 | 14 | 18 | 0 | 2 | 19 | 8 | 18 | 26 | 21 | 37 |
| LTR33B_dup361 | 13.3295994461853 | -3.17713199703532 | 0.756980717822835 | -4.19711086720032 | 2.7034161050583e-05 | 0.00254860161859826 | 2638 | 0 | 0 | 16 | 19 | 0 | 1 | 19 | 19 | 23 | 13 | 28 | 25 |
| THE1C-int_dup384 | 15.6268241764652 | -2.94632995063902 | 0.647973874657287 | -4.54698879981439 | 5.44188929962295e-06 | 0.000768016483563513 | 2639 | 0 | 0 | 21 | 20 | 0 | 2 | 17 | 13 | 18 | 19 | 47 | 40 |
| GSATX_dup53 | 8.70909807492629 | -2.64566462244501 | 0.776717771451071 | -3.40621100699468 | 0.000658712471239089 | 0.0261972250846492 | 2640 | 1 | 0 | 22 | 16 | 1 | 0 | 14 | 8 | 11 | 12 | 5 | 21 |
| L1PA2_dup4059 | 12.7424576047826 | -2.68057785894564 | 0.640463780899555 | -4.18536994423115 | 2.84702040825026e-05 | 0.00265557474739912 | 2641 | 2 | 1 | 28 | 30 | 1 | 0 | 23 | 8 | 9 | 19 | 17 | 29 |
| L1M1_dup9379 | 11.3872860161728 | -2.71923353033386 | 0.837024309115059 | -3.24869122762845 | 0.00115937251882873 | 0.0389094574563077 | 2642 | 1 | 0 | 14 | 25 | 0 | 0 | 3 | 3 | 15 | 28 | 19 | 38 |
| LOR1-int_dup1312 | 7.5032180817586 | -2.41769796750179 | 0.751022144305904 | -3.21920996049488 | 0.00128544326733052 | 0.0417678661324905 | 2643 | 0 | 1 | 15 | 11 | 0 | 0 | 6 | 6 | 10 | 14 | 12 | 19 |
| AluYm1_dup986 | 8.28578500413203 | -2.50183849131581 | 0.737810645143085 | -3.39089508640883 | 0.000696647634929149 | 0.0272370223695122 | 2644 | 0 | 0 | 8 | 7 | 0 | 0 | 7 | 8 | 13 | 15 | 21 | 23 |
| MER57C1_dup175 | 9.15597771821002 | -2.48410053825431 | 0.763813379416688 | -3.2522349112965 | 0.00114501346576659 | 0.0385689940720975 | 2645 | 1 | 0 | 8 | 19 | 0 | 0 | 5 | 9 | 17 | 15 | 17 | 22 |
| AluSq2_dup11329 | 7.53536814929433 | -2.33260604337511 | 0.631020772345264 | -3.69655983701726 | 0.000218540767508667 | 0.0118642257507682 | 2646 | 2 | 2 | 7 | 13 | 0 | 0 | 15 | 12 | 3 | 11 | 18 | 13 |
| L1MD2_dup2108 | 7.82319485929988 | -2.14870611330377 | 0.558766814928689 | -3.84544331534432 | 0.00012033468133905 | 0.00777019161082808 | 2647 | 2 | 6 | 7 | 8 | 0 | 0 | 15 | 7 | 3 | 6 | 24 | 24 |
| L1PB4_dup7311 | 10.9532940784743 | -2.49792244987594 | 0.55403687770347 | -4.50858516896933 | 6.52613704919452e-06 | 0.000881810452190595 | 2648 | 4 | 3 | 6 | 20 | 0 | 0 | 9 | 4 | 3 | 9 | 39 | 50 |
| MIR3_dup66178 | 11.6611230087294 | -2.57625110339394 | 0.488263773483957 | -5.27635111040771 | 1.31781487735978e-07 | 4.12588740705812e-05 | 2649 | 2 | 1 | 12 | 16 | 0 | 0 | 15 | 6 | 9 | 11 | 39 | 41 |
| MLT1D-int_dup506 | 11.3111897355812 | -2.49261901957202 | 0.468851382536732 | -5.31643738808157 | 1.05818611150773e-07 | 3.44919386970984e-05 | 2650 | 2 | 1 | 10 | 12 | 0 | 1 | 10 | 5 | 7 | 10 | 43 | 47 |
| L1MB1_dup5919 | 11.3903693976293 | -2.65206307858928 | 0.582004332982702 | -4.55677548824728 | 5.19448953379463e-06 | 0.000743687127923164 | 2651 | 1 | 2 | 11 | 10 | 0 | 0 | 4 | 12 | 11 | 7 | 30 | 58 |
| L1MC5a_dup16544 | 11.753721936 | -2.61798425289254 | 0.578705359880387 | -4.52386384227313 | 6.07207951509044e-06 | 0.000836610904630985 | 2652 | 2 | 1 | 4 | 16 | 0 | 0 | 11 | 11 | 14 | 6 | 40 | 45 |
| LTR8A_dup1384 | 9.54123302947042 | -3.12254056704268 | 0.567723591913113 | -5.50010711466183 | 3.79560597425933e-08 | 1.5438448874378e-05 | 2653 | 1 | 1 | 7 | 10 | 0 | 0 | 11 | 7 | 6 | 4 | 40 | 37 |
| L2a_dup91381 | 6.5477186186468 | -2.54266966065669 | 0.676991511781332 | -3.7558368405038 | 0.000172763223795403 | 0.0100656998161658 | 2654 | 1 | 1 | 14 | 23 | 0 | 1 | 6 | 6 | 6 | 4 | 11 | 12 |
| THE1D-int_dup1446 | 6.56869668851996 | -2.34632305151514 | 0.628136775064097 | -3.73536965937986 | 0.000187439644370107 | 0.0106343618548616 | 2655 | 2 | 1 | 11 | 23 | 1 | 0 | 9 | 8 | 5 | 5 | 11 | 9 |
| MLT1D_dup4082 | 6.26004639547645 | -2.7522659587827 | 0.678727822776776 | -4.05503629941494 | 5.01265179380479e-05 | 0.00408752722319628 | 2656 | 0 | 0 | 7 | 9 | 0 | 0 | 8 | 6 | 5 | 8 | 19 | 18 |
| MER67C_dup1466 | 4.83084804313033 | -2.92990351367594 | 0.793209513807361 | -3.69373218887979 | 0.000220986541008762 | 0.0119588045228271 | 2657 | 0 | 0 | 10 | 8 | 0 | 0 | 4 | 6 | 4 | 5 | 12 | 12 |
| THE1C_dup2045 | 4.93095162392492 | -2.53717199950444 | 0.786460338239483 | -3.226064781835 | 0.00125504946434618 | 0.0411002382040865 | 2658 | 0 | 0 | 6 | 8 | 0 | 0 | 3 | 6 | 5 | 8 | 11 | 15 |
| MLT2A2_dup3045 | 6.03244837606751 | -2.97224633856532 | 0.938467402879529 | -3.16712794652801 | 0.00153952567074476 | 0.0473194107654694 | 2659 | 0 | 0 | 13 | 13 | 0 | 0 | 10 | 4 | 6 | 15 | 7 | 9 |
| THE1D-int_dup468 | 6.19402889570541 | -3.06084533769529 | 0.808547220102672 | -3.78561110791601 | 0.000153331305652736 | 0.00924980426695434 | 2660 | 0 | 0 | 13 | 9 | 0 | 0 | 13 | 3 | 7 | 6 | 14 | 15 |
| AluSp_dup49092 | 9.09237692403508 | -3.78376017402972 | 0.681364110399428 | -5.55321320316037 | 2.80465650200782e-08 | 1.21117385848775e-05 | 2661 | 1 | 0 | 11 | 17 | 0 | 0 | 10 | 3 | 3 | 4 | 29 | 45 |
| Charlie4a_dup2050 | 8.9127069721716 | -4.71527342137901 | 0.821123048345505 | -5.74246872119824 | 9.33061181347376e-09 | 4.80558822201045e-06 | 2662 | 0 | 0 | 10 | 20 | 0 | 0 | 6 | 4 | 3 | 1 | 33 | 43 |
| Kanga1_dup687 | 8.24515114019912 | -4.00536193699848 | 0.706516873999694 | -5.66916670273357 | 1.43493699378108e-08 | 6.98236232969799e-06 | 2663 | 0 | 0 | 11 | 19 | 0 | 1 | 15 | 8 | 4 | 1 | 24 | 26 |
| L1PA7_dup2715 | 7.68083509458806 | -3.23244333138066 | 0.660822613579313 | -4.8915446671419 | 1.0004768210377e-06 | 0.000205400739998824 | 2664 | 0 | 0 | 12 | 10 | 0 | 1 | 5 | 8 | 4 | 5 | 20 | 35 |
| THE1B_dup14440 | 8.43448291266638 | -4.10617760893059 | 0.769225154215978 | -5.33806985695333 | 9.39412662916455e-08 | 3.13945906587526e-05 | 2665 | 0 | 0 | 23 | 15 | 0 | 1 | 6 | 8 | 3 | 2 | 19 | 34 |
| L1MD3_dup4072 | 9.56317796544186 | -3.05041125302414 | 0.683771339474944 | -4.46115693495796 | 8.15183641706509e-06 | 0.0010439703272847 | 2666 | 3 | 0 | 10 | 31 | 0 | 1 | 8 | 3 | 2 | 8 | 23 | 42 |
| L1PA11_dup3581 | 7.0673970889055 | -3.32544618810281 | 0.714929386924803 | -4.65143306307058 | 3.29636273327138e-06 | 0.000517044381628868 | 2667 | 3 | 0 | 11 | 24 | 0 | 1 | 13 | 2 | 1 | 3 | 19 | 20 |
| L1P1_dup2815 | 7.23615593607627 | -2.35177707433175 | 0.644415082326155 | -3.64947553034064 | 0.000262776268432704 | 0.0136193774708099 | 2668 | 1 | 0 | 7 | 18 | 1 | 1 | 4 | 3 | 8 | 5 | 23 | 23 |
| L1MA6_dup1086 | 7.63956516448611 | -2.21279196761462 | 0.585619948713754 | -3.77854609030098 | 0.000157746658893608 | 0.00944782328866408 | 2669 | 2 | 0 | 12 | 12 | 0 | 1 | 8 | 6 | 10 | 6 | 19 | 21 |
| L1ME4a_dup26456 | 8.06538258241429 | -1.86754476254488 | 0.530896822735684 | -3.51771696978994 | 0.000435276348205598 | 0.0196391699884357 | 2670 | 2 | 0 | 6 | 10 | 0 | 3 | 6 | 4 | 9 | 7 | 31 | 25 |
| L1M2_dup849 | 11.9167606067793 | -2.96788946509743 | 0.761631156884473 | -3.89675427307606 | 9.74904449804718e-05 | 0.00669093205114604 | 2671 | 2 | 4 | 37 | 48 | 0 | 3 | 5 | 4 | 7 | 0 | 20 | 30 |
| L1PB_dup1449 | 16.4643836294396 | -3.64182951524021 | 0.611778138890822 | -5.95285984203847 | 2.63496944912492e-09 | 1.64561795417711e-06 | 2672 | 2 | 1 | 30 | 40 | 0 | 4 | 16 | 8 | 7 | 0 | 37 | 78 |
| L1MC1_dup4044 | 13.7121294491946 | -5.8684802416182 | 1.37545476663373 | -4.26657450610363 | 1.98497223319714e-05 | 0.00203409217485068 | 2673 | 0 | 0 | 43 | 57 | 0 | 0 | 1 | 4 | 4 | 0 | 34 | 43 |
| AluSx_dup86067 | 8.24113933814984 | -4.23638083250856 | 0.934010893533156 | -4.53568674823831 | 5.7416347220935e-06 | 0.000798479845460701 | 2674 | 0 | 0 | 20 | 30 | 2 | 0 | 4 | 2 | 3 | 0 | 21 | 30 |
| ERV3-16A3_I-int_dup1[illegible] | 10.0616050269566 | -4.8995931453914 | 0.868134424970875 | -5.6438185198747 | 1.6631938977893e-08 | 7.88347281281621e-06 | 2675 | 0 | 0 | 27 | 25 | 1 | 0 | 9 | 7 | 3 | 0 | 22 | 41 |
| HERVL-int_dup499 | 9.63811271825331 | -3.64585377614935 | 0.710065877973299 | -5.13452890674807 | 2.82851619519417e-07 | 7.62213040296345e-05 | 2676 | 0 | 2 | 26 | 24 | 1 | 0 | 7 | 3 | 3 | 3 | 27 | 33 |
| MER21B_dup569 | 8.94688376528874 | -3.73891181644119 | 0.712945124220775 | -5.24431921815539 | 1.56860420018391e-07 | 4.74260603579367e-05 | 2677 | 0 | 0 | 17 | 27 | 2 | 0 | 9 | 4 | 4 | 2 | 25 | 30 |
| MER52A_dup1158 | 7.134450608924 | -3.68523552696242 | 0.725996452963678 | -5.07610679352285 | 3.85246819891139e-07 | 9.76229390903269e-05 | 2678 | 1 | 1 | 17 | 14 | 2 | 0 | 5 | 7 | 1 | 1 | 20 | 25 |
| Tigger19b_dup2906 | 7.55800566366721 | -3.36415161887346 | 0.753468918678637 | -4.46488439731953 | 8.01119955885562e-06 | 0.0010326288679477 | 2679 | 0 | 2 | 10 | 24 | 1 | 0 | 3 | 5 | 4 | 1 | 25 | 25 |
| L1MC3_dup10659 | 13.8753212533646 | -3.71276437009262 | 0.619337799213453 | -5.99473239774443 | 2.03820794518528e-09 | 1.31076602087853e-06 | 2680 | 0 | 0 | 25 | 41 | 5 | 0 | 14 | 9 | 4 | 3 | 32 | 54 |
| MLT1F2_dup4803 | 14.9841107993572 | -3.38499049502165 | 0.52520207703235 | -6.44512016050757 | 1.15508864981759e-10 | 1.15482657088023e-07 | 2681 | 1 | 2 | 40 | 34 | 3 | 1 | 12 | 14 | 5 | 4 | 44 | 35 |

| | | | | | | | | | | | | | | | | | | |
|---|---|---|---|---|---|---|---|---|---|---|---|---|---|---|---|---|---|---|
| Tigger3b_dup5415 | 16.7250867883532 | -3.43665591608264 | 0.578358779559747 | -5.94208307635385 | 2.81422723636238e-09 | 1.73032070796766e-06 | 2682 | 1 | 1 | 34 | 57 | 3 | 0 | 12 | 17 | 6 | 9 | 45 | 34 |
| ERVL47-int_dup131 | 13.653401360869 | -5.19271665342762 | 0.822636579780832 | -6.31228513423397 | 2.74945048921688e-10 | 2.45026496669363e-07 | 2683 | 0 | 0 | 24 | 53 | 0 | 0 | 13 | 9 | 3 | 2 | 37 | 44 |
| L1MC3_dup8112 | 15.6245717759221 | -4.7308191883871 | 0.67521992516033 | -7.00633824936931 | 2.44636080268293e-12 | 3.95987596976321e-09 | 2684 | 0 | 0 | 32 | 35 | 0 | 0 | 20 | 8 | 4 | 4 | 47 | 60 |
| L1MC3_dup10664 | 10.8127312709186 | -4.56645249384322 | 0.743539688727855 | -6.14150470118967 | 8.17434059797915e-10 | 6.11651461612184e-07 | 2685 | 0 | 0 | 16 | 38 | 0 | 0 | 21 | 8 | 2 | 4 | 25 | 33 |
| L1MC3_dup10665 | 12.5019167289293 | -4.29634433559731 | 0.651526374438788 | -6.5942753880042 | 4.27337840519851e-11 | 4.86523109092519e-08 | 2686 | 1 | 0 | 35 | 26 | 0 | 0 | 21 | 9 | 3 | 4 | 33 | 34 |
| MER66C_dup450 | 12.8628635045072 | -3.55153009223478 | 0.563570009723994 | -6.30184365909414 | 2.94125773675186e-10 | 2.56359162428263e-07 | 2687 | 0 | 1 | 17 | 33 | 1 | 1 | 10 | 10 | 5 | 5 | 37 | 51 |
| L1MC3_dup10694 | 13.8396494277886 | -3.47375638175857 | 0.594908779025393 | -5.8391412334668 | 5.24705761013189e-09 | 2.92392592529378e-06 | 2688 | 0 | 3 | 30 | 36 | 0 | 1 | 13 | 11 | 8 | 3 | 40 | 36 |
| MLT1A0_dup13174 | 13.9298131402901 | -3.88621905660405 | 0.671393958116614 | -5.78828422511519 | 7.11090166104305e-09 | 3.78525862782673e-06 | 2689 | 0 | 2 | 36 | 42 | 0 | 0 | 10 | 13 | 7 | 4 | 30 | 39 |
| AluJr_dup63191 | 13.16373135596 | -2.72098104167061 | 0.543349231935084 | -5.00779403327783 | 5.50574048117679e-07 | 0.000128943791784852 | 2690 | 4 | 1 | 25 | 34 | 0 | 4 | 16 | 11 | 1 | 13 | 31 | 35 |
| L1MC3_dup10657 | 13.2682724761084 | -3.82896949487884 | 0.548419366828831 | -6.9818276422647 | 2.91364458624064e-12 | 4.56111891261589e-09 | 2691 | 1 | 1 | 29 | 27 | 0 | 1 | 20 | 12 | 3 | 5 | 28 | 50 |
| L1MC3_dup10667 | 10.9764101450956 | -3.34245521813462 | 0.540656832676525 | -6.18221210964406 | 6.32095153280556e-10 | 4.91518017459135e-07 | 2692 | 3 | 1 | 23 | 27 | 0 | 1 | 14 | 13 | 5 | 2 | 23 | 32 |
| AluSx_dup86077 | 12.2583062825863 | -2.8780016976855 | 0.594104729855009 | -4.84426659654414 | 1.27080206235937e-06 | 0.000247246334857043 | 2693 | 3 | 3 | 30 | 32 | 0 | 1 | 5 | 9 | 5 | 7 | 36 | 29 |
| LTR12F_dup493 | 14.6217005448684 | -2.33042367294736 | 0.436405209560768 | -5.34004549417016 | 9.29232598612274e-08 | 3.11857799590121e-05 | 2694 | 5 | 6 | 28 | 29 | 0 | 4 | 14 | 9 | 6 | 8 | 32 | 53 |
| L1PA4_dup2487 | 14.0409905468758 | -3.66762230719867 | 0.687184113298975 | -5.33717563636834 | 9.44055818076421e-08 | 3.15055127192163e-05 | 2695 | 2 | 2 | 34 | 42 | 0 | 1 | 8 | 2 | 3 | 5 | 49 | 41 |
| L1PA4_dup1499 | 14.0531607128549 | -3.89471544868047 | 0.792851865481791 | -4.91228641596722 | 9.00204218282592e-07 | 0.000190399993709751 | 2696 | 6 | 2 | 42 | 59 | 0 | 0 | 6 | 3 | 1 | 2 | 31 | 41 |
| L1MB1_dup509 | 11.9276135171328 | -3.26264893513452 | 0.706499752949267 | -4.61804681673938 | 3.87368870081817e-06 | 0.000584853256094467 | 2697 | 3 | 3 | 37 | 37 | 0 | 0 | 7 | 5 | 1 | 8 | 28 | 31 |
| L1MC3_dup1150 | 11.2564911749304 | -3.86923057918652 | 0.73310539925647 | -5.27786397850946 | 1.30698420550927e-07 | 4.10279239794337e-05 | 2698 | 2 | 2 | 36 | 32 | 0 | 0 | 4 | 6 | 2 | 3 | 30 | 33 |
| L1ME1_dup3529 | 13.101505868906 | -2.40986836926687 | 0.648578240234565 | -3.7156170524551 | 0.000202708396524191 | 0.0112356853074643 | 2699 | 6 | 6 | 35 | 41 | 0 | 2 | 8 | 2 | 7 | 4 | 27 | 38 |
| L1M6B_dup74 | 10.6800909573992 | -2.91750559149278 | 0.714057786685806 | -4.08581160501582 | 4.39230081525615e-05 | 0.00370350960236336 | 2700 | 3 | 3 | 32 | 48 | 0 | 1 | 5 | 3 | 4 | 5 | 15 | 26 |
| MER68_dup117 | 13.2206698618954 | -2.90042884395628 | 0.719129483045114 | -4.03324980040393 | 5.50107474259994e-05 | 0.00438218523837524 | 2701 | 5 | 2 | 42 | 60 | 0 | 3 | 6 | 2 | 3 | 7 | 24 | 26 |
| L1MC4a_dup971 | 9.2419698822971 | -2.53285176891688 | 0.699775228268094 | -3.61952190732094 | 0.000295147824100814 | 0.0148006836993661 | 2702 | 8 | 2 | 28 | 30 | 0 | 3 | 4 | 3 | 2 | 2 | 23 | 19 |
| MLT1A_dup904 | 12.9475539683767 | -3.10627011768403 | 0.677737534301274 | -4.58329362101288 | 4.57708614446714e-06 | 0.000671868809211214 | 2703 | 6 | 5 | 41 | 55 | 0 | 2 | 8 | 4 | 1 | 3 | 27 | 24 |
| L1M8_dup165 | 10.9689262931391 | -3.31780063650142 | 0.839850848214114 | -3.95046411342741 | 7.79997930375985e-05 | 0.00566960116496832 | 2704 | 3 | 1 | 33 | 47 | 0 | 0 | 1 | 5 | 5 | 6 | 24 | 21 |
| L1MEf_dup1148 | 13.6510006751958 | -3.54697342053906 | 0.785086717328464 | -4.51793839107213 | 6.24446463200344e-06 | 0.000852953720623817 | 2705 | 3 | 3 | 39 | 59 | 0 | 0 | 3 | 3 | 3 | 5 | 29 | 39 |
| MLT1E3_dup170 | 12.288561714611 | -2.44518455508511 | 0.765868666317513 | -3.1926943386288 | 0.00140952061896674 | 0.0445168294665064 | 2706 | 7 | 4 | 32 | 45 | 0 | 0 | 2 | 4 | 5 | 9 | 22 | 35 |
| HAL1_dup2713 | 11.1689121868502 | -3.14764403727399 | 0.744055952271276 | -4.23038620639431 | 2.33290435070964e-05 | 0.0022947716355269 | 2707 | 2 | 4 | 31 | 44 | 0 | 0 | 4 | 8 | 2 | 7 | 11 | 38 |
| L1ME1_dup3531 | 10.953221568951 | -3.12756414950881 | 0.733796886566894 | -4.26216601182554 | 2.02454947982981e-05 | 0.00205957011768954 | 2708 | 5 | 5 | 27 | 50 | 0 | 0 | 4 | 7 | 1 | 3 | 18 | 29 |
| HAL1_dup2716 | 11.333783314657 | -3.02290210178239 | 0.73853725119961 | -4.09309360749546 | 4.25655780958189e-05 | 0.00361338175012048 | 2709 | 3 | 8 | 34 | 39 | 0 | 0 | 5 | 5 | 2 | 2 | 26 | 28 |
| MLT1A-int_dup49 | 11.0248806143543 | -3.45941792358723 | 0.749244677767538 | -4.61720720378685 | 3.88938878281651e-06 | 0.000586851302039352 | 2710 | 1 | 6 | 33 | 39 | 0 | 0 | 4 | 7 | 2 | 2 | 26 | 27 |
| L1MB1_dup507 | 9.44029191069824 | -2.62196281818149 | 0.782147601539515 | -3.35226089426169 | 0.000801544453470806 | 0.0300921894170976 | 2711 | 3 | 4 | 24 | 40 | 0 | 0 | 3 | 2 | 6 | 4 | 22 | 18 |
| L1MA6_dup2888 | 7.98532657250538 | -2.63237196569743 | 0.787940397135845 | -3.34082625445537 | 0.000835294772540737 | 0.0310022186104502 | 2712 | 2 | 2 | 22 | 34 | 0 | 0 | 5 | 3 | 8 | 4 | 14 | 11 |
| LTR107_Mam_dup45 | 9.69454145496481 | -2.5906347544983 | 0.714536172780604 | -3.62561736296273 | 0.000288271935886478 | 0.0145355274542157 | 2713 | 5 | 2 | 29 | 37 | 0 | 0 | 6 | 5 | 7 | 4 | 17 | 16 |
| L2a_dup20681 | 11.1728091803877 | -3.07520984520467 | 0.697527585790302 | -4.40872864077546 | 1.03979209426228e-05 | 0.0012591062069279 | 2714 | 2 | 3 | 29 | 36 | 0 | 0 | 6 | 3 | 6 | 5 | 28 | 31 |
| MER4A_dup76 | 10.6682104129508 | -3.09078794244083 | 0.694037266615887 | -4.45334579440014 | 8.4542432672747e-06 | 0.0010734543054829 | 2715 | 2 | 2 | 28 | 41 | 1 | 0 | 6 | 3 | 5 | 5 | 25 | 25 |
| L1PA8A_dup243 | 11.923793799097 | -3.14333669740715 | 0.730742916248619 | -4.30156300870346 | 1.69597531973757e-05 | 0.00180919355428757 | 2716 | 3 | 3 | 44 | 49 | 0 | 0 | 9 | 4 | 4 | 6 | 20 | 18 |
| MLT1B_dup1713 | 9.93173308787889 | -3.15625231715255 | 0.757945815819263 | -4.16421893396292 | 3.12420223130429e-05 | 0.00284606211382056 | 2717 | 1 | 3 | 37 | 35 | 0 | 0 | 5 | 5 | 4 | 6 | 14 | 22 |
| HAL1_dup2711 | 13.60483022926 | -2.48702249090807 | 0.760593821726939 | -3.26984314079918 | 0.00107607136737034 | 0.0369476013824392 | 2718 | 7 | 5 | 42 | 61 | 0 | 0 | 6 | 6 | 9 | 5 | 26 | 13 |
| L1M3b_dup43 | 14.8149584984098 | -3.55861086674097 | 0.801069227250866 | -4.44232626305409 | 8.89914829759924e-06 | 0.00111624498725385 | 2719 | 4 | 1 | 41 | 79 | 0 | 0 | 7 | 3 | 6 | 6 | 28 | 27 |
| Tigger1_dup1273 | 13.7144543166936 | -2.93921937483371 | 0.716840003902925 | -4.10024462757486 | 4.12713639935437e-05 | 0.00353377329140257 | 2720 | 2 | 6 | 34 | 63 | 0 | 0 | 10 | 4 | 6 | 7 | 22 | 32 |
| ERVL47-int_dup135 | 11.5845865123945 | -3.04387847286318 | 0.603994397010941 | -5.03958064499734 | 4.66553005530102e-07 | 0.000113808977408996 | 2721 | 3 | 1 | 30 | 17 | 2 | 0 | 7 | 5 | 6 | 3 | 32 | 47 |
| L1ME3F_dup3795 | 10.3687889753011 | -2.97390574409921 | 0.691186745128611 | -4.30260818086991 | 1.68799206411343e-05 | 0.0018043619033582 | 2722 | 1 | 3 | 25 | 20 | 0 | 0 | 11 | 1 | 7 | 5 | 33 | 31 |
| L1MCc_dup338 | 9.7837699186214 | -3.16402439619263 | 0.786425970797579 | -4.02329591555037 | 5.73893153417946e-05 | 0.00451572686121649 | 2723 | 0 | 5 | 33 | 41 | 0 | 1 | 3 | 5 | 3 | 3 | 16 | 21 |
| L1PA3_dup9510 | 10.7454338503914 | -4.37158401468693 | 0.753365938451542 | -5.80273648112128 | 6.5241284028099e-09 | 3.52815967485228e-06 | 2724 | 0 | 2 | 29 | 32 | 0 | 1 | 7 | 7 | 1 | 2 | 29 | 34 |
| L1MA4_dup6617 | 11.2142385320537 | -3.83891097276226 | 0.74949008115934 | -5.12203039008079 | 3.02263160064256e-07 | 8.04546599502012e-05 | 2725 | 0 | 0 | 26 | 35 | 0 | 1 | 8 | 8 | 8 | 3 | 24 | 35 |
| LTR82B_dup75 | 9.50586444725773 | -4.07221239910744 | 0.816969697823847 | -4.98453297589195 | 6.21116527122177e-07 | 0.000141835118198286 | 2726 | 0 | 0 | 26 | 39 | 0 | 1 | 5 | 6 | 3 | 4 | 22 | 21 |
| L1PA14_dup320 | 12.1668801736557 | -3.19490317455568 | 0.765105818329189 | -4.17576640775337 | 2.96984456428943e-05 | 0.00273794124252 | 2727 | 4 | 0 | 32 | 51 | 0 | 1 | 7 | 4 | 9 | 3 | 18 | 35 |
| THE1D-int_dup464 | 9.04114252118171 | -2.69272248281223 | 0.682892488468701 | -3.94311334255604 | 8.04306194856982e-05 | 0.00580471464487229 | 2728 | 3 | 0 | 22 | 22 | 0 | 2 | 3 | 5 | 7 | 4 | 18 | 33 |
| THE1B_dup14451 | 50.9710805822221 | -3.59674283770736 | 1.13911645940496 | -3.15748474004685 | 0.00159136565611626 | 0.0483880993382834 | 2729 | 5 | 2 | 5 | 3 | 5 | 5 | 262 | 245 | 22 | 16 | 18 | 14 |
| MST-int_dup131 | 28.5419032864562 | -3.99225248303946 | 1.18545392580824 | -3.3676994070584 | 0.000757981895410359 | 0.0289542450226924 | 2730 | 1 | 0 | 8 | 4 | 2 | 3 | 170 | 103 | 12 | 21 | 18 | 9 |
| MSTA-int_dup2074 | 33.1182007979827 | -3.31270032488823 | 1.05231937025678 | -3.1479989996572 | 0.00164392255929975 | 0.0495396146523733 | 2731 | 0 | 3 | 4 | 11 | 6 | 6 | 186 | 131 | 7 | 15 | 19 | 19 |
| L1PREC2_dup3173 | 37.6649147300938 | -3.99587256965464 | 1.16970670597094 | -3.41613205195551 | 0.000635174375625954 | 0.025534245984574 | 2732 | 0 | 2 | 23 | 33 | 2 | 4 | 214 | 137 | 16 | 23 | 3 | 9 |
| THE1A-int_dup1311 | 36.7106175380519 | -3.5220261492752 | 1.11486102156156 | -3.15916161849662 | 0.00158223720233006 | 0.0481687581046096 | 2733 | 3 | 0 | 16 | 24 | 2 | 5 | 163 | 128 | 44 | 24 | 18 | 11 |
| MER126_dup203 | 35.6688347077098 | -5.1196576849923 | 1.27966071578698 | -4.00079303977362 | 6.3130554618624e-05 | 0.00485823510649519 | 2734 | 2 | 5 | 0 | 1 | 0 | 1 | 142 | 120 | 2 | 2 | 68 | 105 |
| MER58A_dup5209 | 75.7191837735987 | -3.85764823015848 | 1.1319277054297 | -3.40803411000001 | 0.000654327148041085 | 0.0260751176633368 | 2735 | 5 | 10 | 2 | 2 | 5 | 12 | 385 | 426 | 3 | 5 | 12 | 18 |
| MLT1J_dup6287 | 59.9521068273588 | -3.84587930720676 | 1.13693350986766 | -3.38267741589781 | 0.000717828953793392 | 0.0278455376979658 | 2736 | 11 | 6 | 1 | 1 | 8 | 16 | 363 | 282 | 2 | 1 | 9 | 30 |
| THE1A_dup3088 | 46.7121390912454 | -2.59498771084381 | 0.81816241966279 | -3.17172684601347 | 0.00151535466222656 | 0.0468397740267811 | 2737 | 19 | 12 | 9 | 20 | 2 | 9 | 208 | 168 | 20 | 17 | 42 | 47 |
| MIRb_dup148188 | 29.946172748025 | -2.9926340099111 | 0.635462218947943 | -4.70938148748739 | 2.48469635761308e-06 | 0.000417236104099225 | 2738 | 14 | 6 | 13 | 7 | 4 | 5 | 147 | 94 | 6 | 5 | 35 | 41 |
| MER5B_dup16930 | 21.2026004423608 | -2.09102380712299 | 0.64667734784269 | -3.23348856133375 | 0.00122288233176704 | 0.0403522339917682 | 2739 | 13 | 7 | 6 | 8 | 5 | 3 | 75 | 66 | 14 | 6 | 32 | 24 |
| MIRb_dup148189 | 25.4007089050168 | -2.12111598416429 | 0.632924598264069 | -3.3512933293822 | 0.000804350528231445 | 0.0301594690813992 | 2740 | 16 | 6 | 6 | 8 | 6 | 7 | 95 | 84 | 10 | 12 | 31 | 31 |
| MLT1C_dup42 | 9.26061278233208 | -4.35335393397359 | 1.11729898137026 | -3.89631961235177 | 9.76654876460684e-05 | 0.0066990810387522 | 2741 | 0 | 0 | 1 | 1 | 0 | 0 | 37 | 19 | 7 | 7 | 16 | 30 |
| THE1B_dup16428 | 11.2101562428427 | -5.05137332932144 | 1.15048747808327 | -4.39063738245733 | 1.13018881498733e-05 | 0.00133707506599419 | 2742 | 0 | 1 | 0 | 1 | 0 | 1 | 41 | 44 | 1 | 1 | 28 | 18 |
| THE1D_dup9311 | 15.9607113347927 | -5.84721389797242 | 1.60180940323993 | -3.65038055473107 | 0.000261852058835142 | 0.0135834810402734 | 2743 | 1 | 0 | 0 | 0 | 0 | 0 | 45 | 52 | 1 | 2 | 48 | 49 |
| Harlequin-int_dup377 | 10.2922413274194 | -5.59332620050486 | 1.05516118774351 | -5.30092109667752 | 1.15219880754131e-07 | 3.69987985774932e-05 | 2744 | 0 | 0 | 0 | 2 | 0 | 0 | 17 | 7 | 1 | 0 | 45 | 67 |
| LTR78_dup623 | 8.27865009982484 | -4.19880325405226 | 1.03129613252143 | -4.07138466018149 | 4.67345014839771e-05 | 0.00388142676792545 | 2745 | 0 | 0 | 0 | 0 | 0 | 0 | 9 | 7 | 1 | 4 | 36 | 53 |
| L2c_dup100858 | 6.22474957802797 | -3.52032066247144 | 1.02967846494501 | -3.41885431454511 | 0.000628853860171847 | 0.0253661558676379 | 2746 | 1 | 0 | 1 | 0 | 0 | 1 | 30 | 10 | 2 | 3 | 19 | 14 |
| MIRc_dup71233 | 4.75899339316483 | -3.36519561846534 | 1.05181068072248 | -3.19943092435019 | 0.00137699180092646 | 0.0438444655510837 | 2747 | 0 | 1 | 1 | 0 | 0 | 2 | 20 | 10 | 1 | 1 | 14 | 11 |
| MIR_dup120595 | 5.68919295288361 | -4.19196893265566 | 1.1998947826395 | -3.49361376789578 | 0.000476529892942685 | 0.0208511184086319 | 2748 | 1 | 0 | 0 | 0 | 0 | 0 | 12 | 9 | 2 | 0 | 21 | 29 |
| L2c_dup100859 | 8.10704345168391 | -4.33115256553172 | 1.16488547981404 | -3.71809301479414 | 0.000200732364761329 | 0.0111492678024041 | 2749 | 1 | 0 | 0 | 0 | 0 | 0 | 30 | 14 | 2 | 3 | 29 | 26 |
| MIR_dup120597 | 4.70209117989265 | -3.86142962275893 | 1.17829192583967 | -3.27714171512058 | 0.00104863722846046 | 0.0362725736245463 | 2750 | 0 | 0 | 0 | 0 | 1 | 0 | 16 | 9 | 1 | 2 | 13 | 19 |
| THE1D-int_dup5 | 4.7778800070341 | -4.08636761330825 | 1.27908501089775 | -3.19475842378933 | 0.00139947929909262 | 0.0443232398911078 | 2751 | 0 | 0 | 0 | 0 | 0 | 0 | 17 | 13 | 3 | 2 | 8 | 17 |
| L1M2_dup6117 | 4.87146365833929 | -4.43129777092046 | 1.38176566129927 | -3.20698212079878 | 0.001341353321551 | 0.0430437838772453 | 2752 | 0 | 0 | 1 | 0 | 0 | 0 | 32 | 19 | 2 | 1 | 3 | 2 |
| L1ME3A_dup5180 | 3.0998407371129 | -4.42761758796031 | 1.37305868439021 | -3.22463827533101 | 0.00126131927987639 | 0.0412468231858415 | 2753 | 0 | 0 | 1 | 0 | 0 | 0 | 22 | 11 | 1 | 0 | 2 | 2 |
| L1PA4_dup2045 | 5.69221304192513 | -5.71412633466733 | 1.35949187650003 | -4.20313385717181 | 2.6324480997721e-05 | 0.0025065245920303 | 2754 | 0 | 0 | 2 | 0 | 0 | 0 | 36 | 25 | 0 | 0 | 4 | 3 |
| MLT1J2_dup1955 | 7.14202917664261 | -6.09995050385813 | 1.89725262835486 | -3.21514932312809 | 0.00130376683119816 | 0.0421904124053144 | 2755 | 0 | 0 | 1 | 0 | 0 | 0 | 37 | 37 | 0 | 0 | 5 | 5 |
| L1MCa_dup6729 | 3.96295805958556 | -4.90550314251068 | 1.43471603039736 | -3.41914569753016 | 0.000628180808607601 | 0.0253473140045691 | 2756 | 0 | 0 | 0 | 0 | 1 | 0 | 25 | 12 | 0 | 0 | 6 | 7 |
| L1MD1_dup4959 | 4.13997663302429 | -4.87455809375436 | 1.33313890704361 | -3.65645175307669 | 0.00025573051638333 | 0.013362576887903 | 2757 | 0 | 0 | 0 | 1 | 1 | 0 | 22 | 18 | 0 | 0 | 3 | 6 |
| AluJo_dup18840 | 7.36612447769091 | -4.23998847594945 | 1.28486677015829 | -3.299944067685 | 0.000967040997089025 | 0.0343489382136655 | 2758 | 1 | 1 | 0 | 0 | 2 | 0 | 53 | 29 | 0 | 0 | 1 | 6 |
| L1M5_dup33675 | 6.06289930459637 | -4.1738227472557 | 1.28969975624929 | -3.23627474304099 | 0.00121100763955538 | 0.0400932257458232 | 2759 | 0 | 0 | 0 | 1 | 1 | 0 | 34 | 31 | 1 | 1 | 2 | 1 |
| L1ME3G_dup9946 | 9.77713626757758 | -3.77972589818576 | 1.05287784697656 | -3.58989972962163 | 0.00033080518733894 | 0.0160378506737065 | 2760 | 1 | 1 | 0 | 2 | 3 | 0 | 69 | 34 | 1 | 1 | 7 | 5 |

| | | | | | | | | | | | | | | | | | | | |
|---|---|---|---|---|---|---|---|---|---|---|---|---|---|---|---|---|---|---|---|
| L1PB3_dup291 | 5.18661570326379 | -3.04759067852727 | 0.918051543252374 | -3.31962916562462 | 0.000901370956272977 | 0.0326798131283452 | 2761 | 1 | 1 | 1 | 1 | 2 | 2 | 23 | 14 | 1 | 0 | 9 | 11 |
| L1MB1_dup5493 | 5.34883479210871 | -4.0539490000973 | 1.20452838363386 | -3.36559026352474 | 0.000763800724622473 | 0.0290882405923526 | 2762 | 0 | 0 | 1 | 0 | 2 | 2 | 19 | 18 | 0 | 0 | 9 | 16 |
| L1PA3_dup10043 | 5.14043112730405 | -3.96032818706291 | 1.162136719357 | -3.40779885971946 | 0.000654891493476378 | 0.0260932369904103 | 2763 | 0 | 1 | 1 | 0 | 2 | 1 | 25 | 11 | 0 | 0 | 7 | 20 |
| MER4-int_dup273 | 5.71789051429168 | -4.85356314626144 | 1.27639206306999 | -3.80256449933384 | 0.00014320596657533 | 0.00880708577964302 | 2764 | 1 | 0 | 0 | 1 | 0 | 1 | 38 | 20 | 0 | 0 | 5 | 7 |
| THE1A-int_dup559 | 6.45493331177521 | -3.10050522512176 | 0.978879285521805 | -3.16740304037488 | 0.00153806989979362 | 0.0473055531490971 | 2765 | 3 | 0 | 0 | 4 | 1 | 4 | 35 | 20 | 0 | 1 | 5 | 10 |
| MIR_dup115840 | 8.74284965639785 | -3.01268098762252 | 0.897889520522331 | -3.35529140140755 | 0.000792814233179907 | 0.0298823024755625 | 2766 | 6 | 3 | 5 | 2 | 0 | 0 | 34 | 33 | 0 | 3 | 8 | 14 |
| MSTA-int_dup196 | 7.15822215320122 | -2.91502491979638 | 0.912764434165633 | -3.19362237471603 | 0.00140499773674136 | 0.0444447741912604 | 2767 | 4 | 1 | 2 | 5 | 0 | 0 | 48 | 16 | 1 | 4 | 8 | 5 |
| L1MB4_dup4632 | 6.74179570462412 | -4.19649009076238 | 0.907091393490182 | -4.62631452671566 | 3.72229894320225e-06 | 0.000567760348935386 | 2768 | 1 | 1 | 7 | 7 | 0 | 0 | 30 | 23 | 1 | 1 | 3 | 11 |
| MIRb_dup202698 | 6.33902549250984 | -4.47506037024098 | 0.937058351263282 | -4.77564749752029 | 1.79129937581218e-06 | 0.000323638968319669 | 2769 | 1 | 1 | 5 | 9 | 0 | 0 | 36 | 19 | 0 | 1 | 4 | 6 |
| L1MB3_dup1728 | 5.39197094430495 | -2.41864370406152 | 0.676871699718734 | -3.57326758537336 | 0.000352554164403447 | 0.0168113934274835 | 2770 | 3 | 2 | 8 | 5 | 2 | 3 | 17 | 16 | 0 | 1 | 3 | 8 |
| LTR25-int_dup41 | 4.60235462291706 | -2.48300226260214 | 0.754916867666518 | -3.28910687911534 | 0.00100505850179374 | 0.035195933052556 | 2771 | 2 | 1 | 5 | 2 | 3 | 2 | 17 | 11 | 0 | 1 | 6 | 9 |
| L2a_dup116389 | 4.81690019950295 | -2.65609225430582 | 0.841756210918404 | -3.15541747106074 | 0.00160268600777234 | 0.048598537632904 | 2772 | 1 | 2 | 4 | 2 | 0 | 2 | 23 | 12 | 0 | 3 | 5 | 8 |
| Zaphod_dup207 | 6.0307419053311 | -2.26467010516972 | 0.719663331517334 | -3.14684659616449 | 0.00165041526300675 | 0.0496546844590019 | 2773 | 1 | 2 | 8 | 9 | 0 | 5 | 26 | 14 | 0 | 4 | 4 | 5 |
| L1PA6_dup377 | 6.19398179489352 | -2.99634415202456 | 0.822526088216574 | -3.64285606857933 | 0.000269629620706983 | 0.0138443827676099 | 2774 | 5 | 1 | 3 | 11 | 0 | 4 | 32 | 15 | 0 | 0 | 3 | 8 |
| MARNA_dup516 | 4.95397228660056 | -2.69057168767077 | 0.819172309540305 | -3.28450028929889 | 0.00102163393282609 | 0.0355972628174312 | 2775 | 2 | 1 | 2 | 8 | 1 | 5 | 22 | 16 | 0 | 0 | 3 | 3 |
| MLT1D_dup16724 | 4.71134288746256 | -2.89887921718466 | 0.795187962926874 | -3.64552703553845 | 0.00026684435825563 | 0.0137523386765203 | 2776 | 4 | 1 | 5 | 7 | 1 | 1 | 19 | 17 | 0 | 0 | 2 | 2 |
| L1MB2_dup3845 | 4.65803744079662 | -3.99993066743185 | 0.964384377235166 | -4.14765187185987 | 3.3590264353779e-05 | 0.00300815545424325 | 2777 | 1 | 0 | 6 | 4 | 0 | 1 | 18 | 19 | 0 | 1 | 3 | 4 |
| L1MB3_dup13935 | 3.90437015681135 | -3.45182182747895 | 0.909765910644921 | -3.79418682002715 | 0.000148128116006805 | 0.00902155378692947 | 2778 | 1 | 0 | 6 | 4 | 0 | 2 | 11 | 12 | 0 | 1 | 5 | 7 |
| LTR48B_dup291 | 4.17707777444077 | -3.22876049230773 | 0.880508256264408 | -3.66692812854007 | 0.000245481755351582 | 0.0129457893969165 | 2779 | 1 | 0 | 6 | 4 | 0 | 3 | 12 | 13 | 0 | 1 | 7 | 5 |
| L1M2_dup3033 | 5.12080061614065 | -4.80070041676088 | 0.974875702160996 | -4.92442308913765 | 8.46095977328341e-07 | 0.000180885133352533 | 2780 | 1 | 0 | 6 | 8 | 0 | 1 | 23 | 10 | 0 | 0 | 8 | 11 |
| L1MA6_dup1877 | 6.63659739472536 | -3.6924970424178 | 0.771769564362093 | -4.78445537751911 | 1.71451580220182e-06 | 0.000314499766555464 | 2781 | 2 | 0 | 8 | 13 | 0 | 3 | 33 | 14 | 0 | 1 | 6 | 8 |
| L1M4c_dup599 | 5.94768201018266 | -3.12862144545715 | 0.765235350257123 | -4.088443436919 | 4.34277405655746e-05 | 0.00366694718119808 | 2782 | 5 | 1 | 6 | 12 | 2 | 0 | 28 | 17 | 0 | 0 | 3 | 3 |
| L1PA5_dup2036 | 4.49095811153405 | -3.35871336081229 | 0.83955242600011 | -4.00059991109102 | 6.31821037067568e-05 | 0.00486063008361072 | 2783 | 3 | 1 | 3 | 17 | 1 | 0 | 19 | 10 | 0 | 0 | 3 | 3 |
| MER20_dup7185 | 4.39976329264323 | -2.69505378521665 | 0.776630852618719 | -3.46572384074126 | 0.000528806134667996 | 0.0224011580060016 | 2784 | 4 | 4 | 6 | 15 | 0 | 0 | 13 | 9 | 0 | 0 | 4 | 3 |
| L1PA2_dup4639 | 3.49453266156231 | -3.57758025184639 | 0.915889538538256 | -3.90612634090804 | 9.37874886955547e-05 | 0.00651002269111798 | 2785 | 1 | 2 | 5 | 8 | 0 | 0 | 14 | 10 | 0 | 0 | 2 | 3 |
| L1PA5_dup2388 | 4.07521345157839 | -2.89400602641581 | 0.848673849501667 | -3.41003322785914 | 0.000649549652662014 | 0.0259238077243065 | 2786 | 3 | 3 | 8 | 6 | 0 | 0 | 14 | 13 | 0 | 0 | 3 | 1 |
| MLT1K_dup15528 | 4.84779288581144 | -3.5742650731608 | 0.911920500657803 | -3.91949196293158 | 8.87358201300703e-05 | 0.0062542456921415 | 2787 | 3 | 2 | 7 | 6 | 0 | 0 | 20 | 12 | 0 | 0 | 10 | 2 |
| Tigger1_dup6132 | 5.13381172309601 | -3.50107532900977 | 0.828793836851392 | -4.22430183881487 | 2.39682638852732e-05 | 0.00233927502807433 | 2788 | 3 | 1 | 8 | 8 | 0 | 0 | 18 | 17 | 1 | 0 | 5 | 3 |
| L1PA3_dup642 | 3.50650096042879 | -3.75869905051385 | 1.1378442107167 | -3.30335121022089 | 0.000955366530392646 | 0.0340373227927919 | 2789 | 2 | 1 | 2 | 1 | 0 | 0 | 14 | 13 | 0 | 0 | 2 | 9 |
| L1MC3_dup4247 | 3.75717007944558 | -4.35155267303842 | 1.00240282656145 | -4.34112171048596 | 1.41757169137755e-05 | 0.00158285083846327 | 2790 | 2 | 0 | 7 | 6 | 0 | 0 | 12 | 7 | 0 | 0 | 4 | 12 |
| MER97a_dup89 | 3.47609346686622 | -3.41172902674056 | 1.02735093022425 | -3.3208993405942 | 0.000897278923294731 | 0.0325761261340079 | 2791 | 4 | 0 | 2 | 6 | 0 | 0 | 13 | 7 | 0 | 0 | 7 | 7 |
| AluSz_dup3697 | 3.2040752484207 | -2.88553462800312 | 0.893410797427942 | -3.22979600908154 | 0.00123878570512907 | 0.0407340794673237 | 2792 | 2 | 1 | 8 | 2 | 0 | 2 | 11 | 8 | 0 | 0 | 2 | 5 |
| L1MEf_dup3783 | 3.62623616286348 | -3.79248128637433 | 1.01505542848173 | -3.73623073180048 | 0.000186799277408837 | 0.0106030870377679 | 2793 | 1 | 2 | 5 | 2 | 0 | 0 | 12 | 10 | 0 | 0 | 6 | 8 |
| L2a_dup59494 | 4.54466606180961 | -3.75508514889627 | 0.899606788895059 | -4.17414051922446 | 2.99113104503191e-05 | 0.00275010690742335 | 2794 | 1 | 2 | 6 | 4 | 0 | 1 | 19 | 10 | 0 | 0 | 8 | 8 |
| ERVL-E-int_dup7394 | 5.39826509968575 | -2.93466676671012 | 0.785837392520026 | -3.73444531227919 | 0.000188129363926302 | 0.0106470863513563 | 2795 | 3 | 2 | 7 | 3 | 0 | 2 | 26 | 12 | 1 | 0 | 6 | 8 |
| L1MA4A_dup3441 | 4.39488162366256 | -2.85342789143675 | 0.8109037626935 | -3.51882433244458 | 0.000433463622691082 | 0.019575049376514 | 2796 | 2 | 3 | 4 | 7 | 0 | 2 | 19 | 13 | 0 | 0 | 4 | 2 |
| L1PB1_dup9512 | 5.08779848297498 | -2.44164508551009 | 0.713085299458974 | -3.42405752490283 | 0.000616935602172576 | 0.0250336560018001 | 2797 | 5 | 4 | 10 | 6 | 0 | 2 | 20 | 11 | 0 | 0 | 4 | 4 |
| Tigger1_dup732 | 25.5344180571573 | -2.35099225316179 | 0.668488750597423 | -3.51687631401534 | 0.00043665720396148 | 0.0196819160927862 | 2798 | 4 | 5 | 12 | 17 | 23 | 25 | 105 | 80 | 3 | 0 | 23 | 22 |
| L2c_dup12450 | 14.3619489153793 | -1.93166996757313 | 0.604904624450381 | -3.19334633840542 | 0.00140634162880083 | 0.0444619624779882 | 2799 | 1 | 8 | 18 | 10 | 13 | 12 | 62 | 29 | 4 | 0 | 14 | 13 |
| MER5B_dup1185 | 11.8532973973983 | -2.16919233084071 | 0.686880472840928 | -3.15803464592459 | 0.00158836679450639 | 0.0483368110097988 | 2800 | 3 | 4 | 9 | 5 | 13 | 10 | 35 | 43 | 1 | 0 | 10 | 11 |
| L1PA16_dup3037 | 14.2122901267566 | -1.95132571179897 | 0.607994024629791 | -3.2094488313222 | 0.0013298971509379 | 0.0427800064184222 | 2801 | 7 | 3 | 10 | 7 | 9 | 12 | 60 | 45 | 1 | 4 | 7 | 13 |
| LTR16A1_dup112 | 15.131907749124 | -1.85190015865594 | 0.570036790506389 | -3.24873795779183 | 0.0011591820896181 | 0.0389085543089673 | 2802 | 4 | 6 | 10 | 20 | 21 | 14 | 51 | 37 | 0 | 2 | 13 | 15 |
| L1PA15_dup7742 | 14.2464205313997 | -3.73313223035109 | 0.947819107338765 | -3.93865475115054 | 8.1939737395256e-05 | 0.00588791570060197 | 2803 | 0 | 1 | 1 | 3 | 6 | 6 | 82 | 34 | 0 | 1 | 18 | 35 |
| L1PA16_dup12849 | 11.6398222266693 | -4.19176915032605 | 1.04883065789032 | -3.9966119590341 | 6.4255499194455e-05 | 0.00492093305803791 | 2804 | 0 | 0 | 1 | 3 | 5 | 4 | 49 | 38 | 0 | 0 | 18 | 29 |
| MIRc_dup67618 | 9.42802089270626 | -2.99255548211194 | 0.886047701417661 | -3.37742028710633 | 0.000731691571342529 | 0.0282333231298448 | 2805 | 3 | 0 | 3 | 2 | 2 | 4 | 42 | 38 | 3 | 0 | 6 | 12 |
| Charlie30a_dup30 | 9.99130702270531 | -2.54718181935468 | 0.772559764395239 | -3.29706766614829 | 0.000976999601096047 | 0.0345890711326291 | 2806 | 4 | 1 | 5 | 7 | 5 | 13 | 30 | 38 | 0 | 0 | 10 | 9 |
| AluSz6_dup3325 | 7.78851974296303 | -2.53405504048699 | 0.744994483519517 | -3.40144134828429 | 0.000670315108126833 | 0.0265010156440533 | 2807 | 2 | 3 | 6 | 5 | 4 | 8 | 32 | 22 | 0 | 0 | 12 | 4 |
| L1PA5_dup584 | 7.07115216109329 | -3.05098486609566 | 0.852316432868566 | -3.57963867460261 | 0.000344069623238305 | 0.0164993935048492 | 2808 | 1 | 2 | 3 | 4 | 2 | 6 | 23 | 31 | 0 | 0 | 6 | 7 |
| L1PA5_dup587 | 9.21210885254417 | -2.29541937749621 | 0.686583806414143 | -3.34324718417788 | 0.000828041060508305 | 0.030802109016452 | 2809 | 12 | 6 | 4 | 12 | 4 | 2 | 32 | 28 | 0 | 0 | 7 | 10 |
| HERVP71A-int_dup58 | 9.25887800940481 | -2.74234103449264 | 0.820338294189548 | -3.34293919218038 | 0.000828960626430136 | 0.030821620780411 | 2810 | 8 | 3 | 3 | 3 | 2 | 2 | 47 | 35 | 1 | 0 | 5 | 6 |
| L1PREC2_dup2607 | 10.9378597908436 | -2.96775281155186 | 0.843596970963433 | -3.51797471269074 | 0.000434853798597303 | 0.0196284076040474 | 2811 | 9 | 4 | 3 | 2 | 4 | 3 | 46 | 43 | 0 | 0 | 12 | 8 |
| L1PBa_dup672 | 31.2703204167291 | -4.24126705910648 | 1.33570465459039 | -3.17530304662083 | 0.00149680092647563 | 0.0464351751305307 | 2812 | 9 | 4 | 1 | 1 | 0 | 3 | 202 | 149 | 0 | 1 | 2 | 10 |
| L1PA16_dup3883 | 19.7174566735377 | -4.17553285408025 | 1.14834270302797 | -3.63613827394045 | 0.000276755812954541 | 0.0141133601948738 | 2813 | 2 | 3 | 2 | 2 | 7 | 1 | 139 | 81 | 0 | 0 | 5 | 4 |
| L1PBa_dup671 | 12.8105920553676 | -6.01887383305364 | 1.76411686963544 | -3.41183395309715 | 0.000645274067908069 | 0.0258137833494374 | 2814 | 0 | 1 | 2 | 0 | 0 | 0 | 83 | 64 | 0 | 0 | 2 | 3 |
| MSTA-int_dup819 | 13.6282360308597 | -4.21049229789674 | 1.31306619515495 | -3.20661084218977 | 0.00134308551693528 | 0.0430761324184771 | 2815 | 4 | 1 | 3 | 0 | 0 | 1 | 85 | 64 | 0 | 1 | 4 | 3 |
| L2a_dup78444 | 14.2761153836258 | -3.37677862064196 | 0.895520823791501 | -3.77074271304912 | 0.000162762414002106 | 0.00965561340367618 | 2816 | 8 | 4 | 7 | 4 | 0 | 1 | 88 | 59 | 1 | 0 | 2 | 3 |
| L1M4_dup14436 | 13.2988645223534 | -3.42239406478784 | 0.876772355772229 | -3.9034009709094 | 9.48503694657698e-05 | 0.00655702092181931 | 2817 | 1 | 5 | 4 | 5 | 3 | 3 | 78 | 61 | 0 | 0 | 1 | 1 |
| L1PB1_dup9690 | 16.7049259179689 | -3.14695767823986 | 0.809623996642443 | -3.88693725888866 | 0.000101516979238563 | 0.00689174297914383 | 2818 | 4 | 8 | 10 | 12 | 2 | 2 | 108 | 64 | 0 | 0 | 0 | 0 |
| L1PB1_dup9010 | 20.1553849409051 | -3.52386784938488 | 0.833089820376459 | -4.22987745522146 | 2.33818648588187e-05 | 0.00229901703127953 | 2819 | 2 | 0 | 16 | 14 | 6 | 3 | 112 | 90 | 1 | 1 | 1 | 0 |
| L1PA3_dup8663 | 12.8044259039832 | -3.48141880313872 | 0.795891380075542 | -4.37423860880146 | 1.2185703668716e-05 | 0.0014117108197428 | 2820 | 2 | 0 | 14 | 21 | 1 | 4 | 61 | 47 | 3 | 0 | 2 | 5 |
| L1PB1_dup10190 | 9.09974420012603 | -3.46558404073238 | 0.753208983320153 | -4.60109228312181 | 4.20281160800738e-06 | 0.000626985712149795 | 2821 | 1 | 0 | 9 | 16 | 4 | 2 | 50 | 30 | 1 | 0 | 1 | 2 |
| L1MC1_dup8639 | 11.0427606578789 | -3.67957959485083 | 1.04995390845576 | -3.50451535559561 | 0.00045743918661571 | 0.0203184291018222 | 2822 | 2 | 0 | 8 | 7 | 1 | 0 | 64 | 47 | 1 | 3 | 0 | 3 |
| L1PA10_dup4707 | 9.32650576793557 | -3.85440654266296 | 0.953913782899455 | -4.04062359907135 | 5.33092603501844e-05 | 0.00429263122276988 | 2823 | 0 | 1 | 6 | 10 | 2 | 0 | 52 | 40 | 0 | 2 | 1 | 1 |
| L1MEg1_dup567 | 7.67949133473793 | -3.89212625833243 | 1.19298010251308 | -3.26252403550859 | 0.00110424808390947 | 0.0375768767625014 | 2824 | 0 | 0 | 2 | 5 | 2 | 0 | 46 | 37 | 1 | 0 | 0 | 0 |
| L1PA11_dup3101 | 8.57929195988043 | -5.07375050623578 | 1.17139409249347 | -4.33137791862652 | 1.48179098316965e-05 | 0.00163385652122931 | 2825 | 0 | 0 | 3 | 7 | 0 | 1 | 72 | 28 | 0 | 0 | 0 | 1 |
| LTR17_dup578 | 6.18256692832174 | -4.34461575043682 | 1.15772829907753 | -3.75270756869172 | 0.000174934879133965 | 0.0101400864191012 | 2826 | 0 | 0 | 3 | 4 | 0 | 1 | 44 | 24 | 0 | 1 | 1 | 0 |
| L1PA16_dup9473 | 18.4562983263371 | -4.08609824765156 | 1.20125273446838 | -3.40153086058104 | 0.000670095622646 78 | 0.0265010156440533 | 2827 | 3 | 2 | 9 | 21 | 0 | 0 | 115 | 74 | 1 | 2 | 4 | 0 |
| L1PA16_dup9475 | 13.9926302587629 | -3.73877659964628 | 0.961955664860915 | -3.88664128318933 | 0.000101640780297419 | 0.00689425397400665 | 2828 | 0 | 2 | 10 | 10 | 0 | 2 | 80 | 60 | 1 | 3 | 1 | 3 |
| L2c_dup86004 | 10.4801333511784 | -3.22324044712593 | 0.979734300434321 | -3.28991283218017 | 0.00100218423711806 | 0.0351511984795541 | 2829 | 2 | 1 | 4 | 2 | 0 | 1 | 63 | 40 | 5 | 2 | 2 | 5 |
| L2a_dup28865 | 14.9763323706493 | -2.31568126579581 | 0.725846400687268 | -3.19031859027366 | 0.00142116032066893 | 0.0447182509470893 | 2830 | 10 | 7 | 2 | 7 | 6 | 4 | 100 | 36 | 1 | 3 | 5 | 16 |
| MLT1F2_dup3654 | 13.8026970385666 | -3.24370895326166 | 0.854857873537581 | -3.79444239056782 | 0.000147975633365647 | 0.00901900872357126 | 2831 | 1 | 1 | 2 | 6 | 1 | 7 | 76 | 55 | 2 | 2 | 11 | 6 |
| L1ME1_dup7135 | 12.9265698439669 | -2.28247210482523 | 0.710858138655392 | -3.21086863989851 | 0.00132334412308067 | 0.04260953189588 | 2832 | 5 | 2 | 9 | 12 | 2 | 6 | 69 | 45 | 2 | 4 | 3 | 3 |
| L1PA16_dup3038 | 12.4790797912597 | -2.41784636011832 | 0.686276068079573 | -3.52313955356807 | 0.00042646674212505 | 0.0193582707786507 | 2833 | 6 | 2 | 6 | 7 | 5 | 5 | 60 | 46 | 3 | 1 | 7 | 6 |
| LTR27_dup57 | 9.61406808043783 | -2.88746847790857 | 0.806995622636389 | -3.57804726186177 | 0.000346170865692731 | 0.0165754515831735 | 2834 | 3 | 1 | 3 | 5 | 3 | 2 | 56 | 36 | 1 | 2 | 4 | 4 |
| L1PB1_dup9011 | 24.6592152692005 | -3.48613003676549 | 0.751255061738296 | -4.6404080508943 | 3.47721872168222e-06 | 0.000539367852639763 | 2835 | 3 | 3 | 13 | 18 | 5 | 2 | 158 | 92 | 1 | 4 | 3 | 9 |
| MER68_dup877 | 16.8617659398158 | -3.32847219653307 | 0.891793198639802 | -3.73233637754783 | 0.000189711926914025 | 0.0106989826347 56 | 2836 | 1 | 2 | 5 | 8 | 3 | 0 | 98 | 62 | 8 | 5 | 8 | 8 |
| L2a_dup78432 | 15.2655758769994 | -3.56776416102253 | 0.731881534292958 | -4.8747836826751 | 1.08927749629887e-06 | 0.00022021174437921 | 2837 | 9 | 6 | 15 | 17 | 2 | 0 | 69 | 59 | 0 | 0 | 4 | 9 |
| L1MA4_dup4716 | 10.2537222601867 | -2.61958611488975 | 0.796048714407671 | -3.2907359404995 | 0.000999256647442012 | 0.0350974833005743 | 2838 | 10 | 4 | 12 | 16 | 0 | 0 | 41 | 38 | 0 | 2 | 2 | 3 |
| L1MA7_dup3917 | 7.08630188074239 | -3.30974002874002 | 0.680291943375009 | -4.86517598947301 | 1.1435500654034e-06 | 0.000228466132546161 | 2839 | 3 | 3 | 12 | 14 | 1 | 0 | 29 | 20 | 0 | 1 | 3 | 5 |

| | | | | | | | | | | | | | | | | | | | |
|---|---|---|---|---|---|---|---|---|---|---|---|---|---|---|---|---|---|---|---|
| LTR12C_dup1417 | 8.95227458054918 | -3.12673529133399 | 0.752520770659957 | -4.15501526767408 | 3.25265822087297e-05 | 0.00293721826574513 | 2840 | 6 | 7 | 12 | 16 | 0 | 0 | 44 | 22 | 0 | 0 | 4 | 6 |
| MER44B_dup1022 | 6.30423435451834 | -4.28611559190882 | 0.952673086563405 | -4.49904133155499 | 6.82605866038754e-06 | 0.000911466528622095 | 2841 | 0 | 2 | 7 | 6 | 1 | 0 | 37 | 22 | 0 | 0 | 1 | 4 |
| MER90a_dup520 | 5.86708729736552 | -3.70369130722575 | 0.903871431342631 | -4.09758642523329 | 4.17480317372191e-05 | 0.00355794310879096 | 2842 | 1 | 2 | 5 | 7 | 1 | 0 | 40 | 18 | 0 | 0 | 1 | 1 |
| L1MD1_dup2177 | 9.37816293991315 | -4.39279280574593 | 0.822443080205133 | -5.34115115255184 | 9.2358205670974e-08 | 3.11131293982015e-05 | 2843 | 2 | 2 | 9 | 11 | 1 | 0 | 52 | 33 | 0 | 0 | 3 | 6 |
| L1PA2_dup761 | 8.29793823969398 | -4.27153323326554 | 0.946162220118878 | -4.51458866401245 | 6.34397770440536e-06 | 0.000865056801634634 | 2844 | 1 | 3 | 5 | 3 | 1 | 0 | 49 | 31 | 0 | 0 | 5 | 6 |
| MLT1H2_dup2076 | 9.21695775619112 | -3.52659813466219 | 0.726851293722737 | -4.85188396184852 | 1.22294172149705e-06 | 0.000239896200217096 | 2845 | 1 | 1 | 13 | 15 | 1 | 5 | 45 | 33 | 0 | 0 | 1 | 1 |
| L1MA4_dup7681 | 7.59254934308692 | -4.22703506646538 | 0.900540499873933 | -4.69388669033444 | 2.68062549942489e-06 | 0.000442947302143162 | 2846 | 1 | 2 | 9 | 9 | 0 | 1 | 47 | 24 | 0 | 0 | 0 | 5 |
| L1MEf_dup7433 | 10.7094571525802 | -3.44245900787648 | 0.820070296065755 | -4.19776088024588 | 2.69567040698692e-05 | 0.00254634375014255 | 2847 | 3 | 4 | 6 | 16 | 0 | 2 | 56 | 45 | 0 | 0 | 0 | 1 |
| L2_dup26571 | 9.96956529366746 | -3.44649081933773 | 0.825861542345606 | -4.17320657594617 | 3.0034239184269e-05 | 0.00275927686368343 | 2848 | 3 | 5 | 5 | 11 | 0 | 2 | 59 | 37 | 0 | 0 | 1 | 3 |
| MLT2B4_dup3973 | 4.90226902290998 | -4.76280458679341 | 1.10549104396581 | -4.3083158500384 | 1.64502357343233e-05 | 0.00176956048464706 | 2849 | 0 | 0 | 19 | 32 | 0 | 0 | 8 | 5 | 1 | 0 | 0 | 2 |
| MER21B_dup2403 | 6.05175016383601 | -4.47238067779204 | 1.06816729075698 | -4.18696651403974 | 2.82707515656466e-05 | 0.00264214935272559 | 2850 | 0 | 0 | 22 | 42 | 0 | 0 | 10 | 6 | 1 | 1 | 0 | 1 |
| MIR_dup28626 | 5.22065409321722 | -4.37836337752439 | 1.01394716239649 | -4.31813761101321 | 1.57351317395408e-05 | 0.00171042104015384 | 2851 | 0 | 0 | 24 | 24 | 0 | 0 | 12 | 6 | 1 | 1 | 0 | 2 |
| L1MA8_dup9902 | 6.88353165355627 | -4.508027557167 | 1.25604760133242 | -3.58905789269839 | 0.000331875141971411 | 0.0160700778452441 | 2852 | 0 | 0 | 27 | 42 | 0 | 0 | 6 | 11 | 2 | 2 | 0 | 1 |
| L1MC4a_dup9044 | 8.07596496348715 | -3.96148224656782 | 1.01998601753793 | -3.88385936518045 | 0.000102811389210254 | 0.00695184962120579 | 2853 | 0 | 1 | 36 | 46 | 0 | 0 | 14 | 8 | 2 | 1 | 0 | 1 |
| FordPrefect_dup483 | 8.31202268382145 | -4.75418379000453 | 1.21919406894562 | -3.89944793130106 | 9.64122672589832e-05 | 0.00663796944580474 | 2854 | 0 | 0 | 39 | 56 | 0 | 0 | 10 | 5 | 2 | 1 | 0 | 1 |
| MLT1D_dup18209 | 8.40244351239423 | -4.85844388742359 | 1.07499007556087 | -4.51952441038932 | 6.19787016465703e-06 | 0.000849026145192563 | 2855 | 0 | 0 | 30 | 63 | 0 | 0 | 15 | 6 | 1 | 1 | 0 | 1 |
| MIRb_dup198502 | 5.38472016177178 | -3.64970607515012 | 1.08987828665293 | -3.34872812849457 | 0.000811834155335779 | 0.0303539739040741 | 2856 | 0 | 0 | 21 | 39 | 1 | 0 | 4 | 6 | 1 | 1 | 0 | 0 |
| MER41B_dup2494 | 6.24780434992255 | -3.86285928792474 | 1.03766913945396 | -3.72263098231625 | 0.000197157571040952 | 0.0110046576117547 | 2857 | 1 | 0 | 29 | 37 | 0 | 0 | 5 | 9 | 1 | 1 | 0 | 0 |
| L1PA7_dup12193 | 6.27584798021473 | -3.83811330098748 | 1.09921602698679 | -3.49168244162946 | 0.000479988539093051 | 0.0209523670744882 | 2858 | 0 | 1 | 29 | 43 | 0 | 0 | 4 | 6 | 1 | 1 | 0 | 0 |
| L2a_dup152031 | 5.91674884419365 | -4.48381122290401 | 1.25320530734159 | -3.57787442858462 | 0.000346399789550619 | 0.0165754515831735 | 2859 | 0 | 0 | 31 | 39 | 0 | 0 | 3 | 5 | 1 | 1 | 0 | 0 |
| THE1B_dup20512 | 6.00518442205274 | -4.50871164466716 | 1.36095318860046 | -3.31290722005193 | 0.000923316037348632 | 0.0332877814883269 | 2860 | 0 | 0 | 27 | 43 | 0 | 0 | 5 | 4 | 2 | 1 | 0 | 0 |
| L2b_dup85160 | 6.34237830168222 | -4.50807080717202 | 1.27175817084641 | -3.54475474230427 | 0.000392978560320818 | 0.0182594515202671 | 2861 | 0 | 0 | 30 | 33 | 0 | 0 | 5 | 8 | 3 | 1 | 3 | 0 |
| MER21B_dup2402 | 6.5678570199832 | -4.47901647937018 | 1.26951751215633 | -3.52812500535135 | 0.00041851446509675 | 0.0190845557919799 | 2862 | 0 | 0 | 34 | 34 | 0 | 0 | 7 | 7 | 2 | 2 | 1 | 0 |
| L1MEb_dup1583 | 5.24752840819823 | -4.30520135858762 | 1.03732906118918 | -4.1502754715579 | 3.32075375661986e-05 | 0.00298286173413617 | 2863 | 0 | 0 | 20 | 31 | 0 | 0 | 11 | 6 | 1 | 1 | 1 | 0 |
| LTR101_Mam_dup989 | 5.52712939963512 | -4.45740853415566 | 1.17543184243025 | -3.79214546794972 | 0.00014935138150932 | 0.00906339296725752 | 2864 | 0 | 0 | 22 | 35 | 0 | 0 | 7 | 7 | 2 | 0 | 1 | 0 |
| MSTB_dup7648 | 5.79069939250217 | -4.86674341324089 | 1.15095193633098 | -4.22845060650854 | 2.35306160054802e-05 | 0.0023107783557383 | 2865 | 0 | 0 | 23 | 42 | 0 | 0 | 6 | 6 | 1 | 0 | 1 | 0 |
| L1PA16_dup10752 | 8.63950932673142 | -3.71639280526194 | 1.17792582445608 | -3.15503126606296 | 0.00160480906386938 | 0.0486381222151908 | 2866 | 0 | 0 | 45 | 59 | 0 | 2 | 4 | 2 | 2 | 2 | 0 | 3 |
| L2-3_Crp_dup1596 | 6.01604268505833 | -4.44975065209268 | 1.30876783836955 | -3.39995415660287 | 0.000673971518384392 | 0.0266128156179045 | 2867 | 0 | 0 | 27 | 35 | 0 | 0 | 8 | 2 | 3 | 3 | 0 | 4 |
| L1MC4_dup8811 | 6.69604827419588 | -4.77159977949468 | 1.39607212980846 | -3.41787482008495 | 0.000631121268556592 | 0.0254271556667739 | 2868 | 0 | 0 | 36 | 45 | 0 | 0 | 4 | 2 | 1 | 3 | 0 | 1 |
| MIR_dup154606 | 6.70915150679176 | -4.61648829536568 | 1.3175932183726 | -3.50372803304775 | 0.000458793659717064 | 0.0203482253063127 | 2869 | 0 | 0 | 36 | 38 | 0 | 0 | 8 | 4 | 1 | 3 | 0 | 1 |
| L1PA5_dup10456 | 8.59135207947083 | -5.43755820804674 | 1.70177659366799 | -3.19522446617195 | 0.00139722125240115 | 0.0442635079382031 | 2870 | 0 | 0 | 51 | 45 | 0 | 0 | 8 | 5 | 0 | 5 | 1 | 1 |
| ALR/Alpha_dup1278 | 8.69521053992924 | -4.73103166036688 | 1.08111344939859 | -4.37607326317021 | 1.20836413977712e-05 | 0.00140461853250321 | 2871 | 0 | 0 | 43 | 61 | 1 | 0 | 11 | 3 | 0 | 1 | 1 | 0 |
| HERVL-int_dup2150 | 9.78958916791812 | -5.53097206601772 | 1.23834763004117 | -4.46641309099436 | 7.95419534114423e-06 | 0.00102750758123991 | 2872 | 0 | 0 | 50 | 75 | 0 | 0 | 5 | 4 | 0 | 1 | 1 | 0 |
| AluSx_dup100157 | 6.11317970419359 | -3.55149090457841 | 0.97930414766096 | -3.62654535167756 | 0.000287238389842884 | 0.0144987751187007 | 2873 | 0 | 1 | 36 | 30 | 1 | 0 | 4 | 6 | 1 | 1 | 0 | 1 |
| L1PA4_dup2372 | 4.58434610195838 | -3.60654990855965 | 1.14275091640172 | -3.15602451662512 | 0.001599354166311 | 0.0485408069460507 | 2874 | 0 | 0 | 23 | 28 | 1 | 0 | 3 | 4 | 0 | 2 | 0 | 1 |
| THE1B_dup20515 | 7.3488561875949 | -3.99570287522098 | 1.17763086986384 | -3.39300113259001 | 0.000691313524006663 | 0.0270952541234211 | 2875 | 0 | 0 | 38 | 42 | 1 | 0 | 6 | 7 | 0 | 3 | 0 | 2 |
| MER5B_dup24102 | 6.26788578823324 | -4.34470699411073 | 1.07469887245203 | -4.04272034286021 | 5.28346228199095e-05 | 0.00426451396251906 | 2876 | 0 | 1 | 35 | 37 | 0 | 0 | 8 | 3 | 0 | 1 | 0 | 1 |
| MSTA-int_dup1586 | 6.82432020029837 | -4.44835735841872 | 1.03413644484489 | -4.3015187991812 | 1.69631379372445e-05 | 0.00180919355428757 | 2877 | 0 | 1 | 42 | 29 | 0 | 0 | 16 | 3 | 0 | 1 | 0 | 1 |
| MLT1A1_dup6080 | 5.55363860982837 | -4.95285878948155 | 1.23169880883288 | -4.02116065548097 | 5.79120899177362e-05 | 0.00454184250084563 | 2878 | 0 | 0 | 27 | 39 | 0 | 0 | 8 | 1 | 0 | 1 | 0 | 2 |
| Tigger19b_dup608 | 4.64634577699752 | -4.28216664442975 | 1.21682962206744 | -3.5191176864631 | 0.000432984591544976 | 0.0195645559475429 | 2879 | 0 | 0 | 29 | 20 | 0 | 0 | 6 | 4 | 0 | 2 | 0 | 1 |
| L2a_dup164502 | 5.38375085980121 | -4.44625251491109 | 1.37585919719268 | -3.23161884877702 | 0.00123091123777665 | 0.0405384644130109 | 2880 | 0 | 0 | 32 | 21 | 0 | 0 | 9 | 4 | 0 | 4 | 0 | 2 |
| MIRc_dup89852 | 5.44384903477287 | -4.38861979318274 | 1.33283605743335 | -3.29269287749756 | 0.000992328072781834 | 0.0349187364102553 | 2881 | 0 | 0 | 33 | 22 | 0 | 0 | 10 | 4 | 0 | 3 | 0 | 1 |
| L1MA2_dup3507 | 11.290584595132 | -4.37667660139108 | 1.05698393689353 | -4.14072196239244 | 3.46214409098411e-05 | 0.00307160081235386 | 2882 | 1 | 2 | 58 | 86 | 0 | 0 | 6 | 2 | 0 | 1 | 0 | 2 |
| L1PREC2_dup4110 | 12.6863539040675 | -4.64663800393269 | 0.935912899534825 | -4.96481884825201 | 6.87652252502784e-07 | 0.000152776940125142 | 2883 | 3 | 1 | 55 | 97 | 0 | 0 | 9 | 4 | 0 | 1 | 2 | 6 |
| L1PA13_dup523 | 11.2462455685609 | -5.88351244963997 | 1.71074344268089 | -3.43915534197225 | 0.000583532289375465 | 0.0240932108498585 | 2884 | 0 | 0 | 64 | 79 | 0 | 0 | 5 | 2 | 0 | 3 | 1 | 2 |
| L1PREC2_dup829 | 17.813202026768 | -6.59973076736015 | 1.58504647299716 | -4.16374590890118 | 3.13068503155457e-05 | 0.00284702111675763 | 2885 | 0 | 0 | 85 | 157 | 0 | 0 | 5 | 2 | 0 | 1 | 1 | 1 |
| L1PB1_dup1302 | 9.79377650732811 | -5.64296165572494 | 1.26458890075405 | -4.46228940674724 | 8.10886066475263e-06 | 0.00104033592015077 | 2886 | 0 | 0 | 43 | 78 | 0 | 0 | 3 | 2 | 0 | 2 | 1 | 9 |
| L1PB1_dup5641 | 9.2462072784256 | -6.28002981654478 | 1.23021402423325 | -5.1048270405297 | 3.31097204159769e-07 | 8.68613620077183e-05 | 2887 | 0 | 0 | 52 | 64 | 0 | 0 | 4 | 1 | 0 | 0 | 2 | 6 |
| Charlie1_dup581 | 10.2668640971469 | -3.74450513854655 | 0.768735464551563 | -4.87099309348357 | 1.11038721777169e-06 | 0.000223340825798734 | 2888 | 3 | 2 | 43 | 70 | 1 | 2 | 9 | 8 | 0 | 0 | 2 | 1 |
| THE1B-int_dup3036 | 10.4609023004639 | -4.35719919038517 | 0.848453265797356 | -5.13546162886223 | 2.8145230293843e-07 | 7.59302150510064e-05 | 2889 | 2 | 0 | 51 | 68 | 2 | 1 | 10 | 6 | 0 | 0 | 2 | 2 |
| ALR/Alpha_dup1273 | 12.1036512462181 | -4.03377059621293 | 0.939738257727843 | -4.29244054186538 | 1.76719881786424e-05 | 0.00186556295437233 | 2890 | 0 | 2 | 68 | 79 | 1 | 2 | 8 | 6 | 0 | 0 | 0 | 0 |
| MLT2B3_dup2261 | 8.26558217821925 | -3.97983748479284 | 0.989553631403462 | -4.021851225131 | 5.77425266614549e-05 | 0.00453302647607606 | 2891 | 1 | 0 | 49 | 48 | 0 | 3 | 7 | 4 | 0 | 0 | 0 | 1 |
| L1PA10_dup4823 | 9.57670102377904 | -4.41521101148172 | 1.00619170663766 | -4.38804154551803 | 1.14375930740456e-05 | 0.00134662519623793 | 2892 | 0 | 1 | 51 | 65 | 0 | 2 | 10 | 3 | 0 | 0 | 0 | 1 |
| L1PREC2_dup7594 | 12.0523976016935 | -5.01771135937816 | 1.04616721134906 | -4.7962804654408 | 1.61638924613593e-06 | 0.000299077259378741 | 2893 | 0 | 0 | 70 | 77 | 0 | 2 | 11 | 5 | 0 | 0 | 0 | 1 |
| L1PA13_dup8132 | 8.75771462648594 | -5.81936067814395 | 1.67956118485454 | -3.46481017221646 | 0.000530605869176051 | 0.0224454300705715 | 2894 | 0 | 0 | 36 | 79 | 0 | 0 | 5 | 3 | 0 | 0 | 0 | 1 |
| L1MB8_dup14376 | 9.39527014090264 | -4.96274630646032 | 1.12661181618376 | -4.40501886734238 | 1.057746712022e-05 | 0.00127436252728499 | 2895 | 1 | 0 | 54 | 65 | 1 | 0 | 3 | 3 | 0 | 0 | 0 | 3 |
| L1MCa_dup1990 | 5.94206054348124 | -4.2028816215493 | 1.10754426252369 | -3.79477530945127 | 0.000147777223544972 | 0.00901384240800615 | 2896 | 1 | 0 | 29 | 45 | 1 | 0 | 4 | 2 | 0 | 0 | 0 | 1 |
| L1MA4_dup2691 | 6.86753021814455 | -3.40425242524284 | 1.05003453346926 | -3.24203853943295 | 0.0011867797229823 | 0.0395337376403116 | 2897 | 2 | 2 | 35 | 49 | 0 | 0 | 3 | 2 | 1 | 0 | 0 | 1 |
| L1M3_dup6420 | 7.94607236019129 | -4.48111206857247 | 1.11396555154396 | -4.02266664562619 | 5.75429128014342e-05 | 0.00452525650661386 | 2898 | 1 | 0 | 45 | 53 | 0 | 0 | 5 | 1 | 1 | 1 | 1 | 2 |
| L1PA6_dup1241 | 7.94209395111008 | -4.32820286366398 | 1.03024046188177 | -4.20115790808523 | 2.65553281328854e-05 | 0.00251992299657343 | 2899 | 1 | 0 | 43 | 51 | 0 | 1 | 9 | 2 | 1 | 0 | 0 | 2 |
| L1PB2_dup2770 | 8.78619035193561 | -4.88024711620437 | 1.13451353645369 | -4.30162087925303 | 1.69553235237191e-05 | 0.00180919355428757 | 2900 | 0 | 0 | 46 | 59 | 1 | 0 | 4 | 9 | 0 | 0 | 0 | 0 |
| HERVL-int_dup2268 | 6.14479422050294 | -4.44462884712054 | 1.27063229920876 | -3.49796620933396 | 0.000468820527365262 | 0.0206359044478966 | 2901 | 0 | 0 | 27 | 53 | 1 | 0 | 2 | 3 | 0 | 0 | 0 | 0 |
| MSTA_dup13236 | 4.99844691302859 | -4.2273697413543 | 1.2008136001206 | -3.52042127181915 | 0.000430861879422715 | 0.01951682100916 | 2902 | 0 | 0 | 25 | 36 | 1 | 0 | 4 | 3 | 0 | 0 | 0 | 0 |
| THE1B_dup20508 | 7.67121449822541 | -3.66153550120975 | 1.11343710326169 | -3.2884978329568 | 0.00100723559890559 | 0.0352452473260574 | 2903 | 0 | 0 | 45 | 46 | 1 | 1 | 7 | 3 | 0 | 2 | 0 | 0 |
| MLT2F_dup1717 | 8.49713080242742 | -3.75537527627574 | 1.0407632073368 | -3.6082898105952 | 0.000308222078747503 | 0.015242808147042 | 2904 | 0 | 0 | 46 | 57 | 2 | 1 | 7 | 3 | 1 | 0 | 0 | 0 |
| L1MA2_dup5853 | 8.90970651926925 | -3.53108826016812 | 1.09881887918997 | -3.21353075292189 | 0.00131113756432505 | 0.0423655538947432 | 2905 | 0 | 0 | 41 | 72 | 3 | 2 | 5 | 2 | 0 | 0 | 0 | 0 |
| MER58A_dup11586 | 6.63154429354458 | -4.06878069343602 | 1.1498913836974 | -3.53840436681344 | 0.00040255307066274 | 0.018563157548821 | 2906 | 0 | 0 | 31 | 54 | 1 | 1 | 4 | 2 | 0 | 0 | 0 | 0 |
| AluSz6_dup40270 | 7.05155868371423 | -4.8718936823814 | 1.27310786189398 | -3.82677212843033 | 0.00012983460198128 | 0.00820112137059665 | 2907 | 0 | 1 | 46 | 44 | 0 | 0 | 1 | 3 | 0 | 0 | 1 | 0 |
| L1PA2_dup4853 | 8.78441663883705 | -4.68414516180703 | 1.0385484210676 | -4.51028095251626 | 6.4741822984913e-06 | 0.000877134904411177 | 2908 | 1 | 1 | 51 | 55 | 0 | 0 | 6 | 5 | 0 | 0 | 1 | 0 |
| L1PA7_dup11984 | 7.66149008085115 | -4.52028031451738 | 1.06633308849748 | -4.23908848302429 | 2.24429193095143e-05 | 0.00222694031610579 | 2909 | 1 | 1 | 41 | 53 | 0 | 0 | 6 | 3 | 0 | 0 | 1 | 0 |
| L1PA5_dup9142 | 5.87793189935426 | -3.89691138530757 | 1.20947414494009 | -3.22198816866862 | 0.00127304391857948 | 0.0415066741916022 | 2910 | 0 | 1 | 32 | 44 | 0 | 1 | 3 | 1 | 0 | 0 | 0 | 0 |
| L1MB4_dup4281 | 7.62237186330013 | -3.87782591934951 | 1.17563070425075 | -3.29850684005477 | 0.000972005119270199 | 0.0344532593639009 | 2911 | 2 | 1 | 42 | 57 | 0 | 0 | 2 | 2 | 0 | 0 | 0 | 0 |
| AluSq2_dup51153 | 5.95842302817234 | -4.45503088994551 | 1.22017347566887 | -3.65114549593317 | 0.000261073279402424 | 0.0135547332622058 | 2912 | 1 | 0 | 37 | 37 | 0 | 0 | 5 | 2 | 0 | 0 | 0 | 0 |
| L1PA16_dup7582 | 7.01819575349955 | -4.64856403861342 | 1.18008577452716 | -3.93917470997057 | 8.17623760626716e-05 | 0.00588120626801949 | 2913 | 1 | 0 | 34 | 54 | 0 | 0 | 6 | 3 | 0 | 0 | 0 | 0 |
| L1PREC2_dup5949 | 13.1104413640582 | -4.63011448736907 | 0.978054199400486 | -4.7340060399589 | 2.20131170957556e-06 | 0.000378242066056012 | 2914 | 1 | 1 | 72 | 89 | 1 | 0 | 14 | 4 | 0 | 0 | 0 | 0 |
| MER52-int_dup368 | 24.2845582178746 | -5.14768020441423 | 1.45081619939617 | -3.54812705190133 | 0.000387981020557929 | 0.0180804018640182 | 2915 | 1 | 1 | 143 | 181 | 1 | 0 | 7 | 4 | 0 | 0 | 0 | 0 |
| ALR/Alpha_dup1277 | 16.0788926095883 | -4.6740201641889 | 1.40739737391343 | -3.32103800307106 | 0.000896833248185969 | 0.0325649144014739 | 2916 | 1 | 0 | 86 | 113 | 0 | 2 | 6 | 12 | 0 | 0 | 0 | 0 |
| HERVL-int_dup2149 | 15.3716090177613 | -6.45877674512441 | 1.93192214707595 | -3.34318686438792 | 0.000828221081620452 | 0.030802109016452 | 2917 | 0 | 0 | 91 | 106 | 0 | 0 | 6 | 7 | 1 | 0 | 0 | 0 |
| L1PA16_dup10755 | 25.846221282442 | -4.97015701116671 | 1.32812397191449 | -3.74223876405321 | 0.000182388096277192 | 0.0104323360473012 | 2918 | 1 | 0 | 125 | 222 | 1 | 3 | 9 | 3 | 0 | 0 | 1 | 0 |

| | | | | | | | | | | | | | | | | | | | |
|---|---|---|---|---|---|---|---|---|---|---|---|---|---|---|---|---|---|---|---|
| MER52-int_dup367 | 27.1620903706769 | -6.29078054128467 | 1.6444169960075 | -3.82553850790774 | 0.000130486534573773 | 0.00823137565103151 | 2919 | 0 | 0 | 143 | 226 | 1 | 0 | 4 | 5 | 0 | 1 | 0 | 1 |
| L1MC3_dup11802 | 10.8624844593384 | -3.64385938666357 | 1.05243579505585 | -3.46231038870186 | 0.000535559129520573 | 0.0226147564565931 | 2920 | 0 | 1 | 53 | 54 | 1 | 0 | 16 | 8 | 4 | 6 | 1 | 1 |
| AluSz_dup99536 | 10.0072785834183 | -3.67066523739472 | 1.08238621498504 | -3.39127123625225 | 0.000695692140370923 | 0.0272041350916515 | 2921 | 0 | 1 | 49 | 58 | 1 | 0 | 15 | 6 | 2 | 3 | 1 | 0 |
| L1ME4b_dup19341 | 10.5227190452396 | -3.7240048069681 | 1.11334818458806 | -3.34486987855107 | 0.000823211819694393 | 0.0306820887888357 | 2922 | 0 | 1 | 45 | 67 | 1 | 0 | 15 | 7 | 4 | 2 | 1 | 0 |
| L2_dup51447 | 11.0055637949944 | -4.0054719929185 | 0.89787945735127 | -4.46103534291182 | 8.15646359014146e-06 | 0.00104400101421183 | 2923 | 0 | 1 | 46 | 77 | 2 | 0 | 10 | 11 | 1 | 1 | 0 | 1 |
| MSTA-int_dup1584 | 13.0522679507192 | -3.46069539730279 | 0.782242248432341 | -4.42407119308401 | 9.68580784587175e-06 | 0.00119442839680939 | 2924 | 3 | 2 | 55 | 75 | 1 | 0 | 26 | 12 | 1 | 2 | 0 | 1 |
| LTR82A_dup945 | 7.4803326765305 | -2.94760858770545 | 0.83930910224337 | -3.51194640904865 | 0.000444837689645691 | 0.0199402598648304 | 2925 | 0 | 0 | 27 | 34 | 3 | 1 | 16 | 14 | 1 | 2 | 0 | 0 |
| L1MD3_dup4663 | 7.56577492311073 | -3.44711063133032 | 0.910300852156896 | -3.78678172514354 | 0.000152611047661449 | 0.00921654526569828 | 2926 | 1 | 0 | 38 | 45 | 1 | 1 | 7 | 7 | 1 | 1 | 0 | 0 |
| MLT2B1_dup4003 | 7.88689010991881 | -3.35810443050905 | 0.854566686432291 | -3.9295990398698 | 8.50876256034513e-05 | 0.00605269362087857 | 2927 | 1 | 0 | 29 | 49 | 1 | 1 | 13 | 9 | 1 | 2 | 0 | 1 |
| L1MB8_dup14689 | 11.7083430231182 | -5.96625846223224 | 1.80425836130352 | -3.30676503442767 | 0.000943800212085843 | 0.0337653714125662 | 2928 | 0 | 0 | 53 | 80 | 0 | 0 | 17 | 8 | 2 | 1 | 0 | 0 |
| L1PB1_dup11944 | 8.30303499129028 | -4.77570038073714 | 1.14098082841195 | -4.18560966303361 | 2.8440171943817e-05 | 0.00265381378562489 | 2929 | 0 | 0 | 44 | 50 | 0 | 0 | 11 | 6 | 1 | 1 | 0 | 0 |
| L2_dup51449 | 8.08913427001132 | -4.66143443767075 | 1.03560094156866 | -4.50118790990081 | 6.75747308722497e-06 | 0.000904338409006093 | 2930 | 0 | 0 | 33 | 49 | 0 | 0 | 17 | 9 | 1 | 1 | 0 | 0 |
| L1M6_dup5196 | 7.75797615321604 | -3.89571968235509 | 1.16182830710834 | -3.35309413492523 | 0.000799135221054244 | 0.0300396570788267 | 2931 | 0 | 0 | 38 | 51 | 1 | 0 | 8 | 5 | 1 | 2 | 0 | 0 |
| MIRb_dup198505 | 7.22184514249061 | -4.01785841857258 | 1.01656018647311 | -3.95240584083102 | 7.736937679549e-05 | 0.0056402370499325 | 2932 | 0 | 0 | 33 | 45 | 1 | 0 | 13 | 5 | 1 | 1 | 0 | 0 |
| L2_dup51446 | 8.1351644413905 | -4.86297076314163 | 1.11083884297243 | -4.37774641560885 | 1.19912752092144e-05 | 0.00139524497356075 | 2933 | 0 | 0 | 31 | 61 | 0 | 0 | 7 | 9 | 1 | 1 | 1 | 0 |
| LTR82A_dup944 | 8.24822416624559 | -5.32409962475744 | 1.63751077740408 | -3.25133715040193 | 0.00114863557508981 | 0.0386581669802432 | 2934 | 0 | 0 | 32 | 56 | 0 | 0 | 13 | 6 | 2 | 3 | 1 | 0 |
| MLT1H1_dup508 | 10.1790939684755 | -5.53371485477368 | 1.00045861948938 | -5.53117814867544 | 3.18087067966435e-08 | 1.32785167498844e-05 | 2935 | 0 | 0 | 43 | 59 | 0 | 0 | 20 | 13 | 0 | 1 | 0 | 2 |
| THE1B_dup20516 | 9.02511136496458 | -5.41682564298116 | 1.00125248929523 | -5.4100496137533 | 6.30072909453752e-08 | 2.31006669511375e-05 | 2936 | 0 | 0 | 38 | 53 | 0 | 0 | 19 | 10 | 0 | 1 | 1 | 1 |
| AluJr_dup40174 | 7.0783160563626 | -3.99472104871543 | 1.02348943285401 | -3.90304083313894 | 9.49916692767876e-05 | 0.00656297553360467 | 2937 | 0 | 0 | 30 | 40 | 1 | 0 | 15 | 8 | 0 | 2 | 0 | 0 |
| L2a_dup152050 | 9.16865081471698 | -5.56828095065655 | 1.59291054763559 | -3.49566456127856 | 0.000472882760669709 | 0.0207525933917954 | 2938 | 0 | 0 | 42 | 62 | 0 | 0 | 16 | 6 | 0 | 1 | 0 | 0 |
| L1PA3_dup10599 | 18.8688528356145 | -5.07433603340487 | 1.36591387999666 | -3.71497508570399 | 0.000203223716602328 | 0.0112524156990625 | 2939 | 0 | 0 | 99 | 123 | 3 | 0 | 18 | 14 | 0 | 0 | 1 | 0 |
| SVA_D_dup758 | 14.7588466758036 | -5.03333477513495 | 0.812439849184286 | -6.19533222082665 | 5.81622190351637e-10 | 4.64411656729566e-07 | 2940 | 3 | 0 | 66 | 92 | 1 | 0 | 22 | 14 | 0 | 0 | 1 | 3 |
| AluSx1_dup112284 | 10.5619065290281 | -3.40898996368864 | 0.987550948425014 | -3.45196363704114 | 0.000556522721879302 | 0.0232795407758408 | 2941 | 0 | 0 | 50 | 50 | 1 | 2 | 21 | 10 | 4 | 1 | 1 | 1 |
| L2a_dup152006 | 9.89884746204685 | -4.11547950291923 | 1.04308139668577 | -3.94550177579192 | 7.96330418282791e-05 | 0.00575938108537134 | 2942 | 0 | 0 | 33 | 68 | 1 | 0 | 19 | 9 | 2 | 2 | 1 | 1 |
| MLT2B1_dup4005 | 9.8918102461869 | -4.62779670844823 | 0.953898739520673 | -4.85145489423088 | 1.22559084942786e-06 | 0.000240217825583165 | 2943 | 0 | 0 | 46 | 54 | 1 | 0 | 20 | 9 | 2 | 0 | 1 | 1 |
| AluJo_dup36031 | 14.5094488337296 | -4.7584266412606 | 0.915665947149655 | -5.19668407029108 | 2.02874712222068e-07 | 5.83714948904378e-05 | 2944 | 0 | 0 | 58 | 66 | 1 | 0 | 43 | 21 | 2 | 1 | 2 | 0 |
| LTR8_dup475 | 12.8612463200713 | -5.483038837437 | 0.851700613074258 | -6.43775377552645 | 1.21254486037789e-10 | 1.19717925040447e-07 | 2945 | 0 | 0 | 55 | 64 | 1 | 0 | 28 | 19 | 1 | 0 | 1 | 3 |
| MER74A_dup270 | 14.1127895138423 | -4.82016024730626 | 0.815237732254813 | -5.91258237517355 | 3.36785318730875e-09 | 2.00341798626842e-06 | 2946 | 0 | 0 | 64 | 77 | 2 | 0 | 25 | 17 | 1 | 1 | 1 | 2 |
| L1MCa_dup3375 | 9.61257735546296 | -4.05538452546029 | 0.722789074536767 | -5.61074408610752 | 2.0145849981539e-08 | 9.21850849943706e-06 | 2947 | 1 | 0 | 36 | 34 | 1 | 1 | 33 | 18 | 0 | 2 | 0 | 1 |
| L1PA16_dup7523 | 13.1413135241136 | -4.50329710933011 | 0.815551935814683 | -5.52177845648985 | 3.35585456275469e-08 | 1.38390324053592e-05 | 2948 | 0 | 0 | 50 | 62 | 1 | 1 | 36 | 23 | 0 | 2 | 0 | 0 |
| MLT1H_dup1400 | 11.3994219455071 | -3.99305605775354 | 1.00338461681826 | -3.97958668174082 | 6.90351857789289e-05 | 0.00519009362254439 | 2949 | 0 | 0 | 44 | 60 | 0 | 2 | 25 | 15 | 1 | 3 | 3 | 0 |
| AluJb_dup60101 | 13.3182203481428 | -4.82740555213066 | 0.85171518431623 | -5.66786367206331 | 1.44588932373637e-08 | 6.99777351904828e-06 | 2950 | 0 | 0 | 59 | 63 | 0 | 1 | 28 | 23 | 1 | 1 | 0 | 0 |
| MER102a_dup1376 | 9.34094107360134 | -3.69699206771991 | 0.952092598755286 | -3.88301733734004 | 0.000103168208943947 | 0.00696805638528996 | 2951 | 0 | 0 | 33 | 53 | 0 | 2 | 22 | 13 | 1 | 2 | 0 | 0 |
| Arthur1B_dup397 | 11.9786532549018 | -5.340408537031 | 0.804929369045888 | -6.63462999661843 | 3.25317534213382e-11 | 3.95767715681267e-08 | 2952 | 0 | 1 | 46 | 58 | 0 | 1 | 25 | 24 | 0 | 0 | 1 | 2 |
| L1PA12_dup1703 | 12.4010881095064 | -5.34175948692071 | 0.901841909677352 | -5.92316616648677 | 3.15801450952757e-09 | 1.89756798102032e-06 | 2953 | 0 | 0 | 62 | 68 | 0 | 2 | 15 | 16 | 0 | 0 | 2 | 1 |
| MER47A_dup384 | 8.56667878411533 | -4.33822034505258 | 0.78901754913173 | -5.49825583705527 | 3.83566167939166e-08 | 1.55538429305711e-05 | 2954 | 0 | 0 | 33 | 37 | 2 | 1 | 23 | 13 | 0 | 1 | 4 | 0 |
| L2b_dup16747 | 9.64727761017482 | -5.21973398890763 | 1.07558295276842 | -4.85293484381905 | 1.21647666649447e-06 | 0.000239417499491889 | 2955 | 0 | 0 | 42 | 55 | 0 | 0 | 19 | 10 | 0 | 2 | 3 | 0 |
| L2b_dup85159 | 11.5996239316699 | -5.10565645686701 | 0.899039005270908 | -5.67901551204502 | 1.3547219491959e-08 | 6.61911024483303e-06 | 2956 | 0 | 0 | 51 | 65 | 1 | 0 | 24 | 14 | 0 | 1 | 1 | 0 |
| MSTA-int_dup1618 | 9.43052034404987 | -2.64101589711825 | 0.690236160658946 | -3.82624969198508 | 0.000130110318863193 | 0.00821635613912456 | 2957 | 0 | 4 | 42 | 40 | 3 | 5 | 13 | 12 | 1 | 1 | 3 | 0 |
| L2a_dup159880 | 6.6877851008583 | -2.552397319044 | 0.791659433657281 | -3.22411028092285 | 0.00126364725700847 | 0.0412979148105248 | 2958 | 1 | 3 | 31 | 31 | 1 | 2 | 6 | 7 | 0 | 3 | 1 | 3 |
| MIR_dup167127 | 8.18828302991744 | -3.02590438518446 | 0.774653428367034 | -3.90613953850699 | 9.37823691710233e-05 | 0.00651002269111798 | 2959 | 3 | 2 | 38 | 46 | 1 | 1 | 12 | 5 | 0 | 2 | 1 | 1 |
| AluSq2_dup43462 | 7.45629740009105 | -3.59244293396036 | 0.770133752573151 | -4.66470002380416 | 3.09067568718975e-06 | 0.000491258461632633 | 2960 | 1 | 1 | 27 | 40 | 2 | 2 | 8 | 8 | 0 | 1 | 8 | 2 |
| L1MC_dup4156 | 7.96843174136661 | -3.15961821829744 | 0.782321422087463 | -4.03877246498843 | 5.37316548734895e-05 | 0.00431560351923593 | 2961 | 2 | 3 | 33 | 37 | 1 | 1 | 14 | 7 | 0 | 2 | 7 | 0 |
| AluSq_dup17735 | 5.76790374622439 | -2.94862354405299 | 0.923134837234228 | -3.19414177119266 | 0.00140247224720815 | 0.0444003008693706 | 2962 | 3 | 1 | 28 | 28 | 0 | 1 | 6 | 4 | 2 | 0 | 4 | 0 |
| L1MA4_dup2693 | 5.83260440698904 | -3.06709448935916 | 0.881254690038944 | -3.48037238726483 | 0.000500717294698598 | 0.021591822653924 | 2963 | 2 | 2 | 28 | 33 | 0 | 1 | 4 | 6 | 1 | 0 | 1 | 0 |
| ERVL-E-int_dup7492 | 6.46643615052749 | -2.97385446388657 | 0.818609501144239 | -3.63281205474621 | 0.000280349220669426 | 0.0142416685869785 | 2964 | 3 | 2 | 29 | 24 | 1 | 0 | 9 | 9 | 2 | 0 | 5 | 0 |
| L1PB3_dup1395 | 6.12226360888644 | -2.82819660852319 | 0.66764481199276 | -4.23607966050346 | 2.27456110810761e-05 | 0.0022504063094793 | 2965 | 4 | 2 | 21 | 24 | 1 | 1 | 15 | 9 | 1 | 0 | 3 | 0 |
| LTR1E_dup103 | 9.39326949062 | -2.81073749697927 | 0.800619841786363 | -3.51070177165218 | 0.000446925488776263 | 0.0199900956602322 | 2966 | 5 | 3 | 44 | 59 | 2 | 1 | 6 | 6 | 1 | 0 | 1 | 0 |
| LTR12C_dup699 | 6.61173608884047 | -2.87728511411952 | 0.83815559733479 | -3.43287704964192 | 0.000597212852107327 | 0.0244965366846285 | 2967 | 3 | 3 | 30 | 39 | 0 | 1 | 7 | 5 | 0 | 1 | 1 | 0 |
| THE1A-int_dup446 | 9.98374025873785 | -3.26896322550176 | 0.905256123617206 | -3.61109208788303 | 0.000304910364709289 | 0.0151276483822178 | 2968 | 3 | 2 | 58 | 58 | 0 | 1 | 7 | 4 | 1 | 1 | 1 | 0 |
| MSTA-int_dup1348 | 4.59313312069126 | -4.23963866067884 | 0.961963938047587 | -4.40727400788398 | 1.04679728536368e-05 | 0.00126565664056477 | 2969 | 0 | 0 | 18 | 15 | 1 | 0 | 16 | 10 | 0 | 0 | 0 | 0 |
| AluSg4_dup3392 | 4.72040511912074 | -4.82301517479987 | 1.08405745197194 | -4.44904019249776 | 8.62548755964409e-06 | 0.00108880650549977 | 2970 | 0 | 0 | 20 | 19 | 0 | 0 | 19 | 6 | 0 | 0 | 0 | 0 |
| L2b_dup26246 | 4.99259344671035 | -4.89275618120076 | 1.03884849041969 | -4.70978802618669 | 2.47974555896325e-06 | 0.00041669882540471 | 2971 | 0 | 0 | 13 | 22 | 0 | 0 | 19 | 12 | 0 | 0 | 0 | 0 |
| L1MD2_dup9605 | 5.72457845373765 | -4.66403130130671 | 1.13223292061977 | -4.1193214014248 | 3.799898181154e-05 | 0.00331205458821924 | 2972 | 0 | 0 | 34 | 24 | 0 | 0 | 13 | 5 | 0 | 1 | 0 | 0 |
| THE1B_dup20513 | 6.47023628490792 | -5.16788260283876 | 1.18528875604401 | -4.36001993310639 | 1.3005059686725e-05 | 0.0014855986232441 | 2973 | 0 | 0 | 40 | 31 | 0 | 0 | 13 | 4 | 0 | 0 | 0 | 0 |
| AluJb_dup110111 | 6.05477624091142 | -5.09001798807637 | 1.16484851053648 | -4.3696823595818 | 1.24427394703832e-05 | 0.00143514303733388 | 2974 | 0 | 0 | 26 | 39 | 0 | 0 | 6 | 10 | 0 | 0 | 0 | 0 |
| L1ME3A_dup13965 | 6.47585274942659 | -5.16765158680429 | 1.09993496513232 | -4.69814284536599 | 2.62537795340681e-06 | 0.000436339063657428 | 2975 | 0 | 0 | 30 | 35 | 0 | 0 | 10 | 11 | 0 | 0 | 0 | 0 |
| MER65-int_dup565 | 6.38353181124232 | -5.15083789266415 | 1.12736061009183 | -4.56893548218309 | 4.90207641293922e-06 | 0.00070993881567452 | 2976 | 0 | 0 | 37 | 29 | 0 | 0 | 11 | 8 | 0 | 0 | 0 | 0 |
| MLT1A0_dup3259 | 6.08619771209658 | -4.87193620656456 | 1.04064761039952 | -4.68163877750522 | 2.84590634505878e-06 | 0.000462233468383178 | 2977 | 0 | 0 | 29 | 28 | 0 | 0 | 10 | 11 | 0 | 1 | 0 | 1 |
| MER58B_dup6018 | 6.47706408479066 | -5.35058177237334 | 1.1252309462314 | -4.75509653399901 | 1.98351615475279e-06 | 0.000350385846294586 | 2978 | 0 | 0 | 37 | 34 | 0 | 0 | 8 | 7 | 0 | 0 | 0 | 1 |
| L1ME3C_dup3050 | 5.35105082639632 | -5.14087925726564 | 1.10874680202088 | -4.63665757402459 | 3.54088282652191e-06 | 0.000545685819326155 | 2979 | 0 | 0 | 29 | 27 | 0 | 0 | 9 | 6 | 0 | 0 | 0 | 1 |
| MSTA-int_dup1463 | 5.42422502268631 | -5.15799726237257 | 1.08971568801658 | -4.73334220943519 | 2.20852666375544e-06 | 0.00037888254184135 | 2980 | 0 | 0 | 24 | 32 | 0 | 0 | 11 | 6 | 0 | 0 | 0 | 1 |
| L3_dup42730 | 6.00154355630772 | -4.30535289149106 | 0.934210168531539 | -4.60854852207243 | 4.05489751351368e-06 | 0.000608423903846844 | 2981 | 0 | 0 | 17 | 23 | 0 | 0 | 24 | 13 | 1 | 1 | 0 | 0 |
| MIRb_dup198509 | 5.22908974663291 | -3.98895994993386 | 1.18439443420952 | -3.36793202899179 | 0.000757342650836005 | 0.0289442104715426 | 2982 | 0 | 0 | 29 | 19 | 0 | 0 | 12 | 6 | 1 | 2 | 0 | 0 |
| AluSx_dup54167 | 4.91846209505969 | -3.59538887485434 | 0.969723116976636 | -3.70764480284217 | 0.000209195800204714 | 0.0114906056961013 | 2983 | 1 | 0 | 21 | 26 | 0 | 0 | 10 | 6 | 1 | 1 | 0 | 0 |
| THE1B_dup20507 | 5.40702743916869 | -4.21793952611982 | 1.09453716068962 | -3.85362843547704 | 0.000116380169513376 | 0.00758274803259302 | 2984 | 0 | 0 | 21 | 34 | 0 | 0 | 12 | 5 | 1 | 1 | 0 | 0 |
| L1ME3A_dup13958 | 6.38847738576398 | -3.26732232316993 | 0.870133262352217 | -3.75496773257171 | 0.000173363811385035 | 0.0100883407835225 | 2985 | 1 | 2 | 30 | 32 | 0 | 0 | 10 | 8 | 1 | 1 | 0 | 0 |
| MIRb_dup198459 | 5.72585955343699 | -3.04041036578778 | 0.80788869718072 | -3.76340252858824 | 0.000167617075837853 | 0.009862466055221 | 2986 | 2 | 2 | 24 | 28 | 0 | 0 | 9 | 8 | 1 | 1 | 0 | 1 |
| L1M5_dup48888 | 5.88835930085277 | -3.34358082588736 | 1.04229504332211 | -3.20790245267823 | 0.00133706840829894 | 0.0429468249839498 | 2987 | 0 | 2 | 24 | 35 | 0 | 0 | 8 | 8 | 0 | 2 | 0 | 0 |
| L1MA7_dup3348 | 5.95883231112245 | -3.05772848503991 | 0.772849754561953 | -3.95643327437298 | 7.60771251802895e-05 | 0.00557664385050714 | 2988 | 2 | 3 | 22 | 26 | 0 | 0 | 12 | 12 | 0 | 1 | 0 | 0 |
| MamGypLTR1a_dup18 | 6.71859918994893 | -3.78217442408577 | 0.843504402879291 | -4.48388225500112 | 7.32971175309339e-06 | 0.00096517741715637 | 2989 | 1 | 1 | 28 | 28 | 0 | 0 | 14 | 13 | 0 | 2 | 0 | 1 |
| L1PA6_dup5780 | 5.6323542676294 | -3.5654675887484 | 1.01907607024094 | -3.49872565244852 | 0.000467487327172499 | 0.0205993962132199 | 2990 | 0 | 2 | 28 | 36 | 1 | 0 | 6 | 4 | 0 | 0 | 0 | 0 |
| LTR16A_dup3757 | 6.87598003682379 | -4.64301312788066 | 1.04592849888027 | -4.43913052646647 | 9.0323028415672e-06 | 0.00113175372929939 | 2991 | 0 | 0 | 36 | 39 | 1 | 0 | 10 | 7 | 0 | 0 | 0 | 0 |
| MLT1H_dup4915 | 6.27997401528895 | -4.1560839564473 | 0.918537078452867 | -4.52467739620002 | 6.04876993684396e-06 | 0.000834849542570923 | 2992 | 0 | 1 | 29 | 33 | 1 | 0 | 14 | 7 | 0 | 0 | 0 | 0 |
| ERV3-16A3_I-int_dup3 | 6.38242652242305 | -4.56324875655384 | 1.0407881192547 | -4.38441664747435 | 1.16297000742668e-05 | 0.00136099802516487 | 2993 | 0 | 1 | 22 | 46 | 0 | 0 | 9 | 9 | 0 | 0 | 0 | 0 |
| L1M6_dup3143 | 5.26862281447327 | -3.9072030286495 | 0.967920056450118 | -4.03670014131054 | 5.42082823643397e-05 | 0.00433422175922889 | 2994 | 0 | 2 | 13 | 35 | 0 | 0 | 11 | 10 | 0 | 0 | 0 | 0 |
| L2_dup51444 | 8.29098884039194 | -4.08868032096024 | 1.00296299472087 | -4.07660137261407 | 4.56987373324526e-05 | 0.0038127039808232 | 2995 | 0 | 2 | 34 | 58 | 0 | 0 | 15 | 5 | 1 | 0 | 0 | 0 |
| MER5A_dup30169 | 5.79450174328264 | -3.73525359237473 | 1.07795869958228 | -3.46511753541411 | 0.000529999792310963 | 0.0224357464118883 | 2996 | 0 | 1 | 22 | 36 | 0 | 0 | 10 | 7 | 2 | 0 | 0 | 0 |
| THE1B_dup20492 | 5.10415022638237 | -3.92363245647191 | 1.08124665572382 | -3.62880424711723 | 0.000284737040716497 | 0.0144027665769684 | 2997 | 0 | 1 | 27 | 29 | 0 | 0 | 7 | 4 | 1 | 0 | 0 | 0 |

| | | | | | | | | | | | | | | | | | | | |
|---|---|---|---|---|---|---|---|---|---|---|---|---|---|---|---|---|---|---|---|
| MIRc_dup89853 | 5.7253624041955 | -3.11079452209173 | 0.810499130736181 | -3.83812197215582 | 0.000123978898319715 | 0.00793432885336416 | 2998 | 2 | 0 | 18 | 34 | 0 | 3 | 12 | 7 | 0 | 1 | 1 | 0 |
| L1PB1_dup2838 | 6.850165099491 | -4.19546543427007 | 0.920139083853462 | -4.55959920395928 | 5.12513417048254e-06 | 0.000735527850017876 | 2999 | 0 | 1 | 28 | 46 | 0 | 1 | 9 | 6 | 0 | 1 | 1 | 1 |
| L2c_dup126708 | 7.36988737279642 | -3.9674589818492 | 1.08024781988586 | -3.67273037613574 | 0.000239972643599778 | 0.0127371248391686 | 3000 | 0 | 0 | 41 | 33 | 0 | 1 | 10 | 8 | 0 | 3 | 1 | 1 |
| LTR85b_dup1085 | 6.83760621370815 | -4.09375957811165 | 1.02458803132347 | -3.99551766462048 | 6.45530433828774e-05 | 0.0049341916032066 | 3001 | 0 | 0 | 38 | 32 | 0 | 1 | 9 | 8 | 0 | 2 | 0 | 1 |
| THE1D_dup11515 | 6.2033267460061 | -3.72336297996344 | 1.05681304133376 | -3.52319931183319 | 0.000426370592073144 | 0.0193582707786507 | 3002 | 0 | 0 | 34 | 33 | 0 | 2 | 4 | 8 | 1 | 0 | 0 | 0 |
| THE1D_dup11520 | 7.63611455024957 | -3.76538652645501 | 0.896819130570562 | -4.19860192328797 | 2.68567966742068e-05 | 0.00253811792759312 | 3003 | 0 | 1 | 40 | 36 | 0 | 1 | 11 | 10 | 1 | 1 | 0 | 0 |
| HAL1_dup2407 | 4.85040051031125 | -3.46832103060372 | 0.996135781034403 | -3.48177537303414 | 0.000498101338168102 | 0.0215170420094504 | 3004 | 0 | 1 | 21 | 31 | 0 | 1 | 9 | 3 | 0 | 1 | 0 | 0 |
| Tigger3a_dup4842 | 3.99185616344307 | -3.56234675998274 | 1.0080302728275 | -3.53396803251796 | 0.000409370587923475 | 0.0187937669137641 | 3005 | 0 | 1 | 17 | 23 | 0 | 1 | 10 | 3 | 0 | 0 | 0 | 0 |
| L1M4_dup13364 | 6.62835893546193 | -4.56879084014848 | 1.16126012204696 | -3.93433887327075 | 8.34260114486679e-05 | 0.0059701911940345 | 3006 | 0 | 0 | 34 | 47 | 0 | 1 | 7 | 3 | 0 | 0 | 0 | 0 |
| REP522_dup166 | 6.99642454767393 | -4.63631270257039 | 1.13767085197599 | -4.07526719570754 | 4.5961542978667e-05 | 0.00382925956078498 | 3007 | 0 | 0 | 31 | 54 | 0 | 1 | 6 | 5 | 0 | 0 | 0 | 0 |
| L1ME4b_dup19342 | 5.63667730397456 | -3.30837203410849 | 1.01362060843646 | -3.26391551885646 | 0.00109883925077653 | 0.0374537178577956 | 3008 | 1 | 0 | 23 | 42 | 0 | 2 | 5 | 4 | 0 | 1 | 0 | 0 |
| L2c_dup126699 | 6.1774643723704 | -4.17651990629897 | 1.03108046717371 | -4.05062460134389 | 5.1081100417337e-05 | 0.00415114191253541 | 3009 | 0 | 0 | 29 | 37 | 0 | 1 | 7 | 8 | 0 | 1 | 0 | 0 |
| L1PA4_dup6269 | 8.35313516524596 | -4.15207002692769 | 0.915941929234266 | -4.5331149218148 | 5.81201727799426e-06 | 0.000805994778747697 | 3010 | 1 | 1 | 42 | 52 | 0 | 1 | 10 | 7 | 0 | 0 | 0 | 0 |
| MSTA-int_dup1462 | 10.5498369068127 | -4.1803224766766 | 0.762775166523314 | -5.48041239429736 | 4.24335588919996e-08 | 1.67167145763506e-05 | 3011 | 2 | 2 | 40 | 66 | 0 | 1 | 17 | 14 | 0 | 0 | 1 | 0 |
| L1P4_dup1851 | 11.7477740101322 | -4.27928735535982 | 0.83561439484227 | -5.12112689988733 | 3.03715166185518e-07 | 8.06560367557806e-05 | 3012 | 4 | 1 | 44 | 81 | 0 | 0 | 22 | 10 | 0 | 0 | 0 | 1 |
| LTR16_dup2938 | 9.92254341222918 | -4.05644504400535 | 0.835195952727877 | -4.85687823409151 | 1.19250875519137e-06 | 0.000236067128338407 | 3013 | 3 | 2 | 49 | 56 | 0 | 0 | 15 | 9 | 0 | 0 | 0 | 1 |
| L1MC_dup7881 | 9.96834488043521 | -3.68852084736154 | 0.806821039228226 | -4.57167161987971 | 4.83848773333245e-06 | 0.000702496054992057 | 3014 | 3 | 3 | 51 | 50 | 0 | 0 | 17 | 8 | 1 | 0 | 0 | 2 |
| L3_dup5399 | 7.76088287464777 | -3.41144965994399 | 0.77621240080623 | -4.39499505083997 | 1.10775277548851e-05 | 0.00131792670958195 | 3015 | 5 | 1 | 25 | 43 | 0 | 0 | 17 | 11 | 1 | 0 | 0 | 2 |
| MSTA-int_dup1588 | 8.69015761803722 | -4.61511746878778 | 0.954670816918244 | -4.83425007552421 | 1.33648493048743e-06 | 0.000258756097046185 | 3016 | 1 | 0 | 48 | 37 | 0 | 0 | 22 | 8 | 1 | 0 | 0 | 0 |
| LTR46_dup95 | 6.96061170877219 | -3.29065947553434 | 0.76275753105304 | -4.31416189492269 | 1.60209514849163e-05 | 0.00173514853073732 | 3017 | 3 | 0 | 25 | 38 | 1 | 1 | 13 | 11 | 1 | 0 | 0 | 0 |
| MER103C_dup7720 | 8.45525494375798 | -3.56112928405281 | 0.923206015671516 | -3.85735060604273 | 0.000114622692630374 | 0.00749697944492222 | 3018 | 1 | 1 | 45 | 42 | 1 | 0 | 10 | 10 | 2 | 0 | 0 | 0 |
| L2a_dup152019 | 7.03582225211887 | -4.34382913515963 | 0.989638640441093 | -4.38930833705476 | 1.13711746883602e-05 | 0.00134212576011734 | 3019 | 0 | 0 | 24 | 49 | 0 | 1 | 15 | 7 | 1 | 0 | 0 | 0 |
| MLT1D_dup10306 | 7.24523197403236 | -3.89629738243762 | 0.821064575562965 | -4.74542136928282 | 2.0807273152341e-06 | 0.000362977083394936 | 3020 | 0 | 0 | 32 | 33 | 1 | 2 | 21 | 7 | 1 | 0 | 1 | 0 |
| MER41-int_dup514 | 5.71431405255218 | -3.55907891920832 | 0.853086628678341 | -4.17200176343437 | 3.01935299714944e-05 | 0.00276806887533308 | 3021 | 0 | 1 | 21 | 24 | 0 | 1 | 15 | 10 | 2 | 0 | 0 | 1 |
| MSTA-int_dup1613 | 8.03398608266466 | -3.43643812620395 | 0.842319319892334 | -4.07973323779774 | 4.5087411532313e-05 | 0.00378392824902677 | 3022 | 1 | 1 | 35 | 41 | 0 | 2 | 22 | 5 | 2 | 0 | 0 | 1 |
| MLT1J1_dup808 | 9.40512422160325 | -5.21564724085709 | 1.11086486978254 | -4.69512303677233 | 2.66446300825761e-06 | 0.000441195765457805 | 3023 | 0 | 0 | 38 | 55 | 0 | 0 | 24 | 7 | 2 | 0 | 3 | 0 |
| L2c_dup126703 | 7.12757221602532 | -5.47716891555694 | 1.1562113213369 | -4.73716942091852 | 2.16723977739423e-06 | 0.000373414942847101 | 3024 | 0 | 0 | 36 | 44 | 0 | 0 | 15 | 3 | 0 | 0 | 1 | 0 |
| MSTA-int_dup1590 | 8.37485981235801 | -5.78409393017235 | 1.11807527861651 | -5.17325983392593 | 2.30044756171855e-07 | 6.41714297210646e-05 | 3025 | 0 | 0 | 52 | 39 | 0 | 0 | 14 | 6 | 0 | 0 | 2 | 0 |
| MIRb_dup212387 | 9.35672573105319 | -3.64009382902666 | 0.885201895020982 | -4.112162264339 | 3.91970638810517e-05 | 0.00338663201243309 | 3026 | 2 | 0 | 42 | 52 | 0 | 1 | 20 | 6 | 2 | 1 | 1 | 1 |
| Charlie15a_dup559 | 6.08435553313242 | -3.49411092886964 | 1.07806670762997 | -3.24108972491241 | 0.0011907369887526 | 0.0396156464661251 | 3027 | 1 | 0 | 22 | 28 | 0 | 0 | 19 | 7 | 1 | 3 | 1 | 0 |
| L1ME3G_dup15981 | 7.80784897915315 | -3.59597881067994 | 0.978157614920821 | -3.67627747903493 | 0.000236662100981028 | 0.0126007608592597 | 3028 | 2 | 0 | 26 | 50 | 0 | 0 | 12 | 11 | 2 | 1 | 1 | 0 |
| L1PA7_dup2240 | 7.06083449071289 | -4.89436180807483 | 1.02772641870473 | -4.76231973703988 | 1.91380131598562e-06 | 0.000340854317315505 | 3029 | 0 | 0 | 28 | 48 | 1 | 0 | 10 | 4 | 1 | 0 | 1 | 5 |
| MLT1F_dup689 | 6.26312777346462 | -4.239735906097 | 0.958689737578322 | -4.42242754867356 | 9.75980819228943e-06 | 0.00120070800872378 | 3030 | 0 | 0 | 27 | 31 | 1 | 0 | 13 | 6 | 2 | 0 | 3 | 1 |
| L1PA6_dup5356 | 6.30586192665512 | -4.03034676445645 | 0.85617852639029 | -4.70736726071452 | 2.50936575607048e-06 | 0.000420784738684952 | 3031 | 0 | 0 | 21 | 32 | 1 | 1 | 16 | 5 | 2 | 0 | 3 | 5 |
| L1MB7_dup15544 | 6.33410037535241 | -4.40012496679221 | 0.896543721738966 | -4.90787550021273 | 9.20682660905877e-07 | 0.000193527169992853 | 3032 | 0 | 0 | 19 | 39 | 1 | 1 | 11 | 7 | 1 | 0 | 5 | 2 |
| MIR_dup84426 | 6.30803372857198 | -4.39272089626204 | 0.886669806379821 | -4.95417895664797 | 7.26363342467278e-07 | 0.000159884599340166 | 3033 | 0 | 0 | 22 | 37 | 1 | 1 | 11 | 6 | 1 | 0 | 3 | 4 |
| MIR_dup84427 | 5.8442219815076 | -5.16853288470321 | 1.04049979736321 | -4.9673559743126 | 6.78719313721093e-07 | 0.00015135758746268 | 3034 | 0 | 0 | 16 | 44 | 0 | 1 | 8 | 6 | 0 | 0 | 3 | 3 |
| L1ME4c_dup1408 | 6.31187797793741 | -5.71202108225287 | 1.03216961139318 | -5.5339946256924 | 3.13018568359667e-08 | 1.31592784923868e-05 | 3035 | 0 | 0 | 28 | 36 | 0 | 0 | 11 | 6 | 0 | 0 | 3 | 2 |
| AluYj4_dup1666 | 6.25325416783529 | -5.74710440602087 | 1.03238231972932 | -5.5668373006695 | 2.5940439009603e-08 | 1.14304142603315e-05 | 3036 | 0 | 0 | 26 | 36 | 0 | 0 | 7 | 9 | 0 | 0 | 2 | 4 |
| THE1D-int_dup174 | 5.76316423645548 | -5.60477062178701 | 1.03582419641468 | -5.41092845792453 | 6.26988186872946e-08 | 2.30478783813032e-05 | 3037 | 0 | 0 | 27 | 30 | 0 | 0 | 7 | 8 | 0 | 0 | 2 | 3 |
| MLT1A-int_dup100 | 10.0032455685749 | -6.20965670045064 | 1.06206093605923 | -5.84679888848143 | 5.01122749576963e-09 | 2.80565067696094e-06 | 3038 | 0 | 0 | 43 | 70 | 0 | 0 | 15 | 6 | 0 | 0 | 3 | 2 |
| L1MA3_dup7268 | 7.44712215055039 | -5.8787521780247 | 0.999676889719166 | -5.88065227723349 | 4.08652851612059e-09 | 2.3487273292194e-06 | 3039 | 0 | 0 | 24 | 45 | 0 | 0 | 21 | 8 | 0 | 0 | 1 | 4 |
| MLT1L_dup10543 | 7.38085515194266 | -5.19876696671596 | 0.996519629872273 | -5.2169237924418 | 1.81919061003003e-07 | 5.35066908398277e-05 | 3040 | 0 | 0 | 30 | 46 | 0 | 1 | 16 | 6 | 0 | 0 | 1 | 2 |
| MIRb_dup198503 | 5.69101894564845 | -5.22058388866086 | 1.09163820619562 | -4.78233892788957 | 1.73267212948617e-06 | 0.000316490373337472 | 3041 | 0 | 0 | 28 | 30 | 0 | 0 | 9 | 8 | 0 | 0 | 1 | 0 |
| ERVL-int_dup121 | 4.4014118617245 | -4.93977627380208 | 1.16884732445141 | -4.22619461966132 | 2.37676448993543e-05 | 0.00232350699803688 | 3042 | 0 | 0 | 24 | 24 | 0 | 0 | 5 | 5 | 0 | 0 | 1 | 0 |
| L1MA4_dup9148 | 4.93319944730938 | -5.06613360885163 | 1.14038551996516 | -4.44247451423831 | 8.89301696961862e-06 | 0.00111606424886755 | 3043 | 0 | 0 | 18 | 35 | 0 | 0 | 6 | 7 | 0 | 0 | 1 | 0 |
| MSTA_dup18847 | 4.83444898466843 | -5.03995629197554 | 1.11075757720354 | -4.53740437644747 | 5.69508396891013e-06 | 0.000794793225845331 | 3044 | 0 | 0 | 20 | 28 | 0 | 0 | 6 | 9 | 0 | 0 | 1 | 0 |
| L2c_dup126707 | 5.11546809533588 | -4.71291796369261 | 1.11048058455829 | -4.24403454614852 | 2.19536532576132e-05 | 0.00218787777556704 | 3045 | 0 | 0 | 29 | 23 | 0 | 0 | 8 | 6 | 0 | 1 | 1 | 0 |
| MLT2C2_dup1167 | 6.52534333876939 | -5.01639585546935 | 1.17009136213203 | -4.2871830506713 | 1.80953175695481e-05 | 0.0018967878565655 | 3046 | 0 | 0 | 34 | 42 | 0 | 0 | 6 | 5 | 0 | 1 | 1 | 0 |
| L1M5_dup5177 | 5.42395296117701 | -3.93323407002651 | 1.0449761782971 | -3.76394615658715 | 0.000167252910861269 | 0.00984590824537246 | 3047 | 0 | 1 | 18 | 37 | 0 | 0 | 8 | 6 | 0 | 2 | 2 | 0 |
| L1MC2_dup3842 | 4.59783588611206 | -4.01958550007054 | 1.01192916101212 | -3.97220048096073 | 7.12117469813189e-05 | 0.00530345863762657 | 3048 | 0 | 1 | 21 | 25 | 0 | 0 | 8 | 5 | 0 | 1 | 1 | 0 |
| LTR78_dup1829 | 4.17602989631889 | -3.58500852135742 | 0.963570661541948 | -3.72054553385896 | 0.000198792894457604 | 0.0110673781152572 | 3049 | 0 | 2 | 19 | 18 | 0 | 0 | 7 | 6 | 0 | 1 | 2 | 0 |
| MLT1D_dup1698 | 4.61619554930391 | -4.4237353541773 | 1.08371819630212 | -4.08199785633574 | 4.46502087708611e-05 | 0.00375551027790432 | 3050 | 0 | 1 | 19 | 29 | 0 | 0 | 11 | 3 | 0 | 0 | 1 | 0 |
| MSTD_dup6960 | 5.98709048113218 | -4.71499223203232 | 1.02593638627178 | -4.59579394504804 | 4.31104570357109e-06 | 0.000639575212342854 | 3051 | 0 | 1 | 24 | 38 | 0 | 0 | 9 | 8 | 0 | 0 | 1 | 0 |
| LTR12C_dup1574 | 4.89660600372486 | -4.10842182172602 | 1.18779019174207 | -3.45887838634231 | 0.000542429763777144 | 0.0228238713654671 | 3052 | 1 | 0 | 18 | 43 | 0 | 0 | 3 | 2 | 0 | 1 | 1 | 0 |
| L1M4c_dup3371 | 5.21110836362202 | -3.73087623465561 | 1.02619499157493 | -3.63564065824347 | 0.000277290641400597 | 0.0141315482884357 | 3053 | 1 | 1 | 26 | 33 | 0 | 0 | 7 | 2 | 0 | 1 | 1 | 0 |
| L1MA4A_dup3229 | 3.93199930378635 | -3.96617271267442 | 1.11359524519342 | -3.56159271494155 | 0.000368611893395049 | 0.0173992710941833 | 3054 | 0 | 1 | 18 | 24 | 0 | 0 | 6 | 2 | 0 | 1 | 2 | 0 |
| LTR16C_dup4625 | 4.7852058337944 | -4.01480170442064 | 1.02524827039333 | -3.91593121428078 | 9.00559238096208e-05 | 0.00630805029343657 | 3055 | 2 | 0 | 19 | 31 | 0 | 0 | 6 | 6 | 0 | 0 | 1 | 0 |
| MER103C_dup807 | 3.47460340247742 | -4.27682535368458 | 1.13998824188388 | -3.75163988237014 | 0.000175681688523241 | 0.0101678703936047 | 3056 | 1 | 0 | 14 | 20 | 0 | 0 | 3 | 6 | 0 | 0 | 2 | 0 |
| L1MA10_dup2243 | 5.381974761084 | -3.79378260606845 | 0.982387396160179 | -3.86179894092398 | 0.000112555179477259 | 0.00741679721293155 | 3057 | 3 | 1 | 20 | 34 | 0 | 0 | 8 | 4 | 0 | 0 | 4 | 0 |
| MLT1G3_dup2647 | 4.04794012546593 | -4.05199538729685 | 1.10432508130748 | -3.66920525113805 | 0.000243305680723964 | 0.0128652474456765 | 3058 | 1 | 1 | 22 | 21 | 0 | 0 | 5 | 2 | 0 | 0 | 3 | 0 |
| L1MB7_dup2180 | 4.66404092310539 | -4.80529409137635 | 1.08831264571775 | -4.41536180828556 | 1.00841183910754e-05 | 0.00123021735284818 | 3059 | 0 | 0 | 22 | 19 | 0 | 0 | 6 | 9 | 1 | 0 | 3 | 0 |
| AluJr_dup8501 | 3.8385273586777 | -3.9517104673169 | 0.957225152370324 | -4.12829777563982 | 3.65458647609935e-05 | 0.00321120470325678 | 3060 | 1 | 0 | 12 | 15 | 0 | 0 | 11 | 8 | 1 | 0 | 2 | 0 |
| MER51A_dup154 | 3.6078045898317 | -3.87392583875952 | 1.03910900140074 | -3.72812268350808 | 0.000192911466795485 | 0.0108311731661441 | 3061 | 1 | 0 | 16 | 16 | 0 | 0 | 5 | 6 | 1 | 0 | 2 | 0 |
| L1MEd_dup12001 | 6.28475849405913 | -4.78965449941051 | 0.997013780444446 | -4.80400029904841 | 1.55526596038073e-06 | 0.000291163897882576 | 3062 | 1 | 0 | 35 | 25 | 0 | 0 | 12 | 9 | 0 | 0 | 1 | 0 |
| THE1D_dup822 | 4.99608067795916 | -4.52994317713306 | 1.04470265688198 | -4.33610764488064 | 1.45027941369227e-05 | 0.00161180843237002 | 3063 | 1 | 0 | 28 | 21 | 0 | 0 | 8 | 7 | 0 | 0 | 1 | 0 |
| MER20_dup1376 | 4.3322867509092 | -4.23502028263951 | 1.02679889858223 | -4.12448853274686 | 3.71559502647225e-05 | 0.00325879459701057 | 3064 | 2 | 0 | 16 | 21 | 0 | 0 | 9 | 6 | 0 | 0 | 4 | 0 |
| THE1B-int_dup626 | 4.74253011332283 | -3.7873323926093 | 0.964721826242394 | -3.92582845084061 | 8.64317409616002e-05 | 0.00612632917332407 | 3065 | 3 | 0 | 22 | 21 | 0 | 0 | 11 | 5 | 0 | 0 | 2 | 0 |
| L3_dup8176 | 5.24508441913112 | -5.38376718846154 | 1.12366935945848 | -4.79123787005823 | 1.65755472115683e-06 | 0.000305269748978625 | 3066 | 0 | 0 | 14 | 29 | 0 | 0 | 25 | 3 | 0 | 0 | 3 | 0 |
| MLT1F2_dup2789 | 3.68382323392197 | -3.40878783380396 | 1.03267982800168 | -3.300914515199 | 0.000963702403619455 | 0.0342712796490263 | 3067 | 0 | 2 | 16 | 18 | 0 | 0 | 9 | 2 | 1 | 0 | 2 | 0 |
| AluSg7_dup4056 | 3.49826035685124 | -4.68435157035295 | 1.17771105280098 | -3.97750497391702 | 6.96421665165034e-05 | 0.00522581991609458 | 3068 | 0 | 0 | 21 | 15 | 0 | 0 | 7 | 3 | 0 | 0 | 1 | 0 |
| L1ME4b_dup21263 | 3.56746135820547 | -4.70991384171194 | 1.21989309706926 | -3.86092343093612 | 0.0001129593010787 | 0.00742697260416478 | 3069 | 0 | 0 | 19 | 21 | 0 | 0 | 6 | 2 | 0 | 0 | 1 | 0 |
| LTR45C_dup357 | 3.74754227000067 | -4.76534725653231 | 1.24125787704919 | -3.83912750496362 | 0.000123472300516988 | 0.00791028362999303 | 3070 | 0 | 0 | 23 | 20 | 0 | 0 | 5 | 2 | 0 | 0 | 1 | 0 |
| L1MA10_dup4458 | 4.77675263534576 | -5.03486257230566 | 1.15248873152964 | -4.36868702882945 | 1.2499574469827e-05 | 0.00144030205656051 | 3071 | 0 | 0 | 17 | 34 | 0 | 0 | 12 | 3 | 0 | 0 | 1 | 0 |
| MLT1D_dup1696 | 4.44935320557185 | -4.96173884881755 | 1.23636379672922 | -4.01317060718193 | 5.99085809233213e-05 | 0.00466887125358525 | 3072 | 0 | 0 | 18 | 31 | 0 | 0 | 12 | 1 | 0 | 0 | 1 | 0 |
| L1MB7_dup22773 | 6.53715952845919 | -3.80861866851645 | 0.790808119299334 | -4.81610971810827 | 1.46384055067672e-06 | 0.000276977082058207 | 3073 | 2 | 1 | 21 | 15 | 0 | 0 | 34 | 12 | 0 | 1 | 0 | 1 |
| L1MB7_dup22777 | 8.30264191299679 | -4.4357465824709 | 0.769942756662211 | -5.76113814188 | 8.35486226683751e-09 | 4.33116788005429e-06 | 3074 | 1 | 1 | 24 | 24 | 0 | 0 | 32 | 23 | 0 | 1 | 0 | 1 |
| ERVL-E-int_dup4370 | 6.05987797185954 | -4.21418767125856 | 0.921414062354041 | -4.57360902490689 | 4.7939404120903e-06 | 0.00069981530386211 | 3075 | 0 | 1 | 14 | 27 | 0 | 0 | 31 | 9 | 0 | 1 | 0 | 0 |
| L1ME4a_dup15897 | 5.57477095101167 | -3.88097238666376 | 0.916801459187126 | -4.23316558647801 | 2.30424717573982e-05 | 0.0022758591751057 | 3076 | 0 | 1 | 20 | 23 | 0 | 0 | 20 | 8 | 0 | 2 | 0 | 1 |

| | | | | | | | | | | | | | | | | | | | |
|---|---|---|---|---|---|---|---|---|---|---|---|---|---|---|---|---|---|---|---|
| MLT2B1_dup422 | 6.42902502791714 | -4.45194272855556 | 0.879519705147802 | -5.06178849944859 | 4.15341639236587e-07 | 0.000103485739989308 | 3077 | 0 | 1 | 20 | 31 | 0 | 0 | 23 | 10 | 0 | 1 | 0 | 1 |
| MLT1D_dup15736 | 5.99955358901264 | -3.99501529948578 | 1.08586204162303 | -3.67911865996757 | 0.000234041361870848 | 0.0124947735902447 | 3078 | 0 | 0 | 12 | 22 | 0 | 0 | 23 | 15 | 1 | 3 | 1 | 1 |
| L1PA13_dup1658 | 6.50788312960495 | -5.14426943537061 | 0.978231226577814 | -5.2587458829821 | 1.4504115491514e-07 | 4.45890990276975e-05 | 3079 | 0 | 0 | 13 | 33 | 0 | 0 | 17 | 18 | 0 | 1 | 1 | 2 |
| MER65-int_dup717 | 5.06091159576401 | -4.10152636467235 | 1.06566942516135 | -3.84877924413694 | 0.000118707923684075 | 0.0076969043281792 | 3080 | 0 | 0 | 8 | 18 | 0 | 0 | 17 | 15 | 0 | 3 | 2 | 2 |
| L1ME1_dup30803 | 4.48047856793865 | -4.14325744562682 | 1.08069787946274 | -3.83387209724758 | 0.000126141749131131 | 0.008033973404378 | 3081 | 0 | 0 | 15 | 23 | 0 | 0 | 9 | 9 | 0 | 2 | 1 | 0 |
| MIRc_dup18013 | 5.145899418652 | -4.81893255905376 | 1.01365732142977 | -4.75400557681231 | 1.99425532546655e-06 | 0.000351760620958831 | 3082 | 0 | 0 | 15 | 26 | 0 | 0 | 16 | 9 | 0 | 1 | 2 | 0 |
| MIR_dup28629 | 5.92521978446328 | -5.06274475350413 | 1.02918476094071 | -4.91917967078777 | 8.69076684386352e-07 | 0.000184967192077813 | 3083 | 0 | 0 | 31 | 25 | 0 | 0 | 12 | 7 | 0 | 1 | 2 | 1 |
| MLT1C_dup1642 | 5.06816972423032 | -4.87243586513933 | 0.992678156731857 | -4.90837421181967 | 9.18345008100838e-07 | 0.000193377452475719 | 3084 | 0 | 0 | 23 | 17 | 0 | 0 | 12 | 10 | 0 | 1 | 1 | 2 |
| MLT1D_dup2951 | 7.29980137555306 | -3.82410500739365 | 0.923839379592265 | -4.13936133473917 | 3.48274034127353e-05 | 0.00308642135286656 | 3085 | 1 | 0 | 31 | 47 | 1 | 0 | 14 | 4 | 1 | 1 | 0 | 1 |
| MSTC_dup530 | 7.97170291078398 | -3.80288286860545 | 0.902079069510569 | -4.21568684735002 | 2.49019113633226e-05 | 0.00240475251674398 | 3086 | 1 | 0 | 26 | 46 | 1 | 0 | 25 | 5 | 1 | 2 | 0 | 4 |
| Charlie8_dup431 | 6.73369058931947 | -4.18482390586071 | 0.978189891684466 | -4.2781303931227 | 1.88469613272039e-05 | 0.00195490804706314 | 3087 | 0 | 0 | 23 | 46 | 1 | 0 | 13 | 6 | 0 | 2 | 1 | 1 |
| MLT2B4_dup2448 | 6.06937528307245 | -4.74387573830498 | 0.9230913483846 | -5.13911840535253 | 2.76030431651458e-07 | 7.48920605356189e-05 | 3088 | 0 | 0 | 22 | 28 | 1 | 0 | 19 | 6 | 0 | 1 | 2 | 4 |
| MLT2F_dup864 | 5.46578160248756 | -4.05812542699004 | 0.833766994869359 | -4.86721764229333 | 1.13180366194863e-06 | 0.000226499709121975 | 3089 | 1 | 0 | 22 | 22 | 1 | 0 | 15 | 8 | 0 | 1 | 1 | 2 |
| L1PB_dup215 | 5.86990847126939 | -3.74398385434992 | 0.786474039231039 | -4.76046718339303 | 1.93145339380004e-06 | 0.000343227490097941 | 3090 | 1 | 0 | 15 | 17 | 1 | 0 | 26 | 13 | 1 | 1 | 0 | 2 |
| L2a_dup133117 | 6.91990709557811 | -3.26461317243875 | 0.971581990643814 | -3.36010054105209 | 0.000779141126199651 | 0.029507164477909 | 3091 | 2 | 0 | 14 | 33 | 0 | 0 | 18 | 15 | 2 | 3 | 0 | 4 |
| L2a_dup28592 | 8.83641253836073 | -5.09567799588203 | 0.927170677008017 | -5.49594386691111 | 3.88626122239158e-08 | 1.56732256410709e-05 | 3092 | 0 | 0 | 43 | 34 | 0 | 1 | 17 | 15 | 1 | 0 | 0 | 5 |
| AluSq2_dup58926 | 6.157446610641 | -3.7613576130081 | 0.897485181655729 | -4.19099689876655 | 2.77731456776952e-05 | 0.00260485176248517 | 3093 | 3 | 0 | 27 | 31 | 0 | 0 | 9 | 9 | 1 | 0 | 0 | 2 |
| MLT1K_dup16703 | 8.20144814499443 | -5.21458143736271 | 0.948417481062084 | -5.49819203197643 | 3.8370494987288e-08 | 1.55538429305711e-05 | 3094 | 1 | 0 | 37 | 41 | 0 | 0 | 17 | 12 | 0 | 0 | 0 | 2 |
| MLT1H_dup3382 | 6.75190435271859 | -2.84726449899285 | 0.797020193897483 | -3.57238689909415 | 0.000353742273729995 | 0.0168511629759674 | 3095 | 4 | 4 | 16 | 18 | 0 | 0 | 15 | 26 | 0 | 0 | 0 | 0 |
| L1P2_dup654 | 5.25621040892716 | -2.91692868084707 | 0.744339062212967 | -3.9188171478934 | 8.89845871063367e-05 | 0.00626552243107462 | 3096 | 2 | 3 | 13 | 23 | 0 | 1 | 12 | 14 | 0 | 0 | 0 | 0 |
| THE1C_dup2506 | 4.7910104579997 | -3.87869401654585 | 0.877400697738345 | -4.42066438577479 | 9.83978978649357e-06 | 0.00120749799924549 | 3097 | 1 | 1 | 14 | 20 | 0 | 0 | 13 | 13 | 0 | 0 | 0 | 0 |
| MLT1C_dup7742 | 6.03944117685312 | -2.19158503896096 | 0.656833567455098 | -3.33659110549459 | 0.00084812626100233 | 0.0313512896225665 | 3098 | 5 | 4 | 18 | 23 | 0 | 2 | 20 | 8 | 1 | 0 | 0 | 0 |
| L1MB7_dup22775 | 4.8055384705871 | -2.44542658639872 | 0.677513972961278 | -3.60941129481101 | 0.000306892693784725 | 0.0152018189335905 | 3099 | 3 | 3 | 13 | 15 | 1 | 1 | 14 | 12 | 0 | 0 | 0 | 0 |
| MSTB1_dup270 | 5.13091471765301 | -2.53432411560798 | 0.695345636330058 | -3.64469694378728 | 0.000267707069300571 | 0.0137763201294569 | 3100 | 2 | 3 | 15 | 11 | 1 | 2 | 25 | 9 | 0 | 0 | 0 | 0 |
| L1P2_dup414 | 6.45165044844455 | -3.84896446460326 | 0.847553195810972 | -4.54126594487136 | 5.59174412993865e-06 | 0.000783127221155022 | 3101 | 4 | 1 | 21 | 29 | 0 | 0 | 13 | 12 | 0 | 0 | 0 | 6 |
| SVA_C_dup50 | 7.98377246892943 | -4.34484954508255 | 0.76890098358131 | -5.65072699588124 | 1.59770667275451e-08 | 7.6033582391049e-06 | 3102 | 3 | 1 | 26 | 31 | 0 | 0 | 27 | 14 | 0 | 0 | 0 | 5 |
| LTR16C_dup1633 | 5.73694295808498 | -3.58097505525145 | 0.802616117277495 | -4.46162863935284 | 8.13390949142542e-06 | 0.00104223544849042 | 3103 | 1 | 4 | 14 | 21 | 0 | 0 | 20 | 12 | 0 | 0 | 0 | 4 |
| L1PA10_dup536 | 7.42502373506036 | -4.13952177821592 | 0.786524040705111 | -5.26305816984931 | 1.41678727464837e-07 | 4.37816707601923e-05 | 3104 | 2 | 2 | 16 | 40 | 0 | 0 | 28 | 11 | 0 | 0 | 0 | 3 |
| L2a_dup46293 | 6.54100249371228 | -3.73037711024791 | 0.784099198185494 | -4.75753210675446 | 1.9597409424204e-06 | 0.000347734912964328 | 3105 | 3 | 2 | 17 | 24 | 0 | 0 | 33 | 8 | 0 | 0 | 0 | 3 |
| L2a_dup73384 | 5.18254533255734 | -3.14778936365621 | 0.747006889173079 | -4.21386925512928 | 2.51032638771188e-05 | 0.00241830818886838 | 3106 | 3 | 2 | 17 | 19 | 0 | 0 | 19 | 6 | 1 | 0 | 1 | 2 |
| Arthur1C_dup364 | 4.0152444605552 | -3.03727507748979 | 0.813321375552147 | -3.7344095074691 | 0.00018815612831253 | 0.0106470863513563 | 3107 | 1 | 3 | 10 | 16 | 0 | 0 | 8 | 9 | 1 | 0 | 1 | 3 |
| MLT1F1_dup290 | 3.89083925626383 | -2.64858590183457 | 0.790187798813996 | -3.35184358175344 | 0.000802753602680536 | 0.0301280770888679 | 3108 | 2 | 3 | 12 | 12 | 0 | 0 | 12 | 7 | 1 | 0 | 0 | 2 |
| AluJb_dup47099 | 5.70176973720808 | -2.46210040894835 | 0.731829472838256 | -3.36430890026823 | 0.000767356049138608 | 0.0291769419092898 | 3109 | 8 | 1 | 10 | 23 | 0 | 1 | 17 | 11 | 1 | 0 | 1 | 3 |
| L1ME1_dup19838 | 6.16451409815238 | -3.38846827836532 | 0.695198266057736 | -4.87410346631088 | 1.09303695586594e-06 | 0.00022071625357832 | 3110 | 2 | 1 | 17 | 15 | 0 | 2 | 19 | 18 | 1 | 0 | 1 | 2 |
| L1MC1_dup1302 | 7.54823366239941 | -3.21734273284589 | 0.658766882036212 | -4.88388657745099 | 1.04014921980394e-06 | 0.000211458504041723 | 3111 | 4 | 3 | 21 | 38 | 1 | 0 | 18 | 10 | 1 | 0 | 2 | 4 |
| LTR3B_dup30 | 6.18878275681771 | -2.64336909274467 | 0.670708323583749 | -3.94116041175773 | 8.10883735865798e-05 | 0.00584155438735462 | 3112 | 3 | 4 | 14 | 19 | 1 | 0 | 27 | 8 | 2 | 0 | 2 | 3 |
| OldhAT1_dup420 | 7.61364417132623 | -3.2947359967295 | 0.60707144868322 | -5.42726231628255 | 5.72249957543568e-08 | 2.14095264776198e-05 | 3113 | 3 | 2 | 23 | 26 | 1 | 1 | 23 | 17 | 1 | 0 | 1 | 1 |
| THE1D_dup5554 | 6.51651757808552 | -2.58960644024909 | 0.620313429184997 | -4.17467415408282 | 2.98412864787576e-05 | 0.00274685290231121 | 3114 | 4 | 4 | 23 | 25 | 1 | 1 | 15 | 10 | 1 | 0 | 1 | 1 |
| UCON17_dup16 | 6.6084533274524 | -2.17497602859641 | 0.582357382596184 | -3.73477883786796 | 0.000187880222702124 | 0.0106422634544002 | 3115 | 6 | 5 | 13 | 27 | 2 | 1 | 19 | 11 | 1 | 0 | 1 | 2 |
| L1MEg_dup13058 | 6.56244896570777 | -2.3764022667214 | 0.652594166159033 | -3.6414702888139 | 0.000271085414896421 | 0.0139030810715334 | 3116 | 2 | 2 | 18 | 24 | 4 | 0 | 20 | 12 | 1 | 2 | 1 | 0 |
| L1MDa_dup2656 | 7.53469294904553 | -2.15569906661041 | 0.626690325654788 | -3.43981545328317 | 0.000582110967385381 | 0.0240512200460986 | 3117 | 6 | 7 | 24 | 24 | 3 | 0 | 17 | 13 | 1 | 0 | 3 | 0 |
| MLT1D_dup8027 | 6.47665115794259 | -2.55425540449742 | 0.741562257487217 | -3.44442476502581 | 0.000572275876863141 | 0.0237438109496211 | 3118 | 1 | 7 | 18 | 27 | 1 | 0 | 20 | 10 | 1 | 0 | 1 | 0 |
| L1PA3_dup1147 | 6.62085186662749 | -2.13011632658511 | 0.585459488986165 | -3.63836673016234 | 0.000274372544981953 | 0.0140339315537996 | 3119 | 3 | 5 | 12 | 18 | 1 | 3 | 23 | 17 | 1 | 1 | 0 | 1 |
| L1PB1_dup9599 | 6.3870685775065 | -3.19320954843774 | 0.727395057981753 | -4.38992472302147 | 1.13389906408194e-05 | 0.00134006315515524 | 3120 | 1 | 3 | 7 | 24 | 1 | 1 | 27 | 16 | 1 | 0 | 0 | 3 |
| MLT2A2_dup1686 | 5.2757208085083 | -2.4116473587447 | 0.7389338783582 | -3.26368492415454 | 0.0010997338981434 | 0.037473475888247 | 3121 | 0 | 4 | 15 | 27 | 3 | 1 | 10 | 8 | 1 | 0 | 0 | 1 |
| MLT2A2_dup3203 | 5.64163053056447 | -2.62801616694382 | 0.692584920988825 | -3.79450387570049 | 0.000147938971169375 | 0.00901900872357126 | 3122 | 0 | 2 | 17 | 17 | 4 | 1 | 19 | 12 | 1 | 0 | 0 | 0 |
| L1ME3_dup3691 | 7.25370998988303 | -2.62583903103939 | 0.58131868383546 | -4.51703876729795 | 6.27104278856403e-06 | 0.000856092683515581 | 3123 | 2 | 4 | 19 | 30 | 2 | 4 | 26 | 10 | 0 | 0 | 0 | 1 |
| MLT1C_dup7745 | 5.55870446719234 | -2.46434225802335 | 0.675733038463995 | -3.64691692983524 | 0.000265405679385395 | 0.0137078836090812 | 3124 | 3 | 2 | 17 | 18 | 4 | 1 | 25 | 5 | 0 | 0 | 0 | 1 |
| MER1A_dup2833 | 3.83800653176451 | -2.56889555683187 | 0.742290823559854 | -3.46076695992556 | 0.00053863887340504 | 0.0226969728017563 | 3125 | 1 | 2 | 11 | 10 | 1 | 1 | 12 | 9 | 1 | 0 | 1 | 0 |
| MLT1G1_dup1958 | 3.91667493540642 | -2.61278726079835 | 0.763561870809444 | -3.42184092826501 | 0.000621986916083869 | 0.025155349593716 | 3126 | 1 | 1 | 8 | 8 | 2 | 2 | 16 | 11 | 0 | 0 | 1 | 0 |
| MER21B_dup1776 | 4.78398772126509 | -3.16571945437428 | 0.820569390027721 | -3.85795460182513 | 0.000114339878599717 | 0.00748877422331086 | 3127 | 1 | 1 | 7 | 10 | 0 | 2 | 20 | 16 | 1 | 0 | 2 | 0 |
| SVA_F_dup702 | 5.15514364834451 | -2.27824069447406 | 0.677465813166816 | -3.36288658437894 | 0.000771320456437646 | 0.0292808898097499 | 3128 | 3 | 2 | 7 | 13 | 1 | 2 | 23 | 12 | 1 | 1 | 2 | 0 |
| AluSc_dup13636 | 6.45624485216855 | -2.56214777761928 | 0.607169670427349 | -4.21982174408669 | 2.44495560911084e-05 | 0.00237165677686705 | 3129 | 1 | 5 | 18 | 19 | 2 | 2 | 21 | 13 | 0 | 1 | 2 | 0 |
| L1MB7_dup8332 | 5.12373429561689 | -2.29865950939931 | 0.65494997498783 | -3.50967187905003 | 0.000448659978355194 | 0.0200609779967691 | 3130 | 3 | 3 | 14 | 10 | 1 | 2 | 22 | 10 | 0 | 1 | 1 | 0 |
| L3_dup40221 | 5.42214005145497 | -2.16699957801213 | 0.67792780043544 | -3.19650496206269 | 0.00139103433888656 | 0.0441498008270909 | 3131 | 2 | 4 | 19 | 13 | 0 | 4 | 14 | 7 | 0 | 2 | 5 | 1 |
| L1MA9_dup8581 | 4.01436461272534 | -2.21713304477267 | 0.694549678249667 | -3.1921878509253 | 0.00141199469923898 | 0.0445594815258778 | 3132 | 2 | 3 | 10 | 9 | 1 | 2 | 11 | 7 | 0 | 1 | 5 | 1 |
| L2_dup5989 | 4.51366125002019 | -2.62238581707039 | 0.762611333055148 | -3.43869242876929 | 0.000584530939856131 | 0.024121886752516 | 3133 | 1 | 3 | 11 | 8 | 1 | 2 | 20 | 5 | 0 | 1 | 7 | 1 |
| AluJr_dup71402 | 3.78788891208533 | -2.38469239847807 | 0.73286619180033 | -3.25392605793416 | 0.00113821901964768 | 0.0384162642339129 | 3134 | 3 | 2 | 6 | 9 | 1 | 2 | 18 | 6 | 0 | 0 | 1 | 3 |
| Tigger17a_dup1851 | 3.21506968956878 | -2.52720417718894 | 0.796626325833906 | -3.17238345662689 | 0.00151193229310182 | 0.0467704422015608 | 3135 | 1 | 2 | 7 | 9 | 2 | 1 | 9 | 7 | 0 | 0 | 0 | 4 |
| MIR_dup72106 | 4.19014393772982 | -2.9272656556255 | 0.802301795000116 | -3.64858420343566 | 0.000263689478652865 | 0.0136488702822568 | 3136 | 1 | 2 | 11 | 17 | 0 | 3 | 8 | 9 | 0 | 0 | 0 | 4 |
| L1MC1_dup2192 | 4.57333825646713 | -3.34381930087775 | 0.790899447650368 | -4.22786905568297 | 2.35915014876268e-05 | 0.0023130980917055 | 3137 | 1 | 1 | 12 | 22 | 0 | 2 | 13 | 9 | 0 | 0 | 0 | 1 |
| L2_dup4736 | 4.21869593706964 | -2.63671063099086 | 0.735748371164231 | -3.58371249510018 | 0.0003387449392525 | 0.0163204674240049 | 3138 | 2 | 2 | 11 | 15 | 0 | 3 | 16 | 6 | 0 | 0 | 0 | 2 |
| L2a_dup165179 | 3.60950464796376 | -2.73919015646586 | 0.785559872435167 | -3.48692728916335 | 0.000488604177686752 | 0.0211989812677116 | 3139 | 3 | 2 | 11 | 13 | 0 | 1 | 11 | 4 | 0 | 0 | 1 | 3 |
| THE1B_dup12172 | 4.14891914583227 | -2.93848671773557 | 0.81775544332396 | -3.59335635332661 | 0.000326445649021485 | 0.0158671995536405 | 3140 | 2 | 2 | 19 | 14 | 0 | 1 | 13 | 4 | 0 | 0 | 0 | 1 |
| L1MEc_dup8721 | 4.78219619202619 | -3.36067550607199 | 0.770981870243953 | -4.35895529554879 | 1.30684802623618e-05 | 0.00148998208170002 | 3141 | 2 | 0 | 14 | 13 | 1 | 2 | 17 | 10 | 0 | 0 | 0 | 4 |
| LTR12_dup565 | 6.19509928347225 | -2.60757674164828 | 0.62063294521709 | -4.20147973410626 | 2.65175989011308e-05 | 0.0025188649054405 | 3142 | 3 | 1 | 14 | 19 | 2 | 5 | 17 | 16 | 0 | 0 | 0 | 3 |
| AluSx_dup71585 | 4.03729778140002 | -2.62320539173318 | 0.814171783073924 | -3.22193110381351 | 0.00127329748870366 | 0.0415092539042447 | 3143 | 2 | 1 | 16 | 19 | 2 | 2 | 4 | 5 | 0 | 0 | 0 | 3 |
| L1ME4a_dup14289 | 4.04996670025386 | -2.64923692627865 | 0.803714444757692 | -3.29624152403701 | 0.000979877354652286 | 0.0346270985938937 | 3144 | 2 | 2 | 16 | 13 | 1 | 1 | 4 | 11 | 0 | 0 | 0 | 1 |
| Charlie15b_dup783 | 3.94538955907421 | -2.95969815199481 | 0.788436840377897 | -3.75388109791551 | 0.00017411748089296 | 0.0101124134997696 | 3145 | 1 | 2 | 8 | 19 | 1 | 1 | 9 | 9 | 0 | 0 | 0 | 2 |
| L1MB7_dup4203 | 3.94692811727554 | -2.6232255624128 | 0.773638924847274 | -3.39076212191709 | 0.000696985682536537 | 0.0272457623158927 | 3146 | 1 | 1 | 10 | 18 | 2 | 2 | 6 | 10 | 0 | 0 | 0 | 1 |
| AluSz_dup4247 | 4.86750025767868 | -2.23632305769227 | 0.705790021632218 | -3.16853878511986 | 0.00153207305682585 | 0.0471971330372196 | 3147 | 5 | 3 | 16 | 5 | 2 | 1 | 13 | 13 | 0 | 0 | 2 | 1 |
| L3_dup42627 | 6.14062129340152 | -2.69419799595913 | 0.626271689068833 | -4.30196357744509 | 1.69291144421013e-05 | 0.00180880784240693 | 3148 | 3 | 3 | 21 | 21 | 3 | 2 | 14 | 8 | 0 | 0 | 1 | 6 |
| LTR73_dup82 | 6.84266050593633 | -2.30350337796572 | 0.60685848251252 | -3.79578343937575 | 0.000147177934478963 | 0.00898419722871505 | 3149 | 5 | 5 | 25 | 29 | 3 | 2 | 13 | 7 | 0 | 0 | 1 | 2 |
| L3_dup5400 | 6.22850238016432 | -3.29528884988967 | 0.67471153290737 | -4.8839966254765 | 1.03956854359629e-06 | 0.000211458504041723 | 3150 | 2 | 1 | 17 | 31 | 2 | 2 | 13 | 11 | 0 | 0 | 2 | 2 |
| MIRc_dup48417 | 4.83598083055934 | -3.24937472644992 | 0.762201934760313 | -4.26314153541441 | 2.01572750932432e-05 | 0.00205235044045222 | 3151 | 1 | 1 | 21 | 19 | 2 | 1 | 10 | 7 | 0 | 0 | 1 | 1 |
| LTR1A1_dup336 | 5.23747425512583 | -2.93401427346199 | 0.707940777222741 | -4.14443463049573 | 3.40653110658347e-05 | 0.00303631442785359 | 3152 | 3 | 1 | 18 | 16 | 3 | 0 | 11 | 13 | 0 | 0 | 1 | 1 |
| MamGypLTR2b_dup40 | 5.80860649101905 | -2.77481436954332 | 0.679761846869503 | -4.08203899986757 | 4.46423029964289e-05 | 0.00375551027790432 | 3153 | 5 | 1 | 16 | 22 | 4 | 0 | 12 | 10 | 0 | 0 | 2 | 5 |
| AluSx1_dup40890 | 4.26154258034639 | -2.2817363460973 | 0.702215907135017 | -3.24933730909998 | 0.00115674224602547 | 0.0388485801655296 | 3154 | 3 | 2 | 11 | 13 | 4 | 0 | 16 | 6 | 0 | 0 | 1 | 1 |
| MIRb_dup89994 | 5.44757284913165 | -2.63831960557775 | 0.670971862294949 | -3.93208680398879 | 8.42116447302584e-05 | 0.0060065419715186 | 3155 | 2 | 4 | 10 | 21 | 3 | 0 | 16 | 13 | 0 | 0 | 1 | 1 |

| | | | | | | | | | | | | | | | | | | | |
|---|---|---|---|---|---|---|---|---|---|---|---|---|---|---|---|---|---|---|---|
| L1M5_dup22419 | 6.23635500877897 | -2.57177552858743 | 0.733671147612424 | -3.50535186909928 | 0.000456004176603622 | 0.020277400449659 | 3156 | 8 | 3 | 17 | 21 | 1 | 0 | 15 | 13 | 0 | 0 | 0 | 4 |
| L3_dup19415 | 5.4720118422102 | -2.76494794456034 | 0.673541602478473 | -4.10508858604426 | 4.04159972710076e-05 | 0.00346677176879855 | 3157 | 4 | 2 | 14 | 20 | 2 | 0 | 20 | 10 | 0 | 0 | 0 | 1 |
| MER5C_dup1168 | 4.83905586606546 | -2.30430283548054 | 0.682416446568752 | -3.37668127294818 | 0.000733660107064379 | 0.0282844277115264 | 3158 | 3 | 5 | 10 | 13 | 1 | 1 | 14 | 13 | 0 | 0 | 0 | 2 |
| MLT1A0_dup9050 | 5.01920488488755 | -2.29250628752983 | 0.689573636336122 | -3.32452716683094 | 0.000885686040181034 | 0.0323352332064057 | 3159 | 5 | 1 | 12 | 12 | 3 | 1 | 12 | 16 | 0 | 0 | 0 | 1 |
| MSTB1_dup2312 | 4.52608524381106 | -2.53486335990867 | 0.773842289474355 | -3.27568471559045 | 0.00105406160089922 | 0.0364284301652239 | 3160 | 4 | 1 | 14 | 13 | 2 | 0 | 20 | 4 | 0 | 1 | 0 | 3 |
| L1M5_dup22410 | 5.02148125189685 | -2.42383651860978 | 0.678139044307728 | -3.57424710898947 | 0.000351237100850861 | 0.0167620263134896 | 3161 | 4 | 3 | 14 | 17 | 1 | 0 | 15 | 10 | 0 | 1 | 0 | 1 |
| L1MCb_dup934 | 4.97164311871801 | -2.45297176087814 | 0.743329075579549 | -3.29998091217626 | 0.000966914047023267 | 0.0343489382136655 | 3162 | 1 | 4 | 14 | 18 | 1 | 0 | 14 | 10 | 0 | 2 | 0 | 1 |
| L1PA11_dup1047 | 5.48001809954344 | -3.53296191362874 | 1.09902569008275 | -3.21463087306241 | 0.00130612360416925 | 0.0422494408805948 | 3163 | 0 | 0 | 16 | 43 | 1 | 3 | 2 | 3 | 0 | 1 | 0 | 8 |
| ERV3-16A3_I-int_dup5( | 6.41250312416539 | -4.05679837810257 | 0.949484173006557 | -4.27263401901335 | 1.93177376454827e-05 | 0.00199244837528913 | 3164 | 0 | 0 | 22 | 42 | 2 | 2 | 10 | 5 | 0 | 0 | 0 | 6 |
| AluJo_dup34581 | 5.64522464317934 | -3.33122773899797 | 0.903429782874933 | -3.68731228717875 | 0.000226635160683401 | 0.012180468247185 | 3165 | 0 | 0 | 28 | 28 | 2 | 4 | 8 | 3 | 0 | 0 | 1 | 3 |
| MER3_dup5223 | 4.97915931693685 | -3.27121972487076 | 0.846610282712712 | -3.86390266178803 | 0.000111589703488224 | 0.00736541569659058 | 3166 | 0 | 0 | 18 | 21 | 3 | 3 | 12 | 4 | 0 | 0 | 1 | 6 |
| L1M4_dup13365 | 7.45789816213736 | -3.61260919868147 | 0.971283985025067 | -3.7194160043607 | 0.00019968393737976 | 0.0110970900987931 | 3167 | 0 | 0 | 38 | 43 | 4 | 2 | 6 | 5 | 0 | 0 | 3 | 0 |
| MER57C1_dup111 | 7.13116855210294 | -3.90464857127332 | 0.998349782103427 | -3.91110274301518 | 9.18756757059796e-05 | 0.00641098226731232 | 3168 | 0 | 0 | 32 | 52 | 3 | 1 | 3 | 5 | 0 | 0 | 1 | 1 |
| MIRb_dup198500 | 6.12561858009331 | -2.97855513234171 | 0.905499157684495 | -3.2894068504253 | 0.00100398782586345 | 0.0351729810384137 | 3169 | 0 | 0 | 25 | 34 | 5 | 1 | 7 | 7 | 0 | 1 | 2 | 0 |
| THE1B-int_dup1292 | 6.55949484713218 | -3.00416080651682 | 0.933053029115031 | -3.21971068393205 | 0.00128320029865706 | 0.0417177726826414 | 3170 | 0 | 0 | 34 | 36 | 5 | 2 | 4 | 3 | 0 | 1 | 2 | 2 |
| MIRb_dup206752 | 8.08612502211914 | -3.54590479915376 | 0.966421875576242 | -3.66910651421199 | 0.000243399659642514 | 0.0128656243633447 | 3171 | 0 | 0 | 51 | 39 | 3 | 3 | 4 | 2 | 1 | 0 | 4 | 2 |
| AluJr_dup38516 | 6.26160586084297 | -2.85001158467474 | 0.85702016106348 | -3.32548954407111 | 0.000882634106849942 | 0.0322709365686104 | 3172 | 0 | 1 | 23 | 39 | 5 | 4 | 6 | 3 | 0 | 0 | 3 | 2 |
| HERV9-int_dup329 | 7.60624639223747 | -3.5059604204249 | 1.03482243318858 | -3.38798262192873 | 0.000704087252035619 | 0.0274692154898946 | 3173 | 0 | 1 | 34 | 57 | 5 | 1 | 3 | 1 | 0 | 0 | 3 | 1 |
| L1MEc_dup5168 | 4.55214800879685 | -3.4117244702321 | 0.984456701370311 | -3.46559118901132 | 0.00052906707716135 | 0.0224042346934036 | 3174 | 1 | 2 | 22 | 27 | 0 | 0 | 4 | 2 | 1 | 0 | 2 | 1 |
| L1MCa_dup3470 | 5.17034364233279 | -3.0586830959712 | 0.875736466763028 | -3.49269810274884 | 0.000478166770636832 | 0.0209035220293867 | 3175 | 1 | 3 | 21 | 26 | 0 | 0 | 9 | 3 | 2 | 0 | 2 | 3 |
| MER4B_dup143 | 5.49462849080803 | -3.17873178215227 | 0.820724529405164 | -3.87308002656643 | 0.00010746851991753 | 0.00718242824695875 | 3176 | 2 | 3 | 20 | 31 | 0 | 0 | 8 | 6 | 1 | 0 | 2 | 1 |
| L2a_dup82398 | 4.18693081030703 | -2.81440055936181 | 0.840425765827246 | -3.34877948035249 | 0.000811683711621392 | 0.0303538661041926 | 3177 | 2 | 1 | 12 | 15 | 0 | 0 | 6 | 6 | 3 | 0 | 6 | 3 |
| AluY_dup46583 | 4.29152910317289 | -2.7704072542299 | 0.819645463847567 | -3.38000681566039 | 0.000724840323183504 | 0.0280306932456074 | 3178 | 4 | 2 | 13 | 20 | 0 | 0 | 6 | 4 | 1 | 0 | 3 | 5 |
| L1MB5_dup4672 | 4.28721359859985 | -2.5413618307185 | 0.805831759206037 | -3.15371267226105 | 0.00161207719492316 | 0.0487838382580676 | 3179 | 4 | 3 | 12 | 19 | 0 | 0 | 5 | 6 | 1 | 0 | 2 | 5 |
| L1MB7_dup9671 | 3.63013553805383 | -2.47433136673969 | 0.77220956832189 | -3.20422262070221 | 0.00135427715458285 | 0.0433182997478655 | 3180 | 3 | 3 | 8 | 12 | 0 | 0 | 9 | 5 | 1 | 0 | 4 | 3 |
| MIRb_dup190715 | 4.44788370399319 | -2.41432034790202 | 0.746259361553879 | -3.23522956264812 | 0.00121544965345829 | 0.0401905757701203 | 3181 | 5 | 3 | 12 | 15 | 0 | 0 | 7 | 8 | 1 | 0 | 3 | 4 |
| L3b_dup9285 | 4.73513676864937 | -4.18095740257538 | 0.964563563870091 | -4.334558715654 | 1.46052803172019e-05 | 0.001619465424217 | 3182 | 1 | 2 | 21 | 17 | 0 | 0 | 4 | 5 | 0 | 0 | 5 | 8 |
| AluSq_dup7837 | 4.16446370363545 | -4.47898054707073 | 1.04914099901287 | -4.26918836580114 | 1.96185540195071e-05 | 0.0020148295225497 | 3183 | 1 | 1 | 11 | 24 | 0 | 0 | 4 | 3 | 0 | 0 | 8 | 5 |
| L1MB3_dup4022 | 3.54475213696118 | -3.77203893499127 | 1.00795448641861 | -3.74227109042772 | 0.00018236462889648 | 0.0104323360473012 | 3184 | 1 | 2 | 11 | 12 | 0 | 0 | 4 | 3 | 0 | 0 | 10 | 4 |
| AluSz_dup91660 | 4.23719859170447 | -3.2166054666136 | 0.916056092814346 | -3.51136299604908 | 0.000445815191365928 | 0.0199623525639362 | 3185 | 2 | 2 | 18 | 20 | 0 | 0 | 4 | 3 | 0 | 1 | 5 | 2 |
| L1MEf_dup10309 | 3.39603194219354 | -3.67905898785752 | 1.05638384430401 | -3.48269145509455 | 0.00049640012741096 | 0.021475713584896 | 3186 | 2 | 1 | 11 | 21 | 0 | 0 | 2 | 2 | 0 | 0 | 3 | 5 |
| MIR3_dup79363 | 3.29576887016842 | -3.28891892134276 | 0.979995328760534 | -3.35605571253332 | 0.000790626400684541 | 0.0298279325046136 | 3187 | 3 | 1 | 11 | 16 | 0 | 0 | 4 | 2 | 0 | 0 | 5 | 3 |
| MLT1F2_dup5731 | 5.69087428109336 | -3.07477820153345 | 0.826291130613375 | -3.72118020830137 | 0.000198293864530788 | 0.0110551152506193 | 3188 | 3 | 4 | 22 | 31 | 0 | 1 | 8 | 3 | 0 | 0 | 4 | 2 |
| Tigger1_dup6298 | 5.65989410297102 | -3.56674961896792 | 0.865264904821654 | -4.12214756324029 | 3.75356619186761e-05 | 0.00328241845310595 | 3189 | 3 | 2 | 23 | 28 | 0 | 1 | 7 | 2 | 0 | 0 | 5 | 7 |
| L1MA9_dup7711 | 5.81834412838081 | -2.95869154325688 | 0.883128740095478 | -3.35023808979086 | 0.000807421262850419 | 0.0302555369779852 | 3190 | 2 | 7 | 26 | 24 | 0 | 0 | 7 | 5 | 0 | 0 | 2 | 5 |
| L1PA4_dup7325 | 5.08801636278473 | -2.81785620901397 | 0.821531145203296 | -3.43000533268481 | 0.000603569386984825 | 0.0246880493255903 | 3191 | 4 | 3 | 18 | 25 | 0 | 0 | 9 | 3 | 0 | 1 | 2 | 5 |
| L1MC5_dup14781 | 5.05307836111557 | -2.68820087444607 | 0.827532382293768 | -3.24845399644044 | 0.00116033970121051 | 0.0389345651193219 | 3192 | 6 | 3 | 21 | 21 | 0 | 0 | 8 | 5 | 0 | 0 | 2 | 2 |
| L1PREC2_dup631 | 4.53680859481349 | -3.07681086457447 | 0.819767747673364 | -3.75327142755612 | 0.000174541685187369 | 0.0101271630879283 | 3193 | 4 | 2 | 16 | 21 | 0 | 0 | 11 | 4 | 0 | 0 | 2 | 2 |
| MER50B_dup298 | 5.09327427651974 | -3.41080224740089 | 0.92184714079715 | -3.69996509882481 | 0.000215629119530593 | 0.011736967608678 | 3194 | 2 | 1 | 13 | 33 | 3 | 0 | 3 | 3 | 0 | 0 | 8 | 4 |
| L1MB8_dup15146 | 4.884242642794 | -2.93572054614681 | 0.857068511876832 | -3.42530440153272 | 0.000614110946420011 | 0.0249616062960123 | 3195 | 1 | 3 | 9 | 26 | 1 | 0 | 3 | 3 | 0 | 2 | 13 | 5 |
| AluSz_dup42645 | 4.73935484411555 | -2.74462114143806 | 0.869879835834153 | -3.15517273579075 | 0.0016040310725457 | 0.0486269305119071 | 3196 | 1 | 3 | 13 | 27 | 2 | 1 | 1 | 2 | 0 | 1 | 6 | 8 |
| MER50-int_dup468 | 4.19775842040597 | -3.21610989866322 | 0.978077730010251 | -3.28819458820468 | 0.0010083212050906 | 0.035267675652872 | 3197 | 1 | 2 | 17 | 23 | 1 | 0 | 1 | 2 | 0 | 1 | 5 | 4 |
| FLAM_A_dup7223 | 5.02947484123202 | -2.46478896692153 | 0.723395877028139 | -3.40724773971258 | 0.000656215356302309 | 0.0261284837969728 | 3198 | 3 | 4 | 20 | 17 | 2 | 0 | 7 | 4 | 0 | 1 | 5 | 4 |
| L2a_dup169312 | 5.64607970316554 | -3.22692104631202 | 0.848167476255226 | -3.80458003478195 | 0.000142044980044067 | 0.00874942656525124 | 3199 | 3 | 2 | 21 | 28 | 1 | 0 | 9 | 1 | 0 | 1 | 7 | 5 |
| L1ME3B_dup7553 | 6.0189590166827 | -3.01628152186039 | 0.95112466298692 | -3.17127884412968 | 0.00151769382017204 | 0.0468633757766229 | 3200 | 4 | 5 | 18 | 33 | 1 | 0 | 3 | 1 | 0 | 0 | 5 | 14 |
| MLT1C_dup7868 | 4.22836989930701 | -3.15060785906478 | 0.995696076424978 | -3.16422644787048 | 0.00155495759572726 | 0.0476173361211657 | 3201 | 2 | 3 | 15 | 21 | 1 | 0 | 4 | 1 | 0 | 0 | 1 | 11 |
| AluSq2_dup60845 | 4.5816369311791 | -3.19157690622809 | 0.827616907665578 | -3.8563457037513 | 0.000115094688983206 | 0.00751545578067997 | 3202 | 1 | 3 | 17 | 14 | 1 | 1 | 6 | 2 | 0 | 0 | 7 | 10 |
| L1MA2_dup6390 | 4.68844556422256 | -3.01755779992109 | 0.81032372707025 | -3.72389169798984 | 0.000196175108606617 | 0.0109626778751785 | 3203 | 2 | 1 | 15 | 19 | 2 | 2 | 4 | 2 | 0 | 0 | 7 | 10 |
| L1PA16_dup4089 | 3.92399105754699 | -3.24613819909425 | 0.879590013591064 | -3.690512794525 | 0.000223802431044803 | 0.0120645431031687 | 3204 | 2 | 1 | 13 | 15 | 1 | 1 | 6 | 1 | 0 | 0 | 6 | 8 |
| LTR16B2_dup1473 | 4.9378809189577 | -2.60737830511811 | 0.80062215338416 | -3.25669018037654 | 0.00112719391723828 | 0.0381469611479418 | 3205 | 4 | 2 | 16 | 22 | 3 | 1 | 5 | 1 | 0 | 0 | 6 | 8 |
| THE1C-int_dup928 | 5.13719403157711 | -2.78447282143379 | 0.806642390782494 | -3.45192969431308 | 0.000556592734160409 | 0.0232795407758408 | 3206 | 2 | 3 | 20 | 26 | 3 | 1 | 5 | 2 | 0 | 0 | 5 | 3 |
| L1PA16_dup1119 | 5.58819811922907 | -3.24642805192 | 0.87253027348141 | -3.72070534466007 | 0.000198667127867082 | 0.011065228514403 | 3207 | 0 | 2 | 18 | 36 | 4 | 0 | 10 | 4 | 0 | 0 | 2 | 1 |
| L1MA9_dup13759 | 6.76249768706634 | -3.94149398498829 | 0.795663180202789 | -4.95372172931733 | 7.28073067981221e-07 | 0.000160112822766968 | 3208 | 1 | 1 | 31 | 34 | 1 | 1 | 15 | 6 | 0 | 0 | 1 | 1 |
| MamRTE1_dup1581 | 7.52249284003416 | -4.35735644712613 | 0.813257283532853 | -5.35790645267563 | 8.41918098490472e-08 | 2.89077985372892e-05 | 3209 | 0 | 1 | 33 | 37 | 1 | 1 | 19 | 8 | 0 | 0 | 1 | 1 |
| HERVH-int_dup5474 | 7.87552780289223 | -3.11975748255939 | 0.85890290973983 | -3.63225860243551 | 0.00028095135659762 | 0.0142600792442358 | 3210 | 2 | 5 | 48 | 33 | 2 | 0 | 13 | 2 | 0 | 0 | 1 | 1 |
| L1MA1_dup3734 | 8.36930337075385 | -2.92870848921547 | 0.737032406801735 | -3.97364954673331 | 7.07796833077118e-05 | 0.0052894919988495 | 3211 | 4 | 6 | 35 | 44 | 2 | 0 | 14 | 6 | 0 | 0 | 2 | 1 |
| L1PA13_dup4383 | 6.01918152032525 | -3.1997168520144 | 0.758765891445095 | -4.21700143363118 | 2.47572401810355e-05 | 0.00239403773781171 | 3212 | 3 | 3 | 15 | 38 | 1 | 0 | 11 | 8 | 0 | 0 | 2 | 1 |
| MLT1B-int_dup47 | 6.91017560291446 | -3.92543511769705 | 0.742444237757416 | -5.28717837389916 | 1.24217499947858e-07 | 3.91484201888648e-05 | 3213 | 1 | 3 | 23 | 32 | 1 | 0 | 18 | 9 | 0 | 0 | 4 | 2 |
| L1PA16_dup12587 | 7.49966831680654 | -3.65648870947311 | 0.76594160054182 | -4.77384791071088 | 1.80738894870831e-06 | 0.000325064981700187 | 3214 | 3 | 2 | 25 | 49 | 1 | 1 | 11 | 5 | 0 | 0 | 3 | 4 |
| L1PA8_dup687 | 6.1281271484599 | -4.42310062063068 | 0.890845662270533 | -4.96505826762107 | 6.86804466273188e-07 | 0.000152776940125142 | 3215 | 1 | 1 | 19 | 37 | 1 | 0 | 14 | 4 | 0 | 0 | 5 | 3 |
| BLACKJACK_dup591 | 5.56623654572333 | -2.86482296460716 | 0.901419149642078 | -3.17812525476597 | 0.00148230699338028 | 0.046128510414423 | 3216 | 3 | 1 | 25 | 33 | 3 | 0 | 3 | 6 | 0 | 0 | 0 | 1 |
| L1PA8A_dup701 | 8.09646951703706 | -2.78700587570241 | 0.873520200819479 | -3.19054541965695 | 0.00142004518218582 | 0.0446949829259771 | 3217 | 5 | 1 | 35 | 51 | 3 | 0 | 4 | 5 | 0 | 2 | 1 | 4 |
| L1MEd_dup3282 | 5.74999486274556 | -2.78419758454097 | 0.831186961036596 | -3.34966465435012 | 0.000809094511522531 | 0.0302830526099167 | 3218 | 3 | 0 | 24 | 26 | 6 | 0 | 9 | 6 | 0 | 0 | 2 | 1 |
| MIR_dup170863 | 4.63534385648061 | -3.17331474252752 | 0.829721435871013 | -3.82455436889645 | 0.000131008834245624 | 0.00825457378141597 | 3219 | 2 | 0 | 14 | 19 | 4 | 0 | 9 | 7 | 0 | 0 | 1 | 6 |
| LTR40a_dup124 | 4.77657276890543 | -2.70387554292601 | 0.79282032553787 | -3.41045184618801 | 0.000648553354553571 | 0.0258927309568128 | 3220 | 3 | 1 | 15 | 19 | 3 | 0 | 3 | 9 | 1 | 0 | 2 | 6 |
| MER4B_dup1259 | 5.57597960418782 | -3.38722287045801 | 0.773068898933443 | -4.38152779801536 | 1.17850004417884e-05 | 0.00137595373656614 | 3221 | 2 | 0 | 22 | 23 | 2 | 1 | 7 | 7 | 1 | 0 | 3 | 6 |
| L1MA1_dup3272 | 5.91701456366264 | -3.24051686604016 | 0.809803271133881 | -4.0016099978243 | 6.29129357345358e-05 | 0.0048461901606636 | 3222 | 2 | 0 | 24 | 27 | 4 | 0 | 9 | 4 | 1 | 0 | 4 | 5 |
| L2a_dup147237 | 5.50731743379635 | -2.63673812176295 | 0.812339354881056 | -3.24585791137624 | 0.00117097267893464 | 0.039169394739703 | 3223 | 3 | 0 | 15 | 31 | 4 | 0 | 7 | 4 | 2 | 0 | 3 | 6 |
| L2_dup42714 | 3.75001784583989 | -3.09441510257488 | 0.887424590321927 | -3.48696118669906 | 0.000488542253169072 | 0.0211989812677116 | 3224 | 0 | 0 | 9 | 12 | 0 | 1 | 7 | 3 | 3 | 1 | 3 | 11 |
| L1MB8_dup13178 | 3.22203953724464 | -2.75430522381215 | 0.84648695733866 | -3.25380704325515 | 0.00113869595788737 | 0.0384174933393487 | 3225 | 0 | 0 | 6 | 10 | 0 | 2 | 7 | 4 | 2 | 1 | 6 | 4 |
| MIRc_dup86525 | 3.58557522986374 | -2.9601766217647 | 0.817471630945742 | -3.6211368195616 | 0.00029331133893823 | 0.0147334304105966 | 3226 | 1 | 0 | 10 | 10 | 0 | 2 | 9 | 3 | 1 | 1 | 5 | 6 |
| AluSc_dup33812 | 4.23685822909957 | -3.550146011233 | 0.958871337039031 | -3.70242166399066 | 0.000213551308404634 | 0.0116491700205523 | 3227 | 0 | 1 | 9 | 26 | 0 | 1 | 7 | 1 | 1 | 1 | 6 | 6 |
| L2a_dup83253 | 3.22922035877148 | -3.14845666057801 | 0.981852526033011 | -3.20664924425948 | 0.0013429062569531 | 0.0430761324184771 | 3228 | 0 | 1 | 7 | 17 | 0 | 1 | 3 | 1 | 1 | 1 | 7 | 5 |
| AluSx1_dup51178 | 4.32596453535081 | -3.04834326340046 | 0.921471238957387 | -3.30812632508157 | 0.000939224329974096 | 0.0336421304260148 | 3229 | 0 | 1 | 14 | 25 | 0 | 1 | 8 | 2 | 2 | 1 | 3 | 2 |
| L2a_dup117062 | 3.61844511266031 | -2.63885290316994 | 0.797597782758349 | -3.30850080104779 | 0.000937969165003098 | 0.0336174139080927 | 3230 | 0 | 1 | 12 | 11 | 0 | 2 | 7 | 3 | 1 | 2 | 4 | 5 |
| MLT1A0_dup15431 | 3.48352587380035 | -2.88129094490278 | 0.844091903053913 | -3.41348013703047 | 0.000641388337721087 | 0.0257014946241651 | 3231 | 0 | 1 | 8 | 12 | 0 | 1 | 10 | 2 | 2 | 1 | 4 | 6 |
| L1P2_dup830 | 2.70193255316639 | -3.69098794517454 | 1.04763046780999 | -3.52317735937016 | 0.000426405910867724 | 0.0193582707786507 | 3232 | 0 | 0 | 10 | 9 | 0 | 1 | 10 | 2 | 0 | 1 | 1 | 3 |
| MIRb_dup217337 | 3.30857473943245 | -3.86881177365284 | 1.00392278333621 | -3.85369456483108 | 0.000116348725041788 | 0.00758274803259302 | 3233 | 0 | 0 | 12 | 10 | 0 | 1 | 15 | 3 | 0 | 1 | 2 | 1 |
| AluSx_dup82002 | 2.82943989461468 | -3.75305157717766 | 1.07079566466115 | -3.50491853958458 | 0.000456747013170568 | 0.0202914721426221 | 3234 | 0 | 0 | 9 | 17 | 0 | 1 | 5 | 2 | 0 | 1 | 2 | 2 |

| | | | | | | | | | | | | | | | | | | | |
|---|---|---|---|---|---|---|---|---|---|---|---|---|---|---|---|---|---|---|---|
| L2c_dup70126 | 2.91553890930218 | -3.24366255217344 | 0.993233601866251 | -3.26575998443741 | 0.00109170739055688 | 0.0372961100866398 | 3235 | 0 | 0 | 12 | 10 | 0 | 1 | 5 | 5 | 0 | 2 | 1 | 2 |
| L1PB2_dup2050 | 2.3710350400576 | -3.47765601473049 | 1.08243705329471 | -3.21280207855527 | 0.00131446837790759 | 0.0424270879882798 | 3236 | 0 | 0 | 8 | 10 | 0 | 0 | 8 | 2 | 1 | 1 | 1 | 1 |
| L2a_dup86885 | 3.06624536820528 | -4.00348315786113 | 1.00464144775031 | -3.98498704868897 | 6.74838062587868e-05 | 0.00510897288070419 | 3237 | 0 | 0 | 9 | 9 | 0 | 0 | 11 | 3 | 1 | 1 | 4 | 3 |
| MSTD_dup4083 | 3.06823514406582 | -3.33615935772027 | 1.05348355570048 | -3.16678826135258 | 0.00154132500386728 | 0.0473534046959591 | 3238 | 0 | 0 | 4 | 15 | 0 | 0 | 9 | 2 | 1 | 3 | 4 | 4 |
| L2a_dup124337 | 3.75477276256992 | -3.50023077688015 | 1.08672801355713 | -3.22088943435169 | 0.00127793439300574 | 0.0416296128190765 | 3239 | 0 | 0 | 10 | 15 | 0 | 0 | 10 | 3 | 1 | 4 | 1 | 7 |
| Tigger19a_dup2584 | 3.12129438081196 | -3.62463134879607 | 1.0538756422489 | -3.43933496846114 | 0.000583145206059998 | 0.0240855874329374 | 3240 | 0 | 0 | 9 | 14 | 0 | 0 | 4 | 3 | 1 | 2 | 2 | 7 |
| LTR105_Mam_dup41 | 2.59246566035624 | -4.01251795184796 | 1.0659469695509 | -3.76427539686942 | 0.000167032721735082 | 0.00983781386286531 | 3241 | 0 | 0 | 7 | 11 | 0 | 0 | 5 | 7 | 1 | 0 | 1 | 1 |
| X6A_LINE_dup847 | 2.85801704922551 | -3.74759745035279 | 1.08057428021347 | -3.46815348002957 | 0.000524047895844781 | 0.0222732329504988 | 3242 | 0 | 0 | 7 | 6 | 0 | 0 | 8 | 9 | 2 | 0 | 2 | 1 |
| AluSz_dup68291 | 3.5249481377548 | -4.67516278245651 | 0.985082384153206 | -4.74596120859005 | 2.07518483110624e-06 | 0.000362977083394936 | 3243 | 0 | 0 | 10 | 10 | 0 | 0 | 8 | 8 | 1 | 0 | 5 | 3 |
| L1P3_dup2646 | 4.08655737842312 | -4.7450335695804 | 1.04002124098503 | -4.5624390950768 | 5.05627635514279e-06 | 0.000727400685369291 | 3244 | 0 | 0 | 7 | 15 | 0 | 0 | 6 | 16 | 1 | 0 | 3 | 2 |
| AluJb_dup60259 | 3.08211526491752 | -3.94034959302103 | 1.01148021565659 | -3.89562695545477 | 9.79450415336082e-05 | 0.00671438514916622 | 3245 | 0 | 0 | 5 | 13 | 0 | 0 | 8 | 7 | 1 | 1 | 3 | 2 |
| MER51-int_dup210 | 2.83228197925403 | -3.86973591417975 | 1.03020672935604 | -3.75627124528555 | 0.000172463767099511 | 0.0100531757781137 | 3246 | 0 | 0 | 6 | 13 | 0 | 0 | 5 | 5 | 1 | 1 | 4 | 2 |
| L1M5_dup39966 | 2.82934282236927 | -3.89970626273028 | 1.05660553566831 | -3.69078727215231 | 0.000223561048044268 | 0.0120569939115914 | 3247 | 0 | 0 | 7 | 11 | 0 | 0 | 2 | 6 | 1 | 1 | 5 | 3 |
| L1MA8_dup7702 | 2.36187988110603 | -3.56288045348813 | 1.08178133266165 | -3.29353109165037 | 0.00098937398681892 | 0.0348663481439003 | 3248 | 0 | 0 | 8 | 9 | 0 | 0 | 2 | 5 | 1 | 1 | 2 | 2 |
| AluSc_dup7507 | 3.42927453332385 | -3.12865088299546 | 0.888623080453476 | -3.52078507953999 | 0.000430271204324223 | 0.0195029261160283 | 3249 | 0 | 0 | 9 | 11 | 0 | 2 | 8 | 7 | 1 | 1 | 4 | 1 |
| HERVL-int_dup1783 | 3.08495281310425 | -3.89311354305563 | 1.05345311443394 | -3.69557362327182 | 0.000219390892097687 | 0.0118940955140297 | 3250 | 0 | 0 | 5 | 16 | 0 | 1 | 5 | 7 | 1 | 0 | 4 | 1 |
| LTR1D1_dup279 | 2.39327338756451 | -3.5968499771729 | 1.046577514314 | -3.4367736053746 | 0.000588687440880919 | 0.0242639566616753 | 3251 | 0 | 0 | 3 | 9 | 0 | 1 | 6 | 5 | 1 | 0 | 4 | 2 |
| Tigger3b_dup3936 | 3.97241992223518 | -3.58436437750696 | 1.08489591290711 | -3.30387858859402 | 0.000953571195944024 | 0.0339956542184604 | 3252 | 0 | 0 | 13 | 18 | 0 | 0 | 3 | 8 | 2 | 2 | 1 | 4 |
| L1ME3G_dup12575 | 4.73725282790753 | -3.44249515223003 | 1.03646910265853 | -3.32136784724222 | 0.000895773920347723 | 0.0325512860798808 | 3253 | 0 | 0 | 23 | 16 | 0 | 1 | 2 | 11 | 2 | 1 | 2 | 1 |
| MER4B-int_dup705 | 4.56940987598435 | -3.39966681816401 | 0.98769811402747 | -3.4420100331076 | 0.000577408833364194 | 0.0238859739679549 | 3254 | 0 | 0 | 22 | 21 | 0 | 1 | 4 | 6 | 1 | 2 | 1 | 2 |
| L1MA4_dup8199 | 3.78996822973225 | -3.48674697106849 | 1.06089601771617 | -3.28660576799463 | 0.00101402685403474 | 0.035420380770721 | 3255 | 0 | 0 | 9 | 31 | 1 | 0 | 3 | 4 | 1 | 1 | 1 | 1 |
| Tigger3b_dup4839 | 2.97096830240437 | -3.35430253313276 | 1.01219872074855 | -3.31387746731409 | 0.000920118197368689 | 0.0332087017916547 | 3256 | 0 | 0 | 14 | 7 | 1 | 0 | 5 | 4 | 1 | 1 | 4 | 1 |
| AluSz_dup18840 | 3.0149170600865 | -2.92453530272746 | 0.920250624275177 | -3.17797698320599 | 0.00148306523715343 | 0.0461353694495633 | 3257 | 0 | 0 | 7 | 13 | 2 | 0 | 4 | 6 | 1 | 1 | 2 | 3 |
| L1PA15_dup6301 | 4.11936701843557 | -2.70222005330391 | 0.836315093206824 | -3.23110281669357 | 0.00123313573578788 | 0.0405892538093489 | 3258 | 0 | 0 | 13 | 12 | 3 | 0 | 6 | 9 | 2 | 1 | 2 | 4 |
| AluSq2_dup61003 | 3.0336451757262 | -3.46766973122595 | 1.02452922315803 | -3.38464697037839 | 0.000712698531050579 | 0.0276850532728205 | 3259 | 0 | 0 | 7 | 6 | 2 | 0 | 3 | 7 | 0 | 1 | 8 | 4 |
| L2b_dup44249 | 3.36350623050477 | -2.83641280300431 | 0.90136338270721 | -3.1468027850047 | 0.00165066256239107 | 0.0496546844590019 | 3260 | 0 | 0 | 9 | 6 | 3 | 0 | 3 | 5 | 0 | 2 | 9 | 6 |
| L1MD1_dup843 | 2.51303292846121 | -4.24971930183099 | 1.13314480117466 | -3.75037620737046 | 0.000176569459269268 | 0.0102099627108834 | 3261 | 0 | 0 | 9 | 6 | 1 | 0 | 1 | 4 | 0 | 0 | 8 | 3 |
| LSAU_dup127 | 3.17979203710065 | -3.20615272771145 | 0.980520644541505 | -3.26984724448169 | 0.00107605575747738 | 0.0369476013824392 | 3262 | 1 | 0 | 10 | 9 | 3 | 0 | 2 | 5 | 0 | 0 | 7 | 4 |
| LTR16C_dup4187 | 3.63851449386306 | -4.26897311391431 | 1.04312961845139 | -4.0924665913062 | 4.26808751726019e-05 | 0.00361929562502492 | 3263 | 0 | 0 | 11 | 16 | 1 | 0 | 2 | 6 | 0 | 1 | 4 | 7 |
| MER4-int_dup312 | 3.76491333068018 | -3.74751971191275 | 1.10399998944736 | -3.39449252512103 | 0.000687559169380302 | 0.0269820092735891 | 3264 | 0 | 0 | 17 | 18 | 1 | 0 | 1 | 4 | 0 | 2 | 3 | 4 |
| LTR27D_dup199 | 3.8324800469792 | -3.8795171691513 | 1.02062632476531 | -3.80111415413803 | 0.000144046915431936 | 0.0088406983078069 | 3265 | 1 | 0 | 15 | 11 | 1 | 0 | 4 | 2 | 0 | 1 | 5 | 12 |
| AluY_dup71776 | 4.23310368602316 | -3.59422436536379 | 0.877605034856182 | -4.09549196120188 | 4.21272845737509e-05 | 0.00358641103942244 | 3266 | 1 | 0 | 11 | 16 | 2 | 0 | 5 | 6 | 0 | 1 | 7 | 7 |
| MIR3_dup79070 | 4.141071627247 | -3.56331474579391 | 0.884224691494185 | -4.02987473667189 | 5.58065837090989e-05 | 0.00442818219589901 | 3267 | 1 | 1 | 14 | 14 | 1 | 0 | 5 | 4 | 0 | 1 | 8 | 6 |
| AluSq2_dup29167 | 2.67517193318202 | -3.25551511743888 | 1.0181840734963 | -3.19737383659902 | 0.00138685063515805 | 0.0440463909814892 | 3268 | 1 | 0 | 5 | 8 | 1 | 0 | 3 | 8 | 0 | 1 | 1 | 6 |
| L2_dup42720 | 2.69410839080299 | -3.29393583081483 | 1.0306555233743 | -3.19596194471525 | 0.00139365491945213 | 0.044189139706152 | 3269 | 0 | 0 | 8 | 5 | 2 | 0 | 4 | 5 | 0 | 1 | 2 | 8 |
| L1M1_dup6423 | 2.30437538511674 | -3.4805028528106 | 1.05090509753618 | -3.3119097632798 | 0.000926614290118353 | 0.0333662475600033 | 3270 | 0 | 0 | 4 | 8 | 1 | 0 | 7 | 5 | 0 | 1 | 3 | 1 |
| L1M4a2_dup1560 | 2.53828644149898 | -3.21078712832363 | 1.0167731012021 | -3.15782068243898 | 0.00158953300601803 | 0.048359931038226 | 3271 | 0 | 0 | 4 | 8 | 1 | 0 | 5 | 5 | 0 | 2 | 4 | 4 |
| AluSg4_dup739 | 2.94668773449145 | -2.9304823821636 | 0.868271356037535 | -3.37507665292245 | 0.000737951333780934 | 0.0283992508600741 | 3272 | 1 | 0 | 7 | 8 | 2 | 0 | 11 | 5 | 0 | 1 | 2 | 2 |
| AluSx1_dup70476 | 2.64490259873889 | -3.1597319196898 | 0.962287982960215 | -3.28356165268712 | 0.00102504221262673 | 0.035695111126252 | 3273 | 1 | 0 | 10 | 7 | 1 | 0 | 5 | 5 | 0 | 1 | 3 | 1 |
| L2b_dup86969 | 3.53227308237888 | -4.21405703861036 | 1.04100290564369 | -4.04807423280403 | 5.16407696210807e-05 | 0.00418233988027831 | 3274 | 1 | 0 | 8 | 15 | 1 | 0 | 2 | 8 | 0 | 0 | 4 | 7 |
| L1MCc_dup2490 | 3.44415235433966 | -2.94657569236758 | 0.877265328549667 | -3.35881927220238 | 0.000782762415321912 | 0.0296113172776133 | 3275 | 2 | 0 | 9 | 17 | 3 | 0 | 5 | 5 | 0 | 0 | 3 | 2 |
| L3_dup40253 | 3.19180968468206 | -2.86353632190372 | 0.903506848970244 | -3.16935762597416 | 0.00152776286554806 | 0.0470948390731569 | 3276 | 2 | 0 | 10 | 13 | 3 | 0 | 3 | 5 | 0 | 0 | 3 | 3 |
| MIRc_dup48410 | 2.65048916456653 | -3.72961197631001 | 1.04282883781793 | -3.57643732226859 | 0.000348308779828494 | 0.016638964249161 | 3277 | 1 | 0 | 8 | 13 | 1 | 0 | 4 | 3 | 0 | 0 | 2 | 4 |
| L2b_dup94926 | 2.4329032871092 | -3.50997962587984 | 1.02342863285122 | -3.42962812766066 | 0.000604408991914277 | 0.0247107735378066 | 3278 | 1 | 0 | 6 | 11 | 1 | 0 | 4 | 5 | 0 | 0 | 2 | 2 |
| MIRb_dup140067 | 2.64255048958752 | -3.54147564083235 | 1.00496154828333 | -3.52399118840107 | 0.000425098385650967 | 0.0193256515995615 | 3279 | 1 | 0 | 7 | 13 | 1 | 0 | 5 | 5 | 0 | 0 | 2 | 1 |
| L1MB5_dup9654 | 2.8040116639548 | -3.64985126170508 | 1.00688834857938 | -3.62488181222345 | 0.000289093628550973 | 0.0145676985470542 | 3280 | 0 | 1 | 8 | 7 | 1 | 0 | 5 | 9 | 0 | 0 | 1 | 3 |
| MLT1C_dup17641 | 2.91412949823827 | -3.63743710818737 | 0.986204620280272 | -3.68831886749185 | 0.000225740641249787 | 0.012138775733971 | 3281 | 0 | 0 | 10 | 10 | 2 | 0 | 7 | 6 | 0 | 0 | 1 | 2 |
| MLT1I_dup4924 | 2.70551376752693 | -3.54218913245529 | 0.999267865985723 | -3.54478438968026 | 0.000392934363983897 | 0.0182594515202671 | 3282 | 0 | 0 | 6 | 9 | 2 | 0 | 8 | 7 | 0 | 0 | 1 | 2 |
| L1ME3A_dup15572 | 3.68405805402593 | -3.74290555853572 | 0.894485201682004 | -4.18442423809527 | 2.85889771399538e-05 | 0.0026635210471979 | 3283 | 0 | 0 | 11 | 11 | 2 | 1 | 8 | 7 | 0 | 0 | 4 | 4 |
| Ricksha_0_dup363 | 2.76946979170923 | -3.28950003348383 | 0.950431323653258 | -3.46106020668562 | 0.000538052464703597 | 0.0226838114398232 | 3284 | 0 | 0 | 8 | 5 | 2 | 1 | 6 | 7 | 0 | 0 | 3 | 3 |
| L1MA6_dup5366 | 3.16941825994224 | -4.39803674804349 | 1.04689648534817 | -4.20102351053439 | 2.65710993183442e-05 | 0.00251992299657343 | 3285 | 0 | 0 | 14 | 8 | 1 | 0 | 4 | 8 | 0 | 0 | 2 | 3 |
| MIRc_dup98062 | 3.45725116211449 | -3.59889657498969 | 0.978587144335985 | -3.67764546654831 | 0.000235396834498781 | 0.0125474317163188 | 3286 | 0 | 0 | 15 | 10 | 3 | 0 | 6 | 5 | 0 | 0 | 4 | 2 |
| Tigger2_dup1506 | 2.91834800839663 | -4.96531974091077 | 1.09271784524836 | -4.54400901614473 | 5.51942953919855e-06 | 0.000775739031975274 | 3287 | 0 | 0 | 6 | 14 | 0 | 0 | 2 | 7 | 0 | 0 | 4 | 5 |
| L1PA16_dup11800 | 2.30586241075893 | -4.55889594757673 | 1.11950043900979 | -4.07225918697192 | 4.65593273617724e-05 | 0.00386822825014815 | 3288 | 0 | 0 | 8 | 10 | 0 | 0 | 2 | 5 | 0 | 0 | 2 | 3 |
| MIR_dup84420 | 2.46515547240514 | -4.68457912510561 | 1.12068142771509 | -4.1801166765624 | 2.91359563751696e-05 | 0.00269548377746738 | 3289 | 0 | 0 | 8 | 11 | 0 | 0 | 2 | 5 | 0 | 0 | 4 | 2 |
| AluY_dup48526 | 1.96197158476022 | -4.16078883311213 | 1.22860553602859 | -3.38659456684663 | 0.00070765882570398 | 0.0275633633886003 | 3290 | 0 | 0 | 3 | 14 | 0 | 0 | 2 | 5 | 0 | 0 | 1 | 1 |
| L1ME2_dup11085 | 2.30714362999259 | -4.56365826822855 | 1.15000577819269 | -3.9683785549327 | 7.23633189921338e-05 | 0.00536236758047408 | 3291 | 0 | 0 | 4 | 13 | 0 | 0 | 2 | 6 | 0 | 0 | 3 | 2 |
| X7A_LINE_dup842 | 2.18793943437015 | -4.45260604225691 | 1.21852300820891 | -3.65410091747198 | 0.00025808474958604 | 0.0134465149605868 | 3292 | 0 | 0 | 4 | 15 | 0 | 0 | 1 | 5 | 0 | 0 | 2 | 2 |
| Charlie3_dup339 | 1.72998873603618 | -4.16669898469937 | 1.15129924225194 | -3.61912770527784 | 0.000295597744895534 | 0.01481076037206 | 3293 | 0 | 0 | 6 | 7 | 0 | 0 | 3 | 3 | 0 | 0 | 1 | 3 |
| Zaphod3_dup643 | 1.92321565372393 | -4.2886772741358 | 1.16401877828426 | -3.68437120959269 | 0.000229267912994998 | 0.0122951054373468 | 3294 | 0 | 0 | 8 | 7 | 0 | 0 | 2 | 4 | 0 | 0 | 1 | 3 |
| AluSq4_dup194 | 1.80027889612852 | -4.05382607323712 | 1.18499119877983 | -3.42097568101036 | 0.000623969115208568 | 0.0252148425847771 | 3295 | 0 | 0 | 6 | 8 | 0 | 0 | 2 | 5 | 0 | 0 | 1 | 1 |
| L1ME1_dup20379 | 1.54025815713766 | -3.96875733798114 | 1.24926588398592 | -3.17687162425215 | 0.00148872918933462 | 0.0462607014129047 | 3296 | 0 | 0 | 6 | 7 | 0 | 0 | 1 | 3 | 0 | 0 | 2 | 1 |
| L2_dup40999 | 2.37008446333543 | -4.00603624188749 | 1.16055152641244 | -3.45183832920469 | 0.00055678123000727 | 0.0232795407758408 | 3297 | 1 | 0 | 9 | 9 | 0 | 0 | 1 | 6 | 0 | 0 | 3 | 1 |
| MIRb_dup41088 | 1.98811722665774 | -3.60776908953964 | 1.11956404546013 | -3.22247673473374 | 0.00127087486169552 | 0.0414586769729918 | 3298 | 1 | 0 | 7 | 6 | 0 | 0 | 3 | 6 | 0 | 0 | 1 | 1 |
| MLT1D_dup3488 | 2.54638557615342 | -3.49638149719563 | 1.04726481406355 | -3.33858394767329 | 0.000842065822208636 | 0.0311902543790872 | 3299 | 2 | 0 | 7 | 8 | 0 | 0 | 4 | 8 | 0 | 0 | 2 | 1 |
| MER5B_dup1912 | 2.49669934323357 | -4.10950960621353 | 1.12738514526442 | -3.64516919836626 | 0.000267215936924442 | 0.013762546174767 | 3300 | 0 | 1 | 8 | 4 | 0 | 0 | 3 | 9 | 0 | 0 | 4 | 1 |
| LTR85a_dup1774 | 1.84250181712048 | -4.23549563607147 | 1.16070458115441 | -3.64907290351085 | 0.000263188412516701 | 0.0136367637314779 | 3301 | 0 | 0 | 7 | 4 | 0 | 0 | 3 | 5 | 0 | 0 | 3 | 1 |
| L1ME2_dup1274 | 2.47968198090664 | -4.41686154532502 | 1.12514660718504 | -3.92558757864931 | 8.65182837295484e-05 | 0.00612794935546866 | 3302 | 0 | 0 | 12 | 6 | 0 | 0 | 4 | 7 | 0 | 0 | 1 | 1 |
| L1PA5_dup9419 | 2.30978852243881 | -4.33637789011614 | 1.1319151518683 | -3.83100966795846 | 0.000127618490239315 | 0.00810889880559255 | 3303 | 0 | 0 | 9 | 8 | 0 | 0 | 3 | 7 | 0 | 0 | 1 | 1 |
| L1MD3_dup4687 | 3.41814883461425 | -3.46352582375771 | 1.01451474097437 | -3.41397289154345 | 0.000640229456004488 | 0.0256748927648831 | 3304 | 0 | 0 | 12 | 17 | 1 | 1 | 1 | 7 | 1 | 0 | 2 | 2 |
| L1MC4_dup19894 | 3.09981672918134 | -3.59910166632937 | 1.09406743133038 | -3.28965250519603 | 0.00100311180633827 | 0.0351668451897249 | 3305 | 0 | 0 | 13 | 18 | 1 | 1 | 1 | 5 | 0 | 0 | 1 | 1 |
| L2_dup29485 | 2.80492531325933 | -3.49813190413282 | 1.03256089652987 | -3.38782140200059 | 0.000704501221975604 | 0.0274808602892143 | 3306 | 0 | 0 | 11 | 16 | 1 | 1 | 4 | 3 | 0 | 0 | 1 | 1 |
| L2_dup52566 | 3.06906679070962 | -3.70651224332593 | 1.01369464999276 | -3.65643859652648 | 0.000255743635730419 | 0.013362576887903 | 3307 | 0 | 0 | 7 | 19 | 1 | 1 | 4 | 6 | 0 | 0 | 1 | 2 |
| LSAU_dup95 | 3.16185262929729 | -4.25974866369287 | 1.06342600607228 | -4.00568411847109 | 6.18382434554039e-05 | 0.00478642346165923 | 3308 | 0 | 0 | 14 | 10 | 0 | 1 | 4 | 8 | 0 | 0 | 2 | 1 |
| MIR_dup170848 | 3.23399535382029 | -4.36907092937588 | 1.04162657319032 | -4.19446953623138 | 2.73510905243133e-05 | 0.0025693180362804 | 3309 | 0 | 0 | 12 | 14 | 0 | 1 | 4 | 7 | 0 | 0 | 2 | 2 |
| L1MA3_dup6848 | 2.95654053184731 | -4.71594460335631 | 1.09976861045935 | -4.28812439135406 | 1.80188182022709e-05 | 0.00188960146141805 | 3310 | 0 | 0 | 10 | 14 | 0 | 0 | 3 | 8 | 0 | 0 | 1 | 2 |
| MLT1E1A_dup288 | 3.48346825592238 | -4.90113486085726 | 1.04468203598893 | -4.69150870026944 | 2.71197737050809e-06 | 0.000446577278479528 | 3311 | 0 | 0 | 15 | 13 | 0 | 0 | 6 | 8 | 0 | 0 | 1 | 2 |
| L3_dup23086 | 2.59370482355245 | -3.65658834355761 | 1.06158130292735 | -3.44447319623513 | 0.000572173363417818 | 0.0237438109496211 | 3312 | 0 | 0 | 9 | 8 | 0 | 1 | 3 | 6 | 1 | 0 | 1 | 4 |
| L1M5_dup47246 | 2.98514051948723 | -3.1741388722501 | 0.945850192343182 | -3.35585793389407 | 0.000791192001900873 | 0.0298400653168973 | 3313 | 0 | 0 | 11 | 12 | 0 | 2 | 6 | 5 | 1 | 0 | 1 | 1 |

| | | | | | | | | | | | | | | | | | | | |
|---|---|---|---|---|---|---|---|---|---|---|---|---|---|---|---|---|---|---|---|
| MSTA_dup9747 | 2.73532747368319 | -3.5214554778321 | 1.01564437261084 | -3.46721310411022 | 0.000525884787923865 | 0.0223171360348372 | 3314 | 0 | 0 | 7 | 12 | 0 | 1 | 5 | 7 | 1 | 0 | 1 | 1 |
| MIRb_dup217354 | 3.32809241881602 | -3.81432885443951 | 0.998194165723001 | -3.8212293614006 | 0.00013278808263163 | 0.00832140982614323 | 3315 | 0 | 0 | 8 | 18 | 0 | 2 | 6 | 7 | 0 | 0 | 2 | 1 |
| L2_dup38314 | 2.60541996090736 | -4.04135889369469 | 1.10056866562289 | -3.67206428815362 | 0.000240599134871035 | 0.0127646826635504 | 3316 | 0 | 0 | 4 | 15 | 0 | 1 | 4 | 7 | 0 | 0 | 2 | 1 |
| UCON132b_dup722 | 3.2827083863977 | -4.30409010788146 | 1.09278832319427 | -3.9386311296779 | 8.19478034620452e-05 | 0.00588791570060197 | 3317 | 0 | 0 | 7 | 18 | 0 | 1 | 3 | 10 | 0 | 0 | 2 | 1 |
| Charlie16a_dup824 | 2.21900698310659 | -3.84365708894368 | 1.09703487429974 | -3.50367812271891 | 0.000458879649137509 | 0.0203482253063127 | 3318 | 0 | 0 | 8 | 7 | 0 | 1 | 3 | 6 | 0 | 0 | 2 | 1 |
| MER89_dup750 | 1.84147621122121 | -3.52537080922948 | 1.10928213670921 | -3.17806506799778 | 0.00148261473853817 | 0.046128510414423 | 3319 | 0 | 0 | 6 | 7 | 0 | 1 | 4 | 4 | 0 | 0 | 1 | 1 |
| MamRep137_dup563 | 2.76737977161462 | -3.76429889419764 | 1.01766356043725 | -3.69896205439474 | 0.000216482959822915 | 0.0117766928116194 | 3320 | 0 | 0 | 8 | 8 | 0 | 2 | 6 | 6 | 0 | 0 | 2 | 4 |
| MIR3_dup84361 | 2.68882336759561 | -3.73271228118532 | 1.02992117953212 | -3.62426985226292 | 0.000289778928065576 | 0.0145960492839736 | 3321 | 0 | 0 | 8 | 8 | 0 | 2 | 6 | 5 | 0 | 0 | 4 | 2 |
| MamRTE1_dup6987 | 2.23769790286036 | -3.39537034389644 | 1.04645937487805 | -3.24462700168568 | 0.00117604560969123 | 0.0392751366517318 | 3322 | 0 | 0 | 6 | 7 | 0 | 2 | 5 | 5 | 0 | 0 | 2 | 2 |
| MLT1A0_dup16492 | 2.73095488750654 | -4.76236248919849 | 1.04977834882913 | -4.53654097030119 | 5.71843845797851e-06 | 0.000796184644424901 | 3323 | 0 | 0 | 7 | 13 | 0 | 0 | 5 | 6 | 0 | 0 | 2 | 3 |
| AluSc_dup34473 | 2.55749961615894 | -4.63233923903344 | 1.05952019723314 | -4.37211036762721 | 1.23051289569201e-05 | 0.00142244960655641 | 3324 | 0 | 0 | 8 | 8 | 0 | 0 | 7 | 6 | 0 | 0 | 3 | 1 |
| MIRb_dup217340 | 2.4292701531034 | -4.62795718513008 | 1.06344653893699 | -4.35184752188498 | 1.34995135768496e-05 | 0.00152814236801002 | 3325 | 0 | 0 | 6 | 11 | 0 | 0 | 5 | 5 | 0 | 0 | 3 | 2 |
| MLT2C1_dup1306 | 2.28391237471567 | -4.50511269938027 | 1.08893025177531 | -4.13719124070203 | 3.51583071649862e-05 | 0.00311342707729059 | 3326 | 0 | 0 | 5 | 10 | 0 | 0 | 6 | 5 | 0 | 0 | 3 | 1 |
| AluJb_dup60036 | 2.71530448440148 | -4.71059742254807 | 1.08790130138391 | -4.32998601670552 | 1.4911881053692e-05 | 0.00164297742621084 | 3327 | 0 | 0 | 11 | 12 | 0 | 0 | 5 | 4 | 0 | 0 | 3 | 1 |
| L1PA13_dup640 | 2.87571595379954 | -4.77578260336152 | 1.08864344860833 | -4.3869116279225 | 1.14971474324671e-05 | 0.00135155305247614 | 3328 | 0 | 0 | 12 | 15 | 0 | 0 | 5 | 3 | 0 | 0 | 2 | 2 |
| LTR105_Mam_dup44 | 2.96433665008425 | -4.73804424493963 | 1.11325445215163 | -4.25602990922896 | 2.0808888209289e-05 | 0.00210086847366168 | 3329 | 0 | 0 | 14 | 15 | 0 | 0 | 5 | 3 | 0 | 0 | 2 | 1 |
| MER65C_dup475 | 3.0653427673168 | -4.95361683082908 | 1.03261404398471 | -4.79716197904281 | 1.60929461848168e-06 | 0.00029892678945296 | 3330 | 0 | 0 | 11 | 12 | 0 | 0 | 8 | 4 | 0 | 0 | 4 | 2 |
| L1PB1_dup12909 | 2.61933328773117 | -4.67249857285062 | 1.06888338255901 | -4.37138293016047 | 1.2346204298653e-05 | 0.00142608248934334 | 3331 | 0 | 0 | 7 | 14 | 0 | 0 | 8 | 3 | 0 | 0 | 2 | 2 |
| BSR/Beta_dup1972 | 2.44532165708792 | -4.64023989220949 | 1.06546189405788 | -4.3551439221696 | 1.32979518072943e-05 | 0.0015096525313337 | 3332 | 0 | 0 | 9 | 10 | 0 | 0 | 6 | 3 | 0 | 0 | 2 | 3 |
| L1PB2_dup1089 | 2.49820457759456 | -4.66711839664603 | 1.07980116934078 | -4.3222016507866 | 1.54479899774476e-05 | 0.00168846459493511 | 3333 | 0 | 0 | 8 | 13 | 0 | 0 | 5 | 3 | 0 | 0 | 2 | 3 |
| MIR3_dup84369 | 2.02546676511333 | -4.4432500340227 | 1.07597660869624 | -4.12950430159124 | 3.63546201296556e-05 | 0.00320371357378448 | 3334 | 0 | 0 | 6 | 7 | 0 | 0 | 5 | 3 | 0 | 0 | 3 | 3 |
| MIR_dup170867 | 2.36916744633843 | -4.67049302429034 | 1.07610871494373 | -4.34016838580714 | 1.42373574938992e-05 | 0.00158824297526646 | 3335 | 0 | 0 | 7 | 11 | 0 | 0 | 4 | 3 | 0 | 0 | 3 | 4 |
| AluSg_dup36237 | 2.10977914444901 | -4.41074697729686 | 1.12475092646894 | -3.92153220192849 | 8.79876868381647e-05 | 0.00621440134532322 | 3336 | 0 | 0 | 9 | 9 | 0 | 0 | 3 | 3 | 0 | 0 | 2 | 2 |
| MLT2B4_dup3865 | 1.90149853575094 | -4.32970580583349 | 1.12178658272463 | -3.85965198060881 | 0.00011354862017416 | 0.00744719955235963 | 3337 | 0 | 0 | 7 | 7 | 0 | 0 | 3 | 3 | 0 | 0 | 3 | 2 |
| MADE1_dup5900 | 1.85078987722513 | -4.26369647691185 | 1.16020072268693 | -3.67496450703588 | 0.000237882482160178 | 0.0126629049440908 | 3338 | 0 | 0 | 6 | 9 | 0 | 0 | 4 | 2 | 0 | 0 | 3 | 1 |
| MIR_dup49179 | 1.90332084632211 | -4.29509553230762 | 1.15838964221687 | -3.70781589870562 | 0.000209054546758281 | 0.0114854983105394 | 3339 | 0 | 0 | 6 | 11 | 0 | 0 | 3 | 2 | 0 | 0 | 2 | 2 |
| MIRb_dup217342 | 1.92741920044595 | -4.23815524799197 | 1.13259544773249 | -3.74198506313701 | 0.000182572369442772 | 0.0104403665031074 | 3340 | 0 | 0 | 6 | 10 | 0 | 0 | 4 | 3 | 0 | 0 | 1 | 2 |
| AluSz_dup7467 | 2.15731316173575 | -4.26679471258673 | 1.11599732736479 | -3.82330190938892 | 0.000131676385460115 | 0.0082779046802358 | 3341 | 0 | 0 | 8 | 9 | 0 | 0 | 4 | 5 | 0 | 0 | 1 | 1 |
| L2a_dup152630 | 2.15832780410635 | -4.26751401246632 | 1.10289017237589 | -3.86939163967076 | 0.000109107259867681 | 0.00726047463797237 | 3342 | 0 | 0 | 8 | 8 | 0 | 0 | 5 | 5 | 0 | 0 | 1 | 1 |
| AluJb_dup104249 | 1.98649043544393 | -4.17437498337407 | 1.15022129802265 | -3.62919291318135 | 0.000284308721314531 | 0.0143905813660726 | 3343 | 0 | 0 | 8 | 9 | 0 | 0 | 3 | 4 | 0 | 0 | 1 | 1 |
| L1PA14_dup1663 | 2.06583611682887 | -4.21947971398636 | 1.12911850270624 | -3.73696800103197 | 0.00018625261582177 | 0.0105796287716226 | 3344 | 0 | 0 | 9 | 8 | 0 | 0 | 4 | 4 | 0 | 0 | 1 | 1 |
| L2c_dup136291 | 2.13918106221206 | -4.26875691113481 | 1.14033660888676 | -3.74341828357341 | 0.000181533657844413 | 0.0104037813088031 | 3345 | 0 | 0 | 10 | 7 | 0 | 0 | 8 | 2 | 0 | 0 | 1 | 1 |
| MER45R_dup47 | 2.07399602909026 | -4.23334752674889 | 1.11017882503252 | -3.81321227832363 | 0.000137172196514709 | 0.00852745751931099 | 3346 | 0 | 0 | 7 | 8 | 0 | 0 | 8 | 3 | 0 | 0 | 1 | 1 |
| MIR3_dup84364 | 2.36988525798647 | -4.46999514106415 | 1.10439873640933 | -4.04744680856589 | 5.17793436226043e-05 | 0.00419213599783063 | 3347 | 0 | 0 | 12 | 7 | 0 | 0 | 5 | 4 | 0 | 0 | 1 | 2 |
| MSTB2_dup269 | 2.37113025214332 | -4.38354252513502 | 1.15877163723141 | -3.78292183230199 | 0.000154998102036334 | 0.00932375194602892 | 3348 | 0 | 0 | 13 | 10 | 0 | 0 | 5 | 2 | 0 | 0 | 1 | 1 |
| LTR37A_dup172 | 3.05223058900739 | -4.23179568086861 | 1.09823262080506 | -3.85327807670336 | 0.000116546898672772 | 0.0075866122539867 | 3349 | 1 | 0 | 14 | 16 | 0 | 0 | 6 | 2 | 0 | 0 | 1 | 2 |
| L1PB3_dup3146 | 2.26897788583508 | -3.77961135676794 | 1.09509954524564 | -3.45138610747953 | 0.000557715086864202 | 0.0233012031085966 | 3350 | 1 | 0 | 9 | 10 | 0 | 0 | 4 | 4 | 0 | 0 | 1 | 1 |
| MLT1I_dup991 | 2.70937497469579 | -3.66555360673367 | 1.08911808563492 | -3.36561632304247 | 0.000763728577728971 | 0.0290882405923526 | 3351 | 2 | 0 | 14 | 10 | 0 | 0 | 3 | 3 | 0 | 0 | 2 | 2 |
| L1M5_dup25388 | 2.35756784389469 | -4.11471265761241 | 1.17775068618707 | -3.49370431779045 | 0.000476368306831381 | 0.0208502760922085 | 3352 | 1 | 0 | 7 | 14 | 0 | 0 | 2 | 2 | 0 | 0 | 5 | 1 |
| L1MA2_dup1808 | 2.4340923968885 | -3.97781170693714 | 1.1971907473197 | -3.32262149189071 | 0.000891758286042266 | 0.0324499643876148 | 3353 | 1 | 0 | 6 | 20 | 0 | 0 | 2 | 2 | 0 | 0 | 2 | 1 |
| L1ME1_dup3210 | 2.36279648295419 | -3.50555021731448 | 1.07205523578185 | -3.26993432829781 | 0.00107572455095665 | 0.0369476013824392 | 3354 | 2 | 0 | 6 | 11 | 0 | 0 | 6 | 3 | 0 | 0 | 3 | 1 |
| L2c_dup136293 | 2.20843629167559 | -3.92792550172513 | 1.0723812083962 | -3.66280709785985 | 0.000249466402283157 | 0.0131007575717652 | 3355 | 1 | 0 | 7 | 8 | 0 | 0 | 5 | 4 | 0 | 0 | 3 | 1 |
| L2a_dup146310 | 2.97656932497139 | -4.4545995362078 | 1.03321743241713 | -4.31138635145425 | 1.6223415285565e-05 | 0.00175070148459821 | 3356 | 1 | 0 | 7 | 13 | 0 | 0 | 4 | 6 | 0 | 0 | 5 | 3 |
| MIR_dup19163 | 2.55667820792 | -4.26403534772986 | 1.0703632965577 | -3.98372717136607 | 6.78427564918984e-05 | 0.00512574933197374 | 3357 | 1 | 0 | 9 | 10 | 0 | 0 | 3 | 3 | 0 | 0 | 5 | 3 |
| LSAU_dup97 | 2.42649563275904 | -4.0936960981463 | 1.11747647427711 | -3.66333984864827 | 0.00024894788975543 | 0.0130764138136303 | 3358 | 0 | 0 | 11 | 10 | 1 | 0 | 2 | 3 | 0 | 0 | 3 | 2 |
| MER90a_dup606 | 2.23041236644911 | -3.94362283390033 | 1.18349407938304 | -3.33218636459606 | 0.000861665328913815 | 0.0316942059597661 | 3359 | 0 | 0 | 12 | 9 | 1 | 0 | 3 | 1 | 0 | 0 | 3 | 1 |
| AluSz_dup74175 | 1.59925487576812 | -4.08351513093023 | 1.28924810729843 | -3.16736174194359 | 0.00153828836632689 | 0.0473055531490971 | 3360 | 0 | 0 | 6 | 10 | 0 | 0 | 1 | 1 | 0 | 0 | 2 | 2 |
| 7SK_dup261 | 2.30044654423713 | -4.63469524094958 | 1.15181534147323 | -4.02381794552379 | 5.72621886809731e-05 | 0.00450870574053039 | 3361 | 0 | 0 | 9 | 11 | 0 | 0 | 2 | 2 | 0 | 0 | 4 | 3 |
| L1ME3D_dup2019 | 2.07800528727593 | -4.51598273271444 | 1.24260215433166 | -3.63429494868643 | 0.000278741841410801 | 0.0141872740526919 | 3362 | 0 | 0 | 9 | 9 | 0 | 0 | 1 | 1 | 0 | 0 | 5 | 3 |
| L1MB1_dup3562 | 2.1639070591125 | -4.3854392650249 | 1.26119035170425 | -3.47722233927563 | 0.000506637497211111 | 0.0217800119081111 | 3363 | 0 | 0 | 9 | 15 | 0 | 0 | 2 | 1 | 0 | 0 | 2 | 1 |
| L1MC3_dup3113 | 2.47816048299387 | -4.53890609846884 | 1.21437831929946 | -3.73763762604654 | 0.000185757413760169 | 0.0105540194781703 | 3364 | 0 | 0 | 10 | 17 | 0 | 0 | 2 | 2 | 0 | 0 | 2 | 1 |
| MER45C_dup614 | 2.31746560733636 | -4.52553589587139 | 1.24915581898319 | -3.62287540681286 | 0.000291346177950521 | 0.0146532778817617 | 3365 | 0 | 0 | 16 | 8 | 0 | 0 | 2 | 1 | 0 | 0 | 2 | 2 |
| LTR8_dup369 | 1.84418569867186 | -4.18659067027547 | 1.31084246885699 | -3.19381677794284 | 0.00140405198882018 | 0.044426627592225 | 3366 | 0 | 0 | 11 | 9 | 0 | 0 | 1 | 1 | 0 | 0 | 1 | 2 |
| ORSL-2a_dup274 | 2.14564506807034 | -4.37257579900009 | 1.31041403966191 | -3.3367894929821 | 0.000847521136036171 | 0.0313386580008178 | 3367 | 0 | 0 | 13 | 11 | 0 | 0 | 1 | 1 | 0 | 0 | 2 | 1 |
| LTR33_dup5583 | 2.31334351905178 | -4.10922528192518 | 1.17102989987368 | -3.50906948009478 | 0.000449677416125743 | 0.0200914211337407 | 3368 | 0 | 0 | 8 | 13 | 0 | 0 | 1 | 2 | 1 | 0 | 3 | 3 |
| L1MD_dup1862 | 2.85704760704937 | -4.43562368380196 | 1.21944569864777 | -3.63740975815534 | 0.000275393632053225 | 0.014074057811971 | 3369 | 0 | 0 | 13 | 15 | 0 | 0 | 1 | 1 | 1 | 0 | 2 | 6 |
| L1ME2z_dup39 | 2.68010851634972 | -4.23446107353919 | 1.25726901800674 | -3.36798331374811 | 0.000757201787798979 | 0.0289434757589744 | 3370 | 0 | 0 | 11 | 19 | 0 | 0 | 1 | 1 | 1 | 0 | 1 | 3 |
| AluSx1_dup1053 | 2.36177057908572 | -4.00734140951016 | 1.17255401811938 | -3.41761773665438 | 0.000631717643643081 | 0.025446874290553 | 3371 | 0 | 0 | 9 | 15 | 0 | 0 | 2 | 2 | 1 | 0 | 1 | 2 |
| L1PA7_dup2242 | 2.68631366023745 | -4.08277815563563 | 1.16947925447109 | -3.49110780719416 | 0.000481022113319315 | 0.0209790150581349 | 3372 | 0 | 0 | 11 | 17 | 0 | 0 | 2 | 3 | 1 | 0 | 1 | 1 |
| AluSx_dup12180 | 2.10321365103651 | -3.90416433120933 | 1.08270367428622 | -3.60593985587347 | 0.000311025173402045 | 0.0153329243090912 | 3373 | 0 | 0 | 6 | 9 | 0 | 0 | 5 | 3 | 1 | 0 | 1 | 3 |
| Charlie17a_dup624 | 2.48265210803967 | -4.11207703264537 | 1.07541440230646 | -3.82371393188163 | 0.000131456427694181 | 0.00827110282377745 | 3374 | 0 | 0 | 10 | 10 | 0 | 0 | 5 | 3 | 1 | 0 | 1 | 3 |
| MER8_dup1238 | 1.8220060272311 | -3.67717217244856 | 1.14278106472328 | -3.21773985058018 | 0.00129204948700498 | 0.0418967439749989 | 3375 | 0 | 0 | 7 | 8 | 0 | 0 | 3 | 2 | 1 | 0 | 2 | 1 |
| Tigger1_dup6534 | 2.20631072191722 | -3.84781944935355 | 1.15479359721277 | -3.33204085876533 | 0.000862115978260003 | 0.0317009810791307 | 3376 | 0 | 0 | 10 | 10 | 0 | 0 | 6 | 1 | 1 | 0 | 1 | 1 |
| MER58A_dup6273 | 2.87324669632947 | -3.89496680136239 | 1.11151285119174 | -3.50420312026647 | 0.000457975894678243 | 0.020323182520728 | 3377 | 0 | 0 | 9 | 17 | 1 | 1 | 3 | 1 | 0 | 0 | 3 | 5 |
| MIR_dup170872 | 2.35712634324603 | -3.62645941637203 | 1.07766326506289 | -3.36511369918542 | 0.000765121229973006 | 0.0291245458626071 | 3378 | 0 | 0 | 8 | 9 | 1 | 1 | 3 | 2 | 0 | 0 | 2 | 6 |
| MIRb_dup104838 | 2.33887861531624 | -3.45863727141604 | 1.05650463728925 | -3.27366028443577 | 0.00106164160958405 | 0.0366053288558306 | 3379 | 0 | 0 | 8 | 11 | 1 | 1 | 5 | 2 | 0 | 0 | 1 | 3 |
| Charlie2a_dup2077 | 2.48921948647285 | -3.16741959682246 | 1.00698785053192 | -3.14543973410338 | 0.00165837358353071 | 0.0498110780998231 | 3380 | 0 | 0 | 10 | 7 | 1 | 1 | 4 | 2 | 0 | 1 | 4 | 3 |
| MIR3_dup49875 | 2.69087445472332 | -3.28421077689661 | 1.02978392149342 | -3.1892232033821 | 0.00142655683269829 | 0.0448583972661857 | 3381 | 0 | 0 | 7 | 15 | 1 | 1 | 3 | 2 | 0 | 1 | 2 | 5 |
| MIR_dup79320 | 3.37461180035075 | -3.63191602990338 | 1.04295391986635 | -3.48233604641784 | 0.000497059495082366 | 0.0214964228674789 | 3382 | 0 | 0 | 13 | 17 | 1 | 1 | 2 | 2 | 0 | 1 | 3 | 6 |
| MER11A_dup911 | 2.82528950307048 | -4.27272089767902 | 1.17816258556961 | -3.62659699943981 | 0.000287180969333186 | 0.0144987751187007 | 3383 | 0 | 0 | 18 | 9 | 0 | 1 | 1 | 3 | 0 | 0 | 2 | 3 |
| MIRb_dup98619 | 2.41982902160835 | -4.61567294850369 | 1.17714417058525 | -3.92107701319957 | 8.81540813023277e-05 | 0.006220857796398 | 3384 | 0 | 0 | 12 | 8 | 0 | 0 | 1 | 5 | 0 | 0 | 2 | 3 |
| L1PBa_dup1084 | 2.33346796185108 | -4.57970970439599 | 1.14614537936461 | -3.99574939344505 | 6.44899265096047e-05 | 0.00493095117392494 | 3385 | 0 | 0 | 11 | 10 | 0 | 0 | 2 | 3 | 0 | 0 | 2 | 3 |
| THE1C_dup3535 | 2.43202636399649 | -4.50140498447378 | 1.19087813285198 | -3.77990397194847 | 0.000156888855283106 | 0.00941752662946364 | 3386 | 0 | 0 | 15 | 9 | 0 | 0 | 2 | 3 | 0 | 0 | 1 | 2 |
| LTR78_dup1059 | 2.19097357158935 | -3.38420344920209 | 1.07624068211822 | -3.14446713029042 | 0.00166389602341567 | 0.0499328292581238 | 3387 | 1 | 0 | 10 | 7 | 0 | 1 | 3 | 3 | 0 | 0 | 1 | 3 |
| MIRb_dup190733 | 2.57840375804472 | -3.71755820900039 | 1.04167376680278 | -3.56883155501815 | 0.000358576866178382 | 0.017033725494047 | 3388 | 1 | 0 | 7 | 10 | 0 | 1 | 3 | 5 | 0 | 0 | 2 | 5 |
| L2b_dup45751 | 2.65261204078448 | -3.72232032487272 | 1.06996229947069 | -3.47892661892307 | 0.000503426419053003 | 0.0216889919804429 | 3389 | 0 | 0 | 8 | 13 | 0 | 2 | 4 | 3 | 0 | 0 | 2 | 4 |
| Kanga11a_dup687 | 2.98250743561917 | -4.42533590227775 | 1.06830479966988 | -4.14239073309906 | 3.43704130721013e-05 | 0.00305616678208304 | 3390 | 0 | 0 | 8 | 16 | 0 | 1 | 3 | 5 | 0 | 0 | 2 | 5 |
| L2c_dup135334 | 2.31096999866389 | -4.07032046753975 | 1.10239767322087 | -3.69224334050661 | 0.000222284624183956 | 0.0120017556582881 | 3391 | 0 | 0 | 6 | 11 | 0 | 1 | 3 | 4 | 0 | 0 | 1 | 5 |
| L1M5_dup39964 | 2.48461541478511 | -4.23070153740751 | 1.13372583450195 | -3.73167957248331 | 0.000190207348785359 | 0.0107197838168223 | 3392 | 0 | 0 | 9 | 8 | 0 | 1 | 2 | 4 | 0 | 0 | 1 | 8 |

| | | | | | | | | | | | | | | | | | | | |
|---|---|---|---|---|---|---|---|---|---|---|---|---|---|---|---|---|---|---|---|
| LTR12F_dup488 | 2.06407843616919 | -3.97031758478642 | 1.16874811616404 | -3.39706864967402 | 0.000681118772935685 | 0.0268202241551396 | 3393 | 0 | 0 | 7 | 9 | 0 | 1 | 1 | 2 | 0 | 0 | 2 | 6 |
| AluJr_dup82512 | 4.83813914724766 | -2.63336880529453 | 0.766768581735245 | -3.43437233609005 | 0.00059392776404695 | 0.0244038569752919 | 3394 | 3 | 2 | 16 | 23 | 0 | 1 | 7 | 5 | 0 | 2 | 4 | 2 |
| L2c_dup56080 | 3.77349330517198 | -2.97707974677982 | 0.823872581982083 | -3.61351962899108 | 0.00030206847582075 | 0.0150368170601766 | 3395 | 1 | 2 | 8 | 20 | 0 | 1 | 9 | 5 | 0 | 1 | 3 | 1 |
| L1M3_dup2446 | 4.32326897998333 | -2.21010926184756 | 0.696469252035514 | -3.17330485931469 | 0.00150714178676791 | 0.046657187727371 | 3396 | 4 | 3 | 14 | 15 | 0 | 3 | 8 | 7 | 0 | 0 | 1 | 2 |
| MER9a1_dup55 | 5.45651317119192 | -2.95824009932038 | 0.713155021637295 | -4.14810245958685 | 3.35242366529816e-05 | 0.00300563602661279 | 3397 | 3 | 2 | 19 | 21 | 0 | 3 | 7 | 11 | 0 | 0 | 2 | 3 |
| AluSz_dup84068 | 4.45783336296927 | -2.38746965970756 | 0.703756288427939 | -3.39246653843865 | 0.000692663916379325 | 0.0271303058842461 | 3398 | 3 | 2 | 13 | 20 | 1 | 2 | 6 | 6 | 1 | 0 | 3 | 2 |
| HERVP71A-int_dup181 | 5.42787032436521 | -3.57720645820754 | 0.784959631161267 | -4.55718525666783 | 5.1843694577378e-06 | 0.00074268511438343 | 3399 | 2 | 1 | 18 | 26 | 0 | 1 | 8 | 8 | 1 | 0 | 4 | 3 |
| L1ME3E_dup2493 | 5.30956188542932 | -4.06432616239702 | 0.839154376903081 | -4.84335930820799 | 1.2766212637749e-06 | 0.000248175590874332 | 3400 | 0 | 1 | 16 | 20 | 0 | 2 | 11 | 14 | 0 | 0 | 3 | 1 |
| MER58A_dup1141 | 5.01394317809121 | -3.67632020546984 | 0.814593916140018 | -4.51307103162544 | 6.38956105801555e-06 | 0.000870773479673864 | 3401 | 1 | 2 | 13 | 21 | 0 | 1 | 8 | 14 | 0 | 0 | 3 | 1 |
| L1MA4_dup5792 | 3.73772676888696 | -3.45080461378398 | 0.937147078674756 | -3.6822444334606 | 0.000231189584380818 | 0.0123758463093539 | 3402 | 0 | 2 | 15 | 20 | 0 | 1 | 5 | 5 | 0 | 0 | 1 | 1 |
| AluY_dup51409 | 5.0654788086401 | -3.82821958020703 | 0.940484083315429 | -4.07047779768016 | 4.69168126199923e-05 | 0.00388977975459118 | 3403 | 0 | 2 | 22 | 28 | 0 | 1 | 7 | 6 | 0 | 0 | 2 | 0 |
| L2a_dup147245 | 3.65743306340177 | -3.68315207759018 | 0.983902730381139 | -3.74341077004982 | 0.000181539088676214 | 0.0104037813088031 | 3404 | 0 | 1 | 18 | 16 | 0 | 1 | 8 | 4 | 0 | 0 | 1 | 0 |
| L1ME3A_dup13443 | 3.80576304292389 | -3.30290975810322 | 0.944581995339163 | -3.49668930214711 | 0.000471070132865495 | 0.0206997698679248 | 3405 | 0 | 2 | 13 | 19 | 1 | 1 | 4 | 6 | 0 | 0 | 4 | 0 |
| MLT1F2_dup1091 | 3.54188741821262 | -3.09188844704072 | 0.928701832561368 | -3.32925847525601 | 0.000870775517518358 | 0.031925409490887 | 3406 | 0 | 2 | 12 | 16 | 0 | 1 | 5 | 6 | 1 | 0 | 3 | 0 |
| L1MC3_dup12156 | 4.46685058839658 | -3.64900803485393 | 1.09572536346581 | -3.33022138258443 | 0.00086776960191484 | 0.0318448960051247 | 3407 | 2 | 1 | 24 | 20 | 0 | 1 | 1 | 4 | 0 | 0 | 6 | 0 |
| MER5A_dup17287 | 3.74033114377607 | -3.24753567478465 | 0.998461628128429 | -3.25253928973917 | 0.00114378781791468 | 0.0385549987422476 | 3408 | 1 | 0 | 11 | 22 | 0 | 1 | 4 | 4 | 1 | 1 | 5 | 0 |
| L1ME2z_dup595 | 2.69378605821566 | -3.15251404460599 | 0.996331580102002 | -3.16412137040085 | 0.00155551912480067 | 0.0476173361211657 | 3409 | 2 | 0 | 7 | 11 | 0 | 1 | 5 | 6 | 0 | 0 | 3 | 0 |
| PABL_B-int_dup137 | 4.25365114725834 | -2.9515127218508 | 0.91334941858379 | -3.23152635978827 | 0.00123130966417256 | 0.0405403638233296 | 3410 | 3 | 0 | 13 | 24 | 0 | 2 | 4 | 9 | 0 | 0 | 1 | 0 |
| L1MEf_dup2014 | 3.41311494133844 | -3.59592728108647 | 0.97464276181961 | -3.68948236415674 | 0.000224710804187178 | 0.0120915958871828 | 3411 | 0 | 0 | 16 | 12 | 1 | 1 | 6 | 7 | 0 | 0 | 1 | 0 |
| MIR3_dup41735 | 3.42297302312458 | -3.43167028138412 | 0.973025982782136 | -3.52680230755204 | 0.000420610690446707 | 0.0191619052937071 | 3412 | 0 | 0 | 14 | 14 | 2 | 1 | 6 | 5 | 0 | 0 | 3 | 0 |
| BLACKJACK_dup1230 | 4.99122417905749 | -3.66483205875247 | 0.934183651519919 | -3.92303167882437 | 8.74416485739794e-05 | 0.00618134002126681 | 3413 | 0 | 0 | 17 | 24 | 1 | 3 | 6 | 10 | 0 | 0 | 4 | 0 |
| L2_dup29474 | 4.62748038874623 | -4.20884784247783 | 0.986582033573027 | -4.26609009616259 | 1.98928472069593e-05 | 0.00203588104150845 | 3414 | 0 | 0 | 21 | 18 | 0 | 2 | 10 | 7 | 0 | 0 | 3 | 0 |
| LOR1b_dup104 | 4.210632193427 | -3.30355082892779 | 1.00894816469451 | -3.27425228027254 | 0.00105941981881759 | 0.0365654801367172 | 3415 | 2 | 0 | 22 | 23 | 1 | 1 | 6 | 1 | 0 | 0 | 2 | 0 |
| AluSx4_dup1801 | 3.16644044750238 | -4.16876231371286 | 1.19322413608561 | -3.49369593493854 | 0.000476383263867308 | 0.0208502760922085 | 3416 | 0 | 0 | 22 | 9 | 0 | 1 | 6 | 2 | 0 | 0 | 2 | 0 |
| MER20_dup14320 | 4.61671982830422 | -4.01760505072056 | 1.1494650022532 | -3.49519562826635 | 0.000473714407761685 | 0.0207799022018599 | 3417 | 0 | 0 | 27 | 27 | 0 | 1 | 4 | 2 | 0 | 1 | 1 | 0 |
| MER4D_dup1174 | 3.72322311757255 | -3.4387472616712 | 1.04951956520917 | -3.27649657582691 | 0.00105103587158523 | 0.0363449762389508 | 3418 | 0 | 0 | 19 | 18 | 0 | 2 | 5 | 3 | 0 | 1 | 2 | 0 |
| LTR33_dup4934 | 3.32491811169335 | -3.58665974435379 | 0.976179449210756 | -3.67418075360387 | 0.000238613776995175 | 0.0126791410855056 | 3419 | 1 | 0 | 15 | 5 | 0 | 1 | 5 | 6 | 1 | 0 | 5 | 3 |
| MLT1E2_dup3424 | 3.20035456288413 | -3.98230028056094 | 1.05118666253334 | -3.78838547186631 | 0.000151629462799782 | 0.00916657117768215 | 3420 | 0 | 0 | 14 | 9 | 0 | 1 | 5 | 5 | 1 | 0 | 5 | 1 |
| L3_dup23084 | 3.98011963355889 | -3.80570289090585 | 0.983843919939064 | -3.86819780432404 | 0.000109642708435005 | 0.00728895891770291 | 3421 | 1 | 0 | 21 | 13 | 0 | 1 | 3 | 5 | 1 | 0 | 2 | 5 |
| LTR10C_dup357 | 3.85692509106558 | -3.85762973796264 | 0.941124200856594 | -4.09895923880344 | 4.1501210909985e-05 | 0.00354579789270638 | 3422 | 1 | 0 | 12 | 15 | 0 | 1 | 7 | 4 | 1 | 0 | 3 | 8 |
| AluSx_dup94214 | 4.12116396847746 | -3.11844679010299 | 0.850322881321627 | -3.66736784179687 | 0.000245060137356383 | 0.0129321533473945 | 3423 | 2 | 0 | 12 | 17 | 0 | 1 | 10 | 4 | 2 | 0 | 5 | 2 |
| AluSx1_dup76150 | 3.24605017497058 | -2.83955288410424 | 0.885212867099427 | -3.20776277621065 | 0.0013377179049599 | 0.0429618875170184 | 3424 | 2 | 1 | 6 | 15 | 0 | 1 | 6 | 2 | 1 | 0 | 7 | 3 |
| LTR81B_dup505 | 3.02023552858856 | -3.47255398646581 | 1.01532564082153 | -3.420138177202 | 0.000625893352964582 | 0.0252807366770515 | 3425 | 1 | 0 | 10 | 15 | 0 | 1 | 3 | 2 | 1 | 0 | 3 | 5 |
| MLT1A_dup3943 | 3.40889935786995 | -2.92537121059936 | 0.927856263687014 | -3.15282800266374 | 0.00161697049171735 | 0.0488759731481423 | 3426 | 2 | 0 | 13 | 12 | 0 | 2 | 5 | 1 | 1 | 0 | 6 | 4 |
| AluSz_dup23441 | 3.79348099770284 | -3.39580795028881 | 0.952975577537109 | -3.56337353268277 | 0.000366119121925544 | 0.0172987650090738 | 3427 | 2 | 0 | 12 | 16 | 0 | 0 | 9 | 1 | 1 | 1 | 6 | 4 |
| Tigger1_dup1543 | 2.89498046444746 | -2.98126991778581 | 0.923754047734508 | -3.22734165560338 | 0.00124946173806492 | 0.040996380249078 | 3428 | 2 | 0 | 7 | 9 | 0 | 0 | 8 | 2 | 1 | 1 | 5 | 4 |
| L2a_dup118748 | 2.84653700959539 | -3.4022852504317 | 1.03208262699424 | -3.29652409743615 | 0.000978892166659561 | 0.0346180581465968 | 3429 | 2 | 0 | 7 | 15 | 0 | 0 | 4 | 2 | 1 | 0 | 3 | 5 |
| MIRc_dup12842 | 3.07320327517294 | -3.36402492618606 | 1.06359039454509 | -3.16289517415668 | 0.00156208567814439 | 0.0477290937039548 | 3430 | 2 | 0 | 15 | 14 | 0 | 0 | 5 | 1 | 1 | 0 | 1 | 3 |
| L1MEi_dup522 | 3.91156388963833 | -3.77146251635506 | 0.890029613446652 | -4.23745733779579 | 2.26065357345336e-05 | 0.00224037265801305 | 3431 | 1 | 0 | 9 | 17 | 1 | 0 | 13 | 5 | 1 | 0 | 2 | 4 |
| MIRb_dup104845 | 4.60894850722172 | -3.23978956186261 | 0.846064004186853 | -3.82924878712499 | 0.000128535016004068 | 0.00814931865656914 | 3432 | 2 | 0 | 9 | 30 | 2 | 0 | 9 | 6 | 1 | 0 | 1 | 3 |
| AluSg_dup3892 | 3.92027122989617 | -3.32712515384804 | 0.828959958523763 | -4.01361382975974 | 5.97961456646834e-05 | 0.00466194419276827 | 3433 | 1 | 1 | 7 | 16 | 1 | 0 | 14 | 7 | 1 | 0 | 3 | 1 |
| BLACKJACK_dup1231 | 4.98687021074139 | -3.72817257310352 | 0.869917919365053 | -4.28566016415053 | 1.82197326435642e-05 | 0.0019089883327193 | 3434 | 1 | 0 | 16 | 21 | 1 | 0 | 11 | 6 | 2 | 0 | 6 | 2 |
| MLT1A-int_dup447 | 4.34071163075483 | -3.90078749718837 | 0.886406468279663 | -4.40067580368521 | 1.07914254628553e-05 | 0.00129359018800231 | 3435 | 1 | 0 | 10 | 15 | 1 | 0 | 15 | 8 | 1 | 0 | 5 | 1 |
| L2c_dup82760 | 3.77067992225593 | -2.77671103525895 | 0.809710799487474 | -3.42926269109482 | 0.000605223438452165 | 0.0247268959871324 | 3436 | 1 | 0 | 12 | 12 | 2 | 2 | 7 | 5 | 1 | 0 | 1 | 7 |
| MER58A_dup13242 | 3.05109606217926 | -2.9924162486451 | 0.879300346838149 | -3.40317874251437 | 0.000666066910285018 | 0.0263882712345453 | 3437 | 0 | 0 | 11 | 9 | 2 | 1 | 7 | 4 | 1 | 0 | 2 | 3 |
| L1PA15_dup5237 | 4.16395527172007 | -3.15294670762485 | 0.876510160697891 | -3.59715933596757 | 0.000321711427552569 | 0.0157154480271861 | 3438 | 0 | 0 | 10 | 25 | 2 | 2 | 8 | 3 | 1 | 0 | 4 | 2 |
| AluSq_dup17907 | 3.45283542040621 | -3.59182911467695 | 0.918279257146321 | -3.91147800271461 | 9.17330125729802e-05 | 0.00640478386434576 | 3439 | 0 | 0 | 5 | 16 | 1 | 1 | 10 | 6 | 1 | 0 | 3 | 3 |
| MLT1B_dup17808 | 3.51668612589263 | -3.13668887552455 | 0.836214482734723 | -3.75105782103468 | 0.000176090081887027 | 0.0101847181878198 | 3440 | 1 | 0 | 9 | 16 | 1 | 1 | 10 | 5 | 1 | 0 | 2 | 1 |
| Ricksha_0_dup361 | 3.92436091861742 | -4.39374343327955 | 1.04462837858426 | -4.20603491476483 | 2.59890100084971e-05 | 0.00247854948917109 | 3441 | 0 | 0 | 9 | 19 | 1 | 0 | 12 | 1 | 1 | 0 | 4 | 8 |
| FLAM_A_dup6860 | 4.70674872854296 | -4.07051327048982 | 0.965117773370828 | -4.21763372595751 | 2.46879410580763e-05 | 0.00239088189784494 | 3442 | 0 | 0 | 18 | 21 | 1 | 0 | 8 | 4 | 2 | 0 | 4 | 5 |
| L2_dup34408 | 3.81805662753382 | -3.19740768303079 | 0.979006778849634 | -3.26597093310002 | 0.00109089446450722 | 0.0372843973357705 | 3443 | 0 | 0 | 14 | 19 | 2 | 0 | 7 | 3 | 2 | 0 | 3 | 1 |
| THE1B_dup22282 | 4.15979122028124 | -3.86252576901015 | 0.949200569045755 | -4.06924089067203 | 4.71665616595838e-05 | 0.003906402603777 | 3444 | 0 | 0 | 8 | 25 | 2 | 0 | 10 | 4 | 1 | 0 | 4 | 3 |
| L1PB4_dup6688 | 4.37847418252387 | -3.10156687212458 | 0.801568016694321 | -3.86937453532077 | 0.000109114913910674 | 0.00726047463797237 | 3445 | 3 | 0 | 13 | 11 | 0 | 0 | 13 | 11 | 1 | 1 | 1 | 2 |
| MSTA_dup17269 | 3.70705444344499 | -3.1887441689365 | 0.862698650741313 | -3.69624337095743 | 0.000218813226352285 | 0.0118735990781347 | 3446 | 2 | 0 | 11 | 16 | 0 | 0 | 9 | 6 | 1 | 1 | 2 | 1 |
| L1MA3_dup1350 | 3.96411730612981 | -4.07687187407765 | 0.940808751941323 | -4.33336941824275 | 1.46844394929828e-05 | 0.00162440894448968 | 3447 | 1 | 0 | 8 | 17 | 0 | 0 | 12 | 10 | 0 | 1 | 1 | 2 |
| L2b_dup11885 | 3.18208791620822 | -3.04865257660427 | 0.907220789458927 | -3.36043068239486 | 0.000778210560801342 | 0.0294860016083497 | 3448 | 3 | 0 | 7 | 10 | 0 | 0 | 8 | 8 | 0 | 1 | 3 | 1 |
| MER58A_dup4406 | 4.5249273129169 | -3.99686079426682 | 0.934513025828689 | -4.27694497968347 | 1.89475608482202e-05 | 0.00196358986320675 | 3449 | 1 | 0 | 15 | 19 | 0 | 0 | 14 | 4 | 0 | 2 | 2 | 5 |
| MSTA-int_dup1630 | 5.40192890557995 | -3.87890825639614 | 0.838636581157527 | -4.62525525781655 | 3.74137351591513e-06 | 0.000569267112838839 | 3450 | 1 | 1 | 16 | 23 | 0 | 0 | 17 | 5 | 0 | 2 | 3 | 6 |
| AluJb_dup60297 | 4.56690945961059 | -3.5326060421835 | 0.884987731485413 | -3.99170058126589 | 6.56011682213126e-05 | 0.00500145324370024 | 3451 | 1 | 0 | 15 | 25 | 1 | 0 | 8 | 5 | 0 | 2 | 2 | 3 |
| L1MEi_dup520 | 3.84349499017369 | -3.67235385189273 | 0.955708763199564 | -3.84254491880797 | 0.00012176511982093 | 0.00783068194619217 | 3452 | 2 | 0 | 13 | 22 | 0 | 0 | 8 | 3 | 0 | 1 | 2 | 2 |
| MER63D_dup1435 | 3.89163971687965 | -2.9576536802932 | 0.903999070338975 | -3.2717441614007 | 0.00106886250803209 | 0.0367850098837438 | 3453 | 3 | 0 | 13 | 18 | 0 | 0 | 9 | 4 | 0 | 2 | 1 | 3 |
| AluSx_dup93130 | 2.48013059856933 | -3.12421192165636 | 0.979773006057846 | -3.18870993826085 | 0.0014290919689158 | 0.0449263124928002 | 3454 | 1 | 1 | 8 | 10 | 0 | 0 | 4 | 3 | 0 | 1 | 3 | 2 |
| MIRb_dup190717 | 3.04607701739826 | -3.03277518624413 | 0.893394282316386 | -3.39466599045247 | 0.000687123729155045 | 0.0269754731657361 | 3455 | 2 | 1 | 9 | 9 | 0 | 0 | 4 | 7 | 0 | 1 | 3 | 3 |
| AluSx_dup54337 | 2.99194327410787 | -3.87964821149113 | 1.04928756214971 | -3.69741179771803 | 0.000217808861104631 | 0.0118325906996353 | 3456 | 1 | 0 | 9 | 15 | 0 | 0 | 5 | 4 | 0 | 1 | 4 | 1 |
| L1PA2_dup1227 | 2.7836214535514 | -3.81429672835275 | 1.01429056530706 | -3.76055625361953 | 0.000169535943028996 | 0.00994831060418679 | 3457 | 1 | 0 | 10 | 9 | 0 | 0 | 6 | 4 | 0 | 1 | 2 | 4 |
| THE1B-int_dup1877 | 2.71672739381458 | -3.22505370169355 | 1.01033495905769 | -3.19206385246876 | 0.00141260101403947 | 0.0445727040030017 | 3458 | 1 | 0 | 9 | 12 | 0 | 0 | 4 | 4 | 0 | 2 | 2 | 2 |
| L1MC5_dup15127 | 4.03441914701004 | -4.2687240897026 | 1.02539647662947 | -4.16299859322133 | 3.14095303558664e-05 | 0.00285367396336655 | 3459 | 1 | 0 | 13 | 18 | 0 | 0 | 3 | 10 | 0 | 1 | 2 | 4 |
| L3_dup44317 | 3.52032668601988 | -3.57688565969094 | 1.0200161790146 | -3.50669502433426 | 0.000453708824922388 | 0.0202244661024696 | 3460 | 2 | 0 | 18 | 5 | 0 | 0 | 4 | 9 | 0 | 1 | 1 | 4 |
| L1PA2_dup1632 | 3.62582283416094 | -2.97980363901861 | 0.837851486978804 | -3.55648188888872 | 0.000375854378622619 | 0.0176535819138843 | 3461 | 3 | 2 | 10 | 13 | 0 | 0 | 7 | 8 | 0 | 0 | 3 | 1 |
| L1ME3Cz_dup1169 | 4.23091826595091 | -3.82345512246325 | 0.879104974406095 | -4.34925888691085 | 1.36598378254677e-05 | 0.00153969861260954 | 3462 | 1 | 2 | 11 | 16 | 0 | 0 | 7 | 12 | 0 | 0 | 2 | 3 |
| LTR43_dup314 | 4.52650504719091 | -3.9219354596709 | 0.887687588143585 | -4.41814835765904 | 9.95500816458508e-06 | 0.00121912216815767 | 3463 | 1 | 2 | 14 | 23 | 0 | 0 | 7 | 8 | 0 | 0 | 3 | 2 |
| THE1D_dup1078 | 4.14033233691133 | -3.99343370848893 | 0.891306279139058 | -4.48042811091415 | 7.4493468018185e-06 | 0.000975532346783437 | 3464 | 1 | 1 | 12 | 16 | 0 | 0 | 12 | 10 | 0 | 0 | 1 | 1 |
| L1M5_dup5181 | 3.75438705378815 | -3.18153655017938 | 0.835012321692802 | -3.81016718858653 | 0.000138872845972876 | 0.00860527036663075 | 3465 | 3 | 1 | 13 | 12 | 0 | 0 | 10 | 8 | 0 | 0 | 1 | 1 |
| MIR_dup19185 | 3.93684995497797 | -3.23702623839346 | 0.82802098317129 | -3.90935290793691 | 9.2543684773359e-05 | 0.00645191902053375 | 3466 | 2 | 2 | 11 | 18 | 0 | 0 | 9 | 8 | 0 | 0 | 1 | 1 |
| THE1B-int_dup3401 | 4.30709052250348 | -4.26714460411496 | 0.940606993131959 | -4.5365860930999 | 5.71721565319511e-06 | 0.000796184644424901 | 3467 | 1 | 1 | 16 | 19 | 0 | 0 | 10 | 6 | 0 | 0 | 1 | 4 |
| L3_dup5398 | 5.31479805768936 | -3.82589449752351 | 0.807394958795319 | -4.738566244248 | 2.15235668607967e-06 | 0.000372198156995576 | 3468 | 3 | 1 | 20 | 22 | 0 | 0 | 12 | 8 | 0 | 0 | 2 | 3 |
| THE1D_dup3189 | 4.86754740825031 | -3.59100202437942 | 0.819781528558502 | -4.38043783530206 | 1.18441079745173e-05 | 0.00137991314652921 | 3469 | 2 | 2 | 22 | 17 | 0 | 0 | 10 | 8 | 0 | 0 | 1 | 2 |
| L1ME4a_dup3401 | 4.22496215905666 | -4.91927261931405 | 1.00280097809815 | -4.90553233069601 | 9.31742907971854e-07 | 0.000194990763395137 | 3470 | 1 | 0 | 14 | 21 | 0 | 0 | 9 | 4 | 0 | 0 | 3 | 6 |
| L1MC4_dup2390 | 4.56533097097605 | -4.34476182576883 | 0.94028070021682 | -4.62070722579648 | 3.82434140033878e-06 | 0.00057865326537906 | 3471 | 2 | 0 | 20 | 16 | 0 | 0 | 9 | 8 | 0 | 0 | 2 | 3 |

| | | | | | | | | | | | | | | | | | | | |
|---|---|---|---|---|---|---|---|---|---|---|---|---|---|---|---|---|---|---|---|
| L1PA13_dup642 | 3.84828588416391 | -4.56756629737663 | 1.03989499762522 | -4.39233413739603 | 1.12140175412212e-05 | 0.00133017478457797 | 3472 | 1 | 0 | 15 | 19 | 0 | 0 | 5 | 7 | 0 | 0 | 1 | 3 |
| MLT1K_dup1451 | 3.36878686559806 | -3.56896512315055 | 0.946452691265933 | -3.77088591546701 | 0.000162669031612399 | 0.00965488785134543 | 3473 | 3 | 0 | 6 | 16 | 0 | 0 | 8 | 7 | 0 | 0 | 2 | 3 |
| HAL1_dup2406 | 2.9987484819799 | -3.60278916843681 | 0.976294515959872 | -3.6902687760103 | 0.000224017232897109 | 0.0120679204208593 | 3474 | 2 | 0 | 7 | 12 | 0 | 0 | 8 | 8 | 0 | 0 | 1 | 1 |
| L2c_dup39586 | 3.22032299112148 | -3.30762066541537 | 0.922119262790363 | -3.58697708516184 | 0.000334533709288563 | 0.016160554047936 4 | 3475 | 3 | 0 | 9 | 12 | 0 | 0 | 11 | 6 | 0 | 0 | 1 | 1 |
| LTR9B_dup360 | 4.53195642831633 | -4.82705363937528 | 0.975638588874969 | -4.94758376146383 | 7.51403731471218e-07 | 0.000165071391477692 | 3476 | 1 | 0 | 12 | 23 | 0 | 0 | 12 | 8 | 0 | 0 | 2 | 3 |
| MIR_dup19184 | 4.43958634066226 | -4.857595904219 | 0.964485395560339 | -5.03646392841114 | 4.74210423631845e-07 | 0.000114671212867381 | 3477 | 1 | 0 | 11 | 20 | 0 | 0 | 12 | 9 | 0 | 0 | 3 | 3 |
| MLT1I-int_dup61 | 3.80045332096798 | -4.13307066265787 | 0.943468046004483 | -4.38072140350817 | 1.18287031958175e-05 | 0.0013797022601137 2 | 3478 | 2 | 0 | 12 | 14 | 0 | 0 | 12 | 6 | 0 | 0 | 2 | 3 |
| L1ME3A_dup2601 | 3.77514965662592 | -4.61042935141873 | 0.976748360113828 | -4.72018130737526 | 2.35634518837626e-06 | 0.000399631441335265 | 3479 | 1 | 0 | 12 | 11 | 0 | 0 | 11 | 9 | 0 | 0 | 2 | 3 |
| L2_dup29781 | 3.36943601723406 | -4.41951825319802 | 0.991740944367305 | -4.45632327504388 | 8.3377272999499e-06 | 0.00106277752445032 | 3480 | 1 | 0 | 9 | 10 | 0 | 0 | 12 | 8 | 0 | 0 | 2 | 2 |
| AluY_dup71772 | 3.46392654410427 | -3.36602690283629 | 0.907294590777742 | -3.70996028969036 | 0.000207291758996291 | 0.0114176492791971 | 3481 | 2 | 1 | 9 | 14 | 1 | 0 | 8 | 2 | 0 | 0 | 3 | 8 |
| L1MCa_dup6563 | 3.13831192676333 | -4.07501780337049 | 1.01118563310368 | -4.02994036897343 | 5.57910042322239e-05 | 0.00442818219589901 | 3482 | 1 | 0 | 9 | 12 | 1 | 0 | 7 | 2 | 0 | 0 | 5 | 6 |
| L1MD_dup8121 | 3.14258746347184 | -2.86910927268782 | 0.899906388138241 | -3.18823080989961 | 0.00143146224303576 | 0.0449842969018768 | 3483 | 3 | 0 | 7 | 9 | 2 | 0 | 6 | 2 | 0 | 0 | 5 | 9 |
| MER5A_dup7689 | 4.01982263447349 | -2.48362287579276 | 0.765172344547326 | -3.24583460640107 | 0.00117106853730782 | 0.039169394739703 | 3484 | 5 | 1 | 8 | 11 | 1 | 0 | 11 | 2 | 0 | 1 | 8 | 7 |
| OldhAT1_dup3769 | 3.92624920425021 | -2.86178823399319 | 0.841893592021563 | -3.39922795601928 | 0.000675763685321098 | 0.0266703045061228 | 3485 | 3 | 0 | 12 | 17 | 2 | 0 | 10 | 2 | 0 | 1 | 4 | 3 |
| MIRb_dup90027 | 2.80838648284405 | -3.10789554748402 | 0.954995278821365 | -3.25435697579543 | 0.00113649370798097 | 0.0383742441707402 | 3486 | 1 | 1 | 8 | 17 | 1 | 0 | 7 | 2 | 0 | 0 | 1 | 1 |
| MIRc_dup51489 | 2.71475200535271 | -3.06794534689572 | 0.919484192911667 | -3.3365938974771 | 0.000848117742069428 | 0.0313512896225665 | 3487 | 1 | 1 | 8 | 13 | 1 | 0 | 8 | 3 | 0 | 0 | 1 | 1 |
| L1ME4a_dup13243 | 2.54996026892213 | -3.21597308121453 | 0.971944828437879 | -3.3088020915583 | 0.000936960430640319 | 0.0335964416258501 | 3488 | 1 | 1 | 7 | 12 | 1 | 0 | 5 | 2 | 0 | 0 | 2 | 4 |
| L2c_dup91498 | 2.62741370193777 | -2.85574347161508 | 0.900685726858566 | -3.17063253747276 | 0.00152107425010566 | 0.0469431335566253 | 3489 | 2 | 1 | 9 | 9 | 1 | 0 | 5 | 3 | 0 | 0 | 3 | 2 |
| HERV16-int_dup153 | 4.10638500445177 | -3.36922492281587 | 0.925538964040701 | -3.64028426000195 | 0.000272337213718703 | 0.013950818225083 | 3490 | 2 | 1 | 18 | 20 | 1 | 0 | 11 | 1 | 0 | 0 | 2 | 1 |
| L1ME4b_dup1932 | 4.38578907396226 | -3.32096384844342 | 0.83146202495827 | -3.99412570719642 | 6.49334083521019e-05 | 0.00496007858226 3 | 3491 | 2 | 2 | 17 | 20 | 1 | 0 | 9 | 3 | 0 | 0 | 2 | 4 |
| L1PA13_dup8816 | 4.19301147859393 | -3.38367968072314 | 0.839979662081072 | -4.0282876282266 | 5.6184580985192e-05 | 0.00444579061744705 | 3492 | 1 | 2 | 20 | 14 | 1 | 0 | 8 | 6 | 0 | 0 | 2 | 1 |
| L1ME3_dup3692 | 4.4847330967568 | -3.10672756812029 | 0.765031455141813 | -4.06091481237774 | 4.8880805976361e-05 | 0.0040024061455097 | 3493 | 3 | 1 | 16 | 12 | 2 | 0 | 15 | 5 | 0 | 0 | 4 | 2 |
| MIRc_dup12844 | 4.26231352070172 | -3.47428808585266 | 0.808701585455255 | -4.29613116672304 | 1.73804766990395e-05 | 0.00184296564555689 | 3494 | 1 | 1 | 13 | 14 | 2 | 0 | 16 | 6 | 0 | 0 | 3 | 1 |
| MLT1A-int_dup448 | 4.76143128579865 | -3.69692333788734 | 0.783156182341283 | -4.7205441535751 | 2.35214541341989e-06 | 0.000399203703667338 | 3495 | 1 | 1 | 15 | 15 | 2 | 0 | 15 | 8 | 0 | 0 | 3 | 3 |
| L1HS_dup33 | 3.22744058286128 | -3.5792820789299 | 0.944281720073388 | -3.79048116980568 | 0.000150355731571855 | 0.00910309530964414 | 3496 | 1 | 1 | 11 | 4 | 1 | 0 | 13 | 4 | 0 | 0 | 3 | 5 |
| AluSc_dup33630 | 4.16164230918812 | -3.02806978920581 | 0.801164575950064 | -3.77958521894825 | 0.000157089823322624 | 0.00942008445068676 | 3497 | 1 | 1 | 15 | 11 | 4 | 0 | 8 | 5 | 0 | 0 | 4 | 6 |
| L2b_dup94908 | 4.61311507913205 | -3.16349667006237 | 0.753901065001201 | -4.19616952001166 | 2.71467093109898e-05 | 0.00255578285850938 | 3498 | 1 | 2 | 15 | 15 | 3 | 0 | 13 | 5 | 0 | 0 | 3 | 5 |
| AluSp_dup8132 | 3.59341143171048 | -4.57111810194023 | 1.34008992486021 | -3.41105325630819 | 0.000647124505271032 | 0.0258757590363231 | 3499 | 0 | 0 | 16 | 22 | 0 | 0 | 8 | 0 | 1 | 0 | 3 | 0 |
| L1PA8_dup7222 | 3.03418086535994 | -3.89758360346958 | 1.18445672355124 | -3.29060870352766 | 0.000999708679469843 | 0.0351057676276758 | 3500 | 0 | 0 | 11 | 16 | 1 | 0 | 8 | 0 | 1 | 0 | 4 | 1 |
| AluSx_dup29571 | 4.53468140091423 | -4.42761365154991 | 1.31920691236485 | -3.35626929335356 | 0.000790016030347446 | 0.029822087382031 | 3501 | 1 | 0 | 28 | 25 | 0 | 0 | 8 | 0 | 0 | 0 | 1 | 0 |
| REP522_dup60 | 4.19334826276669 | -5.05189100530195 | 1.35574352099402 | -3.72628814157853 | 0.000194320255782981 | 0.010892279760315 | 3502 | 0 | 0 | 20 | 18 | 0 | 0 | 19 | 0 | 0 | 0 | 2 | 0 |
| L2b_dup66081 | 2.0527177985118 | -4.31846743857972 | 1.30214259546535 | -3.31643205100468 | 0.000911747557114589 | 0.0329846365085913 | 3503 | 0 | 0 | 9 | 10 | 0 | 0 | 7 | 0 | 0 | 0 | 1 | 2 |
| L1MA5_dup3405 | 2.19935679613318 | -4.47366628702121 | 1.16571064218102 | -3.8377159177779 | 0.000124184027554273 | 0.00794224679894756 | 3504 | 0 | 0 | 6 | 14 | 0 | 0 | 6 | 1 | 0 | 0 | 2 | 2 |
| MSTA1_dup300 | 2.49752249095322 | -4.55153506801015 | 1.19536854553024 | -3.80764165581352 | 0.000140298375164253 | 0.00867102264333332 | 3505 | 0 | 0 | 9 | 17 | 0 | 0 | 5 | 1 | 0 | 0 | 2 | 1 |
| LTR79_dup457 | 1.61030942794241 | -3.94253664067775 | 1.23356616228027 | -3.19604797961539 | 0.00139323941509546 | 0.044189139706152 | 3506 | 0 | 0 | 8 | 7 | 0 | 0 | 4 | 1 | 0 | 0 | 1 | 1 |
| MIR_dup19164 | 2.04820351223851 | -4.38909370879194 | 1.2018808487686 | -3.651854269322 | 0.000260353622666479 | 0.0135262233840607 | 3507 | 0 | 0 | 9 | 10 | 0 | 0 | 4 | 1 | 0 | 0 | 3 | 1 |
| MIR_dup106477 | 2.19001242463232 | -3.84265397985213 | 1.12549717357346 | -3.41418358933027 | 0.00063973452238726 | 0.0256674329159602 | 3508 | 0 | 1 | 7 | 4 | 0 | 0 | 13 | 2 | 0 | 0 | 1 | 2 |
| MIRc_dup73302 | 2.30687815740058 | -3.98332680741198 | 1.08660823960265 | -3.66583526816306 | 0.000246532590917374 | 0.0129689832345436 | 3509 | 0 | 1 | 6 | 11 | 0 | 0 | 8 | 2 | 0 | 0 | 1 | 3 |
| L1MEg_dup5227 | 1.84215096998187 | -3.63480408685516 | 1.11287183191104 | -3.26614798095251 | 0.00109021261307035 | 0.0372664459746642 | 3510 | 0 | 1 | 6 | 5 | 0 | 0 | 8 | 2 | 0 | 0 | 1 | 2 |
| LTR14B_dup346 | 1.86039613162764 | -3.72147691426239 | 1.14084050975275 | -3.26204836035225 | 0.0011061027196236 3 | 0.0376113970273667 | 3511 | 0 | 1 | 5 | 7 | 0 | 0 | 8 | 1 | 0 | 0 | 1 | 3 |
| L2c_dup136304 | 2.067351997808 | -4.39654096515729 | 1.1189651299484 | -3.92911347055117 | 8.52596034782047e-05 | 0.00606311464710845 | 3512 | 0 | 0 | 4 | 8 | 0 | 0 | 9 | 3 | 0 | 0 | 3 | 1 |
| MER5A_dup9215 | 1.86855237333396 | -4.11937102186455 | 1.16434629162201 | -3.53792600320476 | 0.000403283059124997 | 0.0185860140977255 | 3513 | 0 | 0 | 4 | 9 | 0 | 0 | 9 | 2 | 0 | 0 | 1 | 1 |
| AluJb_dup75929 | 2.11373114111078 | -3.80755062639523 | 1.15107495262464 | -3.307821630306 | 0.000940246750935868 | 0.0336686160117661 | 3514 | 0 | 0 | 4 | 10 | 1 | 0 | 11 | 1 | 0 | 0 | 1 | 2 |
| MIRb_dup27266 | 2.2334399393736 | -4.53775760831294 | 1.18717534868045 | -3.82231454970545 | 0.00013220489850792 | 0.00829797593784371 | 3515 | 0 | 0 | 4 | 13 | 0 | 0 | 9 | 1 | 0 | 0 | 1 | 4 |
| MLT1A_dup8915 | 3.15817874758873 | -4.3420380944267 | 1.14710974869107 | -3.78519849507098 | 0.000153585940105316 | 0.00926131781659897 | 3516 | 0 | 0 | 6 | 16 | 0 | 1 | 16 | 2 | 0 | 0 | 0 | 4 |
| MLT2A2_dup3829 | 2.61271478161737 | -3.41762517997011 | 1.064291267296 | -3.21117468966283 | 0.00132193548077731 | 0.0425872271742536 | 3517 | 0 | 0 | 6 | 12 | 1 | 1 | 14 | 1 | 0 | 0 | 1 | 1 |
| AluSz_dup10744 | 2.79515039954921 | -4.67310033367056 | 1.17373282483318 | -3.98140039606946 | 6.8510429622285e-05 | 0.00516060375523508 | 3518 | 0 | 0 | 7 | 20 | 0 | 0 | 9 | 1 | 0 | 0 | 1 | 2 |
| L2a_dup155187 | 2.92539231794954 | -4.72188581796627 | 1.13891569794545 | -4.14594848985254 | 3.38409904226905e-05 | 0.00302605347881154 | 3519 | 0 | 0 | 12 | 13 | 0 | 0 | 12 | 1 | 0 | 0 | 1 | 2 |
| L2a_dup168155 | 3.02019741348687 | -4.3614422639979 | 1.12546821475004 | -3.87522473477097 | 0.00010652633923468 | 0.00714205686121825 | 3520 | 0 | 0 | 8 | 19 | 0 | 1 | 9 | 1 | 0 | 0 | 1 | 4 |
| MER41A_dup2506 | 2.06620735856935 | -3.85284197092266 | 1.13250548595684 | -3.40205148557619 | 0.00066882036634713 | 0.0264736880896866 | 3521 | 0 | 0 | 6 | 10 | 0 | 1 | 7 | 1 | 0 | 0 | 1 | 3 |
| MLT1J1_dup3074 | 2.20578389456251 | -3.75478184670102 | 1.16111818372463 | -3.23376371099145 | 0.00122170487734072 | 0.0403189720865069 | 3522 | 0 | 0 | 7 | 14 | 0 | 1 | 6 | 1 | 0 | 0 | 1 | 1 |
| L2_dup40447 | 2.59415204927326 | -3.67740888825122 | 1.13550713839336 | -3.23856078391054 | 0.00120134419130214 | 0.0398793310468162 | 3523 | 0 | 0 | 5 | 9 | 1 | 0 | 13 | 2 | 1 | 0 | 0 | 5 |
| L1MA7_dup3116 | 2.64837059331189 | -3.7933430275893 | 1.11422568081975 | -3.40446562387477 | 0.000662936438664444 | 0.0263080510064126 | 3524 | 0 | 0 | 4 | 10 | 1 | 0 | 15 | 5 | 0 | 0 | 0 | 1 |
| L1PA10_dup3461 | 2.991587764821 | -3.57850597664884 | 0.996505539440663 | -3.59105477592975 | 0.000329342403187544 | 0.0159823001464637 | 3525 | 0 | 0 | 7 | 10 | 1 | 1 | 16 | 4 | 0 | 0 | 0 | 2 |
| L2a_dup168117 | 2.8323071072163 | -3.61437405342538 | 1.0472070853489 | -3.45144155725528 | 0.000557600502325909 | 0.0233012031085966 | 3526 | 0 | 0 | 9 | 14 | 2 | 0 | 9 | 2 | 0 | 0 | 2 | 1 |
| MIR_dup170849 | 3.04630041315872 | -3.7737704971681 | 1.03677844582952 | -3.63990060976696 | 0.000272743297227584 | 0.013968613130029 | 3527 | 0 | 0 | 5 | 18 | 2 | 0 | 9 | 4 | 0 | 0 | 1 | 3 |
| THE1D_dup1073 | 2.6517906720584 | -3.96186760592988 | 1.09483500382268 | -3.61868920165761 | 0.000296098983825575 | 0.014827915358559 | 3528 | 0 | 0 | 7 | 17 | 1 | 0 | 5 | 4 | 0 | 0 | 1 | 1 |
| Eulor2A_dup30 | 2.67307976081595 | -4.08027861871851 | 1.07468948526374 | -3.79670469904818 | 0.000146632287707981 | 0.00896008380353451 | 3529 | 0 | 0 | 6 | 16 | 1 | 0 | 8 | 3 | 0 | 0 | 1 | 2 |
| MSTC_dup1372 | 2.97101510384973 | -4.19772117189061 | 1.03269489661544 | -4.06482222934212 | 4.80690861054494e-05 | 0.00395457915796686 | 3530 | 0 | 0 | 8 | 15 | 1 | 0 | 8 | 5 | 0 | 0 | 1 | 2 |
| L1MC5a_dup13890 | 2.24908011336153 | -3.32487211326355 | 1.02188183013939 | -3.25367573353372 | 0.00113922238150237 | 0.0384228786376984 | 3531 | 0 | 1 | 8 | 6 | 1 | 0 | 8 | 3 | 0 | 0 | 1 | 2 |
| L1MB1_dup6016 | 1.83908474032505 | -3.63325765685562 | 1.11235614413148 | -3.26627193639716 | 0.00108973546714409 | 0.0372664459746642 | 3532 | 0 | 0 | 6 | 8 | 1 | 0 | 5 | 2 | 0 | 0 | 1 | 2 |
| L1ME1_dup27569 | 1.80953527903832 | -3.60534118921373 | 1.10430560304536 | -3.26480385435991 | 0.00109539902750458 | 0.0373731974769768 | 3533 | 0 | 0 | 6 | 7 | 1 | 0 | 4 | 3 | 0 | 0 | 1 | 2 |
| MER20B_dup392 | 2.635520025494 | -3.6273060630428 | 1.05996709618292 | -3.42209307827121 | 0.000621410367827413 | 0.025142581719664 | 3534 | 1 | 0 | 7 | 9 | 1 | 0 | 14 | 1 | 0 | 0 | 2 | 2 |
| THE1B_dup22283 | 2.70107640570587 | -3.76286645786885 | 1.04422192355378 | -3.60351221612236 | 0.000313945984829492 | 0.0154311487928606 | 3535 | 1 | 0 | 7 | 7 | 1 | 0 | 13 | 2 | 0 | 0 | 4 | 2 |
| AluJo_dup8212 | 2.59433294514662 | -4.04575424499457 | 1.03948958908929 | -3.89205845586113 | 9.93972872930784e-05 | 0.00678656726606566 | 3536 | 1 | 0 | 8 | 10 | 0 | 0 | 9 | 4 | 0 | 0 | 1 | 2 |
| AluJo_dup77454 | 3.41640357628557 | -4.44487260934183 | 1.00290750465706 | -4.43198658769805 | 9.33688270537323e-06 | 0.00116014302465417 | 3537 | 1 | 0 | 11 | 10 | 0 | 0 | 15 | 5 | 0 | 0 | 2 | 2 |
| Charlie1_dup867 | 2.8632860443684 | -4.16652652128478 | 1.05792250873081 | -3.93840426581279 | 8.20253092993914e-05 | 0.00588908258717401 | 3538 | 1 | 0 | 10 | 8 | 0 | 0 | 14 | 3 | 0 | 0 | 2 | 1 |
| L1M4_dup4933 | 4.06967298663051 | -3.2122061281817 | 0.92686583717339 | -3.46566460791972 | 0.000528922638190173 | 0.0224021050314701 | 3539 | 1 | 1 | 16 | 26 | 1 | 0 | 6 | 2 | 1 | 0 | 1 | 1 |
| MER49_dup157 | 4.14238508670007 | -3.06248002570817 | 0.811453445101782 | -3.77406743935142 | 0.000160607313527602 | 0.00956353048664636 | 3540 | 2 | 1 | 15 | 20 | 1 | 0 | 9 | 4 | 1 | 0 | 1 | 2 |
| AluSz_dup72814 | 4.80850584560254 | -3.16846117073153 | 0.939837058509468 | -3.37128775891913 | 0.000748176635339997 | 0.0287045529946164 | 3541 | 1 | 1 | 22 | 24 | 1 | 0 | 7 | 3 | 2 | 0 | 0 | 4 |
| MER3_dup663 | 3.21372294331475 | -2.9251023486268 | 0.869388698091049 | -3.36455069527539 | 0.000766683980612492 | 0.029165369536502 | 3542 | 1 | 1 | 11 | 14 | 1 | 0 | 8 | 4 | 1 | 0 | 0 | 2 |
| L1MB5_dup9400 | 4.23344249005995 | -3.69192220348202 | 1.0428068570545 | -3.54037008723759 | 0.000399566297433734 | 0.0184576203085163 | 3543 | 1 | 2 | 14 | 33 | 0 | 0 | 6 | 1 | 0 | 0 | 1 | 2 |
| L2a_dup19311 | 4.44067682964669 | -4.85110926644323 | 1.08536062746957 | -4.46958286827964 | 7.83722908321147e-06 | 0.0010162601152228 | 3544 | 0 | 1 | 15 | 30 | 0 | 0 | 7 | 3 | 0 | 0 | 1 | 5 |
| THE1B_dup1937 | 3.54068654154939 | -4.47862456176567 | 1.06858498585354 | -4.19117301951267 | 2.77515921996511e-05 | 0.00260385660786206 | 3545 | 0 | 1 | 12 | 22 | 0 | 0 | 7 | 3 | 0 | 0 | 1 | 3 |
| L2_dup30138 | 3.82580843441988 | -4.37309734279743 | 1.10251895745176 | -3.96645999893273 | 7.29480117016157e-05 | 0.005393936479910 7 | 3546 | 0 | 1 | 17 | 23 | 0 | 0 | 8 | 2 | 0 | 0 | 1 | 1 |
| MER44A_dup461 | 2.89302209371329 | -4.06498525499293 | 1.09539740096333 | -3.71096850459755 | 0.000206467793979766 | 0.0113854428056337 | 3547 | 0 | 1 | 11 | 16 | 0 | 0 | 8 | 2 | 0 | 0 | 1 | 1 |
| L1PA6_dup1071 | 3.45181995647467 | -2.97905603857175 | 0.9001054667503 | -3.30967441996236 | 0.000934045500046965 | 0.0335475306496868 | 3548 | 1 | 2 | 14 | 17 | 1 | 1 | 3 | 2 | 0 | 0 | 2 | 4 |
| MIRc_dup86524 | 3.51195215340868 | -3.21329547036882 | 0.959431361236447 | -3.34916660033684 | 0.000810550412244756 | 0.0303250359106904 | 3549 | 1 | 2 | 15 | 20 | 0 | 1 | 3 | 2 | 0 | 0 | 2 | 2 |
| L1MA2_dup3483 | 3.90622827648515 | -4.26298924752528 | 1.07650337625259 | -3.9600333278704 | 7.49393040779757e-05 | 0.00550865234727773 | 3550 | 2 | 0 | 14 | 24 | 0 | 0 | 4 | 3 | 0 | 0 | 2 | 5 |

| | | | | | | | | | | | | | | | | | | | |
|---|---|---|---|---|---|---|---|---|---|---|---|---|---|---|---|---|---|---|---|
| LTR8_dup2803 | 4.05873734990953 | -4.82868180726844 | 1.13936026569622 | -4.23806407213772 | 2.25455434527516e-05 | 0.00223619086386345 | 3551 | 1 | 0 | 20 | 21 | 0 | 0 | 4 | 2 | 0 | 0 | 1 | 7 |
| MER5A_dup33487 | 4.14591276555359 | -3.82336476105102 | 0.925900004615932 | -4.12934954313665 | 3.63790974290514e-05 | 0.00320371357378448 | 3552 | 2 | 1 | 20 | 16 | 0 | 0 | 6 | 5 | 0 | 0 | 3 | 2 |
| LTR89_dup107 | 3.36343289815298 | -3.82842947076587 | 1.00102672973202 | -3.82450274009243 | 0.000131036288862061 | 0.00825457378141597 | 3553 | 1 | 1 | 13 | 18 | 0 | 0 | 4 | 5 | 0 | 0 | 1 | 2 |
| MamRep1527_dup149 | 4.26671686082872 | -4.30721590087257 | 0.996724508308991 | -4.32137051408522 | 1.55063039218054e-05 | 0.00169250596008161 | 3554 | 1 | 1 | 15 | 25 | 0 | 0 | 5 | 5 | 0 | 0 | 2 | 4 |
| AluY_dup8973 | 4.84567930217383 | -4.24064007837086 | 1.0856340918246 | -3.90614122226368 | 9.37817160383436e-05 | 0.00651002269111798 | 3555 | 1 | 1 | 27 | 29 | 0 | 0 | 2 | 3 | 0 | 0 | 1 | 2 |
| L1PA10_dup532 | 5.21542848455644 | -4.20016676352247 | 1.01259232706044 | -4.14793461423469 | 3.35488177586769e-05 | 0.00300670696437142 | 3556 | 1 | 2 | 25 | 27 | 0 | 0 | 4 | 4 | 0 | 0 | 1 | 7 |
| HERVL18-int_dup137 | 2.62822012614448 | -4.49395393484422 | 1.42139751516526 | -3.16164471015115 | 0.00156880847055856 | 0.0478579102994266 | 3557 | 0 | 0 | 14 | 14 | 0 | 0 | 7 | 0 | 0 | 0 | 0 | 2 |
| MLT1B_dup163 | 3.07265603777984 | -4.77161030152997 | 1.45855988224915 | -3.27145313648143 | 0.00106996319964194 | 0.0368069196909066 | 3558 | 0 | 0 | 18 | 18 | 0 | 0 | 4 | 0 | 0 | 0 | 0 | 3 |
| MIRb_dup104849 | 2.35847864818247 | -3.92818640216286 | 1.19557287180409 | -3.28561018303747 | 0.00101761733691222 | 0.035490037228246 | 3559 | 0 | 0 | 7 | 13 | 0 | 0 | 9 | 1 | 1 | 0 | 0 | 2 |
| L1MA3_dup8344 | 2.64848443074112 | -3.9498880169107 | 1.20322010860678 | -3.28276429944669 | 0.00102794574232985 | 0.035754360123435 | 3560 | 0 | 0 | 12 | 16 | 0 | 0 | 4 | 2 | 1 | 0 | 0 | 1 |
| MIRb_dup198429 | 3.76435475277995 | -4.37630407129447 | 1.2190311696689 | -3.58998537542162 | 0.000330696514667318 | 0.0160358493741654 | 3561 | 0 | 0 | 14 | 27 | 0 | 0 | 9 | 1 | 1 | 0 | 0 | 1 |
| L2a_dup23332 | 2.38849016931926 | -4.1800088682867 | 1.28933279405904 | -3.24199375641981 | 0.00118696622806133 | 0.0395344157477998 | 3562 | 0 | 1 | 11 | 8 | 0 | 0 | 3 | 1 | 0 | 0 | 0 | 9 |
| AluJr_dup38506 | 2.65022688721602 | -4.19486304422031 | 1.16097871335822 | -3.61321271092589 | 0.000302426405756811 | 0.0150420471455289 | 3563 | 0 | 1 | 8 | 14 | 0 | 0 | 7 | 2 | 0 | 0 | 0 | 5 |
| AluSz_dup101957 | 2.41224999009632 | -3.53175955597908 | 1.11938704592525 | -3.15508346182427 | 0.00160452198076587 | 0.0486356162083206 | 3564 | 0 | 1 | 7 | 11 | 0 | 0 | 7 | 2 | 1 | 0 | 0 | 4 |
| L1M4_dup10590 | 2.23541859459012 | -3.85773831524686 | 1.17245897135127 | -3.290297067539 | 0.00100081662164098 | 0.0351242695261776 | 3565 | 0 | 0 | 7 | 13 | 1 | 0 | 5 | 2 | 0 | 0 | 0 | 3 |
| Tigger2_dup293 | 2.38584958483081 | -4.01165237988968 | 1.14550086768514 | -3.50209457981167 | 0.000461615721347734 | 0.0204276761078311 | 3566 | 0 | 0 | 8 | 9 | 1 | 0 | 9 | 2 | 0 | 0 | 0 | 4 |
| AluJb_dup21934 | 1.87747543682407 | -3.97309309390923 | 1.21667850423928 | -3.26552419564065 | 0.00109261670530694 | 0.0373111042710915 | 3567 | 0 | 0 | 6 | 9 | 0 | 0 | 8 | 2 | 0 | 0 | 0 | 1 |
| MSTB_dup1371 | 2.04635425292131 | -4.06936622157465 | 1.20423008350115 | -3.3792265093922 | 0.000726900910429595 | 0.0280812305298219 | 3568 | 0 | 0 | 9 | 8 | 0 | 0 | 8 | 2 | 0 | 0 | 0 | 1 |
| L2c_dup82747 | 1.99645582888011 | -4.1810075769989 | 1.27013620464794 | -3.29177891449666 | 0.000995558422756227 | 0.0350012248101338 | 3569 | 0 | 0 | 8 | 11 | 0 | 0 | 6 | 1 | 0 | 0 | 0 | 2 |
| AluSq2_dup60335 | 2.09280167775396 | -4.39719657034863 | 1.21446198626947 | -3.62069510619737 | 0.000293812590951025 | 0.014749268094184 1 | 3570 | 0 | 0 | 7 | 9 | 0 | 0 | 7 | 2 | 0 | 0 | 0 | 4 |
| MSTB1_dup423 | 1.83569240807239 | -4.08167874128178 | 1.21974148214046 | -3.34634740315551 | 0.000818837353990383 | 0.030572583246916 | 3571 | 0 | 0 | 8 | 7 | 0 | 0 | 6 | 2 | 0 | 0 | 0 | 2 |
| AluSx_dup49335 | 2.59487731915693 | -3.67653472001221 | 1.155383909163 | -3.18208925263259 | 0.00146216751678558 | 0.0456523962667711 | 3572 | 1 | 0 | 12 | 10 | 0 | 0 | 4 | 2 | 1 | 0 | 0 | 5 |
| Tigger2_dup1415 | 2.73548532691708 | -3.53727121595409 | 1.0802496699427 | -3.27449414184197 | 0.00105851333831393 | 0.0365504302421195 | 3573 | 1 | 0 | 10 | 15 | 0 | 0 | 5 | 3 | 1 | 0 | 0 | 2 |
| MER20_dup16088 | 2.5119239409695 | -3.74919988818571 | 1.16061536039754 | -3.23035522026998 | 0.00123636504149708 | 0.0406730424670351 | 3574 | 1 | 0 | 9 | 16 | 0 | 0 | 6 | 2 | 0 | 0 | 0 | 1 |
| MER91A_dup3333 | 2.21374141675307 | -3.85040110429865 | 1.20688472812591 | -3.19036359858301 | 0.00142093898666234 | 0.0447171998572089 | 3575 | 1 | 0 | 9 | 11 | 0 | 0 | 6 | 1 | 0 | 0 | 0 | 3 |
| MIR_dup68315 | 2.6866862034043 | -4.07599744196981 | 1.13454357983162 | -3.59263188689034 | 0.00032735487734821 | 0.0158997313013874 | 3576 | 1 | 0 | 12 | 11 | 0 | 0 | 8 | 2 | 0 | 0 | 0 | 3 |
| L1ME3Cz_dup6118 | 2.50956210049907 | -3.99120597191168 | 1.16254291839774 | -3.43316871037546 | 0.000596570759736872 | 0.0244800517309606 | 3577 | 1 | 0 | 8 | 15 | 0 | 0 | 3 | 4 | 0 | 0 | 0 | 3 |
| MLT2B1_dup421 | 2.84632191971935 | -4.02896076381129 | 1.14100794385456 | -3.53105408731918 | 0.000413907101101916 | 0.0189654417636812 | 3578 | 1 | 0 | 11 | 16 | 0 | 0 | 3 | 5 | 0 | 0 | 0 | 2 |
| AluSq2_dup29096 | 2.61342663256771 | -4.03802992131291 | 1.14425985535052 | -3.52894484800042 | 0.00041722007187059 | 0.019068334365099 | 3579 | 1 | 0 | 14 | 9 | 0 | 0 | 5 | 3 | 0 | 0 | 0 | 3 |
| L2a_dup19310 | 2.49548682678253 | -3.98431031545225 | 1.20559423950715 | -3.30485181903411 | 0.000950266266160683 | 0.0339252898676474 | 3580 | 1 | 0 | 13 | 12 | 0 | 0 | 3 | 2 | 0 | 0 | 0 | 3 |
| L1PB1_dup26 | 5.03057974827997 | -4.33970297011717 | 1.260368886715 | -3.44320065011133 | 0.000574872624325886 | 0.0238183251728795 | 3581 | 0 | 0 | 31 | 20 | 0 | 0 | 3 | 5 | 0 | 3 | 3 | 1 |
| MIR_dup1049 | 5.19989135854018 | -4.71329681260234 | 1.22758272653197 | -3.83949424403993 | 0.000123288019724732 | 0.00790724828609733 | 3582 | 0 | 0 | 38 | 18 | 0 | 0 | 4 | 2 | 0 | 2 | 4 | 1 |
| MIRc_dup89851 | 4.63324720758007 | -3.81439606467414 | 0.972326008883649 | -3.92296002557161 | 8.74676682820216e-05 | 0.00618134297506205 | 3583 | 0 | 0 | 21 | 25 | 1 | 0 | 8 | 3 | 1 | 1 | 1 | 2 |
| HAL1_dup21905 | 5.11934361147549 | -4.00026820609251 | 1.19756566509853 | -3.34033307957554 | 0.0008367796560607 | 0.0310476176580414 | 3584 | 0 | 0 | 26 | 26 | 0 | 0 | 4 | 4 | 2 | 3 | 1 | 2 |
| MER102c_dup3023 | 4.67807905840639 | -3.85438738397859 | 1.17609136578883 | -3.27728567362907 | 0.00104810267900216 | 0.0362593544719176 | 3585 | 0 | 0 | 23 | 21 | 0 | 0 | 10 | 3 | 1 | 3 | 1 | 1 |
| HAL1ME_dup6390 | 3.26795192309649 | -3.5569074568167 | 1.03565080478848 | -3.43446598059003 | 0.00059372259132914 | 0.024401638956145 4 | 3586 | 0 | 1 | 16 | 10 | 0 | 0 | 4 | 3 | 0 | 2 | 4 | 3 |
| L1MC2_dup2835 | 3.60198828607533 | -3.94896716503911 | 1.07565363478052 | -3.67122560399738 | 0.000241390144523113 | 0.0127980880857167 | 3587 | 0 | 1 | 19 | 18 | 0 | 0 | 5 | 2 | 0 | 1 | 1 | 2 |
| MIR_dup167827 | 2.98778963009942 | -3.87134665010772 | 1.15610650767357 | -3.34860726447948 | 0.000812188348467122 | 0.0303624459959714 | 3588 | 0 | 1 | 14 | 17 | 0 | 0 | 2 | 1 | 0 | 1 | 2 | 3 |
| SVA_A_dup747 | 4.35526628686357 | -3.86615893950248 | 1.1282481545652 | -3.42669201262059 | 0.000610981623827217 | 0.0248896821542871 | 3589 | 0 | 1 | 23 | 24 | 0 | 0 | 2 | 2 | 0 | 2 | 3 | 2 |
| MER44C_dup363 | 4.10487391727673 | -3.93412146962599 | 1.05899985690336 | -3.71494051106846 | 0.000203251505276704 | 0.0112524156990625 | 3590 | 0 | 0 | 19 | 16 | 1 | 0 | 5 | 2 | 0 | 2 | 7 | 3 |
| HERVL-int_dup2332 | 3.72672349025861 | -4.07915598512092 | 1.0979254996935 | -3.71533039924811 | 0.000202938347149557 | 0.0112429261818041 | 3591 | 0 | 0 | 19 | 17 | 1 | 0 | 8 | 1 | 0 | 1 | 3 | 1 |
| MER34A_dup1308 | 3.03143466380756 | -3.88276363426619 | 1.0414471900247 | -3.72823861973655 | 0.000192822759801743 | 0.0108287477941434 | 3592 | 0 | 0 | 14 | 10 | 1 | 0 | 8 | 2 | 0 | 1 | 2 | 3 |
| L1MA4A_dup554 | 3.56637021249711 | -3.69117915542663 | 1.05482621585886 | -3.4993244384064 | 0.00046643865487771 | 0.0205627678101205 | 3593 | 1 | 1 | 9 | 23 | 0 | 0 | 2 | 4 | 0 | 1 | 1 | 7 |
| L1P3_dup243 | 2.96057892955595 | -3.24794730460873 | 1.00452311987337 | -3.23332259890462 | 0.00122359304537479 | 0.0403644906106683 | 3594 | 1 | 0 | 7 | 18 | 0 | 1 | 4 | 5 | 0 | 1 | 0 | 3 |
| LTR72_dup62 | 3.48423832997992 | -3.0898645137747 | 0.924475371093945 | -3.34228970331402 | 0.000830902899256123 | 0.0308873646721446 | 3595 | 1 | 1 | 8 | 23 | 1 | 0 | 6 | 4 | 0 | 1 | 0 | 3 |
| L1ME1_dup28677 | 2.93297097145791 | -4.27112441625182 | 1.22860211519301 | -3.47640978591416 | 0.000508175163043803 | 0.0218079496676943 | 3596 | 0 | 0 | 14 | 13 | 0 | 0 | 1 | 6 | 0 | 1 | 0 | 3 |
| L2a_dup102460 | 2.92919499678413 | -3.45830728924656 | 1.08194559283158 | -3.19637818404135 | 0.00139164575774964 | 0.0441561342197164 | 3597 | 0 | 1 | 15 | 9 | 0 | 0 | 3 | 7 | 0 | 1 | 0 | 1 |
| MIRc_dup46443 | 2.65059055804299 | -4.21568344742152 | 1.10624656218716 | -3.81079913964812 | 0.000138518283295186 | 0.00859225011390934 | 3598 | 0 | 0 | 7 | 17 | 0 | 0 | 6 | 2 | 0 | 1 | 1 | 3 |
| AluSq_dup15200 | 2.16967343761996 | -3.9196223034719 | 1.18346267332159 | -3.31199487050216 | 0.000926332443855305 | 0.0333611472204623 | 3599 | 0 | 0 | 9 | 13 | 0 | 0 | 3 | 1 | 0 | 1 | 1 | 2 |
| LTR12__dup314 | 2.0111890490201 | -3.86825194890421 | 1.11893483922881 | -3.45708419586817 | 0.000546054211854329 | 0.022931771177883 9 | 3600 | 0 | 0 | 8 | 9 | 0 | 0 | 3 | 2 | 0 | 1 | 2 | 2 |
| MER58A_dup10011 | 1.99963500191247 | -3.85792912851638 | 1.11599859758674 | -3.45693008652416 | 0.000546366578338273 | 0.0229367928456737 | 3601 | 0 | 0 | 8 | 8 | 0 | 0 | 4 | 2 | 0 | 1 | 1 | 3 |
| Charlie7_dup2021 | 2.50179484422593 | -3.62320696476981 | 1.1370342927398 | -3.18654150354545 | 0.00143984827753609 | 0.0451510461579603 | 3602 | 0 | 0 | 9 | 13 | 0 | 0 | 5 | 2 | 1 | 1 | 0 | 3 |
| AluSx1_dup93580 | 2.23928994875246 | -4.05440441312175 | 1.2699290533002 | -3.19262277100084 | 0.00140986996832661 | 0.0445219534815452 | 3603 | 0 | 0 | 10 | 11 | 0 | 0 | 3 | 1 | 0 | 1 | 0 | 5 |
| L3_dup23082 | 2.12757211625141 | -3.80698799039629 | 1.19490406094858 | -3.18601979423693 | 0.00144244727847502 | 0.0452146700202895 | 3604 | 0 | 0 | 6 | 14 | 0 | 0 | 3 | 3 | 0 | 1 | 0 | 2 |
| THE1D_dup7887 | 2.19252679758694 | -3.90845437396373 | 1.14089649311698 | -3.42577472850816 | 0.000613048604068474 | 0.0249419854621203 | 3605 | 0 | 0 | 8 | 9 | 0 | 0 | 4 | 4 | 0 | 1 | 0 | 3 |
| L1PREC2_dup7702 | 4.26701167726445 | -4.81127072242316 | 1.05615229250098 | -4.55547060455648 | 5.22684250657792e-06 | 0.00074687488215709 | 3606 | 0 | 0 | 13 | 29 | 1 | 0 | 6 | 4 | 0 | 0 | 2 | 4 |
| MIRc_dup86526 | 3.55788390010044 | -4.39768783710853 | 1.08100868591475 | -4.06813367404832 | 4.73911931454696e-05 | 0.00391295798829718 | 3607 | 0 | 0 | 10 | 24 | 1 | 0 | 4 | 6 | 0 | 0 | 1 | 2 |
| L2a_dup147244 | 3.00290064491129 | -3.57270557098397 | 1.01250987644081 | -3.52856367539129 | 0.000417821413369411 | 0.0190797204147751 | 3608 | 0 | 1 | 13 | 15 | 1 | 0 | 4 | 4 | 0 | 0 | 1 | 1 |
| MER61-int_dup639 | 3.22479116876259 | -4.292321184674 | 1.10812091979735 | -3.87351335760266 | 0.000107277523942018 | 0.0071743276312275 | 3609 | 0 | 0 | 15 | 17 | 1 | 0 | 3 | 4 | 0 | 0 | 2 | 1 |
| MIR_dup28628 | 3.64292168478063 | -4.5901005799842 | 1.05641630518601 | -4.34497324345634 | 1.39292640396178e-05 | 0.00156097022104272 | 3610 | 0 | 0 | 12 | 21 | 0 | 0 | 6 | 5 | 0 | 1 | 3 | 1 |
| MIR_dup1040 | 3.35061573065851 | -4.34995027422241 | 1.13614196262283 | -3.82870311750511 | 0.000128820289790968 | 0.00815873437773858 | 3611 | 0 | 0 | 13 | 23 | 0 | 0 | 4 | 3 | 0 | 1 | 1 | 1 |
| MLT1F2_dup538 | 3.43532714465974 | -4.47718443495037 | 1.16192847227348 | -3.8532358417813 | 0.000116567012685968 | 0.007586612253986 7 | 3612 | 0 | 0 | 11 | 27 | 0 | 0 | 4 | 2 | 0 | 1 | 2 | 1 |
| MSTA_dup1658 | 3.98412271333068 | -4.96393897524286 | 1.08008400948077 | -4.59588229403486 | 4.30921921619939e-06 | 0.000639575212342854 | 3613 | 0 | 0 | 11 | 28 | 0 | 0 | 9 | 5 | 0 | 0 | 1 | 1 |
| MLT1N2_dup541 | 3.49426570002115 | -4.92440553384895 | 1.110493654461 | -4.4344292415062 | 9.23165330734863e-06 | 0.00114946885288874 | 3614 | 0 | 0 | 14 | 22 | 0 | 0 | 6 | 3 | 0 | 0 | 2 | 1 |
| MLT1C-int_dup42 | 3.50815977142122 | -4.92044872726094 | 1.08227942823749 | -4.54637554672363 | 5.45776171253513e-06 | 0.000769042202955318 | 3615 | 0 | 0 | 16 | 18 | 0 | 0 | 8 | 3 | 0 | 0 | 1 | 2 |
| MamRep1879_dup167 | 3.15735833727719 | -4.7097936099187 | 1.11532815016408 | -4.22278735565476 | 2.41299457432052e-05 | 0.0023502349855966 9 | 3616 | 0 | 0 | 12 | 18 | 0 | 0 | 10 | 2 | 0 | 0 | 1 | 1 |
| Mam_R4_dup462 | 3.40946891147333 | -4.78863936393068 | 1.10106362906671 | -4.34910321031096 | 1.36695371556086e-05 | 0.001540062352286 2 | 3617 | 0 | 0 | 13 | 21 | 0 | 0 | 5 | 5 | 0 | 0 | 1 | 1 |
| THE1D_dup2139 | 4.07678340602616 | -4.98582988638286 | 1.12754953445082 | -4.42182780804502 | 9.7869440191338e-06 | 0.00120287406073182 | 3618 | 0 | 0 | 19 | 25 | 0 | 0 | 4 | 5 | 0 | 0 | 1 | 1 |
| AluSz_dup10736 | 3.02783936309293 | -4.82224003368467 | 1.08931781569656 | -4.42684399740688 | 9.56218357471983e-06 | 0.00118319466087898 | 3619 | 0 | 0 | 14 | 13 | 0 | 0 | 4 | 5 | 0 | 0 | 1 | 3 |
| LSAU_dup141 | 3.72957212289946 | -5.16821652825585 | 1.10031839703513 | -4.69701910118191 | 2.63985772980765e-06 | 0.000438035974460816 | 3620 | 0 | 0 | 20 | 15 | 0 | 0 | 5 | 4 | 0 | 0 | 1 | 5 |
| L2c_dup456 | 3.43500945281759 | -5.1537547792564 | 1.12918136198012 | -4.56415147538287 | 5.01518597264414e-06 | 0.000723238449361687 | 3621 | 0 | 0 | 10 | 23 | 0 | 0 | 2 | 4 | 0 | 0 | 3 | 5 |
| LTR13_dup186 | 2.95587186966907 | -4.90531320014333 | 1.11476047720999 | -4.40032930878596 | 1.08086721614997e-05 | 0.00129500518939588 | 3622 | 0 | 0 | 13 | 15 | 0 | 0 | 3 | 3 | 0 | 0 | 2 | 4 |
| SATR1_dup142 | 3.22281664126835 | -5.06970931548579 | 1.18327721313113 | -4.284464586342 | 1.83179782026702e-05 | 0.00191803784609089 | 3623 | 0 | 0 | 14 | 17 | 0 | 0 | 2 | 3 | 0 | 0 | 1 | 7 |
| MER39_dup2150 | 4.44823063364722 | -5.37573983840913 | 1.08022336417055 | -4.97650765268986 | 6.47417179140501e-07 | 0.000146157806553857 | 3624 | 0 | 0 | 19 | 24 | 0 | 0 | 3 | 7 | 0 | 0 | 3 | 3 |
| Tigger2_dup568 | 4.70695523972997 | -5.43533207626854 | 1.17514733671544 | -4.62523456119162 | 3.74174713792181e-06 | 0.000569267112838839 | 3625 | 0 | 0 | 16 | 36 | 0 | 0 | 2 | 5 | 0 | 0 | 1 | 5 |
| MIR3_dup41742 | 2.91350397143745 | -4.83095577893104 | 1.191590956681 | -4.05420648070958 | 5.03047692306003e-05 | 0.0040978495786869 | 3626 | 0 | 0 | 14 | 12 | 0 | 0 | 4 | 4 | 0 | 0 | 0 | 5 |
| Charlie1a_dup1189 | 4.17480658823153 | -5.17663183092226 | 1.10385577083997 | -4.68959076690166 | 2.73751983287013e-06 | 0.000448325344332882 | 3627 | 0 | 0 | 19 | 20 | 0 | 0 | 7 | 6 | 0 | 0 | 0 | 4 |
| L1PA7_dup2243 | 3.80137972540919 | -5.18693344490615 | 1.16236434502816 | -4.46239896043998 | 8.10471476056943e-06 | 0.00104033592015077 | 3628 | 0 | 0 | 16 | 20 | 0 | 0 | 6 | 4 | 0 | 0 | 0 | 6 |
| MLT1E2_dup3788 | 3.81826206364962 | -4.47505902928032 | 1.06944032395512 | -4.18448690314032 | 2.85810923662946e-05 | 0.0026635210471979 | 3629 | 0 | 1 | 12 | 24 | 0 | 0 | 4 | 7 | 0 | 0 | 1 | 2 |

| | | | | | | | | | | | | | | | | | | | |
|---|---|---|---|---|---|---|---|---|---|---|---|---|---|---|---|---|---|---|---|
| HERVL-int_dup2153 | 3.05626222415805 | -4.10899952198492 | 1.09860357516233 | -3.74020221204702 | 0.000183872271165519 | 0.0104894920725847 | 3630 | 0 | 1 | 11 | 17 | 0 | 0 | 5 | 5 | 0 | 0 | 0 | 2 |
| L1PA15_dup6000 | 3.44975912548135 | -4.35112664550749 | 1.1104299923485 | -3.91841599694645 | 8.91327813520909e-05 | 0.00627107888574944 | 3631 | 0 | 1 | 13 | 19 | 0 | 0 | 4 | 6 | 0 | 0 | 0 | 3 |
| L1PREC2_dup4353 | 2.86270652790199 | -4.23257192441737 | 1.18208303652466 | -3.58060457145311 | 0.00034280011357075 | 0.0164541464870985 | 3632 | 0 | 0 | 12 | 19 | 0 | 1 | 3 | 1 | 0 | 0 | 1 | 3 |
| MER58A_dup3509 | 3.0164268827619 | -4.21221911897904 | 1.21605637435729 | -3.46383540089191 | 0.000532532255357317 | 0.0225149111644016 | 3633 | 0 | 0 | 14 | 21 | 0 | 1 | 2 | 1 | 0 | 0 | 1 | 2 |
| MLT1A0_dup18237 | 3.2044289228805 | -4.90158993004472 | 1.17215456268075 | -4.18169248843313 | 2.89347140138491e-05 | 0.00268676952305871 | 3634 | 0 | 0 | 18 | 16 | 0 | 0 | 5 | 1 | 0 | 0 | 2 | 2 |
| LTR12B_dup156 | 3.38624677070028 | -4.88428918499539 | 1.23244227824374 | -3.96309772166826 | 7.39834798646631e-05 | 0.00545524422060029 | 3635 | 0 | 0 | 18 | 22 | 0 | 0 | 3 | 1 | 0 | 0 | 1 | 2 |
| MER41G_dup98 | 3.38533433227269 | -4.78698940789228 | 1.2300805631525 | -3.89160641285478 | 9.95826957970483e-05 | 0.00679532668027658 | 3636 | 0 | 0 | 18 | 22 | 0 | 0 | 4 | 1 | 0 | 0 | 1 | 1 |
| MLT2B5_dup327 | 3.64593994074349 | -4.97565256038469 | 1.22337048723173 | -4.06716739721562 | 4.75880593750778e-05 | 0.00392355799586357 | 3637 | 0 | 0 | 19 | 24 | 0 | 0 | 2 | 2 | 0 | 0 | 2 | 1 |
| L1M4b_dup4240 | 3.02921250509305 | -4.760828740058 | 1.30553691008929 | -3.64664430646577 | 0.000265687297167688 | 0.0137105247477473 | 3638 | 0 | 0 | 16 | 21 | 0 | 0 | 1 | 1 | 0 | 0 | 1 | 2 |
| MLT1G-int_dup98 | 2.67329508579939 | -4.52263494724249 | 1.27565247101561 | -3.54535035991567 | 0.000392091543299573 | 0.0182326782024546 | 3639 | 0 | 0 | 14 | 18 | 0 | 0 | 2 | 1 | 0 | 0 | 1 | 1 |
| HAL1_dup19119 | 2.79295499821275 | -4.56033189157288 | 1.35038878932225 | -3.37705105939281 | 0.000732674480942626 | 0.0282597361859763 | 3640 | 0 | 0 | 14 | 20 | 0 | 0 | 2 | 1 | 0 | 0 | 0 | 2 |
| Tigger1_dup11787 | 2.5926850145772 | -4.3337336706488 | 1.36206992687943 | -3.18172627199661 | 0.00146400113225096 | 0.0456976536028582 | 3641 | 0 | 0 | 14 | 18 | 0 | 0 | 2 | 1 | 0 | 0 | 0 | 1 |
| L2a_dup60579 | 2.68927804463371 | -4.3771113478505 | 1.33611157003192 | -3.27600736796698 | 0.00105285814112359 | 0.0364027002684967 | 3642 | 0 | 0 | 11 | 22 | 0 | 0 | 3 | 1 | 0 | 0 | 0 | 1 |
| MER74B_dup90 | 2.95464768684446 | -4.62597149407146 | 1.36261231549812 | -3.39492858053346 | 0.000686465051784314 | 0.0269585101851577 | 3643 | 0 | 0 | 11 | 26 | 0 | 0 | 2 | 1 | 0 | 0 | 0 | 2 |
| L1MA4A_dup5608 | 2.33634320608357 | -4.20949947962207 | 1.3260066894101 | -3.17456881118356 | 0.00150059306271334 | 0.0465164300287113 | 3644 | 0 | 0 | 17 | 9 | 0 | 0 | 2 | 2 | 0 | 0 | 0 | 1 |
| L2a_dup163163 | 2.63523550451681 | -4.4916097599525 | 1.21237227831242 | -3.70481067597871 | 0.000211548680101188 | 0.0115757540091628 | 3645 | 0 | 0 | 14 | 13 | 0 | 0 | 5 | 2 | 0 | 0 | 0 | 2 |
| LTR16A_dup778 | 2.4227487828127 | -4.25731761261644 | 1.22431351986367 | -3.4773099729312 | 0.000506471919756361 | 0.0217768282288303 | 3646 | 0 | 0 | 13 | 12 | 0 | 0 | 5 | 2 | 0 | 0 | 0 | 1 |
| MLT2E_dup368 | 2.34441774379836 | -4.22065451734863 | 1.22336333883393 | -3.45004168701967 | 0.00056049999415145 | 0.0233939188928891 | 3647 | 0 | 0 | 12 | 12 | 0 | 0 | 5 | 2 | 0 | 0 | 0 | 1 |
| L2a_dup165189 | 2.40757349217897 | -4.24796547295632 | 1.33084059437657 | -3.19194161262136 | 0.00141319896472616 | 0.0445856590905238 | 3648 | 0 | 0 | 15 | 13 | 0 | 0 | 3 | 1 | 0 | 0 | 0 | 1 |
| MLT1L_dup9729 | 2.39535675411042 | -4.24277456534022 | 1.32860546269277 | -3.193404426278 | 0.00140605872761505 | 0.0444619624779882 | 3649 | 0 | 0 | 14 | 14 | 0 | 0 | 3 | 1 | 0 | 0 | 0 | 1 |
| MLT1D_dup1695 | 2.89876344047598 | -4.59991052977918 | 1.27392119742136 | -3.6108281572598 | 0.000305220849133245 | 0.015139895803181 | 3650 | 0 | 0 | 17 | 15 | 0 | 0 | 5 | 1 | 0 | 0 | 0 | 2 |
| MLT1F1_dup3043 | 3.03099204236756 | -4.65042137495122 | 1.27273900642013 | -3.65386882266742 | 0.00025831827879641 | 0.0134527908002825 | 3651 | 0 | 0 | 17 | 17 | 0 | 0 | 5 | 1 | 0 | 0 | 0 | 2 |
| MLT1O_dup4712 | 3.01566457571222 | -4.50966861472689 | 1.20965625305064 | -3.72805795312009 | 0.000192961011034325 | 0.01083139908742 | 3652 | 0 | 0 | 7 | 23 | 0 | 0 | 10 | 2 | 0 | 0 | 0 | 1 |
| MIRb_dup169271 | 3.28913278381611 | -4.84155314490936 | 1.17939577484433 | -4.10511318437476 | 4.0411696837375e-05 | 0.00346677176879855 | 3653 | 0 | 0 | 13 | 20 | 0 | 0 | 8 | 2 | 0 | 0 | 0 | 3 |
| Charlie7_dup2024 | 2.83663916386234 | -4.44170552776761 | 1.13934795654909 | -3.89846271478018 | 9.6805302637837e-05 | 0.00665927566969146 | 3654 | 0 | 0 | 11 | 15 | 0 | 0 | 9 | 3 | 0 | 0 | 0 | 1 |
| MSTA_dup10398 | 2.96172249586312 | -4.485759600591 | 1.14476183063235 | -3.91850905625771 | 8.90983823797333e-05 | 0.00627051277542976 | 3655 | 0 | 0 | 13 | 16 | 0 | 0 | 6 | 4 | 0 | 0 | 0 | 1 |
| MIRb_dup198463 | 3.44492519671387 | -4.79663821749667 | 1.28584530796353 | -3.73033846901336 | 0.000191222709058346 | 0.0107490382068502 | 3656 | 0 | 0 | 13 | 28 | 0 | 0 | 5 | 1 | 0 | 0 | 0 | 2 |
| L1MEb_dup1581 | 4.0471568620879 | -4.836634424907 | 1.2306420213207 | -3.93017168365209 | 8.48852291332819e-05 | 0.00604010189334566 | 3657 | 0 | 0 | 16 | 32 | 0 | 0 | 6 | 2 | 0 | 0 | 0 | 1 |
| MLT2B4_dup807 | 3.49177969364329 | -4.6710760683587 | 1.25954408384824 | -3.70854512220591 | 0.000208453517792367 | 0.011457687097765 | 3658 | 0 | 0 | 14 | 28 | 0 | 0 | 4 | 2 | 0 | 0 | 0 | 1 |
| MER52D_dup145 | 3.25987136128677 | -4.58966765580653 | 1.21499807650854 | -3.77751022371619 | 0.000158404004408152 | 0.00947977847910013 | 3659 | 0 | 0 | 19 | 17 | 0 | 0 | 4 | 3 | 0 | 0 | 0 | 1 |
| MER20_dup14323 | 4.10598091590323 | -4.9839722977201 | 1.16624549351279 | -4.27351901931737 | 1.92411863527951e-05 | 0.00198541341192636 | 3660 | 0 | 0 | 20 | 25 | 0 | 0 | 5 | 4 | 0 | 0 | 0 | 2 |
| L1MB5_dup9403 | 4.19971759733756 | -4.8753650939007 | 1.18910652195573 | -4.1000238446949 | 4.13107571992049e-05 | 0.00353460289811194 | 3661 | 0 | 0 | 20 | 28 | 0 | 0 | 6 | 3 | 0 | 0 | 0 | 1 |
| L1ME3A_dup13959 | 3.90879763267611 | -4.79552404968869 | 1.22345220348124 | -3.91966603684509 | 8.86717553874398e-05 | 0.00625158207627252 | 3662 | 0 | 0 | 18 | 28 | 0 | 0 | 4 | 3 | 0 | 0 | 0 | 1 |
| HAL1ME_dup2378 | 2.91960093463511 | -3.89889609203719 | 1.18805661500123 | -3.2817426735452 | 0.0010316770761259 | 0.0358422300417366 | 3663 | 0 | 0 | 15 | 17 | 0 | 1 | 4 | 2 | 0 | 0 | 0 | 1 |
| L1PA7_dup9474 | 3.11189858449887 | -3.9656583591108 | 1.21813169819602 | -3.25552513327065 | 0.0011318287840044 | 0.038271157004222 | 3664 | 0 | 0 | 20 | 15 | 0 | 1 | 3 | 2 | 0 | 0 | 0 | 1 |
| THE1D_dup1076 | 2.33955993826714 | -3.7997245955147 | 1.15381740137101 | -3.29317671149674 | 0.000990621920284437 | 0.0348896571113086 | 3665 | 0 | 0 | 10 | 11 | 0 | 1 | 3 | 4 | 0 | 0 | 0 | 2 |
| Tigger3b_dup599 | 2.60445446800656 | -3.93984358787141 | 1.11099855699367 | -3.54621845642401 | 0.000390802089948988 | 0.0182011732423179 | 3666 | 0 | 0 | 12 | 13 | 0 | 1 | 4 | 3 | 0 | 0 | 1 | 1 |
| AluY_dup10647 | 2.53078135843075 | -4.02023547289857 | 1.16547513736384 | -3.44943906910947 | 0.000561752489477228 | 0.0234297559791672 | 3667 | 0 | 0 | 11 | 15 | 0 | 0 | 2 | 3 | 0 | 1 | 1 | 1 |
| AluSc8_dup2189 | 2.09234165100382 | -4.2384203094387 | 1.19816242390701 | -3.5374338444181 | 0.000404035389700479 | 0.0186062714994523 | 3668 | 0 | 0 | 9 | 12 | 0 | 0 | 2 | 3 | 0 | 0 | 1 | 1 |
| MLT1N2_dup1097 | 2.6832690167369 | -4.5203056252184 | 1.1819480891867 | -3.82445360043588 | 0.000131062424861899 | 0.00825457378141597 | 3669 | 0 | 0 | 11 | 17 | 0 | 0 | 2 | 4 | 0 | 0 | 1 | 1 |
| HAL1ME_dup1650 | 3.19768765402681 | -4.56933331044926 | 1.26795049470599 | -3.60371586235217 | 0.000313699984913119 | 0.0154236755256952 | 3670 | 0 | 0 | 11 | 26 | 0 | 0 | 2 | 4 | 0 | 0 | 0 | 1 |
| LTR2_dup823 | 2.45719896318398 | -4.26745924345885 | 1.31702911849667 | -3.2402163198411 | 0.00119439051351915 | 0.0397094376316651 | 3671 | 0 | 0 | 10 | 17 | 0 | 0 | 1 | 4 | 0 | 0 | 0 | 1 |
| LTR33A_dup1656 | 2.32691427148698 | -4.20507520090962 | 1.2537177094021 | -3.35408455138999 | 0.000796280272421157 | 0.0299654761508026 | 3672 | 0 | 0 | 7 | 16 | 0 | 0 | 2 | 5 | 0 | 0 | 0 | 1 |
| MER5A_dup6878 | 2.17342426841333 | -4.13171711333793 | 1.30407597422527 | -3.16831012533032 | 0.0015332786686386 | 0.047222045830686 | 3673 | 0 | 0 | 6 | 18 | 0 | 0 | 2 | 3 | 0 | 0 | 0 | 1 |
| MIR_dup167824 | 2.12155599027728 | -4.10756739219514 | 1.22074721089321 | -3.36479768746689 | 0.00076599803079802 | 0.0291439346716127 | 3674 | 0 | 0 | 7 | 14 | 0 | 0 | 4 | 3 | 0 | 0 | 0 | 1 |
| THE1B_dup19177 | 2.51505289160058 | -4.43660837931121 | 1.20583623752016 | -3.67927935922315 | 0.00023389394773092 | 0.0124897058543046 | 3675 | 0 | 0 | 9 | 16 | 0 | 0 | 3 | 4 | 0 | 0 | 0 | 2 |
| L1MA2_dup2418 | 4.9364511474278 | -4.63596270654822 | 1.41441703078297 | -3.27764909899447 | 0.00104675432253299 | 0.0362293535155833 | 3676 | 0 | 0 | 26 | 39 | 0 | 0 | 2 | 1 | 0 | 1 | 0 | 0 |
| L1MC2_dup5531 | 4.5111605382627 | -4.38392680743682 | 1.36723945449341 | -3.2064074753176 | 0.00134403519690113 | 0.0430861676902808 | 3677 | 0 | 0 | 22 | 34 | 0 | 0 | 3 | 1 | 0 | 2 | 0 | 1 |
| LTR27E_dup116 | 3.69100590790878 | -4.57240144786481 | 1.30701969767531 | -3.49834165161962 | 0.000468160996817199 | 0.0206176266053425 | 3678 | 0 | 0 | 18 | 26 | 0 | 0 | 1 | 2 | 0 | 1 | 0 | 3 |
| MER49_dup388 | 2.90887005105005 | -4.06929338029096 | 1.21370089273142 | -3.35279755058353 | 0.000799991993781176 | 0.030057618024989 | 3679 | 0 | 0 | 13 | 18 | 0 | 0 | 2 | 4 | 0 | 1 | 0 | 1 |
| MIR_dup154608 | 3.10535441823944 | -4.16202570094771 | 1.26994161159563 | -3.27733626723066 | 0.00104791487376631 | 0.0362581289156897 | 3680 | 0 | 0 | 19 | 17 | 0 | 0 | 2 | 2 | 0 | 1 | 0 | 1 |
| L3_dup41591 | 3.58835248522894 | -3.75352676133883 | 1.01208799417617 | -3.70869606490506 | 0.000208329313109384 | 0.011453587508578 | 3681 | 1 | 0 | 15 | 18 | 0 | 0 | 7 | 5 | 0 | 1 | 1 | 0 |
| MLT1C_dup7746 | 3.29364829676923 | -4.20982130665049 | 1.11038404643982 | -3.79132005736958 | 0.000149848697972258 | 0.00908197123483002 | 3682 | 0 | 0 | 13 | 18 | 0 | 0 | 6 | 5 | 0 | 1 | 1 | 0 |
| AluSc_dup30607 | 3.8275979474365 | -4.25577147468507 | 1.28682630565983 | -3.30718408223937 | 0.000942389417082278 | 0.0337301131523857 | 3683 | 0 | 0 | 21 | 24 | 0 | 0 | 3 | 3 | 0 | 1 | 0 | 0 |
| MER58B_dup6014 | 4.36471312483894 | -4.40886012996402 | 1.27171295440008 | -3.46686735769226 | 0.00052656166224187 | 0.0223355444587407 | 3684 | 0 | 0 | 27 | 24 | 0 | 0 | 4 | 3 | 0 | 1 | 0 | 0 |
| AluJr_dup73361 | 3.19786293491331 | -4.01597836530923 | 1.22254274709217 | -3.28493901326662 | 0.00102004448357652 | 0.0355522930553829 | 3685 | 0 | 0 | 15 | 19 | 0 | 0 | 4 | 4 | 0 | 1 | 0 | 0 |
| FordPrefect_a_dup245 | 3.3291622689809 | -4.07340846748796 | 1.25189419859682 | -3.25379610517697 | 0.00113873980030102 | 0.0384174933393487 | 3686 | 0 | 0 | 18 | 19 | 0 | 0 | 4 | 3 | 0 | 1 | 0 | 0 |
| MER72_dup329 | 2.92740499512841 | -3.91459964123666 | 1.23200793041803 | -3.1774143206273 | 0.00148594588167225 | 0.0462008204312536 | 3687 | 0 | 0 | 15 | 15 | 0 | 0 | 7 | 2 | 0 | 1 | 0 | 0 |
| MIR3_dup77113 | 3.25286051470774 | -4.03461966246297 | 1.21049651822397 | -3.333028721456 | 0.000859060736560134 | 0.031617952980903 | 3688 | 0 | 0 | 17 | 17 | 0 | 0 | 6 | 3 | 0 | 1 | 0 | 0 |
| MER121_dup822 | 5.07096721506755 | -3.61071955380745 | 1.05475371240582 | -3.42328214761308 | 0.000618698221714247 | 0.0250923414119683 | 3689 | 1 | 0 | 21 | 27 | 0 | 0 | 7 | 9 | 0 | 2 | 0 | 0 |
| MER102c_dup3022 | 3.92216218403578 | -3.2592158152993 | 0.952550329632525 | -3.42156809347453 | 0.000622611321291358 | 0.0251652579337512 | 3690 | 1 | 0 | 18 | 19 | 0 | 1 | 6 | 6 | 0 | 1 | 0 | 0 |
| MIRb_dup198450 | 3.43249807540084 | -3.49527814912027 | 1.08128064214319 | -3.23253558131987 | 0.00122696854797778 | 0.0404366008472466 | 3691 | 1 | 0 | 12 | 23 | 0 | 0 | 6 | 4 | 0 | 1 | 0 | 0 |
| L2a_dup152011 | 5.34295396022458 | -3.68085240496408 | 1.14005955180686 | -3.22864923953345 | 0.00124376343513556 | 0.0408430215755955 | 3692 | 1 | 0 | 29 | 28 | 0 | 0 | 10 | 3 | 0 | 2 | 0 | 0 |
| MIRb_dup198433 | 4.64411387482165 | -3.83960809392858 | 1.10197231319439 | -3.48430541126605 | 0.000493416102942659 | 0.0213621521162289 | 3693 | 1 | 0 | 30 | 20 | 0 | 0 | 6 | 4 | 0 | 1 | 0 | 0 |
| MIRb_dup198498 | 4.88329963363153 | -4.2018432359724 | 1.21807242032386 | -3.44958408536598 | 0.000561450846318505 | 0.0234242918003361 | 3694 | 0 | 0 | 26 | 23 | 0 | 0 | 8 | 6 | 0 | 2 | 0 | 0 |
| L1ME2_dup11438 | 5.06220745156603 | -4.5454378814502 | 1.18207259519822 | -3.84531195454032 | 0.000120399167273205 | 0.00777224640694249 | 3695 | 0 | 0 | 23 | 32 | 0 | 0 | 5 | 7 | 0 | 1 | 0 | 0 |
| MSTB_dup7642 | 5.54046944815366 | -4.47679719956135 | 1.30671866685447 | -3.42598396511613 | 0.000612576544584189 | 0.0249333114056841 | 3696 | 0 | 0 | 29 | 34 | 0 | 0 | 5 | 5 | 0 | 2 | 0 | 0 |
| AluSg4_dup6198 | 3.93442103207031 | -4.26197679518649 | 1.21570068840844 | -3.50577805525975 | 0.000455274685329126 | 0.0202601066533242 | 3697 | 0 | 0 | 24 | 18 | 0 | 0 | 7 | 3 | 0 | 1 | 0 | 0 |
| MIRb_dup198494 | 4.85693454490293 | -4.49920485971027 | 1.18286302658277 | -3.8036566860224 | 0.0001425757417588 | 0.00877530456506454 | 3698 | 0 | 0 | 26 | 27 | 0 | 0 | 8 | 4 | 0 | 1 | 0 | 0 |
| AluJb_dup110117 | 3.80713766623515 | -4.21809686051719 | 1.1951778805366 | -3.5292628228891 | 0.000416719050153746 | 0.0190539260391782 | 3699 | 0 | 0 | 16 | 25 | 0 | 0 | 6 | 4 | 0 | 1 | 0 | 0 |
| AluSz_dup88905 | 4.45297723721605 | -4.39581641346475 | 1.17464034961261 | -3.74226580494571 | 0.000182368465706038 | 0.0104323360473012 | 3700 | 0 | 0 | 17 | 31 | 0 | 0 | 7 | 5 | 0 | 1 | 0 | 0 |
| AluSx_dup100155 | 4.46743678241566 | -4.43272335816312 | 1.25631505341411 | -3.52835329491352 | 0.000418153658152938 | 0.0190797204147751 | 3701 | 0 | 0 | 22 | 30 | 0 | 0 | 7 | 2 | 0 | 1 | 0 | 0 |
| L2a_dup151995 | 4.67076752255938 | -4.45628526138129 | 1.18295226554278 | -3.76708798079572 | 0.000165162807460315 | 0.00975423911242298 | 3702 | 0 | 0 | 19 | 31 | 0 | 0 | 11 | 3 | 0 | 1 | 0 | 0 |
| L1M5_dup48885 | 5.97909802864474 | -3.78190478347271 | 1.13554355357394 | -3.33047972626834 | 0.00086696476686182 | 0.0318251771699634 | 3703 | 0 | 1 | 33 | 37 | 0 | 0 | 7 | 2 | 1 | 1 | 0 | 0 |
| AluSz6_dup41830 | 3.87526513891451 | -4.04776470493228 | 1.2072597048431 | -3.35285331622853 | 0.00079983083296364 | 0.0300563088092809 | 3704 | 0 | 0 | 19 | 25 | 0 | 0 | 4 | 2 | 1 | 1 | 0 | 1 |
| MSTA_dup17622 | 4.62077214275727 | -4.04569852670065 | 1.2690383765097 | -3.18800329571414 | 0.00143258903664578 | 0.0450005057311837 | 3705 | 0 | 0 | 16 | 37 | 0 | 0 | 4 | 3 | 1 | 2 | 0 | 1 |
| L1MEg_dup13138 | 4.2365438088185 | -4.07441670827431 | 1.26848089053401 | -3.21204421657392 | 0.00131794089222004 | 0.0424997994280759 | 3706 | 0 | 0 | 27 | 21 | 0 | 0 | 5 | 2 | 1 | 1 | 0 | 0 |
| THE1B_dup20493 | 5.05637818104626 | -4.27172042674784 | 1.24317295120189 | -3.43614331587409 | 0.000590058748414741 | 0.0242994321478529 | 3707 | 0 | 0 | 29 | 29 | 0 | 0 | 7 | 2 | 1 | 1 | 0 | 0 |
| MLT2D_dup4009 | 4.49681429527603 | -4.09730806096854 | 1.22210516570131 | -3.35266405540253 | 0.000800377912739555 | 0.0300658759721802 | 3708 | 0 | 0 | 26 | 24 | 0 | 0 | 4 | 4 | 1 | 1 | 0 | 0 |

| | | | | | | | | | | | | | | | | | | |
|---|---|---|---|---|---|---|---|---|---|---|---|---|---|---|---|---|---|---|
| L1MEc_dup11580 | 4.02978661375514 | -3.88921328791059 | 1.13837219271411 | -3.41646898334533 | 0.000634388900585405 | 0.0255199156954683 | 3709 | 0 | 0 | 17 | 24 | 0 | 0 | 6 | 5 | 1 | 1 | 0 | 0 |
| MSTA_dup17624 | 5.14869911923054 | -4.21566882449762 | 1.16979532997264 | -3.6037661601848 | 0.000313639254095281 | 0.0154236755256952 | 3710 | 0 | 0 | 22 | 35 | 0 | 0 | 6 | 5 | 1 | 1 | 0 | 0 |
| L1ME2z_dup3471 | 4.11081605554764 | -4.02048863597697 | 1.19394496267814 | -3.36739863365108 | 0.000758809162781057 | 0.0289689233821965 | 3711 | 0 | 0 | 23 | 25 | 0 | 1 | 4 | 3 | 0 | 0 | 0 | 0 |
| L1PA10_dup4820 | 5.31207556826982 | -4.30119202342352 | 1.23206683565874 | -3.49103790390059 | 0.000481147987375582 | 0.0209798862019553 | 3712 | 0 | 0 | 24 | 43 | 0 | 1 | 3 | 3 | 0 | 0 | 0 | 0 |
| AluSx1_dup99459 | 4.8844225020797 | -4.84626147229902 | 1.27654000019185 | -3.79640392903527 | 0.000146810219011973 | 0.00896635122510855 | 3713 | 0 | 0 | 30 | 28 | 0 | 0 | 4 | 4 | 0 | 0 | 0 | 0 |
| L1ME3A_dup13962 | 5.12647551446693 | -4.90188221881872 | 1.23636822050453 | -3.96474297666627 | 7.34750720127187e-05 | 0.00542448013811305 | 3714 | 0 | 0 | 28 | 32 | 0 | 0 | 6 | 4 | 0 | 0 | 0 | 0 |
| L1PA3_dup10030 | 4.16109578536184 | -4.65913851408721 | 1.31662634097554 | -3.53869459320939 | 0.000402110783637516 | 0.0185499520208244 | 3715 | 0 | 0 | 22 | 27 | 0 | 0 | 2 | 5 | 0 | 0 | 0 | 0 |
| THE1B-int_dup3354 | 3.95472286178155 | -4.60087288608734 | 1.32178494821102 | -3.48080290391749 | 0.000499913209816693 | 0.0215649653051202 | 3716 | 0 | 0 | 22 | 26 | 0 | 0 | 3 | 3 | 0 | 0 | 0 | 0 |
| THE1B_dup20489 | 4.53343348944607 | -4.76078338828198 | 1.30060208584599 | -3.6604457582315 | 0.000251776851234346 | 0.0131958798774907 | 3717 | 0 | 0 | 23 | 32 | 0 | 0 | 3 | 4 | 0 | 0 | 0 | 0 |
| L1ME3A_dup13964 | 3.53339077396861 | -4.14374691612117 | 1.27739557206261 | -3.24390267724998 | 0.00117904024090162 | 0.0393199592377824 | 3718 | 0 | 0 | 16 | 23 | 0 | 0 | 2 | 5 | 1 | 0 | 0 | 0 |
| L3b_dup9033 | 3.79784797775177 | -4.23656146805637 | 1.28437171059273 | -3.29854779042216 | 0.00097186335246592 | 0.0344532593639009 | 3719 | 0 | 0 | 16 | 27 | 0 | 0 | 2 | 5 | 1 | 0 | 0 | 0 |
| L2a_dup152051 | 4.33390765520889 | -4.38451764359344 | 1.24446751220816 | -3.52320779817998 | 0.00042635693932414 | 0.0193582707786507 | 3720 | 0 | 0 | 14 | 36 | 0 | 0 | 5 | 4 | 1 | 0 | 0 | 0 |
| L2a_dup152020 | 4.51711786866246 | -4.41911712029706 | 1.20630038442113 | -3.66336376690923 | 0.000248924634458832 | 0.0130764138136303 | 3721 | 0 | 0 | 21 | 29 | 0 | 0 | 8 | 3 | 1 | 0 | 0 | 0 |
| MIR3_dup77115 | 4.56192625145432 | -4.45708594455316 | 1.27137956703751 | -3.50570833456039 | 0.000455393949804104 | 0.0202601922294915 | 3722 | 0 | 0 | 25 | 29 | 0 | 0 | 4 | 3 | 1 | 0 | 0 | 0 |
| MER85_dup801 | 5.9379623959437 | -5.27281106844896 | 1.29618834898433 | -4.06793586177551 | 4.74314318358105e-05 | 0.00391339286021726 | 3723 | 0 | 0 | 29 | 44 | 0 | 0 | 1 | 6 | 0 | 0 | 0 | 1 |
| MIRb_dup198493 | 6.31714527364911 | -4.94777591568626 | 1.15040789513689 | -4.30088835151597 | 1.70114759499976e-05 | 0.00181353613637909 | 3724 | 0 | 0 | 31 | 38 | 0 | 0 | 4 | 9 | 1 | 0 | 0 | 1 |
| MLT1K_dup15256 | 4.91596053078954 | -4.49187725738577 | 1.03675610649766 | -4.3326267665402 | 1.47340774231938e-05 | 0.00162687461092309 | 3725 | 0 | 0 | 19 | 30 | 0 | 1 | 7 | 8 | 0 | 0 | 0 | 1 |
| MLT1C_dup16509 | 3.55907952630955 | -4.27534707882842 | 1.10724845476086 | -3.86123553430634 | 0.000112815082357426 | 0.00742343515143953 | 3726 | 0 | 0 | 17 | 17 | 0 | 1 | 4 | 6 | 0 | 0 | 0 | 2 |
| THE1B-int_dup627 | 3.72704590397919 | -4.17632420352278 | 1.08755953259021 | -3.84008790174101 | 0.000122990265237409 | 0.00789240605506488 | 3727 | 0 | 0 | 20 | 16 | 0 | 1 | 5 | 6 | 0 | 0 | 0 | 1 |
| L1M4b_dup770 | 5.34265389779869 | -4.12201825423497 | 0.991975230950739 | -4.15536409138392 | 3.24769951425347e-05 | 0.00293496812996033 | 3728 | 1 | 1 | 24 | 33 | 0 | 0 | 5 | 7 | 0 | 0 | 0 | 1 |
| L2a_dup139143 | 4.87645188751089 | -3.65528598338144 | 0.974679281428286 | -3.7502448785256 | 0.000176661963445808 | 0.0102128298236337 | 3729 | 2 | 1 | 18 | 33 | 0 | 0 | 4 | 7 | 0 | 0 | 0 | 1 |
| L2a_dup165192 | 4.08533166356438 | -4.44349625268164 | 1.14231258450679 | -3.88991271999354 | 0.00010028028274558 | 0.00683117439398216 | 3730 | 1 | 0 | 18 | 27 | 0 | 0 | 4 | 4 | 0 | 0 | 0 | 2 |
| MLT1B-int_dup46 | 4.5881055046163 | -4.27285264099816 | 1.04273896830792 | -4.09772030284033 | 4.17239004700023e-05 | 0.00355748909288445 | 3731 | 2 | 0 | 19 | 23 | 0 | 0 | 5 | 8 | 0 | 0 | 0 | 4 |
| AluSc8_dup4079 | 3.56446410613738 | -4.1394981163153 | 1.09434148197766 | -3.78263840353976 | 0.000155174759402577 | 0.00932875525538293 | 3732 | 1 | 0 | 15 | 19 | 0 | 0 | 4 | 7 | 0 | 0 | 0 | 1 |
| MER5A_dup33492 | 4.16304529401357 | -3.45165493487409 | 1.05403619738838 | -3.2747024660314 | 0.00105773312888113 | 0.0365340930591885 | 3733 | 3 | 0 | 20 | 21 | 0 | 0 | 3 | 7 | 0 | 0 | 0 | 1 |
| AluSx_dup54168 | 4.03298055112537 | -3.41203416837414 | 0.964921280936029 | -3.53607515533731 | 0.000406119139250346 | 0.0186841501755728 | 3734 | 0 | 0 | 19 | 18 | 2 | 0 | 9 | 4 | 1 | 0 | 0 | 1 |
| L1ME3Cz_dup11092 | 5.12596323097066 | -3.51572351236304 | 1.011166292335 | -3.47689943683198 | 0.000507248033669372 | 0.0217865777291502 | 3735 | 0 | 0 | 28 | 28 | 1 | 1 | 6 | 4 | 1 | 0 | 0 | 0 |
| L1PA13_dup4460 | 4.42446158664083 | -3.17808690910989 | 0.964692986043261 | -3.2944024213807 | 0.000986311825644289 | 0.0347841935178236 | 3736 | 0 | 0 | 20 | 26 | 2 | 1 | 6 | 3 | 1 | 0 | 1 | 0 |
| L2_dup51456 | 3.94613352158813 | -3.36291653836493 | 0.98511445065635 | -3.41373181169389 | 0.000640796194628812 | 0.0256907988756777 | 3737 | 0 | 0 | 23 | 16 | 2 | 1 | 7 | 3 | 0 | 0 | 0 | 1 |
| REP522_dup82 | 5.53200516343422 | -3.53935880860914 | 1.03504000690446 | -3.41953816760617 | 0.000627275320028984 | 0.025327957457936 | 3738 | 0 | 0 | 31 | 32 | 2 | 1 | 5 | 4 | 0 | 0 | 0 | 0 |
| L2c_dup126705 | 5.58839373969172 | -3.86529871829387 | 0.888542745508647 | -4.35015505762886 | 1.36041300305386e-05 | 0.00153487355345971 | 3739 | 0 | 0 | 17 | 36 | 1 | 1 | 11 | 8 | 0 | 1 | 0 | 1 |
| MIR_dup28627 | 5.14027634243306 | -3.29603026261359 | 0.866609126516146 | -3.80336435627439 | 0.000142744168396628 | 0.00878339898973471 | 3740 | 0 | 1 | 21 | 29 | 1 | 1 | 11 | 5 | 0 | 1 | 0 | 0 |
| LTR67B_dup2052 | 5.90333462988251 | -4.0606506533902 | 0.971739105663688 | -4.17874574535808 | 2.93121155680918e-05 | 0.00270757012848027 | 3741 | 1 | 1 | 27 | 36 | 0 | 0 | 11 | 5 | 0 | 0 | 0 | 0 |
| L2a_dup88270 | 4.0489712317913 | -4.05045446385052 | 1.11420292529259 | -3.63529333113793 | 0.000277664516153551 | 0.0141445431300948 | 3742 | 1 | 0 | 18 | 25 | 0 | 0 | 9 | 3 | 0 | 0 | 0 | 0 |
| MIR_dup154609 | 5.07027606483961 | -4.29110264980925 | 1.14666649456294 | -3.7422412446479 | 0.00018238629538514 | 0.0104323360473012 | 3743 | 1 | 0 | 26 | 33 | 0 | 0 | 7 | 3 | 0 | 0 | 0 | 0 |
| AluSc_dup30610 | 4.75686653423563 | -4.22193134984907 | 1.13986035297593 | -3.70390227085846 | 0.000212308078371025 | 0.0115999673974907 | 3744 | 1 | 0 | 19 | 36 | 0 | 0 | 6 | 4 | 0 | 0 | 0 | 0 |
| L1ME3B_dup7283 | 4.75686653423563 | -4.22193134984907 | 1.13986035297593 | -3.70390227085846 | 0.000212308078371025 | 0.0115999673974907 | 3745 | 1 | 0 | 19 | 36 | 0 | 0 | 6 | 4 | 0 | 0 | 0 | 0 |
| L2a_dup165195 | 5.07053023712732 | -3.49190119808759 | 1.03761107839871 | -3.36532759796326 | 0.000764528278927527 | 0.0291112891434933 | 3746 | 3 | 0 | 19 | 35 | 0 | 0 | 8 | 5 | 0 | 0 | 0 | 0 |
| MLT1I_dup9780 | 5.05171517686097 | -4.31408195921999 | 1.06349405591144 | -4.05651722756712 | 4.98098901035461e-05 | 0.00406449386508175 | 3747 | 1 | 0 | 23 | 30 | 0 | 0 | 7 | 7 | 0 | 0 | 0 | 0 |
| L1PA3_dup5643 | 4.66513319364221 | -4.2557090719332 | 1.00189233867531 | -4.24767103974468 | 2.16004246230287e-05 | 0.00215955236863495 | 3748 | 1 | 0 | 18 | 25 | 0 | 0 | 12 | 7 | 0 | 0 | 0 | 0 |
| L2a_dup46289 | 3.65687076989555 | -3.51043915435513 | 0.972295033758671 | -3.61046702129555 | 0.000305646164317655 | 0.0151578328917734 | 3749 | 2 | 0 | 14 | 16 | 0 | 0 | 11 | 6 | 0 | 0 | 0 | 0 |
| THE1B-int_dup380 | 5.30713505966296 | -3.50213762386423 | 0.927730089179537 | -3.77495315147257 | 0.000160037737454703 | 0.00954155336416859 | 3750 | 4 | 0 | 21 | 27 | 0 | 0 | 15 | 5 | 0 | 0 | 0 | 1 |
| THE1B_dup21736 | 5.11975690963224 | -3.89050900763757 | 1.00628390930216 | -3.86621406908469 | 0.000110537921872179 | 0.00732413496475379 | 3751 | 2 | 0 | 26 | 24 | 0 | 0 | 14 | 4 | 0 | 0 | 0 | 0 |
| LTR79_dup2216 | 3.21383342108655 | -3.79159759391941 | 1.10045401506398 | -3.44548481082963 | 0.000570036002429061 | 0.0236880521540317 | 3752 | 1 | 0 | 17 | 13 | 0 | 0 | 8 | 4 | 0 | 0 | 0 | 0 |
| MIRb_dup107893 | 3.02660723940054 | -3.71942238542679 | 1.105083738845 | -3.36573804742998 | 0.000763391662333993 | 0.029086628100196 | 3753 | 1 | 0 | 15 | 13 | 0 | 0 | 6 | 5 | 0 | 0 | 0 | 0 |
| L1MA7_dup4280 | 2.80793220320509 | -3.63122830769344 | 1.12529767565566 | -3.22690465487517 | 0.00125137150584625 | 0.0410249165514042 | 3754 | 1 | 0 | 11 | 16 | 0 | 0 | 6 | 4 | 0 | 0 | 0 | 0 |
| L1MC4_dup12454 | 3.09069225560515 | -3.72955989944957 | 1.14434221479443 | -3.25912987499073 | 0.0011175449555764 | 0.037880480066617 | 3755 | 1 | 0 | 15 | 16 | 0 | 0 | 4 | 5 | 0 | 0 | 0 | 0 |
| AluSc_dup25588 | 3.15558785732165 | -3.49713150647237 | 1.10788831547711 | -3.15657404958399 | 0.00159634348943371 | 0.0484679785551605 | 3756 | 2 | 0 | 18 | 14 | 0 | 0 | 6 | 2 | 0 | 0 | 0 | 1 |
| AluSx1_dup20485 | 2.80588608906752 | -3.86779129562129 | 1.17956363547802 | -3.27900180989715 | 0.00104174967777967 | 0.0361078177463892 | 3757 | 1 | 0 | 14 | 16 | 0 | 0 | 3 | 3 | 0 | 0 | 0 | 1 |
| MIR3_dup77119 | 3.40217290281736 | -3.37583655044027 | 1.01862412143297 | -3.31411408723686 | 0.00091933987943473 | 0.0331941792996004 | 3758 | 1 | 0 | 18 | 14 | 1 | 0 | 9 | 3 | 0 | 0 | 0 | 0 |
| AluSp_dup43614 | 4.12977376738605 | -3.47060502447953 | 0.924465315910926 | -3.75417548365214 | 0.000173912996693887 | 0.0101073040447031 | 3759 | 1 | 0 | 17 | 24 | 1 | 1 | 9 | 3 | 0 | 0 | 0 | 1 |
| MLT1G1_dup1571 | 4.32849707463246 | -2.99430974960058 | 0.844835480694902 | -3.54425189048366 | 0.000393728886264478 | 0.0182688201316474 | 3760 | 3 | 0 | 21 | 18 | 1 | 1 | 8 | 5 | 0 | 0 | 0 | 1 |
| L1M2_dup4683 | 4.35430622468049 | -4.10593234288135 | 1.19187674599255 | -3.4449303224404 | 0.000571206614869716 | 0.0237201272568568 | 3761 | 1 | 0 | 30 | 19 | 0 | 0 | 7 | 2 | 0 | 0 | 0 | 0 |
| AluSx_dup112487 | 4.180713916961 | -4.66780369314554 | 1.34830233023005 | -3.46198592740628 | 0.000536205191404907 | 0.0226299894420064 | 3762 | 0 | 0 | 30 | 18 | 0 | 0 | 8 | 1 | 0 | 0 | 0 | 0 |
| L1M3_dup6419 | 3.91219814369548 | -4.59157261225364 | 1.40805764924061 | -3.26092657834709 | 0.00111048790838607 | 0.0377049309145024 | 3763 | 0 | 0 | 30 | 18 | 0 | 0 | 4 | 1 | 0 | 0 | 0 | 0 |
| Tigger3b_dup5798 | 4.50710062008625 | -4.76124477235111 | 1.36852881654214 | -3.47909719897703 | 0.000503106071432674 | 0.0216791157684026 | 3764 | 0 | 0 | 19 | 38 | 0 | 0 | 6 | 1 | 0 | 0 | 0 | 0 |
| MLT1A0_dup18674 | 3.76652655469682 | -4.5463099274432 | 1.34374038055514 | -3.38332463117987 | 0.000716139272639064 | 0.0277968015431705 | 3765 | 0 | 0 | 15 | 32 | 0 | 0 | 4 | 2 | 0 | 0 | 0 | 0 |
| L2a_dup122530 | 3.51428608898221 | -4.4638793277596 | 1.33462852246896 | -3.34466051984395 | 0.000823833412925082 | 0.0306820887888357 | 3766 | 0 | 0 | 16 | 27 | 0 | 0 | 4 | 2 | 0 | 0 | 0 | 0 |
| MIRb_dup198490 | 3.38104284472005 | -4.417795128844 | 1.36168656487455 | -3.24435537722368 | 0.00117716778337703 | 0.0392895283207195 | 3767 | 0 | 0 | 16 | 26 | 0 | 0 | 3 | 2 | 0 | 0 | 0 | 0 |
| HUERS-P2-int_dup242 | 4.3749128757694 | -4.72277637306265 | 1.36126884079266 | -3.46939284257224 | 0.000521636108332117 | 0.0222001834078329 | 3768 | 0 | 0 | 27 | 28 | 0 | 0 | 3 | 2 | 0 | 0 | 0 | 0 |
| LTR33_dup3338 | 4.16836920309591 | -4.66864789656486 | 1.39884272879615 | -3.33750735551431 | 0.000845334843608679 | 0.0312869878203936 | 3769 | 0 | 0 | 21 | 33 | 0 | 0 | 2 | 2 | 0 | 0 | 0 | 0 |
| ALR/Alpha_dup1274 | 3.2559149957474 | -4.37046815770404 | 1.36016620495564 | -3.21318684568153 | 0.00131270861289862 | 0.0423875374684185 | 3770 | 0 | 0 | 19 | 20 | 0 | 0 | 2 | 3 | 0 | 0 | 0 | 0 |
| L1PA13_dup8215 | 3.02092187870437 | -4.28215114027584 | 1.36078760482527 | -3.14681815522981 | 0.00165057579871441 | 0.0496546844590019 | 3771 | 0 | 0 | 16 | 20 | 0 | 0 | 2 | 3 | 0 | 0 | 0 | 0 |
| L1MA10_dup4459 | 2.80017478447107 | -4.19793752438973 | 1.2867903937264 | -3.26233203547082 | 0.00110499633691731 | 0.0375828270989318 | 3772 | 0 | 0 | 14 | 17 | 0 | 0 | 4 | 3 | 0 | 0 | 0 | 0 |
| MIR_dup154597 | 2.88970791918338 | -4.2313836317549 | 1.31863385699661 | -3.20891474862669 | 0.00133236991249351 | 0.0428363857854588 | 3773 | 0 | 0 | 16 | 17 | 0 | 0 | 3 | 3 | 0 | 0 | 0 | 0 |
| L1MA9_dup11068 | 3.03322399131952 | -4.28687995539582 | 1.36349380498348 | -3.14404065477053 | 0.00166632288140445 | 0.0499676762030735 | 3774 | 0 | 0 | 20 | 16 | 0 | 0 | 3 | 2 | 0 | 0 | 0 | 0 |
| MLT2D_dup4253 | 3.28237775662569 | -4.38121710459544 | 1.29535176391259 | -3.3822604999294 | 0.000718919354537964 | 0.0278770358800805 | 3775 | 0 | 0 | 21 | 16 | 0 | 0 | 4 | 3 | 0 | 0 | 0 | 0 |
| L1MA9_dup4090 | 3.69299438185132 | -3.91110392271951 | 1.24162424743919 | -3.14998996740442 | 0.00163276069387234 | 0.0492593985120005 | 3776 | 1 | 0 | 14 | 31 | 0 | 0 | 4 | 2 | 0 | 0 | 0 | 0 |
| SVA_A_dup1034 | 4.05006950503329 | -4.01676928024996 | 1.22145764100978 | -3.28850477117594 | 0.00100721077297603 | 0.0352452473260574 | 3777 | 1 | 0 | 24 | 24 | 0 | 0 | 3 | 3 | 0 | 0 | 0 | 0 |
| L1PA10_dup4822 | 2.94965073503423 | -4.25928206155841 | 1.30780058260411 | -3.25682838669277 | 0.00112664526157787 | 0.0381338169859754 | 3778 | 0 | 0 | 10 | 24 | 0 | 0 | 4 | 3 | 0 | 0 | 0 | 0 |
| L2b_dup59139 | 2.78178656130758 | -4.19179659084249 | 1.30908864149613 | -3.20207238682613 | 0.00136442706988285 | 0.0435538777972854 | 3779 | 0 | 0 | 7 | 24 | 0 | 0 | 5 | 3 | 0 | 0 | 0 | 0 |
| MSTA-int_dup1589 | 2.79510331629332 | -4.20036571437156 | 1.27971101173367 | -3.28227676081429 | 0.0010297248425853 | 0.0358057734025723 | 3780 | 0 | 0 | 11 | 19 | 0 | 0 | 7 | 2 | 0 | 0 | 0 | 0 |
| AluJo_dup66907 | 3.01585041052662 | -4.28553052755487 | 1.31132324270487 | -3.26809621609016 | 0.00108273550602804 | 0.0370961242213603 | 3781 | 0 | 0 | 13 | 22 | 0 | 0 | 5 | 2 | 0 | 0 | 0 | 0 |
| MLT1K_dup15257 | 2.61916460485188 | -4.12000557759048 | 1.3071258120377 | -3.15195793675571 | 0.00162179634312449 | 0.0489907265026153 | 3782 | 0 | 0 | 13 | 16 | 0 | 0 | 5 | 2 | 0 | 0 | 0 | 0 |
| LOR1-int_dup1309 | 2.61607790444344 | -4.12134168909999 | 1.25379470830524 | -3.28709449944227 | 0.00101226858469436 | 0.03536934811361 | 3783 | 0 | 0 | 15 | 11 | 0 | 0 | 6 | 3 | 0 | 0 | 0 | 0 |
| MSTA_dup10411 | 2.53356014810347 | -4.08401488741106 | 1.22341066742361 | -3.33822075951947 | 0.000843167313075768 | 0.0312164757238411 | 3784 | 0 | 0 | 13 | 10 | 0 | 0 | 5 | 5 | 0 | 0 | 0 | 0 |
| AluY_dup91706 | 2.61497788752625 | -4.11963206580102 | 1.25868748143554 | -3.27295863871036 | 0.00106428050398785 | 0.0366750599278628 | 3785 | 0 | 0 | 12 | 15 | 0 | 0 | 4 | 4 | 0 | 0 | 0 | 0 |
| AluJr_dup15108 | 2.24877081096223 | -3.94714274820814 | 1.24216317151655 | -3.17763627091689 | 0.00148480895513752 | 0.0461715043961255 | 3786 | 0 | 0 | 9 | 12 | 0 | 0 | 5 | 4 | 0 | 0 | 0 | 0 |
| L2c_dup126681 | 2.24877081096223 | -3.94714274820814 | 1.24216317151655 | -3.17763627091689 | 0.00148480895513752 | 0.0461715043961255 | 3787 | 0 | 0 | 9 | 12 | 0 | 0 | 5 | 4 | 0 | 0 | 0 | 0 |

| | | | | | | | | | | | | | | | | | | |
|---|---|---|---|---|---|---|---|---|---|---|---|---|---|---|---|---|---|---|
| MIR_dup157994 | 2.37601331922266 | -4.01408376395542 | 1.2656991162386 | -3.17143601702468 | 0.00151687279283931 | 0.0468535664540892 | 3788 | 0 | 0 | 9 | 13 | 0 | 0 | 9 | 2 | 0 | 0 | 0 | 0 |
| L4_C_Mam_dup2816 | 2.48896528770148 | -4.06340402169677 | 1.27932261502683 | -3.17621526733625 | 0.00149210184140414 | 0.0463437755847473 | 3789 | 0 | 0 | 13 | 12 | 0 | 0 | 7 | 2 | 0 | 0 | 0 | 0 |
| MIR3_dup77134 | 2.68730819053885 | -4.15396198191304 | 1.27368542439266 | -3.26137200156295 | 0.0011087447715694 | 0.0376726236492723 | 3790 | 0 | 0 | 13 | 15 | 0 | 0 | 7 | 2 | 0 | 0 | 0 | 0 |
| LTR39-int_dup96 | 2.45015764091137 | -4.05709562418416 | 1.1939436886832 | -3.39806279193847 | 0.000678648415162639 | 0.026765549806863 | 3791 | 0 | 0 | 9 | 12 | 0 | 0 | 8 | 4 | 0 | 0 | 0 | 0 |
| MER102b_dup2769 | 2.30469765858065 | -3.98272360253652 | 1.21218085415789 | -3.28558530591819 | 0.0010177072043916 | 0.035490037228246 | 3792 | 0 | 0 | 8 | 12 | 0 | 0 | 7 | 4 | 0 | 0 | 0 | 0 |
| L2a_dup152054 | 2.952694662146 | -4.26728546392059 | 1.24262856518485 | -3.43407964654813 | 0.000594569464973402 | 0.0244176717615115 | 3793 | 0 | 0 | 10 | 21 | 0 | 0 | 7 | 3 | 0 | 0 | 0 | 0 |
| MLT1A0_dup19893 | 2.88860964594139 | -4.24626359434475 | 1.21371092644714 | -3.49857902884231 | 0.00046774444864714 | 0.0206069099384915 | 3794 | 0 | 0 | 10 | 18 | 0 | 0 | 9 | 3 | 0 | 0 | 0 | 0 |
| REP522_dup214 | 3.87543306477416 | -3.94964246462724 | 1.16738313068449 | -3.38333008316763 | 0.000716125054874714 | 0.0277968015431705 | 3795 | 0 | 0 | 22 | 21 | 1 | 0 | 4 | 4 | 0 | 0 | 0 | 0 |
| MIR_dup154637 | 4.37585717039621 | -4.10197781747932 | 1.13634105494076 | -3.60981221231434 | 0.000306418758311521 | 0.0151898162219152 | 3796 | 0 | 0 | 17 | 32 | 1 | 0 | 8 | 3 | 0 | 0 | 0 | 0 |
| MIRb_dup198478 | 4.14703553844812 | -4.03940385408428 | 1.13906490885604 | -3.5462455411264 | 0.0003907619226859 | 0.0182011732423179 | 3797 | 0 | 0 | 20 | 26 | 1 | 0 | 7 | 3 | 0 | 0 | 0 | 0 |
| MER21A_dup1713 | 3.67379185486042 | -3.91240507506197 | 1.11374490162337 | -3.51283769681847 | 0.000443348209140151 | 0.0198884888235821 | 3798 | 0 | 0 | 10 | 27 | 1 | 0 | 6 | 6 | 0 | 0 | 0 | 0 |
| MIRb_dup198501 | 3.90876814250706 | -4.22234551751791 | 1.09978720087059 | -3.83923864014376 | 0.000123416429380747 | 0.00791028362999303 | 3799 | 0 | 0 | 16 | 25 | 1 | 0 | 5 | 5 | 0 | 0 | 0 | 1 |
| LTR16C_dup734 | 4.71831640373323 | -5.00610695214551 | 1.09946265839427 | -4.55323053850393 | 5.28283256360889e-06 | 0.000751590954835126 | 3800 | 0 | 0 | 18 | 30 | 0 | 0 | 11 | 5 | 0 | 0 | 0 | 1 |
| L2c_dup72011 | 3.69810984392692 | -4.73406337593574 | 1.1196637483674 | -4.22811168338579 | 2.35660813706729e-05 | 0.0023130980917055 | 3801 | 0 | 0 | 16 | 21 | 0 | 0 | 7 | 5 | 0 | 0 | 0 | 1 |
| ERV3-16A3_I-int_dup4[illegible] | 3.63914081287193 | -4.71989016155231 | 1.11017682791651 | -4.25147601973484 | 2.12366252160144e-05 | 0.00213034149395016 | 3802 | 0 | 0 | 14 | 21 | 0 | 0 | 10 | 4 | 0 | 0 | 0 | 1 |
| L1ME1_dup3209 | 3.86191719184641 | -4.78503717567768 | 1.11124726913415 | -4.30600579059782 | 1.66228706754251e-05 | 0.00178168720078139 | 3803 | 0 | 0 | 16 | 22 | 0 | 0 | 10 | 4 | 0 | 0 | 0 | 1 |
| AluSz_dup99534 | 4.56479871308411 | -4.77199319682253 | 1.15774063617364 | -4.12181541160554 | 3.75898353127857e-05 | 0.00328474144448626 | 3804 | 0 | 0 | 19 | 29 | 0 | 0 | 8 | 6 | 0 | 0 | 0 | 0 |
| L1PB1_dup11942 | 4.07655048795589 | -4.64207366430329 | 1.1646391233057 | -3.98584726496847 | 6.72397562218512e-05 | 0.0050979692586502 | 3805 | 0 | 0 | 17 | 25 | 0 | 0 | 7 | 6 | 0 | 0 | 0 | 0 |
| L2a_dup151999 | 4.19770514199277 | -4.67147221001533 | 1.25261719809901 | -3.72936936927325 | 0.000191959590372501 | 0.0107828179156693 | 3806 | 0 | 0 | 17 | 32 | 0 | 0 | 5 | 4 | 0 | 0 | 0 | 0 |
| MIRb_dup198489 | 3.85601691611246 | -4.57421481832686 | 1.23356760087767 | -3.70811848095909 | 0.000208804959949079 | 0.011474435334883 | 3807 | 0 | 0 | 19 | 24 | 0 | 0 | 7 | 3 | 0 | 0 | 0 | 0 |
| L1M5_dup48894 | 3.97501413756489 | -4.60871726172585 | 1.25428397445521 | -3.67438104574973 | 0.000238426690525925 | 0.0126761398088187 | 3808 | 0 | 0 | 18 | 28 | 0 | 0 | 6 | 3 | 0 | 0 | 0 | 0 |
| MIRb_dup198488 | 3.89668309855055 | -4.5858085752105 | 1.25493862488946 | -3.65420944439768 | 0.000257975619873876 | 0.0134437728529368 | 3809 | 0 | 0 | 17 | 28 | 0 | 0 | 6 | 3 | 0 | 0 | 0 | 0 |
| L1MA4_dup3658 | 3.64537190065994 | -4.51008657499447 | 1.22212776432914 | -3.69035603856868 | 0.000223940396182174 | 0.0120670218130389 | 3810 | 0 | 0 | 15 | 25 | 0 | 0 | 6 | 4 | 0 | 0 | 0 | 0 |
| MIRb_dup198477 | 3.97091279478586 | -4.60760577008564 | 1.22047866635558 | -3.77524482574055 | 0.000159850586516886 | 0.0095375646086627 | 3811 | 0 | 0 | 20 | 24 | 0 | 0 | 6 | 4 | 0 | 0 | 0 | 0 |
| MLT1K_dup7799 | 3.88036501770297 | -4.58152903759274 | 1.22061136087763 | -3.75347074788702 | 0.000174402892700072 | 0.0101215782210759 | 3812 | 0 | 0 | 18 | 25 | 0 | 0 | 6 | 4 | 0 | 0 | 0 | 0 |
| LTR56_dup876 | 4.23955671589465 | -4.68369739324649 | 1.2807109940299 | -3.65710719676788 | 0.000255077724427416 | 0.0133394998278255 | 3813 | 0 | 0 | 21 | 29 | 0 | 0 | 7 | 2 | 0 | 0 | 0 | 0 |
| MIR_dup154636 | 4.44601501402154 | -4.73786396767061 | 1.24069333878873 | -3.81872282178617 | 0.000134144384764592 | 0.0083843235558696 | 3814 | 0 | 0 | 24 | 27 | 0 | 0 | 7 | 3 | 0 | 0 | 0 | 0 |
| THE1D_dup11525 | 4.04827370840148 | -4.6313502536694 | 1.26312714793961 | -3.66657486637348 | 0.000245820972745499 | 0.0129532246034842 | 3815 | 0 | 0 | 16 | 30 | 0 | 0 | 9 | 2 | 0 | 0 | 0 | 0 |
| MLT1K_dup8491 | 3.90065629348631 | -4.59235528403253 | 1.18783489946372 | -3.86615621927413 | 0.000110564131329779 | 0.00732413496475379 | 3816 | 0 | 0 | 14 | 27 | 0 | 0 | 9 | 4 | 0 | 0 | 0 | 0 |
| MLT1C_dup9549 | 3.59760496530291 | -4.4964158095198 | 1.22503113108391 | -3.67045024034726 | 0.000242123602906295 | 0.0128198359628708 | 3817 | 0 | 0 | 14 | 25 | 0 | 0 | 8 | 3 | 0 | 0 | 0 | 0 |
| MSTA_dup10397 | 3.88649564522318 | -4.58492162556459 | 1.22596155722908 | -3.73985758242487 | 0.000184124546845025 | 0.0104963343132694 | 3818 | 0 | 0 | 16 | 27 | 0 | 0 | 8 | 3 | 0 | 0 | 0 | 0 |
| AluJr4_dup9722 | 3.10311322699333 | -3.38580294341139 | 1.06040114350067 | -3.19294539067916 | 0.00140829576746078 | 0.0444958630572598 | 3819 | 0 | 0 | 15 | 15 | 0 | 1 | 4 | 1 | 1 | 1 | 2 | 2 |
| MIR1_Amn_dup1931 | 3.50566452274922 | -3.8269446840681 | 1.06498104030409 | -3.59343926251998 | 0.000326341746207655 | 0.015866717029654 | 3820 | 0 | 0 | 15 | 21 | 0 | 1 | 6 | 2 | 1 | 0 | 1 | 1 |
| HAL1ME_dup7592 | 3.0436173326121 | -3.64878098679451 | 1.13854291651193 | -3.20478124616768 | 0.001351651653707 | 0.043257579609007 | 3821 | 0 | 0 | 18 | 14 | 1 | 0 | 4 | 1 | 1 | 0 | 1 | 1 |
| L1PA8_dup4017 | 3.87318271725227 | -4.15128581750031 | 1.1050406664697 | -3.75668149006907 | 0.000172181413359416 | 0.0100435294435387 | 3822 | 0 | 0 | 20 | 15 | 0 | 0 | 7 | 4 | 2 | 0 | 1 | 2 |
| L1M1_dup4731 | 5.80762647640113 | -4.68028714245172 | 1.3957587060248 | -3.35322081262989 | 0.000798769534150325 | 0.0300306550123137 | 3823 | 0 | 0 | 19 | 51 | 0 | 0 | 1 | 2 | 3 | 0 | 1 | 4 |
| L1MA5A_dup950 | 5.58435241282215 | -4.97242952249211 | 1.24538150117349 | -3.9926958267862 | 6.53263442595825e-05 | 0.00498369423250741 | 3824 | 0 | 0 | 24 | 47 | 0 | 0 | 2 | 2 | 1 | 0 | 1 | 1 |
| AluSg4_dup3417 | 6.15154766055805 | -4.26349110121174 | 1.02344145517314 | -4.1658378011378 | 3.10211207979695e-05 | 0.00282918804499565 | 3825 | 1 | 0 | 35 | 33 | 1 | 0 | 4 | 3 | 1 | 0 | 4 | 1 |
| L1MC4_dup22006 | 5.79117002405637 | -3.90522797206149 | 1.14999188262028 | -3.39587438057681 | 0.000684097480609856 | 0.0268921624188325 | 3826 | 0 | 0 | 33 | 37 | 1 | 0 | 3 | 2 | 1 | 1 | 1 | 0 |
| L1PREC2_dup5948 | 4.61118612584129 | -4.00115268906905 | 1.19623540347842 | -3.34478705230968 | 0.000823457681628241 | 0.0306820887888357 | 3827 | 0 | 0 | 22 | 34 | 1 | 0 | 4 | 1 | 1 | 0 | 1 | 0 |
| L1MA9_dup15191 | 4.3398234943214 | -4.89967754254012 | 1.32237774416235 | -3.70520266555437 | 0.000211221777575676 | 0.0115618534821766 | 3828 | 1 | 0 | 24 | 24 | 0 | 0 | 2 | 1 | 0 | 0 | 7 | 0 |
| L1PA15-16_dup409 | 2.41602693004946 | -4.64560803149602 | 1.38254172189597 | -3.36019373442504 | 0.000778878338586904 | 0.0295066048644164 | 3829 | 0 | 0 | 10 | 15 | 0 | 0 | 2 | 1 | 0 | 0 | 5 | 0 |
| L1M5_dup44662 | 2.57797608735254 | -3.92948231994769 | 1.19469871467686 | -3.28909897673284 | 0.00100508672174408 | 0.035195933052556 | 3830 | 0 | 1 | 11 | 14 | 0 | 0 | 2 | 4 | 0 | 0 | 2 | 0 |
| MER91A_dup3386 | 2.24834103807158 | -4.33490549254 | 1.33616096269855 | -3.24429886335333 | 0.00117740138584055 | 0.0392895283207195 | 3831 | 0 | 0 | 10 | 14 | 0 | 0 | 1 | 3 | 0 | 0 | 2 | 0 |
| MIR_dup154632 | 2.82075666529582 | -4.43997735685834 | 1.26140012732871 | -3.51988021934083 | 0.000431741727898522 | 0.0195280259354976 | 3832 | 0 | 0 | 17 | 13 | 0 | 0 | 2 | 4 | 0 | 0 | 1 | 0 |
| AluSz6_dup28563 | 2.85449284390818 | -4.46073117540869 | 1.31148993757798 | -3.40126984401164 | 0.000670735825873469 | 0.0265026414518911 | 3833 | 0 | 0 | 18 | 15 | 0 | 0 | 4 | 1 | 0 | 0 | 1 | 0 |
| L1PA15_dup8103 | 2.29294419036863 | -4.21368611213655 | 1.33861256472952 | -3.14780110628048 | 0.00164503582712211 | 0.0495495987569662 | 3834 | 0 | 0 | 10 | 17 | 0 | 0 | 3 | 1 | 0 | 0 | 1 | 0 |
| L1MD_dup7921 | 3.24492178994137 | -4.60861246946098 | 1.28700139397029 | -3.58089159114569 | 0.00034242371960175 | 0.0164436638515354 | 3835 | 0 | 0 | 9 | 30 | 0 | 0 | 4 | 2 | 0 | 0 | 1 | 0 |
| AluSg_dup31689 | 3.47784284894656 | -4.68095915106278 | 1.18780434945703 | -3.94085032034319 | 8.1193280086508e-05 | 0.0058473414719928 | 3836 | 0 | 0 | 14 | 24 | 0 | 0 | 6 | 3 | 0 | 0 | 1 | 0 |
| L2a_dup152042 | 3.62740417405631 | -4.73099862644223 | 1.22076525456202 | -3.87543682846675 | 0.000106433590156991 | 0.00714197604159484 | 3837 | 0 | 0 | 13 | 28 | 0 | 0 | 7 | 2 | 0 | 0 | 1 | 0 |
| L1PB2_dup2771 | 4.77821129214915 | -4.46593947298177 | 1.10278364950363 | -4.04969685122909 | 5.12840232857083e-05 | 0.0041605278570546 | 3838 | 0 | 0 | 19 | 34 | 0 | 1 | 7 | 4 | 0 | 0 | 1 | 0 |
| MIR_dup154643 | 4.23801013072705 | -3.91559614028614 | 1.10282149213927 | -3.5505257815484 | 0.000384462477555676 | 0.0179586393177195 | 3839 | 0 | 0 | 15 | 32 | 0 | 1 | 4 | 4 | 1 | 0 | 1 | 0 |
| AluJb_dup110119 | 3.87038717065631 | -4.43650217466803 | 1.25404429828882 | -3.5377555487647 | 0.000403543473290904 | 0.0185944142710452 | 3840 | 0 | 0 | 8 | 37 | 0 | 0 | 6 | 2 | 1 | 0 | 1 | 0 |
| ERVL-int_dup122 | 4.01228736204923 | -5.10477817243231 | 1.3427918715913 | -3.80161533624924 | 0.000143755792205936 | 0.00883194312735186 | 3841 | 0 | 0 | 7 | 42 | 0 | 0 | 2 | 3 | 0 | 0 | 3 | 0 |
| AluSz_dup99925 | 4.1378381292739 | -3.80690494577921 | 1.09431868584791 | -3.478790040791 | 0.000503683048484191 | 0.0216921930596596 | 3842 | 1 | 0 | 18 | 30 | 1 | 0 | 3 | 3 | 0 | 0 | 1 | 0 |
| MLT2B4_dup3315 | 4.56024465546524 | -4.3944343766275 | 1.2509276740659 | -3.51294041033102 | 0.000443176858349725 | 0.0198853782268308 | 3843 | 0 | 0 | 23 | 32 | 1 | 0 | 1 | 4 | 0 | 0 | 1 | 0 |
| L2a_dup91798 | 5.51955947229598 | -5.34614884370192 | 1.25347540859231 | -4.26506081176797 | 1.99847762295674e-05 | 0.00204002469528985 | 3844 | 0 | 0 | 28 | 38 | 0 | 0 | 2 | 5 | 0 | 0 | 2 | 0 |
| L1MEd_dup5726 | 3.35182850544659 | -4.64742873924289 | 1.36735225840246 | -3.39885257122601 | 0.000676691821295978 | 0.0267025061542443 | 3845 | 0 | 0 | 18 | 24 | 0 | 0 | 1 | 2 | 0 | 0 | 1 | 0 |
| MER103C_dup2495 | 3.22483748832803 | -4.74357645838326 | 1.23927898724355 | -3.82769054200951 | 0.000129351241665714 | 0.0081813510193853 | 3846 | 0 | 0 | 14 | 22 | 0 | 0 | 3 | 3 | 0 | 0 | 2 | 0 |
| MER77_dup112 | 3.84193526622279 | -4.79628239758573 | 1.27605545303264 | -3.75867865788044 | 0.000170813048600484 | 0.00999859327485627 | 3847 | 0 | 0 | 14 | 32 | 0 | 0 | 2 | 4 | 0 | 0 | 1 | 0 |
| Zaphod2_dup678 | 5.77814408792066 | -5.41647836492231 | 1.36665606041878 | -3.96330761030143 | 7.391843655573e-05 | 0.00545213772619024 | 3848 | 0 | 0 | 29 | 47 | 0 | 0 | 2 | 1 | 0 | 0 | 2 | 0 |
| L1PA7_dup12752 | 5.18485231543313 | -5.15882799596903 | 1.41972692951934 | -3.63367622935461 | 0.000279411447918786 | 0.0142047047096345 | 3849 | 0 | 0 | 25 | 45 | 0 | 0 | 1 | 1 | 0 | 0 | 1 | 0 |
| L1PBa_dup221 | 4.3693262601544 | -5.10346694827206 | 1.40943503653867 | -3.6209309517417 | 0.000293544855980139 | 0.0147420481851098 | 3850 | 0 | 0 | 22 | 35 | 0 | 0 | 1 | 1 | 0 | 0 | 2 | 0 |
| MLT1C_dup3256 | 4.68018554918953 | -5.14790664149122 | 1.2192019650233 | -4.22235756599428 | 2.41760175358606e-05 | 0.00235262235269915 | 3851 | 0 | 0 | 21 | 36 | 0 | 0 | 6 | 1 | 0 | 0 | 1 | 1 |
| L2_dup37893 | 4.89828326897065 | -5.21771873859575 | 1.27969655837752 | -4.0773093468432 | 4.55598602221601e-05 | 0.00380645593413698 | 3852 | 0 | 0 | 27 | 32 | 0 | 0 | 6 | 1 | 0 | 0 | 2 | 0 |
| L2a_dup91799 | 6.35998401724355 | -5.51284747194267 | 1.29161252890882 | -4.26818983909984 | 1.97065588943615e-05 | 0.00202203400719178 | 3853 | 0 | 0 | 33 | 47 | 0 | 0 | 6 | 1 | 0 | 0 | 2 | 0 |
| LTR46_dup19 | 13.2831445357937 | -3.26360038966146 | 0.98214061514104 | -3.32294616407121 | 0.00089072103031106 | 0.0324376384242147 | 3854 | 0 | 3 | 69 | 87 | 2 | 1 | 1 | 3 | 4 | 2 | 5 | 4 |
| MER61-int_dup746 | 19.3362145890489 | -4.58538682379271 | 1.39979297898735 | -3.27576069649236 | 0.00105377808553308 | 0.0364239221877184 | 3855 | 0 | 2 | 115 | 137 | 0 | 1 | 1 | 2 | 2 | 2 | 3 | 2 |
| MSTA-int_dup242 | 12.0048798653045 | -3.04635310540671 | 0.924100334368491 | -3.29656098164765 | 0.000978763638097028 | 0.0346180581465968 | 3856 | 2 | 0 | 65 | 66 | 1 | 3 | 4 | 2 | 2 | 7 | 6 | 5 |
| MSTA-int_dup250 | 15.9951473824906 | -3.14546167183674 | 0.956977987962243 | -3.28686940703263 | 0.00101307803047654 | 0.0353924335691925 | 3857 | 2 | 1 | 82 | 106 | 2 | 2 | 3 | 1 | 5 | 6 | 6 | 3 |
| L1PB1_dup1301 | 17.4224861342159 | -3.60396058415415 | 1.1457007904844 | -3.14563855946228 | 0.00165724673227257 | 0.0497898018885517 | 3858 | 2 | 2 | 88 | 130 | 1 | 1 | 6 | 1 | 2 | 4 | 1 | 5 |
| MER52-int_dup239 | 19.8648203762566 | -2.73880762568896 | 0.85703197900964 | -3.19568895066651 | 0.00139497409706957 | 0.0442040892930238 | 3859 | 2 | 6 | 88 | 143 | 3 | 3 | 3 | 5 | 4 | 6 | 2 | 9 |
| HERVL-int_dup1187 | 9.46304404630453 | -3.10487953463205 | 0.813891923992586 | -3.81485482667148 | 0.000136263015996029 | 0.00848775905868878 | 3860 | 2 | 1 | 42 | 53 | 4 | 0 | 5 | 3 | 1 | 3 | 8 | 7 |
| L1PA6_dup31 | 9.13790623120594 | -2.76024522939921 | 0.838252023082183 | -3.29285841655356 | 0.000991744023502553 | 0.0349136741258083 | 3861 | 2 | 0 | 34 | 54 | 6 | 0 | 5 | 4 | 2 | 3 | 9 | 5 |
| SATR1_dup143 | 12.1265521765743 | -3.72485436489735 | 0.770966238113877 | -4.83141048304525 | 1.3556917041301e-06 | 0.000261411197918104 | 3862 | 3 | 2 | 43 | 81 | 3 | 2 | 3 | 10 | 0 | 1 | 7 | 11 |
| AluSz_dup96925 | 10.7031806833154 | -2.61641547144862 | 0.698472410068001 | -3.74591098192969 | 0.000179740323780513 | 0.0103380447382833 | 3863 | 2 | 2 | 58 | 46 | 4 | 5 | 7 | 8 | 2 | 1 | 3 | 4 |
| ERVL-B4-int_dup941 | 13.8562232737296 | -3.21703678592244 | 0.86364310181146 | -3.72496090013899 | 0.000195345497615791 | 0.010934293054737 | 3864 | 1 | 0 | 71 | 83 | 7 | 5 | 3 | 5 | 1 | 1 | 5 | 7 |
| L1MC3_dup3116 | 20.6656110165006 | -3.13506610943066 | 0.757156631214077 | -4.1405780259808 | 3.46431741820831e-05 | 0.00307238342300781 | 3865 | 4 | 4 | 95 | 133 | 8 | 6 | 6 | 2 | 2 | 1 | 9 | 16 |
| L1MA1_dup1056 | 13.3357075466083 | -3.90096662235147 | 0.980569373698147 | -3.97826683862179 | 6.94194398897771e-05 | 0.00521403978662023 | 3866 | 1 | 0 | 59 | 106 | 3 | 6 | 3 | 3 | 0 | 0 | 1 | 5 |

| | | | | | | | | | | | | | | | | | | | |
|---|---|---|---|---|---|---|---|---|---|---|---|---|---|---|---|---|---|---|---|
| L1PA5_dup9437 | 11.2166057128891 | -3.76835778930929 | 1.00002616574196 | -3.76825919001169 | 0.000164389965506525 | 0.00971341811085561 | 3867 | 1 | 0 | 55 | 61 | 4 | 7 | 3 | 1 | 0 | 0 | 6 | 17 |
| L1MB7_dup16447 | 14.8600701174345 | -6.72921756723603 | 1.12408791332365 | -5.98638014649527 | 2.14562562388944e-09 | 1.3761267781725e-06 | 3868 | 0 | 0 | 89 | 74 | 0 | 1 | 5 | 3 | 0 | 0 | 13 | 18 |
| L1PB1_dup2802 | 13.5061221489542 | -7.21152995263418 | 1.4879878141941 | -4.84649799134274 | 1.25659859746396e-06 | 0.000245486543408998 | 3869 | 0 | 0 | 49 | 83 | 0 | 0 | 3 | 1 | 0 | 0 | 16 | 38 |
| L1PREC2_dup1641 | 7.83127566824003 | -6.3922081026364 | 1.69883576066725 | -3.7626992853775 | 0.000168089270313999 | 0.00988047665863016 | 3870 | 0 | 0 | 28 | 55 | 0 | 0 | 0 | 3 | 0 | 0 | 10 | 13 |
| HERVL-int_dup1947 | 6.59429727535009 | -6.14170236985709 | 1.79669589334157 | -3.41833161227663 | 0.000630062906475345 | 0.025402619872667 | 3871 | 0 | 0 | 33 | 38 | 0 | 0 | 0 | 1 | 0 | 0 | 8 | 11 |
| L1M4_dup15152 | 7.01010786095469 | -6.15756128472579 | 1.83831637412505 | -3.34956559784574 | 0.000809383877898627 | 0.0302861544598941 | 3872 | 0 | 0 | 32 | 51 | 0 | 0 | 0 | 1 | 0 | 0 | 6 | 8 |
| L1PA16_dup2974 | 8.14487725933954 | -6.45707123381858 | 1.84292947222678 | -3.50369958868616 | 0.000458842664046339 | 0.0203482253063127 | 3873 | 0 | 0 | 29 | 60 | 0 | 0 | 1 | 0 | 0 | 0 | 11 | 14 |
| L1PB3_dup2790 | 11.8950264195159 | -6.41470289916327 | 1.71979560706314 | -3.72992166791119 | 0.000191539308692148 | 0.0107642920042659 | 3874 | 1 | 0 | 60 | 79 | 0 | 0 | 1 | 0 | 0 | 0 | 16 | 8 |
| L1PA11_dup1046 | 6.56532986162061 | -3.58128480681013 | 1.07969066727907 | -3.31695449015534 | 0.000910044388930201 | 0.032945670034383 3 | 3875 | 1 | 2 | 36 | 37 | 2 | 0 | 1 | 0 | 0 | 1 | 6 | 4 |
| MIRb_dup138700 | 7.32479562595391 | -4.21485339771273 | 1.24645067172124 | -3.38148431649715 | 0.0007209534826698 | 0.0279394132552685 | 3876 | 1 | 2 | 44 | 47 | 1 | 0 | 0 | 0 | 0 | 0 | 4 | 2 |
| L1PA3_dup10554 | 11.0814302416814 | -4.51021996423618 | 1.43350055395725 | -3.14629802673287 | 0.00165351421491312 | 0.0497029240600507 | 3877 | 4 | 1 | 63 | 61 | 1 | 1 | 0 | 0 | 0 | 0 | 13 | 8 |
| MIR3_dup55382 | 9.45396150804953 | -3.41352097366632 | 1.04697869974954 | -3.26035379180389 | 0.00111273319839579 | 0.0377634132088756 | 3878 | 6 | 4 | 49 | 59 | 1 | 1 | 1 | 0 | 0 | 0 | 5 | 5 |
| L1PA3_dup2241 | 10.1950852103985 | -4.85390612912575 | 1.53607604364378 | -3.15993869522997 | 0.00157802338326466 | 0.0480650732148352 | 3879 | 1 | 0 | 66 | 63 | 2 | 0 | 0 | 2 | 0 | 0 | 1 | 5 |
| L1M2_dup2263 | 8.08109698183726 | -3.80173329494415 | 1.02575123331184 | -3.70629171233799 | 0.000210316047326457 | 0.0115335013130079 | 3880 | 3 | 1 | 45 | 49 | 1 | 2 | 0 | 2 | 0 | 0 | 3 | 5 |
| L1PA7_dup13343 | 9.86061809570441 | -4.23702516638352 | 1.09720052983519 | -3.86166890296706 | 0.000112615116589207 | 0.00741804303051745 | 3881 | 1 | 0 | 44 | 77 | 3 | 2 | 0 | 2 | 0 | 0 | 3 | 6 |
| MLT2F_dup1322 | 13.6990958637803 | -3.22738994587254 | 0.941157346645206 | -3.42917149547491 | 0.000605426844834681 | 0.0247268959871324 | 3882 | 5 | 6 | 75 | 79 | 4 | 1 | 0 | 3 | 1 | 0 | 5 | 8 |
| Harlequin-int_dup476 | 15.3809453994437 | -5.10540134029295 | 1.4184704185381 | -3.59923003932269 | 0.000319160768212909 | 0.0156236295198325 | 3883 | 3 | 2 | 90 | 90 | 0 | 1 | 0 | 1 | 0 | 0 | 10 | 15 |
| L1M1_dup9296 | 10.4383819952188 | -4.18706668094173 | 1.11386112593417 | -3.75905629836048 | 0.00017055545986171 | 0.00998842959691227 | 3884 | 4 | 3 | 61 | 54 | 1 | 0 | 0 | 1 | 0 | 0 | 7 | 12 |
| L1PA16_dup9281 | 17.5748094530924 | -6.50881794017006 | 1.66507907216997 | -3.90901432187698 | 9.26734703951538e-05 | 0.00645718348071604 | 3885 | 0 | 0 | 86 | 142 | 0 | 0 | 1 | 1 | 1 | 3 | 3 | 10 |
| LTR12C_dup1783 | 18.1447769093417 | -6.74739912892967 | 2.09152644122412 | -3.22606446465988 | 0.00125505085519779 | 0.041100238204086 5 | 3886 | 0 | 0 | 114 | 124 | 0 | 0 | 0 | 0 | 1 | 2 | 5 | 5 |
| L1MB7_dup4899 | 14.5781885373557 | -5.15349465689537 | 1.3242024982841 | -3.89177234114363 | 9.95146012907241e-05 | 0.00679262803176209 | 3887 | 0 | 0 | 78 | 101 | 2 | 0 | 2 | 1 | 1 | 1 | 9 | 7 |
| MLT1E2_dup2963 | 10.4181149463412 | -5.68625948602661 | 1.2599901166562 | -4.51293975314425 | 6.39351880920268e-06 | 0.000870814096493727 | 3888 | 0 | 0 | 56 | 68 | 1 | 0 | 2 | 0 | 1 | 0 | 10 | 6 |
| L1PA6_dup1301 | 16.8814609062853 | -4.9201568134953 | 1.43649931641489 | -3.42510209178148 | 0.000614568436521792 | 0.0249716702863071 | 3889 | 1 | 1 | 82 | 137 | 0 | 1 | 1 | 0 | 2 | 0 | 6 | 6 |
| L1PA6_dup1302 | 15.0714865809906 | -6.7365677655288 | 1.80053806382218 | -3.74141924621601 | 0.000182983975679953 | 0.0104538547604183 | 3890 | 0 | 0 | 80 | 115 | 0 | 0 | 1 | 0 | 1 | 0 | 6 | 8 |
| L1PA7_dup6499 | 14.9676785194342 | -6.8687045379484 | 2.06442746778436 | -3.3271716469265 | 0.000877323148984613 | 0.0321280058024636 | 3891 | 0 | 0 | 90 | 112 | 0 | 0 | 1 | 0 | 0 | 0 | 3 | 3 |
| LTR108d_Mam_dup59 | 13.7824570862335 | -6.48729202411952 | 2.00127956538008 | -3.24157211033505 | 0.00118872356001717 | 0.03956525614937 | 3892 | 0 | 0 | 90 | 96 | 0 | 0 | 3 | 1 | 0 | 0 | 1 | 0 |
| L1MC1_dup11731 | 14.769734291449 | -6.03485339216591 | 1.63160704571568 | -3.69871741361525 | 0.000216691691001565 | 0.0117851716192173 | 3893 | 0 | 0 | 93 | 106 | 0 | 1 | 1 | 1 | 0 | 0 | 2 | 1 |
| L1ME1_dup27363 | 9.89921614194588 | -5.52130768715555 | 1.17961781394862 | -4.68059029108224 | 2.8605009384422e-06 | 0.000463970522357579 | 3894 | 0 | 0 | 59 | 68 | 0 | 1 | 4 | 2 | 0 | 0 | 2 | 1 |
| L1ME3A_dup13689 | 11.2981177257458 | -5.64641717217504 | 1.69467198268022 | -3.3318643548027 | 0.000862662925956956 | 0.0317112919171565 | 3895 | 0 | 0 | 74 | 75 | 0 | 1 | 3 | 1 | 0 | 0 | 2 | 0 |
| L1M3d_dup375 | 13.1264888317985 | -4.56922635604843 | 1.33487414429149 | -3.42296416151925 | 0.000619422434721162 | 0.02509604812739 | 3896 | 0 | 0 | 62 | 113 | 2 | 1 | 2 | 1 | 1 | 0 | 1 | 2 |
| MLT1A1_dup6012 | 15.1072814496359 | -5.54806946321313 | 1.43566586191318 | -3.86445732979939 | 0.000111336450145585 | 0.00735484190475015 | 3897 | 0 | 0 | 90 | 107 | 2 | 0 | 3 | 3 | 0 | 0 | 3 | 1 |
| L1MA8_dup1067 | 6.59107054505911 | -4.20208526017348 | 1.25369566423001 | -3.35175862856181 | 0.000802999959020265 | 0.0301325702963312 | 3898 | 0 | 1 | 41 | 42 | 0 | 0 | 1 | 1 | 1 | 1 | 2 | 0 |
| L1PA7_dup2510 | 9.6254676953138 | -3.86784606349467 | 1.1921161640684 | -3.24452111302196 | 0.00117648295441417 | 0.0392817819048018 | 3899 | 2 | 1 | 57 | 68 | 0 | 0 | 0 | 1 | 1 | 1 | 1 | 1 |
| Tigger7_dup2752 | 7.59095355102703 | -5.45232263800116 | 1.20895717422092 | -4.50993861012053 | 6.48463886645592e-06 | 0.000877134904411177 | 3900 | 1 | 0 | 35 | 60 | 0 | 0 | 1 | 3 | 0 | 0 | 2 | 4 |
| AluSc_dup29728 | 5.30987604474085 | -5.29470267457962 | 1.29491260343446 | -4.08884944090948 | 4.33518105421763e-05 | 0.00366313562189939 | 3901 | 0 | 0 | 32 | 36 | 0 | 0 | 1 | 2 | 0 | 0 | 1 | 1 |
| L1P3_dup3340 | 6.62104534250527 | -5.54145063635675 | 1.28016474836414 | -4.32870116400088 | 1.49991292204209e-05 | 0.00165001516480919 | 3902 | 0 | 0 | 34 | 53 | 0 | 0 | 3 | 1 | 0 | 0 | 1 | 1 |
| MLT1L_dup10355 | 7.72385137380103 | -5.71947621301916 | 1.30999809032575 | -4.36601874098681 | 1.26531628189299e-05 | 0.00145319890648804 | 3903 | 0 | 0 | 54 | 47 | 0 | 0 | 2 | 1 | 0 | 0 | 1 | 1 |
| L1ME3_dup5374 | 7.42459167500078 | -5.25490578591133 | 1.27877301703444 | -4.10933427270603 | 3.96801358414602e-05 | 0.00341720217261385 | 3904 | 0 | 0 | 38 | 53 | 0 | 0 | 2 | 1 | 0 | 2 | 5 | 2 |
| MER11A_dup101 | 5.4747285107245 | -5.01445075063976 | 1.40595682381483 | -3.56657520750451 | 0.000361677045847135 | 0.0171501806199965 | 3905 | 0 | 0 | 31 | 41 | 0 | 0 | 0 | 1 | 0 | 1 | 1 | 1 |
| L2a_dup164505 | 6.46118627275987 | -3.68132835416507 | 1.14398270355252 | -3.21799301924155 | 0.00129090959629634 | 0.0418825708754197 | 3906 | 0 | 1 | 42 | 34 | 0 | 2 | 7 | 0 | 0 | 1 | 0 | 2 |
| L2c_dup67989 | 6.31830815148788 | -3.76746436056923 | 1.17499001081559 | -3.20637990611863 | 0.00134416398685939 | 0.0430861676902808 | 3907 | 3 | 1 | 37 | 42 | 0 | 0 | 3 | 0 | 0 | 0 | 1 | 1 |
| MSTA-int_dup249 | 5.11417093522147 | -3.89951646506425 | 1.23308644702634 | -3.16240315062108 | 0.00156472774213921 | 0.0477763001836334 | 3908 | 1 | 1 | 33 | 28 | 0 | 0 | 3 | 0 | 0 | 1 | 3 | 0 |
| L1PA3_dup2349 | 7.98276269412195 | -3.30837860882639 | 0.954912050764723 | -3.46458986058134 | 0.000531040689942079 | 0.0224598307872303 | 3909 | 2 | 1 | 38 | 52 | 0 | 3 | 2 | 2 | 0 | 2 | 5 | 3 |
| LTR12D_dup181 | 10.4254694238396 | -3.55029583415911 | 0.940670801728581 | -3.77421710935969 | 0.000160510931291586 | 0.00956137837151469 | 3910 | 1 | 4 | 62 | 65 | 1 | 1 | 3 | 2 | 0 | 1 | 2 | 1 |
| MLT1G_dup341 | 6.24267112807569 | -3.42338248460367 | 1.02606015264201 | -3.33643449245036 | 0.00084860424783453 | 0.031359215150681 | 3911 | 2 | 2 | 34 | 31 | 0 | 0 | 4 | 0 | 1 | 1 | 7 | 3 |
| MLT1H-int_dup72 | 8.53770850784196 | -3.55776437686742 | 0.978199627187064 | -3.63705349908813 | 0.000275774668569576 | 0.0140885772047527 | 3912 | 2 | 1 | 48 | 49 | 0 | 2 | 6 | 0 | 1 | 1 | 2 | 6 |
| L1MA3_dup2580 | 10.6546919597818 | -3.26550963080463 | 1.02055939500846 | -3.19972521616693 | 0.00137558666981547 | 0.043805586956091 | 3913 | 5 | 3 | 60 | 70 | 1 | 2 | 5 | 0 | 0 | 0 | 2 | 0 |
| L1MC1_dup1603 | 17.2434274659961 | -4.26995059661861 | 1.27279229793906 | -3.3547897826948 | 0.000794253164015397 | 0.0299175816629425 | 3914 | 4 | 1 | 91 | 138 | 0 | 1 | 2 | 1 | 0 | 1 | 2 | 1 |
| L1ME3D_dup567 | 13.8072642557777 | -4.04954983013821 | 0.91663367053477 | -4.41784974773584 | 9.96876786826404e-06 | 0.0012201792382623 2 | 3915 | 3 | 1 | 83 | 88 | 1 | 2 | 4 | 2 | 1 | 0 | 3 | 2 |
| L2b_dup70616 | 7.50474221172292 | -3.8335316355808 | 1.07545301300265 | -3.56457380213909 | 0.000364447897484023 | 0.0172471995653805 | 3916 | 2 | 4 | 42 | 45 | 0 | 0 | 1 | 2 | 0 | 0 | 2 | 5 |
| L2_dup50633 | 11.2793754707218 | -4.78957493420542 | 1.41034808595573 | -3.39602328099005 | 0.000683725438360866 | 0.0268908649844983 | 3917 | 2 | 1 | 67 | 82 | 0 | 0 | 1 | 1 | 0 | 0 | 1 | 2 |
| MLT2A2_dup430 | 8.19541141498547 | -4.0777364626314 | 1.06943049290561 | -3.81299812347065 | 0.000137291155370189 | 0.00853170051076391 | 3918 | 2 | 1 | 54 | 49 | 1 | 0 | 1 | 2 | 0 | 0 | 1 | 1 |
| L1PA7_dup7427 | 6.57826989616564 | -3.37323362978816 | 0.98338578474442 | -3.43022411155237 | 0.000603082913729303 | 0.0246779112658 3 | 3919 | 2 | 3 | 24 | 53 | 0 | 1 | 1 | 5 | 0 | 0 | 1 | 1 |
| L1PA16_dup2992 | 10.2636136957239 | -3.55976796702345 | 1.04788225056306 | -3.39710684584139 | 0.000681023704546006 | 0.0268202241551396 | 3920 | 2 | 3 | 54 | 76 | 0 | 1 | 2 | 4 | 0 | 0 | 0 | 0 |
| L1MA3_dup2026 | 10.4852069611417 | -3.44090167367325 | 1.08080250359399 | -3.18365442551367 | 0.00145428517869875 | 0.045522272517261 | 3921 | 3 | 4 | 55 | 79 | 0 | 1 | 1 | 2 | 0 | 0 | 0 | 1 |
| Tigger17a_dup481 | 8.60630277579723 | -3.51849236980455 | 1.08240510693127 | -3.25062432473165 | 0.00115151908381722 | 0.0387442533820661 | 3922 | 1 | 3 | 43 | 67 | 0 | 2 | 1 | 2 | 0 | 0 | 0 | 1 |
| HERV17-int_dup249 | 7.74544108432621 | -3.61428496784492 | 1.1233771834809 | -3.21733877186796 | 0.00129385724391098 | 0.0419312287919356 | 3923 | 2 | 2 | 52 | 46 | 0 | 1 | 1 | 1 | 0 | 0 | 1 | 0 |
| L1P1_dup857 | 9.23855840980573 | -4.50472498772445 | 1.06681634467773 | -4.22258714932349 | 2.4151396686683e-05 | 0.00235136179869455 | 3924 | 1 | 1 | 48 | 72 | 0 | 1 | 2 | 2 | 0 | 0 | 1 | 1 |
| HAL1b_dup1102 | 2.87331001148778 | -4.42268732953528 | 1.19169563113394 | -3.71125580558431 | 0.000206233560672784 | 0.0113751624542991 | 3925 | 0 | 0 | 11 | 7 | 0 | 1 | 2 | 0 | 0 | 0 | 9 | 9 |
| AluSx_dup86069 | 2.21726855787066 | -4.09233166019698 | 1.23366865457747 | -3.3172048629205 | 0.000909229211308517 | 0.0329245858946913 | 3926 | 0 | 0 | 7 | 11 | 0 | 1 | 2 | 0 | 0 | 0 | 4 | 6 |
| MLT1F_dup3272 | 2.35370327276828 | -4.17727216161044 | 1.22394383209817 | -3.41296067030255 | 0.00064261215484851 | 0.0257375848842928 | 3927 | 0 | 0 | 6 | 12 | 0 | 1 | 2 | 0 | 0 | 0 | 5 | 7 |
| AluSx_dup96731 | 3.59842625354038 | -3.8006180396839 | 1.17398011260418 | -3.2373785542697 | 0.00120633272925493 | 0.0400058602920271 | 3928 | 0 | 0 | 15 | 15 | 0 | 3 | 6 | 0 | 0 | 0 | 4 | 7 |
| MIR1_Amn_dup4837 | 2.86471934868839 | -3.85533474408738 | 1.22198513039066 | -3.15497680635023 | 0.00160510864938525 | 0.0486403851240753 | 3929 | 0 | 1 | 16 | 12 | 0 | 1 | 2 | 0 | 0 | 0 | 4 | 3 |
| L1MB7_dup20750 | 1.97442214039521 | -3.90397788188939 | 1.11254562891183 | -3.50904967889526 | 0.000449710896414745 | 0.0200914211337407 | 3930 | 0 | 0 | 6 | 8 | 0 | 1 | 4 | 1 | 0 | 0 | 4 | 3 |
| MIR3_dup3666 | 2.47894823935837 | -4.24381908683295 | 1.07756371837955 | -3.93834630328388 | 8.20451227441598e-05 | 0.00588908258717401 | 3931 | 0 | 0 | 10 | 3 | 0 | 1 | 7 | 2 | 0 | 0 | 5 | 5 |
| L1PA15_dup3037 | 3.11908219629196 | -3.5869831078418 | 1.06677031918651 | -3.36246991815178 | 0.00077248542219989 | 0.0293204364764356 | 3932 | 0 | 0 | 6 | 13 | 0 | 3 | 4 | 2 | 0 | 0 | 5 | 10 |
| L1MCa_dup6159 | 3.02514318118138 | -3.97887761991035 | 1.03084804716266 | -3.85981001842313 | 0.000113475211905449 | 0.00744719955235963 | 3933 | 0 | 1 | 7 | 8 | 0 | 1 | 4 | 1 | 0 | 0 | 10 | 9 |
| MLT1L_dup6323 | 2.93686603906168 | -3.34265696044849 | 1.01175861627677 | -3.30380874120877 | 0.000953808795115034 | 0.0339956542184604 | 3934 | 0 | 1 | 4 | 7 | 0 | 2 | 4 | 1 | 0 | 0 | 10 | 11 |
| L1MEc_dup14916 | 1.9480252225774 | -3.85772970961 | 1.21971093049855 | -3.16282293873776 | 0.00156247330986503 | 0.0477290937039548 | 3935 | 0 | 0 | 6 | 12 | 1 | 0 | 1 | 1 | 0 | 0 | 2 | 4 |
| L1MA9_dup15684 | 2.06849891574414 | -4.49751680618186 | 1.25184821002655 | -3.59270139155806 | 0.000327267544116617 | 0.0158987348004027 | 3936 | 0 | 0 | 7 | 12 | 0 | 0 | 2 | 1 | 0 | 0 | 1 | 6 |
| MLT1A0_dup5527 | 1.5821777714141 | -4.13283363545763 | 1.30622759364462 | -3.1639460501108 | 0.00155645644392763 | 0.0476399003095966 | 3937 | 0 | 0 | 5 | 7 | 0 | 0 | 1 | 1 | 0 | 0 | 1 | 7 |
| MIRb_dup57579 | 2.36318281594563 | -3.67042576855512 | 1.16046840428084 | -3.16288298329823 | 0.00156215109085737 | 0.0477290937039548 | 3938 | 2 | 0 | 6 | 11 | 0 | 0 | 2 | 1 | 0 | 0 | 3 | 8 |
| MamGypsy2-I_dup276 | 2.70129239954977 | -3.86380073995201 | 1.15086206149967 | -3.35731002802991 | 0.000787048076248135 | 0.0297261811985617 | 3939 | 2 | 0 | 11 | 10 | 0 | 0 | 2 | 1 | 0 | 0 | 4 | 7 |
| L2b_dup42967 | 2.41874982847151 | -4.18958888029537 | 1.25750010492782 | -3.33168074012672 | 0.000863232249674503 | 0.0317242822802378 | 3940 | 1 | 0 | 6 | 11 | 0 | 0 | 1 | 0 | 0 | 0 | 5 | 10 |
| MIRb_dup57566 | 2.32448838502595 | -4.15465170465959 | 1.24312556319927 | -3.34210141569875 | 0.000831466755326543 | 0.0309034971201077 | 3941 | 1 | 0 | 11 | 9 | 0 | 0 | 2 | 0 | 0 | 0 | 4 | 5 |
| AluSx1_dup85188 | 2.44563035083531 | -4.73940889149537 | 1.19229862286527 | -3.97501833903477 | 7.03738331447106e-05 | 0.00526757127197664 | 3942 | 0 | 0 | 5 | 6 | 0 | 0 | 9 | 0 | 0 | 0 | 8 | 6 |
| MIR_dup139104 | 2.07552359085729 | -4.52777382148183 | 1.231114296786 | -3.6777851035458 | 0.00023526804036624 | 0.01254618761384 7 | 3943 | 0 | 0 | 7 | 8 | 0 | 0 | 5 | 0 | 0 | 0 | 4 | 5 |
| MLT1G_dup684 | 2.00297379423751 | -4.45128734359531 | 1.22470203245808 | -3.63458802682085 | 0.000278425183910238 | 0.0141795725386253 | 3944 | 0 | 0 | 6 | 5 | 0 | 0 | 5 | 0 | 0 | 0 | 4 | 8 |
| L1MD3_dup4046 | 2.93728270992953 | -4.96286587560117 | 1.22458027950131 | -4.05270765720825 | 5.06282514058174e-05 | 0.00411996918912744 | 3945 | 0 | 0 | 10 | 8 | 0 | 0 | 3 | 0 | 0 | 0 | 5 | 15 |

| | | | | | | | | | | | | | | | | | | | |
|---|---|---|---|---|---|---|---|---|---|---|---|---|---|---|---|---|---|---|---|
| L1MCa_dup6175 | 2.69288554552757 | -4.84858975441599 | 1.20404435550527 | -4.02691955013676 | 5.65123574798433e-05 | 0.00446315851674773 | 3946 | 0 | 0 | 8 | 9 | 0 | 0 | 3 | 0 | 0 | 0 | 9 | 8 |
| MER9a3_dup81 | 3.18828160683062 | -5.09629717616409 | 1.1560800873996 | -4.40825616815813 | 1.04206249455697e-05 | 0.00126121364359029 | 3947 | 0 | 0 | 9 | 10 | 0 | 0 | 5 | 0 | 0 | 0 | 10 | 10 |
| L1MC2_dup4203 | 1.97752604117142 | -3.8157426606046 | 1.18294513980126 | -3.22562943303157 | 0.00125695986215666 | 0.0411457657670559 | 3948 | 0 | 1 | 5 | 5 | 0 | 0 | 2 | 0 | 0 | 0 | 7 | 7 |
| L1MA4_dup2497 | 2.62664245733438 | -4.30031425866237 | 1.15762863768334 | -3.71476146898732 | 0.000203395463893183 | 0.011255148151518 | 3949 | 0 | 1 | 8 | 8 | 0 | 0 | 3 | 0 | 0 | 0 | 8 | 8 |
| MIR_dup104342 | 2.55444461212426 | -4.17258233214164 | 1.1303574667232 | -3.69138299606898 | 0.000223037992996675 | 0.0120356100644547 | 3950 | 0 | 1 | 6 | 6 | 0 | 0 | 3 | 0 | 0 | 0 | 9 | 10 |
| L1MCc_dup2762 | 2.36891978079017 | -4.5904340783696 | 1.19461276215955 | -3.84261262207786 | 0.000121731524304988 | 0.00783068194619217 | 3951 | 0 | 0 | 4 | 8 | 0 | 0 | 3 | 0 | 0 | 0 | 8 | 10 |
| L1ME3E_dup1416 | 1.89669280578697 | -4.28359496286523 | 1.25442049053961 | -3.41479989777795 | 0.000638288842895428 | 0.0256291388814707 | 3952 | 0 | 0 | 6 | 4 | 0 | 0 | 2 | 0 | 0 | 0 | 6 | 8 |
| MER68_dup185 | 1.87335347516957 | -4.33122478459656 | 1.37707924344596 | -3.14522552366576 | 0.00165958841927335 | 0.0498412755285139 | 3953 | 0 | 0 | 8 | 5 | 0 | 0 | 1 | 0 | 0 | 0 | 8 | 3 |
| L1MEc_dup6955 | 2.13824201328164 | -4.56707404232506 | 1.31234321712206 | -3.48009117031181 | 0.0005012431803478 | 0.0216105834011665 | 3954 | 0 | 0 | 8 | 11 | 0 | 0 | 2 | 0 | 0 | 0 | 3 | 6 |
| MSTA_dup4300 | 2.27183720895348 | -4.65301110028851 | 1.27468304857873 | -3.65032790345538 | 0.000261905742641409 | 0.0135834810402734 | 3955 | 0 | 0 | 7 | 13 | 0 | 0 | 3 | 0 | 0 | 0 | 4 | 5 |
| MIR_dup170851 | 1.72583725935164 | -4.25801745782897 | 1.33310899018565 | -3.19405051588166 | 0.00140291565950766 | 0.0444057698186863 | 3956 | 0 | 0 | 6 | 9 | 0 | 0 | 2 | 0 | 0 | 0 | 4 | 3 |
| MLT1D_dup1852 | 1.92316551981842 | -4.40591208147665 | 1.3976528708267 | -3.15236506391647 | 0.00161953655012052 | 0.0489473191864573 | 3957 | 0 | 0 | 6 | 13 | 0 | 0 | 1 | 0 | 0 | 0 | 4 | 3 |
| L1PA3_dup4149 | 2.70805180304082 | -4.38146014414376 | 1.21895616423994 | -3.59443618456596 | 0.000325094812456405 | 0.0158335192013821 | 3958 | 0 | 0 | 14 | 7 | 1 | 0 | 3 | 0 | 0 | 0 | 4 | 8 |
| L1M3c_dup1013 | 2.14746814233196 | -3.98557491414291 | 1.24382685436386 | -3.20428434243871 | 0.00135398683570346 | 0.0433148361936403 | 3959 | 0 | 0 | 9 | 11 | 1 | 0 | 3 | 0 | 0 | 0 | 2 | 4 |
| L1PA15_dup7203 | 2.12610554205265 | -4.02062727202652 | 1.23239534100294 | -3.262449263038 | 0.00110453942774563 | 0.0375780295502375 | 3960 | 0 | 0 | 8 | 8 | 1 | 0 | 2 | 0 | 0 | 0 | 6 | 4 |
| L1MCc_dup2761 | 2.77483171551522 | -4.41364022309389 | 1.17286921699559 | -3.76311370367433 | 0.000167810856869365 | 0.00987006263192797 | 3961 | 0 | 0 | 5 | 14 | 1 | 0 | 4 | 0 | 0 | 0 | 7 | 8 |
| LTR27D_dup229 | 3.25855726265527 | -4.57174898405856 | 1.17304401632914 | -3.89733796892392 | 9.72558501426411e-05 | 0.00667675721813067 | 3962 | 0 | 0 | 10 | 9 | 1 | 0 | 2 | 0 | 0 | 0 | 8 | 15 |
| L1PB2_dup1301 | 3.5243384724641 | -4.12439607040965 | 1.24124902634619 | -3.32277889679435 | 0.000891255272478493 | 0.0324465463685997 | 3963 | 0 | 2 | 18 | 13 | 0 | 0 | 8 | 0 | 0 | 0 | 1 | 7 |
| MamGypsy2-I_dup196 | 4.03285200227699 | -4.97084008794885 | 1.15432229843165 | -4.30628438409497 | 1.66019596835799e-05 | 0.00178105045034676 | 3964 | 0 | 1 | 13 | 18 | 0 | 0 | 8 | 0 | 0 | 0 | 4 | 13 |
| THE1C_dup9204 | 4.77825230467763 | -3.36383256783822 | 1.05821392856848 | -3.17878311466633 | 0.00147894707964172 | 0.0460554301547478 | 3965 | 2 | 4 | 18 | 19 | 0 | 0 | 5 | 0 | 0 | 0 | 7 | 11 |
| L1M5_dup5785 | 3.85630691550747 | -3.49363401034941 | 1.1005061461744 | -3.17457019435471 | 0.00150058591067811 | 0.0465164300287113 | 3966 | 1 | 1 | 13 | 17 | 0 | 0 | 3 | 0 | 0 | 2 | 3 | 14 |
| L1ME1_dup3532 | 4.07225729946483 | -3.54954835592791 | 1.03599980704957 | -3.42620561488006 | 0.000612076848696165 | 0.0249172348742271 | 3967 | 1 | 2 | 17 | 14 | 0 | 0 | 5 | 0 | 0 | 1 | 6 | 10 |
| MIR_dup79322 | 3.57650048362901 | -3.59517619106071 | 1.10960687211057 | -3.24004499379348 | 0.00119510839827829 | 0.0397222046286809 | 3968 | 1 | 1 | 16 | 22 | 0 | 0 | 4 | 0 | 0 | 1 | 2 | 3 |
| L1MDb_dup505 | 3.24889803861702 | -4.08418221522148 | 1.28445198820842 | -3.17970796317438 | 0.00147423543125578 | 0.0459450318658827 | 3969 | 0 | 1 | 11 | 24 | 0 | 0 | 1 | 0 | 0 | 1 | 2 | 6 |
| MLT1L_dup1621 | 2.71106056756059 | -3.85575630182983 | 1.2116499548035 | -3.18223616197398 | 0.00146142599847299 | 0.0456523962667711 | 3970 | 0 | 1 | 9 | 16 | 0 | 0 | 2 | 0 | 0 | 1 | 3 | 6 |
| Ricksha_0_dup268 | 4.29578408737965 | -5.04238050131087 | 1.18123827059938 | -4.26872429281542 | 1.96594083100235e-05 | 0.00201806625096499 | 3971 | 0 | 0 | 19 | 22 | 0 | 0 | 6 | 0 | 0 | 1 | 4 | 8 |
| MLT1I_dup7084 | 2.32996431224036 | -4.11810240137356 | 1.25865516385236 | -3.27182735958377 | 0.00106854803497376 | 0.0367813476597136 | 3972 | 0 | 0 | 12 | 11 | 0 | 0 | 3 | 0 | 0 | 1 | 3 | 2 |
| L1MA8_dup10706 | 2.50321948566119 | -4.28354393991517 | 1.21535796663242 | -3.52451216639015 | 0.00042426333122286 | 0.019304049155399 | 3973 | 0 | 0 | 9 | 11 | 0 | 0 | 3 | 0 | 0 | 1 | 3 | 8 |
| L2a_dup42730 | 2.47673833394825 | -4.25269475223711 | 1.18598289185781 | -3.58579772223812 | 0.000336049370378267 | 0.0162226827924583 | 3974 | 0 | 0 | 9 | 9 | 0 | 0 | 3 | 0 | 0 | 1 | 6 | 6 |
| L1MD2_dup843 | 2.63886878311448 | -4.812971481744 | 1.35845372604728 | -3.54297786480252 | 0.000395635908949611 | 0.0183223008935236 | 3975 | 0 | 0 | 6 | 23 | 0 | 0 | 2 | 0 | 0 | 0 | 3 | 4 |
| L1PB1_dup2801 | 3.38938312064862 | -5.16642826847039 | 1.32313054701756 | -3.90470031858602 | 9.4342219384205e-05 | 0.00653230131034083 | 3976 | 0 | 0 | 13 | 24 | 0 | 0 | 2 | 0 | 0 | 0 | 3 | 6 |
| MLT1J_dup2088 | 3.6631465980368 | -5.3387458755518 | 1.24740417690017 | -4.27988455900373 | 1.86990278939852e-05 | 0.00194464986506215 | 3977 | 0 | 0 | 13 | 20 | 0 | 0 | 3 | 0 | 0 | 0 | 8 | 7 |
| MSTA_dup18179 | 3.83328161150155 | -4.8715196332697 | 1.24770202617335 | -3.90439346180311 | 9.44619928180589e-05 | 0.00653586895698919 | 3978 | 0 | 1 | 13 | 25 | 0 | 0 | 2 | 0 | 0 | 0 | 5 | 8 |
| AluJr_dup75592 | 2.75142623858022 | -3.9012046352002 | 1.12196279559778 | -3.47712477678159 | 0.000506821893851455 | 0.0217800690544299 | 3979 | 1 | 1 | 6 | 12 | 0 | 0 | 7 | 0 | 0 | 0 | 4 | 8 |
| L1PA15_dup3036 | 3.72073149816815 | -4.2741919630299 | 1.21422702488827 | -3.5200929277811 | 0.000431395625869968 | 0.0195280259354976 | 3980 | 2 | 0 | 11 | 26 | 0 | 0 | 5 | 0 | 0 | 0 | 4 | 5 |
| MSTA_dup2426 | 3.49608957894894 | -4.21972754461579 | 1.15880612931908 | -3.64144392910254 | 0.000271113177661801 | 0.0139030810715334 | 3981 | 1 | 0 | 9 | 22 | 0 | 1 | 6 | 0 | 0 | 0 | 3 | 8 |
| L1PA15_dup7306 | 4.58173117150387 | -3.31558028378615 | 0.974360010406882 | -3.40282877824758 | 0.000666920608487222 | 0.0264089018317691 | 3982 | 0 | 0 | 10 | 16 | 0 | 1 | 3 | 0 | 3 | 2 | 11 | 16 |
| AluSz6_dup36182 | 4.87796379744024 | -3.20078172790079 | 0.979687376168681 | -3.26714603633893 | 0.00108637624355229 | 0.0371846020672193 | 3983 | 1 | 0 | 8 | 22 | 0 | 0 | 4 | 0 | 2 | 5 | 10 | 15 |
| MER46C_dup1665 | 4.61965501784095 | -4.06307079367702 | 1.08549251950826 | -3.74306659940655 | 0.000181788021644248 | 0.0104155387907927 | 3984 | 0 | 0 | 13 | 17 | 0 | 0 | 2 | 0 | 2 | 2 | 10 | 17 |
| Tigger9b_dup223 | 5.05887523197826 | -3.95372821480594 | 1.18400558738376 | -3.33928172040328 | 0.000839953334913403 | 0.0311410932111724 | 3985 | 0 | 0 | 15 | 23 | 0 | 0 | 1 | 0 | 2 | 5 | 8 | 15 |
| MLT1F2_dup4801 | 4.92173493472332 | -3.9167489189614 | 1.00392327035492 | -3.90144250523918 | 9.56211742272087e-05 | 0.00659306749425309 | 3986 | 0 | 0 | 11 | 12 | 1 | 0 | 5 | 0 | 0 | 3 | 12 | 24 |
| L1MD2_dup7175 | 4.90446192782385 | -4.7241994963686 | 1.08357442017419 | -4.35982929129053 | 1.30163946154894e-05 | 0.00148617995833841 | 3987 | 0 | 0 | 18 | 11 | 0 | 0 | 6 | 0 | 1 | 1 | 16 | 13 |
| LTR38C_dup97 | 4.33142374620881 | -4.20051532171345 | 1.03865608636036 | -4.04418303312776 | 5.25058893876004e-05 | 0.00424373857208627 | 3988 | 0 | 0 | 11 | 14 | 0 | 0 | 8 | 0 | 1 | 2 | 8 | 16 |
| L1PA13_dup7215 | 3.68704363664395 | -4.41257679995828 | 1.28149753809932 | -3.44329713383833 | 0.000574667553209852 | 0.0238139752030776 | 3989 | 0 | 0 | 13 | 18 | 0 | 0 | 1 | 0 | 0 | 2 | 7 | 10 |
| MIRc_dup79955 | 2.74687457618298 | -4.37412500978866 | 1.24952290824288 | -3.50063610753621 | 0.000464149153057182 | 0.0204978534471686 | 3990 | 0 | 0 | 9 | 11 | 0 | 0 | 1 | 0 | 0 | 1 | 6 | 10 |
| AluY_dup28542 | 2.92668207443828 | -4.02437775968834 | 1.27043867676417 | -3.16770721270742 | 0.00153646172419734 | 0.0472955913875628 | 3991 | 0 | 0 | 14 | 9 | 0 | 0 | 1 | 0 | 1 | 1 | 2 | 12 |
| MLT1A_dup905 | 4.00250887003402 | -4.51768802507585 | 1.28380307803144 | -3.51898831088893 | 0.000433195793864126 | 0.0195703828302247 | 3992 | 0 | 0 | 15 | 21 | 0 | 0 | 1 | 0 | 1 | 1 | 2 | 15 |
| FRAM_dup5553 | 3.2292976916799 | -3.48593380252354 | 1.09630174735322 | -3.17972110410255 | 0.00147416858446363 | 0.0459450318658827 | 3993 | 0 | 0 | 8 | 12 | 0 | 0 | 6 | 0 | 3 | 1 | 4 | 10 |
| L1P1_dup1205 | 3.62913351406243 | -3.30026083613522 | 1.04701482733881 | -3.15206695259843 | 0.00162119095733537 | 0.0489786571272375 | 3994 | 0 | 0 | 11 | 14 | 1 | 1 | 3 | 0 | 2 | 0 | 9 | 8 |
| MIRc_dup13578 | 2.75397858918575 | -3.86961613436402 | 1.22296219992 | -3.1641338829746 | 0.00155545224840022 | 0.0476173361211657 | 3995 | 0 | 1 | 8 | 14 | 0 | 0 | 1 | 0 | 1 | 0 | 5 | 8 |
| MLT1E2_dup2071 | 3.54153593875202 | -4.78473229277073 | 1.23642120782207 | -3.86982386139988 | 0.000108914012574212 | 0.00725319161376528 | 3996 | 0 | 0 | 13 | 20 | 0 | 0 | 2 | 0 | 1 | 0 | 6 | 7 |
| AluSz6_dup36171 | 2.78462680937026 | -3.84741169029486 | 1.12556873926122 | -3.41819344842516 | 0.000630382850123877 | 0.0254060090880041 | 3997 | 1 | 0 | 5 | 11 | 0 | 0 | 4 | 0 | 1 | 0 | 5 | 12 |
| L1PA7_dup11657 | 2.49541655115746 | -3.7180666633641 | 1.14876707786795 | -3.23657139466831 | 0.00120974960576271 | 0.0400689142111379 | 3998 | 0 | 0 | 8 | 8 | 0 | 0 | 2 | 0 | 1 | 1 | 5 | 9 |
| THE1C-int_dup1643 | 2.20887232173525 | -4.06119420771975 | 1.188066615263 | -3.41832196574326 | 0.000630085239879968 | 0.0254026198726667 | 3999 | 0 | 0 | 8 | 7 | 0 | 0 | 3 | 0 | 1 | 0 | 5 | 6 |
| THE1D_dup2537 | 2.270678305732 | -4.11971061043986 | 1.28195901206065 | -3.21360556124 | 0.00131079605230989 | 0.0423602712838421 | 4000 | 0 | 0 | 9 | 10 | 0 | 0 | 1 | 0 | 1 | 0 | 4 | 6 |
| L1M2_dup7677 | 6.31581495350832 | -5.07348914349347 | 0.976144159483714 | -5.19747938273467 | 2.02008936664965e-07 | 5.82881131502447e-05 | 4001 | 1 | 0 | 15 | 21 | 0 | 0 | 5 | 4 | 1 | 0 | 19 | 19 |
| MER54A_dup508 | 5.01495118509321 | -4.74766089335805 | 1.02415808363417 | -4.63567194286182 | 3.55779840722876e-06 | 0.000547937799227478 | 4002 | 0 | 0 | 15 | 16 | 0 | 0 | 5 | 2 | 2 | 0 | 15 | 12 |
| L1PA17_dup3887 | 4.74020262766615 | -4.57067436183747 | 1.00535396784921 | -4.5463334387745 | 5.45885318850299e-06 | 0.000769042202955318 | 4003 | 0 | 0 | 14 | 8 | 1 | 0 | 2 | 4 | 1 | 0 | 16 | 16 |
| LOR1b_dup921 | 5.32857011681717 | -4.2642988464536 | 0.951261314153204 | -4.48278383973766 | 7.36755494260897e-06 | 0.000968276006047014 | 4004 | 0 | 1 | 14 | 8 | 0 | 0 | 2 | 5 | 2 | 0 | 19 | 18 |
| Tigger17c_dup64 | 4.77397058517542 | -3.13505797738398 | 0.926787480012917 | -3.38271507221946 | 0.000717730543261193 | 0.0278455376979658 | 4005 | 3 | 0 | 13 | 16 | 0 | 0 | 2 | 1 | 2 | 1 | 9 | 18 |
| LTR39_dup972 | 4.54504842813019 | -4.06981886800572 | 1.01746688534994 | -3.99995216218361 | 6.33552891829076e-05 | 0.00487080376087759 | 4006 | 1 | 0 | 7 | 18 | 0 | 0 | 3 | 1 | 2 | 0 | 14 | 16 |
| MIR3_dup11282 | 4.43527741708712 | -4.58788002152925 | 1.07683731286277 | -4.26051360472676 | 2.03957666125508e-05 | 0.00207397054802992 | 4007 | 1 | 0 | 13 | 19 | 0 | 0 | 3 | 1 | 1 | 0 | 9 | 14 |
| ERVL47-int_dup126 | 4.43012944120294 | -3.85367848138269 | 0.971653459046899 | -3.96610380532457 | 7.30570547089167e-05 | 0.00540032119905806 | 4008 | 0 | 0 | 10 | 16 | 0 | 0 | 3 | 3 | 3 | 1 | 14 | 8 |
| MER4CL34_dup700 | 4.17151895390488 | -3.7365733096772 | 0.9636020138491 | -3.877714301106 | 0.000105442436555432 | 0.00709838099959188 | 4009 | 0 | 0 | 4 | 16 | 0 | 0 | 3 | 2 | 3 | 1 | 12 | 15 |
| AluY_dup80280 | 3.70342833029864 | -3.89570448676843 | 1.00210536373535 | -3.88751984346953 | 0.000101273710780444 | 0.00687915340718343 | 4010 | 0 | 0 | 10 | 13 | 0 | 0 | 6 | 1 | 2 | 1 | 8 | 9 |
| L1MCa_dup6171 | 3.69806161876205 | -3.9240163056568 | 1.01105621931298 | -3.88110594712842 | 0.000103982526890058 | 0.00701465888816122 | 4011 | 0 | 0 | 8 | 14 | 0 | 0 | 3 | 1 | 1 | 2 | 11 | 10 |
| L1MCb_dup421 | 4.74879715799371 | -4.59966570252051 | 0.998616711400044 | -4.6060371812443 | 4.10415038229128e-06 | 0.00061457908550326 | 4012 | 0 | 0 | 8 | 16 | 0 | 0 | 4 | 1 | 1 | 1 | 15 | 19 |
| MSTC-int_dup148 | 4.98849967245519 | -4.7455796722445 | 0.974730984054382 | -4.86860451742831 | 1.12389075675397e-06 | 0.00022586597128934 | 4013 | 0 | 0 | 11 | 17 | 0 | 0 | 6 | 3 | 1 | 1 | 10 | 19 |
| L1MC3_dup3044 | 3.50940180239893 | -2.95778944464296 | 0.869178593688965 | -3.40297088092048 | 0.000666573842503121 | 0.0264022533136353 | 4014 | 2 | 0 | 5 | 11 | 0 | 0 | 3 | 6 | 1 | 2 | 5 | 11 |
| L1PA16_dup1252 | 4.13526011849994 | -3.99815739599264 | 1.03172628440605 | -3.8752113389205 | 0.000106532199831835 | 0.00714205686121825 | 4015 | 1 | 0 | 9 | 24 | 0 | 0 | 4 | 3 | 1 | 1 | 4 | 10 |
| L1M6_dup575 | 4.88624177116505 | -4.23032961845229 | 1.07494743459363 | -3.93538277529961 | 8.30642009274819e-05 | 0.00595145870336965 | 4016 | 1 | 0 | 14 | 19 | 0 | 0 | 1 | 6 | 2 | 0 | 5 | 17 |
| AluSc_dup3789 | 3.40370850998484 | -3.66804518010076 | 1.06260779926953 | -3.45192759042638 | 0.000556597074031708 | 0.0232795407758408 | 4017 | 1 | 0 | 9 | 13 | 0 | 0 | 1 | 3 | 2 | 0 | 7 | 9 |
| L1M4_dup16437 | 3.57964722142628 | -3.74909702728017 | 1.02177299363822 | -3.66920739794733 | 0.000243303637744862 | 0.0128652474456765 | 4018 | 1 | 0 | 7 | 16 | 0 | 0 | 3 | 3 | 2 | 0 | 7 | 9 |
| MER1A_dup380 | 3.19215939048296 | -3.12897138310327 | 0.989072188753379 | -3.16354197265116 | 0.00155861875418829 | 0.0476815502808032 | 4019 | 0 | 2 | 9 | 16 | 0 | 1 | 2 | 2 | 1 | 0 | 5 | 5 |
| Kanga1_dup690 | 3.23528695715492 | -4.1024876416354 | 1.02927549067092 | -3.98580135135759 | 6.72527611428768e-05 | 0.0050979692586502 | 4020 | 0 | 1 | 12 | 7 | 0 | 0 | 4 | 3 | 1 | 0 | 9 | 5 |
| L1HS_dup174 | 3.73215007250601 | -4.31992943341405 | 1.09472395789837 | -3.9461358292618 | 7.94225694452657e-05 | 0.00575290189017449 | 4021 | 0 | 1 | 15 | 10 | 0 | 0 | 1 | 4 | 1 | 0 | 11 | 5 |
| MER45B_dup184 | 3.92922210951316 | -4.38449186353089 | 1.13145520753596 | -3.87509097516927 | 0.000106584871837249 | 0.00714205686121825 | 4022 | 0 | 1 | 9 | 19 | 0 | 0 | 1 | 1 | 1 | 0 | 9 | 13 |
| L1MEf_dup2411 | 4.53783376207192 | -4.0908313942632 | 1.06601896183677 | -3.83748464212554 | 0.000124301005603291 | 0.00794224679894756 | 4023 | 1 | 1 | 15 | 16 | 0 | 0 | 2 | 1 | 1 | 0 | 14 | 10 |
| L1PREC2_dup4094 | 3.96511274621375 | -3.92759889182886 | 1.06569767599249 | -3.685471949792 | 0.00022827922463275 | 0.0122558953936066 | 4024 | 1 | 1 | 10 | 22 | 0 | 0 | 3 | 1 | 1 | 0 | 6 | 10 |

| | | | | | | | | | | | | | | | | | | | |
|---|---|---|---|---|---|---|---|---|---|---|---|---|---|---|---|---|---|---|---|
| MIRb_dup203194 | 3.79936302008163 | -4.8544829881505 | 1.11646368631301 | -4.34808856540769 | 1.37329150782786e-05 | 0.00154573898354593 | 4025 | 0 | 0 | 17 | 18 | 0 | 0 | 4 | 1 | 1 | 0 | 4 | 7 |
| L1PA3_dup2238 | 3.82344163994588 | -4.82975470493082 | 1.11770071023547 | -4.32115204070447 | 1.55216671769333e-05 | 0.00169316887240008 | 4026 | 0 | 0 | 14 | 10 | 0 | 0 | 2 | 1 | 1 | 0 | 7 | 17 |
| MER57D_dup135 | 2.76882851171423 | -4.39036305882038 | 1.16581907983931 | -3.7659042768673 | 0.000165947366538115 | 0.00979085347837299 | 4027 | 0 | 0 | 7 | 13 | 0 | 0 | 1 | 1 | 1 | 0 | 5 | 10 |
| MER101-int_dup954 | 3.74225137897675 | -3.92432894141506 | 1.1632301389719 | -3.37364792222759 | 0.000741791788630335 | 0.028516755121186 | 4028 | 0 | 0 | 9 | 8 | 0 | 0 | 0 | 1 | 0 | 3 | 9 | 21 |
| MLT1D_dup1854 | 4.28894701761853 | -4.49265690142597 | 1.20788829016861 | -3.71943079338805 | 0.000199672246673057 | 0.0110970900987931 | 4029 | 0 | 1 | 18 | 10 | 0 | 0 | 0 | 1 | 0 | 1 | 9 | 18 |
| FLAM_C_dup2924 | 3.70498428584083 | -3.7657761514449 | 1.14372284627139 | -3.29256004959729 | 0.000992796941929521 | 0.0349300696649951 | 4030 | 0 | 0 | 13 | 19 | 0 | 0 | 1 | 1 | 2 | 2 | 5 | 7 |
| LTR82A_dup86 | 3.95572804442263 | -4.06491051560428 | 1.10850091709119 | -3.66703396716258 | 0.000245380210109459 | 0.0129433029205731 | 4031 | 0 | 0 | 15 | 16 | 0 | 0 | 1 | 2 | 2 | 1 | 5 | 11 |
| LTR13_dup301 | 2.94061224998971 | -3.46707938417228 | 1.07356713484039 | -3.22949471128113 | 0.00124009174982332 | 0.0407564922569379 | 4032 | 0 | 0 | 11 | 8 | 1 | 0 | 1 | 1 | 1 | 1 | 7 | 8 |
| L1PA5_dup1025 | 2.65866010694555 | -3.82441449738494 | 1.1367512992082 | -3.36433703664894 | 0.000767277816038389 | 0.0291769419092898 | 4033 | 0 | 0 | 9 | 9 | 0 | 0 | 1 | 1 | 1 | 1 | 9 | 4 |
| MIR_dup39131 | 2.60816451582132 | -3.40070774361745 | 1.06582348661315 | -3.19068568701167 | 0.00141935600361776 | 0.0446792014336328 | 4034 | 0 | 0 | 8 | 8 | 0 | 0 | 1 | 2 | 1 | 2 | 7 | 5 |
| LTR18A_dup27 | 2.14117143884733 | -4.03497115525426 | 1.14918304086786 | -3.51116489868061 | 0.000446147556897051 | 0.0199734763073238 | 4035 | 0 | 0 | 9 | 6 | 0 | 0 | 2 | 1 | 0 | 1 | 3 | 7 |
| MIR_dup20515 | 3.07116790986804 | -4.5758326643194 | 1.10482594219409 | -4.14167742588683 | 3.44775013275024e-05 | 0.00306111325778877 | 4036 | 0 | 0 | 12 | 11 | 0 | 0 | 3 | 1 | 0 | 1 | 5 | 9 |
| ERVL47-int_dup133 | 3.16692611023376 | -3.70779287526692 | 1.0422309757744 | -3.55755390259049 | 0.000374324306041284 | 0.0175990854229005 | 4037 | 0 | 0 | 11 | 8 | 0 | 0 | 5 | 3 | 0 | 3 | 5 | 7 |
| MER63A_dup3178 | 2.78454283769252 | -4.41584535184462 | 1.04462904379774 | -4.22718990828634 | 2.36627945965038e-05 | 0.00231611160965022 | 4038 | 0 | 0 | 7 | 12 | 0 | 0 | 3 | 4 | 0 | 1 | 5 | 5 |
| L1MD3_dup4035 | 3.18276124170325 | -4.15338292931334 | 1.19667563541866 | -3.4707675216102 | 0.000518973098929668 | 0.0221309412627807 | 4039 | 0 | 0 | 16 | 9 | 0 | 0 | 2 | 1 | 0 | 2 | 9 | 3 |
| LTR88a_dup111 | 2.79732504956061 | -4.43967517119293 | 1.19216266543393 | -3.72405150733097 | 0.000196050899911245 | 0.0109608852044834 | 4040 | 0 | 0 | 11 | 14 | 0 | 0 | 1 | 1 | 0 | 1 | 6 | 4 |
| L1M4_dup1869 | 3.18340151270019 | -4.06293349605652 | 1.14620081506917 | -3.54469604509166 | 0.000393066075924921 | 0.0182594515202671 | 4041 | 1 | 0 | 11 | 20 | 0 | 0 | 2 | 1 | 0 | 1 | 5 | 3 |
| L1PA4_dup4561 | 4.07548940482381 | -4.95941700665255 | 1.16527275792901 | -4.25601385847782 | 2.08103813151748e-05 | 0.00210086847366168 | 4042 | 0 | 0 | 19 | 22 | 0 | 0 | 2 | 1 | 0 | 1 | 5 | 6 |
| L2a_dup166068 | 2.19752290434769 | -3.51907806461996 | 1.11100673413046 | -3.16746780781143 | 0.00153772734100122 | 0.0473055531490971 | 4043 | 0 | 0 | 8 | 10 | 0 | 0 | 2 | 2 | 1 | 1 | 3 | 2 |
| AluY_dup21862 | 2.46110048984676 | -3.71528546905129 | 1.09510137368332 | -3.39264067997203 | 0.000692223763410567 | 0.0271219949956349 | 4044 | 0 | 0 | 8 | 9 | 0 | 0 | 2 | 1 | 1 | 1 | 5 | 6 |
| L1P4a_dup205 | 2.42972615170484 | -3.68975041997511 | 1.05813335399325 | -3.48703724918082 | 0.000488403327742878 | 0.0211989812677116 | 4045 | 0 | 0 | 8 | 8 | 0 | 0 | 3 | 2 | 1 | 1 | 5 | 4 |
| L2a_dup74607 | 2.92725562857193 | -3.46012769812647 | 1.02948284800703 | -3.36103481939976 | 0.000776510358076343 | 0.0294403335982845 | 4046 | 0 | 1 | 11 | 7 | 0 | 0 | 4 | 1 | 1 | 1 | 6 | 7 |
| MamGypsy2-I_dup170 | 2.86478891912354 | -3.00615251382066 | 0.950002485854354 | -3.164362787027 | 0.00155422928389342 | 0.0476084759507342 | 4047 | 0 | 1 | 6 | 9 | 0 | 0 | 7 | 2 | 1 | 2 | 7 | 3 |
| MIR_dup38568 | 2.83057445290991 | -3.40027911460745 | 1.05188549767307 | -3.23255632112942 | 0.00122687948512932 | 0.0404366008472466 | 4048 | 0 | 0 | 12 | 3 | 0 | 1 | 4 | 1 | 1 | 1 | 8 | 6 |
| L2a_dup20680 | 2.97730379697922 | -3.97833524704744 | 1.05342126190845 | -3.77658529488956 | 0.000158993128180604 | 0.00950069785988498 | 4049 | 0 | 0 | 11 | 6 | 0 | 0 | 2 | 3 | 1 | 1 | 5 | 10 |
| AluSz_dup91737 | 2.36057379504476 | -3.63788330603162 | 1.08314592494806 | -3.35862714546617 | 0.000783306774010771 | 0.0296271997534203 | 4050 | 0 | 0 | 7 | 7 | 0 | 0 | 3 | 1 | 1 | 1 | 3 | 9 |
| L4_C_Mam_dup4749 | 2.4597591631444 | -3.62307389954996 | 1.0512778835146 | -3.44635225030838 | 0.000568209181109177 | 0.0236493093944734 | 4051 | 0 | 0 | 6 | 4 | 0 | 0 | 3 | 2 | 1 | 1 | 3 | 13 |
| ERVL47-int_dup130 | 3.82777228336397 | -4.84038106467642 | 1.04241809576659 | -4.64341619196166 | 3.42695039536472e-06 | 0.000533657813334721 | 4052 | 0 | 0 | 12 | 9 | 0 | 0 | 2 | 4 | 0 | 1 | 13 | 9 |
| L1MA6_dup2889 | 3.39959406687146 | -4.23048539436548 | 1.11837760541805 | -3.78269859291766 | 0.000155137228324651 | 0.00932875525538293 | 4053 | 0 | 0 | 12 | 12 | 0 | 0 | 1 | 3 | 0 | 2 | 8 | 7 |
| AluJo_dup10181 | 2.56895411410336 | -3.76295508107603 | 1.09236231861612 | -3.44478660326109 | 0.000571510393811902 | 0.0237286009712034 | 4054 | 1 | 0 | 7 | 9 | 0 | 0 | 2 | 2 | 0 | 1 | 9 | 3 |
| MER20_dup1815 | 2.30576130015746 | -3.54645703127352 | 1.12156467199413 | -3.16206200126467 | 0.0015665620580942 | 0.0478077690746872 | 4055 | 1 | 0 | 6 | 6 | 0 | 0 | 1 | 2 | 0 | 1 | 10 | 3 |
| LTR10C_dup388 | 2.90561935886765 | -3.87478094541073 | 1.02175664099809 | -3.79227380565466 | 0.000149274196745301 | 0.00906339296725752 | 4056 | 0 | 1 | 4 | 8 | 0 | 0 | 2 | 4 | 0 | 1 | 10 | 8 |
| MamGypsy2-I_dup279 | 2.4012640089977 | -3.58905492025047 | 1.10587030072887 | -3.24545737224788 | 0.00117262119150312 | 0.039203977823506 7 | 4057 | 0 | 1 | 6 | 7 | 0 | 0 | 1 | 1 | 0 | 1 | 9 | 6 |
| MER4A1_dup1791 | 3.16954251081711 | -3.06775247265638 | 0.918261669912182 | -3.34082601199042 | 0.000835295501969338 | 0.0310022186104502 | 4058 | 1 | 1 | 4 | 8 | 0 | 1 | 2 | 5 | 0 | 1 | 11 | 7 |
| L1ME2z_dup6926 | 3.71978895223925 | -2.76880627880055 | 0.862320059902653 | -3.21088005202281 | 0.00132329157212837 | 0.04260953189588 | 4059 | 1 | 1 | 11 | 12 | 0 | 1 | 2 | 2 | 0 | 3 | 7 | 10 |
| THE1C_dup1138 | 2.88011143548587 | -3.43720783944028 | 1.03879715332329 | -3.30883448076851 | 0.00093685204993797 | 0.0335964416258501 | 4060 | 1 | 0 | 10 | 12 | 0 | 1 | 2 | 2 | 0 | 1 | 4 | 6 |
| L1M2_dup850 | 5.63791003175139 | -2.99994023139236 | 0.84816413274266 | -3.53698077480785 | 0.000404729126077326 | 0.0186346123517019 | 4061 | 0 | 2 | 15 | 22 | 0 | 0 | 4 | 2 | 2 | 5 | 14 | 9 |
| LTR39_dup977 | 4.65043667196778 | -2.81628382633962 | 0.831921997312907 | -3.38527390240451 | 0.000711072619904192 | 0.0276555877109714 | 4062 | 0 | 3 | 17 | 13 | 1 | 0 | 2 | 3 | 1 | 2 | 10 | 9 |
| AluJr_dup63195 | 4.53480762242619 | -4.07958268042109 | 0.94864242550523 | -4.30044300227073 | 1.70457010821975e-05 | 0.00181555791839954 | 4063 | 0 | 1 | 8 | 14 | 0 | 0 | 3 | 3 | 1 | 1 | 18 | 11 |
| L1M3d_dup32 | 4.89441424764673 | -2.88059464100829 | 0.861972634471065 | -3.34186321677824 | 0.000832180586993778 | 0.0309251978686275 | 4064 | 1 | 4 | 14 | 16 | 0 | 0 | 3 | 1 | 1 | 1 | 12 | 13 |
| L1MC_dup798 | 5.16747111232403 | -3.01031596070525 | 0.857749148771863 | -3.50955283956325 | 0.000448860862778944 | 0.0200609779967691 | 4065 | 1 | 3 | 14 | 23 | 0 | 0 | 2 | 2 | 1 | 2 | 10 | 12 |
| AluY_dup21859 | 3.89318486759188 | -3.41004452050538 | 1.05822894728918 | -3.22240714473059 | 0.0012711836076623 5 | 0.041463064387776 | 4066 | 0 | 2 | 13 | 10 | 0 | 0 | 2 | 0 | 1 | 1 | 13 | 10 |
| L1M5_dup5786 | 4.4063293578 | -3.29125186555737 | 0.991013286955474 | -3.32109761683271 | 0.000896641706651308 | 0.0325629284998248 | 4067 | 0 | 2 | 21 | 13 | 0 | 0 | 3 | 1 | 2 | 1 | 4 | 12 |
| AluYm1_dup3684 | 4.36537073624544 | -3.96572267481314 | 0.914677949810663 | -4.33564914911749 | 1.45330592115893e-05 | 0.00161366463236623 | 4068 | 1 | 0 | 8 | 9 | 1 | 0 | 5 | 2 | 0 | 1 | 15 | 17 |
| L1P1_dup668 | 3.97403937302353 | -3.12830216898125 | 0.904695776801086 | -3.45784986423017 | 0.000544504732597408 | 0.0228828546631266 | 4069 | 2 | 0 | 7 | 15 | 1 | 0 | 2 | 2 | 0 | 2 | 11 | 12 |
| L1MC3_dup10668 | 3.83518860333764 | -3.88040364680937 | 0.998387805991465 | -3.88666971243291 | 0.00010162888269662 | 0.00689425397400665 | 4070 | 1 | 0 | 8 | 15 | 0 | 1 | 6 | 2 | 0 | 1 | 12 | 6 |
| LTR1_dup200 | 4.97510400130976 | -3.49071689633452 | 0.923548007653739 | -3.77968104246431 | 0.000157029382862203 | 0.00941939353153349 | 4071 | 3 | 0 | 9 | 21 | 0 | 1 | 4 | 1 | 0 | 1 | 13 | 16 |
| L1ME3A_dup15374 | 4.23462091542579 | -3.50505892903882 | 1.05394263097639 | -3.32566387014034 | 0.000882082320332181 | 0.0322557184253513 | 4072 | 1 | 0 | 10 | 14 | 0 | 0 | 1 | 1 | 0 | 4 | 9 | 18 |
| L1M5_dup44380 | 3.30615121529741 | -3.72756520548201 | 1.00494659473208 | -3.70921721116513 | 0.000207901016858485 | 0.0114406140974582 | 4073 | 0 | 0 | 8 | 6 | 0 | 0 | 4 | 1 | 0 | 3 | 13 | 9 |
| LTR8A_dup1349 | 3.61734332233272 | -3.15908073126191 | 0.934291081229097 | -3.38125964673238 | 0.000721543267231888 | 0.0279577160502783 | 4074 | 0 | 0 | 8 | 7 | 0 | 1 | 5 | 1 | 0 | 4 | 9 | 14 |
| L1PA4_dup4559 | 3.58687167237821 | -3.76723576882493 | 1.15211292088725 | -3.26984942233246 | 0.00107604747329162 | 0.0369476013824392 | 4075 | 0 | 1 | 16 | 16 | 0 | 2 | 0 | 3 | 0 | 0 | 4 | 6 |
| L1PREC2_dup931 | 3.74089711261492 | -3.49833515252531 | 1.0991593794866 | -3.18273693316371 | 0.00145890098166422 | 0.0456012384201789 | 4076 | 1 | 2 | 17 | 10 | 0 | 1 | 0 | 2 | 0 | 0 | 6 | 11 |
| L1MA4A_dup5831 | 5.14331277913723 | -3.42539989869529 | 0.987792022265739 | -3.46773391714412 | 0.000524866712896911 | 0.0222938120076702 | 4077 | 2 | 3 | 15 | 28 | 1 | 0 | 0 | 8 | 0 | 0 | 6 | 5 |
| L2b_dup90031 | 4.76785126290554 | -3.18612421958809 | 1.00737356798355 | -3.16280307608799 | 0.0015625799128954 | 0.0477290937039548 | 4078 | 3 | 1 | 20 | 19 | 1 | 0 | 0 | 2 | 1 | 0 | 8 | 9 |
| ERVL-E-int_dup7014 | 2.7085189641977 | -4.34610037052214 | 1.25743235358196 | -3.45632936685757 | 0.000547585775598837 | 0.0229758146641934 | 4079 | 0 | 1 | 13 | 10 | 0 | 0 | 0 | 3 | 0 | 0 | 2 | 7 |
| L3_dup7304 | 2.50365262485573 | -4.2437238861837 | 1.18918257025804 | -3.56860585777241 | 0.000358885847809568 | 0.0170415988710628 | 4080 | 0 | 1 | 8 | 12 | 0 | 0 | 1 | 3 | 0 | 0 | 1 | 8 |
| LSAU_dup122 | 2.48111733145023 | -4.21637080596958 | 1.27367149329077 | -3.31040682639116 | 0.000931604616710755 | 0.0334838602028075 | 4081 | 0 | 0 | 6 | 12 | 0 | 0 | 0 | 6 | 1 | 0 | 1 | 6 |
| AluSc_dup17376 | 2.40494557491558 | -4.71614129930288 | 1.29262515387356 | -3.64849878185505 | 0.000263777153511848 | 0.0136504390103371 | 4082 | 0 | 0 | 7 | 8 | 0 | 0 | 0 | 6 | 0 | 0 | 2 | 8 |
| MLT1H2_dup4018 | 2.93854171273492 | -4.97562288686458 | 1.27200841589789 | -3.91162733255375 | 9.16762998062265e-05 | 0.00640270291567137 | 4083 | 0 | 0 | 15 | 9 | 0 | 0 | 0 | 5 | 0 | 0 | 3 | 6 |
| AluJr_dup40130 | 3.20885138823295 | -3.54218313285002 | 1.10283715346517 | -3.21188229986656 | 0.0013186838892158 | 0.0425170131034339 | 4084 | 0 | 1 | 13 | 10 | 1 | 0 | 0 | 5 | 1 | 0 | 5 | 5 |
| L1MC5a_dup8133 | 3.39847719690446 | -3.62292264716351 | 1.005540373436 | -3.60296089831155 | 0.000314612870529203 | 0.0154511608030839 | 4085 | 0 | 1 | 12 | 15 | 1 | 0 | 2 | 4 | 1 | 0 | 3 | 6 |
| MLT1B_dup12838 | 3.42931667854651 | -3.16680546617831 | 0.909519298658902 | -3.48184526798696 | 0.000497971348390249 | 0.0215163047873736 | 4086 | 0 | 1 | 13 | 12 | 2 | 0 | 3 | 6 | 1 | 0 | 3 | 3 |
| LTR62_dup228 | 3.1018887113899 | -3.51370924547375 | 1.08222740187682 | -3.24673838361531 | 0.00116735641466569 | 0.0391001328500764 | 4087 | 0 | 0 | 10 | 12 | 2 | 0 | 1 | 3 | 1 | 0 | 5 | 7 |
| UCON55_dup203 | 3.40786160337123 | -4.10258152417812 | 1.11110703796803 | -3.69233690723519 | 0.000222202835768932 | 0.012000062632745 | 4088 | 0 | 0 | 15 | 18 | 1 | 0 | 2 | 2 | 1 | 0 | 3 | 4 |
| L1ME1_dup8233 | 3.17237151811248 | -4.36162269544605 | 1.365069091145 | -3.19516625476268 | 0.0013975031118682 2 | 0.0442665435911336 | 4089 | 0 | 0 | 19 | 18 | 1 | 0 | 0 | 1 | 0 | 0 | 3 | 1 |
| LTR19B_dup110 | 2.33719541611701 | -4.11790006351737 | 1.26927896126969 | -3.24428292689743 | 0.00117746726759454 | 0.0392895283207195 | 4090 | 0 | 0 | 11 | 10 | 1 | 0 | 0 | 2 | 0 | 0 | 4 | 3 |
| L1MCb_dup1271 | 3.13497115848233 | -4.25078253358721 | 1.30032373199004 | -3.26901865205653 | 0.00107921187112005 | 0.037007371074439 | 4091 | 0 | 1 | 12 | 24 | 0 | 0 | 0 | 3 | 0 | 0 | 1 | 2 |
| MER39_dup960 | 2.70977455091186 | -4.53934409201579 | 1.38820384447691 | -3.26994058550981 | 0.00107570075648766 | 0.0369476013824392 | 4092 | 0 | 0 | 11 | 21 | 0 | 0 | 0 | 3 | 0 | 0 | 1 | 1 |
| L1MB7_dup16448 | 2.88064899295749 | -4.77644105508539 | 1.43696381751503 | -3.32398143701724 | 0.00088742102635886 | 0.0323598422407214 | 4093 | 0 | 0 | 20 | 14 | 0 | 0 | 0 | 1 | 0 | 0 | 1 | 3 |
| L1PA16_dup10795 | 3.36262352904778 | -5.01329180928612 | 1.42751730783166 | -3.51189563992123 | 0.000444922673197382 | 0.0199402598648304 | 4094 | 0 | 0 | 16 | 25 | 0 | 0 | 0 | 1 | 0 | 0 | 1 | 4 |
| L2b_dup22455 | 3.35851998542635 | -4.4285788352698 | 1.36399447200413 | -3.24677183534537 | 0.0011672192258680 4 | 0.0391001328500764 | 4095 | 0 | 1 | 18 | 22 | 0 | 0 | 0 | 1 | 0 | 0 | 3 | 1 |
| MER4E_dup710 | 3.22834725158036 | -5.02557982471163 | 1.39931371019923 | -3.59146043384089 | 0.000328830104086276 | 0.0159616091282972 | 4096 | 0 | 0 | 18 | 19 | 0 | 0 | 0 | 1 | 0 | 0 | 4 | 2 |
| MamGyp-int_dup146 | 3.80467407681002 | -5.03870883466952 | 1.43659561480862 | -3.50739538860472 | 0.00045251623610113 | 0.02018264823155 | 4097 | 0 | 0 | 19 | 30 | 0 | 0 | 0 | 1 | 0 | 0 | 2 | 1 |
| L1PA3_dup615 | 3.92983711531246 | -3.64187513089251 | 1.05702251033212 | -3.44540924653365 | 0.00057019539899326 | 0.0236887000417292 | 4098 | 0 | 0 | 18 | 25 | 1 | 1 | 2 | 2 | 0 | 1 | 1 | 3 |
| AluSx1_dup47607 | 3.60192109758849 | -4.13401953018958 | 1.12428458855253 | -3.67702232360222 | 0.000235972396974597 | 0.01256966374983 6 | 4099 | 0 | 0 | 16 | 20 | 0 | 1 | 2 | 3 | 0 | 1 | 1 | 5 |
| L1M2_dup7040 | 3.00263115538323 | -3.96983507473514 | 1.1413479479321 | -3.47819881038705 | 0.000504795373985988 | 0.0217243696976795 | 4100 | 0 | 0 | 13 | 14 | 0 | 1 | 2 | 2 | 0 | 1 | 1 | 7 |
| L1MA8_dup4762 | 3.74503045084706 | -3.75699118965351 | 1.09381411018142 | -3.43476204474121 | 0.000593074356244446 | 0.0243856351772837 | 4101 | 0 | 0 | 11 | 25 | 1 | 1 | 2 | 2 | 1 | 0 | 1 | 8 |
| AluJb_dup57367 | 3.08535121276442 | -3.84945511070772 | 1.09504170398817 | -3.5153502343225 | 0.000439174388185686 | 0.0197616847525021 | 4102 | 0 | 0 | 12 | 18 | 1 | 1 | 2 | 3 | 0 | 0 | 1 | 4 |
| MSTA-int_dup2375 | 3.08259963436926 | -3.89953562765561 | 1.08610879532956 | -3.59037293908697 | 0.000330205167498315 | 0.0160152871556368 | 4103 | 0 | 0 | 16 | 12 | 1 | 1 | 2 | 3 | 0 | 0 | 2 | 4 |

| | | | | | | | | | | | | | | | | | | | |
|---|---|---|---|---|---|---|---|---|---|---|---|---|---|---|---|---|---|---|---|
| L2b_dup45761 | 3.26208176785581 | -4.05238852051092 | 1.05220077549064 | -3.85134530871384 | 0.000117470722913213 | 0.00763707339735174 | 4104 | 0 | 0 | 12 | 16 | 1 | 1 | 3 | 3 | 0 | 0 | 4 | 4 |
| Tigger10_dup504 | 3.3942081658044 | -4.12843041130459 | 1.11391667221037 | -3.70622912314657 | 0.000210368002088987 | 0.011533692309923 | 4105 | 0 | 0 | 12 | 19 | 1 | 1 | 1 | 3 | 0 | 0 | 4 | 5 |
| L1PA7_dup1424 | 4.46505906760723 | -5.52590678265655 | 1.21332999944354 | -4.55433129090259 | 5.25524806867909e-06 | 0.000749230231845366 | 4106 | 0 | 0 | 17 | 32 | 0 | 0 | 1 | 2 | 0 | 0 | 5 | 5 |
| SATR1_dup141 | 5.62946294201198 | -5.38882385798923 | 1.17213238048915 | -4.59745328061015 | 4.27686489088522e-06 | 0.000636039308329109 | 4107 | 0 | 0 | 25 | 34 | 0 | 1 | 2 | 1 | 0 | 0 | 7 | 8 |
| L1MB3_dup12946 | 2.89064770107752 | -4.47981566666581 | 1.15838044408216 | -3.86730947466519 | 0.00011004274189315 | 0.00730179315741984 | 4108 | 0 | 0 | 6 | 18 | 0 | 1 | 1 | 2 | 0 | 0 | 5 | 7 |
| MER1A_dup335 | 2.81209129705851 | -4.44097403275587 | 1.19783923529897 | -3.70748753412428 | 0.000209325717286406 | 0.0114950881895756 | 4109 | 0 | 0 | 6 | 18 | 0 | 1 | 1 | 1 | 0 | 0 | 7 | 5 |
| L1PA16_dup11798 | 3.43118570574742 | -4.72739971082801 | 1.13531904077276 | -4.16393942235857 | 3.12803138953344e-05 | 0.00284702111675763 | 4110 | 0 | 0 | 12 | 14 | 0 | 1 | 1 | 2 | 0 | 0 | 10 | 6 |
| MIRb_dup225021 | 3.96071315473476 | -4.87017073759679 | 1.14474287237288 | -4.25437961234182 | 2.09629405734737e-05 | 0.00211178994822006 | 4111 | 0 | 0 | 16 | 23 | 0 | 1 | 2 | 2 | 0 | 0 | 7 | 3 |
| L1MA9_dup8728 | 4.92386360911537 | -4.66678595770267 | 1.30082982454945 | -3.58754532655263 | 0.000333805718118779 | 0.0161438486592461 | 4112 | 1 | 0 | 21 | 40 | 0 | 0 | 0 | 4 | 0 | 0 | 1 | 1 |
| MSTA-int_dup758 | 4.80020203148494 | -3.7768013001945 | 1.15342079420127 | -3.27443489763845 | 0.00105873531512606 | 0.0365527906693245 | 4113 | 3 | 0 | 21 | 31 | 1 | 0 | 0 | 3 | 0 | 0 | 2 | 5 |
| AluJo_dup31932 | 4.11834476978457 | -3.9091331003804 | 1.15112653710295 | -3.39591954001732 | 0.000683984625453423 | 0.0268921624188325 | 4114 | 1 | 2 | 14 | 31 | 0 | 0 | 1 | 1 | 0 | 0 | 2 | 6 |
| L1MC2_dup3299 | 5.55081844765359 | -5.19078064584617 | 1.24476624022303 | -4.17008469390694 | 3.0448646047311e-05 | 0.00278658982014364 | 4115 | 0 | 1 | 30 | 36 | 0 | 0 | 1 | 1 | 0 | 0 | 2 | 6 |
| AluJb_dup1168 | 4.00100116915666 | -4.73674354448435 | 1.26595645872252 | -3.74163227482897 | 0.000182828904702304 | 0.0104475039765356 | 4116 | 0 | 0 | 28 | 19 | 0 | 0 | 1 | 1 | 0 | 1 | 2 | 2 |
| ERVL-B4-int_dup942 | 3.21703400622588 | -4.36397160896285 | 1.25494638654592 | -3.47741676915307 | 0.000506270204218872 | 0.021772089284848 | 4117 | 1 | 0 | 20 | 17 | 0 | 0 | 1 | 1 | 0 | 0 | 1 | 3 |
| MIRb_dup206751 | 4.74003890603061 | -3.46723039912895 | 1.06667651189565 | -3.25049849740023 | 0.00115202877232526 | 0.038750443632981 | 4118 | 2 | 0 | 23 | 33 | 0 | 1 | 2 | 1 | 0 | 1 | 1 | 2 |
| HERV17-int_dup250 | 5.40693676947494 | -4.26668010041389 | 1.23935117911921 | -3.44267240173699 | 0.000575996596945982 | 0.0238399871728837 | 4119 | 2 | 0 | 26 | 44 | 0 | 0 | 1 | 1 | 0 | 0 | 1 | 1 |
| L1MA3_dup2582 | 5.52702171024147 | -3.9088052130562 | 1.19197029013924 | -3.27928073828047 | 0.00104072047964448 | 0.036087915367165 | 4120 | 2 | 0 | 29 | 41 | 0 | 0 | 1 | 1 | 0 | 1 | 1 | 1 |
| MLT1I-int_dup394 | 4.6419577962557 | -4.93750401675613 | 1.17238730338111 | -4.21149564014948 | 2.53685451142137e-05 | 0.0024389268023219 | 4121 | 0 | 1 | 28 | 18 | 0 | 0 | 1 | 6 | 0 | 0 | 1 | 6 |
| L1MC1_dup10417 | 4.41564433781063 | -4.87614679892511 | 1.18422928286203 | -4.11756985703015 | 3.82888487414263e-05 | 0.00333236225407001 | 4122 | 0 | 0 | 29 | 18 | 0 | 0 | 1 | 4 | 1 | 0 | 2 | 3 |
| MER103C_dup8383 | 5.5546916442465 | -4.4310982836608 | 1.11548657855689 | -3.97234567303655 | 7.1168343278022e-05 | 0.00530345863762657 | 4123 | 0 | 1 | 32 | 28 | 0 | 0 | 1 | 7 | 1 | 0 | 1 | 2 |
| MADE1_dup4086 | 3.51258950575672 | -5.05333954934209 | 1.45547142170858 | -3.4719606815845 | 0.000516672003123019 | 0.0220559807059759 | 4124 | 0 | 0 | 17 | 20 | 0 | 0 | 0 | 5 | 0 | 0 | 0 | 5 |
| L1PA13_dup4596 | 3.46560416690974 | -4.09033350430603 | 1.27112173468289 | -3.21789282072692 | 0.00129136062860583 | 0.0418857819157909 | 4125 | 0 | 0 | 17 | 20 | 0 | 1 | 0 | 2 | 1 | 0 | 1 | 5 |
| L1PB2_dup739 | 2.5381675736021 | -4.20751260437453 | 1.31299215346164 | -3.2045222762997 | 0.00135286820972781 | 0.0432906911016532 | 4126 | 0 | 0 | 8 | 18 | 0 | 0 | 1 | 2 | 1 | 0 | 0 | 5 |
| LTR9D_dup204 | 3.81607631000115 | -4.51744863117714 | 1.28342756922866 | -3.51983137925901 | 0.000431821233130296 | 0.0195280259354976 | 4127 | 0 | 0 | 18 | 27 | 0 | 0 | 1 | 3 | 1 | 0 | 0 | 2 |
| L2a_dup86184 | 3.83690020625358 | -3.74424781126288 | 1.09372528332313 | -3.42338964670041 | 0.000618453570897358 | 0.0250866950870683 | 4128 | 0 | 0 | 14 | 24 | 2 | 1 | 7 | 0 | 0 | 0 | 3 | 3 |
| L1PA3_dup2912 | 4.08102909708738 | -4.13518837633831 | 1.11473967564756 | -3.70955521425769 | 0.000207623676358096 | 0.0114306393555538 | 4129 | 1 | 0 | 18 | 29 | 1 | 0 | 3 | 1 | 0 | 0 | 2 | 2 |
| L2b_dup1340 | 3.09517647993682 | -3.89249244454485 | 1.18149632815477 | -3.29454468184766 | 0.000985812706119705 | 0.0347820566682027 | 4130 | 0 | 0 | 13 | 18 | 2 | 0 | 3 | 1 | 0 | 0 | 1 | 5 |
| L1ME5_dup778 | 3.3433051930108 | -4.59213465145651 | 1.3866499942202 | -3.31167538354836 | 0.000927390887082785 | 0.0333879521680516 | 4131 | 0 | 0 | 11 | 29 | 0 | 0 | 1 | 0 | 1 | 0 | 4 | 1 |
| L2c_dup33692 | 2.51696812978398 | -4.56442216867389 | 1.44407157335901 | -3.16080051216349 | 0.001573362132654 | 0.047959931593196 | 4132 | 0 | 0 | 14 | 17 | 0 | 0 | 1 | 0 | 0 | 0 | 2 | 1 |
| L1PA7_dup9092 | 2.33150465597609 | -4.46259161358555 | 1.39635854986318 | -3.19587803148611 | 0.00139406028730044 | 0.0441927883189437 | 4133 | 0 | 0 | 16 | 11 | 0 | 0 | 2 | 0 | 0 | 0 | 1 | 2 |
| MER20_dup132 | 2.51570373994672 | -4.45792363978652 | 1.4091902768544 | -3.16346465981691 | 0.00155903278785296 | 0.0476880853243938 | 4134 | 0 | 0 | 15 | 16 | 0 | 0 | 2 | 0 | 0 | 0 | 1 | 1 |
| AluJo_dup685 | 2.9509668406253 | -4.73416364191991 | 1.38387784233838 | -3.42094041618619 | 0.00062405002808096 | 0.0252148425847771 | 4135 | 0 | 0 | 18 | 18 | 0 | 0 | 2 | 0 | 0 | 0 | 1 | 2 |
| L1PA16_dup3587 | 2.97347323596422 | -4.64821583054484 | 1.40782096912103 | -3.30170947336218 | 0.000960975496743047 | 0.0341947473527772 | 4136 | 0 | 0 | 20 | 17 | 0 | 0 | 2 | 0 | 0 | 0 | 1 | 1 |
| L1MB8_dup6906 | 4.45023024153147 | -5.27699260773858 | 1.42224790053497 | -3.71031843728065 | 0.000206998709327962 | 0.0114073429934795 | 4137 | 0 | 0 | 27 | 30 | 0 | 0 | 1 | 0 | 0 | 0 | 1 | 3 |
| MSTA-int_dup240 | 3.26327635431209 | -4.85074110116139 | 1.43485106644305 | -3.3806582540906 | 0.000723124200150621 | 0.0279899139833741 | 4138 | 0 | 0 | 22 | 19 | 0 | 0 | 1 | 0 | 0 | 0 | 1 | 2 |
| THE1D-int_dup1486 | 3.52498197970008 | -5.01870847606068 | 1.41549481818226 | -3.54555058174326 | 0.00039179378552619 | 0.0182295197674648 | 4139 | 0 | 0 | 26 | 17 | 0 | 0 | 1 | 0 | 0 | 0 | 2 | 2 |
| UCON80_AMi_dup134 | 4.1226608263548 | -4.84751981166849 | 1.32964455094212 | -3.64572607636663 | 0.000266637883651102 | 0.0137476528414399 | 4140 | 0 | 0 | 19 | 29 | 1 | 0 | 1 | 0 | 0 | 0 | 2 | 6 |
| L1MC5_dup3177 | 4.25436723251916 | -4.69026544133408 | 1.34370931450558 | -3.49053578084324 | 0.000482053057909084 | 0.0210029742615467 | 4141 | 0 | 0 | 25 | 28 | 1 | 0 | 1 | 0 | 0 | 0 | 2 | 2 |
| L2a_dup47115 | 4.1029259098671 | -4.76966533740884 | 1.28135921091335 | -3.72234834446543 | 0.000197378461235588 | 0.0110144032216612 | 4142 | 0 | 0 | 22 | 26 | 1 | 0 | 2 | 0 | 0 | 0 | 3 | 3 |
| MIRb_dup98627 | 4.39088744374355 | -3.4836384603275 | 1.08028683126037 | -3.2247347274086 | 0.00126089444154765 | 0.0412429434654491 | 4143 | 3 | 0 | 18 | 25 | 1 | 1 | 2 | 0 | 0 | 0 | 6 | 5 |
| THE1C-int_dup190 | 5.58610712786323 | -3.44175592952788 | 1.08940812724811 | -3.15928974958347 | 0.00158154167915861 | 0.0481537448994338 | 4144 | 2 | 0 | 27 | 40 | 1 | 1 | 2 | 0 | 0 | 1 | 3 | 1 |
| L1MB7_dup4902 | 5.99101790077688 | -4.76120257814578 | 1.35959783423396 | -3.50191980176872 | 0.000461918636760698 | 0.0204334805619374 | 4145 | 0 | 0 | 24 | 51 | 1 | 1 | 0 | 0 | 0 | 0 | 3 | 5 |
| MLT1J1_dup3796 | 7.86127842499587 | -5.63509460840966 | 1.28979433249959 | -4.36898695119012 | 1.24824223921283e-05 | 0.00143902251866151 | 4146 | 0 | 0 | 46 | 51 | 0 | 1 | 2 | 0 | 0 | 0 | 5 | 4 |
| L1PA3_dup2320 | 5.35918146898769 | -3.84876676518827 | 1.21912446862927 | -3.15699246814039 | 0.00159405463669708 | 0.0484293826077021 | 4147 | 0 | 0 | 24 | 34 | 0 | 3 | 3 | 0 | 1 | 0 | 1 | 9 |
| L1PA7_dup13362 | 3.97390231846975 | -4.75757174906811 | 1.40242915440704 | -3.39237938267166 | 0.000692884305364794 | 0.0271344715148669 | 4148 | 0 | 0 | 20 | 25 | 0 | 1 | 3 | 0 | 0 | 0 | 0 | 7 |
| L1PB3_dup2354 | 4.1574238998982 | -4.76656587537051 | 1.47233974774504 | -3.23740894903553 | 0.00120620423615682 | 0.0400058602920271 | 4149 | 1 | 0 | 25 | 25 | 0 | 0 | 1 | 0 | 0 | 0 | 0 | 6 |
| ERVL47-int_dup132 | 3.08942282769387 | -4.53500930023926 | 1.13743768051821 | -3.98703979823591 | 6.69028064369484e-05 | 0.00507953260384369 | 4150 | 0 | 0 | 9 | 11 | 1 | 0 | 1 | 1 | 0 | 0 | 9 | 10 |
| L1PA7_dup11371 | 2.39945008431922 | -4.15690901914349 | 1.26776334120923 | -3.27893139359787 | 0.00104200965075924 | 0.0361115683599705 | 4151 | 0 | 0 | 4 | 12 | 1 | 0 | 0 | 1 | 0 | 0 | 7 | 8 |
| L3_dup1625 | 2.84307576146998 | -4.44961071498508 | 1.27083653106479 | -3.50132421142859 | 0.000462952273664144 | 0.0204601860529882 | 4152 | 0 | 0 | 8 | 15 | 1 | 0 | 0 | 1 | 0 | 0 | 7 | 7 |
| MER65A_dup1202 | 2.55503168244661 | -4.58929464532614 | 1.42237784796141 | -3.22649474041207 | 0.00125316535155375 | 0.0410554430319163 | 4153 | 0 | 0 | 7 | 6 | 0 | 0 | 0 | 0 | 0 | 0 | 9 | 13 |
| HAL1ME_dup1924 | 2.61378961160931 | -4.76416118721945 | 1.47345297806095 | -3.23333099742961 | 0.00122355707056634 | 0.0403644906106683 | 4154 | 0 | 0 | 6 | 12 | 0 | 0 | 0 | 0 | 0 | 0 | 10 | 8 |
| MER101-int_dup951 | 2.84293054102003 | -4.76096013156589 | 1.43177966030704 | -3.32520447353256 | 0.000883537117411356 | 0.0322782225708344 | 4155 | 0 | 0 | 8 | 7 | 0 | 0 | 0 | 0 | 0 | 0 | 15 | 8 |
| L1PBa_dup1804 | 2.97224715312218 | -5.03465423708449 | 1.49839294616559 | -3.3600359972117 | 0.0007793231758093 | 0.0295093622519287 | 4156 | 0 | 0 | 12 | 15 | 0 | 0 | 0 | 0 | 0 | 0 | 7 | 7 |
| MER58B_dup5019 | 3.31503539591968 | -5.19290083947388 | 1.49757993108595 | -3.46752833133142 | 0.000525268368360628 | 0.0223029099175478 | 4157 | 0 | 0 | 13 | 19 | 0 | 0 | 0 | 0 | 0 | 0 | 7 | 7 |
| L1MC1_dup4043 | 3.1927941494287 | -5.10879214885525 | 1.46587518265736 | -3.48514812809229 | 0.000491864668985235 | 0.0213104935408523 | 4158 | 0 | 0 | 12 | 13 | 0 | 0 | 0 | 0 | 0 | 0 | 8 | 11 |
| L1PA5_dup4158 | 3.27808287984425 | -5.14330503013144 | 1.4717629493846 | -3.49465586987501 | 0.000474673351765216 | 0.0208112789538306 | 4159 | 0 | 0 | 8 | 18 | 0 | 0 | 0 | 0 | 0 | 0 | 8 | 12 |
| ERV3-16A3_I-int_dup1 | 3.00077851317649 | -4.51797865040671 | 1.40370626962628 | -3.21860687536115 | 0.00128814956481301 | 0.0418215358642374 | 4160 | 1 | 0 | 6 | 18 | 0 | 0 | 0 | 0 | 0 | 0 | 9 | 8 |
| L1MCa_dup5771 | 3.02720057022841 | -5.06056208517886 | 1.53233545807026 | -3.3025158156634 | 0.000958216841581057 | 0.0341067860264542 | 4161 | 0 | 0 | 8 | 20 | 0 | 0 | 0 | 0 | 0 | 0 | 10 | 4 |
| MER1B_dup134 | 2.38686677210921 | -4.18314997061245 | 1.23586023159746 | -3.38480830085888 | 0.000712279800226435 | 0.0276753966924689 | 4162 | 0 | 0 | 7 | 9 | 0 | 0 | 0 | 3 | 0 | 1 | 8 | 3 |
| L2a_dup42728 | 2.02008905054821 | -3.92113511136891 | 1.24003491207878 | -3.1621167058882 | 0.00156626778517957 | 0.0478049197553025 | 4163 | 0 | 1 | 4 | 9 | 0 | 0 | 0 | 2 | 0 | 0 | 6 | 5 |
| AluSq2_dup34127 | 2.31559366292978 | -4.60834527760418 | 1.28187180350274 | -3.5950125940923 | 0.000324375884778887 | 0.0158170015902206 | 4164 | 0 | 0 | 4 | 10 | 0 | 0 | 0 | 2 | 0 | 0 | 9 | 6 |
| L1MD3_dup4056 | 2.03015554365632 | -4.46636000040122 | 1.29463160694533 | -3.4499080483131 | 0.000560777527355327 | 0.02340139767171 | 4165 | 0 | 0 | 6 | 8 | 0 | 0 | 0 | 2 | 0 | 0 | 6 | 5 |
| MSTA_dup3758 | 1.79230607799111 | -4.28278607326379 | 1.36175804767705 | -3.14504186743714 | 0.00166063062709225 | 0.0498662815742165 | 4166 | 0 | 0 | 6 | 7 | 0 | 0 | 0 | 1 | 0 | 0 | 6 | 4 |
| MER58A_dup12733 | 3.24135177459929 | -3.59117310346342 | 1.13657563118922 | -3.15964288245905 | 0.00157962625441329 | 0.0481015803343518 | 4167 | 1 | 1 | 12 | 11 | 0 | 0 | 0 | 2 | 1 | 0 | 7 | 8 |
| MLT1A1_dup5506 | 2.88307678522803 | -3.94274138996702 | 1.18075826995863 | -3.33916051259599 | 0.000840319935006087 | 0.0311467709990085 | 4168 | 1 | 1 | 11 | 8 | 0 | 0 | 0 | 2 | 0 | 0 | 9 | 6 |
| L1M5_dup29231 | 3.51786269661245 | -4.73397206658142 | 1.30198982098737 | -3.63595167202721 | 0.000276956255596371 | 0.0141175306944175 | 4169 | 0 | 1 | 15 | 20 | 0 | 0 | 0 | 1 | 0 | 0 | 7 | 4 |
| L1MCc_dup335 | 2.56995236165581 | -4.27436798402404 | 1.27364326777509 | -3.35601662739589 | 0.000790738145385968 | 0.0298279325046136 | 4170 | 1 | 0 | 8 | 16 | 0 | 0 | 0 | 2 | 0 | 0 | 5 | 3 |
| AluSx_dup29636 | 2.60069273477652 | -4.84550000744356 | 1.36504790462632 | -3.54969227894608 | 0.000385681694427462 | 0.0180014547787832 | 4171 | 0 | 0 | 10 | 13 | 0 | 0 | 0 | 1 | 0 | 0 | 3 | 9 |
| LTR26C_dup147 | 2.68018439824687 | -4.87518351438095 | 1.27276537608296 | -3.83038665726808 | 0.00012794205640351 | 0.00812254550506657 | 4172 | 0 | 0 | 9 | 11 | 0 | 0 | 0 | 2 | 0 | 0 | 7 | 7 |
| MIRc_dup16265 | 2.99491036831555 | -5.04331269903049 | 1.28304810766419 | -3.93072767022892 | 8.46891553698381e-05 | 0.00603092803958492 | 4173 | 0 | 0 | 7 | 19 | 0 | 0 | 0 | 3 | 0 | 0 | 5 | 7 |
| MER102b_dup2724 | 4.06907285153623 | -4.76198295765968 | 1.46024664250053 | -3.26108125782453 | 0.00110988229261281 | 0.0377005070661025 | 4174 | 1 | 0 | 31 | 17 | 0 | 0 | 0 | 0 | 0 | 0 | 3 | 3 |
| L1MCa_dup5588 | 3.30552058670639 | -5.09215989252381 | 1.55800325523186 | -3.2683884808482 | 0.0010816179270099 | 0.0370631709764255 | 4175 | 0 | 0 | 11 | 29 | 0 | 0 | 0 | 0 | 0 | 0 | 4 | 3 |
| L1PA4_dup5700 | 3.14776232651062 | -4.57363491229217 | 1.44474296156052 | -3.1657083882603 | 0.00154705803087459 | 0.0474562322899501 | 4176 | 0 | 0 | 15 | 17 | 1 | 0 | 0 | 0 | 0 | 0 | 7 | 3 |
| L1MEd_dup16929 | 3.09750987114279 | -4.54473390545388 | 1.43899674923326 | -3.15826558181972 | 0.00158710895853901 | 0.0483056906423011 | 4177 | 0 | 1 | 15 | 17 | 0 | 0 | 0 | 0 | 0 | 0 | 3 | 7 |
| L1PA16_dup11229 | 2.74806296850904 | -4.91995794646867 | 1.53854941927495 | -3.19778999935356 | 0.00138485088935608 | 0.0439887504630519 | 4178 | 0 | 0 | 14 | 10 | 0 | 0 | 0 | 0 | 0 | 0 | 3 | 11 |
| L1MEd_dup16928 | 3.20700891353319 | -5.13588807840932 | 1.53147456658297 | -3.35355753890744 | 0.000797798244368556 | 0.0300131068860902 | 4179 | 0 | 0 | 14 | 19 | 0 | 0 | 0 | 0 | 0 | 0 | 3 | 9 |
| L1PA15_dup6839 | 2.94915744974158 | -5.02606328588187 | 1.56377604364935 | -3.21405568674186 | 0.00130874288932577 | 0.0423054117026911 | 4180 | 0 | 0 | 10 | 19 | 0 | 0 | 0 | 0 | 0 | 0 | 2 | 11 |
| L1PB1_dup10569 | 4.61780661078293 | -4.66369905713594 | 1.34956666487571 | -3.45570113616095 | 0.000548863519017993 | 0.0230213077553773 | 4181 | 1 | 0 | 22 | 23 | 0 | 0 | 0 | 0 | 0 | 1 | 5 | 12 |
| AluJb_dup34104 | 4.37535982874777 | -5.07830391612848 | 1.37671861558992 | -3.68870142280486 | 0.000225401545094952 | 0.012123281204603 | 4182 | 1 | 0 | 20 | 15 | 0 | 0 | 0 | 0 | 0 | 0 | 9 | 15 |

| | | | | | | | | | | | | | | | | | | | |
|---|---|---|---|---|---|---|---|---|---|---|---|---|---|---|---|---|---|---|---|
| L1M5_dup12195 | 4.17428956378077 | -4.99609528221599 | 1.40932583657783 | -3.54502497048353 | 0.000392575893841323 | 0.0182480686923164 | 4183 | 1 | 0 | 18 | 25 | 0 | 0 | 0 | 0 | 0 | 0 | 7 | 7 |
| L1PB1_dup6230 | 3.70083041724462 | -3.72425087147245 | 1.15416076857273 | -3.22680424849128 | 0.00125181067949666 | 0.0410336608270441 | 4184 | 1 | 0 | 22 | 15 | 1 | 0 | 1 | 1 | 1 | 0 | 1 | 7 |
| Tigger2a_dup755 | 2.61188503971147 | -3.78576127465608 | 1.16655421740205 | -3.24525102921241 | 0.00117347128131395 | 0.0392219554150969 | 4185 | 1 | 0 | 13 | 9 | 0 | 0 | 1 | 1 | 1 | 0 | 3 | 6 |
| L1MA6_dup4278 | 3.79193903562912 | -3.79068378379433 | 1.15534696460787 | -3.28099168467622 | 0.00103442793842233 | 0.0359115830516252 | 4186 | 1 | 0 | 14 | 25 | 1 | 0 | 2 | 0 | 1 | 0 | 3 | 6 |
| L1MB1_dup510 | 3.70707420870819 | -3.81847973918531 | 1.14358056815889 | -3.33905615879157 | 0.000840635678066656 | 0.0311518294741082 | 4187 | 1 | 1 | 14 | 18 | 0 | 0 | 2 | 0 | 1 | 0 | 6 | 8 |
| L1PA5_dup10730 | 3.96707555201863 | -3.88902502686869 | 1.10490785743096 | -3.51977316543945 | 0.000431916015430559 | 0.0195280259354976 | 4188 | 1 | 1 | 16 | 21 | 0 | 0 | 4 | 0 | 1 | 0 | 4 | 7 |
| LTR27E_dup117 | 6.03633282674094 | -4.87047189220561 | 1.35466305718722 | -3.59533824028427 | 0.000323970380133214 | 0.0158030865254567 | 4189 | 0 | 0 | 26 | 41 | 0 | 0 | 2 | 0 | 0 | 3 | 7 | 5 |
| AluJr_dup578 | 3.44532205312596 | -4.74064258661302 | 1.30806098495965 | -3.62417550949221 | 0.00028988471251813 | 0.0145982873661035 | 4190 | 0 | 0 | 13 | 23 | 0 | 0 | 1 | 0 | 0 | 1 | 6 | 4 |
| L1PB1_dup7091 | 3.63637523491815 | -4.09243767654753 | 1.29942260936872 | -3.14942778972863 | 0.00163590531590952 | 0.0493355039669718 | 4191 | 0 | 0 | 16 | 19 | 0 | 0 | 3 | 0 | 0 | 3 | 5 | 4 |
| L1MDa_dup5329 | 4.62697520221172 | -4.20058647332329 | 1.31203710953031 | -3.20157596367609 | 0.00136678032236415 | 0.04361158348817 | 4192 | 0 | 1 | 15 | 32 | 0 | 0 | 0 | 0 | 1 | 1 | 9 | 5 |
| MER92B_dup533 | 3.6060459833098 | -4.37164365889333 | 1.34518308905754 | -3.2498502950674 | 0.00115465774959952 | 0.0388006947890204 | 4193 | 0 | 0 | 16 | 18 | 0 | 0 | 0 | 0 | 1 | 1 | 7 | 6 |
| MIRb_dup167772 | 3.58782002402969 | -4.29015749956777 | 1.35868581905275 | -3.1575787716389 | 0.00159085249423852 | 0.0483880993382834 | 4194 | 1 | 0 | 11 | 24 | 0 | 0 | 0 | 0 | 0 | 1 | 8 | 5 |
| HSMAR2_dup1428 | 4.55229943308459 | -4.73923339022703 | 1.46343197320251 | -3.23843777982784 | 0.00120186232985366 | 0.0398909658166213 | 4195 | 0 | 0 | 23 | 27 | 0 | 0 | 0 | 0 | 0 | 2 | 4 | 7 |
| L1MB5_dup3888 | 3.71414425598352 | -4.79480083680016 | 1.45440665754959 | -3.29674015992097 | 0.000978139486865521 | 0.0346088294931901 | 4196 | 0 | 0 | 17 | 27 | 0 | 0 | 0 | 0 | 0 | 1 | 3 | 4 |
| L1MA9_dup12876 | 5.34661575328002 | -3.97018280950924 | 1.24225812737924 | -3.19594029775842 | 0.00139375948112039 | 0.044189139706152 | 4197 | 1 | 2 | 14 | 36 | 0 | 0 | 0 | 0 | 0 | 1 | 10 | 11 |
| L1PA14_dup319 | 6.17623504211353 | -3.31894052271331 | 1.03640396378682 | -3.20236185761633 | 0.00136305658332946 | 0.0435289037413651 | 4198 | 2 | 1 | 17 | 31 | 0 | 1 | 0 | 0 | 1 | 2 | 9 | 22 |
| AluY_dup11505 | 4.54267808062455 | -3.16525142487846 | 1.00166821731442 | -3.1599798917099 | 0.00157780027762654 | 0.0480644302727082 | 4199 | 1 | 0 | 13 | 17 | 1 | 0 | 1 | 0 | 2 | 2 | 8 | 17 |
| MLT2C1_dup224 | 5.66257727175932 | -4.02231141160417 | 1.24660621507315 | -3.22660946413469 | 0.00125266306462725 | 0.0410447487177319 | 4200 | 1 | 0 | 23 | 25 | 0 | 0 | 0 | 0 | 3 | 1 | 8 | 16 |
| L1MEd_dup16927 | 5.87994609716784 | -5.01692046805358 | 1.323942184779 | -3.78938032621949 | 0.000151023545193551 | 0.00913690528467448 | 4201 | 0 | 1 | 22 | 31 | 0 | 0 | 0 | 0 | 1 | 0 | 12 | 14 |
| ERVL-E-int_dup6619 | 5.17399002951726 | -5.3516942041929 | 1.39420192132019 | -3.83853595548435 | 0.000123770092420009 | 0.00792321722113842 | 4202 | 0 | 0 | 19 | 31 | 0 | 0 | 0 | 0 | 1 | 0 | 8 | 13 |
| LTR26D_dup108 | 3.85370495720509 | -4.91538833398072 | 1.4087254617897 | -3.48924504263309 | 0.000484386889718142 | 0.0210747710478832 | 4203 | 0 | 0 | 10 | 26 | 0 | 0 | 0 | 0 | 1 | 0 | 7 | 10 |
| MLT1D_dup5580 | 3.81359086454028 | -4.36309954363055 | 1.33614395650686 | -3.26544121416169 | 0.00109293688831069 | 0.037316682569375 | 4204 | 0 | 0 | 13 | 19 | 0 | 1 | 0 | 0 | 1 | 0 | 5 | 14 |
| LTR16C_dup4400 | 3.87726428620532 | -4.92172238407465 | 1.41379023307519 | -3.48122533946866 | 0.000499125388550293 | 0.0215466055341052 | 4205 | 0 | 0 | 19 | 20 | 0 | 0 | 0 | 0 | 1 | 0 | 6 | 7 |
| L1MB1_dup4721 | 3.29457291053942 | -4.68015693838722 | 1.44154403913852 | -3.24662779028528 | 0.00116781007473783 | 0.0391043816554417 | 4206 | 0 | 0 | 18 | 15 | 0 | 0 | 0 | 0 | 1 | 0 | 3 | 8 |
| MER77_dup900 | 2.92963022381266 | -4.50961218948476 | 1.42428428090545 | -3.16623040073004 | 0.00154428422837508 | 0.0474137103232176 | 4207 | 0 | 0 | 14 | 13 | 0 | 0 | 0 | 0 | 1 | 0 | 4 | 8 |
| BSR/Beta_dup2019 | 8.47330692131734 | -3.26763898260483 | 0.760345870206613 | -4.29756918613486 | 1.72681370628063e-05 | 0.00183186988031498 | 4208 | 0 | 1 | 30 | 21 | 7 | 3 | 6 | 10 | 0 | 0 | 14 | 19 |
| LSAU_dup132 | 8.1657935745269 | -2.68030029864829 | 0.828117384215569 | -3.23661880517965 | 0.00120954866021404 | 0.0400678359325199 | 4209 | 0 | 1 | 27 | 17 | 9 | 6 | 5 | 7 | 0 | 0 | 12 | 24 |
| L1MC1_dup4540 | 7.92471315963059 | -2.56075698265188 | 0.684952390856708 | -3.73859120259292 | 0.000185054359247451 | 0.0105351330167045 | 4210 | 3 | 2 | 17 | 29 | 4 | 7 | 6 | 4 | 0 | 0 | 14 | 22 |
| LTR53-int_dup339 | 6.76243952187151 | -2.80568661408474 | 0.732803529450669 | -3.82870237563943 | 0.000128820678041084 | 0.00815873437773858 | 4211 | 1 | 2 | 14 | 14 | 3 | 4 | 5 | 3 | 0 | 0 | 20 | 25 |
| L1MC1_dup1664 | 9.67997564855862 | -3.85054930710158 | 0.772698514069361 | -4.9832492712105 | 6.25253174536243e-07 | 0.00014250621826456 | 4212 | 1 | 0 | 24 | 47 | 1 | 5 | 11 | 4 | 1 | 0 | 14 | 26 |
| L1PB1_dup1590 | 9.2006079103541 | -3.88395926943332 | 0.733143539997048 | -5.29767918223621 | 1.17283794616585e-07 | 3.73770101522728e-05 | 4213 | 1 | 2 | 23 | 32 | 1 | 2 | 6 | 5 | 1 | 0 | 20 | 32 |
| BSR/Beta_dup1966 | 8.98658290231478 | -2.61212378678342 | 0.706234541244408 | -3.69866331117249 | 0.000216737877525552 | 0.0117851716192173 | 4214 | 3 | 2 | 28 | 30 | 2 | 12 | 6 | 9 | 0 | 0 | 9 | 19 |
| BSR/Beta_dup1971 | 11.9217112939001 | -3.75925302280756 | 0.783756652983366 | -4.79645436947549 | 1.61498724952015e-06 | 0.000299077259378741 | 4215 | 1 | 1 | 39 | 65 | 4 | 6 | 4 | 9 | 0 | 0 | 15 | 18 |
| AluSg_dup35568 | 7.74219186553575 | -2.31141554725028 | 0.72220174595624 | -3.20051226709487 | 0.00137183529697644 | 0.0437446692005302 | 4216 | 4 | 4 | 18 | 40 | 5 | 4 | 5 | 1 | 1 | 0 | 10 | 15 |
| L1PA16_dup12336 | 7.37126461231042 | -2.00410043428358 | 0.611284549766525 | -3.27850660555551 | 0.00104357921794302 | 0.0361396449705531 | 4217 | 4 | 8 | 16 | 36 | 4 | 3 | 8 | 5 | 1 | 0 | 6 | 9 |
| MER58A_dup12187 | 6.87347394406121 | -2.51524993087987 | 0.773968647351077 | -3.24980855424617 | 0.00115482723179142 | 0.0388006947890204 | 4218 | 6 | 5 | 24 | 22 | 2 | 2 | 3 | 2 | 0 | 0 | 12 | 15 |
| MLT1H_dup4654 | 6.85442190433305 | -2.90222338471444 | 0.791424757526169 | -3.66708693039398 | 0.000245329410143425 | 0.0129433029205731 | 4219 | 6 | 3 | 25 | 34 | 2 | 1 | 4 | 3 | 0 | 0 | 6 | 10 |
| L1ME4a_dup33291 | 6.28337429387307 | -2.18640359822305 | 0.655454619916132 | -3.33570552680339 | 0.000850832358351423 | 0.0314006179973985 | 4220 | 4 | 2 | 19 | 19 | 5 | 3 | 3 | 5 | 1 | 0 | 11 | 11 |
| THE1A-int_dup49 | 5.70943833489056 | -2.45536064763703 | 0.67418428428329 | -3.64197253611047 | 0.000270556943197922 | 0.0138826329534458 | 4221 | 3 | 2 | 12 | 21 | 5 | 2 | 5 | 5 | 0 | 0 | 8 | 14 |
| THE1B-int_dup2190 | 6.1212890257362 | -2.95521803673664 | 0.798312696382447 | -3.70183018524972 | 0.000214049867272541 | 0.0116683412870635 | 4222 | 3 | 1 | 18 | 31 | 4 | 2 | 3 | 3 | 0 | 0 | 7 | 12 |
| L1PB1_dup4620 | 5.16917400158624 | -2.41326350376681 | 0.72175316561109 | -3.34361332758902 | 0.000826949104210054 | 0.0307779653713189 | 4223 | 1 | 1 | 7 | 23 | 2 | 2 | 2 | 4 | 1 | 3 | 13 | 10 |
| L2c_dup60367 | 5.85684502613072 | -2.32829639840487 | 0.706186904224967 | -3.29699741594636 | 0.000977244003318961 | 0.0345925768541704 | 4224 | 3 | 1 | 12 | 27 | 2 | 1 | 3 | 2 | 2 | 3 | 13 | 10 |
| AluJb_dup53286 | 5.12705945426186 | -2.43734989645273 | 0.737023188971681 | -3.30701928097187 | 0.000942944015056177 | 0.0337448874427067 | 4225 | 3 | 2 | 11 | 28 | 2 | 1 | 2 | 5 | 1 | 1 | 6 | 7 |
| L1PA4_dup5223 | 4.93617205812245 | -2.6820969026527 | 0.783014638230719 | -3.42534707743536 | 0.000614014482395436 | 0.0249616062960123 | 4226 | 1 | 1 | 17 | 17 | 2 | 1 | 1 | 4 | 2 | 1 | 7 | 11 |
| L2_dup27556 | 7.52567428278852 | -2.59797431321073 | 0.757892039086529 | -3.42789497609978 | 0.000608280731244742 | 0.0248178955550002 | 4227 | 1 | 3 | 25 | 37 | 4 | 2 | 3 | 2 | 2 | 1 | 5 | 18 |
| MIRc_dup96581 | 6.47062085276679 | -2.35047054399206 | 0.740127067426825 | -3.1757662264182 | 0.00149441326619432 | 0.0464034528302068 | 4228 | 1 | 1 | 19 | 32 | 4 | 4 | 3 | 2 | 2 | 1 | 7 | 12 |
| MLT1F_dup4136 | 7.02966573804301 | -2.5159517803876 | 0.705693582914604 | -3.5652184479224 | 0.000363553256116999 | 0.0172203754692705 | 4229 | 1 | 1 | 18 | 34 | 5 | 3 | 5 | 2 | 1 | 2 | 11 | 13 |
| L1PA7_dup5925 | 5.48303926982349 | -2.23604484047964 | 0.710353419021841 | -3.1477920435136 | 0.00164508682717941 | 0.0495495987569662 | 4230 | 3 | 1 | 15 | 22 | 4 | 1 | 5 | 1 | 1 | 2 | 7 | 13 |
| MSTB_dup8313 | 8.07831894759436 | -2.61743689438187 | 0.733283724402391 | -3.56947359838788 | 0.000357699262653397 | 0.0170019006814755 | 4231 | 3 | 2 | 27 | 38 | 2 | 2 | 4 | 1 | 1 | 4 | 9 | 18 |
| MSTA-int_dup3049 | 5.83228468712 | -2.36437273059286 | 0.739329092006041 | -3.19799769298614 | 0.00138385387467927 | 0.0439746920492032 | 4232 | 6 | 1 | 22 | 23 | 2 | 3 | 6 | 2 | 0 | 1 | 9 | 4 |
| AluJb_dup55597 | 5.37648542812046 | -2.35391801287817 | 0.748046784860949 | -3.14675239639687 | 0.00165094703143502 | 0.0496569640174234 | 4233 | 8 | 1 | 19 | 20 | 2 | 2 | 6 | 4 | 0 | 0 | 3 | 8 |
| L1M1_dup4464 | 5.39226437277284 | -3.05879420653753 | 0.761507906317121 | -4.0167596175472 | 5.90038555943123e-05 | 0.0046107492358766 | 4234 | 3 | 1 | 18 | 26 | 1 | 3 | 8 | 3 | 0 | 0 | 5 | 6 |
| THE1B_dup18257 | 6.18048006436652 | -3.46864085898723 | 0.786065247297745 | -4.41266277947203 | 1.02106971304263e-05 | 0.00124085522951809 | 4235 | 2 | 1 | 20 | 30 | 2 | 2 | 7 | 3 | 0 | 0 | 7 | 11 |
| Cheshire_dup804 | 7.98370977108724 | -2.91861846203275 | 0.772354441018285 | -3.77885891118175 | 0.000157548652673132 | 0.00944099406045314 | 4236 | 6 | 0 | 22 | 44 | 6 | 2 | 8 | 4 | 0 | 0 | 9 | 9 |
| L1MA7_dup8420 | 5.58872807122041 | -3.51722600811509 | 0.814943648918313 | -4.31591314661273 | 1.58944469903428e-05 | 0.0017238013142954 | 4237 | 2 | 0 | 21 | 24 | 1 | 3 | 8 | 5 | 0 | 0 | 8 | 3 |
| THE1C-int_dup929 | 8.21506463681077 | -3.33879259024777 | 0.811967466744781 | -4.11197828360434 | 3.92283205896711e-05 | 0.00338810234883117 | 4238 | 7 | 0 | 34 | 38 | 1 | 3 | 10 | 4 | 0 | 0 | 10 | 5 |
| L1PA4_dup5707 | 10.7093815581733 | -4.04972271445959 | 0.902584045578122 | -4.48680954898296 | 7.22976470349544e-06 | 0.000955188002297572 | 4239 | 3 | 0 | 49 | 60 | 1 | 3 | 4 | 2 | 0 | 1 | 15 | 9 |
| Tigger3b_dup3098 | 7.87206272901694 | -3.17173018774029 | 0.805204274843767 | -3.93903793960321 | 8.18089941211699e-05 | 0.00588278117713989 | 4240 | 3 | 0 | 37 | 30 | 3 | 2 | 5 | 2 | 0 | 2 | 14 | 8 |
| L2_dup27551 | 6.84476698036669 | -3.90339280960864 | 0.95247064375766 | -4.09817650044214 | 4.16417707815021e-05 | 0.00355373142098866 | 4241 | 2 | 0 | 25 | 38 | 3 | 1 | 4 | 1 | 0 | 0 | 9 | 12 |
| Tigger1_dup11311 | 7.28534487935865 | -4.51160967050906 | 0.989256849775836 | -4.56060493443273 | 5.10064652521493e-06 | 0.000732897607541541 | 4242 | 1 | 1 | 31 | 36 | 1 | 1 | 2 | 2 | 0 | 0 | 10 | 15 |
| L1P4a_dup201 | 9.11521771689078 | -4.45019106956235 | 0.87667785693368 | -5.07619878198772 | 3.85060452706433e-07 | 9.76229390903269e-05 | 4243 | 2 | 0 | 37 | 45 | 1 | 1 | 8 | 3 | 1 | 0 | 12 | 15 |
| L1MA3_dup972 | 7.53315947975661 | -4.25065577265256 | 0.859164906278342 | -4.94742713720139 | 7.52008389978475e-07 | 0.000165071391477692 | 4244 | 2 | 0 | 22 | 38 | 1 | 2 | 6 | 5 | 0 | 0 | 13 | 14 |
| MLT1A_dup8335 | 7.93062717852201 | -4.11829655648829 | 0.879711877373877 | -4.68141520242089 | 2.84901243225077e-06 | 0.000462422313918377 | 4245 | 3 | 0 | 27 | 44 | 1 | 2 | 6 | 3 | 0 | 0 | 9 | 15 |
| L1M5_dup52603 | 3.99256919766535 | -3.53539668142406 | 0.941689952616558 | -3.7543107172384 | 0.000173819137393222 | 0.0101073040447031 | 4246 | 1 | 1 | 11 | 10 | 1 | 1 | 1 | 7 | 0 | 0 | 6 | 13 |
| L2c_dup129459 | 5.55239279066359 | -3.71604299202019 | 0.831044417683142 | -4.47153354616123 | 7.76606744976013e-06 | 0.0010086815968344 | 4247 | 1 | 1 | 16 | 14 | 2 | 1 | 3 | 10 | 0 | 0 | 9 | 15 |
| L3_dup44835 | 4.03056273987319 | -3.23219719454505 | 0.918975967858954 | -3.51717270917919 | 0.000436169881112666 | 0.019667401654583 | 4248 | 0 | 2 | 13 | 13 | 2 | 1 | 1 | 10 | 0 | 0 | 3 | 6 |
| LTR40b_dup1043 | 4.77522767424088 | -2.68814406472537 | 0.844523912441739 | -3.1830289529082 | 0.00145743039974869 | 0.0455822468320982 | 4249 | 0 | 4 | 13 | 23 | 3 | 2 | 2 | 5 | 0 | 0 | 4 | 8 |
| MIRb_dup170215 | 6.14919795199126 | -2.4602927113204 | 0.766024982534577 | -3.21176563090663 | 0.0013192194947045 | 0.0425285194264947 | 4250 | 2 | 2 | 23 | 29 | 1 | 5 | 2 | 4 | 0 | 2 | 5 | 8 |
| MLT2C1_dup547 | 5.30996893502749 | -3.24812485851607 | 0.92271792917306 | -3.52017095996719 | 0.000431268723143227 | 0.0195280259354976 | 4251 | 2 | 2 | 26 | 25 | 1 | 2 | 1 | 3 | 0 | 0 | 2 | 8 |
| ALR/Alpha_dup1731 | 5.06613625627179 | -3.16354327975869 | 0.924184539116367 | -3.42306449184209 | 0.000619193847382983 | 0.025094098408732 | 4252 | 1 | 0 | 20 | 26 | 5 | 1 | 3 | 3 | 0 | 0 | 3 | 7 |
| L1MCa_dup3565 | 4.78951221462752 | -2.74574914077118 | 0.872921105759759 | -3.14547227997355 | 0.00165818908017252 | 0.0498110780998231 | 4253 | 3 | 0 | 15 | 26 | 4 | 2 | 3 | 2 | 0 | 0 | 4 | 7 |
| AluSz_dup42643 | 3.73476616164336 | -2.79489273428551 | 0.854407785649477 | -3.27114614500028 | 0.00107112541504532 | 0.0368362491701652 | 4254 | 1 | 1 | 11 | 17 | 2 | 1 | 4 | 1 | 0 | 1 | 7 | 5 |
| L2a_dup52073 | 4.10012727871607 | -2.93410497429357 | 0.811253947966351 | -3.61675278332854 | 0.000298321974121674 | 0.0149011992442251 | 4255 | 1 | 1 | 8 | 19 | 2 | 1 | 5 | 2 | 0 | 1 | 9 | 7 |
| L1ME1_dup18249 | 3.55562766281041 | -2.88261215420473 | 0.855949448195971 | -3.36773644784774 | 0.000757880073874993 | 0.0289542450226924 | 4256 | 1 | 1 | 14 | 18 | 1 | 1 | 4 | 3 | 0 | 1 | 2 | 2 |
| MER54A_dup47 | 4.3444617322846 | -2.93625860375593 | 0.893418950325187 | -3.28654166411759 | 0.00101425768492807 | 0.035423243666211 | 4257 | 1 | 1 | 17 | 22 | 1 | 1 | 2 | 3 | 0 | 2 | 2 | 7 |
| L1MEc_dup2716 | 5.85529131464089 | -4.47524280909738 | 0.972941147094414 | -4.59970556540056 | 4.23088508616648e-06 | 0.000629988850258429 | 4258 | 0 | 1 | 21 | 30 | 1 | 0 | 5 | 4 | 0 | 1 | 6 | 11 |
| MLT1D_dup1856 | 6.11836593589812 | -4.55415820374055 | 1.01396720737127 | -4.49142553194328 | 7.07480379213596e-06 | 0.000937839143801439 | 4259 | 0 | 2 | 15 | 38 | 0 | 0 | 5 | 4 | 0 | 1 | 6 | 14 |
| L1PA2_dup4270 | 4.83657527536839 | -3.69270692979228 | 0.976106237183141 | -3.7830994098027 | 0.000154887516821172 | 0.00932090669639116 | 4260 | 0 | 2 | 10 | 15 | 1 | 0 | 1 | 3 | 0 | 1 | 9 | 24 |
| MSTB-int_dup25 | 6.34659283087 | -3.95597366010736 | 1.0461855178279 | -3.78133093289303 | 0.000155992144716678 | 0.00937079193808502 | 4261 | 0 | 4 | 16 | 29 | 1 | 0 | 4 | 1 | 0 | 0 | 7 | 27 |

| | | | | | | | | | | | | | | | | | | |
|---|---|---|---|---|---|---|---|---|---|---|---|---|---|---|---|---|---|---|
| L2c_dup129941 | 5.68916200730935 | -3.43656560325463 | 0.933100430982114 | -3.68295361265405 | 0.000230547124326241 | 0.0123497897444691 | 4262 | 0 | 3 | 15 | 34 | 0 | 2 | 5 | 1 | 1 | 0 | 8 | 10 |
| L1MA6_dup509 | 4.96404616592473 | -3.78032334715989 | 1.04770469087705 | -3.60819549638108 | 0.000308334122374128 | 0.0152434803828037 | 4263 | 0 | 1 | 20 | 28 | 0 | 1 | 3 | 1 | 0 | 2 | 8 | 4 |
| MER39_dup1597 | 5.67880437175323 | -3.40710753919769 | 0.887101149551349 | -3.84072046453872 | 0.000122673743629336 | 0.00787846871838757 | 4264 | 0 | 1 | 23 | 24 | 0 | 3 | 5 | 2 | 0 | 2 | 9 | 8 |
| L1M1_dup887 | 5.85459280008285 | -3.43454150264177 | 0.833116176300948 | -4.12252408528568 | 3.74743414132846e-05 | 0.0032782608977667 | 4265 | 0 | 3 | 15 | 25 | 0 | 1 | 5 | 2 | 1 | 1 | 11 | 16 |
| AluSx_dup86074 | 4.30894881444845 | -2.95929204770976 | 0.775712213215399 | -3.81493548418327 | 0.000136218517145971 | 0.00848720902875497 | 4266 | 0 | 2 | 12 | 12 | 0 | 1 | 6 | 3 | 1 | 2 | 10 | 8 |
| MSTA-int_dup1636 | 5.40089929391557 | -3.58196610337775 | 0.842565954443503 | -4.25125900766257 | 2.12572162157879e-05 | 0.00213061060222488 | 4267 | 0 | 1 | 17 | 26 | 0 | 2 | 6 | 4 | 1 | 1 | 7 | 8 |
| L1PREC2_dup4091 | 4.05695377839331 | -4.39829479269282 | 1.0112025932538 | -4.34956834766433 | 1.36405765646242e-05 | 0.00153825622333937 | 4268 | 0 | 0 | 10 | 21 | 0 | 1 | 4 | 7 | 0 | 1 | 4 | 6 |
| L1MB7_dup15545 | 2.77576806218371 | -3.29875659732724 | 0.98959360080278 | -3.33344576465654 | 0.000857773930435149 | 0.0315803613881049 | 4269 | 0 | 1 | 10 | 12 | 0 | 1 | 3 | 3 | 0 | 1 | 3 | 3 |
| THE1B_dup16962 | 4.18948360817225 | -3.46897791748187 | 0.980870051950317 | -3.53663353324358 | 0.00040526157248527 | 0.0186519091153927 | 4270 | 0 | 1 | 15 | 22 | 0 | 1 | 3 | 4 | 0 | 2 | 6 | 2 |
| Tigger12A | 5.01319446017827 | -2.77654023816501 | 0.793777311201832 | -3.49788309514811 | 0.00046896664921905 | 0.0206359044478966 | 4271 | 0 | 0 | 14 | 14 | 0 | 4 | 6 | 6 | 0 | 4 | 8 | 10 |
| Kanga1a_dup701 | 3.14611106240466 | -3.13999244569144 | 0.977298081809501 | -3.2129321689424 | 0.00131387315507146 | 0.0424136293252792 | 4272 | 0 | 0 | 10 | 10 | 0 | 2 | 3 | 2 | 0 | 2 | 7 | 6 |
| L2c_dup104406 | 3.1714578211484 | -3.57862363314982 | 0.99008802704772 | -3.61444996342465 | 0.000300985935794249 | 0.0149954772777426 | 4273 | 0 | 0 | 9 | 10 | 0 | 1 | 5 | 4 | 0 | 2 | 5 | 6 |
| L1ME2_dup6016 | 3.91487652100449 | -3.22496577118105 | 0.99547222905617 | -3.23963409229278 | 0.00119683176902578 | 0.0397623617991143 | 4274 | 2 | 0 | 20 | 15 | 0 | 3 | 2 | 3 | 0 | 0 | 5 | 2 |
| MIRb_dup101994 | 4.7345406060457 | -3.8379046655621 | 0.913256000542767 | -4.20244122489329 | 2.640518207607e-05 | 0.00251219810246803 | 4275 | 3 | 0 | 22 | 13 | 0 | 1 | 5 | 6 | 0 | 0 | 7 | 5 |
| L1MC1_dup10765 | 3.62479049646727 | -3.1242860465666 | 0.944813119063472 | -3.30677674084743 | 0.000943760773902653 | 0.0337653714125662 | 4276 | 2 | 0 | 14 | 7 | 0 | 3 | 2 | 5 | 0 | 0 | 7 | 7 |
| L1P2_dup1361 | 3.78481168825787 | -3.90073265013817 | 1.02907511302626 | -3.79052277211046 | 0.000150330548628853 | 0.00910309530964414 | 4277 | 2 | 0 | 12 | 15 | 0 | 1 | 2 | 3 | 0 | 0 | 8 | 8 |
| MSTD-int_dup645 | 5.47774920739504 | -3.56768848360771 | 0.977933582767377 | -3.64819098809532 | 0.000264093293059464 | 0.0136578880048527 | 4278 | 1 | 0 | 16 | 28 | 0 | 5 | 5 | 2 | 0 | 0 | 12 | 6 |
| L1MA2_dup6387 | 4.5486820083395 | -4.62896580103477 | 1.17294566840349 | -3.94644519838273 | 7.93200661748259e-05 | 0.00574897729699517 | 4279 | 0 | 0 | 21 | 19 | 0 | 2 | 1 | 2 | 0 | 0 | 5 | 12 |
| LSAU_dup128 | 4.3870196029446 | -3.48127570688112 | 1.08717953959025 | -3.20211665148995 | 0.00136421741851994 | 0.0435538777972854 | 4280 | 0 | 0 | 17 | 17 | 0 | 5 | 3 | 2 | 0 | 0 | 4 | 12 |
| L1M2_dup1018 | 3.93356472915629 | -3.88125582110439 | 1.01642256674834 | -3.81854550270466 | 0.000134240825756803 | 0.00838814798464504 | 4281 | 0 | 0 | 18 | 10 | 1 | 2 | 2 | 7 | 0 | 0 | 7 | 3 |
| AluYj4_dup1654 | 2.83281443680741 | -3.91829844624632 | 1.09661802786055 | -3.57307498755125 | 0.000352813673463316 | 0.0168160792937214 | 4282 | 0 | 0 | 13 | 5 | 1 | 1 | 2 | 2 | 0 | 0 | 8 | 5 |
| MLT1F2_dup1977 | 4.05135650705219 | -3.96481480580229 | 0.976655049431767 | -4.05958563170188 | 4.91598772280736e-05 | 0.00402211393683941 | 4283 | 0 | 0 | 15 | 7 | 1 | 2 | 7 | 4 | 0 | 0 | 11 | 6 |
| L1PA10_dup7019 | 3.15581608780024 | -3.55535816313877 | 0.990639053661984 | -3.58895417053879 | 0.000332007194206692 | 0.0160732011867153 | 4284 | 0 | 0 | 8 | 4 | 1 | 2 | 5 | 5 | 0 | 0 | 7 | 9 |
| MIRb_dup202464 | 2.81688939899802 | -3.91751412561417 | 1.01837469068077 | -3.84682981761394 | 0.000119656020644543 | 0.00773686725225173 | 4285 | 0 | 0 | 7 | 8 | 1 | 1 | 4 | 3 | 0 | 0 | 4 | 10 |
| MSTB1_dup3638 | 3.16203694084293 | -3.56607065708079 | 0.99185210485 | -3.59536531670727 | 0.000323936685112589 | 0.0158030865254567 | 4286 | 0 | 0 | 8 | 8 | 2 | 1 | 4 | 3 | 0 | 0 | 8 | 8 |
| L1MC2_dup6272 | 4.48740424000187 | -2.55385675057018 | 0.787916880014046 | -3.2412768597173 | 0.00118995553293573 | 0.0395951837840758 | 4287 | 1 | 0 | 7 | 10 | 3 | 4 | 4 | 4 | 0 | 0 | 11 | 16 |
| L1ME3G_dup8346 | 3.70913562350721 | -2.38071167620523 | 0.755103281051895 | -3.15282920356111 | 0.00161696384004093 | 0.0488759731481423 | 4288 | 2 | 0 | 9 | 6 | 4 | 2 | 8 | 5 | 0 | 0 | 4 | 9 |
| L1MA1_dup1862 | 4.25412426375793 | -3.13466678190225 | 0.993331509329454 | -3.15571060865501 | 0.00160107629046447 | 0.0485682941115323 | 4289 | 0 | 1 | 17 | 18 | 4 | 1 | 2 | 1 | 0 | 0 | 5 | 9 |
| MLT1J_dup2087 | 3.95729001944738 | -3.23480686338659 | 0.920530503033419 | -3.51406808652939 | 0.000441299684209252 | 0.0198348119869389 | 4290 | 0 | 2 | 11 | 17 | 2 | 1 | 5 | 1 | 0 | 0 | 9 | 6 |
| MIRb_dup98628 | 4.61501095993677 | -3.69307150011913 | 0.992586738779653 | -3.72065367774268 | 0.000198707780093147 | 0.011065228514403 | 4291 | 0 | 1 | 19 | 16 | 1 | 1 | 2 | 1 | 1 | 0 | 10 | 10 |
| L1PA16_dup1518 | 5.80288769191702 | -3.54633236529073 | 0.913691091163202 | -3.88132531835893 | 0.000103888759984755 | 0.00701465888816122 | 4292 | 0 | 2 | 20 | 21 | 2 | 2 | 2 | 2 | 0 | 0 | 10 | 18 |
| MSTA-int_dup320 | 5.39175535962732 | -4.02398331696847 | 0.968893984980836 | -4.15317194589464 | 3.27898163530784e-05 | 0.00295650081165199 | 4293 | 0 | 1 | 16 | 25 | 2 | 1 | 3 | 2 | 0 | 0 | 10 | 14 |
| L1MEd_dup19963 | 5.31363511837515 | -3.97570694158562 | 0.95104908150844 | -4.18033834308513 | 2.91075677924972e-05 | 0.00269390483312078 | 4294 | 0 | 0 | 13 | 18 | 3 | 1 | 11 | 1 | 0 | 0 | 14 | 12 |
| MIR_dup157823 | 4.45659108588191 | -3.7375722114153 | 0.967406884991175 | -3.86349556675875 | 0.000111775922831724 | 0.00737413164866463 | 4295 | 0 | 0 | 13 | 23 | 3 | 1 | 7 | 2 | 0 | 0 | 3 | 10 |
| Charlie29a_dup1871 | 4.21149833405727 | -4.51658389018539 | 0.992921710255941 | -4.54878148350807 | 5.39574349832231e-06 | 0.000763314852825089 | 4296 | 0 | 0 | 11 | 11 | 1 | 1 | 8 | 2 | 0 | 0 | 7 | 17 |
| L1MEc_dup2521 | 3.71314372018906 | -4.32748185298738 | 1.06116102934831 | -4.0780633035921 | 4.54124032502724e-05 | 0.00379947246968682 | 4297 | 0 | 0 | 13 | 10 | 1 | 1 | 4 | 1 | 0 | 0 | 6 | 15 |
| L1P3_dup891 | 4.19972176548097 | -2.98536648315899 | 0.852735786552259 | -3.50092787266415 | 0.000463641308293592 | 0.0204792267947331 | 4298 | 3 | 0 | 9 | 22 | 1 | 1 | 3 | 4 | 1 | 0 | 5 | 8 |
| L1PA16_dup7507 | 3.52717078091394 | -3.28780645636224 | 0.875115777070411 | -3.75699597985617 | 0.000171965258100167 | 0.0100418153174827 | 4299 | 1 | 1 | 9 | 10 | 1 | 0 | 6 | 3 | 1 | 0 | 7 | 8 |
| AluSg4_dup6993 | 4.21615015646535 | -3.18828564047169 | 0.905366655963014 | -3.52154082489419 | 0.000429046598351964 | 0.0194605264756875 | 4300 | 2 | 0 | 7 | 19 | 1 | 0 | 3 | 2 | 2 | 0 | 11 | 10 |
| Tigger3a_dup2397 | 4.44366453787255 | -3.27476346762209 | 0.872540744055564 | -3.75313530048008 | 0.000174636534080125 | 0.0101301961819185 | 4301 | 1 | 0 | 12 | 18 | 2 | 0 | 5 | 2 | 2 | 0 | 8 | 10 |
| Eulor11_dup62 | 5.40043579988754 | -3.85156647644889 | 0.896592704529453 | -4.29578163751651 | 1.74078873164631e-05 | 0.00184422847524627 | 4302 | 0 | 0 | 12 | 12 | 1 | 2 | 2 | 6 | 1 | 0 | 15 | 20 |
| Plat_L3_dup2448 | 4.66745124318974 | -2.80059381281233 | 0.805070964667605 | -3.47869186161574 | 0.000503867601866441 | 0.0216962143311097 | 4303 | 0 | 0 | 11 | 19 | 4 | 1 | 5 | 5 | 2 | 0 | 5 | 10 |
| Tigger1_dup1431 | 4.43356264774721 | -2.96358989347585 | 0.84883224759768 | -3.49137288535308 | 0.00048054506914121 | 0.0209689669946588 | 4304 | 0 | 0 | 19 | 10 | 2 | 2 | 4 | 3 | 2 | 0 | 7 | 9 |
| AluJb_dup112925 | 6.93219778816885 | -4.39796619208466 | 0.857505158504021 | -5.12879269409554 | 2.91606241862181e-07 | 7.81379941735794e-05 | 4305 | 1 | 0 | 21 | 22 | 0 | 3 | 8 | 10 | 0 | 0 | 15 | 11 |
| MIRb_dup202452 | 6.91664200103409 | -5.21860146586639 | 0.956064666688914 | -5.45841892048965 | 4.80393143706076e-08 | 1.84963797689783e-05 | 4306 | 1 | 0 | 25 | 22 | 0 | 1 | 12 | 9 | 0 | 0 | 15 | 6 |
| L1MEc_dup17109 | 5.92500451596605 | -4.5581397714013 | 0.938355044681089 | -4.85758540675873 | 1.18825888355312e-06 | 0.000235421688847569 | 4307 | 1 | 0 | 18 | 22 | 0 | 2 | 7 | 7 | 0 | 0 | 6 | 17 |
| AluSx1_dup76146 | 5.54517642638775 | -4.91885140017116 | 0.960470821726687 | -5.12129185905751 | 3.03449557406777e-07 | 8.06560367557806e-05 | 4308 | 1 | 0 | 19 | 13 | 0 | 1 | 6 | 10 | 0 | 0 | 9 | 13 |
| L1MC5a_dup15443 | 4.61152042055023 | -4.64974469226383 | 0.988336602065433 | -4.70461650671113 | 2.54343588870383e-06 | 0.000425598028110775 | 4309 | 1 | 0 | 11 | 14 | 0 | 1 | 3 | 9 | 0 | 0 | 9 | 12 |
| L1PB1_dup1411 | 5.89120060568127 | -3.3250761353544 | 0.769536157788216 | -4.32088356304305 | 1.55405666543192e-05 | 0.00169316887240008 | 4310 | 1 | 0 | 8 | 21 | 1 | 5 | 9 | 8 | 0 | 0 | 11 | 15 |
| Tigger9a_dup967 | 5.95814970442203 | -4.14595068311311 | 0.836812456998242 | -4.95445622067481 | 7.25328444504885e-07 | 0.000159804631533481 | 4311 | 0 | 0 | 12 | 19 | 1 | 3 | 12 | 7 | 0 | 0 | 12 | 14 |
| AluSc_dup17335 | 5.41672193748581 | -3.63633163919943 | 0.795346477945928 | -4.57200948269824 | 4.83069071840723e-06 | 0.000702496054992057 | 4312 | 1 | 0 | 14 | 9 | 0 | 3 | 14 | 5 | 0 | 1 | 9 | 17 |
| AluSq_dup18246 | 4.07289819513503 | -2.75424008213534 | 0.833230456306874 | -3.30549617010275 | 0.000948083997584335 | 0.033880845984224 | 4313 | 2 | 0 | 10 | 8 | 0 | 5 | 7 | 4 | 0 | 0 | 10 | 8 |
| AluSx1_dup85187 | 6.63088542438065 | -4.21324737413593 | 0.820368278471344 | -5.13579996289812 | 2.80946371496476e-07 | 7.58797563133945e-05 | 4314 | 0 | 2 | 15 | 14 | 0 | 1 | 10 | 6 | 1 | 0 | 19 | 20 |
| MLT1G-int_dup39 | 6.95357091881014 | -4.015980659142 | 0.7958220377783 | -5.04633004428154 | 4.50377149209393e-07 | 0.000110479836232761 | 4315 | 0 | 3 | 16 | 19 | 1 | 1 | 8 | 9 | 0 | 0 | 10 | 26 |
| MamRTE1_dup8550 | 5.09232004342574 | -3.03423367501181 | 0.81110658853461 | -3.74085689587804 | 0.000183393924407496 | 0.0104710288659977 | 4316 | 0 | 5 | 12 | 11 | 1 | 1 | 9 | 3 | 0 | 0 | 9 | 18 |
| L2a_dup5596 | 6.20234838966956 | -5.03494412415515 | 0.96171193154589 | -5.23539737732255 | 1.64630276980446e-07 | 4.9212457143705e-05 | 4317 | 1 | 0 | 17 | 12 | 1 | 0 | 9 | 3 | 0 | 0 | 23 | 17 |
| AluJo_dup78138 | 5.94996625887358 | -4.12444111809045 | 0.84804720558251 | -4.8634569997285 | 1.15353094553577e-06 | 0.000229688765160212 | 4318 | 2 | 0 | 13 | 15 | 2 | 0 | 8 | 9 | 0 | 0 | 10 | 20 |
| HAL1_dup12678 | 4.16290098130702 | -4.44834617050732 | 0.952413514700977 | -4.67060378905263 | 3.00315698072969e-06 | 0.000481529104404788 | 4319 | 1 | 0 | 7 | 6 | 1 | 0 | 8 | 6 | 0 | 0 | 11 | 15 |
| L2a_dup134835 | 4.32320590381823 | -5.04963084431698 | 0.972350853441354 | -5.19321891521489 | 2.06688905140632e-07 | 5.92539737621601e-05 | 4320 | 0 | 0 | 6 | 15 | 0 | 1 | 7 | 5 | 0 | 0 | 13 | 11 |
| MSTB1_dup2322 | 4.45774427614276 | -5.10830005777347 | 0.945797298597506 | -5.40105164748135 | 6.62513462177037e-08 | 2.40675463009431e-05 | 4321 | 0 | 0 | 9 | 13 | 0 | 1 | 9 | 7 | 0 | 0 | 10 | 10 |
| L2c_dup87255 | 6.65702291774555 | -6.08661637514539 | 0.933673605099516 | -6.51899801161954 | 7.07785922471043e-11 | 7.4519835888626e-08 | 4322 | 0 | 0 | 13 | 14 | 0 | 0 | 7 | 7 | 0 | 0 | 20 | 28 |
| MIR_dup104345 | 5.31974266472606 | -5.72251461762761 | 0.97103786349489 | -5.89319410988934 | 3.78800987463211e-09 | 2.21459901137399e-06 | 4323 | 0 | 0 | 8 | 14 | 0 | 0 | 6 | 3 | 0 | 0 | 21 | 20 |
| MLT1D_dup14575 | 5.51166117101309 | -2.82234161822793 | 0.823500062947594 | -3.42725124771185 | 0.00060972464453947 | 0.0248554977333543 | 4324 | 0 | 2 | 20 | 25 | 1 | 1 | 5 | 5 | 0 | 4 | 3 | 8 |
| L1PA16_dup6811 | 7.34192984364841 | -2.71889341949178 | 0.793190724105157 | -3.42779275761088 | 0.000608509799430507 | 0.0248229852109191 | 4325 | 0 | 6 | 25 | 39 | 2 | 2 | 3 | 5 | 0 | 2 | 8 | 7 |
| MER65-int_dup107 | 6.69861907527062 | -3.28769750987054 | 0.815089189803763 | -4.03354326250122 | 5.49420598264357e-05 | 0.00437817929251878 | 4326 | 0 | 2 | 22 | 35 | 3 | 2 | 3 | 4 | 0 | 1 | 9 | 10 |
| AluJr_dup30354 | 8.5453417389369 | -2.11304300337707 | 0.655389937935622 | -3.2241004645766 | 0.00126369057574169 | 0.0412979148105248 | 4327 | 0 | 4 | 28 | 31 | 3 | 5 | 3 | 10 | 2 | 5 | 10 | 10 |
| MIR_dup44938 | 6.86975967400476 | -2.03527540251532 | 0.637863446040083 | -3.19076977235567 | 0.00141894301315935 | 0.0446792014336328 | 4328 | 0 | 2 | 22 | 15 | 3 | 3 | 6 | 5 | 1 | 8 | 10 | 15 |
| L1PA4_dup5703 | 6.0352344695586 | -3.13413266540805 | 0.901546860314515 | -3.476394631683 | 0.000508203882004173 | 0.0218079496676943 | 4329 | 0 | 1 | 21 | 34 | 3 | 3 | 3 | 1 | 0 | 1 | 9 | 7 |
| L1MB4_dup932 | 6.79893458348478 | -2.80193671856651 | 0.783122300045813 | -3.57790439424671 | 0.000346360088814262 | 0.0165754515831735 | 4330 | 0 | 1 | 26 | 32 | 4 | 2 | 4 | 3 | 2 | 1 | 10 | 6 |
| L1PA4_dup10384 | 5.3723487229463 | -3.34218888834104 | 0.878220706503552 | -3.80563662823125 | 0.000141439909452806 | 0.00872344756211957 | 4331 | 0 | 1 | 15 | 31 | 2 | 2 | 3 | 3 | 1 | 0 | 9 | 6 |
| L1MC_dup797 | 6.12230124971909 | -2.64444064302394 | 0.802754716433743 | -3.29420754420718 | 0.000986995930198368 | 0.0348031614710997 | 4332 | 0 | 2 | 16 | 32 | 2 | 1 | 6 | 1 | 4 | 1 | 9 | 9 |
| L1MB8_dup6725 | 7.13059634627223 | -2.27114443711868 | 0.614285394116841 | -3.69721380138609 | 0.000217978751158026 | 0.0118391170789883 | 4333 | 0 | 2 | 14 | 25 | 4 | 5 | 15 | 2 | 2 | 2 | 12 | 14 |
| MLT1A0_dup16534 | 7.71981705729509 | -2.2052920464272 | 0.61645934154458 | -3.57735198058918 | 0.000347092651926979 | 0.0165996279655287 | 4334 | 1 | 2 | 20 | 31 | 5 | 5 | 10 | 2 | 2 | 2 | 10 | 15 |
| MER21A_dup1917 | 8.9369241438876 | -2.673739324293 | 0.705711853849282 | -3.78871250314017 | 0.000151430031846909 | 0.00915750092617911 | 4335 | 3 | 1 | 29 | 30 | 2 | 1 | 6 | 4 | 0 | 9 | 16 | 19 |
| AluY_dup40857 | 7.71052806135217 | -2.46284911932337 | 0.656465787311909 | -3.75167932118478 | 0.000175654049199887 | 0.0101678703936047 | 4336 | 2 | 2 | 22 | 31 | 5 | 3 | 6 | 3 | 0 | 3 | 16 | 11 |
| MIR_dup168623 | 7.47921692644639 | -2.49335671041219 | 0.710167731279431 | -3.51094058571232 | 0.000446524185862922 | 0.0199865771123662 | 4337 | 2 | 1 | 25 | 22 | 6 | 1 | 6 | 2 | 0 | 4 | 13 | 19 |
| L1PB4_dup7336 | 7.36908565064831 | -2.74170978606796 | 0.757900826072317 | -3.6175046810232 | 0.000297456952633211 | 0.0148788505468283 | 4338 | 6 | 1 | 21 | 28 | 1 | 4 | 3 | 3 | 0 | 1 | 11 | 22 |
| Tigger1_dup11310 | 9.51246835569162 | -3.88086004549371 | 0.869441657491235 | -4.46362330589483 | 8.05851878995298e-06 | 0.00103536301943529 | 4339 | 3 | 2 | 37 | 48 | 0 | 2 | 3 | 2 | 0 | 1 | 14 | 19 |
| L1MA9_dup7554 | 9.5832491334839 | -2.45320792612988 | 0.70376226924057 | -3.48584747059137 | 0.000490580640966418 | 0.0212664786291484 | 4340 | 4 | 6 | 33 | 47 | 1 | 5 | 6 | 2 | 1 | 2 | 6 | 19 |

| | | | | | | | | | | | | | | | | | | | |
|---|---|---|---|---|---|---|---|---|---|---|---|---|---|---|---|---|---|---|---|
| L1MD_dup8165 | 8.49562109050922 | -3.09622093113348 | 0.73346612631467 | -4.22135504292552 | 2.42838096467846e-05 | 0.00235942644761691 | 4341 | 3 | 4 | 25 | 48 | 1 | 3 | 8 | 2 | 0 | 1 | 10 | 13 |
| L1PA4_dup11265 | 9.41409696750868 | -2.32261058494903 | 0.73281981279342 | -3.16941565225362 | 0.00152745785264046 | 0.0470915374714159 | 4342 | 10 | 6 | 26 | 32 | 1 | 2 | 4 | 2 | 1 | 1 | 14 | 30 |
| L1M5_dup50375 | 12.4276524746196 | -2.90251204912903 | 0.706872700169649 | -4.10613120075571 | 4.02341012285307e-05 | 0.00345490561202597 | 4343 | 6 | 8 | 42 | 52 | 3 | 3 | 8 | 2 | 0 | 1 | 16 | 30 |
| MLT1E2_dup3613 | 10.6343780081105 | -2.87411246627641 | 0.675563886461334 | -4.25439033061888 | 2.09619365467842e-05 | 0.0021178994822006 | 4344 | 5 | 7 | 33 | 48 | 1 | 2 | 6 | 6 | 0 | 2 | 12 | 23 |
| L1M2_dup853 | 11.354852125837 | -3.6225031573997 | 0.843698345982495 | -4.29359992780507 | 1.75799135604336e-05 | 0.00185914227202264 | 4345 | 0 | 4 | 36 | 43 | 1 | 0 | 5 | 3 | 0 | 6 | 21 | 36 |
| L2a_dup38865 | 9.21836110027692 | -3.20216867192563 | 0.828155554474739 | -3.86662705408844 | 0.000110350984934616 | 0.00732020503519714 | 4346 | 1 | 6 | 24 | 35 | 2 | 0 | 3 | 2 | 0 | 2 | 21 | 30 |
| L1M1_dup888 | 7.6377616919612 | -2.96575989942749 | 0.797054477322466 | -3.72089986796175 | 0.000198514144395317 | 0.011062212319037 | 4347 | 3 | 2 | 21 | 31 | 1 | 0 | 3 | 2 | 0 | 5 | 17 | 19 |
| L1MEd_dup2512 | 10.4315532690797 | -3.30250055291715 | 0.817093718593157 | -4.0417647055264 | 5.3050450579649e-05 | 0.00427468422405186 | 4348 | 3 | 2 | 27 | 56 | 3 | 0 | 2 | 3 | 0 | 4 | 18 | 26 |
| L1M6B_dup73 | 9.21281755453807 | -2.68134535453636 | 0.703238021101353 | -3.81285606591216 | 0.000137370119187588 | 0.0085343786893498 | 4349 | 2 | 5 | 31 | 24 | 3 | 1 | 2 | 4 | 1 | 3 | 13 | 35 |
| L1PA3_dup4145 | 7.42080863282758 | -2.85269775195331 | 0.789672009156345 | -3.61250964815255 | 0.000303247819887533 | 0.0150703019529989 | 4350 | 2 | 2 | 31 | 29 | 2 | 0 | 5 | 1 | 2 | 3 | 6 | 18 |
| L1MEc_dup2301 | 9.31782064563463 | -3.93714525009611 | 0.845892355063284 | -4.65442822190013 | 3.24880985094838e-06 | 0.000511270706874182 | 4351 | 2 | 2 | 36 | 38 | 1 | 0 | 3 | 2 | 1 | 1 | 17 | 24 |
| MER52A_dup1282 | 11.208193566801 | -3.01472266164045 | 0.729199994234528 | -4.13428782978136 | 3.56057014769849e-05 | 0.00314718954072907 | 4352 | 4 | 5 | 44 | 43 | 2 | 0 | 7 | 2 | 3 | 1 | 16 | 25 |
| MSTA_dup2235 | 5.19582938387223 | -2.12891815857496 | 0.616999017214528 | -3.45044011283204 | 0.000559673325312361 | 0.0233717144725825 | 4353 | 4 | 2 | 12 | 13 | 1 | 3 | 6 | 4 | 2 | 0 | 9 | 13 |
| L1PA10_dup6056 | 7.20940801578939 | -2.40069734764412 | 0.585546990244793 | -4.09992261533184 | 4.13288309843375e-05 | 0.00353487779201983 | 4354 | 3 | 3 | 17 | 20 | 1 | 4 | 7 | 4 | 2 | 1 | 12 | 23 |
| MER20_dup14687 | 6.45086067941947 | -2.59264086800687 | 0.607180020825064 | -4.26997064969935 | 1.95498692488983e-05 | 0.00201142945604017 | 4355 | 2 | 1 | 17 | 13 | 1 | 3 | 10 | 3 | 3 | 1 | 13 | 19 |
| MLT1D_dup16707 | 6.14283893520235 | -1.87920035575328 | 0.574115633557086 | -3.27320882051269 | 0.00106333887318938 | 0.0366532278023933 | 4356 | 8 | 2 | 8 | 7 | 1 | 2 | 11 | 7 | 1 | 2 | 14 | 18 |
| L1HS_dup63 | 8.23005874258135 | -2.98256985529089 | 0.5764561700877 | -5.17397507400629 | 2.29165409478713e-07 | 6.40011649340632e-05 | 4357 | 3 | 2 | 12 | 21 | 2 | 2 | 10 | 8 | 1 | 1 | 15 | 34 |
| L1PA8_dup4424 | 7.65386308622081 | -2.51970056730363 | 0.553676378698501 | -4.5508543695264 | 5.34285197966011e-06 | 0.000757634241449467 | 4358 | 2 | 3 | 12 | 10 | 3 | 3 | 10 | 7 | 1 | 1 | 22 | 27 |
| L1PA3_dup3650 | 6.29408508480531 | -2.16162104101452 | 0.657157314435054 | -3.28935095681439 | 0.00100418724468031 | 0.0351747884767704 | 4359 | 6 | 2 | 8 | 22 | 3 | 3 | 7 | 3 | 0 | 1 | 9 | 23 |
| MER92B_dup454 | 5.86089224157819 | -2.49190257270956 | 0.63145162411172 | -3.94630796336138 | 7.93655208108909e-05 | 0.00575052016287096 | 4360 | 4 | 1 | 8 | 14 | 2 | 3 | 12 | 3 | 0 | 1 | 14 | 18 |
| L1PA16_dup13714 | 5.72448760412532 | -2.3148515477772 | 0.681479409797289 | -3.39680335824933 | 0.000681779410772193 | 0.0268276308377047 | 4361 | 4 | 2 | 12 | 19 | 1 | 3 | 3 | 3 | 0 | 2 | 15 | 13 |
| MIR3_dup70847 | 4.64274935974732 | -2.37776920761597 | 0.728841810157431 | -3.26239408123742 | 0.00110475448397878 | 0.0375799729013317 | 4362 | 3 | 1 | 9 | 12 | 2 | 2 | 4 | 1 | 0 | 1 | 14 | 14 |
| HERVK14-int_dup305 | 5.29751446013756 | -2.61302524537885 | 0.693238710361409 | -3.76930082859423 | 0.000163705482747737 | 0.00969222811592264 | 4363 | 1 | 1 | 8 | 10 | 2 | 4 | 5 | 8 | 0 | 1 | 8 | 22 |
| L1MDa_dup3425 | 4.70260621034597 | -2.45946192128236 | 0.653350067853198 | -3.76438611135949 | 0.000166958739452056 | 0.00983589111603341 | 4364 | 3 | 2 | 5 | 9 | 1 | 2 | 8 | 9 | 0 | 1 | 10 | 11 |
| L2_dup48336 | 5.56343594766249 | -1.87906811011866 | 0.549525021776598 | -3.41944049070538 | 0.000627500562032534 | 0.025328456104053 | 4365 | 4 | 2 | 7 | 15 | 2 | 4 | 10 | 6 | 0 | 3 | 11 | 10 |
| L1MA9_dup8682 | 5.2063945432705 | -2.46295904268735 | 0.685386017625038 | -3.59353558338097 | 0.000326221074593711 | 0.015866717029654 | 4366 | 4 | 2 | 14 | 13 | 0 | 0 | 8 | 6 | 0 | 4 | 7 | 11 |
| MER94_dup1300 | 5.47474540830513 | -2.22406257983831 | 0.672082230070974 | -3.30921199866189 | 0.00093558965675228 | 0.0335776495423195 | 4367 | 4 | 5 | 7 | 13 | 0 | 0 | 8 | 9 | 0 | 3 | 13 | 9 |
| L1PA3_dup9308 | 5.5216990821678 | -2.90976153566678 | 0.716093405063974 | -4.06338267478784 | 4.83666383742929e-05 | 0.00397232565273664 | 4368 | 3 | 3 | 12 | 16 | 0 | 0 | 5 | 6 | 0 | 2 | 14 | 12 |
| MLT1L_dup6057 | 4.93963165914573 | -2.72810226341424 | 0.731189627868976 | -3.73104617384302 | 0.00019068626692492 | 0.0107298041547547 | 4369 | 2 | 3 | 9 | 18 | 0 | 0 | 6 | 7 | 0 | 3 | 10 | 7 |
| L2a_dup124330 | 5.72798557603523 | -4.46464157747998 | 0.907534397612881 | -4.91952876852214 | 8.67528174663836e-07 | 0.000184802917680001 | 4370 | 0 | 1 | 13 | 19 | 1 | 0 | 8 | 10 | 0 | 1 | 11 | 11 |
| MIRb_dup202454 | 5.69414833565247 | -3.20683020688034 | 0.71330214801979 | -4.49575290889404 | 6.93242035033664e-06 | 0.000923078731214999 | 4371 | 2 | 1 | 13 | 17 | 3 | 0 | 6 | 11 | 0 | 1 | 8 | 12 |
| L1PA10_dup6054 | 6.09983523347524 | -2.85873327745718 | 0.760007002199543 | -3.76145649866868 | 0.0001689268037805 | 0.00992016766590737 | 4372 | 3 | 2 | 7 | 18 | 4 | 0 | 5 | 5 | 0 | 0 | 18 | 20 |
| MER34D_dup65 | 4.09935231876983 | -3.16470789112848 | 0.91003108419765 | -3.47758219041355 | 0.000505957906194837 | 0.021762592181387 | 4373 | 2 | 2 | 6 | 15 | 1 | 0 | 4 | 1 | 0 | 0 | 9 | 17 |
| MER70C_dup229 | 4.21882875095956 | -4.00483390753512 | 0.9896624641992 | -4.04666646701175 | 5.19521829320212e-05 | 0.00420326899692034 | 4374 | 1 | 1 | 8 | 15 | 1 | 0 | 4 | 3 | 0 | 0 | 7 | 18 |
| Tigger17c_dup599 | 3.8553764387122 | -3.17331707891653 | 0.905971864800629 | -3.50266625511032 | 0.000460626220169327 | 0.0204028604959811 | 4375 | 3 | 1 | 9 | 13 | 1 | 0 | 2 | 4 | 0 | 0 | 6 | 13 |
| L1PA2_dup1112 | 4.82494831480971 | -3.02153337409491 | 0.757510603582481 | -3.98876710082371 | 6.64175899818505e-05 | 0.00505559816564984 | 4376 | 5 | 1 | 10 | 11 | 1 | 0 | 11 | 5 | 0 | 0 | 9 | 12 |
| L1MA4_dup2292 | 5.12406335602015 | -3.95455271607259 | 0.8868199226603 | -4.45925110050487 | 8.22465215196481e-06 | 0.00105100198467755 | 4377 | 3 | 1 | 16 | 15 | 0 | 0 | 7 | 5 | 0 | 0 | 8 | 14 |
| L1MEc_dup17110 | 4.2322252113632 | -3.33027960198935 | 0.881071183352409 | -3.77980765335882 | 0.000156949556897156 | 0.00941879426619184 | 4378 | 4 | 1 | 7 | 15 | 0 | 0 | 5 | 5 | 0 | 0 | 9 | 11 |
| L1PA10_dup1563 | 4.73591423834148 | -3.36706811454392 | 0.83540801224557 | -4.03044747618983 | 5.56707684079854e-05 | 0.00442587258924373 | 4379 | 2 | 3 | 5 | 6 | 0 | 0 | 12 | 6 | 0 | 0 | 13 | 16 |
| L1ME3_dup2267 | 4.32523497678137 | -3.24054559699015 | 0.893504466754308 | -3.62678164191117 | 0.000286975777031344 | 0.01449166770829 | 4380 | 1 | 4 | 10 | 9 | 0 | 0 | 5 | 3 | 0 | 0 | 11 | 15 |
| L1PB4_dup3835 | 4.52707449237049 | -4.11253578789315 | 0.93667679932992 | -4.3905601065759 | 1.13059056793571e-05 | 0.00133707506599419 | 4381 | 1 | 2 | 10 | 10 | 0 | 0 | 7 | 6 | 0 | 0 | 10 | 14 |
| MER4E1_dup838 | 5.00020614590709 | -3.89767828344156 | 0.885352700483427 | -4.40240175617392 | 1.07059073041664e-05 | 0.00128600010060026 | 4382 | 1 | 0 | 11 | 14 | 0 | 0 | 8 | 2 | 1 | 2 | 12 | 17 |
| MER5B_dup19916 | 4.71599015120626 | -4.24593727199555 | 0.944628369603349 | -4.49482294691025 | 6.96278568190674e-06 | 0.000926085523681935 | 4383 | 0 | 0 | 11 | 15 | 0 | 0 | 8 | 3 | 1 | 2 | 14 | 9 |
| Kanga1d_dup551 | 6.97276827829679 | -5.25346156040962 | 0.969310089856806 | -5.41979456871815 | 5.96675596611112e-08 | 2.20871875281316e-05 | 4384 | 0 | 0 | 20 | 23 | 0 | 0 | 6 | 5 | 1 | 1 | 15 | 23 |
| L1PBa_dup1859 | 6.77989383810399 | -4.51366210873876 | 0.927894490779058 | -4.86441309178277 | 1.14796933948747e-06 | 0.000229156637964502 | 4385 | 0 | 0 | 16 | 25 | 0 | 0 | 10 | 2 | 1 | 3 | 15 | 21 |
| L1M4_dup16436 | 7.5590555914236 | -4.46249456202473 | 0.934587105330425 | -4.77483001485132 | 1.79859111177248e-06 | 0.000324217849001375 | 4386 | 0 | 0 | 17 | 23 | 0 | 2 | 9 | 1 | 0 | 2 | 22 | 28 |
| L1MC3_dup10256 | 5.86478357809884 | -4.9913020785078 | 1.05197253625466 | -4.74470759120608 | 2.08807744537548e-06 | 0.000363992436496201 | 4387 | 0 | 0 | 21 | 19 | 1 | 0 | 3 | 2 | 0 | 1 | 17 | 15 |
| L2_dup12457 | 8.0718926630733 | -4.63470896946433 | 0.905878437115372 | -5.11625928995819 | 3.11654565559434e-07 | 8.22139215705157e-05 | 4388 | 0 | 0 | 19 | 21 | 1 | 0 | 5 | 3 | 0 | 3 | 27 | 30 |
| L1MD3_dup4032 | 8.37677556672764 | -4.26434766731874 | 0.756548301242742 | -5.63658349415883 | 1.73456955013506e-08 | 8.12468279874876e-06 | 4389 | 0 | 2 | 19 | 22 | 1 | 0 | 12 | 6 | 0 | 2 | 21 | 28 |
| Tigger3b_dup5417 | 8.4076339612332 | -4.06339210871093 | 0.750346504570236 | -5.41535421829979 | 6.1167491456642e-08 | 2.25652091816157e-05 | 4390 | 0 | 1 | 22 | 24 | 1 | 0 | 12 | 8 | 0 | 4 | 25 | 14 |
| AluJr4_dup15642 | 5.3436082590822 | -3.69782293285786 | 0.831928026043562 | -4.44488323159849 | 8.79396178012031e-06 | 0.00110538194914554 | 4391 | 0 | 1 | 13 | 12 | 0 | 0 | 7 | 6 | 0 | 4 | 15 | 12 |
| L1MC3_dup10658 | 4.8836507142694 | -3.39091550336124 | 0.894353303929516 | -3.79147199262594 | 0.000149757038848881 | 0.00908106227470335 | 4392 | 0 | 1 | 6 | 14 | 0 | 0 | 4 | 2 | 0 | 5 | 16 | 18 |
| L1M5_dup5792 | 6.4795085066823 | -2.99782451608949 | 0.893208191925454 | -3.3562438669838 | 0.000790088670775399 | 0.029822087382031 | 4393 | 1 | 4 | 21 | 30 | 0 | 0 | 3 | 2 | 0 | 4 | 12 | 11 |
| L1M6_dup578 | 7.80028332458343 | -4.11263854581061 | 1.00280700469995 | -4.10112666398969 | 4.11143419189359e-05 | 0.00352286395471485 | 4394 | 0 | 1 | 26 | 33 | 0 | 0 | 2 | 5 | 0 | 6 | 19 | 12 |
| LTR10C_dup389 | 6.66214102779584 | -3.46843964589733 | 0.898800681771573 | -3.85896419110507 | 0.000113868619083243 | 0.00746201664455557 | 4395 | 0 | 1 | 18 | 24 | 1 | 0 | 3 | 2 | 0 | 6 | 17 | 18 |
| L1PA16_dup11799 | 6.30289832907752 | -4.15114578686542 | 0.921597610792396 | -4.50429312994447 | 6.6594232948854e-06 | 0.000894234275013996 | 4396 | 1 | 2 | 14 | 19 | 0 | 1 | 2 | 6 | 0 | 0 | 13 | 27 |
| L1MA9_dup13442 | 6.04201759547891 | -5.0333663739971 | 0.991512404239673 | -5.0764532571399 | 3.84545343184984e-07 | 9.76229390903269e-05 | 4397 | 1 | 0 | 7 | 25 | 0 | 1 | 4 | 8 | 0 | 0 | 17 | 18 |
| LTR12E_dup114 | 5.32592737482927 | -3.64089773867744 | 0.863386958857659 | -4.2169941314549 | 2.47580415780564e-05 | 0.00239403773781171 | 4398 | 3 | 1 | 7 | 23 | 0 | 1 | 5 | 4 | 0 | 0 | 13 | 16 |
| ERV3-16A3_I-int_dup14 | 7.72987595845684 | -4.99254783077785 | 1.07293046697941 | -4.65318861233685 | 3.26841026317648e-06 | 0.000513787894657152 | 4399 | 3 | 0 | 28 | 33 | 0 | 0 | 2 | 5 | 0 | 0 | 14 | 20 |
| L1PA13_dup1560 | 5.77645490141855 | -5.99255006368363 | 1.10994503827275 | -5.39896108100001 | 6.70279111946598e-08 | 2.4238637320858e-05 | 4400 | 0 | 0 | 24 | 24 | 0 | 0 | 2 | 3 | 0 | 0 | 13 | 12 |
| L2_dup12458 | 4.5616943603345 | -5.63543849912828 | 1.04840292253795 | -5.37526019622889 | 7.64721679720517e-08 | 2.67985956999673e-05 | 4401 | 0 | 0 | 14 | 16 | 0 | 0 | 3 | 3 | 0 | 0 | 10 | 16 |
| L1PA10_dup740 | 5.5052108480457 | -5.4119645319351 | 1.02137328896577 | -5.29871359511973 | 1.16621396202952e-07 | 3.72978423936932e-05 | 4402 | 0 | 0 | 15 | 30 | 1 | 0 | 6 | 4 | 0 | 0 | 7 | 13 |
| THE1B-int_dup3680 | 6.31428594854141 | -5.12528890099007 | 1.00526743338443 | -5.09843324351487 | 3.42476267129024e-07 | 8.92561422325114e-05 | 4403 | 0 | 1 | 21 | 25 | 1 | 0 | 4 | 5 | 0 | 0 | 13 | 15 |
| L1PB1_dup2932 | 5.93532572256747 | -4.13123488177193 | 0.946074884772348 | -4.3667102343236 | 1.26131880538546e-05 | 0.00145058368519211 | 4404 | 1 | 3 | 10 | 27 | 0 | 0 | 6 | 3 | 0 | 0 | 13 | 19 |
| MLT1D_dup4548 | 4.7255710020556 | -2.78720975352701 | 0.885441213962651 | -3.14782021615337 | 0.00164492829247939 | 0.0495495987569662 | 4405 | 4 | 4 | 9 | 16 | 0 | 0 | 5 | 2 | 0 | 0 | 9 | 16 |
| L1PB1_dup1589 | 8.9538488475215 | -3.38429414567443 | 0.927688439179582 | -3.64809347917216 | 0.000264193519992582 | 0.0136592019467221 | 4406 | 6 | 5 | 24 | 31 | 0 | 0 | 5 | 2 | 0 | 0 | 20 | 30 |
| MLT1A1-int_dup56 | 6.50023860405888 | -3.36485666299805 | 1.01573485032448 | -3.3127313313343 | 0.000923896850827913 | 0.0332986307281276 | 4407 | 4 | 4 | 16 | 28 | 0 | 0 | 2 | 1 | 0 | 0 | 19 | 15 |
| HERVL-int_dup2030 | 5.89225668429077 | -3.446341212505 | 0.795338191303037 | -4.33317706881234 | 1.46972805693011e-05 | 0.00162454507892565 | 4408 | 3 | 1 | 16 | 14 | 0 | 0 | 4 | 4 | 0 | 2 | 15 | 20 |
| HAL1_dup2714 | 5.53264201843005 | -3.41855116738472 | 0.848810076486349 | -4.02746298858258 | 5.6381939333538e-05 | 0.00445709532779337 | 4409 | 3 | 1 | 19 | 18 | 0 | 1 | 2 | 4 | 0 | 1 | 12 | 13 |
| L1MC3_dup10261 | 5.94447080181754 | -3.7714076487287 | 0.822967580267534 | -4.582692853469 | 4.59026032374189e-06 | 0.000673387227492655 | 4410 | 1 | 1 | 13 | 23 | 0 | 1 | 5 | 4 | 0 | 2 | 12 | 19 |
| L1PREC2_dup930 | 7.79192792130034 | -4.17730241208352 | 0.848832162235598 | -4.92123484232927 | 8.5999860285099e-07 | 0.000183363107127224 | 4411 | 2 | 1 | 23 | 27 | 0 | 1 | 4 | 4 | 0 | 1 | 17 | 26 |
| AluJb_dup113372 | 5.68168865056066 | -2.89314050078245 | 0.868666252639273 | -3.33055473490792 | 0.000866731217204821 | 0.0318215130703623 | 4412 | 6 | 3 | 22 | 21 | 0 | 1 | 4 | 2 | 0 | 0 | 10 | 8 |
| THE1B-int_dup1021 | 6.7907129644434 | -3.41671076502223 | 0.766152531451653 | -4.4595698960212 | 8.21242886508616e-06 | 0.00105021314277552 | 4413 | 4 | 3 | 20 | 28 | 0 | 1 | 6 | 6 | 0 | 0 | 9 | 15 |
| X6B_LINE_dup596 | 5.1039536280508 | -3.39401998801471 | 0.795647866670881 | -4.26573127408213 | 1.99248490565202e-05 | 0.00203772930184352 | 4414 | 4 | 1 | 13 | 21 | 0 | 1 | 9 | 5 | 0 | 0 | 9 | 6 |
| L1M1_dup8382 | 6.19334593893455 | -3.11261802887053 | 0.853724311301572 | -3.64592877075867 | 0.00026642777295147 | 0.013745032323694 | 4415 | 1 | 0 | 13 | 16 | 1 | 0 | 0 | 8 | 4 | 3 | 17 | 16 |
| MER47A_dup260 | 5.54736416277083 | -3.51029576699229 | 0.937593485309655 | -3.74394214762804 | 0.000181155381306563 | 0.0103934874016178 | 4416 | 1 | 0 | 11 | 22 | 0 | 0 | 1 | 5 | 1 | 5 | 17 | 10 |
| AluY_dup11507 | 4.39152654614715 | -3.73374722065008 | 1.12976247862188 | -3.30489575579163 | 0.000950117314798758 | 0.033925062212951 | 4417 | 0 | 0 | 9 | 22 | 0 | 0 | 0 | 3 | 3 | 3 | 10 | 8 |
| FLAM_A_dup13112 | 3.49678409083322 | -3.54242727041354 | 1.02858439130251 | -3.44398310957034 | 0.000573211510231578 | 0.0237701962379859 | 4418 | 0 | 0 | 7 | 8 | 0 | 0 | 0 | 4 | 1 | 3 | 12 | 10 |
| MLT1D_dup1855 | 6.3674969580922 | -3.51410185945104 | 0.935366610113073 | -3.75692463410281 | 0.000172014273100412 | 0.0100418153174827 | 4419 | 1 | 2 | 19 | 32 | 0 | 0 | 1 | 7 | 3 | 0 | 10 | 9 |

| | | | | | | | | | | | | | | | | | | | |
|---|---|---|---|---|---|---|---|---|---|---|---|---|---|---|---|---|---|---|---|
| L1ME2_dup1491 | 6.44441886205279 | -3.36968948829307 | 1.02776201283769 | -3.27866709043784 | 0.00104298597866378 | 0.0361296173648466 | 4420 | 1 | 2 | 18 | 30 | 0 | 0 | 0 | 3 | 4 | 0 | 14 | 14 |
| L2_dup6896 | 8.4418929814179 | -3.39285576333148 | 1.0211365501373 | -3.32262689341136 | 0.000891741020224769 | 0.0324499643876148 | 4421 | 1 | 3 | 28 | 36 | 0 | 0 | 0 | 4 | 5 | 0 | 18 | 17 |
| THE1B-int_dup407 | 6.39045324826649 | -3.46885541337567 | 1.04117792408796 | -3.33166439003619 | 0.000863282962371882 | 0.0317242822802378 | 4422 | 2 | 1 | 25 | 29 | 0 | 0 | 0 | 3 | 0 | 4 | 12 | 10 |
| THE1C_dup997 | 9.10853522101875 | -2.89224422211807 | 0.844600491886378 | -3.42439324852673 | 0.000616173871914676 | 0.0250155448091809 | 4423 | 2 | 3 | 29 | 35 | 1 | 0 | 0 | 3 | 3 | 5 | 23 | 17 |
| MLT1A1_dup659 | 8.64466549631121 | -3.40065627025275 | 0.863908274582934 | -3.93636265597117 | 8.27259296093723e-05 | 0.00593079362664408 | 4424 | 3 | 3 | 26 | 31 | 0 | 0 | 2 | 1 | 1 | 2 | 26 | 22 |
| AluSp_dup6018 | 7.64223266810602 | -3.21940886694893 | 0.854278837478828 | -3.76856914359501 | 0.000164186007551877 | 0.0097061872678109 | 4425 | 4 | 2 | 20 | 33 | 0 | 0 | 1 | 3 | 2 | 1 | 15 | 23 |
| AluSz_dup22554 | 7.16982659783515 | -2.72108066582869 | 0.808261683733698 | -3.36658376932936 | 0.000761054634639052 | 0.0290208182842666 | 4426 | 4 | 2 | 17 | 33 | 0 | 0 | 2 | 2 | 2 | 4 | 17 | 14 |
| THE1D_dup11822 | 6.2391287687411 | -3.07056900420989 | 0.81263385606241 | -3.77853935238218 | 0.00015775092636927 | 0.00944782328866408 | 4427 | 3 | 2 | 17 | 35 | 0 | 0 | 6 | 2 | 2 | 1 | 7 | 11 |
| L1MEd_dup2511 | 6.97068174848055 | -2.90229186418437 | 0.722433309819092 | -4.01738378440932 | 5.8847841026467e-05 | 0.00460006845626929 | 4428 | 3 | 4 | 19 | 25 | 0 | 0 | 5 | 4 | 2 | 1 | 11 | 20 |
| MIRb_dup28761 | 7.80256246599887 | -3.43618301693346 | 0.815410024739858 | -4.21405539872988 | 2.50825719604238e-05 | 0.00241827296097852 | 4429 | 1 | 3 | 22 | 36 | 0 | 0 | 4 | 3 | 3 | 1 | 18 | 14 |
| L1M5_dup5787 | 5.42962029335495 | -2.68836704038893 | 0.753828920181233 | -3.56628270475827 | 0.000362080769361792 | 0.0171659037151945 | 4430 | 3 | 4 | 13 | 18 | 0 | 0 | 3 | 6 | 1 | 1 | 14 | 8 |
| L3b_dup9286 | 7.77139954480677 | -2.86330922247482 | 0.749460057107215 | -3.82049609625187 | 0.00013318351335198 | 0.00834179633273238 | 4431 | 4 | 5 | 24 | 37 | 0 | 1 | 4 | 4 | 1 | 1 | 14 | 10 |
| LTR89_dup444 | 5.73119156635067 | -2.27271793252973 | 0.654189926602407 | -3.47409496861758 | 0.000512579578804043 | 0.0219561224947087 | 4432 | 3 | 2 | 13 | 19 | 0 | 3 | 4 | 5 | 2 | 2 | 15 | 7 |
| MLT1F2_dup719 | 5.37965557604957 | -2.67678314160315 | 0.760945909152987 | -3.51770488467794 | 0.000435296170219336 | 0.0196391699884357 | 4433 | 3 | 1 | 18 | 22 | 0 | 2 | 3 | 3 | 1 | 2 | 10 | 7 |
| THE1B_dup2082 | 8.20995158996859 | -2.78994890074259 | 0.701419016136142 | -3.97757807609977 | 6.96207662395399e-05 | 0.00522581991609458 | 4434 | 3 | 3 | 26 | 30 | 0 | 1 | 4 | 5 | 1 | 5 | 19 | 12 |
| THE1D-int_dup1405 | 7.12517103178124 | -2.45221096910855 | 0.767075580414477 | -3.19683096649164 | 0.00138946323793244 | 0.0441117037509102 | 4435 | 2 | 3 | 28 | 27 | 0 | 1 | 4 | 4 | 1 | 7 | 7 | 11 |
| L1MEf_dup1149 | 6.28354584575057 | -3.17500449580239 | 0.835644687899106 | -3.79946709621845 | 0.000145007562343872 | 0.00888267567018937 | 4436 | 1 | 1 | 17 | 30 | 0 | 1 | 3 | 2 | 3 | 2 | 6 | 20 |
| AluSx1_dup12438 | 8.0166596242368 | -2.72487356122555 | 0.708766222617586 | -3.84453078359485 | 0.000120783323041053 | 0.00779493045411622 | 4437 | 1 | 4 | 18 | 31 | 0 | 1 | 5 | 2 | 4 | 3 | 17 | 22 |
| L1MEf_dup1150 | 8.25077105890855 | -3.13634856877297 | 0.789977136909681 | -3.97017637882798 | 7.18194456600127e-05 | 0.00533202802403038 | 4438 | 1 | 2 | 25 | 26 | 0 | 1 | 3 | 1 | 3 | 4 | 14 | 32 |
| MER2_dup963 | 7.33863970015737 | -2.69670775631033 | 0.69299713634738 | -3.89136926383855 | 9.96800945946382e-05 | 0.00679807388604637 | 4439 | 2 | 3 | 25 | 16 | 0 | 1 | 5 | 2 | 3 | 3 | 13 | 25 |
| HAL1ME_dup855 | 9.52394570535601 | -2.30853197303736 | 0.717444577838039 | -3.21771471183731 | 0.00129216272476684 | 0.0418967439749989 | 4440 | 5 | 4 | 23 | 21 | 0 | 0 | 3 | 3 | 7 | 3 | 16 | 42 |
| L1MC_dup799 | 8.77666072226008 | -3.21093597044847 | 0.743562727285 | -4.31831216469482 | 1.57226948977086e-05 | 0.001709850256001 | 4441 | 3 | 1 | 28 | 24 | 0 | 0 | 5 | 3 | 5 | 2 | 18 | 28 |
| AluJo_dup8922 | 6.28267530984046 | -2.77658363456135 | 0.701952746855488 | -3.95551359689026 | 7.63704042397855e-05 | 0.00558623799792192 | 4442 | 3 | 1 | 15 | 16 | 0 | 0 | 3 | 4 | 3 | 3 | 15 | 20 |
| L1M3_dup603 | 7.44650300466986 | -3.32675424719791 | 0.855126854452359 | -3.89036343540916 | 0.000100094195979915 | 0.00682118436943262 | 4443 | 1 | 1 | 20 | 31 | 0 | 0 | 2 | 4 | 5 | 3 | 8 | 25 |
| L1MD_dup863 | 7.99132631670803 | -2.95853867705217 | 0.77451593019163 | -3.81985516594884 | 0.000133530058792734 | 0.00835250877221235 | 4444 | 2 | 2 | 27 | 30 | 0 | 0 | 3 | 5 | 5 | 3 | 9 | 20 |
| L1M4_dup12136 | 5.50628376628804 | -2.06120241489354 | 0.626967311629305 | -3.28757556679801 | 0.0010105406442150З | 0.0353193454947694 | 4445 | 2 | 4 | 12 | 12 | 0 | 0 | 8 | 3 | 4 | 3 | 12 | 12 |
| AluSc_dup26971 | 5.92751542785314 | -2.61539638666475 | 0.69450724157312 | -3.7658302608057 | 0.000165996540679534 | 0.00979132693815875 | 4446 | 2 | 1 | 13 | 12 | 0 | 0 | 7 | 1 | 3 | 5 | 16 | 19 |
| AluJo_dup8923 | 7.17157357791393 | -2.87656850247092 | 0.763279667014126 | -3.76869531154127 | 0.000164103053784916 | 0.00970557867204541 | 4447 | 2 | 2 | 17 | 21 | 0 | 0 | 3 | 1 | 3 | 4 | 21 | 22 |
| LTR8_dup2740 | 7.14263023774029 | -2.79040033417505 | 0.775394254335629 | -3.59868585377372 | 0.000319829247354781 | 0.0156492062700145 | 4448 | 2 | 1 | 19 | 20 | 0 | 0 | 4 | 1 | 5 | 5 | 16 | 22 |
| MLT2A1_dup356 | 10.7071239616939 | -2.72882678920717 | 0.820958463275983 | -3.32395232070324 | 0.000887513681519957 | 0.0323598422407214 | 4449 | 3 | 3 | 32 | 47 | 2 | 0 | 3 | 0 | 5 | 5 | 21 | 24 |
| L1M4_dup20226 | 6.93418449423784 | -2.70917489712647 | 0.778592567288345 | -3.47957970696007 | 0.000502200955113689 | 0.0216440334116069 | 4450 | 1 | 2 | 24 | 25 | 1 | 3 | 2 | 1 | 3 | 1 | 14 | 16 |
| L2a_dup20683 | 5.90586486799304 | -2.63958117713386 | 0.804321101688237 | -3.28175049938822 | 0.00103164844584122 | 0.0358422300417366 | 4451 | 1 | 1 | 15 | 25 | 2 | 1 | 3 | 0 | 3 | 2 | 13 | 14 |
| MIRb_dup214043 | 8.42567880200658 | -2.64746162995475 | 0.742154065747535 | -3.5672668953017 | 0.000360724025779982 | 0.017115222141225 | 4452 | 3 | 2 | 25 | 33 | 2 | 3 | 5 | 0 | 3 | 1 | 21 | 16 |
| L1ME1_dup3528 | 7.93210563635712 | -2.97431767224216 | 0.934989176771272 | -3.18112524308906 | 0.00146704192085723 | 0.0457805582820058 | 4453 | 2 | 6 | 24 | 37 | 0 | 0 | 4 | 0 | 1 | 2 | 10 | 24 |
| HAL1_dup2712 | 7.40920740102728 | -2.99026423071102 | 0.863575277719797 | -3.46265613185057 | 0.000534871489846049 | 0.0225897287048115 | 4454 | 2 | 1 | 23 | 30 | 0 | 1 | 3 | 0 | 4 | 3 | 16 | 17 |
| L1M4_dup1868 | 5.99928021124655 | -3.51021277683031 | 0.970645291523432 | -3.61637027190542 | 0.000298762938402213 | 0.0149134742001672 | 4455 | 0 | 2 | 17 | 32 | 0 | 1 | 3 | 0 | 1 | 2 | 11 | 14 |
| MIR_dup20519 | 6.76327890092927 | -2.71658481443305 | 0.767243267462751 | -3.54070857267566 | 0.000399054086930159 | 0.0184375385958609 | 4456 | 5 | 0 | 17 | 26 | 2 | 0 | 6 | 1 | 2 | 2 | 16 | 15 |
| L1MEc_dup2302 | 9.36258163892273 | -3.39058104324598 | 0.88851907871087 | -3.81599126511193 | 0.000135637302481229 | 0.00845542404406562 | 4457 | 6 | 0 | 35 | 44 | 0 | 0 | 6 | 2 | 3 | 1 | 13 | 18 |
| MER5B_dup2396 | 6.24821503104801 | -4.26281128959749 | 0.963423882941067 | -4.42464772264552 | 9.65997846872323e-06 | 0.00119219566219856 | 4458 | 1 | 0 | 23 | 25 | 0 | 0 | 4 | 3 | 1 | 2 | 8 | 18 |
| AluJb_dup15383 | 8.68391894633928 | -4.77855786009484 | 1.00944276347895 | -4.73385716652817 | 2.20292779403106e-06 | 0.000378242066056012 | 4459 | 1 | 0 | 28 | 41 | 0 | 0 | 3 | 2 | 2 | 1 | 17 | 24 |
| MER45C_dup134 | 6.53435600597683 | -3.52380756550997 | 0.912889751311515 | -3.86005819481208 | 0.00011336002477267 | 0.00744714645349469 | 4460 | 2 | 0 | 18 | 30 | 0 | 0 | 2 | 3 | 4 | 1 | 11 | 17 |
| L1PBa_dup189 | 8.90394209962368 | -3.64730530394632 | 0.766850245562053 | -4.75621586490212 | 1.97255542909534e-06 | 0.000349178663663536 | 4461 | 3 | 0 | 26 | 34 | 1 | 1 | 5 | 3 | 2 | 1 | 14 | 32 |
| AluYj4_dup3518 | 6.8905447121726 | -2.56021279688842 | 0.737012414437107 | -3.47377160375756 | 0.00051319767185834 | 0.021970732858583 | 4462 | 3 | 0 | 16 | 23 | 3 | 3 | 2 | 2 | 1 | 2 | 13 | 26 |
| L1PA8A_dup244 | 8.92300686088437 | -2.71536292007582 | 0.806687720308017 | -3.36606452747169 | 0.00076248869439061 | 0.0290615304942284 | 4463 | 5 | 0 | 30 | 33 | 1 | 2 | 1 | 2 | 2 | 5 | 14 | 26 |
| L1ME3E_dup1413 | 8.69796914939028 | -4.30341342828487 | 1.01434951216834 | -4.2425351189706 | 2.21008942821897e-05 | 0.00219850308982856 | 4464 | 1 | 0 | 21 | 28 | 0 | 1 | 1 | 1 | 3 | 0 | 22 | 41 |
| FLAM_C_dup11421 | 6.13682521387895 | -2.92148813468234 | 0.855251987504205 | -3.41593843378 | 0.000635626159361876 | 0.0255437767464146 | 4465 | 3 | 0 | 15 | 14 | 0 | 2 | 1 | 1 | 2 | 1 | 22 | 21 |
| MIRc_dup29737 | 6.39087766779409 | -3.49703692989268 | 0.921742408302236 | -3.7939416678613 | 0.00014827452167517 | 0.00902566624060374 | 4466 | 2 | 0 | 18 | 19 | 1 | 2 | 1 | 1 | 1 | 0 | 17 | 25 |
| AluSx_dup29637 | 8.98663979280657 | -4.37664583746738 | 1.00346510714187 | -4.36153265949944 | 1.29154512432261e-05 | 0.00147703445175066 | 4467 | 0 | 0 | 18 | 34 | 1 | 0 | 1 | 1 | 3 | 2 | 35 | 26 |
| MER101-int_dup956 | 8.64379792599799 | -4.86644117183699 | 0.980159203010621 | -4.9649497315226 | 6.87188668744042e-07 | 0.000152776940125142 | 4468 | 0 | 0 | 20 | 20 | 0 | 0 | 2 | 1 | 1 | 3 | 37 | 32 |
| AluJb_dup15384 | 9.1521640341884 | -3.75374454265416 | 0.830798357871509 | -4.51823779752188 | 6.23564300783693e-06 | 0.000852237973086023 | 4469 | 1 | 0 | 23 | 36 | 1 | 0 | 2 | 4 | 4 | 3 | 26 | 22 |
| Tigger1_dup1430 | 11.1576335949534 | -3.81116277394441 | 0.876537157772286 | -4.34797628389246 | 1.37399457355688e-05 | 0.00154579911489156 | 4470 | 1 | 0 | 41 | 43 | 2 | 0 | 3 | 2 | 1 | 7 | 24 | 27 |
| L1MA3_dup859 | 7.49604647574356 | -2.99628911228876 | 0.839622293128312 | -3.56861547961646 | 0.000358872670336187 | 0.0170415988710628 | 4471 | 1 | 1 | 26 | 23 | 1 | 0 | 3 | 1 | 5 | 4 | 23 | 10 |
| MLT1B_dup11334 | 6.81282938518029 | -3.92611926686478 | 1.02857709301421 | -3.81703937753409 | 0.000135062620920183 | 0.008424013210347 | 4472 | 0 | 0 | 20 | 29 | 0 | 0 | 2 | 2 | 6 | 4 | 14 | 13 |
| L1PA3_dup5133 | 8.26641246321856 | -3.19554567819374 | 0.930960507429351 | -3.43252549672334 | 0.000597987652328567 | 0.0245198638499006 | 4473 | 0 | 0 | 33 | 41 | 2 | 3 | 1 | 1 | 2 | 3 | 14 | 12 |
| L1MB8_dup16157 | 6.37581675300347 | -3.87630470346168 | 0.901578902009367 | -4.29946252604457 | 1.7121282173631e-05 | 0.001821977016139 | 4474 | 0 | 0 | 23 | 22 | 2 | 0 | 5 | 2 | 2 | 1 | 17 | 11 |
| AluSz_dup100183 | 7.41761214741397 | -3.50951960345552 | 0.846938025650801 | -4.14377380299904 | 3.41636733238325e-05 | 0.00304307669457255 | 4475 | 0 | 0 | 27 | 26 | 1 | 1 | 4 | 3 | 5 | 1 | 15 | 15 |
| ERV3-16A3_I-int_dup1 | 5.15166983214155 | -3.54114025106957 | 0.872289605618946 | -4.05959239713387 | 4.91584529592735e-05 | 0.00402211393683941 | 4476 | 0 | 0 | 15 | 16 | 1 | 1 | 3 | 2 | 2 | 1 | 12 | 16 |
| LTR89_dup443 | 4.36265507323584 | -3.21251918769502 | 0.887734682928421 | -3.61878300969126 | 0.00029599168808758 | 0.0148273764660394 | 4477 | 1 | 0 | 20 | 18 | 1 | 1 | 3 | 3 | 1 | 1 | 5 | 4 |
| L1PA7_dup5097 | 5.74867058322292 | -4.00795151331829 | 0.96257027260979 | -4.16380146714031 | 3.12992294489459e-05 | 0.00284702111675763 | 4478 | 0 | 0 | 19 | 34 | 1 | 1 | 4 | 2 | 1 | 1 | 8 | 8 |
| L2c_dup104404 | 6.48263117936616 | -3.57236779483392 | 0.83027400562692 | -4.30263716631296 | 1.68777117807231e-05 | 0.0018043619033582 | 4479 | 1 | 0 | 19 | 36 | 1 | 1 | 4 | 4 | 2 | 1 | 9 | 10 |
| L1MC4a_dup4208 | 6.62376403598525 | -2.87262314490954 | 0.780875705482498 | -3.67872008917805 | 0.000234407358912117 | 0.0125058953864804 | 4480 | 1 | 0 | 14 | 37 | 1 | 3 | 5 | 2 | 1 | 4 | 12 | 11 |
| L1PA3_dup1010 | 7.29480533737834 | -2.70305123975599 | 0.780025935634058 | -3.46533508217104 | 0.000529571210748402 | 0.0224215927602739 | 4481 | 4 | 0 | 32 | 28 | 1 | 1 | 5 | 2 | 2 | 4 | 9 | 10 |
| L1PA8_dup932 | 8.05292629949022 | -3.55536264814199 | 0.805703581186851 | -4.4127427644106 | 1.0206924285257e-05 | 0.00124085522951809 | 4482 | 3 | 0 | 27 | 36 | 1 | 2 | 3 | 4 | 0 | 2 | 12 | 20 |
| MIR_dup168626 | 6.85390249636562 | -3.11105804175644 | 0.869411977378521 | -3.57834734591189 | 0.000345773729768562 | 0.0165677559209646 | 4483 | 1 | 0 | 26 | 28 | 2 | 4 | 1 | 3 | 0 | 2 | 9 | 17 |
| MSTB-int_dup101 | 6.9952754817251 | -3.19629631316654 | 0.859577923640952 | -3.71844858419333 | 0.000200450080220961 | 0.0111361883698942 | 4484 | 2 | 0 | 28 | 25 | 0 | 2 | 4 | 3 | 0 | 5 | 7 | 19 |
| L1PREC2_dup6012 | 5.32869525012128 | -2.66864908393177 | 0.774169422805448 | -3.44711248638711 | 0.000566612613764759 | 0.023591111985104 | 4485 | 1 | 0 | 18 | 18 | 1 | 4 | 3 | 5 | 0 | 3 | 6 | 12 |
| MSTA-int_dup1476 | 5.02586235081477 | -2.85806032046121 | 0.837294195643417 | -3.41344814681886 | 0.000641463641112414 | 0.0257014946241651 | 4486 | 1 | 0 | 22 | 11 | 1 | 1 | 4 | 3 | 0 | 5 | 10 | 8 |
| L2a_dup91394 | 5.91867966247151 | -3.25517580227489 | 0.996969572667473 | -3.2650703607387 | 0.00109436888237319 | 0.0373494977173224 | 4487 | 3 | 0 | 19 | 20 | 2 | 1 | 1 | 0 | 0 | 1 | 10 | 25 |
| AluYf1_dup738 | 4.70360875415093 | -3.45112378675441 | 0.931568282860886 | -3.70463856514721 | 0.000211692363221782 | 0.0115796209331426 | 4488 | 2 | 1 | 13 | 17 | 1 | 1 | 1 | 2 | 0 | 0 | 11 | 15 |
| L1MB7_dup10054 | 4.82269340287218 | -2.98643988515805 | 0.833174316082767 | -3.58441184216891 | 0.000337838648209285 | 0.0162892310003661 | 4489 | 3 | 1 | 16 | 11 | 1 | 2 | 2 | 3 | 0 | 0 | 8 | 18 |
| L1M4_dup15998 | 3.84956361797567 | -3.91082042377103 | 1.01461751181141 | -3.85447755261883 | 0.000115977023321545 | 0.00756684090794309 | 4490 | 1 | 0 | 16 | 11 | 1 | 1 | 3 | 2 | 0 | 0 | 6 | 11 |
| L1PA4_dup10385 | 4.80039147613248 | -3.86639958921446 | 1.0606745181887 | -3.64522718601467 | 0.000267155689547976 | 0.0137624221054736 | 4491 | 1 | 0 | 18 | 18 | 2 | 1 | 1 | 1 | 0 | 0 | 7 | 17 |
| L1P3_dup2992 | 6.45618794005766 | -4.57489594439063 | 1.03831915870059 | -4.40605945296812 | 1.0526808163713e-05 | 0.00127147811945323 | 4492 | 0 | 2 | 12 | 18 | 0 | 1 | 12 | 0 | 0 | 0 | 23 | 21 |
| L1MA7_dup3843 | 5.42392423797916 | -3.76308562916149 | 1.07634811881341 | -3.49616036242065 | 0.00047200494467895 | 0.020725537657608 | 4493 | 0 | 1 | 19 | 23 | 1 | 3 | 2 | 0 | 0 | 0 | 15 | 10 |
| L1PA3_dup5136 | 6.05265680275818 | -4.16181752221167 | 1.15623051530084 | -3.59947040588945 | 0.000318865917826714 | 0.0156149145262801 | 4494 | 0 | 0 | 14 | 40 | 0 | 3 | 3 | 0 | 0 | 1 | 13 | 11 |
| L1MC3_dup12813 | 7.68516061318101 | -3.13840345202178 | 0.989846392618716 | -3.1705964434733 | 0.00152126323979028 | 0.0469431335566253 | 4495 | 4 | 1 | 22 | 41 | 0 | 2 | 1 | 0 | 0 | 3 | 19 | 13 |
| THE1B_dup2083 | 8.61934042260788 | -3.15428582945164 | 0.992895027145686 | -3.17685731443272 | 0.00148880264455495 | 0.0462607014129047 | 4496 | 3 | 2 | 26 | 54 | 1 | 0 | 3 | 0 | 0 | 5 | 19 | 6 |
| L1PA7_dup12453 | 8.59741546640383 | -3.16354854065138 | 1.00536810247218 | -3.14665696362585 | 0.00165148592106907 | 0.0496668944609075 | 4497 | 7 | 6 | 26 | 44 | 0 | 1 | 5 | 0 | 0 | 0 | 12 | 19 |
| L2c_dup129943 | 6.53458748663937 | -3.65801732152083 | 1.00417454302363 | -3.64281025339112 | 0.000269677633277601 | 0.0138443827676099 | 4498 | 3 | 3 | 22 | 40 | 0 | 1 | 5 | 0 | 0 | 0 | 6 | 12 |

| | | | | | | | | | | | | | | | | | | |
|---|---|---|---|---|---|---|---|---|---|---|---|---|---|---|---|---|---|---|
| L1M2_dup4451 | 7.00945305850874 | -3.41802621449108 | 0.999629163710322 | -3.4192942128703 | 0.000627838018050939 | 0.025337779010032 | 4499 | 4 | 4 | 27 | 42 | 1 | 0 | 5 | 0 | 0 | 0 | 6 | 9 |
| L1PB3_dup2788 | 7.11854330017491 | -4.16661128340138 | 1.16924798930614 | -3.56349664186632 | 0.000365947378653757 | 0.0172940844014194 | 4500 | 4 | 1 | 38 | 42 | 0 | 0 | 2 | 0 | 0 | 0 | 4 | 8 |
| MIRb_dup105687 | 5.54137469280129 | -2.67507043615997 | 0.787242459808454 | -3.3980261135951 | 0.000678739409507246 | 0.026765549806863 | 4501 | 0 | 1 | 6 | 3 | 0 | 5 | 16 | 8 | 0 | 3 | 17 | 13 |
| MER52D_dup259 | 5.19415798962315 | -1.96206831279997 | 0.614906888212327 | -3.19083807713416 | 0.0014186076114947 | 0.0446792014336328 | 4502 | 0 | 3 | 6 | 8 | 3 | 4 | 13 | 6 | 0 | 3 | 13 | 9 |
| MER63B_dup1469 | 6.10250783301241 | -2.8333914439374 | 0.696403115887944 | -4.0686082231621 | 4.72947929621186e-05 | 0.00390985380164496 | 4503 | 0 | 1 | 6 | 8 | 2 | 5 | 16 | 7 | 0 | 1 | 17 | 18 |
| Tigger1_dup601 | 4.02331505551863 | -3.46109008184205 | 1.01646241643991 | -3.4050349780411 | 0.000661555795900923 | 0.0262620212564967 | 4504 | 0 | 0 | 2 | 6 | 0 | 0 | 8 | 9 | 0 | 6 | 11 | 9 |
| Tigger19a_dup80 | 4.31000813680871 | -3.23434970789235 | 0.85431417168749 | -3.78590197269429 | 0.000153152044058054 | 0.00924218013579451 | 4505 | 0 | 0 | 2 | 6 | 2 | 0 | 8 | 9 | 1 | 2 | 12 | 13 |
| MIRb_dup202465 | 3.56977680526827 | -3.73938348978088 | 0.991612688206051 | -3.77101214441486 | 0.000162586759438364 | 0.00965241244044932 | 4506 | 0 | 1 | 2 | 12 | 1 | 0 | 6 | 6 | 0 | 1 | 10 | 8 |
| L1P1_dup2225 | 3.13619239458456 | -3.4595534214604 | 0.948269246389446 | -3.64828178772297 | 0.000263999994492864 | 0.0136573927036814 | 4507 | 0 | 0 | 3 | 4 | 1 | 1 | 5 | 5 | 0 | 1 | 9 | 12 |
| L2c_dup129737 | 2.98292032367983 | -3.39816896653216 | 1.01767651340927 | -3.33914453341183 | 0.000840368276052695 | 0.0311467709990085 | 4508 | 0 | 0 | 1 | 5 | 1 | 1 | 5 | 6 | 0 | 1 | 12 | 6 |
| L1MD1_dup844 | 4.28469229146928 | -3.97995781561417 | 0.935423406500554 | -4.25471266589673 | 2.09317634101197e-05 | 0.00211043617643404 | 4509 | 0 | 1 | 5 | 11 | 1 | 1 | 7 | 4 | 0 | 0 | 11 | 17 |
| SST1_dup94 | 4.32336998782255 | -3.52828495087071 | 0.891917391957031 | -3.95584275257712 | 7.62653160466087e-05 | 0.00558026657196382 | 4510 | 0 | 1 | 4 | 6 | 2 | 1 | 13 | 3 | 0 | 0 | 11 | 18 |
| L1MD2_dup671 | 3.97377363389331 | -2.87824671190144 | 0.821381107544551 | -3.5041549963399 | 0.000458058668179605 | 0.020323182520728 | 4511 | 0 | 2 | 3 | 12 | 1 | 3 | 6 | 6 | 0 | 0 | 7 | 13 |
| MLT2B4_dup2952 | 4.48889283836001 | -2.82491827315065 | 0.831518041007311 | -3.39730244424824 | 0.000680537062976303 | 0.0268101135786938 | 4512 | 0 | 1 | 3 | 6 | 4 | 2 | 9 | 10 | 0 | 0 | 13 | 9 |
| MER1B_dup2922 | 3.6162859581531 | -3.02298053387189 | 0.909218147843406 | -3.32481323766157 | 0.000884777819469463 | 0.0323145576410562 | 4513 | 1 | 0 | 6 | 2 | 1 | 3 | 11 | 5 | 0 | 0 | 12 | 6 |
| L2b_dup85542 | 2.89057207357995 | -3.44163339292285 | 1.09136195480459 | -3.15352150381591 | 0.00161313343261099 | 0.0488033881444444 | 4514 | 0 | 0 | 1 | 3 | 2 | 1 | 13 | 5 | 0 | 0 | 8 | 5 |
| L1PA5_dup10474 | 4.30807814674861 | -3.99170104571273 | 1.06373283305256 | -3.75254097803662 | 0.000175051206589791 | 0.0101418880942816 | 4515 | 0 | 0 | 2 | 2 | 2 | 1 | 15 | 8 | 0 | 0 | 14 | 12 |
| Tigger2_dup1554 | 3.68182432734137 | -3.04598007164064 | 0.942691981219463 | -3.23115092980887 | 0.00123292817415964 | 0.0405880364317363 | 4516 | 0 | 0 | 4 | 3 | 4 | 1 | 9 | 8 | 0 | 0 | 9 | 9 |
| AluY_dup42599 | 3.33107615797048 | -3.26933940076012 | 0.973533136198465 | -3.35822097800031 | 0.000784458737859905 | 0.0296599206088916 | 4517 | 0 | 0 | 2 | 7 | 1 | 3 | 9 | 7 | 0 | 0 | 4 | 11 |
| MLT2A2_dup2122 | 3.15693513142791 | -3.48913600163257 | 1.08762292779563 | -3.20803829384535 | 0.00133643702489578 | 0.0429381372300638 | 4518 | 0 | 0 | 1 | 5 | 0 | 3 | 7 | 5 | 0 | 0 | 8 | 13 |
| L1MC3_dup6648 | 4.46076528755492 | -2.91887486435542 | 0.861880971078768 | -3.38663337781089 | 0.000707558733718423 | 0.0275633633886003 | 4519 | 1 | 0 | 2 | 5 | 2 | 3 | 7 | 8 | 0 | 0 | 19 | 10 |
| L3_dup24279 | 3.90842016333217 | -2.76484740008471 | 0.874275713590853 | -3.16244333121052 | 0.00156451182654177 | 0.0477758381774009 | 4520 | 3 | 0 | 1 | 10 | 2 | 1 | 5 | 5 | 0 | 0 | 13 | 12 |
| AluSz_dup15567 | 4.46942968672697 | -3.55198069052911 | 0.971765079267264 | -3.65518453617144 | 0.000256997051203138 | 0.013407458966363 | 4521 | 0 | 2 | 2 | 3 | 0 | 0 | 9 | 10 | 2 | 0 | 10 | 19 |
| L1PB_dup866 | 2.87518139459631 | -4.31691030037627 | 1.2459260931719 | -3.46482052509729 | 0.000530585444249408 | 0.0224454300705715 | 4522 | 0 | 0 | 0 | 1 | 0 | 1 | 5 | 8 | 0 | 0 | 11 | 10 |
| L1MC4a_dup7167 | 2.44451951617587 | -4.0654672254658 | 1.24104715774542 | -3.27583621628968 | 0.00105349636064989 | 0.0364194747975009 | 4523 | 0 | 0 | 1 | 0 | 0 | 1 | 6 | 5 | 0 | 0 | 6 | 13 |
| MLT1L_dup6321 | 2.50880894026647 | -4.17996058608666 | 1.15301757171032 | -3.62523580615198 | 0.00028869790412831 | 0.0145508391221595 | 4524 | 0 | 0 | 1 | 1 | 0 | 1 | 8 | 5 | 0 | 0 | 6 | 11 |
| L1MD1_dup3585 | 3.37013228850062 | -3.65796857667664 | 1.11800676237504 | -3.27186623532206 | 0.00106840112153586 | 0.0367813476597136 | 4525 | 1 | 1 | 0 | 3 | 0 | 1 | 13 | 6 | 0 | 0 | 7 | 13 |
| AluSz_dup89776 | 2.50773553385209 | -4.23885605205019 | 1.31491615816067 | -3.22367021330057 | 0.00126559058694259 | 0.041342973338913 | 4526 | 0 | 0 | 0 | 1 | 0 | 1 | 13 | 8 | 0 | 0 | 4 | 5 |
| L1MA10_dup3590 | 2.58896072841372 | -4.25885772518564 | 1.26510615842351 | -3.36640344118849 | 0.000761552386926897 | 0.0290351456913487 | 4527 | 0 | 0 | 0 | 1 | 0 | 1 | 11 | 5 | 0 | 0 | 8 | 8 |
| L1MEf_dup3782 | 3.10630040589369 | -4.56673787826267 | 1.01261780925774 | -4.5098336573896 | 6.48784779640621e-06 | 0.000877134904411177 | 4528 | 1 | 0 | 8 | 4 | 0 | 0 | 8 | 6 | 0 | 0 | 8 | 5 |
| MLT1B_dup394 | 3.55243457446076 | -4.24873634555402 | 1.00388820466059 | -4.2322803732817 | 2.31333773098341e-05 | 0.00228026474908339 | 4529 | 2 | 0 | 7 | 8 | 0 | 0 | 5 | 6 | 0 | 0 | 12 | 6 |
| L1PA2_dup67 | 2.46378888554324 | -3.64817074377101 | 1.02454820262277 | -3.56076047416017 | 0.000369782288531609 | 0.0174406771906724 | 4530 | 1 | 0 | 4 | 3 | 0 | 1 | 4 | 4 | 0 | 0 | 8 | 7 |
| L1M4_dup9570 | 2.47822898666366 | -3.65839785897263 | 1.04918564761933 | -3.48689277943687 | 0.000488667228082834 | 0.0211989812677116 | 4531 | 2 | 0 | 3 | 3 | 0 | 0 | 6 | 3 | 0 | 0 | 8 | 8 |
| THE1D_dup7069 | 3.08171217141049 | -4.46489978931047 | 1.01364392667895 | -4.40480100733109 | 1.058810265669e-05 | 0.00127499831718054 | 4532 | 1 | 0 | 3 | 5 | 0 | 0 | 4 | 6 | 0 | 0 | 10 | 11 |
| THE1B_dup395 | 2.14109181368396 | -4.54231211233372 | 1.09527277756572 | -4.14719712328572 | 3.36570270020362e-05 | 0.00301300035629289 | 4533 | 0 | 0 | 2 | 4 | 0 | 0 | 6 | 5 | 0 | 0 | 4 | 7 |
| L1PB2_dup1434 | 2.30834294598958 | -4.53253747044397 | 1.11851043449562 | -4.05229788713403 | 5.07170322477594e-05 | 0.00412578289067533 | 4534 | 0 | 0 | 1 | 3 | 0 | 0 | 4 | 5 | 0 | 0 | 7 | 10 |
| L1MDa_dup6661 | 2.3162816209686 | -4.4359419650681 | 1.07870377479159 | -4.11228927601107 | 3.91754995085252e-05 | 0.00338599833129514 | 4535 | 0 | 0 | 2 | 3 | 0 | 0 | 3 | 3 | 0 | 0 | 7 | 13 |
| Tigger1_dup5520 | 1.82935034008072 | -4.17871173802675 | 1.12217125678561 | -3.72377363326557 | 0.000196266919601261 | 0.0109652332593195 | 4536 | 0 | 0 | 2 | 2 | 0 | 0 | 3 | 3 | 0 | 0 | 6 | 8 |
| L1MB2_dup7830 | 3.30980362602173 | -4.99191133535702 | 1.01187655563693 | -4.93332048019913 | 8.08433451219503e-07 | 0.000174558535375568 | 4537 | 0 | 0 | 4 | 4 | 0 | 0 | 4 | 5 | 0 | 0 | 14 | 12 |
| L1MC3_dup1269 | 3.35174554084177 | -5.16224947648638 | 0.990495556496859 | -5.21178458865984 | 1.87032649332948e-07 | 5.46057310161689e-05 | 4538 | 0 | 0 | 7 | 7 | 0 | 0 | 5 | 5 | 0 | 0 | 10 | 10 |
| MER5A_dup21611 | 3.38973405665673 | -5.00765237357735 | 1.02232662185164 | -4.89829010273397 | 9.66742332953937e-07 | 0.000200202324766804 | 4539 | 0 | 0 | 3 | 5 | 0 | 0 | 5 | 4 | 0 | 0 | 8 | 21 |
| AluJr_dup48403 | 3.46423118881443 | -5.14960697373925 | 1.03045142584674 | -4.99742816067989 | 5.80999722565138e-07 | 0.000135006406235825 | 4540 | 0 | 0 | 2 | 9 | 0 | 0 | 8 | 3 | 0 | 0 | 13 | 12 |
| MER21B_dup1657 | 3.20247848017056 | -5.07605599009622 | 1.03862263068731 | -4.88729577049283 | 1.02230454132683e-06 | 0.000209255520081629 | 4541 | 0 | 0 | 2 | 10 | 0 | 0 | 6 | 4 | 0 | 0 | 11 | 10 |
| Charlie29a_dup1872 | 3.51685215855573 | -4.75288882159694 | 1.04318922123292 | -4.5561138141167 | 5.21087088460436e-06 | 0.000745574132701276 | 4542 | 0 | 0 | 11 | 8 | 0 | 0 | 8 | 3 | 1 | 0 | 12 | 3 |
| THE1D-int_dup2420 | 2.37140523316939 | -4.18691110683801 | 1.09408571000163 | -3.82685841571934 | 0.000129789116705418 | 0.00820042516292813 | 4543 | 1 | 0 | 4 | 8 | 0 | 0 | 7 | 2 | 0 | 0 | 8 | 2 |
| L1M4_dup5965 | 2.17979312880879 | -4.58390732202358 | 1.08187314903682 | -4.2370099730311 | 2.26516078526906e-05 | 0.00224390486349555 | 4544 | 0 | 0 | 4 | 5 | 0 | 0 | 8 | 3 | 0 | 0 | 6 | 3 |
| L2b_dup31912 | 2.84464275633939 | -4.95069613243143 | 1.0409164097616 | -4.75609384769454 | 1.97374741591561e-06 | 0.000349178663663536 | 4545 | 0 | 0 | 5 | 10 | 0 | 0 | 12 | 3 | 0 | 0 | 5 | 4 |
| Kanga1d_dup553 | 1.98929765500636 | -4.4454260341084 | 1.20097173022265 | -3.70152429256954 | 0.000214308133542653 | 0.0116780102951009 | 4546 | 0 | 0 | 3 | 8 | 0 | 0 | 3 | 3 | 0 | 0 | 8 | 1 |
| AluY_dup80284 | 2.75898642201027 | -4.92716430342922 | 1.0921328018233 | -4.51150656330748 | 6.43687912163187e-06 | 0.000874717098501323 | 4547 | 0 | 0 | 6 | 13 | 0 | 0 | 3 | 3 | 0 | 0 | 8 | 4 |
| MIRb_dup217359 | 1.98081347891077 | -4.44724583964289 | 1.20325931257788 | -3.69599951827097 | 0.00021902338682401 | 0.0118822933883324 | 4548 | 0 | 0 | 4 | 10 | 0 | 0 | 2 | 1 | 0 | 0 | 7 | 3 |
| AluSx_dup103320 | 2.33648191442819 | -4.13442597792377 | 1.0644312532097 | -3.88416439808278 | 0.000102682415384853 | 0.00694510233404326 | 4549 | 0 | 0 | 4 | 7 | 1 | 0 | 5 | 1 | 0 | 0 | 7 | 7 |
| L2b_dup28523 | 2.17840359307935 | -4.05368906747589 | 1.10425788494095 | -3.67096230215524 | 0.000241638981910163 | 0.012805574429754 | 4550 | 0 | 0 | 6 | 10 | 1 | 0 | 4 | 1 | 0 | 0 | 4 | 4 |
| ERVL47-int_dup127 | 2.8260690457612 | -4.42991553327603 | 1.0202385640842 | -4.34203890072757 | 1.41166530267755e-05 | 0.00157773655289297 | 4551 | 0 | 0 | 7 | 8 | 1 | 0 | 4 | 4 | 0 | 0 | 8 | 5 |
| MIRc_dup92011 | 3.09781906295884 | -4.57640391693669 | 1.07741991648178 | -4.24755830751724 | 2.16112930726899e-05 | 0.00215973151678886 | 4552 | 0 | 0 | 4 | 16 | 1 | 0 | 3 | 3 | 0 | 0 | 9 | 6 |
| L2b_dup46034 | 2.40305250393747 | -4.12399339208613 | 1.04070488214616 | -3.96269246242176 | 7.41092206148155e-05 | 0.00546282298897549 | 4553 | 1 | 0 | 4 | 7 | 0 | 0 | 9 | 5 | 0 | 0 | 2 | 4 |
| MLT1A0_dup5023 | 2.3239131573184 | -3.55308298776136 | 1.02203698574038 | -3.47647202335585 | 0.000508057232006785 | 0.0218079496676943 | 4554 | 1 | 1 | 5 | 7 | 0 | 0 | 7 | 4 | 0 | 0 | 2 | 4 |
| MLT1F_dup418 | 2.33151303972906 | -4.08133577128738 | 1.0823951365686 | -3.770652355503 | 0.000162821362168248 | 0.00965670285206529 | 4555 | 1 | 0 | 5 | 10 | 0 | 0 | 4 | 5 | 0 | 0 | 1 | 5 |
| AluSz_dup23442 | 2.9095013737396 | -4.39035074275648 | 1.01740775988458 | -4.31523221648571 | 1.5943521521148e-05 | 0.00172833584139913 | 4556 | 1 | 0 | 10 | 8 | 0 | 0 | 6 | 6 | 0 | 0 | 2 | 5 |
| MIRb_dup159981 | 2.72488934627471 | -4.35057617875641 | 1.03992762785662 | -4.18353745223918 | 2.87007780990142e-05 | 0.00267184481437717 | 4557 | 1 | 0 | 9 | 7 | 0 | 0 | 3 | 6 | 0 | 0 | 4 | 5 |
| AluSp_dup6017 | 2.11196184411135 | -4.54662870642358 | 1.161254054216 | -3.91527477550393 | 9.03013067983662e-05 | 0.00632151654823302 | 4558 | 0 | 0 | 7 | 7 | 0 | 0 | 4 | 2 | 0 | 0 | 1 | 8 |
| MIR_dup124602 | 2.48791748660821 | -4.76141042555693 | 1.07289191468442 | -4.43792180776889 | 9.0831601555465e-06 | 0.00113573390770976 | 4559 | 0 | 0 | 8 | 9 | 0 | 0 | 3 | 4 | 0 | 0 | 3 | 6 |
| MIRb_dup159977 | 2.3415282364535 | -4.67999160662816 | 1.12334722580657 | -4.16611311188124 | 3.09837014213493e-05 | 0.0028268588260753 | 4560 | 0 | 0 | 10 | 7 | 0 | 0 | 2 | 3 | 0 | 0 | 3 | 6 |
| L1PREC2_dup780 | 3.07567864878373 | -5.0716292997575 | 1.08784574458614 | -4.6620849739013 | 3.13021887743285e-06 | 0.000495887523974459 | 4561 | 0 | 0 | 8 | 9 | 0 | 0 | 6 | 6 | 0 | 0 | 1 | 11 |
| Charlie1_dup1110 | 2.7437598001685 | -4.85174995169268 | 1.09556196910124 | -4.42854908122897 | 9.48691314117213e-06 | 0.00117510308916676 | 4562 | 0 | 0 | 7 | 14 | 0 | 0 | 7 | 3 | 0 | 0 | 1 | 6 |
| AluSq2_dup60329 | 2.23542470714292 | -4.52667002463852 | 1.09990530216775 | -4.11550886764263 | 3.86326143378618e-05 | 0.00334880730464002 | 4563 | 0 | 0 | 9 | 6 | 0 | 0 | 7 | 3 | 0 | 0 | 1 | 4 |
| Tigger17a_dup1466 | 2.64863556783524 | -4.77801629124448 | 1.04614562753659 | -4.5672573353822 | 4.94147221352364e-06 | 0.000714774192899147 | 4564 | 0 | 0 | 9 | 9 | 0 | 0 | 9 | 3 | 0 | 0 | 2 | 4 |
| L1PA5_dup6255 | 3.57279518215657 | -4.78702588003432 | 1.02077927208535 | -4.68957982488705 | 2.73766621550973e-06 | 0.000448325344332882 | 4565 | 0 | 1 | 9 | 12 | 0 | 0 | 4 | 5 | 0 | 0 | 4 | 13 |
| MLT1C_dup4337 | 3.02385350938595 | -4.53510146815533 | 1.05674685157323 | -4.29156846921636 | 1.77415479132465e-05 | 0.00187041664145563 | 4566 | 0 | 1 | 8 | 10 | 0 | 0 | 2 | 5 | 0 | 0 | 4 | 10 |
| L1MB8_dup16313 | 3.50653589844135 | -4.21727193714765 | 0.978055050416718 | -4.31189628370182 | 1.61860360048178e-05 | 0.00174825352846227 | 4567 | 0 | 1 | 8 | 11 | 0 | 1 | 6 | 7 | 0 | 0 | 4 | 8 |
| L1ME1_dup22153 | 2.7589296252832 | -3.87086455989348 | 1.01549925901001 | -3.81178472120905 | 0.000137967014088372 | 0.0085669882918246 | 4568 | 0 | 1 | 7 | 4 | 0 | 1 | 5 | 5 | 0 | 0 | 4 | 9 |
| AluY_dup71775 | 2.74303340824855 | -3.87802974446431 | 1.03728926471731 | -3.73861937684391 | 0.000185033624982447 | 0.0105351330167045 | 4569 | 1 | 1 | 6 | 12 | 0 | 0 | 6 | 2 | 0 | 0 | 3 | 7 |
| MER58B_dup222 | 2.55476322982862 | -3.27440524047762 | 0.969989203302354 | -3.37571307941348 | 0.000736246559878406 | 0.0283611662517124 | 4570 | 1 | 1 | 5 | 9 | 0 | 1 | 7 | 2 | 0 | 0 | 4 | 5 |
| L4_A_Mam_dup2469 | 2.16007013279097 | -3.50947840974372 | 1.08186720443653 | -3.24390867506847 | 0.00117901541474683 | 0.0393199592377824 | 4571 | 0 | 1 | 7 | 7 | 0 | 1 | 2 | 2 | 0 | 0 | 3 | 6 |
| MER57A1_dup1037 | 2.06249355840241 | -3.97635309737624 | 1.09462804699041 | -3.63260662679793 | 0.000280572577521582 | 0.0142463905013437 | 4572 | 0 | 1 | 6 | 5 | 0 | 0 | 4 | 2 | 0 | 0 | 2 | 8 |
| L1M1_dup3177 | 3.64144560188559 | -3.7987548967951 | 0.955307872694146 | -3.97647188448464 | 6.9945264200501e-05 | 0.00524525554221633 | 4573 | 0 | 2 | 9 | 14 | 1 | 0 | 9 | 3 | 0 | 0 | 3 | 9 |
| L1MC1_dup1662 | 3.72459164040631 | -4.34573363196818 | 1.05561749721104 | -4.11676923075802 | 3.84220439901675e-05 | 0.00334067954751153 | 4574 | 0 | 1 | 9 | 19 | 1 | 0 | 6 | 1 | 0 | 0 | 6 | 9 |
| L1MEd_dup5083 | 4.31211317446479 | -4.56151970544917 | 1.0101056211161 | -4.515883893814 | 6.30532064663395e-06 | 0.000860278570288969 | 4575 | 0 | 1 | 14 | 13 | 1 | 0 | 6 | 2 | 0 | 0 | 7 | 15 |
| L1MC4a_dup8477 | 2.57866759502099 | -3.26687107024558 | 1.02127977511067 | -3.1988012980004 | 0.00138000247320256 | 0.0438933389237609 | 4576 | 0 | 2 | 8 | 8 | 1 | 0 | 6 | 1 | 0 | 0 | 5 | 4 |
| MIRb_dup159994 | 2.95384114777593 | -3.95375267152006 | 1.06731466670423 | -3.70439271084778 | 0.000211897768497869 | 0.0115881926966201 | 4577 | 0 | 1 | 10 | 14 | 1 | 0 | 3 | 2 | 0 | 0 | 5 | 4 |

| | | | | | | | | | | | | | | | | | | | |
|---|---|---|---|---|---|---|---|---|---|---|---|---|---|---|---|---|---|---|---|
| MLT1I_dup1402 | 3.07965108338991 | -4.04205594293013 | 1.03618096351491 | -3.90091700702431 | 9.58290018788662e-05 | 0.0066054831057557 | 4578 | 0 | 1 | 6 | 16 | 1 | 0 | 4 | 3 | 0 | 0 | 5 | 6 |
| LTR16_dup2998 | 2.76364729902087 | -3.80199971783732 | 1.02019504331901 | -3.72673808085582 | 0.000193973844521808 | 0.0108754242244548 | 4579 | 0 | 1 | 7 | 10 | 1 | 0 | 5 | 5 | 0 | 0 | 5 | 2 |
| MLT1K_dup17035 | 3.34890379002135 | -3.27143733858981 | 0.954118055365697 | -3.42875529940152 | 0.00060635595330563 | 0.0247605927004053 | 4580 | 0 | 3 | 11 | 12 | 1 | 0 | 3 | 4 | 0 | 0 | 6 | 4 |
| MER1A_dup785 | 3.52648357837792 | -2.94967812154447 | 0.866017782059731 | -3.40602489076954 | 0.000659161692497016 | 0.0261979648476901 | 4581 | 1 | 2 | 9 | 10 | 1 | 0 | 2 | 7 | 0 | 1 | 7 | 5 |
| MLT1A0_dup1864 | 3.43372148993394 | -2.92084642770312 | 0.826392905879077 | -3.53445244619574 | 0.000408620956029118 | 0.0187738408965639 | 4582 | 2 | 1 | 7 | 10 | 1 | 0 | 4 | 6 | 0 | 1 | 5 | 8 |
| L1MEg_dup11620 | 3.31508888293588 | -2.88097810630687 | 0.848117866993677 | -3.39690769222806 | 0.000681519523676546 | 0.026821839990198 4 | 4583 | 2 | 2 | 7 | 7 | 1 | 0 | 4 | 5 | 0 | 0 | 9 | 6 |
| MLT1F1-int_dup157 | 3.41233134828056 | -2.75087862302531 | 0.868516847098361 | -3.16732903019183 | 0.00153846143005964 | 0.0473055531490971 | 4584 | 3 | 2 | 10 | 13 | 1 | 0 | 2 | 5 | 0 | 0 | 4 | 5 |
| L1MA9_dup13443 | 3.38036841934506 | -3.63186793118419 | 0.984244163962821 | -3.69000707767609 | 0.000224247812798705 | 0.0120748744205025 | 4585 | 0 | 1 | 6 | 10 | 1 | 1 | 3 | 3 | 0 | 0 | 11 | 9 |
| L2a_dup169302 | 3.0357651102551 | -2.69413223326962 | 0.848051638047776 | -3.17684927709301 | 0.0014888439033227 | 0.0462607014129047 | 4586 | 1 | 2 | 7 | 6 | 1 | 1 | 3 | 3 | 0 | 0 | 8 | 8 |
| AluSq2_dup46544 | 3.23187188002363 | -2.59142316208051 | 0.793825944023924 | -3.26447274946006 | 0.00109668011883102 | 0.0373961948345322 | 4587 | 2 | 1 | 5 | 9 | 1 | 2 | 5 | 5 | 0 | 0 | 4 | 9 |
| Charlie4z_dup1859 | 3.11993837821884 | -2.51792690359453 | 0.791744270733505 | -3.18022750106145 | 0.00147159471259809 | 0.0458865254205039 | 4588 | 1 | 2 | 8 | 5 | 1 | 2 | 6 | 4 | 0 | 0 | 5 | 7 |
| L1PA15_dup4086 | 2.77979171712592 | -2.93217003830885 | 0.905803487993771 | -3.23709289837601 | 0.00120754095024701 | 0.0400403482368276 | 4589 | 1 | 1 | 5 | 3 | 1 | 1 | 6 | 5 | 0 | 0 | 6 | 7 |
| LTR32_dup618 | 3.57106780032505 | -4.7605360711228 | 1.05536716931148 | -4.51078658646219 | 6.45876762823456e-06 | 0.000876633285858476 | 4590 | 1 | 0 | 5 | 22 | 0 | 0 | 6 | 4 | 0 | 0 | 7 | 4 |
| LTR83_dup173 | 2.81596173958241 | -4.41959112121298 | 1.08780653097707 | -4.06284665090518 | 4.84778786432931e-05 | 0.00397762665919207 | 4591 | 1 | 0 | 6 | 17 | 0 | 0 | 4 | 2 | 0 | 0 | 5 | 4 |
| AluJb_dup15381 | 4.24474584046838 | -5.01325089593518 | 1.09621986067686 | -4.57321662904351 | 4.80293104094874e-06 | 0.000700697872145671 | 4592 | 1 | 0 | 11 | 28 | 0 | 0 | 4 | 2 | 0 | 0 | 8 | 5 |
| L1MEd_dup5084 | 3.55157167330512 | -4.78666440941248 | 1.02378697702837 | -4.67544959724549 | 2.93310311922839e-06 | 0.000472845633338699 | 4593 | 1 | 0 | 9 | 16 | 0 | 0 | 7 | 2 | 0 | 0 | 7 | 7 |
| L1MC5_dup11824 | 3.88485635839466 | -3.40115936877889 | 1.00931879943203 | -3.36975727658379 | 0.000752344220850086 | 0.0288215668172295 | 4594 | 4 | 0 | 6 | 10 | 0 | 0 | 3 | 3 | 0 | 0 | 15 | 11 |
| L1P2_dup1362 | 4.3245235224488 | -4.98211260931791 | 0.98989277999218 | -5.03298206635801 | 4.8290831737518e-07 | 0.000116398173455619 | 4595 | 1 | 0 | 7 | 12 | 0 | 0 | 3 | 5 | 0 | 0 | 12 | 18 |
| MLT1G_dup682 | 4.11112895592849 | -3.55653170030469 | 1.0078726319293 | -3.52875114139836 | 0.000417525564152099 | 0.0190724779972615 | 4596 | 4 | 0 | 7 | 12 | 0 | 0 | 3 | 4 | 0 | 0 | 8 | 18 |
| AluSg_dup32583 | 3.93366842274539 | -3.59808761027713 | 1.05445837111752 | -3.41226141195489 | 0.000644262974751287 | 0.0257820043373982 | 4597 | 3 | 1 | 6 | 21 | 0 | 0 | 1 | 4 | 0 | 0 | 6 | 12 |
| MLT1A0_dup5022 | 3.66944603445607 | -3.84522918391485 | 1.06620617890331 | -3.60645929464602 | 0.000310403524592645 | 0.0153139699487293 | 4598 | 2 | 1 | 13 | 12 | 0 | 0 | 1 | 3 | 0 | 0 | 8 | 9 |
| L1M2_dup1019 | 2.53497248171468 | -4.77589659438138 | 1.12120663487846 | -4.25960429220888 | 2.04789129563935e-05 | 0.00207975903641571 | 4599 | 0 | 0 | 3 | 10 | 0 | 0 | 3 | 3 | 0 | 0 | 3 | 13 |
| Tigger2a_dup2657 | 1.86784125569222 | -4.33343984992635 | 1.2553818215164 | -3.45188991560503 | 0.000556674794520223 | 0.0232795407758408 | 4600 | 0 | 0 | 2 | 9 | 0 | 0 | 1 | 2 | 0 | 0 | 2 | 10 |
| L1MB7_dup17293 | 2.35573022086133 | -3.6394633102689 | 1.12780917004918 | -3.22702049860989 | 0.00125086498657835 | 0.041024916551 4042 | 4601 | 0 | 1 | 3 | 12 | 1 | 0 | 1 | 2 | 0 | 0 | 6 | 6 |
| THE1B_dup5130 | 2.36322425797378 | -4.15847352593803 | 1.14107373191567 | -3.64435128916398 | 0.000268067077226495 | 0.0137854957332474 | 4602 | 0 | 1 | 2 | 10 | 0 | 0 | 1 | 5 | 0 | 0 | 5 | 7 |
| L1MB1_dup1215 | 2.73977526268805 | -4.91814441964665 | 1.19629302539658 | -4.11115363480134 | 3.93687120493064e-05 | 0.00339776117420539 | 4603 | 0 | 0 | 4 | 17 | 0 | 0 | 1 | 2 | 0 | 0 | 6 | 8 |
| L1MC3_dup10255 | 2.04999266408701 | -4.47803964915241 | 1.18649152464602 | -3.7741859559329 | 0.000160530988519249 | 0.00956137837151469 | 4604 | 0 | 0 | 3 | 10 | 0 | 0 | 1 | 2 | 0 | 0 | 5 | 7 |
| MLT2B3_dup2716 | 2.41235145045213 | -4.73973180608852 | 1.16503051734808 | -4.06833274795018 | 4.73507304947408e-05 | 0.00391212392996583 | 4605 | 0 | 0 | 5 | 12 | 0 | 0 | 3 | 1 | 0 | 0 | 3 | 10 |
| AluSg_dup26535 | 2.78027707220607 | -4.94931410434483 | 1.14978927708352 | -4.30454014747722 | 1.67332946467864e-05 | 0.00179190842845375 | 4606 | 0 | 0 | 6 | 16 | 0 | 0 | 3 | 1 | 0 | 0 | 5 | 8 |
| L1MCc_dup391 | 2.80562298677078 | -4.95727905460093 | 1.1621887820167 | -4.26546799565451 | 1.99483609402943e-05 | 0.00203805611519934 | 4607 | 0 | 0 | 8 | 14 | 0 | 0 | 2 | 1 | 0 | 0 | 5 | 9 |
| L1ME2z_dup169 | 2.69470066847475 | -4.34854888555452 | 1.06667283188001 | -4.07674101710288 | 4.56713127508274e-05 | 0.00381175243206187 | 4608 | 0 | 1 | 5 | 10 | 0 | 0 | 4 | 1 | 0 | 0 | 8 | 8 |
| L1MCc_dup2763 | 2.60259539959421 | -4.32323606626703 | 1.07951908170758 | -4.00477966487498 | 6.20753131543213e-05 | 0.00479252837891568 | 4609 | 1 | 0 | 5 | 11 | 0 | 0 | 4 | 1 | 0 | 0 | 7 | 7 |
| L1MD3_dup4040 | 2.23924813051419 | -4.62974155903652 | 1.09868977641027 | -4.21387516152483 | 2.51026070666705e-05 | 0.00241830818886838 | 4610 | 0 | 0 | 5 | 8 | 0 | 0 | 5 | 1 | 0 | 0 | 5 | 7 |
| AluSx_dup76849 | 2.26352409346418 | -3.59039301106433 | 1.14044822292387 | -3.14822973888222 | 0.00164262538951444 | 0.0495130657377 | 4611 | 1 | 1 | 4 | 12 | 0 | 0 | 3 | 1 | 0 | 0 | 7 | 2 |
| AluJb_dup97920 | 2.73550544872299 | -3.87373844241709 | 1.14065326597852 | -3.39607009242548 | 0.000683608514273519 | 0.026890711115445 | 4612 | 1 | 1 | 5 | 14 | 0 | 0 | 1 | 1 | 0 | 0 | 6 | 9 |
| MLT2B1_dup579 | 2.86996343328252 | -3.93881939768616 | 1.17095063959952 | -3.36377919314626 | 0.00076883028065419 | 0.0292143228937308 | 4613 | 1 | 1 | 7 | 19 | 0 | 0 | 1 | 1 | 0 | 0 | 5 | 5 |
| L1MA9_dup12369 | 2.57468519075136 | -3.26751783865378 | 1.01743653379134 | -3.21152006059563 | 0.00132034751832368 | 0.042559118205777 | 4614 | 1 | 2 | 6 | 10 | 0 | 0 | 2 | 4 | 0 | 0 | 4 | 5 |
| L1MC1_dup310 | 2.87522387259777 | -4.45181314042103 | 1.09396986853848 | -4.06941111309451 | 4.71321166943489e-05 | 0.00390490899685012 | 4615 | 0 | 1 | 7 | 16 | 0 | 0 | 2 | 3 | 0 | 0 | 5 | 5 |
| MLT1G3_dup676 | 2.79573321397318 | -4.41532241782356 | 1.03492229734976 | -4.26633229289807 | 1.9871274362072e-05 | 0.00203542412800585 | 4616 | 0 | 1 | 8 | 10 | 0 | 0 | 3 | 4 | 0 | 0 | 5 | 6 |
| L1ME2_dup10019 | 2.47580704668644 | -4.23375486954082 | 1.08315590415122 | -3.90872159152239 | 9.27858173079322e-05 | 0.0064612274173 6998 | 4617 | 1 | 0 | 4 | 10 | 0 | 0 | 2 | 3 | 0 | 0 | 8 | 5 |
| AluJb_dup91876 | 2.31636836839644 | -3.61243920055599 | 1.09485496093407 | -3.29946826698766 | 0.000968681786392622 | 0.034386686012976 1 | 4618 | 1 | 1 | 7 | 7 | 0 | 0 | 1 | 2 | 0 | 0 | 5 | 7 |
| MER54B_dup73 | 2.89658061392772 | -3.91724038324427 | 1.01917736088864 | -3.84353159083986 | 0.000121276379812885 | 0.00781615099430032 | 4619 | 1 | 1 | 7 | 7 | 0 | 0 | 3 | 2 | 0 | 0 | 8 | 10 |
| LTR39_dup979 | 4.62820313536064 | -2.79004247551715 | 0.800254084083212 | -3.48644578142139 | 0.000489484593971415 | 0.0212266996896523 | 4620 | 0 | 3 | 13 | 6 | 1 | 0 | 6 | 2 | 0 | 3 | 10 | 18 |
| Tigger9a_dup987 | 4.00080208822631 | -3.14737236513005 | 0.850655953530237 | -3.69993574025833 | 0.00021565406598615 | 0.011736967608678 | 4621 | 0 | 3 | 16 | 7 | 1 | 0 | 8 | 5 | 0 | 1 | 6 | 5 |
| L2b_dup31913 | 4.82819737247959 | -4.21188411847205 | 0.98307493025167 | -4.28439785092861 | 1.83234769716395e-05 | 0.00191803784609089 | 4622 | 0 | 1 | 12 | 12 | 0 | 0 | 14 | 7 | 0 | 2 | 2 | 15 |
| AluSz_dup43114 | 3.22350839997074 | -3.16549860785726 | 0.960301065454997 | -3.29636061203101 | 0.000979462044235923 | 0.034624732666 4331 | 4623 | 0 | 3 | 9 | 9 | 0 | 0 | 8 | 2 | 0 | 1 | 3 | 9 |
| L1MB2_dup7375 | 3.41733408480919 | -3.62350857294908 | 1.03754361079544 | -3.4923915826257 | 0.000478715888243815 | 0.0209198403570363 | 4624 | 0 | 2 | 7 | 19 | 0 | 0 | 9 | 2 | 0 | 1 | 1 | 7 |
| MLT1E1A_dup1910 | 2.113671640239 | -3.98980619040897 | 1.17036110390259 | -3.40903860962645 | 0.000651922523325294 | 0.0260054244317117 | 4625 | 0 | 1 | 2 | 1 | 0 | 0 | 9 | 5 | 0 | 0 | 6 | 3 |
| MLT1J_dup8037 | 2.53824363773619 | -4.7715240837424 | 1.11551400747028 | -4.2774219344525 | 1.89070229785782e-05 | 0.00196028343776069 | 4626 | 0 | 0 | 2 | 3 | 0 | 0 | 11 | 3 | 0 | 0 | 9 | 6 |
| L1PA5_dup6206 | 3.62086538437136 | -3.64539924667779 | 0.986445929526162 | -3.69548815354619 | 0.000219464713735476 | 0.0118953874201598 | 4627 | 0 | 1 | 3 | 1 | 0 | 1 | 8 | 6 | 0 | 1 | 12 | 14 |
| L1PA2_dup612 | 3.09983119912105 | -4.04414371547549 | 0.982432774302009 | -4.11645847050323 | 3.84738616452487e-05 | 0.00334235176452732 | 4628 | 0 | 0 | 6 | 6 | 0 | 1 | 8 | 4 | 0 | 1 | 8 | 7 |
| L4_B_Mam_dup485 | 2.71939605600369 | -3.27246892330911 | 0.990385305786086 | -3.30423816285491 | 0.000952348903010289 | 0.0339741547339859 | 4629 | 0 | 0 | 2 | 2 | 0 | 1 | 7 | 3 | 0 | 2 | 9 | 10 |
| MLT1L_dup2932 | 2.48897545213293 | -4.20716259997858 | 1.15394775575213 | -3.64588654816215 | 0.000266471527612 15 | 0.01374503232369 4 | 4630 | 0 | 0 | 2 | 3 | 0 | 0 | 2 | 10 | 0 | 1 | 5 | 7 |
| MLT1L_dup9862 | 2.43364466279078 | -3.67064032341893 | 1.13002970564682 | -3.24826887742555 | 0.00116109494218472 | 0.0389398022710481 | 4631 | 1 | 0 | 2 | 4 | 0 | 0 | 5 | 8 | 0 | 1 | 1 | 9 |
| L1MC3_dup11378 | 3.01004214228746 | -3.48696037095234 | 0.987715180722658 | -3.53032983496428 | 0.000415041900534411 | 0.0190101174330242 | 4632 | 1 | 1 | 7 | 4 | 0 | 0 | 11 | 5 | 0 | 1 | 2 | 8 |
| MLT1A0_dup1859 | 2.88880199050649 | -3.42952233299767 | 1.0101652011247 | -3.39501135970562 | 0.000686257531339072 | 0.0269585101851577 | 4633 | 1 | 0 | 9 | 4 | 0 | 0 | 9 | 4 | 0 | 2 | 4 | 5 |
| MLT1I_dup3799 | 3.36315848868458 | -4.1798460482812 | 1.04343163271062 | -4.00586479961587 | 6.17909872826748e-05 | 0.00478610620441515 | 4634 | 1 | 0 | 10 | 2 | 0 | 0 | 12 | 3 | 0 | 1 | 7 | 9 |
| AluY_dup94139 | 2.16581663369192 | -4.57317878588324 | 1.23379631151971 | -3.70659139047862 | 0.000210067453953533 | 0.0115251810925588 | 4635 | 0 | 0 | 8 | 1 | 0 | 0 | 8 | 3 | 0 | 0 | 1 | 8 |
| HERVK22-int_dup185 | 1.98169102772457 | -3.89631528679848 | 1.14120197998935 | -3.41422058068532 | 0.000639647665635837 | 0.0256674329159602 | 4636 | 1 | 0 | 6 | 1 | 0 | 0 | 8 | 3 | 0 | 0 | 4 | 3 |
| MLT1B_dup11335 | 2.1759883780827 | -3.50219256279605 | 1.08276981140297 | -3.23447562530229 | 0.00121866322352024 | 0.0402632656739444 | 4637 | 0 | 0 | 8 | 2 | 0 | 0 | 7 | 3 | 1 | 1 | 3 | 3 |
| MLT1I_dup271 | 2.23893060407345 | -3.5374632406611 | 1.10493979255332 | -3.20149863775532 | 0.00136714721622948 | 0.0436127110219786 | 4638 | 0 | 0 | 7 | 1 | 0 | 0 | 7 | 5 | 1 | 1 | 3 | 3 |
| L2c_dup103678 | 2.47509801431033 | -3.71059039675019 | 1.0729605676363 | -3.45827284680601 | 0.000543650502301218 | 0.0228590674007008 | 4639 | 0 | 0 | 7 | 4 | 0 | 0 | 11 | 3 | 0 | 2 | 3 | 3 |
| FLAM_C_dup21341 | 2.11576932652286 | -3.91076620295542 | 1.08349745369324 | -3.60939122618615 | 0.000306916435481029 | 0.0152018189335905 | 4640 | 0 | 0 | 7 | 3 | 0 | 0 | 7 | 5 | 0 | 1 | 2 | 2 |
| L1MB7_dup11700 | 2.34956196638091 | -4.09251339606394 | 1.06120247611156 | -3.85648685165121 | 0.00011502828221643 | 0.00751475711202635 | 4641 | 0 | 0 | 4 | 5 | 0 | 0 | 11 | 5 | 0 | 1 | 2 | 3 |
| L2a_dup117069 | 3.56879024461344 | -2.52092587487901 | 0.782288096044037 | -3.22250317706113 | 0.00127075756469259 | 0.0414586769729918 | 4642 | 0 | 0 | 8 | 9 | 2 | 1 | 8 | 8 | 2 | 1 | 3 | 3 |
| L2a_dup99865 | 4.35763092340062 | -2.95682119721983 | 0.749832417903214 | -3.94330936702911 | 8.03648769874552e-05 | 0.00580172967829 4 | 4643 | 0 | 0 | 11 | 11 | 2 | 2 | 12 | 6 | 1 | 1 | 7 | 4 |
| L1PB1_dup9948 | 4.31148291669014 | -2.26455873954068 | 0.707621577842247 | -3.20023980394437 | 0.00137313288693904 | 0.04376296429348 6 | 4644 | 0 | 0 | 6 | 10 | 3 | 3 | 7 | 6 | 1 | 2 | 4 | 15 |
| AluSx_dup113359 | 3.88174087284159 | -2.76460813236304 | 0.775654577002061 | -3.56422589943113 | 0.000364931572730467 | 0.0172666550019733 | 4645 | 0 | 0 | 10 | 9 | 3 | 1 | 5 | 4 | 1 | 1 | 7 | 10 |
| L1ME1_dup338 | 3.86997456048006 | -2.29137209187097 | 0.720082578654428 | -3.18209627589201 | 0.00146213205930813 | 0.0456523962667711 | 4646 | 0 | 0 | 9 | 6 | 2 | 3 | 10 | 3 | 1 | 2 | 8 | 7 |
| L1MCc_dup2583 | 5.982031789242 | -3.21884339021126 | 0.686167501475598 | -4.69104611234015 | 2.71811696636393e-06 | 0.000446909959820496 | 4647 | 1 | 0 | 9 | 19 | 2 | 1 | 9 | 13 | 1 | 2 | 10 | 10 |
| MLT1L_dup2590 | 4.20896021333581 | -2.42176475473687 | 0.679698060634739 | -3.56300083080315 | 0.000366639518621559 | 0.0173164761607633 | 4648 | 2 | 0 | 6 | 10 | 1 | 1 | 6 | 7 | 2 | 2 | 9 | 8 |
| L1ME3_dup6532 | 5.14017712153722 | -2.5538066793094 | 0.641147589354727 | -3.98318066184985 | 6.79990230537702e-05 | 0.00512746329078604 | 4649 | 1 | 0 | 9 | 14 | 2 | 3 | 8 | 10 | 2 | 1 | 7 | 9 |
| L2c_dup117349 | 4.92998975933172 | -2.11004533920301 | 0.630497432435338 | -3.34663589517378 | 0.000817985744393671 | 0.0305503744993716 | 4650 | 2 | 0 | 5 | 9 | 3 | 2 | 8 | 11 | 3 | 1 | 8 | 10 |
| Tigger19a_dup1895 | 5.20595082740121 | -2.44746253191662 | 0.659130174361087 | -3.71317021601843 | 0.000204679124535547 | 0.0113051483209692 | 4651 | 2 | 0 | 8 | 6 | 3 | 3 | 18 | 10 | 1 | 1 | 9 | 6 |
| L1M4a2_dup1728 | 4.61650035124381 | -2.7859475605308 | 0.764390924003137 | -3.64466331695923 | 0.000267742072571176 | 0.0137763201294569 | 4652 | 2 | 0 | 3 | 6 | 1 | 1 | 13 | 11 | 2 | 1 | 12 | 6 |
| L1ME2_dup12223 | 3.66410470912726 | -2.63699404652835 | 0.789092625783217 | -3.34180546157176 | 0.000832353752713489 | 0.0309268029424053 | 4653 | 1 | 0 | 5 | 4 | 1 | 1 | 11 | 9 | 2 | 1 | 5 | 6 |
| L2a_dup124343 | 4.26030026021043 | -2.52464158776785 | 0.696486857159296 | -3.62482301254743 | 0.000289159409006599 | 0.0145679281675808 | 4654 | 3 | 1 | 8 | 11 | 3 | 0 | 11 | 6 | 1 | 0 | 7 | 5 |
| MIRb_dup25939 | 4.08655933381593 | -2.82046431566073 | 0.771618723837922 | -3.65525644794121 | 0.000256925021004689 | 0.0134066405806978 | 4655 | 1 | 1 | 5 | 9 | 2 | 0 | 11 | 10 | 2 | 0 | 4 | 7 |
| L1PA12_dup1726 | 4.54914984788314 | -2.53051020358155 | 0.77205192003663 | -3.27764252365501 | 0.00104677870365392 | 0.0362293535155833 | 4656 | 1 | 0 | 10 | 7 | 5 | 0 | 6 | 10 | 2 | 0 | 7 | 9 |

| | | | | | | | | | | | | | | | | | | | |
|---|---|---|---|---|---|---|---|---|---|---|---|---|---|---|---|---|---|---|---|
| THE1C_dup9739 | 3.20305649066664 | -2.7685989282951 | 0.85600702607755 | -3.23431799500706 | 0.00121933609541427 | 0.0402706929491742 | 4657 | 1 | 0 | 10 | 8 | 3 | 0 | 5 | 6 | 1 | 0 | 3 | 4 |
| MIR_dup136599 | 3.11543437903767 | -2.71802991274631 | 0.856069834250921 | -3.17500956580796 | 0.00149831561853794 | 0.0464700479886115 | 4658 | 1 | 0 | 6 | 3 | 1 | 2 | 10 | 3 | 1 | 0 | 8 | 6 |
| Tigger2_dup1357 | 3.21827240571089 | -2.80397915715606 | 0.838620945821638 | -3.34355965126636 | 0.000827109100496763 | 0.0307779653713189 | 4659 | 1 | 0 | 11 | 5 | 1 | 2 | 8 | 4 | 1 | 0 | 4 | 5 |
| Charlie1b_dup1982 | 5.7588411024282 | -2.95943857790113 | 0.670025152339563 | -4.41690669009589 | 1.00123425746252e-05 | 0.00122425327146032 | 4660 | 2 | 0 | 16 | 9 | 1 | 3 | 14 | 11 | 2 | 0 | 6 | 10 |
| L2_dup32904 | 4.45375491445803 | -2.43271932553771 | 0.72074163846689 | -3.37530010159038 | 0.000737352372251772 | 0.028386577244525 | 4661 | 3 | 0 | 9 | 7 | 2 | 3 | 14 | 7 | 1 | 0 | 2 | 11 |
| L1PA4_dup10778 | 3.1493795770657 | -3.1863819218381 | 0.936645770776615 | -3.40190712567476 | 0.000669173745594197 | 0.0264804949391579 | 4662 | 3 | 0 | 5 | 5 | 0 | 1 | 13 | 7 | 0 | 0 | 5 | 2 |
| AluJo_dup34693 | 2.72808512133962 | -3.81352697964794 | 1.04513581102915 | -3.64883390216316 | 0.000263433349919327 | 0.0136415477431783 | 4663 | 1 | 0 | 3 | 7 | 0 | 1 | 8 | 7 | 0 | 0 | 6 | 2 |
| L2c_dup104392 | 3.20157010675057 | -3.60961965112472 | 0.97834628710267 | -3.68951126887234 | 0.000224685276202249 | 0.0120915958871828 | 4664 | 2 | 0 | 4 | 5 | 0 | 1 | 8 | 8 | 0 | 0 | 7 | 6 |
| L2b_dup60235 | 3.17191266954867 | -3.19206005655401 | 0.92426577851875 | -3.45361705555049 | 0.00055312219783245 | 0.0231713405784226 | 4665 | 2 | 1 | 3 | 18 | 0 | 1 | 6 | 5 | 0 | 0 | 4 | 3 |
| MIRb_dup22138 | 2.74116275087231 | -3.32517328984175 | 1.02154856005093 | -3.25503203653477 | 0.00113379575498374 | 0.0383129580379935 | 4666 | 2 | 0 | 2 | 14 | 0 | 1 | 4 | 7 | 0 | 0 | 3 | 3 |
| L2_dup9326 | 4.5720309751708 | -2.44036305231224 | 0.731691995972572 | -3.33523267405499 | 0.000852280550099027 | 0.031436482370774 | 4667 | 4 | 0 | 6 | 13 | 0 | 3 | 5 | 10 | 0 | 2 | 8 | 8 |
| LTR103b_Mam_dup43 | 3.78955040953242 | -3.41316254512939 | 0.89075324035244 | -3.83177112415433 | 0.000127224068508145 | 0.00808775265969516 | 4668 | 2 | 0 | 9 | 9 | 0 | 0 | 7 | 7 | 0 | 2 | 8 | 5 |
| L1PA5_dup8931 | 4.28353311490435 | -3.61460923886991 | 0.858427703551207 | -4.21073227706507 | 2.54544256943999e-05 | 0.00244619498234235 | 4669 | 1 | 0 | 9 | 11 | 0 | 2 | 10 | 7 | 0 | 1 | 10 | 5 |
| MLT1E3_dup1878 | 3.83836116265829 | -3.82820853366276 | 0.956503581663044 | -4.00229398723921 | 6.27312830048014e-05 | 0.00483376226226052 | 4670 | 1 | 0 | 14 | 14 | 0 | 1 | 7 | 4 | 0 | 1 | 7 | 2 |
| L1MD2_dup8010 | 3.26680889363296 | -2.93457257008347 | 0.881395028381298 | -3.32946349319996 | 0.000870134703565847 | 0.0319169218706149 | 4671 | 2 | 0 | 8 | 17 | 2 | 1 | 6 | 2 | 0 | 0 | 4 | 3 |
| MER5A_dup34109 | 3.10741717431254 | -3.14383419901623 | 0.896366792792223 | -3.50730774979185 | 0.000452665308246231 | 0.0201855133875483 | 4672 | 2 | 0 | 9 | 14 | 1 | 1 | 8 | 2 | 0 | 0 | 2 | 4 |
| L2_dup31236 | 3.24437027368515 | -2.59911937366873 | 0.824243401592474 | -3.15333961866983 | 0.00161413896981472 | 0.0488213945991526 | 4673 | 1 | 0 | 10 | 11 | 3 | 2 | 9 | 3 | 0 | 0 | 1 | 4 |
| L2a_dup124357 | 3.04221442663957 | -2.81682213282618 | 0.87515281540214 | -3.21866316745131 | 0.00128789673609444 | 0.0418195153023083 | 4674 | 1 | 0 | 10 | 7 | 3 | 1 | 8 | 2 | 0 | 0 | 4 | 5 |
| Tigger20a_dup99 | 3.09202747613417 | -2.81984469720048 | 0.837597200760209 | -3.36658801467002 | 0.000761042920041615 | 0.0290208182842666 | 4675 | 1 | 0 | 8 | 7 | 3 | 1 | 9 | 4 | 0 | 0 | 4 | 4 |
| L2a_dup19324 | 4.15871234532357 | -3.54712701725767 | 0.840951382875104 | -4.21799296545598 | 2.46486507215102e-05 | 0.00238804879665329 | 4676 | 2 | 0 | 14 | 14 | 1 | 1 | 6 | 9 | 0 | 0 | 2 | 5 |
| L2a_dup90816 | 4.14163111324727 | -3.55799520881922 | 0.812259443335445 | -4.38036792063475 | 1.18479090144788e-05 | 0.00137991314652921 | 4677 | 2 | 0 | 11 | 11 | 1 | 1 | 11 | 9 | 0 | 0 | 3 | 5 |
| MER91C_dup320 | 3.83816684156961 | -3.46783767827222 | 0.839861510448555 | -4.12905894022945 | 3.64251027886747e-05 | 0.00320414325624916 | 4678 | 2 | 0 | 9 | 13 | 1 | 1 | 7 | 8 | 0 | 0 | 5 | 4 |
| L1M5_dup28719 | 3.72777969061533 | -3.45533878445505 | 0.901073785621368 | -3.83469016588058 | 0.000125722672142419 | 0.00801410773025061 | 4679 | 1 | 0 | 6 | 14 | 2 | 1 | 7 | 4 | 0 | 0 | 4 | 12 |
| L1PA8_dup155 | 2.74798724065173 | -3.36500166336478 | 0.957036268714581 | -3.51606493229812 | 0.000437993851651414 | 0.0197298777117623 | 4680 | 1 | 0 | 5 | 7 | 1 | 1 | 5 | 5 | 0 | 0 | 5 | 6 |
| MLT1I_dup9486 | 3.45505547921507 | -3.71045369923793 | 0.939936520310031 | -3.94755775423436 | 7.8952475667145e-05 | 0.00572756883387027 | 4681 | 1 | 0 | 9 | 15 | 1 | 1 | 4 | 5 | 0 | 0 | 5 | 5 |
| MIRc_dup92012 | 6.0193622242626 | -2.75992346596536 | 0.618504851635763 | -4.46225031002777 | 8.11034071545509e-06 | 0.00104033592015077 | 4682 | 2 | 0 | 12 | 20 | 3 | 4 | 12 | 9 | 0 | 1 | 7 | 10 |
| AluSp_dup50086 | 5.11527377265739 | -2.65383175538233 | 0.648014533494585 | -4.09532752463261 | 4.21571977290931e-05 | 0.00358767402390801 | 4683 | 1 | 1 | 10 | 17 | 3 | 3 | 11 | 5 | 0 | 1 | 7 | 10 |
| L1ME3A_dup10511 | 5.40530503198573 | -2.42441331244562 | 0.606442872067859 | -3.9977604224761 | 6.3944622541813e-05 | 0.00490185797530098 | 4684 | 2 | 1 | 9 | 17 | 4 | 3 | 12 | 8 | 0 | 1 | 5 | 10 |
| HAL1_dup15226 | 4.26060692368154 | -2.6876473395909 | 0.719661820510105 | -3.73459764432947 | 0.000188015534475522 | 0.0106441932824917 | 4685 | 2 | 0 | 10 | 13 | 2 | 1 | 11 | 7 | 0 | 2 | 5 | 3 |
| L1MC5_dup12304 | 4.78714637176008 | -2.02942299066852 | 0.62248406180473 | -3.26020072672179 | 0.00111333391497194 | 0.0377638420145279 | 4686 | 3 | 0 | 7 | 9 | 3 | 4 | 15 | 8 | 0 | 2 | 5 | 7 |
| MSTA-int_dup1642 | 4.20645583146514 | -2.27492008322256 | 0.697026999339184 | -3.26374743787442 | 0.00109949129466762 | 0.0374705751362049 | 4687 | 2 | 1 | 9 | 14 | 1 | 1 | 9 | 3 | 0 | 4 | 5 | 8 |
| MSTB1_dup392 | 3.47541963704564 | -2.36916664520187 | 0.730300324168478 | -3.24409912853238 | 0.00117822734140093 | 0.0393093778711422 | 4688 | 1 | 1 | 6 | 6 | 1 | 2 | 10 | 5 | 0 | 2 | 4 | 8 |
| LTR1F2_dup128 | 5.28057081278313 | -2.39357537682087 | 0.61761682862548 | -3.8755021979369 | 0.000106405019297848 | 0.00714197604159484 | 4689 | 4 | 2 | 11 | 15 | 3 | 1 | 10 | 6 | 0 | 1 | 5 | 13 |
| MER1B_dup3996 | 5.1010727182795 | -3.05081132311879 | 0.715745397580527 | -4.2624253448665 | 2.02220066760355e-05 | 0.00205806056481434 | 4690 | 1 | 2 | 8 | 21 | 2 | 1 | 12 | 4 | 0 | 1 | 7 | 11 |
| AluY_dup51848 | 4.54835751283283 | -3.08183978661035 | 0.732365836579323 | -4.20806055209342 | 2.57571853911964e-05 | 0.00246335178259389 | 4691 | 1 | 2 | 11 | 10 | 1 | 1 | 11 | 8 | 0 | 1 | 8 | 5 |
| L4_C_Mam_dup1365 | 4.23832642985129 | -2.15279676955754 | 0.649136990260036 | -3.31639823621075 | 0.000911857896134054 | 0.0329846365085913 | 4692 | 3 | 2 | 10 | 11 | 3 | 1 | 6 | 7 | 0 | 1 | 6 | 5 |
| AluSc_dup4939 | 4.0604717949646 | -3.01832378408117 | 0.919595859569582 | -3.2822285492824 | 0.0010299009283058 | 0.0358066636888739 | 4693 | 0 | 2 | 3 | 9 | 0 | 4 | 11 | 9 | 0 | 0 | 11 | 3 |
| L1PREC2_dup28 | 3.31413776586913 | -3.58728904420703 | 1.00309433838788 | -3.57622300009422 | 0.000348594317968008 | 0.0166492620600593 | 4694 | 0 | 1 | 3 | 9 | 0 | 1 | 3 | 8 | 1 | 0 | 8 | 8 |
| HSAT5_dup224 | 4.4994783171037 | -3.00206970611107 | 0.784225006731893 | -3.82807189306755 | 0.000129151035620092 | 0.00817530521991442 | 4695 | 0 | 1 | 4 | 11 | 0 | 3 | 8 | 12 | 1 | 1 | 7 | 9 |
| MER5A_dup1497 | 3.63689900494294 | -2.63721612400794 | 0.807194898413397 | -3.2671367586584 | 0.00108641184794579 | 0.0371846020672193 | 4696 | 1 | 2 | 3 | 5 | 0 | 1 | 7 | 10 | 1 | 1 | 9 | 5 |
| L1PA8_dup2784 | 3.71724446586227 | -3.63934687023137 | 0.885934904032871 | -4.10791679350782 | 3.99243874143857e-05 | 0.00343326645742806 | 4697 | 0 | 1 | 9 | 9 | 0 | 1 | 11 | 10 | 1 | 0 | 3 | 2 |
| LTR16B2_dup534 | 3.61868973427194 | -3.30845808263972 | 0.883341392471154 | -3.74539007323577 | 0.000180113701257688 | 0.0103520132269231 | 4698 | 0 | 2 | 4 | 8 | 0 | 1 | 14 | 8 | 1 | 0 | 4 | 5 |
| AluSz_dup52027 | 4.25194286434411 | -3.9612342017897 | 0.987092676771703 | -4.0130316990548 | 5.99438598661165e-05 | 0.00466887125358525 | 4699 | 0 | 1 | 6 | 3 | 0 | 1 | 20 | 11 | 1 | 0 | 7 | 4 |
| L1PA3_dup5076 | 4.00790198190108 | -2.75286241783745 | 0.840474676134329 | -3.27536628527457 | 0.00105525056057069 | 0.0364589298512491 | 4700 | 1 | 1 | 5 | 5 | 0 | 2 | 15 | 11 | 2 | 0 | 6 | 2 |
| L1M4_dup17081 | 4.96577092178916 | -2.98198033037032 | 0.758610609964451 | -3.9308444822701 | 8.46480149741077e-05 | 0.00603092803958492 | 4701 | 0 | 2 | 10 | 6 | 1 | 3 | 20 | 7 | 1 | 0 | 4 | 12 |
| L1PA5_dup174 | 5.12572432672851 | -3.83089866779938 | 0.873374874591527 | -4.38631655117291 | 1.15286309135535e-05 | 0.00135399388517098 | 4702 | 0 | 1 | 10 | 4 | 0 | 1 | 16 | 7 | 2 | 0 | 11 | 15 |
| L2b_dup45753 | 5.22798105493403 | -4.20186388963856 | 0.886807544979943 | -4.73819140739673 | 2.15634089178577e-06 | 0.000372616332488639 | 4703 | 0 | 2 | 15 | 24 | 0 | 1 | 11 | 9 | 0 | 0 | 1 | 7 |
| L2c_dup129463 | 4.43528241152752 | -4.05535827066874 | 0.961377452211776 | -4.21827895103932 | 2.46174147643577e-05 | 0.00238599404216695 | 4704 | 1 | 1 | 15 | 22 | 0 | 1 | 7 | 4 | 0 | 0 | 1 | 9 |
| MIR_dup124597 | 3.53537548016167 | -3.7303230478787 | 0.980596164730376 | -3.80413791329113 | 0.000142298888843402 | 0.00876279798259113 | 4705 | 0 | 1 | 11 | 16 | 0 | 2 | 8 | 2 | 0 | 0 | 2 | 7 |
| L1ME3G_dup7525 | 2.85797399254926 | -3.41388472853991 | 0.986096610009867 | -3.46201852220722 | 0.00053614025638865 | 0.0226299894420064 | 4706 | 0 | 1 | 7 | 13 | 0 | 2 | 5 | 3 | 0 | 0 | 2 | 6 |
| MLT2B1_dup1074 | 2.86375523494382 | -3.44676934991511 | 1.0066295301348 | -3.42406937878481 | 0.000616908691739123 | 0.0250336560018001 | 4707 | 0 | 1 | 7 | 10 | 0 | 2 | 5 | 3 | 0 | 0 | 2 | 9 |
| L4_B_Mam_dup1772 | 3.4454882985469 | -3.65542668925474 | 0.934846854612405 | -3.91018771814803 | 9.22244212678925e-05 | 0.0064334307074201 | 4708 | 0 | 2 | 10 | 13 | 0 | 1 | 5 | 6 | 0 | 0 | 5 | 3 |
| MLT1B_dup3159 | 3.34087111361518 | -3.18280729865017 | 0.876660246050007 | -3.63060525784195 | 0.000282757360553667 | 0.0143272961912911 | 4709 | 0 | 2 | 9 | 15 | 0 | 2 | 8 | 4 | 0 | 0 | 3 | 2 |
| SATR2_dup187 | 3.23792513605525 | -3.7740723614788 | 0.946952531138991 | -3.98549265921425 | 6.73402592745172e-05 | 0.00510297622080709 | 4710 | 0 | 1 | 12 | 10 | 0 | 1 | 8 | 7 | 0 | 0 | 1 | 2 |
| THE1D-int_dup2241 | 3.92354870490613 | -4.20365530455534 | 0.947724999104616 | -4.43552223327109 | 9.18493510252026e-06 | 0.00114544998317835 | 4711 | 0 | 1 | 10 | 19 | 0 | 1 | 10 | 6 | 0 | 0 | 2 | 4 |
| L2c_dup41651 | 3.71070104639851 | -2.52470451693144 | 0.793382645215991 | -3.18220285275349 | 0.00146159409488243 | 0.0456523962667711 | 4712 | 2 | 3 | 13 | 7 | 0 | 1 | 4 | 4 | 0 | 1 | 4 | 10 |
| AluJb_dup112926 | 4.46333062767385 | -3.04722109356892 | 0.773855232722708 | -3.93771465865479 | 8.22613325097116e-05 | 0.00590282117773095 | 4713 | 0 | 4 | 11 | 10 | 0 | 1 | 9 | 6 | 0 | 1 | 6 | 11 |
| MIR3_dup79068 | 4.31344855579884 | -3.62603701350387 | 0.88238988406197 | -4.10933656312091 | 3.9679742320873e-05 | 0.00341720217261385 | 4714 | 0 | 2 | 14 | 13 | 0 | 1 | 7 | 4 | 0 | 1 | 5 | 11 |
| MSTA_dup4950 | 3.99494375695175 | -2.31266507063336 | 0.731116794534835 | -3.16319511180805 | 0.00156047709136442 | 0.0477138630148804 | 4715 | 3 | 3 | 8 | 12 | 0 | 3 | 6 | 3 | 0 | 0 | 7 | 9 |
| Tigger7_dup3091 | 4.25968399204052 | -3.3150251232433 | 0.83494345695954 | -3.97035882563236 | 7.17644689204989e-05 | 0.00533127328184734 | 4716 | 1 | 2 | 4 | 15 | 0 | 2 | 7 | 8 | 0 | 0 | 9 | 8 |
| AluSx_dup11473 | 4.11454176373643 | -2.48767221328711 | 0.721480925193695 | -3.44800829296942 | 0.000564736697503367 | 0.0235294761380032 | 4717 | 2 | 4 | 6 | 13 | 0 | 1 | 9 | 6 | 0 | 1 | 4 | 9 |
| L1MEi_dup4573 | 4.22758988234641 | -2.97477400984735 | 0.803849079355376 | -3.70066233357247 | 0.000215037463441458 | 0.0117114452451456 | 4718 | 1 | 2 | 6 | 21 | 0 | 2 | 8 | 7 | 0 | 1 | 2 | 7 |
| L2a_dup165402 | 2.8036389206885 | -2.96400496799761 | 0.900173367414789 | -3.29270457813026 | 0.000992286780550147 | 0.0349187364102553 | 4719 | 1 | 2 | 10 | 9 | 0 | 1 | 4 | 4 | 0 | 0 | 2 | 4 |
| L1PA2_dup3991 | 3.25583912177136 | -2.61254126832145 | 0.785432157194746 | -3.32624688764006 | 0.000880239243435888 | 0.0322180290753109 | 4720 | 2 | 2 | 7 | 8 | 0 | 2 | 7 | 7 | 0 | 0 | 4 | 3 |
| MIRb_dup27278 | 3.00611024089327 | -2.77792702056755 | 0.837569258724293 | -3.31665350850941 | 0.000911025238265496 | 0.032974568199623 | 4721 | 2 | 1 | 8 | 6 | 0 | 2 | 6 | 6 | 0 | 0 | 3 | 5 |
| AluSc8_dup10057 | 4.24411903915828 | -3.30684858359561 | 0.904767499185319 | -3.65491530870992 | 0.000257266890234221 | 0.0134156536190384 | 4722 | 0 | 0 | 11 | 21 | 1 | 4 | 13 | 2 | 0 | 0 | 4 | 3 |
| FLAM_A_dup6859 | 4.17131771648955 | -3.92895993484155 | 0.985972812157639 | -3.98485626215561 | 6.75209847393632e-05 | 0.00511016165228769 | 4723 | 0 | 0 | 17 | 21 | 1 | 2 | 6 | 2 | 0 | 0 | 5 | 3 |
| L1MA8_dup5166 | 3.77937404235841 | -2.92352866922042 | 0.900080324166831 | -3.24807530030902 | 0.00116188517606282 | 0.0389568319081735 | 4724 | 0 | 0 | 14 | 12 | 2 | 4 | 6 | 2 | 0 | 0 | 5 | 6 |
| MIR3_dup83435 | 3.94666900843407 | -3.79927414884117 | 1.03891334975075 | -3.65696922631004 | 0.000255215006712968 | 0.0133437464265708 | 4725 | 0 | 0 | 11 | 26 | 0 | 3 | 7 | 2 | 0 | 0 | 4 | 2 |
| L1PA10_dup6995 | 3.66707600930038 | -4.65093113067175 | 1.1187338602047 | -4.15731685266123 | 3.22007239010518e-05 | 0.00291221339770417 | 4726 | 0 | 0 | 9 | 26 | 0 | 1 | 9 | 1 | 0 | 0 | 4 | 2 |
| L1PA8_dup3056 | 3.71710514746886 | -4.7418289485346 | 1.11530541441591 | -4.25159681576359 | 2.12251718187372e-05 | 0.00213009056667282 | 4727 | 0 | 0 | 12 | 24 | 0 | 1 | 6 | 1 | 0 | 0 | 5 | 3 |
| L1MA4_dup5279 | 4.32110208700503 | -4.97352627134486 | 1.04391289844936 | -4.76431154240223 | 1.89499530986083e-06 | 0.000338180653495034 | 4728 | 0 | 0 | 20 | 18 | 0 | 1 | 8 | 2 | 0 | 0 | 5 | 5 |
| MLT1H2_dup4694 | 4.17078017142081 | -4.36929315263188 | 1.04509458108774 | -4.18076337941041 | 2.90532073931745e-05 | 0.00269390483312078 | 4729 | 0 | 0 | 15 | 17 | 0 | 2 | 10 | 4 | 0 | 0 | 7 | 1 |
| L2b_dup94898 | 4.95484595920289 | -4.19451356058022 | 0.920883399145865 | -4.55488019924206 | 5.24154419016234e-06 | 0.000747724504719646 | 4730 | 0 | 0 | 14 | 20 | 0 | 2 | 15 | 5 | 0 | 1 | 2 | 9 |
| MER41B_dup2792 | 5.42357467416516 | -3.92735480703933 | 0.829183864202708 | -4.73641007331424 | 2.17537201030309e-06 | 0.000374274091369183 | 4731 | 0 | 0 | 17 | 15 | 0 | 3 | 20 | 7 | 0 | 1 | 3 | 7 |
| MIR1_Amn_dup5136 | 4.79457813297009 | -4.16423310800501 | 0.911051251193112 | -4.57080005384059 | 4.8586569528591e-06 | 0.000704508218954913 | 4732 | 0 | 0 | 13 | 19 | 0 | 1 | 10 | 7 | 0 | 2 | 5 | 7 |
| MLT1G_dup2341 | 4.91503324092295 | -4.20829700665305 | 0.886625809690575 | -4.746418343181 | 2.07050255997494e-06 | 0.00036291185593887 | 4733 | 1 | 0 | 17 | 16 | 0 | 1 | 12 | 6 | 0 | 1 | 4 | 8 |
| THE1D_dup9602 | 5.3138172758872 | -3.49472431022376 | 0.830649583940224 | -4.20721851643671 | 2.58533125182575e-05 | 0.00246907119251075 | 4734 | 0 | 0 | 13 | 23 | 0 | 2 | 8 | 9 | 2 | 1 | 3 | 9 |
| L1PA7_dup1606 | 4.86443877248234 | -4.69850674424783 | 0.968113942469252 | -4.85325800831244 | 1.2144951686909e-06 | 0.00023922538692825 | 4735 | 0 | 0 | 8 | 18 | 0 | 1 | 13 | 9 | 1 | 0 | 3 | 12 |

| | | | | | | | | | | | | | | | | | | | |
|---|---|---|---|---|---|---|---|---|---|---|---|---|---|---|---|---|---|---|---|
| L2c_dup70111 | 5.07859097937341 | -4.20878940458661 | 0.892732613776869 | -4.71450167680169 | 2.42303009139623e-06 | 0.000408900934842105 | 4736 | 0 | 0 | 12 | 25 | 0 | 2 | 13 | 6 | 1 | 0 | 4 | 6 |
| Tigger4b_dup252 | 5.59114363329931 | -4.68409322472588 | 0.926532378980614 | -5.05550947920395 | 4.29243306192327e-07 | 0.000106171234652016 | 4737 | 0 | 0 | 15 | 27 | 0 | 1 | 12 | 11 | 1 | 0 | 2 | 5 |
| AluY_dup48533 | 5.1183096387502 | -3.60202866671616 | 0.893230275993121 | -4.03258685193056 | 5.51662164272484e-05 | 0.00439309924163255 | 4738 | 0 | 0 | 13 | 22 | 0 | 5 | 8 | 10 | 0 | 0 | 6 | 3 |
| L2a_dup168106 | 3.77480081443938 | -4.14847409953438 | 0.985605526668331 | -4.20906132046315 | 2.56433793734608e-05 | 0.00245839627252116 | 4739 | 0 | 0 | 8 | 14 | 0 | 2 | 7 | 11 | 0 | 0 | 4 | 2 |
| MIR_dup170874 | 3.5440868287724 | -3.26789465218195 | 0.858454212557465 | -3.80671980448019 | 0.000140822136343192 | 0.00869210481305247 | 4740 | 1 | 0 | 12 | 9 | 0 | 3 | 9 | 7 | 0 | 0 | 3 | 2 |
| AluJb_dup57360 | 4.05008036958577 | -3.76996991715929 | 0.930914352932688 | -4.04974948047867 | 5.12724915010189e-05 | 0.0041605278570546 | 4741 | 0 | 0 | 12 | 17 | 0 | 3 | 7 | 9 | 0 | 0 | 1 | 4 |
| L1MB5_dup9653 | 4.93916930794637 | -4.52058534314361 | 0.922223621762296 | -4.9018320898191 | 9.49469761930974e-07 | 0.000197830588417187 | 4742 | 0 | 0 | 18 | 18 | 0 | 2 | 11 | 9 | 0 | 0 | 3 | 4 |
| MLT1D_dup9578 | 4.24022301289242 | -4.33443506786254 | 0.985928066859315 | -4.39629950050001 | 1.10111976242075e-05 | 0.00131069056022495 | 4743 | 0 | 0 | 20 | 13 | 0 | 2 | 8 | 6 | 0 | 0 | 2 | 5 |
| L1ME3A_dup1499 | 4.91207216884986 | -4.08696611068985 | 0.890026696543362 | -4.59195901264826 | 4.3910461952342e-06 | 0.000649771068390048 | 4744 | 2 | 0 | 23 | 19 | 0 | 1 | 10 | 5 | 0 | 0 | 3 | 3 |
| AluSz6_dup7608 | 4.01466963469653 | -3.82467033304404 | 0.898823997999781 | -4.25519383278078 | 2.08867992973974e-05 | 0.00210679539873613 | 4745 | 1 | 0 | 17 | 13 | 0 | 2 | 11 | 4 | 0 | 0 | 3 | 3 |
| L2a_dup168109 | 4.29025626683493 | -4.40698974961764 | 0.948050428410018 | -4.64847609109638 | 3.34396372982027e-06 | 0.000524103328910962 | 4746 | 1 | 0 | 13 | 15 | 0 | 1 | 15 | 5 | 0 | 0 | 5 | 3 |
| MIR3_dup41743 | 4.18680459085358 | -4.22910528288796 | 0.943069045833441 | -4.48440684335098 | 7.31170401002251e-06 | 0.000963872976381616 | 4747 | 0 | 0 | 14 | 19 | 1 | 1 | 12 | 5 | 0 | 0 | 1 | 4 |
| MER113_dup3735 | 4.96599474260768 | -4.62308425479203 | 0.932899894549379 | -4.95560593564558 | 7.2105223037146e-07 | 0.000159009725679303 | 4748 | 0 | 0 | 17 | 17 | 1 | 1 | 12 | 8 | 0 | 0 | 3 | 7 |
| THE1D_dup12526 | 4.20299709014192 | -4.36087110653991 | 0.942552713635342 | -4.62666017873995 | 3.71609488284343e-06 | 0.000567760348935386 | 4749 | 0 | 0 | 13 | 13 | 1 | 1 | 13 | 7 | 0 | 0 | 2 | 6 |
| L1MEg_dup14373 | 4.91772484792573 | -3.71553452076407 | 0.791641332409776 | -4.69345695916847 | 2.68626528646902e-06 | 0.000443571186574711 | 4750 | 1 | 0 | 9 | 21 | 1 | 2 | 18 | 9 | 0 | 0 | 3 | 2 |
| THE1A_dup425 | 4.91581383836326 | -4.43418092974317 | 0.890594960039297 | -4.9788973985969 | 6.39475293830927e-07 | 0.000145052991673874 | 4751 | 1 | 0 | 12 | 21 | 0 | 1 | 17 | 9 | 0 | 0 | 2 | 3 |
| L1P4_dup1904 | 2.54481085443819 | -3.75761595604059 | 1.07821786138361 | -3.48502477154168 | 0.000492091482713329 | 0.0213164390944303 | 4752 | 0 | 0 | 4 | 3 | 2 | 0 | 10 | 4 | 0 | 0 | 3 | 8 |
| L1PREC2_dup7669 | 2.56199976360615 | -3.76058884425447 | 1.07255300324342 | -3.50620326723471 | 0.000454547946940477 | 0.0202504897552329 | 4753 | 0 | 1 | 10 | 3 | 1 | 0 | 6 | 3 | 0 | 0 | 2 | 8 |
| MER58A_dup1877 | 2.30806064199974 | -4.07180370920842 | 1.1322481963712 | -3.59621125673537 | 0.000322885612929501 | 0.0157630980825037 | 4754 | 0 | 0 | 10 | 2 | 1 | 0 | 10 | 2 | 0 | 0 | 2 | 4 |
| LTR16_dup2999 | 3.21617542552831 | -3.57569408494362 | 0.973064036640229 | -3.6746749959948 | 0.000238152370225908 | 0.0126659373906513 | 4755 | 1 | 0 | 5 | 7 | 1 | 0 | 9 | 7 | 1 | 0 | 2 | 9 |
| L1MB7_dup4357 | 3.51887907269487 | -4.19950774668756 | 0.972568312727641 | -4.31795658128088 | 1.57480399021664e-05 | 0.00171104251258488 | 4756 | 1 | 0 | 8 | 9 | 1 | 0 | 12 | 6 | 0 | 0 | 3 | 7 |
| L1MEc_dup13720 | 3.17083801806767 | -3.45663117961959 | 0.938045621854592 | -3.68492864215447 | 0.000228766724092125 | 0.0122709934469849 | 4757 | 1 | 0 | 5 | 8 | 2 | 0 | 14 | 7 | 0 | 0 | 1 | 4 |
| MIR_dup76076 | 5.03903189584423 | -3.28577575786741 | 0.766931368742537 | -4.28431524877484 | 1.83302852832604e-05 | 0.00191803784609089 | 4758 | 0 | 3 | 15 | 9 | 3 | 0 | 16 | 10 | 0 | 0 | 5 | 4 |
| L1PB1_dup4483 | 3.98742716926024 | -3.54209114363781 | 0.884304342692355 | -4.00551142025781 | 6.18834437154181e-05 | 0.00478642346165923 | 4759 | 1 | 1 | 5 | 7 | 2 | 0 | 16 | 8 | 0 | 0 | 8 | 4 |
| L2a_dup59493 | 4.3186517667919 | -3.64948641958614 | 0.89482109938292 | -4.07845369549608 | 4.53362293454506e-05 | 0.00379844169994105 | 4760 | 0 | 3 | 7 | 6 | 1 | 0 | 14 | 8 | 0 | 0 | 10 | 7 |
| AluJo_dup9374 | 3.50088926161985 | -2.59375951105439 | 0.824075453394121 | -3.1474781834254 | 0.00164685394980368 | 0.0495839946779686 | 4761 | 0 | 1 | 8 | 6 | 2 | 1 | 16 | 2 | 1 | 1 | 6 | 3 |
| L1M5_dup28720 | 3.49232628296971 | -2.89403597870822 | 0.813074186348165 | -3.55937505740586 | 0.000371738333598128 | 0.0175086400487609 | 4762 | 0 | 1 | 6 | 5 | 2 | 1 | 9 | 6 | 1 | 0 | 7 | 7 |
| LTR8_dup73 | 3.67586610685251 | -3.30365157657783 | 0.915961829100198 | -3.60675682285058 | 0.000310047975920519 | 0.0153027744613947 | 4763 | 0 | 1 | 8 | 5 | 2 | 0 | 10 | 3 | 1 | 0 | 13 | 5 |
| MSTA_dup17192 | 4.46827090477683 | -4.96402822476669 | 1.00355814040397 | -4.94642813895016 | 7.55876134021033e-07 | 0.000165767467820985 | 4764 | 0 | 0 | 14 | 15 | 0 | 0 | 20 | 4 | 0 | 1 | 5 | 2 |
| Tigger1_dup5997 | 3.39935627876466 | -4.3519449281259 | 1.04063199441172 | -4.1820210713261 | 2.88929184552324e-05 | 0.00268461998497634 | 4765 | 0 | 0 | 13 | 11 | 0 | 0 | 15 | 4 | 0 | 1 | 1 | 1 |
| HERVH-int_dup2275 | 3.4523028900895 | -5.16725699770535 | 1.09953782329205 | -4.69948089846915 | 2.60823615343815e-06 | 0.000434607394794114 | 4766 | 0 | 0 | 5 | 11 | 0 | 0 | 23 | 2 | 0 | 0 | 3 | 5 |
| MLT1F2_dup2939 | 3.56864059529946 | -4.64990354575919 | 1.01878427831176 | -4.56416892638411 | 5.0147688680311e-06 | 0.000723238449361687 | 4767 | 0 | 0 | 7 | 10 | 0 | 0 | 19 | 5 | 1 | 0 | 4 | 2 |
| MIRc_dup92142 | 3.89070126364377 | -5.22799311545533 | 1.04723482385178 | -4.9921879948821 | 5.96991039395026e-07 | 0.000137914203747465 | 4768 | 0 | 0 | 7 | 8 | 0 | 0 | 26 | 6 | 0 | 0 | 3 | 3 |
| THE1C_dup4799 | 3.50419860133719 | -4.92281233985781 | 1.03090785403769 | -4.77522052099703 | 1.7951043459177e-06 | 0.00032408034802256 | 4769 | 0 | 0 | 10 | 10 | 0 | 0 | 18 | 6 | 0 | 0 | 2 | 1 |
| L1M4_dup9191 | 5.40304395274684 | -5.78268721269499 | 0.949231527053206 | -6.09196707851325 | 1.11531624519781e-09 | 7.87492698159756e-07 | 4770 | 0 | 0 | 12 | 17 | 0 | 0 | 16 | 13 | 0 | 0 | 8 | 4 |
| LTR26_dup291 | 4.8721013647104 | -5.1343333177066 | 0.95415964819163 | -5.38100026283593 | 7.40730864327561e-08 | 2.62945776336554e-05 | 4771 | 0 | 1 | 10 | 13 | 0 | 0 | 19 | 10 | 0 | 0 | 7 | 4 |
| L1MA4A_dup2140 | 4.54818987085289 | -5.62603719644924 | 1.00338364023699 | -5.60706490602184 | 2.05786504308114e-08 | 9.38047424408016e-06 | 4772 | 0 | 0 | 4 | 16 | 0 | 0 | 20 | 7 | 0 | 0 | 7 | 8 |
| MIR_dup124601 | 4.45364057185461 | -5.0656344130237 | 0.964176613728899 | -5.25384492933575 | 1.48956310090372e-07 | 4.55572727002105e-05 | 4773 | 0 | 1 | 8 | 20 | 0 | 0 | 11 | 7 | 0 | 0 | 6 | 7 |
| ERVL-E-int_dup4748 | 3.16842948008853 | -5.06913779370231 | 1.09758382628403 | -4.61845161372715 | 3.86614104645363e-06 | 0.000584084315834575 | 4774 | 0 | 0 | 3 | 12 | 0 | 0 | 6 | 10 | 0 | 0 | 7 | 2 |
| MIR_dup157826 | 3.09223904426224 | -5.05743454618294 | 1.05748837000925 | -4.78249661141775 | 1.73131306834308e-06 | 0.000316490373337472 | 4775 | 0 | 0 | 6 | 13 | 0 | 0 | 4 | 7 | 0 | 0 | 7 | 3 |
| MIRc_dup92010 | 3.33270009786455 | -5.16248344438097 | 1.08026578624192 | -4.77890118351381 | 1.76255786997501e-06 | 0.000320019705297638 | 4776 | 0 | 0 | 6 | 17 | 0 | 0 | 5 | 6 | 0 | 0 | 8 | 2 |
| L1MA8_dup10811 | 3.50088172118922 | -4.993696028505 | 1.0560803921157 | -4.72851883794648 | 2.26163645397756e-06 | 0.000385975727989599 | 4777 | 0 | 0 | 4 | 20 | 0 | 0 | 11 | 8 | 0 | 0 | 2 | 2 |
| MER4-int_dup2213 | 3.37319178803294 | -4.96195826307253 | 1.08232336606806 | -4.58454323230467 | 4.54979945623712e-06 | 0.000668275667539382 | 4778 | 0 | 0 | 4 | 20 | 0 | 0 | 12 | 6 | 0 | 0 | 3 | 1 |
| L1MC5_dup14050 | 3.86951089147882 | -5.10310619521071 | 1.01159830627835 | -5.04459740940546 | 4.54477257782591e-07 | 0.00011137079874391 | 4779 | 0 | 0 | 14 | 14 | 0 | 0 | 9 | 9 | 0 | 0 | 3 | 1 |
| L1PA7_dup10514 | 4.30526593402513 | -5.27767243151323 | 0.997421766384821 | -5.29131467688165 | 1.21440207753718e-07 | 3.84258133964977e-05 | 4780 | 0 | 0 | 12 | 20 | 0 | 0 | 8 | 11 | 0 | 0 | 2 | 3 |
| L1PA14_dup1660 | 3.17330962660018 | -4.86773504056659 | 1.02682652916876 | -4.74056221015944 | 2.13125999229453e-06 | 0.000369355273216689 | 4781 | 0 | 0 | 9 | 7 | 0 | 0 | 10 | 10 | 0 | 0 | 2 | 2 |
| L3_dup44825 | 3.7177176174171 | -5.1625291476942 | 0.983303656694024 | -5.25018809047369 | 1.51943981772686e-07 | 4.60566403654566e-05 | 4782 | 0 | 0 | 12 | 11 | 0 | 0 | 10 | 9 | 0 | 0 | 3 | 3 |
| MIR_dup19187 | 3.7592677280985 | -4.99298349051555 | 1.01159046230658 | -4.93577556981981 | 7.98328694959253e-07 | 0.000173320364644867 | 4783 | 0 | 0 | 8 | 13 | 0 | 0 | 15 | 10 | 0 | 0 | 2 | 1 |
| L1MA7_dup4439 | 2.82365098267001 | -4.6746479300627 | 1.0449368622169 | -4.47361759268892 | 7.69072317184461e-06 | 0.00100272702238232 | 4784 | 0 | 0 | 6 | 11 | 0 | 0 | 10 | 7 | 0 | 0 | 2 | 1 |
| MIR_dup170878 | 3.31256189875916 | -4.85259777119902 | 1.01426943211597 | -4.78432812578759 | 1.71560226685463e-06 | 0.000314499766555464 | 4785 | 0 | 0 | 9 | 13 | 0 | 0 | 12 | 7 | 0 | 0 | 1 | 2 |
| AluSc_dup34474 | 3.90338165751323 | -2.5672324853517 | 0.786105567774066 | -3.26576046601612 | 0.00109170553407468 | 0.0372961100866398 | 4786 | 1 | 0 | 5 | 11 | 2 | 3 | 20 | 5 | 0 | 1 | 3 | 2 |
| L1PA17_dup1379 | 3.30459988215978 | -3.11545408628712 | 0.865591872196894 | -3.59921827636854 | 0.000319175204020685 | 0.0156236295198325 | 4787 | 0 | 0 | 8 | 6 | 1 | 2 | 13 | 7 | 0 | 1 | 2 | 3 |
| MER2_dup811 | 3.62590553292405 | -3.32554109285704 | 0.888781894961999 | -3.74168410912468 | 0.000182791191398259 | 0.0104475039765356 | 4788 | 0 | 0 | 4 | 8 | 2 | 1 | 16 | 7 | 0 | 1 | 4 | 5 |
| L1PA5_dup5826 | 3.44843275108286 | -3.57821144050411 | 0.942880782748851 | -3.79497758992636 | 0.000147656792717787 | 0.00900880594872477 | 4789 | 0 | 0 | 6 | 15 | 0 | 3 | 13 | 5 | 0 | 0 | 2 | 3 |
| MLT1L_dup11662 | 3.56572193370684 | -3.6642818576575 | 0.943098257614191 | -3.88536595002014 | 0.000102175859918596 | 0.00691673888028172 | 4790 | 0 | 1 | 5 | 14 | 0 | 2 | 17 | 4 | 0 | 0 | 4 | 2 |
| L2a_dup116376 | 3.16712663804871 | -3.51509271423625 | 1.01558379100848 | -3.46115480116687 | 0.00053786343002362 | 0.0226838114398232 | 4791 | 0 | 0 | 4 | 8 | 0 | 2 | 19 | 3 | 0 | 1 | 2 | 5 |
| L2c_dup62512 | 4.10330773594649 | -4.85118757741091 | 1.01828922384076 | -4.76405667842904 | 1.89739171278255e-06 | 0.000338184845310679 | 4792 | 0 | 0 | 9 | 12 | 0 | 1 | 23 | 4 | 0 | 0 | 2 | 6 |
| L2c_dup62515 | 3.33700470654416 | -4.00770959775516 | 1.01855953550531 | -3.93468369599712 | 8.33063334344085e-05 | 0.00596342022123458 | 4793 | 0 | 0 | 8 | 6 | 0 | 2 | 18 | 5 | 0 | 0 | 2 | 4 |
| THE1C_dup6622 | 4.288296588567 | -3.94505726469158 | 0.955410573235089 | -4.12917480213071 | 3.64067540682055e-05 | 0.00320371357378448 | 4794 | 1 | 0 | 9 | 5 | 0 | 2 | 28 | 6 | 0 | 0 | 4 | 3 |
| L1PA16_dup4862 | 3.75547511397393 | -4.32535040791276 | 1.08855668228825 | -3.97347283636206 | 7.08322394379459e-05 | 0.00529009668716305 | 4795 | 0 | 0 | 3 | 3 | 0 | 2 | 19 | 9 | 0 | 0 | 6 | 7 |
| MER58B_dup2879 | 2.77292653450064 | -3.79984677665841 | 1.14011636789248 | -3.33285871834512 | 0.000859585802447613 | 0.0316323852999072 | 4796 | 0 | 0 | 5 | 1 | 0 | 2 | 12 | 8 | 0 | 0 | 4 | 3 |
| MIR_dup158053 | 2.56103638463459 | -4.23705642182429 | 1.08242748191725 | -3.91440211248093 | 9.06284944426683e-05 | 0.00633882649578341 | 4797 | 0 | 0 | 3 | 4 | 0 | 1 | 14 | 5 | 0 | 0 | 4 | 3 |
| L1MB7_dup16451 | 3.09297614624109 | -4.50360681791512 | 0.990236772678857 | -4.54801007412767 | 5.41555438343947e-06 | 0.000765207543540314 | 4798 | 0 | 0 | 9 | 10 | 0 | 1 | 8 | 5 | 0 | 0 | 4 | 4 |
| L1MA1_dup1378 | 2.74217727838107 | -4.37755047877557 | 1.00311358957195 | -4.36396288943065 | 1.27727254233815e-05 | 0.00146482330168208 | 4799 | 0 | 0 | 6 | 6 | 0 | 1 | 8 | 5 | 0 | 0 | 5 | 5 |
| MIRb_dup217336 | 3.29345310404665 | -4.62140589228636 | 0.989380015081093 | -4.67101196895267 | 2.99719472457874e-06 | 0.000480897165161573 | 4800 | 0 | 0 | 10 | 6 | 0 | 1 | 10 | 6 | 0 | 0 | 5 | 5 |
| LTR88c_dup465 | 3.62194755842638 | -4.04590772266784 | 0.976977256705913 | -4.14125067384831 | 3.45417207020394e-05 | 0.0030656711205399 | 4801 | 0 | 0 | 7 | 8 | 0 | 2 | 13 | 11 | 0 | 0 | 3 | 2 |
| MIR_dup158057 | 3.60340592587149 | -4.60779457981165 | 1.02365345078555 | -4.50132276335868 | 6.75318644885201e-06 | 0.000904273327382409 | 4802 | 0 | 0 | 5 | 8 | 0 | 1 | 15 | 11 | 0 | 0 | 4 | 2 |
| L1PA15_dup2777 | 4.15824613070085 | -4.90464373982536 | 0.982193530220263 | -4.99356144071267 | 5.9275916548766e-07 | 0.000137202988707322 | 4803 | 0 | 0 | 6 | 11 | 0 | 1 | 12 | 13 | 0 | 0 | 4 | 6 |
| MIRb_dup159986 | 3.47495265924955 | -4.61076090831114 | 0.999238582194781 | -4.6142742989205 | 3.94471176952037e-06 | 0.00059406859918373 | 4804 | 0 | 0 | 5 | 15 | 0 | 1 | 10 | 8 | 0 | 0 | 3 | 4 |
| MLT1A_dup8914 | 4.15913660716402 | -4.7761352357543 | 0.979560543378587 | -4.87579381186691 | 1.08371760685405e-06 | 0.000219274038843957 | 4805 | 0 | 0 | 7 | 17 | 0 | 1 | 14 | 10 | 0 | 0 | 2 | 4 |
| ALR/Alpha_dup1616 | 3.33248188722049 | -3.62568654949662 | 0.971075756819083 | -3.73368043022014 | 0.000188701898502899 | 0.0106678217964292 | 4806 | 0 | 0 | 5 | 5 | 1 | 2 | 12 | 9 | 0 | 0 | 2 | 7 |
| AluSx1_dup32599 | 2.45108184146516 | -3.66226815485297 | 1.07378043465134 | -3.41063036415085 | 0.000648128919986042 | 0.0258894554666196 | 4807 | 0 | 0 | 4 | 4 | 1 | 1 | 8 | 6 | 0 | 0 | 1 | 7 |
| MLT1H1_dup1565 | 2.59574500824461 | -3.19460804801839 | 0.964448068019252 | -3.31236917149863 | 0.000925093828492798 | 0.0333215828208519 | 4808 | 0 | 0 | 4 | 7 | 1 | 2 | 12 | 4 | 0 | 0 | 2 | 3 |
| L1MB8_dup6579 | 2.53333558169632 | -3.14734180111595 | 0.979495869641299 | -3.21322621020193 | 0.0013125286985774 | 0.042387480145453 | 4809 | 0 | 0 | 8 | 4 | 1 | 2 | 8 | 5 | 0 | 0 | 1 | 4 |
| AluSq2_dup24346 | 2.85166440445644 | -3.25658042708178 | 0.964888261582756 | -3.37508554797821 | 0.000737927481656256 | 0.0283992508600741 | 4810 | 0 | 0 | 6 | 4 | 1 | 2 | 13 | 7 | 0 | 0 | 1 | 3 |
| MLT1A1_dup3053 | 2.83341569654189 | -2.886093299134 | 0.910006204274887 | -3.17150947496419 | 0.00151648920937713 | 0.0468535664540892 | 4811 | 0 | 0 | 6 | 5 | 2 | 2 | 9 | 8 | 0 | 0 | 1 | 3 |
| MIR_dup507 | 2.29383256882783 | -3.77432226417834 | 1.14735678905709 | -3.28958027718659 | 0.00100336930224943 | 0.0351668451897249 | 4812 | 0 | 0 | 3 | 3 | 0 | 1 | 12 | 8 | 0 | 0 | 1 | 1 |
| THE1D-int_dup1568 | 2.18053620200496 | -3.90308768640177 | 1.13165499338772 | -3.44900849570548 | 0.000562648999504579 | 0.0234589239243239 | 4813 | 0 | 0 | 3 | 3 | 0 | 1 | 11 | 6 | 0 | 0 | 3 | 1 |
| L1PA15_dup7864 | 2.77462724544841 | -3.73160008673928 | 1.00769722254266 | -3.70309652866123 | 0.000212983795273551 | 0.0116262083391054 | 4814 | 0 | 0 | 6 | 8 | 0 | 2 | 11 | 5 | 0 | 0 | 2 | 3 |

| | | | | | | | | | | | | | | | | | | | |
|---|---|---|---|---|---|---|---|---|---|---|---|---|---|---|---|---|---|---|---|
| L2c_dup91109 | 2.33993302903242 | -3.99311889168953 | 1.05305034659897 | -3.79195439665928 | 0.000149466365074762 | 0.00906574654705056 | 4815 | 0 | 0 | 4 | 7 | 0 | 1 | 10 | 5 | 0 | 0 | 2 | 2 |
| MIRb_dup169929 | 2.54537668478872 | -4.0874378496062 | 1.03154454478985 | -3.96244434644255 | 7.41863038704001e-05 | 0.00546681147746864 | 4816 | 0 | 0 | 7 | 6 | 0 | 1 | 9 | 6 | 0 | 0 | 2 | 2 |
| AluSc8_dup21730 | 2.91078195901987 | -3.54830266514462 | 0.980994912522197 | -3.6170449202654 | 0.000297985605390842 | 0.0148958997605734 | 4817 | 0 | 0 | 7 | 8 | 0 | 2 | 12 | 7 | 0 | 0 | 1 | 1 |
| Tigger2a_dup3304 | 2.9521133987029 | -3.79235973750269 | 1.02096430068596 | -3.71448809224249 | 0.000203615456996512 | 0.0112620835728712 | 4818 | 0 | 0 | 7 | 6 | 0 | 2 | 10 | 8 | 0 | 0 | 1 | 4 |
| L1PA11_dup882 | 5.91716717611549 | -4.2856495468513 | 0.951352651967247 | -4.5047959218795 | 6.64367582243495e-06 | 0.000892623425886566 | 4819 | 1 | 0 | 4 | 6 | 1 | 1 | 32 | 23 | 0 | 0 | 3 | 3 |
| AluSq_dup16683 | 3.92431011594286 | -4.12805914116325 | 1.0510383547326 | -3.92760085545449 | 8.57974458812547e-05 | 0.00609044124631893 | 4820 | 0 | 0 | 2 | 7 | 1 | 1 | 15 | 17 | 0 | 0 | 3 | 2 |
| L1MEg_dup6688 | 4.63168322957833 | -3.75228811115653 | 0.92940943980776 | -4.03728211748383 | 5.4074026829509e-05 | 0.00432646279909354 | 4821 | 1 | 0 | 4 | 7 | 1 | 2 | 15 | 17 | 0 | 0 | 8 | 2 |
| ERVL-int_dup360 | 3.3530172542946 | -3.24556359608314 | 0.958410081599424 | -3.38640385613102 | 0.000708150852505409 | 0.0275735006955084 | 4822 | 1 | 0 | 1 | 7 | 2 | 1 | 14 | 10 | 0 | 0 | 4 | 3 |
| MIR_dup13279 | 3.32380559670084 | -3.16680232282862 | 1.00459966739528 | -3.15230277851823 | 0.00161988208250554 | 0.0489515445775798 | 4823 | 1 | 0 | 1 | 5 | 1 | 2 | 12 | 14 | 0 | 0 | 1 | 4 |
| L1MA8_dup7648 | 3.83788603151006 | -3.16862954987503 | 0.937849997171755 | -3.3786101822579 | 0.000728532316239376 | 0.028127754424776 | 4824 | 0 | 1 | 3 | 3 | 3 | 1 | 14 | 13 | 0 | 0 | 3 | 7 |
| THE1D_dup11560 | 3.71537184350231 | -3.82341456024479 | 1.10963776783924 | -3.4456420564073 | 0.000569704439036013 | 0.0236825458509544 | 4825 | 0 | 0 | 1 | 2 | 1 | 2 | 14 | 14 | 0 | 0 | 5 | 7 |
| MIR_dup14487 | 2.59790451721771 | -3.72947198601504 | 1.04108273995307 | -3.58230123590673 | 0.000340580735225947 | 0.0163816097885735 | 4826 | 0 | 0 | 5 | 3 | 1 | 0 | 9 | 7 | 0 | 1 | 3 | 4 |
| Tigger3c_dup2044 | 3.38093360351151 | -3.61537258332318 | 1.08929202635563 | -3.31901133566441 | 0.000903367621930061 | 0.0327422257681295 | 4827 | 0 | 0 | 2 | 2 | 2 | 0 | 13 | 12 | 0 | 1 | 4 | 6 |
| THE1B_dup7859 | 2.43908598658933 | -4.0599875978146 | 1.24080561845022 | -3.27205771592617 | 0.00106767777900325 | 0.0367747975742252 | 4828 | 0 | 0 | 2 | 1 | 0 | 0 | 16 | 8 | 0 | 1 | 2 | 1 |
| FRAM_dup6563 | 3.46325147821903 | -4.64496752716525 | 1.07755853078427 | -4.31064057725435 | 1.62782305477504e-05 | 0.00175502485089036 | 4829 | 0 | 0 | 5 | 3 | 0 | 0 | 20 | 9 | 0 | 1 | 3 | 4 |
| LTR67B_dup3372 | 3.16974174179847 | -4.50985321849173 | 1.09852114946067 | -4.10538588238004 | 4.0364051151656e-05 | 0.00346481403872004 | 4830 | 0 | 0 | 2 | 6 | 0 | 0 | 17 | 9 | 0 | 1 | 4 | 2 |
| MIR_dup158056 | 4.79759637535697 | -5.11158482084486 | 1.05744480848108 | -4.83390223286184 | 1.33882356721552e-06 | 0.000258998140263955 | 4831 | 0 | 0 | 3 | 8 | 1 | 0 | 30 | 11 | 0 | 0 | 4 | 7 |
| AluJo_dup68811 | 4.12408303067243 | -4.75791887472317 | 1.01754254131716 | -4.67589184877152 | 2.92678832907758e-06 | 0.000472467826153795 | 4832 | 0 | 0 | 8 | 6 | 1 | 0 | 20 | 12 | 0 | 0 | 2 | 4 |
| LTR39_dup817 | 4.01989293695166 | -4.10366476181021 | 1.00855710364012 | -4.06884721449994 | 4.72463145471677e-05 | 0.00390985380164496 | 4833 | 0 | 0 | 8 | 6 | 2 | 0 | 27 | 7 | 0 | 0 | 2 | 2 |
| L1ME3Cz_dup11359 | 4.15421848168053 | -4.71631003144711 | 1.07062556695032 | -4.40519092485485 | 1.05690747904003e-05 | 0.0012739964894005 | 4834 | 0 | 0 | 6 | 4 | 0 | 1 | 21 | 15 | 0 | 0 | 3 | 2 |
| MLT1J2_dup6210 | 3.52626955465171 | -4.3765416705005 | 1.13044653509859 | -3.87151584317856 | 0.000108160625745634 | 0.00721094694314257 | 4835 | 0 | 0 | 4 | 3 | 0 | 1 | 22 | 12 | 0 | 0 | 1 | 2 |
| MamRep1151_dup116( | 3.9127704732831 | -4.69707948881585 | 1.08687558141276 | -4.32163494068972 | 1.54877285837211e-05 | 0.00169125427516388 | 4836 | 0 | 0 | 4 | 4 | 0 | 1 | 22 | 13 | 0 | 0 | 2 | 4 |
| AluSx1_dup102248 | 3.19731681719704 | -5.00248480220292 | 1.19122884395741 | -4.19943223132848 | 2.67585099502735e-05 | 0.00253365714629041 | 4837 | 0 | 0 | 6 | 1 | 0 | 0 | 22 | 7 | 0 | 0 | 4 | 2 |
| MamTip2_dup6442 | 3.09928864821007 | -4.91315251521062 | 1.16078614501478 | -4.23260782040782 | 2.30997109511366e-05 | 0.00227975272583125 | 4838 | 0 | 0 | 4 | 2 | 0 | 0 | 20 | 9 | 0 | 0 | 3 | 2 |
| MER6_dup753 | 3.29923106289797 | -5.07238695882195 | 1.07679598363368 | -4.71062953049381 | 2.4695278507365e-06 | 0.000415568793473372 | 4839 | 0 | 0 | 5 | 5 | 0 | 0 | 18 | 8 | 0 | 0 | 5 | 2 |
| MER45A_dup3073 | 2.69129423293516 | -4.61280062853826 | 1.11121384364119 | -4.15113675458107 | 3.30828006343673e-05 | 0.00297615125888286 | 4840 | 0 | 0 | 5 | 4 | 0 | 0 | 13 | 9 | 0 | 0 | 2 | 1 |
| UCON132b_dup1153 | 2.64060497008977 | -4.66780273924857 | 1.17698509574358 | -3.96589791674433 | 7.31201545144313e-05 | 0.00540330692114623 | 4841 | 0 | 0 | 2 | 4 | 0 | 0 | 13 | 10 | 0 | 0 | 3 | 1 |
| MIR_dup76075 | 3.84534236259604 | -3.84134641321861 | 1.03135456162166 | -3.72456433137661 | 0.00019565281627534 | 0.0109463449380325 | 4842 | 0 | 1 | 6 | 3 | 1 | 0 | 18 | 16 | 0 | 0 | 1 | 1 |
| MLT1A_dup5882 | 3.72831854150724 | -4.32405327274373 | 1.11841488555718 | -3.86623365674317 | 0.000110529048808401 | 0.00732413496475379 | 4843 | 0 | 1 | 6 | 3 | 0 | 0 | 19 | 15 | 0 | 0 | 1 | 1 |
| L1MC_dup3510 | 4.45418531799761 | -3.74089263535348 | 0.992275915085551 | -3.77001253228135 | 0.000163239350160336 | 0.00966944247280339 | 4844 | 0 | 2 | 4 | 5 | 0 | 1 | 33 | 11 | 0 | 0 | 1 | 2 |
| L1P4_dup3418 | 3.65634265083872 | -4.03807892012265 | 1.05411615855941 | -3.83077224206604 | 0.000127741708436763 | 0.0081141560479696 | 4845 | 0 | 1 | 5 | 3 | 0 | 1 | 18 | 13 | 0 | 0 | 1 | 4 |
| L1PA10_dup2325 | 4.39792152333167 | -3.19163000957231 | 0.937265096141294 | -3.40525857914927 | 0.00066101431090708 | 0.0262492842495154 | 4846 | 0 | 2 | 2 | 4 | 1 | 1 | 26 | 13 | 1 | 0 | 4 | 2 |
| L1M2c_dup178 | 4.11422823925879 | -3.33600241074372 | 0.922613712055792 | -3.61581707181693 | 0.000299401754820795 | 0.0149259270799474 | 4847 | 0 | 3 | 7 | 8 | 0 | 0 | 25 | 7 | 1 | 0 | 1 | 3 |
| L1M3c_dup445 | 5.38437802615304 | -3.6907513643211 | 0.930781232266341 | -3.96521893263207 | 7.33286120028447e-05 | 0.0054153475827526 | 4848 | 0 | 2 | 6 | 6 | 0 | 0 | 35 | 14 | 1 | 1 | 2 | 3 |
| L2a_dup19698 | 2.3757939650017 | -3.81418862747229 | 1.19567150668289 | -3.18999709046664 | 0.00142274226093012 | 0.0447621089540234 | 4849 | 0 | 0 | 2 | 6 | 0 | 0 | 12 | 8 | 1 | 0 | 1 | 0 |
| THE1C_dup6552 | 2.95018972749733 | -4.08840627829765 | 1.26062606410726 | -3.24315544054133 | 0.00118213698263465 | 0.0393956255560202 | 4850 | 0 | 0 | 1 | 6 | 0 | 0 | 14 | 13 | 1 | 0 | 1 | 0 |
| MLT1G1_dup280 | 3.48397455491554 | -3.59892737968502 | 1.07281815247478 | -3.35464810264723 | 0.000794660022602009 | 0.0299281693159319 | 4851 | 0 | 0 | 1 | 9 | 0 | 1 | 17 | 9 | 2 | 0 | 3 | 3 |
| L1PREC2_dup7138 | 3.09829155162465 | -4.39639345499007 | 1.15261198927582 | -3.81428745830788 | 0.00013657642102352 | 0.00850060504233914 | 4852 | 0 | 0 | 1 | 8 | 0 | 0 | 17 | 9 | 1 | 0 | 3 | 1 |
| MamRTE1_dup8572 | 3.24735594808508 | -4.48710987627943 | 1.1047490979568 | -4.06165516186273 | 4.87260160693849e-05 | 0.00399110520469737 | 4853 | 0 | 0 | 3 | 4 | 0 | 0 | 20 | 9 | 1 | 0 | 2 | 3 |
| L1ME3B_dup798 | 2.57808093577535 | -3.57657067976398 | 1.13244984747948 | -3.15825966838571 | 0.00158714115573075 | 0.0483056906423011 | 4854 | 0 | 0 | 3 | 3 | 0 | 0 | 13 | 9 | 1 | 1 | 1 | 1 |
| L1PA4_dup1479 | 1.94642342516455 | -3.7908788790964 | 1.10961569503466 | -3.4163890219469 | 0.000634575229666337 | 0.0255230962809646 | 4855 | 0 | 0 | 3 | 4 | 0 | 0 | 8 | 5 | 1 | 0 | 1 | 3 |
| L2a_dup106964 | 2.43589070054936 | -3.95023720731872 | 1.09502792003862 | -3.60743058239046 | 0.00030924423663465 | 0.0152726087858589 | 4856 | 0 | 0 | 5 | 5 | 0 | 0 | 14 | 5 | 1 | 0 | 1 | 1 |
| L1ME2z_dup5438 | 3.65179079319181 | -4.0087786323745 | 1.02837924342769 | -3.89815202707986 | 9.69295596417533e-05 | 0.00666397023938648 | 4857 | 0 | 0 | 5 | 6 | 1 | 0 | 14 | 14 | 1 | 0 | 1 | 3 |
| L1MA1_dup1379 | 4.06537129241446 | -4.39328126719859 | 1.00793072831687 | -4.35871349466135 | 1.30829254299622e-05 | 0.00149091464289166 | 4858 | 0 | 0 | 7 | 7 | 0 | 0 | 17 | 10 | 2 | 0 | 4 | 5 |
| MER47B_dup341 | 4.05122501348374 | -4.84020006274759 | 0.975212218574194 | -4.96322746019753 | 6.93313053424498e-07 | 0.000153488724156045 | 4859 | 0 | 0 | 9 | 9 | 0 | 0 | 12 | 12 | 1 | 0 | 4 | 4 |
| L1MA8_dup3244 | 2.88387397283185 | -4.28042756290691 | 1.33077799295753 | -3.21648508283043 | 0.00129771279811808 | 0.0420230769544102 | 4860 | 0 | 1 | 0 | 2 | 0 | 0 | 16 | 12 | 0 | 0 | 1 | 4 |
| L2_dup54748 | 3.86714266461542 | -4.93480531042214 | 1.39960291642273 | -3.52586097993787 | 0.000422108483718898 | 0.0192191016584341 | 4861 | 0 | 0 | 0 | 3 | 0 | 0 | 20 | 21 | 0 | 0 | 1 | 1 |
| L1PA11_dup3100 | 3.81322570605777 | -4.54566661711679 | 1.35055912415321 | -3.36576647095468 | 0.000763313010029585 | 0.029086628100196 | 4862 | 0 | 0 | 3 | 3 | 0 | 0 | 21 | 19 | 0 | 0 | 0 | 0 |
| L1MA4_dup4713 | 2.70319267838684 | -4.3851636341245 | 1.30157702899357 | -3.36911572380412 | 0.000754097607447823 | 0.0288572707143422 | 4863 | 0 | 0 | 1 | 6 | 0 | 0 | 15 | 11 | 0 | 0 | 1 | 0 |
| L1PB1_dup9012 | 3.26727109603662 | -4.60128073897503 | 1.36974708090699 | -3.35921923332609 | 0.00078163031919118 | 0.02957789566321 | 4864 | 0 | 0 | 1 | 3 | 0 | 0 | 14 | 19 | 0 | 0 | 1 | 0 |
| MLT1E2_dup3373 | 3.30468790185633 | -3.5469951302207 | 1.0931796780027 | -3.24465886221123 | 0.00117591404774049 | 0.0392751366517318 | 4865 | 2 | 0 | 8 | 3 | 0 | 0 | 12 | 14 | 0 | 0 | 1 | 0 |
| MamRep605_dup1196 | 2.84753442175606 | -3.89028677166196 | 1.09358925280264 | -3.55735644044779 | 0.000374605703237253 | 0.0176088361640639 | 4866 | 1 | 0 | 8 | 5 | 0 | 0 | 12 | 9 | 0 | 0 | 1 | 0 |
| MER91C_dup169 | 2.54792431167478 | -4.08593077072961 | 1.06424881500672 | -3.83926269225285 | 0.000123404340769666 | 0.00791028362999303 | 4867 | 1 | 0 | 6 | 5 | 0 | 0 | 8 | 8 | 0 | 0 | 3 | 1 |
| MIRc_dup61857 | 3.46397381864912 | -4.37001907624731 | 1.03478273819929 | -4.22312715019961 | 2.40935802192256e-05 | 0.0023476539880 6054 | 4868 | 1 | 0 | 5 | 10 | 0 | 0 | 16 | 10 | 0 | 0 | 2 | 1 |
| Tigger2a_dup1587 | 3.0607200594979 | -4.21129311532533 | 1.08884338676415 | -3.86767570664183 | 0.000109877653427853 | 0.00729898049205579 | 4869 | 1 | 0 | 5 | 4 | 0 | 0 | 13 | 12 | 0 | 0 | 1 | 2 |
| L2a_dup81489 | 3.16371492643562 | -4.15374343710122 | 1.04074826998767 | -3.99111250711032 | 6.57640710851603e-05 | 0.00501066207442509 | 4870 | 1 | 0 | 9 | 6 | 0 | 0 | 12 | 10 | 0 | 0 | 1 | 1 |
| MIR_dup85038 | 3.2561044097701 | -4.28600068590815 | 1.02839944401761 | -4.16764197106544 | 3.07766828714764e-05 | 0.00281173169539001 | 4871 | 1 | 0 | 8 | 7 | 0 | 0 | 11 | 11 | 0 | 0 | 1 | 2 |
| HERVK22-int_dup186 | 3.917989333188 | -2.97076563222939 | 0.905499464291902 | -3.28080330180264 | 0.0010351190464962 | 0.0359250928584574 | 4872 | 3 | 0 | 7 | 4 | 0 | 0 | 13 | 14 | 1 | 1 | 3 | 2 |
| L1ME3_dup8254 | 3.0380696679929 | -3.76880702935624 | 1.03941925388095 | -3.62587763819499 | 0.000287981703728246 | 0.01452999452003 | 4873 | 1 | 0 | 8 | 4 | 0 | 0 | 17 | 6 | 0 | 1 | 2 | 1 |
| AluJb_dup13338 | 3.7211606970675 | -3.61396124872699 | 0.99253799436453 | -3.64113139169128 | 0.000271442553756024 | 0.0139169726128056 | 4874 | 1 | 0 | 6 | 5 | 0 | 0 | 17 | 12 | 0 | 2 | 2 | 2 |
| Tigger1_dup11636 | 4.0161543169325 | -3.65111835113806 | 0.978426957877831 | -3.73162076304315 | 0.000190251767453291 | 0.0107197838168223 | 4875 | 1 | 0 | 8 | 6 | 0 | 0 | 16 | 14 | 0 | 2 | 2 | 1 |
| L1ME4c_dup7945 | 3.27014842218751 | -5.1006809287489 | 1.20222556644674 | -4.24269876727402 | 2.20847787374814e-05 | 0.00219781880446204 | 4876 | 0 | 0 | 3 | 1 | 0 | 0 | 11 | 15 | 0 | 0 | 5 | 4 |
| AluJr4_dup18320 | 3.1406351736143 | -4.99371790310265 | 1.07121340730421 | -4.66173954606272 | 3.13547837146266e-06 | 0.000496390243896243 | 4877 | 0 | 0 | 4 | 5 | 0 | 0 | 11 | 12 | 0 | 0 | 3 | 4 |
| L1PA16_dup4869 | 2.98838187280244 | -4.93099414403733 | 1.09335812092909 | -4.50995337177099 | 6.48418765079132e-06 | 0.000877134904411177 | 4878 | 0 | 0 | 4 | 5 | 0 | 0 | 7 | 13 | 0 | 0 | 4 | 3 |
| LTR30_dup53 | 2.03362169059312 | -4.46126451662333 | 1.19466279581095 | -3.73432949637891 | 0.000188215950187357 | 0.0106479392494724 | 4879 | 0 | 0 | 2 | 3 | 0 | 0 | 10 | 5 | 0 | 0 | 1 | 6 |
| AluSq2_dup54302 | 2.06755141891239 | -4.31019121690471 | 1.232966614014 | -3.49578907321149 | 0.000472662169185923 | 0.0207467390719634 | 4880 | 0 | 0 | 2 | 2 | 0 | 0 | 14 | 6 | 0 | 0 | 1 | 2 |
| AluSz_dup59346 | 2.38271728122135 | -4.54120042001258 | 1.13910290127771 | -3.98664634680397 | 6.701379895725e-05 | 0.00508633665284906 | 4881 | 0 | 0 | 3 | 4 | 0 | 0 | 13 | 7 | 0 | 0 | 1 | 3 |
| L1PA5_dup695 | 2.45505612173528 | -4.57822197270261 | 1.30229689463314 | -3.51549788037567 | 0.000438930263607329 | 0.0197544355029903 | 4882 | 0 | 0 | 1 | 1 | 0 | 0 | 13 | 11 | 0 | 0 | 2 | 2 |
| LTR82B_dup1114 | 1.71619032035924 | -4.14729953532314 | 1.28205331985577 | -3.2348884957372 | 0.00121690244064596 | 0.040227433751312 | 4883 | 0 | 0 | 1 | 2 | 0 | 0 | 7 | 7 | 0 | 0 | 3 | 1 |
| AluSc_dup10797 | 1.85757664748641 | -4.29645909115876 | 1.24885415585298 | -3.44032093020842 | 0.000581024776939372 | 0.0240188536437831 | 4884 | 0 | 0 | 2 | 1 | 0 | 0 | 11 | 5 | 0 | 0 | 3 | 2 |
| L1M1_dup8807 | 2.13437218896152 | -4.50219083802173 | 1.2893183110794 | -3.49191568857233 | 0.000479569597577502 | 0.0209426867719755 | 4885 | 0 | 0 | 1 | 1 | 0 | 0 | 12 | 7 | 0 | 0 | 4 | 2 |
| L1MEf_dup991 | 2.77087381682608 | -4.64389538696739 | 1.42251547445997 | -3.26456581340906 | 0.00109631990142999 | 0.0373892697815194 | 4886 | 0 | 0 | 2 | 0 | 0 | 0 | 11 | 16 | 0 | 0 | 1 | 2 |
| L2_dup26027 | 2.74093936324091 | -3.97710283799511 | 1.13788668611623 | -3.49516598315209 | 0.000473767028843057 | 0.0207799022018599 | 4887 | 0 | 0 | 6 | 2 | 1 | 0 | 13 | 10 | 0 | 0 | 1 | 1 |
| MIRb_dup96117 | 2.40196483441072 | -3.8286776393018 | 1.1764615098996 | -3.25440110627038 | 0.00113631715464078 | 0.0383738463920174 | 4888 | 1 | 0 | 7 | 1 | 0 | 0 | 11 | 8 | 0 | 0 | 1 | 1 |
| L1M5_dup25636 | 2.11474103785509 | -4.34127909118317 | 1.22336127809534 | -3.54864844009298 | 0.000387213679694801 | 0.018065871809541 | 4889 | 0 | 0 | 5 | 1 | 0 | 0 | 12 | 6 | 0 | 0 | 2 | 1 |
| MER113_dup3233 | 2.65549428818002 | -4.50099714178294 | 1.22729052111521 | -3.66742597970445 | 0.000245004442834997 | 0.0129320823324792 | 4890 | 0 | 0 | 6 | 1 | 0 | 0 | 17 | 8 | 0 | 0 | 1 | 1 |
| FordPrefect_dup551 | 3.47012918464343 | -4.3559453400389 | 1.27793752600547 | -3.40857455970837 | 0.000653032367592105 | 0.0260356957050735 | 4891 | 0 | 0 | 1 | 1 | 0 | 1 | 22 | 15 | 0 | 0 | 1 | 2 |
| Tigger1_dup7897 | 4.30620937425306 | -4.07205633264762 | 1.16300348849246 | -3.5013277027449 | 0.000462946208260891 | 0.0204601860529882 | 4892 | 0 | 0 | 2 | 1 | 1 | 1 | 29 | 17 | 0 | 0 | 1 | 2 |
| MER5B_dup23086 | 4.58675210398309 | -4.11761694502805 | 1.06330590607509 | -3.87246691803598 | 0.000107739303233776 | 0.00719106206094366 | 4893 | 0 | 0 | 5 | 6 | 0 | 1 | 29 | 15 | 0 | 1 | 0 | 2 |

| | | | | | | | | | | | | | | | | | | | |
|---|---|---|---|---|---|---|---|---|---|---|---|---|---|---|---|---|---|---|---|
| MLT1C_dup11919 | 3.57426018822554 | -3.85663172776233 | 1.21539564578266 | -3.17314920548267 | 0.00150795007082453 | 0.0466714374640312 | 4894 | 0 | 0 | 3 | 2 | 0 | 2 | 27 | 10 | 0 | 0 | 0 | 3 |
| L1ME3A_dup9578 | 5.08201131594701 | -4.35088594834088 | 1.09262810430151 | -3.98203737503375 | 6.83270317116033e-05 | 0.00514987136130794 | 4895 | 0 | 0 | 6 | 5 | 0 | 2 | 25 | 21 | 0 | 0 | 0 | 4 |
| LTR12_dup566 | 3.72153244555718 | -3.90751088890792 | 1.09378175724766 | -3.57247765654876 | 0.000353619662532267 | 0.0168486952785148 | 4896 | 0 | 0 | 6 | 4 | 0 | 2 | 16 | 15 | 0 | 0 | 0 | 3 |
| L2a_dup78433 | 5.62419146380003 | -3.64903936298498 | 1.05239960961673 | -3.46735149808153 | 0.000525614078735323 | 0.0223112209597291 | 4897 | 3 | 1 | 1 | 7 | 1 | 0 | 27 | 24 | 0 | 0 | 0 | 6 |
| LTR34_dup131 | 6.68381604629713 | -4.63362970060269 | 1.20363667242375 | -3.84969136182265 | 0.000118266755038095 | 0.00767205596900067 | 4898 | 0 | 0 | 2 | 3 | 1 | 0 | 44 | 29 | 0 | 1 | 0 | 3 |
| HERV17-int_dup904 | 6.9357499761587 | -4.31351279165861 | 1.13775256450927 | -3.79125736668333 | 0.000149886533243583 | 0.00908197123483002 | 4899 | 1 | 2 | 3 | 2 | 0 | 0 | 44 | 30 | 0 | 0 | 0 | 4 |
| L1PA4_dup8636 | 5.76870926793932 | -4.78550866450098 | 1.12185892454176 | -4.26569558775467 | 1.99280344506311e-05 | 0.00203772930184352 | 4900 | 0 | 1 | 4 | 8 | 0 | 0 | 30 | 26 | 0 | 0 | 0 | 2 |
| L1MCc_dup2256 | 6.75039223777706 | -5.17970259510015 | 1.1315411775664 | -4.57756438545181 | 4.70421023709884e-06 | 0.000688478574665287 | 4901 | 0 | 1 | 6 | 3 | 0 | 0 | 34 | 33 | 0 | 0 | 2 | 2 |
| L1PA5_dup7747 | 7.80831759371466 | -4.86784657536918 | 1.0839641636245 | -4.49078183460635 | 7.09622082968141e-06 | 0.000939631252943446 | 4902 | 1 | 0 | 5 | 2 | 0 | 1 | 54 | 31 | 0 | 0 | 2 | 2 |
| L1MEg1_dup564 | 4.75477827603788 | -4.05083824874544 | 1.16770111869672 | -3.46907113805509 | 0.000522261146144837 | 0.0222148642618125 | 4903 | 1 | 0 | 3 | 3 | 0 | 0 | 25 | 23 | 1 | 0 | 1 | 0 |
| L3_dup20799 | 4.96325250674989 | -3.89572578321078 | 0.984622407693881 | -3.95656827710747 | 7.6034163447819e-05 | 0.00557535440855308 | 4904 | 2 | 1 | 4 | 5 | 0 | 0 | 30 | 18 | 0 | 0 | 1 | 2 |
| L1PA4_dup9389 | 5.23642343839502 | -4.02856905723458 | 1.02659343843884 | -3.92421079893213 | 8.70145193007339e-05 | 0.0061548028565792 | 4905 | 2 | 0 | 5 | 4 | 0 | 0 | 34 | 17 | 0 | 1 | 3 | 1 |
| MER58A_dup12148 | 5.90278637590433 | -5.11780586137252 | 1.09441560325753 | -4.67629102339126 | 2.92109982408773e-06 | 0.000472190230123898 | 4906 | 1 | 0 | 9 | 4 | 0 | 0 | 28 | 25 | 0 | 0 | 4 | 1 |
| Tigger3c_dup2215 | 4.93486398845236 | -4.9712165952628 | 0.979887638395995 | -5.07325166730373 | 3.91074733232035e-07 | 9.87295062566641e-05 | 4907 | 0 | 0 | 12 | 11 | 0 | 0 | 25 | 11 | 0 | 1 | 2 | 3 |
| L1PB1_dup10189 | 6.11703817920804 | -5.22994484327349 | 1.04325866305994 | -5.0130854681175 | 5.35640898863263e-07 | 0.000126191692396948 | 4908 | 0 | 0 | 11 | 8 | 0 | 0 | 34 | 19 | 0 | 1 | 4 | 1 |
| L1PB2_dup2069 | 6.54495334578504 | -5.42004110608684 | 1.00005572049846 | -5.41973911552181 | 5.96860698670649e-08 | 2.20871875281316e-05 | 4909 | 0 | 0 | 13 | 8 | 0 | 0 | 35 | 19 | 0 | 1 | 4 | 4 |
| L1MB8_dup1951 | 6.43438304803532 | -5.57209351976389 | 1.04525560725785 | -5.33084298335588 | 9.77579153447973e-08 | 3.24422662826125e-05 | 4910 | 0 | 0 | 6 | 18 | 0 | 0 | 37 | 20 | 0 | 0 | 1 | 2 |
| MER21C_dup2546 | 6.6319138831662 | -4.64224162047082 | 0.971648922564966 | -4.77769440449354 | 1.77316589157405e-06 | 0.000320930149633559 | 4911 | 0 | 1 | 6 | 15 | 0 | 0 | 31 | 26 | 0 | 1 | 2 | 1 |
| L3b_dup3514 | 4.01801824542101 | -4.52005371809343 | 1.06314120348321 | -4.25160242429153 | 2.12246401833851e-05 | 0.00213009056667282 | 4912 | 0 | 0 | 6 | 8 | 0 | 0 | 18 | 15 | 1 | 0 | 1 | 1 |
| MER20_dup14697 | 3.93155426533197 | -5.03108275457169 | 1.07322524423936 | -4.68781626371213 | 2.76135762251747e-06 | 0.000451273352230455 | 4913 | 0 | 0 | 5 | 8 | 0 | 0 | 20 | 14 | 0 | 0 | 1 | 2 |
| MER4CL34_dup534 | 5.88679117921496 | -5.1899131535809 | 1.14338472992476 | -4.53907859511332 | 5.65005751459992e-06 | 0.000789899286635076 | 4914 | 0 | 0 | 3 | 6 | 0 | 0 | 39 | 21 | 1 | 0 | 1 | 4 |
| MLT1H_dup8786 | 5.1386917435879 | -4.49762206238445 | 0.975456460933683 | -4.61078709559158 | 4.01147215271528e-06 | 0.000602977733954511 | 4915 | 1 | 0 | 6 | 11 | 0 | 0 | 29 | 14 | 1 | 0 | 2 | 3 |
| MIRb_dup202697 | 5.48048211084089 | -5.70119784617631 | 1.15364789989064 | -4.9418872488883 | 7.73699579250517e-07 | 0.000169364047915679 | 4916 | 0 | 0 | 1 | 13 | 0 | 0 | 26 | 21 | 0 | 0 | 6 | 2 |
| L2b_dup57283 | 5.43171618561718 | -5.47695140788164 | 1.13593947023595 | -4.82151694820852 | 1.42470613518243e-06 | 0.000271637120226056 | 4917 | 0 | 0 | 4 | 5 | 0 | 0 | 36 | 20 | 0 | 0 | 2 | 2 |
| Tigger17a_dup2054 | 4.97529471030834 | -5.44462240087085 | 1.12081946265513 | -4.85771578945727 | 1.1874769197003e-06 | 0.000235421688847569 | 4918 | 0 | 0 | 4 | 5 | 0 | 0 | 28 | 20 | 0 | 0 | 3 | 2 |
| L1MD1_dup4960 | 4.55626091099744 | -5.53324840854109 | 1.14179225475511 | -4.84610784974002 | 1.25907089483291e-06 | 0.000245566297657304 | 4919 | 0 | 0 | 2 | 4 | 0 | 0 | 26 | 16 | 0 | 0 | 4 | 6 |
| LTR67B_dup3371 | 5.59307034730232 | -5.7388838501447 | 1.09985042269537 | -5.21787666006492 | 1.80985913181162e-07 | 5.33641562550246e-05 | 4920 | 0 | 0 | 4 | 7 | 0 | 0 | 36 | 17 | 0 | 0 | 2 | 7 |
| L1PA17_dup510 | 3.83297941616703 | -2.89340386343353 | 0.857728547967991 | -3.37333282224099 | 0.000742641277917026 | 0.0285381979191287 | 4921 | 0 | 1 | 7 | 11 | 2 | 1 | 21 | 8 | 0 | 0 | 0 | 0 |
| Tigger1_dup1606 | 4.35021407078017 | -3.68914462669568 | 0.956351922365979 | -3.85751786598481 | 0.000114544309115047 | 0.00749492781368615 | 4922 | 0 | 0 | 9 | 8 | 1 | 1 | 24 | 13 | 0 | 0 | 0 | 0 |
| FordPrefect_a_dup82 | 3.80525756591507 | -3.48663612293614 | 1.03665723485124 | -3.36334518847636 | 0.000770040121376811 | 0.0292509525416869 | 4923 | 1 | 1 | 3 | 10 | 0 | 0 | 21 | 13 | 0 | 0 | 0 | 0 |
| MSTA_dup12948 | 4.28956392923272 | -3.55085745729212 | 1.1205132006917 | -3.16895638096913 | 0.00152987353015508 | 0.0471476864403937 | 4924 | 0 | 1 | 2 | 6 | 1 | 0 | 27 | 17 | 0 | 0 | 0 | 0 |
| L1MEg_dup4932 | 4.39642806951556 | -3.072272685962 | 0.935102285406367 | -3.28549371968103 | 0.00101803811887897 | 0.0354927616461208 | 4925 | 4 | 0 | 9 | 15 | 0 | 0 | 19 | 11 | 0 | 0 | 0 | 0 |
| MER61B_dup421 | 5.48205715927233 | -3.67841198324942 | 0.946013306064992 | -3.88833006858015 | 0.000100936301659087 | 0.0068660346582542 | 4926 | 2 | 0 | 8 | 13 | 0 | 0 | 33 | 15 | 0 | 1 | 0 | 0 |
| L1MA4_dup4710 | 5.16765116039266 | -3.01395666468355 | 0.811379566296172 | -3.71460755222345 | 0.000203519297080244 | 0.0112593821583482 | 4927 | 4 | 1 | 7 | 12 | 1 | 0 | 26 | 15 | 0 | 0 | 1 | 0 |
| THE1B-int_dup2551 | 5.52486119099442 | -2.95092833067481 | 0.787969493200449 | -3.74497789081808 | 0.000180409662498242 | 0.0103615152191182 | 4928 | 3 | 2 | 11 | 15 | 0 | 0 | 29 | 12 | 1 | 0 | 0 | 0 |
| THE1B-int_dup2550 | 4.60665289564272 | -2.67798305059999 | 0.83220439369877 | -3.21793909149839 | 0.00129115232787578 | 0.0418847350795816 | 4929 | 3 | 3 | 12 | 9 | 0 | 0 | 22 | 11 | 0 | 0 | 0 | 0 |
| L1PB1_dup7302 | 4.77231703553499 | -2.74988028675054 | 0.798582440482451 | -3.44345198109947 | 0.000574338575833097 | 0.0238044882015645 | 4930 | 3 | 3 | 9 | 17 | 0 | 0 | 17 | 13 | 0 | 0 | 0 | 0 |
| MER110A_dup207 | 4.87059261587436 | -2.93577159561855 | 0.861523310606317 | -3.40765195726676 | 0.000655244129519415 | 0.0260985469773396 | 4931 | 1 | 4 | 8 | 17 | 0 | 0 | 19 | 14 | 0 | 0 | 0 | 0 |
| MER21C_dup1742 | 4.62119693012967 | -3.4519520262466 | 0.87091026864729 | -3.9636138767869 | 7.38236233176679e-05 | 0.00544852229340751 | 4932 | 1 | 2 | 9 | 13 | 0 | 0 | 22 | 13 | 0 | 0 | 0 | 0 |
| L2b_dup87080 | 4.36097563389318 | -4.14390766717069 | 1.0398044057648 | -3.98527611942821 | 6.74017012898474e-05 | 0.00510600611751481 | 4933 | 0 | 0 | 13 | 4 | 0 | 1 | 9 | 20 | 1 | 0 | 1 | 2 |
| Charlie15a_dup2094 | 7.25445347276235 | -5.84294047493104 | 0.994198634958032 | -5.87703530208296 | 4.17679161345717e-09 | 2.39482134278477e-06 | 4934 | 0 | 0 | 23 | 13 | 0 | 0 | 18 | 29 | 0 | 0 | 3 | 2 |
| MLT2B3_dup1169 | 5.47558924695063 | -5.16766135434217 | 1.03421229996351 | -4.99671233316845 | 5.83159592801001e-07 | 0.000135244144706264 | 4935 | 0 | 0 | 16 | 13 | 0 | 0 | 18 | 20 | 0 | 0 | 1 | 0 |
| L1M4_dup9189 | 3.95785795790495 | -5.29930569685663 | 1.02237365663947 | -5.18333552751682 | 2.17952431683071e-07 | 6.15998251128511e-05 | 4936 | 0 | 0 | 6 | 10 | 0 | 0 | 12 | 14 | 0 | 0 | 2 | 6 |
| L1MD2_dup5174 | 4.14220010339331 | -5.27328295441221 | 1.02857767309531 | -5.12677174738129 | 2.94752501498172e-07 | 7.87151276335399e-05 | 4937 | 0 | 0 | 13 | 6 | 0 | 0 | 11 | 15 | 0 | 0 | 2 | 4 |
| Tigger1_dup6536 | 4.63819786357269 | -5.54412452318074 | 0.975155727131406 | -5.68537349361601 | 1.30520770174205e-08 | 6.39061644141837e-06 | 4938 | 0 | 0 | 12 | 10 | 0 | 0 | 11 | 15 | 0 | 0 | 5 | 5 |
| MIR_dup158058 | 6.28659265272976 | -5.42206849323036 | 0.99477970437039 | -5.45052182851083 | 5.02222377294355e-08 | 1.91509304146927e-05 | 4939 | 1 | 0 | 10 | 9 | 0 | 0 | 31 | 19 | 0 | 0 | 3 | 8 |
| THE1C_dup3509 | 6.43046032524827 | -5.97476240639442 | 0.964887067534432 | -6.19218829584034 | 5.93346082433986e-10 | 4.67497771294163e-07 | 4940 | 0 | 0 | 10 | 15 | 0 | 0 | 24 | 21 | 0 | 0 | 4 | 8 |
| L1MC1_dup2189 | 5.53154960657693 | -4.61030689980337 | 0.958195345154697 | -4.81144781501632 | 1.498408607088e-06 | 0.000282744119287995 | 4941 | 2 | 0 | 18 | 22 | 0 | 0 | 13 | 12 | 0 | 0 | 0 | 6 |
| L1PA8_dup4243 | 4.64578990454088 | -4.88793011883185 | 1.02249915018466 | -4.78037572740194 | 1.74967887077883e-06 | 0.000318781844246814 | 4942 | 1 | 0 | 13 | 12 | 0 | 0 | 15 | 13 | 0 | 0 | 0 | 6 |
| L1MCa_dup1079 | 6.12519795600859 | -4.8985964385036 | 0.938610930158118 | -5.21898507795811 | 1.79906255684649e-07 | 5.31116301676669e-05 | 4943 | 1 | 0 | 17 | 14 | 0 | 0 | 25 | 19 | 0 | 0 | 1 | 1 |
| L1PB1_dup9601 | 4.42356849221415 | -4.16359799625969 | 0.941828530640133 | -4.4207601073943 | 9.83543158386163e-06 | 0.00120749799924549 | 4944 | 2 | 0 | 16 | 7 | 0 | 0 | 18 | 11 | 0 | 0 | 1 | 2 |
| L1M3_dup6108 | 4.18380774729747 | -4.70702682025589 | 1.14870001696284 | -4.09769892116939 | 4.17277536005669e-05 | 0.00355748909288445 | 4945 | 1 | 0 | 6 | 4 | 0 | 0 | 19 | 17 | 0 | 0 | 0 | 5 |
| L1MD3_dup2447 | 5.97605178964982 | -5.62182991123072 | 1.08277103553747 | -5.19207637322895 | 2.07961657072022e-07 | 5.94755341991097e-05 | 4946 | 0 | 0 | 18 | 8 | 0 | 0 | 18 | 24 | 0 | 0 | 0 | 5 |
| L1MA5_dup3407 | 4.98017661071503 | -5.43823077888102 | 0.981347172691129 | -5.54159723512309 | 2.99725065670816e-08 | 1.27127237925326e-05 | 4947 | 0 | 0 | 10 | 18 | 0 | 0 | 21 | 12 | 0 | 0 | 1 | 4 |
| L1ME2z_dup5140 | 5.16812508605502 | -5.41095864318558 | 1.04551498278573 | -5.17540038380732 | 2.27422747118765e-07 | 6.36639211599078e-05 | 4948 | 0 | 0 | 8 | 26 | 0 | 0 | 21 | 11 | 0 | 0 | 0 | 4 |
| L1M4_dup9194 | 3.84484318104491 | -4.61700682441167 | 1.05695875850634 | -4.36819959838003 | 1.25274978189699e-05 | 0.00144212288148651 | 4949 | 0 | 1 | 7 | 10 | 0 | 0 | 16 | 11 | 0 | 0 | 0 | 5 |
| L2a_dup116009 | 4.81314291696062 | -4.6020710489494 | 1.00575447509194 | -4.57574006670832 | 4.74539448283201e-06 | 0.000694005922318344 | 4950 | 0 | 1 | 11 | 11 | 0 | 0 | 17 | 18 | 0 | 0 | 0 | 2 |
| MLT1A1-int_dup449 | 4.50525221606092 | -4.38552921723903 | 1.03536447408768 | -4.23573468763583 | 2.27805630564145e-05 | 0.00225199163150046 | 4951 | 0 | 1 | 6 | 12 | 0 | 0 | 26 | 13 | 0 | 0 | 0 | 1 |
| L1MB2_dup3844 | 2.677899583847 | -3.79316441301043 | 1.18268351608031 | -3.20725228806931 | 0.00134009415557025 | 0.0430207830465893 | 4952 | 0 | 1 | 6 | 3 | 0 | 0 | 20 | 5 | 0 | 0 | 0 | 1 |
| L2b_dup41767 | 2.74570694324964 | -4.38663748657773 | 1.30440945585554 | -3.36292984299215 | 0.000771199602285641 | 0.0292808898097499 | 4953 | 0 | 0 | 6 | 1 | 0 | 0 | 18 | 9 | 0 | 0 | 0 | 1 |
| MIR3_dup52858 | 2.67950726775725 | -4.35708379708901 | 1.2598967425967 | -3.45828642124184 | 0.000543623108870817 | 0.0228590674007008 | 4954 | 0 | 0 | 3 | 3 | 0 | 0 | 17 | 10 | 0 | 0 | 0 | 1 |
| L1MC4a_dup8147 | 2.46060013610035 | -4.39578894036032 | 1.30994438146954 | -3.35570654948646 | 0.000791625180042956 | 0.0298469410696405 | 4955 | 0 | 0 | 1 | 5 | 0 | 0 | 10 | 12 | 0 | 0 | 0 | 2 |
| L2c_dup67795 | 2.27176627050409 | -3.75173158222613 | 1.18568256205468 | -3.16419563067936 | 0.00155512226200452 | 0.0476173361211657 | 4956 | 0 | 1 | 2 | 5 | 0 | 0 | 11 | 8 | 0 | 0 | 0 | 2 |
| AluSx_dup102988 | 2.2482250370943 | -4.00857907014254 | 1.25262431529929 | -3.2001447051463 | 0.00137358605589524 | 0.0437711674911008 | 4957 | 0 | 0 | 3 | 2 | 0 | 0 | 15 | 5 | 0 | 1 | 0 | 4 |
| L1MA4_dup9367 | 2.37592983413769 | -4.65599583121244 | 1.37106219251413 | -3.39590418044763 | 0.000684023007659086 | 0.0268921624188325 | 4958 | 0 | 0 | 3 | 1 | 0 | 0 | 13 | 7 | 0 | 0 | 0 | 7 |
| MIRb_dup72913 | 1.77321874633311 | -4.18357454898927 | 1.29138710799424 | -3.23959757929372 | 0.00119698501999216 | 0.0397623617991143 | 4959 | 0 | 0 | 4 | 2 | 0 | 0 | 8 | 5 | 0 | 0 | 0 | 4 |
| AluJb_dup113123 | 2.76400587662783 | -4.2296264175643 | 1.23109314238419 | -3.4356672715868 | 0.000591096438370315 | 0.0243211193367565 | 4960 | 0 | 0 | 1 | 7 | 0 | 0 | 13 | 9 | 0 | 0 | 0 | 5 |
| L1PB1_dup9629 | 3.32368705206225 | -4.30933762748034 | 1.08491116982802 | -3.97206494626049 | 7.12522863062204e-05 | 0.00530481743348558 | 4961 | 0 | 0 | 4 | 16 | 0 | 1 | 11 | 9 | 0 | 0 | 0 | 3 |
| L1MB2_dup7823 | 3.14793766278561 | -4.14061697873567 | 1.10339703473753 | -3.75260839786525 | 0.000175004119695829 | 0.0101416293875167 | 4962 | 0 | 0 | 6 | 6 | 0 | 1 | 20 | 7 | 0 | 0 | 0 | 2 |
| THE1C-int_dup123 | 3.88106756134776 | -4.37460572096208 | 1.05827733203215 | -4.13370445397502 | 3.56962452750755e-05 | 0.00315285032599225 | 4963 | 0 | 0 | 5 | 12 | 0 | 1 | 20 | 11 | 0 | 0 | 0 | 2 |
| HAL1_dup16860 | 3.11897391515786 | -4.53648960640615 | 1.15325394125789 | -3.93364327154005 | 8.36679293964343e-05 | 0.00598390415634624 | 4964 | 0 | 0 | 4 | 8 | 0 | 0 | 19 | 9 | 0 | 0 | 0 | 1 |
| L1MA8_dup8017 | 3.10053520380519 | -4.84568189065551 | 1.11847848358006 | -4.33238722227835 | 1.47501223666375e-05 | 0.00162789082405006 | 4965 | 0 | 0 | 6 | 9 | 0 | 0 | 17 | 6 | 0 | 0 | 0 | 4 |
| AluSx1_dup115294 | 2.06538370515141 | -4.31001847475255 | 1.19163112030215 | -3.61690660920279 | 0.000298144813517679 | 0.0149007279557399 | 4966 | 0 | 0 | 7 | 5 | 0 | 0 | 10 | 3 | 0 | 0 | 0 | 3 |
| THE1C_dup7933 | 2.02260286342554 | -4.28261132443391 | 1.16802294050073 | -3.666547270551 | 0.000245847489937944 | 0.0129532246034842 | 4967 | 0 | 0 | 6 | 6 | 0 | 0 | 8 | 4 | 0 | 0 | 0 | 3 |
| MER20B_dup390 | 2.42224722166544 | -4.49240749376467 | 1.17593963979784 | -3.82027048134628 | 0.000133305404608448 | 0.00834284266306204 | 4968 | 0 | 0 | 3 | 12 | 0 | 0 | 10 | 5 | 0 | 0 | 0 | 3 |
| MER5A_dup11602 | 1.74493110859963 | -4.01291302671649 | 1.2483678713857 | -3.21452763940657 | 0.00130659335337877 | 0.0422531478748389 | 4969 | 0 | 0 | 2 | 6 | 0 | 0 | 8 | 5 | 0 | 0 | 0 | 2 |
| L1MB8_dup9837 | 1.86175406807122 | -4.17593877233525 | 1.223170536926 | -3.41402825384427 | 0.000640099374287652 | 0.0256748927646831 | 4970 | 0 | 0 | 3 | 5 | 0 | 0 | 7 | 6 | 0 | 0 | 0 | 3 |
| MSTA_dup10002 | 1.92594128821882 | -4.12436593828938 | 1.19694852657335 | -3.44573375272595 | 0.000569511173786502 | 0.0236796912027431 | 4971 | 0 | 0 | 3 | 7 | 0 | 0 | 7 | 6 | 0 | 0 | 0 | 2 |
| MIRc_dup45721 | 2.37099109237261 | -3.91743736203068 | 1.14369234348523 | -3.42525451389566 | 0.000614223729628079 | 0.0249619264850697 | 4972 | 1 | 0 | 3 | 6 | 0 | 0 | 11 | 7 | 0 | 0 | 0 | 3 |

| | | | | | | | | | | | | | | | | | | | |
|---|---|---|---|---|---|---|---|---|---|---|---|---|---|---|---|---|---|---|---|
| MER2B_dup797 | 2.26944388380253 | -3.61201953047312 | 1.14032116338765 | -3.16754581642824 | 0.00153731484210964 | 0.0473055531490971 | 4973 | 1 | 0 | 3 | 8 | 0 | 0 | 8 | 8 | 0 | 0 | 0 | 1 |
| MER5B_dup10092 | 1.97149171137236 | -3.60230372661827 | 1.13038386828859 | -3.18679682864918 | 0.00143857789745153 | 0.0451257104465528 | 4974 | 1 | 0 | 4 | 7 | 0 | 0 | 7 | 5 | 0 | 0 | 0 | 2 |
| L3_dup29520 | 2.7799835424558 | -3.5534109753018 | 1.0373113304679 | -3.425597379428 | 0.000613448987018761 | 0.0249474846462427 | 4975 | 1 | 0 | 5 | 7 | 0 | 0 | 12 | 8 | 0 | 1 | 0 | 2 |
| L1MC4a_dup8146 | 2.10379478285536 | -3.85217447858392 | 1.18659227676241 | -3.2464179600886 | 0.00116867125786293 | 0.0391222075300299 | 4976 | 0 | 0 | 2 | 6 | 0 | 0 | 8 | 7 | 0 | 1 | 0 | 3 |
| MER33_dup4673 | 2.02772338036846 | -3.73947171533918 | 1.17753107094863 | -3.17568835982106 | 0.0014948144176608 | 0.0464098527433086 | 4977 | 0 | 0 | 2 | 8 | 0 | 0 | 9 | 5 | 0 | 1 | 0 | 2 |
| L1M7_dup1546 | 2.62200246174076 | -3.57637614677167 | 1.05585208365958 | -3.38719428802561 | 0.000706113635953775 | 0.0275257069496489 | 4978 | 0 | 0 | 6 | 10 | 1 | 1 | 5 | 7 | 0 | 0 | 0 | 4 |
| L1MA10_dup4891 | 2.54297250668868 | -3.46935250042257 | 1.06263813320222 | -3.26484848606721 | 0.00109522644705556 | 0.0373731974769768 | 4979 | 0 | 0 | 9 | 12 | 1 | 1 | 4 | 4 | 0 | 0 | 0 | 3 |
| MIR_dup170873 | 2.77376027441611 | -3.2667738890053 | 1.01909238524617 | -3.20557187581788 | 0.00134794377421067 | 0.0431679447238672 | 4980 | 0 | 0 | 10 | 9 | 2 | 1 | 6 | 4 | 0 | 0 | 0 | 5 |
| MamTip2_dup4060 | 2.90666176710686 | -4.15160531199388 | 1.14804365538574 | -3.61624341767643 | 0.000298909312397886 | 0.0149138761266151 | 4981 | 0 | 0 | 15 | 11 | 1 | 0 | 3 | 5 | 0 | 0 | 0 | 3 |
| AluJb_dup57349 | 1.92693555492424 | -3.73477434845281 | 1.17921371865136 | -3.16717342190029 | 0.0015392849318048 | 0.0473194107654694 | 4982 | 0 | 0 | 5 | 7 | 1 | 0 | 3 | 5 | 0 | 0 | 0 | 4 |
| L2_dup19406 | 2.05641001213663 | -3.64800196980013 | 1.12698029932524 | -3.23697048829008 | 0.00120805904264893 | 0.0400519460724737 | 4983 | 0 | 0 | 8 | 7 | 1 | 0 | 5 | 4 | 0 | 0 | 0 | 2 |
| MLT1G1_dup2066 | 2.63349805818682 | -3.7794198466255 | 1.18714884655911 | -3.18361076421036 | 0.00145450452853464 | 0.045522272517261 | 4984 | 0 | 0 | 4 | 21 | 0 | 1 | 4 | 5 | 0 | 0 | 0 | 1 |
| L1MA3_dup3731 | 3.1589653817833 | -4.55552314583896 | 1.18540352511894 | -3.84301467753936 | 0.000121532197505052 | 0.00782416132779684 | 4985 | 0 | 0 | 10 | 23 | 0 | 0 | 4 | 5 | 0 | 0 | 0 | 1 |
| MIRb_dup213201 | 2.9128555435889 | -4.4682155419803 | 1.16161009300794 | -3.8465708664859 | 0.00011978249651634 | 0.00774083810865751 | 4986 | 0 | 0 | 9 | 20 | 0 | 0 | 6 | 4 | 0 | 0 | 0 | 1 |
| L1ME3Cz_dup2356 | 2.51892388259335 | -4.53522326141677 | 1.18609827145439 | -3.8236488245242 | 0.000131491162139249 | 0.00827110282377745 | 4987 | 0 | 0 | 6 | 16 | 0 | 0 | 4 | 5 | 0 | 0 | 0 | 3 |
| LTR16_dup1616 | 2.5058800806438 | -4.43577669879721 | 1.14147457900473 | -3.88600568105934 | 0.000101907122470127 | 0.00690247428502043 | 4988 | 0 | 0 | 8 | 14 | 0 | 0 | 6 | 4 | 0 | 0 | 0 | 2 |
| MLT2A2_dup354 | 2.34820336024452 | -4.36159632397396 | 1.17586703953706 | -3.70925978645606 | 0.000207866063559121 | 0.0114406140974582 | 4989 | 0 | 0 | 6 | 15 | 0 | 0 | 5 | 4 | 0 | 0 | 0 | 2 |
| AluJb_dup59479 | 2.33355201464139 | -3.26720890118752 | 1.0215091205498 | -3.19841383249623 | 0.00138185822584366 | 0.0439230079356926 | 4990 | 0 | 1 | 8 | 8 | 0 | 1 | 7 | 4 | 0 | 0 | 0 | 2 |
| Tigger15a_dup4277 | 2.74559894576425 | -3.31403778204336 | 1.02232696619416 | -3.24166131935324 | 0.00118835155527507 | 0.0395584078303696 | 4991 | 0 | 1 | 12 | 12 | 0 | 1 | 4 | 5 | 0 | 0 | 0 | 1 |
| L1ME3A_dup13840 | 3.00913595327929 | -3.57601705207097 | 1.00253418696325 | -3.56697766377722 | 0.000361122248233982 | 0.0171307006535652 | 4992 | 0 | 0 | 11 | 9 | 0 | 2 | 8 | 7 | 0 | 0 | 0 | 2 |
| MER65A_dup1083 | 2.23381092665024 | -3.59397496526644 | 1.11077536882138 | -3.23555515016499 | 0.00121406429609031 | 0.040161510217921 | 4993 | 0 | 0 | 9 | 7 | 0 | 1 | 6 | 5 | 0 | 0 | 0 | 1 |
| Charlie7a_dup805 | 2.07297298947501 | -3.76783714778029 | 1.1304411247654 | -3.33306800790907 | 0.000858939440057876 | 0.031617952980903 | 4994 | 0 | 0 | 7 | 7 | 0 | 1 | 7 | 3 | 0 | 0 | 0 | 3 |
| MLT1L_dup11664 | 2.67975175573607 | -3.441328950885 | 1.05950149126559 | -3.2480642823582 | 0.00116193016924978 | 0.0389568319081735 | 4995 | 0 | 0 | 7 | 13 | 0 | 2 | 10 | 3 | 0 | 0 | 0 | 2 |
| L1MD2_dup53 | 2.12569235879996 | -3.77294585608894 | 1.15644577997449 | -3.26253588488356 | 0.0011042019204679 | 0.0375768767625014 | 4996 | 0 | 0 | 10 | 5 | 0 | 0 | 2 | 6 | 1 | 0 | 1 | 1 |
| L2_dup40072 | 1.8438447448275 | -3.66288264180559 | 1.11613546762929 | -3.28175454327751 | 0.00103163365185084 | 0.0358422300417366 | 4997 | 0 | 0 | 6 | 5 | 0 | 0 | 3 | 5 | 1 | 0 | 1 | 2 |
| MLT1D_dup4162 | 2.28997481919078 | -3.94104446933828 | 1.14965091720031 | -3.42803577188084 | 0.000607965344037965 | 0.024809579982641 | 4998 | 0 | 0 | 8 | 8 | 0 | 0 | 3 | 6 | 1 | 0 | 0 | 3 |
| AluSz_dup10740 | 2.55066323592416 | -3.88544508842914 | 1.1333490838315 | -3.42828625695241 | 0.000607404625363462 | 0.024794904955693 | 4999 | 0 | 0 | 11 | 11 | 0 | 0 | 4 | 5 | 1 | 0 | 0 | 1 |
| MLT1F1_dup1513 | 2.489562716185 | -3.97703390446472 | 1.16073890645015 | -3.42629499395997 | 0.000611875456344256 | 0.0249132986542249 | 5000 | 0 | 0 | 9 | 14 | 0 | 0 | 3 | 4 | 1 | 0 | 0 | 2 |
| L4_A_Mam_dup582 | 2.38488916012285 | -3.81741750879817 | 1.20914084768093 | -3.15713220351442 | 0.00159329092291674 | 0.0484247288216051 | 5001 | 0 | 1 | 5 | 17 | 0 | 0 | 2 | 5 | 0 | 0 | 0 | 2 |
| MIR1_Amn_dup2630 | 2.25479204691544 | -3.60662299938605 | 1.12227202354005 | -3.21367985990549 | 0.00131045694819516 | 0.0423550650631956 | 5002 | 0 | 1 | 5 | 12 | 0 | 0 | 6 | 5 | 0 | 0 | 0 | 1 |
| MLT1A0_dup5499 | 2.07275039552927 | -3.6636879478718 | 1.13881435110388 | -3.21710728734714 | 0.00129490166177649 | 0.041949172336701 | 5003 | 0 | 1 | 7 | 10 | 0 | 0 | 5 | 3 | 0 | 0 | 0 | 2 |
| AluJr4_dup9719 | 1.822026267961 | -4.26446840601624 | 1.28817907302846 | -3.31046241574989 | 0.000931419596136599 | 0.0334838602028075 | 5004 | 0 | 0 | 6 | 7 | 0 | 0 | 2 | 4 | 0 | 0 | 0 | 5 |
| L2a_dup57688 | 1.77866554353678 | -4.04035964959968 | 1.28351621943872 | -3.14788359384079 | 0.00164457170129424 | 0.0495495987569662 | 5005 | 0 | 0 | 6 | 11 | 0 | 0 | 2 | 3 | 0 | 0 | 0 | 2 |
| MIR_dup19155 | 1.79382400477412 | -4.14195215611387 | 1.24412172304338 | -3.32921777620103 | 0.000870902780485269 | 0.031925409490887 | 5006 | 0 | 0 | 7 | 8 | 0 | 0 | 3 | 3 | 0 | 0 | 0 | 3 |
| L1MEg_dup14462 | 2.13161529576138 | -4.10926471088254 | 1.16037701111 | -3.54131861587956 | 0.000398132492497882 | 0.0184059043502628 | 5007 | 0 | 0 | 7 | 10 | 0 | 0 | 5 | 5 | 0 | 0 | 0 | 1 |
| L1MEg_dup1458 | 1.79199912791892 | -3.91360862434543 | 1.20625205953772 | -3.24443684336196 | 0.00117683111502695 | 0.0392848283524414 | 5008 | 0 | 0 | 7 | 8 | 0 | 0 | 5 | 3 | 0 | 0 | 0 | 1 |
| MLT1M_dup529 | 1.80320122361689 | -3.91809077412175 | 1.22210907075681 | -3.20600744064145 | 0.00134590508167581 | 0.0431316808840987 | 5009 | 0 | 0 | 8 | 8 | 0 | 0 | 4 | 3 | 0 | 0 | 0 | 1 |
| L2a_dup123378 | 2.19058432681637 | -4.13018491861501 | 1.2348330689123 | -3.34473138320881 | 0.000823622968414905 | 0.0306820887888357 | 5010 | 0 | 0 | 9 | 10 | 0 | 0 | 2 | 6 | 0 | 0 | 0 | 1 |
| L1ME2_dup9208 | 1.73482728614365 | -3.99914738514929 | 1.23514914160659 | -3.23778501756275 | 0.0012046154616476 | 0.039971856507557 | 5011 | 0 | 0 | 7 | 5 | 0 | 0 | 3 | 5 | 0 | 0 | 0 | 2 |
| L1ME3A_dup1500 | 1.91492502733525 | -3.97282813178876 | 1.22326967216074 | -3.24771244003075 | 0.0011633678086098 | 0.0389995374172255 | 5012 | 0 | 0 | 8 | 6 | 0 | 0 | 3 | 6 | 0 | 0 | 0 | 1 |
| L3_dup5044 | 2.96075236726021 | -3.2732730459937 | 1.00138934458911 | -3.26873165136063 | 0.00108030705353934 | 0.0370342518601747 | 5013 | 1 | 1 | 5 | 5 | 0 | 0 | 10 | 10 | 1 | 0 | 0 | 4 |
| MLT1J1_dup1377 | 3.51540221570283 | -3.76021771212509 | 0.985752505850917 | -3.8145657148284 | 0.000136422631905297 | 0.00849527341110128 | 5014 | 1 | 1 | 13 | 5 | 0 | 0 | 11 | 11 | 0 | 0 | 0 | 2 |
| THE1C-int_dup390 | 3.0675307416074 | -3.2943727835267 | 0.982964095544235 | -3.35146807341189 | 0.000803843072310355 | 0.0301535474269569 | 5015 | 2 | 1 | 9 | 4 | 0 | 0 | 11 | 9 | 0 | 0 | 0 | 3 |
| L2c_dup39576 | 2.57503112107736 | -3.91489146624905 | 1.07449849869104 | -3.6434592240177 | 0.000268998283429078 | 0.01382143501378 | 5016 | 0 | 1 | 6 | 9 | 0 | 0 | 10 | 6 | 0 | 0 | 0 | 2 |
| AluJr4_dup9112 | 2.19986475594643 | -3.73558163042174 | 1.13467843948665 | -3.29219407051726 | 0.000994089872656412 | 0.0349600515576563 | 5017 | 0 | 1 | 6 | 6 | 0 | 0 | 12 | 3 | 0 | 0 | 0 | 2 |
| L2_dup19393 | 2.24570461050526 | -3.60670701771422 | 1.10801842407652 | -3.25509661152092 | 0.00113353798460707 | 0.0383126177962095 | 5018 | 0 | 1 | 7 | 7 | 0 | 0 | 10 | 4 | 0 | 0 | 0 | 1 |
| AluSz_dup869 | 2.59456213445471 | -3.91846268555176 | 1.12264537368728 | -3.49038331907229 | 0.000482328182453663 | 0.0210082485268386 | 5019 | 0 | 1 | 14 | 6 | 0 | 0 | 7 | 4 | 0 | 0 | 0 | 2 |
| HERV9N-int_dup166 | 2.23538726565942 | -3.5800820718207 | 1.13401032612138 | -3.15701011653532 | 0.00159395816205601 | 0.0484293826077021 | 5020 | 0 | 1 | 8 | 4 | 0 | 0 | 7 | 7 | 0 | 0 | 0 | 1 |
| MER102c_dup3322 | 2.06015465787971 | -4.18995765404344 | 1.26133634240235 | -3.321840109723 | 0.00089425922068023 | 0.0325144464958025 | 5021 | 0 | 0 | 8 | 2 | 0 | 0 | 5 | 8 | 0 | 0 | 0 | 2 |
| MLT1F2_dup5878 | 2.2657819768869 | -3.98651513820409 | 1.15913421492594 | -3.43921789804023 | 0.000583397458008128 | 0.0240918242872617 | 5022 | 0 | 0 | 5 | 7 | 0 | 1 | 4 | 7 | 0 | 0 | 0 | 5 |
| L1M5_dup54037 | 3.17156621468106 | -3.22825929584347 | 0.974097720988656 | -3.31410209292653 | 0.000919379317840553 | 0.0331941792996004 | 5023 | 0 | 1 | 12 | 6 | 0 | 2 | 6 | 10 | 0 | 0 | 2 | 0 |
| L1PA10_dup3003 | 3.19029321925496 | -3.35144377662435 | 0.938318849720607 | -3.57175365028878 | 0.000354598885239653 | 0.0168750772694469 | 5024 | 0 | 1 | 8 | 11 | 0 | 2 | 11 | 6 | 0 | 0 | 3 | 0 |
| LTR104_Mam_dup182 | 2.69111079704812 | -4.27216623394824 | 1.14072059664641 | -3.74514692424064 | 0.000180288235385736 | 0.0103570411533304 | 5025 | 0 | 0 | 7 | 10 | 0 | 1 | 6 | 5 | 0 | 0 | 6 | 0 |
| L1MB2_dup4597 | 2.96625963806723 | -3.57874387732916 | 1.02668890763311 | -3.48571397891056 | 0.000490825496765947 | 0.0212732174232186 | 5026 | 0 | 0 | 9 | 12 | 0 | 2 | 11 | 4 | 0 | 0 | 2 | 0 |
| L2c_dup103826 | 2.31781687639388 | -3.80358581731365 | 1.11406085700492 | -3.41416341252608 | 0.000639781902719775 | 0.0256674329159602 | 5027 | 0 | 0 | 8 | 5 | 0 | 1 | 9 | 5 | 0 | 0 | 2 | 0 |
| MIR_dup18074 | 2.52704725286741 | -3.37252285108642 | 1.02045575628839 | -3.3049182488352 | 0.00095004106880553 | 0.033925062212951 | 5028 | 0 | 1 | 3 | 9 | 0 | 1 | 13 | 5 | 0 | 0 | 1 | 1 |
| L1PB1_dup11672 | 2.51347147408421 | -3.55506027856739 | 1.02302065945587 | -3.47506205833445 | 0.000510735181747609 | 0.0218928830041644 | 5029 | 0 | 1 | 6 | 10 | 0 | 1 | 9 | 3 | 0 | 0 | 3 | 1 |
| L1PB1_dup10383 | 2.09889401961163 | -3.2440791268173 | 1.03077983101263 | -3.1472085786063 | 0.00164837329697367 | 0.0496109072251071 | 5030 | 0 | 1 | 7 | 7 | 0 | 1 | 6 | 3 | 0 | 0 | 1 | 2 |
| MER94_dup1079 | 2.50800353752493 | -3.54053741311846 | 1.00345232664166 | -3.52835637440584 | 0.000418148793065151 | 0.0190797204147751 | 5031 | 0 | 1 | 9 | 6 | 0 | 1 | 8 | 4 | 0 | 0 | 2 | 2 |
| L1M5_dup13708 | 3.53065309424697 | -3.48062462070125 | 0.888406771056802 | -3.9178276596889 | 8.93505482365491e-05 | 0.00627712003268199 | 5032 | 1 | 1 | 6 | 14 | 0 | 1 | 8 | 11 | 0 | 0 | 2 | 1 |
| L1PB3_dup1596 | 2.92857278228572 | -3.15867060614899 | 0.897457654767478 | -3.51957620436963 | 0.000432236846619955 | 0.0195344783866727 | 5033 | 1 | 1 | 11 | 6 | 0 | 1 | 10 | 6 | 0 | 0 | 1 | 1 |
| THE1B_dup9486 | 2.92360177437576 | -3.03650743046687 | 0.901035665422416 | -3.37001913131077 | 0.000751629651127342 | 0.0288091605939347 | 5034 | 1 | 1 | 11 | 10 | 0 | 1 | 7 | 6 | 0 | 0 | 1 | 0 |
| L2a_dup62305 | 2.2701840656272 | -4.53323796012731 | 1.07364640259562 | -4.22228207458982 | 2.41841185592619e-05 | 0.00235262235269915 | 5035 | 0 | 0 | 7 | 4 | 0 | 0 | 7 | 6 | 0 | 0 | 2 | 3 |
| MLT1L_dup4094 | 2.42562883707451 | -4.68949872621471 | 1.07051848661388 | -4.38058640262057 | 1.18360347020546e-05 | 0.00137988099618574 | 5036 | 0 | 0 | 6 | 4 | 0 | 0 | 10 | 5 | 0 | 0 | 2 | 5 |
| AluSx4_dup5092 | 2.61863038975759 | -4.78614653529264 | 1.05332711590711 | -4.54383682240145 | 5.52394255878031e-06 | 0.000775915015402326 | 5037 | 0 | 0 | 9 | 4 | 0 | 0 | 9 | 5 | 0 | 0 | 4 | 3 |
| L1M4_dup20640 | 2.03844839808581 | -4.41481982698145 | 1.11622740815428 | -3.9551258056649 | 7.64943884430261e-05 | 0.00559015164387724 | 5038 | 0 | 0 | 8 | 3 | 0 | 0 | 8 | 3 | 0 | 0 | 2 | 3 |
| MIR_dup58589 | 2.0937125547433 | -4.45399036698434 | 1.08854681855306 | -4.09168470392919 | 4.28250654049199e-05 | 0.00362506332722842 | 5039 | 0 | 0 | 6 | 5 | 0 | 0 | 9 | 3 | 0 | 0 | 3 | 2 |
| MER4D1_dup741 | 2.11867626403885 | -4.24206357985724 | 1.14807046635547 | -3.69495053149795 | 0.000219929601906758 | 0.0119070236758374 | 5040 | 0 | 0 | 10 | 3 | 0 | 0 | 7 | 5 | 0 | 0 | 1 | 1 |
| HAL1_dup21329 | 2.00063599798587 | -4.28168200053593 | 1.13431564127479 | -3.77468302890013 | 0.000160211243768176 | 0.00954950516274107 | 5041 | 0 | 0 | 6 | 4 | 0 | 0 | 11 | 3 | 0 | 0 | 1 | 2 |
| MLT1L_dup6049 | 1.92932033734386 | -4.22737193479809 | 1.13858363316725 | -3.71283392071834 | 0.000204951385613145 | 0.0113123086061483 | 5042 | 0 | 0 | 5 | 3 | 0 | 0 | 9 | 5 | 0 | 0 | 1 | 2 |
| L1MA10_dup3961 | 2.18969302774439 | -4.27862855726116 | 1.12438179080982 | -3.80531647900447 | 0.000141622989835232 | 0.00873005915322085 | 5043 | 0 | 0 | 4 | 5 | 0 | 0 | 10 | 7 | 0 | 0 | 1 | 1 |
| L1MEg_dup6767 | 2.2782115200861 | -4.32318915335781 | 1.09222069593423 | -3.95816447120147 | 7.55279448657057e-05 | 0.00554335976835756 | 5044 | 0 | 0 | 6 | 6 | 0 | 0 | 8 | 7 | 0 | 0 | 1 | 1 |
| MLT1J1_dup3256 | 2.1379528974516 | -4.25909562354735 | 1.09799992239347 | -3.87895803695793 | 0.000104904847718468 | 0.00707129997031691 | 5045 | 0 | 0 | 6 | 6 | 0 | 0 | 9 | 5 | 0 | 0 | 1 | 1 |
| AluSx1_dup51356 | 1.8457879386333 | -4.16413588116229 | 1.17113609822382 | -3.55563788655968 | 0.000377063124412314 | 0.0176987900832067 | 5046 | 0 | 0 | 3 | 3 | 0 | 0 | 7 | 7 | 0 | 0 | 1 | 2 |
| HSMAR1_dup411 | 1.76687958320463 | -4.12022272568025 | 1.18776332876007 | -3.46889201401886 | 0.000522609467629322 | 0.0222257073073327 | 5047 | 0 | 0 | 4 | 2 | 0 | 0 | 7 | 6 | 0 | 0 | 2 | 1 |
| MIR3_dup79197 | 2.07094555375862 | -4.30908913159912 | 1.14917052357372 | -3.74973865340594 | 0.000177018960101623 | 0.0102284976882809 | 5048 | 0 | 0 | 4 | 3 | 0 | 0 | 9 | 7 | 0 | 0 | 2 | 1 |
| L2c_dup49373 | 1.82980241351497 | -3.93500136917095 | 1.23609973235443 | -3.18340119828021 | 0.00145555779040288 | 0.0455416376062071 | 5049 | 0 | 0 | 5 | 3 | 0 | 0 | 8 | 6 | 0 | 0 | 1 | 0 |
| L3_dup44817 | 2.08495691482288 | -4.07653088826039 | 1.23835899611912 | -3.29188135349747 | 0.00099519587412425 | 0.0349937753013697 | 5050 | 0 | 0 | 6 | 3 | 0 | 0 | 6 | 9 | 0 | 0 | 1 | 0 |
| AluSz6_dup42940 | 2.6465518290095 | -4.36685301030533 | 1.22738114331304 | -3.55786222893893 | 0.000373885314815186 | 0.0175819199839949 | 5051 | 0 | 0 | 6 | 3 | 0 | 0 | 19 | 6 | 0 | 0 | 1 | 0 |

| | | | | | | | | | | | | | | | | | | | |
|---|---|---|---|---|---|---|---|---|---|---|---|---|---|---|---|---|---|---|---|
| THE1B_dup14865 | 2.04864356630569 | -4.21830971598336 | 1.33464030576413 | -3.1606341407232 | 0.00157426098592596 | 0.0479811841369461 | 5052 | 0 | 0 | 2 | 2 | 0 | 0 | 13 | 7 | 0 | 0 | 2 | 0 |
| AluSx_dup102992 | 2.55680342823212 | -4.31814373169535 | 1.23697915635522 | -3.49087833009153 | 0.000481435445219266 | 0.0209885742851124 | 5053 | 0 | 0 | 3 | 4 | 0 | 0 | 14 | 10 | 0 | 0 | 1 | 0 |
| L1ME3_dup8253 | 2.52568969573595 | -4.45782886598158 | 1.19982979677239 | -3.71538436366008 | 0.000202895038668479 | 0.0112429261818041 | 5054 | 0 | 0 | 3 | 6 | 0 | 0 | 15 | 7 | 0 | 0 | 2 | 0 |
| Tigger2_dup1549 | 2.41451552085644 | -4.25663231915675 | 1.20173252623667 | -3.5420796443671 | 0.000396985591691733 | 0.0183705043953094 | 5055 | 0 | 0 | 3 | 6 | 0 | 0 | 13 | 8 | 0 | 0 | 1 | 0 |
| AluJr_dup75402 | 1.94633002340769 | -3.77534465023288 | 1.1820005913847 | -3.19402940891097 | 0.00140301823731806 | 0.0444057698186863 | 5056 | 0 | 0 | 3 | 5 | 0 | 0 | 8 | 5 | 0 | 1 | 3 | 0 |
| HAL1_dup13390 | 2.49164133814832 | -4.15002990161721 | 1.21235351120986 | -3.4231186392785 | 0.000619070513321053 | 0.025094098408732 | 5057 | 0 | 0 | 2 | 5 | 0 | 0 | 13 | 7 | 0 | 1 | 4 | 0 |
| L1ME3Cz_dup4131 | 2.14884856355752 | -4.48407476630861 | 1.20854576246315 | -3.71030614278898 | 0.000207008762684997 | 0.0114073429934795 | 5058 | 0 | 0 | 3 | 2 | 0 | 0 | 13 | 5 | 0 | 0 | 4 | 1 |
| AluSz_dup91408 | 2.23873364967949 | -4.54065195258746 | 1.27304314107045 | -3.56676989655622 | 0.000361408561912778 | 0.0171408653922459 | 5059 | 0 | 0 | 4 | 3 | 0 | 0 | 12 | 5 | 0 | 0 | 5 | 0 |
| L1PA15_dup4083 | 1.9798204602614 | -4.27070708969009 | 1.24922300929414 | -3.41869070447496 | 0.000629232069257047 | 0.0253768227036334 | 5060 | 0 | 0 | 4 | 3 | 0 | 0 | 9 | 6 | 0 | 0 | 3 | 0 |
| LTR79_dup1987 | 2.06144517385419 | -4.38791245626227 | 1.2958163742503 | -3.38621470098409 | 0.000708639179981555 | 0.0275834873744301 | 5061 | 0 | 0 | 4 | 2 | 0 | 0 | 10 | 6 | 0 | 0 | 4 | 0 |
| L2a_dup90811 | 2.90008942228777 | -4.7142635908482 | 1.25596802026261 | -3.75349014846931 | 0.000174389389060196 | 0.0101215782210759 | 5062 | 0 | 0 | 5 | 15 | 0 | 0 | 2 | 11 | 0 | 0 | 3 | 0 |
| MLT1C_dup1645 | 3.58753081970388 | -4.84438133981541 | 1.08020105985573 | -4.48470337592748 | 7.30154354904209e-06 | 0.000963601265291387 | 5063 | 0 | 0 | 11 | 16 | 0 | 0 | 7 | 10 | 0 | 0 | 2 | 0 |
| MamRTE1_dup1582 | 2.95813747741383 | -4.63854706818409 | 1.18849097392747 | -3.90288792253559 | 9.50517237436621e-05 | 0.00656521842029301 | 5064 | 0 | 0 | 10 | 18 | 0 | 0 | 3 | 6 | 0 | 0 | 2 | 0 |
| L2a_dup19351 | 2.22953040262628 | -4.16237620025817 | 1.19970048864225 | -3.46951279895609 | 0.000521403224278905 | 0.0221958935541952 | 5065 | 0 | 0 | 8 | 9 | 0 | 0 | 3 | 7 | 0 | 0 | 1 | 0 |
| MLT1F1_dup2685 | 2.3251941648588 | -4.21613086339804 | 1.16927528784009 | -3.60576410640317 | 0.000311235768896079 | 0.0153391272298564 | 5066 | 0 | 0 | 8 | 11 | 0 | 0 | 4 | 6 | 0 | 0 | 1 | 0 |
| MER65-int_dup716 | 2.21655051332901 | -4.30850603128473 | 1.18485571937482 | -3.63631281077673 | 0.00027656845299949 | 0.0141109311366803 | 5067 | 0 | 0 | 3 | 10 | 0 | 0 | 8 | 6 | 0 | 0 | 2 | 0 |
| MIR3_dup23712 | 1.97859884372081 | -4.1796910323574 | 1.21106581380536 | -3.45125011763328 | 0.000557996196793628 | 0.0233057684820531 | 5068 | 0 | 0 | 3 | 10 | 0 | 0 | 6 | 5 | 0 | 0 | 2 | 0 |
| MER103C_dup3539 | 1.98333518988529 | -4.0279120690356 | 1.21673277669578 | -3.31043278046146 | 0.000931518228368601 | 0.0334838602028075 | 5069 | 0 | 0 | 4 | 9 | 0 | 0 | 4 | 7 | 0 | 0 | 1 | 0 |
| MIR_dup170854 | 2.31593597935542 | -4.21133036935494 | 1.17520084811013 | -3.58349840891221 | 0.000339022830158606 | 0.0163231740884095 | 5070 | 0 | 0 | 4 | 12 | 0 | 0 | 6 | 7 | 0 | 0 | 1 | 0 |
| ERVL-E-int_dup306 | 2.19600949007899 | -4.14915428536235 | 1.15105551533722 | -3.60465175665033 | 0.000312571764924545 | 0.0153849969913431 | 5071 | 0 | 0 | 8 | 6 | 0 | 0 | 7 | 6 | 0 | 0 | 1 | 0 |
| L1MC2_dup6133 | 1.88492814115267 | -3.98327561267961 | 1.19876826245842 | -3.32280703237068 | 0.000891165388274388 | 0.0324465463685997 | 5072 | 0 | 0 | 8 | 6 | 0 | 0 | 7 | 3 | 0 | 0 | 1 | 0 |
| MIRb_dup90021 | 2.0547362250739 | -4.08056648175892 | 1.16385177162122 | -3.506087786484 | 0.00045474521027515 | 0.020255485549257 | 5073 | 0 | 0 | 8 | 7 | 0 | 0 | 7 | 4 | 0 | 0 | 1 | 0 |
| L1ME1_dup22784 | 1.9443984108162 | -4.26057206188003 | 1.20754089996317 | -3.52830455847084 | 0.000418230660698446 | 0.0190797204147751 | 5074 | 0 | 0 | 5 | 7 | 0 | 0 | 8 | 3 | 0 | 0 | 3 | 0 |
| MLT1H_dup7571 | 1.95332147430631 | -4.16887185456588 | 1.18619481024226 | -3.51449173320397 | 0.000440596385180777 | 0.0198144296859242 | 5075 | 0 | 0 | 7 | 7 | 0 | 0 | 7 | 3 | 0 | 0 | 2 | 0 |
| L1M5_dup24010 | 1.91324943767894 | -3.99550545018994 | 1.19368956063525 | -3.34718974007249 | 0.000816353133098243 | 0.0304989735607151 | 5076 | 0 | 0 | 4 | 6 | 0 | 0 | 9 | 5 | 0 | 0 | 1 | 0 |
| L3_dup41757 | 1.90010343178639 | -3.99011270030688 | 1.19181125116059 | -3.34794011754905 | 0.000814146010241683 | 0.0304260698214179 | 5077 | 0 | 0 | 6 | 5 | 0 | 0 | 9 | 4 | 0 | 0 | 1 | 0 |
| MIRb_dup217372 | 2.03233203367797 | -4.07020394535918 | 1.16561166983033 | -3.49190390823013 | 0.000479590748407178 | 0.0209426867719755 | 5078 | 0 | 0 | 6 | 7 | 0 | 0 | 9 | 4 | 0 | 0 | 1 | 0 |
| L2c_dup50547 | 1.77009543630209 | -4.25812551532984 | 1.31575712631055 | -3.23625495175531 | 0.00121109161299627 | 0.0400932257458232 | 5079 | 0 | 0 | 3 | 4 | 0 | 0 | 4 | 5 | 0 | 0 | 6 | 0 |
| AluSx1_dup51355 | 2.17382555598068 | -3.71498946601951 | 1.16633717268327 | -3.18517625351236 | 0.00144665869776759 | 0.0453347360067176 | 5080 | 0 | 0 | 3 | 5 | 1 | 0 | 8 | 8 | 0 | 0 | 2 | 0 |
| MIRb_dup202218 | 2.10691013505417 | -3.68656516043521 | 1.12615869009123 | -3.27357520114377 | 0.00106196128560514 | 0.0366110460829615 | 5081 | 0 | 0 | 7 | 7 | 1 | 0 | 5 | 5 | 0 | 0 | 2 | 0 |
| MLT1A_dup8918 | 2.26813027749856 | -3.95590847316017 | 1.14227192101881 | -3.46319330832524 | 0.000533804745180733 | 0.0225606934097291 | 5082 | 0 | 0 | 7 | 7 | 1 | 0 | 5 | 5 | 0 | 0 | 4 | 0 |
| L1PA5_dup870 | 2.08921109215838 | -3.947918237764 | 1.2492400278409 | -3.1602559554446 | 0.00157630596279919 | 0.0480250574422811 | 5083 | 0 | 0 | 4 | 9 | 0 | 0 | 10 | 1 | 0 | 1 | 4 | 0 |
| LTR78_dup1153 | 2.52278856761039 | -4.22988902334803 | 1.1720261348958 | -3.60903984766865 | 0.000307332404020884 | 0.0152129220318605 | 5084 | 0 | 0 | 6 | 9 | 0 | 0 | 4 | 6 | 0 | 1 | 6 | 0 |
| MLT1B_dup13200 | 2.55196668180704 | -4.17961516641068 | 1.14479855622416 | -3.65096124875989 | 0.000261260660925567 | 0.0135615028849465 | 5085 | 0 | 0 | 9 | 11 | 0 | 0 | 6 | 3 | 0 | 1 | 4 | 0 |
| L3b_dup2732 | 2.40571996580545 | -3.69321164686603 | 1.16817816066568 | -3.16151403203899 | 0.00156951256254714 | 0.0478671156380211 | 5086 | 0 | 1 | 3 | 16 | 0 | 0 | 8 | 4 | 0 | 0 | 1 | 0 |
| MLT1I_dup994 | 2.33271014243553 | -3.81333470381255 | 1.19603932672097 | -3.18830210563987 | 0.00143110930976404 | 0.0449837167531194 | 5087 | 0 | 1 | 4 | 16 | 0 | 0 | 3 | 5 | 0 | 0 | 2 | 0 |
| MIR_dup19179 | 3.0484335178809 | -3.42155517275058 | 1.03716386377342 | -3.29895332093644 | 0.000970460472527566 | 0.0344162667346355 | 5088 | 1 | 2 | 6 | 15 | 0 | 0 | 8 | 4 | 0 | 0 | 5 | 0 |
| MER44B_dup564 | 3.13111219234129 | -4.15618794006125 | 1.06740489674551 | -3.89373137853626 | 9.87139574748577e-05 | 0.00675736229151683 | 5089 | 0 | 1 | 6 | 15 | 0 | 0 | 12 | 6 | 0 | 0 | 2 | 0 |
| MLT2D_dup439 | 3.97608650980951 | -4.68832094383059 | 1.06328368938217 | -4.4092851142622 | 1.03712409787233e-05 | 0.00125651492154954 | 5090 | 0 | 1 | 9 | 19 | 0 | 0 | 12 | 7 | 0 | 0 | 5 | 0 |
| MIR_dup170890 | 3.18617110046207 | -4.17273198412201 | 1.07787569405969 | -3.87125529142039 | 0.000108276320163893 | 0.00721436221615462 | 5091 | 0 | 0 | 14 | 13 | 1 | 0 | 7 | 5 | 0 | 0 | 2 | 0 |
| LTR13_dup380 | 2.44031859272489 | -3.71128987162512 | 1.13117213394333 | -3.28092406121015 | 0.00103467597511764 | 0.0359149539861915 | 5092 | 0 | 0 | 6 | 15 | 1 | 0 | 6 | 4 | 0 | 0 | 1 | 0 |
| MIRb_dup217381 | 2.46906038581103 | -3.88439166831929 | 1.09728778995283 | -3.53999352210623 | 0.000400136853484726 | 0.018480388924549 | 5093 | 0 | 0 | 7 | 11 | 1 | 0 | 8 | 4 | 0 | 0 | 2 | 0 |
| OldhAT1_dup1146 | 3.0323682779202 | -3.96051464637557 | 1.08234181358063 | -3.65920876074564 | 0.000252995184797135 | 0.0132509778212061 | 5094 | 0 | 1 | 11 | 16 | 0 | 0 | 8 | 4 | 0 | 0 | 1 | 0 |
| L1ME3A_dup1502 | 3.26841945003463 | -4.75118960648535 | 1.10527681603521 | -4.29864223835668 | 1.71847602023523e-05 | 0.001827097833382 | 5095 | 0 | 0 | 11 | 18 | 0 | 0 | 8 | 5 | 0 | 0 | 2 | 0 |
| L1PA13_dup2347 | 3.43628362376128 | -4.8061702066151 | 1.11351738600816 | -4.31620580604019 | 1.58733993415627e-05 | 0.00172230363872662 | 5096 | 0 | 0 | 14 | 18 | 0 | 0 | 7 | 5 | 0 | 0 | 2 | 0 |
| MER65-int_dup715 | 3.12203019987992 | -4.7008954252321 | 1.12971454811687 | -4.16113560108443 | 3.16668976375179e-05 | 0.00287048061914546 | 5097 | 0 | 0 | 13 | 16 | 0 | 0 | 7 | 4 | 0 | 0 | 2 | 0 |
| Charlie1a_dup641 | 2.58373600718681 | -4.34669565197254 | 1.15247911587684 | -3.77160470163093 | 0.000162201071749961 | 0.00963501924333564 | 5098 | 0 | 0 | 11 | 12 | 0 | 0 | 8 | 3 | 0 | 0 | 1 | 0 |
| L1ME3A_dup1501 | 2.67656281647756 | -4.52916229979974 | 1.14808566919961 | -3.94496893507718 | 7.98103245270442e-05 | 0.00576869607530743 | 5099 | 0 | 0 | 12 | 11 | 0 | 0 | 8 | 3 | 0 | 0 | 2 | 0 |
| HERVI-int | 2.35890748892892 | -3.84183370866807 | 1.18124366301978 | -3.25236344451291 | 0.00114449575002717 | 0.0385660757328165 | 5100 | 1 | 0 | 5 | 16 | 0 | 0 | 7 | 2 | 0 | 0 | 2 | 0 |
| L2c_dup88665 | 2.14196747303612 | -3.57302736965327 | 1.12990389129356 | -3.16224007828022 | 0.0015656043141437 | 0.047790799760518 | 5101 | 1 | 0 | 7 | 10 | 0 | 0 | 7 | 3 | 0 | 0 | 1 | 0 |
| L1ME3A_dup4076 | 2.43901806521778 | -3.72997033690354 | 1.14198140333908 | -3.2662268632373 | 0.00108990894644697 | 0.0372664459746642 | 5102 | 1 | 0 | 7 | 12 | 0 | 0 | 11 | 2 | 0 | 0 | 1 | 0 |
| L2c_dup74403 | 2.26535838598419 | -3.79276604570267 | 1.14461616498375 | -3.31357022706081 | 0.000921129719082566 | 0.0332341367910266 | 5103 | 1 | 0 | 8 | 9 | 0 | 0 | 9 | 2 | 0 | 0 | 2 | 0 |
| L2b_dup94938 | 1.91087575459371 | -4.23661679354604 | 1.23290485830645 | -3.43628850596433 | 0.000589742598064077 | 0.024294822063531 | 5104 | 0 | 0 | 8 | 7 | 0 | 0 | 4 | 3 | 0 | 0 | 3 | 0 |
| LTR40b_dup653 | 1.8963799843173 | -4.13322198172644 | 1.21905747547308 | -3.39050624345866 | 0.000697636654408971 | 0.0272667298570954 | 5105 | 0 | 0 | 8 | 8 | 0 | 0 | 4 | 3 | 0 | 0 | 2 | 0 |
| MIRb_dup213209 | 2.09789496210971 | -4.26018210524561 | 1.30795399954431 | -3.2571345068174 | 0.0011254308954907 | 0.0381035543338688 | 5106 | 0 | 0 | 9 | 13 | 0 | 0 | 4 | 1 | 0 | 0 | 2 | 0 |
| MIRc_dup48425 | 1.86391648743727 | -4.12540671121155 | 1.30809249363556 | -3.15375765191182 | 0.00161182876734347 | 0.048782524532474 | 5107 | 0 | 0 | 6 | 12 | 0 | 0 | 5 | 1 | 0 | 0 | 2 | 0 |
| AluJb_dup60099 | 2.50362543089796 | -4.45619695184233 | 1.21084757974809 | -3.68022947427405 | 0.000233024158257571 | 0.0124544398833684 | 5108 | 0 | 0 | 9 | 16 | 0 | 0 | 4 | 3 | 0 | 0 | 2 | 0 |
| AluSq_dup1948 | 2.13741835433394 | -4.27591038498933 | 1.20580974675504 | -3.5460904147576 | 0.000390992031048873 | 0.018206455933455 | 5109 | 0 | 0 | 6 | 13 | 0 | 0 | 5 | 3 | 0 | 0 | 2 | 0 |
| AluSp_dup10975 | 1.95412914250689 | -4.03123254002531 | 1.23505613980831 | -3.26400752977186 | 0.00109848246029272 | 0.0374469208448155 | 5110 | 0 | 0 | 6 | 12 | 0 | 0 | 6 | 2 | 0 | 0 | 1 | 0 |
| SST1_dup439 | 2.17690552148137 | -4.15392573724336 | 1.21974850718558 | -3.40555918926931 | 0.000660286986012891 | 0.0262291564563979 | 5111 | 0 | 0 | 8 | 13 | 0 | 0 | 6 | 2 | 0 | 0 | 1 | 0 |
| THE1B_dup20503 | 2.25422191812512 | -4.19268436524581 | 1.23139658846458 | -3.40482051397724 | 0.000662075541603642 | 0.0262782696951493 | 5112 | 0 | 0 | 9 | 14 | 0 | 0 | 5 | 2 | 0 | 0 | 1 | 0 |
| MER54B_dup45 | 1.98773542960078 | -4.04648473918319 | 1.27938046646083 | -3.1628470539159 | 0.0015623438924753 | 0.0477290937039548 | 5113 | 0 | 0 | 9 | 12 | 0 | 0 | 3 | 2 | 0 | 0 | 1 | 0 |
| Charlie4z_dup3295 | 2.0182141587115 | -4.06721125529212 | 1.27347937543146 | -3.19377866164039 | 0.00140423737369455 | 0.044426627592225 | 5114 | 0 | 0 | 6 | 15 | 0 | 0 | 4 | 2 | 0 | 0 | 1 | 0 |
| L1PA15_dup1545 | 2.06699573643912 | -4.09040646339195 | 1.25376501451265 | -3.26249848739155 | 0.00110434762173003 | 0.0375768767625014 | 5115 | 0 | 0 | 7 | 14 | 0 | 0 | 3 | 3 | 0 | 0 | 1 | 0 |
| MLT1E_dup288 | 3.71179911576565 | -4.32176805737306 | 0.978026662100927 | -4.41886527724034 | 9.92204710838496e-06 | 0.00121571133947053 | 5116 | 0 | 1 | 6 | 7 | 0 | 0 | 14 | 6 | 0 | 1 | 8 | 6 |
| MLT1K_dup5703 | 3.26204923730028 | -4.63196164992509 | 1.05479571632106 | -4.39133528725407 | 1.1265666155996e-05 | 0.00133430572382012 | 5117 | 0 | 1 | 4 | 8 | 0 | 0 | 10 | 7 | 0 | 0 | 10 | 2 |
| Tigger2a_dup1197 | 4.0679173955877 | -4.9196508437542 | 1.02400014382654 | -4.80434585230641 | 1.55258259607662e-06 | 0.000291119636253781 | 5118 | 0 | 1 | 4 | 6 | 0 | 0 | 17 | 12 | 0 | 0 | 6 | 6 |
| L1MA1_dup39 | 3.1210979150571 | -4.04620167826348 | 1.04483299060186 | -3.87258223530321 | 0.000107688323518658 | 0.00718967615824088 | 5119 | 0 | 2 | 2 | 9 | 0 | 0 | 10 | 7 | 0 | 0 | 5 | 6 |
| L1PB1_dup2534 | 2.26209173657316 | -4.09864124478083 | 1.11254872921381 | -3.68401054008407 | 0.00022959274068067 | 0.0123042059175682 | 5120 | 0 | 1 | 1 | 5 | 0 | 0 | 9 | 5 | 0 | 0 | 3 | 6 |
| AluJr4_dup14032 | 2.88140283142991 | -3.88970590213187 | 1.03216830459331 | -3.76848028061127 | 0.000164244457473836 | 0.00970723091854679 | 5121 | 0 | 1 | 3 | 12 | 1 | 0 | 9 | 5 | 0 | 0 | 2 | 6 |
| L1M3_dup112 | 2.81145633085218 | -3.88613574378197 | 1.04045322182637 | -3.73504128994902 | 0.000187684390949074 | 0.0106381014980392 | 5122 | 0 | 1 | 2 | 7 | 1 | 0 | 6 | 8 | 0 | 0 | 4 | 7 |
| Tigger19b_dup1974 | 3.37765665904852 | -4.12813370863859 | 0.99450105650451 | -4.15095960093619 | 3.3108420828317e-05 | 0.00297733042419302 | 5123 | 1 | 0 | 5 | 14 | 0 | 0 | 12 | 5 | 0 | 1 | 3 | 5 |
| AluY_dup10038 | 2.90137995631548 | -3.3767110451712 | 1.02117118238259 | -3.30670420731291 | 0.000944005159341662 | 0.0337676265250618 | 5124 | 1 | 0 | 3 | 6 | 0 | 0 | 13 | 7 | 0 | 2 | 2 | 4 |
| L1PA3_dup877 | 3.22592303746721 | -3.48207152242176 | 0.961953184678415 | -3.61979312286993 | 0.000294838645969168 | 0.0147914139687497 | 5125 | 1 | 0 | 5 | 8 | 0 | 0 | 13 | 8 | 0 | 2 | 2 | 3 |
| L1MB7_dup10217 | 2.79147422538002 | -3.70505490752693 | 1.01843912906983 | -3.63797383836858 | 0.000274791329594436 | 0.0140493119277348 | 5126 | 0 | 0 | 9 | 6 | 0 | 1 | 6 | 8 | 0 | 1 | 2 | 2 |
| THE1D_dup6694 | 2.8160375930356 | -3.73322978457736 | 1.00801862584256 | -3.70353254282072 | 0.000212617891834656 | 0.0116142279362922 | 5127 | 0 | 0 | 9 | 6 | 0 | 1 | 11 | 5 | 0 | 1 | 2 | 2 |
| L1MA9_dup2613 | 2.22051056648264 | -3.46746321514216 | 1.07702611294091 | -3.21947924333418 | 0.00128423657727845 | 0.0417400569071301 | 5128 | 0 | 0 | 4 | 4 | 0 | 1 | 7 | 6 | 0 | 1 | 4 | 1 |
| THE1B_dup19667 | 3.1055524598819 | -3.10069539479224 | 0.951185226345529 | -3.25982291241545 | 0.00111481796223788 | 0.0378087905990101 | 5129 | 0 | 0 | 7 | 6 | 0 | 1 | 9 | 6 | 0 | 3 | 5 | 3 |
| L1PA2_dup613 | 3.32703016343643 | -3.63773318781478 | 1.03450386141518 | -3.51640368247486 | 0.000437435340255739 | 0.0197132555819114 | 5130 | 0 | 0 | 2 | 8 | 0 | 1 | 8 | 10 | 0 | 2 | 6 | 5 |

| | | | | | | | | | | | | | | | | | | |
|---|---|---|---|---|---|---|---|---|---|---|---|---|---|---|---|---|---|---|
| LTR84a_dup11 | 2.89665286491155 | -2.97488837850877 | 0.942164176413848 | -3.15750529788987 | 0.00159125345207027 | 0.0483880993382834 | 5131 | 1 | 0 | 3 | 6 | 0 | 1 | 6 | 8 | 0 | 2 | 3 | 7 |
| MIRc_dup92140 | 3.17683638216461 | -3.87332932371726 | 1.04389281428575 | -3.71046650643672 | 0.000206877667372387 | 0.0114054016312766 | 5132 | 0 | 0 | 3 | 9 | 1 | 0 | 14 | 9 | 0 | 1 | 3 | 1 |
| MIRc_dup68733 | 2.68132639750126 | -3.52519649505663 | 1.04845754666948 | -3.36226917938139 | 0.00077304725475756 | 0.0293328379960391 | 5133 | 0 | 1 | 2 | 7 | 1 | 0 | 11 | 9 | 0 | 0 | 2 | 1 |
| LTR53_dup167 | 2.51245733531205 | -4.16388468596451 | 1.07436341866541 | -3.8756761572699 | 0.000106329022609821 | 0.00713898578270779 | 5134 | 0 | 0 | 2 | 7 | 1 | 0 | 8 | 8 | 0 | 0 | 3 | 3 |
| MLT1D_dup8545 | 2.2640684649098 | -3.99245449148745 | 1.10820393235031 | -3.60263519641203 | 0.00031500746936318 | 0.0154673477724084 | 5135 | 0 | 0 | 2 | 4 | 1 | 0 | 10 | 7 | 0 | 0 | 2 | 3 |
| MSTA_dup6684 | 2.8818533162275 | -4.29935982339031 | 1.05425378009839 | -4.07810709769432 | 4.54038520331047e-05 | 0.00379947246968682 | 5136 | 0 | 0 | 3 | 14 | 1 | 0 | 8 | 7 | 0 | 0 | 3 | 2 |
| L1PA12_dup446 | 2.21812209644875 | -3.91596669948926 | 1.10624695125088 | -3.53986665912283 | 0.000400329242423167 | 0.0184856863802878 | 5137 | 0 | 0 | 3 | 11 | 1 | 0 | 4 | 6 | 0 | 0 | 1 | 3 |
| MER5A_dup34112 | 2.29591204412557 | -4.01149345051613 | 1.15279989246115 | -3.47978298467038 | 0.000501820089769063 | 0.0216315367898894 | 5138 | 0 | 0 | 1 | 10 | 1 | 0 | 4 | 8 | 0 | 0 | 3 | 2 |
| L2b_dup64689 | 1.95867136196192 | -4.14243651866466 | 1.24344365180684 | -3.33142278915922 | 0.00086403265230766 | 0.0317409449090447 | 5139 | 0 | 0 | 1 | 7 | 0 | 0 | 5 | 9 | 0 | 0 | 1 | 1 |
| L4_C_Mam_dup4085 | 1.84500364804897 | -4.08663794656186 | 1.13533142514799 | -3.59951099391895 | 0.000318816154890158 | 0.0156149145262801 | 5140 | 0 | 0 | 4 | 8 | 0 | 0 | 5 | 5 | 0 | 0 | 1 | 1 |
| LTR33B_dup834 | 1.93739313138346 | -4.23007981040496 | 1.14403237555412 | -3.69751757100064 | 0.00021771815366789 | 0.0118303639626992 | 5141 | 0 | 0 | 3 | 9 | 0 | 0 | 4 | 6 | 0 | 0 | 1 | 2 |
| AluJb_dup57588 | 1.99897806033477 | -4.38642399105998 | 1.1041620024713 | -3.97262718807787 | 7.10842584931455e-05 | 0.0053022617465235 1 | 5142 | 0 | 0 | 3 | 5 | 0 | 0 | 8 | 5 | 0 | 0 | 3 | 2 |
| L1ME3A_dup7930 | 2.08751338207288 | -4.39470610506034 | 1.1460430013363 | -3.83467819264729 | 0.000125728796269009 | 0.00801410773025061 | 5143 | 0 | 0 | 2 | 8 | 0 | 0 | 7 | 6 | 0 | 0 | 3 | 1 |
| MIR_dup46784 | 2.33565250500845 | -4.52539490461573 | 1.10977217007741 | -4.07776931755289 | 4.54698463877544e-05 | 0.00380026978172835 | 5144 | 0 | 0 | 3 | 9 | 0 | 0 | 7 | 7 | 0 | 0 | 3 | 1 |
| MER103C_dup4486 | 3.49037471902564 | -4.09029320823232 | 1.1005602603861 | -3.71655542677631 | 0.000201957352058472 | 0.0112016186696749 | 5145 | 0 | 1 | 5 | 24 | 0 | 0 | 3 | 7 | 0 | 1 | 1 | 5 |
| MIR_dup40556 | 2.91989851999003 | -3.76279575415732 | 1.04024005699146 | -3.61723789510655 | 0.000297763607520855 | 0.0148879301019452 | 5146 | 0 | 1 | 4 | 14 | 0 | 0 | 6 | 8 | 0 | 1 | 1 | 3 |
| L1MEj_dup201 | 2.80246215928398 | -3.15316103437638 | 0.95150439215419 | -3.31386913226715 | 0.000920145625109812 | 0.0332087017916547 | 5147 | 0 | 1 | 7 | 9 | 0 | 1 | 4 | 9 | 0 | 1 | 1 | 2 |
| L1ME4c_dup824 | 2.64256691439583 | -3.27874792610087 | 1.00241675529202 | -3.27084309873265 | 0.00107227383985027 | 0.0368704148984123 | 5148 | 0 | 1 | 4 | 8 | 0 | 1 | 5 | 5 | 0 | 1 | 2 | 8 |
| L1PB1_dup12631 | 2.72190892522569 | -3.81680810773777 | 1.044471309324 | -3.65429674675132 | 0.000257887864025366 | 0.0134421436349134 | 5149 | 0 | 0 | 6 | 9 | 0 | 1 | 3 | 7 | 0 | 1 | 2 | 6 |
| L1M4_dup9188 | 4.07992361094542 | -4.9301422985976 | 0.991761017360599 | -4.97109909776281 | 6.65744081912344e-07 | 0.00014944447293841 | 5150 | 0 | 0 | 8 | 15 | 0 | 0 | 10 | 9 | 0 | 1 | 2 | 9 |
| LTR33C_dup339 | 4.03385582509791 | -4.889277028702 | 0.983725950143268 | -4.97016168780537 | 6.68970918431119e-07 | 0.000149886020863475 | 5151 | 0 | 0 | 7 | 18 | 0 | 0 | 14 | 6 | 0 | 1 | 3 | 6 |
| L1MC5a_dup12192 | 4.28701252220397 | -4.52197237179315 | 1.00341563744385 | -4.50657953000677 | 6.58809994920808e-06 | 0.000887662531435032 | 5152 | 0 | 0 | 7 | 17 | 0 | 0 | 6 | 11 | 0 | 2 | 7 | 5 |
| L1MEc_dup9514 | 3.230001106591 | -4.57925218477378 | 1.02026454731111 | -4.48829884057258 | 7.17941714072961e-06 | 0.00094959065645806 | 5153 | 0 | 0 | 4 | 13 | 0 | 0 | 8 | 8 | 0 | 1 | 5 | 3 |
| MLT2B3_dup1752 | 3.04734433186538 | -4.29451516700764 | 1.02937717122913 | -4.17195493259266 | 3.01997377571578e-05 | 0.00276806887533308 | 5154 | 0 | 0 | 6 | 11 | 0 | 0 | 9 | 9 | 0 | 1 | 2 | 1 |
| MIR_dup133231 | 3.03123977340767 | -4.29093166528528 | 1.02722035637081 | -4.17722608267345 | 2.95085691211377e-05 | 0.00272149069306133 | 5155 | 0 | 0 | 11 | 10 | 0 | 0 | 7 | 7 | 0 | 1 | 2 | 1 |
| L1MC1_dup12814 | 2.92006559852815 | -4.15972318186567 | 1.05873528488854 | -3.9289548966941 | 8.53158379301716e-05 | 0.0060653009776315 | 5156 | 0 | 0 | 11 | 10 | 0 | 0 | 5 | 8 | 0 | 1 | 1 | 1 |
| MLT1A0_dup16491 | 2.77663490093861 | -4.09824130751868 | 1.05459709728367 | -3.88607300178859 | 0.000101878881288634 | 0.00690247428502043 | 5157 | 0 | 0 | 10 | 8 | 0 | 0 | 6 | 8 | 0 | 1 | 1 | 1 |
| MER117_dup1990 | 2.91157487421271 | -4.29187186066396 | 1.06642772336728 | -4.0245314020084 | 5.70888764462904e-05 | 0.00450096777238542 | 5158 | 0 | 0 | 9 | 4 | 0 | 0 | 8 | 10 | 0 | 1 | 1 | 3 |
| L1PA15_dup2775 | 2.114419562139 | -3.85129699322457 | 1.07759708351843 | -3.57396753585285 | 0.000351612543721097 | 0.0167765787704552 | 5159 | 0 | 0 | 5 | 5 | 0 | 0 | 7 | 6 | 0 | 1 | 1 | 2 |
| MER4CL34_dup428 | 2.28549203589749 | -4.00255878404098 | 1.06145579102406 | -3.77082005476594 | 0.000162711973155704 | 0.00965502822057533 | 5160 | 0 | 0 | 4 | 7 | 0 | 0 | 6 | 7 | 0 | 1 | 2 | 2 |
| AluJr_dup38655 | 3.18897367600242 | -4.49317571992259 | 1.06821625486702 | -4.20624166637675 | 2.59652576358428e-05 | 0.00247727714250932 | 5161 | 0 | 0 | 7 | 11 | 0 | 0 | 4 | 11 | 0 | 1 | 1 | 5 |
| L3_dup44827 | 2.41253013627193 | -4.15727365349534 | 1.04651653064377 | -3.97248732510509 | 7.11260219424042e-05 | 0.00530345863762657 | 5162 | 0 | 0 | 4 | 10 | 0 | 0 | 6 | 5 | 0 | 1 | 2 | 4 |
| L1PA16_dup1031 | 2.67672076319202 | -4.29392310950825 | 1.0460626011067 | -4.10484334777425 | 4.04588951095012e-05 | 0.00346920081287401 | 5163 | 0 | 0 | 8 | 10 | 0 | 0 | 7 | 4 | 0 | 1 | 1 | 5 |
| MIR_dup124589 | 2.93325724825257 | -3.94563694277832 | 1.01881735059867 | -3.87276182571864 | 0.000107608975108938 | 0.00718648085006103 | 5164 | 0 | 0 | 7 | 10 | 0 | 0 | 8 | 5 | 0 | 2 | 2 | 5 |
| L1MB3_dup12790 | 2.49593460897331 | -3.54171948433656 | 1.10591103187692 | -3.20253563103142 | 0.00136223447087102 | 0.0435084890477686 | 5165 | 0 | 0 | 5 | 11 | 0 | 0 | 5 | 7 | 0 | 2 | 1 | 1 |
| AluSx_dup51629 | 1.95369891462795 | -3.75813642702604 | 1.1315357415499 | -3.32127063160938 | 0.000896086017559474 | 0.0325544037700857 | 5166 | 0 | 0 | 3 | 9 | 0 | 0 | 3 | 6 | 0 | 1 | 1 | 2 |
| L1PA16_dup7477 | 2.16233147473568 | -3.81478042283294 | 1.13289568868624 | -3.36728302608047 | 0.00075912736059431 | 0.0289689233821965 | 5167 | 0 | 0 | 4 | 12 | 0 | 0 | 3 | 6 | 0 | 1 | 1 | 1 |
| L2a_dup124315 | 2.50757206511826 | -4.20135327009113 | 1.07210956531514 | -3.91877230276943 | 8.90011423475588e-05 | 0.00626552243107462 | 5168 | 0 | 0 | 11 | 6 | 0 | 0 | 3 | 5 | 0 | 1 | 3 | 3 |
| THE1D_dup10833 | 2.50669585176746 | -4.18963079440673 | 1.06428856043477 | -3.93655531982344 | 8.26595721459993e-05 | 0.00592782234896081 | 5169 | 0 | 0 | 6 | 6 | 0 | 0 | 4 | 8 | 0 | 1 | 4 | 2 |
| L1ME3C_dup2770 | 3.06813256477003 | -2.79666775632315 | 0.835258871272924 | -3.34826465483816 | 0.000813193148687914 | 0.030395233578023 | 5170 | 2 | 2 | 5 | 8 | 1 | 0 | 8 | 7 | 0 | 0 | 2 | 5 |
| L1PA2_dup242 | 3.15195640917273 | -2.86317939033414 | 0.874876795776436 | -3.27266582466977 | 0.00106538357317413 | 0.0367077555317828 | 5171 | 2 | 3 | 4 | 8 | 0 | 0 | 7 | 7 | 0 | 0 | 4 | 6 |
| L1MA2_dup608 | 3.06739869201711 | -3.72610294680351 | 0.953723672025683 | -3.90689992929437 | 9.34878486889589e-05 | 0.00650060188315109 | 5172 | 1 | 1 | 10 | 12 | 0 | 0 | 9 | 5 | 0 | 0 | 1 | 2 |
| LTR16E1_dup635 | 3.21480258454242 | -3.78362233464066 | 0.942471465938014 | -4.01457494617615 | 5.95530188513472e-05 | 0.00464755743640625 | 5173 | 1 | 1 | 8 | 13 | 0 | 0 | 11 | 6 | 0 | 0 | 1 | 2 |
| AluSz6_dup11624 | 2.96607831816437 | -2.83741513105401 | 0.89565115241557 | -3.16799138079765 | 0.00153496071148123 | 0.04726773151988 | 5174 | 1 | 3 | 8 | 13 | 0 | 0 | 9 | 4 | 0 | 0 | 1 | 1 |
| L2b_dup26249 | 2.59975028138109 | -3.00218628130739 | 0.940306414490494 | -3.19277443505916 | 0.0014091297318616 | 0.0445115856757183 | 5175 | 1 | 2 | 6 | 8 | 0 | 0 | 12 | 4 | 0 | 0 | 1 | 1 |
| L2a_dup19337 | 3.01683932069019 | -3.79119927135996 | 1.00443115169926 | -3.77447400446127 | 0.000160345626728757 | 0.00955512158717775 | 5176 | 1 | 1 | 5 | 14 | 0 | 0 | 14 | 3 | 0 | 0 | 2 | 2 |
| L1ME2z_dup2134 | 3.09643731582581 | -3.99716818099975 | 0.988907667288193 | -4.04200342784366 | 5.29964577322373e-05 | 0.00427178019361617 | 5177 | 1 | 1 | 6 | 11 | 0 | 0 | 11 | 4 | 0 | 0 | 4 | 4 |
| MSTA-int_dup1069 | 3.74804698264534 | -3.41755311591172 | 0.851355731741657 | -4.0142480851339 | 5.96355973399793e-05 | 0.004652476014642 2 | 5178 | 2 | 2 | 10 | 12 | 0 | 0 | 14 | 4 | 0 | 0 | 3 | 4 |
| L1MB8_dup2579 | 3.6700469858548 | -3.67383181043929 | 0.928644506998517 | -3.956122911138 | 7.61759786993354e-05 | 0.00557967784924931 | 5179 | 0 | 3 | 12 | 13 | 0 | 0 | 7 | 7 | 0 | 0 | 2 | 4 |
| MamGypsy2-I_dup2187 | 3.80805878577161 | -4.71894355053919 | 1.00915718941435 | -4.67612340281475 | 2.9234872361326e-06 | 0.000472255325111207 | 5180 | 0 | 1 | 14 | 13 | 0 | 0 | 6 | 8 | 0 | 0 | 5 | 2 |
| MER102c_dup3278 | 2.90999457362674 | -4.1648426098883 | 1.04554390992166 | -3.98342199726489 | 6.79299747060435e-05 | 0.00512746329078604 | 5181 | 0 | 1 | 11 | 12 | 0 | 0 | 5 | 6 | 0 | 0 | 1 | 2 |
| MSTA_dup10340 | 2.1992003213103 | -3.73700581336104 | 1.11560142473068 | -3.34976787454642 | 0.000808793084198461 | 0.0302825264335606 | 5182 | 0 | 1 | 8 | 11 | 0 | 0 | 3 | 4 | 0 | 0 | 1 | 1 |
| Eulor2B_dup45 | 2.26417183204223 | -3.76431513204907 | 1.07450260832352 | -3.50330944093499 | 0.00045951530895897 | 0.02036502695204 9 | 5183 | 0 | 1 | 7 | 8 | 0 | 0 | 5 | 6 | 0 | 0 | 1 | 1 |
| Tigger3d_dup56 | 2.30164911012889 | -3.87844075417522 | 1.09086529797867 | -3.55538008346386 | 0.000377433063360224 | 0.0177032697997855 | 5184 | 0 | 1 | 7 | 8 | 0 | 0 | 3 | 7 | 0 | 0 | 1 | 2 |
| MIR_dup19180 | 2.54573001460939 | -3.38688654726832 | 0.995626770660032 | -3.40176323806867 | 0.000669526141443189 | 0.0264900344615632 | 5185 | 1 | 1 | 11 | 7 | 0 | 0 | 6 | 5 | 0 | 0 | 1 | 1 |
| L1MC1_dup12813 | 2.9616950199809 | -3.81198809481605 | 0.981256544597668 | -3.88480272137092 | 0.000102413013899138 | 0.00693082110501828 | 5186 | 1 | 1 | 8 | 12 | 0 | 0 | 6 | 6 | 0 | 0 | 2 | 3 |
| MIR_dup149092 | 2.41331383422822 | -3.41397989896142 | 1.03090210135376 | -3.31164316619225 | 0.000927497683637551 | 0.0333879521680516 | 5187 | 1 | 1 | 8 | 9 | 0 | 0 | 3 | 6 | 0 | 0 | 1 | 2 |
| Charlie5_dup1230 | 3.04227922428746 | -2.95852345001113 | 0.88953485136513 | -3.32592190791717 | 0.000881266151872025 | 0.0322412843617357 | 5188 | 1 | 2 | 8 | 14 | 0 | 0 | 5 | 5 | 1 | 0 | 2 | 2 |
| THE1D_dup982 | 3.11210733092995 | -3.38090465526801 | 0.935796715173618 | -3.61286228135643 | 0.000302835564725474 | 0.0150561034860359 | 5189 | 1 | 1 | 11 | 13 | 0 | 0 | 8 | 3 | 1 | 0 | 1 | 3 |
| MIR_dup80876 | 3.12065232020969 | -3.39186856690276 | 0.967459359169609 | -3.50595457551217 | 0.000454972859464254 | 0.0202580411710482 | 5190 | 1 | 2 | 7 | 19 | 0 | 0 | 4 | 5 | 0 | 0 | 2 | 2 |
| L1ME3A_dup13442 | 2.86749941606749 | -3.21504296368209 | 0.94522105362832 | -3.40136622152125 | 0.000670499371652109 | 0.0265017036927055 | 5191 | 1 | 2 | 8 | 14 | 0 | 0 | 5 | 5 | 0 | 0 | 2 | 1 |
| MER5A_dup30817 | 3.079073699344 | -3.31051219294713 | 0.928393699039344 | -3.56584948430034 | 0.000362679491169178 | 0.0171908633875979 | 5192 | 1 | 2 | 9 | 15 | 0 | 0 | 6 | 5 | 0 | 0 | 2 | 1 |
| L1ME1_dup3215 | 2.97801246142434 | -3.32881620718806 | 1.02381227123582 | -3.25139315156861 | 0.00114840932323532 | 0.0386560199217078 | 5193 | 1 | 2 | 8 | 20 | 0 | 0 | 3 | 3 | 0 | 0 | 1 | 3 |
| Tigger3b_dup1716 | 2.52008115877078 | -3.19140131185505 | 1.00871828545259 | -3.16381824130722 | 0.0015571400789907 | 0.0476485665169136 | 5194 | 1 | 2 | 8 | 10 | 0 | 0 | 4 | 3 | 0 | 0 | 1 | 5 |
| L2_dup16033 | 2.29987786474944 | -3.90068070988989 | 1.15854911754099 | -3.36686692936165 | 0.000760273649286574 | 0.0290024180302933 | 5195 | 0 | 1 | 6 | 17 | 0 | 0 | 3 | 2 | 0 | 0 | 1 | 2 |
| MER68_dup1388 | 2.44757603287123 | -3.61167444121975 | 1.11162418871673 | -3.24900670377558 | 0.00115808748896064 | 0.0388827833566008 | 5196 | 0 | 2 | 10 | 10 | 0 | 0 | 3 | 2 | 0 | 0 | 2 | 4 |
| L2c_dup70106 | 1.78318019525168 | -3.59104341024469 | 1.13958325698582 | -3.1511900409479 | 0.001626066513044 | 0.0490867014268497 | 5197 | 0 | 1 | 7 | 8 | 0 | 0 | 3 | 2 | 0 | 0 | 1 | 2 |
| AluSz_dup10745 | 2.0535208453724 | -3.88829662493437 | 1.10186945322188 | -3.52881787725846 | 0.00041742029227462 | 0.0190713304273381 | 5198 | 0 | 1 | 7 | 8 | 0 | 0 | 5 | 2 | 0 | 0 | 1 | 4 |
| HERVL74-int_dup139 | 2.10549132745144 | -3.87020319999227 | 1.10666913115012 | -3.49716377827409 | 0.0004702330461041 | 0.0206782613912929 | 5199 | 0 | 1 | 6 | 11 | 0 | 0 | 5 | 2 | 0 | 0 | 1 | 3 |
| BSR/Beta_dup2005 | 3.13555278485587 | -4.44857854652588 | 1.06310391204182 | -4.18451902597352 | 2.85770513408261e-05 | 0.0026635210471979 | 5200 | 0 | 1 | 8 | 18 | 0 | 0 | 6 | 4 | 0 | 0 | 1 | 5 |
| L1MEc_dup9513 | 2.7227888918922 | -3.62951477730514 | 1.02412427289585 | -3.54401792181159 | 0.00039407845669614 | 0.0182750716150886 | 5201 | 0 | 2 | 5 | 12 | 0 | 0 | 7 | 6 | 0 | 0 | 1 | 3 |
| MER20B_dup391 | 2.84206882028885 | -4.33085495822972 | 1.04494763770935 | -4.14456648538242 | 3.40457170642992e-05 | 0.00303631442785359 | 5202 | 0 | 1 | 13 | 8 | 0 | 0 | 7 | 3 | 0 | 0 | 2 | 4 |
| LTR88b_dup130 | 3.08449061348214 | -4.39100024266391 | 1.02161706164541 | -4.29808820497947 | 1.72277609409835e-05 | 0.00183085163236412 | 5203 | 0 | 1 | 7 | 15 | 0 | 0 | 10 | 4 | 0 | 0 | 3 | 2 |
| AluSx1_dup11539 | 3.02367094487636 | -3.76881867990285 | 1.00814829393633 | -3.73835744460514 | 0.000185226472922795 | 0.0105389522539664 | 5204 | 0 | 2 | 9 | 13 | 0 | 0 | 7 | 5 | 0 | 0 | 3 | 1 |
| AluYc_dup3233 | 2.70701752856788 | -4.09322626873853 | 1.04074736885567 | -3.93296816425215 | 8.39033535719352e-05 | 0.00599353568568829 | 5205 | 0 | 1 | 10 | 9 | 0 | 0 | 9 | 4 | 0 | 0 | 2 | 1 |
| MLT1L_dup8433 | 3.25070761864245 | -3.17422543608827 | 0.967218659706174 | -3.2818074840001 | 0.00103143999390002 | 0.0358422300417366 | 5206 | 0 | 1 | 3 | 11 | 0 | 0 | 9 | 3 | 3 | 0 | 7 | 6 |
| AluSx3_dup9814 | 2.27540664166366 | -3.54388053716437 | 1.05956146697339 | -3.34466724926055 | 0.000823813426283506 | 0.0306820887888357 | 5207 | 0 | 1 | 4 | 7 | 0 | 0 | 10 | 2 | 1 | 0 | 2 | 4 |
| L3_dup34601 | 2.45835145514666 | -3.69548090211615 | 1.0571046257397 | -3.49585160459425 | 0.000472551421716179 | 0.0207457049062136 | 5208 | 0 | 1 | 8 | 7 | 0 | 0 | 6 | 1 | 1 | 0 | 5 | 4 |
| FLAM_A_dup10035 | 2.5143791612817 | -3.73462532277548 | 1.03127002119605 | -3.62138455110342 | 0.000293030566321414 | 0.0147224347833436 | 5209 | 0 | 0 | 6 | 8 | 1 | 0 | 6 | 2 | 1 | 0 | 3 | 7 |

| | | | | | | | | | | | | | | | | | | | |
|---|---|---|---|---|---|---|---|---|---|---|---|---|---|---|---|---|---|---|---|
| L1MC4a_dup10261 | 2.82970306460954 | -3.31064398345812 | 1.02888506913168 | -3.21770048257392 | 0.0012922268247103 | 0.0418967439749989 | 5210 | 0 | 0 | 11 | 10 | 1 | 0 | 4 | 3 | 2 | 0 | 2 | 4 |
| L1M5_dup28723 | 2.15894111711693 | -3.49986645651942 | 1.10688477694076 | -3.16190675798474 | 0.00156739743646555 | 0.0478271288044668 | 5211 | 0 | 0 | 3 | 7 | 0 | 0 | 6 | 2 | 2 | 0 | 2 | 7 |
| MIRb_dup159972 | 2.66665366036755 | -4.33922567094248 | 1.03138745624145 | -4.20717320603779 | 2.58584948338564e-05 | 0.00246907119251075 | 5212 | 0 | 0 | 8 | 6 | 0 | 0 | 5 | 4 | 1 | 0 | 3 | 8 |
| MSTA_dup18218 | 2.75250886942838 | -4.3516777971761 | 1.01918387430139 | -4.26976712142252 | 1.95677170221719e-05 | 0.00201212618606643 | 5213 | 0 | 0 | 7 | 6 | 0 | 0 | 4 | 3 | 1 | 0 | 8 | 7 |
| MIR_dup167807 | 2.4179837502749 | -3.5593889079909 | 1.07193318471174 | -3.32053243500252 | 0.000898459185086657 | 0.0326068405930723 | 5214 | 1 | 0 | 5 | 11 | 0 | 0 | 9 | 2 | 1 | 0 | 3 | 1 |
| L1MA7_dup5207 | 3.62907221669492 | -4.10671759019378 | 1.0110018159873 | -4.06202790662975 | 4.86482597671019e-05 | 0.00398748150828206 | 5215 | 1 | 0 | 5 | 21 | 0 | 0 | 10 | 6 | 1 | 0 | 3 | 2 |
| L2a_dup18278 | 3.46760987984947 | -3.99986874535475 | 1.01147964142444 | -3.95447281541114 | 7.67035919711683e-05 | 0.00559856223778271 | 5216 | 1 | 0 | 9 | 19 | 0 | 0 | 9 | 4 | 1 | 0 | 3 | 1 |
| L1MEc_dup9512 | 3.33808802800042 | -3.5964323686379 | 0.951576738685242 | -3.77944544294657 | 0.000157178025973868 | 0.00942299888344116 | 5217 | 1 | 1 | 7 | 4 | 0 | 0 | 10 | 9 | 1 | 0 | 4 | 5 |
| MLT2C1_dup1307 | 2.44841297563215 | -3.58046009901002 | 1.04860492398785 | -3.41449865159273 | 0.000638995098313221 | 0.025653168156443 | 5218 | 1 | 0 | 5 | 5 | 0 | 0 | 7 | 7 | 1 | 0 | 1 | 4 |
| L2a_dup124341 | 3.00065330976281 | -3.9619926778568 | 0.989067501818355 | -4.00578592520011 | 6.1811612251627e-05 | 0.0047861457496992 | 5219 | 1 | 0 | 7 | 10 | 0 | 0 | 9 | 4 | 1 | 0 | 4 | 4 |
| THE1D-int_dup1538 | 2.95017962115906 | -2.93650854804878 | 0.932038260986448 | -3.15063090322156 | 0.00162918231380325 | 0.0491576737053295 | 5220 | 2 | 0 | 5 | 6 | 0 | 0 | 11 | 6 | 2 | 0 | 3 | 3 |
| L1PB1_dup12905 | 4.18980277898297 | -3.45389396901268 | 0.963573862384633 | -3.58446207794082 | 0.00033777363456707 | 0.0162892310003661 | 5221 | 0 | 0 | 12 | 15 | 3 | 0 | 9 | 8 | 1 | 0 | 0 | 7 |
| MIRb_dup104839 | 4.13677385392589 | -4.30799132226578 | 1.00217281561554 | -4.29865114593016 | 1.71840696861883e-05 | 0.001827097833382 | 5222 | 0 | 0 | 10 | 13 | 1 | 0 | 7 | 13 | 1 | 0 | 1 | 6 |
| LTR33B_dup1368 | 4.01934604433647 | -3.00682833037272 | 0.792000500693149 | -3.79649801703557 | 0.000146754536036482 | 0.00896525156142148 | 5223 | 0 | 1 | 10 | 14 | 1 | 1 | 14 | 6 | 2 | 0 | 2 | 2 |
| UCON31_dup15 | 3.62361409459308 | -2.89932430587359 | 0.8217351780816 | -3.52829522601464 | 0.000418245407291852 | 0.0190797204147751 | 5224 | 0 | 2 | 9 | 18 | 1 | 1 | 9 | 4 | 1 | 0 | 1 | 3 |
| L1PREC2_dup528 | 3.34490175349631 | -2.66819841589711 | 0.827458327963756 | -3.22457134785642 | 0.00126161415011689 | 0.0412470514651984 | 5225 | 0 | 2 | 4 | 13 | 1 | 1 | 13 | 7 | 1 | 0 | 1 | 1 |
| LTR16B1_dup1178 | 4.17604008478441 | -3.15636325729686 | 0.783414511684793 | -4.02898237168069 | 5.60188183431644e-05 | 0.00443575831929537 | 5226 | 0 | 1 | 6 | 14 | 2 | 1 | 13 | 10 | 1 | 0 | 3 | 3 |
| L1ME1_dup10300 | 3.51139808891497 | -3.39088548049731 | 0.904927413713068 | -3.74713532722359 | 0.000178865599793792 | 0.0103002532427594 | 5227 | 0 | 1 | 13 | 16 | 1 | 0 | 9 | 4 | 1 | 0 | 1 | 1 |
| L1PA8_dup7221 | 4.96728104128655 | -2.74059563678901 | 0.799796564621901 | -3.4266159146165 | 0.000611152853757154 | 0.0248923959157994 | 5228 | 0 | 3 | 13 | 29 | 3 | 0 | 10 | 6 | 1 | 0 | 1 | 1 |
| L1MC4_dup24223 | 3.2439963793847 | -2.77338763710014 | 0.861282088076921 | -3.22006886651107 | 0.00128159805253866 | 0.0417098372636057 | 5229 | 0 | 1 | 9 | 13 | 3 | 0 | 6 | 4 | 1 | 0 | 2 | 4 |
| L2b_dup23281 | 3.01735255568623 | -2.9388413650681 | 0.885216649952363 | -3.3199119845139 | 0.000900458324622297 | 0.0326574534107386 | 5230 | 0 | 1 | 11 | 9 | 2 | 0 | 9 | 3 | 1 | 0 | 2 | 2 |
| MER77_dup108 | 3.77272382976375 | -2.90499147702087 | 0.855837096625074 | -3.39432759864754 | 0.000687973412838191 | 0.026982119942508 | 5231 | 0 | 2 | 6 | 19 | 1 | 0 | 9 | 6 | 2 | 0 | 2 | 3 |
| MIRc_dup34861 | 4.07630042473403 | -3.09246907257962 | 0.805210483695874 | -3.84057229158934 | 0.000122747817422899 | 0.00788109880315953 | 5232 | 0 | 2 | 12 | 12 | 1 | 0 | 10 | 6 | 2 | 0 | 4 | 4 |
| L1ME3Cz_dup10670 | 3.6694639551365 | -3.32851287264716 | 0.956029874041398 | -3.48159922929671 | 0.000498429068483441 | 0.0215204505769118 | 5233 | 0 | 2 | 8 | 18 | 2 | 0 | 9 | 5 | 0 | 0 | 0 | 6 |
| L1PA7_dup1989 | 4.57278782849187 | -4.18824269420141 | 0.921100457598713 | -4.54699882043275 | 5.44163031008765e-06 | 0.000768016483563513 | 5234 | 0 | 1 | 16 | 17 | 1 | 0 | 18 | 6 | 0 | 0 | 0 | 3 |
| THE1D_dup2100 | 3.8061600012008 | -3.35833620029197 | 0.883905371610727 | -3.7994295635653 | 0.000145029523415757 | 0.00888267567018937 | 5235 | 0 | 2 | 15 | 15 | 1 | 0 | 9 | 7 | 0 | 0 | 0 | 1 |
| AluJr_dup56476 | 3.34627051408234 | -3.95307029646257 | 0.965096731806058 | -4.09603531561535 | 4.2028584466634e-05 | 0.00357928903346374 | 5236 | 0 | 1 | 10 | 13 | 1 | 0 | 10 | 5 | 0 | 0 | 1 | 4 |
| MER46C_dup977 | 3.92210970740349 | -4.14518712192041 | 0.974801594340801 | -4.25233929241114 | 2.11549021977714e-05 | 0.00212662626039329 | 5237 | 0 | 1 | 14 | 13 | 1 | 0 | 17 | 3 | 0 | 0 | 1 | 4 |
| CR1_Mam_dup899 | 3.71416664566722 | -3.65850209653785 | 0.875616969530706 | -4.17819917137815 | 2.93826300254288e-05 | 0.00271092643816622 | 5238 | 0 | 1 | 9 | 14 | 2 | 0 | 11 | 7 | 0 | 0 | 3 | 2 |
| AluSg_dup16690 | 4.14663847356535 | -3.80080866816623 | 0.915071264378853 | -4.15356575615584 | 3.27334091509439e-05 | 0.00295360591946033 | 5239 | 0 | 2 | 13 | 23 | 1 | 0 | 7 | 5 | 0 | 0 | 3 | 2 |
| L1PA6_dup5357 | 3.92301529444885 | -3.80360878769554 | 0.910926135084761 | -4.17554030035774 | 2.97279616751796e-05 | 0.00273960091741374 | 5240 | 0 | 1 | 15 | 15 | 2 | 0 | 7 | 5 | 0 | 0 | 4 | 3 |
| MLT1L-int_dup52 | 4.16177817852641 | -4.2238476225856 | 0.933540908454367 | -4.52454475677867 | 6.05256440998991e-06 | 0.000834888980347513 | 5241 | 0 | 1 | 15 | 17 | 1 | 0 | 12 | 5 | 0 | 0 | 3 | 2 |
| L1M5_dup28502 | 5.03642189063759 | -2.87457621463736 | 0.783458785548789 | -3.66908415306595 | 0.00024342094796205 | 0.0128656243633447 | 5242 | 2 | 3 | 5 | 17 | 0 | 0 | 22 | 9 | 2 | 0 | 3 | 4 |
| L1MA4_dup2916 | 5.35110542418522 | -3.3826657833832 | 0.796027045916022 | -4.24943574560412 | 2.14309670689348e-05 | 0.0021471212169199 | 5243 | 0 | 2 | 9 | 19 | 0 | 1 | 21 | 12 | 2 | 0 | 1 | 3 |
| L1MB7_dup16638 | 6.49602820100637 | -4.2158953810087 | 0.944380375997252 | -4.4641920651483 | 8.0371445702514e-06 | 0.00103485194908714 | 5244 | 0 | 1 | 9 | 16 | 0 | 0 | 38 | 16 | 2 | 0 | 2 | 1 |
| HAL1_dup12457 | 4.80305522225556 | -2.95762871625285 | 0.869138470770014 | -3.40294304730584 | 0.000666641750264014 | 0.0264022533136353 | 5245 | 1 | 2 | 7 | 10 | 0 | 0 | 20 | 13 | 0 | 3 | 1 | 5 |
| MER4E_dup468 | 4.44110178645282 | -2.81439252688493 | 0.828916544630784 | -3.39526644161568 | 0.000685618429006116 | 0.0269474996214551 | 5246 | 3 | 1 | 6 | 11 | 0 | 0 | 18 | 13 | 1 | 1 | 0 | 3 |
| L2a_dup81494 | 3.44181654785111 | -3.06362817070825 | 0.955621523065717 | -3.20590118238429 | 0.00134640216860876 | 0.0431350699295524 | 5247 | 0 | 0 | 4 | 6 | 1 | 0 | 14 | 11 | 1 | 2 | 2 | 2 |
| L2b_dup23282 | 3.68575686969095 | -2.84311445395777 | 0.859325480373092 | -3.30854201218772 | 0.00093783112885625 | 0.0336174139080927 | 5248 | 0 | 0 | 6 | 12 | 2 | 0 | 12 | 8 | 1 | 2 | 1 | 4 |
| ALR/Alpha_dup1437 | 5.79213271157258 | -3.04312926135571 | 0.808001666336038 | -3.76624131872782 | 0.000165723618862812 | 0.00978007743401057 | 5249 | 0 | 2 | 8 | 19 | 1 | 0 | 19 | 19 | 1 | 2 | 1 | 1 |
| L1PA3_dup9562 | 5.14401930579422 | -2.33883063146868 | 0.737460970482434 | -3.17146360971301 | 0.00151672869856195 | 0.0468535664540892 | 5250 | 1 | 3 | 8 | 9 | 2 | 0 | 18 | 16 | 2 | 1 | 1 | 3 |
| L1PA2_dup760 | 5.31437043488487 | -4.137012581731 | 0.922584076826738 | -4.4841577972605 | 7.32024782813182e-06 | 0.000964464944469908 | 5251 | 0 | 3 | 5 | 17 | 0 | 0 | 17 | 19 | 0 | 0 | 3 | 3 |
| L1M4_dup14164 | 3.89213779204871 | -4.02229701158155 | 0.969150739125378 | -4.15033167617611 | 3.31993840197566e-05 | 0.00298286173413617 | 5252 | 0 | 1 | 11 | 10 | 0 | 0 | 8 | 14 | 0 | 1 | 1 | 2 |
| L1PA13_dup6978 | 5.16346797125604 | -4.43374422383584 | 0.941308393684979 | -4.71019301812329 | 2.4748230129993e-06 | 0.000416165538269351 | 5253 | 0 | 1 | 12 | 26 | 0 | 0 | 9 | 14 | 0 | 1 | 1 | 3 |
| HERVL-int_dup376 | 4.87811964649732 | -4.61786915545068 | 0.966376997047704 | -4.77853795108777 | 1.76574438106613e-06 | 0.000320122812288222 | 5254 | 0 | 1 | 7 | 18 | 0 | 0 | 14 | 12 | 1 | 0 | 7 | 3 |
| MSTA_dup14710 | 5.13903414462414 | -4.15679794796879 | 0.917478204016321 | -4.53067760059273 | 5.87948044376696e-06 | 0.000814318308307668 | 5255 | 0 | 1 | 9 | 24 | 0 | 0 | 12 | 11 | 2 | 0 | 5 | 3 |
| AluJb_dup122075 | 3.31506481151804 | -3.85253246143177 | 0.986648346091611 | -3.90466621334007 | 9.43555243779741e-05 | 0.00653230131034083 | 5256 | 0 | 1 | 7 | 17 | 0 | 0 | 9 | 6 | 1 | 0 | 2 | 1 |
| L1PA7_dup2459 | 3.74670342556413 | -4.11978475249102 | 0.974460094613356 | -4.2277613780846 | 2.36027912320749e-05 | 0.0023130980917055 | 5257 | 0 | 1 | 7 | 14 | 0 | 0 | 15 | 7 | 1 | 0 | 1 | 4 |
| L1P1_dup1629 | 3.12704173026385 | -3.02074082790246 | 0.92400209386315 | -3.26919262192695 | 0.00107854850952659 | 0.0369899544029713 | 5258 | 0 | 2 | 4 | 5 | 0 | 0 | 9 | 9 | 1 | 1 | 4 | 4 |
| L1P4b_dup56 | 3.65977214746986 | -3.69788575951129 | 0.914763568579519 | -4.04244975043498 | 5.28956506973072e-05 | 0.00426686322936619 | 5259 | 0 | 1 | 7 | 10 | 0 | 0 | 14 | 7 | 1 | 1 | 3 | 4 |
| AluSg4_dup3423 | 4.52479862642631 | -3.28044348373394 | 0.801455093955121 | -4.09310953099717 | 4.25626538991166e-05 | 0.00361338175012048 | 5260 | 1 | 2 | 13 | 13 | 0 | 0 | 7 | 10 | 1 | 1 | 3 | 7 |
| L1M5_dup28506 | 3.95344229787856 | -3.36183842376953 | 0.832476563503181 | -4.03835804052241 | 5.38266523296848e-05 | 0.00431729839865828 | 5261 | 1 | 1 | 11 | 13 | 0 | 0 | 15 | 5 | 1 | 1 | 2 | 3 |
| L2a_dup34009 | 3.59910449560756 | -2.58455718355856 | 0.759189693295372 | -3.40436284420553 | 0.000663185957238111 | 0.0263135644123694 | 5262 | 1 | 2 | 10 | 12 | 0 | 1 | 9 | 6 | 1 | 1 | 2 | 2 |
| AluJb_dup43991 | 3.14720495713922 | -3.73458780840107 | 1.15438075245309 | -3.23514386433157 | 0.00121581453785258 | 0.0401970551651898 | 5263 | 1 | 0 | 9 | 14 | 0 | 0 | 3 | 12 | 0 | 0 | 0 | 0 |
| THE1C_dup3385 | 3.88116227060967 | -3.54940954184249 | 1.00307404497826 | -3.53853193551573 | 0.000402358607924661 | 0.0185577866488159 | 5264 | 2 | 0 | 8 | 17 | 0 | 0 | 8 | 14 | 0 | 0 | 0 | 0 |
| L1ME3F_dup4607 | 2.80568939607404 | -3.63074475897959 | 1.10743273439818 | -3.27852396466559 | 0.00104351503418844 | 0.0361396449705531 | 5265 | 1 | 0 | 7 | 16 | 0 | 0 | 6 | 7 | 0 | 0 | 0 | 0 |
| L1PA6_dup524 | 3.07319052696897 | -3.75155596751856 | 1.06990019292808 | -3.50645414620534 | 0.000454119672549613 | 0.0202389885005601 | 5266 | 1 | 0 | 7 | 17 | 0 | 0 | 9 | 7 | 0 | 0 | 0 | 0 |
| L1PA8A_dup901 | 3.35789448956361 | -3.85342356031685 | 1.05670385126704 | -3.64664475831748 | 0.000265686830176768 | 0.0137105247477473 | 5267 | 1 | 0 | 8 | 18 | 0 | 0 | 8 | 9 | 0 | 0 | 0 | 0 |
| X6A_LINE_dup961 | 3.63291251320203 | -4.30566707956503 | 1.08883876383816 | -3.95436608482558 | 7.67378374690312e-05 | 0.00559934421171607 | 5268 | 0 | 0 | 14 | 19 | 0 | 0 | 6 | 7 | 0 | 1 | 0 | 1 |
| AluJr_dup73369 | 3.94096676441298 | -4.2459064037755 | 1.17860581156727 | -3.60248215485162 | 0.000315193044333585 | 0.0154700751087045 | 5269 | 0 | 0 | 11 | 24 | 0 | 0 | 4 | 11 | 0 | 1 | 0 | 0 |
| L2a_dup152008 | 3.90554471496777 | -4.25366420045727 | 1.2256994831088 | -3.47039731930741 | 0.000519689000324911 | 0.0221450427778136 | 5270 | 0 | 0 | 12 | 28 | 0 | 0 | 3 | 8 | 0 | 1 | 0 | 0 |
| L1PA6_dup5477 | 3.71537402199489 | -4.53528227443185 | 1.18781650701152 | -3.81816740856916 | 0.0001344466832598 | 0.00839413174973455 | 5271 | 0 | 0 | 9 | 28 | 0 | 0 | 8 | 6 | 0 | 0 | 0 | 0 |
| MLT1H-int_dup443 | 2.85897481010806 | -4.23154191241463 | 1.21245928597936 | -3.4900486650128 | 0.000482932595298869 | 0.0210153400367565 | 5272 | 0 | 0 | 7 | 20 | 0 | 0 | 7 | 5 | 0 | 0 | 0 | 0 |
| AluJb_dup121854 | 3.07764984630351 | -4.32172566800382 | 1.15996194014761 | -3.72574781846193 | 0.000194737022408464 | 0.0109079320936922 | 5273 | 0 | 0 | 11 | 17 | 0 | 0 | 7 | 6 | 0 | 0 | 0 | 0 |
| MIRb_dup107900 | 2.89461038194313 | -4.24639941509729 | 1.19875037218125 | -3.54235503374278 | 0.000396571329895498 | 0.0183620474145387 | 5274 | 0 | 0 | 10 | 18 | 0 | 0 | 6 | 5 | 0 | 0 | 0 | 0 |
| L1ME3_dup8112 | 2.94034803255898 | -4.24925574236224 | 1.31005957478149 | -3.24355916643791 | 0.00118046290627379 | 0.0393508583211298 | 5275 | 0 | 0 | 11 | 20 | 0 | 0 | 2 | 6 | 0 | 0 | 0 | 0 |
| MER126_dup254 | 3.36248195674141 | -4.41012085207074 | 1.26194316861152 | -3.49470638754919 | 0.000474583524451499 | 0.0208112789538306 | 5276 | 0 | 0 | 13 | 23 | 0 | 0 | 3 | 6 | 0 | 0 | 0 | 0 |
| MLT1B_dup17201 | 3.27100491183453 | -4.37810293053264 | 1.27366630134999 | -3.43740187354583 | 0.000587323484117692 | 0.0242196281150699 | 5277 | 0 | 0 | 14 | 22 | 0 | 0 | 3 | 5 | 0 | 0 | 0 | 0 |
| L1MEd_dup12027 | 3.80494801428193 | -4.55487523280093 | 1.21144690217989 | -3.75986370067466 | 0.000170005955553683 | 0.00996605989164245 | 5278 | 0 | 0 | 19 | 20 | 0 | 0 | 4 | 7 | 0 | 0 | 0 | 0 |
| MIRc_dup28165 | 3.75485289724725 | -4.54134059866123 | 1.17851460491361 | -3.85344447979424 | 0.000116467682500352 | 0.00758595305672931 | 5279 | 0 | 0 | 11 | 23 | 0 | 0 | 5 | 10 | 0 | 0 | 0 | 0 |
| L1MEc_dup6193 | 3.94726112446478 | -2.9322611318831 | 0.860826639234519 | -3.4063317725513 | 0.000658421136573499 | 0.0261900160085453 | 5280 | 1 | 2 | 15 | 17 | 1 | 0 | 6 | 9 | 0 | 0 | 0 | 0 |
| L1MA6_dup1824 | 3.11716605090374 | -3.73595660479766 | 1.071590082286 | -3.48636728405308 | 0.000489628263220016 | 0.021229060991281 | 5281 | 0 | 1 | 6 | 14 | 0 | 0 | 9 | 10 | 0 | 0 | 0 | 0 |
| MLT1I_dup8470 | 2.96464617322881 | -3.66673967689658 | 1.08693497341378 | -3.37346737991169 | 0.000742278408622689 | 0.0285288609623864 | 5282 | 0 | 1 | 10 | 10 | 0 | 0 | 6 | 10 | 0 | 0 | 0 | 0 |
| L2_dup19395 | 3.35858150477944 | -3.3613426297889 | 0.976655363467901 | -3.44168757529111 | 0.00057809751197446 | 0.0239061506055396 | 5283 | 0 | 2 | 12 | 12 | 0 | 0 | 11 | 7 | 0 | 0 | 0 | 0 |
| MSTA_dup9921 | 3.43886364161237 | -3.87893461888399 | 1.0295084384503 | -3.76775407953225 | 0.000164722852889435 | 0.00973067128938173 | 5284 | 0 | 1 | 12 | 16 | 0 | 0 | 11 | 6 | 0 | 0 | 0 | 0 |
| L1ME3F_dup1522 | 3.51947307343043 | -3.36954794695816 | 1.05438076066454 | -3.1957600827565 | 0.00139463025759461 | 0.0442040892930238 | 5285 | 0 | 2 | 13 | 21 | 0 | 0 | 8 | 4 | 0 | 0 | 0 | 0 |
| AluYc_dup469 | 2.9700610553149 | -3.67047454142067 | 1.10874151460176 | -3.31048715420298 | 0.000931337268972603 | 0.0334838602028075 | 5286 | 0 | 1 | 15 | 12 | 0 | 0 | 6 | 5 | 0 | 0 | 0 | 0 |
| THE1D_dup3190 | 2.71164910450535 | -3.5759328850686 | 1.09449534027022 | -3.26719790710643 | 0.00108617720212786 | 0.0371846020672193 | 5287 | 0 | 1 | 10 | 13 | 0 | 0 | 7 | 5 | 0 | 0 | 0 | 0 |
| AluSq2_dup52827 | 3.05493115060959 | -3.6887481937934 | 1.16862228781434 | -3.15649310496417 | 0.00159678662557148 | 0.0484752475642039 | 5288 | 0 | 0 | 20 | 10 | 0 | 1 | 5 | 4 | 0 | 0 | 0 | 0 |

| | | | | | | | | | | | | | | | | | | | |
|---|---|---|---|---|---|---|---|---|---|---|---|---|---|---|---|---|---|---|---|
| L1MC4_dup24101 | 3.42850896741867 | -4.42577575980344 | 1.25310382468021 | -3.53185081127087 | 0.000412662096769835 | 0.0189193247163767 | 5289 | 0 | 0 | 20 | 11 | 0 | 0 | 3 | 9 | 0 | 0 | 0 | 0 |
| L1M4_dup13367 | 2.98930035937979 | -4.2694679967608 | 1.27972976051006 | -3.33622623190315 | 0.000849240251119563 | 0.0313681031966618 | 5290 | 0 | 0 | 18 | 13 | 0 | 0 | 3 | 5 | 0 | 0 | 0 | 0 |
| L1MC4a_dup868 | 2.85605711511762 | -4.21196877253722 | 1.32881281702519 | -3.16972316835906 | 0.00152584234211604 | 0.0470478271267518 | 5291 | 0 | 0 | 18 | 12 | 0 | 0 | 2 | 5 | 0 | 0 | 0 | 0 |
| LTR2B_dup249 | 2.54230591428052 | -4.0741279980649 | 1.22718197481459 | -3.319905345481 | 0.000900479738377009 | 0.0326574534107386 | 5292 | 0 | 0 | 6 | 8 | 0 | 0 | 6 | 11 | 0 | 0 | 0 | 0 |
| MLT1A1_dup6666 | 2.71342753750881 | -4.1594802025457 | 1.20154553465194 | -3.46177492453551 | 0.000536625726376972 | 0.0226437214201977 | 5293 | 0 | 0 | 13 | 7 | 0 | 0 | 6 | 8 | 0 | 0 | 0 | 0 |
| L1PB4_dup5654 | 2.39093057049465 | -4.01056822470912 | 1.2143498937261 | -3.3026463340011 | 0.000957771003075569 | 0.0340960172199012 | 5294 | 0 | 0 | 8 | 5 | 0 | 0 | 9 | 8 | 0 | 0 | 0 | 0 |
| L4_A_Mam_dup2132 | 2.21807855946165 | -3.92454264166344 | 1.21625138677266 | -3.22675286075297 | 0.00125203550253507 | 0.0410353773672464 | 5295 | 0 | 0 | 8 | 7 | 0 | 0 | 6 | 7 | 0 | 0 | 0 | 0 |
| AluSz6_dup20295 | 3.56683870666141 | -3.88917100864067 | 1.066262904841 | -3.64747848863842 | 0.00026482647522925 | 0.0136839088979151 | 5296 | 0 | 0 | 16 | 12 | 1 | 0 | 6 | 10 | 0 | 0 | 0 | 0 |
| LTR16C_dup342 | 3.37304290471107 | -3.385818537614 | 0.951107661835487 | -3.55986884921094 | 0.000371040051061721 | 0.0174792112433047 | 5297 | 0 | 0 | 13 | 11 | 1 | 1 | 8 | 9 | 0 | 0 | 0 | 0 |
| MIRb_dup71040 | 3.01758497811301 | -3.19714297265985 | 1.00191849621577 | -3.19102101092596 | 0.00141770969905688 | 0.0446746592572889 | 5298 | 0 | 1 | 10 | 7 | 1 | 0 | 7 | 11 | 0 | 0 | 0 | 0 |
| L1MA5_dup3408 | 2.46234488881087 | -3.46046958878163 | 1.09594166792948 | -3.15753081577724 | 0.00159111418639265 | 0.0483880993382834 | 5299 | 0 | 0 | 8 | 8 | 1 | 0 | 9 | 6 | 0 | 0 | 0 | 0 |
| L2c_dup49372 | 2.5680679565275 | -3.5018121731968 | 1.10417971915108 | -3.17141504454461 | 0.00151698232356208 | 0.0468535664540892 | 5300 | 0 | 0 | 8 | 6 | 1 | 0 | 11 | 7 | 0 | 0 | 0 | 0 |
| L1PA14_dup1659 | 2.62893896394789 | -3.6973572310663 | 1.13635150845076 | -3.25370908875464 | 0.00113908863836394 | 0.0384228786376984 | 5301 | 0 | 0 | 7 | 11 | 0 | 0 | 10 | 3 | 0 | 2 | 0 | 3 |
| L1PA4_dup11805 | 3.33633711037905 | -3.77635502569027 | 1.00659993458655 | -3.75159474577294 | 0.000175713325958422 | 0.0101678703936047 | 5302 | 0 | 0 | 8 | 15 | 0 | 1 | 15 | 4 | 0 | 1 | 0 | 2 |
| L2_dup30936 | 3.61585882058102 | -3.11389911062726 | 0.836177476870723 | -3.72396912947307 | 0.000196114917264435 | 0.0109618886782718 | 5303 | 1 | 1 | 10 | 13 | 0 | 1 | 15 | 5 | 0 | 1 | 0 | 2 |
| THE1A_dup424 | 4.60931670553907 | -3.80987422742591 | 0.956477321061117 | -3.98323529845876 | 6.79833851894655e-05 | 0.00512746329078604 | 5304 | 0 | 1 | 12 | 22 | 0 | 0 | 17 | 6 | 0 | 2 | 0 | 3 |
| L2a_dup79132 | 3.01059572360584 | -3.92789324314286 | 1.13038102851902 | -3.4748400265431 | 0.000511158084457063 | 0.0219070644027774 | 5305 | 0 | 0 | 12 | 8 | 0 | 0 | 14 | 5 | 0 | 1 | 0 | 0 |
| MSTB-int_dup462 | 3.79398110641472 | -3.89868395566548 | 1.22728759036227 | -3.17666697380576 | 0.00148978001947091 | 0.0462837441589013 | 5306 | 0 | 0 | 16 | 18 | 0 | 0 | 13 | 3 | 0 | 2 | 0 | 0 |
| L1MA3_dup4736 | 3.31155285559794 | -3.26450429047926 | 0.941503154272776 | -3.46733229268976 | 0.000525651638187829 | 0.0223112209597291 | 5307 | 1 | 0 | 10 | 10 | 0 | 0 | 15 | 5 | 1 | 1 | 1 | 0 |
| Helitron1Nb_Mam_dup | 3.41050783234492 | -3.82017291193666 | 0.981512276257714 | -3.89212952740859 | 9.93681664786837e-05 | 0.00678652633207143 | 5308 | 1 | 0 | 12 | 11 | 0 | 0 | 12 | 6 | 0 | 1 | 2 | 0 |
| L1MC1_dup7001 | 3.0780960140123 | -4.23798456634744 | 1.07234178853825 | -3.95208375878401 | 7.74736120938882e-05 | 0.00564103777374032 | 5309 | 0 | 0 | 9 | 7 | 0 | 0 | 14 | 7 | 0 | 1 | 2 | 0 |
| AluJb_dup110104 | 3.18763342467805 | -3.59756341468882 | 1.13180717059964 | -3.17860100920091 | 0.00147987645093261 | 0.0460723121802446 | 5310 | 0 | 0 | 16 | 13 | 0 | 0 | 7 | 4 | 1 | 1 | 0 | 0 |
| MIRb_dup53866 | 3.27403070951212 | -3.7430517346915 | 1.05237421169385 | -3.55676877397714 | 0.000375444338982203 | 0.0176441355553167 | 5311 | 0 | 0 | 12 | 11 | 0 | 0 | 6 | 9 | 1 | 1 | 1 | 0 |
| L1MC1_dup9423 | 2.83031493571795 | -3.60264152401033 | 1.11630529835629 | -3.22729053540734 | 0.00124968500297013 | 0.0409978695321565 | 5312 | 0 | 0 | 6 | 10 | 0 | 0 | 11 | 7 | 0 | 2 | 0 | 1 |
| MLT1I_dup8469 | 3.85546481585875 | -3.93726312726574 | 1.05053876254846 | -3.74785135744492 | 0.000178355894142796 | 0.0102782929493102 | 5313 | 0 | 0 | 10 | 12 | 0 | 0 | 15 | 10 | 0 | 2 | 0 | 1 |
| MSTA_dup10403 | 2.72969246317856 | -3.8122797013494 | 1.16049160879751 | -3.28505580949408 | 0.00101961729468 | 0.0355427647289763 | 5314 | 0 | 0 | 5 | 15 | 0 | 0 | 8 | 7 | 0 | 1 | 0 | 0 |
| L1PA16_dup7913 | 2.88305431840895 | -3.86579483789409 | 1.1625420581122 | -3.32529460841321 | 0.000883251506939119 | 0.0322782225708344 | 5315 | 0 | 0 | 5 | 8 | 0 | 0 | 11 | 11 | 0 | 1 | 0 | 0 |
| L2b_dup92684 | 2.57501515291919 | -3.73056371405736 | 1.16120079512461 | -3.21267754011229 | 0.00131503843119224 | 0.0424397307132062 | 5316 | 0 | 0 | 8 | 8 | 0 | 0 | 6 | 9 | 0 | 1 | 0 | 0 |
| THE1D_dup6202 | 3.34764930811719 | -3.07039288106482 | 0.917769679314974 | -3.3454939188627 | 0.000821361602909491 | 0.0306475941611889 | 5317 | 0 | 1 | 7 | 16 | 0 | 1 | 10 | 8 | 0 | 1 | 0 | 0 |
| L1MC3_dup8509 | 3.45812165900553 | -3.86710810895388 | 1.05638485538677 | -3.66070006516521 | 0.000251527064153902 | 0.0131856926210981 | 5318 | 0 | 0 | 6 | 11 | 0 | 1 | 14 | 12 | 0 | 0 | 0 | 0 |
| LTR33_dup975 | 2.70068643214953 | -3.58475918788496 | 1.0847881594514 | -3.30457071885616 | 0.0009512197445731 | 0.0339493383905337 | 5319 | 0 | 0 | 6 | 12 | 0 | 1 | 11 | 6 | 0 | 0 | 0 | 0 |
| MLT1D_dup7699 | 2.74333738235693 | -3.59680473729342 | 1.07530517654845 | -3.34491530008119 | 0.000823077019055946 | 0.0306820887888357 | 5320 | 0 | 0 | 9 | 9 | 0 | 1 | 8 | 8 | 0 | 0 | 0 | 0 |
| L1MC4_dup7964 | 2.86776317753504 | -4.22192471852957 | 1.20355128159468 | -3.50788934638133 | 0.000451676878706055 | 0.0201641100526437 | 5321 | 0 | 0 | 5 | 7 | 0 | 0 | 13 | 11 | 0 | 0 | 0 | 0 |
| L1MB7_dup22776 | 2.82303842561459 | -4.20519360402245 | 1.18148155744058 | -3.559254545735 | 0.000371908938370198 | 0.0175132088362231 | 5322 | 0 | 0 | 7 | 7 | 0 | 0 | 10 | 11 | 0 | 0 | 0 | 0 |
| L1MEc_dup452 | 2.95729631224735 | -4.26183775709761 | 1.17202810807748 | -3.63629312959777 | 0.000276589574195308 | 0.0141109311366803 | 5323 | 0 | 0 | 7 | 7 | 0 | 0 | 12 | 11 | 0 | 0 | 0 | 0 |
| L1PB1_dup2272 | 3.23295561172847 | -4.37517368024404 | 1.12462578599921 | -3.89033733239255 | 0.000100104964227424 | 0.00682118436943262 | 5324 | 0 | 0 | 8 | 11 | 0 | 0 | 11 | 11 | 0 | 0 | 0 | 0 |
| MLT1L_dup3721 | 3.55828298346453 | -4.47674239794978 | 1.14178444872705 | -3.92082971785157 | 8.82446048198301e-05 | 0.00622515586672377 | 5325 | 0 | 0 | 9 | 8 | 0 | 0 | 13 | 14 | 0 | 0 | 0 | 0 |
| L2a_dup57692 | 3.30314723843289 | -4.03779403702541 | 1.03873394274023 | -3.88722643102769 | 0.000101396161455145 | 0.0068855054319652 | 5326 | 0 | 1 | 7 | 11 | 0 | 0 | 11 | 11 | 0 | 0 | 0 | 1 |
| MER4D_dup1215 | 3.59398901617174 | -4.70122338082418 | 1.06463599755557 | -4.41580351558498 | 1.00635464842497e-05 | 0.00122862012916433 | 5327 | 0 | 0 | 8 | 17 | 0 | 0 | 11 | 10 | 0 | 0 | 0 | 1 |
| L2b_dup26235 | 2.75198080433431 | -4.18684907622277 | 1.17504647021639 | -3.56313489070074 | 0.000366452253693485 | 0.0173110676905599 | 5328 | 0 | 0 | 5 | 13 | 0 | 0 | 10 | 8 | 0 | 0 | 0 | 0 |
| MIR_dup56972 | 2.58614591534884 | -4.09928166589956 | 1.28262582159495 | -3.19600743793081 | 0.00139343519636474 | 0.044189139706152 | 5329 | 0 | 0 | 2 | 11 | 0 | 0 | 13 | 8 | 0 | 0 | 0 | 0 |
| MSTA-int_dup2164 | 2.59417593609177 | -4.09685583622425 | 1.29226602347507 | -3.17028828569467 | 0.00152287764760249 | 0.0469746752315819 | 5330 | 0 | 0 | 2 | 9 | 0 | 0 | 12 | 10 | 0 | 0 | 0 | 0 |
| L2a_dup106921 | 2.93535649497902 | -3.67278279109857 | 1.08929237162725 | -3.37171441457168 | 0.000747018656402161 | 0.0286739960019791 | 5331 | 0 | 1 | 6 | 13 | 0 | 0 | 15 | 5 | 0 | 0 | 0 | 0 |
| L2c_dup91372 | 3.13345028975386 | -4.33896134493155 | 1.17898848016753 | -3.68024066216066 | 0.000233013934333324 | 0.0124544398833684 | 5332 | 0 | 0 | 6 | 9 | 0 | 0 | 20 | 7 | 0 | 0 | 0 | 0 |
| MSTB1_dup1486 | 2.87512196981571 | -4.24562321114118 | 1.1456770217676 | -3.70577669838473 | 0.000210743914709747 | 0.0115410061057596 | 5333 | 0 | 0 | 7 | 10 | 0 | 0 | 14 | 7 | 0 | 0 | 0 | 0 |
| MLT1A_dup8954 | 4.32039322924747 | -3.9420875374609 | 0.980034471285505 | -4.02239681660389 | 5.76088941243684e-05 | 0.00452552192846384 | 5334 | 0 | 1 | 7 | 11 | 0 | 0 | 17 | 16 | 0 | 1 | 1 | 0 |
| L1MA6_dup4492 | 4.2657068439926 | -4.34851723977439 | 0.977270474546423 | -4.44965580464575 | 8.60080165169877e-06 | 0.00108626664002925 | 5335 | 0 | 1 | 11 | 15 | 0 | 0 | 15 | 12 | 0 | 0 | 1 | 0 |
| LTR18B_dup98 | 4.75177478199415 | -4.15319204684131 | 0.912396431499606 | -4.55195998521735 | 5.31484457496717e-06 | 0.000755012541632919 | 5336 | 1 | 1 | 8 | 19 | 0 | 0 | 18 | 13 | 0 | 0 | 2 | 0 |
| L1MA3_dup1349 | 3.35616824292776 | -4.95666152405162 | 1.10946168486051 | -4.46762749150261 | 7.90918754263348e-06 | 0.00102336027135371 | 5337 | 0 | 0 | 6 | 11 | 0 | 0 | 15 | 8 | 0 | 0 | 4 | 0 |
| L1M4_dup9195 | 2.49330982972732 | -4.44451090832298 | 1.1152863910245 | -3.98508485720897 | 6.74560150469723e-05 | 0.00510849425727781 | 5338 | 0 | 0 | 7 | 10 | 0 | 0 | 9 | 5 | 0 | 0 | 2 | 0 |
| LTR79_dup2215 | 2.95167347155554 | -4.4926699042236 | 1.093445281008 | -4.10872860513148 | 3.97843268426681e-05 | 0.00342369672148481 | 5339 | 0 | 0 | 7 | 14 | 0 | 0 | 10 | 7 | 0 | 0 | 1 | 0 |
| THE1D_dup3191 | 3.32523279218037 | -4.76627393127226 | 1.06325323607011 | -4.48272694554769 | 7.36952017243889e-06 | 0.000968276006047014 | 5340 | 0 | 0 | 9 | 12 | 0 | 0 | 14 | 7 | 0 | 0 | 2 | 0 |
| L2b_dup11883 | 3.36880582258776 | -4.22194602420806 | 1.06828189196865 | -3.95208985189086 | 7.7471638952326e-05 | 0.00564103777374032 | 5341 | 0 | 1 | 7 | 8 | 0 | 0 | 12 | 12 | 0 | 0 | 2 | 0 |
| L3_dup12584 | 2.78510846188794 | -3.32446256474761 | 1.03355975025134 | -3.21651705567984 | 0.0012975682067737 | 0.0420230769544102 | 5342 | 0 | 2 | 7 | 6 | 0 | 0 | 13 | 7 | 0 | 0 | 1 | 0 |
| L1MA8_dup5256 | 2.50190997926056 | -3.86990039149746 | 1.10927676058156 | -3.48866985139811 | 0.000485430305081599 | 0.0211085890666934 | 5343 | 0 | 1 | 3 | 6 | 0 | 0 | 9 | 10 | 0 | 0 | 1 | 1 |
| MSTA-int_dup415 | 2.78162784822405 | -3.99745750758779 | 1.08589097526018 | -3.68126966579677 | 0.000232075388716612 | 0.0124120949524754 | 5344 | 0 | 1 | 4 | 6 | 0 | 0 | 12 | 10 | 0 | 0 | 1 | 1 |
| L1PA16_dup5759 | 2.98811383392642 | -4.19362247303234 | 1.04949548030119 | -3.99584614869312 | 6.44635901996783e-05 | 0.0049305218365968 | 5345 | 0 | 1 | 5 | 7 | 0 | 0 | 15 | 8 | 0 | 0 | 1 | 2 |
| MIRb_dup60701 | 3.11127008512901 | -4.13293449134944 | 1.02745885562566 | -4.02248174583372 | 5.7588118646038e-05 | 0.00452538390995712 | 5346 | 0 | 1 | 7 | 9 | 0 | 0 | 12 | 9 | 0 | 0 | 1 | 1 |
| L1MA10_dup2570 | 2.96512033530106 | -3.05955266158717 | 0.923562798935556 | -3.31277165463294 | 0.000923763666761205 | 0.0332986307281276 | 5347 | 0 | 1 | 7 | 14 | 0 | 1 | 8 | 6 | 1 | 0 | 0 | 1 |
| L1PA7_dup1085 | 3.12790074324687 | -2.78676490010931 | 0.873386910464516 | -3.19075642961852 | 0.00141900853945652 | 0.0446792014336328 | 5348 | 0 | 1 | 8 | 13 | 0 | 1 | 9 | 6 | 1 | 1 | 0 | 1 |
| L1PB1_dup4394 | 3.85884452015421 | -3.49447555877595 | 0.919051319815542 | -3.80226379466743 | 0.00014337994232378 | 0.00881255146167725 | 5349 | 0 | 2 | 12 | 17 | 0 | 0 | 13 | 5 | 1 | 0 | 0 | 2 |
| MIRc_dup77320 | 3.04400342918161 | -3.64163599358129 | 1.02469394593851 | -3.55387675316646 | 0.000379597058837999 | 0.0177767372096568 | 5350 | 0 | 1 | 12 | 13 | 0 | 0 | 6 | 5 | 1 | 0 | 0 | 2 |
| L1M5_dup25215 | 3.34802469955905 | -4.04026941910988 | 1.14965261060849 | -3.51433935941 | 0.000440849221493516 | 0.0198220538279471 | 5351 | 0 | 0 | 12 | 5 | 0 | 0 | 17 | 8 | 1 | 0 | 0 | 0 |
| L2a_dup79133 | 3.30148418528729 | -4.02698347082121 | 1.14741412502284 | -3.50961643490405 | 0.000448753532373807 | 0.0200609779967691 | 5352 | 0 | 0 | 18 | 9 | 0 | 0 | 10 | 5 | 1 | 0 | 0 | 0 |
| L1PB3_dup3506 | 2.74403687452677 | -3.81545417681475 | 1.15327454102606 | -3.30836591035831 | 0.000938421111156109 | 0.0336247186288139 | 5353 | 0 | 0 | 8 | 15 | 0 | 0 | 9 | 4 | 1 | 0 | 0 | 0 |
| L1ME1_dup25452 | 2.60453485490112 | -3.73844636753215 | 1.13664874600857 | -3.2890076029732 | 0.0010054130769907 | 0.0352021806971202 | 5354 | 0 | 0 | 9 | 7 | 0 | 0 | 9 | 7 | 1 | 0 | 0 | 0 |
| L1M5_dup42397 | 2.43168284386812 | -3.65848267734469 | 1.15514868357733 | -3.16710976635051 | 0.00153962192323579 | 0.0473194107654694 | 5355 | 0 | 0 | 9 | 9 | 0 | 0 | 6 | 6 | 1 | 0 | 0 | 0 |
| MER34_dup593 | 2.72158816615897 | -3.7944404149134 | 1.13470877105737 | -3.34397733735465 | 0.00082586483547075 | 0.0307481198780978 | 5356 | 0 | 0 | 11 | 10 | 0 | 0 | 7 | 6 | 1 | 0 | 0 | 0 |
| LTR16B1_dup453 | 2.873391582011 | -4.23973547048003 | 1.20100252071484 | -3.53016367355875 | 0.000415302661071811 | 0.0190154726476381 | 5357 | 0 | 0 | 14 | 11 | 0 | 0 | 11 | 3 | 0 | 0 | 0 | 0 |
| L2a_dup57676 | 2.8661626812488 | -4.24280391357953 | 1.1707495490755 | -3.62400644692105 | 0.000290074369128201 | 0.0145987912737432 | 5358 | 0 | 0 | 10 | 10 | 0 | 0 | 15 | 4 | 0 | 0 | 0 | 0 |
| THE1B_dup22338 | 3.162154114088 | -4.35234937155954 | 1.16167318964679 | -3.74662117568788 | 0.000179232442731175 | 0.0103163141795143 | 5359 | 0 | 0 | 15 | 8 | 0 | 0 | 14 | 5 | 0 | 0 | 0 | 0 |
| AluSx1_dup11538 | 2.96994009509564 | -4.27563073670667 | 1.17985516571035 | -3.62386067457057 | 0.00029023799177988 | 0.01460371520238 | 5360 | 0 | 0 | 16 | 10 | 0 | 0 | 8 | 5 | 0 | 0 | 0 | 0 |
| MIRc_dup49649 | 3.6777926242433 | -4.52655362867389 | 1.13141105074558 | -4.00080379778063 | 6.31276843080214e-05 | 0.00485823510649519 | 5361 | 0 | 0 | 19 | 13 | 0 | 0 | 9 | 7 | 0 | 0 | 0 | 0 |
| LTR22C0_dup95 | 3.10195517459735 | -4.33323755877613 | 1.12103936752897 | -3.86537501205473 | 0.000110918640007232 | 0.0073435299708294 | 5362 | 0 | 0 | 12 | 10 | 0 | 0 | 10 | 8 | 0 | 0 | 0 | 0 |
| MIRb_dup202632 | 3.61873821864171 | -4.51839980123449 | 1.09844014078304 | -4.11346930385617 | 3.89756888645987e-05 | 0.00337363014280676 | 5363 | 0 | 0 | 14 | 16 | 0 | 0 | 11 | 7 | 0 | 0 | 0 | 0 |
| MSTA_dup10399 | 3.39494719729665 | -4.44349277875233 | 1.10931839250521 | -4.00560633337868 | 6.18585982581393e-05 | 0.00478642346165923 | 5364 | 0 | 0 | 12 | 16 | 0 | 0 | 10 | 7 | 0 | 0 | 0 | 0 |
| AluSx_dup34103 | 3.35924349686314 | -3.338799514998 | 1.0118288931095 | -3.29976692475877 | 0.000967651569985405 | 0.0343649490556445 | 5365 | 0 | 0 | 13 | 8 | 0 | 1 | 11 | 6 | 2 | 0 | 2 | 0 |
| AmnSINE1_dup483 | 2.29201083438876 | -3.9019760313866 | 1.159486854974 | -3.36526111930274 | 0.000764712519409192 | 0.0291136456229343 | 5366 | 0 | 0 | 7 | 7 | 0 | 0 | 12 | 2 | 1 | 0 | 2 | 0 |
| L2c_dup56054 | 2.92347449104146 | -4.17276400909548 | 1.08815350333936 | -3.83472000622152 | 0.000125707410485806 | 0.00801410773025061 | 5367 | 0 | 0 | 9 | 14 | 0 | 0 | 6 | 6 | 1 | 0 | 2 | 0 |

| | | | | | | | | | | | | | | | | | | |
|---|---|---|---|---|---|---|---|---|---|---|---|---|---|---|---|---|---|---|
| MLT1C_dup13677 | 2.61374290649899 | -4.14377925824067 | 1.16383480814686 | -3.56045310660427 | 0.00037021542285535 | 0.0174576454630874 | 5368 | 0 | 0 | 11 | 12 | 0 | 0 | 6 | 2 | 1 | 0 | 3 | 0 |
| Tigger2_dup1721 | 3.46862798108189 | -4.26703207447895 | 1.11415687422591 | -3.82983058596985 | 0.000128231509884308 | 0.00813441078403935 | 5369 | 0 | 0 | 15 | 17 | 0 | 0 | 10 | 3 | 1 | 0 | 1 | 0 |
| L1PBa_dup1691 | 3.91192172485835 | -3.39199035784704 | 0.938735402028809 | -3.61336149730395 | 0.000302252840497673 | 0.0150387799455394 | 5370 | 0 | 0 | 9 | 21 | 2 | 0 | 8 | 9 | 1 | 0 | 1 | 0 |
| MLT1F2_dup1307 | 3.7512673369352 | -3.46665983875541 | 0.983783880273073 | -3.52380223773656 | 0.000425401626044159 | 0.0193320502885224 | 5371 | 0 | 0 | 11 | 12 | 1 | 0 | 9 | 10 | 2 | 0 | 2 | 0 |
| LTR9_dup632 | 3.71511015080206 | -4.32539406900819 | 1.10055547505345 | -3.93019176865949 | 8.48781385091762e-05 | 0.00604010189334566 | 5372 | 0 | 0 | 7 | 16 | 0 | 0 | 7 | 14 | 1 | 0 | 1 | 0 |
| MIR1_Amn_dup2279 | 3.04788757901569 | -3.93054458119696 | 1.17128379542228 | -3.35575767082128 | 0.000791478874550312 | 0.0298461540861725 | 5373 | 0 | 0 | 7 | 17 | 0 | 0 | 5 | 9 | 1 | 0 | 0 | 0 |
| MLT1F2_dup541 | 4.36935154040515 | -3.18623142242258 | 0.838673670558958 | -3.79913133590927 | 0.000145204133456612 | 0.00888650790624151 | 5374 | 3 | 1 | 9 | 25 | 1 | 0 | 9 | 8 | 0 | 0 | 0 | 3 |
| MLT1J_dup8042 | 3.83801416056099 | -3.98612529388398 | 0.982184678961189 | -4.05842748239559 | 4.94042699043222e-05 | 0.00403970735623844 | 5375 | 1 | 1 | 9 | 23 | 0 | 0 | 11 | 5 | 0 | 0 | 0 | 3 |
| L2c_dup39573 | 4.32411320222925 | -3.26205959773979 | 0.815675093737424 | -3.99921442101785 | 6.35530827808557e-05 | 0.00488127883646659 | 5376 | 2 | 2 | 11 | 20 | 0 | 0 | 14 | 8 | 0 | 0 | 0 | 1 |
| THE1B_dup6490 | 4.48864401743883 | -3.15233243976905 | 0.824864218117543 | -3.82163799875223 | 0.00013256819548887 | 0.00830981871543592 | 5377 | 3 | 2 | 14 | 22 | 0 | 0 | 12 | 6 | 0 | 0 | 0 | 2 |
| L2_dup32067 | 3.49732791627862 | -3.38898214658369 | 0.887743525454582 | -3.8175239237575 | 0.000134797719623821 | 0.00841190091623648 | 5378 | 1 | 1 | 6 | 13 | 1 | 0 | 16 | 7 | 0 | 0 | 0 | 2 |
| HAL1_dup11199 | 2.72592980599562 | -3.15896870212487 | 0.992527226144512 | -3.18275269323939 | 0.0014588215806184 | 0.0456012384201789 | 5379 | 2 | 1 | 4 | 9 | 0 | 0 | 13 | 5 | 0 | 0 | 0 | 3 |
| L1M3_dup5823 | 3.11558937511894 | -3.10532122829677 | 0.970463728415554 | -3.19983234547749 | 0.00137507549664291 | 0.0438010326806149 | 5380 | 2 | 1 | 3 | 15 | 0 | 0 | 13 | 7 | 0 | 0 | 0 | 1 |
| MIR3_dup43945 | 2.92973773923891 | -3.22540144812691 | 0.967304954240277 | -3.3344204782453 | 0.000854773374175574 | 0.0314893799801178 | 5381 | 1 | 2 | 9 | 11 | 0 | 0 | 5 | 7 | 0 | 0 | 0 | 3 |
| MLT1H2_dup1263 | 2.86779139934248 | -3.5340246111868 | 0.982142646957649 | -3.59828037417378 | 0.000320328191527566 | 0.0156639564038673 | 5382 | 1 | 1 | 11 | 8 | 0 | 0 | 10 | 5 | 0 | 0 | 0 | 2 |
| L1M6_dup3230 | 2.68375412200791 | -3.32027455984713 | 0.978169965014995 | -3.39437385996229 | 0.000687857195555259 | 0.0269820092735891 | 5383 | 1 | 1 | 7 | 12 | 0 | 0 | 9 | 5 | 0 | 0 | 0 | 1 |
| MIR3_dup10600 | 2.86985626975633 | -3.12210803009758 | 0.928680618729369 | -3.36187486540775 | 0.000774151974914873 | 0.0293602750754055 | 5384 | 1 | 2 | 7 | 11 | 0 | 0 | 9 | 6 | 0 | 0 | 0 | 2 |
| L3_dup19411 | 3.51692013842366 | -2.79479085003074 | 0.878605943279033 | -3.18093779288624 | 0.00146799147866019 | 0.0458041831079565 | 5385 | 3 | 2 | 6 | 13 | 0 | 0 | 8 | 11 | 0 | 0 | 0 | 2 |
| LTR79_dup459 | 4.27415289758782 | -4.02304434635791 | 0.904808876304085 | -4.44629186529538 | 8.73652317407363e-06 | 0.00110107030888672 | 5386 | 1 | 1 | 10 | 18 | 0 | 0 | 13 | 11 | 0 | 0 | 0 | 2 |
| MER53_dup607 | 3.24813862421817 | -4.03682806928188 | 1.07061705357663 | -3.77056208454365 | 0.000162880273904837 | 0.00965778959744839 | 5387 | 1 | 0 | 16 | 11 | 0 | 0 | 6 | 7 | 0 | 0 | 0 | 1 |
| MIRc_dup21330 | 2.3855679272364 | -3.68345273093988 | 1.09224382037102 | -3.37237223250087 | 0.000745236544832871 | 0.0286194407515821 | 5388 | 1 | 0 | 8 | 9 | 0 | 0 | 6 | 6 | 0 | 0 | 0 | 1 |
| MLT1J1_dup1378 | 2.92564165720401 | -3.92416346695111 | 1.05011433980761 | -3.73689161093644 | 0.000186309186725811 | 0.01058031640684 | 5389 | 1 | 0 | 11 | 9 | 0 | 0 | 9 | 7 | 0 | 0 | 0 | 1 |
| MIRb_dup27279 | 3.17090899476892 | -4.69201466301115 | 1.06695539510186 | -4.3975733986173 | 1.09467871052614e-05 | 0.00130498206640708 | 5390 | 0 | 0 | 12 | 10 | 0 | 0 | 9 | 8 | 0 | 0 | 0 | 2 |
| X7C_LINE_dup310 | 3.0518095693735 | -4.74319817883406 | 1.10318825607142 | -4.29953650497071 | 1.71155683058781e-05 | 0.001821977016139 | 5391 | 0 | 0 | 13 | 7 | 0 | 0 | 8 | 8 | 0 | 0 | 0 | 3 |
| L1MB8_dup1953 | 2.74481870504712 | -4.3915039804243 | 1.11288710171998 | -3.94604625539929 | 7.94522713221421e-05 | 0.0057533019330549 | 5392 | 0 | 0 | 10 | 10 | 0 | 0 | 6 | 8 | 0 | 0 | 0 | 1 |
| L2_dup6368 | 2.62787671223614 | -4.47948686983087 | 1.15003996705089 | -3.89507060464851 | 9.81701296810321e-05 | 0.00672400393698413 | 5393 | 0 | 0 | 12 | 10 | 0 | 0 | 4 | 6 | 0 | 0 | 0 | 2 |
| MER65-int_dup710 | 2.89742395726864 | -4.46207248212478 | 1.07886663856408 | -4.13588883243586 | 3.53583344275462e-05 | 0.00312923477336041 | 5394 | 0 | 0 | 9 | 11 | 0 | 0 | 10 | 7 | 0 | 0 | 0 | 1 |
| MLT1F_dup3505 | 2.47937454646885 | -4.42189889113939 | 1.10601061059964 | -3.9980619071475 | 6.3863250128239e-05 | 0.00489719784563775 | 5395 | 0 | 0 | 8 | 10 | 0 | 0 | 8 | 5 | 0 | 0 | 0 | 2 |
| L1PA16_dup3935 | 3.41826382712017 | -4.64506928555172 | 1.09048280221627 | -4.25964469692799 | 2.0475211560201e-05 | 0.00207975903641571 | 5396 | 0 | 0 | 14 | 16 | 0 | 0 | 7 | 7 | 0 | 0 | 0 | 1 |
| MER67D_dup42 | 3.51991323963324 | -4.81551647138362 | 1.05418931081165 | -4.56798074311343 | 4.92445261520325e-06 | 0.000712745621640604 | 5397 | 0 | 0 | 14 | 13 | 0 | 0 | 12 | 6 | 0 | 0 | 0 | 2 |
| L1PA14_dup1641 | 2.32161280255026 | -3.66157934433344 | 1.12675921239975 | -3.24965556441742 | 0.0011554486200784 | 0.0388160916777037 | 5398 | 1 | 0 | 6 | 8 | 0 | 0 | 13 | 3 | 0 | 0 | 0 | 1 |
| Tigger15a_dup2431 | 2.46187142518476 | -3.72822151645409 | 1.09135056131379 | -3.41615393679369 | 0.000635123328894114 | 0.025534245984574 | 5399 | 1 | 0 | 6 | 8 | 0 | 0 | 12 | 5 | 0 | 0 | 0 | 1 |
| L2a_dup19155 | 2.52200007585 | -3.51146829363541 | 1.11623817563891 | -3.14580559084134 | 0.00165630061933809 | 0.049773946346176 | 5400 | 1 | 0 | 6 | 14 | 0 | 0 | 8 | 5 | 0 | 0 | 0 | 0 |
| MER5A_dup34115 | 2.59318758864874 | -3.5489760009534 | 1.10220366524622 | -3.21989130762017 | 0.0012823920898023 | 0.0417142950376073 | 5401 | 1 | 0 | 6 | 10 | 0 | 0 | 13 | 5 | 0 | 0 | 0 | 0 |
| MIRc_dup12843 | 3.19807119475897 | -4.17184614715825 | 1.06066348762706 | -3.93324197148675 | 8.38077958417382e-05 | 0.00598850744425172 | 5402 | 1 | 0 | 6 | 17 | 0 | 0 | 10 | 7 | 0 | 0 | 0 | 2 |
| AluSz6_dup5157 | 2.57772251558243 | -3.92068363014911 | 1.12022765482723 | -3.4998989832596 | 0.000465434500365099 | 0.0205355604716992 | 5403 | 1 | 0 | 5 | 16 | 0 | 0 | 6 | 5 | 0 | 0 | 0 | 2 |
| MER20_dup1358 | 2.88911959084162 | -3.91198968055457 | 1.09231626421816 | -3.58137089843172 | 0.000341796023929454 | 0.0164201487401409 | 5404 | 1 | 0 | 9 | 17 | 0 | 0 | 6 | 5 | 0 | 0 | 0 | 1 |
| MLT1D_dup6433 | 2.85388526014268 | -3.38719449302302 | 0.988333174887037 | -3.42717878858025 | 0.000609887373376555 | 0.0248578727210445 | 5405 | 1 | 0 | 10 | 12 | 0 | 1 | 5 | 7 | 0 | 0 | 0 | 1 |
| Tigger9a_dup985 | 2.88140718036342 | -3.4185236792713 | 0.967069695246573 | -3.53493000150283 | 0.000407883192766232 | 0.0187471845056894 | 5406 | 1 | 0 | 7 | 12 | 0 | 1 | 12 | 5 | 0 | 0 | 0 | 1 |
| L1ME2z_dup6853 | 3.60765761739498 | -3.25527765780442 | 1.00311913787235 | -3.24515557016385 | 0.00117386474510518 | 0.0392243253249259 | 5407 | 1 | 2 | 20 | 16 | 0 | 0 | 4 | 4 | 0 | 0 | 0 | 1 |
| L2c_dup126679 | 4.08693936633228 | -3.10031131390781 | 0.972081758983422 | -3.18935242355543 | 0.00142591923770587 | 0.0448502021064322 | 5408 | 1 | 3 | 23 | 17 | 0 | 0 | 7 | 3 | 0 | 0 | 0 | 1 |
| L1MCa_dup4763 | 3.54898520536929 | -3.84263226060564 | 1.19206555162735 | -3.22350751211615 | 0.00126630977052321 | 0.0413519595461451 | 5409 | 0 | 1 | 20 | 20 | 0 | 0 | 4 | 3 | 0 | 0 | 0 | 0 |
| LTR12C_dup457 | 3.5251638780754 | -3.38125583878831 | 1.07207137941771 | -3.15394655962631 | 0.00161078579389292 | 0.0487571602230815 | 5410 | 1 | 1 | 17 | 20 | 0 | 0 | 3 | 5 | 0 | 0 | 0 | 0 |
| THE1D_dup12207 | 5.60291552984435 | -3.59082311107315 | 0.81901190390463 | -4.38433567809448 | 1.16340261369611e-05 | 0.00136099802516487 | 5411 | 1 | 2 | 20 | 32 | 0 | 1 | 12 | 7 | 0 | 0 | 1 | 0 |
| Tigger3b_dup5881 | 4.34359165567375 | -3.10707759742587 | 0.805389526531404 | -3.85785696867356 | 0.000114385549553507 | 0.00748970390040144 | 5412 | 2 | 2 | 13 | 22 | 0 | 1 | 9 | 7 | 0 | 0 | 2 | 0 |
| MSTB2_dup430 | 4.10740696425636 | -3.23477431485174 | 0.981898444074679 | -3.2944082296618 | 0.00098629144276339 | 0.0347841935178236 | 5413 | 1 | 1 | 15 | 29 | 0 | 1 | 4 | 5 | 0 | 0 | 0 | 0 |
| THE1B_dup11735 | 3.41870292067819 | -3.03346569645332 | 0.945991535307178 | -3.20665205050519 | 0.00134289315832781 | 0.0430761324184771 | 5414 | 1 | 1 | 7 | 24 | 0 | 1 | 8 | 5 | 0 | 0 | 0 | 0 |
| LTR33A__dup451 | 3.68031974538592 | -2.85608456556696 | 0.868578997608741 | -3.28822660164471 | 0.00100820654691864 | 0.035267675652872 | 5415 | 2 | 2 | 7 | 14 | 0 | 0 | 16 | 8 | 0 | 0 | 0 | 0 |
| MIRb_dup71042 | 3.80242716119317 | -3.56296674555156 | 0.930855010122552 | -3.8276280481989 | 0.000129384078261231 | 0.0081813510193853 | 5416 | 1 | 1 | 7 | 15 | 0 | 0 | 16 | 10 | 0 | 0 | 0 | 0 |
| L1MA2_dup2220 | 4.09725949749614 | -3.32442728324192 | 0.87131000833207 | -3.81543566750232 | 0.000135942870921559 | 0.00847225310746496 | 5417 | 2 | 1 | 10 | 20 | 0 | 0 | 14 | 8 | 0 | 0 | 0 | 0 |
| MER52A_dup550 | 3.86570386979305 | -3.22262518567361 | 0.881584107305334 | -3.65549374015367 | 0.000256687472279321 | 0.0134001222639261 | 5418 | 1 | 2 | 9 | 18 | 0 | 0 | 12 | 9 | 0 | 0 | 0 | 0 |
| AluJb_dup31103 | 3.76095267115035 | -3.18933514002063 | 0.876089684279349 | -3.64042083504739 | 0.000272192789512165 | 0.0139494261238987 | 5419 | 1 | 2 | 11 | 11 | 0 | 0 | 15 | 9 | 0 | 0 | 0 | 0 |
| L1ME3A_dup4081 | 3.63790406783956 | -3.51619272090973 | 0.925517078540813 | -3.79916567985263 | 0.000145184015255841 | 0.00888650790624151 | 5420 | 1 | 1 | 14 | 14 | 0 | 0 | 11 | 7 | 0 | 0 | 0 | 0 |
| MER11B_dup165 | 3.2501184711127 | -3.00001851457864 | 0.907551637430989 | -3.30561743359398 | 0.000947673825479788 | 0.0338736920948435 | 5421 | 2 | 1 | 12 | 11 | 0 | 0 | 11 | 6 | 0 | 0 | 0 | 0 |
| L1M4_dup9611 | 4.62960955495382 | -3.09548689384026 | 0.768187553563029 | -4.02959782345169 | 5.58723614110146e-05 | 0.00443005828334064 | 5422 | 2 | 1 | 13 | 21 | 1 | 0 | 15 | 7 | 1 | 0 | 1 | 0 |
| L1MB8_dup6088 | 3.97339675474953 | -3.13574612031745 | 0.844262310489408 | -3.71418465725386 | 0.000203859900449267 | 0.0112703642825979 | 5423 | 2 | 1 | 12 | 18 | 0 | 0 | 9 | 8 | 1 | 0 | 1 | 0 |
| MLT1J-int_dup385 | 3.42829998968519 | -3.04305666154824 | 0.954824533476739 | -3.18703233406432 | 0.001437407047076 | 0.0451136031879259 | 5424 | 2 | 1 | 14 | 14 | 0 | 0 | 13 | 3 | 0 | 0 | 0 | 0 |
| MLT2A1_dup346 | 3.6995508773911 | -3.14500083674877 | 0.94300278091691 | -3.3350917944174 | 0.000852712459679607 | 0.0314457951614279 | 5425 | 2 | 2 | 14 | 15 | 0 | 0 | 14 | 2 | 0 | 0 | 2 | 0 |
| L1MC4_dup2388 | 3.08899264892844 | -3.50040329536586 | 0.958470556114532 | -3.65207180652044 | 0.000260133118454041 | 0.0135192207904764 | 5426 | 1 | 1 | 12 | 11 | 0 | 0 | 10 | 5 | 0 | 0 | 1 | 0 |
| L2b_dup26245 | 3.12650586371505 | -2.80722273331421 | 0.891831893442789 | -3.14770390468694 | 0.0016455828978622 | 0.0495582670819793 | 5427 | 2 | 2 | 11 | 10 | 0 | 0 | 12 | 4 | 0 | 0 | 1 | 0 |
| L1PA7_dup9990 | 3.99645183134659 | -3.42003386405107 | 0.937836474382402 | -3.64672728932118 | 0.000265601546892022 | 0.0137105247477473 | 5428 | 1 | 2 | 15 | 23 | 0 | 0 | 7 | 5 | 0 | 0 | 1 | 0 |
| L1MB8_dup15925 | 4.45209457073547 | -2.67876170497336 | 0.835722635735482 | -3.20532386036894 | 0.00134910590091453 | 0.0431935350779076 | 5429 | 2 | 5 | 18 | 18 | 0 | 0 | 8 | 6 | 0 | 0 | 1 | 1 |
| L1ME1_dup12949 | 4.29031351917681 | -3.65110223051415 | 0.883610317005755 | -4.13202761471421 | 3.59577208536488e-05 | 0.00317241225296341 | 5430 | 1 | 2 | 17 | 21 | 0 | 0 | 10 | 5 | 0 | 0 | 1 | 1 |
| L2c_dup59406 | 5.61242722430476 | -3.59645401397765 | 0.826992896768394 | -4.34883301662126 | 1.36863869966991e-05 | 0.00154123097033439 | 5431 | 2 | 2 | 24 | 25 | 0 | 0 | 16 | 6 | 0 | 0 | 1 | 0 |
| L1MC_dup7879 | 5.2410519195688 | -3.89858728623392 | 0.921247690243379 | -4.23185569692335 | 2.31771098082082e-05 | 0.00228265752087082 | 5432 | 2 | 1 | 23 | 31 | 0 | 0 | 9 | 4 | 0 | 0 | 1 | 1 |
| L2c_dup18343 | 6.47384021739821 | -3.88365037382497 | 0.821343848335992 | -4.72840988788445 | 2.2628501447618e-06 | 0.000385975727989599 | 5433 | 2 | 2 | 24 | 37 | 0 | 0 | 16 | 6 | 0 | 0 | 1 | 1 |
| Tigger3b_dup2786 | 3.98706648352446 | -2.83445165607273 | 0.828083856402066 | -3.42290413483981 | 0.000619559233893798 | 0.0250973172400568 | 5434 | 1 | 2 | 7 | 21 | 1 | 0 | 8 | 10 | 0 | 1 | 1 | 0 |
| L1PB2_dup674 | 4.48856920210342 | -2.90775177327967 | 0.763739979470334 | -3.80725358294879 | 0.000140518639519896 | 0.00868012673914882 | 5435 | 2 | 3 | 11 | 18 | 1 | 0 | 10 | 11 | 0 | 0 | 2 | 0 |
| MER1A_dup1654 | 3.3943291048233 | -3.01208160108946 | 0.877482126600687 | -3.43264154309112 | 0.000597731790144683 | 0.0245135967834827 | 5436 | 1 | 2 | 5 | 15 | 1 | 0 | 9 | 9 | 0 | 0 | 2 | 0 |
| MLT1A1-int_dup187 | 4.693845597917 | -2.49128387899922 | 0.774323373200393 | -3.21736882189462 | 0.00129372172045288 | 0.0419312287919356 | 5437 | 1 | 5 | 15 | 19 | 1 | 0 | 11 | 7 | 0 | 1 | 2 | 0 |
| MSTB_dup4365 | 5.45165777245996 | -2.56131199513285 | 0.707026995874982 | -3.62265091725825 | 0.000291599227841402 | 0.0146598077050391 | 5438 | 3 | 4 | 21 | 20 | 2 | 0 | 10 | 10 | 0 | 0 | 1 | 0 |
| L1MC5a_dup14617 | 4.63443392666883 | -2.95053269837755 | 0.82490653454068 | -3.57680849263783 | 0.000347814792702332 | 0.016618702543042 | 5439 | 2 | 2 | 20 | 22 | 2 | 0 | 7 | 5 | 0 | 0 | 2 | 0 |
| MIRb_dup90020 | 4.79313022835538 | -3.49713934478433 | 0.801566946780404 | -4.36287868259917 | 1.28362132575555e-05 | 0.00147066114457063 | 5440 | 1 | 2 | 17 | 20 | 1 | 0 | 14 | 7 | 0 | 0 | 2 | 0 |
| BLACKJACK_dup1902 | 4.8933101787357 | -3.23840808806114 | 0.809936657856135 | -3.99834734809146 | 6.37862983467361e-05 | 0.00489287380606462 | 5441 | 0 | 2 | 16 | 13 | 2 | 0 | 15 | 14 | 0 | 0 | 0 | 0 |
| L1MEi_dup1846 | 5.49328275985616 | -2.96482933709246 | 0.745880785456224 | -3.97493727537035 | 7.03978072541068e-05 | 0.00526757127197664 | 5442 | 1 | 4 | 12 | 20 | 1 | 0 | 21 | 13 | 0 | 0 | 0 | 0 |
| L2c_dup74406 | 4.43897977299489 | -2.42738042852662 | 0.743873644330147 | -3.26316229514014 | 0.00110176405650379 | 0.0375265316617307 | 5443 | 2 | 2 | 14 | 8 | 3 | 0 | 19 | 7 | 1 | 0 | 2 | 0 |
| MIR_dup117029 | 4.78049394036561 | -3.26574083101288 | 0.857739277775967 | -3.80738170167585 | 0.000140445885256336 | 0.0086778853838494 | 5444 | 0 | 4 | 14 | 11 | 1 | 0 | 19 | 10 | 0 | 0 | 3 | 0 |
| MER94_dup2035 | 2.99127549681611 | -3.19033834634318 | 0.900097921670373 | -3.54443474374728 | 0.000393455888621689 | 0.0182673636422175 | 5445 | 1 | 0 | 8 | 11 | 2 | 0 | 11 | 5 | 0 | 0 | 1 | 1 |
| L1PA16_dup1421 | 3.5020006271142 | -2.94592442901723 | 0.865933171999507 | -3.40202283995526 | 0.000668890474289106 | 0.0264736880896866 | 5446 | 1 | 0 | 10 | 11 | 3 | 0 | 13 | 7 | 0 | 0 | 1 | 0 |

| | | | | | | | | | | | | | | | | | | | |
|---|---|---|---|---|---|---|---|---|---|---|---|---|---|---|---|---|---|---|---|
| MLT1A0_dup1311 | 3.4175684507487 | -3.2372912104319 | 0.895568466884269 | -3.61478918713452 | 0.000300592118749492 | 0.0149789931493124 | 5447 | 1 | 0 | 13 | 12 | 2 | 0 | 10 | 6 | 0 | 0 | 1 | 0 |
| L1MC4_dup12451 | 2.816577787149 | -3.35291744248561 | 0.978820214906811 | -3.42546812113482 | 0.000613740953585256 | 0.0249550931206079 | 5448 | 0 | 1 | 7 | 11 | 1 | 0 | 7 | 8 | 0 | 0 | 1 | 0 |
| L1MA4_dup4996 | 2.50651108059327 | -3.2158736093203 | 0.992552256444877 | -3.24000433069279 | 0.00119527884198654 | 0.0397223211362191 | 5449 | 0 | 1 | 7 | 10 | 1 | 0 | 8 | 5 | 0 | 0 | 1 | 0 |
| THE1B-int_dup3371 | 2.82198548161109 | -2.95948097114244 | 0.933386348725496 | -3.17069236676056 | 0.00152076102909722 | 0.0469425684457881 | 5450 | 0 | 2 | 10 | 10 | 1 | 0 | 8 | 5 | 0 | 0 | 1 | 0 |
| L1M2_dup3805 | 3.3717587968321 | -2.88909372251765 | 0.857684270651004 | -3.36848164456223 | 0.000755834296517956 | 0.0289097809868609 | 5451 | 1 | 2 | 10 | 13 | 1 | 0 | 6 | 9 | 0 | 0 | 1 | 0 |
| MLT1A0_dup9052 | 3.08909130660265 | -3.11138424760468 | 0.894867784143387 | -3.47692061636016 | 0.000507207966898936 | 0.0217865777291502 | 5452 | 1 | 1 | 11 | 13 | 1 | 0 | 8 | 5 | 0 | 0 | 1 | 0 |
| HERVK9-int_dup187 | 2.66515446752945 | -3.16045076963276 | 0.956138240773135 | -3.30543287033182 | 0.000948298173516723 | 0.033880845984224 | 5453 | 1 | 1 | 8 | 4 | 1 | 0 | 6 | 8 | 0 | 0 | 3 | 1 |
| L2a_dup106865 | 2.88465574499708 | -3.12238017210378 | 0.957590848293906 | -3.26066208513456 | 0.00111152418532942 | 0.0377347316025674 | 5454 | 1 | 1 | 5 | 5 | 1 | 0 | 14 | 8 | 0 | 0 | 2 | 0 |
| ERVL-E-int_dup4369 | 4.78822336591231 | -3.41291487155941 | 0.777664174322438 | -4.38867442303486 | 1.14043648964385e-05 | 0.00134404309541751 | 5455 | 0 | 1 | 13 | 18 | 1 | 1 | 17 | 10 | 0 | 1 | 1 | 0 |
| LTR1D_dup273 | 3.91327677208737 | -3.07713377145904 | 0.883411991883887 | -3.48323749250564 | 0.00049538868694912 | 0.0214397520015997 | 5456 | 0 | 1 | 13 | 15 | 1 | 0 | 13 | 6 | 0 | 2 | 1 | 0 |
| L1PA5_dup6270 | 3.64455650382868 | -2.78795381673971 | 0.826729699967068 | -3.37226764304073 | 0.000745519626613334 | 0.028621073422739 | 5457 | 0 | 1 | 7 | 13 | 1 | 2 | 8 | 11 | 0 | 1 | 2 | 0 |
| LTR40A1_dup501 | 4.29092882140233 | -3.40083888435448 | 0.82858958812907 | -4.1043707682032 | 4.0541682077747e-05 | 0.00347504722034279 | 5458 | 1 | 1 | 9 | 18 | 0 | 1 | 12 | 10 | 0 | 1 | 3 | 0 |
| L2a_dup86172 | 4.57262194743373 | -3.77488804119073 | 0.983119677545768 | -3.83970347395981 | 0.000123183001070386 | 0.0079026428613357 | 5459 | 0 | 0 | 11 | 27 | 2 | 0 | 6 | 10 | 0 | 1 | 3 | 0 |
| MLT1A1_dup596 | 5.20943830064139 | -3.28293273890753 | 0.797499230493473 | -4.11653405217223 | 3.84612526940339e-05 | 0.00334235176452732 | 5460 | 2 | 0 | 16 | 17 | 1 | 0 | 17 | 11 | 0 | 2 | 2 | 0 |
| AluSg_dup35345 | 4.26437130782538 | -3.221054234698 | 0.862807611842254 | -3.7332241747618 | 0.000189044197241126 | 0.0106795609109062 | 5461 | 1 | 0 | 17 | 22 | 1 | 1 | 10 | 4 | 0 | 1 | 1 | 0 |
| AluJb_dup47100 | 4.62262121767709 | -2.84646700190024 | 0.742333522414906 | -3.83448533031395 | 0.000125827481168471 | 0.00801825008947859 | 5462 | 1 | 0 | 15 | 18 | 2 | 2 | 10 | 10 | 0 | 1 | 1 | 0 |
| MLT1O_dup801 | 4.10236621730551 | -2.79144253356466 | 0.769536775485507 | -3.62743227158119 | 0.000286253829604672 | 0.014461348989196 | 5463 | 1 | 0 | 11 | 14 | 2 | 2 | 10 | 9 | 0 | 1 | 3 | 0 |
| MER67D_dup43 | 4.8107393697516 | -4.09854852859891 | 0.906929710179504 | -4.5191468341992 | 6.20893242321361e-06 | 0.000850052148661672 | 5464 | 2 | 1 | 10 | 23 | 0 | 0 | 15 | 7 | 0 | 0 | 6 | 1 |
| Tigger3c_dup212 | 5.95065342400709 | -4.20592722209591 | 0.905600101564879 | -4.64435374380817 | 3.41142615120388e-06 | 0.000532634650245643 | 5465 | 1 | 2 | 11 | 35 | 0 | 0 | 13 | 13 | 0 | 0 | 4 | 0 |
| MIRb_dup213168 | 4.20516609153038 | -3.06202803613027 | 0.861199813816305 | -3.55553727138102 | 0.000377207463534057 | 0.0176987900832067 | 5466 | 3 | 0 | 7 | 11 | 1 | 0 | 14 | 13 | 1 | 0 | 3 | 0 |
| MIR_dup19186 | 3.413818795759 | -3.92303635977027 | 1.03593693833024 | -3.78694514561239 | 0.000152510751939144 | 0.00921654526569828 | 5467 | 1 | 0 | 6 | 13 | 1 | 0 | 7 | 11 | 0 | 0 | 4 | 0 |
| MLT1F2_dup4043 | 4.95650832295302 | -4.36216922944941 | 0.964668968046876 | -4.52193381765074 | 6.12772205820838e-06 | 0.000841760032295107 | 5468 | 1 | 0 | 13 | 13 | 1 | 0 | 11 | 18 | 0 | 0 | 4 | 0 |
| L1MEc_dup102 | 7.38087390886815 | -4.46764732161638 | 1.26575684808337 | -3.52962524230571 | 0.000416148683988354 | 0.0190351623914442 | 5469 | 0 | 0 | 35 | 35 | 0 | 0 | 3 | 4 | 7 | 7 | 3 | 2 |
| L1PA13_dup5293 | 7.65015476734109 | -4.45896097631902 | 1.22779543660572 | -3.63168068831234 | 0.000281581399316811 | 0.014279130225429 | 5470 | 0 | 0 | 37 | 34 | 0 | 0 | 4 | 5 | 8 | 5 | 3 | 3 |
| L1PA14_dup1953 | 8.25621844091253 | -4.59542863235667 | 1.26181006597039 | -3.64193372385454 | 0.000270597747548106 | 0.0138826329534458 | 5471 | 0 | 0 | 31 | 47 | 0 | 0 | 5 | 4 | 8 | 9 | 2 | 3 |
| LTR76_dup85 | 7.00110803809777 | -4.33391849120657 | 1.28137801904718 | -3.38223258615691 | 0.000718992415069193 | 0.0278770358800805 | 5472 | 0 | 0 | 24 | 37 | 0 | 0 | 6 | 4 | 9 | 8 | 2 | 1 |
| THE1A-int_dup3 | 10.9452882871589 | -5.41186925924194 | 1.42743067103622 | -3.79133597802916 | 0.000149839090919715 | 0.00908197123483002 | 5473 | 0 | 0 | 50 | 54 | 0 | 0 | 9 | 9 | 8 | 5 | 3 | 6 |
| L1MA4A_dup3627 | 9.38045772111208 | -4.7175032183792 | 1.21321811016154 | -3.88842136369946 | 0.00010089834942373 | 0.0068654156854386 | 5474 | 0 | 0 | 33 | 51 | 0 | 0 | 11 | 7 | 7 | 10 | 3 | 2 |
| THE1C-int_dup1139 | 10.7782480176781 | -5.45613403907887 | 1.51658680919188 | -3.59764044234711 | 0.00032111711256008 | 0.015692859409575 | 5475 | 0 | 0 | 43 | 53 | 0 | 0 | 18 | 8 | 6 | 11 | 3 | 1 |
| L1PA13_dup3472 | 10.4350405808646 | -3.96435437121427 | 1.03807079068557 | -3.81896341442775 | 0.000134013634673691 | 0.00837835215871417 | 5476 | 0 | 0 | 36 | 54 | 0 | 1 | 21 | 7 | 7 | 7 | 2 | 5 |
| THE1D-int_dup1009 | 8.70153945524023 | -4.5237064629973 | 1.13789922188651 | -3.97548954774529 | 7.02346287631615e-05 | 0.00526198015607029 | 5477 | 0 | 0 | 22 | 54 | 0 | 0 | 15 | 8 | 6 | 5 | 3 | 4 |
| HERVL40-int_dup602 | 13.799072485127 | -3.27691182215106 | 0.974406824122134 | -3.36298119125274 | 0.000771056170424869 | 0.0292801997331497 | 5478 | 0 | 0 | 29 | 62 | 1 | 1 | 10 | 5 | 20 | 30 | 12 | 7 |
| HERVL-int_dup1388 | 14.7296092023171 | -4.66824082028493 | 1.46077966636925 | -3.19571864789697 | 0.00139483053660087 | 0.0442040892930238 | 5479 | 0 | 0 | 51 | 66 | 1 | 0 | 13 | 11 | 24 | 17 | 2 | 2 |
| MLT2A2_dup2308 | 9.6970209688086 | -3.28663799104887 | 1.03424167710193 | -3.17782396882169 | 0.00148384810988988 | 0.0461536891968441 | 5480 | 0 | 0 | 41 | 36 | 1 | 1 | 16 | 9 | 9 | 9 | 2 | 1 |
| HERV4_I-int_dup189 | 11.4020219720734 | -3.02100990434114 | 0.886939948316803 | -3.40610422393792 | 0.000658970174391399 | 0.0261979648476901 | 5481 | 1 | 0 | 39 | 46 | 1 | 1 | 11 | 13 | 10 | 14 | 6 | 4 |
| L1M1_dup5533 | 13.1718069288932 | -3.27535894271379 | 0.920891467543803 | -3.5567263441476 | 0.000375504956805845 | 0.0176441355553167 | 5482 | 1 | 0 | 36 | 67 | 1 | 1 | 23 | 11 | 10 | 16 | 5 | 3 |
| Arthur1_dup434 | 22.7691050147165 | -4.7588752619431 | 0.778186113831398 | -6.115343331575 | 9.63492990577769e-10 | 6.94725161018236e-07 | 5483 | 0 | 1 | 91 | 103 | 0 | 2 | 59 | 34 | 2 | 2 | 4 | 6 |
| Arthur1_dup435 | 20.5551178228781 | -6.29417138453112 | 1.30614987587675 | -4.81887377610946 | 1.44370850619517e-06 | 0.000274474461862497 | 5484 | 0 | 0 | 77 | 84 | 0 | 0 | 45 | 39 | 5 | 3 | 8 | 7 |
| THE1B_dup3264 | 14.6504784804265 | -5.56623705679066 | 0.864180643800256 | -6.4410573144904 | 1.18644055132104e-10 | 1.18120829884785e-07 | 5485 | 0 | 0 | 57 | 79 | 0 | 1 | 21 | 22 | 0 | 2 | 6 | 8 |
| MIRb_dup38502 | 12.6319100349812 | -4.50626519704104 | 0.756853035952051 | -5.95395008407752 | 2.6174666540138e-09 | 1.63898873804202e-06 | 5486 | 0 | 0 | 59 | 40 | 1 | 1 | 31 | 24 | 1 | 2 | 2 | 3 |
| MLT1D_dup2952 | 13.9398986301042 | -4.63012196337176 | 0.900175782486705 | -5.14357534767399 | 2.69558469339886e-07 | 7.33032680520555e-05 | 5487 | 1 | 0 | 47 | 67 | 0 | 0 | 28 | 29 | 1 | 3 | 3 | 4 |
| L1MC1_dup10413 | 20.0171776128991 | -3.41227213652312 | 0.72109134555221 | -4.73209414808606 | 2.2221529581987e-06 | 0.000380397415677373 | 5488 | 7 | 0 | 87 | 100 | 0 | 3 | 22 | 15 | 4 | 5 | 12 | 15 |
| MLT1K_dup122 | 16.3890400332211 | -5.09222511970987 | 1.31032260105327 | -3.88623772162412 | 0.000101809812132959 | 0.00690177708199119 | 5489 | 0 | 1 | 92 | 83 | 0 | 0 | 10 | 11 | 5 | 6 | 7 | 3 |
| THE1D_dup10197 | 14.8757313889369 | -3.65926968160107 | 0.890115421243221 | -4.11100582494142 | 3.9393926124692e-05 | 0.0033987045488274 | 5490 | 2 | 1 | 56 | 77 | 0 | 0 | 21 | 7 | 7 | 11 | 12 | 6 |
| MER77_dup917 | 10.5760003313299 | -3.07228648937885 | 0.852967811508642 | -3.60187857962061 | 0.000315925925155857 | 0.0154996515849764 | 5491 | 1 | 4 | 52 | 59 | 0 | 1 | 5 | 7 | 2 | 3 | 5 | 3 |
| THE1D-int_dup673 | 14.4458871367267 | -3.12401360434135 | 0.823369516156696 | -3.7941817653434 | 0.000148131133305596 | 0.00902155378692947 | 5492 | 5 | 4 | 64 | 99 | 0 | 1 | 8 | 6 | 2 | 2 | 4 | 4 |
| L2a_dup20678 | 10.9548540692005 | -3.75401795657577 | 0.793265054420011 | -4.73236270230066 | 2.21921409198119e-06 | 0.000380167830331575 | 5493 | 1 | 3 | 41 | 55 | 0 | 0 | 11 | 4 | 3 | 2 | 9 | 21 |
| L2a_dup82413 | 14.8226622874595 | -3.63855367827885 | 0.785794298266262 | -4.6304149652228 | 3.64933631831765e-06 | 0.000559861366601167 | 5494 | 4 | 4 | 72 | 76 | 0 | 0 | 14 | 5 | 3 | 1 | 10 | 13 |
| AluJo_dup44013 | 10.7214326625749 | -1.70158274006216 | 0.514187870193768 | -3.30926270085082 | 0.0009354202323043 | 0.0335776495423195 | 5495 | 10 | 4 | 18 | 10 | 6 | 2 | 20 | 28 | 1 | 8 | 10 | 17 |
| LTR1B_dup17 | 12.1655967869833 | -2.25315306789315 | 0.571288639405992 | -3.94398367563534 | 8.01391157118657e-05 | 0.00578718726166179 | 5496 | 5 | 2 | 13 | 14 | 7 | 2 | 50 | 34 | 2 | 6 | 8 | 11 |
| L2a_dup139684 | 9.97012067022679 | -1.47615608103972 | 0.468415884033188 | -3.15137921525978 | 0.00162501357948727 | 0.0490771125227367 | 5497 | 9 | 3 | 14 | 17 | 9 | 5 | 23 | 24 | 3 | 3 | 11 | 4 |
| AluSx1_dup103889 | 8.63407495965161 | -1.91030615427989 | 0.515887339747676 | -3.70295218955021 | 0.00021310505499506 | 0.0116274926429384 | 5498 | 3 | 3 | 10 | 8 | 6 | 5 | 17 | 23 | 3 | 2 | 12 | 15 |
| L1PA8A_dup194 | 8.79035272816267 | -1.43030049590567 | 0.44682592774069 | -3.20102394938847 | 0.00136940149552336 | 0.0436729135844795 | 5499 | 5 | 3 | 11 | 12 | 5 | 9 | 22 | 17 | 4 | 3 | 8 | 13 |
| L1M4_dup17080 | 12.5827047669366 | -2.95661341106993 | 0.529367744002056 | -5.58517862973238 | 2.3346012552259e-08 | 1.04813024580374e-05 | 5500 | 5 | 4 | 13 | 18 | 3 | 3 | 59 | 26 | 2 | 1 | 13 | 18 |
| L1MA5A_dup2459 | 11.867783916322 | -2.14076280734131 | 0.508647696926152 | -4.2087339041901 | 2.56805599620525e-05 | 0.00245997847050344 | 5501 | 2 | 3 | 15 | 14 | 8 | 7 | 43 | 30 | 3 | 3 | 11 | 11 |
| L2a_dup18238 | 11.6960154341843 | -2.2985335459281 | 0.452133919926588 | -5.08374498047239 | 3.70064732666855e-07 | 9.48873091370124e-05 | 5502 | 2 | 4 | 18 | 21 | 4 | 7 | 44 | 18 | 2 | 5 | 12 | 17 |
| L1PA3_dup8230 | 16.7220350182736 | -2.05277252409543 | 0.485994713878402 | -4.22385772000195 | 2.40155696042443e-05 | 0.00234101135807108 | 5503 | 3 | 5 | 12 | 25 | 10 | 8 | 56 | 43 | 6 | 6 | 12 | 26 |
| L2b_dup63331 | 12.4143896734842 | -1.52291989012814 | 0.478322474557088 | -3.1838769264152 | 0.00145316783085566 | 0.0454967727211555 | 5504 | 5 | 10 | 10 | 15 | 4 | 7 | 31 | 34 | 4 | 8 | 12 | 14 |
| L1PA12_dup188 | 9.10192254927634 | -2.50193635027433 | 0.787214315014322 | -3.17821500772482 | 0.0014818481806039 | 0.0461276618500752 | 5505 | 1 | 0 | 12 | 24 | 3 | 0 | 18 | 23 | 6 | 12 | 6 | 8 |
| MLT2B3_dup299 | 8.87998601122934 | -2.46553420322942 | 0.777580786035219 | -3.17077562551519 | 0.00152032524794367 | 0.0469385378807843 | 5506 | 3 | 0 | 6 | 22 | 3 | 0 | 38 | 21 | 6 | 4 | 4 | 7 |
| MIR_dup164139 | 14.1470476868284 | -2.80297735485506 | 0.595644526234909 | -4.70578882437279 | 2.52886191885075e-06 | 0.000423457127475623 | 5507 | 1 | 1 | 19 | 12 | 3 | 5 | 50 | 40 | 6 | 5 | 17 | 17 |
| MER21A_dup165 | 15.5986320065074 | -2.78504633211168 | 0.819199525818798 | -3.39971672875177 | 0.000674556972156264 | 0.0266270957691928 | 5508 | 0 | 0 | 12 | 18 | 3 | 4 | 49 | 56 | 12 | 12 | 9 | 12 |
| MER21B_dup260 | 15.6183386064652 | -2.80900839607212 | 0.752055528481357 | -3.73510770108215 | 0.000187634867915048 | 0.0106378285158242 | 5509 | 0 | 1 | 12 | 17 | 3 | 3 | 56 | 50 | 13 | 9 | 12 | 14 |
| LTR33C_dup504 | 11.8684958901757 | -3.79987614505421 | 0.861554495585947 | -4.41048844213842 | 1.03137711226185e-05 | 0.00125082598549571 | 5510 | 0 | 0 | 21 | 18 | 0 | 2 | 40 | 39 | 2 | 8 | 9 | 8 |
| L1PA3_dup5677 | 9.30670295045976 | -3.79274102769921 | 0.715767428954509 | -5.29884551080944 | 1.16537183156627e-07 | 3.72978423936932e-05 | 5511 | 1 | 0 | 14 | 15 | 1 | 2 | 40 | 28 | 2 | 1 | 6 | 8 |
| L1PA7_dup13356 | 9.04113538917707 | -2.3024238250905 | 0.671392897129583 | -3.42932407377869 | 0.00060508656385597 | 0.0247268959871324 | 5512 | 0 | 0 | 17 | 18 | 2 | 8 | 30 | 20 | 3 | 4 | 4 | 10 |
| AluJo_dup60286 | 13.6360407832693 | -1.3651278041773 | 0.369273158865933 | -3.69679672459736 | 0.000218337029584376 | 0.0118558701144418 | 5513 | 19 | 9 | 29 | 29 | 8 | 9 | 19 | 20 | 3 | 2 | 13 | 17 |
| MIR_dup168043 | 15.7155810654699 | -1.29153502846313 | 0.347176960953182 | -3.72010580689799 | 0.000199139333359581 | 0.0110814798446162 | 5514 | 18 | 12 | 32 | 25 | 17 | 12 | 35 | 20 | 1 | 2 | 12 | 20 |
| AluSg_dup35425 | 14.24135454086 | -1.65204164486062 | 0.427305376978125 | -3.86618501396764 | 0.000110551084862046 | 0.00732413496475379 | 5515 | 13 | 7 | 29 | 43 | 14 | 12 | 20 | 13 | 1 | 1 | 21 | 15 |
| BSR/Beta_dup2035 | 11.0811314697214 | -1.62119480419226 | 0.418573795269765 | -3.87313974862527 | 0.000107442177636958 | 0.00718242824695875 | 5516 | 7 | 7 | 30 | 21 | 8 | 11 | 22 | 9 | 1 | 2 | 11 | 18 |
| AluSz_dup102969 | 12.6913370276328 | -1.39260142576956 | 0.437514951162917 | -3.18298019774642 | 0.00145767583077073 | 0.0455839312956463 | 5517 | 11 | 7 | 21 | 22 | 9 | 17 | 19 | 11 | 1 | 2 | 21 | 27 |
| MIRb_dup1008 | 12.3003603476159 | -1.42635716206818 | 0.39422234212386 | -3.61815404571879 | 0.00029671178144037 | 0.0148478194630095 | 5518 | 8 | 8 | 27 | 20 | 13 | 9 | 17 | 13 | 1 | 4 | 21 | 19 |
| Arthur1B_dup2027 | 13.1934744313968 | -1.43769648839149 | 0.432505767660228 | -3.32410940128995 | 0.000887013919353066 | 0.0323598422407214 | 5519 | 18 | 7 | 26 | 32 | 10 | 3 | 15 | 17 | 1 | 7 | 15 | 22 |
| L1PA16_dup5441 | 12.967182948866 | -1.9586488436883 | 0.469353432716659 | -4.17307876572132 | 3.00510992964688e-05 | 0.00275976027525958 | 5520 | 10 | 6 | 31 | 51 | 7 | 3 | 12 | 16 | 2 | 5 | 9 | 20 |
| L2c_dup119824 | 12.6504082277787 | -1.2752207324717 | 0.39348522895932 | -3.24083507745482 | 0.00119180113209212 | 0.0396344251819416 | 5521 | 11 | 15 | 31 | 24 | 3 | 7 | 26 | 11 | 4 | 6 | 12 | 16 |
| MER34_dup439 | 14.1819196316118 | -1.42324782105481 | 0.447175274678732 | -3.1827516002027 | 0.00145882708733251 | 0.0456012384201789 | 5522 | 12 | 22 | 36 | 28 | 4 | 4 | 22 | 21 | 2 | 5 | 11 | 16 |
| L1MB7_dup18739 | 14.0826676269815 | -2.13884524247839 | 0.51874501542483 | -4.12311478448957 | 3.7378331539399e-05 | 0.00327239977457619 | 5523 | 17 | 11 | 38 | 41 | 3 | 4 | 19 | 12 | 1 | 1 | 15 | 27 |
| L2a_dup143044 | 13.5601437068631 | -2.0814419272131 | 0.494770045879922 | -4.20688751177603 | 2.5891193490209e-05 | 0.00247120173534748 | 5524 | 12 | 13 | 33 | 44 | 2 | 5 | 20 | 10 | 2 | 1 | 18 | 22 |
| HERV9NC-int_dup132 | 14.2038474300688 | -1.6384182644672 | 0.428622581204247 | -3.82251970921351 | 0.000132094916657161 | 0.00829544920530608 | 5525 | 12 | 15 | 33 | 30 | 6 | 4 | 22 | 8 | 3 | 4 | 23 | 29 |

| | | | | | | | | | | | | | | | | | | | |
|---|---|---|---|---|---|---|---|---|---|---|---|---|---|---|---|---|---|---|---|
| MIR_dup44945 | 14.7981065563411 | -1.69219473574647 | 0.437047043341633 | -3.87188235574839 | 0.000107998078311114 | 0.00720631260286491 | 5526 | 10 | 18 | 35 | 38 | 6 | 2 | 22 | 13 | 4 | 4 | 19 | 24 |
| Tigger7_dup3032 | 11.9827554561306 | -1.94450135929336 | 0.454414250173553 | -4.27913816204203 | 1.87618378229022e-05 | 0.00194777672888669 | 5527 | 4 | 11 | 24 | 47 | 7 | 9 | 21 | 12 | 1 | 1 | 13 | 10 |
| AluSc8_dup21210 | 10.9148166389282 | -1.98342604285812 | 0.399957596458997 | -4.95909081467204 | 7.0823855473871e-07 | 0.000156474030776098 | 5528 | 6 | 6 | 24 | 30 | 6 | 6 | 18 | 16 | 1 | 3 | 11 | 16 |
| AluSx3_dup31281 | 13.4005398556288 | -1.75285357921086 | 0.384539480000132 | -4.55831889929809 | 5.15647008302842e-06 | 0.000739578921263581 | 5529 | 10 | 6 | 23 | 41 | 11 | 9 | 19 | 21 | 1 | 3 | 16 | 15 |
| L1MCa_dup6565 | 13.1019125464796 | -1.91375318651307 | 0.409216174312766 | -4.67663134216777 | 2.91625843255398e-06 | 0.000471728095848055 | 5530 | 6 | 9 | 21 | 51 | 9 | 6 | 20 | 16 | 2 | 3 | 14 | 17 |
| L1PA11_dup2178 | 14.815995762856 | -2.23397825950607 | 0.446286423075913 | -5.00570517944274 | 5.56578902956164e-07 | 0.000130094031083308 | 5531 | 9 | 8 | 35 | 68 | 4 | 4 | 27 | 19 | 2 | 4 | 9 | 9 |
| LTR16A1_dup2229 | 12.9257462178679 | -2.15666458233592 | 0.4826338514956 | -4.46853152892774 | 7.87584053386173e-06 | 0.00102015500907472 | 5532 | 3 | 9 | 29 | 58 | 4 | 4 | 23 | 18 | 3 | 4 | 8 | 8 |
| LTR16E2_dup1976 | 13.9007126599977 | -2.36815346391084 | 0.462347608276089 | -5.12201949684728 | 3.02280626677817e-07 | 8.04546599502012e-05 | 5533 | 4 | 7 | 39 | 61 | 4 | 6 | 25 | 13 | 2 | 4 | 9 | 13 |
| L2a_dup147227 | 12.7913303468226 | -1.71123235644145 | 0.443161945380111 | -3.86141539065065 | 0.000112732052088547 | 0.00742228579586646 | 5534 | 10 | 8 | 39 | 41 | 3 | 7 | 25 | 15 | 5 | 2 | 6 | 7 |
| THE1D_dup11291 | 13.4925039394666 | -1.94124377612308 | 0.423025236530256 | -4.58895500430561 | 4.45470428981915e-06 | 0.000655927640436452 | 5535 | 7 | 5 | 32 | 47 | 6 | 5 | 33 | 18 | 5 | 4 | 6 | 10 |
| L1PA16_dup7551 | 10.4866775543563 | -1.92839590683392 | 0.46020241343504 | -4.19032115116477 | 2.78559907567163e-05 | 0.00261070479052471 | 5536 | 4 | 4 | 28 | 28 | 3 | 8 | 13 | 20 | 3 | 4 | 7 | 12 |
| MIR_dup149089 | 11.3194615132976 | -2.04818725847781 | 0.464938349884191 | -4.40528783867363 | 1.05643504920832e-05 | 0.0012739964894005 | 5537 | 7 | 4 | 32 | 36 | 3 | 4 | 14 | 18 | 4 | 4 | 11 | 9 |
| L1PA3_dup9880 | 10.8482734218316 | -1.62340601706496 | 0.434862088690181 | -3.73315140428708 | 0.000189098846119967 | 0.0106801125181252 | 5538 | 6 | 7 | 28 | 30 | 9 | 3 | 18 | 16 | 3 | 4 | 4 | 13 |
| Tigger13a_dup2707 | 10.4379233053945 | -1.65472723573951 | 0.433372976561493 | -3.81825200285583 | 0.000134400599160042 | 0.00839372308864447 | 5539 | 8 | 6 | 25 | 33 | 5 | 3 | 20 | 14 | 3 | 5 | 5 | 10 |
| AluJr_dup68011 | 9.9406890672075 | -1.49727961750754 | 0.428935327217587 | -3.49068850826551 | 0.000481777600741725 | 0.0209957972501997 | 5540 | 6 | 9 | 13 | 35 | 5 | 6 | 21 | 15 | 2 | 4 | 5 | 10 |
| L1PA16_dup6686 | 11.0073094369362 | -1.83407288832853 | 0.425421873600143 | -4.31118614754725 | 1.62381132024515e-05 | 0.00175149323847258 | 5541 | 4 | 5 | 20 | 41 | 7 | 8 | 21 | 12 | 2 | 4 | 8 | 15 |
| L1PA7_dup6599 | 11.1390094597893 | -1.84378924798996 | 0.43294804626754 | -4.25868476341522 | 2.05633216021169e-05 | 0.00208570290670831 | 5542 | 6 | 5 | 23 | 46 | 7 | 7 | 19 | 13 | 3 | 2 | 8 | 9 |
| ERV3-16A3_I-int_dup44 | 9.97782536060614 | -1.78165530906317 | 0.496311040957423 | -3.58979583775975 | 0.000330937056594717 | 0.0160409755283126 | 5543 | 5 | 11 | 11 | 26 | 2 | 6 | 19 | 24 | 3 | 1 | 6 | 13 |
| AluJr_dup68012 | 8.11017704865857 | -1.43770009676514 | 0.446490666706235 | -3.22000033588845 | 0.00128190446533562 | 0.0417098372636057 | 5544 | 3 | 9 | 14 | 18 | 5 | 6 | 18 | 14 | 3 | 1 | 6 | 7 |
| Tigger1_dup8713 | 8.25176542227352 | -1.48300058774387 | 0.446183701678485 | -3.3237444177477 | 0.000888175539740465 | 0.0323785569898267 | 5545 | 4 | 8 | 13 | 17 | 2 | 8 | 17 | 14 | 2 | 3 | 10 | 8 |
| L1PB1_dup12602 | 9.65359579739998 | -1.71842783758941 | 0.440857996560013 | -3.89791690521255 | 9.70236947971926e-05 | 0.00666432509233256 | 5546 | 7 | 5 | 17 | 16 | 2 | 3 | 27 | 12 | 7 | 3 | 12 | 14 |
| L1PA4_dup6333 | 11.6013445298201 | -1.35867742257755 | 0.414962943729317 | -3.27421386200649 | 0.00105956387391536 | 0.0365654801367172 | 5547 | 4 | 9 | 18 | 27 | 4 | 6 | 25 | 18 | 9 | 6 | 9 | 13 |
| THE1B_dup9645 | 9.91419058044599 | -1.31907224687841 | 0.407002882495625 | -3.24094079823277 | 0.00119135923012382 | 0.03962931596808 | 5548 | 9 | 8 | 18 | 22 | 3 | 5 | 24 | 15 | 5 | 4 | 7 | 8 |
| MIRb_dup91829 | 9.43844156986759 | -1.40350385476517 | 0.442859822689272 | -3.16918307522768 | 0.00152868072306691 | 0.0471170289288703 | 5549 | 12 | 5 | 23 | 14 | 4 | 6 | 27 | 9 | 1 | 5 | 9 | 10 |
| AluJb_dup30098 | 9.4832747207299 | -1.79467649706348 | 0.493116233030356 | -3.63945937458725 | 0.000273211034532282 | 0.0139835390025421 | 5550 | 7 | 8 | 25 | 28 | 3 | 4 | 14 | 6 | 1 | 4 | 17 | 9 |
| AluY_dup101623 | 7.95714685633564 | -1.62795652933695 | 0.490169220748588 | -3.32121328803701 | 0.000896270158346142 | 0.0325544037700857 | 5551 | 4 | 10 | 17 | 21 | 4 | 4 | 21 | 6 | 1 | 2 | 10 | 6 |
| MIR1_Amn_dup4797 | 8.12971125263279 | -1.63073501877242 | 0.480565079896169 | -3.39336977860471 | 0.000690383744850291 | 0.0270721902689762 | 5552 | 6 | 6 | 18 | 21 | 3 | 5 | 23 | 5 | 1 | 4 | 11 | 6 |
| MIR_dup44936 | 9.98105162399635 | -1.70338553467148 | 0.520881588145464 | -3.27019724528214 | 0.0010747251688196 | 0.0369351624374795 | 5553 | 13 | 9 | 19 | 27 | 1 | 2 | 12 | 13 | 3 | 2 | 17 | 12 |
| FLAM_C_dup5372 | 9.8296804733208 | -2.07818162836827 | 0.501151264773751 | -4.14681509245812 | 3.37132111976965e-05 | 0.00301620717569165 | 5554 | 7 | 9 | 18 | 34 | 2 | 3 | 20 | 14 | 2 | 1 | 14 | 5 |
| THE1-int_dup182 | 11.1824313941341 | -1.89392393989909 | 0.487992416092954 | -3.88105199474726 | 0.000104005600251455 | 0.00701465888816122 | 5555 | 9 | 11 | 35 | 30 | 2 | 4 | 16 | 15 | 2 | 2 | 13 | 6 |
| L1MB7_dup55 | 10.8263255439791 | -1.77950406088252 | 0.445632832594589 | -3.99320680776995 | 6.51856672777063e-05 | 0.00497769755429928 | 5556 | 6 | 12 | 26 | 23 | 2 | 9 | 17 | 15 | 1 | 2 | 12 | 16 |
| L1MD3_dup394 | 8.65416413558203 | -1.97525509314414 | 0.519879846116234 | -3.7994454062805 | 0.00014502025315868 | 0.00888267567018937 | 5557 | 3 | 8 | 17 | 30 | 1 | 6 | 11 | 9 | 2 | 2 | 15 | 10 |
| MER5A_dup29020 | 7.69383980321176 | -1.79111899060304 | 0.476071083414265 | -3.76229318058465 | 0.000168362520088524 | 0.00989165140863799 | 5558 | 6 | 4 | 13 | 21 | 3 | 6 | 9 | 12 | 1 | 2 | 12 | 11 |
| MER4B-int_dup90 | 13.5380828959461 | -1.83743804486286 | 0.573625292643135 | -3.20320262796706 | 0.00135908319017531 | 0.0434369924472067 | 5559 | 9 | 8 | 11 | 23 | 2 | 3 | 51 | 38 | 7 | 3 | 8 | 7 |
| FRAM_dup6547 | 15.1637768171769 | -1.54309386338931 | 0.433643438412073 | -3.55843932296048 | 0.000373064948621575 | 0.0175559726861849 | 5560 | 14 | 4 | 25 | 34 | 6 | 8 | 43 | 30 | 9 | 3 | 9 | 9 |
| LTR40b_dup578 | 16.6868628623759 | -1.43426144451176 | 0.429230159678345 | -3.34147406041216 | 0.000833348029306772 | 0.0309589118159921 | 5561 | 15 | 9 | 18 | 35 | 5 | 8 | 42 | 40 | 8 | 7 | 15 | 8 |
| L1MC1_dup85 | 19.1887872803645 | -1.47897319084973 | 0.437552859849618 | -3.38010175812366 | 0.000724589975296802 | 0.0280255666875769 | 5562 | 19 | 10 | 19 | 27 | 9 | 9 | 68 | 49 | 5 | 7 | 7 | 15 |
| L1PB_dup871 | 17.5626615718498 | -1.73921463432061 | 0.442759761352998 | -3.92812262118369 | 8.56115598287894e-05 | 0.00608268982234133 | 5563 | 16 | 13 | 32 | 46 | 1 | 9 | 43 | 41 | 4 | 4 | 6 | 10 |
| LTR8_dup905 | 18.406733106706 | -1.94113518284062 | 0.592893988534957 | -3.27400044590969 | 0.00106036443983986 | 0.0365771929547892 | 5564 | 10 | 10 | 59 | 48 | 1 | 2 | 40 | 35 | 8 | 8 | 6 | 8 |
| MER52A_dup725 | 20.1426087871613 | -2.21675069252216 | 0.481154079136728 | -4.60715348501126 | 4.08218686014034e-06 | 0.000612060513309989 | 5565 | 9 | 9 | 58 | 65 | 3 | 4 | 52 | 35 | 4 | 8 | 7 | 9 |
| L1MA8_dup4860 | 15.8019981134355 | -2.12318079403476 | 0.531764008704344 | -3.9927124801243 | 6.53217549418054e-05 | 0.00498369423250741 | 5566 | 11 | 6 | 39 | 57 | 2 | 1 | 41 | 20 | 4 | 9 | 9 | 11 |
| L1ME1_dup27018 | 16.573965110417 | -2.12358368878966 | 0.496066944863502 | -4.28084094450999 | 1.86188399624321e-05 | 0.00194055124559828 | 5567 | 10 | 8 | 49 | 72 | 5 | 2 | 29 | 19 | 5 | 4 | 10 | 7 |
| AluJb_dup59480 | 13.1057868691245 | -2.02289901092323 | 0.602521835197223 | -3.35738705678786 | 0.000786828818461963 | 0.0297261811985617 | 5568 | 1 | 8 | 30 | 38 | 2 | 4 | 37 | 28 | 4 | 7 | 4 | 5 |
| MLT1D_dup10168 | 17.0440232784824 | -1.9689964551195 | 0.583088262366689 | -3.3768411786023 | 0.000733233744647242 | 0.0282725711560254 | 5569 | 3 | 9 | 43 | 51 | 4 | 4 | 54 | 28 | 5 | 10 | 2 | 10 |
| L1PA12_dup130 | 16.3120110021726 | -2.02154848137569 | 0.487329380505948 | -4.1482179450722 | 3.35073336337734e-05 | 0.00300525292454649 | 5570 | 7 | 6 | 25 | 36 | 3 | 5 | 61 | 30 | 3 | 13 | 10 | 14 |
| L1PBa1_dup36 | 14.5240746814437 | -2.35442615170225 | 0.547811269809028 | -4.29787826840989 | 1.7244081725879e-05 | 0.00183176797783304 | 5571 | 3 | 10 | 18 | 36 | 1 | 2 | 47 | 34 | 3 | 8 | 9 | 16 |
| AluSx1_dup114049 | 17.4240398847771 | -2.28205890151396 | 0.429959999821598 | -5.30760745757943 | 1.11073525248841e-07 | 3.59585048147765e-05 | 5572 | 7 | 9 | 53 | 46 | 15 | 3 | 38 | 21 | 3 | 1 | 21 | 11 |
| L1MB1_dup5373 | 18.4240103643161 | -2.57008833274199 | 0.465601507018749 | -5.51993130176552 | 3.3913222034993e-08 | 1.39503003945546e-05 | 5573 | 9 | 3 | 49 | 75 | 11 | 2 | 42 | 24 | 5 | 1 | 12 | 12 |
| L1MC3_dup11473 | 20.9375670506796 | -2.45963326238447 | 0.460344675533684 | -5.34302533103697 | 9.14079837624607e-08 | 3.09390670047546e-05 | 5574 | 6 | 9 | 70 | 96 | 8 | 5 | 37 | 20 | 3 | 5 | 11 | 11 |
| MLT1E2_dup3422 | 17.2805072787126 | -2.35122611181281 | 0.532592685392454 | -4.41467969106683 | 1.01159659804669e-05 | 0.00123312174241198 | 5575 | 2 | 8 | 56 | 72 | 5 | 4 | 39 | 15 | 5 | 6 | 11 | 8 |
| L1M4_dup17744 | 17.0974872448661 | -2.48220497654343 | 0.408829946345719 | -6.07148521954017 | 1.26732627764509e-09 | 8.79169733121102e-07 | 5576 | 4 | 10 | 44 | 62 | 4 | 5 | 34 | 24 | 2 | 6 | 13 | 18 |
| L1PA2_dup4555 | 16.2836863956492 | -2.02923183477549 | 0.413252324643359 | -4.91039424043588 | 9.08934677639917e-07 | 0.000191905386695393 | 5577 | 10 | 8 | 60 | 42 | 6 | 5 | 29 | 20 | 3 | 6 | 9 | 15 |
| LTR78_dup4185 | 16.7573025579663 | -2.40576452772961 | 0.446936290956434 | -5.38279073865612 | 7.33397826635919e-08 | 2.60915319258269e-05 | 5578 | 6 | 7 | 56 | 61 | 8 | 3 | 32 | 21 | 1 | 6 | 9 | 12 |
| HAL1ME_dup7590 | 18.6549031670661 | -2.35620634366286 | 0.395133543815044 | -5.96306332515713 | 2.47552427050797e-09 | 1.56244323095568e-06 | 5579 | 6 | 11 | 49 | 74 | 9 | 6 | 38 | 18 | 2 | 4 | 12 | 22 |
| L1MEc_dup16318 | 20.5985847026903 | -2.01606524001952 | 0.389412644571416 | -5.17719511198302 | 2.2524661932497e-07 | 6.32034576437493e-05 | 5580 | 9 | 14 | 56 | 86 | 8 | 11 | 34 | 19 | 4 | 4 | 12 | 19 |
| L2b_dup44447 | 16.4416504055598 | -2.19201260293271 | 0.416100101981661 | -5.26799342872865 | 1.37923043238399e-07 | 4.28996554854955e-05 | 5581 | 8 | 16 | 45 | 51 | 6 | 6 | 24 | 22 | 2 | 1 | 14 | 21 |
| MER102a_dup1380 | 16.0559582103825 | -2.33160644226591 | 0.416702089598923 | -5.59537977001769 | 2.20139359096038e-08 | 9.97738665513634e-06 | 5582 | 8 | 7 | 40 | 62 | 9 | 5 | 20 | 17 | 2 | 3 | 18 | 23 |
| L1PA13_dup5751 | 21.883378524121 | -2.48689866986083 | 0.394771079286016 | -6.29959690653794 | 2.98420683019568e-10 | 2.59153313291147e-07 | 5583 | 10 | 4 | 68 | 83 | 14 | 10 | 49 | 24 | 3 | 1 | 10 | 17 |
| L1PA8_dup5175 | 16.5936631285241 | -2.37286482751567 | 0.506558272624637 | -4.68428797978388 | 2.8093479483923e-06 | 0.000458172108929509 | 5584 | 2 | 4 | 40 | 84 | 10 | 12 | 33 | 18 | 3 | 1 | 6 | 11 |
| L1MA4A_dup5681 | 14.7848871369694 | -2.2979786616599 | 0.524358390475556 | -4.38245807333377 | 1.1734775380943e-05 | 0.00137076232832295 | 5585 | 1 | 6 | 18 | 56 | 5 | 11 | 42 | 36 | 1 | 4 | 5 | 7 |
| L1PREC2_dup6949 | 16.0866345727394 | -1.40954684485991 | 0.406413414683904 | -3.46825865961196 | 0.000523842814550562 | 0.0222701987405839 | 5586 | 8 | 5 | 37 | 40 | 17 | 14 | 36 | 33 | 2 | 5 | 4 | 5 |
| L1ME2_dup11350 | 14.3236358145496 | -1.74212997545407 | 0.522059610366725 | -3.33703266994798 | 0.000846779939662838 | 0.0313209854691456 | 5587 | 5 | 2 | 32 | 36 | 12 | 8 | 41 | 34 | 2 | 5 | 2 | 4 |
| MER61-int_dup1050 | 12.6652025161964 | -1.90330561530041 | 0.425667026353474 | -4.4713484894644 | 7.77279178998628e-06 | 0.00100900420908897 | 5588 | 8 | 3 | 28 | 30 | 10 | 7 | 38 | 28 | 1 | 3 | 4 | 3 |
| L1MA5_dup2424 | 13.8083510389667 | -2.0101336748762 | 0.51114791267705 | -3.93258707513539 | 8.40365236855839e-05 | 0.00599654610756935 | 5589 | 1 | 2 | 29 | 34 | 16 | 9 | 38 | 26 | 2 | 3 | 6 | 13 |
| L2_dup51125 | 15.246984207456 | -1.53366142869158 | 0.476164318776321 | -3.22086592425256 | 0.00127803922588887 | 0.0416296128190765 | 5590 | 4 | 6 | 27 | 40 | 20 | 8 | 53 | 26 | 4 | 2 | 3 | 6 |
| L1PA16_dup12584 | 19.724641357186 | -1.95658173765426 | 0.372528085108434 | -5.25217242905257 | 1.50315640041599e-07 | 4.57964216201516e-05 | 5591 | 13 | 5 | 46 | 74 | 15 | 9 | 33 | 28 | 2 | 6 | 16 | 13 |
| LTR43_dup80 | 16.1666767922484 | -1.44865867257574 | 0.388874532250445 | -3.72525982658789 | 0.000195114146180915 | 0.0109264840261624 | 5592 | 12 | 9 | 44 | 46 | 10 | 9 | 36 | 20 | 1 | 10 | 7 | 9 |
| L1MB1_dup4349 | 13.180028488434 | -1.52523382108003 | 0.383604721562142 | -3.97605591210887 | 7.00676582302498e-05 | 0.00525112409614331 | 5593 | 7 | 9 | 27 | 40 | 6 | 8 | 17 | 20 | 4 | 7 | 14 | 11 |
| AluJr_dup37707 | 15.4154418064116 | -1.28509540716175 | 0.374317644824978 | -3.43316812586442 | 0.000596572045897627 | 0.0244800517309606 | 5594 | 7 | 13 | 42 | 28 | 12 | 7 | 22 | 25 | 6 | 8 | 12 | 13 |
| Ricksha_c_dup832 | 13.5248143467611 | -1.24866865829235 | 0.390037036561482 | -3.20141048475923 | 0.00136756559290828 | 0.0436202094598062 | 5595 | 8 | 9 | 34 | 33 | 9 | 6 | 26 | 16 | 5 | 10 | 11 | 8 |
| L1MB5_dup4320 | 14.8507593961383 | -1.33417454939293 | 0.392986140761316 | -3.39496590594337 | 0.000686371472890885 | 0.0269585101851577 | 5596 | 13 | 8 | 39 | 43 | 4 | 13 | 26 | 17 | 6 | 6 | 10 | 9 |
| MLT1H1_dup3109 | 15.6474182190735 | -1.7630372812175 | 0.457460251870868 | -3.85396823878629 | 0.000116218678397779 | 0.00757997716800584 | 5597 | 4 | 9 | 40 | 59 | 5 | 8 | 26 | 17 | 9 | 5 | 11 | 12 |
| L1MA9_dup3206 | 16.068121280558 | -1.96008144462298 | 0.55353091248238 | -3.54105145787205 | 0.00039853584404205 | 0.0184171703139308 | 5598 | 1 | 2 | 38 | 47 | 6 | 13 | 36 | 33 | 5 | 8 | 6 | 11 |
| L2a_dup82515 | 13.8649052832183 | -1.57175761035128 | 0.457699825788671 | -3.43403585011849 | 0.000594665541009133 | 0.0244176717615115 | 5599 | 6 | 2 | 29 | 45 | 8 | 10 | 24 | 22 | 5 | 9 | 12 | 7 |
| L1M1_dup4231 | 10.8445088708471 | -1.65228804208948 | 0.459170113536865 | -3.59842244383536 | 0.000320153291416537 | 0.0156586218046042 | 5600 | 8 | 2 | 31 | 34 | 6 | 7 | 20 | 13 | 2 | 6 | 5 | 9 |
| L1PBa_dup665 | 11.5460831550833 | -1.59936490998248 | 0.424635411164561 | -3.76644261861304 | 0.000165590119887986 | 0.00977462333586373 | 5601 | 5 | 3 | 26 | 23 | 6 | 10 | 23 | 17 | 1 | 10 | 10 | 16 |
| L1MA1_dup1891 | 12.7303491549087 | -1.87202642165128 | 0.477691123407905 | -3.91890560640111 | 8.89519398673871e-05 | 0.00626552243107462 | 5602 | 5 | 4 | 24 | 43 | 5 | 10 | 30 | 14 | 3 | 6 | 3 | 24 |
| L1MA2_dup3323 | 12.8395514546798 | -2.67449398951699 | 0.47812010500223 | -5.59377018773247 | 2.22191015906772e-08 | 1.00512287967591e-05 | 5603 | 3 | 3 | 38 | 49 | 4 | 7 | 28 | 18 | 1 | 3 | 4 | 13 |
| LTR12D_dup37 | 15.0026524297838 | -2.73255645039001 | 0.456973496958831 | -5.97968256053192 | 2.23572872563478e-09 | 1.43006104126315e-06 | 5604 | 9 | 10 | 43 | 46 | 3 | 1 | 44 | 26 | 0 | 2 | 5 | 9 |

| | | | | | | | | | | | | | | | | | | | |
|---|---|---|---|---|---|---|---|---|---|---|---|---|---|---|---|---|---|---|---|
| L1ME2_dup6728 | 17.4883708443897 | -3.09065596605374 | 0.455171570474911 | -6.79009008148082 | 1.12063550247282e-11 | 1.53247549006551e-08 | 5605 | 5 | 9 | 53 | 68 | 4 | 2 | 44 | 22 | 0 | 3 | 8 | 17 |
| THE1A-int_dup137 | 19.0257245933881 | -3.46410531921193 | 0.434795483497931 | -7.96720630891377 | 1.62301586179649e-15 | 5.94138665001581e-12 | 5606 | 5 | 7 | 50 | 76 | 3 | 3 | 43 | 39 | 0 | 2 | 8 | 14 |
| L1MB3_dup8238 | 15.7826239655389 | -3.04336230325815 | 0.447807728636049 | -6.79613617328077 | 1.07462210827303e-11 | 1.47804642875801e-08 | 5607 | 8 | 6 | 32 | 62 | 2 | 2 | 45 | 33 | 1 | 2 | 6 | 9 |
| L1PA6_dup3339 | 12.669306746751 | -2.81686402797761 | 0.516578184493008 | -5.45292873864234 | 4.95469283301543e-08 | 1.89237454389196e-05 | 5608 | 4 | 7 | 28 | 50 | 2 | 1 | 30 | 28 | 1 | 3 | 5 | 6 |
| L1MA7_dup4440 | 15.3943309798501 | -3.10303410273809 | 0.477994847707387 | -6.49177311768362 | 8.48319492985134e-11 | 8.85325570516845e-08 | 5609 | 6 | 2 | 33 | 68 | 5 | 2 | 39 | 29 | 1 | 3 | 8 | 8 |
| L1MEg_dup1338 | 13.7374186015245 | -3.05902237103147 | 0.466492300385925 | -6.5574980948255 | 5.47180095543708e-11 | 6.08408014085248e-08 | 5610 | 6 | 3 | 34 | 57 | 3 | 2 | 33 | 19 | 2 | 2 | 9 | 14 |
| L1MA10_dup432 | 18.4734376249467 | -3.23245393983864 | 0.545867865010331 | -5.92167838965473 | 3.18672371311335e-09 | 1.90999536685257e-06 | 5611 | 9 | 10 | 57 | 81 | 0 | 2 | 42 | 21 | 1 | 1 | 8 | 18 |
| MER4B-int_dup92 | 14.9750310781189 | -3.59485678036353 | 0.579976753392505 | -6.19827736083532 | 5.70844809735535e-10 | 4.59276639579826e-07 | 5612 | 4 | 7 | 51 | 61 | 0 | 1 | 43 | 16 | 1 | 1 | 5 | 13 |
| HAL1_dup6239 | 16.9861700715923 | -3.63012075301883 | 0.502472904547192 | -7.22451045651934 | 5.02910168914099e-13 | 1.0141140936647e-09 | 5613 | 8 | 6 | 54 | 55 | 0 | 1 | 52 | 36 | 1 | 0 | 4 | 5 |
| L1MA7_dup685 | 19.0205067580436 | -3.83726362496765 | 0.504260832641944 | -7.60968010317855 | 2.7477516692187e-14 | 7.69195904334015e-11 | 5614 | 6 | 7 | 54 | 84 | 0 | 2 | 45 | 35 | 1 | 0 | 9 | 9 |
| L1PA13_dup4415 | 16.7853680564574 | -3.56232150641789 | 0.640559336722156 | -5.56126700868472 | 2.67823063924289e-08 | 1.16762961513692e-05 | 5615 | 2 | 3 | 36 | 54 | 0 | 1 | 48 | 43 | 3 | 5 | 7 | 14 |
| L1PA2_dup1462 | 17.3173933993422 | -3.26178002987935 | 0.600428866745205 | -5.43241707808079 | 5.55957908939118e-08 | 2.08656089275122e-05 | 5616 | 7 | 6 | 30 | 44 | 0 | 1 | 70 | 47 | 0 | 4 | 6 | 9 |
| L1PA10_dup1447 | 9.22083391853567 | -2.91837693321932 | 0.623223096146719 | -4.6827162716901 | 2.8309824268079e-06 | 0.000460655673537113 | 5617 | 3 | 5 | 17 | 23 | 0 | 2 | 36 | 27 | 0 | 2 | 1 | 2 |
| L1PREC2_dup6836 | 12.9177509070136 | -2.07900954439615 | 0.593476832525111 | -3.50310143624382 | 0.000459874301067815 | 0.0203771416092891 | 5618 | 13 | 12 | 23 | 45 | 0 | 3 | 37 | 29 | 0 | 2 | 2 | 3 |
| L2b_dup93081 | 9.31667682114768 | -2.04353627958637 | 0.605975269906613 | -3.37230969821805 | 0.000745405788089125 | 0.028621073422739 | 5619 | 9 | 10 | 20 | 23 | 0 | 5 | 13 | 25 | 0 | 1 | 3 | 9 |
| LTR3B_dup31 | 9.2450754070997 | -2.40688591483488 | 0.573108703589203 | -4.1997022550195 | 2.67266199923503e-05 | 0.00253265325396248 | 5620 | 5 | 7 | 18 | 28 | 0 | 2 | 24 | 24 | 1 | 2 | 5 | 2 |
| L1PA5_dup4886 | 9.76194740060888 | -2.15803589719681 | 0.490574466676575 | -4.39899759116397 | 1.08752033527745e-05 | 0.00130023601977413 | 5621 | 7 | 7 | 26 | 28 | 1 | 5 | 21 | 23 | 1 | 1 | 2 | 3 |
| L3_dup19409 | 9.20257677980582 | -1.94458047212185 | 0.539273999916071 | -3.60592291196032 | 0.000311045470989146 | 0.0153329243090912 | 5622 | 5 | 10 | 25 | 19 | 1 | 3 | 25 | 21 | 2 | 1 | 2 | 3 |
| L1PA2_dup4733 | 11.5634507092914 | -2.20438892149561 | 0.669521764660739 | -3.29248284051321 | 0.000993069576496302 | 0.0349344963703414 | 5623 | 11 | 15 | 13 | 27 | 0 | 1 | 29 | 22 | 1 | 1 | 8 | 24 |
| L1MA4_dup4714 | 14.6951973552988 | -2.67791548371544 | 0.594362472836365 | -4.50552584677178 | 6.62087791621136e-06 | 0.00089056609194507 | 5624 | 9 | 12 | 20 | 30 | 0 | 2 | 52 | 45 | 1 | 1 | 7 | 7 |
| THE1B_dup20031 | 13.3090873775604 | -2.21103897407438 | 0.670508036939651 | -3.29755775063646 | 0.000975296158944807 | 0.0345441827681276 | 5625 | 17 | 10 | 21 | 38 | 0 | 1 | 41 | 33 | 1 | 1 | 7 | 2 |
| LTR16E1_dup1989 | 12.6859755957297 | -1.61726768367054 | 0.450225521168701 | -3.59212796172109 | 0.000327988716409317 | 0.0159240161425692 | 5626 | 14 | 11 | 17 | 38 | 5 | 2 | 24 | 17 | 6 | 1 | 16 | 15 |
| LTR84a_dup572 | 15.2750720365516 | -2.044634978118 | 0.51869055344343 | -3.94191674505018 | 8.08330368442042e-05 | 0.00582668821112724 | 5627 | 17 | 15 | 29 | 43 | 4 | 2 | 26 | 17 | 3 | 0 | 25 | 21 |
| MLT1E2_dup3787 | 20.564956421101 | -1.90638907499779 | 0.551752123151249 | -3.45515494187813 | 0.000549976667041725 | 0.0230598674917904 | 5628 | 24 | 20 | 51 | 79 | 3 | 1 | 45 | 18 | 2 | 5 | 18 | 11 |
| L2_dup49994 | 16.28304114905 | -1.96815275201702 | 0.478085003351648 | -4.11674229105526 | 3.84265334216127e-05 | 0.00334067954751153 | 5629 | 13 | 12 | 38 | 63 | 3 | 4 | 34 | 7 | 5 | 4 | 14 | 25 |
| LTR8B_dup753 | 16.9208240429155 | -1.6462736465473 | 0.494020848044424 | -3.33239711049453 | 0.000861013010395202 | 0.0316751084989945 | 5630 | 12 | 20 | 38 | 52 | 1 | 3 | 33 | 16 | 7 | 6 | 13 | 23 |
| THE1B_dup10980 | 13.7585937147105 | -1.8519252401568 | 0.573394286695089 | -3.22975879447782 | 0.0012389469516132 | 0.0407340794673237 | 5631 | 9 | 9 | 32 | 40 | 0 | 1 | 34 | 15 | 8 | 8 | 14 | 10 |
| L2a_dup82386 | 14.0595776609539 | -2.14551971142932 | 0.657938274491919 | -3.26097415914306 | 0.00111030158300476 | 0.0377039846538676 | 5632 | 11 | 7 | 45 | 49 | 0 | 0 | 31 | 9 | 6 | 6 | 13 | 11 |
| THE1C_dup4936 | 13.6230970479685 | -2.19339569641324 | 0.647951084836925 | -3.38512543267872 | 0.000711457355606894 | 0.0276615084864768 | 5633 | 10 | 5 | 33 | 53 | 0 | 0 | 29 | 12 | 8 | 6 | 14 | 11 |
| L1M5_dup25974 | 11.1420520236879 | -2.29404669323805 | 0.668204024880295 | -3.43315306077215 | 0.000596605196085838 | 0.0244800517309606 | 5634 | 4 | 3 | 26 | 31 | 1 | 0 | 25 | 23 | 7 | 8 | 7 | 6 |
| MIR3_dup40247 | 12.4830478225803 | -2.2364323742952 | 0.619908058321082 | -3.60768398519002 | 0.00030894245324827 | 0.0152672110032426 | 5635 | 7 | 3 | 33 | 44 | 1 | 0 | 20 | 16 | 8 | 8 | 12 | 10 |
| MLT1A0-int_dup750 | 12.7415680145213 | -2.68545871046241 | 0.689959321951647 | -3.89219860508038 | 9.93398703522664e-05 | 0.00678652633207143 | 5636 | 4 | 2 | 37 | 61 | 2 | 0 | 16 | 17 | 6 | 6 | 11 | 5 |
| MIRb_dup100619 | 10.922178194901 | -1.89720050939276 | 0.59283119292441 | -3.2002373222535 | 0.00137314471104858 | 0.043762964293486 | 5637 | 5 | 7 | 29 | 43 | 2 | 0 | 16 | 10 | 6 | 7 | 8 | 11 |
| THE1B-int_dup2211 | 12.8724939511183 | -2.4157178822664 | 0.634530525923416 | -3.80709482613286 | 0.0001406088413576 | 0.00868344442400094 | 5638 | 6 | 5 | 44 | 54 | 1 | 0 | 19 | 12 | 5 | 6 | 8 | 11 |
| LTR33_dup4385 | 9.29901688625522 | -2.0233731699622 | 0.610630189231581 | -3.31358194475845 | 0.000921091122219491 | 0.0332341367910266 | 5639 | 7 | 5 | 27 | 27 | 1 | 0 | 17 | 12 | 6 | 2 | 11 | 5 |
| MIR_dup80863 | 9.76795136718226 | -2.42911681311022 | 0.61913307217245 | -3.92341634179999 | 8.7302089461006e-05 | 0.00617330846326554 | 5640 | 5 | 5 | 29 | 34 | 1 | 0 | 13 | 19 | 3 | 3 | 9 | 4 |
| Arthur1B_dup1318 | 14.3640413482952 | -2.54001203510601 | 0.60163172786423 | -4.22187181537608 | 2.42281888716908e-05 | 0.00235498391718273 | 5641 | 8 | 5 | 33 | 47 | 1 | 0 | 34 | 35 | 3 | 5 | 7 | 6 |
| LTR50_dup234 | 12.1017226983947 | -2.02560615858643 | 0.608718665378318 | -3.32765573621357 | 0.000875800217363535 | 0.0320802276048004 | 5642 | 8 | 5 | 21 | 21 | 2 | 0 | 41 | 31 | 4 | 7 | 6 | 7 |
| L1MEg_dup1334 | 9.85625181736808 | -1.83234985584903 | 0.566497702942377 | -3.23452301100577 | 0.00121846101659074 | 0.0402621753997639 | 5643 | 7 | 9 | 17 | 31 | 2 | 0 | 28 | 16 | 1 | 6 | 5 | 8 |
| LTR50_dup1282 | 11.3694712378857 | -1.83151829673897 | 0.541412330417934 | -3.38285294560093 | 0.000717370333746448 | 0.0278368234562349 | 5644 | 10 | 8 | 30 | 36 | 1 | 1 | 25 | 14 | 2 | 7 | 9 | 7 |
| L1MC4_dup21206 | 9.78350183621519 | -2.58805037604631 | 0.593251223876582 | -4.36248636645834 | 1.28592601889769e-05 | 0.00147247811594143 | 5645 | 4 | 3 | 24 | 37 | 0 | 3 | 10 | 20 | 4 | 2 | 11 | 7 |
| MIRb_dup100618 | 11.3583297365636 | -1.92384621347257 | 0.540116670125992 | -3.56190860212442 | 0.000368168563032384 | 0.0173852424884508 | 5646 | 10 | 6 | 22 | 31 | 1 | 1 | 14 | 26 | 4 | 6 | 14 | 8 |
| Tigger2a_dup3239 | 11.5613868275343 | -1.92373367435658 | 0.609631855634831 | -3.15556619388488 | 0.0016018691345444 | 0.0485799583339667 | 5647 | 8 | 13 | 36 | 45 | 1 | 2 | 32 | 7 | 1 | 3 | 6 | 3 |
| MLT1C_dup18807 | 14.3041190916943 | -2.47441950895388 | 0.567287604940922 | -4.36184307113774 | 1.28971365661652e-05 | 0.00147610488570118 | 5648 | 11 | 9 | 53 | 58 | 1 | 1 | 23 | 15 | 2 | 2 | 7 | 9 |
| THE1D_dup12206 | 15.7454724040037 | -1.9335654095967 | 0.612302411695057 | -3.15786018912444 | 0.00158931761485722 | 0.0483595612768308 | 5649 | 16 | 11 | 49 | 68 | 1 | 1 | 29 | 11 | 5 | 3 | 11 | 7 |
| L1MC5_dup7019 | 12.7874422802382 | -2.05956690681442 | 0.526510076077534 | -3.91173312799264 | 9.16361406594759e-05 | 0.00640177719476208 | 5650 | 9 | 13 | 28 | 36 | 1 | 1 | 38 | 22 | 3 | 2 | 8 | 6 |
| L1PA13_dup8625 | 13.0514263825104 | -1.76044792920028 | 0.487191091059628 | -3.61346494528737 | 0.000302132219269693 | 0.0150368443937139 | 5651 | 10 | 13 | 31 | 39 | 2 | 2 | 26 | 25 | 4 | 3 | 6 | 7 |
| Kanga2_a_dup1465 | 14.1026980313893 | -1.80908588839082 | 0.509664346442163 | -3.5495633567849 | 0.000385870599368718 | 0.0180067396817786 | 5652 | 10 | 12 | 38 | 46 | 1 | 2 | 26 | 16 | 7 | 4 | 10 | 13 |
| MER2_dup4521 | 11.2251965914559 | -1.65901461630935 | 0.511743784431593 | -3.24188523003178 | 0.00118741831361644 | 0.0395439381458049 | 5653 | 8 | 9 | 27 | 36 | 2 | 1 | 26 | 11 | 6 | 5 | 8 | 9 |
| MIR_dup168036 | 11.6430905359225 | -2.50038477371913 | 0.596929187998563 | -4.18874604222762 | 2.80500084725817e-05 | 0.00262667741677171 | 5654 | 3 | 7 | 23 | 55 | 8 | 8 | 20 | 5 | 0 | 0 | 16 | 15 |
| L1M5_dup25389 | 13.4775355070144 | -2.79824865930108 | 0.696526812214255 | -4.01743136119263 | 5.88359649378177e-05 | 0.00460006845626929 | 5655 | 8 | 6 | 46 | 65 | 7 | 6 | 9 | 3 | 0 | 0 | 11 | 25 |
| L1MCa_dup3567 | 12.3971643419137 | -2.41756294708418 | 0.589705936148215 | -4.09960761608571 | 4.13851198774513e-05 | 0.0035371493298707 | 5656 | 12 | 8 | 35 | 52 | 7 | 3 | 18 | 6 | 0 | 0 | 10 | 19 |
| Tigger1_dup6297 | 11.5176610427069 | -2.09990464779611 | 0.619127885416609 | -3.39171388861461 | 0.000694569276376262 | 0.0271825626704977 | 5657 | 12 | 9 | 41 | 44 | 8 | 5 | 8 | 6 | 0 | 0 | 11 | 11 |
| LTR18B_dup349 | 15.3139930177166 | -3.57979728679371 | 0.64324824723811 | -5.56518778273264 | 2.61870255877903e-08 | 1.15177412024258e-05 | 5658 | 4 | 7 | 49 | 76 | 2 | 2 | 12 | 9 | 1 | 0 | 25 | 21 |
| MER67C_dup1373 | 15.8122141269712 | -3.1852780757919 | 0.605347365964371 | -5.26190127335811 | 1.42573321965653e-07 | 4.40009749266398e-05 | 5659 | 8 | 5 | 46 | 91 | 3 | 5 | 12 | 11 | 1 | 0 | 13 | 22 |
| L1PREC2_dup929 | 17.6670808636462 | -3.39395370519873 | 0.753309457006709 | -4.50539107617828 | 6.62508159478366e-06 | 0.000890628059408131 | 5660 | 10 | 7 | 50 | 81 | 3 | 1 | 9 | 3 | 1 | 0 | 27 | 53 |
| Tigger17a_dup2067 | 15.5801956799992 | -3.02473565669589 | 0.714726239176816 | -4.23201988523553 | 2.31601926357868e-05 | 0.00228193589934365 | 5661 | 12 | 6 | 50 | 84 | 2 | 2 | 11 | 2 | 1 | 1 | 21 | 24 |
| HSMAR2_dup1691 | 13.6633495176148 | -3.38249559487115 | 0.697889956108569 | -4.84674634627489 | 1.25502722461776e-06 | 0.000245381025463351 | 5662 | 5 | 2 | 38 | 60 | 6 | 2 | 6 | 5 | 1 | 0 | 31 | 30 |
| L1PA7_dup1608 | 12.8271082342879 | -3.26124985948816 | 0.720854592949262 | -4.52414383065145 | 6.06404773060937e-06 | 0.000835988355334634 | 5663 | 7 | 6 | 33 | 47 | 1 | 1 | 5 | 7 | 1 | 0 | 27 | 39 |
| L1PB1_dup2933 | 11.9487905472612 | -3.27140947700898 | 0.870413153060039 | -3.75845593039116 | 0.000170965142623822 | 0.010003191049461 | 5664 | 12 | 5 | 28 | 39 | 0 | 0 | 11 | 6 | 0 | 0 | 22 | 41 |
| MLT1C_dup17809 | 13.1079965181375 | -2.86667850904185 | 0.874227531131058 | -3.27909886952771 | 0.00104139143751107 | 0.0361028256680255 | 5665 | 17 | 8 | 39 | 56 | 1 | 1 | 4 | 4 | 0 | 0 | 26 | 23 |
| L1MA9_dup12784 | 8.03607380336969 | -2.33279685029978 | 0.727555277132083 | -3.20634998277427 | 0.00134430378713088 | 0.0430861676902808 | 5666 | 8 | 7 | 18 | 19 | 0 | 0 | 11 | 5 | 2 | 0 | 20 | 17 |
| L1PA2_dup4273 | 8.74603334627771 | -2.49687956433789 | 0.637575165615049 | -3.91621207819353 | 8.9951126755707e-05 | 0.00630256501973305 | 5667 | 8 | 2 | 27 | 23 | 1 | 4 | 6 | 6 | 2 | 0 | 19 | 18 |
| THE1C-int_dup635 | 7.28836904700184 | -2.70968620872001 | 0.720421947419928 | -3.76124883261024 | 0.00016906713539736 | 0.00992324829779486 | 5668 | 5 | 3 | 12 | 17 | 0 | 2 | 5 | 3 | 1 | 0 | 20 | 31 |
| L1PA4_dup10304 | 11.0999247559639 | -3.93210238186672 | 0.727652460683697 | -5.40381926032675 | 6.52366807758342e-08 | 2.37715271728739e-05 | 5669 | 3 | 4 | 33 | 36 | 0 | 2 | 14 | 5 | 0 | 0 | 25 | 29 |
| L1PA5_dup9083 | 10.9472133092117 | -4.45320791634955 | 0.77587315887932 | -5.73960816324903 | 9.48958584676615e-09 | 4.8663986937384e-06 | 5670 | 0 | 4 | 27 | 41 | 1 | 1 | 13 | 7 | 0 | 0 | 24 | 31 |
| L1MA5_dup4134 | 8.93898229083016 | -3.71087928335266 | 0.853850214822405 | -4.34605416609809 | 1.38608354734764e-05 | 0.00155597019613423 | 5671 | 2 | 3 | 27 | 48 | 4 | 0 | 4 | 3 | 0 | 0 | 15 | 17 |
| L1ME3B_dup7554 | 8.3487751281547 | -3.46940111710067 | 0.837130061987455 | -4.14439914971381 | 3.4070585436902e-05 | 0.00303631442785359 | 5672 | 5 | 4 | 29 | 34 | 1 | 0 | 6 | 3 | 0 | 0 | 9 | 24 |
| L1M3_dup6141 | 10.1930706044019 | -2.95526491625824 | 0.786961781264903 | -3.75528391163822 | 0.000173145092400488 | 0.0100830108067368 | 5673 | 6 | 8 | 34 | 61 | 1 | 0 | 9 | 4 | 0 | 1 | 7 | 10 |
| L1MC4a_dup9363 | 11.4478577507176 | -3.5011468706405 | 0.834886962597471 | -4.19355796352102 | 2.74612864798139e-05 | 0.00257763442711076 | 5674 | 6 | 5 | 39 | 60 | 1 | 0 | 4 | 3 | 0 | 1 | 16 | 23 |
| L1PA6_dup1400 | 7.405092517077 | -2.63412099283458 | 0.685403746058031 | -3.84316690415573 | 0.000121456808545896 | 0.00782354135524138 | 5675 | 9 | 4 | 24 | 17 | 1 | 1 | 13 | 6 | 0 | 0 | 7 | 18 |
| L2b_dup71300 | 8.61928140564474 | -2.61914877495603 | 0.77462947095794 | -3.38116334731892 | 0.00072179620173256 | 0.0279629630441966 | 5676 | 12 | 6 | 25 | 29 | 1 | 0 | 8 | 8 | 0 | 0 | 10 | 17 |
| L1PA3_dup5335 | 7.11226296021957 | -2.49012065833954 | 0.76800411574116 | -3.2423272314582 | 0.00118557807466461 | 0.0394992383861865 | 5677 | 4 | 10 | 26 | 29 | 1 | 1 | 7 | 4 | 0 | 0 | 6 | 8 |
| L1M5_dup50373 | 5.5568271263102 | -2.61589217523538 | 0.762398260315099 | -3.43113607598505 | 0.000601059012282968 | 0.0246151831313543 | 5678 | 4 | 4 | 14 | 22 | 2 | 1 | 4 | 4 | 0 | 0 | 4 | 17 |
| L1MCa_dup3570 | 6.61000757544965 | -2.81190103551856 | 0.768501368790028 | -3.65894082914358 | 0.000253259801273224 | 0.0132619182820771 | 5679 | 3 | 5 | 22 | 31 | 1 | 1 | 4 | 5 | 0 | 1 | 3 | 14 |
| L1PA15-16_dup461 | 11.2387745934296 | -5.10235109798281 | 0.763767191777162 | -6.68050572597975 | 2.38119077229109e-11 | 2.96568807215841e-08 | 5680 | 2 | 0 | 27 | 33 | 1 | 1 | 31 | 17 | 0 | 0 | 17 | 21 |
| L1PB3_dup3334 | 11.3391536325035 | -3.72278951644327 | 0.561413676510074 | -6.63109872845513 | 3.33197268236869e-11 | 3.99059507548812e-08 | 5681 | 2 | 0 | 26 | 41 | 2 | 4 | 24 | 15 | 1 | 1 | 14 | 22 |
| MLT1J_dup13658 | 14.0072371522929 | -4.35819176581256 | 0.581104824219806 | -7.49983752357225 | 6.38969856152657e-14 | 1.53576082214243e-10 | 5682 | 2 | 0 | 34 | 39 | 3 | 1 | 29 | 26 | 2 | 0 | 20 | 27 |
| L1PA12_dup1609 | 11.9292743527591 | -3.19675132953378 | 0.541224730316142 | -5.90651378340839 | 3.49423073869416e-09 | 2.06312215223156e-06 | 5683 | 5 | 1 | 33 | 46 | 3 | 4 | 17 | 11 | 2 | 0 | 19 | 19 |

| | | | | | | | | | | | | | | | | | | |
|---|---|---|---|---|---|---|---|---|---|---|---|---|---|---|---|---|---|---|
| L1PA3_dup10176 | 11.3385402601316 | -3.09315450013564 | 0.497904914887921 | -6.21233976136169 | 5.22014025743246e-10 | 4.25380648525694e-07 | 5684 | 4 | 1 | 28 | 31 | 6 | 2 | 24 | 13 | 2 | 0 | 16 | 24 |
| MLT1H1_dup3286 | 10.9546997925013 | -3.31406197304709 | 0.527334621568458 | -6.28455223211029 | 3.28799977446277e-10 | 2.80418062485418e-07 | 5685 | 4 | 3 | 22 | 41 | 1 | 4 | 29 | 12 | 1 | 0 | 16 | 15 |
| Ricksha_0_dup362 | 10.6909036027115 | -3.81987183498119 | 0.641030684863249 | -5.95895317522293 | 2.53858817824546e-09 | 1.59379129989655e-06 | 5686 | 2 | 2 | 23 | 49 | 3 | 1 | 24 | 14 | 1 | 0 | 18 | 6 |
| REP522_dup117 | 8.48194131080587 | -3.74693516902391 | 0.738554426265124 | -5.07333655553619 | 3.90900237877233e-07 | 9.87295062566641e-05 | 5687 | 0 | 0 | 28 | 20 | 1 | 0 | 14 | 7 | 5 | 2 | 11 | 24 |
| LTR49-int_dup1685 | 9.42700618705897 | -4.8118737314007 | 0.864088279425849 | -5.56872931385898 | 2.56603759130999e-08 | 1.13701746871852e-05 | 5688 | 0 | 0 | 28 | 27 | 1 | 0 | 6 | 12 | 2 | 1 | 18 | 29 |
| MLT1E1A_dup2907 | 10.1214013428698 | -4.58356454869257 | 0.777442047498342 | -5.89569931732095 | 3.73097837063044e-09 | 2.18662901324638e-06 | 5689 | 1 | 0 | 29 | 32 | 1 | 0 | 9 | 9 | 1 | 2 | 24 | 27 |
| L1ME4b_dup19809 | 11.1854384654117 | -4.71120495775168 | 0.721157337855317 | -6.53283924387784 | 6.45344021565548e-11 | 6.93629549999411e-08 | 5690 | 0 | 1 | 22 | 41 | 1 | 0 | 28 | 14 | 1 | 2 | 21 | 19 |
| MLT1A_dup8281 | 8.12920225654585 | -3.00874647906104 | 0.651011306384666 | -4.62165011506459 | 3.80699704038267e-06 | 0.000576980711955983 | 5691 | 0 | 0 | 14 | 26 | 5 | 0 | 18 | 10 | 2 | 4 | 17 | 11 |
| L2_dup42716 | 6.76534568984439 | -3.2358731563814 | 0.721117422536186 | -4.48730408565191 | 7.21300887810505e-06 | 0.000953503672505325 | 5692 | 3 | 0 | 17 | 25 | 2 | 1 | 9 | 8 | 2 | 0 | 4 | 20 |
| MIRb_dup202457 | 6.34751955834239 | -3.85006313639198 | 0.7970388128959 | -4.8304587858193 | 1.36218809643308e-06 | 0.000262451181209608 | 5693 | 1 | 0 | 10 | 29 | 1 | 1 | 13 | 5 | 2 | 0 | 9 | 16 |
| L1MD1_dup4620 | 7.15104939764579 | -3.59726076145791 | 0.745441656246674 | -4.82567714228669 | 1.39528383895697e-06 | 0.000267454897421193 | 5694 | 5 | 0 | 18 | 31 | 1 | 0 | 10 | 11 | 1 | 0 | 9 | 9 |
| L2c_dup104390 | 6.5887993322755 | -3.61874728646635 | 0.720254444565475 | -5.02426234752291 | 5.05371159666763e-07 | 0.000120371417375443 | 5695 | 3 | 0 | 13 | 12 | 1 | 0 | 14 | 12 | 2 | 0 | 9 | 20 |
| MER20_dup10195 | 6.42857271546163 | -3.71329416662169 | 0.773411220275225 | -4.80118993528473 | 1.57725579898765e-06 | 0.000294123595882378 | 5696 | 3 | 0 | 22 | 19 | 2 | 1 | 10 | 5 | 0 | 0 | 14 | 10 |
| MIR_dup157825 | 5.6434477362664 | -3.70406571872375 | 0.805264122912112 | -4.59981466121771 | 4.22866999416381e-06 | 0.000629988850258429 | 5697 | 2 | 0 | 20 | 14 | 1 | 1 | 11 | 4 | 1 | 0 | 13 | 8 |
| MIR_dup44933 | 6.80715191907868 | -3.44677393232626 | 0.704531157426064 | -4.89229453658049 | 9.96671406702999e-07 | 0.000205021688222428 | 5698 | 1 | 1 | 19 | 19 | 3 | 0 | 15 | 3 | 1 | 1 | 15 | 14 |
| L1PA11_dup3485 | 6.04356536261216 | -3.77506335613722 | 0.816997681717343 | -4.62065369414755 | 3.82532837888118e-06 | 0.00057865326537906 | 5699 | 2 | 0 | 17 | 22 | 1 | 0 | 10 | 3 | 2 | 0 | 11 | 14 |
| L1PA5_dup3081 | 6.18014355436466 | -3.36053033209444 | 0.763380216416001 | -4.4021711066496 | 1.0717298040271e-05 | 0.00128600010060026 | 5700 | 3 | 1 | 23 | 22 | 2 | 0 | 10 | 3 | 1 | 0 | 7 | 12 |
| MIR1_Amn_dup5134 | 7.56721760995078 | -4.09549351297127 | 0.772574225897729 | -5.30110036768612 | 1.15106782095205e-07 | 3.69987985774932e-05 | 5701 | 0 | 1 | 34 | 26 | 1 | 2 | 13 | 10 | 1 | 0 | 3 | 9 |
| MIRc_dup48415 | 10.1844235845107 | -3.86953693160329 | 0.679890073786218 | -5.69141554024219 | 1.25990473213325e-08 | 6.21969484216137e-06 | 5702 | 0 | 1 | 38 | 44 | 5 | 1 | 18 | 14 | 1 | 0 | 7 | 6 |
| L1ME1_dup22780 | 10.0253477767237 | -2.99900580773667 | 0.765598568667266 | -3.91720404200502 | 8.95819224641319e-05 | 0.00628896961191906 | 5703 | 0 | 0 | 24 | 34 | 2 | 3 | 18 | 19 | 8 | 0 | 9 | 11 |
| MIR1_Amn_dup5135 | 7.57036844036204 | -2.72953545779134 | 0.693051552243257 | -3.93843062461433 | 8.20163005146481e-05 | 0.00588908258717401 | 5704 | 0 | 0 | 13 | 34 | 7 | 3 | 17 | 11 | 2 | 0 | 5 | 9 |
| PRIMA41-int_dup442 | 8.63083506477303 | -3.25946652318808 | 0.593982096650709 | -5.487482773584 | 4.07701761306372e-08 | 1.61954930376972e-05 | 5705 | 1 | 0 | 19 | 31 | 4 | 3 | 27 | 13 | 2 | 0 | 6 | 9 |
| AluSz6_dup21886 | 8.24625692500607 | -2.30285667761318 | 0.704069254601 | -3.27078147861779 | 0.00107250749487431 | 0.0368731207015405 | 5706 | 1 | 0 | 28 | 31 | 5 | 3 | 9 | 17 | 3 | 2 | 3 | 3 |
| L1MA2_dup4477 | 9.22472997552697 | -3.09638356764816 | 0.796735288205864 | -3.88633918126155 | 0.000101767290794681 | 0.00690086058006017 | 5707 | 0 | 0 | 28 | 40 | 3 | 1 | 11 | 20 | 4 | 2 | 4 | 5 |
| AluSz_dup73269 | 5.8058588863971 | -2.2378713461644 | 0.631336909764197 | -3.54465470266935 | 0.000393127727040287 | 0.0182594515202671 | 5708 | 0 | 1 | 19 | 16 | 4 | 3 | 7 | 8 | 2 | 2 | 5 | 8 |
| L2a_dup99867 | 6.54038868225942 | -2.42275650116292 | 0.758826665316572 | -3.192766690865 | 0.00140916752076179 | 0.0445115856757183 | 5709 | 0 | 0 | 21 | 28 | 3 | 2 | 8 | 7 | 2 | 5 | 6 | 4 |
| L1M5_dup27226 | 8.2082118947747 | -2.81971469723463 | 0.67637648042602 | -4.1688538540823 | 3.06135197836799e-05 | 0.00279844970359105 | 5710 | 0 | 0 | 29 | 34 | 5 | 5 | 12 | 12 | 1 | 1 | 4 | 5 |
| L1PA8_dup6656 | 8.54876904168517 | -2.79717863230372 | 0.761452695133652 | -3.67347656680466 | 0.000239272629262967 | 0.01271018377112 | 5711 | 0 | 0 | 34 | 40 | 4 | 7 | 11 | 5 | 1 | 1 | 4 | 9 |
| AluSc_dup16511 | 7.32170828261664 | -2.58647434223581 | 0.794643667175857 | -3.25488574196793 | 0.00113437993397031 | 0.0383247434003263 | 5712 | 0 | 0 | 28 | 36 | 5 | 3 | 9 | 7 | 2 | 1 | 1 | 6 |
| MamTip2_dup3480 | 9.48214310807854 | -3.17715475023415 | 0.752149914389764 | -4.22409773563805 | 2.3989993100288e-05 | 0.00234043579263679 | 5713 | 0 | 1 | 39 | 51 | 3 | 3 | 10 | 8 | 2 | 1 | 2 | 8 |
| THE1B-int_dup93 | 9.37705925420471 | -3.93554310937427 | 0.774748252391161 | -5.07977023146257 | 3.77891636181479e-07 | 9.64783296811568e-05 | 5714 | 0 | 0 | 34 | 37 | 0 | 2 | 11 | 11 | 2 | 3 | 11 | 13 |
| L1PA10_dup6261 | 8.34989473817411 | -3.06667652656507 | 0.671656340652391 | -4.5658416975359 | 4.97494101896835e-06 | 0.000719178199088362 | 5715 | 0 | 1 | 20 | 38 | 2 | 3 | 8 | 18 | 2 | 2 | 5 | 9 |
| MER103C_dup4483 | 9.36646087142237 | -4.06834578416632 | 0.797117675897328 | -5.10382081238698 | 3.3286347016588e-07 | 8.72285586697031e-05 | 5716 | 0 | 1 | 39 | 41 | 1 | 1 | 5 | 13 | 1 | 2 | 9 | 10 |
| L2_dup9558 | 7.51186233665631 | -4.66521879496792 | 0.898633010195937 | -5.19146163343223 | 2.08649586326972e-07 | 5.95291780193737e-05 | 5717 | 0 | 0 | 24 | 38 | 0 | 1 | 18 | 9 | 2 | 0 | 6 | 3 |
| L2a_dup86165 | 8.10933571057479 | -4.40340472402997 | 0.801008592160604 | -5.49732520615344 | 3.85595207400823e-08 | 1.56038838809857e-05 | 5718 | 0 | 1 | 28 | 37 | 0 | 1 | 16 | 13 | 1 | 1 | 7 | 2 |
| HERVK9-int_dup557 | 8.15527602863838 | -3.87465494041297 | 0.714026620785789 | -5.42648527046358 | 5.74745658085745e-08 | 2.14691413436218e-05 | 5719 | 0 | 0 | 26 | 21 | 1 | 1 | 18 | 16 | 2 | 2 | 8 | 10 |
| L1PA7_dup9970 | 8.69945511573484 | -3.44596670385652 | 0.690428910730222 | -4.99105215656735 | 6.00512782200839e-07 | 0.000138191116512148 | 5720 | 2 | 0 | 18 | 34 | 1 | 1 | 22 | 19 | 3 | 1 | 6 | 6 |
| MIRb_dup159974 | 6.8529711158048 | -3.39293192390787 | 0.683487966257709 | -4.96414288386837 | 6.90051294835129e-07 | 0.000153166926679888 | 5721 | 2 | 0 | 20 | 18 | 1 | 1 | 16 | 13 | 2 | 1 | 10 | 4 |
| L1ME3Cz_dup7237 | 13.2316003254513 | -2.46117449740451 | 0.702908339658781 | -3.50141598632797 | 0.00046279285936337 | 0.0204601860529882 | 5722 | 3 | 0 | 46 | 54 | 2 | 4 | 16 | 16 | 7 | 9 | 10 | 5 |
| LTR40a_dup1661 | 10.1659340821039 | -2.32796924449554 | 0.708063638559696 | -3.28779662973651 | 0.00100974752665608 | 0.0353026550351607 | 5723 | 1 | 0 | 38 | 49 | 3 | 8 | 15 | 6 | 3 | 3 | 6 | 5 |
| L1M3_dup3890 | 13.3118731487622 | -2.41069618404913 | 0.707560119169935 | -3.40705491835409 | 0.000656679125963878 | 0.0261382019582805 | 5724 | 1 | 0 | 41 | 71 | 5 | 5 | 19 | 11 | 5 | 7 | 5 | 9 |
| L1PA16_dup8754 | 11.785399771604 | -2.26093459349588 | 0.626345624360714 | -3.60972361833537 | 0.000306523428634218 | 0.0151918399812117 | 5725 | 3 | 1 | 41 | 56 | 4 | 4 | 17 | 9 | 4 | 6 | 5 | 8 |
| L1PA8_dup5043 | 10.282712460222 | -2.14423102486167 | 0.582934833506447 | -3.67833744290724 | 0.000234759238152268 | 0.0125218609462855 | 5726 | 1 | 1 | 31 | 38 | 4 | 8 | 22 | 10 | 5 | 2 | 7 | 7 |
| MLT1E2_dup3122 | 11.3738125136867 | -2.20818235643975 | 0.549405994799044 | -4.01921780494485 | 5.83916729817255e-05 | 0.00457192004584062 | 5727 | 1 | 1 | 32 | 44 | 7 | 8 | 16 | 9 | 4 | 3 | 14 | 12 |
| MER63A_dup2835 | 10.3034680058554 | -2.72718467598274 | 0.595774871744876 | -4.57754229881414 | 4.7047067923216e-06 | 0.000688478574665287 | 5728 | 1 | 2 | 37 | 47 | 4 | 6 | 12 | 11 | 2 | 1 | 7 | 7 |
| MIR3_dup66146 | 10.0418236426708 | -2.37473777877065 | 0.593469313887351 | -4.00144998772659 | 6.29555027523965e-05 | 0.0048478996951203 | 5729 | 3 | 1 | 25 | 55 | 5 | 5 | 13 | 7 | 3 | 2 | 10 | 7 |
| HERV9-int_dup48 | 8.83031840217299 | -2.06481558425607 | 0.601280277038527 | -3.43403178701597 | 0.000594674454951793 | 0.0244176717615115 | 5730 | 2 | 2 | 24 | 36 | 5 | 1 | 9 | 9 | 5 | 5 | 6 | 12 |
| MSTA_dup10457 | 10.2552646860106 | -2.54079721597711 | 0.576560268698357 | -4.40681981384742 | 1.04899380089301e-05 | 0.00126766825265255 | 5731 | 5 | 2 | 32 | 36 | 1 | 1 | 16 | 11 | 3 | 6 | 13 | 9 |
| THE1B_dup131 | 10.5000214622513 | -1.91288109721411 | 0.529973265156021 | -3.60939168629756 | 0.000306915891138213 | 0.0152018189335905 | 5732 | 5 | 2 | 30 | 37 | 4 | 3 | 16 | 11 | 3 | 9 | 10 | 8 |
| AluJb_dup125013 | 8.52274751206632 | -2.91985407089941 | 0.616006920347636 | -4.73996959198385 | 2.13750295048905e-06 | 0.000370167596111403 | 5733 | 1 | 0 | 14 | 28 | 3 | 6 | 18 | 25 | 1 | 1 | 5 | 6 |
| L1PA10_dup5955 | 6.27076957351379 | -2.77978052455496 | 0.666387101009281 | -4.17142006552171 | 3.02707245666147e-05 | 0.00277243950258957 | 5734 | 1 | 0 | 15 | 16 | 2 | 3 | 12 | 19 | 1 | 2 | 4 | 3 |
| L1PA13_dup4370 | 7.02775999828035 | -2.63637315297502 | 0.701742229624165 | -3.75689682290745 | 0.00017203338313305 | 0.0100418153174827 | 5735 | 1 | 0 | 21 | 37 | 2 | 6 | 14 | 8 | 1 | 1 | 1 | 3 |
| AluJr_dup40299 | 7.79356436108194 | -3.35173134828533 | 0.686420554957784 | -4.88291226723458 | 1.04530387187258e-06 | 0.000212223442915182 | 5736 | 1 | 0 | 22 | 39 | 1 | 3 | 18 | 8 | 1 | 2 | 4 | 7 |
| L1MA8_dup5168 | 6.98716403350411 | -3.16909122618663 | 0.685816305606044 | -4.62090387802336 | 3.82071776120019e-06 | 0.00057865326537906 | 5737 | 1 | 0 | 24 | 30 | 1 | 3 | 13 | 9 | 1 | 2 | 4 | 5 |
| AluSx_dup34105 | 8.74483971256993 | -2.08347131219933 | 0.525852841762227 | -3.96208054180566 | 7.42994657301577e-05 | 0.00547220503773886 | 5738 | 3 | 0 | 22 | 30 | 5 | 8 | 22 | 11 | 2 | 2 | 5 | 6 |
| L3b_dup3562 | 7.20935052250454 | -2.96831738336108 | 0.633172299723362 | -4.68800891109412 | 2.75876008671667e-06 | 0.000451158713122944 | 5739 | 2 | 0 | 27 | 19 | 2 | 4 | 14 | 12 | 1 | 1 | 3 | 9 |
| L1PA10_dup6262 | 7.22155471487063 | -2.74577052600591 | 0.60709826286245 | -4.52277776757206 | 6.10333124132949e-06 | 0.0008394585292193 | 5740 | 4 | 0 | 24 | 22 | 2 | 2 | 22 | 9 | 2 | 1 | 3 | 5 |
| Tigger3a_dup2812 | 7.17418822011038 | -2.93084590721336 | 0.617988786884777 | -4.74255515538959 | 2.11039345166825e-06 | 0.000367305488203946 | 5741 | 2 | 0 | 18 | 32 | 3 | 3 | 16 | 12 | 1 | 1 | 4 | 3 |
| MER47C_dup666 | 6.78022363486658 | -1.84524967560303 | 0.54513691035929 | -3.3849288876566 | 0.000711966968309671 | 0.0276693103837291 | 5742 | 8 | 0 | 14 | 17 | 4 | 4 | 9 | 10 | 1 | 2 | 7 | 13 |
| L1MA7_dup7990 | 7.09482156481647 | -1.91202915990396 | 0.520066463072322 | -3.67650924577704 | 0.000236447289390514 | 0.0125921411641261 | 5743 | 4 | 0 | 15 | 22 | 4 | 4 | 13 | 10 | 2 | 4 | 6 | 9 |
| L1ME4b_dup4151 | 5.8232146710726 | -2.02955032920266 | 0.554852233004112 | -3.65782132337138 | 0.000254368265897662 | 0.0133082478885851 | 5744 | 3 | 0 | 13 | 13 | 2 | 4 | 10 | 8 | 2 | 3 | 6 | 12 |
| L2c_dup110865 | 7.16666765266319 | -2.55797587723467 | 0.677264519179709 | -3.77692290795456 | 0.000158777850511381 | 0.00949498728770571 | 5745 | 1 | 0 | 14 | 37 | 6 | 1 | 14 | 7 | 2 | 2 | 5 | 8 |
| MamGypLTR1a_dup85 | 9.08384963434981 | -2.99523463722116 | 0.609105011517126 | -4.91743554984188 | 8.76853118885362e-07 | 0.000185791355499819 | 5746 | 3 | 0 | 22 | 43 | 3 | 2 | 16 | 9 | 3 | 1 | 11 | 9 |
| HAL1b_dup5235 | 9.34677058861849 | -2.99311297204363 | 0.671824937650113 | -4.45519778189961 | 8.38158895806353e-06 | 0.00106650565038256 | 5747 | 3 | 0 | 26 | 27 | 3 | 5 | 7 | 4 | 0 | 2 | 24 | 25 |
| L1MB7_dup20749 | 9.89025756036312 | -3.11755208363556 | 0.571108520946212 | -5.45877354179623 | 4.79434788469415e-08 | 1.8489398732114e-05 | 5748 | 2 | 0 | 18 | 28 | 5 | 5 | 13 | 15 | 0 | 1 | 21 | 22 |
| L1PA7_dup1413 | 10.9090307155215 | -3.0031699955472 | 0.5800076203736 | -5.17781127360494 | 2.24504166554223e-07 | 6.31440524998949e-05 | 5749 | 3 | 0 | 18 | 19 | 3 | 6 | 12 | 10 | 0 | 2 | 28 | 45 |
| L1PA17_dup4724 | 9.05808167827259 | -2.23119668577728 | 0.654199016004072 | -3.41057786880467 | 0.00064825370323569 | 0.0258894554666196 | 5750 | 1 | 0 | 25 | 19 | 6 | 10 | 9 | 8 | 2 | 1 | 11 | 28 |
| L1MA9_dup1741 | 11.1658874318757 | -2.61679404429945 | 0.50477057049617 | -5.18412561518244 | 2.17030627822371e-07 | 6.1478059247407e-05 | 5751 | 4 | 0 | 25 | 36 | 5 | 4 | 13 | 11 | 4 | 2 | 24 | 19 |
| LTR31_dup131 | 10.8237733278891 | -1.80590763572199 | 0.484196288456098 | -3.72970152555336 | 0.00019170672625984 | 0.0107711567146691 | 5752 | 7 | 0 | 14 | 22 | 4 | 11 | 16 | 9 | 4 | 3 | 20 | 34 |
| LSAU | 12.4014953757675 | -2.4779783967461 | 0.758608275976128 | -3.26647951942997 | 0.00108893684321061 | 0.0372495925955421 | 5753 | 0 | 1 | 33 | 26 | 8 | 17 | 13 | 12 | 1 | 0 | 19 | 34 |
| AluY_dup101594 | 11.4134435036541 | -2.89908582403945 | 0.724807727306548 | -3.99979982941506 | 6.33960828701339e-05 | 0.00487236574115532 | 5754 | 2 | 0 | 42 | 40 | 2 | 14 | 14 | 10 | 1 | 0 | 9 | 19 |
| BSR/Beta_dup1954 | 13.92353046444 | -3.36567493587939 | 0.708612956212802 | -4.74966609962555 | 2.03752785871854e-06 | 0.000358066177157048 | 5755 | 1 | 0 | 60 | 45 | 4 | 11 | 14 | 11 | 1 | 0 | 19 | 19 |
| BSR/Beta_dup2010 | 15.4493070325096 | -3.75325992335056 | 0.662982397416505 | -5.66117582906602 | 1.50339287591303e-08 | 7.19903305860017e-06 | 5756 | 3 | 0 | 69 | 49 | 4 | 3 | 9 | 15 | 3 | 0 | 18 | 31 |
| L2_dup42718 | 8.72283560253338 | -2.98333388115352 | 0.618230395988012 | -4.8256020741034 | 1.39580953185992e-06 | 0.000267454897421193 | 5757 | 3 | 1 | 25 | 33 | 2 | 3 | 14 | 7 | 3 | 0 | 6 | 21 |
| MLT1J2_dup6575 | 8.24708148796989 | -2.74579050734587 | 0.658089850565788 | -4.17236416119348 | 3.01455323685372e-05 | 0.00276629727919936 | 5758 | 2 | 1 | 26 | 26 | 4 | 3 | 5 | 6 | 3 | 0 | 10 | 24 |
| L1ME2z_dup6928 | 9.99559946108029 | -2.72292247552182 | 0.646873187230009 | -4.20936055053033 | 2.56094443245345e-05 | 0.00245613254302527 | 5759 | 4 | 1 | 34 | 39 | 8 | 4 | 7 | 7 | 1 | 0 | 13 | 16 |
| LSAU_dup133 | 9.21053848460707 | -2.15742030835918 | 0.528822070165666 | -4.07967146243219 | 4.50993944602553e-05 | 0.00378392824902677 | 5760 | 2 | 4 | 22 | 24 | 9 | 3 | 12 | 8 | 3 | 0 | 17 | 17 |
| AluSc_dup31361 | 8.64316325392092 | -2.20048538470595 | 0.505253588791669 | -4.35520980656166 | 1.3293952653688e-05 | 0.0015096525313337 | 5761 | 2 | 6 | 19 | 19 | 4 | 3 | 12 | 6 | 0 | 4 | 19 | 21 |
| L2c_dup38253 | 9.61711670999116 | -1.6169474192654 | 0.500301790565392 | -3.23194409805746 | 0.00122951106611008 | 0.0404979568295443 | 5762 | 4 | 10 | 12 | 31 | 5 | 5 | 15 | 7 | 0 | 6 | 11 | 24 |

| | | | | | | | | | | | | | | | | | | | |
|---|---|---|---|---|---|---|---|---|---|---|---|---|---|---|---|---|---|---|---|
| AluSq2_dup54210 | 8.58541207523984 | -3.05697875362249 | 0.597580765952248 | -5.11559094234095 | 3.12760209504677e-07 | 8.24142201669988e-05 | 5763 | 2 | 2 | 25 | 29 | 2 | 5 | 12 | 7 | 0 | 1 | 11 | 20 |
| L2a_dup131066 | 9.93764469060877 | -2.61502635492374 | 0.625383594951151 | -4.18147577908243 | 2.89623107657982e-05 | 0.00268697648734055 | 5764 | 1 | 8 | 28 | 38 | 4 | 5 | 9 | 6 | 0 | 1 | 15 | 19 |
| MSTB-int_dup394 | 7.79539196533391 | -2.06860926961659 | 0.620261548300214 | -3.3350596619853 | 0.000852810999894318 | 0.0314457951614279 | 5765 | 2 | 4 | 19 | 24 | 1 | 8 | 5 | 7 | 0 | 4 | 10 | 20 |
| L1MC4_dup11944 | 8.98728149073686 | -1.90592397625257 | 0.544653983917311 | -3.49932990950439 | 0.000466429083301277 | 0.0205627678101205 | 5766 | 6 | 8 | 26 | 31 | 2 | 6 | 11 | 5 | 0 | 3 | 11 | 12 |
| L1ME2z_dup5006 | 7.96143374950562 | -2.37703238708453 | 0.597089555547845 | -3.9810315973515 | 6.86168263284889e-05 | 0.00516516904699735 | 5767 | 6 | 5 | 19 | 33 | 1 | 3 | 8 | 6 | 0 | 2 | 10 | 15 |
| MLT2D_dup3616 | 10.365882149213 | -1.90516229381749 | 0.594166464129989 | -3.20644534626696 | 0.00134385830070933 | 0.0430861676902808 | 5768 | 5 | 4 | 29 | 52 | 10 | 3 | 10 | 4 | 1 | 4 | 10 | 9 |
| L1MB4_dup7649 | 9.13317095350204 | -2.18499068057029 | 0.592613846885519 | -3.68703953181908 | 0.000226878122949227 | 0.0121885974162253 | 5769 | 6 | 3 | 23 | 47 | 5 | 6 | 8 | 5 | 1 | 1 | 7 | 13 |
| L1MEf_dup5574 | 9.29683052629094 | -2.23377747306447 | 0.554261953260322 | -4.03018367023891 | 5.57332864982555e-05 | 0.00442750622676088 | 5770 | 4 | 3 | 23 | 35 | 5 | 6 | 8 | 5 | 1 | 2 | 12 | 22 |
| L1P1_dup1535 | 9.58311388652583 | -1.88541552436234 | 0.577252497979871 | -3.26618859331135 | 0.00109005626125222 | 0.0372664459746642 | 5771 | 9 | 10 | 23 | 42 | 2 | 1 | 15 | 6 | 1 | 3 | 9 | 9 |
| L1M1_dup4298 | 9.57348163295584 | -2.53439685682583 | 0.670298894445426 | -3.7809951319145 | 0.000156202728945223 | 0.00937870667211707 | 5772 | 9 | 6 | 32 | 44 | 3 | 1 | 8 | 5 | 0 | 1 | 12 | 9 |
| L1MB2_dup3999 | 9.44266380179438 | -1.94523149869791 | 0.616221231502121 | -3.15670963487601 | 0.00159560147330715 | 0.0484578159754362 | 5773 | 12 | 7 | 30 | 35 | 5 | 1 | 13 | 4 | 0 | 2 | 11 | 8 |
| L1ME3Cz_dup55 | 8.88633812715684 | -1.48674077979753 | 0.44108600860499 | -3.37063690707308 | 0.000749946312867799 | 0.028753614264793 | 5774 | 4 | 5 | 12 | 15 | 5 | 10 | 18 | 7 | 2 | 3 | 20 | 16 |
| MER20_dup10746 | 7.55957433217576 | -1.5568912882808 | 0.476315951346906 | -3.2686104336382 | 0.00108076992390536 | 0.0370447825286062 | 5775 | 3 | 4 | 18 | 19 | 4 | 8 | 12 | 5 | 2 | 3 | 11 | 11 |
| MIRb_dup22556 | 7.51750968795431 | -2.03738813863265 | 0.525025715231096 | -3.88054923697569 | 0.00010422084287017 | 0.00702718409679387 | 5776 | 1 | 4 | 16 | 15 | 3 | 6 | 12 | 5 | 1 | 3 | 20 | 13 |
| L1PA16_dup3621 | 9.01220398711427 | -1.73232513027292 | 0.51555545154557 | -3.36011407711747 | 0.000779102951957648 | 0.029507164477909 | 5777 | 6 | 9 | 22 | 29 | 6 | 5 | 10 | 5 | 1 | 1 | 11 | 16 |
| LTR1E_dup171 | 8.83175444667965 | -1.69531915873959 | 0.470234889315221 | -3.60526025877918 | 0.000311840255127418 | 0.0153537974260931 | 5778 | 6 | 7 | 16 | 29 | 7 | 4 | 16 | 6 | 2 | 1 | 14 | 10 |
| LTR5_Hs_dup633 | 7.2190524612321 | -1.83664189538792 | 0.532151680526936 | -3.45135036230513 | 0.00055778896433042 | 0.0233012031085966 | 5779 | 4 | 2 | 9 | 23 | 6 | 5 | 14 | 4 | 2 | 1 | 10 | 18 |
| MER31A_dup871 | 8.18093485908128 | -1.55357739579347 | 0.478953859855002 | -3.24368906070368 | 0.00117992475698115 | 0.0393439428565914 | 5780 | 5 | 5 | 15 | 15 | 7 | 5 | 16 | 3 | 2 | 2 | 13 | 22 |
| AluYh3_dup534 | 7.82889870194046 | -1.78337731067917 | 0.494661604055299 | -3.60524709429399 | 0.000311856063814092 | 0.0153537974260931 | 5781 | 5 | 1 | 14 | 16 | 3 | 7 | 11 | 11 | 0 | 6 | 14 | 14 |
| L1ME1_dup21949 | 6.49414676627948 | -2.08077033283955 | 0.544068947520776 | -3.82446074586916 | 0.000131058624101918 | 0.00825457378141597 | 5782 | 3 | 2 | 16 | 13 | 2 | 7 | 12 | 10 | 0 | 2 | 6 | 12 |
| SAR_dup11 | 7.24017233884065 | -2.00383092898305 | 0.498609331543098 | -4.01883960490989 | 5.84854660917836e-05 | 0.00457775747193274 | 5783 | 5 | 3 | 15 | 14 | 2 | 4 | 14 | 14 | 0 | 4 | 10 | 8 |
| L1ME1_dup22779 | 9.28815708438834 | -2.50866642573282 | 0.46964972791982 | -5.34156899620542 | 9.21455311319004e-08 | 3.11002277315052e-05 | 5784 | 2 | 4 | 14 | 24 | 3 | 4 | 17 | 18 | 2 | 2 | 13 | 17 |
| MER103C_dup5679 | 9.37038667609373 | -2.5689232757203 | 0.511291073001576 | -5.02438515235407 | 5.05047917751989e-07 | 0.000120371417375443 | 5785 | 2 | 3 | 14 | 29 | 5 | 5 | 15 | 10 | 1 | 1 | 13 | 28 |
| L1MC3_dup9445 | 8.04802373206005 | -3.50996263546154 | 0.619994820549999 | -5.6612773512331 | 1.50250358597523e-08 | 7.19903305860017e-06 | 5786 | 1 | 3 | 20 | 25 | 1 | 1 | 14 | 10 | 1 | 1 | 12 | 18 |
| MLT2B4_dup4071 | 8.34634088500302 | -2.88587856638865 | 0.584430332668945 | -4.93793426704869 | 7.89544444428825e-07 | 0.000171883753315701 | 5787 | 2 | 1 | 21 | 25 | 1 | 4 | 7 | 13 | 3 | 1 | 14 | 16 |
| L1ME3G_dup12635 | 10.2410478883879 | -1.80566027524859 | 0.543189459607575 | -3.32418135755632 | 0.000886785072915653 | 0.0323598422407214 | 5788 | 3 | 7 | 34 | 32 | 11 | 7 | 22 | 5 | 1 | 1 | 10 | 4 |
| MIR_dup133436 | 8.11485337436588 | -1.9128972586007 | 0.570565469858775 | -3.35263411414325 | 0.000800464493136097 | 0.0300658759721802 | 5789 | 2 | 4 | 27 | 26 | 6 | 10 | 17 | 5 | 0 | 1 | 6 | 5 |
| AluSz_dup73267 | 9.82041261487319 | -2.07039820454867 | 0.631718573722147 | -3.27740593782082 | 0.00104765630503693 | 0.0362544542115061 | 5790 | 1 | 2 | 38 | 34 | 8 | 10 | 10 | 8 | 0 | 3 | 9 | 7 |
| MLT1A0_dup16538 | 11.1834852406962 | -2.16741020881621 | 0.566671446433954 | -3.82480928314926 | 0.000130873358089781 | 0.00825218678151476 | 5791 | 3 | 3 | 35 | 46 | 10 | 9 | 14 | 10 | 0 | 2 | 12 | 5 |
| L1MB3_dup5953 | 7.09776680128604 | -1.77142585461148 | 0.558690400601605 | -3.17067530192749 | 0.00152085036163071 | 0.0469425684457881 | 5792 | 10 | 4 | 16 | 17 | 4 | 3 | 8 | 6 | 0 | 1 | 10 | 16 |
| MER5B_dup24695 | 6.99511534361035 | -1.93933891703077 | 0.596882715800343 | -3.24911220528536 | 0.00115765804316729 | 0.0388738499491228 | 5793 | 9 | 5 | 14 | 26 | 5 | 2 | 10 | 5 | 0 | 0 | 8 | 11 |
| AluSx3_dup25474 | 5.71435401734722 | -1.97590488254971 | 0.571547341383286 | -3.45711499202766 | 0.000545991810616263 | 0.0229317711778839 | 5794 | 7 | 3 | 14 | 14 | 3 | 3 | 10 | 6 | 0 | 0 | 8 | 8 |
| L1MEc_dup7092 | 6.4170702987294 | -1.91191138816146 | 0.580643333512758 | -3.29274664464816 | 0.000992138338436376 | 0.0349187364102553 | 5795 | 7 | 6 | 17 | 17 | 4 | 2 | 11 | 6 | 0 | 0 | 10 | 5 |
| AluJr_dup34446 | 7.87976142851383 | -1.81902786135104 | 0.542103760876071 | -3.35549758668228 | 0.000792223477225359 | 0.029864766716075 | 5796 | 7 | 7 | 20 | 9 | 3 | 6 | 18 | 10 | 0 | 1 | 5 | 18 |
| ALR/Alpha_dup1622 | 7.26751730943161 | -2.35826126314001 | 0.541983125944038 | -4.3511710056145 | 1.35412387379446e-05 | 0.00153140855168202 | 5797 | 3 | 6 | 15 | 16 | 4 | 3 | 16 | 11 | 0 | 0 | 6 | 16 |
| AluJb_dup122347 | 8.28623798161055 | -1.58508747510504 | 0.479473033832725 | -3.3058949372699 | 0.000946735791514137 | 0.033850036761476 | 5798 | 7 | 8 | 21 | 20 | 8 | 6 | 12 | 10 | 0 | 0 | 6 | 11 |
| L1PA8_dup7220 | 7.96935596795291 | -2.39332842976592 | 0.539494601904954 | -4.43624166268778 | 9.15430759872996e-06 | 0.00114282837139948 | 5799 | 3 | 2 | 24 | 25 | 4 | 4 | 14 | 10 | 0 | 3 | 4 | 13 |
| MER34A1_dup130 | 7.45275478554773 | -1.9774010397494 | 0.540080535426212 | -3.66130773105702 | 0.00025093113931074 | 0.0131573514487513 | 5800 | 2 | 4 | 21 | 24 | 6 | 5 | 11 | 13 | 0 | 2 | 2 | 7 |
| L2_dup49993 | 6.89013865309849 | -2.15447328139289 | 0.587795561531687 | -3.66534458984129 | 0.000247005771951493 | 0.0129829987657985 | 5801 | 4 | 4 | 12 | 33 | 6 | 2 | 12 | 6 | 0 | 1 | 5 | 9 |
| MIRb_dup101990 | 7.15204742326396 | -2.18157420782408 | 0.620873822913603 | -3.51371587480771 | 0.00044188519114916 | 0.019842866920714 | 5802 | 4 | 5 | 18 | 31 | 3 | 3 | 6 | 6 | 0 | 2 | 4 | 15 |
| Tigger1_dup11052 | 6.9858726657263 | -1.91588604959947 | 0.585993195779828 | -3.26946808153607 | 0.00107749893044187 | 0.0369752755268131 | 5803 | 4 | 4 | 21 | 30 | 4 | 3 | 7 | 8 | 0 | 3 | 3 | 6 |
| LTR10B_dup13 | 8.48445840310541 | -2.0329066099874 | 0.470468676614961 | -4.32102435514779 | 1.55306528639891e-05 | 0.00169316887240008 | 5804 | 5 | 7 | 23 | 27 | 4 | 5 | 20 | 10 | 0 | 1 | 5 | 6 |
| MLT1C_dup16505 | 9.01632581230745 | -2.40749564905878 | 0.495348299698144 | -4.86020775790663 | 1.17262627562394e-06 | 0.00023290628696128 | 5805 | 6 | 5 | 24 | 33 | 2 | 5 | 20 | 11 | 0 | 1 | 5 | 9 |
| AluSx3_dup27403 | 10.2630471392965 | -1.77016313401444 | 0.43017744074556 | -4.11496040086734 | 3.87245889682799e-05 | 0.00335539634683104 | 5806 | 11 | 6 | 29 | 24 | 5 | 7 | 16 | 15 | 0 | 2 | 9 | 10 |
| MER45C_dup813 | 9.47162087708838 | -1.85743928228611 | 0.437850003188328 | -4.24218172607205 | 2.21357334403012e-05 | 0.0022010485704914 | 5807 | 9 | 4 | 26 | 21 | 6 | 6 | 22 | 13 | 0 | 2 | 5 | 11 |
| L2a_dup154985 | 6.9533447280735 | -1.61985904499651 | 0.498334508526946 | -3.25054560195869 | 0.00115183794157549 | 0.0387495024524421 | 5808 | 5 | 2 | 18 | 19 | 6 | 7 | 13 | 10 | 0 | 2 | 5 | 4 |
| L2a_dup165404 | 7.60413859970959 | -1.77235281622914 | 0.521040701851882 | -3.40156308313314 | 0.000670016628827759 | 0.0265010156440533 | 5809 | 9 | 4 | 19 | 18 | 3 | 5 | 18 | 7 | 0 | 2 | 12 | 4 |
| AluJo_dup75781 | 6.9660117110584 | -2.1398427743841 | 0.504227388966155 | -4.24380511889989 | 2.19761219338278e-05 | 0.00218883646281565 | 5810 | 6 | 4 | 15 | 16 | 2 | 4 | 18 | 10 | 0 | 1 | 8 | 8 |
| MER121_dup844 | 7.42800440348197 | -2.93301246231244 | 0.564932689542242 | -5.1917910161811 | 2.08280715020807e-07 | 5.94952737291008e-05 | 5811 | 3 | 3 | 16 | 18 | 1 | 3 | 19 | 16 | 0 | 1 | 9 | 7 |
| MIR_dup89665 | 9.42119275892818 | -2.19894511038531 | 0.555197981363439 | -3.96065040615819 | 7.47458952539119e-05 | 0.00549613312487247 | 5812 | 13 | 3 | 32 | 26 | 2 | 2 | 19 | 14 | 0 | 2 | 6 | 5 |
| MLT1A0_dup1996 | 9.72516871207068 | -2.81903551542351 | 0.527551027479693 | -5.34362624387462 | 9.11053255129519e-08 | 3.08805523995795e-05 | 5813 | 8 | 3 | 22 | 39 | 1 | 3 | 23 | 15 | 0 | 1 | 8 | 7 |
| L2a_dup165171 | 7.68923633018196 | -2.12684251765684 | 0.597695283788575 | -3.55840605630272 | 0.000373112192993524 | 0.0175559726861849 | 5814 | 10 | 5 | 24 | 23 | 1 | 1 | 16 | 7 | 0 | 2 | 8 | 6 |
| LTR16D_dup986 | 7.53151996069718 | -2.72353377394016 | 0.605828785928143 | -4.49555028945621 | 6.93902551480166e-06 | 0.000923441479387582 | 5815 | 5 | 3 | 24 | 34 | 2 | 2 | 13 | 7 | 0 | 1 | 5 | 6 |
| MamRep564_dup713 | 9.72412992740741 | -3.63253386195116 | 0.661443357983948 | -5.49182907062968 | 3.97792296034856e-08 | 1.58597375599916e-05 | 5816 | 0 | 1 | 18 | 29 | 0 | 6 | 18 | 22 | 0 | 2 | 18 | 11 |
| L1MC4_dup21207 | 8.29744453527922 | -2.60222273350868 | 0.597219742969202 | -4.35722824662693 | 1.31719887510102e-05 | 0.00149891058601046 | 5817 | 0 | 6 | 22 | 28 | 3 | 1 | 16 | 12 | 0 | 4 | 9 | 8 |
| LTR10B1_dup26 | 6.47123104456386 | -3.35872415905501 | 0.689292128022297 | -4.87271509786684 | 1.1007490618008e-06 | 0.000221777168720791 | 5818 | 0 | 1 | 21 | 20 | 3 | 1 | 12 | 10 | 0 | 2 | 6 | 9 |
| LTR3A_dup95 | 7.36987095857448 | -3.18576895560276 | 0.632648742755324 | -5.03560465753561 | 4.76342781155576e-07 | 0.000115069908025223 | 5819 | 1 | 1 | 14 | 27 | 2 | 1 | 11 | 9 | 0 | 4 | 15 | 13 |
| L1MEd_dup12043 | 8.2254266096409 | -3.3566574449842 | 0.661650862334949 | -5.07315509744617 | 3.91273332605021e-07 | 9.87295062566641e-05 | 5820 | 0 | 2 | 19 | 25 | 1 | 2 | 18 | 8 | 0 | 4 | 22 | 8 |
| L2c_dup129460 | 8.13872637462046 | -3.74560408664938 | 0.665418042321744 | -5.62894879372433 | 1.81311209925254e-08 | 8.4427156862768e-06 | 5821 | 0 | 1 | 23 | 26 | 2 | 2 | 20 | 7 | 0 | 2 | 12 | 15 |
| L1M4b_dup3993 | 7.76856357489387 | -3.72531126449589 | 0.68697582759059 | -5.42276906243057 | 5.86827872770957e-08 | 2.18434989596504e-05 | 5822 | 3 | 1 | 12 | 34 | 1 | 1 | 18 | 10 | 0 | 1 | 8 | 17 |
| MIR_dup157827 | 7.3231406841693 | -3.8324853492416 | 0.70347530889302 | -5.44793157740299 | 5.09589530131737e-08 | 1.94007664538762e-05 | 5823 | 1 | 2 | 13 | 21 | 1 | 1 | 17 | 18 | 0 | 1 | 6 | 14 |
| L1PBa1_dup436 | 6.04926228445864 | -3.23253592559879 | 0.712910370571559 | -4.53428096859803 | 5.78000463258091e-06 | 0.000802408974506475 | 5824 | 1 | 2 | 17 | 24 | 0 | 1 | 12 | 7 | 0 | 3 | 7 | 7 |
| LTR40b_dup597 | 6.84864402258987 | -2.17374789772581 | 0.56197092494408 | -3.86807893654306 | 0.000109696157465535 | 0.00728895891770291 | 5825 | 3 | 4 | 15 | 23 | 1 | 2 | 16 | 9 | 0 | 5 | 6 | 7 |
| LSAU_dup3 | 6.89717075409387 | -2.94729860128403 | 0.717713105813534 | -4.10651356009897 | 4.01675891786419e-05 | 0.00345043941324951 | 5826 | 2 | 1 | 22 | 18 | 1 | 7 | 5 | 10 | 0 | 0 | 11 | 13 |
| MER4A1_dup265 | 5.06796467952335 | -2.73821302971318 | 0.712388609997443 | -3.84370691963058 | 0.000121189725874249 | 0.00781268233889843 | 5827 | 2 | 1 | 17 | 14 | 1 | 5 | 6 | 8 | 0 | 0 | 6 | 6 |
| AluSz_dup58123 | 5.86526699746493 | -2.47500555973007 | 0.642397145766897 | -3.8527654987872 | 0.000116791230771556 | 0.00759704926111771 | 5828 | 2 | 2 | 17 | 8 | 2 | 6 | 10 | 8 | 0 | 0 | 12 | 9 |
| MLT1G_dup1717 | 7.26466586029805 | -3.1086010294344 | 0.613638721848364 | -5.06584887614468 | 4.0658438832111e-07 | 0.000101623034520436 | 5829 | 1 | 1 | 18 | 18 | 2 | 6 | 17 | 11 | 0 | 0 | 9 | 13 |
| L1M5_dup27224 | 5.91892181597918 | -3.83409821673145 | 0.802788477158062 | -4.77597564716484 | 1.78838036243504e-06 | 0.000323357107689945 | 5830 | 1 | 1 | 19 | 22 | 0 | 3 | 6 | 12 | 0 | 0 | 5 | 8 |
| L2b_dup60231 | 7.2228254035835 | -3.27848541661273 | 0.632730111602022 | -5.18149105992739 | 2.20119141413522e-07 | 6.19839872458958e-05 | 5831 | 4 | 2 | 21 | 20 | 0 | 3 | 14 | 14 | 0 | 0 | 8 | 8 |
| LTR84b_dup790 | 6.90524265789306 | -2.79832874140356 | 0.642693352606465 | -4.35406516973428 | 1.33635950667425e-05 | 0.00151636337231812 | 5832 | 3 | 2 | 17 | 25 | 0 | 7 | 12 | 12 | 0 | 0 | 6 | 7 |
| L1MA7_dup6577 | 7.71916856637988 | -3.23009685983379 | 0.653438344756956 | -4.94323127155205 | 7.68382404750705e-07 | 0.000168354990497985 | 5833 | 2 | 1 | 33 | 14 | 2 | 5 | 15 | 15 | 0 | 0 | 5 | 7 |
| MIRb_dup104834 | 7.29210304216372 | -3.43521361639167 | 0.684436622856135 | -5.01903828882888 | 5.19307995724783e-07 | 0.000122586567808263 | 5834 | 1 | 1 | 32 | 20 | 3 | 3 | 11 | 14 | 0 | 0 | 3 | 6 |
| CR1_Mam_dup1562 | 6.44353435865712 | -3.48955405946571 | 0.714009197076631 | -4.88726766231162 | 1.02245045550067e-06 | 0.000209255520081629 | 5835 | 2 | 2 | 20 | 21 | 1 | 2 | 9 | 11 | 0 | 0 | 4 | 13 |
| MIRc_dup64206 | 6.20035471659197 | -3.60132452171072 | 0.701090297073587 | -5.13674848552743 | 2.79532666278716e-07 | 7.56697153701425e-05 | 5836 | 1 | 2 | 22 | 14 | 1 | 2 | 15 | 13 | 0 | 0 | 4 | 6 |
| L1PA15-16_dup1354 | 6.088541902679 | -3.3596056326798 | 0.696239641182461 | -4.82535815825425 | 1.39751895862816e-06 | 0.000267525379026335 | 5837 | 1 | 1 | 17 | 19 | 1 | 4 | 13 | 14 | 0 | 0 | 2 | 7 |
| THE1B-int_dup4174 | 5.47273530270082 | -3.41831053615971 | 0.728015473162624 | -4.6953817084546 | 2.66109329994905e-06 | 0.000441033325652405 | 5838 | 1 | 1 | 14 | 18 | 1 | 3 | 11 | 13 | 0 | 0 | 2 | 7 |
| L1PA2_dup2288 | 7.07539344076475 | -2.06343305192209 | 0.594397164153896 | -3.47147189852314 | 0.000517613504481315 | 0.0220878143334192 | 5839 | 0 | 3 | 11 | 17 | 6 | 8 | 16 | 15 | 1 | 0 | 5 | 9 |
| LSAU_dup137 | 8.50324521064062 | -2.86810952792291 | 0.632537302461822 | -4.53429310296846 | 5.77967238457318e-06 | 0.000802408974506475 | 5840 | 0 | 2 | 22 | 21 | 5 | 7 | 11 | 13 | 0 | 0 | 13 | 17 |
| MLT1E2_dup3911 | 10.3748061112519 | -3.66024838466865 | 0.597105121270667 | -6.12998993691342 | 8.78846315312153e-10 | 6.49434674979085e-07 | 5841 | 0 | 1 | 30 | 25 | 4 | 5 | 23 | 21 | 0 | 0 | 13 | 12 |

| | | | | | | | | | | | | | | | | | | | |
|---|---|---|---|---|---|---|---|---|---|---|---|---|---|---|---|---|---|---|---|
| BSR/Beta_dup2023 | 8.53761741730942 | -2.92926694764871 | 0.675584325384446 | -4.3359012895715 | 1.45164081014948e-05 | 0.00161256827363132 | 5842 | 1 | 0 | 22 | 30 | 7 | 6 | 10 | 16 | 0 | 0 | 9 | 10 |
| L4_C_Mam_dup2665 | 7.70619414122304 | -3.04747358289623 | 0.747333512531372 | -4.07779596632006 | 4.54646365371581e-05 | 0.00380026978172835 | 5843 | 1 | 0 | 21 | 46 | 4 | 6 | 10 | 10 | 0 | 0 | 4 | 2 |
| AluSz_dup45938 | 5.74556640711405 | -3.09969817814203 | 0.737667911130949 | -4.20202388008142 | 2.64539219568293e-05 | 0.00251582930213418 | 5844 | 0 | 0 | 17 | 22 | 3 | 5 | 11 | 9 | 0 | 0 | 4 | 5 |
| L1MA3_dup8616 | 7.30446964653833 | -3.03451333955617 | 0.722215304146575 | -4.20167410207678 | 2.64948368944805e-05 | 0.00251871372740473 | 5845 | 0 | 0 | 16 | 33 | 6 | 5 | 16 | 10 | 0 | 0 | 6 | 6 |
| L1MC1_dup12812 | 7.83559077641556 | -3.03143688134345 | 0.743831006812898 | -4.07543763782084 | 4.5927889682275e-05 | 0.00382779602043384 | 5846 | 0 | 0 | 22 | 36 | 6 | 6 | 13 | 10 | 0 | 0 | 5 | 7 |
| L1PA8_dup8021 | 5.49522178853741 | -3.9838057224659 | 0.818424380842105 | -4.86765279202316 | 1.12931513365966e-06 | 0.000226245953966858 | 5847 | 0 | 0 | 16 | 16 | 1 | 3 | 19 | 8 | 0 | 0 | 7 | 3 |
| MIR_dup157820 | 6.61128344340028 | -4.00712371779592 | 0.804414948412837 | -4.98141379110649 | 6.31214035279797e-07 | 0.000143727133816925 | 5848 | 0 | 1 | 27 | 15 | 1 | 3 | 13 | 9 | 0 | 0 | 12 | 5 |
| MLT1L_dup10797 | 7.88745438696915 | -3.60434382092292 | 0.704270964557063 | -5.11783674510819 | 3.09059934083115e-07 | 8.16676507162132e-05 | 5849 | 0 | 2 | 15 | 37 | 5 | 1 | 20 | 9 | 0 | 0 | 10 | 8 |
| MIRb_dup217364 | 5.47012296667565 | -3.23031342035289 | 0.74760550222373 | -4.32087967617202 | 1.55408404315339e-05 | 0.00169316887240008 | 5850 | 0 | 0 | 14 | 16 | 5 | 2 | 16 | 7 | 0 | 0 | 6 | 7 |
| L1ME3D_dup4121 | 6.61264416220985 | -3.96546347589603 | 0.765216292273065 | -5.18214721241321 | 2.19345979800739e-07 | 6.1912809620008e-05 | 5851 | 0 | 0 | 17 | 31 | 3 | 2 | 18 | 8 | 0 | 0 | 4 | 7 |
| MIRb_dup104846 | 6.81920368429498 | -3.78405825912846 | 0.742295699229091 | -5.09777742624453 | 3.4366454794519e-07 | 8.94678386491971e-05 | 5852 | 0 | 0 | 20 | 27 | 4 | 2 | 14 | 10 | 0 | 0 | 5 | 9 |
| L2c_dup119844 | 7.48435335243761 | -1.79123636658732 | 0.550028932107975 | -3.25662208299197 | 0.00112746434314675 | 0.0381506869445955 | 5853 | 9 | 7 | 19 | 9 | 2 | 2 | 13 | 12 | 2 | 0 | 10 | 11 |
| AluJr_dup75256 | 8.54119408886309 | -3.44418688151861 | 0.617895711680828 | -5.57405856103059 | 2.48872013763951e-08 | 1.11103377461683e-05 | 5854 | 3 | 1 | 20 | 21 | 3 | 1 | 12 | 15 | 1 | 0 | 10 | 25 |
| AluSp_dup23917 | 8.70365263880795 | -2.56480431731033 | 0.536670318897506 | -4.77910595573697 | 1.76076391142226e-06 | 0.000320019705297638 | 5855 | 3 | 1 | 17 | 9 | 3 | 5 | 16 | 12 | 3 | 0 | 18 | 26 |
| L1MA1_dup2056 | 8.86027630128539 | -1.98800032878229 | 0.60081858980454 | -3.30881960464811 | 0.000936901826936319 | 0.0335964416258501 | 5856 | 3 | 7 | 6 | 11 | 6 | 4 | 16 | 8 | 1 | 0 | 22 | 34 |
| L1MD2_dup5373 | 6.64894057763449 | -2.21027642750062 | 0.613769635479872 | -3.60114984471744 | 0.000316812904684194 | 0.015536760803377 | 5857 | 4 | 3 | 4 | 12 | 2 | 4 | 11 | 7 | 1 | 0 | 15 | 26 |
| L1PA2_dup3791 | 5.98340653975159 | -2.15953955938929 | 0.666582848262012 | -3.23971666090701 | 0.00119648528202867 | 0.0397568617396443 | 5858 | 6 | 6 | 6 | 7 | 1 | 2 | 15 | 10 | 0 | 0 | 13 | 12 |
| ERV3-16A3_I-int_dup3( | 8.23046477288878 | -2.9136355152994 | 0.698899575564738 | -4.16889009117665 | 3.06086536463551e-05 | 0.00279844970359105 | 5859 | 4 | 1 | 6 | 7 | 2 | 5 | 18 | 14 | 0 | 0 | 26 | 24 |
| L1PA15_dup4253 | 7.55713902063462 | -2.59700188512721 | 0.653762062843671 | -3.97239612502295 | 7.11532670263638e-05 | 0.00530345863762657 | 5860 | 2 | 2 | 10 | 9 | 2 | 8 | 19 | 9 | 0 | 0 | 19 | 20 |
| L1MA9_dup8547 | 10.63705059536 | -2.31913785590609 | 0.608768605528869 | -3.80955560921433 | 0.000139216792755402 | 0.00861760639151325 | 5861 | 3 | 5 | 12 | 16 | 12 | 4 | 23 | 12 | 0 | 0 | 23 | 31 |
| LSAU_dup119 | 8.89461357486586 | -2.66292174976483 | 0.631890917950571 | -4.21421114644527 | 2.50652713495264e-05 | 0.00241758453852226 | 5862 | 3 | 1 | 18 | 18 | 6 | 6 | 11 | 9 | 0 | 0 | 15 | 32 |
| MER121_dup843 | 12.8232405103919 | -3.69487342979841 | 0.558055224724072 | -6.62098169876525 | 3.56821151487079e-11 | 4.16196890743845e-08 | 5863 | 4 | 5 | 23 | 42 | 2 | 1 | 19 | 22 | 0 | 0 | 18 | 34 |
| THE1B_dup20912 | 12.9907565062843 | -3.4382856035765 | 0.493850397336645 | -6.96220074362461 | 3.34997676778915e-12 | 5.08001000040174e-09 | 5864 | 2 | 2 | 34 | 35 | 4 | 5 | 24 | 18 | 0 | 1 | 25 | 21 |
| L1MD_dup8120 | 12.9283295743123 | -3.52521611321926 | 0.553242372098377 | -6.37191995950811 | 1.86676310577605e-10 | 1.73511255455854e-07 | 5865 | 1 | 2 | 28 | 28 | 7 | 2 | 18 | 24 | 0 | 1 | 22 | 35 |
| MIRb_dup202453 | 10.8607058464117 | -3.27282490066511 | 0.517305634216403 | -6.32667553606423 | 2.5049930795634e-10 | 2.25777683071891e-07 | 5866 | 3 | 3 | 22 | 31 | 5 | 1 | 19 | 22 | 0 | 1 | 16 | 18 |
| ERV3-16A3_I-int_dup3( | 15.6273677560995 | -2.69626282544106 | 0.570526536243894 | -4.72592010039027 | 2.29075714720873e-06 | 0.000389062455495881 | 5867 | 4 | 4 | 19 | 13 | 12 | 6 | 36 | 24 | 1 | 0 | 36 | 49 |
| L1P4_dup1903 | 12.8806747021293 | -2.55184384081056 | 0.633645448961157 | -4.02724243501509 | 5.64348349053358e-05 | 0.00445979515987547 | 5868 | 4 | 7 | 9 | 19 | 9 | 1 | 15 | 18 | 1 | 0 | 35 | 51 |
| L1MB4_dup5465 | 12.5840852381519 | -2.32124333209638 | 0.422676415523257 | -5.49177395957324 | 3.97916473624898e-08 | 1.58597375599916e-05 | 5869 | 7 | 5 | 19 | 33 | 9 | 6 | 33 | 16 | 1 | 0 | 21 | 17 |
| L2a_dup913 | 12.253360231088 | -1.60510103190309 | 0.449941580812923 | -3.5673542974248 | 0.000360603768787282 | 0.017115222141225 | 5870 | 8 | 6 | 16 | 26 | 12 | 11 | 28 | 9 | 3 | 0 | 18 | 27 |
| L1PA8_dup4423 | 13.3553233265045 | -2.51581071116332 | 0.57156403607408 | -4.40162528147108 | 1.07442999889755e-05 | 0.0012885903251899 | 5871 | 9 | 7 | 15 | 14 | 2 | 6 | 33 | 12 | 1 | 0 | 39 | 40 |
| MER68_dup1155 | 11.2713686718775 | -2.01969117092427 | 0.468820485812868 | -4.30802670114216 | 1.64717504227598e-05 | 0.00177107416452508 | 5872 | 8 | 4 | 14 | 19 | 5 | 7 | 20 | 8 | 2 | 1 | 32 | 30 |
| L1ME4a_dup31833 | 12.4810729925198 | -2.21970517418218 | 0.440570257931473 | -5.0382547033564 | 4.69796003521412e-07 | 0.000114300695147143 | 5873 | 8 | 5 | 22 | 21 | 2 | 0 | 22 | 16 | 4 | 8 | 32 | 21 |
| AluSz_dup83738 | 9.519720754784 | -2.84873214328572 | 0.512251508022259 | -5.5611981588582 | 2.6792876109065e-08 | 1.16762961513692e-05 | 5874 | 2 | 2 | 16 | 13 | 5 | 1 | 23 | 13 | 1 | 3 | 25 | 20 |
| L2a_dup124323 | 10.8568531520824 | -2.39702596291535 | 0.452015047172487 | -5.30297824797999 | 1.13928501308662e-07 | 3.67328335669252e-05 | 5875 | 5 | 2 | 11 | 28 | 3 | 2 | 20 | 20 | 4 | 5 | 20 | 20 |
| L1MA8_dup7703 | 8.51010398512092 | -2.30433840160153 | 0.602806179034017 | -3.82268543646016 | 0.000132006136597492 | 0.0082942519616654 | 5876 | 2 | 5 | 16 | 27 | 2 | 0 | 30 | 11 | 4 | 3 | 4 | 9 |
| MLT1J_dup7268 | 8.47178784618353 | -2.02450334949814 | 0.60695985703248 | -3.33548145901485 | 0.000851518319235655 | 0.031415106915987 | 5877 | 3 | 5 | 19 | 32 | 2 | 0 | 21 | 8 | 5 | 4 | 8 | 5 |
| HAL1_dup16904 | 10.0459914117609 | -3.21373120680363 | 0.554274059070354 | -5.79809059113068 | 6.70742181309736e-09 | 3.61086920981734e-06 | 5878 | 2 | 2 | 23 | 36 | 4 | 0 | 21 | 9 | 1 | 3 | 18 | 16 |
| L1MEd_dup18894 | 7.74640607456311 | -2.7501952355937 | 0.584975353116146 | -4.70138651302742 | 2.58400865116995e-06 | 0.000431069250145492 | 5879 | 2 | 2 | 10 | 29 | 4 | 0 | 20 | 14 | 1 | 3 | 9 | 8 |
| L1P4_dup976 | 7.76810325195428 | -2.22838283911347 | 0.532099466654513 | -4.18790654522557 | 2.81539396314618e-05 | 0.00263330083315559 | 5880 | 5 | 5 | 13 | 26 | 3 | 0 | 11 | 11 | 1 | 3 | 13 | 11 |
| AluSz_dup91342 | 6.18195975329246 | -1.80206782826807 | 0.562099399987504 | -3.20595935222156 | 0.00134613002321789 | 0.0431330803265019 | 5881 | 4 | 3 | 13 | 11 | 3 | 0 | 11 | 3 | 1 | 6 | 14 | 13 |
| Charlie25_dup1041 | 8.98642981571784 | -2.09899249183981 | 0.587453279587351 | -3.57303732020054 | 0.00035286444787284 | 0.0168160792937214 | 5882 | 5 | 8 | 25 | 22 | 2 | 0 | 8 | 6 | 1 | 5 | 17 | 20 |
| L1M5_dup28836 | 9.96413749028492 | -1.91593709595049 | 0.556571364983933 | -3.44239250613585 | 0.000576592969638195 | 0.0238563711969275 | 5883 | 5 | 6 | 27 | 33 | 3 | 0 | 11 | 7 | 7 | 4 | 11 | 17 |
| L1MB5_dup5131 | 9.06555665337893 | -1.66806269656838 | 0.470599622185726 | -3.54454746228009 | 0.000393287689387575 | 0.018263316264445 | 5884 | 7 | 5 | 18 | 18 | 5 | 0 | 14 | 14 | 6 | 3 | 9 | 17 |
| AluSx3_dup24724 | 7.70167424331732 | -1.74738841127492 | 0.553557335062354 | -3.15665297990874 | 0.00159591148969386 | 0.0484610458558825 | 5885 | 2 | 9 | 8 | 17 | 1 | 0 | 12 | 8 | 5 | 4 | 15 | 19 |
| L3_dup12172 | 6.43425377354375 | -1.93177839812535 | 0.549427503797154 | -3.51598415582515 | 0.000438127129412442 | 0.0197298777117623 | 5886 | 5 | 2 | 12 | 12 | 3 | 0 | 13 | 7 | 5 | 1 | 13 | 10 |
| AluSc8_dup3116 | 8.52232638258996 | -1.77165835669007 | 0.497010474770993 | -3.56462981490762 | 0.000364370081210895 | 0.0172469471541292 | 5887 | 9 | 5 | 9 | 17 | 1 | 4 | 20 | 17 | 3 | 2 | 14 | 8 |
| MLT1D_dup1598 | 8.45726794715608 | -1.69623840870194 | 0.529574322563988 | -3.20302238312732 | 0.0013599341077141 | 0.0434564319787437 | 5888 | 7 | 7 | 6 | 17 | 0 | 6 | 25 | 16 | 2 | 3 | 9 | 12 |
| L1MEi_dup16 | 7.86966298344021 | -1.80703572066064 | 0.513900640366984 | -3.51631342465386 | 0.00043758408704276 | 0.0197162248012643 | 5889 | 3 | 6 | 14 | 14 | 0 | 9 | 14 | 12 | 1 | 3 | 12 | 14 |
| MLT1A0_dup1857 | 8.43956333931908 | -2.20376748668258 | 0.578499708946397 | -3.80945306039347 | 0.000139274543790909 | 0.00861893903689768 | 5890 | 2 | 5 | 12 | 13 | 0 | 5 | 34 | 15 | 1 | 5 | 10 | 8 |
| MLT2A2_dup3403 | 9.74344273762761 | -1.41683315562637 | 0.44889015829926 | -3.15630255961597 | 0.0015978302244896 | 0.0485007412420465 | 5891 | 9 | 9 | 21 | 19 | 0 | 6 | 17 | 9 | 3 | 6 | 12 | 17 |
| MSTA_dup5677 | 8.00242165388823 | -1.69709076645025 | 0.513685832048862 | -3.30375233375878 | 0.000954000715811892 | 0.0339974021753147 | 5892 | 7 | 6 | 14 | 23 | 0 | 2 | 15 | 6 | 5 | 3 | 11 | 14 |
| L2a_dup155354 | 8.92601035308941 | -1.88514147440282 | 0.477419436798518 | -3.94860646446281 | 7.86074548488943e-05 | 0.00570601874663667 | 5893 | 8 | 3 | 15 | 13 | 0 | 2 | 12 | 9 | 4 | 6 | 16 | 29 |
| MIRb_dup202622 | 9.25881286905784 | -2.34123589594918 | 0.499303606574067 | -4.68900257303044 | 2.74539937034407e-06 | 0.000449282529969663 | 5894 | 3 | 2 | 18 | 22 | 0 | 3 | 15 | 7 | 5 | 6 | 24 | 16 |
| AluSc_dup15791 | 7.51634359351025 | -2.02191002471296 | 0.578578646652227 | -3.49461570421262 | 0.0004747447831079 | 0.0208112789538306 | 5895 | 7 | 3 | 23 | 9 | 0 | 1 | 18 | 10 | 3 | 4 | 9 | 10 |
| L1PA6_dup2500 | 7.30954445972324 | -2.05928671702407 | 0.625327155374951 | -3.29313496035415 | 0.000990769040944286 | 0.0348896744437663 | 5896 | 10 | 2 | 18 | 27 | 0 | 2 | 9 | 8 | 1 | 3 | 6 | 12 |
| L1PB1_dup3500 | 6.81136893627095 | -2.10356931654859 | 0.598005874125278 | -3.51763988878127 | 0.000435402791013285 | 0.0196402564000848 | 5897 | 5 | 4 | 18 | 18 | 0 | 2 | 9 | 3 | 1 | 4 | 15 | 12 |
| THE1B_dup1465 | 6.65314110042592 | -1.91969077927356 | 0.555065092061409 | -3.45849668215341 | 0.000543198963149025 | 0.0228481563523878 | 5898 | 5 | 5 | 18 | 13 | 0 | 2 | 10 | 5 | 3 | 2 | 13 | 11 |
| L1MB8_dup5156 | 6.84993978131998 | -1.82725258771123 | 0.538704921312369 | -3.3919359475309 | 0.000694006620310502 | 0.0271650105735158 | 5899 | 7 | 3 | 21 | 19 | 3 | 3 | 15 | 5 | 2 | 1 | 8 | 4 |
| L1PA5_dup10154 | 7.0542512638919 | -1.88726449158607 | 0.53805211751801 | -3.50758677484973 | 0.000452190850575935 | 0.0201808046916367 | 5900 | 5 | 2 | 17 | 15 | 2 | 2 | 16 | 8 | 4 | 3 | 13 | 4 |
| AluSg7_dup4067 | 8.21734893702195 | -2.27697193743171 | 0.51843906114227 | -4.39197604519784 | 1.12325077500805e-05 | 0.00133104098062782 | 5901 | 6 | 3 | 23 | 23 | 2 | 1 | 19 | 10 | 2 | 3 | 10 | 6 |
| L2a_dup68331 | 8.17943737729443 | -2.05684480584822 | 0.556518156630571 | -3.69591680225018 | 0.000219094717314587 | 0.0118834537511139 | 5902 | 8 | 3 | 24 | 26 | 1 | 1 | 12 | 11 | 2 | 4 | 8 | 7 |
| L1MEh_dup935 | 7.35891849962997 | -1.78167738053719 | 0.506785862393488 | -3.51564144296083 | 0.000438693012290781 | 0.0197474929062698 | 5903 | 5 | 5 | 15 | 21 | 2 | 1 | 10 | 7 | 5 | 2 | 11 | 12 |
| Tigger1_dup2855 | 7.69289518749772 | -1.64107935210491 | 0.506910723941495 | -3.23741296957512 | 0.00120618724037496 | 0.0400058602920271 | 5904 | 10 | 4 | 14 | 15 | 1 | 1 | 18 | 9 | 4 | 3 | 8 | 14 |
| MLT1D_dup9009 | 7.23411600847276 | -1.73441673459733 | 0.503643906515562 | -3.44373616390361 | 0.00057373527763273 | 0.0237836261971424 | 5905 | 1 | 2 | 16 | 11 | 3 | 7 | 15 | 12 | 3 | 4 | 9 | 9 |
| ERVL-E-int_dup891 | 6.75225009123377 | -2.16732194383597 | 0.514734290693292 | -4.21056452430403 | 2.5473335397916e-05 | 0.0024470239259558 | 5906 | 2 | 2 | 12 | 15 | 2 | 5 | 11 | 12 | 2 | 2 | 9 | 13 |
| MSTA-int_dup1933 | 5.97162661748416 | -1.8284042393751 | 0.526426774022375 | -3.47323565137929 | 0.000514223643765095 | 0.0220027799207569 | 5907 | 1 | 2 | 9 | 11 | 3 | 6 | 12 | 9 | 2 | 2 | 10 | 10 |
| AluSq2_dup19535 | 7.0008077326857 | -1.60648408527383 | 0.478029682821561 | -3.36063667802299 | 0.000777630446252235 | 0.0294734077992887 | 5908 | 5 | 3 | 15 | 12 | 4 | 4 | 17 | 11 | 4 | 1 | 5 | 9 |
| MLT1C_dup15383 | 6.53469226639158 | -1.74102414010696 | 0.518587922601348 | -3.35724004402881 | 0.000787247330718775 | 0.029728987994796 | 5909 | 4 | 4 | 12 | 12 | 1 | 3 | 20 | 9 | 4 | 2 | 5 | 9 |
| ERV24_Prim-int_dup57 | 7.5122135196486 | -2.17448957233077 | 0.630621781438728 | -3.44816756466894 | 0.000564403771483972 | 0.0235197232591112 | 5910 | 2 | 2 | 9 | 18 | 3 | 1 | 26 | 19 | 2 | 5 | 3 | 6 |
| HSMAR2_dup154 | 7.6702128792253 | -1.93209433997382 | 0.500547407347992 | -3.85996273601829 | 0.000113404317398826 | 0.00744719955235963 | 5911 | 3 | 3 | 13 | 14 | 2 | 2 | 23 | 12 | 4 | 5 | 8 | 10 |
| Arthur1B_dup213 | 8.14334234109813 | -1.6596617540113 | 0.50546509836313 | -3.28343491842633 | 0.00102550320241745 | 0.0356954922472825 | 5912 | 5 | 4 | 9 | 14 | 3 | 3 | 26 | 18 | 3 | 5 | 5 | 9 |
| MLT1G3_dup222 | 7.34695701114263 | -1.72299152211856 | 0.528667085024855 | -3.25912388140744 | 0.00111756856626147 | 0.037880480066617 | 5913 | 4 | 5 | 9 | 18 | 2 | 1 | 19 | 15 | 3 | 5 | 6 | 7 |
| THE1B_dup4511 | 7.81855518595863 | -1.7932625335227 | 0.539490733465398 | -3.32399135385282 | 0.000887389470634079 | 0.0323598422407214 | 5914 | 7 | 2 | 17 | 16 | 4 | 2 | 24 | 16 | 3 | 2 | 5 | 2 |
| LTR37A_dup11 | 6.37051997383024 | -1.7167302586872 | 0.519373378801387 | -3.30538747028021 | 0.00094845181302025 | 0.033880845984224 | 5915 | 4 | 1 | 9 | 15 | 4 | 4 | 19 | 9 | 3 | 2 | 8 | 5 |
| MER67C_dup1384 | 7.4312283132323 | -2.22917126356829 | 0.555473275117218 | -4.01310263414182 | 5.99258417972714e-05 | 0.00466887125358525 | 5916 | 2 | 1 | 11 | 18 | 4 | 3 | 23 | 17 | 3 | 2 | 5 | 6 |
| L1MB2_dup5253 | 7.71229240918848 | -1.91405267753024 | 0.473390181056371 | -4.04328766021092 | 5.27068898001507e-05 | 0.00425709219293505 | 5917 | 3 | 1 | 15 | 15 | 6 | 5 | 18 | 7 | 2 | 3 | 12 | 15 |
| MIR_dup35766 | 8.44205982307643 | -1.50605426491116 | 0.458696401073207 | -3.28333569085664 | 0.00102586427185834 | 0.0357028375064507 | 5918 | 3 | 2 | 16 | 17 | 8 | 6 | 17 | 8 | 2 | 6 | 16 | 9 |
| L2c_dup30778 | 7.74356957988978 | -1.64665615024407 | 0.498345633889653 | -3.30424516292379 | 0.000952325122236465 | 0.0339741547339859 | 5919 | 3 | 1 | 10 | 22 | 5 | 5 | 10 | 6 | 2 | 7 | 18 | 13 |
| Charlie1a_dup2166 | 7.41098893885221 | -1.82999918628128 | 0.483996477321305 | -3.78101757353581 | 0.000156188647236014 | 0.00937870667211707 | 5920 | 4 | 2 | 8 | 16 | 5 | 4 | 13 | 11 | 1 | 4 | 17 | 11 |

| | | | | | | | | | | | | | | | | | | |
|---|---|---|---|---|---|---|---|---|---|---|---|---|---|---|---|---|---|---|
| MER4A_dup141 | 8.45249396544822 | -1.65526844683802 | 0.456267820036793 | -3.62784394197368 | 0.000285797913912752 | 0.0144444499625921 | 5921 | 4 | 1 | 9 | 14 | 4 | 6 | 19 | 11 | 2 | 8 | 16 | 16 |
| LTR41B_dup166 | 8.57012039611416 | -1.71335117305642 | 0.473764205129451 | -3.61646395929862 | 0.000298654877909319 | 0.0149134742001672 | 5922 | 5 | 3 | 11 | 18 | 4 | 5 | 12 | 12 | 5 | 2 | 23 | 9 |
| MER77_dup1295 | 7.99776036643316 | -1.56870869914882 | 0.449422432459014 | -3.49049932947412 | 0.000482118822827733 | 0.0210029742615467 | 5923 | 3 | 3 | 16 | 13 | 5 | 3 | 11 | 11 | 6 | 4 | 14 | 12 |
| MIR_dup35786 | 7.59427848737705 | -1.4959224299655 | 0.46272803724109 | -3.23283291603551 | 0.00122569227582348 | 0.0404057320951917 | 5924 | 5 | 2 | 12 | 10 | 5 | 5 | 16 | 13 | 4 | 3 | 13 | 8 |
| THE1B-int_dup2373 | 7.70881374208596 | -1.51445876039868 | 0.455912363660408 | -3.32181989591039 | 0.000894324004301034 | 0.0325144464958025 | 5925 | 3 | 2 | 14 | 16 | 6 | 4 | 19 | 8 | 5 | 4 | 10 | 9 |
| AluSx3_dup28578 | 10.7149081825143 | -1.65127134107371 | 0.396462255555405 | -4.16501525160431 | 3.1133175147573e-05 | 0.00283831972937334 | 5926 | 7 | 2 | 13 | 27 | 7 | 8 | 22 | 12 | 4 | 4 | 17 | 18 |
| MLT1A_dup8677 | 10.3477747177397 | -1.32281257254453 | 0.414736017503842 | -3.18952904188572 | 0.00142504819681502 | 0.044828730597481 | 5927 | 8 | 3 | 13 | 32 | 7 | 11 | 17 | 14 | 2 | 6 | 10 | 13 |
| AluSx_dup54389 | 10.958071267518 | -1.55401606502633 | 0.434408029171931 | -3.57731892752672 | 0.000347136529954153 | 0.0165996279655287 | 5928 | 6 | 3 | 20 | 21 | 10 | 6 | 12 | 14 | 3 | 6 | 27 | 12 |
| MER103C_dup3841 | 9.91585281714882 | -1.53593673154505 | 0.451046913982883 | -3.40527045841474 | 0.000660985554943113 | 0.0262492842495154 | 5929 | 6 | 5 | 23 | 26 | 5 | 5 | 6 | 11 | 4 | 6 | 16 | 15 |
| Tigger10_dup564 | 9.93321806807215 | -1.41122672314587 | 0.447343446528206 | -3.15468290437309 | 0.00160672630713913 | 0.0486714319387049 | 5930 | 3 | 5 | 18 | 27 | 8 | 3 | 16 | 15 | 5 | 8 | 10 | 9 |
| L1MB1_dup405 | 9.40104932815036 | -1.54139902105542 | 0.431636834798028 | -3.57105533353443 | 0.000355545764838329 | 0.0169099925165343 | 5931 | 3 | 7 | 9 | 18 | 8 | 4 | 15 | 15 | 4 | 3 | 18 | 16 |
| MIR_dup53682 | 10.702822424333 | -1.25926483058106 | 0.397790210494294 | -3.16565063030661 | 0.00154736521939002 | 0.0474593148353929 | 5932 | 3 | 6 | 14 | 19 | 8 | 10 | 15 | 18 | 5 | 6 | 18 | 13 |
| MER5A1_dup13562 | 9.93763034565515 | -1.85722597786011 | 0.466208611265868 | -3.98368012297605 | 6.78561959256188e-05 | 0.00512574933197374 | 5933 | 1 | 3 | 8 | 12 | 3 | 6 | 27 | 11 | 6 | 7 | 21 | 24 |
| AluJr_dup67118 | 8.66363306330969 | -1.4455537932399 | 0.422690733065353 | -3.41988522614807 | 0.000626475614629248 | 0.0252999603867229 | 5934 | 2 | 4 | 11 | 15 | 4 | 7 | 18 | 12 | 5 | 6 | 13 | 14 |
| MIR_dup35772 | 9.34760098547031 | -1.69511833472992 | 0.438056720447799 | -3.86963207184017 | 0.000108999722223043 | 0.00725686846784672 | 5935 | 1 | 3 | 15 | 16 | 4 | 5 | 19 | 11 | 5 | 9 | 12 | 21 |
| AluSx_dup21622 | 8.32935827077744 | -1.73810245782283 | 0.54537858119748 | -3.18696501429613 | 0.00143774164767517 | 0.0451136031879259 | 5936 | 2 | 1 | 11 | 16 | 3 | 6 | 23 | 16 | 3 | 8 | 11 | 6 |
| MamGypsy2-I_dup733 | 9.13206928947969 | -1.64673191634214 | 0.477076550611475 | -3.45171422538271 | 0.000557037364505034 | 0.0232861582456983 | 5937 | 3 | 3 | 12 | 18 | 2 | 6 | 28 | 11 | 4 | 9 | 13 | 10 |
| L1PA12_dup191 | 8.51602937904876 | -1.52941332363683 | 0.441258566239629 | -3.46602523021904 | 0.0005282137097202 | 0.0223800461846657 | 5938 | 5 | 4 | 17 | 13 | 2 | 4 | 17 | 7 | 3 | 9 | 16 | 14 |
| MER47C_dup128 | 8.68838306298283 | -1.56713771178228 | 0.490896441597445 | -3.19239981997546 | 0.0014109587905801 | 0.0445445137831197 | 5939 | 5 | 5 | 9 | 16 | 1 | 3 | 16 | 16 | 3 | 10 | 16 | 10 |
| L3_dup9753 | 9.34125962910801 | -1.98891119436371 | 0.549309288897092 | -3.62074924739951 | 0.000293751108924821 | 0.0147492680941841 | 5940 | 1 | 0 | 12 | 18 | 2 | 6 | 17 | 13 | 7 | 8 | 24 | 10 |
| L4_A_Mam_dup2324 | 7.97227631471646 | -1.92816339342851 | 0.487584375555062 | -3.95452251978646 | 7.66876487937417e-05 | 0.00559856223778271 | 5941 | 3 | 0 | 15 | 14 | 3 | 5 | 12 | 14 | 4 | 5 | 13 | 13 |
| L2c_dup116145 | 8.14305063336432 | -1.80271248349707 | 0.522593616888686 | -3.44954937304764 | 0.000561523036441534 | 0.0234242918003361 | 5942 | 1 | 1 | 14 | 13 | 5 | 1 | 16 | 10 | 7 | 8 | 10 | 18 |
| AluSq_dup13961 | 11.0371672668147 | -1.42140428622539 | 0.444835696353243 | -3.19534672661849 | 0.00139662943834704 | 0.0442506509103761 | 5943 | 3 | 2 | 19 | 21 | 6 | 5 | 13 | 17 | 9 | 12 | 17 | 14 |
| AluSz_dup77059 | 9.6798116575869 | -1.71131932739756 | 0.515065668784496 | -3.32252648761487 | 0.000892062015350355 | 0.0324560532575773 | 5944 | 1 | 1 | 19 | 14 | 3 | 5 | 12 | 16 | 9 | 10 | 15 | 15 |
| L2a_dup155355 | 12.8879491681812 | -1.87573827906187 | 0.469884463100771 | -3.99191381362954 | 6.55421949634789e-05 | 0.0049985586931987 | 5945 | 5 | 11 | 21 | 22 | 4 | 3 | 12 | 6 | 5 | 3 | 38 | 40 |
| L2a_dup44319 | 10.4870146579556 | -1.56512299328386 | 0.483526399754954 | -3.23689253384521 | 0.00120838908657045 | 0.0400573069926293 | 5946 | 7 | 12 | 20 | 23 | 6 | 3 | 16 | 5 | 4 | 1 | 21 | 21 |
| L1M2_dup9208 | 11.173199027258 | -1.90704825059038 | 0.453596366534543 | -4.20428467088516 | 2.61909139467764e-05 | 0.00249580424909077 | 5947 | 9 | 5 | 32 | 25 | 2 | 5 | 11 | 10 | 4 | 4 | 19 | 20 |
| MLT1G1_dup970 | 10.2011067572995 | -2.32509824495433 | 0.502615554738731 | -4.62599739111329 | 3.72799989979389e-06 | 0.000568234748367321 | 5948 | 6 | 2 | 27 | 28 | 1 | 3 | 12 | 7 | 3 | 6 | 21 | 19 |
| AmnSINE1_dup2384 | 8.67491607018444 | -1.90689297607158 | 0.533642448714559 | -3.57335324553906 | 0.000352438802038505 | 0.0168105948577582 | 5949 | 6 | 5 | 21 | 13 | 1 | 5 | 9 | 4 | 1 | 4 | 23 | 23 |
| L1PA15_dup131 | 10.6157624699664 | -2.24013328914523 | 0.51434854251677 | -4.35528266141085 | 1.32895317322454e-05 | 0.0015096525313337 | 5950 | 4 | 6 | 20 | 25 | 4 | 4 | 8 | 6 | 1 | 3 | 32 | 28 |
| L1ME2_dup1611 | 10.7692308380635 | -2.08923422880072 | 0.489264546801145 | -4.27015250228188 | 1.95339353731748e-05 | 0.00201142945604017 | 5951 | 5 | 2 | 26 | 29 | 4 | 2 | 10 | 7 | 7 | 5 | 16 | 29 |
| L2b_dup87935 | 9.47331696914082 | -1.54157893277776 | 0.480980415958649 | -3.20507630171432 | 0.00135026680888667 | 0.0432248871394253 | 5952 | 6 | 5 | 23 | 19 | 5 | 1 | 10 | 5 | 8 | 4 | 17 | 20 |
| AluSx3_dup10815 | 9.14113765126023 | -1.52945944724804 | 0.432526897288546 | -3.53610251023929 | 0.000406077087645352 | 0.0186841501755728 | 5953 | 6 | 3 | 13 | 12 | 5 | 2 | 13 | 7 | 4 | 8 | 28 | 17 |
| MLT1D_dup18652 | 8.91892249301478 | -1.35315669066036 | 0.429836942052569 | -3.14806978711214 | 0.00164352450618439 | 0.0495338926090628 | 5954 | 7 | 4 | 14 | 18 | 4 | 3 | 9 | 5 | 6 | 6 | 22 | 18 |
| L1M3de_dup338 | 7.73309775385133 | -1.63471091220595 | 0.506246897761256 | -3.22907837941335 | 0.00124189852867664 | 0.040793040765391 | 5955 | 10 | 2 | 15 | 10 | 4 | 2 | 9 | 9 | 3 | 2 | 17 | 17 |
| MIR_dup59084 | 7.22899328731581 | -1.59320751827151 | 0.492356194383066 | -3.2358839727158 | 0.001212666654279 | 0.0401320138691335 | 5956 | 5 | 2 | 14 | 9 | 4 | 2 | 7 | 5 | 5 | 3 | 15 | 23 |
| L2a_dup20688 | 10.9688111469514 | -2.81076513197526 | 0.638479406643816 | -4.40228001518502 | 1.07119180993324e-05 | 0.00128600010060026 | 5957 | 4 | 1 | 37 | 36 | 2 | 1 | 7 | 4 | 6 | 3 | 20 | 25 |
| LTR40a_dup154 | 11.2108169165036 | -3.02252284496648 | 0.676866780294307 | -4.46546193868794 | 7.98961758218775e-06 | 0.00103040517355623 | 5958 | 4 | 2 | 36 | 42 | 1 | 1 | 4 | 5 | 5 | 2 | 26 | 21 |
| AluSg_dup35566 | 8.63728610599455 | -2.68015790605986 | 0.725309890437495 | -3.69519007171298 | 0.000219722354557285 | 0.0119009725769799 | 5959 | 3 | 2 | 30 | 31 | 2 | 1 | 5 | 1 | 5 | 1 | 15 | 20 |
| MIRb_dup214045 | 9.14309842903621 | -2.10534221303111 | 0.648297241495718 | -3.24749525105764 | 0.00116425607029907 | 0.0390238167211413 | 5960 | 4 | 2 | 28 | 32 | 4 | 2 | 3 | 2 | 5 | 4 | 15 | 21 |
| L1MC3_dup12812 | 8.34866423495111 | -2.56199795429613 | 0.621871880184372 | -4.11981637365007 | 3.79174460406802e-05 | 0.00330729641334153 | 5961 | 3 | 4 | 21 | 36 | 3 | 1 | 10 | 3 | 1 | 3 | 18 | 10 |
| L1PB1_dup5898 | 9.00443013063301 | -3.0265863173759 | 0.637470449663663 | -4.74780645749581 | 2.05634669418528e-06 | 0.000360840317473901 | 5962 | 2 | 1 | 31 | 31 | 4 | 1 | 12 | 3 | 2 | 2 | 13 | 20 |
| MER121_dup888 | 6.32539723141956 | -2.15175090262974 | 0.630899987358501 | -3.41060539823253 | 0.000648188262049543 | 0.0258894554666196 | 5963 | 3 | 2 | 19 | 22 | 1 | 3 | 6 | 3 | 4 | 1 | 9 | 11 |
| MIR_dup73406 | 7.30164134286772 | -2.43050538806111 | 0.611921396710263 | -3.97192417380352 | 7.1294415391022e-05 | 0.00530629370179297 | 5964 | 2 | 1 | 20 | 26 | 1 | 2 | 11 | 4 | 5 | 3 | 10 | 12 |
| L1MCc_dup3259 | 9.75620512318224 | -2.06453441477305 | 0.529910426064403 | -3.89600640641506 | 9.77918030273196e-05 | 0.00670581220839729 | 5965 | 4 | 5 | 16 | 43 | 3 | 4 | 11 | 4 | 4 | 3 | 17 | 18 |
| L1M5_dup22820 | 8.63925852729848 | -1.99081588340939 | 0.545051349078125 | -3.6525290447892 | 0.00025967021455778 | 0.0134995606015218 | 5966 | 3 | 2 | 20 | 33 | 4 | 4 | 10 | 4 | 5 | 3 | 9 | 19 |
| L1MC4_dup10260 | 9.34062069605664 | -2.11570075517506 | 0.616874346953729 | -3.42971103535573 | 0.000604224357987317 | 0.0247074701985994 | 5967 | 4 | 3 | 29 | 42 | 2 | 4 | 6 | 4 | 4 | 4 | 7 | 17 |
| MLT1E3_dup1858 | 10.4469990127198 | -2.40426698692809 | 0.624908267751813 | -3.84739186693392 | 0.000119381939887794 | 0.00772334286801005 | 5968 | 2 | 3 | 27 | 35 | 4 | 9 | 7 | 3 | 1 | 2 | 24 | 24 |
| L2c_dup134810 | 8.78211560891363 | -2.17436948696938 | 0.639794487518127 | -3.39854364079337 | 0.00067745653817677 | 0.0267282496158199 | 5969 | 2 | 4 | 22 | 37 | 6 | 4 | 5 | 2 | 1 | 3 | 17 | 16 |
| MER5A1_dup15497 | 9.99966304275519 | -2.20388163246478 | 0.612501982204117 | -3.598162449261 | 0.000320473435276604 | 0.0156678389213739 | 5970 | 4 | 4 | 22 | 41 | 6 | 6 | 5 | 3 | 1 | 2 | 19 | 23 |
| L1MC_dup7974 | 8.42377417385389 | -2.16023674664792 | 0.603897838697732 | -3.57715594959958 | 0.000347352959456196 | 0.0166020943228561 | 5971 | 4 | 3 | 20 | 25 | 3 | 6 | 5 | 3 | 2 | 1 | 14 | 28 |
| L2a_dup141148 | 8.33903664771699 | -1.82173755009761 | 0.57350398686487 | -3.1765037241613 | 0.00149061875655318 | 0.0463037559591127 | 5972 | 5 | 3 | 16 | 22 | 5 | 5 | 5 | 2 | 2 | 2 | 21 | 24 |
| L1MC4_dup10258 | 7.55417374227831 | -2.18317472482817 | 0.614509589212441 | -3.55271058931096 | 0.000381283695211355 | 0.0178416775104742 | 5973 | 5 | 1 | 17 | 30 | 3 | 4 | 4 | 4 | 2 | 2 | 12 | 18 |
| MER5A1_dup14497 | 7.42387112029151 | -1.87861794647376 | 0.53035417916386 | -3.54219504678087 | 0.000396811945349383 | 0.0183660406823778 | 5974 | 2 | 2 | 14 | 18 | 3 | 6 | 5 | 5 | 2 | 4 | 17 | 20 |
| AluJr4_dup2453 | 8.63365642855589 | -2.87303901220758 | 0.674631334844642 | -4.25868005800412 | 2.05637543885639e-05 | 0.00208570290670831 | 5975 | 2 | 1 | 19 | 38 | 2 | 1 | 3 | 5 | 3 | 4 | 17 | 21 |
| MIR_dup73408 | 9.12082998013755 | -2.2694135874938 | 0.58473223680187 | -3.88111591026025 | 0.000103978266569657 | 0.00701465888816122 | 5976 | 3 | 4 | 22 | 37 | 2 | 2 | 5 | 7 | 4 | 4 | 19 | 11 |
| MIRb_dup91543 | 9.30606313863477 | -2.42626797713173 | 0.597897054473702 | -4.05800289360424 | 4.94941548020113e-05 | 0.00404427752696407 | 5977 | 5 | 2 | 23 | 38 | 1 | 3 | 4 | 10 | 2 | 5 | 15 | 15 |
| L3_dup19694 | 8.96668890419717 | -2.02261740667861 | 0.627206031749815 | -3.22480541367849 | 0.00126058317666073 | 0.0412416780632083 | 5978 | 4 | 5 | 20 | 32 | 2 | 2 | 4 | 3 | 1 | 8 | 22 | 17 |
| THE1B_dup9618 | 10.2103877133717 | -2.42102503863794 | 0.653620047563622 | -3.704025064198 | 0.000212205277555602 | 0.0115996776873984 | 5979 | 2 | 7 | 20 | 53 | 1 | 5 | 5 | 5 | 1 | 4 | 18 | 18 |
| MSTA-int_dup1631 | 11.8838213865226 | -3.25183933537596 | 0.607645527983335 | -5.35153997786871 | 8.72088657684606e-08 | 2.96866963864694e-05 | 5980 | 0 | 2 | 46 | 29 | 1 | 3 | 16 | 7 | 4 | 4 | 16 | 30 |
| ERVL-E-int_dup1231 | 9.03321830585856 | -2.89173750515592 | 0.539266334487627 | -5.36235496306931 | 8.21439182712463e-08 | 2.83683842916836e-05 | 5981 | 0 | 2 | 24 | 17 | 4 | 4 | 16 | 9 | 2 | 1 | 17 | 23 |
| L1PA5_dup36 | 9.2703828632119 | -2.80658765578761 | 0.555352030216563 | -5.05370918459264 | 4.33311239549857e-07 | 0.000106955058304907 | 5982 | 0 | 2 | 28 | 23 | 3 | 2 | 12 | 8 | 4 | 3 | 19 | 17 |
| L2c_dup60371 | 9.33983658193798 | -2.40273271338609 | 0.687680096156101 | -3.49396867353956 | 0.00047589685687457 | 0.0208387474247104 | 5983 | 0 | 2 | 27 | 30 | 8 | 4 | 4 | 4 | 2 | 2 | 22 | 19 |
| AluY_dup102226 | 7.86397755808455 | -2.76599526275041 | 0.7170308230833 | -3.85756814589416 | 0.000114520756178958 | 0.00749492781368615 | 5984 | 0 | 2 | 21 | 24 | 2 | 5 | 4 | 2 | 1 | 2 | 20 | 23 |
| Kanga1c_dup707 | 8.66260361107008 | -2.82372626981067 | 0.707145515452511 | -3.99313324924889 | 6.52059008209882e-05 | 0.00497769755429928 | 5985 | 0 | 1 | 17 | 30 | 3 | 2 | 3 | 2 | 3 | 4 | 27 | 24 |
| L1MA9_dup15238 | 9.58046496284148 | -3.09640618433192 | 0.740411126713465 | -4.18200925488011 | 2.88944205051752e-05 | 0.00268461998497634 | 5986 | 0 | 0 | 16 | 37 | 2 | 5 | 4 | 4 | 1 | 4 | 32 | 24 |
| L1PREC2_dup4060 | 8.93218585230049 | -4.11318086909411 | 0.833619183465465 | -4.93412453873131 | 8.05110585023103e-07 | 0.000174157130239916 | 5987 | 0 | 0 | 28 | 43 | 1 | 2 | 5 | 4 | 1 | 2 | 15 | 21 |
| MLT1G1_dup3269 | 7.1468769892145 | -2.47663522873892 | 0.668139320298365 | -3.70676467242334 | 0.000209923836537034 | 0.0115199578433213 | 5988 | 0 | 0 | 21 | 13 | 5 | 2 | 4 | 6 | 3 | 3 | 14 | 22 |
| HSMAR1_dup539 | 6.09009242168564 | -2.56496233318993 | 0.679792113590737 | -3.77315694299764 | 0.000161194815611762 | 0.00958172910206254 | 5989 | 1 | 0 | 18 | 17 | 1 | 2 | 4 | 7 | 4 | 3 | 12 | 9 |
| L1ME2z_dup2734 | 6.54629888301732 | -2.00118577765257 | 0.628591996596564 | -3.1836004729423 | 0.00145455623523293 | 0.045522272517261 | 5990 | 2 | 0 | 21 | 15 | 3 | 4 | 4 | 4 | 3 | 4 | 9 | 17 |
| L1PA5_dup110 | 8.39314774948821 | -2.71145019520471 | 0.590384424280396 | -4.59268585635475 | 4.37577507974545e-06 | 0.000648317669442784 | 5991 | 3 | 1 | 19 | 12 | 1 | 2 | 6 | 4 | 1 | 5 | 29 | 28 |
| MLT1D_dup2309 | 8.5928590546491 | -2.54860288433155 | 0.595449454221066 | -4.2801330428046 | 1.86781622504007e-05 | 0.00194332925003666 | 5992 | 1 | 2 | 20 | 21 | 3 | 3 | 8 | 2 | 1 | 5 | 21 | 29 |
| MER5A_dup31892 | 9.31288414686633 | -2.96203887906603 | 0.582368986079218 | -5.08618925435551 | 3.65329512615081e-07 | 9.37849614490391e-05 | 5993 | 2 | 0 | 18 | 11 | 3 | 2 | 6 | 8 | 1 | 4 | 27 | 41 |
| MLT1F1-int_dup15 | 8.56337321350859 | -2.7210684113995 | 0.548249569939069 | -4.96319297013203 | 6.93436235396891e-07 | 0.000153488724156045 | 5994 | 2 | 0 | 10 | 19 | 2 | 4 | 10 | 5 | 2 | 3 | 25 | 33 |
| L3_dup1368 | 7.64529245691952 | -2.62182755901751 | 0.571747066416474 | -4.58564234609943 | 4.52592792745738e-06 | 0.000665180016446433 | 5995 | 3 | 0 | 9 | 14 | 1 | 6 | 20 | 9 | 2 | 1 | 19 | 17 |
| L1PA7_dup1881 | 5.98043703214291 | -2.31726706193236 | 0.599599448069009 | -3.86469178614998 | 0.000111229563901071 | 0.0073538843601012 | 5996 | 3 | 0 | 5 | 8 | 2 | 1 | 12 | 5 | 1 | 5 | 14 | 24 |
| THE1B-int_dup2885 | 6.20494342085883 | -1.94485563719272 | 0.573431199039007 | -3.39161113042339 | 0.000694829790013977 | 0.0271882863368962 | 5997 | 5 | 0 | 9 | 16 | 1 | 2 | 13 | 3 | 3 | 5 | 13 | 13 |
| MIR3_dup71126 | 6.00379639345892 | -1.90860136682015 | 0.601460845169234 | -3.17327616942899 | 0.00150729073848815 | 0.046657187727371 | 5998 | 0 | 1 | 7 | 9 | 1 | 3 | 12 | 9 | 6 | 5 | 14 | 8 |
| AluJb_dup61918 | 5.78262839168168 | -1.92084242439221 | 0.580540005195534 | -3.30871672443184 | 0.000937246141583047 | 0.0336016227821485 | 5999 | 1 | 1 | 8 | 13 | 3 | 2 | 16 | 5 | 2 | 6 | 12 | 7 |

| | | | | | | | | | | | | | | | | | | | |
|---|---|---|---|---|---|---|---|---|---|---|---|---|---|---|---|---|---|---|---|
| MamGypsy2-I_dup732 | 6.63385199094137 | -1.68705422030243 | 0.519528721863326 | -3.24727806049238 | 0.0011651449652367 | 0.039048110408199 | 6000 | 1 | 2 | 8 | 16 | 4 | 2 | 14 | 10 | 4 | 6 | 9 | 9 |
| L2c_dup104393 | 6.98142258315049 | -3.46809759916063 | 0.706554749952679 | -4.90846264835514 | 9.17931069087667e-07 | 0.000193377452475719 | 6001 | 0 | 0 | 5 | 13 | 2 | 2 | 20 | 13 | 1 | 2 | 11 | 23 |
| HAL1_dup980 | 6.43156131462221 | -2.07767102803193 | 0.598392726330259 | -3.47208603415617 | 0.000516430804398907 | 0.0220559807059759 | 6002 | 0 | 2 | 6 | 15 | 1 | 7 | 11 | 10 | 1 | 4 | 11 | 16 |
| L1PA15-16_dup666 | 7.80642042984474 | -1.94384038856097 | 0.598987040132053 | -3.2452127647577 | 0.00117362898538109 | 0.0392219554150969 | 6003 | 0 | 0 | 6 | 15 | 3 | 7 | 15 | 12 | 2 | 8 | 15 | 18 |
| MER44C_dup108 | 5.38144294982889 | -2.6023633056755 | 0.632809794678716 | -4.11239416260418 | 3.9157700078667e-05 | 0.00338568971763027 | 6004 | 1 | 1 | 5 | 11 | 1 | 2 | 7 | 9 | 2 | 2 | 15 | 13 |
| THE1B-int_dup453 | 5.66880590628885 | -1.83503010846478 | 0.561304555578197 | -3.26922361528765 | 0.00107843036885152 | 0.0369899544029713 | 6005 | 2 | 2 | 7 | 16 | 3 | 1 | 5 | 8 | 3 | 4 | 10 | 12 |
| L1MEd_dup18469 | 5.24077436884962 | -2.43008145990972 | 0.624556556061333 | -3.89089096307729 | 9.98768097997583e-05 | 0.00680953793255395 | 6006 | 2 | 1 | 10 | 15 | 3 | 1 | 5 | 8 | 1 | 2 | 10 | 10 |
| MSTB1_dup1230 | 4.56242561038071 | -2.359308220691 | 0.681692926678777 | -3.46095452711472 | 0.00053826372468924 | 0.022688698004412 | 6007 | 1 | 1 | 5 | 15 | 3 | 1 | 4 | 8 | 1 | 2 | 9 | 9 |
| L2a_dup60612 | 5.64711885082806 | -1.81727475847428 | 0.56718396893762 | -3.20403054035216 | 0.00135518100523855 | 0.0433355596657024 | 6008 | 5 | 1 | 10 | 12 | 1 | 2 | 5 | 7 | 4 | 2 | 9 | 15 |
| AluJo_dup15398 | 5.43558654759365 | -2.20644701091921 | 0.581862085995076 | -3.7920446511751 | 0.000149412041006821 | 0.00906476224105279 | 6009 | 2 | 2 | 9 | 15 | 3 | 3 | 6 | 8 | 1 | 1 | 9 | 12 |
| L1PA4_dup10479 | 5.90227316920321 | -2.18286744723406 | 0.58944535946821 | -3.7032566499521 | 0.000212849352704748 | 0.0116220194979671 | 6010 | 3 | 4 | 10 | 17 | 1 | 2 | 6 | 12 | 2 | 1 | 10 | 7 |
| L2c_dup38256 | 5.90999636460953 | -2.21417479516544 | 0.61512903312712 | -3.59952900273506 | 0.000318794077517134 | 0.0156149145262801 | 6011 | 4 | 1 | 19 | 16 | 2 | 2 | 7 | 4 | 1 | 3 | 12 | 7 |
| L1PA8_dup4483 | 7.14517465838499 | -2.51113050739529 | 0.55844687400668 | -4.49663275824023 | 6.90380794882438e-06 | 0.000919783586893038 | 6012 | 2 | 1 | 19 | 20 | 3 | 1 | 10 | 8 | 2 | 4 | 9 | 15 |
| MIR_dup157992 | 7.65979934461905 | -2.3905749936 | 0.519634097701733 | -4.60049678066388 | 4.21484535050034e-06 | 0.000628386962262001 | 6013 | 4 | 1 | 14 | 20 | 3 | 2 | 9 | 8 | 1 | 4 | 18 | 17 |
| L2c_dup129573 | 6.69504902214018 | -1.90255257889978 | 0.5248074709634 | -3.62523912894614 | 0.000288694192032515 | 0.0145508391221595 | 6014 | 3 | 4 | 15 | 12 | 1 | 2 | 6 | 10 | 3 | 4 | 13 | 12 |
| MER58B_dup2201 | 7.02038815615426 | -1.71102645572789 | 0.500885197353881 | -3.41600523386806 | 0.000635470255968451 | 0.0255418252578833 | 6015 | 3 | 3 | 13 | 16 | 3 | 3 | 8 | 9 | 2 | 6 | 16 | 8 |
| AluJb_dup32127 | 5.76652132712422 | -1.65684664606308 | 0.512998670538397 | -3.22972892760951 | 0.00123907637524686 | 0.0407340794673237 | 6016 | 3 | 5 | 10 | 13 | 1 | 3 | 11 | 7 | 2 | 3 | 9 | 8 |
| MIR_dup141121 | 4.70933295829045 | -1.93045579273751 | 0.596785502155152 | -3.23475651765353 | 0.00121746503647614 | 0.0402382513060916 | 6017 | 3 | 2 | 11 | 9 | 1 | 1 | 7 | 6 | 1 | 4 | 8 | 8 |
| L1MC4a_dup6064 | 6.72350524771691 | -1.88589358945603 | 0.559076729362218 | -3.3732285577463 | 0.000742922567170918 | 0.0285443974112791 | 6018 | 4 | 2 | 9 | 7 | 1 | 1 | 11 | 11 | 1 | 9 | 14 | 16 |
| L2b_dup21862 | 6.33322827032776 | -1.92256063058291 | 0.559779275241887 | -3.43449769509982 | 0.000593653120601572 | 0.0244016389561454 | 6019 | 2 | 3 | 15 | 8 | 3 | 1 | 11 | 12 | 1 | 6 | 9 | 9 |
| AluSz6_dup36677 | 6.39729556208442 | -1.68342302066687 | 0.52127144872099 | -3.22945564119687 | 0.00124026120120227 | 0.0407564922569379 | 6020 | 3 | 5 | 7 | 14 | 1 | 6 | 17 | 10 | 1 | 3 | 6 | 11 |
| L1PB1_dup6286 | 6.5628568862976 | -2.05711311376865 | 0.530893156075808 | -3.87481565777599 | 0.000106705445027645 | 0.00714611140516407 | 6021 | 4 | 3 | 8 | 10 | 2 | 4 | 21 | 9 | 1 | 2 | 8 | 15 |
| MER44A_dup1756 | 6.76557299902369 | -1.85752857380243 | 0.521234949069079 | -3.56370687944076 | 0.000365654261200878 | 0.0172836648461867 | 6022 | 5 | 3 | 12 | 19 | 1 | 2 | 10 | 9 | 2 | 5 | 6 | 15 |
| THE1B_dup6142 | 6.28061331993182 | -1.84881364227807 | 0.541120218257764 | -3.4166412192667 | 0.000633987722780731 | 0.0255080897336462 | 6023 | 4 | 5 | 9 | 20 | 3 | 1 | 10 | 9 | 1 | 3 | 5 | 13 |
| L1PA7_dup8218 | 9.15285974282011 | -1.64441075935889 | 0.488070173769268 | -3.36920969101519 | 0.000753840554738456 | 0.0288520740932599 | 6024 | 3 | 4 | 9 | 20 | 1 | 5 | 8 | 10 | 5 | 9 | 11 | 35 |
| MIR_dup157989 | 8.32812076768321 | -1.9037355390606 | 0.500938583571628 | -3.80033720997733 | 0.000144499319117958 | 0.00886389146348072 | 6025 | 6 | 2 | 11 | 17 | 2 | 4 | 9 | 6 | 1 | 6 | 15 | 33 |
| L3_dup36495 | 6.81519717493476 | -2.15861526295014 | 0.558991763499472 | -3.8616226640559 | 0.000112636436289156 | 0.00741804303051745 | 6026 | 0 | 1 | 11 | 17 | 3 | 3 | 13 | 6 | 6 | 2 | 11 | 16 |
| MLT1F_dup507 | 6.20423755785223 | -1.87897728843293 | 0.557135165807417 | -3.372569896409 | 0.000744701820388113 | 0.0286035220910525 | 6027 | 0 | 2 | 11 | 17 | 4 | 3 | 6 | 6 | 3 | 4 | 10 | 15 |
| AluJo_dup36119 | 6.27312521934189 | -2.21333171074385 | 0.61152870627157 | -3.61934229422902 | 0.000295352745118356 | 0.0148048568047797 | 6028 | 1 | 1 | 5 | 18 | 4 | 3 | 8 | 4 | 2 | 2 | 11 | 26 |
| L2b_dup20545 | 5.57094933663774 | -2.06865834511003 | 0.609962299992873 | -3.39145279164663 | 0.000695231389910309 | 0.0271950564365606 | 6029 | 0 | 2 | 5 | 16 | 2 | 3 | 12 | 3 | 3 | 3 | 9 | 17 |
| L3_dup3326 | 7.23155648772803 | -1.80611740443114 | 0.560085395238627 | -3.22471790870682 | 0.00126096851265053 | 0.0412429434654491 | 6030 | 2 | 2 | 17 | 22 | 3 | 2 | 7 | 3 | 5 | 6 | 13 | 13 |
| LTR81C_dup610 | 7.67563713383044 | -1.72500029417515 | 0.501019230819714 | -3.4429822012079 | 0.000575337177859143 | 0.0238284593385645 | 6031 | 3 | 4 | 9 | 15 | 2 | 1 | 10 | 2 | 6 | 5 | 22 | 22 |
| L3_dup9932 | 5.80585061811161 | -1.63402015135892 | 0.51937725159132 | -3.14611420957011 | 0.00165455382297015 | 0.0497277332272901 | 6032 | 1 | 3 | 6 | 11 | 4 | 2 | 10 | 5 | 2 | 5 | 13 | 14 |
| LTR41B_dup165 | 6.42784360603784 | -1.63193898794409 | 0.505329822404344 | -3.22945315235774 | 0.00124027199630373 | 0.0407564922569379 | 6033 | 3 | 2 | 13 | 9 | 5 | 3 | 9 | 5 | 1 | 5 | 15 | 14 |
| HAL1ME_dup937 | 8.05945116134164 | -1.76095167339995 | 0.458099309861786 | -3.84403913188878 | 0.00012102569442598 | 0.00780422219129656 | 6034 | 1 | 2 | 11 | 14 | 3 | 5 | 12 | 7 | 5 | 6 | 21 | 17 |
| L2a_dup2482 | 7.11481386801462 | -2.36312696723604 | 0.529081933872191 | -4.46646694197442 | 7.95219435261843e-06 | 0.00102750758123991 | 6035 | 1 | 1 | 12 | 8 | 2 | 2 | 13 | 6 | 3 | 5 | 18 | 22 |
| X9_LINE_dup411 | 6.24860551243581 | -3.69556263749229 | 1.07692020504387 | -3.43160302888157 | 0.000600025163385775 | 0.0245864624637492 | 6036 | 0 | 0 | 2 | 3 | 0 | 1 | 34 | 24 | 2 | 3 | 3 | 5 |
| L1PA8_dup5933 | 5.61411547641577 | -2.74693829791345 | 0.819367350128305 | -3.35251129726283 | 0.00080081973061777 | 0.0300697256780142 | 6037 | 0 | 0 | 4 | 8 | 1 | 2 | 28 | 12 | 2 | 4 | 4 | 8 |
| MIRb_dup121955 | 5.23392671926004 | -3.10196160685665 | 0.874412450343847 | -3.54748106072467 | 0.000388933713987524 | 0.0181212495610878 | 6038 | 0 | 0 | 2 | 10 | 0 | 2 | 25 | 9 | 3 | 2 | 6 | 10 |
| L1MA3_dup2020 | 9.6246865075546 | -3.50081550024772 | 0.884780085936796 | -3.95670693304663 | 7.59900630423843e-05 | 0.00557535440855308 | 6039 | 0 | 1 | 5 | 4 | 3 | 1 | 47 | 42 | 1 | 2 | 3 | 8 |
| MER102a_dup126 | 8.00154801631609 | -2.51945510404985 | 0.68299864507482 | -3.68881420514949 | 0.000225301666376392 | 0.01212064895 0395 | 6040 | 0 | 2 | 8 | 5 | 6 | 2 | 33 | 21 | 2 | 2 | 9 | 11 |
| MLT1A0-int_dup533 | 9.11829024235742 | -2.17495117532011 | 0.632666471603762 | -3.43775318108257 | 0.000586562086003868 | 0.024197313126955 | 6041 | 0 | 3 | 7 | 7 | 5 | 3 | 27 | 25 | 2 | 7 | 10 | 18 |
| MamRep564_dup714 | 5.66182554290147 | -2.77880592674831 | 0.709654457154103 | -3.9157168657716 | 9.01359799573786e-05 | 0.00631179984901071 | 6042 | 1 | 2 | 6 | 17 | 5 | 0 | 15 | 5 | 1 | 0 | 8 | 17 |
| Charlie1_dup1247 | 6.19009942508476 | -3.1523018095162 | 0.676051580413507 | -4.66281257354372 | 3.11916809697443e-06 | 0.000495071274840737 | 6043 | 4 | 1 | 19 | 14 | 2 | 0 | 18 | 9 | 1 | 0 | 4 | 10 |
| L1PA4_dup6885 | 4.91170278070875 | -3.18927416536745 | 0.759540935671162 | -4.19894967550271 | 2.68155902116235e-05 | 0.00253805188086514 | 6044 | 3 | 1 | 6 | 13 | 1 | 0 | 19 | 12 | 1 | 0 | 3 | 5 |
| MER41A_dup2192 | 5.92928967979917 | -3.04828216766463 | 0.718133307281776 | -4.24473024263804 | 2.18856546790492e-05 | 0.00218256663380596 | 6045 | 2 | 2 | 7 | 10 | 3 | 0 | 24 | 16 | 1 | 0 | 4 | 7 |
| L2_dup40222 | 5.04137283237229 | -3.15094494263133 | 0.879717977833444 | -3.58176713677198 | 0.000341277927392667 | 0.0164018819857353 | 6046 | 0 | 0 | 5 | 4 | 3 | 0 | 18 | 16 | 1 | 2 | 4 | 10 |
| L2c_dup120446 | 4.56955838202702 | -2.9837145829871 | 0.911914264211162 | -3.27192445615281 | 0.00106818113693267 | 0.0367813476597136 | 6047 | 0 | 3 | 4 | 2 | 1 | 0 | 21 | 13 | 1 | 1 | 3 | 9 |
| MER53_dup553 | 4.79985770790647 | -2.60880265910209 | 0.758351211809802 | -3.44009822688381 | 0.000581503098620558 | 0.0240302772324362 | 6048 | 0 | 2 | 5 | 9 | 4 | 0 | 16 | 13 | 1 | 1 | 3 | 7 |
| MER97c_dup220 | 5.85041424481547 | -3.16770806348209 | 0.727744872512542 | -4.35277276849175 | 1.34426461297453e-05 | 0.00152315422666114 | 6049 | 0 | 2 | 6 | 9 | 1 | 0 | 21 | 13 | 2 | 2 | 9 | 10 |
| MER1B_dup4975 | 6.49373118948772 | -3.80633623957112 | 0.882501819482479 | -4.31311999084971 | 1.60966698540715e-05 | 0.00174097191140769 | 6050 | 0 | 0 | 6 | 11 | 4 | 0 | 35 | 18 | 0 | 1 | 5 | 4 |
| MER94_dup4213 | 5.15034261122247 | -3.81105265659164 | 0.932830290258349 | -4.08547266999249 | 4.39871782019245e-05 | 0.00370629359222207 | 6051 | 0 | 1 | 4 | 7 | 1 | 0 | 30 | 11 | 0 | 2 | 6 | 6 |
| L1PA16_dup6247 | 6.79245053779134 | -3.55025985171656 | 0.803702597243305 | -4.41738009046373 | 9.99044605488666e-06 | 0.00122220394703139 | 6052 | 0 | 1 | 6 | 14 | 5 | 1 | 31 | 19 | 0 | 0 | 3 | 8 |
| THE1D-int_dup2225 | 6.29458325569969 | -3.85674359376569 | 0.861375512334818 | -4.47742423430603 | 7.55490265205189e-06 | 0.000986640431638386 | 6053 | 0 | 1 | 8 | 6 | 1 | 2 | 27 | 18 | 0 | 1 | 3 | 14 |
| L1MEb_dup167 | 7.70133612928018 | -2.2921179543503 | 0.552746298440259 | -4.14678119205539 | 3.37182011352987e-05 | 0.00301620717569165 | 6054 | 1 | 8 | 13 | 14 | 3 | 1 | 22 | 14 | 2 | 1 | 11 | 9 |
| BSR/Beta_dup2037 | 6.25867663377559 | -2.52565583571625 | 0.629700134243896 | -4.01088660835192 | 6.04911591618187e-05 | 0.00470635401913803 | 6055 | 2 | 4 | 11 | 21 | 1 | 0 | 20 | 6 | 3 | 1 | 6 | 9 |
| L1PA7_dup1030 | 5.52774538848164 | -2.89896923528014 | 0.669045933142139 | -4.33298984669899 | 1.47097896334105e-05 | 0.00162494689141666 | 6056 | 2 | 3 | 7 | 15 | 1 | 0 | 16 | 12 | 1 | 1 | 5 | 9 |
| LTR12F_dup48 | 6.02486904758257 | -2.25812984505425 | 0.666230918184334 | -3.38941016308323 | 0.000700431555344889 | 0.0273535018657632 | 6057 | 2 | 4 | 7 | 10 | 1 | 0 | 23 | 12 | 4 | 1 | 7 | 6 |
| L1PA4_dup1160 | 8.52185072513993 | -2.46075922339484 | 0.609115683535046 | -4.03988813604938 | 5.34767013077135e-05 | 0.00430320161290673 | 6058 | 3 | 5 | 19 | 12 | 6 | 0 | 37 | 17 | 1 | 1 | 2 | 8 |
| AluJb_dup110799 | 8.25073537872514 | -1.91079610613912 | 0.490175072267973 | -3.89819110404398 | 9.69139228545794e-05 | 0.00666397023938648 | 6059 | 4 | 7 | 17 | 17 | 5 | 1 | 26 | 15 | 1 | 3 | 5 | 6 |
| L1PA2_dup4074 | 9.01736907305877 | -2.65302405257219 | 0.504063099291901 | -5.26327766563176 | 1.41509611609979e-07 | 4.3781670760192 3e-05 | 6060 | 3 | 4 | 19 | 20 | 3 | 2 | 29 | 21 | 1 | 2 | 5 | 7 |
| L1M5_dup28499 | 6.83385255090343 | -2.27792847920433 | 0.593354351023717 | -3.83906931039471 | 0.000123501566260513 | 0.00791028362999303 | 6061 | 2 | 4 | 19 | 20 | 2 | 1 | 20 | 12 | 1 | 3 | 3 | 2 |
| L1PA7_dup4740 | 8.5042826407784 | -2.75337388923598 | 0.605282391907475 | -4.54890795775349 | 5.39250208810566e-06 | 0.000763310114132296 | 6062 | 2 | 4 | 23 | 18 | 1 | 1 | 29 | 20 | 1 | 3 | 3 | 4 |
| L1ME1_dup27656 | 5.86946779606021 | -2.00634025096193 | 0.552155982881178 | -3.6336475799696 | 0.000279442490085142 | 0.0142047047096345 | 6063 | 3 | 3 | 13 | 17 | 4 | 4 | 20 | 8 | 1 | 0 | 2 | 3 |
| LTR16C_dup5273 | 6.12086791996937 | -1.85702880521089 | 0.587237938912967 | -3.16231067878282 | 0.00156522475576831 | 0.0477853438203807 | 6064 | 1 | 3 | 13 | 13 | 4 | 4 | 23 | 12 | 2 | 1 | 1 | 2 |
| AluSq2_dup60327 | 5.19099316395409 | -2.72842194586994 | 0.712019332601438 | -3.83194924764383 | 0.000127131969444798 | 0.00808405761 6652 | 6065 | 0 | 1 | 8 | 15 | 2 | 3 | 23 | 8 | 2 | 0 | 3 | 4 |
| MER1B_dup1213 | 6.36503353203299 | -2.41377533573871 | 0.680283888800833 | -3.5481883012011 | 0.000387890804729206 | 0.0180797385743576 | 6066 | 0 | 3 | 8 | 19 | 1 | 4 | 29 | 12 | 2 | 1 | 2 | 3 |
| MER2_dup4226 | 8.02528672176489 | -2.35363457547269 | 0.652756450471635 | -3.60568566388294 | 0.000311329807378491 | 0.0153405844577516 | 6067 | 5 | 3 | 15 | 7 | 3 | 1 | 25 | 27 | 3 | 0 | 5 | 4 |
| THE1D_dup8578 | 7.57013582926312 | -2.78016294223541 | 0.628502760472954 | -4.42346973964492 | 9.71282415596416e-06 | 0.00119685533744953 | 6068 | 2 | 2 | 11 | 13 | 2 | 5 | 28 | 24 | 1 | 0 | 4 | 3 |
| MSTA_dup12687 | 5.54647829403084 | -2.44788820478061 | 0.699844526540549 | -3.49776001947309 | 0.000469183104835217 | 0.0206359044478966 | 6069 | 2 | 2 | 4 | 8 | 2 | 4 | 12 | 20 | 1 | 0 | 5 | 8 |
| Arthur1C_dup81 | 7.3557428912094 | -2.40141138902169 | 0.638149405279289 | -3.76308646400869 | 0.000167829143624102 | 0.00987006263192797 | 6070 | 2 | 3 | 5 | 10 | 4 | 2 | 31 | 18 | 3 | 0 | 7 | 9 |
| L1PA2_dup66 | 6.79663869778318 | -2.38953977677512 | 0.597992168917226 | -3.99593824297369 | 6.44385320389418e-05 | 0.00493019001432092 | 6071 | 1 | 3 | 8 | 8 | 4 | 5 | 21 | 13 | 1 | 0 | 10 | 14 |
| L1MB7_dup16450 | 12.5930582374691 | -4.4066619601925 | 0.659649826297657 | -6.68030488983114 | 2.38445646309957e-11 | 2.96568807215841e-08 | 6072 | 1 | 4 | 21 | 32 | 1 | 0 | 43 | 38 | 0 | 1 | 7 | 13 |
| L1PA6_dup3214 | 11.3466066353239 | -5.39726553084147 | 0.826781787214654 | -6.52804115221783 | 6.66353909980941e-11 | 7.11014562620291e-08 | 6073 | 1 | 1 | 23 | 24 | 0 | 0 | 41 | 32 | 0 | 1 | 7 | 16 |
| MER1B_dup521 | 9.42066077924811 | -3.69169803884545 | 0.602158461151893 | -6.13077499863317 | 8.74520122562087e-10 | 6.48250981567472e-07 | 6074 | 1 | 3 | 22 | 34 | 2 | 0 | 30 | 21 | 1 | 1 | 4 | 4 |
| THE1D-int_dup159 | 9.89767296520447 | -3.24802069552524 | 0.675012055761825 | -4.81179657133604 | 1.49579564774289e-06 | 0.000282475072379229 | 6075 | 2 | 2 | 25 | 39 | 1 | 0 | 31 | 15 | 3 | 2 | 3 | 9 |
| L1PA16_dup7137 | 8.7993964337035 | -3.58774053096517 | 0.65605366092599 | -5.46866932485559 | 4.53426772883147e-08 | 1.76292625654335e-05 | 6076 | 3 | 2 | 15 | 21 | 0 | 0 | 27 | 24 | 1 | 2 | 5 | 13 |
| L1PA13_dup4416 | 9.17772260391497 | -4.30108237997587 | 0.799681221638332 | -5.37849616021257 | 7.51105839002582e-08 | 2.65167106776422e-05 | 6077 | 0 | 1 | 17 | 29 | 0 | 0 | 26 | 24 | 1 | 3 | 8 | 9 |
| L1PB4_dup3665 | 9.36213556130276 | -4.04096294608901 | 0.730802079725512 | -5.52949021109353 | 3.21162720724455e-08 | 1.33367163604714e-05 | 6078 | 1 | 1 | 23 | 30 | 0 | 0 | 31 | 18 | 1 | 3 | 8 | 7 |

| | | | | | | | | | | | | | | | | | | | |
|---|---|---|---|---|---|---|---|---|---|---|---|---|---|---|---|---|---|---|---|
| L1MA4A_dup5809 | 10.0026056921917 | -4.86367806346906 | 0.784842644204565 | -6.19701044455653 | 5.75456838479449e-10 | 4.61035868312562e-07 | 6079 | 0 | 2 | 18 | 19 | 1 | 0 | 35 | 28 | 0 | 1 | 10 | 14 |
| L1P3_dup3210 | 8.14113007758276 | -4.52626465458482 | 0.791750584132275 | -5.71678094755738 | 1.08560914853718e-08 | 5.44971211936346e-06 | 6080 | 0 | 1 | 16 | 14 | 1 | 1 | 37 | 20 | 0 | 1 | 5 | 10 |
| MLT1A0_dup19118 | 9.57481725824903 | -4.32463514665229 | 0.8111464761804 | -5.33150950370459 | 9.73997229301988e-08 | 3.23685397658507e-05 | 6081 | 1 | 0 | 16 | 21 | 1 | 0 | 48 | 22 | 0 | 3 | 6 | 8 |
| L2b_dup87079 | 7.75437744616944 | -5.2455869290597 | 0.950909652804697 | -5.51638834834404 | 3.46037155900112e-08 | 1.41474496731629e-05 | 6082 | 0 | 1 | 11 | 16 | 0 | 0 | 42 | 18 | 0 | 1 | 8 | 5 |
| MLT1H1_dup3290 | 8.53094173571611 | -5.98491606288046 | 0.980830851764789 | -6.10188398143464 | 1.04825455724582e-09 | 7.4903296960485e-07 | 6083 | 0 | 0 | 6 | 20 | 0 | 0 | 42 | 22 | 0 | 1 | 10 | 11 |
| L1MA4A_dup3440 | 8.70998934234186 | -3.43684458487621 | 0.666348111923505 | -5.15773140701981 | 2.49959832698144e-07 | 6.88390536472137e-05 | 6084 | 4 | 4 | 10 | 18 | 2 | 0 | 29 | 25 | 0 | 0 | 11 | 8 |
| L1P2_dup916 | 9.06534153543245 | -3.75307144580135 | 0.700058552416252 | -5.36108220212098 | 8.27248607764393e-08 | 2.85276082932038e-05 | 6085 | 3 | 2 | 9 | 12 | 2 | 0 | 38 | 23 | 0 | 1 | 14 | 13 |
| Tigger17a_dup2055 | 9.98091369846779 | -4.79230904661851 | 0.88078087506953 | -5.44097763957488 | 5.29889656461625e-08 | 2.01413936416003e-05 | 6086 | 1 | 1 | 6 | 17 | 1 | 0 | 52 | 32 | 1 | 0 | 12 | 4 |
| L1PA5_dup8564 | 7.31523166253564 | -4.25727003989998 | 0.767854257908585 | -5.54437251086627 | 2.95010186469475e-08 | 1.25800168153523e-05 | 6087 | 1 | 2 | 12 | 21 | 1 | 0 | 35 | 18 | 0 | 0 | 4 | 2 |
| Tigger1_dup6537 | 7.97571957366462 | -4.17733199097602 | 0.745419531015922 | -5.60400125991171 | 2.09459183311764e-08 | 9.52963189910153e-06 | 6088 | 1 | 2 | 10 | 25 | 1 | 0 | 35 | 18 | 1 | 0 | 7 | 5 |
| L1MA6_dup1876 | 16.6312734040195 | -6.04730771976629 | 0.824331251487609 | -7.33601656961709 | 2.20044790682638e-13 | 4.67489087176526e-10 | 6089 | 1 | 1 | 30 | 46 | 0 | 1 | 71 | 25 | 0 | 0 | 20 | 30 |
| L1MA4A_dup1787 | 11.0550241388812 | -6.90995411881903 | 0.898408443495165 | -7.69132811345424 | 1.45615465470772e-14 | 4.38590095530486e-11 | 6090 | 0 | 0 | 18 | 31 | 0 | 0 | 36 | 22 | 0 | 0 | 14 | 26 |
| L1MA6_dup1879 | 11.5712002364036 | -6.37593046575306 | 0.907033154008491 | -7.02943485315353 | 2.0737174717148e-12 | 3.52451983946178e-09 | 6091 | 0 | 0 | 21 | 30 | 0 | 0 | 37 | 34 | 1 | 0 | 10 | 15 |
| L1MA6_dup1875 | 8.53665440034506 | -5.00987178007009 | 0.84257262360054 | -5.94592280800859 | 2.74903681356252e-09 | 1.6990189964674e-06 | 6092 | 0 | 0 | 15 | 23 | 0 | 2 | 33 | 18 | 1 | 0 | 11 | 9 |
| L1PA16_dup4861 | 7.31211664177922 | -4.03805759844795 | 0.826250743729506 | -4.88720600749003 | 1.02277058599403e-06 | 0.000209255520081629 | 6093 | 0 | 1 | 4 | 12 | 0 | 2 | 28 | 15 | 2 | 0 | 17 | 14 |
| L1M2_dup3031 | 7.72414922641732 | -5.40541739164493 | 0.936527391319529 | -5.7717664659321 | 7.84447861008585e-09 | 4.11137072104733e-06 | 6094 | 1 | 0 | 10 | 20 | 0 | 0 | 22 | 17 | 0 | 1 | 20 | 9 |
| LTR1A2_dup439 | 6.60095682528688 | -5.68907567002835 | 0.929176068925963 | -6.1227100657084 | 9.19970079308865e-10 | 6.69428747679594e-07 | 6095 | 0 | 1 | 11 | 16 | 0 | 0 | 21 | 10 | 0 | 0 | 9 | 21 |
| L1M2_dup3032 | 7.07933081953817 | -6.26191716180553 | 0.976765577944282 | -6.41087002163249 | 1.44691518737645e-10 | 1.38825677893337e-07 | 6096 | 0 | 0 | 7 | 12 | 0 | 0 | 32 | 17 | 0 | 0 | 10 | 15 |
| L1ME1_dup28703 | 8.10451554652218 | -5.95161458606771 | 0.945250198669112 | -6.29633783145185 | 3.04759794062042e-10 | 2.63695905301406e-07 | 6097 | 0 | 1 | 10 | 14 | 0 | 0 | 26 | 24 | 0 | 0 | 14 | 14 |
| MLT1H1_dup1564 | 9.64082385706514 | -4.63613413313564 | 0.779395229079514 | -5.94837376488824 | 2.70819627306792e-09 | 1.6815482144846e-06 | 6098 | 0 | 0 | 21 | 25 | 0 | 4 | 34 | 31 | 0 | 0 | 5 | 2 |
| HERVH-int_dup2874 | 6.4420778696325 | -3.88790244053759 | 0.833012026546562 | -4.66728248409061 | 3.05209582905652e-06 | 0.000486752006796704 | 6099 | 0 | 0 | 11 | 17 | 0 | 5 | 30 | 14 | 0 | 0 | 5 | 3 |
| THE1B_dup22278 | 7.30594828756194 | -4.73574241568923 | 0.835663399454205 | -5.66704539026392 | 1.45280867020662e-08 | 6.99777351904828e-06 | 6100 | 0 | 0 | 16 | 18 | 0 | 3 | 24 | 20 | 0 | 0 | 5 | 8 |
| L1PA3_dup8337 | 11.1819360652802 | -5.72013074478036 | 1.0268155622778 | -5.57074800472571 | 2.53647974195895e-08 | 1.12601718037344e-05 | 6101 | 0 | 0 | 11 | 11 | 0 | 1 | 74 | 41 | 0 | 0 | 2 | 2 |
| SVA_D_dup900 | 9.76633590204842 | -3.95259300268095 | 0.774210146435769 | -5.10532317469294 | 3.3022965696309e-07 | 8.67293884721184e-05 | 6102 | 2 | 0 | 15 | 11 | 0 | 4 | 59 | 26 | 0 | 1 | 5 | 4 |
| L1M2_dup4660 | 8.06284319397183 | -4.14810438128864 | 0.84531349596635 | -4.90717869888803 | 9.23958433027355e-07 | 0.000193876565593509 | 6103 | 1 | 0 | 10 | 28 | 0 | 0 | 27 | 19 | 2 | 2 | 3 | 14 |
| L1MEc_dup9515 | 6.43734754913088 | -4.57161434440096 | 0.950482441273618 | -4.80978305951148 | 1.51094191318065e-06 | 0.000284883188964884 | 6104 | 0 | 0 | 11 | 13 | 0 | 0 | 20 | 20 | 2 | 1 | 4 | 10 |
| L1P4b_dup57 | 6.96975226670539 | -3.86932461845832 | 0.936796709816518 | -4.13037810435539 | 3.62167086611075e-05 | 0.00319289401966502 | 6105 | 0 | 0 | 11 | 18 | 0 | 0 | 19 | 18 | 4 | 4 | 5 | 9 |
| L1MD3_dup2446 | 11.9536915350344 | -4.95572017015193 | 0.977083490971665 | -5.07195159466229 | 3.93756557968426e-07 | 9.91458179284182e-05 | 6106 | 0 | 0 | 14 | 33 | 0 | 0 | 36 | 43 | 3 | 2 | 7 | 11 |
| Tigger2_dup1508 | 10.2434428682635 | -5.59980774020167 | 0.907630133614493 | -6.16970231905074 | 6.841867940956e-10 | 5.26821348908516e-07 | 6107 | 0 | 0 | 22 | 25 | 0 | 0 | 49 | 23 | 1 | 1 | 4 | 10 |
| L1MEf_dup9156 | 8.68574692372695 | -3.6788873761296 | 0.787955620043446 | -4.66890175353627 | 3.02814136852576e-06 | 0.000483904718653614 | 6108 | 0 | 1 | 19 | 20 | 2 | 0 | 27 | 31 | 2 | 1 | 3 | 1 |
| L1MA3_dup6847 | 8.22530581739936 | -4.65990544750701 | 0.863823192196371 | -5.39451300868486 | 6.8709649291546e-08 | 2.47340184725056e-05 | 6109 | 0 | 0 | 17 | 26 | 1 | 0 | 34 | 21 | 2 | 0 | 1 | 5 |
| L1MB3_dup9914 | 6.59677739494848 | -3.38958211263627 | 0.801892501240402 | -4.22697818896313 | 2.36850615935936e-05 | 0.00231733785606465 | 6110 | 1 | 0 | 17 | 18 | 1 | 1 | 26 | 13 | 3 | 0 | 2 | 4 |
| AluJr_dup12741 | 8.31741509027913 | -4.73725279682057 | 1.1383476227685 | -4.16151683551586 | 3.16140686223697e-05 | 0.00286826105032305 | 6111 | 0 | 0 | 31 | 35 | 0 | 0 | 19 | 17 | 3 | 0 | 1 | 2 |
| MER4-int_dup2823 | 6.00826910992925 | -3.55948509795516 | 1.07923724014964 | -3.29814888287365 | 0.000973245153188454 | 0.0344766698259014 | 6112 | 1 | 0 | 13 | 20 | 0 | 0 | 16 | 18 | 4 | 0 | 2 | 1 |
| MamRTE1_dup841 | 5.96936792218197 | -2.9935186286733 | 0.738035233357439 | -4.05606466110745 | 4.99064487510798e-05 | 0.0040709769813248 | 6113 | 0 | 3 | 14 | 11 | 0 | 0 | 15 | 13 | 1 | 4 | 7 | 8 |
| L2b_dup11357 | 5.43605425401339 | -3.8607207877627 | 0.804842049429479 | -4.79686764688725 | 1.61166013726911e-06 | 0.000299077259378741 | 6114 | 0 | 2 | 14 | 23 | 0 | 0 | 14 | 9 | 1 | 1 | 4 | 4 |
| MER47B_dup63 | 7.70927447793132 | -4.10994104345617 | 0.729449210875383 | -5.63430734063581 | 1.75763326210188e-08 | 8.20042753302145e-06 | 6115 | 1 | 2 | 17 | 32 | 0 | 0 | 18 | 16 | 1 | 1 | 7 | 6 |
| MLT1H1_dup3090 | 7.03712486969216 | -3.90813705019082 | 0.811159392831312 | -4.81796436647261 | 1.45030263156319e-06 | 0.000275179052885578 | 6116 | 0 | 2 | 19 | 27 | 0 | 0 | 9 | 15 | 2 | 1 | 10 | 5 |
| Arthur1_dup1248 | 7.24955107144421 | -2.22993432138859 | 0.69611337231631 | -3.20340681571524 | 0.00135811983392945 | 0.0434120341220012 | 6117 | 2 | 4 | 16 | 28 | 0 | 0 | 13 | 11 | 4 | 5 | 6 | 5 |
| MIR1_Amn_dup4971 | 7.17419318140504 | -2.18771824825453 | 0.693819085935007 | -3.15315374368277 | 0.0016151671605389 | 0.0488462843392971 | 6118 | 5 | 3 | 20 | 25 | 0 | 0 | 11 | 11 | 2 | 5 | 8 | 3 |
| L1MA4A_dup2911 | 9.11246000455826 | -2.31092173116812 | 0.693967399599019 | -3.33001482851124 | 0.000868413591709771 | 0.0318636145710059 | 6119 | 4 | 6 | 26 | 38 | 0 | 0 | 11 | 10 | 2 | 6 | 12 | 5 |
| LTR12B_dup64 | 8.95868042337644 | -2.19694956587929 | 0.67333169117477 | -3.2628043424575 | 0.00110315652690362 | 0.0375632059173738 | 6120 | 3 | 8 | 30 | 30 | 0 | 0 | 17 | 10 | 2 | 5 | 6 | 7 |
| LTR16E1_dup1125 | 7.27685815323842 | -2.26288480915639 | 0.646173431823185 | -3.50197748423614 | 0.000461818644274153 | 0.0204328559186422 | 6121 | 7 | 5 | 25 | 18 | 0 | 0 | 16 | 11 | 2 | 1 | 3 | 7 |
| MIR3_dup82741 | 6.5270489279925 | -2.2908977397041 | 0.678301843120187 | -3.37740161395697 | 0.000731741251131572 | 0.0282333231298448 | 6122 | 4 | 8 | 18 | 23 | 0 | 0 | 12 | 7 | 1 | 1 | 5 | 8 |
| MIRc_dup46431 | 6.80473304163986 | -2.43627264685981 | 0.623257355579198 | -3.90893524970238 | 9.27038048148101e-05 | 0.00645740618866066 | 6123 | 4 | 6 | 21 | 20 | 0 | 0 | 14 | 8 | 1 | 2 | 6 | 8 |
| L1MA5A_dup1665 | 7.58127279731383 | -2.1784886890411 | 0.658256350839937 | -3.30948373876125 | 0.00093468195270047 | 0.0335653232594727 | 6124 | 7 | 3 | 18 | 32 | 0 | 0 | 16 | 7 | 3 | 3 | 5 | 8 |
| MLT1F1_dup1512 | 7.32430706315273 | -2.72472808019034 | 0.636520018829289 | -4.28066360772401 | 1.86336839453825e-05 | 0.0019412480560718 | 6125 | 7 | 2 | 23 | 19 | 0 | 0 | 13 | 8 | 2 | 1 | 9 | 13 |
| Charlie15b_dup790 | 5.29919743687247 | -2.49137229401907 | 0.696821684180258 | -3.57533691987488 | 0.000349777154458355 | 0.0167020254247992 | 6126 | 1 | 2 | 5 | 4 | 0 | 1 | 18 | 10 | 2 | 4 | 9 | 12 |
| L1MC1_dup12193 | 6.4360165289272 | -2.81888129709537 | 0.68315167902245 | -4.12628905652259 | 3.68663844458576e-05 | 0.00323697738500149 | 6127 | 1 | 2 | 11 | 17 | 0 | 1 | 19 | 15 | 3 | 2 | 6 | 5 |
| L1PA13_dup583 | 6.1599388155307 | -2.06321460932242 | 0.599252536307154 | -3.44298018667859 | 0.000575341463584224 | 0.0238284593385645 | 6128 | 4 | 2 | 8 | 15 | 0 | 3 | 23 | 10 | 3 | 2 | 4 | 7 |
| L2a_dup155332 | 5.18541735652992 | -2.08486333759241 | 0.620077026169408 | -3.36226508901947 | 0.000773058706904765 | 0.0293328379960391 | 6129 | 4 | 1 | 6 | 9 | 0 | 2 | 16 | 10 | 2 | 3 | 6 | 8 |
| L2_dup5993 | 4.51791554958959 | -2.49333301679768 | 0.777871905925795 | -3.20532596408686 | 0.00134909603963186 | 0.0431935350779076 | 6130 | 1 | 2 | 4 | 9 | 0 | 1 | 22 | 8 | 1 | 3 | 2 | 7 |
| L1MC4_dup17878 | 4.69315227521296 | -3.07622631293331 | 0.738853206855535 | -4.16351486924627 | 3.13385607032907e-05 | 0.00284831366123194 | 6131 | 1 | 1 | 11 | 9 | 0 | 1 | 16 | 7 | 1 | 2 | 4 | 9 |
| L1PA7_dup1032 | 4.57582870641442 | -2.50662785271641 | 0.698943388802226 | -3.58631026900763 | 0.000335389881973055 | 0.0161941316671998 | 6132 | 1 | 1 | 9 | 8 | 0 | 3 | 13 | 11 | 2 | 1 | 3 | 6 |
| L2c_dup17419 | 9.47826839230232 | -2.34841314512766 | 0.631949692747672 | -3.71613938906581 | 0.000202290012041698 | 0.0112174549534547 | 6133 | 6 | 3 | 12 | 14 | 0 | 1 | 41 | 21 | 3 | 5 | 9 | 7 |
| LTR108d_Mam_dup2 | 9.60390273258013 | -2.50139604308328 | 0.766584063782529 | -3.26304206056767 | 0.00110223159917778 | 0.0375370831684495 | 6134 | 4 | 1 | 8 | 18 | 0 | 1 | 51 | 18 | 7 | 5 | 6 | 6 |
| L1MC4a_dup4724 | 7.23632759367671 | -2.73369305532105 | 0.701583638097224 | -3.89646067393354 | 9.760864809475e-05 | 0.0066971128545425 | 6135 | 6 | 2 | 17 | 12 | 0 | 0 | 33 | 12 | 2 | 1 | 7 | 3 |
| MER61-int_dup628 | 8.23364809630942 | -2.56041555222734 | 0.602076233590281 | -4.25264345174223 | 2.11261798859265e-05 | 0.00212463651696393 | 6136 | 6 | 3 | 10 | 32 | 0 | 1 | 28 | 14 | 2 | 2 | 5 | 7 |
| MST-int_dup85 | 8.35402588029276 | -3.96107308022549 | 0.83272024317429 | -4.75678730365203 | 1.96698223763395e-06 | 0.000348759728402421 | 6137 | 2 | 0 | 30 | 37 | 0 | 1 | 21 | 3 | 0 | 3 | 6 | 13 |
| L2a_dup19338 | 8.15686586952147 | -5.23796346987383 | 0.913269615334791 | -5.73539662540253 | 9.72844142077194e-09 | 4.94685954022896e-06 | 6138 | 1 | 0 | 27 | 42 | 0 | 0 | 15 | 13 | 0 | 1 | 6 | 4 |
| LTR10C_dup412 | 6.50398437162692 | -5.52475672141928 | 0.945176593360637 | -5.84521110682147 | 5.05926243335808e-09 | 2.82589497410287e-06 | 6139 | 0 | 0 | 27 | 21 | 0 | 0 | 18 | 8 | 0 | 1 | 5 | 7 |
| L1PA5_dup11211 | 7.43399981632437 | -4.14492682342036 | 0.788498999419198 | -5.25673060647317 | 1.46638869782161e-07 | 4.4906219445542e-05 | 6140 | 1 | 0 | 21 | 34 | 1 | 2 | 11 | 8 | 0 | 1 | 7 | 15 |
| MLT1A_dup6670 | 6.43872150462126 | -3.89454939942822 | 0.773184944521664 | -5.03702177211639 | 4.7283101649278e-07 | 0.000114642320552505 | 6141 | 1 | 0 | 16 | 27 | 1 | 1 | 12 | 7 | 0 | 2 | 9 | 11 |
| L1MCc_dup2582 | 7.51572125718318 | -4.4405221379365 | 0.81335075880442 | -5.45954139695378 | 4.77366025751791e-08 | 1.84395026564181e-05 | 6142 | 1 | 0 | 18 | 25 | 1 | 0 | 15 | 15 | 0 | 2 | 6 | 16 |
| L2a_dup124345 | 7.75466456863928 | -3.93834290200923 | 0.741930468647464 | -5.30823718452865 | 1.10690543902994e-07 | 3.59322948970557e-05 | 6143 | 2 | 0 | 23 | 31 | 2 | 1 | 17 | 10 | 0 | 1 | 3 | 15 |
| AluSq2_dup6846 | 5.84237992316344 | -3.02653292112704 | 0.808208462964513 | -3.74474292192599 | 0.000180578582857355 | 0.0103662126602114 | 6144 | 4 | 0 | 20 | 22 | 0 | 0 | 6 | 4 | 1 | 3 | 11 | 7 |
| AluY_dup71778 | 4.37241955665604 | -3.62437264475165 | 0.886964607178817 | -4.08626524149566 | 4.38372607980102e-05 | 0.00369758981135886 | 6145 | 2 | 0 | 13 | 18 | 0 | 0 | 7 | 4 | 1 | 1 | 5 | 8 |
| THE1C_dup8595 | 5.08415851354495 | -3.50340022408618 | 0.8129563679966 | -4.30945664737179 | 1.63656134839405e-05 | 0.00176284846810761 | 6146 | 3 | 0 | 16 | 20 | 0 | 0 | 9 | 6 | 1 | 1 | 5 | 7 |
| L1PA15_dup3870 | 4.72425831419787 | -2.78816773831962 | 0.876934190119193 | -3.17944923317525 | 0.00147555213794325 | 0.0459737762229991 | 6147 | 4 | 1 | 21 | 20 | 0 | 0 | 6 | 3 | 1 | 1 | 1 | 6 |
| L1PREC2_dup5491 | 5.16052889735163 | -3.75415661534591 | 0.879807018948925 | -4.26702280669557 | 1.9809891445438e-05 | 0.00203088514858948 | 6148 | 1 | 1 | 22 | 24 | 0 | 0 | 10 | 3 | 1 | 1 | 4 | 3 |
| MIRb_dup159983 | 4.99539987763176 | -3.15349154160534 | 0.757238944394577 | -4.16446032649126 | 3.12089886859684e-05 | 0.00284414171653445 | 6149 | 1 | 2 | 15 | 17 | 0 | 0 | 12 | 4 | 2 | 1 | 5 | 8 |
| MIRb_dup100635 | 4.77177250749358 | -2.60681911493612 | 0.784311071657334 | -3.32370561775652 | 0.000888299110006423 | 0.0323785569898267 | 6150 | 2 | 1 | 13 | 18 | 0 | 0 | 10 | 6 | 1 | 4 | 5 | 3 |
| THE1C_dup4940 | 6.33250269037403 | -2.82217723260625 | 0.710835346034023 | -3.97022636585544 | 7.18043790960654e-05 | 0.00533202802403038 | 6151 | 3 | 2 | 24 | 23 | 0 | 0 | 8 | 8 | 2 | 2 | 5 | 6 |
| MER113A_dup697 | 8.02268407400802 | -2.81149190275928 | 0.780169922950448 | -3.60369173439392 | 0.000313729121499688 | 0.0154236755256952 | 6152 | 4 | 0 | 21 | 23 | 0 | 0 | 20 | 17 | 3 | 5 | 4 | 6 |
| MIR1_Amn_dup4973 | 8.76488369578167 | -2.32140214909533 | 0.668905423520049 | -3.47044898646385 | 0.000519589030558416 | 0.0221450427778136 | 6153 | 6 | 2 | 23 | 25 | 0 | 0 | 19 | 13 | 2 | 8 | 5 | 12 |
| AluSg_dup15953 | 8.20319917563707 | -2.639003776249 | 0.79141181026004 | -3.33455192611023 | 0.000854369471023769 | 0.0314805513293488 | 6154 | 11 | 0 | 24 | 35 | 0 | 0 | 15 | 9 | 1 | 2 | 5 | 9 |
| LTR16A1_dup2228 | 6.89499141863299 | -2.54252061340143 | 0.752313219903027 | -3.37960379551639 | 0.00072590391672904 | 0.0280627024159233 | 6155 | 5 | 0 | 23 | 26 | 0 | 1 | 15 | 9 | 2 | 3 | 5 | 2 |
| L2c_dup91497 | 7.782836394217 | -3.75341357450517 | 0.658923813267444 | -5.69627853619814 | 1.22450769911424e-08 | 6.0575199370777e-06 | 6156 | 1 | 3 | 16 | 27 | 1 | 2 | 20 | 15 | 0 | 0 | 6 | 12 |
| THE1B_dup1942 | 8.15499346067151 | -3.4379802040382 | 0.62558994622671 | -5.49558097084938 | 3.89426207772071e-08 | 1.56789185168415e-05 | 6157 | 1 | 5 | 20 | 36 | 1 | 2 | 21 | 13 | 0 | 0 | 5 | 5 |

| | | | | | | | | | | | | | | | | | | | |
|---|---|---|---|---|---|---|---|---|---|---|---|---|---|---|---|---|---|---|---|
| L1MC4a_dup10262 | 7.08328509897826 | -3.26151086938109 | 0.655483550429343 | -4.975732598087 | 6.50013284159321e-07 | 0.000146465966773271 | 6158 | 3 | 5 | 19 | 22 | 0 | 1 | 22 | 10 | 0 | 0 | 4 | 9 |
| L1MEi_dup517 | 7.65585135355445 | -3.09353822994187 | 0.66551110011064 | -4.64836458689805 | 3.34577154739206e-06 | 0.000524103328910962 | 6159 | 4 | 5 | 19 | 40 | 0 | 2 | 15 | 9 | 0 | 0 | 5 | 5 |
| SST1_dup438 | 6.43681037112576 | -3.68834941144188 | 0.716982417317148 | -5.14426758921536 | 2.68566501566917e-07 | 7.31170764094297e-05 | 6160 | 3 | 2 | 19 | 28 | 0 | 1 | 16 | 8 | 0 | 0 | 5 | 5 |
| L1MB3_dup1800 | 7.75879068208063 | -3.40412954359375 | 0.672725069873037 | -5.06020913452237 | 4.18796872851958e-07 | 0.000104128569182479 | 6161 | 3 | 5 | 23 | 28 | 0 | 1 | 12 | 15 | 0 | 0 | 8 | 6 |
| L2a_dup19335 | 8.21500529019173 | -3.56085092488031 | 0.677837013921942 | -5.25325535747502 | 1.49434124144942e-07 | 4.56447395427373e-05 | 6162 | 1 | 5 | 26 | 33 | 0 | 1 | 15 | 14 | 0 | 1 | 7 | 5 |
| L1PA3_dup8342 | 7.93424218564834 | -2.83528424243955 | 0.661166352147706 | -4.28830691887076 | 1.80040205559438e-05 | 0.00188960146141805 | 6163 | 1 | 1 | 25 | 28 | 0 | 8 | 14 | 12 | 1 | 1 | 8 | 5 |
| MLT2C2_dup800 | 8.09799235503553 | -2.32109598139286 | 0.544397928692483 | -4.26360178659678 | 2.01157803186002e-05 | 0.00205075812497414 | 6164 | 3 | 4 | 20 | 24 | 0 | 8 | 20 | 13 | 2 | 0 | 6 | 6 |
| L1MEi_dup515 | 8.99335205641679 | -3.51561312871145 | 0.622073823548598 | -5.651440384771 | 1.59108870703292e-08 | 7.58703794556423e-06 | 6165 | 3 | 2 | 26 | 44 | 0 | 4 | 20 | 16 | 0 | 0 | 2 | 3 |
| MLT2D_dup4369 | 8.48155849555134 | -3.59677399431966 | 0.640609503069065 | -5.61461229826915 | 1.97003453321703e-08 | 9.07667809109646e-06 | 6166 | 1 | 1 | 24 | 25 | 0 | 5 | 28 | 17 | 0 | 1 | 6 | 3 |
| L1PA13_dup648 | 8.32152901888677 | -4.06675467365988 | 0.765900179173726 | -5.30977114804597 | 1.09762972801546e-07 | 3.56798638336566e-05 | 6167 | 1 | 3 | 24 | 47 | 0 | 0 | 15 | 15 | 1 | 0 | 1 | 4 |
| L2b_dup11886 | 6.42575702007225 | -3.20856281052006 | 0.720072430520711 | -4.45588898355659 | 8.35462604049319e-06 | 0.00106364357829384 | 6168 | 1 | 4 | 21 | 23 | 0 | 0 | 13 | 14 | 1 | 1 | 1 | 4 |
| MIR_dup19182 | 6.00380022366031 | -3.54131515748464 | 0.757534568478224 | -4.67479017439247 | 2.94254312241919e-06 | 0.000473725551966953 | 6169 | 2 | 4 | 21 | 18 | 0 | 0 | 10 | 14 | 0 | 0 | 3 | 5 |
| L1PA13_dup643 | 5.85048352480593 | -3.77110675187504 | 0.755795226525767 | -4.98958794594408 | 6.05082221506519e-07 | 0.000138973836176245 | 6170 | 2 | 3 | 15 | 25 | 0 | 0 | 14 | 10 | 0 | 0 | 4 | 5 |
| MIR3_dup10598 | 6.08011809685668 | -4.08529269486509 | 0.789147934862786 | -5.17684012640218 | 2.25675440845954e-07 | 6.324919722913e-05 | 6171 | 2 | 2 | 23 | 23 | 0 | 0 | 14 | 9 | 0 | 0 | 3 | 5 |
| L1MC4a_dup869 | 5.86076126239675 | -4.76621046218313 | 0.918696260810736 | -5.18801552319046 | 2.12546889854649e-07 | 6.04237541856086e-05 | 6172 | 1 | 1 | 16 | 31 | 0 | 0 | 10 | 11 | 0 | 0 | 4 | 4 |
| L1MC1_dup2193 | 7.09178429722196 | -4.24686630092509 | 0.790430758300308 | -5.37285050756031 | 7.75015802589403e-08 | 2.70795756480085e-05 | 6173 | 1 | 3 | 24 | 32 | 0 | 0 | 12 | 14 | 0 | 0 | 4 | 3 |
| MIRb_dup140065 | 7.39635986764461 | -4.40367397632042 | 0.796055306411409 | -5.53186938250815 | 3.16835809037033e-08 | 1.32785167498844e-05 | 6174 | 2 | 2 | 28 | 30 | 0 | 0 | 14 | 11 | 0 | 0 | 7 | 4 |
| L3_dup5394 | 7.30370811405542 | -4.01120192629916 | 0.722041603969229 | -5.55536122052892 | 2.77038249764072e-08 | 1.20292232442266e-05 | 6175 | 2 | 1 | 27 | 22 | 0 | 1 | 15 | 17 | 0 | 1 | 3 | 5 |
| MSTA_dup1663 | 7.69750112884206 | -4.40380848760987 | 0.755530552792796 | -5.82876294192382 | 5.58397593048557e-09 | 3.07565911285953e-06 | 6176 | 2 | 1 | 20 | 25 | 0 | 0 | 20 | 20 | 0 | 1 | 6 | 4 |
| L1M5_dup28508 | 8.94286872065389 | -3.62668502163571 | 0.675408005388193 | -5.36962101826327 | 7.89022752141204e-08 | 2.74882588259138e-05 | 6177 | 4 | 1 | 31 | 28 | 1 | 0 | 17 | 18 | 2 | 0 | 9 | 4 |
| L2_dup6360 | 9.28900565165464 | -4.59992160568921 | 0.854424208115354 | -5.38365083994458 | 7.29900261649618e-08 | 2.60385078948246e-05 | 6178 | 2 | 1 | 39 | 54 | 0 | 0 | 17 | 5 | 1 | 0 | 5 | 4 |
| MER50_dup226 | 8.92241371399076 | -3.85090675404616 | 0.742943060492276 | -5.18331344463267 | 2.17978250296372e-07 | 6.15998251128511e-05 | 6179 | 2 | 2 | 34 | 47 | 0 | 0 | 18 | 11 | 1 | 1 | 3 | 1 |
| L2a_dup19334 | 6.64713319360297 | -3.62723878716852 | 0.722303795826571 | -5.02176342991203 | 5.11992209565031e-07 | 0.000121220396315583 | 6180 | 3 | 2 | 19 | 29 | 0 | 0 | 18 | 10 | 0 | 1 | 5 | 2 |
| MSTA_dup18846 | 7.46236193379354 | -3.84517604673131 | 0.758749733952651 | -5.0677791037898 | 4.02484093093532e-07 | 0.000100703974779425 | 6181 | 3 | 2 | 22 | 39 | 0 | 0 | 17 | 7 | 0 | 1 | 7 | 4 |
| L1PB4_dup6658 | 8.8451253624823 | -3.54679541054202 | 0.641366123737767 | -5.53006353044019 | 3.20114833269645e-08 | 1.33164412792271e-05 | 6182 | 4 | 4 | 28 | 24 | 0 | 0 | 30 | 10 | 0 | 1 | 8 | 10 |
| LTR8B_dup1165 | 6.76599654875352 | -3.93689090560177 | 0.771718525189924 | -5.10145963469372 | 3.37043911875188e-07 | 8.8129945774784e-05 | 6183 | 1 | 2 | 25 | 14 | 0 | 0 | 15 | 12 | 0 | 2 | 6 | 11 |
| L1MA4_dup9720 | 6.88172288435903 | -2.86305296046242 | 0.76464366735795 | -3.74429696168809 | 0.000180899595026423 | 0.0103821358027394 | 6184 | 5 | 7 | 23 | 18 | 0 | 0 | 12 | 12 | 0 | 0 | 3 | 10 |
| MLT1G1_dup931 | 6.04553919389749 | -3.88966834773655 | 0.826863742677955 | -4.70412251375194 | 2.54960120327868e-06 | 0.000426030483086621 | 6185 | 3 | 2 | 13 | 17 | 0 | 0 | 7 | 15 | 0 | 0 | 6 | 15 |
| MamGypsy2-I_dup218 | 7.22277200858496 | -4.1337815572891 | 0.792798281948305 | -5.21416563508477 | 1.84646428923828e-07 | 5.39752815438687e-05 | 6186 | 0 | 4 | 22 | 26 | 1 | 0 | 11 | 11 | 0 | 0 | 11 | 9 |
| MIR_dup124598 | 5.75969635569134 | -4.43670475678789 | 0.893088631950583 | -4.96782133157125 | 6.77093024226334e-07 | 0.000151136563548367 | 6187 | 0 | 2 | 21 | 19 | 0 | 0 | 13 | 6 | 1 | 0 | 6 | 9 |
| Tigger19b_dup1976 | 5.28507529720906 | -3.31269486133145 | 0.757161241983578 | -4.37515112719319 | 1.21348376443903e-05 | 0.0014078186631558 | 6188 | 1 | 4 | 16 | 15 | 0 | 0 | 11 | 8 | 1 | 0 | 7 | 6 |
| L1MB7_dup9541 | 4.86307301743149 | -4.09642342443007 | 0.85865664136015 | -4.77073515432334 | 1.83554756499174e-06 | 0.00032963109501851 | 6189 | 1 | 1 | 7 | 15 | 0 | 1 | 23 | 10 | 0 | 0 | 3 | 4 |
| L2a_dup19343 | 5.51588957032183 | -3.4655077783507 | 0.754713829573817 | -4.59181698089149 | 4.39403626562027e-06 | 0.000649809417811859 | 6190 | 2 | 3 | 10 | 27 | 0 | 1 | 14 | 8 | 0 | 0 | 3 | 7 |
| L2b_dup60233 | 5.22232347966695 | -4.34382162991338 | 0.885901652687443 | -4.90327748767157 | 9.42506945192563e-07 | 0.000196722507389869 | 6191 | 1 | 1 | 11 | 20 | 0 | 1 | 18 | 5 | 0 | 0 | 5 | 10 |
| LTR12C_dup1416 | 5.17210257972435 | -3.51412843820141 | 0.81060469628202 | -4.33519378103727 | 1.45631774374737e-05 | 0.00161625457954156 | 6192 | 1 | 1 | 10 | 16 | 0 | 2 | 30 | 4 | 0 | 1 | 3 | 4 |
| MSTA_dup10461 | 5.30114964809318 | -3.59690234075158 | 0.810456670350958 | -4.43811800474661 | 9.07488653045713e-06 | 0.00113529597811575 | 6193 | 1 | 1 | 19 | 21 | 0 | 2 | 15 | 3 | 0 | 1 | 3 | 7 |
| MIRb_dup213177 | 4.69247900716039 | -2.51988703901817 | 0.70569667283424 | -3.57077925406355 | 0.000355920765415965 | 0.0169244449335866 | 6194 | 4 | 4 | 12 | 10 | 0 | 0 | 11 | 9 | 0 | 1 | 4 | 6 |
| MIRc_dup51484 | 4.89687216822946 | -3.10153813360448 | 0.782809247347224 | -3.96206118427311 | 7.43054914851868e-05 | 0.0054722050377388 6 | 6195 | 1 | 4 | 15 | 18 | 0 | 0 | 8 | 11 | 0 | 1 | 3 | 2 |
| L2a_dup19342 | 4.91055450650774 | -2.74207165189397 | 0.786244326667365 | -3.48755667785448 | 0.000487455594096553 | 0.0211734408165203 | 6196 | 2 | 3 | 22 | 14 | 0 | 0 | 13 | 6 | 0 | 2 | 1 | 2 |
| LTR8B_dup147 | 6.55054997410921 | -3.15707986126161 | 0.690569852986336 | -4.57170241013125 | 4.83777667372965e-06 | 0.000702496054992057 | 6197 | 4 | 3 | 24 | 25 | 0 | 0 | 15 | 10 | 0 | 1 | 2 | 3 |
| MER30_dup435 | 4.76317240820742 | -4.22806775246475 | 0.918191015375016 | -4.60478014015187 | 4.12901850298717e-06 | 0.000617137837130519 | 6198 | 0 | 2 | 21 | 11 | 0 | 0 | 13 | 11 | 0 | 0 | 1 | 2 |
| Tigger1_dup4283 | 4.9051023390276 | -3.90462281251661 | 0.824113136162262 | -4.73796938937255 | 2.15870409881729e-06 | 0.000372644923299795 | 6199 | 1 | 2 | 15 | 13 | 0 | 0 | 16 | 13 | 0 | 0 | 1 | 2 |
| MLT1D_dup4948 | 4.87827653736862 | -3.87552332387132 | 0.888066919833948 | -4.3639991956867 | 1.27706046332412e-05 | 0.00146482330168208 | 6200 | 0 | 3 | 14 | 26 | 0 | 0 | 10 | 9 | 0 | 0 | 1 | 2 |
| L3_dup5401 | 3.89658765871516 | -3.99777240130105 | 0.936867697823434 | -4.26716857736564 | 1.97969563298337e-05 | 0.0020304338667494 | 6201 | 0 | 2 | 8 | 18 | 0 | 0 | 13 | 8 | 0 | 0 | 1 | 2 |
| MLT2F_dup324 | 3.63538071680963 | -3.50438508068443 | 0.907970768499079 | -3.85957918719933 | 0.000113582447641771 | 0.00744736541376934 | 6202 | 0 | 3 | 10 | 10 | 0 | 0 | 15 | 7 | 0 | 0 | 1 | 2 |
| THE1B_dup22284 | 4.94535286084105 | -4.55731189506442 | 0.938631486424554 | -4.85527276782946 | 1.2022114262583e-06 | 0.00023739535272403 | 6203 | 0 | 1 | 23 | 12 | 0 | 1 | 13 | 7 | 0 | 0 | 4 | 4 |
| L1P2_dup161 | 4.96287848873381 | -5.03498461595716 | 0.967226733282675 | -5.20558876497231 | 1.93382479190231e-07 | 5.62525518256707e-05 | 6204 | 0 | 1 | 13 | 24 | 0 | 0 | 17 | 6 | 0 | 0 | 3 | 4 |
| MER20_dup16085 | 5.54216342051938 | -4.16858643079549 | 0.869741194820077 | -4.79290443596598 | 1.64383939444015e-06 | 0.00030316187519907 | 6205 | 0 | 3 | 20 | 18 | 0 | 0 | 23 | 6 | 0 | 0 | 3 | 2 |
| L2a_dup124329 | 6.25013485593157 | -3.90300528896244 | 0.753839816955958 | -5.17749951803152 | 2.24879524868002e-07 | 6.31749509140986e-05 | 6206 | 3 | 0 | 12 | 20 | 1 | 1 | 18 | 15 | 0 | 0 | 3 | 9 |
| MLT2B4_dup4392 | 5.80355118817619 | -4.45015083342469 | 0.849588328660487 | -5.23800843691093 | 1.62318644089384e-07 | 4.85824806119404e-05 | 6207 | 1 | 0 | 19 | 21 | 1 | 1 | 12 | 9 | 0 | 0 | 6 | 7 |
| L1MCa_dup605 | 6.28097770594065 | -4.99582615665738 | 0.961440836914654 | -5.19618676973345 | 2.03417893724726e-07 | 5.84110420366877e-05 | 6208 | 2 | 0 | 13 | 36 | 0 | 0 | 13 | 9 | 0 | 0 | 4 | 9 |
| L1M3_dup2033 | 6.49217509875298 | -5.06540692149037 | 0.910663858504623 | -5.56232343491499 | 2.66206319719827e-08 | 1.16438839985393e-05 | 6209 | 2 | 0 | 14 | 22 | 0 | 0 | 20 | 13 | 0 | 0 | 7 | 8 |
| MSTC_dup3143 | 6.14485663286361 | -5.40902468301501 | 0.927054927467883 | -5.83463236400564 | 5.39092977970815e-09 | 2.99009365352549e-06 | 6210 | 1 | 0 | 21 | 19 | 0 | 0 | 18 | 11 | 0 | 0 | 6 | 5 |
| L1PA11_dup3215 | 6.22326535202345 | -4.64568544917842 | 0.848126146437711 | -5.47758781956101 | 4.31163034998924e-08 | 1.69017330355608e-05 | 6211 | 1 | 0 | 16 | 24 | 1 | 0 | 19 | 16 | 0 | 0 | 2 | 2 |
| L1PA16_dup1122 | 7.61818867272257 | -5.5025279828566 | 0.927485670655265 | -5.93273638283715 | 2.9792700075386e-09 | 1.81305724095596e-06 | 6212 | 1 | 0 | 20 | 25 | 0 | 0 | 24 | 21 | 0 | 0 | 5 | 2 |
| L1PA3_dup940 | 6.71477889185969 | -3.62080673785864 | 0.751259496299759 | -4.81964854446766 | 1.43811341574036e-06 | 0.000273754667857405 | 6213 | 5 | 0 | 16 | 30 | 1 | 0 | 27 | 9 | 0 | 0 | 2 | 2 |
| MER119_dup119 | 5.68179835613691 | -4.27496046559621 | 0.855269413915631 | -4.99837875181856 | 5.78143415962746e-07 | 0.000134474011001634 | 6214 | 3 | 0 | 21 | 20 | 0 | 0 | 20 | 7 | 0 | 0 | 2 | 4 |
| MLT1F2_dup543 | 6.2001299617019 | -3.77536039686228 | 0.794719006771813 | -4.75056009066396 | 2.02854001301652e-06 | 0.0003570140398944 | 6215 | 5 | 0 | 22 | 20 | 0 | 0 | 16 | 13 | 0 | 0 | 3 | 2 |
| MSTB_dup1295 | 7.92725856103109 | -4.71310033230294 | 0.861454256672436 | -5.47109761870394 | 4.47256841622646e-08 | 1.74463895797938e-05 | 6216 | 2 | 0 | 20 | 52 | 0 | 1 | 22 | 7 | 0 | 0 | 2 | 5 |
| L1ME1_dup3213 | 5.22999804170572 | -3.83905533356948 | 0.846918240671313 | -4.53297042052895 | 5.81599621388801e-06 | 0.000805994778747697 | 6217 | 2 | 0 | 17 | 27 | 0 | 2 | 10 | 7 | 0 | 0 | 1 | 5 |
| MER5B_dup24170 | 6.49411949474687 | -4.03263713823286 | 0.842245222375857 | -4.78796083503728 | 1.68484517869062e-06 | 0.000309576965937233 | 6218 | 3 | 0 | 30 | 30 | 0 | 1 | 8 | 10 | 0 | 0 | 2 | 2 |
| AluSx1_dup67054 | 6.71976893687809 | -4.92927390825375 | 0.906004630618794 | -5.4406718703933 | 5.3080002354933e-08 | 2.01438185651466e-05 | 6219 | 1 | 0 | 30 | 27 | 0 | 1 | 12 | 10 | 0 | 0 | 4 | 4 |
| MER77_dup113 | 6.16626574082995 | -4.36718335954902 | 0.853216926371793 | -5.11849123542352 | 3.07989547529168e-07 | 8.15182212195498e-05 | 6220 | 2 | 0 | 25 | 26 | 0 | 1 | 10 | 11 | 0 | 0 | 4 | 2 |
| L1PA3_dup2692 | 6.59936088633577 | -4.34841336930295 | 0.824091954444129 | -5.27661184635162 | 1.3159420881314e-07 | 4.12546977737172e-05 | 6221 | 1 | 1 | 31 | 21 | 0 | 1 | 23 | 7 | 0 | 0 | 1 | 3 |
| Ricksha_c_dup103 | 7.89608527298435 | -4.7359644847996 | 0.817125490462955 | -5.79588391266116 | 6.79622645570937e-09 | 3.64219572123924e-06 | 6222 | 1 | 2 | 28 | 35 | 0 | 0 | 26 | 8 | 0 | 0 | 3 | 5 |
| MLT1F2_dup544 | 8.99978885715428 | -4.07938596246904 | 0.67789967052852 | -6.01768394324277 | 1.76930213370981e-09 | 1.16297891024231e-06 | 6223 | 1 | 4 | 30 | 38 | 0 | 1 | 24 | 15 | 0 | 0 | 2 | 5 |
| THE1C-int_dup166 | 7.41209326203951 | -4.73334874161073 | 0.821272125564401 | -5.76343527835897 | 8.24188678297256e-09 | 4.30071050978331e-06 | 6224 | 1 | 1 | 22 | 36 | 0 | 1 | 17 | 11 | 0 | 0 | 4 | 7 |
| L1MB8_dup1418 | 6.38312835577209 | -4.60255099013295 | 0.871223134947905 | -5.28286130786478 | 1.27181675021041e-07 | 3.99839815483212e-05 | 6225 | 1 | 1 | 22 | 34 | 0 | 0 | 15 | 10 | 0 | 0 | 1 | 2 |
| Tigger20a_dup110 | 6.55885471845977 | -4.40120229452333 | 0.80331053712562 | -5.47883052831795 | 4.28146189445221e-08 | 1.68389542468979e-05 | 6226 | 2 | 1 | 23 | 28 | 0 | 0 | 16 | 12 | 0 | 0 | 2 | 3 |
| L1MA6_dup455 | 6.12897124074969 | -3.5379936729562 | 0.773542939581736 | -4.573752136978 | 4.79066542164227e-06 | 0.000699766528187902 | 6227 | 5 | 1 | 18 | 29 | 0 | 0 | 19 | 7 | 0 | 0 | 2 | 3 |
| MER67D_dup986 | 6.23202379465853 | -3.4622415697139 | 0.798284619396801 | -4.33710168727795 | 1.44373841793401e-05 | 0.0016060391846364 | 6228 | 5 | 1 | 27 | 25 | 0 | 0 | 16 | 7 | 0 | 0 | 2 | 1 |
| L2a_dup73380 | 5.86690167321706 | -2.85427763868607 | 0.759446025099649 | -3.75836800029542 | 0.000171025222552012 | 0.0100036301881418 | 6229 | 6 | 2 | 21 | 26 | 0 | 1 | 18 | 4 | 0 | 0 | 1 | 2 |
| L1P2_dup413 | 7.6102783366065 | -2.87829129531485 | 0.684489840000817 | -4.20501682729347 | 2.61062733091347e-05 | 0.00248873530000616 | 6230 | 9 | 2 | 23 | 32 | 0 | 1 | 19 | 12 | 0 | 0 | 2 | 2 |
| LTR9B_dup291 | 6.52056931120328 | -2.74810427198145 | 0.661365593034003 | -4.15519691518056 | 3.25007511223603e-05 | 0.00293599989618874 | 6231 | 6 | 4 | 15 | 28 | 0 | 1 | 17 | 12 | 0 | 0 | 1 | 3 |
| L1PA16_dup6248 | 8.26822649526254 | -3.70184672339484 | 0.792951112192407 | -4.66844256408156 | 3.03491594685829e-06 | 0.000484661818718887 | 6232 | 0 | 0 | 6 | 20 | 5 | 3 | 28 | 27 | 0 | 0 | 6 | 10 |
| MamGypsy2-I_dup111 | 7.19205086354892 | -3.67807826090315 | 0.831591210938454 | -4.42294027705322 | 9.73666631458903e-06 | 0.0011985518897523 | 6233 | 0 | 0 | 6 | 17 | 3 | 4 | 19 | 26 | 0 | 0 | 10 | 4 |
| L1PA5_dup5154 | 7.3283441350336 | -3.87882847888475 | 0.773590462144626 | -5.01405933590685 | 5.32935399995632e-07 | 0.000125678738044956 | 6234 | 0 | 0 | 16 | 10 | 4 | 2 | 28 | 20 | 0 | 0 | 8 | 5 |
| MLT1E3_dup906 | 6.66823680891443 | -3.51218269584672 | 0.769617381736475 | -4.56354388452382 | 5.02972902145758e-06 | 0.000724457567639071 | 6235 | 0 | 1 | 13 | 11 | 5 | 1 | 23 | 19 | 0 | 0 | 8 | 3 |
| L1PA4_dup5452 | 7.36563907261366 | -3.80345106053337 | 0.755479356097975 | -5.03448708403902 | 4.7912997925624e-07 | 0.000115625823574143 | 6236 | 0 | 0 | 15 | 21 | 2 | 4 | 27 | 19 | 0 | 0 | 6 | 1 |

| | | | | | | | | | | | | | | | | | | | |
|---|---|---|---|---|---|---|---|---|---|---|---|---|---|---|---|---|---|---|---|
| MLT1F_dup1847 | 6.19393960221609 | -3.29822336963007 | 0.834554031761314 | -3.95207888777341 | 7.74751895160388e-05 | 0.00564103777374032 | 6237 | 0 | 0 | 6 | 14 | 4 | 3 | 25 | 21 | 0 | 0 | 4 | 1 |
| LTR16C_dup2532 | 5.37180381589858 | -3.66274741250344 | 0.789604376938084 | -4.63871214431053 | 3.50586948744337e-06 | 0.000541341739817781 | 6238 | 0 | 0 | 11 | 11 | 3 | 2 | 13 | 15 | 0 | 0 | 6 | 7 |
| LTR32_dup573 | 5.54644788364327 | -3.65280063480771 | 0.772725104688408 | -4.72716702569235 | 2.27673981857233e-06 | 0.000387511539964243 | 6239 | 0 | 0 | 11 | 14 | 3 | 2 | 15 | 17 | 0 | 0 | 4 | 4 |
| AluYc_dup1983 | 5.74677505500407 | -4.10228106225532 | 0.921034833939436 | -4.45399121845274 | 8.4288549220994e-06 | 0.00107080208931867 | 6240 | 0 | 0 | 8 | 11 | 3 | 1 | 11 | 20 | 0 | 0 | 3 | 15 |
| ERVL-int_dup361 | 4.50708744760887 | -4.05644680269825 | 0.928857079352782 | -4.36713773611409 | 1.25885347728753e-05 | 0.00144844849858152 | 6241 | 0 | 0 | 7 | 12 | 2 | 1 | 7 | 16 | 0 | 0 | 4 | 7 |
| L1MC1_dup816 | 4.1250411085967 | -2.83986218102685 | 0.877928011953448 | -3.23473239532245 | 0.00121756789105032 | 0.0402382513060916 | 6242 | 1 | 0 | 4 | 3 | 2 | 4 | 20 | 9 | 0 | 0 | 4 | 7 |
| L1MEg_dup6687 | 6.13339463803038 | -3.11793114196477 | 0.872555565994923 | -3.57333247700916 | 0.000352466768701115 | 0.0168105948577582 | 6243 | 0 | 0 | 6 | 5 | 4 | 5 | 26 | 18 | 0 | 0 | 5 | 9 |
| MLT1K_dup15616 | 5.76441432209439 | -2.68612173909376 | 0.795594349666248 | -3.37624536954114 | 0.000734823542400476 | 0.0283201040848759 | 6244 | 0 | 0 | 7 | 18 | 2 | 7 | 20 | 10 | 1 | 0 | 2 | 10 |
| L1MA2_dup6892 | 4.58510649367332 | -2.51849428807359 | 0.721488628281661 | -3.49069159145555 | 0.000481772041452834 | 0.0209957972501997 | 6245 | 2 | 0 | 6 | 15 | 3 | 4 | 14 | 11 | 0 | 0 | 1 | 4 |
| L1ME3A_dup15568 | 6.19828516609722 | -3.01186823357632 | 0.668183726475236 | -4.50754502727019 | 6.55820161543367e-06 | 0.000885137743383994 | 6246 | 1 | 0 | 14 | 14 | 3 | 5 | 20 | 15 | 0 | 0 | 3 | 5 |
| L1PB3_dup277 | 5.06616858650386 | -4.9953863400731 | 1.00177626536125 | -4.98652894144156 | 6.14737015071628e-07 | 0.000140648282488686 | 6247 | 0 | 0 | 19 | 23 | 1 | 0 | 6 | 11 | 0 | 0 | 2 | 4 |
| MSTA_dup19402 | 5.30412199978726 | -5.05177916140546 | 0.944490869876611 | -5.34867971996958 | 8.85981695139739e-08 | 3.01165429188172e-05 | 6248 | 0 | 0 | 16 | 20 | 1 | 0 | 16 | 11 | 0 | 0 | 2 | 4 |
| MSTA_dup19404 | 4.71993607495462 | -5.04821557015454 | 0.976927272324191 | -5.16744256524277 | 2.37318859693234e-07 | 6.57381529552576e-05 | 6249 | 0 | 0 | 14 | 24 | 1 | 0 | 10 | 6 | 0 | 0 | 4 | 5 |
| L1PA2_dup615 | 5.4067387326557 | -5.30211161702877 | 0.966851562766365 | -5.48389413764644 | 4.16063807539765e-08 | 1.64453021177503e-05 | 6250 | 0 | 0 | 12 | 10 | 1 | 0 | 12 | 18 | 0 | 0 | 8 | 6 |
| L1PA7_dup4684 | 4.21990601721652 | -4.98311280374549 | 0.963492108952174 | -5.17192902510097 | 2.31689593410548e-07 | 6.44790783551653e-05 | 6251 | 0 | 0 | 12 | 11 | 1 | 0 | 9 | 9 | 0 | 0 | 4 | 9 |
| MamRTE1_dup8551 | 4.39653216752251 | -4.56115727130721 | 0.951058883242344 | -4.79587263383456 | 1.61968173345524e-06 | 0.000299453605088376 | 6252 | 0 | 1 | 12 | 11 | 1 | 0 | 8 | 8 | 0 | 0 | 9 | 7 |
| L1MEd_dup21118 | 6.43896545519303 | -3.337058569391 | 0.823286742806297 | -4.05333694311186 | 5.04921969809873e-05 | 0.00411101067512712 | 6253 | 1 | 0 | 25 | 13 | 7 | 0 | 16 | 8 | 0 | 0 | 5 | 10 |
| L1MCc_dup2584 | 4.4089858904638 | -4.04070273016455 | 0.903588065956708 | -4.47184163049597 | 7.75488503294715e-06 | 0.00100812484955221 | 6254 | 0 | 0 | 14 | 10 | 3 | 0 | 13 | 7 | 0 | 0 | 5 | 6 |
| MIR_dup170841 | 4.88197008637986 | -3.83862093492747 | 0.886107359414965 | -4.33200434929447 | 1.47758022303348e-05 | 0.00162996895572519 | 6255 | 0 | 0 | 10 | 14 | 4 | 0 | 18 | 7 | 0 | 0 | 8 | 4 |
| L1MA5_dup3406 | 4.27304055114989 | -3.87813314692546 | 0.888148337370086 | -4.36653764213428 | 1.26231541920289e-05 | 0.00145102852530266 | 6256 | 0 | 0 | 9 | 17 | 3 | 0 | 14 | 8 | 0 | 0 | 2 | 4 |
| MLT1J_dup14763 | 5.41658341244376 | -3.92547232663743 | 0.823103510708982 | -4.7691113882581 | 1.85040329679591e-06 | 0.000331298768141008 | 6257 | 0 | 0 | 13 | 15 | 4 | 0 | 16 | 13 | 0 | 0 | 4 | 5 |
| AluSg_dup35830 | 4.3611120219389 | -2.84773941962949 | 0.811912555165936 | -3.50744597002504 | 0.000452430218809788 | 0.0201825948340671 | 6258 | 2 | 0 | 15 | 13 | 5 | 0 | 12 | 5 | 0 | 0 | 4 | 2 |
| MLT1E1A_dup3403 | 4.20260617804723 | -3.91341715248229 | 0.897265072766396 | -4.36149502667775 | 1.29176733128236e-05 | 0.00147703445175066 | 6259 | 1 | 0 | 10 | 18 | 2 | 0 | 14 | 5 | 0 | 0 | 5 | 2 |
| L2_dup29486 | 5.98880970644974 | -3.35168845752081 | 0.787070695312281 | -4.25843380713213 | 2.0586415747325e-05 | 0.00208622456618952 | 6260 | 1 | 0 | 20 | 22 | 6 | 0 | 12 | 9 | 0 | 0 | 5 | 4 |
| HERVL-int_dup2333 | 5.77581477609758 | -3.53692361004781 | 0.753991645589588 | -4.6909320955169 | 2.71963227715565e-06 | 0.000446909959820496 | 6261 | 2 | 0 | 11 | 22 | 4 | 0 | 20 | 8 | 0 | 0 | 6 | 5 |
| THE1B_dup22279 | 5.54633964211238 | -3.38559676502924 | 0.768676769485331 | -4.40444787644106 | 1.06053635694357e-05 | 0.0012757855611878 | 6262 | 1 | 0 | 12 | 22 | 5 | 0 | 18 | 10 | 0 | 0 | 2 | 4 |
| L1PA11_dup3099 | 6.33667572786351 | -4.14475323599394 | 0.803331083825859 | -5.15945830983482 | 2.47665363674015e-07 | 6.82861907587222e-05 | 6263 | 1 | 1 | 19 | 24 | 2 | 0 | 13 | 14 | 0 | 0 | 1 | 8 |
| THE1C_dup7434 | 5.77805161261811 | -4.79895839185047 | 0.959039273627405 | -5.00392270036994 | 5.61752903739986e-07 | 0.000131174540170083 | 6264 | 0 | 1 | 17 | 32 | 1 | 0 | 11 | 7 | 0 | 0 | 2 | 8 |
| L1MC_dup3937 | 6.51702863184087 | -3.42803597804479 | 0.710389699688708 | -4.82557106268145 | 1.39602675647887e-06 | 0.000267454897421193 | 6265 | 1 | 2 | 18 | 27 | 3 | 0 | 18 | 6 | 0 | 1 | 4 | 9 |
| Tigger3a_dup543 | 7.03294323407329 | -3.89890728507487 | 0.76459645590733 | -5.09930075525674 | 3.40910506989027e-07 | 8.89454950614156e-05 | 6266 | 1 | 1 | 20 | 35 | 1 | 0 | 14 | 12 | 0 | 2 | 2 | 6 |
| L1M4_dup7960 | 4.9508646466179 | -3.77813916041055 | 0.892667181918444 | -4.23241633269288 | 2.31193930270202e-05 | 0.00228026474908339 | 6267 | 0 | 0 | 15 | 7 | 3 | 0 | 16 | 11 | 0 | 1 | 2 | 9 |
| AluY_dup71779 | 3.84831798206835 | -3.42311909729945 | 0.865325898291747 | -3.95587269958877 | 7.62557617754729e-05 | 0.00558026657196382 | 6268 | 0 | 0 | 9 | 11 | 3 | 0 | 11 | 6 | 0 | 1 | 4 | 6 |
| MIR3_dup21260 | 3.13673723496431 | -3.86098045637312 | 0.985452964808067 | -3.91797538213821 | 8.92958231105297e-05 | 0.00627712003268199 | 6269 | 0 | 0 | 9 | 12 | 1 | 0 | 10 | 5 | 0 | 1 | 1 | 3 |
| LTR33A_dup1986 | 5.10573874969058 | -3.73721979559846 | 0.825960089181824 | -4.52469779659748 | 6.04818653590489e-06 | 0.000834849542570923 | 6270 | 0 | 0 | 16 | 18 | 3 | 0 | 11 | 11 | 0 | 1 | 4 | 2 |
| MIRb_dup190731 | 5.07172076135812 | -3.39420437143783 | 0.780847033217363 | -4.3468236761463 | 1.3812316606528e-05 | 0.00155174007896927 | 6271 | 0 | 0 | 11 | 20 | 2 | 1 | 12 | 12 | 0 | 2 | 2 | 4 |
| MLT1E2_dup1944 | 7.31705213188545 | -4.35490584517963 | 0.822845472721977 | -5.29249535854354 | 1.20658533072113e-07 | 3.82293146609547e-05 | 6272 | 0 | 0 | 19 | 34 | 2 | 1 | 8 | 17 | 0 | 1 | 6 | 7 |
| L2a_dup133365 | 6.62157647249124 | -3.11377018622969 | 0.778943238505046 | -3.99742886555594 | 6.40342249346829e-05 | 0.00490610748948006 | 6273 | 0 | 1 | 20 | 33 | 3 | 2 | 4 | 12 | 0 | 2 | 3 | 7 |
| X7A_LINE_dup469 | 6.4982441781003 | -3.26172758424228 | 0.73627628061538 | -4.43003213619231 | 9.4219046829366e-06 | 0.00116826708269204 | 6274 | 1 | 1 | 16 | 28 | 2 | 1 | 7 | 18 | 0 | 2 | 2 | 5 |
| L1ME2z_dup3268 | 6.99627778331006 | -3.21130463842369 | 0.72897534380997 | -4.40523080196361 | 1.0567130640143e-05 | 0.0012739964894005 | 6275 | 0 | 0 | 24 | 28 | 3 | 2 | 10 | 9 | 0 | 3 | 5 | 9 |
| L2a_dup86188 | 5.39998800832697 | -2.75210128233784 | 0.822782580487917 | -3.34487062269333 | 0.000823209611086554 | 0.0306820887888357 | 6276 | 0 | 0 | 18 | 26 | 3 | 2 | 7 | 5 | 0 | 3 | 2 | 7 |
| AluSp_dup22706 | 4.90795219462471 | -3.07463540633363 | 0.779019445771351 | -3.94680187127969 | 7.92020449537983e-05 | 0.00574244960288883 | 6277 | 0 | 0 | 19 | 18 | 2 | 3 | 8 | 9 | 0 | 1 | 1 | 3 |
| MIR_dup84424 | 5.59586246414475 | -3.25197019191151 | 0.804850378502469 | -4.04046550610094 | 5.33452109972074e-05 | 0.00429407292826167 | 6278 | 0 | 0 | 16 | 31 | 1 | 4 | 9 | 9 | 0 | 1 | 1 | 3 |
| Charlie3_dup338 | 5.05608691938415 | -3.76785895620284 | 0.829873595722215 | -4.54028056275701 | 5.61794213717206e-06 | 0.000786260221373366 | 6279 | 0 | 0 | 19 | 21 | 1 | 2 | 11 | 6 | 0 | 1 | 2 | 5 |
| OldhAT1_dup2264 | 4.16746309990488 | -3.47594812515247 | 0.842421607672204 | -4.12613837714501 | 3.68905347838397e-05 | 0.00323790305779252 | 6280 | 0 | 0 | 13 | 18 | 1 | 2 | 7 | 7 | 0 | 1 | 3 | 3 |
| BSR/Beta_dup2027 | 5.53212656028417 | -4.42168104768121 | 0.972832844239005 | -4.54516012063723 | 5.48935082504942e-06 | 0.000771967536298587 | 6281 | 0 | 0 | 27 | 19 | 1 | 2 | 5 | 4 | 0 | 0 | 7 | 9 |
| MER103C_dup8109 | 5.46322493552647 | -4.40905742235966 | 0.978994481545748 | -4.50365911705461 | 6.67933152209806e-06 | 0.000896044185789028 | 6282 | 0 | 0 | 19 | 17 | 1 | 2 | 3 | 11 | 0 | 0 | 4 | 14 |
| L1PA4_dup12247 | 4.53296763258498 | -3.43189516567125 | 0.863287044020998 | -3.97538129344123 | 7.02665861600433e-05 | 0.00526271737816735 | 6283 | 0 | 0 | 16 | 15 | 2 | 3 | 4 | 8 | 0 | 0 | 4 | 7 |
| L2c_dup70119 | 5.31155580387059 | -3.62909845451554 | 0.830290645078387 | -4.37087720550305 | 1.23748375148735e-05 | 0.00142869630631445 | 6284 | 0 | 0 | 18 | 23 | 3 | 2 | 6 | 9 | 0 | 0 | 3 | 6 |
| MLT1H_dup9870 | 5.70196941079756 | -4.31848596072457 | 0.881116563400309 | -4.9011517205614 | 9.52764374563781e-07 | 0.000198230688463463 | 6285 | 0 | 0 | 26 | 22 | 2 | 1 | 13 | 5 | 0 | 0 | 3 | 5 |
| AluJb_dup57365 | 6.27325175150257 | -3.88718988547035 | 0.822221293241412 | -4.72766871573714 | 2.27112332934316e-06 | 0.000387109392352354 | 6286 | 0 | 0 | 22 | 31 | 2 | 3 | 13 | 5 | 0 | 0 | 3 | 7 |
| MER41A_dup2505 | 5.14993455609578 | -3.79983142685549 | 0.86415867280375 | -4.39714550862169 | 1.09683817450721e-05 | 0.00130690163380717 | 6287 | 0 | 0 | 17 | 29 | 2 | 2 | 11 | 4 | 0 | 0 | 2 | 4 |
| L1ME3_dup9071 | 5.86049222178832 | -4.4769669323168 | 0.890252368402218 | -5.02887393644551 | 4.93368604816691e-07 | 0.000118461237176299 | 6288 | 0 | 0 | 15 | 29 | 1 | 2 | 12 | 7 | 0 | 0 | 4 | 10 |
| L1PA15_dup4046 | 6.73910939553616 | -4.92399507562234 | 0.906012543252015 | -5.43479790903148 | 5.48585714580886e-08 | 2.06540785508961e-05 | 6289 | 0 | 0 | 24 | 35 | 1 | 1 | 13 | 9 | 0 | 0 | 3 | 5 |
| Tigger10_dup554 | 6.89678256966666 | -4.58096300260171 | 0.872298336223089 | -5.25160121528667 | 1.50782638277885e-07 | 4.58258476452757e-05 | 6290 | 0 | 0 | 25 | 37 | 2 | 1 | 10 | 9 | 0 | 0 | 3 | 6 |
| L1PA12_dup872 | 8.19602083231676 | -3.67855126197249 | 0.797162476913213 | -4.61455646559863 | 3.93935668071609e-06 | 0.000593637849746466 | 6291 | 0 | 0 | 28 | 43 | 2 | 6 | 11 | 9 | 0 | 0 | 4 | 8 |
| MIRc_dup48419 | 7.65497031560221 | -3.7484367040063 | 0.773781210339066 | -4.84431083867203 | 1.27051895390017e-06 | 0.000247246334857043 | 6292 | 0 | 0 | 31 | 25 | 1 | 6 | 12 | 13 | 0 | 0 | 5 | 7 |
| MER63B_dup1148 | 5.33484733732901 | -3.1112310296976 | 0.873949279713115 | -3.55996749687682 | 0.000370900698128466 | 0.0174761064389853 | 6293 | 0 | 0 | 23 | 25 | 2 | 5 | 5 | 6 | 0 | 0 | 3 | 2 |
| Tigger10_dup553 | 6.26845352023048 | -3.00063175179574 | 0.886065139092262 | -3.38646857822413 | 0.000707983836205012 | 0.0275715093109063 | 6294 | 0 | 0 | 26 | 22 | 1 | 9 | 8 | 7 | 0 | 0 | 5 | 5 |
| LTR9_dup355 | 5.06906904496205 | -2.98797234286259 | 0.791276335825482 | -3.77614267934027 | 0.000159275776407989 | 0.00951288585068369 | 6295 | 0 | 0 | 8 | 18 | 2 | 2 | 14 | 15 | 2 | 0 | 2 | 1 |
| MIR_dup153546 | 3.59618555797565 | -2.93093582856067 | 0.884303244772075 | -3.31440130508184 | 0.000918395948603054 | 0.0331941792996004 | 6296 | 0 | 0 | 7 | 8 | 2 | 3 | 7 | 13 | 0 | 0 | 3 | 1 |
| LTR79_dup1914 | 4.13711171980079 | -3.89234674137963 | 0.903148146485514 | -4.3097544478458 | 1.63435915772802e-05 | 0.00176127366007587 | 6297 | 0 | 0 | 10 | 7 | 1 | 2 | 12 | 12 | 0 | 0 | 5 | 3 |
| MLT1E3_dup905 | 4.24162160312855 | -3.55579521550211 | 0.856533163346947 | -4.15138066762956 | 3.30475564220403e-05 | 0.00297410509092239 | 6298 | 0 | 0 | 6 | 12 | 1 | 3 | 12 | 13 | 0 | 0 | 3 | 4 |
| L2c_dup62506 | 4.79252455517287 | -3.18589287450647 | 0.850728512756551 | -3.74489960867006 | 0.000180465923431559 | 0.0103622453225979 | 6299 | 0 | 0 | 9 | 7 | 1 | 5 | 19 | 14 | 0 | 0 | 2 | 4 |
| L1M4_dup7959 | 4.37419721144444 | -2.66679793191628 | 0.840206625329487 | -3.17397869943062 | 0.00150364725074216 | 0.0465807640574267 | 6300 | 0 | 0 | 5 | 9 | 4 | 4 | 12 | 14 | 0 | 0 | 2 | 5 |
| MIRb_dup94819 | 4.52056204423007 | -3.05080724030126 | 0.841833642736641 | -3.62400251715252 | 0.000290078778976145 | 0.0145987912737432 | 6301 | 0 | 0 | 6 | 11 | 4 | 2 | 14 | 15 | 0 | 0 | 1 | 4 |
| Charlie7_dup2023 | 3.49277462787389 | -3.20558897665548 | 0.964390462081331 | -3.32395342207889 | 0.000887510176512782 | 0.0323598422407214 | 6302 | 0 | 1 | 4 | 18 | 1 | 1 | 5 | 13 | 0 | 0 | 0 | 1 |
| LTR33_dup4275 | 3.94725964209277 | -2.82840414072524 | 0.807481846145826 | -3.5027464137125 | 0.000460487633583864 | 0.0204005195421236 | 6303 | 1 | 1 | 9 | 6 | 1 | 1 | 17 | 12 | 0 | 1 | 0 | 1 |
| MLT1F2_dup498 | 3.67884221058961 | -2.8578778517704 | 0.825794632005957 | -3.46076099432647 | 0.00053865080905549 | 0.0226969728017563 | 6304 | 0 | 1 | 7 | 10 | 1 | 1 | 14 | 8 | 1 | 1 | 0 | 4 |
| L1MCa_dup3374 | 3.70157017923431 | -3.42739232303071 | 0.913488180795458 | -3.7519832167354 | 0.000175441211670852 | 0.010162009384672 | 6305 | 0 | 0 | 10 | 8 | 1 | 2 | 10 | 13 | 0 | 0 | 0 | 2 |
| MLT1K_dup1452 | 4.43385604426241 | -3.94257109080668 | 0.889566441305924 | -4.43201418998991 | 9.33568722389946e-06 | 0.00116014302465417 | 6306 | 0 | 0 | 18 | 14 | 1 | 1 | 14 | 9 | 0 | 0 | 0 | 1 |
| L1P3_dup1548 | 3.52607771352891 | -3.26648453853639 | 0.996256499766954 | -3.27875857201483 | 0.00104264795272902 | 0.0361231668258677 | 6307 | 0 | 0 | 8 | 5 | 3 | 1 | 12 | 10 | 0 | 0 | 0 | 6 |
| L2a_dup77187 | 3.3525886002568 | -4.30241449144517 | 1.09855020044738 | -3.91644777789219 | 8.98632704633367e-05 | 0.0062992343273615 | 6308 | 0 | 0 | 7 | 6 | 1 | 0 | 10 | 14 | 0 | 0 | 0 | 3 |
| L2_dup4735 | 4.2569594572749 | -2.60967951413876 | 0.738417842635882 | -3.53415012944861 | 0.000409088641721522 | 0.0187880729478999 | 6309 | 0 | 1 | 13 | 10 | 4 | 2 | 14 | 9 | 0 | 0 | 1 | 1 |
| MER112_dup2273 | 4.13150874318463 | -3.42436452816084 | 0.803035744468687 | -4.2642741020532 | 2.00553126684985e-05 | 0.00204632287707526 | 6310 | 0 | 1 | 14 | 15 | 2 | 1 | 12 | 6 | 0 | 0 | 2 | 2 |
| L1MA7_dup8419 | 2.92587955598722 | -3.14096357027575 | 0.909379963081536 | -3.45396170774671 | 0.000552415806615511 | 0.023152740137846 | 6311 | 0 | 0 | 10 | 7 | 2 | 1 | 10 | 6 | 0 | 0 | 1 | 1 |
| MIR3_dup84354 | 3.68553473543204 | -3.21598378284684 | 0.817230655555541 | -3.93522166720565 | 8.31199431384189e-05 | 0.00595232743482503 | 6312 | 1 | 0 | 10 | 9 | 2 | 1 | 14 | 8 | 0 | 0 | 2 | 1 |
| L1ME1_dup15516 | 3.72055303919271 | -2.75588574149191 | 0.868154362723644 | -3.17441904322858 | 0.00150136766151295 | 0.0465343792609563 | 6313 | 0 | 0 | 16 | 12 | 2 | 4 | 6 | 5 | 0 | 0 | 1 | 3 |
| L1M5_dup54042 | 2.90272313046004 | -3.72739828598391 | 0.996809186693513 | -3.73932978923274 | 0.000184511532697292 | 0.0105158757267521 | 6314 | 0 | 0 | 12 | 11 | 1 | 1 | 7 | 3 | 0 | 0 | 2 | 2 |
| MLT1F2_dup5879 | 3.05901718584334 | -2.95771820811202 | 0.90084081665059 | -3.28328618491011 | 0.0010260444581671 | 0.0357038863180798 | 6315 | 0 | 0 | 12 | 11 | 2 | 2 | 8 | 3 | 0 | 0 | 2 | 1 |

| | | | | | | | | | | | | | | | | | | | |
|---|---|---|---|---|---|---|---|---|---|---|---|---|---|---|---|---|---|---|---|
| L1ME3D_dup4123 | 3.6976239170742 | -3.39372286810844 | 0.881786046193717 | -3.84869196190806 | 0.000118750221182193 | 0.0076969043281792 | 6316 | 0 | 1 | 15 | 13 | 2 | 1 | 5 | 4 | 0 | 0 | 4 | 4 |
| MER2_dup6107 | 3.01015772453293 | -2.94244461730111 | 0.876288954226691 | -3.35784743503672 | 0.000785519560697291 | 0.0296882070323522 | 6317 | 1 | 1 | 15 | 8 | 1 | 1 | 7 | 3 | 0 | 0 | 1 | 2 |
| AluSx3_dup15643 | 4.31480602372325 | -3.68442635911428 | 0.891350377957341 | -4.1335331764347 | 3.57228701779177e-05 | 0.00315403119382368 | 6318 | 0 | 1 | 6 | 16 | 1 | 2 | 13 | 6 | 0 | 0 | 2 | 12 |
| MER5A_dup34108 | 2.94809103832697 | -3.45703836849577 | 0.988079442878418 | -3.49874536244263 | 0.000467452773471196 | 0.020599396213219[9] | 6319 | 0 | 1 | 3 | 16 | 1 | 1 | 8 | 3 | 0 | 0 | 2 | 6 |
| MER1B_dup4960 | 2.76177537381686 | -2.79548982345791 | 0.872801014447492 | -3.2028947918072 | 0.00136053675037599 | 0.043465934157487[3] | 6320 | 0 | 1 | 8 | 7 | 1 | 2 | 6 | 7 | 0 | 0 | 2 | 1 |
| L1M4b_dup3010 | 3.12395601868988 | -2.82015039836078 | 0.883585549543841 | -3.19171177009029 | 0.00141432389862876 | 0.0446152345728647 | 6321 | 0 | 1 | 4 | 5 | 1 | 3 | 7 | 10 | 0 | 0 | 3 | 5 |
| HERVL-int_dup280 | 2.77356394686425 | -2.91205190669725 | 0.894371427995157 | -3.2559760023025 | 0.00113003302145348 | 0.0382212988376362 | 6322 | 0 | 1 | 4 | 6 | 2 | 1 | 7 | 8 | 0 | 0 | 3 | 3 |
| L1MB7_dup10759 | 3.02344270791414 | -3.07093336089739 | 0.871057214505308 | -3.52552428216949 | 0.000422645428582411 | 0.0192398678303942 | 6323 | 0 | 1 | 5 | 9 | 1 | 2 | 7 | 7 | 0 | 0 | 4 | 3 |
| MIRc_dup57645 | 2.8612805951012 | -2.89802706094715 | 0.913314360286381 | -3.17308824536432 | 0.0015082667352308 | 0.0466714374640312 | 6324 | 0 | 1 | 9 | 6 | 1 | 1 | 4 | 7 | 1 | 0 | 1 | 5 |
| L3b_dup6721 | 2.50617988152738 | -3.66003995962671 | 1.07642323747383 | -3.4001866851334 | 0.000673398603337109 | 0.0265946064845895 | 6325 | 0 | 0 | 9 | 4 | 1 | 1 | 4 | 6 | 0 | 0 | 1 | 6 |
| MIR_dup170856 | 2.63060162683829 | -3.23588848651907 | 0.956119443207675 | -3.38439774392938 | 0.000713345844133796 | 0.0277032463241815 | 6326 | 0 | 1 | 8 | 9 | 1 | 1 | 5 | 4 | 0 | 0 | 1 | 5 |
| AluSc8_dup10055 | 4.17807764444321 | -3.50159253948325 | 0.873028414279856 | -4.01085747291701 | 6.04986252544815e-05 | 0.00470635401913803 | 6327 | 0 | 0 | 15 | 17 | 2 | 1 | 9 | 4 | 0 | 1 | 5 | 2 |
| MIR_dup170850 | 4.04448599504015 | -3.8089683157995 | 0.907575403329729 | -4.19686155202652 | 2.70639261099709e-05 | 0.00255039721273788 | 6328 | 0 | 0 | 17 | 12 | 1 | 1 | 12 | 4 | 0 | 1 | 4 | 2 |
| L1MB5_dup3648 | 4.56710749994104 | -2.58013618794912 | 0.779413995401633 | -3.31035393663874 | 0.000931780683674135 | 0.0334838602028075 | 6329 | 0 | 0 | 12 | 22 | 4 | 3 | 8 | 6 | 0 | 1 | 3 | 2 |
| L1ME1_dup22782 | 4.23483524355688 | -2.91489146980194 | 0.81790415049801 | -3.56385460084425 | 0.000365448436375248 | 0.0172824167455316 | 6330 | 0 | 0 | 12 | 17 | 2 | 2 | 6 | 5 | 0 | 2 | 7 | 3 |
| L1ME4a_dup33789 | 4.03816087872973 | -3.15616271589428 | 0.83996637922352 | -3.75748695895649 | 0.000171628308323118 | 0.0100290445609658 | 6331 | 0 | 0 | 10 | 13 | 3 | 1 | 10 | 6 | 0 | 1 | 7 | 2 |
| L1P4a_dup304 | 3.05423289755767 | -3.30144567522472 | 0.922933362180442 | -3.57712247764564 | 0.000347397424766546 | 0.016602094322856[1] | 6332 | 0 | 0 | 6 | 15 | 1 | 1 | 9 | 5 | 0 | 1 | 2 | 1 |
| L1PA4_dup2366 | 3.52811350025276 | -3.38942138389667 | 0.96634301027172 | -3.5074723445701 | 0.000452385373078827 | 0.0201825948340671 | 6333 | 0 | 0 | 9 | 25 | 1 | 1 | 7 | 3 | 0 | 1 | 1 | 1 |
| L1ME3D_dup4118 | 3.63140977594353 | -2.9755313099046 | 0.863238775117721 | -3.4469388953234 | 0.000566976802853713 | 0.0236021452644908 | 6334 | 0 | 0 | 8 | 15 | 3 | 1 | 9 | 5 | 0 | 1 | 1 | 6 |
| L1P5_dup618 | 4.32495523463462 | -2.58905999557579 | 0.782414553979331 | -3.30906420695771 | 0.000936083673946102 | 0.0335817797964463 | 6335 | 0 | 0 | 14 | 16 | 3 | 2 | 12 | 5 | 0 | 2 | 1 | 3 |
| AluSx3_dup14943 | 3.52206209940797 | -2.63357807729771 | 0.779317756075614 | -3.37933796165448 | 0.00072660626105865 | 0.0280807298024787 | 6336 | 0 | 1 | 9 | 7 | 1 | 3 | 9 | 8 | 0 | 1 | 3 | 3 |
| L2a_dup86152 | 3.70362678929802 | -2.44797060136909 | 0.769448066539172 | -3.18146306141178 | 0.00146533207932783 | 0.0457331987077308 | 6337 | 0 | 1 | 12 | 13 | 1 | 4 | 8 | 5 | 0 | 1 | 2 | 2 |
| L1MEd_dup19218 | 4.62534094162803 | -2.52641722963434 | 0.748104398909906 | -3.3770917980374 | 0.000732565971700499 | 0.0282597361859763 | 6338 | 0 | 2 | 12 | 11 | 1 | 3 | 9 | 12 | 0 | 2 | 5 | 1 |
| MIRc_dup77443 | 5.39644311540133 | -3.00676361389545 | 0.741535932680972 | -4.05477803755875 | 5.01819302559272e-05 | 0.00409064391116028 | 6339 | 0 | 1 | 21 | 20 | 1 | 4 | 11 | 8 | 0 | 1 | 3 | 1 |
| ERV3-16A3_I-int_dup4 | 3.50135970982394 | -3.60172786124761 | 0.919650214158587 | -3.91641061546746 | 8.98771172509406e-05 | 0.0062992343273615 | 6340 | 0 | 1 | 11 | 5 | 0 | 1 | 12 | 8 | 0 | 1 | 2 | 4 |
| L1PA10_dup1265 | 5.10476193455432 | -4.02066000222231 | 0.837419222764384 | -4.80125114509529 | 1.57677369137546e-06 | 0.000294123595882378 | 6341 | 0 | 1 | 15 | 13 | 0 | 1 | 19 | 12 | 0 | 1 | 3 | 1 |
| LTR1C_dup31 | 3.83520485404678 | -3.28973504813372 | 0.854350669776912 | -3.85056764688056 | 0.00011784437426418 | 0.00765136822309103 | 6342 | 0 | 2 | 13 | 16 | 0 | 1 | 11 | 4 | 0 | 1 | 1 | 3 |
| MIRb_dup112370 | 4.28923962215599 | -2.6312162880579 | 0.754120159591575 | -3.48912073837537 | 0.00048461220429912 | 0.0210807194815646 | 6343 | 0 | 3 | 13 | 15 | 0 | 3 | 13 | 6 | 0 | 1 | 2 | 1 |
| MER44A_dup2056 | 4.12685544508864 | -3.47955540314173 | 0.833987606174312 | -4.17219078243049 | 3.01684863838118e-05 | 0.0027673364152207 | 6344 | 0 | 1 | 10 | 14 | 0 | 2 | 8 | 10 | 0 | 1 | 4 | 3 |
| L2_dup49992 | 5.23417305703962 | -3.54142897166088 | 0.788999615497618 | -4.48850532002772 | 7.17246334082015e-06 | 0.000949198532866959 | 6345 | 0 | 2 | 19 | 17 | 0 | 2 | 8 | 10 | 0 | 1 | 3 | 6 |
| L2a_dup19336 | 5.35201648984245 | -3.81337120762546 | 0.801841344695973 | -4.75576774987992 | 1.97693646417789e-06 | 0.000349483004387276 | 6346 | 0 | 2 | 19 | 21 | 0 | 1 | 10 | 10 | 0 | 1 | 3 | 3 |
| L1PA3_dup9975 | 5.63984343774257 | -1.99726655803209 | 0.625930342533059 | -3.19087671952347 | 0.00141841789547876 | 0.0446792014336328 | 6347 | 3 | 1 | 14 | 17 | 5 | 2 | 13 | 13 | 2 | 0 | 0 | 2 |
| L1PA5_dup10415 | 6.64835828827622 | -2.11317557950557 | 0.617340371371368 | -3.42303156816286 | 0.000619268850310888 | 0.025094098408732 | 6348 | 3 | 3 | 22 | 12 | 4 | 1 | 16 | 16 | 1 | 2 | 0 | 4 |
| L3_dup40944 | 5.45050394456942 | -1.95885099254701 | 0.620500582985874 | -3.15688823872 | 0.00159462451462007 | 0.0484405112671756 | 6349 | 4 | 0 | 15 | 14 | 2 | 3 | 13 | 9 | 2 | 2 | 1 | 6 |
| MLT1C_dup5386 | 5.53001220868372 | -2.37245746680331 | 0.648263021143276 | -3.65971432801958 | 0.000252496579011663 | 0.0132306872912374 | 6350 | 3 | 0 | 16 | 16 | 2 | 2 | 17 | 10 | 2 | 1 | 1 | 2 |
| L2a_dup90817 | 5.78226166032298 | -2.46472495181794 | 0.70249591205854 | -3.50852568607197 | 0.000450597720417077 | 0.0201257241515209 | 6351 | 1 | 1 | 21 | 22 | 1 | 3 | 11 | 11 | 1 | 2 | 0 | 1 |
| HERVK9-int_dup103 | 4.92845637122597 | -2.07845785485254 | 0.631721583056762 | -3.29014855689328 | 0.00100134501314952 | 0.0351299650698384 | 6352 | 3 | 2 | 9 | 16 | 3 | 1 | 15 | 11 | 1 | 1 | 0 | 2 |
| Tigger6a_dup436 | 5.12845059639319 | -2.8174699579822 | 0.697525224851762 | -4.03923737464973 | 5.36252738933296e-05 | 0.00431078358845345 | 6353 | 1 | 1 | 8 | 21 | 2 | 1 | 17 | 12 | 1 | 1 | 1 | 1 |
| L1MEf_dup6281 | 6.75559159348771 | -3.32237426021974 | 0.774773669933126 | -4.28818684623925 | 1.80137536330558e-05 | 0.00188960146141805 | 6354 | 0 | 0 | 13 | 38 | 3 | 2 | 26 | 9 | 0 | 1 | 0 | 1 |
| L2c_dup62513 | 4.74930273495444 | -3.07909684324623 | 0.927095723778247 | -3.32122861132161 | 0.000896220949019597 | 0.0325544037700857 | 6355 | 0 | 0 | 9 | 14 | 5 | 1 | 26 | 6 | 0 | 0 | 0 | 4 |
| L1ME3A_dup13841 | 3.9984712905818 | -2.74808405268389 | 0.861963405234562 | -3.18816789204185 | 0.00143177377015908 | 0.0449867742657167 | 6356 | 0 | 0 | 7 | 13 | 4 | 1 | 15 | 8 | 0 | 1 | 0 | 4 |
| L1PA13_dup5037 | 3.59152886407453 | -3.17152772953481 | 0.913318473910244 | -3.47253211243649 | 0.000515573328287015 | 0.0220407134697417 | 6357 | 0 | 0 | 9 | 7 | 2 | 1 | 18 | 6 | 0 | 1 | 0 | 4 |
| L1ME3D_dup3749 | 4.41199799939545 | -2.28305942794816 | 0.721359684144245 | -3.16493904238158 | 0.00155115444670655 | 0.0475632726773244 | 6358 | 2 | 0 | 10 | 19 | 2 | 3 | 15 | 7 | 1 | 0 | 0 | 0 |
| L2_dup4739 | 4.07330536129953 | -2.77596318386992 | 0.790454400656818 | -3.51185746016882 | 0.000444986593102003 | 0.0199402598648304 | 6359 | 2 | 0 | 8 | 14 | 2 | 1 | 18 | 9 | 0 | 0 | 0 | 0 |
| AluSx_dup20521 | 6.51900773981039 | -2.28371104976172 | 0.713586652230678 | -3.20032758827932 | 0.00137271469557823 | 0.0437609821749809 | 6360 | 1 | 0 | 19 | 26 | 5 | 7 | 13 | 13 | 0 | 0 | 1 | 0 |
| Charlie19a_dup1397 | 5.61562640893742 | -2.69922169542995 | 0.686582633358639 | -3.93138649928537 | 8.44573673170928e-05 | 0.0060222633274297[2] | 6361 | 3 | 0 | 9 | 26 | 4 | 2 | 15 | 13 | 0 | 0 | 1 | 1 |
| MIR_dup170843 | 6.25413274426963 | -3.06713228253982 | 0.654911040259115 | -4.68328077249441 | 2.82319376989215e-06 | 0.000459799907440628 | 6362 | 2 | 0 | 16 | 21 | 3 | 3 | 15 | 17 | 0 | 0 | 2 | 1 |
| L1MA3_dup8108 | 6.06471598814346 | -3.03287799293307 | 0.777784903611951 | -3.89937883706499 | 9.64397820267917e-05 | 0.0066379694458047[4] | 6363 | 1 | 0 | 10 | 30 | 0 | 4 | 16 | 14 | 1 | 1 | 0 | 3 |
| L1M5_dup28503 | 6.23841973258786 | -4.31573371043162 | 0.823076311180244 | -5.24341868646798 | 1.57628293251801e-07 | 4.75374168138062e-05 | 6364 | 0 | 0 | 15 | 23 | 0 | 2 | 16 | 18 | 0 | 1 | 2 | 3 |
| L1MC3_dup5606 | 5.3225182806028 | -4.00193704205173 | 0.868377168222237 | -4.60852402446807 | 4.05537521749092e-06 | 0.000608423903846844 | 6365 | 0 | 0 | 9 | 27 | 0 | 3 | 17 | 12 | 0 | 0 | 1 | 2 |
| AluJr_dup29924 | 5.74259649905507 | -2.36997050299107 | 0.696778627594445 | -3.40132491028484 | 0.000670600715850569 | 0.026501703692705[5] | 6366 | 3 | 2 | 8 | 20 | 0 | 7 | 15 | 16 | 0 | 0 | 1 | 2 |
| LTR12C_dup1843 | 5.91241167748612 | -2.55432785773682 | 0.648928616803574 | -3.93622317092235 | 8.27740025957923e-05 | 0.00593245265712602 | 6367 | 2 | 3 | 16 | 20 | 0 | 5 | 17 | 13 | 0 | 0 | 1 | 0 |
| L1ME1_dup30805 | 4.81954240050733 | -3.70329591281771 | 0.851226352476202 | -4.35054189998335 | 1.35801501455874e-05 | 0.00153362240462361 | 6368 | 1 | 0 | 15 | 22 | 0 | 2 | 19 | 6 | 0 | 0 | 0 | 1 |
| L1MB8_dup10312 | 3.79759628999366 | -3.51011789533582 | 0.882229526188518 | -3.97869011537227 | 6.9295988834665e-05 | 0.00520641091229972 | 6369 | 2 | 0 | 8 | 9 | 0 | 1 | 19 | 9 | 0 | 0 | 1 | 1 |
| LTR1D_dup141 | 3.48672603742985 | -2.7899989190778 | 0.873551959328731 | -3.19385571663274 | 0.00140386262743222 | 0.0444265855495394 | 6370 | 2 | 0 | 8 | 10 | 0 | 2 | 16 | 8 | 0 | 0 | 0 | 0 |
| L1PA5_dup8933 | 7.59817752246556 | -4.2603586051086 | 0.97266410178324 | -4.38009236415515 | 1.18629014915832e-05 | 0.0013809833455559 | 6371 | 0 | 0 | 39 | 39 | 0 | 1 | 8 | 4 | 1 | 2 | 4 | 5 |
| L1M1_dup3589 | 8.50126809537257 | -3.39704224352804 | 0.950998187180107 | -3.57208067199468 | 0.000354156272848587 | 0.0168668370030781 | 6372 | 0 | 1 | 39 | 44 | 0 | 1 | 11 | 5 | 4 | 3 | 3 | 3 |
| MIRb_dup212393 | 6.93957062405563 | -3.23408708924338 | 0.979572426096653 | -3.30152932349311 | 0.000961592829210616 | 0.0342064833815741 | 6373 | 0 | 1 | 38 | 29 | 0 | 1 | 12 | 3 | 3 | 2 | 1 | 3 |
| LTR16A_dup3574 | 5.81959399472937 | -2.4800403648389 | 0.768742093943351 | -3.22610194547465 | 0.00125488650710431 | 0.041100238204086[5] | 6374 | 0 | 3 | 24 | 20 | 1 | 2 | 14 | 1 | 2 | 2 | 3 | 7 |
| L1PA4_dup5929 | 5.80491906357717 | -2.71994886094033 | 0.823859228820178 | -3.3014728315121 | 0.000961786489965899 | 0.034208258317104 | 6375 | 0 | 1 | 24 | 29 | 1 | 5 | 6 | 2 | 1 | 1 | 4 | 5 |
| SVA_B_dup275 | 5.99886025789772 | -3.20359864476709 | 0.798703082661306 | -4.01100072644341 | 6.04619242622735e-05 | 0.00470635401913803 | 6376 | 0 | 1 | 27 | 22 | 1 | 2 | 7 | 2 | 1 | 2 | 6 | 10 |
| AluSx1_dup81355 | 5.92896117915504 | -3.00564768165176 | 0.816287086300708 | -3.6820963262728 | 0.000231323969652738 | 0.012380254899458 | 6377 | 0 | 1 | 27 | 27 | 1 | 2 | 10 | 3 | 1 | 2 | 4 | 2 |
| X1_LINE_dup55 | 6.00330769766638 | -2.93824263147095 | 0.73061374189327 | -4.02160876943945 | 5.78020058724458e-05 | 0.00453470362325255 | 6378 | 0 | 2 | 16 | 30 | 1 | 2 | 14 | 3 | 2 | 1 | 6 | 5 |
| L1PA7_dup10171 | 5.1517684187698 | -4.19583560821558 | 0.999630050396578 | -4.19738843040081 | 2.70010601553186e-05 | 0.00254767794676055 | 6379 | 0 | 0 | 12 | 37 | 0 | 2 | 6 | 4 | 1 | 0 | 6 | 3 |
| MER5A_dup17291 | 5.87378470214734 | -3.70035690153333 | 0.858732287061662 | -4.30909255106141 | 1.63925762979897e-05 | 0.00176415556752666 | 6380 | 0 | 0 | 22 | 26 | 0 | 4 | 9 | 7 | 1 | 0 | 6 | 3 |
| MER30_dup3180 | 6.5921581514997 | -3.09770431335337 | 0.883728549732632 | -3.50526676352209 | 0.000456149980099803 | 0.020280094014355 | 6381 | 1 | 0 | 21 | 40 | 0 | 2 | 6 | 4 | 3 | 2 | 2 | 9 |
| AluSx4_dup1189 | 4.87485831856369 | -3.05894779965793 | 0.781976025322731 | -3.91181788264603 | 9.16039804235391e-05 | 0.00640140990480453 | 6382 | 2 | 0 | 17 | 23 | 0 | 2 | 8 | 6 | 1 | 1 | 3 | 2 |
| HAL1_dup13799 | 4.87818726643127 | -3.19322759698794 | 0.884142062191587 | -3.61166800397739 | 0.000304233891290054 | 0.0151035337986445 | 6383 | 1 | 0 | 18 | 25 | 0 | 1 | 10 | 4 | 2 | 1 | 1 | 3 |
| L1M4_dup1623 | 4.21544667074644 | -2.42787715672278 | 0.737073185499272 | -3.29394313141131 | 0.000987924837817016 | 0.0348204360034377 | 6384 | 1 | 3 | 7 | 19 | 0 | 2 | 7 | 5 | 2 | 0 | 5 | 5 |
| L2a_dup124338 | 4.31103876443408 | -3.25294474743369 | 0.817037009169882 | -3.98139216574622 | 6.85128023282074e-05 | 0.00516060375523508 | 6385 | 0 | 3 | 11 | 18 | 0 | 1 | 8 | 6 | 1 | 0 | 5 | 4 |
| MSTA_dup16927 | 5.50516018511894 | -2.3068864893768 | 0.658847574746553 | -3.50139634385726 | 0.000462826974304856 | 0.0204601860529882 | 6386 | 2 | 4 | 19 | 13 | 0 | 1 | 7 | 6 | 3 | 1 | 8 | 7 |
| AluJo_dup58251 | 3.97574138056672 | -2.53834447908363 | 0.754202895555557 | -3.36559895757739 | 0.000763776654061117 | 0.0290882405923526 | 6387 | 1 | 2 | 13 | 8 | 0 | 2 | 5 | 8 | 1 | 1 | 6 | 3 |
| MLT1B_dup13198 | 4.30211315443054 | -2.4066765261835 | 0.71180359499857 | -3.3810963348511 | 0.000721972261427988 | 0.0279652306231066 | 6388 | 1 | 1 | 11 | 8 | 0 | 2 | 8 | 8 | 3 | 1 | 7 | 4 |
| L1ME4b_dup7251 | 7.81177430158527 | -3.42086226436619 | 0.680131800729131 | -5.02970491998591 | 4.91235244102002e-07 | 0.000118068142821308 | 6389 | 2 | 2 | 32 | 28 | 0 | 2 | 10 | 12 | 1 | 1 | 6 | 6 |
| MLT1B_dup13202 | 7.94721790676709 | -3.69482658606104 | 0.73357982912032 | -5.03670689867758 | 4.73609142843806e-07 | 0.000114642320552505 | 6390 | 2 | 1 | 25 | 38 | 0 | 2 | 8 | 7 | 1 | 1 | 12 | 10 |
| AluJb_dup18988 | 6.46142238545411 | -2.18223532901439 | 0.651134615871918 | -3.35143498106335 | 0.000803939149540712 | 0.0301535474269569 | 6391 | 2 | 3 | 16 | 27 | 0 | 3 | 12 | 10 | 1 | 4 | 4 | 3 |
| LTR22C0_dup99 | 5.08093748266937 | -2.35867739020765 | 0.709339990514257 | -3.32517188055006 | 0.000883640415897887 | 0.0322782225708344 | 6392 | 4 | 2 | 12 | 26 | 0 | 1 | 8 | 4 | 1 | 2 | 5 | 4 |
| L1PB1_dup3752 | 4.82987062003214 | -2.92390182821031 | 0.913914433321823 | -3.19931683055136 | 0.00137753691126727 | 0.0438559536912501 | 6393 | 0 | 0 | 14 | 16 | 0 | 1 | 10 | 5 | 1 | 7 | 3 | 7 |
| L1PB3_dup266 | 4.51328104646219 | -3.64246780205906 | 0.878824118582616 | -4.14470623306709 | 3.40249618561848e-05 | 0.00303631442785359 | 6394 | 0 | 0 | 5 | 13 | 0 | 1 | 16 | 9 | 1 | 2 | 7 | 5 |

| | | | | | | | | | | | | | | | | | | | |
|---|---|---|---|---|---|---|---|---|---|---|---|---|---|---|---|---|---|---|---|
| MER45B_dup154 | 4.66485166988488 | -2.5437383622236 | 0.798690821606312 | -3.18488493095198 | 0.00144811577109144 | 0.0453590776452584 | 6395 | 0 | 1 | 5 | 16 | 0 | 1 | 11 | 9 | 2 | 5 | 6 | 4 |
| L2a_dup99862 | 4.71377828534566 | -2.70760381812449 | 0.747302778390259 | -3.62316840833491 | 0.000291016209217343 | 0.0146397765155243 | 6396 | 1 | 0 | 10 | 17 | 1 | 0 | 9 | 4 | 3 | 3 | 9 | 5 |
| L1M5_dup4794 | 3.22681043977563 | -2.74719613932722 | 0.853394336375626 | -3.21914034606157 | 0.00128575538851224 | 0.0417723036148189 | 6397 | 1 | 0 | 8 | 12 | 1 | 0 | 4 | 4 | 2 | 1 | 5 | 4 |
| L2c_dup104395 | 3.7678923408786 | -2.40556932233025 | 0.753827208762716 | -3.19114154327037 | 0.00141711836434313 | 0.04467326233182 | 6398 | 1 | 1 | 9 | 11 | 1 | 0 | 6 | 6 | 3 | 1 | 5 | 4 |
| L2b_dup64694 | 4.29256836602327 | -2.31948703926224 | 0.721240487117045 | -3.21596898772798 | 0.0013004880178662 | 0.0420929939025606 | 6399 | 1 | 1 | 10 | 11 | 1 | 1 | 6 | 12 | 2 | 2 | 3 | 3 |
| L1MD1_dup4876 | 4.43005488544186 | -2.44027941421946 | 0.727730675725309 | -3.35327270873569 | 0.00079861976789766 | 0.0300297691675374 | 6400 | 2 | 0 | 10 | 11 | 2 | 0 | 8 | 11 | 1 | 3 | 3 | 5 |
| MER57A-int_dup137 | 4.28884047269195 | -2.57306072746353 | 0.749662822106994 | -3.43229069334359 | 0.00059850566503288 | 0.0245326492630342 | 6401 | 1 | 0 | 6 | 10 | 2 | 1 | 13 | 12 | 1 | 2 | 4 | 2 |
| L1MA3_dup708 | 3.46946652998406 | -3.15044007952221 | 0.929234121853771 | -3.39036202548961 | 0.000698003803480251 | 0.0272721203650101 | 6402 | 0 | 1 | 10 | 16 | 0 | 0 | 6 | 4 | 2 | 1 | 2 | 4 |
| LTR12__dup310 | 4.57583701532945 | -2.81207051154706 | 0.831531036137497 | -3.38179862126285 | 0.000720129147205557 | 0.0279165607789139 | 6403 | 0 | 1 | 10 | 17 | 0 | 0 | 9 | 4 | 4 | 3 | 6 | 6 |
| L1M1_dup3588 | 6.18059549539852 | -3.26733385474283 | 0.993028588745622 | -3.29027169184532 | 0.00100090688846999 | 0.0351242695261776 | 6404 | 1 | 0 | 18 | 31 | 0 | 0 | 10 | 8 | 5 | 3 | 2 | 3 |
| MSTA_dup14133 | 5.23798677599739 | -3.47553751503502 | 1.02066764123159 | -3.40516087180079 | 0.000661250873790476 | 0.0262542967486982 | 6405 | 0 | 0 | 16 | 14 | 0 | 0 | 8 | 11 | 5 | 4 | 3 | 4 |
| L1MB7_dup22102 | 4.34354852017748 | -2.69921697820378 | 0.814885915928488 | -3.31238634199275 | 0.000925037045709016 | 0.0333215828208519 | 6406 | 5 | 0 | 9 | 14 | 0 | 0 | 15 | 6 | 1 | 1 | 5 | 2 |
| MSTA_dup1574 | 5.02298199982399 | -2.65512131417609 | 0.727630012425533 | -3.64899917380444 | 0.00026326395065815 | 0.0136367637314779 | 6407 | 3 | 0 | 11 | 9 | 0 | 1 | 19 | 9 | 2 | 2 | 6 | 3 |
| L1MC4_dup568 | 4.87289921796083 | -4.23367916705886 | 0.946697595380018 | -4.47205019609182 | 7.74732354918948e-06 | 0.00100812484955221 | 6408 | 0 | 0 | 12 | 7 | 0 | 0 | 16 | 9 | 1 | 2 | 8 | 8 |
| L2a_dup86881 | 5.81969133217534 | -3.97735377691229 | 0.939587422168733 | -4.23308537669848 | 2.30506947458866e-05 | 0.0022758591751057 | 6409 | 0 | 0 | 14 | 17 | 0 | 0 | 18 | 14 | 1 | 3 | 3 | 5 |
| MLT1B_dup9158 | 4.78351555056181 | -4.53329518453009 | 0.973923862198419 | -4.65467102766861 | 3.24498388990314e-06 | 0.000511006576218989 | 6410 | 0 | 0 | 12 | 9 | 0 | 0 | 13 | 16 | 1 | 1 | 4 | 3 |
| LTR12__dup311 | 5.23434487001929 | -2.55166863701969 | 0.773833305993329 | -3.29743966466298 | 0.000975706352041585 | 0.0345484261373344 | 6411 | 0 | 1 | 12 | 17 | 1 | 0 | 13 | 2 | 3 | 6 | 7 | 8 |
| L1MC1_dup9424 | 5.3238158077379 | -2.71585783252854 | 0.777474237266292 | -3.49318048412495 | 0.000477303794862675 | 0.0208734660489605 | 6412 | 0 | 0 | 12 | 13 | 2 | 0 | 14 | 9 | 3 | 4 | 7 | 4 |
| LTR87_dup48 | 4.80740175014495 | -2.42346938367465 | 0.740963164795933 | -3.27070156630808 | 0.00107281058153707 | 0.0368782124580207 | 6413 | 0 | 1 | 8 | 16 | 2 | 0 | 11 | 9 | 2 | 4 | 4 | 5 |
| ERVL-E-int_dup1786 | 5.59028523932045 | -2.89361868461375 | 0.835924872286618 | -3.46157744618657 | 0.000537019584949055 | 0.022656323078065 2 | 6414 | 1 | 0 | 27 | 16 | 3 | 0 | 7 | 2 | 1 | 3 | 4 | 11 |
| L2a_dup124344 | 4.94400272436083 | -3.18136712611694 | 0.806304383936764 | -3.94561556342281 | 7.95952315434627e-05 | 0.00575839676644596 | 6415 | 1 | 0 | 14 | 18 | 2 | 0 | 12 | 2 | 1 | 2 | 4 | 12 |
| L2c_dup82766 | 4.20503845309556 | -2.90793916639388 | 0.83320685939643 | -3.49005668112283 | 0.000482918109288725 | 0.0210153400367565 | 6416 | 0 | 0 | 16 | 14 | 1 | 1 | 7 | 2 | 2 | 2 | 5 | 6 |
| L2_dup34395 | 4.70859742752227 | -3.33343937929003 | 0.851483233002052 | -3.91486203144296 | 9.04559176646093e-05 | 0.00632861622599916 | 6417 | 0 | 0 | 14 | 21 | 1 | 1 | 9 | 3 | 2 | 1 | 3 | 9 |
| THE1D-int_dup1463 | 4.97395455029585 | -3.46505314602172 | 0.846509090488283 | -4.09334428295745 | 4.25195660870237e-05 | 0.00361205307823739 | 6418 | 0 | 0 | 15 | 16 | 1 | 1 | 13 | 2 | 2 | 1 | 4 | 13 |
| L1MA2_dup3481 | 4.44991583901095 | -3.08913440047286 | 0.925033076347396 | -3.33948534323841 | 0.000839337799676775 | 0.0311231217207246 | 6419 | 0 | 0 | 12 | 25 | 1 | 1 | 10 | 4 | 1 | 2 | 0 | 5 |
| L1PA15-16_dup838 | 6.90564940141629 | -2.49183898469033 | 0.731011210495776 | -3.40875618446447 | 0.000652597775917798 | 0.0260279970480282 | 6420 | 1 | 0 | 19 | 27 | 2 | 3 | 19 | 5 | 2 | 4 | 1 | 11 |
| AluJo_dup14234 | 4.45035298313453 | -2.1693803182513 | 0.685876261745425 | -3.16293249853937 | 0.00156188542170081 | 0.0477290937039548 | 6421 | 2 | 1 | 19 | 10 | 2 | 3 | 9 | 3 | 1 | 1 | 3 | 5 |
| MIRb_dup101576 | 3.99591411328359 | -2.35836550178013 | 0.710267379569039 | -3.32039112258132 | 0.000898914142544307 | 0.0326155877648441 | 6422 | 2 | 0 | 13 | 7 | 2 | 1 | 10 | 3 | 1 | 2 | 5 | 7 |
| MIRb_dup75801 | 4.75265006755916 | -2.76019194182004 | 0.816024263148896 | -3.38248758825987 | 0.000718325237743041 | 0.0278602472730245 | 6423 | 1 | 0 | 25 | 11 | 2 | 1 | 9 | 2 | 2 | 1 | 3 | 6 |
| MLT1E2_dup1548 | 5.2876789406557 | -2.58605111760278 | 0.77397436500844 | -3.34126197781056 | 0.000833984901919015 | 0.0309650765041459 | 6424 | 1 | 0 | 17 | 22 | 1 | 2 | 12 | 2 | 1 | 4 | 5 | 5 |
| MLT1D-int_dup366 | 5.51971781099271 | -3.35736730021282 | 0.742973628104954 | -4.51882432055657 | 6.2183964137272e-06 | 0.000850858282381099 | 6425 | 2 | 0 | 15 | 22 | 0 | 2 | 15 | 5 | 1 | 1 | 5 | 7 |
| MamRep1527_dup107 | 6.45318258585181 | -3.08100487585105 | 0.712207650283096 | -4.3259923908798 | 1.51846664142995e-05 | 0.00166657317094875 | 6426 | 2 | 0 | 15 | 32 | 1 | 2 | 18 | 4 | 2 | 1 | 4 | 8 |
| MER44A_dup219 | 4.10264405508361 | -2.59949850902946 | 0.715415519400305 | -3.63355062692585 | 0.00027954756457072 | 0.0142070108499241 | 6427 | 1 | 0 | 10 | 9 | 1 | 2 | 12 | 6 | 1 | 2 | 3 | 7 |
| MER67D_dup25 | 5.200907630414 | -2.72533266756619 | 0.702753102743162 | -3.87807987887636 | 0.000105284151094321 | 0.00709107997151219 | 6428 | 1 | 0 | 17 | 16 | 2 | 1 | 12 | 6 | 1 | 3 | 3 | 7 |
| AluJo_dup34580 | 4.65096104659187 | -2.65369731865641 | 0.76212080908685 | -3.48199037083896 | 0.000497701588868846 | 0.0215085547157622 | 6429 | 1 | 0 | 14 | 22 | 2 | 1 | 12 | 4 | 1 | 2 | 2 | 2 |
| L1M7_dup1837 | 4.82746684662896 | -2.71806958187441 | 0.777417745928189 | -3.49627931200516 | 0.000471794570636332 | 0.0207219823447359 | 6430 | 1 | 0 | 12 | 21 | 1 | 2 | 18 | 3 | 2 | 1 | 3 | 2 |
| AluSq_dup5570 | 4.75919242323139 | -2.7730240998539 | 0.709709218041386 | -3.90726797589966 | 9.33456079345543e-05 | 0.00649260860146024 | 6431 | 1 | 1 | 14 | 14 | 1 | 1 | 16 | 6 | 1 | 2 | 4 | 2 |
| MIRb_dup75803 | 5.22490391553012 | -2.64254601475376 | 0.766536896708034 | -3.44738267147012 | 0.000566046206572212 | 0.023571653949778 | 6432 | 1 | 0 | 16 | 16 | 1 | 1 | 17 | 6 | 2 | 3 | 4 | 2 |
| AluSz6_dup34538 | 6.59053056486539 | -2.21603265470574 | 0.623727532525316 | -3.55288573799778 | 0.000381029929410566 | 0.0178364961027393 | 6433 | 2 | 0 | 14 | 24 | 6 | 5 | 10 | 4 | 1 | 1 | 10 | 12 |
| THE1D_dup10297 | 7.48269886587578 | -1.99712988396746 | 0.611759501335268 | -3.26456700649257 | 0.00109631528413911 | 0.0373892697815194 | 6434 | 4 | 0 | 22 | 23 | 6 | 8 | 15 | 4 | 0 | 2 | 9 | 8 |
| L1P4_dup3901 | 6.07347253111311 | -2.34669007778868 | 0.66051128378811 | -3.55283874081627 | 0.000381098006212866 | 0.0178364961027393 | 6435 | 2 | 0 | 8 | 14 | 5 | 5 | 10 | 11 | 0 | 1 | 16 | 6 |
| MIR1_Amn_dup11093 | 7.51489021772166 | -3.11229600301629 | 0.652581043850688 | -4.76920994310768 | 1.84949834133477e-06 | 0.000331298768141008 | 6436 | 3 | 0 | 16 | 26 | 5 | 1 | 11 | 8 | 0 | 1 | 18 | 11 |
| HAL1_dup15223 | 4.35642448665898 | -3.09842451433247 | 0.807408779550217 | -3.83749173009799 | 0.000124297419000477 | 0.00794224679894756 | 6437 | 1 | 0 | 11 | 15 | 3 | 2 | 5 | 9 | 0 | 0 | 7 | 3 |
| MER1B_dup5503 | 6.12506209556619 | -2.56505635958227 | 0.791269890342123 | -3.24169590033713 | 0.00118820738028734 | 0.0395584078303696 | 6438 | 2 | 0 | 14 | 28 | 4 | 7 | 4 | 8 | 0 | 0 | 10 | 4 |
| MER58B_dup5108 | 4.76178221607563 | -2.67389951355586 | 0.818859503321948 | -3.26539473830174 | 0.00109311625274543 | 0.0373174519190478 | 6439 | 0 | 2 | 12 | 21 | 3 | 4 | 6 | 3 | 0 | 0 | 10 | 3 |
| MLT2B2_dup63 | 3.77833622661261 | -2.43502639179937 | 0.767785652982147 | -3.17149243716852 | 0.00151657816956302 | 0.0468535664540892 | 6440 | 2 | 1 | 11 | 13 | 3 | 2 | 3 | 4 | 0 | 0 | 6 | 5 |
| AluSx1_dup114061 | 4.63404167869983 | -2.40378055877852 | 0.688326988016325 | -3.49220733841328 | 0.00047904623679085 | 0.0209304325852801 | 6441 | 2 | 3 | 14 | 14 | 2 | 3 | 7 | 5 | 0 | 0 | 8 | 3 |
| MLT1F_dup3612 | 4.50666001192497 | -2.23685849729007 | 0.694947332811774 | -3.21874535188111 | 0.00128752769749684 | 0.0418195153023083 | 6442 | 3 | 1 | 9 | 19 | 2 | 5 | 8 | 4 | 0 | 0 | 6 | 4 |
| AluSx1_dup81353 | 6.67040004425897 | -2.51897460425579 | 0.667484599260736 | -3.77383179633754 | 0.000160759169740404 | 0.00957017798425071 | 6443 | 2 | 3 | 24 | 30 | 4 | 3 | 8 | 4 | 0 | 1 | 7 | 4 |
| MIR_dup151753 | 6.95835950299662 | -2.36029980857686 | 0.639174474894217 | -3.69273164258866 | 0.000221858101159912 | 0.0119896088436512 | 6444 | 2 | 6 | 22 | 28 | 5 | 3 | 10 | 4 | 0 | 0 | 7 | 7 |
| AluSx1_dup115290 | 7.17990536433972 | -3.10907101337672 | 0.612920000894419 | -5.07255597604863 | 3.92507623822672e-07 | 9.89360371378278e-05 | 6445 | 3 | 1 | 19 | 28 | 4 | 2 | 15 | 10 | 0 | 0 | 6 | 8 |
| MER65-int_dup106 | 6.52720009217213 | -3.72247872435827 | 0.748226285073999 | -4.97507077553432 | 6.52238048198083e-07 | 0.000146828225748856 | 6446 | 1 | 2 | 26 | 22 | 2 | 1 | 10 | 8 | 0 | 0 | 9 | 5 |
| MLT1A_dup8917 | 6.79230037985839 | -3.02653197636968 | 0.658546284360622 | -4.59577716592558 | 4.3113926714378e-06 | 0.000639575212342854 | 6447 | 2 | 1 | 25 | 16 | 5 | 2 | 12 | 8 | 0 | 0 | 11 | 7 |
| L1ME3G_dup10796 | 5.77874861098641 | -2.37312306794522 | 0.616619323730516 | -3.84860314397536 | 0.00011879327748005 | 0.0076969043281792 | 6448 | 6 | 2 | 14 | 19 | 1 | 4 | 16 | 6 | 0 | 0 | 6 | 4 |
| L2a_dup154992 | 6.79876269321041 | -2.03777582274164 | 0.588275411618033 | -3.46398265590738 | 0.000532240826202199 | 0.0225065888806661 | 6449 | 8 | 1 | 14 | 25 | 3 | 7 | 17 | 7 | 0 | 0 | 6 | 4 |
| LTR85b_dup231 | 6.11469902355451 | -2.7062895799872 | 0.647380799688534 | -4.18036738390951 | 2.91038505152678e-05 | 0.00269390483312078 | 6450 | 1 | 3 | 12 | 25 | 4 | 3 | 8 | 10 | 0 | 0 | 5 | 10 |
| MIRb_dup215002 | 5.18727558230348 | -2.40451861670258 | 0.624380590680506 | -3.85104638515736 | 0.000117614217473755 | 0.00764431373695988 | 6451 | 2 | 3 | 8 | 13 | 3 | 3 | 11 | 10 | 0 | 0 | 4 | 11 |
| MER51B_dup622 | 6.30839572227552 | -2.163972818713 | 0.599170925525118 | -3.61161185652737 | 0.000304299780434569 | 0.0151036536410688 | 6452 | 3 | 6 | 11 | 28 | 2 | 5 | 12 | 7 | 0 | 0 | 6 | 5 |
| LTR43B_dup198 | 5.99025073874559 | -1.72343287731462 | 0.540711671434196 | -3.18734173564878 | 0.0014358701459312 | 0.04507382028281 6 | 6453 | 5 | 6 | 15 | 16 | 4 | 4 | 9 | 10 | 0 | 0 | 3 | 6 |
| MLT1E1-int_dup38 | 5.05039383540648 | -1.97882756566982 | 0.613209826322342 | -3.22699911307295 | 0.00125095847921047 | 0.0410249165514042 | 6454 | 3 | 4 | 11 | 19 | 4 | 3 | 7 | 7 | 0 | 0 | 3 | 6 |
| L2a_dup86156 | 4.12190179045961 | -3.14460069197994 | 0.846866886558736 | -3.71321720318774 | 0.000204641111287344 | 0.0113051483209692 | 6455 | 0 | 1 | 9 | 19 | 3 | 1 | 17 | 3 | 0 | 0 | 3 | 1 |
| L1ME1_dup25579 | 5.5452376016681 | -3.50564141562024 | 0.768277420572831 | -4.56298899557197 | 5.04304589456955e-06 | 0.000725936236194342 | 6456 | 0 | 1 | 15 | 27 | 1 | 3 | 20 | 6 | 0 | 0 | 2 | 1 |
| L2b_dup94897 | 5.36552206927397 | -3.70875727277369 | 0.829343579413647 | -4.47191895474226 | 7.7520808426687e-06 | 0.00100812484955221 | 6457 | 0 | 0 | 19 | 22 | 2 | 2 | 22 | 4 | 0 | 0 | 1 | 2 |
| MIR_dup110266 | 6.21414255911293 | -2.8643425952263 | 0.683563589994277 | -4.19030890052274 | 2.78574948229524e-05 | 0.00261070479052471 | 6458 | 1 | 4 | 18 | 25 | 3 | 2 | 21 | 4 | 0 | 0 | 5 | 2 |
| AluSq2_dup49946 | 6.32192616326449 | -2.97640969599459 | 0.718613665212276 | -4.14187739543662 | 3.44474481319217e-05 | 0.00306038517169789 | 6459 | 0 | 2 | 19 | 31 | 2 | 4 | 15 | 4 | 0 | 1 | 3 | 6 |
| HERV16-int_dup1770 | 6.89767061860108 | -3.08052552749028 | 0.71319825152858 | -4.31931166528783 | 1.56516613527398e-05 | 0.00170290361785049 | 6460 | 0 | 2 | 20 | 28 | 2 | 6 | 20 | 5 | 0 | 0 | 7 | 4 |
| MER44A_dup2381 | 5.97908499928155 | -2.35671544190738 | 0.707422718758821 | -3.3314104557488 | 0.000864070939194004 | 0.0317409449090447 | 6461 | 3 | 0 | 18 | 25 | 5 | 6 | 9 | 5 | 0 | 0 | 3 | 7 |
| L1ME3A_dup15569 | 7.55342940767819 | -3.72903474471579 | 0.722053243299674 | -5.16448721658622 | 2.41098952490555e-07 | 6.66301177111704e-05 | 6462 | 1 | 0 | 22 | 36 | 2 | 4 | 21 | 6 | 0 | 0 | 5 | 7 |
| L2a_dup168121 | 5.25692239958403 | -3.15519433506225 | 0.805341852436824 | -3.91783231619614 | 8.93488227075534e-05 | 0.00627712003268199 | 6463 | 1 | 0 | 16 | 20 | 2 | 4 | 20 | 3 | 0 | 0 | 1 | 6 |
| LOR1-int_dup385 | 6.65414624292901 | -2.81994118175289 | 0.784443379067488 | -3.59483075133493 | 0.000324602527615055 | 0.0158209490036649 | 6464 | 2 | 0 | 33 | 29 | 3 | 4 | 12 | 2 | 0 | 1 | 2 | 3 |
| AluY_dup76548 | 5.59836941414831 | -2.52180486582414 | 0.698897602518858 | -3.60826086215698 | 0.000308256464927194 | 0.015242808147042 | 6465 | 1 | 1 | 18 | 24 | 3 | 2 | 9 | 4 | 0 | 3 | 3 | 8 |
| LTR79_dup1837 | 5.88001457360178 | -2.24081841739405 | 0.652239720944581 | -3.43557490511138 | 0.000591297977164831 | 0.0243252054762211 | 6466 | 2 | 1 | 16 | 25 | 4 | 2 | 16 | 5 | 0 | 3 | 2 | 4 |
| AluSc_dup16510 | 5.04542946075863 | -3.48193285946803 | 0.818016505185854 | -4.25655575088541 | 2.07600286556642e-05 | 0.00209756508641218 | 6467 | 2 | 0 | 21 | 22 | 1 | 1 | 11 | 3 | 0 | 1 | 3 | 4 |
| AluY_dup86855 | 5.80367604342948 | -3.1531214407518 | 0.720678448814796 | -4.37521261519242 | 1.21314174487728e-05 | 0.00140781866315558 | 6468 | 2 | 0 | 21 | 23 | 1 | 2 | 10 | 8 | 0 | 2 | 4 | 4 |
| AluJr_dup38511 | 7.45441762543734 | -2.7607236861656 | 0.765996884729498 | -3.60409257687845 | 0.000313245398119241 | 0.0154126322375684 | 6469 | 0 | 0 | 22 | 38 | 6 | 3 | 10 | 9 | 2 | 0 | 6 | 3 |
| ERVL-E-int_dup7590 | 5.20256728796327 | -2.89600982281139 | 0.763591290563642 | -3.79261767204509 | 0.00014906757396091 | 0.00905788374908434 | 6470 | 0 | 0 | 21 | 20 | 3 | 3 | 12 | 6 | 1 | 0 | 2 | 1 |
| L2a_dup168157 | 5.64613429815815 | -2.93200476698994 | 0.792006817254635 | -3.70199435549465 | 0.000213911378303011 | 0.0116634639829717 | 6471 | 0 | 0 | 20 | 28 | 4 | 3 | 8 | 5 | 1 | 0 | 3 | 4 |
| (CATTC)n | 5.15980780370476 | -2.46091509118237 | 0.777184472538478 | -3.16644912261874 | 0.00154312337438759 | 0.0473964159599885 | 6472 | 0 | 1 | 17 | 18 | 3 | 5 | 5 | 9 | 1 | 0 | 1 | 7 |
| L1P3_dup3407 | 5.19725909735589 | -3.28447006039321 | 0.768118627420449 | -4.27599324263669 | 1.9028699491324e-05 | 0.00196689962170398 | 6473 | 0 | 0 | 10 | 22 | 3 | 2 | 11 | 9 | 1 | 0 | 3 | 8 |

| | | | | | | | | | | | | | | | | | | | |
|---|---|---|---|---|---|---|---|---|---|---|---|---|---|---|---|---|---|---|---|
| MER58A_dup6275 | 5.09262029009984 | -3.0645172031489 | 0.809525781481832 | -3.78557085302376 | 0.000153356130521777 | 0.00924980426695434 | 6474 | 0 | 0 | 16 | 21 | 1 | 3 | 8 | 10 | 2 | 0 | 1 | 4 |
| AluJo_dup34577 | 6.02707811659407 | -2.34255240961144 | 0.74090076832625 | -3.16176269448801 | 0.00156817302230679 | 0.0478446593109133 | 6475 | 1 | 0 | 21 | 27 | 3 | 5 | 14 | 5 | 2 | 0 | 2 | 1 |
| MSTA_dup3997 | 4.5529133068665 | -2.89022635264055 | 0.78389885501691 | -3.68698886870833 | 0.00022692327903759 | 0.0121885974162253 | 6476 | 2 | 0 | 18 | 18 | 2 | 1 | 15 | 3 | 1 | 0 | 1 | 1 |
| L1MA2_dup4476 | 8.39933846137825 | -3.35661703483382 | 0.672256781901408 | -4.99305789870944 | 5.94307315351208e-07 | 0.000137427646704139 | 6477 | 3 | 0 | 30 | 38 | 2 | 2 | 15 | 8 | 2 | 0 | 6 | 7 |
| Cheshire_dup504 | 7.25832722151288 | -3.55336926334121 | 0.676689069373767 | -5.25111077474568 | 1.51184719305248e-07 | 4.58849479844473e-05 | 6478 | 1 | 0 | 25 | 29 | 3 | 2 | 15 | 11 | 1 | 0 | 4 | 5 |
| MIR_dup84431 | 7.59534149202987 | -4.37727817645723 | 0.79972175031883 | -5.4735014706204 | 4.41229219870075e-08 | 1.72395283992126e-05 | 6479 | 1 | 0 | 24 | 42 | 1 | 1 | 14 | 9 | 1 | 0 | 4 | 6 |
| AluJb_dup57362 | 6.89487609936663 | -3.05321112623098 | 0.820049336387196 | -3.72320419120414 | 0.000196710303641396 | 0.0109822688667892 | 6480 | 1 | 0 | 25 | 29 | 1 | 3 | 10 | 9 | 3 | 0 | 8 | 1 |
| MLT2A2_dup2387 | 6.52674609355596 | -2.79980984144598 | 0.674371941742617 | -4.1517294361493 | 3.2997223077585e-05 | 0.00297144635917868 | 6481 | 2 | 0 | 13 | 28 | 1 | 5 | 19 | 7 | 2 | 0 | 7 | 4 |
| L1PA2_dup726 | 5.61868604552661 | -2.67560269770439 | 0.796720683729181 | -3.3582694065137 | 0.000784321303378937 | 0.0296599206088916 | 6482 | 0 | 2 | 16 | 28 | 2 | 0 | 13 | 5 | 2 | 2 | 1 | 5 |
| Charlie1a_dup601 | 5.69090193289626 | -2.63092958345059 | 0.722182751647293 | -3.64302467408624 | 0.000269452997555714 | 0.0138418097919671 | 6483 | 0 | 4 | 12 | 29 | 3 | 0 | 13 | 8 | 1 | 1 | 3 | 2 |
| ERV3-16A3_I-int_dup1: | 5.54856811572338 | -2.7977374012962 | 0.687753935956235 | -4.06793368242422 | 4.74318753367007e-05 | 0.00391339286021726 | 6484 | 0 | 2 | 14 | 23 | 3 | 1 | 20 | 6 | 1 | 1 | 2 | 2 |
| L1MA9_dup3204 | 5.77287163780602 | -2.83598028106989 | 0.701491073347543 | -4.0427888376918 | 5.28191854701527e-05 | 0.00426451396251906 | 6485 | 0 | 3 | 14 | 30 | 2 | 1 | 12 | 9 | 1 | 1 | 1 | 3 |
| L1MA4A_dup2100 | 5.50304336872296 | -2.88922261643813 | 0.727170548318664 | -3.97323932207991 | 7.090174647045e-05 | 0.00529362628354493 | 6486 | 0 | 2 | 16 | 16 | 2 | 0 | 20 | 9 | 2 | 1 | 2 | 2 |
| LTR16A1_dup653 | 6.1597215901848 | -2.63306548261837 | 0.72554815047274 | -3.62907062874154 | 0.000284443416593374 | 0.0143943371341401 | 6487 | 1 | 2 | 15 | 19 | 1 | 0 | 20 | 11 | 3 | 2 | 3 | 3 |
| L1MCa_dup2787 | 4.37337763175069 | -2.77424191961193 | 0.863111636260067 | -3.21423301814462 | 0.0013079348406546 | 0.0422850358144565 | 6488 | 0 | 1 | 11 | 19 | 1 | 0 | 12 | 6 | 1 | 3 | 3 | 1 |
| MLT1J_dup1330 | 4.22001003876699 | -2.5635278768872 | 0.747303322217448 | -3.43037131064871 | 0.000602755809517743 | 0.02467285977146 2 | 6489 | 1 | 1 | 9 | 13 | 2 | 0 | 13 | 7 | 1 | 2 | 5 | 1 |
| AluSx_dup94212 | 4.72884464843871 | -2.55701892030878 | 0.760525224635625 | -3.36217503046512 | 0.000773310891698534 | 0.0293330517194483 | 6490 | 3 | 1 | 10 | 22 | 4 | 0 | 9 | 4 | 0 | 1 | 2 | 9 |
| AluJb_dup59481 | 4.52634846029707 | -2.38430811085752 | 0.709349424394338 | -3.36126037304296 | 0.000775876474450198 | 0.0294209886198449 | 6491 | 5 | 1 | 7 | 13 | 1 | 0 | 11 | 10 | 1 | 1 | 2 | 7 |
| LTR16A_dup320 | 4.3485624441469 | -2.9374178745943 | 0.741894975357822 | -3.95934461367333 | 7.51557243498698e-05 | 0.00552285484439596 | 6492 | 2 | 1 | 11 | 15 | 1 | 0 | 10 | 8 | 1 | 1 | 2 | 5 |
| L2a_dup87261 | 5.50157310797139 | -2.82922866787248 | 0.686830209693815 | -4.11925484339678 | 3.80099584669324e-05 | 0.00331205458821924 | 6493 | 1 | 1 | 14 | 14 | 3 | 0 | 14 | 11 | 0 | 3 | 6 | 4 |
| Helitron1Nb_Mam_dup | 5.61927639691268 | -2.85080747719655 | 0.749984413118558 | -3.80115563381168 | 0.000144022799946052 | 0.0088406983078069 | 6494 | 3 | 1 | 21 | 22 | 2 | 0 | 12 | 6 | 0 | 2 | 5 | 1 |
| LOR1-int_dup1336 | 4.73500866279637 | -3.14454815459407 | 0.82101081301247 | -3.83009347106652 | 0.00012809459267416 | 0.0081278922529188 8 | 6495 | 1 | 2 | 13 | 18 | 2 | 0 | 11 | 5 | 0 | 1 | 9 | 1 |
| L1PA13_dup4165 | 5.93452991020777 | -2.35509102905299 | 0.695699571244561 | -3.38521270731831 | 0.000711231173821361 | 0.0276572336797022 | 6496 | 2 | 2 | 18 | 21 | 1 | 0 | 13 | 7 | 2 | 4 | 5 | 3 |
| L2c_dup64097 | 5.5675862598282 | -1.87955802335487 | 0.595345451234987 | -3.15708807290944 | 0.00159353207921089 | 0.0484258728122495 | 6497 | 2 | 3 | 15 | 13 | 3 | 0 | 9 | 9 | 2 | 4 | 6 | 5 |
| LTR81A_dup826 | 6.85589529317532 | -2.54174301582427 | 0.657409091681004 | -3.86630341440061 | 0.00011049745456499 | 0.00732413496475379 | 6498 | 3 | 3 | 18 | 33 | 2 | 0 | 9 | 6 | 1 | 3 | 9 | 5 |
| L2a_dup83623 | 6.30566651180909 | -2.48423172235073 | 0.66380361967351 | -3.74241966859505 | 0.000182256804853418 | 0.0104323360473012 | 6499 | 4 | 2 | 19 | 27 | 4 | 0 | 13 | 3 | 1 | 1 | 6 | 6 |
| MADE1_dup6961 | 6.03699945883469 | -2.41414972515815 | 0.627960669910269 | -3.84442822749251 | 0.000120833842702031 | 0.00779604741127 8 | 6500 | 5 | 3 | 18 | 23 | 2 | 0 | 11 | 6 | 1 | 1 | 4 | 7 |
| L2c_dup34029 | 4.22942621790895 | -2.49959498298404 | 0.698573336242492 | -3.5781425561258 | 0.000346044705564108 | 0.0165754515831735 | 6501 | 0 | 3 | 6 | 9 | 1 | 3 | 10 | 8 | 0 | 1 | 6 | 8 |
| MER5A_dup12060 | 5.13860942219399 | -2.82503120307903 | 0.708739045405512 | -3.9859962864931 | 6.71975626242008e-05 | 0.00509703259042974 | 6502 | 0 | 4 | 17 | 16 | 1 | 2 | 8 | 8 | 0 | 1 | 5 | 5 |
| L2c_dup70109 | 5.05597591659153 | -3.89863793572583 | 0.85166643233281 | -4.57765832691917 | 4.70209879120586e-06 | 0.000688478574665287 | 6503 | 0 | 1 | 9 | 28 | 1 | 2 | 10 | 6 | 0 | 0 | 6 | 6 |
| LTR33A_dup1985 | 4.13893658068562 | -3.60114503656702 | 0.841321246954047 | -4.28034481430815 | 1.86603969134588e-05 | 0.0019423301854636 3 | 6504 | 0 | 1 | 11 | 13 | 2 | 1 | 12 | 4 | 0 | 0 | 5 | 7 |
| MLT1B_dup17807 | 4.99191299016597 | -3.56663138258956 | 0.790487615490664 | -4.51193834374713 | 6.42378633361741e-06 | 0.000873436721679388 | 6505 | 0 | 1 | 13 | 16 | 3 | 1 | 9 | 8 | 0 | 0 | 5 | 10 |
| MER112_dup4185 | 5.34350690617318 | -3.61562497909415 | 0.746083087120152 | -4.84614252958113 | 1.25885094200605e-06 | 0.000245566297657304 | 6506 | 0 | 2 | 13 | 14 | 1 | 2 | 22 | 9 | 0 | 0 | 4 | 4 |
| MIRc_dup64205 | 4.55007679983551 | -2.92575500941194 | 0.792263262844593 | -3.69290758088053 | 0.000221704610622525 | 0.0119867587546438 | 6507 | 0 | 5 | 11 | 12 | 1 | 1 | 12 | 8 | 0 | 0 | 7 | 2 |
| THE1D_dup12527 | 4.41544834702976 | -2.86388224818833 | 0.762849535874586 | -3.75419019545573 | 0.000173902783609646 | 0.0101073040447031 | 6508 | 0 | 2 | 15 | 13 | 1 | 4 | 10 | 6 | 0 | 0 | 5 | 2 |
| MIR_dup44935 | 5.10845907150826 | -2.51161642553839 | 0.78694507671358 | -3.1916032006037 | 0.00141485556468393 | 0.0446260898984997 | 6509 | 1 | 3 | 19 | 21 | 1 | 1 | 9 | 2 | 0 | 3 | 7 | 2 |
| L1PREC2_dup7666 | 4.96900559929956 | -2.27653046596246 | 0.68094609887224 | -3.3431874706864 | 0.000828219271975181 | 0.030802109016452 | 6510 | 1 | 4 | 14 | 15 | 4 | 1 | 15 | 3 | 0 | 1 | 6 | 3 |
| MLT1I_dup8493 | 5.42866857552958 | -2.60685054572225 | 0.721602818349899 | -3.61258365326702 | 0.000303161258704472 | 0.0150691474542917 | 6511 | 2 | 2 | 22 | 17 | 4 | 1 | 9 | 2 | 0 | 1 | 8 | 5 |
| L2a_dup143049 | 4.74596142088858 | -2.07458941830518 | 0.651932054346987 | -3.18221723333302 | 0.00146152152048551 | 0.0456523962667711 | 6512 | 3 | 3 | 9 | 21 | 2 | 3 | 12 | 3 | 0 | 1 | 5 | 3 |
| L1M5_dup28504 | 5.46948991276773 | -2.29193846383494 | 0.647656270523627 | -3.53881922888188 | 0.000401920985977492 | 0.0185447917256933 | 6513 | 4 | 2 | 22 | 16 | 3 | 2 | 9 | 6 | 0 | 1 | 5 | 2 |
| MIRc_dup21182 | 5.6896888860383 | -2.14477881006222 | 0.600874960003115 | -3.56942617487522 | 0.000357764016676674 | 0.0170019006814755 | 6514 | 3 | 4 | 13 | 23 | 3 | 1 | 10 | 9 | 0 | 2 | 4 | 3 |
| L2c_dup67458 | 4.99492565693132 | -1.98487274252017 | 0.618265771368026 | -3.21038756217779 | 0.00132556115909278 | 0.0426751421406429 | 6515 | 1 | 2 | 10 | 17 | 1 | 2 | 8 | 5 | 4 | 2 | 7 | 6 |
| MER21B_dup2753 | 4.4939599538211 | -2.29115766763302 | 0.683928297755582 | -3.34999688586042 | 0.000808124689158104 | 0.0302723610337554 | 6516 | 1 | 2 | 16 | 11 | 1 | 1 | 7 | 6 | 1 | 3 | 5 | 4 |
| MIR1_Amn_dup11095 | 4.8306527226323 | -2.26995609343401 | 0.666903285007339 | -3.40372606413091 | 0.000664733818036248 | 0.0263530078417949 | 6517 | 1 | 2 | 14 | 12 | 1 | 3 | 8 | 7 | 1 | 2 | 9 | 2 |
| MIRb_dup104840 | 4.27122845831102 | -2.85109065382122 | 0.771843348771715 | -3.69387215470386 | 0.000220864875569873 | 0.0119549394181868 | 6518 | 0 | 2 | 15 | 17 | 1 | 1 | 10 | 4 | 1 | 1 | 3 | 2 |
| MLT1D_dup11763 | 4.19662047888349 | -2.74715237661463 | 0.780272184816241 | -3.5207616394292 | 0.000430309238689234 | 0.019502926116028 3 | 6519 | 0 | 1 | 14 | 15 | 1 | 1 | 7 | 7 | 2 | 1 | 3 | 2 |
| L2b_dup33056 | 5.18285982702784 | -2.42446828360381 | 0.683800886053606 | -3.54557639958036 | 0.000391755406185688 | 0.0182295197674648 | 6520 | 0 | 1 | 10 | 19 | 2 | 2 | 13 | 10 | 2 | 2 | 4 | 2 |
| MLT1C_dup1567 | 4.51031125447091 | -2.39974094400444 | 0.734225860606661 | -3.26839610637199 | 0.00108158878237491 | 0.0370631709764255 | 6521 | 0 | 2 | 10 | 11 | 1 | 1 | 12 | 9 | 3 | 1 | 5 | 2 |
| MER63B_dup1691 | 6.18675418360938 | -2.21529423877229 | 0.555184612944054 | -3.99019386907167 | 6.60193100434628e-05 | 0.00502688983923257 | 6522 | 2 | 3 | 12 | 15 | 2 | 1 | 13 | 11 | 3 | 2 | 9 | 6 |
| AluSz_dup80704 | 6.77913770978158 | -2.19108983536532 | 0.530636195456133 | -4.12917523178355 | 3.64066860416394e-05 | 0.00320371357378448 | 6523 | 3 | 5 | 18 | 19 | 3 | 1 | 12 | 11 | 1 | 2 | 6 | 7 |
| L1MA8_dup8085 | 6.16583755061174 | -2.22252645384488 | 0.559924509195696 | -3.96933232488328 | 7.2074303516826e-05 | 0.00534710773564649 | 6524 | 2 | 3 | 19 | 12 | 3 | 1 | 12 | 10 | 2 | 2 | 6 | 7 |
| MER117_dup3550 | 5.86421191368056 | -2.02532047272428 | 0.614629498743366 | -3.29518917797654 | 0.000983554425827477 | 0.0347126730062214 | 6525 | 4 | 4 | 13 | 22 | 2 | 1 | 8 | 9 | 2 | 1 | 8 | 2 |
| HAL1_dup11378 | 4.7885178402736 | -2.14192728308575 | 0.614060998037734 | -3.48813438718693 | 0.000486403538006519 | 0.021140517384827 | 6526 | 3 | 3 | 11 | 17 | 1 | 2 | 9 | 7 | 1 | 1 | 4 | 4 |
| MIRb_dup190727 | 5.6194053803734 | -1.98688837696492 | 0.56194548869601 | -3.5357315201079 | 0.000406647742680464 | 0.0187012376845469 | 6527 | 2 | 4 | 14 | 16 | 3 | 3 | 9 | 11 | 1 | 1 | 4 | 4 |
| L2c_dup104399 | 4.52170248153441 | -2.1803169953107 | 0.634031044531485 | -3.43881741141215 | 0.000584261156000495 | 0.0241149358240579 | 6528 | 3 | 1 | 9 | 13 | 2 | 1 | 8 | 8 | 1 | 2 | 3 | 8 |
| AluJb_dup60303 | 4.81114311714668 | -2.10855848302943 | 0.598823515438768 | -3.52116847229095 | 0.000429649549377318 | 0.0194841607920974 | 6529 | 2 | 1 | 7 | 14 | 3 | 1 | 8 | 6 | 2 | 2 | 8 | 9 |
| L2_dup42367 | 4.47226127368458 | -2.15570052227534 | 0.620885333975195 | -3.47197848670954 | 0.000516637736771441 | 0.0220559807059759 | 6530 | 3 | 1 | 8 | 9 | 2 | 1 | 9 | 5 | 2 | 1 | 6 | 12 |
| MIRc_dup48869 | 5.78749295010806 | -2.22465222640551 | 0.60677151358653 | -3.6663755245462 | 0.000246012583279589 | 0.0129565538160791 | 6531 | 1 | 3 | 9 | 10 | 3 | 1 | 18 | 13 | 2 | 2 | 3 | 9 |
| MER41A_dup217 | 4.6291510550675 | -2.37211365757272 | 0.673149301800195 | -3.52390420851511 | 0.000425237951585043 | 0.0193283036538404 | 6532 | 1 | 2 | 5 | 8 | 2 | 1 | 18 | 9 | 1 | 2 | 6 | 5 |
| MLT2B3_dup301 | 4.75598779467726 | -2.25159502229539 | 0.647832677619756 | -3.47558111851986 | 0.000509747802883667 | 0.0218584340791056 | 6533 | 1 | 3 | 4 | 10 | 1 | 2 | 12 | 11 | 1 | 2 | 5 | 9 |
| MER3_dup9090 | 5.60562446887719 | -2.09249994801612 | 0.654661202686567 | -3.19630969336356 | 0.00139197617435087 | 0.0441561342197164 | 6534 | 2 | 2 | 14 | 21 | 1 | 4 | 9 | 11 | 3 | 0 | 2 | 3 |
| MIRb_dup196941 | 4.51165224988651 | -2.27985444591576 | 0.685799866080987 | -3.32437283626785 | 0.000886176369304127 | 0.0323457773232766 | 6535 | 2 | 1 | 8 | 14 | 1 | 3 | 10 | 12 | 2 | 0 | 2 | 2 |
| L1MB5_dup696 | 4.58472783598137 | -2.25634757979587 | 0.672642302009862 | -3.35445387995058 | 0.000795218080762606 | 0.0299444471348535 | 6536 | 3 | 3 | 10 | 10 | 1 | 2 | 6 | 13 | 1 | 0 | 4 | 4 |
| L1MC2_dup4283 | 4.12355637336184 | -2.38709933127177 | 0.733165120839555 | -3.2558822882058 | 0.00113040605757424 | 0.0382284820601989 | 6537 | 1 | 2 | 18 | 6 | 2 | 2 | 8 | 8 | 1 | 0 | 2 | 2 |
| AluSz6_dup10070 | 4.02275626835644 | -2.19251437617353 | 0.674153207111502 | -3.25224941904177 | 0.00114495501954516 | 0.0385689940720975 | 6538 | 1 | 3 | 8 | 9 | 2 | 2 | 13 | 8 | 1 | 0 | 2 | 3 |
| L1PA2_dup765 | 4.06471970440891 | -2.33832678555104 | 0.68881726077247 | -3.39469830202678 | 0.00068704264756002 | 0.0269754731657361 | 6539 | 2 | 2 | 10 | 10 | 1 | 1 | 10 | 8 | 2 | 0 | 3 | 3 |
| L1ME1_dup21940 | 4.92816996801134 | -1.99322702709037 | 0.616077843863991 | -3.23534931006285 | 0.00121493996498077 | 0.0401793057550123 | 6540 | 5 | 2 | 9 | 8 | 3 | 2 | 17 | 4 | 1 | 0 | 8 | 7 |
| L2_dup42719 | 5.67224307614752 | -2.83575989935782 | 0.703788583140533 | -4.02927806345442 | 5.59484082632436e-05 | 0.00443313351735124 | 6541 | 3 | 2 | 13 | 28 | 2 | 1 | 9 | 4 | 1 | 0 | 9 | 5 |
| L2c_dup110866 | 5.05317565464003 | -2.46882957960191 | 0.679090845434102 | -3.63549235894019 | 0.000277450218082248 | 0.0141366530175158 | 6542 | 3 | 1 | 12 | 24 | 3 | 2 | 7 | 5 | 1 | 0 | 5 | 5 |
| L4_C_Mam_dup2666 | 5.48098612850597 | -2.94942258606492 | 0.842860991076019 | -3.4992989559282 | 0.000466483238377743 | 0.0205627678101205 | 6543 | 0 | 0 | 20 | 22 | 3 | 2 | 13 | 5 | 1 | 1 | 6 | 0 |
| L1PA5_dup5825 | 5.02509746840944 | -3.01502147392097 | 0.801096309010338 | -3.76361923030913 | 0.000167471822593459 | 0.00985635673975343 | 6544 | 0 | 0 | 14 | 26 | 1 | 2 | 12 | 5 | 1 | 2 | 3 | 2 |
| MLT1H1_dup2113 | 4.87560387138085 | -2.36373283936226 | 0.730047325551578 | -3.23778028715655 | 0.00120463543415603 | 0.03997185650755 7 | 6545 | 0 | 0 | 14 | 19 | 3 | 2 | 11 | 5 | 1 | 3 | 3 | 4 |
| L1M3_dup3891 | 5.60879082460881 | -2.73134297072859 | 0.712740260397503 | -3.83217158127883 | 0.000127017099616596 | 0.00807891226553612 | 6546 | 0 | 0 | 15 | 22 | 3 | 2 | 13 | 10 | 2 | 1 | 3 | 2 |
| MER5A_dup17290 | 5.98968353550114 | -2.87731711548507 | 0.762170290943572 | -3.77516304384278 | 0.000159903040563087 | 0.00953788423004232 | 6547 | 0 | 0 | 23 | 28 | 4 | 2 | 10 | 6 | 1 | 1 | 3 | 2 |
| L1ME4b_dup12381 | 5.73146885824378 | -3.26700090786413 | 0.801369020765633 | -4.07677464839209 | 4.56647102824617e-05 | 0.00381175243206187 | 6548 | 0 | 0 | 14 | 35 | 3 | 1 | 8 | 8 | 1 | 1 | 3 | 3 |
| MIR_dup84423 | 6.64877274994123 | -3.26205974109217 | 0.766258960859585 | -4.25712442883905 | 2.07073118304286e-05 | 0.00209532321696719 | 6549 | 0 | 0 | 18 | 41 | 2 | 3 | 10 | 8 | 1 | 1 | 3 | 3 |
| L1PREC2_dup7699 | 5.7014701119015 | -3.95927653989209 | 0.902375746169796 | -4.38761409168802 | 1.14600880921027e-05 | 0.00134860639028856 | 6550 | 0 | 0 | 17 | 29 | 3 | 0 | 6 | 9 | 1 | 0 | 7 | 3 |
| MIRb_dup104852 | 5.34259997187376 | -4.22806223943084 | 1.02838790962456 | -4.11134961803898 | 3.93353039582705e-05 | 0.00339610966460618 | 6551 | 0 | 0 | 21 | 31 | 3 | 0 | 4 | 5 | 0 | 0 | 6 | 2 |
| L1MC3_dup1338 | 5.61585412232925 | -2.87523810143951 | 0.844204912595954 | -3.40585331658172 | 0.0006595760665624 | 0.0262052908222169 | 6552 | 0 | 0 | 18 | 21 | 5 | 0 | 3 | 7 | 1 | 2 | 10 | 6 |

| | | | | | | | | | | | | | | | | | | | |
|---|---|---|---|---|---|---|---|---|---|---|---|---|---|---|---|---|---|---|---|
| AluSz6_dup21888 | 5.88995775618628 | -3.93124924640237 | 0.904337524249644 | -4.34710397499456 | 1.37946835900003e-05 | 0.00155049115800955 | 6553 | 0 | 0 | 22 | 28 | 1 | 0 | 8 | 7 | 2 | 1 | 5 | 4 |
| L1MA9_dup9885 | 4.77599138269295 | -3.00863083969249 | 0.889880527284109 | -3.38093794329307 | 0.000722388557064506 | 0.0279768013670689 | 6554 | 0 | 0 | 20 | 19 | 2 | 0 | 6 | 5 | 1 | 3 | 3 | 4 |
| LTR33_dup7906 | 7.02184144048874 | -2.30958612636411 | 0.687214317579275 | -3.36079454004926 | 0.000777186154860564 | 0.0294612612401548 | 6555 | 1 | 2 | 21 | 25 | 4 | 4 | 6 | 10 | 3 | 0 | 12 | 2 |
| GSATII_dup151 | 6.84769421133693 | -1.9259064768383 | 0.549896109074005 | -3.50230970006575 | 0.00046124314155385 | 0.0204149852232464 | 6556 | 1 | 6 | 14 | 19 | 6 | 3 | 9 | 11 | 2 | 0 | 9 | 8 |
| MER39B_dup534 | 6.16119167419108 | -1.8202850503011 | 0.54938913476417 | -3.3132891335437 | 0.000922056060854858 | 0.0332525825530458 | 6557 | 2 | 5 | 14 | 20 | 3 | 5 | 11 | 9 | 2 | 0 | 6 | 3 |
| AluSx1_dup76149 | 4.60932269487003 | -2.36648259923759 | 0.727069913979268 | -3.25482124034782 | 0.00113463758823967 | 0.0383280061855872 | 6558 | 0 | 3 | 15 | 12 | 3 | 1 | 9 | 4 | 2 | 0 | 8 | 3 |
| HERVK14-int_dup169 | 5.0566318090645 | -2.28626218959805 | 0.678569254880338 | -3.36923928273353 | 0.000753759621739295 | 0.0288520740932599 | 6559 | 0 | 5 | 12 | 17 | 2 | 3 | 12 | 5 | 1 | 0 | 7 | 3 |
| L1ME3_dup857 | 5.89833869257821 | -3.2486459316513 | 0.701407807540365 | -4.63160788449642 | 3.62836846473324e-06 | 0.000557003717877042 | 6560 | 0 | 1 | 13 | 21 | 4 | 1 | 11 | 8 | 1 | 0 | 9 | 9 |
| MLT1N2_dup5118 | 6.29363569496278 | -2.9643768551792 | 0.64565080606289 | -4.59130047905563 | 4.40492618144311e-06 | 0.000651015257869355 | 6561 | 1 | 1 | 18 | 17 | 4 | 2 | 10 | 11 | 1 | 0 | 10 | 6 |
| AluSg_dup26186 | 6.11298708455944 | -2.48632384327709 | 0.672300261948906 | -3.69823423252816 | 0.000217104503647259 | 0.0118024099211448 | 6562 | 1 | 2 | 23 | 22 | 5 | 3 | 9 | 4 | 1 | 0 | 4 | 8 |
| AluY_dup48532 | 6.69760949908029 | -2.66600537968251 | 0.723154003417612 | -3.68663571947747 | 0.000227238275769455 | 0.0122027620776887 | 6563 | 0 | 1 | 24 | 28 | 2 | 6 | 8 | 6 | 2 | 0 | 6 | 6 |
| MIR_dup132697 | 4.82783715781997 | -3.05593842202329 | 0.861932936480377 | -3.54544801884695 | 0.000391946284447119 | 0.0182326782024546 | 6564 | 0 | 3 | 21 | 25 | 1 | 1 | 5 | 3 | 1 | 0 | 3 | 2 |
| MIRb_dup170213 | 5.36518176296736 | -2.49521830265148 | 0.759818675101055 | -3.2839654833696 | 0.00102357457681346 | 0.0356596597737123 | 6565 | 2 | 3 | 17 | 32 | 3 | 1 | 8 | 2 | 1 | 0 | 2 | 3 |
| MIRb_dup170214 | 4.21240166869345 | -2.48092494856908 | 0.766328416773383 | -3.23741739738034 | 0.0012061685232399 | 0.0400058602920271 | 6566 | 1 | 2 | 11 | 22 | 3 | 1 | 7 | 3 | 1 | 0 | 4 | 2 |
| L1MA10_dup2242 | 6.88273205029003 | -3.29503528978697 | 0.816152599815989 | -4.03727843362856 | 5.40748756629832e-05 | 0.00432646279909354 | 6567 | 2 | 1 | 25 | 36 | 1 | 0 | 9 | 5 | 3 | 0 | 4 | 7 |
| L2a_dup82414 | 6.81323932959339 | -3.30998137880342 | 0.822356955556952 | -4.02499347325602 | 5.69768957147498e-05 | 0.00449516725057754 | 6568 | 4 | 2 | 27 | 36 | 1 | 0 | 6 | 3 | 1 | 0 | 7 | 6 |
| AluJb_dup123581 | 5.02377603096289 | -2.55939576442508 | 0.744773514292778 | -3.4364752710835 | 0.000589336150275128 | 0.0242822821841326 | 6569 | 4 | 1 | 20 | 13 | 1 | 1 | 11 | 2 | 2 | 0 | 8 | 4 |
| MER47C_dup672 | 6.444651961229 | -3.31185327915933 | 0.811787290300125 | -4.07970575387415 | 4.50927423763503e-05 | 0.00378392824902677 | 6570 | 1 | 1 | 24 | 25 | 1 | 1 | 6 | 2 | 3 | 0 | 14 | 8 |
| L1P2_dup780 | 10.2103706535943 | -4.26538011562889 | 0.963329589212139 | -4.42774743285664 | 9.52223090297692e-06 | 0.00117886407619134 | 6571 | 5 | 2 | 41 | 67 | 0 | 0 | 3 | 7 | 0 | 0 | 3 | 13 |
| THE1D-int_dup173 | 8.54514109067047 | -3.91882965393067 | 0.893576143249947 | -4.38555760864188 | 1.15689033947663e-05 | 0.0013560464468823 | 6572 | 1 | 5 | 40 | 46 | 0 | 0 | 7 | 10 | 0 | 0 | 4 | 1 |
| L1PA15-16_dup70 | 6.66202945225095 | -4.87927892143542 | 0.994541697756048 | -4.90605766700821 | 9.29252141312593e-07 | 0.000194812185036798 | 6573 | 1 | 1 | 23 | 46 | 0 | 0 | 6 | 7 | 0 | 0 | 5 | 2 |
| MER67D_dup38 | 6.2377739662772 | -4.82862380148763 | 0.971838409440707 | -4.96854595844437 | 6.74568135624098e-07 | 0.000150997920601328 | 6574 | 1 | 1 | 26 | 28 | 0 | 0 | 4 | 13 | 0 | 0 | 3 | 5 |
| L1P4_dup283 | 10.0369552507645 | -3.45665544605627 | 0.717738153446144 | -4.81603970676423 | 1.46435396782978e-06 | 0.000276977082058207 | 6575 | 2 | 7 | 40 | 50 | 1 | 1 | 23 | 6 | 0 | 0 | 2 | 6 |
| L1ME2z_dup597 | 11.7036428903729 | -4.90020742725043 | 0.791285352025261 | -6.19271848607909 | 5.91352932983121e-10 | 4.67475298975421e-07 | 6576 | 1 | 2 | 47 | 65 | 1 | 0 | 24 | 12 | 0 | 0 | 1 | 7 |
| L1P4_dup285 | 10.3192895030764 | -4.33007008431222 | 0.729464099509882 | -5.93596050473429 | 2.92130124947773e-09 | 1.78692017251472e-06 | 6577 | 1 | 4 | 37 | 49 | 1 | 0 | 19 | 16 | 0 | 0 | 2 | 9 |
| MLT1D_dup10293 | 10.1052154109878 | -4.59685877345053 | 1.01503167695905 | -4.52878356193014 | 5.93242281364574e-06 | 0.000821059877208587 | 6578 | 0 | 3 | 57 | 64 | 0 | 0 | 9 | 4 | 0 | 0 | 1 | 1 |
| MLT1D_dup13027 | 14.9360689906808 | -4.95385924041658 | 0.950057093636904 | -5.2142753036586 | 1.84537234006309e-07 | 5.39752815438687e-05 | 6579 | 0 | 3 | 80 | 103 | 0 | 1 | 11 | 5 | 0 | 0 | 1 | 3 |
| L1M4c_dup3372 | 8.83873110414047 | -5.45292903648589 | 1.084930209106 | -5.02606434102261 | 5.00647977689319e-07 | 0.000119846261266864 | 6580 | 0 | 1 | 47 | 52 | 0 | 0 | 8 | 8 | 0 | 0 | 0 | 4 |
| MLT1F1_dup2598 | 9.51792673038298 | -5.92442577104515 | 1.06861436075534 | -5.54402597290341 | 2.95594959742026e-08 | 1.25824039876165e-05 | 6581 | 0 | 1 | 47 | 59 | 0 | 0 | 7 | 5 | 0 | 0 | 4 | 8 |
| THE1C_dup1489 | 11.6687333850376 | -5.02317235127719 | 1.08131669854019 | -4.64542197309876 | 3.39382030559373e-06 | 0.000530582106724576 | 6582 | 0 | 0 | 59 | 51 | 0 | 0 | 21 | 13 | 3 | 1 | 3 | 4 |
| HERV16-int_dup300 | 12.284720044734 | -5.16887036881665 | 0.950472486227719 | -5.43821146189209 | 5.38180779627036e-08 | 2.03651646353014e-05 | 6583 | 1 | 0 | 44 | 87 | 0 | 0 | 14 | 13 | 2 | 0 | 3 | 4 |
| LOR1b_dup590 | 11.2131507567806 | -4.76995526881366 | 1.40199914304285 | -3.40225262795891 | 0.000668328278296841 | 0.0264602396851281 | 6584 | 1 | 0 | 51 | 55 | 0 | 0 | 18 | 16 | 4 | 0 | 1 | 2 |
| LTR12C_dup2225 | 11.6009743272715 | -3.85971898382826 | 0.784143579385501 | -4.92220951021872 | 8.55725299946664e-07 | 0.000182615616341802 | 6585 | 4 | 2 | 49 | 60 | 0 | 0 | 23 | 10 | 2 | 0 | 1 | 7 |
| L1MB5_dup9402 | 14.2630756432611 | -4.40641684678188 | 0.71179041749937 | -6.19060995828282 | 5.99318422959926e-10 | 4.70644955345289e-07 | 6586 | 4 | 2 | 70 | 64 | 0 | 0 | 28 | 16 | 1 | 0 | 4 | 2 |
| MLT1F1-int_dup174 | 17.932979102599 | -5.18086573011084 | 0.766668751100238 | -6.75763257948865 | 1.40264675800356e-11 | 1.86454852223416e-08 | 6587 | 3 | 1 | 88 | 90 | 0 | 0 | 30 | 19 | 1 | 0 | 2 | 8 |
| AluJr_dup40173 | 15.8072973026736 | -5.30429336931479 | 0.800245224803853 | -6.62833492147975 | 3.39494497434852e-11 | 4.03906788433166e-08 | 6588 | 0 | 2 | 75 | 76 | 0 | 0 | 35 | 20 | 1 | 0 | 1 | 2 |
| L1PA8_dup1259 | 12.6075172956437 | -4.94532835879203 | 0.702520202064631 | -7.03941088705812 | 1.93054301707179e-12 | 3.35302911489171e-09 | 6589 | 1 | 1 | 45 | 54 | 1 | 0 | 32 | 25 | 1 | 0 | 2 | 4 |
| L1MB7_dup2182 | 11.6250711551385 | -4.78460883978535 | 0.806776205983158 | -5.93052795100062 | 3.01962172817852e-09 | 1.82826186191645e-06 | 6590 | 1 | 0 | 46 | 63 | 4 | 0 | 19 | 10 | 0 | 0 | 5 | 11 |
| L1ME2z_dup596 | 14.1308489873048 | -5.03721808205662 | 0.716167573587688 | -7.0335746378775 | 2.01308208579949e-12 | 3.47104949266409e-09 | 6591 | 3 | 0 | 54 | 62 | 1 | 0 | 35 | 16 | 1 | 0 | 8 | 11 |
| AluSg7_dup4075 | 8.26826447174603 | -5.25082801461545 | 0.966432141179443 | -5.43320921446931 | 5.53494480612602e-08 | 2.080597119808e-05 | 6592 | 2 | 0 | 36 | 44 | 0 | 0 | 15 | 6 | 0 | 0 | 7 | 3 |
| THE1B_dup21628 | 12.1605282698875 | -5.48982000683335 | 0.912043230517943 | -6.01925415719134 | 1.75222560156775e-09 | 1.15494480052809e-06 | 6593 | 3 | 0 | 63 | 55 | 0 | 0 | 17 | 8 | 0 | 0 | 9 | 10 |
| L1MA3_dup779 | 10.1611620822089 | -5.4284586535665 | 0.908925813013692 | -5.97238913874341 | 2.33804021737265e-09 | 1.48353951350121e-06 | 6594 | 0 | 0 | 37 | 61 | 1 | 1 | 11 | 16 | 0 | 0 | 5 | 4 |
| L1ME2z_dup3469 | 10.4222016117027 | -5.63270320304728 | 0.953208750030312 | -5.90920215836055 | 3.43768619589095e-09 | 2.03478527255589e-06 | 6595 | 0 | 0 | 45 | 56 | 1 | 1 | 9 | 12 | 0 | 0 | 8 | 8 |
| L1PA10_dup533 | 11.6409633609983 | -6.13856379685283 | 0.990387510919816 | -6.19814338243389 | 5.71330828488937e-10 | 4.59276639579826e-07 | 6596 | 0 | 1 | 48 | 70 | 0 | 0 | 16 | 12 | 0 | 0 | 3 | 9 |
| L1PA15_dup669 | 13.1166000319338 | -6.80257912823815 | 0.97682201164367 | -6.96399041703784 | 3.30767117805034e-12 | 5.08001000040174e-09 | 6597 | 0 | 0 | 52 | 80 | 0 | 0 | 20 | 14 | 0 | 0 | 6 | 7 |
| L1PA5_dup6227 | 9.9725986900855 | -2.63572280716484 | 0.660488961116006 | -3.99056299549859 | 6.59166376574784e-05 | 0.00502067870167937 | 6598 | 4 | 0 | 34 | 50 | 3 | 6 | 25 | 10 | 0 | 2 | 0 | 2 |
| L1ME1_dup3212 | 8.16879760270501 | -3.56382090816129 | 0.685722716922163 | -5.19717492247792 | 2.0233994882075e-07 | 5.82881131502447e-05 | 6599 | 2 | 0 | 34 | 36 | 3 | 2 | 18 | 12 | 0 | 0 | 0 | 2 |
| MER41-int_dup233 | 10.6216771272403 | -3.34175737693862 | 0.771060887484711 | -4.33397340103686 | 1.46441876430367e-05 | 0.00162146387757562 | 6600 | 3 | 0 | 57 | 46 | 1 | 6 | 11 | 15 | 0 | 0 | 0 | 1 |
| HERV9NC-int_dup102 | 13.8834905977036 | -3.32278692090272 | 0.940162942053742 | -3.5342670640094 | 0.00040890768404019 | 0.018783387603596 | 6601 | 4 | 0 | 81 | 73 | 0 | 2 | 9 | 9 | 1 | 4 | 2 | 2 |
| MER3_dup10448 | 10.5065458921024 | -3.43288045994654 | 1.04600421644688 | -3.28189925620713 | 0.00103110436950776 | 0.0358422300417366 | 6602 | 3 | 0 | 42 | 60 | 0 | 0 | 13 | 12 | 1 | 7 | 2 | 1 |
| L2a_dup79131 | 13.5162247513662 | -2.78751892163125 | 0.848401477791421 | -3.28561299644101 | 0.00101760717408023 | 0.035490037228246 | 6603 | 3 | 2 | 45 | 59 | 0 | 3 | 42 | 18 | 0 | 7 | 1 | 2 |
| Tigger17c_dup119 | 18.8934466352546 | -5.23711988900852 | 0.765034543723363 | -6.84559923728395 | 7.61564042380591e-12 | 1.07863760493031e-08 | 6604 | 2 | 0 | 80 | 100 | 0 | 2 | 44 | 19 | 0 | 1 | 8 | 1 |
| ERV3-16A3_I-int_dup1 | 14.277474553291 | -2.73709739206392 | 0.859042538010707 | -3.186218692269 | 0.00144145591790077 | 0.0451895480686189 | 6605 | 12 | 6 | 74 | 79 | 0 | 1 | 10 | 9 | 1 | 0 | 0 | 2 |
| L1MB3_dup16146 | 12.9836733783324 | -2.66153850173476 | 0.669134631170378 | -3.97758295229689 | 6.96193389800129e-05 | 0.00522581991609458 | 6606 | 6 | 6 | 66 | 61 | 2 | 5 | 11 | 7 | 2 | 0 | 3 | 6 |
| L1PA16_dup3769 | 18.5329412687817 | -3.23454452632875 | 0.732576278918721 | -4.41530065797779 | 1.00869695452195e-05 | 0.00123021735284818 | 6607 | 8 | 6 | 82 | 122 | 1 | 3 | 12 | 9 | 2 | 0 | 4 | 6 |
| L1PREC2_dup6610 | 13.4226364570491 | -2.8036295287872 | 0.736198216096918 | -3.80825362991387 | 0.000139951688470054 | 0.00865409159685431 | 6608 | 11 | 8 | 65 | 61 | 5 | 1 | 13 | 8 | 0 | 0 | 1 | 9 |
| L1MB5_dup9401 | 11.7858258203644 | -3.50378428150039 | 0.693894234536889 | -5.0494500560865 | 4.43083755690235e-07 | 0.000109027928988075 | 6609 | 7 | 3 | 53 | 62 | 2 | 1 | 13 | 10 | 0 | 0 | 3 | 6 |
| L1MC4_dup6651 | 11.0265924584336 | -3.24213315401842 | 0.771930586295863 | -4.20003198678253 | 2.66877274869149e-05 | 0.00252997530063803 | 6610 | 8 | 5 | 45 | 66 | 1 | 1 | 6 | 9 | 0 | 0 | 5 | 4 |
| L1ME1_dup27013 | 8.17552269711516 | -2.09876529468195 | 0.652308343709719 | -3.21744358311613 | 0.00129338460956713 | 0.0419285687068202 | 6611 | 12 | 7 | 20 | 30 | 1 | 0 | 18 | 12 | 0 | 1 | 4 | 4 |
| MER4E1_dup987 | 10.2950116932689 | -2.49142556831037 | 0.729494584638729 | -3.41527630331103 | 0.000637173417797644 | 0.0255929888709112 | 6612 | 10 | 10 | 32 | 36 | 0 | 0 | 20 | 17 | 0 | 1 | 2 | 8 |
| MER41C_dup333 | 8.23795514129839 | -2.24928395163553 | 0.621010272698421 | -3.62197543345283 | 0.000292361891385472 | 0.0146948506446909 | 6613 | 7 | 7 | 26 | 34 | 2 | 0 | 10 | 11 | 1 | 1 | 4 | 6 |
| MLT1A0_dup3769 | 9.10462070584196 | -2.01959500850352 | 0.520346170707163 | -3.88125275479369 | 0.000103919767344158 | 0.00701465888816122 | 6614 | 9 | 6 | 20 | 35 | 3 | 1 | 17 | 17 | 1 | 2 | 5 | 3 |
| MLT1K_dup16894 | 8.97364843806888 | -2.58386469778297 | 0.805803736232251 | -3.20656827661847 | 0.00134328423784309 | 0.0430766996304327 | 6615 | 4 | 4 | 29 | 46 | 0 | 0 | 15 | 8 | 0 | 6 | 3 | 7 |
| L1MC1_dup6612 | 7.62354269499972 | -2.8015414257518 | 0.785861362552576 | -3.56493086344396 | 0.000363952112724073 | 0.0172334696425477 | 6616 | 3 | 3 | 29 | 41 | 0 | 1 | 7 | 4 | 0 | 4 | 6 | 6 |
| L2b_dup43842 | 7.38640657113965 | -2.41161974533532 | 0.660056263100559 | -3.65365784729765 | 0.000258530729948821 | 0.0134579639178303 | 6617 | 7 | 5 | 26 | 28 | 0 | 1 | 11 | 7 | 0 | 2 | 6 | 6 |
| MLT1I_dup10538 | 8.73546879610305 | -2.72330000751787 | 0.707638899239684 | -3.84843175021031 | 0.000118876405705604 | 0.00769901067828675 | 6618 | 7 | 4 | 41 | 32 | 0 | 1 | 16 | 6 | 0 | 2 | 3 | 6 |
| HERVL-int_dup1267 | 9.07360308031848 | -3.25190554943064 | 0.777886951013485 | -4.18043463152819 | 2.90952444325625e-05 | 0.00269390483312078 | 6619 | 0 | 6 | 40 | 42 | 0 | 3 | 12 | 11 | 0 | 1 | 1 | 5 |
| LTR12C_dup763 | 10.0147687743004 | -3.43908391087057 | 0.850166276863755 | -4.0451897522415 | 5.22807606315955e-05 | 0.00422841531925412 | 6620 | 2 | 5 | 41 | 69 | 0 | 2 | 4 | 6 | 0 | 1 | 2 | 6 |
| THE1D-int_dup672 | 10.0949133478473 | -2.53846789150605 | 0.76845742547815 | -3.30332925070841 | 0.000955441354191694 | 0.0340373227927919 | 6621 | 3 | 7 | 38 | 63 | 0 | 2 | 10 | 7 | 1 | 2 | 1 | 4 |
| Tigger3_dup566 | 12.1356115754835 | -3.05658850885371 | 0.616660466896049 | -4.95667984724073 | 7.17079904295384e-07 | 0.000158280421992087 | 6622 | 6 | 1 | 22 | 39 | 3 | 1 | 41 | 35 | 0 | 3 | 3 | 2 |
| L2c_dup14109 | 9.62545000060403 | -2.40969126368903 | 0.485271922677303 | -4.96565152666256 | 6.84708067963279e-07 | 0.000152550136647557 | 6623 | 6 | 4 | 22 | 24 | 3 | 5 | 29 | 26 | 0 | 1 | 1 | 2 |
| THE1D_dup5549 | 11.6133249370281 | -2.39216226189816 | 0.467426747566943 | -5.11772651939556 | 3.09240555820682e-07 | 8.16676507162132e-05 | 6624 | 9 | 5 | 20 | 39 | 6 | 3 | 39 | 25 | 0 | 1 | 1 | 5 |
| L1PA13_dup7681 | 11.5679518283906 | -2.72386967343907 | 0.598429758714761 | -4.55169488778279 | 5.32154717096837e-06 | 0.000755320671976583 | 6625 | 3 | 1 | 19 | 27 | 2 | 6 | 47 | 33 | 1 | 3 | 3 | 3 |
| Tigger5b_dup1422 | 14.5519797969754 | -3.0182725722312 | 0.543482158041788 | -5.55358170929896 | 2.79874748200468e-08 | 1.21081957882379e-05 | 6626 | 3 | 3 | 34 | 53 | 2 | 5 | 51 | 30 | 2 | 2 | 5 | 1 |
| L1MA3_dup1348 | 11.8986445434875 | -4.24510790986621 | 0.602317084006992 | -7.04796198312205 | 1.81557223335324e-12 | 3.17653051939316e-09 | 6627 | 2 | 1 | 31 | 47 | 1 | 1 | 38 | 25 | 1 | 1 | 3 | 6 |
| L1ME3Cz_dup1167 | 11.0782417008375 | -3.77202771054715 | 0.568731206868484 | -6.6323557859898 | 3.30371068669665e-11 | 3.9903794063793e-08 | 6628 | 1 | 2 | 37 | 36 | 1 | 3 | 34 | 21 | 1 | 1 | 5 | 3 |
| Helitron1Na_Mam_dup | 10.5212284195729 | -2.48686309723031 | 0.650396717387429 | -3.82360954590878 | 0.000131512121263742 | 0.00827110282377745 | 6629 | 4 | 1 | 39 | 34 | 2 | 3 | 27 | 19 | 2 | 3 | 1 | 2 |
| Tigger3b_dup3522 | 10.2640049660288 | -2.22086727713614 | 0.578643241028733 | -3.83805965345383 | 0.000124010359517123 | 0.00793432885336416 | 6630 | 7 | 2 | 27 | 40 | 3 | 2 | 32 | 14 | 3 | 2 | 2 | 3 |
| MER41-int_dup811 | 17.4320725262858 | -2.67947603046715 | 0.57683483630036 | -4.64513559488272 | 3.39853163385502e-06 | 0.000530969801805165 | 6631 | 8 | 14 | 53 | 55 | 1 | 2 | 37 | 45 | 2 | 0 | 1 | 4 |

| | | | | | | | | | | | | | | | | | | | |
|---|---|---|---|---|---|---|---|---|---|---|---|---|---|---|---|---|---|---|---|
| L1PA13_dup1660 | 11.4196086680074 | -2.97223069747368 | 0.591777093680867 | -5.02255110786113 | 5.09896227838661e-07 | 0.000120844390268225 | 6632 | 5 | 6 | 21 | 24 | 1 | 1 | 45 | 34 | 2 | 0 | 2 | 4 |
| L1PA2_dup526 | 12.0271537838699 | -3.32920864336171 | 0.514963513869195 | -6.46494082337522 | 1.01338458792166e-10 | 1.0348961766421e-07 | 6633 | 3 | 4 | 29 | 36 | 2 | 3 | 54 | 24 | 1 | 0 | 2 | 1 |
| LTR62_dup116 | 12.8369412220545 | -3.25065055876269 | 0.639985240662407 | -5.07925863321185 | 3.78910583866222e-07 | 9.66347886266154e-05 | 6634 | 4 | 4 | 38 | 30 | 0 | 1 | 42 | 38 | 1 | 2 | 2 | 0 |
| L1PA5_dup3579 | 10.1857645414155 | -2.66414976009278 | 0.655858261546411 | -4.06208157508168 | 4.86370739694044e-05 | 0.00398748150828206 | 6635 | 2 | 3 | 22 | 30 | 2 | 2 | 29 | 32 | 2 | 2 | 0 | 2 |
| L1PB1_dup9598 | 10.6187282725164 | -3.57986144969316 | 0.668929587781244 | -5.35162671091752 | 8.71670684550495e-08 | 2.96866963864694e-05 | 6636 | 1 | 2 | 25 | 48 | 0 | 2 | 39 | 20 | 1 | 2 | 0 | 2 |
| L1PA16_dup1121 | 10.9868407231347 | -3.69418104961834 | 0.603427616331076 | -6.12199533073987 | 9.24107325940384e-10 | 6.70389151762837e-07 | 6637 | 4 | 4 | 30 | 45 | 0 | 1 | 19 | 24 | 1 | 0 | 7 | 8 |
| HERVL-int_dup1870 | 11.4047286086596 | -4.67837561783576 | 0.662260914638333 | -7.06424841694102 | 1.61487628001043e-12 | 2.84632112091379e-09 | 6638 | 2 | 2 | 32 | 34 | 0 | 1 | 36 | 30 | 0 | 0 | 5 | 5 |
| L1ME3Cz_dup1168 | 13.8027486811892 | -3.53337581789567 | 0.59592041972856 | -5.92927461607224 | 3.0427584268856e-09 | 1.83759440772518e-06 | 6639 | 11 | 3 | 34 | 55 | 0 | 1 | 38 | 30 | 0 | 0 | 5 | 5 |
| L1P4_dup284 | 14.5077916756004 | -4.32627943714188 | 0.595971800424285 | -7.25920158313181 | 3.89381819611093e-13 | 7.98722814217079e-10 | 6640 | 6 | 2 | 44 | 64 | 0 | 1 | 35 | 26 | 0 | 0 | 8 | 7 |
| L1MA7_dup688 | 11.9259405292238 | -5.01456961263783 | 0.747765123456776 | -6.70607581891013 | 1.99927872836435e-11 | 2.59956489781083e-08 | 6641 | 1 | 2 | 38 | 64 | 1 | 0 | 30 | 15 | 0 | 0 | 7 | 4 |
| L1MC3_dup1030 | 12.0305085579011 | -4.85694586306357 | 0.702457180256049 | -6.91422338553532 | 4.70433376482183e-12 | 6.99610876252684e-09 | 6642 | 2 | 3 | 38 | 51 | 0 | 0 | 30 | 18 | 0 | 0 | 10 | 9 |
| L1MC4_dup2389 | 14.2555671187349 | -3.69268508651926 | 0.668993349574121 | -5.51976352062395 | 3.39456176125973e-08 | 1.39503003945546e-05 | 6643 | 6 | 8 | 38 | 71 | 0 | 0 | 37 | 21 | 0 | 0 | 3 | 9 |
| L1MEi_dup519 | 11.9611319296872 | -3.11356170067327 | 0.553287227015792 | -5.62738763637572 | 1.82959265447143e-08 | 8.48624276336959e-06 | 6644 | 5 | 9 | 31 | 40 | 1 | 0 | 36 | 21 | 1 | 0 | 6 | 8 |
| L1PA4_dup1024 | 13.6717467115195 | -4.18269873999142 | 0.607536649049423 | -6.88468546965165 | 5.79153533781474e-12 | 8.301642575251e-09 | 6645 | 5 | 3 | 50 | 49 | 0 | 0 | 37 | 21 | 1 | 0 | 5 | 11 |
| L1MC3_dup3059 | 10.1611917954216 | -5.16312547604539 | 0.803383381982971 | -6.4267267556635 | 1.30380876083158e-10 | 1.27669168479355e-07 | 6646 | 1 | 1 | 22 | 45 | 0 | 0 | 45 | 20 | 0 | 0 | 1 | 2 |
| THE1D-int_dup632 | 11.7734523593608 | -4.17016775684542 | 0.702139242599506 | -5.9392318557874 | 2.86360635852881e-09 | 1.75614350151159e-06 | 6647 | 5 | 2 | 22 | 61 | 0 | 0 | 53 | 16 | 0 | 0 | 1 | 3 |
| ERVL-E-int_dup2276 | 8.97408366432143 | -3.71715972878679 | 0.651866809758412 | -5.7023300974081 | 1.18180684719965e-08 | 5.92013078028985e-06 | 6648 | 4 | 3 | 30 | 35 | 0 | 0 | 31 | 15 | 0 | 0 | 1 | 1 |
| L2a_dup46290 | 9.36033877291793 | -3.69258297568063 | 0.645852212513654 | -5.71738070124296 | 1.081785721855e-08 | 5.44199969075076e-06 | 6649 | 3 | 4 | 24 | 46 | 1 | 0 | 34 | 12 | 0 | 0 | 2 | 2 |
| MLT2A2_dup2221 | 14.8338748162907 | -5.46743543010564 | 0.731891740437614 | -7.47027890605315 | 8.00250222971581e-14 | 1.8853102926257e-10 | 6650 | 1 | 2 | 46 | 45 | 0 | 0 | 41 | 46 | 0 | 0 | 2 | 5 |
| L1MC3_dup3058 | 12.9000604045492 | -3.57556203259973 | 0.580310559808117 | -6.16146298247962 | 7.20759210162108e-10 | 5.47928980898508e-07 | 6651 | 4 | 7 | 39 | 47 | 0 | 0 | 38 | 27 | 1 | 0 | 2 | 4 |
| L2b_dup26240 | 13.0362143022265 | -3.98365682836768 | 0.604534255297606 | -6.5896296090063 | 4.40925173354573e-11 | 4.97234034592546e-08 | 6652 | 3 | 5 | 38 | 46 | 0 | 0 | 30 | 37 | 1 | 0 | 3 | 3 |
| L1M1_dup3085 | 10.6245812795923 | -3.55486267333773 | 0.648922748682213 | -5.47809840317155 | 4.29921037258865e-08 | 1.68808566223759e-05 | 6653 | 6 | 3 | 26 | 36 | 0 | 0 | 35 | 30 | 0 | 0 | 1 | 0 |
| MIR_dup149071 | 10.2499862209599 | -3.05601312345739 | 0.690497108157788 | -4.42581596266302 | 9.60784119136905e-06 | 0.00118760902863455 | 6654 | 6 | 7 | 37 | 25 | 1 | 0 | 24 | 26 | 0 | 0 | 4 | 0 |
| AluSg4_dup2976 | 8.30072909567192 | -3.3920576968362 | 0.639943437501078 | -5.30055860887001 | 1.15448896028031e-07 | 3.70223760300347e-05 | 6655 | 3 | 1 | 31 | 31 | 3 | 0 | 21 | 16 | 0 | 1 | 0 | 2 |
| L1MEc_dup8722 | 11.9999219468983 | -3.87600209338496 | 0.610650412432348 | -6.34733394831592 | 2.19078236457756e-10 | 1.9972716495087e-07 | 6656 | 2 | 3 | 53 | 38 | 3 | 0 | 35 | 22 | 0 | 0 | 1 | 0 |
| MER11A_dup252 | 8.3925403661785 | -3.09989542078181 | 0.663929128856809 | -4.66901554104033 | 3.02646486695803e-06 | 0.000483904718653614 | 6657 | 3 | 5 | 18 | 40 | 1 | 0 | 28 | 17 | 0 | 0 | 0 | 0 |
| L1M5_dup27223 | 8.23504718968636 | -2.78016893553919 | 0.773470020445789 | -3.59441072316783 | 0.000325126603590533 | 0.0158335192013821 | 6658 | 0 | 1 | 16 | 32 | 7 | 0 | 15 | 23 | 1 | 2 | 1 | 7 |
| Tigger10_dup555 | 7.99944322301738 | -4.38212720683613 | 0.900609124185459 | -4.8657370763365 | 1.14031028251643e-06 | 0.000228010311330803 | 6659 | 0 | 1 | 19 | 46 | 2 | 0 | 12 | 16 | 0 | 1 | 0 | 10 |
| AluJo_dup17330 | 10.4030652251652 | -2.40722402948606 | 0.660091628988852 | -3.64680284337725 | 0.000265523495726558 | 0.0137105247477473 | 6660 | 5 | 4 | 30 | 41 | 5 | 2 | 18 | 9 | 3 | 1 | 0 | 23 |
| L1PA6_dup3780 | 13.1417691453781 | -2.40980686122357 | 0.763745153863688 | -3.1552499535122 | 0.00160360657187025 | 0.0486202560334147 | 6661 | 3 | 1 | 36 | 79 | 8 | 1 | 25 | 11 | 3 | 4 | 0 | 9 |
| L1MB3_dup6407 | 13.00676666318 | -2.54021094669427 | 0.535287342978004 | -4.74550908034202 | 2.07982582689612e-06 | 0.000362977083394936 | 6662 | 6 | 13 | 33 | 62 | 3 | 4 | 35 | 14 | 0 | 0 | 2 | 5 |
| L1ME3A_dup6486 | 11.8749282516558 | -2.41108527267711 | 0.455959112175012 | -5.28794185332928 | 1.23700284215235e-07 | 3.90371191351171e-05 | 6663 | 8 | 8 | 35 | 41 | 5 | 4 | 40 | 14 | 0 | 0 | 2 | 3 |
| L1MA7_dup3347 | 12.7344913752978 | -3.19002969895041 | 0.50591746880143 | -6.30543496848987 | 2.87385652712865e-10 | 2.53634202212811e-07 | 6664 | 2 | 8 | 40 | 51 | 3 | 3 | 36 | 21 | 0 | 0 | 3 | 2 |
| MLT1C_dup9855 | 9.78257887131069 | -3.42535278736007 | 0.565810774636908 | -6.05388398543343 | 1.41394459309238e-09 | 9.70323082590349e-07 | 6665 | 2 | 3 | 29 | 45 | 2 | 3 | 25 | 18 | 0 | 0 | 1 | 2 |
| MER4B_dup250 | 11.6267473745737 | -2.49830411391178 | 0.510019781816133 | -4.89844551718279 | 9.65978147088089e-07 | 0.000200202324766804 | 6666 | 10 | 6 | 41 | 36 | 6 | 2 | 19 | 22 | 0 | 0 | 3 | 6 |
| MER89_dup1151 | 10.3597727428468 | -2.45338766334059 | 0.530203464952895 | -4.62725694099067 | 3.7054070041611e-06 | 0.000567001141486893 | 6667 | 6 | 5 | 30 | 44 | 7 | 4 | 12 | 18 | 0 | 0 | 4 | 6 |
| L1M5_dup5184 | 11.987495854113 | -2.7675421935541 | 0.497342140905379 | -5.56466457581086 | 2.62657139198925e-08 | 1.15310360966471e-05 | 6668 | 9 | 5 | 30 | 46 | 1 | 6 | 24 | 25 | 0 | 0 | 5 | 6 |
| L1MC3_dup5226 | 8.94014275596513 | -2.21590402745799 | 0.548803331380377 | -4.03770148749724 | 5.39774781214224e-05 | 0.00432303096906091 | 6669 | 12 | 4 | 27 | 29 | 1 | 5 | 22 | 13 | 0 | 0 | 3 | 3 |
| L1MEi_dup523 | 7.59638983661382 | -2.9046077060842 | 0.605228088732508 | -4.79919514668783 | 1.59304516573549e-06 | 0.000296371168886706 | 6670 | 5 | 5 | 19 | 31 | 1 | 1 | 15 | 15 | 0 | 0 | 3 | 5 |
| Tigger3b_dup2755 | 7.00170945274909 | -2.99723640509154 | 0.660476352389587 | -4.53799200266234 | 5.67924136382632e-06 | 0.000793047397627357 | 6671 | 7 | 1 | 18 | 28 | 1 | 1 | 17 | 15 | 0 | 0 | 3 | 1 |
| L2a_dup14886 | 8.14961591659992 | -2.36866468527255 | 0.593871725069167 | -3.98851230877624 | 6.64889536101274e-05 | 0.00505779429530543 | 6672 | 8 | 4 | 10 | 21 | 1 | 5 | 37 | 17 | 0 | 0 | 3 | 2 |
| L1MA5_dup1879 | 8.84077367684709 | -1.82248281863191 | 0.559355022484046 | -3.25818620621019 | 0.00112126806106197 | 0.0379842326391587 | 6673 | 13 | 8 | 11 | 28 | 4 | 2 | 23 | 21 | 0 | 0 | 2 | 3 |
| MIR_dup72105 | 7.82452785330315 | -2.05468009008418 | 0.573024303054597 | -3.58567704568791 | 0.000336204819963412 | 0.0162268949474673 | 6674 | 7 | 9 | 14 | 33 | 2 | 2 | 20 | 14 | 0 | 0 | 1 | 2 |
| AluJr_dup5793 | 8.62344252364388 | -2.05425509212412 | 0.513091960537481 | -4.00367819049868 | 6.236518674597e-05 | 0.00481022584293568 | 6675 | 9 | 5 | 25 | 30 | 2 | 4 | 22 | 14 | 0 | 1 | 2 | 0 |
| L1PA12_dup778 | 10.2306280150255 | -2.11203706102673 | 0.55284779283249 | -3.82028668362011 | 0.000133296647626713 | 0.00834284266306204 | 6676 | 11 | 6 | 20 | 45 | 2 | 5 | 18 | 25 | 0 | 0 | 1 | 0 |
| L1ME2z_dup2690 | 9.19888689876266 | -2.35652555769551 | 0.629651682745095 | -3.74258597614121 | 0.000182136185588286 | 0.0104323360473012 | 6677 | 10 | 5 | 24 | 48 | 2 | 1 | 13 | 15 | 0 | 1 | 1 | 3 |
| L1ME3_dup3690 | 8.92652619318237 | -2.09927790841562 | 0.597946889807055 | -3.51080998028606 | 0.000446743613404791 | 0.0199888784947755 | 6678 | 9 | 8 | 26 | 44 | 2 | 4 | 12 | 11 | 0 | 0 | 1 | 3 |
| L1ME3F_dup1936 | 9.49226149342623 | -2.15067143801458 | 0.59021192685373 | -3.64389694643967 | 0.000268540976858092 | 0.0138038996067133 | 6679 | 8 | 11 | 21 | 42 | 2 | 2 | 23 | 15 | 0 | 0 | 0 | 3 |
| L1MA7_dup3346 | 10.3221552027802 | -2.64006560877301 | 0.600865920564953 | -4.39376825746871 | 1.11402570799289e-05 | 0.0013227443169365 | 6680 | 10 | 5 | 32 | 48 | 1 | 2 | 27 | 13 | 0 | 0 | 0 | 2 |
| L1PA14_dup1397 | 8.87692807250339 | -3.24674724139303 | 0.621728611234782 | -5.22212937079547 | 1.76877323431061e-07 | 5.23473278620985e-05 | 6681 | 6 | 2 | 30 | 39 | 1 | 1 | 21 | 16 | 0 | 0 | 0 | 2 |
| L1PREC2_dup4559 | 8.25022440764798 | -1.89718477746496 | 0.572389507203601 | -3.31449957343492 | 0.000918073199546587 | 0.0331892806957326 | 6682 | 6 | 6 | 25 | 35 | 3 | 2 | 8 | 8 | 3 | 1 | 7 | 5 |
| L1PA3_dup9589 | 9.58060269673861 | -2.2712271332952 | 0.579556351844802 | -3.91890646365207 | 8.89516235385241e-05 | 0.00626552243107462 | 6683 | 8 | 8 | 38 | 33 | 3 | 2 | 14 | 9 | 1 | 0 | 7 | 4 |
| MER41A_dup2049 | 6.70500412460258 | -2.38482662374059 | 0.667677422066836 | -3.57182457414573 | 0.00035450284838101 | 0.0168738817285192 | 6684 | 4 | 5 | 27 | 24 | 2 | 2 | 7 | 7 | 1 | 0 | 7 | 2 |
| AluSg4_dup7042 | 6.54610170091112 | -2.46112166876042 | 0.607426424375616 | -4.05171979683013 | 5.08425323288708e-05 | 0.00413316610779826 | 6685 | 2 | 5 | 23 | 20 | 1 | 4 | 14 | 7 | 1 | 0 | 3 | 7 |
| LTR12_dup72 | 7.65043196528828 | -2.9840066085206 | 0.618784971555785 | -4.82236438454224 | 1.41866480382476e-06 | 0.000270919434519171 | 6686 | 2 | 5 | 16 | 33 | 1 | 2 | 14 | 12 | 1 | 0 | 10 | 5 |
| MLT1A1_dup5445 | 8.37042333435958 | -2.34014196880956 | 0.600090514505993 | -3.8996483234467 | 9.63325088381646e-05 | 0.00663634486045336 | 6687 | 3 | 2 | 28 | 36 | 1 | 7 | 16 | 9 | 2 | 1 | 2 | 5 |
| L1PA7_dup2209 | 6.92064062662303 | -2.05264614087447 | 0.528535288500603 | -3.88365012806921 | 0.000102899947112301 | 0.00695354856729831 | 6688 | 5 | 5 | 18 | 24 | 1 | 4 | 16 | 9 | 1 | 1 | 3 | 5 |
| MER39_dup2777 | 8.20134189636726 | -2.08576964698763 | 0.516646822818755 | -4.03712856610238 | 5.41094188257265e-05 | 0.00432777135190129 | 6689 | 2 | 6 | 24 | 28 | 2 | 6 | 14 | 13 | 1 | 2 | 4 | 5 |
| MER87_dup61 | 6.29576213435858 | -1.67658593799458 | 0.520511968793711 | -3.2210324421167 | 0.00127729688414186 | 0.0416168265636066 | 6690 | 5 | 2 | 11 | 22 | 3 | 6 | 13 | 7 | 2 | 1 | 4 | 8 |
| L1MC4_dup2227 | 5.90355006462153 | -1.98746861513616 | 0.595796977931836 | -3.33581520006223 | 0.000850496791598685 | 0.0313998928741258 | 6691 | 2 | 3 | 19 | 19 | 2 | 3 | 8 | 7 | 3 | 1 | 3 | 7 |
| L1PA2_dup493 | 6.5655789292026 | -2.01939598297553 | 0.613664503624806 | -3.29071662292234 | 0.000999325264393564 | 0.0350974833005743 | 6692 | 2 | 4 | 19 | 28 | 2 | 3 | 9 | 7 | 3 | 1 | 2 | 7 |
| L1ME3Cz_dup7235 | 7.71636748969132 | -2.49888206999101 | 0.629007273788496 | -3.97273954391704 | 7.10507255497284e-05 | 0.00530142237116829 | 6693 | 3 | 1 | 25 | 30 | 1 | 3 | 15 | 8 | 4 | 1 | 6 | 5 |
| LTR33_dup7905 | 8.09213961566427 | -2.57465190968988 | 0.60011291461454 | -4.29027912412717 | 1.78448689811064e-05 | 0.00187773358988088 | 6694 | 3 | 2 | 27 | 29 | 2 | 2 | 23 | 8 | 3 | 1 | 4 | 4 |
| L2a_dup147235 | 6.77245161429368 | -2.06069126575993 | 0.610579809517598 | -3.37497446466174 | 0.000738225403349736 | 0.0284052040468073 | 6695 | 5 | 2 | 24 | 29 | 2 | 4 | 8 | 5 | 1 | 2 | 4 | 5 |
| MER41A_dup2152 | 7.69683306259264 | -2.43074992334071 | 0.644749280953073 | -3.77007000263351 | 0.000163201764314118 | 0.00966944247280339 | 6696 | 5 | 1 | 25 | 37 | 1 | 3 | 9 | 8 | 1 | 3 | 6 | 4 |
| AluJo_dup63342 | 6.37853546203006 | -2.10111799821977 | 0.6409208542181 | -3.27827996919067 | 0.00104441752084206 | 0.0361581509405333 | 6697 | 4 | 4 | 20 | 26 | 2 | 1 | 18 | 3 | 2 | 1 | 2 | 4 |
| MSTA_dup17274 | 6.87682944091459 | -2.06883388799661 | 0.54285633569642 | -3.81101546018163 | 0.000138397110451373 | 0.00858996949817145 | 6698 | 3 | 4 | 16 | 24 | 3 | 2 | 21 | 5 | 2 | 2 | 4 | 7 |
| AluSg_dup29865 | 5.07412545112552 | -2.45138410856528 | 0.678844722232113 | -3.61111168472352 | 0.00030488732304971 | 0.0151276483822178 | 6699 | 1 | 2 | 20 | 18 | 3 | 1 | 11 | 6 | 1 | 1 | 1 | 2 |
| MIRb_dup75804 | 5.70272122567141 | -2.58118981248138 | 0.678322118002629 | -3.80525674156385 | 0.000141657175958351 | 0.00873005915322085 | 6700 | 1 | 1 | 13 | 21 | 2 | 1 | 14 | 10 | 3 | 1 | 4 | 3 |
| L1PA3_dup3025 | 5.82029286350938 | -2.44687882286041 | 0.587856250917056 | -4.16237612348815 | 3.14953009458796e-05 | 0.00286037438124743 | 6701 | 1 | 3 | 15 | 18 | 3 | 2 | 15 | 10 | 1 | 1 | 3 | 4 |
| MER67D_dup909 | 6.39877519232823 | -2.00047099568345 | 0.535643906953712 | -3.73470316699844 | 0.000187936721085413 | 0.0106422634544002 | 6702 | 2 | 3 | 15 | 19 | 4 | 2 | 16 | 11 | 2 | 2 | 3 | 4 |
| MLT1L_dup10176 | 7.38663512073613 | -1.90041092467546 | 0.603411275499885 | -3.14944549735352 | 0.00163580618079903 | 0.0493355039669718 | 6703 | 3 | 4 | 16 | 36 | 5 | 1 | 17 | 9 | 2 | 2 | 1 | 3 |
| MIRc_dup51486 | 6.737216208035 | -2.90253298871991 | 0.621365687005073 | -4.6712154362914 | 2.99422693375631e-06 | 0.000480897165161573 | 6704 | 2 | 2 | 18 | 23 | 1 | 1 | 23 | 12 | 1 | 2 | 1 | 3 |
| MSTC_dup1692 | 6.62060169161373 | -2.80237880457147 | 0.625910977584934 | -4.47728016432049 | 7.56000102661928e-06 | 0.000986764675962672 | 6705 | 3 | 3 | 13 | 26 | 1 | 1 | 17 | 14 | 1 | 1 | 1 | 6 |
| LTR16E2_dup1018 | 5.59762639589519 | -2.71009887062177 | 0.655565854243288 | -4.13398418035364 | 3.56528026342748e-05 | 0.0031501826125669 | 6706 | 3 | 2 | 17 | 16 | 1 | 2 | 25 | 6 | 0 | 1 | 1 | 2 |
| L4_C_Mam_dup4498 | 6.00055171605891 | -2.72068205864156 | 0.709874807925693 | -3.83262235575256 | 0.000126784504804412 | 0.00806627427277826 | 6707 | 4 | 3 | 15 | 34 | 1 | 1 | 13 | 5 | 0 | 1 | 1 | 5 |
| L1PA8_dup4439 | 6.8925210196216 | -2.62476681866902 | 0.572114283744672 | -4.5878365446307 | 4.47863074618088e-06 | 0.00065904283953664 5 | 6708 | 3 | 3 | 21 | 27 | 3 | 2 | 18 | 10 | 0 | 1 | 1 | 3 |
| LTR78_dup4186 | 7.82443870514347 | -2.23404029110851 | 0.593510289634639 | -3.7641138327758 | 0.000167140738378399 | 0.00984173970160519 | 6709 | 6 | 3 | 18 | 35 | 1 | 3 | 19 | 12 | 0 | 3 | 1 | 4 |
| L1MB7_dup10215 | 5.74147684613084 | -2.75216146708014 | 0.640393004498748 | -4.2976132589617 | 1.72647049975656e-05 | 0.00183186988031498 | 6710 | 1 | 1 | 12 | 22 | 2 | 3 | 16 | 10 | 0 | 2 | 4 | 3 |

| | | | | | | | | | | | | | | | | | | | |
|---|---|---|---|---|---|---|---|---|---|---|---|---|---|---|---|---|---|---|---|
| L1PA3_dup1808 | 5.50290113094208 | -2.56734615753044 | 0.663829212769744 | -3.86747993029489 | 0.000109965875552539 | 0.00730155533608211 | 6711 | 1 | 1 | 16 | 24 | 3 | 4 | 9 | 8 | 0 | 1 | 3 | 3 |
| ALR/Alpha_dup1617 | 5.98870369563255 | -2.27383566056692 | 0.595539512289723 | -3.81811049249193 | 0.000134477697586797 | 0.00839413174973455 | 6712 | 1 | 2 | 10 | 17 | 5 | 4 | 19 | 13 | 0 | 1 | 2 | 4 |
| AluSx_dup98449 | 5.94045607394615 | -2.0796004206065 | 0.567894907767666 | -3.66194588498986 | 0.000250306741161521 | 0.0131361905232508 | 6713 | 2 | 4 | 15 | 13 | 2 | 4 | 18 | 11 | 0 | 2 | 4 | 2 |
| AluSz_dup99899 | 4.61959166867317 | -2.00798881231472 | 0.630165657554737 | -3.18644595788736 | 0.001440323934538 | 0.0451600104002607 | 6714 | 5 | 1 | 9 | 12 | 1 | 4 | 14 | 8 | 0 | 1 | 3 | 3 |
| L1PB1_dup5250 | 4.81906511143598 | -2.18366713787534 | 0.653073482609391 | -3.34367754322282 | 0.000826757730478211 | 0.0307765480188311 | 6715 | 4 | 2 | 7 | 10 | 3 | 1 | 17 | 12 | 0 | 1 | 3 | 2 |
| LTR1A1_dup292 | 6.0308187884096 | -3.30734076576154 | 0.693938315884861 | -4.76604431554446 | 1.87877953263609e-06 | 0.000336126371934306 | 6716 | 2 | 1 | 14 | 20 | 2 | 1 | 13 | 10 | 0 | 1 | 11 | 4 |
| L1M5_dup47247 | 4.58081066019348 | -2.86124834515751 | 0.71128550905008 | -4.02264394360951 | 5.75484613694432e-05 | 0.00452525650661386 | 6717 | 3 | 1 | 10 | 15 | 1 | 1 | 15 | 7 | 0 | 1 | 5 | 2 |
| HAL1ME_dup7591 | 5.01629912587672 | -3.01478696925619 | 0.686358615068743 | -4.39243699003361 | 1.12087120852214e-05 | 0.00133017478457797 | 6718 | 3 | 1 | 12 | 17 | 1 | 1 | 15 | 8 | 0 | 1 | 3 | 5 |
| MLT1C_dup19524 | 5.42927264367773 | -3.61073315230723 | 0.737976372488151 | -4.89274899158808 | 9.94371937228179e-07 | 0.000204854393918352 | 6719 | 1 | 1 | 16 | 20 | 1 | 1 | 17 | 7 | 0 | 1 | 4 | 4 |
| L2b_dup84584 | 7.28104809644153 | -1.95429983356297 | 0.606672697216591 | -3.22134133038998 | 0.00127592090463942 | 0.0415833824921698 | 6720 | 2 | 1 | 21 | 17 | 3 | 6 | 20 | 16 | 1 | 3 | 2 | 1 |
| L2b_dup84591 | 8.10054931716122 | -2.01554326347124 | 0.545753684352297 | -3.69313725451674 | 0.000221504390776389 | 0.0119786554017451 | 6721 | 4 | 1 | 20 | 25 | 4 | 4 | 20 | 16 | 1 | 4 | 3 | 3 |
| MLT1A1_dup1241 | 9.87212057543256 | -2.41636068023631 | 0.484364561659871 | -4.98872310549657 | 6.07796899319702e-07 | 0.000139194264682893 | 6722 | 4 | 3 | 22 | 28 | 2 | 7 | 21 | 25 | 1 | 2 | 4 | 7 |
| MLT1D_dup17478 | 9.27324490804928 | -1.79762750029467 | 0.551376280480946 | -3.26025540802491 | 0.00111311927925631 | 0.0377638420145279 | 6723 | 5 | 4 | 28 | 31 | 2 | 5 | 15 | 18 | 3 | 3 | 2 | 3 |
| MER2_dup4954 | 8.1800767121198 | -1.97141521401809 | 0.506949819312374 | -3.88877782162367 | 0.000100750295462042 | 0.00685906291437348 | 6724 | 5 | 5 | 16 | 26 | 4 | 3 | 17 | 20 | 0 | 3 | 2 | 4 |
| MER5A1_dup6553 | 8.07857205727963 | -2.06541031573351 | 0.525458651300301 | -3.93068095961964 | 8.47056118123137e-05 | 0.00603092803958492 | 6725 | 5 | 5 | 18 | 31 | 2 | 7 | 14 | 17 | 0 | 1 | 3 | 2 |
| MIRb_dup190728 | 6.90368625667816 | -2.0876773973266 | 0.539813109317807 | -3.86740774036577 | 0.000109998423150478 | 0.00730155533608211 | 6726 | 4 | 4 | 14 | 27 | 4 | 4 | 11 | 14 | 0 | 1 | 3 | 4 |
| L1MC1_dup5770 | 10.2404285321421 | -1.87425006990241 | 0.505181367250959 | -3.71005383690515 | 0.000207215177613249 | 0.0114160737791994 | 6727 | 9 | 11 | 25 | 33 | 2 | 3 | 23 | 21 | 0 | 2 | 2 | 2 |
| MER34A_dup1200 | 9.6698687298949 | -1.70930526584936 | 0.473614663042457 | -3.6090632305785 | 0.000307304706502291 | 0.0152129220318605 | 6728 | 6 | 8 | 23 | 38 | 3 | 6 | 20 | 14 | 1 | 3 | 2 | 4 |
| MER4A1_dup816 | 8.53857829210152 | -1.51914165533546 | 0.468860497708376 | -3.24007175430749 | 0.00119499624099108 | 0.0397222046286809 | 6729 | 10 | 7 | 16 | 24 | 3 | 4 | 23 | 17 | 1 | 2 | 2 | 2 |
| MLT1F1_dup2066 | 10.3633790889486 | -2.94564651371298 | 0.503742939251948 | -5.8475192090776 | 4.98958233653869e-09 | 2.80012065719348e-06 | 6730 | 3 | 3 | 25 | 34 | 7 | 3 | 26 | 12 | 0 | 0 | 12 | 14 |
| L1MA2_dup7367 | 10.1757050278462 | -1.90027160823771 | 0.46076657652647 | -4.12415245602898 | 3.72102377481392e-05 | 0.00326115183470303 | 6731 | 6 | 7 | 23 | 26 | 12 | 5 | 26 | 11 | 0 | 0 | 11 | 8 |
| MIRb_dup215003 | 11.1725668651155 | -2.14513125858663 | 0.426905783014617 | -5.02483532417546 | 5.0386469786148e-07 | 0.000120253349445685 | 6732 | 6 | 8 | 24 | 33 | 8 | 6 | 28 | 17 | 0 | 0 | 11 | 6 |
| MLT1H2_dup4617 | 7.56663684468619 | -1.80753555243061 | 0.565026560318583 | -3.19902758449347 | 0.00137891974525832 | 0.0438809714567221 | 6733 | 3 | 3 | 11 | 17 | 7 | 11 | 20 | 9 | 0 | 0 | 12 | 7 |
| L1MA1_dup1778 | 8.26547906854168 | -2.62823197268596 | 0.650897590044438 | -4.03785789482878 | 5.39415113596222e-05 | 0.00432303096906091 | 6734 | 1 | 1 | 15 | 12 | 7 | 7 | 26 | 24 | 0 | 0 | 5 | 6 |
| L2a_dup63141 | 7.76715895547772 | -2.14624302439676 | 0.61847702317254 | -3.47020656222181 | 0.000520058247761265 | 0.0221528427894382 | 6735 | 2 | 1 | 18 | 17 | 8 | 9 | 27 | 12 | 0 | 0 | 6 | 2 |
| L1PA2_dup4707 | 13.5420517610742 | -1.26461025548227 | 0.370604811714056 | -3.41228774023039 | 0.000644200747158758 | 0.0257820043373982 | 6736 | 18 | 13 | 23 | 24 | 9 | 12 | 26 | 27 | 0 | 2 | 9 | 11 |
| MLT2A2_dup3549 | 11.5575212803917 | -1.25773385896041 | 0.391927922351275 | -3.20909480349076 | 0.00133153579879906 | 0.0428211403136966 | 6737 | 14 | 14 | 27 | 23 | 6 | 7 | 24 | 20 | 1 | 2 | 6 | 5 |
| ALR/Alpha_dup1618 | 10.6760929984433 | -1.97385543291214 | 0.545465144486616 | -3.61866464404414 | 0.00029612707834497 | 0.014827915358559 | 6738 | 5 | 8 | 8 | 23 | 9 | 9 | 25 | 33 | 0 | 0 | 5 | 9 |
| L1PA5_dup5369 | 10.6106346574988 | -2.02306730752281 | 0.559450753970165 | -3.6161669157938 | 0.000298997618514438 | 0.0149151545775758 | 6739 | 6 | 6 | 5 | 20 | 9 | 7 | 36 | 28 | 1 | 0 | 6 | 12 |
| L2_dup40333 | 12.2560379774071 | -2.38689425542058 | 0.495464540442478 | -4.81748755075176 | 1.45377158351577e-06 | 0.000275413322620417 | 6740 | 9 | 6 | 24 | 16 | 5 | 5 | 36 | 38 | 1 | 0 | 5 | 8 |
| MER20_dup7510 | 12.1001210673103 | -1.79446198617951 | 0.454786663872656 | -3.94572253042578 | 7.95597031489315e-05 | 0.00575757698463372 | 6741 | 10 | 6 | 15 | 37 | 9 | 7 | 20 | 29 | 3 | 0 | 5 | 14 |
| L1ME5_dup1462 | 12.1061457685157 | -1.82040478876946 | 0.558446801948003 | -3.25976401408233 | 0.00111504947847419 | 0.0378112531288326 | 6742 | 13 | 10 | 12 | 17 | 8 | 6 | 46 | 36 | 0 | 0 | 4 | 1 |
| L2a_dup78627 | 15.9421290387821 | -2.08456529462085 | 0.558672638438372 | -3.7312822415068 | 0.000190507641813701 | 0.010729119843551 | 6743 | 19 | 10 | 16 | 24 | 6 | 8 | 71 | 45 | 0 | 0 | 1 | 5 |
| MIR3_dup7894 | 9.73622411132327 | -2.73285539635304 | 0.569197658488904 | -4.80124145908853 | 1.57684997195827e-06 | 0.000294123595882378 | 6744 | 3 | 5 | 22 | 17 | 2 | 6 | 41 | 26 | 0 | 0 | 0 | 3 |
| MamRTE1_dup8792 | 13.0835025223701 | -2.65287963769292 | 0.506162268035541 | -5.24116435622309 | 1.59566508424125e-07 | 4.800026853791e-05 | 6745 | 3 | 9 | 27 | 33 | 4 | 8 | 36 | 38 | 0 | 0 | 1 | 7 |
| MER11C_dup368 | 8.37907854026951 | -1.92961965751496 | 0.552238556835205 | -3.49417771293137 | 0.000475524365858846 | 0.0208262692359008 | 6746 | 10 | 6 | 23 | 17 | 5 | 3 | 32 | 11 | 0 | 0 | 0 | 5 |
| MER5B_dup1409 | 10.1440668965233 | -1.45591435145622 | 0.457019756704611 | -3.18567048819561 | 0.00144418983361814 | 0.0452633290503296 | 6747 | 7 | 8 | 17 | 21 | 10 | 10 | 28 | 26 | 0 | 0 | 0 | 2 |
| Tigger1_dup5448 | 11.3433701685632 | -1.63942330661465 | 0.488833382843583 | -3.35374662237262 | 0.000797253312194906 | 0.0299973492447073 | 6748 | 15 | 10 | 25 | 26 | 6 | 5 | 22 | 32 | 0 | 0 | 0 | 2 |
| L1ME3_dup3693 | 9.42931146520005 | -2.21707898827031 | 0.478481799552981 | -4.63357015949532 | 3.59412874114285e-06 | 0.000552470703322225 | 6749 | 5 | 6 | 23 | 39 | 5 | 4 | 24 | 15 | 1 | 0 | 2 | 1 |
| MER6A_dup60 | 8.11812223079383 | -2.24794717827321 | 0.631642817133999 | -3.55888979862543 | 0.000372425746090275 | 0.0175306066427689 | 6750 | 1 | 9 | 16 | 26 | 4 | 2 | 34 | 12 | 1 | 0 | 3 | 0 |
| L1MDa_dup2998 | 10.1607769999559 | -3.57696179357347 | 0.594474337830039 | -6.01701632173084 | 1.77661174403161e-09 | 1.16456655108911e-06 | 6751 | 3 | 1 | 20 | 39 | 1 | 4 | 40 | 24 | 0 | 0 | 2 | 0 |
| L2a_dup80477 | 11.6587364808589 | -3.34588264608756 | 0.557442241916118 | -6.00220506179552 | 1.94655639463969e-09 | 1.25701083793171e-06 | 6752 | 3 | 1 | 45 | 47 | 3 | 4 | 29 | 19 | 1 | 0 | 2 | 0 |
| ERVL-E-int_dup7594 | 11.1648858364669 | -2.03131290392496 | 0.452180000770796 | -4.49226613397837 | 7.04692842665926e-06 | 0.00093518596286105 | 6753 | 3 | 5 | 28 | 30 | 9 | 6 | 36 | 21 | 2 | 1 | 2 | 2 |
| L1M1_dup8761 | 10.4234104395547 | -1.89647620586371 | 0.576132118937146 | -3.29173837654173 | 0.000995701927075033 | 0.0350012248101338 | 6754 | 2 | 2 | 24 | 23 | 16 | 7 | 35 | 20 | 1 | 0 | 2 | 3 |
| L2b_dup9247 | 11.5238445894605 | -2.0727809404123 | 0.523967860139804 | -3.95593145705395 | 7.6237019128506e-05 | 0.00558026657196382 | 6755 | 8 | 3 | 20 | 31 | 8 | 8 | 59 | 15 | 0 | 1 | 3 | 0 |
| MER52D_dup54 | 14.1062553392678 | -2.33086529932114 | 0.533587633622137 | -4.36828957878689 | 1.25223386501523e-05 | 0.00144212288148651 | 6756 | 1 | 5 | 31 | 42 | 10 | 12 | 49 | 29 | 0 | 1 | 4 | 0 |
| L1ME2z_dup3266 | 13.289737906995 | -2.87166031531249 | 0.563780635579849 | -5.09357742015915 | 3.51369469816875e-07 | 9.10751196786996e-05 | 6757 | 3 | 0 | 47 | 53 | 7 | 7 | 22 | 13 | 0 | 3 | 12 | 11 |
| THE1B-int_dup3600 | 11.6718390538752 | -2.63476807400712 | 0.609622659729697 | -4.32196545183436 | 1.54645407920785e-05 | 0.00168949753136452 | 6758 | 1 | 0 | 34 | 53 | 6 | 6 | 20 | 9 | 1 | 5 | 10 | 13 |
| L1M5_dup42466 | 15.8575199348241 | -2.39480156710611 | 0.59021718670466 | -4.05749209113501 | 4.96024964559973e-05 | 0.00405173897072391 | 6759 | 2 | 2 | 53 | 72 | 15 | 9 | 17 | 10 | 2 | 2 | 17 | 12 |
| THE1B_dup18259 | 17.5709812105764 | -2.51611573986076 | 0.53879274691711 | -4.66991390336561 | 3.01325997059818e-06 | 0.000482174281751146 | 6760 | 4 | 1 | 64 | 74 | 11 | 9 | 21 | 17 | 2 | 4 | 8 | 20 |
| L1PA3_dup5608 | 13.7024766583653 | -2.17223368129456 | 0.428314463826498 | -5.07158610028752 | 3.94513699810927e-07 | 9.92314554071997e-05 | 6761 | 5 | 3 | 38 | 47 | 7 | 10 | 21 | 15 | 2 | 4 | 11 | 19 |
| THE1C_dup7977 | 14.1928824650187 | -2.16058179773378 | 0.483111656853016 | -4.4722203802901 | 7.74115879477045e-06 | 0.00100812484955221 | 6762 | 3 | 2 | 40 | 54 | 12 | 9 | 23 | 12 | 2 | 4 | 12 | 17 |
| L2a_dup86167 | 18.0072411044939 | -3.256206443667 | 0.626662476339035 | -5.19610885701946 | 2.03503121920159e-07 | 5.84110420366877e-05 | 6763 | 1 | 1 | 53 | 106 | 4 | 6 | 17 | 26 | 2 | 4 | 8 | 13 |
| L1MCa_dup3643 | 12.5374337513383 | -3.39294905451719 | 0.630337930337483 | -5.38274612904956 | 7.33579671790162e-08 | 2.60915319258269e-05 | 6764 | 0 | 1 | 46 | 54 | 6 | 6 | 20 | 17 | 0 | 1 | 5 | 11 |
| L1PB4_dup3637 | 14.6841863078229 | -2.79928243200493 | 0.704013362888333 | -3.97617798122525 | 7.00317201177885e-05 | 0.0052500843336948 | 6765 | 0 | 1 | 49 | 77 | 5 | 8 | 19 | 20 | 0 | 5 | 6 | 6 |
| MIR_dup60667 | 16.3850457326334 | -2.58662601970426 | 0.641865920382818 | -4.02985411370892 | 5.58114799387303e-05 | 0.00442818219589901 | 6766 | 3 | 2 | 56 | 68 | 4 | 4 | 28 | 29 | 1 | 9 | 6 | 4 |
| L1MB8_dup10310 | 13.3473175819332 | -2.52594486868113 | 0.626940796426746 | -4.02900063782382 | 5.60144663806409e-05 | 0.00443575831929537 | 6767 | 5 | 2 | 35 | 46 | 1 | 3 | 35 | 28 | 2 | 7 | 6 | 3 |
| L1MEf_dup6279 | 15.2390649274785 | -2.48750875628946 | 0.643689140313736 | -3.8644566150006 | 0.000111336776163121 | 0.00735484190475015 | 6768 | 4 | 4 | 45 | 58 | 2 | 2 | 43 | 23 | 3 | 8 | 7 | 2 |
| MLT1L_dup11391 | 13.4291656236637 | -2.39389079220708 | 0.667272167345429 | -3.58757776715094 | 0.000333764202252533 | 0.0161438486592461 | 6769 | 5 | 5 | 48 | 70 | 4 | 1 | 16 | 20 | 1 | 4 | 2 | 2 |
| MIR_dup167126 | 13.0783736425506 | -2.60280536001255 | 0.741206551840121 | -3.51157899717808 | 0.000445453050487573 | 0.0199536467528833 | 6770 | 5 | 1 | 59 | 64 | 1 | 4 | 22 | 10 | 1 | 5 | 2 | 3 |
| MIRc_dup95963 | 14.9612413429605 | -2.34297126104006 | 0.692461174165054 | -3.38354170378597 | 0.000715573390492237 | 0.0277852196432875 | 6771 | 4 | 2 | 67 | 69 | 2 | 7 | 20 | 16 | 3 | 4 | 1 | 4 |
| L1MB8_dup14134 | 13.3036512785205 | -2.50384192787386 | 0.485800669745872 | -5.15405202957759 | 2.54917163719082e-07 | 7.00421702520792e-05 | 6772 | 5 | 2 | 36 | 51 | 3 | 5 | 33 | 15 | 2 | 6 | 10 | 10 |
| LTR50_dup2279 | 10.0262506159844 | -2.62995794201579 | 0.577610753910403 | -4.55316651258841 | 5.28444129104573e-06 | 0.000751590954835126 | 6773 | 3 | 1 | 29 | 41 | 1 | 5 | 21 | 12 | 1 | 5 | 7 | 8 |
| THE1D-int_dup1431 | 14.3429298713349 | -3.03859434701088 | 0.622903438684355 | -4.87811458133663 | 1.07104703662771e-06 | 0.000217079521445841 | 6774 | 4 | 3 | 56 | 64 | 1 | 2 | 23 | 14 | 1 | 6 | 11 | 7 |
| AluJb_dup60261 | 10.781188533144 | -3.70406142813484 | 0.631300991458213 | -5.86734612847511 | 4.42825633165481e-09 | 2.52077961983716e-06 | 6775 | 1 | 2 | 43 | 34 | 1 | 1 | 20 | 13 | 1 | 3 | 13 | 10 |
| L1MA2_dup4468 | 12.3911196348008 | -3.38569661411183 | 0.72160584554956 | -4.69189188944742 | 2.70690165738956e-06 | 0.000446359266645334 | 6776 | 1 | 1 | 53 | 59 | 2 | 1 | 17 | 9 | 2 | 5 | 7 | 10 |
| L1PA3_dup9882 | 13.2605459796391 | -2.42727444520006 | 0.596530675656913 | -4.06898512390346 | 4.72183616313715e-05 | 0.00390933203783519 | 6777 | 7 | 2 | 39 | 50 | 2 | 1 | 31 | 15 | 7 | 4 | 5 | 13 |
| L1PA3_dup6584 | 10.7395433701108 | -1.98257412479188 | 0.552384245794699 | -3.58912141301136 | 0.000331794296351898 | 0.0160700778452441 | 6778 | 10 | 3 | 25 | 48 | 1 | 3 | 23 | 9 | 4 | 4 | 7 | 8 |
| MLT1G_dup1432 | 10.4615373895096 | -2.2614446691049 | 0.52730431166315 | -4.28868988757585 | 1.7973010608717e-05 | 0.00188812626150189 | 6779 | 5 | 2 | 22 | 38 | 2 | 3 | 27 | 17 | 5 | 3 | 6 | 7 |
| AluSg_dup31156 | 11.8607219292122 | -3.02749019604135 | 0.590453721531495 | -5.12739624739559 | 2.93776781351276e-07 | 7.86310573851638e-05 | 6780 | 2 | 2 | 46 | 32 | 1 | 3 | 22 | 23 | 3 | 3 | 6 | 9 |
| L1MA4_dup7723 | 9.97084950146249 | -3.16290764909571 | 0.568142107490039 | -5.56710655203666 | 2.59004031126628e-08 | 1.14304142603315e-05 | 6781 | 3 | 4 | 31 | 40 | 1 | 1 | 21 | 13 | 2 | 1 | 5 | 11 |
| MSTB_dup7480 | 11.4929088088747 | -2.75363102389955 | 0.534722513595037 | -5.14964482304362 | 2.60980194610654e-07 | 7.13783832032492e-05 | 6782 | 4 | 5 | 34 | 45 | 1 | 2 | 19 | 19 | 3 | 2 | 9 | 7 |
| L2_dup55055 | 12.6853830776436 | -2.13661623223924 | 0.623568118629803 | -3.42643597131639 | 0.000611557926119589 | 0.0249046315742686 | 6783 | 3 | 6 | 54 | 53 | 3 | 3 | 18 | 10 | 3 | 6 | 3 | 7 |
| MER11A_dup303 | 10.2422475690664 | -2.00158122467579 | 0.567756931147388 | -3.52541926812018 | 0.000422813028962702 | 0.0192438157975438 | 6784 | 6 | 5 | 30 | 30 | 1 | 2 | 33 | 12 | 3 | 5 | 3 | 6 |
| AluSp_dup40884 | 10.0564034850083 | -2.2228259129812 | 0.54579092486877 | -4.07266924329248 | 4.64774046583704e-05 | 0.00386412038044745 | 6785 | 5 | 4 | 27 | 39 | 2 | 2 | 20 | 15 | 1 | 6 | 4 | 8 |
| MLT1E1_dup931 | 10.7678281838461 | -2.04385892942162 | 0.500043608595736 | -4.08736137066395 | 4.36307229811996e-05 | 0.00368147376258316 | 6786 | 8 | 4 | 31 | 40 | 3 | 3 | 21 | 16 | 2 | 4 | 3 | 7 |
| L1MC1_dup10415 | 13.0141825460568 | -3.39440706358515 | 0.621013601857595 | -5.46591419806538 | 4.60527074731283e-08 | 1.78469992384381e-05 | 6787 | 3 | 5 | 52 | 48 | 1 | 0 | 14 | 21 | 1 | 3 | 8 | 14 |
| Tigger3b_dup3523 | 13.3177560890191 | -3.13482152869143 | 0.652711745155418 | -4.80276561891036 | 1.56489023603864e-06 | 0.000292674451890849 | 6788 | 2 | 5 | 39 | 51 | 1 | 0 | 33 | 25 | 1 | 5 | 4 | 9 |
| L1PA8_dup686 | 15.1534251980195 | -4.3880338698276 | 0.769933056230089 | -5.69924077726086 | 1.20342159100873e-08 | 5.97806584330194e-06 | 6789 | 1 | 1 | 63 | 85 | 0 | 2 | 18 | 18 | 2 | 1 | 3 | 10 |

| | | | | | | | | | | | | | | | | | | | |
|---|---|---|---|---|---|---|---|---|---|---|---|---|---|---|---|---|---|---|---|
| L2a_dup19341 | 12.1512012364564 | -2.95415194959945 | 0.636365534206492 | -4.6422249333209 | 3.44677332761175e-06 | 0.000535692962907842 | 6790 | 6 | 3 | 39 | 56 | 0 | 1 | 27 | 14 | 2 | 3 | 3 | 10 |
| L1PA5_dup1219 | 14.6892913454276 | -1.68126015016374 | 0.509226995223939 | -3.30159273945088 | 0.000961375475324829 | 0.0342038649598747 | 6791 | 11 | 3 | 33 | 48 | 5 | 10 | 32 | 30 | 7 | 1 | 3 | 6 |
| L1PA16_dup7469 | 13.7365031884452 | -2.33713803630001 | 0.544677687326956 | -4.29086428667507 | 1.77979064035e-05 | 0.00187402678208973 | 6792 | 7 | 7 | 35 | 52 | 1 | 5 | 41 | 25 | 3 | 1 | 1 | 3 |
| THE1B-int_dup2450 | 14.0932849931955 | -2.15268681919372 | 0.455460429430494 | -4.72639702615973 | 2.28538602101975e-06 | 0.00038862122532163 | 6793 | 12 | 6 | 30 | 40 | 3 | 5 | 52 | 28 | 2 | 2 | 2 | 3 |
| L1ME1_dup12948 | 14.1563089482404 | -1.65125565653942 | 0.438403011579061 | -3.76652443739346 | 0.000165535887860152 | 0.00977384674262325 | 6794 | 9 | 17 | 37 | 53 | 6 | 8 | 25 | 19 | 2 | 1 | 2 | 8 |
| HAL1_dup19929 | 13.4495680284235 | -2.20420744424146 | 0.451240933889125 | -4.88476837693775 | 1.03550509979359e-06 | 0.000210954020955039 | 6795 | 9 | 6 | 42 | 44 | 6 | 5 | 16 | 23 | 2 | 2 | 5 | 15 |
| THE1B_dup3842 | 11.0679654398349 | -1.82762493970072 | 0.451069578733066 | -4.05175836693318 | 5.08341498257014e-05 | 0.00413316610779826 | 6796 | 7 | 6 | 34 | 31 | 10 | 3 | 19 | 16 | 3 | 1 | 4 | 10 |
| L1MC5a_dup13074 | 11.2981595503965 | -1.89596166683411 | 0.485451587670761 | -3.90556280994177 | 9.40063362691956e-05 | 0.00652140865595044 | 6797 | 6 | 9 | 34 | 28 | 4 | 5 | 24 | 22 | 3 | 1 | 7 | 1 |
| MIR_dup63985 | 10.8176187463057 | -1.74194814877707 | 0.419972453462936 | -4.14776763193305 | 3.35732894576076e-05 | 0.00300776729415659 | 6798 | 8 | 6 | 24 | 21 | 5 | 6 | 26 | 24 | 4 | 1 | 6 | 6 |
| MER57C2_dup385 | 12.666748371329 | -2.20186204250609 | 0.503597303693023 | -4.37226733812753 | 1.22962826139142e-05 | 0.00142238758524571 | 6799 | 8 | 10 | 32 | 42 | 1 | 4 | 20 | 29 | 1 | 3 | 8 | 4 |
| MIRb_dup213181 | 11.1422892419827 | -1.91243254026299 | 0.504251934563612 | -3.79261319427172 | 0.000149070262838687 | 0.00905788374908434 | 6800 | 10 | 12 | 27 | 42 | 2 | 2 | 25 | 15 | 2 | 1 | 4 | 6 |
| MSTA_dup17270 | 9.60632496328409 | -1.50442517566594 | 0.437317775979548 | -3.44011896679978 | 0.000581458538001031 | 0.0240302772324362 | 6801 | 10 | 11 | 20 | 25 | 2 | 6 | 24 | 14 | 2 | 1 | 5 | 6 |
| L2a_dup82516 | 8.39610039516147 | -1.78119953922281 | 0.537436106774926 | -3.31425357687917 | 0.000918881339655287 | 0.0331941792996004 | 6802 | 2 | 3 | 19 | 30 | 7 | 6 | 22 | 5 | 2 | 3 | 3 | 12 |
| L2a_dup168132 | 7.6420055155106 | -2.79691952947684 | 0.566048031784432 | -4.9411346253775 | 7.76692543922527e-07 | 0.000169551269776321 | 6803 | 1 | 2 | 20 | 28 | 3 | 4 | 22 | 8 | 1 | 1 | 5 | 8 |
| MLT1B_dup13201 | 7.79303560208712 | -2.93342855140608 | 0.578437098722564 | -5.07130085169908 | 3.95105583926059e-07 | 9.92753888837065e-05 | 6804 | 1 | 2 | 13 | 34 | 2 | 3 | 17 | 12 | 1 | 2 | 7 | 10 |
| L1MC_dup661 | 8.39087484921982 | -2.14581558802438 | 0.595549573230693 | -3.60308475478191 | 0.000314462935720977 | 0.0154479128898742 | 6805 | 2 | 2 | 20 | 28 | 2 | 4 | 24 | 12 | 3 | 4 | 1 | 9 |
| L1M5_dup47243 | 6.15496952166807 | -1.96258891344316 | 0.564964431929723 | -3.47382738191081 | 0.000513091005614466 | 0.0219701192049558 | 6806 | 4 | 1 | 17 | 20 | 3 | 3 | 12 | 9 | 1 | 3 | 2 | 6 |
| L1ME1_dup27651 | 7.6241315427656 | -2.42977668931093 | 0.544215411234431 | -4.464733337484 | 8.01685365833265e-06 | 0.00103279819197922 | 6807 | 2 | 2 | 17 | 26 | 4 | 2 | 23 | 12 | 1 | 3 | 3 | 6 |
| MIRb_dup190725 | 7.84844753761379 | -2.01918827314099 | 0.557888699368499 | -3.61933890295072 | 0.00029535661551762 | 0.0148048568047797 | 6808 | 2 | 4 | 24 | 29 | 5 | 2 | 14 | 9 | 1 | 4 | 3 | 7 |
| MSTA_dup17271 | 9.34583632414661 | -2.20062040944693 | 0.515237218717685 | -4.27108199777144 | 1.94526861993075e-05 | 0.00200462922060651 | 6809 | 7 | 2 | 20 | 43 | 3 | 6 | 16 | 12 | 1 | 2 | 8 | 5 |
| MSTB_dup7479 | 10.5315205606919 | -2.91607832883608 | 0.537449878684952 | -5.42576795434623 | 5.77058879846676e-08 | 2.15217636714729e-05 | 6810 | 5 | 1 | 33 | 43 | 2 | 4 | 21 | 14 | 1 | 2 | 9 | 5 |
| MLT1C_dup16507 | 9.70157856362478 | -2.47993956138546 | 0.522291914185723 | -4.74818677836856 | 2.0524844692999e-06 | 0.000360428390798549 | 6811 | 5 | 4 | 32 | 38 | 3 | 5 | 12 | 11 | 1 | 1 | 7 | 10 |
| LTR16C_dup5050 | 9.40304483070712 | -2.03058027232465 | 0.435543974851015 | -4.66217050303415 | 3.12891791612243e-06 | 0.000495887523974459 | 6812 | 6 | 5 | 23 | 28 | 5 | 3 | 23 | 12 | 2 | 2 | 8 | 7 |
| MER51C_dup144 | 11.2524664511489 | -2.26397152038956 | 0.493301476255589 | -4.58942782327403 | 4.44462646163368e-06 | 0.000655660934928634 | 6813 | 7 | 3 | 35 | 46 | 7 | 3 | 18 | 10 | 2 | 2 | 7 | 11 |
| ERVL-E-int_dup7491 | 7.81362978796276 | -2.0124243661026 | 0.53300971587869 | -3.77558664720592 | 0.000159631521061336 | 0.00952927293288023 | 6814 | 4 | 4 | 25 | 27 | 6 | 2 | 17 | 7 | 1 | 2 | 6 | 3 |
| MIR3_dup73999 | 8.25551419157039 | -2.70388565077289 | 0.578303227487988 | -4.6755499922038 | 2.93166846084608e-06 | 0.000472845633338699 | 6815 | 2 | 3 | 22 | 35 | 3 | 2 | 17 | 12 | 1 | 2 | 8 | 2 |
| L1MEf_dup6278 | 10.8295994308164 | -3.21139528626893 | 0.523698068442129 | -6.13215033582619 | 8.66991146277128e-10 | 6.44678360287679e-07 | 6816 | 3 | 2 | 36 | 40 | 3 | 3 | 33 | 14 | 1 | 1 | 4 | 5 |
| MER31A_dup730 | 9.43407362790743 | -2.77750265983068 | 0.502823845152961 | -5.5238085596075 | 3.31728910574607e-08 | 1.37037443325669e-05 | 6817 | 2 | 3 | 28 | 31 | 5 | 3 | 26 | 13 | 1 | 1 | 6 | 6 |
| L1MB7_dup2181 | 10.1911583888111 | -3.35895639469778 | 0.615721672081525 | -5.4553161712538 | 4.88857827867964e-08 | 1.87313630772738e-05 | 6818 | 4 | 1 | 35 | 50 | 3 | 1 | 20 | 10 | 1 | 1 | 6 | 6 |
| LTR86B1_dup161 | 9.57692960292543 | -3.14363048765554 | 0.609624920614794 | -5.1566633537311 | 2.51389176768468e-07 | 6.91526578674565e-05 | 6819 | 2 | 2 | 26 | 49 | 3 | 1 | 23 | 12 | 2 | 1 | 4 | 4 |
| L1PA15_dup3421 | 11.988884900842 | -2.03514120767135 | 0.570451039988526 | -3.56760013569663 | 0.000360265719738843 | 0.0171037084894212 | 6820 | 5 | 5 | 43 | 46 | 10 | 8 | 7 | 19 | 1 | 1 | 2 | 9 |
| LTR37-int_dup1170 | 8.3168286761092 | -1.69141765178651 | 0.50710101847923 | -3.3354649076805 | 0.000851569009809436 | 0.031415106915987 | 6821 | 4 | 6 | 21 | 25 | 9 | 2 | 13 | 16 | 1 | 2 | 2 | 6 |
| MER94B_dup510 | 8.32247411258286 | -1.5659538524721 | 0.490413578024224 | -3.19312907032674 | 0.00140740023922113 | 0.0444793807041719 | 6822 | 4 | 4 | 25 | 20 | 5 | 6 | 9 | 15 | 2 | 4 | 4 | 8 |
| L1M1_dup4232 | 8.26046070817469 | -1.81522578051219 | 0.486478643794155 | -3.73135759127029 | 0.00019045066009132 | 0.0107284499919719 | 6823 | 5 | 3 | 11 | 22 | 5 | 4 | 15 | 19 | 4 | 1 | 5 | 11 |
| MLT1K_dup3426 | 8.27105069840721 | -1.86956940566809 | 0.54282196464221 | -3.44416683083261 | 0.000572822129553709 | 0.0237581897226402 | 6824 | 5 | 2 | 14 | 37 | 8 | 4 | 12 | 12 | 3 | 0 | 6 | 6 |
| L1MB8_dup7971 | 8.73474377747788 | -2.18260054046193 | 0.496163839927736 | -4.39895124316156 | 1.08775258759254e-05 | 0.00130023601977413 | 6825 | 2 | 2 | 18 | 21 | 8 | 6 | 26 | 16 | 1 | 1 | 4 | 9 |
| AluSc_dup16512 | 10.4285725109926 | -2.63347755990146 | 0.533950051679203 | -4.93206724415423 | 8.13638949377963e-07 | 0.000175523239754024 | 6826 | 1 | 2 | 30 | 41 | 8 | 5 | 18 | 17 | 1 | 1 | 5 | 8 |
| L1ME2_dup7506 | 9.61866876053147 | -2.30752659497216 | 0.589417324961116 | -3.91492834915294 | 9.04310586926049e-05 | 0.00632861622599916 | 6827 | 1 | 1 | 31 | 29 | 7 | 4 | 14 | 18 | 4 | 1 | 4 | 9 |
| HERVL18-int_dup496 | 13.2733612891477 | -4.05290946265544 | 0.604820328078767 | -6.70101396150101 | 2.06978262803913e-11 | 2.66214322816973e-08 | 6828 | 1 | 0 | 37 | 62 | 3 | 6 | 31 | 21 | 0 | 0 | 7 | 10 |
| MSTA_dup19401 | 13.7163155657443 | -3.97739565178154 | 0.597888146467997 | -6.65240760379323 | 2.8833683400813e-11 | 3.53652558272672e-08 | 6829 | 1 | 0 | 28 | 40 | 3 | 7 | 59 | 24 | 0 | 0 | 12 | 9 |
| Tigger1_dup5772 | 14.1089040678083 | -3.63678544401531 | 0.695783383464769 | -5.22689321194382 | 1.72381992622917e-07 | 5.12720070208157e-05 | 6830 | 0 | 0 | 22 | 31 | 10 | 5 | 62 | 31 | 0 | 0 | 11 | 13 |
| L1M1_dup3993 | 12.3110883280346 | -3.42005413736394 | 0.678065512490086 | -5.04384026965823 | 4.56280237153663e-07 | 0.000111697589824862 | 6831 | 1 | 0 | 12 | 32 | 6 | 8 | 39 | 38 | 0 | 0 | 8 | 13 |
| MLT1F2_dup2599 | 11.5232086824955 | -3.61080620938846 | 0.709286984804121 | -5.09075492254469 | 3.56640740733209e-07 | 9.21403232296462e-05 | 6832 | 0 | 0 | 23 | 24 | 3 | 9 | 36 | 36 | 0 | 0 | 8 | 6 |
| HAL1M8_dup1424 | 11.1028144879233 | -3.50395271211154 | 0.587837656035167 | -5.96074898594437 | 2.51084407708469e-09 | 1.58054313430157e-06 | 6833 | 0 | 3 | 31 | 50 | 3 | 6 | 30 | 14 | 0 | 0 | 6 | 7 |
| L2c_dup70124 | 9.85196180103855 | -4.53894567422664 | 0.737999444426321 | -6.15033752194077 | 7.73182293833639e-10 | 5.80965556768678e-07 | 6834 | 0 | 1 | 32 | 49 | 3 | 1 | 21 | 12 | 0 | 0 | 8 | 6 |
| MLT1A-int_dup445 | 8.00593538943672 | -3.80175454998514 | 0.705043200490093 | -5.39222922417015 | 6.95889162252907e-08 | 2.49749687181644e-05 | 6835 | 0 | 1 | 28 | 34 | 2 | 4 | 15 | 10 | 0 | 0 | 7 | 6 |
| L1MB1_dup6018 | 10.0420760505937 | -3.03109799357408 | 0.566263721202681 | -5.35280273144882 | 8.66022461515902e-08 | 2.9564789186207e-05 | 6836 | 1 | 3 | 28 | 41 | 6 | 3 | 16 | 19 | 0 | 1 | 8 | 5 |
| MER47A_dup1240 | 9.75810876790342 | -3.05171879523359 | 0.684641349189946 | -4.45739772925536 | 8.2960594366553e-06 | 0.00105845263201844 | 6837 | 0 | 1 | 28 | 44 | 3 | 5 | 21 | 13 | 0 | 3 | 10 | 2 |
| MER66-int_dup747 | 11.1658503360351 | -3.22832049982162 | 0.630173210729871 | -5.12290977282667 | 3.00856335464022e-07 | 8.02551138994644e-05 | 6838 | 0 | 2 | 33 | 38 | 7 | 4 | 38 | 11 | 0 | 1 | 3 | 15 |
| L1PA5_dup7294 | 9.84775389972991 | -2.37843943960396 | 0.618150594684859 | -3.84766990447776 | 0.000119246575113417 | 0.00772088314610534 | 6839 | 0 | 2 | 23 | 40 | 11 | 3 | 21 | 14 | 1 | 2 | 3 | 11 |
| L1MB7_dup10218 | 11.8278856950641 | -3.0032635727119 | 0.599655490064633 | -5.00831497830229 | 5.49086240336038e-07 | 0.000128788430838433 | 6840 | 0 | 2 | 41 | 54 | 5 | 5 | 18 | 16 | 2 | 1 | 4 | 9 |
| L1PA7_dup8445 | 11.4725217688567 | -2.64389169469983 | 0.573031518734718 | -4.61386783843526 | 3.95243802915952e-06 | 0.000594855673172924 | 6841 | 0 | 1 | 35 | 45 | 7 | 5 | 22 | 14 | 3 | 2 | 7 | 11 |
| Tigger3a_dup2811 | 13.9350274788782 | -3.19538887155098 | 0.627119051316436 | -5.09534651330282 | 3.48103952604895e-07 | 9.04257020813584e-05 | 6842 | 2 | 0 | 40 | 68 | 4 | 11 | 20 | 24 | 0 | 1 | 5 | 10 |
| L1PA5_dup10951 | 10.2848152161973 | -2.33441849909222 | 0.554850458866328 | -4.2072930855314 | 2.58447859293851e-05 | 0.00246907119251075 | 6843 | 5 | 0 | 30 | 37 | 6 | 11 | 18 | 15 | 0 | 1 | 6 | 7 |
| L2a_dup168108 | 9.67959829906943 | -2.77722413492748 | 0.570098517573555 | -4.87148106742648 | 1.10764778518557e-06 | 0.000222978307862746 | 6844 | 2 | 0 | 26 | 37 | 7 | 6 | 28 | 12 | 0 | 1 | 6 | 5 |
| L1MEc_dup18239 | 11.5899460032999 | -3.52643117336457 | 0.527446517089274 | -6.68585545473929 | 2.29579546085007e-11 | 2.89237561814592e-08 | 6845 | 3 | 2 | 28 | 45 | 3 | 3 | 34 | 14 | 0 | 1 | 10 | 14 |
| L1MEd_dup21119 | 11.9438514346905 | -3.3100344377938 | 0.525146542440479 | -6.30306813487011 | 2.91810530903511e-10 | 2.55741922875659e-07 | 6846 | 4 | 1 | 39 | 39 | 2 | 5 | 22 | 16 | 0 | 2 | 10 | 19 |
| L2b_dup60232 | 11.1957859967299 | -3.53497932135952 | 0.554749015215363 | -6.37221378389862 | 1.86318916584369e-10 | 1.73511255455854e-07 | 6847 | 6 | 2 | 27 | 41 | 2 | 2 | 38 | 15 | 0 | 0 | 11 | 7 |
| MER41-int_dup513 | 10.075268275498 | -3.77374804285868 | 0.605086820323443 | -6.23670507455684 | 4.46883360111234e-10 | 3.70502118484417e-07 | 6848 | 2 | 1 | 39 | 32 | 4 | 2 | 24 | 15 | 0 | 0 | 7 | 7 |
| MSTA_dup13430 | 14.0581994255567 | -3.67127143208237 | 0.554127643613373 | -6.62531724305004 | 3.46503193703497e-11 | 4.08163608559269e-08 | 6849 | 3 | 2 | 41 | 52 | 2 | 7 | 34 | 20 | 0 | 0 | 6 | 22 |
| L1MA5_dup3295 | 14.0211618360763 | -3.18261397634471 | 0.506266021726714 | -6.28644593901406 | 3.24816054121345e-10 | 2.78016837460279e-07 | 6850 | 3 | 5 | 43 | 63 | 6 | 5 | 25 | 18 | 0 | 0 | 8 | 12 |
| L1ME4c_dup4308 | 12.1782497875272 | -3.43906837681363 | 0.564782001492378 | -6.08919612828711 | 1.13479032229317e-09 | 7.94172994201091e-07 | 6851 | 5 | 1 | 36 | 55 | 4 | 4 | 21 | 21 | 0 | 0 | 4 | 11 |
| MSTA_dup17272 | 8.07688279129578 | -3.1218858187753 | 0.623460436387165 | -5.00735192896286 | 5.5183973681999e-07 | 0.00012911313472701 | 6852 | 2 | 2 | 31 | 27 | 2 | 1 | 14 | 10 | 0 | 3 | 6 | 9 |
| Tigger3a_dup4712 | 7.62577204767246 | -2.86410832819766 | 0.703938504788838 | -4.06869109831804 | 4.7277976736515e-05 | 0.00390985380164496 | 6853 | 1 | 2 | 32 | 25 | 2 | 1 | 20 | 7 | 0 | 4 | 4 | 4 |
| L2a_dup147242 | 7.29887704767287 | -3.30485638141465 | 0.675327330384108 | -4.89371040193936 | 9.89524172588801e-07 | 0.000204209296418747 | 6854 | 1 | 2 | 28 | 31 | 2 | 1 | 14 | 7 | 1 | 1 | 4 | 6 |
| LTR71A_dup139 | 8.85589597853998 | -2.68335885592175 | 0.646567888466856 | -4.15015793977108 | 3.32245939304989e-05 | 0.00298326763297603 | 6855 | 2 | 2 | 36 | 35 | 2 | 2 | 12 | 11 | 1 | 4 | 3 | 7 |
| L1ME3Cz_dup5052 | 8.57320753332686 | -2.72768416553439 | 0.558268052650329 | -4.88597574692829 | 1.02917867713911e-06 | 0.00021016339570728 | 6856 | 3 | 6 | 18 | 37 | 2 | 2 | 19 | 19 | 0 | 1 | 3 | 2 |
| MER84_dup28 | 8.6204730623638 | -3.14087779289538 | 0.59241458439354 | -5.30182388421569 | 1.14651420028929e-07 | 3.69158955212498e-05 | 6857 | 2 | 6 | 26 | 32 | 1 | 1 | 23 | 14 | 0 | 1 | 5 | 3 |
| L1MC5a_dup1465 | 10.5217499681133 | -3.84670923249237 | 0.631716108490109 | -6.08930052723612 | 1.13405063944518e-09 | 7.94172994201091e-07 | 6858 | 1 | 3 | 37 | 33 | 1 | 1 | 24 | 24 | 0 | 2 | 2 | 8 |
| L1ME3A_dup15573 | 8.43903609479688 | -3.37751030932954 | 0.595911853200651 | -5.66780185893079 | 1.44641089249293e-08 | 6.99777351904828e-06 | 6859 | 1 | 1 | 23 | 23 | 3 | 2 | 30 | 16 | 0 | 2 | 4 | 6 |
| LTR32_dup1057 | 9.64372413794682 | -2.40333739654495 | 0.563424841468782 | -4.26558649824479 | 1.99377748869938e-05 | 0.00203784956325629 | 6860 | 7 | 1 | 33 | 35 | 7 | 3 | 22 | 6 | 0 | 2 | 7 | 8 |
| L2a_dup19314 | 11.0568630437885 | -2.79894236837315 | 0.59002569657351 | -4.74376350831431 | 2.09783745338077e-06 | 0.000365426083954715 | 6861 | 5 | 1 | 42 | 47 | 5 | 4 | 24 | 10 | 0 | 2 | 7 | 2 |
| MER2_dup8081 | 9.05325018807624 | -2.71825767615508 | 0.633264667458818 | -4.29245119116305 | 1.76711403594554e-05 | 0.00186556295437233 | 6862 | 4 | 1 | 33 | 31 | 5 | 1 | 24 | 9 | 0 | 3 | 8 | 2 |
| L2a_dup40240 | 6.99032262842552 | -2.5128051080921 | 0.647027153661233 | -3.88361615717869 | 0.000102914331806387 | 0.00695354856729831 | 6863 | 2 | 0 | 22 | 22 | 3 | 3 | 22 | 7 | 0 | 4 | 4 | 5 |
| L2a_dup86179 | 9.1573450721665 | -3.26012545915056 | 0.694970898341657 | -4.69102442552614 | 2.71840512688366e-06 | 0.000446909959820496 | 6864 | 1 | 0 | 31 | 45 | 3 | 6 | 22 | 6 | 0 | 1 | 4 | 7 |
| L1PA13_dup4373 | 9.57003917419597 | -3.81155504607243 | 0.666348559505862 | -5.72006195811232 | 1.06485215713305e-08 | 5.36816337671991e-06 | 6865 | 1 | 0 | 28 | 49 | 3 | 3 | 20 | 11 | 0 | 1 | 6 | 8 |
| MIRb_dup104843 | 9.19604415925054 | -3.49769668714853 | 0.686382619414619 | -5.09584099045447 | 3.47196462590545e-07 | 9.02885349590928e-05 | 6866 | 1 | 0 | 38 | 39 | 3 | 3 | 16 | 8 | 0 | 2 | 6 | 8 |
| MIR_dup84432 | 9.37474881754911 | -3.64139061781348 | 0.695794881582654 | -5.23342541631059 | 1.66397166438495e-07 | 4.96782185261908e-05 | 6867 | 1 | 0 | 28 | 51 | 3 | 2 | 17 | 13 | 0 | 2 | 6 | 3 |
| AluJr4_dup9721 | 7.02146114610724 | -3.94776090513565 | 0.769217556670261 | -5.13217732864089 | 2.86409462898356e-07 | 7.70056339647595e-05 | 6868 | 1 | 0 | 24 | 36 | 2 | 1 | 16 | 7 | 0 | 1 | 2 | 6 |

| | | | | | | | | | | | | | | | | | | | |
|---|---|---|---|---|---|---|---|---|---|---|---|---|---|---|---|---|---|---|---|
| AluJr_dup38509 | 7.77467548208846 | -4.03252666449287 | 0.7717677568799 | -5.22505200372138 | 1.74106170030572e-07 | 5.16556941821626e-05 | 6869 | 1 | 0 | 21 | 49 | 1 | 2 | 20 | 7 | 0 | 1 | 2 | 4 |
| L1PA6_dup5433 | 8.33491435316859 | -2.90997889235297 | 0.640678184804276 | -4.54202899579287 | 5.57153754986273e-06 | 0.000781216896782344 | 6870 | 1 | 1 | 17 | 33 | 6 | 1 | 32 | 14 | 2 | 0 | 2 | 2 |
| L2a_dup168116 | 7.94275147077023 | -3.1620111910673 | 0.758467324945731 | -4.1689484662949 | 3.06008162263907e-05 | 0.00279844970359105 | 6871 | 1 | 1 | 13 | 36 | 7 | 0 | 32 | 7 | 1 | 0 | 8 | 3 |
| L1MA8_dup8018 | 7.81172020635532 | -4.06315274490977 | 0.714661955640361 | -5.68541911716714 | 1.30492233807463e-08 | 6.39061644141837e-06 | 6872 | 2 | 0 | 22 | 30 | 3 | 0 | 28 | 14 | 0 | 0 | 4 | 1 |
| MSTB1_dup4474 | 7.6693318910277 | -3.48899405190124 | 0.685174220927843 | -5.09212685669427 | 3.54069065506167e-07 | 9.16749922316979e-05 | 6873 | 3 | 0 | 20 | 30 | 5 | 0 | 21 | 15 | 0 | 0 | 5 | 2 |
| HAL1ME_dup4649 | 7.36420141220061 | -3.72109454620658 | 0.720346139887251 | -5.16570345860257 | 2.39536304745299e-07 | 6.62752390336335e-05 | 6874 | 2 | 0 | 20 | 37 | 2 | 0 | 15 | 12 | 1 | 1 | 5 | 3 |
| Tigger3b_dup3524 | 7.89054699599546 | -3.31136970142756 | 0.663769752954067 | -4.98873244327648 | 6.0776752604231e-07 | 0.000139194264682893 | 6875 | 2 | 0 | 18 | 34 | 3 | 0 | 25 | 14 | 1 | 2 | 3 | 3 |
| MER44B_dup2130 | 7.7288722937473 | -4.35998605952103 | 0.756171134807591 | -5.76587211389712 | 8.1236644974612e-09 | 4.24833730222616e-06 | 6876 | 0 | 0 | 28 | 28 | 2 | 2 | 19 | 15 | 0 | 0 | 5 | 2 |
| AluSq_dup8355 | 7.6967918416805 | -3.63829530230071 | 0.753411633751639 | -4.82909360475853 | 1.37155927581227e-06 | 0.000263829461149899 | 6877 | 0 | 0 | 30 | 38 | 4 | 3 | 15 | 8 | 0 | 0 | 3 | 3 |
| Cheshire_dup502 | 7.18725883775395 | -3.588508824867 | 0.717864882350533 | -4.99886387131379 | 5.76690965802854e-07 | 0.000134398930997969 | 6878 | 0 | 0 | 27 | 28 | 4 | 3 | 17 | 9 | 0 | 0 | 4 | 4 |
| ERV3-16A3_I-int_dup6 | 6.04652952132946 | -3.75365795795131 | 0.76588202978348 | -4.90109156760408 | 9.530561875476e-07 | 0.000198230688463463 | 6879 | 0 | 0 | 19 | 21 | 4 | 1 | 12 | 14 | 0 | 0 | 3 | 4 |
| L1MC2_dup6215 | 5.48020286393523 | -2.68885618183757 | 0.748772982690177 | -3.59101656175828 | 0.000329390701748539 | 0.0159823001464637 | 6880 | 0 | 0 | 18 | 16 | 5 | 4 | 11 | 13 | 0 | 0 | 1 | 2 |
| LTR1B_dup178 | 4.46957496622642 | -3.00397013997631 | 0.769439001657617 | -3.90410433251343 | 9.45749781729597e-05 | 0.00653988310268616 | 6881 | 0 | 0 | 14 | 11 | 4 | 2 | 14 | 9 | 0 | 0 | 2 | 2 |
| MLT2A2_dup1776 | 7.9936112210745 | -2.97693053488161 | 0.702751198068609 | -4.23610880075792 | 2.27426609863078e-05 | 0.002250406309479З | 6882 | 1 | 0 | 23 | 10 | 7 | 4 | 36 | 16 | 0 | 0 | 3 | 4 |
| THE1D-int_dup2261 | 8.21823251794345 | -2.5184658951052 | 0.629145232395972 | -4.00299607375889 | 6.25453405471094e-05 | 0.00482255787486147 | 6883 | 2 | 0 | 21 | 26 | 9 | 4 | 31 | 14 | 0 | 0 | 1 | 1 |
| MLT2B1_dup3987 | 8.68454508709376 | -2.67459727203162 | 0.743654452241781 | -3.59655921371669 | 0.000322454207007304 | 0.0157452675437225 | 6884 | 0 | 0 | 17 | 24 | 10 | 3 | 30 | 22 | 0 | 1 | 1 | 4 |
| THE1D_dup11443 | 7.26192723693284 | -2.96252432631611 | 0.656773377929941 | -4.51072535195257 | 6.46063254260085e-06 | 0.000876633285858476 | 6885 | 0 | 1 | 17 | 20 | 6 | 2 | 19 | 20 | 0 | 1 | 4 | 2 |
| L1PA5_dup10355 | 37.8619646721327 | -5.79320209099383 | 1.2890456703476 | -4.49417908477335 | 6.98388370190932e-06 | 0.000928372732589114 | 6886 | 0 | 0 | 215 | 286 | 3 | 1 | 13 | 2 | 1 | 0 | 5 | 3 |
| L1PA12_dup1706 | 38.5657936822046 | -5.70031734542663 | 1.05953884748386 | -5.37999843890902 | 7.4486484629958e-08 | 2.63746444520238e-05 | 6887 | 0 | 0 | 209 | 259 | 1 | 3 | 34 | 20 | 1 | 0 | 2 | 2 |
| L1PA8_dup6180 | 58.5109754896935 | -4.71397640638622 | 1.09007272300708 | -4.32446047579487 | 1.5290561381687e-05 | 0.00167664881038106 | 6888 | 3 | 1 | 316 | 452 | 4 | 6 | 18 | 13 | 1 | 0 | 0 | 2 |
| L1MA3_dup2025 | 30.4422397414132 | -4.29846095900465 | 1.18105280330503 | -3.63951632558335 | 0.000273150620632759 | 0.0139835390025421 | 6889 | 6 | 7 | 158 | 239 | 2 | 0 | 6 | 4 | 0 | 0 | 2 | 2 |
| LTR1A2_dup110 | 33.7991739800622 | -4.62742202722188 | 1.38538852528864 | -3.34016194212218 | 0.000837295499999905 | 0.0310619137890517 | 6890 | 9 | 5 | 200 | 242 | 0 | 0 | 5 | 3 | 0 | 0 | 3 | 3 |
| L1PA16_dup2993 | 45.8578454870208 | -3.98734326659025 | 1.18535760904245 | -3.36383150213317 | 0.000768684582228305 | 0.0292134517813283 | 6891 | 17 | 15 | 245 | 342 | 2 | 1 | 10 | 2 | 0 | 0 | 4 | 3 |
| L1MA3_dup2024 | 54.2013180107203 | -4.72154356274893 | 1.47318914610693 | -3.20498123083933 | 0.00135071288156717 | 0.0432333504596962 | 6892 | 11 | 12 | 282 | 437 | 0 | 0 | 9 | 2 | 0 | 0 | 4 | 5 |
| L1PA7_dup2815 | 57.2821995395335 | -4.68312211495279 | 1.4081023523154 | -3.32583928096714 | 0.000881527423076092 | 0.0322412843617357 | 6893 | 17 | 12 | 307 | 434 | 0 | 0 | 10 | 7 | 0 | 0 | 7 | 6 |
| L1PREC2_dup1646 | 47.3082658427876 | -4.64290876051937 | 1.27842022211472 | -3.63175478626208 | 0.000281500543710119 | 0.014279130225429 | 6894 | 14 | 8 | 246 | 371 | 1 | 0 | 6 | 6 | 0 | 0 | 7 | 3 |
| MER57-int_dup685 | 36.5789934855054 | -5.56366902701385 | 1.53899784337202 | -3.61512464164575 | 0.000300203152102847 | 0.0149627438689284 | 6895 | 3 | 0 | 240 | 227 | 0 | 0 | 4 | 1 | 3 | 7 | 10 | 7 |
| HERVIP10FH-int_dup14 | 19.8405854427811 | -3.96556002160029 | 1.1255154291313 | -3.52332799618839 | 0.000426163609812255 | 0.0193582707786507 | 6896 | 7 | 2 | 110 | 129 | 0 | 1 | 3 | 5 | 2 | 0 | 6 | 8 |
| L1PA5_dup8505 | 32.0053854401685 | -4.80373061742651 | 1.4297848738876 | -3.35975761469984 | 0.000780108822191765 | 0.0295297126638946 | 6897 | 6 | 5 | 202 | 208 | 0 | 0 | 1 | 1 | 1 | 0 | 8 | 10 |
| BSR/Beta_dup1970 | 59.1244038726683 | -3.86971586775424 | 0.891267977628199 | -4.34180960708602 | 1.41313967037588e-05 | 0.00157864287327352 | 6898 | 6 | 0 | 214 | 279 | 31 | 25 | 58 | 32 | 1 | 1 | 79 | 78 |
| BSR/Beta_dup2013 | 38.1498790088699 | -3.30327681418537 | 0.993916083414162 | -3.32349669082566 | 0.000888964774783376 | 0.0323928962175506 | 6899 | 0 | 0 | 129 | 157 | 31 | 18 | 47 | 38 | 2 | 1 | 38 | 49 |
| BSR/Beta_dup2006 | 37.1594281942905 | -4.6581704593467 | 1.03269819260592 | -4.51067939568312 | 6.46203249222835e-06 | 0.000876633285858476 | 6900 | 0 | 0 | 155 | 178 | 7 | 14 | 30 | 21 | 0 | 1 | 37 | 64 |
| BSR/Beta_dup2017 | 53.672454322998 | -4.65797841048209 | 0.912848843433289 | -5.10268314846421 | 3.34871405781502e-07 | 8.76582084929432e-05 | 6901 | 1 | 1 | 232 | 237 | 10 | 21 | 34 | 29 | 0 | 1 | 69 | 93 |
| BSR/Beta_dup13 | 49.0961684520086 | -3.77191531030775 | 0.644044991579386 | -5.85660219336217 | 4.72432926791656e-09 | 2.65753014653351e-06 | 6902 | 2 | 3 | 194 | 217 | 26 | 9 | 19 | 30 | 2 | 3 | 68 | 89 |
| BSR/Beta_dup2009 | 32.4814243752506 | -3.64879381538672 | 0.70097162983047 | -5.20533736332407 | 1.93644480177071e-07 | 5.62599871553276e-05 | 6903 | 1 | 3 | 130 | 148 | 13 | 16 | 24 | 7 | 1 | 1 | 42 | 59 |
| BSR/Beta_dup1489 | 33.1523077659081 | -3.86003405877225 | 1.0165305995796 | -3.79726302422043 | 0.000146302528598555 | 0.00894592725757062 | 6904 | 0 | 1 | 111 | 149 | 17 | 16 | 19 | 22 | 1 | 0 | 46 | 67 |
| BSR/Beta_dup2008 | 39.2378718279482 | -3.97467386718341 | 0.869575185999766 | -4.57082254780955 | 4.85813540113174e-06 | 0.000704508218954913 | 6905 | 0 | 2 | 152 | 159 | 20 | 11 | 27 | 26 | 1 | 1 | 58 | 71 |
| L1ME2_dup12727 | 29.4710370252614 | -2.2249944019054 | 0.551550477919317 | -4.03407211303492 | 5.48184824215038e-05 | 0.00437391430415276 | 6906 | 22 | 25 | 73 | 78 | 20 | 16 | 40 | 13 | 0 | 1 | 52 | 57 |
| L1MC4_dup11945 | 26.4131665539326 | -1.55892942095784 | 0.433055677919269 | -3.5998360036477 | 0.000318417938939977 | 0.015602610152185З | 6907 | 35 | 27 | 62 | 75 | 16 | 15 | 24 | 24 | 2 | 3 | 25 | 43 |
| L1MEc_dup18387 | 26.4386926990695 | -1.83029307642378 | 0.548810056008521 | -3.33502102664709 | 0.000852929496490194 | 0.0314457951614279 | 6908 | 25 | 25 | 71 | 70 | 23 | 17 | 27 | 11 | 1 | 1 | 25 | 61 |
| L1MCa_dup3569 | 16.2257648478822 | -1.98535554706026 | 0.560540126751972 | -3.54186159439527 | 0.00039731388593194 | 0.0183785478036473 | 6909 | 18 | 15 | 53 | 61 | 7 | 9 | 13 | 9 | 1 | 0 | 17 | 15 |
| LTR76_dup155 | 22.4269959203509 | -2.36075981029231 | 0.558167635177316 | -4.22948172110043 | 2.34230308270622e-05 | 0.00230211338008102 | 6910 | 13 | 13 | 76 | 96 | 12 | 11 | 10 | 15 | 3 | 0 | 21 | 31 |
| L1MCa_dup3568 | 24.8760393084005 | -1.82903012929921 | 0.518298847555915 | -3.52891027623188 | 0.000417274579270384 | 0.019068334365099 | 6911 | 25 | 24 | 84 | 88 | 19 | 4 | 20 | 16 | 3 | 2 | 20 | 27 |
| AluJo_dup33604 | 19.8513213614935 | -1.54841028417593 | 0.430912960794524 | -3.5933249288231 | 0.000326485038683667 | 0.0158671995536405 | 6912 | 18 | 24 | 54 | 56 | 8 | 7 | 26 | 12 | 4 | 5 | 18 | 33 |
| L1P3_dup842 | 20.8828302845236 | -1.53570193276561 | 0.442513684592395 | -3.47040551792237 | 0.000519673135789254 | 0.0221450427778136 | 6913 | 27 | 21 | 46 | 56 | 10 | 11 | 24 | 14 | 1 | 5 | 25 | 40 |
| LOR1a_dup412 | 21.5346694437197 | -1.48703191031207 | 0.4457958135949 | -3.33567939618059 | 0.000850912328205666 | 0.0314006179973985 | 6914 | 22 | 28 | 39 | 76 | 10 | 11 | 23 | 14 | 2 | 5 | 28 | 31 |
| AluSc_dup16090 | 20.8449592867121 | -1.34193482342996 | 0.416961941941923 | -3.21836284908916 | 0.00128924610743382 | 0.0418455865662082 | 6915 | 27 | 26 | 47 | 56 | 7 | 12 | 21 | 19 | 3 | 5 | 25 | 27 |
| LTR49_dup715 | 20.0055817717755 | -1.53395420445034 | 0.458476935479635 | -3.34576090037245 | 0.000820571208574835 | 0.0306229041398287 | 6916 | 24 | 21 | 46 | 69 | 8 | 10 | 16 | 14 | 3 | 4 | 20 | 33 |
| REP522_dup224 | 30.5723027114488 | -2.01449350414188 | 0.392193187509999 | -5.13648265267362 | 2.79928176752734e-07 | 7.56906703927341e-05 | 6917 | 27 | 25 | 92 | 96 | 10 | 23 | 62 | 36 | 2 | 0 | 8 | 26 |
| L1PREC2_dup3226 | 34.2249201238605 | -2.07655953984112 | 0.375408824776861 | -5.53146170997819 | 3.1757319162664e-08 | 1.32785167498844e-05 | 6918 | 45 | 19 | 74 | 112 | 13 | 11 | 99 | 60 | 1 | 1 | 5 | 14 |
| L1MA3_dup3848 | 30.6615519808897 | -2.36968998206625 | 0.370205413491294 | -6.40101385800501 | 1.5434856152079e-10 | 1.46906491278504e-07 | 6919 | 20 | 22 | 77 | 144 | 9 | 13 | 71 | 37 | 1 | 1 | 9 | 10 |
| L1PREC2_dup3228 | 34.9274690331052 | -2.22393746629018 | 0.404645880564979 | -5.49600916036771 | 3.88482337514926e-08 | 1.56732256410709e-05 | 6920 | 19 | 30 | 86 | 151 | 12 | 11 | 97 | 44 | 2 | 2 | 2 | 15 |
| AluJr_dup83492 | 27.314506471586 | -2.12703030298061 | 0.321874923590558 | -6.60825105371149 | 3.88886804136227e-11 | 4.47024442014486e-08 | 6921 | 20 | 12 | 71 | 88 | 11 | 25 | 63 | 37 | 0 | 2 | 16 | 18 |
| AluJb_dup123575 | 29.2400216938139 | -1.90653361980612 | 0.351242357862864 | -5.42797181810998 | 5.69980367598027e-08 | 2.13581966218079e-05 | 6922 | 21 | 15 | 71 | 85 | 25 | 24 | 44 | 39 | 1 | 2 | 34 | 22 |
| L1PA16_dup13067 | 26.8317219999142 | -1.86984101294281 | 0.306472068809686 | -6.10117920437294 | 1.05288780274269e-09 | 7.5009114105213e-07 | 6923 | 17 | 17 | 63 | 75 | 26 | 15 | 69 | 33 | 1 | 2 | 22 | 15 |
| Cheshire_dup1051 | 19.9095333599643 | -1.7998658443777 | 0.428964507861799 | -4.1958386099336 | 2.71863789294427e-05 | 0.00255788458708785 | 6924 | 13 | 18 | 42 | 59 | 26 | 6 | 38 | 30 | 0 | 1 | 17 | 10 |
| AluSg7_dup8350 | 20.4430399293767 | -1.29897870832668 | 0.365637450038073 | -3.55264130682299 | 0.000381384119380628 | 0.0178428678077355 | 6925 | 12 | 25 | 45 | 55 | 20 | 21 | 35 | 26 | 0 | 3 | 16 | 9 |
| L1MA5A_dup3135 | 18.9511528802459 | -1.0338141003664 | 0.309402609339618 | -3.34132314712196 | 0.000833801167461447 | 0.0309650765041459 | 6926 | 15 | 19 | 40 | 49 | 20 | 20 | 35 | 25 | 1 | 5 | 12 | 6 |
| MSTA-int_dup3086 | 17.2364650279455 | -3.43942761821212 | 0.482076440635416 | -7.13461046484385 | 9.70616392151318e-13 | 1.86253458102288e-09 | 6927 | 3 | 2 | 40 | 67 | 10 | 3 | 35 | 35 | 1 | 0 | 11 | 19 |
| L2a_dup168107 | 15.3785440266792 | -3.17933120685818 | 0.472031460565619 | -6.73542225988178 | 1.63454539678936e-11 | 2.13699746694747e-08 | 6928 | 1 | 3 | 31 | 47 | 5 | 10 | 49 | 29 | 1 | 0 | 15 | 11 |
| L1MA8_dup10812 | 15.5647497811356 | -3.13292828533194 | 0.473118722744991 | -6.62186494576874 | 3.54695056031094e-11 | 4.15755023657018e-08 | 6929 | 4 | 1 | 42 | 52 | 8 | 9 | 39 | 20 | 0 | 0 | 16 | 17 |
| L1PA16_dup13858 | 16.3596196633182 | -3.48468226101605 | 0.458743988263617 | -7.59613717055096 | 3.05101120004777e-14 | 8.44158036054146e-11 | 6930 | 4 | 2 | 48 | 54 | 7 | 5 | 48 | 23 | 0 | 0 | 10 | 18 |
| PRIMA41-int_dup441 | 23.2101957674641 | -3.98937610944393 | 0.489011394223644 | -8.15804326150207 | 3.40495570816622e-16 | 1.44677784095593e-12 | 6931 | 1 | 0 | 67 | 59 | 8 | 8 | 80 | 37 | 1 | 0 | 23 | 23 |
| AluSx1_dup102096 | 21.168563487517 | -3.92077946290911 | 0.469477346504657 | -8.35137092790529 | 6.74751543906134e-17 | 3.27662103808753e-13 | 6932 | 1 | 1 | 46 | 68 | 5 | 9 | 43 | 36 | 0 | 1 | 34 | 35 |
| L1PA16_dup13857 | 26.6485002276535 | -3.88986113154781 | 0.402125170565069 | -9.67325951290676 | 3.91699635469941e-22 | 4.66016807307653e-18 | 6933 | 3 | 3 | 58 | 84 | 6 | 9 | 77 | 55 | 0 | 1 | 18 | 37 |
| L1PA5_dup6256 | 23.2426679499039 | -4.94934193293965 | 0.558906143278988 | -8.85540800089058 | 8.33782643106753e-19 | 5.83515425872587e-15 | 6934 | 2 | 2 | 38 | 68 | 2 | 3 | 48 | 32 | 0 | 0 | 62 | 52 |
| MSTB1_dup4192 | 18.8640607892716 | -3.68810057999609 | 0.473863983985025 | -7.78303628180496 | 7.08043167861003e-15 | 2.24634719493139e-11 | 6935 | 2 | 6 | 45 | 51 | 7 | 3 | 35 | 34 | 0 | 0 | 20 | 45 |
| L1M4_dup18817 | 15.9842570016353 | -3.92641692217406 | 0.555487855407315 | -7.06841181846359 | 1.56716886425505e-12 | 2.78284748189577e-09 | 6936 | 2 | 2 | 46 | 44 | 8 | 1 | 30 | 19 | 0 | 0 | 25 | 36 |
| LTR2_dup769 | 14.4599566788737 | -3.99217745252128 | 0.543364182924728 | -7.34714870426842 | 2.02479566956767e-13 | 4.37992754900862e-10 | 6937 | 2 | 1 | 26 | 51 | 5 | 3 | 32 | 17 | 0 | 0 | 27 | 31 |
| BSR/Beta_dup2012 | 26.0879040996498 | -4.36598520795528 | 0.679266019385028 | -6.4275042227315 | 1.29715993784415e-10 | 1.27542982880275e-07 | 6938 | 1 | 0 | 87 | 128 | 9 | 8 | 27 | 26 | 0 | 0 | 32 | 34 |
| L1PA3_dup10791 | 20.1717406347755 | -3.17647121548459 | 0.581380363309488 | -5.46367131735004 | 4.6638673413255e-08 | 1.80446923316917e-05 | 6939 | 0 | 1 | 53 | 55 | 9 | 13 | 25 | 25 | 2 | 1 | 40 | 41 |
| BSR/Beta_dup2014 | 15.8939985996729 | -3.24456192143401 | 0.674601186455359 | -4.8096000816161 | 1.51232561558798e-06 | 0.000284918314272919 | 6940 | 3 | 1 | 55 | 54 | 9 | 10 | 11 | 12 | 0 | 0 | 24 | 34 |
| BSR/Beta_dup2016 | 20.3809104987064 | -2.84681870226318 | 0.663134774590114 | -4.2929715215475 | 1.76297626344374e-05 | 0.00186358573958856 | 6941 | 5 | 3 | 67 | 87 | 18 | 11 | 10 | 12 | 1 | 0 | 20 | 42 |
| L1MB8_dup6723 | 18.1367690794847 | -2.05150127599294 | 0.527970879650611 | -3.88563338445964 | 0.000102063435114899 | 0.00691109451674722 | 6942 | 8 | 13 | 36 | 50 | 17 | 17 | 18 | 22 | 0 | 0 | 23 | 35 |
| L1PA4_dup11834 | 23.4996608091291 | -2.93021123533173 | 0.522093882108083 | -5.61242208680991 | 1.99514051034216e-08 | 9.14712338870667e-06 | 6943 | 5 | 7 | 73 | 79 | 17 | 13 | 31 | 21 | 0 | 0 | 31 | 37 |
| LSAU_dup93 | 20.5318237862207 | -2.54984422702668 | 0.688867099709313 | -3.70150385771458 | 0.00021432539718552 | 0.0116780102951009 | 6944 | 2 | 5 | 57 | 40 | 20 | 22 | 22 | 21 | 0 | 0 | 33 | 48 |
| L1M5_dup52918 | 18.3149226589404 | -2.14752861683108 | 0.479370032603373 | -4.47989751292595 | 7.46788881962566e-06 | 0.000976885251827734 | 6945 | 17 | 11 | 47 | 47 | 15 | 9 | 37 | 12 | 0 | 0 | 27 | 24 |
| MIR3_dup82835 | 17.391235796332 | -2.20325538450033 | 0.401171048149014 | -5.49205979510749 | 3.97272829807764e-08 | 1.58597375599916e-05 | 6946 | 7 | 9 | 52 | 30 | 13 | 13 | 39 | 23 | 1 | 0 | 19 | 21 |
| MLT1A1_dup6595 | 18.8364058443141 | -1.90754579109699 | 0.374781659377269 | -5.08975224205618 | 3.58531647311607e-07 | 9.25283854134573e-05 | 6947 | 14 | 13 | 44 | 35 | 11 | 19 | 37 | 36 | 0 | 0 | 13 | 21 |

| | | | | | | | | | | | | | | | | | | | |
|---|---|---|---|---|---|---|---|---|---|---|---|---|---|---|---|---|---|---|---|
| MSTA-int_dup1607 | 18.4157164369457 | -3.24224953061236 | 0.421521488008352 | -7.69177757919691 | 1.45104714369001e-14 | 4.38590095530486e-11 | 6948 | 7 | 4 | 48 | 68 | 4 | 6 | 38 | 23 | 0 | 2 | 26 | 20 |
| MER21B_dup2755 | 14.6793624201693 | -2.48805189508098 | 0.45624009906007 | -5.45338276974509 | 4.94205308790305e-08 | 1.89058161423501e-05 | 6949 | 7 | 10 | 42 | 35 | 5 | 4 | 18 | 27 | 0 | 3 | 23 | 14 |
| MIRb_dup213465 | 15.737043362142 | -2.39109644866791 | 0.430214984339701 | -5.55791066259068 | 2.73023056700783e-08 | 1.18765528792915e-05 | 6950 | 11 | 12 | 45 | 51 | 6 | 5 | 30 | 16 | 0 | 1 | 18 | 15 |
| MER41A_dup2266 | 16.5322393495056 | -2.94141265340084 | 0.474462985204641 | -6.19945653322605 | 5.66584619703672e-10 | 4.58559673197313e-07 | 6951 | 4 | 6 | 22 | 49 | 8 | 6 | 20 | 28 | 0 | 1 | 32 | 41 |
| L1M5_dup52926 | 13.6873728189396 | -1.92364232013574 | 0.494184803002709 | -3.89255660726014 | 9.91933445756644e-05 | 0.00678042504119536 | 6952 | 4 | 5 | 19 | 27 | 14 | 14 | 22 | 20 | 0 | 2 | 27 | 24 |
| Tigger13a_dup3017 | 15.7503815914211 | -1.94032584134128 | 0.489150484496957 | -3.96672578856119 | 7.28667447444709e-05 | 0.00538960225208824 | 6953 | 9 | 7 | 39 | 22 | 13 | 14 | 19 | 16 | 0 | 2 | 26 | 40 |
| MER21A_dup1036 | 15.2440566678741 | -2.83747117585711 | 0.49004000190535 | -5.79028480292342 | 7.02671736824145e-09 | 3.7488324908152e-06 | 6954 | 4 | 2 | 44 | 44 | 3 | 3 | 32 | 21 | 5 | 5 | 25 | 10 |
| MSTA-int_dup2374 | 16.2239092608506 | -3.57457696364053 | 0.509532272280066 | -7.015408361957 | 2.29278300921957e-12 | 3.73670236925862e-09 | 6955 | 2 | 1 | 41 | 61 | 3 | 3 | 27 | 24 | 3 | 3 | 28 | 17 |
| THE1A_dup3201 | 16.3362264378017 | -2.87986339702558 | 0.411792483547843 | -6.99348218358384 | 2.6814592880525e-12 | 4.25361674503293e-09 | 6956 | 6 | 2 | 25 | 44 | 7 | 2 | 40 | 32 | 1 | 6 | 23 | 25 |
| HERVL-int_dup1832 | 16.2691194990742 | -3.17195305604052 | 0.404745599649114 | -7.83690559895001 | 4.61784951754442e-15 | 1.56971260185946e-11 | 6957 | 4 | 4 | 33 | 36 | 4 | 4 | 36 | 23 | 2 | 2 | 37 | 28 |
| LTR2_dup770 | 14.3805100745958 | -3.3426575007041 | 0.451185077473257 | -7.40861714537141 | 1.27623151709986e-13 | 2.91359388978609e-10 | 6958 | 2 | 2 | 23 | 43 | 4 | 5 | 33 | 25 | 2 | 1 | 23 | 26 |
| L1PA15-16_dup460 | 24.8459001543403 | -4.69762803854656 | 0.534484017601531 | -8.78908982092096 | 1.50776486860865e-18 | 9.96573942849872e-15 | 6959 | 2 | 1 | 46 | 80 | 2 | 1 | 67 | 50 | 3 | 2 | 23 | 51 |
| L1PB3_dup3333 | 26.7052212340154 | -4.14427089696182 | 0.460874813818247 | -8.99218350125805 | 2.42366630707602e-19 | 1.9886265624259e-15 | 6960 | 4 | 3 | 70 | 97 | 2 | 6 | 48 | 30 | 2 | 1 | 29 | 69 |
| MIRb_dup113170 | 22.5889085019202 | -3.38443595769585 | 0.5316964060943 | -6.36535421135722 | 1.94839564375257e-10 | 1.7912288271436e-07 | 6961 | 2 | 2 | 88 | 71 | 3 | 6 | 27 | 25 | 1 | 10 | 29 | 34 |
| MSTA-int_dup1634 | 19.4929493552568 | -3.30719181479597 | 0.559559181401764 | -5.91035215705164 | 3.41377123764844e-09 | 2.02566885514587e-06 | 6962 | 2 | 2 | 60 | 70 | 1 | 4 | 37 | 17 | 2 | 12 | 27 | 27 |
| AluJb_dup60201 | 23.1541382899009 | -3.50121115998665 | 0.519416792946901 | -6.74065838365101 | 1.5767052457304e-11 | 2.07276633370478e-08 | 6963 | 3 | 5 | 102 | 82 | 2 | 5 | 35 | 15 | 4 | 3 | 26 | 28 |
| AluJr_dup40235 | 21.4489600316225 | -3.32308663427917 | 0.442560854816643 | -7.50876766011299 | 5.96865481838756e-14 | 1.4492015708327e-10 | 6964 | 5 | 5 | 68 | 79 | 3 | 5 | 27 | 25 | 2 | 4 | 33 | 28 |
| L1MEc_dup13778 | 20.9621767563005 | -3.31121055323205 | 0.468385918600154 | -7.06940670447167 | 1.55597500221088e-12 | 2.78284748189577e-09 | 6965 | 5 | 3 | 47 | 98 | 5 | 2 | 30 | 22 | 5 | 3 | 28 | 34 |
| L2c_dup70123 | 21.2686938070755 | -3.36885202338514 | 0.515712567042873 | -6.53242181531922 | 6.47145823421572e-11 | 6.93629549999411e-08 | 6966 | 1 | 2 | 66 | 96 | 7 | 6 | 29 | 24 | 4 | 2 | 16 | 32 |
| ERV3-16A3_I-int_dup1[illegible] | 25.0178719195133 | -2.75301743635999 | 0.502831398467994 | -5.47503088460221 | 4.37435311676163e-08 | 1.71194050447527e-05 | 6967 | 5 | 1 | 58 | 85 | 5 | 12 | 73 | 54 | 6 | 4 | 10 | 12 |
| L1PA10_dup3641 | 20.4729870791376 | -2.58262261549558 | 0.62877568712825 | -4.10738307533957 | 4.00167237790862e-05 | 0.00343996364029568 | 6968 | 6 | 0 | 50 | 68 | 4 | 7 | 62 | 40 | 10 | 2 | 6 | 11 |
| HSMAR1_dup538 | 28.8691734999604 | -3.56894470211334 | 0.473992999591202 | -7.52953040486125 | 5.09231277098768e-14 | 1.24917057175818e-10 | 6969 | 3 | 1 | 83 | 128 | 8 | 5 | 39 | 44 | 3 | 6 | 29 | 33 |
| L1MA2_dup3321 | 33.3006168253947 | -3.1277624787697 | 0.504953365554599 | -6.19416106937802 | 5.85962827000966e-10 | 4.66312745262782e-07 | 6970 | 4 | 0 | 89 | 140 | 20 | 12 | 50 | 57 | 3 | 3 | 27 | 33 |
| L1MC5_dup15126 | 27.7056995288594 | -2.76152455519408 | 0.376935212429713 | -7.32625783989078 | 2.36668250392507e-13 | 4.98356314229163e-10 | 6971 | 7 | 6 | 66 | 82 | 13 | 18 | 74 | 53 | 1 | 2 | 9 | 33 |
| L1PA7_dup6921 | 28.1802990982546 | -2.87093251621924 | 0.380590875325567 | -7.54335614001092 | 4.58028061004821e-14 | 1.17189188176186e-10 | 6972 | 4 | 5 | 66 | 111 | 14 | 14 | 64 | 51 | 2 | 3 | 12 | 26 |
| L1PA6_dup3196 | 27.1983740366468 | -2.88759661843279 | 0.423660162283275 | -6.81583230028137 | 9.37195718236994e-12 | 1.30410510158842e-08 | 6973 | 6 | 5 | 83 | 119 | 12 | 10 | 38 | 42 | 3 | 3 | 11 | 28 |
| L1MA2_dup3322 | 27.4723204847092 | -2.71116595547755 | 0.448170591128873 | -6.04940620634778 | 1.45380674280029e-09 | 9.91196272843432e-07 | 6974 | 2 | 3 | 80 | 104 | 15 | 14 | 50 | 38 | 3 | 6 | 17 | 32 |
| L1PA5_dup5827 | 31.7551823346335 | -2.79808580107409 | 0.375715348951458 | -7.44735558151392 | 9.52297018564181e-14 | 2.22152221940463e-10 | 6975 | 5 | 5 | 94 | 122 | 12 | 18 | 69 | 42 | 4 | 4 | 22 | 25 |
| L1ME2z_dup3479 | 30.1697517727915 | -3.05951396498034 | 0.343602746351263 | -8.90421859973324 | 5.37634117943687e-19 | 3.95022716806549e-15 | 6976 | 5 | 4 | 67 | 95 | 12 | 14 | 59 | 44 | 3 | 3 | 40 | 53 |
| L1MEc_dup17108 | 37.6661591743813 | -3.47125288800376 | 0.344493348321223 | -10.0764003279593 | 7.02528340195964e-24 | 1.44265534334244e-19 | 6977 | 6 | 7 | 78 | 125 | 12 | 8 | 63 | 44 | 4 | 2 | 63 | 92 |
| AluJr_dup40282 | 34.9153088936003 | -3.38721370840762 | 0.363991927750278 | -9.30573853476077 | 1.33064321193138e-20 | 1.31925512377593e-16 | 6978 | 4 | 6 | 101 | 115 | 7 | 10 | 49 | 48 | 6 | 4 | 46 | 64 |
| REP522_dup54 | 31.8433266334376 | -3.05980901739055 | 0.323732283280806 | -9.45166477183392 | 3.33483931835691e-21 | 3.6068712565716e-17 | 6979 | 6 | 9 | 68 | 67 | 10 | 9 | 48 | 60 | 5 | 3 | 51 | 75 |
| L1PREC2_dup6013 | 33.0644744073658 | -3.16582124738646 | 0.386931774251125 | -8.1818590719608 | 2.79497821063233e-16 | 1.23158126908726e-12 | 6980 | 5 | 11 | 86 | 120 | 7 | 3 | 55 | 32 | 6 | 8 | 39 | 72 |
| MLT1D_dup10232 | 27.9894066145066 | -2.82528841758652 | 0.382463936876872 | -7.38707142079144 | 1.50097984444414e-13 | 3.36935990628402e-10 | 6981 | 9 | 6 | 78 | 70 | 8 | 6 | 22 | 42 | 7 | 6 | 53 | 54 |
| LTR28B_dup259 | 28.6984244736748 | -3.24499287124894 | 0.339152011959501 | -9.56795996137694 | 1.09035852420638e-21 | 1.23545928286101e-17 | 6982 | 10 | 8 | 51 | 96 | 9 | 7 | 63 | 47 | 0 | 2 | 44 | 43 |
| Tigger18a_dup816 | 25.1192889568887 | -3.31892927068668 | 0.371785896094115 | -8.92699079108295 | 4.3775384686621e-19 | 3.36005731762669e-15 | 6983 | 9 | 9 | 72 | 77 | 6 | 4 | 50 | 40 | 0 | 2 | 33 | 28 |
| L1PA3_dup9727 | 22.4113538125628 | -3.17624018042916 | 0.445513549820138 | -7.12939074852263 | 1.00814160689141e-12 | 1.91906610234706e-09 | 6984 | 6 | 5 | 33 | 22 | 5 | 7 | 56 | 53 | 1 | 4 | 30 | 66 |
| L1PA4_dup9689 | 21.3872351407362 | -2.64853267410013 | 0.319908205683385 | -8.27903950898152 | 1.24172737703388e-16 | 5.68200120107124e-13 | 6985 | 8 | 5 | 37 | 44 | 8 | 5 | 43 | 35 | 3 | 7 | 41 | 42 |
| L1PBa_dup1725 | 22.38912020627 | -2.7788035261544 | 0.321546013490157 | -8.64200894917785 | 5.52331018046205e-18 | 3.36987067743647e-14 | 6986 | 8 | 6 | 44 | 44 | 3 | 8 | 56 | 33 | 4 | 6 | 35 | 47 |
| L1PB3_dup2923 | 16.6887039364103 | -1.86562680429463 | 0.455677046809208 | -4.09418648000448 | 4.23653243793656e-05 | 0.0036015217844865 | 6987 | 9 | 1 | 40 | 50 | 11 | 14 | 15 | 15 | 4 | 6 | 27 | 28 |
| L3_dup36489 | 24.1504277495597 | -1.79887655520201 | 0.511775598611012 | -3.51497132744168 | 0.000439801469872646 | 0.0197847965455045 | 6988 | 9 | 4 | 71 | 118 | 21 | 20 | 22 | 14 | 4 | 7 | 17 | 19 |
| MIR3_dup66147 | 18.9136166790691 | -2.06020844512398 | 0.48408827733475 | -4.25585278880722 | 2.08253703170699e-05 | 0.0021014900616902 1 | 6989 | 3 | 4 | 59 | 72 | 13 | 17 | 23 | 16 | 3 | 5 | 15 | 22 |
| THE1B-int_dup15 | 20.1377489591638 | -1.23854538284536 | 0.320112411031386 | -3.86909516833423 | 0.000109239999858533 | 0.00726676572724027 | 6990 | 14 | 10 | 26 | 36 | 16 | 14 | 55 | 36 | 7 | 11 | 18 | 15 |
| L1M1_dup4229 | 16.3475529279887 | -1.47036821839291 | 0.323145971948383 | -4.55016724957901 | 5.36032920934789e-06 | 0.000759207675028269 | 6991 | 8 | 10 | 34 | 46 | 11 | 12 | 33 | 19 | 5 | 7 | 15 | 14 |
| MER68_dup1196 | 18.5282944130786 | -1.18883378824561 | 0.335986708897153 | -3.53833576378022 | 0.000402657683818825 | 0.018564383885672 2 | 6992 | 11 | 12 | 35 | 47 | 9 | 15 | 42 | 27 | 7 | 12 | 10 | 13 |
| AluSx_dup112500 | 24.3263895279147 | -1.51287895575262 | 0.398817105097462 | -3.79341541878677 | 0.000148589259585635 | 0.00903557882989102 | 6993 | 7 | 5 | 59 | 64 | 22 | 15 | 36 | 27 | 11 | 15 | 16 | 40 |
| Tigger2a_dup3232 | 26.2370341755409 | -1.42328584071511 | 0.289367461157644 | -4.91861052732434 | 8.71606970226269e-07 | 0.000185077886391031 | 6994 | 13 | 13 | 61 | 58 | 17 | 20 | 40 | 30 | 15 | 8 | 32 | 31 |
| AluSq2_dup53192 | 20.7261695696255 | -1.10483255238001 | 0.320434173194145 | -3.44792361366094 | 0.000564913777325251 | 0.0235306142639945 | 6995 | 8 | 18 | 48 | 46 | 15 | 14 | 34 | 18 | 11 | 13 | 18 | 26 |
| MIRc_dup98469 | 20.3859511686406 | -1.14520086214547 | 0.334438456694094 | -3.42424993066204 | 0.000616498941658098 | 0.0250244723936844 | 6996 | 12 | 14 | 55 | 43 | 16 | 21 | 27 | 17 | 6 | 10 | 25 | 19 |
| L2a_dup44323 | 20.5979089740622 | -1.62034160906091 | 0.441373006235018 | -3.67113889198319 | 0.000241472066461863 | 0.0127995794890476 | 6997 | 17 | 20 | 51 | 42 | 2 | 8 | 25 | 16 | 10 | 5 | 41 | 31 |
| MER121_dup230 | 24.0174510055698 | -1.81467837366597 | 0.391751879797118 | -4.63221356999172 | 3.61776663488817e-06 | 0.000555734731894836 | 6998 | 21 | 17 | 54 | 51 | 6 | 9 | 29 | 16 | 8 | 5 | 56 | 44 |
| AluSx_dup26359 | 17.8816998593095 | -1.68938518466066 | 0.381932797503068 | -4.42325245620492 | 9.72260186237175e-06 | 0.00119744007388401 | 6999 | 12 | 13 | 36 | 53 | 7 | 12 | 26 | 12 | 2 | 8 | 34 | 23 |
| MIR_dup44940 | 19.282490137602 | -1.78514455897713 | 0.428897202778131 | -4.16217347050544 | 3.15232726458282e-05 | 0.00286182244677003 | 7000 | 15 | 15 | 42 | 61 | 4 | 10 | 25 | 12 | 4 | 7 | 41 | 20 |
| MER4E1_dup1049 | 17.3268145871306 | -1.60504649472049 | 0.446343044816125 | -3.59599306713001 | 0.000323156405472739 | 0.0157698470173536 | 7001 | 18 | 11 | 53 | 50 | 9 | 4 | 17 | 12 | 3 | 9 | 18 | 26 |
| LTR81C_dup348 | 19.3976391105358 | -1.33244554756767 | 0.391918953596582 | -3.39979869649073 | 0.000674354801248134 | 0.02662353202883 5 | 7002 | 22 | 14 | 33 | 70 | 13 | 5 | 26 | 15 | 6 | 9 | 25 | 20 |
| MER104_dup530 | 19.9194700523964 | -1.96758335826116 | 0.433689780957277 | -4.53684510139516 | 5.71020150089556e-06 | 0.00079596930658002 | 7003 | 16 | 15 | 45 | 68 | 6 | 3 | 29 | 15 | 4 | 7 | 36 | 21 |
| L1MB8_dup16403 | 23.3798128909358 | -1.58681900754231 | 0.309109207442148 | -5.1335222935386 | 2.84369329007518e-07 | 7.65436022172204e-05 | 7004 | 15 | 11 | 60 | 65 | 18 | 12 | 38 | 22 | 7 | 9 | 21 | 30 |
| L1MCa_dup6564 | 21.7807582146101 | -1.67935702498804 | 0.337328901484871 | -4.97839650737236 | 6.41132097746766e-07 | 0.000145290302981383 | 7005 | 19 | 9 | 47 | 73 | 13 | 12 | 44 | 21 | 4 | 8 | 12 | 30 |
| L1PA7_dup10167 | 19.0445141303218 | -1.37975977041574 | 0.344403170350188 | -4.00623423127263 | 6.16944708371592e-05 | 0.00478151274018318 | 7006 | 13 | 16 | 40 | 55 | 12 | 9 | 35 | 13 | 9 | 6 | 15 | 30 |
| L1ME2_dup12449 | 18.0219445474388 | -1.31041838724709 | 0.398272842513771 | -3.29025293056926 | 0.0010009736312315 | 0.0351242695261776 | 7007 | 9 | 12 | 44 | 56 | 15 | 9 | 22 | 12 | 6 | 12 | 12 | 30 |
| L1PA16_dup13728 | 17.9899308652811 | -1.3556215865153 | 0.370251423676504 | -3.66135414971351 | 0.000250885672016573 | 0.0131573514487513 | 7008 | 13 | 15 | 35 | 64 | 14 | 6 | 27 | 16 | 5 | 9 | 13 | 22 |
| MER3_dup2795 | 24.7637604161195 | -1.59548962926707 | 0.336856110807866 | -4.73641290175999 | 2.17534166475956e-06 | 0.000374274091369183 | 7009 | 17 | 19 | 59 | 55 | 8 | 11 | 24 | 23 | 7 | 14 | 36 | 51 |
| MLT1A1_dup3339 | 26.8492566901866 | -1.53003267623108 | 0.372585320715222 | -4.10652967565657 | 4.01647881432171e-05 | 0.00345043941324951 | 7010 | 25 | 18 | 63 | 76 | 10 | 9 | 28 | 19 | 7 | 17 | 46 | 36 |
| MIRb_dup101985 | 24.3768793942337 | -1.4779963138943 | 0.427246389298623 | -3.45935355081785 | 0.00054147364380806 | 0.0227876702599138 | 7011 | 25 | 25 | 63 | 75 | 6 | 8 | 24 | 23 | 4 | 13 | 28 | 27 |
| MIRb_dup58991 | 21.919514241568 | -1.43547881873289 | 0.380900329144701 | -3.76864683198413 | 0.000164134923790737 | 0.00970557867204541 | 7012 | 20 | 24 | 40 | 57 | 5 | 11 | 27 | 18 | 5 | 10 | 27 | 47 |
| Tigger15a_dup4217 | 24.663974697863 | -1.3787174554968 | 0.411858794462245 | -3.34754890276645 | 0.000815296018110796 | 0.030464262927982 3 | 7013 | 25 | 27 | 45 | 82 | 17 | 3 | 33 | 21 | 7 | 8 | 28 | 31 |
| HERV9-int_dup136 | 23.2855904158233 | -1.99522025946611 | 0.400247011813231 | -4.98497228106015 | 6.1970696507668e-07 | 0.000141649177245087 | 7014 | 10 | 20 | 73 | 73 | 13 | 5 | 30 | 22 | 5 | 5 | 22 | 29 |
| L1PA7_dup12029 | 26.4959505009984 | -2.00823736391046 | 0.369481342564978 | -5.43528761146385 | 5.47081151826408e-08 | 2.06300747626761e-05 | 7015 | 14 | 20 | 64 | 101 | 8 | 8 | 32 | 33 | 7 | 7 | 24 | 31 |
| AluY_dup47378 | 16.8636477164315 | -1.52909092439717 | 0.43304850122394 | -3.53099230242213 | 0.000414003795984173 | 0.0189662201076723 | 7016 | 14 | 17 | 39 | 50 | 12 | 11 | 19 | 12 | 2 | 3 | 12 | 35 |
| L2_dup55260 | 18.6243040927448 | -1.19590926975344 | 0.34075792979053 | -3.50955668291735 | 0.000448854375636937 | 0.0200609779967691 | 7017 | 16 | 22 | 36 | 37 | 15 | 19 | 38 | 17 | 1 | 4 | 14 | 28 |
| MIR_dup168039 | 19.537985702307 | -1.40902287584902 | 0.364920762410612 | -3.86117486585644 | 0.000112843102816424 | 0.00742343515143953 | 7018 | 21 | 14 | 49 | 55 | 15 | 20 | 28 | 16 | 2 | 2 | 16 | 22 |
| AluJo_dup53084 | 21.0460733723483 | -1.59687202279868 | 0.439182290798638 | -3.63601187082209 | 0.000276891577095218 | 0.0141172580251765 | 7019 | 20 | 19 | 42 | 44 | 17 | 19 | 22 | 24 | 1 | 1 | 34 | 32 |
| MIR3_dup75706 | 19.1024248965066 | -1.57216302015756 | 0.448372129789613 | -3.50637989229005 | 0.000454246391833889 | 0.0202408449347016 | 7020 | 12 | 8 | 38 | 44 | 17 | 23 | 21 | 14 | 3 | 2 | 38 | 32 |
| L1ME3B_dup31 | 14.9055528094648 | -1.4084388087204 | 0.356018016629029 | -3.95608857679803 | 7.61869219579695e-05 | 0.00557967784924931 | 7021 | 9 | 6 | 22 | 35 | 14 | 11 | 18 | 20 | 2 | 9 | 21 | 27 |
| MIRb_dup219409 | 15.1220660103645 | -1.21049984466741 | 0.364109927025095 | -3.32454502011966 | 0.000885629334095043 | 0.0323352332064057 | 7022 | 11 | 11 | 30 | 38 | 11 | 14 | 18 | 15 | 1 | 10 | 22 | 17 |
| MER6B_dup2 | 14.7206765038465 | -1.35177431270523 | 0.368670930031256 | -3.66661486597446 | 0.000245782541360271 | 0.0129532246034842 | 7023 | 6 | 11 | 26 | 27 | 16 | 7 | 22 | 13 | 4 | 7 | 31 | 21 |
| MIRb_dup984 | 16.6467506839783 | -1.30667993693992 | 0.339619250253038 | -3.84748489953489 | 0.000119336630109992 | 0.00772334286801005 | 7024 | 15 | 7 | 25 | 35 | 13 | 8 | 25 | 16 | 6 | 10 | 37 | 19 |
| L2a_dup44314 | 17.0471949085347 | -1.63369632861154 | 0.343158883616064 | -4.76075779066633 | 1.92867403249162e-06 | 0.000342989739413491 | 7025 | 16 | 10 | 29 | 32 | 6 | 8 | 26 | 19 | 2 | 10 | 24 | 43 |
| MLT1F1_dup2341 | 17.3583982790466 | -1.15171104444722 | 0.355820998048282 | -3.23677087851612 | 0.0012089043179395 | 0.0400654707787832 | 7026 | 21 | 14 | 29 | 30 | 7 | 21 | 26 | 18 | 3 | 5 | 23 | 31 |

| | | | | | | | | | | | | | | | | | | | |
|---|---|---|---|---|---|---|---|---|---|---|---|---|---|---|---|---|---|---|---|
| L1PA6_dup2501 | 18.7246566059241 | -1.72356049397508 | 0.349590933148253 | -4.93022081108769 | 8.21367238532969e-07 | 0.00017686972754748 | 7027 | 18 | 8 | 36 | 61 | 8 | 11 | 25 | 22 | 6 | 4 | 21 | 27 |
| AluSx_dup26358 | 18.0002094327893 | -1.70411007801005 | 0.361961809097231 | -4.70798309429458 | 2.50179844165977e-06 | 0.000419811658673607 | 7028 | 10 | 13 | 35 | 54 | 8 | 7 | 19 | 15 | 6 | 8 | 26 | 37 |
| L3_dup4390 | 16.1240594360074 | -1.05259820573866 | 0.324048592291277 | -3.2482727306296 | 0.00116107921741211 | 0.0389398022710481 | 7029 | 17 | 17 | 26 | 34 | 11 | 9 | 18 | 19 | 5 | 7 | 21 | 25 |
| LTR16E1_dup1248 | 15.6561772507506 | -1.09253535375267 | 0.312321713356667 | -3.49810886348784 | 0.000468569828161407 | 0.0206318127926895 | 7030 | 13 | 14 | 25 | 25 | 8 | 10 | 35 | 17 | 10 | 6 | 20 | 19 |
| MIR_dup102543 | 17.112005212049 | -1.06379910098801 | 0.296475705374108 | -3.58814932119195 | 0.000333033547742988 | 0.0161130541991161 | 7031 | 13 | 15 | 21 | 28 | 9 | 14 | 30 | 21 | 7 | 10 | 30 | 22 |
| MIR1_Amn_dup3269 | 18.6409728861968 | -1.27156963181494 | 0.299816231290692 | -4.24116341647315 | 2.22364158229025e-05 | 0.00221013625705779 | 7032 | 13 | 13 | 34 | 38 | 9 | 17 | 27 | 16 | 5 | 11 | 26 | 36 |
| MLT1D_dup6635 | 17.0937144361753 | -1.035722607208 | 0.296518747007215 | -3.49294139969775 | 0.000477731332926456 | 0.0208883240985885 | 7033 | 10 | 15 | 29 | 25 | 11 | 13 | 24 | 18 | 10 | 9 | 21 | 35 |
| L1PA3_dup5238 | 54.7851730863444 | -2.09373603706092 | 0.535838757444543 | -3.907399395755 | 9.32948670051425e-05 | 0.00649097673228235 | 7034 | 42 | 46 | 153 | 218 | 4 | 3 | 72 | 52 | 10 | 20 | 51 | 60 |
| L1PBa_dup1082 | 50.3792422416995 | -2.03829514525888 | 0.554271379803182 | -3.6774317049938 | 0.000235594125556315 | 0.012555135453443 | 7035 | 44 | 35 | 127 | 193 | 4 | 1 | 95 | 56 | 15 | 15 | 41 | 44 |
| L1PA16_dup6671 | 34.7668687538601 | -1.58989516750534 | 0.480429847278929 | -3.30931805446775 | 0.000935235297282616 | 0.0335776495423195 | 7036 | 29 | 26 | 80 | 96 | 5 | 3 | 77 | 50 | 18 | 17 | 22 | 28 |
| L1MDa_dup3067 | 36.249856034866 | -1.82514908562957 | 0.535729925733606 | -3.40684549800029 | 0.000657183164426286 | 0.0261538894869672 | 7037 | 39 | 26 | 96 | 120 | 1 | 4 | 64 | 45 | 13 | 10 | 28 | 31 |
| PRIMA41-int_dup161 | 44.8680507729476 | -1.59439497633581 | 0.502996138885237 | -3.16979565662151 | 0.00152546176028607 | 0.0470421881820929 | 7038 | 47 | 45 | 105 | 146 | 3 | 6 | 92 | 51 | 17 | 12 | 34 | 34 |
| L1MC1_dup6611 | 33.2836165106168 | -2.04367546916515 | 0.632258961532608 | -3.23233926840869 | 0.00122781186922409 | 0.0404531879028517 | 7039 | 29 | 31 | 97 | 133 | 2 | 0 | 39 | 30 | 6 | 11 | 32 | 34 |
| L1P3_dup1649 | 38.5840645671083 | -2.22187454749138 | 0.570405124216945 | -3.89525699044444 | 9.80946674606123e-05 | 0.00672076986570195 | 7040 | 25 | 29 | 128 | 184 | 2 | 3 | 42 | 33 | 7 | 11 | 25 | 29 |
| L1MA8_dup4858 | 21.4930528061281 | -1.88779336151065 | 0.489384530602992 | -3.85748474555299 | 0.000114559826424384 | 0.00749492781368615 | 7041 | 17 | 13 | 52 | 66 | 2 | 2 | 33 | 33 | 7 | 13 | 13 | 29 |
| L1PA17_dup2209 | 21.3279085467685 | -1.83048991589154 | 0.556571997267558 | -3.28886455818505 | 0.00100592418015079 | 0.0352148940665233 | 7042 | 19 | 17 | 58 | 69 | 1 | 1 | 29 | 20 | 9 | 9 | 28 | 20 |
| L1PBa_dup1081 | 19.4143850065623 | -1.79871708031409 | 0.51329102956364 | -3.50428310006357 | 0.000457838359722517 | 0.020323182520728 | 7043 | 19 | 13 | 64 | 55 | 1 | 3 | 27 | 22 | 7 | 8 | 19 | 15 |
| L1ME4a_dup16372 | 22.5248072425041 | -1.90664238046456 | 0.593592555026912 | -3.2120389049996 | 0.001317965259633 | 0.0424997994280759 | 7044 | 17 | 16 | 50 | 77 | 1 | 0 | 48 | 21 | 7 | 16 | 21 | 26 |
| MLT2B1_dup2240 | 25.3948208538557 | -1.8119157412139 | 0.501400986124765 | -3.61370597855792 | 0.000301851347287469 | 0.0150304376367637 | 7045 | 19 | 16 | 54 | 79 | 2 | 2 | 59 | 30 | 12 | 15 | 26 | 19 |
| L1MA4A_dup5679 | 31.3692941412707 | -1.72479397357368 | 0.480121713150866 | -3.59240985427314 | 0.000327634009447357 | 0.0159100412269308 | 7046 | 8 | 5 | 69 | 99 | 23 | 22 | 88 | 66 | 6 | 7 | 4 | 9 |
| L1PA5_dup5368 | 31.0514387646069 | -1.52057174287667 | 0.366422235451571 | -4.14978021462795 | 3.32794662568939e-05 | 0.00298706747565555 | 7047 | 13 | 23 | 38 | 50 | 21 | 20 | 105 | 71 | 7 | 10 | 10 | 29 |
| SAR_dup12 | 29.3289793717251 | -1.95669605407305 | 0.354086346996105 | -5.52604208174841 | 3.27535600941796e-08 | 1.35776630839193e-05 | 7048 | 12 | 23 | 65 | 59 | 13 | 11 | 75 | 69 | 2 | 10 | 10 | 25 |
| L1PA13_dup522 | 21.3488586935799 | -1.61847916506485 | 0.330836746168092 | -4.89207799257744 | 9.97768884375327e-07 | 0.000205021688222428 | 7049 | 12 | 12 | 32 | 57 | 9 | 8 | 54 | 33 | 9 | 12 | 15 | 25 |
| Tigger1_dup1102 | 23.5644428888927 | -1.94468235582444 | 0.376212409175683 | -5.16910742015507 | 2.35214677353551e-07 | 6.53836350672524e-05 | 7050 | 10 | 9 | 43 | 52 | 8 | 7 | 70 | 42 | 10 | 12 | 13 | 29 |
| HERV35I-int_dup136 | 22.4739148841712 | -1.69468971720483 | 0.538483550699332 | -3.14715224820504 | 0.00164869090754904 | 0.0496141908040552 | 7051 | 14 | 7 | 53 | 56 | 4 | 6 | 66 | 37 | 8 | 20 | 5 | 16 |
| Tigger1_dup1100 | 24.0439466205613 | -2.16741404723533 | 0.48978088946709 | -4.42527279819676 | 9.63204850446977e-06 | 0.00118936554297264 | 7052 | 10 | 5 | 43 | 60 | 5 | 4 | 85 | 40 | 12 | 15 | 15 | 19 |
| L1M4_dup17743 | 24.9235266077092 | -1.8467177900192 | 0.370968556787113 | -4.97809789059554 | 6.4212180359409e-07 | 0.000145310429698672 | 7053 | 14 | 12 | 68 | 97 | 17 | 11 | 34 | 36 | 7 | 3 | 13 | 14 |
| L1MC1_dup11598 | 27.8027961756159 | -1.75857224318671 | 0.359421610066198 | -4.89278383362319 | 9.94195853046534e-07 | 0.000204854393918352 | 7054 | 28 | 16 | 75 | 98 | 12 | 10 | 46 | 34 | 5 | 7 | 10 | 28 |
| L1PB1_dup11673 | 24.7137083343036 | -1.77768502271506 | 0.399224503898945 | -4.45284546753432 | 8.47397435383401e-06 | 0.00107538576085194 | 7055 | 19 | 13 | 64 | 97 | 19 | 4 | 46 | 29 | 3 | 8 | 11 | 16 |
| L1PA4_dup6507 | 23.9417850109435 | -1.7846439141193 | 0.491029275465484 | -3.63449595225763 | 0.000278524629923079 | 0.0141795725386253 | 7056 | 9 | 17 | 61 | 95 | 8 | 7 | 61 | 30 | 6 | 8 | 2 | 15 |
| L1MA9_dup8577 | 26.6370855843567 | -2.1255459486832 | 0.417262789528653 | -5.09402228529474 | 3.5054553443123e-07 | 9.09606409332318e-05 | 7057 | 13 | 11 | 58 | 82 | 7 | 7 | 68 | 62 | 5 | 9 | 10 | 10 |
| MLT2C2_dup712 | 21.9475048356635 | -2.04642015261633 | 0.368812260105099 | -5.54867712920707 | 2.87839173184154e-08 | 1.23852043223285e-05 | 7058 | 16 | 13 | 55 | 63 | 8 | 4 | 61 | 38 | 3 | 6 | 10 | 10 |
| L1PA8_dup4438 | 23.9904267170679 | -2.4528719671527 | 0.467783492751065 | -5.24360522584327 | 1.57468935327121e-07 | 4.75374168138062e-05 | 7059 | 14 | 8 | 61 | 64 | 4 | 4 | 71 | 56 | 5 | 4 | 4 | 14 |
| L1PB3_dup1798 | 25.2858092911239 | -2.40479280758672 | 0.438518588975306 | -5.48390163620211 | 4.16046162641256e-08 | 1.64453021177503e-05 | 7060 | 11 | 8 | 67 | 99 | 8 | 4 | 65 | 36 | 7 | 5 | 7 | 19 |
| L1PA4_dup939 | 33.2725205991317 | -2.19635994718946 | 0.411721811441947 | -5.33457272884641 | 9.57697971355785e-08 | 3.19160226179585e-05 | 7061 | 15 | 11 | 78 | 101 | 10 | 3 | 74 | 38 | 11 | 22 | 22 | 56 |
| L1PA7_dup889 | 37.9996494720327 | -2.48261193673322 | 0.302103117900626 | -8.21776337161094 | 2.07333547011069e-16 | 9.30833739190488e-13 | 7062 | 14 | 15 | 87 | 119 | 8 | 9 | 94 | 53 | 10 | 12 | 37 | 43 |
| Arthur1_dup2110 | 27.2543230599907 | -2.18086489582933 | 0.326036305625325 | -6.68902468283866 | 2.24662841066477e-11 | 2.85869649093069e-08 | 7063 | 8 | 15 | 72 | 90 | 10 | 12 | 66 | 35 | 6 | 7 | 18 | 21 |
| MSTB-int_dup792 | 31.8301594278196 | -2.21901043072988 | 0.457015496462288 | -4.85543804949071 | 1.20120904649485e-06 | 0.000237394425064173 | 7064 | 6 | 10 | 93 | 131 | 8 | 12 | 58 | 36 | 9 | 17 | 18 | 24 |
| L1PB1_dup11675 | 36.4361721033278 | -1.60752685284279 | 0.305178569456878 | -5.26749586546551 | 1.3829727502598e-07 | 4.29599000043498e-05 | 7065 | 26 | 21 | 96 | 110 | 18 | 15 | 68 | 45 | 9 | 17 | 18 | 36 |
| L1PBa_dup1935 | 38.21593822297 | -2.00214568904468 | 0.355080702256822 | -5.63856519467107 | 1.71472900645928e-08 | 8.04759187713925e-06 | 7066 | 16 | 22 | 105 | 148 | 15 | 11 | 75 | 33 | 10 | 14 | 27 | 35 |
| HERVP71A-int_dup182 | 34.832648273858 | -2.04244286696917 | 0.438396012605091 | -4.6588992788331 | 3.17904693188893e-06 | 0.000502618937711124 | 7067 | 17 | 14 | 115 | 135 | 12 | 7 | 49 | 50 | 9 | 13 | 13 | 22 |
| L1M4b_dup3776 | 37.453327414041 | -2.00477549000416 | 0.399335389264694 | -5.02023498016434 | 5.16083085198238e-07 | 0.000121946281817855 | 7068 | 23 | 19 | 118 | 140 | 7 | 12 | 60 | 48 | 8 | 14 | 23 | 21 |
| L1PA6_dup3254 | 45.3721992234071 | -3.31955718504456 | 0.504008714130902 | -6.58630910929528 | 4.50894611272146e-11 | 5.0607815648001e-08 | 7069 | 4 | 4 | 73 | 107 | 6 | 7 | 159 | 119 | 15 | 12 | 35 | 37 |
| HERV9NC-int_dup150 | 37.8391828635962 | -1.68337792550727 | 0.488129160756682 | -3.44863216714558 | 0.00056343365576944 | 0.0234834094685085 | 7070 | 8 | 3 | 85 | 130 | 26 | 21 | 90 | 33 | 21 | 24 | 28 | 31 |
| L1PA15-16_dup840 | 31.0017073851737 | -1.87361491471722 | 0.552860849685382 | -3.38894482360877 | 0.000701621270476294 | 0.0273954668238845 | 7071 | 1 | 3 | 89 | 95 | 25 | 16 | 54 | 34 | 18 | 12 | 25 | 31 |
| HERVL-int_dup1389 | 56.8634586030389 | -3.74651530781033 | 1.14781656276799 | -3.26403663210393 | 0.00109836963256017 | 0.0374469208448155 | 7072 | 3 | 1 | 219 | 250 | 4 | 4 | 36 | 37 | 73 | 85 | 8 | 7 |
| HERVIP10FH-int_dup14 | 32.3274825896526 | -3.25707827110934 | 0.91732425404371 | -3.55062918782712 | 0.00038431147007492 | 0.0179568234586598 | 7073 | 13 | 9 | 158 | 212 | 4 | 0 | 16 | 7 | 2 | 3 | 11 | 12 |
| L1PA3_dup2913 | 38.2372271641099 | -2.37857705139148 | 0.668098537843947 | -3.56021891481383 | 0.000370545758121047 | 0.0174697604441985 | 7074 | 22 | 19 | 172 | 233 | 5 | 6 | 31 | 12 | 6 | 2 | 4 | 14 |
| L1MA4_dup8197 | 30.2501615691679 | -2.89673896390916 | 0.568973382368419 | -5.09116780094552 | 3.55864914390822e-07 | 9.20398183909115e-05 | 7075 | 3 | 9 | 124 | 149 | 6 | 5 | 26 | 35 | 7 | 6 | 12 | 19 |
| L1MA5_dup2751 | 24.3610122344663 | -2.39514582417311 | 0.601907929189279 | -3.97925614204681 | 6.91312285248287e-05 | 0.00519567255292764 | 7076 | 1 | 1 | 72 | 114 | 11 | 11 | 30 | 25 | 8 | 12 | 15 | 24 |
| L1MA1_dup2704 | 21.400653108268 | -2.43554694630831 | 0.493633630667812 | -4.93391615764385 | 8.05970479262317e-07 | 0.000174184788064079 | 7077 | 3 | 3 | 70 | 77 | 6 | 10 | 38 | 31 | 7 | 7 | 15 | 12 |
| L1ME3Cz_dup7236 | 22.8267094093893 | -2.31577229250745 | 0.536955597765054 | -4.3127817312014 | 1.61213254831919e-05 | 0.0017428463941043 | 7078 | 2 | 2 | 75 | 86 | 13 | 9 | 32 | 35 | 5 | 10 | 14 | 14 |
| L1MA8_dup799 | 19.1838656484282 | -1.98625707945432 | 0.377550250393768 | -5.26090785897439 | 1.43345858406212e-07 | 4.41820901869489e-05 | 7079 | 8 | 8 | 34 | 61 | 3 | 10 | 41 | 22 | 5 | 13 | 17 | 33 |
| L1PREC2_dup605 | 19.4340560848713 | -2.22743003012591 | 0.452864257982149 | -4.91853793021951 | 8.71930230574778e-07 | 0.000185077886391031 | 7080 | 7 | 4 | 45 | 65 | 3 | 6 | 41 | 28 | 9 | 11 | 14 | 21 |
| L1PA5_dup876 | 16.1090234887519 | -2.58463363715916 | 0.432611537290642 | -5.97449077143481 | 2.30809999119755e-09 | 1.46845764841041e-06 | 7081 | 5 | 4 | 40 | 50 | 1 | 7 | 30 | 21 | 2 | 9 | 18 | 26 |
| L2_dup42715 | 20.7972165540315 | -3.11262579171749 | 0.405149277139467 | -7.68266406321641 | 1.55813799905385e-14 | 4.63440880403584e-11 | 7082 | 2 | 5 | 49 | 72 | 2 | 9 | 42 | 30 | 4 | 4 | 24 | 32 |
| MLT2B3_dup2932 | 19.8127832402123 | -1.74319307071255 | 0.468874760900405 | -3.71782236127406 | 0.000200947485352415 | 0.0111560080143877 | 7083 | 10 | 15 | 49 | 78 | 2 | 10 | 32 | 21 | 3 | 13 | 16 | 14 |
| L1PA3_dup5862 | 29.2714765331662 | -2.74901956779339 | 0.540844800177101 | -5.0828251781162 | 3.71861932885776e-07 | 9.52454892168344e-05 | 7084 | 8 | 7 | 102 | 119 | 1 | 8 | 59 | 31 | 3 | 15 | 15 | 24 |
| MLT1A0-int_dup740 | 29.6941734520394 | -2.62747267268171 | 0.568410106567989 | -4.62249464307763 | 3.79152606210521e-06 | 0.000575368916054646 | 7085 | 10 | 7 | 114 | 146 | 3 | 5 | 36 | 27 | 6 | 13 | 13 | 18 |
| L1MA3_dup3648 | 26.3106201841730 | -2.78570840434844 | 0.753835839213866 | -3.69537803781458 | 0.000219559856920599 | 0.0118978341413866 | 7086 | 6 | 4 | 92 | 113 | 3 | 0 | 47 | 25 | 18 | 18 | 9 | 11 |
| L1MEf_dup4421 | 23.8384376040462 | -2.7351258641623 | 0.744382525548339 | -3.67435528144285 | 0.000238450748423055 | 0.0126761398088187 | 7087 | 2 | 3 | 76 | 108 | 3 | 3 | 36 | 26 | 19 | 19 | 6 | 10 |
| L1MA3_dup3647 | 19.1808809924931 | -2.89757842569952 | 0.814364173088534 | -3.55808681355695 | 0.000373565857023469 | 0.0175738436961855 | 7088 | 2 | 3 | 65 | 102 | 1 | 2 | 35 | 13 | 8 | 18 | 5 | 4 |
| MLT1D_dup18210 | 15.6164872751858 | -2.58905849484999 | 0.807626718042393 | -3.20576132142533 | 0.0013470567105763 | 0.0431453433914105 | 7089 | 6 | 2 | 61 | 88 | 1 | 1 | 27 | 8 | 5 | 8 | 2 | 4 |
| THE1D_dup10196 | 16.2265065630513 | -2.83032450426091 | 0.702362822515972 | -4.02971856357979 | 5.58436718912341e-05 | 0.00442925945061053 | 7090 | 2 | 4 | 61 | 73 | 1 | 1 | 20 | 11 | 8 | 11 | 9 | 15 |
| L1MC3_dup10032 | 16.0492558991905 | -2.53592026198139 | 0.652100722131877 | -3.88884749842147 | 0.000100721379338357 | 0.00685906291437348 | 7091 | 4 | 6 | 61 | 86 | 2 | 2 | 21 | 10 | 6 | 4 | 5 | 10 |
| MLT1E1A_dup2766 | 19.3422136959543 | -2.63080986874374 | 0.624161784819435 | -4.21494864429224 | 2.49835033774247e-05 | 0.00241165302013983 | 7092 | 3 | 5 | 82 | 95 | 2 | 6 | 23 | 15 | 6 | 5 | 7 | 10 |
| L1PA8_dup2471 | 14.4918823100053 | -1.74424782632495 | 0.554000004030191 | -3.14846175746579 | 0.00164132197722667 | 0.0494800455078184 | 7093 | 7 | 3 | 46 | 50 | 7 | 3 | 22 | 20 | 7 | 9 | 4 | 10 |
| MLT2B4_dup4271 | 15.7340001227339 | -2.51944850452553 | 0.526823272892484 | -4.78234093701421 | 1.73265480656212e-06 | 0.000316490373337472 | 7094 | 4 | 2 | 45 | 53 | 6 | 2 | 34 | 28 | 6 | 5 | 7 | 12 |
| L1MA4_dup8198 | 19.3714758914064 | -2.57141947691956 | 0.630269237667675 | -4.07987463648892 | 4.50599949683793e-05 | 0.00378392824902677 | 7095 | 7 | 2 | 79 | 96 | 6 | 2 | 16 | 19 | 5 | 7 | 7 | 12 |
| L1PB2_dup1088 | 19.6748204845513 | -2.29858975835176 | 0.729547915968175 | -3.15070430336481 | 0.00162877297767729 | 0.0491515554783008 | 7096 | 3 | 2 | 69 | 87 | 9 | 1 | 25 | 23 | 13 | 10 | 3 | 12 |
| L1MC3_dup4879 | 14.6917838489317 | -2.42595611070444 | 0.762933136330615 | -3.17977552052891 | 0.00147389180242673 | 0.0459450318658827 | 7097 | 2 | 3 | 40 | 72 | 3 | 1 | 24 | 20 | 13 | 5 | 5 | 3 |
| L1ME3Cz_dup7234 | 14.9669462546636 | -3.08037007870575 | 0.701345750757539 | -4.39208489589988 | 1.12268841314252e-05 | 0.00133103745467668 | 7098 | 0 | 1 | 42 | 73 | 3 | 2 | 29 | 15 | 7 | 7 | 9 | 12 |
| L1MA2_dup1202 | 34.8356285286904 | -1.84650626412376 | 0.572473524588252 | -3.2254876161336 | 0.00125758276266672 | 0.0411604935413335 | 7099 | 6 | 9 | 74 | 91 | 52 | 50 | 102 | 58 | 0 | 0 | 3 | 11 |
| AluJo_dup76472 | 18.1895490882472 | -1.69572338396277 | 0.462008523005046 | -3.67032922452005 | 0.000242238266765004 | 0.0128230537538744 | 7100 | 12 | 7 | 30 | 52 | 26 | 21 | 36 | 24 | 0 | 0 | 16 | 16 |
| L1PA2_dup2773 | 15.3870731302446 | -2.09015260035997 | 0.589591059475623 | -3.54508869625448 | 0.000392480992389208 | 0.0182472220037207 | 7101 | 3 | 2 | 28 | 36 | 19 | 21 | 43 | 29 | 0 | 0 | 9 | 10 |
| L1PA3_dup1626 | 28.3349747431205 | -2.84497172633538 | 0.396537058181618 | -7.17454186849884 | 7.25497177246323e-13 | 1.42668720113267e-09 | 7102 | 12 | 18 | 67 | 62 | 3 | 6 | 97 | 69 | 1 | 3 | 10 | 17 |
| L1MC5_dup13211 | 24.6812236895945 | -1.71635374257752 | 0.534475601873145 | -3.21125180749575 | 0.00132158075209768 | 0.0425815644737489 | 7103 | 30 | 25 | 60 | 70 | 4 | 2 | 57 | 46 | 6 | 1 | 10 | 9 |
| Tigger1_dup10677 | 19.9380728182471 | -2.02978271028169 | 0.579636103823048 | -3.50182243116682 | 0.000462087474411857 | 0.0204371498487739 | 7104 | 29 | 20 | 31 | 74 | 1 | 3 | 50 | 37 | 1 | 1 | 8 | 9 |
| MLT2B4_dup302 | 19.7101992236926 | -1.70982302953818 | 0.432819317004296 | -3.95043142106619 | 7.80104486484592e-05 | 0.00566960116496832 | 7105 | 14 | 13 | 27 | 31 | 10 | 16 | 72 | 57 | 2 | 1 | 2 | 4 |

| | | | | | | | | | | | | | | | | | | | |
|---|---|---|---|---|---|---|---|---|---|---|---|---|---|---|---|---|---|---|---|
| L1MB7_dup10650 | 17.9948127345959 | -1.96507611040759 | 0.432707818891125 | -4.54134643428301 | 5.58960936056033e-06 | 0.000783127221155022 | 7106 | 8 | 12 | 35 | 62 | 7 | 5 | 43 | 41 | 3 | 5 | 4 | 7 |
| LTR1A2_dup338 | 20.9334342845745 | -1.30402870448334 | 0.355367856317547 | -3.66951788492119 | 0.000243008336847498 | 0.0128552382657881 | 7107 | 20 | 26 | 43 | 61 | 11 | 9 | 52 | 34 | 2 | 5 | 4 | 7 |
| MER41-int_dup657 | 18.8225047271898 | -2.3112242968044 | 0.392897956586497 | -5.88250526137715 | 4.04102487849208e-09 | 2.32819783471593e-06 | 7108 | 13 | 7 | 42 | 64 | 8 | 6 | 61 | 34 | 3 | 1 | 6 | 3 |
| L1PA7_dup5842 | 23.4817309203332 | -1.94585682718544 | 0.326108824531648 | -5.96689411879622 | 2.41812210378655e-09 | 1.53027255879679e-06 | 7109 | 15 | 17 | 63 | 86 | 15 | 11 | 52 | 38 | 2 | 1 | 3 | 6 |
| L1MA4_dup4178 | 18.023111726733 | -1.73801280230307 | 0.375878274529041 | -4.62387139687902 | 3.76643432465813e-06 | 0.00057192596159228 | 7110 | 15 | 12 | 50 | 61 | 11 | 7 | 40 | 28 | 2 | 3 | 2 | 6 |
| L1P3_dup1465 | 19.7037559774048 | -2.11692613203844 | 0.433277309835707 | -4.88584581740768 | 1.02985769626171e-06 | 0.00021016339570728 | 7111 | 17 | 10 | 44 | 87 | 11 | 5 | 33 | 38 | 2 | 1 | 2 | 9 |
| L1PA15_dup462 | 26.5649098645704 | -1.61009377978742 | 0.428580402221256 | -3.75680682420986 | 0.000172095238032931 | 0.0100418153174827 | 7112 | 32 | 30 | 84 | 88 | 9 | 20 | 39 | 40 | 0 | 0 | 4 | 2 |
| MER41-int_dup656 | 23.4972762050275 | -1.95932438028544 | 0.429320698366198 | -4.5637780515632 | 5.02411931742947e-06 | 0.000724087883164793 | 7113 | 20 | 21 | 50 | 81 | 10 | 5 | 62 | 52 | 2 | 0 | 2 | 0 |
| L1M4a2_dup804 | 24.4430761937064 | -2.23160969703535 | 0.417556904794811 | -5.34444448507437 | 9.06947663083448e-08 | 3.07852451697652e-05 | 7114 | 25 | 16 | 65 | 105 | 6 | 9 | 54 | 37 | 1 | 0 | 5 | 3 |
| L1MA3_dup3847 | 21.7746381156597 | -1.87679451832821 | 0.473086224260463 | -3.96712992702768 | 7.27433407760147e-05 | 0.00538454815896774 | 7115 | 20 | 28 | 40 | 81 | 7 | 4 | 47 | 44 | 1 | 1 | 7 | 4 |
| L1ME2_dup5464 | 26.1238247663036 | -2.14995373142091 | 0.444791192185365 | -4.83362478662779 | 1.34069173158324e-06 | 0.000259148850335747 | 7116 | 32 | 17 | 63 | 83 | 7 | 4 | 66 | 57 | 1 | 1 | 4 | 5 |
| L1PREC2_dup3229 | 23.6582898849347 | -2.42923220811421 | 0.412875377252264 | -5.88369358396 | 4.0121032480936e-09 | 2.31714543560893e-06 | 7117 | 10 | 20 | 70 | 95 | 7 | 7 | 57 | 30 | 0 | 3 | 5 | 13 |
| THE1B-int_dup2005 | 21.1195387960532 | -2.4795714117255 | 0.470000095504381 | -5.27568278271209 | 1.3226270598633e-07 | 4.13010260349387e-05 | 7118 | 7 | 17 | 68 | 94 | 7 | 7 | 33 | 34 | 0 | 2 | 2 | 8 |
| L1MB5_dup2904 | 13.4311205471823 | -1.86693391870322 | 0.449486382911298 | -4.15348270755432 | 3.2745296897348e-05 | 0.00295360591946033 | 7119 | 14 | 14 | 26 | 41 | 9 | 3 | 37 | 24 | 0 | 0 | 2 | 7 |
| L2a_dup73367 | 18.0370169627912 | -1.88299909864924 | 0.41447556808524 | -4.54308828708087 | 5.54360198212684e-06 | 0.000777758205919313 | 7120 | 24 | 16 | 46 | 53 | 6 | 7 | 46 | 32 | 0 | 0 | 3 | 4 |
| L1MD3_dup1979 | 14.8799776413106 | -1.98678156298895 | 0.451728595912701 | -4.3981753224516 | 1.09164780756859e-05 | 0.0013026741686044 | 7121 | 14 | 12 | 37 | 57 | 5 | 4 | 46 | 20 | 0 | 2 | 1 | 2 |
| MER57-int_dup491 | 16.0169326260735 | -2.23928831123364 | 0.440985443434914 | -5.07791888501223 | 3.81591532910847e-07 | 9.72143242933666e-05 | 7122 | 9 | 11 | 48 | 62 | 10 | 5 | 24 | 32 | 0 | 1 | 2 | 4 |
| L1MA5A_dup3053 | 17.4869970453854 | -2.10481862369312 | 0.567497625787065 | -3.70894701237549 | 0.000208122972346038 | 0.0114495269098621 | 7123 | 23 | 21 | 58 | 48 | 2 | 5 | 32 | 25 | 0 | 0 | 6 | 10 |
| L1PA3_dup4639 | 13.7493443639576 | -1.96176667658986 | 0.498810951170716 | -3.93288614050186 | 8.39319995995231e-05 | 0.00599378258604686 | 7124 | 16 | 13 | 34 | 39 | 3 | 5 | 36 | 24 | 0 | 1 | 8 | 1 |
| L1PA2_dup3773 | 25.4332080127126 | -5.59313813091872 | 0.657114332914301 | -8.51166661684757 | 1.71450938406173e-17 | 9.71334880714172e-14 | 7125 | 2 | 3 | 42 | 55 | 0 | 1 | 119 | 85 | 0 | 0 | 7 | 10 |
| L1MA9_dup7820 | 35.5578309829605 | -5.68957279746538 | 0.838379467708252 | -6.78639329397949 | 1.14971466734963e-11 | 1.56325717849815e-08 | 7126 | 1 | 1 | 13 | 30 | 1 | 2 | 221 | 160 | 1 | 0 | 2 | 7 |
| L1P3_dup353 | 22.0539164067497 | -5.44027848367329 | 0.755096021728005 | -7.20475055771508 | 5.81502022084697e-13 | 1.16274016930223e-09 | 7127 | 1 | 0 | 21 | 41 | 1 | 3 | 101 | 96 | 0 | 0 | 3 | 5 |
| L1ME3A_dup7931 | 23.8084723726837 | -4.6586282708057 | 0.77121007882656 | -6.04067348016777 | 1.53472274958596e-09 | 1.03248149604292e-06 | 7128 | 0 | 0 | 53 | 58 | 1 | 1 | 71 | 69 | 7 | 4 | 17 | 19 |
| L1PA10_dup6519 | 20.9996929867219 | -4.36637771918083 | 0.659974148990527 | -6.61598295305882 | 3.69090903103301e-11 | 4.26328660338923e-08 | 7129 | 2 | 0 | 24 | 35 | 1 | 2 | 104 | 57 | 2 | 4 | 16 | 25 |
| L1PA10_dup6521 | 37.0353623607461 | -4.54322087541994 | 0.565847161164054 | -8.02906011947411 | 9.82222201122884e-16 | 3.76961038497396e-12 | 7130 | 2 | 2 | 39 | 50 | 5 | 2 | 194 | 107 | 3 | 3 | 29 | 41 |
| THE1C-int_dup663 | 27.7920078902186 | -4.65867149790913 | 0.5973858177713 | -7.79843002515442 | 6.26820944104674e-15 | 2.02375955837588e-11 | 7131 | 3 | 2 | 47 | 52 | 0 | 1 | 110 | 73 | 5 | 1 | 29 | 34 |
| L1ME2z_dup3460 | 29.3602880259673 | -5.51159025505537 | 1.44987331209312 | -3.80142886215247 | 0.000143864045213348 | 0.00883512601993136 | 7132 | 0 | 0 | 138 | 150 | 2 | 0 | 44 | 48 | 2 | 2 | 2 | 0 |
| L1MC3_dup5877 | 23.9707352118582 | -3.91582023539288 | 1.07185721234913 | -3.65330399448521 | 0.000258887425854407 | 0.0134735843027893 | 7133 | 1 | 1 | 45 | 85 | 1 | 3 | 94 | 73 | 3 | 1 | 1 | 0 |
| L1MDa_dup2999 | 17.8183631025881 | -3.74780261618994 | 0.476776681134018 | -7.86070872274154 | 3.81966067235095e-15 | 1.31720721499017e-11 | 7134 | 2 | 2 | 52 | 58 | 3 | 6 | 65 | 42 | 1 | 0 | 1 | 0 |
| L1PA3_dup5641 | 19.5504370493723 | -2.79693888747781 | 0.560646832220776 | -4.98877140962139 | 6.07644966786045e-07 | 0.000139194264682893 | 7135 | 6 | 6 | 77 | 93 | 5 | 3 | 32 | 24 | 3 | 2 | 2 | 8 |
| HERV-Fc1-int_dup2 | 26.1327434603562 | -3.3700156518148 | 0.488001539942748 | -6.9057479863899 | 4.99395857720227e-12 | 7.33513868895661e-09 | 7136 | 4 | 7 | 88 | 91 | 5 | 4 | 87 | 49 | 2 | 2 | 2 | 3 |
| L1PA5_dup8420 | 21.0057134040178 | -3.32809492744892 | 0.659464562194058 | -5.04666227458265 | 4.4959504657284e-07 | 0.00011040179871189 | 7137 | 4 | 3 | 74 | 68 | 2 | 2 | 79 | 33 | 1 | 5 | 1 | 8 |
| L1MCa_dup3376 | 23.6089370240019 | -3.75142614196221 | 0.431488628455985 | -8.69414833801322 | 3.49440913392593e-18 | 2.18810704152931e-14 | 7138 | 6 | 2 | 76 | 77 | 9 | 2 | 77 | 53 | 0 | 1 | 1 | 4 |
| MLT1C_dup19335 | 18.4541212967639 | -3.72168930710718 | 0.502412514797843 | -7.40763654863312 | 1.28570078264623e-13 | 2.91359388978609e-10 | 7139 | 5 | 3 | 81 | 67 | 2 | 7 | 42 | 28 | 0 | 0 | 4 | 5 |
| L1PA15-16_dup68 | 26.0231173934498 | -4.71227503800813 | 0.665134837259861 | -7.08469136486847 | 1.39354523134163e-12 | 2.5506808739755e-09 | 7140 | 6 | 6 | 96 | 161 | 1 | 0 | 33 | 29 | 0 | 0 | 12 | 10 |
| L1M2_dup780 | 26.4370541929101 | -3.80583333177195 | 0.496077061591143 | -7.67185912520309 | 1.69521080836634e-14 | 4.97985964206836e-11 | 7141 | 8 | 6 | 78 | 145 | 2 | 2 | 56 | 35 | 2 | 1 | 10 | 13 |
| HERVL-int_dup220 | 36.4075632269072 | -4.97236231734849 | 0.599338415469501 | -8.29641849914345 | 1.07297072079317e-16 | 5.00606061038924e-13 | 7142 | 4 | 3 | 144 | 226 | 1 | 3 | 60 | 32 | 1 | 1 | 9 | 16 |
| L1M2_dup781 | 36.6990560355874 | -4.53367474509134 | 0.540669107959709 | -8.38530383620361 | 5.05949014984496e-17 | 2.61714226781524e-13 | 7143 | 4 | 6 | 147 | 175 | 1 | 2 | 71 | 43 | 2 | 2 | 12 | 30 |
| ERVL-E-int_dup8117 | 475.831481347102 | -1.17939444381276 | 0.362906376222396 | -3.24985869934125 | 0.00115462362811716 | 0.0388006947890204 | 7144 | 277 | 235 | 698 | 854 | 728 | 671 | 660 | 583 | 69 | 105 | 622 | 662 |
| HSATI_dup10 | 884.647365056593 | -1.18274259510046 | 0.367186746723167 | -3.22109379125322 | 0.00127702348816105 | 0.0416136169424773 | 7145 | 505 | 411 | 1192 | 1571 | 1371 | 1247 | 1273 | 1164 | 156 | 167 | 1252 | 1100 |
| Harlequin-int_dup136 | 750.008466063126 | 1.23666626894912 | 0.337332712553501 | 3.6660134725386 | 0.000246360951774386 | 0.012966642019494 | 7146 | 1942 | 1756 | 351 | 420 | 1006 | 1111 | 363 | 335 | 455 | 435 | 574 | 871 |
| L1PA7_dup3105 | 1058.90566713177 | -1.39997288476112 | 0.281788466173176 | -4.96816956737307 | 6.75878497476342e-07 | 0.000151007122028644 | 7147 | 1022 | 911 | 1700 | 2389 | 480 | 527 | 1227 | 712 | 347 | 368 | 1826 | 2525 |
| THE1A-int_dup1058 | 276.205819219632 | -2.36895403204581 | 0.70972365823787 | -3.3378541134271 | 0.00084428064671973 | 0.0312528317928396 | 7148 | 134 | 129 | 76 | 86 | 31 | 66 | 1123 | 949 | 87 | 79 | 273 | 374 |
| L1PA11_dup2177 | 105.755736817602 | -2.11413399952737 | 0.347659001173101 | -6.08105640616209 | 1.19393278422449e-09 | 8.32824296068398e-07 | 7149 | 66 | 53 | 218 | 298 | 23 | 26 | 349 | 213 | 13 | 30 | 36 | 57 |
| HERVIP10F-int_dup195 | 222.438422163457 | -1.66243591147998 | 0.298283210311812 | -5.57334725525499 | 2.49890757259224e-08 | 1.11349262409744e-05 | 7150 | 141 | 92 | 575 | 627 | 100 | 103 | 566 | 398 | 54 | 61 | 72 | 98 |
| L1PA6_dup3338 | 192.756273831246 | -1.9666309523951 | 0.308461766131849 | -6.37560686063918 | 1.82239866258146e-10 | 1.72076377843892e-07 | 7151 | 108 | 109 | 453 | 625 | 52 | 57 | 507 | 339 | 45 | 43 | 85 | 96 |
| L1PA3_dup3690 | 391.861516270849 | 1.60016173299289 | 0.227617221134063 | 7.03005565668698 | 2.0645116990773e-12 | 3.52451983946178e-09 | 7152 | 1123 | 975 | 194 | 238 | 543 | 556 | 241 | 166 | 268 | 310 | 159 | 232 |
| HUERS-P3-int_dup162 | 200.326476095577 | 1.70726090356131 | 0.209644936928698 | 8.14358280515958 | 3.83749856368669e-16 | 1.6019604091842e-12 | 7153 | 593 | 467 | 105 | 153 | 261 | 263 | 140 | 76 | 166 | 183 | 66 | 82 |
| L1PA7_dup4534 | 200.654262801682 | 1.15049768210163 | 0.234518369816219 | 4.90578918403375 | 9.30524292313412e-07 | 0.000194907159558809 | 7154 | 533 | 403 | 149 | 233 | 223 | 273 | 119 | 86 | 144 | 180 | 93 | 142 |
| Harlequin-int_dup243 | 236.978642956105 | 1.20672583283459 | 0.289108236126163 | 4.17395868413789 | 2.99352066411997e-05 | 0.00275124089588525 | 7155 | 681 | 579 | 127 | 129 | 320 | 360 | 181 | 144 | 124 | 114 | 124 | 153 |
| L1PREC2_dup6400 | 214.952992435434 | 1.02008916834818 | 0.251873184381825 | 4.05001100395742 | 5.12152245866408e-05 | 0.00415920062440028 | 7156 | 515 | 404 | 122 | 138 | 298 | 308 | 215 | 87 | 121 | 193 | 168 | 200 |
| L1MCc_dup813 | 200.085866993351 | -1.44473357704836 | 0.311201266222028 | -4.64244119115377 | 3.44316657720523e-06 | 0.000535482166261225 | 7157 | 196 | 200 | 343 | 482 | 93 | 99 | 224 | 148 | 51 | 59 | 363 | 386 |
| L1PA5_dup3084 | 164.61994024006 | -1.47536732980904 | 0.307796852885478 | -4.793315188177 | 1.64047582284109e-06 | 0.000302827509807405 | 7158 | 166 | 152 | 279 | 340 | 79 | 65 | 186 | 145 | 41 | 55 | 244 | 423 |
| L1MD1_dup3272 | 173.100157130905 | -1.16614055785432 | 0.319861152812643 | -3.64577113413138 | 0.000266591163862739 | 0.0137476528414399 | 7159 | 237 | 213 | 321 | 426 | 129 | 126 | 204 | 148 | 30 | 31 | 179 | 243 |
| L1PA7_dup6706 | 189.518325863232 | -1.23225956709038 | 0.359812393146145 | -3.42472797091755 | 0.000615415284684482 | 0.0249922467841018 | 7160 | 231 | 186 | 378 | 533 | 147 | 133 | 168 | 98 | 37 | 48 | 231 | 342 |
| THE1A-int_dup399 | 273.983755418067 | -1.17142641791601 | 0.303070150161066 | -3.86519892273607 | 0.000110998696587084 | 0.00734678605232555 | 7161 | 309 | 236 | 351 | 393 | 165 | 206 | 289 | 281 | 81 | 78 | 482 | 705 |
| AluSc5_dup6526 | 227.404298110092 | -1.22700178823318 | 0.382439040316222 | -3.20835913409527 | 0.00133494686633859 | 0.0429018459019181 | 7162 | 100 | 130 | 371 | 432 | 360 | 307 | 332 | 263 | 45 | 29 | 275 | 305 |
| AluSc8_dup23025 | 269.2545589973 | -1.22975414833278 | 0.328778405004533 | -3.74037384941943 | 0.000183746750474065 | 0.01048484515307 | 7163 | 173 | 162 | 357 | 561 | 372 | 341 | 369 | 358 | 44 | 52 | 337 | 363 |
| L1PA5_dup4270 | 111.270276147156 | 1.10382892385433 | 0.218039532400656 | 5.06251738710394 | 4.13756316710471e-07 | 0.000103198805593281 | 7164 | 185 | 199 | 79 | 79 | 191 | 216 | 141 | 91 | 77 | 72 | 22 | 57 |
| HAL1M8_dup478 | 85.0400828762559 | 1.02252480890367 | 0.269799730619602 | 3.78994006612022 | 0.000150683637492788 | 0.00911865941171388 | 7165 | 187 | 156 | 48 | 38 | 108 | 96 | 66 | 56 | 85 | 71 | 67 | 92 |
| L1PA7_dup9491 | 102.292620095424 | 1.46300799472276 | 0.322364428202592 | 4.53836672637878 | 5.66916072065804e-06 | 0.000792104590039753 | 7166 | 275 | 261 | 42 | 36 | 102 | 99 | 84 | 55 | 86 | 117 | 67 | 68 |
| Harlequin-int_dup135 | 131.901047476828 | 1.3428455117713 | 0.3463431953996 | 3.87721060961505 | 0.000105660889467929 | 0.007108166809538 | 7167 | 347 | 283 | 74 | 69 | 181 | 198 | 59 | 38 | 93 | 84 | 122 | 143 |
| MLT1D_dup6489 | 136.98612734297 | 1.66791565877033 | 0.272284034690691 | 6.12564618658251 | 9.03163036451736e-10 | 6.61243174989369e-07 | 7168 | 381 | 364 | 62 | 76 | 213 | 194 | 63 | 68 | 93 | 92 | 50 | 86 |
| L1PA16_dup12620 | 107.658899722374 | 1.18281596603957 | 0.264435752703412 | 4.47298050262592 | 7.71368132417328e-06 | 0.00100516955989142 | 7169 | 254 | 177 | 49 | 47 | 155 | 169 | 94 | 66 | 99 | 82 | 65 | 106 |
| L1PA16_dup12806 | 132.923054383471 | 1.2353984971704 | 0.264953571464852 | 4.66269803550953 | 3.12090521488533e-06 | 0.000495071274840737 | 7170 | 269 | 220 | 55 | 69 | 217 | 269 | 108 | 64 | 97 | 105 | 79 | 144 |
| L1PA3_dup9280 | 131.006601079725 | 1.09229324521973 | 0.311410306070114 | 3.50756935120123 | 0.000452220464373457 | 0.0201808046916667 | 7171 | 310 | 228 | 54 | 69 | 243 | 219 | 104 | 71 | 75 | 74 | 102 | 129 |
| L1PB1_dup313 | 243.431909232024 | 1.66846931869969 | 0.332111135401645 | 5.02382829374838 | 5.06515258012257e-07 | 0.000120504515706563 | 7172 | 304 | 268 | 87 | 87 | 484 | 599 | 80 | 78 | 277 | 250 | 202 | 333 |
| LTR16C_dup4795 | 205.52343273169 | 1.35526206712408 | 0.4221916237938 | 3.21006384481467 | 0.00132705493182276 | 0.0427058984051257 | 7173 | 372 | 356 | 53 | 51 | 357 | 388 | 107 | 84 | 148 | 125 | 224 | 357 |
| HERVI-int_dup36 | 161.441877609244 | 1.48923346299639 | 0.317547222062561 | 4.68980157761541 | 2.73470106599558e-06 | 0.000448325344332882 | 7174 | 208 | 175 | 27 | 34 | 172 | 181 | 58 | 43 | 245 | 275 | 265 | 319 |
| L2b_dup82791 | 131.389138945754 | 1.04444777857948 | 0.220470696298102 | 4.73735419770828 | 2.16526533152463e-06 | 0.000373345090271709 | 7175 | 193 | 144 | 54 | 70 | 163 | 200 | 105 | 55 | 137 | 222 | 100 | 218 |
| Tigger2_dup586 | 90.1907026823156 | -1.31050135391643 | 0.236365022176977 | -5.54439629792261 | 2.94970087659062e-08 | 1.25800168153523e-05 | 7176 | 35 | 28 | 113 | 198 | 67 | 59 | 112 | 90 | 52 | 72 | 147 | 196 |
| L1PREC2_dup1391 | 112.459593793981 | -1.00933295206545 | 0.152323547751204 | -6.626243722435 | 3.44336474523298e-11 | 4.07629287397616e-08 | 7177 | 71 | 53 | 136 | 169 | 87 | 83 | 188 | 123 | 72 | 85 | 161 | 218 |
| THE1A_dup3715 | 121.188021469263 | -1.19761164230457 | 0.208995411584787 | -5.73032504983351 | 1.00238364300883e-08 | 5.07474847488042e-06 | 7178 | 78 | 63 | 123 | 149 | 76 | 97 | 192 | 160 | 72 | 54 | 223 | 263 |
| L1M3f_dup115 | 130.289748908571 | -1.25338389932745 | 0.329344267217632 | -3.80569520737766 | 0.000141406434544274 | 0.00872344756211957 | 7179 | 39 | 19 | 75 | 106 | 56 | 69 | 155 | 159 | 161 | 128 | 236 | 434 |
| L1MA4_dup1393 | 85.1175439817247 | -1.24093934587873 | 0.303265970844432 | -4.09191754163312 | 4.27820788492083e-05 | 0.00362271335724331 | 7180 | 22 | 21 | 62 | 57 | 41 | 49 | 114 | 110 | 81 | 92 | 177 | 243 |
| L1PA8_dup1199 | 81.701073309614 | -1.14960427670409 | 0.227768690399716 | -5.04724452990719 | 4.48227519567205e-07 | 0.000110179695631135 | 7181 | 25 | 30 | 63 | 77 | 34 | 40 | 123 | 77 | 91 | 88 | 158 | 230 |
| L1PB1_dup12761 | 207.979123853446 | -1.80924077727292 | 0.471799819232786 | -3.83476360846216 | 0.000125685113531234 | 0.00801410773025061 | 7182 | 109 | 106 | 445 | 690 | 144 | 133 | 94 | 48 | 46 | 49 | 387 | 558 |
| L1PB1_dup12758 | 143.466841064762 | -2.00146717834681 | 0.490782426690953 | -4.07811500473128 | 4.54023082712961e-05 | 0.00379947246968682 | 7183 | 70 | 57 | 358 | 459 | 70 | 82 | 37 | 39 | 32 | 43 | 270 | 416 |
| HERVL-int_dup2286 | 173.310485021956 | -2.0842709268815 | 0.490946339947634 | -4.24541494107851 | 2.18189268487215e-05 | 0.00217682447293328 | 7184 | 67 | 64 | 454 | 600 | 112 | 100 | 66 | 41 | 37 | 38 | 349 | 405 |

| | | | | | | | | | | | | | | | | | | |
|---|---|---|---|---|---|---|---|---|---|---|---|---|---|---|---|---|---|---|
| L1PB1_dup12760 | 161.620239539819 | -1.87088852353233 | 0.49509535185049 | -3.77884485592445 | 0.000157557544207759 | 0.00944099406045314 | 7185 | 95 | 72 | 367 | 532 | 96 | 81 | 51 | 32 | 37 | 49 | 339 | 428 |
| MLT1A0-int_dup440 | 131.629203027166 | -1.39711516755377 | 0.343220637964301 | -4.07060360892134 | 4.68914799257041e-05 | 0.00388903453551815 | 7186 | 97 | 79 | 292 | 325 | 92 | 72 | 97 | 53 | 41 | 66 | 240 | 292 |
| L1PA6_dup2742 | 140.859070260529 | -1.76054108255758 | 0.387316963987781 | -4.54547888744972 | 5.48104913468188e-06 | 0.000771602701100299 | 7187 | 70 | 72 | 322 | 462 | 65 | 92 | 97 | 47 | 45 | 51 | 224 | 346 |
| L1PA7_dup5928 | 139.047739341535 | -1.86380817760316 | 0.413639873940541 | -4.50587164106688 | 6.6101037902823e-06 | 0.000890123235134415 | 7188 | 63 | 59 | 371 | 487 | 82 | 59 | 76 | 48 | 49 | 53 | 202 | 315 |
| L1MB3_dup4108 | 131.43237044327 | -1.5103059864007 | 0.261349744916982 | -5.77886918114442 | 7.52043709308826e-09 | 3.98542967606232e-06 | 7189 | 99 | 95 | 234 | 331 | 65 | 62 | 152 | 85 | 39 | 63 | 251 | 263 |
| L1MB7_dup5537 | 113.887033076139 | -1.79838975766113 | 0.244167638088026 | -7.36538949937657 | 1.76630383058509e-13 | 3.92789655394767e-10 | 7190 | 64 | 57 | 242 | 278 | 42 | 52 | 142 | 73 | 44 | 50 | 200 | 264 |
| L2a_dup44324 | 102.565432966824 | -1.17916214265779 | 0.313238988009024 | -3.76441690784616 | 0.00016693816592452 | 0.00983589111603341 | 7191 | 132 | 95 | 161 | 248 | 66 | 54 | 134 | 60 | 26 | 38 | 147 | 207 |
| L2_dup28580 | 90.9319034570417 | -1.40264295065195 | 0.325981311254089 | -4.30283240856913 | 1.68628403532096e-05 | 0.00180415710912087 | 7192 | 114 | 84 | 197 | 262 | 54 | 47 | 102 | 67 | 18 | 22 | 114 | 127 |
| L2b_dup44441 | 87.1286259924438 | -1.33535175082606 | 0.365645105207935 | -3.65204328406549 | 0.00026016201995239 | 0.0135192207904764 | 7193 | 112 | 79 | 151 | 278 | 58 | 49 | 99 | 49 | 20 | 19 | 84 | 176 |
| L1PA10_dup1637 | 91.916370414565 | -1.44401558176533 | 0.286593612394059 | -5.03854768326045 | 4.69077565083841e-07 | 0.000114242712693388 | 7194 | 73 | 57 | 183 | 208 | 49 | 54 | 101 | 50 | 40 | 30 | 159 | 212 |
| MLT1F_dup1079 | 94.3625234104152 | -1.30655079701707 | 0.299808732566924 | -4.35794776833399 | 1.31287704349435e-05 | 0.00149542288650697 | 7195 | 93 | 74 | 171 | 210 | 35 | 43 | 94 | 56 | 42 | 45 | 181 | 198 |
| L3b_dup2649 | 80.8030825455831 | -1.21582851108533 | 0.295415345134203 | -4.11565794096782 | 3.86076514842978e-05 | 0.00334856591289842 | 7196 | 87 | 61 | 106 | 142 | 42 | 60 | 102 | 58 | 27 | 26 | 136 | 223 |
| LTR14B_dup304 | 86.9189426365603 | -1.23408726559527 | 0.223105202995159 | -5.5314140998408 | 3.17659415523261e-08 | 1.32785167498844e-05 | 7197 | 65 | 42 | 113 | 99 | 48 | 75 | 122 | 97 | 42 | 39 | 165 | 215 |
| L1PA7_dup10205 | 191.972643557466 | 4.05651969399786 | 0.447558817350414 | 9.06365719261838 | 1.26148416097652e-19 | 1.072018250599e-15 | 7198 | 597 | 514 | 59 | 88 | 422 | 432 | 15 | 5 | 147 | 152 | 6 | 5 |
| MLT2B5_dup347 | 213.784556742855 | 2.92492882308891 | 0.387352775607837 | 7.55107232289503 | 4.31689023662141e-14 | 1.15414243173384e-10 | 7199 | 648 | 530 | 83 | 67 | 291 | 308 | 16 | 14 | 269 | 286 | 69 | 96 |
| L1MA5A_dup2326 | 53.8676971062912 | 1.23555074201988 | 0.314112102074363 | 3.93347067451537 | 8.37280581559348e-05 | 0.00598640520612141 | 7200 | 138 | 114 | 21 | 31 | 42 | 60 | 36 | 36 | 58 | 55 | 41 | 44 |
| L1PA3_dup7753 | 55.9211695327298 | 1.53455142272933 | 0.346373971738202 | 4.43033122560715 | 9.40884602042117e-06 | 0.00116725613929882 | 7201 | 147 | 141 | 16 | 22 | 56 | 51 | 42 | 26 | 49 | 72 | 44 | 40 |
| Harlequin-int_dup134 | 60.5761119833512 | 1.2457756387161 | 0.278046562824459 | 4.48045689204437 | 7.448342290278e-06 | 0.000975532346783437 | 7202 | 155 | 113 | 26 | 35 | 85 | 85 | 45 | 28 | 50 | 48 | 40 | 63 |
| MIR_dup57369 | 66.632143644566 | 1.74792279794671 | 0.280177254440322 | 6.23863204541118 | 4.41413926416791e-10 | 3.68535712754982e-07 | 7203 | 200 | 153 | 25 | 30 | 94 | 105 | 49 | 22 | 49 | 55 | 28 | 43 |
| L1PA16_dup4748 | 60.66357197871 | 1.66084063292928 | 0.280878441112614 | 5.9130228235046 | 3.35885615002183e-09 | 2.00307364780224e-06 | 7204 | 172 | 151 | 30 | 38 | 97 | 99 | 29 | 23 | 40 | 35 | 30 | 31 |
| L1PA7_dup11334 | 50.2207381105332 | 1.50811064308214 | 0.308799101875976 | 4.88379219343664 | 1.04064749242625e-06 | 0.000211458504041723 | 7205 | 119 | 116 | 21 | 34 | 95 | 80 | 27 | 15 | 32 | 33 | 32 | 39 |
| MIRc_dup18444 | 61.3574758650803 | 1.39596531704758 | 0.319039126372771 | 4.37553015179871 | 1.21137694968566e-05 | 0.00140674621605615 | 7206 | 122 | 120 | 39 | 42 | 78 | 112 | 17 | 16 | 47 | 70 | 47 | 72 |
| THE1D-int_dup491 | 49.3141087922325 | 1.3122464974693 | 0.329428969725629 | 3.9833973877957 | 6.79370126694032e-05 | 0.00512746329078604 | 7207 | 113 | 109 | 32 | 26 | 42 | 51 | 20 | 16 | 40 | 75 | 43 | 59 |
| L1PA3_dup3504 | 46.3286644446552 | 1.04701655339215 | 0.245420299211094 | 4.26621822545974 | 1.98814317516714e-05 | 0.00203558827865026 | 7208 | 72 | 69 | 23 | 27 | 59 | 84 | 32 | 20 | 45 | 50 | 49 | 55 |
| LTR40a_dup679 | 41.2751554875038 | 1.06789863560218 | 0.212318055985071 | 5.02971181912699 | 4.91217569424124e-07 | 0.000118068142821308 | 7209 | 61 | 67 | 33 | 27 | 58 | 67 | 30 | 18 | 41 | 48 | 39 | 29 |
| L1P3_dup2922 | 43.4325401264932 | 1.09376066095538 | 0.344582795495603 | 3.17415923038832 | 0.0015027122855364 | 0.0465578616528964 | 7210 | 102 | 86 | 31 | 17 | 64 | 75 | 37 | 15 | 19 | 35 | 31 | 49 |
| MIRb_dup145913 | 51.1443268144477 | 1.09789203057389 | 0.345213858547651 | 3.18032432183585 | 0.00147110307194179 | 0.0458832208095769 | 7211 | 134 | 100 | 26 | 31 | 64 | 63 | 29 | 21 | 36 | 39 | 38 | 77 |
| AluJo_dup46970 | 45.8978142696908 | 1.09432741927451 | 0.274972073139829 | 3.97977658886842 | 6.89800628291978e-05 | 0.00518758850504307 | 7212 | 119 | 86 | 24 | 28 | 50 | 71 | 41 | 27 | 33 | 38 | 26 | 44 |
| MamTip2_dup4766 | 43.5915781057799 | 1.06887908498529 | 0.313237142642945 | 3.41236379557864 | 0.000644021019679887 | 0.0257820043373982 | 7213 | 99 | 95 | 22 | 34 | 46 | 67 | 29 | 20 | 37 | 27 | 30 | 51 |
| L2_dup19568 | 82.1402822435663 | 1.6022978303338 | 0.240688917078877 | 6.65713174407905 | 2.7922288848007e-11 | 3.44248546229424e-08 | 7214 | 259 | 190 | 47 | 69 | 117 | 115 | 48 | 34 | 49 | 64 | 26 | 38 |
| Tigger4_dup540 | 74.1498228172761 | 1.53739046971553 | 0.2347942718261 | 6.54781932181973 | 5.83832914023874e-11 | 6.37250947524426e-08 | 7215 | 207 | 155 | 59 | 58 | 93 | 96 | 47 | 35 | 64 | 73 | 30 | 23 |
| L1PB1_dup1567 | 77.8968345197853 | 1.15904917752158 | 0.218912879507921 | 5.29456823247188 | 1.19297940086437e-07 | 3.78991557434009e-05 | 7216 | 150 | 149 | 35 | 51 | 134 | 131 | 66 | 54 | 55 | 60 | 43 | 58 |
| L2c_dup107622 | 80.9138440292744 | 1.18635256738901 | 0.20664187779132 | 5.74110427213144 | 9.40611475458219e-09 | 4.83401162288081e-06 | 7217 | 170 | 151 | 49 | 79 | 141 | 142 | 61 | 32 | 55 | 57 | 39 | 62 |
| L2b_dup71189 | 53.0247097130348 | 1.40553779781117 | 0.195953722109696 | 7.17280479635058 | 7.34765872218403e-13 | 1.43307049369574e-09 | 7218 | 121 | 93 | 43 | 45 | 87 | 112 | 48 | 24 | 34 | 39 | 9 | 25 |
| L1P2_dup1496 | 48.9552168501987 | 1.03530450763836 | 0.194281242401977 | 5.32889585653495 | 9.8811632835621e-08 | 3.27007410107158e-05 | 7219 | 84 | 77 | 48 | 51 | 70 | 66 | 37 | 27 | 43 | 64 | 23 | 28 |
| MIR1_Amn_dup8493 | 48.7199849307805 | 1.22237291714926 | 0.209671207537091 | 5.82995124370151 | 5.54435729592307e-09 | 3.06092260124295e-06 | 7220 | 83 | 72 | 31 | 28 | 66 | 82 | 28 | 33 | 55 | 58 | 25 | 46 |
| L1M1_dup8111 | 57.8262913540514 | 1.09168290744033 | 0.240449457566405 | 4.54017621203756 | 5.62072334292695e-06 | 0.000786260221373366 | 7221 | 117 | 91 | 44 | 49 | 107 | 101 | 31 | 25 | 39 | 44 | 42 | 48 |
| L1PA6_dup4874 | 64.7543383060331 | 1.14501982150983 | 0.19522848036568 | 5.86502450546714 | 4.49066370400102e-09 | 2.55020397544684e-06 | 7222 | 121 | 96 | 48 | 64 | 115 | 125 | 52 | 23 | 43 | 63 | 30 | 51 |
| L1PA3_dup3692 | 69.4328653517214 | 1.73304762438664 | 0.268583718693642 | 6.45254162395236 | 1.09989659273777e-10 | 1.11055469205789e-07 | 7223 | 182 | 178 | 39 | 50 | 88 | 90 | 35 | 21 | 70 | 60 | 23 | 44 |
| L4_B_Mam_dup1569 | 57.4362667468797 | 1.12369104168449 | 0.29256951301195 | 3.84076601186603 | 0.000122650982333042 | 0.00787846871838757 | 7224 | 143 | 124 | 49 | 61 | 61 | 85 | 45 | 16 | 48 | 38 | 22 | 48 |
| L1MB7_dup20863 | 96.8984831671461 | 2.96466136311078 | 0.335909159964563 | 8.82578302843406 | 1.08696581263519e-18 | 7.38969049295125e-15 | 7225 | 257 | 238 | 31 | 33 | 130 | 139 | 7 | 9 | 145 | 140 | 31 | 40 |
| L1PA16_dup2969 | 86.4452803894846 | 2.15496125803856 | 0.420325707153528 | 5.12688427417893 | 2.94576458567936e-07 | 7.87151276335399e-05 | 7226 | 193 | 197 | 29 | 32 | 110 | 143 | 12 | 8 | 78 | 111 | 81 | 102 |
| L2a_dup112240 | 67.7065022700435 | 2.01299737761248 | 0.31382318808267 | 6.41443161007655 | 1.41349225783345e-10 | 1.36167946875481e-07 | 7227 | 121 | 111 | 22 | 22 | 96 | 104 | 12 | 12 | 91 | 109 | 52 | 92 |
| L1PA6_dup4481 | 70.9709910584055 | 2.36091397197562 | 0.518270362084787 | 4.55537137504572 | 5.22931065734962e-06 | 0.00074687488215709 | 7228 | 232 | 196 | 8 | 8 | 70 | 92 | 27 | 12 | 52 | 69 | 53 | 92 |
| L1PA7_dup7319 | 63.7420007145768 | 1.56706430143014 | 0.456574291706359 | 3.43222194042844 | 0.000598657423671345 | 0.024534643357372 | 7229 | 122 | 97 | 4 | 10 | 81 | 73 | 30 | 27 | 75 | 85 | 75 | 121 |
| Tigger1_dup6809 | 64.4430216833842 | 1.32868149607921 | 0.378173744260522 | 3.51341550344089 | 0.00044238509293038 | 0.0198610874193231 | 7230 | 127 | 106 | 10 | 17 | 62 | 76 | 37 | 27 | 72 | 84 | 85 | 108 |
| AluSq2_dup47275 | 105.515655204328 | 1.69088517194527 | 0.495762254103128 | 3.41067751316447 | 0.000648016864196143 | 0.0258894554666196 | 7231 | 108 | 90 | 29 | 34 | 278 | 314 | 33 | 24 | 70 | 73 | 108 | 186 |
| L2b_dup20194 | 76.7590600706887 | 1.153898292748 | 0.337162721546581 | 3.42237803590805 | 0.000620759402496916 | 0.0251288221821251 | 7232 | 186 | 139 | 30 | 41 | 90 | 115 | 42 | 31 | 53 | 67 | 94 | 94 |
| LTR33_dup2036 | 66.0699147520848 | 1.56724495912629 | 0.42332772586149 | 3.70220248611613 | 0.00021373592666096 | 0.011656568600795 | 7233 | 197 | 161 | 24 | 26 | 74 | 71 | 22 | 21 | 58 | 41 | 64 | 84 |
| L1PA7_dup11039 | 60.3143996803966 | 1.25120078104359 | 0.32988130032288 | 3.7928818026937 | 0.000148909045753548 | 0.00905271124191972 | 7234 | 101 | 82 | 20 | 21 | 89 | 76 | 34 | 16 | 70 | 66 | 71 | 117 |
| L1PA5_dup6584 | 60.9202910842431 | 1.0491614312683 | 0.200574833450055 | 5.23077303977695 | 1.68802652081543e-07 | 5.03332278849558e-05 | 7235 | 76 | 63 | 36 | 46 | 69 | 92 | 25 | 24 | 94 | 91 | 64 | 74 |
| L1PA7_dup539 | 74.8858981998522 | 1.05378398991513 | 0.197810598214048 | 5.32723726346982 | 9.97178696727934e-08 | 3.29548169682812e-05 | 7236 | 81 | 86 | 40 | 47 | 100 | 94 | 35 | 24 | 103 | 127 | 76 | 120 |
| L2_dup40203 | 58.1967343635411 | 1.16536034163695 | 0.272149546220533 | 4.28205873506256 | 1.8517208606866e-05 | 0.00193249812244269 | 7237 | 105 | 58 | 31 | 43 | 79 | 66 | 26 | 12 | 84 | 82 | 47 | 100 |
| L2a_dup46234 | 69.5823156971057 | 1.10824260015804 | 0.297634824162085 | 3.72349775695102 | 0.00019648160747039 | 0.0109742294291344 | 7238 | 99 | 77 | 39 | 50 | 81 | 109 | 30 | 9 | 73 | 115 | 83 | 119 |
| AluSx_dup24300 | 32.6876569584122 | 1.58771542731645 | 0.423220356992356 | 3.7515100610936 | 0.000175772698147315 | 0.0101688330739998 | 7239 | 81 | 79 | 10 | 12 | 35 | 27 | 16 | 6 | 34 | 34 | 41 | 39 |
| L1M4b_dup2130 | 29.0749633548267 | 1.76893153455548 | 0.436902435138197 | 4.04880218622712 | 5.14804330195686e-05 | 0.00417219452155118 | 7240 | 61 | 61 | 5 | 6 | 30 | 38 | 16 | 5 | 35 | 35 | 24 | 52 |
| AluY_dup97495 | 34.9074982747564 | 1.45130768099958 | 0.377216092565189 | 3.84741719561916 | 0.000119369602421529 | 0.00772334286801005 | 7241 | 72 | 57 | 13 | 23 | 38 | 37 | 12 | 3 | 45 | 48 | 31 | 64 |
| L1MA9_dup608 | 29.6401335035717 | 1.22808880047873 | 0.313202661301724 | 3.92106757769741 | 8.81575335977661e-05 | 0.006220857796398 | 7242 | 49 | 44 | 14 | 14 | 22 | 23 | 12 | 6 | 45 | 57 | 29 | 55 |
| L1MA9_dup610 | 35.8285302788585 | 1.34836460878729 | 0.322084838071219 | 4.18636473812885 | 2.83457723183797e-05 | 0.00264708129516059 | 7243 | 45 | 57 | 12 | 13 | 33 | 40 | 11 | 8 | 49 | 63 | 46 | 70 |
| LTR104_Mam_dup105 | 27.0531703241951 | 1.16528500192735 | 0.350678825747148 | 3.32294086888373 | 0.000890737938287758 | 0.0324376384242147 | 7244 | 43 | 38 | 8 | 20 | 31 | 47 | 14 | 5 | 29 | 32 | 29 | 49 |
| L2a_dup50572 | 26.9927505004617 | 1.27586787539187 | 0.266983415526194 | 4.77882820128464 | 1.76319767361168e-06 | 0.000320019705297638 | 7245 | 39 | 43 | 15 | 19 | 29 | 41 | 16 | 4 | 38 | 39 | 25 | 30 |
| MER11C_dup821 | 29.1020258178014 | 1.40768076617271 | 0.281545856186499 | 4.99982768434085 | 5.73815734649032e-07 | 0.000133859959604704 | 7246 | 49 | 46 | 9 | 17 | 33 | 29 | 14 | 6 | 42 | 50 | 35 | 32 |
| L2a_dup113760 | 28.1045259564077 | 1.2686686961549 | 0.32869872972741 | 3.85967021292417 | 0.000113540149012709 | 0.00744719955235963 | 7247 | 44 | 17 | 17 | 22 | 50 | 43 | 11 | 6 | 41 | 42 | 20 | 39 |
| L2a_dup116342 | 30.836273229255 | 1.05430058924505 | 0.273343851197912 | 3.85704885851517 | 0.000114764229430936 | 0.00750417404126781 | 7248 | 30 | 37 | 23 | 29 | 43 | 35 | 13 | 6 | 40 | 64 | 27 | 38 |
| L1MEc_dup6582 | 34.0629072988748 | 1.40091804449138 | 0.255004423838897 | 5.49370094605271 | 3.93596782841449e-08 | 1.57933861871824e-05 | 7249 | 38 | 48 | 17 | 18 | 56 | 44 | 13 | 10 | 44 | 63 | 27 | 45 |
| L1P3_dup912 | 32.5839163599797 | 1.01836060610069 | 0.289536745075364 | 3.51720679126798 | 0.000436113877060507 | 0.019667401654583 | 7250 | 45 | 30 | 21 | 26 | 44 | 49 | 10 | 11 | 36 | 56 | 27 | 56 |
| L2c_dup29943 | 31.5108204923254 | 1.09061948696687 | 0.284941272046019 | 3.82752375300244 | 0.000129438896370532 | 0.00818264283628655 | 7251 | 39 | 36 | 15 | 26 | 53 | 49 | 14 | 11 | 33 | 47 | 24 | 51 |
| L1ME1_dup3330 | 26.014682977547 | 1.00903279249253 | 0.300500648532984 | 3.35783898443657 | 0.00078554357491065 | 0.0296882070323522 | 7252 | 21 | 28 | 13 | 15 | 47 | 45 | 22 | 10 | 32 | 34 | 25 | 33 |
| L1MEc_dup9647 | 29.7440389835702 | 1.20060849012538 | 0.2796027415291 | 4.29397967830877 | 1.75498545320103e-05 | 0.00185678865561304 | 7253 | 47 | 30 | 15 | 23 | 56 | 54 | 33 | 8 | 36 | 31 | 20 | 23 |
| L1MA3_dup1447 | 32.2748359125512 | 1.22462533679807 | 0.324697652878791 | 3.77158666205457 | 0.000162212800756485 | 0.00963501924333564 | 7254 | 53 | 40 | 18 | 17 | 43 | 59 | 16 | 8 | 36 | 39 | 21 | 60 |
| MIR3_dup12982 | 30.13546839479 | 1.20216689960151 | 0.346632989518532 | 3.46812604671962 | 0.000524101398173715 | 0.0222732329504988 | 7255 | 48 | 31 | 11 | 13 | 45 | 61 | 14 | 13 | 31 | 33 | 28 | 53 |
| L1ME3_dup1073 | 31.2772330010975 | 1.39579056162375 | 0.297961869940173 | 4.6844603368344 | 2.80698513854283e-06 | 0.000458100744702134 | 7256 | 50 | 52 | 14 | 15 | 59 | 39 | 15 | 9 | 29 | 44 | 36 | 31 |
| L1PA6_dup5534 | 30.0811231415314 | 1.09020171455582 | 0.283128602489635 | 3.85055308778183 | 0.000117851380303214 | 0.00765136822309103 | 7257 | 59 | 53 | 15 | 16 | 33 | 38 | 20 | 14 | 34 | 31 | 26 | 40 |
| L2b_dup59690 | 28.7650244537325 | 1.15412646165648 | 0.268000334887685 | 4.30643663986521 | 1.65905420916312e-05 | 0.00178062838454456 | 7258 | 44 | 46 | 15 | 22 | 36 | 45 | 14 | 9 | 35 | 33 | 25 | 38 |
| L1MC2_dup867 | 22.9733631766913 | 1.62705283669424 | 0.459448722870594 | 3.54131539756725 | 0.000398137349191844 | 0.0184059043502628 | 7259 | 27 | 27 | 8 | 9 | 65 | 61 | 8 | 7 | 16 | 17 | 24 | 21 |
| AluSq2_dup7131 | 26.5851293844868 | 1.49586379637132 | 0.473227334158524 | 3.16098350284693 | 0.00157237403647647 | 0.0479420953976716 | 7260 | 56 | 46 | 9 | 5 | 33 | 63 | 10 | 10 | 19 | 19 | 24 | 46 |
| L2_dup55532 | 28.798877778022 | 1.65179728910259 | 0.456700849091378 | 3.61680363062364 | 0.000298263402602601 | 0.0149011992442251 | 7261 | 41 | 29 | 8 | 5 | 53 | 69 | 10 | 8 | 29 | 29 | 30 | 53 |
| MER117_dup424 | 25.3699308042036 | 1.1639897412215 | 0.317527611742742 | 3.66579062158713 | 0.000246575610235412 | 0.0129689832345436 | 7262 | 42 | 32 | 15 | 17 | 43 | 37 | 7 | 9 | 28 | 28 | 27 | 34 |
| MLT1C_dup5302 | 25.0742756411917 | 1.36269683432232 | 0.368210893556412 | 3.70085963823759 | 0.000214870312529788 | 0.0117050209215231 | 7263 | 40 | 33 | 14 | 15 | 30 | 49 | 4 | 9 | 38 | 23 | 21 | 36 |

| | | | | | | | | | | | | | | | | | | | |
|---|---|---|---|---|---|---|---|---|---|---|---|---|---|---|---|---|---|---|---|
| MIR3_dup45882 | 25.158184401657 | 1.22278038949701 | 0.385556967214951 | 3.17146490265677 | 0.00151672194687246 | 0.0468535664540892 | 7264 | 29 | 27 | 11 | 7 | 33 | 39 | 9 | 4 | 29 | 36 | 46 | 46 |
| L1MB7_dup10259 | 27.6658887806059 | 1.26214185546558 | 0.394863018456088 | 3.19640431357829 | 0.00139151972122073 | 0.0441561342197164 | 7265 | 36 | 23 | 14 | 14 | 56 | 54 | 9 | 6 | 25 | 34 | 33 | 47 |
| MIR3_dup79766 | 30.0000689787043 | 1.2048631191423 | 0.362759746778555 | 3.32138041732012 | 0.000895733573225038 | 0.0325512860798808 | 7266 | 34 | 26 | 18 | 12 | 38 | 59 | 8 | 8 | 47 | 33 | 41 | 48 |
| X7A_LINE_dup507 | 29.6580620344896 | 1.47662642176841 | 0.411660977228105 | 3.58699634760424 | 0.000334509007286765 | 0.0161605540479364 | 7267 | 38 | 23 | 10 | 7 | 50 | 49 | 6 | 8 | 36 | 43 | 49 | 51 |
| MER51A_dup449 | 32.3646514047192 | 1.27711419581275 | 0.365233847817467 | 3.49670273838096 | 0.000471046409096097 | 0.0206997698679248 | 7268 | 55 | 49 | 7 | 8 | 29 | 39 | 13 | 16 | 34 | 56 | 49 | 49 |
| L1MCc_dup1255 | 33.3915556331116 | 1.17120262415396 | 0.331162335150006 | 3.53664200254972 | 0.000405248578206568 | 0.0186519091153927 | 7269 | 31 | 29 | 5 | 14 | 43 | 48 | 15 | 11 | 51 | 53 | 50 | 64 |
| L1M5_dup25451 | 31.2146537494163 | 1.66611764584595 | 0.341856236054119 | 4.87373775911518 | 1.09506332580533e-06 | 0.000220818591628876 | 7270 | 52 | 29 | 4 | 8 | 35 | 32 | 11 | 6 | 48 | 69 | 46 | 46 |
| MER21B_dup1166 | 28.5920768478543 | 1.24595454618003 | 0.325395478683559 | 3.82904689155715 | 0.000128640496697403 | 0.00815383367798621 | 7271 | 32 | 26 | 4 | 7 | 29 | 30 | 11 | 13 | 47 | 60 | 52 | 37 |
| L1MC4a_dup3331 | 27.514278780946 | 1.26512144652857 | 0.298676520104108 | 4.23575795676068 | 2.27782038675901e-05 | 0.00225199163150046 | 7272 | 28 | 24 | 9 | 11 | 35 | 46 | 18 | 7 | 39 | 55 | 24 | 47 |
| L1ME4b_dup7265 | 29.3576417586832 | 1.33669513188096 | 0.354257371099652 | 3.77323166976518 | 0.000161146521740943 | 0.00958125193957281 | 7273 | 37 | 20 | 10 | 6 | 54 | 34 | 19 | 7 | 44 | 51 | 31 | 52 |
| L1MB1_dup848 | 25.9000738836694 | 1.03631058050215 | 0.302862893054707 | 3.42171525223778 | 0.000622274463757297 | 0.025155915656336 | 7274 | 36 | 18 | 8 | 10 | 22 | 38 | 18 | 13 | 41 | 49 | 22 | 46 |
| L1ME4a_dup16605 | 22.6954235115085 | 1.10264445653528 | 0.34621205834206 | 3.18488172195855 | 0.00144813182866063 | 0.0453590776452584 | 7275 | 25 | 21 | 12 | 5 | 35 | 23 | 13 | 14 | 37 | 40 | 13 | 40 |
| L1PA7_dup8747 | 43.7564426637676 | 1.28402441130145 | 0.320359394155669 | 4.00807478952065 | 6.12157361203548e-05 | 0.00474927927841342 | 7276 | 51 | 40 | 10 | 12 | 56 | 64 | 45 | 25 | 63 | 98 | 30 | 44 |
| L1P3_dup1599 | 34.9285337875486 | 1.29005919330415 | 0.250458660482934 | 5.15078692354521 | 2.59395759520067e-07 | 7.10266782448354e-05 | 7277 | 37 | 34 | 16 | 21 | 52 | 62 | 24 | 9 | 45 | 67 | 28 | 41 |
| L1PA11_dup2632 | 37.1729172989345 | 1.1493581467983 | 0.248050439700047 | 4.63356626622915 | 3.59419636700261e-06 | 0.000552470703322225 | 7278 | 33 | 37 | 11 | 15 | 43 | 49 | 26 | 19 | 70 | 66 | 35 | 48 |
| L1PA7_dup538 | 35.5143015312162 | 1.02387459634657 | 0.244443769821492 | 4.18858945390289 | 2.80693666593235e-05 | 0.00262745614442148 | 7279 | 34 | 38 | 16 | 30 | 48 | 41 | 18 | 18 | 56 | 65 | 23 | 51 |
| L1ME2z_dup1143 | 35.5271969062205 | 1.18882195360226 | 0.22308268054246 | 5.32906432140523 | 9.87200327609899e-08 | 3.27007410107158e-05 | 7280 | 68 | 43 | 23 | 25 | 63 | 48 | 26 | 14 | 31 | 52 | 25 | 32 |
| LTR33_dup2870 | 32.2508727021586 | 1.11713535188615 | 0.261244510370671 | 4.27620603510896 | 1.90105295749435e-05 | 0.00196642906849961 | 7281 | 37 | 36 | 19 | 17 | 62 | 41 | 23 | 19 | 38 | 51 | 33 | 22 |
| L1MD2_dup9738 | 29.5099075104441 | 1.03436118906379 | 0.237617047242163 | 4.35305968603188 | 1.34250580918669e-05 | 0.00152261147413125 | 7282 | 34 | 36 | 14 | 27 | 34 | 39 | 26 | 14 | 36 | 61 | 21 | 26 |
| THE1C_dup4209 | 34.9278204317609 | 1.03489324491218 | 0.232989107085232 | 4.44180956723269 | 8.92054923664083e-06 | 0.00111833983596509 | 7283 | 65 | 47 | 21 | 35 | 34 | 39 | 24 | 19 | 49 | 50 | 26 | 26 |
| AluSp_dup34924 | 29.501110485907 | 1.3651219713761 | 0.375659865987367 | 3.6339308373762 | 0.00027913571710098 | 0.0142016820250244 | 7284 | 31 | 40 | 30 | 28 | 56 | 79 | 10 | 6 | 31 | 25 | 14 | 22 |
| L1PA5_dup2639 | 28.7195695663499 | 1.13460224456849 | 0.253294936525657 | 4.47937199271082 | 7.48629688758142e-06 | 0.000978755164402444 | 7285 | 51 | 42 | 24 | 25 | 56 | 55 | 15 | 18 | 22 | 21 | 14 | 20 |
| L1PA14_dup2535 | 29.9029965726795 | 1.01168629645193 | 0.252505802692351 | 4.00658632659051 | 6.16026164779252e-05 | 0.00477617992194734 | 7286 | 59 | 51 | 23 | 40 | 47 | 51 | 25 | 10 | 15 | 31 | 15 | 20 |
| L1PB_dup1355 | 33.3288906991953 | 1.09974954524151 | 0.233174383527315 | 4.71642522907188 | 2.40024518518278e-06 | 0.000406497324436657 | 7287 | 61 | 62 | 30 | 26 | 53 | 65 | 31 | 14 | 24 | 22 | 14 | 24 |
| MER112_dup3344 | 26.6649397819452 | 1.24776648868762 | 0.315476283589168 | 3.95518317412581 | 7.64760346712511e-05 | 0.00559015164387724 | 7288 | 26 | 26 | 24 | 25 | 52 | 58 | 21 | 10 | 32 | 33 | 14 | 12 |
| MER30_dup3119 | 32.7408532507904 | 1.29567696507736 | 0.310597744489401 | 4.17155947866702 | 3.02522065843678e-05 | 0.0027718103765591 | 7289 | 33 | 35 | 22 | 32 | 51 | 79 | 25 | 15 | 36 | 49 | 8 | 26 |
| L1M6_dup2822 | 44.7876220399474 | 1.07362429895932 | 0.316940467627197 | 3.38746360474921 | 0.000705420758353483 | 0.0275040126812403 | 7290 | 64 | 42 | 24 | 41 | 76 | 98 | 22 | 12 | 34 | 57 | 42 | 61 |
| LTR9A1_dup34 | 39.8125769103048 | 1.10469617490699 | 0.304448497335303 | 3.62851577385299 | 0.000285055336040205 | 0.0144099802399453 | 7291 | 76 | 61 | 25 | 44 | 63 | 69 | 25 | 8 | 36 | 34 | 21 | 50 |
| MIR3_dup15594 | 36.2821894861655 | 1.09566317634354 | 0.292763535329683 | 3.74248512578489 | 0.000182209321210373 | 0.0104323360473012 | 7292 | 64 | 53 | 29 | 35 | 27 | 62 | 19 | 9 | 52 | 41 | 34 | 30 |
| L1PA11_dup3919 | 40.7545759330587 | 1.1936307307796 | 0.343349890017564 | 3.47642671654428 | 0.000508143079398369 | 0.0218079496676943 | 7293 | 42 | 46 | 29 | 42 | 63 | 63 | 7 | 12 | 68 | 53 | 28 | 51 |
| L2c_dup97460 | 40.1635919664842 | 1.04750786633459 | 0.222983789501555 | 4.69768618013051 | 2.6312530019405e-06 | 0.000436912858897233 | 7294 | 55 | 41 | 24 | 40 | 52 | 44 | 20 | 12 | 59 | 73 | 42 | 37 |
| MER4B-int_dup1363 | 45.3333134393235 | 1.24077312846208 | 0.248292065731035 | 4.9972322909672 | 5.81589953725684e-07 | 0.000135011710369962 | 7295 | 49 | 60 | 23 | 26 | 62 | 68 | 23 | 10 | 71 | 62 | 44 | 65 |
| L1PA10_dup5116 | 56.1017994364039 | 2.45255912782283 | 0.492697694591534 | 4.97781734062324 | 6.43052971988189e-07 | 0.000145310429698672 | 7296 | 209 | 184 | 14 | 14 | 78 | 69 | 15 | 6 | 35 | 28 | 33 | 37 |
| L1PA10_dup5117 | 59.8388854783214 | 2.40875631093164 | 0.456540129880015 | 5.27611080227425 | 1.31954322273593e-07 | 4.12588740705812e-05 | 7297 | 207 | 170 | 11 | 18 | 74 | 71 | 14 | 8 | 47 | 52 | 40 | 54 |
| L1PB1_dup6025 | 37.2812728631202 | 1.888118522698 | 0.576502012456614 | 3.27512910952776 | 0.00105613693691078 | 0.0364842566187824 | 7298 | 59 | 45 | 6 | 7 | 76 | 92 | 13 | 4 | 19 | 28 | 53 | 80 |
| L1MB2_dup1708 | 47.5054209136178 | 2.40979013172767 | 0.398567375053454 | 6.04612992070531 | 1.48366470292068e-09 | 1.00293204943513e-06 | 7299 | 95 | 91 | 9 | 13 | 69 | 80 | 6 | 5 | 54 | 75 | 46 | 52 |
| L1PA4_dup6348 | 42.5430828130251 | 1.62900729076269 | 0.468289687599949 | 3.47863156908618 | 0.000503980968654187 | 0.0216971694531191 | 7300 | 89 | 58 | 17 | 35 | 77 | 77 | 4 | 4 | 38 | 45 | 41 | 59 |
| L1MB3_dup15527 | 40.4848400464184 | 1.4963672358329 | 0.433715524195338 | 3.45011223337941 | 0.000560353539110795 | 0.0233929062572123 | 7301 | 63 | 48 | 11 | 12 | 86 | 76 | 12 | 14 | 34 | 37 | 43 | 78 |
| L2a_dup43448 | 41.8954767829755 | 1.37129900327522 | 0.424037044696489 | 3.23391321684348 | 0.00122106553256382 | 0.0403034622293565 | 7302 | 72 | 54 | 21 | 21 | 90 | 84 | 9 | 12 | 33 | 29 | 41 | 68 |
| MIR3_dup4570 | 38.4638644416444 | 1.64087418510948 | 0.453474024456456 | 3.61845242861763 | 0.000296369962094064 | 0.0148338340345886 | 7303 | 62 | 49 | 19 | 15 | 87 | 86 | 7 | 7 | 23 | 41 | 35 | 61 |
| L1MEc_dup5873 | 49.1675599337068 | 1.26175848292132 | 0.376434048498967 | 3.35187129844548 | 0.000802673242032053 | 0.0301280770888679 | 7304 | 97 | 96 | 26 | 26 | 82 | 83 | 16 | 16 | 27 | 42 | 55 | 63 |
| AluSz_dup31508 | 46.6198481492432 | 1.40669624527746 | 0.40533431209507 | 3.47045933024175 | 0.000519569018739118 | 0.0221450427778136 | 7305 | 104 | 70 | 19 | 22 | 85 | 105 | 18 | 14 | 28 | 29 | 39 | 67 |
| MER58A_dup3976 | 46.1381021003166 | 1.43432305168382 | 0.409871763559184 | 3.49944343379173 | 0.000466230516178518 | 0.0205627678101205 | 7306 | 86 | 77 | 14 | 23 | 88 | 103 | 19 | 13 | 26 | 33 | 48 | 62 |
| L1MEf_dup1901 | 40.2206590689482 | 1.19878512769329 | 0.319686146413018 | 3.74988137942181 | 0.000176918239216534 | 0.0102251608813742 | 7307 | 79 | 64 | 17 | 22 | 66 | 46 | 19 | 14 | 28 | 55 | 42 | 61 |
| L2c_dup42629 | 39.1166577703483 | 1.09812161036114 | 0.277913475192526 | 3.95130754131447 | 7.77253649214451e-05 | 0.00565406288034185 | 7308 | 61 | 56 | 19 | 28 | 53 | 73 | 16 | 20 | 37 | 44 | 31 | 56 |
| AluSx_dup107084 | 60.2001896369823 | 1.42212617646843 | 0.349132912507204 | 4.07330883317706 | 4.63498976277929e-05 | 0.00385486638970388 | 7309 | 73 | 69 | 29 | 38 | 60 | 65 | 5 | 9 | 122 | 102 | 67 | 98 |
| L1PA7_dup730 | 55.1065048438186 | 2.02125764516616 | 0.31468601712379 | 6.42309329038616 | 1.33532598347978e-10 | 1.29687949577583e-07 | 7310 | 50 | 42 | 16 | 13 | 63 | 58 | 3 | 6 | 135 | 135 | 58 | 79 |
| L1ME1_dup26091 | 63.3398188709712 | 1.68696984461506 | 0.470386833672881 | 3.58634579850554 | 0.000335344211461754 | 0.0161941316671998 | 7311 | 41 | 29 | 21 | 24 | 142 | 153 | 11 | 17 | 111 | 61 | 66 | 102 |
| Tigger1_dup9386 | 64.585001472566 | 1.68468225860991 | 0.393720062589143 | 4.27888344711538 | 1.87833182515559e-05 | 0.00194915632125755 | 7312 | 78 | 51 | 19 | 17 | 144 | 165 | 18 | 34 | 71 | 79 | 54 | 71 |
| L1PA3_dup5692 | 54.8885921587941 | 1.53504009318473 | 0.465660163735747 | 3.29648145306205 | 0.000979040786852401 | 0.0346180581465968 | 7313 | 40 | 42 | 10 | 16 | 114 | 129 | 21 | 14 | 53 | 68 | 67 | 120 |
| L1PA6_dup3264 | 47.8280849139947 | 1.197618452596 | 0.374263761695286 | 3.1999316395774 | 0.00137460186603981 | 0.0437918082228824 | 7314 | 51 | 42 | 15 | 26 | 83 | 101 | 24 | 11 | 53 | 49 | 64 | 86 |
| Tigger16a_dup991 | 45.9225503225644 | 1.30797339061583 | 0.369665660929256 | 3.53826045764674 | 0.000402772547811291 | 0.018566082266855 | 7315 | 49 | 57 | 17 | 16 | 59 | 70 | 8 | 15 | 61 | 61 | 56 | 105 |
| L1ME2_dup4575 | 45.255085666881 | 1.21189959131239 | 0.283547896125616 | 4.2743734228783 | 1.91675558089849e-05 | 0.00197953265387358 | 7316 | 38 | 34 | 13 | 18 | 55 | 62 | 19 | 23 | 72 | 94 | 41 | 76 |
| L1PA15_dup2317 | 50.8436992563856 | 1.61751019333729 | 0.298925791855166 | 5.411076051012 | 6.26471575599277e-08 | 2.30478783813032e-05 | 7317 | 48 | 51 | 9 | 14 | 70 | 80 | 15 | 21 | 79 | 109 | 57 | 68 |
| L1MA3_dup4343 | 113.037761211608 | 3.3231195924464 | 0.850073967855626 | 3.90921227811412 | 9.25975696474709e-05 | 0.00645378480004016 | 7318 | 320 | 297 | 2 | 2 | 61 | 63 | 53 | 45 | 241 | 246 | 8 | 19 |
| Charlie8_dup1995 | 51.4696396235175 | 3.54733174177425 | 0.486045090465551 | 7.29835937315299 | 2.91297742103003e-13 | 6.08009934582818e-10 | 7319 | 135 | 119 | 18 | 35 | 135 | 108 | 1 | 5 | 38 | 54 | 2 | 2 |
| L1PA13_dup6826 | 61.2571452597168 | 3.90394961996604 | 0.393852636281243 | 9.91220893384678 | 3.68409352959796e-23 | 5.84410212662478e-19 | 7320 | 196 | 138 | 23 | 22 | 153 | 132 | 4 | 7 | 53 | 45 | 1 | 1 |
| L1PA13_dup6827 | 65.2183400413548 | 3.96015883255697 | 0.390309470158427 | 10.1462022711095 | 3.44509876137027e-24 | 8.19747469873011e-20 | 7321 | 204 | 168 | 14 | 31 | 147 | 125 | 6 | 8 | 54 | 66 | 2 | 0 |
| L1MC2_dup1942 | 29.1488823560377 | 1.66962737324024 | 0.328227955415511 | 5.08679210802329 | 3.64170642986745e-07 | 9.3678862503918e-05 | 7322 | 87 | 79 | 10 | 20 | 38 | 39 | 19 | 13 | 15 | 27 | 12 | 15 |
| LTR9A1_dup125 | 27.5626657364364 | 1.58816696623624 | 0.385325164519125 | 4.12162794562931 | 3.76204434691281e-05 | 0.00328620926641158 | 7323 | 92 | 78 | 12 | 17 | 42 | 31 | 18 | 9 | 15 | 13 | 11 | 18 |
| L1PA17_dup1856 | 33.0576230918687 | 1.10884974591729 | 0.344981055407545 | 3.2142337341027 | 0.00130793157917102 | 0.0422850358144565 | 7324 | 92 | 89 | 19 | 36 | 37 | 39 | 26 | 14 | 21 | 15 | 15 | 24 |
| Plat_L3_dup1086 | 36.3614956005378 | 1.66893685175458 | 0.399410290249929 | 4.17850238838426 | 2.93434915851863e-05 | 0.00270941654975893 | 7325 | 124 | 99 | 15 | 23 | 50 | 42 | 16 | 11 | 26 | 17 | 15 | 30 |
| L1PA7_dup9490 | 28.7580475311526 | 1.30251398447285 | 0.39032696624984 | 3.33698180524669 | 0.000846934924062918 | 0.0313218491515504 | 7326 | 88 | 63 | 10 | 12 | 33 | 22 | 27 | 14 | 22 | 30 | 17 | 30 |
| MER92A_dup98 | 28.2696317155907 | 1.72340206918232 | 0.41581823332429 | 4.14460437534077 | 3.40400884917 95e-05 | 0.00303631442785359 | 7327 | 96 | 64 | 6 | 8 | 33 | 31 | 29 | 9 | 21 | 31 | 15 | 20 |
| L1PA16_dup11749 | 24.0558796932518 | 1.04887002345342 | 0.315181118683434 | 3.32783266914824 | 0.000875244203092725 | 0.0320725613067184 | 7328 | 49 | 51 | 9 | 18 | 39 | 34 | 32 | 12 | 16 | 17 | 11 | 21 |
| L1PB1_dup10953 | 25.327998267327 | 1.09547545097365 | 0.341521528192393 | 3.20763220044075 | 0.00133832534654547 | 0.0429755955342927 | 7329 | 65 | 38 | 7 | 19 | 41 | 36 | 29 | 10 | 14 | 26 | 23 | 19 |
| MLT1B_dup5970 | 37.7485474260335 | 1.57739837114248 | 0.349741867752712 | 4.51017883926265 | 6.47729957468196e-06 | 0.000877134904411177 | 7330 | 114 | 93 | 13 | 25 | 58 | 52 | 16 | 32 | 21 | 25 | 18 | 10 |
| L1PREC2_dup2563 | 31.7995677738609 | 1.50480672827175 | 0.339168816790286 | 4.43674846795307 | 9.13279052532946e-06 | 0.0011407406689449 | 7331 | 90 | 78 | 19 | 37 | 33 | 32 | 15 | 15 | 30 | 30 | 10 | 15 |
| MER61E_dup93 | 31.0005493843465 | 1.43233823889822 | 0.303746293856019 | 4.71557437200263 | 2.41029829142889e-06 | 0.000407619642681121 | 7332 | 76 | 68 | 16 | 28 | 27 | 41 | 20 | 21 | 31 | 36 | 9 | 17 |
| L1PA8A_dup919 | 38.8051353305854 | 1.78060321285994 | 0.28464985608887 | 6.25541581972211 | 3.96458145956298e-10 | 3.32167711259568e-07 | 7333 | 119 | 100 | 19 | 29 | 51 | 52 | 26 | 14 | 28 | 32 | 10 | 17 |
| L2b_dup31120 | 37.8465391668318 | 1.78999106365036 | 0.325105056234616 | 5.5058850341552 | 3.6731819172317e-08 | 1.49660778163975e-05 | 7334 | 123 | 102 | 24 | 23 | 48 | 50 | 20 | 15 | 25 | 28 | 6 | 21 |
| MSTA-int_dup1036 | 37.7961813040913 | 1.25829362527748 | 0.298570739789531 | 4.21439028541268 | 2.50453864515996e-05 | 0.00241664619813662 | 7335 | 101 | 89 | 31 | 38 | 44 | 42 | 20 | 19 | 30 | 30 | 13 | 25 |
| L3_dup15436 | 38.8557750325348 | 1.51995104003146 | 0.292987546566226 | 5.18776670832969 | 2.12830988422136e-07 | 6.04323178652667e-05 | 7336 | 106 | 94 | 17 | 37 | 54 | 50 | 19 | 20 | 21 | 42 | 16 | 21 |
| MER45R_dup198 | 41.3548894574259 | 1.61411255655724 | 0.248848362509868 | 6.48632982864508 | 8.7952514699974e-11 | 9.13884238550219e-08 | 7337 | 100 | 107 | 25 | 27 | 54 | 67 | 26 | 17 | 29 | 37 | 17 | 20 |
| HAL1ME_dup2899 | 44.9075977531126 | 1.77700003243122 | 0.315355837870456 | 5.63490450797104 | 1.75155365558811e-08 | 8.18811760574793e-06 | 7338 | 134 | 131 | 26 | 26 | 75 | 63 | 18 | 17 | 21 | 28 | 19 | 17 |
| L1M3c_dup294 | 48.7273083803275 | 1.75356089141255 | 0.334640038281255 | 5.24014072081456 | 1.60454193061513e-07 | 4.81455654756808e-05 | 7339 | 132 | 133 | 29 | 20 | 103 | 64 | 22 | 15 | 25 | 30 | 21 | 29 |
| LTR36_dup149 | 39.1138721051591 | 1.39964287873369 | 0.349168361236484 | 4.00850430370391 | 6.1104525458076e-05 | 0.00474219746074604 | 7340 | 105 | 101 | 25 | 19 | 63 | 45 | 18 | 26 | 28 | 21 | 13 | 31 |
| MLT1L_dup3967 | 43.8742076343424 | 1.24768954363333 | 0.323512219610782 | 3.8566998957085 | 0.000114928118429675 | 0.00751076244654418 | 7341 | 131 | 116 | 28 | 26 | 43 | 53 | 33 | 25 | 25 | 31 | 22 | 29 |
| MamRTE1_dup3302 | 44.6878122238471 | 1.7498380170073 | 0.315958405063501 | 5.53819106871241 | 3.05611747504401e-08 | 1.28934561829224e-05 | 7342 | 134 | 119 | 25 | 13 | 54 | 69 | 29 | 24 | 30 | 34 | 18 | 18 |

| | | | | | | | | | | | | | | | | | | | |
|---|---|---|---|---|---|---|---|---|---|---|---|---|---|---|---|---|---|---|---|
| AluSx_dup81816 | 38.9940473019126 | 1.26244501606429 | 0.350671332288301 | 3.60008047371944 | 0.000318118710530978 | 0.0155911585367671 | 7343 | 97 | 67 | 18 | 20 | 75 | 78 | 21 | 22 | 22 | 18 | 33 | 26 |
| L1MA7_dup6976 | 38.6960937192628 | 1.27087845163755 | 0.32532760381344 | 3.90645748083013 | 9.36591148396082e-05 | 0.00650870669965695 | 7344 | 102 | 76 | 16 | 19 | 67 | 54 | 37 | 14 | 22 | 30 | 29 | 32 |
| L1PA5_dup9587 | 40.8786168018995 | 1.13279362641659 | 0.317034936506362 | 3.57308768206957 | 0.000352796563189594 | 0.0168160792937214 | 7345 | 105 | 76 | 14 | 22 | 70 | 57 | 40 | 26 | 27 | 25 | 26 | 34 |
| L2a_dup34410 | 36.3680922143706 | 1.28160888174406 | 0.400767927703961 | 3.19788284727902 | 0.00138440509936055 | 0.0439863340596135 | 7346 | 99 | 60 | 8 | 17 | 49 | 69 | 25 | 17 | 18 | 32 | 26 | 50 |
| AluSx_dup12026 | 35.1883538507961 | 1.43688701146407 | 0.393265670770928 | 3.65373109899804 | 0.00025845694728926 | 0.0134570671290351 | 7347 | 109 | 71 | 10 | 9 | 41 | 65 | 32 | 22 | 23 | 23 | 17 | 28 |
| MSTB1_dup4194 | 36.0895718775969 | 1.62090850624175 | 0.382626991240665 | 4.23626284435913 | 2.27270720029698e-05 | 0.0022504063094793 | 7348 | 105 | 71 | 11 | 9 | 48 | 73 | 33 | 13 | 28 | 23 | 27 | 20 |
| L1M1_dup6855 | 36.3625660213896 | 1.41197392163795 | 0.368565619221515 | 3.83099738011462 | 0.000127624864578283 | 0.00810889880559255 | 7349 | 78 | 78 | 10 | 8 | 44 | 45 | 30 | 19 | 33 | 46 | 22 | 46 |
| L1P1_dup2290 | 42.6513482498929 | 1.62194527997373 | 0.338917566133915 | 4.78566306985946 | 1.70423749033916e-06 | 0.000312898529225495 | 7350 | 95 | 93 | 10 | 13 | 48 | 58 | 27 | 19 | 55 | 41 | 29 | 45 |
| HERVL-int_dup1698 | 30.7670556051381 | 1.05587155474072 | 0.326685813014764 | 3.23207042569983 | 0.00122896763261747 | 0.0404856614025747 | 7351 | 61 | 59 | 20 | 9 | 32 | 44 | 34 | 20 | 36 | 23 | 21 | 26 |
| L2_dup10160 | 32.8815527536693 | 1.06228394185199 | 0.300973542794761 | 3.52949276533711 | 0.000416357088167344 | 0.0190410347301685 | 7352 | 61 | 53 | 21 | 17 | 53 | 52 | 25 | 12 | 32 | 23 | 25 | 44 |
| L1MA4_dup3692 | 34.5960928701454 | 1.35412642601423 | 0.328877826630986 | 4.11741478556295 | 3.83146127505748e-05 | 0.00333338531829919 | 7353 | 68 | 57 | 24 | 26 | 56 | 75 | 37 | 35 | 27 | 18 | 1 | 11 |
| HERVL-int_dup859 | 47.1047819158221 | 1.56990833930639 | 0.409554663865642 | 3.83320830603803 | 0.000126482760765662 | 0.00804922893638571 | 7354 | 136 | 112 | 52 | 60 | 50 | 34 | 22 | 15 | 46 | 50 | 13 | 9 |
| L1PB2_dup871 | 39.2249732741923 | 2.41054525190052 | 0.427508673938777 | 5.63858793715548 | 1.71450259560627e-08 | 8.04759187713925e-06 | 7355 | 149 | 129 | 20 | 21 | 33 | 30 | 9 | 9 | 34 | 43 | 9 | 13 |
| L1PA8_dup688 | 63.0887050215275 | -3.77975848226479 | 0.387380981156175 | -9.75721232101726 | 1.71812998799143e-22 | 2.40483622425064e-18 | 7356 | 15 | 20 | 206 | 304 | 10 | 6 | 111 | 95 | 1 | 3 | 23 | 50 |
| L1PA8_dup535 | 72.453512219252 | -3.25678629665891 | 0.43240943145413 | -7.53171892136305 | 5.00766649820503e-14 | 1.24120230477281e-10 | 7357 | 38 | 29 | 225 | 296 | 4 | 7 | 175 | 114 | 5 | 2 | 27 | 43 |
| L1PB1_dup1068 | 93.8373004268277 | -3.53543476616608 | 0.361634712413931 | -9.77625942644408 | 1.42375412127141e-22 | 2.1173537383753e-18 | 7358 | 41 | 28 | 252 | 429 | 7 | 11 | 219 | 135 | 4 | 5 | 51 | 81 |
| HERVIP10F-int_dup249 | 101.296237095206 | -2.55110892080642 | 0.81137704357335 | -3.14417192477026 | 0.00166557554305962 | 0.0499580282577669 | 7359 | 155 | 114 | 380 | 384 | 14 | 13 | 194 | 83 | 0 | 0 | 9 | 22 |
| L1M4_dup7343 | 70.6262205234589 | -1.89931903288766 | 0.301715409950569 | -6.2950680351356 | 3.07265062462074e-10 | 2.6490033533551e-07 | 7360 | 74 | 59 | 177 | 258 | 25 | 31 | 157 | 117 | 2 | 4 | 10 | 19 |
| L1PB1_dup5583 | 78.1416066963819 | -1.80253953960079 | 0.37754383038973 | -4.77438483828505 | 1.80257395940504e-06 | 0.000324444223407407 | 7361 | 85 | 65 | 208 | 341 | 29 | 24 | 194 | 78 | 3 | 6 | 8 | 19 |
| BSR/Beta_dup1545 | 147.425687358471 | -2.59719974202395 | 0.653913711292696 | -3.97177746416977 | 7.13383463737536e-05 | 0.00530790311640062 | 7362 | 19 | 23 | 416 | 467 | 108 | 153 | 141 | 111 | 8 | 12 | 223 | 287 |
| BSR/Beta_dup2004 | 261.295474323361 | -3.5728373442495 | 0.752868264468788 | -4.7456341472574 | 2.07854104639616e-06 | 0.000362977083394936 | 7363 | 20 | 21 | 885 | 1094 | 127 | 168 | 191 | 167 | 8 | 5 | 392 | 447 |
| ERVL-E-int_dup6527 | 105.392891160974 | -1.12906535511757 | 0.35906292411835 | -3.14447769256573 | 0.00166383596009961 | 0.0499328292581238 | 7364 | 97 | 91 | 147 | 191 | 128 | 136 | 150 | 104 | 12 | 19 | 168 | 133 |
| MER1B_dup5191 | 151.303688746829 | -1.18448356630749 | 0.373425359616587 | -3.1719419578886 | 0.00151423267640487 | 0.0468294266207219 | 7365 | 107 | 85 | 230 | 288 | 235 | 215 | 213 | 178 | 16 | 27 | 157 | 221 |
| MLT1A1_dup6676 | 95.2080990888843 | -2.16930302838348 | 0.514921326984264 | -4.21288246320739 | 2.52132281739877e-05 | 0.00242693640416168 | 7366 | 67 | 60 | 241 | 270 | 68 | 93 | 78 | 73 | 5 | 2 | 140 | 169 |
| HSMAR2_dup1786 | 114.193707854955 | -2.23439560730771 | 0.509460947902161 | -4.3858034978116 | 1.15558408635481e-05 | 0.0013560464468823 | 7367 | 58 | 76 | 303 | 414 | 93 | 74 | 76 | 75 | 12 | 2 | 140 | 207 |
| AluY_dup105517 | 105.76590340779 | -1.95748851995954 | 0.463751678472038 | -4.22098422675049 | 2.43237958283826e-05 | 0.00236234690701238 | 7368 | 62 | 60 | 255 | 315 | 83 | 105 | 85 | 65 | 12 | 6 | 149 | 217 |
| L2c_dup137285 | 102.503788896044 | -2.10683639470286 | 0.448374779455152 | -4.69882895122471 | 2.61657478766387e-06 | 0.000435387066031795 | 7369 | 83 | 73 | 270 | 328 | 62 | 67 | 89 | 60 | 9 | 6 | 138 | 191 |
| Cheshire_dup1041 | 92.5300891868491 | -1.94624948412301 | 0.406352375453309 | -4.78956098620529 | 1.67146594044934e-06 | 0.00030759368496996 | 7370 | 87 | 66 | 199 | 249 | 84 | 81 | 158 | 103 | 1 | 6 | 101 | 88 |
| REP522_dup205 | 129.367359238224 | -2.15810263499682 | 0.350633757003086 | -6.1548627075796 | 7.5142562236944e-10 | 5.69422678790824e-07 | 7371 | 80 | 50 | 288 | 348 | 96 | 125 | 208 | 197 | 11 | 5 | 134 | 147 |
| HERVE_a-int_dup169 | 127.534192630409 | -3.69725105314911 | 0.322386714710039 | -11.4683728716132 | 1.90201983271463e-30 | 1.13144502778779e-25 | 7372 | 25 | 18 | 280 | 405 | 43 | 32 | 223 | 202 | 4 | 6 | 215 | 228 |
| HERVE_a-int_dup171 | 67.2807465315076 | -4.03714836666375 | 0.3909649231174 | -10.3261140014125 | 5.36918836927075e-25 | 1.59697111964312e-20 | 7373 | 9 | 11 | 162 | 218 | 16 | 19 | 130 | 81 | 0 | 1 | 129 | 122 |
| L1MA4_dup9912 | 76.7059856513465 | -3.42090989017865 | 0.351979218342864 | -9.71906780827705 | 2.50060212713347e-22 | 3.30560152079389e-18 | 7374 | 14 | 12 | 197 | 243 | 33 | 32 | 224 | 129 | 3 | 0 | 50 | 79 |
| HERVIP10B3-int_dup36 | 73.7519614486322 | -2.90222906957996 | 0.799131986341788 | -3.63172682258106 | 0.000281531055093908 | 0.014279130225429 | 7375 | 0 | 2 | 225 | 232 | 42 | 37 | 178 | 105 | 6 | 14 | 46 | 89 |
| BSR/Beta_dup1967 | 145.021300255215 | -4.14506814822885 | 0.890422260310076 | -4.65517129680181 | 3.23711462346212e-06 | 0.000510104951122064 | 7376 | 12 | 0 | 552 | 710 | 48 | 46 | 79 | 67 | 5 | 4 | 206 | 248 |
| BSR/Beta_dup2015 | 153.768273314594 | -4.35533449231702 | 0.740026172902871 | -5.88537899305983 | 3.97142867642012e-09 | 2.29923495824687e-06 | 7377 | 16 | 4 | 597 | 702 | 40 | 56 | 120 | 95 | 1 | 4 | 222 | 226 |
| BSR/Beta_dup2011 | 75.8511636521864 | -4.15990395845489 | 0.790323627580499 | -5.2635449748479 | 1.41303920408337e-07 | 4.37795607363049e-05 | 7378 | 4 | 5 | 271 | 345 | 28 | 30 | 75 | 38 | 0 | 2 | 93 | 146 |
| BSR/Beta_dup2020 | 93.5424721079071 | -3.85296001383993 | 0.875953380425055 | -4.39859026740703 | 1.08956307781241e-05 | 0.00130149184795759 | 7379 | 5 | 4 | 323 | 421 | 42 | 49 | 67 | 49 | 0 | 4 | 137 | 171 |
| BSR/Beta_dup2007 | 99.3727738753997 | -3.56339136298869 | 0.786848195219236 | -4.52868975825234 | 5.93505664646084e-06 | 0.000821059877208587 | 7380 | 6 | 2 | 313 | 378 | 61 | 45 | 90 | 77 | 3 | 4 | 171 | 183 |
| BSR/Beta_dup2018 | 70.7339442570266 | -3.01117024943151 | 0.538415325021523 | -5.59265330961956 | 2.2362554183879e-08 | 1.00969265993117e-05 | 7381 | 12 | 7 | 224 | 259 | 54 | 41 | 61 | 67 | 3 | 3 | 100 | 110 |
| L1M5_dup8263 | 62.612797987996 | -1.95920183200365 | 0.552789725761763 | -3.54420811512696 | 0.000393794268578733 | 0.0182688201316474 | 7382 | 34 | 56 | 93 | 93 | 51 | 71 | 82 | 88 | 1 | 2 | 112 | 129 |
| MIR3_dup14769 | 62.3980509483507 | -1.85811414731409 | 0.53945557035484 | -3.44442480423712 | 0.000572275793858382 | 0.0237438109496211 | 7383 | 45 | 51 | 90 | 104 | 70 | 66 | 93 | 89 | 0 | 3 | 86 | 115 |
| AluJb_dup122348 | 43.7133137485681 | -2.11470148138625 | 0.474239515460396 | -4.45914229507695 | 8.22882795867244e-06 | 0.00105100198467755 | 7384 | 33 | 28 | 101 | 101 | 32 | 53 | 90 | 51 | 0 | 0 | 40 | 49 |
| Charlie4a_dup3298 | 35.7577426516374 | -1.4233252043149 | 0.4501947829078 | -3.16157640726459 | 0.0015691764493462 | 0.0478629995405886 | 7385 | 23 | 19 | 67 | 46 | 51 | 48 | 73 | 39 | 1 | 5 | 36 | 61 |
| HSATI_dup17 | 41.9223485188593 | -1.81508332822842 | 0.390526775054962 | -4.64778203228951 | 3.35523175569007e-06 | 0.000524894132373063 | 7386 | 19 | 22 | 78 | 100 | 46 | 44 | 58 | 66 | 3 | 3 | 44 | 61 |
| MER4D1_dup716 | 79.2872274248372 | -1.2317651687248 | 0.324139000140565 | -3.80011405042478 | 0.000144629508937 | 0.00886959101379471 | 7387 | 78 | 87 | 161 | 247 | 55 | 63 | 78 | 55 | 18 | 18 | 95 | 96 |
| MLT1D_dup9282 | 70.5911233357535 | -1.19505349717564 | 0.344604536477618 | -3.46789833178315 | 0.000524545700055721 | 0.0222861020129775 | 7388 | 97 | 79 | 143 | 187 | 43 | 33 | 80 | 52 | 17 | 19 | 105 | 76 |
| MER72_dup416 | 73.3866274822049 | -1.28118972645807 | 0.390636707871482 | -3.2797473986484 | 0.00103900068618348 | 0.0360387838592733 | 7389 | 87 | 77 | 155 | 208 | 47 | 59 | 59 | 55 | 20 | 7 | 82 | 115 |
| L1PA5_dup2963 | 73.8235314660789 | -1.42885951497538 | 0.412381716720775 | -3.46489540403864 | 0.000530437739363977 | 0.0224454300705715 | 7390 | 96 | 76 | 147 | 199 | 36 | 52 | 68 | 42 | 12 | 12 | 106 | 141 |
| L2b_dup44445 | 70.3421388333203 | -1.3635241805005 | 0.357905899212155 | -3.8097281534112 | 0.000139119674270338 | 0.00861383554877174 | 7391 | 71 | 83 | 155 | 178 | 46 | 50 | 84 | 42 | 11 | 14 | 82 | 121 |
| L1MCc_dup1781 | 64.1951293688125 | -1.13370471632057 | 0.348381605950301 | -3.25420371499837 | 0.00113710705733285 | 0.0383841787294826 | 7392 | 80 | 60 | 136 | 186 | 55 | 48 | 67 | 32 | 15 | 15 | 75 | 87 |
| L1MEf_dup5818 | 64.0955891196437 | -1.28090001090571 | 0.345024017761665 | -3.71249520313259 | 0.000205225951592095 | 0.0113222105906637 | 7393 | 77 | 67 | 142 | 192 | 42 | 40 | 70 | 38 | 13 | 15 | 70 | 89 |
| HAL1ME_dup4372 | 61.9933395066858 | -1.36399322080909 | 0.380115455151751 | -3.58836559346041 | 0.000332757463437022 | 0.0161029707941805 | 7394 | 64 | 58 | 122 | 208 | 47 | 39 | 48 | 37 | 13 | 14 | 67 | 113 |
| L1MB7_dup9946 | 67.1315085730641 | -1.18830596159822 | 0.368635276350053 | -3.22352752933447 | 0.00126622126861242 | 0.0413519595461451 | 7395 | 83 | 69 | 134 | 202 | 45 | 41 | 47 | 48 | 19 | 16 | 68 | 119 |
| MLT1C_dup4483 | 71.3545060734437 | -1.60260421280341 | 0.329210519104648 | -4.86802249564199 | 1.12720501528702e-06 | 0.000226245953966858 | 7396 | 73 | 55 | 139 | 172 | 24 | 34 | 77 | 52 | 19 | 21 | 124 | 156 |
| MSTB2_dup388 | 63.1596504565743 | -1.40106740727449 | 0.302133467099798 | -4.63724664706442 | 3.53080989309896e-06 | 0.00054448612496651 | 7397 | 58 | 57 | 122 | 150 | 17 | 28 | 85 | 45 | 24 | 31 | 105 | 110 |
| MLT1J2-int_dup267 | 54.6186274047955 | -1.06214655000844 | 0.321986525903522 | -3.29872980562763 | 0.00097123346246009 | 0.0344361671075143 | 7398 | 46 | 45 | 122 | 118 | 37 | 44 | 51 | 27 | 29 | 18 | 76 | 104 |
| AluSx_dup26360 | 52.3530685469332 | -1.31125431566608 | 0.30454064642095 | -4.30567916327862 | 1.66474190123734e-05 | 0.00178351497718064 | 7399 | 48 | 46 | 94 | 110 | 38 | 26 | 63 | 35 | 18 | 14 | 96 | 101 |
| L1MA8_dup4964 | 51.8993528596651 | -1.3275066484744 | 0.311724824044999 | -4.25858496364975 | 2.05725026772646e-05 | 0.00208570290670831 | 7400 | 47 | 47 | 129 | 146 | 33 | 32 | 56 | 34 | 14 | 17 | 64 | 67 |
| MSTA-int_dup3047 | 97.9043150912204 | -2.03434262702012 | 0.466487490699142 | -4.36098002107446 | 1.2948119036186e-05 | 0.00147980457837863 | 7401 | 58 | 32 | 249 | 367 | 56 | 60 | 54 | 25 | 17 | 26 | 176 | 205 |
| L1M4_dup20223 | 83.2840969207975 | -1.8570056523979 | 0.489396095590031 | -3.79448399595227 | 0.00014795082407868 | 0.00901900872357126 | 7402 | 40 | 43 | 211 | 267 | 62 | 52 | 34 | 22 | 18 | 17 | 157 | 195 |
| L1MC3_dup12704 | 85.4796308950788 | -1.96789333246594 | 0.503935333460829 | -3.90505130678421 | 9.42053962579169e-05 | 0.00652570515807461 | 7403 | 39 | 29 | 225 | 323 | 58 | 56 | 33 | 20 | 15 | 26 | 153 | 177 |
| MER52A_dup1348 | 83.5485541080818 | -1.84808846537626 | 0.503822139810432 | -3.66813666837195 | 0.000244324581592582 | 0.0129047851035801 | 7404 | 42 | 45 | 204 | 269 | 58 | 50 | 38 | 15 | 13 | 25 | 164 | 204 |
| MER21A_dup1916 | 50.2549808657572 | -1.81193491427733 | 0.450710668754536 | -4.02017311745541 | 5.81553906938062e-05 | 0.00455942095355137 | 7405 | 23 | 31 | 107 | 155 | 27 | 28 | 32 | 12 | 12 | 16 | 121 | 109 |
| THE1D-int_dup2520 | 65.2398148690056 | -2.05552592721461 | 0.503961564070508 | -4.07873551032759 | 4.52813165033045e-05 | 0.00379651449495959 | 7406 | 37 | 30 | 179 | 236 | 28 | 34 | 26 | 13 | 15 | 16 | 110 | 159 |
| L1PREC2_dup3278 | 57.764925391123 | -1.82929699575719 | 0.455118008915777 | -4.01939048756851 | 5.83488953047963e-05 | 0.0045713089602932 | 7407 | 20 | 19 | 165 | 177 | 39 | 28 | 30 | 17 | 23 | 24 | 92 | 136 |
| HSMAR1_dup704 | 65.8514429157817 | -1.93342856057929 | 0.452880688294469 | -4.26917863921403 | 1.96194094615907e-05 | 0.0020148295225497 | 7408 | 38 | 20 | 174 | 207 | 43 | 33 | 33 | 21 | 14 | 23 | 137 | 138 |
| L1PREC2_dup7637 | 58.0106815824003 | -1.81995148391595 | 0.444560523600811 | -4.09382162225037 | 4.24320798994178e-05 | 0.00360590845848102 | 7409 | 31 | 28 | 141 | 191 | 36 | 37 | 23 | 27 | 10 | 21 | 101 | 130 |
| L1MB4_dup3283 | 64.1205867981279 | -1.60521735032216 | 0.345640768254041 | -4.64417828495956 | 3.41432630880601e-06 | 0.000532737893688625 | 7410 | 38 | 30 | 158 | 208 | 36 | 33 | 46 | 40 | 29 | 27 | 95 | 106 |
| L1PA5_dup9438 | 60.9661541566658 | -1.6142022284996 | 0.406105845855645 | -3.97483130315092 | 7.04291595768716e-05 | 0.0052682605484685 | 7411 | 25 | 15 | 164 | 228 | 56 | 51 | 59 | 36 | 17 | 21 | 63 | 78 |
| L1MA1_dup1892 | 39.7347939110143 | -1.55728799528684 | 0.325636599957674 | -4.78228797220354 | 1.73311153143455e-06 | 0.000316490373337472 | 7412 | 30 | 11 | 108 | 123 | 25 | 28 | 65 | 37 | 10 | 18 | 24 | 48 |
| Charlie1a_dup5590 | 44.1393792495337 | -1.59753894135582 | 0.282193641528371 | -5.66114435712827 | 1.50366866016839e-08 | 7.19903305860017e-06 | 7413 | 15 | 15 | 104 | 82 | 26 | 35 | 56 | 55 | 18 | 24 | 54 | 85 |
| MIRb_dup212421 | 40.9814672046664 | -1.54761776341712 | 0.347665468536765 | -4.45145665438286 | 8.52897504312344e-06 | 0.00108006146653063 | 7414 | 16 | 13 | 84 | 108 | 33 | 38 | 47 | 42 | 20 | 8 | 55 | 70 |
| MamRep605_dup1139 | 42.7685878163045 | -1.4722676879855 | 0.286014424073592 | -5.1475295092344 | 2.63939493066217e-07 | 7.19396868466599e-05 | 7415 | 47 | 28 | 81 | 107 | 18 | 15 | 67 | 34 | 14 | 20 | 63 | 72 |
| L1ME1_dup7859 | 48.8059825170102 | -1.48804465875858 | 0.292517170996765 | -5.08703353614423 | 3.63707539737888e-07 | 9.36609894485622e-05 | 7416 | 51 | 34 | 76 | 116 | 22 | 22 | 58 | 41 | 15 | 17 | 91 | 101 |
| L1ME3_dup2321 | 44.2817915001965 | -1.47383861673336 | 0.323146162061624 | -4.56090398020045 | 5.09338695940584e-06 | 0.000732296709028871 | 7417 | 40 | 41 | 77 | 99 | 20 | 26 | 53 | 32 | 11 | 12 | 80 | 95 |
| MER33_dup2390 | 43.8529473507 | -1.57722524565085 | 0.302963293052796 | -5.20599452744921 | 1.92960332015453e-07 | 5.61984567463268e-05 | 7418 | 32 | 36 | 77 | 85 | 24 | 16 | 52 | 34 | 14 | 13 | 86 | 108 |
| L1MCb_dup1136 | 48.8499305104569 | -1.34195920776871 | 0.348690755348448 | -3.84856549015097 | 0.000118811535379465 | 0.0076969043281792 | 7419 | 48 | 48 | 100 | 163 | 35 | 28 | 42 | 40 | 9 | 15 | 42 | 80 |
| L1MEf_dup5817 | 45.3117986753112 | -1.29489525011246 | 0.343903605939392 | -3.76528546880275 | 0.000166358906823688 | 0.00980825995433598 | 7420 | 48 | 43 | 100 | 146 | 39 | 27 | 39 | 39 | 9 | 10 | 39 | 61 |
| L1MB7_dup9947 | 55.4196400251403 | -1.26703036870096 | 0.370758030541157 | -3.4174050575563 | 0.000632211406976222 | 0.0254505833552071 | 7421 | 67 | 51 | 106 | 178 | 35 | 50 | 50 | 35 | 11 | 11 | 55 | 93 |

| | | | | | | | | | | | | | | | | | | |
|---|---|---|---|---|---|---|---|---|---|---|---|---|---|---|---|---|---|---|
| LTR16_dup1513 | 46.4714070720071 | -1.25788571025077 | 0.394956874381189 | -3.18486850550863 | 0.00144819796454249 | 0.0453590776452584 | 7422 | 71 | 40 | 113 | 123 | 36 | 31 | 43 | 27 | 8 | 8 | 52 | 67 |
| MIRb_dup101982 | 45.8942306863637 | -1.24054175896428 | 0.334491497026614 | -3.70873929529387 | 0.000208293753365635 | 0.011453587508578 | 7423 | 62 | 43 | 97 | 127 | 34 | 35 | 44 | 43 | 7 | 9 | 45 | 60 |
| L1MEd_dup10967 | 40.9680388517886 | -1.38010493657987 | 0.318824168310557 | -4.32873374654444 | 1.49969106940149e-05 | 0.00165001516480919 | 7424 | 47 | 28 | 64 | 138 | 39 | 23 | 48 | 49 | 6 | 10 | 35 | 55 |
| MLT1C_dup9080 | 42.6392989410298 | -1.46437261173426 | 0.362894568145105 | -4.03525635343412 | 5.45427133228892e-05 | 0.00435656947443041 | 7425 | 33 | 44 | 108 | 149 | 31 | 25 | 47 | 26 | 8 | 9 | 47 | 41 |
| L2b_dup44442 | 47.1060278489057 | -1.35844946856094 | 0.33647647465038 | -4.03727918860434 | 5.40747017005281e-05 | 0.00432646279909354 | 7426 | 45 | 42 | 103 | 162 | 19 | 38 | 53 | 37 | 7 | 20 | 50 | 51 |
| L2b_dup44446 | 42.8049593085276 | -1.27072985325767 | 0.366967171404094 | -3.46278891486555 | 0.000534607619827707 | 0.0225825927050459 | 7427 | 56 | 44 | 79 | 134 | 27 | 28 | 49 | 29 | 4 | 14 | 49 | 60 |
| HERVIP10F-int_dup269 | 53.9857990101931 | -1.6517298622526 | 0.525358572599828 | -3.14400477768685 | 0.00166652718830151 | 0.0499676762030735 | 7428 | 8 | 19 | 78 | 78 | 81 | 69 | 98 | 70 | 5 | 7 | 90 | 97 |
| HSATI_dup15 | 51.7322882840326 | -1.14255867758662 | 0.346303189796857 | -3.29930162715754 | 0.000969257049894975 | 0.0343992137530909 | 7429 | 44 | 36 | 71 | 114 | 63 | 63 | 68 | 47 | 9 | 9 | 70 | 87 |
| SST1_dup643 | 54.4951707677946 | -1.52757617318993 | 0.433725693479678 | -3.52198681368066 | 0.000428325448012553 | 0.0194389332543954 | 7430 | 22 | 34 | 68 | 126 | 55 | 65 | 79 | 38 | 7 | 9 | 129 | 88 |
| AluY_dup105516 | 59.3708540105595 | -1.77696856285233 | 0.512191742068857 | -3.46934246865198 | 0.00052173393346715 | 0.0222003759894089 | 7431 | 48 | 41 | 128 | 196 | 50 | 55 | 38 | 24 | 5 | 5 | 76 | 137 |
| LTR12C_dup281 | 50.1484057240174 | -2.13760981461216 | 0.435544490206048 | -4.90790232153069 | 9.20556793446578e-07 | 0.000193527169992853 | 7432 | 34 | 17 | 103 | 118 | 26 | 32 | 51 | 29 | 7 | 6 | 68 | 188 |
| LTR37B_dup2008 | 40.3658261539165 | -2.0070311800542 | 0.458508023788306 | -4.37730874036099 | 1.20153717288219e-05 | 0.00139736541612231 | 7433 | 28 | 20 | 103 | 116 | 36 | 29 | 35 | 30 | 4 | 2 | 48 | 88 |
| LSAU_dup124 | 47.7744148783248 | -2.37691337063045 | 0.587343642246576 | -4.04688703454558 | 5.19032736670582e-05 | 0.00420074025713667 | 7434 | 5 | 10 | 110 | 115 | 42 | 48 | 36 | 49 | 2 | 5 | 62 | 150 |
| LTR108d_Mam_dup67 | 36.286605950664 | -2.27739729155754 | 0.390119625324084 | -5.83768963088087 | 5.29296371848247e-09 | 2.94261575924774e-06 | 7435 | 24 | 13 | 83 | 116 | 24 | 21 | 42 | 39 | 1 | 5 | 65 | 47 |
| MLT1D_dup10278 | 51.6033488997626 | -3.08636740837314 | 0.332201529076354 | -9.29064780934154 | 1.53352093597123e-20 | 1.45958069052244e-16 | 7436 | 16 | 9 | 137 | 193 | 19 | 16 | 75 | 56 | 6 | 4 | 74 | 84 |
| L2b_dup87045 | 28.415204261793 | -1.00368437844363 | 0.282547855888261 | -3.55226329815278 | 0.000381932474139227 | 0.0178650097290215 | 7437 | 22 | 14 | 35 | 44 | 11 | 34 | 38 | 23 | 16 | 17 | 50 | 66 |
| MIR_dup44942 | 30.6282279001979 | -1.32043673572841 | 0.262843240725137 | -5.02366631945931 | 5.06942836703578e-07 | 0.000120504515706563 | 7438 | 25 | 18 | 61 | 63 | 13 | 23 | 44 | 26 | 14 | 15 | 36 | 64 |
| L1PA2_dup5103 | 32.9129489859123 | -1.0663510964787 | 0.251308329789629 | -4.24319837456777 | 2.2035648229152e-05 | 0.00219384700984678 | 7439 | 27 | 22 | 50 | 48 | 17 | 20 | 71 | 40 | 26 | 16 | 34 | 51 |
| L2c_dup30793 | 29.6928288802787 | -1.19290694905363 | 0.295660239539536 | -4.03472225724863 | 5.46669229533249e-05 | 0.00436502538558786 | 7440 | 14 | 16 | 34 | 49 | 13 | 17 | 66 | 31 | 19 | 32 | 28 | 67 |
| MLT1H1_dup709 | 28.2828661393687 | -1.21173198110531 | 0.348764130208419 | -3.47435953456907 | 0.00051207439262891 | 0.0219384323781921 | 7441 | 7 | 9 | 42 | 54 | 30 | 26 | 43 | 34 | 19 | 11 | 41 | 46 |
| MER67B_dup70 | 30.4610993028702 | -1.36647324402663 | 0.314857612074447 | -4.33997207507099 | 1.42500823587896e-05 | 0.00158891757120176 | 7442 | 6 | 10 | 43 | 59 | 24 | 20 | 53 | 37 | 17 | 26 | 52 | 43 |
| MLT1B_dup11746 | 28.8812472619451 | -1.07893202806164 | 0.24759344952269 | -4.35767598109561 | 1.31450795044151e-05 | 0.00149656415682179 | 7443 | 14 | 12 | 46 | 50 | 22 | 25 | 44 | 31 | 19 | 20 | 43 | 44 |
| MER39_dup749 | 29.9583184097331 | -1.48082721234595 | 0.27846516876396 | -5.3178184507563 | 1.05018776313529e-07 | 3.43252716328284e-05 | 7444 | 15 | 9 | 50 | 59 | 22 | 15 | 37 | 41 | 13 | 22 | 37 | 66 |
| MIR_dup35793 | 30.2090317028554 | -1.01835861018144 | 0.256722629697781 | -3.96676604388274 | 7.28544438798659e-05 | 0.00538960225208824 | 7445 | 14 | 20 | 31 | 37 | 28 | 19 | 51 | 39 | 16 | 23 | 46 | 62 |
| L1M5_dup10629 | 25.4157000769434 | -1.03105845843127 | 0.298173917812029 | -3.45790961864497 | 0.000544383980356479 | 0.0228818213372024 | 7446 | 9 | 10 | 25 | 40 | 24 | 20 | 39 | 30 | 12 | 26 | 37 | 55 |
| MLT1A_dup1702 | 26.2054590868667 | -1.4682765816382 | 0.256168013130572 | -5.73169367906132 | 9.94327238354782e-09 | 5.04469486263469e-06 | 7447 | 9 | 10 | 36 | 41 | 17 | 13 | 43 | 29 | 12 | 23 | 50 | 55 |
| LOR1-int_dup301 | 31.2642582038798 | -1.19830673563592 | 0.35272336086715 | -3.39729904106706 | 0.000680545527200753 | 0.0268101135786938 | 7448 | 7 | 9 | 54 | 60 | 20 | 17 | 51 | 23 | 30 | 36 | 47 | 46 |
| L2a_dup35500 | 30.9134763836121 | -1.07932291082962 | 0.331749736167736 | -3.25342507667861 | 0.00114022789509241 | 0.0384458929751536 | 7449 | 15 | 7 | 24 | 55 | 21 | 26 | 61 | 40 | 26 | 25 | 52 | 41 |
| L1MD_dup1694 | 33.1216396978707 | -1.24180058611617 | 0.258704545852851 | -4.80007253843344 | 1.58608171112873e-06 | 0.000295537822111383 | 7450 | 17 | 8 | 44 | 56 | 21 | 19 | 52 | 38 | 22 | 33 | 60 | 53 |
| MIR_dup35776 | 32.673073547143 | -1.09115228267258 | 0.318712677315127 | -3.42362372235889 | 0.000617921163313417 | 0.0250693723999615 | 7451 | 11 | 7 | 40 | 52 | 31 | 22 | 63 | 31 | 27 | 28 | 45 | 63 |
| MER5A_dup7100 | 18.7334745127223 | -1.49099293092726 | 0.337922506107804 | -4.41223329011299 | 1.02309786925655e-05 | 0.00124145147070941 | 7452 | 9 | 6 | 27 | 32 | 18 | 5 | 38 | 23 | 6 | 16 | 28 | 35 |
| LOR1-int_dup304 | 21.1051157707153 | -1.04321530901559 | 0.323117909976846 | -3.22859017344581 | 0.00124402032007284 | 0.040845820212509 | 7453 | 10 | 12 | 32 | 38 | 14 | 9 | 44 | 17 | 12 | 28 | 25 | 33 |
| MamGypsy2-I_dup444 | 17.4455510388057 | -1.22422281281439 | 0.326674365410871 | -3.7475325352653 | 0.000178582678599373 | 0.0102888702280887 | 7454 | 8 | 9 | 23 | 28 | 11 | 7 | 36 | 16 | 9 | 20 | 34 | 24 |
| AluSx1_dup20913 | 19.3693369710292 | -1.19625684337424 | 0.328196857704108 | -3.64493691908759 | 0.000267457395635202 | 0.0137720011819549 | 7455 | 10 | 10 | 12 | 33 | 11 | 18 | 37 | 26 | 11 | 11 | 29 | 42 |
| L1MB8_dup11364 | 19.9652293856634 | -1.14045550345169 | 0.348427280717437 | -3.27315215129944 | 0.0010635520964456 | 0.0366552675464723 | 7456 | 15 | 13 | 14 | 21 | 10 | 15 | 40 | 37 | 8 | 15 | 29 | 36 |
| L2a_dup35508 | 21.0180697679827 | -1.06639905498141 | 0.292303776544408 | -3.64825616551485 | 0.000264026318731705 | 0.0136573927036814 | 7457 | 17 | 10 | 27 | 19 | 17 | 12 | 34 | 28 | 10 | 16 | 36 | 42 |
| AluSz6_dup18129 | 21.3830266982125 | -1.15460285963144 | 0.303282866786283 | -3.80701643935944 | 0.000140653398962362 | 0.00868394231175354 | 7458 | 9 | 11 | 26 | 29 | 23 | 18 | 40 | 26 | 8 | 12 | 32 | 42 |
| Charlie16a_dup1528 | 22.1176760634896 | -1.29872933768394 | 0.322408024352885 | -4.02821654420874 | 5.620156748708e-05 | 0.00444579061744705 | 7459 | 8 | 9 | 29 | 24 | 22 | 15 | 32 | 25 | 8 | 14 | 49 | 49 |
| MIRc_dup69070 | 26.1190832650495 | -1.14904275242922 | 0.343025534108891 | -3.34972950458103 | 0.000808905121379267 | 0.0302825264335606 | 7460 | 12 | 7 | 37 | 48 | 17 | 15 | 48 | 29 | 19 | 30 | 47 | 23 |
| L1MB1_dup966 | 22.8145216121319 | -1.47080701421018 | 0.295322003830969 | -4.9803502452598 | 6.34693004162829e-07 | 0.000144381129606624 | 7461 | 12 | 9 | 29 | 42 | 8 | 13 | 46 | 24 | 11 | 21 | 48 | 32 |
| AluSz6_dup9264 | 22.2184752863635 | -1.42489015955417 | 0.335742477406365 | -4.2439972760121 | 2.19573017745795e-05 | 0.00218787777556704 | 7462 | 11 | 5 | 35 | 47 | 14 | 8 | 40 | 21 | 11 | 26 | 38 | 32 |
| AluSz_dup19286 | 22.8029612658712 | -1.498524871999 | 0.385860442635548 | -3.88359289115932 | 0.000102924184705877 | 0.00695354856729831 | 7463 | 8 | 6 | 30 | 55 | 12 | 11 | 39 | 25 | 9 | 29 | 48 | 22 |
| Tigger4_dup306 | 23.5233645120608 | -1.22435548763256 | 0.290589866213087 | -4.2133454397021 | 2.51615789186803e-05 | 0.00242294498477714 | 7464 | 15 | 6 | 38 | 50 | 15 | 14 | 30 | 24 | 14 | 22 | 38 | 37 |
| L1MCc_dup665 | 23.0443807114849 | -1.26104800100273 | 0.291433602009346 | -4.32705080096527 | 1.51119117269334e-05 | 0.00166088635001242 | 7465 | 14 | 4 | 35 | 34 | 14 | 13 | 43 | 21 | 15 | 23 | 36 | 46 |
| L2a_dup35506 | 21.3520450604331 | -1.10675961906967 | 0.279856520113971 | -3.95473944512333 | 7.66181044856483e-05 | 0.00559747359224503 | 7466 | 14 | 6 | 28 | 26 | 15 | 12 | 38 | 23 | 14 | 21 | 33 | 44 |
| L1PA7_dup7119 | 24.3369025857959 | -1.06621962456187 | 0.301594470619977 | -3.53527577070653 | 0.000407349799340155 | 0.0187299044161918 | 7467 | 16 | 8 | 33 | 49 | 19 | 16 | 34 | 38 | 19 | 16 | 30 | 29 |
| L2a_dup35507 | 20.3762540670331 | -1.10706178738736 | 0.309642485475353 | -3.57529034069027 | 0.000349839437171455 | 0.0167020254247992 | 7468 | 9 | 9 | 23 | 42 | 14 | 10 | 33 | 26 | 17 | 19 | 28 | 29 |
| MER58B_dup1369 | 21.6405872800555 | -1.27606679476373 | 0.305407630728748 | -4.17824136128769 | 2.93771812945657e-05 | 0.00271092643816622 | 7469 | 10 | 7 | 29 | 34 | 12 | 12 | 41 | 27 | 20 | 15 | 37 | 30 |
| L1MEf_dup5543 | 27.7081709964341 | -2.25860718688145 | 0.291990572609005 | -7.7352058551078 | 1.03235922943675e-14 | 3.23218091062576e-11 | 7470 | 5 | 8 | 49 | 59 | 6 | 12 | 53 | 38 | 13 | 14 | 51 | 50 |
| MSTB_dup7172 | 26.4600587341849 | -1.96673525315949 | 0.293616429247023 | -6.69831473055907 | 2.10836801830335e-11 | 2.69719213163015e-08 | 7471 | 6 | 11 | 31 | 36 | 9 | 13 | 43 | 35 | 8 | 17 | 73 | 58 |
| L1PA16_dup10858 | 24.4354071091017 | -2.06137567972918 | 0.318246485010753 | -6.47729284318577 | 9.33828484513272e-11 | 9.61498323146392e-08 | 7472 | 10 | 5 | 36 | 48 | 16 | 6 | 63 | 25 | 12 | 8 | 44 | 48 |
| THE1D_dup2284 | 25.4571669594014 | -1.44029440890155 | 0.340692860961319 | -4.22754502350747 | 2.36254910473941e-05 | 0.00231436438565798 | 7473 | 11 | 5 | 38 | 25 | 16 | 16 | 53 | 32 | 20 | 14 | 55 | 37 |
| MIR3_dup70139 | 27.6681302243631 | -1.62690217041114 | 0.270089322687345 | -6.02357084768746 | 1.70610376040087e-09 | 1.13080937429622e-06 | 7474 | 18 | 8 | 37 | 34 | 12 | 13 | 39 | 36 | 10 | 20 | 67 | 61 |
| MIRb_dup179587 | 24.8329880293936 | -1.58450888761639 | 0.295341224128157 | -5.36501090321486 | 8.09443295577612e-08 | 2.79947375595219e-05 | 7475 | 14 | 6 | 25 | 30 | 14 | 15 | 39 | 23 | 11 | 12 | 67 | 67 |
| L2a_dup155351 | 26.9446681237776 | -1.08860248298109 | 0.282728094127764 | -3.85035129366789 | 0.000117948526871659 | 0.00765558651800427 | 7476 | 17 | 19 | 23 | 46 | 7 | 18 | 43 | 16 | 16 | 26 | 61 | 60 |
| MER21B_dup1884 | 25.2907737725434 | -1.02073823983495 | 0.319862258160977 | -3.19118062163259 | 0.00141692669373468 | 0.04467326233182 | 7477 | 12 | 19 | 42 | 49 | 8 | 14 | 41 | 20 | 27 | 21 | 35 | 35 |
| MIRb_dup179590 | 18.6200189910853 | -1.55754935497543 | 0.37379465124287 | -4.16685832661481 | 3.08826292236297e-05 | 0.00281871810251086 | 7478 | 5 | 10 | 26 | 29 | 5 | 16 | 27 | 12 | 10 | 9 | 59 | 33 |
| MER72_dup175 | 19.1053089259981 | -1.45455966321584 | 0.331665160924977 | -4.3856269351874 | 1.15652190893171e-05 | 0.0013560464468823 | 7479 | 8 | 7 | 23 | 21 | 5 | 9 | 43 | 20 | 14 | 20 | 37 | 38 |
| THE1C_dup1857 | 16.8672175522677 | -1.28153238574314 | 0.322516136042655 | -3.97354501845345 | 7.08107670182931e-05 | 0.00529009668716305 | 7480 | 8 | 5 | 28 | 26 | 6 | 12 | 29 | 18 | 12 | 17 | 26 | 29 |
| L1MDa_dup783 | 22.1236094963136 | -1.20039808971028 | 0.325191806316529 | -3.69135404519345 | 0.000223063385744015 | 0.0120356100644547 | 7481 | 9 | 12 | 29 | 55 | 14 | 12 | 18 | 22 | 15 | 19 | 30 | 51 |
| MER39B_dup226 | 21.7781280470884 | -1.23980663998009 | 0.354390062738828 | -3.49842382824875 | 0.000468016754596251 | 0.0206150897240206 | 7482 | 4 | 9 | 31 | 51 | 15 | 17 | 21 | 24 | 15 | 18 | 39 | 34 |
| LTR24B_dup86 | 21.7522852924142 | -1.13884158201641 | 0.334608741225687 | -3.40350218540251 | 0.000665278811557782 | 0.0263658281304011 | 7483 | 7 | 8 | 30 | 49 | 14 | 19 | 38 | 12 | 13 | 22 | 36 | 37 |
| MER67D_dup124 | 21.985278671735 | -1.34408086060972 | 0.384644774987635 | -3.49434321746064 | 0.000475229643309987 | 0.0208248605353661 | 7484 | 2 | 10 | 30 | 54 | 19 | 17 | 32 | 17 | 10 | 17 | 30 | 51 |
| CER_dup71 | 22.3848935004307 | -1.30079691148551 | 0.386152996574187 | -3.36860499083451 | 0.000755496170887843 | 0.0289061411606494 | 7485 | 9 | 4 | 46 | 36 | 18 | 23 | 21 | 18 | 9 | 15 | 49 | 41 |
| AluSp_dup53766 | 27.8082576116601 | -1.01350213800579 | 0.314776228076599 | -3.21975437662073 | 0.001283004750647 | 0.0417177726826414 | 7486 | 7 | 15 | 45 | 51 | 23 | 21 | 26 | 19 | 19 | 25 | 56 | 50 |
| LTR16E1_dup2220 | 24.7476336578519 | -1.16735896944889 | 0.314364567258896 | -3.71339231907616 | 0.000204499498597828 | 0.0113004732218669 | 7487 | 13 | 18 | 44 | 44 | 14 | 15 | 21 | 13 | 11 | 20 | 47 | 64 |
| L1ME3A_dup12784 | 22.4292605343247 | -2.02246285501906 | 0.365240321808538 | -5.53734824513502 | 3.07085572659575e-08 | 1.29327050747001e-05 | 7488 | 11 | 5 | 34 | 56 | 11 | 8 | 18 | 16 | 9 | 8 | 52 | 69 |
| L2a_dup155352 | 24.6714932925123 | -1.37588646326283 | 0.365414627190728 | -3.76527473418487 | 0.000166366054380826 | 0.00980825995433598 | 7489 | 20 | 7 | 40 | 44 | 15 | 19 | 21 | 16 | 13 | 7 | 51 | 71 |
| AluJb_dup97228 | 25.3308531064685 | -1.52512463850018 | 0.316298747698243 | -4.82178525713034 | 1.42279070291326e-06 | 0.000271489459980271 | 7490 | 8 | 8 | 32 | 38 | 17 | 23 | 50 | 23 | 13 | 9 | 48 | 62 |
| AluY_dup35528 | 25.0131690197687 | -1.18031021254187 | 0.306834740165857 | -3.84672938893381 | 0.000119705056663026 | 0.00773793518411854 | 7491 | 17 | 15 | 36 | 42 | 25 | 19 | 37 | 16 | 9 | 9 | 48 | 56 |
| LTR79_dup792 | 21.4733365693233 | -1.24998508154182 | 0.306450103626122 | -4.07891877585019 | 4.5245640276148e-05 | 0.00379486045863529 | 7492 | 9 | 11 | 28 | 49 | 19 | 19 | 33 | 17 | 8 | 12 | 31 | 47 |
| LTR39_dup1133 | 35.0525874854453 | -1.89327150285609 | 0.383595972109786 | -4.9355875465612 | 7.99098247083657e-07 | 0.000173329290337801 | 7493 | 6 | 7 | 23 | 35 | 15 | 21 | 32 | 34 | 9 | 19 | 97 | 163 |
| AluSq2_dup95 | 18.0497775934438 | -1.09944909216898 | 0.337230184465437 | -3.26023334450854 | 0.00111320587863419 | 0.0377638420145279 | 7494 | 12 | 14 | 16 | 26 | 9 | 8 | 17 | 18 | 10 | 14 | 26 | 65 |
| L1PA4_dup3783 | 20.447741931568 | -1.09352850874596 | 0.303545853508441 | -3.60251506026765 | 0.000315153135284856 | 0.0154700751087045 | 7495 | 16 | 9 | 25 | 27 | 6 | 11 | 20 | 15 | 21 | 13 | 39 | 61 |
| L1MB4_dup8117 | 26.2608190571334 | -1.85633638123624 | 0.328579940860313 | -5.64957305785569 | 1.60846816152367e-08 | 7.63929271780261e-06 | 7496 | 13 | 8 | 28 | 23 | 11 | 7 | 25 | 21 | 8 | 14 | 87 | 98 |
| L1PA3_dup6649 | 28.6015670092003 | -1.57874290651262 | 0.273386603624257 | -5.77476323120223 | 7.70613693384608e-09 | 4.06313162923373e-06 | 7497 | 20 | 12 | 35 | 37 | 8 | 8 | 34 | 30 | 14 | 23 | 55 | 96 |
| L1PA16_dup13718 | 50.8145246884707 | -1.79548831866722 | 0.537773708475197 | -3.33874321182072 | 0.000841583220923533 | 0.0311772319921954 | 7498 | 15 | 12 | 114 | 173 | 35 | 43 | 23 | 9 | 16 | 17 | 78 | 153 |
| L1MCc_dup3261 | 46.0297436284002 | -1.70273772842972 | 0.496252599152703 | -3.43119155715649 | 0.000600936088431543 | 0.0246151831313543 | 7499 | 24 | 18 | 91 | 137 | 27 | 27 | 20 | 7 | 16 | 16 | 91 | 147 |
| L1PA10_dup6932 | 42.9348292687582 | -1.80317754394361 | 0.510435052587817 | -3.53262875423977 | 0.000411449816372221 | 0.0188681401546048 | 7500 | 16 | 18 | 93 | 118 | 31 | 21 | 17 | 8 | 15 | 11 | 92 | 136 |

| | | | | | | | | | | | | | | | | | | | |
|---|---|---|---|---|---|---|---|---|---|---|---|---|---|---|---|---|---|---|---|
| ERVL-B4-int_dup3877 | 38.734736238146 | -2.03957212036114 | 0.440605332732917 | -4.62902277580347 | 3.67395363376537e-06 | 0.000562912151538915 | 7501 | 18 | 20 | 111 | 127 | 23 | 14 | 25 | 15 | 8 | 12 | 59 | 88 |
| MLT1J2-int_dup572 | 44.4631779335804 | -2.06923031548936 | 0.451666205606355 | -4.5813264083184 | 4.62036038536603e-06 | 0.000677385257089528 | 7502 | 25 | 16 | 109 | 154 | 24 | 17 | 22 | 18 | 7 | 18 | 90 | 97 |
| L1MB1_dup5943 | 52.0061163928309 | -2.27266027494433 | 0.500889723691838 | -4.53724675801601 | 5.69934060592991e-06 | 0.000794921043269988 | 7503 | 22 | 19 | 140 | 202 | 20 | 23 | 22 | 12 | 9 | 18 | 85 | 135 |
| HSMAR2_dup1765 | 51.1124674517141 | -2.0090500337351 | 0.518524318358441 | -3.87455315518355 | 0.000106820525920775 | 0.00715180553200468 | 7504 | 29 | 21 | 139 | 185 | 34 | 27 | 20 | 11 | 9 | 12 | 94 | 109 |
| L4_C_Mam_dup4863 | 46.0867572382499 | -1.90212663787276 | 0.513204175627951 | -3.70637404800018 | 0.000210247719537234 | 0.011532412142233 | 7505 | 25 | 13 | 117 | 173 | 31 | 24 | 22 | 8 | 12 | 16 | 76 | 107 |
| L1MA3_dup8737 | 35.7033615848573 | -1.9960883777512 | 0.552590999335257 | -3.61223469103262 | 0.000303569630359688 | 0.0150799956713082 | 7506 | 22 | 16 | 84 | 114 | 18 | 23 | 16 | 5 | 5 | 8 | 71 | 103 |
| Tigger6a_dup775 | 36.1579689646954 | -2.11369306280071 | 0.562110438912468 | -3.76028074997172 | 0.000169722771547493 | 0.00995681819493088 | 7507 | 12 | 11 | 94 | 115 | 26 | 19 | 13 | 6 | 6 | 10 | 71 | 106 |
| L1PA15-16_dup1341 | 37.9057607607267 | -1.68062992605901 | 0.517642701581103 | -3.24669877683129 | 0.00116751886520384 | 0.0391001328500764 | 7508 | 14 | 15 | 99 | 139 | 27 | 35 | 13 | 12 | 8 | 15 | 61 | 70 |
| L1PA10_dup6931 | 37.947044360605 | -1.99323595295624 | 0.52539476260759 | -3.79378725258623 | 0.000148366808900936 | 0.00902666548472053 | 7509 | 12 | 19 | 91 | 136 | 19 | 30 | 17 | 10 | 10 | 6 | 62 | 101 |
| L1MDa_dup6679 | 37.7782785616709 | -2.06393429842032 | 0.498316082734295 | -4.14181755301851 | 3.44564391709827e-05 | 0.00306038517169789 | 7510 | 15 | 17 | 100 | 120 | 22 | 20 | 13 | 13 | 8 | 9 | 81 | 87 |
| MLT2A2_dup3784 | 36.0118497543022 | -2.02717040374686 | 0.487151850890107 | -4.16127004350468 | 3.16482579046201e-05 | 0.0028698842893951 | 7511 | 10 | 16 | 87 | 127 | 21 | 19 | 16 | 12 | 8 | 13 | 57 | 100 |
| L1PA3_dup400 | 52.4302055694384 | -2.57065420187509 | 0.369029028963512 | -6.96599454274713 | 3.26091784714753e-12 | 5.07138796115926e-09 | 7512 | 18 | 20 | 70 | 97 | 11 | 9 | 48 | 26 | 13 | 9 | 161 | 221 |
| MER103C_dup8118 | 33.5060894090808 | -1.70041215919993 | 0.281981406307778 | -6.03022795532823 | 1.63728567264115e-09 | 1.0912761251044e-06 | 7513 | 18 | 9 | 45 | 62 | 18 | 13 | 34 | 22 | 18 | 15 | 83 | 103 |
| LTR41B_dup765 | 36.1224218134288 | -1.19480003558583 | 0.261348590473404 | -4.57167200872054 | 4.8384787529504e-06 | 0.000702496054992057 | 7514 | 12 | 16 | 49 | 55 | 25 | 20 | 41 | 17 | 26 | 29 | 80 | 100 |
| MER31B_dup266 | 30.9886843166755 | -1.10034316850578 | 0.260359663916327 | -4.22624285173224 | 2.37625536375497e-05 | 0.0023235069980368 | 7515 | 17 | 9 | 43 | 54 | 17 | 23 | 44 | 17 | 24 | 29 | 57 | 69 |
| L1PA4_dup11224 | 40.7139037303749 | -1.84108624552738 | 0.3210053462156 | -5.73537564788978 | 9.72964565417008e-09 | 4.94685954022896e-06 | 7516 | 14 | 12 | 78 | 91 | 11 | 21 | 48 | 17 | 24 | 25 | 98 | 97 |
| Tigger19b_dup3296 | 35.0929882793878 | -1.76649233693495 | 0.347588690368095 | -5.08213410241929 | 3.73217763326891e-07 | 9.54899719490111e-05 | 7517 | 15 | 14 | 70 | 73 | 11 | 20 | 25 | 20 | 10 | 25 | 92 | 86 |
| LTR9_dup322 | 37.0419063668167 | -1.38554812717164 | 0.41097958639663 | -3.37133077416276 | 0.000748059812874006 | 0.0287045529946164 | 7518 | 19 | 16 | 78 | 123 | 32 | 20 | 24 | 16 | 20 | 17 | 47 | 80 |
| L1MDa_dup2697 | 29.3253085865268 | -1.3233282019005 | 0.364183815403765 | -3.63368207462306 | 0.00027940511485355 | 0.0142047047096345 | 7519 | 15 | 23 | 66 | 76 | 25 | 15 | 24 | 17 | 11 | 14 | 37 | 64 |
| MLT1A0_dup9181 | 30.7477137267857 | -1.2610407728096 | 0.371843572176233 | -3.39132061750939 | 0.00069556679275092 | 0.027203704153174 | 7520 | 22 | 20 | 80 | 68 | 23 | 22 | 20 | 20 | 14 | 11 | 35 | 68 |
| MLT1A0_dup9182 | 28.9596462945262 | -1.26677483445208 | 0.366618206142445 | -3.45529712716964 | 0.000549686690062584 | 0.0230517710880563 | 7521 | 27 | 25 | 64 | 75 | 19 | 14 | 24 | 15 | 9 | 13 | 46 | 52 |
| SATR1_dup587 | 32.4474784413747 | -1.31833482466428 | 0.362421322764975 | -3.63757522489703 | 0.000275216825123382 | 0.0140680435383047 | 7522 | 27 | 22 | 75 | 87 | 22 | 27 | 30 | 15 | 8 | 13 | 51 | 54 |
| L1MEg1_dup415 | 46.6598637414706 | -1.32195679533103 | 0.383245597253357 | -3.4493724254244 | 0.000561891163250282 | 0.0234314326552316 | 7523 | 31 | 39 | 107 | 118 | 23 | 38 | 27 | 23 | 16 | 19 | 83 | 91 |
| LTR49-int_dup827 | 43.8524748790464 | -1.51242247682103 | 0.458744541163738 | -3.29687296765284 | 0.000977677101054539 | 0.034602760001119 | 7524 | 20 | 21 | 111 | 161 | 15 | 33 | 29 | 13 | 15 | 36 | 57 | 76 |
| L1M1_dup4295 | 48.1979973697657 | -2.25579615747446 | 0.472427466227376 | -4.77490476048817 | 1.79792321691236e-06 | 0.000324217849001375 | 7525 | 49 | 30 | 142 | 208 | 19 | 16 | 42 | 27 | 5 | 3 | 52 | 60 |
| L1PA16_dup6813 | 43.7207068873038 | -1.96645902803771 | 0.546447062519777 | -3.5986267708531 | 0.000319901903806569 | 0.0156495432572282 | 7526 | 55 | 35 | 150 | 194 | 11 | 17 | 25 | 18 | 6 | 3 | 34 | 45 |
| L1PA3_dup2926 | 30.0829879448684 | -1.66974753015564 | 0.495878075695035 | -3.36725419411874 | 0.000759206736870456 | 0.0289689233821965 | 7527 | 42 | 29 | 43 | 65 | 5 | 11 | 42 | 31 | 8 | 2 | 40 | 82 |
| SVA_D_dup1152 | 40.3958157882258 | -1.59622848437408 | 0.455299395953599 | -3.50588755126914 | 0.000455087439899458 | 0.0202593519128749 | 7528 | 48 | 38 | 69 | 91 | 14 | 44 | 73 | 34 | 6 | 1 | 39 | 84 |
| HSMAR2_dup1462 | 27.7233116954628 | -1.47173131796757 | 0.441028452012629 | -3.33704392823487 | 0.000846745639294545 | 0.0313209854691456 | 7529 | 5 | 10 | 63 | 98 | 24 | 29 | 33 | 17 | 7 | 15 | 31 | 37 |
| L1PA7_dup11172 | 31.2259143654055 | -1.38141825009362 | 0.382229065635899 | -3.61411094626048 | 0.000301379995769377 | 0.0150119666052628 | 7530 | 16 | 13 | 80 | 93 | 27 | 35 | 32 | 19 | 9 | 10 | 33 | 47 |
| L2a_dup148463 | 25.9876057761619 | -1.01744789531603 | 0.31919624659339 | -3.18753088789329 | 0.00143493130971862 | 0.0450562371895363 | 7531 | 13 | 20 | 49 | 44 | 24 | 23 | 57 | 15 | 6 | 21 | 27 | 45 |
| MLT1E3_dup13 | 32.3161364643623 | -1.15909617193285 | 0.317020087301938 | -3.65622311758719 | 0.000255958595191535 | 0.0133708724240274 | 7532 | 24 | 14 | 60 | 74 | 23 | 32 | 38 | 21 | 7 | 25 | 54 | 54 |
| FLAM_A_dup7237 | 25.8796529712885 | -2.34447222861066 | 0.322512594926524 | -7.26939742971832 | 3.6109458093112e-13 | 7.47139227428141e-10 | 7533 | 16 | 6 | 51 | 71 | 13 | 10 | 41 | 30 | 4 | 5 | 36 | 61 |
| MSTA_dup4584 | 23.2590891810563 | -2.03368965664506 | 0.368616792749812 | -5.51708358556896 | 3.44671527258776e-08 | 1.41158711230838e-05 | 7534 | 14 | 15 | 57 | 72 | 12 | 9 | 28 | 19 | 2 | 7 | 34 | 41 |
| L1MA2_dup1656 | 25.8410188946885 | -1.77258857123162 | 0.350318657794788 | -5.05993195563675 | 4.19406120856806e-07 | 0.000104171199199784 | 7535 | 20 | 15 | 56 | 62 | 17 | 9 | 29 | 22 | 6 | 7 | 33 | 67 |
| MER102a_dup700 | 20.6342496647446 | -1.52600888161289 | 0.33510677794104 | -4.55379891445042 | 5.26857198361066e-06 | 0.000750680017492349 | 7536 | 22 | 14 | 42 | 45 | 11 | 7 | 27 | 18 | 6 | 7 | 31 | 42 |
| L1MB8_dup13661 | 22.7331809349812 | -1.08765546710024 | 0.305102364600554 | -3.56488704544855 | 0.000364012920677148 | 0.0172334696425477 | 7537 | 20 | 23 | 29 | 61 | 15 | 10 | 43 | 28 | 11 | 10 | 15 | 32 |
| L1PA5_dup10371 | 25.8293866335857 | -1.21862886800577 | 0.287440893395894 | -4.23958071382469 | 2.23937665548057e-05 | 0.00222391785335968 | 7538 | 33 | 16 | 51 | 52 | 17 | 13 | 64 | 25 | 6 | 13 | 20 | 32 |
| AluSp_dup50642 | 21.3658995938113 | -1.04249291801932 | 0.29896125163123 | -3.48705028605259 | 0.000488379520057565 | 0.0211989812677116 | 7539 | 22 | 15 | 24 | 53 | 12 | 16 | 36 | 17 | 8 | 14 | 37 | 27 |
| L1MEd_dup7876 | 21.8388678521746 | -1.04684790802736 | 0.271225755339327 | -3.85969211042538 | 0.000113529975707819 | 0.00744719955235963 | 7540 | 17 | 18 | 28 | 37 | 14 | 15 | 29 | 19 | 7 | 16 | 44 | 40 |
| MER20_dup95 | 20.0633868981711 | -1.52499937968262 | 0.322149044643167 | -4.73383176216402 | 2.20320368299376e-06 | 0.000378242066056012 | 7541 | 14 | 11 | 37 | 48 | 17 | 14 | 29 | 17 | 4 | 6 | 27 | 42 |
| MER3_dup44 | 19.3165554661035 | -1.3452926835942 | 0.342800628889858 | -3.92441719827312 | 8.69399553905226e-05 | 0.00615135730756863 | 7542 | 13 | 7 | 28 | 36 | 24 | 16 | 32 | 24 | 3 | 7 | 27 | 35 |
| L2a_dup906 | 21.6221703002861 | -1.0191947700065 | 0.296861490557167 | -3.43323335099348 | 0.000596428540150685 | 0.0244800517309606 | 7543 | 22 | 17 | 43 | 44 | 17 | 23 | 35 | 18 | 7 | 6 | 17 | 36 |
| L3_dup160 | 22.5813794030039 | -1.04440412674509 | 0.304867255373663 | -3.42576681600328 | 0.000613066462140824 | 0.0249419854621203 | 7544 | 23 | 10 | 33 | 36 | 25 | 22 | 38 | 23 | 8 | 6 | 34 | 36 |
| Charlie7a_dup393 | 32.5876653824054 | -1.90412035470789 | 0.386647292429769 | -4.92469594896687 | 8.44916255127913e-07 | 0.000180885133352533 | 7545 | 27 | 31 | 76 | 86 | 9 | 11 | 41 | 28 | 7 | 5 | 53 | 57 |
| HAL1b_dup1280 | 31.9693750723559 | -1.73623423681968 | 0.326545289715096 | -5.31697835339935 | 1.05504614155841e-07 | 3.44367639505154e-05 | 7546 | 30 | 21 | 71 | 73 | 8 | 10 | 40 | 31 | 9 | 14 | 56 | 56 |
| L1ME3G_dup4059 | 32.5648381196795 | -1.56840129501117 | 0.35338312766999 | -4.43824611931059 | 9.06948782699052e-06 | 0.0011352174384435 | 7547 | 31 | 26 | 72 | 67 | 9 | 9 | 35 | 26 | 13 | 13 | 61 | 64 |
| L1MA9_dup3580 | 30.2762000810438 | -1.7324642509175 | 0.328792006980397 | -5.26917995004908 | 1.37034574939136e-07 | 4.26790955084655e-05 | 7548 | 23 | 18 | 54 | 78 | 16 | 9 | 37 | 19 | 10 | 10 | 61 | 67 |
| MIR3_dup23130 | 27.9325937490096 | -1.75553757409375 | 0.302723830413273 | -5.79913900962842 | 6.6656261737211e-09 | 3.5965058628849e-06 | 7549 | 18 | 21 | 56 | 56 | 11 | 9 | 47 | 24 | 12 | 7 | 57 | 47 |
| MIR_dup44941 | 34.6606270083592 | -1.38677446822589 | 0.320748408899627 | -4.32355837082221 | 1.53532491703778e-05 | 0.00168119844781165 | 7550 | 27 | 29 | 60 | 100 | 18 | 20 | 31 | 33 | 16 | 9 | 48 | 63 |
| MLT2D_dup1182 | 35.341490855122 | -1.5108066110725 | 0.342118298593482 | -4.41603567328535 | 1.00527501363517e-05 | 0.00122793207594679 | 7551 | 24 | 28 | 75 | 93 | 11 | 20 | 34 | 24 | 18 | 11 | 55 | 72 |
| L1ME3F_dup1170 | 37.484166097341 | -1.52311662096518 | 0.319699703878466 | -4.76421029637298 | 1.89594694494707e-06 | 0.000338180653495034 | 7552 | 43 | 22 | 64 | 83 | 12 | 22 | 50 | 32 | 14 | 9 | 63 | 81 |
| AluJr_dup18562 | 36.2400124764944 | -1.70042826242975 | 0.316173932652833 | -5.37814186059692 | 7.52585087820739e-08 | 2.65295720454213e-05 | 7553 | 31 | 25 | 79 | 95 | 11 | 18 | 47 | 28 | 7 | 16 | 57 | 67 |
| L3_dup12174 | 35.1722564377667 | -1.61939151039648 | 0.364487281206428 | -4.44293009357256 | 8.87420050848485e-06 | 0.00111429050880841 | 7554 | 38 | 27 | 70 | 84 | 12 | 13 | 39 | 24 | 8 | 12 | 65 | 75 |
| L1PA14_dup2120 | 33.2683985426903 | -1.2875079541857 | 0.382119193604837 | -3.36938833676373 | 0.000753352083396646 | 0.0288426572543682 | 7555 | 35 | 37 | 53 | 59 | 15 | 8 | 33 | 26 | 10 | 14 | 68 | 78 |
| AluSq2_dup13952 | 31.9552817553726 | -1.33729554411663 | 0.307745283847446 | -4.3454623492445 | 1.38982610370292e-05 | 0.0015591869970377 | 7556 | 26 | 31 | 49 | 84 | 13 | 18 | 42 | 28 | 6 | 20 | 44 | 62 |
| L1MA4_dup5044 | 31.011565931129 | -1.47840861075534 | 0.334505458613026 | -4.41968456026196 | 9.88450742358598e-06 | 0.00121236031103742 | 7557 | 28 | 26 | 51 | 93 | 9 | 14 | 40 | 23 | 11 | 14 | 40 | 64 |
| L1PA3_dup4886 | 29.0270092234713 | -1.64662634843267 | 0.328646067386388 | -5.01033333983801 | 5.43358362562769e-07 | 0.000127883233371277 | 7558 | 19 | 18 | 69 | 64 | 8 | 21 | 36 | 20 | 10 | 11 | 41 | 67 |
| LTR16B2_dup354 | 28.3727050419671 | -1.58737114790601 | 0.361674195705315 | -4.38895328103353 | 1.13897531879513e-05 | 0.00134365206349046 | 7559 | 25 | 19 | 61 | 64 | 9 | 14 | 34 | 15 | 13 | 7 | 50 | 65 |
| L1PA2_dup2565 | 23.9042196944953 | -1.27428056478085 | 0.404713102911204 | -3.14860219650569 | 0.00164053349438653 | 0.0494625421762922 | 7560 | 27 | 23 | 54 | 46 | 5 | 18 | 30 | 12 | 8 | 8 | 36 | 50 |
| MER102a_dup699 | 23.9480267846839 | -1.4792369722619 | 0.367265599810224 | -4.02770358298261 | 5.63242909083635e-05 | 0.00445401785459669 | 7561 | 22 | 18 | 42 | 57 | 10 | 17 | 28 | 13 | 6 | 7 | 41 | 59 |
| L1PA5_dup7606 | 29.0481814400566 | -1.49224199407797 | 0.36508215666137 | -4.08741420759735 | 4.36207905537807e-05 | 0.00368147376258316 | 7562 | 15 | 23 | 51 | 60 | 21 | 25 | 35 | 18 | 5 | 8 | 51 | 74 |
| MIR3_dup23132 | 31.042295627603 | -1.38313438474424 | 0.324902133411441 | -4.25708003275159 | 2.07114227862911e-05 | 0.00209532321696719 | 7563 | 20 | 27 | 51 | 63 | 23 | 16 | 48 | 16 | 12 | 8 | 59 | 68 |
| LTR16D_dup588 | 29.37003570961 | -1.15149568027396 | 0.359974990518215 | -3.1988213364942 | 0.0013799065620644 | 0.0438933389237609 | 7564 | 25 | 33 | 45 | 73 | 19 | 20 | 22 | 23 | 7 | 12 | 56 | 50 |
| MIRb_dup58994 | 26.6387566395594 | -1.04596291934026 | 0.305885853642236 | -3.41945502508794 | 0.000627467041120108 | 0.0253284561040531 | 7565 | 30 | 23 | 38 | 39 | 17 | 21 | 28 | 28 | 8 | 10 | 46 | 58 |
| MSTC_dup2707 | 26.3155216172745 | -1.02594988297244 | 0.278759939927206 | -3.68040645739971 | 0.000232862473553005 | 0.01245139193785 | 7566 | 28 | 19 | 36 | 39 | 16 | 22 | 30 | 33 | 12 | 10 | 49 | 43 |
| L1MB7_dup9945 | 21.9013171770116 | -1.33190672113124 | 0.388654294191784 | -3.42697029477309 | 0.000610355834598516 | 0.0248727058433599 | 7567 | 21 | 21 | 43 | 62 | 16 | 10 | 16 | 19 | 7 | 5 | 29 | 39 |
| L1MEf_dup5816 | 21.5568764626721 | -1.23512733783696 | 0.372410398519794 | -3.31657586025035 | 0.000911278440008853 | 0.0329787163021059 | 7568 | 22 | 19 | 42 | 64 | 16 | 15 | 17 | 23 | 8 | 3 | 27 | 25 |
| MIRb_dup91530 | 22.5548464453899 | -1.64696946571279 | 0.365834324540567 | -4.50195445105141 | 6.73314135783662e-06 | 0.000902096876988622 | 7569 | 10 | 15 | 46 | 60 | 14 | 10 | 24 | 13 | 11 | 6 | 44 | 44 |
| MIR3_dup69389 | 22.9502073472438 | -1.28650969923403 | 0.345270840544084 | -3.7260884736363 | 0.000194474167626484 | 0.0108983396820658 | 7570 | 11 | 19 | 45 | 39 | 16 | 14 | 16 | 21 | 11 | 9 | 48 | 46 |
| MamTip1_dup252 | 25.4908937045544 | -1.18205455573244 | 0.373375319245168 | -3.16586152004405 | 0.00154624386377293 | 0.0474432678799888 | 7571 | 19 | 31 | 40 | 55 | 17 | 11 | 21 | 21 | 11 | 7 | 52 | 46 |
| L1MB8_dup16409 | 29.6303921660244 | -1.29756667481325 | 0.339445766282119 | -3.82260379625658 | 0.000132049864212729 | 0.00829480913145773 | 7572 | 30 | 16 | 72 | 89 | 27 | 23 | 28 | 34 | 9 | 5 | 29 | 24 |
| L2a_dup83616 | 27.8517298085777 | -1.11136417690055 | 0.337488855304483 | -3.29303963503588 | 0.000991105019521448 | 0.0348963413694955 | 7573 | 27 | 28 | 59 | 60 | 31 | 14 | 25 | 29 | 6 | 9 | 33 | 41 |
| L2c_dup67756 | 32.05696456305 | -1.46655753186678 | 0.330951601133128 | -4.43133535793607 | 9.36513055354131e-06 | 0.00116243889133695 | 7574 | 34 | 26 | 65 | 96 | 25 | 15 | 33 | 35 | 6 | 7 | 35 | 45 |
| L2b_dup44444 | 26.8827078288947 | -1.46771489407112 | 0.391586304815923 | -3.74812621386507 | 0.000178160600780228 | 0.0102699739702481 | 7575 | 27 | 25 | 65 | 82 | 17 | 18 | 28 | 16 | 3 | 7 | 28 | 44 |
| MSTB1_dup2178 | 26.3432314419238 | -1.30380988532077 | 0.404447987239757 | -3.22367752209351 | 0.00126555828893775 | 0.041342973338913 | 7576 | 31 | 30 | 44 | 83 | 17 | 16 | 36 | 14 | 4 | 6 | 24 | 51 |
| AluSz_dup44718 | 26.1759001922032 | -1.40067837853366 | 0.41063055977304 | -3.41104271271927 | 0.000647149529830965 | 0.0258757590363231 | 7577 | 29 | 20 | 73 | 73 | 20 | 14 | 28 | 11 | 7 | 5 | 32 | 37 |
| L1ME3Cz_dup5948 | 26.283272438781 | -1.34646278453643 | 0.416881592521772 | -3.2298446577876 | 0.00123857494525548 | 0.0407340794673237 | 7578 | 33 | 25 | 62 | 71 | 18 | 17 | 34 | 11 | 6 | 3 | 33 | 39 |
| AluSx1_dup49930 | 25.0806754332402 | -1.32038134776704 | 0.408418726383254 | -3.23291088892925 | 0.00122535778989546 | 0.040404293276638 | 7579 | 36 | 23 | 60 | 76 | 14 | 16 | 27 | 16 | 6 | 3 | 23 | 35 |

| | | | | | | | | | | | | | | | | | | | |
|---|---|---|---|---|---|---|---|---|---|---|---|---|---|---|---|---|---|---|---|
| L2a_dup83621 | 23.9233550598164 | -1.32570011778388 | 0.371310484268456 | -3.57032772827769 | 0.000356534873160376 | 0.0169502591266771 | 7580 | 33 | 15 | 58 | 66 | 12 | 20 | 29 | 16 | 5 | 6 | 25 | 34 |
| MIRb_dup101984 | 30.4293827623352 | -1.28161301200739 | 0.388895773503018 | -3.29551797506856 | 0.00098240418433528 | 0.0346825666932116 | 7581 | 36 | 34 | 71 | 89 | 14 | 19 | 39 | 14 | 6 | 9 | 38 | 38 |
| MSTD_dup3921 | 30.9786018429422 | -1.19242231918058 | 0.36673115034612 | -3.25148904873549 | 0.00114802198211405 | 0.0386524401139303 | 7582 | 38 | 34 | 69 | 99 | 15 | 22 | 33 | 22 | 9 | 6 | 31 | 33 |
| L1MDa_dup3194 | 30.6446859391371 | -1.66142958494207 | 0.426224178768864 | -3.89801815031015 | 9.69831489730531e-05 | 0.00666432509233256 | 7583 | 26 | 30 | 72 | 122 | 16 | 20 | 31 | 16 | 5 | 4 | 28 | 44 |
| MER5A_dup16817 | 31.2570974599593 | -1.5528837200873 | 0.396357755885501 | -3.91788402529935 | 8.93296633460352e-05 | 0.00627712003268199 | 7584 | 31 | 30 | 62 | 103 | 20 | 15 | 42 | 18 | 8 | 3 | 35 | 53 |
| L1P4a_dup518 | 67.2053211162326 | -2.22442203200844 | 0.312859043675525 | -7.10998156190573 | 1.16058458353531e-12 | 2.17446030955821e-09 | 7585 | 35 | 37 | 189 | 244 | 15 | 19 | 128 | 84 | 17 | 14 | 46 | 61 |
| L1PBa_dup1936 | 69.4746516753633 | -1.90806862189826 | 0.302734169728137 | -6.30278578599752 | 2.92342813168447e-10 | 2.55741922875659e-07 | 7586 | 34 | 47 | 165 | 260 | 25 | 23 | 115 | 84 | 24 | 17 | 54 | 68 |
| L1MC3_dup11469 | 49.0418840000239 | -1.6190270745738 | 0.313524378565378 | -5.16395912171845 | 2.41780519773889e-07 | 6.67409600442201e-05 | 7587 | 42 | 35 | 125 | 178 | 25 | 16 | 69 | 62 | 11 | 15 | 32 | 35 |
| L1MC3_dup11472 | 58.6798186264731 | -1.86712453656893 | 0.356267005021225 | -5.24080116949839 | 1.59880914782941e-07 | 4.80341214001788e-05 | 7588 | 37 | 42 | 158 | 230 | 17 | 15 | 105 | 72 | 13 | 18 | 27 | 44 |
| L1P2_dup868 | 78.2757026612886 | -2.71470059521048 | 0.359705629345316 | -7.54700614541779 | 4.45377579048798e-14 | 1.16456937828951e-10 | 7589 | 22 | 18 | 220 | 322 | 20 | 20 | 188 | 124 | 7 | 21 | 36 | 40 |
| LTR8_dup904 | 73.9555961532612 | -1.46732845130529 | 0.37846433030353 | -3.87705877097714 | 0.00010572682619495 | 0.00711059225149336 | 7590 | 50 | 38 | 187 | 173 | 26 | 32 | 176 | 153 | 24 | 31 | 28 | 24 |
| L1PA15_dup4275 | 62.8374356633359 | -2.02310600825993 | 0.370719440820149 | -5.45724282434225 | 4.83584824413672e-08 | 1.85592055854735e-05 | 7591 | 47 | 28 | 133 | 208 | 12 | 17 | 188 | 113 | 12 | 16 | 20 | 33 |
| L1PA15_dup4276 | 58.1737110904116 | -1.68032234509496 | 0.38648807811237 | -4.3476692820688 | 1.37591865968308e-05 | 0.00154723223722567 | 7592 | 39 | 32 | 130 | 196 | 25 | 14 | 158 | 92 | 10 | 25 | 14 | 32 |
| L1PA10_dup3642 | 52.1348319549984 | -2.47231626801037 | 0.533101394321726 | -4.63760983247088 | 3.52461325337567e-06 | 0.000543883025413571 | 7593 | 10 | 9 | 169 | 244 | 15 | 12 | 113 | 68 | 14 | 13 | 12 | 15 |
| HAL1ME_dup6764 | 41.0756551536438 | -1.70200652273032 | 0.483521730201144 | -3.52002074864823 | 0.000431513040890499 | 0.0195280259354976 | 7594 | 14 | 18 | 118 | 153 | 13 | 15 | 45 | 54 | 21 | 28 | 11 | 44 |
| HAL1ME_dup6763 | 58.4644582470452 | -1.63495240809889 | 0.441977565346341 | -3.69917510817028 | 0.000216301332962146 | 0.0117695030809538 | 7595 | 30 | 24 | 186 | 201 | 19 | 20 | 74 | 72 | 38 | 24 | 29 | 39 |
| HERVP71A-int_dup176 | 69.3747237296716 | -2.24060015376998 | 0.530321431633553 | -4.22498511302522 | 2.38956573988127e-05 | 0.00233314571006068 | 7596 | 28 | 15 | 251 | 274 | 14 | 12 | 98 | 97 | 21 | 40 | 21 | 36 |
| THE1B-int_dup4153 | 59.3119263831122 | -1.08805406843857 | 0.245673991678062 | -4.42885329866086 | 9.47354319232118e-06 | 0.0011740581814792 | 7597 | 27 | 25 | 90 | 83 | 47 | 60 | 120 | 71 | 28 | 44 | 65 | 105 |
| MLT1F2_dup3032 | 60.7057032948681 | -1.072111479959 | 0.211455040785363 | -5.07016279194425 | 3.97475571655026e-07 | 9.97655299293532e-05 | 7598 | 50 | 41 | 108 | 119 | 29 | 34 | 75 | 60 | 43 | 40 | 76 | 107 |
| L1PA16_dup2520 | 65.3977994552509 | -1.14472528293756 | 0.227238180213178 | -5.03755698916295 | 4.71511197732633e-07 | 0.000114600820690183 | 7599 | 28 | 27 | 94 | 119 | 57 | 42 | 85 | 71 | 38 | 54 | 84 | 144 |
| L1PA6_dup302 | 66.3497784484172 | -1.3741648257008 | 0.182143399242963 | -7.54441188323156 | 4.54333102013687e-14 | 1.17189188176186e-10 | 7600 | 40 | 20 | 82 | 100 | 40 | 38 | 95 | 71 | 42 | 40 | 131 | 156 |
| MIRb_dup202619 | 43.761403044387 | -1.36846026320297 | 0.30408783473057 | -4.50021377677099 | 6.78851563042674e-06 | 0.000907471988875012 | 7601 | 17 | 29 | 79 | 70 | 23 | 12 | 39 | 26 | 25 | 39 | 72 | 141 |
| LTR41B_dup764 | 57.5578694117639 | -1.42368376057071 | 0.281809057063226 | -5.05194465858239 | 4.3733442722687e-07 | 0.0001078362462393 | 7602 | 31 | 31 | 87 | 123 | 26 | 24 | 43 | 31 | 33 | 38 | 116 | 174 |
| MER5A1_dup14494 | 50.3015441846439 | -1.51240820276292 | 0.322333189682702 | -4.69206476767626 | 2.70461470270203e-06 | 0.000446291435540317 | 7603 | 19 | 29 | 84 | 70 | 29 | 22 | 32 | 34 | 25 | 22 | 107 | 185 |
| L1PA6_dup26 | 68.6641148121592 | -1.807509307417 | 0.245670228607127 | -7.35746173911672 | 1.87440106345908e-13 | 4.1296873652392e-10 | 7604 | 16 | 14 | 78 | 83 | 39 | 30 | 90 | 54 | 28 | 46 | 190 | 229 |
| BLACKJACK_dup2105 | 49.3359132758601 | -1.44172585904048 | 0.321489608294245 | -4.48451776307785 | 7.30790184890575e-06 | 0.000963872976381616 | 7605 | 13 | 18 | 93 | 91 | 23 | 38 | 35 | 32 | 29 | 36 | 107 | 125 |
| L1M2_dup3211 | 40.6280538298076 | -1.34222857389418 | 0.31335759387712 | -4.28337656441325 | 1.84078238902637e-05 | 0.0019233484536458 | 7606 | 10 | 12 | 75 | 71 | 19 | 38 | 52 | 29 | 26 | 34 | 75 | 85 |
| L2c_dup30781 | 50.4048719889811 | -1.04825254886267 | 0.243026820225037 | -4.31332043060931 | 1.60820768328865e-05 | 0.00174018456301865 | 7607 | 23 | 20 | 64 | 78 | 32 | 39 | 91 | 46 | 25 | 63 | 84 | 86 |
| L2c_dup30783 | 46.9600606936047 | -1.235001934873 | 0.298608414913962 | -4.13585777624132 | 3.53631172852922e-05 | 0.00312923477336041 | 7608 | 19 | 14 | 56 | 103 | 34 | 32 | 95 | 35 | 20 | 54 | 72 | 83 |
| L2c_dup30782 | 45.7651516050068 | -1.07461694459962 | 0.308620692519511 | -3.48199900605072 | 0.000497685539513941 | 0.0215085547157622 | 7609 | 15 | 16 | 60 | 92 | 40 | 22 | 65 | 42 | 32 | 57 | 50 | 100 |
| ERV3-16A3_I-int_dup1[illegible] | 51.4210638300729 | -1.25852125566638 | 0.304023791001443 | -4.13954859098643 | 3.47989889507558e-05 | 0.0030850522447379 | 7610 | 15 | 15 | 80 | 116 | 46 | 32 | 62 | 56 | 34 | 41 | 70 | 94 |
| LTR28C_dup79 | 51.0515462604552 | -1.15167201498741 | 0.344251287291145 | -3.34543996639699 | 0.000821521414056961 | 0.0306487510801502 | 7611 | 14 | 10 | 72 | 106 | 45 | 30 | 76 | 48 | 47 | 48 | 77 | 79 |
| L1MA2_dup5618 | 47.0645688320924 | -2.09214620063895 | 0.413140786689744 | -5.06400304216414 | 4.10543091203548e-07 | 0.000102504812570325 | 7612 | 12 | 11 | 37 | 63 | 24 | 22 | 180 | 112 | 13 | 23 | 45 | 59 |
| MIRc_dup68986 | 33.2344290643722 | -1.16576888603771 | 0.338630229008829 | -3.44260135738596 | 0.000576147916511849 | 0.0238421029812745 | 7613 | 19 | 26 | 41 | 25 | 25 | 31 | 79 | 69 | 10 | 16 | 35 | 42 |
| FLAM_A_dup11469 | 36.7725820934458 | -1.10571350603792 | 0.240452209282031 | -4.59847513707396 | 4.2559449317822e-06 | 0.00063332399795988 | 7614 | 32 | 23 | 42 | 41 | 24 | 25 | 63 | 36 | 15 | 22 | 79 | 74 |
| MLT2B1_dup865 | 32.9048781745102 | -1.12572102159125 | 0.271148842163006 | -4.15167187368813 | 3.30055253179682e-05 | 0.00297144635917868 | 7615 | 24 | 11 | 34 | 35 | 19 | 18 | 64 | 37 | 27 | 25 | 71 | 53 |
| L1MD2_dup3242 | 40.4018834038218 | -1.13621002494617 | 0.262407299762361 | -4.32994823686361 | 1.49144395813143e-05 | 0.00164297742621084 | 7616 | 31 | 31 | 29 | 58 | 18 | 14 | 50 | 40 | 25 | 28 | 82 | 117 |
| L1MC_dup1538 | 40.4206150565744 | -1.31685711272596 | 0.215976849395466 | -6.09721419870664 | 1.07932852362227e-09 | 7.66632551886043e-07 | 7617 | 17 | 18 | 52 | 70 | 28 | 22 | 77 | 43 | 24 | 31 | 60 | 81 |
| L3_dup38594 | 45.7244330238791 | -1.2756887759082 | 0.215331791044464 | -5.92429371306714 | 3.13642446741687e-09 | 1.88936621854171e-06 | 7618 | 25 | 24 | 62 | 51 | 35 | 25 | 73 | 50 | 20 | 31 | 89 | 104 |
| MER5A_dup28412 | 40.2101752910413 | -1.24756327353158 | 0.24003515467867 | -5.19741899973636 | 2.02074544015592e-07 | 5.82881131502447e-05 | 7619 | 20 | 22 | 41 | 56 | 32 | 26 | 57 | 43 | 14 | 28 | 88 | 93 |
| AluSc8_dup6995 | 46.9123572739741 | -1.37523633953337 | 0.423867101503218 | -3.24449888810945 | 0.00117657476753348 | 0.0392817819048018 | 7620 | 23 | 12 | 41 | 34 | 28 | 58 | 80 | 51 | 19 | 12 | 94 | 160 |
| AluJb_dup131614 | 46.9152109623388 | -1.02436063110159 | 0.248337968232338 | -4.12486515208672 | 3.70952031575479e-05 | 0.00325466637556264 | 7621 | 23 | 32 | 74 | 84 | 37 | 46 | 59 | 34 | 17 | 34 | 75 | 98 |
| LTR10E_dup219 | 47.6213494947179 | -1.06741522389085 | 0.271806548704762 | -3.92711371001691 | 8.59713421864028e-05 | 0.00609914638875546 | 7622 | 36 | 22 | 47 | 49 | 30 | 40 | 71 | 31 | 20 | 30 | 106 | 143 |
| AluSz6_dup6828 | 47.2008279262313 | -1.08150952898282 | 0.213003108370471 | -5.0774354292604 | 3.82563455912516e-07 | 9.7357694203807e-05 | 7623 | 37 | 25 | 66 | 67 | 40 | 47 | 94 | 63 | 16 | 24 | 59 | 71 |
| MLT1A0_dup4061 | 50.1123790424041 | -1.00987538896243 | 0.260554253551158 | -3.8758737391487 | 0.000106242768189975 | 0.00713520793670103 | 7624 | 31 | 21 | 49 | 59 | 36 | 45 | 100 | 73 | 27 | 42 | 84 | 69 |
| MLT1A0_dup4062 | 43.1257159460196 | -1.05123487510042 | 0.277841639435355 | -3.78357569886499 | 0.000154591277551645 | 0.00931007316711119 | 7625 | 21 | 21 | 35 | 60 | 43 | 29 | 81 | 64 | 25 | 30 | 73 | 65 |
| L1PA10_dup6908 | 263.368410949215 | -3.64576990713183 | 0.341425399989056 | -10.6780863616142 | 1.2890138111656e-26 | 5.11192800519351e-22 | 7626 | 17 | 16 | 206 | 268 | 2 | 6 | 176 | 137 | 129 | 158 | 914 | 1466 |
| L1MA4_dup932 | 105.453998285956 | -2.59971599584102 | 0.647104197761385 | -4.01746118296647 | 5.88285220014461e-05 | 0.00460006845626929 | 7627 | 53 | 39 | 250 | 380 | 6 | 6 | 58 | 40 | 38 | 42 | 191 | 322 |
| L1PA4_dup10537 | 152.248254985836 | -3.43140146911549 | 0.289490572339376 | -11.8532408201977 | 2.07026428853205e-32 | 1.64203702133016e-27 | 7628 | 18 | 17 | 259 | 340 | 15 | 15 | 136 | 91 | 54 | 38 | 435 | 616 |
| L1PA5_dup6275 | 44.7115607166852 | -1.99503757157294 | 0.476955090759646 | -4.1828625172979 | 2.87861485032543e-05 | 0.00267874418918864 | 7629 | 23 | 13 | 117 | 134 | 5 | 6 | 43 | 32 | 35 | 40 | 49 | 87 |
| L1M1_dup886 | 40.524208303658 | -2.61880047216498 | 0.550063139955798 | -4.76090885198274 | 1.92723080370925e-06 | 0.000342989424696635 | 7630 | 16 | 14 | 109 | 179 | 5 | 2 | 22 | 17 | 17 | 17 | 63 | 85 |
| L1M1_dup889 | 41.7613632333671 | -2.61084634231042 | 0.590509386945127 | -4.42134604467012 | 9.80879403618939e-06 | 0.00120493717384364 | 7631 | 19 | 15 | 118 | 165 | 2 | 2 | 17 | 19 | 15 | 22 | 66 | 101 |
| L1M2c_dup33 | 70.8745150877275 | -2.68795168983131 | 0.793501690851049 | -3.38745552885769 | 0.000705441526201783 | 0.0275040126812403 | 7632 | 29 | 31 | 189 | 253 | 0 | 5 | 33 | 30 | 25 | 28 | 140 | 188 |
| L1M1_dup693 | 73.6906364115489 | -2.89118843694319 | 0.762759308425707 | -3.79043350242483 | 0.000150384590743306 | 0.00910309530964414 | 7633 | 27 | 27 | 212 | 270 | 5 | 1 | 37 | 19 | 31 | 15 | 130 | 223 |
| L1PA13_dup717 | 58.184080161523 | -3.10610182418768 | 0.583472116830943 | -5.32347945101146 | 1.01800957825874e-07 | 3.34112147735661e-05 | 7634 | 18 | 12 | 169 | 233 | 1 | 4 | 28 | 23 | 25 | 20 | 98 | 153 |
| L1PA16_dup1254 | 65.7950399211968 | -2.89294361560893 | 0.57194560423227 | -5.05807474382493 | 4.23510447231545e-07 | 0.000105080935221019 | 7635 | 22 | 21 | 185 | 261 | 1 | 8 | 27 | 25 | 21 | 25 | 125 | 166 |
| HERVL18-int_dup69 | 59.8682693978022 | -3.09916145197717 | 0.573471556340805 | -5.40421127728153 | 6.5094181920917e-08 | 2.37559819192554e-05 | 7636 | 14 | 16 | 211 | 234 | 4 | 4 | 29 | 15 | 19 | 24 | 109 | 127 |
| L1PA10_dup580 | 56.0080810383213 | -2.81317071514077 | 0.535680501744861 | -5.25158318433747 | 1.50797402378064e-07 | 4.58258476452757e-05 | 7637 | 21 | 16 | 138 | 204 | 4 | 6 | 22 | 16 | 20 | 18 | 104 | 191 |
| L1PA4_dup8359 | 74.1954449305104 | -4.32451590160264 | 0.380298798787006 | -11.3713635578025 | 5.80732038509297e-30 | 2.76365731270266e-25 | 7638 | 0 | 2 | 91 | 111 | 8 | 8 | 60 | 43 | 5 | 15 | 287 | 366 |
| L1M4_dup12829 | 110.165051307909 | -4.83976444154904 | 0.339074504068532 | -14.2734543101207 | 3.20311215039694e-46 | 7.6216772373835e-41 | 7639 | 0 | 0 | 124 | 157 | 8 | 5 | 88 | 65 | 14 | 17 | 406 | 599 |
| L1PA4_dup8363 | 94.5787751168963 | -4.55240709589614 | 0.357795832299546 | -12.7234771479531 | 4.37929737183191e-37 | 5.21018146218958e-32 | 7640 | 1 | 0 | 97 | 148 | 4 | 8 | 67 | 70 | 20 | 14 | 365 | 466 |
| L1MA4A_dup6036 | 89.3860885884918 | -3.3204527859817 | 0.330944037456775 | -10.0332757510864 | 1.08844395554164e-23 | 1.99223758034855e-19 | 7641 | 5 | 4 | 47 | 73 | 1 | 4 | 74 | 46 | 54 | 51 | 323 | 500 |
| HSMAR2_dup1463 | 57.9134514914616 | -2.54762782417748 | 0.292558070971559 | -8.70810986590469 | 3.08983753109818e-18 | 1.98706616533699e-14 | 7642 | 3 | 4 | 49 | 47 | 10 | 4 | 41 | 36 | 33 | 57 | 216 | 252 |
| L1MC2_dup4897 | 63.9671531112043 | -2.65637718119144 | 0.281376990054908 | -9.44063400732619 | 3.70530539200541e-21 | 3.83331563828747e-17 | 7643 | 4 | 2 | 35 | 53 | 5 | 7 | 46 | 34 | 41 | 55 | 260 | 291 |

**Table S6**. A list of candidate insertions detected using L1-targeted amplicon-based NGS protocol and validated using site-specific PCR in paired primary samples from CLL patients.

| Patient ID | Timepoint | Chromosome | Coordinate | Sample reads | UMI | Forward Primer | Reverse primer | Product size |
|---|---|---|---|---|---|---|---|---|
| BRNO0859 | T2 | chr13 | 67009364 | 3 | 3 | GAATTTGCCTTCCCAACAAG | TGAGGGGACTTGCAACTTTT | 214 bp |
| BRNO1077 | T1 | chr4 | 70295573 | 7 | 2 | TATTGCTTCCCTGCCTTGTT | AAGCCAAAGAGCAGCTCAAC | 188 bp |

**Table S7**. Sums of normalized TE read counts per individual patient samples with RNAseq data.

| Patient ID | Time point | Category based on TP53 status development [stable / expanding mutation] | Group based on TE expression | Sum of normalized TE read counts |
|---|---|---|---|---|
| BRNO0209 | T1 | stable | 1 | 6641899 |
| BRNO0311 | T1 | stable | 1 | 6557419 |
| BRNO0311 | T2 | stable | 1 | 6586871 |
| BRNO1340 | T1 | stable | 1 | 6542651 |
| BRNO1340 | T2 | stable | 1 | 6516788 |
| BRNO0209 | T2 | stable | 1 | 6521365 |
| BRNO0513 | T1 | stable | 1 | 6625208 |
| BRNO0513 | T2 | stable | 1 | 6646966 |
| BRNO0823 | T1 | stable | 1 | 6547047 |
| BRNO0823 | T2 | stable | 1 | 6627341 |
| BRNO1530 | T1 | stable | 1 | 6586443 |
| BRNO1530 | T2 | stable | 1 | 6561265 |
| BRNO0178 | T1 | stable | 2 | 6545214 |
| BRNO0467 | T1 | stable | 2 | 6483173 |
| BRNO1876 | T1 | stable | 2 | 6525658 |
| BRNO1876 | T2 | stable | 2 | 6535172 |
| BRNO0874 | T1 | stable | 2 | 6585117 |
| BRNO0178 | T2 | stable | 2 | 6531296 |
| BRNO0475 | T1 | stable | 2 | 6501588 |
| BRNO0467 | T2 | stable | 2 | 6520556 |
| BRNO0475 | T2 | stable | 2 | 6497409 |
| BRNO1266 | T1 | stable | 2 | 6496685 |
| BRNO1266 | T2 | stable | 2 | 6512707 |
| BRNO0874 | T2 | stable | 2 | 6606541 |
| BRNO1794 | T1 | stable | 2 | 6580581 |
| BRNO1794 | T2 | stable | 2 | 6572945 |
| BRNO1921 | T1 | stable | 2 | 6574804 |
| BRNO1921 | T2 | stable | 2 | 6550605 |
| BRNO0462 | T2 | expanding mutation | 3 | 6533181 |
| BRNO0047 | T1 | expanding mutation | 3 | 6573130 |
| BRNO0332 | T1 | expanding mutation | 3 | 6562445 |
| BRNO0488 | T1 | expanding mutation | 3 | 6537840 |
| BRNO0837 | T1 | expanding mutation | 3 | 6580027 |
| BRNO0047 | T2 | expanding mutation | 3 | 6547309 |
| BRNO0462 | T1 | expanding mutation | 3 | 6561954 |
| BRNO1208 | T1 | expanding mutation | 3 | 6532786 |
| BRNO1503 | T1 | expanding mutation | 3 | 6646705 |
| BRNO0332 | T2 | expanding mutation | 3 | 6503698 |
| BRNO1503 | T2 | expanding mutation | 3 | 6745086 |
| BRNO0488 | T2 | expanding mutation | 3 | 6562406 |
| BRNO1208 | T2 | expanding mutation | 3 | 6535357 |
| BRNO0837 | T2 | expanding mutation | 3 | 6559725 |

**Table S8.** Differentially expressed TE subfamilies between the groups of patients with different *TP53* mutation evolution.

| TE families | baseMean | log2FoldChange | lfcSE | stat | pvalue | FDR | Heatmap_Order |
|---|---|---|---|---|---|---|---|
| L1HS_dup251 | 16.1647009000338 | 1.57529052558829 | 0.283419369349435 | 5.55816114192983 | 2.72631629609187e-08 | 0.00093111015129125 | 1 |
| L3_dup8732 | 22.6058352640787 | 2.0113578664643 | 0.283215352369411 | 7.10186735866207 | 1.23082334625224e-12 | 2.94251706565176e-07 | 2 |
| L1MC3_dup8780 | 20.5698323802536 | 1.14784246838939 | 0.248260182292509 | 4.62354638504601 | 3.77234336626016e-06 | 0.0300616785409484 | 3 |
| MER53_dup2282 | 18.2457327896288 | 1.13642535294297 | 0.248519982866688 | 4.57277253858726 | 4.81312558448926e-06 | 0.0328762605816646 | 4 |
| AluJb_dup61131 | 25.5263948626659 | -1.00725806046082 | 0.215946078099975 | -4.66439617391195 | 3.09524559634737e-06 | 0.0286524198590821 | 5 |
| L1M4_dup15411 | 29.1909471031418 | -1.04289052679523 | 0.220596348161724 | -4.72759651501875 | 2.27193080598875e-06 | 0.028088723591278 | 6 |
| L1PA3_dup1629 | 52.1244956173228 | 1.51005654986627 | 0.290025877590065 | 5.20662694796031 | 1.92304149857539e-07 | 0.00510821786692133 | 7 |
| L1PA8_dup4285 | 41.6128983826216 | 1.63450304919457 | 0.335260228604975 | 4.87532641731997 | 1.08628680712877e-06 | 0.0185498214781048 | 8 |
| L1PA4_dup5403 | 14.5009396861925 | -5.69958659491847 | 1.27495628155832 | -4.47041728203586 | 7.80671331529391e-06 | 0.039709428629234 | 9 |
| AluY_dup11877 | 12.1153428324468 | -6.46457862294425 | 1.3947356707356 | -4.63498479216121 | 3.56963722081582e-06 | 0.0300616785409484 | 10 |
| MIR_dup21066 | 19.8149045173946 | -7.37913786610729 | 1.51257434279929 | -4.87852904634815 | 1.06879926173679e-06 | 0.0185498214781048 | 11 |
| L1PA12_dup153 | 23.8451546302626 | -4.57396092307523 | 0.99419598835127 | -4.600663226031 | 4.21147854996893e-06 | 0.030510120165531 | 12 |
| Tigger1_dup1341 | 8.41818444126574 | -5.25703275320783 | 1.1288729528947 | -4.65688609132457 | 3.21027945185344e-06 | 0.0286524198590821 | 13 |
| L1HS_dup649 | 16.5474214466003 | -3.0177308928174 | 0.68099473745618 | -4.43135714101106 | 9.3641843689238e-06 | 0.0430516575556585 | 14 |
| AluSq2_dup14427 | 16.1146746417253 | -1.43997696925327 | 0.317786647535627 | -4.53126958108531 | 5.86302630481326e-06 | 0.0373037432020931 | 15 |
| LTR21B_dup47 | 12.8114162346999 | -1.70682437984889 | 0.302061126951605 | -5.65059263690806 | 1.59895608230201e-08 | 0.000637101386066434 | 16 |
| Tigger1_dup4137 | 8.36273951154196 | -1.46063281233911 | 0.327361019034102 | -4.46184098720365 | 8.12585160795286e-06 | 0.040471650376285 | 17 |
| AluSg_dup13034 | 9.95519709143086 | -1.14306281140376 | 0.252653176108287 | -4.52423685706544 | 6.06138141319755e-06 | 0.0373037432020931 | 18 |
| AluSg_dup1925 | 7.55487038083914 | -1.4397374476042 | 0.312525532707402 | -4.60678343664207 | 4.08945513208721e-06 | 0.030510120165531 | 19 |
| L1M4b_dup2618 | 2.56755410416446 | 2.99037157535726 | 0.521244713818902 | 5.73698206634708 | 9.63784576277168e-09 | 0.000460822029732012 | 20 |
| MIRb_dup107183 | 2.92919295270719 | 2.04756074643943 | 0.46311134427363 | 4.4213141650653 | 9.81024155429304e-06 | 0.0439569592615176 | 21 |
| FRAM_dup2158 | 5.97996904426257 | 1.41023133968924 | 0.281898125208354 | 5.00262759337943 | 5.6554126332086e-07 | 0.0135203384280855 | 22 |
| AluJo_dup65197 | 10.0521746125609 | 1.0333759524775 | 0.236141874896744 | 4.37608091715777 | 1.20832173184071e-05 | 0.0477755538317194 | 23 |
| L2a_dup87945 | 12.5255295563094 | 1.73131026260904 | 0.38374929668881 | 4.51156595607522 | 6.43507665391174e-06 | 0.0382278401487876 | 24 |
| MIR_dup114518 | 4.99270368230988 | 1.75637068815041 | 0.398760478205087 | 4.40457563913114 | 1.05991154746308e-05 | 0.0452485703107949 | 25 |
| AluJb_dup77912 | 5.01751074798377 | 1.61646990514876 | 0.337971594909523 | 4.78285728592516 | 1.72820829549061e-06 | 0.0229533904997025 | 26 |
| MLT1L_dup1387 | 6.42081557318024 | 1.1822004384498 | 0.26429150538523 | 4.47309283257754 | 7.70962865403032e-06 | 0.039709428629234 | 27 |
| MIRb_dup145987 | 6.00100744936283 | 1.25227658424008 | 0.279313901602257 | 4.48340228343995 | 7.34622506710135e-06 | 0.0390278817903745 | 28 |
| MIRb_dup172272 | 5.16437344582101 | 1.13474675019009 | 0.25908215521447 | 4.37987228124892 | 1.18748887670725e-05 | 0.0477755538317194 | 29 |
| L1M5_dup9143 | 10.3564290730048 | 1.27804710207316 | 0.289234850073738 | 4.41871752918894 | 9.92883142574718e-06 | 0.0439569592615176 | 30 |
| X1_LINE_dup20 | 10.4636551774124 | 1.63974767463158 | 0.260242840641361 | 6.30083682836564 | 2.9604292268226e-10 | 3.53873427413626e-05 | 31 |
| MIR_dup30573 | 8.60004940650453 | 1.43811099772494 | 0.317161823589122 | 4.53431305650453 | 5.77912608178979e-06 | 0.0373037432020931 | 32 |
| Tigger1_dup2076 | 10.4229041666772 | 1.52744580935623 | 0.318076766488626 | 4.80212945515731 | 1.56987142758602e-06 | 0.0229533904997025 | 33 |
| L2a_dup30312 | 5.82825183995955 | 1.47174656425525 | 0.295305912958526 | 4.98380323478978 | 6.23464819510763e-07 | 0.0135501009941471 | 34 |
| MLT1J2_dup1154 | 4.4245178440637 | 1.73323880538318 | 0.29377667054315 | 5.89985175534419 | 3.63828321109472e-09 | 0.000217450182248301 | 35 |
| L1PREC2_dup1129 | 5.23576024798689 | 1.71322590220838 | 0.371995852841915 | 4.60549731702638 | 4.11481291366062e-06 | 0.030510120165531 | 36 |
| MIRb_dup41134 | 3.99443861820542 | 1.81071414049786 | 0.305509421998745 | 5.92686840442291 | 3.08766213540964e-09 | 0.000217450182248301 | 37 |
| HERVL74-int_dup285 | 4.21981521306723 | -4.30311340037787 | 0.981488696009984 | -4.38427199199663 | 1.16374298610925e-05 | 0.0477755538317194 | 38 |
| MLT1H_dup4071 | 5.47964162460913 | -4.22225227223704 | 0.789560590081481 | -5.34759754384564 | 8.91293789500015e-08 | 0.00266350893702474 | 39 |
| Tigger17a_dup1745 | 5.84327246906994 | -2.70762171630944 | 0.598581355451488 | -4.52339801707852 | 6.08546480255337e-06 | 0.0373037432020931 | 40 |
| Tigger1_dup4135 | 3.69345225961644 | -2.45576065973853 | 0.52556803190035 | -4.67258377732561 | 2.9743413120312e-06 | 0.0286524198590821 | 41 |
| L1MA4_dup8077 | 2.42002854952993 | -3.27031287447274 | 0.681379943436991 | -4.7995437875332 | 1.59027465220531e-06 | 0.0229533904997025 | 42 |
| HAL1_dup14920 | 2.78390144004605 | -2.94854616572787 | 0.654125468184372 | -4.50761560150229 | 6.55602125788074e-06 | 0.0382278401487876 | 43 |
| L2a_dup38602 | 2.58542469740183 | -3.14727741482641 | 0.657042387361421 | -4.79006754414336 | 1.66725181316626e-06 | 0.0229533904997025 | 44 |
| AluSg_dup33413 | 6.07371798099363 | -2.49113696956358 | 0.535124733105468 | -4.65524543241883 | 3.23595002361333e-06 | 0.0286524198590821 | 45 |
| L2_dup24256 | 11.0284494537574 | -1.64021642655521 | 0.354541054891754 | -4.62630886867527 | 3.72240058113862e-06 | 0.0300616785409484 | 46 |
| L1MD1_dup1202 | 8.35707324281766 | -1.86766372176405 | 0.399223452150349 | -4.67824150035323 | 2.89345643488336e-06 | 0.0286524198590821 | 47 |
| AluY_dup2197 | 5.72105137944561 | -1.65637084942337 | 0.338530583509624 | -4.89282484392222 | 9.93988634236293e-07 | 0.0185498214781048 | 48 |
| Charlie1a_dup3640 | 5.89485228121248 | -1.3364588980667 | 0.297096893192787 | -4.49839405489664 | 6.84687020007202e-06 | 0.0389732002824052 | 49 |
| MIRb_dup206602 | 5.48924815734764 | -1.62704696970217 | 0.3446590579272 | -4.72074339054752 | 2.34984239623523e-06 | 0.028088723591278 | 50 |
| Tigger15a_dup1664 | 5.34510038459067 | -1.54320885569191 | 0.344109044655285 | -4.48465066426205 | 7.30334868647254e-06 | 0.0390278817903745 | 51 |

| TE families | RNA3043 | RNA3335 | RNA3336 | RNA3337 | RNA3338 | RNA3346 | RNA3347 | RNA3480 | RNA3482 | RNA3484 | RNA3485 | RNA3487 | RNA3494 | RNA3496 | RNA3497 |
|---|---|---|---|---|---|---|---|---|---|---|---|---|---|---|---|
| L1HS_dup251 | 48 | 26 | 51 | 47 | 35 | 49 | 43 | 8 | 41 | 19 | 13 | 13 | 32 | 13 | 3 |
| L3_dup8732 | 63 | 27 | 94 | 54 | 46 | 51 | 74 | 9 | 30 | 35 | 19 | 11 | 38 | 14 | 3 |
| L1MC3_dup8780 | 36 | 51 | 47 | 12 | 16 | 81 | 29 | 44 | 18 | 22 | 27 | 14 | 52 | 18 | 9 |
| MER53_dup2282 | 21 | 43 | 32 | 23 | 37 | 35 | 21 | 8 | 25 | 8 | 20 | 5 | 43 | 11 | 12 |
| AluJb_dup61131 | 14 | 15 | 21 | 23 | 13 | 21 | 14 | 35 | 20 | 31 | 18 | 20 | 3 | 47 | 47 |
| L1M4_dup15411 | 8 | 21 | 17 | 21 | 28 | 9 | 22 | 48 | 41 | 58 | 34 | 26 | 12 | 20 | 6 |
| L1PA3_dup1629 | 108 | 70 | 167 | 134 | 136 | 112 | 145 | 30 | 78 | 91 | 35 | 31 | 90 | 34 | 21 |
| L1PA8_dup4285 | 124 | 16 | 126 | 26 | 37 | 19 | 54 | 12 | 12 | 12 | 23 | 13 | 21 | 22 | 27 |
| L1PA4_dup5403 | 1 | 0 | 0 | 2 | 0 | 0 | 0 | 1 | 159 | 68 | 47 | 0 | 2 | 30 | 27 |
| AluY_dup11877 | 0 | 0 | 0 | 1 | 0 | 0 | 0 | 0 | 22 | 91 | 49 | 0 | 0 | 31 | 26 |
| MIR_dup21066 | 0 | 0 | 0 | 0 | 0 | 0 | 0 | 0 | 45 | 136 | 68 | 0 | 0 | 31 | 50 |
| L1PA12_dup153 | 1 | 0 | 0 | 12 | 1 | 0 | 0 | 2 | 76 | 116 | 62 | 20 | 1 | 109 | 29 |
| Tigger1_dup1341 | 0 | 0 | 0 | 4 | 0 | 0 | 0 | 0 | 31 | 44 | 31 | 6 | 0 | 32 | 6 |
| L1HS_dup649 | 1 | 4 | 5 | 2 | 3 | 6 | 2 | 3 | 11 | 6 | 1 | 25 | 3 | 90 | 49 |
| AluSq2_dup14427 | 15 | 6 | 10 | 9 | 9 | 10 | 15 | 15 | 11 | 8 | 5 | 10 | 2 | 24 | 36 |
| LTR21B_dup47 | 10 | 4 | 8 | 6 | 4 | 4 | 1 | 10 | 22 | 8 | 13 | 33 | 3 | 19 | 8 |
| Tigger1_dup4137 | 5 | 9 | 4 | 6 | 6 | 6 | 1 | 6 | 5 | 6 | 11 | 9 | 1 | 16 | 26 |
| AluSg_dup13034 | 3 | 8 | 5 | 2 | 3 | 13 | 3 | 10 | 8 | 11 | 11 | 8 | 7 | 20 | 13 |
| AluSg_dup1925 | 5 | 6 | 6 | 5 | 3 | 1 | 1 | 6 | 4 | 8 | 3 | 2 | 3 | 10 | 5 |
| L1M4b_dup2618 | 9 | 11 | 14 | 6 | 5 | 8 | 6 | 1 | 0 | 2 | 0 | 0 | 10 | 1 | 0 |
| MIRb_dup107183 | 16 | 3 | 10 | 0 | 4 | 2 | 7 | 1 | 0 | 1 | 0 | 0 | 3 | 0 | 1 |
| FRAM_dup2158 | 13 | 10 | 13 | 19 | 10 | 10 | 13 | 0 | 5 | 7 | 0 | 1 | 9 | 6 | 2 |
| AluJo_dup65197 | 24 | 5 | 20 | 17 | 23 | 6 | 28 | 2 | 12 | 11 | 7 | 1 | 16 | 9 | 5 |
| L2a_dup87945 | 43 | 4 | 39 | 9 | 3 | 5 | 16 | 2 | 7 | 4 | 9 | 4 | 8 | 10 | 7 |
| MIR_dup114518 | 12 | 26 | 14 | 9 | 1 | 23 | 3 | 7 | 8 | 2 | 0 | 0 | 12 | 5 | 5 |
| AluJb_dup77912 | 5 | 9 | 16 | 8 | 6 | 24 | 6 | 6 | 7 | 5 | 3 | 2 | 21 | 3 | 2 |
| MLT1L_dup1387 | 7 | 11 | 9 | 23 | 3 | 12 | 4 | 8 | 8 | 6 | 5 | 6 | 16 | 5 | 4 |
| MIRb_dup145987 | 9 | 15 | 19 | 6 | 7 | 26 | 11 | 3 | 13 | 6 | 7 | 3 | 9 | 2 | 4 |
| MIRb_dup172272 | 7 | 9 | 12 | 12 | 4 | 8 | 9 | 4 | 12 | 6 | 6 | 4 | 16 | 4 | 1 |
| L1M5_dup9143 | 9 | 21 | 30 | 23 | 34 | 15 | 25 | 9 | 19 | 21 | 11 | 4 | 21 | 4 | 6 |
| X1_LINE_dup20 | 19 | 12 | 42 | 25 | 24 | 20 | 27 | 6 | 19 | 19 | 5 | 8 | 25 | 7 | 2 |
| MIR_dup30573 | 21 | 13 | 37 | 9 | 19 | 17 | 27 | 10 | 17 | 11 | 10 | 10 | 15 | 3 | 1 |
| Tigger1_dup2076 | 9 | 18 | 37 | 22 | 30 | 22 | 35 | 11 | 25 | 20 | 14 | 6 | 17 | 4 | 2 |
| L2a_dup30312 | 9 | 12 | 16 | 11 | 11 | 15 | 16 | 3 | 7 | 2 | 5 | 2 | 8 | 2 | 1 |
| MLT1J2_dup1154 | 11 | 7 | 11 | 8 | 16 | 11 | 12 | 1 | 7 | 6 | 1 | 1 | 11 | 1 | 2 |
| L1PREC2_dup1129 | 15 | 8 | 18 | 15 | 13 | 10 | 15 | 2 | 9 | 5 | 1 | 7 | 12 | 0 | 1 |
| MIRb_dup41134 | 3 | 11 | 16 | 8 | 13 | 12 | 9 | 3 | 7 | 5 | 4 | 4 | 10 | 1 | 0 |
| HERVL74-int_dup285 | 2 | 0 | 0 | 0 | 0 | 1 | 1 | 7 | 0 | 0 | 0 | 16 | 0 | 1 | 0 |
| MLT1H_dup4071 | 2 | 0 | 0 | 0 | 0 | 0 | 0 | 7 | 1 | 33 | 16 | 14 | 1 | 15 | 10 |
| Tigger17a_dup1745 | 0 | 0 | 0 | 1 | 1 | 1 | 1 | 2 | 5 | 0 | 5 | 1 | 0 | 25 | 14 |
| Tigger1_dup4135 | 1 | 2 | 0 | 0 | 0 | 6 | 0 | 9 | 1 | 7 | 3 | 1 | 0 | 9 | 8 |
| L1MA4_dup8077 | 1 | 1 | 0 | 1 | 0 | 0 | 0 | 5 | 0 | 3 | 1 | 1 | 1 | 1 | 2 |
| HAL1_dup14920 | 0 | 3 | 0 | 0 | 0 | 0 | 0 | 4 | 1 | 8 | 3 | 7 | 0 | 0 | 1 |
| L2a_dup38602 | 1 | 1 | 1 | 0 | 0 | 0 | 0 | 3 | 2 | 3 | 5 | 2 | 0 | 2 | 0 |
| AluSg_dup33413 | 2 | 2 | 2 | 3 | 0 | 0 | 1 | 2 | 16 | 5 | 9 | 33 | 1 | 0 | 1 |
| L2_dup24256 | 0 | 7 | 16 | 2 | 6 | 3 | 0 | 7 | 28 | 9 | 4 | 16 | 10 | 3 | 4 |
| L1MD1_dup1202 | 4 | 2 | 4 | 2 | 5 | 3 | 1 | 5 | 4 | 12 | 11 | 0 | 0 | 1 | 3 |
| AluY_dup2197 | 2 | 2 | 2 | 1 | 3 | 1 | 3 | 4 | 10 | 6 | 5 | 3 | 2 | 7 | 6 |
| Charlie1a_dup3640 | 3 | 5 | 4 | 3 | 2 | 2 | 1 | 5 | 8 | 7 | 4 | 9 | 3 | 3 | 1 |
| MIRb_dup206602 | 1 | 2 | 0 | 7 | 0 | 3 | 1 | 2 | 5 | 14 | 8 | 4 | 2 | 3 | 5 |
| Tigger15a_dup1664 | 2 | 2 | 3 | 1 | 4 | 3 | 1 | 5 | 9 | 6 | 12 | 7 | 3 | 3 | 1 |

| TE families | RNA3498 | RNA3500 | RNA3501 | RNA3531 | RNA3537 | RNA354 | RNA3542 | RNA3638 | RNA3639 | RNA3640 | RNA3652 | RNA3658 | RNA3659 | RNA3661 | RNA3662 |
|---|---|---|---|---|---|---|---|---|---|---|---|---|---|---|---|
| L1HS_dup251 | 37 | 1 | 1 | 7 | 12 | 8 | 5 | 3 | 7 | 34 | 11 | 10 | 3 | 18 | 5 |
| L3_dup8732 | 39 | 3 | 2 | 40 | 25 | 10 | 10 | 6 | 5 | 34 | 16 | 8 | 5 | 58 | 5 |
| L1MC3_dup8780 | 43 | 8 | 16 | 27 | 18 | 14 | 35 | 15 | 10 | 16 | 32 | 10 | 12 | 20 | 7 |
| MER53_dup2282 | 42 | 12 | 10 | 13 | 10 | 23 | 14 | 8 | 9 | 8 | 61 | 23 | 5 | 30 | 28 |
| AluJb_dup61131 | 27 | 44 | 35 | 7 | 8 | 25 | 34 | 17 | 21 | 6 | 33 | 42 | 9 | 23 | 17 |
| L1M4_dup15411 | 30 | 14 | 12 | 10 | 17 | 60 | 47 | 14 | 33 | 23 | 13 | 23 | 13 | 19 | 15 |
| L1PA3_dup1629 | 148 | 1 | 5 | 30 | 40 | 20 | 18 | 19 | 22 | 138 | 27 | 42 | 9 | 66 | 13 |
| L1PA8_dup4285 | 28 | 28 | 21 | 80 | 91 | 9 | 34 | 32 | 16 | 385 | 25 | 22 | 17 | 17 | 27 |
| L1PA4_dup5403 | 1 | 0 | 0 | 0 | 0 | 1 | 0 | 0 | 88 | 0 | 0 | 2 | 0 | 0 | 1 |
| AluY_dup11877 | 0 | 0 | 0 | 0 | 0 | 0 | 0 | 0 | 25 | 0 | 0 | 0 | 0 | 0 | 0 |
| MIR_dup21066 | 0 | 0 | 0 | 0 | 0 | 0 | 0 | 0 | 52 | 0 | 0 | 0 | 0 | 0 | 0 |
| L1PA12_dup153 | 0 | 0 | 0 | 0 | 0 | 0 | 0 | 0 | 48 | 6 | 1 | 27 | 0 | 0 | 0 |
| Tigger1_dup1341 | 0 | 0 | 0 | 0 | 0 | 0 | 0 | 0 | 10 | 0 | 0 | 5 | 0 | 0 | 0 |
| L1HS_dup649 | 9 | 2 | 7 | 1 | 2 | 1 | 1 | 24 | 0 | 2 | 3 | 68 | 77 | 1 | 36 |
| AluSq2_dup14427 | 17 | 13 | 12 | 2 | 3 | 10 | 4 | 12 | 5 | 11 | 6 | 16 | 9 | 3 | 16 |
| LTR21B_dup47 | 6 | 10 | 11 | 3 | 6 | 17 | 2 | 19 | 17 | 3 | 12 | 44 | 7 | 4 | 9 |
| Tigger1_dup4137 | 5 | 8 | 13 | 0 | 2 | 12 | 0 | 4 | 6 | 3 | 7 | 4 | 4 | 2 | 12 |
| AluSg_dup13034 | 8 | 8 | 10 | 7 | 2 | 14 | 10 | 1 | 7 | 5 | 8 | 19 | 4 | 6 | 8 |
| AluSg_dup1925 | 7 | 9 | 5 | 2 | 1 | 24 | 8 | 5 | 3 | 3 | 9 | 8 | 8 | 1 | 6 |
| L1M4b_dup2618 | 2 | 2 | 1 | 6 | 0 | 1 | 0 | 0 | 1 | 11 | 2 | 2 | 1 | 1 | 0 |
| MIRb_dup107183 | 3 | 0 | 1 | 9 | 5 | 1 | 6 | 1 | 0 | 14 | 1 | 1 | 3 | 3 | 2 |
| FRAM_dup2158 | 7 | 2 | 6 | 11 | 7 | 12 | 12 | 5 | 1 | 18 | 5 | 2 | 1 | 4 | 4 |
| AluJo_dup65197 | 12 | 5 | 12 | 11 | 14 | 15 | 4 | 4 | 20 | 11 | 13 | 5 | 4 | 16 | 4 |
| L2a_dup87945 | 5 | 6 | 11 | 24 | 30 | 2 | 5 | 8 | 7 | 124 | 9 | 3 | 8 | 5 | 5 |
| MIR_dup114518 | 6 | 0 | 1 | 12 | 2 | 1 | 5 | 1 | 5 | 7 | 12 | 6 | 0 | 1 | 0 |
| AluJb_dup77912 | 8 | 9 | 0 | 4 | 1 | 1 | 5 | 2 | 1 | 8 | 10 | 2 | 0 | 8 | 5 |
| MLT1L_dup1387 | 14 | 4 | 1 | 10 | 3 | 9 | 4 | 4 | 2 | 12 | 17 | 1 | 4 | 7 | 5 |
| MIRb_dup145987 | 12 | 4 | 2 | 12 | 3 | 7 | 7 | 3 | 4 | 2 | 12 | 1 | 5 | 3 | 6 |
| MIRb_dup172272 | 6 | 3 | 4 | 5 | 4 | 7 | 6 | 5 | 6 | 13 | 5 | 3 | 2 | 7 | 6 |
| L1M5_dup9143 | 33 | 2 | 1 | 8 | 3 | 10 | 3 | 1 | 4 | 24 | 5 | 8 | 3 | 14 | 3 |
| X1_LINE_dup20 | 22 | 2 | 3 | 11 | 9 | 7 | 4 | 2 | 11 | 17 | 4 | 4 | 1 | 27 | 2 |
| MIR_dup30573 | 13 | 0 | 0 | 16 | 5 | 5 | 5 | 1 | 2 | 6 | 8 | 5 | 1 | 12 | 1 |
| Tigger1_dup2076 | 26 | 0 | 1 | 13 | 4 | 19 | 5 | 1 | 2 | 19 | 5 | 4 | 0 | 21 | 4 |
| L2a_dup30312 | 18 | 1 | 2 | 10 | 2 | 4 | 3 | 0 | 3 | 12 | 4 | 3 | 4 | 10 | 1 |
| MLT1J2_dup1154 | 18 | 0 | 0 | 5 | 0 | 4 | 3 | 1 | 1 | 7 | 4 | 3 | 3 | 8 | 0 |
| L1PREC2_dup1129 | 12 | 0 | 0 | 4 | 1 | 6 | 0 | 2 | 1 | 14 | 2 | 1 | 0 | 7 | 0 |
| MIRb_dup41134 | 16 | 0 | 0 | 2 | 4 | 2 | 3 | 1 | 0 | 5 | 2 | 4 | 1 | 8 | 2 |
| HERVL74-int_dup285 | 0 | 10 | 15 | 0 | 0 | 0 | 16 | 2 | 0 | 0 | 0 | 53 | 0 | 0 | 13 |
| MLT1H_dup4071 | 0 | 18 | 10 | 0 | 1 | 0 | 9 | 10 | 0 | 1 | 0 | 17 | 7 | 1 | 3 |
| Tigger17a_dup1745 | 1 | 0 | 2 | 1 | 1 | 9 | 1 | 2 | 4 | 4 | 7 | 0 | 3 | 0 | 1 |
| Tigger1_dup4135 | 5 | 3 | 4 | 0 | 0 | 15 | 4 | 2 | 2 | 1 | 0 | 0 | 2 | 0 | 4 |
| L1MA4_dup8077 | 1 | 2 | 0 | 0 | 0 | 9 | 8 | 2 | 0 | 0 | 0 | 2 | 5 | 0 | 2 |
| HAL1_dup14920 | 3 | 0 | 6 | 0 | 0 | 18 | 3 | 1 | 0 | 0 | 0 | 2 | 1 | 2 | 4 |
| L2a_dup38602 | 1 | 0 | 1 | 1 | 0 | 6 | 8 | 1 | 0 | 0 | 1 | 1 | 3 | 0 | 1 |
| AluSg_dup33413 | 3 | 2 | 4 | 1 | 0 | 3 | 1 | 16 | 14 | 4 | 2 | 38 | 10 | 1 | 10 |
| L2_dup24256 | 3 | 3 | 4 | 5 | 2 | 21 | 14 | 21 | 12 | 2 | 0 | 24 | 5 | 10 | 23 |
| L1MD1_dup1202 | 5 | 7 | 14 | 6 | 2 | 3 | 7 | 9 | 10 | 1 | 0 | 2 | 9 | 7 | 9 |
| AluY_dup2197 | 5 | 2 | 8 | 2 | 1 | 9 | 1 | 3 | 3 | 1 | 7 | 8 | 2 | 3 | 9 |
| Charlie1a_dup3640 | 6 | 4 | 7 | 1 | 4 | 13 | 1 | 7 | 1 | 6 | 2 | 13 | 4 | 2 | 11 |
| MIRb_dup206602 | 2 | 3 | 6 | 3 | 2 | 20 | 4 | 5 | 9 | 3 | 5 | 3 | 10 | 2 | 5 |
| Tigger15a_dup1664 | 4 | 4 | 5 | 2 | 3 | 21 | 6 | 6 | 3 | 1 | 1 | 1 | 15 | 4 | 4 |

| TE families | RNA3663 | RNA3744 | RNA3745 | RNA3750 | RNA3751 | RNA3752 | RNA3753 | RNA3754 | RNA3755 | RNA3756 | RNA3757 | RNA3758 |
|---|---|---|---|---|---|---|---|---|---|---|---|---|
| L1HS_dup251 | 4 | 16 | 17 | 18 | 27 | 1 | 19 | 18 | 16 | 2 | 3 | 16 |
| L3_dup8732 | 2 | 20 | 20 | 20 | 21 | 0 | 29 | 25 | 12 | 3 | 13 | 12 |
| L1MC3_dup8780 | 16 | 23 | 14 | 12 | 12 | 5 | 11 | 10 | 12 | 2 | 16 | 17 |
| MER53_dup2282 | 28 | 11 | 15 | 13 | 18 | 42 | 7 | 11 | 3 | 9 | 10 | 7 |
| AluJb_dup61131 | 17 | 31 | 55 | 29 | 58 | 22 | 33 | 47 | 40 | 27 | 39 | 45 |
| L1M4_dup15411 | 24 | 46 | 55 | 41 | 73 | 70 | 50 | 48 | 85 | 31 | 57 | 55 |
| L1PA3_dup1629 | 22 | 49 | 89 | 60 | 77 | 2 | 57 | 56 | 54 | 6 | 30 | 39 |
| L1PA8_dup4285 | 29 | 16 | 35 | 17 | 44 | 4 | 13 | 25 | 19 | 83 | 20 | 13 |
| L1PA4_dup5403 | 0 | 0 | 0 | 0 | 1 | 0 | 3 | 0 | 1 | 44 | 68 | 95 |
| AluY_dup11877 | 1 | 62 | 60 | 39 | 42 | 0 | 14 | 85 | 0 | 0 | 0 | 5 |
| MIR_dup21066 | 2 | 78 | 91 | 80 | 69 | 0 | 39 | 161 | 0 | 0 | 0 | 3 |
| L1PA12_dup153 | 1 | 101 | 96 | 71 | 104 | 0 | 44 | 73 | 121 | 1 | 0 | 6 |
| Tigger1_dup1341 | 0 | 40 | 35 | 24 | 46 | 0 | 21 | 20 | 49 | 1 | 0 | 3 |
| L1HS_dup649 | 113 | 4 | 2 | 1 | 3 | 0 | 2 | 4 | 3 | 0 | 53 | 26 |
| AluSq2_dup14427 | 18 | 15 | 39 | 28 | 75 | 16 | 46 | 35 | 100 | 15 | 16 | 32 |
| LTR21B_dup47 | 35 | 28 | 23 | 10 | 18 | 14 | 10 | 9 | 12 | 27 | 21 | 29 |
| Tigger1_dup4137 | 17 | 13 | 16 | 9 | 16 | 24 | 10 | 22 | 14 | 8 | 5 | 13 |
| AluSg_dup13034 | 7 | 17 | 22 | 12 | 15 | 21 | 16 | 18 | 17 | 21 | 14 | 17 |
| AluSg_dup1925 | 7 | 15 | 24 | 9 | 21 | 22 | 13 | 13 | 8 | 12 | 20 | 16 |
| L1M4b_dup2618 | 0 | 0 | 0 | 0 | 0 | 2 | 3 | 6 | 0 | 0 | 0 | 0 |
| MIRb_dup107183 | 2 | 3 | 2 | 0 | 2 | 0 | 2 | 2 | 1 | 8 | 0 | 2 |
| FRAM_dup2158 | 0 | 3 | 6 | 5 | 9 | 2 | 2 | 7 | 4 | 4 | 6 | 5 |
| AluJo_dup65197 | 11 | 7 | 13 | 6 | 10 | 12 | 5 | 6 | 8 | 10 | 8 | 8 |
| L2a_dup87945 | 8 | 4 | 6 | 1 | 9 | 2 | 4 | 4 | 11 | 25 | 6 | 3 |
| MIR_dup114518 | 3 | 4 | 8 | 5 | 3 | 3 | 4 | 2 | 1 | 0 | 1 | 5 |
| AluJb_dup77912 | 4 | 5 | 5 | 0 | 3 | 3 | 8 | 5 | 2 | 1 | 1 | 0 |
| MLT1L_dup1387 | 6 | 4 | 6 | 4 | 6 | 10 | 4 | 2 | 9 | 2 | 0 | 6 |
| MIRb_dup145987 | 1 | 4 | 5 | 7 | 5 | 2 | 5 | 4 | 2 | 1 | 2 | 8 |
| MIRb_dup172272 | 1 | 4 | 2 | 3 | 1 | 1 | 3 | 2 | 5 | 4 | 2 | 3 |
| L1M5_dup9143 | 3 | 17 | 26 | 9 | 14 | 3 | 5 | 15 | 8 | 1 | 6 | 3 |
| X1_LINE_dup20 | 5 | 11 | 8 | 12 | 12 | 2 | 5 | 14 | 11 | 0 | 8 | 2 |
| MIR_dup30573 | 1 | 14 | 11 | 4 | 9 | 2 | 12 | 8 | 12 | 0 | 12 | 7 |
| Tigger1_dup2076 | 2 | 12 | 13 | 11 | 14 | 0 | 8 | 6 | 9 | 0 | 7 | 7 |
| L2a_dup30312 | 2 | 11 | 11 | 14 | 5 | 0 | 7 | 9 | 2 | 1 | 5 | 2 |
| MLT1J2_dup1154 | 3 | 3 | 5 | 7 | 7 | 0 | 3 | 5 | 3 | 0 | 3 | 5 |
| L1PREC2_dup1129 | 0 | 8 | 4 | 7 | 10 | 1 | 13 | 5 | 7 | 0 | 7 | 2 |
| MIRb_dup41134 | 0 | 1 | 6 | 2 | 4 | 0 | 6 | 2 | 3 | 0 | 4 | 2 |
| HERVL74-int_dup285 | 5 | 1 | 1 | 0 | 0 | 0 | 17 | 12 | 0 | 1 | 8 | 4 |
| MLT1H_dup4071 | 6 | 1 | 0 | 17 | 14 | 2 | 0 | 1 | 2 | 0 | 0 | 4 |
| Tigger17a_dup1745 | 2 | 18 | 16 | 18 | 11 | 16 | 9 | 6 | 11 | 12 | 21 | 21 |
| Tigger1_dup4135 | 10 | 13 | 10 | 3 | 6 | 7 | 6 | 10 | 9 | 3 | 1 | 3 |
| L1MA4_dup8077 | 1 | 6 | 13 | 1 | 19 | 9 | 3 | 1 | 9 | 1 | 1 | 8 |
| HAL1_dup14920 | 1 | 3 | 9 | 3 | 16 | 7 | 1 | 10 | 6 | 3 | 3 | 7 |
| L2a_dup38602 | 1 | 0 | 10 | 7 | 9 | 3 | 4 | 9 | 12 | 8 | 10 | 5 |
| AluSg_dup33413 | 22 | 6 | 15 | 2 | 4 | 3 | 1 | 3 | 5 | 10 | 2 | 3 |
| L2_dup24256 | 6 | 19 | 20 | 34 | 38 | 14 | 13 | 14 | 15 | 22 | 26 | 19 |
| L1MD1_dup1202 | 11 | 35 | 23 | 10 | 11 | 24 | 6 | 25 | 9 | 37 | 9 | 21 |
| AluY_dup2197 | 4 | 19 | 25 | 15 | 15 | 8 | 8 | 13 | 14 | 4 | 11 | 11 |
| Charlie1a_dup3640 | 7 | 8 | 13 | 10 | 11 | 11 | 9 | 11 | 21 | 5 | 9 | 17 |
| MIRb_dup206602 | 7 | 14 | 8 | 11 | 7 | 11 | 7 | 4 | 13 | 4 | 10 | 8 |
| Tigger15a_dup1664 | 4 | 8 | 11 | 8 | 4 | 10 | 9 | 13 | 14 | 4 | 7 | 4 |

**Table S9.** Differentially expressed genes between Group 1 and Group 2 of the *TP53* non-exp cases.

| Genes | Gene_Symbol | logFC | AveExpr | t | P.Value | adj.P.Val | B |
|---|---|---|---|---|---|---|---|
| lnc-SYTL2-4 | lnc-SYTL2-4 | 0.657176682800122 | 2.63314834749958 | 5.93012256940611 | 4.80803399444587e-07 | 0.00620348476706947 | 6.15756759083015 |
| lnc-C5orf63-1 | lnc-C5orf63-1 | -0.987910622860897 | 1.63197391162526 | -5.80469723933686 | 7.29809061306218e-07 | 0.00620348476706947 | 5.6802558661708 |
| ENSG00000112320.11 | SOBP | -1.59770329279989 | 2.42207084596341 | -5.68237450993697 | 1.09577602882676e-06 | 0.00620348476706947 | 5.40110551885141 |
| ENSG00000198211.8 | | -1.18087593519111 | 0.426136217878717 | -5.52582051210337 | 1.8414157256307e-06 | 0.00620348476706947 | 4.52592708360408 |
| ENSG00000258839.3 | MC1R | -1.00764611741808 | 2.41081552952392 | -5.52083592952634 | 1.872056284967e-06 | 0.00620348476706947 | 4.91401762335156 |
| ENSG00000238923.1 | RNU7-1 | -1.44472861670299 | 1.29292527126279 | -5.49149850776797 | 2.06294176841142e-06 | 0.00620348476706947 | 4.71469770478951 |
| lnc-ZNF582-1 | lnc-ZNF582-1 | -1.22045580153832 | 0.838059292098076 | -5.48472092633233 | 2.1097212927895e-06 | 0.00620348476706947 | 4.52113048714727 |
| lnc-KCMF1-1 | lnc-KCMF1-1 | -1.79357179111004 | -0.646071526192307 | -5.30288881862142 | 3.84518767461777e-06 | 0.00945325848277165 | 3.12655759947701 |
| ENSG00000112293.14 | GPLD1 | -0.899560732886413 | 3.09239780923581 | -5.26157641754556 | 4.40532329109597e-06 | 0.00945325848277165 | 4.14614716253528 |
| lnc-IL4R-4 | lnc-IL4R-4 | -1.69529984803517 | 0.930194880895377 | -5.24890927435024 | 4.59275056248926e-06 | 0.00945325848277165 | 3.92332079116568 |
| UGDH-AS1 | UGDH-AS1 | -1.09982712575706 | 2.1153756495925 | -5.16883728031364 | 5.97443516034966e-06 | 0.0102198465896668 | 3.83858102046386 |
| ENSG00000089234.15 | BRAP | 0.415268982652549 | 5.09084270648892 | 5.15087263086098 | 6.3370082977068e-06 | 0.0102198465896668 | 3.79013175712751 |
| ENSG00000168411.13 | RFWD3 | 0.35177694375801 | 5.13701349328695 | 5.12637963940646 | 6.86666338315485e-06 | 0.0102198465896668 | 3.71428080879294 |
| ENSG00000170190.15 | SLC16A5 | -1.09952824048582 | 3.3794265325149 | -5.11662576742722 | 7.08958308524543e-06 | 0.0102198465896668 | 3.70547229091352 |
| lnc-ZNF227-1 | lnc-ZNF227-1 | -0.969210656727036 | 2.68805313006282 | -5.10157109267752 | 7.44778209420407e-06 | 0.0102198465896668 | 3.65149940084286 |
| ENSG00000228439.5 | NA | -0.842959348173902 | 1.79668661179422 | -5.02910596432561 | 9.43854985302104e-06 | 0.0111915615620049 | 3.39959468542134 |
| ENSG00000049089.14 | COL9A2 | -0.997088823587443 | 7.02888584787175 | -5.01971928988141 | 9.73220906470956e-06 | 0.0111915615620049 | 3.37853715759121 |
| ENSG00000086062.12 | B4GALT1 | 0.978593490624934 | 6.8364452207291 | 5.00645394920663 | 1.01626675361257e-05 | 0.0111915615620049 | 3.33573786877463 |
| ENSG00000204228.3 | HSD17B8 | -0.670381644606016 | 1.92798725874921 | -5.00142172057467 | 1.03308395121262e-05 | 0.0111915615620049 | 3.33333388124427 |
| ENSG00000115306.15 | SPTBN1 | 0.982268444104982 | 8.82458068289533 | 4.93849618034885 | 1.26808059576141e-05 | 0.0117282169370032 | 3.13673328628087 |
| lnc-CYP4F22-5 | lnc-CYP4F22-5 | -0.652378076867728 | 3.45553011486175 | -4.93793535063965 | 1.2703961223094e-05 | 0.0117282169370032 | 3.16167209196135 |
| ENSG00000174437.16 | ATP2A2 | 0.384580757259219 | 6.79336935852534 | 4.92108011799073 | 1.3419689876622e-05 | 0.0117282169370032 | 3.0724661204159 |
| ENSG00000138439.11 | FAM117B | 0.571458104868288 | 5.89288923010934 | 4.91755619326356 | 1.35742836162035e-05 | 0.0117282169370032 | 3.06252139997602 |
| ENSG00000156239.11 | N6AMT1 | -0.952828065962562 | 2.69127854974296 | -4.91302844589529 | 1.37754973456242e-05 | 0.0117282169370032 | 3.08804198899855 |
| ENSG00000132481.6 | TRIM47 | -1.61676050297039 | -0.343051616981124 | -4.9027118515429 | 1.42450285879162e-05 | 0.0117282169370032 | 2.68681420872411 |
| ENSG00000143622.10 | RIT1 | 0.463725954844726 | 5.18186675755355 | 4.80408122878693 | 1.9610605122417e-05 | 0.0151312384983376 | 2.72189487191551 |
| ENSG00000200169.1 | RNU5D-1 | -1.51915662598192 | 4.0617449256703 | -4.80811982487766 | 1.99310910500202e-05 | 0.0151312384983376 | 2.73570957110854 |
| ENSG00000121644.18 | DESI2 | 0.448049767260489 | 5.06813947605526 | 4.77585018446093 | 2.14838283671998e-05 | 0.0151312384983376 | 2.63753943138209 |
| ENSG00000023734.10 | STRAP | 0.521738699085643 | 5.16368124285931 | 4.77065366790415 | 2.18473538647369e-05 | 0.0151312384983376 | 2.61995299423547 |
| URB1-AS1 | URB1-AS1 | -1.59783032119997 | 0.235089911368046 | -4.76773776780543 | 2.20539841106801e-05 | 0.0151312384983376 | 2.36232746796873 |
| lnc-TMCC1-10 | lnc-TMCC1-10 | -0.872521608517218 | 0.164505103636654 | -4.73082601948976 | 2.48420932583128e-05 | 0.0156234554115578 | 2.28053769671165 |
| ENSG00000162885.12 | B3GALNT2 | 0.825219660907792 | 1.72738157895089 | 4.72375362805574 | 2.54146067229725e-05 | 0.0156234554115578 | 2.50565963594477 |
| ENSG00000160908.14 | ZNF394 | 0.480075487636931 | 6.0141501230803 | 4.71044327849398 | 2.65275221492427e-05 | 0.0156234554115578 | 2.42837306008358 |
| ENSG00000150316.11 | CWC15 | 0.520405930625372 | 5.07342129506469 | 4.70513261916885 | 2.69848254365203e-05 | 0.0156234554115578 | 2.42437862568846 |
| lnc-GPLD1-2 | lnc-GPLD1-2 | -0.906746245654968 | -0.291902370707993 | -4.70410397215254 | 2.7074295248373e-05 | 0.0156234554115578 | 2.01818650231204 |
| lnc-LTC4S-2 | lnc-LTC4S-2 | -0.59239782232529 | 1.44711511504079 | -4.70123161076631 | 2.73256762773202e-05 | 0.0156234554115578 | 2.43098425750447 |
| ENSG00000135108.14 | FBXO21 | 0.513308143139258 | 6.46971353574817 | 4.64614221350067 | 3.26162643539293e-05 | 0.0181443397080251 | 2.23212027133994 |
| ENSG00000067596.11 | DHX8 | 0.41466530033156 | 6.53056110428398 | 4.62677627190643 | 3.47056725669692e-05 | 0.0182880594649202 | 2.17346247953486 |
| ENSG00000066117.14 | SMARCD1 | 0.341336649192562 | 6.02242784595084 | 4.62400940312778 | 3.50147370137714e-05 | 0.0182880594649202 | 2.16561968535252 |
| ENSG00000270722.1 | RNVU1-31 | -1.55759666532933 | 1.61456732818864 | -4.61366263895421 | 3.63157855661666e-05 | 0.0182880594649202 | 2.18448985537078 |
| lnc-GK2-2 | lnc-GK2-2 | 0.582324481640608 | 2.98927288103303 | 4.61156701244508 | 3.64383349028923e-05 | 0.0182880594649202 | 2.19352707332587 |
| lnc-NMU-3 | lnc-NMU-3 | -2.18875118088092 | 1.97596151593234 | -4.6165511669503 | 3.7317130521627e-05 | 0.0182880594649202 | 2.17220947534447 |
| LINC01572 | LINC01572 | -1.32860291398357 | 3.61653239678559 | -4.58064308662678 | 4.08239314822656e-05 | 0.0195413716674296 | 2.0712930616717 |
| ENSG00000107951.14 | MTPAP | 0.360284784802166 | 5.07717818363729 | 4.56812585671301 | 4.18688421453362e-05 | 0.0195860540426694 | 2.00894199841667 |
| LINC01829 | LINC01829 | -0.979747180077188 | 1.15252075427702 | -4.55203578962168 | 4.40756820325409e-05 | 0.0201602169616842 | 1.97594380983401 |
| ENSG00000137502.9 | RAB30 | 0.87017171341454 | 7.18196815292742 | 4.52596394403652 | 4.78962420813693e-05 | 0.0214314858861049 | 1.8714409344239 |
| ENSG00000204410.14 | MSH5 | -0.411742790040192 | 3.10652216803362 | -4.50552100379749 | 5.11175267812486e-05 | 0.0218006934561435 | 1.87647790985414 |
| ENSG00000141569.11 | TRIM65 | -0.707766084886708 | 4.03570805549053 | -4.49287376591219 | 5.32158149807279e-05 | 0.0218006934561435 | 1.81141420447043 |
| ENSG00000143924.18 | EML4 | 0.804834112372551 | 7.21299787415873 | 4.48622424096258 | 5.43526720500254e-05 | 0.0218006934561435 | 1.7532007882164 |
| ENSG00000136305.11 | CIDEB | -0.916919290594023 | 2.36435221352156 | -4.4795067540479 | 5.55253183738391e-05 | 0.0218006934561435 | 1.80962418470688 |
| LINC00494 | LINC00494 | -0.63475873065477 | 6.49456003247466 | -4.47854815679317 | 5.56946666175799e-05 | 0.0218006934561435 | 1.72801868399547 |
| ENSG00000109436.7 | TBC1D9 | 1.03402349352899 | 7.64555324554722 | 4.47957345540004 | 5.5786061715469e-05 | 0.0218006934561435 | 1.7313549675237 |
| ENSG00000129566.12 | TEP1 | 0.455190642059581 | 7.3043006926135 | 4.47606621916569 | 5.61354881783805e-05 | 0.0218006934561435 | 1.7229808111257 |
| ENSG00000099797.14 | TECR | -0.491641211079155 | 3.26929931837439 | -4.44678953494745 | 6.16006048869656e-05 | 0.0234006004417492 | 1.70067557243372 |
| ENSG00000073921.17 | PICALM | 0.534383373992648 | 6.23685157570667 | 4.44207080903465 | 6.25289328230194e-05 | 0.0234006004417492 | 1.61850825266301 |
| ENSG00000282413.1 | | -0.913309216537963 | 0.421564162190687 | -4.42066943873534 | 6.69142114643366e-05 | 0.0235623762091295 | 1.55392035628229 |
| lnc-FAM126B-1 | lnc-FAM126B-1 | -0.749372062404783 | 3.30071596852529 | -4.42039356385744 | 6.69726616999622e-05 | 0.0235623762091295 | 1.62132360479585 |
| lnc-ZNF395-2 | lnc-ZNF395-2 | -1.12362717698716 | 4.40414084731466 | -4.42071389153633 | 6.73691140225039e-05 | 0.0235623762091295 | 1.58163705722357 |
| lnc-ANTXR2-3 | lnc-ANTXR2-3 | 0.620273948560346 | 2.84323450227505 | 4.4158563329829 | 6.79411792905043e-05 | 0.0235623762091295 | 1.62286016531636 |
| ENSG00000199879.1 | RNVU1-22 | -1.80379652996679 | 2.76196541748931 | -4.4196178896769 | 6.94073612411029e-05 | 0.0235623762091295 | 1.60523982700113 |
| lnc-DHX38-4 | lnc-DHX38-4 | -0.817669149147343 | 0.961775397405172 | -4.40718840557876 | 6.98297113519 36e-05 | 0.0235623762091295 | 1.56927968387144 |
| ENSG00000140853.15 | NLRC5 | 0.815841061813994 | 7.66305024865437 | 4.39866149463671 | 7.17376312741482e-05 | 0.0238157365244483 | 1.49402366428897 |
| ENSG00000270103.3 | | -1.21792063730726 | 3.2295369199639 | -4.38833122220587 | 7.44617472754721e-05 | 0.0240671228321693 | 1.52616359994459 |

| | | | | | | | |
|---|---|---|---|---|---|---|---|
| lnc-RASGRP3-9 | lnc-RASGRP3-9 | -1.0018794579535 | 3.27682347336197 | -4.38166262474896 | 7.56946646894745e-05 | 0.0240671228321693 | 1.50872907038708 |
| MANEA-DT | MANEA-DT | -0.727382439061935 | 4.54238221567341 | -4.38037618772983 | 7.60026713351312e-05 | 0.0240671228321693 | 1.46127134731333 |
| lnc-TSTD3-1 | lnc-TSTD3-1 | -0.954031197818699 | 0.97833007000932 | -4.37046648355018 | 7.84167795962885e-05 | 0.0244431403201365 | 1.46027019739299 |
| ENSG00000060642.10 | PIGV | -0.579608239771347 | 3.34077679126839 | -4.35819210223227 | 8.15111381486372e-05 | 0.0244431403201365 | 1.43991797050721 |
| ENSG00000199377.1 | RNU5F-1 | -1.28841952801713 | 3.58850483956739 | -4.35706638205582 | 8.28875678771338e-05 | 0.0244431403201365 | 1.41638417083364 |
| ENSG00000284057.1 | | 0.51597359632868 | 4.31926676657624 | 4.3488057506764 | 8.39577156464426e-05 | 0.0244431403201365 | 1.3820628612818 |
| ENSG00000007944.14 | MYLIP | 0.496396526018679 | 5.81633627230815 | 4.34727970910574 | 8.4362212093407e-05 | 0.0244431403201365 | 1.33868007693876 |
| ENSG00000161267.11 | BDH1 | -0.624597125196617 | 2.83553335995156 | -4.34689136411002 | 8.44654516377299e-05 | 0.0244431403201365 | 1.41881679960967 |
| lnc-RPRML-3 | lnc-RPRML-3 | -0.494595518105833 | 5.03041011786169 | -4.33389487578374 | 8.799266322169e-05 | 0.0244431403201365 | 1.31200666351226 |
| lnc-ASPH-5 | lnc-ASPH-5 | 4.30472147101757 | -0.944857532242974 | 4.33278886008731 | 8.85449712688348e-05 | 0.0244431403201365 | -0.326100356797097 |
| LINC01703 | LINC01703 | -0.929101953433374 | 0.736399012539551 | -4.33135135568002 | 8.86996265855014e-05 | 0.0244431403201365 | 1.3348812673925 |
| ENSG00000119231.10 | SENP5 | 0.361304087721003 | 6.28590579754783 | 4.32817815291187 | 8.95893971987845e-05 | 0.0244431403201365 | 1.27958374463638 |
| lnc-ABCC6-11 | lnc-ABCC6-11 | 0.882451861735986 | 3.83846260255495 | 4.32581054842361 | 9.02589550868885e-05 | 0.0244431403201365 | 1.33422965178248 |
| lnc-ZSCAN1-3 | lnc-ZSCAN1-3 | -0.608621926677699 | 4.31514036673844 | -4.32167376043388 | 9.14405968347915e-05 | 0.0244431403201365 | 1.29934081049556 |
| ENSG00000184441.4 | | -0.672389268320023 | 2.89302342950751 | -4.3101443834308 | 9.4814242913382e-05 | 0.0249299299794387 | 1.31336488856435 |
| LINC00662 | LINC00662 | -0.439860424582122 | 6.13819265139827 | -4.30723690442655 | 9.56840338325635e-05 | 0.0249299299794387 | 1.21836553054292 |
| lnc-ZNF667-2 | lnc-ZNF667-2 | -1.57713278729656 | 0.456848361589311 | -4.29463445698441 | 9.95451389190348e-05 | 0.0256117199296312 | 1.15736257210778 |
| lnc-ZEB1-9 | lnc-ZEB1-9 | -1.98513467564804 | -0.749576233233451 | -4.2835829562597 | 0.000103056030228004 | 0.0260371136013675 | 0.417935776168973 |
| lnc-CHSY1-5 | lnc-CHSY1-5 | -1.10819530019363 | 0.304867156753758 | -4.28150826582653 | 0.00010372848055736 | 0.0260371136013675 | 1.09409309026257 |
| ENSG00000142512.14 | SIGLEC10 | -0.896631778180827 | 5.12754460022112 | -4.26667761372714 | 0.000108661790835092 | 0.0269468149489001 | 1.11335023867523 |
| ENSG00000101367.8 | MAPRE1 | 0.445517255307188 | 6.12022786287108 | 4.26093708909143 | 0.00011063221353033 | 0.026996950798481 | 1.08174862449786 |
| ENSG00000163297.16 | ANTXR2 | 0.550299388882131 | 5.517312364456 | 4.25692067829205 | 0.000112031535637415 | 0.026996950798481 | 1.07611294662354 |
| ENSG00000197863.8 | ZNF790 | -0.570739882333271 | 3.2241097083584 | -4.25324578483019 | 0.000113327000360612 | 0.026996950798481 | 1.14166967762983 |
| lnc-SIRT2-1 | lnc-SIRT2-1 | -0.958331699591974 | 1.24268630168124 | -4.25104328304103 | 0.00011411041730884 | 0.026996950798481 | 1.14634616632999 |
| CCDC144NL-AS1 | CCDC144NL-AS1 | -1.28542764748481 | 4.26490049240231 | -4.25134876365397 | 0.000116116611388274 | 0.0271594115023278 | 1.08342879888819 |
| ENSG00000266009.1 | | -1.26232893668956 | 0.446046810240421 | -4.23723349072017 | 0.000119144337458437 | 0.0275453814859681 | 1.02528169498676 |
| ENSG00000284697.1 | | 0.570323618920743 | 2.40314071835309 | 4.23325855204091 | 0.000120633007368107 | 0.0275453814859681 | 1.10518198013815 |
| ENSG00000107819.13 | SFXN3 | -0.668362602857183 | 4.28063888175022 | -4.23022436235771 | 0.000121781553477292 | 0.0275453814859681 | 1.03114604662749 |
| lnc-SH3BGRL-5 | lnc-SH3BGRL-5 | -0.899586312364968 | 0.632907798216022 | -4.22522631056986 | 0.000123696792694135 | 0.027674468304602 | 1.03407487560753 |
| lnc-GCNT3-1 | lnc-GCNT3-1 | -0.785266815832141 | 0.775184384837642 | -4.21931779757105 | 0.000125998840775529 | 0.0278499774675373 | 1.03098983581711 |
| ENSG00000135655.15 | USP15 | 0.419542603867463 | 7.85128159459362 | 4.21630834319695 | 0.000127187381914614 | 0.0278499774675373 | 0.958851499070531 |
| SNAI3-AS1 | SNAI3-AS1 | -0.825326083170053 | 1.97027169998409 | -4.21048366995628 | 0.000129518898874163 | 0.0278771601200499 | 1.04371137350213 |
| ENSG00000198840.2 | MT-ND3 | -1.25316098724654 | 7.14015830275038 | -4.2154760056469 | 0.000130597386117255 | 0.0278771601200499 | 0.934652538863119 |
| ENSG00000068650.18 | ATP11A | 0.726421515291145 | 6.00165182555064 | 4.20592019069358 | 0.000131374655377974 | 0.0278771601200499 | 0.921784416851467 |
| ENSG00000259768.5 | ZFHX3-AS1 | -0.74933008749457 | 2.02328853858617 | -4.19314396213127 | 0.00013670915193575 | 0.0284380813405135 | 0.994287938057556 |
| lnc-GRM1-1 | lnc-GRM1-1 | -0.717601169931875 | 2.59164171109452 | -4.19297445588862 | 0.000136781326954809 | 0.0284380813405135 | 0.984131451604568 |
| ENSG00000163155.11 | LYSMD1 | 2.54571515463834 | -0.619110930500723 | 4.18091776006152 | 0.000142011429984246 | 0.0292302126336574 | 0.119540950458802 |
| ENSG00000134324.11 | LPIN1 | 1.54495883298453 | 2.76123490439383 | 4.18023730359476 | 0.000145094935713522 | 0.0293009789728854 | 0.931103345647588 |
| lnc-SGSH-1 | lnc-SGSH-1 | -0.890832679642947 | 2.28284609169734 | -4.17377273117954 | 0.00014520234442182 | 0.0293009789728854 | 0.937925462185954 |
| lnc-RIMS3-4 | lnc-RIMS3-4 | -0.68890512883576 | 2.30553687869119 | -4.16923055185124 | 0.000147267062245982 | 0.0294291062350392 | 0.923922476102237 |
| lnc-SOD1-9 | lnc-SOD1-9 | -1.13937541324959 | 1.48623959928074 | -4.16592699261677 | 0.000148786688940072 | 0.0294468886389759 | 0.917414027176622 |
| ENSG00000244627.5 | TPTEP2 | -0.395434925635753 | 3.77237417659237 | -4.15223087127747 | 0.000155251472020667 | 0.0304337247723941 | 0.824424102538196 |
| LINC01341 | LINC01341 | -1.24357028853209 | 0.790709261047993 | -4.14085196362267 | 0.000160829888007995 | 0.0312298262723449 | 0.823767948644798 |
| ENSG00000226174.6 | TEX22 | -0.531365997025796 | 1.96408452495382 | -4.13472111608269 | 0.000163915916368419 | 0.031531600996366 | 0.831606024643489 |
| ENSG00000223509.8 | WHAMMP1 | 0.619092870786583 | 4.17560391400895 | 4.1216236591049 | 0.000170703053200615 | 0.0322660630247605 | 0.723613775473158 |
| lnc-KCNMB1-4 | lnc-KCNMB1-4 | -2.26482009282174 | 0.172468862325609 | -4.12140968309389 | 0.000170869206126361 | 0.0322660630247605 | 0.483461015266373 |
| ENSG00000058335.15 | RASGRF1 | -0.912005196665256 | 6.05992355338566 | -4.11852488144377 | 0.000172752542046123 | 0.0323251415721395 | 0.665596418400944 |
| lnc-RIOK2-7 | lnc-RIOK2-7 | 0.655728795172585 | 1.95004074307131 | 4.1134865528557 | 0.000175056685839761 | 0.0324007531588585 | 0.773353492758838 |
| ENSG00000044090.8 | CUL7 | -0.415672191082972 | 4.75729644127997 | -4.1058072827035 | 0.000179264369473595 | 0.0324007531588585 | 0.651787879776609 |
| ENSG00000143315.7 | PIGM | -0.4964897484855 | 2.85895963361603 | -4.1015937080654 | 0.000181614751800764 | 0.0324007531588585 | 0.717130895985704 |
| ENSG00000273768.1 | RNVU1-29 | -1.21060591073111 | 2.82757896726101 | -4.09770690995819 | 0.000184662930726863 | 0.0324007531588585 | 0.70081613905546 |
| lnc-BRF1-12 | lnc-BRF1-12 | -0.703089800238605 | 8.36739814374266 | -4.09616869866354 | 0.000184685079890865 | 0.0324007531588585 | 0.615458108141749 |
| ENSG00000274210.1 | RNVU1-27 | -1.22627937700761 | 4.53043122501367 | -4.1006100748155 | 0.00018549482341393 | 0.0324007531588585 | 0.631185604914131 |
| ENSG00000082153.17 | BZW1 | 0.391624233209161 | 6.87164056450234 | 4.09241034926187 | 0.000186841698655644 | 0.0324007531588585 | 0.590761089945555 |
| lnc-SPATA21-4 | lnc-SPATA21-4 | -0.640212845946332 | 4.65735059944617 | -4.0919293792511 | 0.000187119451316462 | 0.0324007531588585 | 0.617009799219237 |
| ENSG00000121073.14 | SLC35B1 | 0.389657644229405 | 4.03783762255989 | 4.09157563902915 | 0.000187323987072057 | 0.0324007531588585 | 0.640468367350461 |
| ENSG00000134508.12 | CABLES1 | -2.58645862586162 | 2.72226276312976 | -4.10139561008549 | 0.000189684470431813 | 0.0324928237909104 | 0.698569860754175 |
| ENSG00000068878.14 | PSME4 | 0.707955090076009 | 6.32263789190855 | 4.08001254581043 | 0.000194130928200306 | 0.0324928237909104 | 0.553858770262972 |
| ENSG00000090520.11 | DNAJB11 | 0.519400629043736 | 5.14636270514775 | 4.07940078101517 | 0.000194497683846378 | 0.0324928237909104 | 0.565971764522473 |
| ENSG00000119760.15 | SUPT7L | 0.378340024308367 | 6.00217904762888 | 4.07737609561072 | 0.000195716303478868 | 0.0324928237909104 | 0.546570207373478 |
| ENSG00000146856.14 | AGBL3 | -0.984053189443649 | 3.10848934251055 | -4.07532088417922 | 0.000196960889188055 | 0.0324928237909104 | 0.628724581328389 |
| lnc-DTNBP1-14 | lnc-DTNBP1-14 | -1.98935597677936 | 0.890247648736039 | -4.07787483663071 | 0.00019732803643122 | 0.0324928237909104 | 0.61055056743156 |
| ENSG00000139620.12 | KANSL2 | 0.366785800928317 | 5.64564448624562 | 4.07053202338947 | 0.000199890840507509 | 0.0326535965886194 | 0.530275983027166 |
| ENSG00000042429.11 | MED17 | 0.510285674255664 | 5.57434412382252 | 4.06294178355155 | 0.000204621769035841 | 0.0330161239084921 | 0.509811813322623 |
| ENSG00000135763.9 | URB2 | 0.487977846029041 | 4.29633388556581 | 4.05948573103984 | 0.00020681179514181 | 0.0330161239084921 | 0.53983873096289 |
| ENSG00000204310.12 | AGPAT1 | -0.542875006833597 | 3.31511677330388 | -4.05931241987874 | 0.000206922216596 | 0.0330161239084921 | 0.578952622805621 |
| ENSG00000170185.9 | USP38 | 0.365109023688041 | 5.95947026524371 | 4.04857916219653 | 0.000213873502981077 | 0.0332808680822343 | 0.463875319360281 |
| ENSG00000144747.16 | TMF1 | 0.42873032677393 | 6.76546140908432 | 4.0484183989204 | 0.00021397932568253 | 0.0332808680822343 | 0.462951380400285 |

| | | | | | | | |
|---|---|---|---|---|---|---|---|
| ENSG00000213760.10 | ATP6V1G2 | -0.811472001706718 | -0.11821623381711 | -4.04839762322393 | 0.000213993005032769 | 0.0332808680822343 | 0.443966799676126 |
| lnc-IDE-3 | lnc-IDE-3 | -0.583317463391261 | 4.41085803113249 | -4.04679756116314 | 0.000215049091723129 | 0.0332808680822343 | 0.494706921316669 |
| ENSG00000246273.7 | SBF2-AS1 | -0.711185448847298 | 4.2830339322032 | -4.04181457377554 | 0.000218370556373216 | 0.0332959264945589 | 0.485749188260028 |
| lnc-ID3-4 | lnc-ID3-4 | -1.2260509155619 | -0.599657068955431 | -4.04179801284026 | 0.000218381677926709 | 0.0332959264945589 | 0.186762715179324 |
| ENSG00000160602.13 | NEK8 | -0.470001874067321 | 4.930010130558 | -4.03077000974969 | 0.000225911087664903 | 0.034106308750048 | 0.430792796368492 |
| lnc-PKDCC-6 | lnc-PKDCC-6 | 1.08155573058532 | 0.567364321123898 | 4.02820066197662 | 0.000227701213669309 | 0.034106308750048 | 0.492177829107338 |
| ENSG00000164366.3 | CCDC127 | -0.568443510972216 | 3.10341806612437 | -4.02682138925367 | 0.000228667862192422 | 0.034106308750048 | 0.495261835312522 |
| ENSG00000106351.12 | AGFG2 | -0.664505824281932 | 4.05856996056795 | -4.01910048446332 | 0.000234153028494656 | 0.0344969588266439 | 0.433617947934479 |
| lnc-PRR7-4 | lnc-PRR7-4 | -0.751521178620907 | 0.0928456909510764 | -4.00778235587932 | 0.00024242537586797 | 0.0344969588266439 | 0.375314154667637 |
| ENSG00000120705.12 | ETF1 | 0.546257436348975 | 6.20002608481568 | 4.00664352625342 | 0.000243273282973321 | 0.0344969588266439 | 0.342591128863223 |
| ENSG00000284360.1 | MIR6125 | 0.487273680404242 | 1.90281783255366 | 4.00274814246781 | 0.00024619538619147 | 0.0344969588266439 | 0.467467769183636 |
| lnc-SLC10A5-2 | lnc-SLC10A5-2 | 0.556677621266408 | 1.19627528374714 | 4.00200332557839 | 0.000246757974540482 | 0.0344969588266439 | 0.463208080421392 |
| ENSG00000004139.13 | SARM1 | -0.430651900071561 | 5.18758905607734 | -4.0016104865017 | 0.000247055203273991 | 0.0344969588266439 | 0.340562166650959 |
| ENSG00000020256.19 | ZFP64 | -0.708603124976179 | 4.22035135350874 | -4.00151814415155 | 0.00024712512150548 | 0.0344969588266439 | 0.372469735759096 |
| ENSG00000132967.9 | HMGB1P5 | 1.23324236492458 | 1.65883091940399 | 4.00144526553995 | 0.000247180316072546 | 0.0344969588266439 | 0.465126656602774 |
| ENSG00000115816.14 | CEBPZ | 0.430446479156958 | 6.3280003042627 | 4.00093053784338 | 0.000247570485641393 | 0.0344969588266439 | 0.325773033952904 |
| ENSG00000132199.19 | ENOSF1 | -3.92002479976903 | 2.43544584413282 | -4.011394900243 | 0.000250620754887648 | 0.0344969588266439 | 0.458559035080246 |
| lnc-ARID2-8 | lnc-ARID2-8 | -1.03722137959199 | 0.510855970753705 | -3.99267217019144 | 0.000253912776682256 | 0.0344969588266439 | 0.412663222523753 |
| LINC00565 | LINC00565 | 1.3281270731862 | 0.611573361232409 | 3.99118394097241 | 0.000255072361848979 | 0.0344969588266439 | 0.331556623834202 |
| ENSG00000164241.13 | C5orf63 | -0.609173919347509 | 3.57572555548565 | -3.99046094530472 | 0.000255637552575656 | 0.0344969588266439 | 0.369091593565408 |
| ENSG00000110876.9 | SELPLG | -1.17535264631296 | 3.76207056128618 | -3.99296534325698 | 0.000256324105158045 | 0.0344969588266439 | 0.365293336459351 |
| ENSG00000154124.4 | OTULIN | 0.428601214331884 | 6.04878990404835 | 3.98945381029099 | 0.000256426891146894 | 0.0344969588266439 | 0.293844349353055 |
| lnc-CLEC19A-3 | lnc-CLEC19A-3 | -1.57721827556122 | 0.854253951443914 | -3.98451584770816 | 0.000260331358086097 | 0.0345439867144076 | 0.379140167007696 |
| ENSG00000156469.8 | MTERF3 | 0.423536892250374 | 3.17226844735102 | 3.98190071918788 | 0.000262422455931794 | 0.0345439867144076 | 0.373970725437558 |
| ENSG00000135925.8 | WNT10A | -1.06842457628976 | 2.69591885221662 | -3.97576446773039 | 0.000267393289628402 | 0.0345439867144076 | 0.366926605585552 |
| ENSG00000184465.16 | WDR27 | -0.543989702104854 | 5.05824725898527 | -3.97457489271127 | 0.000268367452351407 | 0.0345439867144076 | 0.265602514500268 |
| ENSG00000107949.16 | BCCIP | 0.273274869653901 | 4.83467680725465 | 3.97353137222194 | 0.000269224842950815 | 0.0345439867144076 | 0.270699605560897 |
| ENSG00000049656.13 | CLPTM1L | 0.508161333567702 | 5.18660422677654 | 3.96923190116442 | 0.00027278554442632 | 0.0345439867144076 | 0.250496357378008 |
| lnc-UPRT-6 | lnc-UPRT-6 | -4.27105748081297 | -1.48898288463873 | -3.97196563821471 | 0.000273130233962464 | 0.0345439867144076 | -0.817619869851515 |
| lnc-VN1R2-5 | lnc-VN1R2-5 | -0.791472071050003 | 4.04760264816444 | -3.96794646031709 | 0.000273858952420944 | 0.0345439867144076 | 0.288145397781149 |
| ENSG00000132912.12 | DCTN4 | 0.367265771197707 | 5.41016850874201 | 3.96764233748184 | 0.000274113509068688 | 0.0345439867144076 | 0.239199037504473 |
| ENSG00000165219.21 | GAPVD1 | 0.347441166051289 | 6.34011115961661 | 3.9653739879808 | 0.000276019414733792 | 0.0345439867144076 | 0.223944671046582 |
| lnc-TMEM234-2 | lnc-TMEM234-2 | -0.54589725585406 | 1.12063482887853 | -3.96218761801236 | 0.000278718380617217 | 0.0345439867144076 | 0.358389769823725 |
| ENSG00000137692.11 | DCUN1D5 | 0.556907061441144 | 2.6037502634128 | 3.95936999040565 | 0.000281126319804676 | 0.0345439867144076 | 0.330870942802149 |
| ENSG00000121104.7 | FAM117A | -0.63641206719168 | 5.12172480953337 | -3.95785541172273 | 0.000282429000653134 | 0.0345439867144076 | 0.217792779887679 |
| ENSG00000198873.11 | GRK5 | -0.901836449434946 | 4.00473324584691 | -3.95339834352637 | 0.000286296552070356 | 0.0345439867144076 | 0.247639865810381 |
| ENSG00000198585.11 | NUDT16 | -0.393907618287199 | 4.059358161921 | -3.95001376909822 | 0.000289267721140754 | 0.0345439867144076 | 0.23408621904789 |
| ENSG00000137714.2 | FDX1 | 0.804864632834813 | 3.08688676410929 | 3.949467330059 | 0.00028975021065372 | 0.0345439867144076 | 0.290374394357942 |
| lnc-PRCP-1 | lnc-PRCP-1 | 0.74336912986509 | 3.37992199072367 | 3.94793421018282 | 0.000291108079659885 | 0.0345439867144076 | 0.273239002522899 |
| SGMS1-AS1 | SGMS1-AS1 | -0.397293066965089 | 5.63474713347394 | -3.94409047132343 | 0.000294539621213666 | 0.0345439867144076 | 0.168456340740338 |
| ENSG00000177042.14 | TMEM80 | -0.536182360284792 | 4.13083111263063 | -3.94327125109435 | 0.000295276042741445 | 0.0345439867144076 | 0.211317540992185 |
| lnc-FUBP1-3 | lnc-FUBP1-3 | -1.57689269843052 | 3.79193840194057 | -3.95104208090954 | 0.000296461343629235 | 0.0345439867144076 | 0.230923673833347 |
| lnc-PCDHGA1-2 | lnc-PCDHGA1-2 | -1.82431952774734 | -0.211363600891305 | -3.94105970096383 | 0.000297272995828328 | 0.0345439867144076 | 0.126965897641439 |
| ENSG00000143753.12 | DEGS1 | 0.467919164476992 | 4.8140530018628 | 3.94087767044136 | 0.000297437944983446 | 0.0345439867144076 | 0.180121423864797 |
| KCTD21-AS1 | KCTD21-AS1 | -1.64643723246241 | 0.375364599317114 | -3.9388364842403 | 0.000299293676679171 | 0.0345439867144076 | 0.178620613153526 |
| ENSG00000230124.7 | ACBD6 | -0.349550559353789 | 5.16740498515301 | -3.93735600351525 | 0.000300646663504521 | 0.0345439867144076 | 0.157330955854794 |
| lnc-RAB23-53 | lnc-RAB23-53 | -0.414114679501763 | 3.51722707985307 | -3.9370926713758 | 0.000300887938624602 | 0.0345439867144076 | 0.226982427142971 |
| ENSG00000224775.2 | BRAFP1 | -2.61629423080843 | -1.48482757449348 | -3.9361874291321 | 0.000301718789073184 | 0.0345439867144076 | -0.538762718447757 |
| ENSG00000261147.1 | | 0.399553915768968 | 3.80232914086975 | 3.93578380726149 | 0.000302089958149607 | 0.0345439867144076 | 0.213902590665488 |
| ENSG00000207205.1 | RNVU1-15 | -1.28965285686134 | 4.10526119345364 | -3.9243291101691 | 0.000318881324211105 | 0.0357015471634708 | 0.151010240245228 |
| lnc-SIKE1-2 | lnc-SIKE1-2 | 0.358462600089296 | 5.19038958046756 | 3.91734570092051 | 0.000319525580841233 | 0.0357015471634708 | 0.10149392672878 |
| ENSG00000129315.10 | CCNT1 | 0.367359336262398 | 6.39694796762872 | 3.91630258443796 | 0.000320540592499715 | 0.0357015471634708 | 0.084180801390473 |
| ENSG00000111786.8 | SRSF9 | 0.330444781816564 | 5.96214310867485 | 3.91565098060259 | 0.000321176224440363 | 0.0357015471634708 | 0.0833098619782007 |
| ENSG00000169129.14 | AFAP1L2 | -0.889315846018799 | 1.41906014010146 | -3.91463512801696 | 0.000322169613274964 | 0.0357015471634708 | 0.230464674278348 |
| lnc-AMZ2-7 | lnc-AMZ2-7 | -0.917277285843607 | 1.43906972876289 | -3.91384779929118 | 0.000322941578457312 | 0.0357015471634708 | 0.228878858011696 |
| lnc-SBDS-4 | lnc-SBDS-4 | -0.48626044371595 | 3.01310304010014 | -3.91241143687483 | 0.000324354531388478 | 0.0357015471634708 | 0.182591881246664 |
| ENSG00000115946.7 | PNO1 | 0.535139196802167 | 4.31883669576871 | 3.90965622808 | 0.000327081615820428 | 0.0358102175448503 | 0.113343587506884 |
| lnc-LGMN-2 | lnc-LGMN-2 | -1.38173104080445 | 2.06566831528036 | -3.90849702994894 | 0.000331135681722804 | 0.0358929932364605 | 0.195273546978449 |
| lnc-C12orf49-8 | lnc-C12orf49-8 | 0.741451260501217 | 2.34146014223373 | 3.90517347497713 | 0.000331566138622098 | 0.0358929932364605 | 0.189472316648804 |
| lnc-SNX10-2 | lnc-SNX10-2 | -1.03015452208526 | 1.4101193238266 | -3.89966292366292 | 0.00033716050593673 | 0.0358929932364605 | 0.190623231303694 |
| ENSG00000062650.18 | WAPL | 0.238380987544858 | 6.86666320805909 | 3.8984881200552 | 0.000338364944907244 | 0.0358929932364605 | 0.034958708957233 |
| TNFRSF14-AS1 | TNFRSF14-AS1 | -0.816042972937792 | 2.85132864310883 | -3.89797140572856 | 0.000338896012083005 | 0.0358929932364605 | 0.13979452724209 |
| ENSG00000160055.19 | TMEM234 | -0.453445043065769 | 3.42179137620621 | -3.8975714591709 | 0.000339307622073517 | 0.0358929932364605 | 0.121146474005546 |
| ENSG00000090447.11 | TFAP4 | -0.801116090114414 | 2.59435683162579 | -3.89607104531653 | 0.000340856110667456 | 0.0358929932364605 | 0.150997931937133 |
| ENSG00000253251.2 | SHLD3 | 0.491627153315623 | 3.20663300981076 | 3.89381135866326 | 0.000343201120317513 | 0.0358929932364605 | 0.123764041539641 |
| lnc-GLB1-2 | lnc-GLB1-2 | -1.33707425450172 | -0.652202321930147 | -3.89131367244641 | 0.000345811291560188 | 0.0358929932364605 | -0.197508657693574 |
| ENSG00000101452.14 | DHX35 | 0.430346262926999 | 3.96862159081529 | 3.89054117644179 | 0.000346622461655303 | 0.0358929932364605 | 0.0765885576530874 |
| lnc-HK3-2 | lnc-HK3-2 | 0.396014616062585 | 3.3231555747853 | 3.89016355376984 | 0.000347019659624721 | 0.0358929932364605 | 0.1120166357743 |

| | | | | | | | |
|---|---|---|---|---|---|---|---|
| ENSG00000134248.13 | LAMTOR5 | 0.316304109329468 | 5.32689570902695 | 3.88433034430506 | 0.000353211515593064 | 0.0362072128027118 | 0.00385303453365182 |
| ENSG00000285646.1 | | -1.27966384642092 | 3.62710545398964 | -3.8891233949 | 0.000353575755397419 | 0.0362072128027118 | 0.0753422207058172 |
| ENSG00000178878.12 | APOLD1 | 0.385184418410676 | 4.45681518936817 | 3.87971884847946 | 0.000358182106648974 | 0.0362212946242184 | 0.0207288986132745 |
| ENSG00000170634.12 | ACYP2 | 0.413932718786715 | 3.16061071574253 | 3.87818099918868 | 0.000359854711057478 | 0.0362212946242184 | 0.0855989525139371 |
| ENSG00000254685.6 | FPGT | -0.6177901979635 | 3.15962787394387 | -3.87780679098565 | 0.000360262851639598 | 0.0362212946242184 | 0.0804214887613792 |
| ENSG00000243477.5 | NAA80 | -0.669112725151508 | 2.6330877939023 | -3.87652614811692 | 0.000361663013396535 | 0.0362212946242184 | 0.094488873934961 |
| ENSG00000132274.15 | TRIM22 | 0.511791421572402 | 8.27005145631771 | 3.87439937740161 | 0.00036399991321343 | 0.0362212946242184 | -0.0196790990597977 |
| ENSG00000123213.22 | NLN | -0.864711283436146 | 3.90665066191112 | -3.87415272946799 | 0.000364271874226945 | 0.0362212946242184 | 0.0292011066137325 |
| ENSG00000141452.9 | RMC1 | 0.469430238269108 | 5.54296528797955 | 3.86990845104023 | 0.000368982656861688 | 0.0363459393522763 | -0.0406694440626003 |
| ENSG00000112304.10 | ACOT13 | -0.712131460303322 | 2.40555123985413 | -3.86984184362448 | 0.000369057053132475 | 0.0363459393522763 | 0.0888639860928091 |
| ENSG00000124226.10 | RNF114 | 0.455426561976379 | 5.93236327849367 | 3.86579485938226 | 0.000373604612887706 | 0.0366173030507164 | -0.0572731271716425 |
| lnc-ACO2-1 | lnc-ACO2-1 | -0.828431503606183 | 1.37311264663264 | -3.86423619078816 | 0.000375370497191914 | 0.0366173030507164 | 0.0954068489828908 |
| ENSG00000107937.18 | GTPBP4 | 0.379703331225155 | 5.46796585878247 | 3.86240076246994 | 0.000377460289499943 | 0.0366474770697044 | -0.0604303423834178 |
| ENSG00000176531.10 | PHLDB3 | -0.595115063701531 | 2.86486993870911 | -3.85708063138885 | 0.000383581508363269 | 0.0370118931794641 | 0.0326419429355003 |
| ENSG00000174574.15 | AKIRIN1 | 0.424309159336628 | 5.51062052590243 | 3.85602277712056 | 0.000384810044230935 | 0.0370118931794641 | -0.0798734893122957 |
| lnc-CTSH-1 | lnc-CTSH-1 | -1.08111613745004 | 1.87171293873705 | -3.85234852831848 | 0.000389106759305447 | 0.0372510903571349 | 0.0549818876756252 |
| ENSG00000178409.13 | BEND3 | -0.965032198197109 | 2.22928650179989 | -3.8481788147848 | 0.000394039094104185 | 0.0375486420090113 | 0.0330742274292595 |
| ENSG00000144711.15 | IQSEC1 | 1.06222874107772 | 4.88984656543097 | 3.84681996061567 | 0.000399864341513921 | 0.0379281462736453 | -0.0925593402784237 |
| ENSG00000259332.3 | ST20-MTHFS | -0.48086967633494 | 1.11666598618573 | -3.83730367431026 | 0.000407189424402237 | 0.0384457794608773 | 0.0245203311712059 |
| lnc-ZNF66-2 | lnc-ZNF66-2 | -0.851455048385196 | 1.3853712961899 | -3.83524824751887 | 0.000409722100037536 | 0.038508264772021 | 0.0179904169000373 |
| ENSG00000255152.8 | MSH5-SAPCD1 | -0.374213713701145 | 2.96716111010127 | -3.83269259203541 | 0.000412892398091163 | 0.0386121933181995 | -0.0380649829779234 |
| ENSG00000026559.13 | KCNG1 | -3.05692430319295 | -0.862042835735792 | -3.83235559005119 | 0.000414579736837297 | 0.0386121933181995 | -0.632642077369097 |
| ENSG00000122068.12 | FYTTD1 | 0.635983210607995 | 6.17764492145398 | 3.82422509596722 | 0.000423566691116682 | 0.0392715009155616 | -0.175310411147797 |
| ENSG00000197780.9 | TAF13 | 0.625538746717999 | 2.5194736262004 | 3.81569651725184 | 0.000434587372565243 | 0.0400092666200733 | -0.0615605576203491 |
| ENSG00000025434.18 | NR1H3 | -0.849911133212726 | -0.200513643286592 | -3.81401065349806 | 0.000436798417400725 | 0.0400092666200733 | -0.179195507231669 |
| ENSG00000151779.12 | NBAS | 0.352096323536193 | 6.43057302407761 | 3.8135873137242 | 0.000437355341277583 | 0.0400092666200733 | -0.205814049670487 |
| ENSG00000076351.12 | SLC46A1 | -1.13341195902574 | 1.41837181397252 | -3.80444273209838 | 0.000449554312166362 | 0.04094325843947 | -0.0640995056186258 |
| ENSG00000072840.12 | EVC | -4.47743242865904 | 0.9337592009348 | -3.81229703686788 | 0.000455378123869377 | 0.0412909600158739 | -0.114280473241024 |
| ENSG00000275993.2 | | 1.38606275054895 | 2.58280003135904 | 3.80194819610858 | 0.000458418967544161 | 0.0413061083349645 | -0.10975331470515 |
| ENSG00000089505.17 | CMTM1 | -1.06118460045059 | 0.4246139313593 | -3.79712034306124 | 0.000459558801375255 | 0.0413061083349645 | -0.0994839453801779 |
| ENSG00000123091.4 | RNF11 | 0.645332074842892 | 4.10530148325909 | 3.79333317400608 | 0.000464817189126204 | 0.0413964912241166 | -0.201678777180916 |
| ENSG00000132664.11 | POLR3F | 0.380676245652717 | 3.42602463903377 | 3.79312805531394 | 0.000465103643777735 | 0.0413964912241166 | -0.167271296481402 |
| ENSG00000150991.14 | UBC | 0.517119564243542 | 9.42763548989742 | 3.79122780236775 | 0.000467765524045774 | 0.0413964912241166 | -0.234691150119343 |
| ENSG00000138074.14 | SLC5A6 | 0.603286417185927 | 4.72314778509338 | 3.79062772551182 | 0.00046860916558418 | 0.0413964912241166 | -0.236007192958841 |
| ENSG00000188906.15 | LRRK2 | 0.668088347195527 | 7.49312377296229 | 3.78534859455178 | 0.000476094627581219 | 0.0418360177141106 | -0.279881431500588 |
| LINC01762 | LINC01762 | -0.869717240494186 | 1.81983753926011 | -3.7838654096674 | 0.000478218375762483 | 0.0418360177141106 | -0.124806983459569 |
| ENSG00000280128.1 | | -0.686562781981386 | 2.65752049719342 | -3.78284675132786 | 0.000479682270831759 | 0.0418360177141106 | -0.157703467724911 |
| ENSG00000114423.20 | CBLB | 0.890045728649546 | 6.69502010516067 | 3.78217672339826 | 0.000482717192849694 | 0.0419230716473639 | -0.295998398266039 |
| ENSG00000188419.13 | CHM | -0.338371492096316 | 4.36152666249099 | -3.77610581888255 | 0.000489479144655012 | 0.0420993941274255 | -0.266338881736957 |
| ENSG00000145476.15 | CYP4V2 | -0.678273529577146 | 4.50804034009828 | -3.77609289852536 | 0.000489498106504396 | 0.0420993941274255 | -0.274505993382228 |
| lnc-PRPF18-14 | lnc-PRPF18-14 | -1.94888403310734 | 1.65030918598332 | -3.7828617003859 | 0.000490883476197936 | 0.0420993941274255 | -0.143618619390455 |
| lnc-UBTF-4 | lnc-UBTF-4 | 0.457129986929972 | 2.05119852113441 | 3.77374736163241 | 0.000492952177240578 | 0.0421013886478955 | -0.157950873857237 |
| lnc-TMEM234-1 | lnc-TMEM234-1 | -0.617946820639276 | 2.48435680581931 | -3.7713502492031 | 0.000496506499919081 | 0.0422297243298944 | -0.183313931267425 |
| lnc-CSNK1E-1 | lnc-CSNK1E-1 | -0.356845584315175 | 3.62415329979452 | -3.76783007238311 | 0.000501770913153812 | 0.0423585478870216 | -0.253341666100104 |
| lnc-ZNF583-4 | lnc-ZNF583-4 | -1.50098942120748 | 1.14863953512373 | -3.7671845162641 | 0.000504378021248837 | 0.0423585478870216 | -0.164368198505963 |
| ENSG00000102383.13 | ZDHHC15 | -3.56872661991742 | -1.67103035040913 | -3.76615566846594 | 0.000506056551975168 | 0.0423585478870216 | -1.08328140888645 |
| ENSG00000271503.5 | CCL5 | -1.89827299781982 | -0.512858363179263 | -3.76486108100654 | 0.000506252867910767 | 0.0423585478870216 | -0.41121058743455 |
| lnc-DTNBP1-15 | lnc-DTNBP1-15 | -1.26230126810254 | 3.92626596946562 | -3.7647767445819 | 0.00051416030555137 | 0.0426201609084801 | -0.271786950475932 |
| ENSG00000124788.18 | ATXN1 | -1.49423656790565 | 6.75008544579054 | -3.76716294483202 | 0.000516486031243619 | 0.0426201609084801 | -0.350901392619944 |
| ENSG00000089818.17 | NECAP1 | 0.422577401577937 | 5.57221890055213 | 3.75809864388429 | 0.000516606021969449 | 0.0426201609084801 | -0.35542501714465 |
| lnc-NIT2-1 | lnc-NIT2-1 | -0.491726876211152 | 0.271929193889761 | -3.75546692565582 | 0.000520690094710449 | 0.0426201609084801 | -0.221886816456402 |
| lnc-CTSH-2 | lnc-CTSH-2 | -1.02924908046612 | 1.79055389353759 | -3.75453416358251 | 0.000522145078279619 | 0.0426201609084801 | -0.205088030749689 |
| lnc-ACTR8-3 | lnc-ACTR8-3 | -1.67723358435111 | 0.0358917737562435 | -3.75412445215199 | 0.000522785412088637 | 0.0426201609084801 | -0.297223849209551 |
| ENSG00000187601.4 | MAGEH1 | -0.732955694751745 | 2.56450470602186 | -3.75342897217829 | 0.000523874105322133 | 0.0426201609084801 | -0.239062043741952 |
| ENSG00000139180.10 | NDUFA9 | 0.388355998174702 | 4.64058261279927 | 3.74588939323219 | 0.000535817682822992 | 0.0434202179745892 | -0.361708279133905 |
| ENSG00000135776.4 | ABCB10 | -0.597808577682076 | 5.68547037496473 | -3.7428010322879 | 0.000540785454460838 | 0.0436509294477154 | -0.399187107387824 |
| ENSG00000135778.11 | NTPCR | -0.475791193803872 | 4.18508707728211 | -3.7407441111775 | 0.000544118718009424 | 0.043748420206203 | -0.357562367259737 |
| lnc-SYTL2-3 | lnc-SYTL2-3 | 0.536695901093741 | 2.13668592246615 | 3.73800731990533 | 0.000548584447351236 | 0.0437507428387015 | -0.254816110176178 |
| ENSG00000119684.15 | MLH3 | -0.427112432960772 | 4.88372633934298 | -3.73793530274136 | 0.000548702436479088 | 0.0437507428387015 | -0.39378008096399 |
| ENSG00000002586.19 | CD99 | -0.787509911806033 | 5.54065432582003 | -3.73620358019336 | 0.000551546978826992 | 0.0437507428387015 | -0.413755208102026 |
| ENSG00000212443.1 | SNORA53 | 0.815879913844455 | 0.65339034469715 | 3.73553444377161 | 0.000552649911969217 | 0.0437507428387015 | -0.248131959190494 |
| ENSG00000249456.1 | | 0.472867133683776 | 2.69353059484805 | 3.73384254864397 | 0.000555448145512329 | 0.043803789958162 | -0.290679024223052 |
| lnc-CNOT1-2 | lnc-CNOT1-2 | -0.659384368384683 | 0.183397530770468 | -3.72619114096084 | 0.000568274318598303 | 0.044565585061545 | -0.303701698801767 |
| lnc-MARCH3-3 | lnc-MARCH3-3 | -0.598080268009475 | 2.56516760798351 | -3.72394753328737 | 0.00057208912918713 | 0.044565585061545 | -0.31987625709753 |
| ENSG00000103489.11 | XYLT1 | 0.752332162756186 | 6.06586137378163 | 3.72347698114619 | 0.000572892335266138 | 0.044565585061545 | -0.455176780660915 |
| ENSG00000108469.14 | RECQL5 | -0.400464224508342 | 4.46617806334914 | -3.72158390702344 | 0.000576134702252683 | 0.044565585061545 | -0.42324637596017 |
| ENSG00000179456.10 | ZBTB18 | 0.380990883548786 | 6.34335888238827 | 3.72018500694403 | 0.000578542027214336 | 0.044565585061545 | -0.466299485372208 |
| ENSG00000121058.4 | COIL | 0.387379178592775 | 4.79200299663097 | 3.71902382799182 | 0.000580547625200643 | 0.044565585061545 | -0.44065319139046 |

| | | | | | | | |
|---|---|---|---|---|---|---|---|
| lnc-SELENBP1-2 | lnc-SELENBP1-2 | 0.450047253233891 | 1.30403655366516 | 3.71775789707275 | 0.000582741785589564 | 0.044565585061545 | -0.291128408302426 |
| lnc-MINOS1-1 | lnc-MINOS1-1 | -0.841180860889042 | 0.0542178802642651 | -3.71652100909082 | 0.000584893324826287 | 0.044565585061545 | -0.35368818158652 |
| ENSG00000100225.17 | FBXO7 | 0.318990770744808 | 7.07377395990972 | 3.71597582291652 | 0.000585844092808868 | 0.044565585061545 | -0.474794252800229 |
| lnc-PPP2R5B-5 | lnc-PPP2R5B-5 | -0.594459484810184 | 1.53785417193191 | -3.71545164141724 | 0.000586759634245673 | 0.044565585061545 | -0.301275447631013 |
| ENSG00000215251.3 | FASTKD5 | 0.366943047552577 | 4.80840083049983 | 3.71401098947393 | 0.00058928300066755 | 0.0445773813815522 | -0.454102704520533 |
| ENSG00000186162.10 | CIDECP1 | 0.350552457148061 | 2.16983125913626 | 3.71178705197707 | 0.000593198867621918 | 0.0445773813815522 | -0.330804987007792 |
| ENSG00000115128.6 | SF3B6 | 0.452762655891179 | 4.54270901958586 | 3.71166632321203 | 0.000593412160450143 | 0.0445773813815522 | -0.449741943016412 |
| ENSG00000074266.19 | EED | 0.558666144653653 | 5.06020687020444 | 3.71041789536097 | 0.000595622108711976 | 0.0445806904131586 | -0.471363324028912 |
| ENSG00000079950.13 | STX7 | -0.531680363876045 | 8.796882170352 | -3.70867328090233 | 0.000598723686173363 | 0.0445913803804753 | -0.472452713990542 |
| ENSG00000136051.14 | WASHC4 | 0.504171052062081 | 7.31836459619799 | 3.70546434694713 | 0.000604469209293541 | 0.0445913803804753 | -0.50203258536397 |
| ENSG00000212195.2 | | -1.1843272497017 | 5.37269825231854 | -3.71097790101981 | 0.000606228476781255 | 0.0445913803804753 | -0.494592997221043 |
| ENSG00000234345.2 | ELF2P1 | -0.647008480581019 | 0.775003678936202 | -3.70320185834346 | 0.000608552030196993 | 0.0445913803804753 | -0.329905809893439 |
| ENSG00000173040.12 | EVC2 | -4.4662857439622 | 1.33067774072071 | -3.71481973265609 | 0.000608840204217804 | 0.0445913803804753 | -0.328020443023583 |
| ENSG00000143554.13 | SLC27A3 | -0.775670842667143 | 2.43953361965482 | -3.70236536649426 | 0.00061006825469069 | 0.0445913803804753 | -0.372817337372859 |
| ENSG00000255767.1 | NA | 0.473675875822429 | 1.53021021490394 | 3.70189091703 | 0.000610929858003889 | 0.0445913803804753 | -0.337095985816914 |
| ENSG00000184588.17 | PDE4B | 0.761209147676436 | 8.4937175680099 | 3.69728379505193 | 0.000619357598400254 | 0.0449216674734901 | -0.512186710090831 |
| lnc-GK2-4 | lnc-GK2-4 | 0.573506404552782 | 2.43874944691428 | 3.69614315238151 | 0.000621461388390963 | 0.0449216674734901 | -0.379190582612868 |
| ENSG00000108021.20 | TASOR2 | 0.329953033957236 | 6.87580341023042 | 3.69559321430208 | 0.000622478145947428 | 0.0449216674734901 | -0.532989697509555 |
| lnc-RIOK2-6 | lnc-RIOK2-6 | 0.481748236999453 | 3.38876329432079 | 3.69383239968186 | 0.000625744430197888 | 0.0449216674734901 | -0.432317677731342 |
| lnc-VPS54-3 | lnc-VPS54-3 | -1.24590959924609 | 1.36872515351376 | -3.69338547911078 | 0.000626576083579784 | 0.0449216674734901 | -0.355036687024859 |
| LINC01715 | LINC01715 | -1.21646077926463 | 4.41636487547743 | -3.69833543920968 | 0.000628549785374588 | 0.0449216674734901 | -0.497064736394283 |
| EDRF1-AS1 | EDRF1-AS1 | -0.968501000546209 | 0.469688952946604 | -3.68945041667474 | 0.000633944743598428 | 0.0450456679184402 | -0.390674445992698 |
| lnc-RPS25-1 | lnc-RPS25-1 | -0.870908976432626 | 1.30318537737561 | -3.68906984716899 | 0.000634661793535813 | 0.0450456679184402 | -0.366883281135951 |
| FRG1-DT | FRG1-DT | -1.13428629214226 | 1.29100469172447 | -3.68702451364638 | 0.000638528904496046 | 0.0451644001417255 | -0.373307724832267 |
| ENSG00000063601.16 | MTMR1 | 0.349175039995671 | 6.56613025408852 | 3.68566600509891 | 0.00064110996352373 | 0.0451916656822224 | -0.561632880877959 |
| ENSG00000145287.10 | PLAC8 | 2.16413277283104 | 5.54388975598284 | 3.6947528142784 | 0.000645068228827129 | 0.0452494155801636 | -0.524930627816113 |
| ENSG00000211584.13 | SLC48A1 | -0.664394232866698 | 3.73700936155611 | -3.68230518897469 | 0.000647538483033879 | 0.0452494155801636 | -0.491377240099315 |
| ENSG00000106052.13 | TAX1BP1 | 0.323488370795562 | 6.78783359283363 | 3.68179259985577 | 0.000648524393730178 | 0.0452494155801636 | -0.571310971046668 |
| ENSG00000170852.10 | KBTBD2 | 0.382225316620278 | 6.27840889648314 | 3.6718977867944 | 0.000667841597797939 | 0.0462208171938653 | -0.599647786952702 |
| ENSG00000139239.7 | RPL14P1 | 1.05970264559979 | 2.53759237895521 | 3.67146169275824 | 0.000668705581971452 | 0.0462208171938653 | -0.44066519946823 |
| lnc-PSD4-2 | lnc-PSD4-2 | -0.745298131907249 | 0.349421666896271 | -3.6703716426244 | 0.000670869883158608 | 0.0462208171938653 | -0.435440772664924 |
| lnc-B3GNT5-11 | lnc-B3GNT5-11 | -0.795044189169478 | -0.0198698827611598 | -3.66957562644556 | 0.000672454635196041 | 0.0462208171938653 | -0.459488541538605 |
| ENSG00000101782.14 | RIOK3 | 0.406156223893169 | 6.27703239965481 | 3.66896404950708 | 0.000673674642091026 | 0.0462208171938653 | -0.607722718074379 |
| ENSG00000149313.10 | AASDHPPT | 0.556484515360408 | 4.39526600722763 | 3.66285248230768 | 0.000685983896416045 | 0.0468393103326162 | -0.573606074551617 |
| lnc-HMGA1-2 | lnc-HMGA1-2 | -0.700175098997306 | 0.31504506004354 | -3.66211183530371 | 0.000687490246649685 | 0.0468393103326162 | -0.454294542391391 |
| ENSG00000198522.13 | GPN1 | 0.391297588017487 | 4.44716858825932 | 3.66111818816573 | 0.000689516155603299 | 0.0468393103326162 | -0.586077993517221 |
| ENSG00000240225.10 | ZNF542P | -1.62404017802257 | 2.43304706092051 | -3.66505234322467 | 0.000694582193953926 | 0.0470282411123476 | -0.476470785156184 |
| ENSG00000166575.16 | TMEM135 | 0.588613114788434 | 3.39956933033078 | 3.65496014882144 | 0.000702200225136 | 0.0473881548654895 | -0.53695692762907 |
| ENSG00000270802.2 | | -0.657843451945735 | -0.256888252275067 | -3.65168493738769 | 0.000709037494004742 | 0.0475846469101408 | -0.562351981837369 |
| ENSG00000231259.5 | ANAPC1P2 | 0.505510039952964 | 2.65402711269615 | 3.65135226822082 | 0.00070973553910573 | 0.0475846469101408 | -0.512090772499477 |
| ENSG00000049618.23 | ARID1B | 0.599871458180623 | 8.04283325101479 | 3.64740227563255 | 0.000718074551523399 | 0.0479147931078001 | -0.654347197125299 |
| ENSG00000171202.6 | TMEM126A | 0.629190474137017 | 2.7111838765057 | 3.64681823888176 | 0.000719315506501006 | 0.0479147931078001 | -0.523145313056191 |
| lnc-ATL1-8 | lnc-ATL1-8 | -0.689500535920342 | 2.3684968953387 | -3.64324131774298 | 0.000726960821992327 | 0.0482407352263624 | -0.527167687458299 |
| ENSG00000006634.7 | DBF4 | 0.499962067220674 | 5.04277778143683 | 3.64234161506978 | 0.000728896111130482 | 0.0482407352263624 | -0.660474212400518 |
| ENSG00000263264.1 | NA | 0.642411509686818 | 5.78913971732943 | 3.63971391944402 | 0.000734576771639778 | 0.0484608772136588 | -0.681462582394544 |
| CEBPB-AS1 | CEBPB-AS1 | -0.904124096691479 | -0.19529517793382 | -3.63620412342373 | 0.000742230822790284 | 0.0487585957516335 | -0.603692606442457 |
| lnc-HPDL-1 | lnc-HPDL-1 | -1.21239861439538 | -0.30890321814626 | -3.63547640105212 | 0.000743827385027106 | 0.0487585957516335 | -0.597900549678567 |
| ENSG00000101335.9 | MYL9 | -2.21664864342579 | 1.76957084660398 | -3.6362116515055 | 0.000759670283737715 | 0.0493571844305309 | -0.522416835879304 |
| ENSG00000167674.14 | HDGFL2 | -0.520009685973962 | 3.6008764011424 | -3.62830481979938 | 0.000759738828548515 | 0.0493571844305309 | -0.631383356239807 |
| ENSG00000160285.14 | LSS | -0.500994746034785 | 5.1966349863673 | -3.6260655976908 | 0.000764773610353415 | 0.0493571844305309 | -0.712177429491563 |
| LINC00205 | LINC00205 | -0.614400735994259 | 3.42981982650833 | -3.62540279554758 | 0.000766270028758741 | 0.0493571844305309 | -0.619225067193597 |
| lnc-PUS7L-2 | lnc-PUS7L-2 | 0.442239483012135 | 2.15051014315892 | 3.62521751531818 | 0.000766688841717877 | 0.0493571844305309 | -0.562583879615791 |
| ENSG00000143977.13 | SNRPG | 0.428417057976635 | 4.34393286551477 | 3.62472718605855 | 0.000767798258290544 | 0.0493571844305309 | -0.67786722032956 |
| ENSG00000233426.3 | EIF3FP3 | 1.34961706564309 | 1.09752965528963 | 3.62386853705055 | 0.000769744750629181 | 0.0493571844305309 | -0.537642357410923 |
| ENSG00000108061.11 | SHOC2 | 0.299935852956939 | 6.5187207337432 | 3.61894905107832 | 0.000780988592821419 | 0.0496535132686836 | -0.74492445895546 |
| ENSG00000169592.14 | INO80E | -0.509560190839379 | 4.81365029653167 | -3.61861215876438 | 0.000781764331182552 | 0.0496535132686836 | -0.722033133376951 |
| ENSG00000125037.12 | EMC3 | 0.318461691544664 | 4.702985903305 | 3.61708760061781 | 0.000785284089828031 | 0.0496535132686836 | -0.714867762013371 |
| ENSG00000057468.6 | MSH4 | -0.877654161962214 | -0.316834820350176 | -3.6163907330445 | 0.000786898018830406 | 0.0496535132686836 | -0.629224439936385 |
| ENSG00000094975.13 | SUCO | 0.381656468634474 | 5.98865864206783 | 3.61630108471272 | 0.000787105873479009 | 0.0496535132686836 | -0.751163031278726 |
| ENSG00000163349.21 | HIPK1 | 0.378828771845375 | 6.21019403807853 | 3.61555392895188 | 0.000788840248693559 | 0.0496535132686836 | -0.754261139248546 |
| ENSG00000285802.1 | | -1.45193621495949 | -0.438412326952695 | -3.61199972907125 | 0.000797141022003262 | 0.0500230294386986 | -0.785281499261218 |
| lnc-ZNF394-1 | lnc-ZNF394-1 | 0.371068547302669 | 3.36180409033785 | 3.60941417735782 | 0.000803232188708317 | 0.0502520612163626 | -0.657466854532265 |
| lnc-RFC3-7 | lnc-RFC3-7 | -0.697731716317689 | 0.966936550887784 | -3.59909806025092 | 0.000827983404449863 | 0.051619661431245 | -0.596350824709844 |
| lnc-TAOK1-8 | lnc-TAOK1-8 | -0.744248466965766 | 3.09340299516889 | -3.59731518006582 | 0.000832334595088479 | 0.051619661431245 | -0.68674614747014 |
| lnc-MRNIP-4 | lnc-MRNIP-4 | -0.67826172034689 | 2.97797243937535 | -3.59719957103462 | 0.000832617500298238 | 0.051619661431245 | -0.677649989428299 |
| ENSG00000183508.4 | TENT5C | 0.838447938799645 | 6.96819841192321 | 3.59723427548444 | 0.000835123512442529 | 0.051619661431245 | -0.804352811804527 |
| ENSG00000156127.6 | BATF | -0.789844530063579 | 1.52809252772648 | -3.59496094394101 | 0.000838113850583462 | 0.0516493933729323 | -0.624516289728321 |
| ENSG00000228253.1 | MT-ATP8 | -1.23248111529862 | 5.76724759458838 | -3.59871130017575 | 0.000845522987671373 | 0.0519504467320593 | -0.80735944502582 |

| | | | | | | | |
|---|---|---|---|---|---|---|---|
| lnc-UGDH-1 | lnc-UGDH-1 | -0.823347235381463 | 2.09687306776901 | -3.59070726090052 | 0.000848653763659754 | 0.0519876202899069 | -0.653124363456619 |
| ENSG00000173757.9 | STAT5B | -0.671085564102648 | 5.55369308976375 | -3.58816634063724 | 0.000855010320986935 | 0.0522215947681724 | -0.82200731245594 |
| lnc-TMTC3-20 | lnc-TMTC3-20 | -1.0802774676399 | 0.273765582747886 | -3.58496016269708 | 0.000863096374566935 | 0.0525201205341393 | -0.665160162310736 |
| ENSG00000136045.11 | PWP1 | 0.342515778462461 | 5.85355707381213 | 3.5842090602486 | 0.000865001256428762 | 0.0525201205341393 | -0.837403521209329 |
| lnc-ZNF721-2 | lnc-ZNF721-2 | -1.17769631918687 | -0.360614652540022 | -3.58174618759299 | 0.000871275732470629 | 0.0526065282589297 | -0.773351343439222 |
| ENSG00000115548.16 | KDM3A | 0.520680176247598 | 6.93850188301158 | 3.58121487479644 | 0.000872635034690623 | 0.0526065282589297 | -0.846166288316265 |
| ENSG00000226800.5 | CACTIN-AS1 | -1.53625940519358 | -0.113402711422583 | -3.5793934871238 | 0.000877310308056258 | 0.0526065282589297 | -0.751406920832829 |
| ENSG00000119801.12 | YPEL5 | 0.584469985477348 | 7.38228895948837 | 3.57899404921478 | 0.000878338824074016 | 0.0526065282589297 | -0.848308989960123 |
| ENSG00000171448.8 | ZBTB26 | 0.457665457563929 | 4.29681490545136 | 3.57865858641713 | 0.000879203503914484 | 0.0526065282589297 | -0.798674386779997 |
| ENSG00000106636.7 | YKT6 | 0.32867199903408 | 5.23835463627015 | 3.57708205732963 | 0.000883278074407734 | 0.0526971379870562 | -0.846026183765164 |
| ENSG00000152782.16 | PANK1 | -1.15679623847237 | 1.56786403506162 | -3.57228882777504 | 0.000895777862864942 | 0.0531532647326408 | -0.672100980064182 |
| ENSG00000248487.8 | ABHD14A | -0.682694394458153 | 2.06214700757866 | -3.57217065295502 | 0.000896088172872097 | 0.0531532647326408 | -0.696379062602081 |
| lnc-RBM22-4 | lnc-RBM22-4 | 0.443958613067604 | 0.933169687193922 | 3.57054048634225 | 0.000900379292964004 | 0.0532543304226382 | -0.667889778131575 |
| ENSG00000204524.6 | ZNF805 | 0.437274842452589 | 4.26817473764677 | 3.56838275966139 | 0.000906089444961106 | 0.0534057978765588 | -0.822723889944698 |
| lnc-SYTL2-2 | lnc-SYTL2-2 | 0.45925284114364 | 2.36597340221084 | 3.56761503881811 | 0.000908129488257085 | 0.0534057978765588 | -0.718597013072954 |
| ENSG00000115761.15 | NOL10 | 0.337380770929242 | 4.76011421523133 | 3.56590117807787 | 0.000912699592132189 | 0.053410448494608 | -0.858367205447005 |
| ENSG00000125484.11 | GTF3C4 | 0.339746582803137 | 4.86466386960492 | 3.56563986540562 | 0.000913398332123695 | 0.053410448494608 | -0.863417006411931 |
| ENSG00000171865.9 | RNASEH1 | 0.372370473038654 | 4.75522495291784 | 3.5634270067059 | 0.000919336036967341 | 0.0535480086869068 | -0.864990338941745 |
| lnc-BCKDK-4 | lnc-BCKDK-4 | 0.365334847942765 | 3.38876087490508 | 3.56282642221724 | 0.00092095394622577 | 0.0535480086869068 | -0.787018972828974 |
| ENSG00000186792.16 | HYAL3 | -0.917597742828311 | 0.563760108945675 | -3.56061432957835 | 0.000926936668976405 | 0.0537440491761728 | -0.697813545645134 |
| lnc-CLLU1OS-1 | lnc-CLLU1OS-1 | 0.547520811783614 | 6.50932935923742 | 3.55800220203275 | 0.000934049307048749 | 0.0539431038502263 | -0.910813914809111 |
| ENSG00000174738.12 | NR1D2 | 0.419526360908369 | 5.94574292840704 | 3.55693838977409 | 0.000936960965472477 | 0.0539431038502263 | -0.912135735982836 |
| lnc-GLMP-1 | lnc-GLMP-1 | -0.596605747863227 | 0.184251763381061 | -3.55562284570546 | 0.000940573653020447 | 0.0539431038502263 | -0.731513293281425 |
| ENSG00000161036.12 | LRWD1 | -0.871806906526882 | 4.48622374412922 | -3.55600814665285 | 0.000942093478622898 | 0.0539431038502263 | -0.877357191483473 |
| lnc-GIPC3-3 | lnc-GIPC3-3 | -1.63752399038676 | 0.0171072308132068 | -3.55433115221551 | 0.000944133837249255 | 0.0539431038502263 | -0.781850119680842 |
| LINC01771 | LINC01771 | -2.1417796018775 | 0.607913783495738 | -3.55572207609054 | 0.000948128216025368 | 0.0539431038502263 | -0.769994672157114 |
| ENSG00000156521.13 | TYSND1 | -0.428247829766059 | 3.9464741350369 | -3.55180942470793 | 0.000951121531384084 | 0.0539431038502263 | -0.858917613277308 |
| ENSG00000222011.8 | FAM185A | -0.452025284954605 | 2.21727573158926 | -3.5517323465673 | 0.000951335893583645 | 0.0539431038502263 | -0.75937664752882 |
| ENSG00000079785.14 | DDX1 | 0.363330159968372 | 5.4885687035001 | 3.5496721222897 | 0.000957082784521653 | 0.0540491202084845 | -0.925849180746093 |
| ENSG00000171747.8 | LGALS4 | -0.676952955006151 | 0.586370292413219 | -3.54918108557003 | 0.000958457410294751 | 0.0540491202084845 | -0.722422491079516 |
| ENSG00000119906.12 | SLF2 | -0.314550775948939 | 5.85890110257989 | -3.54768396428985 | 0.000962660198148169 | 0.0541378001597917 | -0.936616486263047 |
| ENSG00000170759.10 | KIF5B | 0.270867818233111 | 7.3868569593285 | 3.53811177458179 | 0.000989952211128404 | 0.0550952647448409 | -0.958715763528097 |
| ENSG00000139146.13 | SINHCAF | 0.502178332770475 | 6.51350593858186 | 3.53743180161599 | 0.000991918890020494 | 0.0550952647448409 | -0.966433054997331 |
| lnc-STMND1-4 | lnc-STMND1-4 | -1.24550678119928 | 2.35030737014207 | -3.53913381810259 | 0.000993392698180652 | 0.0550952647448409 | -0.793426556559373 |
| ENSG00000155545.19 | MIER3 | 0.401315898337007 | 5.12222470240604 | 3.53684010128786 | 0.000993633305182063 | 0.0550952647448409 | -0.951877914616063 |
| ENSG00000162434.11 | JAK1 | 0.312724354916229 | 8.66276000820909 | 3.53533418958772 | 0.000998009411275305 | 0.0550952647448409 | -0.949276433777856 |
| ENSG00000135974.9 | C2orf49 | 0.277763390352326 | 4.97428776611305 | 3.53488411277167 | 0.000999320893463834 | 0.0550952647448409 | -0.952064561470606 |
| ENSG00000188649.13 | CC2D2B | 0.921745279020074 | 0.961889395880565 | 3.53471778892802 | 0.000999805963547872 | 0.0550952647448409 | -0.75829368756384 |
| ENSG00000116783.14 | TNNI3K | -0.905650232479366 | 0.860425646376542 | -3.53427466404645 | 0.00100109940312736 | 0.0550952647448409 | -0.759241513610696 |
| ENSG00000285531.1 | | 0.572495918303894 | 1.58888048174762 | 3.53132467085932 | 0.0010097511115284 | 0.0553415248941435 | -0.778644900400991 |
| ENSG00000138095.18 | LRPPRC | 0.344052841516882 | 6.59284888143895 | 3.53033169451122 | 0.00101267938440499 | 0.0553415248941435 | -0.985447663214088 |
| ENSG00000206562.11 | METTL6 | -0.308788533423788 | 2.93950996178114 | -3.53000650384907 | 0.00101364013433863 | 0.0553415248941435 | -0.850198487942574 |
| lnc-CXCR5-1 | lnc-CXCR5-1 | -0.77683737175816 | 0.405064235894789 | -3.52639236358813 | 0.00102437678091443 | 0.0557708581184998 | -0.78741273917035 |
| lnc-VIP-4 | lnc-VIP-4 | -0.771562878143757 | 1.00771626967267 | -3.52554060155061 | 0.0010269229571448 | 0.0557708581184998 | -0.78443727970376 |
| lnc-DHX38-26 | lnc-DHX38-26 | -0.783409379324057 | 0.233411262582292 | -3.52352888710873 | 0.00103296067624549 | 0.0558855548343324 | -0.800064643909506 |
| ENSG00000172115.8 | CYCS | 0.605562907503041 | 5.76130129808998 | 3.52302935961597 | 0.00103446516017493 | 0.0558855548343324 | -1.00178754979355 |
| lnc-APAF1-3 | lnc-APAF1-3 | 0.928924368759934 | 0.0808291119835157 | 3.52134262164007 | 0.00103956082676547 | 0.0560138232913973 | -0.812844519655576 |
| lnc-CDKL2-2 | lnc-CDKL2-2 | 0.465995473027657 | 3.73880232215181 | 3.51708646477664 | 0.00105252584991446 | 0.0565643330777791 | -0.930361725001331 |
| ENSG00000078237.6 | TIGAR | 0.706132529825301 | 2.17516845250944 | 3.51466614595943 | 0.00105996750092506 | 0.0568159142488035 | -0.84143213961828 |
| ENSG00000158615.8 | PPP1R15B | 0.436816602948644 | 7.07286310239181 | 3.51181611597243 | 0.00106879497026274 | 0.0570690745016795 | -1.03298431796876 |
| lnc-ERLIN1-2 | lnc-ERLIN1-2 | 0.551477425970571 | 0.510027274423664 | 3.51135308293005 | 0.00107023576532324 | 0.0570690745016795 | -0.81975187300334 |
| ENSG00000131174.5 | COX7B | 0.389575269590366 | 4.78516362383683 | 3.50729979126207 | 0.00108292781854399 | 0.0575966493258166 | -1.01839819081788 |
| ENSG00000136937.12 | NCBP1 | 0.418713020882735 | 5.98110321834338 | 3.50447337011157 | 0.00109186332245461 | 0.0579222236239257 | -1.05405420162197 |
| ENSG00000118557.15 | PMFBP1 | -1.31046816901883 | 2.40436215233879 | -3.50617240846332 | 0.00109870849607557 | 0.0581355192152273 | -0.90447008370504 |
| TMEM9B-AS1 | TMEM9B-AS1 | 0.544199890959419 | 4.26447006445673 | 3.50122662397884 | 0.0011022147590495 | 0.0581417540452699 | -1.00343599303552 |
| ENSG00000152700.13 | SAR1B | 0.402574239828382 | 5.32517071689854 | 3.50052132307409 | 0.00110447582139147 | 0.0581417540452699 | -1.05395825082977 |
| lnc-MTA3-7 | lnc-MTA3-7 | 0.87494569013752 | 2.22060382969618 | 3.49588947909346 | 0.00111943533114137 | 0.0586696491023175 | -0.900893752858615 |
| ENSG00000205323.8 | SARNP | 0.281833778477956 | 4.21471762776735 | 3.49438331675467 | 0.00112434143968802 | 0.0586696491023175 | -1.02332409229413 |
| ENSG00000213782.7 | DDX47 | 0.419334514753911 | 5.05642945049742 | 3.49434389612116 | 0.00112447012269151 | 0.0586696491023175 | -1.06318278031324 |
| ENSG00000158683.8 | PKD1L1 | -1.31004043406499 | -0.433948480634899 | -3.49373785716028 | 0.00112645022478753 | 0.0586696491023175 | -1.01624688928336 |
| EIF3J-DT | EIF3J-DT | -0.641994297747565 | 3.15464938307595 | -3.49212025829589 | 0.0011317517339897 | 0.0586696491023175 | -0.96885681282608 |
| ENSG00000073050.11 | XRCC1 | -0.430413878378806 | 3.52568339148207 | -3.49160646167129 | 0.00113344063855673 | 0.0586696491023175 | -0.991948309091355 |
| lnc-AK5-1 | lnc-AK5-1 | -1.01010797606568 | 0.522748731603166 | -3.49090032156129 | 0.00113576573218044 | 0.0586696491023175 | -0.874907577639987 |
| ENSG00000224672.4 | RPL17P10 | -0.626811442887327 | -0.178653708233854 | -3.48965861151254 | 0.0011398653595162 | 0.0586696491023175 | -0.924969343764657 |
| ENSG00000262488.1 | | -1.86227836592241 | 0.546497477365375 | -3.49275449458902 | 0.00114015739401093 | 0.0586696491023175 | -0.876191298293897 |
| lnc-GOLGA8Q-4 | lnc-GOLGA8Q-4 | -0.686990467537051 | 1.71012372558565 | -3.48690206048583 | 0.00114901704853042 | 0.0589780995259395 | -0.904546628758004 |
| lnc-RIPOR1-1 | lnc-RIPOR1-1 | -0.952439594141433 | 3.90197714465212 | -3.48572023735789 | 0.0011575685096761 | 0.0592072850818097 | -1.02684620471001 |
| ENSG00000175110.11 | MRPS22 | 0.342143861826918 | 4.55471078743545 | 3.48384902743764 | 0.00115923509148177 | 0.0592072850818097 | -1.07002090233398 |

| | | | | | | | |
|---|---|---|---|---|---|---|---|
| ENSG00000198899.2 | MT-ATP6 | -1.00415718991963 | 8.08557570540721 | -3.48575057914415 | 0.00116554707338012 | 0.0593317439591504 | -1.09596542200656 |
| lnc-SCARB1-1 | lnc-SCARB1-1 | 0.492688613392597 | 6.36466846433291 | 3.48097177453354 | 0.00116894434755902 | 0.0593317439591504 | -1.11836814262287 |
| ENSG00000285733.1 | | -0.412773098976244 | 2.57006500727737 | -3.4805660841822 | 0.00117031958642642 | 0.0593317439591504 | -0.964735452907473 |
| ENSG00000175224.16 | ATG13 | 0.281171513028568 | 5.72896971115681 | 3.47905447564844 | 0.00117545738655764 | 0.0594457970209234 | -1.12022004737025 |
| ENSG00000239557.1 | PPP2R5CP | 0.740967745702922 | -0.186627491482409 | 3.47813600129736 | 0.0011785897148136 | 0.0594581178921774 | -1.0174936099444 |
| lnc-CYP4F22-4 | lnc-CYP4F22-4 | -0.62034224548451 | -0.230031154444614 | -3.47656469837609 | 0.00118396694405837 | 0.059579162685926 | -0.966294132255866 |
| ENSG00000196636.7 | SDHAF3 | -0.988425542353462 | 1.14110230048957 | -3.4757459100304 | 0.00118677824910021 | 0.059579162685926 | -0.913655715311892 |
| ENSG00000115760.14 | BIRC6 | 0.299021036979957 | 8.88398650707569 | 3.4737576120845 | 0.00119363165812806 | 0.0597774219446467 | -1.1102964915505 |
| ENSG00000136522.13 | MRPL47 | 0.370308543183998 | 3.94709774285095 | 3.4721550381578 | 0.00119918305666393 | 0.0599096719789168 | -1.06600347969729 |
| ENSG00000167981.6 | ZNF597 | 0.45955792347905 | 3.31943958751923 | 3.47124394676883 | 0.00120235012358333 | 0.0599224518007643 | -1.02501477557356 |
| ENSG00000138185.19 | ENTPD1 | 0.649586026457212 | 6.35437302567025 | 3.46713100479886 | 0.00121674703937612 | 0.0604934886750691 | -1.15509004201039 |
| ENSG00000102901.12 | CENPT | -0.404404503214376 | 4.97694981611333 | -3.46426756372021 | 0.00122686737306924 | 0.0608496653973113 | -1.14271821938376 |
| lnc-ANKLE2-2 | lnc-ANKLE2-2 | -0.491788144395695 | 2.12453234299702 | -3.46013835416633 | 0.00124160314526995 | 0.0614324940843541 | -0.992407640186563 |
| ENSG00000279253.1 | | -0.472266526912502 | 1.27986942364862 | -3.45564737343593 | 0.00125782198143653 | 0.0620857310405471 | -0.964823762181998 |
| ENSG00000136021.18 | SCYL2 | 0.293982081902131 | 5.69998208922614 | 3.45234715077235 | 0.0012698693485809 | 0.0623146684055667 | -1.1912378416305 |
| ENSG00000119787.13 | ATL2 | 0.4264379491932 | 5.01654404486517 | 3.45211817196139 | 0.00127070930728204 | 0.0623146684055667 | -1.17460207740707 |
| ENSG00000189376.11 | C8orf76 | 0.385073798280683 | 2.86680870169304 | 3.45189116123608 | 0.00127154257058437 | 0.0623146684055667 | -1.04842909413431 |
| ENSG00000130475.14 | FCHO1 | -0.605496716396683 | 5.12439555023913 | -3.45053379611565 | 0.00127653580690263 | 0.0624107755664532 | -1.18135253982207 |
| ENSG00000215717.5 | TMEM167B | 0.390754186680174 | 5.33219704159684 | 3.44790811070545 | 0.00128624794386612 | 0.0627365910630243 | -1.19506348043805 |
| ENSG00000166866.12 | MYO1A | -0.935545422076692 | -0.444866773451934 | -3.44623199840898 | 0.00129248458536959 | 0.0628917499306439 | -1.11186973421005 |
| lnc-TYSND1-1 | lnc-TYSND1-1 | -0.382810318847087 | 3.05218712492584 | -3.44267420621338 | 0.00130581865052121 | 0.0633907200086747 | -1.09033886061024 |
| ENSG00000103528.16 | SYT17 | -0.928234390000756 | 4.56720782918954 | -3.44312357819476 | 0.00131148750316275 | 0.063516111241409 | -1.17238435231934 |
| ENSG00000257315.2 | ZBED6 | 0.353003809286609 | 8.00565093269543 | 3.43920047028767 | 0.00131896441142188 | 0.0637282734279262 | -1.2162716141502 |
| ENSG00000215302.8 | WHAMMP4 | -0.730838073684482 | 1.60879890030693 | -3.43810066575646 | 0.00132315271460111 | 0.0637809187930555 | -1.02497035823609 |
| ENSG00000168883.19 | USP39 | 0.304361529010037 | 5.21768730228087 | 3.4361075341596 | 0.00133077540007717 | 0.0638803914538237 | -1.22350731386168 |
| DGUOK-AS1 | DGUOK-AS1 | -0.724771819428606 | 0.719985876220126 | -3.43368479739894 | 0.00134009758347404 | 0.0638803914538237 | -1.009340882282 |
| ENSG00000058673.16 | ZC3H11A | 0.35422367194625 | 8.00982791439347 | 3.43266946329536 | 0.00134402289258777 | 0.0638803914538237 | -1.23356042440044 |
| lnc-AQP4-5 | lnc-AQP4-5 | -0.918404579424925 | 0.128774425499432 | -3.43255485414793 | 0.00134446666367653 | 0.0638803914538237 | -1.03481296633302 |
| lnc-TRIM49D1-2 | lnc-TRIM49D1-2 | 0.58517921549351 | 1.12892485532484 | 3.43209098921362 | 0.00134626419644607 | 0.0638803914538237 | -1.01725643707965 |
| lnc-ARID3B-1 | lnc-ARID3B-1 | -0.575545450875441 | 0.534841508918865 | -3.43201313978693 | 0.00134656609721355 | 0.0638803914538237 | -1.01399395662652 |
| ENSG00000105641.3 | SLC5A5 | -3.58934340691748 | -0.786313999174726 | -3.43656411648227 | 0.00134971668369688 | 0.0638803914538237 | -1.33996053170015 |
| ENSG00000068697.6 | LAPTM4A | 0.553721882206644 | 4.83639573557017 | 3.43077781464612 | 0.00135136536010408 | 0.0638803914538237 | -1.22193524384344 |
| lnc-WDR73-10 | lnc-WDR73-10 | -0.820056292499623 | 0.507025970151379 | -3.43031996462968 | 0.00135314826186013 | 0.0638803914538237 | -1.01795313580158 |
| ENSG00000008018.8 | PSMB1 | 0.254054162265358 | 5.9769802474315 | 3.42827482380239 | 0.00136113965180202 | 0.0639444419238957 | -1.25786815334185 |
| lnc-ZNF26-12 | lnc-ZNF26-12 | -1.19652151734113 | -0.422612605332722 | -3.42712087868811 | 0.00136566855939757 | 0.0639444419238957 | -1.11845844719252 |
| ENSG00000187653.11 | TMSB4XP8 | 1.51002526674832 | 1.9318885722797 | 3.43059788527383 | 0.00136644521090566 | 0.0639444419238957 | -1.04563059856216 |
| ENSG00000025800.13 | KPNA6 | 0.23893329462964 | 6.46033184386666 | 3.42679969688939 | 0.00136693166431104 | 0.0639444419238957 | -1.26300913207949 |
| ENSG00000279339.1 | | -0.807606626939147 | 1.7639980740908 | -3.42575602115132 | 0.00137104381156672 | 0.0639913713684305 | -1.05652081383846 |
| ENSG00000092096.16 | SLC22A17 | -2.45807074859252 | -1.37828236893627 | -3.4247786615656 | 0.00137490537581212 | 0.0640264193446627 | -1.48018309136139 |
| lnc-LANCL2-1 | lnc-LANCL2-1 | -1.09883995059812 | 2.52948928121195 | -3.42382437445547 | 0.00138366976760447 | 0.0642891079607289 | -1.10771249088207 |
| lnc-NEK11-5 | lnc-NEK11-5 | -0.430559883291867 | 1.24679968880792 | -3.42127604700197 | 0.00138882972509618 | 0.0643023728707037 | -1.05470795894487 |
| ENSG00000145882.10 | PCYOX1L | -0.38679693028858 | 3.58915008788623 | -3.4209313662246 | 0.00139020722669943 | 0.0643023728707037 | -1.17469252142 |
| ENSG00000138448.11 | ITGAV | -0.982827198157498 | 3.01002915116235 | -3.42090027214244 | 0.0013933274207032 | 0.0643023728707037 | -1.14383381016925 |
| ENSG00000088888.17 | MAVS | -0.290325737162059 | 6.75285174149707 | -3.41824105221387 | 0.00140100385401077 | 0.0645119962575029 | -1.28426309109635 |
| ENSG00000138050.14 | THUMPD2 | 0.573481659137533 | 4.04612234176035 | 3.41318720067382 | 0.00142150245512095 | 0.0652454586086827 | -1.22100955844916 |
| lnc-LINC00854-5 | lnc-LINC00854-5 | -1.30467117826093 | 1.5980414357421 | -3.41413853175738 | 0.00142584979670556 | 0.0652454586086827 | -1.09036928749721 |
| ENSG00000120727.12 | PAIP2 | 0.367519885849278 | 5.9659480013992 | 3.41197976869169 | 0.00142644203342113 | 0.0652454586086827 | -1.30081809960112 |
| lnc-SPICE1-2 | lnc-SPICE1-2 | -1.33916555877511 | 0.498124758031627 | -3.41041742768004 | 0.00143285784955782 | 0.0652627546397466 | -1.0675001590129 |
| ENSG00000173264.14 | GPR137 | -0.565782767165025 | 2.83142086516765 | -3.40963593329197 | 0.00143607741023242 | 0.0652627546397466 | -1.16596304130596 |
| ENSG00000165030.3 | NFIL3 | 1.01096256981902 | 0.887916667483387 | 3.40854418789335 | 0.00144058668641912 | 0.0652627546397466 | -1.07108635820445 |
| ENSG00000076258.9 | FMO4 | -0.574034626181576 | 2.51661655404401 | -3.40825077974351 | 0.00144180086229228 | 0.0652627546397466 | -1.14317351918484 |
| ENSG00000220842.6 | RPL21P16 | 1.25638635022253 | 2.32033946055418 | 3.41015225540022 | 0.00144267372885802 | 0.0652627546397466 | -1.11382004490549 |
| lnc-HIST4H4-4 | lnc-HIST4H4-4 | -0.737861626572648 | 0.551391837394311 | -3.4069354812186 | 0.0014472558145423 | 0.0653264614708863 | -1.07496781040458 |
| lnc-ZNF12-2 | lnc-ZNF12-2 | 0.672628860082967 | 0.081954704904172 | 3.40506321683739 | 0.00145505465171008 | 0.0655347700134542 | -1.09926365546463 |
| lnc-PUS10-4 | lnc-PUS10-4 | 0.321091335075963 | 4.85306350442007 | 3.40031221005182 | 0.00147502512615468 | 0.066262361924405 | -1.30378008132952 |
| lnc-SQSTM1-3 | lnc-SQSTM1-3 | -0.735060878379617 | 0.165373188000825 | -3.39969284193822 | 0.00147764777356566 | 0.066262361924405 | -1.09879747832738 |
| ENSG00000135426.15 | TESPA1 | -0.529060067057133 | 5.0217547892065 | -3.39771070530017 | 0.00148607084411928 | 0.0664952090967546 | -1.31933183286516 |
| ENSG00000125746.16 | EML2 | -0.365615667775539 | 4.62825828189187 | -3.39587536032115 | 0.0014939109425667 | 0.0665800380914317 | -1.31138495481784 |
| lnc-CYBA-2 | lnc-CYBA-2 | -0.680807233581616 | 1.59848251157929 | -3.39456885462959 | 0.00149951600118779 | 0.0665800380914317 | -1.14093900703939 |
| ENSG00000043462.11 | LCP2 | -2.31667225070331 | 3.44316244337684 | -3.40131870978656 | 0.00150896167486542 | 0.0665800380914317 | -1.21318537016546 |
| ENSG00000167508.11 | MVD | -0.777292290057174 | 3.5093038862631 | -3.39180742160102 | 0.00151142892707414 | 0.0665800380914317 | -1.26107648163008 |
| ENSG00000177981.10 | ASB8 | 0.35504399698469 | 4.79774500901902 | 3.39178282634801 | 0.00151153543631317 | 0.0665800380914317 | -1.32507695666679 |
| ENSG00000141837.19 | CACNA1A | -1.18864168185427 | 0.173288700121657 | -3.39175373664533 | 0.00151166141791287 | 0.0665800380914317 | -1.1185258314815 |
| lnc-PANO1-1 | lnc-PANO1-1 | -0.642452625170607 | 1.54456605133264 | -3.39010007493633 | 0.00151883957706827 | 0.0665800380914317 | -1.14446241778283 |
| lnc-DHX57-3 | lnc-DHX57-3 | 0.453721306021198 | 5.51244803858258 | 3.38886959689195 | 0.00152420187211913 | 0.0665800380914317 | -1.35532534174135 |
| ENSG00000128394.16 | APOBEC3F | -0.523601677447348 | 4.3290519882755 | -3.38821657373872 | 0.00152705500603264 | 0.0665800380914317 | -1.30827317828092 |
| ENSG00000182134.15 | TDRKH | -0.492196843297019 | 1.94194550929639 | -3.38760970432307 | 0.00152971105336418 | 0.0665800380914317 | -1.1695852615018 |
| ENSG00000238286.1 | SLC35E1P1 | -0.805162400199942 | 0.0591042021437426 | -3.38759012953745 | 0.00152979679838614 | 0.0665800380914317 | -1.13932213732422 |

| | | | | | | | |
|---|---|---|---|---|---|---|---|
| lnc-GPR75-ASB3-5 | lnc-GPR75-ASB3-5 | 0.704025279840282 | 1.45134709816125 | 3.38732873450574 | 0.00153094224751451 | 0.0665800380914317 | -1.14564846005121 |
| lnc-FAM181B-5 | lnc-FAM181B-5 | 0.654906806922826 | 1.94195051441313 | 3.38668418823627 | 0.00153377018407154 | 0.0665800380914317 | -1.16354695446585 |
| ENSG00000075399.13 | VPS9D1 | -0.508151811445158 | 2.68195331600235 | -3.38533152629059 | 0.00153972116848322 | 0.0665800380914317 | -1.21791135585372 |
| ENSG00000133460.19 | SLC2A11 | -0.434565187496547 | 4.45130230069106 | -3.3836022756105 | 0.00154736100350063 | 0.0665800380914317 | -1.33380014488659 |
| ENSG00000156273.15 | BACH1 | 0.756295103431227 | 7.49047502962729 | 3.38339948901778 | 0.00155027535991717 | 0.0665800380914317 | -1.37244520981309 |
| lnc-CCDC90B-4 | lnc-CCDC90B-4 | -1.02928732679527 | -0.156861099888651 | -3.38234139247494 | 0.00155295435582894 | 0.0665800380914317 | -1.16597650949245 |
| ENSG00000225032.5 | | -0.555900438122459 | 3.3436958768553 | -3.38173006821719 | 0.00155567315900507 | 0.0665800380914317 | -1.26619304033896 |
| lnc-C12orf49-6 | lnc-C12orf49-6 | 0.637442129215852 | 3.23160301716379 | 3.38006613290818 | 0.00156309634480296 | 0.0665800380914317 | -1.25627268916223 |
| NIFK-AS1 | NIFK-AS1 | -0.642980921936368 | 3.1129095830574 | -3.37940323845139 | 0.00156606305899431 | 0.0665800380914317 | -1.2535016836123 |
| lnc-C12orf49-17 | lnc-C12orf49-17 | 0.665694938340724 | 2.14542560789808 | 3.37881095191604 | 0.00156871831410226 | 0.0665800380914317 | -1.19698199237562 |
| lnc-GSN-2 | lnc-GSN-2 | -1.45982789652703 | 1.24864659960357 | -3.3800714365181 | 0.00157236401516343 | 0.0665800380914317 | -1.15116694708879 |
| lnc-RALGAPA2-5 | lnc-RALGAPA2-5 | -0.330890717396962 | 2.03328803496864 | -3.37778109486881 | 0.00157334544343959 | 0.0665800380914317 | -1.20066841607145 |
| ENSG00000182149.20 | IST1 | 0.330786430093148 | 7.39106711176913 | 3.37714592578107 | 0.00157620572918484 | 0.0665800380914317 | -1.38840603676721 |
| lnc-MITD1-2 | lnc-MITD1-2 | -0.44218291880019 | 1.98322067577794 | -3.37681057020011 | 0.00157771789674262 | 0.0665800380914317 | -1.20238119883188 |
| lnc-NFE2L3-7 | lnc-NFE2L3-7 | -1.5600117517791 | 2.09408737008133 | -3.38116623905138 | 0.00158055308356465 | 0.0665800380914317 | -1.18916988700968 |
| lnc-CDR2-8 | lnc-CDR2-8 | -0.763151019058944 | 1.12262124570838 | -3.3758472260009 | 0.00158206945147933 | 0.0665800380914317 | -1.1614744622586 |
| lnc-RAB33B-4 | lnc-RAB33B-4 | -1.43392440239546 | -0.815877135429146 | -3.37546775541597 | 0.00158378670920934 | 0.0665800380914317 | -1.33866714256519 |
| ENSG00000176788.8 | BASP1 | -1.01979230853932 | 2.51902971392503 | -3.37653019668235 | 0.0015838350755831 | 0.0665800380914317 | -1.25350284526131 |
| ENSG00000101343.14 | CRNKL1 | 0.254713900639268 | 5.59933585470346 | 3.37519813023491 | 0.00158500795145516 | 0.0665800380914317 | -1.39365235627059 |
| ENSG00000127903.13 | ZNF835 | -2.07829662021008 | -1.58126234882433 | -3.3721282548376 | 0.00159897609398809 | 0.0669509446019147 | -1.53309320961669 |
| lnc-PPFIA2-5 | lnc-PPFIA2-5 | 0.374703190599771 | 2.90575858968054 | 3.37170980297835 | 0.00160088913835442 | 0.0669509446019147 | -1.25830624459354 |
| ENSG00000008294.20 | SPAG9 | 0.398693853800053 | 6.70765238653462 | 3.37111853624046 | 0.00160359596214079 | 0.0669509446019147 | -1.40931943941138 |
| lnc-KANSL2-1 | lnc-KANSL2-1 | 0.371531636098693 | 4.78914315735321 | 3.37018355558292 | 0.00160788521343064 | 0.0669941322834876 | -1.38137599877106 |
| lnc-TTC30B-1 | lnc-TTC30B-1 | -0.580436125467099 | 3.88269531648899 | -3.36809247214245 | 0.00161751771783481 | 0.0672593276488767 | -1.33892432852228 |
| ENSG00000279995.1 | | -0.908497944960198 | -0.577170045429492 | -3.36548179757633 | 0.00162962088250328 | 0.0676259810979133 | -1.27857672694837 |
| ENSG00000274114.2 | ALOX15P1 | -0.941819012810505 | 3.31090459929869 | -3.36461849566571 | 0.00163766091367964 | 0.0678228864914851 | -1.31268793278864 |
| ENSG00000138166.5 | DUSP5 | 0.959261157290609 | 4.88186595309909 | 3.36566300263461 | 0.00164131477346464 | 0.0678377148237403 | -1.38728467496052 |
| ENSG00000159915.12 | ZNF233 | -1.07755751938022 | 1.01842040793098 | -3.36212241856323 | 0.00164532203637044 | 0.0678670610713685 | -1.18907630231934 |
| lnc-SLC38A1-9 | lnc-SLC38A1-9 | 0.354118613215264 | 3.83810611436579 | 3.35903223620567 | 0.00165989216573511 | 0.0683099547982599 | -1.35406799095485 |
| lnc-PFKL-1 | lnc-PFKL-1 | -0.664722678015273 | -0.0281942851796789 | -3.35844037614928 | 0.00166269675722335 | 0.0683099547982599 | -1.21472834557581 |
| ENSG00000140950.15 | MEAK7 | -0.403464666795694 | 2.8522838393048 | -3.3569373239352 | 0.00166983942733195 | 0.0684604652826314 | -1.29754581432031 |
| ENSG00000278318.4 | ZNF229 | -4.26565102649634 | -0.064916752285573 | -3.36271581543138 | 0.00167330672287196 | 0.0684604652826314 | -1.4958637246618 |
| ENSG00000132952.11 | USPL1 | 0.446937271806298 | 6.25415341951274 | 3.3547289726893 | 0.00168038678471647 | 0.0684604652826314 | -1.45302746395416 |
| ENSG00000127364.3 | TAS2R4 | -1.15911168080041 | -0.0930711703794721 | -3.35438461545607 | 0.00168203718421072 | 0.0684604652826314 | -1.23214448764271 |
| lnc-NPBWR1-12 | lnc-NPBWR1-12 | -0.563475777728828 | 3.62686194542766 | -3.35381803817468 | 0.00168475597641873 | 0.0684604652826314 | -1.35599493854786 |
| ENSG00000167807.15 | | -0.545830221514657 | 0.804269970539944 | -3.35349319498591 | 0.0016863166641546 | 0.0684604652826314 | -1.21012326475368 |
| ENSG00000177106.15 | EPS8L2 | -0.839184477279987 | 2.32559027624042 | -3.35235002969029 | 0.00169181988087565 | 0.0685486783623295 | -1.28148252885109 |
| ENSG00000226849.1 | NA | -0.838295013446692 | -0.159988339024291 | -3.35103249647568 | 0.00169818372003205 | 0.0686713467768559 | -1.26444866322181 |
| lnc-PDCD1-17 | lnc-PDCD1-17 | -0.785777772207406 | 0.363821320242451 | -3.34987297577855 | 0.00170380319042236 | 0.0687634922911046 | -1.21408971826142 |
| ENSG00000153922.10 | CHD1 | 0.464397239173294 | 7.14965941622549 | 3.34912421436853 | 0.00170744137100524 | 0.0687754711142875 | -1.46439526288213 |
| ENSG00000128487.16 | SPECC1 | -1.29387801974213 | 2.92528733964461 | -3.35248172383063 | 0.001712504145871 | 0.0688446735048099 | -1.3167652544609 |
| ENSG00000198727.2 | MT-CYB | -0.817725261662266 | 8.43813124917075 | -3.34809526265498 | 0.00171757686510784 | 0.0689140050965199 | -1.45005862327353 |
| ENSG00000162413.16 | KLHL21 | -0.762055419601971 | 4.37092712335722 | -3.34599827767752 | 0.00172299116887166 | 0.068996745581489 | -1.43109597613671 |
| lnc-CHMP7-3 | lnc-CHMP7-3 | -0.832141996061355 | 1.38448117555698 | -3.34490961761533 | 0.00172805818747944 | 0.0690425755241231 | -1.25619190269635 |
| lnc-N6AMT1-1 | lnc-N6AMT1-1 | -1.11076404839 | 0.82389010024703 | -3.34434373399972 | 0.00173084433612435 | 0.0690425755241231 | -1.23016518690784 |
| ENSG00000123815.11 | COQ8B | -0.430464763184052 | 3.31994067356949 | -3.34177063758442 | 0.00174356708485257 | 0.0694155537863065 | -1.37236374493876 |
| ENSG00000160094.14 | ZNF362 | -0.562821706584998 | 3.18617863076587 | -3.33889429246595 | 0.0017578946134088 | 0.0696242533218931 | -1.36889660720856 |
| ASB16-AS1 | ASB16-AS1 | -0.376949453177828 | 3.18999659164539 | -3.33817791373453 | 0.00176148039889 | 0.0696242533218931 | -1.36863757801753 |
| ENSG00000161958.10 | FGF11 | -0.631043755147155 | -0.126860493233138 | -3.33762463606631 | 0.00176425455601598 | 0.0696242533218931 | -1.26158403357831 |
| lnc-GPR75-ASB3-6 | lnc-GPR75-ASB3-6 | 0.759911477558844 | 1.24486953419264 | 3.33660834693007 | 0.0017693611082996 | 0.0696242533218931 | -1.26298884706036 |
| ENSG00000263465.4 | SRSF8 | 0.38864556212036 | 5.45144378627518 | 3.33626798496486 | 0.00177107446845403 | 0.0696242533218931 | -1.4917841597222 |
| lnc-FAM117A-1 | lnc-FAM117A-1 | -0.522112443650927 | 3.33091770809337 | -3.33608164360621 | 0.00177201316752204 | 0.0696242533218931 | -1.38407485447681 |
| ENSG00000278768.2 | BACE1-AS | -0.562764026008156 | 2.272159315516 | -3.33598753367221 | 0.00177248742849303 | 0.0696242533218931 | -1.32107574512978 |
| ENSG00000110330.8 | BIRC2 | 0.500441184144037 | 5.86413638776922 | 3.33332286800139 | 0.00178596607696657 | 0.0700200757375292 | -1.50601097383137 |
| lnc-COX7A2L-4 | lnc-COX7A2L-4 | 1.42536324823185 | -0.112943321990036 | 3.33174112274317 | 0.00179401307061618 | 0.0702018460693782 | -1.35852824267791 |
| ENSG00000111647.12 | BLTP3B | 0.213120573508918 | 5.38527114074384 | 3.33030053744247 | 0.00180137193159175 | 0.0703560502238199 | -1.50721601574245 |
| lnc-CDC20-2 | lnc-CDC20-2 | -0.852407091389772 | 1.55761726698405 | -3.32840958795147 | 0.00181107495970913 | 0.070401504229572 | -1.29359817278982 |
| ENSG00000167554.14 | ZNF610 | -0.768301842424708 | 1.60372463885627 | -3.32783531842863 | 0.00181403153813046 | 0.070401504229572 | -1.29795869092901 |
| lnc-HECA-1 | lnc-HECA-1 | -0.51349615352895 | 5.69174210137963 | -3.32743982109446 | 0.00181607039237495 | 0.070401504229572 | -1.51947597385853 |
| ENSG00000126106.13 | TMEM53 | -0.704341768132553 | 0.930226451396527 | -3.32601121961313 | 0.0018234532432681 | 0.070401504229572 | -1.27601418267696 |
| ENSG00000283149.1 | | 0.420774975532251 | 2.74803583946204 | 3.32502317004163 | 0.00182857606357756 | 0.070401504229572 | -1.36773129622752 |
| ENSG00000114735.9 | HEMK1 | -0.408411183047936 | 5.23687068016098 | -3.32393704435868 | 0.00183422318194228 | 0.070401504229572 | -1.51829612184504 |
| ENSG00000108829.9 | LRRC59 | 0.417279542470613 | 5.34080214542506 | 3.32283847020394 | 0.00183995189616545 | 0.070401504229572 | -1.52444584324415 |
| lnc-APRT-1 | lnc-APRT-1 | -0.625317758664299 | 0.449721580935508 | -3.32201836373594 | 0.00184423957987014 | 0.070401504229572 | -1.28066854848447 |
| ENSG00000253540.5 | FAM86HP | -0.6070285136355 | 0.150377597760299 | -3.32121521055268 | 0.0018484478377022 | 0.070401504229572 | -1.29645949887478 |
| ENSG00000183960.8 | KCNH8 | -1.23629677602209 | 4.37171113636641 | -3.3263377501654 | 0.00184856498069838 | 0.070401504229572 | -1.45318493739018 |
| ENSG00000134900.11 | TPP2 | 0.359511836118545 | 6.85568559389467 | 3.32100629285552 | 0.00184954399342627 | 0.070401504229572 | -1.53997557346025 |
| ENSG00000075415.12 | SLC25A3 | 0.419317772190317 | 7.59834588055311 | 3.32032980992471 | 0.00185309762813674 | 0.070401504229572 | -1.53377857102491 |

| | | | | | | | |
|---|---|---|---|---|---|---|---|
| lnc-PDZRN4-6 | lnc-PDZRN4-6 | 0.762589071688008 | 1.01068545033784 | 3.31993665147927 | 0.0018551659126992 | 0.070401504229572 | -1.29275678988458 |
| ENSG00000144843.11 | ADPRH | -0.622943755989345 | 4.15978410294107 | -3.31975534010742 | 0.00185612047506479 | 0.070401504229572 | -1.48253885325537 |
| ENSG00000070010.18 | UFD1 | 0.391551954482796 | 5.56223964773431 | 3.31874733308595 | 0.00186143592145681 | 0.070401504229572 | -1.54146780311229 |
| lnc-SCARB2-2 | lnc-SCARB2-2 | 0.350740680740033 | 3.36171385821794 | 3.31853932941819 | 0.00186253457161178 | 0.070401504229572 | -1.42611093937227 |
| ENSG00000172824.15 | CES4A | -1.73807111544804 | 2.05409313685078 | -3.32374817718088 | 0.00186381240454806 | 0.070401504229572 | -1.32246286562924 |
| ENSG00000148204.11 | CRB2 | -0.666555165204457 | 0.348776775475722 | -3.31824269884333 | 0.00186410240514584 | 0.070401504229572 | -1.29052533927048 |
| lnc-CFL2-1 | lnc-CFL2-1 | 0.418586853575667 | 1.93338047038846 | 3.31759097389705 | 0.00186755149024018 | 0.070402586673285 | -1.34142234773242 |
| ENSG00000080845.17 | DLGAP4 | -0.617231326930368 | 5.28467848449011 | -3.31433659547427 | 0.00188486544675854 | 0.0709253848091975 | -1.54379631489137 |
| ENSG00000173960.13 | UBXN2A | 0.426177062954336 | 4.06608322080592 | 3.31201285529899 | 0.00189732149927666 | 0.0712638109846924 | -1.49539305552707 |
| ENSG00000264070.1 | DND1P1 | -2.05462125729777 | -0.894219386053784 | -3.31083639682846 | 0.00190365752653149 | 0.0713715534947136 | -1.55774633771766 |
| ENSG00000136052.9 | SLC41A2 | 0.874886158860552 | 1.75907808213563 | 3.30789438445518 | 0.00191959042679096 | 0.0718380540993423 | -1.35202637825956 |
| ENSG00000270099.1 | | 0.672923991426235 | 2.18190621271345 | 3.3029027934545 | 0.00194691349268472 | 0.072684688777991 | -1.3869936972157 |
| ENSG00000141428.16 | C18orf21 | 0.301617598252414 | 3.59489903842939 | 3.30247432224319 | 0.0019492760144513 | 0.072684688777991 | -1.48417946686965 |
| ZRANB2-AS2 | ZRANB2-AS2 | -0.637076848739499 | 1.87310704647245 | -3.30151067941431 | 0.00195459934548475 | 0.0727513893817587 | -1.38209617915254 |
| ENSG00000272356.1 | | -0.590978224450451 | 1.83229416715448 | -3.29986669458594 | 0.00196371290877763 | 0.0729112649001618 | -1.38770436870139 |
| ENSG00000256771.3 | ZNF253 | -0.418224535276154 | 5.05836336456589 | -3.29924150438526 | 0.00196718928811813 | 0.0729112649001618 | -1.57928018389735 |
| ENSG00000162851.7 | TFB2M | 0.407528232020113 | 3.37526295998158 | 3.2988226491911 | 0.00196952160931302 | 0.0729112649001618 | -1.4774928094713 |
| LENG8-AS1 | LENG8-AS1 | -0.421104883287951 | 3.38276008892067 | -3.29520855281318 | 0.00198975550615471 | 0.0733225222332165 | -1.48957391635621 |
| ENSG00000064703.11 | DDX20 | 0.384439374645324 | 4.84449335499857 | 3.29434796427181 | 0.00199460263371115 | 0.0733225222332165 | -1.58224458663528 |
| lnc-NCOA1-2 | lnc-NCOA1-2 | 0.434468287568393 | 1.88569365353159 | 3.29382153361966 | 0.00199757319825934 | 0.0733225222332165 | -1.39892136134461 |
| lnc-LGI2-1 | lnc-LGI2-1 | 0.546530810479711 | 1.34420688042026 | 3.29374298281323 | 0.00199801680819994 | 0.0733225222332165 | -1.37419837849975 |
| ENSG00000160703.15 | NLRX1 | -0.421786869860668 | 2.16577490669642 | -3.29366768062289 | 0.00199844215968685 | 0.0733225222332165 | -1.42970467373985 |
| ENSG00000185650.9 | ZFP36L1 | -0.630043425865707 | 6.71039849909311 | -3.29160388578228 | 0.00201013324289874 | 0.0734354339557044 | -1.61598443225004 |
| ENSG00000275674.1 | | 0.445434142984154 | 1.25500330592386 | 3.29124201322124 | 0.00201218987950598 | 0.0734354339557044 | -1.37350504198059 |
| ENSG00000115875.18 | SRSF7 | 0.430579753619513 | 7.48916893019311 | 3.2912361991757 | 0.00201222293888244 | 0.0734354339557044 | -1.61198077096338 |
| ENSG00000079332.14 | SAR1A | 0.327440763620053 | 6.03965361129961 | 3.28799546760504 | 0.00203073065277113 | 0.0739796973911295 | -1.62645710963341 |
| ENSG00000121577.13 | POPDC2 | -0.570060976536485 | 1.23039633763935 | -3.28692351163829 | 0.00203688806554897 | 0.074072909987976 | -1.38882386322028 |
| ENSG00000162607.12 | USP1 | 0.280346306120051 | 5.89389506148445 | 3.28600271222821 | 0.00204219137593432 | 0.0741347885200283 | -1.63065937852358 |
| ENSG00000264187.1 | | -0.397377292674743 | 2.55739386530989 | -3.28535114380994 | 0.0020459519807916 | 0.0741405451067491 | -1.46842885573851 |
| ENSG00000198431.15 | TXNRD1 | 0.393296625416631 | 4.81700482911462 | 3.28456724034263 | 0.00205048508978298 | 0.0741742260158226 | -1.60466921799863 |
| ENSG00000156990.14 | RPUSD3 | -0.493511917463134 | 2.99242916181071 | -3.2797147877333 | 0.00207875853017022 | 0.0749912226488355 | -1.5058024325085 |
| ENSG00000166689.15 | PLEKHA7 | -1.38737176600619 | 1.72216642851324 | -3.28263615827648 | 0.00208035700007215 | 0.0749912226488355 | -1.42599847544666 |
| lnc-PKDCC-4 | lnc-PKDCC-4 | 1.11137587182154 | -0.0326395088137472 | 3.2785427298998 | 0.00208564297313385 | 0.0750503309720527 | -1.43115025308351 |
| ENSG00000269001.2 | ZNF818P | -0.652145686763901 | 4.79552023442584 | -3.27745527444612 | 0.00209204983440815 | 0.0750949323900093 | -1.62165358260109 |
| SEC24B-AS1 | SEC24B-AS1 | -0.424547306072505 | 3.06257462671166 | -3.27709454505543 | 0.0020941792389605 | 0.0750949323900093 | -1.5124468492872 |
| ENSG00000065621.14 | GSTO2 | -0.692465548323574 | -0.110126621499661 | -3.27596745741265 | 0.00210084574713465 | 0.0752029704578653 | -1.41353283167701 |
| ENSG00000071462.11 | BUD23 | 0.337972154225638 | 5.06082547055289 | 3.27442234895495 | 0.00211001752331097 | 0.0754001574345655 | -1.64078657949285 |
| lnc-RBM22-3 | lnc-RBM22-3 | 0.362308318481281 | 1.84701980154417 | 3.27245635386592 | 0.00212174264931425 | 0.0756877451487611 | -1.45001797635296 |
| ENSG00000101146.12 | RAE1 | 0.320291838613701 | 4.71379387362688 | 3.2718153026971 | 0.00212557920935006 | 0.0756934201834816 | -1.63320852266059 |
| lnc-HSD11B1-4 | lnc-HSD11B1-4 | -0.961406585791799 | -0.336600428828121 | -3.27015521126669 | 0.00213554514778764 | 0.0758309759739826 | -1.43257055442828 |
| ENSG00000134504.13 | KCTD1 | -0.80835979599974 | 0.625505219697142 | -3.26956317100848 | 0.00213911002855802 | 0.0758309759739826 | -1.40783839437609 |
| lnc-PHF10-3 | lnc-PHF10-3 | -0.454012449473738 | 1.70864690533858 | -3.26844796105538 | 0.00214584044251384 | 0.0758309759739826 | -1.46190387802198 |
| lnc-NIPA1-1 | lnc-NIPA1-1 | -1.21798708394934 | 0.253872465798118 | -3.26819565783725 | 0.00214736590552019 | 0.0758309759739826 | -1.40990948551324 |
| ENSG00000067248.9 | DHX29 | 0.228535675405065 | 4.95110084695343 | 3.26811351913222 | 0.00214786275046552 | 0.0758309759739826 | -1.65307621308749 |
| ENSG00000136802.11 | LRRC8A | 0.470833926036057 | 4.45392003767695 | 3.26701454335657 | 0.0021545207976518 | 0.0759153264250179 | -1.63118680505459 |
| ENSG00000137055.14 | PLAA | 0.312045024146046 | 4.35472098838868 | 3.26650270125466 | 0.00215762842922001 | 0.0759153264250179 | -1.62772260329563 |
| ENSG00000132823.10 | OSER1 | 0.426058348775361 | 5.8441543729553 | 3.26397187497261 | 0.00217305687270621 | 0.0761954403056255 | -1.68678123921769 |
| ENSG00000108671.10 | PSMD11 | 0.302769557930414 | 5.54854864174444 | 3.26392812980192 | 0.00217332447057126 | 0.0761954403056255 | -1.68338849935328 |
| lnc-TRIM26-2 | lnc-TRIM26-2 | -0.432157374526878 | 5.94727061134656 | -3.26289766927991 | 0.00217963705299584 | 0.0761954403056255 | -1.69031896881566 |
| ENSG00000129292.20 | PHF20L1 | 0.31579490716942 | 6.404953077177 | 3.26277378435159 | 0.0021803971403592 | 0.0761954403056255 | -1.69280201445081 |
| ENSG00000160447.6 | PKN3 | -0.792335014260627 | 1.91559680013941 | -3.26147087574344 | 0.00218840629284619 | 0.0763457063146663 | -1.4930889943658 |
| ENSG00000162642.13 | C1orf52 | 0.319643832106972 | 4.24276299598217 | 3.26017627728291 | 0.00219639200505679 | 0.0764946474451504 | -1.63741361950999 |
| ENSG00000111252.10 | SH2B3 | 0.731336625557935 | 5.28224467145761 | 3.25871694548115 | 0.00220733099048542 | 0.0765647937761798 | -1.68609123509751 |
| lnc-ZNF585A-4 | lnc-ZNF585A-4 | -0.893896423469122 | 0.769459357871427 | -3.2575801530231 | 0.00221248955243244 | 0.0765647937761798 | -1.43819923520576 |
| lnc-H2AFV-4 | lnc-H2AFV-4 | -0.710224897791988 | 0.0964955217875777 | -3.25728011534712 | 0.00221435716729112 | 0.0765647937761798 | -1.4430770715532 |
| ENSG00000204262.12 | COL5A2 | -1.21173376315285 | 1.78443171272856 | -3.25877107091904 | 0.00221704718468049 | 0.0765647937761798 | -1.49412823878126 |
| ENSG00000100154.14 | TTC28 | -0.925660128340005 | 4.44070621350144 | -3.25886091922525 | 0.00221848420128184 | 0.0765647937761798 | -1.65419043534347 |
| lnc-C1QTNF1-5 | lnc-C1QTNF1-5 | -1.19064062069151 | -0.0500579587020136 | -3.25625891258278 | 0.00222072496158866 | 0.0765647937761798 | -1.46950090708367 |
| TIPARP-AS1 | TIPARP-AS1 | -0.596002510522879 | 0.0296326353232139 | -3.25530384283905 | 0.00222669609544638 | 0.0766422838337337 | -1.44613540328924 |
| ENSG00000140836.16 | ZFHX3 | -0.983012831756493 | 3.18830427275507 | -3.25619305781496 | 0.00223122107553332 | 0.0766698220328921 | -1.58894916341477 |
| ENSG00000085733.15 | CTTN | -1.53411889714812 | 1.91766394555821 | -3.25876500308717 | 0.00223584454807182 | 0.0767006472216038 | -1.50767653632155 |
| ENSG00000205758.11 | CRYZL1 | -0.406044764282144 | 4.41497500078052 | -3.25322016498423 | 0.00223977623554829 | 0.0767076776310988 | -1.66703786178897 |
| ENSG00000082515.17 | MRPL22 | 0.309602749242135 | 2.98461729650415 | 3.24917769558233 | 0.00226536068321016 | 0.0773089270401371 | -1.57890478724571 |
| lnc-TVP23A-4 | lnc-TVP23A-4 | -1.67726814443854 | -0.696210416334066 | -3.24895644999722 | 0.00226676889762592 | 0.0773089270401371 | -1.60290574300878 |
| lnc-SOWAHB-7 | lnc-SOWAHB-7 | 0.350645911110128 | 5.00438795897853 | 3.24866897328789 | 0.00226859990925729 | 0.0773089270401371 | -1.70522189621253 |
| lnc-DHX38-25 | lnc-DHX38-25 | -0.856534231005925 | 0.231860625945648 | -3.24718630439451 | 0.00227806569497088 | 0.0775031838009681 | -1.45956298075066 |
| ENSG00000226742.3 | HSBP1L1 | -1.98631754300255 | -0.279964730216915 | -3.2495990661631 | 0.00228212143722446 | 0.0775130454494902 | -1.46950950647137 |
| ENSG00000171517.5 | LPAR3 | 1.86373347780013 | -0.615563036248398 | 3.24462230017739 | 0.00229452348809193 | 0.0777213886683829 | -1.67948978669814 |

| | | | | | | | |
|---|---|---|---|---|---|---|---|
| ENSG00000257390.5 | | 0.306035145392458 | 4.02423798567471 | 3.24442300551707 | 0.00229580742896452 | 0.0777213886683829 | -1.66434379385253 |
| ENSG00000115464.14 | USP34 | 0.362000637667705 | 8.46208578667695 | 3.24244171082294 | 0.00230860887600128 | 0.0780264310258365 | -1.72408529650575 |
| ENSG00000188375.4 | H3-5 | -1.04829711290552 | 1.34531403190354 | -3.24122400853403 | 0.002316510185188 | 0.0781651297405321 | -1.51276599877677 |
| ENSG00000180530.10 | NRIP1 | -2.29774160971248 | 1.98741839027467 | -3.24764277558471 | 0.0023238784092641 | 0.0782854161994811 | -1.52228353936607 |
| ENSG00000149809.14 | TM7SF2 | -0.335476825807763 | 3.23211490954692 | -3.2386642691629 | 0.00233320324751477 | 0.0782948080687914 | -1.62756827331665 |
| ENSG00000120063.9 | GNA13 | 0.40346189732664 | 7.33593180339637 | 3.23828519560517 | 0.00233568500778808 | 0.0782948080687914 | -1.74989346701337 |
| ENSG00000255154.7 | HTD2 | 0.337609126153601 | 2.12940852271194 | 3.2380217087116 | 0.00233741150662363 | 0.0782948080687914 | -1.55272686283895 |
| ENSG00000139372.14 | TDG | 0.360719393452117 | 5.09374040258882 | 3.23772263779561 | 0.00233937263578228 | 0.0782948080687914 | -1.73751331337133 |
| ENSG00000120690.15 | ELF1 | 0.381236895449119 | 7.93674535260839 | 3.23644841229345 | 0.002347745741975 | 0.0784474847517393 | -1.74852669085821 |
| ENSG00000071564.14 | TCF3 | -0.658119591573461 | 6.87964989408498 | -3.2358552144663 | 0.00235165340775874 | 0.078450700311018 | -1.75857621018782 |
| ENSG00000119403.14 | PHF19 | -0.525470160051199 | 4.77455616054241 | -3.23443219346567 | 0.00236105264764848 | 0.0785658028646867 | -1.73865991991933 |
| lnc-ASMT-2 | lnc-ASMT-2 | 1.9557022669705 | 1.11657349086688 | 3.23757367643541 | 0.00236613799295685 | 0.0785658028646867 | -1.48937383457367 |
| ENSG00000259660.2 | DNM1P47 | -0.574764264244788 | 2.61408955836664 | -3.23360166875606 | 0.00236655481592119 | 0.0785658028646867 | -1.59951870411209 |
| ENSG00000207031.1 | SNORD59A | 0.656093960608538 | 1.46809980663588 | 3.23191546262001 | 0.0023777632200102 | 0.0787524061028223 | -1.53123049214262 |
| ENSG00000068912.13 | ERLEC1 | 0.371537492141369 | 4.76866907241192 | 3.23122158095129 | 0.00238239012596907 | 0.0787524061028223 | -1.74019855631663 |
| lnc-C19orf71-3 | lnc-C19orf71-3 | -0.718250147426515 | -0.196917431895812 | -3.23103227336287 | 0.00238365393781559 | 0.0787524061028223 | -1.52373538306614 |
| ENSG00000273544.1 | SNORA44 | -0.46404810826739 | 1.11479579404958 | -3.22859666446383 | 0.00239997076705692 | 0.079164420349892 | -1.52815158336815 |
| lnc-YIPF2-1 | lnc-YIPF2-1 | -0.561154290520735 | 0.512413470319511 | -3.2262449023572 | 0.00241582627835438 | 0.0795014479638548 | -1.51021925523606 |
| ENSG00000125814.17 | NAPB | -0.326536197412967 | 3.96653660918683 | -3.22587420620395 | 0.00241833453835058 | 0.0795014479638548 | -1.71285869041376 |
| ENSG00000131100.12 | ATP6V1E1 | 0.329756569535476 | 5.16110210005123 | 3.22536623839709 | 0.00242177563393757 | 0.0795014479638548 | -1.76970473605465 |
| lnc-EIF2AK3-1 | lnc-EIF2AK3-1 | -0.655366174405698 | 1.25803775892302 | -3.22425329980828 | 0.00242933117039024 | 0.079622489618061 | -1.53665470464203 |
| ENSG00000143093.14 | STRIP1 | 0.22996347354445 | 5.95539722728631 | 3.22135736137071 | 0.00244909596373513 | 0.0800429491123823 | -1.79801883367906 |
| ENSG00000115827.13 | DCAF17 | -0.282823891221804 | 4.94954832624301 | -3.22123459129268 | 0.00244993722687659 | 0.0800429491123823 | -1.77638001004921 |
| lnc-CARD16-1 | lnc-CARD16-1 | 1.19426328921125 | 0.696279600162152 | 3.21931758637926 | 0.00246310877020696 | 0.0802721867448538 | -1.52482825944929 |
| TMSB15B-AS1 | TMSB15B-AS1 | -0.555769938500403 | 0.594391103959549 | -3.2187558562445 | 0.00246698104986305 | 0.0802721867448538 | -1.5299055035847 |
| LINC00884 | LINC00884 | -0.779866121153074 | 0.21966430094798 | -3.21683502284271 | 0.00248026590752521 | 0.0802721867448538 | -1.53015727925995 |
| lnc-C1GALT1C1-1 | lnc-C1GALT1C1-1 | -0.507279932203396 | 1.05190835159781 | -3.21676482546794 | 0.00248075268729252 | 0.0802721867448538 | -1.54985369797468 |
| ENSG00000198961.9 | PJA2 | 0.315632631965531 | 7.40506255609764 | 3.2167497070829 | 0.00248085753672945 | 0.0802721867448538 | -1.80500047844861 |
| ENSG00000155508.13 | CNOT8 | 0.308608682719816 | 6.13906737184007 | 3.21615496529524 | 0.00248498553807299 | 0.0802721867448538 | -1.81215747939977 |
| LAMTOR5-AS1 | LAMTOR5-AS1 | 0.356621223312282 | 3.83292324389478 | 3.21565049103179 | 0.00248849210645799 | 0.0802721867448538 | -1.72158675041748 |
| ENSG00000104812.14 | GYS1 | -0.461865089678855 | 4.40365930064527 | -3.21546517411796 | 0.00248978140904763 | 0.0802721867448538 | -1.76421795134056 |
| ENSG00000176055.9 | MBLAC2 | -0.558628118225415 | 2.73144142652821 | -3.21464630783433 | 0.0024954860729832 | 0.0802721867448538 | -1.6571893103134 |
| ENSG00000077238.13 | IL4R | -0.903305506736086 | 9.49800197844513 | -3.21617926372058 | 0.00249989777208242 | 0.0802721867448538 | -1.77238802513557 |
| ENSG00000136371.10 | MTHFS | -0.445291813627244 | 0.443724118572427 | -3.21314436669559 | 0.00250598159309464 | 0.0802721867448538 | -1.53985998513389 |
| ENSG00000204837.4 | FGF7P3 | -1.52256445185377 | -0.658007291749202 | -3.21276186238163 | 0.0025086611928831 | 0.0802721867448538 | -1.66108344570369 |
| ENSG00000086666.18 | ZFAND6 | 0.584304128223534 | 5.88708220342706 | 3.21237941119797 | 0.00251134313309012 | 0.0802721867448538 | -1.81911673447005 |
| lnc-CTTNBP2-3 | lnc-CTTNBP2-3 | -0.845378245020851 | 0.485879373403554 | -3.21234954939212 | 0.00251155265333945 | 0.0802721867448538 | -1.54357348745371 |
| ENSG00000170903.10 | MSANTD4 | 0.543651435347858 | 3.92518748096611 | 3.21143105424996 | 0.00251800520789125 | 0.0803536452620552 | -1.73248501594849 |
| ENSG00000110172.11 | CHORDC1 | 0.600699485078771 | 5.7730546380909 | 3.21035265648023 | 0.00252560113524088 | 0.0804712819917385 | -1.82215674763383 |
| lnc-MRPS30-8 | lnc-MRPS30-8 | -0.565114666313166 | 2.8627780779881 | -3.20947174147263 | 0.00253182214597929 | 0.0805448148851495 | -1.6745678679425 |
| ENSG00000248527.1 | MTATP6P1 | -1.14860830120271 | 5.07697715715045 | -3.21188942041985 | 0.00255130909698878 | 0.081039498677963 | -1.80887580987833 |
| ENSG00000188295.14 | ZNF669 | 0.41490427633941 | 4.38864945737292 | 3.2058490749488 | 0.00255755821471184 | 0.0810756811844498 | -1.77788285717155 |
| lnc-TRIM38-2 | lnc-TRIM38-2 | -0.391644826911679 | 3.23847660127672 | -3.20485359074115 | 0.00256467357479685 | 0.0810756811844498 | -1.71050951342192 |
| ENSG00000205089.7 | CCNI2 | -0.864502529747071 | 1.17562320570248 | -3.20472526871663 | 0.00256559213643987 | 0.0810756811844498 | -1.59528424370242 |
| ENSG00000114026.21 | OGG1 | -0.512260449878525 | 4.1201357042557 | -3.20436062370242 | 0.00256820405831323 | 0.0810756811844498 | -1.77796285755015 |
| ENSG00000166707.10 | ZCCHC18 | -0.569874987408441 | 3.91428977784279 | -3.20360354087849 | 0.00257363502032976 | 0.0811227099899656 | -1.75644475290841 |
| ENSG00000280163.1 | | -0.792243404500611 | -0.0210071324126012 | -3.20192817568918 | 0.00258569198880818 | 0.0813781318129033 | -1.57732366540907 |
| ENSG00000113811.10 | SELENOK | 0.548112427690454 | 4.21457724582398 | 3.20106734029994 | 0.00259190786950886 | 0.0814492208825966 | -1.78255313755636 |
| ENSG00000074319.12 | TSG101 | 0.313713177402724 | 4.40473504877581 | 3.19921636504657 | 0.00260532124075208 | 0.0817459254548782 | -1.80310976084645 |
| lnc-SNRPN-14 | lnc-SNRPN-14 | -0.60049942561727 | 2.24953078853446 | -3.194555025931 | 0.00263939197574292 | 0.0818723941769493 | -1.67522613736865 |
| ENSG00000103995.13 | CEP152 | 0.443279690346316 | 4.08954120618604 | 3.19389814608527 | 0.00264422701145942 | 0.0818723941769493 | -1.79083577577774 |
| ENSG00000259030.7 | FPGT-TNNI3K | -0.729499154465353 | 2.39600452523767 | -3.19317088020475 | 0.00264958990701139 | 0.0818723941769493 | -1.6778757434599 |
| lnc-EPSTI1-3 | lnc-EPSTI1-3 | -2.61054535208844 | 0.0156118716729636 | -3.19862160914677 | 0.00265020966190916 | 0.0818723941769493 | -1.59434661128357 |
| ENSG00000087206.16 | UIMC1 | 0.371426896586123 | 5.43272946740588 | 3.1927568228574 | 0.00265264778293293 | 0.0818723941769493 | -1.86065922607883 |
| lnc-MFSD8-12 | lnc-MFSD8-12 | -0.594670130015436 | 0.990774390154143 | -3.19207970334421 | 0.00265765560899959 | 0.0818723941769493 | -1.60411675070366 |
| ENSG00000269069.1 | | -0.52503107155918 | 2.81561793811046 | -3.19207678764718 | 0.00265767719217884 | 0.0818723941769493 | -1.71063011671649 |
| ENSG00000119778.14 | ATAD2B | 0.36713329661811 | 6.35406773949921 | 3.19188978982573 | 0.00265906177281234 | 0.0818723941769493 | -1.87467451807659 |
| ENSG00000137841.11 | PLCB2 | -0.607699925171961 | 6.04182281493944 | -3.19148498145967 | 0.00266206141774923 | 0.0818723941769493 | -1.87330652958569 |
| MRPS30-DT | MRPS30-DT | -0.722118960430005 | 1.35353342496618 | -3.19135237164481 | 0.00266304475736284 | 0.0818723941769493 | -1.61852443555781 |
| ENSG00000180773.14 | SLC36A4 | 0.5546853525138 | 3.87968018943772 | 3.19076220219526 | 0.00266742520183562 | 0.0818723941769493 | -1.78210500082783 |
| ENSG00000125482.12 | TTF1 | 0.291120086889571 | 4.21581913584849 | 3.19053422549405 | 0.00266911914941889 | 0.0818723941769493 | -1.8094350868879 |
| lnc-ZNF732-10 | lnc-ZNF732-10 | -1.30234431578875 | 0.119531176587131 | -3.18993907144743 | 0.00267354615250484 | 0.0818723941769493 | -1.59857783594786 |
| lnc-ACOX3-5 | lnc-ACOX3-5 | -0.556267190491002 | 1.05057847517204 | -3.18958438686809 | 0.00267618774639408 | 0.0818723941769493 | -1.61629945145441 |
| ENSG00000255639.3 | | 0.402048180026717 | 3.57276993406656 | 3.18916703052665 | 0.00267929926470575 | 0.0818723941769493 | -1.77203680035142 |
| ENSG00000213398.7 | LCAT | -0.811745705871142 | 0.686784569711042 | -3.18912352648669 | 0.00267962379743536 | 0.0818723941769493 | -1.60618761548288 |
| ENSG00000240024.5 | SNHG33 | -0.720351228064683 | 0.220310029982179 | -3.18903753382623 | 0.00268026539744657 | 0.0818723941769493 | -1.59441137767221 |
| ENSG00000108406.9 | DHX40 | 0.490034649272401 | 5.58107193822132 | 3.188936102459 | 0.00268102237403813 | 0.0818723941769493 | -1.87514732890248 |
| ENSG00000178202.12 | POGLUT3 | 0.70851018151064 | 2.51540650724958 | 3.18835940357503 | 0.00268533009081707 | 0.0818723941769493 | -1.68783879834399 |

| | | | | | | | |
|---|---|---|---|---|---|---|---|
| LINC01806 | LINC01806 | -2.11998284364144 | 2.35145738940754 | -3.19510955676985 | 0.00268890533273175 | 0.0818723941769493 | -1.70163768742383 |
| ENSG00000132963.7 | POMP | 0.437266687551467 | 4.14806728979152 | 3.18733258369495 | 0.00269301624931732 | 0.08187644528759 | -1.81697749126661 |
| lnc-PKD2L1-3 | lnc-PKD2L1-3 | 0.654185816599143 | 2.57067575100112 | 3.18625322651604 | 0.00270111810385741 | 0.0818928712329593 | -1.69643038480197 |
| ENSG00000183317.16 | EPHA10 | -0.979507349410119 | 3.34704465255418 | -3.1880132871166 | 0.00270151384964191 | 0.0818928712329593 | -1.76270078327512 |
| ENSG00000275405.1 | | -1.12957306946773 | 1.26908252857967 | -3.1857330038188 | 0.00270794448233916 | 0.0819670901176278 | -1.63913643257987 |
| lnc-GBP7-1 | lnc-GBP7-1 | 1.22140707664844 | 0.406688807353641 | 3.18409703005179 | 0.00271737199687774 | 0.0820926756650435 | -1.605880961231 |
| lnc-ZNF100-9 | lnc-ZNF100-9 | -2.04212732877438 | 1.00966458255367 | -3.19039244172439 | 0.00272007019402223 | 0.0820926756650435 | -1.6647502256687 |
| ENSG00000108039.17 | XPNPEP1 | 0.316998062061806 | 5.49613193475078 | 3.18230164156004 | 0.00273097657994043 | 0.0823011580452619 | -1.89094373930583 |
| ENSG00000066135.12 | KDM4A | 0.350107291396753 | 5.04086987721127 | 3.18047829818203 | 0.00274485890559962 | 0.0825827261938957 | -1.88044441604978 |
| ENSG00000117036.11 | ETV3 | 0.508013461929297 | 4.45985665106469 | 3.18002190738951 | 0.0027483441404469 | 0.0825827261938957 | -1.85138483341029 |
| lnc-ADAMTS15-1 | lnc-ADAMTS15-1 | -0.639706616567603 | -0.130905333712496 | -3.17758280961168 | 0.00276704140242444 | 0.0830233428368837 | -1.64462058827774 |
| lnc-PABPN1L-2 | lnc-PABPN1L-2 | -0.504071277286798 | 0.922304024557182 | -3.17630595588306 | 0.00277687721215224 | 0.0830291334414125 | -1.6434812407768 |
| ENSG00000079387.13 | SENP1 | 0.228786476742879 | 5.04609005956369 | 3.17603514429972 | 0.00277896756231363 | 0.0830291334414125 | -1.89415106642356 |
| lnc-RPL35A-2 | lnc-RPL35A-2 | -0.586922669948568 | 1.44944199029478 | -3.17575917251063 | 0.0027810992738203 | 0.0830291334414125 | -1.67604193777697 |
| ENSG00000173214.5 | MFSD4B | -0.42603905701219 | 4.55215741297472 | -3.17546544032025 | 0.00278336987196107 | 0.0830291334414125 | -1.87317075107938 |
| ENSG00000224631.4 | RPS27AP16 | 1.18269110372784 | 3.2185564544787 | 3.17811600640121 | 0.00278775175491171 | 0.0830394998138171 | -1.75485562038035 |
| lnc-HLA-C-2 | lnc-HLA-C-2 | 0.430786759931573 | 8.56098132879394 | 3.17331472424664 | 0.00280004874074057 | 0.0831903897184396 | -1.89874647802316 |
| ENSG00000089327.14 | FXYD5 | -0.656997699899888 | 6.57847415442313 | -3.17320518471155 | 0.00280090074696976 | 0.0831903897184396 | -1.92061227136636 |
| ENSG00000122218.15 | COPA | 0.246180540904722 | 7.13110071575461 | 3.17117654420307 | 0.00281672402536476 | 0.0835398135649609 | -1.92363636573217 |
| ENSG00000187522.15 | HSPA14 | 0.379430941317481 | 3.9201028451368 | 3.16802531717338 | 0.0028414710988852 | 0.0841525174508692 | -1.8497893826165 |
| ENSG00000259305.6 | ZHX1-C8orf76 | 0.35132646655119 | 2.38051548931454 | 3.16545712529734 | 0.00286179135223141 | 0.0846325451192229 | -1.7452806746504 |
| lnc-ELF2-4 | lnc-ELF2-4 | 0.368045279334018 | 2.48900626804923 | 3.16454235727571 | 0.00286906236697313 | 0.0846948360178199 | -1.75658861435204 |
| ENSG00000080503.23 | SMARCA2 | -0.234696752843265 | 7.50507455390013 | -3.16415741772384 | 0.00287212726718352 | 0.0846948360178199 | -1.9370423128522 |
| ENSG00000101974.14 | ATP11C | 0.356792416670798 | 5.81950766964031 | 3.16335024422761 | 0.0028785640559318 | 0.084763210247846 | -1.94354511306878 |
| ENSG00000168811.6 | IL12A | 0.593521121727678 | 2.63018558985483 | 3.16245417225055 | 0.00288572573007801 | 0.0848527038602796 | -1.77666755276647 |
| ENSG00000100292.16 | HMOX1 | -0.873106960784195 | 4.04111016652785 | -3.16325898058391 | 0.00289092301159987 | 0.0848842629782597 | -1.87581519474709 |
| lnc-GATA5-10 | lnc-GATA5-10 | -0.632099380828371 | 1.92989086033116 | -3.16110839007613 | 0.00289651326200071 | 0.0849272542332772 | -1.73662492066152 |
| lnc-MVD-3 | lnc-MVD-3 | -0.663547412471017 | 0.870518988035083 | -3.16002965539117 | 0.00290518769407571 | 0.0850604243345097 | -1.68363932763756 |
| ENSG00000207501.1 | RNVU1-14 | -0.911971454277013 | 5.49002981136128 | -3.16249038463653 | 0.00291010832024032 | 0.0850834652777083 | -1.94413460827829 |
| lnc-SKP1-1 | lnc-SKP1-1 | 0.408125166168576 | 3.51807756149453 | 3.15853332785929 | 0.0029172607617488 | 0.0851057415637785 | -1.84308402911712 |
| ENSG00000137876.9 | RSL24D1 | 0.380516954635553 | 5.77166945612745 | 3.15796328117796 | 0.00292187262104663 | 0.0851057415637785 | -1.95753823786328 |
| ENSG00000106346.11 | USP42 | 0.449239310890489 | 3.87458724354329 | 3.15779016955148 | 0.00292327451224755 | 0.0851057415637785 | -1.87483441293004 |
| lnc-GRIK5-3 | lnc-GRIK5-3 | -0.717342875657388 | 1.59958068587776 | -3.1566069448233 | 0.00293287352247438 | 0.0852532920764424 | -1.72850523868283 |
| lnc-RAB27A-3 | lnc-RAB27A-3 | 0.322019062877295 | 3.62253260636194 | 3.15614532259165 | 0.0029366265404556 | 0.0852532920764424 | -1.8551241952232 |
| lnc-DHX38-27 | lnc-DHX38-27 | -0.774003560146564 | 0.0809509912643559 | -3.15474115317555 | 0.00294807043584253 | 0.0854649771562631 | -1.67602388576913 |
| ENSG00000217555.12 | CKLF | -0.552252378885719 | 2.98783078963349 | -3.15152198342472 | 0.00297446567430376 | 0.0861088986978823 | -1.83267447965506 |
| lnc-MOB1A-2 | lnc-MOB1A-2 | 0.394327650316111 | 3.78945767287498 | 3.1507644734014 | 0.00298070915901612 | 0.0861177735310239 | -1.88139282257886 |
| ENSG00000213199.7 | ASIC3 | -0.642590423096769 | 0.673766736306304 | -3.1502799893449 | 0.00298470881998525 | 0.0861177735310239 | -1.69453160591814 |
| ENSG00000276027.1 | RNU12 | -1.01269839750121 | 2.52528934102875 | -3.15073803828051 | 0.0029888607063351 | 0.0861177735310239 | -1.79445511009927 |
| ENSG00000198964.13 | SGMS1 | -0.29879912014743 | 4.97394730026565 | -3.14934017023665 | 0.00299248195986149 | 0.0861177735310239 | -1.95750423678492 |
| ENSG00000188157.14 | AGRN | 1.21772078719772 | 0.0231643685892261 | 3.14895274485403 | 0.00299569187427552 | 0.0861177735310239 | -1.69705498858088 |
| ENSG00000103855.17 | CD276 | -1.60519360927033 | 0.932172417430983 | -3.15123441507678 | 0.00300200153333089 | 0.0861270987143033 | -1.69883329067092 |
| ENSG00000172893.15 | DHCR7 | -1.01575825609142 | 1.30289746486017 | -3.14788588449162 | 0.00300454788871327 | 0.0861270987143033 | -1.73417597759221 |
| lnc-FAM71F2-5 | lnc-FAM71F2-5 | -0.915464302198537 | 0.397391197123257 | -3.14740240381347 | 0.00300856940074742 | 0.0861270987143033 | -1.69075737435322 |
| ENSG00000139218.17 | SCAF11 | 0.263347873951068 | 8.19143223814378 | 3.14676811487402 | 0.00301385301999101 | 0.0861585232089929 | -1.97168767350955 |
| ENSG00000196422.10 | PPP1R26 | -0.651493533331142 | 3.23234633826151 | -3.14517252242895 | 0.00302718305785646 | 0.0861982766818986 | -1.86852335235352 |
| lnc-CCDC57-4 | lnc-CCDC57-4 | 0.490818696679444 | 2.25893855796692 | 3.14493824527429 | 0.00302914495739494 | 0.0861982766818986 | -1.78821026918125 |
| lnc-RGS6-3 | lnc-RGS6-3 | -0.678343640562573 | 1.04084217654508 | -3.14464817508407 | 0.00303157574629022 | 0.0861982766818986 | -1.71911593285975 |
| ENSG00000213341.10 | CHUK | 0.410066552491687 | 4.37619187716306 | 3.14411088855437 | 0.00303608307485701 | 0.0861982766818986 | -1.93862817539758 |
| lnc-RGS10-2 | lnc-RGS10-2 | 0.294048491187487 | 3.55707985313059 | 3.14402972557899 | 0.00303676450574947 | 0.0861982766818986 | -1.88527675327494 |
| lnc-DEXI-1 | lnc-DEXI-1 | -1.63437897226407 | -0.363372214709452 | -3.14356111200197 | 0.00304070173247312 | 0.0861982766818986 | -1.76457730898299 |
| ENSG00000207306.1 | RNU6-1152P | -1.34374691444193 | -0.0640003504094735 | -3.14266486441081 | 0.00304824530909041 | 0.0861982766818986 | -1.71725657980768 |
| ENSG00000103222.18 | ABCC1 | 0.384824102141206 | 6.42610848742371 | 3.14237844438648 | 0.0030506597852293 | 0.0861982766818986 | -2.00028831951826 |
| ENSG00000204839.8 | MROH6 | -0.992784099978929 | 0.627012486435043 | -3.14210245384917 | 0.00305298805055442 | 0.0861982766818986 | -1.71453024203215 |
| ENSG00000141076.17 | UTP4 | 0.410523135884556 | 3.82159688626894 | 3.14116179763195 | 0.00306093606700907 | 0.0861982766818986 | -1.90557097935321 |
| ENSG00000183666.16 | GUSBP1 | -0.33998556480701 | 3.63741984775058 | -3.14093610084405 | 0.00306284598178519 | 0.0861982766818986 | -1.89991971661568 |
| ENSG00000130733.10 | YIPF2 | -0.463132216247802 | 2.6913160176773 | -3.14062296810701 | 0.0030654976694918 | 0.0861982766818986 | -1.83849152533296 |
| lnc-TARDBP-5 | lnc-TARDBP-5 | -0.687467869333166 | 0.712529159228307 | -3.13732735632621 | 0.00309353749857794 | 0.0867622115110865 | -1.71955221595614 |
| ENSG00000156502.13 | SUPV3L1 | 0.386618694775222 | 5.26804071637719 | 3.13727515956838 | 0.00309398354220169 | 0.0867622115110865 | -1.998806662345 |
| ENSG00000135299.16 | ANKRD6 | -0.886007044970207 | 1.9504491980042 | -3.13539646421871 | 0.00311007827113785 | 0.087094885788885 | -1.81093799502254 |
| lnc-UBA3-1 | lnc-UBA3-1 | 0.366542450543791 | 1.99553647434642 | 3.13461317716797 | 0.00311681198338901 | 0.0871648655626304 | -1.79771291165091 |
| lnc-ZHX1-1 | lnc-ZHX1-1 | 0.512837739359208 | 1.29738231504025 | 3.13259529264535 | 0.00313422264551803 | 0.0875328422153292 | -1.76499080906399 |
| ENSG00000122557.9 | HERPUD2 | 0.302297104183562 | 5.69323519556874 | 3.13177765014668 | 0.00314130350566564 | 0.0876117209446014 | -2.02182780256017 |
| lnc-ZNF843-2 | lnc-ZNF843-2 | -0.361209387035876 | 5.42960576581485 | -3.13104522044815 | 0.00314765923107449 | 0.0876623099707299 | -2.01816962975797 |
| ENSG00000137040.9 | RANBP6 | 0.296574615392095 | 6.18909791175265 | 3.13040662970024 | 0.00315321056111126 | 0.0876623099707299 | -2.03030236558795 |
| lnc-TMEM86A-5 | lnc-TMEM86A-5 | -0.732372952923633 | 0.193804792133186 | -3.12962386543297 | 0.00316002780845586 | 0.0876623099707299 | -1.73178201999448 |
| ENSG00000213780.10 | GTF2H4 | -0.450654820195104 | 1.30670771602892 | -3.12960947895534 | 0.00316015323316725 | 0.0876623099707299 | -1.7744993115987 |
| lnc-FAAP100-2 | lnc-FAAP100-2 | -0.812389237935183 | 3.10133468825093 | -3.12863402670852 | 0.00316866842199762 | 0.0877218383548763 | -1.89091955356888 |

| | | | | | | | |
|---|---|---|---|---|---|---|---|
| ENSG00000133935.6 | ERG28 | -0.421943398962352 | 2.89448090315383 | -3.1283876139764 | 0.00317082289928718 | 0.0877218383548763 | -1.88175482558388 |
| ENSG00000110944.8 | IL23A | -0.792452843474053 | 0.795583138033614 | -3.12780820685248 | 0.00317589431661605 | 0.0877442049918229 | -1.75656372137104 |
| ENSG00000102606.18 | ARHGEF7 | 0.332487471760814 | 6.25344905355377 | 3.12575920758951 | 0.00319389019069155 | 0.0880613777424457 | -2.04217683907879 |
| MAPKAPK5-AS1 | MAPKAPK5-AS1 | 0.36102401129624 | 3.59491107406368 | 3.12551753263993 | 0.00319601909599553 | 0.0880613777424457 | -1.93251193140784 |
| ENSG00000151239.13 | TWF1 | 0.354771886007863 | 5.43066685570783 | 3.12504228689061 | 0.00320020942289022 | 0.0880613777424457 | -2.03541533860068 |
| ENSG00000173818.16 | ENDOV | -0.471402217300231 | 2.70080802747051 | -3.1241421138112 | 0.0032081606204404 | 0.0881208469076791 | -1.87822756317574 |
| lnc-RHOG-3 | lnc-RHOG-3 | 0.428780019754943 | 2.21578699174211 | 3.12322123090814 | 0.00321631402981332 | 0.0881208469076791 | -1.83860791113859 |
| ENSG00000263740.2 | RN7SL4P | 1.39531455932223 | 4.48476747274375 | 3.12828629804118 | 0.00321893465631641 | 0.0881208469076791 | -1.95084394471855 |
| lnc-BTD-2 | lnc-BTD-2 | 1.39531455932224 | 4.48478677043073 | 3.12819316027804 | 0.00321975722012 | 0.0881208469076791 | -1.95107492743518 |
| ENSG00000185090.14 | MANEAL | -0.769917895542466 | 1.44117352677479 | -3.12225732620341 | 0.00322486928658218 | 0.0881208469076791 | -1.79858524911394 |
| ENSG00000010810.17 | FYN | 1.86769546272773 | 4.08776797695425 | 3.12870406218937 | 0.0032306575148786 | 0.0881208469076791 | -1.93999385114949 |
| lnc-GPD1-1 | lnc-GPD1-1 | 0.315082154211575 | 2.44647111674111 | 3.121417666556 | 0.00323233928073156 | 0.0881208469076791 | -1.86428070619864 |
| lnc-PEA15-1 | lnc-PEA15-1 | -0.953294592804992 | -0.42079377812478 | -3.12086300474464 | 0.00323728275326073 | 0.0881388768655629 | -1.78959251476083 |
| ENSG00000174977.8 | PAIP1P2 | 0.871055909806282 | 0.758019408916165 | 3.12005090497428 | 0.00324453353029717 | 0.0882195953158608 | -1.7596377929921 |
| ENSG00000137411.17 | VARS2 | -0.442950522942904 | 4.39729250036121 | -3.11933378148737 | 0.00325094905096606 | 0.0882774199419979 | -2.00745305530914 |
| ENSG00000037474.14 | NSUN2 | 0.216896839755457 | 5.81035683303104 | 3.11789698461728 | 0.00326383892180203 | 0.0885106673615956 | -2.059235804799 |
| ENSG00000088356.5 | PDRG1 | 0.51238228973001 | 2.91927067135901 | 3.11550435018146 | 0.00328541087835378 | 0.0888640865066996 | -1.90525246461828 |
| ENSG00000137947.11 | GTF2B | 0.395242633309937 | 5.07690637440336 | 3.11549383358802 | 0.00328550599191558 | 0.0888640865066996 | -2.04333570624415 |
| ENSG00000254788.7 | CKLF-CMTM1 | -0.652248454744072 | 2.52090970014812 | -3.11329839957045 | 0.00330541876039199 | 0.089251228195378 | -1.89680423481417 |
| ENSG00000279267.1 | | -0.787567883821565 | 0.168365955368678 | -3.11225699069028 | 0.0033149041682919 | 0.089251228195378 | -1.77086435222612 |
| lnc-SLC35F5-15 | lnc-SLC35F5-15 | -0.787567883821565 | 0.168365955368678 | -3.11225699069028 | 0.0033149041682919 | 0.089251228195378 | -1.77086435222612 |
| lnc-MLKL-4 | lnc-MLKL-4 | 0.366391877979018 | 0.915851799218465 | 3.11166063033554 | 0.0033203475155675 | 0.089251228195378 | -1.790193892804 |
| ENSG00000105227.14 | PRX | -0.65168032742322 | 0.587517829314278 | -3.11153445161432 | 0.00332150030596412 | 0.089251228195378 | -1.78652616753245 |
| lnc-PRICKLE3-4 | lnc-PRICKLE3-4 | -0.788197755465893 | -0.397866124043884 | -3.10988763210378 | 0.00333658057517587 | 0.0895395540793285 | -1.80539502298631 |
| ENSG00000198406.7 | BZW1P2 | 1.195650302673 | -0.0276664972322578 | 3.1082979104394 | 0.00335119920048745 | 0.0897813389319884 | -1.83608915833284 |
| lnc-PTPN14-2 | lnc-PTPN14-2 | -0.67001110575577 | 1.6786976634667 | -3.10796000837847 | 0.00335431422235335 | 0.0897813389319884 | -1.84056394563937 |
| ENSG00000135537.16 | AFG1L | -0.428632498291351 | 2.77371777108802 | -3.10712033656431 | 0.00336206673832213 | 0.0898719736037461 | -1.9166182793992 |
| ENSG00000105982.16 | RNF32 | -0.490480593731108 | 1.4358552286312 | -3.1063463527522 | 0.00336922772785344 | 0.0899465814817215 | -1.83316422790608 |
| PXN-AS1 | PXN-AS1 | -0.495595392697839 | 1.90777307528101 | -3.10490075832972 | 0.00338264105784502 | 0.090079984773528 | -1.87193453881534 |
| ENSG00000112739.16 | PRP4K | 0.325856362811928 | 7.34144926403 | 3.10486455619919 | 0.00338297761404738 | 0.090079984773528 | -2.0887928750839 |
| ENSG00000165119.20 | HNRNPK | 0.289169323355129 | 8.84806007073815 | 3.10176073713568 | 0.00341195017521601 | 0.0907340703571979 | -2.07223151242105 |
| ENSG00000160785.13 | SLC25A44 | 0.338932611162786 | 4.17192442969208 | 3.10090172455587 | 0.0034200098387555 | 0.0908310484014251 | -2.0355568846063 |
| ENSG00000184860.9 | SDR42E1 | 0.603764680479943 | 3.30260699724933 | 3.1003147284659 | 0.00342552763005769 | 0.0908603546513883 | -1.94898829009786 |
| lnc-PLCB1-1 | lnc-PLCB1-1 | -0.62938708963196 | -0.386746049296992 | -3.09882202804947 | 0.00343959690100312 | 0.0908685775150019 | -1.83810425872017 |
| ENSG00000089335.20 | ZNF302 | -0.386667181412433 | 5.73020894462364 | -3.09857697087215 | 0.00344191185546373 | 0.0908685775150019 | -2.10598214477162 |
| lnc-TSTD1-1 | lnc-TSTD1-1 | 0.359958553218755 | 4.95756297054792 | 3.09844340986581 | 0.00344317416946728 | 0.0908685775150019 | -2.0847965193593 |
| ENSG00000066422.4 | ZBTB11 | 0.276387444859788 | 5.87500461866684 | 3.09800634766532 | 0.00344730798915286 | 0.0908685775150019 | -2.10986842735518 |
| ENSG00000099840.13 | IZUMO4 | -0.707084737915026 | 2.57081898546713 | -3.09794259801077 | 0.0034479113365018 | 0.0908685775150019 | -1.93020296624701 |
| LINC00665 | LINC00665 | -0.306063008601633 | 3.64216530041893 | -3.09715823414545 | 0.00345534295801889 | 0.0909479847888782 | -2.00686236858672 |
| ENSG00000124789.11 | NUP153 | 0.272903821841123 | 6.6376254422547 | 3.09597098618319 | 0.00346662050831723 | 0.0910678331678368 | -2.11635348985203 |
| ENSG00000198648.10 | STK39 | -0.608376904579861 | 3.07109599180134 | -3.09574772237051 | 0.00346874513936666 | 0.0910678331678368 | -1.97336976008018 |
| ENSG00000143751.9 | SDE2 | 0.355289121653187 | 5.69470603000682 | 3.09198595133811 | 0.00350472803300975 | 0.091895308412025 | -2.12185778975666 |
| lnc-RPL11-1 | lnc-RPL11-1 | -1.04668211584476 | 1.9179517883555 | -3.09187704626682 | 0.0035105002757175 | 0.091907622989904 | -1.88875922050121 |
| ENSG00000111581.9 | NUP107 | 0.264307698508794 | 5.708322785035 | 3.09100175270979 | 0.00351420016939589 | 0.091907622989904 | -2.12511868519401 |
| lnc-INO80E-2 | lnc-INO80E-2 | -0.60172822197621 | 1.79732498947193 | -3.09054612901297 | 0.00351859334965964 | 0.091907622989904 | -1.90229199917191 |
| lnc-CELA3A-5 | lnc-CELA3A-5 | -0.438259856512538 | 2.83528042766374 | -3.08952499510271 | 0.00352845804605264 | 0.0920484815740197 | -1.96891923813759 |
| ENSG00000127081.13 | ZNF484 | 0.433970840333311 | 5.02056992162834 | 3.08819734631241 | 0.00354132279516611 | 0.0921650876263871 | -2.10914328620157 |
| ENSG00000257108.1 | NHLRC4 | -0.665890250261145 | 1.24948508252113 | -3.08813960350536 | 0.00354188331693496 | 0.0921650876263871 | -1.87535244334376 |
| ENSG00000082269.16 | FAM135A | 0.652376454417613 | 3.28654538778605 | 3.08736830476311 | 0.00354937849577221 | 0.0922435070435347 | -1.99666602556936 |
| ENSG00000273559.4 | CWC25 | 0.338746524103886 | 4.6435696861302 | 3.08643218487732 | 0.00355849540956298 | 0.0923638222131585 | -2.097715834764 |
| ENSG00000149527.17 | PLCH2 | -0.884516922937574 | 3.83779418284219 | -3.08606281301667 | 0.00357834931678908 | 0.0927621712688533 | -2.06084845363568 |
| ENSG00000198689.11 | SLC9A6 | 0.217345863101191 | 4.23128120161551 | 3.08369753215787 | 0.003585254793321 | 0.0928242759885863 | -2.08558213966254 |
| ENSG00000223459.6 | TCAF1P1 | -0.771887278808559 | -0.0775491312917358 | -3.0813613689512 | 0.00360826480810342 | 0.09330265646381 | -1.84936902191321 |
| ENSG00000140543.14 | DET1 | -0.481537247151993 | 2.40215393359296 | -3.08075282669089 | 0.00361428142642623 | 0.0933409718947694 | -1.95796902927097 |
| ENSG00000202444.1 | RNU5E-6P | -1.28988152750763 | -0.150118485275163 | -3.07882721620967 | 0.0036333821073033 | 0.0937166715722104 | -1.85582485031645 |
| lnc-TNFRSF13B-1 | lnc-TNFRSF13B-1 | 3.71033297994516 | -1.9541601274927 | 3.07856566060911 | 0.00363931720525582 | 0.0937522728858329 | -2.69240243350191 |
| ENSG00000143442.21 | POGZ | 0.262096882838506 | 6.18108520476397 | 3.07460700120724 | 0.00367557649701785 | 0.094567988797648 | -2.16991538658256 |
| lnc-ACTR8-2 | lnc-ACTR8-2 | 0.568512000733706 | 1.97318861094355 | 3.07382677901965 | 0.00368342762980522 | 0.094573050267529 | -1.94170547422291 |
| ENSG00000171566.11 | PLRG1 | 0.197900428964362 | 5.34628923637075 | 3.07367441581244 | 0.00368496265435351 | 0.094573050267529 | -2.16006878450507 |
| lnc-TRIM69-2 | lnc-TRIM69-2 | 0.447982412774862 | 4.67793635772936 | 3.07270120721119 | 0.00369478172348417 | 0.0947069641525215 | -2.13411184918151 |
| lnc-NT5C3A-1 | lnc-NT5C3A-1 | -0.536610300104876 | 0.269830608793926 | -3.07101422074199 | 0.00371186073070368 | 0.0948658128929439 | -1.86655892739575 |
| ENSG00000100106.20 | TRIOBP | -0.363341152813526 | 4.91614076362073 | -3.07075944979429 | 0.00371444647368463 | 0.0948658128929439 | -2.15625716760378 |
| lnc-ZFAND1-1 | lnc-ZFAND1-1 | 0.338839032241173 | 2.0129616667322 | 3.07030236043423 | 0.00371908985335708 | 0.0948658128929439 | -1.95542732074032 |
| ENSG00000278053.4 | DDX52 | 0.267320242761852 | 5.38934120537358 | 3.07027041141773 | 0.00371941461422561 | 0.0948658128929439 | -2.17075417554256 |
| ENSG00000196072.11 | BLOC1S2 | 0.523632114941676 | 4.98314075012428 | 3.06932515538358 | 0.00372903519983321 | 0.0949334866132123 | -2.15221184303706 |
| lnc-DTNBP1-5 | lnc-DTNBP1-5 | -1.92032293713484 | 2.00193333768945 | -3.0751320246817 | 0.00373129236117615 | 0.0949334866132123 | -1.95372014254039 |
| ENSG00000184508.11 | HDDC3 | -0.374741216037688 | 2.30373995291654 | -3.06829775392243 | 0.00373951839742206 | 0.0950253174989361 | -1.98172480660901 |
| lnc-DNASE1L2-2 | lnc-DNASE1L2-2 | -0.591570162294663 | 1.37365043400869 | -3.06728519080131 | 0.0037498773195019 | 0.0951371310958135 | -1.92451721619712 |

| | | | | | | | |
|---|---|---|---|---|---|---|---|
| ENSG00000071967.11 | CYBRD1 | -1.38573050156641 | 2.97595170435275 | -3.07152086196461 | 0.00375316282610895 | 0.0951371310958135 | -2.01259356852082 |
| ENSG00000171307.18 | ZDHHC16 | -0.389883690003521 | 3.7261892806113 | -3.06531215146573 | 0.00377013990620421 | 0.0954499258172217 | -2.09688241516291 |
| lnc-ATP11A-2 | lnc-ATP11A-2 | -0.823816537565965 | 1.18311336051309 | -3.06472324963532 | 0.00377620770472855 | 0.0954860972806237 | -1.91913225046645 |
| lnc-IGFBP4-1 | lnc-IGFBP4-1 | -1.47930677840335 | 2.21555301927908 | -3.06733407847415 | 0.00379687232181576 | 0.0958908257667899 | -1.99322110577324 |
| lnc-CTBP2-6 | lnc-CTBP2-6 | -0.636998508368925 | 1.12556061779667 | -3.06136391716317 | 0.00381099692527554 | 0.0960217909872329 | -1.92634743034578 |
| ENSG00000163636.10 | PSMD6 | 0.244617024034851 | 5.80026953405903 | 3.06132629941895 | 0.00381138819591747 | 0.0960217909872329 | -2.20060451876223 |
| ENSG00000111275.12 | ALDH2 | 0.986094026030228 | 0.646552644030914 | 3.0594298658121 | 0.00383116247327864 | 0.0963108836477135 | -1.89325853270815 |
| ENSG00000177628.15 | GBA1 | -0.360043398918667 | 2.87463735555844 | -3.05880539192433 | 0.00383769500876915 | 0.0963108836477135 | -2.04363271046281 |
| ENSG00000188993.3 | LRRC66 | -1.19300052224408 | -0.254714703613039 | -3.05858790340395 | 0.00383997258503077 | 0.0963108836477135 | -1.92361359134839 |
| ENSG00000133321.10 | PLAAT4 | 1.33988290337414 | 0.843740251895317 | 3.05956091289044 | 0.00384157972476183 | 0.0963108836477135 | -1.90826299063995 |
| RFPL3S | RFPL3S | 0.527254046447022 | 1.21683942581934 | 3.05501356760809 | 0.00387758624380831 | 0.0970953256159446 | -1.94134804788373 |
| ENSG00000119446.13 | RBM18 | 0.295236557954256 | 4.36735101896048 | 3.05394039597834 | 0.00388894696522271 | 0.0972614767742152 | -2.16446060505058 |
| lnc-TOB2-2 | lnc-TOB2-2 | -0.664748696635873 | 1.28875275435978 | -3.05349130105447 | 0.003893710420648 | 0.0972624291118905 | -1.95809674474095 |
| ENSG00000125122.15 | FBXL9P | -0.5514428862982 | 0.841274401553681 | -3.05110461609131 | 0.00391911767698698 | 0.0977784232065734 | -1.93432022871517 |
| ENSG00000005156.11 | LIG3 | 0.322517798054966 | 5.12118013456159 | 3.05058464483968 | 0.00392467362895505 | 0.0977984955263702 | -2.21128151840303 |
| ENSG00000164134.12 | NAA15 | 0.346899880725407 | 6.84464868360304 | 3.05013148631982 | 0.00392952171775887 | 0.0978009014711376 | -2.22962589839739 |
| lnc-ULK4-14 | lnc-ULK4-14 | -0.597124377856722 | 1.8713394920087 | -3.0489521284998 | 0.00394216541391701 | 0.0979970902350891 | -1.99052606334171 |
| ENSG00000137752.23 | CASP1 | 0.938113112067431 | 3.19450783175948 | 3.04906396568339 | 0.00395222011615711 | 0.0981285243074327 | -2.07370232676042 |
| ENSG00000117425.13 | PTCH2 | -0.682540933993606 | 1.28385472896978 | -3.04750956517632 | 0.00395768289626595 | 0.0981457675347496 | -1.96627624724505 |
| ENSG00000250067.11 | YJEFN3 | -0.621713033531747 | 0.449084810370967 | -3.0464755823163 | 0.00396884063638813 | 0.0983040274594187 | -1.92581791493149 |
| lnc-ASMT-3 | lnc-ASMT-3 | 1.790923534806 | 1.95429394405016 | 3.05065209645878 | 0.00397688037761284 | 0.0983503887595041 | -1.9679913780633 |
| ENSG00000114098.17 | ARMC8 | 0.240634091580805 | 6.00469018558635 | 3.04505558606685 | 0.00398421205475164 | 0.0983503887595041 | -2.2426961321652 |
| ENSG00000179833.4 | SERTAD2 | 0.500637504794153 | 5.93268900303334 | 3.04497859402629 | 0.00398504708863754 | 0.0983503887595041 | -2.24150897527246 |
| ENSG00000170876.7 | TMEM43 | -0.289521748102919 | 5.71464414968705 | -3.04421985546037 | 0.00399328495560789 | 0.0984218114506606 | -2.2414787599238 |
| ENSG00000060688.12 | SNRNP40 | 0.225298833294165 | 5.14068872481767 | 3.0437652031034 | 0.00399822892802339 | 0.0984218114506606 | -2.22837505855399 |
| ENSG00000102181.20 | CD99L2 | -0.332279423991424 | 3.56770321082842 | -3.04339249012748 | 0.00400228616742958 | 0.0984218114506606 | -2.13669568908382 |
| lnc-LINC02210-CRHR1-7 | lnc-LINC02210-CR | -1.48086395754868 | 0.124825279360171 | -3.04303789735853 | 0.00401253815763501 | 0.0985561729100256 | -1.92762288666292 |
| ENSG00000185043.11 | CIB1 | 0.488815095903487 | 5.18660535157266 | 3.04129901882929 | 0.0040251470572217 | 0.0986566654622935 | -2.23180474007841 |
| lnc-GBP5-11 | lnc-GBP5-11 | -0.748895792620163 | 1.85371058178722 | -3.04102255848282 | 0.00402817518535625 | 0.0986566654622935 | -2.03005691861652 |
| ENSG00000204576.11 | PRR3 | -0.369872940087099 | 3.68257897764945 | -3.0407640268086 | 0.00403100887401199 | 0.0986566654622935 | -2.15036464986053 |
| ENSG00000149781.12 | FERMT3 | -0.537959735615662 | 5.18679219949568 | -3.03785573509286 | 0.00406301512492815 | 0.0993219006132971 | -2.24446824080146 |
| ENSG00000268043.7 | NBPF12 | 0.33286363601595 | 5.58644971942596 | 3.03535060719879 | 0.00409077576901443 | 0.0998818952000285 | -2.26014889889824 |
| ENSG00000091009.7 | RBM27 | 0.257448189148858 | 6.12135499860201 | 3.03381089834713 | 0.00410792641658587 | 0.100085052905639 | -2.27109854727904 |
| ENSG00000166333.13 | ILK | -0.435917311261386 | 4.7763669123426 | -3.03339045885179 | 0.00411262137407902 | 0.100085052905639 | -2.2429405768497 |
| ENSG00000196220.15 | SRGAP3 | -0.681769344454521 | 3.35854927066141 | -3.03329537501257 | 0.00411368385357676 | 0.100085052905639 | -2.15031524993013 |
| LINC02498 | LINC02498 | -1.48719138271497 | 0.328490423365803 | -3.03091362382423 | 0.00415101783814866 | 0.100582348660653 | -1.95591467456076 |
| ENSG00000270441.1 | LAMB2P1 | -0.477334531190436 | 2.37076993930122 | -3.02977711505602 | 0.0041531791363862 | 0.100582348660653 | -2.07647302105827 |
| lnc-TMEM259-1 | lnc-TMEM259-1 | -0.741368451869265 | 0.506791226245446 | -3.02966044301453 | 0.00415449495513032 | 0.100582348660653 | -1.97390137227044 |
| ENSG00000112294.12 | ALDH5A1 | -0.541458182991129 | 5.88190697453101 | -3.02935276534436 | 0.0041579667994704 | 0.100582348660653 | -2.27916025729098 |
| lnc-TXNDC17-1 | lnc-TXNDC17-1 | -1.00248143409696 | 1.20454130430348 | -3.02930048538107 | 0.00415855699899024 | 0.100582348660653 | -2.00619461386366 |
| ENSG00000076321.10 | KLHL20 | 0.229303367544446 | 4.5911918604336 | 3.02766805571616 | 0.00417702547768005 | 0.100595938756443 | -2.24292353529729 |
| lnc-MAP1S-1 | lnc-MAP1S-1 | -0.580109025502367 | 1.02556030165438 | -3.02742665992861 | 0.00417976303576001 | 0.100595938756443 | -1.99376128982766 |
| TMCC1-AS1 | TMCC1-AS1 | -1.04405692842525 | -0.325008371810649 | -3.02682271434602 | 0.00418661948687082 | 0.100595938756443 | -2.00158704227452 |
| lnc-ALG1L-2 | lnc-ALG1L-2 | -0.732320743834316 | 1.32659009364395 | -3.02658549011613 | 0.00418931552557675 | 0.100595938756443 | -2.01333680876472 |
| ENSG00000165689.16 | ENTR1 | -0.42394822533286 | 4.83236153317969 | -3.02598447883908 | 0.00419615328161737 | 0.100595938756443 | -2.25896832964006 |
| lnc-ORMDL1-3 | lnc-ORMDL1-3 | -0.571566166323134 | 0.445824022298716 | -3.02572485259058 | 0.00419911031189771 | 0.100595938756443 | -1.97535327815557 |
| ENSG00000243323.7 | PTPRVP | -0.803021516435892 | 2.36630784373827 | -3.02564868882917 | 0.0041999781552578 | 0.100595938756443 | -2.09524720276433 |
| ENSG00000133398.3 | MED10 | 0.327179935315552 | 4.09411531213122 | 3.02552226653732 | 0.00420141903823239 | 0.100595938756443 | -2.21741085029486 |
| lnc-SAMSN1-3 | lnc-SAMSN1-3 | -0.79312426680503 | 2.30496490531011 | -3.02537440513648 | 0.00420310485986205 | 0.100595938756443 | -2.07503219101968 |
| ENSG00000214784.4 | RPS3AP21 | 1.29819810738534 | 0.892406885457271 | 3.02347292413467 | 0.00422534116130003 | 0.100998768393796 | -1.97207379868522 |
| ENSG00000162654.8 | GBP4 | 1.22248005838713 | 3.81629692678088 | 3.027551845675 | 0.00422974971362058 | 0.100998768393796 | -2.1985225066548 |
| ENSG00000163536.12 | SERPINI1 | -0.607995225192963 | 2.8309852442581 | -3.02248022677594 | 0.00423623061440291 | 0.101036309080249 | -2.12518684778536 |
| ENSG00000152242.10 | ARK2N | 0.252112673011321 | 5.85954664990556 | 3.02185879132073 | 0.00424337524416475 | 0.101089574827133 | -2.29868365215728 |
| ENSG00000083168.10 | KAT6A | 0.313584053923348 | 7.59375080698012 | 3.02106870384305 | 0.00425247519920432 | 0.101096186227974 | -2.29388180769025 |
| lnc-BAG1-1 | lnc-BAG1-1 | 0.333711384308121 | 1.98129579932814 | 3.02074179755909 | 0.00425624574928145 | 0.101096186227974 | -2.07582783828474 |
| lnc-ELOVL1-1 | lnc-ELOVL1-1 | -0.719102014435052 | -0.403394140296455 | -3.02055621414139 | 0.00425838767233413 | 0.101096186227974 | -2.01686514317441 |
| lnc-ZNF519-1 | lnc-ZNF519-1 | -0.440688914510342 | 1.23660343388905 | -3.01653392822629 | 0.00430506054440957 | 0.101904139094646 | -2.03682642318738 |
| ENSG00000166233.14 | ARIH1 | 0.308173353741386 | 6.62822539623688 | 3.0165200804306 | 0.00430522205454427 | 0.101904139094646 | -2.31361099219866 |
| ENSG00000138380.17 | CARF | -0.315673169158749 | 5.27771190322974 | -3.01634427227187 | 0.00430727304145858 | 0.101904139094646 | -2.30046871718706 |
| ENSG00000261427.6 | | -0.861715152711427 | 0.46994398194076 | -3.01552666131864 | 0.00431682338885524 | 0.101985436052996 | -1.98891774116065 |
| ENSG00000175073.7 | VCPIP1 | 0.288288530753153 | 6.58472218222279 | 3.0150927112714 | 0.00432190034191797 | 0.101985436052996 | -2.31729506419776 |
| ENSG00000206053.12 | JPT2 | -0.436746581138117 | 3.76542279742362 | -3.01455462860245 | 0.00432820336680454 | 0.101985436052996 | -2.22355975775923 |
| ENSG00000169193.11 | CCDC126 | 0.512710302261559 | 2.2060609045353 | 3.01435630767883 | 0.00433052864549962 | 0.101985436052996 | -2.1116382772275 |
| lnc-MDM4-5 | lnc-MDM4-5 | 0.774509621344094 | 2.83061725211888 | 3.01344777859898 | 0.00434119598946751 | 0.102003380790051 | -2.14256461386627 |
| ENSG00000177025.3 | C19orf18 | -0.529518378579589 | 1.13919758481311 | -3.01344726394607 | 0.00434120203916265 | 0.102003380790051 | -2.03609801902096 |
| ZNF561-AS1 | ZNF561-AS1 | -0.445603886857934 | 0.945965399502337 | -3.0126209772679 | 0.00435092516988651 | 0.10201239115676 | -2.02401681450548 |
| lnc-DHRS7C-1 | lnc-DHRS7C-1 | -0.634395951456444 | 0.401436023484244 | -3.01257236742908 | 0.00435149781060271 | 0.10201239115676 | -2.00553019455023 |
| ENSG00000207357.1 | RNU6-2 | -0.938692396368249 | 1.40103889123919 | -3.01116259828189 | 0.00436813618324494 | 0.102285946598101 | -2.05046431195531 |

| | | | | | | | |
|---|---|---|---|---|---|---|---|
| ENSG00000006638.11 | TBXA2R | -1.48249440597073 | 0.79553601737084 | -3.0117620071778 | 0.00438253819286545 | 0.102506572299715 | -2.01110902876732 |
| lnc-SP6-3 | lnc-SP6-3 | -0.403430165840496 | 1.16472372816778 | -3.00803723026259 | 0.00440523551384262 | 0.102877345205323 | -2.05579373981315 |
| ENSG00000166262.15 | FAM227B | -0.393139275136525 | 1.99715230973846 | -3.0062847325964 | 0.00442616750464727 | 0.102877345205323 | -2.10463708787897 |
| ENSG00000165661.16 | QSOX2 | 0.35089334298092 | 7.19318938986915 | 3.00592201762041 | 0.00443051144465457 | 0.102877345205323 | -2.3363087925424 |
| lnc-ZNF717-4 | lnc-ZNF717-4 | -0.459680254145099 | 1.65905104829399 | -3.005751296125 | 0.00443255742014698 | 0.102877345205323 | -2.09350101921723 |
| lnc-GPR75-ASB3-8 | lnc-GPR75-ASB3-8 | 0.710370219858074 | 0.90932515145197 | 3.00573355401278 | 0.00443277009756104 | 0.102877345205323 | -2.03657977150081 |
| ENSG00000239382.10 | ALKBH6 | -0.31559199062391 | 3.338428918305 | -3.00568598536981 | 0.00443334035740776 | 0.102877345205323 | -2.20850782532626 |
| ENSG00000187735.13 | TCEA1 | 0.335331882701434 | 6.68036618469778 | 3.00568290303741 | 0.00443337731123361 | 0.102877345205323 | -2.33990522565039 |
| ENSG00000168421.12 | RHOH | 0.34604778097505 | 7.55257276002432 | 3.00358753839184 | 0.00445856544359216 | 0.10334532942056 | -2.33722462047287 |
| ENSG00000131467.10 | PSME3 | 0.250624624597592 | 5.37097118340486 | 3.0020191821293 | 0.00447750624899492 | 0.103553954138482 | -2.33778916775091 |
| ENSG00000136048.13 | DRAM1 | 0.674336511338556 | 2.56680917452783 | 3.0017071234297 | 0.00448128392733464 | 0.103553954138482 | -2.16364148310259 |
| ENSG00000177200.17 | CHD9 | 0.288121571197435 | 7.75219701110988 | 3.00159358370951 | 0.00448265914285514 | 0.103553954138482 | -2.34030980092371 |
| lnc-WDCP-2 | lnc-WDCP-2 | 0.432792816086838 | 1.32684026359415 | 3.00074814716385 | 0.00449291169226739 | 0.103674440988722 | -2.07881947778262 |
| ENSG00000171943.11 | SRGAP2C | 0.437301695709445 | 2.4034981662276 | 3.00006026003371 | 0.00450126987498631 | 0.103724974866797 | -2.14640326644782 |
| lnc-DDX5-1 | lnc-DDX5-1 | -0.611920226626468 | 0.00059842751205227 | -2.9997388336098 | 0.00450518036879545 | 0.103724974866797 | -2.02245974339815 |
| lnc-GPN2-2 | lnc-GPN2-2 | -0.54901932991112 | 1.85901837966173 | -2.99883248452891 | 0.00451622423456654 | 0.103863065273836 | -2.11822527672307 |
| lnc-ETHE1-2 | lnc-ETHE1-2 | -0.459390274589881 | 1.40517291768429 | -2.99786524892106 | 0.00452803800633629 | 0.104018533799576 | -2.09637310772171 |
| ENSG00000186283.13 | TOR3A | 0.430871177380279 | 4.72716823732852 | 2.99701039506481 | 0.00453850327283356 | 0.104142712223783 | -2.32028919423879 |
| ENSG00000059728.10 | MXD1 | 0.306750465615818 | 4.51543260019595 | 2.99643642918333 | 0.00454554258387245 | 0.104188087977557 | -2.31818545732599 |
| lnc-FNDC1-10 | lnc-FNDC1-10 | -0.584467630134636 | 3.22542631542962 | -2.99459581300419 | 0.00456818564463407 | 0.104298148938844 | -2.2230187550058 |
| ENSG00000182446.13 | NPLOC4 | -0.763700152732796 | 6.83387862330147 | -2.99573611131749 | 0.00457304733200402 | 0.104298148938844 | -2.36472476646388 |
| lnc-KDSR-3 | lnc-KDSR-3 | 0.247878433086656 | 2.8704038252162 | 2.99414240513934 | 0.00457377963223731 | 0.104298148938844 | -2.19823186826304 |
| ENSG00000162241.12 | SLC25A45 | -0.490250833751943 | 5.08484455846454 | -2.99406510445819 | 0.00457473398163722 | 0.104298148938844 | -2.34802653297107 |
| ENSG00000157106.16 | SMG1 | 0.32188641594882 | 7.92315927940477 | 2.99385064154232 | 0.00457738270515864 | 0.104298148938844 | -2.35599005833879 |
| ENSG00000226979.8 | LTA | -0.816374515404011 | 1.62521215717221 | -2.9935783638788 | 0.00458074754120947 | 0.104298148938844 | -2.12154319437223 |
| ENSG00000198060.9 | MARCHF5 | 0.407718262576861 | 3.62466963500079 | 2.99187799514937 | 0.00460181339876941 | 0.104613764897395 | -2.26372812216231 |
| ENSG00000176853.15 | FAM91A1 | 0.471855400990393 | 6.3339955686782 | 2.99163957967694 | 0.00460477437676917 | 0.104613764897395 | -2.37514507431959 |
| ENSG00000198924.7 | DCLRE1A | -0.43011867299887 | 2.65785678649497 | -2.98877857942561 | 0.00464044589542556 | 0.105143377434986 | -2.19954577210448 |
| lnc-CDKL2-1 | lnc-CDKL2-1 | 0.412371379955979 | 3.56882347234444 | 2.98857306308506 | 0.00464301826086921 | 0.105143377434986 | -2.25957107056341 |
| ENSG00000080947.14 | CROCCP3 | -0.422518481994114 | 3.36761946850098 | -2.98854168928656 | 0.00464341107168063 | 0.105143377434986 | -2.25155133576813 |
| ENSG00000070495.14 | JMJD6 | 0.346824320361444 | 3.78314767850995 | 2.98774996348913 | 0.00465333407078056 | 0.105211273051754 | -2.2852336589212 |
| ENSG00000113734.17 | BNIP1 | 0.466790753946006 | 2.84999987593534 | 2.98708827998934 | 0.0046616424449899 | 0.105211273051754 | -2.21122860424005 |
| ENSG00000189144.13 | ZNF573 | -0.383106717775701 | 3.74728109182242 | -2.98708018242071 | 0.00466174420751104 | 0.105211273051754 | -2.28433352164337 |
| ENSG00000140025.15 | EFCAB11 | -0.387906987103037 | 1.87197143920911 | -2.98631034295406 | 0.00467142833038605 | 0.105271824348129 | -2.14858477560588 |
| ENSG00000197535.14 | MYO5A | 0.258346641445999 | 7.26379076963014 | 2.98605408006507 | 0.00467465614605209 | 0.105271824348129 | -2.38425203774357 |
| ENSG00000143952.19 | VPS54 | 0.43321287792587 | 4.9852238595466 | 2.98488095181752 | 0.0046894592479743 | 0.105489770165088 | -2.36598511180267 |
| lnc-ERMP1-5 | lnc-ERMP1-5 | -0.722482776824854 | 0.164772514579704 | -2.98333676147541 | 0.00470901157356758 | 0.105730996103707 | -2.05991671748644 |
| ENSG00000048707.15 | VPS13D | 0.316879579201692 | 7.98930512800432 | 2.98322289400194 | 0.0047104563682213 | 0.105730996103707 | -2.38178689574488 |
| ENSG00000123066.8 | MED13L | 0.528527344428804 | 8.19011816440973 | 2.98236459927143 | 0.00472136011902332 | 0.105860299923592 | -2.37988207034849 |
| ENSG00000116815.15 | CD58 | -0.496050671015131 | 3.07584053129362 | -2.98134350948349 | 0.0047343628201847 | 0.105930938228142 | -2.24360796265231 |
| lnc-MED26-2 | lnc-MED26-2 | -0.338498253854893 | 2.53173281024071 | -2.9810228440265 | 0.00473845313799997 | 0.105930938228142 | -2.21131646031606 |
| ENSG00000110042.7 | DTX4 | 0.675436930669159 | 4.3166770820169 | 2.98016010901371 | 0.00474947438708065 | 0.105930938228142 | -2.33269551962827 |
| ENSG00000065548.17 | ZC3H15 | 0.276785585356095 | 6.49695793504952 | 2.97992520900461 | 0.00475247934044198 | 0.105930938228142 | -2.40378554230518 |
| lnc-N4BP2-3 | lnc-N4BP2-3 | 0.37439502775877 | 5.86932237535231 | 2.97971253427987 | 0.00475520151380806 | 0.105930938228142 | -2.40173946950542 |
| ENSG00000137713.15 | PPP2R1B | 0.493033236593972 | 4.65661522241287 | 2.9796978333475 | 0.00475538973535456 | 0.105930938228142 | -2.35571704144733 |
| lnc-SDE2-1 | lnc-SDE2-1 | -0.57673249710763 | 3.54839798918928 | -2.97916078564999 | 0.00476227054579084 | 0.105969529344879 | -2.28447990969599 |
| ENSG00000152492.14 | CCDC50 | 0.415231177395771 | 7.50345426956978 | 2.97874123941376 | 0.00476765238264477 | 0.105974718133885 | -2.39872508751354 |
| ENSG00000187609.15 | EXD3 | -0.614282702909222 | 2.77720744323795 | -2.97822258144233 | 0.00477431347746333 | 0.106008300222899 | -2.24076018886364 |
| ENSG00000284989.1 | | 0.295271236269835 | 3.92274384268951 | 2.97609924251972 | 0.00480167441967547 | 0.106500931659677 | -2.32033803429588 |
| ENSG00000197471.11 | SPN | -1.1195918147414 | 5.39505036695696 | -2.97998745341624 | 0.0048111519705782 | 0.106596276652757 | -2.3949127540663 |
| lnc-FYB1-3 | lnc-FYB1-3 | 0.442337265804607 | 2.86853175841976 | 2.9742431140614 | 0.00482571239905305 | 0.106803912160977 | -2.24674528230299 |
| ENSG00000285793.1 | | 0.322732963244794 | 2.65376263886174 | 2.97287266098137 | 0.00484353288025901 | 0.106990997072082 | -2.23657779312455 |
| ENSG00000127993.15 | RBM48 | 0.255239666765172 | 4.9047430512632 | 2.97279370133147 | 0.00484456149595204 | 0.106990997072082 | -2.39068697520059 |
| ENSG00000106588.10 | PSMA2 | 0.372766835256363 | 4.5625641468256 | 2.97083573082593 | 0.00487013372145663 | 0.107412783823797 | -2.38158601750648 |
| ENSG00000138134.11 | STAMBPL1 | -0.403178069364845 | 5.95336253079788 | -2.97053313374004 | 0.00487409707483974 | 0.107412783823797 | -2.42450034119932 |
| ENSG00000083828.15 | ZNF586 | 0.326337088082494 | 5.21654052282794 | 2.96812356778151 | 0.00490576503300153 | 0.107995039223819 | -2.41520248984898 |
| ENSG00000080561.13 | MID2 | -2.54996881466836 | 1.46861857197412 | -2.97311229281492 | 0.00492225450918859 | 0.108224257640235 | -2.18482776761483 |
| ENSG00000180667.10 | YOD1 | 0.481635845220921 | 5.13326118230299 | 2.96653914488858 | 0.00492669335902932 | 0.108224257640235 | -2.420900096664 |
| ENSG00000263335.1 | | -0.431141100095926 | 1.87444097083935 | -2.96570736327187 | 0.00493771363219302 | 0.108350703295766 | -2.19840269579239 |
| lnc-ERCC4-3 | lnc-ERCC4-3 | -0.637994402820657 | 0.530035755138789 | -2.96449356645246 | 0.00495383664641302 | 0.108588732367539 | -2.11139676172867 |
| TMEM147-AS1 | TMEM147-AS1 | -0.313027860888334 | 3.75595481877325 | -2.96357215601771 | 0.00496610875244013 | 0.108593638920944 | -2.34378135080634 |
| ENSG00000113790.10 | EHHADH | -2.61207635612977 | -0.328441450439006 | -2.96674716088119 | 0.00496970020742348 | 0.108593638920944 | -2.16161537070529 |
| ENSG00000185728.16 | YTHDF3 | 0.319259170376498 | 6.0608737693285 | 2.96286882856525 | 0.00497549539771563 | 0.108593638920944 | -2.4446329411391 |
| ENSG00000054965.10 | FAM168A | 0.259805502999956 | 5.03883268893015 | 2.96282850043891 | 0.0049760341222059 | 0.108593638920944 | -2.41993097423007 |
| ENSG00000135829.16 | DHX9 | 0.265238838859088 | 8.0730898636393 | 2.96203988308653 | 0.00498657987211187 | 0.108593638920944 | -2.43122397307471 |
| ENSG00000134285.10 | FKBP11 | 0.591110214960607 | 2.56810566804452 | 2.96081935077223 | 0.00500294264265378 | 0.108593638920944 | -2.25548417424389 |
| ENSG00000161057.11 | PSMC2 | 0.296874013206015 | 5.24458228427902 | 2.96030674213638 | 0.00500982978203008 | 0.108593638920944 | -2.43699763891841 |
| lnc-AHR-73 | lnc-AHR-73 | -1.40287678978241 | -0.284727792853644 | -2.95985229736204 | 0.00501594288291344 | 0.108593638920944 | -2.13631913077471 |

| | | | | | | | |
|---|---|---|---|---|---|---|---|
| ENSG00000239713.8 | APOBEC3G | -0.624248045484044 | 6.5720561771566 | -2.95984073352215 | 0.00501609852845059 | 0.108593638920944 | -2.45116481783627 |
| lnc-FASTKD1-4 | lnc-FASTKD1-4 | -0.490262329615321 | 0.888909502405608 | -2.95983190988988 | 0.00501621729472201 | 0.108593638920944 | -2.14393447589894 |
| ENSG00000204397.7 | CARD16 | 0.680143094271915 | 2.55392215044958 | 2.95852338952043 | 0.00503385917347744 | 0.108593638920944 | -2.25002935966061 |
| ENSG00000117758.13 | STX12 | 0.254587132676879 | 4.80904939923643 | 2.95816618764227 | 0.0050386851611189 | 0.108593638920944 | -2.42303255206944 |
| ENSG00000166860.2 | ZBTB39 | 0.299160952284462 | 4.66503442768992 | 2.95809599064127 | 0.00503963406897838 | 0.108593638920944 | -2.41848957116227 |
| lnc-MRPL3-2 | lnc-MRPL3-2 | -0.648960935830418 | 1.21157121148613 | -2.95794539353931 | 0.00504167037234286 | 0.108593638920944 | -2.17068959284912 |
| FAM66C | FAM66C | -0.685339168298335 | 1.06729625685786 | -2.95701992586727 | 0.0050542010290589 | 0.108593638920944 | -2.16843565701326 |
| lnc-WNT10A-4 | lnc-WNT10A-4 | -0.919750032065331 | 1.68947895971725 | -2.95617597175192 | 0.00506565341214749 | 0.108593638920944 | -2.20605741418635 |
| ENSG00000189298.13 | ZKSCAN3 | -0.528529166742266 | 2.16648258032442 | -2.95536647890598 | 0.00507666097365989 | 0.108593638920944 | -2.23961044250031 |
| ENSG00000204482.10 | LST1 | -0.858981984613028 | 0.0504087345475999 | -2.95513467302049 | 0.00507981721274992 | 0.108593638920944 | -2.12000693558168 |
| ENSG00000220201.7 | ZGLP1 | -0.665546774279423 | -0.508761301928467 | -2.95508387124661 | 0.00508050916844176 | 0.108593638920944 | -2.14451356058634 |
| lnc-ZNF585A-3 | lnc-ZNF585A-3 | -0.443124272505128 | 2.46548214649996 | -2.95498307117519 | 0.00508188239706665 | 0.108593638920944 | -2.26263188625023 |
| ENSG00000164022.16 | AIMP1 | 0.249765078908037 | 5.19472495827848 | 2.95435491404547 | 0.005090447793801 | 0.108593638920944 | -2.44773852963201 |
| lnc-SHF-2 | lnc-SHF-2 | -0.5346150895834 | 0.348148943605025 | -2.95316933721984 | 0.00510665085210097 | 0.108593638920944 | -2.13012488160421 |
| ENSG00000108094.14 | CUL2 | 0.222231657667396 | 4.96754260342575 | 2.95297965927143 | 0.00510924761923538 | 0.108593638920944 | -2.44307698999549 |
| ENSG00000158301.18 | GPRASP2 | 0.547354666463103 | 1.10984293125109 | 2.95292268856297 | 0.00511002781232683 | 0.108593638920944 | -2.17741754548074 |
| ENSG00000135249.7 | RINT1 | 0.358089407630495 | 4.49695838495972 | 2.9525789941904 | 0.00511473694690421 | 0.108593638920944 | -2.41957705040153 |
| ENSG00000129460.15 | NGDN | 0.251829992331634 | 5.53566834339421 | 2.9520899345282 | 0.00512144480697782 | 0.108593638920944 | -2.46376852117644 |
| ENSG00000170464.9 | DNAJC18 | 0.493792140427876 | 2.89677374445243 | 2.95158731414265 | 0.00512834723988745 | 0.108593638920944 | -2.29820510017118 |
| ENSG00000095485.17 | CWF19L1 | 0.269045792919602 | 5.82830736759123 | 2.95156692184004 | 0.00512862746888825 | 0.108593638920944 | -2.46944968954321 |
| lnc-CEACAM8-3 | lnc-CEACAM8-3 | -0.796189025391008 | 1.73256782167643 | -2.95152541077303 | 0.00512919795412307 | 0.108593638920944 | -2.21664625604542 |
| lnc-MC5R-1 | lnc-MC5R-1 | -2.36924725936959 | -0.632198599682217 | -2.95286964468949 | 0.00513341664737944 | 0.108593638920944 | -2.23002848220558 |
| ENSG00000164904.17 | ALDH7A1 | -1.963129725469 | -0.572319576591108 | -2.95220709992951 | 0.00513390034231506 | 0.108593638920944 | -2.17028090372539 |
| ENSG00000236296.8 | GUSBP5 | -0.984916668100842 | 0.00098795413161576 | -2.9511587115252 | 0.00513424006941772 | 0.108593638920944 | -2.12906841295474 |
| lnc-CASP4-1 | lnc-CASP4-1 | 0.686893646962227 | 2.99204244554214 | 2.95052420568582 | 0.00514297548179235 | 0.108593638920944 | -2.30686014304468 |
| ENSG00000151553.14 | FHIP2A | 0.35288710742635 | 5.08945496604945 | 2.95021563351459 | 0.00514722870155456 | 0.108593638920944 | -2.4570699968546 |
| ENSG00000198804.2 | MT-CO1 | -0.465272934280121 | 11.6232093557089 | -2.9497389479169 | 0.00515380559005969 | 0.108593638920944 | -2.37459969819424 |
| ENSG00000146021.14 | KLHL3 | 1.83075244422267 | 2.71954578055127 | 2.95480610808161 | 0.00516113906579792 | 0.108593638920944 | -2.25966719489229 |
| ENSG00000142178.8 | SIK1 | 2.22508101367614 | -0.980187347954314 | 2.94971951730849 | 0.00516332177053614 | 0.108593638920944 | -2.26126305888667 |
| ENSG00000196449.3 | YRDC | 0.39308008844832 | 3.71112449873463 | 2.9488897355485 | 0.00516554175844971 | 0.108593638920944 | -2.37265284215044 |
| lnc-MMP13-1 | lnc-MMP13-1 | 0.495717417627208 | 0.12866864677422 | 2.9484073939301 | 0.00517221887871015 | 0.108593638920944 | -2.13453974795937 |
| lnc-INTS2-4 | lnc-INTS2-4 | 0.36147662836148 | 3.82305707359943 | 2.94840292194543 | 0.00517228082275914 | 0.108593638920944 | -2.38000487507396 |
| lnc-MOB2-1 | lnc-MOB2-1 | -0.390290436598088 | 0.607648019585155 | -2.94796257234421 | 0.00517838375992906 | 0.108593638920944 | -2.16106760603808 |
| ENSG00000228727.8 | SAPCD1 | -0.428236339409955 | 0.705967537343566 | -2.94783277349377 | 0.00518018396619374 | 0.108593638920944 | -2.16253228504278 |
| ENSG00000237489.4 | C10orf143 | -0.444169039723089 | 0.515110098363081 | -2.94769299622887 | 0.00518212321975667 | 0.108593638920944 | -2.15082337037302 |
| ENSG00000143256.4 | PFDN2 | 0.361651612485685 | 2.55809588795605 | 2.94703366348901 | 0.00519127988627731 | 0.108593638920944 | -2.28661612608194 |
| ENSG00000130749.9 | ZC3H4 | -0.413981465713172 | 5.0328188730549 | -2.94701958836234 | 0.00519147552340325 | 0.108593638920944 | -2.4610027389985 |
| ENSG00000113716.12 | HMGXB3 | 0.246510779549589 | 5.51435660252847 | 2.94592208326904 | 0.00520675154041598 | 0.108802606047089 | -2.47893614732847 |
| lnc-PSMB1-1 | lnc-PSMB1-1 | -0.484716285286855 | 0.912202564266486 | -2.94414873338952 | 0.00523152342830772 | 0.108995493641977 | -2.18620412407281 |
| ENSG00000185813.10 | PCYT2 | -0.451930810447867 | 3.68390549911322 | -2.9435160010483 | 0.00524038870364282 | 0.108995493641977 | -2.39001311538695 |
| ENSG00000236439.4 | RPS27P8 | 0.960580063965791 | -0.319875419907347 | 2.94279556341071 | 0.00525049993266885 | 0.108995493641977 | -2.18030201940548 |
| lnc-SYT2-4 | lnc-SYT2-4 | 0.960580063965791 | -0.319875419907347 | 2.94279556341071 | 0.00525049993266885 | 0.108995493641977 | -2.18030201940548 |
| ENSG00000140525.17 | FANCI | 0.693337356004803 | 2.83165421938854 | 2.94254849855173 | 0.00525397164859819 | 0.108995493641977 | -2.3141957680687 |
| ENSG00000119396.10 | RAB14 | 0.330136513264288 | 5.56481596347466 | 2.94251848066964 | 0.00525439360131602 | 0.108995493641977 | -2.48861340449142 |
| ENSG00000207332.1 | RNU6-146P | 0.607629884089744 | -0.256534952674109 | 2.94234514160973 | 0.00525683079848677 | 0.108995493641977 | -2.15480009935612 |
| ENSG00000114439.18 | BBX | 0.286955068156348 | 7.26837670962302 | 2.94179347015078 | 0.00526459449681461 | 0.108995493641977 | -2.49128511795527 |
| ENSG00000141098.12 | GFOD2 | -0.711151759253745 | 3.84272175402171 | -2.94144101857598 | 0.0052717371343999 | 0.108995493641977 | -2.41781473939927 |
| lnc-ZNF256-2 | lnc-ZNF256-2 | -0.832293134113164 | 0.993904283989152 | -2.94102542405622 | 0.00527542110289385 | 0.108995493641977 | -2.19462280391281 |
| ENSG00000183397.5 | TEKTIP1 | -0.630785071167178 | 1.34524082113901 | -2.94041859431568 | 0.00528398987552902 | 0.108995493641977 | -2.21964682527204 |
| lnc-MCF2L-4 | lnc-MCF2L-4 | 0.788115363254731 | 1.47103299359799 | 2.94037507315813 | 0.00528460491833976 | 0.108995493641977 | -2.21960208297015 |
| lnc-MCF2L-2 | lnc-MCF2L-2 | 0.596826767422166 | 1.74707740795669 | 2.94019079679895 | 0.00528720986141937 | 0.108995493641977 | -2.24769337506524 |
| lnc-RGS5-1 | lnc-RGS5-1 | -0.520456002533368 | 4.74718263351984 | -2.93998517456436 | 0.00529011796863117 | 0.108995493641977 | -2.46258234224994 |
| ENSG00000182472.8 | CAPN12 | -0.674931382950316 | 3.01563208615354 | -2.93831566086513 | 0.00531378526870213 | 0.109373642185696 | -2.3465055545903 |
| ENSG00000169896.17 | ITGAM | -1.42725453718574 | 4.25429990760486 | -2.94176826927846 | 0.00533993953406504 | 0.109802173256404 | -2.41174514236159 |
| ENSG00000144504.15 | ANKMY1 | -0.282143419704716 | 4.75554257241936 | -2.93594718779821 | 0.00534753102555548 | 0.109842349738724 | -2.47793884102526 |
| ENSG00000136149.6 | RPL13AP25 | 0.960117851974584 | 1.41973585169215 | 2.93505977207199 | 0.00536022635054935 | 0.109842349738724 | -2.2220640079521 |
| ENSG00000214106.8 | PAXIP1-AS2 | -0.437982831716326 | 4.31031163629525 | -2.93446416347522 | 0.00536876289315049 | 0.109842349738724 | -2.45011748210259 |
| lnc-ST8SIA4-3 | lnc-ST8SIA4-3 | -0.661075257709594 | 0.0258774098648326 | -2.93424726957507 | 0.00537187467402764 | 0.109842349738724 | -2.16526918336038 |
| ENSG00000214026.10 | MRPL23 | -0.637477839622463 | 1.00197319106192 | -2.93412262089328 | 0.00537366377385471 | 0.109842349738724 | -2.21705415174 |
| ENSG00000215105.4 | TTC3P1 | -1.16095369063377 | 2.05818952109957 | -2.93640019612053 | 0.0053776020060384 | 0.109842349738724 | -2.3003333128283 |
| ENSG00000189043.9 | NDUFA4 | 0.331252896322463 | 4.35967703227063 | 2.9337337195495 | 0.00537924930946088 | 0.109842349738724 | -2.46128437058011 |
| ENSG00000133466.13 | C1QTNF6 | -0.511792283504358 | 4.57208634552154 | -2.93306421787184 | 0.00538887763955855 | 0.109929899360787 | -2.46883363279183 |
| ENSG00000178761.14 | FAM219B | -0.344453659663139 | 5.31319988038421 | -2.93247495978722 | 0.00539736528598744 | 0.109994029387604 | -2.5061896032263 |
| ENSG00000136874.10 | STX17 | 0.2952082987636 | 4.47318151464975 | 2.93168433823266 | 0.00540877299412451 | 0.110050142737694 | -2.46561107668472 |
| LINC01135 | LINC01135 | -0.715033866484649 | 3.01334457116121 | -2.93143266012504 | 0.00541240912960302 | 0.110050142737694 | -2.3608399349247 |
| ENSG00000198182.12 | ZNF607 | -0.501202785200738 | 2.67966688872759 | -2.93117329574213 | 0.00541615870345837 | 0.110050142737694 | -2.33575556648885 |
| ENSG00000047365.11 | ARAP2 | 0.498076182270754 | 6.69232187237674 | 2.92967034807453 | 0.00543793432797846 | 0.110383631432722 | -2.52451857168618 |
| SPON1-AS1 | SPON1-AS1 | -0.97022809846593 | 0.180703463342077 | -2.92887052041492 | 0.00544955605184522 | 0.110489321347095 | -2.1776422926784 |

| | | | | | | | |
|---|---|---|---|---|---|---|---|
| ENSG00000239697.10 | TNFSF12 | -0.533838963743757 | 1.97085738511967 | -2.92847728547026 | 0.0054552783753587 | 0.110489321347095 | -2.29753058073912 |
| LINC02352 | LINC02352 | -0.565660445785299 | 0.956774765074398 | -2.92767656601085 | 0.00546694774374674 | 0.110489321347095 | -2.22167221659729 |
| ENSG00000188739.14 | RBM34 | 0.327515274697915 | 4.66869823706446 | 2.92755002708419 | 0.00546879400505383 | 0.110489321347095 | -2.48682506100364 |
| ENSG00000177868.11 | SVBP | 0.425823354957911 | 2.58996383324354 | 2.92689263550707 | 0.00547839503144922 | 0.110489321347095 | -2.33739973066227 |
| ENSG00000204389.9 | HSPA1A | -1.05381855902231 | 2.13925132435065 | -2.92792958678728 | 0.0054843367955244 | 0.110489321347095 | -2.30671960828367 |
| ENSG00000203414.2 | BTBD7P1 | -2.8902031670862 | -2.50991435549951 | -2.92687226846323 | 0.00548589726439078 | 0.110489321347095 | -2.58151111393228 |
| ENSG00000163933.9 | RFT1 | -0.197120895143363 | 4.39514821621591 | -2.92624291673414 | 0.00548789947294412 | 0.110489321347095 | -2.474091364217 |
| ENSG00000096872.16 | IFT74 | -0.348396529656798 | 3.79190845642641 | -2.92600027397294 | 0.00549145293388128 | 0.110489321347095 | -2.43338693685569 |
| ENSG00000157593.18 | SLC35B2 | -0.585058092331717 | 4.87589079908476 | -2.92541704110887 | 0.00550000307426742 | 0.110553284450827 | -2.50731848987052 |
| ENSG00000134815.18 | DHX34 | -0.368499493493924 | 3.89067093311446 | -2.92476905027659 | 0.00550951714719408 | 0.110636479454337 | -2.44901631490521 |
| ENSG00000107443.15 | CCNJ | 0.350530640532745 | 3.1414674182519 | 2.92363588544762 | 0.00552619171554749 | 0.110863161872431 | -2.38754534042094 |
| ENSG00000110851.11 | PRDM4 | 0.431188334421083 | 6.26159621076653 | 2.92283558354513 | 0.00553799658817909 | 0.110991805038452 | -2.54222943470885 |
| lnc-TEX35-1 | lnc-TEX35-1 | -0.711560020113893 | 2.93082469384271 | -2.9222182317823 | 0.00554711894233643 | 0.111066487539018 | -2.37544954432468 |
| ENSG00000112039.4 | FANCE | -0.574636661686894 | 2.05509812202178 | -2.92025423812697 | 0.00557623352696473 | 0.111540927779898 | -2.32172433200742 |
| ENSG00000165494.11 | PCF11 | 0.285077748189511 | 8.02452535995588 | 2.91980802355732 | 0.00558286815992617 | 0.111565218772583 | -2.53514291785631 |
| lnc-NAGLU-6 | lnc-NAGLU-6 | -0.655431566401767 | 0.642359617570347 | -2.91821684756119 | 0.00560658696408502 | 0.111837272273714 | -2.23092315866849 |
| ENSG00000130518.16 | IQCN | -0.790666256534248 | 6.11481649133739 | -2.91994116779352 | 0.00561027878682569 | 0.111837272273714 | -2.55053635333973 |
| lnc-UBE2D2-2 | lnc-UBE2D2-2 | -0.737497340563707 | 0.790559086552363 | -2.91780225555824 | 0.00561278250297557 | 0.111837272273714 | -2.23508134094151 |
| ENSG00000165185.14 | KIAA1958 | 0.530633191716318 | 2.94071447700192 | 2.9172387898649 | 0.00562121301735944 | 0.111878052975975 | -2.37909172979355 |
| ENSG00000198113.2 | TOR4A | -0.505384261386292 | 1.90595669067354 | -2.91687166263366 | 0.00562671229807963 | 0.111878052975975 | -2.32665799113926 |
| lnc-NAA50-2 | lnc-NAA50-2 | -0.619965839420548 | -0.114410122720115 | -2.91617064572169 | 0.00563722694702332 | 0.111878052975975 | -2.20624300385739 |
| ENSG00000079385.21 | CEACAM1 | -0.830237442032309 | 2.78919260380019 | -2.91549048209691 | 0.00565023837073527 | 0.111878052975975 | -2.37882408641783 |
| ENSG00000104714.13 | ERICH1 | -0.472630459902161 | 4.55295983624852 | -2.91529313505472 | 0.00565041471273423 | 0.111878052975975 | -2.51677667419895 |
| ENSG00000159905.14 | ZNF221 | -1.04439661106934 | 1.2854230849061 | -2.91525247904471 | 0.00565440894224046 | 0.111878052975975 | -2.26807130359162 |
| ENSG00000104983.8 | CCDC61 | -0.450912782582196 | 1.57652363645851 | -2.91491634725072 | 0.0056560861467736 | 0.111878052975975 | -2.2966104649621 |
| ENSG00000144597.13 | EAF1 | 0.391816726281139 | 4.88675156067084 | 2.91476851106888 | 0.00565831283816693 | 0.111878052975975 | -2.53106676342614 |
| ENSG00000137700.18 | SLC37A4 | -0.555632496418296 | 2.3320600987798 | -2.911930976782 | 0.00570121041621702 | 0.112618055659304 | -2.36135132387633 |
| lnc-SNAPC5-4 | lnc-SNAPC5-4 | 0.586986658652647 | 3.61350322784728 | 2.91100994078492 | 0.00571519970633451 | 0.112749913305772 | -2.44419264512671 |
| lnc-RUBCN-9 | lnc-RUBCN-9 | -0.508705986113429 | 0.732856881059135 | -2.91077053162195 | 0.00571884125206366 | 0.112749913305772 | -2.25088501799801 |
| ENSG00000205213.13 | LGR4 | -0.842209381407493 | 1.32812145647681 | -2.91020240346895 | 0.00572749146738151 | 0.11281239892164 | -2.30187188189663 |
| lnc-TIGD2-2 | lnc-TIGD2-2 | -0.571873685153159 | 1.35798445583272 | -2.90926885254683 | 0.00574173207642447 | 0.112934745281565 | -2.30078987696676 |
| lnc-RAPGEF1-3 | lnc-RAPGEF1-3 | 0.344521221669796 | 0.853281366279588 | 2.90907609288493 | 0.00574467659280955 | 0.112934745281565 | -2.25685893224464 |
| ENSG00000147526.19 | TACC1 | 0.317116226651905 | 7.35831469098124 | 2.90792386391218 | 0.0057623069824816 | 0.113173248683606 | -2.57217488109075 |
| ENSG00000028277.21 | POU2F2 | -0.421149010834488 | 6.44019909263943 | -2.90743224991304 | 0.00576984458792845 | 0.113213261347313 | -2.57881391093434 |
| ENSG00000198786.2 | MT-ND5 | -0.878575883166879 | 7.68266393084886 | -2.90933579963171 | 0.00578587565746884 | 0.113419693599696 | -2.56724659944845 |
| ENSG00000125449.6 | ARMC7 | -0.583629985353224 | 2.43977467575838 | -2.90539374590182 | 0.00580119799302375 | 0.113556925140927 | -2.37729874755483 |
| ENSG00000153879.8 | CEBPG | 0.283047664678085 | 4.91527152037662 | 2.90521788492857 | 0.00580391027781446 | 0.113556925140927 | -2.56066870573912 |
| ENSG00000173581.7 | CCDC106 | -0.684081388014243 | 3.22710902798838 | -2.90474036538625 | 0.00581128098173982 | 0.113593159019136 | -2.44440619107684 |
| lnc-KDSR-5 | lnc-KDSR-5 | -0.702637255999366 | 1.19740652248589 | -2.90331476208822 | 0.00583333774411611 | 0.113816790773353 | -2.29417734805601 |
| lnc-PEX6-3 | lnc-PEX6-3 | -0.370918352291143 | 1.18102730061328 | -2.90317481316668 | 0.0058355072233971 | 0.113816790773353 | -2.29473197578538 |
| ENSG00000170340.10 | B3GNT2 | 0.598572975382027 | 3.87824679669661 | 2.90270559332587 | 0.00584278653561619 | 0.113816790773353 | -2.49592812053 |
| lnc-TERB1-3 | lnc-TERB1-3 | 0.384625154603371 | 2.43819406286333 | 2.90257330780488 | 0.00584484029769391 | 0.113816790773353 | -2.38202941734752 |
| ENSG00000277610.1 | RNVU1-4 | -1.125986792581 | 2.47263926625586 | -2.90338731560133 | 0.00587139317953764 | 0.114225789994729 | -2.39285211789931 |
| ENSG00000177301.14 | KCNA2 | -1.09929363034955 | 1.30143287375038 | -2.90049432953669 | 0.00588623588329674 | 0.114406414717561 | -2.30207406443594 |
| ENSG00000170417.15 | TMEM182 | -0.450310000180273 | 2.05048729200497 | -2.89925951657629 | 0.0058965079886991 | 0.114497947105088 | -2.3639550451333 |
| ENSG00000088930.7 | XRN2 | 0.174313520164668 | 6.5452686979028 | 2.89886754602514 | 0.00590264759052401 | 0.114509137941334 | -2.59987124863513 |
| lnc-ZNF814-4 | lnc-ZNF814-4 | -0.39275163739841 | 1.35263466278345 | -2.89839367922353 | 0.00591007793269159 | 0.114545324000557 | -2.32433557435556 |
| lnc-SINHCAF-9 | lnc-SINHCAF-9 | 0.594837665854485 | 1.67903679775349 | 2.89786413165714 | 0.0059183916875655 | 0.114598547605984 | -2.33987479844195 |
| ENSG00000151694.13 | ADAM17 | 0.444373826114223 | 6.5258777230368 | 2.89726649949213 | 0.00592778745824138 | 0.114672602681374 | -2.60380037895655 |
| ENSG00000161013.16 | MGAT4B | -0.534647559452185 | 5.46257764814569 | -2.89639627419018 | 0.00594149373471814 | 0.114829826799722 | -2.59634616323452 |
| LINC01184 | LINC01184 | -0.35075943649804 | 4.68132834599353 | -2.89602447087344 | 0.00594735874446501 | 0.114835351817376 | -2.56759937151302 |
| ENSG00000103423.13 | DNAJA3 | -0.352754789931996 | 4.59850614992609 | -2.89566948047468 | 0.00595296357892667 | 0.11483584755862 | -2.56347012983936 |
| ENSG00000131089.15 | ARHGEF9 | -0.260695796809328 | 5.25111211565714 | -2.89502022705763 | 0.00596322719170602 | 0.114869780954059 | -2.59396813322616 |
| ENSG00000111790.13 | FGFR1OP2 | 0.273687951266787 | 5.30928755706925 | 2.89485231632018 | 0.00596588426565071 | 0.114869780954059 | -2.59754050659988 |
| ENSG00000032742.17 | IFT88 | -0.420471350454874 | 3.98792630259695 | -2.8936825504262 | 0.00598442569132532 | 0.115119097200513 | -2.53122256724515 |
| ENSG00000166170.9 | BAG5 | 0.273083340614258 | 5.84981465412872 | 2.89220717617804 | 0.00600788783079618 | 0.115315165498152 | -2.6134198686648 |
| ENSG00000164654.15 | MIOS | 0.337598361758921 | 5.43979351476734 | 2.89204499060273 | 0.00601047221466463 | 0.115315165498152 | -2.60776264875501 |
| ENSG00000113391.18 | ARB2A | -0.222769930790268 | 5.46542168187341 | -2.89177295950793 | 0.00601480928591539 | 0.115315165498152 | -2.60783005526645 |
| ENSG00000138785.14 | INTS12 | 0.288195263224966 | 3.54470632827333 | 2.89163386609995 | 0.00601702802045449 | 0.115315165498152 | -2.49005379993832 |
| lnc-REPS1-2 | lnc-REPS1-2 | -0.73221402843863 | -0.401735874861231 | -2.89092574746117 | 0.00602833534431251 | 0.115424396643706 | -2.27260873823905 |
| ENSG00000126581.12 | BECN1 | 0.198814710649272 | 6.03825919194116 | 2.88820001341559 | 0.00607204541728285 | 0.116084130461517 | -2.62481785328608 |
| lnc-VTCN1-3 | lnc-VTCN1-3 | -0.667642298990973 | 0.662435479003528 | -2.88807412796664 | 0.00607407124846006 | 0.116084130461517 | -2.29774548342628 |
| ENSG00000170855.3 | TRIAP1 | 0.48403378072041 | 3.09668344207677 | 2.88721995434344 | 0.00608783382686827 | 0.11623922417294 | -2.46612103530479 |
| lnc-CCNT1-3 | lnc-CCNT1-3 | 0.426799440578515 | 2.03379988431129 | 2.88665278473174 | 0.00609698821017587 | 0.116299618274441 | -2.38970272925438 |
| lnc-ONECUT2-5 | lnc-ONECUT2-5 | -0.400085865914201 | 3.35809507475594 | -2.88632421084394 | 0.00610229741711103 | 0.116299618274441 | -2.49280528533193 |
| ENSG00000284934.1 | | 0.324952104263587 | 2.48746441575196 | 2.88501989620631 | 0.00612341552031706 | 0.116522894013172 | -2.42923946301274 |
| lnc-DET1-1 | lnc-DET1-1 | -0.598887114995581 | 1.20435098671695 | -2.8848409789488 | 0.00612631767107674 | 0.116522894013172 | -2.33369443512025 |
| ENSG00000179119.14 | SPTY2D1 | 0.437139917428086 | 5.01826823192518 | 2.88455271620038 | 0.00613099617238814 | 0.116522894013172 | -2.60729185035543 |

| | | | | | | | |
|---|---|---|---|---|---|---|---|
| ENSG00000111142.13 | METAP2 | 0.237283047322404 | 5.49872474942165 | 2.88398903202897 | 0.00614015438779139 | 0.116589296830175 | -2.62816200608334 |
| ENSG00000171793.14 | CTPS1 | 0.469004099222682 | 3.88761233685625 | 2.88311770766455 | 0.00615433595318424 | 0.11668932287697 | -2.54269319867933 |
| lnc-XPO1-9 | lnc-XPO1-9 | 0.351823395610139 | 3.78268533716086 | 2.88283923548097 | 0.00615887476667149 | 0.11668932287697 | -2.5320525590776 |
| ENSG00000167637.16 | ZNF283 | -0.379245433616861 | 3.2621035078141 | -2.88239997349294 | 0.0061660406346363 | 0.11668932287697 | -2.48950497200029 |
| ENSG00000125347.13 | IRF1 | 0.715990254576577 | 6.80599488582149 | 2.88327875115787 | 0.00617035087592485 | 0.11668932287697 | -2.63787672145178 |
| lnc-EIF2AK3-4 | lnc-EIF2AK3-4 | -4.339183464653 | 6.50799649287914 | -2.88716807003019 | 0.00618617689622701 | 0.11668932287697 | -2.59954063663642 |
| lnc-MTR-1 | lnc-MTR-1 | -0.818091152484674 | 0.424701437161128 | -2.88053097967722 | 0.00619661730588224 | 0.11668932287697 | -2.29072480897208 |
| ENSG00000198886.2 | MT-ND4 | -0.768306222574964 | 8.61304161516072 | -2.8819163821618 | 0.00619726759783887 | 0.11668932287697 | -2.61423832467146 |
| lnc-BICDL1-1 | lnc-BICDL1-1 | -0.561469490828718 | 1.36219576171935 | -2.88026481233872 | 0.00620098327385429 | 0.11668932287697 | -2.36299242575138 |
| ENSG00000100711.13 | ZFYVE21 | -0.498262396179331 | 2.40343778292804 | -2.8801620346643 | 0.00620266991299887 | 0.11668932287697 | -2.4407058028154 |
| lnc-TRIAP1-2 | lnc-TRIAP1-2 | 0.247823308723371 | 3.95336504802366 | 2.87998268748176 | 0.00620561412482807 | 0.11668932287697 | -2.56011943081626 |
| ENSG00000138821.12 | SLC39A8 | 1.07683069777624 | 1.57295178049889 | 2.8804502072061 | 0.00620778353739891 | 0.11668932287697 | -2.35780162377604 |
| ENSG00000163788.13 | SNRK | 0.254615182622885 | 6.36382301074911 | 2.87877462901522 | 0.00622547990386858 | 0.116915194216539 | -2.64829401345688 |
| ENSG00000179889.18 | PDXDC1 | 0.307594443796329 | 6.47346418699667 | 2.87754576941598 | 0.0062457485477512 | 0.117052737840619 | -2.65107206819098 |
| ENSG00000066827.15 | ZFAT | -0.609555845673011 | 4.4323406164034 | -2.87738880738667 | 0.00624834188329898 | 0.117052737840619 | -2.60421104929104 |
| ENSG00000165678.20 | GHITM | 0.227045507424442 | 6.20713884933422 | 2.87729668528699 | 0.00624986439716465 | 0.117052737840619 | -2.65170719614816 |
| lnc-PLCD3-1 | lnc-PLCD3-1 | -0.503314367565844 | 2.56488429241261 | -2.87666660476298 | 0.00626028709287502 | 0.11714135384786 | -2.45259442546099 |
| ENSG00000009307.15 | CSDE1 | 0.258163801598819 | 9.0623609802187 | 2.87582101736568 | 0.00627430012891401 | 0.117212626314998 | -2.61994449378686 |
| ENSG00000125971.16 | DYNLRB1 | -0.414944909475365 | 4.53412971388785 | -2.87560514662209 | 0.00627788220999973 | 0.117212626314998 | -2.61134382218022 |
| lnc-PCBD2-1 | lnc-PCBD2-1 | -0.356164883165122 | 0.952919975556562 | -2.87523643413004 | 0.00628400489967906 | 0.117212626314998 | -2.34599943666805 |
| ENSG00000069248.11 | NUP133 | 0.288838386452934 | 5.89328734612421 | 2.87506373534623 | 0.00628687457862111 | 0.117212626314998 | -2.65512601233023 |
| lnc-SERHL2-4 | lnc-SERHL2-4 | -0.996711929215822 | 3.16136694371032 | -2.87600653369485 | 0.00630834265510286 | 0.117506440606319 | -2.50692706802056 |
| ENSG00000103591.12 | AAGAB | 0.268317225076644 | 4.65791231959666 | 2.87124845364385 | 0.00635058419904158 | 0.118186324203321 | -2.623167402739 |
| ENSG00000165732.12 | DDX21 | 0.471883424722447 | 7.04093198237644 | 2.87058746291017 | 0.00636168272445217 | 0.118285921876603 | -2.66459350846037 |
| ENSG00000113810.15 | SMC4 | 0.532871142442366 | 5.36555890993902 | 2.87007809073407 | 0.00637024778720904 | 0.118320238886822 | -2.65934217984632 |
| ENSG00000200488.1 | RN7SKP203 | 1.57211936512604 | 2.59050236873844 | 2.87364688721676 | 0.00638077370472584 | 0.118320238886822 | -2.41726812381967 |
| lnc-LRRTM4-6 | lnc-LRRTM4-6 | 1.57211936512604 | 2.59050236873844 | 2.87364688721676 | 0.00638077370472584 | 0.118320238886822 | -2.41726812381967 |
| ENSG00000103494.13 | RPGRIP1L | -0.438701465090517 | 2.11386224313958 | -2.86880793298414 | 0.00639165223061509 | 0.118415281604636 | -2.43928524770894 |
| ENSG00000178921.13 | PFAS | -0.306604450222051 | 4.35168541763771 | -2.86836648528143 | 0.00639910709467606 | 0.118446781771328 | -2.61747539858175 |
| lnc-ZSCAN10-3 | lnc-ZSCAN10-3 | -0.304091006792659 | 4.26374453309218 | -2.86689112213946 | 0.00642408081969031 | 0.118633544069493 | -2.61250486136727 |
| ENSG00000166341.7 | DCHS1 | 1.58145064504827 | 0.191622177584516 | 2.86741189328054 | 0.00642500745104935 | 0.118633544069493 | -2.31032404862725 |
| ENSG00000177311.11 | ZBTB38 | -0.502889900387782 | 6.94273674822369 | -2.86635895969019 | 0.00643311108205141 | 0.118633544069493 | -2.67415357504548 |
| ENSG00000234664.1 | HMGN2P5 | 0.473935525218067 | 2.47288397835238 | 2.86615928865367 | 0.00643650234835927 | 0.118633544069493 | -2.46705169165452 |
| ENSG00000171103.10 | TRMT61B | 0.31927227016471 | 3.4910587458124 | 2.86607024122395 | 0.00643801529056133 | 0.118633544069493 | -2.54922040831533 |
| lnc-MTERF4-3 | lnc-MTERF4-3 | -0.376094583913937 | 4.42761028015051 | -2.86414230830606 | 0.00647085277128171 | 0.119131988006522 | -2.63317204087597 |
| ENSG00000109790.16 | KLHL5 | 0.388641502600629 | 5.00680570442852 | 2.86098831873671 | 0.006524909555145 | 0.119586691455907 | -2.66342010931666 |
| ENSG00000265354.3 | TIMM23 | 0.261535714981266 | 4.20242191079465 | 2.8607692037638 | 0.00652868058375209 | 0.119586691455907 | -2.61991459353987 |
| ENSG00000134516.16 | DOCK2 | 0.264034736221593 | 8.33219175437576 | 2.86057970664589 | 0.0065319435173362 | 0.119586691455907 | -2.67104778441265 |
| ENSG00000211879.1 | TRAJ10 | -1.2147608502361 | 0.0827159850725781 | -2.86053033562065 | 0.00653279388182706 | 0.119586691455907 | -2.32228019424575 |
| ENSG00000156958.14 | GALK2 | -0.328304942283971 | 4.0628067019481 | -2.86035954725006 | 0.00653573632877046 | 0.119586691455907 | -2.60865160918861 |
| lnc-ZBTB46-2 | lnc-ZBTB46-2 | -0.388454259695633 | 2.46547788478033 | -2.86035466707561 | 0.00653582042554527 | 0.119586691455907 | -2.48444963905132 |
| ENSG00000160766.14 | GBA1LP | -0.666731774885169 | 0.720527659118018 | -2.85968179019504 | 0.00654742531332606 | 0.119586691455907 | -2.35868831277469 |
| ENSG00000136205.16 | TNS3 | 0.951616980704761 | 5.7757347465946 | 2.86291458129545 | 0.00655180039038358 | 0.119586691455907 | -2.68373124907255 |
| lnc-CD47-8 | lnc-CD47-8 | 1.04597541093316 | 5.4521282962383 | 2.86341762490464 | 0.00655181546779776 | 0.119586691455907 | -2.67376983134849 |
| ENSG00000172590.18 | MRPL52 | -0.289159390033571 | 4.31001369068788 | -2.85903939826731 | 0.00655852233602177 | 0.119586691455907 | -2.63449956350984 |
| lnc-SEPT14-6 | lnc-SEPT14-6 | -0.570463716312263 | 9.09668772973069 | -2.85868919460405 | 0.00656457931486111 | 0.119586691455907 | -2.65812621695012 |
| lnc-PPP2CA-1 | lnc-PPP2CA-1 | 0.27323034569378 | 3.82212830828053 | 2.85858894067087 | 0.00656631422438154 | 0.119586691455907 | -2.59266615506277 |
| LINC01772 | LINC01772 | 0.495826125539705 | 1.99280904207223 | 2.85811564031638 | 0.00657451052522265 | 0.119586691455907 | -2.45269193759285 |
| ENSG00000167380.16 | ZNF226 | -0.364749186991327 | 5.47240997077514 | -2.85797831696935 | 0.0065768903817756 | 0.119586691455907 | -2.68727609096362 |
| ENSG00000163811.11 | WDR43 | 0.321448867149794 | 6.06271431725074 | 2.85708185639575 | 0.0065924460420943 | 0.119763739527297 | -2.69917913590983 |
| ENSG00000251441.2 | RTEL1P1 | -0.809181611158203 | 0.431566360986646 | -2.85608501049659 | 0.00660978383627708 | 0.119939816905981 | -2.34636654853387 |
| ENSG00000107643.16 | MAPK8 | 0.355912376389907 | 5.63611824899321 | 2.85585487345069 | 0.00661379255639549 | 0.119939816905981 | -2.69702653016886 |
| ENSG00000268861.6 | NA | 0.446390309490373 | 6.32613105149045 | 2.85545493042724 | 0.00662076448408724 | 0.119960559309831 | -2.70331521525098 |
| lnc-NOL4L-2 | lnc-NOL4L-2 | -0.696353222818637 | 1.09523017665911 | -2.85380985224742 | 0.00664951399121413 | 0.120375502621953 | -2.40264702971268 |
| lnc-KHDC3L-2 | lnc-KHDC3L-2 | -0.439208116608393 | 3.08039934512378 | -2.853011211077 | 0.00666351296351044 | 0.120522923838256 | -2.54966418725345 |
| ENSG00000277117.4 | | 1.14375557970844 | 2.74851141653068 | 2.85508817791972 | 0.00667748720029866 | 0.120669639195564 | -2.51144653667665 |
| lnc-CDKN2AIPNL-3 | lnc-CDKN2AIPNL-3 | 0.489433708498198 | 0.251187405274277 | 2.85171530015631 | 0.00668628670300996 | 0.120711099248464 | -2.34879424653064 |
| lnc-SULF1-3 | lnc-SULF1-3 | -2.55711315933015 | -0.473562063482003 | -2.85432111230999 | 0.00669822697293396 | 0.120711099248464 | -2.37430036600415 |
| ENSG00000162402.13 | USP24 | 0.273262447636708 | 7.90677148643871 | 2.85085424694879 | 0.00670145848718088 | 0.120711099248464 | -2.70060479771496 |
| lnc-FAM24A-1 | lnc-FAM24A-1 | 1.21509886105642 | -0.551717395446421 | 2.85061949128913 | 0.0067056004439022 | 0.120711099248464 | -2.41238032269899 |
| ENSG00000230673.3 | PABPC1P3 | 0.52320292309954 | 2.09587237861982 | 2.85042099663076 | 0.00670910448137992 | 0.120711099248464 | -2.47330411644369 |
| lnc-RXFP4-2 | lnc-RXFP4-2 | -0.79158514581116 | 1.62906021754465 | -2.84996934425845 | 0.0067170838771186 | 0.120749115670508 | -2.45393104699655 |
| ENSG00000187741.14 | FANCA | -0.56214423817414 | 4.32862806589507 | -2.8480952793282 | 0.00675028764402461 | 0.1212401139415 | -2.66616454833744 |
| lnc-ZNF574-1 | lnc-ZNF574-1 | -0.801231217669639 | 3.72936185857062 | -2.84720368282336 | 0.0067826953948518 | 0.121715971501512 | -2.61741382700927 |
| lnc-LMNB1-3 | lnc-LMNB1-3 | -0.611622319000831 | 0.791965577574465 | -2.8443156760982 | 0.00681771790961382 | 0.122032376921407 | -2.40318512971416 |
| lnc-GRAP-3 | lnc-GRAP-3 | -1.02134829571569 | 3.59382412647868 | -2.84733708529643 | 0.00682165481351215 | 0.122032376921407 | -2.63718393167888 |
| ENSG00000100577.18 | GSTZ1 | -0.381387701365574 | 4.37395913073995 | -2.84399315747386 | 0.0068235007479242 | 0.122032376921407 | -2.66918791737428 |
| ENSG00000162631.18 | NTNG1 | -1.61706274815879 | 0.794572825673841 | -2.8468623030237 | 0.00682459058067319 | 0.122032376921407 | -2.38207557881355 |

| | | | | | | | |
|---|---|---|---|---|---|---|---|
| ENSG00000171100.14 | MTM1 | 0.256334038108503 | 4.85798594527071 | 2.84363258804339 | 0.00682997124877135 | 0.122032376921407 | -2.69772742989892 |
| ATP2B1-AS1 | ATP2B1-AS1 | -0.599363611953273 | 2.69200946236321 | -2.84281775087737 | 0.00684461471819132 | 0.122099782972137 | -2.5456576454545 |
| ENSG00000089775.11 | ZBTB25 | -0.234072830044101 | 5.65012689903388 | -2.84253061208599 | 0.00684978185553382 | 0.122099782972137 | -2.72875108223454 |
| lnc-SLC39A8-1 | lnc-SLC39A8-1 | 0.940378712064997 | -0.220500492388108 | 2.84224411234586 | 0.0068549411092269 | 0.122099782972137 | -2.36919943852803 |
| ENSG00000272772.1 | | 0.268911022255038 | 4.4484416265734 | 2.84210363424771 | 0.00685747214282611 | 0.122099782972137 | -2.67839130214058 |
| ENSG00000152689.17 | RASGRP3 | -0.565357865680236 | 7.67041251691793 | -2.83989093289971 | 0.00689745386179052 | 0.122705525356296 | -2.72803519745251 |
| ENSG00000117410.13 | ATP6V0B | 0.418655300571475 | 5.01705729117119 | 2.8392555732705 | 0.00690897429766501 | 0.122804333306424 | -2.71672227383382 |
| ENSG00000112096.17 | NA | 0.334982413458016 | 5.19210621264951 | 2.83851573467079 | 0.00692241171287414 | 0.122937014914658 | -2.72765773270055 |
| ENSG00000187994.13 | RINL | -0.598804905534431 | 4.35941069742371 | -2.83770181754786 | 0.00693722266424994 | 0.123041982154843 | -2.68589413532218 |
| lnc-VIP-1 | lnc-VIP-1 | -0.397417184963874 | 2.19718737730115 | -2.83741303414314 | 0.00694248477098706 | 0.123041982154843 | -2.52445746710866 |
| ENSG00000273885.1 | | 0.612655748234341 | 3.5025658309423 | 2.83687444347178 | 0.00695230869357134 | 0.123041982154843 | -2.62249460850304 |
| ENSG00000100410.7 | PHF5A | 0.352386633090853 | 3.34320207262737 | 2.83620867975667 | 0.00696447012505307 | 0.123041982154843 | -2.60837621628585 |
| ENSG00000136925.14 | TSTD2 | 0.446924167489527 | 4.68861975027914 | 2.83611667061497 | 0.00696615240077771 | 0.123041982154843 | -2.7077873359031 |
| ENSG00000081377.16 | CDC14B | -0.643021939380022 | 2.99454917517263 | -2.83590239418418 | 0.00697007165058384 | 0.123041982154843 | -2.5884402485798 |
| ENSG00000170836.11 | PPM1D | 0.493327021302637 | 4.39771419901141 | 2.83589717238444 | 0.00697016718615107 | 0.123041982154843 | -2.68348147310996 |
| ENSG00000179761.11 | PIPOX | -0.482898679925268 | 0.983908835562131 | -2.8354218155607 | 0.00697886919115896 | 0.123090029615788 | -2.43686428228104 |
| ENSG00000271982.1 | SNORD58B | 0.479141349661166 | 2.65166785281933 | 2.83150444217879 | 0.00705096722846664 | 0.124090741507612 | -2.56453142398956 |
| ENSG00000102580.14 | DNAJC3 | 0.357149363141628 | 5.25476569567181 | 2.83039912717168 | 0.0070714351076608 | 0.124090741507612 | -2.74604102740372 |
| ENSG00000199347.1 | RNU5E-1 | -0.989800640223737 | 2.23026781508492 | -2.83116932252953 | 0.00707831335231706 | 0.124090741507612 | -2.53481153076368 |
| ENSG00000157500.11 | APPL1 | -0.388884254797251 | 7.52024593333022 | -2.82996336462762 | 0.00707951959161821 | 0.124090741507612 | -2.7543743027273 |
| ENSG00000174780.16 | VIII.72 | 0.216728048055235 | 6.55870228386544 | 2.82979548917685 | 0.00708263639251349 | 0.124090741507612 | -2.76415443500529 |
| lnc-SAG-4 | lnc-SAG-4 | 0.80725298860718 | 4.30794550966502 | 2.83108795408564 | 0.00708375729233292 | 0.124090741507612 | -2.68328915926299 |
| ENSG00000135966.12 | TGFBRAP1 | 0.217149803116281 | 4.95150339547601 | 2.82955109673755 | 0.00708717609945891 | 0.124090741507612 | -2.73881525105031 |
| ENSG00000166788.9 | SAAL1 | 0.291207508528387 | 3.4863857582203 | 2.82942864041792 | 0.00708945180143192 | 0.124090741507612 | -2.63903660001706 |
| ENSG00000257379.1 | | 0.334608367525943 | 5.81981922914629 | 2.82940635582568 | 0.00708986600655648 | 0.124090741507612 | -2.76176340491651 |
| ENSG00000213064.9 | SFT2D2 | 0.423376590548425 | 7.49553436451651 | 2.82854905559199 | 0.00710581778906147 | 0.124201208697421 | -2.75998629259551 |
| ENSG00000252010.1 | SCARNA5 | 0.807009059964684 | 4.3077334261376 | 2.82965153194562 | 0.00711044814583239 | 0.124201208697421 | -2.68664519303218 |
| lnc-STRN3-7 | lnc-STRN3-7 | 0.323252896652977 | 2.3613341405963 | 2.82809499548661 | 0.00711427999097599 | 0.124201208697421 | -2.54887234428978 |
| lnc-RASGRF1-2 | lnc-RASGRF1-2 | -1.08014206827843 | 0.00022618329025118 | -2.82753925814301 | 0.00712464987299962 | 0.124276577389735 | -2.39427032588828 |
| lnc-ACTN2-1 | lnc-ACTN2-1 | -0.355351695431879 | 2.00163119474052 | -2.82721682648703 | 0.00713067278323265 | 0.124276577389735 | -2.53318353488621 |
| ENSG00000178386.12 | ZNF223 | -1.49102400765145 | 1.01017144098356 | -2.82877143991083 | 0.00714915779756012 | 0.124328349372891 | -2.43168594927372 |
| ENSG00000172728.15 | FUT10 | 1.40310631517416 | 1.81968121433659 | 2.82936065003693 | 0.00715018679450931 | 0.124328349372891 | -2.49115589127892 |
| ENSG00000198625.12 | MDM4 | 0.342802409330474 | 8.64696721167717 | 2.82598008502391 | 0.00715381859242349 | 0.124328349372891 | -2.74630468735167 |
| ENSG00000113194.12 | FAF2 | 0.311831798454826 | 5.69373901362926 | 2.82575402606587 | 0.00715805686134278 | 0.124328349372891 | -2.76847766959547 |
| lnc-TMEM234-4 | lnc-TMEM234-4 | -0.468370655827946 | 3.09286060813426 | -2.8254454998124 | 0.00716384503504101 | 0.124328349372891 | -2.61323363621052 |
| ENSG00000115275.12 | MOGS | -0.329043989216522 | 5.07107466513696 | -2.82457796076308 | 0.00718014400851831 | 0.124328366524949 | -2.75795310767273 |
| ENSG00000022840.15 | RNF10 | 0.417103071445618 | 6.31841927715244 | 2.8244274766141 | 0.00718297474872721 | 0.124328366524949 | -2.77710461821187 |
| lnc-CCDC71-1 | lnc-CCDC71-1 | -0.437640719746307 | 1.5152331234334 | -2.82435743296208 | 0.00718429268555885 | 0.124328366524949 | -2.49366998892183 |
| ENSG00000123989.13 | CHPF | -1.08397009492022 | 0.679084919977257 | -2.824305869393 | 0.00718800739273622 | 0.124328366524949 | -2.44433442322722 |
| ENSG00000188779.10 | SKOR1 | -0.516534701640781 | 1.49823073725098 | -2.82376962325117 | 0.00719536173396149 | 0.124351075205818 | -2.49762581665393 |
| ENSG00000075856.11 | SART3 | 0.231447403904497 | 6.77982485867191 | 2.82324304447065 | 0.00720529119743993 | 0.124418212010827 | -2.77875024157587 |
| DPH6-DT | DPH6-DT | -0.834354223602944 | 0.902539765739953 | -2.82274938318656 | 0.00721461151212117 | 0.124474726533101 | -2.44730545588218 |
| ENSG00000277406.2 | SEC22B4P | -0.573954559904756 | 0.155899154412279 | -2.82200701245529 | 0.00722864854742732 | 0.124612456492208 | -2.40562247749548 |
| ENSG00000106367.14 | AP1S1 | -0.795914267147159 | 2.45513706453189 | -2.82034641232103 | 0.00726022213700817 | 0.124846940177381 | -2.5876559722048 |
| ENSG00000117000.8 | RLF | 0.351703891127242 | 6.43444511035072 | 2.82027653802192 | 0.00726146760621235 | 0.124846940177381 | -2.78698500007893 |
| ENSG00000198712.1 | MT-CO2 | -0.705595167321084 | 9.37939574001382 | -2.82082316769169 | 0.00726347576414889 | 0.124846940177381 | -2.73964948088942 |
| ENSG00000110713.16 | NUP98 | 0.222070282267595 | 6.95837175443757 | 2.82001115751963 | 0.00726651286656474 | 0.124846940177381 | -2.78516765854219 |
| lnc-PSMD5-4 | lnc-PSMD5-4 | -0.503829618341073 | 0.630955530355302 | -2.81874874672045 | 0.00729055771261104 | 0.125040390603423 | -2.45243390148037 |
| ENSG00000148057.15 | IDNK | -0.325499643514688 | 3.09460851047676 | -2.81838622040217 | 0.00729747631551834 | 0.125040390603423 | -2.63405985684991 |
| lnc-ANKRD34B-4 | lnc-ANKRD34B-4 | 1.30055396983816 | 2.57864458263477 | 2.82165103833636 | 0.0073018922414238 | 0.125040390603423 | -2.57740141437403 |
| lnc-AKNA-3 | lnc-AKNA-3 | -1.45570176501637 | -0.0103047566708685 | -2.81869527491933 | 0.00730207207429987 | 0.125040390603423 | -2.41166448472833 |
| ENSG00000224464.2 | PGAM1P6 | 0.824726947113444 | -0.250813281055786 | 2.8174796540423 | 0.00731480430954896 | 0.125154295181585 | -2.41987881795464 |
| ENSG00000134759.13 | ELP2 | 0.348469410109685 | 6.54949980142375 | 2.81654930179293 | 0.00733262667523732 | 0.125355028950506 | -2.79557871156313 |
| ENSG00000179562.2 | GCC1 | 0.348002476741741 | 5.20834552125536 | 2.81565626280252 | 0.00734977217881021 | 0.125476561375866 | -2.77865045305923 |
| ENSG00000162972.10 | MAIP1 | 0.25022768291119 | 3.36262763849127 | 2.81554411442902 | 0.00735192795118761 | 0.125476561375866 | -2.66271907295521 |
| ENSG00000163596.16 | ICA1L | -0.525811235985005 | 2.30172721312491 | -2.81314076670632 | 0.00739826768899713 | 0.126076375038561 | -2.58434503353532 |
| TAPT1-AS1 | TAPT1-AS1 | -0.926849655435352 | 3.35099967198796 | -2.81509762887942 | 0.00739958748896821 | 0.126076375038561 | -2.67675648924296 |
| lnc-GAS2-1 | lnc-GAS2-1 | -0.802449090822072 | 1.16166637151035 | -2.81276961806079 | 0.00740544806012829 | 0.126076375038561 | -2.49498881858468 |
| ENSG00000099326.8 | MZF1 | -0.3554338486781 | 4.8141163100113 | -2.81173145740847 | 0.00742556706068388 | 0.126314418851286 | -2.77451511062768 |
| lnc-FAM199X-1 | lnc-FAM199X-1 | -0.577268403372376 | 1.43335681869332 | -2.81042931262699 | 0.00745087368864109 | 0.126640242058877 | -2.51268078327454 |
| lnc-UCK2-1 | lnc-UCK2-1 | -0.619748474494274 | 1.59784808806955 | -2.80990919041842 | 0.00746100440548202 | 0.126698064182805 | -2.5354530234246 |
| ENSG00000114544.16 | SLC41A3 | 0.286091972665887 | 4.11644225990969 | 2.80955784821772 | 0.00746785493378389 | 0.126698064182805 | -2.73469679953306 |
| ENSG00000257103.8 | LSM14A | 0.215976667893506 | 6.75235617713714 | 2.80930738537706 | 0.00747274206470995 | 0.126698064182805 | -2.81157248262875 |
| lnc-MDM4-1 | lnc-MDM4-1 | 0.730417892427362 | 3.64488792761553 | 2.80899665153891 | 0.00748227280616948 | 0.126755243760812 | -2.68622567743529 |
| ENSG00000122882.10 | ECD | 0.272555444635861 | 5.55330718405894 | 2.80788878492641 | 0.00750047844691887 | 0.12695916765866 | -2.80748501877571 |
| ENSG00000117614.9 | SYF2 | 0.298296666091812 | 5.70249412959794 | 2.80738479689038 | 0.00751035537550724 | 0.127014556474167 | -2.81187307977138 |
| ENSG00000107854.5 | TNKS2 | 0.290517714054969 | 6.59931780840317 | 2.80709233754036 | 0.00751609239593525 | 0.127014556474167 | -2.81755598894489 |
| ENSG00000101460.12 | MAP1LC3A | -0.902415658814197 | 0.781626878584789 | -2.80576974187857 | 0.00754208789190289 | 0.127349298670252 | -2.49952129599081 |

| | | | | | | | |
|---|---|---|---|---|---|---|---|
| ENSG00000174485.15 | DENND4A | 0.34685963385841 | 7.96016261821974 | 2.80485326093889 | 0.00756015016750286 | 0.127549648276813 | -2.80906976441559 |
| ENSG00000108510.9 | MED13 | 0.279167524483466 | 7.77074510920409 | 2.80367812878016 | 0.00758336876645171 | 0.127836592399571 | -2.81387270445533 |
| ENSG00000166716.9 | ZNF592 | 0.306350659920214 | 6.33794540692067 | 2.80323850362386 | 0.00759207200554881 | 0.127878574541908 | -2.82706323825836 |
| ENSG00000087152.15 | ATXN7L3 | 0.315854557402402 | 5.68645243106249 | 2.80165132661131 | 0.00762357048675657 | 0.128270385026254 | -2.8254091952133 |
| ENSG00000105401.8 | CDC37 | -0.3491687859859 | 6.4820556554543 | -2.80143880638814 | 0.007627797273096 | 0.128270385026254 | -2.83057755614166 |
| lnc-SLCO2A1-4 | lnc-SLCO2A1-4 | -0.846660807961778 | -0.492636725654094 | -2.79967525724048 | 0.00766295626222297 | 0.128756431628845 | -2.45501463312396 |
| ENSG00000235552.4 | RPL6P27 | 1.0566363987877 | 2.80176887481506 | 2.80128488068214 | 0.00767134032420856 | 0.128792167938976 | -2.62435120009166 |
| lnc-DTNBP1-16 | lnc-DTNBP1-16 | -1.905349433843 | 1.90918973215851 | -2.80173317014003 | 0.00772374484216226 | 0.129461077711595 | -2.59825799279771 |
| lnc-SP9-13 | lnc-SP9-13 | -0.648822897536957 | 0.392591781960392 | -2.79664290706139 | 0.00772376249476943 | 0.129461077711595 | -2.4804455238225 |
| lnc-C19orf18-2 | lnc-C19orf18-2 | -0.389198043439918 | 0.77181076371944 | -2.79506206220487 | 0.0077556396333004 | 0.129889609904168 | -2.50599762686214 |
| ENSG00000198162.12 | MAN1A2 | 0.226340763227738 | 6.7945889851155 | 2.79446713799085 | 0.0077676676311963 | 0.129985286872287 | -2.84634504187759 |
| lnc-ZNF527-1 | lnc-ZNF527-1 | -0.238237658013067 | 3.46626007959133 | -2.79210185240839 | 0.00781565962317906 | 0.130662045940943 | -2.72419801694866 |
| lnc-MCM9-2 | lnc-MCM9-2 | -0.505931482246242 | 1.13149957375074 | -2.79135594506616 | 0.00783085114182817 | 0.130662045940943 | -2.54486449175092 |
| ENSG00000122085.16 | MTERF4 | -0.329442206996397 | 5.57995051841736 | -2.7912411477521 | 0.00783319159119913 | 0.130662045940943 | -2.84861144843969 |
| lnc-SLTM-7 | lnc-SLTM-7 | 0.385452303682153 | 2.04014785825775 | 2.7909642360216 | 0.00783883984548154 | 0.130662045940943 | -2.60760957190197 |
| lnc-UBE2H-2 | lnc-UBE2H-2 | -0.526875404408621 | 4.01069426833591 | -2.79047972431732 | 0.00784873166828947 | 0.130662045940943 | -2.76470144856079 |
| lnc-CEP97-8 | lnc-CEP97-8 | -0.303717478463343 | 4.79322390315681 | -2.79037711817205 | 0.00785082796696669 | 0.130662045940943 | -2.82322001247798 |
| ENSG00000184497.12 | TMEM255B | 1.36463582472677 | 0.771923145639463 | 2.79190777038063 | 0.00785850112249073 | 0.130662045940943 | -2.51569281767073 |
| lnc-CHORDC1-1 | lnc-CHORDC1-1 | 0.401089788944602 | 4.01306885943703 | 2.78983952734278 | 0.00786181972645506 | 0.130662045940943 | -2.76256123156144 |
| ENSG00000278867.1 | | -0.593903944904239 | -0.406774923414946 | -2.78967229402665 | 0.00786524194339158 | 0.130662045940943 | -2.46990340546913 |
| lnc-ZBED5-5 | lnc-ZBED5-5 | 0.347706166097398 | 5.91497837094959 | 2.78920800487196 | 0.00787475027328107 | 0.130714503931407 | -2.85818950638864 |
| ENSG00000109220.10 | CHIC2 | 0.317012259662997 | 3.5058058278638 | 2.78841753488507 | 0.00789096309495247 | 0.130878076860118 | -2.73379420287122 |
| ENSG00000044574.7 | HSPA5 | 0.657640213354334 | 5.85586575531475 | 2.78743900196876 | 0.00792364705465472 | 0.131218529702553 | -2.86060101311924 |
| ENSG00000016391.10 | CHDH | -1.17733058028545 | 4.84221617860929 | -2.79090558918135 | 0.0079255260446722 | 0.131218529702553 | -2.82851298113973 |
| lnc-RASA2-2 | lnc-RASA2-2 | -0.512186605759896 | 2.8990161502412 | -2.78632215497965 | 0.00793408998262478 | 0.131218529702553 | -2.69530303147175 |
| ENSG00000106992.18 | AK1 | -0.506662972793348 | 2.77459690971581 | -2.78618162269621 | 0.00793699020938048 | 0.131218529702553 | -2.67830693499957 |
| lnc-HK3-1 | lnc-HK3-1 | 0.327228081770567 | 1.52436273932955 | 2.78545010663507 | 0.0079521027109916 | 0.131270618186943 | -2.58321546085848 |
| ENSG00000162461.7 | SLC25A34 | -0.510253864004854 | 0.780216291144616 | -2.785303535999 | 0.00795513394262026 | 0.131270618186943 | -2.54155654979202 |
| lnc-ERICH1-7 | lnc-ERICH1-7 | -0.633441715705285 | 1.27984212343909 | -2.78503209754757 | 0.00796075039845634 | 0.131270618186943 | -2.56869489282004 |
| lnc-IL5-1 | lnc-IL5-1 | 0.70780259417696 | 3.08317389781463 | 2.78465047886168 | 0.00796865286131106 | 0.131270618186943 | -2.7009941380632 |
| lnc-SLC38A7-2 | lnc-SLC38A7-2 | 0.372440491036576 | 2.39235287705825 | 2.78448754875183 | 0.00797202899157939 | 0.131270618186943 | -2.64836508818175 |
| ENSG00000184489.11 | PTP4A3 | 4.76146186043377 | -1.81574407607292 | 2.78664024496595 | 0.00799085819312653 | 0.131310997015984 | -3.05162067268353 |
| ENSG00000158435.7 | CNOT11 | 0.214039641306249 | 4.6911100171025 | 2.78346092899675 | 0.0079933324503728 | 0.131310997015984 | -2.83539964653528 |
| ENSG00000111615.13 | KRR1 | 0.1908422695588 | 6.64483727681001 | 2.78344709205981 | 0.00799361994174941 | 0.131310997015984 | -2.87243944528592 |
| lnc-VPS18-1 | lnc-VPS18-1 | -0.541300230484815 | 0.0228914045875549 | -2.78273903187379 | 0.00800834417878855 | 0.131404737714726 | -2.49091538892481 |
| ENSG00000095397.13 | WHRN | -1.42578231019924 | 4.55353392059316 | -2.78716211085951 | 0.00802076183153405 | 0.131404737714726 | -2.82582291318486 |
| lnc-LYRM7-3 | lnc-LYRM7-3 | -0.749071720405432 | 0.608573080494529 | -2.78207803603873 | 0.0080221123918861 | 0.131404737714726 | -2.53186148174253 |
| ENSG00000267673.6 | FDX2 | -0.43642176698983 | 0.657661182424879 | -2.78194610818394 | 0.00802486300866786 | 0.131404737714726 | -2.53192436527401 |
| ENSG00000198399.14 | ITSN2 | 0.328659439259061 | 7.79045597658466 | 2.78119070074415 | 0.00804062964338144 | 0.131506720629704 | -2.86648649361883 |
| lnc-SMNDC1-10 | lnc-SMNDC1-10 | 0.42557020713425 | 1.07288464252074 | 2.78080529921155 | 0.00804868468437043 | 0.131506720629704 | -2.55841335745289 |
| CHKB-DT | CHKB-DT | -0.577003658639427 | 1.77434021273157 | -2.78073004647386 | 0.00805025836823725 | 0.131506720629704 | -2.61075162514634 |
| ENSG00000172086.7 | KRCC1 | -0.496519524376158 | 4.16709657945114 | -2.78003213579524 | 0.00806486665707047 | 0.131640880572943 | -2.80510973654835 |
| ENSG00000113712.17 | CSNK1A1 | 0.191608186815642 | 6.54245781029146 | 2.77962893318238 | 0.00807331747006627 | 0.13166921577906 | -2.88192167881809 |
| THAP7-AS1 | THAP7-AS1 | -0.660040394017262 | 0.696713016084899 | -2.77933913102467 | 0.00807939656653319 | 0.13166921577906 | -2.53519929350984 |
| ENSG00000128694.11 | OSGEPL1 | -0.434886654399353 | 2.68922647780251 | -2.77786545007805 | 0.00811037524020345 | 0.131967283864841 | -2.69669596247087 |
| lnc-POLR2B-1 | lnc-POLR2B-1 | -0.593202644405731 | -0.527665075698216 | -2.77746643818195 | 0.00811878190247782 | 0.131967283864841 | -2.50288326759948 |
| ENSG00000168890.13 | TMEM150A | -0.466535058746875 | 1.69555498756312 | -2.77742235364898 | 0.00811971120153652 | 0.131967283864841 | -2.63218304369533 |
| ENSG00000095015.5 | MAP3K1 | 0.278715068028724 | 9.18807507373401 | 2.77725061901496 | 0.00812333229639771 | 0.131967283864841 | -2.84900869379884 |
| ENSG00000137200.12 | CMTR1 | 0.182791144982698 | 6.11334994936162 | 2.77688230059108 | 0.00813110348789977 | 0.131989355750348 | -2.88821206292968 |
| ENSG00000165684.4 | SNAPC4 | -0.316961918091786 | 3.76597942801673 | -2.77426716951772 | 0.0081864788506237 | 0.132783525754443 | -2.79387906861203 |
| ENSG00000130598.15 | TNNI2 | -0.911377239815976 | 1.25541130925343 | -2.77357459084905 | 0.00820120260807263 | 0.132917601009416 | -2.58908796181124 |
| lnc-PRDX2-3 | lnc-PRDX2-3 | -0.495611879794366 | -0.145448900510926 | -2.77183316111265 | 0.00823833270848366 | 0.133414321116223 | -2.50644423296317 |
| ENSG00000204396.10 | VWA7 | -1.22874796742759 | 0.0593145439246351 | -2.77092169539014 | 0.00825782866245993 | 0.133552902423388 | -2.50825536009204 |
| ENSG00000235863.3 | B3GALT4 | -0.360964774149486 | 3.49895135178053 | -2.77019139863937 | 0.00827348030335922 | 0.133552902423388 | -2.78390061682106 |
| ENSG00000140854.12 | KATNB1 | -0.29790363903354 | 2.78032728752822 | -2.76964370065467 | 0.00828523653514725 | 0.133552902423388 | -2.72192118221692 |
| lnc-PDE6G-3 | lnc-PDE6G-3 | -0.778290669476514 | 1.9415708790013 | -2.76960626463332 | 0.00828604065685191 | 0.133552902423388 | -2.64997228591676 |
| lnc-C2orf73-5 | lnc-C2orf73-5 | 0.587992329482094 | 0.152740325360672 | 2.76957077402564 | 0.00828680305803467 | 0.133552902423388 | -2.51179196379725 |
| ENSG00000182919.14 | C11orf54 | 0.375966866637063 | 4.2198591323479 | 2.76889667094642 | 0.00830129633116558 | 0.133552902423388 | -2.82643681571622 |
| ENSG00000176396.10 | EID2 | 0.509446978864633 | 1.9212021723661 | 2.76836068698649 | 0.00831283678353739 | 0.133552902423388 | -2.65952542888369 |
| ENSG00000213699.8 | SLC35F6 | 0.404006314175937 | 4.03966559231906 | 2.76789161806689 | 0.00832294865666518 | 0.133552902423388 | -2.82214558319308 |
| ENSG00000223773.7 | CD99P1 | -0.40609949492704 | 2.42756819474241 | -2.76776253614643 | 0.0083257333163682 | 0.133552902423388 | -2.69498854962601 |
| lnc-ASMT-1 | lnc-ASMT-1 | 1.64884304117939 | 1.1925047456214 | 2.77063357462106 | 0.00832771922719636 | 0.133552902423388 | -2.54356711337986 |
| ENSG00000236085.1 | ACTG1P4 | -0.629309930039677 | -0.303519074623843 | -2.76754246535678 | 0.00833048285141037 | 0.133552902423388 | -2.51552452162609 |
| ENSG00000114742.13 | WDR48 | 0.26230065481388 | 5.97550973297291 | 2.76747817848734 | 0.00833187075458807 | 0.133552902423388 | -2.90919830307799 |
| ENSG00000165115.14 | KIF27 | 0.505661741190029 | 3.51975189340543 | 2.76743491120474 | 0.00833280498191492 | 0.133552902423388 | -2.78746828334595 |
| ENSG00000110047.17 | EHD1 | -0.687199496168597 | 5.92470794867479 | -2.76783652978417 | 0.00834593536159647 | 0.133552902423388 | -2.90758061387471 |
| lnc-LYPLA1-2 | lnc-LYPLA1-2 | 0.361577077384434 | 3.93634338694013 | 2.76676884701955 | 0.0083471988902494 | 0.133552902423388 | -2.81539108263237 |
| lnc-TOMM22-2 | lnc-TOMM22-2 | 0.345705662623719 | 2.08130549718094 | 2.76660671150165 | 0.00835070618563382 | 0.133552902423388 | -2.67099373252534 |

| | | | | | | | |
|---|---|---|---|---|---|---|---|
| ENSG00000154803.12 | FLCN | -0.307759609369917 | 4.70023130270422 | -2.76542631947376 | 0.00837628144588524 | 0.133857920031565 | -2.87977431079142 |
| lnc-FAM24B-1 | lnc-FAM24B-1 | 1.21218921642657 | -0.269616253650229 | 2.76441708510238 | 0.00839820577847376 | 0.134043008474056 | -2.53843729712083 |
| ENSG00000119640.8 | ACYP1 | -0.341957022310859 | 2.45465384605612 | -2.76429377576866 | 0.00840088815680575 | 0.134043008474056 | -2.70776342330205 |
| SNHG12 | SNHG12 | -0.398669667293922 | 3.29854233671391 | -2.76196267872959 | 0.00845174658083775 | 0.134715107697292 | -2.78147980164521 |
| ENSG00000272897.5 | | -0.249321879075717 | 3.59174772509602 | -2.76176371507336 | 0.00845610062405389 | 0.134715107697292 | -2.80629336782495 |
| ENSG00000196684.12 | HSH2D | -0.399081955548083 | 7.0400510854235 | -2.76055450007588 | 0.00848260731034263 | 0.134996244128005 | -2.92228883659629 |
| ENSG00000236493.2 | EIF2S2P3 | -0.545549022597223 | 0.870593494619951 | -2.76036059895213 | 0.00848686488372143 | 0.134996244128005 | -2.59213021562107 |
| ENSG00000137492.7 | THAP12 | 0.29833793774413 | 6.25959405456663 | 2.75916414582745 | 0.00851317973385904 | 0.135030009691568 | -2.92980300241142 |
| ENSG00000196531.10 | NACA | 0.454636575461777 | 7.68530775708133 | 2.75897560568337 | 0.00851733337785112 | 0.135030009691568 | -2.91978173680697 |
| lnc-ABCC10-2 | lnc-ABCC10-2 | -0.530911479363773 | 0.926131067820933 | -2.75895979376773 | 0.00851768180837731 | 0.135030009691568 | -2.59820869593572 |
| ENSG00000115421.12 | PAPOLG | 0.303656059886744 | 5.30165211385518 | 2.75878163623214 | 0.00852160859028122 | 0.135030009691568 | -2.91646317181182 |
| lnc-GLYCTK-1 | lnc-GLYCTK-1 | -0.452266456283194 | 0.836464861461159 | -2.75877345377131 | 0.00852178898068051 | 0.135030009691568 | -2.58469409191583 |
| lnc-NT5M-1 | lnc-NT5M-1 | 0.489425533700188 | 2.9752958264344 | 2.75845959939966 | 0.00852871087781933 | 0.135035735383196 | -2.76038506113901 |
| lnc-CORO2B-1 | lnc-CORO2B-1 | -0.547751573551783 | 0.549758314659766 | -2.75729474824823 | 0.00855444660096027 | 0.135243201097254 | -2.57139759362959 |
| lnc-ASAH2-7 | lnc-ASAH2-7 | -0.265563738816093 | 3.17097808885594 | -2.75727174986149 | 0.0085549554403452 | 0.135243201097254 | -2.77317741810943 |
| lnc-MAP1B-2 | lnc-MAP1B-2 | -0.401529463744587 | 1.13606896093938 | -2.75691683617711 | 0.00856281145977763 | 0.135263505968229 | -2.61457875412942 |
| ENSG00000212464.1 | SNORA12 | 0.784191456886778 | 4.01046901789406 | 2.75728057045978 | 0.0085741682228893 | 0.135339037217585 | -2.82066068481272 |
| ENSG00000136758.18 | YME1L1 | 0.202226094120315 | 7.19679120138678 | 2.75513945858844 | 0.00860225428526456 | 0.135389827500595 | -2.93425354416324 |
| ENSG00000135070.14 | ISCA1 | 0.348636213212193 | 4.75103696211774 | 2.75471797263945 | 0.00861163235943819 | 0.135389827500595 | -2.90261175879344 |
| lnc-ASB8-1 | lnc-ASB8-1 | 0.2425664026029 | 2.56767304080595 | 2.75434097920897 | 0.0086200284893357 | 0.135389827500595 | -2.73763336252744 |
| ENSG00000263874.2 | LASP1NB | -0.53104750941194 | 1.85314797476236 | -2.75419402194298 | 0.00862330346884852 | 0.135389827500595 | -2.68924122847841 |
| ENSG00000114786.16 | ABHD14A-ACY1 | -0.493144177412511 | 2.15487603752248 | -2.7539456920524 | 0.00862884017973323 | 0.135389827500595 | -2.70699025295041 |
| ENSG00000184164.14 | CRELD2 | 0.525959437855974 | 2.67127490336232 | 2.753917998851 | 0.00862945782536521 | 0.135389827500595 | -2.73977366224095 |
| lnc-CFL1-1 | lnc-CFL1-1 | -0.9651037575829 | 0.0171791151031095 | -2.75368070555991 | 0.0086347518907982 | 0.135389827500595 | -2.54315960196413 |
| lnc-PPARA-1 | lnc-PPARA-1 | -0.709260254006324 | -0.0885517845784278 | -2.75351774823852 | 0.00863838924500227 | 0.135389827500595 | -2.54600701841082 |
| ENSG00000122958.14 | VPS26A | 0.39249592745782 | 4.22114862982518 | 2.75320403080889 | 0.00864539569335316 | 0.135389827500595 | -2.87119759559576 |
| lnc-TEX29-9 | lnc-TEX29-9 | 0.660425005499469 | 0.519052169994066 | 2.75305195448015 | 0.00864879400166116 | 0.135389827500595 | -2.57956204407938 |
| ZNF529-AS1 | ZNF529-AS1 | -0.29668439992767 | 3.49558408442847 | -2.75300957654725 | 0.00864974120212227 | 0.135389827500595 | -2.81671126120112 |
| lnc-PYDC2-5 | lnc-PYDC2-5 | 0.426805693825687 | 5.28850551663308 | 2.75259390766001 | 0.00865903701796462 | 0.135432339620643 | -2.92935634930073 |
| ENSG00000100211.10 | CBY1 | -0.363694118219974 | 1.60870601143114 | -2.75130250659439 | 0.00868797628724846 | 0.135781671858927 | -2.66978942751059 |
| ENSG00000176208.8 | ATAD5 | 0.386377603787845 | 3.75626825686193 | 2.75100901855174 | 0.00869456558859575 | 0.135781671858927 | -2.83707115844075 |
| ENSG00000143514.16 | TP53BP2 | 0.271066735705332 | 5.49855204849145 | 2.75058171732554 | 0.00870416748902139 | 0.135827765283718 | -2.9406694798303 |
| ENSG00000083799.17 | CYLD | 0.271339002908619 | 8.06982200550554 | 2.75029058456452 | 0.00871071516176008 | 0.135827765283718 | -2.9333246418982 |
| lnc-ARAP2-1 | lnc-ARAP2-1 | 0.570572469402483 | 1.62155040843129 | 2.7494990462292 | 0.00872854016771057 | 0.135927537944252 | -2.68256270729053 |
| ENSG00000109814.11 | UGDH | -0.567635950725218 | 3.03970525849055 | -2.74942003138628 | 0.00873032138960798 | 0.135927537944252 | -2.79645423925853 |
| lnc-THSD7A-3 | lnc-THSD7A-3 | 0.367694767802592 | 3.9399498083261 | 2.74830116207776 | 0.00875558004571875 | 0.136217765745298 | -2.86556644321292 |
| lnc-PCDHGA1-4 | lnc-PCDHGA1-4 | -0.763417357768731 | -0.415201028042672 | -2.7473344891086 | 0.00877745726421876 | 0.136454987061492 | -2.5593859415074 |
| lnc-MXRA5-8 | lnc-MXRA5-8 | 0.5239767837054 | 1.55565225056719 | 2.74673364358013 | 0.00879108074630693 | 0.136559702771115 | -2.68393631459706 |
| lnc-METAP1-11 | lnc-METAP1-11 | -0.768448906467205 | 0.0888384330752369 | -2.74645249121045 | 0.00879746226859534 | 0.136559702771115 | -2.5611289935485 |
| ENSG00000092148.12 | HECTD1 | 0.187440905563449 | 7.84879100686362 | 2.74556803196607 | 0.00881756544443188 | 0.136768613069134 | -2.94826695771496 |
| ENSG00000072609.17 | CHFR | 0.360735808186464 | 5.35911911442068 | 2.74508057474106 | 0.00882866318012764 | 0.136810059496383 | -2.94930607079457 |
| SDCBP2-AS1 | SDCBP2-AS1 | -0.399388813285209 | 1.49620592379544 | -2.7443585423002 | 0.00884512513410191 | 0.136810059496383 | -2.67333960822225 |
| ENSG00000120675.5 | DNAJC15 | -0.340777534068139 | 5.51266493319811 | -2.74355588305036 | 0.00886345865530682 | 0.136810059496383 | -2.95868225403297 |
| lnc-PKN2-1 | lnc-PKN2-1 | -0.677957935567528 | 0.819824267868349 | -2.74333545864596 | 0.00886849951134568 | 0.136810059496383 | -2.61327666893467 |
| lnc-KMT5A-1 | lnc-KMT5A-1 | -0.455877851154689 | 1.10200690219424 | -2.74329758984704 | 0.00886936579464471 | 0.136810059496383 | -2.64991858321276 |
| TTC28-AS1 | TTC28-AS1 | -0.272909918805234 | 3.60461041049397 | -2.74309579314827 | 0.0088739833978954 | 0.136810059496383 | -2.85250004278973 |
| lnc-ERAL1-5 | lnc-ERAL1-5 | -0.615753640988231 | 1.14147155172366 | -2.74309209692052 | 0.00887406799737992 | 0.136810059496383 | -2.64461900068535 |
| ENSG00000172059.10 | KLF11 | 0.700311983193284 | 5.46108316829393 | 2.74416305678376 | 0.00887645558095275 | 0.136810059496383 | -2.95518059324756 |
| ENSG00000108312.14 | UBTF | -0.381620276281855 | 7.44902665323967 | -2.74256820823486 | 0.00888606634109935 | 0.136810059496383 | -2.95942552396461 |
| lnc-DSCC1-2 | lnc-DSCC1-2 | 0.39517575394109 | 1.34498366244974 | 2.74252921317437 | 0.00888696002385304 | 0.136810059496383 | -2.66120859206707 |
| ENSG00000234797.5 | RPS3AP6 | 1.09899837344511 | 3.7764868538686 | 2.74536836359161 | 0.00889335177603653 | 0.136810059496383 | -2.82351838804269 |
| lnc-A3GALT2-4 | lnc-A3GALT2-4 | 0.533189553280923 | 0.110424465066121 | 2.74092989495852 | 0.00892368464117425 | 0.137174160544652 | -2.57468810817413 |
| ENSG00000185158.12 | LRRC37B | -0.27010903794986 | 3.84298769191206 | -2.74061209184735 | 0.00893099894250466 | 0.137184142711622 | -2.87647028534985 |
| ENSG00000212907.2 | MT-ND4L | -0.74456984667211 | 6.22549477523529 | -2.74163372072737 | 0.00894639658697331 | 0.137318181170523 | -2.97099476473068 |
| ENSG00000160326.13 | SLC2A6 | 0.719944278643503 | 3.40811116624011 | 2.73949179139297 | 0.0089599307180856 | 0.137326449690636 | -2.83519497267131 |
| lnc-ZNF567-4 | lnc-ZNF567-4 | -0.459988408704679 | 1.11527126627683 | -2.73909654149948 | 0.00896595594716861 | 0.137326449690636 | -2.65491949601729 |
| lnc-RARRES2-6 | lnc-RARRES2-6 | -1.04152410659 | 0.349246311903573 | -2.73885586453802 | 0.00897151892037716 | 0.137326449690636 | -2.58584447341652 |
| ENSG00000279689.1 | | -0.953544662361007 | 0.112187714373768 | -2.73876488490922 | 0.00897362264169001 | 0.137326449690636 | -2.57602639836223 |
| ENSG00000104517.12 | UBR5 | 0.285830449021067 | 8.11576510476014 | 2.73776947379704 | 0.00899666933799646 | 0.137576853628515 | -2.96164604295717 |
| ENSG00000186409.15 | CCDC30 | -0.38421694610918 | 2.83755068259675 | -2.73638942537501 | 0.00902871205285752 | 0.13796435054489 | -2.79427686150885 |
| lnc-PHIP-3 | lnc-PHIP-3 | -0.592672233406059 | 2.37897716924279 | -2.73601555738039 | 0.00903741086751754 | 0.13799482780869 | -2.76269692443991 |
| lnc-NUDT21-1 | lnc-NUDT21-1 | 0.495647605329284 | 1.40929835170409 | 2.73503605286554 | 0.00906023782405451 | 0.138064757124905 | -2.68541938583928 |
| lnc-SLC35G3-1 | lnc-SLC35G3-1 | -0.789579933276458 | 1.66563180798595 | -2.73499378107181 | 0.0090612241497467 | 0.138064757124905 | -2.70264010705125 |
| lnc-SUMF1-5 | lnc-SUMF1-5 | -0.905451918915354 | 0.139689435956531 | -2.73495565928698 | 0.00906211372859865 | 0.138064757124905 | -2.58735778945906 |
| lnc-RAP1GDS1-3 | lnc-RAP1GDS1-3 | -0.868323899017878 | -0.373849966067842 | -2.73368766294181 | 0.00909174869485569 | 0.138413804279745 | -2.58445581243434 |
| ENSG00000100226.15 | GTPBP1 | 0.315235033876793 | 6.43707035901305 | 2.73102610821149 | 0.0091542447543528 | 0.139182585661212 | -2.99493131686557 |
| ENSG00000196843.15 | ARID5A | 0.684031605685364 | 5.18523934590409 | 2.73182027487934 | 0.00915577034374395 | 0.139182585661212 | -2.97273213159258 |
| ENSG00000206656.1 | SNORD116-17 | -0.872428349985522 | 0.233043938100943 | -2.73046484281138 | 0.00916747443226156 | 0.139257657741136 | -2.59525517238284 |

| | | | | | | | |
|---|---|---|---|---|---|---|---|
| ENSG00000185989.10 | RASA3 | 0.566985285671278 | 5.41477395293604 | 2.73003041057873 | 0.0091777266331026 | 0.139310580596719 | -2.98032738881907 |
| lnc-TMEM154-4 | lnc-TMEM154-4 | 0.586513436827595 | 2.51976099633009 | 2.72933261367278 | 0.00919421617608645 | 0.13945803356845 | -2.77640162963656 |
| ENSG00000130313.6 | PGLS | -0.298765134894938 | 4.56523797755034 | -2.72808103670038 | 0.0092238606236971 | 0.139804656272134 | -2.95770727030365 |
| ENSG00000106397.11 | PLOD3 | -0.360476502802838 | 3.40204230389804 | -2.72650467256667 | 0.00926132353713338 | 0.140104653685161 | -2.87037283706133 |
| ATP1A1-AS1 | ATP1A1-AS1 | -0.570820809645855 | 0.935359704161559 | -2.72640527438101 | 0.00926369047916483 | 0.140104653685161 | -2.65975448488342 |
| ENSG00000241014.2 | GPR199P | -0.402980865256119 | 1.48386404724896 | -2.72605366800196 | 0.00927206767222967 | 0.140104653685161 | -2.71900076642678 |
| lnc-PPM1M-2 | lnc-PPM1M-2 | -0.59800593551213 | 0.24584410680798 | -2.72593865063401 | 0.00927480953578457 | 0.140104653685161 | -2.6042612943845 |
| ENSG00000135951.14 | TSGA10 | -0.36957229488117 | 3.30071989810612 | -2.72581795545378 | 0.00927768755637539 | 0.140104653685161 | -2.86754515507898 |
| ENSG00000196954.13 | CASP4 | 0.608981450508735 | 5.61578923366624 | 2.72477846453318 | 0.0093085997816233 | 0.14046840858149 | -3.00133295003194 |
| ENSG00000137970.7 | RPL7P9 | 0.988454940048443 | 2.92877421670757 | 2.72542967960982 | 0.00932618800318937 | 0.14055416445872 | -2.80289816725021 |
| ENSG00000160218.12 | TRAPPC10 | 0.236765084865364 | 7.10598748433315 | 2.72333845647953 | 0.00933699498772027 | 0.14055416445872 | -3.00888231156074 |
| lnc-MSN-1 | lnc-MSN-1 | -1.28329480287034 | -0.312250920764464 | -2.72323493169048 | 0.0093394788089525 | 0.14055416445872 | -2.60715003537077 |
| ENSG00000138764.14 | CCNG2 | -0.508869280640978 | 5.58921703028138 | -2.72314665432648 | 0.00934159728803038 | 0.14055416445872 | -3.005810020999 |
| ENSG00000262074.6 | SNORD3B-2 | -1.64836703241923 | 3.27347531398703 | -2.72728195210432 | 0.00935606804928417 | 0.140634903570137 | -2.86449055211754 |
| ENSG00000140688.16 | RUSF1 | -0.296856290055839 | 5.96165968664388 | -2.72209893269851 | 0.00936677441668102 | 0.140634903570137 | -3.01373634838163 |
| ENSG00000166478.9 | ZNF143 | 0.173115152018939 | 4.61305533617877 | 2.72178878467903 | 0.0093742393991899 | 0.140634903570137 | -2.96803731741736 |
| ENSG00000162894.11 | FCMR | -0.554508161900554 | 9.0289162511921 | -2.72170250675807 | 0.00937631700745168 | 0.140634903570137 | -2.98003490028613 |
| ENSG00000115525.17 | ST3GAL5 | 0.3273063517384 | 6.74398097751149 | 2.72130125468843 | 0.00938598491363712 | 0.140634903570137 | -3.01607012175657 |
| lnc-TMEM160-2 | lnc-TMEM160-2 | -0.520423785237403 | 1.16557634890769 | -2.72121937532362 | 0.00938795887408869 | 0.140634903570137 | -2.69684145536674 |
| ENSG00000185361.8 | TNFAIP8L1 | -0.527351969468042 | 2.77402409465922 | -2.72050734345793 | 0.00940514082037627 | 0.140789828004222 | -2.83599812477754 |
| ENSG00000113387.11 | SUB1 | 0.287494332460916 | 6.62857005415881 | 2.72012536844003 | 0.00941437017316181 | 0.140825567786475 | -3.01941652249197 |
| lnc-STX16-1 | lnc-STX16-1 | -0.510763252008154 | 0.503662922986572 | -2.71791180344206 | 0.00946801963028117 | 0.141301611290883 | -2.64778985783631 |
| ENSG00000251143.1 | | -0.575017439034032 | 0.0156946766159553 | -2.71776126284487 | 0.00947167846704489 | 0.141301611290883 | -2.61968508258116 |
| ENSG00000133895.14 | MEN1 | -0.389593953882025 | 3.99823856177409 | -2.71749076797101 | 0.00947825603143799 | 0.141301611290883 | -2.93825844842235 |
| ENSG00000198945.7 | L3MBTL3 | 0.460079089271681 | 4.7222212201915 | 2.71745007467662 | 0.0094792459262383 | 0.141301611290883 | -2.97448712206459 |
| ENSG00000105771.13 | SMG9 | -0.274277324795187 | 4.60991987220188 | -2.717397740806 | 0.00948051912708103 | 0.141301611290883 | -2.98101851482658 |
| lnc-MTMR2-3 | lnc-MTMR2-3 | 0.525713214102657 | 3.98738670318918 | 2.71706888632374 | 0.0094885232541641 | 0.141318577525658 | -2.93129363235955 |
| ENSG00000143727.15 | ACP1 | 0.311699025848758 | 5.5905546554843 | 2.71675654582162 | 0.00949613121758388 | 0.141321541467271 | -3.0217021858434 |
| ENSG00000275740.1 | | 0.243809696382375 | 5.29999804249859 | 2.71649713802882 | 0.00950245413159905 | 0.141321541467271 | -3.01272181790332 |
| ENSG00000274512.5 | TBC1D3L | 1.24923989615934 | 0.438829098195991 | 2.71594607694394 | 0.00951691032736074 | 0.141434343153838 | -2.62805153088937 |
| LINC02099 | LINC02099 | -1.39040815693988 | -0.209455453457056 | -2.71554492430639 | 0.00952569712412892 | 0.141462787810927 | -2.62450888325191 |
| ENSG00000065268.10 | WDR18 | -0.594780135233537 | 2.2397901442533 | -2.71522616435655 | 0.00953348959133993 | 0.141476435658652 | -2.80203903689872 |
| ENSG00000200831.1 | SNORD36B | 0.458267851519805 | 0.436874729908831 | 2.71406824682466 | 0.00956184581148034 | 0.141795008888833 | -2.65736251777869 |
| ENSG00000283088.1 | NA | -0.439892927979823 | -0.119391316211933 | -2.71369985283535 | 0.00957088371854789 | 0.141826853548503 | -2.62656668048681 |
| ENSG00000284488.1 | MIR6890 | 0.549192812425496 | -0.299308858566156 | 2.7132381030227 | 0.0095822230856091 | 0.141888391719807 | -2.62520068918225 |
| ENSG00000164898.12 | FMC1 | -0.392178925945695 | 1.00180823346939 | -2.71295831111898 | 0.00958910007532131 | 0.141888391719807 | -2.70409538727287 |
| lnc-DONSON-3 | lnc-DONSON-3 | -0.335328647728571 | 1.43151459067363 | -2.71249859635322 | 0.00960040927238377 | 0.141888391719807 | -2.74082611549044 |
| lnc-HINT1-4 | lnc-HINT1-4 | -1.06961000615512 | -0.223181769516221 | -2.71198683635048 | 0.00961301328718108 | 0.141888391719807 | -2.6274973233291 |
| lnc-TOB2-1 | lnc-TOB2-1 | 0.330116052391046 | 2.57582959000419 | 2.71196221349271 | 0.00961362010261404 | 0.141888391719807 | -2.82953862563472 |
| lnc-ZNF701-1 | lnc-ZNF701-1 | -0.504381408781522 | -0.0379322497347879 | -2.71178489751059 | 0.00961799099191117 | 0.141888391719807 | -2.62934813282411 |
| ENSG00000099817.11 | POLR2E | -0.386218326169118 | 4.1047370817571 | -2.71156999677304 | 0.00962329081479134 | 0.141888391719807 | -2.96700193376739 |
| ENSG00000066379.14 | POLR1H | 0.314684830700155 | 3.25030965617139 | 2.71101520755704 | 0.00963698533827609 | 0.141988596433598 | -2.88637769754566 |
| ENSG00000163904.12 | SENP2 | 0.221005767363605 | 5.30495560405082 | 2.71067881753868 | 0.00964529761143012 | 0.142009413974296 | -3.02604716551112 |
| lnc-GHRH-3 | lnc-GHRH-3 | 0.418673194087991 | 0.46040422263493 | 2.71020941869222 | 0.00965690762241441 | 0.142023145550519 | -2.66205685278904 |
| ENSG00000143162.8 | CREG1 | 0.693490452866173 | 2.86907358874003 | 2.71008325587401 | 0.00966003030514146 | 0.142023145550519 | -2.85188927362691 |
| ENSG00000131876.16 | SNRPA1 | 0.363964428907925 | 5.72678379701084 | 2.70888861761183 | 0.00968964527643178 | 0.142192077838326 | -3.04130403779222 |
| ENSG00000185946.15 | RNPC3 | -0.24547909569096 | 6.68799407691848 | -2.70879867636396 | 0.00969187829972122 | 0.142192077838326 | -3.0444354355388 |
| ENSG00000146587.17 | RBAK | 0.265310394187266 | 5.21760377880536 | 2.7087838958469 | 0.00969224530958416 | 0.142192077838326 | -3.02894737476317 |
| ENSG00000106336.12 | FBXO24 | -0.488430272898746 | 0.0327011563931332 | -2.70772386512523 | 0.00971860003151264 | 0.142250399087243 | -2.63945350946145 |
| ENSG00000042286.14 | AIFM2 | -0.278130820637752 | 2.75856500015828 | -2.70731589131179 | 0.00972876077982853 | 0.142250399087243 | -2.85944008902289 |
| ENSG00000166548.15 | TK2 | -0.534363401443348 | 3.32447727575067 | -2.70724173147031 | 0.00973060881241621 | 0.142250399087243 | -2.91324234816198 |
| ENSG00000109381.19 | ELF2 | 0.272346625956488 | 6.16797760745366 | 2.70699456534004 | 0.00973677042939481 | 0.142250399087243 | -3.0498575977226 |
| lnc-HPS1-1 | lnc-HPS1-1 | -0.377455730768562 | 1.06936621985161 | -2.70643052345128 | 0.00975084494306787 | 0.142250399087243 | -2.72689370448041 |
| ENSG00000028116.17 | VRK2 | 0.28937696769435 | 5.48309020203432 | 2.70640714448151 | 0.00975142872250132 | 0.142250399087243 | -3.04217046480888 |
| ENSG00000285130.1 | | 1.02271050639246 | 1.99136275869914 | 2.70739354523621 | 0.00975612520307978 | 0.142250399087243 | -2.78303639104567 |
| ENSG00000146574.15 | CCZ1B | 0.431989631847402 | 3.30851500383688 | 2.70620300104163 | 0.00975652761189536 | 0.142250399087243 | -2.90265240744765 |
| lnc-VEPH1-2 | lnc-VEPH1-2 | 0.459741646440754 | 5.21185499561805 | 2.70612725179754 | 0.00975842022597224 | 0.142250399087243 | -3.03417956291308 |
| ENSG00000168386.18 | FILIP1L | -0.833442463321891 | 4.16635838228872 | -2.70722866803287 | 0.00977631632421989 | 0.142410416773827 | -2.97067704895368 |
| ENSG00000108515.17 | ENO3 | -0.413799930827115 | 2.70939408130661 | -2.7034721059321 | 0.00982497414452229 | 0.142970857576458 | -2.86201246217526 |
| ENSG00000234851.4 | RPL23AP42 | 0.889767401482783 | 4.18795840830083 | 2.70541287704728 | 0.00982868209059363 | 0.142970857576458 | -2.95254680381522 |
| ENSG00000183283.15 | DAZAP2 | 0.309226157572646 | 8.00169446368163 | 2.70219750386606 | 0.00985707187453933 | 0.143282563837319 | -3.04529240079077 |
| lnc-FIG4-1 | lnc-FIG4-1 | -0.841517754728647 | -0.131003902500314 | -2.70190739152133 | 0.00986439114470349 | 0.143287764948082 | -2.64885756493216 |
| ENSG00000204859.12 | ZBTB48 | -0.535767208357645 | 4.70255989513659 | -2.70153679241405 | 0.00987374831525404 | 0.143322539896244 | -3.02935026385398 |
| ENSG00000113558.18 | SKP1 | 0.291663140690106 | 6.99116766005644 | 2.70054645216035 | 0.0098987933736767 | 0.14358482312219 | -3.06204824026954 |
| ENSG00000102218.5 | RP2 | 0.417875141526604 | 4.07854050973293 | 2.69870025267256 | 0.00994563905348722 | 0.143958470104217 | -2.98078050085808 |
| lnc-MYH11-2 | lnc-MYH11-2 | -0.350231936571576 | 2.63933254534948 | -2.69849730540413 | 0.00995080111549413 | 0.143958470104217 | -2.86676648454858 |
| lnc-SLC35F5-3 | lnc-SLC35F5-3 | -0.473289148838568 | 2.96669032876839 | -2.69812030566428 | 0.00996039684532274 | 0.143958470104217 | -2.89502757480369 |
| lnc-SCARB2-4 | lnc-SCARB2-4 | 0.473950142675809 | 1.80600764151948 | 2.698106559996 | 0.00996074687338588 | 0.143958470104217 | -2.79652674044414 |

| | | | | | | | |
|---|---|---|---|---|---|---|---|
| lnc-ZBTB7C-5 | lnc-ZBTB7C-5 | 0.253581343612721 | 2.58672555385428 | 2.69803850468101 | 0.00996248004247443 | 0.143958470104217 | -2.86004124619288 |
| STK24-AS1 | STK24-AS1 | -0.693998048692198 | 1.87565508613095 | -2.69758071811986 | 0.00997414575178025 | 0.143958470104217 | -2.81686935670977 |
| ENSG00000049769.12 | PPP1R3F | -0.475875947461236 | 1.93335829259021 | -2.69744150163582 | 0.0099776958827091 | 0.143958470104217 | -2.822818531444 |
| ENSG00000162337.11 | LRP5 | 2.00665683843992 | 1.28986678858421 | 2.70195328722538 | 0.00998050511775336 | 0.143958470104217 | -2.77827508696051 |
| lnc-CPSF6-2 | lnc-CPSF6-2 | -0.880099211538717 | 0.109710370697053 | -2.69650806557394 | 0.0100015293598286 | 0.144103167149177 | -2.67555773210913 |
| ENSG00000145016.15 | RUBCN | 0.433344167921542 | 6.80799242712328 | 2.69603098064859 | 0.0100137310597237 | 0.144103167149177 | -3.07413744515873 |
| ENSG00000037757.13 | MRI1 | -0.423907297809864 | 4.41355764181815 | -2.69585016277511 | 0.0100183591586979 | 0.144103167149177 | -3.023544586016 |
| ENSG00000146425.10 | DYNLT1 | 0.395046573444229 | 4.29420087979503 | 2.69565324781635 | 0.0100234015119192 | 0.144103167149177 | -3.01173826468182 |
| ENSG00000155970.11 | MICU3 | -2.48163635163985 | 1.57387306062812 | -2.70039796955009 | 0.0100255422123899 | 0.144103167149177 | -2.75726835995021 |
| ENSG00000172687.13 | ZNF738 | -0.328460603345426 | 3.83116765606934 | -2.69503174829543 | 0.0100393314508915 | 0.144200669402443 | -2.9856181227922 |
| lnc-MYOC-1 | lnc-MYOC-1 | -1.48981572401833 | 0.918889957643369 | -2.69671407118707 | 0.0100577464250986 | 0.144319526567427 | -2.70715244803404 |
| ENSG00000112406.4 | HECA | 0.365352599718332 | 6.63746162203667 | 2.69405945124735 | 0.0100642996740938 | 0.144319526567427 | -3.07907710829134 |
| lnc-ABAT-2 | lnc-ABAT-2 | -0.580463489564395 | 1.44401864525602 | -2.69389061641301 | 0.0100686411189246 | 0.144319526567427 | -2.78192242487595 |
| ENSG00000206652.1 | RNU1-1 | -1.12430753581386 | 0.469399122870968 | -2.69332857736674 | 0.0100871885492913 | 0.14440437358479 | -2.70206845788381 |
| lnc-RELL1-6 | lnc-RELL1-6 | 0.509979959179244 | 0.62957840533294 | 2.69303830081875 | 0.0100905840646541 | 0.14440437358479 | -2.72542520672375 |
| lnc-ZNF468-2 | lnc-ZNF468-2 | -0.445204427911842 | 0.671894660965826 | -2.69284341085324 | 0.0100956077145466 | 0.14440437358479 | -2.7215708934621 |
| ENSG00000133731.9 | IMPA1 | 0.339185149937629 | 4.18683378679466 | 2.69191143964386 | 0.0101196628915264 | 0.144548052515068 | -3.01169250035919 |
| ENSG00000101290.13 | CDS2 | 0.252559345621373 | 6.22835667590639 | 2.69191008141575 | 0.0101196979873786 | 0.144548052515068 | -3.08454588372396 |
| ENSG00000148700.14 | ADD3 | 1.07371653498982 | 7.48088955663447 | 2.69470500740586 | 0.0101432385289065 | 0.14473149218363 | -3.07900039047911 |
| lnc-DHCR24-2 | lnc-DHCR24-2 | 0.292053226830022 | 1.16328866712303 | 2.69060913650608 | 0.0101533652447064 | 0.14473149218363 | -2.77113487131277 |
| lnc-ROGDI-1 | lnc-ROGDI-1 | -0.28106468510371 | 2.93510336103996 | -2.69059871856741 | 0.0101536352676074 | 0.14473149218363 | -2.91621503808635 |
| ENSG00000123473.15 | STIL | 0.519177318417312 | 1.89468607221214 | 2.69028877728148 | 0.0101616716747405 | 0.144745804900474 | -2.81935149473779 |
| ENSG00000262180.1 | NA | 0.619220302446101 | 0.292103286683018 | 2.69001401748648 | 0.0101688007700405 | 0.144747182745328 | -2.690328396331 |
| ENSG00000176542.9 | USF3 | 0.332815047035415 | 6.96581181108282 | 2.68892094360628 | 0.0101972080481857 | 0.145051232381346 | -3.08913456595863 |
| ENSG00000119866.21 | BCL11A | 0.474811983685441 | 7.40668523448441 | 2.68823679132688 | 0.0102150252929134 | 0.14520432707461 | -3.08712939515887 |
| ENSG00000170871.11 | KIAA0232 | 0.209672844204318 | 5.14579631178054 | 2.68686346670716 | 0.0102508771533131 | 0.14561339161259 | -3.07812704207577 |
| RPL34-AS1 | RPL34-AS1 | -0.532312712475574 | 1.52959667332458 | -2.68585604836741 | 0.0102772503987118 | 0.145879733121816 | -2.80461832911816 |
| lnc-RBM12B-4 | lnc-RBM12B-4 | -0.444360472382243 | 2.93647438602542 | -2.68560616353542 | 0.0102838018150782 | 0.145879733121816 | -2.91468843522944 |
| ENSG00000106799.12 | TGFBR1 | 0.499755561478348 | 5.08883726224197 | 2.6851827490168 | 0.0102949115740791 | 0.145936752706108 | -3.08189882099626 |
| ENSG00000239650.4 | GUSBP4 | -0.306868165220547 | 1.35624015449356 | -2.68487013792748 | 0.0103031211139125 | 0.145952609695568 | -2.79333880159046 |
| ENSG00000073614.12 | KDM5A | 0.263796058446663 | 7.98069859149715 | 2.68434989805248 | 0.0103167966070581 | 0.145963230687819 | -3.08652197742399 |
| lnc-TSSK3-2 | lnc-TSSK3-2 | -0.513512914412952 | 1.81090003538065 | -2.68408959860719 | 0.0103236453483696 | 0.145963230687819 | -2.8270010848823 |
| ENSG00000174405.13 | LIG4 | 0.463364273499491 | 4.24802613707041 | 2.68403261470652 | 0.0103251452111677 | 0.145963230687819 | -3.0200482476593 |
| ENSG00000164758.7 | MED30 | 0.475225748225341 | 3.68752895951287 | 2.68344748436974 | 0.0103405579394752 | 0.146080785221838 | -2.98100813989901 |
| lnc-NXNL1-7 | lnc-NXNL1-7 | -0.396927339876069 | 1.8164668277131 | -2.68296436451131 | 0.0103532996167361 | 0.14616047051528 | -2.83953889810952 |
| ENSG00000100246.12 | DNAL4 | -0.453814609812397 | 1.27207242262944 | -2.68154962142654 | 0.0103906949301449 | 0.146425584292174 | -2.80052248964371 |
| ENSG00000166595.11 | CIAO2B | 0.261962695481159 | 3.3753954001468 | 2.681354764022 | 0.0103958552531927 | 0.146425584292174 | -2.96501513627843 |
| ENSG00000164751.14 | PEX2 | 0.228768054797604 | 5.46039435938486 | 2.68118559949446 | 0.0104003370774755 | 0.146425584292174 | -3.09889813848283 |
| ENSG00000184983.10 | NDUFA6 | 0.297572913074745 | 4.30358472058925 | 2.68106214771691 | 0.0104036089171736 | 0.146425584292174 | -3.04217941182901 |
| ENSG00000163444.11 | TMEM183A | 0.207543037431833 | 5.46377452659093 | 2.68083435702045 | 0.0104096485380204 | 0.146425584292174 | -3.10068735840494 |
| ENSG00000164754.14 | RAD21 | 0.248730358883726 | 7.7164313863892 | 2.68064157577537 | 0.0104147624449178 | 0.146425584292174 | -3.09893406798722 |
| ENSG00000130479.10 | MAP1S | -0.450467672307261 | 2.58723574554699 | -2.67912226820286 | 0.0104551461705113 | 0.146893019541047 | -2.91814568092312 |
| lnc-PRIMPOL-7 | lnc-PRIMPOL-7 | -0.48982947711248 | 3.30514624730998 | -2.67841010376261 | 0.0104741253580415 | 0.146900487558996 | -2.96417551129986 |
| lnc-RNASE1-2 | lnc-RNASE1-2 | 1.80425622658573 | -1.45575410634763 | 2.67820458719848 | 0.010479608273279 | 0.146900487558996 | -2.86368878989449 |
| lnc-EBLN1-1 | lnc-EBLN1-1 | -1.91992223926402 | 2.89855543431804 | -2.68288617516293 | 0.0104813192975356 | 0.146900487558996 | -2.9496860037164 |
| lnc-POLG2-1 | lnc-POLG2-1 | 0.392844244148443 | 7.58655781390373 | 2.67803159109257 | 0.0104842256339778 | 0.146900487558996 | -3.10723304389232 |
| ENSG00000218510.8 | LINC00339 | -0.398820679291356 | 2.49277495681762 | -2.67616788957578 | 0.0105340878125368 | 0.14749285959155 | -2.90632803664624 |
| ENSG00000134590.13 | RTL8C | -0.399106526546129 | 2.53420234462862 | -2.67591634264799 | 0.0105408344973605 | 0.14749285959155 | -2.91615650839708 |
| ENSG00000119139.19 | TJP2 | 1.23358108549071 | 1.97620704685155 | 2.67809053156755 | 0.0105504768246536 | 0.147527489457776 | -2.84184275102411 |
| ENSG00000123836.14 | PFKFB2 | 0.620886089739663 | 5.65666228639041 | 2.6756125129822 | 0.0105608724044733 | 0.147572597896316 | -3.11599170030545 |
| lnc-PITPNB-6 | lnc-PITPNB-6 | -0.985717113688669 | -0.328907780651621 | -2.67472669787726 | 0.0105727956417572 | 0.147638977404538 | -2.70246428798913 |
| ENSG00000212452.1 | SNORD69 | 0.615507184871618 | -0.234475864958991 | 2.67389710043896 | 0.0105951364983327 | 0.147850640369615 | -2.70329700523405 |
| ENSG00000243989.8 | ACY1 | -0.540925308807573 | 1.50858609354123 | -2.67248645373017 | 0.01063322469263 | 0.148185010408037 | -2.83762795520017 |
| lnc-FMOD-1 | lnc-FMOD-1 | -1.39335350481123 | 1.33699679615141 | -2.67511401405146 | 0.0106368998065723 | 0.148185010408037 | -2.81121991904413 |
| ENSG00000239763.2 | RPL21P118 | -0.431955642783721 | 0.271495740052207 | -2.67181657920459 | 0.0106513557648637 | 0.148185010408037 | -2.73096510654765 |
| ENSG00000067334.13 | DNTTIP2 | 0.246492501928285 | 5.95682831252248 | 2.67177881332099 | 0.0106523787973767 | 0.148185010408037 | -3.12867508238489 |
| lnc-PRC1-4 | lnc-PRC1-4 | -0.391450562681245 | 1.64730366757534 | -2.67167856813764 | 0.0106550947579991 | 0.148185010408037 | -2.84749131589356 |
| ENSG00000188559.14 | RALGAPA2 | -0.245565141629841 | 5.70562144708142 | -2.67091904334035 | 0.0106756934222096 | 0.148288061195426 | -3.12725686694491 |
| ENSG00000155465.18 | SLC7A7 | 0.678433156265027 | 5.53285194510566 | 2.67169849708314 | 0.0106823514124996 | 0.148288061195426 | -3.11789800174116 |
| ENSG00000127080.9 | IPPK | 0.308790380507061 | 3.08828817270445 | 2.67060880735412 | 0.0106841177064965 | 0.148288061195426 | -2.9676662163008 |
| ENSG00000133639.4 | BTG1 | 0.405641662476734 | 9.16745438550204 | 2.669171869988 | 0.0107232168111246 | 0.148730439099311 | -3.09778624928202 |
| ENSG00000214575.9 | CPEB1 | -1.32095188973472 | 1.44994595296597 | -2.67135363385364 | 0.010738311134613 | 0.148821476603121 | -2.8377813724279 |
| ENSG00000285441.1 | NA | 0.314425160641079 | 3.02459697276859 | 2.66840119640954 | 0.0107442410840129 | 0.148821476603121 | -2.96677713052615 |
| ENSG00000179029.14 | TMEM107 | -0.733375885030284 | 3.9917080262924 | -2.66856693264229 | 0.0107631671780048 | 0.1489373263496 | -3.05204497802568 |
| ENSG00000153207.15 | AHCTF1 | 0.303797025173417 | 6.49745523946902 | 2.66756569328742 | 0.0107670767919256 | 0.1489373263496 | -3.13990608456666 |
| lnc-SLC6A9-2 | lnc-SLC6A9-2 | -0.491831482669111 | 2.04031043780909 | -2.66719515043616 | 0.0107772186447453 | 0.148977495879646 | -2.88482846084249 |
| ENSG00000104880.17 | ARHGEF18 | 0.411907350451552 | 6.29579855500957 | 2.66619702930006 | 0.0108045812328263 | 0.149147675736492 | -3.14258497528907 |
| ENSG00000182481.8 | KPNA2 | 0.406034366635101 | 3.70884082888032 | 2.66581935527215 | 0.0108149514879029 | 0.149147675736492 | -3.03223889165464 |

| | | | | | | | |
|---|---|---|---|---|---|---|---|
| ENSG00000184517.11 | ZFP1 | 0.267294779307155 | 3.04742565375106 | 2.66553776167922 | 0.0108226895059514 | 0.149147675736492 | -2.97314396214652 |
| ENSG00000214534.5 | ZNF705EP | -0.545813538492323 | 0.735217356719412 | -2.66525467292505 | 0.0108304737457435 | 0.149147675736492 | -2.77717047338504 |
| ENSG00000110921.13 | MVK | -0.419115045279415 | 1.97683203851455 | -2.66521012287308 | 0.010831699230994 | 0.149147675736492 | -2.89976094409591 |
| ENSG00000197343.10 | ZNF655 | 0.220400633161643 | 7.36333403375859 | 2.66511855365267 | 0.0108342185228082 | 0.149147675736492 | -3.1388986528059 |
| ENSG00000136869.14 | TLR4 | 3.31362290929031 | 0.501219190736019 | 2.66817985691916 | 0.0108486924633864 | 0.149147675736492 | -2.76712054444547 |
| ENSG00000024862.17 | CCDC28A | 0.340524677320996 | 3.82335429293844 | 2.6643073734371 | 0.0108565596204248 | 0.149147675736492 | -3.03818753251423 |
| ENSG00000170946.14 | DNAJC24 | -0.205992053054608 | 4.25934395668593 | -2.66425044911851 | 0.010858128990785 | 0.149147675736492 | -3.08422201331873 |
| lnc-PSKH1-1 | lnc-PSKH1-1 | -0.706516959149776 | -0.37420791842885 | -2.663707069469 | 0.0108731201627301 | 0.149147675736492 | -2.72264226783437 |
| ENSG00000135643.4 | KCNMB4 | -0.966142720627031 | -0.708815725863376 | -2.66356603354696 | 0.0108770142812586 | 0.149147675736492 | -2.73993884146936 |
| ENSG00000111530.12 | CAND1 | 0.258781122030807 | 6.78104591412925 | 2.66335864887875 | 0.0108827426745611 | 0.149147675736492 | -3.14787676429306 |
| ENSG00000233476.3 | EEF1A1P6 | 1.27201135486131 | 3.33176412932304 | 2.6665709969948 | 0.0108907452957735 | 0.149147675736492 | -2.98710824601298 |
| lnc-STX4-1 | lnc-STX4-1 | -0.448977073550675 | 1.13923223559057 | -2.66295127217958 | 0.01089400334452 | 0.149147675736492 | -2.83337312974418 |
| ENSG00000118690.12 | ARMC2 | -0.577123180817481 | 2.25015373670351 | -2.66279875230521 | 0.0108982220428355 | 0.149147675736492 | -2.91196587999707 |
| ENSG00000139154.14 | AEBP2 | 0.209365397791644 | 4.78951417471959 | 2.66212198628833 | 0.0109169595241987 | 0.149304835805038 | -3.11728677812758 |
| ENSG00000255624.3 | C10orf88B | 1.25325631003296 | 0.77403458229795 | 2.66223654739739 | 0.0109270717279912 | 0.149330529468793 | -2.75280369157997 |
| ENSG00000225190.10 | PLEKHM1 | 0.256852385396778 | 4.71218448158097 | 2.66114649406424 | 0.0109440199352295 | 0.149330529468793 | -3.11471028790752 |
| ENSG00000141030.12 | COPS3 | 0.314589851239705 | 4.42453908003491 | 2.66111484423129 | 0.0109448989426648 | 0.149330529468793 | -3.09906061557351 |
| ENSG00000134897.13 | BIVM | -1.37402628825639 | -0.179321297027738 | -2.66100830305599 | 0.0109478583767385 | 0.149330529468793 | -2.73030808746811 |
| ENSG00000067900.7 | ROCK1 | 0.184247388658679 | 7.4964530491135 | 2.66053565040713 | 0.0109609963060905 | 0.14941071984653 | -3.14750087054857 |
| ENSG00000249884.8 | RNF103-CHMP3 | 0.261925218356352 | 4.13424990110617 | 2.65986356670555 | 0.0109797026306046 | 0.149566657343305 | -3.07602999908663 |
| ENSG00000183207.13 | RUVBL2 | -0.466401313720106 | 3.63273792472073 | -2.65948994232694 | 0.0109901145371003 | 0.149574489336568 | -3.04534257453978 |
| ENSG00000165644.10 | COMTD1 | -0.895307289061945 | -0.408157772469777 | -2.65932150626071 | 0.0109948113669643 | 0.149574489336568 | -2.73241429756628 |
| lnc-RBM23-4 | lnc-RBM23-4 | 0.672586813238535 | 2.49416539364528 | 2.65899213338992 | 0.0110040012465983 | 0.14960063253549 | -2.92463939206908 |
| lnc-ZNF780B-1 | lnc-ZNF780B-1 | -0.706875708948462 | 0.678791658392812 | -2.65684062246669 | 0.011064204896569 | 0.150248477482199 | -2.78851816360753 |
| ENSG00000007129.17 | CEACAM21 | -0.597397034973091 | 3.20704812922876 | -2.65676760749228 | 0.0110662533091878 | 0.150248477482199 | -3.00728153604794 |
| ENSG00000093072.16 | ADA2 | 0.534279912759759 | 5.92358132282362 | 2.65617170357026 | 0.0110829842551521 | 0.150376443588527 | -3.16121085734715 |
| ENSG00000173599.13 | PC | -1.18308209576934 | -0.412134066653952 | -2.65549248948408 | 0.0111020826354386 | 0.150526403236731 | -2.74260325441418 |
| ENSG00000163877.10 | SNIP1 | 0.345720837111999 | 4.7627915167192 | 2.65525872146419 | 0.0111086628050622 | 0.150526403236731 | -3.13044851558694 |
| ENSG00000083896.12 | YTHDC1 | 0.317460496901981 | 7.27689802758811 | 2.65386460315836 | 0.0111479793837918 | 0.15095977608986 | -3.16506974551068 |
| ENSG00000118939.17 | UCHL3 | 0.329201749320353 | 2.12288301062517 | 2.65328778486218 | 0.0111642840645425 | 0.151067910116877 | -2.92831200318025 |
| ENSG00000167543.15 | TP53I13 | -0.532696948594162 | 5.61559984876189 | -2.65306300719072 | 0.0111706436961516 | 0.151067910116877 | -3.16675936353216 |
| ENSG00000084676.15 | NCOA1 | 0.300526243671403 | 6.75143089989188 | 2.65219877446152 | 0.0111951264599814 | 0.151299598112801 | -3.17373296051379 |
| ENSG00000138757.14 | G3BP2 | 0.29796749247793 | 7.14146201169425 | 2.65159671880664 | 0.0112122111830186 | 0.15143106481632 | -3.17160878280206 |
| ENSG00000276550.4 | HERC2P2 | 0.720605025275524 | 3.92956830260475 | 2.65111201566615 | 0.0112392893078003 | 0.151616572432137 | -3.05831486533248 |
| ENSG00000107104.18 | KANK1 | -0.970675415870325 | 3.75310421083998 | -2.65328268736252 | 0.0112452529120145 | 0.151616572432137 | -3.09484296193155 |
| ENSG00000267426.5 | | -0.294910813958183 | 3.64989586892486 | -2.65033669815924 | 0.0112480447992942 | 0.151616572432137 | -3.06410341800561 |
| ENSG00000227855.3 | DPY19L2P3 | -1.14898448987186 | 0.312959438524097 | -2.65060363846733 | 0.0112564423307339 | 0.151630466291555 | -2.8063012594162 |
| ENSG00000105993.14 | DNAJB6 | 0.359314297545387 | 5.77121502469423 | 2.6496671848216 | 0.0112671278383736 | 0.15167514211723 | -3.17603457955413 |
| lnc-CD200R1-2 | lnc-CD200R1-2 | 0.347470988048465 | 1.48238541543341 | 2.64903153647387 | 0.0112852731684692 | 0.151774529409589 | -2.88459811905526 |
| lnc-PIGP-4 | lnc-PIGP-4 | -0.764015228955179 | 0.0858285804500451 | -2.64886072322393 | 0.0112901538111504 | 0.151774529409589 | -2.76099596798706 |
| ENSG00000172922.9 | RNASEH2C | -0.426689461340064 | 4.34900998593614 | -2.64843877470079 | 0.0113022184537583 | 0.151774529409589 | -3.12084705642967 |
| ENSG00000285791.1 | | -0.938650837841127 | 1.87702685304799 | -2.64898061755449 | 0.0113040059070544 | 0.151774529409589 | -2.90746847360433 |
| lnc-ESR2-3 | lnc-ESR2-3 | -0.377856054453806 | 3.15140181307804 | -2.64764288851489 | 0.01132500724482 | 0.151957382086134 | -3.02417432952697 |
| lnc-LRIG2-1 | lnc-LRIG2-1 | -0.539441648024021 | 0.542854468327842 | -2.64704532309395 | 0.0113421452254508 | 0.152088192296713 | -2.81130685253115 |
| lnc-APOL1-1 | lnc-APOL1-1 | -0.582479240788745 | 0.335498468338924 | -2.64583475276838 | 0.0113769370720254 | 0.1524166179325 | -2.77638773638038 |
| ENSG00000104375.16 | STK3 | 1.19515767657338 | 1.36570985643771 | 2.64721667760746 | 0.0113887037796369 | 0.1524166179325 | -2.87064703486593 |
| ENSG00000198841.3 | KTI12 | 0.313395386977571 | 2.65475619261896 | 2.64542092924545 | 0.0113888528581929 | 0.1524166179325 | -2.99010784330126 |
| lnc-CMTM6-3 | lnc-CMTM6-3 | 0.576935086687358 | 2.92018681701834 | 2.64460603807882 | 0.0114123506720409 | 0.152533951790289 | -3.00233191204629 |
| ENSG00000267858.5 | MZF1-AS1 | -0.674607790253238 | 0.128525925179126 | -2.64460288723164 | 0.0114124416147813 | 0.152533951790289 | -2.77907388177039 |
| lnc-ZNF705E-1 | lnc-ZNF705E-1 | -0.453121837759511 | 0.642910882903696 | -2.64338527469856 | 0.0114476354040572 | 0.152824021695418 | -2.8145431584271 |
| ENSG00000103876.12 | FAH | 0.752793237820786 | -0.283273623847068 | 2.64333834328477 | 0.0114489939005167 | 0.152824021695418 | -2.76252020539397 |
| ENSG00000258864.1 | | 0.289201215262802 | 3.44428719267556 | 2.64246133998776 | 0.0114744072733965 | 0.153063982442203 | -3.05326021093629 |
| ENSG00000154743.17 | TSEN2 | -0.305478453899136 | 3.47468563949683 | -2.64168965885973 | 0.0114968115329359 | 0.153263517993795 | -3.06659532244047 |
| lnc-KIAA0319L-1 | lnc-KIAA0319L-1 | -0.514907251253236 | 1.15395579591703 | -2.6402956275248 | 0.0115373864403659 | 0.153621369379741 | -2.87746469627066 |
| ENSG00000166908.17 | PIP4K2C | 0.240964368398685 | 4.33704162677461 | 2.64015029672757 | 0.0115416240270758 | 0.153621369379741 | -3.13877723451147 |
| ENSG00000129084.17 | PSMA1 | 0.285576623618641 | 4.49245579403247 | 2.63999870518381 | 0.0115460456896691 | 0.153621369379741 | -3.15146698634413 |
| ENSG00000211771.1 | TRBJ2-7 | -0.663371491139698 | -0.289876692731124 | -2.63949212663948 | 0.011560833000593 | 0.153705711788614 | -2.77003171895982 |
| ENSG00000160284.14 | SPATC1L | -0.6918531463456 | 0.142182440575473 | -2.63927008510584 | 0.0115673200000274 | 0.153705711788614 | -2.79015013058915 |
| lnc-SRD5A1-2 | lnc-SRD5A1-2 | -0.69605930224359 | -0.471785535351332 | -2.63881361039765 | 0.0115806665290476 | 0.153755944318262 | -2.77138536185013 |
| ENSG00000166780.10 | BMERB1 | -1.75355561404068 | 0.777385331348487 | -2.64198410041428 | 0.0115860404041017 | 0.153755944318262 | -2.84183639903634 |
| ENSG00000178201.4 | VN1R1 | -0.592300541699611 | 1.3825134474734 | -2.63759944537172 | 0.0116162354983855 | 0.154057329422209 | -2.89657677528091 |
| ENSG00000141905.18 | NFIC | -0.62919479037557 | 2.86151617108129 | -2.63713864472215 | 0.0116297609059861 | 0.154095853997314 | -3.02484959696177 |
| lnc-DNER-3 | lnc-DNER-3 | 0.321292767524621 | 2.99770539678726 | 2.63699046150641 | 0.0116341134485656 | 0.154095853997314 | -3.03470859764237 |
| lnc-UBE3C-8 | lnc-UBE3C-8 | 0.588968215612021 | 1.8710761695466 | 2.63651807754792 | 0.0116479986354775 | 0.154180550427031 | -2.93588461802379 |
| ENSG00000220785.7 | MTMR9LP | -0.936833028437112 | 2.00149562198522 | -2.636348006952 | 0.0116844653639016 | 0.154479228279298 | -2.96949903053655 |
| lnc-PLA2G4F-3 | lnc-PLA2G4F-3 | 0.321218116754842 | 1.93252313655431 | 2.63498772568434 | 0.0116930861814016 | 0.154479228279298 | -2.94473786080113 |
| ENSG00000198938.2 | MT-CO3 | -0.533433553121295 | 11.4501035789989 | -2.63488935576934 | 0.0116959898522145 | 0.154479228279298 | -3.10645029185855 |
| ENSG00000196205.8 | EEF1A1P5 | 1.00208344627622 | 6.92549978979001 | 2.63807478679539 | 0.0117005838258478 | 0.154479228279298 | -3.20970311163526 |

| | | | | | | | |
|---|---|---|---|---|---|---|---|
| ENSG00000117118.9 | SDHB | 0.242130998505417 | 4.70692449266767 | 2.63342962320815 | 0.0117391559528557 | 0.154889132677967 | -3.17628139981212 |
| ENSG00000159461.14 | AMFR | 0.296761581810924 | 7.61164991478938 | 2.63150885231506 | 0.0117961784455387 | 0.155479803303116 | -3.21181627080388 |
| lnc-PLCD3-4 | lnc-PLCD3-4 | -0.342630475602329 | 5.51705024636534 | -2.63141299347341 | 0.0117990308875998 | 0.155479803303116 | -3.21214995670442 |
| lnc-PAPPA2-1 | lnc-PAPPA2-1 | -0.572562386067388 | 0.799688131110062 | -2.63051468296774 | 0.0118257924307874 | 0.155732748306396 | -2.85353400608745 |
| ENSG00000154832.14 | CXXC1 | -0.363199061060572 | 4.78661565499805 | -2.62960717366639 | 0.0118528845952458 | 0.155937480204166 | -3.19210508374093 |
| ENSG00000150787.7 | PTS | 0.466466920442927 | 3.6090553899151 | 2.62948650949741 | 0.0118564911101161 | 0.155937480204166 | -3.09304421769442 |
| ENSG00000172113.9 | NME6 | -0.307196885363783 | 4.37166720980114 | -2.62915669935689 | 0.0118663539013244 | 0.155942008122369 | -3.16630073917912 |
| ENSG00000162775.14 | RBM15 | 0.268743263567721 | 6.75696428653317 | 2.62896841263356 | 0.0118719878894113 | 0.155942008122369 | -3.22579506513501 |
| ENSG00000275377.1 | MIR1299 | -0.862456685567819 | -0.538435842432778 | -2.62770428218642 | 0.0119098773258448 | 0.156339926656801 | -2.79458440131406 |
| lnc-GLCCI1-6 | lnc-GLCCI1-6 | -0.754508213143574 | 1.16154944609859 | -2.62703612439507 | 0.0119299486495169 | 0.156503590218615 | -2.90995619935511 |
| ENSG00000158480.10 | SPATA2 | 0.396934264106027 | 2.75844355745516 | 2.6262487081283 | 0.0119536423293379 | 0.156714535073097 | -3.03676209164552 |
| ENSG00000100722.19 | ZC3H14 | -0.279512099621999 | 5.46058771555763 | -2.62550511840885 | 0.0119760568975498 | 0.156863957399739 | -3.22483309486642 |
| ENSG00000144028.14 | SNRNP200 | 0.261554169803751 | 8.00703654226261 | 2.62509743300325 | 0.0119883624239796 | 0.156863957399739 | -3.21996932922254 |
| ENSG00000116752.5 | BCAS2 | 0.349623615064163 | 4.70199781393064 | 2.62481564963487 | 0.0119968745202392 | 0.156863957399739 | -3.19387373275729 |
| ENSG00000259316.11 | | 0.278111543924664 | 2.88480522860572 | 2.62467449587657 | 0.0120011405701116 | 0.156863957399739 | -3.04500791221785 |
| ENSG00000161010.14 | MRNIP | -0.547225336451246 | 6.36729980233309 | -2.62460819082584 | 0.0120031449693723 | 0.156863957399739 | -3.23570849934845 |
| ENSG00000196345.12 | ZKSCAN7 | -0.503344898016601 | 2.51460012901884 | -2.62338015879349 | 0.012040323919055 | 0.157113492465226 | -3.02287934881665 |
| MIR600HG | MIR600HG | -0.396127350645209 | 6.44287200627801 | -2.62331032673553 | 0.0120424412729872 | 0.157113492465226 | -3.23901161619773 |
| ENSG00000068354.15 | TBC1D25 | 0.246994525282916 | 4.2559131504325 | 2.6231415849906 | 0.012047559044244 | 0.157113492465226 | -3.16779830102527 |
| lnc-PSMD5-2 | lnc-PSMD5-2 | -0.555150020270681 | 0.0179210118237656 | -2.62296977482322 | 0.0120527719284163 | 0.157113492465226 | -2.82464399031601 |
| lnc-NFATC1-2 | lnc-NFATC1-2 | 0.708049597556919 | -0.355674850662819 | 2.62229246120954 | 0.0120733424397129 | 0.157280533539548 | -2.80326521141119 |
| ENSG00000205085.11 | GARIN1A | -0.568050151906218 | 1.82466829023514 | -2.62204490796285 | 0.0120808688493429 | 0.157280533539548 | -2.970809050473 |
| ENSG00000165996.13 | HACD1 | -1.0615961274975 | 2.07795916326304 | -2.62274058787118 | 0.0121112058716955 | 0.157537158776854 | -2.9787019285843 |
| lnc-ROBO3-2 | lnc-ROBO3-2 | -0.442372729270174 | 2.39051398762848 | -2.62070392951293 | 0.0121217137599904 | 0.157537158776854 | -3.02394545072747 |
| ENSG00000265929.1 | MIR5195 | 2.01177243415514 | -0.591672993120863 | 2.62179442082129 | 0.0121235417335926 | 0.157537158776854 | -2.82788999950428 |
| ENSG00000137574.10 | TGS1 | 0.283312085604795 | 5.47685174316458 | 2.61950840482622 | 0.0121582350889254 | 0.157827412320955 | -3.23795390596645 |
| lnc-TSHR-9 | lnc-TSHR-9 | -0.627469335212073 | -0.462481205981797 | -2.61941102266242 | 0.0121612143973685 | 0.157827412320955 | -2.80928718761831 |
| ENSG00000224578.5 | HNRNPA1L3 | 0.793561322218462 | 2.31551679345282 | 2.61825441337802 | 0.0121975632526417 | 0.158107651693438 | -3.00265084248963 |
| PIK3CD-AS2 | PIK3CD-AS2 | -0.826801261087063 | 1.09233588180441 | -2.61808643873128 | 0.0122018052698594 | 0.158107651693438 | -2.92070403954359 |
| lnc-ABCG5-4 | lnc-ABCG5-4 | 0.492124846075284 | 2.38327392594089 | 2.61795459446925 | 0.0122058523315782 | 0.158107651693438 | -3.02178105659685 |
| lnc-HIST4H4-1 | lnc-HIST4H4-1 | -0.430741149983192 | 3.92010871620599 | -2.61603785132235 | 0.0122648274947781 | 0.158771663097495 | -3.1646714776705 |
| ENSG00000125841.12 | NRSN2 | -0.471347992540644 | 1.65932608310206 | -2.61544903942135 | 0.012282996719692 | 0.15885233799326 | -2.97436888161949 |
| lnc-TLK1-2 | lnc-TLK1-2 | -0.603848688907559 | 3.61570712442952 | -2.61533576910296 | 0.0122864947813861 | 0.15885233799326 | -3.14061356089645 |
| lnc-ERMN-4 | lnc-ERMN-4 | -0.504950241252852 | 2.7572066599423 | -2.61475897147096 | 0.0123043218665893 | 0.158982961067174 | -3.06825729219461 |
| ENSG00000198492.15 | YTHDF2 | 0.224783572411419 | 5.86922726889782 | 2.61351598130649 | 0.0123428196418983 | 0.159321347466763 | -3.25925837197128 |
| ENSG00000201801.1 | RNU5E-4P | -1.41444092240483 | 0.0157446598065182 | -2.6142897825403 | 0.0123459917995184 | 0.159321347466763 | -2.83310534773738 |
| lnc-TMEM99-5 | lnc-TMEM99-5 | -0.446993999836459 | 2.38728277224472 | -2.61266200228118 | 0.0123693329863165 | 0.159522544396837 | -3.03675200067697 |
| ENSG00000267318.1 | | 0.448895301682171 | 1.42910790662763 | 2.61113721154221 | 0.0124168028283726 | 0.1600313690493 | -2.96024397678836 |
| ENSG00000115896.15 | PLCL1 | -2.46757043036024 | -0.0352535905746507 | -2.61370596983558 | 0.0124413240901039 | 0.1600313690493 | -2.82562782427679 |
| ENSG00000275395.5 | FCGBP | -1.25379401645903 | 2.82016718796121 | -2.61356987522954 | 0.0124445211112357 | 0.1600313690493 | -3.10147158111747 |
| ENSG00000112232.8 | KHDRBS2 | -0.428195077463605 | 5.09667629284341 | -2.61023365499226 | 0.0124450111501089 | 0.1600313690493 | -3.24546435048776 |
| lnc-TBX1-3 | lnc-TBX1-3 | -0.661282140126823 | 1.40998323299312 | -2.61014884861374 | 0.0124476617523164 | 0.1600313690493 | -2.96314981010338 |
| ENSG00000280239.1 | | -0.597762709969248 | 0.279674801233029 | -2.6098448508135 | 0.0124571673816405 | 0.160053605628156 | -2.86189191954459 |
| MEF2C-AS1 | MEF2C-AS1 | -0.601851896900075 | 5.75146165699622 | -2.60969400977748 | 0.0124732476183474 | 0.160073957094098 | -3.26345100150454 |
| lnc-FKBP11-1 | lnc-FKBP11-1 | 0.591915131208853 | 0.785778153531583 | 2.60929730133011 | 0.0124743053577677 | 0.160073957094098 | -2.90127613102952 |
| ENSG00000166266.13 | CUL5 | 0.412132096753642 | 5.57890003523562 | 2.60839210707283 | 0.0125026848185133 | 0.160239344567555 | -3.26326945846256 |
| lnc-DDT-3 | lnc-DDT-3 | -0.547221087247484 | 1.30164850072976 | -2.60838959037671 | 0.0125027638038912 | 0.160239344567555 | -2.95780385604332 |
| lnc-GPR75-ASB3-7 | lnc-GPR75-ASB3-7 | 0.630708469101966 | 0.568275121349965 | 2.60776678294605 | 0.0125223244059314 | 0.160390170035648 | -2.88491801191323 |
| lnc-ZNF8-5 | lnc-ZNF8-5 | -0.573866318825919 | 0.346189963202666 | -2.60745240148266 | 0.0125322088809851 | 0.160416949873952 | -2.88027004832924 |
| ENSG00000099999.14 | RNF215 | -0.430588890745252 | 2.07516724443971 | -2.60696366793522 | 0.0125475893674801 | 0.160500255146457 | -3.02989453336928 |
| ENSG00000214389.2 | RPS3AP26 | 1.06838027989586 | 1.96748733037485 | 2.60785442945634 | 0.0125612321784562 | 0.160500255146457 | -2.97872322128383 |
| ENSG00000164032.11 | H2AZ1 | 0.344775604116004 | 4.80980127416796 | 2.60624811947292 | 0.0125701389531438 | 0.160500255146457 | -3.24523713997346 |
| ENSG00000001497.16 | LAS1L | 0.192237177603563 | 5.26634682817289 | 2.60621414365976 | 0.0125712105802378 | 0.160500255146457 | -3.26140559117962 |
| lnc-AAMP-2 | lnc-AAMP-2 | -0.459843485267827 | 0.659058380493257 | -2.60600828093028 | 0.0125777054633063 | 0.160500255146457 | -2.90865049887333 |
| ENSG00000129083.12 | COPB1 | 0.200030107021202 | 6.45920352634434 | 2.60553866744913 | 0.0125925330853453 | 0.160589906131142 | -3.27939821553989 |
| ENSG00000076108.11 | BAZ2A | 0.453417523240184 | 8.26426944147338 | 2.60462393661984 | 0.0126214608948959 | 0.160859151454886 | -3.26134161628469 |
| ENSG00000130066.16 | SAT1 | 0.540658112513729 | 6.66643061878568 | 2.60436488472244 | 0.0126296643146496 | 0.160864096898782 | -3.28136322615098 |
| ENSG00000168477.19 | TNXB | -0.66332629407098 | 0.823930030046093 | -2.60317102409553 | 0.012667533599698 | 0.161246656822872 | -2.93342936642375 |
| ENSG00000125657.4 | TNFSF9 | 0.708687492247237 | 3.64477328986157 | 2.60311127161633 | 0.0126808892037165 | 0.161256410966635 | -3.14770641057988 |
| ENSG00000182986.13 | ZNF320 | -0.294450044963106 | 4.1363458623789 | -2.60243184029118 | 0.0126910326589553 | 0.161256410966635 | -3.20579262128228 |
| ENSG00000232940.5 | HCG25 | -0.549459535583862 | -0.492078974378479 | -2.60210207834885 | 0.012701528852853 | 0.161256410966635 | -2.84545453265694 |
| lnc-HNRNPA0-4 | lnc-HNRNPA0-4 | 2.02204827737825 | 2.18770930147494 | 2.60616915930734 | 0.0127109636876221 | 0.161256410966635 | -2.99378507245175 |
| ENSG00000104549.11 | SQLE | -0.596309085659718 | 3.63697678955291 | -2.6017846118522 | 0.0127116412027008 | 0.161256410966635 | -3.18440813138613 |
| ENSG00000124587.13 | PEX6 | -0.378490449359258 | 4.23247135015413 | -2.60141259996046 | 0.0127235003901084 | 0.161256410966635 | -3.21528692566639 |
| lnc-KMT2B-2 | lnc-KMT2B-2 | -0.663792675649872 | 1.06773969369255 | -2.60115831903368 | 0.0127316123213738 | 0.161256410966635 | -2.94687878100373 |
| ENSG00000156671.13 | SAMD8 | 0.38340919284464 | 6.25318725138364 | 2.60111664349606 | 0.0127329422835366 | 0.161256410966635 | -3.2890952968644 |
| ENSG00000109519.12 | GRPEL1 | 0.32207499553803 | 3.73595042460631 | 2.60072383763535 | 0.0127454838771924 | 0.161256410966635 | -3.18148811870363 |
| lnc-XRCC3-6 | lnc-XRCC3-6 | -0.583979973626044 | -0.23282845339775 | -2.60068750926032 | 0.0127466443493522 | 0.161256410966635 | -2.84686749671332 |

| | | | | | | | |
|---|---|---|---|---|---|---|---|
| ENSG00000101457.12 | DNTTIP1 | 0.312778816449751 | 2.3705848470991 | 2.60018599972154 | 0.0127626744467964 | 0.161360029569048 | -3.06504059036939 |
| ENSG00000185129.5 | PURA | -0.361337185664088 | 4.70024450825171 | -2.59956205775797 | 0.0127826437221967 | 0.161405483650298 | -3.25766676396636 |
| ENSG00000279050.1 | PWAR1 | -0.858254661407587 | -0.204794778980715 | -2.59928395618036 | 0.0127915535891851 | 0.161405483650298 | -2.85008088503832 |
| lnc-SNRPN-11 | lnc-SNRPN-11 | -0.858254661407587 | -0.204794778980715 | -2.59928395618036 | 0.0127915535891851 | 0.161405483650298 | -2.85008088503832 |
| SNHG26 | SNHG26 | -0.42804221295874 | 4.55989512882701 | -2.59895472445292 | 0.0128021089281636 | 0.161405483650298 | -3.25238086233979 |
| ENSG00000249353.2 | NPM1P27 | 0.435519242936405 | 3.05019626384287 | 2.59884969033735 | 0.0128054780547509 | 0.161405483650298 | -3.10955450286806 |
| ENSG00000123684.12 | LPGAT1 | 0.259714096585422 | 6.70113544941902 | 2.59830782237294 | 0.0128228721904974 | 0.161525812911265 | -3.29460116218609 |
| ENSG00000243667.6 | DNAAF10 | -0.424308487165151 | 2.43340081580624 | -2.59802089963777 | 0.0128320912653622 | 0.161543079214037 | -3.0744867604279 |
| ENSG00000183763.8 | TRAIP | 0.585311010366419 | 1.00455366581626 | 2.59735378514753 | 0.0128535497059458 | 0.161677440466239 | -2.94471957434928 |
| ENSG00000163374.19 | YY1AP1 | 0.201725273048737 | 6.23887090168626 | 2.59720083975997 | 0.0128584739854848 | 0.161677440466239 | -3.29791749831159 |
| ENSG00000269343.6 | ZNF587B | 0.225517442568044 | 5.33090475182374 | 2.59609266841735 | 0.0128942046872126 | 0.162003363051464 | -3.2863535798261 |
| ENSG00000186026.6 | ZNF284 | -0.558106161220263 | 1.71309735763423 | -2.59548743072517 | 0.0129137577082991 | 0.162003363051464 | -3.01420084634945 |
| ENSG00000140548.9 | ZNF710 | -0.375944617953627 | 3.91727279120776 | -2.59528374992644 | 0.0129203439970342 | 0.162003363051464 | -3.20973254115253 |
| ENSG00000158793.13 | NIT1 | -0.318048192868793 | 3.90584925656813 | -2.59521130792292 | 0.0129226872469574 | 0.162003363051464 | -3.20229213012689 |
| ENSG00000198960.10 | ARMCX6 | -0.324012914765154 | 2.73424550178186 | -2.59517849290361 | 0.0129237488281836 | 0.162003363051464 | -3.10854435295934 |
| lnc-CHN1-6 | lnc-CHN1-6 | 0.234074256042976 | 2.31848842484286 | 2.59416516780123 | 0.012956569712856 | 0.16220621051771 | -3.06877713525146 |
| lnc-DNAJB7-2 | lnc-DNAJB7-2 | -0.706969318150991 | -0.0178877241082668 | -2.59408612501939 | 0.0129591330597187 | 0.16220621051771 | -2.87014867375042 |
| lnc-CCDC189-1 | lnc-CCDC189-1 | -0.696557554067895 | -0.408073479768057 | -2.59394926002396 | 0.0129635726717015 | 0.16220621051771 | -2.8575407590283 |
| lnc-ZNF8-4 | lnc-ZNF8-4 | 0.430516496426723 | 1.56075405560421 | 2.59354061485021 | 0.0129768365497905 | 0.162264346243428 | -3.00682904647243 |
| ACTA2-AS1 | ACTA2-AS1 | -0.644080543075386 | 2.55009483868042 | -2.59310513748256 | 0.0129909850311849 | 0.162264346243428 | -3.09118969913406 |
| lnc-DOK4-2 | lnc-DOK4-2 | 0.333154799000165 | 0.682469678128241 | 2.5929302650212 | 0.0129966705403893 | 0.162264346243428 | -2.93388995055661 |
| ENSG00000160679.12 | CHTOP | 0.184405717299522 | 6.30022262204238 | 2.59283549884081 | 0.0129997525606283 | 0.162264346243428 | -3.30776434357089 |
| lnc-RAB3IL1-2 | lnc-RAB3IL1-2 | -0.455069334858274 | 1.77274216241436 | -2.59227511951866 | 0.013017991106296 | 0.162393521782358 | -3.04100734795837 |
| ENSG00000101236.16 | RNF24 | -0.775702032206241 | 2.27759622215706 | -2.59183169175437 | 0.0130333836050764 | 0.162462101405705 | -3.06808873122633 |
| ENSG00000074071.14 | MRPS34 | -0.634528304744182 | 3.05317054209369 | -2.59162208887359 | 0.0130392747180793 | 0.162462101405705 | -3.14357259906268 |
| ENSG00000139116.18 | KIF21A | -0.722010371959546 | 2.47209627134735 | -2.59014472994684 | 0.0130884488047833 | 0.162973515743787 | -3.11347667976728 |
| PTOV1-AS2 | PTOV1-AS2 | -0.436977107129222 | 3.15091476493647 | -2.58988159332097 | 0.0130961567818211 | 0.162973515743787 | -3.15668421893231 |
| ENSG00000138069.17 | RAB1A | 0.346284765540293 | 5.13418853754947 | 2.58942937991288 | 0.0131109729480079 | 0.162982467215824 | -3.29524442607821 |
| ENSG00000092871.16 | RFFL | -0.262958308095866 | 4.76740982524505 | -2.58937418795721 | 0.0131127822889308 | 0.162982467215824 | -3.28211362240837 |
| ENSG00000130382.8 | MLLT1 | -0.338334008456858 | 4.18424039510752 | -2.5891348519026 | 0.013120631014751 | 0.162982467215824 | -3.24302038414917 |
| LINC00910 | LINC00910 | -0.851248791667855 | 5.16779322812368 | -2.59137175187494 | 0.0131305816664412 | 0.16300769749117 | -3.29693524714891 |
| lnc-ENDOU-4 | lnc-ENDOU-4 | -1.00441774927077 | 2.05863691704591 | -2.58985967090176 | 0.0131479460925326 | 0.163124879097408 | -3.06626307925948 |
| ENSG00000100403.11 | ZC3H7B | -0.259030617654335 | 5.89980025371007 | -2.58774204626463 | 0.0131663917395048 | 0.163246856911314 | -3.31680214420123 |
| lnc-ZNF394-2 | lnc-ZNF394-2 | 0.387760377134372 | 2.296585479342 | 2.58752184391379 | 0.0131736398644363 | 0.163246856911314 | -3.07496116414795 |
| lnc-SCP2D1-1 | lnc-SCP2D1-1 | -0.37765733892148 | 3.41960606231689 | -2.58657377456439 | 0.013204888013175 | 0.163535625737172 | -3.18116509227357 |
| ENSG00000197989.14 | SNHG12 | -0.378372896023541 | 3.96715448263368 | -2.58559080671454 | 0.0132373580390976 | 0.163839170486317 | -3.23499098090564 |
| ENSG00000099974.7 | DDTL | 1.15271024098916 | -0.394883340386148 | 2.58523533317635 | 0.0132491182515735 | 0.163886178468833 | -2.87476623280941 |
| ENSG00000183696.13 | UPP1 | -1.4311495360294 | -0.716313327261636 | -2.58474560960069 | 0.0132653355516813 | 0.163914906967256 | -2.88013146980068 |
| ENSG00000213185.6 | FAM24B | 1.27426141366612 | 0.888231320606965 | 2.58575171999433 | 0.0132673679739323 | 0.163914906967256 | -2.92543220073214 |
| lnc-FBXO3-3 | lnc-FBXO3-3 | 0.45192050725517 | 2.8066813619861 | 2.58399300860787 | 0.0132902935114376 | 0.164099646878176 | -3.12913889129877 |
| lnc-PRTFDC1-3 | lnc-PRTFDC1-3 | -0.455987126422755 | 1.62719879094713 | -2.58331610480961 | 0.0133127778826838 | 0.164278721318513 | -3.04740362601312 |
| ENSG00000164344.15 | KLKB1 | -0.640599593775098 | 1.152375723227 | -2.58294554166124 | 0.0133251014318012 | 0.164308400830013 | -2.99725732978911 |
| ENSG00000177731.15 | FLII | -0.253704452865166 | 6.33734846982245 | -2.58264536996814 | 0.0133350916762263 | 0.164308400830013 | -3.32993249735666 |
| ENSG00000103496.14 | STX4 | -0.280081386709129 | 4.21688951874577 | -2.58252405448611 | 0.0133391312144465 | 0.164308400830013 | -3.25803317127396 |
| ENSG00000087053.18 | MTMR2 | 0.373410007839446 | 4.45702161113825 | 2.58182609097687 | 0.0133623936240866 | 0.164441232224836 | -3.27424233244557 |
| ENSG00000133142.17 | TCEAL4 | -0.34225226115251 | 3.6725285931173 | -2.58172118362505 | 0.0133658932863115 | 0.164441232224836 | -3.21992574895118 |
| ENSG00000159363.17 | ATP13A2 | -0.391256404901704 | 3.64464418704506 | -2.58126986806557 | 0.0133809585388225 | 0.164528237517673 | -3.2182613529384 |
| ENSG00000173692.12 | PSMD1 | 0.194125668489959 | 5.36701869317173 | 2.58092061889861 | 0.0133926274013745 | 0.164573402867159 | -3.32230908549393 |
| ENSG00000141551.14 | CSNK1D | 0.419360022910819 | 5.16804920311754 | 2.58044567074913 | 0.0134085109695305 | 0.164670275230219 | -3.31328110499079 |
| lnc-HLA-C-5 | lnc-HLA-C-5 | 0.345715852459842 | 8.53961210995827 | 2.57997373996809 | 0.0134243106997373 | 0.16476600306064 | -3.31059267207889 |
| ENSG00000226571.2 | | -0.361373677762717 | 5.3495803378597 | -2.579063706335 | 0.0134548257418445 | 0.165042120527048 | -3.32533242563875 |
| ENSG00000124333.15 | VAMP7 | 0.298023232351763 | 4.83680491055923 | 2.57788154632134 | 0.0134945604291992 | 0.165209649352126 | -3.30774644761355 |
| lnc-CENPB-3 | lnc-CENPB-3 | -0.892521833841163 | 0.134532477373451 | -2.57787031147374 | 0.0134949385686175 | 0.165209649352126 | -2.90375281151288 |
| ENSG00000029639.10 | TFB1M | -0.312687058014899 | 2.65804982074454 | -2.57722960351819 | 0.0135165193876649 | 0.165209649352126 | -3.13747742806552 |
| ENSG00000133030.20 | MPRIP | 0.386292993962077 | 6.41294114719717 | 2.57720773084946 | 0.0135172566767331 | 0.165209649352126 | -3.34226509413067 |
| ENSG00000077458.12 | FAM76B | 0.484193693458668 | 5.18099891323435 | 2.57704178237848 | 0.0135228517054347 | 0.165209649352126 | -3.32339763132859 |
| lnc-IDI2-4 | lnc-IDI2-4 | -0.65860813197161 | -0.298642690263575 | -2.57703598743616 | 0.0135230471228454 | 0.165209649352126 | -2.89113484900347 |
| ENSG00000231767.4 | RPS27AP5 | 1.03322832553415 | 1.23802740179098 | 2.57718072882393 | 0.0135246688606292 | 0.165209649352126 | -2.96194050182629 |
| ENSG00000107864.14 | CPEB3 | -0.368448635479654 | 3.36510519511276 | -2.57488754223929 | 0.0135956754018281 | 0.165839397297616 | -3.21145877418269 |
| ENSG00000240087.3 | RPSAP12 | 1.52270396292696 | -1.1120672556372 | 2.57485826527101 | 0.0135966675703638 | 0.165839397297616 | -3.03633181401229 |
| ENSG00000128271.21 | ADORA2A | -0.433621040977093 | 3.60097667646943 | -2.57457763388375 | 0.0136061812607898 | 0.165839397297616 | -3.22468320109892 |
| lnc-EGR2-4 | lnc-EGR2-4 | 0.344960997324969 | 4.18536114409603 | 2.57434145688018 | 0.0136141926200186 | 0.165839397297616 | -3.26642802400971 |
| NNT-AS1 | NNT-AS1 | -0.329686613480052 | 3.21746530283883 | -2.57427322644904 | 0.0136165078673163 | 0.165839397297616 | -3.19583820343676 |
| lnc-CASP5-2 | lnc-CASP5-2 | 0.98903798382665 | 0.0117112714111863 | 2.57380609303273 | 0.0136323686715496 | 0.165876257952643 | -2.89645362381658 |
| lnc-TBC1D7-1 | lnc-TBC1D7-1 | -0.425622525827854 | -0.176600818156775 | -2.5737094495621 | 0.0136356521622636 | 0.165876257952643 | -2.90013878366494 |
| lnc-SMYD3-4 | lnc-SMYD3-4 | 0.335121129330369 | 1.15919577096423 | 2.57279657102417 | 0.0136667031852478 | 0.166155789522714 | -3.01813724978528 |
| ENSG00000164896.19 | FASTK | -0.345299214563012 | 5.02496043532112 | -2.57226334515179 | 0.013684870451489 | 0.166195728939499 | -3.33085909224322 |
| lnc-DNAJC10-2 | lnc-DNAJC10-2 | -0.786648886821131 | 1.10204200420916 | -2.57222619111699 | 0.013686137130275 | 0.166195728939499 | -3.01803046293373 |

| | | | | | | | |
|---|---|---|---|---|---|---|---|
| ENSG00000134369.15 | NAV1 | 0.859164652846006 | 3.18349087851814 | 2.57290491869422 | 0.0137075846305591 | 0.166358027388442 | -3.18310972516239 |
| lnc-QPRT-3 | lnc-QPRT-3 | -1.08898225584262 | 2.09133075820207 | -2.57270954470751 | 0.0137389004884066 | 0.166509202775663 | -3.09319616262995 |
| lnc-PPIL1-2 | lnc-PPIL1-2 | 0.308742544352234 | 3.07072839711503 | 2.57029434651939 | 0.0137521467003983 | 0.166509202775663 | -3.18717588032245 |
| lnc-GLRX2-1 | lnc-GLRX2-1 | -0.349569891037688 | 1.95694379750005 | -2.57020761588078 | 0.013755117033056 | 0.166509202775663 | -3.10427094292055 |
| lnc-LASP1-5 | lnc-LASP1-5 | -0.49820518757084 | 1.55198822943516 | -2.57012175746582 | 0.0137580580717631 | 0.166509202775663 | -3.06519622623261 |
| ENSG00000161298.17 | ZNF382 | -0.361017128998373 | 4.28483535621342 | -2.57005079096273 | 0.0137604894292087 | 0.166509202775663 | -3.2867373009639 |
| ENSG00000033178.12 | UBA6 | 0.238991621534378 | 6.43536563673202 | 2.56821006503971 | 0.0138236912787366 | 0.167175697761595 | -3.36219721285121 |
| lnc-ASMT-5 | lnc-ASMT-5 | 1.6570370280605 | 1.01945012124512 | 2.56990311182905 | 0.0138549481223509 | 0.16745531250872 | -2.95152587223802 |
| lnc-POLR1A-7 | lnc-POLR1A-7 | 0.325904553529973 | 2.88067619570019 | 2.56643075410872 | 0.0138850365677935 | 0.167706470894091 | -3.17382933108472 |
| lnc-ZNF670-ZNF695-2 | lnc-ZNF670-ZNF69 | 0.371800127294822 | 1.52651923530489 | 2.56604027139685 | 0.0138985324919549 | 0.167706470894091 | -3.05886476552984 |
| ENSG00000180964.16 | TCEAL8 | -0.482797362846396 | 2.82800499986916 | -2.56598042018798 | 0.0139006021381919 | 0.167706470894091 | -3.17957589363942 |
| ENSG00000141012.12 | GALNS | -0.34238813482016 | 4.41239466268339 | -2.56575730912386 | 0.0139083197695289 | 0.167706470894091 | -3.30780509479364 |
| GLIDR | GLIDR | -0.514658212687284 | 1.68399414245321 | -2.56416093138571 | 0.0139636544125981 | 0.168089926921578 | -3.09060440970238 |
| ENSG00000160194.17 | NDUFV3 | -0.276765538762215 | 4.1204106111964 | -2.56408337706912 | 0.0139663477667997 | 0.168089926921578 | -3.29214459005241 |
| lnc-MGAT1-3 | lnc-MGAT1-3 | 0.26891317856395 | 2.13792102404692 | 2.56391401115005 | 0.0139722312617372 | 0.168089926921578 | -3.11547522492882 |
| ENSG00000000460.16 | FIRRM | -0.306905177303643 | 3.59675144948458 | -2.56374380972515 | 0.013978146062676 | 0.168089926921578 | -3.24130101063947 |
| lnc-ZNF573-2 | lnc-ZNF573-2 | -0.561916602057069 | 2.51434145225247 | -2.56346115162579 | 0.0139879739882535 | 0.168089926921578 | -3.16196883300339 |
| ENSG00000055917.15 | PUM2 | 0.363000618021696 | 7.04537779183468 | 2.56342822029452 | 0.0139891194100308 | 0.168089926921578 | -3.36950874818432 |
| AZIN1-AS1 | AZIN1-AS1 | -0.439709590554408 | 3.11449282204413 | -2.56277846662381 | 0.0140117367630147 | 0.168214782936647 | -3.20170522480775 |
| lnc-PSMG1-11 | lnc-PSMG1-11 | 0.357515635828754 | 2.24851811197159 | 2.5626602477175 | 0.0140158554504373 | 0.168214782936647 | -3.12391234471741 |
| UFL1-AS1 | UFL1-AS1 | -0.523118069503988 | 4.74169810111342 | -2.56072807570087 | 0.0140833283520862 | 0.168926076614796 | -3.3311729733874 |
| lnc-EARS2-1 | lnc-EARS2-1 | 0.34931794413275 | 4.1434077675278 | 2.56033306655956 | 0.014097158856722 | 0.168993489078572 | -3.29573983965625 |
| ENSG00000188878.19 | FBF1 | -0.404135353526737 | 3.06574543771224 | -2.55980405948925 | 0.014115700486674 | 0.16911726607521 | -3.2069988512685 |
| lnc-KIF1C-1 | lnc-KIF1C-1 | -0.488562872587065 | 1.05120048966548 | -2.55918839751541 | 0.0141373074140609 | 0.169277602387211 | -3.04375145864222 |
| lnc-CYYR1-5 | lnc-CYYR1-5 | -0.734853037679001 | 0.282530615160432 | -2.55810644550073 | 0.0141753522206228 | 0.169634462068069 | -2.97412209973646 |
| ENSG00000164610.8 | RP9 | -0.42148118320681 | 3.9161643249701 | -2.55703234634096 | 0.0142132133903642 | 0.169892100658665 | -3.29212396045382 |
| ENSG00000164323.13 | CFAP97 | -0.378386151615918 | 5.85380938611248 | -2.55702735459533 | 0.0142133895610077 | 0.169892100658665 | -3.38369005484051 |
| ENSG00000177000.12 | MTHFR | -0.257889857493203 | 6.21074591405457 | -2.55567664493093 | 0.014261132694959 | 0.170327717059993 | -3.38922108568429 |
| XXYLT1-AS2 | XXYLT1-AS2 | -1.19438255361267 | 2.29699364801806 | -2.55816599451747 | 0.0142690063154574 | 0.170327717059993 | -3.15698674520557 |
| ENSG00000135245.9 | HILPDA | -0.803853642855419 | 0.0504053994998405 | -2.5550876523218 | 0.0142819974699406 | 0.170327717059993 | -2.96731022932599 |
| lnc-C1D-5 | lnc-C1D-5 | 0.435245117481482 | 0.668268944130332 | 2.55506122090606 | 0.0142829344432564 | 0.170327717059993 | -3.00122573292279 |
| ENSG00000201882.1 | | 0.844561218264581 | -0.521817996941874 | 2.55419651757254 | 0.0143136184900849 | 0.170513607742591 | -2.93479336800656 |
| ENSG00000121989.14 | ACVR2A | -1.19353620704342 | 1.36106641935888 | -2.55638467461591 | 0.014315090811796 | 0.170513607742591 | -3.09194365941527 |
| ENSG00000198826.10 | ARHGAP11A | 0.645106928599626 | 1.93950940894848 | 2.55378127968035 | 0.0143283746164613 | 0.170531883293441 | -3.11883109653803 |
| ENSG00000183044.11 | ABAT | -0.366171417570869 | 4.01416592102975 | -2.55364571115774 | 0.0143331952629672 | 0.170531883293441 | -3.29830826209719 |
| lnc-PFKP-37 | lnc-PFKP-37 | -1.19478345294251 | 1.41162474711349 | -2.55535151086826 | 0.0143518016782472 | 0.170654612330076 | -3.09457973241945 |
| lnc-C21orf58-1 | lnc-C21orf58-1 | -0.540636147292675 | 1.43906344598478 | -2.5521491293142 | 0.0143865103492965 | 0.170968558036703 | -3.08724714163649 |
| ENSG00000269896.2 | | -0.43828617171292 | 0.63033015161957 | -2.55154036431103 | 0.0144082491085825 | 0.171106263790317 | -3.01841256076238 |
| ENSG00000047315.15 | POLR2B | 0.18844446726733 | 6.93993922410455 | 2.55135920934563 | 0.0144147238698153 | 0.171106263790317 | -3.39671230361761 |
| ENSG00000106404.13 | CLDN15 | -0.453981288944069 | 3.206372789651 | -2.55089649650863 | 0.0144312740103854 | 0.171203984412543 | -3.24402396683241 |
| ENSG00000171148.13 | TADA3 | -0.321362016416087 | 4.68540292295892 | -2.55023695386497 | 0.0144548942779438 | 0.171216929582764 | -3.35989642047297 |
| ENSG00000134375.10 | TIMM17A | 0.260346338949083 | 4.63490932758822 | 2.5502144567241 | 0.014455700592532 | 0.171216929582764 | -3.35720579701001 |
| ENSG00000278873.1 | | -0.91498266431475 | -0.341394238710976 | -2.55016926857443 | 0.0144573202941672 | 0.171216929582764 | -2.94281853441998 |
| ENSG00000158092.6 | NCK1 | 0.187417486842933 | 3.98626766989142 | 2.54960019072335 | 0.0144777322262599 | 0.171297130003565 | -3.31203949878518 |
| ENSG00000136986.9 | DERL1 | 0.252650007583609 | 6.41662883942533 | 2.54950598774486 | 0.0144811136768664 | 0.171297130003565 | -3.40340029774507 |
| lnc-TFAP4-3 | lnc-TFAP4-3 | -0.552223963492548 | 1.20390285306446 | -2.54902818782787 | 0.0144982755868458 | 0.171297130003565 | -3.06872107242473 |
| ENSG00000164144.15 | ARFIP1 | -0.386174381116214 | 3.6163943644008 | -2.54901149935549 | 0.0144988753490857 | 0.171297130003565 | -3.28449676818371 |
| ENSG00000118900.14 | UBN1 | 0.191597130405819 | 5.73305619733455 | 2.54882154615405 | 0.0145057036193078 | 0.171297130003565 | -3.39993187580682 |
| ENSG00000138758.11 | SEPTIN11 | -0.482699204864633 | 4.50350286724215 | -2.54827944630702 | 0.0145252066944749 | 0.171351713188581 | -3.35665896486025 |
| lnc-PPP4C-10 | lnc-PPP4C-10 | -0.619021628251647 | 0.592000374186187 | -2.54817933209329 | 0.0145288111117256 | 0.171351713188581 | -3.00782802501306 |
| ENSG00000133835.15 | HSD17B4 | -0.520996830121989 | 6.89642163623033 | -2.54786636732141 | 0.0145400840653067 | 0.171351713188581 | -3.40360061860507 |
| ENSG00000155961.4 | RAB39B | 0.31629454444016 | 3.04673806626196 | 2.54776809513148 | 0.0145436254647258 | 0.171351713188581 | -3.22784620611403 |
| ENSG00000116005.11 | PCYOX1 | -0.613620381330004 | 3.49117923848987 | -2.54693960348748 | 0.0145735128344108 | 0.171603703713701 | -3.28308977689732 |
| lnc-CNPY1-2 | lnc-CNPY1-2 | -0.315965070447411 | 3.33665487581123 | -2.54653598920848 | 0.0145880932923632 | 0.171603703713701 | -3.2618015253932 |
| ENSG00000154719.13 | MRPL39 | 0.282293801898957 | 4.19204500165165 | 2.54624856229487 | 0.0145984846288996 | 0.171603703713701 | -3.331280200868 |
| ENSG00000231345.3 | BEND3P1 | -0.632504382237881 | -0.437921315471224 | -2.54617844255784 | 0.0146010206905965 | 0.171603703713701 | -2.94775376698473 |
| lnc-WRB-2 | lnc-WRB-2 | -0.611792232601986 | 0.0574264493006569 | -2.54602147835784 | 0.01460669916467 | 0.171603703713701 | -2.96236390501069 |
| lnc-UACA-6 | lnc-UACA-6 | 0.753808459023078 | 1.26454227673936 | 2.54523261637347 | 0.0146352682591688 | 0.171841258744137 | -3.06228324091078 |
| ENSG00000205763.13 | RP9P | -0.477472833554566 | 0.746719422229745 | -2.54389115488997 | 0.0146839671895668 | 0.172246524234624 | -3.03589075296284 |
| ENSG00000128829.11 | EIF2AK4 | 0.390750282837241 | 4.95267707814149 | 2.54368776972907 | 0.0146913635561122 | 0.172246524234624 | -3.38720438797237 |
| ENSG00000196724.12 | ZNF418 | -0.71039912979761 | 4.31629191239837 | -2.54405278074701 | 0.0147148929369357 | 0.172246524234624 | -3.34459257677196 |
| ENSG00000173442.12 | EHBP1L1 | -0.304317536061656 | 6.78586277970264 | -2.54292091220918 | 0.0147192819171295 | 0.172246524234624 | -3.41547588496266 |
| lnc-IRF2BP2-1 | lnc-IRF2BP2-1 | -1.08420558928665 | -0.329465027417558 | -2.54286455503047 | 0.01472133557587 | 0.172246524234624 | -2.95389880550271 |
| ENSG00000124196.5 | GTSF1L | -1.35141016467303 | 1.43119141077551 | -2.54539139228372 | 0.014725536416534 | 0.172246524234624 | -3.09488711165183 |
| ENSG00000172269.18 | DPAGT1 | -0.589995515284029 | 2.99549998251304 | -2.54267177770148 | 0.0147283623695738 | 0.172246524234624 | -3.24338066154359 |
| ENSG00000162222.13 | TTC9C | -0.313985536768037 | 3.25268961177485 | -2.54228641324495 | 0.0147424181993352 | 0.172313000452537 | -3.26308918031182 |
| lnc-H2AFJ-4 | lnc-H2AFJ-4 | -0.417166982063581 | 5.1585797209837 | -2.54181786260438 | 0.0147595246605526 | 0.172415037507466 | -3.40106893707133 |
| ENSG00000164327.12 | RICTOR | 0.327153637030469 | 7.43554748101536 | 2.54123778893357 | 0.0147807278420594 | 0.172518944705558 | -3.41301177139626 |

| | | | | | | | |
|---|---|---|---|---|---|---|---|
| CPEB1-AS1 | CPEB1-AS1 | -1.25764258147338 | -0.0496501332315983 | -2.54134451165424 | 0.0147851828431523 | 0.172518944705558 | -2.97322364952507 |
| ENSG00000258653.3 | | 0.268534574056839 | 4.42987322047228 | 2.54072329660554 | 0.0147995571105194 | 0.172588829465054 | -3.36054433168802 |
| ENSG00000136630.12 | HLX | 2.32358124698322 | -0.917283319495298 | 2.54119709228462 | 0.0148119086594136 | 0.172600464466439 | -3.04933142221912 |
| ENSG00000204525.16 | HLA-C | 0.399944202640622 | 8.98218366759662 | 2.54023832605767 | 0.0148173259831996 | 0.172600464466439 | -3.38819919780276 |
| ENSG00000124574.14 | ABCC10 | -0.288833007168378 | 4.82518720733692 | -2.53922620217662 | 0.0148544719817167 | 0.172935292307509 | -3.39153916500005 |
| lnc-GRHPR-4 | lnc-GRHPR-4 | -0.713923647503166 | -0.278720234418014 | -2.53863555555471 | 0.0148761885623987 | 0.173090214347005 | -2.96386443168441 |
| ENSG00000135018.13 | UBQLN1 | 0.350260760646218 | 6.5437789790597 | 2.53747713663233 | 0.0149188648207686 | 0.173488697517446 | -3.42950050077713 |
| ENSG00000186265.9 | BTLA | -0.524151516058285 | 7.11589268795651 | -2.53649361515822 | 0.0149551854680668 | 0.173728229879098 | -3.42727122445631 |
| ENSG00000273294.1 | C1QTNF3-AMACR | -0.275945810057685 | 2.17092901293204 | -2.53646228615412 | 0.0149563437470612 | 0.173728229879098 | -3.18441084628692 |
| lnc-ZNF595-1 | lnc-ZNF595-1 | -1.03479019873921 | 1.78812090416424 | -2.53760101915877 | 0.0149689194329894 | 0.173776237275364 | -3.14794413579726 |
| ENSG00000023287.12 | RB1CC1 | 0.324149056875488 | 6.87844027566344 | 2.53562352703836 | 0.0149873843316934 | 0.173892520687286 | -3.4318359004306 |
| lnc-ALB-1 | lnc-ALB-1 | -0.504643921289091 | 4.22967935079794 | -2.53536246610841 | 0.0149970575916687 | 0.173906724737643 | -3.35722899094407 |
| ENSG00000279456.1 | NA | -0.543943107945143 | 0.297758187961273 | -2.53490212566808 | 0.015014128758937 | 0.174006651038964 | -3.02147924878488 |
| lnc-ATP8B4-1 | lnc-ATP8B4-1 | -0.437972521405171 | 0.388232623688645 | -2.53415550694773 | 0.015041853893896 | 0.174172089197894 | -3.01272693637827 |
| lnc-MRPS30-7 | lnc-MRPS30-7 | -0.408294223011987 | 2.85540729502769 | -2.53406205537284 | 0.0150453274349636 | 0.174172089197894 | -3.25260034001782 |
| ENSG00000143815.14 | LBR | 0.314408668300071 | 5.07643646144869 | 2.5335224523963 | 0.0150653984763077 | 0.174306406316943 | -3.41898664830507 |
| ENSG00000115942.8 | ORC2 | -0.264010198201428 | 5.05246278525437 | -2.53271970109197 | 0.0150953027523599 | 0.174554278961699 | -3.41858041509381 |
| ENSG00000139178.10 | C1RL | -0.404980342998136 | 3.47748815553758 | -2.53231653096654 | 0.0151103421486271 | 0.174587511280356 | -3.30256280801286 |
| lnc-ATP5ME-1 | lnc-ATP5ME-1 | -0.520385121050891 | 2.56966552044895 | -2.53218796501453 | 0.0151151408979058 | 0.174587511280356 | -3.23218674630448 |
| ENSG00000283761.1 | | 0.287170599119841 | 4.98641065900384 | 2.53174307864679 | 0.0151317570945711 | 0.174681411260548 | -3.41700295049664 |
| ENSG00000062194.15 | GPBP1 | 0.208611511222524 | 7.27859866386348 | 2.53148571627408 | 0.0151413770030046 | 0.174688373473504 | -3.43677903139214 |
| ENSG00000277734.7 | TRAC | -1.10690198743325 | 4.90980631703643 | -2.53457697827774 | 0.0151552430695945 | 0.174688373473504 | -3.40661755922284 |
| lnc-BZW1-1 | lnc-BZW1-1 | 0.332945091786229 | 2.78439360782051 | 2.53104290197501 | 0.0151579419425981 | 0.174688373473504 | -3.25373718360284 |
| ENSG00000143771.11 | CNIH4 | -0.283382526013455 | 3.02212141261554 | -2.53069161535055 | 0.0151710947369483 | 0.174688373473504 | -3.27680348164785 |
| ENSG00000129562.10 | DAD1 | -0.397427394797196 | 4.26785540349345 | -2.53059283011671 | 0.0151747953053794 | 0.174688373473504 | -3.37529458530777 |
| ENSG00000189079.15 | ARID2 | 0.325490233435108 | 6.45593682162228 | 2.52898717420432 | 0.0152350598828537 | 0.175284090312341 | -3.44819104339884 |
| ENSG00000186625.13 | KATNA1 | 0.250043228540035 | 3.8224872842498 | 2.52844936456496 | 0.0152552940496765 | 0.175411366258516 | -3.3403232941842 |
| ENSG00000107036.11 | RIC1 | 0.331762896913496 | 7.17759399279499 | 2.52824029502288 | 0.0152631665437012 | 0.175411366258516 | -3.44502303848938 |
| lnc-SENP1-1 | lnc-SENP1-1 | 0.352214961740888 | 0.136240460574894 | 2.52738297947584 | 0.0152954874240384 | 0.175684719670192 | -3.0053374264053 |
| lnc-CIPC-1 | lnc-CIPC-1 | -0.431902613778419 | 1.87660175348429 | -2.5267724776292 | 0.0153185414208647 | 0.175826179081271 | -3.18551746130116 |
| lnc-DIP2C-2 | lnc-DIP2C-2 | -0.307798366964528 | 1.6774451926931 | -2.52660456410552 | 0.015324887784667 | 0.175826179081271 | -3.16416711702342 |
| lnc-ATG4C-2 | lnc-ATG4C-2 | -0.478944873380732 | 2.61457308767111 | -2.5244487694265 | 0.0154065801306922 | 0.176610377939037 | -3.24829828911745 |
| lnc-PLEKHM1-7 | lnc-PLEKHM1-7 | -1.27578265307015 | 0.562219720734386 | -2.52543776261277 | 0.0154103988135116 | 0.176610377939037 | -3.04355089872151 |
| lnc-CDIPT-1 | lnc-CDIPT-1 | -0.484738714159307 | 1.8384777599407 | -2.5238154713028 | 0.0154306538293926 | 0.17674039980813 | -3.18616608483488 |
| ENSG00000200156.1 | RNU5B-1 | -0.936803504248802 | 1.5545639091956 | -2.52430986405756 | 0.0154389174976931 | 0.17674039980813 | -3.16302915429108 |
| ENSG00000187535.13 | IFT140 | 0.430299386723977 | 2.05845928390049 | 2.52327730163986 | 0.0154511383259425 | 0.17678197896769 | -3.19282104147127 |
| ENSG00000162065.13 | TBC1D24 | -0.401538875739193 | 2.24821862975786 | -2.52285283776857 | 0.0154673122845634 | 0.176868715973982 | -3.22163452027404 |
| lnc-ZNF665-1 | lnc-ZNF665-1 | -0.554453322870566 | 1.21682080523198 | -2.52219745690685 | 0.0154923155070984 | 0.177056263232986 | -3.11801479274702 |
| ENSG00000039650.11 | PNKP | -0.345636648944235 | 4.80792965099173 | -2.52126695543518 | 0.0155278780310136 | 0.177364213935823 | -3.43116384424932 |
| ENSG00000280046.1 | | 0.587232097947182 | 1.57021709159481 | 2.52066468311846 | 0.0155509357081153 | 0.177461660013028 | -3.16885575397774 |
| lnc-STAT3-1 | lnc-STAT3-1 | -0.81380497679912 | 0.888339679483623 | -2.52030299962249 | 0.0155647975562922 | 0.177461660013028 | -3.09587679720692 |
| ENSG00000225217.1 | HSPA7 | 1.32113719526337 | 2.66767569814046 | 2.52322625961045 | 0.0155723155601785 | 0.177461660013028 | -3.26293619125335 |
| lnc-KRBA2-2 | lnc-KRBA2-2 | -0.461995527776296 | 1.86206887272343 | -2.52002381345945 | 0.0155755053166296 | 0.177461660013028 | -3.19003159595574 |
| lnc-PLEKHM1-4 | lnc-PLEKHM1-4 | -1.79280949737925 | -0.0300125843012198 | -2.5220591118765 | 0.0155875528763139 | 0.177461660013028 | -3.01271701083154 |
| ENSG00000155368.16 | DBI | -0.685306924914166 | 5.89676454489412 | -2.52086018202086 | 0.0155945739935742 | 0.177461660013028 | -3.46087805290898 |
| ENSG00000207112.1 | SNORA25 | 0.790821469356558 | 3.06558139157573 | 2.52011311517407 | 0.0155967615490243 | 0.177461660013028 | -3.27193970193621 |
| lnc-RPEL1-1 | lnc-RPEL1-1 | -0.516126009564185 | 1.38026507578763 | -2.51896072477575 | 0.0156163399213758 | 0.17758625668601 | -3.14461474511197 |
| ENSG00000100567.12 | PSMA3 | 0.234776271913049 | 4.56458052884574 | 2.51840592823274 | 0.015637689076265 | 0.177599882398032 | -3.42378942364443 |
| ENSG00000089486.16 | CDIP1 | -0.425286171284647 | 4.63878337484339 | -2.51826632933826 | 0.0156430651734665 | 0.177599882398032 | -3.42721565809475 |
| ENSG00000137074.18 | APTX | 0.196837845835975 | 3.85625272698155 | 2.51812114532622 | 0.0156486581451972 | 0.177599882398032 | -3.36285548764628 |
| lnc-CNPY1-10 | lnc-CNPY1-10 | -0.692432434493736 | 1.69382868886574 | -2.51790945046976 | 0.0156568166005352 | 0.177599882398032 | -3.17906854293446 |
| lnc-C9orf57-4 | lnc-C9orf57-4 | 0.358824795942142 | 4.39973051331933 | 2.51780922564408 | 0.0156606804913 | 0.177599882398032 | -3.40991962090477 |
| ENSG00000244734.4 | HBB | 3.47548220529227 | -0.168673484763371 | 2.52042486945457 | 0.0156771835314746 | 0.177617478099817 | -3.06333993858749 |
| lnc-MAP4K3-3 | lnc-MAP4K3-3 | 0.450308916270844 | 0.0942197603009897 | 2.51690115477999 | 0.015695728254789 | 0.177617478099817 | -3.02167712236037 |
| ENSG00000102053.12 | ZC3H12B | -1.54518611259403 | 3.15596919732334 | -2.52040968814056 | 0.015708747116161 | 0.177617478099817 | -3.29906901591071 |
| SNHG17 | SNHG17 | 0.356945740945643 | 3.87810921758136 | 2.51650894499158 | 0.0157108880029373 | 0.177617478099817 | -3.36955624944829 |
| CKMT2-AS1 | CKMT2-AS1 | -0.355454029702091 | 2.72278576940683 | -2.51647613362849 | 0.0157121568366656 | 0.177617478099817 | -3.2657310174498 |
| lnc-ARL8A-1 | lnc-ARL8A-1 | -0.455320336809272 | 2.93513722185463 | -2.51637758598355 | 0.0157159682902397 | 0.177617478099817 | -3.2947082376215 |
| ENSG00000167842.15 | MIS12 | 0.29598852614936 | 4.83800877302469 | 2.51612179904837 | 0.0157258650999451 | 0.177617478099817 | -3.44242295836511 |
| ENSG00000263934.4 | SNORD3A | 1.40246472628104 | 8.57333727749245 | 2.51950359280162 | 0.0157505628417969 | 0.177617478099817 | -3.4487468867434 |
| ENSG00000159459.11 | UBR1 | 0.200257690615643 | 6.28286011738739 | 2.51533712465194 | 0.0157562608441896 | 0.177617478099817 | -3.47797230159252 |
| ENSG00000174327.6 | SLC16A13 | -0.745771190068406 | 0.080059872933752 | -2.51518206860194 | 0.0157622735405307 | 0.177617478099817 | -3.03120475447788 |
| lnc-PRICKLE2-5 | lnc-PRICKLE2-5 | 0.225340253203598 | 2.75822885591068 | 2.51506874166557 | 0.0157666694050031 | 0.177617478099817 | -3.27975894084259 |
| ENSG00000196656.7 | | 1.95541675881405 | -0.537945541437218 | 2.51530268004253 | 0.015770367085363 | 0.177617478099817 | -3.05513404808213 |
| lnc-EPN2-3 | lnc-EPN2-3 | 1.402472638861 | 8.57240065934933 | 2.51860404031556 | 0.0157853161065105 | 0.177617478099817 | -3.45070720980683 |
| lnc-IGSF3-2 | lnc-IGSF3-2 | -0.507961590599094 | 1.95792531859236 | -2.51452700163231 | 0.0157876985116786 | 0.177617478099817 | -3.20668912054351 |
| lnc-EPHB6-4 | lnc-EPHB6-4 | -0.522412203234588 | 0.835469227463955 | -2.5143830612014 | 0.0157932902425289 | 0.177617478099817 | -3.10044925047337 |
| ENSG00000102349.17 | KLF8 | -0.661947339198168 | 6.11618044513478 | -2.51516355305054 | 0.0158066536771457 | 0.177617478099817 | -3.47678216044472 |

| | | | | | | | |
|---|---|---|---|---|---|---|---|
| LINC02362 | LINC02362 | -0.761837724335605 | 3.13843080463509 | -2.51467666058505 | 0.0158129483297069 | 0.177617478099817 | -3.32695450832042 |
| ENSG00000219023.1 | RPS15AP19 | 1.44398507636898 | 0.200874468871855 | 2.51375885486739 | 0.0158175599940225 | 0.177617478099817 | -3.00948353200919 |
| lnc-FER-2 | lnc-FER-2 | -0.707165060661621 | 3.2260693883603 | -2.51328366279963 | 0.0158493679428381 | 0.177877611977883 | -3.32490256305368 |
| ENSG00000138658.15 | ZGRF1 | 0.406687531376839 | 3.93166776936717 | 2.51242300536456 | 0.0158696132724665 | 0.178007765660588 | -3.37899408798825 |
| lnc-TLK1-1 | lnc-TLK1-1 | -0.613783864779766 | 0.989093998738818 | -2.51216206348657 | 0.0158797994351851 | 0.17802500641308 | -3.11767110773028 |
| lnc-PXYLP1-3 | lnc-PXYLP1-3 | -0.497378824875427 | 2.03243138422647 | -2.51172854919815 | 0.0158967353187526 | 0.178117856867656 | -3.22427663004477 |
| lnc-FAM103A1-2 | lnc-FAM103A1-2 | -0.579597181258594 | 0.36716365802864 | -2.51126764307362 | 0.015914759340095 | 0.178208301354595 | -3.06199459202363 |
| ENSG00000213523.10 | SRA1 | 0.251092704481043 | 3.99884334987219 | 2.51099440819 | 0.0159254531427 | 0.178208301354595 | -3.39346322497766 |
| ITGB1-DT | ITGB1-DT | -0.927530845689876 | 0.218991997973636 | -2.51073550478302 | 0.0159355920790231 | 0.178208301354595 | -3.05428187334677 |
| ZKSCAN2-DT | ZKSCAN2-DT | -0.525121554924496 | 3.66156542376799 | -2.51055018450204 | 0.0159428530290618 | 0.178208301354595 | -3.37034529281356 |
| lnc-C19orf25-2 | lnc-C19orf25-2 | -0.509698694733675 | 0.68748230327115 | -2.51027985164143 | 0.0159534502198191 | 0.178208301354595 | -3.10805822168057 |
| ENSG00000099991.17 | CABIN1 | -0.240262254792967 | 6.24710559880808 | -2.51019556668469 | 0.0159567555456696 | 0.178208301354595 | -3.48881446542034 |
| ENSG00000153140.8 | CETN3 | -0.470170710271202 | 2.1482146841605 | -2.50959371513631 | 0.0159803759312682 | 0.178272319730426 | -3.24340693308107 |
| ENSG00000171311.12 | EXOSC1 | 0.177157294785331 | 4.19865655723535 | 2.50934423684497 | 0.0159901763337578 | 0.178272319730426 | -3.4133020451037 |
| ENSG00000204623.9 | POLR1HASP | -0.430555824936445 | 3.26878830894247 | -2.50930655962155 | 0.0159916569057318 | 0.178272319730426 | -3.33408870900718 |
| lnc-TSPAN3-4 | lnc-TSPAN3-4 | -0.683071600638319 | 0.265077475898949 | -2.50911431756483 | 0.0159992132323413 | 0.178272319730426 | -3.05461278755288 |
| lnc-ABCA5-111 | lnc-ABCA5-111 | -0.899013846192952 | -0.356417171326987 | -2.50894697586103 | 0.0160057934636267 | 0.178272319730426 | -3.02089450763282 |
| lnc-STOML3-4 | lnc-STOML3-4 | 0.398102675106575 | 0.986515204367366 | 2.50865922616027 | 0.0160171141533998 | 0.178301925700069 | -3.12250420049181 |
| ENSG00000161939.19 | RNASEK-C17orf49 | -0.301556440255739 | 4.15732140938877 | -2.50799885179382 | 0.0160431222570662 | 0.178433642671571 | -3.42392583254757 |
| ENSG00000120913.23 | PDLIM2 | -0.51697747086812 | 3.33980636463251 | -2.50791862699506 | 0.0160462844378894 | 0.178433642671571 | -3.35032472709449 |
| ENSG00000162419.12 | GMEB1 | 0.201038450475778 | 4.70680946486295 | 2.50681378171947 | 0.0160898913146427 | 0.178821940026615 | -3.45466501396077 |
| ENSG00000136153.19 | LMO7 | -0.645050677877149 | 6.40702119881442 | -2.50695484984701 | 0.0161233125926153 | 0.178983842345234 | -3.49593780798842 |
| ENSG00000243335.9 | KCTD7 | -0.307455307328227 | 5.13223595947109 | -2.50580965574537 | 0.0161296164490236 | 0.178983842345234 | -3.47594757738055 |
| ENSG00000156026.14 | MCU | -0.22081413752751 | 3.81776409751246 | -2.50575167030562 | 0.0161319131870765 | 0.178983842345234 | -3.39380644161066 |
| ENSG00000268279.4 | | -0.231721705133388 | 3.03208171895069 | -2.50556669916992 | 0.0161392416748168 | 0.178983842345234 | -3.32607056494072 |
| ENSG00000104164.10 | BLOC1S6 | 0.221441655240305 | 5.64921717094867 | 2.5053378862118 | 0.0161483113511658 | 0.178987987367284 | -3.49401233011317 |
| lnc-ZNF484-3 | lnc-ZNF484-3 | 0.415817969677867 | 1.93409550602319 | 2.5050917389393 | 0.0161580733025359 | 0.178999796978523 | -3.22133369459776 |
| ENSG00000185716.11 | MOSMO | -0.36529712715848 | 3.56092509411788 | -2.50477259575181 | 0.016170738192933 | 0.17904373546269 | -3.3752882981954 |
| ENSG00000137409.19 | MTCH1 | 0.232122806409305 | 6.67805220629619 | 2.5041980924125 | 0.0161935595741118 | 0.179076016519098 | -3.50113250839761 |
| ENSG00000118620.12 | ZNF430 | 0.222070945226092 | 5.60624457719682 | 2.504011034616 | 0.0162009965184877 | 0.179076016519098 | -3.49455637575784 |
| lnc-ESCO1-7 | lnc-ESCO1-7 | 0.210555883096853 | 2.86801348908392 | 2.50383627897041 | 0.0162079471682727 | 0.179076016519098 | -3.30960788755286 |
| lnc-DHRSX-1 | lnc-DHRSX-1 | -0.422266243517955 | 3.63682243027137 | -2.5037877773253 | 0.0162098767311287 | 0.179076016519098 | -3.37770503730246 |
| lnc-SNRNP200-1 | lnc-SNRNP200-1 | -0.437117934025519 | 1.81334582571732 | -2.50343929168074 | 0.0162237468402393 | 0.179076016519098 | -3.21872697315506 |
| NRSN2-AS1 | NRSN2-AS1 | -0.257965231248782 | 4.05601577249115 | -2.5033863512713 | 0.0162258548709186 | 0.179076016519098 | -3.41855925881245 |
| ENSG00000172716.16 | SLFN11 | 1.69370660044556 | 3.25011714753112 | 2.50676493196325 | 0.0162422764956672 | 0.179161188162014 | -3.27628497741792 |
| ENSG00000083750.12 | RRAGB | -0.195858220733108 | 4.42839047844097 | -2.50266192131878 | 0.0162547259484313 | 0.179202476805871 | -3.44781818536209 |
| lnc-TMEM71-3 | lnc-TMEM71-3 | 1.02816678947297 | 1.87668265621024 | 2.50334667404654 | 0.0162717121842661 | 0.179293710861214 | -3.19020578113406 |
| ENSG00000155380.11 | SLC16A1 | 0.368028601216266 | 3.50679871482987 | 2.50183869694128 | 0.0162875910597152 | 0.179372652103862 | -3.37815786120325 |
| lnc-TEX35-6 | lnc-TEX35-6 | -0.797994142077485 | 0.189510333600842 | -2.50155883672279 | 0.0162987775193472 | 0.179399859722312 | -3.05887021327634 |
| ENSG00000137075.17 | RNF38 | 0.246213682155791 | 6.19867612471428 | 2.50066515498307 | 0.0163345462178585 | 0.179501197217074 | -3.50964534982888 |
| ENSG00000198742.9 | SMURF1 | 0.414340814653679 | 5.25557175009025 | 2.5005194424729 | 0.0163403849781775 | 0.179501197217074 | -3.49207137341065 |
| lnc-FAM173B-1 | lnc-FAM173B-1 | -0.825921925699433 | -0.748501153114431 | -2.50009763828054 | 0.0163572975501102 | 0.179501197217074 | -3.04243999413142 |
| ENSG00000114125.13 | RNF7 | 0.240846309893861 | 4.60701240857924 | 2.49989490400388 | 0.0163654320054487 | 0.179501197217074 | -3.46564541290396 |
| lnc-ZNF404-1 | lnc-ZNF404-1 | -0.794244137976819 | 0.353746778175309 | -2.49975186248109 | 0.0163711735787181 | 0.179501197217074 | -3.08510832013443 |
| ENSG00000162385.10 | MAGOH | 0.256255801765433 | 3.63722607350767 | 2.49962741652168 | 0.016376170231244 | 0.179501197217074 | -3.38454506895396 |
| ENSG00000110628.14 | SLC22A18 | -0.602478084341285 | 1.32133263002528 | -2.49951192077347 | 0.0163808087632908 | 0.179501197217074 | -3.19130350513159 |
| ENSG00000126522.16 | ASL | -0.327490867714618 | 3.71906403630994 | -2.49927462629547 | 0.0163903427176145 | 0.179501197217074 | -3.40621429183061 |
| ENSG00000230409.3 | TCEA1P2 | 0.86660676606489 | 1.76550796546114 | 2.49916640778876 | 0.0164027242597226 | 0.179501197217074 | -3.19575326685123 |
| ENSG00000204152.10 | TIMM23B | 0.266810325461032 | 3.72157489864704 | 2.49851280627915 | 0.0164209850421145 | 0.179501197217074 | -3.39501612074989 |
| ENSG00000006015.17 | REX1BD | -0.522379495296168 | 3.4765682653519 | -2.49847705208397 | 0.0164224244453949 | 0.179501197217074 | -3.38507505055171 |
| ENSG00000158161.15 | EYA3 | -0.279446862981878 | 4.88569919486137 | -2.49798809777496 | 0.0164421204358584 | 0.179501197217074 | -3.48362763839549 |
| lnc-RASA3-8 | lnc-RASA3-8 | -0.554739490280407 | 0.181626717743617 | -2.49784630049805 | 0.0164478363116803 | 0.179501197217074 | -3.0809708824017 |
| ENSG00000244026.6 | FAM86DP | -0.354106684814201 | 1.77352464126985 | -2.49726921932169 | 0.0164711172146505 | 0.179501197217074 | -3.23655784037391 |
| ENSG00000160219.11 | GAB3 | 0.426200429644917 | 3.42293535591714 | 2.497216767919 | 0.016473234720926 | 0.179501197217074 | -3.37651993261558 |
| ENSG00000143303.11 | METTL25B | -0.252255881772809 | 4.83803900553368 | -2.49714640609979 | 0.0164760756742667 | 0.179501197217074 | -3.48356490500174 |
| ENSG00000198105.13 | ZNF248 | -0.362220949545065 | 3.92609496603788 | -2.49680743536079 | 0.0164897683199501 | 0.179501197217074 | -3.41750407297429 |
| lnc-WDCP-8 | lnc-WDCP-8 | 0.391475217381749 | 0.732716347104477 | 2.4962834715487 | 0.0165109540811289 | 0.179501197217074 | -3.12931596747621 |
| lnc-PLPPR5-4 | lnc-PLPPR5-4 | -1.75735111354769 | -1.30746291056181 | -2.49624526877832 | 0.0165124997257278 | 0.179501197217074 | -3.09728461882925 |
| lnc-AKAP8-4 | lnc-AKAP8-4 | -0.517808632504575 | 2.56720058700972 | -2.49608967260906 | 0.0165187963453186 | 0.179501197217074 | -3.31369339065558 |
| ENSG00000113643.8 | RARS1 | 0.23470425617454 | 5.50000412786598 | 2.49599046301792 | 0.0165228122694577 | 0.179501197217074 | -3.5108034982725 |
| ENSG00000104228.12 | TRIM35 | -0.380229944921581 | 3.58636804064403 | -2.4959036421145 | 0.0165263274379551 | 0.179501197217074 | -3.39856764508705 |
| lnc-SMARCC2-9 | lnc-SMARCC2-9 | -0.581641400140174 | 1.56802294653055 | -2.49583890781752 | 0.0165289488156952 | 0.179501197217074 | -3.21297318461152 |
| ENSG00000164615.4 | CAMLG | 0.291615335092232 | 4.10349550986188 | 2.49583291718028 | 0.0165291914221828 | 0.179501197217074 | -3.43760362714638 |
| ENSG00000117308.14 | GALE | -0.389719784409954 | 1.31980732429671 | -2.49575217277619 | 0.0165324616934968 | 0.179501197217074 | -3.2032641936803 |
| ENSG00000121797.9 | CCRL2 | 1.28176759511297 | -0.528666777001607 | 2.49569626898941 | 0.0165347262266712 | 0.179501197217074 | -3.04582476928724 |
| ENSG00000214078.12 | CPNE1 | -0.216394449592024 | 6.34809539639365 | -2.49518578062683 | 0.0165554179817267 | 0.179631085038418 | -3.52134116944175 |
| lnc-CLK1-3 | lnc-CLK1-3 | 0.460799754179121 | 4.37053019331577 | 2.49485518488581 | 0.0165688306619883 | 0.179678321791716 | -3.45381996734176 |
| ENSG00000178502.5 | KLHL11 | 0.266030985525801 | 4.21095655501617 | 2.4946482723284 | 0.0165772303883044 | 0.179678321791716 | -3.4477370000344 |

| | | | | | | | |
|---|---|---|---|---|---|---|---|
| ENSG00000198677.11 | SKIC3 | 0.175457874681128 | 6.62743625887359 | 2.49441079067622 | 0.0165868758556091 | 0.179688245124211 | -3.52285621088434 |
| ENSG00000198939.7 | ZFP2 | -0.472569828482944 | 0.661541622186265 | -2.49329845903076 | 0.016632121869168 | 0.180077069655063 | -3.13036429255591 |
| ENSG00000138413.13 | IDH1 | -0.504877694210609 | 2.79601877509708 | -2.49309854553743 | 0.0166402655824675 | 0.180077069655063 | -3.34486802450102 |
| ENSG00000229638.1 | RPL4P4 | 1.16339871439399 | 3.25979283145077 | 2.49539923993838 | 0.0166576488695503 | 0.180109095465473 | -3.31887574598392 |
| ENSG00000091947.9 | TMEM101 | -0.359361805262728 | 3.11712733844355 | -2.49259667788448 | 0.0166607257331906 | 0.180109095465473 | -3.36287938250954 |
| lnc-ICA1-1 | lnc-ICA1-1 | -0.634474426321748 | 1.15101473279926 | -2.49236405158399 | 0.0166702172022749 | 0.180117102716234 | -3.1895409820929 |
| lnc-DAZL-6 | lnc-DAZL-6 | -0.691353423032166 | -0.414978058098734 | -2.49200007379654 | 0.0166850778537169 | 0.180183083663722 | -3.04831495345727 |
| ENSG00000189057.10 | FAM111B | 1.24579880459864 | 6.41624654949665 | 2.49548625113091 | 0.016695694628723 | 0.180203189587313 | -3.52266244837586 |
| ENSG00000277985.1 | SNORA67 | 0.3439325828878 | 1.83929727973438 | 2.49134320839717 | 0.0167119271364569 | 0.180283855476778 | -3.24718721380579 |
| ENSG00000011198.9 | ABHD5 | 0.41051132059762 | 2.75284684162116 | 2.49028821130381 | 0.0167551322041087 | 0.180626729009334 | -3.32482777511606 |
| ENSG00000004975.11 | DVL2 | -0.289600392409416 | 4.75709476666828 | -2.49013873602297 | 0.016761261837819 | 0.180626729009334 | -3.4986835319473 |
| lnc-MARCH10-8 | lnc-MARCH10-8 | -0.42134465974701 | 4.63627115007797 | -2.48983690810363 | 0.0167736453127372 | 0.180665589467331 | -3.48962228921059 |
| ENSG00000163126.14 | ANKRD23 | -0.25652842480239 | 2.38774083125842 | -2.48940245849097 | 0.0167914846252465 | 0.180712243494033 | -3.29883802791485 |
| ENSG00000187912.11 | CLEC17A | -0.549332505249296 | 6.69446048800829 | -2.48949408828584 | 0.0167955362096918 | 0.180712243494033 | -3.53259470707951 |
| ENSG00000155229.20 | MMS19 | -0.231520423215605 | 6.19349982417239 | -2.48787019782215 | 0.0168545398556949 | 0.180831168440769 | -3.53688407672086 |
| ENSG00000197013.9 | ZNF429 | -0.386745699391104 | 3.32605643818323 | -2.48785897794727 | 0.0168550023659282 | 0.180831168440769 | -3.384288384968 |
| LINC02100 | LINC02100 | -0.643759096943192 | 4.17902150543468 | -2.48811073557778 | 0.0168644500595898 | 0.180831168440769 | -3.4670859277738 |
| ENSG00000176700.20 | SCAND2P | -0.26803441863552 | 4.53134658230609 | -2.48747919865777 | 0.0168706645863802 | 0.180831168440769 | -3.4853404384674 |
| ENSG00000011021.22 | CLCN6 | -0.232075016713947 | 5.71193150852993 | -2.48737303194902 | 0.0168750453005697 | 0.180831168440769 | -3.53321659644096 |
| lnc-BCOR-11 | lnc-BCOR-11 | -1.898655726989 | 2.13557155814523 | -2.49096716595353 | 0.0168870768178116 | 0.180831168440769 | -3.2585862111908 |
| lnc-RIOK2-9 | lnc-RIOK2-9 | 0.529733129028507 | 1.79710579317773 | 2.48680038734483 | 0.01689869193476 | 0.180831168440769 | -3.2540976674142 |
| lnc-ZNF132-1 | lnc-ZNF132-1 | -0.43187785035104 | 1.63473587505376 | -2.486725971717 | 0.0169017670436769 | 0.180831168440769 | -3.23413638289202 |
| ENSG00000204186.8 | ZDBF2 | 1.38515788474208 | 3.86386609267839 | 2.49032114175212 | 0.0169071604682917 | 0.180831168440769 | -3.42563097496923 |
| lnc-CYB561D2-2 | lnc-CYB561D2-2 | -0.388839057470618 | 0.524284053615668 | -2.4865054533766 | 0.0169108826040455 | 0.180831168440769 | -3.13615688334578 |
| ENSG00000218565.2 | | -0.705608051486513 | 2.6466452615961 | -2.48648117727002 | 0.0169120147329583 | 0.180831168440769 | -3.32048110087479 |
| lnc-HECA-4 | lnc-HECA-4 | -0.705608051486513 | 2.6466452615961 | -2.48648117727002 | 0.0169120147329583 | 0.180831168440769 | -3.32048110087479 |
| ENSG00000138744.14 | NAAA | -0.274392519823195 | 4.23030881017015 | -2.48621644570914 | 0.0169228360699273 | 0.180852925663195 | -3.46806867785556 |
| ENSG00000130775.15 | THEMIS2 | 0.759075207423603 | 7.0216621612856 | 2.48766204217287 | 0.0169488939085929 | 0.181037407016382 | -3.53749293371159 |
| ENSG00000107372.12 | ZFAND5 | 0.365242418619865 | 6.14179422073363 | 2.48537474152121 | 0.016957693026577 | 0.181037445833006 | -3.54239129379558 |
| ENSG00000284024.2 | MSANTD7 | 0.228349749097609 | 4.33545942135879 | 2.48442383959644 | 0.0169971505908433 | 0.181188587914693 | -3.48127148788305 |
| ENSG00000243279.3 | PRAF2 | -0.518493688397051 | 0.80364282002165 | -2.4844237648737 | 0.0169971536947294 | 0.181188587914693 | -3.17726141778469 |
| lnc-FBXL12-3 | lnc-FBXL12-3 | -0.424293654832206 | 1.51071921647568 | -2.48432802211684 | 0.0170011311486264 | 0.181188587914693 | -3.22832324255612 |
| lnc-USP12-9 | lnc-USP12-9 | -0.666761463309754 | 2.59642889180166 | -2.48413446138524 | 0.0170091748505189 | 0.181188587914693 | -3.3168434128028 |
| ENSG00000140992.18 | PDPK1 | 0.174023425464759 | 5.5600922985957 | 2.48397354573608 | 0.017015864569747 | 0.181188587914693 | -3.53838600817748 |
| ENSG00000157350.12 | ST3GAL2 | -0.501158811535341 | 4.88611115882075 | -2.48343144419467 | 0.0170384188607216 | 0.181221431046039 | -3.5177369529892 |
| ENSG00000139531.12 | SUOX | -0.772555686244466 | 2.15104501128718 | -2.48365727829468 | 0.0170406151866247 | 0.181221431046039 | -3.30772045788575 |
| ENSG00000198420.9 | TCAF1 | -0.504416243834796 | 2.96920284195431 | -2.48326468905613 | 0.0170453622166415 | 0.181221431046039 | -3.36021010836881 |
| ENSG00000025293.16 | PHF20 | 0.225344023637154 | 5.64231617339991 | 2.48304635085073 | 0.0170544572762425 | 0.181224519420186 | -3.54128711993321 |
| ENSG00000112200.16 | ZNF451 | 0.285707336223029 | 7.27073971610262 | 2.48254784744498 | 0.0170752393765292 | 0.181347188941342 | -3.54288066152045 |
| lnc-DENND5B-3 | lnc-DENND5B-3 | 0.4450562345598 | 3.43422213380576 | 2.48234691916026 | 0.017083622375614 | 0.181347188941342 | -3.39852523914691 |
| lnc-RWDD4-2 | lnc-RWDD4-2 | -0.560774765944245 | -0.266936353163038 | -2.4812663976293 | 0.017128767336822 | 0.181666146267913 | -3.07284745651139 |
| lnc-PEX19-2 | lnc-PEX19-2 | 0.33094603949148 | 2.23117848019423 | 2.48120534300421 | 0.0171313214743244 | 0.181666146267913 | -3.29823919058062 |
| ENSG00000198736.11 | MSRB1 | -0.674662607411771 | 1.70487548516046 | -2.4805696594484 | 0.0171579349936567 | 0.181854673519277 | -3.26949078637285 |
| ENSG00000055950.16 | MRPL43 | -0.247067206786548 | 3.04181622164719 | -2.48003439061629 | 0.0171803736686669 | 0.181917641834042 | -3.38410261110534 |
| lnc-INPP5F-2 | lnc-INPP5F-2 | -0.856620629324532 | 1.15507705664254 | -2.47982391834984 | 0.017189204044913 | 0.181917641834042 | -3.19687333027509 |
| ENSG00000213380.15 | COG8 | -0.337171856277709 | 4.65173069225806 | -2.47962677429621 | 0.0171974789738884 | 0.181917641834042 | -3.51322967579748 |
| ENSG00000240021.9 | TEX35 | -0.574963789928121 | 0.578084848059069 | -2.47950043957126 | 0.0172027836524484 | 0.181917641834042 | -3.14861456688212 |
| lnc-CDK19-7 | lnc-CDK19-7 | -0.638798598877488 | 0.586172993026734 | -2.47932143326965 | 0.0172103025065484 | 0.181917641834042 | -3.15154051909438 |
| ENSG00000236552.2 | RPL13AP5 | 1.2227971723178 | 4.94636406698718 | 2.48211877692718 | 0.0172409726646525 | 0.181917641834042 | -3.47751480231012 |
| ENSG00000242110.7 | AMACR | -0.309330583672945 | 1.32101980191159 | -2.47858630306142 | 0.0172412117024028 | 0.181917641834042 | -3.23380296265169 |
| ENSG00000202408.2 | RNVU1-21 | -1.15069135303313 | 0.452355852425018 | -2.47902488899941 | 0.0172422427193649 | 0.181917641834042 | -3.12979914915459 |
| lnc-TEC-4 | lnc-TEC-4 | 0.371507388378227 | 1.25783124686519 | 2.47841402556829 | 0.0172484625487585 | 0.181917641834042 | -3.22093647362052 |
| lnc-KBTBD2-2 | lnc-KBTBD2-2 | -0.461198445192057 | 1.9637561389519 | -2.47807286064243 | 0.0172628297326304 | 0.181917641834042 | -3.28794023994665 |
| ENSG00000103145.10 | HCFC1R1 | -0.780288155818024 | -0.339783995982279 | -2.47779868701756 | 0.0172743836449491 | 0.181917641834042 | -3.07849868848792 |
| ENSG00000170085.17 | SIMC1 | 0.419819091334579 | 5.19213580859075 | 2.47724676521762 | 0.0172976634030326 | 0.181917641834042 | -3.5374316328971 |
| lnc-ABCG5-5 | lnc-ABCG5-5 | 0.282257524755759 | 2.80066198189426 | 2.47707731407426 | 0.0173048164729384 | 0.181917641834042 | -3.35861618444025 |
| lnc-BBOX1-11 | lnc-BBOX1-11 | -0.592879253776837 | -0.475039488880237 | -2.47701323941287 | 0.0173075219672885 | 0.181917641834042 | -3.07849997691914 |
| ENSG00000101216.10 | GMEB2 | 0.325314500034494 | 5.05834622120189 | 2.47700565803468 | 0.0173078421095031 | 0.181917641834042 | -3.53642889144413 |
| ENSG00000102471.14 | NDFIP2 | -0.962228606417749 | 2.33028827482415 | -2.47838916881355 | 0.0173193559089325 | 0.181917641834042 | -3.33497120588394 |
| ENSG00000181035.13 | SLC25A42 | -0.431716322325768 | 3.65232551229454 | -2.47667831592436 | 0.0173216700591785 | 0.181917641834042 | -3.43789667226535 |
| ENSG00000256646.7 | | 0.382137867871795 | 4.04950520535856 | 2.47664768577861 | 0.0173229644849984 | 0.181917641834042 | -3.47164724865609 |
| ENSG00000085840.12 | ORC1 | 0.589039838880794 | 2.15961625993814 | 2.47615093948132 | 0.017343969182647 | 0.182021377560609 | -3.28962265632274 |
| ENSG00000135720.12 | DYNC1LI2 | 0.245860124219234 | 6.62831238556733 | 2.47599591134268 | 0.0173505292121612 | 0.182021377560609 | -3.56232346585077 |
| ENSG00000137261.13 | KIAA0319 | -0.625561212223833 | -0.271727204315578 | -2.47548444820704 | 0.0173721878003602 | 0.182132079501985 | -3.09571138544121 |
| lnc-BCAT1-4 | lnc-BCAT1-4 | -0.810674695199891 | 0.291720403141279 | -2.47525413923167 | 0.0173819485539391 | 0.182132079501985 | -3.1290949032897 |
| lnc-RNF186-1 | lnc-RNF186-1 | -0.491872467268802 | 1.82552661662114 | -2.4750416230693 | 0.0173909596448449 | 0.182132079501985 | -3.27489908603993 |
| VIM-AS1 | VIM-AS1 | -0.317018797292604 | 5.29871487889423 | -2.47491157233973 | 0.0173964761356898 | 0.182132079501985 | -3.55151178833855 |
| ZNF426-DT | ZNF426-DT | -0.768521367160331 | -0.11320843945418 | -2.47445323900131 | 0.0174159303835545 | 0.182218272632544 | -3.09149711363439 |

| | | | | | | | |
|---|---|---|---|---|---|---|---|
| ENSG00000164062.12 | APEH | -0.258439140867332 | 5.35714407441803 | -2.47430057610367 | 0.0174224146402783 | 0.182218272632544 | -3.55416448473097 |
| ENSG00000166037.10 | CEP57 | 0.373315234047507 | 6.01311471804472 | 2.47395258181668 | 0.0174372036630113 | 0.182280326559554 | -3.56584369466594 |
| ENSG00000180304.14 | OAZ2 | 0.299013639230298 | 4.35548559685927 | 2.473535153145 | 0.0174549585343435 | 0.182344159492706 | -3.49844575083639 |
| lnc-ZNF708-14 | lnc-ZNF708-14 | -2.0194890165328 | 2.80311018259598 | -2.47739220924339 | 0.0174610279531712 | 0.182344159492706 | -3.40270996464086 |
| ENSG00000151065.13 | DCP1B | -0.385240894303353 | 3.0983051613671 | -2.47233993554229 | 0.0175058866116789 | 0.182685860521002 | -3.40283094292196 |
| ENSG00000285816.1 | | -0.292909113963722 | 2.52217845749643 | -2.4722083915234 | 0.0175114999177932 | 0.182685860521002 | -3.35236431806932 |
| ENSG00000165244.6 | ZNF367 | 0.584954829387412 | 2.07661130605564 | 2.4717620885433 | 0.017530556938303 | 0.182700534323768 | -3.30704561595236 |
| ENSG00000233762.3 | RPS15P4 | 1.10755823183714 | 1.51951588076165 | 2.4729406483681 | 0.017530659053075 | 0.182700534323768 | -3.21524070206284 |
| LINC01480 | LINC01480 | -0.656999610247222 | 5.51323796303412 | -2.47240785874509 | 0.0175488867468928 | 0.182797943274946 | -3.56360528820152 |
| ENSG00000232346.1 | RPS17P16 | 0.871642903013322 | 1.52482396857269 | 2.47107203628761 | 0.0175611509745656 | 0.182833166671463 | -3.22561029573215 |
| lnc-FAM109A-5 | lnc-FAM109A-5 | -0.606659293111889 | 0.38003378872399 | -2.46999086873181 | 0.0176063732941382 | 0.183029126064352 | -3.15726822716167 |
| ENSG00000117010.16 | ZNF684 | 0.301810441199005 | 1.78072444487249 | 2.46985822048274 | 0.0176120632095505 | 0.183029126064352 | -3.28100049598989 |
| TMEM161B-AS1 | TMEM161B-AS1 | -0.300535063742477 | 6.45484369521853 | -2.46942405447335 | 0.0176306983301682 | 0.183029126064352 | -3.57655226053346 |
| ENSG00000131051.22 | RBM39 | 0.204361450478213 | 8.94249616489917 | 2.46874364930626 | 0.0176599384380015 | 0.183029126064352 | -3.54230745049615 |
| ENSG00000163214.20 | DHX57 | 0.317574255051705 | 5.16600897748561 | 2.46832173714711 | 0.0176780920217829 | 0.183029126064352 | -3.55817003554588 |
| ENSG00000128604.19 | IRF5 | -0.452540429520585 | 5.12636649215802 | -2.4683191436098 | 0.0176782036660714 | 0.183029126064352 | -3.56127081392312 |
| ENSG00000135999.11 | EPC2 | 0.197525139766468 | 6.27697058685044 | 2.46814254726758 | 0.0176858071335538 | 0.183029126064352 | -3.5798724059578 |
| ENSG00000088808.17 | PPP1R13B | -0.442645315355035 | 3.05564426931148 | -2.46802923926721 | 0.017690687244063 | 0.183029126064352 | -3.40917216824983 |
| ENSG00000131378.13 | RFTN1 | -1.19079251374206 | 4.0009778562747 | -2.47121221691409 | 0.0176930851395694 | 0.183029126064352 | -3.46871538320966 |
| ENSG00000234287.1 | RPS27P4 | 0.774733261874594 | 2.58824179219681 | 2.46810326398213 | 0.017699407473208 | 0.183029126064352 | -3.33545056311067 |
| lnc-FBXO25-2 | lnc-FBXO25-2 | -0.385624619882737 | 1.05850913615523 | -2.4677790668703 | 0.0177014663529982 | 0.183029126064352 | -3.23339780811427 |
| ENSG00000234911.2 | TEX21P | -0.390084377882629 | 0.807778804386223 | -2.46768367596094 | 0.0177055780039343 | 0.183029126064352 | -3.19686876930352 |
| lnc-HJURP-9 | lnc-HJURP-9 | -0.615385732182521 | 0.333240003818822 | -2.46767866603164 | 0.0177057939717375 | 0.183029126064352 | -3.15546717534826 |
| ENSG00000136044.11 | APPL2 | 0.277200636199511 | 5.56879348256519 | 2.46740310239059 | 0.017717676638766 | 0.183029126064352 | -3.5732709529363 |
| ENSG00000168569.7 | TMEM223 | -0.532458699483427 | 1.4309735523319 | -2.46730245045979 | 0.0177220186864119 | 0.183029126064352 | -3.26986312021218 |
| ENSG00000277370.1 | SNORD49A | 0.525022348803446 | 0.69124221907591 | 2.46729711553147 | 0.017722248858099 | 0.183029126064352 | -3.17239951207976 |
| lnc-ZNF621-5 | lnc-ZNF621-5 | 0.774434623199899 | 2.58827295650452 | 2.46715431150051 | 0.0177403325216837 | 0.183124004159386 | -3.33741840321047 |
| ENSG00000106635.7 | BCL7B | 0.293757843784536 | 4.46967032618778 | 2.46626379881595 | 0.0177668817488549 | 0.183287178419816 | -3.527907227056 |
| ENSG00000134686.18 | PHC2 | 0.303066626410593 | 4.17914768582985 | 2.46610037983549 | 0.0177739497704879 | 0.183287178419816 | -3.50280322830913 |
| ENSG00000215271.7 | NA | -0.407271520135485 | 2.93200135547676 | -2.46488645044261 | 0.0178265332457448 | 0.183737372958019 | -3.40180705644002 |
| lnc-VAT1-1 | lnc-VAT1-1 | -0.529013573882253 | 1.79793849563383 | -2.46444469428549 | 0.0178457036827872 | 0.183842902353758 | -3.30933644566471 |
| lnc-ASF1A-7 | lnc-ASF1A-7 | -0.893319883837043 | 0.0140845679702374 | -2.46353317889169 | 0.0178853188954243 | 0.184047335634391 | -3.12625019222102 |
| ENSG00000164164.16 | OTUD4 | 0.220943710065563 | 6.02885284195182 | 2.46339117260773 | 0.0178914977827479 | 0.184047335634391 | -3.58902352992128 |
| ENSG00000284310.1 | MIR939 | -0.468516768366044 | 0.422948988084638 | -2.46337105577926 | 0.0178923732499838 | 0.184047335634391 | -3.17128407225055 |
| lnc-PLEC-3 | lnc-PLEC-3 | -0.493415303721501 | 2.16879169720623 | -2.46269650996808 | 0.0179217514146668 | 0.184257447236807 | -3.34183823164456 |
| ENSG00000170049.9 | KCNAB3 | -0.651904384800525 | 1.14551004934074 | -2.46210410892951 | 0.0179475880455294 | 0.184430955936661 | -3.24733073476564 |
| ENSG00000163873.9 | GRIK3 | 3.68698516944851 | 2.50646712062253 | 2.46576711406111 | 0.0179601393009854 | 0.184467837940211 | -3.21242656633534 |
| lnc-VKORC1L1-7 | lnc-VKORC1L1-7 | 0.411570454001065 | 1.42618564454838 | 2.46123893158865 | 0.0179853820812322 | 0.184479974890192 | -3.2688212896825 |
| ENSG00000240489.1 | SETP14 | 0.967902081006701 | 1.1302430013557 | 2.46132149374443 | 0.0179861984194688 | 0.184479974890192 | -3.19934561585649 |
| ENSG00000276141.4 | WHAMMP3 | -0.948116222112084 | 1.27309091580896 | -2.46186782362988 | 0.0179882091825591 | 0.184479974890192 | -3.27368698344038 |
| ENSG00000163882.9 | POLR2H | -0.24308608808778 | 3.35918773088723 | -2.46041155476085 | 0.018021592358409 | 0.184661065250848 | -3.4515691081959 |
| ENSG00000255432.1 | | -0.34568800197964 | 2.18370940010853 | -2.46022630739276 | 0.0180297087916592 | 0.184661065250848 | -3.35051019116048 |
| ENSG00000185669.5 | SNAI3 | -1.05662821417954 | -0.477469266404838 | -2.4601561969384 | 0.0180327814776371 | 0.184661065250848 | -3.11290160195563 |
| ENSG00000196152.10 | ZNF79 | 0.246166020301477 | 3.38087307096988 | 2.45895095440735 | 0.0180856771714514 | 0.185105604478881 | -3.44460706820037 |
| lnc-UBP1-1 | lnc-UBP1-1 | 0.225276669383638 | 1.7125975120147 | 2.45875754741684 | 0.0180941785070936 | 0.185105604478881 | -3.30190470057968 |
| lnc-GAST-2 | lnc-GAST-2 | 0.462930220354248 | 5.56432153927969 | 2.45852800173195 | 0.0181042730458239 | 0.18511686641937 | -3.59103872233313 |
| ENSG00000164010.14 | ERMAP | -0.326796166305422 | 2.89894311632691 | -2.45808395972143 | 0.0181238148045575 | 0.18514297953186 | -3.4127833469521 |
| ENSG00000047346.12 | ATOSA | 0.438873116387755 | 6.44734020923686 | 2.45718307220894 | 0.0181635205621777 | 0.18514297953186 | -3.60317838870727 |
| ENSG00000283795.1 | MIR4426 | 1.07187016547514 | 1.09948063153449 | 2.45722929419145 | 0.0181751429239238 | 0.18514297953186 | -3.18562057089849 |
| ENSG00000124486.12 | USP9X | 0.294890867681641 | 7.83980199929057 | 2.45658766442207 | 0.0181898058614639 | 0.18514297953186 | -3.59012084169692 |
| lnc-ADGRD2-7 | lnc-ADGRD2-7 | -0.587306389318439 | -0.138458762527381 | -2.45632891597952 | 0.0182012395017353 | 0.18514297953186 | -3.12762114957563 |
| ENSG00000006607.13 | FARP2 | 0.284261609936164 | 6.50691154128045 | 2.45627298498914 | 0.0182037118504386 | 0.18514297953186 | -3.60507647316108 |
| ENSG00000111215.12 | PRR4 | -0.540131099043467 | 0.116783115530786 | -2.45626284789079 | 0.0182041599788295 | 0.18514297953186 | -3.16226474779273 |
| lnc-MBP-13 | lnc-MBP-13 | -0.459660830825511 | 1.61770865727139 | -2.45626003544941 | 0.018204284309555 | 0.18514297953186 | -3.30105712413415 |
| lnc-ZNF311-2 | lnc-ZNF311-2 | -0.377259165080716 | 1.88180112132166 | -2.4561632212936 | 0.0182085646812133 | 0.18514297953186 | -3.33338794330647 |
| ENSG00000133704.9 | IPO8 | 0.227711691801574 | 5.65366583161864 | 2.4560540498282 | 0.0182133924918411 | 0.18514297953186 | -3.60035656115025 |
| ENSG00000204842.15 | ATXN2 | -0.370668010021878 | 4.87243323559398 | -2.45592807261437 | 0.0182189649330019 | 0.18514297953186 | -3.57401615419945 |
| ENSG00000101464.10 | PIGU | -0.32427974834165 | 2.75341628096621 | -2.45572257610606 | 0.01822805812559 | 0.18514297953186 | -3.4088143993645 |
| ENSG00000205643.10 | CDPF1 | -0.565100608721949 | 1.70959894394487 | -2.45569160466984 | 0.0182294289641468 | 0.18514297953186 | -3.31615592335272 |
| ENSG00000143612.20 | C1orf43 | 0.2822220971895 | 5.75437175043576 | 2.45561644250009 | 0.0182327561342409 | 0.18514297953186 | -3.60211902396545 |
| ENSG00000176142.12 | TMEM39A | 0.322679158294391 | 4.76584474964096 | 2.4549879280713 | 0.0182605998564755 | 0.185300835964257 | -3.56810824352753 |
| ENSG00000198673.10 | TAFA2 | -0.412348093894934 | 1.31413927655976 | -2.4547567108892 | 0.0182708526679199 | 0.185300835964257 | -3.27391640702038 |
| ENSG00000175324.9 | LSM1 | 0.325003130586783 | 3.4350812883997 | 2.45450442370742 | 0.0182820457345184 | 0.185300835964257 | -3.4659833082647 |
| ENSG00000168036.17 | CTNNB1 | 0.225540163556888 | 7.83662836326449 | 2.45437051886259 | 0.0182879891311414 | 0.185300835964257 | -3.59474277794761 |
| ENSG00000197980.12 | LEKR1 | -0.475735927787778 | 2.07807203929112 | -2.4542253651039 | 0.018294433791541 | 0.185300835964257 | -3.35854922667207 |
| ENSG00000162909.17 | CAPN2 | 0.544319281842765 | 6.27535187025018 | 2.45424097856358 | 0.0183023174228895 | 0.185300835964257 | -3.60914804135949 |
| ENSG00000143443.9 | C1orf56 | -0.593551530571462 | 1.64971277946436 | -2.45380306292624 | 0.01831319523222 | 0.185319811929589 | -3.30692120932744 |
| ADORA2A-AS1 | ADORA2A-AS1 | -0.7011271657061 | -0.317390874296186 | -2.45307941601257 | 0.0183453849379398 | 0.185554328342808 | -3.12105308409129 |

| | | | | | | | |
|---|---|---|---|---|---|---|---|
| ENSG00000166012.16 | TAF1D | 0.437139613873671 | 6.72342913439596 | 2.45188933130862 | 0.0183984345459252 | 0.185999498162464 | -3.61382207397793 |
| lnc-TBX19-2 | lnc-TBX19-2 | 0.511113501570397 | 3.89704399173957 | 2.4514708033996 | 0.0184171240018352 | 0.186038292666162 | -3.50500919654781 |
| ENSG00000038002.8 | AGA | -0.365009217422464 | 3.64098007671467 | -2.45139862565822 | 0.0184203488535995 | 0.186038292666162 | -3.49198166389006 |
| lnc-LENG9-5 | lnc-LENG9-5 | -0.554978090710937 | 0.141352028295087 | -2.45081797138858 | 0.0184463107092301 | 0.186149009966627 | -3.16234230943148 |
| lnc-TMEM230-5 | lnc-TMEM230-5 | -0.547247104692513 | 1.18095073917172 | -2.45067093510551 | 0.0184528901640178 | 0.186149009966627 | -3.26611371516512 |
| ENSG00000085274.15 | MYNN | 0.233908978340895 | 5.32685372816369 | 2.45054688162378 | 0.0184584428577898 | 0.186149009966627 | -3.60371524854224 |
| lnc-NPIPA8-8 | lnc-NPIPA8-8 | 0.948123364118455 | 0.0865259433603556 | 2.45018244042012 | 0.0184747641808427 | 0.186152397101242 | -3.13791528190087 |
| ENSG00000179859.9 | RNF227 | -0.467674418742227 | 1.29333028711401 | -2.45013551418583 | 0.0184768666995986 | 0.186152397101242 | -3.29201780948169 |
| ENSG00000118816.9 | CCNI | 0.233224664392584 | 7.19268585382138 | 2.44921206953167 | 0.0185182855555894 | 0.186340421305838 | -3.61507123634522 |
| ENSG00000244682.7 | FCGR2C | 1.16018101062273 | 4.26181455244885 | 2.45237087481262 | 0.0185248842413601 | 0.186340421305838 | -3.54825581055292 |
| ENSG00000263917.1 | | 0.454851454977954 | 1.69333828210744 | 2.44889693670241 | 0.0185324393107235 | 0.186340421305838 | -3.30520253560737 |
| ENSG00000025156.12 | HSF2 | 0.475753467709799 | 3.79252349642392 | 2.44879832526768 | 0.0185368703199059 | 0.186340421305838 | -3.51316633257943 |
| lnc-MPLKIP-1 | lnc-MPLKIP-1 | -0.493421952167308 | 1.1081450186952 | -2.44871100064702 | 0.0185407949683893 | 0.186340421305838 | -3.27373606932611 |
| ENSG00000153561.12 | RMND5A | 0.330631142256793 | 6.59193281812049 | 2.44798870782641 | 0.0185732860015828 | 0.186575864212093 | -3.62231138297437 |
| ENSG00000207442.1 | SNORD116-6 | -0.706871849116683 | 0.894962174735626 | -2.44776309040016 | 0.0185834455604646 | 0.186583700276692 | -3.23761093898842 |
| ENSG00000167257.10 | RNF214 | -0.318595062913872 | 3.95655347154401 | -2.44755411910066 | 0.0185928600346919 | 0.186583700276692 | -3.53076691983066 |
| lnc-ADAM17-2 | lnc-ADAM17-2 | 0.285440156451775 | 3.30345225925293 | 2.44736772775252 | 0.0186012608933475 | 0.186583700276692 | -3.46600741616522 |
| lnc-ANKRD30B-13 | lnc-ANKRD30B-13 | -1.63608700487459 | -0.750988948961199 | -2.44704287935877 | 0.0186293856383297 | 0.186593756896704 | -3.13415445781808 |
| ENSG00000147576.16 | ADHFE1 | -1.12325020262229 | 2.27887362489024 | -2.44887145124629 | 0.0186354025740965 | 0.186593756896704 | -3.36700937294941 |
| ENSG00000254522.1 | | 0.872975087303165 | 0.220162823392868 | 2.44656202318826 | 0.0186376144377698 | 0.186593756896704 | -3.14523612224683 |
| ENSG00000169047.5 | IRS1 | -0.729492826861857 | 4.19866280521051 | -2.44766750460688 | 0.018648445707597 | 0.186593756896704 | -3.55803476694897 |
| NRAV | NRAV | -0.431910188602766 | 1.87662052706293 | -2.44629945632731 | 0.0186494754021518 | 0.186593756896704 | -3.34537937967055 |
| lnc-OPA3-2 | lnc-OPA3-2 | -0.278109190940306 | 1.70463597770485 | -2.44614057117533 | 0.018656656060507 | 0.186593756896704 | -3.3392447574513 |
| ENSG00000138668.18 | HNRNPD | 0.34677746604597 | 6.75442072801377 | 2.44584783615559 | 0.0186698924903672 | 0.186635452709678 | -3.6261101619737 |
| ENSG00000116771.5 | AGMAT | -0.989861099098569 | 2.63176114515348 | -2.44716640753741 | 0.0186987110702119 | 0.186832800950569 | -3.40550357258173 |
| ENSG00000138029.13 | HADHB | 0.321234602025201 | 6.11244103647261 | 2.44384205843649 | 0.0187608156346552 | 0.187276867128233 | -3.63104710026239 |
| ENSG00000124784.8 | RIOK1 | 0.310875468001 | 4.1778755399614 | 2.44383026105073 | 0.0187613516017304 | 0.187276867128233 | -3.54886139434887 |
| ENSG00000168288.12 | MMADHC | 0.294615286134848 | 5.41927310352817 | 2.44242563724002 | 0.0188252642745076 | 0.187823758876486 | -3.62289124410059 |
| lnc-CHD1L-5 | lnc-CHD1L-5 | -0.340701631121311 | 2.55275232782546 | -2.44205237932713 | 0.0188422812442966 | 0.187902458745813 | -3.42236810860234 |
| FLG-AS1 | FLG-AS1 | -2.65047338106217 | -1.86432325596833 | -2.44097600877929 | 0.0189071334231307 | 0.188253876616128 | -3.35149981261097 |
| lnc-ANXA3-8 | lnc-ANXA3-8 | -0.622116371048603 | 3.34255816378649 | -2.44018215636259 | 0.0189296336730203 | 0.188253876616128 | -3.50423175088993 |
| ENSG00000075539.14 | FRYL | 0.252316437160669 | 7.60169699099639 | 2.43984040871135 | 0.0189434119961964 | 0.188253876616128 | -3.62986240295722 |
| ENSG00000115350.11 | POLE4 | 0.292033959538367 | 3.72983489861261 | 2.43981862057023 | 0.0189444105842331 | 0.188253876616128 | -3.51761717706589 |
| ENSG00000188785.11 | ZNF548 | 0.333458500506143 | 5.19150269808685 | 2.43960951145148 | 0.0189539968392422 | 0.188253876616128 | -3.61971470051131 |
| lnc-IRF2BP2-9 | lnc-IRF2BP2-9 | 0.351158110241422 | 2.43209917314651 | 2.43953605442813 | 0.0189573653948615 | 0.188253876616128 | -3.4080684807991 |
| ENSG00000126217.20 | MCF2L | -0.450584640297146 | 3.20474800618171 | -2.43940910711721 | 0.0189631881592313 | 0.188253876616128 | -3.48916566965107 |
| ENSG00000263266.2 | RPS7P1 | 1.04955338767741 | 2.88736565979913 | 2.44153374766428 | 0.0189646664916234 | 0.188253876616128 | -3.4084692334423 |
| ENSG00000099338.22 | CATSPERG | -0.438482603013773 | 2.52258513459503 | -2.43923772417251 | 0.0189710516465776 | 0.188253876616128 | -3.41727671334415 |
| ENSG00000169967.16 | MAP3K2 | 0.302953202924219 | 7.4047679519991 | 2.43917446582445 | 0.0189739548470303 | 0.188253876616128 | -3.63424865534835 |
| lnc-C6orf106-2 | lnc-C6orf106-2 | -0.702347602588498 | 1.35109768858423 | -2.43908356710533 | 0.0189781272884645 | 0.188253876616128 | -3.31669344288811 |
| ENSG00000158805.11 | ZNF276 | -0.255604038276997 | 5.25449427594831 | -2.43850501900509 | 0.0190047033313753 | 0.188426690110644 | -3.62821966802879 |
| ENSG00000240038.6 | AMY2B | -0.288993743854278 | 5.82406007238636 | -2.43824389106314 | 0.0190167094714323 | 0.18845494995209 | -3.64069836870586 |
| FAM182A | FAM182A | -0.634358019791849 | 0.0760224827844613 | -2.43803191675214 | 0.0190264606747007 | 0.188460847000656 | -3.18408336975902 |
| lnc-CXorf40B-2 | lnc-CXorf40B-2 | -0.395169767838125 | 0.00056155427526403 | -2.43731023259906 | 0.0190596934038688 | 0.188561246871067 | -3.17892678345697 |
| ENSG00000139428.11 | MMAB | -0.298563887524118 | 3.07295752420918 | -2.43719186040047 | 0.0190651493213387 | 0.188561246871067 | -3.47670328318572 |
| ENSG00000036054.12 | TBC1D23 | 0.232379404313587 | 5.07774714868936 | 2.43704519144105 | 0.0190719114316649 | 0.188561246871067 | -3.623139339133 |
| ENSG00000167645.16 | YIF1B | -0.468573345557846 | 1.91999610371876 | -2.43694194327079 | 0.0190766729468598 | 0.188561246871067 | -3.37348428328052 |
| lnc-ZNF805-1 | lnc-ZNF805-1 | 0.361654222730304 | 1.6035594768801 | 2.43681774932835 | 0.0190824018477594 | 0.188561246871067 | -3.33370817829532 |
| ENSG00000069956.11 | MAPK6 | 0.365999544129155 | 5.1760024714782 | 2.43619255218743 | 0.0191112650163628 | 0.188613470142812 | -3.62582049962641 |
| ENSG00000004961.14 | HCCS | 0.250113255617574 | 2.94576935895612 | 2.43614505510837 | 0.0191134594048453 | 0.188613470142812 | -3.45637843591576 |
| ENSG00000128989.10 | ARPP19 | 0.294685314992053 | 7.58879258621469 | 2.43610787049598 | 0.0191151775114369 | 0.188613470142812 | -3.63727568203187 |
| lnc-LGALS7-1 | lnc-LGALS7-1 | -0.663657340253924 | 1.11190950290828 | -2.4353239190708 | 0.0191514323639392 | 0.188880657569219 | -3.29719467694759 |
| lnc-DPP6-2 | lnc-DPP6-2 | -0.459709122658262 | 0.138111032315838 | -2.43489750572373 | 0.0191711785245403 | 0.188907323852375 | -3.19612495695481 |
| lnc-ZNF117-5 | lnc-ZNF117-5 | -0.245936385574213 | 1.83432270598586 | -2.43485210643365 | 0.0191732819397175 | 0.188907323852375 | -3.36043755835504 |
| ENSG00000186765.11 | FSCN2 | -0.713333522332359 | 0.10153049372861 | -2.43453956319953 | 0.0191877681918103 | 0.188907323852375 | -3.18195977953376 |
| ENSG00000177173.5 | NAP1L4P1 | -0.523908997852893 | 1.35760106599405 | -2.43447315406214 | 0.0191908475040236 | 0.188907323852375 | -3.30716196384401 |
| ENSG00000100138.14 | SNU13 | 0.257181972343865 | 5.41068651964079 | 2.4337006811333 | 0.0192266990201005 | 0.18899701619064 | -3.64068970788119 |
| ENSG00000213326.4 | RPS7P11 | 1.20216739025064 | 1.15356552741794 | 2.43460194640504 | 0.0192327697790796 | 0.18899701619064 | -3.23643059738555 |
| ENSG00000230629.2 | RPS23P8 | 1.1156017233429 | 2.29357513594125 | 2.43553005385331 | 0.0192365204966638 | 0.18899701619064 | -3.36300513905794 |
| ENSG00000234745.10 | HLA-B | 0.295970906650417 | 10.1219566423176 | 2.43348568645076 | 0.019236687991031 | 0.18899701619064 | -3.58700463171397 |
| ENSG00000072310.16 | SREBF1 | 0.506410743787066 | 3.96822811237473 | 2.43324969044847 | 0.0192476581208731 | 0.189014573999013 | -3.55682823214207 |
| ENSG00000100319.12 | ZMAT5 | -0.406703965200728 | 2.63360238923126 | -2.43253926979156 | 0.0192807157697069 | 0.189248914014248 | -3.44421865854914 |
| ZEB1-AS1 | ZEB1-AS1 | -0.325157968328376 | 2.60663643927341 | -2.43215893850358 | 0.0192984346413741 | 0.189332545387704 | -3.44545344563743 |
| ENSG00000100629.16 | CEP128 | 0.65386894043306 | 2.62997648012678 | 2.43132365267519 | 0.01933740065452 | 0.189624448628864 | -3.42104421997178 |
| ENSG00000198000.11 | NOL8 | 0.248128027983041 | 5.82447125758044 | 2.43063506699415 | 0.0193695766322895 | 0.189849521820198 | -3.65617271019236 |
| ENSG00000197258.5 | EIF4BP6 | 0.975671508007294 | 2.32668691937766 | 2.43161458904542 | 0.0193858390515095 | 0.189918479389443 | -3.3797307426483 |
| ENSG00000114021.11 | NIT2 | -0.228411671189715 | 5.43926228073644 | -2.42989226462256 | 0.0194043403284893 | 0.190009294472548 | -3.65112706144249 |
| ENSG00000157181.15 | ODR4 | 0.265104171989512 | 4.71514900608242 | 2.4295608296863 | 0.0194198699301917 | 0.190070938075671 | -3.61898889318214 |

| | | | | | | | |
|---|---|---|---|---|---|---|---|
| lnc-C1QTNF6-1 | lnc-C1QTNF6-1 | -0.38485622083087 | 2.05771497904136 | -2.42934158921815 | 0.0194301487639975 | 0.190081155897985 | -3.39053847327323 |
| ENSG00000265794.5 | | 0.148913824001967 | 4.77915764812806 | 2.42893236640405 | 0.0194493478661996 | 0.190158782930145 | -3.62576842227043 |
| lnc-C10orf67-1 | lnc-C10orf67-1 | -0.647782785695569 | -0.173714421132943 | -2.42877871330839 | 0.0194565610868623 | 0.190158782930145 | -3.18440667557153 |
| ENSG00000211877.1 | TRAJ12 | -1.12702553260078 | 0.244312557376212 | -2.42819595993876 | 0.0194850861350056 | 0.190239019538095 | -3.19691808635795 |
| ENSG00000243710.7 | CFAP57 | -0.737293983977348 | 0.104106147321346 | -2.42810354266094 | 0.0194882855550355 | 0.190239019538095 | -3.19130829975597 |
| ENSG00000151612.15 | ZNF827 | 0.405767662941895 | 5.03289218311081 | 2.42792023708984 | 0.0194969066671568 | 0.190239019538095 | -3.63683993974417 |
| ENSG00000155287.10 | SLC25A28 | 0.371902815953947 | 5.07895158080026 | 2.42781748353126 | 0.019501740816469 | 0.190239019538095 | -3.64278384901094 |
| ENSG00000283676.1 | MIR5087 | -1.18583351053455 | 0.686820519708421 | -2.42775540300569 | 0.0195529186000694 | 0.190560786596809 | -3.25557581589761 |
| ENSG00000180957.17 | PITPNB | 0.247118073463427 | 4.18254703985599 | 2.42672421284423 | 0.0195532420586144 | 0.190560786596809 | -3.58931658417056 |
| lnc-SDHAF4-4 | lnc-SDHAF4-4 | 0.750896210912019 | -0.508585202263896 | 2.42647265626729 | 0.0195651096531209 | 0.190566171537556 | -3.16630402489911 |
| ENSG00000151576.10 | QTRT2 | 0.19215467367184 | 5.54141049585819 | 2.4260342660731 | 0.0195858069996418 | 0.190566171537556 | -3.66148631327431 |
| ENSG00000171533.11 | MAP6 | -2.20763386259143 | -0.262138663956733 | -2.42834138331354 | 0.0195967448197988 | 0.190566171537556 | -3.16822560046299 |
| ENSG00000285796.1 | | -0.511947707632751 | 2.80110885771761 | -2.42530316474628 | 0.0196203679208655 | 0.190566171537556 | -3.47332344793721 |
| ENSG00000167207.13 | NOD2 | -0.886018934746446 | 4.81373736210972 | -2.42779125625486 | 0.0196213507137123 | 0.190566171537556 | -3.64020269511027 |
| ENSG00000064726.9 | BTBD1 | 0.232657040295488 | 5.79664968344189 | 2.42511327831347 | 0.0196293533277434 | 0.190566171537556 | -3.66832708730526 |
| ENSG00000100426.6 | ZBED4 | 0.295177915504079 | 4.30132541627938 | 2.42499499878956 | 0.0196349521836178 | 0.190566171537556 | -3.60231792503092 |
| ENSG00000279453.1 | | -0.488524972153374 | 1.8804145891121 | -2.42494920132597 | 0.0196371204309943 | 0.190566171537556 | -3.39412778263593 |
| lnc-SERINC1-3 | lnc-SERINC1-3 | -0.488524972153374 | 1.8804145891121 | -2.42494920132597 | 0.0196371204309943 | 0.190566171537556 | -3.39412778263593 |
| LINC00847 | LINC00847 | -0.334691711764019 | 4.19609100666323 | -2.42472009403132 | 0.0196479705993478 | 0.190581611143438 | -3.59126620922511 |
| ENSG00000134146.11 | DPH6 | -0.348433357651207 | 3.52256660788684 | -2.42425625957059 | 0.0196699536838189 | 0.190682448365576 | -3.53831380406723 |
| ENSG00000165782.10 | PIP4P1 | 0.354791365291083 | 5.84038496311083 | 2.42407484531764 | 0.0196785577242727 | 0.190682448365576 | -3.67012943678056 |
| ENSG00000171492.14 | LRRC8D | -0.347527017508469 | 5.09000919996774 | -2.42375609258434 | 0.0196936836408056 | 0.190682448365576 | -3.65403473737899 |
| lnc-ZFP90-4 | lnc-ZFP90-4 | -0.648337601580406 | 0.195700733381228 | -2.42353946236698 | 0.0197039694911078 | 0.190682448365576 | -3.22133915601376 |
| lnc-AKAP10-2 | lnc-AKAP10-2 | -0.558200065547771 | 1.41198950794297 | -2.42352435969157 | 0.0197046867644941 | 0.190682448365576 | -3.34630395421084 |
| lnc-TUT1-1 | lnc-TUT1-1 | -0.315334463149258 | 0.734719523277074 | -2.42319163213994 | 0.0197204950293841 | 0.19073945851181 | -3.28555144741857 |
| ENSG00000161091.12 | MFSD12 | -0.348921508965657 | 4.30570577659264 | -2.42301037154069 | 0.0197291117510394 | 0.19073945851181 | -3.61148477118779 |
| lnc-CREG1-1 | lnc-CREG1-1 | 0.706928420154091 | -0.536984673930307 | 2.42215634212185 | 0.0197697562124682 | 0.191042672357387 | -3.17333353968913 |
| ENSG00000111186.12 | WNT5B | -1.32741553843145 | -1.02128338284671 | -2.4216666060915 | 0.019793097576657 | 0.191166590691618 | -3.17670099179319 |
| ENSG00000166224.16 | SGPL1 | 0.220571685712351 | 5.25014623863345 | 2.42149767280997 | 0.0198011549023238 | 0.191166590691618 | -3.66334975194652 |
| ENSG00000186812.12 | ZNF397 | -0.224591194135684 | 5.02221282419711 | -2.42080918833669 | 0.0198340230609996 | 0.191299092327739 | -3.65371698110577 |
| ENSG00000124688.13 | MAD2L1BP | 0.311844501546918 | 3.54103038769311 | 2.42067005908077 | 0.0198406710608245 | 0.191299092327739 | -3.54159792395872 |
| ENSG00000270898.5 | NA | 0.294578601222642 | 3.77064223534125 | 2.42059668085574 | 0.0198441780967619 | 0.191299092327739 | -3.56872476732235 |
| ENSG00000233558.1 | | 1.18120675327968 | -0.86367444299621 | 2.42026698425203 | 0.019859942519358 | 0.191299092327739 | -3.23034413610845 |
| ENSG00000100614.17 | PPM1A | 0.269671942947837 | 6.18768876502224 | 2.42023756660529 | 0.0198613496722722 | 0.191299092327739 | -3.68166918803057 |
| ENSG00000146677.7 | RPL32P18 | 1.42668235901583 | 1.96343659355809 | 2.42180303159453 | 0.0199096571062689 | 0.191585583272559 | -3.33298871653787 |
| ENSG00000106105.13 | GARS1 | 0.273401885532911 | 5.47817124217224 | 2.41912362203833 | 0.0199147000528143 | 0.191585583272559 | -3.67566780320029 |
| ENO1-AS1 | ENO1-AS1 | -0.588392283076097 | -0.268434656375496 | -2.41903358153734 | 0.0199190180344593 | 0.191585583272559 | -3.18925051729941 |
| ENSG00000107968.9 | MAP3K8 | 0.909576410745793 | 3.91425068080558 | 2.42059211263925 | 0.0199512927359668 | 0.191652523802486 | -3.57549410342945 |
| ENSG00000278619.4 | MRM1 | -0.490877224548146 | 1.71301478881496 | -2.41769931914314 | 0.0199831031678143 | 0.191652523802486 | -3.39317450983994 |
| lnc-FAM46C-2 | lnc-FAM46C-2 | 0.289405189621606 | 2.49388805233557 | 2.4176592379045 | 0.0199850311647009 | 0.191652523802486 | -3.45631975552242 |
| ENSG00000151062.14 | CACNA2D4 | -0.394333077550987 | 1.34281641980938 | -2.41764350804061 | 0.0199857878521357 | 0.191652523802486 | -3.35515379457998 |
| ENSG00000123352.17 | SPATS2 | 0.340074270315546 | 3.33433152521564 | 2.41761118284793 | 0.0199873429416034 | 0.191652523802486 | -3.52966097983887 |
| ENSG00000144724.19 | PTPRG | 1.01041113901728 | 4.42829594263782 | 2.42026631358075 | 0.0199876702978239 | 0.191652523802486 | -3.59608425946892 |
| ENSG00000095066.11 | HOOK2 | -0.376994884519902 | 2.21204821705752 | -2.41753192682886 | 0.019991156226203 | 0.191652523802486 | -3.43824953942494 |
| DHRS4-AS1 | DHRS4-AS1 | -0.529292371338855 | 1.97694278654523 | -2.41718350393621 | 0.0200079278714416 | 0.191724012745755 | -3.415057363196 |
| ENSG00000137571.10 | SLCO5A1 | -2.62965146478349 | -0.892936946399116 | -2.41909593305808 | 0.0200438862765451 | 0.19197920485348 | -3.18574155658042 |
| ENSG00000144645.13 | OSBPL10 | 0.553878758002689 | 7.35321006511039 | 2.41622074422602 | 0.0200686343131756 | 0.192010280576591 | -3.68407505963278 |
| ENSG00000108100.17 | CCNY | 0.207196647699016 | 6.03991317126579 | 2.41591286246722 | 0.020069199123321 | 0.192010280576591 | -3.68993060475547 |
| ENSG00000198836.9 | OPA1 | 0.206716405246734 | 6.77340406325266 | 2.41566911627888 | 0.0200809721050678 | 0.192010280576591 | -3.68994845944849 |
| ENSG00000068654.15 | POLR1A | 0.218817494687102 | 5.2881198511027 | 2.41559723594492 | 0.0200844451285722 | 0.192010280576591 | -3.67660922906414 |
| ENSG00000198763.3 | MT-ND2 | -0.742280385957481 | 6.99704956652658 | -2.41702179266482 | 0.0201090037419363 | 0.192155814308391 | -3.68532818246746 |
| ENSG00000272980.4 | | 0.687102355637311 | 3.03335064460983 | 2.41454144406936 | 0.020145728885959 | 0.192331369370783 | -3.50604503787823 |
| ENSG00000214226.8 | C17orf67 | -0.460794504445105 | 1.60615038096952 | -2.41401465400829 | 0.0201610479824643 | 0.192331369370783 | -3.38925210826771 |
| ENSG00000226084.5 | RPL17P6 | 0.858390330836707 | 1.35456593953668 | 2.41400857590545 | 0.0201616391578799 | 0.192331369370783 | -3.33235888218991 |
| ENSG00000183943.5 | PRKX | 0.375708660201997 | 5.04467552719511 | 2.41393826369107 | 0.0201647522276709 | 0.192331369370783 | -3.67422144649091 |
| ENSG00000165338.16 | HECTD2 | -0.377097483784594 | 4.86844129991728 | -2.41367257857849 | 0.020177640361437 | 0.192365155886734 | -3.6685119519708 |
| ENSG00000114503.10 | NCBP2 | 0.219063212484681 | 5.80704075225112 | 2.41169470758789 | 0.0202738191682568 | 0.193066084799468 | -3.69595507749286 |
| ENSG00000196652.11 | ZKSCAN5 | 0.240786026254502 | 4.63745852962922 | 2.41168044314737 | 0.020274514313086 | 0.193066084799468 | -3.65003234879093 |
| lnc-SOWAHB-5 | lnc-SOWAHB-5 | -0.522371655150771 | 0.694579022356982 | -2.41135940852103 | 0.020290164896248 | 0.193066084799468 | -3.3158577850125 |
| lnc-FSCN2-1 | lnc-FSCN2-1 | -0.499911346220543 | 2.27092512422601 | -2.41064366413046 | 0.0203250970524577 | 0.193066084799468 | -3.46117891621151 |
| lnc-KLF8-1 | lnc-KLF8-1 | -0.5794931457253 | 1.59567425772143 | -2.41023350738295 | 0.0203451393711429 | 0.193066084799468 | -3.3774069471686 |
| ENSG00000284773.1 | | -0.337721094822409 | 2.31337247992918 | -2.41010615356337 | 0.0203513661491968 | 0.193066084799468 | -3.46221760552596 |
| ENSG00000112855.15 | HARS2 | 0.177983198463097 | 5.15997446873754 | 2.41002358205035 | 0.0203554042818508 | 0.193066084799468 | -3.68315721577785 |
| ENSG00000264801.1 | ERVFRD-3 | -0.939925629346726 | -0.0022047302170277 | -2.41001220337389 | 0.0203559608090068 | 0.193066084799468 | -3.2250067346593 |
| ENSG00000143653.9 | SCCPDH | -0.580412751609829 | 3.64065325388475 | -2.40995129299107 | 0.0203589401485896 | 0.193066084799468 | -3.58041853860109 |
| lnc-MRNIP-1 | lnc-MRNIP-1 | -0.534677236890452 | 4.97693574852564 | -2.40960568017639 | 0.0203772568625312 | 0.193066084799468 | -3.67520832873668 |
| ENSG00000166387.12 | PPFIBP2 | 0.910801232855378 | 4.74994146916176 | 2.41204472517113 | 0.020377952069249 | 0.193066084799468 | -3.64809225515875 |
| ENSG00000181074.4 | OR52N4 | 0.528130675369499 | 1.80103421703519 | 2.40956261371156 | 0.0203779610866846 | 0.193066084799468 | -3.41036650587204 |

| | | | | | | | |
|---|---|---|---|---|---|---|---|
| lnc-TMEM106C-6 | lnc-TMEM106C-6 | -0.937105883389985 | 0.121638322640145 | -2.40945022565217 | 0.020383464052307 | 0.193066084799468 | -3.22811035936041 |
| ENSG00000267022.1 | | -0.569111674578573 | 1.81460693985326 | -2.40940802793297 | 0.0203855305667215 | 0.193066084799468 | -3.40587329691624 |
| lnc-MELTF-2 | lnc-MELTF-2 | 0.322034176896246 | 1.2862603945264 | 2.40927879262712 | 0.0203918606789119 | 0.193066084799468 | -3.36450326087196 |
| ENSG00000125531.6 | FNDC11 | -0.608211036828435 | 1.68982036090951 | -2.40880743125111 | 0.020414963615813 | 0.193195952231852 | -3.41096638892097 |
| ENSG00000185475.10 | TMEM179B | -0.361574815238418 | 3.59585494539521 | -2.40779009601725 | 0.020464907070496 | 0.193579587422803 | -3.58431621951475 |
| ENSG00000101193.7 | GID8 | 0.214278599089443 | 5.39139101704402 | 2.40755594651247 | 0.0204764176460202 | 0.193599496742321 | -3.69642510757416 |
| lnc-HECA-11 | lnc-HECA-11 | -0.740225167368906 | 2.08502075066952 | -2.40715770203178 | 0.0204971807397286 | 0.193706827899832 | -3.43639178290269 |
| ENSG00000157014.10 | TATDN2 | 0.279926473679924 | 5.03608500330586 | 2.40643947947547 | 0.0205313825803848 | 0.193765337685828 | -3.68264223452034 |
| ENSG00000246705.4 | H2AJ | -0.540192838907065 | 3.44990492123872 | -2.40622316918355 | 0.0205420471737064 | 0.193765337685828 | -3.59472685185762 |
| ENSG00000278949.1 | | 0.532829760528817 | 2.0634653992399 | 2.4062127290481 | 0.0205425620228245 | 0.193765337685828 | -3.42991517750855 |
| ENSG00000148835.10 | TAF5 | 0.27379203292764 | 2.3766548808282 | 2.40611783150417 | 0.0205472423734856 | 0.193765337685828 | -3.47048939215689 |
| ENSG00000042980.12 | ADAM28 | -0.416411016828389 | 8.71773545363738 | -2.40587423512806 | 0.0205592609646152 | 0.193765337685828 | -3.67999488538159 |
| lnc-HJURP-12 | lnc-HJURP-12 | 0.604361255403562 | 3.89880307489509 | 2.40601826631985 | 0.0205598550991521 | 0.193765337685828 | -3.60427636329476 |
| ENSG00000277986.1 | | 0.461134568789174 | 1.38321836866851 | 2.4053619186035 | 0.0205845584212464 | 0.193901610371897 | -3.38480525748294 |
| ENSG00000207221.1 | SNORA70E | 0.921530591155403 | -0.502196896330821 | 2.40518794194059 | 0.0205931555299503 | 0.193901610371897 | -3.20784879852448 |
| ENSG00000180011.6 | PTGR3 | -0.378661967040576 | 4.54290933772443 | -2.40483383901794 | 0.0206106636457262 | 0.193953569070416 | -3.66092104467337 |
| ENSG00000131507.10 | NDFIP1 | -0.230433634636145 | 5.8688662122901 | -2.40447749171411 | 0.0206282962971092 | 0.193953569070416 | -3.71260765242611 |
| ENSG00000103248.18 | MTHFSD | -0.233212975033373 | 3.57641468121297 | -2.40430172871205 | 0.0206369983539533 | 0.193953569070416 | -3.5808985262094 |
| ENSG00000106683.14 | LIMK1 | -0.380726243578917 | 4.37826972572466 | -2.40424176680734 | 0.0206399678363554 | 0.193953569070416 | -3.65626750555246 |
| ENSG00000235173.6 | HGH1 | -0.602125888480273 | 1.91668879534686 | -2.40400357048131 | 0.0206517677993286 | 0.193953569070416 | -3.44128020120022 |
| ENSG00000188747.8 | NOXA1 | -0.735170265398539 | 1.01089496961098 | -2.40376164452254 | 0.0206637587572952 | 0.193953569070416 | -3.36047519505389 |
| lnc-STPG2-3 | lnc-STPG2-3 | -0.525267494431578 | 1.3929692847371 | -2.40372028688665 | 0.0206658092599899 | 0.193953569070416 | -3.39176789803415 |
| ENSG00000188522.14 | FAM83G | -0.458326456800626 | 3.45647259045153 | -2.40355395641861 | 0.0206740577437931 | 0.193953569070416 | -3.58868311339879 |
| ENSG00000114988.11 | LMAN2L | -0.457968735567032 | 1.98070350538681 | -2.40331682496627 | 0.020685822453579 | 0.193975527818687 | -3.44290795338217 |
| ENSG00000064313.11 | TAF2 | 0.206795707778372 | 5.88718680554086 | 2.40238003118327 | 0.0207323583535554 | 0.19420341708613 | -3.71675032995278 |
| ENSG00000271601.3 | LIX1L | -0.30673938970834 | 3.38152121289387 | -2.40228944227624 | 0.0207368634291626 | 0.19420341708613 | -3.57673588770612 |
| lnc-FAM153A-3 | lnc-FAM153A-3 | 0.538258079020745 | 1.18402745198358 | 2.40223888401105 | 0.0207393781256751 | 0.19420341708613 | -3.35266201789422 |
| ENSG00000160505.15 | NLRP4 | -3.74222612108652 | -1.36877306751924 | -2.40450777809823 | 0.0207521848361078 | 0.19420341708613 | -3.34470108837272 |
| ENSG00000116793.15 | PHTF1 | -0.374348228250524 | 5.47611252130046 | -2.40187871139418 | 0.0207573005679195 | 0.19420341708613 | -3.70983331103674 |
| ENSG00000143436.10 | MRPL9 | 0.181823204785424 | 4.64899175551413 | 2.40141744712994 | 0.0207802738164364 | 0.194330020883104 | -3.67792760216956 |
| ENSG00000110108.9 | TMEM109 | -0.401568310774875 | 4.64706685065716 | -2.40087805758317 | 0.0208071671970277 | 0.194345477197947 | -3.67780435275465 |
| ENSG00000121671.11 | CRY2 | 0.240697863176988 | 5.55438278839434 | 2.40084700836735 | 0.0208087162331191 | 0.194345477197947 | -3.71435432417083 |
| ENSG00000184182.18 | UBE2F | 0.356654350515922 | 3.1449784730219 | 2.40081621274298 | 0.0208102527204137 | 0.194345477197947 | -3.54595118853099 |
| ENSG00000168944.15 | CEP120 | 0.182773628832237 | 6.23123380866014 | 2.40044741582236 | 0.0208286610765539 | 0.194347386291194 | -3.72334721952008 |
| lnc-ST8SIA6-4 | lnc-ST8SIA6-4 | -0.882176957680103 | 0.381456481343202 | -2.40043379159625 | 0.0208293414059357 | 0.194347386291194 | -3.26401919804969 |
| ENSG00000167721.10 | TSR1 | 0.202334111564629 | 5.35125418814906 | 2.39972909395551 | 0.0208645580281059 | 0.194587765243545 | -3.71273426749486 |
| ENSG00000284268.1 | MIR1181 | -0.363621545403124 | 2.21796601009674 | -2.39938242626906 | 0.0208819021399669 | 0.194615456439431 | -3.47092715580123 |
| ENSG00000141456.14 | PELP1 | -0.24224164794082 | 4.05423908074201 | -2.39912088815298 | 0.0208949957481769 | 0.194615456439431 | -3.64199134703321 |
| lnc-ZNF705D-2 | lnc-ZNF705D-2 | -0.600606399709361 | 2.45901843160937 | -2.39891838981163 | 0.0209051386854653 | 0.194615456439431 | -3.48975915874734 |
| lnc-TEX35-4 | lnc-TEX35-4 | -0.674229377945839 | 1.3552367793629 | -2.39858151645751 | 0.0209220221751256 | 0.194615456439431 | -3.380760233968 |
| ENSG00000144026.11 | ZNF514 | -0.214864516976131 | 4.83928234287008 | -2.39826605199411 | 0.0209378438441991 | 0.194615456439431 | -3.69249145342995 |
| ENSG00000257065.1 | | 0.3069410646569 | 1.7212188992201 | 2.39822855328655 | 0.0209397252559933 | 0.194615456439431 | -3.43086761955341 |
| lnc-DHRS4L2-2 | lnc-DHRS4L2-2 | -0.618417630582544 | 2.18923519387656 | -2.39809564887921 | 0.0209463946588045 | 0.194615456439431 | -3.46419544539641 |
| ENSG00000205060.10 | SLC35B4 | -0.202576805815078 | 4.15042385902221 | -2.39800291613701 | 0.0209510493041528 | 0.194615456439431 | -3.64957023603216 |
| ENSG00000115520.8 | COQ10B | 0.329783326339966 | 4.95793852569787 | 2.39776488647278 | 0.0209630012841238 | 0.194615456439431 | -3.69954256399366 |
| BEAN1-AS1 | BEAN1-AS1 | -0.522154929434726 | -0.163975606155307 | -2.39724161477666 | 0.0209892975082284 | 0.194615456439431 | -3.24096175706015 |
| ENSG00000121289.17 | CEP89 | -0.345089297404663 | 3.45048452457053 | -2.39719142123109 | 0.0209918214722654 | 0.194615456439431 | -3.59673139722874 |
| lnc-MARCH4-2 | lnc-MARCH4-2 | -0.728346720525743 | -0.574008011455064 | -2.39712718277649 | 0.0209950520788888 | 0.194615456439431 | -3.21708771670779 |
| ENSG00000138018.17 | SELENOI | 0.365839630221382 | 5.22653401041992 | 2.3970653671416 | 0.0209981612633506 | 0.194615456439431 | -3.71355162540433 |
| lnc-DHX33-3 | lnc-DHX33-3 | 0.311861535890959 | 2.68729248752068 | 2.39703081433944 | 0.0209998993709361 | 0.194615456439431 | -3.51410917458722 |
| lnc-ZNF331-1 | lnc-ZNF331-1 | -0.669759581198744 | 0.912496773459013 | -2.39637429116281 | 0.0210329490910246 | 0.194834019415193 | -3.35982081331725 |
| lnc-OSGEP-1 | lnc-OSGEP-1 | 0.382547616332366 | 3.1533765948971 | 2.39562317355278 | 0.0210708182927715 | 0.194837134191189 | -3.55822723611085 |
| lnc-ZNF280D-3 | lnc-ZNF280D-3 | -0.328389697030945 | 0.812100642100774 | -2.39543469719961 | 0.0210803303798904 | 0.194837134191189 | -3.34888434761022 |
| lnc-ZNF708-2 | lnc-ZNF708-2 | -1.85327965031963 | -0.150272398818434 | -2.39793277469783 | 0.0210805723094215 | 0.194837134191189 | -3.25865462184599 |
| lnc-SIPA1L2-10 | lnc-SIPA1L2-10 | -0.734563876714216 | -0.561173920796653 | -2.3953836437959 | 0.0210829076266702 | 0.194837134191189 | -3.22027482334873 |
| ENSG00000100918.12 | REC8 | 0.43048994151317 | 4.20527935146121 | 2.39536905666942 | 0.0210836440572977 | 0.194837134191189 | -3.65219036068437 |
| ENSG00000117385.15 | P3H1 | -0.264237870104722 | 3.62817946660839 | -2.39524157622991 | 0.021090080890928 | 0.194837134191189 | -3.6128807723127 |
| ENSG00000095370.19 | SH2D3C | -0.549318601679586 | 5.3741838455597 | -2.39500170704971 | 0.0211151061485276 | 0.194980811958342 | -3.72076692519244 |
| lnc-CCDC112-2 | lnc-CCDC112-2 | 0.236293596779306 | 2.08017909471083 | 2.3942117050622 | 0.0211421469267585 | 0.195142964212319 | -3.46511824842051 |
| ENSG00000065150.18 | IPO5 | 0.226489117067278 | 6.59068558419191 | 2.39371935133867 | 0.0211670792393084 | 0.195211937448717 | -3.73704205466733 |
| ENSG00000204590.12 | GNL1 | 0.26382911893885 | 5.1389902781489 | 2.39368957601924 | 0.0211685878825019 | 0.195211937448717 | -3.714888896332 |
| ENSG00000285710.1 | | -0.682338610450359 | -0.353696425850854 | -2.39326256483239 | 0.0211902341745268 | 0.195324043893545 | -3.23084697879742 |
| ENSG00000275778.2 | | -0.374243712394942 | 1.79035402310465 | -2.39287075793322 | 0.0212101134318428 | 0.19541976936778 | -3.45202214242075 |
| lnc-CAPZA1-1 | lnc-CAPZA1-1 | 0.478801629900105 | 3.03139150847766 | 2.39134458741339 | 0.021287707789375 | 0.196000470939089 | -3.55313117356931 |
| ENSG00000197016.11 | ZNF470 | -0.537287459349062 | 3.15526652128995 | -2.39103260264747 | 0.0213036013605831 | 0.196000470939089 | -3.57796269051265 |
| ENSG00000138286.14 | FAM149B1 | -0.210089865751229 | 4.15945324027506 | -2.39092885851584 | 0.0213088888117956 | 0.196000470939089 | -3.66203014520942 |
| lnc-FCGR2B-2 | lnc-FCGR2B-2 | 1.29049601646865 | 0.362074048425767 | 2.39218236359153 | 0.0213112303338523 | 0.196000470939089 | -3.30373942185625 |
| ENSG00000143248.12 | RGS5 | -0.46217546324264 | 0.949812590514586 | -2.39026847485862 | 0.0213425738527459 | 0.196201070840138 | -3.35828925476336 |

| | | | | | | | |
|---|---|---|---|---|---|---|---|
| lnc-ZDHHC20-5 | lnc-ZDHHC20-5 | -0.630730283431734 | 0.147278495500668 | -2.3895163504804 | 0.0213809969090079 | 0.1964111641926 | -3.26053813845181 |
| ENSG00000126777.17 | KTN1 | 0.19321161045192 | 7.64015423318789 | 2.38936996527753 | 0.02138848239618 | 0.1964111641926 | -3.73563661205341 |
| ENSG00000090006.17 | LTBP4 | -0.431968650774024 | 4.22090915349257 | -2.38903278805256 | 0.0214057331275705 | 0.1964111641926 | -3.67895435825297 |
| lnc-KCNE4-7 | lnc-KCNE4-7 | -0.63459326837303 | 0.42181948122053 | -2.38890269003108 | 0.0214123925828531 | 0.1964111641926 | -3.33249852565564 |
| ENSG00000104852.14 | SNRNP70 | -0.366580864940637 | 8.32149445144032 | -2.38888809864456 | 0.0214131396029828 | 0.1964111641926 | -3.72388817360909 |
| ENSG00000165905.17 | LARGE2 | -0.61857118613687 | 3.49537818004028 | -2.38871679119682 | 0.0214339626770897 | 0.196514589658145 | -3.63216828771903 |
| ENSG00000169895.5 | SYAP1 | 0.421357702570216 | 5.42882999882597 | 2.38794424620844 | 0.0214615109869721 | 0.196643894637894 | -3.73844171264194 |
| ENSG00000247746.4 | USP51 | -0.451799101430618 | 2.72403282997896 | -2.3878338820234 | 0.0214671734563158 | 0.196643894637894 | -3.53765229407091 |
| ENSG00000110321.16 | EIF4G2 | 0.210563375160849 | 8.8517409694773 | 2.38687856938674 | 0.0215162441189272 | 0.196892666605462 | -3.71718806842533 |
| lnc-PDE6G-2 | lnc-PDE6G-2 | -0.725639690301655 | -0.083668689575887 | -2.38663853698206 | 0.0215285895260887 | 0.196892666605462 | -3.26125236169866 |
| lnc-CEP19-1 | lnc-CEP19-1 | 0.56350735438138 | 0.923674614173036 | 2.38661708470209 | 0.0215296931759343 | 0.196892666605462 | -3.35409414611541 |
| ENSG00000132600.16 | PRMT7 | -0.236559426084447 | 4.79305658914064 | -2.38656069477813 | 0.0215325944968613 | 0.196892666605462 | -3.71711535567952 |
| ENSG00000156050.8 | FAM161B | -0.480366102692341 | 2.19205538127155 | -2.3863073240109 | 0.0215456350393526 | 0.196924425406303 | -3.49212836822703 |
| ENSG00000111727.11 | HCFC2 | 0.229972597453665 | 4.03456603236187 | 2.38601146473999 | 0.0215608713989241 | 0.196976216601889 | -3.6618819652192 |
| lnc-LYZL1-15 | lnc-LYZL1-15 | -2.42040026456859 | -1.66582184013106 | -2.38690346242508 | 0.0215921270108015 | 0.19713242004688 | -3.2754114644726 |
| ENSG00000050555.17 | LAMC3 | -2.15783261459169 | -1.85004987347959 | -2.3853968512659 | 0.021597124190143 | 0.19713242004688 | -3.34765765602299 |
| ENSG00000163781.13 | TOPBP1 | 0.197734001906548 | 6.13221603972248 | 2.38441059287958 | 0.021643482747626 | 0.197311925705273 | -3.75661384805401 |
| ENSG00000132589.15 | FLOT2 | -0.323382445437424 | 6.56972001934343 | -2.38438729449576 | 0.0216446871400686 | 0.197311925705273 | -3.75621257840385 |
| ENSG00000138593.8 | SECISBP2L | 0.337616623354505 | 7.81296474451758 | 2.38393421333409 | 0.0216681208422771 | 0.197311925705273 | -3.74470177937214 |
| ENSG00000073464.11 | CLCN4 | -0.997623914817263 | 4.59394705014879 | -2.38623947670282 | 0.0216876311490658 | 0.197311925705273 | -3.70967572919605 |
| ENSG00000197223.11 | C1D | 0.271861959689775 | 3.49229908397553 | 2.38348223907501 | 0.0216915200730312 | 0.197311925705273 | -3.62404960225668 |
| ENSG00000075711.20 | DLG1 | 0.210864368593363 | 5.96938011036341 | 2.38347816675019 | 0.0216917310054756 | 0.197311925705273 | -3.75726703635168 |
| ENSG00000225569.1 | CCT4P2 | -1.41675190552905 | 0.705442336718677 | -2.38547375819614 | 0.021693479467086 | 0.197311925705273 | -3.33855210453944 |
| lnc-ZC3H12B-8 | lnc-ZC3H12B-8 | -1.41675190552905 | 0.705442336718677 | -2.38547375819614 | 0.021693479467086 | 0.197311925705273 | -3.33855210453944 |
| lnc-VSTM5-1 | lnc-VSTM5-1 | 0.414659264919148 | 5.63440979191609 | 2.38302829617872 | 0.0217150441408065 | 0.197378020173565 | -3.7514762797017 |
| PSMG3-AS1 | PSMG3-AS1 | -0.503279224290888 | 1.68544524755554 | -2.38270660442564 | 0.0217317286528254 | 0.197378020173565 | -3.46058836358527 |
| lnc-ACOX1-1 | lnc-ACOX1-1 | -0.433247122491167 | 0.0882212368000476 | -2.38215739128682 | 0.0217602402380042 | 0.197378020173565 | -3.283185747195 |
| lnc-C1QTNF3-AMACR-2 | lnc-C1QTNF3-AMA | -0.373955543624521 | 3.53808338797097 | -2.38211835230678 | 0.0217622681712763 | 0.197378020173565 | -3.63034365326192 |
| lnc-DCUN1D5-1 | lnc-DCUN1D5-1 | 0.393287687633671 | 2.70406135140144 | 2.38210683217659 | 0.0217628666327534 | 0.197378020173565 | -3.54650751424976 |
| ENSG00000134153.9 | EMC7 | 0.294731908241393 | 3.35672774017482 | 2.38208320159467 | 0.0217640942688448 | 0.197378020173565 | -3.60556836040118 |
| ENSG00000125505.16 | MBOAT7 | -0.329471239648008 | 3.0486044423378 | -2.38186910515639 | 0.0217752196708049 | 0.197378020173565 | -3.58336870908758 |
| ENSG00000156253.6 | RWDD2B | -0.653201567175992 | 1.62463663033356 | -2.38163699152753 | 0.0217872871152548 | 0.197378020173565 | -3.44548301618562 |
| lnc-C1R-1 | lnc-C1R-1 | -0.472738662040366 | -0.269238377354916 | -2.38156014812521 | 0.0217912834838312 | 0.197378020173565 | -3.25725234265317 |
| ENSG00000131584.18 | ACAP3 | 0.604839532763624 | 2.59291208467905 | 2.38105382847025 | 0.0218176319949098 | 0.197378020173565 | -3.53204856376991 |
| lnc-SERPINB1-5 | lnc-SERPINB1-5 | 0.335512639851485 | 2.32960256605248 | 2.38099607423553 | 0.0218206393069531 | 0.197378020173565 | -3.51535678676127 |
| ENSG00000221539.1 | SNORD99 | -0.457427011559382 | 0.0124896990305236 | -2.38092179501117 | 0.021824507638306 | 0.197378020173565 | -3.28954072024612 |
| ENSG00000256206.2 | | 0.253142438741383 | 4.46038092741638 | 2.38090450709773 | 0.0218254080510632 | 0.197378020173565 | -3.7067925968896 |
| lnc-GIMD1-5 | lnc-GIMD1-5 | -0.243309115498739 | 2.93955861035459 | -2.38067672889537 | 0.0218372746330251 | 0.197398605081931 | -3.57129450839923 |
| ENSG00000059758.7 | CDK17 | 0.242617826881621 | 5.94539327332184 | 2.37996183495979 | 0.0218745562901474 | 0.197526085521951 | -3.76490449191556 |
| ENSG00000178464.6 | RPL10P16 | 0.980095284982645 | 2.32875865941038 | 2.3808512538843 | 0.0219030619440044 | 0.197526085521951 | -3.48691877537488 |
| ENSG00000175166.16 | PSMD2 | 0.197492858971161 | 5.92192304244797 | 2.37940169103289 | 0.0219038078041689 | 0.197526085521951 | -3.76550263604117 |
| ENSG00000124256.14 | ZBP1 | -0.536910346262144 | 3.89608938867809 | -2.37936137840484 | 0.0219059143457405 | 0.197526085521951 | -3.66495828290539 |
| ENSG00000101977.20 | MCF2 | 0.70478554828921 | 0.505664208140038 | 2.37923471611 | 0.0219125342879257 | 0.197526085521951 | -3.32103424431887 |
| ENSG00000106327.12 | TFR2 | 1.13083539922725 | 0.230579335906612 | 2.37881436051646 | 0.0219345168821773 | 0.197526085521951 | -3.26081939003133 |
| ENSG00000160179.18 | ABCG1 | 0.599143290468137 | 5.49522860351082 | 2.37944309308464 | 0.0219367588405334 | 0.197526085521951 | -3.75156123479541 |
| lnc-ABCA10-1 | lnc-ABCA10-1 | -0.57071176938281 | 1.26100304268486 | -2.37865469624509 | 0.0219428717672149 | 0.197526085521951 | -3.42938799522762 |
| ENSG00000184014.7 | DENND5A | 0.42421910399761 | 5.06517282630678 | 2.37864859072163 | 0.0219431913129162 | 0.197526085521951 | -3.74407550728447 |
| ENSG00000130669.17 | PAK4 | -0.396823120807829 | 1.41606593297302 | -2.37856927532435 | 0.0219473428357723 | 0.197526085521951 | -3.45164000513466 |
| ENSG00000196189.12 | SEMA4A | -0.886127858994265 | 4.79572635931409 | -2.3804503162424 | 0.0219736129842927 | 0.197635757313349 | -3.7234143132229 |
| ENSG00000249661.1 | TNRC18P1 | 0.834560798375861 | 3.78087436052336 | 2.37958916735988 | 0.0219854502403488 | 0.197635757313349 | -3.63938477117327 |
| lnc-TRABD2A-2 | lnc-TRABD2A-2 | -0.893523391209805 | -0.100742858548697 | -2.37778268169412 | 0.0219885530014977 | 0.197635757313349 | -3.27475494415615 |
| lnc-ZC3H12B-2 | lnc-ZC3H12B-2 | -1.17973648049704 | -0.694427825370327 | -2.37760376747794 | 0.0219979361611467 | 0.197635757313349 | -3.25181359820259 |
| lnc-PSTPIP1-1 | lnc-PSTPIP1-1 | -0.937181959333915 | -0.783982888187211 | -2.37719449984437 | 0.0220194137729643 | 0.19774240562344 | -3.2516957331279 |
| lnc-MFSD12-1 | lnc-MFSD12-1 | -0.436335127908197 | 0.764763821708429 | -2.37693819018769 | 0.0220328740481437 | 0.197776966616908 | -3.38540829370149 |
| ENSG00000100364.18 | KIAA0930 | -0.443535951194655 | 3.84982611905752 | -2.37650897727241 | 0.0220554310520621 | 0.197776966616908 | -3.66366855129906 |
| MIR583HG | MIR583HG | -0.621850324961735 | 1.64425593552558 | -2.37643889388738 | 0.0220591162132404 | 0.197776966616908 | -3.47470258887803 |
| ENSG00000172974.11 | VDAC2P5 | 1.22088271670408 | -0.270159064902662 | 2.3763898139262 | 0.0220616972915717 | 0.197776966616908 | -3.25385718015912 |
| lnc-ZSCAN2-5 | lnc-ZSCAN2-5 | -0.257061812092793 | 3.04088039732379 | -2.37600495676188 | 0.0220819460716783 | 0.197872309966632 | -3.59252958661239 |
| ENSG00000065883.15 | CDK13 | -0.216629100056873 | 7.37526474479226 | -2.37576251347637 | 0.0220947105147002 | 0.197900533735454 | -3.76675487071358 |
| lnc-SNN-6 | lnc-SNN-6 | -0.626564376619353 | 2.6490626896744 | -2.37531100850326 | 0.0221184995985634 | 0.197942528258473 | -3.55689152712047 |
| ENSG00000214189.9 | ZNF788P | -0.796418492902578 | 0.236812877035612 | -2.37521409706972 | 0.0221236087148006 | 0.197942528258473 | -3.30713477752974 |
| lnc-RNF24-2 | lnc-RNF24-2 | -0.469939162108889 | 1.91866721120794 | -2.37506167772045 | 0.0221316463273565 | 0.197942528258473 | -3.49479540010264 |
| lnc-ERICH1-9 | lnc-ERICH1-9 | -0.535959887239356 | 3.79021674248226 | -2.37494376280647 | 0.0221378662027403 | 0.197942528258473 | -3.66869796218659 |
| ENSG00000140043.11 | PTGR2 | -0.422187717525818 | 2.27808820045418 | -2.37448475226741 | 0.0221620934714826 | 0.19807310895507 | -3.52057378816458 |
| ENSG00000172239.13 | PAIP1 | 0.204327613293195 | 5.08494140908499 | 2.37333769251881 | 0.0222227414313477 | 0.198403529095474 | -3.75792121615858 |
| lnc-LHFPL4-6 | lnc-LHFPL4-6 | -0.661376894189615 | 0.271011905531377 | -2.37332403352091 | 0.0222234645167046 | 0.198403529095474 | -3.31743855602208 |
| lnc-ZNF706-1 | lnc-ZNF706-1 | -0.447237314385739 | 0.998379149875908 | -2.37323872198894 | 0.0222279812512347 | 0.198403529095474 | -3.39979565861114 |
| lnc-WDR73-9 | lnc-WDR73-9 | -0.597432192651193 | -0.0395670431188046 | -2.37304561479942 | 0.0222382081731781 | 0.198408772790865 | -3.29292544390521 |

| | | | | | | | |
|---|---|---|---|---|---|---|---|
| lnc-MYOCD-1 | lnc-MYOCD-1 | -0.975940363484843 | 2.56350359679243 | -2.37388003821646 | 0.0222913254696168 | 0.19877372696133 | -3.55087882298624 |
| ENSG00000278991.1 | | 0.506000294555859 | 2.28536254719513 | 2.37191015585475 | 0.0222984276127732 | 0.19877372696133 | -3.53549027049057 |
| ENSG00000115137.11 | DNAJC27 | 0.355970197626469 | 3.14493814150751 | 2.37112190554941 | 0.022340319022242 | 0.198938371238063 | -3.61181632611995 |
| ENSG00000196510.12 | ANAPC7 | 0.226388015250535 | 4.95932835851744 | 2.37106225722429 | 0.0223434919025757 | 0.198938371238063 | -3.75515872502714 |
| ENSG00000172986.12 | GXYLT2 | 0.276206537298324 | 1.10731772321458 | 2.37101712424469 | 0.0223458929360347 | 0.198938371238063 | -3.42211053151742 |
| ENSG00000164081.12 | TEX264 | -0.270230554793538 | 4.29430859787007 | -2.36952572385863 | 0.0224253648458688 | 0.199559569659541 | -3.72231642988003 |
| ENSG00000058063.15 | ATP11B | 0.225479835268673 | 7.30766441568743 | 2.36928197173767 | 0.0224383777700719 | 0.199589079361016 | -3.7817911856153 |
| lnc-RHEX-1 | lnc-RHEX-1 | -0.871701344619592 | 4.24729505308374 | -2.37026076521876 | 0.0225031781055557 | 0.200079012935919 | -3.71095347473426 |
| ENSG00000224470.7 | ATXN1L | 0.22704305270584 | 6.91317197080873 | 2.36727081994263 | 0.0225460045498873 | 0.200373234736757 | -3.79037362558403 |
| lnc-CLDN19-3 | lnc-CLDN19-3 | -0.426881641340638 | 0.849816296501022 | -2.36707183090996 | 0.0225566786650023 | 0.200381578317541 | -3.40327807695689 |
| ENSG00000178440.5 | TIMM23B-AGAP6 | 0.385117000881669 | 1.34305391353017 | 2.36625034387023 | 0.0226007928045632 | 0.200686849998414 | -3.44604825962456 |
| lnc-PINX1-85 | lnc-PINX1-85 | -0.655990753979758 | 1.64397479674802 | -2.36582206363806 | 0.0226238223766498 | 0.200804715816551 | -3.48994701758132 |
| lnc-ACTR8-1 | lnc-ACTR8-1 | -0.433313240128234 | 1.58126542771737 | -2.36486392839745 | 0.0226754198986928 | 0.201175934385687 | -3.48818984432998 |
| ENSG00000177352.9 | CCDC71 | -0.281485559728504 | 3.77838337398085 | -2.36419556284949 | 0.0227114754014114 | 0.201409003958316 | -3.69079449973202 |
| ENSG00000166200.14 | COPS2 | 0.285593643616552 | 6.09094153573297 | 2.36327634273846 | 0.0227611477024503 | 0.201762576726759 | -3.80036481354303 |
| lnc-PRCP-2 | lnc-PRCP-2 | 0.857991116260026 | 0.21586348421092 | 2.36304053318749 | 0.0227739059887537 | 0.201783339929513 | -3.29999942695567 |
| ENSG00000253729.7 | PRKDC | 0.235279895273608 | 7.96294234964394 | 2.36287073388665 | 0.0227830968272938 | 0.201783339929513 | -3.78588479801465 |
| ENSG00000225067.4 | RPL23AP2 | 1.08759235518488 | -0.424803818021594 | 2.36227581403319 | 0.022815324777357 | 0.201981862319286 | -3.27930088060491 |
| lnc-MECP2-2 | lnc-MECP2-2 | 0.352831346499053 | 3.22423739726323 | 2.36181068351057 | 0.0228405503490685 | 0.202118249284126 | -3.6331780946271 |
| ENSG00000083290.19 | ULK2 | -1.65382284454599 | 1.77883679906343 | -2.36406923526549 | 0.022894420803344 | 0.202464934117283 | -3.52730883730714 |
| LINC01366 | LINC01366 | -1.050856445412 | 0.0877860264629462 | -2.36050326478368 | 0.0229115904276095 | 0.202464934117283 | -3.30121918212007 |
| ENSG00000148297.15 | MED22 | -0.232198786254388 | 4.7708595804266 | -2.36030271565685 | 0.0229225050543603 | 0.202464934117283 | -3.77316843144208 |
| ENSG00000164924.17 | YWHAZ | 0.334699478440406 | 9.19921761946925 | 2.3601713333167 | 0.0229296579033141 | 0.202464934117283 | -3.76560482294866 |
| ENSG00000149761.8 | NUDT22 | -0.307490071510707 | 3.8858500509103 | -2.35999926935231 | 0.0229390286180572 | 0.202464934117283 | -3.7107989224806 |
| ENSG00000218227.3 | | 1.2315379817741 | 2.43723128669711 | 2.3621261689712 | 0.0229517950384727 | 0.202464934117283 | -3.51754001776334 |
| lnc-DTWD2-14 | lnc-DTWD2-14 | -0.528899186268687 | 0.574254726868384 | -2.35974465906227 | 0.0229529011781836 | 0.202464934117283 | -3.3781070205291 |
| lnc-FAM117A-3 | lnc-FAM117A-3 | 0.324687073171766 | 1.00609659713014 | 2.35964341008673 | 0.022958419872212 | 0.202464934117283 | -3.44450553339278 |
| lnc-CTAGE6-1 | lnc-CTAGE6-1 | -0.539463474318916 | 0.469148380009301 | -2.35937763165767 | 0.0229729121149649 | 0.202488424916586 | -3.37109999212328 |
| ENSG00000205485.13 | NA | -0.355183253688486 | 1.94092559809898 | -2.35923378942646 | 0.0229807589080865 | 0.202488424916586 | -3.53056394917455 |
| lnc-TEC-3 | lnc-TEC-3 | 0.266272410600766 | 2.5993091417363 | 2.35875291555457 | 0.0230070087455779 | 0.20262462573574 | -3.58567457584836 |
| ENSG00000283515.1 | | 0.197182999714625 | 4.18249506748657 | 2.35859005386463 | 0.0230159051144226 | 0.20262462573574 | -3.73071820016567 |
| ENSG00000277203.1 | NA | -0.609402130200212 | 1.85842517160421 | -2.35827416723454 | 0.0230331693362867 | 0.202689920670709 | -3.53274914567427 |
| lnc-DHH-1 | lnc-DHH-1 | 0.401782029658443 | 1.94982085142324 | 2.35773364025434 | 0.0230627378902122 | 0.202791820637256 | -3.52715784436216 |
| lnc-NUP107-3 | lnc-NUP107-3 | 0.608672561966397 | 1.40604427890733 | 2.35742986532735 | 0.0230793703271037 | 0.202791820637256 | -3.46857111576007 |
| lnc-POF1B-2 | lnc-POF1B-2 | -0.276586964127487 | 1.76074239462446 | -2.35714118980749 | 0.0230951860276935 | 0.202791820637256 | -3.51876657713206 |
| ENSG00000197579.7 | TOPORS | 0.302101004725529 | 5.5494666910319 | 2.35687363221375 | 0.0231098534390577 | 0.202791820637256 | -3.80536332315422 |
| ENSG00000160953.15 | PWWP3A | -0.297888328919534 | 4.03606971067825 | -2.35674246531764 | 0.0231170470202757 | 0.202791820637256 | -3.72126567635609 |
| ENSG00000196646.11 | ZNF136 | 0.25595207855246 | 5.58601396874894 | 2.35670748623698 | 0.0231189657167104 | 0.202791820637256 | -3.80638624952711 |
| ENSG00000196781.14 | TLE1 | -0.87421259618572 | 5.78684797889131 | -2.35890629015863 | 0.0231348764320547 | 0.202791820637256 | -3.80714304632084 |
| lnc-SINHCAF-3 | lnc-SINHCAF-3 | 0.804658513704503 | 2.07867974104814 | 2.35690631887123 | 0.0231362821692886 | 0.202791820637256 | -3.54837425036086 |
| ENSG00000020922.12 | MRE11 | 0.309561225912464 | 5.62228414897528 | 2.3563364515143 | 0.0231393267832639 | 0.202791820637256 | -3.80707828115499 |
| ENSG00000172009.14 | THOP1 | -0.339661281053337 | 4.17019328010139 | -2.35623095044307 | 0.0231451192502141 | 0.202791820637256 | -3.73978703674974 |
| ENSG00000175130.6 | MARCKSL1 | -0.763440083392951 | 3.46635462708211 | -2.35740973389349 | 0.0231531253217486 | 0.202791820637256 | -3.69718699711196 |
| ENSG00000146707.14 | POMZP3 | 0.381514905732959 | 2.17917203806614 | 2.35564536913493 | 0.0231772939132813 | 0.202917158918362 | -3.5426671795303 |
| lnc-TUBB2B-3 | lnc-TUBB2B-3 | 1.9437513509558 | -1.65460053826889 | 2.35537224775909 | 0.0231923142664439 | 0.202931732928217 | -3.40675301849002 |
| ENSG00000164934.13 | DCAF13 | 0.315340796118556 | 5.44675803004835 | 2.35525660043645 | 0.023198676946028 | 0.202931732928217 | -3.80614104141584 |
| ENSG00000166704.11 | ZNF606 | -0.224695357150363 | 4.58189098905728 | -2.35443986244907 | 0.0232436568964955 | 0.203181796288544 | -3.76761404491285 |
| lnc-SEC24C-5 | lnc-SEC24C-5 | -0.374668376746623 | 2.18079792094298 | -2.35437910179128 | 0.0232470062799165 | 0.203181796288544 | -3.55652771816123 |
| lnc-ZNF311-3 | lnc-ZNF311-3 | -0.287461578447678 | 1.8397740533418 | -2.3530490549959 | 0.0233204328400666 | 0.203725295513112 | -3.53553229416304 |
| ENSG00000132953.16 | XPO4 | 0.220171703810387 | 6.35205780359074 | 2.3528943657843 | 0.0233289861305158 | 0.203725295513112 | -3.82252848692899 |
| ENSG00000201183.1 | RNVU1-3 | -1.09799082106863 | 2.27266140590073 | -2.35467314968364 | 0.0233503384765488 | 0.203751303797202 | -3.5705117496256 |
| ENSG00000116209.11 | TMEM59 | 0.212806456269337 | 6.17082401613338 | 2.3524826973961 | 0.0233517624086674 | 0.203751303797202 | -3.82300047285735 |
| lnc-ARID3B-2 | lnc-ARID3B-2 | -0.403956521771289 | 0.474593698898511 | -2.3517229034369 | 0.023393851836638 | 0.203908541044619 | -3.37752140897596 |
| ENSG00000176915.14 | ANKLE2 | 0.219385851000767 | 6.75978590854979 | 2.35139310585307 | 0.0234121424629571 | 0.203908541044619 | -3.82422197116545 |
| lnc-PRDM4-6 | lnc-PRDM4-6 | -0.926117080976749 | 1.17826296101354 | -2.35186040344957 | 0.0234293685329679 | 0.203908541044619 | -3.4969739277733 |
| ENSG00000167220.11 | HDHD2 | -0.225086231829767 | 4.26683013174481 | -2.35106233339433 | 0.0234305000640838 | 0.203908541044619 | -3.75595081167063 |
| lnc-SLC27A4-3 | lnc-SLC27A4-3 | 0.428227467112649 | 1.18708564239108 | 2.35091751654143 | 0.0234385413518197 | 0.203908541044619 | -3.46578398603086 |
| lnc-RBM8A-1 | lnc-RBM8A-1 | -0.341878719520367 | 2.13004953528311 | -2.35088732240691 | 0.0234402182627793 | 0.203908541044619 | -3.56794720510382 |
| lnc-TRPC4AP-1 | lnc-TRPC4AP-1 | -0.585114343539703 | 0.974466762508881 | -2.35044831323466 | 0.0234646119788523 | 0.203908541044619 | -3.43876105013142 |
| lnc-ZNFX1-2 | lnc-ZNFX1-2 | 0.215429407249552 | 2.77824050878639 | 2.35040828698799 | 0.0234668371872643 | 0.203908541044619 | -3.61809069538765 |
| ENSG00000166004.14 | CEP295 | 0.364528888150526 | 5.86871535731906 | 2.35019073610683 | 0.0234789349691904 | 0.203908541044619 | -3.8229360845933 |
| ENSG00000137078.8 | SIT1 | -0.48599370483071 | 4.27627095479369 | -2.35018244224237 | 0.0234793962933914 | 0.203908541044619 | -3.76512066367538 |
| ENSG00000138190.16 | EXOC6 | -0.225874153304659 | 5.05328844975346 | -2.35001688325804 | 0.0234886067768036 | 0.203908541044619 | -3.80406080996876 |
| ENSG00000168394.11 | TAP1 | 0.376886208890363 | 6.20968620087631 | 2.34987336899145 | 0.0234965934812493 | 0.203908541044619 | -3.82836740536924 |
| ENSG00000137218.10 | FRS3 | -0.43351053229989 | 1.31627333153406 | -2.34983786546971 | 0.0234985696622375 | 0.203908541044619 | -3.49460846333404 |
| lnc-RNASE6-1 | lnc-RNASE6-1 | 1.36261622900384 | 1.05766919148521 | 2.35068720774243 | 0.0235600648445048 | 0.204356011249238 | -3.42069550220826 |
| ENSG00000163608.14 | NEPRO | 0.217534111116123 | 5.44208517149811 | 2.34794138919342 | 0.0236043477709087 | 0.204628510976639 | -3.82293296161236 |
| lnc-SLC25A25-5 | lnc-SLC25A25-5 | -0.891455268194801 | 1.5731494482142 | -2.34842864840486 | 0.0236113644060398 | 0.204628510976639 | -3.51094555795997 |

| | | | | | | | |
|---|---|---|---|---|---|---|---|
| ENSG00000138756.17 | BMP2K | -0.462738744441132 | 6.29753959676407 | -2.34711995732328 | 0.0236502966915314 | 0.204879653536107 | -3.83346963155243 |
| ENSG00000008324.11 | SS18L2 | 0.431437703889503 | 2.78602823105679 | 2.3462044904512 | 0.0237016003701978 | 0.205237711577527 | -3.62583659831711 |
| ENSG00000120925.15 | RNF170 | -0.335381399909113 | 3.74428286078168 | -2.34524599502292 | 0.0237554225225615 | 0.205604247794748 | -3.71896522054562 |
| ENSG00000196418.12 | ZNF124 | 0.431222302596703 | 3.95505065326482 | 2.34509506998313 | 0.0237639073751983 | 0.205604247794748 | -3.73240909911476 |
| ENSG00000165071.14 | TMEM71 | 0.96699010211713 | 3.49455823401387 | 2.34697531554961 | 0.0237741145843357 | 0.205606134659404 | -3.65966823825625 |
| ENSG00000274349.4 | ZNF658 | -1.38035668867624 | 0.761583365294742 | -2.34635379238566 | 0.0238024341546279 | 0.205673077974898 | -3.42114823049895 |
| lnc-MYOM1-1 | lnc-MYOM1-1 | -0.597960254252558 | 0.0527584689531573 | -2.34438531771742 | 0.0238038454389374 | 0.205673077974898 | -3.36415021913925 |
| lnc-RCCD1-4 | lnc-RCCD1-4 | -0.515128340604359 | -0.00864070594822068 | -2.34415072652544 | 0.0238170592354396 | 0.205673077974898 | -3.35682675092789 |
| lnc-CNDP1-9 | lnc-CNDP1-9 | 0.457021309103028 | 0.74055102164841 | 2.34406615143067 | 0.0238218247044725 | 0.205673077974898 | -3.42041992994952 |
| lnc-H1FNT-1 | lnc-H1FNT-1 | -0.630856770664665 | 0.873668112368484 | -2.34344007859604 | 0.0238571280513747 | 0.205891516428279 | -3.46346840606185 |
| ENSG00000250151.9 | ARPC4-TTLL3 | 0.253995212332939 | 5.1588971724074 | 2.34270440123859 | 0.023898671879538 | 0.206163605740374 | -3.81959230270104 |
| ENSG00000174579.3 | MSL2 | 0.18013181761162 | 6.08679078369959 | 2.34235479893248 | 0.023918436744738 | 0.206247668000394 | -3.84335403026246 |
| ENSG00000174004.5 | NRROS | 0.551403775369686 | 4.18890053610228 | 2.34126383701823 | 0.0239802089924841 | 0.206624664110993 | -3.74411287919064 |
| ENSG00000137343.17 | ATAT1 | -0.308045917231432 | 2.40577310534241 | -2.34110031835651 | 0.0239894800507406 | 0.206624664110993 | -3.61003314605362 |
| lnc-ACTB-3 | lnc-ACTB-3 | 0.2503713339395 | 2.7049707648212 | 2.34105107552008 | 0.0239922726145495 | 0.206624664110993 | -3.63076376453415 |
| ENSG00000182866.16 | LCK | -0.649121580646129 | 5.1961935861222 | -2.34123291652195 | 0.0240436665033962 | 0.206980672371143 | -3.82325766544048 |
| lnc-TRIM32-2 | lnc-TRIM32-2 | -0.464150175574268 | 1.1026656867205 | -2.33884937270861 | 0.0241174299501607 | 0.207460370316477 | -3.48430621659092 |
| ENSG00000135862.5 | LAMC1 | 1.16430305352951 | 5.0012961797204 | 2.34198375975432 | 0.0241195484704528 | 0.207460370316477 | -3.80929057921376 |
| ENSG00000106333.12 | PCOLCE | -0.554961934330299 | 0.139278582752483 | -2.33811088153392 | 0.0241595410792076 | 0.207717558075743 | -3.38506819173578 |
| lnc-SEPT14-4 | lnc-SEPT14-4 | -0.451432907565199 | 6.81941893518022 | -2.33790587293578 | 0.0241712430242985 | 0.207731396730328 | -3.85116467517351 |
| ENSG00000135074.15 | ADAM19 | 0.37951592590482 | 7.7682978717818 | 2.33698821450474 | 0.0242236855914537 | 0.20804499654528 | -3.84408477826907 |
| ENSG00000117899.10 | MESD | -0.215362654664593 | 4.67095563621046 | -2.33679155372671 | 0.0242349376891128 | 0.20804499654528 | -3.81168221128526 |
| ENSG00000187815.9 | ZFP69 | -0.387046757921517 | 2.41547281803184 | -2.33673707151721 | 0.0242380557603645 | 0.20804499654528 | -3.61890169372086 |
| ENSG00000121350.15 | PYROXD1 | 0.218735764834732 | 5.19747400911452 | 2.33628037885848 | 0.0242642068997563 | 0.208182647193699 | -3.84039207484313 |
| ENSG00000165355.7 | FBXO33 | 0.265005261352958 | 5.88313413024158 | 2.33579169340022 | 0.0242922180551866 | 0.208267993976036 | -3.85531088037111 |
| lnc-TLK1-7 | lnc-TLK1-7 | -0.422800163761734 | 3.03987579241044 | -2.33572883189981 | 0.0242958233443197 | 0.208267993976036 | -3.67463778252998 |
| ENSG00000152223.13 | EPG5 | 0.238380332234748 | 6.38279310773774 | 2.33552867996743 | 0.0243073058362907 | 0.208267993976036 | -3.85831765058257 |
| ENSG00000137193.13 | PIM1 | -0.408976099317347 | 7.15402062746383 | -2.33540109005689 | 0.0243146280680374 | 0.208267993976036 | -3.85254003702717 |
| ENSG00000143198.12 | MGST3 | -0.567055678108258 | 3.32919791371657 | -2.33501792776339 | 0.0243366291917251 | 0.208369733216837 | -3.70301176465417 |
| lnc-CAPN12-3 | lnc-CAPN12-3 | 0.360107053598663 | 2.28106495972446 | 2.3342547470217 | 0.0243805041575035 | 0.208568305181493 | -3.60284864451842 |
| ENSG00000284368.1 | MIR5047 | 0.394957172283584 | 5.25794894641095 | 2.3342169610864 | 0.0243826782993217 | 0.208568305181493 | -3.84345719446874 |
| ENSG00000163682.16 | RPL9 | 0.260011903571563 | 9.44464291531046 | 2.33408590161861 | 0.0243902205981564 | 0.208568305181493 | -3.81136917166924 |
| ENSG00000160593.18 | JAML | -2.6006817486286 | 0.964492701876198 | -2.33664470209895 | 0.0244328368180615 | 0.208776917684847 | -3.41330158965068 |
| ENSG00000128294.15 | TPST2 | -0.645848318212452 | 4.63518071455318 | -2.33411923007468 | 0.024441534197271 | 0.208776917684847 | -3.81934188183426 |
| lnc-CCNB1IP1-5 | lnc-CCNB1IP1-5 | -0.426121801343386 | 2.91437240951686 | -2.33294104695099 | 0.0244561945390942 | 0.208776917684847 | -3.66801144432932 |
| ENSG00000135407.10 | AVIL | -0.560591731860372 | 1.06094913215775 | -2.33293808671674 | 0.0244563653343084 | 0.208776917684847 | -3.504633752762 |
| ENSG00000118260.14 | CREB1 | -0.189057051163511 | 6.68283541049585 | -2.33274288404693 | 0.0244676302114175 | 0.208776917684847 | -3.86280886657899 |
| lnc-DUSP19-4 | lnc-DUSP19-4 | -0.624980106416398 | 0.660310869253953 | -2.33260699339894 | 0.0244754750217916 | 0.208776917684847 | -3.44863576866252 |
| ENSG00000137845.14 | ADAM10 | 0.239616431701744 | 6.23367032213382 | 2.33228971693382 | 0.024493799812179 | 0.208811814936448 | -3.86481309155858 |
| lnc-HTD2-1 | lnc-HTD2-1 | 0.304393333609217 | 0.580339180147977 | 2.33209973869929 | 0.0245047781884311 | 0.208811814936448 | -3.44340605920394 |
| ENSG00000204713.10 | TRIM27 | -0.315242546282268 | 5.02908795269523 | -2.33200939075708 | 0.0245100007232405 | 0.208811814936448 | -3.84299261335486 |
| lnc-TFB1M-1 | lnc-TFB1M-1 | -0.49020914971732 | 1.47912739463149 | -2.33062221124197 | 0.0245903116680135 | 0.209409344254333 | -3.5439254355914 |
| ENSG00000087191.12 | PSMC5 | 0.275537872747782 | 4.54385002339232 | 2.3300453516225 | 0.024623778471322 | 0.209587920772446 | -3.81846756909167 |
| ENSG00000135116.9 | HRK | -1.53448121807415 | 3.1587045796414 | -2.3331950171236 | 0.0246316465213306 | 0.209587920772446 | -3.7012516337452 |
| ENSG00000204514.9 | ZNF814 | 0.205516656608015 | 5.17035203804131 | 2.32944521471505 | 0.0246586390757899 | 0.209730895907844 | -3.84879375634328 |
| ENSG00000139826.5 | ABHD13 | 0.290661512826829 | 5.11381096811616 | 2.3277892351821 | 0.0247550609192582 | 0.210248748337144 | -3.85273548713667 |
| lnc-GOLGA8A-2 | lnc-GOLGA8A-2 | -0.38640585059482 | 1.22949603434173 | -2.32768616116679 | 0.0247610737190272 | 0.210248748337144 | -3.52350353609692 |
| ENSG00000186260.16 | MRTFB | -0.279775731238409 | 5.09161090161438 | -2.32746669856923 | 0.0247738803842941 | 0.210248748337144 | -3.8490709219859 |
| lnc-CRABP2-3 | lnc-CRABP2-3 | 0.440942650275569 | -0.0658936939892408 | 2.3273817075578 | 0.0247788416013359 | 0.210248748337144 | -3.36354386844955 |
| ENSG00000125534.9 | PPDPF | -0.4574614449604 | 4.03412637904802 | -2.32681584803474 | 0.0248118954837779 | 0.210248748337144 | -3.79684578713389 |
| ENSG00000204116.11 | CHIC1 | -0.471593069741366 | 5.26066035085752 | -2.32677934643987 | 0.0248140290296825 | 0.210248748337144 | -3.85968756196448 |
| lnc-MRPS25-6 | lnc-MRPS25-6 | 0.358608820815207 | 0.554633228206265 | 2.32671876402511 | 0.0248175704809018 | 0.210248748337144 | -3.43779077327027 |
| ENSG00000136950.13 | ARPC5L | 0.397091879362992 | 3.89839122516439 | 2.32640876080053 | 0.024835699357197 | 0.210248748337144 | -3.76994200033989 |
| ENSG00000177383.4 | MAGEF1 | -0.458239288570495 | 3.99497042641432 | -2.32632970862347 | 0.0248403241996956 | 0.210248748337144 | -3.7886167750047 |
| ENSG00000281348.1 | | -0.424067312287663 | 2.20167899195215 | -2.32622005141562 | 0.0248467408274544 | 0.210248748337144 | -3.63094564988994 |
| ENSG00000153896.17 | ZNF599 | -0.36553090325458 | 1.44721569140658 | -2.3260138322626 | 0.0248588118352434 | 0.210248748337144 | -3.54951520627271 |
| ENSG00000090097.21 | PCBP4 | -0.400400995429052 | 3.52272180063075 | -2.32599464243976 | 0.0248599353760906 | 0.210248748337144 | -3.74698541474377 |
| lnc-H2BFWT-18 | lnc-H2BFWT-18 | -0.237830473207111 | 3.31828296431363 | -2.32597949575216 | 0.0248608222284281 | 0.210248748337144 | -3.71920360486664 |
| ENSG00000109618.11 | SEPSECS | 0.370994365297229 | 4.86117350935384 | 2.32595033109892 | 0.0248625299253077 | 0.210248748337144 | -3.8451419660921 |
| lnc-HSPB7-4 | lnc-HSPB7-4 | -0.334730540032107 | 2.2239727366439 | -2.32563996992932 | 0.0248807092208774 | 0.210316073056805 | -3.62523602298708 |
| LINC02413 | LINC02413 | -0.594763697085362 | 1.47625003102908 | -2.32478923051464 | 0.0249306021094384 | 0.210544572615257 | -3.53240411163038 |
| lnc-GTF2B-2 | lnc-GTF2B-2 | 0.302933463434622 | 3.04502543954528 | 2.32452330168621 | 0.0249462162965995 | 0.210544572615257 | -3.6885075687134 |
| lnc-ZNF431-5 | lnc-ZNF431-5 | -0.399218935112104 | 0.394673140336504 | -2.32427082485066 | 0.0249610487542518 | 0.210544572615257 | -3.43146565621986 |
| ENSG00000242247.10 | ARFGAP3 | 0.228168430946433 | 4.51629348953565 | 2.32410995896966 | 0.0249705033935513 | 0.210544572615257 | -3.8259665691278 |
| lnc-TMEM240-1 | lnc-TMEM240-1 | 0.300532646220534 | 3.74632673827272 | 2.32397779436062 | 0.0249782735630854 | 0.210544572615257 | -3.75848337900575 |
| ENSG00000179010.14 | MRFAP1 | 0.327831853248186 | 5.75223553743484 | 2.32392852159392 | 0.0249811709428104 | 0.210544572615257 | -3.87794607506589 |
| ENSG00000173914.11 | RBM4B | -0.327236006265763 | 3.6403023953744 | -2.32379850353447 | 0.024988817823465 | 0.210544572615257 | -3.75656763712832 |
| ENSG00000135093.12 | USP30 | 0.301863262929844 | 3.86215766434793 | 2.32378565725895 | 0.024989573478068 | 0.210544572615257 | -3.77503963789197 |

| | | | | | | | |
|---|---|---|---|---|---|---|---|
| ENSG00000074855.10 | ANO8 | -0.760806602623234 | -0.527446756710486 | -2.32356466770214 | 0.0250025759225435 | 0.210567929711011 | -3.35538138390686 |
| lnc-RGPD8-5 | lnc-RGPD8-5 | -0.679727322557406 | 0.683097924151816 | -2.32259166859439 | 0.0250598968002977 | 0.210831172627775 | -3.45725368987365 |
| ENSG00000171132.13 | PRKCE | 0.336249082052519 | 7.38603701735619 | 2.3225650272484 | 0.0250614679386461 | 0.210831172627775 | -3.8780631112986 |
| lnc-HUS1-3 | lnc-HUS1-3 | 1.09305896622145 | 0.534008883997061 | 2.32301809036201 | 0.0250647772079653 | 0.210831172627775 | -3.43577436634009 |
| ENSG00000175643.9 | RMI2 | -0.740990780424668 | 0.866164182534667 | -2.32233893536559 | 0.0250748049649124 | 0.210831172627775 | -3.48783099337593 |
| ENSG00000169554.19 | ZEB2 | 0.26114039485859 | 7.52719396738614 | 2.32186914417005 | 0.025102538015882 | 0.210889135479288 | -3.8777909979893 |
| ENSG00000264229.1 | RNU4ATAC | -0.775007659833711 | 1.69797237012154 | -2.32190213379007 | 0.0251092466072228 | 0.210889135479288 | -3.57519006649156 |
| lnc-GPBP1L1-3 | lnc-GPBP1L1-3 | -0.244920468315877 | 1.76499184717033 | -2.32122011582103 | 0.0251408971523763 | 0.210889135479288 | -3.58657518682942 |
| lnc-CETP-2 | lnc-CETP-2 | 0.60088754645896 | 2.36088214195518 | 2.3210338475895 | 0.0251519157515145 | 0.210889135479288 | -3.62540318705174 |
| ENSG00000124198.8 | ARFGEF2 | 0.19401014458455 | 7.31412216100313 | 2.32100257197495 | 0.0251537662683544 | 0.210889135479288 | -3.88118719893374 |
| ENSG00000130414.11 | NDUFA10 | 0.17698614831043 | 6.25374503446873 | 2.3209953479155 | 0.0251541937191472 | 0.210889135479288 | -3.8880490625324 |
| ENSG00000099337.4 | KCNK6 | -0.543128337610436 | 1.44253121334881 | -2.32090833591007 | 0.0251593427694874 | 0.210889135479288 | -3.54653211635755 |
| lnc-ZNF708-12 | lnc-ZNF708-12 | -0.481662392514256 | 1.68665778476329 | -2.32083530833107 | 0.0251636650020468 | 0.210889135479288 | -3.58945975078087 |
| lnc-FAM43A-10 | lnc-FAM43A-10 | -1.01221223766092 | -0.267986801033426 | -2.3197817940945 | 0.0252260927114422 | 0.21132453688795 | -3.38704213108569 |
| ENSG00000163950.12 | SLBP | 0.368588872819011 | 4.37808206454388 | 2.31937540046358 | 0.0252502113043794 | 0.21132453688795 | -3.82911412890829 |
| ENSG00000148719.14 | DNAJB12 | 0.2052939709593 | 5.21615476960292 | 2.31926775578645 | 0.0252566032466203 | 0.21132453688795 | -3.87361883087008 |
| ENSG00000138642.14 | HERC6 | -0.447233338002137 | 3.8804532989842 | -2.31926636820809 | 0.0252566856505056 | 0.21132453688795 | -3.79020949395496 |
| ENSG00000135049.15 | AGTPBP1 | 0.391895340250215 | 5.97195641897468 | 2.31884655293427 | 0.0252816282169112 | 0.211447278987681 | -3.89054178626486 |
| ENSG00000137266.14 | SLC22A23 | 2.62737879491139 | -0.281377342898283 | 2.31955068298812 | 0.025384185063179 | 0.212218798194725 | -3.3524949205846 |
| ENSG00000168785.7 | TSPAN5 | -0.461198991698794 | 2.83801399939604 | -2.31636480335765 | 0.025429528324083 | 0.21251156374121 | -3.7093439243989 |
| ENSG00000212127.5 | TAS2R14 | -0.515926294635185 | 2.1155059224725 | -2.31561608851841 | 0.0254743000161334 | 0.212799317058471 | -3.63975931633167 |
| lnc-CHRM4-2 | lnc-CHRM4-2 | 1.25311791629138 | 2.22301800767355 | 2.31767329616565 | 0.0254856539082033 | 0.212807794885415 | -3.58231648805326 |
| ENSG00000138161.13 | CUZD1 | 1.20599914639839 | 1.14360788414257 | 2.31599033121019 | 0.0255413314442107 | 0.213186222674853 | -3.49313501141221 |
| lnc-SMDT1-1 | lnc-SMDT1-1 | -0.485359316316753 | -0.43227017045105 | -2.31365942885758 | 0.0255916380521461 | 0.21351953223645 | -3.36488985196052 |
| lnc-RHBDD1-6 | lnc-RHBDD1-6 | -0.801273618296169 | 0.910591441368285 | -2.31338938449179 | 0.0256078701807675 | 0.21355377049711 | -3.5159974275621 |
| lnc-AMMECR1L-1 | lnc-AMMECR1L-1 | 0.367257692552243 | 2.15554530907446 | 2.31324600704873 | 0.025616492219665 | 0.21355377049711 | -3.6337070273511 |
| lnc-VSTM2B-9 | lnc-VSTM2B-9 | -0.434053746484191 | 1.31369146756292 | -2.31199638853237 | 0.0256917484983131 | 0.214094436980072 | -3.56761360772953 |
| ENSG00000258388.7 | PPT2-EGFL8 | 0.674242909871953 | -0.453843808132064 | 2.31177008683657 | 0.0257053983122588 | 0.214121494723279 | -3.37816503780149 |
| ENSG00000158773.14 | USF1 | 0.271739774033971 | 6.45852502474355 | 2.31076507343695 | 0.0257660960899668 | 0.214367460419133 | -3.90891541437753 |
| ENSG00000215158.9 | GUSBP18 | -0.361293400966048 | 3.45204272738848 | -2.31071187557872 | 0.0257693125478955 | 0.214367460419133 | -3.76717588521888 |
| ENSG00000184602.5 | SNN | 0.364024050255808 | 5.85954577291411 | 2.31065127603386 | 0.025772976965914 | 0.214367460419133 | -3.90567643864681 |
| ENSG00000206828.1 | RNVU1-30 | -0.855665695793349 | 2.80672256762854 | -2.31191777481632 | 0.0257798505339327 | 0.214367460419133 | -3.7078382899052 |
| ENSG00000164162.13 | ANAPC10 | 0.250621561406903 | 2.7146062935039 | 2.31041943544837 | 0.0257870005343134 | 0.214367460419133 | -3.69725098505408 |
| lnc-GAREM2-4 | lnc-GAREM2-4 | 0.37712200113149 | 2.48709041805614 | 2.30951616428738 | 0.0258417027708142 | 0.214467292461101 | -3.66950224341656 |
| ENSG00000182768.8 | NGRN | 0.236198670584406 | 4.60790299502028 | 2.30950917258555 | 0.0258421265942372 | 0.214467292461101 | -3.86360029077896 |
| ENSG00000214960.9 | CRPPA | -1.01977198246387 | 0.695885711899739 | -2.30982229405221 | 0.0258468446836648 | 0.214467292461101 | -3.47984059363108 |
| lnc-C6orf201-6 | lnc-C6orf201-6 | 0.337930713013102 | 3.0388152659328 | 2.30939015675659 | 0.025849342057993 | 0.214467292461101 | -3.73108762228021 |
| ENSG00000162194.12 | LBHD1 | -0.250612022703821 | 4.18465771469332 | -2.30936103556689 | 0.0258511078363694 | 0.214467292461101 | -3.83466187281412 |
| ENSG00000139405.15 | RITA1 | -0.424974554645717 | 2.75148845140032 | -2.30825963829664 | 0.025917970962277 | 0.21490233334762 | -3.70654705331839 |
| ENSG00000118507.16 | AKAP7 | -1.0415625755768 | 1.80065752412297 | -2.30995744841482 | 0.0259281757637932 | 0.21490233334762 | -3.64077179631246 |
| ENSG00000013288.8 | MAN2B2 | -0.525234314945157 | 6.16919618895464 | -2.30808861737141 | 0.0259443449890865 | 0.21490233334762 | -3.91314188434135 |
| ENSG00000077380.15 | DYNC1I2 | -0.196645149208375 | 4.82449202658412 | -2.30762352145331 | 0.0259566585490194 | 0.21490233334762 | -3.88144644532844 |
| ENSG00000196151.10 | WDSUB1 | 0.361365974982885 | 3.20575220801237 | 2.30756438007767 | 0.0259602580561476 | 0.21490233334762 | -3.74022628665338 |
| ENSG00000201662.1 | RNU6-60P | -0.677630757474407 | -0.109231131320121 | -2.30743954782944 | 0.0259678571578605 | 0.21490233334762 | -3.41245762728043 |
| ENSG00000142794.18 | NBPF3 | -1.66763055560157 | 2.42157145303051 | -2.31047561317976 | 0.0259766314613457 | 0.21490233334762 | -3.64523370452483 |
| ENSG00000125726.10 | CD70 | 1.4193829653106 | 2.02088801605587 | 2.30983492067432 | 0.0259872426933903 | 0.214903743012476 | -3.62505511560208 |
| ENSG00000131381.12 | RBSN | 0.200301418102355 | 5.39267850412578 | 2.30634381197702 | 0.0260346450354745 | 0.215209276612518 | -3.90532109629159 |
| lnc-PPM1D-1 | lnc-PPM1D-1 | -0.555631639514513 | 0.847188915729607 | -2.30566720200959 | 0.0260759628732217 | 0.215464288968094 | -3.51947305672548 |
| lnc-DRD5-29 | lnc-DRD5-29 | -1.29124181811334 | 4.91286669281402 | -2.30829525528829 | 0.0261106008019839 | 0.215663923076739 | -3.88247103954649 |
| ENSG00000101294.17 | HM13 | -0.187077157074606 | 4.69875604792226 | -2.30489140770918 | 0.0261234097159651 | 0.215683169748781 | -3.87774775020968 |
| lnc-BRCA1-3 | lnc-BRCA1-3 | -0.411967497183984 | 3.40053898403117 | -2.30432074837997 | 0.0261583600044074 | 0.215885133909671 | -3.77614176174852 |
| lnc-SH3KBP1-3 | lnc-SH3KBP1-3 | -0.328722777865263 | 0.150961081808444 | -2.30393537193482 | 0.0261819862251631 | 0.21590312920769 | -3.45012329045137 |
| ENSG00000198718.12 | TOGARAM1 | -0.26011556877148 | 5.42838686359195 | -2.30379587728694 | 0.0261905429104493 | 0.21590312920769 | -3.91313115602305 |
| ENSG00000199568.1 | RNU5A-1 | -0.922141907771441 | 2.38698205749029 | -2.30536530425162 | 0.0261920086300152 | 0.21590312920769 | -3.69024922024669 |
| ENSG00000161677.11 | JOSD2 | -0.570190196239342 | 0.452846537758636 | -2.30355924471799 | 0.0262050638173967 | 0.215924270838061 | -3.48768980319505 |
| ENSG00000141873.10 | SLC39A3 | 0.510866724991565 | 3.01363278277914 | 2.30324552786988 | 0.0262243260915919 | 0.215996520185369 | -3.73504987089891 |
| ENSG00000023572.9 | GLRX2 | -0.295136277045085 | 2.64065414202419 | -2.3017916910491 | 0.0263137574564279 | 0.216585815152666 | -3.71736279452501 |
| ENSG00000067221.13 | STOML1 | -0.319557065503235 | 1.94773134158667 | -2.30153299565432 | 0.0263296994207101 | 0.216585815152666 | -3.64254721327845 |
| ENSG00000113966.9 | ARL6 | -0.385153265832248 | 1.60492015518292 | -2.30142696613499 | 0.0263362359290078 | 0.216585815152666 | -3.61305566437303 |
| lnc-MRPS2-2 | lnc-MRPS2-2 | -0.62148215198538 | -0.331173668519713 | -2.30139895274902 | 0.0263379631408018 | 0.216585815152666 | -3.41305537828124 |
| lnc-CD83-8 | lnc-CD83-8 | -0.583062775535974 | -0.276271738347055 | -2.30100413723882 | 0.0263623169282111 | 0.216699508519716 | -3.4051160333597 |
| ENSG00000078687.17 | TNRC6C | -0.36601497085386 | 4.78740717547304 | -2.30065073385935 | 0.0263841333369574 | 0.216734953755342 | -3.89638802053183 |
| lnc-TNFSF4-3 | lnc-TNFSF4-3 | -0.373686859086181 | 2.96768597799432 | -2.30036528850307 | 0.026401766329835 | 0.216734953755342 | -3.73927807739087 |
| lnc-INTS12-7 | lnc-INTS12-7 | 0.274391313192846 | 2.10631823441836 | 2.30027092459296 | 0.0264075978483725 | 0.216734953755342 | -3.65982091412951 |
| ENSG00000175768.12 | TOMM5 | 0.254360467422723 | 4.00126129271602 | 2.30020661925657 | 0.0264115724617045 | 0.216734953755342 | -3.83477862199527 |
| ENSG00000256861.1 | | 0.174450993861301 | 4.22905443837968 | 2.30003197105251 | 0.0264223698975495 | 0.216734953755342 | -3.85505136002559 |
| lnc-TAGLN-1 | lnc-TAGLN-1 | -0.978604577901391 | 3.81419511858799 | -2.30199017196565 | 0.0264456806833029 | 0.216734953755342 | -3.80507689686395 |
| lnc-CCNL2-3 | lnc-CCNL2-3 | 0.215852129474086 | 2.82899283718309 | 2.29962392314826 | 0.0264476124090538 | 0.216734953755342 | -3.72330622738517 |

| | | | | | | | |
|---|---|---|---|---|---|---|---|
| ENSG00000139636.15 | LMBR1L | 0.428111493373112 | 5.86598907755069 | 2.29953843453836 | 0.0264529036068579 | 0.216734953755342 | -3.92830731995196 |
| ENSG00000186481.16 | ANKRD20A5P | -1.86370472789914 | 1.10634794551139 | -2.30240878242218 | 0.0264626629243138 | 0.216734953755342 | -3.53225094634746 |
| ENSG00000185261.14 | KIAA0825 | -0.399228316807988 | 2.41339701310824 | -2.29916691406284 | 0.0264759093478318 | 0.216734953755342 | -3.69249099533491 |
| ENSG00000100852.12 | ARHGAP5 | 0.398855151026945 | 7.49935262809554 | 2.29897424642499 | 0.0264878469926163 | 0.216734953755342 | -3.92421483197565 |
| ENSG00000121486.11 | TRMT1L | -0.272689434787011 | 3.96922297682077 | -2.29830310729819 | 0.0265294682315113 | 0.216734953755342 | -3.83403075139863 |
| ENSG00000105856.13 | HBP1 | 0.271716832751655 | 7.36411459409351 | 2.29814431185597 | 0.0265393246138345 | 0.216734953755342 | -3.92777264643332 |
| LINC01006 | LINC01006 | -0.412404453204705 | 1.03865204295127 | -2.29810787299656 | 0.0265415868241608 | 0.216734953755342 | -3.55728010403776 |
| ENSG00000135749.18 | PCNX2 | -0.6736273310616 | 4.80295384623175 | -2.29924795933887 | 0.0265463472356599 | 0.216734953755342 | -3.89055721449388 |
| ENSG00000161243.8 | FBXO27 | -1.58196393188718 | 0.148154550637931 | -2.29952933505097 | 0.0265678194538518 | 0.216734953755342 | -3.44261303947297 |
| lnc-PIGBOS1-5 | lnc-PIGBOS1-5 | -0.61647606665155 | -0.118539425772135 | -2.29750644263363 | 0.0265789499633707 | 0.216734953755342 | -3.42670465022788 |
| ENSG00000242615.1 | RPL17P47 | -0.497609729518451 | -0.0117263585218072 | -2.29742773327157 | 0.0265838431673993 | 0.216734953755342 | -3.43681463102831 |
| ENSG00000138172.10 | CALHM2 | -0.772898264979045 | 1.15322081066347 | -2.29738551867343 | 0.0265864678966801 | 0.216734953755342 | -3.56131276255626 |
| lnc-ERV3-1-2 | lnc-ERV3-1-2 | -0.411695368950126 | 3.19320153830228 | -2.29733133086687 | 0.0265898374102893 | 0.216734953755342 | -3.77487539921855 |
| ENSG00000162959.13 | MEMO1 | 0.213901167219531 | 3.7569817582778 | 2.29708889717549 | 0.0266049171325371 | 0.216734953755342 | -3.82360465368386 |
| ENSG00000112137.17 | PHACTR1 | -0.850371004073089 | 4.72684570802882 | -2.29920432638911 | 0.026606954753555 | 0.216734953755342 | -3.88111388730304 |
| ENSG00000101417.11 | PXMP4 | -0.376813402950372 | 2.44123631503615 | -2.29702626051792 | 0.0266088144653233 | 0.216734953755342 | -3.69585067681122 |
| lnc-ASAH2-5 | lnc-ASAH2-5 | -0.470345362202823 | 0.505723198586225 | -2.29603406207116 | 0.0266706184585374 | 0.217061203460804 | -3.49153361979755 |
| ENSG00000143319.16 | ISG20L2 | 0.240470663420544 | 4.51974962676714 | 2.2959495442185 | 0.026675889000109 | 0.217061203460804 | -3.87865324002585 |
| ENSG00000101407.12 | TTI1 | 0.252990157932834 | 4.76285696589441 | 2.29565197768063 | 0.0266944526915917 | 0.217061203460804 | -3.89742848609083 |
| lnc-UBE3C-11 | lnc-UBE3C-11 | 0.514811863927918 | -0.318232596427206 | 2.29562042964592 | 0.0266964214931937 | 0.217061203460804 | -3.41212642123935 |
| ENSG00000282164.3 | PEG13 | -0.597585480329949 | 2.27229868023622 | -2.29553751038403 | 0.0267015968110944 | 0.217061203460804 | -3.68389454016963 |
| lnc-TBC1D19-3 | lnc-TBC1D19-3 | -0.6983583518458 | -0.525631059704552 | -2.29521164453025 | 0.0267219440749417 | 0.217140850728198 | -3.40620864914457 |
| lnc-FZD5-4 | lnc-FZD5-4 | 0.39658783423967 | 1.13290170275751 | 2.29458369103723 | 0.0267611929810911 | 0.21737396808595 | -3.57994417146502 |
| ENSG00000149639.14 | MTCL2 | -0.354827866455214 | 5.54205515343154 | -2.29432869246568 | 0.0267771458381194 | 0.217396794571905 | -3.9321095370218 |
| lnc-FAM167B-2 | lnc-FAM167B-2 | -0.715697991801571 | -0.504205133409614 | -2.29420116694592 | 0.0267851270968446 | 0.217396794571905 | -3.39462865410601 |
| ENSG00000064933.17 | PMS1 | -0.310949350043718 | 5.31695302070449 | -2.29396436568495 | 0.026799953083266 | 0.217431389165496 | -3.92767456421437 |
| ENSG00000175455.14 | CCDC14 | -0.307686389514459 | 6.51236156735849 | -2.29356932373396 | 0.0268247027559036 | 0.21754643688919 | -3.94313046705396 |
| lnc-ARMCX1-1 | lnc-ARMCX1-1 | 1.07748144331666 | -0.452001944339903 | 2.29307487209985 | 0.0268557093130948 | 0.217624733540243 | -3.39397125706923 |
| ENSG00000119725.18 | ZNF410 | 0.259031646006864 | 4.83869797595746 | 2.29277157862837 | 0.0268747443882496 | 0.217624733540243 | -3.90944074231437 |
| ENSG00000159346.12 | ADIPOR1 | 0.212002241841396 | 5.52137322409446 | 2.29276836740897 | 0.0268749459928819 | 0.217624733540243 | -3.93626594445395 |
| ENSG00000231793.5 | DOC2GP | -0.890949887035001 | -0.554741354385549 | -2.29252312129062 | 0.0268903468660848 | 0.217624733540243 | -3.40541989167682 |
| ENSG00000169570.9 | DTWD2 | -0.357739619227257 | 3.36824851745228 | -2.29222285330257 | 0.0269092137250316 | 0.217624733540243 | -3.80389358388587 |
| ENSG00000116985.11 | BMP8B | -0.563382093329907 | 2.04551818574514 | -2.29213016225389 | 0.0269150402086816 | 0.217624733540243 | -3.7114202016787 |
| ENSG00000196839.12 | ADA | -1.00584958263927 | 1.13030472287425 | -2.29332090056045 | 0.0269175873806157 | 0.217624733540243 | -3.61075650880525 |
| lnc-LRRC40-5 | lnc-LRRC40-5 | 0.263982137209942 | 1.87775206596938 | 2.29188462367827 | 0.0269304800140211 | 0.217624733540243 | -3.65043187722285 |
| ENSG00000121749.15 | TBC1D15 | 0.401246964659236 | 5.3938575940459 | 2.29167940574334 | 0.0269433904751151 | 0.217624733540243 | -3.93384332618656 |
| ENSG00000075413.17 | MARK3 | 0.150004090287903 | 6.14607744488364 | 2.29145146795587 | 0.0269577367493412 | 0.217624733540243 | -3.94784907699894 |
| lnc-MRPS22-1 | lnc-MRPS22-1 | -0.355224674734259 | 2.62891468075213 | -2.2913959654934 | 0.0269612310762287 | 0.217624733540243 | -3.72108916506009 |
| ENSG00000144476.5 | ACKR3 | 1.38549468685103 | -0.816488113388527 | 2.29132321239799 | 0.0269671305128383 | 0.217624733540243 | -3.39764690248325 |
| lnc-CNPY3-3 | lnc-CNPY3-3 | -0.279784852440361 | 3.35337260797861 | -2.29122802734803 | 0.026971806600649 | 0.217624733540243 | -3.80211913584398 |
| ENSG00000211455.7 | STK38L | 0.27215420386098 | 4.0771611829833 | 2.29080642379071 | 0.0269983724705687 | 0.217753722790641 | -3.86309948438989 |
| ENSG00000186020.12 | ZNF529 | 0.284767038268947 | 4.97757498077412 | 2.29037189239872 | 0.0270257774147635 | 0.217837417381183 | -3.92195405525531 |
| ENSG00000244313.3 | | 1.24039302282849 | 2.22345667602295 | 2.29249208683198 | 0.0270299161439801 | 0.217837417381183 | -3.63236314456719 |
| lnc-ALKBH6-1 | lnc-ALKBH6-1 | -0.21810054716097 | 1.52766727973255 | -2.2899377461681 | 0.0270531828949285 | 0.217928799494963 | -3.62265441751362 |
| ENSG00000071242.11 | RPS6KA2 | -0.401822931427789 | 4.00719294845181 | -2.28979133466319 | 0.0270624307199691 | 0.217928799494963 | -3.86425536231469 |
| ENSG00000166526.16 | ZNF3 | 0.208530970592482 | 5.06192808273996 | 2.28950011077587 | 0.0270808337637311 | 0.217950913921309 | -3.92641258340728 |
| ENSG00000175322.11 | ZNF519 | -0.358374777004552 | 3.1145532700231 | -2.28940022122962 | 0.0270871485676733 | 0.217950913921309 | -3.77190153369485 |
| ENSG00000109670.14 | FBXW7 | 0.32281363153255 | 6.4268180032635 | 2.28905121757792 | 0.0271092221662656 | 0.217950913921309 | -3.95307851709888 |
| ENSG00000182397.14 | DNM1P46 | -0.419046225646861 | 1.38880418373461 | -2.288992293181 | 0.0271129505711658 | 0.217950913921309 | -3.61479689641388 |
| LPP-AS2 | LPP-AS2 | -0.62234193838563 | 1.42253030280676 | -2.28874871524848 | 0.0271283676776041 | 0.217950913921309 | -3.62429160249722 |
| ENSG00000168610.14 | STAT3 | 0.190182809529069 | 6.84580737299971 | 2.28859269030527 | 0.0271382472905651 | 0.217950913921309 | -3.95200923968494 |
| ENSG00000121417.13 | ZNF211 | 0.205156866435352 | 5.01137590019416 | 2.28842383919065 | 0.0271489426885279 | 0.217950913921309 | -3.92726376543399 |
| lnc-CLEC18B-5 | lnc-CLEC18B-5 | -0.268351644782101 | 4.59647963136105 | -2.2884089192454 | 0.0271498879315083 | 0.217950913921309 | -3.90543266722481 |
| ENSG00000164182.10 | NDUFAF2 | -0.358314909628404 | 1.85912588356138 | -2.28820368233438 | 0.0271628935614598 | 0.217970307280907 | -3.66207422368377 |
| ENSG00000117226.11 | GBP3 | 0.342576298615272 | 3.61209340242674 | 2.28782332018663 | 0.0271870113991494 | 0.218078821367378 | -3.82156059391846 |
| lnc-RFX2-2 | lnc-RFX2-2 | -0.317099581405309 | 3.27152386831363 | -2.28739394414196 | 0.0272142600798274 | 0.218135722568488 | -3.79901786661598 |
| ENSG00000141738.13 | GRB7 | 1.65571508519434 | -0.0247152445628566 | 2.28885482498688 | 0.027231649900647 | 0.218135722568488 | -3.43487029829181 |
| ENSG00000256349.1 | | -0.353235072696358 | 2.30983606545269 | -2.28701432485298 | 0.0272383714637981 | 0.218135722568488 | -3.71414202114372 |
| lnc-SEC22A-14 | lnc-SEC22A-14 | -0.579847050282651 | 1.96822809952011 | -2.28695949086438 | 0.0272418558022669 | 0.218135722568488 | -3.66499658664295 |
| ENSG00000010072.15 | SPRTN | 0.266331264385671 | 4.15774352850123 | 2.28687706241213 | 0.0272470943362766 | 0.218135722568488 | -3.87410180242312 |
| lnc-COX5A-1 | lnc-COX5A-1 | -0.258603308608558 | 2.99251428675715 | -2.28659625912125 | 0.0272649468431699 | 0.218185972591675 | -3.77136905115464 |
| ENSG00000136111.13 | TBC1D4 | -2.49544635516334 | -1.75159169718941 | -2.28711205760442 | 0.0272813531969799 | 0.218185972591675 | -3.49102748383432 |
| ENSG00000213213.13 | CCDC183 | -0.451737369035434 | 0.488896521304939 | -2.28613408662565 | 0.0272943529371438 | 0.218185972591675 | -3.50904381113576 |
| ENSG00000113282.13 | CLINT1 | 0.544571010898122 | 7.38948952653385 | 2.28646146946324 | 0.0272977483196182 | 0.218185972591675 | -3.95005443827903 |
| ENSG00000196263.7 | ZNF471 | -2.21698076907247 | 1.89767655765277 | -2.28919799545558 | 0.0273063725111089 | 0.218185972591675 | -3.70423660965767 |
| ENSG00000163472.18 | TMEM79 | -0.524297462966383 | 1.40213971187052 | -2.2855740546921 | 0.0273300233841072 | 0.218208966445083 | -3.62769730722409 |
| ENSG00000152749.7 | GPR180 | -0.374632434423738 | 3.58058130307762 | -2.2855673125676 | 0.0273304530678436 | 0.218208966445083 | -3.8361108120997 |
| lnc-ZNF821-3 | lnc-ZNF821-3 | 0.284817483650353 | 3.14451776796649 | 2.2853135262623 | 0.0273466315655038 | 0.218253477127865 | -3.78153903246213 |

| | | | | | | | |
|---|---|---|---|---|---|---|---|
| ENSG00000126882.12 | FAM78A | -0.382007470048965 | 4.9423689491277 | -2.28469353486651 | 0.0273861910771913 | 0.218415809143981 | -3.93203740877347 |
| ENSG00000272419.6 | LINC01145 | -0.345409607893684 | 1.99252457561859 | -2.28466216080566 | 0.0273881943060106 | 0.218415809143981 | -3.68690073199925 |
| lnc-MDM4-12 | lnc-MDM4-12 | -0.452574824199214 | 0.735428980229189 | -2.28442790055032 | 0.0274031559242416 | 0.218441707327066 | -3.55716555687661 |
| ENSG00000064012.21 | CASP8 | 0.306088318385804 | 6.74795778779967 | 2.28427903716414 | 0.0274126672509261 | 0.218441707327066 | -3.96127845238303 |
| ENSG00000155592.15 | ZKSCAN2 | 0.311047167666555 | 4.26290361291316 | 2.2829288648237 | 0.0274990685540975 | 0.218857230818992 | -3.88721361281468 |
| ENSG00000143851.15 | PTPN7 | -0.399633752703834 | 5.8625623513082 | -2.28273434056556 | 0.0275115367312089 | 0.218857230818992 | -3.96251140025123 |
| ENSG00000115415.18 | STAT1 | 0.509619775016636 | 7.33891824200872 | 2.28265171450656 | 0.0275270773725573 | 0.218857230818992 | -3.95851254962199 |
| ENSG00000137124.7 | ALDH1B1 | -0.644149914907987 | 1.73407815093692 | -2.28239618452462 | 0.0275332231123867 | 0.218857230818992 | -3.68175934688383 |
| lnc-STAT3-2 | lnc-STAT3-2 | -0.605892193487658 | 0.454011510356245 | -2.28235080513676 | 0.0275361345124161 | 0.218857230818992 | -3.51685442530939 |
| ENSG00000056586.15 | RC3H2 | 0.251309767992042 | 6.67481086327972 | 2.28233605338706 | 0.0275370809978961 | 0.218857230818992 | -3.96580507188732 |
| lnc-SLC39A4-1 | lnc-SLC39A4-1 | -0.439264054023124 | 2.56834361256867 | -2.28230236672941 | 0.0275392424729723 | 0.218857230818992 | -3.7448326548913 |
| ENSG00000141469.17 | SLC14A1 | -1.05818552536808 | 3.69018462980787 | -2.28393985454591 | 0.027585056668057 | 0.219082802179445 | -3.8291360737232 |
| ENSG00000236567.2 | TCF3P1 | -0.664617482908088 | 0.134484558910338 | -2.28137106586326 | 0.0275990586139705 | 0.219082802179445 | -3.47171484234707 |
| lnc-TP53TG3-73 | lnc-TP53TG3-73 | -0.406526942375911 | 3.75552962514387 | -2.28136329549327 | 0.0275995581815722 | 0.219082802179445 | -3.85406168212171 |
| ENSG00000186532.11 | SMYD4 | -0.286423612861816 | 4.70221754537926 | -2.28086466800292 | 0.0276316325081109 | 0.219252849620064 | -3.92756262582484 |
| lnc-IFFO1-4 | lnc-IFFO1-4 | 0.306588075201681 | 1.174707501051 | 2.27958139897117 | 0.0277143321966696 | 0.219824315839711 | -3.61131627795904 |
| ENSG00000139505.11 | MTMR6 | 0.391156599747708 | 5.04946506799558 | 2.27885072675389 | 0.0277615187137786 | 0.219926896367048 | -3.9470271167892 |
| ENSG00000284388.1 | MIR4496 | 0.439632145123414 | 0.431049279706854 | 2.27873749395533 | 0.0277688376580937 | 0.219926896367048 | -3.49965604121016 |
| ENSG00000229023.1 | RAB1AP1 | 0.70352715125911 | -0.275161423739638 | 2.27855279086614 | 0.0277807798714624 | 0.219926896367048 | -3.43589911878679 |
| ENSG00000138750.14 | NUP54 | 0.313729792570264 | 4.56703634909885 | 2.27851095459071 | 0.0277834854859481 | 0.219926896367048 | -3.92238466683365 |
| ENSG00000131069.19 | ACSS2 | 0.431867858255925 | 3.15871754056531 | 2.27803506107417 | 0.0278142787935532 | 0.219926896367048 | -3.78900260460476 |
| ENSG00000236254.1 | MTND4P14 | -0.754430402247596 | 0.243421490064914 | -2.27780608319724 | 0.0278291059584917 | 0.219926896367048 | -3.49031966944159 |
| lnc-CYGB-1 | lnc-CYGB-1 | 0.569567462754069 | 2.79697688474615 | 2.27780381201365 | 0.0278292530613697 | 0.219926896367048 | -3.77889097573946 |
| ENSG00000152926.14 | ZNF117 | -0.472841736127294 | 5.7826635926025 | -2.27764564882551 | 0.0278394988813914 | 0.219926896367048 | -3.97274031970967 |
| ENSG00000111863.12 | ADTRP | -0.991827420576082 | 6.13249643638333 | -2.28037207113287 | 0.0278399447881444 | 0.219926896367048 | -3.96978810377788 |
| lnc-TUBB2B-2 | lnc-TUBB2B-2 | 2.65624807601893 | -1.48487364970662 | 2.27847497782449 | 0.0278419819141306 | 0.219926896367048 | -3.52274170211881 |
| ENSG00000164708.5 | PGAM2 | -0.651331513256095 | -0.387745820046436 | -2.27745261354863 | 0.0278520082802281 | 0.219926896367048 | -3.4428590428298 |
| ENSG00000184619.5 | KRBA2 | -0.391760004067501 | 2.26792404386394 | -2.27724231271955 | 0.0278656422615715 | 0.219926896367048 | -3.72184977368776 |
| ENSG00000186687.15 | LYRM7 | -0.315146944537855 | 4.33854978990223 | -2.27723419784396 | 0.0278661684752106 | 0.219926896367048 | -3.91454405314148 |
| lnc-PCP4L1-46 | lnc-PCP4L1-46 | -0.400676369111327 | 0.353899291324903 | -2.27680375774752 | 0.02789409332119 | 0.220062906412439 | -3.51140867565865 |
| lnc-ANXA2-2 | lnc-ANXA2-2 | -0.229838758112438 | 3.38319971834104 | -2.2764395987907 | 0.0279177376746329 | 0.220137379443479 | -3.82660768173512 |
| ENSG00000164338.9 | UTP15 | 0.292558017998198 | 5.1969979412065 | 2.27618326355426 | 0.0279343919052948 | 0.220137379443479 | -3.96071829757676 |
| ENSG00000115423.18 | DNAH6 | -1.31129473599098 | 0.873119516249481 | -2.27791074551436 | 0.0279356184767219 | 0.220137379443479 | -3.5482581375991 |
| ENSG00000213859.5 | KCTD11 | -0.524247237398725 | 2.24245611540306 | -2.27586984773906 | 0.0279547667397858 | 0.220203966247612 | -3.73094771478971 |
| ENSG00000070950.9 | RAD18 | 0.215141216800463 | 3.6861018298427 | 2.27546688445558 | 0.0279809824453875 | 0.220326152132139 | -3.85105869671516 |
| ENSG00000007520.3 | TSR3 | -0.42634938611576 | 3.22531847589463 | -2.27517693777184 | 0.0279998591791187 | 0.220390478578891 | -3.82284670063581 |
| lnc-ANTXR2-1 | lnc-ANTXR2-1 | -0.651921286356156 | 1.12674225775413 | -2.27469712176643 | 0.0280311221520335 | 0.220534082385023 | -3.61859586177972 |
| ENSG00000148384.12 | INPP5E | -0.342387682600856 | 3.43132626490209 | -2.27456812631708 | 0.0280395323131519 | 0.220534082385023 | -3.84749970786172 |
| ENSG00000005810.17 | MYCBP2 | 0.191249259050737 | 8.86199181802194 | 2.27430943647034 | 0.0280564049916433 | 0.220582499596255 | -3.94816162788175 |
| lnc-AMZ2-8 | lnc-AMZ2-8 | -0.606345268474105 | 5.14552281894709 | -2.27478167194078 | 0.0280772819179554 | 0.220662349643862 | -3.95671235103534 |
| ENSG00000182224.11 | CYB5D1 | 0.509157080255068 | 1.81035211898522 | 2.27374330518405 | 0.0280933617187861 | 0.220704452006784 | -3.69201104854115 |
| lnc-TMEM154-25 | lnc-TMEM154-25 | 0.40537068670185 | 2.5855986496043 | 2.27344045530462 | 0.0281131494163222 | 0.220775640761603 | -3.74692964324587 |
| ENSG00000163960.11 | UBXN7 | 0.256060459449246 | 6.75916062685771 | 2.27279371598929 | 0.0281554479117148 | 0.22102348755409 | -3.98473138405147 |
| ENSG00000164463.12 | CREBRF | 0.252064896005164 | 7.52260087882224 | 2.27250459195641 | 0.0281743757646773 | 0.221087753093539 | -3.97705954831575 |
| ENSG00000197111.15 | PCBP2 | 0.305549563764052 | 7.28919377265824 | 2.27225663097925 | 0.0281906178720757 | 0.221130902309807 | -3.98085281935104 |
| ENSG00000213465.7 | ARL2 | -0.551721411001192 | 2.50329607607493 | -2.27197317789663 | 0.028209195040906 | 0.22119232820075 | -3.76179396963249 |
| lnc-ANKDD1B-2 | lnc-ANKDD1B-2 | -0.60950252779714 | -0.206250140697986 | -2.27093245050746 | 0.0282774967538644 | 0.221618355488433 | -3.47595820547601 |
| ENSG00000169021.5 | UQCRFS1 | 0.273799705648959 | 4.83696356607819 | 2.27081732362062 | 0.0282850614520776 | 0.221618355488433 | -3.95393472214997 |
| ENSG00000119699.7 | TGFB3 | -0.364624466662149 | 2.14224502692111 | -2.2702351044433 | 0.0283233452690307 | 0.221771086623735 | -3.72196757128144 |
| ENSG00000170271.10 | FAXDC2 | -1.26197804995258 | 2.72268558267014 | -2.27270534633771 | 0.0283261034219404 | 0.221771086623735 | -3.75100121639342 |
| ENSG00000099899.14 | TRMT2A | -0.2900541424939 | 4.52495689215978 | -2.26874768185034 | 0.0284213607083351 | 0.222346504578244 | -3.94396599647552 |
| lnc-USP45-4 | lnc-USP45-4 | 0.346337341537827 | 1.7643059591235 | 2.26873290956449 | 0.028422335660721 | 0.222346504578244 | -3.68977395339301 |
| lnc-PASK-4 | lnc-PASK-4 | 0.233626000008275 | 1.42730123254227 | 2.26858639584103 | 0.0284320069984909 | 0.222346504578244 | -3.65564936934589 |
| lnc-BCOR-9 | lnc-BCOR-9 | -1.67382261811971 | 2.32615954146207 | -2.27129613475893 | 0.0284545980051145 | 0.222381957981418 | -3.7205023114103 |
| lnc-TFCP2-3 | lnc-TFCP2-3 | 0.53819149772596 | 0.209713459410871 | 2.26801344714823 | 0.0284698554017223 | 0.222381957981418 | -3.49766202559527 |
| lnc-MICALL2-2 | lnc-MICALL2-2 | -0.488699865464007 | 0.531276880883009 | -2.26791879934454 | 0.028476112061661 | 0.222381957981418 | -3.56240424009942 |
| ENSG00000216937.13 | CCDC7 | -0.361359186469686 | 5.25143312766034 | -2.26786366684766 | 0.0284797571412826 | 0.222381957981418 | -3.97802957129647 |
| ENSG00000175395.15 | ZNF25 | -0.284635370423119 | 3.52976185259225 | -2.26760357313818 | 0.0284969588244568 | 0.222431893622979 | -3.85790449903325 |
| ENSG00000119574.12 | ZBTB45 | -0.350303135023358 | 1.84640582837415 | -2.26703975184666 | 0.0285342798011728 | 0.222624976399849 | -3.70805334185781 |
| lnc-ARHGEF2-1 | lnc-ARHGEF2-1 | -0.381864668974383 | 2.12145407849916 | -2.26690316204502 | 0.0285433276353885 | 0.222624976399849 | -3.73837531465745 |
| LINC01374 | LINC01374 | 1.72943363289406 | 1.92294000866995 | 2.26842576747509 | 0.0286359415475027 | 0.223197085483924 | -3.63566262618777 |
| ENSG00000213793.5 | ZNF888 | 0.234962492805819 | 4.07566514733853 | 2.2654707409377 | 0.0286383667474636 | 0.223197085483924 | -3.90736613236051 |
| ENSG00000250731.1 | TPM3P6 | 0.400405232415157 | 0.882485578556054 | 2.26441896386636 | 0.0287083302250033 | 0.223594128177139 | -3.590299819253 |
| ENSG00000146457.15 | WTAP | 0.25526191499604 | 7.16508061222524 | 2.26426074761971 | 0.0287188678355705 | 0.223594128177139 | -3.99785858790197 |
| ENSG00000129467.13 | ADCY4 | -0.427477033269466 | 2.04513905119048 | -2.2639604835356 | 0.0287388756820976 | 0.223594128177139 | -3.72740192684515 |
| ENSG00000262560.1 | | -0.682245674267089 | 1.9446175573112 | -2.26391212925704 | 0.0287422607408502 | 0.223594128177139 | -3.73019011461628 |
| lnc-MARCH3-2 | lnc-MARCH3-2 | -0.438008613353921 | 1.19815499933222 | -2.2636450893329 | 0.0287599050933132 | 0.223594128177139 | -3.63896999135949 |
| ENSG00000104976.11 | SNAPC2 | -0.489523176004462 | 1.62102825164286 | -2.26359102702738 | 0.0287635111607565 | 0.223594128177139 | -3.7047372492757 |

| | | | | | | | |
|---|---|---|---|---|---|---|---|
| ENSG00000124203.6 | ZNF831 | -0.562828729374145 | 6.38973142144564 | -2.2641232545226 | 0.0287689929719708 | 0.223594128177139 | -4.00262923383987 |
| ENSG00000108883.12 | EFTUD2 | 0.193716821982413 | 5.98790581897905 | 2.26340060981693 | 0.0287762155925396 | 0.223594128177139 | -4.00351751735947 |
| ENSG00000101082.13 | SLA2 | -0.902153273489337 | 1.01563695103888 | -2.26351764559201 | 0.0288001700989291 | 0.223695811753305 | -3.63469646951396 |
| ENSG00000142669.14 | SH3BGRL3 | -0.485912940777151 | 5.68914998511047 | -2.26276178573823 | 0.0288202278416106 | 0.223767163207798 | -3.99995939975123 |
| ENSG00000175985.9 | PLEKHD1 | 0.820713379766951 | -0.464579995697574 | 2.26177056153984 | 0.0288851755335885 | 0.224087625894501 | -3.44761944312645 |
| ENSG00000085721.12 | RRN3 | 0.287764418293074 | 5.72764105517492 | 2.26176287134871 | 0.0288856904518433 | 0.224087625894501 | -4.00327413709477 |
| lnc-AIDA-2 | lnc-AIDA-2 | -1.59153664582052 | 1.94722612682161 | -2.26447417813521 | 0.0288985005632001 | 0.224087625894501 | -3.69014683223102 |
| ENSG00000157895.11 | C12orf43 | 0.20054848529659 | 3.70323913187929 | 2.2614664571707 | 0.0289055439288357 | 0.224087625894501 | -3.88942764835646 |
| lnc-HACL1-7 | lnc-HACL1-7 | 0.312818905780539 | 1.85452212442787 | 2.2611385264442 | 0.0289275225184925 | 0.224087625894501 | -3.71543362588348 |
| ENSG00000136875.12 | PRPF4 | 0.26583563679312 | 4.29366268964667 | 2.26110778128525 | 0.028929583885624 | 0.224087625894501 | -3.93600287781335 |
| ENSG00000183718.5 | TRIM52 | -0.179736091264644 | 5.59860721760496 | -2.26098658193554 | 0.0289377111999021 | 0.224087625894501 | -4.00226722659049 |
| ENSG00000100749.7 | VRK1 | 0.26719561039931 | 4.65400488523206 | 2.26076263205028 | 0.0289527340533595 | 0.224119640850056 | -3.95836129054651 |
| lnc-ZNF674-1 | lnc-ZNF674-1 | 0.28210390090428 | 6.69850860585785 | 2.26017860509801 | 0.0289919440571762 | 0.224294285664298 | -4.01057751752778 |
| ENSG00000127481.14 | UBR4 | 0.262465861861456 | 8.63470532441129 | 2.26010201607667 | 0.0289970895473302 | 0.224294285664298 | -3.98111327189695 |
| ENSG00000184470.20 | TXNRD2 | -0.280439515622306 | 3.64471440296192 | -2.25965911376789 | 0.0290268610759218 | 0.224306636108491 | -3.88418866336761 |
| lnc-AUNIP-2 | lnc-AUNIP-2 | -0.484694367870559 | -0.156688192532616 | -2.25962270664757 | 0.0290293095438251 | 0.224306636108491 | -3.48291182580588 |
| lnc-PML-1 | lnc-PML-1 | -0.509470690649496 | 0.26070611021475 | -2.25958815800374 | 0.0290316331948447 | 0.224306636108491 | -3.53402609607227 |
| ENSG00000105926.15 | PALS2 | 0.384873798595796 | 4.26724338966027 | 2.25939795623419 | 0.0290444286346804 | 0.224306636108491 | -3.94244731996222 |
| ENSG00000221988.12 | PPT2 | 0.786883461814669 | -0.422935263276884 | 2.25926799275929 | 0.029053174554984 | 0.224306636108491 | -3.47076479853002 |
| ENSG00000163541.11 | SUCLG1 | 0.202224186856722 | 4.02021842693727 | 2.25882089005902 | 0.0290832803565284 | 0.224340720823181 | -3.92203906709279 |
| ENSG00000103510.19 | KAT8 | 0.163571475470681 | 5.25356893528124 | 2.25879144224221 | 0.0290852642100459 | 0.224340720823181 | -3.99686552654631 |
| lnc-PGC-2 | lnc-PGC-2 | -0.35691318779883 | 2.81367010297798 | -2.258716888501 | 0.0290902873185187 | 0.224340720823181 | -3.81503381984969 |
| lnc-KLF8-2 | lnc-KLF8-2 | -0.619115405351045 | 3.51634442172672 | -2.2587676526875 | 0.0291036031717283 | 0.224359349844077 | -3.86605580807101 |
| ENSG00000114796.15 | KLHL24 | 0.314116963956252 | 6.86330224475199 | 2.25797869842646 | 0.0291400651083531 | 0.224556331009072 | -4.01397919970904 |
| lnc-PPA2-8 | lnc-PPA2-8 | 0.28512477302574 | 1.06321224997461 | 2.25721365513093 | 0.029191733746378 | 0.224868548681271 | -3.63049891751479 |
| ENSG00000278571.1 | MIR7161 | 0.476626234415222 | -0.331508308825699 | 2.25705541938918 | 0.0292024306770169 | 0.224868548681271 | -3.47087022688633 |
| ENSG00000141858.11 | SAMD1 | -0.395287394598772 | 2.70843658342532 | -2.2567658589364 | 0.0292220143627719 | 0.224935198814112 | -3.81065650845928 |
| lnc-HOXB1-3 | lnc-HOXB1-3 | -1.08097646989913 | 1.78251598224654 | -2.25808342154712 | 0.0292529596708846 | 0.225056451660827 | -3.71502031846029 |
| lnc-TMEM199-1 | lnc-TMEM199-1 | -0.474070649866269 | -0.299355086963242 | -2.25621009431756 | 0.0292596348755951 | 0.225056451660827 | -3.47808949547911 |
| ENSG00000185112.5 | FAM43A | -1.00439579910299 | 0.969419757197121 | -2.25697131892075 | 0.0292756337016369 | 0.225095393530367 | -3.64950053393932 |
| lnc-SLFN12-6 | lnc-SLFN12-6 | 1.69010022270194 | -0.294474576429701 | 2.25560275582983 | 0.0293175803238143 | 0.225333740031766 | -3.46078246604298 |
| ENSG00000116525.13 | TRIM62 | 0.337946211850391 | 3.07756244894819 | 2.25512894203611 | 0.0293329432769638 | 0.225367663855822 | -3.83285248853529 |
| ENSG00000008282.8 | SYPL1 | 0.350404133190902 | 5.62572906380481 | 2.2541377941813 | 0.0294002925614339 | 0.225715463702321 | -4.01785459412264 |
| ENSG00000197275.13 | RAD54B | -0.45193000245952 | 1.11448321495463 | -2.25411741073095 | 0.0294016790773899 | 0.225715463702321 | -3.64623582267243 |
| ENSG00000139899.10 | CBLN3 | -0.743279575032449 | 2.95635242576882 | -2.25485698857264 | 0.0294111098309102 | 0.225715463702321 | -3.84610730257253 |
| lnc-MTIF3-2 | lnc-MTIF3-2 | -0.421606741068397 | 0.392283869424436 | -2.25330779853899 | 0.0294567973558438 | 0.225877387631442 | -3.5747277908518 |
| ENSG00000116731.22 | PRDM2 | 0.401410158036377 | 8.85094005446149 | 2.25325347470566 | 0.0294604990070337 | 0.225877387631442 | -3.99390662133706 |
| ENSG00000099994.10 | SUSD2 | -0.749200125811792 | -0.524775406417309 | -2.25305875299342 | 0.0294737708390851 | 0.225877387631442 | -3.46811421130659 |
| ENSG00000126351.12 | THRA | -0.420722577617084 | 4.73255412939523 | -2.25297321820172 | 0.0294796023979667 | 0.225877387631442 | -3.9869062484616 |
| ENSG00000165156.14 | ZHX1 | 0.287641960162351 | 5.08485341075035 | 2.25281073084526 | 0.0294906832302177 | 0.225877387631442 | -4.00243950340671 |
| lnc-MELTF-4 | lnc-MELTF-4 | 0.33229399014877 | 1.13809230639919 | 2.25270269690212 | 0.0294980526625524 | 0.225877387631442 | -3.64989458622241 |
| ENSG00000204519.10 | ZNF551 | 0.271485826743834 | 5.04977558307563 | 2.25185985390197 | 0.0295556027263532 | 0.226026084594483 | -4.00193075389612 |
| ENSG00000152944.8 | MED21 | 0.271059210306628 | 5.19397489045115 | 2.25183118788562 | 0.0295575618242491 | 0.226026084594483 | -4.00963615538703 |
| ENSG00000283701.1 | MIR555 | 0.268904206822819 | 0.781142909831973 | 2.25155494303466 | 0.029576446925775 | 0.226026084594483 | -3.60696757800988 |
| ENSG00000179152.19 | TCAIM | -0.380670075325977 | 3.0296356964442 | -2.25146596766022 | 0.0295825318929791 | 0.226026084594483 | -3.84044047150053 |
| ENSG00000143033.17 | MTF2 | 0.19334648893058 | 5.62272663813952 | 2.25140696912557 | 0.0295865673785471 | 0.226026084594483 | -4.02323664492042 |
| ENSG00000109606.12 | DHX15 | 0.327058601276 | 7.4494035169788 | 2.25133108735726 | 0.0295917583939447 | 0.226026084594483 | -4.01920513224798 |
| ENSG00000133318.13 | RTN3 | 0.362072750506134 | 3.89181661635548 | 2.25110167459056 | 0.0296074572837818 | 0.226026084594483 | -3.91830857413724 |
| lnc-C6orf163-2 | lnc-C6orf163-2 | -0.413344005927397 | 2.67040279403005 | -2.25108337879152 | 0.0296087095984589 | 0.226026084594483 | -3.80636022168325 |
| ENSG00000011590.13 | ZBTB32 | -0.507044580934813 | 5.10536162473215 | -2.25099177924779 | 0.0296227510182049 | 0.226026084594483 | -4.00469438666107 |
| ENSG00000041357.15 | PSMA4 | 0.242440687284867 | 5.54177076101936 | 2.25081210267882 | 0.0296272834978338 | 0.226026084594483 | -4.02164501492195 |
| LINC01358 | LINC01358 | -0.77803668915098 | -0.542435224043532 | -2.25042480832489 | 0.0296538189851521 | 0.22606527179995 | -3.48086879193314 |
| ENSG00000110324.10 | IL10RA | 0.547451915615637 | 9.01782444212995 | 2.25063775376547 | 0.0296600341714736 | 0.22606527179995 | -3.99246078341518 |
| lnc-AMZ2-1 | lnc-AMZ2-1 | -0.495832720548297 | 6.25366157010143 | -2.25027253309469 | 0.0296735199699675 | 0.22606527179995 | -4.02949304127315 |
| lnc-RBM19-2 | lnc-RBM19-2 | -0.27729007927223 | 2.77889532910913 | -2.24997589230119 | 0.0296846029621643 | 0.22606527179995 | -3.82621237809593 |
| ENSG00000125827.8 | TMX4 | -0.203462958382877 | 6.68312557037626 | -2.24993606215056 | 0.029687335649578 | 0.22606527179995 | -4.03044923221722 |
| ENSG00000183049.12 | CAMK1D | 0.333963008107841 | 5.27312962869127 | 2.24946082519252 | 0.0297199582346647 | 0.226228129426628 | -4.01635116603088 |
| ENSG00000173402.11 | DAG1 | -0.298848189710852 | 4.35006727157699 | -2.24924360299834 | 0.0297348800521421 | 0.226228129426628 | -3.97458564398998 |
| ENSG00000225031.1 | EIF4BP7 | 1.12114184079811 | 0.965193070590362 | 2.24981380790308 | 0.0297451499713675 | 0.226228129426628 | -3.5706571362064 |
| ENSG00000135241.16 | PNPLA8 | 0.323188495963236 | 5.95007420537545 | 2.24887940408956 | 0.0297599132255749 | 0.226228129426628 | -4.03152593899313 |
| lnc-ZC3HAV1L-2 | lnc-ZC3HAV1L-2 | 0.253206070975234 | 2.58763450509849 | 2.24882466220911 | 0.0297636775245255 | 0.226228129426628 | -3.79321021622854 |
| ENSG00000124275.14 | MTRR | 0.243799259452146 | 5.53805748958008 | 2.24814131097111 | 0.0298107035280636 | 0.226346688529776 | -4.02549630188755 |
| ENSG00000126246.9 | IGFLR1 | -0.464799296180426 | 4.8682211069475 | -2.2480418123621 | 0.029817556211551 | 0.226346688529776 | -3.99969384157569 |
| ENSG00000139350.11 | NEDD1 | 0.17578100056797 | 4.84752305537952 | 2.24795901259458 | 0.0298232598788543 | 0.226346688529776 | -4.00393704443516 |
| ENSG00000165671.20 | NSD1 | 0.138745260265482 | 7.74786647981018 | 2.24778253262005 | 0.029835419954506 | 0.226346688529776 | -4.02387027324086 |
| ENSG00000099810.19 | MTAP | -0.32894581027725 | 4.97532098197467 | -2.24760450244627 | 0.0298476913159968 | 0.226346688529776 | -4.00866055799451 |
| ENSG00000017797.12 | RALBP1 | -0.402063811834271 | 5.17752811389502 | -2.2474821247798 | 0.029856129236429 | 0.226346688529776 | -4.01676922945444 |
| lnc-TDRP-5 | lnc-TDRP-5 | -0.480735288982824 | 1.63081384359591 | -2.24736284809158 | 0.0298643553882035 | 0.226346688529776 | -3.72837044978309 |

| | | | | | | | |
|---|---|---|---|---|---|---|---|
| ENSG00000015285.10 | WAS | -0.337967897644483 | 4.58659442576315 | -2.24720479714577 | 0.0298752587909565 | 0.226346688529776 | -3.98800755105986 |
| ENSG00000173726.10 | TOMM20 | 0.264065446406622 | 6.34868457087826 | 2.24680263857415 | 0.0299030183378221 | 0.226346688529776 | -4.03801317608895 |
| ENSG00000204305.13 | AGER | -0.419897042342471 | 2.53044232156201 | -2.2467351726447 | 0.0299076775143277 | 0.226346688529776 | -3.80882901818357 |
| ENSG00000101474.11 | APMAP | -0.281031350087271 | 4.38999785316437 | -2.24673057235195 | 0.0299079952326724 | 0.226346688529776 | -3.97412786001899 |
| ENSG00000104823.8 | ECH1 | -0.312424033559509 | 4.25004101277528 | -2.24668363564402 | 0.0299112370791911 | 0.226346688529776 | -3.96287585829638 |
| ENSG00000164808.16 | SPIDR | -0.234830522384551 | 5.55062529429075 | -2.24632454238043 | 0.0299360494526325 | 0.226441448147565 | -4.02981328368961 |
| MORF4L2-AS1 | MORF4L2-AS1 | -0.424637592781419 | 1.93906766814786 | -2.2460993305381 | 0.0299516203281456 | 0.226441448147565 | -3.75114156016067 |
| ENSG00000231130.1 | HLA-T | -0.67274061071162 | -0.764432396612631 | -2.24598134107024 | 0.0299597808550438 | 0.226441448147565 | -3.4777568317313 |
| lnc-ZNF726-2 | lnc-ZNF726-2 | -0.372307185685002 | 3.08366312965635 | -2.24586593108512 | 0.0299677648911221 | 0.226441448147565 | -3.86085822537636 |
| ENSG00000176973.7 | FAM89B | -0.477874954035015 | 2.44538574002701 | -2.24562634822978 | 0.029984345223867 | 0.226483588162515 | -3.80712663908773 |
| ENSG00000100220.11 | RTCB | 0.221090777510927 | 5.28816407481433 | 2.24460748072946 | 0.0300549471769118 | 0.226875002417962 | -4.02768954940383 |
| ENSG00000095209.11 | TMEM38B | 0.36709906376202 | 3.48717171200125 | 2.24456044933299 | 0.0300582097650382 | 0.226875002417962 | -3.89609129206669 |
| ENSG00000069869.16 | NEDD4 | -1.79267317595784 | -1.42410556202037 | -2.24433709903708 | 0.0300750848119917 | 0.226919160808367 | -3.50361710831994 |
| ENSG00000139160.13 | ETFBKMT | -0.432163414335937 | 1.09486269211747 | -2.24413582081845 | 0.0300876806987332 | 0.226931012027126 | -3.67475270839112 |
| lnc-COL4A4-2 | lnc-COL4A4-2 | -1.66191051557188 | -1.45142759758258 | -2.24393420075531 | 0.0301186449584099 | 0.227063432723669 | -3.47763691629327 |
| lnc-SLFN12-5 | lnc-SLFN12-5 | 1.48177960719885 | 0.104425030664493 | 2.24397320658823 | 0.0301273009166953 | 0.227063432723669 | -3.47861728255822 |
| lnc-NDUFS5-2 | lnc-NDUFS5-2 | -0.436466801177443 | -0.00707967004558348 | -2.24300322063051 | 0.0301664135200369 | 0.227132402029147 | -3.5375383755827 |
| ENSG00000164296.6 | TIGD6 | -0.432591300798243 | 2.15815242934233 | -2.24292206018454 | 0.0301720624295458 | 0.227132402029147 | -3.77418475324168 |
| ENSG00000151474.22 | FRMD4A | -2.00386424979598 | 0.710115911950348 | -2.24560310146456 | 0.0301743939050056 | 0.227132402029147 | -3.5741057683529 |
| ENSG00000183963.18 | SMTN | 0.549699805833946 | 0.709625881818595 | 2.24279954149007 | 0.0301805917285973 | 0.227132402029147 | -3.62132530420844 |
| ENSG00000198887.8 | SMC5 | 0.29228846818893 | 6.47620492138122 | 2.24236317700346 | 0.0302109872425128 | 0.227226672723474 | -4.04670080465757 |
| ENSG00000181061.13 | HIGD1A | 0.345080509846839 | 3.19207034733113 | 2.24227125784218 | 0.0302173934584345 | 0.227226672723474 | -3.87921322335095 |
| ENSG00000196227.10 | FAM217B | -0.313364813523677 | 3.97804638706421 | -2.24213292026967 | 0.030227037038307 | 0.227226672723474 | -3.9530472935494 |
| lnc-OSBPL3-2 | lnc-OSBPL3-2 | 1.37916174918713 | -1.07335646658553 | 2.24198608162927 | 0.0302372762274496 | 0.227226672723474 | -3.47704678659618 |
| lnc-PRTFDC1-4 | lnc-PRTFDC1-4 | -0.428308956019264 | 1.79012851968289 | -2.2408543294758 | 0.0303162979157084 | 0.22766889345839 | -3.76130278253788 |
| ENSG00000114127.10 | XRN1 | 0.247024650286376 | 7.35744243504689 | 2.24082647629759 | 0.030318245006532 | 0.22766889345839 | -4.04270732058934 |
| ENSG00000115339.13 | GALNT3 | 0.603383266430819 | 3.31568203193369 | 2.24060325536671 | 0.0303390788307002 | 0.22768981534715 | -3.87350558986171 |
| ENSG00000140859.15 | KIFC3 | -3.651553020091 | -2.03413338141984 | -2.2422081804896 | 0.0303558779332401 | 0.22768981534715 | -3.61105138604316 |
| ENSG00000113575.9 | PPP2CA | 0.24324439048535 | 6.42429424078469 | 2.24017287084789 | 0.0303639675790623 | 0.22768981534715 | -4.05113290497358 |
| ENSG00000183520.11 | UTP11 | 0.26908860004797 | 3.04801527052233 | 2.24000108802786 | 0.0303759947162672 | 0.22768981534715 | -3.86648394148689 |
| ENSG00000277916.1 | | 0.561481705413462 | 1.07064046527892 | 2.23988942122287 | 0.0303838151860593 | 0.22768981534715 | -3.66149192858979 |
| ENSG00000107438.8 | PDLIM1 | 0.404744002885386 | 5.28243231970968 | 2.23978844209358 | 0.0303908886974647 | 0.22768981534715 | -4.03547626748121 |
| lnc-ENTPD1-1 | lnc-ENTPD1-1 | 0.560793795105893 | 1.21730013598102 | 2.23955747671648 | 0.0304070731538905 | 0.22768981534715 | -3.67304596721933 |
| lnc-PPP1R10-2 | lnc-PPP1R10-2 | 0.259563062692406 | 1.57477913426206 | 2.23952246223263 | 0.030409527395876 | 0.22768981534715 | -3.72658303165914 |
| ENSG00000006695.10 | COX10 | -0.272626500640064 | 3.43730428585394 | -2.23858141285113 | 0.0304755536469801 | 0.228101207533015 | -3.91315639297585 |
| ENSG00000285542.1 | | 0.336254619976645 | 4.30058118390186 | 2.23802359933002 | 0.0305147513243915 | 0.228311569069411 | -3.97969973698442 |
| ENSG00000127366.5 | TAS2R5 | -0.582643783953896 | -0.42508972329469 | -2.23774620271927 | 0.0305342607322742 | 0.228374523492878 | -3.50355378900751 |
| ENSG00000175221.14 | MED16 | -0.353726416316495 | 4.06154312610633 | -2.23752820040354 | 0.0305496007005723 | 0.228406259070062 | -3.97108507134941 |
| ENSG00000177485.6 | ZBTB33 | 0.237867501162926 | 5.77079859968116 | 2.23703016510353 | 0.0305846712078648 | 0.228585434811721 | -4.05383088027855 |
| ENSG00000157379.13 | DHRS1 | -0.317160376615967 | 2.60453995638494 | -2.23671596136454 | 0.0306068151310059 | 0.228660541886964 | -3.82877867887252 |
| lnc-PABPC1-4 | lnc-PABPC1-4 | 0.319817983018027 | 5.66895250250728 | 2.23657238084089 | 0.0306169389030011 | 0.228660541886964 | -4.04987089157114 |
| ENSG00000100441.9 | KHNYN | 0.253726180900748 | 7.0780383721313 | 2.23626428615312 | 0.0306386725152841 | 0.228739860856762 | -4.05453327481537 |
| lnc-POLDIP3-3 | lnc-POLDIP3-3 | -0.296920827501874 | 2.03314411596647 | -2.23592650997821 | 0.0306625156820176 | 0.228834865947414 | -3.78122962651149 |
| ENSG00000083520.14 | DIS3 | 0.213254552873716 | 6.79102099707748 | 2.2354044912271 | 0.0306993967367341 | 0.22898282076699 | -4.05924727830963 |
| ENSG00000133706.17 | LARS1 | -0.264798734726563 | 6.35302115665956 | -2.23518322317638 | 0.030715041405447 | 0.22898282076699 | -4.06068579908087 |
| ENSG00000170456.15 | DENND5B | 0.753090345354829 | 6.27482063729381 | 2.23706828228335 | 0.0307157153057212 | 0.22898282076699 | -4.05847333167464 |
| lnc-FAM173B-6 | lnc-FAM173B-6 | -0.577883134472513 | -0.366388086946106 | -2.23486763661598 | 0.0307373670912801 | 0.229061269674083 | -3.5036322072526 |
| ENSG00000170915.8 | PAQR8 | -0.67117218645173 | 2.74395945562869 | -2.23468480755796 | 0.0307696970507492 | 0.229219208974148 | -3.85000424642028 |
| lnc-PPIE-1 | lnc-PPIE-1 | -0.491894019470839 | 2.29014394721141 | -2.23405304122626 | 0.0307950610860862 | 0.229325160034339 | -3.81702435178714 |
| ENSG00000081014.10 | AP4E1 | 0.192908759228172 | 4.71798171549691 | 2.23354976708605 | 0.0308307537419465 | 0.229507921978476 | -4.02301215109555 |
| lnc-ANKRD44-1 | lnc-ANKRD44-1 | -0.740515583219529 | -0.110474078948783 | -2.23309100513168 | 0.0308633215887894 | 0.229602901694471 | -3.54124545725388 |
| ENSG00000143224.17 | PPOX | -0.359120952195271 | 4.06185047341198 | -2.23297935951504 | 0.0308712520181185 | 0.229602901694471 | -3.97474809240025 |
| ENSG00000279670.1 | | -1.04536834549908 | 1.42640603019733 | -2.23430646727824 | 0.0308813113522768 | 0.229602901694471 | -3.72269036407979 |
| ENSG00000154760.13 | SLFN13 | 0.477900162744693 | 4.46154133216368 | 2.23252034405835 | 0.0309038759133899 | 0.229602901694471 | -3.99778257250344 |
| lnc-C12orf49-2 | lnc-C12orf49-2 | -1.48222980095988 | 0.816875271674233 | -2.23459857159071 | 0.0309097922656378 | 0.229602901694471 | -3.64033687896117 |
| ENSG00000127311.9 | HELB | 0.38791289634942 | 4.10734190256554 | 2.23242800302583 | 0.0309104426271865 | 0.229602901694471 | -3.9731483619423 |
| ENSG00000146281.5 | PM20D2 | -0.390289180689669 | 3.96022007200687 | -2.23190943739188 | 0.0309473428000457 | 0.229731988089897 | -3.98068924651218 |
| ENSG00000245849.7 | RAD51-AS1 | -0.401635984825808 | 4.63666026547275 | -2.2318343458845 | 0.030952689417064 | 0.229731988089897 | -4.02039132580566 |
| ENSG00000108588.13 | CCDC47 | 0.238575019128302 | 4.99391035578788 | 2.23161305392306 | 0.0309684504794439 | 0.229731988089897 | -4.04068360881132 |
| lnc-EMC3-2 | lnc-EMC3-2 | 0.323793878537125 | 1.08643058571524 | 2.23155669081844 | 0.0309724659646049 | 0.229731988089897 | -3.69223771596448 |
| ENSG00000233560.2 | KRT8P39 | 0.440422572885255 | 1.7467789986788 | 2.23093095870704 | 0.0310170761873177 | 0.229804040991651 | -3.75254451215666 |
| lnc-ATP5PF-2 | lnc-ATP5PF-2 | -0.540179800467216 | 1.63214685529499 | -2.23075510798085 | 0.0310296233606189 | 0.229804040991651 | -3.75550643755823 |
| lnc-PPP2R3C-5 | lnc-PPP2R3C-5 | -0.489631162124941 | -0.419888028999662 | -2.23069263114005 | 0.0310340822486629 | 0.229804040991651 | -3.52025242183814 |
| lnc-TUBA3E-1 | lnc-TUBA3E-1 | 0.322239599331597 | 1.19451865963912 | 2.23066065852874 | 0.031036364311159 | 0.229804040991651 | -3.71357585319488 |
| ENSG00000105808.17 | RASA4 | -0.579135100646067 | 0.152478368548346 | -2.23053752128088 | 0.0310451546923661 | 0.229804040991651 | -3.59822249682764 |
| ENSG00000173992.8 | CCS | -0.367830806387533 | 4.44001392436987 | -2.23048130356302 | 0.031049168634202 | 0.229804040991651 | -4.00686080667564 |
| ENSG00000251201.8 | TMED7-TICAM2 | 0.227863003541302 | 3.08271726362761 | 2.23018857696425 | 0.0310700767506177 | 0.229876128597399 | -3.88802821158793 |
| ENSG00000110011.13 | DNAJC4 | -0.30004234630493 | 3.69385156333815 | -2.22971183012272 | 0.0311041553705381 | 0.230045573119578 | -3.95323987940588 |

| | | | | | | | |
|---|---|---|---|---|---|---|---|
| ENSG00000015171.19 | ZMYND11 | -0.317205736671617 | 6.13510173377085 | -2.22887214806603 | 0.0311642579505186 | 0.230407299351841 | -4.07293747206948 |
| lnc-NLRC4-3 | lnc-NLRC4-3 | -0.72729064677637 | -0.286708429063495 | -2.22862167136709 | 0.0311822064886752 | 0.230457219445746 | -3.53605199149939 |
| ENSG00000284952.1 | | -0.170751677236827 | 3.45055878333236 | -2.22769452929341 | 0.0312487231326275 | 0.230865925426731 | -3.93143859727922 |
| lnc-COPS3-1 | lnc-COPS3-1 | -0.643892617545852 | -0.225840886168212 | -2.22735975721836 | 0.0312727718941798 | 0.230960697487586 | -3.54404631711582 |
| ENSG00000233024.7 | NPIPA9 | 1.12885682884382 | -0.846231290214619 | 2.2263336769748 | 0.0313465840859998 | 0.231370062855408 | -3.50254918791016 |
| ENSG00000262468.6 | LINC01569 | -0.439508021293935 | 0.502551156926177 | -2.22603254001953 | 0.0313682760535817 | 0.231370062855408 | -3.61752709700049 |
| lnc-MAPKAP1-1 | lnc-MAPKAP1-1 | -0.710419913576783 | -1.00835277420751 | -2.22601087067241 | 0.0313698374879989 | 0.231370062855408 | -3.50419855404082 |
| lnc-CPNE9-2 | lnc-CPNE9-2 | 0.753725235943643 | 0.189637218215211 | 2.22596470175158 | 0.0313731645255523 | 0.231370062855408 | -3.5500656396296 |
| lnc-SMIM2-6 | lnc-SMIM2-6 | -0.468998440349416 | -0.47928265306144 | -2.22534898874015 | 0.0314175641794512 | 0.231614514149586 | -3.52301067591511 |
| ENSG00000096654.15 | ZNF184 | 0.220553188324119 | 3.80079731713229 | 2.2243131081559 | 0.0314923884663714 | 0.23190268048718 | -3.96335565713026 |
| lnc-ANXA11-1 | lnc-ANXA11-1 | -0.353220489023676 | 2.51458716349483 | -2.22424498053069 | 0.0314973150401872 | 0.23190268048718 | -3.83935992773098 |
| ENSG00000233448.2 | PMS2P9 | 0.476492549145366 | -0.162611805482584 | 2.22421754169561 | 0.0314992994425271 | 0.23190268048718 | -3.52513091175467 |
| lnc-NR1I2-1 | lnc-NR1I2-1 | -0.475353242223497 | 2.36102136973565 | -2.22418407956402 | 0.0315017196056044 | 0.23190268048718 | -3.84240092877633 |
| lnc-ACTL7B-11 | lnc-ACTL7B-11 | -2.20267672375647 | -1.60510938223206 | -2.22348215686877 | 0.031573839542434 | 0.232350496711447 | -3.57556981149644 |
| ENSG00000168461.12 | RAB31 | -1.09469750328278 | 5.90373790107725 | -2.22568216788089 | 0.0315944082977121 | 0.232418765543892 | -4.07161487093189 |
| lnc-C18orf32-2 | lnc-C18orf32-2 | 0.29931148324363 | 1.9987588199575 | 2.22249225128008 | 0.0316242972661456 | 0.232555523625965 | -3.79640229356178 |
| ENSG00000105223.19 | PLD3 | -0.339964289646018 | 3.86448297442674 | -2.22226563225665 | 0.0316407485698192 | 0.232579755441316 | -3.98048106002983 |
| lnc-ZNF761-1 | lnc-ZNF761-1 | -0.539363479261042 | -0.0981371587355973 | -2.22213559958514 | 0.0316501916625917 | 0.232579755441316 | -3.55408064961571 |
| ENSG00000005194.14 | CIAPIN1 | 0.214633942222579 | 3.47622702403355 | 2.22172300036963 | 0.0316801715690399 | 0.232716977660795 | -3.94536547980641 |
| ENSG00000164609.9 | SLU7 | 0.22401638244204 | 6.30099234985814 | 2.221535551451 | 0.031693800138711 | 0.232732975972323 | -4.08804827742275 |
| ENSG00000172409.5 | CLP1 | 0.395722308622836 | 3.62357705645599 | 2.22120925939858 | 0.031717535787396 | 0.232732975972323 | -3.95312014797828 |
| RORA-AS1 | RORA-AS1 | -0.300113663336476 | 4.22887980878376 | -2.22120833072895 | 0.0317176033646575 | 0.232732975972323 | -4.00540054500753 |
| lnc-BDP1-6 | lnc-BDP1-6 | -0.388719456218865 | 2.29193722696023 | -2.22073753642971 | 0.031751878501947 | 0.232732975972323 | -3.84821785202807 |
| lnc-ARPP19-2 | lnc-ARPP19-2 | 0.445967272348307 | 3.75844261776661 | 2.22073387724931 | 0.0317521450292185 | 0.232732975972323 | -3.96058858618746 |
| ENSG00000239305.6 | RNF103 | 0.338995723084343 | 6.06990463065769 | 2.22063522325353 | 0.0317593315346864 | 0.232732975972323 | -4.08934489876268 |
| lnc-SNX14-3 | lnc-SNX14-3 | -0.493096303195647 | 1.45986693833824 | -2.2203801557614 | 0.0317779187625301 | 0.232732975972323 | -3.76470306422787 |
| lnc-ARHGAP27-1 | lnc-ARHGAP27-1 | 0.318708449359775 | 0.60097400123357 | 2.22031431853617 | 0.0317827180076425 | 0.232732975972323 | -3.65148487762837 |
| lnc-IFITM2-1 | lnc-IFITM2-1 | -1.19248985097158 | -1.16762230276557 | -2.22029518495663 | 0.0317841128823883 | 0.232732975972323 | -3.5117557296398 |
| ENSG00000280893.1 | | -0.373474771716183 | 1.82334454754101 | -2.21990027935776 | 0.0318129143992894 | 0.23286103025625 | -3.79651292731273 |
| ENSG00000144134.18 | RABL2A | -0.221832259442055 | 3.43484391000397 | -2.21940647295224 | 0.0318489616137752 | 0.233042010983411 | -3.93829082057996 |
| ENSG00000163806.15 | SPDYA | 0.311051402430474 | 3.83924521006804 | 2.2189239654315 | 0.0318842190437441 | 0.233217086203762 | -3.97761724134565 |
| ENSG00000011260.13 | UTP18 | 0.170768179415196 | 4.87063817506545 | 2.21863093001971 | 0.0319056484151772 | 0.233224363712527 | -4.06074231836749 |
| ENSG00000011454.16 | RABGAP1 | 0.267691482363382 | 6.07079605516233 | 2.2185795729113 | 0.0319094054228563 | 0.233224363712527 | -4.09293497574775 |
| LINC01281 | LINC01281 | -1.78310822134803 | 3.39288769524802 | -2.22145423380375 | 0.0319192067520861 | 0.233224363712527 | -3.91353125115146 |
| ENSG00000174944.8 | P2RY14 | -2.02254824099034 | 3.0230181866035 | -2.22106001024869 | 0.0319479656718168 | 0.23322690448131 | -3.89592336463679 |
| ENSG00000185436.11 | IFNLR1 | -0.302203892812193 | 5.76316018293685 | -2.21801793390379 | 0.0319505175247339 | 0.23322690448131 | -4.09069553125931 |
| ENSG00000280194.1 | | -0.610676844315603 | -0.067001216143781 | -2.21783638163372 | 0.0319638172434312 | 0.23322690448131 | -3.57621168851214 |
| NDUFV2-AS1 | NDUFV2-AS1 | -0.334771544825672 | 3.77654925999563 | -2.21782189524508 | 0.0319648786640322 | 0.23322690448131 | -3.97853781590228 |
| ENSG00000106392.10 | C1GALT1 | 0.326702447777567 | 5.28102288075928 | 2.21712497576486 | 0.0320159790628682 | 0.23343199629663 | -4.08317993651822 |
| lnc-C9orf64-1 | lnc-C9orf64-1 | 0.211879868330938 | 6.01840281585387 | 2.21701018293624 | 0.0320244029968181 | 0.23343199629663 | -4.09612047144192 |
| ENSG00000169871.12 | TRIM56 | -0.24628660509998 | 5.90492135306395 | -2.21697465568063 | 0.0320270105204141 | 0.23343199629663 | -4.09461285097414 |
| lnc-PTX3-4 | lnc-PTX3-4 | -0.461127336853867 | 0.53294237774437 | -2.2161929541723 | 0.0320844312210439 | 0.233767733742565 | -3.63407186630953 |
| lnc-LINC00672-1 | lnc-LINC00672-1 | -0.457305411917641 | 0.266849346450557 | -2.21598231144236 | 0.0320999198207492 | 0.23379782366259 | -3.62141322641439 |
| ENSG00000108175.16 | ZMIZ1 | 1.28446092321479 | 0.344578617392133 | 2.21612099628982 | 0.0321415676418137 | 0.234018354004758 | -3.5716999191915 |
| ENSG00000133134.11 | BEX2 | -1.71342073473758 | 2.53479086438682 | -2.21800022932838 | 0.0321634225458797 | 0.234094669823848 | -3.85186831147787 |
| ENSG00000280789.1 | PAGR1 | -0.36937830981467 | 1.96341121695483 | -2.21449756238934 | 0.0322092820381174 | 0.234204342933883 | -3.82024661436219 |
| ENSG00000100490.9 | CDKL1 | -0.610167554052125 | 4.90521305971055 | -2.21526461941819 | 0.0322115280110502 | 0.234204342933883 | -4.05625319924063 |
| ENSG00000141564.14 | RPTOR | -0.29066976583364 | 5.09470782030152 | -2.21438593820686 | 0.032217517290677 | 0.234204342933883 | -4.08177271523106 |
| lnc-SEPT11-2 | lnc-SEPT11-2 | -0.360648599794996 | 2.72458865765627 | -2.21412971571857 | 0.032236427580718 | 0.234204342933883 | -3.88354270459468 |
| ENSG00000236801.1 | RPL24P8 | 1.55048986045745 | -1.07915894434657 | 2.21370405514544 | 0.0322678648798567 | 0.234204342933883 | -3.59484652347735 |
| ENSG00000196230.12 | TUBB | -0.47805048247928 | 6.66435992887897 | -2.21375295786048 | 0.0322681432157811 | 0.234204342933883 | -4.10115517422318 |
| lnc-ASF1A-3 | lnc-ASF1A-3 | -0.685588388526445 | 1.11228998055312 | -2.21359835536231 | 0.0322756755860326 | 0.234204342933883 | -3.72982626038926 |
| ENSG00000204264.9 | PSMB8 | 0.283294625894284 | 6.02897785307956 | 2.21358457382979 | 0.0322766940986764 | 0.234204342933883 | -4.10222105879998 |
| LINC01372 | LINC01372 | -0.485934562978415 | 2.61246325712239 | -2.21352769649599 | 0.0322808978721967 | 0.234204342933883 | -3.87413346552223 |
| ENSG00000131263.12 | RLIM | 0.282712074033281 | 7.07693091655549 | 2.21329845657847 | 0.0322978457972758 | 0.234244735745359 | -4.10046301176732 |
| lnc-TRIM56-1 | lnc-TRIM56-1 | -0.221322772927433 | 3.92046517627587 | -2.21299363312104 | 0.0323203939123123 | 0.234325701971512 | -3.99858024093081 |
| lnc-LDAH-3 | lnc-LDAH-3 | -0.518313285832501 | 0.845223045242638 | -2.21253436394436 | 0.03235439291263 | 0.234487768082636 | -3.70433030755604 |
| lnc-TMEM119-1 | lnc-TMEM119-1 | 0.234671855030594 | 2.37111178448037 | 2.21238398416265 | 0.0323655321927207 | 0.234487768082636 | -3.85250497168021 |
| PPP3CB-AS1 | PPP3CB-AS1 | -0.325075121235015 | 3.36424094145392 | -2.21210181754662 | 0.0323864426740535 | 0.234556702871233 | -3.93673571575649 |
| lnc-IDH3B-2 | lnc-IDH3B-2 | -0.523322593220347 | 1.78148264638881 | -2.21164683405301 | 0.0324201853130948 | 0.234718492542888 | -3.79907060467538 |
| ENSG00000110651.11 | CD81 | 0.395238686342921 | 5.35525205838054 | 2.21088804751683 | 0.0324765280683747 | 0.234965999659493 | -4.09240905855643 |
| ENSG00000136868.10 | SLC31A1 | 0.427488156589571 | 1.08579580188601 | 2.21044230882233 | 0.0325096663225987 | 0.234965999659493 | -3.72404409538442 |
| ENSG00000164306.10 | PRIMPOL | -0.271519201890984 | 3.39010463738027 | -2.210275882443 | 0.0325220469081838 | 0.234965999659493 | -3.95433154844309 |
| ENSG00000152518.7 | ZFP36L2 | 0.616119834899654 | 6.48225539868841 | 2.21137783800524 | 0.0325221157872821 | 0.234965999659493 | -4.10884248931648 |
| ENSG00000054523.17 | KIF1B | 0.449111210292173 | 4.39530420997175 | 2.21021876095342 | 0.0325262971832841 | 0.234965999659493 | -4.03783379759612 |
| ENSG00000081320.10 | STK17B | 0.388712414931518 | 7.37039073636599 | 2.2098545877341 | 0.0325534060363696 | 0.234965999659493 | -4.10397754206615 |
| ENSG00000103035.10 | PSMD7 | 0.21382926829561 | 5.79310145424645 | 2.20969359604258 | 0.0325653965545817 | 0.234965999659493 | -4.1081881612794 |
| ENSG00000133059.16 | DSTYK | 0.215889531416104 | 4.8430067978969 | 2.20967929980996 | 0.0325664615146187 | 0.234965999659493 | -4.07363216025997 |

| | | | | | | | |
|---|---|---|---|---|---|---|---|
| ENSG00000198846.5 | TOX | -3.04878042516508 | 0.795001620919037 | -2.2125586592622 | 0.0325737741864794 | 0.234965999659493 | -3.59526218313231 |
| lnc-GIP-5 | lnc-GIP-5 | 0.216979595515746 | 2.34417811593785 | 2.20953802450225 | 0.0325769871057944 | 0.234965999659493 | -3.85517592974756 |
| MCPH1-AS1 | MCPH1-AS1 | -0.532391179254645 | 3.24416754132262 | -2.20946381192477 | 0.0325825174564616 | 0.234965999659493 | -3.93506898185058 |
| lnc-RIPK1-8 | lnc-RIPK1-8 | -0.507388617615443 | 1.066982794936 | -2.20925019108613 | 0.0325984412139104 | 0.234965999659493 | -3.72713890314249 |
| ENSG00000149150.8 | SLC43A1 | -0.617354119454268 | 1.4981447658548 | -2.20919208323072 | 0.0326027738924119 | 0.234965999659493 | -3.7815376900147 |
| ENSG00000224295.3 | OLFM5P | -0.584339370160822 | -0.0499678729425031 | -2.20820384858713 | 0.0326765375686352 | 0.235326440155404 | -3.59731026587287 |
| lnc-SIX2-8 | lnc-SIX2-8 | 0.336010118735008 | 1.14597710922818 | 2.20807671063769 | 0.0326860381182203 | 0.235326440155404 | -3.73960386862436 |
| ENSG00000210156.1 | MT-TK | 0.984222011547383 | 1.62024294627701 | 2.20893721089883 | 0.0326872493420416 | 0.235326440155404 | -3.73250957708006 |
| ENSG00000067141.16 | NEO1 | 1.52302934174768 | 2.17144870171853 | 2.21063928779962 | 0.0326985191101616 | 0.235326440155404 | -3.82616126332907 |
| lnc-TRAP1-2 | lnc-TRAP1-2 | 0.377753290559818 | 2.24724383381993 | 2.20734867631216 | 0.0327404886199305 | 0.235513716712999 | -3.84245823058549 |
| lnc-RIOK2-8 | lnc-RIOK2-8 | 0.385823593883686 | 2.51143882862517 | 2.20725239288473 | 0.0327476957834138 | 0.235513716712999 | -3.87117959907226 |
| ENSG00000087302.8 | RTRAF | 0.205686195840084 | 5.98993814565152 | 2.20702642714097 | 0.0327646156629765 | 0.235513716712999 | -4.11544759232474 |
| ENSG00000100991.11 | TRPC4AP | 0.180502606653648 | 6.3757980284726 | 2.20693345057062 | 0.0327715798198517 | 0.235513716712999 | -4.11698592576545 |
| lnc-SLC39A4-2 | lnc-SLC39A4-2 | -0.462477937492471 | 1.90820334636201 | -2.20679767799352 | 0.0327817518526329 | 0.235513716712999 | -3.82742794906951 |
| lnc-ECE1-4 | lnc-ECE1-4 | -0.338425797115113 | 1.0750833892634 | -2.20661383196464 | 0.032795529992583 | 0.235530493313795 | -3.73719065134377 |
| lnc-SFPQ-2 | lnc-SFPQ-2 | 0.315894750302236 | 6.78058931535349 | 2.20632016732797 | 0.0328175490186684 | 0.23555013013159 | -4.11658748803654 |
| lnc-SMC5-6 | lnc-SMC5-6 | 1.54475429370139 | -1.07772749214567 | 2.20620321348802 | 0.0328263218867775 | 0.23555013013159 | -3.60400360104269 |
| ENSG00000214595.11 | EML6 | 1.2613434986355 | 3.72827207014947 | 2.20884340175263 | 0.0328325959941472 | 0.23555013013159 | -3.98482999817882 |
| ENSG00000178199.13 | ZC3H12D | -0.348259135104496 | 5.84389457576143 | -2.20585882395925 | 0.0328521670944408 | 0.235608416482535 | -4.11641849897842 |
| ENSG00000166912.16 | MTMR10 | 0.198634055862127 | 5.96412352740925 | 2.2054616548606 | 0.032881995617698 | 0.235674485312262 | -4.11789130548934 |
| lnc-DPP6-3 | lnc-DPP6-3 | -0.601432321599095 | -0.649649842399425 | -2.20543125998092 | 0.032884279347851 | 0.235674485312262 | -3.54790310723548 |
| lnc-APRT-3 | lnc-APRT-3 | -0.33149526690169 | 0.489062537202784 | -2.20522558651395 | 0.0328997363923871 | 0.235703193235121 | -3.65938444905695 |
| lnc-CAMK1D-1 | lnc-CAMK1D-1 | 0.356879749367311 | 2.8207812306294 | 2.20473131631948 | 0.0329369087795912 | 0.235887402021686 | -3.91279741320216 |
| ENSG00000168070.11 | MAJIN | 2.64825445559481 | -3.06312331338853 | 2.20437974307175 | 0.0329633720664882 | 0.235957889623146 | -3.76543113117148 |
| ENSG00000103642.11 | LACTB | -0.335723315089548 | 2.86288769738992 | -2.20429599829258 | 0.0329696784023791 | 0.235957889623146 | -3.93758119439088 |
| ENSG00000258359.1 | PCNPP1 | 0.38905760459686 | 1.2072124724498 | 2.20410060983626 | 0.0329843961374617 | 0.235981169863529 | -3.74581548286898 |
| ENSG00000137404.14 | NRM | 0.800190231355728 | -0.173465967856969 | 2.20371421693328 | 0.0330135185257194 | 0.236107453723031 | -3.58431356466421 |
| ENSG00000120370.12 | GORAB | 0.2755948982189 | 4.21030536794185 | 2.20274960431492 | 0.0330863207507104 | 0.236444634413235 | -4.04588573293431 |
| ENSG00000165240.19 | ATP7A | 0.191581426329583 | 4.76481872267479 | 2.20270291983669 | 0.0330898477760207 | 0.236444634413235 | -4.08189213811064 |
| lnc-IL17RA-33 | lnc-IL17RA-33 | 2.5596825391955 | 2.62664290242598 | 2.20558634585643 | 0.0330951266455098 | 0.236444634413235 | -3.82678751947746 |
| lnc-NEDD9-3 | lnc-NEDD9-3 | -0.903027562410463 | 5.60290964513616 | -2.20460092382374 | 0.0331170631438853 | 0.23644757003192 | -4.11447514088572 |
| ENSG00000250673.2 | REELD1 | 0.574222746519386 | 1.51591070776269 | 2.20232366971717 | 0.0331185125784397 | 0.23644757003192 | -3.79573596141199 |
| ENSG00000224546.2 | EIF4BP3 | 0.541655434890756 | 2.67167056935914 | 2.20212338203791 | 0.0331336597684516 | 0.236473688978516 | -3.88793565991883 |
| ENSG00000102007.10 | PLP2 | -0.37758213127641 | 5.66990158981529 | -2.20192131936743 | 0.0331489474167783 | 0.236500791916655 | -4.12155974332002 |
| ENSG00000146247.13 | PHIP | 0.283161102906844 | 7.86754952501357 | 2.20128352881309 | 0.0331972423550003 | 0.236763284612949 | -4.11311161439258 |
| ENSG00000143761.15 | ARF1 | 0.212281633198121 | 6.84388279176091 | 2.20085366260116 | 0.0332298279553699 | 0.236867443822939 | -4.12693202076636 |
| ENSG00000237438.7 | CECR7 | 2.64884161295848 | 2.96905304164897 | 2.2037341559862 | 0.0332348626420175 | 0.236867443822939 | -3.86344023454121 |
| ENSG00000147166.10 | ITGB1BP2 | -0.435332859722964 | 1.18939242137549 | -2.20032048642579 | 0.0332702842434872 | 0.237037819516683 | -3.77107375262991 |
| lnc-FAM129A-1 | lnc-FAM129A-1 | -0.352963595424078 | 0.719079627318649 | -2.19952691624738 | 0.033330579495924 | 0.237385231060417 | -3.70136201393794 |
| ENSG00000171204.12 | TMEM126B | 0.29648969259359 | 3.82311482059274 | 2.19859870592011 | 0.0334012274947936 | 0.23773770697008 | -4.0199927326811 |
| lnc-F5-1 | lnc-F5-1 | -0.91693565025591 | 0.925008638951057 | -2.19880479953997 | 0.0334031700217398 | 0.23773770697008 | -3.70050478844311 |
| ENSG00000125650.4 | PSPN | -0.400709308907827 | 0.366103946452863 | -2.19825632320286 | 0.0334273204194568 | 0.237827354370439 | -3.66295659770995 |
| ENSG00000258315.5 | C17orf49 | -0.311862268420189 | 3.33702527664761 | -2.19791500483571 | 0.0334533502009319 | 0.237930306560395 | -3.99174893858859 |
| lnc-ZDHHC2-1 | lnc-ZDHHC2-1 | -2.08400687368272 | -0.253337476918226 | -2.19882926193827 | 0.0335218845541263 | 0.238335388524208 | -3.55289110255785 |
| lnc-SRRM5-2 | lnc-SRRM5-2 | -0.593429901471752 | -0.557795340560849 | -2.19597226609765 | 0.0336018504634977 | 0.238821439257657 | -3.56810925163664 |
| ENSG00000250741.6 | NT5C1B-RDH14 | 0.320165579413522 | 1.12476914092671 | 2.19563136705849 | 0.0336279684393557 | 0.238924568307649 | -3.76025383008002 |
| ENSG00000218283.2 | MORF4L1P1 | 0.618328579075273 | 2.52085289055196 | 2.19531002322788 | 0.0336526046524601 | 0.239001330632372 | -3.87897324254509 |
| lnc-VEZT-2 | lnc-VEZT-2 | -0.341864288927324 | 2.91790038445253 | -2.19518758540443 | 0.0336619957004929 | 0.239001330632372 | -3.95645832697206 |
| ENSG00000174483.19 | BBS1 | -0.368149480566639 | 2.50231021554615 | -2.19483373638848 | 0.0336891491599646 | 0.239111640399846 | -3.91128932400168 |
| ENSG00000158482.10 | SNX29P1 | 1.00813641807502 | 2.20434398137435 | 2.19602146418737 | 0.0337135197602406 | 0.239168725162064 | -3.82601724765005 |
| ENSG00000189195.12 | BTBD8 | -0.529568725507504 | 2.72593615399347 | -2.19436685225371 | 0.0337250063209025 | 0.239168725162064 | -3.91516092227379 |
| ENSG00000280227.1 | | 0.331280759032724 | 1.69028853643472 | 2.19416820119577 | 0.0337402731631774 | 0.239168725162064 | -3.81970331837162 |
| ENSG00000115239.21 | ASB3 | 0.23199549583568 | 4.32883721499831 | 2.19412400108261 | 0.0337436708871707 | 0.239168725162064 | -4.07521349943883 |
| lnc-SIGLEC12-2 | lnc-SIGLEC12-2 | 2.51217924499193 | -1.48527581144013 | 2.1941871725654 | 0.0337730340957991 | 0.239224386038224 | -3.66147557628553 |
| ENSG00000100243.20 | CYB5R3 | 0.383517887720651 | 5.85679208392954 | 2.19361490206949 | 0.0337828278810232 | 0.239224386038224 | -4.14030283707897 |
| ENSG00000169957.9 | ZNF768 | -0.338568631730915 | 3.37482375511836 | -2.1935685994858 | 0.0337863912069726 | 0.239224386038224 | -3.99695091215995 |
| RTCA-AS1 | RTCA-AS1 | -0.395515059062043 | 1.72739143010355 | -2.19341741317283 | 0.0337980284282355 | 0.239224490762851 | -3.81669215995046 |
| ENSG00000213918.10 | DNASE1 | -0.447481101813252 | 4.89579395800755 | -2.19294136850422 | 0.0338346940629202 | 0.239401687142347 | -4.10929999784016 |
| ENSG00000132300.18 | PTCD3 | 0.25743640575771 | 7.21553406311222 | 2.19262256226771 | 0.0338592686559042 | 0.239493239431091 | -4.13904657301072 |
| lnc-KCTD7-2 | lnc-KCTD7-2 | -0.38361695714411 | 0.812251413251648 | -2.19154337184792 | 0.0339425733879081 | 0.23995125728896 | -3.73208830622933 |
| lnc-DOCK4-7 | lnc-DOCK4-7 | -0.643284157774035 | 1.46772985949897 | -2.19135311471883 | 0.0339572784902963 | 0.23995125728896 | -3.81651995404471 |
| ENSG00000243199.1 | RPL6P10 | 0.859683783787206 | 2.13483363169889 | 2.19209728159353 | 0.0339604211935028 | 0.23995125728896 | -3.83780012966942 |
| lnc-VCL-1 | lnc-VCL-1 | -0.761612637348916 | -0.111750658927228 | -2.19118012815386 | 0.0339706536335825 | 0.23995125728896 | -3.61818600144842 |
| lnc-SLC7A10-1 | lnc-SLC7A10-1 | -0.319332028531183 | 1.77563779432285 | -2.19073941450962 | 0.034004750235974 | 0.240109699522145 | -3.83626271943889 |
| ENSG00000107816.17 | LZTS2 | -0.316840012613644 | 3.95485008661463 | -2.19045057375818 | 0.0340271133487234 | 0.240115958009362 | -4.0537185990329 |
| ENSG00000146828.17 | SLC12A9 | 0.309380120733712 | 5.21218484810013 | 2.19042662599532 | 0.0340289680568095 | 0.240115958009362 | -4.12868151164605 |
| ENSG00000130244.12 | FAM98C | -0.320470487722449 | 3.34656604328754 | -2.18968163379292 | 0.034086710974987 | 0.240366082411509 | -4.00184250922648 |
| ENSG00000116691.10 | MIIP | -0.306464602189052 | 4.35018990500564 | -2.18958720955318 | 0.0340940358056759 | 0.240366082411509 | -4.08594544991007 |

| | | | | | | | |
|---|---|---|---|---|---|---|---|
| lnc-SCGB3A1-3 | lnc-SCGB3A1-3 | -0.345082143794301 | 1.13681497021874 | -2.18880319314909 | 0.034154908624489 | 0.240366082411509 | -3.77898244467992 |
| LINC00324 | LINC00324 | -0.745871882313785 | 4.01430347570686 | -2.19005120171465 | 0.0341736094533871 | 0.240366082411509 | -4.05788675708373 |
| ENSG00000124733.4 | MEA1 | -0.28810515753456 | 4.14714810468105 | -2.18855005384833 | 0.0341745834670058 | 0.240366082411509 | -4.07024048634804 |
| ENSG00000112640.14 | PPP2R5D | -0.252841903689791 | 4.20941271519369 | -2.1884249441589 | 0.0341843111181187 | 0.240366082411509 | -4.07612013575142 |
| ENSG00000184574.9 | LPAR5 | -0.352662096953697 | 4.19889417941879 | -2.18819949452563 | 0.0342018466851845 | 0.240366082411509 | -4.07343179384951 |
| ENSG00000070882.12 | OSBPL3 | 0.68718769040739 | 4.07032919813217 | 2.18939398540372 | 0.0342020032231281 | 0.240366082411509 | -4.07485521615386 |
| lnc-JUN-11 | lnc-JUN-11 | 0.289500141315578 | 1.86655106709846 | 2.18812379042219 | 0.0342077367670931 | 0.240366082411509 | -3.84735861786551 |
| lnc-FCMR-1 | lnc-FCMR-1 | -1.40204862329016 | -1.2857704829685 | -2.18801337695308 | 0.0342163289857035 | 0.240366082411509 | -3.56420723006876 |
| lnc-ANGPT2-2 | lnc-ANGPT2-2 | -0.316183041891712 | 1.13050750735699 | -2.1878664927462 | 0.03422776226524 | 0.240366082411509 | -3.78642540161696 |
| ENSG00000282826.1 | FRG1CP | -0.837621628657999 | 3.31113932295268 | -2.1892288564946 | 0.0342480484291541 | 0.240366082411509 | -3.99976159015608 |
| ENSG00000138032.20 | PPM1B | 0.291623543481241 | 5.53093729439179 | 2.18749283486531 | 0.0342568625586234 | 0.240366082411509 | -4.14655858833881 |
| lnc-C22orf23-2 | lnc-C22orf23-2 | -0.39848555900185 | 0.122352288860486 | -2.18725816855432 | 0.034275149446125 | 0.240366082411509 | -3.64503017305257 |
| LINC00467 | LINC00467 | -0.239561124945683 | 3.29534331090407 | -2.18724375496774 | 0.0342762729380145 | 0.240366082411509 | -3.99301758062219 |
| ENSG00000115020.16 | PIKFYVE | 0.319010128737917 | 7.15512290302086 | 2.18713790905429 | 0.0342845242787697 | 0.240366082411509 | -4.15028360028786 |
| ENSG00000251184.1 | | 0.322906079756278 | 1.14334973854099 | 2.18707379112392 | 0.0342895235210484 | 0.240366082411509 | -3.77183795307928 |
| ENSG00000212916.4 | MAP10 | -1.24774889711172 | 1.79085926648625 | -2.18906298890491 | 0.0342942014920812 | 0.240366082411509 | -3.82820490088594 |
| lnc-CD99-2 | lnc-CD99-2 | -0.30312568033553 | 3.28380061576532 | -2.18701183708797 | 0.034294354658294 | 0.240366082411509 | -3.99183331187986 |
| ENSG00000176222.8 | ZNF404 | -0.808726206791631 | 1.03056091362103 | -2.18698398622026 | 0.0342979732809893 | 0.240366082411509 | -3.75113246350742 |
| ENSG00000133997.11 | MED6 | 0.262099071461166 | 4.98479393108494 | 2.18653785162759 | 0.034331335683936 | 0.24051799944944 | -4.12764133162283 |
| ENSG00000214022.11 | REPIN1 | -0.346856234075187 | 5.57782218145258 | -2.18572521651916 | 0.0343948207474458 | 0.240828387360691 | -4.150104630426 |
| LINC02422 | LINC02422 | -0.402459328598467 | 6.0405848421576 | -2.18566013309037 | 0.0343999097150948 | 0.240828387360691 | -4.15729676958897 |
| ENSG00000183137.14 | CEP57L1 | -0.380523853277783 | 3.01936255153591 | -2.18552163557256 | 0.0344107412538402 | 0.240828387360691 | -3.96181625420546 |
| lnc-SHISA9-8 | lnc-SHISA9-8 | -0.555967470919495 | -0.308135083093592 | -2.18509807307184 | 0.034443885726146 | 0.240978416009947 | -3.60418120681567 |
| ENSG00000120800.4 | UTP20 | 0.209155989511132 | 5.62428048940163 | 2.18470709232946 | 0.0344745056849999 | 0.24111068654922 | -4.1539361863645 |
| ENSG00000103365.15 | GGA2 | 0.293331953353483 | 8.58542669283988 | 2.18405278720704 | 0.0345258019648858 | 0.241270328166014 | -4.13336325928989 |
| ENSG00000091651.8 | ORC6 | 0.531214541641294 | 1.93602514519924 | 2.18404135100321 | 0.0345266991415763 | 0.241270328166014 | -3.86286062488995 |
| ENSG00000143314.12 | MRPL24 | -0.413612555407997 | 2.65851668963247 | -2.18388633666184 | 0.0345388621391144 | 0.241270328166014 | -3.942033258605 |
| ENSG00000263961.7 | RHEX | -0.944848659107435 | 3.97787599729897 | -2.18568816838744 | 0.0345581980801933 | 0.241270328166014 | -4.05705537919863 |
| ENSG00000171316.11 | CHD7 | 0.263688950486809 | 8.1566690621971 | 2.18363911496524 | 0.0345582679083213 | 0.241270328166014 | -4.14267902205505 |
| ENSG00000176102.12 | CSTF3 | -0.211587509573087 | 4.38250371192493 | -2.18351947542913 | 0.0345676625254616 | 0.241270328166014 | -4.0969799262537 |
| ENSG00000183011.13 | NAA38 | -0.377017135814346 | 2.86270555201453 | -2.18299821384359 | 0.0346086206204267 | 0.24147431804415 | -3.96115243794737 |
| lnc-PCSK9-1 | lnc-PCSK9-1 | -0.480538988789163 | 0.184916644226913 | -2.18253627668941 | 0.0346449531686291 | 0.241609412832137 | -3.6642777263113 |
| lnc-SPZ1-1 | lnc-SPZ1-1 | -0.300283793748974 | 2.44346491027046 | -2.18245360210586 | 0.0346514592955579 | 0.241609412832137 | -3.91524244846156 |
| ENSG00000273112.1 | | 1.14491615143848 | 0.494776349606809 | 2.18326290714083 | 0.0346812634094763 | 0.241658661368113 | -3.72713485766647 |
| lnc-PFDN6-3 | lnc-PFDN6-3 | -0.467858582284892 | 0.776438831739903 | -2.18206564983397 | 0.0346820038712241 | 0.241658661368113 | -3.74367066630779 |
| ENSG00000116560.10 | SFPQ | 0.249851600642616 | 8.52917888153014 | 2.18182015973155 | 0.0347013442973099 | 0.241711597181567 | -4.13793411826612 |
| ENSG00000227730.2 | MTND6P5 | -0.700252888687358 | -0.189513632324911 | -2.18147820913875 | 0.0347283000574504 | 0.24181752370856 | -3.61258404212854 |
| ENSG00000169826.7 | CSGALNACT2 | 0.429444132710061 | 5.03224700846964 | 2.18129380445064 | 0.0347428442522373 | 0.241836984526141 | -4.13917927745258 |
| ENSG00000163719.19 | MTMR14 | 0.187235245913928 | 6.10901242348582 | 2.18084585875684 | 0.0347781966435103 | 0.241904345546845 | -4.1673548958449 |
| ENSG00000240853.3 | RN7SL328P | 0.695954439238484 | 0.791499421343322 | 2.18072450657253 | 0.0347877793722325 | 0.241904345546845 | -3.7178418507202 |
| lnc-MED27-3 | lnc-MED27-3 | 0.695954439238484 | 0.791499421343322 | 2.18072450657253 | 0.0347877793722325 | 0.241904345546845 | -3.7178418507202 |
| lnc-TUBGCP3-1 | lnc-TUBGCP3-1 | -0.501151364003741 | 1.11822111475651 | -2.18035609097516 | 0.0348168860808617 | 0.242024980142647 | -3.78133816343662 |
| lnc-ARL6IP1-6 | lnc-ARL6IP1-6 | 0.270463525698102 | 1.21608657647308 | 2.17958122358723 | 0.0348781748326681 | 0.242300320077604 | -3.80466809167697 |
| lnc-ZNF408-5 | lnc-ZNF408-5 | 0.214668351990811 | 2.20380235128713 | 2.17955767051302 | 0.0348800392746413 | 0.242300320077604 | -3.90198790772575 |
| ENSG00000122692.8 | SMU1 | 0.153500256945173 | 6.62693366292524 | 2.1793825704396 | 0.0348939028151699 | 0.242314845359191 | -4.17040790326902 |
| ENSG00000118894.14 | EEF2KMT | -0.368159401397972 | 1.70713271451738 | -2.17901778547825 | 0.0349228002956048 | 0.242372975273082 | -3.86176481284655 |
| ENSG00000227507.2 | LTB | -0.640556755093947 | 4.78372712212213 | -2.18012834761005 | 0.0349258244502727 | 0.242372975273082 | -4.14222107861497 |
| ENSG00000108773.10 | KAT2A | -0.288549344003813 | 5.0557317734429 | -2.17825346470952 | 0.0349834167592654 | 0.242632745690388 | -4.1501292109467 |
| lnc-KRTAP5-7-1 | lnc-KRTAP5-7-1 | -0.461241137423763 | 1.11800452223964 | -2.17821042034223 | 0.0349868332706151 | 0.242632745690388 | -3.79896717908164 |
| ENSG00000121039.9 | RDH10 | 0.554091264636777 | 0.965744284458702 | 2.17759769238207 | 0.0350354986188764 | 0.242756791922978 | -3.77425535003864 |
| ENSG00000204267.13 | TAP2 | 0.212079845592948 | 5.7179209414469 | 2.17724972692126 | 0.0350631620828147 | 0.242756791922978 | -4.1699684541259 |
| lnc-ANXA3-7 | lnc-ANXA3-7 | -0.55697675449822 | 2.6795839351369 | -2.17723910683035 | 0.0350640066899857 | 0.242756791922978 | -3.96444098057272 |
| lnc-SLC45A4-3 | lnc-SLC45A4-3 | 0.368816218783645 | 2.13934167141157 | 2.17710758131641 | 0.0350744682974477 | 0.242756791922978 | -3.88863160125336 |
| ENSG00000186132.14 | C2orf76 | -0.540944519419958 | 2.26150653997024 | -2.17703008298559 | 0.0350806338464674 | 0.242756791922978 | -3.90026586500884 |
| ENSG00000141295.13 | SCRN2 | -0.274340082635917 | 3.80005680511154 | -2.17696720336653 | 0.0350856370758769 | 0.242756791922978 | -4.06601703320657 |
| ENSG00000158411.11 | MITD1 | -0.319016659540331 | 3.36048550667064 | -2.17694657440174 | 0.0350872786265782 | 0.242756791922978 | -4.020328985751 |
| ENSG00000158406.4 | H4C8 | 0.386903669571113 | 4.17856552237803 | 2.17652912197617 | 0.0351205120098058 | 0.242905073487175 | -4.08800462373429 |
| ENSG00000165813.19 | CCDC186 | 0.22277194968469 | 5.94537620195315 | 2.17593104536858 | 0.035168173364167 | 0.242992791265599 | -4.17572994552297 |
| ENSG00000170949.17 | ZNF160 | 0.268574476776013 | 6.01395842160241 | 2.17583398138572 | 0.0351759138843316 | 0.242992791265599 | -4.17584532288664 |
| NIPBL-DT | NIPBL-DT | -0.438955558042252 | 4.03697512927588 | -2.17582264884577 | 0.0351768177137292 | 0.242992791265599 | -4.08884251237709 |
| lnc-IGLL5-8 | lnc-IGLL5-8 | -0.917231877230935 | -0.652359300289466 | -2.17577752537565 | 0.0351804167503029 | 0.242992791265599 | -3.62073297166603 |
| ENSG00000134152.10 | KATNBL1 | -0.30499059535474 | 4.05133000883293 | -2.17556963865419 | 0.0351970019463076 | 0.243025793713804 | -4.09151834911708 |
| lnc-BAIAP2-4 | lnc-BAIAP2-4 | -0.305317618421056 | 1.1024520812465 | -2.17539726277455 | 0.0352107593283634 | 0.243039255283603 | -3.7960895273741 |
| ENSG00000242071.3 | RPL7AP6 | 0.916403124541516 | 3.50288299467129 | 2.17653612258083 | 0.0352607462046129 | 0.24327701920021 | -3.99761157149637 |
| ENSG00000119943.12 | PYROXD2 | -0.294847333608957 | 2.41916881278751 | -2.17466548267994 | 0.0352692158890885 | 0.24327701920021 | -3.93153296179081 |
| ENSG00000265355.1 | MIR3136 | 0.466839875923941 | 0.565864596296973 | 2.17452229957646 | 0.0352806637668283 | 0.24327701920021 | -3.72678608711254 |
| lnc-DLGAP3-2 | lnc-DLGAP3-2 | -0.715232939969103 | -0.672193469444934 | -2.17395821890073 | 0.035325795502615 | 0.24348501068657 | -3.59661792325091 |
| ENSG00000100104.13 | SRRD | 0.187425213227695 | 4.60974023731836 | 2.17384967255848 | 0.0353344860768977 | 0.24348501068657 | -4.12922180706184 |

| | | | | | | | |
|---|---|---|---|---|---|---|---|
| ENSG00000143816.7 | WNT9A | -1.59891374483287 | 1.66077195393019 | -2.17614046604047 | 0.0353696810060313 | 0.24357901287571 | -3.89560244910468 |
| THAP9-AS1 | THAP9-AS1 | 0.198347463231822 | 6.24785184869257 | 2.17332607655635 | 0.0353764334209799 | 0.24357901287571 | -4.1826661025902 |
| ENSG00000143061.17 | IGSF3 | -1.08836599329377 | 5.48288054877158 | -2.17585619252453 | 0.0353836296214532 | 0.24357901287571 | -4.16208047334042 |
| ENSG00000272054.1 | | -0.554309631701013 | 1.14691091609588 | -2.17293128760223 | 0.0354080906121638 | 0.243665907412293 | -3.81049355626133 |
| ENSG00000116641.17 | DOCK7 | -0.371401940303443 | 5.51878028675059 | -2.17262364762011 | 0.0354327768824073 | 0.243672179818049 | -4.17453961783834 |
| ENSG00000135604.9 | STX11 | 0.420012318675523 | 2.80993459534183 | 2.17253370059636 | 0.0354399974658706 | 0.243672179818049 | -3.96503297312522 |
| ENSG00000103089.8 | FA2H | 1.4695643514471 | 0.569158560679087 | 2.17394368384956 | 0.0354445176298517 | 0.243672179818049 | -3.65629933114886 |
| lnc-ZNF217-5 | lnc-ZNF217-5 | 0.317495474144776 | 1.79424089597823 | 2.17219296127756 | 0.0354673624324292 | 0.243691958080865 | -3.87166254618994 |
| ENSG00000074706.13 | IPCEF1 | -2.51622948599883 | 0.823352490253819 | -2.17488479166842 | 0.0354710735271958 | 0.243691958080865 | -3.6776680299247 |
| ENSG00000183298.5 | RPSAP19 | 1.56590988066459 | -1.49921066167698 | 2.17172118489707 | 0.035505281874422 | 0.243845584524934 | -3.68129535732023 |
| ENSG00000119414.11 | PPP6C | 0.227696481445585 | 6.08989896141132 | 2.17111072990679 | 0.0355544008822985 | 0.243958847251899 | -4.18644466365273 |
| ENSG00000122026.10 | RPL21 | 0.289252223530841 | 9.25708735427435 | 2.17101699485821 | 0.0355619483943757 | 0.243958847251899 | -4.14228429128045 |
| lnc-LAX1-4 | lnc-LAX1-4 | -0.893689308028294 | -0.531658959406292 | -2.17082753906091 | 0.0355772076276383 | 0.243958847251899 | -3.59914537415472 |
| ENSG00000238228.1 | OR7E7P | -0.354806691436096 | 0.0779143301945963 | -2.17078002070274 | 0.0355810357795366 | 0.243958847251899 | -3.68003031057381 |
| lnc-OCM2-1 | lnc-OCM2-1 | -0.354806691436096 | 0.0779143301945963 | -2.17078002070274 | 0.0355810357795366 | 0.243958847251899 | -3.68003031057381 |
| lnc-FBXL3-1 | lnc-FBXL3-1 | 0.242267514648228 | 3.49219999429594 | 2.17055691714707 | 0.03559901420951 | 0.244000835655792 | -4.04468135931413 |
| ENSG00000116212.14 | LRRC42 | 0.285687664565703 | 2.07515137357828 | 2.17032032333559 | 0.0356180884954332 | 0.244044187168053 | -3.90435233269339 |
| ENSG00000188732.10 | FAM221A | 1.14968112688024 | 1.68511903502242 | 2.17168738284574 | 0.0356503741909138 | 0.244044187168053 | -3.84688539070851 |
| ENSG00000096401.7 | CDC5L | 0.165976694916689 | 5.99089352426721 | 2.16975010397562 | 0.0356640969051369 | 0.244044187168053 | -4.18802103856195 |
| ENSG00000123219.12 | CENPK | 0.483941809580238 | 1.92000375110952 | 2.16957436836842 | 0.0356782867902763 | 0.244044187168053 | -3.89379112356278 |
| ENSG00000143575.14 | HAX1 | -0.235853499110405 | 5.0850274591406 | -2.16948534895894 | 0.0356854766225306 | 0.244044187168053 | -4.16694074197582 |
| WAC-AS1 | WAC-AS1 | -0.285260055486113 | 5.11408061292744 | -2.16948179076684 | 0.0356857640336916 | 0.244044187168053 | -4.16563881058598 |
| ENSG00000250182.3 | EEF1A1P13 | 0.658944268224518 | 0.62234503386092 | 2.16944996050631 | 0.0356883351977768 | 0.244044187168053 | -3.71407283461344 |
| ENSG00000162231.13 | NXF1 | 0.233596185627522 | 6.75450657466362 | 2.1689503856593 | 0.0357287109709589 | 0.244170579440223 | -4.18993198743982 |
| ENSG00000185838.13 | GNB1L | -0.454691189370514 | 1.18301386676012 | -2.16892771830776 | 0.0357305439087574 | 0.244170579440223 | -3.81891236327212 |
| ENSG00000064995.17 | TAF11 | 0.181470961797876 | 5.41813565210821 | 2.16847669770762 | 0.0357670318129098 | 0.244312142636768 | -4.17993657355356 |
| ENSG00000071575.11 | TRIB2 | 1.85393315739484 | 2.35593620827501 | 2.17110263150468 | 0.0357839919299623 | 0.244312142636768 | -3.91085434061258 |
| lnc-MAP4K2-1 | lnc-MAP4K2-1 | -0.405465275246292 | 0.165732219818658 | -2.16823167293124 | 0.0357868682917872 | 0.244312142636768 | -3.69200352677768 |
| lnc-USE1-4 | lnc-USE1-4 | 0.483947892514783 | 1.27588121559526 | 2.1679008704206 | 0.035813664493315 | 0.244414010698244 | -3.82574313084982 |
| ENSG00000197417.7 | SHPK | -0.252908657389744 | 2.40767360405607 | -2.16700882383225 | 0.0358860118260343 | 0.244826576538039 | -3.95228467080036 |
| ENSG00000255423.1 | EBLN2 | 0.403008169026159 | 2.92657017819131 | 2.16599081761096 | 0.0359687324559385 | 0.245253993043624 | -3.98868743126575 |
| ENSG00000123106.10 | CCDC91 | 0.220311282853102 | 5.80129665304729 | 2.1659445983982 | 0.0359724921050722 | 0.245253993043624 | -4.19336605478389 |
| lnc-CHAF1B-1 | lnc-CHAF1B-1 | -0.611068493956613 | -0.21989051809967 | -2.16569357643585 | 0.0359929172600199 | 0.245311992040725 | -3.65182302512303 |
| ENSG00000214485.6 | RPL7P1 | 1.03825536528839 | 2.01296176532168 | 2.16684709829958 | 0.03602153016542 | 0.24536867959759 | -3.8597593231854 |
| ENSG00000139193.3 | CD27 | -0.623102440560383 | 5.47253510170802 | -2.16613214061983 | 0.036047531709478 | 0.24536867959759 | -4.18331393299265 |
| ENSG00000119812.18 | FAM98A | 0.256933479769771 | 3.58733698665466 | 2.16483752971747 | 0.0360626490850509 | 0.24536867959759 | -4.06474480500779 |
| lnc-RND3-7 | lnc-RND3-7 | 0.241619095654988 | 2.32615292125295 | 2.16470251430094 | 0.036073658047758 | 0.24536867959759 | -3.94011049563605 |
| ENSG00000213442.5 | RPL18AP3 | 0.767392915015705 | 3.764644432854 | 2.16597738962869 | 0.0360756027787126 | 0.24536867959759 | -4.04385464792242 |
| CRTC3-AS1 | CRTC3-AS1 | -0.347538785553976 | 1.15424441890776 | -2.16465723971724 | 0.0360773503362963 | 0.24536867959759 | -3.82305393742775 |
| ENSG00000037749.12 | MFAP3 | 0.290821376774933 | 3.86102767716917 | 2.16456736129937 | 0.036084681200112 | 0.24536867959759 | -4.08052722983319 |
| lnc-DLGAP1-11 | lnc-DLGAP1-11 | -0.677957582252573 | 2.13613840027657 | -2.16446601170964 | 0.0361009739797535 | 0.245398397432386 | -3.91592619768786 |
| ENSG00000106133.17 | NSUN5P2 | -0.305187374051897 | 4.77797562094734 | -2.16386099680155 | 0.036142341066516 | 0.245560245089707 | -4.16383314010067 |
| ENSG00000107863.17 | ARHGAP21 | -0.401295134720862 | 6.52183809480811 | -2.16364582812449 | 0.0361599212345767 | 0.245560245089707 | -4.19975742840034 |
| ENSG00000120738.7 | EGR1 | -1.54354344500495 | 1.31446428612915 | -2.16563895552505 | 0.0361902646311954 | 0.245560245089707 | -3.81359487211577 |
| lnc-SNX10-6 | lnc-SNX10-6 | -0.670774307811646 | -0.429443389354439 | -2.16318559336278 | 0.0361975496524754 | 0.245560245089707 | -3.63521580765688 |
| ENSG00000207005.1 | RNU1-2 | -0.740372073145819 | 0.866817544512928 | -2.16311706208248 | 0.0362031556691979 | 0.245560245089707 | -3.79723033270297 |
| ENSG00000266127.1 | ZNF415P1 | -0.916441407292219 | 0.630229353478881 | -2.16302174969805 | 0.0362124288525478 | 0.245560245089707 | -3.72148362541758 |
| lnc-NABP1-1 | lnc-NABP1-1 | -0.533914714197138 | 3.39142475509073 | -2.16260682422702 | 0.0362449183992823 | 0.245560245089707 | -4.0536444307303 |
| ENSG00000198888.2 | MT-ND1 | -0.765600074365912 | 7.97209741573713 | -2.16441202814392 | 0.0362460972197627 | 0.245560245089707 | -4.1814366089999 |
| lnc-TRIM65-1 | lnc-TRIM65-1 | 0.351722388186318 | 0.577547090143319 | 2.16253453555455 | 0.0362508386307293 | 0.245560245089707 | -3.746050737032 |
| lnc-TMEM138-4 | lnc-TMEM138-4 | -0.355926739705661 | -0.0021713084636123[illegible] | -2.16249817568405 | 0.0362538167206575 | 0.245560245089707 | -3.67760892169823 |
| ENSG00000100564.8 | PIGH | 0.219682941647096 | 3.61753663930757 | 2.16247132165286 | 0.0362560163643599 | 0.245560245089707 | -4.07626605099409 |
| lnc-TMEM250-1 | lnc-TMEM250-1 | -0.405161374767764 | 0.465245642052524 | -2.16216880718426 | 0.0362808038008149 | 0.245647297576373 | -3.74889232391644 |
| ENSG00000196199.13 | MPHOSPH8 | -0.198637341315917 | 6.576977105604 | -2.16120030944049 | 0.0363602612018062 | 0.246104326312653 | -4.20550816249068 |
| ENSG00000125304.9 | TM9SF2 | 0.173350458380318 | 6.44574169881727 | 2.16102263863333 | 0.0363748543111666 | 0.246122165117272 | -4.20645160860019 |
| ENSG00000224790.2 | HLCS-AS1 | -0.435148205069116 | 0.251568577169963 | -2.16083851075522 | 0.0363899832294895 | 0.246143616435288 | -3.72212560321582 |
| lnc-USP34-5 | lnc-USP34-5 | 0.264012939399919 | 3.11334640701617 | 2.16045942798412 | 0.0364211481548348 | 0.246216357812651 | -4.02580698999518 |
| ENSG00000169093.15 | ASMTL | -0.222753700435369 | 4.33788804213409 | -2.16015595053807 | 0.0364461144315368 | 0.246216357812651 | -4.13946484414854 |
| HCP5 | HCP5 | 0.405031909676532 | 5.71869522937253 | 2.16007565295729 | 0.036452722824245 | 0.246216357812651 | -4.20172573900637 |
| ENSG00000106682.14 | EIF4H | 0.22188309751156 | 7.23655549603469 | 2.16003818171703 | 0.0364558070232785 | 0.246216357812651 | -4.20236644731671 |
| lnc-STRN3-10 | lnc-STRN3-10 | 0.245971880436056 | 2.26649417249741 | 2.15998058755111 | 0.0364605479576815 | 0.246216357812651 | -3.94041828706991 |
| ENSG00000123737.12 | EXOSC9 | 0.234691436188562 | 5.49302558276478 | 2.15972518185359 | 0.036481578546559 | 0.246277576655895 | -4.19992843837638 |
| ENSG00000011132.11 | APBA3 | -0.301484348680366 | 3.17397863719469 | -2.15901145972697 | 0.0365404045316684 | 0.24647413876734 | -4.04384147210599 |
| ENSG00000198715.12 | GLMP | -0.413750106932072 | 3.03906393446826 | -2.15899026790999 | 0.0365421524696891 | 0.24647413876734 | -4.02146635104172 |
| ENSG00000285713.1 | | 0.228687280110726 | 1.8504874175327 | 2.15893611278279 | 0.0365466196141438 | 0.24647413876734 | -3.90916711011115 |
| ENSG00000137936.17 | BCAR3 | -0.459072818896323 | 3.12387419124778 | -2.15855979907065 | 0.0365776742615834 | 0.24660277409963 | -4.02344903358504 |
| lnc-CYYR1-6 | lnc-CYYR1-6 | -0.444839351996307 | 1.14037491927786 | -2.15737175201346 | 0.0366758686303212 | 0.247183825808088 | -3.83881617377141 |
| ENSG00000134283.17 | PPHLN1 | 0.193637253551151 | 6.13246982153579 | 2.1567662036362 | 0.0367260077714368 | 0.247440726009651 | -4.21434995799646 |

| | | | | | | | |
|---|---|---|---|---|---|---|---|
| ENSG00000280778.1 | | -0.441199028317871 | 2.45711331469644 | -2.1564720332018 | 0.0367503867665063 | 0.247523956418521 | -3.96621073271277 |
| ENSG00000108840.15 | HDAC5 | -0.245059189040152 | 5.67897438638163 | -2.15628370148482 | 0.0367660020075356 | 0.247548125391268 | -4.21013332634009 |
| ENSG00000088205.12 | DDX18 | 0.191614356348964 | 6.5055549825597 | 2.15527832447803 | 0.0368494603701438 | 0.247707839851692 | -4.21751938733503 |
| ENSG00000265303.1 | | 0.393484313431077 | 0.520794793854712 | 2.15526177494843 | 0.0368508355764067 | 0.247707839851692 | -3.75917463615142 |
| ENSG00000178295.14 | GEN1 | 0.650193025832474 | 5.45334185064414 | 2.15654187742079 | 0.0368543158646281 | 0.247707839851692 | -4.19964644077702 |
| ENSG00000138246.16 | DNAJC13 | -0.189296007290919 | 6.72112644804527 | -2.15502679925602 | 0.0368703660934599 | 0.247707839851692 | -4.21680806963539 |
| ENSG00000159200.17 | RCAN1 | -0.716269420810218 | 1.52199231719786 | -2.15495691296299 | 0.0368786428922365 | 0.247707839851692 | -3.88094990802092 |
| ENSG00000164631.18 | ZNF12 | 0.268485757827264 | 5.43236528330827 | 2.1543723121396 | 0.0369248133458639 | 0.247707839851692 | -4.20880781080945 |
| ENSG00000224420.3 | ADM5 | -0.406792385338998 | 1.19477438449106 | -2.15428786405818 | 0.0369318438040589 | 0.247707839851692 | -3.85520918330453 |
| lnc-NOP16-2 | lnc-NOP16-2 | 0.28931727903598 | 1.20144302567035 | 2.15408190537265 | 0.0369489951937226 | 0.247707839851692 | -3.84683498973443 |
| ENSG00000162430.16 | SELENON | -0.682006350367163 | 3.90642197207328 | -2.15494554900793 | 0.0369675993841117 | 0.247707839851692 | -4.12416585534874 |
| ENSG00000105492.15 | SIGLEC6 | 2.46541635829336 | 0.589134846779827 | 2.15636194727979 | 0.0369722992809007 | 0.247707839851692 | -3.66110122625674 |
| lnc-ARF4-4 | lnc-ARF4-4 | 0.33827166828302 | 0.871480013037002 | 2.15364175080991 | 0.0369856729953587 | 0.247707839851692 | -3.80809707270372 |
| lnc-MBNL1-3 | lnc-MBNL1-3 | -0.282774361092504 | 5.62942533950457 | -2.15357929609495 | 0.0369908799071891 | 0.247707839851692 | -4.21355178686871 |
| lnc-SLTM-9 | lnc-SLTM-9 | -0.710101820565505 | -0.382804222573181 | -2.15349017768745 | 0.0369983109167882 | 0.247707839851692 | -3.65352407301877 |
| lnc-BCOR-10 | lnc-BCOR-10 | -1.62116901830352 | 2.60210640825265 | -2.15604846091202 | 0.0370082336966438 | 0.247707839851692 | -3.9737325747188 |
| lnc-ZC3H12B-3 | lnc-ZC3H12B-3 | -1.50412515280312 | 4.42808526403171 | -2.15610511255051 | 0.0370125110641842 | 0.247707839851692 | -4.13563977838862 |
| ENSG00000130545.15 | CRB3 | -0.952653458307183 | -0.697515714354532 | -2.1533189468829 | 0.0370125924440513 | 0.247707839851692 | -3.63323150381996 |
| lnc-H1FOO-2 | lnc-H1FOO-2 | -0.727658176221677 | -0.534360929698498 | -2.15316802516927 | 0.0370251841187647 | 0.247707839851692 | -3.64091813378182 |
| ENSG00000232457.4 | SLC16A6P1 | -0.672923211029395 | -0.500708964849897 | -2.15312593844512 | 0.0370286961645721 | 0.247707839851692 | -3.6422999464153 |
| lnc-ABCB10-1 | lnc-ABCB10-1 | 0.375845440504978 | 0.763879820424515 | 2.15309471080996 | 0.0370313022327821 | 0.247707839851692 | -3.79297860693939 |
| lnc-APOL2-5 | lnc-APOL2-5 | 0.554865199134294 | 2.04080630008882 | 2.15287993783447 | 0.0370492302582154 | 0.247707839851692 | -3.91742533476192 |
| ENSG00000164151.11 | ICE1 | 0.177919177067024 | 6.85136945504443 | 2.15284948993257 | 0.037051772494978 | 0.247707839851692 | -4.22030977095779 |
| lnc-SBDS-15 | lnc-SBDS-15 | -0.293428084952468 | 2.51181730008563 | -2.15281701983115 | 0.0370544837440296 | 0.247707839851692 | -3.975358101236 |
| ENSG00000131042.14 | LILRB2 | -1.19258566359501 | 2.15795150167042 | -2.15450821880404 | 0.0370837204865818 | 0.247822798303673 | -3.95059751353757 |
| ENSG00000116350.16 | SRSF4 | 0.248074905789023 | 6.67539837295515 | 2.1520115406058 | 0.0371217971614218 | 0.247996738388038 | -4.22314047471151 |
| ENSG00000186314.11 | PRELID2 | -2.75008374576371 | -1.00266996603701 | -2.15363523110085 | 0.0371377129062681 | 0.248022564811719 | -3.62965669722023 |
| ENSG00000182944.17 | EWSR1 | 0.186452436812798 | 8.03493034818596 | 2.15122860077164 | 0.037187330270109 | 0.248273376240562 | -4.20667424304149 |
| lnc-ZIC5-3 | lnc-ZIC5-3 | -0.487796517383604 | -0.13417243448253 | -2.15071896606195 | 0.0372300421510898 | 0.248477936963645 | -3.68237268469083 |
| lnc-LEKR1-44 | lnc-LEKR1-44 | -0.375147176066557 | 1.33522205956189 | -2.15042402872955 | 0.0372547802480961 | 0.24856244468284 | -3.86763210097648 |
| ENSG00000071243.15 | ING3 | 0.232043767920352 | 4.90136710790804 | 2.14953893210172 | 0.0373291054137359 | 0.248977633418965 | -4.19526735581204 |
| ENSG00000134371.12 | CDC73 | 0.214727309957969 | 5.77573124259989 | 2.14907880065149 | 0.0373677961299687 | 0.249154955537138 | -4.22550367415508 |
| ENSG00000159720.11 | ATP6V0D1 | 0.329661089315728 | 4.70713851433588 | 2.14804949437544 | 0.0374544745881198 | 0.249508180668509 | -4.18410640711847 |
| lnc-MCM6-2 | lnc-MCM6-2 | 0.245802204956797 | 3.05587000689764 | 2.14798199459935 | 0.0374601649675102 | 0.249508180668509 | -4.03725144593131 |
| lnc-MTMR2-1 | lnc-MTMR2-1 | 0.384515535155651 | 1.25657492725931 | 2.14792197023924 | 0.0374652257928359 | 0.249508180668509 | -3.86303969454184 |
| lnc-ZDHHC22-2 | lnc-ZDHHC22-2 | -0.521621236037217 | 0.543096524637319 | -2.1478445238795 | 0.0374717564068913 | 0.249508180668509 | -3.77752638701571 |
| ENSG00000177479.19 | ARIH2 | 0.155822631951374 | 6.34775986954131 | 2.14773039079752 | 0.0374813824334174 | 0.249508180668509 | -4.23221077464995 |
| ENSG00000077232.17 | DNAJC10 | -0.273960988069495 | 7.85905396692379 | -2.14650496352349 | 0.0375848727953306 | 0.250116209746618 | -4.21808584399115 |
| ENSG00000136908.17 | DPM2 | -0.35197492655169 | 3.36981170931249 | -2.14582134746376 | 0.0376427151919234 | 0.250404125833983 | -4.08328666022482 |
| ENSG00000198455.4 | ZXDB | 0.222329216747074 | 5.15298019981692 | 2.1457061623041 | 0.0376524690014175 | 0.250404125833983 | -4.21587521908778 |
| ENSG00000275400.1 | | 0.412320632156649 | 0.336268130443445 | 2.1455120927967 | 0.0376689077257078 | 0.250432534792715 | -3.7580513952794 |
| lnc-BOD1-1 | lnc-BOD1-1 | -0.872749861942328 | -0.285624246024937 | -2.14466724438607 | 0.0377405445966943 | 0.250698015973297 | -3.66675285844152 |
| ENSG00000079337.15 | RAPGEF3 | -0.799438408201564 | 6.46689255363747 | -2.14658707117912 | 0.0377429452314003 | 0.250698015973297 | -4.23377154966894 |
| ENSG00000140553.17 | UNC45A | -0.361889670011077 | 4.53411475134367 | -2.14438602000981 | 0.0377644169432308 | 0.250698015973297 | -4.18682547298968 |
| ENSG00000122515.14 | ZMIZ2 | -0.308647417095743 | 6.35823890888321 | -2.14437085504893 | 0.0377657046320034 | 0.250698015973297 | -4.23810290661841 |
| lnc-ARF5-18 | lnc-ARF5-18 | -0.509172059613116 | -0.00551665369330039 | -2.1442563103588 | 0.0377754321123253 | 0.250698015973297 | -3.69994510655816 |
| lnc-POLR1A-3 | lnc-POLR1A-3 | 0.266784824104926 | 1.70675376112356 | 2.14384732649089 | 0.0378101822654922 | 0.250698015973297 | -3.91955612391269 |
| ENSG00000162601.9 | MYSM1 | 0.234039081995832 | 6.22452025853909 | 2.1437414298541 | 0.0378191845783155 | 0.250698015973297 | -4.23961827074383 |
| ENSG00000104921.14 | FCER2 | -0.647442273468827 | 8.09449601287579 | -2.14475200538214 | 0.0378430621375605 | 0.250698015973297 | -4.21945471654471 |
| lnc-MLYCD-2 | lnc-MLYCD-2 | -0.538031323605761 | -0.0736926642592908 | -2.14337434782009 | 0.0378504049759132 | 0.250698015973297 | -3.70145374726099 |
| ENSG00000172586.7 | CHCHD1 | 0.310824358947364 | 2.76434690552836 | 2.14313865928219 | 0.0378704622955981 | 0.250698015973297 | -4.02194056253356 |
| ENSG00000251369.8 | ZNF550 | -0.204681492010311 | 4.34527412946538 | -2.14313167189272 | 0.0378710570719648 | 0.250698015973297 | -4.17421748267759 |
| ENSG00000226049.3 | TLK2P1 | 0.986616916613735 | -0.162967135905167 | 2.14308028211445 | 0.0378754316945492 | 0.250698015973297 | -3.64368388414763 |
| ENSG00000130193.7 | THEM6 | -0.630399483503286 | 1.47927048720577 | -2.14304435435897 | 0.037878490356408 | 0.250698015973297 | -3.90333865553005 |
| ENSG00000130021.13 | PUDP | -0.362240210042884 | 2.0687442840687 | -2.14289892837629 | 0.0378908732272849 | 0.250698015973297 | -3.96439973814951 |
| lnc-ZNF613-7 | lnc-ZNF613-7 | -0.412663370962466 | 1.11813021836047 | -2.1428664530825 | 0.0378936389516329 | 0.250698015973297 | -3.86767161736563 |
| lnc-LRRC9-4 | lnc-LRRC9-4 | -0.370541656631269 | 0.61034398479731 | -2.14274812305689 | 0.0379037179084148 | 0.250698015973297 | -3.79996651219173 |
| lnc-ZEB2-1 | lnc-ZEB2-1 | 0.294459200558502 | 4.98187254016037 | 2.14253035760044 | 0.0379222726141436 | 0.250737433814236 | -4.20908864415422 |
| ENSG00000175573.6 | C11orf68 | -0.333389253013546 | 2.91853830566296 | -2.14229883980774 | 0.0379420078601034 | 0.250737433814236 | -4.04988400155437 |
| ENSG00000242299.1 | RPS18P5 | 1.35241177132255 | 3.09054696051754 | 2.14460956009548 | 0.0379582671225264 | 0.250737433814236 | -3.98903032505407 |
| ENSG00000139697.13 | SBNO1 | 0.172597623348734 | 7.24866042242766 | 2.14210656621403 | 0.0379584046915008 | 0.250737433814236 | -4.23711308591581 |
| ENSG00000144161.12 | ZC3H8 | -0.2145620588418 | 3.62633798403357 | -2.14196344811997 | 0.0379706136593435 | 0.250737613394375 | -4.11149196573432 |
| ENSG00000120458.11 | MSANTD2 | -0.320153824135331 | 3.28408682364275 | -2.14181929993503 | 0.0379829139965241 | 0.250738396020031 | -4.086171814636 |
| lnc-TIAF1-2 | lnc-TIAF1-2 | -0.292346077608175 | 1.09949963324577 | -2.14156935651723 | 0.0380042502846394 | 0.250798808467051 | -3.87193043587834 |
| ENSG00000163510.13 | CWC22 | 0.225874341304936 | 4.99012501246414 | 2.14123621117408 | 0.0380327054697387 | 0.250839922852675 | -4.21249695116307 |
| ENSG00000072849.10 | DERL2 | 0.23558263340956 | 4.69359987970799 | 2.14121106607039 | 0.0380348539679929 | 0.250839922852675 | -4.20306432485377 |
| lnc-COMMD3-1 | lnc-COMMD3-1 | -1.0397382291428 | -0.352399742271734 | -2.1398981181725 | 0.0381471860696702 | 0.25148667361967 | -3.68125127804735 |
| ENSG00000136930.12 | PSMB7 | 0.225275808284866 | 4.88137923854095 | 2.13977940606979 | 0.0381573571255031 | 0.25148667361967 | -4.21439013231137 |

| | | | | | | | |
|---|---|---|---|---|---|---|---|
| lnc-TMEM150A-1 | lnc-TMEM150A-1 | -0.4514023423331 | 0.19452535306849 | -2.13945913600928 | 0.0381848092431209 | 0.251587045022777 | -3.76638840623739 |
| ENSG00000117155.16 | SSX2IP | 0.653571339025876 | 1.52283616900904 | 2.13927522081594 | 0.0382005814894099 | 0.251610422014888 | -3.91944858337058 |
| ENSG00000267508.5 | ZNF285 | -2.69299009138198 | 0.251391801615572 | -2.14163211217162 | 0.0382141397072729 | 0.251619205884452 | -3.66424506369521 |
| LINC02580 | LINC02580 | 0.798995152474731 | 0.329170764934505 | 2.13876675171217 | 0.0382442167644283 | 0.251735088864523 | -3.73087513683678 |
| LINC00167 | LINC00167 | -0.561711979422113 | -0.432245757440832 | -2.13836163489606 | 0.0382790140602483 | 0.251735088864523 | -3.67409590019707 |
| ENSG00000170468.7 | RIOX1 | -0.356455287414822 | 3.47805639691914 | -2.13831322668819 | 0.0382831739213823 | 0.251735088864523 | -4.10872794825733 |
| lnc-ADRB2-1 | lnc-ADRB2-1 | -0.553949544395593 | 0.155302041060456 | -2.1382299584625 | 0.0382903303384024 | 0.251735088864523 | -3.71502626201858 |
| ENSG00000141965.4 | FEM1A | 0.309005909967308 | 4.93850574945807 | 2.13820017421292 | 0.0382928904063947 | 0.251735088864523 | -4.22097834905302 |
| lnc-ZC3H12B-18 | lnc-ZC3H12B-18 | -1.43063982286591 | 4.01875250221796 | -2.14045110583104 | 0.0383252833119853 | 0.251838684631226 | -4.13474094644437 |
| lnc-ROR1-2 | lnc-ROR1-2 | -0.764679866776465 | 2.56480461253558 | -2.13856042100168 | 0.0383331195136584 | 0.251838684631226 | -4.01785189507795 |
| lnc-TNFAIP8L1-2 | lnc-TNFAIP8L1-2 | -0.446895966733494 | 1.55639949312704 | -2.13749519970321 | 0.0383535296148809 | 0.251892373983119 | -3.92507552781678 |
| ENSG00000180198.16 | RCC1 | -0.393016736231568 | 4.30293738548369 | -2.13666997438192 | 0.0384246197024107 | 0.252029433375851 | -4.18563114324541 |
| ENSG00000004700.15 | RECQL | 0.31764409396171 | 5.76032888035154 | 2.1365812438725 | 0.0384322704062993 | 0.252029433375851 | -4.2497226522824 |
| ENSG00000067992.13 | PDK3 | 0.213314958559451 | 4.23336718461938 | 2.13638140559697 | 0.0384495061819936 | 0.252029433375851 | -4.17614174153551 |
| lnc-C12orf49-16 | lnc-C12orf49-16 | 0.692711465845715 | 0.055072804564751 | 2.13635157732214 | 0.0384520794129731 | 0.252029433375851 | -3.71559060238356 |
| ENSG00000238825.1 | RNVU1-2 | -0.972606600414011 | 1.71217336558422 | -2.13765843572899 | 0.038458317070873 | 0.252029433375851 | -3.93538597295139 |
| ENSG00000099953.9 | MMP11 | -0.694638346068672 | 2.5675908257714 | -2.1366939789266 | 0.0384639031045612 | 0.252029433375851 | -4.02612965420696 |
| ENSG00000215883.10 | CYB5RL | -0.264555529927792 | 2.95578032691326 | -2.13593199357922 | 0.0384882922030099 | 0.252029433375851 | -4.04639386705912 |
| ENSG00000123154.11 | WDR83 | -0.359854699197037 | 2.50741840274491 | -2.13586736332362 | 0.0384938728787526 | 0.252029433375851 | -4.02201632957177 |
| ENSG00000254783.1 | | 0.408327169103475 | 1.57222482103463 | 2.13578794270441 | 0.038500731643995 | 0.252029433375851 | -3.90712129807209 |
| lnc-SH3BP5L-3 | lnc-SH3BP5L-3 | -0.406226019212059 | 1.64226277814172 | -2.13575873578438 | 0.0385032542242826 | 0.252029433375851 | -3.93065893490289 |
| ENSG00000215492.6 | HNRNPA1P7 | 1.12692480278926 | 1.37137387795082 | 2.13703896872754 | 0.0385166857173088 | 0.252029433375851 | -3.84956507260654 |
| ENSG00000213553.4 | RPLP0P6 | 1.01271960551486 | 3.44938974529818 | 2.13734334746194 | 0.0385328002641575 | 0.252029433375851 | -4.05609050612345 |
| ENSG00000120889.12 | TNFRSF10B | 0.341775726973543 | 6.1070360062065 | 2.135407778137 | 0.0385335775559346 | 0.252029433375851 | -4.2546593528775 |
| lnc-FGFR1OP2-6 | lnc-FGFR1OP2-6 | 0.281416743455042 | 4.48619025781455 | 2.13503175922018 | 0.0385660895417438 | 0.252161950774369 | -4.19815583267125 |
| ENSG00000130812.10 | ANGPTL6 | -0.597886713448963 | 1.87793638899521 | -2.13459496246628 | 0.0386038869085368 | 0.252184598285921 | -3.96300678901259 |
| ENSG00000069188.16 | SDK2 | -1.60803248994221 | 5.19674942962041 | -2.137225282593 | 0.0386118936784481 | 0.252184598285921 | -4.22170420650222 |
| ENSG00000108960.8 | MMD | 0.425477517073011 | 4.16579516519588 | 2.13447838723163 | 0.0386139800312098 | 0.252184598285921 | -4.17253822670367 |
| ENSG00000258427.3 | RMB8AP1 | 1.65046571528222 | -1.11759175759405 | 2.13440592040289 | 0.038620255403683 | 0.252184598285921 | -3.70793255057052 |
| ENSG00000132326.11 | PER2 | 0.309263455254537 | 3.76978494297836 | 2.13428401854293 | 0.0386308137004085 | 0.252184598285921 | -4.13400022154697 |
| ENSG00000245680.9 | ZNF585B | -0.288069892707801 | 4.5489873694606 | -2.13399215990564 | 0.0386561027992142 | 0.252186324813274 | -4.19861639997488 |
| ENSG00000250299.6 | MRPS31P4 | 0.369443909930929 | 2.38067448876741 | 2.13395153033943 | 0.038659624442974 | 0.252186324813274 | -4.00950815323625 |
| lnc-CNIH3-2 | lnc-CNIH3-2 | -0.359760568610401 | 1.52404922100291 | -2.13362780631378 | 0.0386876939156399 | 0.252186324813274 | -3.92187523962953 |
| ENSG00000104783.12 | KCNN4 | -0.388126426456657 | 7.73263345396079 | -2.13362061002758 | 0.0386883180951248 | 0.252186324813274 | -4.24642467644009 |
| ENSG00000270757.1 | HSPE1-MOB4 | 0.238178797742129 | 3.81640537846061 | 2.13343085074078 | 0.0387047803175549 | 0.252186324813274 | -4.14531302689217 |
| lnc-CCDC96-2 | lnc-CCDC96-2 | -0.330499596563947 | 1.29238944636054 | -2.13312707614321 | 0.0387311465612362 | 0.252186324813274 | -3.90887522158887 |
| ENSG00000144228.8 | SPOPL | 0.231743900814782 | 6.32135202194223 | 2.13311139588838 | 0.0387325079643168 | 0.252186324813274 | -4.26025418503668 |
| ENSG00000188042.7 | ARL4C | 0.572739119645176 | 5.18108580341146 | 2.1337513509714 | 0.0387397930715711 | 0.252186324813274 | -4.23659941449438 |
| ENSG00000131043.11 | AAR2 | 0.243211355257555 | 4.57315118801975 | 2.13300959447564 | 0.0387413476684435 | 0.252186324813274 | -4.20599643811255 |
| lnc-SEH1L-5 | lnc-SEH1L-5 | -0.44845025548309 | 1.56745556618753 | -2.13285066945322 | 0.0387551511251366 | 0.252196419730853 | -3.92683749599086 |
| lnc-JPT2-4 | lnc-JPT2-4 | 0.625401087928693 | 1.61616030906929 | 2.13262470609823 | 0.038774784655655 | 0.252244435071855 | -3.91701973843781 |
| MIR17HG | MIR17HG | -0.408490329152027 | 4.92861249345769 | -2.13233855873064 | 0.0387996600178585 | 0.252326509367324 | -4.23482788814692 |
| ENSG00000169413.2 | RNASE6 | 0.947014859942608 | 4.58599948185971 | 2.13435719255123 | 0.0388139764743297 | 0.252339885587848 | -4.20359811006342 |
| lnc-SAG-9 | lnc-SAG-9 | -0.784120289417711 | 0.780780333337646 | -2.13165605016931 | 0.0388651083309668 | 0.252592524400471 | -3.86562051668121 |
| ENSG00000132155.11 | RAF1 | 0.166190453672362 | 6.75420386109191 | 2.13019418973433 | 0.0389865215093777 | 0.253301632647576 | -4.26446580953627 |
| lnc-LSM14B-6 | lnc-LSM14B-6 | -0.521352085920899 | 0.013768828628105 | -2.12917856171247 | 0.0390752997793586 | 0.253798326083477 | -3.73667369216216 |
| lnc-MAPK1IP1L-1 | lnc-MAPK1IP1L-1 | 0.265662913815709 | 1.02062289114383 | 2.12880070938539 | 0.039108374025728 | 0.253933016584088 | -3.87351611361246 |
| ENSG00000204390.9 | HSPA1L | -0.463532073530609 | 1.65178785823606 | -2.12855251557038 | 0.0391301123706315 | 0.253943102176716 | -3.94837740430232 |
| ENSG00000111254.7 | AKAP3 | -0.626847147454 | 0.37641229257711 | -2.12850126714764 | 0.0391346023468174 | 0.253943102176716 | -3.77723663849771 |
| lnc-CMYA5-4 | lnc-CMYA5-4 | -0.741591877103372 | 0.533676381681355 | -2.12778918099964 | 0.0391970365259229 | 0.254217462271558 | -3.80202069489989 |
| ENSG00000207145.1 | SNORA18 | 0.511266826434412 | 0.968639225936761 | 2.12773734220243 | 0.0392015850580539 | 0.254217462271558 | -3.84909915615594 |
| lnc-GBP4-1 | lnc-GBP4-1 | 0.576892438916723 | 0.068183491046818 | 2.12729585463426 | 0.0392403416579059 | 0.254362025831152 | -3.74209642315006 |
| lnc-CDH1-1 | lnc-CDH1-1 | -0.551007261606767 | -0.444899789667365 | -2.1272019083386 | 0.0392485932099178 | 0.254362025831152 | -3.68688515016886 |
| ENSG00000197872.11 | CYRIA | 2.26994424161786 | 2.10217391486077 | 2.12930488746155 | 0.0393007952129607 | 0.254620166153091 | -3.95502775491075 |
| ENSG00000128708.12 | HAT1 | 0.226237531011488 | 4.79720089963541 | 2.12602094821181 | 0.0393524502624415 | 0.254805638872914 | -4.23599508797627 |
| ENSG00000014216.15 | CAPN1 | -0.292579557940163 | 6.11065334440846 | -2.12600128100826 | 0.0393541818965648 | 0.254805638872914 | -4.27289249255266 |
| lnc-LIX1-2 | lnc-LIX1-2 | 0.316952588441539 | 0.698726264114181 | 2.12492436390976 | 0.0394491033019907 | 0.255339903542413 | -3.83466164141365 |
| ENSG00000285417.1 | | -0.832865079513517 | 4.69005563222788 | -2.1262565916826 | 0.0394993133596038 | 0.255548155851419 | -4.22579550681138 |
| ENSG00000186815.12 | TPCN1 | -0.234740557023724 | 5.44505334782748 | -2.12427871350338 | 0.0395061085322458 | 0.255548155851419 | -4.26495576134015 |
| ENSG00000218052.5 | ADAMTS7P4 | -0.540753728148941 | 0.621492425571718 | -2.12393268076077 | 0.039536689958205 | 0.255615964655788 | -3.8245613561047 |
| lnc-BRCC3-1 | lnc-BRCC3-1 | -0.384280324296221 | 0.340473535919754 | -2.12387907985347 | 0.0395414289201782 | 0.255615964655788 | -3.79883806290894 |
| ENSG00000140598.14 | EFL1 | -0.289437123319515 | 4.90062807020985 | -2.12334507165159 | 0.0395886689080545 | 0.255752091322828 | -4.24532746934029 |
| ENSG00000100228.12 | RAB36 | 1.01400009227616 | -0.644716322670487 | 2.12332552241427 | 0.0395903992337112 | 0.255752091322828 | -3.69142764349716 |
| ENSG00000154582.16 | ELOC | 0.232669841069958 | 3.38761751251863 | 2.12317333265345 | 0.0396038719988177 | 0.255752091322828 | -4.12664808345842 |
| ENSG00000134987.11 | WDR36 | 0.18349565917813 | 5.85517841528745 | 2.1230794161033 | 0.0396121880744874 | 0.255752091322828 | -4.27708988443274 |
| ENSG00000169084.13 | DHRSX | -0.318439750611871 | 3.54173942924122 | -2.12286360791472 | 0.0396313031648947 | 0.255795269063352 | -4.14184623368208 |
| ENSG00000079999.13 | KEAP1 | -0.414187598900127 | 2.63199695752542 | -2.1223911581885 | 0.0396731784370615 | 0.255985276416939 | -4.06415400137454 |
| lnc-SERTAD2-1 | lnc-SERTAD2-1 | -0.60863135248267 | 2.65954934982712 | -2.12225513837543 | 0.039691344716047 | 0.25600723796593 | -4.05614228767823 |

| | | | | | | | |
|---|---|---|---|---|---|---|---|
| ENSG00000212493.1 | SNORD19 | 0.470483763723582 | 0.579129836896656 | 2.12190665112656 | 0.0397161627811284 | 0.25600723796593 | -3.81964443904664 |
| ENSG00000226763.4 | SRRM5 | -0.623462385343618 | 0.288476046606214 | -2.12179228442222 | 0.0397263150982126 | 0.25600723796593 | -3.78096585408538 |
| lnc-PUS7-1 | lnc-PUS7-1 | -0.518371308422843 | -0.268737776279232 | -2.12179207953782 | 0.039726333287819 | 0.25600723796593 | -3.71737426573427 |
| ENSG00000139514.12 | SLC7A1 | -0.315596164173741 | 4.49288616493828 | -2.12134401223378 | 0.0397661301496776 | 0.256183491978346 | -4.22138575701065 |
| ENSG00000073008.14 | PVR | -2.53625112716985 | -0.287534539187953 | -2.12364293853184 | 0.0397805281190021 | 0.256196060786427 | -3.70695895887585 |
| ENSG00000226314.8 | ZKSCAN8P1 | -0.480541129699253 | 1.00331664982833 | -2.11753527711086 | 0.0401058347341836 | 0.258210321030247 | -3.88768613937649 |
| ENSG00000154263.17 | ABCA10 | -0.565180718538785 | 3.1393684973212 | -2.11694243277351 | 0.0401647216275621 | 0.258508588261448 | -4.11709691409862 |
| lnc-RPL3L-3 | lnc-RPL3L-3 | -0.51628886293965 3 | 0.656458601323543 | -2.11668313542617 | 0.0401821860390321 | 0.258540148559361 | -3.83905358578844 |
| lnc-LEF1-1 | lnc-LEF1-1 | -0.8406669902476 | 4.90439007638893 | -2.11833163066084 | 0.0402032009136325 | 0.258594526376656 | -4.25256705999649 |
| ENSG00000140474.13 | ULK3 | -0.276394371378221 | 5.40314997903209 | -2.11616622456197 | 0.0402285630708183 | 0.258676824019573 | -4.28131328867111 |
| lnc-NAGK-2 | lnc-NAGK-2 | -0.369851503144128 | 1.67058390952832 | -2.11552638504261 | 0.0402860343080191 | 0.258922195484914 | -3.96521673034993 |
| lnc-TMEM234-3 | lnc-TMEM234-3 | -0.852984102847209 | 0.54199244377849 | -2.11557424813946 | 0.0402918812679483 | 0.258922195484914 | -3.84755888725513 |
| lnc-NAGLU-3 | lnc-NAGLU-3 | -0.507708254028858 | 1.196317291338 | -2.11525795275385 | 0.0403101667131621 | 0.258958851890454 | -3.92021253779374 |
| ENSG00000129219.13 | PLD2 | -0.294330624034894 | 3.64832193876676 | -2.11490474116158 | 0.0403419402249865 | 0.259082107847394 | -4.1757513404413 |
| lnc-RIMKLB-1 | lnc-RIMKLB-1 | 1.24986867101007 | 1.82040936647186 | 2.11595267220191 | 0.0404277910752003 | 0.259483026841751 | -3.99073019537578 |
| ENSG00000069667.15 | RORA | -1.68581909839493 | 0.793743058767284 | -2.11629270693587 | 0.04043662603373 | 0.259483026841751 | -3.92098727947379 |
| ENSG00000239528.1 | RPS14P8 | -1.06733041128369 | -0.683288500717658 | -2.11379193425445 | 0.040442187733 | 0.259483026841751 | -3.68597231982738 |
| ENSG00000178685.13 | PARP10 | -0.371081581725397 | 4.98559288926966 | -2.11332799347002 | 0.0404840465164781 | 0.259670654237665 | -4.2694422152857 |
| lnc-CCDC90B-3 | lnc-CCDC90B-3 | -0.777844263362496 | 0.579027829693022 | -2.11316867589142 | 0.0404984296214836 | 0.259681986572896 | -3.85277902142607 |
| lnc-TFRC-2 | lnc-TFRC-2 | 0.466689438425166 | 0.776684467085707 | 2.11265404610484 | 0.040544920773829 | 0.25982461545589 | -3.88980856342984 |
| ENSG00000188070.9 | ZFTA | 0.595374405215206 | 0.310942381862098 | 2.11264299444701 | 0.0405459196834436 | 0.25982461545589 | -3.8442961099922 |
| lnc-PRIM1-1 | lnc-PRIM1-1 | 0.335067529562576 | 6.35031348770709 | 2.11246686586547 | 0.0405618420690803 | 0.259845750173632 | -4.29955942296327 |
| lnc-TIGIT-1 | lnc-TIGIT-1 | -0.3577850798128 | 3.47590523493324 | -2.11231302691282 | 0.0405757539168492 | 0.259853995914906 | -4.14558453243861 |
| ENSG00000076053.10 | RBM7 | 0.40673832302067 | 4.40417978758016 | 2.11122310541472 | 0.0406744368784891 | 0.26035725937242 | -4.23064132184936 |
| ENSG00000069943.9 | PIGB | 0.234242827291402 | 4.63651834257978 | 2.11116574806083 | 0.0406796359200167 | 0.26035725937242 | -4.25082976394549 |
| ENSG00000112486.16 | CCR6 | 1.27938616449496 | 1.65543632743323 | 2.11262568725389 | 0.0407168020875181 | 0.260514124142799 | -3.93230682184956 |
| OIP5-AS1 | OIP5-AS1 | -0.269976682474908 | 7.50099674178989 | -2.11035231496644 | 0.0407534307329597 | 0.260532888032778 | -4.29369743656377 |
| ENSG00000056736.9 | IL17RB | -1.24213985052472 | 2.44363358440471 | -2.11256672008428 | 0.0407560345766424 | 0.260532888032778 | -4.06011681137404 |
| ENSG00000111860.13 | CEP85L | -0.332791858224176 | 7.08608872244709 | -2.11030520900964 | 0.0407577077911649 | 0.260532888032778 | -4.29817185721134 |
| LINC00667 | LINC00667 | -0.335760061393624 | 4.78755408627425 | -2.10966348094627 | 0.0408160137321047 | 0.260824591942847 | -4.26458690810049 |
| ENSG00000198825.13 | INPP5F | -0.435549458956634 | 8.53522963409738 | -2.10845053496248 | 0.0409264191656211 | 0.26141138563399 | -4.27649981820557 |
| ENSG00000283199.2 | C13orf46 | 0.708700770679745 | 0.499759200123447 | 2.10837568445967 | 0.0409332408248724 | 0.26141138563399 | -3.7913316868957 |
| ENSG00000112234.8 | FBXL4 | -0.252654006476765 | 5.33229206934894 | -2.10816108302523 | 0.0409528045122799 | 0.261455203249459 | -4.29346027968227 |
| ENSG00000232713.2 | RPS12P3 | -0.372744161004783 | 0.115289808896469 | -2.10760891013006 | 0.0410031799047769 | 0.261670946153865 | -3.77833153795388 |
| ENSG00000065989.15 | PDE4A | -0.520277932306076 | 2.40520882463751 | -2.10742149786096 | 0.0410202901027228 | 0.261670946153865 | -4.0563722216124 |
| ENSG00000197217.12 | ENTPD4 | 0.373115718221317 | 7.44046861129872 | 2.10737281091954 | 0.0410247361044805 | 0.261670946153865 | -4.30088555808347 |
| ENSG00000105287.12 | PRKD2 | -0.291239065294963 | 4.99749283179035 | -2.10691392885253 | 0.0410666611428363 | 0.261819348297846 | -4.28270634107523 |
| lnc-SDC2-3 | lnc-SDC2-3 | 0.486760662719409 | 0.272086576612811 | 2.10683809976297 | 0.0410735927658242 | 0.261819348297846 | -3.79466606321235 |
| RAD51-AS1 | RAD51-AS1 | -0.352577306530907 | 6.87309728231261 | -2.10670061418372 | 0.0410861630958579 | 0.261819348297846 | -4.30742886881177 |
| lnc-GLT8D1-1 | lnc-GLT8D1-1 | -0.356519377227516 | 0.229096711092085 | -2.10645275812028 | 0.0411088331443069 | 0.261849053115058 | -3.79143132371889 |
| ENSG00000149016.15 | TUT1 | -0.256878392312699 | 3.31686399101577 | -2.10634267554273 | 0.0411189053184303 | 0.261849053115058 | -4.15006065374065 |
| ENSG00000262730.1 | | -0.46374949723747 | 0.796291399322882 | -2.10623248656982 | 0.0411289893951796 | 0.261849053115058 | -3.89212115345803 |
| ENSG00000102921.7 | N4BP1 | 0.153284679927895 | 5.18763178888201 | 2.10584664519169 | 0.0411643172336628 | 0.261921560164916 | -4.29164593573505 |
| ENSG00000176903.4 | PNMA1 | -0.312413717735323 | 3.70189014910447 | -2.10583014648459 | 0.0411658284571493 | 0.261921560164916 | -4.19255540711479 |
| ENSG00000169223.14 | LMAN2 | -0.229809735658055 | 5.22940405467582 | -2.10519228543604 | 0.041224291605092 | 0.262204596635865 | -4.29679265513469 |
| ENSG00000269971.1 | | 0.491758792720697 | -0.183589473324106 | 2.10499959726935 | 0.0412419667532956 | 0.262204596635865 | -3.73921497523374 |
| ENSG00000187860.10 | CCDC157 | -0.36771409515358 | 1.87739331263596 | -2.10492807034233 | 0.0412485295587101 | 0.262204596635865 | -4.01308299682851 |
| ENSG00000172992.11 | DCAKD | -0.243961104601117 | 3.65381950920412 | -2.10470316149759 | 0.041269171573543 | 0.262247031680039 | -4.18731202435792 |
| ENSG00000005889.15 | ZFX | 0.228677661067664 | 6.56387444483528 | 2.10457773481029 | 0.0412806871031106 | 0.262247031680039 | -4.31402261896891 |
| lnc-C15orf40-4 | lnc-C15orf40-4 | 0.248065674982161 | 2.61430415086149 | 2.10439257568521 | 0.0412976918687043 | 0.262274110377519 | -4.08062216678577 |
| lnc-FECH-2 | lnc-FECH-2 | 0.223158383757915 | 2.56608624957151 | 2.10374224455897 | 0.0413574660961036 | 0.262526251890434 | -4.08004926299117 |
| ENSG00000071894.16 | CPSF1 | -0.379565902735886 | 5.89000132561908 | -2.10362489766781 | 0.0413682599392627 | 0.262526251890434 | -4.31355465239229 |
| ENSG00000066697.14 | MSANTD3 | -0.313650767388629 | 2.74540599408581 | -2.10354448799421 | 0.0413756576365237 | 0.262526251890434 | -4.10834719250552 |
| ENSG00000109911.18 | ELP4 | -0.229801252486866 | 5.3010852393448 | -2.1033664954107 | 0.0413920370965363 | 0.262530509434855 | -4.30142854613691 |
| ENSG00000171806.11 | METTL18 | -0.314372568945921 | 2.73815808265888 | -2.10317484177626 | 0.0414096800482276 | 0.262530509434855 | -4.09647103713395 |
| ENSG00000148290.9 | SURF1 | 0.172222738814642 | 3.62047065262969 | 2.10303196029745 | 0.0414228374990962 | 0.262530509434855 | -4.18213523058865 |
| ENSG00000185641.6 | | 1.11244369801039 | 2.0258932437658 | 2.10454328244906 | 0.0414339801572942 | 0.262530509434855 | -3.9746028839126 |
| ENSG00000064886.13 | CHI3L2 | 1.48204242851284 | 2.15141062626396 | 2.10525108035473 | 0.04144010227633 7 | 0.262530509434855 | -4.00657000302843 |
| lnc-VTI1B-2 | lnc-VTI1B-2 | 0.240795119931713 | 1.86389422538261 | 2.10223304557186 | 0.0414964743097966 | 0.262806747913398 | -4.01439498829241 |
| ENSG00000284564.1 | MIR3064 | 0.3383014502059 | 5.09324465535286 | 2.10180676389937 | 0.0415358120794487 | 0.262974967711871 | -4.29663632975439 |
| ENSG00000188786.9 | MTF1 | 0.257320019674992 | 4.14615996750875 | 2.10125392136536 | 0.0415868777271277 | 0.263217313732309 | -4.23392336780445 |
| ENSG00000161638.10 | ITGA5 | 2.05491135713991 | 1.2597401469026 | 2.10338219099867 | 0.041622323348358 | 0.263360676753536 | -3.87621035750801 |
| lnc-TPH1-1 | lnc-TPH1-1 | 0.26250761526122 | 0.685973806964046 | 2.10046869404835 | 0.0416595031862089 | 0.263408464436388 | -3.88790833249203 |
| ENSG00000170248.14 | PDCD6IP | -0.154697373781455 | 7.15170968928025 | -2.1004295359657 | 0.0416631278152355 | 0.263408464436388 | -4.31675881733698 |
| ENSG00000204196.5 | RPL12P16 | 0.976005093448111 | 2.10392585128856 | 2.10164337654468 | 0.0416795380229068 | 0.263408464436388 | -4.00027286888759 |
| ENSG00000253846.2 | PCDHGA10 | -1.38959186618011 | -0.250203654993154 | -2.10141586973088 | 0.0416852509178842 | 0.263408464436388 | -3.77506268650658 |
| lnc-MARVELD2-2 | lnc-MARVELD2-2 | -0.248842601629424 | 1.04936401728196 | -2.09966951081605 | 0.0417335335631008 | 0.263408464436388 | -3.93240821611236 |
| lnc-TMOD1-2 | lnc-TMOD1-2 | 0.437644681936984 | 1.07145568748252 | 2.09948332377803 | 0.0417507970950256 | 0.263408464436388 | -3.92904428961965 |

| | | | | | | | |
|---|---|---|---|---|---|---|---|
| ENSG00000166068.12 | SPRED1 | -2.49331836647667 | -1.22519732536089 | -2.1011373671713 | 0.0417528731569214 | 0.263408464436388 | -3.69675741810268 |
| ENSG00000179115.10 | FARSA | -0.274625758339816 | 3.87208186375471 | -2.09945151881019 | 0.0417537467235961 | 0.263408464436388 | -4.22284820959333 |
| ENSG00000171246.5 | NPTX1 | -3.43190572073633 | 0.834758839409493 | -2.10181567054139 | 0.0417764009942783 | 0.263408464436388 | -3.81301195396813 |
| lnc-MCM9-3 | lnc-MCM9-3 | -0.354067688566742 | 3.33716167283162 | -2.09905445225247 | 0.0417905865010435 | 0.263408464436388 | -4.16616796129031 |
| lnc-ZNF772-1 | lnc-ZNF772-1 | -0.350847993142478 | 0.986553466948376 | -2.09893491700202 | 0.0418016825420082 | 0.263408464436388 | -3.92131701529537 |
| ENSG00000070371.15 | CLTCL1 | -0.697334658753827 | -0.0998789325235017 | -2.09892188761981 | 0.0418028921702352 | 0.263408464436388 | -3.81082849213678 |
| ENSG00000171861.10 | MRM3 | 0.231348071276452 | 3.73829367475419 | 2.09877732259208 | 0.0418163154284771 | 0.263408464436388 | -4.20566980712785 |
| ENSG00000113621.14 | TXNDC15 | -0.179576116615332 | 4.48937655558981 | -2.09867069336849 | 0.0418262186642392 | 0.263408464436388 | -4.26907084343933 |
| ENSG00000130813.17 | SHFL | -0.258787267522141 | 4.72607144566089 | -2.09859519887127 | 0.0418332314915015 | 0.263408464436388 | -4.28596313169545 |
| ENSG00000171467.15 | ZNF318 | -0.445404888089274 | 6.98301148389287 | -2.09858209813681 | 0.0418346339329812 | 0.263408464436388 | -4.32266686008571 |
| ENSG00000169251.12 | NMD3 | 0.25689051208481 | 5.069937792529 | 2.09810139316107 | 0.0418791274621568 | 0.263607975704457 | -4.30262909034545 |
| ENSG00000178449.8 | COX14 | 0.288679259904595 | 2.93413808982788 | 2.097561406178 | 0.0419293661828751 | 0.263843517010736 | -4.12624104508587 |
| ENSG00000257591.5 | ZNF625 | -0.376328542960431 | 1.17709196296078 | -2.09718657668527 | 0.0419642702077036 | 0.263966594479141 | -3.94955070619478 |
| lnc-SUDS3-3 | lnc-SUDS3-3 | -0.494437106559613 | 0.148794885609551 | -2.09704401013601 | 0.0419775526500083 | 0.263966594479141 | -3.82152787122747 |
| ENSG00000023892.10 | DEF6 | -0.210446304909931 | 5.74174126709759 | -2.09693835196969 | 0.041987398840048 | 0.263966594479141 | -4.32476039199983 |
| ENSG00000113013.13 | HSPA9 | 0.182052796815852 | 7.47081680054948 | 2.09667492514635 | 0.04201195616328 | 0.264040333956883 | -4.32065258411625 |
| lnc-COX7B-2 | lnc-COX7B-2 | -0.4574367689605 | -0.311241848784679 | -2.0964703941543 | 0.0420310317446365 | 0.264079586813142 | -3.77792101659243 |
| lnc-ZNF780B-3 | lnc-ZNF780B-3 | -0.239526185691758 | 3.74955556426664 | -2.09569103359767 | 0.042103788373589 | 0.264445618693669 | -4.21914445911756 |
| lnc-ADCY3-3 | lnc-ADCY3-3 | 0.370760031842954 | 0.318744386992746 | 2.09557120047256 | 0.0421149850885607 | 0.264445618693669 | -3.82353488715105 |
| POT1-AS1 | POT1-AS1 | -0.660229376863924 | -0.238755427947508 | -2.09542172893864 | 0.0421289547510026 | 0.264452661067364 | -3.7359389974953 |
| ENSG00000104957.13 | YJU2B | -0.201266616209463 | 4.90783760501079 | -2.09464853830982 | 0.0422012822547373 | 0.264825912393067 | -4.3018176644965 |
| ENSG00000205744.9 | DENND1C | -0.332580672541548 | 5.69767407937645 | -2.09418076116743 | 0.0422450928904848 | 0.264934034478766 | -4.32829975677716 |
| lnc-POPDC2-1 | lnc-POPDC2-1 | -0.305714793281109 | 2.71645003063747 | -2.09415360785355 | 0.0422476372132928 | 0.264934034478766 | -4.12018290383346 |
| ENSG00000065923.9 | SLC9A7 | 0.245671686944323 | 8.70734817627921 | 2.09405234946588 | 0.0422571265215853 | 0.264934034478766 | -4.30319117804812 |
| ENSG00000170448.11 | NFXL1 | 0.2621264139762 | 3.40885120284948 | 2.09381799106785 | 0.0422790962992697 | 0.264991059417743 | -4.17564515611518 |
| lnc-C17orf62-6 | lnc-C17orf62-6 | -0.260506757259896 | 3.63292559913761 | -2.0934769986532 | 0.0423110802841605 | 0.265110796191439 | -4.20475497469911 |
| lnc-WNT7B-2 | lnc-WNT7B-2 | -0.812487042786679 | -0.221867900463268 | -2.0928741660568 | 0.0423676758860305 | 0.265384623482096 | -3.75330717672731 |
| lnc-OCIAD2-2 | lnc-OCIAD2-2 | 0.34118143491857 | 0.844960700095303 | 2.09226670701541 | 0.0424247728911027 | 0.265661423917726 | -3.90964888633498 |
| ENSG00000171617.13 | ENC1 | 0.79649860664063 | 4.40743948057321 | 2.09342039730238 | 0.0424748442972815 | 0.265810763709849 | -4.25839576154344 |
| ENSG00000104853.15 | CLPTM1 | -0.235206574380193 | 5.56164491667717 | -2.09172538760523 | 0.0424757099946371 | 0.265810763709849 | -4.33061747761498 |
| lnc-ZNF487-7 | lnc-ZNF487-7 | -0.522866264240655 | -0.203818087834962 | -2.09160161824829 | 0.0424873639705292 | 0.265810763709849 | -3.78708316308463 |
| ENSG00000141298.18 | SSH2 | 0.230082558845041 | 8.22834151674804 | 2.09141012185643 | 0.0425054005615006 | 0.265842801506341 | -4.31765277444842 |
| ENSG00000147168.12 | IL2RG | -0.261679541340679 | 5.97589832764338 | -2.09126271190553 | 0.0425192893222248 | 0.265848885819974 | -4.33796006055761 |
| ENSG00000182700.4 | IGIP | -0.382640282465656 | 1.96828750978689 | -2.09107528294418 | 0.0425369543573052 | 0.265878570159858 | -4.05014203031033 |
| lnc-HENMT1-2 | lnc-HENMT1-2 | -0.550235262296097 | -0.417859447369918 | -2.09056373794162 | 0.0425851997895976 | 0.265928258242303 | -3.76091569753057 |
| ENSG00000073803.13 | MAP3K13 | -1.13604248152657 | 1.46700988639567 | -2.09233252752236 | 0.0425904108077055 | 0.265928258242303 | -4.02380810528725 |
| ENSG00000236965.4 | OR52N3P | 1.88176565577892 | -1.54755462093068 | 2.09041525434602 | 0.0425992127181766 | 0.265928258242303 | -3.77379610501786 |
| ENSG00000160325.14 | CACFD1 | -0.363109958515804 | 3.53334873248759 | -2.09039166432887 | 0.0426014393642036 | 0.265928258242303 | -4.21236508858021 |
| lnc-TMEM126B-2 | lnc-TMEM126B-2 | -1.25069684862631 | -0.205013317928588 | -2.09131045675262 | 0.0426095027781721 | 0.265928258242303 | -3.81646689140806 |
| ENSG00000167325.14 | RRM1 | 0.240637159330334 | 4.32359708455985 | 2.09014558520427 | 0.0426246726860029 | 0.265942297028189 | -4.26701805183964 |
| ENSG00000188641.13 | DPYD | -2.16183758803887 | 3.39506348845579 | -2.09253706727546 | 0.0426423772654446 | 0.2659703273023 | -4.19686817488321 |
| ENSG00000284172.1 | MIR4315-2 | 0.344197232248017 | 0.268593713788549 | 2.08982445033656 | 0.0426550090086427 | 0.2659703273023 | -3.82662860449055 |
| ENSG00000224114.1 | | -1.86309672923375 | 0.0388829347076459 | -2.09075798766438 | 0.0427388943433659 | 0.266330412895141 | -3.76969059080904 |
| lnc-IL10-1 | lnc-IL10-1 | -1.86309672923375 | 0.0388829347076459 | -2.09075798766438 | 0.0427388943433659 | 0.266330412895141 | -3.76969059080904 |
| ENSG00000182973.18 | CNOT10 | -0.202896356208671 | 4.84229353454594 | -2.08880354153202 | 0.0427515757763954 | 0.266330412895141 | -4.30887741340231 |
| ENSG00000197961.11 | ZNF121 | 0.238005829851141 | 6.51332307587845 | 2.08821542632726 | 0.0428072918606092 | 0.266596819475619 | -4.34509735621431 |
| lnc-SRSF12-3 | lnc-SRSF12-3 | -0.560978164532926 | -0.457368619272726 | -2.08769525102295 | 0.0428566245078223 | 0.266823321913039 | -3.74523471359531 |
| ENSG00000207280.1 | SNORD20 | 0.325691601689805 | 1.65402140621925 | 2.08659956291625 | 0.0429607007680645 | 0.267366788537046 | -4.02427671078249 |
| ENSG00000166401.14 | SERPINB8 | -0.401541157607397 | 4.26224712722602 | -2.08645950441788 | 0.0429740204549733 | 0.267366788537046 | -4.27347124996077 |
| ENSG00000100612.13 | DHRS7 | -0.201673693154564 | 4.29117835375591 | -2.08636632300093 | 0.0429828840921676 | 0.267366788537046 | -4.27666894719149 |
| ENSG00000138071.13 | ACTR2 | 0.307704673936716 | 7.69852223061283 | 2.0860945601545 | 0.0430087439569648 | 0.267446820805501 | -4.33757420064097 |
| lnc-B4GALT1-3 | lnc-B4GALT1-3 | 0.297476545733681 | 1.61633404621097 | 2.08577567020659 | 0.043039105610161 | 0.267450894600632 | -4.01873278331849 |
| ENSG00000172766.18 | NAA16 | 0.290997864797586 | 5.17298477132722 | 2.08573328940044 | 0.0430431421172822 | 0.267450894600632 | -4.33146320002783 |
| ENSG00000168701.18 | TMEM208 | 0.190308467444744 | 3.85597854390396 | 2.08557883061731 | 0.0430578561559965 | 0.267450894600632 | -4.23657426332688 |
| ENSG00000172748.13 | ZNF596 | -0.260546132891132 | 3.4067024355065 | -2.08543110828158 | 0.0430719325857601 | 0.267450894600632 | -4.19415949405974 |
| ENSG00000072858.10 | SIDT1 | 0.311901282192157 | 5.71104740348869 | 2.08521985445155 | 0.0430920699155876 | 0.267450894600632 | -4.34415997920322 |
| ENSG00000189042.13 | ZNF567 | 0.201938906291494 | 4.16739463600374 | 2.08515927814273 | 0.0430978457453672 | 0.267450894600632 | -4.26160822455772 |
| ENSG00000071082.10 | RPL31 | 0.39886447958051 | 8.53396037648305 | 2.08513295824021 | 0.0431003555065004 | 0.267450894600632 | -4.32366688973615 |
| ENSG00000169738.7 | DCXR | -0.441631541979925 | 3.30002407503886 | -2.08481888905313 | 0.0431303137642163 | 0.267480444969293 | -4.19893801727908 |
| lnc-MGAT2-1 | lnc-MGAT2-1 | -0.459195412289083 | 2.71191894802922 | -2.08481056477998 | 0.043131108043195 | 0.267480444969293 | -4.12466424393601 |
| ENSG00000104221.12 | BRF2 | 0.234742701254569 | 2.86232430518987 | 2.08432586073215 | 0.0431773792312816 | 0.267672994780452 | -4.14657045817422 |
| ENSG00000183475.12 | ASB7 | 0.197194995616829 | 4.69024641504746 | 2.08410597318492 | 0.0431983846223183 | 0.267672994780452 | -4.30825363435794 |
| lnc-MBTD1-2 | lnc-MBTD1-2 | 0.366426862206514 | 0.249288172351139 | 2.08406240332846 | 0.043202547819529 | 0.267672994780452 | -3.84216176529214 |
| lnc-SORBS1-2 | lnc-SORBS1-2 | 0.42273641763836 | 2.70156603041814 | 2.08394073996477 | 0.0432141748848779 | 0.267672994780452 | -4.12448698259926 |
| ENSG00000187942.11 | LDLRAD2 | 1.63644282658586 | 1.80875675826768 | 2.08585615475549 | 0.0432557372274196 | 0.267849831333327 | -3.979487020615 |
| ENSG00000105700.10 | KXD1 | -0.177668649622751 | 5.15176320596693 | -2.08326835030587 | 0.0432784828857351 | 0.267866673032206 | -4.33361903203616 |
| lnc-ABHD4-12 | lnc-ABHD4-12 | -0.829030835530773 | 3.58243310345334 | -2.08470755005194 | 0.0432918982198472 | 0.267866673032206 | -4.21411013772517 |
| LINC00271 | LINC00271 | -0.49800374844068 | 1.38006673736847 | -2.08306969002886 | 0.0432974989641038 | 0.267866673032206 | -4.00250275255433 |

| | | | | | | | |
|---|---|---|---|---|---|---|---|
| ENSG00000197951.9 | ZNF71 | -0.449533048711805 | 2.48058449385217 | -2.08286659943195 | 0.0433169466757961 | 0.267906464371368 | -4.1244361797548 |
| ENSG00000140284.10 | SLC27A2 | -1.4270429003981 | -0.766737058995051 | -2.08252485705263 | 0.04337350559364 | 0.268120867826978 | -3.72914925297727 |
| ENSG00000149925.18 | ALDOA | -0.265638334058856 | 5.96429316246212 | -2.08223303101267 | 0.0433776655426244 | 0.268120867826978 | -4.35476418244241 |
| ENSG00000186660.14 | ZFP91 | 0.177523046550382 | 6.46747746264218 | 2.08197295277661 | 0.0434026120314612 | 0.268158701984857 | -4.3570410641616 |
| ENSG00000167549.18 | CORO6 | -0.776057584596645 | 1.06478257314369 | -2.08195504428942 | 0.0434146472670099 | 0.268158701984857 | -3.95423718200112 |
| lnc-C19orf47-2 | lnc-C19orf47-2 | 0.41197859032839 | 0.0534156012279794 | 2.08176184026292 | 0.0434228709962361 | 0.268158701984857 | -3.8137380783945 |
| lnc-UBR4-3 | lnc-UBR4-3 | 0.239962531244525 | 1.57927160687108 | 2.08160242651369 | 0.0434381742796473 | 0.268172747809832 | -4.01686222177554 |
| ENSG00000100902.10 | PSMA6 | 0.167816337210005 | 4.58353034085497 | 2.08116255482568 | 0.0434804252282347 | 0.268318671808816 | -4.30833808477742 |
| ENSG00000244045.12 | TMEM199 | 0.169351547376803 | 4.55812927553594 | 2.08108495466067 | 0.0434878826776569 | 0.268318671808816 | -4.30370734811979 |
| ENSG00000188171.15 | ZNF626 | -0.702568132769382 | 4.40314809220949 | -2.0822525044834 | 0.0435087564842867 | 0.268367016696456 | -4.28983747224894 |
| ENSG00000166169.16 | POLL | -0.167359477246889 | 4.51861459503087 | -2.08049600912691 | 0.0435445173767721 | 0.268507130367316 | -4.30205622955358 |
| ENSG00000105968.18 | H2AZ2 | 0.210936459333291 | 6.1581377607965 | 2.08032265840924 | 0.0435611995647996 | 0.268529550956055 | -4.36002883707161 |
| lnc-MECR-1 | lnc-MECR-1 | 0.276180220608068 | 3.77876404576477 | 2.07993292467305 | 0.0435987255116987 | 0.268680409343501 | -4.2360012716543 |
| ENSG00000188223.9 | | -0.293838525706761 | 2.05755861565511 | -2.07936622540677 | 0.0436533412239529 | 0.268882124667951 | -4.07877051902823 |
| lnc-OR2B2-4 | lnc-OR2B2-4 | -0.578607507498684 | 0.870676539826655 | -2.07932222318776 | 0.0436575844454303 | 0.268882124667951 | -3.92456130803956 |
| lnc-HNRNPA1L2-2 | lnc-HNRNPA1L2-2 | 0.359037584637012 | 2.30144722950585 | 2.07906997533907 | 0.043681916168335 | 0.268951504783978 | -4.10370725746707 |
| ENSG00000143947.13 | RPS27A | 0.299123043987368 | 9.45987844523131 | 2.07795517030639 | 0.0437895918690337 | 0.269533842535981 | -4.31245170046111 |
| ENSG00000142207.6 | URB1 | -0.26561365607537 | 4.88669431715731 | -2.07735690602722 | 0.0438474720716392 | 0.269809422317055 | -4.33210517049082 |
| ENSG00000204592.8 | HLA-E | 0.256424840792633 | 9.83747478636757 | 2.0770900544135 | 0.0438733106972592 | 0.269887732839715 | -4.30584514013597 |
| lnc-CARF-1 | lnc-CARF-1 | -0.537969775788182 | -0.528800299935835 | -2.0746331602457 | 0.0441118322497413 | 0.271194961609713 | -3.78298288915816 |
| ENSG00000240616.1 | RPS6P25 | 0.637588961566458 | 0.112131630848661 | 2.07462972125951 | 0.044112166908095 | 0.271194961609713 | -3.79816115040269 |
| lnc-SMIM14-5 | lnc-SMIM14-5 | -0.512451600572622 | -0.237299426796051 | -2.07448004884338 | 0.0441267341427311 | 0.271203514141486 | -3.80142299781962 |
| ENSG00000113845.9 | TIMMDC1 | -0.239161306451363 | 3.9833752959609 | -2.07431791941497 | 0.0441425185304644 | 0.271219539973895 | -4.26800928275714 |
| ENSG00000263276.1 | | -0.220778266678633 | 3.76119919078669 | -2.07398119481908 | 0.044175316697514 | 0.271340060753486 | -4.24583902971067 |
| ENSG00000100147.13 | CCDC134 | 0.211250060638777 | 2.38479698875191 | 2.0729525226163 | 0.0442756449599416 | 0.271875179060405 | -4.11359348626505 |
| lnc-PLEKHM1-3 | lnc-PLEKHM1-3 | -2.22317312897117 | 2.83468422051023 | -2.07500141060266 | 0.044322042052878 | 0.272078911892153 | -4.17056502543182 |
| ENSG00000115216.13 | NRBP1 | 0.215435791748575 | 5.80147980106808 | 2.07184060049341 | 0.0443843166268709 | 0.272379960981182 | -4.37297939272238 |
| ENSG00000008853.16 | RHOBTB2 | -0.661798462794748 | 6.31683135432638 | -2.07207016174308 | 0.0444998061496478 | 0.273007305507661 | -4.37509222633534 |
| ENSG00000197061.4 | H4C3 | 0.434206725092602 | 5.38957672956193 | 2.07030084384792 | 0.0445351866962636 | 0.273142952255421 | -4.36132781626205 |
| ENSG00000196182.10 | STK40 | -0.346448940914089 | 5.0169967506057 | -2.0699170533909 | 0.0445728613404179 | 0.273237029800078 | -4.35338154011139 |
| ENSG00000157388.16 | CACNA1D | -1.4312946221945 | 2.84693561936826 | -2.07228289141509 | 0.0445770755511181 | 0.273237029800078 | -4.13626752012145 |
| ENSG00000025772.7 | TOMM34 | -0.314344689035341 | 3.7578288922953 | -2.06969991031026 | 0.0445941894382071 | 0.27326055409545 | -4.26244617186746 |
| ENSG00000115970.18 | THADA | 0.25600501845659 | 6.12121575581448 | 2.06943979833364 | 0.0446197497363641 | 0.273335806197495 | -4.37983361406015 |
| ENSG00000151320.10 | AKAP6 | 1.82475472310812 | 4.6038768166313 | 2.07097802683906 | 0.0447154938112646 | 0.273804704920703 | -4.27337082013262 |
| ENSG00000172081.13 | MOB3A | -0.264092886440126 | 5.731430389614 | -2.06839143619453 | 0.0447228984085607 | 0.273804704920703 | -4.37806956486738 |
| lnc-ASF1A-6 | lnc-ASF1A-6 | -0.408123786075858 | 3.03073830289683 | -2.06817177522328 | 0.0447445372903293 | 0.273855727340722 | -4.19271984061564 |
| ENSG00000138279.15 | ANXA7 | 0.204948279810838 | 6.1019057017231 | 2.06803231699424 | 0.0447582801160747 | 0.273858406548504 | -4.38272368852136 |
| ENSG00000104626.14 | ERI1 | -0.258991960605417 | 3.42734017426512 | -2.06741929597366 | 0.0448187336241092 | 0.274059285131432 | -4.24475632653659 |
| ENSG00000104047.14 | DTWD1 | -0.228556778594386 | 3.87768445403651 | -2.06729317375789 | 0.0448311800978391 | 0.274059285131432 | -4.27850112439302 |
| lnc-ARL13B-1 | lnc-ARL13B-1 | 0.384255843794917 | 0.270438944142153 | 2.06719181062645 | 0.044841185388872 | 0.274059285131432 | -3.87367422952612 |
| lnc-BCL11A-2 | lnc-BCL11A-2 | 0.763686017538589 | -0.119630873575858 | 2.06688866291144 | 0.0448711199488405 | 0.274059285131432 | -3.79853485413029 |
| ENSG00000125637.15 | PSD4 | -0.24336661326539 | 7.64647560772767 | -2.06683265736321 | 0.0448766521702011 | 0.274059285131432 | -4.37225649782158 |
| ENSG00000182247.9 | UBE2E2 | 1.14038703453143 | 2.491961642135 | 2.06872736435877 | 0.044883316274429 | 0.274059285131432 | -4.140310409226 |
| ENSG00000133794.17 | BMAL1 | 0.397899020513092 | 3.92919899944629 | 2.0665558205667 | 0.0449040068253726 | 0.274059285131432 | -4.27339255284127 |
| ENSG00000070770.8 | CSNK2A2 | 0.224791471462932 | 5.62367982776347 | 2.06653202637424 | 0.0449063586442483 | 0.274059285131432 | -4.37921781345844 |
| ENSG00000123144.10 | TRIR | -0.44367991454568 | 5.47746169485262 | -2.06638803330951 | 0.0449221779204693 | 0.274059285131432 | -4.37833479145601 |
| ENSG00000184640.17 | SEPTIN9 | -0.256662670538191 | 8.70967081128232 | -2.06635095525176 | 0.0449242592446899 | 0.274059285131432 | -4.35261751458921 |
| ENSG00000207155.1 | RNY1P14 | 0.879201353543678 | -0.844301317831342 | 2.06597459778455 | 0.0449614856906898 | 0.274109137744507 | -3.74840944839684 |
| ENSG00000225178.5 | NA | 0.873289443047277 | 2.96786793922133 | 2.06716778098392 | 0.0449806780648883 | 0.274109137744507 | -4.14485757410805 |
| ENSG00000152443.12 | ZNF776 | 0.201017647975614 | 5.99920056841514 | 2.06562867402842 | 0.044995725616334 | 0.274109137744507 | -4.38618176521262 |
| ENSG00000124067.16 | SLC12A4 | -0.388557614897551 | 3.34270976110885 | -2.06561533969361 | 0.0449970459195041 | 0.274109137744507 | -4.23491086030196 |
| lnc-TMEM173-3 | lnc-TMEM173-3 | -0.462711875200698 | 2.69548600339337 | -2.06559542882722 | 0.0449990174628912 | 0.274109137744507 | -4.16305680157133 |
| ENSG00000196700.8 | ZNF512B | -0.456703025492948 | 3.60568245550102 | -2.06520015172422 | 0.045038172808346 | 0.27411005437547 | -4.25536046491178 |
| ENSG00000000457.13 | SCYL3 | -0.316495769953999 | 3.94535986192618 | -2.06491742568579 | 0.0450661973186501 | 0.27411005437547 | -4.29127495506692 |
| ENSG00000278769.1 | | -0.724325055373138 | -0.682225354107614 | -2.06489695215478 | 0.0450682272971461 | 0.27411005437547 | -3.77413708530796 |
| ENSG00000197265.8 | GTF2E2 | 0.271754238566608 | 4.56128291448492 | 2.06479152717568 | 0.0450786815914454 | 0.27411005437547 | -4.33191009187951 |
| ENSG00000142039.3 | CCDC97 | -0.244357621663617 | 4.39656571644625 | -2.06472620660268 | 0.0450851600606325 | 0.27411005437547 | -4.32392926832328 |
| ENSG00000187838.16 | PLSCR3 | -0.3058688495152 | 2.09810245772314 | -2.06467979074269 | 0.0450897640616856 | 0.27411005437547 | -4.10554815503287 |
| ENSG00000101577.9 | LPIN2 | -0.3902481870245 | 5.59429086797825 | -2.0646533282731 | 0.0450923890645359 | 0.27411005437547 | -4.38202064448932 |
| ENSG00000174840.8 | PDE12 | 0.229695066606836 | 5.80365199080988 | 2.06448218704363 | 0.0451093690168446 | 0.274125584567067 | -4.38642316127261 |
| ENSG00000085644.13 | ZNF213 | -0.437180670206706 | 1.14168002432777 | -2.06394402596599 | 0.0451627995984345 | 0.274125584567067 | -4.01010258737282 |
| ENSG00000163399.15 | ATP1A1 | 0.270202582970135 | 6.9468525744439 | 2.0638810488794 | 0.0451690558011834 | 0.274125584567067 | -4.38861573087341 |
| ENSG00000054267.21 | ARID4B | 0.197676338664951 | 7.51587610209246 | 2.06386770392328 | 0.0451703815987564 | 0.274125584567067 | -4.38170501975751 |
| ENSG00000155256.17 | ZFYVE27 | 0.197039422233795 | 5.16027845718292 | 2.06379125056648 | 0.045177977758112 | 0.274125584567067 | -4.37015222504365 |
| ENSG00000243566.6 | UPK3B | -0.564148095880443 | -0.116750047303505 | -2.06378185318424 | 0.045178911528704 | 0.274125584567067 | -3.82587974646566 |
| ENSG00000112851.14 | ERBIN | 0.14259454297877 | 8.14295452798374 | 2.06368868334189 | 0.04518817025876 | 0.274125584567067 | -4.37122913101063 |
| lnc-C17orf62-1 | lnc-C17orf62-1 | -0.452358963009635 | 2.09191298463125 | -2.06313543532353 | 0.0452431832701355 | 0.274287142924416 | -4.10755199245922 |
| ENSG00000197971.14 | MBP | -0.302273296854779 | 6.160208013855 | -2.06307531766378 | 0.0452491646759071 | 0.274287142924416 | -4.39181335546969 |

| | | | | | | | |
|---|---|---|---|---|---|---|---|
| lnc-VMP1-8 | lnc-VMP1-8 | -0.486793226990808 | 2.82130122171228 | -2.06277626044908 | 0.045278929627827 | 0.274287142924416 | -4.18992030095446 |
| ENSG00000167077.12 | MEI1 | 0.547395398264331 | 4.4868451230738 | 2.06322345116639 | 0.0452813309184528 | 0.274287142924416 | -4.33336678413397 |
| ENSG00000232573.1 | RPL3P4 | 0.811168946597007 | 3.17957162261551 | 2.06403426504718 | 0.0452814318445885 | 0.274287142924416 | -4.18284813483894 |
| ENSG00000164687.10 | FABP5 | -0.764899266580107 | 0.960431365353365 | -2.06261141124039 | 0.045301712887839 | 0.274329260479668 | -3.98427247623303 |
| ENSG00000040531.14 | CTNS | -0.309475012234254 | 2.31185865881208 | -2.06195692173897 | 0.0453605653922459 | 0.274471546399128 | -4.13593978312385 |
| lnc-C12orf49-15 | lnc-C12orf49-15 | -1.29587479982807 | 0.429370212253513 | -2.06324531115601 | 0.0453710305957886 | 0.274471546399128 | -3.89504535696292 |
| lnc-PLEKHG7-7 | lnc-PLEKHG7-7 | -0.686114906134327 | 0.444066822635646 | -2.06173999187301 | 0.0453822009475506 | 0.274471546399128 | -3.88850967633155 |
| lnc-RBP4-2 | lnc-RBP4-2 | -0.480343018413951 | 1.22873250414944 | -2.06160388185157 | 0.0453957805128078 | 0.274471546399128 | -4.02926773336833 |
| ENSG00000152359.14 | POC5 | -0.171795044875952 | 3.42538048845788 | -2.06141246714205 | 0.0454148837735151 | 0.274471546399128 | -4.24861002125244 |
| lnc-FERMT2-5 | lnc-FERMT2-5 | 0.293398541433327 | 1.95597869142288 | 2.0611610730588 | 0.0454399836557173 | 0.274471546399128 | -4.08916178149349 |
| ENSG00000259984.1 | | -0.376806286972964 | 0.686882361307982 | -2.06114712546042 | 0.0454413765769931 | 0.274471546399128 | -3.96211227011427 |
| ENSG00000250312.7 | ZNF718 | -0.695392622172645 | 4.2684414910568 | -2.06232673818921 | 0.0454559328972563 | 0.274471546399128 | -4.32071732925454 |
| ENSG00000207279.1 | SNORD116-24 | -0.630784749031815 | -0.0943700641191502 | -2.06099717609043 | 0.0454563541016356 | 0.274471546399128 | -3.8166419188809 |
| ENSG00000169359.14 | SLC33A1 | -0.205383751064127 | 4.30780724122399 | -2.06097511397608 | 0.0454585581146882 | 0.274471546399128 | -4.32860607951754 |
| ENSG00000267618.5 | | -0.198354040264196 | 2.73190385604892 | -2.05984421391533 | 0.0455716604095111 | 0.275008656435807 | -4.17801090810112 |
| lnc-RAB33B-5 | lnc-RAB33B-5 | 0.52260337168168 | 0.491970532969318 | 2.0598184759907 | 0.0455742373367604 | 0.275008656435807 | -3.91941447491589 |
| ENSG00000100307.12 | CBX7 | -0.267133277186235 | 6.54319659079942 | -2.05952865779204 | 0.045603263232968 | 0.275076607920263 | -4.39801128271862 |
| ENSG00000200816.1 | SNORA38 | 0.4677820955466 | -0.194858949286071 | 2.05932391295618 | 0.045623778575888 | 0.275076607920263 | -3.82038414488827 |
| lnc-ASAH2-6 | lnc-ASAH2-6 | -0.250161371632952 | 1.53327122975106 | -2.05929601406923 | 0.0456265746554188 | 0.275076607920263 | -4.0525896925015 |
| ENSG00000203896.9 | LIME1 | -0.533280478351197 | 1.55660709216321 | -2.05917250009967 | 0.0456389552566534 | 0.275076607920263 | -4.06219447966633 |
| lnc-ZNF221-2 | lnc-ZNF221-2 | -1.1525979648146 | 0.0757523891715414 | -2.05916319402856 | 0.0456640905574359 | 0.275119969509756 | -3.8168472780502 |
| ENSG00000077312.8 | SNRPA | -0.316698352298729 | 5.14648230266277 | -2.05883417998909 | 0.0456728822724984 | 0.275119969509756 | -4.3782551395537 |
| lnc-RETREG2-1 | lnc-RETREG2-1 | -0.585183991826735 | -0.531361907907268 | -2.05852414307833 | 0.0457039923449732 | 0.275226821075654 | -3.79361670234952 |
| ENSG00000119559.15 | C19orf25 | -0.251400769615118 | 3.54979206454749 | -2.05822404195825 | 0.045734123035104 | 0.275327714077668 | -4.26081114932556 |
| lnc-MRPL11-1 | lnc-MRPL11-1 | -0.332877176068225 | 0.50421284765168 | -2.05773876982858 | 0.0457828818925405 | 0.275540660232211 | -3.93587909881352 |
| lnc-FGL2-4 | lnc-FGL2-4 | -0.356569929293817 | 0.044998374146753 | -2.05675031455819 | 0.0458823395294131 | 0.275980925091466 | -3.86870250481433 |
| lnc-IARS-2 | lnc-IARS-2 | 0.233826377635483 | 2.47860890899449 | 2.05674523446007 | 0.0458828511715005 | 0.275980925091466 | -4.15222610165072 |
| ENSG00000238227.7 | TMEM250 | -0.401085808385896 | 2.11212562251236 | -2.05628428223553 | 0.0459292966914084 | 0.276179583347724 | -4.13276022727394 |
| ENSG00000124191.17 | TOX2 | -0.90340947591214 | 2.78385690165727 | -2.05715981641955 | 0.0459962953619962 | 0.276471918552672 | -4.19636193699304 |
| ENSG00000116761.11 | CTH | -0.655595707033902 | -0.75284873247937 | -2.05511864079689 | 0.0460469296518159 | 0.276471918552672 | -3.77007103333156 |
| ENSG00000077721.15 | UBE2A | 0.211572525875727 | 4.98955079574516 | 2.05500986981411 | 0.046057919876526 | 0.276471918552672 | -4.38238580439005 |
| ENSG00000078070.12 | MCCC1 | 0.294663468929747 | 4.83522655290699 | 2.05486061977908 | 0.0460730038290951 | 0.276471918552672 | -4.36614638884653 |
| lnc-ARPP19-1 | lnc-ARPP19-1 | 0.558758918042062 | 1.31993355199325 | 2.0546846525011 | 0.0460907934927961 | 0.276471918552672 | -4.03790790057614 |
| DICER1-AS1 | DICER1-AS1 | -0.242959729307045 | 3.21594490352348 | -2.05449883443539 | 0.0461095855377953 | 0.276471918552672 | -4.22758865255188 |
| lnc-DPP4-2 | lnc-DPP4-2 | -0.627538401544109 | 0.517090742756426 | -2.05427918622463 | 0.0461318074894371 | 0.276471918552672 | -3.92508284693308 |
| ENSG00000205038.11 | PKHD1L1 | -0.732378096783668 | 0.759359828140817 | -2.05417025783476 | 0.0461428313086317 | 0.276471918552672 | -3.97905755774237 |
| ENSG00000103066.12 | PLA2G15 | -0.393214561066334 | 3.53663209394505 | -2.0541451369583 | 0.0461453739286271 | 0.276471918552672 | -4.26492025214155 |
| ENSG00000131127.13 | ZNF141 | -0.268859220717331 | 6.26772633197041 | -2.05401698368705 | 0.0461583469168499 | 0.276471918552672 | -4.40891663051787 |
| ENSG00000254087.7 | LYN | 0.197260398857935 | 8.60874937118982 | 2.05399619368755 | 0.0461604517935517 | 0.276471918552672 | -4.37963914702071 |
| ENSG00000189045.13 | ANKDD1B | -0.505332142849521 | 0.637583344104904 | -2.05379398408241 | 0.0461809288049921 | 0.276471918552672 | -3.97534922702712 |
| lnc-MSRB3-6 | lnc-MSRB3-6 | 0.363955574586171 | 1.71290094099028 | 2.05369590128021 | 0.0461908641363137 | 0.276471918552672 | -4.09785980019183 |
| lnc-DPM2-1 | lnc-DPM2-1 | -0.300252802913288 | 1.76800944906865 | -2.05363866621757 | 0.0461966626427804 | 0.276471918552672 | -4.09765937599354 |
| ENSG00000146063.19 | TRIM41 | -0.189850213873541 | 4.96880932369197 | -2.05363458902941 | 0.0461970757285144 | 0.276471918552672 | -4.38239996952168 |
| lnc-BORCS8-1 | lnc-BORCS8-1 | -0.312585661813061 | 0.489944190197261 | -2.05358701161624 | 0.0462018963353897 | 0.276471918552672 | -3.9355747927673 |
| lnc-SLC17A9-5 | lnc-SLC17A9-5 | 0.261669109941765 | 0.101000809679384 | 2.05347621597445 | 0.046213123997335 | 0.276471918552672 | -3.87188922774316 |
| ENSG00000101255.10 | TRIB3 | -0.499033049911132 | 2.48061816395936 | -2.05341143651703 | 0.0462196896341518 | 0.276471918552672 | -4.17017990132258 |
| lnc-IL9R-1 | lnc-IL9R-1 | -0.362816616876584 | 0.35534830203045 | -2.05312983200441 | 0.0462482407427646 | 0.276562329810669 | -3.9372738830203 |
| lnc-DICER1-2 | lnc-DICER1-2 | 0.215804052137311 | 2.01716377613773 | 2.05273188590186 | 0.0462886136231292 | 0.276670372464463 | -4.11622920504289 |
| ENSG00000232811.1 | | 0.561866154292025 | 2.54754057394461 | 2.05247521585059 | 0.0463146699019678 | 0.276670372464463 | -4.1562395490073 |
| ENSG00000092020.10 | PPP2R3C | 0.18267351015733 | 4.35836446606324 | 2.0523298476645 | 0.0463294328614313 | 0.276670372464463 | -4.34226004652781 |
| ENSG00000126775.8 | ATG14 | 0.195499339364034 | 5.6288383071581 | 2.0523142256973 | 0.0463310196044279 | 0.276670372464463 | -4.40654879239426 |
| lnc-CCDC188-3 | lnc-CCDC188-3 | -0.257531640309594 | 1.57108458478232 | -2.0522896421447 | 0.0463335166829425 | 0.276670372464463 | -4.07956131153737 |
| lnc-VOPP1-3 | lnc-VOPP1-3 | -0.346993778992813 | 4.34567503464621 | -2.0520060580526 | 0.0463623302689306 | 0.276762135709222 | -4.3441256144832 |
| ENSG00000258725.1 | PRC1-AS1 | -0.403566980083999 | 0.717079036073284 | -2.05147237122046 | 0.0464165979257718 | 0.277005750973082 | -3.96379931640379 |
| lnc-ARF1-3 | lnc-ARF1-3 | -0.567780842387959 | 1.56226760023821 | -2.05090825421528 | 0.0464740200455902 | 0.277268044811125 | -4.07075898277801 |
| ENSG00000279283.1 | | 0.436174221482032 | 1.39002371504414 | 2.05052169248853 | 0.0465134043518345 | 0.277422602658305 | -4.04412410139659 |
| lnc-TP53BP2-2 | lnc-TP53BP2-2 | 0.231971649068816 | 1.99405210605015 | 2.04990885360232 | 0.0465759022091902 | 0.277471423202126 | -4.12097499082055 |
| lnc-SLC41A1-1 | lnc-SLC41A1-1 | 0.254773779789506 | 1.03206167258505 | 2.04984043049274 | 0.0465828845987643 | 0.277471423202126 | -4.01183104538879 |
| lnc-LTBP3-7 | lnc-LTBP3-7 | -0.378751914276044 | 1.31932908434959 | -2.04975835879583 | 0.0465912609923754 | 0.277471423202126 | -4.04461862127042 |
| LRRC8C-DT | LRRC8C-DT | -0.52573284869675 | 2.73211695919658 | -2.04967254256085 | 0.0466000209645106 | 0.277471423202126 | -4.20091553138025 |
| ENSG00000221823.10 | PPP3R1 | 0.235596681861581 | 5.17477378838318 | 2.04965897048981 | 0.0466014065093838 | 0.277471423202126 | -4.39719372269314 |
| ENSG00000258728.1 | | -0.21166702180853 | 2.84862821262162 | -2.04964852002007 | 0.0466024734008527 | 0.277471423202126 | -4.21382605864084 |
| lnc-RCHY1-5 | lnc-RCHY1-5 | 0.28807253825819 | 0.910028680308582 | 2.04884804012447 | 0.0466842579100911 | 0.277599398130378 | -4.00622657121674 |
| lnc-GLRX2-2 | lnc-GLRX2-2 | -0.306344927436714 | -0.120562217672276 | -2.04872644998992 | 0.0466966916303966 | 0.277599398130378 | -3.85696417567516 |
| ENSG00000087095.12 | NLK | -0.220940547757704 | 4.39678182506549 | -2.04865575855479 | 0.0467039218125271 | 0.277599398130378 | -4.35536468116466 |
| ENSG00000140740.10 | UQCRC2 | 0.167315782206624 | 6.17705811925168 | 2.04865561532412 | 0.0467039364628713 | 0.277599398130378 | -4.41906555462311 |
| ENSG00000083817.8 | ZNF416 | 0.31501277230455 | 2.88918063709038 | 2.04860491155492 | 0.0467091229476682 | 0.277599398130378 | -4.20857577803052 |
| ENSG00000216809.1 | | -0.498059658601179 | 2.46565365393042 | -2.04851940134292 | 0.0467178709178453 | 0.277599398130378 | -4.16780503774873 |

| | | | | | | | |
|---|---|---|---|---|---|---|---|
| ENSG00000150756.13 | ATPSCKMT | -0.239976310287648 | 2.36698284800171 | -2.04851447324748 | 0.0467183751213929 | 0.277599398130378 | -4.1599854164503 |
| lnc-NDFIP1-1 | lnc-NDFIP1-1 | -0.496325398114395 | -0.398482601101291 | -2.04823654635039 | 0.0467468180673077 | 0.277646579899839 | -3.81455458478649 |
| ENSG00000173198.5 | CYSLTR1 | -0.489428030410526 | 4.37251397570149 | -2.04813938309655 | 0.046768493801698 | 0.277646579899839 | -4.35079219421582 |
| lnc-TOM1L2-2 | lnc-TOM1L2-2 | 0.587755560206935 | 0.209433073536147 | 2.04801914921524 | 0.0467690769336943 | 0.277646579899839 | -3.8946633826511 |
| ENSG00000084693.15 | AGBL5 | 0.354317579368207 | 3.59529493732784 | 2.04785114082731 | 0.0467862853087519 | 0.277646579899839 | -4.28679058813369 |
| ENSG00000112759.17 | SLC29A1 | -0.789300511519867 | 1.7229640004122 | -2.04843004513993 | 0.0467937611462149 | 0.277646579899839 | -4.10874706990344 |
| ENSG00000218426.5 | | 0.886729745913527 | 3.20366036989843 | 2.04905289507408 | 0.0468166508531456 | 0.277702341357434 | -4.20265738547594 |
| ENSG00000233820.2 | | 1.59888016523463 | -0.919905671161847 | 2.04719518625529 | 0.0468535247365412 | 0.277840996730691 | -3.7853638558505 |
| ENSG00000076662.9 | ICAM3 | -0.37549729742517 | 6.40885893370499 | -2.04670110642342 | 0.0469042265705784 | 0.27787990283809 | -4.42220260344329 |
| ENSG00000230532.1 | | -0.311413563832536 | 1.15287662567792 | -2.04663453967026 | 0.0469110612172308 | 0.27787990283809 | -4.03031930537335 |
| ENSG00000093010.13 | COMT | -0.282216371512922 | 3.41694919196606 | -2.04656427364322 | 0.0469182766228246 | 0.27787990283809 | -4.27292636965965 |
| ENSG00000072954.6 | TMEM38A | -0.700970175838925 | 1.51677414046721 | -2.04659812624784 | 0.0469271743076777 | 0.27787990283809 | -4.08155903321449 |
| lnc-MYO5C-1 | lnc-MYO5C-1 | 0.241449957191096 | 1.46020794587485 | 2.04634309159003 | 0.0469409954502103 | 0.27787990283809 | -4.062197233935 |
| DLGAP1-AS2 | DLGAP1-AS2 | -0.460740462259885 | 3.45351865300115 | -2.04623326046535 | 0.0469522803706858 | 0.27787990283809 | -4.26499821536194 |
| ENSG00000196937.10 | FAM3C | -0.474532238457676 | 6.58698302892855 | -2.04644229389086 | 0.0469545888388902 | 0.27787990283809 | -4.42212795650784 |
| ENSG00000137098.13 | SPAG8 | -0.390809195869979 | 0.653982890468176 | -2.04582341588335 | 0.0469944119188903 | 0.277963518399147 | -3.96512763551553 |
| lnc-GPX7-1 | lnc-GPX7-1 | -0.469105704030574 | 0.755861454972381 | -2.04581063054668 | 0.0469957267662164 | 0.277963518399147 | -3.98004492355133 |
| ENSG00000204371.11 | EHMT2 | -0.252646047983735 | 5.13957206623441 | -2.04556952956389 | 0.0470205276542493 | 0.278030313331633 | -4.40451871181397 |
| ENSG00000185252.18 | ZNF74 | -0.521301345541389 | 1.29324807875503 | -2.04530456636504 | 0.047047796280839 | 0.278111657337309 | -4.05706783162761 |
| ENSG00000145604.15 | SKP2 | -0.257829595794865 | 3.26741721453686 | -2.04470991735791 | 0.0471090445972735 | 0.278329180145794 | -4.25897043376412 |
| ENSG00000152804.10 | HHEX | -0.279395126083051 | 4.87551178423983 | -2.04464898298839 | 0.0471153247039687 | 0.278329180145794 | -4.39473419598432 |
| ENSG00000164930.11 | FZD6 | -1.21061080848523 | 0.376998348081813 | -2.04585827753015 | 0.047131221200391 | 0.278329180145794 | -3.9648147634639 |
| ENSG00000126524.9 | SBDS | 0.324098938934371 | 6.33265225494202 | 2.04428717554105 | 0.0471526288504698 | 0.278329180145794 | -4.42737732698893 |
| ENSG00000165280.15 | VCP | 0.180978242065359 | 7.17218354403 | 2.04416499958742 | 0.0471652316072718 | 0.278329180145794 | -4.42244238158486 |
| ENSG00000143486.15 | EIF2D | 0.12824092548108 | 5.26847215356487 | 2.04397953524607 | 0.0471843683270574 | 0.278329180145794 | -4.41010317945349 |
| ENSG00000181315.10 | ZNF322 | -0.288734489343533 | 3.04835450494148 | -2.04395122293816 | 0.0471872902635495 | 0.278329180145794 | -4.2388870700288 |
| ENSG00000213860.4 | RPL21P75 | 1.17880300092517 | 0.288513797838365 | 2.04386681172562 | 0.0472065572293581 | 0.278329180145794 | -3.81355954731828 |
| lnc-DENND1A-1 | lnc-DENND1A-1 | -0.51502622667714 | 0.690223315050156 | -2.04371744388298 | 0.0472114231631434 | 0.278329180145794 | -3.96044989277332 |
| ENSG00000164405.10 | UQCRQ | 0.281498244512846 | 3.80273314264903 | 2.04363615413876 | 0.0472198171825833 | 0.278329180145794 | -4.30814366707011 |
| ENSG00000160392.13 | C19orf47 | -0.2292717784991 | 2.46762725734215 | -2.04316921900926 | 0.0472680583150145 | 0.278533765902646 | -4.186984270054 |
| ENSG00000165521.15 | EML5 | 0.638345671984927 | 4.04502212423482 | 2.04373774434163 | 0.0472988251278759 | 0.278554313715112 | -4.32455790871832 |
| lnc-INTS14-4 | lnc-INTS14-4 | 0.273443206230632 | 3.5976882137003 | 2.04282600583641 | 0.047303544546585 | 0.278554313715112 | -4.28517849706488 |
| ENSG00000172795.15 | DCP2 | 0.253749990874508 | 6.66252272144637 | 2.0427428582466 | 0.0473121450103499 | 0.278554313715112 | -4.42938128316026 |
| ENSG00000198393.7 | ZNF26 | 0.188176590920762 | 5.6647851583469 | 2.04227951348373 | 0.0473600965325201 | 0.278636916841459 | -4.42456837163571 |
| ENSG00000125508.3 | SRMS | -0.795829114887185 | 1.12887359181257 | -2.04247541009112 | 0.047373727947875 | 0.278636916841459 | -4.04893898355345 |
| ENSG00000087111.20 | PIGS | 0.201849066116059 | 4.39356054347543 | 2.04213927974119 | 0.04737461765363 6 | 0.278636916841459 | -4.36601221361246 |
| ENSG00000124098.9 | FAM210B | -0.352210044063038 | 2.80897584602417 | -2.04191845950673 | 0.0473974913167955 | 0.278636916841459 | -4.21557979044238 |
| ENSG00000111596.12 | CNOT2 | 0.173024199052216 | 6.75761075118768 | 2.04189893754379 | 0.0473995139618072 | 0.278636916841459 | -4.43020325973984 |
| lnc-ZC2HC1A-2 | lnc-ZC2HC1A-2 | 0.749048842879727 | -0.165380107768101 | 2.04182284552635 | 0.0474073984734387 | 0.278636916841459 | -3.83533508723023 |
| ENSG00000001167.14 | NFYA | 0.356884373715673 | 4.66629270519437 | 2.04137401509511 | 0.0474539286746122 | 0.278830777593361 | -4.38243221676549 |
| ENSG00000133773.11 | CCDC59 | 0.199194525494344 | 5.6954125675248 | 2.04118683016475 | 0.047473345861625 | 0.278865261949152 | -4.42848392283933 |
| ENSG00000116205.13 | TCEANC2 | -0.368513997863729 | 4.34704223933758 | -2.04094660445479 | 0.0474982752433237 | 0.278868590755879 | -4.36133667995844 |
| lnc-AGPS-6 | lnc-AGPS-6 | 0.281631176734412 | 3.37872151661103 | 2.040826076497 | 0.0475107873043735 | 0.278868590755879 | -4.27282457284175 |
| ENSG00000170037.13 | CNTROB | -0.247309066176286 | 4.2387021399688 | -2.04078975880513 | 0.0475145580226823 | 0.278868590755879 | -4.36234089448045 |
| lnc-SERP1-4 | lnc-SERP1-4 | -0.438903403723792 | 2.42033093818656 | -2.04064549512602 | 0.0475295389070363 | 0.27887699524616 | -4.18069230457889 |
| ENSG00000206560.11 | ANKRD28 | 0.199892661471834 | 5.778616704966 | 2.04038836985667 | 0.0475562499601399 | 0.278914145928009 | -4.43191986000971 |
| lnc-CCDC57-6 | lnc-CCDC57-6 | 0.332186676750512 | 3.1387146691059 | 2.04023545245189 | 0.047572141739598 | 0.278914145928009 | -4.24693735095312 |
| lnc-CAMK1-1 | lnc-CAMK1-1 | -0.266332437295243 | 2.5351599894735 | -2.04019330500031 | 0.0475765226814963 | 0.278914145928009 | -4.19157288026459 |
| ENSG00000234043.3 | NUDT9P1 | -0.4746827291402 | -0.319557596279172 | -2.03974586902392 | 0.0476230522765296 | 0.279085147462561 | -3.83346411778645 |
| lnc-TSHR-2 | lnc-TSHR-2 | -0.621742154039575 | 1.03836084938289 | -2.03951324820679 | 0.047647258540136 | 0.279085147462561 | -4.04593494352667 |
| lnc-REL-3 | lnc-REL-3 | 0.475459401072978 | -0.20202580499547 | 2.03939095738253 | 0.047659988281818 | 0.279085147462561 | -3.84207177107136 |
| ENSG00000225648.5 | SBDSP1 | 0.244408728118872 | 5.60628711681164 | 2.039380481775 | 0.0476610788674014 | 0.279085147462561 | -4.42904326288143 |
| ENSG00000214776.12 | | 2.92540859386976 | 0.582944219240847 | 2.04158839770552 | 0.0476734867841599 | 0.279085147462561 | -3.81318872486725 |
| lnc-DNA2-1 | lnc-DNA2-1 | 0.28202141466063 | 1.94955120274299 | 2.03910959256136 | 0.0476892879141863 | 0.279098269302729 | -4.13206813754285 |
| ENSG00000167393.17 | PPP2R3B | -0.457769964214608 | 2.7213031377931 | -2.03803160288007 | 0.0478016883910297 | 0.279575511172568 | -4.222372801341 |
| ENSG00000100385.13 | IL2RB | -1.12786511147203 | 1.81648314057364 | -2.03975471226812 | 0.0478035597353923 | 0.279575511172568 | -4.14808313929185 |
| ENSG00000188549.12 | CCDC9B | -0.534286837052534 | 0.206343886419294 | -2.03793681948812 | 0.0478115823411281 | 0.279575511172568 | -3.90971723750068 |
| ENSG00000156030.13 | MIDEAS | -0.23842724172165 | 7.06047252595655 | -2.03774230106532 | 0.0478318926992584 | 0.279614838803986 | -4.43558092315377 |
| lnc-SLC9C1-2 | lnc-SLC9C1-2 | -0.844351995481871 | 1.38339747660416 | -2.03802495977773 | 0.047864684427271 | 0.279687185141779 | -4.06932004566923 |
| lnc-CMTM6-4 | lnc-CMTM6-4 | 0.544614564648931 | 3.00000775012897 | 2.0373637025701 | 0.0478714450398139 | 0.279687185141779 | -4.23046018716718 |
| ENSG00000105298.13 | CACTIN | -0.262510933400184 | 4.07926085066523 | -2.03703657151754 | 0.0479056434729075 | 0.279740788800724 | -4.35330465563768 |
| lnc-KIAA0100-4 | lnc-KIAA0100-4 | -0.304152816892525 | 1.38396143959019 | -2.03701593430717 | 0.0479078016092188 | 0.279740788800724 | -4.07999628048849 |
| ENSG00000240376.1 | | 1.2371123790249 | 1.07943718664175 | 2.0378776495422 | 0.0479521432134157 | 0.279920296018643 | -3.9451010469091 |
| lnc-UBE2K-2 | lnc-UBE2K-2 | -0.442843057174667 | -0.664390167304084 | -2.03639485250501 | 0.0479727908372222 | 0.279961427219321 | -3.81910704611363 |
| ENSG00000178927.17 | CYBC1 | -0.312453842530906 | 6.24311180135211 | -2.03598716057104 | 0.0480154929065582 | 0.280056743651573 | -4.44214737891155 |
| ENSG00000135605.12 | TEC | -0.249829099754231 | 4.23514826154903 | -2.03597911236538 | 0.048016336216606 | 0.280056743651573 | -4.36658128234308 |
| lnc-COPZ1-3 | lnc-COPZ1-3 | 0.290651557558132 | 3.74294543685744 | 2.03562589455566 | 0.048053359910981 | 0.280193288115502 | -4.3230725487479 |
| ENSG00000099785.10 | MARCHF2 | -0.285810078957164 | 3.81877069429111 | -2.03531539537294 | 0.0480859264216866 | 0.280227235929618 | -4.32964087811275 |

| | | | | | | | |
|---|---|---|---|---|---|---|---|
| ENSG00000080822.16 | CLDND1 | 0.202783943269795 | 5.19936864778292 | 2.0351242443809 | 0.0481059847204856 | 0.280227235929618 | -4.42773854476969 |
| ENSG00000171135.14 | JAGN1 | 0.278430993708803 | 3.33768179176365 | 2.03510746445175 | 0.048107745858382 | 0.280227235929618 | -4.27711697598102 |
| ENSG00000105185.11 | PDCD5 | -0.267560858587599 | 3.88551103598924 | -2.03503073739139 | 0.0481157994636168 | 0.280227235929618 | -4.3450487798911 |
| ENSG00000180423.4 | HARBI1 | 0.264684933203043 | 1.67012232338597 | 2.03479448025737 | 0.0481406054044262 | 0.280227235929618 | -4.10698838086347 |
| ENSG00000183337.16 | BCOR | -0.368404219897086 | 6.1437902116979 | -2.03479197046071 | 0.0481408689815444 | 0.280227235929618 | -4.44372942618547 |
| ENSG00000178381.11 | ZFAND2A | 0.259564464113282 | 4.08962074633099 | 2.03452844699787 | 0.0481685510213069 | 0.280309099709234 | -4.35977304117662 |
| ENSG00000163507.13 | CIP2A | 0.347793887161459 | 2.91601345210924 | 2.03414026925092 | 0.0482093526943769 | 0.280467243218869 | -4.24556258028323 |
| ENSG00000105708.8 | ZNF14 | 0.281603319792621 | 4.54156203266096 | 2.03361182668268 | 0.048264946034893 | 0.280679640834081 | -4.38619751480945 |
| ENSG00000110697.12 | PITPNM1 | -0.446290908578734 | 5.52195321832563 | -2.033590855333 | 0.0482736450394635 | 0.280679640834081 | -4.43848232044511 |
| ENSG00000213598.4 | RPL31P4 | 1.36746250988594 | 0.0162722609385075 | 2.03363830182284 | 0.048286771034032 | 0.280679640834081 | -3.80719664801519 |
| ENSG00000184675.10 | AMER1 | -0.275446431259626 | 4.25372286036767 | -2.03305826740517 | 0.0483232415023353 | 0.280812331971363 | -4.37853546712906 |
| ENSG00000105552.14 | BCAT2 | -0.310187123037192 | 3.19366614995373 | -2.0326223158131 | 0.0483691947714265 | 0.280900583607054 | -4.27803187547312 |
| ENSG00000090861.15 | AARS1 | -0.279411960139909 | 4.63723364808645 | -2.03260934812347 | 0.0483705622659215 | 0.280900583607054 | -4.39953571677447 |
| ENSG00000090013.10 | BLVRB | -0.620706077094482 | 1.00179803362995 | -2.03252583546054 | 0.0483793698142645 | 0.280900583607054 | -4.03512232754147 |
| ENSG00000159079.18 | CFAP298 | -0.374956725116169 | 2.95332627058739 | -2.03226318363569 | 0.0484070791203379 | 0.280982207990388 | -4.25094784083029 |
| ENSG00000108528.13 | SLC25A11 | -0.264211220989854 | 3.3969977365927 | -2.03202207179577 | 0.0484325281455732 | 0.281007468471617 | -4.29895929649345 |
| ENSG00000130303.12 | BST2 | -0.454807817859331 | 3.83991551335052 | -2.03190960082772 | 0.0484444032816726 | 0.281007468471617 | -4.34039487739513 |
| lnc-SFRP4-1 | lnc-SFRP4-1 | 0.389675788181475 | 1.78293319407379 | 2.03169304797871 | 0.0484672749316913 | 0.281007468471617 | -4.12343642775354 |
| lnc-GEMIN5-1 | lnc-GEMIN5-1 | -1.26483185565443 | 1.75548191737241 | -2.03332646240746 | 0.0484799854931815 | 0.281007468471617 | -4.08741867688138 |
| lnc-ZNF131-4 | lnc-ZNF131-4 | -0.371300038541805 | 3.45465793467037 | -2.03147744157194 | 0.0484900559496245 | 0.281007468471617 | -4.31101251561682 |
| ENSG00000151116.16 | UEVLD | -0.29728058453802 | 3.23530472729373 | -2.03144631666626 | 0.0484933453826548 | 0.281007468471617 | -4.28042828493349 |
| lnc-KCNJ5-2 | lnc-KCNJ5-2 | -0.646759435885347 | 1.50223486835046 | -2.03106126711208 | 0.0485340553640718 | 0.281164216594058 | -4.10456136907195 |
| ENSG00000128590.4 | DNAJB9 | 0.325503860379707 | 4.83856579181694 | 2.0309015636638 | 0.0485509489842143 | 0.281182943990456 | -4.4147458539726 |
| ENSG00000267681.1 | | -0.503852040445671 | -0.450217339310679 | -2.03044356943908 | 0.0485994245164915 | 0.281288679366853 | -3.83683071142343 |
| ENSG00000162755.13 | KLHDC9 | -0.627717793869418 | -0.500154494608162 | -2.03037757087356 | 0.0486064134777979 | 0.281288679366853 | -3.84118574455464 |
| ENSG00000228532.4 | SUMO2P21 | 1.40601561404378 | -1.25871759732586 | 2.03034177780468 | 0.0486102041737306 | 0.281288679366853 | -3.82401139055503 |
| ENSG00000185485.14 | SDHAP1 | 0.387596293864354 | 3.45081122497396 | 2.02959709196976 | 0.0486891290966727 | 0.281526976419577 | -4.30689703709588 |
| ENSG00000275302.1 | CCL4 | -1.78433364424773 | -1.88939183606619 | -2.02957789570124 | 0.0486911650696895 | 0.281526976419577 | -3.83701192710399 |
| lnc-PAK1IP1-5 | lnc-PAK1IP1-5 | -0.586719388131998 | -0.120552552526432 | -2.02948845248266 | 0.048700652471543 | 0.281526976419577 | -3.88632886712371 |
| ENSG00000168152.12 | THAP9 | 0.31350731502907 | 2.66779590517138 | 2.02943714504301 | 0.0487060954685962 | 0.281526976419577 | -4.2165107872755 |
| ENSG00000267278.5 | MAP3K14-AS1 | -0.368845247352408 | 1.63971792813754 | -2.02906722665774 | 0.0487453542562969 | 0.281674796927951 | -4.11906167187774 |
| ENSG00000166225.8 | FRS2 | 0.228048719516764 | 5.80465819542223 | 2.0286781019248 | 0.0487866810659473 | 0.281834481162053 | -4.45283073626035 |
| SAP30L-AS1 | SAP30L-AS1 | -0.260272808050362 | 3.63335156542898 | -2.0284915459495 | 0.0488065049549545 | 0.281869890989851 | -4.31886675231195 |
| ENSG00000213866.3 | YBX1P10 | 0.770260508135977 | 0.135894245562107 | 2.02812838536705 | 0.0488451153600699 | 0.282012980922386 | -3.88324766365636 |
| lnc-JADE2-2 | lnc-JADE2-2 | -1.04953280870609 | -0.795645438080798 | -2.02800082239632 | 0.0488586838638308 | 0.282012980922386 | -3.81083022065513 |
| lnc-PFKFB3-12 | lnc-PFKFB3-12 | 0.847997867205629 | -0.828954225990848 | 2.02780004443486 | 0.0488800466706083 | 0.282057191090869 | -3.80090093053951 |
| ENSG00000175470.19 | PPP2R2D | 0.154486089287325 | 4.54523809571827 | 2.02723179192812 | 0.0489405528393989 | 0.282145581171596 | -4.40333329958751 |
| lnc-CD200R1-3 | lnc-CD200R1-3 | 0.24451657940407 | 1.95552553359637 | 2.02722449369129 | 0.0489413303614265 | 0.282145581171596 | -4.16231698631716 |
| lnc-SLC28A1-4 | lnc-SLC28A1-4 | -0.735626268633028 | 1.01692878548721 | -2.02717457418492 | 0.0489466488537759 | 0.282145581171596 | -4.00788145685411 |
| ENSG00000134352.19 | IL6ST | 0.632646465109978 | 5.81558549886125 | 2.02843629848565 | 0.0489501953244799 | 0.282145581171596 | -4.45497673814157 |
| lnc-SDHC-3 | lnc-SDHC-3 | -0.482413773522358 | -0.441950560170651 | -2.02694475223649 | 0.0489711408782707 | 0.282187287989207 | -3.83881857518835 |
| lnc-TFRC-7 | lnc-TFRC-7 | 0.377866443182836 | 1.21138651531445 | 2.02659391899643 | 0.049008549563539 | 0.282283348680412 | -4.08570473598821 |
| ENSG00000122642.10 | FKBP9 | -2.33640156112679 | -2.52216947012714 | -2.02650444935398 | 0.0490353401450223 | 0.282283348680412 | -3.88552221338425 |
| lnc-GTF2F2-6 | lnc-GTF2F2-6 | -0.463618386081268 | 0.668790436094386 | -2.02633051769323 | 0.0490366518667376 | 0.282283348680412 | -4.03196568142431 |
| lnc-BCL2L10-5 | lnc-BCL2L10-5 | 0.265301369106109 | 1.83087666966142 | 2.02627413701625 | 0.0490426689443304 | 0.282283348680412 | -4.14921131450882 |
| lnc-DIP2A-6 | lnc-DIP2A-6 | -0.66673967107383 | 0.853465160295117 | -2.02601906393485 | 0.0490698989581702 | 0.282361121122734 | -4.02770055174764 |
| ENSG00000101544.8 | ADNP2 | 0.401063574965113 | 5.21025216402042 | 2.02558055314532 | 0.0491167423494 | 0.282551679088233 | -4.44595810031169 |
| lnc-DHX33-1 | lnc-DHX33-1 | 0.295840622437732 | 1.91165587635022 | 2.02527981785027 | 0.0491488904968267 | 0.282571137915546 | -4.16469671908989 |
| ENSG00000229320.3 | KRT8P12 | -0.37201741343751 | 1.01743589034455 | -2.02517867718968 | 0.0491597063880363 | 0.282571137915546 | -4.05370598368309 |
| lnc-WARS-1 | lnc-WARS-1 | -0.525907464064392 | 0.11148781993749 | -2.02511800058751 | 0.0491661960817795 | 0.282571137915546 | -3.91173268900483 |
| ENSG00000188636.3 | RTL6 | -0.505869506210554 | 3.12631765010076 | -2.0250353385993 | 0.0491750384304272 | 0.282571137915546 | -4.29246258980846 |
| lnc-ALG1L-3 | lnc-ALG1L-3 | 0.286683241609985 | 0.572947713557129 | 2.02437860361456 | 0.0492453384498495 | 0.282832246517167 | -3.99588808611845 |
| ENSG00000256294.7 | ZNF225 | -0.287758956276954 | 2.75061110546633 | -2.02435412422664 | 0.049247960526528 | 0.282832246517167 | -4.24137377624237 |
| lnc-ESRP1-2 | lnc-ESRP1-2 | -0.399741885101464 | 1.55167012152576 | -2.02415629092163 | 0.049269155626413 | 0.282869550850828 | -4.1201863371233 |
| ENSG00000136381.12 | IREB2 | 0.280340592367476 | 6.28866003542424 | 2.02403698101149 | 0.0492819418622683 | 0.282869550850828 | -4.46470243173172 |
| ENSG00000234616.8 | JRK | -0.311121368343541 | 4.66226842374665 | -2.02390061712664 | 0.0492965592733932 | 0.282874569145317 | -4.42053102152062 |
| ENSG00000141562.17 | NARF | 0.318483497195428 | 4.73698628475083 | 2.02323853301274 | 0.0493675843738184 | 0.283175047331591 | -4.4223841207812 |
| ENSG00000168769.13 | TET2 | 0.207444748997668 | 7.46511617031333 | 2.02315604926669 | 0.0493764390454783 | 0.283175047331591 | -4.45840994035373 |
| ENSG00000118965.14 | WDR35 | 0.234559805817137 | 5.13369134588105 | 2.02287652528622 | 0.0494064563463348 | 0.283268270467022 | -4.44552155094731 |
| ENSG00000126453.9 | BCL2L12 | 0.337310757827594 | 1.38019644007362 | 2.02227777172548 | 0.0494708081714139 | 0.283439417455844 | -4.10599562980953 |
| ENSG00000185730.7 | ZNF696 | -0.372092040651526 | 1.25827590726339 | -2.0222735912285 | 0.0494712577316388 | 0.283439417455844 | -4.08967562308402 |
| lnc-KHDRBS2-1 | lnc-KHDRBS2-1 | -0.376921081548759 | 2.47149842541159 | -2.02214272199885 | 0.0494853328768209 | 0.283439417455844 | -4.21692106118353 |
| ENSG00000158887.15 | MPZ | -0.468346202734885 | -0.0942690971378645 | -2.02208641978887 | 0.0494913893181899 | 0.283439417455844 | -3.89552001237437 |
| ENSG00000212802.4 | RPL15P3 | 1.03870555957243 | 2.08249481848918 | 2.02336966861802 | 0.0495098910803809 | 0.283466505732261 | -4.12542381278369 |
| lnc-ACTA1-4 | lnc-ACTA1-4 | 0.267878122013193 | 1.29978039457484 | 2.02095373997255 | 0.0496133687777932 | 0.28397996928624 | -4.10428694517195 |
| ENSG00000113273.16 | ARSB | 0.22194199147263 | 3.76868618221758 | 2.02025293806832 | 0.0496889695358677 | 0.284333628011333 | -4.3460110646334 |
| ENSG00000100266.18 | PACSIN2 | 0.274731932052357 | 4.59210373360218 | 2.01955212711644 | 0.049764671292726 | 0.284687667931678 | -4.42264877218151 |
| lnc-DCTN5-2 | lnc-DCTN5-2 | 0.248673671268183 | 1.01423148692864 | 2.01939235551244 | 0.0497819438692431 | 0.284707349447244 | -4.07405816856794 |

| | | | | | | | |
|---|---|---|---|---|---|---|---|
| ENSG00000151502.10 | VPS26B | 0.428428584354906 | 3.5169967524356 | 2.01899061463309 | 0.0498253982446595 | 0.284876714463841 | -4.31830561842787 |
| ENSG00000213058.3 | | -2.95943212639169 | 0.514046019272586 | -2.02118048494477 | 0.0498438188749949 | 0.284902894724804 | -3.92335687386843 |
| ENSG00000156875.13 | MFSD14A | 0.260893732506945 | 4.75269116217718 | 2.01859840840075 | 0.0498678530708111 | 0.284942417438226 | -4.43587654445326 |
| lnc-ZNF415-2 | lnc-ZNF415-2 | -0.724151767689728 | 0.670408487291874 | -2.01838674590281 | 0.0498907777735743 | 0.284942417438226 | -4.01494189728352 |
| ENSG00000138182.14 | KIF20B | 0.226882235169906 | 5.31023198098668 | 2.01828654293538 | 0.049901633728817 | 0.284942417438226 | -4.45882294768223 |
| lnc-TMEM138-3 | lnc-TMEM138-3 | -0.390017179615511 | 0.71811445776289 | -2.01812056720668 | 0.0499196199944742 | 0.284942417438226 | -4.03421340060466 |
| ENSG00000214425.7 | LRRC37A4P | -1.57154900846418 | 4.89851985928084 | -2.02040854491392 | 0.0499271863761074 | 0.284942417438226 | -4.43272712924119 |
| ENSG00000254901.7 | BORCS8 | -0.264147118196537 | 2.76400238396336 | -2.01795485969161 | 0.0499375828090707 | 0.284942417438226 | -4.25848401184435 |
| ENSG00000114978.17 | MOB1A | 0.2805749027515 | 8.29470547635806 | 2.01786211873171 | 0.0499476384451791 | 0.284942417438226 | -4.45152154370877 |
| ENSG00000171163.15 | ZNF692 | -0.352688725917357 | 3.44366917239001 | -2.01724896932241 | 0.0500141647161682 | 0.285242879565777 | -4.32253444872103 |
| ENSG00000172139.14 | SLC9C1 | -2.1294381809931 | -1.13517432316206 | -2.01755465001449 | 0.0500426878915224 | 0.285326494424157 | -3.82985794430059 |
| ENSG00000227347.1 | HNRNPKP2 | 1.36806270932212 | -0.921814936488535 | 2.0158991675179 | 0.0501608883458561 | 0.285906094883898 | -3.82503580359882 |
| lnc-STXBP4-1 | lnc-STXBP4-1 | -0.393247393707413 | -0.223407837372015 | -2.0157959511361 | 0.0501721233406519 | 0.285906094883898 | -3.89370612765953 |
| lnc-RPLP0-2 | lnc-RPLP0-2 | 0.192908675243239 | 2.6175365620433 | 2.01537932652647 | 0.0502174946948812 | 0.285907879148004 | -4.24377434749162 |
| ENSG00000184007.20 | PTP4A2 | 0.214413596305983 | 6.90337247448728 | 2.01534965770276 | 0.0502207270541 | 0.285907879148004 | -4.47803054967848 |
| ENSG00000169635.9 | HIC2 | 0.403926488829972 | 3.06633603034168 | 2.0153000000992 | 0.0502261375550853 | 0.285907879148004 | -4.28647155080301 |
| ENSG00000124508.16 | BTN2A2 | -0.319262438714901 | 4.97241439971077 | -2.01528292235906 | 0.0502279983966955 | 0.285907879148004 | -4.44336505011845 |
| lnc-USP28-4 | lnc-USP28-4 | 0.448442419893924 | -0.255276654579415 | 2.01472191054056 | 0.0502891611978202 | 0.286059419179827 | -3.87875316078791 |
| ENSG00000285959.1 | | -0.559720802909774 | 1.51120570836359 | -2.01468480867398 | 0.0502932084064266 | 0.286059419179827 | -4.13545605259865 |
| ENSG00000105325.13 | FZR1 | -0.250864327744357 | 5.53404780717151 | -2.0146563375114 | 0.0502963143376472 | 0.286059419179827 | -4.47366296999465 |
| ENSG00000100982.11 | PCIF1 | 0.209705060091425 | 5.42647503652537 | 2.01449094517204 | 0.0503143603504856 | 0.286083005274598 | -4.47107392556533 |
| ENSG00000163159.12 | VPS72 | -0.328923090174761 | 3.57390018581089 | -2.01430417825146 | 0.0503347453056366 | 0.286119873688461 | -4.34553680618165 |
| ENSG00000001630.16 | CYP51A1 | -0.422198480963992 | 4.01151409204417 | -2.01405743339084 | 0.050361687631459 | 0.286193985786394 | -4.39974949670943 |
| ENSG00000033011.12 | ALG1 | -0.243348214188998 | 3.1540421464414 | -2.0137096612213 | 0.0503996824814109 | 0.286209866766145 | -4.30971456473013 |
| ENSG00000237350.1 | CDC42P6 | 1.09013339902801 | 0.777584796373913 | 2.01415613362606 | 0.0504191306174282 | 0.286209866766145 | -3.95778918650518 |
| ENSG00000265735.2 | RN7SL5P | 0.850599548546662 | 0.0834718822809938 | 2.01343000064702 | 0.0504302540349197 | 0.286209866766145 | -3.89583455780879 |
| lnc-PTPN9-3 | lnc-PTPN9-3 | -0.408552265626392 | 0.0418687190482067 | -2.01341270921229 | 0.0504321448044972 | 0.286209866766145 | -3.94669356815801 |
| ENSG00000207475.1 | SNORA80E | 0.393010647387927 | -0.245195386610689 | 2.01332513687435 | 0.0504417215370717 | 0.286209866766145 | -3.89796217596291 |
| ENSG00000137054.15 | POLR1E | 0.366029294415314 | 4.69806236108928 | 2.01326852658534 | 0.0504479131626864 | 0.286209866766145 | -4.43138332864804 |
| TMEM72-AS1 | TMEM72-AS1 | -0.409801238034031 | 3.48261215839074 | -2.013018963754 | 0.050475216409392 | 0.286285858185317 | -4.3278072342671 |
| ENSG00000162928.8 | PEX13 | 0.255956580405127 | 4.56577279706398 | 2.01274260665042 | 0.0505054660235331 | 0.286378514369802 | -4.43468875354031 |
| KCNQ1OT1 | KCNQ1OT1 | -0.831359355152658 | 3.77332979585289 | -2.01333919778309 | 0.0506212584087334 | 0.286938403932045 | -4.38570526849836 |
| ENSG00000169410.9 | PTPN9 | -0.326344434504372 | 3.97819434494725 | -2.01149833831418 | 0.0506418565839225 | 0.286938403932045 | -4.3946717783018 |
| ENSG00000130590.13 | SAMD10 | -0.512159620265762 | 1.76629247468695 | -2.01146031699472 | 0.05064602931959 | 0.286938403932045 | -4.18132132994748 |
| ENSG00000261359.2 | PYCARD-AS1 | -0.479373321164774 | 1.26494707726989 | -2.01094569457749 | 0.0507025370652063 | 0.287112547146174 | -4.11030885518713 |
| ENSG00000128886.11 | ELL3 | -0.601230500146755 | 2.28126417239685 | -2.01102442792382 | 0.0507046644743887 | 0.287112547146174 | -4.23525446071595 |
| ENSG00000183647.10 | ZNF530 | 0.290667912212577 | 2.7516953334462 | 2.01042303928404 | 0.0507599828227813 | 0.287346734444804 | -4.26147548088002 |
| ENSG00000051523.10 | CYBA | -0.429220178043573 | 6.71083109757411 | -2.00986687371647 | 0.0508228174145927 | 0.287599820413235 | -4.48823836064822 |
| lnc-KDM4C-18 | lnc-KDM4C-18 | 0.84730572709027 | 0.082338753108285 | 2.00964053601707 | 0.05084609435325 | 0.287599820413235 | -3.90183247719631 |
| lnc-CCNJL-3 | lnc-CCNJL-3 | -0.510673254529094 | -0.242225269495458 | -2.00963586572714 | 0.050846608681133 | 0.287599820413235 | -3.88764394659459 |
| lnc-SETBP1-8 | lnc-SETBP1-8 | -0.26803385727047 | 1.65376644777695 | -2.00937430259733 | 0.0508754212080378 | 0.287661786855646 | -4.16050105336994 |
| ENSG00000166532.15 | RIMKLB | 1.26378719266405 | 5.2046197274767 | 2.01161443916135 | 0.0508855155196719 | 0.287661786855646 | -4.4673624753711 |
| LINC02175 | LINC02175 | 0.569636530278809 | 2.26123809610203 | 2.00907943207231 | 0.0509079196939105 | 0.287709420938979 | -4.21109907039452 |
| ENSG00000225630.1 | MTND2P28 | -1.26105955438628 | 0.679818080634672 | -2.0101494321562 | 0.0509404677343119 | 0.287814341854335 | -4.00457183086124 |
| ENSG00000143578.15 | CREB3L4 | -0.269195093187026 | 3.20280987950156 | -2.00833361079824 | 0.0509901990585713 | 0.287959809122858 | -4.31281864395106 |
| ENSG00000119720.17 | NRDE2 | 0.27631408370785 | 5.00906442561691 | 2.00817146764464 | 0.0510081020221531 | 0.287959809122858 | -4.46294495219321 |
| lnc-CSF1R-4 | lnc-CSF1R-4 | -0.41626068712003 | -0.206776819875654 | -2.00814930384837 | 0.0510105496511769 | 0.287959809122858 | -3.90186433319613 |
| ENSG00000137817.16 | PARP6 | 0.145217192596491 | 5.7399349403669 | 2.00804404828334 | 0.0510221747981861 | 0.287959809122858 | -4.48903419570541 |
| ENSG00000083807.9 | SLC27A5 | -0.286709728727368 | 2.0091257152526 | -2.00780038390766 | 0.0510490955791791 | 0.288032767079562 | -4.20206301297151 |
| lnc-RARS2-4 | lnc-RARS2-4 | 0.405984626085429 | 3.47188289712761 | 2.00753220293753 | 0.0510787392644999 | 0.288115478294121 | -4.34381292631601 |
| ENSG00000170471.14 | RALGAPB | 0.133827740572061 | 6.79619131557177 | 2.00735598950964 | 0.051098225338389 | 0.288115478294121 | -4.49346735929955 |
| lnc-LUZP1-3 | lnc-LUZP1-3 | -0.549186918711699 | 0.759272868349753 | -2.007287977262 | 0.0511057480081541 | 0.288115478294121 | -4.05499342535567 |
| ENSG00000156170.12 | NDUFAF6 | 0.263325872630578 | 3.58265238948671 | 2.00710030231451 | 0.0511265112597026 | 0.288153609325975 | -4.35764542648565 |
| lnc-PTGIR-4 | lnc-PTGIR-4 | -0.472169676072164 | -0.39786045351529 | -2.00664901455498 | 0.0511764690217224 | 0.288268252725635 | -3.86910753598746 |
| ENSG00000256664.1 | | 0.316198475852103 | 2.57870820793989 | 2.00657219161082 | 0.0511849775709223 | 0.288268252725635 | -4.24518404256751 |
| ENSG00000124228.14 | DDX27 | 0.226659187589523 | 6.34620688546729 | 2.0065370717335 | 0.0511888676923771 | 0.288268252725635 | -4.49672526461043 |
| lnc-METAP1-3 | lnc-METAP1-3 | -0.378767713039281 | 1.92493576839328 | -2.00622853862858 | 0.0512230539795267 | 0.28838187091373 | -4.18220892749638 |
| ENSG00000120280.5 | TASL | -0.313721380116593 | 4.08162170301728 | -2.00576563052607 | 0.0512743825506619 | 0.288591910320009 | -4.40566215425409 |
| ENSG00000256223.5 | ZNF10 | 0.502150491055544 | 4.73369261946576 | 2.00580607984203 | 0.0513000344160434 | 0.288633619243487 | -4.44642497061781 |
| ENSG00000242028.6 | HYPK | -0.279070900696992 | 5.1766711175947 | -2.00538840468719 | 0.051316243393101 | 0.288633619243487 | -4.47945815284268 |
| lnc-DRC3-3 | lnc-DRC3-3 | -0.400637256817522 | 0.726212537555889 | -2.00531978111388 | 0.0513238617515018 | 0.288633619243487 | -4.05210780807559 |
| ENSG00000204370.10 | SDHD | 0.367417564991224 | 4.204459276999 | 2.00490914854954 | 0.051369469286296 | 0.288779078645017 | -4.4117912880687 |
| ENSG00000263528.7 | IKBKE | -0.417267550738459 | 3.92856441445269 | -2.00472527957151 | 0.0513899023596364 | 0.288779078645017 | -4.40927585206086 |
| ENSG00000135521.8 | LTV1 | 0.228921991584166 | 4.29504723176496 | 2.0047080555938 | 0.0513918167942816 | 0.288779078645017 | -4.42676486404553 |
| lnc-SLC2A13-2 | lnc-SLC2A13-2 | 0.462333863250757 | 0.711978914151235 | 2.00441454866975 | 0.0514244494132439 | 0.288841113616955 | -4.04106018152814 |
| ENSG00000268133.1 | | 0.269354725242562 | 0.956445400203607 | 2.00428170024027 | 0.0514392256396504 | 0.288841113616955 | -4.08100612777871 |
| ENSG00000159322.17 | ADPGK | 0.239605583470211 | 7.09197779720714 | 2.00416414977068 | 0.0514523034005135 | 0.288841113616955 | -4.49711293910497 |
| ENSG00000115902.10 | SLC1A4 | 0.753516651481092 | 3.83233977917933 | 2.00541182852016 | 0.0514589886621665 | 0.288841113616955 | -4.36938660517549 |

| | | | | | | | |
|---|---|---|---|---|---|---|---|
| ENSG00000118369.12 | USP35 | 0.58850847199358 | 0.27019980798142 | 2.00395703633539 | 0.0514753522592284 | 0.288854191807988 | -3.96150991992802 |
| lnc-FOXN3-8 | lnc-FOXN3-8 | -0.308195055430176 | -0.231932821587392 | -2.0036070229337 | 0.0515143242458669 | 0.288994095380943 | -3.90148919661168 |
| ENSG00000116852.14 | KIF21B | 0.376800468795637 | 5.10598839663518 | 2.0034502398751 | 0.0515317894316409 | 0.289013302962252 | -4.47316523442426 |
| ENSG00000205695.3 | | -0.564434916373506 | 0.0952639741663932 | -2.00304811571509 | 0.0515766083776224 | 0.289107116077506 | -3.96338052185853 |
| lnc-HECA-6 | lnc-HECA-6 | -0.564434916373506 | 0.0952639741663932 | -2.00304811571509 | 0.0515766083776224 | 0.289107116077506 | -3.96338052185853 |
| ENSG00000197619.13 | ZNF615 | 0.340300248065999 | 4.13569346342411 | 2.00274728725581 | 0.0516101594451762 | 0.289216420326725 | -4.41249936813835 |
| ENSG00000198633.10 | ZNF534 | -0.803870729260002 | -0.201423397069887 | -2.00254206619607 | 0.051633058374603 | 0.289265988166699 | -3.92083673286323 |
| ENSG00000151748.14 | SAV1 | -0.280136094634522 | 4.79699801048934 | -2.00219975909243 | 0.0516712731970525 | 0.28940080134519 | -4.47119944503916 |
| ENSG00000178252.17 | WDR6 | -0.195193507263914 | 7.20715497743271 | -2.00186755833859 | 0.0517083831866927 | 0.28940080134519 | -4.49911093493128 |
| ENSG00000268083.5 | | 0.200657794468697 | 5.17255325959506 | 2.00186732793498 | 0.0517084089329821 | 0.28940080134519 | -4.48406249324826 |
| lnc-CALB1-4 | lnc-CALB1-4 | 0.324659416653256 | 0.901188933209964 | 2.00159346783536 | 0.0517390190920266 | 0.28940080134519 | -4.08417213418973 |
| ENSG00000147050.14 | KDM6A | 0.388394103484306 | 6.48949451872887 | 2.00133334034518 | 0.0517681088537475 | 0.28940080134519 | -4.50586391169909 |
| lnc-SLC38A8-7 | lnc-SLC38A8-7 | 0.21388060250919 | 2.21586888349323 | 2.0013222484953 | 0.0517693495576442 | 0.28940080134519 | -4.22818810927876 |
| lnc-TMEM168-3 | lnc-TMEM168-3 | 0.262667234815436 | 1.44529811773513 | 2.00125014204482 | 0.0517774158154902 | 0.28940080134519 | -4.15193173878624 |
| ENSG00000204237.4 | OXLD1 | -0.293839772904662 | 3.58884748659152 | -2.00119940471795 | 0.0517830922482748 | 0.28940080134519 | -4.37208149238357 |
| ENSG00000170989.9 | S1PR1 | 0.958072091935076 | 4.9960370933116 | 2.00326101442394 | 0.0517836637688546 | 0.28940080134519 | -4.46008770452691 |
| lnc-NCR3-1 | lnc-NCR3-1 | -0.585573990133714 | 3.7239215342114 | -2.00144011972667 | 0.0518230454207012 | 0.289475886402022 | -4.4049218505576 |
| ENSG00000214193.10 | SH3D21 | -0.732746057058024 | 4.69974794930561 | -2.0022990656547 | 0.0518284715206563 | 0.289475886402022 | -4.46032056741674 |
| ENSG00000149428.18 | HYOU1 | -0.30690314099445 | 5.33441419781079 | -2.00068325539271 | 0.0518408690781876 | 0.289475886402022 | -4.49599247659932 |
| ENSG00000179909.15 | ZNF154 | -0.305971237439741 | 5.87168776350364 | -2.00057178344841 | 0.0518533543780914 | 0.289475886402022 | -4.50348014107745 |
| ENSG00000172159.15 | FRMD3 | 0.58144090818891 | 0.691139450261084 | 2.00030035159622 | 0.0518837667164986 | 0.289567128613257 | -4.06328133478719 |
| ENSG00000103657.13 | HERC1 | 0.210185591734497 | 8.49036629400152 | 1.99983038418854 | 0.0519364603316558 | 0.289782641096902 | -4.4815913413316 |
| lnc-MYO1G-1 | lnc-MYO1G-1 | -0.590105836701026 | 1.31256179737262 | -1.99968675115957 | 0.0519525739786578 | 0.289793991924855 | -4.13460276849941 |
| lnc-TRMT112-1 | lnc-TRMT112-1 | -0.400171796012818 | 0.555924974492653 | -1.99944871650518 | 0.0519792877224967 | 0.289864448440029 | -4.00758767596448 |
| ENSG00000206585.1 | RNVU1-7 | -0.865472857027883 | 0.493760202007483 | -1.99944419218493 | 0.0520030357354656 | 0.28991833275815 | -4.03544562656478 |
| ENSG00000122779.17 | TRIM24 | -0.233275953930155 | 4.31674882381684 | -1.99884432146795 | 0.0520471701931413 | 0.290085812100034 | -4.44348392672527 |
| lnc-SOX30-4 | lnc-SOX30-4 | 0.380508627651967 | 1.9730217436567 | 1.99859681674728 | 0.0520749907728518 | 0.290125503162436 | -4.19541042059569 |
| ENSG00000121089.4 | NACA3P | 1.2511865587636 | -0.729261715495927 | 1.99842789844354 | 0.0520939852998889 | 0.290125503162436 | -3.84002851753605 |
| ENSG00000123595.7 | RAB9A | 0.265667435189523 | 3.04029521571522 | 1.99835892162601 | 0.0521017433309323 | 0.290125503162436 | -4.31333728991111 |
| ENSG00000086848.14 | ALG9 | 0.352401766204771 | 3.17144467436932 | 1.99827953753751 | 0.0521106731376149 | 0.290125503162436 | -4.32169886009201 |
| lnc-CDHR3-3 | lnc-CDHR3-3 | -0.623120665646359 | 0.451779412034198 | -1.99797416411208 | 0.0521450365308497 | 0.290238314471195 | -4.01250118857267 |
| lnc-SERPINB9-2 | lnc-SERPINB9-2 | 0.956529024551055 | 0.149716258921481 | 1.99778709167103 | 0.0521660973139154 | 0.290277042717578 | -3.93593895963729 |
| lnc-SLC7A7-2 | lnc-SLC7A7-2 | 0.512355855132541 | 1.51463120677238 | 1.99723314391611 | 0.0522285044556438 | 0.290507063979773 | -4.14427740153813 |
| ENSG00000151729.10 | SLC25A4 | -0.416435998881567 | 1.13879059912924 | -1.99716964214237 | 0.0522356626239684 | 0.290507063979773 | -4.12183117742535 |
| ENSG00000013275.7 | PSMC4 | 0.201199639598415 | 4.96214218378777 | 1.99621234227058 | 0.0523436761883373 | 0.290667396336409 | -4.48603471976858 |
| ENSG00000279672.1 | NA | -0.55555135622649 | 1.04211822916801 | -1.9962042361132 | 0.0523445916431803 | 0.290667396336409 | -4.12509264404685 |
| ENSG00000243746.1 | EEF1A1P10 | -0.518613707838228 | 0.999059560006222 | -1.99601584233456 | 0.0523658714734808 | 0.290667396336409 | -4.09907598389091 |
| ENSG00000187257.15 | RSBN1L | -0.223439692867037 | 5.84610703382075 | -1.99593226921867 | 0.052375313789003 | 0.290667396336409 | -4.51311965604848 |
| ENSG00000163939.18 | PBRM1 | 0.12098036598571 | 6.56450450734705 | 1.99591857097721 | 0.0523768615937837 | 0.290667396336409 | -4.51571128199649 |
| ENSG00000124257.6 | NEURL2 | -0.438020437353263 | -0.0379522423858411 | -1.99566750720407 | 0.0524052370467273 | 0.290667396336409 | -3.94933724769661 |
| ENSG00000090263.15 | MRPS33 | -0.519865508754471 | 2.16009396943997 | -1.99544206631905 | 0.0524307279178018 | 0.290667396336409 | -4.24785317344656 |
| ENSG00000235174.1 | RPL39P3 | 1.29835228447637 | 3.22558400605253 | 1.99750617851229 | 0.0524352723875218 | 0.290667396336409 | -4.28729262071797 |
| lnc-GTF3C1-4 | lnc-GTF3C1-4 | -0.491574094781744 | -0.0914333435370321 | -1.99540178115762 | 0.0524352841396016 | 0.290667396336409 | -3.94812645817333 |
| ENSG00000158863.21 | FHIP2B | -0.293547963683968 | 4.73063534137597 | -1.99538072287739 | 0.0524376659518536 | 0.290667396336409 | -4.47526996631198 |
| ENSG00000160991.15 | ORAI2 | -0.35032919800364 | 7.70609854088445 | -1.99533149254366 | 0.0524432345503527 | 0.290667396336409 | -4.50260445571783 |
| ENSG00000116044.15 | NFE2L2 | 0.298402724922664 | 4.96102503616385 | 1.99532910938192 | 0.0524435041302935 | 0.290667396336409 | -4.48210390708438 |
| ENSG00000157193.16 | LRP8 | -0.502917107563516 | 2.6285087615697 | -1.99528871085023 | 0.0524480741385329 | 0.290667396336409 | -4.29086552032629 |
| ENSG00000103150.5 | MLYCD | -0.268367661987844 | 4.51138835281467 | -1.99515296449165 | 0.0524634327167753 | 0.290674249154612 | -4.46058696002105 |
| ENSG00000104904.12 | OAZ1 | -0.255675659676774 | 7.3716970602174 | -1.99486504318683 | 0.0524960215197333 | 0.290776536851634 | -4.50970609619654 |
| ENSG00000155324.9 | GRAMD2B | -0.967505252256583 | 1.10171996318791 | -1.99537822424424 | 0.0525395579378581 | 0.290939392261214 | -4.09978730101514 |
| ENSG00000229473.2 | RGS17P1 | 0.370374933821207 | 2.94993519849293 | 1.99432328070883 | 0.0525573892263794 | 0.290959855418657 | -4.30576267513361 |
| ENSG00000033170.16 | FUT8 | 1.67638445432901 | 3.70837470081201 | 1.99640113267527 | 0.052581115200451 | 0.290998027051326 | -4.35280746433903 |
| ENSG00000284691.1 | | -0.307684665314443 | 3.67024731766551 | -1.99394953525787 | 0.0525997611283061 | 0.290998027051326 | -4.38050150589412 |
| ENSG00000152601.17 | MBNL1 | -0.250749968324076 | 8.61159221333243 | -1.9938883754262 | 0.0526066976950874 | 0.290998027051326 | -4.48819522248432 |
| lnc-SYNCRIP-6 | lnc-SYNCRIP-6 | -0.248180111105583 | 2.66379459658789 | -1.99369132812004 | 0.0526290516015874 | 0.291043462954184 | -4.29072240473096 |
| ENSG00000118960.12 | HS1BP3 | -0.256073617750185 | 3.59067722211712 | -1.9933609612939 | 0.052666548303886 | 0.291172592999969 | -4.3845991389984 |
| ENSG00000086061.15 | DNAJA1 | 0.309345307709553 | 5.91142218057409 | 1.99291802229581 | 0.0527168582525126 | 0.291297739632887 | -4.51921385820186 |
| ENSG00000146232.16 | NFKBIE | -0.456805321250669 | 5.00806595715917 | -1.99296818339431 | 0.0527239533993529 | 0.291297739632887 | -4.49450695696213 |
| ENSG00000077348.8 | EXOSC5 | -0.518473302486817 | 1.80557390127925 | -1.99277464686811 | 0.052733152045191 | 0.291297739632887 | -4.21112225737541 |
| ENSG00000136731.12 | UGGT1 | 0.126064601571783 | 6.62032982848685 | 1.99244686434218 | 0.0527704190169631 | 0.291297739632887 | -4.52173604233724 |
| ENSG00000228929.1 | RPS13P2 | 1.15039902255253 | 0.450546011725869 | 1.99278621435992 | 0.0527786371444303 | 0.291297739632887 | -3.93437360379771 |
| lnc-MCUB-1 | lnc-MCUB-1 | -0.448706818604744 | 0.850784440008437 | -1.99233201054306 | 0.0527834826189661 | 0.291297739632887 | -4.09541730102813 |
| ENSG00000259315.1 | ACTG1P17 | -0.296604223400039 | 0.304820060138502 | -1.99229009421416 | 0.052788250927011 | 0.291297739632887 | -4.00686377304009 |
| ENSG00000144445.16 | KANSL1L | -0.78201632702179 | 3.9920307384891 | -1.99287656666899 | 0.0528929172608994 | 0.291797082814552 | -4.42663656452241 |
| ENSG00000068745.14 | IP6K2 | 0.193964351216928 | 5.94836864163148 | 1.99085799143285 | 0.0529513878289193 | 0.292041376120752 | -4.52322739573655 |
| lnc-METTL22-5 | lnc-METTL22-5 | -0.372412580017387 | -0.164704491831291 | -1.99027761355298 | 0.0530176252454042 | 0.292257121932432 | -3.92468940473323 |
| ENSG00000107929.14 | LARP4B | -0.199430012293423 | 6.14202950695763 | -1.99026641846181 | 0.0530189036241413 | 0.292257121932432 | -4.52565940537181 |
| ENSG00000170802.15 | FOXN2 | 0.298296297317242 | 5.69518545925161 | 1.99013372489009 | 0.0530340580682584 | 0.292262387474957 | -4.52228384551611 |

| | | | | | | | |
|---|---|---|---|---|---|---|---|
| ENSG00000141510.17 | TP53 | -0.792143521331647 | 4.69288878134381 | -1.99150772465052 | 0.0530607821045406 | 0.292331391343083 | -4.48044175301603 |
| ENSG00000115661.13 | STK16 | -0.2181783616184 | 4.15336827735171 | -1.98941529325417 | 0.0531161725874843 | 0.292558250031627 | -4.44733843260223 |
| ENSG00000250135.1 | | 0.177116453853047 | 2.34258968323496 | 1.98925141799031 | 0.0531349184226286 | 0.292583206498921 | -4.26700545879488 |
| lnc-POLR3E-1 | lnc-POLR3E-1 | -0.39897426322206 | -0.214633620725298 | -1.98865972106942 | 0.0532026507974532 | 0.292846741193106 | -3.94071195340702 |
| ENSG00000174010.9 | KLHL15 | 0.349933633664388 | 4.7401302961719 | 1.98858479376073 | 0.053211233156596 | 0.292846741193106 | -4.48405900473375 |
| ENSG00000284146.1 | MIR922 | 0.381400610266769 | 1.41630709837063 | 1.98793836546733 | 0.0532853263975654 | 0.293176122224295 | -4.15817442081695 |
| ENSG00000115687.13 | PASK | -0.365328270820746 | 5.87131415362824 | -1.98776775966923 | 0.0533048960133837 | 0.293205418130272 | -4.52753638092555 |
| ENSG00000173065.13 | FAM222B | -0.255903918421868 | 3.32938405855609 | -1.9866928259338 | 0.0534283410628705 | 0.293805916136004 | -4.36908700926157 |
| lnc-MED4-2 | lnc-MED4-2 | 0.339658778897874 | 1.81381369009463 | 1.98642340271756 | 0.0534593202600512 | 0.293897753448887 | -4.21444930019905 |
| ENSG00000149257.14 | SERPINH1 | -0.544300978310716 | 2.67009373189961 | -1.98610817922745 | 0.0534955854457722 | 0.294018594187004 | -4.31024843066711 |
| ENSG00000141367.11 | CLTC | 0.276283980991672 | 7.32789788900959 | 1.98562422497191 | 0.0535513038233294 | 0.294246259101866 | -4.5277168236649 |
| ENSG00000171681.12 | ATF7IP | -0.299884528126029 | 8.83308463883841 | -1.98507690876652 | 0.053614377614748 | 0.294514207217603 | -4.49969380810452 |
| ENSG00000158234.12 | FAIM | -0.328741101827813 | 2.48647390839637 | -1.98479265367044 | 0.0536471610638814 | 0.294615666002634 | -4.29586249025231 |
| ENSG00000111696.11 | NT5DC3 | -0.601720610975486 | 4.31106338713766 | -1.98531125501458 | 0.0536880713771076 | 0.294748747140521 | -4.47296989753312 |
| ENSG00000100473.16 | COCH | -1.45800352864002 | 3.54266118983383 | -1.98659325634829 | 0.053700034094979 | 0.294748747140521 | -4.43249605391201 |
| ENSG00000170035.15 | UBE2E3 | -0.24162176208254 | 3.97695447712813 | -1.98370005080709 | 0.0537733335264654 | 0.295050150088286 | -4.44277620559843 |
| ENSG00000172602.10 | RND1 | 0.766694401007246 | -0.0613760986010459 | 1.98361110851379 | 0.0537836157572389 | 0.295050150088286 | -3.97327313222815 |
| ENSG00000170684.8 | ZNF296 | -0.625745877061121 | 0.2169692544131 | -1.98341229196078 | 0.0538066062181558 | 0.295097622112524 | -4.03071666782624 |
| LINC01058 | LINC01058 | -0.641609714151115 | 0.085044049968939 | -1.98319206692907 | 0.0538320821978394 | 0.295119335545084 | -3.98506414415314 |
| lnc-CD38-7 | lnc-CD38-7 | -0.792347864953265 | -0.428026327628806 | -1.98313019632565 | 0.0538392413628621 | 0.295119335545084 | -3.9293585929221 |
| ENSG00000175287.18 | PHYHD1 | 2.98816696019198 | -0.517601625808638 | 1.98476901576731 | 0.0538548015959021 | 0.295126033346234 | -3.86058978878303 |
| ENSG00000185022.11 | MAFF | 0.635595216380838 | 1.64988554559244 | 1.98279266814181 | 0.0538783118831611 | 0.295176282536892 | -4.18987272352255 |
| ENSG00000204177.10 | BMS1P1 | -0.714967788018272 | 2.01438559210632 | -1.98286941944029 | 0.0539164962476677 | 0.295180743545057 | -4.22966449742062 |
| ENSG00000219201.4 | | 1.11011936321945 | -1.02325558883061 | 1.9822773023111 | 0.0539380152481533 | 0.295180743545057 | -3.86888491944761 |
| lnc-ZNF787-2 | lnc-ZNF787-2 | -0.355681455258239 | 1.08604313177637 | -1.98213471640545 | 0.0539545434317664 | 0.295180743545057 | -4.15317033449754 |
| lnc-TLE4-10 | lnc-TLE4-10 | -1.36480932260091 | 2.41852859288512 | -1.98415946115617 | 0.0539596471708634 | 0.295180743545057 | -4.27253606725652 |
| lnc-DNAJB7-1 | lnc-DNAJB7-1 | -0.470264614147537 | 0.500470158556959 | -1.9820609519697 | 0.0539630957292905 | 0.295180743545057 | -4.07520096271033 |
| ENSG00000119318.12 | RAD23B | 0.231076564289092 | 6.2313601982841 | 1.98204304432334 | 0.0539651721304013 | 0.295180743545057 | -4.54111095082496 |
| ENSG00000169372.12 | CRADD | 0.348237401747459 | 1.78189540737125 | 1.98132240298275 | 0.0540487881975269 | 0.295559566277815 | -4.21635396939536 |
| ENSG00000132471.11 | WBP2 | -0.268091305283306 | 5.10540299899205 | -1.9808557663808 | 0.0541029917792852 | 0.295560851426731 | -4.51893031785706 |
| lnc-SLC25A47-6 | lnc-SLC25A47-6 | 0.192022000014449 | 3.03494883801483 | 1.98084834433187 | 0.0541038542891363 | 0.295560851426731 | -4.34810324610736 |
| lnc-ZNF77-2 | lnc-ZNF77-2 | -0.358868365353876 | 1.23881377798926 | -1.98075694053299 | 0.0541144772193531 | 0.295560851426731 | -4.16691673219637 |
| ZNF460-AS1 | ZNF460-AS1 | -0.284435370711678 | 4.09111295950055 | -1.98071024174513 | 0.0541199052382359 | 0.295560851426731 | -4.44862463689632 |
| lnc-CLEC18B-2 | lnc-CLEC18B-2 | 0.319911572191128 | 0.436829700924527 | 1.9804074827791 | 0.0541551077453628 | 0.295560851426731 | -4.04431196434962 |
| ENSG00000076513.16 | ANKRD13A | 0.35147667783208 | 7.64528803620359 | 1.98040275700128 | 0.0541556573796144 | 0.295560851426731 | -4.53298803326865 |
| lnc-ATF7IP2-3 | lnc-ATF7IP2-3 | 0.418262734678376 | 2.749931115867 | 1.98033701908582 | 0.0541633035647956 | 0.295560851426731 | -4.30780967190683 |
| lnc-SHCBP1L-1 | lnc-SHCBP1L-1 | 0.362093327146502 | 0.4010168971061 | 1.98022892265703 | 0.0541758786309321 | 0.295560851426731 | -4.0274869526762 |
| lnc-SPATA32-2 | lnc-SPATA32-2 | -0.453127922766517 | 5.78189175736345 | -1.98038553177745 | 0.0541782583895961 | 0.295560851426731 | -4.53944850375679 |
| ENSG00000168259.14 | DNAJC7 | 0.200222977676385 | 5.67259554014347 | 1.98003594410989 | 0.0541983344696001 | 0.295592029249544 | -4.53878521649966 |
| ENSG00000250318.1 | RPL13AP26 | -0.411664673027215 | -0.0124774278430469 | -1.97988223780441 | 0.0542162261698275 | 0.295611280332069 | -3.97355150650498 |
| lnc-NAA35-6 | lnc-NAA35-6 | 0.289632068617521 | 2.09344747662521 | 1.97935101966805 | 0.0542781002518015 | 0.295783783528533 | -4.25208447254627 |
| ENSG00000182934.11 | SRPRA | 0.240797087840049 | 6.74368607320993 | 1.97908760135481 | 0.054308804743424 | 0.295783783528533 | -4.54487779536554 |
| ENSG00000188868.13 | ZNF563 | -0.38048385285195 | 2.11693194434676 | -1.97902930401971 | 0.0543156020086952 | 0.295783783528533 | -4.24671132050219 |
| lnc-CDT1-1 | lnc-CDT1-1 | -0.743372235135868 | 2.05012690216995 | -1.9795970930279 | 0.054318974799677 | 0.295783783528533 | -4.24783655708851 |
| ENSG00000199785.1 | SNORA52 | 0.588259914469999 | 0.940623506840982 | 1.97889079041986 | 0.0543317551575861 | 0.295783783528533 | -4.09919881481344 |
| lnc-LIMA1-1 | lnc-LIMA1-1 | 0.221934676201521 | 1.69828670584752 | 1.9788242937286 | 0.0543395113286073 | 0.295783783528533 | -4.21288450308001 |
| lnc-GSC2-1 | lnc-GSC2-1 | 0.27947284899684 | 1.05533176207152 | 1.9786279407203 | 0.0543624195172291 | 0.295783783528533 | -4.13449822062505 |
| ENSG00000120896.13 | SORBS3 | -0.469876606536728 | 3.14276346508665 | -1.97851699908208 | 0.0543753665874559 | 0.295783783528533 | -4.36746512169694 |
| ENSG00000197021.8 | EOLA2 | 0.221102153072678 | 3.01257474720781 | 1.97819334187423 | 0.0544131531342335 | 0.295783783528533 | -4.34960105396636 |
| lnc-SRPX-2 | lnc-SRPX-2 | 0.399971542105056 | 0.232143847702899 | 1.97817334783205 | 0.0544154881556994 | 0.295783783528533 | -4.01631771103542 |
| ENSG00000126870.15 | DYNC2I1 | -0.239337225527686 | 3.54207213524259 | -1.97810775998348 | 0.0544231484968916 | 0.295783783528533 | -4.40886906731819 |
| ENSG00000273269.3 | | 0.282836269160274 | 3.56638279309295 | 1.97797712001731 | 0.0544384093841991 | 0.295783783528533 | -4.40607028582816 |
| ENSG00000149591.16 | TAGLN | -0.857101190161442 | 4.75743738139628 | -1.97954194189789 | 0.0544582698940404 | 0.295783783528533 | -4.49321252155427 |
| lnc-SCAP-1 | lnc-SCAP-1 | -0.45959771288729 | 0.589710079016629 | -1.97776952403876 | 0.0544626675981873 | 0.295783783528533 | -4.06754248977053 |
| ENSG00000138433.15 | CIR1 | 0.230686761997012 | 5.06993157984679 | 1.97776310069413 | 0.0544634183342147 | 0.295783783528533 | -4.52092079771186 |
| lnc-ANKRD55-3 | lnc-ANKRD55-3 | 0.614118588169204 | 1.48026818713115 | 1.97749361242732 | 0.0544949231508972 | 0.295827191830581 | -4.20880859076961 |
| ENSG00000285043.1 | | -0.262622791504905 | 5.89887980185748 | -1.97744886412615 | 0.0545001560230075 | 0.295827191830581 | -4.546759648724 |
| lnc-ZNF852-2 | lnc-ZNF852-2 | -0.459266748280437 | 1.63379195292966 | -1.9771410985469 | 0.0545361579288612 | 0.295920264828676 | -4.21269010101933 |
| ENSG00000273047.1 | | -0.562803084466609 | 0.715208673356311 | -1.97705650887971 | 0.0545460566856142 | 0.295920264828676 | -4.09363416247856 |
| ENSG00000106803.9 | SEC61B | 0.280272700984979 | 4.20666124811075 | 1.9759780345568 | 0.0546723967732404 | 0.2965275211551 | -4.4704337149518 |
| lnc-PANK1-2 | lnc-PANK1-2 | -0.710390073295441 | -0.505769079136089 | -1.97562030129189 | 0.0547143599499724 | 0.296676941741381 | -3.91812063405941 |
| ENSG00000137509.10 | PRCP | 0.375087857156981 | 5.00679795578351 | 1.97544924418543 | 0.0547344352854529 | 0.296707632731229 | -4.51996926334039 |
| ENSG00000137038.7 | DMAC1 | -0.353411317203912 | 3.4335956088903 | -1.97498546256364 | 0.0547888969003687 | 0.296847038032249 | -4.4016362361399 |
| ENSG00000196214.10 | ZNF766 | 0.211932366038439 | 5.42569250272302 | 1.97498462198936 | 0.0547889956509991 | 0.296847038032249 | -4.53878547207637 |
| ENSG00000117877.10 | POLR1G | -0.843865326132094 | -0.254340189018956 | -1.97444811520418 | 0.054852055807554 | 0.297012027982338 | -3.95145420332681 |
| ENSG00000185344.13 | ATP6V0A2 | 0.210769200781137 | 5.63910335880006 | 1.97438330065582 | 0.0548596782478589 | 0.297012027982338 | -4.54815076287891 |
| ENSG00000138081.20 | FBXO11 | 0.23466258569205 | 6.66154874840324 | 1.97435728781645 | 0.0548627377150488 | 0.297012027982338 | -4.554354168505 |
| ENSG00000101413.11 | RPRD1B | 0.316401728510011 | 4.56994206295226 | 1.9739244898645 | 0.0549136623185873 | 0.297044156327606 | -4.49755193076737 |

| | | | | | | | |
|---|---|---|---|---|---|---|---|
| lnc-CPNE1-2 | lnc-CPNE1-2 | -1.01720065865126 | -0.0496117582621012 | -1.97402232768174 | 0.0549296603946278 | 0.297044156327606 | -3.993170534747 |
| FAM66B | FAM66B | -0.86212528220976 | 0.0951055367474153 | -1.97347122353506 | 0.0549670390535577 | 0.297044156327606 | -4.0124678623553 |
| lnc-MICU2-7 | lnc-MICU2-7 | 0.317718989440657 | 2.2388208156473 | 1.97345387721435 | 0.0549690826493838 | 0.297044156327606 | -4.27210098656797 |
| ENSG00000117222.13 | RBBP5 | 0.178066305582036 | 4.56920801866329 | 1.97341955564823 | 0.0549731263173902 | 0.297044156327606 | -4.50160710082429 |
| ENSG00000108654.14 | DDX5 | 0.26424740978309 | 10.600960071072 | 1.97325837707305 | 0.0549921193475205 | 0.297044156327606 | -4.46992236192924 |
| ENSG00000185267.9 | CDNF | -0.358550890363174 | 0.56447495782776 | -1.97324930494799 | 0.054993188560871 | 0.297044156327606 | -4.06784748831011 |
| ENSG00000257511.1 | | 0.318052806836958 | 1.5922746361066 | 1.97313141890764 | 0.055007083886608 | 0.297044156327606 | -4.20289371089814 |
| ENSG00000121621.6 | KIF18A | 0.409615475496912 | 1.76795554635718 | 1.97305944266661 | 0.0550155692762298 | 0.297044156327606 | -4.23325332266875 |
| lnc-ADORA2A-1 | lnc-ADORA2A-1 | -0.434053872314656 | -0.22651744729986 | -1.97300083161614 | 0.0550224798577155 | 0.297044156327606 | -3.9523241511857 |
| ENSG00000142192.20 | APP | -0.414175723096574 | 5.29927229386481 | -1.972958943769 | 0.0550274191360425 | 0.297044156327606 | -4.54241639693091 |
| ENSG00000279891.1 | | -0.419544298934755 | 1.23301695122065 | -1.97187766484935 | 0.0551550525509565 | 0.297655072537058 | -4.17634990385508 |
| ENSG00000144320.13 | LNPK | -0.293628494382738 | 4.06889323704344 | -1.97166750841658 | 0.0551798889072276 | 0.297674572175496 | -4.47401243208002 |
| LINC01963 | LINC01963 | -0.36628426384645 | 3.398297646984 | -1.97151176732413 | 0.0551983006676163 | 0.297674572175496 | -4.40994531837999 |
| lnc-TCHP-4 | lnc-TCHP-4 | 0.49462393814329 | 1.28391327864187 | 1.97141395372406 | 0.0552098669325092 | 0.297674572175496 | -4.16731982671409 |
| ENSG00000173451.6 | THAP2 | -0.293777620542597 | 2.50446647323681 | -1.97135774514214 | 0.0552165144325922 | 0.297674572175496 | -4.30852764157774 |
| lnc-MAMDC2-4 | lnc-MAMDC2-4 | 0.500049183878277 | 0.815210510972509 | 1.97077382196182 | 0.0552856128998964 | 0.297951966533277 | -4.11610456306664 |
| ENSG00000126970.15 | ZC4H2 | -1.01461420834408 | 3.99011367528814 | -1.97224442793184 | 0.0553448498997788 | 0.297951966533277 | -4.45012026583141 |
| lnc-SIPA1L2-12 | lnc-SIPA1L2-12 | -0.933568612134529 | -0.593070599542606 | -1.97027104722958 | 0.0553451684549036 | 0.297951966533277 | -3.89727783388748 |
| ENSG00000198814.12 | GK | -0.342870523310531 | 2.93319810248972 | -1.97004864313584 | 0.0553715307119333 | 0.297951966533277 | -4.36147280170012 |
| ENSG00000164187.6 | LMBRD2 | -0.173322324493124 | 4.5885010000401 | -1.96983328207488 | 0.0553970684620942 | 0.297951966533277 | -4.51280164295282 |
| ENSG00000135269.17 | TES | -1.14256322090555 | 3.64145511418641 | -1.97186697863054 | 0.0553973905172548 | 0.297951966533277 | -4.41171183713887 |
| lnc-CYLD-1 | lnc-CYLD-1 | -0.798073228699017 | 0.683487500488282 | -1.96995346636794 | 0.055406199331536 | 0.297951966533277 | -4.12457025737611 |
| ENSG00000145780.7 | FEM1C | 0.37683387740182 | 4.59185709669516 | 1.9696634725248 | 0.0554172118300457 | 0.297951966533277 | -4.51337853716402 |
| ENSG00000164989.16 | CCDC171 | -0.562853177490817 | 3.06074348487386 | -1.96973223359901 | 0.0554265860471384 | 0.297951966533277 | -4.36151979392923 |
| lnc-COL9A2-3 | lnc-COL9A2-3 | -0.574462866588001 | 0.645505847844043 | -1.96944236829839 | 0.0554434494135976 | 0.297951966533277 | -4.11685662960371 |
| lnc-GDPGP1-5 | lnc-GDPGP1-5 | -0.33591544753516 | 0.748016532943105 | -1.9693786869833 | 0.0554510082169155 | 0.297951966533277 | -4.10384910867981 |
| ENSG00000160949.16 | TONSL | -0.424314481947332 | 2.60030126003647 | -1.96916845381252 | 0.0554759686507951 | 0.297951966533277 | -4.32969725769891 |
| lnc-ZNF705A-7 | lnc-ZNF705A-7 | -0.602309806601722 | -0.155433118224112 | -1.96912613569205 | 0.055480994142382 | 0.297951966533277 | -3.98807250279766 |
| ENSG00000173486.12 | FKBP2 | -0.4253397710511 | 2.48281088385611 | -1.96907150962899 | 0.0554874818454086 | 0.297951966533277 | -4.32155659483229 |
| ENSG00000126067.11 | PSMB2 | 0.207289508767476 | 4.81714822255512 | 1.96898953150602 | 0.0554972192634575 | 0.297951966533277 | -4.52651133073603 |
| ENSG00000170004.16 | CHD3 | -0.478433959171294 | 7.06498523161352 | -1.96917135067108 | 0.0555165191753519 | 0.297951966533277 | -4.5606115678419 |
| ENSG00000171262.11 | FAM98B | 0.331629019732647 | 4.6087043408896 | 1.96861796406154 | 0.0555413727893817 | 0.297951966533277 | -4.50928037895734 |
| ENSG00000120533.12 | ENY2 | 0.229369075689631 | 4.97447160765682 | 1.96852440551306 | 0.0555524951689753 | 0.297951966533277 | -4.53647326373571 |
| ENSG00000104219.12 | ZDHHC2 | -0.259223301601498 | 5.77671378968413 | -1.96845723211189 | 0.0555604820280727 | 0.297951966533277 | -4.56220046975604 |
| lnc-FILIP1L-7 | lnc-FILIP1L-7 | 0.213203074078029 | 1.63728491447221 | 1.96836429081459 | 0.0555715343012849 | 0.297951966533277 | -4.22707717737929 |
| ENSG00000148926.9 | ADM | -1.67003206706161 | -0.816344440058002 | -1.96935820123732 | 0.0555719574173469 | 0.297951966533277 | -3.90328337643388 |
| ENSG00000056558.10 | TRAF1 | -0.385434159460647 | 6.79780609983899 | -1.9682149185645 | 0.0555893011316646 | 0.297967339894024 | -4.56326473865091 |
| lnc-PLSCR2-3 | lnc-PLSCR2-3 | 0.624587070162221 | -0.15939635471983 | 1.96798011066348 | 0.0556172398680308 | 0.298039481438083 | -3.96291062417495 |
| lnc-LYRM2-2 | lnc-LYRM2-2 | 0.30182414256174 | 1.36800734949319 | 1.96749086233339 | 0.0556754922455902 | 0.298162794484662 | -4.19151677152307 |
| lnc-TUFM-3 | lnc-TUFM-3 | -0.39911736839947 | 1.21526697750885 | -1.96746647880673 | 0.0556783968478871 | 0.298162794484662 | -4.18435962271269 |
| lnc-METTL21C-3 | lnc-METTL21C-3 | -0.347795887835756 | 1.46470958722864 | -1.96742188738501 | 0.0556837089830948 | 0.298162794484662 | -4.22141270160488 |
| ENSG00000151348.13 | EXT2 | 0.210113628343787 | 4.43117917718495 | 1.96707064721342 | 0.0557255671899908 | 0.298309323659709 | -4.50202155606897 |
| ENSG00000154429.10 | CCSAP | -0.309166757038419 | 5.00560880875453 | -1.96657041858895 | 0.0557852275917159 | 0.298551050317288 | -4.54805894098919 |
| lnc-DGAT2-1 | lnc-DGAT2-1 | -0.719068942304795 | 0.488413940262786 | -1.96644540456635 | 0.0558001461498595 | 0.298553264414494 | -4.10731405253693 |
| ENSG00000171044.10 | XKR6 | -0.280421748819911 | 2.56874197394811 | -1.96522578333948 | 0.0559458703806336 | 0.299255158535494 | -4.32985154085494 |
| ENSG00000173812.10 | EIF1 | 0.281857894883688 | 8.1530452629228 | 1.96473197854064 | 0.0560049650286466 | 0.299429891263247 | -4.55149843691199 |
| ENSG00000164933.11 | SLC25A32 | 0.251621704819934 | 4.18232778402063 | 1.96470970676275 | 0.0560076316068359 | 0.299429891263247 | -4.48948565140488 |
| ENSG00000110429.13 | FBXO3 | 0.250327781151213 | 6.15491028497168 | 1.96429249047083 | 0.0560576047455459 | 0.299619236166598 | -4.57270833454187 |
| lnc-ARFGAP1-4 | lnc-ARFGAP1-4 | 0.330215708904863 | 0.362333929228782 | 1.96411459951845 | 0.0560789237742221 | 0.299635681066632 | -4.05944080145868 |
| lnc-PRTFDC1-1 | lnc-PRTFDC1-1 | -0.353802340677456 | 0.904065272198938 | -1.96402389830448 | 0.0560897963926412 | 0.299635681066632 | -4.17120429743395 |
| ENSG00000196704.11 | AMZ2 | 0.196277510008529 | 5.18530895745461 | 1.96388806421788 | 0.0561060826191841 | 0.299644913998616 | -4.55349444999198 |
| FMR1-AS1 | FMR1-AS1 | -0.548281249016983 | -0.47888438519632 | -1.96317930888911 | 0.0561911270785716 | 0.299947285766681 | -3.95856312265742 |
| lnc-RGS10-4 | lnc-RGS10-4 | 0.273135100047439 | 2.17660768184886 | 1.96307555164791 | 0.0562035863643511 | 0.299947285766681 | -4.2876930085136 |
| lnc-BAZ2B-1 | lnc-BAZ2B-1 | -0.376672476305508 | 2.41919772958224 | -1.96305198168039 | 0.0562064170044255 | 0.299947285766681 | -4.32468927946259 |
| ENSG00000165121.11 | | 0.293379703154883 | 2.43926060131748 | 1.96290279092401 | 0.0562243369502309 | 0.299965144491084 | -4.31322155907431 |
| ENSG00000224411.3 | HSP90AA2P | 1.40378460012488 | -1.06589003678793 | 1.962670790339 | 0.0562522133275536 | 0.300036099228048 | -3.90803794074017 |
| lnc-USP34-1 | lnc-USP34-1 | -0.32369958269877 | 2.36371610034227 | -1.96242675737433 | 0.0562815483312232 | 0.300036455335176 | -4.32250801603122 |
| ENSG00000030066.13 | NUP160 | 0.247981149870498 | 5.78934837859857 | 1.96235384974721 | 0.0562903150558721 | 0.300036455335176 | -4.5719931976917 |
| ENSG00000279095.1 | | -2.26795326069841 | -1.75877223728097 | -1.96309911446525 | 0.056307716000266 | 0.300036455335176 | -3.89622651644348 |
| LINC00304 | LINC00304 | -0.634813043250731 | 0.866989120816618 | -1.9621852915904 | 0.0563105877160659 | 0.300036455335176 | -4.15168466745565 |
| lnc-TREM2-1 | lnc-TREM2-1 | 1.4169132183248 | -1.12746713063578 | 1.96189175333777 | 0.0563459068833905 | 0.300146946527129 | -3.89911488181406 |
| ENSG00000168310.10 | IRF2 | -0.392766618743538 | 5.34240084678826 | -1.96174893168989 | 0.056363098402789 | 0.300160842024478 | -4.56233030021266 |
| lnc-FGFR1OP-2 | lnc-FGFR1OP-2 | -0.473475354465925 | 0.105367066938059 | -1.96125563366761 | 0.0564225117366911 | 0.300301456604251 | -4.0253230124131 |
| ENSG00000174013.7 | FBXO45 | 0.361253623344404 | 2.57364247468703 | 1.96115964508937 | 0.0564340789743482 | 0.300301456604251 | -4.34238510986701 |
| ENSG00000162923.15 | WDR26 | 0.251021551629772 | 7.22176304539511 | 1.96112820696084 | 0.0564378679142617 | 0.300301456604251 | -4.57314086150969 |
| ENSG00000177119.15 | ANO6 | 1.0562599593168 | 4.81423933682529 | 1.96305493773649 | 0.0564483185225165 | 0.300301456604251 | -4.50847208753454 |
| ENSG00000064102.14 | INTS13 | 0.250780874906521 | 3.33344881747297 | 1.96084488672603 | 0.0564720237122263 | 0.300301456604251 | -4.4188004707124 |
| lnc-TLE4-9 | lnc-TLE4-9 | -1.27259676886236 | 2.1384603272254 | -1.96261709782585 | 0.0564900180132083 | 0.300301456604251 | -4.27133678371633 |

| | | | | | | | |
|---|---|---|---|---|---|---|---|
| ENSG00000182208.13 | MOB2 | -0.240622755554222 | 3.74252443162075 | -1.96056045282815 | 0.0565063316885073 | 0.300301456604251 | -4.46731809001627 |
| ENSG00000166710.18 | B2M | 0.232314041702598 | 10.6010385834147 | 1.96038632434456 | 0.0565273436603489 | 0.300301456604251 | -4.49364000424544 |
| lnc-VTI1A-1 | lnc-VTI1A-1 | -0.170904804440138 | 4.82770264538784 | -1.96032168640491 | 0.0565351451924936 | 0.300301456604251 | -4.54363491682946 |
| ENSG00000088543.14 | C3orf18 | -0.349595753958322 | 1.71121549053602 | -1.96031957290903 | 0.056535400298376 | 0.300301456604251 | -4.24970631308242 |
| ENSG00000122591.11 | HYCC1 | -0.543544802761418 | 5.23030717670729 | -1.96056858491232 | 0.0565954894336146 | 0.300543075080518 | -4.56555919785606 |
| ENSG00000125319.14 | HROB | 0.505438202127687 | 0.0809950145607334 | 1.95969879657444 | 0.0566103729908641 | 0.300544572419643 | -4.00348600994348 |
| ENSG00000274049.4 | INO80B-WBP1 | -0.206508620473994 | 4.47701102344247 | -1.95902749203842 | 0.056691544509523 | 0.300649406525904 | -4.52361726990367 |
| ENSG00000100271.16 | TTLL1 | -0.61416373333755 | 1.40033472640397 | -1.95883923101551 | 0.0567143262953735 | 0.300649406525904 | -4.20954566138042 |
| lnc-TENM1-12 | lnc-TENM1-12 | 0.173425657352861 | 2.05972999995992 | 1.95883426289481 | 0.0567149276029119 | 0.300649406525904 | -4.28667295268689 |
| ENSG00000168005.8 | SPINDOC | -0.238998447712231 | 2.50435112411076 | -1.95872771901765 | 0.0567278242713547 | 0.300649406525904 | -4.34077198928596 |
| lnc-ENC1-5 | lnc-ENC1-5 | 0.73242331773349 | 2.93607431930115 | 1.95958560176582 | 0.0567334170648099 | 0.300649406525904 | -4.36415429874833 |
| ENSG00000162302.12 | RPS6KA4 | -0.416651633025498 | 2.79958774654379 | -1.95864634352751 | 0.0567376761190794 | 0.300649406525904 | -4.36828180833998 |
| USP46-AS1 | USP46-AS1 | -0.310793248377934 | 2.99287324816365 | -1.95861505790052 | 0.0567414641535237 | 0.300649406525904 | -4.38274364322521 |
| ENSG00000127922.9 | SEM1 | 0.24592475561828 | 3.71398827132102 | 1.95856956354816 | 0.0567469729559897 | 0.300649406525904 | -4.45486815742011 |
| lnc-CCDC90B-7 | lnc-CCDC90B-7 | -1.16232562393851 | -0.508041049416098 | -1.9586652335068 | 0.0567964564410202 | 0.30083413868387 | -3.9895197954001 |
| ENSG00000088876.11 | ZNF343 | -0.207573616038739 | 3.32096996167164 | -1.95775012686671 | 0.0568462755184174 | 0.300968213450106 | -4.41818678663734 |
| ENSG00000106479.10 | ZNF862 | -0.280166975966432 | 5.08644829066069 | -1.95771105905119 | 0.0568510136468937 | 0.300968213450106 | -4.55766291149639 |
| ENSG00000118197.13 | DDX59 | 0.159468693038272 | 4.4666102180642 | 1.95729895495905 | 0.0569010141880211 | 0.301155457709447 | -4.52172528768339 |
| ENSG00000230797.2 | YY2 | -0.494159460726643 | -0.077243100390254 | -1.95715472733392 | 0.0569185222563953 | 0.301166918805993 | -4.00554764390264 |
| ENSG00000197056.10 | ZMYM1 | -0.320076877101061 | 3.84664940349434 | -1.95685603335149 | 0.0569547960512566 | 0.301166918805993 | -4.47244074912046 |
| ENSG00000107611.15 | CUBN | -0.465635057678084 | 3.7002499685592 | -1.95677476987136 | 0.0569646682418182 | 0.301166918805993 | -4.45464806291671 |
| lnc-MNX1-7 | lnc-MNX1-7 | 0.287186762431129 | 0.973242378599137 | 1.95671812975757 | 0.0569715499657788 | 0.301166918805993 | -4.16820355698393 |
| lnc-ZFYVE21-2 | lnc-ZFYVE21-2 | 0.268018564729414 | 2.33748959256847 | 1.95667871855212 | 0.0569763388150677 | 0.301166918805993 | -4.30861601636647 |
| ENSG00000187122.16 | SLIT1 | 0.755769741684311 | -0.296525835503421 | 1.95609777920554 | 0.0570469689003021 | 0.301453108195479 | -3.94380505287774 |
| ENSG00000213563.6 | C8orf82 | -0.363408076199615 | 3.34377504981042 | -1.95599252983467 | 0.0570597730908802 | 0.301453108195479 | -4.43153948032302 |
| ENSG00000244716.3 | RPL17P7 | 0.621652238785481 | 2.2106204126166 | 1.955728528816 | 0.057103026621021 | 0.301558259247251 | -4.28475945580018 |
| ENSG00000104381.12 | GDAP1 | -0.348080528868787 | 3.58181284829381 | -1.95539283126156 | 0.0571327771135986 | 0.301558259247251 | -4.45781772815676 |
| ENSG00000173715.16 | TOP6BL | -0.299147276853032 | 6.06561193242396 | -1.95532725829911 | 0.057140764484296 | 0.301558259247251 | -4.58696235522502 |
| ENSG00000285641.1 | | 0.224654995398909 | 3.91860540513691 | 1.95527360137055 | 0.0571473010914347 | 0.301558259247251 | -4.48164467925774 |
| lnc-PHLDA1-1 | lnc-PHLDA1-1 | 0.225041772025909 | 4.73615626484095 | 1.95516099522052 | 0.0571610211187286 | 0.301558259247251 | -4.54963222044078 |
| ENSG00000265590.9 | CFAP298-TCP10L | -0.406394091662896 | 2.30287002603705 | -1.95510716055287 | 0.0571675813818576 | 0.301558259247251 | -4.32270323294471 |
| CAMTA1-DT | CAMTA1-DT | -0.481693787072804 | -0.279560578099967 | -1.95482475303613 | 0.0572020060313869 | 0.301572275742168 | -4.00209260491932 |
| lnc-CENPJ-9 | lnc-CENPJ-9 | 0.323402535153786 | 2.47294037942761 | 1.95481724792407 | 0.0572029211260534 | 0.301572275742168 | -4.32657848326614 |
| lnc-CS-2 | lnc-CS-2 | 0.266135558977035 | 1.57383547932818 | 1.95472480968586 | 0.0572141931095159 | 0.301572275742168 | -4.24264605252394 |
| ENSG00000159899.14 | NPR2 | -0.412533465905168 | 1.1998214783196 | -1.95422494441973 | 0.057275180176046 | 0.30181644484474 | -4.21524569307129 |
| ENSG00000105699.16 | LSR | -0.515431109705814 | 4.82827409479387 | -1.95430478827324 | 0.0573260742694906 | 0.301964695486902 | -4.5532058966414 |
| lnc-STARD8-3 | lnc-STARD8-3 | -0.216363808698322 | 2.01197995929098 | -1.95375428886876 | 0.0573326546160818 | 0.301964695486902 | -4.29813806906471 |
| lnc-KLHL25-9 | lnc-KLHL25-9 | -0.485717295324338 | 0.458677731370624 | -1.95359196798463 | 0.0573524880617216 | 0.301991880730216 | -4.09120744954331 |
| ENSG00000162415.6 | ZSWIM5 | -0.55077585166875 | 3.28240482712166 | -1.95330870308376 | 0.0574129185183576 | 0.302232762624899 | -4.43202688188775 |
| ENSG00000109062.11 | NHERF1 | -0.431508297505206 | 3.21635242125669 | -1.95287541332275 | 0.0574401120991343 | 0.302283643925263 | -4.42196415011922 |
| ENSG00000137497.17 | NUMA1 | -0.212555931749101 | 8.71326464850983 | -1.95277862895193 | 0.0574519562277428 | 0.302283643925263 | -4.56043373236587 |
| ENSG00000275342.4 | PRAG1 | -0.310904978765689 | 5.57198747637957 | -1.95251047972753 | 0.057484782364544 | 0.30237906348311 | -4.58467902600405 |
| lnc-FAM19A2-2 | lnc-FAM19A2-2 | -0.313587136213256 | 0.683996348526805 | -1.95225992619782 | 0.0575154690739808 | 0.302463183431208 | -4.12818179304444 |
| ENSG00000158158.11 | CNNM4 | 0.351280706775716 | 4.07564444694514 | 1.95188192752528 | 0.0575617913996937 | 0.302629464209424 | -4.48789146892051 |
| lnc-MYO18B-2 | lnc-MYO18B-2 | -0.311962463724249 | 3.32762176744273 | -1.95163349559919 | 0.0575922532926287 | 0.302654939881043 | -4.42905695337844 |
| ENSG00000259917.1 | HNRNPLP2 | 1.34485082047472 | -0.14438771141302 | 1.95177322535144 | 0.057596045256476 | 0.302654939881043 | -3.91923599946544 |
| ENSG00000134748.12 | PRPF38A | 0.174524072181499 | 5.41430933722409 | 1.95137347951383 | 0.0576241504589296 | 0.302725341729492 | -4.58333770504694 |
| lnc-AIM2-2 | lnc-AIM2-2 | -0.590194309381873 | 5.00581985120296 | -1.95197510993948 | 0.0576700182650075 | 0.302888998711061 | -4.56597072860285 |
| ENSG00000228409.6 | CCT6P1 | 0.22222753117576 | 3.53191451572367 | 1.95056246144805 | 0.0577237388542119 | 0.303040279226675 | -4.44974669150754 |
| PTOV1-AS1 | PTOV1-AS1 | -0.388317407844534 | 0.774269307720003 | -1.95052560821563 | 0.0577282677378318 | 0.303040279226675 | -4.13553802666393 |
| ENSG00000124214.19 | STAU1 | 0.158309976376574 | 5.40439071513783 | 1.94991434365016 | 0.0578034304912735 | 0.303357473177431 | -4.58618653171846 |
| lnc-CNOT8-2 | lnc-CNOT8-2 | 0.2262585096571 | 2.56194499312856 | 1.94950900385974 | 0.0578533186064973 | 0.303454577917535 | -4.34906185315354 |
| ENSG00000089902.9 | RCOR1 | 0.227190884950432 | 4.73074568459843 | 1.9492273335967 | 0.0578880076318252 | 0.303454577917535 | -4.5534128310861 |
| lnc-PPIL2-8 | lnc-PPIL2-8 | -0.581976706366594 | -0.340550811514454 | -1.94921542462089 | 0.0578894746728554 | 0.303454577917535 | -3.97166925563492 |
| ENSG00000212128.2 | TAS2R13 | -0.574374937654098 | 0.604001090163384 | -1.94904096715168 | 0.0579109693798057 | 0.303454577917535 | -4.13885572769335 |
| ENSG00000156802.12 | ATAD2 | 0.290470533264179 | 5.21748879562495 | 1.94902461789565 | 0.0579129841050513 | 0.303454577917535 | -4.57953964187415 |
| ENSG00000252542.1 | SNORD36C | 0.32696205055959 | 0.681057096645553 | 1.94894133317928 | 0.0579232482496881 | 0.303454577917535 | -4.15459300882995 |
| lnc-COX7B-4 | lnc-COX7B-4 | 0.318066006497694 | 0.322300478020148 | 1.94881872236293 | 0.0579383618560229 | 0.303454577917535 | -4.07491948499927 |
| lnc-C20orf173-4 | lnc-C20orf173-4 | -1.00201898573054 | 0.0116105891891184 | -1.94909605060967 | 0.0579409663648961 | 0.303454577917535 | -4.04541221144723 |
| ENSG00000137815.14 | RTF1 | 0.287054653501574 | 5.84977029477001 | 1.94868685616319 | 0.0579546201133863 | 0.303454577917535 | -4.59741037022144 |
| FAM111A-DT | FAM111A-DT | -0.340648211301938 | 3.06098740050681 | -1.94839739995757 | 0.057990321976431 | 0.303564292278962 | -4.40609655806627 |
| ENSG00000139344.7 | AMDHD1 | -0.323783417099693 | 1.03352079398827 | -1.94818029174724 | 0.0580171127737611 | 0.303627315591743 | -4.18655590812704 |
| KCNQ5-IT1 | KCNQ5-IT1 | -0.593516957521797 | 1.13710297812966 | -1.94797563737527 | 0.0580423765384702 | 0.303682317308422 | -4.21055089191823 |
| ENSG00000108395.13 | TRIM37 | 0.230263750739155 | 5.04814918780718 | 1.9474905614623 | 0.0581022950330786 | 0.303881940902345 | -4.57671338225183 |
| ENSG00000144744.16 | UBA3 | -0.168750946671752 | 5.37480041799056 | -1.94733311312876 | 0.0581217551111212 | 0.303881940902345 | -4.59179399973926 |
| ENSG00000126264.9 | HCST | -0.467191853114832 | 3.10957592741917 | -1.94715521553114 | 0.0581437493958683 | 0.303881940902345 | -4.41321598318119 |
| ENSG00000181029.8 | TRAPPC5 | -0.413921047414318 | 2.56286229271733 | -1.94707779799666 | 0.0581533231118331 | 0.303881940902345 | -4.37210254055494 |
| ENSG00000164669.13 | INTS4P1 | 0.915072533611635 | -0.684292601569218 | 1.94706950293599 | 0.0581543489877246 | 0.303881940902345 | -3.91972706477158 |

| | | | | | | | |
|---|---|---|---|---|---|---|---|
| lnc-LRP5-1 | lnc-LRP5-1 | -0.456928090829925 | 0.808754500814581 | -1.94677886302105 | 0.0581903031590972 | 0.303943620272002 | -4.14933221326539 |
| ENSG00000110852.4 | CLEC2B | -3.1376695982045 | 1.75222963389571 | -1.94882856874114 | 0.0582029045377155 | 0.303943620272002 | -4.17456622832448 |
| lnc-ABHD12-3 | lnc-ABHD12-3 | -1.70430451384508 | -1.31968506201067 | -1.94687092286181 | 0.0582104528548916 | 0.303943620272002 | -3.90947797159535 |
| lnc-ARMCX6-3 | lnc-ARMCX6-3 | -0.458137395989513 | 1.0768479289763 | -1.94644791717237 | 0.0582312667381273 | 0.303975187235829 | -4.22056152333774 |
| ENSG00000102804.14 | TSC22D1 | -0.290596546369851 | 4.45949572528527 | -1.94621292571393 | 0.0582603684022839 | 0.304049990574089 | -4.54809827322194 |
| lnc-BRD2-4 | lnc-BRD2-4 | 0.517293796516545 | 2.92539580659051 | 1.94597106574412 | 0.0582903337367356 | 0.304129262180793 | -4.37538576473563 |
| lnc-SLTM-3 | lnc-SLTM-3 | -0.54491847955485 | -0.359664416595854 | -1.9457522595034 | 0.0583174542422079 | 0.304193654502627 | -3.98309577621928 |
| ENSG00000206680.1 | SNORD21 | 0.395067761164898 | -0.173812602263583 | 1.9453374521304 | 0.0583688984092747 | 0.304384858362833 | -4.00814279942113 |
| lnc-DTWD2-1 | lnc-DTWD2-1 | -0.516793353864392 | -0.247663500869071 | -1.94519435146022 | 0.0583866547240891 | 0.304400332873841 | -3.9938166223144 |
| lnc-ARMC1-3 | lnc-ARMC1-3 | -0.495485695004545 | -0.615617424700659 | -1.94467904235587 | 0.0584506341539759 | 0.304633713115052 | -3.96297073323825 |
| lnc-ST3GAL2-1 | lnc-ST3GAL2-1 | -0.411539179089189 | 1.65527310816049 | -1.94459544054914 | 0.0584610196183479 | 0.304633713115052 | -4.280433025697 |
| ENSG00000108443.13 | RPS6KB1 | 0.325539776719747 | 5.88536519925015 | 1.94422346022075 | 0.0585072482506818 | 0.304739838327481 | -4.60648695452318 |
| ENSG00000111707.11 | SUDS3 | 0.134962472586724 | 6.23809027070359 | 1.94409566772507 | 0.0585231371840483 | 0.304739838327481 | -4.60897833120451 |
| ENSG00000129968.15 | ABHD17A | -0.285970679204166 | 5.00604766466801 | -1.94388085256074 | 0.0585498543430521 | 0.304739838327481 | -4.58306696389836 |
| ENSG00000160796.16 | NBEAL2 | -0.262773547457319 | 5.74582622414197 | -1.94385997479813 | 0.0585524515275883 | 0.304739838327481 | -4.60364121801599 |
| ENSG00000141665.12 | FBXO15 | -0.378283416062612 | 0.44093479819075 | -1.94383617166617 | 0.058555412747665 | 0.304739838327481 | -4.0997688678676 |
| ENSG00000136504.11 | KAT7 | 0.193309091229342 | 7.01330341471663 | 1.9433048232674 | 0.0586215484877424 | 0.304941626011271 | -4.60711663880528 |
| lnc-CHFR-2 | lnc-CHFR-2 | 0.298832629344519 | 2.0370633352853 | 1.9432866110738 | 0.0586238164566194 | 0.304941626011271 | -4.30925321892121 |
| lnc-TCF7-4 | lnc-TCF7-4 | -0.379870434691866 | 0.706591229176261 | -1.94293308730316 | 0.0586678558072951 | 0.305093601839706 | -4.17107522572067 |
| ENSG00000188554.13 | NBR1 | -0.244484936062271 | 6.29066658704898 | -1.9425110885372 | 0.0587204625050649 | 0.305201992545974 | -4.61138650311681 |
| lnc-PMEPA1-3 | lnc-PMEPA1-3 | -0.5295672302232 | 2.19717689175133 | -1.94244082077512 | 0.0587292260774861 | 0.305201992545974 | -4.32838563590574 |
| lnc-CUL1-1 | lnc-CUL1-1 | -0.649405228014592 | -0.0029852138033124 | -1.94233210037675 | 0.0587427875567709 | 0.305201992545974 | -4.05062902049899 |
| ENSG00000142945.12 | KIF2C | 0.877676762209257 | 0.384951707635872 | 1.94225023281507 | 0.0587530012651443 | 0.305201992545974 | -4.04582591632357 |
| ENSG00000135119.14 | RNFT2 | -0.445714170150791 | 3.42008839010038 | -1.94196731810639 | 0.0587883091543992 | 0.305201992545974 | -4.4564274398697 |
| ENSG00000168081.8 | PNOC | -0.422283534405935 | 4.58882082755587 | -1.94194423395858 | 0.0587911908717689 | 0.305201992545974 | -4.56050185783294 |
| lnc-AHNAK2-7 | lnc-AHNAK2-7 | -0.250307379759382 | 0.752368865861308 | -1.94193379688545 | 0.0587924938271772 | 0.305201992545974 | -4.17634478429932 |
| ENSG00000252620.1 | RNU6ATAC24P | 0.383655847773199 | 1.4587339614679 | 1.94103180678103 | 0.0589051913362074 | 0.305709922660907 | -4.25672350214288 |
| ENSG00000158169.13 | FANCC | 0.354487983916307 | 3.03467691236695 | 1.94090623857701 | 0.0589208949433348 | 0.305714338446852 | -4.41501232961501 |
| ENSG00000042445.13 | RETSAT | -0.27877611181388 | 3.55989764431088 | -1.9401296896299 | 0.0590180904179694 | 0.306141470532526 | -4.4769945778866 |
| lnc-STIM2-4 | lnc-STIM2-4 | -0.46965828052469 | 2.56104509498688 | -1.93991783273434 | 0.0590446310549605 | 0.306157935920851 | -4.37928344659077 |
| ENSG00000196437.10 | ZNF569 | -0.224359182456818 | 3.50517837954416 | -1.93968746607974 | 0.0590735021670108 | 0.306157935920851 | -4.47009946576959 |
| lnc-MCM5-1 | lnc-MCM5-1 | -0.451264176420158 | 2.18362800063467 | -1.9396682053292 | 0.059075916604508 | 0.306157935920851 | -4.32939598355823 |
| ENSG00000121578.12 | B4GALT4 | 0.197454580781443 | 4.16869892248682 | 1.93955897097813 | 0.0590896113158802 | 0.306157935920851 | -4.52606684220267 |
| ENSG00000285967.1 | NIPBL-DT | -0.356309903604029 | 3.6869581336707 | -1.9395109212384 | 0.0590956361761424 | 0.306157935920851 | -4.49215099544171 |
| lnc-BTG1-3 | lnc-BTG1-3 | 0.619614989985799 | 3.23082724736022 | 1.93944518087618 | 0.0591671442775985 | 0.306335168459308 | -4.42937718172447 |
| ENSG00000138138.13 | ATAD1 | 0.184414294635543 | 5.40204284903902 | 1.93894039197594 | 0.0591672140562738 | 0.306335168459308 | -4.60618398961492 |
| lnc-CARHSP1-3 | lnc-CARHSP1-3 | 0.228499739661851 | 3.81958369730626 | 1.93888091819336 | 0.0591746798484791 | 0.306335168459308 | -4.49936329145606 |
| ENSG00000229419.1 | RALGAPA1P1 | 0.485945490864856 | 1.98323226623268 | 1.9387638499702 | 0.0591893778828483 | 0.306335168459308 | -4.29610702810325 |
| ENSG00000226360.5 | RPL10AP6 | 1.051370234564 | 1.64414115390671 | 1.9398502924464 | 0.0592078971016328 | 0.306353983419534 | -4.21127723965275 |
| lnc-PDPK1-3 | lnc-PDPK1-3 | -0.600676027668728 | 2.10225157264513 | -1.93853130883754 | 0.059232546637869 | 0.306367160355545 | -4.34525500285877 |
| SUCLG2-AS1 | SUCLG2-AS1 | -0.456599630539138 | 2.13418299004295 | -1.93835914870872 | 0.0592402127102498 | 0.306367160355545 | -4.34320696913335 |
| ENSG00000183386.9 | FHL3 | 0.574386029871576 | -0.32324786683503 | 1.93805149862125 | 0.0592788819648729 | 0.306490135012052 | -3.97467705360021 |
| ENSG00000105750.14 | ZNF85 | -0.314688074180088 | 3.96617707890695 | -1.937782261624 | 0.0593127407841432 | 0.306547468009073 | -4.51346742730537 |
| ENSG00000005483.20 | KMT2E | 0.230090388907636 | 8.16470621316171 | 1.93772648431573 | 0.0593197573278987 | 0.306547468009073 | -4.5997447282047 |
| lnc-PHLPP1-1 | lnc-PHLPP1-1 | -0.281767113946519 | -0.201322753413516 | -1.93709649593126 | 0.0593990566719173 | 0.306775129899943 | -4.02313154918544 |
| lnc-MCPH1-2 | lnc-MCPH1-2 | -0.578361524182495 | 0.196710776527882 | -1.93702783613181 | 0.0594077046776731 | 0.306775129899943 | -4.08805000186506 |
| ENSG00000176225.13 | RTTN | 0.294985032478155 | 5.6175659604879 | 1.93702132461222 | 0.0594085248885571 | 0.306775129899943 | -4.61349845903752 |
| ENSG00000243811.9 | APOBEC3D | -0.422064390584441 | 3.52116508568245 | -1.9368748162414 | 0.0594269821026161 | 0.306793471938336 | -4.47132738533869 |
| ENSG00000152778.8 | IFIT5 | 0.311875533498577 | 4.69035222785569 | 1.936321707706 | 0.0594967074707647 | 0.307062904217715 | -4.57808127837275 |
| ENSG00000147162.13 | OGT | 0.290042508946849 | 10.3424331357019 | 1.93622418431761 | 0.0595090086442436 | 0.307062904217715 | -4.545049838852 |
| ENSG00000215012.8 | RTL10 | -0.277695792132701 | 3.69607653069688 | -1.93586415198059 | 0.0595544404724588 | 0.307082546921286 | -4.50126695024505 |
| ENSG00000213281.4 | NRAS | 0.272826181364871 | 6.36599953710632 | 1.93572171385834 | 0.0595724227053018 | 0.307082546921286 | -4.62386789226162 |
| lnc-IL7-4 | lnc-IL7-4 | 0.235222018421181 | 2.26583204883518 | 1.93563268405154 | 0.059583664724514 | 0.307082546921286 | -4.34130233146468 |
| ENSG00000131152.4 | | -0.266013604208602 | 0.215977923235894 | -1.93561890742393 | 0.0595854044969925 | 0.307082546921286 | -4.09949474333075 |
| ENSG00000174173.6 | TRMT10C | 0.276964852728249 | 3.78949226859218 | 1.93525422224435 | 0.059631474397801 | 0.307082546921286 | -4.50558540050776 |
| lnc-C7orf77-5 | lnc-C7orf77-5 | -0.25133357932579 | 1.07146585250306 | -1.93522750540185 | 0.0596348506821017 | 0.307082546921286 | -4.22003458069375 |
| ENSG00000139668.8 | WDFY2 | 0.39426303071055 | 5.01609307925722 | 1.93517238208745 | 0.0596418172929523 | 0.307082546921286 | -4.60106185125687 |
| ENSG00000252699.1 | SNORA21B | 0.502643259890025 | 0.361654792127808 | 1.93511989724277 | 0.0596484510976338 | 0.307082546921286 | -4.08841570149565 |
| lnc-ZNF212-2 | lnc-ZNF212-2 | 0.20458143386512 | 2.89759470972914 | 1.93499198159361 | 0.0596646216088324 | 0.307082546921286 | -4.409166526352 |
| ENSG00000215041.9 | NEURL4 | -0.232677877346148 | 4.54092498605947 | -1.93494180110985 | 0.0596709662242966 | 0.307082546921286 | -4.57434876857633 |
| lnc-CREB1-2 | lnc-CREB1-2 | -0.216764476028187 | 2.49891581492607 | -1.93489466093616 | 0.0596769269632776 | 0.307082546921286 | -4.37417433151947 |
| ENSG00000164961.15 | WASHC5 | 0.208612383379171 | 5.48114832444558 | 1.93474314822225 | 0.0596960887727092 | 0.307104372708991 | -4.61428691607996 |
| lnc-PAX8-6 | lnc-PAX8-6 | -0.224778511333638 | 4.42374958753376 | -1.93423466153591 | 0.0597604357067706 | 0.307358582746742 | -4.56767087957189 |
| ENSG00000100478.14 | AP4S1 | -0.187578750494979 | 4.89220112750489 | -1.93367845631907 | 0.0598308894391813 | 0.307479009211239 | -4.59515533200281 |
| lnc-CCDC69-2 | lnc-CCDC69-2 | -0.431668579742755 | 2.04734045058736 | -1.9336570961148 | 0.0598335965279529 | 0.307479009211239 | -4.34388354147156 |
| lnc-TEF-1 | lnc-TEF-1 | -0.397798008365082 | 1.42728240221462 | -1.93361392761598 | 0.0598390678154023 | 0.307479009211239 | -4.26173692623514 |
| ENSG00000256407.2 | | -0.275497817758304 | 0.398056832896259 | -1.93357813591179 | 0.0598436044745772 | 0.307479009211239 | -4.10422214743107 |
| lnc-HSPB7-5 | lnc-HSPB7-5 | -0.348737983524589 | 1.55665584973917 | -1.93300701246073 | 0.0599160353113441 | 0.307773401049087 | -4.29092451394901 |

| | | | | | | | |
|---|---|---|---|---|---|---|---|
| lnc-ALDH1A2-5 | lnc-ALDH1A2-5 | 0.223158305477591 | 2.66240138740181 | 1.93287449298822 | 0.0599328524102476 | 0.307773401049087 | -4.38681017507395 |
| ENSG00000278902.1 | | -0.984192201108975 | 1.446172036101 | -1.9339525531629 | 0.0599607121511364 | 0.307773401049087 | -4.28624608096174 |
| lnc-FBXO15-7 | lnc-FBXO15-7 | -0.984192201108975 | 1.446172036101 | -1.9339525531629 | 0.0599607121511364 | 0.307773401049087 | -4.28624608096174 |
| ENSG00000279800.2 | BCLAF1P2 | 0.99694801122103 | 0.759631647825184 | 1.93287540790725 | 0.0599899578333331 | 0.307846746966715 | -4.09373396067102 |
| lnc-BRD1-3 | lnc-BRD1-3 | -0.69256641114274 | 1.3132576000985 | -1.93229202329696 | 0.0600134172230791 | 0.30785224109484 | -4.22949869282785 |
| lnc-NPEPL1-3 | lnc-NPEPL1-3 | 0.51388914309639 | 1.67973693835212 | 1.93217065683864 | 0.0600222391734082 | 0.30785224109484 | -4.28494676946047 |
| lnc-COG7-1 | lnc-COG7-1 | 0.250558102304621 | 5.19030547389137 | 1.93205961959993 | 0.0600363512840939 | 0.30785224109484 | -4.6087491304773 |
| ENSG00000251667.1 | BRCC3P1 | 0.490151839132267 | 1.07694812843754 | 1.93188283425889 | 0.0600588254295444 | 0.30785224109484 | -4.20292158245319 |
| ENSG00000198690.9 | FAN1 | 0.235860588238711 | 4.69332186412009 | 1.93178455171014 | 0.0600713228942641 | 0.30785224109484 | -4.58912969204948 |
| lnc-TMUB2-2 | lnc-TMUB2-2 | -0.58811552065498 | -0.0191732615561554 | -1.93171028335843 | 0.0600807682299942 | 0.30785224109484 | -4.04756351649616 |
| ENSG00000088340.15 | FER1L4 | -1.0212831435965 | 4.60247914987235 | -1.93345399497234 | 0.060105426781916 | 0.307901941127969 | -4.57886367426314 |
| lnc-UBP1-5 | lnc-UBP1-5 | 0.225722330884593 | 2.13945008307948 | 1.9313529595118 | 0.0601262299597856 | 0.307931871426292 | -4.34005969537332 |
| ENSG00000109787.12 | KLF3 | 1.63729912306195 | 2.604586212758 | 1.9331550268771 | 0.0601547999414034 | 0.308001554028335 | -4.3206608463768 |
| lnc-TRIM28-14 | lnc-TRIM28-14 | -0.325486160575313 | 1.20584343591889 | -1.93092303507043 | 0.0601809676863732 | 0.308007899415928 | -4.25548951268714 |
| ENSG00000204392.10 | LSM2 | -0.283320592219542 | 3.36146182424277 | -1.93075677765016 | 0.0602021469511179 | 0.308007899415928 | -4.47191737467474 |
| lnc-GADL1-10 | lnc-GADL1-10 | 0.478096646171565 | 3.066285201422 | 1.93066602120864 | 0.0602137109674825 | 0.308007899415928 | -4.42853774970745 |
| lnc-CRK-4 | lnc-CRK-4 | -0.383820545333711 | 0.771297177837191 | -1.93064887441364 | 0.0602158959942522 | 0.308007899415928 | -4.18584904628749 |
| ENSG00000091972.18 | CD200 | -0.934574535161145 | 6.02489377338788 | -1.93215664722519 | 0.0602702672028446 | 0.308110479517156 | -4.62523966575194 |
| ENSG00000117394.21 | SLC2A1 | 0.265229316880625 | 6.03709472918089 | 1.93016469845329 | 0.0602776229418338 | 0.308110479517156 | -4.63282352795704 |
| ENSG00000216316.1 | RPL13AP15 | 0.885552339137081 | 1.78164864858446 | 1.93117408173309 | 0.0602808580389442 | 0.308110479517156 | -4.28732107624323 |
| ENSG00000171791.12 | BCL2 | -0.272715000978778 | 9.38093493943821 | -1.9295822548838 | 0.0603519498601383 | 0.308363635404571 | -4.58174972532051 |
| ENSG00000244187.7 | TMEM141 | -0.299542221929499 | 1.58942638054099 | -1.92951646696312 | 0.0603603501455093 | 0.308363635404571 | -4.29243347337241 |
| ENSG00000186446.11 | ZNF501 | -0.458864600151057 | 1.94313418955049 | -1.92914196393871 | 0.0604081885645676 | 0.30848855344304 | -4.3179238941113 |
| ENSG00000172819.16 | RARG | -0.431043314028658 | 2.51894023190226 | -1.92905868620425 | 0.0604188307511123 | 0.30848855344304 | -4.38789959563022 |
| ENSG00000228903.7 | RASA4CP | -0.505368476435705 | 1.39166936505966 | -1.92897313821697 | 0.0604297647321739 | 0.30848855344304 | -4.26994709097173 |
| ENSG00000273654.1 | | 0.87170136397552 | -0.749829745558761 | 1.92863053364233 | 0.060473570415795 | 0.308635631011235 | -3.94279835263394 |
| ENSG00000169155.9 | ZBTB43 | 0.308906352354684 | 5.55910706120546 | 1.92820064052085 | 0.0605285754303621 | 0.308839778900135 | -4.62665938071702 |
| ENSG00000189060.5 | H1-0 | -1.82625826694458 | -0.980291004805846 | -1.92895498499906 | 0.0605823417768315 | 0.309037507011778 | -3.95462083074576 |
| ENSG00000278637.1 | H4C1 | 0.503470659959742 | 2.36141406529607 | 1.9275547057168 | 0.0606113037944723 | 0.309039932310576 | -4.36300621867391 |
| lnc-CENPC-2 | lnc-CENPC-2 | -0.380916566996173 | 2.16984548932512 | -1.92744407097328 | 0.0606254831126701 | 0.309039932310576 | -4.35579158275575 |
| ENSG00000171827.10 | ZNF570 | 0.259191109204292 | 4.02145421294661 | 1.92742552575405 | 0.0606278602084296 | 0.309039932310576 | -4.5360107945716 |
| ENSG00000005486.16 | RHBDD2 | 0.237727706963678 | 4.5152734634715 | 1.92710609448903 | 0.0606688169381569 | 0.309109172626599 | -4.58511880118829 |
| ENSG00000204899.5 | MZT1 | 0.437943547402851 | 2.78517434971394 | 1.92701410996764 | 0.0606806153800281 | 0.309109172626599 | -4.42153642447973 |
| ENSG00000144451.18 | SPAG16 | -0.328148970515031 | 2.03976577845676 | -1.92696826110356 | 0.060686496943307 | 0.309109172626599 | -4.34931960400556 |
| ENSG00000116161.17 | CACYBP | -0.293775511028124 | 4.53567159371719 | -1.92651640470072 | 0.0607444879523181 | 0.309219678807309 | -4.5856672583454 |
| lnc-CCKAR-10 | lnc-CCKAR-10 | -0.738379678150546 | 0.29993784493784 | -1.92650662542273 | 0.0607457435450814 | 0.309219678807309 | -4.11844356018369 |
| lnc-UBE2E2-1 | lnc-UBE2E2-1 | 1.17404823591232 | 1.02464590611749 | 1.92773460494622 | 0.0607658028224797 | 0.309219678807309 | -4.2070049392169 |
| PLCG1-AS1 | PLCG1-AS1 | -0.436159782817857 | 0.356710328284327 | -1.92633109266423 | 0.0607682845443115 | 0.309219678807309 | -4.13506583877425 |
| ENSG00000268154.2 | | 0.544260150936657 | -0.0760666391436348 | 1.92618730692628 | 0.0607867541035594 | 0.30922964781759 | -4.02244154163501 |
| ENSG00000204253.4 | HNRNPCP2 | 0.953180722687978 | 1.43051889263749 | 1.92687802548156 | 0.0608002907602288 | 0.30922964781759 | -4.19818501386603 |
| ENSG00000244398.1 | RPL36AP37 | 0.894025393929299 | 2.92245980319235 | 1.92715241062157 | 0.0608303579253482 | 0.309306140607075 | -4.38314934752054 |
| ENSG00000284308.1 | C2orf81 | -0.530252512268772 | 0.913042622546994 | -1.92555232424095 | 0.0608683764796564 | 0.309386302618814 | -4.20219974154668 |
| ENSG00000224796.1 | RPL32P1 | 0.514655322012747 | 2.93323378106562 | 1.9253529991492 | 0.0608940176431102 | 0.309386302618814 | -4.41191300903594 |
| lnc-PEX19-4 | lnc-PEX19-4 | -0.237133602874927 | 3.09484609127971 | -1.92534955525444 | 0.060894460746869 | 0.309386302618814 | -4.45586538619068 |
| ENSG00000112695.11 | COX7A2 | 0.205208151192699 | 5.05684940937395 | 1.92496213347738 | 0.0609443254724159 | 0.309386302618814 | -4.6151248651489 |
| ENSG00000163684.11 | RPP14 | 0.172612219206183 | 3.48192838506035 | 1.92491984506965 | 0.0609497704969196 | 0.309386302618814 | -4.49228608745731 |
| ENSG00000157833.12 | GAREM2 | 0.737115617193844 | 1.34418615987138 | 1.92509680190874 | 0.0609509727594108 | 0.309386302618814 | -4.23799333169175 |
| ENSG00000112941.13 | TENT4A | 0.234620199748623 | 5.13540629667214 | 1.92490764646197 | 0.0609513412582855 | 0.309386302618814 | -4.61828203458485 |
| lnc-CCNB1IP1-4 | lnc-CCNB1IP1-4 | 0.289807249356299 | 1.65911243392631 | 1.92435419304464 | 0.061022643570532 | 0.309530459954914 | -4.30581921223103 |
| ENSG00000137878.17 | GCOM1 | 0.169910233799013 | 3.91812678657643 | 1.9243439850971 | 0.061023959348847 | 0.309530459954914 | -4.53245783226911 |
| lnc-ZNF569-1 | lnc-ZNF569-1 | -0.252418046258489 | 0.526661839183727 | -1.92433703060595 | 0.0610248557788973 | 0.309530459954914 | -4.15713780796434 |
| ENSG00000134077.15 | THUMPD3 | 0.198248714757989 | 4.86763870932621 | 1.92403932139609 | 0.0610632409232115 | 0.309606379201477 | -4.60785071837423 |
| ENSG00000244055.1 | | -0.2587985704592 | 2.55456419162464 | -1.92398763527756 | 0.0610699071835008 | 0.309606379201477 | -4.39944072570001 |
| ENSG00000238311.1 | SNORD13E | 0.372107376790399 | 1.29311363433574 | 1.92379198466645 | 0.0610951470380776 | 0.309658067344189 | -4.26056148641305 |
| ENSG00000113597.17 | TRAPPC13 | 0.246762111935521 | 4.27510018642708 | 1.92361121588752 | 0.0611184750172376 | 0.309700042166371 | -4.57219771185532 |
| ENSG00000205352.10 | PRR13 | 0.217680828161134 | 5.50333896006315 | 1.92345330460337 | 0.0611388595170759 | 0.30972708477479 | -4.63507284585644 |
| ENSG00000048052.21 | HDAC9 | -1.12784861719771 | 6.81422520804511 | -1.92517455196859 | 0.0611755634987563 | 0.309836767592249 | -4.63927757985878 |
| ENSG00000108375.12 | RNF43 | -0.480410381415921 | 1.95475882931314 | -1.92301832876934 | 0.0611950399635875 | 0.309859165454003 | -4.3287920884778 |
| lnc-ZNF761-4 | lnc-ZNF761-4 | -0.444604070232031 | 0.473602965130029 | -1.92273005213957 | 0.0612322974772352 | 0.309971563938498 | -4.14100141163812 |
| ENSG00000201428.1 | RN7SKP71 | 0.821847757652554 | -0.574659751544659 | 1.92250619647572 | 0.0612612424976726 | 0.309971829889339 | -4.01333615580669 |
| lnc-MGARP-2 | lnc-MGARP-2 | 0.217048466738799 | 3.35458759333813 | 1.92242742613842 | 0.0612714304626102 | 0.309971829889339 | -4.47825455608259 |
| ENSG00000271949.1 | | -0.303494958347878 | 1.53954998129067 | -1.92238028076861 | 0.0612775288257163 | 0.309971829889339 | -4.30862249901028 |
| lnc-RRM2B-2 | lnc-RRM2B-2 | 0.263830367148408 | 3.14203192166589 | 1.92201753898842 | 0.0613244677283188 | 0.310133051413264 | -4.46109081300547 |
| ENSG00000285982.1 | | 0.244947627924279 | 3.9724058807772 | 1.92124798675045 | 0.0614241501250245 | 0.310471633400868 | -4.54622327589076 |
| ENSG00000240857.1 | RDH14 | 0.260587682011623 | 3.32084919381655 | 1.92117418322479 | 0.0614337174123511 | 0.310471633400868 | -4.48224200552014 |
| ENSG00000196576.14 | PLXNB2 | 1.19994855181728 | 1.30091050622586 | 1.9225834971353 | 0.0614469631417265 | 0.310471633400868 | -4.27202992239824 |
| ENSG00000207304.1 | SNORA8 | 0.493657219954258 | 2.36218811928248 | 1.92102304451226 | 0.0614533137900048 | 0.310471633400868 | -4.37339486616065 |
| ENSG00000006555.10 | TTC22 | -0.342837066656103 | 1.66883523494439 | -1.92079504509549 | 0.0614828859305705 | 0.310471633400868 | -4.31923713548002 |

| | | | | | | | |
|---|---|---|---|---|---|---|---|
| ENSG00000106080.10 | FKBP14 | 0.388148440497672 | 2.02349673198209 | 1.92077889518677 | 0.0614849810799941 | 0.310471633400868 | -4.35014291958128 |
| ENSG00000219200.11 | RNASEK | -0.20941717658841 | 3.59272239320903 | -1.92065217788435 | 0.0615014224108111 | 0.310471633400868 | -4.51870246352106 |
| lnc-IGSF10-6 | lnc-IGSF10-6 | -0.639721688744225 | 2.28065652935627 | -1.92077222553861 | 0.0615120886658281 | 0.310471633400868 | -4.35423153811798 |
| ENSG00000172123.12 | SLFN12 | 0.542291501883325 | 1.73135186998939 | 1.91973641392484 | 0.0616203531363871 | 0.310941830989521 | -4.30117710361728 |
| ENSG00000263753.7 | LINC00667 | -0.309329309344077 | 4.66353331693018 | -1.9194497965425 | 0.0616576167640843 | 0.311053609278222 | -4.60273072386047 |
| ENSG00000278324.1 | | 0.619013611995949 | 0.652345779805427 | 1.91926663997189 | 0.06168143938461 | 0.311097541498022 | -4.16242205511024 |
| lnc-ROPN1B-15 | lnc-ROPN1B-15 | -1.4493932755026 | 0.529332610127245 | -1.92038833295222 | 0.0617263125846169 | 0.311238419533174 | -4.11556657132793 |
| ENSG00000149716.12 | LTO1 | -0.156210389011177 | 4.4549504920833 | -1.91881963102336 | 0.0617396136109386 | 0.311238419533174 | -4.59061197682918 |
| ENSG00000004864.13 | SLC25A13 | 0.226490845861852 | 3.90283238867343 | 1.91856067824392 | 0.0617733355115436 | 0.311332165728232 | -4.54546419437298 |
| lnc-FER-17 | lnc-FER-17 | -0.388807979510783 | 0.125589111934828 | -1.9184121728935 | 0.0617926816131651 | 0.311353431002149 | -4.10202199273346 |
| lnc-PIP4K2B-1 | lnc-PIP4K2B-1 | 0.489009491115902 | 0.799382635411269 | 1.9181774836684 | 0.0618232656550777 | 0.311396763976226 | -4.20057791647268 |
| ENSG00000186222.4 | BLOC1S4 | 0.273049572133656 | 2.92240935815188 | 1.9181140118985 | 0.0618315393465888 | 0.311396763976226 | -4.4549468086306 |
| ENSG00000239559.2 | RPL37P2 | 1.16296511999911 | -0.605703529236082 | 1.91784500338542 | 0.0618666157605898 | 0.311464545084623 | -3.955143611477 |
| ENSG00000235821.1 | IFITM4P | -0.527458454919907 | 0.233595273935044 | -1.91777871096246 | 0.0618752623451888 | 0.311464545084623 | -4.10944032452941 |
| ENSG00000151445.15 | VIPAS39 | -0.263928938327066 | 3.94474178227398 | -1.91723876075872 | 0.0619457271471175 | 0.311693178003153 | -4.55473731318768 |
| ENSG00000255185.5 | PDXDC2P | 0.320008272536578 | 4.30972239304833 | 1.91719861646401 | 0.0619509688194578 | 0.311693178003153 | -4.57546015040294 |
| lnc-IGSF10-11 | lnc-IGSF10-11 | -2.63700698603619 | -1.40665924030813 | -1.91797322419517 | 0.061995246928467 | 0.311725759850973 | -3.96379514616664 |
| lnc-ZNF595-2 | lnc-ZNF595-2 | -1.08961132190157 | -1.20850524313043 | -1.9167498775995 | 0.0620095868817003 | 0.311725759850973 | -3.96659534422526 |
| lnc-ASF1A-2 | lnc-ASF1A-2 | -0.515800629956029 | -0.126538143587881 | -1.91667337599184 | 0.0620195849083127 | 0.311725759850973 | -4.04589717732954 |
| ENSG00000121406.8 | ZNF549 | 0.247284419176292 | 4.54210517450883 | 1.91654761701028 | 0.0620360234029655 | 0.311725759850973 | -4.59589269461609 |
| lnc-P4HA3-2 | lnc-P4HA3-2 | 0.295251435890719 | 0.855368039643467 | 1.91641703269346 | 0.0620530965885494 | 0.311725759850973 | -4.21522270755786 |
| ENSG00000145949.10 | MYLK4 | 0.412843622986243 | 0.663712874589787 | 1.91639750049649 | 0.0620556506626256 | 0.311725759850973 | -4.18047115752035 |
| lnc-RSBN1-2 | lnc-RSBN1-2 | -0.330658732696847 | 3.03666204422635 | -1.91633779549461 | 0.0620634583816396 | 0.311725759850973 | -4.46883299178495 |
| ENSG00000047249.17 | ATP6V1H | 0.234957933582261 | 4.10515266457612 | 1.91615334395131 | 0.0620875847232169 | 0.311770860297139 | -4.56610135479262 |
| lnc-BTLA-3 | lnc-BTLA-3 | 0.201896776309625 | 1.72119487923748 | 1.91559615936162 | 0.0621605135127443 | 0.312000471043835 | -4.33152368320939 |
| ENSG00000116353.15 | MECR | -0.23070726732645 | 2.01091918929739 | -1.91552510797696 | 0.0621698185618494 | 0.312000471043835 | -4.356313421726 |
| lnc-OPTC-2 | lnc-OPTC-2 | -1.03670467519254 | -0.539290205767575 | -1.91540101610232 | 0.0621860727748294 | 0.312000471043835 | -3.97262400010296 |
| ENSG00000163517.14 | HDAC11 | -0.480066669132929 | 0.473279487981238 | -1.91534093984002 | 0.0621939431906357 | 0.312000471043835 | -4.1603851233712 |
| ENSG00000031823.14 | RANBP3 | -0.208634402682263 | 5.45689558485752 | -1.91486407207618 | 0.0622564465199317 | 0.312217901622005 | -4.65079663740742 |
| ENSG00000110756.17 | HPS5 | 0.1845996047534 | 5.65644695532063 | 1.91477884387557 | 0.0622676230947058 | 0.312217901622005 | -4.65336074749045 |
| ENSG00000256742.1 | KDM2B-DT | -0.949563012812241 | -0.342618572861856 | -1.91456314988653 | 0.0622959162474743 | 0.312283693161657 | -4.00444247105752 |
| ENSG00000176454.13 | LPCAT4 | -0.297609976657644 | 3.32665983855096 | -1.91431672927526 | 0.0623282533691105 | 0.312369719770246 | -4.49780752075358 |
| ENSG00000061273.17 | HDAC7 | -0.30961627645529 | 6.43032965312006 | -1.9141084551229 | 0.0623555958430525 | 0.312430678977008 | -4.66123642262903 |
| lnc-GVQW2-1 | lnc-GVQW2-1 | -0.442039518725444 | 0.736564966549096 | -1.91357827222442 | 0.0624252452415285 | 0.312633908595375 | -4.21344266775681 |
| ZNF252P-AS1 | ZNF252P-AS1 | -0.538830296326769 | -0.481222175263716 | -1.91356846120742 | 0.0624265347289992 | 0.312633908595375 | -4.00036420325261 |
| ENSG00000275538.1 | RNVU1-19 | -1.25837583619849 | 4.23338979734136 | -1.91548459025232 | 0.0624454261675279 | 0.31265244631628 | -4.61934361620658 |
| ENSG00000196968.10 | FUT11 | -0.274909247328685 | 4.00241210811017 | -1.91316921988608 | 0.062479027409975 | 0.312727023901203 | -4.56935979988755 |
| lnc-RTKN2-6 | lnc-RTKN2-6 | 0.706878730038502 | 1.4607019620879 | 1.91319029075166 | 0.0625079566649799 | 0.312727023901203 | -4.30004546996207 |
| ENSG00000186432.8 | KPNA4 | 0.184995479110876 | 6.55314543128169 | 1.91274498402459 | 0.0625348478218341 | 0.312727023901203 | -4.66401309594609 |
| ENSG00000282458.1 | WASH5P | -0.496746279422302 | 0.975411725557644 | -1.91267167852624 | 0.0625444975917662 | 0.312727023901203 | -4.26138530512083 |
| ENSG00000275183.1 | LENG9 | -0.561035542919008 | -0.365929862775799 | -1.91255541378948 | 0.0625598050337161 | 0.312727023901203 | -4.04978469663046 |
| lnc-SLC39A1-1 | lnc-SLC39A1-1 | 0.217570070344888 | 1.85146620317134 | 1.91246479719101 | 0.062571737860484 | 0.312727023901203 | -4.34564231774853 |
| lnc-PGAP1-2 | lnc-PGAP1-2 | -0.403126064389674 | 4.84771769616886 | -1.9124083367719 | 0.062579173823827 | 0.312727023901203 | -4.63371004439123 |
| ENSG00000284135.1 | MIR3654 | 1.05770813093472 | 1.20822107691007 | 1.91335582127113 | 0.0625910290356407 | 0.312727023901203 | -4.1960197092359 |
| ENSG00000278384.1 | | 1.98139527515424 | 0.804967257578859 | 1.91411099716919 | 0.0626037270460815 | 0.312727023901203 | -4.08968099182183 |
| ENSG00000154451.14 | GBP5 | 0.425983554097044 | 4.32982881954406 | 1.91215721711477 | 0.0626122560120905 | 0.312727023901203 | -4.59971610735871 |
| lnc-JPT1-1 | lnc-JPT1-1 | -0.390105404251718 | 3.25964320133158 | -1.91174813577242 | 0.062666179942979 | 0.312920422553696 | -4.49598345376901 |
| ENSG00000256590.2 | TRDV3 | -0.722141945783433 | 1.88186110936888 | -1.91204138829263 | 0.0626903544130599 | 0.312965210983267 | -4.34240410897745 |
| lnc-HARBI1-1 | lnc-HARBI1-1 | 0.214622115389384 | 0.801745375213354 | 1.91106978601841 | 0.0627556857068771 | 0.313215392556899 | -4.22205433007129 |
| lnc-GPR149-6 | lnc-GPR149-6 | 0.365594169656686 | 2.16856088617716 | 1.9108175784335 | 0.0627889914745848 | 0.31330565127791 | -4.3772426809591 |
| ENSG00000172732.11 | MUS81 | -0.184454050313665 | 4.673388008304 | -1.9104804660735 | 0.0628335331408647 | 0.313400486757847 | -4.62363283089281 |
| lnc-SLC25A24-2 | lnc-SLC25A24-2 | -0.323048104740858 | 1.39848597179875 | -1.91044326825877 | 0.0628384496356039 | 0.313400486757847 | -4.3049746838601 |
| ENSG00000170917.13 | NUDT6 | -0.319295928991342 | 2.29905263245779 | -1.91029987069967 | 0.0628574058027046 | 0.313419085183399 | -4.40671269027781 |
| ENSG00000167925.15 | GHDC | -0.25266260018372 | 4.12494077721793 | -1.9100850112999 | 0.0628858178943795 | 0.313428508447617 | -4.58673964232941 |
| HEXA-AS1 | HEXA-AS1 | -0.465744230834289 | 0.155611634860272 | -1.91000498764267 | 0.0628964026913935 | 0.313428508447617 | -4.1314075559233 |
| ENSG00000285772.1 | | -0.412271126962521 | 3.80159743167986 | -1.90982965159271 | 0.0629195998712729 | 0.313428508447617 | -4.56615690048958 |
| ENSG00000158545.15 | ZC3H18 | -0.258208448495076 | 4.81226304009752 | -1.9098248533726 | 0.0629202347847674 | 0.313428508447617 | -4.63107565473661 |
| lnc-RNF39-4 | lnc-RNF39-4 | -0.380283766413736 | 3.69035234171695 | -1.90971000562037 | 0.0629354333874557 | 0.313428508447617 | -4.5378396927657 |
| ENSG00000072778.19 | ACADVL | -0.320850047389519 | 6.42181320050454 | -1.90946104695441 | 0.0629683906114005 | 0.31351678373354 | -4.66922893185248 |
| ENSG00000143621.16 | ILF2 | 0.198240939862666 | 6.80498325130098 | 1.90931119816158 | 0.0629882347613833 | 0.313539742707026 | -4.66849570644251 |
| ENSG00000130363.11 | RSPH3 | -0.273829859197453 | 1.55438530962464 | -1.90913920388126 | 0.0630110181880651 | 0.313577318028275 | -4.32031043054342 |
| ENSG00000100697.14 | DICER1 | 0.131267894425967 | 7.71153924854199 | 1.90898420768489 | 0.0630315559856104 | 0.313603702405564 | -4.65763950668425 |
| lnc-CYTIP-1 | lnc-CYTIP-1 | 0.393908702985425 | 4.39814871094015 | 1.90885913148356 | 0.0630481334062039 | 0.313610374552899 | -4.5930097316202 |
| ENSG00000142347.17 | MYO1F | -0.872838085425492 | 3.65291924213397 | -1.90997744393557 | 0.0630952569284751 | 0.313698324456133 | -4.52574657496827 |
| lnc-RNF113A-5 | lnc-RNF113A-5 | -0.342411584616727 | -0.345617374340674 | -1.90834092300213 | 0.0631168557807231 | 0.313698324456133 | -4.05478598603969 |
| lnc-ARPC2-1 | lnc-ARPC2-1 | -0.280697114796164 | 1.80501530301091 | -1.90831608590508 | 0.0631201511692348 | 0.313698324456133 | -4.3486066718088 |
| ENSG00000146433.8 | TMEM181 | 0.203671411153533 | 3.92967717480827 | 1.90826614761141 | 0.0631267774327018 | 0.313698324456133 | -4.56685305117344 |
| lnc-PHLDA1-2 | lnc-PHLDA1-2 | 0.229593643968869 | 5.2685440432896 | 1.90806809297296 | 0.0631530629723776 | 0.31374434639535 | -4.65484115507687 |

| | | | | | | | |
|---|---|---|---|---|---|---|---|
| ENSG00000182487.12 | NCF1B | -0.409937113384746 | 4.64642649487065 | -1.90796669206222 | 0.0631665243872287 | 0.31374434639535 | -4.62662023748794 |
| ENSG00000172575.11 | RASGRP1 | 1.62029273149842 | 2.88428276057152 | 1.90978278536989 | 0.0631968473953942 | 0.313753599276182 | -4.40330121805633 |
| ENSG00000196421.8 | C20orf204 | -0.396079084888145 | 1.17382640048719 | -1.90772308781313 | 0.063198873954188 | 0.313753599276182 | -4.28730939914797 |
| ENSG00000181894.14 | ZNF329 | 0.173942104576614 | 4.36320041936323 | 1.9075640534765 | 0.0632200006591764 | 0.313782800474519 | -4.60086320443166 |
| lnc-ZNF316-3 | lnc-ZNF316-3 | 0.411535083460136 | 0.507335841592562 | 1.90737018864311 | 0.0632457625491294 | 0.313834988078286 | -4.18638512840962 |
| ENSG00000121851.12 | POLR3GL | -0.353573957889564 | 4.13632939073355 | -1.90706606176509 | 0.0632861948031475 | 0.313959929533185 | -4.58264803176479 |
| ENSG00000256682.2 | | -0.546067249830228 | -0.253559705755664 | -1.9067724894851 | 0.0633252448504276 | 0.314077955362976 | -4.07003176621463 |
| ENSG00000278420.1 | MIR6819 | 0.47801004674392 | 0.42973639310522 | 1.90661362068568 | 0.0633463856672157 | 0.31410712025736 | -4.12298249144716 |
| lnc-RESP18-5 | lnc-RESP18-5 | 0.29022335801307 | 1.05798954266313 | 1.90618172968861 | 0.063403888324384 | 0.314316530197687 | -4.27063798746138 |
| LINC00526 | LINC00526 | -0.359095347169843 | 0.780015208695586 | -1.9056782861603 | 0.0634709739662844 | 0.314573334251874 | -4.23436604687831 |
| lnc-NOBOX-1 | lnc-NOBOX-1 | -0.456830746874855 | 2.41300573196671 | -1.90541013431685 | 0.063506730934237 | 0.314674781612759 | -4.41064299893534 |
| lnc-COX18-3 | lnc-COX18-3 | 0.246300275852779 | 1.59634327263843 | 1.90482761124361 | 0.0635844674247155 | 0.314887080616962 | -4.32641314054509 |
| ENSG00000137710.15 | RDX | 0.462900895901089 | 3.84472458589824 | 1.90473916004743 | 0.0635962781679449 | 0.314887080616962 | -4.55032642750999 |
| ENSG00000197381.15 | ADARB1 | -0.961050891992923 | 3.96288692463629 | -1.90646631175994 | 0.0635963358974076 | 0.314887080616962 | -4.55260878355873 |
| ENSG00000163626.16 | COX18 | -0.249662470939531 | 3.40571705590665 | -1.90458593555832 | 0.0636167424241127 | 0.314887080616962 | -4.52913315883923 |
| ENSG00000196689.12 | TRPV1 | -0.269834369756384 | 2.34052730706501 | -1.90451612116519 | 0.063626068517026 | 0.314887080616962 | -4.4172908177058 |
| lnc-MTFR2-4 | lnc-MTFR2-4 | 0.214325988890506 | 4.73657595168719 | 1.90417533753487 | 0.0636716085804821 | 0.315036711397131 | -4.63500230240539 |
| ENSG00000100865.14 | CINP | -0.214884122973586 | 3.32161883858213 | -1.90380662995369 | 0.0637209116135792 | 0.315204884340856 | -4.51556871914361 |
| lnc-FUT8-1 | lnc-FUT8-1 | 1.3646009681105 | -0.35171769026491 | 1.90439445629616 | 0.063745160999506 | 0.315219450821266 | -4.03800454802681 |
| ADPGK-AS1 | ADPGK-AS1 | -0.35953152205374 | 1.2940408939746 | -1.90340274346495 | 0.063774956189909 | 0.315219450821266 | -4.29130524177263 |
| ENSG00000105939.12 | ZC3HAV1 | 0.188445057327661 | 7.66607226097049 | 1.90338273077024 | 0.0637776351337131 | 0.315219450821266 | -4.66971062225756 |
| lnc-ZNF780B-2 | lnc-ZNF780B-2 | -0.444924607849658 | 0.95183718259036 | -1.90332686062808 | 0.063785114544555 | 0.315219450821266 | -4.25036844924346 |
| ENSG00000159210.9 | SNF8 | 0.15251906821566 | 4.10458318681846 | 1.90287987521936 | 0.0638449800940162 | 0.31524807044706 | -4.58698835301491 |
| ENSG00000083937.8 | CHMP2B | 0.25263510736796 | 4.86909472036925 | 1.90274545768836 | 0.0638629922677254 | 0.31524807044706 | -4.6447453860896 |
| ENSG00000157353.16 | FCSK | -0.217152535031788 | 3.85095731949704 | -1.90268486447208 | 0.0638711132808757 | 0.31524807044706 | -4.56659549693033 |
| lnc-RMND1-1 | lnc-RMND1-1 | -0.349050938607962 | 1.83858909961876 | -1.90267872993961 | 0.0638719355115618 | 0.31524807044706 | -4.36834773255298 |
| ENSG00000008300.16 | CELSR3 | -0.648881012735525 | 4.52695822281788 | -1.9037346933606 | 0.0638826532891823 | 0.31524807044706 | -4.62693988114016 |
| ENSG00000105821.14 | DNAJC2 | 0.20336248676218 | 5.34540238007151 | 1.90234302260566 | 0.0639169452254688 | 0.31524807044706 | -4.66822609825789 |
| lnc-SRRM2-1 | lnc-SRRM2-1 | -0.408364964822429 | 1.77579911042792 | -1.90234109063625 | 0.0639172043313219 | 0.31524807044706 | -4.36693042668035 |
| lnc-XPO1-3 | lnc-XPO1-3 | 0.660968864856028 | -0.561372663194944 | 1.90227577354259 | 0.0639259648542849 | 0.31524807044706 | -4.00749084292131 |
| ENSG00000140450.8 | ARRDC4 | 3.21924178591346 | -2.08064092049119 | 1.90328069126683 | 0.0639378542275447 | 0.31524807044706 | -4.03265153616275 |
| lnc-CHST7-4 | lnc-CHST7-4 | 0.331151275584948 | 0.552875550640507 | 1.9020269951571 | 0.0639593411552389 | 0.31524807044706 | -4.18035830540785 |
| lnc-VAX1-4 | lnc-VAX1-4 | 0.241396739540778 | 1.74152252566725 | 1.90202669712153 | 0.063959381148857 | 0.31524807044706 | -4.34643817649758 |
| ENSG00000198258.10 | UBL5 | 0.19571057425619 | 4.90924665070509 | 1.90162419156502 | 0.0640134131925763 | 0.315373675008296 | -4.64842971322287 |
| lnc-TTLL7-10 | lnc-TTLL7-10 | -1.28906173247095 | -1.13637316153324 | -1.90160858767565 | 0.0640155086325929 | 0.315373675008296 | -3.97469863447058 |
| ERICD | ERICD | -0.606136985284442 | -0.685160916428431 | -1.90129405899938 | 0.0640577590696559 | 0.315437199595923 | -4.01275811580936 |
| ENSG00000157326.18 | DHRS4 | -0.277966034988415 | 2.1285437313958 | -1.90128442696021 | 0.0640590533115171 | 0.315437199595923 | -4.39037747382479 |
| ENSG00000197629.5 | MPEG1 | 0.845649393539216 | 6.18332736661902 | 1.90271507658074 | 0.0641041760411248 | 0.315549226618623 | -4.67867335806247 |
| FAM53B-AS1 | FAM53B-AS1 | -0.70223390127109 | 0.777715442074539 | -1.90091405694621 | 0.0641124649331527 | 0.315549226618623 | -4.2567107993542 |
| lnc-OTOGL-1 | lnc-OTOGL-1 | -0.413115582402945 | 0.902443701425268 | -1.90031039079122 | 0.0641900485092233 | 0.315753217798559 | -4.26414310027437 |
| ENSG00000134910.13 | STT3A | -0.190413892730882 | 5.12828749806689 | -1.90028751268973 | 0.0641931280669597 | 0.315753217798559 | -4.66631132855594 |
| lnc-MDH1B-8 | lnc-MDH1B-8 | -1.08920024611771 | 0.609039295793941 | -1.90122135286475 | 0.0642070772893299 | 0.315753217798559 | -4.19831355560321 |
| ENSG00000105229.6 | PIAS4 | -0.267031532960748 | 3.23448178049696 | -1.9001230231258 | 0.0642152732694344 | 0.315753217798559 | -4.51572923776943 |
| ENSG00000005469.11 | CROT | 0.252922438176927 | 2.6797451709925 | 1.89974860629747 | 0.0642657052976028 | 0.315925725373909 | -4.44235871151629 |
| ENSG00000148396.18 | SEC16A | 0.222610178441054 | 6.76665621328787 | 1.89954836157188 | 0.064292691129319 | 0.315950795841148 | -4.68620199530045 |
| lnc-PNPLA8-2 | lnc-PNPLA8-2 | 0.301772049756736 | 3.67365632549504 | 1.89948297270449 | 0.0643015053091663 | 0.315950795841148 | -4.54452510823425 |
| lnc-COG7-3 | lnc-COG7-3 | 0.280738865962847 | 5.50101442860276 | 1.89911714845873 | 0.0643508361342709 | 0.316117723186563 | -4.67775096790378 |
| lnc-BRD3OS-7 | lnc-BRD3OS-7 | -0.655756571941857 | -0.273770779797139 | -1.89894925079395 | 0.0643734876989278 | 0.316153542664526 | -4.06803340240238 |
| ENSG00000112365.4 | ZBTB24 | -0.260386548125133 | 6.40233147399341 | -1.89874500506542 | 0.0644010522747372 | 0.316213468265963 | -4.688211286269 |
| ENSG00000107566.13 | ERLIN1 | 0.305763799679799 | 1.82922563204004 | 1.89858227706151 | 0.0644230209254596 | 0.316232777881094 | -4.36313767083151 |
| lnc-TMEM189-UBE2V1-4 | lnc-TMEM189-UBE | 0.240038709320174 | 2.56546757675535 | 1.89848828857932 | 0.0644357125022255 | 0.316232777881094 | -4.43822067632624 |
| lnc-RPRML-6 | lnc-RPRML-6 | 0.343927305175244 | 1.78087870545207 | 1.8982887333259 | 0.0644626661958864 | 0.316289644412379 | -4.369152935335 |
| lnc-PDSS1-9 | lnc-PDSS1-9 | -0.463933548784057 | 1.45245255049213 | -1.89805568134965 | 0.0644941564501162 | 0.316368737419624 | -4.32479972247636 |
| ENSG00000169495.4 | HTRA4 | -0.509267752483344 | -0.13711510002961 | -1.89789613295869 | 0.0645157223946434 | 0.316399121765295 | -4.11664285793651 |
| lnc-DCTN6-1 | lnc-DCTN6-1 | -0.325893188838397 | 0.966418939107675 | -1.89714002296125 | 0.0646180086078564 | 0.316825267073727 | -4.27835670982816 |
| lnc-WDFY1-1 | lnc-WDFY1-1 | 0.291639342623486 | 2.39781791673341 | 1.89690226973119 | 0.0646502003805053 | 0.316907614773027 | -4.43290148927097 |
| ENSG00000166887.15 | VPS39 | 0.162444638568131 | 6.6616323157588 | 1.89677123405343 | 0.0646679484601962 | 0.316919138846719 | -4.6914669679832 |
| ENSG00000203950.6 | RTL8A | -0.342676971993301 | 0.324357277485136 | -1.89664632726104 | 0.0646848702907957 | 0.316926609187205 | -4.14691963361463 |
| ENSG00000169105.7 | CHST14 | -0.357939360528877 | 2.64862342164861 | -1.89575193114625 | 0.0648061497745617 | 0.317431604752076 | -4.47310473873166 |
| lnc-GATAD1-6 | lnc-GATAD1-6 | 0.522403024000588 | -0.460685847622159 | 1.89555251544276 | 0.0648332168745316 | 0.317431604752076 | -4.04317588512113 |
| ENSG00000113742.12 | CPEB4 | 0.329617688741528 | 6.46852623234969 | 1.89554522791871 | 0.0648342062079253 | 0.317431604752076 | -4.69394177357675 |
| ENSG00000169288.17 | MRPL1 | -0.241787107077702 | 3.75370385379831 | -1.8953032439881 | 0.0648670645786384 | 0.317516953679456 | -4.57456480861827 |
| ENSG00000163660.11 | CCNL1 | 0.293711708721892 | 9.13074116027477 | 1.8949438981478 | 0.064915885467821 | 0.317604381337954 | -4.64992556616621 |
| lnc-SETD9-6 | lnc-SETD9-6 | 0.364992375164249 | 1.8625367063671 | 1.89494148792377 | 0.0649162130278975 | 0.317604381337954 | -4.37589121027354 |
| ENSG00000107959.15 | PITRM1 | 0.135993750660952 | 6.13382889127202 | 1.89482413216136 | 0.0649321639018123 | 0.317604381337954 | -4.69506421487425 |
| lnc-ZDHHC19-2 | lnc-ZDHHC19-2 | -0.208486728721407 | 1.7681955169684 | -1.89464897106679 | 0.0649559778446424 | 0.317604381337954 | -4.36828035384594 |
| ENSG00000151702.16 | FLI1 | -0.363686780304864 | 6.21266538333861 | -1.89459412013118 | 0.0649634366111148 | 0.317604381337954 | -4.69506995145509 |
| ENSG00000165359.15 | INTS6L | 0.196348407620514 | 6.30393957727736 | 1.89411942283832 | 0.0650280176971406 | 0.317604381337954 | -4.69670694449043 |

| | | | | | | | |
|---|---|---|---|---|---|---|---|
| lnc-DOLPP1-3 | lnc-DOLPP1-3 | -0.312031342452607 | 1.11380957525075 | -1.89411923557392 | 0.065028043184695 | 0.317604381337954 | -4.30733362802425 |
| ENSG00000197114.11 | ZGPAT | -0.20654796928787 | 3.732754251528 | -1.89410547038667 | 0.0650299167139689 | 0.317604381337954 | -4.57650836753583 |
| ENSG00000117228.9 | GBP1 | 1.33600285643754 | 0.348101048055841 | 1.8952463080021 | 0.0650483287770598 | 0.317604381337954 | -4.15632620278211 |
| ENSG00000116396.14 | KCNC4 | -0.392481003709103 | 4.31280494894484 | -1.89393246753532 | 0.0650534674305268 | 0.317604381337954 | -4.61858963570356 |
| lnc-CLN6-1 | lnc-CLN6-1 | -0.611733828149901 | 0.686817283122148 | -1.89392370661613 | 0.0650546602400434 | 0.317604381337954 | -4.23315906194313 |
| ENSG00000100647.7 | SUSD6 | 0.183073797233 | 6.27670668612086 | 1.89357650156507 | 0.0651019476614755 | 0.317659562085778 | -4.69762238147912 |
| lnc-AMIGO1-2 | lnc-AMIGO1-2 | -0.542405897057554 | 0.225912861371203 | -1.89354424599673 | 0.0651063421827472 | 0.317659562085778 | -4.18326615471591 |
| lnc-RPUSD3-2 | lnc-RPUSD3-2 | -0.284330478532815 | 0.522331998951515 | -1.89350079645388 | 0.0651122621794636 | 0.317659562085778 | -4.20857294230673 |
| lnc-PLEKHO2-3 | lnc-PLEKHO2-3 | -0.522815574206064 | 0.803923640588285 | -1.89321715944921 | 0.0651509189961374 | 0.31774931195265 | -4.23634775296158 |
| lnc-FBF1-3 | lnc-FBF1-3 | -0.278341664884483 | 0.209016667764655 | -1.8930511140092 | 0.0651735583825044 | 0.31774931195265 | -4.17892805336823 |
| lnc-OXGR1-6 | lnc-OXGR1-6 | -0.416675430778698 | 2.3484751416899 | -1.89294599656269 | 0.0651878940433616 | 0.31774931195265 | -4.42188907693993 |
| lnc-OTUD7A-3 | lnc-OTUD7A-3 | -0.470778120547294 | 1.65127816542734 | -1.89279451120747 | 0.0652085579831985 | 0.31774931195265 | -4.35963580869923 |
| lnc-GCSH-4 | lnc-GCSH-4 | -0.547516264684278 | -0.367060318998183 | -1.8926976789434 | 0.0652217696894063 | 0.31774931195265 | -4.06123098366323 |
| lnc-PRR14L-5 | lnc-PRR14L-5 | 0.675694057548621 | -0.356398615255571 | 1.89268198982078 | 0.0652239105142333 | 0.31774931195265 | -4.04078578899709 |
| NBR2 | NBR2 | -0.251751183098897 | 2.99840528238613 | -1.89257346333422 | 0.0652387209013214 | 0.31774931195265 | -4.49301345653688 |
| lnc-FGD4-8 | lnc-FGD4-8 | -0.49838985515095 | 1.25502491615276 | -1.89178943783565 | 0.0653458005928353 | 0.318137663198752 | -4.3207396918308 |
| ENSG00000110218.8 | PANX1 | 0.361439225105824 | 3.35800088897533 | 1.89176333711623 | 0.0653493679251967 | 0.318137663198752 | -4.5315617042081 |
| lnc-XRN1-4 | lnc-XRN1-4 | 0.248068349053236 | 1.76612877810966 | 1.89149413554024 | 0.0653861709243984 | 0.318143443595055 | -4.37216009469476 |
| ENSG00000239002.3 | SCARNA10 | 0.705167545415788 | 7.19756377778641 | 1.89286291675972 | 0.0653923809442287 | 0.318143443595055 | -4.69597337309308 |
| lnc-SUSD1-1 | lnc-SUSD1-1 | -0.591593337953657 | 2.11996015249687 | -1.89142347686129 | 0.0654140092351714 | 0.318143443595055 | -4.425074016919 |
| lnc-GAPDH-2 | lnc-GAPDH-2 | 0.705162423440481 | 7.1972093818566 | 1.89261704704355 | 0.0654259492430223 | 0.318143443595055 | -4.69640346653131 |
| ENSG00000156650.13 | KAT6B | 0.2228225358144 | 6.72887552806092 | 1.89109681677364 | 0.06544052137376 | 0.318143443595055 | -4.70095421577365 |
| ENSG00000197497.10 | ZNF665 | -0.507266997168301 | 3.10688286686632 | -1.89107654854142 | 0.0654432949609612 | 0.318143443595055 | -4.49588461090069 |
| ENSG00000137770.13 | CTDSPL2 | 0.218122184729498 | 6.08113832692419 | 1.89064595605962 | 0.0655022427046446 | 0.318354819737827 | -4.702025017546 |
| ENSG00000144224.16 | UBXN4 | 0.1851573165709 | 6.756815614878 | 1.89026788659323 | 0.065554037460666 | 0.318531339247613 | -4.70175340321631 |
| ENSG00000015532.9 | XYLT2 | -0.316489715762842 | 3.57050544899071 | -1.89013147118923 | 0.065572734678397 | 0.318546990296305 | -4.56709937426709 |
| MIR155HG | MIR155HG | -0.550010406686686 | 7.23209253733057 | -1.89052642676558 | 0.0656340421837543 | 0.318712166705582 | -4.69835297763456 |
| ENSG00000285625.1 | | 0.293528242159289 | 6.04900085639013 | 1.88950785741146 | 0.0656582657274989 | 0.318712166705582 | -4.70323356171922 |
| lnc-GTF2F2-8 | lnc-GTF2F2-8 | -0.39475891135213 | 0.648550294082581 | -1.88943844684391 | 0.0656677915432654 | 0.318712166705582 | -4.24896888556144 |
| lnc-SELENOT-45 | lnc-SELENOT-45 | -0.602288112136616 | -0.302476771744894 | -1.88943202354378 | 0.0656686731282308 | 0.318712166705582 | -4.06735784144733 |
| lnc-CHRNA1-4 | lnc-CHRNA1-4 | 0.294946675994295 | 2.68179286494584 | 1.88922524098761 | 0.0656970590142944 | 0.318774767961156 | -4.46466119558671 |
| lnc-SYF2-1 | lnc-SYF2-1 | -0.267241986987178 | 3.53249657505564 | -1.8888590996635 | 0.0657473464200994 | 0.318876337565952 | -4.562519199682 |
| lnc-VSIG10-2 | lnc-VSIG10-2 | -0.485333973998335 | -0.375580036938956 | -1.88880386990411 | 0.0657549347609787 | 0.318876337565952 | -4.0597690964504 |
| ENSG00000197976.11 | AKAP17A | -0.323771967450635 | 6.37243226278329 | -1.88873448954868 | 0.0657644683946687 | 0.318876337565952 | -4.70552132392418 |
| lnc-NBN-2 | lnc-NBN-2 | -0.418007086146783 | 0.261325210966628 | -1.88856187979827 | 0.0657881920159415 | 0.318916240288301 | -4.17105409502655 |
| ENSG00000274430.1 | | 0.294187245666419 | 0.422752944379076 | 1.88816158867974 | 0.0658432364569774 | 0.319052288027825 | -4.19957524413259 |
| ENSG00000100519.11 | PSMC6 | 0.145561308583567 | 6.0268414534928 | 1.88813235176456 | 0.0658472583948986 | 0.319052288027825 | -4.70641881800934 |
| ENSG00000262664.2 | OVCA2 | 0.178349322748773 | 3.23649279868935 | 1.88752008942704 | 0.065931531642594 | 0.319385435584729 | -4.53225090678109 |
| ENSG00000127870.16 | RNF6 | 0.187524286743865 | 5.74289167999483 | 1.88740626538567 | 0.0659472087968123 | 0.319386211450538 | -4.7034068243953 |
| ENSG00000204271.12 | SPIN3 | -0.774867590591996 | 4.62191819751769 | -1.88837374658898 | 0.066012971720634 | 0.319552953547379 | -4.66364408900557 |
| ENSG00000001461.16 | NIPAL3 | -0.312490149128566 | 4.69739954227162 | -1.88688383053163 | 0.0660192053381636 | 0.319552953547379 | -4.66228832764781 |
| ENSG00000196605.7 | ZNF846 | -0.318642652525931 | 2.78972796683242 | -1.88681850051402 | 0.066028213158286 | 0.319552953547379 | -4.47927163702446 |
| ENSG00000100575.13 | TIMM9 | 0.215325510862017 | 3.06323351931735 | 1.8864437123631 | 0.0660799098894591 | 0.319727970205627 | -4.50910417760515 |
| ENSG00000196118.11 | CFAP119 | 0.471989113531546 | 1.05515165084191 | 1.88622221792976 | 0.0661104781507184 | 0.319800698419797 | -4.29560220714267 |
| ENSG00000173230.15 | GOLGB1 | 0.239448408384609 | 7.37049653201015 | 1.88586276523424 | 0.0661601115933634 | 0.319840008762119 | -4.70393875767729 |
| ENSG00000110700.6 | RPS13 | 0.265588296377191 | 8.07460272467335 | 1.88575780079898 | 0.0661746111410617 | 0.319840008762119 | -4.69206888625133 |
| lnc-COL4A4-1 | lnc-COL4A4-1 | -1.46122905687762 | -1.2808982951615 | -1.88628539691841 | 0.0661756141214157 | 0.319840008762119 | -4.01845660634388 |
| lnc-ADH5-5 | lnc-ADH5-5 | 0.189171332734754 | 2.64574986995745 | 1.88570630523754 | 0.0661817256117029 | 0.319840008762119 | -4.47064145027055 |
| ENSG00000248503.2 | C4orf46P2 | 0.317357572002053 | 0.354379965557177 | 1.88560083042441 | 0.0661962997292245 | 0.319840008762119 | -4.18071573431567 |
| ENSG00000144959.9 | NCEH1 | -0.360523770749472 | 2.57523421037181 | -1.88512462484319 | 0.066262134137854 | 0.320050490089436 | -4.46644035939235 |
| lnc-SYNE2-3 | lnc-SYNE2-3 | -0.377024773106405 | 3.20878243179031 | -1.8850608079737 | 0.0662709609270358 | 0.320050490089436 | -4.53484043880817 |
| ENSG00000105948.13 | IFT56 | -0.363779005353296 | 2.29328445291423 | -1.8846757708756 | 0.0663242383706386 | 0.32023265268188 | -4.44259253372085 |
| lnc-C12orf73-4 | lnc-C12orf73-4 | -0.592555818826287 | -0.446760284114371 | -1.88436836396564 | 0.0663668003785399 | 0.320345539108046 | -4.07727225050663 |
| ENSG00000169609.13 | C15orf40 | -0.195322165875589 | 4.12046268369472 | -1.88418336726435 | 0.066392425314615 | 0.320345539108046 | -4.6237139932958 |
| lnc-RPGR-5 | lnc-RPGR-5 | -1.32659100511293 | 1.7435869310886 | -1.88587546348874 | 0.0664038055119978 | 0.320345539108046 | -4.35949855768667 |
| lnc-FRYL-3 | lnc-FRYL-3 | 0.22878475536725 | 2.90928736574735 | 1.88405744047233 | 0.06640987297158 | 0.320345539108046 | -4.49968978915294 |
| lnc-TCEA1-3 | lnc-TCEA1-3 | 0.258794896750675 | 1.67714823704716 | 1.88368086405436 | 0.0664620724657307 | 0.320522220609685 | -4.3773900216175 |
| DLGAP1-AS1 | DLGAP1-AS1 | -0.350900093861719 | 4.22014539971877 | -1.88339977590832 | 0.0665010585688488 | 0.320594181739551 | -4.63089137277372 |
| ENSG00000171988.18 | JMJD1C | 0.230325600380631 | 9.23872593354024 | 1.88323511178235 | 0.066523906059403 | 0.320594181739551 | -4.67078247709914 |
| lnc-LETM2-3 | lnc-LETM2-3 | -0.507532443421456 | 0.869473086495801 | -1.88318121787859 | 0.066531385404803 | 0.320594181739551 | -4.28251938355511 |
| lnc-CDC42SE2-2 | lnc-CDC42SE2-2 | -0.265620935035934 | 0.607148828802502 | -1.88269612488587 | 0.0665987384391589 | 0.320594181739551 | -4.23899674185897 |
| ENSG00000129226.13 | CD68 | -0.569030666669672 | 2.77203743272433 | -1.88282275034395 | 0.0666032200227506 | 0.320594181739551 | -4.48164403414409 |
| ENSG00000112343.10 | TRIM38 | 0.218891425298352 | 6.94549801925117 | 1.88254048315277 | 0.0666203609286003 | 0.320594181739551 | -4.71476167342896 |
| lnc-DMAC1-1 | lnc-DMAC1-1 | -0.370954109272773 | 0.794966627694846 | -1.88248146192747 | 0.0666285620031608 | 0.320594181739551 | -4.25595373501708 |
| ENSG00000267221.2 | C17orf113 | -0.599611806682455 | 0.0430925697588714 | -1.88246139466084 | 0.0666313505715709 | 0.320594181739551 | -4.175714875378 |
| ENSG00000089050.15 | RBBP9 | -0.320464355356939 | 3.31160112168393 | -1.88242142677011 | 0.0666369048480818 | 0.320594181739551 | -4.55344990795094 |
| ENSG00000145388.14 | METTL14 | 0.191078656837099 | 5.31243832463362 | 1.88232828573007 | 0.0666498500477821 | 0.320594181739551 | -4.70070356243227 |
| ENSG00000156256.14 | USP16 | 0.251920324056383 | 7.01872683051009 | 1.88222192483584 | 0.0666646352303696 | 0.320594181739551 | -4.71355364583005 |

| | | | | | | | |
|---|---|---|---|---|---|---|---|
| ENSG00000153774.8 | CFDP1 | -0.221473923049145 | 3.31112569928043 | -1.88207798153586 | 0.0666846491808686 | 0.320594181739551 | -4.54439161636207 |
| ENSG00000149582.15 | TMEM25 | -0.388326561779999 | 1.89186871560125 | -1.88203759053564 | 0.0666902660867912 | 0.320594181739551 | -4.41225414183037 |
| ENSG00000067208.14 | EVI5 | 0.170015487553106 | 4.54795708329614 | 1.88195238587575 | 0.0667021162514108 | 0.320594181739551 | -4.66222578654962 |
| lnc-SPATA21-3 | lnc-SPATA21-3 | -0.399239778537053 | 0.12563379297257 | -1.88189118447341 | 0.0667106291789582 | 0.320594181739551 | -4.15475174678649 |
| lnc-MRPS25-5 | lnc-MRPS25-5 | -0.227153325778111 | 4.62957528428806 | -1.88166206912865 | 0.0667425066476994 | 0.320662147740533 | -4.6695817036465 |
| lnc-BLCAP-1 | lnc-BLCAP-1 | -0.242511799800244 | 4.36154419716922 | -1.88156561980792 | 0.0667559297997466 | 0.320662147740533 | -4.64833314217238 |
| ENSG00000171469.10 | ZNF561 | 0.175142851535395 | 4.28884669163409 | 1.88122947396114 | 0.0668027302554515 | 0.320812085125515 | -4.64132536532644 |
| ENSG00000227008.2 | RPS24P19 | 0.951139289506031 | 0.613342776584465 | 1.88121195563051 | 0.0668312202035257 | 0.320835014733382 | -4.1447501887188 |
| lnc-CC2D1B-1 | lnc-CC2D1B-1 | 0.187611710187418 | 2.28507648907795 | 1.88093909575665 | 0.0668431811537181 | 0.320835014733382 | -4.43932947654237 |
| ENSG00000254893.6 | RAP1BL | 1.13567360261888 | -1.00863430992065 | 1.88077948446169 | 0.0668654245673385 | 0.320835014733382 | -3.99436164359745 |
| lnc-DKK4-3 | lnc-DKK4-3 | -0.243933640094957 | 3.32643253687643 | -1.88068845370796 | 0.0668781134294261 | 0.320835014733382 | -4.55378544904812 |
| SMIM15-AS1 | SMIM15-AS1 | -0.386167677103885 | 1.4622434209794 | -1.88063588631142 | 0.0668854417830705 | 0.320835014733382 | -4.35947513819159 |
| ENSG00000137767.13 | SQOR | 0.378406851927471 | 3.46614352969165 | 1.88041200366673 | 0.0669166606439926 | 0.320909978107013 | -4.56330151033686 |
| lnc-SLC38A1-8 | lnc-SLC38A1-8 | 0.29231515510924 | 3.14006260746373 | 1.88028482603398 | 0.0669344002026951 | 0.32092027937854 | -4.53115933532406 |
| ENSG00000117153.15 | KLHL12 | 0.190579375594965 | 5.16606207386476 | 1.88013762189686 | 0.0669549381932757 | 0.320939581977297 | -4.69882952470292 |
| ENSG00000075884.13 | ARHGAP15 | 0.229547309898914 | 6.991844011645 | 1.87999701270353 | 0.0669745610704412 | 0.320939581977297 | -4.7184506947564 |
| ENSG00000123728.9 | RAP2C | 0.241382581431472 | 5.61147472784622 | 1.87992076808065 | 0.0669852035259423 | 0.320939581977297 | -4.71459669894652 |
| lnc-RELT-1 | lnc-RELT-1 | -0.635883254669515 | -0.209784265497461 | -1.87915808608268 | 0.0670917403463721 | 0.321303513853111 | -4.09967135113228 |
| ENSG00000189149.12 | ABCA15P | 0.456083892642141 | 1.3418855573031 | 1.87915349437086 | 0.0670923821863028 | 0.321303513853111 | -4.30714989850069 |
| ENSG00000155903.11 | RASA2 | 0.171302625284413 | 6.17471652645668 | 1.87881979969845 | 0.0671390407998061 | 0.321452169523705 | -4.72307795413611 |
| ENSG00000121101.15 | TEX14 | -0.774578185715579 | 0.38801748964199 | -1.8781517952553 | 0.067234288562115 | 0.321833339877677 | -4.21888852272811 |
| ENSG00000137106.17 | GRHPR | -0.43666720417078 | 5.2322526762144 | -1.87805888818992 | 0.0672672300046266 | 0.321916157913329 | -4.70584506086779 |
| ENSG00000267281.2 | ATF7-NPFF | -0.27650469029209 | 1.51443534090728 | -1.87758435600115 | 0.0673120266196447 | 0.322010678853944 | -4.37047901647176 |
| ENSG00000173409.13 | ARV1 | -0.269964699357068 | 2.7883226498698 | -1.8775021646775 | 0.0673235484442968 | 0.322010678853944 | -4.50405762831855 |
| PAXIP1-AS1 | PAXIP1-AS1 | -0.350379419259965 | 3.43677908941554 | -1.87736289388142 | 0.0673430756760386 | 0.322010678853944 | -4.56860060346952 |
| ENSG00000141759.14 | TXNL4A | 0.238675640990154 | 4.54455443736685 | 1.8773166617172 | 0.0673495589790715 | 0.322010678853944 | -4.66982328321556 |
| ENSG00000239280.1 | DBTP1 | 0.374078020907764 | 0.816467590784326 | 1.87672253231016 | 0.067432923234135 | 0.322259544678012 | -4.28143765905068 |
| lnc-USF3-2 | lnc-USF3-2 | 0.374078020907764 | 0.816467590784326 | 1.87672253231016 | 0.067432923234135 | 0.322259544678012 | -4.28143765905068 |
| ENSG00000276256.1 | | -1.5555177590736 | -1.42279734788107 | -1.87653241386112 | 0.0674598288607028 | 0.322266510657708 | -3.9994686564396 |
| ENSG00000153827.13 | TRIP12 | 0.172159206086342 | 7.82292379375628 | 1.87644252094872 | 0.0674722430241796 | 0.322266510657708 | -4.71206330764049 |
| lnc-UBL5-1 | lnc-UBL5-1 | -0.317444437972022 | 1.85992725463687 | -1.87608431320985 | 0.0675225717930484 | 0.322266510657708 | -4.41663631351068 |
| lnc-NCOA4-6 | lnc-NCOA4-6 | -0.427735351618593 | 1.81651654784461 | -1.87607775255219 | 0.0675234938742859 | 0.322266510657708 | -4.4127858332165 |
| ENSG00000178467.17 | P4HTM | -0.194852740694693 | 4.96758001898694 | -1.87606963921377 | 0.0675246341950963 | 0.322266510657708 | -4.69743287907108 |
| ENSG00000100908.13 | EMC9 | -0.301676076348755 | 1.47257496297676 | -1.87604339840257 | 0.067528322424656 | 0.322266510657708 | -4.37809814280852 |
| ELOA-AS1 | ELOA-AS1 | -0.251666431110352 | 3.26701209363957 | -1.87532088577917 | 0.0676299413158219 | 0.322676653246074 | -4.55267558531063 |
| ENSG00000120805.13 | ARL1 | 0.130581775370058 | 5.15450165339199 | 1.87519516490414 | 0.0676476368197107 | 0.322686282424126 | -4.70915919274899 |
| ENSG00000173208.3 | ABCD2 | -0.519695107643148 | 3.34531111460627 | -1.87480086440128 | 0.0677248837989491 | 0.322979908070846 | -4.55826962414716 |
| lnc-ZMAT5-3 | lnc-ZMAT5-3 | 0.275228855042552 | 0.305659482999432 | 1.8744662977166 | 0.0677503042970313 | 0.32302629449752 | -4.18165063022487 |
| ENSG00000258472.8 | | -0.257419326701769 | 1.69354032409181 | -1.87426276681122 | 0.0677789971440532 | 0.323060526169237 | -4.39860961604082 |
| lnc-SBNO2-1 | lnc-SBNO2-1 | -0.382399543457897 | 2.14532732373925 | -1.87410223427294 | 0.0678016355805568 | 0.323060526169237 | -4.45651407011365 |
| ENSG00000111087.9 | GLI1 | 0.78139583157757 | -0.222592085389241 | 1.87396432237643 | 0.067821089177341 | 0.323060526169237 | -4.09745648590249 |
| ENSG00000231584.8 | FAHD2CP | -0.309625673584576 | 1.52249638752719 | -1.87395199038065 | 0.0678228289373647 | 0.323060526169237 | -4.37474075366246 |
| ENSG00000120539.14 | MASTL | 0.361348778241179 | 4.7105116914412 | 1.87385891161464 | 0.0678359614295021 | 0.323060526169237 | -4.68188452578349 |
| ENSG00000142556.18 | ZNF614 | 0.265311072208728 | 4.80673924705129 | 1.87372196108297 | 0.0678552877307981 | 0.323077813408054 | -4.68942675872468 |
| ENSG00000115484.14 | CCT4 | 0.231943872167209 | 5.76943040740754 | 1.87351641613027 | 0.0678843027984704 | 0.323141212881803 | -4.72830561777435 |
| lnc-DDX20-6 | lnc-DDX20-6 | -0.302437824422178 | 2.07121565607943 | -1.8731683977922 | 0.0679334537374484 | 0.32330041116252 | -4.4379646294179 |
| ENSG00000265787.2 | CYP4F35P | -1.24747358141471 | -1.1256096577231 | -1.87304092833178 | 0.0679514639441775 | 0.323311369015951 | -4.00706023306091 |
| ENSG00000285942.1 | | 0.257362548502124 | 2.31402828788712 | 1.87271248954726 | 0.0679978879173671 | 0.323457482552153 | -4.45725897906742 |
| ENSG00000197912.15 | SPG7 | -0.224022509487207 | 6.89095282331603 | -1.87238490212619 | 0.0680442184535061 | 0.323585761457749 | -4.73139841346899 |
| ENSG00000124193.14 | SRSF6 | 0.164068479174888 | 8.03857080398058 | 1.87223013887961 | 0.0680661158949767 | 0.323585761457749 | -4.71513167843176 |
| lnc-RAB22A-2 | lnc-RAB22A-2 | -0.347858169073056 | 1.13526937727664 | -1.87185232116103 | 0.0681195985050019 | 0.323585761457749 | -4.3482309542952 |
| ENSG00000162739.13 | SLAMF6 | -0.386830624426842 | 6.75243369943996 | -1.8717690597774 | 0.0681313895177569 | 0.323585761457749 | -4.73321661008658 |
| ENSG00000279573.1 | | -0.385662163458302 | 0.35101384237021 | -1.871577205992 | 0.0681585653963091 | 0.323585761457749 | -4.223008582576 |
| ENSG00000277474.1 | MIR6894 | 0.411600159835994 | -0.272929780230227 | 1.87154684008309 | 0.0681628675399217 | 0.323585761457749 | -4.10244470767506 |
| ENSG00000198169.8 | ZNF251 | -0.173847328803885 | 5.17999196203738 | -1.8715380158358 | 0.0681641177740487 | 0.323585761457749 | -4.71573452622486 |
| lnc-TUBB1-1 | lnc-TUBB1-1 | -0.700769226814866 | 3.24518085918992 | -1.87227346160463 | 0.0681992027494556 | 0.323585761457749 | -4.55292662313866 |
| ENSG00000285583.1 | | -0.268257835828882 | 0.642192305774256 | -1.87121429487595 | 0.0682099965996451 | 0.323585761457749 | -4.26604698544317 |
| lnc-MAN2B1-2 | lnc-MAN2B1-2 | -0.268257835828882 | 0.642192305774256 | -1.87121429487595 | 0.0682099965996451 | 0.323585761457749 | -4.26604698544317 |
| ENSG00000198453.12 | ZNF568 | -0.229327578842714 | 3.91982751189738 | -1.8711226755227 | 0.0682229859754677 | 0.323585761457749 | -4.62870675074598 |
| ENSG00000169375.15 | SIN3A | 0.190000014500928 | 6.98635645319831 | 1.87098072384085 | 0.0682431153993482 | 0.323585761457749 | -4.73363253689624 |
| ENSG00000278897.1 | | -0.393265762517136 | 4.11490946434256 | -1.87097355093465 | 0.0682441326857683 | 0.323585761457749 | -4.6392465996222 |
| ENSG00000138795.9 | LEF1 | -0.732706000309566 | 7.25943044988084 | -1.8723696749737 | 0.0682449492536603 | 0.323585761457749 | -4.72774073331469 |
| ENSG00000110665.11 | C11orf21 | -0.832773171829518 | 3.62327894595791 | -1.87220055209976 | 0.0682652338838968 | 0.323607394986699 | -4.57481687167763 |
| ENSG00000170502.12 | NUDT9 | -0.25264956613608 | 2.55725262217053 | -1.87053702993333 | 0.0683060659116426 | 0.32372639987551 | -4.48530917233985 |
| ENSG00000076067.12 | RBMS2 | 1.43553905952744 | -0.207384731850755 | 1.87139503143176 | 0.0683372914094635 | 0.323799831740559 | -4.1069904703856 |
| ENSG00000123159.15 | GIPC1 | -0.435915082710559 | 1.86443922024286 | -1.86936539284284 | 0.0684725333847854 | 0.324365973454324 | -4.43517023874279 |
| lnc-LRRFIP2-3 | lnc-LRRFIP2-3 | -0.353746717848629 | 0.34320753500981 | -1.86923293780067 | 0.0684913744489662 | 0.324380570704802 | -4.23117630741686 |
| lnc-RBM11-2 | lnc-RBM11-2 | -0.418126516438281 | 0.559233556238962 | -1.86893254101965 | 0.0685341207337748 | 0.324480795532095 | -4.2763326943648 |

| | | | | | | | |
|---|---|---|---|---|---|---|---|
| lnc-GRAP-1 | lnc-GRAP-1 | -0.940064322169428 | 1.93299299210882 | -1.87011813713972 | 0.0685440654410702 | 0.324480795532095 | -4.42883641198272 |
| ENSG00000170776.21 | AKAP13 | 0.196967611605302 | 9.45989683736794 | 1.86857325470244 | 0.0685852767820881 | 0.324532010397333 | -4.68928547556659 |
| ENSG00000135924.15 | DNAJB2 | -0.305582177022872 | 5.73487669573833 | -1.86850020627687 | 0.068595681566583 | 0.324532010397333 | -4.73699731203562 |
| lnc-CBS-2 | lnc-CBS-2 | -0.289587543997764 | 2.37463903602728 | -1.86834063448777 | 0.068618415135183 | 0.324532010397333 | -4.48089012266444 |
| lnc-ARF4-1 | lnc-ARF4-1 | 0.256111835081465 | 1.00185150030255 | 1.86824772833363 | 0.0686316540646075 | 0.324532010397333 | -4.32015768640548 |
| lnc-SRPX-1 | lnc-SRPX-1 | 0.356329393142118 | 0.37162521425545 | 1.86816460921904 | 0.0686435002039411 | 0.324532010397333 | -4.21963680763301 |
| ENSG00000157303.10 | SUSD3 | -0.346031959275098 | 3.272016169768 | -1.86786857831964 | 0.0686857046741954 | 0.324532010397333 | -4.56683298533093 |
| ENSG00000205423.11 | CNEP1R1 | 0.1879813266629 | 3.52969736892091 | 1.867854714188 | 0.0686876817939426 | 0.324532010397333 | -4.59745428216938 |
| lnc-DRICH1-2 | lnc-DRICH1-2 | -0.561383122147438 | -0.115773617346307 | -1.86758342328641 | 0.0687263794738947 | 0.324532010397333 | -4.15178340651029 |
| lnc-RC3H2-2 | lnc-RC3H2-2 | 0.336728298896503 | 1.07104769867705 | 1.86754291974747 | 0.0687321586013835 | 0.324532010397333 | -4.33432607002206 |
| lnc-KCNK12-5 | lnc-KCNK12-5 | 0.22479463564294 | 3.3757701928722 | 1.8674858352953 | 0.0687403042294509 | 0.324532010397333 | -4.57504535834906 |
| ENSG00000100139.13 | MICALL1 | 0.204593256449994 | 4.64386782512825 | 1.86737973818687 | 0.0687554458681911 | 0.324532010397333 | -4.69364297706731 |
| lnc-IKZF1-15 | lnc-IKZF1-15 | -0.547489832259768 | -0.360312144573135 | -1.86735147462423 | 0.0687594799785304 | 0.324532010397333 | -4.10073178511207 |
| ENSG00000070756.15 | PABPC1 | 0.30862091040165 | 8.71956552335496 | 1.8671506478253 | 0.0687881501575382 | 0.324532010397333 | -4.71192942275008 |
| ENSG00000137216.18 | TMEM63B | -0.250709665523539 | 2.67878850840965 | -1.86713712456539 | 0.0687900811133814 | 0.324532010397333 | -4.50758005434773 |
| ENSG00000127463.14 | EMC1 | 0.1485444930128 | 5.00450677093732 | 1.86712796455676 | 0.0687913890765953 | 0.324532010397333 | -4.71393604626165 |
| lnc-CTTN-8 | lnc-CTTN-8 | 0.588634220964387 | -0.599394030193217 | 1.86697353945959 | 0.0688134427159763 | 0.324535137625794 | -4.05005672860703 |
| ENSG00000235363.1 | SNRPGP10 | 0.723226739836923 | -0.412121625223775 | 1.86676486917709 | 0.068843252733729 | 0.324535137625794 | -4.05299472872227 |
| lnc-VAMP2-4 | lnc-VAMP2-4 | 0.648739766438754 | 0.2790160872096 | 1.86674216793286 | 0.0688464964287423 | 0.324535137625794 | -4.18531005576919 |
| lnc-ZNF33A-2 | lnc-ZNF33A-2 | -0.317365964470319 | 1.20993586054894 | -1.86668181588693 | 0.0688551205369403 | 0.324535137625794 | -4.3458529383225 |
| ENSG00000119314.15 | PTBP3 | 0.166133247438857 | 7.71365840366766 | 1.86640849324321 | 0.0688941889488142 | 0.324644938446301 | -4.73112456663 |
| ENSG00000164574.15 | GALNT10 | 0.348099160502902 | 5.20114635798073 | 1.86616948633004 | 0.0689283677663038 | 0.32473165340669 | -4.72352116112574 |
| lnc-PNMA1-2 | lnc-PNMA1-2 | -0.289749777705693 | 1.77502166743901 | -1.86588815422318 | 0.0689686177161099 | 0.324846924130593 | -4.40600495239262 |
| lnc-YRDC-3 | lnc-YRDC-3 | 0.272753945539911 | 1.85309117151457 | 1.86543978310847 | 0.069032807124083 | 0.325015397690575 | -4.42572059366223 |
| ENSG00000230383.1 | RPL6P19 | 1.10525294413802 | 0.147827619568337 | 1.86559147264476 | 0.0690359674830293 | 0.325015397690575 | -4.09327627428038 |
| ENSG00000108666.9 | C17orf75 | -0.190238787246416 | 3.7060982525121 | -1.86519097678903 | 0.0690684484890877 | 0.325037021993883 | -4.62333829852609 |
| ENSG00000002587.9 | HS3ST1 | -0.831022791105909 | 3.60981794732833 | -1.86653053593875 | 0.0690721437206064 | 0.325037021993883 | -4.60832342556273 |
| lnc-CKAP2L-1 | lnc-CKAP2L-1 | -0.456626881328703 | 0.0227221307129769 | -1.86414409204806 | 0.0692185856772071 | 0.325651691198618 | -4.16619806884007 |
| ENSG00000132323.8 | ILKAP | 0.256997935688753 | 5.41537267610031 | 1.86358607817234 | 0.0692987256308133 | 0.325941643837579 | -4.73660283130719 |
| lnc-EFCAB11-4 | lnc-EFCAB11-4 | -0.61727749879125 | -0.474012454626448 | -1.86349448645729 | 0.0693118872407852 | 0.325941643837579 | -4.07718839061416 |
| ENSG00000101346.13 | POFUT1 | -0.233657791830994 | 4.43057794756986 | -1.86319011836059 | 0.0693556398101958 | 0.326072894977903 | -4.68604322968614 |
| ENSG00000118503.14 | TNFAIP3 | -0.373125744160256 | 5.34760324670202 | -1.86251432588737 | 0.0694528681843697 | 0.32645544321509 | -4.73797906157093 |
| lnc-ADRA2A-4 | lnc-ADRA2A-4 | 0.610049179409886 | 0.854087032063344 | 1.86233255761506 | 0.0694790395242016 | 0.326490640618689 | -4.29545865165678 |
| ENSG00000161999.11 | JMJD8 | -0.26857109369825 | 3.34782807633405 | -1.86222050284578 | 0.0694951775596489 | 0.326490640618689 | -4.59774747399848 |
| lnc-DENND2D-1 | lnc-DENND2D-1 | -0.320049435466546 | 1.38041869887865 | -1.86213188263557 | 0.0695079428261718 | 0.326490640618689 | -4.38945462151649 |
| ENSG00000140931.19 | CMTM3 | -0.429136758180368 | 4.11126342627039 | -1.86192421274413 | 0.0695378643717008 | 0.326556664924188 | -4.67323589673003 |
| ENSG00000241106.7 | HLA-DOB | -0.333314989704189 | 6.46876271112966 | -1.86156822804532 | 0.0695891809250839 | 0.326713645113494 | -4.75225784913678 |
| lnc-C5-1 | lnc-C5-1 | -0.69984133040909 | 0.0959752545970432 | -1.86146865665617 | 0.0696035402819521 | 0.326713645113494 | -4.1926948504274 |
| lnc-ZNF781-2 | lnc-ZNF781-2 | -0.418977112838001 | 0.962653068325532 | -1.861362090792 | 0.0696189111144044 | 0.326713645113494 | -4.32625231439347 |
| lnc-ZBTB7C-10 | lnc-ZBTB7C-10 | -0.320806054779935 | 2.76493492168682 | -1.86102200519732 | 0.069667983634924 | 0.326773448114849 | -4.53182606856279 |
| ENSG00000178209.15 | PLEC | -0.279924860700261 | 8.04770192917444 | -1.86100853604104 | 0.0696699277668862 | 0.326773448114849 | -4.73572513197985 |
| lnc-ZNF577-1 | lnc-ZNF577-1 | -0.245349029003523 | 1.2671629374285 | -1.8609437325727 | 0.0696792821151471 | 0.326773448114849 | -4.36885074806708 |
| ENSG00000227939.1 | RPL3P2 | 1.08781570767654 | -0.25549132662401 | 1.86063149065865 | 0.0697243690470259 | 0.326784783933503 | -4.05713853080192 |
| ENSG00000198380.12 | GFPT1 | 0.207516955844784 | 4.86555739946529 | 1.86059139796687 | 0.069730160125189 | 0.326784783933503 | -4.71966686198297 |
| ENSG00000171612.6 | SLC25A33 | 0.361016469265986 | 2.86374973662308 | 1.86040982362009 | 0.0697563922495956 | 0.326784783933503 | -4.53470150621374 |
| ENSG00000135392.16 | DNAJC14 | 0.229215385853971 | 5.00917156324463 | 1.86038060746719 | 0.0697606139032171 | 0.326784783933503 | -4.72791687746946 |
| ENSG00000160199.14 | PKNOX1 | -0.274899250107482 | 3.69229105103285 | -1.86037737150403 | 0.0697610815043392 | 0.326784783933503 | -4.62294400430085 |
| TDRKH-AS1 | TDRKH-AS1 | -0.371504627115253 | 0.0184080142806428 | -1.86007506700097 | 0.0698047766724415 | 0.326915066723291 | -4.18461287241488 |
| ENSG00000143252.14 | SDHC | -0.20274139069172 | 4.50447366003773 | -1.85995312634986 | 0.0698224085878365 | 0.326923256588589 | -4.6949825732547 |
| lnc-MPEG1-4 | lnc-MPEG1-4 | 1.14247108776261 | -0.502889905697659 | 1.85977322079644 | 0.0698484288063908 | 0.326970709602443 | -4.04055102563731 |
| lnc-RALB-1 | lnc-RALB-1 | 0.46187267988356 | 0.904486551664821 | 1.85944931705677 | 0.0698952966430935 | 0.327115709596361 | -4.30852334445784 |
| ENSG00000149658.17 | YTHDF1 | 0.167757230109105 | 5.015986488754 | 1.85928408704228 | 0.069919215191834 | 0.327153263535694 | -4.73033164210571 |
| lnc-MIS18BP1-2 | lnc-MIS18BP1-2 | 0.182762730834482 | 2.75236625732784 | 1.85905253611543 | 0.0699527460014073 | 0.32723576612431 | -4.5246490004604 |
| ENSG00000151233.10 | GXYLT1 | 0.254928736274776 | 3.88860674766579 | 1.85878897992489 | 0.0699909281113641 | 0.327339984848036 | -4.6451571480792 |
| ENSG00000167895.14 | TMC8 | 0.300604573291442 | 6.11359828416426 | 1.85851053895248 | 0.070031285859294 | 0.327400011886116 | -4.75766329363508 |
| ENSG00000126003.6 | PLAGL2 | 0.235313352906493 | 4.17481658921186 | 1.85848095076201 | 0.0700355755883286 | 0.327400011886116 | -4.66972933936378 |
| ENSG00000260170.1 | | 0.324843275788617 | 2.7408225667984 | 1.85834820230682 | 0.0700548243566438 | 0.327415633454314 | -4.52762340333235 |
| ENSG00000280088.1 | | -0.311016780570843 | 0.825117657681754 | -1.85789843179155 | 0.0701200753259116 | 0.327646199871337 | -4.32870165507865 |
| ENSG00000128513.14 | POT1 | -0.196982260693216 | 4.06800052961008 | -1.85770500231661 | 0.0701481532040133 | 0.32767098064545 | -4.66367444510868 |
| lnc-CMKLR1-4 | lnc-CMKLR1-4 | 0.33630646530184 | 2.47630366071987 | 1.85756482947362 | 0.0701685064137734 | 0.32767098064545 | -4.50717187395992 |
| lnc-FBXO15-8 | lnc-FBXO15-8 | -1.0527668532803 | 2.8108494336323 | -1.85919666498593 | 0.0701750093550602 | 0.32767098064545 | -4.56025723813047 |
| lnc-INPP4B-2 | lnc-INPP4B-2 | -0.583472835363061 | 2.74012441560127 | -1.85765355744186 | 0.0701961899852844 | 0.32767098064545 | -4.53082665600539 |
| lnc-LYL1-2 | lnc-LYL1-2 | -0.226755668512428 | 2.30238188311264 | -1.85731374809029 | 0.0702049761767689 | 0.32767098064545 | -4.49017019487574 |
| ENSG00000059691.11 | GATB | -0.173242081188762 | 3.31378721924807 | -1.85684472504547 | 0.0702731452755101 | 0.327711551872333 | -4.59395458992715 |
| lnc-SLC46A3-7 | lnc-SLC46A3-7 | -0.423333035341937 | 0.272208405746351 | -1.85672486874974 | 0.0702905745381277 | 0.327711551872333 | -4.24735377771911 |
| ENSG00000173011.11 | TADA2B | 0.21572720876578 | 5.24321106996487 | 1.85659386706371 | 0.0703096287417392 | 0.327711551872333 | -4.74350108334787 |
| lnc-PIGV-3 | lnc-PIGV-3 | -0.410527294454946 | -0.3666958534366 | -1.85655325798703 | 0.0703155362254497 | 0.327711551872333 | -4.11321994635888 |
| ENSG00000257767.2 | | 0.430872527124597 | -0.262542702150487 | 1.85650864372526 | 0.0703220268375717 | 0.327711551872333 | -4.12291704806821 |

| | | | | | | | |
|---|---|---|---|---|---|---|---|
| ENSG00000196114.3 | IFITM3P3 | -0.610527343724506 | 0.121651254044102 | -1.8564171268583 | 0.0703353425695852 | 0.327711551872333 | -4.20915143084036 |
| BDNF-AS | BDNF-AS | -0.332031060492263 | 1.05491063890576 | -1.856346769772 | 0.0703455810034114 | 0.327711551872333 | -4.3326787209349 |
| lnc-TCP11L2-1 | lnc-TCP11L2-1 | -0.462334764836809 | 1.62358387690874 | -1.85634631987189 | 0.0703456464774007 | 0.327711551872333 | -4.40025799353613 |
| lnc-LTBP1-1 | lnc-LTBP1-1 | -0.322649075600866 | 3.70367337521421 | -1.8562685717377 | 0.0703569619454812 | 0.327711551872333 | -4.63549997450708 |
| ENSG00000185842.15 | DNAH14 | -2.50355236497497 | 0.38514584502216 | -1.85767253070982 | 0.0704302602315247 | 0.327838700505348 | -4.25923663991371 |
| ENSG00000164985.14 | PSIP1 | -0.303211730189248 | 6.4060732199399 | -1.85567809485148 | 0.0704429504631274 | 0.327838700505348 | -4.76234854756732 |
| ENSG00000159917.16 | ZNF235 | -0.247601430003429 | 3.7498438419589 | -1.85566140355751 | 0.0704453824387417 | 0.327838700505348 | -4.63367784973628 |
| ENSG00000283829.1 | MIR378I | 0.398473928664259 | 2.3175317961335 | 1.85564364303316 | 0.070447970283008 | 0.327838700505348 | -4.48348590762675 |
| lnc-ATAD5-3 | lnc-ATAD5-3 | -0.565734162510556 | -0.478974615400982 | -1.85516141024845 | 0.0705182661581972 | 0.327911752103734 | -4.08644631702644 |
| lnc-CCDC14-1 | lnc-CCDC14-1 | -0.265120983758583 | 4.15060688366685 | -1.85515088769771 | 0.0705198007119013 | 0.327911752103734 | -4.67830318376703 |
| ENSG00000255663.1 | | 0.405514544358878 | 2.23018708990472 | 1.85514663214599 | 0.0705204213273388 | 0.327911752103734 | -4.47077032765764 |
| lnc-AFAP1L2-4 | lnc-AFAP1L2-4 | 0.258447635562197 | 1.60379826607298 | 1.85488967498814 | 0.0705579036983117 | 0.327911752103734 | -4.41211753983623 |
| ENSG00000172932.14 | ANKRD13D | -0.24044075052755 | 4.75310509513178 | -1.85466011531395 | 0.0705914039048831 | 0.327911752103734 | -4.72481379095765 |
| lnc-B9D2-1 | lnc-B9D2-1 | 0.338506822762134 | 2.05524576893961 | 1.85464500259793 | 0.0705936098144894 | 0.327911752103734 | -4.45856890616502 |
| lnc-DENND2D-2 | lnc-DENND2D-2 | -0.235666791533857 | 3.86424198241378 | -1.85460247393072 | 0.0705998177749781 | 0.327911752103734 | -4.6480561976618 |
| ENSG00000083544.14 | TDRD3 | -0.203890580805444 | 4.76215138019869 | -1.85457103130804 | 0.0706044077903844 | 0.327911752103734 | -4.72567703149231 |
| ENSG00000167004.12 | PDIA3 | 0.257346655972187 | 6.95746048207247 | 1.85450800737398 | 0.0706136088302268 | 0.327911752103734 | -4.76230588509318 |
| ENSG00000136247.14 | ZDHHC4 | 0.208473125464414 | 4.11569860883942 | 1.85441487535867 | 0.0706272072971189 | 0.327911752103734 | -4.67286663696046 |
| lnc-RTL8B-2 | lnc-RTL8B-2 | -0.374616798490074 | -0.279592735045984 | -1.85433473099831 | 0.0706389111804866 | 0.327911752103734 | -4.12459734994198 |
| ENSG00000096384.19 | HSP90AB1 | -0.300326585941814 | 9.26282572685732 | -1.85415447878453 | 0.0706652403349068 | 0.32793723680631 | -4.71619548588434 |
| ENSG00000248235.6 | | -0.316016100139735 | 1.08754287374888 | -1.85407901304372 | 0.0706762659705968 | 0.32793723680631 | -4.3683391003105 |
| ENSG00000160229.11 | ZNF66 | -0.262212846627606 | 3.4870684492236 | -1.85390625257578 | 0.070701511983811 | 0.327980442001979 | -4.60888466420478 |
| ENSG00000139687.15 | RB1 | 0.867590031222326 | 6.09709844762593 | 1.85538712745672 | 0.0707404086285092 | 0.328086937990222 | -4.76147970218723 |
| ENSG00000239523.5 | MYLK-AS1 | -0.401021922691172 | 0.038474446256227 | -1.85328657445237 | 0.0707921304455304 | 0.328252854462796 | -4.1660835978843 |
| lnc-FBXO15-2 | lnc-FBXO15-2 | -1.01117529686306 | 2.0909085844331 | -1.85436152342124 | 0.0708368057115248 | 0.328386029720792 | -4.48444817349222 |
| BACH1-IT2 | BACH1-IT2 | 0.530778850170502 | 1.53626387137892 | 1.85282688097721 | 0.0708594172983153 | 0.328416884992394 | -4.39861666118556 |
| lnc-DOCK11-1 | lnc-DOCK11-1 | -0.465176248905514 | 3.18601881873198 | -1.85220686227047 | 0.0709502574950005 | 0.328758517572817 | -4.5893428658458 |
| ENSG00000213626.12 | LBH | -0.433497601538511 | 6.85022534681553 | -1.85227610249947 | 0.0709671703099284 | 0.328758517572817 | -4.76606854086972 |
| lnc-CFAP20-3 | lnc-CFAP20-3 | 0.280229061768868 | 1.58722439725574 | 1.85198526423256 | 0.0709827482475338 | 0.328758517572817 | -4.4164525805126 |
| ENSG00000101182.14 | PSMA7 | 0.21982472581005 | 5.23948749884578 | 1.85188797046323 | 0.070997017471271 | 0.328758517572817 | -4.75343520689073 |
| lnc-TBCA-6 | lnc-TBCA-6 | 0.532051511438056 | 2.87448187922899 | 1.85174878296972 | 0.07103049455929 | 0.328839556795741 | -4.54433133422289 |
| ENSG00000104863.11 | LIN7B | -0.381802741916659 | 1.00544720331379 | -1.85120968483804 | 0.0710965633577474 | 0.329040942388666 | -4.34984964260505 |
| ENSG00000185627.17 | PSMD13 | 0.166269214855545 | 5.49922389680347 | 1.85108842923477 | 0.0711143714313552 | 0.329040942388666 | -4.76130688726277 |
| lnc-PFKP-38 | lnc-PFKP-38 | -1.01727359495834 | -0.8585050333773 | -1.85101062995004 | 0.0711257993335301 | 0.329040942388666 | -4.07194111791117 |
| ENSG00000226981.2 | ABHD17AP6 | -2.00054387636384 | -2.48596508491528 | -1.85092799830248 | 0.0711379387664366 | 0.329040942388666 | -4.07429412827257 |
| ENSG00000101247.17 | NDUFAF5 | 0.229174575863043 | 4.28213041253365 | 1.85026025632798 | 0.0712361014278892 | 0.329409589290006 | -4.69578492023877 |
| ENSG00000149547.14 | EI24 | 0.250554858231153 | 3.86242408032344 | 1.85009299830807 | 0.071260707505016 | 0.329409589290006 | -4.65749328363702 |
| ENSG00000170473.16 | PYM1 | -0.285623392057857 | 4.0076977318538 | -1.8500593990029 | 0.0712656513194577 | 0.329409589290006 | -4.67089055946258 |
| ENSG00000104611.11 | SH2D4A | -2.74601544919939 | -2.40451864397736 | -1.8507094033799 | 0.071295916982832 | 0.329475496016531 | -4.06360358357896 |
| ENSG00000221963.5 | APOL6 | 0.64737463897552 | 5.01828544987288 | 1.85040746074405 | 0.0713875867558942 | 0.32975201213329 | -4.74507690202589 |
| ENSG00000145220.13 | LYAR | -0.360965553059176 | 1.60871456738497 | -1.84917591660287 | 0.0713957517556786 | 0.32975201213329 | -4.43056212721419 |
| ENSG00000205808.5 | PLPP6 | -0.272974058339356 | 2.52671792842932 | -1.84912120792546 | 0.0714038147052458 | 0.32975201213329 | -4.52273381286709 |
| ENSG00000249141.1 | | 0.269577579688834 | 4.59858815491483 | 1.84877441347099 | 0.0714549431513408 | 0.329914108318539 | -4.71528719142911 |
| lnc-ARMCX6-2 | lnc-ARMCX6-2 | -0.231421565582603 | 2.13586138187165 | -1.84836207586314 | 0.0715157751529358 | 0.330089335327938 | -4.48972891701763 |
| ENSG00000104497.13 | SNX16 | 0.370651554239658 | 2.97336141993787 | 1.84820911560461 | 0.0715383524901609 | 0.330089335327938 | -4.56814983064748 |
| lnc-RNASEH1-1 | lnc-RNASEH1-1 | 0.754632762477163 | -0.789576677528119 | 1.84813460199295 | 0.0715493530861359 | 0.330089335327938 | -4.06861785360393 |
| lnc-BRF1-8 | lnc-BRF1-8 | -1.72027969105449 | 0.863911276107736 | -1.84978568184261 | 0.0715734260296907 | 0.330089335327938 | -4.34208993028689 |
| lnc-ITGAL-2 | lnc-ITGAL-2 | -0.333072803844209 | 0.988114469138729 | -1.84792260085533 | 0.0715806590965712 | 0.330089335327938 | -4.35654013000136 |
| ENSG00000183741.11 | CBX6 | -0.272578015876615 | 5.26347383855783 | -1.84786533904587 | 0.071589116887913 | 0.330089335327938 | -4.7608367775562 |
| lnc-TTC34-5 | lnc-TTC34-5 | -0.735097118523992 | -0.335344874381779 | -1.84754775640141 | 0.071636040470113 | 0.330231718028295 | -4.11241444971016 |
| ENSG00000168795.4 | ZBTB5 | 0.270113966777036 | 5.54519643106261 | 1.84701734269119 | 0.0717144685285274 | 0.330457481875722 | -4.76734179888045 |
| lnc-SMAD9-3 | lnc-SMAD9-3 | 0.274414274979295 | 1.06597958707423 | 1.84699938948348 | 0.0717171244006633 | 0.330457481875722 | -4.3626981609881 |
| ENSG00000019995.6 | ZRANB1 | 0.198198842052738 | 6.00573673672289 | 1.84648122092954 | 0.0717938146039118 | 0.330736814232837 | -4.77729357402987 |
| ENSG00000147144.12 | CCDC120 | 0.285165374683386 | 1.79276574024571 | 1.84629824615378 | 0.0718209119297712 | 0.330787610259674 | -4.4456709086411 |
| lnc-AKNA-4 | lnc-AKNA-4 | -0.897423074000799 | 0.623009927532091 | -1.84672272304133 | 0.0718425310735492 | 0.33081315818498 | -4.30902960572882 |
| lnc-RPL39-1 | lnc-RPL39-1 | 0.361856029150575 | 6.80906501861162 | 1.84563924447234 | 0.0719185775210671 | 0.331044950600576 | -4.77833467580936 |
| ENSG00000111642.15 | CHD4 | 0.175154981296862 | 7.13977824556171 | 1.84554793369945 | 0.0719321188729877 | 0.331044950600576 | -4.77518391302613 |
| ENSG00000117090.14 | SLAMF1 | 2.05720793250748 | 1.07191782028554 | 1.84721037797345 | 0.0719411195664566 | 0.331044950600576 | -4.14903096198727 |
| ENSG00000254986.7 | DPP3 | -0.270776691270443 | 3.89524292604324 | -1.8451240341397 | 0.0719950113388367 | 0.331115672771511 | -4.67006476407225 |
| ENSG00000163931.15 | TKT | -0.332027962342249 | 4.77557676197063 | -1.84498491351227 | 0.0720156623323727 | 0.331115672771511 | -4.74722644151789 |
| lnc-OR6J1-2 | lnc-OR6J1-2 | -0.325110319396632 | 2.69381422031673 | -1.84497762006137 | 0.0720167451070069 | 0.331115672771511 | -4.54935651670292 |
| ENSG00000243156.7 | MICAL3 | -0.327585088305717 | 6.33712785791006 | -1.84476093246363 | 0.0720489205176907 | 0.331115672771511 | -4.78094645479604 |
| ENSG00000197329.11 | PELI1 | 0.518604309652347 | 7.97447883316563 | 1.84535570798905 | 0.0720599529242087 | 0.331115672771511 | -4.76490582973475 |
| lnc-C2orf71-3 | lnc-C2orf71-3 | 0.258841568766048 | 1.28427419541352 | 1.84455060260495 | 0.0720801635426217 | 0.331115672771511 | -4.38753525030116 |
| ENSG00000122477.12 | LRRC39 | 0.372466717418782 | 0.967402848116348 | 1.84450734445015 | 0.0720865906623725 | 0.331115672771511 | -4.33971395334634 |
| lnc-KLHL42-5 | lnc-KLHL42-5 | 0.264026778066761 | 2.28068891532723 | 1.84441892434782 | 0.0720997292715867 | 0.331115672771511 | -4.50984386816979 |
| ENSG00000106009.15 | BRAT1 | -0.338595007712609 | 4.28092195200658 | -1.84433188050225 | 0.0721126653622264 | 0.331115672771511 | -4.71598681706107 |
| ENSG00000234782.3 | TPT1P9 | 1.23806480072325 | 0.412841948653465 | 1.8450753046645 | 0.0721295849794961 | 0.331115672771511 | -4.17636272181606 |

| | | | | | | | |
|---|---|---|---|---|---|---|---|
| ENSG00000065518.7 | NDUFB4 | 0.167178428179835 | 4.78923516755904 | 1.84419209412834 | 0.0721334439443936 | 0.331115672771511 | -4.74193257046514 |
| ENSG00000178631.7 | ACTG1P1 | -0.356092804662842 | 0.015087367374499 | -1.84407448726941 | 0.0721509295792795 | 0.331122092202968 | -4.19326311116447 |
| lnc-SLC2A9-3 | lnc-SLC2A9-3 | 0.309225190944464 | 2.22045838481199 | 1.84386279646903 | 0.0721824125500133 | 0.331167910726899 | -4.50350936428119 |
| ENSG00000108107.14 | RPL28 | -0.374735416236235 | 7.18632329565528 | -1.84370381533128 | 0.0722060641200297 | 0.331167910726899 | -4.77599876046444 |
| lnc-AKAIN1-4 | lnc-AKAIN1-4 | -0.264350690765871 | 0.739543606284664 | -1.84361059094678 | 0.072219936136402 | 0.331167910726899 | -4.31971372137104 |
| lnc-GSR-2 | lnc-GSR-2 | -0.406544616328754 | -0.29646229380461 | -1.8435747435685 | 0.0722252709154667 | 0.331167910726899 | -4.1429045574311 |
| ENSG00000160049.11 | DFFA | -0.234526387150155 | 5.01557308609265 | -1.84328865120898 | 0.0722678589349551 | 0.331289385402713 | -4.75701550771893 |
| ENSG00000167208.14 | SNX20 | -0.305179591828058 | 5.51151047182797 | -1.84309744063003 | 0.0722963346191847 | 0.331346126801754 | -4.77525241425942 |
| lnc-ERCC1-1 | lnc-ERCC1-1 | -0.216168800397817 | 2.52551426450209 | -1.84239455737032 | 0.07240109199 2208 | 0.331752376775516 | -4.53869735312115 |
| lnc-RNF113A-4 | lnc-RNF113A-4 | 0.245546547020143 | 1.17127908742835 | 1.84143553570812 | 0.0725442316602674 | 0.332334280049696 | -4.3814396440536 |
| ENSG00000201098.1 | RNY1 | 0.565729196707336 | 1.3053740692792 | 1.84094939735529 | 0.0726168822651879 | 0.332559878833312 | -4.41338205522671 |
| ENSG00000196235.13 | SUPT5H | 0.194357950341393 | 6.92614069409961 | 1.84088981358313 | 0.0726257909612659 | 0.332559878833312 | -4.78560542870704 |
| ENSG00000119541.9 | VPS4B | 0.14501533885271 | 5.34110900387641 | 1.84075273228937 | 0.0726462902534871 | 0.332579758071069 | -4.77426684222529 |
| lnc-C11orf21-2 | lnc-C11orf21-2 | -0.727897023273457 | 0.469295485948045 | -1.84055671081967 | 0.07267561204 6846 | 0.332640009508613 | -4.25032828623666 |
| ENSG00000114473.13 | IQCG | -0.202928721372327 | 3.84696724324261 | -1.84029381757868 | 0.0727149525744192 | 0.332720677548166 | -4.66560669196789 |
| lnc-CA12-2 | lnc-CA12-2 | 0.193356481657281 | 2.7293167372413 | 1.84020605996878 | 0.0727280890354829 | 0.332720677548166 | -4.55693640713109 |
| ENSG00000284055.1 | MIR4315-1 | 0.417753013762181 | -0.18111229435123 | 1.84005411737294 | 0.0727508381208179 | 0.332720677548166 | -4.1631836172151 |
| ENSG00000153944.10 | MSI2 | 0.462944886147804 | 8.37115269643752 | 1.84022292789206 | 0.0727727476541231 | 0.332720677548166 | -4.76219154959663 |
| ENSG00000143457.10 | GOLPH3L | -0.398136925351541 | 2.82462475188818 | -1.83987640611056 | 0.0727774529828175 | 0.332720677548166 | -4.56805844963828 |
| ENSG00000210195.2 | MT-TT | 0.64768614935089 | 0.183640469786793 | 1.83979114188466 | 0.072790225477306 | 0.332720677548166 | -4.21095547041028 |
| ENSG00000111711.9 | GOLT1B | 0.304550886935462 | 4.96899768830598 | 1.83944455481101 | 0.0728421634257736 | 0.332884158479729 | -4.7651486403244 |
| ENSG00000129667.12 | RHBDF2 | 0.422835787951401 | 4.57749285842336 | 1.83939282477241 | 0.0728601768440587 | 0.332892568253332 | -4.7466607359421 |
| lnc-URGCP-2 | lnc-URGCP-2 | -0.216934380013357 | 1.34671894764814 | -1.83895152977656 | 0.0729161000037745 | 0.332905528420585 | -4.41660524763561 |
| ENSG00000173848.18 | NET1 | 1.15327270598931 | 0.446285296368817 | 1.83961460347665 | 0.0729263376585547 | 0.332905528420585 | -4.20109326667395 |
| ENSG00000147677.10 | EIF3H | 0.254741851557362 | 6.96922473937748 | 1.83886671426742 | 0.0729288257839021 | 0.332905528420585 | -4.78860384398328 |
| lnc-ZNF487-4 | lnc-ZNF487-4 | -0.469764166795748 | 0.644478706559358 | -1.83879227041517 | 0.0729399969439175 | 0.332905528420585 | -4.32811469708222 |
| lnc-NMT1-1 | lnc-NMT1-1 | -0.282122685317927 | 2.44906803485152 | -1.83876637970143 | 0.072943882484421 | 0.332905528420585 | -4.53342211454125 |
| ENSG00000030110.12 | BAK1 | 0.241671291700306 | 4.44130829923479 | 1.83827002126581 | 0.0730184072342889 | 0.333171774795693 | -4.73089711755175 |
| ENSG00000100603.13 | SNW1 | 0.260899353139975 | 5.88937868784146 | 1.83805693092176 | 0.0730504210469609 | 0.333186108998141 | -4.78984044866745 |
| lnc-TSTD3-2 | lnc-TSTD3-2 | -0.229221182498143 | 2.72659846397027 | -1.83803362188434 | 0.0730539236218535 | 0.333186108998141 | -4.55848797435148 |
| lnc-HEATR4-5 | lnc-HEATR4-5 | -0.40494864735509 | 0.318589401347622 | -1.83755560701222 | 0.0731257847806966 | 0.333377155137633 | -4.26679933417394 |
| ENSG00000155366.16 | RHOC | -0.454861256066397 | 4.3241285092607 | -1.8376410398852 | 0.0731349918771753 | 0.333377155137633 | -4.71703926182021 |
| lnc-STK36-2 | lnc-STK36-2 | -0.425292274628814 | 0.0559279188353475 | -1.83743182960375 | 0.0731444023029466 | 0.333377155137633 | -4.22778891091104 |
| lnc-MITD1-1 | lnc-MITD1-1 | -0.300157710165293 | 2.61317910976916 | -1.83729953026398 | 0.0731643060596101 | 0.333394047293548 | -4.55206291707278 |
| LINC01138 | LINC01138 | -0.345661716670187 | 2.22268980469169 | -1.83666672828172 | 0.0732595713393314 | 0.333640654004593 | -4.52160615648579 |
| ENSG00000230457.3 | PA2G4P4 | -0.422219402821016 | 1.69485641822964 | -1.8365569998355 | 0.0732761011114591 | 0.333640654004593 | -4.45019697875545 |
| ENSG00000074582.13 | BCS1L | -0.287488890396779 | 3.04566169335371 | -1.836494341087 | 0.0732855416016578 | 0.333640654004593 | -4.60429074972729 |
| lnc-MYOC-2 | lnc-MYOC-2 | -0.785234771455285 | 4.33303612026959 | -1.83774402183432 | 0.073298911266062 | 0.333640654004593 | -4.71129785384426 |
| BACH1-IT1 | BACH1-IT1 | 0.487605159492571 | 1.83773254058947 | 1.83625667244798 | 0.0733213593500192 | 0.333640654004593 | -4.46082656078926 |
| ENSG00000153201.15 | RANBP2 | 0.197056716124398 | 7.85297978776136 | 1.83623839728539 | 0.0733241141142098 | 0.333640654004593 | -4.78057243428914 |
| ENSG00000146109.4 | ABT1 | 0.199378902986069 | 5.05968084359859 | 1.83612279793294 | 0.0733415413778461 | 0.333640654004593 | -4.77106668769232 |
| ENSG00000181126.13 | HLA-V | 2.96858864768372 | -1.66149713501813 | 1.83706855335002 | 0.0733481008293633 | 0.333640654004593 | -4.08504650633549 |
| ENSG00000178935.5 | ZNF552 | 0.188566297186943 | 2.73704050006542 | 1.83553840387613 | 0.0734296958308018 | 0.333938009121828 | -4.56285535128518 |
| ENSG00000203879.11 | GDI1 | -0.176212508641045 | 5.47374430333492 | -1.83540323327845 | 0.0734500987679149 | 0.333957009706206 | -4.78725154492713 |
| lnc-CD47-1 | lnc-CD47-1 | -0.384640402848455 | 1.74501092543102 | -1.8352160209086 | 0.0734783649302659 | 0.334011745883318 | -4.46302124776599 |
| lnc-HDAC9-1 | lnc-HDAC9-1 | -0.964027302210194 | 0.342535935874268 | -1.83533943579875 | 0.0735243840954995 | 0.334147140171708 | -4.24599578502219 |
| lnc-HUS1-2 | lnc-HUS1-2 | 0.993664688580219 | 0.279890540083334 | 1.83495930679884 | 0.0735863033558994 | 0.334302886485416 | -4.23887707344721 |
| ENSG00000256566.1 | | 0.227563198293585 | 3.6263115710729 | 1.83446971390569 | 0.0735911372815149 | 0.334302886485416 | -4.65794978937593 |
| ENSG00000147649.9 | MTDH | -0.33997486160861 | 7.34850542718143 | -1.83430646710902 | 0.0736158245718262 | 0.334341243857436 | -4.79174593129885 |
| lnc-CMTM4-5 | lnc-CMTM4-5 | -0.454539826859597 | -0.00907571717319061 | -1.83412243280615 | 0.0736436638987911 | 0.334393896763472 | -4.21753666316919 |
| ENSG00000166046.10 | TCP11L2 | 0.385180888053638 | 6.82902429389002 | 1.83361725053263 | 0.073720129904017 | 0.334667276977147 | -4.79916295925896 |
| lnc-KIAA1522-1 | lnc-KIAA1522-1 | -0.340764325749547 | -0.0390672896839711 | -1.83316795348541 | 0.0737881934121013 | 0.334836257692361 | -4.19280093708329 |
| ENSG00000197302.10 | KRBOX5 | -0.160274882915972 | 4.94602341016582 | -1.8330914663303 | 0.0737997856636922 | 0.334836257692361 | -4.77156091797699 |
| ENSG00000259113.1 | | -1.46660334924153 | -0.932103432678621 | -1.83306365479838 | 0.0738061556211555 | 0.334836257692361 | -4.05336756481463 |
| ENSG00000178149.16 | DALRD3 | -0.274362612752274 | 3.55634297302925 | -1.8328043673054 | 0.073843311616291 | 0.334879047535812 | -4.66461376507748 |
| ENSG00000182463.15 | TSHZ2 | -4.19029688569393 | 1.06482124118585 | -1.83455105624666 | 0.0738587659459874 | 0.334879047535812 | -4.17381579252429 |
| lnc-CEACAM21-5 | lnc-CEACAM21-5 | -0.436075383243835 | 5.18986709840393 | -1.83284514155562 | 0.0738643966288969 | 0.334879047535812 | -4.78023629135202 |
| ENSG00000145022.4 | TCTA | -0.222271325720855 | 4.37006212845146 | -1.83240553765814 | 0.0739038126714698 | 0.3349839630515 | -4.73300486571436 |
| ENSG00000215146.5 | | 3.0158509411207 | -0.281942156412209 | 1.83357909795598 | 0.0739831462831287 | 0.335027410404695 | -4.06383376940291 |
| ENSG00000132383.11 | RPA1 | -0.287360419266863 | 5.69930973729756 | -1.83181203702633 | 0.0739939223102209 | 0.335027410404695 | -4.79729392155337 |
| lnc-SHC3-5 | lnc-SHC3-5 | -0.823187184288886 | 0.024565965720873 | -1.83182817928147 | 0.0740104992521476 | 0.335027410404695 | -4.27177088449117 |
| ENSG00000100350.14 | FOXRED2 | -0.5030880123966 | 0.883924502481981 | -1.83166224726696 | 0.0740166791871218 | 0.335027410404695 | -4.34282615667573 |
| ENSG00000166226.12 | CCT2 | 0.213979690577493 | 5.97295929479486 | 1.83152844545709 | 0.0740370120966389 | 0.335027410404695 | -4.80259177234552 |
| ENSG00000177427.12 | MIEF2 | -0.336881057227324 | 2.41284306447173 | -1.83145239652023 | 0.074048570819789 | 0.335027410404695 | -4.54752841570908 |
| lnc-CFLAR-1 | lnc-CFLAR-1 | -0.327522583542917 | 3.09646860726702 | -1.83133814105385 | 0.0740659394472206 | 0.335027410404695 | -4.6163684842772 |
| ENSG00000118495.19 | PLAGL1 | -0.413059004826487 | 3.80097306422568 | -1.83123408037848 | 0.0740817613063444 | 0.335027410404695 | -4.68855716858941 |
| ENSG00000115756.12 | HPCAL1 | -0.310863629312032 | 4.65369343109938 | -1.83113278859102 | 0.0740971649184944 | 0.335027410404695 | -4.75858396699287 |
| ENSG00000077984.5 | CST7 | -1.81429854076305 | -1.55017708592565 | -1.83152613176456 | 0.0741006198122194 | 0.335027410404695 | -4.05405575563837 |

| | | | | | | | |
|---|---|---|---|---|---|---|---|
| ENSG00000102054.17 | RBBP7 | 0.140911165637619 | 6.6433432832578 | 1.83079240413178 | 0.0741489476162147 | 0.335027410404695 | -4.80424063074378 |
| ENSG00000185808.13 | PIGP | -0.315672854968221 | 1.82688791878483 | -1.83077536586963 | 0.0741515404522299 | 0.335027410404695 | -4.46819432387998 |
| lnc-PYGO2-2 | lnc-PYGO2-2 | 0.405974281976766 | 5.28913502994485 | 1.83079630638035 | 0.0741555515502656 | 0.335027410404695 | -4.78462187127104 |
| lnc-AGXT2-1 | lnc-AGXT2-1 | 0.25771440999887 | 0.800677497475436 | 1.83074847777037 | 0.0741556323655426 | 0.335027410404695 | -4.34814438538186 |
| lnc-PINX1-86 | lnc-PINX1-86 | -0.516202957114528 | 1.59073014178745 | -1.83064098919932 | 0.0741719922145759 | 0.335027410404695 | -4.45914112201726 |
| HM13-IT1 | HM13-IT1 | -0.390768560709623 | -0.43073663433529 | -1.83060051486565 | 0.074178153233062 | 0.335027410404695 | -4.15076546383835 |
| lnc-SURF6-2 | lnc-SURF6-2 | -0.41090561994753 | 0.077992320067288 | -1.83052200504422 | 0.0741901052628189 | 0.335027410404695 | -4.29043746109367 |
| ENSG00000159239.13 | | -0.55517851004983 | 0.544679101283056 | -1.83033022173328 | 0.0742193084612256 | 0.335032168042374 | -4.30005768840107 |
| ENSG00000234329.1 | RPL7AP16 | 0.30245705387467 | 1.49587499117758 | 1.83030130127008 | 0.0742237130774536 | 0.335032168042374 | -4.43988356490558 |
| ENSG00000268061.5 | NAPA-AS1 | -0.396524753989647 | 0.179813730530069 | -1.83009534907667 | 0.0742550862050447 | 0.335100293654557 | -4.23475019831659 |
| ENSG00000214826.5 | DDX12P | 0.520492741989248 | 2.69461647005186 | 1.83003326129526 | 0.0742713874012344 | 0.335100387303728 | -4.57353133612414 |
| lnc-IFNA1-5 | lnc-IFNA1-5 | -0.853525960294302 | 1.49865829054831 | -1.83045741371263 | 0.0743386304869393 | 0.33530887909899 | -4.44439742859298 |
| ENSG00000066084.12 | DIP2B | 0.16989550167481 | 7.06946479547667 | 1.82940391540401 | 0.0743604958295744 | 0.33530887909899 | -4.80315766256681 |
| LINC00893 | LINC00893 | 0.289116457718597 | 2.69785546448044 | 1.82921399367844 | 0.0743894717064032 | 0.33530887909899 | -4.55975122614322 |
| ENSG00000166411.13 | IDH3A | 0.153273337176391 | 4.64031991993359 | 1.82903355464561 | 0.0744170096821115 | 0.33530887909899 | -4.76150376710367 |
| ENSG00000225200.2 | | 1.02616439630436 | 0.763220842240791 | 1.82956848230407 | 0.0744223375408473 | 0.33530887909899 | -4.24943977435147 |
| ENSG00000127314.17 | RAP1B | 0.214451297457435 | 6.21963359437288 | 1.82897962240739 | 0.0744252423062702 | 0.33530887909899 | -4.8081526416509 |
| ENSG00000108590.10 | MED31 | -0.312320078346488 | 2.58819882441738 | -1.82893777084668 | 0.0744316313755666 | 0.33530887909899 | -4.56886396001608 |
| ENSG00000226521.7 | | -0.650654332429752 | 0.485136901345764 | -1.82878701499882 | 0.0744546496460372 | 0.335315141941419 | -4.24974057415416 |
| ENSG00000197568.13 | ANKRD13C-DT | -0.493415130705645 | -0.357411142482528 | -1.82871528864355 | 0.0744656033529722 | 0.335315141941419 | -4.14309432128327 |
| ENSG00000173575.20 | CHD2 | 0.231263624172536 | 9.14760801535061 | 1.82818603784297 | 0.0745464701648904 | 0.335600515428659 | -4.76585007186331 |
| lnc-FXYD4-4 | lnc-FXYD4-4 | -0.26806283691841 | 1.19751369419404 | -1.82800734154838 | 0.0745737908151199 | 0.335600515428659 | -4.41066984641595 |
| lnc-CALB1-3 | lnc-CALB1-3 | 0.331298588211221 | 1.15434466430249 | 1.82787513503467 | 0.0745940091527694 | 0.335600515428659 | -4.40482135081488 |
| ENSG00000279555.1 | | 0.465502071269545 | 3.64888327509123 | 1.82783823197345 | 0.074601451168591 | 0.335600515428659 | -4.65815165086708 |
| ENSG00000142539.9 | | -0.459634149286583 | 3.6717816344324 | -1.82780183902516 | 0.0746170140182484 | 0.335600515428659 | -4.6917947153316 |
| ENSG00000110711.9 | AIP | -0.259750463281021 | 5.4139631631739 | -1.82763642744011 | 0.0746305264420581 | 0.335600515428659 | -4.79952159402209 |
| lnc-MED4-4 | lnc-MED4-4 | 0.356354274369807 | 0.212064336534865 | 1.82755418523957 | 0.0746431112875869 | 0.335600515428659 | -4.260237274278 |
| BOLA3-AS1 | BOLA3-AS1 | -0.568557191968871 | -0.301631387689497 | -1.82727657033678 | 0.0746856056640927 | 0.335709812517977 | -4.19102523448442 |
| ENSG00000123600.19 | METTL8 | -0.41800933319961 | 6.36053786630846 | -1.827326899003 | 0.0747000408751075 | 0.335709812517977 | -4.81022979646772 |
| lnc-NEURL3-6 | lnc-NEURL3-6 | 0.263082175478531 | 1.36852877837584 | 1.82690849100931 | 0.0747419789360923 | 0.335824962331715 | -4.43620749297293 |
| ENSG00000033627.16 | ATP6V0A1 | -0.309409888406249 | 5.48904978053292 | -1.82679723875364 | 0.0747590248817227 | 0.335828242937691 | -4.8019117132618 |
| lnc-BTLA-4 | lnc-BTLA-4 | -0.614811260610677 | 5.07293488016134 | -1.82757317493232 | 0.0748011035255101 | 0.335943948039619 | -4.77598523488983 |
| lnc-TMEM268-1 | lnc-TMEM268-1 | -0.600786767951476 | 0.0673384039699249 | -1.82623019429761 | 0.0748459578944737 | 0.33596032495037 | -4.24142977814593 |
| ENSG00000063176.15 | SPHK2 | -0.22268485959991 | 2.73955522807849 | -1.8262278842217 | 0.074846312224663 | 0.33596032495037 | -4.58651471278929 |
| ENSG00000283944.1 | MIR4709 | 0.419776417282691 | 1.50902759241969 | 1.82601052434473 | 0.0748796582409478 | 0.33596032495037 | -4.43666017065605 |
| ENSG00000140379.7 | BCL2A1 | 0.683417334620116 | 2.33438025396149 | 1.82636697756606 | 0.0748928087154074 | 0.33596032495037 | -4.50068211916215 |
| lnc-OR4F29-11 | lnc-OR4F29-11 | 0.323682232761144 | 2.73910726536528 | 1.82581990184149 | 0.0749089127138915 | 0.33596032495037 | -4.5923423015147 |
| lnc-MFSD4B-1 | lnc-MFSD4B-1 | -0.355009846379375 | 2.8631753499605 | -1.82554602671319 | 0.0749509607277975 | 0.33596032495037 | -4.59392582713159 |
| lnc-PRSS16-5 | lnc-PRSS16-5 | -0.782986149232398 | -0.357791254143216 | -1.82553894756887 | 0.0749520478525025 | 0.33596032495037 | -4.18283398550306 |
| lnc-C5orf67-4 | lnc-C5orf67-4 | -1.04373471271887 | 0.130585633869231 | -1.82624276316811 | 0.0749535877172245 | 0.33596032495037 | -4.27276666806928 |
| ENSG00000131503.20 | ANKHD1 | 0.23237438476827 | 6.53136552500262 | 1.82544154266113 | 0.0749670074100485 | 0.33596032495037 | -4.81403253362692 |
| lnc-ELF1-1 | lnc-ELF1-1 | -0.265827205220576 | 3.46326430311618 | -1.82543526104694 | 0.0749679722342248 | 0.33596032495037 | -4.65609512176201 |
| ENSG00000197714.8 | ZNF460 | 0.210672024372611 | 6.34263922832005 | 1.82527444691254 | 0.0749926760477202 | 0.335964128207575 | -4.81449996854731 |
| ENSG00000208772.1 | SNORD94 | 0.544043930592245 | 2.50846554755009 | 1.82527236400554 | 0.0750014657296705 | 0.335964128207575 | -4.54931638674626 |
| ENSG00000110048.11 | OSBP | 0.234723695015051 | 5.4633814164885 | 1.82501025726247 | 0.0750332750596034 | 0.336023510448606 | -4.80289055629897 |
| ENSG00000133985.2 | TTC9 | -0.490377726277794 | 4.37117255915133 | -1.82530888355252 | 0.0750473729549747 | 0.336023510448606 | -4.75264113808188 |
| ENSG00000087995.15 | METTL2A | -0.211992030482284 | 3.16139297932014 | -1.82454101394167 | 0.0751054312570476 | 0.336210328743761 | -4.63469816822949 |
| ENSG00000116685.15 | KIAA2013 | -0.386878109624399 | 4.82846748181366 | -1.82428961302563 | 0.0751441136560717 | 0.336310348202419 | -4.77944706723948 |
| lnc-TNK2-2 | lnc-TNK2-2 | -0.29900441341146 | 3.25768053110183 | -1.82392608972049 | 0.0752000778204851 | 0.336400507365033 | -4.637176918346 |
| lnc-SPOPL-2 | lnc-SPOPL-2 | 0.267136419721103 | 2.75829744260382 | 1.82390512172656 | 0.0752033069034113 | 0.336400507365033 | -4.58765814906373 |
| ENSG00000256806.5 | C17orf100 | -0.442951352294106 | 0.910226177506795 | -1.82354880771383 | 0.0752581973868892 | 0.336400507365033 | -4.40533530100981 |
| ENSG00000185386.14 | MAPK11 | -0.597597977809896 | 2.32507750296108 | -1.82372222743529 | 0.0752737341780244 | 0.336400507365033 | -4.55607246788423 |
| ENSG00000252481.1 | SCARNA13 | 0.343905701239498 | 6.38438632711477 | 1.82331660559069 | 0.0752939865509931 | 0.336400507365033 | -4.81774876876906 |
| ENSG00000099330.8 | OCEL1 | -0.322842480323141 | 0.795646530665313 | -1.82328463545272 | 0.0752989152144813 | 0.336400507365033 | -4.36466812536797 |
| LINC01678 | LINC01678 | -0.842464958733438 | -0.777320485369096 | -1.82321689532182 | 0.0753093592457392 | 0.336400507365033 | -4.09738723693713 |
| lnc-DIP2A-7 | lnc-DIP2A-7 | -0.680611287333333 | 1.77731441635828 | -1.82358851555813 | 0.0753107164365651 | 0.336400507365033 | -4.48252509709174 |
| ENSG00000129465.15 | RIPK3 | -0.236574212204402 | 3.76448358287791 | -1.82312938115009 | 0.0753228538100368 | 0.336400507365033 | -4.69287342195022 |
| lnc-HVCN1-2 | lnc-HVCN1-2 | -0.353970134392523 | 1.11645559666945 | -1.82300985794362 | 0.0753412874254551 | 0.336400507365033 | -4.40866301771416 |
| ENSG00000141096.4 | DPEP3 | -0.437876213423494 | 0.861439071068095 | -1.82293552449332 | 0.0753527535176163 | 0.336400507365033 | -4.36860797027425 |
| PPP4R1-AS1 | PPP4R1-AS1 | -0.527578058112686 | -0.719077174681902 | -1.82288607617583 | 0.0753603818423052 | 0.336400507365033 | -4.14431748882642 |
| ENSG00000115368.9 | WDR75 | 0.200403771278247 | 5.85083949869924 | 1.82275800476144 | 0.0753801422801281 | 0.336415756407605 | -4.81560819813138 |
| ENSG00000230847.4 | OCLNP1 | 2.6318159599294 | -1.34708205602034 | 1.8231748967637 | 0.0754319295776405 | 0.336534278209077 | -4.08852669414718 |
| ENSG00000213079.9 | SCAF8 | 0.18328852731358 | 6.93060524149717 | 1.82237230483692 | 0.0754396792815522 | 0.336534278209077 | -4.81639948013011 |
| ENSG00000234444.9 | ZNF736 | 0.178849310667729 | 6.82767545890486 | 1.82226824024699 | 0.0754557495960206 | 0.336534278209077 | -4.81785742618617 |
| ENSG00000158711.13 | ELK4 | 0.174396838054681 | 7.48673073352634 | 1.82203672633928 | 0.0754915118222752 | 0.336620837919821 | -4.81049756802448 |
| MIRLET7BHG | MIRLET7BHG | -0.56582498602347 | 2.08197854556097 | -1.82178224586775 | 0.0755495386476186 | 0.336761817264081 | -4.54645027826615 |
| ENSG00000160961.11 | ZNF333 | -0.24482428801799 | 4.36751130201055 | -1.82140772755319 | 0.0755887463459908 | 0.336761817264081 | -4.75509286278921 |
| ENSG00000166446.14 | CDYL2 | 0.61936671025965 | 2.98644240354883 | 1.82169991203019 | 0.0756185344273982 | 0.336761817264081 | -4.59643431774886 |

| | | | | | | | |
|---|---|---|---|---|---|---|---|
| ENSG00000213337.8 | ANKRD39 | -0.178904234495163 | 2.43742387909566 | -1.82120617248203 | 0.0756199263591733 | 0.336761817264081 | -4.55724263100984 |
| ENSG00000104320.13 | NBN | 0.238751542717297 | 5.60177347911713 | 1.8211187751389 | 0.0756334498648908 | 0.336761817264081 | -4.81486197303171 |
| ENSG00000159147.17 | DONSON | -0.274083011818473 | 2.50657112049076 | -1.8210884822865 | 0.0756381377333288 | 0.336761817264081 | -4.57259762009524 |
| lnc-BTBD6-2 | lnc-BTBD6-2 | -0.277344919983592 | 0.887794783099774 | -1.82096913060938 | 0.0756566099900863 | 0.336761817264081 | -4.3859351625953 |
| ENSG00000183401.11 | CCDC159 | -0.251278821087383 | 3.60517839992994 | -1.82092411873722 | 0.0756635775420159 | 0.336761817264081 | -4.6724266945747 |
| ENSG00000277108.1 | SNORD49B | 0.341115328498959 | 1.57525320055033 | 1.82088018516339 | 0.0756703787031227 | 0.336761817264081 | -4.4579050045062 |
| lnc-RUNX1-12 | lnc-RUNX1-12 | 0.249666772093893 | 0.752761686949832 | 1.82064415521377 | 0.0757069262932268 | 0.336851635082898 | -4.33826422864418 |
| lnc-TMEM109-2 | lnc-TMEM109-2 | -0.296812758711493 | 2.5404417576085 | -1.82042056549391 | 0.0757415613570645 | 0.336913571402045 | -4.57425963644423 |
| lnc-GTF2F2-9 | lnc-GTF2F2-9 | -0.435719813061387 | -0.243297044674944 | -1.82034297591423 | 0.0757535834644447 | 0.336913571402045 | -4.20899482305308 |
| ENSG00000067182.7 | TNFRSF1A | -1.47055181711252 | 1.25658767044556 | -1.82188383709264 | 0.0757844220903384 | 0.33697791313144 | -4.4537345360134 |
| lnc-SNRPN-8 | lnc-SNRPN-8 | -0.369343282222418 | 7.98711102731759 | -1.81940556959685 | 0.0758989572245138 | 0.337414305950792 | -4.80658389092568 |
| ENSG00000185338.5 | SOCS1 | -0.577614085255342 | 0.511309290592762 | -1.81913536684698 | 0.075940904229336 | 0.337527884204799 | -4.32990326936732 |
| ENSG00000099821.13 | POLRMT | -0.291123863141073 | 4.62446034560221 | -1.81873561067129 | 0.0760029993791101 | 0.337730944779841 | -4.77784949227269 |
| ENSG00000183722.8 | LHFPL6 | -1.91117505241446 | -0.874253404547867 | -1.81920408553065 | 0.0760649603989902 | 0.337927420168673 | -4.07570537953344 |
| ENSG00000213722.8 | DDAH2 | -0.421825287275098 | 3.40881645057886 | -1.81807158422441 | 0.0761062391019201 | 0.337927420168673 | -4.67295210380636 |
| ENSG00000136819.15 | C9orf78 | 0.199130322928958 | 5.58944515744203 | 1.81804232146689 | 0.0761107914622612 | 0.337927420168673 | -4.81861665848123 |
| ENSG00000181284.2 | TMEM102 | -0.416722326750443 | 1.2389980340324 | -1.81800246121513 | 0.0761169928278514 | 0.337927420168673 | -4.44903074727996 |
| ENSG00000133065.10 | SLC41A1 | 0.187558815663343 | 4.31307800410534 | 1.81792334262891 | 0.0761293031784549 | 0.337927420168673 | -4.75031658840632 |
| ENSG00000237441.9 | RGL2 | -0.215037001364685 | 5.15946681251641 | -1.81776842866358 | 0.076153411679652 | 0.337928400057057 | -4.80835294316808 |
| ENSG00000171121.16 | KCNMB3 | -0.472883664685893 | 2.08610151106799 | -1.81757824487151 | 0.0761830178687458 | 0.337928400057057 | -4.55185447770124 |
| ENSG00000159921.15 | GNE | 0.225661956341432 | 5.13522942158872 | 1.8175106197356 | 0.0761935475152679 | 0.337928400057057 | -4.80651923333853 |
| ENSG00000100815.12 | TRIP11 | -0.210466998125733 | 6.10534703338985 | -1.817500037851 | 0.0761951952905214 | 0.337928400057057 | -4.82647911106818 |
| ENSG00000161835.10 | TAMALIN | 1.1754673708008 | 0.650167306621633 | 1.81846653703696 | 0.0762136972676192 | 0.337936147338726 | -4.30539926118443 |
| ENSG00000122188.12 | LAX1 | 0.34067305863462 | 6.34713311663184 | 1.81727799136911 | 0.0762297785596708 | 0.337936147338726 | -4.82773269985231 |
| ENSG00000125458.6 | NT5C | -0.361212442926206 | 3.93505322099995 | -1.81707217853763 | 0.0762618453098498 | 0.338005504309354 | -4.72392379068857 |
| lnc-TARSL2-3 | lnc-TARSL2-3 | 0.314456470963824 | 1.49475083017188 | 1.8168411454384 | 0.0762978550850129 | 0.33809230381374 | -4.46584528632564 |
| ENSG00000233276.4 | GPX1 | -0.423949913561276 | 4.86535853813323 | -1.81567999625504 | 0.0764981952957529 | 0.338855725234519 | -4.79716295212194 |
| lnc-AES-1 | lnc-AES-1 | 0.189701170198223 | 5.66577324609623 | 1.81544741625271 | 0.0765153922140388 | 0.338855725234519 | -4.82460002182399 |
| ENSG00000262481.5 | TMEM256-PLSCR3 | -0.24064774691741 | 2.78978042820064 | -1.81542096184185 | 0.07651952635136 | 0.338855725234519 | -4.60930124953664 |
| ENSG00000136271.10 | DDX56 | 0.165061325520916 | 5.00609920188642 | 1.8148332639406 | 0.0766114169143184 | 0.33904319572732 | -4.80443132112661 |
| ENSG00000108509.20 | CAMTA2 | -0.215028818553168 | 3.80324386833427 | -1.8147696673873 | 0.0766213662518308 | 0.33904319572732 | -4.71334458716913 |
| ENSG00000132294.14 | EFR3A | 0.21072783410219 | 6.39990862359535 | 1.81465286634699 | 0.0766396419856841 | 0.33904319572732 | -4.83235282403055 |
| ENSG00000213463.4 | SYNJ2BP | -0.169930307692921 | 5.10942549202564 | -1.81450709105737 | 0.0766624564519582 | 0.33904319572732 | -4.807530899031 |
| ENSG00000187049.9 | TMEM216 | -0.313928265990407 | 2.4900207851499 | -1.81441485486365 | 0.0766768947784588 | 0.33904319572732 | -4.57794453731738 |
| lnc-NPPA-1 | lnc-NPPA-1 | -0.221683794169834 | 1.1622876715984 | -1.81431599332142 | 0.076692372762327 | 0.33904319572732 | -4.42960232499804 |
| ENSG00000120915.13 | EPHX2 | -1.48228587049328 | -1.57388985455436 | -1.81451119428462 | 0.0767204268665411 | 0.33904319572732 | -4.09344378991282 |
| ENSG00000142396.10 | ERVK3-1 | 0.281144760473373 | 5.73765051983703 | 1.8140315405066 | 0.0767369220435862 | 0.33904319572732 | -4.82857407590235 |
| ENSG00000197162.9 | ZNF785 | -0.253319138580559 | 2.81817779498347 | -1.81390730743763 | 0.0767563855289516 | 0.33904319572732 | -4.61207968693656 |
| ENSG00000198053.11 | SIRPA | 1.99022254475591 | -0.854658791879013 | 1.81491009080489 | 0.0767571844477394 | 0.33904319572732 | -4.08553159655159 |
| ENSG00000186854.10 | TRABD2A | -0.66503342800712 | 3.41063319246879 | -1.81471331489855 | 0.0767725642616892 | 0.33904319572732 | -4.66713124491843 |
| lnc-ZMIZ2-2 | lnc-ZMIZ2-2 | 0.302460453120847 | 2.14185054639503 | 1.81374451243597 | 0.0767818967864741 | 0.33904319572732 | -4.53410863712346 |
| ENSG00000163714.17 | U2SURP | 0.154831795846166 | 7.79010977107054 | 1.81371248928174 | 0.0767869159090223 | 0.33904319572732 | -4.81910050019116 |
| ENSG00000116138.12 | DNAJC16 | -0.225391009813925 | 5.45973079402141 | -1.81367706473787 | 0.0767924684681905 | 0.33904319572732 | -4.82096589346429 |
| ENSG00000152117.17 | SMPD4BP | 0.229909638619226 | 5.08975354387002 | 1.81286808051854 | 0.0769193635834924 | 0.339530615620636 | -4.81050567945463 |
| ENSG00000186908.14 | ZDHHC17 | 0.167599143321646 | 6.43443687143463 | 1.81260498913339 | 0.0769606695170291 | 0.33959369196642 | -4.83560120393156 |
| ENSG00000152818.18 | UTRN | 0.268396307807697 | 8.97532342659958 | 1.81256690233027 | 0.0769665507809041 | 0.33959369196642 | -4.79613674059098 |
| lnc-ZNF708-13 | lnc-ZNF708-13 | -0.351347145611746 | 2.99386974113093 | -1.81230640102864 | 0.0770075711926573 | 0.339698807961932 | -4.63522092950268 |
| lnc-MRPL1-3 | lnc-MRPL1-3 | -0.489884243809969 | 0.1443745879165 | -1.8122051415211 | 0.0770234823280542 | 0.339698807961932 | -4.28448684931354 |
| ENSG00000116954.7 | RRAGC | 0.285199273066345 | 3.99161681251499 | 1.8120355135631 | 0.0770501425692846 | 0.33974359136752 | -4.73423848964485 |
| lnc-CLEC2A-3 | lnc-CLEC2A-3 | -3.2606337996123 | 0.4698586963183 | -1.8136018684851 | 0.0770863215743016 | 0.339778489726032 | -4.1636711224718 |
| ENSG00000284179.1 | MIR7-1 | 0.349283332168612 | 1.29046917341411 | 1.81174493012517 | 0.0770958313480307 | 0.339778489726032 | -4.46190535582273 |
| ENSG00000182670.13 | TTC3 | 0.159256043947181 | 8.03165019435528 | 1.81167022949887 | 0.0771075803094931 | 0.339778489726032 | -4.81978419985275 |
| lnc-CDHR4-5 | lnc-CDHR4-5 | 0.313697214740495 | 1.46476284031614 | 1.81126078883845 | 0.0771720042330875 | 0.339989589710968 | -4.4707567574996 |
| ENSG00000139304.13 | PTPRQ | -1.92528209645883 | 1.03402048270146 | -1.81268831176826 | 0.0772067774129185 | 0.340069997750931 | -4.25173369138082 |
| ENSG00000214279.13 | SCART1 | -0.36834541735849 | 2.33247851058663 | -1.81027053047025 | 0.077328005386422 | 0.340531094323647 | -4.56781141880691 |
| lnc-DOK4-1 | lnc-DOK4-1 | 0.227475791494181 | 1.08806140435516 | 1.81013974535844 | 0.0773486286008471 | 0.340549052939302 | -4.42702757260224 |
| lnc-SEC24A-1 | lnc-SEC24A-1 | -1.05821694334718 | 0.00128536161210392 | -1.81053024091943 | 0.0773999026496276 | 0.340641168766136 | -4.28667623023363 |
| ENSG00000011243.17 | AKAP8L | -0.236217839596666 | 6.14655301574215 | -1.80977600082055 | 0.0774060110717326 | 0.340641168766136 | -4.83978867607121 |
| ENSG00000101842.13 | VSIG1 | -1.67635158030403 | -1.72320651546056 | -1.80970619769854 | 0.0774191997030551 | 0.340641168766136 | -4.08203918781663 |
| ENSG00000166510.13 | CCDC68 | -1.22167707567233 | 1.06054666053605 | -1.81012286477922 | 0.0775509609331975 | 0.341115056971113 | -4.37961990521999 |
| ENSG00000126216.14 | TUBGCP3 | 0.195185695692492 | 5.06890143431302 | 1.80865991685972 | 0.0775823029370709 | 0.341115056971113 | -4.81653688272203 |
| ENSG00000128928.8 | IVD | -0.270584073754196 | 5.39436236405049 | -1.80851234496625 | 0.0776056380942326 | 0.341115056971113 | -4.82814606793989 |
| ENSG00000196670.13 | ZFP62 | -0.187001604756001 | 4.84811795917786 | -1.80846342724708 | 0.0776133746313964 | 0.341115056971113 | -4.80663017150717 |
| ENSG00000215190.9 | LINC00680 | -0.242126046420967 | 3.00520507525597 | -1.808391281672 | 0.0776247859360353 | 0.341115056971113 | -4.64319650068223 |
| lnc-ZC2HC1C-1 | lnc-ZC2HC1C-1 | -0.579237417357946 | -0.0463091405681707 | -1.80837867510012 | 0.0776267800695586 | 0.341115056971113 | -4.25169328034735 |
| ENSG00000164919.10 | COX6C | 0.326357492031191 | 5.11611254086806 | 1.80817505813715 | 0.0776589946056474 | 0.341115056971113 | -4.81738095645098 |
| lnc-KIAA1109-1 | lnc-KIAA1109-1 | 0.294426021737194 | 0.68718820391692 | 1.80817196607086 | 0.077659483892855 | 0.341115056971113 | -4.36231338812095 |
| ENSG00000277947.1 | SNORD3D | -0.661553062614367 | 2.01876300835694 | -1.80850271520033 | 0.0776905805737206 | 0.341178839331959 | -4.57323947951918 |

| | | | | | | | |
|---|---|---|---|---|---|---|---|
| lnc-KIZ-9 | lnc-KIZ-9 | 0.43534941821294 | 1.39057083301632 | 1.80778570918728 | 0.0777206254632033 | 0.341237976516449 | -4.47195884058526 |
| ENSG00000101246.19 | ARFRP1 | -0.202606469223855 | 4.80336314525475 | -1.80757853453105 | 0.0777534363821954 | 0.341309230337967 | -4.80665388428383 |
| ENSG00000106692.14 | FKTN | -0.200691709855212 | 4.43044936586814 | -1.80735485833843 | 0.077788873815981 | 0.34131174791028 | -4.77952100374537 |
| ENSG00000109944.10 | JHY | 0.348882512173232 | 1.73808377939662 | 1.80729329935949 | 0.0777986291088285 | 0.34131174791028 | -4.52453029309855 |
| ENSG00000103932.11 | RPAP1 | -0.203789666325128 | 4.34157834533066 | -1.80720276016491 | 0.0778129787904354 | 0.34131174791028 | -4.76936081806996 |
| lnc-NADSYN1-1 | lnc-NADSYN1-1 | -0.345331619650003 | 1.48279807574733 | -1.80715632788651 | 0.077820338771945 | 0.34131174791028 | -4.48301903549341 |
| ENSG00000065243.19 | PKN2 | 0.185239393859328 | 6.45573863828102 | 1.80667403355359 | 0.0778968219800214 | 0.341528599950576 | -4.84560901965962 |
| ENSG00000176945.16 | MUC20 | -0.590312063879013 | 1.8494997546655 | -1.80671250398229 | 0.0779029673404243 | 0.341528599950576 | -4.51552987755764 |
| ENSG00000174132.8 | FAM174A | -0.339018122607535 | 1.65435288113309 | -1.80646658783815 | 0.0779297386289278 | 0.341573213415507 | -4.51613786588703 |
| lnc-NECAB1-1 | lnc-NECAB1-1 | -0.597914660882829 | -0.367881468114104 | -1.80613781845267 | 0.0779819304503135 | 0.34172920469636 | -4.18273207006324 |
| ENSG00000279166.1 | | 0.389023031262187 | 4.07189027861003 | 1.80590836992466 | 0.0780183726403712 | 0.341793862960344 | -4.7375740875815 |
| ENSG00000225159.1 | NPM1P39 | 0.98355184200815 | -0.889884412295711 | 1.80578075581643 | 0.0780386471792372 | 0.341793862960344 | -4.08686627326577 |
| ENSG00000272162.1 | | 0.301426107079505 | 1.26526300669962 | 1.80573132109563 | 0.0780465022549491 | 0.341793862960344 | -4.45650886528927 |
| ENSG00000181555.20 | SETD2 | 0.175225845239489 | 7.76468294632173 | 1.80555346598366 | 0.0780747685808236 | 0.341838009768044 | -4.83486513962562 |
| ENSG00000213903.8 | LTB4R | -0.298329639447604 | 3.11512499808415 | -1.80544802264727 | 0.078091530654135 | 0.341838009768044 | -4.6517523579633 |
| lnc-SEPSECS-4 | lnc-SEPSECS-4 | -0.433418353293273 | 0.776185169076015 | -1.80522459411559 | 0.0781270585751347 | 0.341838009768044 | -4.37417019834956 |
| lnc-AKR1E2-13 | lnc-AKR1E2-13 | -0.470541480105874 | 0.609572746660135 | -1.80517965272633 | 0.0781342064619086 | 0.341838009768044 | -4.37025118639879 |
| ENSG00000268225.2 | NA | -0.37960986563786 | 0.0443013850894814 | -1.80508011404353 | 0.0781500399609263 | 0.341838009768044 | -4.24456065267115 |
| ENSG00000124243.17 | BCAS4 | -0.223142198595959 | 7.1244024332865 | -1.80501319760939 | 0.0781606857982429 | 0.341838009768044 | -4.84397195297655 |
| ENSG00000084110.10 | HAL | 0.381715660782885 | 1.9363800233733 | 1.80493682586511 | 0.0781728373890193 | 0.341838009768044 | -4.51726187881119 |
| ENSG00000205542.10 | TMSB4X | 0.459420994923319 | 8.79529519737597 | 1.80496574967137 | 0.0782173443237795 | 0.341915595382299 | -4.81311825654293 |
| ENSG00000133246.11 | PRAM1 | -0.776743179222623 | -0.380904862548121 | -1.80461661971489 | 0.0782238030731792 | 0.341915595382299 | -4.1842406907334 |
| ENSG00000281028.1 | NA | 0.193858601859148 | 3.21296571286035 | 1.80419999997025 | 0.0782901563744496 | 0.342132969990509 | -4.67189471216936 |
| ENSG00000166682.11 | TMPRSS5 | 0.441170011720468 | 0.147134436762356 | 1.80377641964524 | 0.078357666888202 | 0.342336602620109 | -4.24820845871128 |
| lnc-FBXL20-1 | lnc-FBXL20-1 | -0.199988873174295 | 2.34974188000663 | -1.8036405489243 | 0.0783793324447472 | 0.342336602620109 | -4.58219780611412 |
| ENSG00000214182.5 | PTMAP5 | 0.662653786916566 | 1.51035705607016 | 1.80348521520746 | 0.0784160030237911 | 0.342336602620109 | -4.43237444345693 |
| ENSG00000176769.9 | TCERG1L | 2.17695371169781 | -3.30243060956679 | 1.80328630147549 | 0.078435843472385 | 0.342336602620109 | -4.17384365221344 |
| ENSG00000174428.16 | GTF2IRD2B | 0.197524413433893 | 3.03583765258979 | 1.80328296394162 | 0.0784363760529173 | 0.342336602620109 | -4.6506045640992 |
| lnc-ALPK1-2 | lnc-ALPK1-2 | -0.415152849641512 | -0.143746163115384 | -1.80328190148036 | 0.0784365455937636 | 0.342336602620109 | -4.23266096986033 |
| ENSG00000242689.2 | CNTF | 0.316336229205717 | 1.0837721455172 | 1.802851468273 | 0.0785052568136969 | 0.342563854355803 | -4.43644293157486 |
| ENSG00000084652.15 | TXLNA | -0.254962931529203 | 5.74679458787438 | -1.80260799281716 | 0.078544145916729 | 0.342637135183685 | -4.84738476568353 |
| ENSG00000112992.16 | NNT | -0.202209060170609 | 5.81916269618505 | -1.80247660337793 | 0.0785651388302764 | 0.342637135183685 | -4.8491212325936 |
| ENSG00000183655.12 | KLHL25 | 0.438264967064736 | -0.13367071504646 | 1.80243372760788 | 0.0785719903836659 | 0.342637135183685 | -4.22723804171284 |
| lnc-ITGB7-4 | lnc-ITGB7-4 | 0.269798033598367 | 0.222010207456556 | 1.80225178554035 | 0.0786010703504111 | 0.34268731105581 | -4.29649855254335 |
| lnc-RELL1-8 | lnc-RELL1-8 | 0.602931803408709 | -0.542420733715819 | 1.80212910133393 | 0.0786206841980005 | 0.34268731105581 | -4.17295317985955 |
| ENSG00000205659.10 | LIN52 | 0.225051734377114 | 3.49589214743007 | 1.80201566716964 | 0.0786388228861328 | 0.34268731105581 | -4.69499743585685 |
| ENSG00000257524.6 | | -0.231980789365877 | 5.18880460162761 | -1.80165726107367 | 0.078696156979268 | 0.34268731105581 | -4.83394596169924 |
| ENSG00000132570.14 | PCBD2 | -0.254363295637053 | 3.62936279455595 | -1.80160573854912 | 0.0787044019134996 | 0.34268731105581 | -4.71790422884658 |
| ENSG00000198081.10 | ZBTB14 | -0.203960073148949 | 4.89814723770404 | -1.80156976672143 | 0.0787101587661545 | 0.34268731105581 | -4.82132646230409 |
| lnc-ZHX1-5 | lnc-ZHX1-5 | 0.271485069346769 | 1.59999661302902 | 1.80148172901386 | 0.0787242496250686 | 0.34268731105581 | -4.49721046765385 |
| ENSG00000108424.9 | KPNB1 | 0.169192691619556 | 7.79726693201285 | 1.80147428678322 | 0.0787254408873284 | 0.34268731105581 | -4.84019988963391 |
| ENSG00000176473.13 | WDR25 | -0.21108315753519 | 1.40038873309678 | -1.80142495379496 | 0.0787333378993794 | 0.34268731105581 | -4.47091579924918 |
| ENSG00000109475.16 | RPL34 | 0.238141408667094 | 8.46939103177496 | 1.80123099103895 | 0.0787643930917712 | 0.342750000635925 | -4.8270762474241 |
| lnc-FAAH2-1 | lnc-FAAH2-1 | -0.376794010749597 | 0.887827492138993 | -1.8008859241204 | 0.0788196669385438 | 0.342918030986271 | -4.41312696580435 |
| ENSG00000132819.16 | RBM38 | 0.310862394114402 | 7.56448347180144 | 1.80035435996795 | 0.0789048784072239 | 0.343132041607754 | -4.84584350935281 |
| ENSG00000105662.15 | CRTC1 | -0.29177776737431 | 4.44654853927957 | -1.80020498173259 | 0.0789288381889034 | 0.343132041607754 | -4.79098503315564 |
| lnc-TRAFD1-4 | lnc-TRAFD1-4 | 0.783698481513716 | -0.606702361453625 | 1.80013117263378 | 0.0789406791895973 | 0.343132041607754 | -4.18968408865553 |
| ENSG00000157259.7 | GATAD1 | -0.198627579476449 | 5.51829571192953 | -1.80003755776068 | 0.0789556997313216 | 0.343132041607754 | -4.84691316277694 |
| lnc-GLB1L-2 | lnc-GLB1L-2 | 0.322397576927608 | 1.51550367792132 | 1.80002934522501 | 0.078957017550624 | 0.343132041607754 | -4.49758534736269 |
| ENSG00000155252.13 | PI4K2A | 0.364556549019682 | 5.37966551594024 | 1.79995541738305 | 0.0789688811687281 | 0.343132041607754 | -4.83961089428561 |
| ENSG00000167323.11 | STIM1 | 0.215588487567859 | 6.00622621804787 | 1.79965082147828 | 0.079017777231699 | 0.343255769105831 | -4.85579253084679 |
| lnc-TRIM23-1 | lnc-TRIM23-1 | -0.594346098558873 | 3.04004820546812 | -1.80004527678039 | 0.0790339003168592 | 0.343255769105831 | -4.66540162708047 |
| ENSG00000244193.1 | | -1.12067340841984 | -0.820387814408453 | -1.79949755356225 | 0.0790473859768567 | 0.343255769105831 | -4.14588379470095 |
| lnc-KLHL36-1 | lnc-KLHL36-1 | -0.487661342380765 | 2.29161998175121 | -1.79927784272918 | 0.0790776853595349 | 0.34331491199226 | -4.59836800352205 |
| lnc-TMC8-1 | lnc-TMC8-1 | -0.270939506968907 | 0.561999206866613 | -1.79899727646021 | 0.0791227753321063 | 0.343438229578394 | -4.37997294166213 |
| ENSG00000134899.19 | ERCC5 | -0.12072845006705 | 6.22157419588053 | -1.79847733223719 | 0.0792063931242702 | 0.343728692742326 | -4.85896471951309 |
| ENSG00000074755.14 | ZZEF1 | 0.209344989285463 | 6.95528407011239 | 1.79820615679337 | 0.0792500332671505 | 0.343845580678279 | -4.85642894177006 |
| ENSG00000114316.12 | USP4 | 0.167583243815631 | 5.63598929495993 | 1.79780578072886 | 0.0793145026225146 | 0.344052772914482 | -4.85388978371999 |
| ENSG00000153250.19 | RBMS1 | -0.600851978297333 | 4.03129201617399 | -1.79827947050188 | 0.0793811906862046 | 0.344269500188401 | -4.77092867827822 |
| ENSG00000113356.11 | POLR3G | -0.389847723978723 | 0.92817145898746 | -1.79703074939885 | 0.0794394252028688 | 0.344283819434013 | -4.42291152999305 |
| ENSG00000256464.1 | YWHABP2 | 0.506220918537293 | -0.327133665564259 | 1.79696107388306 | 0.0794506638862356 | 0.344283819434013 | -4.17416900636956 |
| lnc-METTL14-3 | lnc-METTL14-3 | 0.44465494601413 | 3.36204154660325 | 1.79695935062223 | 0.0794509418657323 | 0.344283819434013 | -4.67434061447293 |
| ENSG00000115165.9 | CYTIP | 0.36087058914471 | 7.40594629645049 | 1.79686009477808 | 0.079466954224589 | 0.344283819434013 | -4.85501205162574 |
| lnc-EIF2AK3-26 | lnc-EIF2AK3-26 | -2.01957340832544 | 1.36770086782578 | -1.79861264200108 | 0.0794681254496914 | 0.344283819434013 | -4.59968273670415 |
| lnc-IL17RA-11 | lnc-IL17RA-11 | 1.99855666474184 | 1.60076570971703 | 1.79841232871882 | 0.0795003709862048 | 0.344351038722444 | -4.41387638783124 |
| ENSG00000254996.5 | ANKHD1-EIF4EBP3 | 0.228454058814435 | 6.47242271309447 | 1.79649693898079 | 0.0795255631609316 | 0.344387684944552 | -4.86257118879969 |
| ENSG00000204569.9 | PPP1R10 | -0.321040469557654 | 6.09708030743356 | -1.79635264232779 | 0.0795488609888474 | 0.344416113953186 | -4.86148365343508 |
| ENSG00000262304.2 | | -0.210970914177353 | 2.54292789202529 | -1.79587902915208 | 0.0796253699388313 | 0.344674769306752 | -4.61831510367554 |

| | | | | | | | |
|---|---|---|---|---|---|---|---|
| lnc-PBDC1-8 | lnc-PBDC1-8 | -0.386546007501483 | 0.609441791845803 | -1.79577555866278 | 0.0796420931265079 | 0.344674769306752 | -4.38688664332192 |
| ENSG00000173825.6 | TIGD3 | 0.474079400381857 | -0.0628171665977778 | 1.79553271465219 | 0.0796813538619726 | 0.344772189729027 | -4.25073092486462 |
| ENSG00000134440.11 | NARS1 | 0.132688256086878 | 6.01993084728007 | 1.79530310031685 | 0.0797184907306744 | 0.344860381401738 | -4.86359948874722 |
| ENSG00000132128.16 | LRRC41 | -0.184197873445243 | 4.89356686471297 | -1.79510286997355 | 0.0797508870447586 | 0.344899112987589 | -4.832654664013 |
| lnc-RAB30-2 | lnc-RAB30-2 | 0.401804812269171 | 0.868352004216703 | 1.7949875249731 | 0.0797695543451713 | 0.344899112987589 | -4.3903657611481 |
| lnc-THAP4-3 | lnc-THAP4-3 | 0.247662155055054 | 2.21156563835462 | 1.79493711681007 | 0.0797777134982224 | 0.344899112987589 | -4.57482574647563 |
| ENSG00000177842.12 | ZNF620 | 0.259958889101225 | 2.04748062624076 | 1.79463909652041 | 0.0798259659345757 | 0.345024041730861 | -4.55562902819074 |
| ENSG00000182195.7 | LDOC1 | -2.35738329927653 | -0.771521319740457 | -1.79582757058578 | 0.0798401355858762 | 0.345024041730861 | -4.11426105098537 |
| ENSG00000262803.1 | | 0.464981471222865 | 0.149872558112429 | 1.7943775466744 | 0.0798683336802008 | 0.345073449231648 | -4.27950360772563 |
| ENSG00000138035.14 | PNPT1 | 0.199322987180152 | 4.48267957609447 | 1.79409942589654 | 0.0799134064650876 | 0.345195728283504 | -4.80763804948534 |
| ENSG00000168724.16 | DNAJC21 | 0.130511508719369 | 5.50878158974326 | 1.79382742025856 | 0.0799575089217546 | 0.345313765450372 | -4.8579173734773 |
| lnc-ZNF708-3 | lnc-ZNF708-3 | -0.238504111359573 | 1.2751827804386 | -1.79326443575393 | 0.0800488551851732 | 0.345595763827636 | -4.4814778376271 |
| lnc-TEX29-5 | lnc-TEX29-5 | 0.265779139017106 | 2.59794868517521 | 1.79321804336427 | 0.0800563864320152 | 0.345595763827636 | -4.6231932255537 |
| ENSG00000133678.13 | TMEM254 | -0.222632303138238 | 2.23943451664332 | -1.79286674364286 | 0.0801134350720847 | 0.345769518575953 | -4.58237712695177 |
| lnc-SELE-1 | lnc-SELE-1 | 0.60797315993022 | 5.76693547807634 | 1.79380624530956 | 0.0801354113774181 | 0.345791860038029 | -4.85339546562816 |
| ENSG00000173928.2 | SWSAP1 | -0.306418947490718 | 1.06329989250025 | -1.79254206893913 | 0.0801661904044654 | 0.345852168747666 | -4.4571097931004 |
| ENSG00000148229.12 | POLE3 | 0.2133563528794 | 5.45898837860169 | 1.79232264422229 | 0.0802018605591018 | 0.345866541583855 | -4.85949020896948 |
| lnc-C9orf78-3 | lnc-C9orf78-3 | 0.355289642642591 | 2.91438732877244 | 1.79231484329518 | 0.080203128940375 | 0.345866541583855 | -4.65388731381323 |
| ENSG00000139496.15 | NUP58 | 0.237134837662417 | 5.25969507395647 | 1.79198354611856 | 0.0802570113393827 | 0.345895122714866 | -4.85428072240715 |
| ENSG00000068028.17 | RASSF1 | 0.232776590956525 | 5.02163359391013 | 1.79195179710633 | 0.0802621766190888 | 0.345895122714866 | -4.83992538359767 |
| ENSG00000182504.10 | CEP97 | -0.235680031331851 | 4.60635516443657 | -1.79191410492285 | 0.0802683091625685 | 0.345895122714866 | -4.82029527412886 |
| ENSG00000273003.1 | ARL2-SNX15 | -0.27718294738589 | 3.09830854090843 | -1.79180156623447 | 0.0802866216279832 | 0.345895122714866 | -4.67887697171223 |
| ENSG00000103653.16 | CSK | -0.281188894025242 | 6.82846145300752 | -1.7917575738074 | 0.0802937810975868 | 0.345895122714866 | -4.86804695650679 |
| ENSG00000144674.16 | GOLGA4 | 0.227891309456113 | 7.23556417080976 | 1.79149969520408 | 0.0803357598963316 | 0.346003545918852 | -4.86549843742013 |
| lnc-OLA1-4 | lnc-OLA1-4 | 0.160288951293592 | 4.43750427583055 | 1.79129264885918 | 0.0803694773203674 | 0.34603888490041 | -4.80439907771516 |
| ENSG00000168389.17 | MFSD2A | 0.572701785125009 | 1.45734538353212 | 1.79122798108875 | 0.0803800108832057 | 0.34603888490041 | -4.48225927031172 |
| ENSG00000118922.17 | KLF12 | 0.24235417767349 | 6.71058177324713 | 1.79113965107989 | 0.0803944006021358 | 0.34603888490041 | -4.87080993138548 |
| lnc-MTRF1-4 | lnc-MTRF1-4 | -0.507381345329727 | 0.12452092138637 | -1.79100875348114 | 0.0804157289327435 | 0.346057514262702 | -4.29352776724387 |
| ENSG00000169710.8 | FASN | -0.419196227582993 | 3.51365287128791 | -1.7908383695176 | 0.0804434982635203 | 0.346057514262702 | -4.74014981469736 |
| ENSG00000160710.16 | ADAR | 0.133135127731004 | 8.45520284016349 | 1.79080359358374 | 0.0804491670673386 | 0.346057514262702 | -4.84399932017937 |
| lnc-S100A13-2 | lnc-S100A13-2 | -0.302183399226069 | 2.0884113961939 | -1.79046036455307 | 0.0805051346482179 | 0.346218462915776 | -4.57312070758141 |
| ENSG00000244005.12 | NFS1 | -0.133785033632787 | 3.76487425392553 | -1.790354781943 | 0.0805223577350096 | 0.346218462915776 | -4.74565805884681 |
| lnc-ENDOU-5 | lnc-ENDOU-5 | -0.638251036484764 | 2.7582932399937 | -1.79088439359732 | 0.0805370451557468 | 0.346218462915776 | -4.65170422959299 |
| ENSG00000005059.15 | MCUB | 0.190034013510987 | 5.6042339845082 | 1.79010446515819 | 0.0805632028623787 | 0.346258593551126 | -4.86568163597186 |
| ENSG00000174282.11 | ZBTB4 | -0.353552943694369 | 6.64676843372894 | -1.78998828335156 | 0.0805821666027541 | 0.346267794401772 | -4.8715541150413 |
| ENSG00000064042.17 | LIMCH1 | -2.28422862844731 | 0.0862298542674112 | -1.79089181589102 | 0.080684054419939 | 0.346518347693855 | -4.16168717607231 |
| ENSG00000167261.13 | DPEP2 | -0.344082410941098 | 6.22188661711179 | -1.7892872403262 | 0.0806966738236596 | 0.346518347693855 | -4.87313546104291 |
| lnc-STPG2-1 | lnc-STPG2-1 | -0.727733083896199 | 0.0280690575069477 | -1.78926670707319 | 0.0807000297530768 | 0.346518347693855 | -4.26207483748404 |
| ENSG00000027697.14 | IFNGR1 | 0.347165471413562 | 4.70637462367303 | 1.78921907500499 | 0.0807078151311441 | 0.346518347693855 | -4.83548459302092 |
| lnc-TPST2-1 | lnc-TPST2-1 | -0.527744014839454 | 2.2757928375835 | -1.78868663801282 | 0.0807981507470975 | 0.346833855438479 | -4.60580343418202 |
| ENSG00000171067.10 | C11orf24 | -0.343819911995566 | 4.5607576555302 | -1.78821236595592 | 0.0808725077513897 | 0.346991428611726 | -4.81940225883175 |
| ENSG00000171649.11 | ZIK1 | -0.330300188968177 | 4.19379384724028 | -1.78816376282469 | 0.0808804661282108 | 0.346991428611726 | -4.79167686562944 |
| lnc-AMMECR1L-2 | lnc-AMMECR1L-2 | 0.294559817593224 | 0.820744921111912 | 1.78813342926146 | 0.0808854333420328 | 0.346991428611726 | -4.42073933954469 |
| lnc-MAT1A-3 | lnc-MAT1A-3 | -0.254446779622451 | 1.74432462013234 | -1.78801270388208 | 0.0809052050341376 | 0.347003924821349 | -4.53552870767601 |
| lnc-ZNF397-4 | lnc-ZNF397-4 | -0.273588675893595 | 1.19014487066893 | -1.78784721640699 | 0.0809323142011114 | 0.347047879833641 | -4.4722699276441 |
| ENSG00000202314.1 | SNORD6 | 0.467461972103083 | 0.613706670353951 | 1.78767495070462 | 0.0809605418485144 | 0.347096611720053 | -4.37315189640106 |
| ENSG00000070718.11 | AP3M2 | 0.269588223332617 | 3.66262822581351 | 1.78740864629428 | 0.0810041950691467 | 0.347211442546491 | -4.73977579606199 |
| ENSG00000175104.14 | TRAF6 | 0.141104970999261 | 5.41915230556462 | 1.78712939240937 | 0.0810499922486603 | 0.34729207414478 | -4.86563272274224 |
| ENSG00000196850.5 | PPTC7 | 0.260592577746986 | 5.62579370590297 | 1.78707205124314 | 0.0810593987989497 | 0.34729207414478 | -4.87108073869797 |
| ENSG00000068308.13 | OTUD5 | -0.170711464999743 | 6.06348185052184 | -1.78695418331557 | 0.0810787373635132 | 0.34729207414478 | -4.8773944165732 |
| lnc-FNBP4-4 | lnc-FNBP4-4 | 0.254239761935731 | 1.45752471698393 | 1.78682616760568 | 0.0810997452657225 | 0.34729207414478 | -4.50598758945416 |
| ENSG00000116741.7 | RGS2 | 0.693771374940073 | 6.31411716619489 | 1.78801117624986 | 0.0811103593459156 | 0.34729207414478 | -4.87610698413927 |
| ENSG00000122203.14 | KIAA1191 | 0.130577427476895 | 5.38619455532826 | 1.78663537397962 | 0.0811310637721992 | 0.34729207414478 | -4.86576360201195 |
| ENSG00000169239.12 | CA5B | 0.288250825397037 | 3.4718136221507 | 1.7865574056179 | 0.0811438650952547 | 0.34729207414478 | -4.71348995481233 |
| ENSG00000148832.15 | PAOX | -0.302774658514416 | 2.24406373452677 | -1.7864187012683 | 0.0811666426214103 | 0.34729207414478 | -4.60807022909035 |
| ENSG00000181666.17 | ZNF875 | -0.177467292635042 | 5.49818496229247 | -1.78636866186145 | 0.0811748612306533 | 0.34729207414478 | -4.86878140445063 |
| ENSG00000112245.11 | PTP4A1 | 0.321700284628299 | 4.9058133866793 | 1.7859660464718 | 0.0812410133429539 | 0.347502863183296 | -4.84466904344099 |
| ENSG00000148690.11 | FRA10AC1 | -0.206646731192464 | 4.10646382084193 | -1.7852736937929 | 0.0813548769705261 | 0.347917604962464 | -4.79089783808522 |
| ENSG00000011028.13 | MRC2 | -1.39353976772831 | 1.28655839903518 | -1.78664972434947 | 0.0813901337276014 | 0.347996078628006 | -4.48963894374075 |
| lnc-C16orf46-3 | lnc-C16orf46-3 | 0.290414204874667 | 1.42817399795659 | 1.78447955139462 | 0.0814856459390636 | 0.348332097687175 | -4.50766940474698 |
| ENSG00000174669.11 | SLC29A2 | -0.762196862544219 | 2.41661865306436 | -1.7849470917458 | 0.0815679911338205 | 0.348611702970811 | -4.62897032196121 |
| ENSG00000163116.9 | STPG2 | -0.554976063678985 | 0.619434348156256 | -1.78382506981472 | 0.081593550111575 | 0.348636834055304 | -4.39247496112335 |
| ENSG00000167604.14 | NFKBID | -0.415682977028138 | 5.99866398204879 | -1.78387958154299 | 0.0816122874685739 | 0.348636834055304 | -4.88151913897225 |
| ENSG00000166821.8 | PEX11A | -0.487841784220298 | -0.0582910600237977 | -1.78363635604467 | 0.0816246855809411 | 0.348636834055304 | -4.25858347825893 |
| ENSG00000129295.8 | DNAAF11 | 1.10260112291381 | -1.11009941688266 | 1.78326010113994 | 0.0816867928402921 | 0.348829721375878 | -4.11044606937787 |
| ENSG00000116191.17 | RALGPS2 | 0.23055578561096 | 9.40463422553783 | 1.78296418368655 | 0.0817356669253054 | 0.34892081071523 | -4.83454760376841 |
| lnc-CDR2-7 | lnc-CDR2-7 | -0.607935689727452 | -0.622091657002167 | -1.7829256841208 | 0.081742027365731 | 0.34892081071523 | -4.20105717013598 |
| lnc-JAML-1 | lnc-JAML-1 | -0.537622144067526 | -0.458293767261775 | -1.78218860191585 | 0.0818638795251758 | 0.349348716976017 | -4.20745441715397 |

| | | | | | | | |
|---|---|---|---|---|---|---|---|
| LINC02387 | LINC02387 | 0.578763020995936 | 0.316357015330441 | 1.78211401344623 | 0.0818762187578248 | 0.349348716976017 | -4.30127699887332 |
| ENSG00000177398.18 | UMODL1 | -0.741650408617875 | -0.661928841629665 | -1.78190557646217 | 0.0819107089444447 | 0.349423445016271 | -4.16661339730736 |
| lnc-WDYHV1-1 | lnc-WDYHV1-1 | -0.442020591461024 | 1.620112073811 | -1.78164316981952 | 0.0819541468721438 | 0.349525245261153 | -4.53658342249005 |
| lnc-ADH5-6 | lnc-ADH5-6 | -0.357126314247452 | 0.080206315253625 | -1.78154401146702 | 0.081970566253851 | 0.349525245261153 | -4.33227888394054 |
| lnc-FAM104A-8 | lnc-FAM104A-8 | -0.454528859209469 | 0.787462878563755 | -1.78145374047159 | 0.0819855164028978 | 0.349525245261153 | -4.41742362435279 |
| ENSG00000166965.12 | RCCD1 | -0.330936053901606 | 2.67924279225694 | -1.78103804122886 | 0.0820543916130084 | 0.349746436647453 | -4.66073512182805 |
| ENSG00000167625.10 | ZNF526 | -0.241015336017768 | 2.74806934757566 | -1.78083002536947 | 0.0820888749986994 | 0.349789555621832 | -4.66466572131579 |
| ENSG00000167536.13 | DHRS13 | -0.369829675498478 | 1.69321074778514 | -1.78077200120122 | 0.0820984960019952 | 0.349789555621832 | -4.56315965911808 |
| lnc-CTSF-2 | lnc-CTSF-2 | -0.29269154196278 | 1.74770598804393 | -1.78063025900182 | 0.0821220022969996 | 0.349817295794525 | -4.55093048672996 |
| lnc-LTBP3-2 | lnc-LTBP3-2 | -0.34050541182988 | 3.77028245525456 | -1.78049626287787 | 0.0821442291921116 | 0.349839575721339 | -4.76862658916884 |
| ENSG00000105402.7 | NAPA | 0.212120742242777 | 4.34220991692608 | 1.78033049424629 | 0.0821717334104646 | 0.349884317084732 | -4.81566679780179 |
| ENSG00000168646.12 | AXIN2 | -0.968615062551419 | 3.49697001089656 | -1.78178748681002 | 0.0821920725519018 | 0.349898537608231 | -4.81127321770343 |
| ENSG00000174652.18 | ZNF266 | 0.782690119640017 | 7.98939363495868 | 1.78152213583402 | 0.0822160438837092 | 0.349913202799907 | -4.8695027707649 |
| ENSG00000199744.1 | SNORD36A | 0.249284098152334 | 1.82162187709304 | 1.77994944615469 | 0.0822349859691266 | 0.349913202799907 | -4.56759229433779 |
| ENSG00000138613.13 | APH1B | 0.283505470660015 | 6.10436630907405 | 1.77988000246869 | 0.0822465177644634 | 0.349913202799907 | -4.88959255518666 |
| ENSG00000136383.6 | ALPK3 | 0.326267332970624 | 0.277912119729406 | 1.77969015027283 | 0.0822780514872181 | 0.349954155645389 | -4.33771974490277 |
| lnc-FAM120B-8 | lnc-FAM120B-8 | 0.214396143333505 | 2.10316404431026 | 1.77941564457722 | 0.082323663781388 | 0.349954155645389 | -4.59262747076266 |
| ENSG00000142089.15 | IFITM3 | -0.983349146776809 | -0.737413346267332 | -1.77929431439999 | 0.0823438309571139 | 0.349954155645389 | -4.19918473152453 |
| ENSG00000186184.17 | POLR1D | 0.143982551913527 | 5.7919924884575 | 1.77926611537734 | 0.0823485187168029 | 0.349954155645389 | -4.88723478539491 |
| lnc-TOMM22-3 | lnc-TOMM22-3 | 0.236507229966495 | 2.11268256654544 | 1.77924710642787 | 0.0823516788600886 | 0.349954155645389 | -4.59177440326027 |
| ENSG00000112531.16 | QKI | 0.274738793398146 | 6.00610318048652 | 1.77920814547526 | 0.0823581562428347 | 0.349954155645389 | -4.8903033918953 |
| ENSG00000142959.4 | BEST4 | -0.394659130197309 | 0.0396413079758095 | -1.77884882316114 | 0.0824179148903352 | 0.35008607444502 | -4.30524214534561 |
| lnc-STARD9-1 | lnc-STARD9-1 | -0.58766686453598 | -0.586813766140881 | -1.77881694066578 | 0.0824232190040601 | 0.35008607444502 | -4.21655174364875 |
| ENSG00000096093.15 | EFHC1 | -0.297936631065852 | 5.25566524887951 | -1.7786672001755 | 0.0824481343316823 | 0.350119651113888 | -4.87482702579611 |
| lnc-FCRL5-1 | lnc-FCRL5-1 | -0.685651492844281 | 2.55860106863741 | -1.77926149362492 | 0.0824753638848509 | 0.350163039365076 | -4.65139351318904 |
| ENSG00000136146.14 | MED4 | 0.311988289640227 | 4.81658233207392 | 1.77834050969842 | 0.0825025143212686 | 0.350206073886301 | -4.85505702290953 |
| ENSG00000138688.15 | BLTP1 | 0.13892061420875 | 8.14949193649869 | 1.77789522612783 | 0.0825766834717256 | 0.350391299225489 | -4.87254860946653 |
| ENSG00000032444.16 | PNPLA6 | -0.241721373626131 | 5.00210124281814 | -1.77787414123347 | 0.0825801968878611 | 0.350391299225489 | -4.8657591334569 |
| ENSG00000277998.1 | | 1.29402251006588 | -0.956479291311444 | 1.77757491724778 | 0.0826300706777742 | 0.350530656381003 | -4.11757224007205 |
| ENSG00000106245.10 | BUD31 | 0.210100654310294 | 4.96826929028036 | 1.77713131672183 | 0.0827040552528394 | 0.350772217034658 | -4.86432450856064 |
| ENSG00000117751.17 | PPP1R8 | 0.187458952211108 | 5.07207417045477 | 1.77613210446962 | 0.0828709095275958 | 0.351405123550246 | -4.87147292553854 |
| lnc-HAO2-2 | lnc-HAO2-2 | -0.225268508110652 | 5.28134863395118 | -1.77603329111403 | 0.0828874252944878 | 0.351405123550246 | -4.8800609790517 |
| lnc-ITGB3BP-11 | lnc-ITGB3BP-11 | 0.371989500325457 | 0.106869709869237 | 1.77565761226519 | 0.0829502418615168 | 0.351599017346705 | -4.31480849326381 |
| ENSG00000185527.11 | PDE6G | -0.857227596932384 | -0.316603907412382 | -1.77551280131999 | 0.082974466096374 | 0.35162928467401 | -4.24284632998547 |
| ENSG00000235297.3 | FAUP1 | 0.727982721318482 | -0.575511415465935 | 1.77485087458003 | 0.0830852699776863 | 0.352026371336088 | -4.15676640719301 |
| ENSG00000115808.11 | STRN | 0.183411194459586 | 5.72173553284989 | 1.77460467894373 | 0.0831265138003452 | 0.352128634194794 | -4.89410156886872 |
| lnc-ANKRD20A4-7 | lnc-ANKRD20A4-7 | -0.492285969768628 | 1.02496879890629 | -1.77442495854094 | 0.0831566322312768 | 0.352176925471129 | -4.47062873463858 |
| ENSG00000222937.1 | SNORD63B | 0.361058634829569 | 0.318351853605999 | 1.77431578516646 | 0.0831749325127131 | 0.352176925471129 | -4.36014689497993 |
| ENSG00000107140.15 | TESK1 | -0.308854095143615 | 3.90422290062946 | -1.77423042095546 | 0.0831892441160486 | 0.352176925471129 | -4.79256089344084 |
| ENSG00000258973.1 | | -0.270245142584344 | 1.15830594001401 | -1.77389809979576 | 0.0832449785767722 | 0.352201056813716 | -4.48974709062169 |
| lnc-CDK12-1 | lnc-CDK12-1 | -0.23359198846547 | 3.84615123801754 | -1.77387022480064 | 0.0832496549894682 | 0.352201056813716 | -4.78049100265737 |
| ENSG00000204642.13 | HLA-F | 0.318378477941602 | 7.04495184928765 | 1.77380867781616 | 0.0832599811205746 | 0.352201056813716 | -4.89643732477977 |
| ENSG00000285976.1 | | 0.317777653373323 | 4.91613709285348 | 1.77378183085684 | 0.0832644857431247 | 0.352201056813716 | -4.8653291319941 |
| lnc-NPAS3-3 | lnc-NPAS3-3 | 2.15224213893397 | -1.59225594228602 | 1.77384199417781 | 0.0832993932652539 | 0.352201056813716 | -4.13650875418245 |
| ENSG00000023608.4 | SNAPC1 | -0.225381487206017 | 1.52724547012463 | -1.77350689199948 | 0.0833106292075892 | 0.352201056813716 | -4.54153772120477 |
| ENSG00000178188.14 | SH2B1 | -0.237439565475039 | 5.36609071580106 | -1.77348250531294 | 0.083314723102851 | 0.352201056813716 | -4.88624038458333 |
| UBA6-AS1 | UBA6-AS1 | -0.236931666897623 | 3.6828337370821 | -1.77323964482427 | 0.0833555023193434 | 0.352269852089885 | -4.7636647362104 |
| ENSG00000184887.13 | BTBD6 | -0.259079903898515 | 3.78268689676739 | -1.77315670135179 | 0.0833694333672214 | 0.352269852089885 | -4.78312732527155 |
| lnc-REL-5 | lnc-REL-5 | 0.423217686481308 | -0.220569936902764 | 1.77307986308921 | 0.0833823407366234 | 0.352269852089885 | -4.26188033531855 |
| lnc-STAT1-2 | lnc-STAT1-2 | -0.328770373400631 | 1.58857673137436 | -1.77294494395384 | 0.08340500864527 | 0.352293308628277 | -4.56222980095609 |
| lnc-CNPY2-2 | lnc-CNPY2-2 | 0.22853626213174 | 2.32679101831964 | 1.77260002094192 | 0.0834629830279496 | 0.352465855491237 | -4.61994830415316 |
| lnc-SLC27A4-2 | lnc-SLC27A4-2 | 0.33194886590385 | 1.1604543833574 | 1.77246529193334 | 0.0834856373483329 | 0.352489204828869 | -4.48379302763817 |
| ENSG00000227534.1 | AP1M2P1 | -0.963481548297628 | -0.753011135696935 | -1.77225795365962 | 0.0835205107957247 | 0.35256412504274 | -4.16369326904279 |
| ENSG00000166342.18 | NETO1 | -1.89675506547864 | -1.94924768480109 | -1.77312600832363 | 0.0835386027733982 | 0.352568189642168 | -4.14569303296605 |
| ENSG00000168496.3 | FEN1 | 0.274785895650508 | 3.07291924295609 | 1.77195383043462 | 0.0835716851755485 | 0.35263550552856 | -4.70207543117506 |
| lnc-BMT2-2 | lnc-BMT2-2 | 0.26354830381879 | 1.08569277506786 | 1.77174282752445 | 0.0836072057811059 | 0.352713079850892 | -4.48262328775799 |
| ENSG00000179094.15 | PER1 | 0.438339775919919 | 5.81591544622436 | 1.77176570630838 | 0.0836550940110065 | 0.352756014370252 | -4.89883761386434 |
| lnc-SEC61B-1 | lnc-SEC61B-1 | 0.45296360651523 | -0.250026724757971 | 1.77129891501463 | 0.0836819761380741 | 0.352756014370252 | -4.2641869462585 |
| ENSG00000258659.6 | TRIM34 | -0.319627519721139 | 3.46391614168277 | -1.77120810668597 | 0.0836972783351926 | 0.352756014370252 | -4.74780264154288 |
| ENSG00000079277.20 | MKNK1 | -0.145183050999328 | 4.58787218687459 | -1.77116731636833 | 0.0837041527137031 | 0.352756014370252 | -4.85036558334566 |
| lnc-H2AFZ-2 | lnc-H2AFZ-2 | 0.189396092065031 | 2.22233938456135 | 1.77113733835263 | 0.083709205200415 | 0.352756014370252 | -4.61786499273321 |
| ENSG00000104899.7 | AMH | -0.585718125697898 | -0.67080464336836 | -1.77104724301096 | 0.0837243913841937 | 0.352756014370252 | -4.21816975349729 |
| ENSG00000102978.12 | POLR2C | 0.16647250695248 | 5.3964619621443 | 1.77097037081818 | 0.0837373505423432 | 0.352756014370252 | -4.89212894066024 |
| lnc-DUSP4-9 | lnc-DUSP4-9 | -0.264214899930767 | 0.766787091796571 | -1.77062554751778 | 0.0837955017448677 | 0.352872228986876 | -4.44552893707416 |
| lnc-DNAH8-2 | lnc-DNAH8-2 | -0.448866428272114 | 3.29909162823701 | -1.77060347425466 | 0.0837992253455691 | 0.352872228986876 | -4.73875952772117 |
| ENSG00000279841.1 | | -0.479487089416539 | 0.316593614632694 | -1.77032495055777 | 0.0838462222096947 | 0.352997911994712 | -4.37197047854518 |
| lnc-TF-3 | lnc-TF-3 | -0.330189585834455 | 0.200503480307228 | -1.77020986156676 | 0.083865648270755 | 0.353007492506534 | -4.3507515396855 |
| ENSG00000165028.11 | NIPSNAP3B | 0.630087377012364 | 2.27985495487419 | 1.77044419665856 | 0.0838921725486823 | 0.353046940823866 | -4.61770603904316 |

| | | | | | | | |
|---|---|---|---|---|---|---|---|
| ENSG00000263002.7 | ZNF234 | -0.330961009593378 | 3.26919229566846 | -1.76989375316247 | 0.0839190241459803 | 0.353087750203744 | -4.72349259722598 |
| lnc-TRPM7-2 | lnc-TRPM7-2 | 0.282778415461942 | 1.74543246809174 | 1.76965032546512 | 0.0839601470668077 | 0.353127841891366 | -4.57263486993717 |
| lnc-MROH1-1 | lnc-MROH1-1 | -0.472564886327369 | 0.0174915330225467 | -1.76959759345192 | 0.0839690574586996 | 0.353127841891366 | -4.32030324563731 |
| lnc-PSMC5-4 | lnc-PSMC5-4 | 0.219002748683954 | 1.57799820449169 | 1.76953271329258 | 0.0839800216711964 | 0.353127841891366 | -4.55272118803661 |
| lnc-RPL12-1 | lnc-RPL12-1 | -0.47599514934915 | -0.0181665781024604 | -1.76933500277384 | 0.0840134405210471 | 0.353168901428358 | -4.29055218241477 |
| ENSG00000112561.17 | TFEB | -0.339095482149399 | 6.02618609406595 | -1.76927193645565 | 0.0840241029147679 | 0.353168901428358 | -4.90612024857138 |
| ENSG00000205307.11 | SAP25 | -0.391462248643917 | 2.03042796677832 | -1.76911562753823 | 0.0840505343861564 | 0.353207870410424 | -4.61836791795623 |
| lnc-AAGAB-3 | lnc-AAGAB-3 | -0.415500003499491 | 0.0331626517830671 | -1.76897988412437 | 0.0840734939328832 | 0.353232236297313 | -4.33170558633617 |
| ENSG00000076003.4 | MCM6 | 0.365314639542065 | 3.63422356392197 | 1.76846023278548 | 0.0841614359856879 | 0.353434101521409 | -4.76641109985865 |
| ENSG00000040199.18 | PHLPP2 | 0.185173462616145 | 5.40498881384594 | 1.7684376644957 | 0.0841652570268186 | 0.353434101521409 | -4.89497369513391 |
| ENSG00000135372.8 | NAT10 | 0.126700119725104 | 5.44809922099442 | 1.76811238126495 | 0.0842203469399616 | 0.353434101521409 | -4.89823682261346 |
| ENSG00000081692.12 | JMJD4 | -0.237478873910296 | 3.85098150980665 | -1.76811087982425 | 0.084220601293753 | 0.353434101521409 | -4.79623960551242 |
| ENSG00000155438.11 | NIFK | 0.170230912158359 | 4.05604399307277 | 1.76807565155383 | 0.0842265693759919 | 0.353434101521409 | -4.81248442434327 |
| lnc-LINC02210-CRHR1-5 | lnc-LINC02210-CRI | -0.282565620387788 | 2.4326251923734 | -1.76799421943692 | 0.0842403662883336 | 0.353434101521409 | -4.64860244134069 |
| lnc-GAB2-1 | lnc-GAB2-1 | 0.51072790222582 | 2.24528293892543 | 1.76798612140701 | 0.0842417384280247 | 0.353434101521409 | -4.60165627988369 |
| ENSG00000187742.14 | SECISBP2 | 0.286949834865798 | 7.39400901591215 | 1.76777464915335 | 0.0842775771663559 | 0.35351240489405 | -4.90151112233583 |
| ENSG00000083720.12 | OXCT1 | -0.207563159115885 | 5.06737815391487 | -1.76741442275269 | 0.0843386550656653 | 0.353663693512921 | -4.88561241876759 |
| ENSG00000226525.5 | RPS7P10 | 1.00883137766765 | 0.0226226150851301 | 1.76743019677533 | 0.0843480091072696 | 0.353663693512921 | -4.22996477371389 |
| lnc-SMNDC1-11 | lnc-SMNDC1-11 | 0.211828835958825 | 1.19467801794452 | 1.76698576117353 | 0.0844113847637867 | 0.353832580557966 | -4.50668221076804 |
| ENSG00000280054.1 | | -0.835100002414425 | 1.82878690472062 | -1.76771489288382 | 0.0844478804150104 | 0.353832580557966 | -4.57899902836978 |
| lnc-SUMF1-4 | lnc-SUMF1-4 | -0.465070654031302 | 1.06936392540826 | -1.76653467705873 | 0.0844879755647324 | 0.353832580557966 | -4.47487571462322 |
| ENSG00000166507.17 | NDST2 | 0.15025734167027 | 4.75272958479535 | 1.76651735244585 | 0.0844909183186503 | 0.353832580557966 | -4.86805798226938 |
| lnc-ATP5S-1 | lnc-ATP5S-1 | -1.17388176423337 | 1.91953318256013 | -1.76787380758735 | 0.0844982951604637 | 0.353832580557966 | -4.53865300024923 |
| ENSG00000160336.14 | ZNF761 | 0.131747182605032 | 4.72963799049186 | 1.7663914862034 | 0.0845123005019739 | 0.353832580557966 | -4.86352901898698 |
| lnc-ERAP2-8 | lnc-ERAP2-8 | 0.481308614432081 | 0.0843531564440974 | 1.76635637708133 | 0.0845182656555135 | 0.353832580557966 | -4.31839441557135 |
| lnc-PPM1G-1 | lnc-PPM1G-1 | 0.23149517444113 | 1.14297350491531 | 1.76612798383504 | 0.0845570790195791 | 0.353832580557966 | -4.51097925533242 |
| lnc-ENDOU-1 | lnc-ENDOU-1 | -0.834150842984694 | 1.82476829168183 | -1.76702443887808 | 0.0845650256733569 | 0.353832580557966 | -4.57959110092248 |
| ENSG00000144554.10 | FANCD2 | 0.198254287659674 | 5.9274722211657 | 1.76592424231224 | 0.084591715647592 | 0.353832580557966 | -4.90958769548161 |
| lnc-PRPF4B-1 | lnc-PRPF4B-1 | 0.249939276563648 | 3.08797771617901 | 1.76592265326225 | 0.0845919858372355 | 0.353832580557966 | -4.72243331801293 |
| ENSG00000124596.16 | OARD1 | -0.123224487032727 | 4.98863449696416 | -1.76590742836297 | 0.0845945745967912 | 0.353832580557966 | -4.88219456525706 |
| ENSG00000080986.12 | NDC80 | 0.524801055242538 | 2.60314273317262 | 1.76551876357046 | 0.0846730902251973 | 0.354089032122153 | -4.65571080397313 |
| lnc-RFTN1-3 | lnc-RFTN1-3 | 0.309939926353046 | 1.46494805237971 | 1.76449736324719 | 0.0848346223262094 | 0.354636372039098 | -4.54445428466218 |
| lnc-DAPP1-2 | lnc-DAPP1-2 | -0.356941894867287 | 0.215615789706516 | -1.76443419308858 | 0.0848453896740039 | 0.354636372039098 | -4.3643001715102 |
| lnc-NEMF-1 | lnc-NEMF-1 | 0.508107757842778 | 11.6463787040807 | 1.76473288439905 | 0.0848556640087722 | 0.354636372039098 | -4.7928168583829 |
| ENSG00000215712.10 | TMEM242 | 0.217922868051283 | 2.94366021549252 | 1.76417583233362 | 0.0848894391557271 | 0.354647185783947 | -4.70252497012934 |
| ENSG00000136940.13 | PDCL | 0.245151193623733 | 3.75776611789285 | 1.76415664271972 | 0.0848927116726186 | 0.354647185783947 | -4.7927811786276 |
| ENSG00000149923.13 | PPP4C | 0.200260106147661 | 4.83973631330241 | 1.76396428654555 | 0.0849255211322004 | 0.354712256790601 | -4.88035577316086 |
| lnc-UNK-1 | lnc-UNK-1 | -0.536047279299074 | 0.305202160698159 | -1.76378882220557 | 0.0849554586770168 | 0.354765308571523 | -4.37023980872208 |
| MIR3681HG | MIR3681HG | 1.44909579510127 | 1.19334727823473 | 1.7651054876986 | 0.0849941268089863 | 0.354854789474516 | -4.48835428391659 |
| lnc-SLX1B-5 | lnc-SLX1B-5 | -0.529026431928412 | -0.553526640473105 | -1.76329290218051 | 0.0850401198336827 | 0.354936602564472 | -4.22503364816243 |
| lnc-CDKL1-1 | lnc-CDKL1-1 | -0.55113408682459 | 2.24376653273528 | -1.76331604468411 | 0.0850482108462311 | 0.354936602564472 | -4.61700092157866 |
| ENSG00000095574.11 | IKZF5 | 0.228522477512717 | 5.07170799217256 | 1.76294456658273 | 0.0850996282953359 | 0.35507919099998 | -4.89489330114958 |
| lnc-PPP1R3B-10 | lnc-PPP1R3B-10 | -0.549658773929788 | 0.718088685670306 | -1.76260304608258 | 0.0851580063374968 | 0.355250758906505 | -4.49896230900516 |
| ENSG00000185483.11 | ROR1 | -0.612950809439695 | 6.45941125558096 | -1.76353912295572 | 0.0851851094023119 | 0.35526728679419 | -4.91583754151611 |
| ENSG00000071189.21 | SNX13 | 0.237806973302803 | 6.39342933681575 | 1.76234552772416 | 0.085202047586899 | 0.35526728679419 | -4.91878341661339 |
| ENSG00000111554.14 | MDM1 | -0.18796780727728 | 4.39962978506579 | -1.76227712608153 | 0.0852137489628778 | 0.35526728679419 | -4.85557823598722 |
| ENSG00000277726.4 | NA | 0.175526511316396 | 5.13555374454247 | 1.76163968871727 | 0.0853228592128874 | 0.355650144021641 | -4.89758194169742 |
| lnc-CHFR-3 | lnc-CHFR-3 | 0.244804222400453 | 2.56080540299244 | 1.7614620202917 | 0.0853532915787309 | 0.355704960632723 | -4.66718800024099 |
| lnc-NFS1-3 | lnc-NFS1-3 | 0.33971651362137 | 3.12502590540416 | 1.76118705068802 | 0.0854004083257705 | 0.35582478719501 | -4.72826544316747 |
| ENSG00000172071.11 | EIF2AK3 | 0.312115958981462 | 5.28993541142757 | 1.7610661985029 | 0.0854211235442857 | 0.35582478719501 | -4.90339778006402 |
| ENSG00000164038.14 | SLC9B2 | -0.157802874845945 | 3.99675844904124 | -1.76099163499109 | 0.0854339065402389 | 0.35582478719501 | -4.8203053521512 |
| ENSG00000105705.15 | SUGP1 | -0.181405004853683 | 4.46529304235443 | -1.76074617861969 | 0.0854759983250662 | 0.355928074757199 | -4.86131415469098 |
| lnc-PHF20L1-8 | lnc-PHF20L1-8 | 0.277903994277473 | 2.51840624157853 | 1.7603970932886 | 0.0855358906923318 | 0.356105428422384 | -4.66382646030617 |
| ENSG00000284554.2 | | -0.356860707478612 | 3.59688112230217 | -1.76020430123702 | 0.0855689829662667 | 0.35611964990884 | -4.77568682301879 |
| ENSG00000213976.4 | | -0.326684141476525 | 2.47370133426408 | -1.76017560075762 | 0.0855739102480667 | 0.35611964990884 | -4.6694522534614 |
| lnc-SPINT2-10 | lnc-SPINT2-10 | 0.262507737423756 | 2.73848052106058 | 1.76007020391283 | 0.0855920067550799 | 0.35611964990884 | -4.67250846915431 |
| lnc-FBXO15-1 | lnc-FBXO15-1 | -0.93735670820519 | 2.75934710117075 | -1.76132530999074 | 0.0856085132268834 | 0.35611964990884 | -4.70951459803923 |
| ENSG00000153066.12 | TXNDC11 | 0.26070008330878 | 5.39370206468754 | 1.75962019289299 | 0.0856693091235143 | 0.356300543481369 | -4.90986417723953 |
| lnc-GRAMD4-2 | lnc-GRAMD4-2 | -0.726460277948863 | 0.553281255633505 | -1.75961685343533 | 0.0856893638778386 | 0.356311954888394 | -4.43767685768865 |
| lnc-TRPM2-1 | lnc-TRPM2-1 | -0.480771648798318 | 0.23128819002072 | -1.75926004062819 | 0.0857312177445834 | 0.356413988050244 | -4.36333750234726 |
| lnc-MRPL49-1 | lnc-MRPL49-1 | -0.50767327236377 | 0.123765828267456 | -1.75908733305449 | 0.0857609187104976 | 0.356431173290516 | -4.35750533028752 |
| lnc-NFS1-1 | lnc-NFS1-1 | 0.145036204433277 | 7.3459974113297 | 1.75903462361894 | 0.0857699850025714 | 0.356431173290516 | -4.91648813025578 |
| lnc-CELF4-17 | lnc-CELF4-17 | -0.335602300538747 | 1.11774905325252 | -1.75869143262437 | 0.0858290352196651 | 0.356604568414689 | -4.50304802729735 |
| ENSG00000213997.3 | PGAM1P7 | 0.339155387006941 | 2.63101639106574 | 1.75845704355216 | 0.0858693842799735 | 0.356648827272433 | -4.66788139333548 |
| ENSG00000164048.13 | ZNF589 | 0.365047853347416 | 5.78313322878449 | 1.75842824848901 | 0.0858743423194956 | 0.356648827272433 | -4.92084480692855 |
| ENSG00000150712.10 | MTMR12 | 0.150711003746511 | 4.79322903156548 | 1.75830934115937 | 0.0858948187562984 | 0.356661903260216 | -4.8887358116653 |
| ENSG00000273749.4 | CYFIP1 | -0.493378136131986 | 4.6727664417016 | -1.75816875490661 | 0.0860055359429079 | 0.356859927967689 | -4.88359464768096 |
| ENSG00000114738.10 | MAPKAPK3 | -0.2755796300061 | 3.79516154865453 | -1.7574874282107 | 0.0860364680169788 | 0.356859927967689 | -4.81157964273747 |

| | | | | | | | |
|---|---|---|---|---|---|---|---|
| ENSG00000100197.21 | CYP2D6 | -0.473222820089627 | -0.21790172933923 | -1.75735946154085 | 0.0860585394870429 | 0.356859927967689 | -4.29510915561584 |
| ENSG00000156515.23 | HK1 | -0.240069666358998 | 5.59359261122746 | -1.75724944761035 | 0.0860775182861069 | 0.356859927967689 | -4.92045747279933 |
| ENSG00000131591.17 | C1orf159 | -0.303720830889108 | 2.09805827864514 | -1.75724192011528 | 0.0860788170025053 | 0.356859927967689 | -4.63946187534269 |
| ENSG00000148634.15 | HERC4 | -0.169518570034404 | 6.2226773625094 | -1.7572396389162 | 0.0860792105803135 | 0.356859927967689 | -4.92682077905127 |
| ENSG00000197622.12 | CDC42SE1 | 0.20966179788978 | 8.69464306273171 | 1.75723771422469 | 0.0860795426506471 | 0.356859927967689 | -4.89487123896567 |
| lnc-DHX58-1 | lnc-DHX58-1 | -0.274600559974029 | 2.01043070741577 | -1.75722805116677 | 0.0860812098508273 | 0.356859927967689 | -4.62135553078981 |
| ENSG00000080824.18 | HSP90AA1 | 0.337612633925475 | 7.54976224618627 | 1.75595740079362 | 0.0863006749087467 | 0.357697702707759 | -4.92083903265899 |
| ENSG00000177463.15 | NR2C2 | 0.126064370806446 | 6.42174193658427 | 1.75558337487672 | 0.0863653652555561 | 0.357893761436503 | -4.9298357365971 |
| lnc-UNC119-1 | lnc-UNC119-1 | 0.204749425471474 | 1.61823873521794 | 1.75542119505957 | 0.0863934279692771 | 0.357937309824576 | -4.58123802935177 |
| ENSG00000228672.3 | PROB1 | -0.473060923342382 | 0.229144467082038 | -1.75528633784069 | 0.0864167687357785 | 0.357937309824576 | -4.41118814524858 |
| lnc-RCL1-3 | lnc-RCL1-3 | -0.425215031392309 | -0.185141841474255 | -1.7552212029941 | 0.0864280440085577 | 0.357937309824576 | -4.29516342656301 |
| ENSG00000126464.13 | PRR12 | -0.359607355289727 | 3.88037569503444 | -1.75498788482512 | 0.0864684430240593 | 0.358032581525691 | -4.81579117368532 |
| ENSG00000089041.16 | P2RX7 | -0.414691891414863 | 4.26649223649266 | -1.75488216855696 | 0.0864941283050759 | 0.358066903238813 | -4.85405834402561 |
| lnc-HPCAL1-1 | lnc-HPCAL1-1 | -0.329361203086042 | 0.948442300188381 | -1.75448227213068 | 0.0865560439988347 | 0.358251167027552 | -4.49718979634216 |
| lnc-CDH24-1 | lnc-CDH24-1 | 0.223068162729687 | 1.74354381778706 | 1.75433517660647 | 0.0865815432696105 | 0.358284661262242 | -4.58689644745301 |
| lnc-EMC10-4 | lnc-EMC10-4 | -0.424144250704697 | -0.090982649256782 | -1.75390440274538 | 0.0866562548197326 | 0.35851545594502 | -4.32484827493573 |
| ENSG00000099958.14 | DERL3 | -0.573484792410055 | 3.52488277693125 | -1.75441443983549 | 0.0866721522024204 | 0.35851545594502 | -4.78795292657949 |
| ENSG00000178694.9 | NSUN3 | 0.128668459044879 | 4.59453612818964 | 1.75346288970658 | 0.0867328848539216 | 0.358591036101241 | -4.87955938147355 |
| ENSG00000100591.7 | AHSA1 | -0.385379259863001 | 4.52016836208155 | -1.75344404550558 | 0.0867361567576856 | 0.358591036101241 | -4.87483960493501 |
| ENSG00000187091.13 | PLCD1 | 0.247423725961504 | 2.13419025129732 | 1.75340642519298 | 0.0867426890515513 | 0.358591036101241 | -4.63011205009981 |
| ENSG00000171443.6 | ZNF524 | -0.340668662399219 | 1.48956447257038 | -1.75309146671723 | 0.0867973937648095 | 0.358745131699011 | -4.57539706307961 |
| lnc-SRM-3 | lnc-SRM-3 | 0.222045261491474 | 2.00096289693481 | 1.7524526501816 | 0.086908437524104 | 0.359119506113488 | -4.63296461630213 |
| ENSG00000115750.16 | TAF1B | 0.228520442832128 | 3.71024795999857 | 1.7523696876385 | 0.0869228673884952 | 0.359119506113488 | -4.80152549209701 |
| ENSG00000204650.14 | LINC02210 | -0.277654222427452 | 3.26075708409323 | -1.75209879579936 | 0.0869699981704462 | 0.359242117668532 | -4.75999554777253 |
| ENSG00000241852.9 | C8orf58 | -0.282358317920905 | 2.26631061434072 | -1.75193294759122 | 0.0869988636065796 | 0.359289247514893 | -4.66400539568601 |
| lnc-RPS3A-1 | lnc-RPS3A-1 | 0.423116769166918 | 1.93212741975696 | 1.75175719475673 | 0.0870294616552516 | 0.35934351238717 | -4.62963265021644 |
| ENSG00000074219.13 | TEAD2 | -0.587560704044863 | 5.53308351313547 | -1.75255136174769 | 0.0870657223836863 | 0.359421131934099 | -4.92392786962083 |
| ENSG00000164087.7 | POC1A | 0.466998164737832 | 0.214762409588805 | 1.75140772552487 | 0.0870903299390354 | 0.359450623849041 | -4.35540109573401 |
| ENSG00000082196.20 | C1QTNF3 | -0.279139101210148 | 2.73773003103223 | -1.75129337852575 | 0.0871102538813737 | 0.359460776992846 | -4.70726026865468 |
| ENSG00000155975.9 | VPS37A | -0.20785707652955 | 4.7087421765868 | -1.75020256365374 | 0.0873005101885767 | 0.360173662299353 | -4.89412384412374 |
| ENSG00000024048.10 | UBR2 | 0.134594812616564 | 7.21826261610378 | 1.75006899941569 | 0.0873238298623308 | 0.360183653248766 | -4.93300095294366 |
| ENSG00000014138.8 | POLA2 | -0.2030921819668 | 2.93493136040271 | -1.74988870520246 | 0.0873553166199012 | 0.360183653248766 | -4.73164471242929 |
| lnc-ZNF814-2 | lnc-ZNF814-2 | -0.281629340677017 | 0.472128332384373 | -1.74986273686 | 0.0873598525380782 | 0.360183653248766 | -4.42731055788395 |
| ENSG00000167186.10 | COQ7 | -0.197503998196531 | 3.35367112091567 | -1.74978788452239 | 0.0873729281771895 | 0.360183653248766 | -4.76898370556508 |
| ENSG00000107021.15 | TBC1D13 | 0.139190530828189 | 4.37703674019715 | 1.74930600789778 | 0.0874571443261929 | 0.360441241231351 | -4.87047489466158 |
| ENSG00000175857.8 | GAPT | -0.823806376266208 | 3.07826398393333 | -1.7503938383612 | 0.0874704367658065 | 0.360441241231351 | -4.73045099562188 |
| ENSG00000172780.16 | RAB43 | -0.343339179061252 | 4.0758636659888 | -1.74905087412676 | 0.0875017607511578 | 0.360452673534432 | -4.85245167071598 |
| ENSG00000122482.20 | ZNF644 | 0.166671513969888 | 7.53594700362333 | 1.74900683438915 | 0.0875094641083174 | 0.360452673534432 | -4.93084507107876 |
| ENSG00000130299.16 | GTPBP3 | -0.269160777630804 | 1.96755304061243 | -1.74888034910733 | 0.0875315918524473 | 0.360452673534432 | -4.63246755522886 |
| ENSG00000202058.1 | RN7SKP80 | 0.826150885934725 | 2.28609252591448 | 1.74973424825859 | 0.0875432597288357 | 0.360452673534432 | -4.6012626915584 |
| ENSG00000285667.1 | | -0.599760238973143 | 0.56060404279052 | -1.74805292615041 | 0.0876764592303738 | 0.360928912067757 | -4.44930763105469 |
| ENSG00000183597.15 | TANGO2 | -0.191064856289363 | 4.44671941868956 | -1.74757373309054 | 0.0877604491465565 | 0.36120242447182 | -4.87843001271593 |
| ENSG00000148154.9 | UGCG | 0.263057356590091 | 7.5704443641956 | 1.74717853476666 | 0.0878297675539445 | 0.361415454930596 | -4.93249773185257 |
| lnc-FAM162A-2 | lnc-FAM162A-2 | -0.23685799949486 | 2.93987047624843 | -1.74684023598817 | 0.0878891420118388 | 0.361583407697111 | -4.72779989181244 |
| ENSG00000072756.16 | TRNT1 | 0.165542288527877 | 4.52562441473101 | 1.74674583074454 | 0.0879057169565343 | 0.361583407697111 | -4.88592496609146 |
| lnc-MESD-4 | lnc-MESD-4 | -0.393569147833046 | 0.157198277573397 | -1.74632935091205 | 0.0879788704378408 | 0.361614121226922 | -4.37001831597653 |
| ENSG00000144655.14 | CSRNP1 | 0.348166286620011 | 5.24049077535647 | 1.74627255486216 | 0.087988850438073 | 0.361614121226922 | -4.9232071738677 |
| ENSG00000121879.4 | PIK3CA | 0.225461372201234 | 6.7457101863595 | 1.74618814367186 | 0.0880036846192827 | 0.361614121226922 | -4.9434805690579 |
| lnc-EVI2A-3 | lnc-EVI2A-3 | -0.726851087344873 | -0.0382277311026267 | -1.74612192526265 | 0.0880153231148382 | 0.361614121226922 | -4.33270687179558 |
| ENSG00000106153.12 | CHCHD2 | 0.273746142086983 | 5.88368656961708 | 1.74602575742036 | 0.0880322277878725 | 0.361614121226922 | -4.9430322739735 |
| lnc-HSD17B6-2 | lnc-HSD17B6-2 | -0.433467701966736 | -0.408352927138677 | -1.74592470384085 | 0.0880499942116584 | 0.361614121226922 | -4.27034228443651 |
| lnc-TRPM1-2 | lnc-TRPM1-2 | 0.180736345234239 | 2.16961903138229 | 1.74590855836158 | 0.088052833056729 | 0.361614121226922 | -4.64805175030282 |
| ENSG00000197429.10 | IPP | -0.170588513831915 | 3.63092228729069 | -1.74590344313847 | 0.0880537324777406 | 0.361614121226922 | -4.80140314369812 |
| ENSG00000253710.3 | ALG11 | 0.30562035774391 | 4.79562147271644 | 1.74573262920728 | 0.0880837714752201 | 0.361665323812977 | -4.91010982494197 |
| ENSG00000135185.11 | TMEM243 | 0.235526099700133 | 5.88848387696888 | 1.74510299858956 | 0.0881945709684888 | 0.361904327169203 | -4.94470783182577 |
| lnc-IL17B-2 | lnc-IL17B-2 | 0.223684017645869 | 5.80796969111291 | 1.74500291280531 | 0.0882121943173818 | 0.361904327169203 | -4.94378691041129 |
| ENSG00000153179.12 | RASSF3 | 0.218556122250091 | 6.52936212137649 | 1.744861873648 | 0.0882370338266725 | 0.361904327169203 | -4.94715462283314 |
| ENSG00000123104.11 | ITPR2 | 0.264101438118526 | 7.78190241968704 | 1.74483295506229 | 0.0882421276263561 | 0.361904327169203 | -4.93316700832011 |
| ENSG00000178700.7 | DHFR2 | -0.346988813304026 | 2.11655190177063 | -1.74467504350174 | 0.0882699469359328 | 0.361904327169203 | -4.64453962593038 |
| ENSG00000250050.1 | MTND4P9 | -0.43661284082739 | -0.319778094902528 | -1.74466454143997 | 0.0882717973457808 | 0.361904327169203 | -4.29082349080497 |
| ENSG00000135845.9 | PIGC | 0.149725561638004 | 4.19781842088911 | 1.74459589355346 | 0.0882838935517623 | 0.361904327169203 | -4.86385343949829 |
| lnc-RBPMS2-1 | lnc-RBPMS2-1 | 0.284421399348658 | 0.437441012069274 | 1.74458085903646 | 0.0882865429165665 | 0.361904327169203 | -4.41604547099304 |
| lnc-SAT2-1 | lnc-SAT2-1 | -0.676924891833407 | 0.140421816104379 | -1.74450322248813 | 0.0883002249936228 | 0.361904327169203 | -4.37701652794588 |
| ENSG00000127184.12 | COX7C | 0.257888405888016 | 6.33677885472207 | 1.74431048408324 | 0.0883341994017886 | 0.361971496374082 | -4.94818183124052 |
| ENSG00000101191.16 | DIDO1 | -0.146898855287673 | 7.26143102757556 | -1.743703409708 | 0.0884412810130201 | 0.362338154277666 | -4.94216830845099 |
| lnc-PEX19-3 | lnc-PEX19-3 | 0.244242324088868 | 2.79626521643439 | 1.74313222767049 | 0.0885421304975872 | 0.362679138712804 | -4.71954755847763 |
| ENSG00000100802.14 | C14orf93 | -0.25153688037065 | 2.4607949896965 | -1.74292463818636 | 0.0885788068448407 | 0.362757178927051 | -4.68889860044455 |
| ENSG00000253873.5 | PCDHGA11 | -1.36319214263819 | -1.50562981741689 | -1.74298188422325 | 0.0885967398583957 | 0.362758443705064 | -4.18038305084677 |

| | | | | | | | |
|---|---|---|---|---|---|---|---|
| lnc-SYPL1-4 | lnc-SYPL1-4 | 0.34653003037083 | 1.60855983473269 | 1.74252566506382 | 0.0886493319363499 | 0.362804634008505 | -4.58052850591239 |
| ENSG00000177084.16 | POLE | 0.235995756873813 | 5.2867386707758 | 1.74250028998871 | 0.0886538189846336 | 0.362804634008505 | -4.93351344366846 |
| ENSG00000254912.2 | MPHOSPH10P2 | -0.739146206990752 | -1.2158954416249 | -1.74246024653902 | 0.0886609002119604 | 0.362804634008505 | -4.20207276175694 |
| lnc-SHF-3 | lnc-SHF-3 | -0.473411590031617 | -0.298671075163759 | -1.7423364709186 | 0.0886827915030953 | 0.362822082589587 | -4.27534632869398 |
| ENSG00000177674.15 | AGTRAP | -0.555010244705639 | 1.30318777353176 | -1.74221416027923 | 0.0887044281244049 | 0.362838482528741 | -4.54251803825126 |
| ENSG00000258588.3 | TRIM6-TRIM34 | -0.314222348487667 | 3.43718467631112 | -1.74157255740235 | 0.088817999161054 | 0.36323085172501 | -4.79251596769715 |
| ENSG00000163576.17 | EFHB | -0.997132207596227 | -1.09390460648013 | -1.7413489174278 | 0.0888576144957987 | 0.363249004312984 | -4.16952908811425 |
| lnc-LIG3-1 | lnc-LIG3-1 | 0.234957578313914 | 2.60702918717204 | 1.741348243523 | 0.0888577338928181 | 0.363249004312984 | -4.707382594309 |
| ENSG00000249967.1 | | 0.379434065117549 | 3.7121307707465 | 1.74083838903709 | 0.0889481041672794 | 0.3635462327393 | -4.80750421079323 |
| ENSG00000100280.16 | AP1B1 | -0.219635289554826 | 6.9791663280231 | -1.7406453057399 | 0.0889823476507781 | 0.363578836436287 | -4.95007666557891 |
| ENSG00000283563.1 | | -1.98591388693024 | -0.173027205828008 | -1.7418018197757 | 0.0889914093167185 | 0.363578836436287 | -4.22967155446901 |
| ENSG00000233170.4 | | 0.664885610254731 | -0.523798979460561 | 1.74009311733323 | 0.0890803394313618 | 0.363869939772915 | -4.26961553581603 |
| ZNF571-AS1 | ZNF571-AS1 | -0.57475572419658 | 0.627793604684343 | -1.73984261065323 | 0.0891248242139091 | 0.363930613532656 | -4.46644243531732 |
| ENSG00000160917.14 | CPSF4 | 0.204442180966439 | 4.23477140613986 | 1.73980737307513 | 0.0891310831621624 | 0.363930613532656 | -4.87210019814377 |
| lnc-NSUN6-1 | lnc-NSUN6-1 | -0.228846063722172 | 2.75220736969287 | -1.73954910874094 | 0.0891769676290803 | 0.363930613532656 | -4.72558570030898 |
| ENSG00000189171.14 | S100A13 | -0.301951822228321 | 1.77114167474891 | -1.73953986537026 | 0.0891786102153054 | 0.363930613532656 | -4.61956589915011 |
| ENSG00000221857.7 | | -0.698805316716585 | -0.578244285775272 | -1.7395117936882 | 0.0891835988271252 | 0.363930613532656 | -4.23718392789078 |
| ENSG00000254415.3 | SIGLEC14 | 0.908929456452292 | 0.915475535039666 | 1.74014053304315 | 0.0892332738519865 | 0.364061144835567 | -4.54857902473472 |
| ENSG00000211460.11 | TSN | 0.143750918723114 | 5.91578469201701 | 1.73884728785787 | 0.0893017559839238 | 0.364234424200407 | -4.95486038290837 |
| lnc-CPLX4-1 | lnc-CPLX4-1 | 0.29575246144164 | 1.97602259574037 | 1.73879455958947 | 0.0893111372948285 | 0.364234424200407 | -4.64549683071233 |
| ENSG00000198408.13 | OGA | 0.182129214336959 | 8.8754497258638 | 1.73852185938527 | 0.0893596687031451 | 0.364329289238605 | -4.92040090471315 |
| ENSG00000213186.7 | TRIM59 | 0.217922166058555 | 3.41885514668267 | 1.73846495003319 | 0.089369799415329 | 0.364329289238605 | -4.8024432302381 |
| ENSG00000274012.1 | RN7SL2 | 0.502242014330763 | 11.6832713912566 | 1.73857419587807 | 0.0894131693269122 | 0.364353480835882 | -4.83350965611697 |
| LINC01857 | LINC01857 | -0.638116471510385 | 5.57262310028017 | -1.73929665164103 | 0.0894238824501289 | 0.364353480835882 | -4.94547936601618 |
| lnc-BIN3-4 | lnc-BIN3-4 | -1.03183040114205 | -0.823507303352195 | -1.73813340388443 | 0.0894288386135585 | 0.364353480835882 | -4.21072757386464 |
| LINC00377 | LINC00377 | -1.62494530245674 | 1.63729094967248 | -1.73902818817796 | 0.089546776731187 | 0.364761786158326 | -4.56495203995355 |
| ENSG00000099246.16 | RAB18 | 0.204971493634866 | 5.42466624681625 | 1.7371308476834 | 0.0896075640460256 | 0.364859336922977 | -4.94854502857504 |
| lnc-SH3D21-3 | lnc-SH3D21-3 | -0.525232789284381 | 3.33302017791843 | -1.73744253171012 | 0.0896135837850849 | 0.364859336922977 | -4.79533886611767 |
| ENSG00000138442.9 | WDR12 | -0.317736416948852 | 3.29401699911889 | -1.73703927462391 | 0.0896239035846363 | 0.364859336922977 | -4.78712189151971 |
| ENSG00000154727.10 | GABPA | 0.167119113019529 | 5.69451739796628 | 1.73690248130298 | 0.0896483164793463 | 0.364886552915638 | -4.95508391942185 |
| lnc-POLDIP3-2 | lnc-POLDIP3-2 | 0.8235725761756 | 2.2783633240883 | 1.73751895772968 | 0.0896997962713579 | 0.365023904043764 | -4.61912366975309 |
| ENSG00000277957.1 | SENP3-EIF4A1 | 0.185873951873847 | 6.22740403435633 | 1.73601500507241 | 0.0898068351080379 | 0.365316984630035 | -4.96152135344907 |
| ENSG00000283710.1 | MIR1204 | 0.746074146778273 | -0.197952491654024 | 1.73595219896251 | 0.0898180622149463 | 0.365316984630035 | -4.27033929253608 |
| ENSG00000137842.6 | TMEM62 | 0.233862393336126 | 5.42502462978748 | 1.73591304224118 | 0.0898250623919062 | 0.365316984630035 | -4.94976889833884 |
| ENSG00000148248.13 | SURF4 | 0.194333594406809 | 5.46582925871497 | 1.73561855287903 | 0.0898777238055125 | 0.365423685982214 | -4.95169490415118 |
| ENSG00000129518.8 | EAPP | 0.180018953923143 | 5.3547792492985 | 1.73551305873882 | 0.0898965948231832 | 0.365423685982214 | -4.94643223192484 |
| lnc-UEVLD-6 | lnc-UEVLD-6 | 0.185928291695205 | 2.96349359223481 | 1.73543564119092 | 0.0899104455409837 | 0.365423685982214 | -4.75455306266434 |
| ENSG00000285581.1 | | -0.472736871645593 | 0.298197136700432 | -1.73536931654928 | 0.0899223130496 | 0.365423685982214 | -4.44120185485553 |
| lnc-POLR2E-1 | lnc-POLR2E-1 | 0.205970605840457 | 3.25310896464813 | 1.73518685159468 | 0.0899549683523524 | 0.365484230871786 | -4.78273114086125 |
| ENSG00000141349.8 | G6PC3 | -0.249777822755363 | 3.05950765747654 | -1.73502416254695 | 0.0899840927446976 | 0.365530408716027 | -4.76974298210257 |
| ENSG00000090432.6 | MUL1 | 0.236352428405371 | 4.13399423212077 | 1.73461913102169 | 0.0900566351289642 | 0.365752904668404 | -4.86913966668723 |
| ENSG00000135837.15 | CEP350 | 0.119682543031844 | 7.8326534774831 | 1.73444224028473 | 0.0900883320885537 | 0.36578211600298 | -4.94937383530793 |
| ENSG00000225663.7 | MCRIP1 | -0.304098342100507 | 3.11288648867064 | -1.73437207355389 | 0.0901009078039616 | 0.36578211600298 | -4.78089689379548 |
| lnc-ZNF674-9 | lnc-ZNF674-9 | 0.338938660038353 | 2.45962127498977 | 1.73428151262313 | 0.090117140856586 | 0.36578211600298 | -4.68675539710039 |
| ENSG00000220583.1 | RPL35P2 | 1.237547400092 | -1.20434723639649 | 1.73412090123074 | 0.0901459364533273 | 0.365826855287625 | -4.16493289772823 |
| ENSG00000108518.7 | PFN1 | -0.385262025688264 | 6.93799235010121 | -1.73401334308013 | 0.0901794771313547 | 0.36586516363054 | -4.96006483952114 |
| ENSG00000138767.13 | CNOT6L | 0.176712830816725 | 6.99800826270769 | 1.73378093054056 | 0.0902069142034933 | 0.36586516363054 | -4.96138602923525 |
| ENSG00000089280.18 | FUS | 0.394660567919943 | 7.05356092883391 | 1.73389793777474 | 0.090208701619054 | 0.36586516363054 | -4.96123370438537 |
| lnc-PEAK1-4 | lnc-PEAK1-4 | 0.32989948038488 | 2.28867699184034 | 1.73342907800022 | 0.0902700592940311 | 0.366041889371364 | -4.68429894751701 |
| lnc-VPS72-2 | lnc-VPS72-2 | -0.334698406101773 | 1.26632082885361 | -1.73324770532633 | 0.0903026236597969 | 0.366101812643214 | -4.57348059860106 |
| ENSG00000182400.14 | TRAPPC6B | -0.181491565272626 | 5.09890431206083 | -1.73309503439297 | 0.0903300423885929 | 0.3661114147045 | -4.9432457827275 |
| ENSG00000239883.8 | PARGP1 | 0.178490298874482 | 3.78572878982641 | 1.73292912912183 | 0.0903598457845419 | 0.3661114147045 | -4.84054323654516 |
| ENSG00000131871.14 | SELENOS | 0.184092910784245 | 3.31183896655659 | 1.732779170226 | 0.090386791606112 | 0.3661114147045 | -4.79849632887629 |
| lnc-GALNT15-8 | lnc-GALNT15-8 | -0.40069965148133 | 2.76654829828711 | -1.73268804345342 | 0.0904031692681765 | 0.3661114147045 | -4.7395135600712 |
| lnc-CCDC17-3 | lnc-CCDC17-3 | 0.2881855383918 | 1.48185913374769 | 1.7326856590403 | 0.090403597837473 | 0.3661114147045 | -4.59309176812032 |
| ENSG00000038219.12 | BOD1L1 | 0.234015103668062 | 8.19474083925141 | 1.73264050210607 | 0.0904117145675058 | 0.3661114147045 | -4.94437458434258 |
| ENSG00000002834.17 | LASP1 | -0.271595673809626 | 6.71640530887372 | -1.73247814436418 | 0.0904409025645643 | 0.366157572282932 | -4.96542526161149 |
| lnc-YWHAZ-7 | lnc-YWHAZ-7 | 0.222024500415575 | 2.52480531081004 | 1.73165403273627 | 0.0905891789923558 | 0.366611252902081 | -4.70826909374141 |
| ENSG00000267980.1 | | -0.510515349300752 | -0.369238231364362 | -1.73146538147404 | 0.0906231501142096 | 0.366611252902081 | -4.29489111534854 |
| lnc-HLA-DMA-3 | lnc-HLA-DMA-3 | -0.362055972793829 | 5.15565850069289 | -1.73144675010895 | 0.0906265057078482 | 0.366611252902081 | -4.94261741202747 |
| ENSG00000215835.2 | | 0.828552198128101 | -0.775868658523528 | 1.73138143668613 | 0.0906382697698836 | 0.366611252902081 | -4.19119182695053 |
| lnc-SERINC4-2 | lnc-SERINC4-2 | -0.556985915323186 | 0.789622589800885 | -1.73128057020913 | 0.090656440042728 | 0.366611252902081 | -4.52167441143351 |
| ENSG00000101150.17 | TPD52L2 | 0.150008824774245 | 4.7594744937319 | 1.73126175478176 | 0.090659829824204 | 0.366611252902081 | -4.92466717415775 |
| ENSG00000153162.8 | BMP6 | -1.22722461697583 | 1.346069564482 | -1.73215790396685 | 0.0907571955508007 | 0.366932892559837 | -4.60053844972623 |
| lnc-ZNF75D-2 | lnc-ZNF75D-2 | 0.208040825655739 | 0.831731981958674 | 1.73061769731465 | 0.0907759266885544 | 0.366936547335137 | -4.51635325292099 |
| ENSG00000114354.13 | TFG | 0.216988924387528 | 5.07108365987007 | 1.73042428118309 | 0.090810815766141 | 0.367005501848514 | -4.94693966580576 |
| lnc-SYPL1-5 | lnc-SYPL1-5 | 0.349317127455852 | 1.60127026070761 | 1.73017306763472 | 0.0908561472150829 | 0.367061253475396 | -4.59543767335606 |
| ENSG00000083097.14 | DOP1A | 0.245963666824075 | 6.08331833370758 | 1.73015018543371 | 0.090860277241274 | 0.367061253475396 | -4.97002110141794 |

| | | | | | | | |
|---|---|---|---|---|---|---|---|
| lnc-ZNF705E-2 | lnc-ZNF705E-2 | -0.387578476518526 | 0.423507295389833 | -1.73003244251414 | 0.0908815312241885 | 0.367075070091733 | -4.42630728479283 |
| ENSG00000012660.13 | ELOVL5 | -0.26779117986436 | 7.90919693554284 | -1.72972318390112 | 0.0909373757637328 | 0.367170951298124 | -4.95516991925439 |
| ENSG00000131462.7 | TUBG1 | -0.328685981511622 | 1.54556352661238 | -1.72927584180638 | 0.0910182053611353 | 0.367170951298124 | -4.62181094645486 |
| ENSG00000143772.9 | ITPKB | -0.351234884754509 | 6.60831362675356 | -1.72919155784866 | 0.0910334412033388 | 0.367170951298124 | -4.97160709970458 |
| lnc-RPS27-2 | lnc-RPS27-2 | -0.241412947709267 | 1.4791699603827 | -1.7291809796701 | 0.0910353535494653 | 0.367170951298124 | -4.606658626693 |
| lnc-DNAH7-4 | lnc-DNAH7-4 | -1.64977951306286 | -1.70210501344878 | -1.72949466542624 | 0.091055706041989 | 0.367170951298124 | -4.18122970775546 |
| lnc-TRABD2A-5 | lnc-TRABD2A-5 | -0.296895529087072 | 5.03952805048908 | -1.72903284501988 | 0.0910621371679732 | 0.367170951298124 | -4.94695413957674 |
| ENSG00000121022.13 | COPS5 | 0.165248076936696 | 5.46733751585637 | 1.72894377456913 | 0.0910782447892138 | 0.367170951298124 | -4.96219973144416 |
| lnc-NADK-1 | lnc-NADK-1 | -0.381277481220903 | 0.327485481792681 | -1.72892308655728 | 0.0910819863768384 | 0.367170951298124 | -4.40232398346434 |
| ENSG00000120029.12 | ARMH3 | 0.154401856203003 | 4.52719100492643 | 1.72884129132881 | 0.0910967809325192 | 0.367170951298124 | -4.91137248156556 |
| ENSG00000103522.15 | IL21R | -0.472003334377638 | 4.98313155095978 | -1.72926509446955 | 0.0911013422021201 | 0.367170951298124 | -4.94499076276641 |
| ENSG00000175348.10 | TMEM9B | 0.184034349069991 | 4.4899491754702 | 1.72881523706569 | 0.0911014938677315 | 0.367170951298124 | -4.91379885590092 |
| lnc-PRELID1-1 | lnc-PRELID1-1 | 0.159751525509649 | 3.3202670044834 | 1.72865984524984 | 0.0911296067875771 | 0.367212352487999 | -4.79589835168821 |
| lnc-PHLPP1-2 | lnc-PHLPP1-2 | -0.861881318458718 | -0.903920534381606 | -1.72835065912187 | 0.0911855650963242 | 0.367365920214844 | -4.23915988074842 |
| ENSG00000154608.14 | CEP170P1 | 0.376256637062795 | 0.234118997375998 | 1.7280238190943 | 0.0912447496115008 | 0.367385977022706 | -4.36988728116924 |
| ENSG00000136932.13 | TRMO | 0.220036770490352 | 5.09289238615322 | 1.72790916382418 | 0.0912655190765097 | 0.367385977022706 | -4.94851262210665 |
| ENSG00000169241.18 | SLC50A1 | -0.319844807266068 | 4.15496162943553 | -1.72789853953018 | 0.0912674438353479 | 0.367385977022706 | -4.88477551612363 |
| lnc-ZNF746-3 | lnc-ZNF746-3 | -0.268061429392452 | 3.80854466613287 | -1.72788426678276 | 0.0912700296223999 | 0.367385977022706 | -4.8499985102258 |
| ENSG00000240036.4 | | 0.802455529257635 | -0.476532524785657 | 1.72779843826692 | 0.0912855804210338 | 0.367385977022706 | -4.22252333108335 |
| lnc-FAM120AOS-1 | lnc-FAM120AOS-1 | -0.405066170863399 | 0.990572283308674 | -1.72767647367961 | 0.0913076823105661 | 0.367385977022706 | -4.53117732165539 |
| lnc-DPH6-3 | lnc-DPH6-3 | -0.526482208661997 | 0.361463978265411 | -1.72763341383582 | 0.09131548649119 | 0.367385977022706 | -4.44661303905753 |
| lnc-TBC1D31-4 | lnc-TBC1D31-4 | 0.331588083732428 | 3.72714223500071 | 1.72744397315909 | 0.0913498273787236 | 0.367452315211309 | -4.82544477754801 |
| ENSG00000108784.9 | NAGLU | -0.313156753867836 | 3.08129045343669 | -1.72717213217824 | 0.0913991241743367 | 0.367485635869751 | -4.79703555545727 |
| ENSG00000266028.7 | SRGAP2 | 0.258860843881974 | 4.44914427290032 | 1.72713379600963 | 0.091406077998886 | 0.367485635869751 | -4.91026648901657 |
| lnc-TNFRSF13B-5 | lnc-TNFRSF13B-5 | 0.980399067894629 | 1.50547156998112 | 1.72811995468639 | 0.091431763037261 | 0.367485635869751 | -4.62293392876638 |
| ENSG00000066044.14 | ELAVL1 | -0.134896901712406 | 4.66951385402315 | -1.72690938574004 | 0.0914467927696534 | 0.367485635869751 | -4.93130984615985 |
| ENSG00000150776.17 | NKAPD1 | 0.333045496008128 | 4.19104672312651 | 1.72688072763116 | 0.0914519932990828 | 0.367485635869751 | -4.88325343382902 |
| ENSG00000172366.19 | MCRIP2 | -0.37777663600205 | 2.39312342718516 | -1.72680776906068 | 0.091465234055324 | 0.367485635869751 | -4.71739898100609 |
| ENSG00000212664.5 | RPL17P39 | 0.901320630884714 | -0.0588391282348791 | 1.72651783097712 | 0.0915180904371908 | 0.36762624033347 | -4.31143010556628 |
| ENSG00000279679.1 | | 0.53495687512515 | -0.27954988463445 | 1.72639475669294 | 0.0915402189549023 | 0.36763149975554 | -4.33180429971221 |
| ENSG00000104824.17 | HNRNPL | 0.141914533687268 | 7.43016136116968 | 1.72625510573722 | 0.0915655850042322 | 0.36763149975554 | -4.96847994504415 |
| ENSG00000133104.13 | SPART | -2.66825738982238 | 0.850611342486268 | -1.72780805107589 | 0.0915729825218215 | 0.36763149975554 | -4.45015616733604 |
| ENSG00000154957.13 | ZNF18 | 0.18836384661769 | 4.13554010325123 | 1.72609158981188 | 0.091595293295411 | 0.367649360744821 | -4.88263702594068 |
| ENSG00000203872.6 | C6orf163 | -0.29591842326755 | 2.55202291520335 | -1.72575282294914 | 0.0916568638348898 | 0.367824766682304 | -4.71823321099849 |
| lnc-SOCS2-1 | lnc-SOCS2-1 | -0.251350350636635 | 1.68438729990235 | -1.72491018769346 | 0.0918101740412284 | 0.368368189530332 | -4.64468404774004 |
| ENSG00000214629.3 | RPSAP6 | -0.723777843985232 | 0.210364080244767 | -1.72479976800315 | 0.0918302792757498 | 0.368377048983192 | -4.40330153904345 |
| lnc-NATD1-4 | lnc-NATD1-4 | 0.245814577212747 | 0.362409262364895 | 1.72465240244921 | 0.0918571173256514 | 0.368412908400991 | -4.46409997537461 |
| ENSG00000172273.12 | HINFP | -0.239234286042155 | 3.85564516619742 | -1.72430556958077 | 0.0919203078944869 | 0.368594525110505 | -4.87006859328364 |
| ENSG00000116670.14 | MAD2L2 | 0.26304378619298 | 2.77871229164633 | 1.72393978390351 | 0.0919869907147001 | 0.36879007204532 | -4.7528785478621 |
| ENSG00000188229.5 | TUBB4B | -0.279937283631818 | 4.87753636037049 | -1.72365785263751 | 0.0920384143129024 | 0.368924378150432 | -4.95107642820349 |
| ENSG00000173531.15 | MST1 | 0.485830575539354 | 1.61454045510422 | 1.72345679780623 | 0.0920751008461024 | 0.368997817514575 | -4.6193978255012 |
| ENSG00000175390.13 | EIF3F | 0.175183994105792 | 7.09822829456657 | 1.72336097238748 | 0.0920925904179358 | 0.368997817514575 | -4.97739720939636 |
| lnc-RNF6-4 | lnc-RNF6-4 | 0.158469744754874 | 2.65778402439041 | 1.72313966199868 | 0.0921329934343717 | 0.369087855947776 | -4.7378059493319 |
| ENSG00000110768.11 | GTF2H1 | -0.207330069221333 | 4.63236335545616 | -1.72288137806373 | 0.0921801650839197 | 0.369204969434193 | -4.93490084409703 |
| ENSG00000181026.14 | AEN | 0.382178264779525 | 5.83377395111724 | 1.72281811188618 | 0.092205280420536 | 0.369210332662192 | -4.97624105165299 |
| ENSG00000106012.17 | IQCE | 0.256248434995385 | 2.52136545596635 | 1.72252980553633 | 0.0922444067626633 | 0.369210332662192 | -4.73457957291586 |
| ENSG00000180921.6 | FAM83H | -1.22363392855395 | -0.195725246554847 | -1.72307774060549 | 0.0922605907521063 | 0.369210332662192 | -4.31137686936793 |
| lnc-SLC45A3-3 | lnc-SLC45A3-3 | 0.202071599755214 | 4.16554435823326 | 1.72243537275891 | 0.0922616684927695 | 0.369210332662192 | -4.89659387231806 |
| lnc-C10orf143-46 | lnc-C10orf143-46 | -0.242598500862914 | 1.97565821411839 | -1.72222200906225 | 0.0923006799549401 | 0.369210332662192 | -4.67131605438222 |
| LINC01252 | LINC01252 | 0.479615428953411 | 0.670252736192757 | 1.72220389999096 | 0.0923039916527575 | 0.369210332662192 | -4.5204443719063 |
| ENSG00000108389.9 | MTMR4 | 0.157588772208586 | 6.56484867516673 | 1.7221870806793 | 0.0923070675741943 | 0.369210332662192 | -4.98334132606086 |
| ENSG00000204947.8 | ZNF425 | -0.386636721490614 | -0.127321443177163 | -1.72184807330319 | 0.0923690835356635 | 0.369386603150294 | -4.34314807172826 |
| ENSG00000165475.14 | CRYL1 | -0.557291607291617 | 1.73251481336238 | -1.72146555898578 | 0.092442802893666 | 0.369573567729476 | -4.63914658310554 |
| lnc-ARPC3-1 | lnc-ARPC3-1 | 0.290589380455279 | 1.66165912343444 | 1.72139649367781 | 0.0924517465986042 | 0.369573567729476 | -4.63972851965734 |
| lnc-NEK7-4 | lnc-NEK7-4 | 0.337509974103456 | 3.87141859283617 | 1.72105085087091 | 0.0925150591490965 | 0.369680465318123 | -4.86308924161067 |
| lnc-KATNBL1-2 | lnc-KATNBL1-2 | 0.259485658265037 | 0.143127606831651 | 1.72098802081341 | 0.0925265718125969 | 0.369680465318123 | -4.41106248032857 |
| ENSG00000277053.4 | GTF2IP1 | 0.312749158073824 | 2.08087673242282 | 1.7209563836351 | 0.0925323693008293 | 0.369680465318123 | -4.69252908351926 |
| ENSG00000184903.9 | IMMP2L | -0.294814847579828 | 5.24173116624719 | -1.7206952972049 | 0.0925802247177063 | 0.369799876841558 | -4.96847909228467 |
| ENSG00000152404.15 | CWF19L2 | 0.393460443841089 | 5.61996347042512 | 1.72068989344643 | 0.0926013786087923 | 0.36981260688878 | -4.97407777390245 |
| lnc-GPR83-3 | lnc-GPR83-3 | 0.321329735248058 | 0.799057820205728 | 1.7204387323711 | 0.092627271400561 | 0.369844253586372 | -4.50003616690115 |
| ENSG00000204778.4 | ZNG1DP | 0.670902357841919 | -0.55180787846877 | 1.7200021290195 | 0.0927073777124825 | 0.370092310988368 | -4.26752810658658 |
| ENSG00000278601.1 | | -0.481610192754834 | 1.49441045614086 | -1.71987744851451 | 0.0927302641834701 | 0.37009725725915 | -4.63359027360407 |
| ENSG00000172375.13 | C2CD2L | -0.304349723503158 | 2.93287331461679 | -1.71979948191453 | 0.0927445781928143 | 0.37009725725915 | -4.77918147139103 |
| PCF11-AS1 | PCF11-AS1 | 0.453078327588881 | 2.94505980781089 | 1.71966894644879 | 0.092768547514018 | 0.370121150122317 | -4.75562750553096 |
| ENSG00000243829.1 | | 1.45946475445762 | -1.17971114278764 | 1.71951409337711 | 0.092796988786173 | 0.370162872129031 | -4.18043329684897 |
| lnc-EFCAB8-1 | lnc-EFCAB8-1 | 0.22503108359935 | 3.97632227667192 | 1.71938151045372 | 0.0928213455446461 | 0.370188288189392 | -4.87567707657738 |
| ENSG00000138639.17 | ARHGAP24 | 0.313330103594211 | 8.45563514111591 | 1.7185843678674 | 0.0929679003575938 | 0.370673128895892 | -4.95951703992225 |
| lnc-ZMYND19-2 | lnc-ZMYND19-2 | -0.290981808301562 | 0.756510942643361 | -1.71852440482973 | 0.0929789323465719 | 0.370673128895892 | -4.52189869888863 |

| | | | | | | | |
|---|---|---|---|---|---|---|---|
| ENSG00000150401.14 | DCUN1D2 | 0.195769190072616 | 3.08710064777575 | 1.71840700567039 | 0.0930005345734509 | 0.370680564246322 | -4.79344204745561 |
| lnc-EEA1-2 | lnc-EEA1-2 | -0.338043870689427 | 0.0955240999907223 | -1.71831854031537 | 0.093016815543519 | 0.370680564246322 | -4.39555712884183 |
| lnc-GOLGA6L22-5 | lnc-GOLGA6L22-5 | 0.386310100505834 | 1.58141492801495 | 1.71810279728968 | 0.093056530349962 | 0.370767046882166 | -4.6127856932605 |
| ENSG00000109321.10 | AREG | 2.01291473821388 | -1.53561123515149 | 1.71806048701605 | 0.0930986838352196 | 0.370863210640667 | -4.18892520433917 |
| ENSG00000075914.12 | EXOSC7 | 0.151344166696818 | 4.04006018856949 | 1.7175094294545 | 0.0931658324693443 | 0.371036679463061 | -4.8906531932787 |
| ENSG00000178741.11 | COX5A | 0.338246071404082 | 3.55547481899411 | 1.71744187678153 | 0.0931782828618063 | 0.371036679463061 | -4.84031922786696 |
| ENSG00000128881.16 | TTBK2 | 0.147215730997647 | 5.79262820664042 | 1.71700418049216 | 0.0932589866107598 | 0.37123961942645 | -4.98785695827605 |
| ENSG00000004799.7 | PDK4 | 1.35778544730087 | 1.84317175765659 | 1.71840907217622 | 0.0932775874838854 | 0.37123961942645 | -4.63923500975942 |
| ENSG00000164944.11 | VIRMA | 0.140008936787849 | 6.1085000376221 | 1.71687207640993 | 0.0932833557631832 | 0.37123961942645 | -4.99161404191832 |
| lnc-RRM2B-4 | lnc-RRM2B-4 | 0.212957070241249 | 3.24833573246879 | 1.71666100737019 | 0.0933223025090773 | 0.371322820905536 | -4.80772552204783 |
| ENSG00000006756.15 | ARSD | -0.912892571116591 | 3.47347236799967 | -1.71777563288987 | 0.0933696194091703 | 0.371439288036133 | -4.83211920893792 |
| lnc-GFAP-4 | lnc-GFAP-4 | 0.191978607175176 | 1.85820351071327 | 1.71624028901791 | 0.0933999743357711 | 0.371488245749406 | -4.66228429912812 |
| lnc-NFE2L1-1 | lnc-NFE2L1-1 | 0.4612605204957 | 0.483580327012875 | 1.71609444472709 | 0.0934269122274781 | 0.371523596286357 | -4.4600555009939 |
| ENSG00000143543.14 | JTB | 0.220947462712012 | 4.76355048563404 | 1.71569070138892 | 0.0935015185396879 | 0.371728116580718 | -4.95060509741153 |
| lnc-ZNF131-1 | lnc-ZNF131-1 | -0.285336426483493 | 5.53550371593867 | -1.71562067885545 | 0.0935144627923509 | 0.371728116580718 | -4.98334672740182 |
| ENSG00000238622.1 | SNORD97 | 0.458415617751717 | 2.58370303590595 | 1.71518051852771 | 0.0935958642144988 | 0.371814382598193 | -4.73728639183652 |
| lnc-NAPB-2 | lnc-NAPB-2 | -0.399499754624381 | 1.30994477592816 | -1.71505958365131 | 0.0936182397125495 | 0.371814382598193 | -4.59565976440716 |
| lnc-CCNG1-1 | lnc-CCNG1-1 | 0.263244137858126 | 3.01282212630866 | 1.71504742014725 | 0.093620490462769 | 0.371814382598193 | -4.79974914293336 |
| ENSG00000182183.14 | SHISAL2A | -0.416485759329789 | 3.61673531009426 | -1.71495397092465 | 0.0936377839246604 | 0.371814382598193 | -4.85001685095248 |
| lnc-MTF2-4 | lnc-MTF2-4 | -0.33244589093752 | 1.43222199611539 | -1.71489771655275 | 0.0936481954868193 | 0.371814382598193 | -4.63032213964767 |
| ENSG00000187630.16 | DHRS4L2 | -0.256620353378608 | 2.35975776202244 | -1.71470482340834 | 0.0936839034620289 | 0.371814382598193 | -4.71739915061706 |
| ENSG00000003756.16 | RBM5 | 0.153275650919414 | 7.97917063661833 | 1.7146534645963 | 0.0936934128039716 | 0.371814382598193 | -4.97827400884238 |
| ENSG00000145425.9 | RPS3A | 0.247066700822707 | 9.85655255144999 | 1.71463133919402 | 0.093697509680381 | 0.371814382598193 | -4.93078374208725 |
| ENSG00000177889.9 | UBE2N | 0.183871199882791 | 5.76437284309426 | 1.71442525972727 | 0.0937356757772178 | 0.371814382598193 | -4.99267259181029 |
| ENSG00000173890.16 | GPR160 | 0.763296618316952 | 2.6943143176339 | 1.71538683311017 | 0.0937381439217272 | 0.371814382598193 | -4.76017860311573 |
| ENSG00000091536.17 | MYO15A | -0.348562765073413 | 0.921445504669174 | -1.71431576901228 | 0.0937559588049984 | 0.371814382598193 | -4.54405486550038 |
| lnc-TACSTD2-2 | lnc-TACSTD2-2 | 0.180122820532482 | 4.4095637759837 | 1.71425568270754 | 0.0937670912721608 | 0.371814382598193 | -4.92892403903564 |
| lnc-DHH-2 | lnc-DHH-2 | 0.326239896030148 | 2.30717950170068 | 1.71423459580613 | 0.0937709984000011 | 0.371814382598193 | -4.71456848833899 |
| ENSG00000160799.11 | CCDC12 | -0.176398686780313 | 4.68728890562938 | -1.71406829931603 | 0.0938018157089659 | 0.37186494082004 | -4.94884738162756 |
| ENSG00000122966.15 | CIT | 0.492819992822635 | 1.61659874170646 | 1.71382034660576 | 0.0938477807971979 | 0.371975519381615 | -4.62039311124382 |
| LINC01535 | LINC01535 | -0.83544678558303 | 1.94397805939108 | -1.71454521562199 | 0.0938939321852091 | 0.372086793640385 | -4.6892598639271 |
| ENSG00000048405.9 | ZNF800 | 0.210333799862829 | 6.86635431173632 | 1.71334169078435 | 0.0939365661778742 | 0.372184088862211 | -4.99566040305373 |
| lnc-ANKRD31-3 | lnc-ANKRD31-3 | 0.178854329844295 | 2.18637243624844 | 1.71279634454649 | 0.0940378068883294 | 0.372507591215709 | -4.70141608508686 |
| ENSG00000138600.9 | SPPL2A | 0.179493575641201 | 5.61105066637551 | 1.71270694828178 | 0.0940544114826818 | 0.372507591215709 | -4.99283135025202 |
| ENSG00000225022.1 | UBE2D3P1 | 0.917103656197959 | -0.695909420628448 | 1.71239548851454 | 0.0941122814969227 | 0.37266508081015 | -4.2190687388485 |
| lnc-DENND1B-6 | lnc-DENND1B-6 | 0.460429563099922 | 0.314169304212208 | 1.71199249003198 | 0.0941872035053393 | 0.372843189470418 | -4.48172177961483 |
| ENSG00000228863.8 | | 1.2783267952258 | -1.23552095811378 | 1.71189568504591 | 0.0942052080289885 | 0.372843189470418 | -4.18850253753752 |
| ENSG00000177169.9 | ULK1 | 0.326972726464088 | 4.24484636571105 | 1.7118613038641 | 0.0942116031888278 | 0.372843189470418 | -4.91814008569776 |
| ENSG00000181467.4 | RAP2B | -0.293916923525252 | 4.08093742533667 | -1.71156798376242 | 0.0942661776011042 | 0.37298745358776 | -4.91124255387937 |
| lnc-DLX2-4 | lnc-DLX2-4 | -0.301080054193544 | 1.44955089064384 | -1.711125587549 | 0.0943485383837184 | 0.373241584768802 | -4.62943396069916 |
| lnc-ZNF585B-6 | lnc-ZNF585B-6 | -0.236940641792563 | 0.937529518654488 | -1.7109834806672 | 0.0943750070549791 | 0.373274552308346 | -4.55627508947007 |
| lnc-USP53-1 | lnc-USP53-1 | -0.547187230178077 | 1.85987937000254 | -1.71063751297081 | 0.0944500287660342 | 0.373499508567009 | -4.67844034686535 |
| ENSG00000115282.19 | TTC31 | -0.197016788598089 | 5.02402375703281 | -1.71036789658669 | 0.0944897362563438 | 0.373513208090185 | -4.97751283157528 |
| ENSG00000159958.6 | TNFRSF13C | 0.304985285737835 | 7.10548986817008 | 1.71027440593955 | 0.0945071706540964 | 0.373513208090185 | -4.99879857167535 |
| ENSG00000095794.19 | CREM | 0.35229484515691 | 2.38218597003203 | 1.71018141694276 | 0.0945245141509716 | 0.373513208090185 | -4.72384341888537 |
| lnc-RNMT-5 | lnc-RNMT-5 | -0.45279644836202 | 2.71164452749851 | -1.71017302313743 | 0.0945260798203262 | 0.373513208090185 | -4.7657014277583 |
| ENSG00000169976.6 | SF3B5 | 0.267389280827105 | 4.62571741278329 | 1.70991380790031 | 0.0945744409962583 | 0.373535120695886 | -4.95504488083656 |
| ENSG00000260027.4 | HOXB7 | 0.977544710573238 | -0.985904921931841 | 1.70988021892059 | 0.0945807091155541 | 0.373535120695886 | -4.25307991878245 |
| ENSG00000138777.19 | PPA2 | 0.150868197232407 | 4.33614974269368 | 1.70979091531263 | 0.0945973759508394 | 0.373535120695886 | -4.93318907696405 |
| lnc-RPS27L-3 | lnc-RPS27L-3 | 0.353253724038975 | 2.37423497878089 | 1.70975426736899 | 0.0946042163041176 | 0.373535120695886 | -4.7173772586942 |
| ENSG00000157837.15 | SPPL3 | 0.202141592055154 | 4.63603960814264 | 1.70937443418268 | 0.0946751364965259 | 0.373741900017548 | -4.95136873772546 |
| lnc-NOC2L-12 | lnc-NOC2L-12 | -0.406792659469771 | 1.93100126614377 | -1.70927932122787 | 0.0946929023267508 | 0.373741900017548 | -4.69109940653809 |
| ENSG00000145439.11 | CBR4 | -0.213834836757912 | 4.36213929670096 | -1.70917232353323 | 0.0947128913741107 | 0.373749126371419 | -4.93699223260367 |
| ENSG00000251485.1 | NRBF2P6 | -0.258379380934131 | 0.613798711791029 | -1.70905725674414 | 0.0947343917771961 | 0.373762312813883 | -4.50620118603767 |
| lnc-GTF2A2-3 | lnc-GTF2A2-3 | 0.264432378462086 | 4.92593044109677 | 1.70890249255775 | 0.0947633160853877 | 0.373804778648052 | -4.97275872798225 |
| lnc-SCGN-1 | lnc-SCGN-1 | -0.715100120144404 | 0.0889918429962745 | -1.70871008960477 | 0.0947992850242478 | 0.373823550530867 | -4.40795787281284 |
| lnc-RAX2-2 | lnc-RAX2-2 | -0.382547946293555 | -0.0966636451691201 | -1.70868274195999 | 0.0948043984730664 | 0.373823550530867 | -4.39482420066618 |
| lnc-ERV3-1-10 | lnc-ERV3-1-10 | -0.243875182994763 | 2.61984748880255 | -1.70844463111531 | 0.0948489299900694 | 0.37392750928665 | -4.75079071786498 |
| ENSG00000166971.16 | AKTIP | -0.218106544576671 | 2.64375538970678 | -1.70816508791003 | 0.0949012323376846 | 0.374005379539343 | -4.77603908084415 |
| ENSG00000039123.15 | MTREX | 0.145474652443446 | 6.52578104741282 | 1.70814483040186 | 0.0949050234336098 | 0.374005379539343 | -5.00599192125713 |
| lnc-FGF1-8 | lnc-FGF1-8 | 0.64672120280533 | -0.461135562233775 | 1.70788488161108 | 0.0949536827604354 | 0.374059131367601 | -4.26433271397317 |
| lnc-TMEM206-5 | lnc-TMEM206-5 | -0.340658262204673 | 0.49504091652175 | -1.70787779520291 | 0.0949550095416468 | 0.374059131367601 | -4.49308999555994 |
| LINC01534 | LINC01534 | -0.208901138412049 | 3.368510631482 | -1.70763525073034 | 0.0950004301766091 | 0.374166447440709 | -4.83907293152943 |
| ENSG00000241975.1 | ELOCP19 | 0.2414355782574 | 1.59212855889206 | 1.70726094212926 | 0.09507056128606 | 0.374244744687568 | -4.64550515206549 |
| ENSG00000222588.1 | | 0.501401916122948 | -0.465271117332817 | 1.70724234281636 | 0.0950740472053305 | 0.374244744687568 | -4.32033877326469 |
| ENSG00000137462.7 | TLR2 | 0.601176293443711 | 2.43097440722517 | 1.70760757868925 | 0.0950748564335273 | 0.374244744687568 | -4.73643063676897 |
| ENSG00000254965.1 | C1DP5 | 0.477737270932823 | 0.290907663397882 | 1.70704911616397 | 0.0951102683917894 | 0.37431255340501 | -4.42063867214191 |
| lnc-ZNF726-1 | lnc-ZNF726-1 | 0.377910879979194 | 2.28007831337342 | 1.70682961630537 | 0.0951514284924878 | 0.374402954054841 | -4.71224315146231 |

| | | | | | | | |
|---|---|---|---|---|---|---|---|
| ENSG00000272410.5 | | 0.206783169335055 | 4.2073271300674 | 1.70669137285988 | 0.0951773591625254 | 0.374433406659453 | -4.91806679848474 |
| ENSG00000174953.13 | DHX36 | 0.264186895102609 | 6.28686380449361 | 1.70652795485707 | 0.0952080194466977 | 0.374448926829678 | -5.00871749880373 |
| lnc-RPS28-4 | lnc-RPS28-4 | -0.474762485782209 | -0.411875762662905 | -1.7064764281064 | 0.095217688530658 | 0.374448926829678 | -4.34290317064814 |
| ENSG00000272647.3 | | 0.177859550019466 | 2.51764649718487 | 1.7059173954788 | 0.0953226443317804 | 0.374725240325527 | -4.75122916320319 |
| lnc-ZNF626-1 | lnc-ZNF626-1 | -0.400066763088156 | 2.87998242095883 | -1.70590824684491 | 0.095324362743257 | 0.374725240325527 | -4.7825646739839 |
| ENSG00000258529.5 | | 0.309218242498131 | 2.41433526176356 | 1.70580689171281 | 0.0953434022578858 | 0.374728517982445 | -4.73239159976097 |
| ENSG00000139644.12 | TMBIM6 | 0.177125766187162 | 8.25656609680596 | 1.70569426360065 | 0.0953645630958514 | 0.374740130240914 | -4.98623848824796 |
| lnc-HNRNPH1-6 | lnc-HNRNPH1-6 | -0.852430409609551 | -0.854696646903205 | -1.70546704601261 | 0.095407265131957 | 0.374788791961193 | -4.28173448132996 |
| lnc-FBXO15-3 | lnc-FBXO15-3 | -1.11093965929411 | 2.93768773405656 | -1.70691414162576 | 0.0954136693956701 | 0.374788791961193 | -4.80772904261766 |
| ENSG00000182220.14 | ATP6AP2 | 0.196575368408397 | 5.36284055063137 | 1.70533774364846 | 0.0954315725923632 | 0.374788791961193 | -4.99508149941286 |
| ENSG00000167700.8 | MFSD3 | -0.426878632784892 | 1.16872061247093 | -1.70515636920508 | 0.0954656777000991 | 0.374851210244399 | -4.60629542510773 |
| lnc-URM1-1 | lnc-URM1-1 | 0.228128096058935 | -0.130814103113437 | 1.70499878692655 | 0.095495317204803 | 0.374896073627019 | -4.38412795984362 |
| ENSG00000163947.11 | ARHGEF3 | 0.291117681482329 | 6.41962486146513 | 1.70432452283156 | 0.0956222250994189 | 0.37532270389423 | -5.0121645208279 |
| lnc-YPEL5-5 | lnc-YPEL5-5 | -0.413878402571292 | 5.69717920111168 | -1.70426765955813 | 0.0956766358483759 | 0.375464670289251 | -5.00561791673348 |
| ENSG00000188389.10 | PDCD1 | -0.533483259537766 | 1.90230732418662 | -1.7037102550881 | 0.0957426914626856 | 0.375652271897914 | -4.69381620042374 |
| lnc-ZNF383-2 | lnc-ZNF383-2 | 0.254060881953989 | 0.197154350161838 | 1.70302102819779 | 0.0958679612663353 | 0.376072088192296 | -4.43141397219216 |
| ENSG00000099364.16 | FBXL19 | -0.283153961788533 | 2.92024630451783 | -1.70276704482256 | 0.0959159033603964 | 0.376159406992311 | -4.80875613293248 |
| ENSG00000272886.5 | DCP1A | 0.200580368533498 | 5.75678027744915 | 1.70254104538958 | 0.0959585798911811 | 0.376159406992311 | -5.01031443510225 |
| ENSG00000174442.11 | ZWILCH | 0.197750745317186 | 3.36541263915831 | 1.70252002681944 | 0.0959625497259506 | 0.376159406992311 | -4.84641342869399 |
| lnc-RBM15-3 | lnc-RBM15-3 | 0.285728892083413 | 3.78457717481392 | 1.70248596459301 | 0.0959689834407641 | 0.376159406992311 | -4.88897107878512 |
| lnc-TDRP-6 | lnc-TDRP-6 | -0.427017258638043 | 0.680319859007746 | -1.70239288461614 | 0.0959865663234573 | 0.376159406992311 | -4.55449163279277 |
| ZNF582-AS1 | ZNF582-AS1 | -0.411172682671114 | 2.07416285813975 | -1.70232245675056 | 0.0959998719783613 | 0.376159406992311 | -4.70754941262563 |
| lnc-MIA3-1 | lnc-MIA3-1 | 0.180745106289673 | 0.731703131917439 | 1.70199368015266 | 0.0960620066522142 | 0.376277844805165 | -4.54417363170781 |
| ENSG00000254598.2 | CSNK2A3 | 0.871251199308748 | -0.38696479256825 | 1.70196906173413 | 0.0960666605670283 | 0.376277844805165 | -4.27173159855161 |
| lnc-HBS1L-1 | lnc-HBS1L-1 | -0.455442358510459 | 0.80178011406642 | -1.70143744009312 | 0.0961672049728545 | 0.376599996186504 | -4.55669003468956 |
| ENSG00000148444.15 | COMMD3 | -0.285938831002431 | 4.38201821900049 | -1.70094782291494 | 0.0962598822928995 | 0.376867159655897 | -4.95350175558076 |
| ENSG00000178035.11 | IMPDH2 | -0.356529654474463 | 5.30873044896318 | -1.70084292233928 | 0.0962797480501687 | 0.376867159655897 | -5.00310799936019 |
| ENSG00000122126.16 | OCRL | 0.384917886136027 | 3.24659291134998 | 1.70078691596322 | 0.0962903557610826 | 0.376867159655897 | -4.84526011102135 |
| ENSG00000163069.12 | SGCB | -0.409827521669313 | 2.15465584576413 | -1.70060928453872 | 0.0963240058868114 | 0.376927188815254 | -4.72463184022942 |
| ENSG00000173867.10 | | -0.178923593449865 | 2.80064770117358 | -1.70002989709175 | 0.0964338315593692 | 0.377268137776174 | -4.79155855229431 |
| ENSG00000160563.13 | MED27 | 0.183729183409831 | 2.9041551398735 | 1.69995627789011 | 0.0964477938579522 | 0.377268137776174 | -4.79921216993173 |
| LINC02018 | LINC02018 | -0.475061388291556 | 1.02323377896712 | -1.69953831565215 | 0.0965270945119092 | 0.377487807563193 | -4.57851004509882 |
| ENSG00000104903.4 | LYL1 | -0.433720064082289 | 2.547989917914 | -1.69946699577488 | 0.0965406315412062 | 0.377487807563193 | -4.77685802675528 |
| ENSG00000065457.10 | ADAT1 | 0.163794265031195 | 5.0228152200456 | 1.69922660312957 | 0.0965862713908177 | 0.377581782351154 | -4.98917192866121 |
| ENSG00000147121.15 | KRBOX4 | 0.131116102256786 | 3.43638231474689 | 1.69910440553519 | 0.096609478187581 | 0.377581782351154 | -4.85816319311604 |
| ENSG00000167468.16 | GPX4 | -0.258349248140806 | 4.86794133036 | -1.69905059838204 | 0.096619698180223 | 0.377581782351154 | -4.98697305046276 |
| lnc-FAM167A-5 | lnc-FAM167A-5 | -0.422692120485349 | 1.37733595856701 | -1.69866634198549 | 0.0966927093823197 | 0.377795375325795 | -4.63549894543617 |
| ENSG00000113758.13 | DBN1 | 0.803019407186597 | 0.782057420904381 | 1.69903669289429 | 0.0967424618292824 | 0.377918028436538 | -4.60758278650236 |
| LINC00402 | LINC00402 | 0.908486572119227 | -0.216066988309544 | 1.698115633504 | 0.0967974272284243 | 0.378017916829369 | -4.27199394039707 |
| ENSG00000282977.1 | PCBP2-OT1 | 0.23606687039116 | 3.27394545465769 | 1.69807707265541 | 0.0968047631349951 | 0.378017916829369 | -4.84759389064311 |
| PCBP1-AS1 | PCBP1-AS1 | 0.258828202799934 | 5.74328794516013 | 1.69756621744288 | 0.0969019928952011 | 0.37832581937821 | -5.01820539135107 |
| ENSG00000117305.14 | HMGCL | -0.201887161744451 | 3.34575821179189 | -1.69714687883829 | 0.0969818649586115 | 0.378511876093166 | -4.85259874066551 |
| ENSG00000189266.12 | PNRC2 | 0.216046123051665 | 7.46427993382015 | 1.69712293423775 | 0.096986427367991 | 0.378511876093166 | -5.01449924925511 |
| ENSG00000197465.13 | GYPE | 1.89123516795521 | -2.06009261797492 | 1.69687784688129 | 0.0970614679452024 | 0.378732927908266 | -4.20661932629158 |
| ENSG00000101846.6 | STS | 0.378324598931508 | 1.79133751999975 | 1.6963321115287 | 0.0971372107919141 | 0.378956635657689 | -4.69072352167626 |
| POC1B-AS1 | POC1B-AS1 | -0.307272720168257 | 4.41126454659983 | -1.69613241650313 | 0.0971753166273289 | 0.379033455019957 | -4.96241003361462 |
| lnc-ZNF347-1 | lnc-ZNF347-1 | -0.266919620740256 | 0.597949475230584 | -1.69591016527722 | 0.0972177411910553 | 0.379090895324344 | -4.50816318128684 |
| lnc-LHFPL3-4 | lnc-LHFPL3-4 | 0.221479653073994 | 3.71759626739429 | 1.69586230876153 | 0.0972268783178939 | 0.379090895324344 | -4.89543853522806 |
| RPS6KB2-AS1 | RPS6KB2-AS1 | -0.566406759002765 | -0.551803653781334 | -1.69567165646893 | 0.0972632861491668 | 0.379161026289451 | -4.3553247850948 |
| lnc-SNU13-3 | lnc-SNU13-3 | -0.473082188103982 | 0.184589406919686 | -1.69556508633005 | 0.0972836421859583 | 0.379168567906378 | -4.45278692179818 |
| lnc-PPP1R9A-1 | lnc-PPP1R9A-1 | -2.52763388478481 | -2.19926321765988 | -1.69596194428175 | 0.0973852758014732 | 0.379449625535327 | -4.20544037898338 |
| ENSG00000278948.1 | | 0.195282978003133 | 2.68159525852155 | 1.69499057429883 | 0.097393440892236 | 0.379449625535327 | -4.78069796584536 |
| lnc-PTDSS1-3 | lnc-PTDSS1-3 | 0.369890828651845 | 0.778903594154981 | 1.69489844056816 | 0.0974110587051775 | 0.379449625535327 | -4.54180596193303 |
| lnc-UBQLN2-5 | lnc-UBQLN2-5 | 0.321244047727333 | 0.255049993596487 | 1.69455037645753 | 0.0974776393360202 | 0.379637133482177 | -4.44609780339926 |
| lnc-MBD2-7 | lnc-MBD2-7 | -0.428033698750001 | -0.0508191780863945 | -1.69401087027868 | 0.0975809150500551 | 0.379967456389573 | -4.4349232696299 |
| ENSG00000229447.2 | | 0.380508004160764 | -0.0331560085606583 | 1.69364863366479 | 0.0976503074914215 | 0.380033947666408 | -4.39796147967487 |
| lnc-NKAIN1-2 | lnc-NKAIN1-2 | 0.380508004160764 | -0.0331560085606583 | 1.69364863366479 | 0.0976503074914215 | 0.380033947666408 | -4.39796147967487 |
| lnc-RUFY3-1 | lnc-RUFY3-1 | -0.427185947987647 | -0.157865481390408 | -1.69363259246996 | 0.0976533813927821 | 0.380033947666408 | -4.37329975762798 |
| ENSG00000219529.2 | | 0.293955854452974 | 0.504496579219054 | 1.69315717179278 | 0.0977445204594871 | 0.380316722990099 | -4.495775662652 |
| ENSG00000277971.1 | | -0.302345286114909 | 0.291012803321803 | -1.69297387074717 | 0.0977796784203308 | 0.380381614236566 | -4.47043115677081 |
| ENSG00000240849.10 | PEDS1 | -0.257733910408421 | 3.79302510375649 | -1.69278118622373 | 0.0978166474544997 | 0.380453525048369 | -4.90854498938369 |
| ENSG00000267359.1 | | -0.680932897473121 | -0.446985680992071 | -1.69263679270846 | 0.097844358810005 | 0.380481938238685 | -4.31122976396236 |
| ENSG00000226232.8 | NPIPB14P | 0.267821640406965 | 3.96661759962339 | 1.69229721699184 | 0.0979095542582361 | 0.380481938238685 | -4.91889259131955 |
| ENSG00000156711.16 | MAPK13 | -0.287531175958284 | 5.54704980573995 | -1.69219418066029 | 0.0979293434021133 | 0.380481938238685 | -5.02131184502561 |
| lnc-DHRS13-1 | lnc-DHRS13-1 | -0.21456205274658 | 1.86505197308444 | -1.69204248729983 | 0.0979584836302097 | 0.380481938238685 | -4.71183311059114 |
| ENSG00000200312.1 | RN7SKP255 | 0.425198527285807 | 0.845636123042898 | 1.69192970241137 | 0.0979801542071296 | 0.380481938238685 | -4.54671054808446 |
| lnc-ANKRD11-1 | lnc-ANKRD11-1 | -0.357140941769555 | -0.294278911651915 | -1.69174670083305 | 0.0980153247120475 | 0.380481938238685 | -4.36221866087155 |
| ENSG00000105254.11 | TBCB | -0.194086048368159 | 4.9615121486483 | -1.69172831287347 | 0.0980188592142274 | 0.380481938238685 | -5.00509452384615 |

| | | | | | | | |
|---|---|---|---|---|---|---|---|
| ENSG00000279491.1 | | -0.417618919490474 | -0.279419841646619 | -1.6917052054244 | 0.0980233010381195 | 0.380481938238685 | -4.36756686124692 |
| ENSG00000196511.14 | TPK1 | -0.329443549369913 | 4.36535250466224 | -1.69165836372703 | 0.0980323056831295 | 0.380481938238685 | -4.95777287114301 |
| ENSG00000103978.15 | TMEM87A | 0.180875093201648 | 6.08833479678142 | 1.69160181957279 | 0.0980431763979759 | 0.380481938238685 | -5.03171471478399 |
| ENSG00000196466.10 | ZNF799 | 0.361074785278625 | 2.46894199724598 | 1.69151723964781 | 0.0980594389012949 | 0.380481938238685 | -4.77021812168461 |
| ENSG00000123143.12 | PKN1 | -0.203562798763871 | 6.18916322803901 | -1.69151221323876 | 0.0980604054182865 | 0.380481938238685 | -5.03197075497155 |
| ENSG00000043143.20 | JADE2 | -0.167588404963662 | 6.9284608486867 | -1.69149216300105 | 0.09806426091222 | 0.380481938238685 | -5.02929190896788 |
| ENSG00000061987.15 | MON2 | 0.214872620461146 | 6.73437574106196 | 1.69127620989213 | 0.0981057948479251 | 0.380571348540302 | -5.03150591427232 |
| ENSG00000133103.16 | COG6 | -0.142615684182551 | 4.45267099547171 | -1.69084395394687 | 0.0981889736945025 | 0.380676867504377 | -4.96983207834123 |
| lnc-RAB30-4 | lnc-RAB30-4 | 0.471581759547789 | 0.296321058716582 | 1.69067152570897 | 0.0982221702656978 | 0.380676867504377 | -4.44542817867628 |
| lnc-CDAN1-1 | lnc-CDAN1-1 | 0.203363646200506 | 2.22933062328839 | 1.69063527022948 | 0.0982291514960158 | 0.380676867504377 | -4.7428972106459 |
| lnc-RPAP2-1 | lnc-RPAP2-1 | -0.35196725608319 | -0.0530756310418914 | -1.69047358367783 | 0.0982602902981719 | 0.380676867504377 | -4.40699283571175 |
| ENSG00000224616.2 | RTCA-AS1 | -0.338306112703986 | 0.192091817586401 | -1.69045268336353 | 0.0982643160323975 | 0.380676867504377 | -4.44244932947589 |
| ENSG00000047932.13 | GOPC | -0.187688446165216 | 5.06543254934029 | -1.69039784713168 | 0.0982748790146321 | 0.380676867504377 | -5.00989438658368 |
| ENSG00000075702.17 | WDR62 | 0.362580699933597 | 1.62294928723005 | 1.69028859345707 | 0.0982959271080775 | 0.380676867504377 | -4.66926156717079 |
| lnc-KIAA0408-4 | lnc-KIAA0408-4 | -0.289301508075589 | 1.74504036072968 | -1.69024138462408 | 0.0983050232044304 | 0.380676867504377 | -4.68248077452723 |
| ENSG00000213983.11 | AP1G2 | -0.2115790689528 | 6.51463139100796 | -1.69020638575922 | 0.098311767159742 | 0.380676867504377 | -5.03398550524 |
| ENSG00000135587.8 | SMPD2 | -0.23351733360342 | 3.66883108509236 | -1.69016447292978 | 0.0983198438751015 | 0.380676867504377 | -4.90555185437136 |
| ENSG00000285238.2 | | 0.166690101809007 | 7.05817305748254 | 1.69007836899867 | 0.0983364380566861 | 0.380676867504377 | -5.03060406374387 |
| ENSG00000277785.1 | SNORD116-21 | -0.775620222005041 | 1.38076562566171 | -1.69050078507544 | 0.0983654273016066 | 0.380717485924966 | -4.62373482360348 |
| ENSG00000130935.9 | NOL11 | 0.204404799019697 | 5.79573405008838 | 1.68968929752397 | 0.0984114498412065 | 0.380787294535876 | -5.03214632314255 |
| ENSG00000166126.10 | AMN | -0.439004660346604 | 1.60122989094963 | -1.68955292898455 | 0.0984377524727731 | 0.380787294535876 | -4.71022057588351 |
| ENSG00000263001.5 | GTF2I | 0.234661089500289 | 5.95478665294371 | 1.68954664888028 | 0.0984389639132001 | 0.380787294535876 | -5.03333142139864 |
| ENSG00000249222.1 | ATP5MGL | 0.42814499417792 | 0.0168852022227294 | 1.68929889595533 | 0.0984867656180831 | 0.380846771656003 | -4.42848079890435 |
| lnc-FAM193A-5 | lnc-FAM193A-5 | 0.330462791775251 | 0.395316477417358 | 1.68922587657588 | 0.0985008577140735 | 0.380846771656003 | -4.51676073629417 |
| ENSG00000197457.9 | STMN3 | -0.363302320084827 | 5.09306331169312 | -1.68917929442032 | 0.0985098485301734 | 0.380846771656003 | -5.01341890213893 |
| ENSG00000260537.2 | | 0.119778186023678 | 3.98054221156586 | 1.68873518555302 | 0.0985956000065375 | 0.381106710785833 | -4.92894285823804 |
| ENSG00000138670.17 | RASGEF1B | 0.61256333139024 | 5.15027381074712 | 1.68948636732776 | 0.0986418942516795 | 0.381183795851128 | -5.00124129793289 |
| ENSG00000162704.15 | ARPC5 | -0.265236117193595 | 5.77161847888989 | -1.68828302841872 | 0.0986829688914865 | 0.381183795851128 | -5.03329200848858 |
| ENSG00000103502.13 | CDIPT | -0.235367285644815 | 4.45172874095922 | -1.68825832618685 | 0.0986877438685381 | 0.381183795851128 | -4.9789559379261 |
| lnc-NUFIP2-1 | lnc-NUFIP2-1 | -0.426987065772177 | -0.428662271383662 | -1.68824862082083 | 0.0986896199820562 | 0.381183795851128 | -4.31174821322667 |
| ENSG00000176845.12 | METRNL | 1.46934601550778 | -0.0348296537975598 | 1.68902773091727 | 0.0987451428741437 | 0.381326693391839 | -4.40375956690711 |
| ENSG00000115758.12 | ODC1 | 0.297694083252379 | 7.09515089047965 | 1.68785592361493 | 0.0987655557701831 | 0.381333977568501 | -5.03357323594363 |
| ENSG00000139624.12 | CERS5 | 0.150212534915292 | 3.84883368614337 | 1.68762656063466 | 0.0988099299885234 | 0.381433756367925 | -4.9180057079948 |
| ENSG00000140332.15 | TLE3 | 0.428899274655048 | 5.25944220293361 | 1.68779226968713 | 0.0988325781392823 | 0.38144964482296 | -5.01326884419235 |
| ENSG00000201487.1 | SNORD45B | 0.447559164878605 | 1.14963641350741 | 1.68733537499379 | 0.0988662886065454 | 0.381508214920983 | -4.6292650435556 |
| lnc-ULK3-1 | lnc-ULK3-1 | -0.325418643988826 | 2.37427675596412 | -1.68704740877727 | 0.0989220502388563 | 0.381512028215248 | -4.76943395556504 |
| ENSG00000166562.8 | SEC11C | -0.372920254582202 | 4.580954604554 | -1.68701859421515 | 0.0989276313065155 | 0.381512028215248 | -4.99054851627083 |
| lnc-ACADL-2 | lnc-ACADL-2 | -0.727887997405864 | -0.289882324802522 | -1.68699116569251 | 0.0989329441551666 | 0.381512028215248 | -4.40580681822345 |
| lnc-PSMB10-1 | lnc-PSMB10-1 | 0.27013838339846 | 1.50326753366061 | 1.68694742050663 | 0.0989414179960644 | 0.381512028215248 | -4.66171487082025 |
| ENSG00000241506.1 | PSMC1P1 | 0.783675519544312 | -0.302157692420672 | 1.68658961861714 | 0.099010750003187 | 0.381596447481247 | -4.28843394013588 |
| lnc-MTSS1-4 | lnc-MTSS1-4 | 0.242691428834195 | 1.41830507567623 | 1.68658797034125 | 0.0990110694858843 | 0.381596447481247 | -4.65227852779491 |
| ENSG00000137070.17 | IL11RA | -0.184541032634521 | 4.59245175863697 | -1.68654742029711 | 0.099018929504802 | 0.381596447481247 | -4.98814768576391 |
| ENSG00000136861.17 | CDK5RAP2 | 0.213408409068641 | 6.46797019889871 | 1.68638205157226 | 0.0990509890962175 | 0.38164031382195 | -5.04058565932229 |
| ENSG00000062524.15 | LTK | 1.1178119326194 | 3.88856425701312 | 1.68771230027065 | 0.099073507472892 | 0.38164031382195 | -4.89878882669588 |
| ENSG00000115211.15 | EIF2B4 | 0.191405382182221 | 4.44942126449843 | 1.68620183588943 | 0.0990859367956323 | 0.38164031382195 | -4.97715278101441 |
| ENSG00000272617.3 | | -0.247871543322666 | 2.47728004665095 | -1.68582498072827 | 0.099159050057137 | 0.381808590398191 | -4.78029411552913 |
| lnc-TOMM5-1 | lnc-TOMM5-1 | 0.466283688537395 | 2.39882252016088 | 1.68577717634693 | 0.0991683277188908 | 0.381808590398191 | -4.75342429746818 |
| ENSG00000066027.11 | PPP2R5A | 0.160562910637448 | 5.43546199693432 | 1.6856898583878 | 0.0991852758518744 | 0.381808590398191 | -5.02998367917857 |
| ENSG00000136699.19 | SMPD4 | 0.15331773949046 | 5.47912760277139 | 1.68550802913617 | 0.0992205760022528 | 0.381873058312335 | -5.03173757772245 |
| ENSG00000133612.18 | AGAP3 | -0.447739877797929 | 3.67251909817771 | -1.68542757033116 | 0.0992673708506025 | 0.38190891207998 | -4.93063941547668 |
| lnc-SMC1B-7 | lnc-SMC1B-7 | -0.354469824817893 | 0.854149615871785 | -1.68518605795421 | 0.0992831086346108 | 0.38190891207998 | -4.58105178972379 |
| ENSG00000136449.13 | MYCBPAP | 0.766779405665668 | 0.00559560531411808 | 1.68506102620137 | 0.099307400841183 | 0.38190891207998 | -4.40637186237812 |
| ENSG00000119685.19 | TTLL5 | -0.202968565637637 | 4.61387087770029 | -1.68501892952161 | 0.0993155808385131 | 0.38190891207998 | -4.9896497333814 |
| ENSG00000249115.8 | HAUS5 | -0.187420366273146 | 4.53285565464776 | -1.68498247648397 | 0.0993226646438388 | 0.38190891207998 | -4.98325410200816 |
| ENSG00000275801.1 | | 0.715608106388926 | -0.489618670766988 | 1.68469375478165 | 0.0993787857824659 | 0.382053333537634 | -4.28350071126326 |
| ENSG00000149743.13 | TRPT1 | -0.234860283151042 | 2.70931192429284 | -1.68452193101361 | 0.0994121969715604 | 0.382110410880603 | -4.81028123160854 |
| lnc-VSIR-1 | lnc-VSIR-1 | 0.417522321822429 | 3.90760106256569 | 1.68431513174648 | 0.0994595375436253 | 0.382175775560064 | -4.92309910087669 |
| ENSG00000162976.12 | SLC66A3 | 0.245274014748786 | 3.41996250504667 | 1.68407121173756 | 0.0994998835881193 | 0.382175775560064 | -4.87628721000879 |
| lnc-ZNF101-3 | lnc-ZNF101-3 | 0.254862149988076 | 3.14511070175274 | 1.68398902883879 | 0.0995158790072462 | 0.382175775560064 | -4.85895273072084 |
| ENSG00000110075.14 | PPP6R3 | 0.118583792009918 | 7.52793175169364 | 1.68392515502547 | 0.0995283123601203 | 0.382175775560064 | -5.03368040859913 |
| lnc-C11orf95-1 | lnc-C11orf95-1 | 0.448160460670324 | 0.0621772310049813 | 1.68386953802528 | 0.0995391395287156 | 0.382175775560064 | -4.47697742146044 |
| lnc-ORC1-1 | lnc-ORC1-1 | 0.189403515355172 | 1.10927659731994 | 1.68379059106202 | 0.099554510098927 | 0.382175775560064 | -4.62042532099209 |
| ENSG00000237651.6 | C2orf74 | -0.323385892838891 | 2.0139367007359 | -1.68370754077976 | 0.0995706816805072 | 0.382175775560064 | -4.72725729692643 |
| lnc-IGIP-2 | lnc-IGIP-2 | -0.263870507069326 | 2.47191152984446 | -1.68357490391242 | 0.0995965132804026 | 0.382175775560064 | -4.78744912665667 |
| STT3A-AS1 | STT3A-AS1 | -0.419681930895669 | 0.526997862248734 | -1.68346447520335 | 0.0996180239688213 | 0.382175775560064 | -4.54509184876851 |
| ENSG00000213402.2 | PTPRCAP | -0.500515242778172 | 4.77994348545378 | -1.68392088390512 | 0.0996495867886968 | 0.382175775560064 | -5.01222550953436 |
| lnc-RERE-3 | lnc-RERE-3 | -0.364524390030731 | 1.77056397707574 | -1.68324551399541 | 0.0996606873267122 | 0.382175775560064 | -4.69965514014814 |
| lnc-SMG1-3 | lnc-SMG1-3 | 0.172615781406335 | 4.07066957557471 | 1.68315081208897 | 0.0996791441344227 | 0.382175775560064 | -4.95296863422463 |

| | | | | | | | |
|---|---|---|---|---|---|---|---|
| ENSG00000215301.10 | DDX3X | 0.315001619561041 | 9.44667978547936 | 1.68306970007079 | 0.0996949546027098 | 0.382175775560064 | -4.98990696393966 |
| ENSG00000140993.10 | TIGD7 | -0.296299885745702 | 3.02095034417669 | -1.6830367366892 | 0.0997013804631645 | 0.382175775560064 | -4.8333222261057 |
| ENSG00000072818.11 | ACAP1 | -0.249447763211954 | 6.40569178094694 | -1.68297715410555 | 0.0997129963218158 | 0.382175775560064 | -5.04548183202574 |
| ENSG00000168116.13 | KIAA1586 | -0.212313322708154 | 3.72116449362412 | -1.68290900634067 | 0.0997262833665212 | 0.382175775560064 | -4.92921961933126 |
| ENSG00000140577.15 | CRTC3 | 0.386013505979144 | 4.40527732401537 | 1.68276956019359 | 0.0997536999238775 | 0.382209680851298 | -4.97057975040519 |
| ENSG00000159214.12 | CCDC24 | 0.549187868312543 | 2.19920672491684 | 1.68280791513696 | 0.0997743971310503 | 0.382217832895665 | -4.754720721031 |
| lnc-BACH1-3 | lnc-BACH1-3 | -1.05649295686862 | -1.09438071704874 | -1.6818230421945 | 0.0999382155741592 | 0.38269428566881 | -4.25804730295989 |
| ENSG00000135365.15 | PHF21A | -0.165351994603157 | 5.75213001285528 | -1.68174330805666 | 0.0999537907994408 | 0.38269428566881 | -5.04275323455575 |
| lnc-APOBEC3G-3 | lnc-APOBEC3G-3 | -0.44155300323388 | 3.4222028623504 | -1.68165465488663 | 0.0999855134829445 | 0.38269428566881 | -4.89735652581892 |
| ENSG00000100058.12 | CRYBB2P1 | -0.175051874033973 | 3.90776624722193 | -1.68155267249907 | 0.0999910375761857 | 0.38269428566881 | -4.9358333168389 |
| ENSG00000094880.10 | CDC23 | 0.141735188738011 | 4.93572433560763 | 1.68154910679693 | 0.0999917343597561 | 0.38269428566881 | -5.01553041659528 |
| ENSG00000038274.16 | MAT2B | -0.195776941012109 | 6.40623730543976 | -1.68118444069981 | 0.100063015930792 | 0.382836096474235 | -5.04832908617168 |
| lnc-EXOSC6-3 | lnc-EXOSC6-3 | 0.309569168449853 | 2.19560305498121 | 1.68109389523626 | 0.100080721425158 | 0.382836096474235 | -4.74706125488062 |
| lnc-TRUB2-6 | lnc-TRUB2-6 | 0.197506925395019 | 1.22446250697357 | 1.68107413328379 | 0.100084586072383 | 0.382836096474235 | -4.63988687605175 |
| ENSG00000244509.3 | APOBEC3C | -0.367135230199191 | 6.11123705047536 | -1.68092740295088 | 0.10012252909385 | 0.382910073641531 | -5.04760216878371 |
| lnc-ARID3B-4 | lnc-ARID3B-4 | -0.486625401608765 | 0.23903990255412 | -1.68072247941732 | 0.100153376105878 | 0.382931241323447 | -4.46223143035029 |
| ENSG00000029363.16 | BCLAF1 | 0.174333368676026 | 8.50619994905217 | 1.68062032253287 | 0.100173367207252 | 0.382931241323447 | -5.01918158814611 |
| ENSG00000164011.17 | ZNF691 | -0.336159967227005 | 1.93172836113519 | -1.68056662443416 | 0.100183876719952 | 0.382931241323447 | -4.72241815697642 |
| ENSG00000204220.11 | PFDN6 | -0.283518177898318 | 3.22434420617159 | -1.68031299672808 | 0.100233527720169 | 0.38299963542998 | -4.87148597574065 |
| ENSG00000115232.13 | ITGA4 | 2.06894285972134 | 3.67911714350172 | 1.6817017960929 | 0.100253656355649 | 0.38299963542998 | -4.85140275292271 |
| lnc-PPP1CC-1 | lnc-PPP1CC-1 | -0.523448592792764 | -0.593493420857774 | -1.68019010381039 | 0.100257592950334 | 0.38299963542998 | -4.33345434022655 |
| ENSG00000080839.11 | RBL1 | -0.183360410729226 | 4.67375265610017 | -1.67968256531564 | 0.100357031071181 | 0.383093459023156 | -5.00120664443428 |
| ENSG00000116809.11 | ZBTB17 | -0.23249864832469 | 4.89112471472109 | -1.67967790750848 | 0.100357944016442 | 0.383093459023156 | -5.0204594426749 |
| ENSG00000021776.10 | AQR | 0.121744809114488 | 6.77487217759725 | 1.6796542164694 | 0.100362587642806 | 0.383093459023156 | -5.049441929196 |
| ENSG00000122912.14 | SLC25A16 | 0.140824908809527 | 4.43271370975813 | 1.67955826092614 | 0.100381397483298 | 0.383093459023156 | -4.98575942659474 |
| ENSG00000116679.15 | IVNS1ABP | 0.286387141202183 | 5.58400988145244 | 1.67942541645015 | 0.100407443339432 | 0.383093459023156 | -5.04437688260038 |
| ENSG00000104312.7 | RIPK2 | 0.26818609003861 | 3.56411523745763 | 1.67941850837947 | 0.100408797907484 | 0.383093459023156 | -4.90935407134541 |
| ENSG00000151304.5 | SRFBP1 | 0.163098192164991 | 4.06708015939001 | 1.67939994487827 | 0.100412438003689 | 0.383093459023156 | -4.94960883179728 |
| LEF1-AS1 | LEF1-AS1 | -0.668248346749773 | 1.20157420949659 | -1.67943610548695 | 0.100448037705566 | 0.383158257986225 | -4.62992641725069 |
| LINC01278 | LINC01278 | -0.394843087734544 | 3.6218039788719 | -1.67890205367637 | 0.100510109579408 | 0.383323992120244 | -4.90567655007307 |
| ENSG00000167173.18 | C15orf39 | 0.61966218926442 | 2.47645050081263 | 1.67927952195144 | 0.100537472374333 | 0.38332629571362 | -4.79505985245643 |
| ENSG00000170954.11 | ZNF415 | -0.596211801941974 | 2.76048451971459 | -1.67918188445033 | 0.100547960479903 | 0.38332629571362 | -4.8133421890941 |
| lnc-EML4-1 | lnc-EML4-1 | 0.623992443473179 | 0.493488013995318 | 1.67834226247971 | 0.100620017737284 | 0.383467644271 | -4.5008643183077 |
| lnc-SH3GLB2-1 | lnc-SH3GLB2-1 | -0.262920987744463 | 0.182910483470025 | -1.67829898930812 | 0.10062851802389 | 0.383467644271 | -4.50138512540098 |
| ENSG00000157349.16 | DDX19B | 0.174497994341642 | 4.55171652546277 | 1.67823581979603 | 0.100640927675846 | 0.383467644271 | -4.99603903659507 |
| ENSG00000169230.9 | PRELID1 | -0.304918910545489 | 4.61535896132209 | -1.67811468437438 | 0.100664728264968 | 0.383487340714018 | -5.00780194481001 |
| ENSG00000171522.5 | PTGER4 | 0.57736713712913 | 4.01574896111597 | 1.6783423803013 | 0.100763532175158 | 0.383792705914375 | -4.93082930855752 |
| ENSG00000165733.7 | BMS1 | 0.129018511342333 | 5.54038706707292 | 1.67741712191788 | 0.100801875172502 | 0.383867714463571 | -5.04557935028538 |
| ENSG00000074621.13 | SLC24A1 | -0.261952172141019 | 2.56915226115962 | -1.6770364019262 | 0.100876793081546 | 0.384081951904821 | -4.79886911197843 |
| ENSG00000179299.16 | NSUN7 | 1.17436679388813 | 1.98277357163449 | 1.67788735772159 | 0.100977582227811 | 0.384394594968565 | -4.75145915157495 |
| ENSG00000135622.12 | SEMA4F | -0.212827723554224 | 3.53970313662874 | -1.67640178133841 | 0.101001775544637 | 0.384415596530188 | -4.90148840821017 |
| lnc-ITGB1BP2-2 | lnc-ITGB1BP2-2 | -0.242161032851411 | 4.90445548305358 | -1.67629031709525 | 0.101023740546384 | 0.384428110864525 | -5.02789332792574 |
| ENSG00000246203.2 | NA | 0.160050182117338 | 3.06486526086136 | 1.67593141287388 | 0.101094492528479 | 0.384626236545966 | -4.85217971582523 |
| ENSG00000235194.8 | PPP1R3E | -0.225029481267142 | 4.98629626708039 | -1.67541056637548 | 0.10119724142641 | 0.384854387114462 | -5.02328845673238 |
| lnc-NAF1-1 | lnc-NAF1-1 | -0.402036502535079 | 1.16728122275701 | -1.67539488629954 | 0.101200336015558 | 0.384854387114462 | -4.63675484168528 |
| ENSG00000219712.1 | MARK2P18 | 0.370741587929983 | 0.160322876261221 | 1.67517728860337 | 0.101243288729738 | 0.384854387114462 | -4.45877721701931 |
| ENSG00000179588.8 | ZFPM1 | -0.408728296959813 | 3.57344721043244 | -1.67510205175526 | 0.10125814361111 | 0.384854387114462 | -4.9181687746431 |
| lnc-HLA-DQA1-9 | lnc-HLA-DQA1-9 | -0.528889647546803 | -0.360835207885018 | -1.67494711390005 | 0.101288740447288 | 0.384854387114462 | -4.39736849409171 |
| lnc-COPRS-2 | lnc-COPRS-2 | -0.246088496262905 | 1.37427408043743 | -1.6749378485245 | 0.101290570397825 | 0.384854387114462 | -4.66645798966878 |
| ENSG00000079459.12 | FDFT1 | -0.228410923554744 | 5.90167550690118 | -1.67487336088381 | 0.101303307732734 | 0.384854387114462 | -5.0566461105745 |
| SRP14-AS1 | SRP14-AS1 | -0.327202519158898 | 3.93670349257216 | -1.67486965024365 | 0.101304040683387 | 0.384854387114462 | -4.94528272851723 |
| ENSG00000196911.10 | KPNA5 | 0.263859128286501 | 6.06329655668161 | 1.67450985316328 | 0.101375131002765 | 0.384993894801102 | -5.05844119080938 |
| ENSG00000213300.5 | HNRNPA3P6 | 0.951495896036815 | -0.174743354809007 | 1.67443804398867 | 0.101389324305182 | 0.384993894801102 | -4.31431697986827 |
| lnc-FAM161A-2 | lnc-FAM161A-2 | 0.246908871526596 | 2.35225870906651 | 1.67435898413207 | 0.101404952622321 | 0.384993894801102 | -4.786979008861 |
| ENSG00000198728.10 | LDB1 | -0.203490958291641 | 6.03398831268829 | -1.67430522499556 | 0.101415580703084 | 0.384993894801102 | -5.05850306331753 |
| lnc-AHNAK2-9 | lnc-AHNAK2-9 | -0.267075739811607 | 1.23202258367755 | -1.67409911228027 | 0.101456337314296 | 0.38507759375625 | -4.66601345707875 |
| ENSG00000168395.15 | ING5 | -0.132490383926505 | 5.26808351058429 | -1.67389523083954 | 0.101496666001859 | 0.385137144802798 | -5.04405497979323 |
| ENSG00000241913.1 | RPS29P21 | 0.595124452576653 | -0.434509589581483 | 1.67373858404307 | 0.101527660438413 | 0.385137144802798 | -4.33487642217167 |
| ENSG00000171109.18 | MFN1 | 0.170854660230782 | 6.64120328903948 | 1.67364686768307 | 0.101545811236102 | 0.385137144802798 | -5.05992804534444 |
| ENSG00000240505.8 | TNFRSF13B | 1.09893520987947 | 4.06616747156718 | 1.67511890614891 | 0.101546872897283 | 0.385137144802798 | -4.97840165501547 |
| lnc-TRMT10B-4 | lnc-TRMT10B-4 | -0.288368561972236 | 1.58702113441559 | -1.67352812251654 | 0.101569315047476 | 0.385151291750589 | -4.70404168540665 |
| ENSG00000165806.19 | CASP7 | 0.235163681857148 | 4.54139893326931 | 1.67301429298269 | 0.101671071545637 | 0.385466138446093 | -5.0006305331024 |
| lnc-LPIN1-10 | lnc-LPIN1-10 | 1.22169116669111 | -1.19277697348094 | 1.6728587012875 | 0.101704148550693 | 0.385520532158181 | -4.26234590950868 |
| ENSG00000134461.15 | ANKRD16 | -0.326288819734513 | 2.32894797516831 | -1.67245613776508 | 0.101781701326849 | 0.385743464999179 | -4.78400861813577 |
| lnc-TLE4-12 | lnc-TLE4-12 | -1.2564779493517 | 7.30134583999864 | -1.67364260461603 | 0.101838777256192 | 0.385888724643631 | -5.05150383421064 |
| lnc-GOLGA8Q-3 | lnc-GOLGA8Q-3 | -0.287306642272174 | 3.28035027250656 | -1.67205851339973 | 0.101860573425012 | 0.385900272926011 | -4.89200810498426 |
| lnc-GPD2-1 | lnc-GPD2-1 | 0.410060836017944 | 1.55185063082703 | 1.67166635308408 | 0.101938411025105 | 0.386124091669072 | -4.66059109441336 |
| ENSG00000136108.14 | CKAP2 | 0.35070196733674 | 4.64032827811694 | 1.67155802366689 | 0.101959921335696 | 0.386134509816491 | -5.01053593455369 |

| | | | | | | | |
|---|---|---|---|---|---|---|---|
| lnc-LDAH-4 | lnc-LDAH-4 | -0.362806653899375 | 0.87568468104433 | -1.67141394605751 | 0.101988535743121 | 0.386171823252512 | -4.60555145177213 |
| lnc-FAM83C-2 | lnc-FAM83C-2 | -0.452281343679503 | 1.76735097553355 | -1.67083089642408 | 0.102104399402174 | 0.386517190967966 | -4.72980397548066 |
| lnc-PHLPP2-3 | lnc-PHLPP2-3 | 0.25383734509155 | 2.28957012219665 | 1.67076598763036 | 0.102117304789574 | 0.386517190967966 | -4.77114351784357 |
| ENSG00000204227.4 | RING1 | -0.213770273451321 | 3.91665582596903 | -1.67063730157582 | 0.102142894563663 | 0.386542967237338 | -4.95783852304566 |
| ENSG00000116406.18 | EDEM3 | -0.166602869472882 | 5.14405907936252 | -1.67036366192079 | 0.102197326549774 | 0.386644427296819 | -5.04259367515877 |
| ENSG00000181847.11 | TIGIT | -0.496326507389492 | 4.33574422188108 | -1.67068760894615 | 0.102224247913458 | 0.386644427296819 | -4.97673463383646 |
| lnc-FNTB-5 | lnc-FNTB-5 | 1.06607112835075 | -0.172699192101842 | 1.67044030997258 | 0.102251641620542 | 0.386644427296819 | -4.38586767614498 |
| ENSG00000140563.14 | MCTP2 | 0.255581602936513 | 7.89698116432409 | 1.67007432750993 | 0.10225490651025 | 0.386644427296819 | -5.05075517987852 |
| lnc-HECA-5 | lnc-HECA-5 | -0.346980937230951 | 0.785333730115037 | -1.66977571395624 | 0.102314361127492 | 0.386644427296819 | -4.59339927347765 |
| ENSG00000177853.14 | ZNF518A | -0.181173454247002 | 5.84477405157385 | -1.66970401906366 | 0.102328639978396 | 0.386644427296819 | -5.06359876509689 |
| lnc-RAB1A-11 | lnc-RAB1A-11 | -0.392163986529325 | 0.259114063279706 | -1.66965488313042 | 0.102338426904962 | 0.386644427296819 | -4.50177654947427 |
| ENSG00000139675.12 | HNRNPA1L2 | 0.2548743880242 | 2.48143809567843 | 1.66960217053199 | 0.102348927091496 | 0.386644427296819 | -4.80959417912127 |
| lnc-ACTL7B-1 | lnc-ACTL7B-1 | 0.416519442311388 | 0.707773790247038 | 1.66954750705418 | 0.102359816824727 | 0.386644427296819 | -4.56927853809852 |
| lnc-BARHL2-5 | lnc-BARHL2-5 | 0.247736580466822 | 3.28603434336994 | 1.66929542033026 | 0.102410048397289 | 0.386644427296819 | -4.88574614216815 |
| ENSG00000196155.12 | PLEKHG4 | 1.43895985227519 | -0.869133984288078 | 1.66946872814769 | 0.102427139374312 | 0.386644427296819 | -4.25985645959223 |
| ENSG00000160299.16 | PCNT | 0.327741662537538 | 5.76607160828951 | 1.66913903389658 | 0.102441220643655 | 0.386644427296819 | -5.06282041127577 |
| lnc-LRRC57-4 | lnc-LRRC57-4 | 0.29222677868244 | 1.45818824693369 | 1.66911453430372 | 0.102446104814738 | 0.386644427296819 | -4.68079511003044 |
| lnc-ARL5A-1 | lnc-ARL5A-1 | -0.327565832821362 | 2.06348590371525 | -1.66908826560743 | 0.102451341882521 | 0.386644427296819 | -4.75649818767835 |
| ENSG00000103121.8 | CMC2 | 0.217841535007795 | 2.87506799604368 | 1.66908759875592 | 0.102451474832476 | 0.386644427296819 | -4.85138303486281 |
| lnc-DYRK1B-1 | lnc-DYRK1B-1 | -0.36090254079611 | 0.552327442011549 | -1.66898463443185 | 0.102472004499451 | 0.386651011661266 | -4.55116184801144 |
| ENSG00000251791.1 | SCARNA6 | 0.55335457244036 | 4.58588352954407 | 1.66962442474396 | 0.102494989182387 | 0.386666855267792 | -4.9930485209515 |
| ENSG00000213995.11 | NAXD | 0.312714540036054 | 4.43297972039272 | 1.66851949233579 | 0.102564789664974 | 0.386859275366349 | -5.00066118012673 |
| ENSG00000122877.15 | EGR2 | -0.790076899900449 | 2.04983856376996 | -1.66907914382848 | 0.102616766175413 | 0.386984407876242 | -4.7343880147998 |
| lnc-PPM1J-6 | lnc-PPM1J-6 | -0.449269863386031 | 0.372345814629434 | -1.66789304254385 | 0.102689861374675 | 0.387137514372657 | -4.51559535380288 |
| ENSG00000087301.8 | TXNDC16 | 0.264151672919339 | 3.30144834581714 | 1.66778505962491 | 0.102711433027103 | 0.387137514372657 | -4.89057103086853 |
| lnc-MRPS16-1 | lnc-MRPS16-1 | -0.245630786925005 | 0.847077627586607 | -1.66773048003613 | 0.102722337763384 | 0.387137514372657 | -4.61351028075406 |
| ENSG00000273154.3 | | -0.204123901839451 | 3.13377134079906 | -1.66767912115681 | 0.102732599888425 | 0.387137514372657 | -4.88209746597586 |
| ENSG00000105379.9 | ETFB | -0.345505121092403 | 2.89156629710326 | -1.66752008138771 | 0.102764383313384 | 0.387156875589726 | -4.84523282340413 |
| ENSG00000255054.3 | | -0.260813970711859 | 1.91991262916714 | -1.66738703231456 | 0.102790978829162 | 0.387156875589726 | -4.75279025022084 |
| ENSG00000164346.9 | NSA2 | 0.203050537667873 | 6.05127496240761 | 1.66733271412871 | 0.102801838258239 | 0.387156875589726 | -5.06923893754243 |
| ENSG00000244119.1 | PDCL3P4 | -0.322005603837769 | 1.42664529652863 | -1.66726991516785 | 0.102814394362989 | 0.387156875589726 | -4.68719359679681 |
| lnc-FTHL17-2 | lnc-FTHL17-2 | -2.40041457013841 | 0.999592555300564 | -1.66864649014527 | 0.102831785398097 | 0.387156875589726 | -4.52618693817641 |
| ENSG00000113615.12 | SEC24A | 0.158229498014085 | 4.60356368132851 | 1.66708043182799 | 0.1028522875602 | 0.387163247046745 | -5.01678818020499 |
| ENSG00000202093.1 | SNORD58C | 0.350066431620395 | 0.503602305433758 | 1.66697044556722 | 0.102874288073101 | 0.387175255331621 | -4.56295652912734 |
| ENSG00000109861.15 | CTSC | 0.329428771770734 | 5.18457228832988 | 1.66606319091802 | 0.103055913777577 | 0.387749091503955 | -5.04394445340407 |
| ENSG00000228782.7 | MRPL45P2 | -0.227665080475056 | 2.10469039250559 | -1.66602065475243 | 0.103064435680811 | 0.387749091503955 | -4.76622844887507 |
| ENSG00000196453.7 | ZNF777 | -0.253845886547698 | 3.1058885475648 | -1.66580316933688 | 0.103108016840019 | 0.387784906317321 | -4.87968782801843 |
| lnc-GOLGA8O-1 | lnc-GOLGA8O-1 | 0.212647382756425 | 4.65354785573448 | 1.66578511464201 | 0.10311163544064 | 0.387784906317321 | -5.02658410571554 |
| ENSG00000215837.7 | SDHAP2 | 0.361116318893516 | 3.20268911194588 | 1.66559659045129 | 0.103149426527252 | 0.387810120819952 | -4.8911801943414 |
| lnc-GTF2F2-1 | lnc-GTF2F2-1 | -0.395677202873505 | 0.170295495139213 | -1.66556369159493 | 0.10315602251806 | 0.387810120819952 | -4.48559333512179 |
| ENSG00000001561.6 | ENPP4 | -0.294360286111348 | 4.96184080227223 | -1.66523142009059 | 0.103222660100583 | 0.387989775904 | -5.04139354039621 |
| ENSG00000138709.18 | LARP1B | -0.226715397608407 | 3.93209865932765 | -1.66495442985999 | 0.103278238018027 | 0.388127802286846 | -4.96980543500868 |
| OSER1-DT | OSER1-DT | -0.263838527441631 | 2.86074609178515 | -1.66484936569032 | 0.103299325521753 | 0.388136184230421 | -4.856920973636 |
| ENSG00000113240.12 | CLK4 | 0.141975509266715 | 6.01900661533226 | 1.66473897941376 | 0.103321485047676 | 0.388148590388085 | -5.07353955194829 |
| ENSG00000184319.15 | RPL23AP82 | -0.410887127589609 | 0.865987460444884 | -1.66463800587808 | 0.103341758436972 | 0.388153907647479 | -4.61789857952245 |
| ENSG00000235944.8 | ZNF815P | -0.280594104640432 | 1.22713011060114 | -1.66430392143563 | 0.103408858998654 | 0.388174877631437 | -4.67414129991839 |
| ENSG00000162961.13 | DPY30 | 0.177003736575561 | 4.05282366299059 | 1.66418926585323 | 0.103431895734546 | 0.388174877631437 | -4.97890010830684 |
| ENSG00000072121.15 | ZFYVE26 | 0.196818096021596 | 6.11446479814131 | 1.66410742648993 | 0.103448341580319 | 0.388174877631437 | -5.07478818114448 |
| ENSG00000230897.1 | RPS18P12 | 1.22372417586318 | 0.381776881892872 | 1.66468925354279 | 0.103448561585027 | 0.388174877631437 | -4.3737362456774 |
| lnc-SYT13-8 | lnc-SYT13-8 | -0.61907679826577 | -0.0199608135660145 | -1.66405123508494 | 0.10345963464437 | 0.388174877631437 | -4.4674063468591 |
| ENSG00000132196.14 | HSD17B7 | 0.160153492999502 | 3.56533446667506 | 1.66404695194013 | 0.103460495490748 | 0.388174877631437 | -4.92493873837693 |
| lnc-DHRS7B-9 | lnc-DHRS7B-9 | -1.83557882374937 | -2.98086990650323 | -1.66393287639276 | 0.103483425094846 | 0.388190147389686 | -4.25686200428656 |
| lnc-LTA4H-5 | lnc-LTA4H-5 | 0.256643253847821 | 1.88693849150857 | 1.66348527350768 | 0.103573435327621 | 0.388457000610136 | -4.74310214763664 |
| ENSG00000213593.9 | TMX2 | 0.142735841556648 | 4.65761650396233 | 1.66337840290638 | 0.103594935885653 | 0.388466854679248 | -5.02553144500336 |
| ENSG00000174165.7 | ZDHHC24 | -0.221964347730883 | 3.99661326840878 | -1.66305657043991 | 0.103659705324426 | 0.38863892799502 | -4.97170999720845 |
| ENSG00000181450.17 | ZNF678 | -0.183960725019691 | 5.54228600174037 | -1.66279773347292 | 0.103711820967253 | 0.38870497782763 | -5.06806657086102 |
| ENSG00000102309.12 | PIN4 | -0.27353401034576 | 2.61218559502911 | -1.6627814912667 | 0.103715091980243 | 0.38870497782763 | -4.82647142320092 |
| ENSG00000154222.14 | CC2D1B | 0.145349476028659 | 5.87765186969728 | 1.66229254714423 | 0.103813600041014 | 0.388978629824224 | -5.07546673162116 |
| ENSG00000259326.1 | | 0.372005903990974 | 2.20316097995509 | 1.66220053492761 | 0.103832146438812 | 0.388978629824224 | -4.77418302962128 |
| lnc-BIRC2-1 | lnc-BIRC2-1 | 0.479039050321188 | 4.42396843853778 | 1.66250583106768 | 0.103857562699275 | 0.388978629824224 | -5.01060784081785 |
| lnc-HYOU1-1 | lnc-HYOU1-1 | -0.364718188825755 | 0.735731148227855 | -1.6619866703973 | 0.103875264472043 | 0.388978629824224 | -4.59614470018892 |
| lnc-KIAA1586-4 | lnc-KIAA1586-4 | -0.280077572487811 | 1.33520767747686 | -1.6619179994612 | 0.103889112602454 | 0.388978629824224 | -4.70443608403033 |
| ENSG00000272896.1 | | -0.308075133481931 | 0.714708570059827 | -1.66185659479017 | 0.103901496709595 | 0.388978629824224 | -4.5953337465894 |
| lnc-ANKRD20A4-18 | lnc-ANKRD20A4-1[illegible] | -0.389968883870123 | 0.136173653655533 | -1.66142068338406 | 0.103989446336496 | 0.389169706390271 | -4.49896490641693 |
| ENSG00000180626.9 | ZNF594 | -0.322499065102036 | 4.43934063300055 | -1.66135088074756 | 0.1040035354267 | 0.389169706390271 | -5.00999683737653 |
| lnc-TRABD2A-9 | lnc-TRABD2A-9 | 0.199891504889227 | 2.13424767692849 | 1.66132253181139 | 0.104009257875571 | 0.389169706390271 | -4.7815864027892 |
| ENSG00000173706.13 | HEG1 | 0.897343011987059 | 2.45142216912127 | 1.66224018697897 | 0.104031036221424 | 0.389180446845796 | -4.76474796005526 |
| ENSG00000197170.9 | PSMD12 | 0.239479214126598 | 4.92463972588528 | 1.66110841129494 | 0.104052488101613 | 0.38918996231065 | -5.04768245267876 |

| | | | | | | | |
|---|---|---|---|---|---|---|---|
| ENSG00000019485.13 | PRDM11 | -0.282566702147989 | 4.02394007688784 | -1.66080175832868 | 0.104114426067308 | 0.389350877860356 | -4.98718259416203 |
| ENSG00000154309.8 | DISP1 | -0.280170204938182 | 3.08994096199108 | -1.66046773146814 | 0.104181927538134 | 0.389532536697078 | -4.88281011778818 |
| ENSG00000129657.15 | SEC14L1 | 0.362292550129356 | 6.01036981163149 | 1.66038517967138 | 0.104207406322427 | 0.389557036748005 | -5.07892593395552 |
| lnc-ARHGAP9-2 | lnc-ARHGAP9-2 | 0.258759164864811 | 0.962855762296667 | 1.65994297145364 | 0.104288045818568 | 0.389787696946356 | -4.61567027180423 |
| lnc-HSP90B1-2 | lnc-HSP90B1-2 | 0.310328510080431 | 1.80667369369464 | 1.65946103781954 | 0.10438558194307 | 0.390048315343633 | -4.74973642082493 |
| ENSG00000177225.16 | GATD1 | -0.244755963007329 | 4.20292225159959 | -1.65941119061294 | 0.10439567454832 | 0.390048315343633 | -5.00569956092935 |
| lnc-TYROBP-1 | lnc-TYROBP-1 | -0.400811073309038 | 2.88108461237251 | -1.65909889648614 | 0.104458923264189 | 0.390213796287986 | -4.87042535401327 |
| ENSG00000121966.6 | CXCR4 | 0.317236123287721 | 10.621966937532 | 1.65880179373914 | 0.104519124527766 | 0.390366089228375 | -4.99382076661116 |
| ENSG00000188277.9 | C15orf62 | -0.383791491466489 | 1.08323301777995 | -1.65866347555907 | 0.104547161362134 | 0.390366089228375 | -4.66969589060324 |
| ENSG00000137393.9 | RNF144B | -0.652598961599392 | 4.66334372496763 | -1.65962647363004 | 0.104559045420511 | 0.390366089228375 | -5.05209552851381 |
| lnc-ANKUB1-3 | lnc-ANKUB1-3 | 0.221591524037676 | 1.77821148039529 | 1.65852343599328 | 0.104575553418125 | 0.390366089228375 | -4.75167223229382 |
| ENSG00000138231.12 | DBR1 | 0.168664015848823 | 3.32255577318887 | 1.65819243394945 | 0.104642687003297 | 0.390545861575497 | -4.91197342933952 |
| ENSG00000113384.13 | GOLPH3 | 0.185783668451374 | 5.37483942494113 | 1.65791777988402 | 0.104698419031094 | 0.390627924900135 | -5.07208044496512 |
| lnc-ZNF749-2 | lnc-ZNF749-2 | 0.383339110001046 | -0.121091278970332 | 1.65789702581872 | 0.104702631379004 | 0.390627924900135 | -4.43042999227523 |
| LINC01032 | LINC01032 | -0.544606082432843 | 0.385655170645173 | -1.65763473835987 | 0.104755878549972 | 0.390755753569061 | -4.53363783880813 |
| ENSG00000084733.10 | RAB10 | -0.245579025740915 | 5.32678443887868 | -1.65746896895305 | 0.104789543000083 | 0.390759591901778 | -5.0731898946106 |
| ENSG00000139746.15 | RBM26 | 0.239393199345141 | 7.48094534181744 | 1.65744270889333 | 0.104794876708828 | 0.390759591901778 | -5.07594597411078 |
| lnc-IRF2-1 | lnc-IRF2-1 | -0.37057258456465 | 2.38623358526966 | -1.65672257562348 | 0.104941230749037 | 0.391234441678578 | -4.8217637605718 |
| ENSG00000186312.10 | CA5BP1 | 0.225246134733721 | 2.5200221841519 | 1.65650014841639 | 0.104986468996162 | 0.39133221502137 | -4.82876725901378 |
| lnc-SASS6-2 | lnc-SASS6-2 | -0.282366887974049 | 1.66030151354736 | -1.65622164762215 | 0.105043134341266 | 0.391472539226194 | -4.74080209347804 |
| lnc-XPNPEP1-3 | lnc-XPNPEP1-3 | -1.48231376060891 | -1.95518261901299 | -1.65609840622733 | 0.105119937744089 | 0.391617465951179 | -4.26751249223268 |
| ENSG00000140575.12 | IQGAP1 | 0.17853833292687 | 8.35698085326365 | 1.65576200484055 | 0.105136710838203 | 0.391617465951179 | -5.06169079989024 |
| ENSG00000112079.8 | STK38 | 0.350818891676073 | 5.3717066825557 | 1.65555741928349 | 0.105178383470947 | 0.391617465951179 | -5.0689228789235 |
| lnc-BPHL-9 | lnc-BPHL-9 | -0.361223257431415 | 0.494610445256871 | -1.65551889109489 | 0.105186232909527 | 0.391617465951179 | -4.56931054271876 |
| ENSG00000175193.13 | PARL | 0.215183698547432 | 2.29185459683679 | 1.65547853732957 | 0.105194454793329 | 0.391617465951179 | -4.80319834531498 |
| ENSG00000237440.8 | ZNF737 | -0.402362002009644 | 6.54618811448082 | -1.65570046374025 | 0.105196179820438 | 0.391617465951179 | -5.08760409188727 |
| lnc-BID-3 | lnc-BID-3 | -0.369141181860977 | 0.885517590370242 | -1.65528247481053 | 0.105234409096988 | 0.391620703199103 | -4.63816500527514 |
| ENSG00000140678.16 | ITGAX | -0.83547953186344 | 5.23450905702819 | -1.6565546896206 | 0.105244356172416 | 0.391620703199103 | -5.05255679545099 |
| lnc-FBXO28-1 | lnc-FBXO28-1 | -0.305757442704387 | 5.96267580663646 | -1.65515744054369 | 0.105259895527569 | 0.391620703199103 | -5.08781560821861 |
| ENSG00000124588.19 | NQO2 | -0.316106362086052 | 2.63736867315715 | -1.65501671846926 | 0.105288585757981 | 0.391620703199103 | -4.84157824146873 |
| ENSG00000250571.6 | GLI4 | -0.23997243193706 | 3.53582419177418 | -1.65499908409021 | 0.105292181484907 | 0.391620703199103 | -4.93982771254886 |
| ENSG00000080819.7 | CPOX | -0.294741973123043 | 1.80631073208552 | -1.6546917115352 | 0.105354872301488 | 0.391783077973177 | -4.75908475529672 |
| ENSG00000240356.6 | RPL23AP7 | -0.280629897983071 | 1.14542135626553 | -1.65458539959347 | 0.105376562514755 | 0.391792952716981 | -4.67492846430727 |
| lnc-HMGB2-5 | lnc-HMGB2-5 | -0.781720771046891 | 0.0502351533192192 | -1.65459813683313 | 0.105402479600963 | 0.391805854077183 | -4.52368218629353 |
| ENSG00000138175.8 | ARL3 | -0.310541942394557 | 1.56672158180948 | -1.6543818427527 | 0.105418103283265 | 0.391805854077183 | -4.74011630970142 |
| ENSG00000141086.17 | CTRL | 0.250556864768922 | 1.88066572410279 | 1.65378526712887 | 0.105539926688021 | 0.391840537235405 | -4.75851223287383 |
| ENSG00000112237.12 | CCNC | 0.238019389347733 | 5.17475291224748 | 1.65376779761602 | 0.105543495782815 | 0.391840537235405 | -5.07200376208349 |
| ENSG00000104129.9 | DNAJC17 | -0.164911160976027 | 3.92821028232769 | -1.65375901414418 | 0.10554529032044 | 0.391840537235405 | -4.98261226664944 |
| ENSG00000140367.11 | UBE2Q2 | 0.136646777575931 | 5.46813960669455 | 1.65373796175309 | 0.105549591604131 | 0.391840537235405 | -5.08270804592554 |
| lnc-CCNYL1-2 | lnc-CCNYL1-2 | -0.170810224235547 | 3.21433925010174 | -1.65371049292378 | 0.105555204068083 | 0.391840537235405 | -4.90737439456528 |
| ENSG00000107897.18 | ACBD5 | 0.146933622801182 | 5.18162928772321 | 1.65366851072471 | 0.105563782394762 | 0.391840537235405 | -5.07227152575365 |
| ENSG00000145495.15 | MARCHF6 | 0.23504085354035 | 7.58875639053035 | 1.65364280897732 | 0.10556903437908 | 0.391840537235405 | -5.08013297421621 |
| ENSG00000082438.15 | COBLL1 | 1.48383377124 32 | 5.82278701583427 | 1.65502514388733 | 0.105579731793594 | 0.391840537235405 | -5.07101556806796 |
| ENSG00000181938.13 | GINS3 | 0.392886549868563 | 0.26034493981074 | 1.65348202588354 | 0.105601894226801 | 0.391852134283442 | -4.51923629851918 |
| lnc-HELZ-7 | lnc-HELZ-7 | 0.280559556354954 | 1.23491797937211 | 1.65331086668889 | 0.105636883910331 | 0.39191131606459 | -4.6821614171784 |
| ENSG00000162869.15 | PPP1R21 | 0.211410364009776 | 4.63084228984671 | 1.65310144163485 | 0.105679709151182 | 0.391999504679557 | -5.04137655601133 |
| ENSG00000147130.14 | ZMYM3 | -0.164703809404942 | 5.35165782674687 | -1.65299454364869 | 0.105701574172324 | 0.391999504679557 | -5.07727447504014 |
| ENSG00000170209.4 | ANKK1 | 0.474150539409766 | 0.00944822330448026 | 1.65291528128248 | 0.105717788975185 | 0.391999504679557 | -4.43474418483452 |
| ENSG00000105974.11 | CAV1 | -1.48565365360568 | 0.244574900690474 | -1.65404280596231 | 0.105744216990359 | 0.392026876497217 | -4.53250897706582 |
| ENSG00000100523.14 | DDHD1 | 0.173156083065941 | 6.85264904830784 | 1.65246884333668 | 0.105809155519596 | 0.392196983263073 | -5.09154330294408 |
| ENSG00000026025.15 | VIM | -0.232703999359126 | 8.47226131382491 | -1.65219829208176 | 0.105864557218859 | 0.392331685494377 | -5.06235920687208 |
| ENSG00000168876.8 | ANKRD49 | 0.246721803992098 | 5.98113715966899 | 1.65201522765491 | 0.105902057455941 | 0.392398362573778 | -5.09208857549883 |
| ENSG00000054118.14 | THRAP3 | -0.270492306706354 | 7.97329384655013 | -1.65192435080217 | 0.105920677377443 | 0.392398362573778 | -5.07475226391752 |
| ENSG00000197324.8 | LRP10 | -0.203880792642943 | 6.59881953234503 | -1.65167269815904 | 0.105972252971069 | 0.392518784038781 | -5.09372877824284 |
| ENSG00000064651.13 | SLC12A2 | 0.14541068046316 | 4.85362817942764 | 1.65153842930873 | 0.105999779479973 | 0.392550100942118 | -5.06045666411469 |
| lnc-SERPINB1-3 | lnc-SERPINB1-3 | 0.863275615468683 | -0.31710014197246 | 1.6511300537444 | 0.106083536802002 | 0.392789609281455 | -4.36932914203688 |
| lnc-FRMD7-4 | lnc-FRMD7-4 | -0.366869429073754 | 1.51083746820811 | -1.65086853888506 | 0.106137201714005 | 0.392917630014276 | -4.7315309332689 |
| ENSG00000103326.11 | CAPN15 | -0.32661728295851 | 4.38898632071562 | -1.65068769203891 | 0.106174325949544 | 0.392922320732324 | -5.03772920389692 |
| lnc-RFLNB-5 | lnc-RFLNB-5 | 0.278639054465546 | 0.154972552103085 | 1.6506763816336 | 0.106176648103444 | 0.392922320732324 | -4.50365993505032 |
| ENSG00000103343.12 | ZNF174 | -0.276092127985101 | 1.94562001074401 | -1.65033263945034 | 0.106247242149124 | 0.393112886060654 | -4.7771661572175 |
| lnc-ZBTB7A-1 | lnc-ZBTB7A-1 | -0.518767306213243 | 0.639489134689489 | -1.65005355047548 | 0.106304586760741 | 0.393254369032412 | -4.63512410382519 |
| lnc-ITGB3BP-13 | lnc-ITGB3BP-13 | 0.198363089411059 | 3.4504638924176 | 1.64958343121481 | 0.106401239951604 | 0.393420410159942 | -4.93675649925934 |
| ENSG00000162407.8 | PLPP3 | 1.04041087641977 | 0.327488429841412 | 1.65030515078522 | 0.106403763224437 | 0.393420410159942 | -4.52156196098123 |
| lnc-B3GNT10-4 | lnc-B3GNT10-4 | -0.365214125565042 | 0.808754036960379 | -1.64936014811037 | 0.10644717063858 | 0.393420410159942 | -4.62869410579952 |
| ENSG00000059588.9 | TARBP1 | 0.233304665124963 | 5.36989563436723 | 1.64931581678992 | 0.106456291795937 | 0.393420410159942 | -5.08492661377706 |
| ENSG00000131016.16 | AKAP12 | -1.31695219242369 | 4.31618351148366 | -1.65058771186604 | 0.10648787342812 | 0.393420410159942 | -5.03701204072317 |
| ENSG00000100335.14 | MIEF1 | 0.158629542315566 | 4.94050567904943 | 1.64905622030357 | 0.10650971658288 | 0.393420410159942 | -5.06670987651548 |
| ENSG00000108828.15 | VAT1 | -0.335485584231431 | 4.76563751216995 | -1.64899362435391 | 0.10652260208091 | 0.393420410159942 | -5.05814624482927 |

| | | | | | | | |
|---|---|---|---|---|---|---|---|
| ENSG00000167523.13 | SPATA33 | -0.219525785445428 | 1.41402844265923 | -1.64885137578244 | 0.106551888984026 | 0.393420410159942 | -4.71837835088094 |
| ENSG00000284627.1 | | -0.518405602763922 | 0.597155774920953 | -1.64874291340183 | 0.106574224244567 | 0.393420410159942 | -4.62631641950445 |
| lnc-SLC52A1-3 | lnc-SLC52A1-3 | -0.320092644600638 | 0.806248696443012 | -1.64870436733536 | 0.106582162820232 | 0.393420410159942 | -4.64146661502999 |
| ENSG00000133027.17 | PEMT | -0.271420483814421 | 1.05001390968467 | -1.64864077418267 | 0.106595260912818 | 0.393420410159942 | -4.68804906642995 |
| lnc-SEMA4B-6 | lnc-SEMA4B-6 | -0.333284585597534 | 0.446796179121292 | -1.64862129422878 | 0.106599273405329 | 0.393420410159942 | -4.55441108202542 |
| ENSG00000055163.19 | CYFIP2 | 0.198665293203039 | 8.75518388114952 | 1.64860042514277 | 0.106603572169641 | 0.393420410159942 | -5.06433475930945 |
| ENSG00000130402.11 | ACTN4 | -0.249994581931918 | 5.59137387020658 | -1.6484705555491 | 0.106630326839732 | 0.393420410159942 | -5.09166182604879 |
| lnc-BRD2-6 | lnc-BRD2-6 | 0.401116846066502 | 3.66167445490533 | 1.64840947459084 | 0.10664291214392 | 0.393420410159942 | -4.93630425121971 |
| ENSG00000162542.13 | TMCO4 | -0.131649027534007 | 4.56679194616857 | -1.64833194693128 | 0.106658887931557 | 0.393420410159942 | -5.04147736423402 |
| ENSG00000261553.5 | | -0.33902539076379 | 4.04818260975608 | -1.64820033733233 | 0.106686012646642 | 0.393420410159942 | -4.99810549478752 |
| ENSG00000285669.1 | | -0.318186651622281 | 0.618344154059942 | -1.64816391463986 | 0.106693520357227 | 0.393420410159942 | -4.59831070084142 |
| ENSG00000105928.14 | GSDME | 1.13503711530298 | 2.57140509005303 | 1.64877300640442 | 0.106849681335992 | 0.393925665581001 | -4.88253650421518 |
| ENSG00000169660.16 | HEXD | -0.401834663920912 | 3.96778570241319 | -1.64701014812147 | 0.106942367209542 | 0.394105970906174 | -5.00442367033704 |
| lnc-CSF3R-1 | lnc-CSF3R-1 | 2.2554937514359 | -1.92718972024694 | 1.64718742547296 | 0.106945854610439 | 0.394105970906174 | -4.26709432153206 |
| ENSG00000184867.13 | ARMCX2 | -0.394950486901577 | 3.48378817582672 | -1.64689170619716 | 0.1069560294166 | 0.394105970906174 | -4.96353628571686 |
| ENSG00000276991.1 | | -0.264334298104726 | 0.260172603658698 | -1.64661855451711 | 0.10701246092274 | 0.394243329724853 | -4.53600174268533 |
| ENSG00000147604.13 | RPL7 | 0.271470910567114 | 8.96956052858739 | 1.64645052644834 | 0.107047186652294 | 0.394300687699387 | -5.06137238949828 |
| lnc-ADIPOQ-4 | lnc-ADIPOQ-4 | 0.354982321578795 | 2.65747967498824 | 1.64616799231927 | 0.107105597760873 | 0.394378990021592 | -4.83448174450078 |
| lnc-BMP6-106 | lnc-BMP6-106 | -0.511863720215742 | 1.75843926469361 | -1.6461623452776 | 0.107106765496803 | 0.394378990021592 | -4.76416299281344 |
| lnc-ADAM10-8 | lnc-ADAM10-8 | 0.137137249236448 | 2.72074618522409 | 1.64570668401791 | 0.107201024841579 | 0.394655463121843 | -4.86448401272138 |
| ENSG00000115486.11 | GGCX | -0.181029279951982 | 4.03302144428878 | -1.64546745849045 | 0.107250538928479 | 0.394713609334369 | -5.00703799634946 |
| BRWD1-IT1 | BRWD1-IT1 | 0.499778584222304 | -0.140671412520972 | 1.64543120063118 | 0.107258045093849 | 0.394713609334369 | -4.45390363290105 |
| ENSG00000131944.9 | FAAP24 | 0.386078722541175 | 0.29086545911225 | 1.64535245203476 | 0.10727434925018 | 0.394713609334369 | -4.52521853546624 |
| ENSG00000119431.9 | HDHD3 | 0.313351157045934 | 3.00792615489241 | 1.64501437086261 | 0.107344368877541 | 0.394738294243401 | -4.89455265357655 |
| ENSG00000128000.15 | ZNF780B | -0.215332907658189 | 5.08588466558781 | -1.64497457732764 | 0.107352612938957 | 0.394738294243401 | -5.0789650346758 |
| ENSG00000099256.18 | PRTFDC1 | -0.593796061369417 | 0.115726334931974 | -1.64495872147602 | 0.107355897954274 | 0.394738294243401 | -4.51611150201037 |
| ENSG00000163605.14 | PPP4R2 | 0.170330583317673 | 5.67977875522505 | 1.64494968786047 | 0.107357769575599 | 0.394738294243401 | -5.09978459927193 |
| ENSG00000160588.9 | MPZL3 | -0.451819880098269 | 1.08687891504957 | -1.64483273354922 | 0.10738200306319 | 0.394756879630226 | -4.6734436318989 |
| ENSG00000163171.7 | CDC42EP3 | -0.346036806371207 | 5.6870202141961 | -1.64418600775716 | 0.107516088722985 | 0.395088436052262 | -5.1006860151282 |
| ENSG00000255274.9 | SMIM35 | -0.78518750889029 | 0.247587331775025 | -1.64418334273829 | 0.107518520194029 | 0.395088436052262 | -4.47982571057641 |
| ENSG00000152291.13 | TGOLN2 | 0.164159703080806 | 7.77730875409784 | 1.64411762067035 | 0.107530275441722 | 0.395088436052262 | -5.09277302300977 |
| ENSG00000284138.1 | ATP6V0CP4 | 0.452370109761994 | 0.412980621364441 | 1.6439849027027 | 0.107557811810519 | 0.395088436052262 | -4.57589368149452 |
| ENSG00000276168.1 | RN7SL1 | 0.433739030077168 | 11.4632126510064 | 1.64407259260826 | 0.107568167693576 | 0.395088436052262 | -4.99235075714371 |
| lnc-LRR1-1 | lnc-LRR1-1 | 0.433711629247833 | 11.463175219885 | 1.64396634734501 | 0.107590212929819 | 0.395098903253251 | -4.99251396823072 |
| lnc-DLX5-3 | lnc-DLX5-3 | 0.219935184493937 | 2.95991309142596 | 1.64368785930832 | 0.10761946342939 | 0.395135821578154 | -4.89414780095768 |
| lnc-ATR-3 | lnc-ATR-3 | 0.218980978559681 | 2.49572008272287 | 1.64340683197728 | 0.107677817573499 | 0.395279564671898 | -4.84095331166618 |
| ENSG00000161204.11 | ABCF3 | 0.132770061365189 | 4.7855469663291 | 1.64321141828079 | 0.107718409696991 | 0.395358064692076 | -5.06580947140267 |
| ENSG00000177879.15 | AP3S1 | 0.26178137237927 | 4.21245309632799 | 1.64302909133163 | 0.107756294702207 | 0.395426602577203 | -5.03423773912116 |
| ENSG00000185345.20 | PRKN | -0.764086664669527 | 0.634884838195817 | -1.6431007074712 | 0.10780917650811 | 0.395550139049276 | -4.6116608249272 |
| ENSG00000103269.13 | RHBDL1 | -0.460426611930281 | -0.190328887484516 | -1.64266188048095 | 0.107832629169947 | 0.395565675673681 | -4.45466975077309 |
| ENSG00000141580.15 | WDR45B | 0.189402887419809 | 5.22277698937625 | 1.64250814676735 | 0.107864599942756 | 0.395591529727762 | -5.09022201062572 |
| lnc-MFSD14B-6 | lnc-MFSD14B-6 | -0.542724764423191 | -0.48004969192504 | -1.64244316631839 | 0.107878115744154 | 0.395591529727762 | -4.40202826393254 |
| ENSG00000166136.15 | NDUFB8 | 0.122455488046077 | 4.52969914547086 | 1.64214861465359 | 0.107939399304177 | 0.395688584439033 | -5.04924839307785 |
| ENSG00000231205.11 | ZNF826P | -0.365509553292624 | 3.33989052755315 | -1.64196806068115 | 0.107976978943726 | 0.395688584439033 | -4.94394086914256 |
| lnc-MIOS-5 | lnc-MIOS-5 | 0.350309884703496 | 3.17310555655738 | 1.6419556706987 | 0.107979558128561 | 0.395688584439033 | -4.93258199792942 |
| lnc-ACADL-1 | lnc-ACADL-1 | -0.521399594052423 | 0.837367549832631 | -1.64192432938391 | 0.107986082579857 | 0.395688584439033 | -4.64989557066831 |
| ENSG00000115524.15 | SF3B1 | 0.130745914206278 | 9.99345874132251 | 1.64185410391434 | 0.108000702879973 | 0.395688584439033 | -5.03788684922975 |
| ENSG00000258643.5 | BCL2L2-PABPN1 | -0.188899599202208 | 5.09369544364889 | -1.64161168165973 | 0.108051185444053 | 0.395774670891298 | -5.08654038333691 |
| lnc-CTSV-5 | lnc-CTSV-5 | -0.436216712396664 | 0.0893508676425561 | -1.6413117959915 | 0.108113661081664 | 0.395774670891298 | -4.50911786469049 |
| lnc-UBA6-3 | lnc-UBA6-3 | 0.239862208245897 | 2.77762278873597 | 1.64119888871893 | 0.108137190896611 | 0.395774670891298 | -4.87029879014663 |
| lnc-CD226-10 | lnc-CD226-10 | 0.237239382835941 | 1.39476203361852 | 1.64113106433387 | 0.108151327479889 | 0.395774670891298 | -4.72238831595778 |
| ENSG00000104231.10 | ZFAND1 | 0.190165035728711 | 4.10004108012686 | 1.64106696838644 | 0.108164688339879 | 0.395774670891298 | -5.01848508918759 |
| lnc-PHYHD1-1 | lnc-PHYHD1-1 | 0.269731194785112 | 1.24111307659147 | 1.64104631345943 | 0.108168994166875 | 0.395774670891298 | -4.70916940735822 |
| ENSG00000122033.14 | MTIF3 | -0.180423726455321 | 4.06368937394503 | -1.64101470882982 | 0.108175582894303 | 0.395774670891298 | -5.0125639801739 |
| ENSG00000127863.15 | TNFRSF19 | 1.19475761871863 | 0.495497364940366 | 1.6415606276158 | 0.108182949630769 | 0.395774670891298 | -4.41419322195918 |
| lnc-NLRP1-2 | lnc-NLRP1-2 | -0.253687212986509 | 2.83796621866253 | -1.64086228846453 | 0.108207363118266 | 0.395774670891298 | -4.8863564592597 |
| ENSG00000115138.10 | POMC | 2.0499332802819 | -2.32293471306399 | 1.64105141221482 | 0.108246126630503 | 0.395774670891298 | -4.26041613089619 |
| ENSG00000122873.11 | CISD1 | -0.366400944501065 | 1.98696062165759 | -1.64065114860277 | 0.108251399221008 | 0.395774670891298 | -4.79817034765062 |
| ENSG00000087274.16 | ADD1 | 0.273893887000592 | 7.78305753406377 | 1.64046552241239 | 0.108290126230033 | 0.395774670891298 | -5.09756477699086 |
| ENSG00000114982.17 | KANSL3 | 0.179177083499624 | 5.68886908130219 | 1.6404560014442 | 0.108292112886311 | 0.395774670891298 | -5.1057453542761 |
| lnc-ANXA6-4 | lnc-ANXA6-4 | -0.298789433711997 | 3.14569895808017 | -1.64044985770108 | 0.108293394862741 | 0.395774670891298 | -4.91953520878999 |
| lnc-NANP-1 | lnc-NANP-1 | -0.18121200652778 | 1.62164410328396 | -1.64029767533555 | 0.108325153774287 | 0.395820458039436 | -4.75063372793435 |
| ENSG00000151923.17 | TIAL1 | 0.14288283352075 | 5.86030416564972 | 1.64006946440514 | 0.108372793378233 | 0.395915641760029 | -5.11025564986725 |
| ENSG00000177483.11 | RBM44 | 0.426297913338278 | 1.08581403319725 | 1.63998862418096 | 0.108389673095164 | 0.395915641760029 | -4.68981171934032 |
| lnc-COL4A2-3 | lnc-COL4A2-3 | -0.714410116956897 | -0.391771094131864 | -1.639670171073 | 0.108456188160144 | 0.396088311018495 | -4.48152041138723 |
| ENSG00000048028.11 | USP28 | 0.356201443493187 | 6.15473696206502 | 1.63950824311148 | 0.108497556926337 | 0.396169099559126 | -5.11186044724173 |
| ENSG00000243749.1 | TMEM35B | -0.231245768554887 | 1.43102121405619 | -1.63873028610084 | 0.108652696284573 | 0.396469658821695 | -4.73388759809936 |
| ENSG00000232472.1 | EEF1B2P3 | 0.710994627798972 | 2.08527993130258 | 1.63928224934157 | 0.108658416349508 | 0.396469658821695 | -4.76137850697509 |

| | | | | | | | |
|---|---|---|---|---|---|---|---|
| lnc-RSL1D1-2 | lnc-RSL1D1-2 | 0.408523001438712 | 0.245602728080923 | 1.63869491834172 | 0.108660096552744 | 0.396469658821695 | -4.52685943573158 |
| lnc-DYNC1H1-1 | lnc-DYNC1H1-1 | -0.300111695568608 | 2.24436228149855 | -1.63865834139726 | 0.108667750262388 | 0.396469658821695 | -4.82515377872972 |
| ENSG00000182841.12 | RRP7BP | -0.206551896406302 | 3.93205660069771 | -1.63861805777599 | 0.108676180103581 | 0.396469658821695 | -5.00909339262389 |
| lnc-RBM12B-2 | lnc-RBM12B-2 | -0.655087076119388 | 0.723780559714944 | -1.63841857696343 | 0.108727895566562 | 0.396588033749168 | -4.61798264621611 |
| ENSG00000119737.5 | GPR75 | 0.295681203095012 | 1.37292777915416 | 1.63777693407699 | 0.108852317946942 | 0.396907600331553 | -4.73020635411037 |
| ENSG00000101898.6 | MCTS2 | -0.276385716137865 | 0.0524058535450427 | -1.63768089922857 | 0.108872443255688 | 0.396907600331553 | -4.5092687770833 |
| ENSG00000197888.2 | UGT2B17 | 2.02648631475008 | 0.114906142305376 | 1.63907794148587 | 0.1088733572109 | 0.396907600331553 | -4.61329494931734 |
| lnc-VPS54-4 | lnc-VPS54-4 | 0.430136047780769 | 5.69529835269825 | 1.63766846276458 | 0.108952829181113 | 0.397126984776844 | -5.10795134650585 |
| ENSG00000178074.5 | C2orf69 | 0.282314041359253 | 3.61715223664118 | 1.63718832197466 | 0.108975716917036 | 0.397139613922437 | -4.97993739182226 |
| ENSG00000149634.4 | SPATA25 | -0.352548072039373 | -0.277994053760705 | -1.6369683458524 | 0.109021862981129 | 0.397139613922437 | -4.47925998336324 |
| ENSG00000154710.17 | RABGEF1 | 0.215045415733319 | 3.90184220120821 | 1.63692497312574 | 0.109030963496331 | 0.397139613922437 | -4.99809371832344 |
| ENSG00000145494.11 | NDUFS6 | 0.198543367225243 | 3.32585964120891 | 1.63691301725991 | 0.109033472199179 | 0.397139613922437 | -4.94464286459556 |
| lnc-ZDHHC2-2 | lnc-ZDHHC2-2 | -0.275729393029282 | 2.77749802725289 | -1.63661308379141 | 0.10909642279695 | 0.397298597032843 | -4.89231629389699 |
| ENSG00000112159.11 | MDN1 | 0.196271143374799 | 7.58026396081938 | 1.63642493404716 | 0.109135927206526 | 0.397372154553675 | -5.10676635975618 |
| ENSG00000265241.6 | RBM8A | -0.195953728409163 | 6.53521544786719 | -1.63622690180095 | 0.109177519224078 | 0.397408373715569 | -5.11753754430537 |
| ENSG00000105617.3 | LENG1 | -0.25472263639293 | 3.19475368458372 | -1.63617646576284 | 0.109188114201676 | 0.397408373715569 | -4.93461502417655 |
| ENSG00000145982.12 | FARS2 | -0.139616269733467 | 3.82906459092328 | -1.63610181528994 | 0.109203797392764 | 0.397408373715569 | -4.9997993064551 |
| lnc-LIPM-1 | lnc-LIPM-1 | -0.327273681347401 | 1.86800296516296 | -1.63566688832087 | 0.109295207143879 | 0.397518191457061 | -4.7879943687976 |
| lnc-ARPC3-2 | lnc-ARPC3-2 | 0.337119704921636 | 4.14046873844601 | 1.63565107124038 | 0.109298532641749 | 0.397518191457061 | -5.0225303923449 |
| ENSG00000009790.14 | TRAF3IP3 | -0.307019963024014 | 7.1048871758243 | -1.63552824997704 | 0.10932435829551 | 0.397518191457061 | -5.11513146689795 |
| lnc-PPIL4-1 | lnc-PPIL4-1 | -0.272464740957686 | 2.47682827178419 | -1.63551279763156 | 0.109327607817355 | 0.397518191457061 | -4.86203277578743 |
| ENSG00000175782.9 | SLC35E3 | -0.253279262309893 | 4.90673420732243 | -1.63549885995082 | 0.109330538883468 | 0.397518191457061 | -5.08751829233842 |
| ENSG00000086827.8 | ZW10 | 0.344479462756532 | 3.05942217398524 | 1.63533708972551 | 0.109364563542376 | 0.397571672799845 | -4.90687073448358 |
| ENSG00000150636.17 | CCDC102B | -0.425510822063342 | 0.168747291245473 | -1.63520990656059 | 0.109391319692551 | 0.397598716798829 | -4.51219999337418 |
| ENSG00000198155.5 | ZNF876P | -0.806097645197186 | 2.29851770528394 | -1.63579928011955 | 0.109461339881993 | 0.397764439300478 | -4.83016742169839 |
| ENSG00000285331.2 | TCF12-DT | -0.306020918080421 | 0.197094508548469 | -1.63480962099546 | 0.109475564720265 | 0.397764439300478 | -4.5592025663697 |
| ENSG00000261915.6 | | -0.246842316972377 | 4.77357360057355 | -1.63460313193993 | 0.109519043646886 | 0.397852184148227 | -5.08276996868891 |
| ENSG00000214999.3 | | -0.499651125233461 | -0.484908328919765 | -1.63442277789842 | 0.109557031074245 | 0.397882341682986 | -4.4811978409906 |
| ENSG00000110955.8 | ATP5F1B | 0.253530774558254 | 7.26387973258316 | 1.63433930294797 | 0.109574616801991 | 0.397882341682986 | -5.11418023695689 |
| ENSG00000142186.16 | SCYL1 | -0.203785892583698 | 4.55074560357235 | -1.63400104713072 | 0.109645901051203 | 0.397882341682986 | -5.06868217007531 |
| lnc-LSM2-1 | lnc-LSM2-1 | -0.245015168471065 | 0.385586370271921 | -1.63399743635653 | 0.109646662192794 | 0.397882341682986 | -4.58273958559471 |
| lnc-FBN1-3 | lnc-FBN1-3 | -0.43606949397462 | 1.40133305538855 | -1.63393739933314 | 0.109659318471326 | 0.397882341682986 | -4.75066127733782 |
| ENSG00000280351.2 | | -0.319711483133135 | 0.436988574534614 | -1.63386536495891 | 0.109674505464126 | 0.397882341682986 | -4.5725812433644 |
| ENSG00000254779.4 | EGLN1P1 | -0.301699994278048 | -0.0683055348108541 | -1.63378248229009 | 0.109691981730579 | 0.397882341682986 | -4.48785126601178 |
| ENSG00000174943.10 | KCTD13 | -0.157247646273319 | 3.66811133760185 | -1.63372089033327 | 0.109704970209509 | 0.397882341682986 | -4.98438194417723 |
| lnc-RCC1L-1 | lnc-RCC1L-1 | -0.226696979654408 | 1.97577828940349 | -1.63366327241055 | 0.109717121785766 | 0.397882341682986 | -4.81081558719248 |
| lnc-ATOH7-3 | lnc-ATOH7-3 | -0.195364317416804 | 2.68432662186769 | -1.63363992271247 | 0.109722046533069 | 0.397882341682986 | -4.88907654852282 |
| lnc-RHCE-1 | lnc-RHCE-1 | -0.285200020915159 | 0.867228109875483 | -1.63351516077758 | 0.109748363472618 | 0.397882341682986 | -4.6696959812137 |
| CAHM | CAHM | -0.439663400383615 | 0.115472821189591 | -1.63346325984222 | 0.109759312834669 | 0.397882341682986 | -4.54774368003536 |
| lnc-SCD5-3 | lnc-SCD5-3 | 0.264077970560951 | 2.82254498882465 | 1.63324892570284 | 0.109804539641767 | 0.397957244255662 | -4.89031510720177 |
| ENSG00000116133.12 | DHCR24 | -1.47406190200988 | 1.07673594132242 | -1.63435297731208 | 0.109852777117539 | 0.397957244255662 | -4.65447176905612 |
| ENSG00000124151.18 | NCOA3 | 0.161917675167338 | 9.53882039413681 | 1.6329823424842 | 0.109860812846294 | 0.397957244255662 | -5.06607607931834 |
| lnc-CLK2-2 | lnc-CLK2-2 | -0.186976269451599 | 2.10964646195574 | -1.63297368632022 | 0.109862640476443 | 0.397957244255662 | -4.82196136708557 |
| lnc-ENO1-3 | lnc-ENO1-3 | -0.564807399925773 | -0.838920022954797 | -1.63290735284907 | 0.10987664670383 | 0.397957244255662 | -4.34153185272599 |
| lnc-OGDHL-1 | lnc-OGDHL-1 | -0.234147086563983 | -0.177830568559916 | -1.63248088435187 | 0.109966729880557 | 0.398213441437633 | -4.47683918071327 |
| lnc-DHRS1-1 | lnc-DHRS1-1 | -0.299454708625259 | 0.444682508267009 | -1.63227662549193 | 0.110009896991774 | 0.398299685097921 | -4.59184205110755 |
| ENSG00000240509.1 | RPL34P18 | 0.727595650664486 | 0.197588380554131 | 1.63201731628723 | 0.11006471815792 | 0.398428085445739 | -4.46565740706866 |
| ENSG00000162543.5 | UBXN10 | 1.57724554026754 | -0.984659575637424 | 1.63253106128995 | 0.110091265551469 | 0.398454109169312 | -4.32071014518011 |
| ENSG00000170144.20 | HNRNPA3 | 0.217017590098112 | 7.34333870688265 | 1.63178161787195 | 0.1101145671019 | 0.398468378104503 | -5.1178150785514 |
| lnc-LAMC2-1 | lnc-LAMC2-1 | 0.791813207058871 | -0.38588033830156 | 1.63160458772381 | 0.110152020169445 | 0.398533842704813 | -4.41854930089668 |
| ENSG00000163930.9 | BAP1 | 0.12940768902681 | 5.77832566839386 | 1.63144369362933 | 0.110186068481342 | 0.398545747223637 | -5.12167827670699 |
| ENSG00000138768.14 | USO1 | -0.137834509864015 | 5.91757654520047 | -1.63135280973094 | 0.110205305082583 | 0.398545747223637 | -5.12358284079914 |
| ENSG00000140948.11 | ZCCHC14 | 1.45880093217558 | -0.109576005257731 | 1.63249236667055 | 0.11021339907676 | 0.398545747223637 | -4.54516915497629 |
| ENSG00000115107.19 | STEAP3 | -0.780260785273023 | -0.449456158355606 | -1.63121531709004 | 0.11023441217156 | 0.398551713635556 | -4.44813171661264 |
| ENSG00000087589.16 | CASS4 | -0.498076146510947 | 3.18584666563309 | -1.6311194578902 | 0.110309821591192 | 0.398596311463574 | -4.93997202661716 |
| ENSG00000172292.14 | CERS6 | -1.91766171235051 | 2.51308849495262 | -1.63224044899259 | 0.110311113658424 | 0.398596311463574 | -4.84627015111299 |
| lnc-ADAMTS10-3 | lnc-ADAMTS10-3 | -0.596657544867939 | 0.155154665960131 | -1.63070623338074 | 0.110342239567964 | 0.398596311463574 | -4.50679175051428 |
| lnc-POLR1A-6 | lnc-POLR1A-6 | 0.395789770274888 | 2.89293654693593 | 1.63063827432927 | 0.110356640283022 | 0.398596311463574 | -4.89567820618535 |
| ENSG00000186567.12 | CEACAM19 | -1.93116572946231 | -1.42787364884687 | -1.6314571514734 | 0.110364625646333 | 0.398596311463574 | -4.30470372914712 |
| lnc-EXD2-4 | lnc-EXD2-4 | -0.525935008666478 | -0.597745709265702 | -1.63051065086996 | 0.110383688209049 | 0.398596311463574 | -4.3855372516742 |
| ENSG00000196268.11 | ZNF493 | -0.23440101134085 | 5.83281104254612 | -1.63046709074148 | 0.110392921382486 | 0.398596311463574 | -5.1233605483363 |
| ENSG00000165259.13 | HDX | -2.9673104400735 | -1.68752488861823 | -1.63162569772436 | 0.110417249369381 | 0.398596311463574 | -4.28904826226716 |
| ENSG00000214029.4 | ZNF891 | 0.164865990236063 | 5.04163953184411 | 1.63033447403791 | 0.110421035221557 | 0.398596311463574 | -5.0993592661993 |
| LMO7-AS1 | LMO7-AS1 | -1.37218631434319 | -1.36687992968956 | -1.63024578094164 | 0.110459367242576 | 0.398606446254782 | -4.29141930844948 |
| ENSG00000131849.11 | ZNF132 | -0.26180987428183 | 2.35207420215239 | -1.63010670093088 | 0.110469335237392 | 0.398606446254782 | -4.83406876704356 |
| ENSG00000179899.8 | PHC1P1 | 0.405777436201209 | 1.02682982564888 | 1.63004727170775 | 0.110481940236289 | 0.398606446254782 | -4.68090961466176 |
| ENSG00000170629.14 | DPY19L2P2 | 0.460574625250342 | 2.33458083862247 | 1.62975644863693 | 0.110543641071002 | 0.398611693810622 | -4.84944917136751 |
| ENSG00000170266.15 | GLB1 | -0.195828740518761 | 4.32382257637522 | -1.62970319580189 | 0.110554942210086 | 0.398611693810622 | -5.05234545790381 |

| | | | | | | | |
|---|---|---|---|---|---|---|---|
| lnc-KCNE1B-157 | lnc-KCNE1B-157 | -1.90365491052329 | -1.00281824608188 | -1.63067950683396 | 0.110559402321448 | 0.398611693810622 | -4.35332116533275 |
| lnc-MRPL28-2 | lnc-MRPL28-2 | 0.291428122020663 | 1.05488228778944 | 1.62967531687605 | 0.110560858959571 | 0.398611693810622 | -4.68996156876512 |
| ENSG00000101574.14 | METTL4 | -0.247775519521383 | 4.06026744504722 | -1.62941480654712 | 0.11061615963093 | 0.398741228315838 | -5.02646578048315 |
| ENSG00000068001.13 | HYAL2 | -0.440969119817141 | 0.786651845078129 | -1.62930783050358 | 0.110638874866682 | 0.398753276375578 | -4.68126831842612 |
| lnc-TCL1B-2 | lnc-TCL1B-2 | -0.824782887347237 | 1.62330516904999 | -1.6298518015022 | 0.110703830399423 | 0.398917531707164 | -4.77063626669567 |
| ENSG00000142197.12 | DOP1B | 0.250645956438177 | 6.19273196344606 | 1.62889823589232 | 0.110725883232901 | 0.398927158162576 | -5.12883281920488 |
| ENSG00000247315.3 | ZCCHC3 | 0.229116925093617 | 4.6138704389827 | 1.6283002896788 | 0.110853002747327 | 0.39931525298359 | -5.08027010075372 |
| ENSG00000075651.15 | PLD1 | -1.82574265173416 | 0.721719003436037 | -1.629505833618 | 0.110879092038622 | 0.399339344082407 | -4.575331424937 |
| ENSG00000156735.10 | BAG4 | 0.164613949779254 | 4.74161294324515 | 1.62796217277506 | 0.110924937020782 | 0.399434565902513 | -5.08916518998156 |
| lnc-HNRNPA2B1-14 | lnc-HNRNPA2B1-1 | -0.370740682595027 | 2.45903049177598 | -1.62782825692422 | 0.110953438128056 | 0.399451288284914 | -4.87942983027038 |
| ENSG00000151849.14 | CENPJ | -0.262475540088297 | 3.90514328548945 | -1.62775013203494 | 0.110970068094611 | 0.399451288284914 | -5.0153951341453 |
| ENSG00000089154.10 | GCN1 | 0.135042350947487 | 7.18289975046812 | 1.6275982651578 | 0.11100240089688 | 0.399451288284914 | -5.12611200680731 |
| lnc-KCNT1-1 | lnc-KCNT1-1 | -0.341740178092456 | 1.8075657562687 | -1.62757568779183 | 0.11100720832676 | 0.399451288284914 | -4.79437690789483 |
| ENSG00000188822.7 | CNR2 | -0.209091082546165 | 7.58172078662701 | -1.62748236306909 | 0.111027081895458 | 0.399452967427759 | -5.12082995454965 |
| ENSG00000221983.7 | UBA52 | 0.25167604202449 | 7.90941730954531 | 1.62723253644681 | 0.111080296974266 | 0.399521421692216 | -5.11542531753598 |
| lnc-TNK2-7 | lnc-TNK2-7 | 0.373200482993617 | 2.31839450183313 | 1.62721079467252 | 0.111084929133001 | 0.399521421692216 | -4.85279170859339 |
| ENSG00000118096.7 | IFT46 | -0.273377832811462 | 2.68740578436904 | -1.62661126710602 | 0.111212722734863 | 0.399658423633564 | -4.89105649993532 |
| ENSG00000184209.14 | SNRNP35 | -0.185103063400716 | 4.73831005954905 | -1.62656974971993 | 0.111221576910533 | 0.399658423633564 | -5.09044148406438 |
| ENSG00000178913.7 | TAF7 | 0.166333833339994 | 6.94165559055457 | 1.62654650279202 | 0.111226534901616 | 0.399658423633564 | -5.1304629426956 |
| ENSG00000048544.5 | MRPS10 | 0.16671479649333 | 4.50163484029665 | 1.62645755701804 | 0.11124550648911 | 0.399658423633564 | -5.06998126598596 |
| SNHG10 | SNHG10 | 0.280659095519156 | 6.50563476101873 | 1.62640113581858 | 0.11125754215543 | 0.399658423633564 | -5.13336000812389 |
| ENSG00000197238.4 | H4C11 | 0.66073717499937 | 0.708413492127895 | 1.62642975716697 | 0.111259499658356 | 0.399658423633564 | -4.62135582031881 |
| ENSG00000148450.12 | MSRB2 | -0.459897022142934 | 0.0477866193734337 | -1.62624309013365 | 0.111291261835813 | 0.399658423633564 | -4.55786575786415 |
| ENSG00000152127.8 | MGAT5 | 0.372884524214707 | 7.11490631130673 | 1.62626274775498 | 0.111310240327918 | 0.399658423633564 | -5.12919501067833 |
| ENSG00000267430.1 | | 0.343094591264653 | 1.91564982669761 | 1.62610322069345 | 0.111321110507875 | 0.399658423633564 | -4.80485768809493 |
| ENSG00000119523.9 | ALG2 | 0.172916540579838 | 4.14523165584111 | 1.62609509858966 | 0.11132284399678 | 0.399658423633564 | -5.04391319869155 |
| lnc-HAL-1 | lnc-HAL-1 | 0.317998582512094 | 3.52433736619698 | 1.62603061258626 | 0.111336607934453 | 0.399658423633564 | -4.97061297281338 |
| ENSG00000126804.13 | ZBTB1 | 0.164555765710659 | 6.78737814381482 | 1.62579063056841 | 0.111387842083152 | 0.399749359095533 | -5.13299801165065 |
| ENSG00000136813.14 | ECPAS | 0.160494965707064 | 6.53385102604188 | 1.62570666888156 | 0.111405771748381 | 0.399749359095533 | -5.13394901800154 |
| lnc-ZFP62-2 | lnc-ZFP62-2 | -0.162900414902737 | 2.74737102739896 | -1.62563908980156 | 0.111420204689845 | 0.399749359095533 | -4.89896421518384 |
| ENSG00000142082.14 | SIRT3 | -0.167691889409162 | 3.29486017625548 | -1.62533646385751 | 0.11148485560232 | 0.399911603844991 | -4.95758602502774 |
| ENSG00000116539.12 | ASH1L | 0.117418074509953 | 8.09361043400044 | 1.62498206988233 | 0.111560604843141 | 0.400078759981586 | -5.11574231490906 |
| ENSG00000132330.16 | SCLY | -0.229734688369482 | 2.86147961664479 | -1.62475723119936 | 0.111608684343783 | 0.400078759981586 | -4.91262360898634 |
| lnc-DUOXA1-2 | lnc-DUOXA1-2 | -0.890882592455978 | 0.741612212781672 | -1.62540915801404 | 0.111613761271894 | 0.400078759981586 | -4.6363390533798 |
| ENSG00000226683.2 | PWWP2AP1 | -1.35303339282406 | 1.82293883685659 | -1.62600082903055 | 0.111625304693569 | 0.400078759981586 | -4.7900578615298 |
| ENSG00000181896.11 | ZNF101 | 0.208178893221346 | 7.55612431227642 | 1.62466392905728 | 0.111628641042329 | 0.400078759981586 | -5.12470321193156 |
| ENSG00000155313.15 | USP25 | 0.129354955941105 | 6.28018494862123 | 1.62438454376782 | 0.111688417107259 | 0.400186490485915 | -5.13648641009293 |
| lnc-ZNF599-10 | lnc-ZNF599-10 | -0.430725507986732 | -0.191038286946428 | -1.62431406320697 | 0.111703500952484 | 0.400186490485915 | -4.46754372051165 |
| ENSG00000106993.11 | CDC37L1 | 0.219119416831915 | 3.47112447841259 | 1.62425086647869 | 0.111717027368803 | 0.400186490485915 | -4.98303512505855 |
| ENSG00000035862.12 | TIMP2 | -0.397157250213611 | 4.50189975196195 | -1.62409966959322 | 0.111772571408072 | 0.400315788636217 | -5.07434064019204 |
| lnc-MAP1LC3C-6 | lnc-MAP1LC3C-6 | -1.52347618960752 | -2.31047858666926 | -1.62385170928379 | 0.111802492560005 | 0.400353288859181 | -4.27655756473574 |
| lnc-MESD-2 | lnc-MESD-2 | -0.372665080100255 | 0.61166071118604 | -1.62361900812379 | 0.111852341804345 | 0.400462124083986 | -4.62991188273912 |
| ENSG00000180089.5 | TMEM86B | -0.26799404797825 | 1.66350025282805 | -1.62347137417401 | 0.111883977362585 | 0.400505722792016 | -4.79014212804109 |
| lnc-PIK3IP1-1 | lnc-PIK3IP1-1 | -0.171633770108262 | 3.14358637046049 | -1.62337982061167 | 0.111903599475773 | 0.400506309860865 | -4.95075948718244 |
| ENSG00000173653.7 | RCE1 | -0.226929528455049 | 4.05190967482492 | -1.62309183310843 | 0.111965340398508 | 0.400657614990001 | -5.04160176432867 |
| ENSG00000058668.14 | ATP2B4 | -0.492375827656859 | 7.00165117809894 | -1.62364848453033 | 0.111994296529109 | 0.400691570564688 | -5.13412031213474 |
| lnc-C19orf48-5 | lnc-C19orf48-5 | -0.252627497526847 | 1.28385721342435 | -1.62259383164446 | 0.11207217134589 | 0.400816252199991 | -4.75286103125427 |
| lnc-RNASET2-2 | lnc-RNASET2-2 | 0.352607807710126 | 0.0292671733782975 | 1.62253556456585 | 0.112084676199569 | 0.400816252199991 | -4.50884176694301 |
| ENSG00000184635.15 | ZNF93 | -0.270096413630755 | 3.63733644033624 | -1.62245272089124 | 0.112102457464117 | 0.400816252199991 | -5.00102348701779 |
| ENSG00000204248.10 | COL11A2 | -0.298318919373695 | -0.0930719337292059 | -1.62240983662323 | 0.112111662892087 | 0.400816252199991 | -4.51254462773249 |
| ENSG00000213551.4 | DNAJC9 | 0.120560408831995 | 4.71169351801404 | 1.62234067054832 | 0.112126511206702 | 0.400816252199991 | -5.09417915560678 |
| lnc-CTAGE5-8 | lnc-CTAGE5-8 | -0.413335282261235 | 3.60325378059853 | -1.62228330785268 | 0.112150660983012 | 0.400832966663194 | -5.00486852917723 |
| lnc-PABPN1L-1 | lnc-PABPN1L-1 | 0.496487087003113 | 1.98503050017285 | 1.62157569257791 | 0.112290840951466 | 0.401264301962504 | -4.80650064489424 |
| ENSG00000243547.1 | HNRNPKP4 | 0.927559529641462 | 0.682472461722473 | 1.62172500599558 | 0.112345504227006 | 0.401384937723212 | -4.5356320085639 |
| ENSG00000065427.14 | KARS1 | 0.136470501316329 | 6.15626537105814 | 1.62123727365768 | 0.112363601572227 | 0.401384937723212 | -5.14088260687996 |
| ENSG00000213261.3 | EEF1B2P6 | 0.490689984368174 | 0.334300165982035 | 1.62114120654001 | 0.112384263172855 | 0.401389083617364 | -4.53882640197724 |
| ENSG00000115364.13 | MRPL19 | 0.189213053432307 | 4.07119628085333 | 1.62079538089451 | 0.112458667182801 | 0.401585139941638 | -5.04527297223429 |
| ENSG00000015475.18 | BID | 0.664946354542451 | 0.991639324585347 | 1.62082435655748 | 0.112486926879363 | 0.401616377442831 | -4.68216218481218 |
| lnc-CDNF-2 | lnc-CDNF-2 | -0.27264910771035 | 2.543225219607 | -1.62046508880629 | 0.112529766707574 | 0.401699651082553 | -4.88668817250221 |
| lnc-TBC1D9B-5 | lnc-TBC1D9B-5 | 0.298836723765452 | 3.0123337235392 | 1.61972973464531 | 0.112688192738552 | 0.402179473009368 | -4.93638025269128 |
| lnc-NUDT6-3 | lnc-NUDT6-3 | -0.394451784578663 | -0.408327666703709 | -1.61955282792555 | 0.112726332997932 | 0.402179473009368 | -4.46014088843076 |
| lnc-PACSIN3-4 | lnc-PACSIN3-4 | 0.171446507416986 | 2.6254984450811 | 1.61945753049321 | 0.112746883043919 | 0.402179473009368 | -4.89575229951333 |
| lnc-MFSD8-8 | lnc-MFSD8-8 | 0.216984834996955 | 0.386182356762089 | 1.61939322315873 | 0.112760752078013 | 0.402179473009368 | -4.59800903477486 |
| ENSG00000102900.12 | NUP93 | 0.127786088273401 | 5.15128469391505 | 1.61933899361449 | 0.11277244873748 | 0.402179473009368 | -5.12193788296762 |
| ENSG00000255302.4 | EID1 | -0.413892043585817 | 5.49373605091678 | -1.61961228353303 | 0.112781417587819 | 0.402179473009368 | -5.13384172061635 |
| ENSG00000275004.3 | ZNF280B | 0.406171851420764 | 2.10883553339229 | 1.61909824067401 | 0.112824388204363 | 0.402263014448364 | -4.84191728625858 |
| ENSG00000182162.10 | P2RY8 | -0.318275383967948 | 6.08827095750421 | -1.61883090387987 | 0.112882085683538 | 0.402399024874308 | -5.14329798114707 |
| lnc-TTC3-4 | lnc-TTC3-4 | 0.429962043063344 | 1.45368146216334 | 1.61851465554125 | 0.112950370497678 | 0.402565813247137 | -4.7598656812319 |

| | | | | | | | |
|---|---|---|---|---|---|---|---|
| ENSG00000218175.2 | | 0.888219372002776 | 1.03489900597052 | 1.61897797106698 | 0.112967989958483 | 0.402565813247137 | -4.60951375891574 |
| lnc-SEC62-2 | lnc-SEC62-2 | 0.576089196074045 | 0.00348882316318596 | 1.61823318499495 | 0.113011174387941 | 0.4026499917651 | -4.52589709008204 |
| ENSG00000079819.18 | EPB41L2 | -1.28468047722839 | 0.826956516362132 | -1.61929935320308 | 0.113061447877134 | 0.402712820462234 | -4.76223987028179 |
| lnc-DERL3-1 | lnc-DERL3-1 | -0.523309292077748 | 1.88686390327895 | -1.61803765017448 | 0.113067939049276 | 0.402712820462234 | -4.82213516390251 |
| ENSG00000138663.8 | COPS4 | 0.154477081845932 | 4.35016718246356 | 1.61786164001237 | 0.113091477253704 | 0.402726968220241 | -5.07526903989454 |
| ENSG00000171984.14 | SHLD1 | -0.205028308553505 | 1.97821695132153 | -1.6175704148512 | 0.113154452957767 | 0.402881526592235 | -4.82289759430035 |
| ENSG00000115109.13 | EPB41L5 | 0.979999202539393 | 2.43109615095874 | 1.61840798020341 | 0.113236622461734 | 0.403104358375973 | -4.89854738034246 |
| lnc-HPSE-1 | lnc-HPSE-1 | 0.393310750123168 | -0.307214277229906 | 1.61673645160643 | 0.113334950835229 | 0.403384626152777 | -4.44074547019439 |
| ENSG00000140398.13 | NEIL1 | -0.664058483223052 | 1.56660652743827 | -1.61695989456258 | 0.113396675880211 | 0.403492701674249 | -4.81528232615725 |
| lnc-ROPN1L-7 | lnc-ROPN1L-7 | 0.374040231261093 | 1.89904116204953 | 1.61637661940306 | 0.113412903208146 | 0.403492701674249 | -4.81131198699443 |
| lnc-RBM34-1 | lnc-RBM34-1 | 0.315119632503876 | 0.751672452820673 | 1.61632483407048 | 0.113424125340679 | 0.403492701674249 | -4.66225147043304 |
| ENSG00000104936.17 | DMPK | -0.219840246962185 | 3.07891987169522 | -1.61600866465183 | 0.113492660434449 | 0.403666740923149 | -4.95299959282343 |
| ENSG00000170892.10 | TSEN34 | -0.237360962621227 | 3.04990979649293 | -1.61576593893728 | 0.113545298274063 | 0.403744737294945 | -4.94636686726583 |
| lnc-SATB1-7 | lnc-SATB1-7 | -0.504581133397576 | -0.358884440527817 | -1.61556946219245 | 0.113587921078239 | 0.403744737294945 | -4.46954116050142 |
| ENSG00000110025.12 | SNX15 | -0.236387994709388 | 2.86278083709808 | -1.61555306248751 | 0.113591479348368 | 0.403744737294945 | -4.92943685967776 |
| lnc-RAB40B-4 | lnc-RAB40B-4 | 0.237187087826901 | 1.28187990573974 | 1.61554581793188 | 0.113593051240102 | 0.403744737294945 | -4.74793329305412 |
| ENSG00000271011.1 | | -0.427859969407714 | -0.307373275276399 | -1.61543186445239 | 0.113617778697906 | 0.403762903822341 | -4.47561484512412 |
| lnc-MGAT1-4 | lnc-MGAT1-4 | -0.201422536907355 | 3.74502422763438 | -1.6151228395255 | 0.113684857989947 | 0.403931543588311 | -5.02716309651883 |
| ENSG00000188486.3 | H2AX | -0.402960608940484 | 1.47128694445599 | -1.61477644564932 | 0.113760087257576 | 0.404069673895566 | -4.78672923596082 |
| ENSG00000154845.15 | PPP4R1 | 0.205593097840328 | 5.49429965864761 | 1.61476305303686 | 0.113762996658641 | 0.404069673895566 | -5.1402782733288 |
| lnc-CHRNA10-3 | lnc-CHRNA10-3 | 0.222712046838363 | 1.53178608314459 | 1.61446251479497 | 0.113828301320607 | 0.404231871304701 | -4.77857176899099 |
| ENSG00000003509.15 | NDUFAF7 | 0.172707560628164 | 4.40364812640849 | 1.61410119207827 | 0.113906854447813 | 0.404393214906352 | -5.08328889537984 |
| STARD7-AS1 | STARD7-AS1 | -0.240697141976845 | 1.9903625236055 | -1.61407280300268 | 0.113913028228491 | 0.404393214906352 | -4.83415673383362 |
| ENSG00000207631.1 | MIR641 | 0.574478350446491 | -0.450975566075084 | 1.6133619086671 | 0.114067715607758 | 0.404872528083202 | -4.41184443699737 |
| ENSG00000246596.7 | SIMC1P1 | 0.29879595435009 | 4.789728956743 | 1.61315252256857 | 0.114113309706026 | 0.404964526496401 | -5.10567107967709 |
| ENSG00000175376.8 | EIF1AD | 0.179714294214953 | 5.17931179278058 | 1.61282943182129 | 0.114183692289115 | 0.405144447230968 | -5.13134451721902 |
| lnc-WDR47-1 | lnc-WDR47-1 | -0.209674641177603 | 0.761722944495432 | -1.61260446947726 | 0.114232719324858 | 0.405248545650389 | -4.67426522966882 |
| lnc-BRAP-2 | lnc-BRAP-2 | -0.323328956164507 | 0.895348448066931 | -1.61237944328458 | 0.114281777440344 | 0.405352718430916 | -4.69305008262277 |
| lnc-ZNF385C-4 | lnc-ZNF385C-4 | 0.217250144997636 | 2.89799586974322 | 1.61179340101188 | 0.114409621522272 | 0.405736257717595 | -4.93483013035633 |
| ENSG00000230207.1 | RPL4P5 | 0.89705842364366 | -0.258218319840757 | 1.61150945120749 | 0.114471606574121 | 0.405882682493421 | -4.39000085388454 |
| ENSG00000239779.6 | WBP1 | -0.168832362493879 | 4.85758418230588 | -1.61139876859651 | 0.114495775536021 | 0.405882682493421 | -5.11938170002306 |
| ENSG00000220749.4 | RPL21P28 | 0.374958943127595 | 0.734152263907936 | 1.61133332302417 | 0.114510068369008 | 0.405882682493421 | -4.68519331945344 |
| ENSG00000138386.16 | NAB1 | 0.298214934370733 | 4.38602074888225 | 1.6110392013737 | 0.114574320331829 | 0.406040501961095 | -5.08975852461269 |
| lnc-HERPUD2-1 | lnc-HERPUD2-1 | 0.409788934978557 | 0.0445308131100386 | 1.61088695288271 | 0.114607591115699 | 0.406088491639601 | -4.53643860688245 |
| ENSG00000201501.1 | | -0.457948260959578 | -0.426675815105371 | -1.61072871659206 | 0.114642178752994 | 0.406141129995332 | -4.46816856649855 |
| ENSG00000264357.1 | MIR4648 | 0.569634997539142 | -0.137421345710102 | 1.61055736169025 | 0.114679643477053 | 0.406203941092441 | -4.50394176096044 |
| ENSG00000155100.10 | OTUD6B | 0.220241295302102 | 4.34446438511158 | 1.6103877674676 | 0.114716733071458 | 0.406265402066384 | -5.08570095583286 |
| ENSG00000185164.14 | NOMO2 | 0.221542891953137 | 2.6011312568086 | 1.60991973628701 | 0.114819140369447 | 0.406558122522677 | -4.90317507262268 |
| ENSG00000178093.13 | TSSK6 | -0.23779011105511 | 1.0111516939034 | -1.60976437523458 | 0.114853150517016 | 0.406608599430986 | -4.7238777461415 |
| ENSG00000256087.6 | ZNF432 | 0.200188347743983 | 5.12444946948425 | 1.60937099912225 | 0.114939301331599 | 0.406843618109768 | -5.13377483458731 |
| lnc-ZNF629-2 | lnc-ZNF629-2 | 0.518730287718752 | -0.504886951654435 | 1.60916259829707 | 0.114984963205192 | 0.40693526438316 | -4.43623721715031 |
| ENSG00000166394.14 | CYB5R2 | -1.25649896840521 | 0.954000727394069 | -1.61012656430877 | 0.115043988973144 | 0.407074166242776 | -4.72964251042546 |
| ENSG00000128438.10 | TBC1D27P | 1.06047841561167 | 4.93943431469247 | 1.60973170548634 | 0.115154348911863 | 0.40739463108506 | -5.13308380325372 |
| ENSG00000137877.9 | SPTBN5 | -0.588006010153197 | 0.730633447535106 | -1.60790249024606 | 0.115261375334904 | 0.407703194452369 | -4.67039797221743 |
| ENSG00000213236.3 | YWHAZP2 | 1.6008712390441 | 0.10196830459283 | 1.60849554317986 | 0.115304610006482 | 0.407786046007461 | -4.37689853874594 |
| lnc-PHKB-5 | lnc-PHKB-5 | -0.294362733550178 | 0.803043390491988 | -1.60757246663224 | 0.1153338573114 | 0.407819409902173 | -4.69199181404893 |
| ENSG00000206535.7 | LNP1 | -0.256087016065643 | 0.453243444255693 | -1.60726699904247 | 0.115400979203762 | 0.40798666350928 | -4.64840665628818 |
| lnc-SRSF7-2 | lnc-SRSF7-2 | 0.286299558428368 | -0.193013408708246 | 1.60705903735889 | 0.115446693841214 | 0.408011401622748 | -4.4927474592431 |
| ENSG00000154642.10 | C21orf91 | 0.246139716288975 | 4.45971371462146 | 1.60705481586851 | 0.115447621972059 | 0.408011401622748 | -5.10047411082112 |
| ENSG00000175582.19 | RAB6A | -0.177651840101078 | 5.60178886553259 | -1.60693985812196 | 0.115472898753497 | 0.408030673827163 | -5.15571504093773 |
| ENSG00000092847.11 | AGO1 | 0.115063859032287 | 5.88633216275148 | 1.60673757947908 | 0.11551738649658 | 0.408117810892396 | -5.16037264338609 |
| lnc-B3GNT7-4 | lnc-B3GNT7-4 | -0.299543083393095 | 0.252079584890944 | -1.60649625089974 | 0.115570480863 | 0.408235319650442 | -4.59041619849636 |
| ENSG00000197746.13 | PSAP | 0.38892465324056 | 7.62259213553586 | 1.60649753518939 | 0.115610924603353 | 0.408243976850994 | -5.15242410731703 |
| lnc-RALGPS2-2 | lnc-RALGPS2-2 | -0.421560930714756 | -0.121756989763349 | -1.60613827109346 | 0.115649276096875 | 0.408243976850994 | -4.49956900321579 |
| lnc-ZNF14-3 | lnc-ZNF14-3 | -0.250245253591345 | 3.58295900391866 | -1.60612624775775 | 0.115651923322265 | 0.408243976850994 | -5.01549006248193 |
| ENSG00000185453.12 | ZSWIM9 | -0.287874013942273 | 0.287023088620477 | -1.60612468300691 | 0.115652267843276 | 0.408243976850994 | -4.62038828469565 |
| ENSG00000005801.17 | ZNF195 | 0.12818067502538 | 4.91912302810297 | 1.60556901603104 | 0.115774665367472 | 0.408597603711421 | -5.13167100887955 |
| lnc-MKLN1-1 | lnc-MKLN1-1 | -0.457975441669504 | -0.500296783687229 | -1.60548967760015 | 0.115792149951354 | 0.408597603711421 | -4.44532265738536 |
| ENSG00000196757.7 | ZNF700 | 0.259703361798056 | 5.37135363136481 | 1.60453643871307 | 0.116002392597742 | 0.409067271161812 | -5.14895962910272 |
| lnc-SNAPC5-1 | lnc-SNAPC5-1 | 0.365940185972091 | -0.191303950390141 | 1.60440643359575 | 0.116031090078136 | 0.409067271161812 | -4.48615436270066 |
| ENSG00000256053.7 | COA8 | -0.174836482241077 | 2.99744484738412 | -1.60435586084582 | 0.116042255126399 | 0.409067271161812 | -4.95833852345227 |
| RBM26-AS1 | RBM26-AS1 | -0.279201487138145 | 4.23518656355656 | -1.60415538283353 | 0.116086523665457 | 0.409067271161812 | -5.07502180602391 |
| ENSG00000130703.16 | OSBPL2 | 0.124185848582884 | 5.44513578372116 | 1.60409358590029 | 0.116100172121868 | 0.409067271161812 | -5.15585014071379 |
| ENSG00000102158.19 | MAGT1 | 0.18532257588118 | 5.62694094735515 | 1.60393742088999 | 0.116134668509508 | 0.409067271161812 | -5.16078023413171 |
| ENSG00000162408.10 | NOL9 | -0.16550387284783 | 5.61702620242689 | -1.60390190393795 | 0.116142515262941 | 0.409067271161812 | -5.16143512317351 |
| lnc-HEATR5B-1 | lnc-HEATR5B-1 | 0.34252180332315 | 1.17934671264435 | 1.60382422481058 | 0.116159678401327 | 0.409067271161812 | -4.75027722268453 |
| ENSG00000207741.1 | MIR590 | 0.299195306057824 | 1.37440716090446 | 1.60368205308039 | 0.116191096471844 | 0.409067271161812 | -4.77550325005778 |
| lnc-CDC14B-1 | lnc-CDC14B-1 | 0.280163892655511 | 1.67858452427577 | 1.60364272827773 | 0.116199787953401 | 0.409067271161812 | -4.81171514740064 |

| | | | | | | | |
|---|---|---|---|---|---|---|---|
| ENSG00000123505.15 | AMD1 | 0.211302969361241 | 6.99675023471074 | 1.60358690595582 | 0.116212126588916 | 0.409067271161812 | -5.16426830304179 |
| ENSG00000162927.13 | PUS10 | -0.197238672852197 | 3.74308208261165 | -1.60357079232889 | 0.116215688447879 | 0.409067271161812 | -5.03395458979632 |
| ENSG00000229677.1 | RPL17P28 | 0.339818433488333 | 0.294254244014664 | 1.60354847705118 | 0.116220621306005 | 0.409067271161812 | -4.56512914078383 |
| lnc-ZC3HAV1L-1 | lnc-ZC3HAV1L-1 | 0.339818433488333 | 0.294254244014664 | 1.60354847705118 | 0.116220621306005 | 0.409067271161812 | -4.56512914078383 |
| lnc-NDRG1-2 | lnc-NDRG1-2 | -0.274820468583339 | 4.67527640533292 | -1.60353609183597 | 0.116223359167968 | 0.409067271161812 | -5.11355034905042 |
| ENSG00000157637.12 | SLC38A10 | -0.244226951619423 | 5.45353233624868 | -1.60325603394248 | 0.116285282467126 | 0.4091953307678 | -5.15647427871842 |
| ENSG00000200354.1 | SNORA71D | 0.54694339408142 | -0.241477050086092 | 1.60315352379043 | 0.116307955086393 | 0.4091953307678 | -4.47851856342868 |
| ENSG00000259845.1 | HERC2P10 | 3.15757852910949 | -1.98866334445714 | 1.60406037081078 | 0.116319383973298 | 0.4091953307678 | -4.2966096744056 |
| ENSG00000076382.16 | SPAG5 | 0.316083316816057 | 2.51587246429995 | 1.60119708405889 | 0.116741359183537 | 0.410609602883586 | -4.90095711375355 |
| ENSG00000141570.10 | CBX8 | 0.673964013359838 | -0.0539953069868538 | 1.60101091940741 | 0.116782668019003 | 0.410684718235972 | -4.50334187614909 |
| EIF1B-AS1 | EIF1B-AS1 | -0.418827823071002 | 4.02082415848378 | -1.60097374762553 | 0.116819560338518 | 0.410693936973925 | -5.0688043067792 |
| ENSG00000211450.9 | SELENOH | -0.194416245528072 | 4.04400688099412 | -1.60081931824137 | 0.116825195597451 | 0.410693936973925 | -5.07439408493357 |
| ENSG00000103507.13 | BCKDK | -0.278015872668756 | 2.43905953377834 | -1.6006699087823 | 0.116858367091262 | 0.410740397855098 | -4.90448822483586 |
| ZNF337-AS1 | ZNF337-AS1 | -0.256674055751666 | 3.06718742998698 | -1.60053714988312 | 0.116887848297927 | 0.410773874255802 | -4.96430396198526 |
| lnc-AMZ2-3 | lnc-AMZ2-3 | 0.328596932877112 | 0.89075742419565 | 1.60036998430735 | 0.116924978651741 | 0.410834215703103 | -4.70710685242259 |
| lnc-NAA38-2 | lnc-NAA38-2 | -0.538209426869437 | 1.99474700050798 | -1.60035909891417 | 0.116956040254651 | 0.410873216685695 | -4.85526623955128 |
| ENSG00000164307.12 | ERAP1 | 0.138589065347421 | 6.14675520025175 | 1.59988538462637 | 0.117032670792594 | 0.411019864725571 | -5.17324022096824 |
| ENSG00000108984.14 | MAP2K6 | -0.646807059266797 | 5.5929883232385 | -1.60084421680071 | 0.117037721768283 | 0.411019864725571 | -5.16100367684855 |
| ENSG00000266929.1 | | -0.355915045282048 | -0.269525907909145 | -1.59959936288435 | 0.117096270979558 | 0.411029141366731 | -4.52859734433105 |
| ENSG00000140368.12 | PSTPIP1 | -0.236745828121955 | 4.46044332238606 | -1.59958443716957 | 0.117099590653292 | 0.411029141366731 | -5.10778324338242 |
| ENSG00000253719.3 | ATXN7L3B | 0.17308965488701 | 6.6731357656823 | 1.59958137675362 | 0.117100271339188 | 0.411029141366731 | -5.17296076536528 |
| ENSG00000253320.6 | MAILR | -0.22617035921696 | 3.26471225253465 | -1.59939883819165 | 0.117140876679651 | 0.411101562608226 | -4.98906032595297 |
| ENSG00000012061.15 | ERCC1 | -0.15897655816128 | 5.35091704598359 | -1.59923701783062 | 0.11717688286885 | 0.411112001718093 | -5.1608634238242 |
| ENSG00000166747.12 | AP1G1 | 0.181380342250876 | 7.43681831352117 | 1.59920594425094 | 0.117183797992521 | 0.411112001718093 | -5.16561766288365 |
| ENSG00000145348.16 | TBCK | -0.175565908309053 | 5.85565558656522 | -1.58899045853085 | 0.117231761376941 | 0.411170158621119 | -5.17142617163112 |
| ENSG00000270878.1 | GCATP1 | -0.310604963779895 | 0.554066671497792 | -1.59895198093876 | 0.117240327500721 | 0.411170158621119 | -4.64824050412784 |
| lnc-ERAP1-2 | lnc-ERAP1-2 | -0.296998592955828 | 1.17792197436193 | -1.59798285230299 | 0.11745624887209 | 0.411857235184706 | -4.76274218520439 |
| lnc-KLF8-4 | lnc-KLF8-4 | -0.358405563131315 | 4.36472180548614 | -1.59727103530568 | 0.117615047131103 | 0.412343811122379 | -5.10262899696497 |
| lnc-ZNF710-1 | lnc-ZNF710-1 | -0.240644372868055 | 0.552607300793435 | -1.5971614021905 | 0.117639520527803 | 0.412359375174349 | -4.66653477362286 |
| lnc-USP32-2 | lnc-USP32-2 | -0.462773946461141 | 0.538181373071336 | -1.59671764949434 | 0.117738621705167 | 0.412636480598919 | -4.67223842939503 |
| ENSG00000133243.8 | BTBD2 | -0.291339712337083 | 4.48383026149017 | -1.59659365559636 | 0.117766324794705 | 0.412663306647841 | -5.11913479358695 |
| ENSG00000166451.13 | CENPN | 0.370305103205469 | 1.17599100646954 | 1.59643882870177 | 0.1178009241206 | 0.412714284455202 | -4.75441636147769 |
| ENSG00000267001.1 | | 0.323901289190607 | 0.75911308912413 | 1.59625545168859 | 0.117841914254873 | 0.412730082488747 | -4.70080702021268 |
| ENSG00000277443.2 | MARCKS | 1.28204992161407 | 3.13108238080042 | 1.59755656709449 | 0.117845537326258 | 0.412730082488747 | -5.00918468461679 |
| ENSG00000175984.14 | DENND2C | -0.815849355779621 | -0.855075610828348 | -1.59602472360992 | 0.117893505202011 | 0.412827835585743 | -4.43891750152385 |
| lnc-MINDY1-3 | lnc-MINDY1-3 | -0.464960740719729 | 0.881477017107542 | -1.59592100333158 | 0.117916703101099 | 0.412838833122967 | -4.70968274031641 |
| ENSG00000188585.9 | CLEC20A | 3.28440515092872 | 0.701262085365695 | 1.59692110011789 | 0.117987394433057 | 0.413016078165919 | -4.37812826148556 |
| lnc-ABCG1-1 | lnc-ABCG1-1 | 0.380713444806196 | 3.73384831728635 | 1.59537644899041 | 0.118038558042394 | 0.413124917562762 | -5.03500380237857 |
| ENSG00000284770.1 | TBCE | -0.220093077436612 | 2.5921029012203 | -1.59506233937904 | 0.118108892867732 | 0.41324527727069 | -4.92660828581622 |
| ENSG00000178665.15 | ZNF713 | -0.257522783607986 | 2.9697767609918 | -1.59504354921993 | 0.118113101403268 | 0.41324527727069 | -4.96047783706065 |
| ENSG00000173295.7 | FAM86B3P | -0.583548368552859 | 0.805049261424173 | -1.59442805058695 | 0.118267572248835 | 0.413641438963387 | -4.76287254821315 |
| ENSG00000196981.3 | WDR5B | -0.214713334923714 | 3.73468201519755 | -1.59431220054485 | 0.118277000258568 | 0.413641438963387 | -5.05076429913994 |
| FRY-AS1 | FRY-AS1 | -0.529460860907008 | 3.44868086054398 | -1.59481070538362 | 0.118286620499368 | 0.413641438963387 | -5.04729614662381 |
| LINC01376 | LINC01376 | -0.332523436529288 | 3.39705275321467 | -1.59380750002569 | 0.118390213824742 | 0.413802258464455 | -5.02030352730302 |
| lnc-GPT2-1 | lnc-GPT2-1 | -0.265970848531056 | 2.75973694485585 | -1.59378143415541 | 0.118396063265747 | 0.413802258464455 | -4.94647761114165 |
| ENSG00000102409.9 | BEX4 | -0.849889969735222 | 3.72876923018586 | -1.59482447073858 | 0.11842456068226 | 0.413802258464455 | -5.03733672291223 |
| ENSG00000103187.7 | COTL1 | 0.407187543234595 | 5.19563893980382 | 1.59384502900178 | 0.118437203107834 | 0.413802258464455 | -5.15313827106958 |
| ENSG00000242737.1 | | 0.508051152082244 | 0.779293494037467 | 1.59354012205255 | 0.118450227250868 | 0.413802258464455 | -4.66871878460862 |
| lnc-FARP1-2 | lnc-FARP1-2 | 0.256326451238553 | 0.780522313895017 | 1.59341029251192 | 0.118479376613616 | 0.413802258464455 | -4.71063657438449 |
| ENSG00000198520.11 | ARMH1 | 0.752914805655471 | 3.00700683798842 | 1.59425412636811 | 0.11849807456219 | 0.413802258464455 | -4.9609392650934 |
| ENSG00000228828.1 | TLK2P2 | -0.415611086369641 | 0.989525763642421 | -1.5932885957866 | 0.118506705283338 | 0.413802258464455 | -4.72536128912997 |
| ENSG00000243406.6 | MRPS31P5 | 0.233561605890893 | 3.27640472527759 | 1.59325813789488 | 0.118513545821696 | 0.413802258464455 | -5.01097722470979 |
| lnc-CMAS-4 | lnc-CMAS-4 | 0.493697148703464 | -0.0420776118978229 | 1.59308530587436 | 0.118552368239632 | 0.413863846041329 | -4.5719007940948 |
| ENSG00000008513.15 | ST3GAL1 | -0.2419330238069 | 6.05897784312644 | -1.59300060154506 | 0.118571398732241 | 0.413863846041329 | -5.18158347674854 |
| ENSG00000138794.9 | CASP6 | -0.24700070156411 | 1.46604245476699 | -1.59271269372389 | 0.118636101428004 | 0.41399059266021 | -4.80802130373069 |
| ENSG00000136143.15 | SUCLA2 | 0.246262895084214 | 3.57705692093038 | 1.59261155690277 | 0.118658837119837 | 0.41399059266021 | -5.04276163806954 |
| ENSG00000246922.8 | UBAP1L | -0.237728100852361 | 3.59806376603444 | -1.59257057397364 | 0.118668051143917 | 0.41399059266021 | -5.03931901995527 |
| ENSG00000157020.17 | SEC13 | 0.141427945711258 | 4.98982134001891 | 1.59219980360447 | 0.118751436329702 | 0.41421128859079 | -5.15593649298832 |
| lnc-POLR3G-1 | lnc-POLR3G-1 | -0.447035420963542 | 1.0038543176585 | -1.59205395724951 | 0.118784249786969 | 0.414255542759264 | -4.75025563653111 |
| ENSG00000224531.5 | SMIM13 | 0.256550005333505 | 1.8921016378853 | 1.59175110732299 | 0.118852410435681 | 0.414402328603533 | -4.86075992120012 |
| lnc-SSTR3-2 | lnc-SSTR3-2 | -0.394819707384426 | 5.05808603035639 | -1.59188466396851 | 0.118866605843427 | 0.414402328603533 | -5.16014531658407 |
| ENSG00000187239.16 | FNBP1 | 0.29513863514169 | 7.00418588221985 | 1.59123140684794 | 0.118969450345947 | 0.414616402240249 | -5.1829037581121 |
| lnc-CSNK2A1-1 | lnc-CSNK2A1-1 | -0.256531445209456 | 0.968761089509563 | -1.59114993137523 | 0.118987807629594 | 0.414616402240249 | -4.75029869062348 |
| lnc-HYAL4-5 | lnc-HYAL4-5 | -0.359258035341335 | -0.0965001453404915 | -1.59114711896343 | 0.118988441336693 | 0.414616402240249 | -4.58164018446856 |
| ENSG00000215210.3 | RBMXP2 | 0.854918031491138 | -0.946649523755325 | 1.59079260731824 | 0.119068343663357 | 0.414824596754041 | -4.33487698974437 |
| ENSG00000217130.1 | RPS10P13 | 1.00445304664612 | -0.893171039145917 | 1.59062319649904 | 0.119106542038919 | 0.414878616543657 | -4.33338353041228 |
| ENSG00000112312.9 | GMNN | 0.462349195568867 | 0.824272181312191 | 1.59050748590837 | 0.119132637921954 | 0.414878616543657 | -4.71993656290977 |
| ENSG00000117906.13 | RCN2 | 0.247134862908351 | 4.71862805421675 | 1.59034783595168 | 0.119168650943684 | 0.414878616543657 | -5.14560900588542 |

| | | | | | | | |
|---|---|---|---|---|---|---|---|
| ENSG00000213089.4 | PDCL3P5 | -0.427833928662905 | 2.0627823536232 | -1.59028219292188 | 0.119183460925893 | 0.414878616543657 | -4.87958392799402 |
| ENSG00000154358.20 | OBSCN | 0.38913227617682 | 5.86487026912373 | 1.59049278263507 | 0.119184630987837 | 0.414878616543657 | -5.18349091091879 |
| ENSG00000124172.9 | ATP5F1E | 0.248382435260734 | 6.50427483960269 | 1.58995485064056 | 0.119257336202708 | 0.414935067416316 | -5.18816977302749 |
| lnc-ROCK1-4 | lnc-ROCK1-4 | 0.174429538271825 | 2.73584767454917 | 1.58991012143314 | 0.119267433666892 | 0.414935067416316 | -4.94656174759521 |
| ENSG00000186480.12 | INSIG1 | -0.2971242076565 | 6.08133241227346 | -1.5898559586661 | 0.119279661653246 | 0.414935067416316 | -5.18735575144007 |
| ENSG00000183621.15 | ZNF438 | -0.247223650262276 | 2.12276219461563 | -1.58984788525199 | 0.119281484424153 | 0.414935067416316 | -4.87712996796103 |
| ENSG00000102886.14 | GDPD3 | -0.349471212516724 | 1.30075033039035 | -1.58942606507702 | 0.119376752102275 | 0.415118763445869 | -4.78450546701422 |
| ENSG00000100288.19 | CHKB | -0.153030804212836 | 5.2579871664719 | -1.5893882832465 | 0.119385288103757 | 0.415118763445869 | -5.17233272495198 |
| lnc-XRCC2-8 | lnc-XRCC2-8 | 0.256547288493627 | 2.82981407503906 | 1.58934620361149 | 0.119394795685738 | 0.415118763445869 | -4.95555507050055 |
| lnc-L2HGDH-1 | lnc-L2HGDH-1 | -0.43438548027531 | 1.4472412468561 | -1.58914387456943 | 0.119440519013529 | 0.41516095466444 | -4.78071875667107 |
| lnc-THYN1-2 | lnc-THYN1-2 | 0.287781807376415 | 0.0885448000352624 | 1.58911400304066 | 0.119447270734238 | 0.41516095466444 | -4.5812302549217 |
| ENSG00000166024.13 | R3HCC1L | 0.158906251271691 | 5.26374250627387 | 1.58866786957967 | 0.119548145027386 | 0.415441409606397 | -5.16967368314928 |
| ENSG00000248483.6 | POU5F2 | -0.366460785659224 | 3.52330493432882 | -1.58845494362813 | 0.119596313621452 | 0.415508243840329 | -5.04162839434557 |
| ENSG00000147324.10 | MFHAS1 | -0.534459341023343 | 4.91887440519604 | -1.58918004859431 | 0.119607751287662 | 0.415508243840329 | -5.17107685123738 |
| ENSG00000164406.7 | LEAP2 | -0.331576197559547 | 1.07269971594845 | -1.58785950716529 | 0.119731098194684 | 0.415866553179411 | -4.76003372133804 |
| ENSG00000059804.15 | SLC2A3 | 1.19213426070922 | 3.40754457602518 | 1.58903874123738 | 0.119758569133971 | 0.415891788170156 | -5.0232084991754 |
| ENSG00000168872.16 | DDX19A | 0.125885146244447 | 4.63072389325542 | 1.58762645299593 | 0.11978388649207 | 0.415909537055714 | -5.13854464045837 |
| lnc-DYNLL2-1 | lnc-DYNLL2-1 | -0.273186398884487 | 2.19848197788347 | -1.58697446908874 | 0.119931665401459 | 0.416296011912608 | -4.91110294598544 |
| LINC01355 | LINC01355 | -0.368637328742963 | 3.27197063451704 | -1.58695692707046 | 0.119935643523382 | 0.416296011912608 | -5.02147486647505 |
| lnc-ZNF121-3 | lnc-ZNF121-3 | -0.28513124661821 | 0.426131158935396 | -1.58674875179984 | 0.119982861005715 | 0.416316042046882 | -4.65370034900171 |
| ENSG00000257921.5 | | 0.247381980488284 | 0.882446611474083 | 1.58662241864085 | 0.120011522736781 | 0.416316042046882 | -4.72973211136899 |
| lnc-SLC9A3-6 | lnc-SLC9A3-6 | -0.250896731134953 | 0.498310370292743 | -1.58658067338264 | 0.120020994878102 | 0.416316042046882 | -4.68661757036522 |
| ENSG00000134480.14 | CCNH | 0.228344753825815 | 5.96734290104349 | 1.58657483764879 | 0.120022319074294 | 0.416316042046882 | -5.19154969701626 |
| ENSG00000116750.13 | UCHL5 | -0.16933020183373 | 3.89133565283903 | -1.58638610404326 | 0.120065151322188 | 0.416389217703995 | -5.08291263569661 |
| lnc-PHB2-3 | lnc-PHB2-3 | 0.395135504773131 | -0.0932248737734375 | 1.58619242313115 | 0.120109119224691 | 0.416389217703995 | -4.53117926761644 |
| ENSG00000188895.11 | MSL1 | -0.168038701184858 | 6.37523758212113 | -1.5861679937982 | 0.120114665904942 | 0.416389217703995 | -5.19375420737828 |
| FSIP2-AS1 | FSIP2-AS1 | -0.464274216812407 | 0.127256138283402 | -1.58609584749005 | 0.120131047937254 | 0.416389217703995 | -4.61037235839045 |
| ENSG00000185019.16 | UBOX5 | 0.198107670358243 | 3.5450507777855 | 1.58603628661 | 0.12014457360163 | 0.416389217703995 | -5.0460169963 |
| lnc-LSR-1 | lnc-LSR-1 | -0.322715349247563 | 0.660753740367922 | -1.58594725567414 | 0.120164793915451 | 0.416389217703995 | -4.66727602290864 |
| ENSG00000145779.7 | TNFAIP8 | -0.252020947936621 | 7.62181667383587 | -1.58585626931783 | 0.120185461186477 | 0.416390733479425 | -5.18153478634198 |
| ENSG00000226121.2 | AHCTF1P1 | 0.828754119930472 | -0.342015012064825 | 1.58566453901682 | 0.120229021570108 | 0.416471550147683 | -4.40967708700111 |
| lnc-C9orf78-2 | lnc-C9orf78-2 | 0.274522068987949 | 2.93547937659552 | 1.58532530833682 | 0.120306124829012 | 0.416657097448918 | -4.97665708899408 |
| LINC00539 | LINC00539 | -0.519145797402794 | 0.50514902002619 | -1.58525077041319 | 0.120323071818314 | 0.416657097448918 | -4.63799256002393 |
| SEPT7-AS1 | SEPT7-AS1 | -0.174559140262855 | 3.59821835488002 | -1.58513021323696 | 0.120350485870322 | 0.416681926100729 | -5.04623930120643 |
| ENSG00000173598.13 | NUDT4 | 0.224739119171394 | 5.74537510439406 | 1.58434670117794 | 0.120528775699008 | 0.416926480538059 | -5.19242938560346 |
| ENSG00000179240.9 | GVQW3 | -0.293384219195407 | 1.42582600623811 | -1.58433893818668 | 0.120530543254616 | 0.416926480538059 | -4.79871883563419 |
| lnc-GRIP2-4 | lnc-GRIP2-4 | -0.34941187586432 | 1.31426224617156 | -1.58420053925767 | 0.120562058835413 | 0.416926480538059 | -4.77177157652195 |
| ENSG00000104131.12 | EIF3J | 0.177329365045184 | 5.01482117915952 | 1.58418572989166 | 0.120565431553095 | 0.416926480538059 | -5.16999116086233 |
| ENSG00000050344.8 | NFE2L3 | -0.404328637752188 | 2.23499800601443 | -1.58417605023297 | 0.120567636061245 | 0.416926480538059 | -4.92594744775967 |
| ENSG00000049883.14 | PTCD2 | -0.198682556958618 | 3.99728536527301 | -1.58407753007447 | 0.120590075538649 | 0.416926480538059 | -5.08562225469256 |
| lnc-WNT11-4 | lnc-WNT11-4 | 0.197478707200605 | 3.18877638744431 | 1.58406380109749 | 0.120593202792456 | 0.416926480538059 | -5.00538442890579 |
| ENSG00000005022.5 | SLC25A5 | 0.329378245397799 | 5.59030720220583 | 1.58404640750311 | 0.120597164885205 | 0.416926480538059 | -5.18981755356238 |
| ENSG00000250566.1 | UGT2B29P | 1.47875318136878 | -2.3669873424309 | 1.58389479227209 | 0.120631705838222 | 0.416926480538059 | -4.31376327762619 |
| lnc-SLC38A1-10 | lnc-SLC38A1-10 | 0.179494904966997 | 4.68424074938309 | 1.58379642820304 | 0.12065411940976 | 0.416926480538059 | -5.14735460175223 |
| ENSG00000205078.5 | SYCE1L | 0.305148466692278 | 0.0832508648979222 | 1.58375117188516 | 0.120664432801974 | 0.416926480538059 | -4.58293607411886 |
| ENSG00000213462.4 | ERV3-1 | -0.36050889717778 | 5.0280402010915 | -1.58371054258679 | 0.120688222533336 | 0.416926480538059 | -5.17168324707667 |
| lnc-NIPBL-3 | lnc-NIPBL-3 | -0.540087739257491 | 1.3364586550241 | -1.58357403081505 | 0.120712366886058 | 0.416926480538059 | -4.81245971536129 |
| ENSG00000105516.10 | DBP | -0.506247381130229 | 1.74033557761241 | -1.58345553683812 | 0.120733062011271 | 0.416926480538059 | -4.84460207690942 |
| ENSG00000129646.14 | QRICH2 | -0.245952762449318 | 2.13732490151073 | -1.58344637514933 | 0.120733911069291 | 0.416926480538059 | -4.90160424710187 |
| ENSG00000023445.13 | BIRC3 | 0.445238291609865 | 8.72418079309845 | 1.58371628845258 | 0.120766722556097 | 0.416926480538059 | -5.16365951733853 |
| ENSG00000183474.15 | GTF2H2C | -0.211830692304193 | 2.62496971242841 | -1.58330010633106 | 0.120767264485447 | 0.416926480538059 | -4.95176355050071 |
| lnc-HLA-DRB1-4 | lnc-HLA-DRB1-4 | -1.05404139282891 | 3.39608642402568 | -1.58446070729735 | 0.120785726252172 | 0.416926480538059 | -5.03412605500488 |
| ENSG00000124207.16 | CSE1L | 0.116379060673856 | 5.28689286358223 | 1.58293604653846 | 0.120850312808237 | 0.417079474938287 | -5.18223904418087 |
| lnc-WNT8A-4 | lnc-WNT8A-4 | -0.302941160961919 | 0.821357726763452 | -1.58276719451766 | 0.120888846574864 | 0.41714251953905 | -4.7298458483896 |
| ENSG00000014824.13 | SLC30A9 | -0.175136119682862 | 5.44728005239355 | -1.5825467018438 | 0.120939180257192 | 0.417246253307707 | -5.18870980403618 |
| ENSG00000055332.17 | EIF2AK2 | 0.292412705020407 | 6.06616349829023 | 1.58229589002884 | 0.120996455782066 | 0.41731057711413 | -5.1991697129393 |
| HYMAI | HYMAI | -0.353678939558228 | 1.2871261637969 | -1.58228749340508 | 0.120998373619838 | 0.41731057711413 | -4.8021437368386 |
| ENSG00000201558.1 | RNVU1-6 | -0.716953406205592 | 2.15629598014827 | -1.58278907373682 | 0.121049733097779 | 0.417417767859202 | -4.90152460837936 |
| ENSG00000204070.9 | SYS1 | 0.233393315132306 | 3.8611082613556 | 1.58181444306264 | 0.121106460858993 | 0.417428282299138 | -5.07684732242679 |
| ENSG00000174839.12 | DENND6A | 0.171339695932693 | 5.01520818773557 | 1.58178791839905 | 0.121112523792721 | 0.417428282299138 | -5.17358009583277 |
| lnc-DOCK7-7 | lnc-DOCK7-7 | -0.31425465772789 | 1.44109190456044 | -1.58172176348334 | 0.121127646370543 | 0.417428282299138 | -4.82313457822803 |
| ENSG00000108604.15 | SMARCD2 | 0.176791789389202 | 5.95338273733124 | 1.58157201229257 | 0.121161884156567 | 0.417428282299138 | -5.19921936546022 |
| ENSG00000182108.10 | DEXI | -0.338849058175206 | 2.06401891713366 | -1.58152196809684 | 0.121173327565837 | 0.417428282299138 | -4.89249153588898 |
| ENSG00000062370.16 | ZNF112 | -2.33197884776872 | -2.32812639177771 | -1.58198477172995 | 0.121174463719446 | 0.417428282299138 | -4.31711480298878 |
| lnc-OSBPL9-6 | lnc-OSBPL9-6 | -0.645944977093616 | -0.0602390583213461 | -1.58109429114478 | 0.121271158487902 | 0.41769147509312 | -4.52361772910773 |
| ENSG00000167515.10 | TRAPPC2L | -0.466760098787362 | 5.77254879255083 | -1.58151802949159 | 0.121309144660753 | 0.41770146278847 | -5.19497979357528 |
| ENSG00000176444.18 | CLK2 | -0.16768574985892 | 5.64923803700733 | -1.58084649233035 | 0.121327871629905 | 0.41770146278847 | -5.19588420035963 |
| lnc-ZNF785-2 | lnc-ZNF785-2 | -0.267325837501014 | 1.07000028375785 | -1.58066514041388 | 0.121369390836325 | 0.41770146278847 | -4.77408902092359 |

| | | | | | | | |
|---|---|---|---|---|---|---|---|
| lnc-CREB3L4-4 | lnc-CREB3L4-4 | -0.396613346592997 | -0.315186801978305 | -1.58061822036928 | 0.121380134713964 | 0.41770146278847 | -4.50770831283299 |
| ENSG00000171453.18 | POLR1C | 0.202454645929927 | 3.7633049671514 | 1.58046412165075 | 0.121415426071276 | 0.41770146278847 | -5.07699963904249 |
| ENSG00000170633.16 | RNF34 | 0.146519432193762 | 5.35514834150743 | 1.58038900760179 | 0.121432631541786 | 0.41770146278847 | -5.18826803747296 |
| ENSG00000185920.15 | PTCH1 | -2.10518657437999 | -2.32150590067424 | -1.58081340894676 | 0.121440104272195 | 0.41770146278847 | -4.31659060372568 |
| ENSG00000099904.15 | ZDHHC8 | -0.337119398904323 | 4.1308554593538 | -1.58030411663002 | 0.121452078871087 | 0.41770146278847 | -5.11311299723483 |
| lnc-ZNF534-1 | lnc-ZNF534-1 | -1.31390082646135 | -0.815165564949625 | -1.58077088491605 | 0.121456699936306 | 0.41770146278847 | -4.42074385300293 |
| ENSG00000284325.1 | MIR3655 | -0.216258954062103 | 2.07905890967783 | -1.58017246321063 | 0.121482243809473 | 0.41771951625967 | -4.89944829820922 |
| lnc-AMY1B-1 | lnc-AMY1B-1 | -0.503881417211456 | 0.985391542790836 | -1.5796858317792 | 0.12159379534313 | 0.417990342891729 | -4.74082676921732 |
| ENSG00000165917.9 | RAPSN | -0.565897379966882 | 0.0729023921066243 | -1.57957759764772 | 0.121618617350585 | 0.417990342891729 | -4.62636720425481 |
| ENSG00000196381.10 | ZNF781 | -0.534874919991202 | 1.67577918495435 | -1.57962563924509 | 0.121621928950035 | 0.417990342891729 | -4.83557441273 |
| lnc-NEIL3-6 | lnc-NEIL3-6 | -0.419084351958246 | 1.66807707351504 | -1.57934697299677 | 0.121671521624392 | 0.418090973221177 | -4.86327380809479 |
| ENSG00000108379.9 | WNT3 | -0.984033314257562 | 4.68531043960181 | -1.5803235476109 | 0.121741048299709 | 0.418260056276566 | -5.15782793887321 |
| ENSG00000171115.3 | GIMAP8 | -1.37808613422394 | -0.537685787682956 | -1.57947076539506 | 0.121803033560223 | 0.418403177531721 | -4.44637903436944 |
| lnc-MMADHC-2 | lnc-MMADHC-2 | -1.42077459242894 | -1.40217483327298 | -1.57885321300588 | 0.121854107354177 | 0.418508775516609 | -4.36319084725861 |
| lnc-TMTC3-13 | lnc-TMTC3-13 | -0.850658911903194 | -0.407856091547922 | -1.57853557466225 | 0.1219303477629 | 0.418648727395137 | -4.61245494949024 |
| ENSG00000049167.14 | ERCC8 | 0.15274395293872 | 3.42505717337031 | 1.57811534403131 | 0.12195436741626 | 0.418648727395137 | -5.04160374704182 |
| ENSG00000234534.1 | CSNK1G2P1 | -0.456918491186593 | 0.158571432564443 | -1.57810878677648 | 0.121955874724833 | 0.418648727395137 | -4.60968841115296 |
| lnc-JPH4-1 | lnc-JPH4-1 | -0.188115950375191 | 4.74412427722623 | -1.57794990849618 | 0.121992400500787 | 0.418699736679847 | -5.16121132452774 |
| ENSG00000116489.12 | CAPZA1 | 0.177669477467858 | 7.1010083523604 | 1.5778151078064 | 0.122023397833933 | 0.418699736679847 | -5.20181221402313 |
| ENSG00000166888.11 | STAT6 | -0.222788685060539 | 8.09372692387707 | -1.57774161082627 | 0.122040301103422 | 0.418699736679847 | -5.18480293018503 |
| ENSG00000151532.13 | VTI1A | 0.103342302238187 | 5.90383473455133 | 1.57769030334576 | 0.122052102224121 | 0.418699736679847 | -5.20419335187368 |
| ENSG00000214367.7 | HAUS3 | 0.220438307597615 | 4.19869101512546 | 1.57719267982145 | 0.122166607446704 | 0.418800583910897 | -5.12566510864102 |
| ENSG00000155980.11 | KIF5A | -0.911531520508902 | 0.444923645953108 | -1.57786626230916 | 0.122176374369761 | 0.418800583910897 | -4.70841377604875 |
| ENSG00000239039.1 | SNORD13 | -0.43904010307789 | 4.46420501845809 | -1.57744664923606 | 0.12218090539101 | 0.418800583910897 | -5.13968214705967 |
| ENSG00000205482.9 | SPDYE18 | -0.318437498696468 | -0.308062950468834 | -1.57710349426671 | 0.122187138595111 | 0.418800583910897 | -4.53597287693076 |
| ENSG00000167862.9 | MRPL58 | -0.22631402447462 | 2.33419111829686 | -1.57704839927442 | 0.122199823249194 | 0.418800583910897 | -4.92703198538827 |
| ENSG00000241962.9 | | 0.137903608301982 | 2.17122633855186 | 1.57703207876944 | 0.12220358096336 | 0.418800583910897 | -4.90321326506467 |
| lnc-PUS7L-1 | lnc-PUS7L-1 | -0.245667364993515 | 3.02366850318528 | -1.57675855115932 | 0.122266573238745 | 0.41894670833579 | -5.0016025310466 |
| ENSG00000197006.13 | METTL9 | 0.154012705169706 | 5.31545903895232 | 1.57648468685206 | 0.12232966937832 | 0.419039879552817 | -5.193784635804 |
| ENSG00000108219.14 | TSPAN14 | 0.525924146280589 | 3.74268572278186 | 1.57698408223111 | 0.122349090565826 | 0.419039879552817 | -5.09145416213105 |
| ENSG00000105372.7 | RPS19 | 0.292422645344779 | 8.83773551277849 | 1.5763754665288 | 0.122354840213401 | 0.419039879552817 | -5.17137554643757 |
| ENSG00000170846.17 | MRFAP1L2 | -0.284376751895874 | 2.23867260028396 | -1.57618525485932 | 0.122398686259604 | 0.419081274779277 | -4.92693886239685 |
| ENSG00000206910.1 | SNORA29 | 0.301079487163006 | -0.0104919843201756 | 1.5761065329841 | 0.122416836305195 | 0.419081274779277 | -4.57891282032673 |
| ENSG00000258724.1 | PINX1 | 0.193028038736244 | 0.610203050598618 | 1.57605807938919 | 0.122428008805703 | 0.419081274779277 | -4.68909100924757 |
| PSORS1C3 | PSORS1C3 | 1.15481133888901 | 0.828453902048143 | 1.57682933472107 | 0.122451184860518 | 0.419090910871971 | -4.63056698306605 |
| lnc-SLC36A1-5 | lnc-SLC36A1-5 | -0.21665887595271 | 5.23352943056111 | -1.57536484880659 | 0.122587945288436 | 0.419489223253844 | -5.19372843461482 |
| ENSG00000143748.17 | NVL | 0.142925121196144 | 4.75776936624574 | 1.57511329538914 | 0.122646023416231 | 0.419618201458825 | -5.16874991582997 |
| lnc-NIPSNAP3B-2 | lnc-NIPSNAP3B-2 | 0.638392276410065 | 2.00934832128839 | 1.57541513894894 | 0.122683456652822 | 0.419647697148119 | -4.88201070969271 |
| ENSG00000114650.19 | SCAP | -0.184430075986427 | 5.08092643092945 | -1.57469776177118 | 0.122742009705312 | 0.419647697148119 | -5.18721141074416 |
| ENSG00000163291.14 | PAQR3 | 0.372003347541807 | 2.90858022877529 | 1.57467466150057 | 0.122747347540501 | 0.419647697148119 | -4.99135708668101 |
| ENSG00000279277.1 | | -0.359337930456085 | 0.888474908665476 | -1.57451603665225 | 0.122784006438584 | 0.419647697148119 | -4.7517906392343 |
| ENSG00000185189.17 | NRBP2 | -0.297832540643966 | 4.02922401596512 | -1.57450957588462 | 0.122785499738002 | 0.419647697148119 | -5.10513533832778 |
| ENSG00000105519.15 | CAPS | -0.193490815901366 | 4.55605942064489 | -1.57443229315803 | 0.122803363499031 | 0.419647697148119 | -5.15848170524619 |
| ENSG00000198046.11 | ZNF667 | -2.01932236128341 | 0.65046531726012 | -1.57565158056325 | 0.122816112260583 | 0.419647697148119 | -4.71852356275562 |
| ENSG00000104635.13 | SLC39A14 | -0.35933968042205 | 3.10581597179891 | -1.57431612842455 | 0.122830218719783 | 0.419647697148119 | -5.01952075436034 |
| ENSG00000165406.15 | MARCHF8 | 0.145125385412549 | 6.18453397913953 | 1.57425583336176 | 0.122844159738799 | 0.419647697148119 | -5.21152750432341 |
| lnc-OST4-4 | lnc-OST4-4 | -0.325648177413333 | 0.730595708460314 | -1.57413582320671 | 0.12287191148884 | 0.419647697148119 | -4.7242876806039 |
| ENSG00000226253.1 | MRPL35P3 | -0.444242235171978 | -0.122618467514594 | -1.57401739977428 | 0.122899301287901 | 0.419647697148119 | -4.54755688015984 |
| lnc-KAT6B-3 | lnc-KAT6B-3 | -0.444242235171978 | -0.122618467514594 | -1.57401739977428 | 0.122899301287901 | 0.419647697148119 | -4.54755688015984 |
| ENSG00000103254.9 | ANTKMT | -0.356053719671886 | 1.53596500642786 | -1.57357781012693 | 0.123001015845271 | 0.419864225877105 | -4.86296470493809 |
| lnc-TTC7B-9 | lnc-TTC7B-9 | 0.223455806928683 | 0.959340604716015 | 1.57356702709552 | 0.123003511734876 | 0.419864225877105 | -4.74855208943439 |
| lnc-CFL2-4 | lnc-CFL2-4 | 0.18148914954723 | 1.76485839363669 | 1.57321111143592 | 0.123085916586754 | 0.420075844985103 | -4.86161002966516 |
| lnc-EZH2-7 | lnc-EZH2-7 | -0.369316294412861 | 0.52525996062141 | -1.57302180029434 | 0.123129765821012 | 0.420155830552702 | -4.69187868191957 |
| ENSG00000154813.9 | DPH3 | 0.258012075128899 | 3.08903267760787 | 1.57285907502443 | 0.123167467192191 | 0.420174385280316 | -5.01089449563274 |
| ENSG00000183617.4 | MRPL54 | 0.239562661226175 | 3.92802405408604 | 1.57265731797207 | 0.123214224691416 | 0.420174385280316 | -5.10613870899649 |
| ENSG00000148841.16 | ITPRIP | 0.318836746998481 | 3.79690162185577 | 1.57264876926964 | 0.123216206182937 | 0.420174385280316 | -5.08231043387067 |
| lnc-KYNU-14 | lnc-KYNU-14 | 0.377224027533974 | 2.85305790388936 | 1.57264595689441 | 0.123216858065004 | 0.420174385280316 | -4.98545925834279 |
| ENSG00000078596.10 | ITM2A | -0.524861031973919 | 3.67153371971397 | -1.57279168269877 | 0.123294608777114 | 0.420338158626834 | -5.05895507131825 |
| lnc-ING1-4 | lnc-ING1-4 | -0.348179999247813 | 0.0157538864767909 | -1.57226266280551 | 0.123305728114892 | 0.420338158626834 | -4.6087316292956 |
| ENSG00000138496.16 | PARP9 | 0.305712226309269 | 5.40209305711827 | 1.57192279724442 | 0.123384572228772 | 0.420537282693297 | -5.20554991037679 |
| lnc-CHAF1B-6 | lnc-CHAF1B-6 | -0.286602777140042 | 2.48471764971438 | -1.57173530538392 | 0.123428085191742 | 0.420615939983712 | -4.9463037924356 |
| ENSG00000120158.11 | RCL1 | 0.18248357150426 | 4.16967242374906 | 1.57159587903484 | 0.1234604511912 | 0.420656591105524 | -5.1255956499512 |
| ENSG00000180346.3 | TIGD2 | -0.244035345338689 | 2.80662051449365 | -1.57140627558691 | 0.123504476155698 | 0.420677709735897 | -4.98908092508631 |
| ENSG00000130023.15 | ERMARD | -0.139502914539904 | 5.15560778978683 | -1.57139314438358 | 0.12350752562474 | 0.420677709735897 | -5.19516183192016 |
| ENSG00000256825.4 | | 0.233346946380984 | 3.08303687023947 | 1.57127222863947 | 0.123535608838173 | 0.420690268949412 | -5.00689166353031 |
| ENSG00000137814.10 | HAUS2 | 0.206757113478389 | 4.26759337997418 | 1.57119742959845 | 0.123552983828598 | 0.420690268949412 | -5.13721054059647 |
| ENSG00000069345.11 | DNAJA2 | 0.1352565826672 | 6.05173663769804 | 1.57104619869761 | 0.12358811912983 | 0.420690268949412 | -5.2154711488838 |
| ENSG00000223705.9 | NSUN5P1 | -0.226569204375315 | 4.89499544946183 | -1.57102533154545 | 0.123592967805329 | 0.420690268949412 | -5.17958763203737 |

| | | | | | | | |
|---|---|---|---|---|---|---|---|
| lnc-DIRAS1-3 | lnc-DIRAS1-3 | 0.875762866608001 | 0.486379573741204 | 1.57139094962583 | 0.123619996955863 | 0.420712697973301 | -4.65683056599713 |
| ENSG00000265817.2 | FSBP | -0.367735403718334 | 0.730997702013323 | -1.57040360564613 | 0.123737502113341 | 0.421001738676904 | -4.72174565930075 |
| ENSG00000090054.14 | SPTLC1 | 0.159438273797399 | 5.06414662235244 | 1.57036777955351 | 0.123745834863493 | 0.421001738676904 | -5.1919086670645 |
| ENTPD3-AS1 | ENTPD3-AS1 | -0.348041362313662 | 2.55736015687791 | -1.57027670010691 | 0.123767020970567 | 0.421004229488873 | -4.95239549247421 |
| lnc-COPG1-3 | lnc-COPG1-3 | -0.273018289324852 | 1.80735035143227 | -1.56960788601185 | 0.123922684491988 | 0.421409339975187 | -4.88693874437148 |
| lnc-SLC25A19-1 | lnc-SLC25A19-1 | 0.374823665713266 | 3.84628663049717 | 1.56943227360676 | 0.123963583716646 | 0.421409339975187 | -5.0827026317336 |
| lnc-EIF2B1-1 | lnc-EIF2B1-1 | -0.266988469770811 | 2.18962457663064 | -1.56936129312715 | 0.123980117799816 | 0.421409339975187 | -4.91670782699859 |
| ENSG00000111962.7 | UST | 2.30345380835516 | -1.55005271088247 | 1.57021782874323 | 0.123998894667899 | 0.421409339975187 | -4.36212414638594 |
| ENSG00000102312.21 | PORCN | -0.230479420744213 | 2.54614149499998 | -1.5691521948053 | 0.124028835196795 | 0.421409339975187 | -4.96868123968558 |
| lnc-SIGLEC11-2 | lnc-SIGLEC11-2 | 0.151331057267758 | 2.52438514557504 | 1.56898887194794 | 0.124066898219021 | 0.421409339975187 | -4.95344821633287 |
| ENSG00000101400.5 | SNTA1 | 0.5832718947001 | -0.0240202522593605 | 1.56894334106734 | 0.124077511048666 | 0.421409339975187 | -4.59709091961237 |
| ENSG00000181830.8 | SLC35C1 | -0.304197569561704 | 3.00592785043038 | -1.56893981709241 | 0.124078332485405 | 0.421409339975187 | -5.01937618123125 |
| ENSG00000189190.10 | ZNF600 | 0.217093954705274 | 4.89959708743864 | 1.56891375822704 | 0.124084406929865 | 0.421409339975187 | -5.18398437743809 |
| ENSG00000126432.13 | PRDX5 | -0.275700048224196 | 4.44191275890987 | -1.56872018551435 | 0.124129537164544 | 0.421409339975187 | -5.1573880859045 |
| ENSG00000168439.16 | STIP1 | -0.209694649413406 | 5.09469168327156 | -1.56859910964672 | 0.124157771966524 | 0.421409339975187 | -5.19580246918398 |
| ENSG00000115649.15 | CNPPD1 | 0.291702068947826 | 5.91627106074227 | 1.56859534132367 | 0.124158650819904 | 0.421409339975187 | -5.21826933615652 |
| ENSG00000090857.13 | PDPR | 0.206957199575688 | 6.2736050954561 | 1.56828443283983 | 0.124231178631315 | 0.421409339975187 | -5.22051907785958 |
| LINC01144 | LINC01144 | -0.402001265309713 | -0.199555078101226 | -1.56814199851601 | 0.124264416719269 | 0.421409339975187 | -4.55882289396153 |
| ENSG00000188021.8 | UBQLN2 | 0.20407522814042 | 4.61253983076938 | 1.56795087006391 | 0.124309029224777 | 0.421409339975187 | -5.16282607866133 |
| ENSG00000253797.2 | UTP14C | 0.296147613787525 | 3.56722043172666 | 1.56792671579693 | 0.124314668146367 | 0.421409339975187 | -5.07756554368155 |
| lnc-CAAP1-2 | lnc-CAAP1-2 | 0.209846644256963 | 0.951357370602372 | 1.56780831664855 | 0.12434231194576 | 0.421409339975187 | -4.77301218547163 |
| ENSG00000180822.11 | PSMG4 | -0.234962818969344 | 3.34734853868868 | -1.56774001850081 | 0.124358260436071 | 0.421409339975187 | -5.05150451626975 |
| lnc-SLC25A6-4 | lnc-SLC25A6-4 | -0.263628870028977 | 3.52500668126299 | -1.56767033316723 | 0.124374534555061 | 0.421409339975187 | -5.07521634149847 |
| ENSG00000133028.11 | SCO1 | 0.161138240038616 | 4.96343899220942 | 1.5675641914831 | 0.124399325894485 | 0.421409339975187 | -5.19212375853239 |
| ENSG00000034510.5 | TMSB10 | -0.290477492328671 | 8.63329103508207 | -1.56751350769818 | 0.124411165431194 | 0.421409339975187 | -5.18685938952621 |
| ENSG00000171757.15 | LRRC34 | -0.694366049766079 | 2.18088947469529 | -1.56820368433853 | 0.124414760857117 | 0.421409339975187 | -4.92795551141126 |
| lnc-RHD-4 | lnc-RHD-4 | -0.403104321111152 | 1.28614293314889 | -1.56745205601142 | 0.12442552152902 | 0.421409339975187 | -4.8210843551823 |
| ENSG00000134709.10 | HOOK1 | -0.834231928947427 | 3.82262756535024 | -1.56859694528647 | 0.124428662612012 | 0.421409339975187 | -5.10953839029504 |
| ENSG00000134058.11 | CDK7 | 0.155198497884179 | 3.61530343467675 | 1.56736783107012 | 0.124445199997458 | 0.421409339975187 | -5.08092979118712 |
| ENSG00000148339.12 | SLC25A25 | -0.342726358974788 | 3.84339318455671 | -1.56727346507959 | 0.124467250825083 | 0.421409339975187 | -5.09991619892012 |
| ENSG00000166147.13 | FBN1 | -0.809817253282776 | 0.785921572849811 | -1.56795145433727 | 0.124468805988847 | 0.421409339975187 | -4.79084017894608 |
| ENSG00000204149.10 | AGAP6 | -0.180035489154291 | 3.56368014715787 | -1.56725173055252 | 0.124472330054376 | 0.421409339975187 | -5.07373810549808 |
| ENSG00000158457.5 | TSPAN33 | -0.331415932638933 | 4.70850851279753 | -1.5670920061157 | 0.124509661838295 | 0.421409339975187 | -5.18487258810794 |
| lnc-MFSD4B-5 | lnc-MFSD4B-5 | -0.349660725036125 | 0.741452167028222 | -1.56703056800397 | 0.124524023942054 | 0.421409339975187 | -4.72882526029127 |
| ENSG00000170540.14 | ARL6IP1 | 0.182145137191263 | 6.28468995341998 | 1.56692872365181 | 0.124547834573483 | 0.421409339975187 | -5.22254770487155 |
| lnc-ADAM19-2 | lnc-ADAM19-2 | 0.500951709790533 | 2.78770152505196 | 1.56706032534331 | 0.124560122107893 | 0.421409339975187 | -4.97026805518989 |
| ENSG00000244563.1 | RPS26P19 | 0.253859257698144 | 0.904688875361026 | 1.56686750115482 | 0.124562149815323 | 0.421409339975187 | -4.76281324156485 |
| ENSG00000138495.6 | COX17 | -0.164486704288789 | 3.16257008340918 | -1.56675710341113 | 0.124587966730639 | 0.421409339975187 | -5.0299038961612 |
| lnc-ATG3-3 | lnc-ATG3-3 | -0.330159328996682 | 4.37829209928405 | -1.56669413681158 | 0.124602693634989 | 0.421409339975187 | -5.14274771605827 |
| ENSG00000124786.11 | SLC35B3 | -0.179482859783401 | 4.03979754870281 | -1.56629029283528 | 0.124697179861274 | 0.421551662238134 | -5.12513271331825 |
| ENSG00000100304.12 | TTLL12 | -0.345560375738008 | 2.88079326554755 | -1.56619409363863 | 0.124719695844544 | 0.421551662238134 | -5.00919756751283 |
| ENSG00000165997.4 | ARL5B | 0.250329466186045 | 5.17657356984409 | 1.5661642352194 | 0.124726685049391 | 0.421551662238134 | -5.20477978829063 |
| ENSG00000137968.16 | SLC44A5 | -3.19298293991174 | -1.92318126573355 | -1.56737393928801 | 0.124726697907508 | 0.421551662238134 | -4.3712186058486 |
| ENSG00000108306.12 | FBXL20 | -0.143299581289355 | 6.26358826863925 | -1.5655654924111 | 0.124866904457962 | 0.421956246011859 | -5.22427748790266 |
| ENSG00000095303.15 | PTGS1 | 1.69610231313309 | -1.21400669509481 | 1.56580513537905 | 0.124887762162042 | 0.421957453805207 | -4.3427152929635 |
| ENSG00000169299.13 | PGM2 | -0.178247680354485 | 3.96345251767224 | -1.56531813073928 | 0.124924871159671 | 0.422013560328166 | -5.11867704556244 |
| lnc-ZNF705B-1 | lnc-ZNF705B-1 | -0.531869819560099 | 0.311934107306287 | -1.56504871435099 | 0.124988030877327 | 0.422157636945851 | -4.72223189338827 |
| lnc-VPS37C-3 | lnc-VPS37C-3 | -0.518779010313993 | 2.81188472058568 | -1.56514252888413 | 0.125025573139529 | 0.422215155361925 | -4.97272288301486 |
| lnc-OR4F21-1 | lnc-OR4F21-1 | -0.241754296707194 | 2.52779256134922 | -1.56479416896761 | 0.125047728044038 | 0.422220699857355 | -4.96204593393536 |
| lnc-TRMT1L-4 | lnc-TRMT1L-4 | 0.330644441177815 | 1.59448791493946 | 1.56451257199519 | 0.125113796290576 | 0.422374490577155 | -4.85845306651267 |
| ENSG00000131778.18 | CHD1L | 0.141136687212463 | 5.19122957861098 | 1.56424821759157 | 0.125175844728256 | 0.422453943666574 | -5.20693059069988 |
| ENSG00000141179.13 | PCTP | -0.336976250428975 | 1.57127703615823 | -1.56422041229767 | 0.12518237254324 | 0.422453943666574 | -4.86316968482897 |
| lnc-PLCXD3-2 | lnc-PLCXD3-2 | -0.244537087672016 | 0.821588077909529 | -1.56414999845236 | 0.125198904744989 | 0.422453943666574 | -4.75724605858212 |
| lnc-DNAH12-1 | lnc-DNAH12-1 | -0.312051280880173 | 0.695211960430571 | -1.56386036037736 | 0.125266926313391 | 0.422575498968049 | -4.7513149523069 |
| ENSG00000181790.11 | ADGRB1 | -0.4347441848325 | 1.85961998267992 | -1.56382177803128 | 0.125275989637227 | 0.422575498968049 | -4.88494516820527 |
| ENSG00000160877.5 | NACC1 | -0.264433180010912 | 3.52626428701234 | -1.56356738818389 | 0.125335761243417 | 0.422707844285312 | -5.08468099540189 |
| LCMT1-AS1 | LCMT1-AS1 | -0.284699108133135 | 0.489685055645027 | -1.56330163486414 | 0.125398227398384 | 0.422849232395305 | -4.70935449475251 |
| ENSG00000252464.1 | RN7SKP70 | 0.202362170857364 | 1.72606030165302 | 1.56310113973742 | 0.125445371035844 | 0.42284938975329 | -4.87800158849308 |
| ENSG00000116649.9 | SRM | -0.259336888902482 | 2.97339139868229 | -1.56302937283204 | 0.125462249500243 | 0.42284938975329 | -5.02311905998071 |
| ENSG00000002919.14 | SNX11 | 0.356373779411788 | 3.485931772269 | 1.56276454335474 | 0.125524549150623 | 0.42284938975329 | -5.07498812218432 |
| ENSG00000251669.5 | FAM86EP | -0.355630173117521 | 0.46813677262544 | -1.56272285830375 | 0.125534357598811 | 0.42284938975329 | -4.69992298773198 |
| lnc-MTRF1-5 | lnc-MTRF1-5 | -0.37618148931295 | 2.62954775786458 | -1.56271693878165 | 0.125535750506216 | 0.42284938975329 | -4.98842081645645 |
| ENSG00000119508.17 | NR4A3 | 1.74745982808025 | -0.572373433371167 | 1.5634063056126 | 0.125538352144212 | 0.42284938975329 | -4.38269734474872 |
| ENSG00000282936.1 | | -0.372293666811435 | -0.582852451712206 | -1.56269004306823 | 0.125542079423911 | 0.42284938975329 | -4.48947065549652 |
| ENSG00000110395.6 | CBL | -0.240085110731467 | 6.42315324318325 | -1.56246536921422 | 0.12559495821775 | 0.422898494986158 | -5.22866565415119 |
| ZNF433-AS1 | ZNF433-AS1 | 0.291669233410451 | 3.76046445127105 | 1.56225259032113 | 0.125645054005675 | 0.422898494986158 | -5.09300643194766 |
| LINC02062 | LINC02062 | -0.333267829140482 | 3.84537199277245 | -1.56225243596884 | 0.125645090351596 | 0.422898494986158 | -5.10814181498447 |
| ENSG00000108946.14 | PRKAR1A | -0.186909877835169 | 7.51982789758568 | -1.56223728742895 | 0.125648657477076 | 0.422898494986158 | -5.21783626283677 |

| | | | | | | | |
|---|---|---|---|---|---|---|---|
| ENSG00000073417.14 | PDE8A | -0.495068596478488 | 3.75022603867179 | -1.56256083899624 | 0.125659388589387 | 0.422898494986158 | -5.07409455982906 |
| lnc-HMGN1-5 | lnc-HMGN1-5 | 0.230677087068802 | 1.56092734020591 | 1.56208396122906 | 0.125684766796577 | 0.422914754777496 | -4.85376193722414 |
| lnc-SSH3-4 | lnc-SSH3-4 | -0.275146615273222 | 0.348573544494011 | -1.56196100847121 | 0.125713729021229 | 0.422943067087932 | -4.68340599531626 |
| lnc-AGBL2-1 | lnc-AGBL2-1 | 0.189020695386536 | 3.8822036673409 | 1.56174969381423 | 0.125763517975765 | 0.422953379989205 | -5.10842658518233 |
| ENSG00000118518.15 | RNF146 | -0.189364761626816 | 5.14133763073847 | -1.56173309931251 | 0.125767428566967 | 0.422953379989205 | -5.20975195923862 |
| ENSG00000167034.9 | NKX3-1 | 0.463288263895552 | -0.0938690679595968 | 1.56168637306156 | 0.125778440407809 | 0.422953379989205 | -4.58042977887136 |
| lnc-BCL7B-1 | lnc-BCL7B-1 | 0.238649567885225 | 2.53372938562856 | 1.56135143301922 | 0.12585739753567 | 0.423144131545432 | -4.96864670858198 |
| lnc-ZFHX3-27 | lnc-ZFHX3-27 | -0.369193565250889 | 0.213538072762068 | -1.56127134826881 | 0.125876282245187 | 0.423144131545432 | -4.6568585201017 |
| lnc-PPRC1-1 | lnc-PPRC1-1 | -0.353958079658942 | 0.913977932792312 | -1.56094418581898 | 0.125953453859013 | 0.423150816067314 | -4.77799152931863 |
| ENSG00000181264.8 | TLCD5 | -1.69927303880441 | -1.13019350762572 | -1.56159983324207 | 0.125980073795899 | 0.423150816067314 | -4.4086537062925 |
| ENSG00000255073.8 | ZFP91-CNTF | 0.161127781361217 | 4.6658732520536 | 1.5608057192956 | 0.125986127061356 | 0.423150816067314 | -5.18141973499113 |
| ENSG00000174917.8 | MICOS13 | -0.284031163694281 | 3.03184275218021 | -1.56078916373111 | 0.125990034046587 | 0.423150816067314 | -5.03160194788847 |
| lnc-LRP8-3 | lnc-LRP8-3 | -0.726112699994093 | 0.332556473163853 | -1.56082959061285 | 0.125995023591048 | 0.423150816067314 | -4.63366956119098 |
| ENSG00000189233.11 | NUGGC | 1.30530260112253 | 2.22369424590329 | 1.56182938696119 | 0.126027643355818 | 0.423150816067314 | -4.86463761872401 |
| lnc-FBXO11-1 | lnc-FBXO11-1 | -0.253994166444783 | 3.00174106981186 | -1.56056661024072 | 0.126042564437699 | 0.423150816067314 | -5.01433918305693 |
| ENSG00000131747.14 | TOP2A | 0.67037529591029 | 2.1461294030764 | 1.56106981630182 | 0.126065764229015 | 0.423150816067314 | -4.91185403498076 |
| lnc-CHST7-5 | lnc-CHST7-5 | 0.233600574392014 | -0.095158867593669 | 1.56040269823102 | 0.126081264688481 | 0.423150816067314 | -4.57536865542349 |
| ENSG00000145391.13 | SETD7 | -0.862688184059009 | 4.62631640753614 | -1.56158620484184 | 0.126083853419853 | 0.423150816067314 | -5.17240329594295 |
| lnc-PARD3-2 | lnc-PARD3-2 | 0.160198610615909 | 1.5602385245294 | 1.56002555252759 | 0.126170346623566 | 0.423372064648329 | -4.8530030295188 |
| lnc-PPIL4-9 | lnc-PPIL4-9 | -0.237131485983442 | 3.52402440066675 | -1.55983463181424 | 0.126215461505386 | 0.42345441632687 | -5.08719104486086 |
| ENSG00000125835.18 | SNRPB | 0.225247241999309 | 5.69867938132681 | 1.55971460603063 | 0.126243830457148 | 0.423480567519471 | -5.22862587157313 |
| ENSG00000186834.3 | HEXIM1 | 0.252844015714166 | 4.16615748479345 | 1.55955703424052 | 0.12628108148225 | 0.423516026940323 | -5.1540778805371 |
| lnc-SUCLG1-5 | lnc-SUCLG1-5 | -0.616307340759873 | 0.0757503472062594 | -1.55949582862298 | 0.126295553289593 | 0.423516026940323 | -4.59936130182132 |
| ENSG00000269556.7 | TMEM185A | -0.256026050501962 | 1.6896705432775 | -1.559404989213 | 0.126317034346238 | 0.423519061402285 | -4.87912629089233 |
| ENSG00000225091.3 | SNORA71A | 0.646922791933483 | 1.17655137936478 | 1.55941400209049 | 0.126362303396551 | 0.423601838894335 | -4.78174692244873 |
| ENSG00000150779.11 | TIMM8B | 0.40508911753915 | 1.46811117832072 | 1.5589373730325 | 0.12642765951341 | 0.423721448539185 | -4.83950795962518 |
| lnc-KANK4-7 | lnc-KANK4-7 | -0.147519419298413 | 3.01448523310448 | -1.5588313234009 | 0.126452758812458 | 0.423721448539185 | -5.02677295751283 |
| ENSG00000215912.12 | TTC34 | -0.529073843654323 | 0.556159564607983 | -1.55879724139823 | 0.126460826024264 | 0.423721448539185 | -4.69589748673904 |
| ENSG00000054219.10 | LY75 | -0.154010482339483 | 6.06615207639929 | -1.5585455713086 | 0.126520409193747 | 0.423721448539185 | -5.23389237597459 |
| ENSG00000015568.12 | RGPD5 | 0.973105463592734 | -0.833273308712346 | 1.55850482837265 | 0.126530057256842 | 0.423721448539185 | -4.38925934419072 |
| lnc-RPL13A-1 | lnc-RPL13A-1 | 0.25001409851409 | 6.54464463556603 | 1.55847114350992 | 0.126538034392593 | 0.423721448539185 | -5.23470729526976 |
| ENSG00000151876.12 | FBXO4 | -0.19256614520489 | 3.41369902129551 | -1.55832283439537 | 0.126573161280408 | 0.423721448539185 | -5.07544633463671 |
| ENSG00000136250.11 | AOAH | -0.773325984548529 | 1.20905327737356 | -1.55873120764825 | 0.126581048229206 | 0.423721448539185 | -4.74225452267606 |
| ENSG00000142065.13 | ZFP14 | -0.203546243403509 | 4.48114183579151 | -1.55818277382444 | 0.126606341725068 | 0.423721448539185 | -5.17031763234112 |
| HIPK1-AS1 | HIPK1-AS1 | -0.421046246613145 | 0.594316426633963 | -1.55816058750958 | 0.12661159832148 | 0.423721448539185 | -4.72070261169891 |
| ENSG00000035403.17 | VCL | -0.387214802661431 | 5.22568526736647 | -1.5583055338089 | 0.126624429381748 | 0.423721448539185 | -5.21381055570793 |
| ENSG00000255292.8 | | 0.340790994617313 | 2.26610693625333 | 1.55729035296332 | 0.126817921547688 | 0.424247584636293 | -4.93741089929049 |
| ENSG00000178096.8 | BOLA1 | -0.415751050370268 | 0.83029626546387 | -1.55719426334798 | 0.126840719963916 | 0.424247584636293 | -4.77186793480105 |
| lnc-VWA8-10 | lnc-VWA8-10 | -0.469876713176164 | -0.593892096904594 | -1.55718257266606 | 0.126843493944116 | 0.424247584636293 | -4.51102683235828 |
| lnc-CFAP206-1 | lnc-CFAP206-1 | -0.402543632979899 | 0.167663727444518 | -1.55707831909336 | 0.126868233529729 | 0.424261389235162 | -4.64680744967197 |
| lnc-TTC38-7 | lnc-TTC38-7 | 0.305301975070982 | -0.272644888778132 | 1.55679626294155 | 0.126935185525039 | 0.424416329379773 | -4.56633468373189 |
| ENSG00000059915.16 | PSD | -0.491819520456447 | -0.168923232208869 | -1.55633315674599 | 0.12704517533673 | 0.424672140831775 | -4.58027976131533 |
| ENSG00000089094.18 | KDM2B | -0.253666450243867 | 6.17557156292181 | -1.55619071071002 | 0.127079022354476 | 0.424672140831775 | -5.23739155429815 |
| PAXBP1-AS1 | PAXBP1-AS1 | -0.3560011891581 | 0.727042230572808 | -1.55615791945621 | 0.127086815008707 | 0.424672140831775 | -4.73814268892272 |
| lnc-EEF1E1-2 | lnc-EEF1E1-2 | -0.55511531846355 | 4.55003753931462 | -1.55682591014744 | 0.127109981511627 | 0.424672140831775 | -5.1842923992365 |
| lnc-LIMK2-5 | lnc-LIMK2-5 | -0.291541520496959 | 0.687014350646935 | -1.55601907814143 | 0.127119814121445 | 0.424672140831775 | -4.73538624945745 |
| ENSG00000143398.19 | PIP5K1A | 0.136459089403422 | 5.49325848293098 | 1.5559182636658 | 0.127143779531004 | 0.424672140831775 | -5.22761826287439 |
| ENSG00000173905.8 | GOLIM4 | -0.251430736091134 | 3.59453996678417 | -1.55585390016871 | 0.127159081793055 | 0.424672140831775 | -5.09750629629438 |
| lnc-CDK2AP1-4 | lnc-CDK2AP1-4 | 0.2343750208834 | 0.621282303545724 | 1.55577958663216 | 0.127176751498181 | 0.424672140831775 | -4.7248129292392 |
| lnc-SLFN12-1 | lnc-SLFN12-1 | 0.433360891292635 | -0.312212697021377 | 1.55559694976871 | 0.127220185902851 | 0.424748270306307 | -4.52055759415791 |
| ENSG00000249471.7 | ZNF324B | -0.300130171088446 | 1.71960352170754 | -1.55539528059937 | 0.127268160424723 | 0.4247773099254 | -4.88575699705482 |
| ENSG00000102984.14 | ZNF821 | -0.190073971063623 | 3.58244339568876 | -1.55538688310228 | 0.12727015839819 | 0.4247773099254 | -5.0968018472551 |
| ENSG00000177051.5 | FBXO46 | -0.254092155368693 | 3.47052834544051 | -1.55515895536733 | 0.127324397729262 | 0.424886331335232 | -5.08599457270314 |
| ENSG00000183199.6 | HSP90AB3P | 0.78824219535283 | 0.871572368243891 | 1.55534025981918 | 0.127344108147843 | 0.424886331335232 | -4.66511506863295 |
| ENSG00000197562.9 | RAB40C | -0.270966224910466 | 2.76474558589793 | -1.55417502436102 | 0.127558754818739 | 0.425460149429917 | -5.01072011056704 |
| ENSG00000163590.13 | PPM1L | -1.58219928688652 | -1.6845645209657 | -1.55439700039514 | 0.127607847583332 | 0.425460149429917 | -4.37671334324825 |
| ENSG00000152464.14 | RPP38 | 0.194864682881716 | 2.64366743340976 | 1.55393342038274 | 0.127616354239375 | 0.425460149429917 | -4.98778298533679 |
| lnc-TLDC1-2 | lnc-TLDC1-2 | 0.389399608516325 | 5.04438685591223 | 1.55406922016038 | 0.127628139456693 | 0.425460149429917 | -5.20385218337629 |
| ENSG00000269028.3 | NA | 0.506958263515333 | 4.26562360172454 | 1.55440205315866 | 0.127640111875321 | 0.425460149429917 | -5.14659360457395 |
| lnc-MTRNR2L12-3 | lnc-MTRNR2L12-3 | 0.506958263515333 | 4.26562360172454 | 1.55440205315866 | 0.127640111875321 | 0.425460149429917 | -5.14659360457395 |
| ENSG00000126773.12 | PCNX4 | -0.16369938604075 | 5.37757608573583 | -1.55355767166594 | 0.12770597596288 | 0.425610768012299 | -5.2289334315773 |
| lnc-DR1-3 | lnc-DR1-3 | -0.324307928876212 | 0.96722899593489 | -1.55317156410396 | 0.127798121226425 | 0.425848911964303 | -4.78680425935367 |
| ENSG00000185104.19 | FAF1 | 0.196437060118555 | 5.46072047482366 | 1.55283719254322 | 0.127877962856404 | 0.425941171376586 | -5.22994216520347 |
| ENSG00000109501.13 | WFS1 | -0.643679891465739 | -0.861064924666251 | -1.5527706818847 | 0.127893849137385 | 0.425941171376586 | -4.50000032831413 |
| lnc-SRPK2-1 | lnc-SRPK2-1 | -0.272399745067651 | 6.04350926517905 | -1.55276773277117 | 0.127894553579266 | 0.425941171376586 | -5.24150828936046 |
| lnc-GJE1-4 | lnc-GJE1-4 | 0.305969251031423 | 0.928398880231556 | 1.55270899863753 | 0.127908583796273 | 0.425941171376586 | -4.77476481290884 |
| lnc-SCRN1-1 | lnc-SCRN1-1 | -0.98819544149154 | 0.516073185430372 | -1.55321934287212 | 0.12799477354666 | 0.42606085930937 | -4.78198065445575 |
| ENSG00000179526.16 | SHARPIN | -0.197486409171517 | 3.72080992760673 | -1.55231130934258 | 0.128003615119118 | 0.42606085930937 | -5.11632365245669 |

| | | | | | | | |
|---|---|---|---|---|---|---|---|
| lnc-TLDC1-3 | lnc-TLDC1-3 | 0.449199478831941 | 1.95458848651236 | 1.55229871911603 | 0.128006624591612 | 0.42606085930937 | -4.89592606790582 |
| lnc-ZNF563-1 | lnc-ZNF563-1 | -0.239087889709064 | 1.36085347434734 | -1.55199626901962 | 0.128078937098626 | 0.42618991946459 | -4.84834084690944 |
| lnc-MAN2A1-7 | lnc-MAN2A1-7 | -0.495724182562712 | -0.550708360580569 | -1.55196334216401 | 0.128086811534177 | 0.42618991946459 | -4.50717242795378 |
| ENSG00000125656.9 | CLPP | -0.277794547325822 | 2.77560407734813 | -1.55180530551513 | 0.128124611318519 | 0.426246787581876 | -5.01647559972887 |
| lnc-GRK3-4 | lnc-GRK3-4 | -0.28984954804919 | 0.337651809391742 | -1.55168733234833 | 0.128152834426582 | 0.426271782644205 | -4.67536229485312 |
| lnc-GINS2-4 | lnc-GINS2-4 | -0.471948637680022 | 1.8922422240515 | -1.55136561500588 | 0.128229825358924 | 0.4264589586949 | -4.91695673033013 |
| ENSG00000132388.12 | UBE2G1 | 0.234221414023108 | 6.57409279708171 | 1.55083396883215 | 0.128357136457617 | 0.426813398983381 | -5.24535482221283 |
| lnc-C8orf89-4 | lnc-C8orf89-4 | 0.259219644828816 | 6.9635565565574 | 1.55065514138774 | 0.128399982382415 | 0.426842883949211 | -5.24321249896406 |
| lnc-TXK-1 | lnc-TXK-1 | -0.284713357027173 | 0.213058218234536 | -1.55062385802239 | 0.128407478861853 | 0.426842883949211 | -4.66613003958197 |
| ENSG00000283881.1 | MIR3916 | -0.194526945941588 | 1.8698711939534 | -1.55040726835771 | 0.128459390229056 | 0.426912707689419 | -4.92210822091951 |
| ENSG00000124201.14 | ZNFX1 | 0.195473455831144 | 6.88770112757137 | 1.55035213786427 | 0.128472606387545 | 0.426912707689419 | -5.2450394949369 |
| ENSG00000105501.12 | SIGLEC5 | 0.919859457427904 | -0.32254445476309 | 1.55047379676056 | 0.128499112581057 | 0.426912707689419 | -4.56501927776487 |
| ENSG00000128789.20 | PSMG2 | 0.205050792765323 | 4.49846744661483 | 1.55019013804706 | 0.128511448129215 | 0.426912707689419 | -5.18730822190551 |
| lnc-KHDRBS1-1 | lnc-KHDRBS1-1 | 0.252482135437826 | 1.47182603932497 | 1.54995233103845 | 0.128568482833963 | 0.426969862209471 | -4.86691261483688 |
| ENSG00000047648.21 | ARHGAP6 | -1.46542823674015 | -0.249551941665251 | -1.55084258738806 | 0.128573167423346 | 0.426969862209471 | -4.56958013023538 |
| ENSG00000101773.18 | RBBP8 | 0.260583895947597 | 4.10976399002333 | 1.54985895016179 | 0.128590884508406 | 0.426969862209471 | -5.15152167047907 |
| ENSG00000171735.18 | CAMTA1 | -0.17750448191365 | 3.26083300426926 | -1.54949936208944 | 0.128677177493996 | 0.427187474896599 | -5.07121162543126 |
| ENSG00000285533.1 | RELA-DT | -0.176831949701944 | 0.831613465861651 | -1.54905801430575 | 0.128783154646946 | 0.427449863033403 | -4.79025565179734 |
| ENSG00000130150.11 | MOSPD2 | -0.4004020004542 | 3.48898440631872 | -1.54903405265381 | 0.128797748167573 | 0.427449863033403 | -5.09506230945514 |
| lnc-PAX1-5 | lnc-PAX1-5 | 0.400959043827172 | 1.56407928404989 | 1.54880348938135 | 0.128844303511162 | 0.427464507623184 | -4.88383502797405 |
| lnc-WIPF1-2 | lnc-WIPF1-2 | 0.400026647151121 | -0.271407650075883 | 1.54869882610764 | 0.128869455332049 | 0.427464507623184 | -4.55684396007454 |
| lnc-FDXACB1-3 | lnc-FDXACB1-3 | 0.361710393949952 | -0.0931089893043557 | 1.54867292841386 | 0.128875679463384 | 0.427464507623184 | -4.58889543523126 |
| ENSG00000169016.16 | E2F6 | 0.265675952955501 | 2.64791762400585 | 1.54858518358707 | 0.128896769445139 | 0.427464507623184 | -5.00299668462387 |
| ENSG00000236698.1 | EIF1AXP1 | 1.06974869631073 | -1.19163545602009 | 1.54853876996875 | 0.128907926352278 | 0.427464507623184 | -4.35919444455355 |
| ENSG00000138674.16 | SEC31A | 0.116875599214416 | 7.14497813907344 | 1.54846039504638 | 0.128926767882463 | 0.427464507623184 | -5.24495720788728 |
| ENSG00000140416.20 | TPM1 | -0.275749142047272 | 2.7175399843159 | -1.54836966926226 | 0.128948581357946 | 0.427467973923434 | -5.01290438570711 |
| lnc-NAA11-1 | lnc-NAA11-1 | 0.310803050625498 | 0.376677729255096 | 1.54800253792158 | 0.129036882179478 | 0.427669885928142 | -4.69332474579299 |
| lnc-NDUFA2-1 | lnc-NDUFA2-1 | 0.283002714345277 | 1.45265777521027 | 1.5479411158517 | 0.12905165989528 | 0.427669885928142 | -4.86287020715369 |
| ENSG00000181191.11 | PJA1 | -0.198959430311023 | 3.75613779378039 | -1.54785731937487 | 0.129071822930847 | 0.427669885928142 | -5.12232973829571 |
| lnc-XPO4-1 | lnc-XPO4-1 | 0.176927703360941 | 0.77941844654498 | 1.54769248202448 | 0.129111493346619 | 0.427732475060914 | -4.76065461692613 |
| lnc-DLL3-1 | lnc-DLL3-1 | -0.212670465054564 | 1.07584864042482 | -1.54749625623108 | 0.129158730617044 | 0.427754311966998 | -4.81840335067699 |
| ENSG00000163521.15 | GLB1L | -0.412307260884844 | -0.0990290137936207 | -1.54749244310819 | 0.129159648684589 | 0.427754311966998 | -4.64473948098386 |
| ENSG00000253767.2 | PCDHGA8 | -1.34662676910819 | -1.49103682492606 | -1.54765245090921 | 0.129265794253931 | 0.428036976050299 | -4.46879710306805 |
| lnc-PSMA1-2 | lnc-PSMA1-2 | 0.277583703132481 | 1.97104093392465 | 1.54624922116938 | 0.129459253294648 | 0.428542729331549 | -4.94312533679382 |
| ENSG00000171475.13 | WIPF2 | 0.128422865981307 | 5.14724806512151 | 1.54619975899029 | 0.129471184757515 | 0.428542729331549 | -5.23055064009949 |
| ENSG00000165916.8 | PSMC3 | 0.157966767034704 | 5.13731936015065 | 1.54601133974978 | 0.129516644093891 | 0.428542729331549 | -5.23076731306972 |
| RNASEH2B-AS1 | RNASEH2B-AS1 | -0.762271897709314 | -0.257230456552679 | -1.54594020564529 | 0.129533809739757 | 0.428542729331549 | -4.58339648746093 |
| ENSG00000171863.14 | RPS7 | 0.258816439355304 | 8.93957257497928 | 1.54590034421772 | 0.129543429654773 | 0.428542729331549 | -5.21361573055857 |
| ENSG00000168010.10 | ATG16L2 | 0.231853718440124 | 5.60199115421558 | 1.54584011374935 | 0.129557966400269 | 0.428542729331549 | -5.24473417394036 |
| lnc-NSMCE4A-4 | lnc-NSMCE4A-4 | 0.161049283163495 | 1.93406721268258 | 1.54581399048369 | 0.129564271711132 | 0.428542729331549 | -4.92774805597563 |
| ENSG00000115207.13 | GTF3C2 | 0.14995416025293 | 5.93943443815165 | 1.54529123268199 | 0.129690500406129 | 0.428782852981318 | -5.25209923316832 |
| ENSG00000210164.1 | MT-TG | 0.474915821315454 | -0.187378521412976 | 1.54524099465608 | 0.129702636431281 | 0.428782852981318 | -4.55276108207036 |
| lnc-RFX3-3 | lnc-RFX3-3 | -0.283482707930246 | 2.028352287037 | -1.54520729404692 | 0.129710778016025 | 0.428782852981318 | -4.93875037473167 |
| lnc-LRIG1-1 | lnc-LRIG1-1 | 0.641457842376092 | -0.644289639499763 | 1.54516830585342 | 0.129720197518081 | 0.428782852981318 | -4.43632703163015 |
| ENSG00000091073.19 | DTX2 | -0.219057021150051 | 3.09039378634687 | -1.54503683602599 | 0.12975196452888 | 0.428795566041258 | -5.05327470705112 |
| lnc-EVX2-1 | lnc-EVX2-1 | -0.333314752518966 | -0.0334698888571477 | -1.54492990119377 | 0.12977780776903 | 0.428795566041258 | -4.65963658655086 |
| lnc-HSD17B11-1 | lnc-HSD17B11-1 | 0.274948129212476 | 2.41236935355171 | 1.54486749133802 | 0.129792892445706 | 0.428795566041258 | -4.97460847637618 |
| lnc-FANCE-1 | lnc-FANCE-1 | -0.278970178283479 | 0.38791448442826 | -1.54478526821009 | 0.129812768210173 | 0.428795566041258 | -4.69877006512002 |
| ENSG00000266618.1 | MIR4742 | 0.343152642045338 | 0.527532947736671 | 1.54466043590833 | 0.129842948550747 | 0.428795566041258 | -4.71421046301673 |
| lnc-SIPA1-2 | lnc-SIPA1-2 | -0.179040568413322 | 0.821510698901839 | -1.54463524867106 | 0.129849038679258 | 0.428795566041258 | -4.79438809317985 |
| lnc-CCDC144A-1 | lnc-CCDC144A-1 | -1.64455176338776 | -2.5291386834659 | -1.5447388545339 | 0.129872647014336 | 0.428804731070913 | -4.35720762156776 |
| ENSG00000111229.15 | ARPC3 | 0.270933745281683 | 7.02000027781249 | 1.54409077580768 | 0.129980745215535 | 0.429049638518934 | -5.25247778114302 |
| lnc-TBXA2R-2 | lnc-TBXA2R-2 | -0.275154960610089 | 0.296404204517858 | -1.54405868122107 | 0.129988512160719 | 0.429049638518934 | -4.70271718508324 |
| lnc-SPATA31D4-12 | lnc-SPATA31D4-12 | -1.02466970644977 | 2.29989831297027 | -1.54491831109029 | 0.130050832426243 | 0.429186513360488 | -4.96742090689163 |
| ENSG00000075420.12 | FNDC3B | -0.350346825704823 | 2.57995413168007 | -1.54351106156823 | 0.130121094552713 | 0.429349099773164 | -5.00596260271978 |
| lnc-RSBN1-3 | lnc-RSBN1-3 | -0.34354626897372 | 0.418711671070184 | -1.54342550704797 | 0.130141817688615 | 0.429349099773164 | -4.73355180483756 |
| lnc-DDX1-3 | lnc-DDX1-3 | 0.272473306614493 | 1.41491552370926 | 1.54311391526763 | 0.130217314308889 | 0.429388449489819 | -4.86637109796685 |
| ENSG00000170631.14 | ZNF16 | 0.229532563638155 | 2.8653211152097 | 1.54309889652497 | 0.130220954139467 | 0.429388449489819 | -5.03203726837194 |
| lnc-PPP1R14B-2 | lnc-PPP1R14B-2 | -0.212081019383782 | 2.64841902202024 | -1.54298849017723 | 0.130247713909113 | 0.429388449489819 | -5.0133840678065 |
| lnc-NRP1-5 | lnc-NRP1-5 | -0.598380151175633 | -0.685030137137584 | -1.54298663012006 | 0.130248164778841 | 0.429388449489819 | -4.50238221819975 |
| ENSG00000211841.1 | TRAJ48 | -0.623680551584766 | -0.610517277237261 | -1.54294020572689 | 0.130259418255159 | 0.429388449489819 | -4.49701657149122 |
| lnc-DPH5-1 | lnc-DPH5-1 | 0.972284300963879 | -0.208506080284942 | 1.54302350125409 | 0.130278913038134 | 0.429388449489819 | -4.52060055972619 |
| ENSG00000061936.9 | SFSWAP | 0.21258058393925 | 6.78628155921671 | 1.54264957701352 | 0.130329885685856 | 0.42944837357116 | -5.25613796399774 |
| ENSG00000116663.10 | FBXO6 | 0.38131353011738 | 1.71617449984583 | 1.54261272670962 | 0.130338822800322 | 0.42944837357116 | -4.91109003161367 |
| ENSG00000163694.14 | RBM47 | 0.235060176352026 | 4.26664947838617 | 1.54199335658886 | 0.130489109154172 | 0.429823679120351 | -5.17903016260693 |
| ENSG00000276045.3 | ORAI1 | -0.368116165414928 | 3.75678577545391 | -1.54185403964859 | 0.13052293272637 | 0.429823679120351 | -5.13000841040556 |
| ENSG00000073536.17 | NLE1 | -0.221703593427016 | 3.1971646894809 | -1.54184251509292 | 0.130525730990392 | 0.429823679120351 | -5.08053165569021 |
| ENSG00000108352.12 | RAPGEFL1 | 0.361742225251021 | 0.798459203807961 | 1.54179915778303 | 0.130536258960371 | 0.429823679120351 | -4.78401889516426 |

| | | | | | | | |
|---|---|---|---|---|---|---|---|
| ENSG00000121940.15 | CLCC1 | -0.151715842925374 | 4.90880992290512 | -1.54152563846342 | 0.130602690321222 | 0.429871944792952 | -5.22438968080859 |
| ENSG00000008405.11 | CRY1 | -2.17559689497108 | 1.27133016930688 | -1.54266099819378 | 0.130621244497664 | 0.429871944792952 | -4.79957000584522 |
| ENSG00000181222.15 | POLR2A | 0.354387587414552 | 6.47545336775844 | 1.54154546123702 | 0.13062410527731 | 0.429871944792952 | -5.25964843542133 |
| lnc-UPK3B-5 | lnc-UPK3B-5 | -0.316836720616394 | -0.383282827776923 | -1.54139488705958 | 0.13063445633192 | 0.429871944792952 | -4.56202644484665 |
| LINC01590 | LINC01590 | -0.27706280753234 | 1.08607307313849 | -1.54120193447698 | 0.13068134543632 | 0.429948246537821 | -4.82082698880371 |
| ENSG00000167447.12 | SMG8 | 0.202451077686923 | 4.57010424100294 | 1.54103553008453 | 0.130721793962996 | 0.429948246537821 | -5.20476412080673 |
| lnc-ZNF284-1 | lnc-ZNF284-1 | -0.160639387646264 | 2.40976775851757 | -1.54100071301772 | 0.130730258347335 | 0.429948246537821 | -4.98399217281702 |
| ENSG00000123810.7 | B9D2 | 0.247477475428045 | 1.5549424170738 | 1.54095571763174 | 0.130741197837061 | 0.429948246537821 | -4.88456445001708 |
| lnc-MOK-3 | lnc-MOK-3 | -0.244305293621901 | -0.0044373981151183 | -1.54070004009476 | 0.130803373322291 | 0.430064983818345 | -4.64111822043516 |
| ENSG00000182307.13 | C8orf33 | 0.168718769792332 | 4.36728127843741 | 1.54063791533013 | 0.13081848436509 | 0.430064983818345 | -5.18605723800583 |
| ENSG00000187650.3 | VMAC | -0.18066767809176 | 3.89498756253403 | -1.54045700632587 | 0.130862496119293 | 0.430140970556279 | -5.14855934418978 |
| lnc-FAN1-12 | lnc-FAN1-12 | 0.298360944061967 | 2.88354258298675 | 1.54036972958918 | 0.130883733158089 | 0.43014208519766 | -5.03434404145168 |
| ENSG00000196074.12 | SYCP2 | -0.551783524289974 | 1.87144873396904 | -1.54044756092687 | 0.130941141883835 | 0.430210297390324 | -4.97944669927355 |
| ENSG00000132676.15 | DAP3 | 0.153274288915355 | 5.7018469860353 | 1.5401127039125 | 0.130946291267083 | 0.430210297390324 | -5.25663732296737 |
| ENSG00000112983.17 | BRD8 | -0.198596202331908 | 4.93784669077289 | -1.54002406427952 | 0.130967871052326 | 0.430212526311847 | -5.23149623912671 |
| ENSG00000164136.16 | IL15 | 1.52996730050417 | 1.33393953056595 | 1.54101172771183 | 0.131010286786361 | 0.430231004188054 | -4.80800164314475 |
| ENSG00000189241.6 | TSPYL1 | 0.169348137084522 | 6.9597635086432 | 1.53976336758808 | 0.131031355586676 | 0.430231004188054 | -5.25899568222516 |
| ENSG00000224660.1 | SH3BP5-AS1 | -0.24661384571865 | 4.40754869556087 | -1.53974182626615 | 0.131036602407776 | 0.430231004188054 | -5.19835350128138 |
| ENSG00000196305.17 | IARS1 | 0.12164084590335 | 6.65747458011358 | 1.53965765676248 | 0.131057105196477 | 0.430231004188054 | -5.26145647068007 |
| ENSG00000131400.7 | NAPSA | 0.552036143194622 | -0.279301696047189 | 1.53944100606867 | 0.131109890832029 | 0.430335653483599 | -4.52373477358205 |
| lnc-TFDP2-5 | lnc-TFDP2-5 | -0.282548208124195 | 2.29441264834749 | -1.53930538222372 | 0.131142943451891 | 0.430375511012481 | -4.98205284820626 |
| lnc-TBC1D28-7 | lnc-TBC1D28-7 | 2.03034862579356 | -1.56721209221589 | 1.53986544968788 | 0.131214437590873 | 0.430541490344802 | -4.40829893307006 |
| ENSG00000197070.13 | ARRDC1 | -0.198302970655682 | 4.13084445790909 | -1.53865013343165 | 0.13130272710329 | 0.430762517049254 | -5.18082809713128 |
| lnc-RGS9-8 | lnc-RGS9-8 | -0.425725721856524 | 0.00417518229309121 | -1.53831245246848 | 0.131385132250159 | 0.43093866848111 | -4.64209282264678 |
| ENSG00000159267.14 | HLCS | -0.211668269429275 | 4.52761922746227 | -1.53825853421524 | 0.131398293902125 | 0.43093866848111 | -5.20543149837446 |
| ENSG00000085998.13 | POMGNT1 | -0.134210375977848 | 4.73634516333371 | -1.53796751463942 | 0.131469351169247 | 0.431103019773236 | -5.22128146502949 |
| ENSG00000170881.4 | RNF139 | 0.222313295808958 | 5.85403626294794 | 1.53779430087807 | 0.131511658829728 | 0.431173060479817 | -5.26235032980226 |
| ENSG00000136114.16 | THSD1 | 0.353629583937303 | 0.927295375258147 | 1.53751849821547 | 0.131579046501091 | 0.431325292902047 | -4.79895638570465 |
| ENSG00000040341.17 | STAU2 | -0.137013736572661 | 4.79644908267055 | -1.53724967740578 | 0.131644754945195 | 0.431471973095057 | -5.22843343306696 |
| lnc-NPM3-1 | lnc-NPM3-1 | 0.170985230633532 | 4.35116832045031 | 1.53688312646291 | 0.131734394210261 | 0.431697028503391 | -5.1907737153543 |
| ENSG00000177030.16 | DEAF1 | 0.256080234513951 | 3.69658176089681 | 1.53677501085243 | 0.131760843009185 | 0.431697097500248 | -5.13219538862315 |
| lnc-DERL2-4 | lnc-DERL2-4 | 0.25184924204859 | 0.888536598596227 | 1.53663429008649 | 0.131795274534152 | 0.431697097500248 | -4.82006158476309 |
| ENSG00000116199.11 | FAM20B | 0.164398165194593 | 5.20217977160263 | 1.53662586709439 | 0.131797335698953 | 0.431697097500248 | -5.24473600720491 |
| lnc-RPLP0-4 | lnc-RPLP0-4 | -0.743263111542701 | -0.660105464734518 | -1.53636047593346 | 0.131862292012382 | 0.431841138662029 | -4.48143986635371 |
| ENSG00000186063.12 | AIDA | 0.19117579730987 | 5.25404221593955 | 1.53601836424688 | 0.131946064102288 | 0.432046744737097 | -5.24618633674722 |
| ENSG00000114416.17 | FXR1 | 0.136722198307753 | 6.73839449975097 | 1.5357078594986 | 0.132022133637963 | 0.432213898538192 | -5.26688526952345 |
| ENSG00000175054.14 | ATR | 0.139144042010063 | 6.41140575498832 | 1.53561339842108 | 0.132045282325517 | 0.432213898538192 | -5.2683024924443 |
| ENSG00000273173.5 | SNURF | -0.263129391037058 | 5.06879597814213 | -1.53555290642556 | 0.132060108240135 | 0.432213898538192 | -5.23976138773914 |
| lnc-ORC1-3 | lnc-ORC1-3 | 0.222274598629487 | 1.62116081704899 | 1.53497417637173 | 0.132202016051692 | 0.432604486464895 | -4.89784751543856 |
| ENSG00000144580.13 | CNOT9 | 0.181253015733621 | 5.62935667162231 | 1.53487044119476 | 0.132227465415182 | 0.432604486464895 | -5.26481694104871 |
| ENSG00000248333.8 | CDK11B | 0.146174729895162 | 4.57530374665304 | 1.53472835401481 | 0.132262330058091 | 0.432604486464895 | -5.21603491990734 |
| lnc-USE1-1 | lnc-USE1-1 | -0.332777652753384 | -0.0965870571355942 | -1.53472350483178 | 0.132263520056531 | 0.432604486464895 | -4.59575534811048 |
| ENSG00000135205.14 | CCDC146 | 0.282851466733755 | 3.09248688088932 | 1.53463343743376 | 0.132285624323041 | 0.432608040267105 | -5.05945986914108 |
| lnc-ZNF320-2 | lnc-ZNF320-2 | -0.329567421364514 | -0.202213465064578 | -1.53444268635098 | 0.13233244807992 | 0.432687710343272 | -4.5746539365409 |
| ENSG00000143368.9 | SF3B4 | -0.210310174842414 | 3.78192194425175 | -1.53436293120056 | 0.132352029554547 | 0.432687710343272 | -5.14799767748532 |
| ENSG00000185220.11 | PGBD2 | -0.307399625018983 | 2.9430097074681 | -1.53406885033169 | 0.132424252337953 | 0.432855071601093 | -5.04383718548482 |
| ENSG00000253341.1 | PCBP2P2 | 0.983749662608223 | -0.458805150646392 | 1.5338663535072 | 0.132474001537234 | 0.432948931984899 | -4.44610943280167 |
| ENSG00000196998.17 | WDR45 | 0.149742885889784 | 5.28807396593778 | 1.53342006058655 | 0.132583699219152 | 0.433181425906918 | -5.25458013823464 |
| ENSG00000099949.19 | LZTR1 | -0.218350591178827 | 4.59719280356229 | -1.53333710508442 | 0.132604097500215 | 0.433181425906918 | -5.22057815463892 |
| lnc-RHEBL1-1 | lnc-RHEBL1-1 | 0.402790947956049 | 0.673408994295864 | 1.5333201093727 | 0.132608276958631 | 0.433181425906918 | -4.74092377300223 |
| ENSG00000163743.13 | RCHY1 | 0.228242876672002 | 4.03744838471323 | 1.53315949062271 | 0.132647780338035 | 0.433233031389124 | -5.17519572280413 |
| FBXL19-AS1 | FBXL19-AS1 | -0.277437508170009 | 2.11953509872184 | -1.53302955264965 | 0.132679744838063 | 0.433233031389124 | -4.95656115865259 |
| lnc-PGC-1 | lnc-PGC-1 | -0.342501521726952 | 1.96428911710809 | -1.53299731815983 | 0.13268767541879 | 0.433233031389124 | -4.95935633058757 |
| lnc-FBXL7-10 | lnc-FBXL7-10 | -0.469790113911461 | -0.338982667306508 | -1.53276984348989 | 0.132743651318493 | 0.433233031389124 | -4.55726886338641 |
| ENSG00000132842.13 | AP3B1 | 0.258038314145693 | 6.65958805218665 | 1.53273517620158 | 0.132752183744642 | 0.433233031389124 | -5.2717711453476 |
| ENSG00000117724.12 | CENPF | 0.640569873560772 | 1.31310398326153 | 1.53297010349042 | 0.132762303250669 | 0.433233031389124 | -4.84608441516535 |
| ENSG00000006740.16 | ARHGAP44 | -0.580621599220458 | 6.13489282685446 | -1.53350204579023 | 0.132771411456182 | 0.433233031389124 | -5.26913962671976 |
| lnc-PTCH1-9 | lnc-PTCH1-9 | 0.321847833823631 | -0.0151205502434205 | 1.53225980545013 | 0.132869227909179 | 0.433483486773598 | -4.62804080812162 |
| ENSG00000240531.1 | RPL21P123 | 0.437680945706426 | 1.79672376863576 | 1.53204965795725 | 0.132920996072991 | 0.433570426702851 | -4.90862921403327 |
| ENSG00000136527.17 | TRA2B | 0.218189095631073 | 8.67491637172599 | 1.53198062511859 | 0.132938005291828 | 0.433570426702851 | -5.23906429933735 |
| ENSG00000183808.11 | RBM12B | -0.174689516566132 | 5.75060720104617 | -1.53173416076267 | 0.132998746658296 | 0.433574332575482 | -5.26883994021735 |
| ENSG00000130348.11 | QRSL1 | -0.227327637283796 | 5.90717563112382 | -1.53172249361662 | 0.133001622588581 | 0.433574332575482 | -5.27019043094165 |
| lnc-NBAS-8 | lnc-NBAS-8 | -0.498607832145013 | -0.418577804028133 | -1.53171935230885 | 0.133002396923752 | 0.433574332575482 | -4.53694880968609 |
| ENSG00000100949.14 | RABGGTA | -0.23334112158627 | 3.53620011140539 | -1.53163077635918 | 0.133024232462142 | 0.433576845093947 | -5.12365355169829 |
| lnc-AIG1-9 | lnc-AIG1-9 | 0.521272002535005 | 0.393566031919968 | 1.53140955053688 | 0.133078781092289 | 0.43362421564828 | -4.71952686550636 |
| lnc-SLC45A3-4 | lnc-SLC45A3-4 | 0.21334528793978 | 3.30284468955557 | 1.53140095753035 | 0.133080900269649 | 0.43362421564828 | -5.09792684089554 |
| lnc-MAN2A1-12 | lnc-MAN2A1-12 | -0.26770494251979 | 1.09766150061015 | -1.53119827496478 | 0.133130892982652 | 0.433718450500464 | -4.85159343786577 |
| ENSG00000267048.1 | | -0.164455539125629 | 1.82542512629525 | -1.53102488977683 | 0.133173671290179 | 0.433789155905326 | -4.93852658912991 |

| | | | | | | | |
|---|---|---|---|---|---|---|---|
| lnc-PCNX1-2 | lnc-PCNX1-2 | -0.249995349115667 | 0.538651213194603 | -1.53061380305236 | 0.133275140327914 | 0.434050983128079 | -4.73967203433962 |
| lnc-TMEM252-2 | lnc-TMEM252-2 | -0.289168370670753 | 0.163908030520015 | -1.53046334977291 | 0.133312292379507 | 0.434103292840909 | -4.67921140267533 |
| ENSG00000162236.11 | STX5 | 0.1658508605972 | 5.55555116358369 | 1.53013469217623 | 0.133393478030073 | 0.434251503880762 | -5.2670852118248 |
| lnc-WASF3-3 | lnc-WASF3-3 | -0.288188255990021 | 0.346240775589056 | -1.53010828513771 | 0.133400002868292 | 0.434251503880762 | -4.73238403142983 |
| ENSG00000100505.13 | TRIM9 | -0.434893247787085 | -0.18270688716786 | -1.52985154927129 | 0.133463452317256 | 0.434345626771571 | -4.62284549532603 |
| lnc-PJVK-2 | lnc-PJVK-2 | -0.585921213233262 | 2.91812600971958 | -1.53042737599743 | 0.133471121281164 | 0.434345626771571 | -5.06947773788719 |
| lnc-SLC25A29-1 | lnc-SLC25A29-1 | -0.303669190082497 | 0.609722127313591 | -1.52966346804477 | 0.133509949871666 | 0.43438084725293 | -4.75982807061794 |
| lnc-CDYL2-1 | lnc-CDYL2-1 | 0.600315897873697 | 1.09057146485494 | 1.52969981738232 | 0.13352415199773 | 0.43438084725293 | -4.81342478051403 |
| lnc-CNOT6-4 | lnc-CNOT6-4 | -0.292496459689916 | 0.804292814378789 | -1.52922976783532 | 0.133617218975447 | 0.434614920697161 | -4.78735828923296 |
| ENSG00000196659.9 | IFT70B | -0.340805464861655 | 2.05744810948214 | -1.52908830288749 | 0.133652223095135 | 0.434660089740428 | -4.97682434619561 |
| ENSG00000100075.9 | SLC25A1 | -0.234468235240104 | 4.11510218129534 | -1.52894998220595 | 0.133686456300339 | 0.434702737761435 | -5.19055382959052 |
| ENSG00000255730.5 | | -0.218143187100408 | 3.11374882347675 | -1.52841220661053 | 0.133819617990477 | 0.435067003174535 | -5.08596389395995 |
| ENSG00000071054.16 | MAP4K4 | -0.785438504725315 | 6.37798414112101 | -1.52935984012917 | 0.133866455975171 | 0.435150546957825 | -5.27515127077497 |
| ENSG00000172936.13 | MYD88 | 0.279195993559443 | 4.47993146272828 | 1.52801440877032 | 0.133918187339407 | 0.435249968420497 | -5.21246547110853 |
| ENSG00000213178.3 | RPL22P1 | 0.955013664267331 | 1.40553075314827 | 1.52844600015527 | 0.134001226286539 | 0.435270854729443 | -4.78131998270363 |
| lnc-DHRS13-2 | lnc-DHRS13-2 | -0.188618650058646 | 3.09089097910824 | -1.52763742914131 | 0.134011651857439 | 0.435270854729443 | -5.07412112816135 |
| lnc-FAM161A-1 | lnc-FAM161A-1 | -0.606184899021967 | 0.66861366182572 | -1.5276895315586 | 0.134013239539578 | 0.435270854729443 | -4.78887008932004 |
| ENSG00000273155.1 | | 0.154468866059926 | 1.81047303617723 | 1.52755392614457 | 0.134032361812722 | 0.435270854729443 | -4.93517203654672 |
| ENSG00000141084.10 | RANBP10 | -0.172406103672257 | 4.87455262925618 | -1.52750140142807 | 0.134045390018082 | 0.435270854729443 | -5.24764619224019 |
| lnc-DHX9-4 | lnc-DHX9-4 | 0.785380487804352 | -0.574899581424153 | 1.52745665266415 | 0.134056490280027 | 0.435270854729443 | -4.56063938003706 |
| ENSG00000235408.5 | SNORA71B | 0.300779460049364 | 0.188350384707447 | 1.52737625987586 | 0.134076434155261 | 0.435270854729443 | -4.6872972614736 |
| lnc-HSP90AA1-15 | lnc-HSP90AA1-15 | -0.370253501309005 | -0.071875208388066 | -1.52703668396743 | 0.134160702502016 | 0.435270854729443 | -4.61030501594727 |
| lnc-STARD10-1 | lnc-STARD10-1 | -0.33326878918251 | 0.660614036941623 | -1.52695071031578 | 0.134182044245471 | 0.435270854729443 | -4.7791122584361 |
| ENSG00000144120.12 | TMEM177 | -0.243532363946305 | 1.21657831125039 | -1.52692911313527 | 0.134187405867416 | 0.435270854729443 | -4.85586749921695 |
| lnc-NOC3L-5 | lnc-NOC3L-5 | -0.369329692849697 | -0.0916560528468024 | -1.52688219126494 | 0.134199055077808 | 0.435270854729443 | -4.61269829659701 |
| ENSG00000130713.15 | EXOSC2 | 0.126771661336986 | 4.99945535912823 | 1.52687580008383 | 0.134200641867791 | 0.435270854729443 | -5.25152601327679 |
| lnc-XRN2-10 | lnc-XRN2-10 | 0.269994579184343 | 2.40506778937813 | 1.52679617869704 | 0.134220411369999 | 0.435270854729443 | -5.00419642980491 |
| ENSG00000055208.18 | TAB2 | 0.213999743207378 | 6.67753681891386 | 1.5267951245385 | 0.134220673126744 | 0.435270854729443 | -5.28028692082487 |
| ENSG00000130311.10 | DDA1 | -0.178248319443452 | 4.18324584802449 | -1.52667801891884 | 0.134249754016978 | 0.435296579541818 | -5.19929409451215 |
| lnc-TMEM35B-5 | lnc-TMEM35B-5 | -0.242458430437801 | 2.11583548209922 | -1.5263100752991 | 0.134341158450555 | 0.435524344682277 | -4.98098687866166 |
| ENSG00000105426.16 | PTPRS | -1.31886352278655 | 1.94353986288165 | -1.52739119395844 | 0.134366593073019 | 0.435538202397156 | -4.9686726336876 |
| lnc-YWHAZ-5 | lnc-YWHAZ-5 | -0.393856083953807 | 0.0537365514750492 | -1.52595656647945 | 0.134429023925454 | 0.435671957086697 | -4.66120044227195 |
| ENSG00000172663.8 | TMEM134 | -0.166048037224662 | 3.99121863133901 | -1.52578015413477 | 0.134472888848402 | 0.435745508684926 | -5.18043649238654 |
| ENSG00000104972.15 | LILRB1 | -0.270951072595471 | 5.38068613427843 | -1.52558313865158 | 0.13452189028527 | 0.435835678851207 | -5.26819456359018 |
| ENSG00000148840.10 | PPRC1 | 0.213636476468838 | 4.89661176732736 | 1.52502398094055 | 0.134661041131959 | 0.436107632656527 | -5.24862424918943 |
| ENSG00000215388.3 | ACTG1P3 | 0.268355629373595 | 1.39456039029131 | 1.52495680578833 | 0.134677765953027 | 0.436107632656527 | -4.88689633499044 |
| ENSG00000188818.12 | ZDHHC11 | -0.44807300381602 | 0.00623620416285046 | -1.52492587292395 | 0.134685467970247 | 0.436107632656527 | -4.64473385012069 |
| ENSG00000125485.17 | DDX31 | 0.184107178072289 | 4.14686759958502 | 1.52490534028705 | 0.134690580614951 | 0.436107632656527 | -5.18969162430469 |
| ENSG00000129675.15 | ARHGEF6 | 0.408556161331496 | 5.84000565193625 | 1.52508783688242 | 0.134732151298904 | 0.43617361909175 | -5.27980487588176 |
| ENSG00000267303.1 | | -0.249482611997231 | 1.57209844649683 | -1.52457682554583 | 0.134772402203659 | 0.436192537529905 | -4.91706736256526 |
| ENSG00000160201.11 | U2AF1 | 1.29605335981355 | 2.31440523118595 | 1.52568881374198 | 0.13478943300385 | 0.436192537529905 | -5.02204701403164 |
| ENSG00000187824.8 | TMEM220 | 0.534284521777545 | 1.76376945862111 | 1.5246242275824 | 0.134801570773343 | 0.436192537529905 | -4.95143473539843 |
| lnc-TFB1M-5 | lnc-TFB1M-5 | -0.233081297935864 | 0.94108787097278 | -1.5241141137547 | 0.134887715081867 | 0.436402678329152 | -4.83292650739827 |
| lnc-IRF1-1 | lnc-IRF1-1 | 0.493912519761827 | 2.55682347676548 | 1.52404125322414 | 0.13494885627295 | 0.436531873120561 | -5.01975290375086 |
| ENSG00000150433.9 | TMEM218 | -0.224581190392143 | 2.44214285911959 | -1.52366078171586 | 0.135000766986263 | 0.43656797884379 | -5.01814202057311 |
| lnc-ZNF415-3 | lnc-ZNF415-3 | 0.236138080835745 | 1.90665777162001 | 1.52365408254328 | 0.13500243819369 | 0.43656797884379 | -4.94091261828587 |
| ENSG00000011566.14 | MAP4K3 | 1.33383780672242 | 1.6972412674396 | 1.52463510552019 | 0.135040235735984 | 0.436621610454566 | -4.85786430750041 |
| ENSG00000132478.9 | UNK | -0.158407211494698 | 4.97614251904262 | -1.52317128798918 | 0.135122922012989 | 0.436816705412838 | -5.2569231803432 |
| ENSG00000141639.11 | MAPK4 | -1.96726368010478 | -1.40057843448132 | -1.52376243828736 | 0.135149398674563 | 0.436816705412838 | -4.39576215310077 |
| ENSG00000226221.1 | RPL26P19 | 0.778915952314541 | 1.77227934256555 | 1.52358503029817 | 0.135185176460228 | 0.436816705412838 | -4.88618912675487 |
| lnc-CHUK-1 | lnc-CHUK-1 | 0.160471085741468 | 2.63237547497431 | 1.52285556685617 | 0.135201758334407 | 0.436816705412838 | -5.0284566291479 |
| ENSG00000266076.1 | | -0.772624086968327 | 1.8500688470763 | -1.52366803425834 | 0.135206686595015 | 0.436816705412838 | -5.0331246066669 |
| ENSG00000110031.12 | LPXN | -0.223533182408365 | 5.66214362515458 | -1.52256058672678 | 0.135275448828119 | 0.436933105052833 | -5.28025788888707 |
| ENSG00000102144.14 | PGK1 | 0.182840770714924 | 6.82675728221522 | 1.52252167811725 | 0.135285171185041 | 0.436933105052833 | -5.28509904120288 |
| ENSG00000120910.14 | PPP3CC | 0.194823730897749 | 6.43769486278912 | 1.52239500088063 | 0.135316828754249 | 0.436943845118298 | -5.28739597007833 |
| lnc-ZBED5-3 | lnc-ZBED5-3 | 0.130445114384136 | 5.89409423875165 | 1.5223384892285 | 0.135330953341551 | 0.436943845118298 | -5.28519420264929 |
| ENSG00000126261.12 | UBA2 | 0.119688865900163 | 6.59684063996911 | 1.52221803081175 | 0.135361064793337 | 0.436972521430561 | -5.28705996309832 |
| ENSG00000155959.10 | VBP1 | 0.194426147917969 | 3.96498942115688 | 1.52205865381551 | 0.135400913108293 | 0.437032616356907 | -5.18379452194293 |
| lnc-FNTB-4 | lnc-FNTB-4 | 0.942855414469363 | -0.196106173014884 | 1.52188061931642 | 0.135523748607329 | 0.437360507617537 | -4.60609423461064 |
| ENSG00000074695.5 | LMAN1 | 0.221348091842803 | 5.79357138964129 | 1.52141976793011 | 0.135560745092767 | 0.437411320935011 | -5.28538250089946 |
| ENSG00000142875.19 | PRKACB | 0.708048414704904 | 3.08046353142963 | 1.52193130923598 | 0.135640460019043 | 0.43756479526687 | -5.10206704976232 |
| ENSG00000128739.22 | SNRPN | -0.248606700894547 | 5.6046428682405 | -1.52105995751206 | 0.135650826342025 | 0.43756479526687 | -5.28028715348154 |
| ENSG00000285708.1 | | -0.190065763683329 | 7.42831098387791 | -1.5208126822572 | 0.135712761316816 | 0.437661532144646 | -5.28068304798722 |
| ENSG00000280388.1 | | -0.223413695408363 | 2.13451180999527 | -1.52077044573849 | 0.13572334254867 | 0.437661532144646 | -4.97421677667544 |
| ENSG00000184792.15 | OSBP2 | -1.10098732890767 | -1.28068255135791 | -1.52060830189542 | 0.135765008822404 | 0.437727314629002 | -4.43761962232451 |
| ENSG00000163539.16 | CLASP2 | 0.170239892515413 | 6.02889608802969 | 1.51991164680243 | 0.135938634815039 | 0.438218468347367 | -5.28977750583832 |
| lnc-TRIM39-2 | lnc-TRIM39-2 | 0.448446278820319 | 0.661027896860523 | 1.51976731412689 | 0.135974844355176 | 0.438266555177354 | -4.75774944397915 |
| lnc-ZDHHC21-3 | lnc-ZDHHC21-3 | -0.310786861198785 | 2.04251245261384 | -1.51956407451364 | 0.13602584528067 | 0.438362294255838 | -4.97952238620103 |

| | | | | | | | |
|---|---|---|---|---|---|---|---|
| ENSG00000090661.11 | CERS4 | -0.393940771402431 | 3.07269182590471 | -1.51942061740955 | 0.136062078759348 | 0.438399505131827 | -5.10654573917881 |
| ENSG00000109189.12 | USP46 | 0.192317974500324 | 4.1627553380345 | 1.51934837269097 | 0.136079990200031 | 0.438399505131827 | -5.20145968523291 |
| ENSG00000185875.12 | THNSL1 | -0.369678626478243 | 1.99390548172684 | -1.51899320802236 | 0.136169180267504 | 0.438577459222038 | -4.97460391090462 |
| lnc-PDIA4-3 | lnc-PDIA4-3 | -0.602175925369254 | -0.624130139823719 | -1.51888496412407 | 0.136196372105042 | 0.438577459222038 | -4.54956568583228 |
| ENSG00000272741.1 | | -0.138883156130974 | 1.87343105459486 | -1.51887390411142 | 0.136199150723766 | 0.438577459222038 | -4.95497863122944 |
| ENSG00000166822.12 | TMEM170A | 0.212716383962914 | 4.43140093803736 | 1.51877848310783 | 0.136223125325895 | 0.438586045453292 | -5.22642341435824 |
| lnc-NVL-7 | lnc-NVL-7 | 0.233082839649119 | 3.84641922510969 | 1.51818870766725 | 0.136371381739763 | 0.438988378070641 | -5.16415025470007 |
| ENSG00000189046.10 | ALKBH2 | -0.262842710398735 | 2.31442965445632 | -1.51811172210246 | 0.136390743708971 | 0.438988378070641 | -5.01322223155941 |
| lnc-TLL2-1 | lnc-TLL2-1 | 0.161064662085413 | 2.8765711377874 | 1.51790856849408 | 0.136441847653656 | 0.439084201103065 | -5.06850603550694 |
| ENSG00000285314.1 | | -0.210483923646752 | 3.82310105207567 | -1.51728467425812 | 0.136598885929135 | 0.439520848691479 | -5.17545203134946 |
| TNRC6C-AS1 | TNRC6C-AS1 | -0.215711214707996 | 4.78401707842004 | -1.51717582477205 | 0.136626298851695 | 0.4395403421795 | -5.25419218536119 |
| lnc-ABCG2-1 | lnc-ABCG2-1 | -0.595026356617582 | 2.76895901438662 | -1.51756359737405 | 0.136689084217263 | 0.439673608445684 | -5.0913299278936 |
| ENSG00000171914.16 | TLN2 | -0.767355552019349 | 0.0319768675602468 | -1.51687664857099 | 0.136724833400259 | 0.439719882168363 | -4.65439588817126 |
| ENSG00000196323.13 | ZBTB44 | 0.122027336287688 | 6.69261387189677 | 1.51669716495267 | 0.136746897892814 | 0.439722137061053 | -5.29450716107934 |
| ENSG00000113812.13 | ACTR8 | -0.233303303938715 | 4.67525209937408 | -1.51658355716363 | 0.136775534027387 | 0.439745519663498 | -5.25194249689527 |
| ENSG00000129473.9 | BCL2L2 | -0.222534308005452 | 3.65641431217661 | -1.51648204221975 | 0.136801126076143 | 0.439759109483875 | -5.15547258083442 |
| ENSG00000109270.12 | LAMTOR3 | 0.150127124869319 | 4.39354684948221 | 1.51637750877869 | 0.136827483090043 | 0.439775153723041 | -5.22830763652375 |
| ENSG00000116260.16 | QSOX1 | -0.206494121769246 | 4.27982876640332 | -1.51621797520701 | 0.136867715632292 | 0.439835783116231 | -5.22104264192285 |
| ENSG00000254402.6 | LRRC24 | -0.362969378918723 | 1.28879436887833 | -1.51561244586418 | 0.137020509231158 | 0.440258061427555 | -4.90348899441658 |
| lnc-SERPINB9-1 | lnc-SERPINB9-1 | 0.505541895584229 | 2.88412073509008 | 1.51572074603209 | 0.137061015711916 | 0.440319476572244 | -5.05292107759575 |
| ENSG00000109572.13 | CLCN3 | -0.181144860856426 | 4.99665224316084 | -1.51526293317371 | 0.137108763990719 | 0.44040413377356 | -5.2688058832744 |
| lnc-MYBPC1-1 | lnc-MYBPC1-1 | -0.439109688248868 | 0.104945523798084 | -1.5149250240836 | 0.137194131911512 | 0.440471909475783 | -4.70029804016827 |
| ENSG00000170265.11 | ZNF282 | -0.16407947184044 | 3.56161250305534 | -1.51492269964689 | 0.13719471929434 | 0.440471909475783 | -5.15446465765656 |
| ENSG00000272398.5 | CD24 | -0.537724998790537 | 7.06167267746698 | -1.51561235450542 | 0.137214576102587 | 0.440471909475783 | -5.29088486601983 |
| lnc-TDP1-3 | lnc-TDP1-3 | 0.379229798974239 | 0.905039295233497 | 1.5148406144896 | 0.137215463419264 | 0.440471909475783 | -4.81184915880997 |
| ENSG00000106477.18 | CEP41 | -0.296894254152804 | 2.38971496504721 | -1.51454274850212 | 0.137290759568723 | 0.440644893841109 | -5.00699205322785 |
| TRAM2-AS1 | TRAM2-AS1 | -0.238580655752066 | 1.89138843930886 | -1.51440587897924 | 0.137325369238287 | 0.440686778755844 | -4.97065215824529 |
| ENSG00000142046.14 | TMEM91 | -0.292263321019055 | 1.07885380305391 | -1.51432181344268 | 0.13734663002083 | 0.440686778755844 | -4.86854098593831 |
| ENSG00000169180.11 | XPO6 | 0.148134885786817 | 7.02193871717568 | 1.51416356223273 | 0.137386660028725 | 0.440746512370829 | -5.29525854556297 |
| ENSG00000100938.17 | GMPR2 | 0.177372505675044 | 4.41924600622376 | 1.51376924797609 | 0.137486443257156 | 0.440997890223164 | -5.22633556931212 |
| ENSG00000214265.11 | | -0.254637798335402 | 3.79798986036813 | -1.51365159616718 | 0.137516226852336 | 0.441024695746593 | -5.17191852201786 |
| ENSG00000163599.15 | CTLA4 | -1.15634981484843 | 7.18288672898804 | -1.51440180578166 | 0.137619954528583 | 0.441288600103105 | -5.29208875244211 |
| ENSG00000144214.9 | LYG1 | 0.249095876236042 | 0.289770876624158 | 1.51313704604149 | 0.137646545930584 | 0.441303740258128 | -4.7187399177307 |
| lnc-PLA2G15-2 | lnc-PLA2G15-2 | -0.36261690715869 | 1.03865813341408 | -1.51302096080727 | 0.137675960232304 | 0.441303740258128 | -4.859326040363 |
| ENSG00000256966.6 | | 0.261511937367416 | 4.18149027348653 | 1.51296951780467 | 0.13768899674186 | 0.441303740258128 | -5.2066692130613 |
| ENSG00000283041.1 | | 0.644083939707159 | 4.51497385676552 | 1.51368188664509 | 0.137720008484883 | 0.441334412991491 | -5.20988034298154 |
| ENSG00000112799.8 | LY86 | 0.527044510014771 | 3.52884255466489 | 1.51268944315127 | 0.137882148879699 | 0.44165429287638 | -5.13176062253718 |
| ENSG00000126821.7 | SGPP1 | -0.253133433686648 | 6.83403814979668 | -1.51205952102966 | 0.137919767952731 | 0.44165429287638 | -5.29936763412978 |
| ENSG00000089737.16 | DDX24 | 0.165512744944277 | 8.22908865902776 | 1.51204400909353 | 0.13792370438657 | 0.44165429287638 | -5.27922881106677 |
| ENSG00000236375.3 | POU5F1P5 | -0.433812560565958 | -0.430854406390321 | -1.5120176510189 | 0.13793039342952 | 0.44165429287638 | -4.57699272065281 |
| lnc-C6orf229-3 | lnc-C6orf229-3 | 0.411604422999416 | 2.04674451981336 | 1.51196463186337 | 0.13794384919536 | 0.44165429287638 | -4.96401179503296 |
| ENSG00000120438.11 | TCP1 | 0.162323638948311 | 6.72132306769671 | 1.51194602542624 | 0.137948571583455 | 0.44165429287638 | -5.30087021314159 |
| ENSG00000081870.11 | IFT25 | -0.147550460206563 | 3.67295865135188 | -1.51175349885245 | 0.137997443172983 | 0.441742048651557 | -5.17281900585379 |
| ENSG00000109680.10 | TBC1D19 | -0.285744795186099 | 2.52073218489228 | -1.5113152105084 | 0.138108751289943 | 0.441873964397832 | -5.05201803322945 |
| lnc-FAM169A-3 | lnc-FAM169A-3 | -0.207433998631959 | -0.135541984630995 | -1.51128540815988 | 0.138116322523103 | 0.441873964397832 | -4.64650995847028 |
| ENSG00000201998.1 | SNORA23 | 0.474517732170024 | 0.486258521781435 | 1.51127343564447 | 0.138119364212423 | 0.441873964397832 | -4.74565477858011 |
| ENSG00000280184.2 | | -0.342963687023709 | 0.805029403353647 | -1.51125312467283 | 0.138124524458808 | 0.441873964397832 | -4.82752019204454 |
| ENSG00000075407.18 | ZNF37A | 0.335262333907959 | 6.06280861324126 | 1.51071529004274 | 0.138275711878914 | 0.442264444890018 | -5.30205197363943 |
| lnc-CCDC68-4 | lnc-CCDC68-4 | -0.327435227003431 | 5.34794691882845 | -1.51061125855189 | 0.138289557756992 | 0.442264444890018 | -5.28790788727854 |
| ENSG00000101654.17 | RNMT | 0.182297989393286 | 6.39378255488351 | 1.51045713043012 | 0.138326877554254 | 0.442315072347244 | -5.30448379021225 |
| ENSG00000163041.9 | H3-3A | 0.153682515922434 | 6.88853227128407 | 1.51008692724489 | 0.138421069039659 | 0.442466597166044 | -5.30249310825422 |
| lnc-WDPCP-3 | lnc-WDPCP-3 | 0.290382190600308 | 0.390934174982946 | 1.51000150895896 | 0.138442809437223 | 0.442466597166044 | -4.74249764608408 |
| ENSG00000068394.10 | GPKOW | 0.14895416250514 | 3.4698169496122 | 1.50998044333534 | 0.13844817141324 | 0.442466597166044 | -5.14302669172025 |
| ENSG00000139351.14 | SYCP3 | -0.425960165958937 | -0.00099140389247188 | -1.50993298743587 | 0.138460251292158 | 0.442466597166044 | -4.71661348564077 |
| ENSG00000013374.16 | NUB1 | -0.174492009904053 | 5.86263057772695 | -1.50978168467579 | 0.138498770959489 | 0.44252098768382 | -5.30182463736067 |
| ENSG00000118217.5 | ATF6 | 0.103202334232866 | 6.31250786688679 | 1.5096874716711 | 0.138522760654332 | 0.442528943434443 | -5.3055364270024 |
| ENSG00000137628.16 | DDX60 | -0.231563983855605 | 5.02726022682708 | -1.50947945322853 | 0.138575740654709 | 0.442613446803741 | -5.27828858244847 |
| ENSG00000179941.7 | BBS10 | -0.238220327411015 | 2.93374235464873 | -1.50937354044758 | 0.138602721680113 | 0.442613446803741 | -5.0948611124543 |
| ENSG00000155858.5 | LSM11 | -0.236938021599336 | 3.04819016172586 | -1.50933035663605 | 0.138613723854487 | 0.442613446803741 | -5.1112732496579 |
| ENSG00000117448.13 | AKR1A1 | -0.233323580956324 | 4.76400000027914 | -1.50890862551935 | 0.138721207210662 | 0.442782615457448 | -5.26535412988812 |
| ENSG00000156171.14 | DRAM2 | -0.152791539547107 | 6.4272063493834 | -1.50890123549936 | 0.138723091240535 | 0.442782615457448 | -5.30635133361991 |
| ENSG00000083312.17 | TNPO1 | 0.143932574120578 | 7.34225021354257 | 1.50886927821241 | 0.138731238744842 | 0.442782615457448 | -5.29849112838007 |
| ENSG00000197557.6 | IFT70A | -0.549875218446375 | 1.24652247396804 | -1.50886793881462 | 0.138752902002098 | 0.442783097970415 | -4.89542763890953 |
| ENSG00000083838.15 | ZNF446 | -0.188015880173484 | 3.29085843780642 | -1.508613079171 | 0.138796570438477 | 0.442814753494189 | -5.13408334255045 |
| lnc-CXorf67-1 | lnc-CXorf67-1 | -2.1659702418071 | -0.0800364901457963 | -1.50963429944518 | 0.138805848979474 | 0.442814753494189 | -4.50580932164181 |
| ENSG00000283204.1 | MIR4434 | 0.422755194993389 | -0.034884382327078 | 1.50829908839373 | 0.138876672699299 | 0.442972036908364 | -4.64721500024598 |
| ENSG00000093144.18 | ECHDC1 | -0.169765002577495 | 4.39888166871673 | -1.5080998430022 | 0.138927521355317 | 0.443048438686859 | -5.23789763247659 |
| ENSG00000244480.1 | | -0.257135777012753 | 1.55313513176254 | -1.50803655697699 | 0.138943675446906 | 0.443048438686859 | -4.93745336381978 |

| | | | | | | | |
|---|---|---|---|---|---|---|---|
| lnc-TSTA3-1 | lnc-TSTA3-1 | -0.277622113924117 | 1.21370750227018 | -1.50772019193573 | 0.13902445175335 | 0.443237343624411 | -4.89115551630749 |
| ENSG00000172548.14 | NIPAL4 | 0.661928020613436 | 0.114588466442227 | 1.507496868348 | 0.139081494675986 | 0.443255949048263 | -4.65177462651221 |
| ENSG00000063978.15 | RNF4 | 0.192695250447074 | 6.37544701481559 | 1.50747701935133 | 0.13908656555294 | 0.443255949048263 | -5.30865754913164 |
| lnc-HMGB2-3 | lnc-HMGB2-3 | -0.289136202737482 | 2.24099810536691 | -1.50737748116432 | 0.139111997066585 | 0.443255949048263 | -5.01179372206249 |
| lnc-PPFIA4-1 | lnc-PPFIA4-1 | 0.172959448089926 | 2.36549286148364 | 1.50736014137578 | 0.139116427675838 | 0.443255949048263 | -5.03292616245344 |
| lnc-BAZ1A-3 | lnc-BAZ1A-3 | 0.180120497504342 | 1.88216612509902 | 1.50719934939107 | 0.139157518106946 | 0.443318247205582 | -4.96892955869457 |
| ENSG00000136636.12 | KCTD3 | -1.11052659374061 | 0.965682814818893 | -1.50773899894095 | 0.139274950824499 | 0.443557808222369 | -4.87131781511801 |
| ENSG00000167535.7 | CACNB3 | 0.657885424288948 | -0.163623368815129 | 1.50667844513555 | 0.139290701775269 | 0.443557808222369 | -4.64786343690465 |
| ENSG00000105953.14 | OGDH | -0.214831693513263 | 6.67439202700574 | -1.5066523929236 | 0.139297365415605 | 0.443557808222369 | -5.30865418792466 |
| LINC02102 | LINC02102 | -0.391996034134255 | 0.260642756384786 | -1.50656440435593 | 0.139319873028915 | 0.443560857935677 | -4.72007497472838 |
| ENSG00000124357.12 | NAGK | -0.167860095564461 | 6.67685002505617 | -1.50640861474118 | 0.139359731367251 | 0.443588694694669 | -5.30923192445601 |
| lnc-ADGRF3-2 | lnc-ADGRF3-2 | 0.221701445053223 | 3.48302855523486 | 1.50636176771531 | 0.139371718825751 | 0.443588694694669 | -5.14800357271592 |
| ENSG00000251660.1 | | 0.333307248129119 | 1.44907881976401 | 1.50615253271737 | 0.139425268990417 | 0.443690524370711 | -4.91355702660768 |
| ENSG00000007314.12 | SCN4A | -0.421936530392091 | 1.82213425353707 | -1.50573615252363 | 0.139531883264791 | 0.443826325064982 | -4.96201951032588 |
| lnc-ITGB7-1 | lnc-ITGB7-1 | 0.183531357799518 | 3.09997844819392 | 1.50569404827398 | 0.1395426676873 | 0.443826325064982 | -5.11016089348475 |
| ENSG00000102317.17 | RBM3 | 0.181080488075366 | 7.40260202481213 | 1.50562140697837 | 0.13956127531459 | 0.443826325064982 | -5.30219689495034 |
| ENSG00000212126.3 | TAS2R50 | -0.443262474905804 | 0.537240867658024 | -1.5055922535012 | 0.139568743758388 | 0.443826325064982 | -4.80373145164848 |
| ENSG00000155561.14 | NUP205 | 0.152949382779608 | 6.65014897365848 | 1.50556487885628 | 0.139575756796659 | 0.443826325064982 | -5.31065908606586 |
| ENSG00000017260.19 | ATP2C1 | 0.16443652154193 | 5.87262568864088 | 1.5054588425429 | 0.139602924611941 | 0.443844145395054 | -5.30849420239801 |
| ENSG00000138660.11 | AP1AR | -0.249995606059213 | 3.19406534962444 | -1.50520930503576 | 0.139666875838376 | 0.443978888861976 | -5.13491304696399 |
| ENSG00000147224.11 | PRPS1 | 0.144283489164081 | 5.12045731621814 | 1.50511773237762 | 0.139690349844199 | 0.443984939907836 | -5.28790474990919 |
| lnc-SLC22A31-1 | lnc-SLC22A31-1 | 0.584922859363172 | -0.221360406140097 | 1.50493596011482 | 0.139736955198361 | 0.444064497274645 | -4.56778693959506 |
| ENSG00000268643.1 | | 0.171633025037884 | 1.50991009234842 | 1.50482725480639 | 0.139764832529632 | 0.444076824178486 | -4.93188294969854 |
| ENSG00000116704.7 | SLC35D1 | 0.180916267141717 | 3.96798071270984 | 1.50475258504711 | 0.139783984057349 | 0.444076824178486 | -5.20164260170098 |
| lnc-ZDHHC21-4 | lnc-ZDHHC21-4 | -0.495345238597809 | -0.474637964312859 | -1.5045573791855 | 0.139834060932174 | 0.444099712149925 | -4.57406326744252 |
| ENSG00000198830.10 | HMGN2 | 0.156225626938027 | 6.99692279973325 | 1.50449534537298 | 0.139849977686304 | 0.444099712149925 | -5.30914076024247 |
| lnc-XPNPEP1-4 | lnc-XPNPEP1-4 | 0.20141839313822 | 1.57748968209903 | 1.50447220002008 | 0.139855916734966 | 0.444099712149925 | -4.93591791512417 |
| lnc-SPCS2-1 | lnc-SPCS2-1 | 0.259317028700906 | 2.96917908437138 | 1.5043810395814 | 0.139879310265595 | 0.444105482523019 | -5.09003764085616 |
| ENSG00000232810.3 | TNF | -0.564228850764779 | 1.30630987791773 | -1.50411628597299 | 0.139991377108455 | 0.444392738282437 | -4.92999979693434 |
| lnc-BAIAP2L1-1 | lnc-BAIAP2L1-1 | 0.268250077348385 | 0.569941727407078 | 1.50356366725298 | 0.14008920314105 | 0.44460402559894 | -4.76291188026652 |
| ENSG00000197555.9 | SIPA1L1 | -0.348472102344579 | 7.3586561122335 | -1.50361931074544 | 0.140101137348041 | 0.44460402559894 | -5.30716447341372 |
| ENSG00000237522.1 | NONOP2 | -0.256137236720773 | 1.41720324444523 | -1.50329205121071 | 0.140159006887507 | 0.444638041828491 | -4.91355447377537 |
| lnc-ZNF582-8 | lnc-ZNF582-8 | -1.62371032249132 | -1.11951098001268 | -1.50398020898562 | 0.140177405363224 | 0.444638041828491 | -4.49927414535691 |
| ENSG00000160113.5 | NR2F6 | -0.769258160087224 | -0.609374208742499 | -1.50316572948243 | 0.14019148027692 | 0.444638041828491 | -4.58353205468024 |
| ENSG00000271425.7 | NBPF10 | 0.315081893236842 | 3.86024536179055 | 1.50313933924103 | 0.140198265144387 | 0.444638041828491 | -5.1862456170575 |
| ENSG00000183506.17 | PI4KAP2 | 0.20897857042478 | 2.76758157983779 | 1.50276631141023 | 0.140294197683603 | 0.444873743787029 | -5.08476373258927 |
| ENSG00000154059.10 | IMPACT | -0.577279015494556 | 2.89337864548374 | -1.5029478075079 | 0.14037909362062 | 0.44507438139144 | -5.07148128997229 |
| ENSG00000124787.13 | RPP40 | -0.443200113179946 | -0.18989042847291 | -1.50235195030778 | 0.140400821298516 | 0.445074711964786 | -4.63070694694956 |
| lnc-ARNTL-5 | lnc-ARNTL-5 | 0.435230179629167 | -0.52680134982205 | 1.50211867175071 | 0.140460877048134 | 0.445196524835501 | -4.57387994211444 |
| ENSG00000226752.9 | CUTALP | -0.83174000040058 | 3.53981557391959 | -1.50286180963975 | 0.14052680940844 | 0.445275183164193 | -5.12814926802398 |
| lnc-MGAT5-3 | lnc-MGAT5-3 | 0.315223727067185 | 5.37133759965729 | 1.50177118619614 | 0.140550372433367 | 0.445275183164193 | -5.30297205312545 |
| ENSG00000145592.13 | RPL37 | 0.219368778278866 | 8.13683493528842 | 1.50177032826164 | 0.140550593451769 | 0.445275183164193 | -5.2957962023749 |
| ENSG00000205583.13 | STAG3L1 | -0.221369425977129 | 1.41110249075521 | -1.50159720005596 | 0.14059519986196 | 0.445296198669638 | -4.93302630507056 |
| ENSG00000130731.15 | METTL26 | -0.274419338952948 | 2.79492778810464 | -1.50155027178572 | 0.140607292853095 | 0.445296198669638 | -5.09171350790821 |
| SEPSECS-AS1 | SEPSECS-AS1 | -0.343777675066246 | 0.655786610510493 | -1.50149270085532 | 0.140622129492914 | 0.445296198669638 | -4.8107869808753 |
| ENSG00000105849.5 | POLR1F | 0.290573154737198 | 4.41811081590268 | 1.50137038680841 | 0.14065365525889 | 0.445327516719539 | -5.25441484271642 |
| ENSG00000283904.1 | MIR155 | -0.522835117693355 | 1.27006059957345 | -1.50120886604111 | 0.140695294961855 | 0.445359426583679 | -4.85991997381354 |
| ENSG00000184056.14 | VPS33B | 0.113852249432374 | 3.82379505845835 | 1.50116343690529 | 0.140707008262822 | 0.445359426583679 | -5.1947406823609 |
| ENSG00000283969.1 | MIR6838 | -0.241269254716236 | 0.356632692255632 | -1.50105958187984 | 0.140733788822123 | 0.44537570346337 | -4.75609540878238 |
| ENSG00000007080.10 | CCDC124 | -0.27187841882356 | 1.9454044788573 | -1.5009322256774 | 0.140766635048365 | 0.445411168209146 | -5.00108461596358 |
| ENSG00000229833.9 | PET100 | 0.172032309019254 | 3.45602460386411 | 1.50080442950574 | 0.140799600881459 | 0.445447000452363 | -5.15251129617908 |
| ENSG00000229036.7 | VDAC1P8 | 0.21171148591154 | 1.67568790780997 | 1.50069336853218 | 0.140828254755781 | 0.445462584250052 | -4.95700556185357 |
| lnc-MSRA-2 | lnc-MSRA-2 | -0.42409582677678 | 0.212633872252073 | -1.50061757899916 | 0.140847811217963 | 0.445462584250052 | -4.7255729521499 |
| lnc-AARS2-3 | lnc-AARS2-3 | 0.264028277980621 | 0.575750455617787 | 1.50045365384116 | 0.140890117279709 | 0.445527928094678 | -4.77598958126591 |
| ENSG00000140968.10 | IRF8 | 0.173511766760422 | 8.91336245562692 | 1.50002548599452 | 0.14100066723104 | 0.445809022060907 | -5.28279940272331 |
| ENSG00000134049.5 | IER3IP1 | -0.258263361980599 | 2.96150393128515 | -1.49984408732692 | 0.141047523921088 | 0.445888678370104 | -5.12106575120667 |
| ENSG00000073969.18 | NSF | -0.264680786596661 | 5.16033668000136 | -1.49959291204417 | 0.141112424917265 | 0.446025344298536 | -5.29646292948208 |
| lnc-PPEF1-7 | lnc-PPEF1-7 | -0.29705126449368 | 0.693634127240288 | -1.49935973072218 | 0.141172697741812 | 0.446106880951699 | -4.8123785943736 |
| ENSG00000165102.14 | HGSNAT | -0.206010641243647 | 5.27380153708338 | -1.49928189416618 | 0.141192821534482 | 0.446106880951699 | -5.29857706162828 |
| lnc-SLC10A5-1 | lnc-SLC10A5-1 | -0.286532751189465 | 0.412680237035705 | -1.49912898474435 | 0.14123236124599 | 0.446106880951699 | -4.76878375332752 |
| lnc-FCRLB-3 | lnc-FCRLB-3 | -0.342883415283129 | 0.240962101300434 | -1.49911481861119 | 0.141236024807123 | 0.446106880951699 | -4.72397131850163 |
| ENSG00000173200.12 | PARP15 | 0.206964360689289 | 8.10870522458589 | 1.49905132283266 | 0.141252446637112 | 0.446106880951699 | -5.2986195417227 |
| ENSG00000072958.8 | AP1M1 | -0.2384365003086 | 5.88217846498379 | -1.49899017504351 | 0.141268262646027 | 0.446106880951699 | -5.31667049432851 |
| ENSG00000214827.9 | MTCP1 | 0.137459264666026 | 2.64299051777883 | 1.4987988742974 | 0.141317752102855 | 0.446194706478459 | -5.06807205923205 |
| ENSG00000101751.10 | POLI | -0.148357005700531 | 5.26786152618186 | -1.49863785411791 | 0.14135941867781 | 0.446257808994687 | -5.30629150191741 |
| ENSG00000145354.11 | CISD2 | -0.25084147541198 | 3.4382936294212 | -1.49854778749276 | 0.141382729136583 | 0.446262952586765 | -5.16318300515729 |
| lnc-ATF2-5 | lnc-ATF2-5 | 0.145327325465037 | 2.50706247765559 | 1.49825638854475 | 0.14145816807775 | 0.446366175716981 | -5.0556692439843 |
| ENSG00000118046.14 | STK11 | -0.250740357962975 | 4.8388610885359 | -1.49825393240973 | 0.141458804071412 | 0.446366175716981 | -5.28845670673948 |

| | | | | | | | |
|---|---|---|---|---|---|---|---|
| ENSG00000188542.9 | DUSP28 | -0.150095167439551 | 2.77293717713114 | -1.49808733533728 | 0.141501948168566 | 0.446433874793623 | -5.08588351112878 |
| ENSG00000138641.16 | HERC3 | 0.227432799990345 | 5.74266209566174 | 1.49798739924672 | 0.141527833913642 | 0.446447111945516 | -5.31643117564144 |
| ENSG00000081052.12 | COL4A4 | -2.00808388857703 | 3.69678117242752 | -1.49896888500565 | 0.141566857941493 | 0.446501783176487 | -5.22248978369642 |
| ENSG00000090686.15 | USP48 | 0.151926776333828 | 6.64884434984879 | 1.49762440387561 | 0.141621889759895 | 0.446606918481372 | -5.32205811141382 |
| ENSG00000271858.5 | | -0.280002469601192 | 1.42841048744711 | -1.49733843080052 | 0.141696023351312 | 0.446772250098049 | -4.9312043009685 |
| ENSG00000131979.18 | GCH1 | 0.249873045669215 | 3.05810362058054 | 1.49723760725815 | 0.141722167476926 | 0.446786241871276 | -5.13056401489925 |
| ENSG00000285755.1 | | 0.280723203627299 | 0.395609775571186 | 1.4965245229142 | 0.141907183922339 | 0.447301005616156 | -4.77870407244076 |
| ENSG00000173567.14 | ADGRF3 | 0.276810078360674 | -0.039344482664886 | 1.49627364205505 | 0.141972322991403 | 0.447402765000784 | -4.6678220975105 |
| lnc-PDZD7-6 | lnc-PDZD7-6 | -0.18034867008909 | 2.46900908592264 | -1.49615265119318 | 0.14200374573275 | 0.447402765000784 | -5.06142947366193 |
| ENSG00000099968.17 | BCL2L13 | -0.212577539248808 | 5.27013032363551 | -1.49604197522849 | 0.142032494418656 | 0.447402765000784 | -5.30686990905366 |
| ENSG00000109756.9 | RAPGEF2 | 0.241625803864502 | 7.00381051699722 | 1.49598473885974 | 0.142047363693659 | 0.447402765000784 | -5.32116672889743 |
| ENSG00000165006.13 | UBAP1 | 0.168417077200389 | 4.50404475400212 | 1.49598171266556 | 0.142048149894579 | 0.447402765000784 | -5.26460097757045 |
| lnc-KCNK7-6 | lnc-KCNK7-6 | -0.469075858300567 | -0.683059532530545 | -1.49571361086383 | 0.142117816094915 | 0.447550017644704 | -4.52162796959287 |
| ENSG00000099889.13 | ARVCF | -0.308510371261118 | -0.124820523994887 | -1.49563445770776 | 0.142138389221369 | 0.447550017644704 | -4.69184138936676 |
| ENSG00000117592.8 | PRDX6 | 0.16493334514337 | 5.38494624591968 | 1.49541956359928 | 0.142194255459156 | 0.447583407983162 | -5.31362518361965 |
| lnc-CCDC169-SOHLH2-2 | lnc-CCDC169-SOH | -1.74010964405181 | -1.28571312598807 | -1.49582610082106 | 0.142267193020329 | 0.447583407983162 | -4.45599209744332 |
| ENSG00000086504.16 | MRPL28 | 0.215678946770486 | 3.21104676125745 | 1.49499775938099 | 0.14230396309866 | 0.447583407983162 | -5.13558873245908 |
| ENSG00000087470.17 | DNM1L | 0.154144038459874 | 5.74012934680471 | 1.49498451241035 | 0.142307409611208 | 0.447583407983162 | -5.32268999194816 |
| ENSG00000228835.1 | | -0.591207907307031 | 2.3683326976394 | -1.49543986367824 | 0.142309988063647 | 0.447583407983162 | -5.04517581549922 |
| ENSG00000266777.1 | SH3GL1P1 | -0.288988307965819 | -0.270719089073527 | -1.49494421626538 | 0.142317894013879 | 0.447583407983162 | -4.631599301845 |
| lnc-KAT2B-5 | lnc-KAT2B-5 | -0.515418887952553 | -0.550840999678995 | -1.494931315955 | 0.14232125059497 | 0.447583407983162 | -4.57955529584472 |
| ENSG00000111850.10 | SMIM8 | -0.22090163119347 | 3.56747638407737 | -1.49492476132859 | 0.142322956092396 | 0.447583407983162 | -5.17472629798303 |
| ENSG00000175575.12 | PAAF1 | 0.145841104498103 | 3.73939932653456 | 1.49466768318949 | 0.142389860000788 | 0.447725403054723 | -5.19030915460279 |
| ENSG00000155868.7 | MED7 | 0.140283483883246 | 3.92477721432113 | 1.49431486902211 | 0.142481719662966 | 0.447936000441088 | -5.21273148928675 |
| lnc-STRN3-6 | lnc-STRN3-6 | 0.147773497689055 | 2.46242266231191 | 1.49423708331597 | 0.142501978499584 | 0.447936000441088 | -5.05198611765029 |
| ENSG00000188404.8 | SELL | 0.496204771862142 | 7.707007457485 | 1.49475112266138 | 0.14254198132847 | 0.447936000441088 | -5.31880376287276 |
| ENSG00000111817.17 | DSE | 0.64891238945265 | 2.71564179536757 | 1.49470585562088 | 0.142543885871308 | 0.447936000441088 | -5.05267633266147 |
| ENSG00000104859.14 | CLASRP | -0.222903951927193 | 6.30420904934828 | -1.49389345790671 | 0.142591501151239 | 0.448017229155236 | -5.32754208646702 |
| lnc-LITAF-1 | lnc-LITAF-1 | 0.265068999471789 | 2.12537919481124 | 1.49374895760428 | 0.142629160292371 | 0.44805774368419 | -5.01791770911705 |
| lnc-B3GNT10-21 | lnc-B3GNT10-21 | -0.828650348042867 | 4.01435193520054 | -1.49468230703093 | 0.142655157122785 | 0.44805774368419 | -5.19013310531577 |
| ENSG00000175938.6 | ORAI3 | -0.24141994253594 | 3.45277794034428 | -1.4935934351251 | 0.142669700826225 | 0.44805774368419 | -5.16664000457006 |
| ENSG00000196440.11 | ARMCX4 | 0.494859777340408 | 2.45512674635053 | 1.49357431002539 | 0.142714206552706 | 0.44812914011813 | -5.03800037457173 |
| ENSG00000186814.13 | ZSCAN30 | -0.164912331047033 | 4.80496255383373 | -1.49312086275523 | 0.14279294395577 | 0.448300730993451 | -5.28573148756951 |
| ENSG00000143793.12 | C1orf35 | -0.186362174582456 | 2.73063708270908 | -1.49304623957835 | 0.142812412822429 | 0.448300730993451 | -5.09663809663763 |
| ENSG00000274134.1 | MIR6774 | 0.347824472933383 | 1.85098649456368 | 1.49295988501116 | 0.142834944991688 | 0.448303091302824 | -4.97312602288521 |
| ENSG00000099940.11 | SNAP29 | -0.17794419024246 | 4.15973855268375 | -1.49281985516461 | 0.14287148845308 | 0.448349420160045 | -5.24150223821419 |
| ENSG00000226415.1 | TPI1P1 | 1.07845709524142 | -0.711750928967911 | 1.49241971461341 | 0.142975953721382 | 0.448608849915734 | -4.45563663820349 |
| lnc-ACRV1-2 | lnc-ACRV1-2 | -0.37437767008109 | -0.361931038322947 | -1.49226712007262 | 0.143015807781377 | 0.448665503972577 | -4.62099608455719 |
| ENSG00000234912.12 | SNHG20 | -0.278791687823419 | 1.58615652737837 | -1.49187765121869 | 0.143117567790186 | 0.44891632091213 | -4.96371316800898 |
| ENSG00000061656.9 | SPAG4 | -0.508440570943424 | 2.97334825930252 | -1.49192417106826 | 0.143199279071999 | 0.449104184235708 | -5.12487636858419 |
| ENSG00000129993.14 | CBFA2T3 | 0.339969204956985 | 3.53693658852401 | 1.49135288414906 | 0.143254769344362 | 0.449209768040067 | -5.17867604106176 |
| lnc-RPL37-1 | lnc-RPL37-1 | 0.236545965512883 | 8.01682852303894 | 1.4910698477207 | 0.14332881323628 | 0.449292731478336 | -5.31348089351699 |
| lnc-SAV1-1 | lnc-SAV1-1 | -0.24236007003582 | 1.39797526306703 | -1.49100679358289 | 0.143345312687104 | 0.449292731478336 | -4.94567169858139 |
| ENSG00000160310.17 | PRMT2 | -0.237514066459686 | 5.82764652647378 | -1.49100144726481 | 0.143346711733869 | 0.449292731478336 | -5.3279221659937 |
| ENSG00000213693.4 | SEC14L1P1 | -0.597684530725776 | 1.03841841207176 | -1.49062292878934 | 0.143472148960045 | 0.44955456058862 | -4.86038320508748 |
| ENSG00000131323.14 | TRAF3 | 0.173226257894891 | 6.0554162849875 | 1.4904529286256 | 0.143490308030364 | 0.44955456058862 | -5.33234615602499 |
| lnc-COL6A3-14 | lnc-COL6A3-14 | -0.387041711196084 | -0.109152301686594 | -1.49043206783327 | 0.143495771416569 | 0.44955456058862 | -4.65550014086918 |
| ENSG00000011405.13 | PIK3C2A | 0.153580066841054 | 6.1591803894769 | 1.49032750743111 | 0.143523157998354 | 0.449571931377283 | -5.33261928976431 |
| ENSG00000121766.15 | ZCCHC17 | -0.1433835388206 | 3.87228314528363 | -1.49023818860343 | 0.14354655577707 | 0.449576804254327 | -5.21637718708448 |
| ENSG00000168000.14 | BSCL2 | -0.132357797790535 | 2.74187248505979 | -1.48983997310998 | 0.143650908365058 | 0.449777847450107 | -5.09619181093639 |
| ENSG00000260274.1 | | -0.256499439357649 | 1.13965655901943 | -1.48982645746434 | 0.143654451204246 | 0.449777847450107 | -4.89836320402809 |
| lnc-PRPF18-1 | lnc-PRPF18-1 | -0.230169970051932 | 4.46008509472203 | -1.48973202797817 | 0.143679205822805 | 0.449786934365139 | -5.26625816836434 |
| lnc-EIF2D-2 | lnc-EIF2D-2 | 0.157364971236307 | 1.54265405310806 | 1.48952943943679 | 0.143732325686489 | 0.449884802251369 | -4.95540898582118 |
| lnc-ZDHHC1-2 | lnc-ZDHHC1-2 | 0.309600032463052 | 1.7536470637318 | 1.48938112462751 | 0.143771224557614 | 0.449938135178557 | -4.98378587287821 |
| lnc-C14orf93-2 | lnc-C14orf93-2 | -0.184625577429717 | 1.37422225350681 | -1.48925992469437 | 0.143803018153601 | 0.449969219011185 | -4.94059021094404 |
| ENSG00000135828.11 | RNASEL | -0.215682860460741 | 4.32435483432836 | -1.4891695129872 | 0.143826738908709 | 0.449975036777316 | -5.25042119631718 |
| lnc-EPAS1-3 | lnc-EPAS1-3 | 0.403084799844356 | 3.98348533185328 | 1.48887160976376 | 0.143938676873354 | 0.450256806395781 | -5.20764958725448 |
| ENSG00000132591.11 | ERAL1 | 0.155218345167916 | 3.4285426593527 | 1.48860479669029 | 0.143974970316363 | 0.450301901538017 | -5.1708521658567 |
| ENSG00000199293.1 | SNORA21 | 0.354847032399913 | 0.170478603520023 | 1.48842339456997 | 0.14402261197794 | 0.450329071573602 | -4.72891870934448 |
| ENSG00000198298.12 | ZNF485 | -0.226682522715514 | 1.48542812218004 | -1.48840510993593 | 0.144027414767965 | 0.450329071573602 | -4.94782993867857 |
| ENSG00000256628.3 | ZBTB11-AS1 | -0.187336745427409 | 2.43489694018339 | -1.48827113670829 | 0.144062609139958 | 0.450334080045993 | -5.06416173549783 |
| ENSG00000282804.1 | | 0.320944999207037 | 3.1961664063476 | 1.4882324454916 | 0.144072774479078 | 0.450334080045993 | -5.12964104679124 |
| ENSG00000198843.12 | SELENOT | 0.15341238419511 | 5.96254235347362 | 1.48814054928187 | 0.144096920642655 | 0.450339850415633 | -5.3347346096354 |
| ENSG00000206651.1 | | 0.439784831911676 | -0.244783358502918 | 1.48805889271628 | 0.14411837898692 | 0.450339850415633 | -4.61132518920329 |
| lnc-SLC26A8-1 | lnc-SLC26A8-1 | -0.297878465144744 | 2.20950444429234 | -1.48791838756588 | 0.144155307955633 | 0.450386870620947 | -5.03505473372153 |
| ENSG00000205155.7 | PSENEN | -0.252296817790427 | 2.31709665456542 | -1.48757002433193 | 0.144246900665936 | 0.450548020112699 | -5.05382795052855 |
| lnc-CCDC141-1 | lnc-CCDC141-1 | -0.793435837689392 | 2.16656747868806 | -1.48834311622751 | 0.144250665721357 | 0.450548020112699 | -5.02281347483613 |
| ENSG00000101213.6 | PTK6 | -0.479468540568133 | 0.0674326999552343 | -1.48718820643579 | 0.144347342432567 | 0.450763716528821 | -4.76551316905399 |

| | | | | | | | |
|---|---|---|---|---|---|---|---|
| lnc-SIGIRR-2 | lnc-SIGIRR-2 | -0.259380223283988 | 1.62889378568796 | -1.48712671288321 | 0.144363524236408 | 0.450763716528821 | -4.97353685573032 |
| ENSG00000180008.8 | SOCS4 | 0.185986161115652 | 5.62651553087508 | 1.48679433640479 | 0.144451012817309 | 0.450968481240509 | -5.33106736343584 |
| lnc-HLA-DOB-1 | lnc-HLA-DOB-1 | 0.169618478940767 | 4.3400473733397 | 1.48661785639554 | 0.144497483221271 | 0.450993450536716 | -5.2625086741468 |
| ENSG00000139324.11 | TMTC3 | 0.191419528219374 | 4.13270949240314 | 1.48655702290958 | 0.14451350453664 | 0.450993450536716 | -5.24635764784478 |
| ENSG00000158106.13 | RHPN1 | 0.703530872869679 | 2.21909665994668 | 1.48716507032203 | 0.144539548656805 | 0.450993450536716 | -5.05387423805103 |
| ENSG00000101868.10 | POLA1 | 0.130573515440686 | 4.67261022688763 | 1.48643116768265 | 0.144546654675738 | 0.450993450536716 | -5.28677219725153 |
| lnc-PNPLA3-1 | lnc-PNPLA3-1 | -0.560466124349305 | -0.684111489249902 | -1.48597041578371 | 0.14466806774734 | 0.451281187580666 | -4.53980605716486 |
| LINC01619 | LINC01619 | 0.402245947278005 | 6.7951006029992 | 1.48626795660513 | 0.144682726368596 | 0.451281187580666 | -5.33705978020807 |
| ENSG00000064201.15 | TSPAN32 | -0.629836788753194 | 3.25279495382462 | -1.48650083866511 | 0.144710434595455 | 0.451299223527007 | -5.13814275861815 |
| ENSG00000127191.17 | TRAF2 | -0.20314232268215 | 3.59994718728871 | -1.48566916577324 | 0.144747494100155 | 0.451346412825857 | -5.20019654297714 |
| lnc-GPR149-4 | lnc-GPR149-4 | 0.406468662346906 | 1.17281459378727 | 1.48554145524199 | 0.144781176180924 | 0.451347252146464 | -4.91866195639835 |
| ENSG00000101972.18 | STAG2 | 0.111149203414312 | 7.67812285385117 | 1.48550186233604 | 0.144791619585245 | 0.451347252146464 | -5.32642215649001 |
| lnc-HLA-DMB-3 | lnc-HLA-DMB-3 | -0.348725475472432 | 0.750116470936481 | -1.48525670716474 | 0.144856297371053 | 0.451432438985771 | -4.85842324517779 |
| ENSG00000103415.11 | HMOX2 | -0.302113061418756 | 4.05196615246048 | -1.48510678911149 | 0.144895860624812 | 0.451432438985771 | -5.2403177648597 |
| ENSG00000168495.12 | POLR3D | -0.12939583288926 | 4.46994168715591 | -1.48507554372915 | 0.144904107335499 | 0.451432438985771 | -5.27507263442437 |
| ENSG00000114648.11 | KLHL18 | 0.186889071107158 | 4.9120917641929 | 1.48506580952624 | 0.144906676596171 | 0.451432438985771 | -5.30493746282567 |
| ENSG00000211751.9 | TRBC1 | -1.47171656559002 | -2.12503867678833 | -1.48430608723136 | 0.145107310384227 | 0.451989069255228 | -4.40020474387341 |
| lnc-SF1-2 | lnc-SF1-2 | -0.331315347741041 | 0.193769109178501 | -1.48408080984824 | 0.145166845847977 | 0.452106095943245 | -4.73960195892366 |
| ENSG00000125503.12 | PPP1R12C | -0.178784570276735 | 5.2551361280261 | -1.48391865094332 | 0.145209712587287 | 0.452171182176116 | -5.32724366309181 |
| C21orf62-AS1 | C21orf62-AS1 | -0.314517826642622 | 1.45736108015818 | -1.48380128162529 | 0.145240745452742 | 0.45219940457628 | -4.94504794225505 |
| ENSG00000221716.1 | SNORA11 | -0.42497513249553 | 0.176025393322812 | -1.48355204672417 | 0.145306661491384 | 0.452336208934841 | -4.7831203361667 |
| ENSG00000155363.18 | MOV10 | -0.194885542727321 | 4.96938528933159 | -1.4834068785812 | 0.145345065557375 | 0.45238734074814 | -5.31281088803284 |
| ENSG00000118007.12 | STAG1 | 0.118758046138133 | 6.44244390356343 | 1.48306412841785 | 0.145435771658559 | 0.452453150601741 | -5.34314797863226 |
| ENSG00000163162.8 | RNF149 | 0.179494415247653 | 5.43290647584624 | 1.48303598980528 | 0.145443220311675 | 0.452453150601741 | -5.33263771089375 |
| ENSG00000081154.11 | PCNP | 0.173662041034448 | 6.13493952993318 | 1.48301707267021 | 0.145448228090983 | 0.452453150601741 | -5.34298322154911 |
| lnc-WRNIP1-35 | lnc-WRNIP1-35 | -0.386685750540368 | 2.32079389967873 | -1.48299475293348 | 0.145454136789181 | 0.452453150601741 | -5.05370698982919 |
| lnc-ROGDI-5 | lnc-ROGDI-5 | 0.198542415120151 | 2.04572641940488 | 1.48287352348735 | 0.145486233153971 | 0.452484608191023 | -5.02710169150926 |
| ENSG00000158122.11 | PRXL2C | 0.203852055966366 | 4.15343375543685 | 1.48251958683974 | 0.145579972551295 | 0.452707746641985 | -5.25581896268348 |
| ENSG00000104695.12 | PPP2CB | 0.197672977054783 | 2.91946670369318 | 1.48218881180725 | 0.145667620925276 | 0.452911879381412 | -5.12505834579389 |
| lnc-CASP3-7 | lnc-CASP3-7 | -0.243264046331357 | 1.56757897849194 | -1.48194039519275 | 0.145733473567616 | 0.453044099852951 | -4.96182326542973 |
| ENSG00000105647.16 | PIK3R2 | -0.247609367226912 | 1.69811411016784 | -1.48186234970862 | 0.145754167479291 | 0.453044099852951 | -4.9870698979751 |
| ENSG00000140943.16 | MBTPS1 | 0.100210583560159 | 6.58260749559193 | 1.4817357750855 | 0.145787733938848 | 0.453056506112535 | -5.34463604454765 |
| lnc-RPLP0-3 | lnc-RPLP0-3 | 0.158440644228964 | 1.95957207063898 | 1.48162308329119 | 0.145817623952477 | 0.453056506112535 | -5.01385821444746 |
| ENSG00000088766.11 | CRLS1 | -0.149435122185423 | 3.24693890583274 | -1.48159832113099 | 0.145824192440147 | 0.453056506112535 | -5.16679811064761 |
| lnc-ATG12-5 | lnc-ATG12-5 | -0.301326112980897 | 1.22618889965881 | -1.48098950201955 | 0.145985763478342 | 0.453286372875648 | -4.93376161476189 |
| ENSG00000270181.3 | BIVM-ERCC5 | -0.107129994646781 | 5.99877549397691 | -1.48093572026696 | 0.146000043129458 | 0.453286372875648 | -5.34449144844448 |
| ENSG00000051825.14 | MPHOSPH9 | 0.194552710569059 | 5.52445319263355 | 1.48080928446656 | 0.146033617595573 | 0.453286372875648 | -5.33582326234855 |
| ENSG00000079435.9 | LIPE | -0.277051140904731 | 1.96231422627427 | -1.48080009298643 | 0.146036058590932 | 0.453286372875648 | -5.02674950349846 |
| ENSG00000204438.10 | GPANK1 | -0.193630607049998 | 3.85767635897197 | -1.4807852595162 | 0.146039998006388 | 0.453286372875648 | -5.2310834199179 |
| ENSG00000105819.13 | PMPCB | 0.107913332762844 | 6.06507492901512 | 1.48075322302288 | 0.146048506421332 | 0.453286372875648 | -5.34549947888143 |
| ENSG00000186635.14 | ARAP1 | -0.151903204058914 | 6.59238364781559 | -1.4807388053779 | 0.146052335661045 | 0.453286372875648 | -5.34577966750171 |
| lnc-RNMT-3 | lnc-RNMT-3 | -0.456179067998 | 0.142305943978439 | -1.48058755219839 | 0.146092512391149 | 0.453331636655721 | -4.73838646435781 |
| ENSG00000158669.11 | GPAT4 | 0.128560526607478 | 5.86773138451737 | 1.48043003858692 | 0.14613436136236 | 0.453331636655721 | -5.34393247768914 |
| ENSG00000141293.15 | SKAP1 | -0.731033450245349 | 5.53944302793481 | -1.48144049892275 | 0.146134694419904 | 0.453331636655721 | -5.33229697442827 |
| ENSG00000033327.12 | GAB2 | -0.432112869069066 | 7.47278342427946 | -1.48081041474819 | 0.146155018260087 | 0.453331636655721 | -5.33559327755991 |
| ENSG00000064961.18 | HMG20B | -0.257531882502456 | 3.61531459441174 | -1.47981142522224 | 0.146298809453103 | 0.453709265477357 | -5.20364198102044 |
| ENSG00000139890.9 | REM2 | -0.273298965369526 | 1.78897024300991 | -1.47969427118616 | 0.14632996941632 | 0.453731471027174 | -4.9880172079583 |
| lnc-NCKIPSD-2 | lnc-NCKIPSD-2 | 0.197642669231441 | 1.51520476969921 | 1.47961875477765 | 0.146350057627625 | 0.453731471027174 | -4.96972969911201 |
| ENSG00000026103.21 | FAS | 0.308146331620674 | 2.96910056601374 | 1.47938907903035 | 0.146411167380155 | 0.45383653626214 | -5.13590286341356 |
| ENSG00000187790.10 | FANCM | 0.140636098294701 | 4.28343328219256 | 1.47931203379501 | 0.146431671309002 | 0.45383653626214 | -5.26846355209428 |
| lnc-PDCD2L-4 | lnc-PDCD2L-4 | -0.263596689965671 | 0.137483186931929 | -1.47921472724643 | 0.146457570600635 | 0.45383653626214 | -4.74014652092525 |
| ENSG00000068903.19 | SIRT2 | -0.211143479608138 | 4.237265579974 | -1.47912924218753 | 0.146480326457098 | 0.45383653626214 | -5.26520537521571 |
| ENSG00000206633.1 | SNORA80B | 0.641357888557801 | 0.729283058361238 | 1.47907277313046 | 0.146514236938204 | 0.45383653626214 | -4.79642344052044 |
| lnc-ANKRD27-2 | lnc-ANKRD27-2 | -0.367554180814835 | 2.07052157991304 | -1.47894944428293 | 0.146528197240741 | 0.45383653626214 | -5.03116239239911 |
| ENSG00000168298.6 | H1-4 | -0.309440811833078 | 8.66790248686062 | -1.47880478628159 | 0.146566721101278 | 0.45383653626214 | -5.31049883887894 |
| ENSG00000164830.18 | OXR1 | -1.00557834759047 | 5.39034335633349 | -1.4798674367305 | 0.146576273140778 | 0.45383653626214 | -5.32430654534319 |
| lnc-FANCG-3 | lnc-FANCG-3 | 0.14235832273824 | 3.48715558780814 | 1.47874596516617 | 0.146582388042108 | 0.45383653626214 | -5.19059915500569 |
| lnc-MRPL53-2 | lnc-MRPL53-2 | -0.175716619784674 | 2.26760764663682 | -1.47842595668248 | 0.146667645199951 | 0.454017307586116 | -5.06440974896759 |
| ENSG00000118655.5 | DCLRE1B | -0.236527218137849 | 1.41220323230642 | -1.47813477466221 | 0.146745256493474 | 0.454017307586116 | -4.96878645575242 |
| lnc-MUC20-71 | lnc-MUC20-71 | -0.306872919813716 | 1.4602770444687 | -1.47795862000098 | 0.146792224327492 | 0.454017307586116 | -4.9557992455866 |
| ENSG00000143158.10 | MPC2 | 0.15738371568379 | 4.22652182157139 | 1.47788087796079 | 0.146812956349055 | 0.454017307586116 | -5.27034969143242 |
| ENSG00000084463.7 | WBP11 | 0.175943362307414 | 6.53528539409047 | 1.47786630657374 | 0.146816842462303 | 0.454017307586116 | -5.35016783770807 |
| ENSG00000145331.13 | TRMT10A | -0.199756176613589 | 1.46336217580482 | -1.47785084524367 | 0.146820966007816 | 0.454017307586116 | -4.9622157378048 |
| lnc-PRPS1L1-3 | lnc-PRPS1L1-3 | 0.245496310304444 | 3.00631663153852 | 1.47784118259724 | 0.146823543087683 | 0.454017307586116 | -5.14139577935065 |
| ENSG00000173614.13 | NMNAT1 | -0.278053566022705 | 2.33428066408539 | -1.47776887936936 | 0.146842827885161 | 0.454017307586116 | -5.05602031597159 |
| lnc-SLC7A1-7 | lnc-SLC7A1-7 | 0.405142472105263 | 1.40915050223443 | 1.47772011813747 | 0.146855834667865 | 0.454017307586116 | -4.93780803769032 |
| ENSG00000063587.14 | ZNF275 | -0.249003107954451 | 5.74461287236196 | -1.47769943052077 | 0.146861353248232 | 0.454017307586116 | -5.34527265940653 |
| lnc-NUTM2G-6 | lnc-NUTM2G-6 | 1.26836403720308 | -0.742271694256568 | 1.47729263850208 | 0.147010352982484 | 0.454409685454044 | -4.4681192986054 |

| | | | | | | | |
|---|---|---|---|---|---|---|---|
| lnc-TM9SF3-1 | lnc-TM9SF3-1 | -0.262644445572752 | 4.16673446364071 | -1.47694190183869 | 0.147063542970051 | 0.454505841584467 | -5.26268623984906 |
| lnc-LNX2-2 | lnc-LNX2-2 | -0.152030611883225 | 2.0653050074586 | -1.47667327594273 | 0.147135294023932 | 0.454659323959556 | -5.04228066806604 |
| ENSG00000230989.6 | HSBP1 | 0.197519404661228 | 4.30315638965397 | 1.47610159981894 | 0.147288083268211 | 0.454809183658131 | -5.27532003051187 |
| ENSG00000099804.8 | CDC34 | -0.232657703978976 | 2.26542719647377 | -1.47605255098937 | 0.147301198171658 | 0.454809183658131 | -5.06697809819495 |
| lnc-SLC22A31-3 | lnc-SLC22A31-3 | 0.657936681801402 | -0.0446933026450686 | 1.47603091415531 | 0.147306983822515 | 0.454809183658131 | -4.60095303932822 |
| ENSG00000119682.16 | AREL1 | 0.143662616552014 | 5.50388832167637 | 1.47602568177791 | 0.147308382977919 | 0.454809183658131 | -5.34344426235544 |
| ENSG00000136280.16 | CCM2 | -0.206224405469195 | 5.73626051819013 | -1.47596072852684 | 0.147325752572747 | 0.454809183658131 | -5.3479871970378 |
| ENSG00000256238.1 | SUPT16HP1 | 0.401865188286488 | -0.468074324614648 | 1.47594822076097 | 0.147329097546461 | 0.454809183658131 | -4.57521681181038 |
| ENSG00000185495.10 | SEPTIN7P13 | -0.280722014514248 | 3.10339268830359 | -1.47591319240658 | 0.147338465560531 | 0.454809183658131 | -5.16188446187582 |
| ENSG00000249264.1 | EEF1A1P9 | 0.644082872647279 | -0.0082018152703308 | 1.47560942371963 | 0.147419725504502 | 0.454852604457064 | -4.64239664809673 |
| lnc-NUP62CL-2 | lnc-NUP62CL-2 | -0.170888871036609 | 1.40935735337923 | -1.4755547356815 | 0.147434358650044 | 0.454852604457064 | -4.95006685138196 |
| lnc-SEPT3-4 | lnc-SEPT3-4 | -0.360539096313381 | 1.40601126788528 | -1.47554494339939 | 0.147436978940172 | 0.454852604457064 | -4.99139701472933 |
| lnc-MAPK1-3 | lnc-MAPK1-3 | -0.237520425892876 | 2.84213096155884 | -1.47551359746213 | 0.14744536696212 | 0.454852604457064 | -5.11082538680672 |
| ENSG00000239264.8 | TXNDC5 | -0.536984170748167 | 7.35746072618276 | -1.47617087767627 | 0.147469896454497 | 0.454852604457064 | -5.34291572377942 |
| lnc-MOB3A-2 | lnc-MOB3A-2 | -0.203098641573874 | 0.642521836278191 | -1.47536347166003 | 0.147485545141228 | 0.454852604457064 | -4.85292832957568 |
| LINC01730 | LINC01730 | -0.469252929170649 | -0.0218723069349049 | -1.47528249234653 | 0.147507221238444 | 0.454852604457064 | -4.69828687007527 |
| ENSG00000151327.12 | FAM177A1 | -0.161705913994396 | 3.71488614829493 | -1.47489189106557 | 0.147611810658064 | 0.455054053740282 | -5.22048330897744 |
| ENSG00000182979.17 | MTA1 | -0.255052565079854 | 5.12124305073235 | -1.47480875680148 | 0.147634078692361 | 0.455054053740282 | -5.32919406037324 |
| ENSG00000172890.12 | NADSYN1 | -0.148171490402943 | 6.86027619657605 | -1.4747908506026 | 0.147638875328067 | 0.455054053740282 | -5.35233221243512 |
| ENSG00000185033.14 | SEMA4B | 0.262013731835126 | 6.65945445013888 | 1.47465585629514 | 0.147675040993879 | 0.455097375172484 | -5.35461597036557 |
| ENSG00000004487.16 | KDM1A | 0.104102390657031 | 5.66471086953081 | 1.47390838850784 | 0.147875418342845 | 0.45564666702856 | -5.34965085574648 |
| ENSG00000005238.19 | ATOSB | -0.301552014193953 | 1.96430152674099 | -1.47373153856347 | 0.147922858831671 | 0.455654739651809 | -5.03215558265253 |
| ENSG00000089876.11 | DHX32 | -0.470449816945572 | 3.17704205896153 | -1.47396360268755 | 0.147949054407474 | 0.455654739651809 | -5.19755597182256 |
| ENSG00000163686.14 | ABHD6 | 0.414378043420503 | 1.0546232489267 | 1.47362572993963 | 0.147951248046205 | 0.455654739651809 | -4.90609901337271 |
| lnc-TMEM218-1 | lnc-TMEM218-1 | -0.236084546504425 | 2.41654864743852 | -1.47356856370055 | 0.14796658795281 | 0.455654739651809 | -5.08406413762134 |
| lnc-TRAPPC3-2 | lnc-TRAPPC3-2 | -0.418293747918402 | -0.579865226132714 | -1.47345965500368 | 0.147995815838689 | 0.455676571040798 | -4.61434887741725 |
| ENSG00000163046.15 | ANKRD30BL | -1.71977711247949 | -0.412868756103028 | -1.47413039287231 | 0.148037367790571 | 0.455736335811145 | -4.55769628373946 |
| ENSG00000163328.13 | GPR155 | 0.32056499766913 | 3.86954982605666 | 1.47321567127423 | 0.148061310463133 | 0.455741880254625 | -5.23421584027465 |
| lnc-HEPH-1 | lnc-HEPH-1 | -0.685930665042466 | 1.29149815716136 | -1.47345915444564 | 0.14811657872173 | 0.455805107605828 | -4.9382331545663 |
| ENSG00000268034.1 | TPM3P5 | 0.992720162753122 | -0.976845687629751 | 1.47297424468498 | 0.148126141222144 | 0.455805107605828 | -4.45454058061605 |
| ENSG00000125826.20 | RBCK1 | -0.166888671301668 | 5.07505321800377 | -1.47286332324183 | 0.148155934704513 | 0.455828640362181 | -5.33337493890368 |
| ENSG00000068024.16 | HDAC4 | 0.218889284079729 | 2.40369045063662 | 1.47277572128295 | 0.148179467927271 | 0.455832908137351 | -5.08597808634922 |
| ENSG00000146872.17 | TLK2 | 0.162450671833606 | 6.22128064669786 | 1.47258324663105 | 0.148231184335749 | 0.455872137008864 | -5.3576660601638 |
| ENSG00000285253.1 | | -0.18333033771167 | 4.30754760134432 | -1.47256340586785 | 0.148236516202707 | 0.455872137008864 | -5.27239113696488 |
| ENSG00000136279.20 | DBNL | -0.177892571475978 | 8.28195587486395 | -1.47232918945592 | 0.148299469338199 | 0.455931253706922 | -5.3353416563139 |
| ENSG00000163166.14 | IWS1 | 0.114776038323656 | 5.24721104457344 | 1.47232706301573 | 0.148300040983717 | 0.455931253706922 | -5.3392863799987 |
| ENSG00000149292.16 | TTC12 | 0.35169478888499 | 4.50469444722624 | 1.4722593631183 | 0.148330246694936 | 0.455956013698008 | -5.27938857623093 |
| lnc-SBNO1-1 | lnc-SBNO1-1 | 0.192140924060628 | 1.03545618267711 | 1.47205817283405 | 0.148372340094282 | 0.455960173459244 | -4.91096103949425 |
| lnc-AGAP6-1 | lnc-AGAP6-1 | 0.318032716765216 | 1.14016321142933 | 1.4720449190965 | 0.148375904476025 | 0.455960173459244 | -4.92465625360154 |
| ENSG00000151131.10 | NOPCHAP1 | 0.210529491153252 | 3.86834032690696 | 1.47185174710194 | 0.148427862696349 | 0.456051753676513 | -5.23018944900575 |
| ENSG00000143570.17 | SLC39A1 | -0.23442566812248 | 3.12977511756331 | -1.47176499100278 | 0.148451202502871 | 0.456055388226358 | -5.16389150451729 |
| ENSG00000124659.6 | TBCC | 0.181984645395943 | 4.714288099147 | 1.47139814133889 | 0.148549927365988 | 0.456290576775725 | -5.31487013640582 |
| ENSG00000136463.7 | TACO1 | 0.141564911806926 | 3.23478671325715 | 1.47115529002434 | 0.148615310931469 | 0.45639613554515 | -5.17275422427664 |
| ENSG00000206775.1 | SNORD37 | 0.481877221407709 | -0.257144200615037 | 1.47110579291779 | 0.148628639972751 | 0.45639613554515 | -4.67065215637037 |
| lnc-HHIPL1-1 | lnc-HHIPL1-1 | -0.650160040840139 | 2.16547090778127 | -1.47156531388798 | 0.148666843178414 | 0.456445351005563 | -5.05750433921573 |
| ENSG00000100664.10 | EIF5 | 0.146084641898074 | 8.48719178897352 | 1.47078914510579 | 0.148713932209708 | 0.456521829481344 | -5.33120946350807 |
| ENSG00000185834.10 | RPL12P4 | 0.87362055591223 | 1.62295604390875 | 1.47100825170025 | 0.148854207902111 | 0.456884306777387 | -4.92518287498936 |
| ENSG00000165704.14 | HPRT1 | 0.28083622237736 | 2.01774689691381 | 1.46999497318953 | 0.148928020569981 | 0.457042708721025 | -5.04103191306435 |
| lnc-KIAA0825-1 | lnc-KIAA0825-1 | -0.275079303503508 | 0.68484742479431 | -1.46988831333762 | 0.148956791877513 | 0.457062857366556 | -4.88072090460884 |
| ENSG00000118263.14 | KLF7 | -0.897159658806751 | 4.1323859068516 | -1.47066706833912 | 0.14902745902319 | 0.45721153511318 | -5.26252159936447 |
| lnc-AKAP8-3 | lnc-AKAP8-3 | -0.204214709837148 | 3.43044667424383 | -1.46933489150301 | 0.149106146997786 | 0.457246925520344 | -5.19724126584297 |
| ENSG00000112773.15 | TENT5A | -0.692670308028775 | 3.84036879378854 | -1.47008456136603 | 0.149129135757371 | 0.457246925520344 | -5.22736279997285 |
| lnc-GPS1-2 | lnc-GPS1-2 | -0.455801079563385 | -0.306635764372011 | -1.46914929611391 | 0.149156261184772 | 0.457246925520344 | -4.64270293991565 |
| ENSG00000124613.8 | ZNF391 | -0.599417165149097 | 1.76568859723933 | -1.46948201032007 | 0.149169586007126 | 0.457246925520344 | -5.01678030590262 |
| ENSG00000186352.8 | ANKRD37 | -0.468399877098834 | 0.812140979572568 | -1.46909592101139 | 0.149170675914001 | 0.457246925520344 | -4.88549408995258 |
| ENSG00000120784.16 | ZFP30 | -0.481758869004029 | 3.92572612656806 | -1.46952710335939 | 0.149177088369314 | 0.457246925520344 | -5.24767507188672 |
| ENSG00000128340.14 | RAC2 | -0.356465826484505 | 6.53047103556337 | -1.46917843454821 | 0.149194497973795 | 0.457246925520344 | -5.36152059625114 |
| ENSG00000196352.15 | CD55 | 0.238185533039885 | 7.12989647104 | 1.46871189107603 | 0.149274421292534 | 0.457387909180802 | -5.35955669349338 |
| ENSG00000130592.15 | LSP1 | -0.277622911548033 | 7.24655163644741 | -1.46867295651235 | 0.149284942616559 | 0.457387909180802 | -5.3562780699022 |
| lnc-ANKS1B-1 | lnc-ANKS1B-1 | -0.233326322392704 | 0.301542691064793 | -1.46827382796343 | 0.149392833303766 | 0.457597201170757 | -4.79796573779654 |
| ENSG00000173621.8 | LRFN4 | -0.454560481486214 | 1.84665504618304 | -1.46825659726802 | 0.149397716167103 | 0.457597201170757 | -5.02899077373521 |
| ENSG00000137073.21 | UBAP2 | -0.176358096524679 | 4.88344963128165 | -1.46796646144065 | 0.149475961269833 | 0.457745697206341 | -5.33120038991591 |
| lnc-ARSJ-3 | lnc-ARSJ-3 | -0.222098445927534 | 3.9673362599632 | -1.4677993630801 | 0.149521168730413 | 0.457745697206341 | -5.25398708922229 |
| lnc-SH3KBP1-1 | lnc-SH3KBP1-1 | -0.247116328702212 | 0.550006537172834 | -1.46770140012985 | 0.149547677042695 | 0.457745697206341 | -4.83968896737027 |
| lnc-TTC5-2 | lnc-TTC5-2 | -0.239247715075354 | 1.91037374582645 | -1.46770105119296 | 0.149547771470007 | 0.457745697206341 | -5.03214262005545 |
| ENSG00000175274.18 | TP53I11 | -0.558495661516358 | 4.86285002426792 | -1.46842843121602 | 0.149557392688755 | 0.457745697206341 | -5.32934291991285 |
| ENSG00000122042.9 | UBL3 | 0.179642210132702 | 5.08702381007856 | 1.46718114018187 | 0.149688519160898 | 0.457939065261671 | -5.34175753745793 |
| lnc-TTC30B-2 | lnc-TTC30B-2 | 0.259129569078214 | 3.18790259128046 | 1.46709612739297 | 0.149711543346858 | 0.457939065261671 | -5.1656455772073 |

| | | | | | | | |
|---|---|---|---|---|---|---|---|
| lnc-TET3-6 | lnc-TET3-6 | -0.43948175364968 | 2.60154288325239 | -1.46711774269102 | 0.149720094678257 | 0.457939065261671 | -5.10006439146308 |
| ENSG00000112146.16 | FBXO9 | -0.128865062682927 | 5.28851328526172 | -1.46704546866057 | 0.14972526468432 | 0.457939065261671 | -5.34920946136481 |
| lnc-ENAH-4 | lnc-ENAH-4 | 0.22031382646814 | 1.58036886407154 | 1.46702129338416 | 0.149731813108441 | 0.457939065261671 | -4.99568748389493 |
| ENSG00000114956.19 | DGUOK | 0.157645798644563 | 4.39063648421982 | 1.46683258537037 | 0.149782936747511 | 0.458026701770344 | -5.29081889349148 |
| ENSG00000145715.14 | RASA1 | -0.260972217614581 | 6.79445223195985 | -1.46675126118634 | 0.149804972857113 | 0.458026701770344 | -5.36310628519588 |
| ENSG00000130429.14 | ARPC1B | -0.309440203130248 | 5.44744912073979 | -1.46648217415724 | 0.149877904618058 | 0.458141087434194 | -5.35473057552894 |
| lnc-MAP1LC3C-5 | lnc-MAP1LC3C-5 | -1.29323199739425 | -2.53535160217788 | -1.46644898860906 | 0.149886900975653 | 0.458141087434194 | -4.41505674994973 |
| lnc-ST3GAL1-7 | lnc-ST3GAL1-7 | -0.364431432933091 | 0.918203146817997 | -1.46634623658778 | 0.149914758996526 | 0.458158201102523 | -4.91284592408593 |
| ENSG00000179085.7 | DPM3 | -0.337663672734534 | 1.07001880460688 | -1.466207592032 | 0.149952354645832 | 0.458194242677675 | -4.92469959160085 |
| lnc-STK10-2 | lnc-STK10-2 | 0.203038306554601 | 2.11976397136292 | 1.46610925920755 | 0.149979023649422 | 0.458194242677675 | -5.05024197072746 |
| ENSG00000106785.14 | TRIM14 | 0.327404823123806 | 3.97372162610626 | 1.46602687986563 | 0.150001368767048 | 0.458194242677675 | -5.24741099673203 |
| lnc-SERPINB1-4 | lnc-SERPINB1-4 | 0.47041036776469 | 1.31361535380759 | 1.46597443561398 | 0.15001559546251 | 0.458194242677675 | -4.92941887439449 |
| ENSG00000182473.21 | EXOC7 | -0.144531840400004 | 6.44239012949603 | -1.46560174100496 | 0.150116728046072 | 0.458435105841587 | -5.36725529930597 |
| ENSG00000095564.13 | BTAF1 | 0.161180609912432 | 8.54880995920111 | 1.46550831621414 | 0.150142087777959 | 0.458444532374089 | -5.33800482238996 |
| ENSG00000047410.13 | TPR | 0.119755451864835 | 8.25784582664873 | 1.46539471645331 | 0.150172928470985 | 0.458468766155002 | -5.34426375050043 |
| lnc-ZNF672-2 | lnc-ZNF672-2 | 0.194156634447294 | 1.85867033654699 | 1.46531500223945 | 0.15019457271453 | 0.458468766155002 | -5.01920390600332 |
| lnc-ONECUT1-5 | lnc-ONECUT1-5 | 0.303889167436157 | 2.0347734319458 | 1.46511860743138 | 0.150247908935796 | 0.458563569042926 | -5.04672406150874 |
| ENSG00000207233.1 | SNORA37 | 0.414391455800725 | 0.775998966692961 | 1.46499498779196 | 0.150281488798117 | 0.458598055438346 | -4.84931927159043 |
| lnc-RAB11FIP2-8 | lnc-RAB11FIP2-8 | 0.207102648025184 | 1.47755994515999 | 1.46465770519273 | 0.150373137903462 | 0.458685400265213 | -4.99067451965907 |
| ENSG00000181109.3 | OR52P1 | 0.438990052770654 | 0.235882539547362 | 1.46465176440774 | 0.150374752576125 | 0.458685400265213 | -4.75281653064853 |
| ENSG00000136213.9 | CHST12 | 0.268349982590776 | 2.90947693642228 | 1.46464362257012 | 0.150376965505012 | 0.458685400265213 | -5.14768516881681 |
| ENSG00000075234.16 | TTC38 | -0.448632642826847 | 1.7483654300767 | -1.46442464348155 | 0.15043671666716 | 0.458799665011136 | -5.04094704783393 |
| ENSG00000162636.15 | EEIG2 | -0.295336649899693 | 3.95712037779706 | -1.46427587176543 | 0.150476945965627 | 0.458854367231186 | -5.26124995296759 |
| lnc-EGR3-2 | lnc-EGR3-2 | -0.596342212757876 | -0.295331551181508 | -1.4640273045481 | 0.150544553686938 | 0.458992526816507 | -4.62408346184453 |
| lnc-NETO2-2 | lnc-NETO2-2 | 0.237650070223624 | 0.732035426283704 | 1.46388407095128 | 0.15058352263783 | 0.459024092260579 | -4.85184408619866 |
| ENSG00000218596.2 | | 0.426701778570605 | 0.963369828457477 | 1.46381635146768 | 0.150601949559719 | 0.459024092260579 | -4.88875257112975 |
| ENSG00000183527.11 | PSMG1 | -0.208111621137496 | 2.97354059928662 | -1.46369657373929 | 0.150634546234701 | 0.459024092260579 | -5.1652177650081 |
| lnc-ERLEC1-5 | lnc-ERLEC1-5 | -0.451812555033898 | -0.641072707872732 | -1.46366143036464 | 0.150644111316145 | 0.459024092260579 | -4.59455540392704 |
| ENSG00000213928.8 | IRF9 | 0.192524931641946 | 6.09264609367383 | 1.46347206610204 | 0.150695659427664 | 0.459113196861992 | -5.36977490572965 |
| ENSG00000136059.14 | VILL | -0.161306073039147 | 4.50744543561844 | -1.46332090954608 | 0.150736816755759 | 0.459170622951574 | -5.30944857605664 |
| ENSG00000196262.13 | PPIA | -0.212390358297579 | 8.68509463341167 | -1.46265534555928 | 0.15091814392051 | 0.459504263627304 | -5.33656708901672 |
| ENSG00000166133.17 | RPUSD2 | -0.252612900454259 | 3.20249777067647 | -1.46259197811024 | 0.150935416806254 | 0.459504263627304 | -5.18874530792222 |
| ENSG00000089682.16 | RBM41 | -0.172866623093013 | 4.03164967716259 | -1.46257315201114 | 0.150940548780342 | 0.459504263627304 | -5.26751833196737 |
| ENSG00000120008.15 | WDR11 | 0.296677047129389 | 8.71195876212206 | 1.46247116568043 | 0.150968352536509 | 0.459504263627304 | -5.33831948428088 |
| lnc-SPATA9-4 | lnc-SPATA9-4 | 0.153895737600442 | 2.8965818097217 | 1.46241531436509 | 0.15098358056764 | 0.459504263627304 | -5.14128781453066 |
| ENSG00000213917.2 | RPL5P8 | -0.340523062467508 | 0.139009922992859 | -1.46231716997847 | 0.151010342869858 | 0.459504263627304 | -4.74171933407863 |
| ENSG00000162441.11 | LZIC | 0.156783961775137 | 4.09381574413346 | 1.46224059860788 | 0.15103122517755 | 0.459504263627304 | -5.27032183871624 |
| ENSG00000214535.3 | RPS15AP1 | 0.912288516248838 | -0.964853108650743 | 1.46209830014932 | 0.151070038425703 | 0.459504263627304 | -4.46732251264901 |
| lnc-ERG28-2 | lnc-ERG28-2 | -0.306862357272357 | 2.211309404663 | -1.46203347148574 | 0.151087723665277 | 0.459504263627304 | -5.07687849463397 |
| lnc-SNRPN-1 | lnc-SNRPN-1 | -0.303392722935294 | 6.87049096154902 | -1.46191677160886 | 0.151119563473152 | 0.459504263627304 | -5.37024477978388 |
| ENSG00000028839.9 | TBPL1 | 0.194866274835624 | 3.95909885458823 | 1.46181272128193 | 0.151147956505801 | 0.459504263627304 | -5.26514762131425 |
| ENSG00000064999.14 | ANKS1A | -0.151968581908867 | 3.7164680735732 | -1.46178606971272 | 0.151155229806614 | 0.459504263627304 | -5.24232552405898 |
| ENSG00000198918.7 | RPL39 | 0.250699885293636 | 7.35137421668666 | 1.46173621436651 | 0.151168836235863 | 0.459504263627304 | -5.36543011675686 |
| ENSG00000198925.11 | ATG9A | 0.22852890527269 | 4.80429562222077 | 1.46171207102485 | 0.151175425739494 | 0.459504263627304 | -5.33362792509024 |
| lnc-SPTBN2-3 | lnc-SPTBN2-3 | -0.295372269702742 | 0.48782647089202 | -1.46162606737685 | 0.151198900775403 | 0.459504263627304 | -4.83417857923693 |
| ENSG00000285269.2 | | 0.276295407537416 | 3.30119160963665 | 1.4616090876148 | 0.151203535808567 | 0.459504263627304 | -5.1857829457596 |
| ENSG00000183935.5 | HTR7P1 | -0.569330662876515 | -0.0812031604015058 | -1.46133711525815 | 0.151277792463709 | 0.459662061157444 | -4.69770217019041 |
| ENSG00000188811.13 | NHLRC3 | -0.156608747070385 | 5.4463268850145 | -1.4611038957413 | 0.15134149133574 | 0.459704032067113 | -5.36250431520651 |
| ENSG00000236086.4 | HMGN2P28 | -0.404705503415631 | 0.207419366086782 | -1.46108462653215 | 0.151346755250094 | 0.459704032067113 | -4.75409152663598 |
| lnc-LMNTD1-1 | lnc-LMNTD1-1 | -0.363529070086983 | 3.00200030631063 | -1.4610412405708 | 0.151358607847195 | 0.459704032067113 | -5.18068288677439 |
| lnc-JAM2-1 | lnc-JAM2-1 | -0.477073534830566 | 0.979464514508878 | -1.46081211686731 | 0.151421214224597 | 0.459826328177172 | -4.88044337952174 |
| ENSG00000065491.8 | TBC1D22B | 0.115556197979183 | 3.80004470532851 | 1.46062616620364 | 0.151472038899427 | 0.459908639981491 | -5.24727430568631 |
| ENSG00000081237.19 | PTPRC | 0.16274272091044 | 9.93319015991368 | 1.46054946236178 | 0.151493007777025 | 0.459908639981491 | -5.30555219982793 |
| lnc-PC-2 | lnc-PC-2 | -0.25976846829503 | 4.07994245196724 | -1.46014482141838 | 0.151603664211504 | 0.460176702619876 | -5.27716204224861 |
| ENSG00000173926.5 | MARCHF3 | -0.236315666644857 | 4.73997764367339 | -1.45979531210112 | 0.15169929520279 | 0.460399084806699 | -5.33016670276777 |
| ENSG00000135414.9 | GDF11 | 0.640239958616829 | 1.0436193013591 | 1.4597876634984 | 0.151779080317247 | 0.460573317141369 | -4.92843041988703 |
| ENSG00000183281.14 | PLGLB1 | 0.421941130854715 | 0.0303517536652355 | 1.45940527104906 | 0.151806072446955 | 0.46058732151764 | -4.73096121295131 |
| ENSG00000112282.17 | MED23 | 0.172554139779675 | 6.50834003250342 | 1.45924585056523 | 0.151849732296446 | 0.460625625593577 | -5.37607049439122 |
| lnc-GDAP2-3 | lnc-GDAP2-3 | 1.01747471763257 | -0.940374747149921 | 1.45913439220504 | 0.151880262831997 | 0.460625625593577 | -4.47611286847764 |
| ENSG00000132825.6 | PPP1R3D | -0.258740493649863 | 2.59690401991231 | -1.45897754685726 | 0.151923233912586 | 0.460625625593577 | -5.13864577359417 |
| ENSG00000126653.17 | NSRP1 | -0.17897225140501 | 4.72834140843466 | -1.45896244132412 | 0.151927372896998 | 0.460625625593577 | -5.32741219117306 |
| ENSG00000087008.15 | ACOX3 | -0.191438226832805 | 3.39337774537508 | -1.45895069371988 | 0.151930591855162 | 0.460625625593577 | -5.20941217892864 |
| SNHG25 | SNHG25 | -0.289857084932488 | 1.76704375062673 | -1.45868248705092 | 0.152004097728858 | 0.46078061024346 | -5.03687758815747 |
| ENSG00000065809.13 | FAM107B | 0.169899765998824 | 6.56716062794673 | 1.45857992575725 | 0.152032213528258 | 0.460797975416307 | -5.3771325720851 |
| ENSG00000166987.14 | MBD6 | -0.297103850495418 | 3.60647714960684 | -1.45836060652053 | 0.152092350706063 | 0.460912375527517 | -5.23963534090869 |
| lnc-CAMKMT-1 | lnc-CAMKMT-1 | -0.602507965198924 | 1.14197822623098 | -1.45842586383103 | 0.1521182693666 | 0.460923058791804 | -4.91387041036759 |
| ENSG00000165118.14 | QNG1 | -0.258821059105732 | 1.72309393619484 | -1.45808485206574 | 0.152167988991277 | 0.461005845953409 | -5.02553529787697 |
| ENSG00000146409.11 | SLC18B1 | 0.230792350672482 | 4.80771623191534 | 1.45783849554855 | 0.152235588623404 | 0.461086357089347 | -5.33418859358772 |

| | | | | | | | |
|---|---|---|---|---|---|---|---|
| lnc-GSX1-2 | lnc-GSX1-2 | 0.251448233481068 | 0.144333447567628 | 1.4578247298263 | 0.152239366602497 | 0.461086357089347 | -4.77218228102823 |
| lnc-GPR160-1 | lnc-GPR160-1 | 0.780672425875516 | -0.984932645719168 | 1.45732260772668 | 0.152377223680735 | 0.461364170652576 | -4.54902363891975 |
| ENSG00000152409.8 | JMY | 0.258766172589007 | 6.11012847343599 | 1.45731309922531 | 0.152379835178411 | 0.461364170652576 | -5.37807225349251 |
| lnc-ZNF100-7 | lnc-ZNF100-7 | 0.259336572559508 | 1.641837071458 | 1.45719727332316 | 0.152411649445662 | 0.461364170652576 | -5.00915340353585 |
| lnc-JUND-1 | lnc-JUND-1 | -0.436862886893805 | 0.542058496034485 | -1.45716413335101 | 0.152420753069889 | 0.461364170652576 | -4.87876239299896 |
| lnc-MUC16-1 | lnc-MUC16-1 | -0.651633987158729 | 0.888410343853458 | -1.4568332399307 | 0.152581005981615 | 0.461750820320257 | -4.90454473662923 |
| lnc-LRRC63-6 | lnc-LRRC63-6 | -0.350395436959694 | 1.93229367506843 | -1.4565360950714 | 0.152593357616401 | 0.461750820320257 | -5.06379894783828 |
| ENSG00000130770.17 | ATP5IF1 | -0.194746358118791 | 4.6566192723238 | -1.45534898452697 | 0.152920033384997 | 0.462671328408555 | -5.33030817671507 |
| ENSG00000130985.16 | UBA1 | 0.164672722029103 | 6.768915127186 | 1.45518598341535 | 0.152964931937561 | 0.4627246813417 | -5.38029660149755 |
| ENSG00000091831.23 | ESR1 | -0.392390405911601 | 2.54095213353931 | -1.45507585656717 | 0.152995272185703 | 0.4627246813417 | -5.12476717475824 |
| ENSG00000244510.2 | | -0.210077969933551 | 0.288203302794319 | -1.45504015130371 | 0.153005110101133 | 0.4627246813417 | -4.79663460699754 |
| ENSG00000223803.1 | RPS20P14 | 0.551144371349508 | 0.686013579429446 | 1.45492452439844 | 0.153036972345341 | 0.462732321948729 | -4.82784519609511 |
| ENSG00000100532.11 | CGRRF1 | 0.197032073686367 | 3.86173660851558 | 1.45486604747309 | 0.153053088284468 | 0.462732321948729 | -5.26271847122859 |
| ENSG00000151500.14 | THYN1 | -0.213542700691363 | 3.34075564653748 | -1.45466720478779 | 0.153107898305016 | 0.462732321948729 | -5.20803994847794 |
| ENSG00000116604.17 | MEF2D | 0.223828704790831 | 5.37073890882522 | 1.45464415890391 | 0.153114251790645 | 0.462732321948729 | -5.36887111780224 |
| ENSG00000230510.6 | PPP5D1P | -0.347470706080884 | -0.416327745201913 | -1.45462315325949 | 0.153120042986581 | 0.462732321948729 | -4.62485488240188 |
| lnc-KANTR-23 | lnc-KANTR-23 | -0.251187606559665 | 3.64077447352838 | -1.45453647435996 | 0.15314394193551 | 0.462736605528275 | -5.23529938672622 |
| ENSG00000120709.10 | FAM53C | 0.205034606279726 | 5.42098060322621 | 1.45435474026306 | 0.153194058868622 | 0.46282009594787 | -5.37059970813051 |
| ENSG00000256043.2 | CTSO | 0.182279928367141 | 4.52382343929195 | 1.45417468949751 | 0.153243724326178 | 0.462899287573727 | -5.32236434824658 |
| ENSG00000183955.12 | KMT5A | -0.147964742626421 | 4.79874545182755 | -1.45409666664674 | 0.153265250197491 | 0.462899287573727 | -5.34661660703675 |
| ENSG00000103152.11 | MPG | -0.272160224035828 | 3.94423697087784 | -1.45363597024127 | 0.153392401143779 | 0.463161881975485 | -5.28087172824771 |
| ENSG00000120948.17 | TARDBP | 0.161120795443447 | 6.35829745322298 | 1.45361859199659 | 0.153397199117081 | 0.463161881975485 | -5.38417240744218 |
| ENSG00000078747.14 | ITCH | 0.092450176079482 | 7.11607919981509 | 1.45334136873555 | 0.153473753887078 | 0.463325062519466 | -5.37981002907661 |
| lnc-SERPINF1-1 | lnc-SERPINF1-1 | -0.241354909753559 | 2.47697795482005 | -1.45306476532033 | 0.153550167467669 | 0.463397120221731 | -5.1180322581995 |
| ENSG00000227032.1 | RPS2P36 | -0.376703066172813 | 0.839308079958087 | -1.45303354949397 | 0.153558792933457 | 0.463397120221731 | -4.90379939922167 |
| ENSG00000178719.16 | GRINA | -0.338110582649033 | 3.99730680797187 | -1.45301049568189 | 0.153565163340253 | 0.463397120221731 | -5.28855251991671 |
| lnc-LYN-2 | lnc-LYN-2 | -0.273348148539328 | 1.03562521626778 | -1.45286217973602 | 0.153606152119654 | 0.463452862661806 | -4.94305910497418 |
| ENSG00000176160.10 | HSF5 | -0.309898732001652 | 3.02970536186217 | -1.4525777219362 | 0.15368478933368 | 0.463622163103494 | -5.16333548211687 |
| ENSG00000213930.11 | GALT | -0.146330684960523 | 4.53605364172404 | -1.45222475184246 | 0.153782410555294 | 0.463848674744962 | -5.32851033506584 |
| lnc-LY75-1 | lnc-LY75-1 | -0.282455602191579 | 1.8214300337732 | -1.45200946257694 | 0.153841977242818 | 0.463960354225482 | -5.04258771492464 |
| ENSG00000241685.9 | ARPC1A | 0.138974302358234 | 3.75860477034463 | 1.45192723567859 | 0.153864732746854 | 0.463961001190814 | -5.25438476870909 |
| ENSG00000227057.9 | WDR46 | -0.251676880434699 | 4.03231944938712 | -1.45151781613153 | 0.153978075113492 | 0.464190022480746 | -5.28728037772991 |
| ENSG00000176658.16 | MYO1D | 0.514428589394223 | 3.99343184763642 | 1.45201351537305 | 0.154003731406899 | 0.464190022480746 | -5.29255137490483 |
| lnc-AOAH-3 | lnc-AOAH-3 | 0.172953536736262 | 2.41483115638897 | 1.45134503614418 | 0.15402592667302 | 0.464190022480746 | -5.11116438135511 |
| ENSG00000248098.11 | BCKDHA | -0.196873195455881 | 3.17092267924522 | -1.45132710939729 | 0.154030892170407 | 0.464190022480746 | -5.1986676726689 |
| ENSG00000157654.17 | PALM2AKAP2 | -0.337072209001 | 6.87085105802447 | -1.45111527540414 | 0.154117184058422 | 0.464340432958149 | -5.38389768591391 |
| lnc-MEX3C-4 | lnc-MEX3C-4 | -0.292395203813818 | 1.6187845775159 | -1.45095008653621 | 0.154135352245475 | 0.464340432958149 | -5.02165848525751 |
| ENSG00000113456.18 | RAD1 | -0.125921947128451 | 4.42824913114595 | -1.45090271681808 | 0.154148480707527 | 0.464340432958149 | -5.32516740479218 |
| ENSG00000189339.11 | SLC35E2B | -0.21228532919337 | 5.81438838657678 | -1.45078488573701 | 0.154181141271004 | 0.464359030103147 | -5.38313412459871 |
| ENSG00000213999.15 | MEF2B | -0.254437116293272 | 0.695329335100029 | -1.45071766840096 | 0.154199775093767 | 0.464359030103147 | -4.89355822193313 |
| ENSG00000258890.6 | CEP95 | 0.255846070341954 | 7.09573368651558 | 1.45051726380286 | 0.154255341276515 | 0.464436613925666 | -5.3839616764206 |
| ENSG00000090238.11 | YPEL3 | -0.249333177973038 | 5.74971941304133 | -1.45046200180831 | 0.154270666540824 | 0.464436613925666 | -5.38296081538399 |
| ENSG00000284482.1 | MIR4680 | -0.340404054739393 | 2.11499434915468 | -1.45023210738777 | 0.154334433754207 | 0.464505644663432 | -5.0966028633193 |
| ENSG00000260075.1 | NSFP1 | -1.34041260437356 | -0.254017126093607 | -1.4508038185532 | 0.154338731178799 | 0.464505644663432 | -4.58537798007984 |
| ENSG00000126457.21 | PRMT1 | -0.231448223435132 | 5.6194569150535 | -1.45003254006343 | 0.154389805770881 | 0.464542512535197 | -5.38230406530429 |
| lnc-USP13-5 | lnc-USP13-5 | -0.230691045216717 | 0.454491248088072 | -1.44995509970324 | 0.154411296608215 | 0.464542512535197 | -4.83200539927833 |
| ENSG00000202335.1 | | -0.249753134083621 | 1.36210062115723 | -1.44992846468505 | 0.15441868876091 | 0.464542512535197 | -4.99275639307998 |
| ENSG00000185811.18 | IKZF1 | -0.220206395909153 | 7.81719624272179 | -1.44981406841716 | 0.154450440907669 | 0.464570133742882 | -5.37405179716597 |
| lnc-NCAPD3-3 | lnc-NCAPD3-3 | -0.238355809353741 | 1.1974164378931 | -1.44926305870071 | 0.154603452662435 | 0.464962429303171 | -4.96618234196474 |
| ENSG00000143179.15 | UCK2 | -0.245653768329234 | 3.21537158263782 | -1.44881541944986 | 0.154727846889544 | 0.465212473263747 | -5.21554453924928 |
| ENSG00000148843.14 | PDCD11 | -0.137857372055542 | 5.78086878850277 | -1.44880120690356 | 0.154731797695361 | 0.465212473263747 | -5.38654944634702 |
| ENSG00000182489.8 | XKRX | -0.529532984196153 | 4.05350569697753 | -1.44924897546102 | 0.154777845873381 | 0.465282956274542 | -5.28554683710789 |
| ENSG00000172164.14 | SNTB1 | 2.71694137259765 | -1.3771773524884 | 1.44887512922101 | 0.154916886348625 | 0.465632925191845 | -4.42749332174254 |
| ENSG00000076984.17 | MAP2K7 | 0.180430375210102 | 4.28683469422583 | 1.44789701372256 | 0.154983308594654 | 0.465764555526903 | -5.31439478980329 |
| ENSG00000110442.11 | COMMD9 | 0.127922931649451 | 3.93529133028815 | 1.44779877854735 | 0.155010653098703 | 0.465778725946074 | -5.28179813594416 |
| ENSG00000169689.14 | CENPX | -0.257997238716643 | 1.50374801714054 | -1.44735541594904 | 0.155134113635417 | 0.466048000334068 | -5.01703735222155 |
| ENSG00000168675.18 | LDLRAD4 | -0.567953690829571 | 4.90869879425657 | -1.4480528132268 | 0.155145552071566 | 0.466048000334068 | -5.35549001406202 |
| ENSG00000118705.16 | RPN2 | 0.122966302837008 | 6.42229110462126 | 1.44697240400388 | 0.15524083092127 | 0.466186970124835 | -5.39329126309481 |
| lnc-ABCA8-4 | lnc-ABCA8-4 | -0.615978097101856 | 2.09165322809812 | -1.44742907170878 | 0.155250109795244 | 0.466186970124835 | -5.069576379322 |
| lnc-RBM4-2 | lnc-RBM4-2 | -0.411125041382825 | 0.480440257445041 | -1.44689225284551 | 0.155263170460907 | 0.466186970124835 | -4.84116782392685 |
| ENSG00000078018.19 | MAP2 | -0.799984406029684 | 4.49936650734824 | -1.44782266568881 | 0.15528241107593 | 0.466186970124835 | -5.33798474391673 |
| ENSG00000040933.15 | INPP4A | 0.309881943423954 | 4.25597447004118 | 1.44654195512294 | 0.155360834268522 | 0.466354389929852 | -5.3069781075771 |
| lnc-INTS13-4 | lnc-INTS13-4 | 0.261952473415668 | 2.61681173138309 | 1.44644998671708 | 0.155386483273275 | 0.466363369089211 | -5.13778453808047 |
| lnc-KCNJ16-4 | lnc-KCNJ16-4 | -0.64760108874867 | 2.13695829109448 | -1.44686282157364 | 0.155429603776233 | 0.466424775408399 | -5.07428700802559 |
| lnc-IAH1-2 | lnc-IAH1-2 | -0.384250150096568 | 1.3022174380652 | -1.4460928087812 | 0.155486127948803 | 0.46650657774469 | -4.97782753754161 |
| ENSG00000138375.12 | SMARCAL1 | -0.191426179842806 | 3.71214504566338 | -1.44603524169137 | 0.155502192581563 | 0.46650657774469 | -5.26156012610391 |
| lnc-CA13-1 | lnc-CA13-1 | -0.287617994590054 | 1.18411690179118 | -1.4459171897817 | 0.155535140147734 | 0.466537421984961 | -4.95973900149233 |
| lnc-TOR1AIP2-7 | lnc-TOR1AIP2-7 | -0.327946818121578 | 0.10001676237313 | -1.44558143614439 | 0.155628876930748 | 0.466750571742035 | -4.78203735894042 |

| | | | | | | | |
|---|---|---|---|---|---|---|---|
| ENSG00000172943.19 | PHF8 | 0.156590891122931 | 5.93607633838365 | 1.44497693264867 | 0.155797756092809 | 0.46718898800383 | -5.39369320164052 |
| ENSG00000185760.15 | KCNQ5 | -0.278571108049554 | 4.03974802447601 | -1.4448537691787 | 0.155832181725095 | 0.46722415097562 | -5.29684905142985 |
| ENSG00000132394.10 | EEFSEC | -0.146768002822407 | 3.0017726835446 | -1.44437892366784 | 0.155964962557357 | 0.467496748899142 | -5.18452374341248 |
| ENSG00000152439.12 | ZNF773 | -0.155234889924752 | 4.11456402741381 | -1.4442690181672 | 0.155995708047399 | 0.467496748899142 | -5.30149799215882 |
| ENSG00000205683.11 | DPF3 | 1.22171693479425 | -1.05295277780155 | 1.44427760580619 | 0.156012629886958 | 0.467496748899142 | -4.48724840775619 |
| ENSG00000126860.11 | EVI2A | -0.470241702211162 | 3.94033548603527 | -1.44461971491374 | 0.156013951716864 | 0.467496748899142 | -5.27308778339934 |
| lnc-HSPG2-3 | lnc-HSPG2-3 | 0.158743884011422 | 1.46409166150965 | 1.44400690439083 | 0.156069052229049 | 0.467572763772757 | -5.00019606481562 |
| lnc-ZNF8-2 | lnc-ZNF8-2 | 0.15020312000535 | 3.07832583830161 | 1.44388361362621 | 0.156103560585395 | 0.467572763772757 | -5.19582244843213 |
| lnc-FAM184B-1 | lnc-FAM184B-1 | -0.339118223639966 | -0.0140991744529788 | -1.44386965137977 | 0.156107468913491 | 0.467572763772757 | -4.7402272770567 |
| ENSG00000145545.11 | SRD5A1 | -0.280460816448418 | 2.82640651539199 | -1.44329170350208 | 0.156269316208976 | 0.467989427546828 | -5.17098156556455 |
| ENSG00000000419.12 | DPM1 | 0.138383962747455 | 4.35903405497542 | 1.44295957347217 | 0.156362384817236 | 0.468109113673313 | -5.32455523308162 |
| lnc-ABCG2-2 | lnc-ABCG2-2 | 0.166694715802146 | 3.72632763451484 | 1.44292503702693 | 0.156372065028002 | 0.468109113673313 | -5.25874974391138 |
| lnc-RGS9-15 | lnc-RGS9-15 | -0.433703167510484 | 2.2168247621912 | -1.44279974397393 | 0.156409422389121 | 0.468109113673313 | -5.09780073827171 |
| lnc-TACR1-1 | lnc-TACR1-1 | 1.44249016521334 | 4.037375740511 | 1.44379334696079 | 0.156419357405455 | 0.468109113673313 | -5.28922337372864 |
| ENSG00000151176.7 | PLBD2 | -0.303098723723263 | 1.54594566222603 | -1.44273573612285 | 0.156425132487591 | 0.468109113673313 | -5.02524675394491 |
| ENSG00000149474.13 | KAT14 | -0.209986980014254 | 4.11831058231588 | -1.44261816278642 | 0.156458099398931 | 0.468109113673313 | -5.30923540816572 |
| ENSG00000011258.15 | MBTD1 | 0.14810898733173 | 5.60866347809856 | 1.44258114916449 | 0.156468478942447 | 0.468109113673313 | -5.39173920757052 |
| ENSG00000248734.2 | | -0.305570983936658 | 1.90565278295229 | -1.44246412017298 | 0.156501300351051 | 0.468115393839819 | -5.06730960429447 |
| ENSG00000229932.3 | YWHAZP3 | 1.02628907485278 | 0.0822231119378219 | 1.44265519469359 | 0.15651606376163 | 0.468115393839819 | -4.61303896863992 |
| lnc-TRIM69-3 | lnc-TRIM69-3 | 0.208617340028484 | 8.93538633846896 | 1.44226890658802 | 0.156556061080113 | 0.468166991894808 | -5.35922168571227 |
| ENSG00000157869.14 | RAB28 | 0.115866245749388 | 3.70856197471833 | 1.44212902096492 | 0.156595310629205 | 0.468216339145979 | -5.26466209771261 |
| ENSG00000198911.11 | SREBF2 | -0.298695805648761 | 5.78194667158582 | -1.44193987939431 | 0.156648392854559 | 0.468228818970513 | -5.39806241870473 |
| lnc-NNT-2 | lnc-NNT-2 | -0.317854706276776 | 0.703234767600686 | -1.44190235559437 | 0.156658925517756 | 0.468228818970513 | -4.87750352717625 |
| ENSG00000100600.14 | LGMN | 1.13389534676394 | -0.0859578023763094 | 1.44222971161622 | 0.15666772949764 | 0.468228818970513 | -4.59898682726071 |
| ENSG00000140694.16 | PARN | -0.112885285260615 | 5.84454196745452 | -1.44164391167786 | 0.156731483988836 | 0.468351355247128 | -5.39641152779677 |
| ENSG00000165271.16 | NOL6 | 0.158229664532861 | 5.23690196518567 | 1.44146804030739 | 0.156780875194877 | 0.46843094123039 | -5.38246303882216 |
| ENSG00000135541.20 | AHI1 | -0.33704691000628 | 5.43848848381014 | -1.44142187985193 | 0.15682139446378 | 0.468448274010476 | -5.38980372700316 |
| ENSG00000168056.15 | LTBP3 | -0.424539767832733 | 2.60788703651981 | -1.44129134820282 | 0.156837967329789 | 0.468448274010476 | -5.12971383258172 |
| ENSG00000174197.16 | MGA | 0.13704781734617 | 7.59692531205484 | 1.4412043456555 | 0.156854953334315 | 0.468448274010476 | -5.39069010667142 |
| ENSG00000232888.4 | RPS11P5 | 1.0303374229779 | 1.78229187150164 | 1.44195841351735 | 0.156881003380304 | 0.46845810134583 | -4.94504638097615 |
| lnc-MALL-6 | lnc-MALL-6 | -1.48205434671691 | -2.46923560055157 | -1.44100597611404 | 0.156910698196374 | 0.468478807800401 | -4.43398025661456 |
| lnc-ZDHHC19-3 | lnc-ZDHHC19-3 | -0.157634385147278 | 3.6473729563982 | -1.44089603471 | 0.156941600103243 | 0.468503111664256 | -5.25999361344667 |
| ENSG00000257923.10 | CUX1 | 0.212705610701779 | 5.3468088567916 | 1.44028338040278 | 0.157113890120011 | 0.468823306514324 | -5.38818997368196 |
| lnc-ZNF207-1 | lnc-ZNF207-1 | -0.267192747231164 | 3.47683199108233 | -1.44027190712959 | 0.157117118037765 | 0.468823306514324 | -5.24569473349235 |
| lnc-SUMO1-7 | lnc-SUMO1-7 | -0.270766518809042 | 0.418580377149833 | -1.44027164330879 | 0.157117192262343 | 0.468823306514324 | -4.84564507976376 |
| ENSG00000162813.17 | BPNT1 | -0.285069923828472 | 2.66373829180238 | -1.44011744295731 | 0.157160580420591 | 0.468826138515169 | -5.14814143750815 |
| ENSG00000130147.15 | SH3BP4 | 1.92857171585339 | -2.12657698365919 | 1.44038457522634 | 0.157163696047936 | 0.468826138515169 | -4.43450492633702 |
| lnc-ABCA8-3 | lnc-ABCA8-3 | -0.603854641351648 | 1.44510028612754 | -1.44023157906576 | 0.157209866600895 | 0.468867174058992 | -5.00099000587706 |
| lnc-SLC22A17-3 | lnc-SLC22A17-3 | -0.236904346903471 | 1.92205516231121 | -1.43989562498013 | 0.15722301099719 | 0.468867174058992 | -5.05239220741391 |
| ENSG00000214439.4 | FAM185BP | -0.254170385671562 | 0.844027276015534 | -1.43975941586995 | 0.157261356638957 | 0.468913588830894 | -4.92535187001936 |
| ENSG00000137942.16 | FNBP1L | -0.910496186333567 | -0.19526741044835 | -1.43985706832292 | 0.157375940599169 | 0.46914281574148 | -4.78318798579083 |
| ENSG00000106484.15 | MEST | 0.709779240386733 | 2.35855794179551 | 1.43996529749962 | 0.157383818816252 | 0.46914281574148 | -5.07023856837485 |
| ENSG00000163161.13 | ERCC3 | 0.112033016732974 | 5.6641304459541 | 1.43901044713544 | 0.157472337766206 | 0.46927696026611 | -5.3978025603004 |
| ENSG00000085719.12 | CPNE3 | 0.226333399721734 | 5.82318057483922 | 1.43900306311584 | 0.157474418916486 | 0.46927696026611 | -5.4000348451757 |
| ENSG00000097046.12 | CDC7 | 0.339624169830719 | 2.44851318439985 | 1.43887250765956 | 0.15751121890596 | 0.469318676714153 | -5.11387806567885 |
| ENSG00000160959.7 | LRRC14 | -0.174041744709005 | 4.8478219788101 | -1.43859913366439 | 0.157588297368149 | 0.469480377005154 | -5.36717962199843 |
| lnc-ELF1-5 | lnc-ELF1-5 | -0.353482472433684 | 0.369504185830315 | -1.43830081839077 | 0.157672441848717 | 0.469663078230412 | -4.84555656931247 |
| lnc-BCL2-2 | lnc-BCL2-2 | -0.428033311693549 | 0.747340993217982 | -1.4381552963006 | 0.157713501415868 | 0.469717406980583 | -4.90625078872952 |
| lnc-ZBED5-2 | lnc-ZBED5-2 | -0.181558798078878 | 1.7174651620143 | -1.43793253340537 | 0.157776370994715 | 0.469812845916259 | -5.04846581345964 |
| ENSG00000151353.14 | TMEM18 | 0.166471548143057 | 4.09260363626265 | 1.43784408265178 | 0.157801339588409 | 0.469812845916259 | -5.31019958781599 |
| ENSG00000277775.1 | H3C7 | 2.95928112362325 | -1.00053925869159 | 1.43849587979237 | 0.157846631337886 | 0.469812845916259 | -4.43803908197796 |
| lnc-SARDH-2 | lnc-SARDH-2 | 0.29767688045809 | 2.40563438145644 | 1.43756888060663 | 0.157879045672141 | 0.469812845916259 | -5.11132929674686 |
| lnc-LRRC52-3 | lnc-LRRC52-3 | -0.42015212880428 | -0.344149948996063 | -1.43746947492347 | 0.15790712125113 | 0.469812845916259 | -4.68647423351604 |
| ENSG00000185591.9 | SP1 | 0.218193629452789 | 7.090600523991 | 1.43720086120435 | 0.157983006580078 | 0.469812845916259 | -5.40209870286501 |
| lnc-AMFR-6 | lnc-AMFR-6 | 0.166516531334047 | 1.92413888090143 | 1.43709339289816 | 0.158013375168085 | 0.469812845916259 | -5.07440759994144 |
| lnc-ISG20L2-1 | lnc-ISG20L2-1 | 0.239083889550035 | 0.874832403753898 | 1.43701944295336 | 0.158034274736142 | 0.469812845916259 | -4.9117335063004 |
| lnc-ANKRD31-2 | lnc-ANKRD31-2 | -0.171093384388278 | 2.73236246949539 | -1.43696951162214 | 0.158048387443432 | 0.469812845916259 | -5.16818662708178 |
| ENSG00000145901.15 | TNIP1 | -0.161859265982685 | 7.08869691404587 | -1.4369397460959 | 0.158056800911057 | 0.469812845916259 | -5.40170545042331 |
| lnc-C5orf67-3 | lnc-C5orf67-3 | -0.713795503224364 | 0.381161391763655 | -1.43681671106208 | 0.158134019830105 | 0.469812845916259 | -4.80345760509539 |
| ENSG00000148572.15 | NRBF2 | 0.278024097883878 | 3.59917648235981 | 1.43659427424178 | 0.158154477034982 | 0.469812845916259 | -5.25239498746869 |
| lnc-TM9SF3-2 | lnc-TM9SF3-2 | -0.372848399499368 | 0.838199923150063 | -1.43657958852659 | 0.158158630212756 | 0.469812845916259 | -4.92481908233233 |
| ENSG00000197147.13 | LRRC8B | 0.198588227254454 | 5.25132538477074 | 1.43656252981002 | 0.158163454591991 | 0.469812845916259 | -5.38767848899309 |
| ENSG00000131470.14 | PSMC3IP | 0.24365418827346 | 0.499128361386012 | 1.43654293960541 | 0.158168995044258 | 0.469812845916259 | -4.88165686138063 |
| ENSG00000068305.17 | MEF2A | 0.157004455279799 | 6.36940711628189 | 1.43653358186392 | 0.158171641630824 | 0.469812845916259 | -5.40748839890782 |
| ENSG00000259781.1 | HMGB1P6 | 0.305343513641818 | 1.01272563342149 | 1.43648303890407 | 0.158185936953759 | 0.469812845916259 | -4.97003537371754 |
| lnc-BCKDHB-1 | lnc-BCKDHB-1 | -0.35056483679213 | 0.515682651696564 | -1.43647460748314 | 0.158188321754113 | 0.469812845916259 | -4.87511418147651 |
| lnc-LYL1-1 | lnc-LYL1-1 | -0.22310067597204 | 1.76394475716429 | -1.43622898121275 | 0.15825780873213 | 0.469812845916259 | -5.06943507096135 |
| ENSG00000134758.13 | RNF138 | 0.226968147080929 | 5.2944853465417 | 1.436220045999 | 0.158260336929974 | 0.469812845916259 | -5.39245761860397 |

| | | | | | | | |
|---|---|---|---|---|---|---|---|
| ENSG00000139192.11 | TAPBPL | -0.234429605477162 | 4.18561855020866 | -1.43615031420974 | 0.158280068464772 | 0.469812845916259 | -5.32504410025768 |
| lnc-MSRA-1 | lnc-MSRA-1 | -0.421445671049957 | 0.532045237734052 | -1.43601587811665 | 0.158318114385082 | 0.469812845916259 | -4.86281435858456 |
| ENSG00000102178.12 | UBL4A | -0.441144407737905 | 2.30070818602201 | -1.43591066578665 | 0.158368630474387 | 0.469812845916259 | -5.13443206021254 |
| ENSG00000219481.10 | NBPF1 | -0.309843270738523 | 3.1791914723016 | -1.43580482059093 | 0.15837785892475 | 0.469812845916259 | -5.21187462256682 |
| ENSG00000166278.14 | C2 | -0.408682771757895 | -0.412353732555435 | -1.43578535251049 | 0.158383370689479 | 0.469812845916259 | -4.66105895898863 |
| lnc-TBXA2R-1 | lnc-TBXA2R-1 | -0.187924684028983 | 0.864465396862962 | -1.43570619012057 | 0.158405784539132 | 0.469812845916259 | -4.94401054182445 |
| lnc-FAM204A-6 | lnc-FAM204A-6 | 0.126173513224196 | 2.71615524913403 | 1.43569863951235 | 0.158407922530186 | 0.469812845916259 | -5.16647083468238 |
| lnc-RRAS-5 | lnc-RRAS-5 | -0.182259607770674 | 3.21923789923458 | -1.435647183436 | 0.158422493168308 | 0.469812845916259 | -5.22057653987469 |
| lnc-GOLGA6B-1 | lnc-GOLGA6B-1 | 0.214652135374053 | 1.12135748797908 | 1.43563165439041 | 0.158426890680529 | 0.469812845916259 | -4.97221715880043 |
| lnc-PINX1-84 | lnc-PINX1-84 | -0.298121383215281 | 4.40525774070598 | -1.43561959829065 | 0.158430304790592 | 0.469812845916259 | -5.33859652762393 |
| ENSG00000131368.7 | MRPS25 | -0.168656417371019 | 5.65746040144879 | -1.43545939862614 | 0.158475676454562 | 0.469879695831785 | -5.40246884502442 |
| ENSG00000145632.14 | PLK2 | 0.470537691783306 | 1.42337445175443 | 1.43524252381392 | 0.158537115853617 | 0.469994160393922 | -4.99847849378537 |
| lnc-BRD2-2 | lnc-BRD2-2 | 0.338838017711175 | 4.17678271108747 | 1.43450001446972 | 0.158747605937162 | 0.470482697642051 | -5.2997389937352 |
| lnc-PELI1-1 | lnc-PELI1-1 | -0.447441815982893 | 0.916077436399979 | -1.43446044326392 | 0.158758829916274 | 0.470482697642051 | -4.92774718870398 |
| ENSG00000244171.4 | PBX2P1 | 0.597235488283819 | -0.826984695807892 | 1.43441936731249 | 0.158770481359456 | 0.470482697642051 | -4.5321578524674 |
| ENSG00000241837.6 | ATP5PO | 0.202499783074884 | 4.86764308626301 | 1.4342733190248 | 0.158811914271791 | 0.470537733043942 | -5.37138003107617 |
| ENSG00000171208.9 | NETO2 | -0.282761803989667 | 2.74754788903401 | -1.43393219670212 | 0.15890872138231 | 0.470756795079459 | -5.18999156118283 |
| ENSG00000100330.15 | MTMR3 | 0.120182976683624 | 6.3762944768035 | 1.4338161059563 | 0.158941677322918 | 0.470786666331504 | -5.41128774162219 |
| ENSG00000213676.12 | ATF6B | -0.173917009884772 | 5.490309447451 | -1.43334553854423 | 0.159075317265183 | 0.47109855820942 | -5.40125528944754 |
| ENSG00000176095.11 | IP6K1 | 0.198117691996059 | 4.95530210459583 | 1.43322955301787 | 0.159108270380424 | 0.47109855820942 | -5.38079667845238 |
| ENSG00000197728.11 | RPS26 | 0.707226460435963 | 6.15083838335237 | 1.43408676898232 | 0.159131192681796 | 0.47109855820942 | -5.40928715080828 |
| lnc-HK3-3 | lnc-HK3-3 | 0.256015166241632 | 2.10636020025598 | 1.4330975423723 | 0.159145782965197 | 0.47109855820942 | -5.08954872180988 |
| ENSG00000171703.16 | TCEA2 | 0.211131606673635 | 3.47060459389367 | 1.43304254415634 | 0.15916141348629 | 0.47109855820942 | -5.25390518095474 |
| CARD8-AS1 | CARD8-AS1 | 0.297899492571291 | 2.90619084514006 | 1.43261392891913 | 0.159283267373575 | 0.471235848703306 | -5.18532389161161 |
| ENSG00000041880.14 | PARP3 | -0.230737395346829 | 4.79621827888977 | -1.43230635769643 | 0.159370753883754 | 0.471235848703306 | -5.37380490390958 |
| ENSG00000010278.13 | CD9 | -1.49073753530226 | 0.146396313623309 | -1.433285311906 | 0.159371075782755 | 0.471235848703306 | -4.89095945843649 |
| ENSG00000168438.14 | CDC40 | 0.185812145296345 | 6.16194435311843 | 1.43224286097958 | 0.159388819773392 | 0.471235848703306 | -5.41295438511954 |
| lnc-SRP14-1 | lnc-SRP14-1 | -0.481338872965578 | 2.80593902523747 | -1.43251789673705 | 0.159390463551143 | 0.471235848703306 | -5.19581496355794 |
| ENSG00000196867.7 | ZFP28 | -0.898182527269389 | 2.93481669323237 | -1.43319791960568 | 0.1593948190946 | 0.471235848703306 | -5.25132446615673 |
| lnc-KCNG3-4 | lnc-KCNG3-4 | 0.253662010226535 | 0.819343671588773 | 1.43221785159763 | 0.159395935805819 | 0.471235848703306 | -4.92584450798181 |
| ENSG00000280148.1 | | 0.165002528993528 | 2.54164448368233 | 1.43214429787919 | 0.159416865819972 | 0.471235848703306 | -5.1507756297697 |
| ENSG00000145860.11 | RNF145 | 0.198313814149701 | 5.30430946551555 | 1.43207536520312 | 0.159436482851666 | 0.471235848703306 | -5.39874382792889 |
| lnc-LYPLAL1-7 | lnc-LYPLAL1-7 | -0.536665984566846 | 0.181780291267482 | -1.43197568115443 | 0.159464854529622 | 0.471235848703306 | -4.78740508420412 |
| lnc-FBXL17-4 | lnc-FBXL17-4 | -0.433229247474617 | -0.127468829140542 | -1.43194733396295 | 0.159472923316842 | 0.471235848703306 | -4.77358660824082 |
| ENSG00000233327.10 | USP32P2 | 1.67493350968039 | -1.25174274378234 | 1.43263390418173 | 0.159482530343839 | 0.471235848703306 | -4.57482603061333 |
| lnc-CCDC12-1 | lnc-CCDC12-1 | -0.212592207993162 | 3.28595990552104 | -1.43174139613808 | 0.159531551370605 | 0.471313036007057 | -5.23010354330113 |
| ENSG00000215421.9 | ZNF407 | 0.163990585486776 | 6.32567892656184 | 1.43162935447937 | 0.159563455391141 | 0.471339638679085 | -5.41425124989593 |
| ENSG00000168374.10 | ARF4 | 0.182393119970667 | 4.74777230787314 | 1.43145696477408 | 0.159612553362979 | 0.471368354263479 | -5.37013834184397 |
| ENSG00000170175.10 | CHRNB1 | -0.385794776420396 | 0.80349438042977 | -1.43139122466566 | 0.159631279789825 | 0.471368354263479 | -4.93613844354409 |
| ENSG00000198270.12 | TMEM116 | -0.205277967271782 | 3.67856383931108 | -1.43122456549411 | 0.159678761270604 | 0.471368354263479 | -5.27787081835936 |
| lnc-DNAJC25-5 | lnc-DNAJC25-5 | -0.726579354982317 | -0.398115399521912 | -1.43118919056822 | 0.159688841069957 | 0.471368354263479 | -4.65638760089401 |
| lnc-ZFP36L2-1 | lnc-ZFP36L2-1 | 0.460831215344395 | -0.530265854386164 | 1.43116538082707 | 0.159695625743928 | 0.471368354263479 | -4.62781305617043 |
| ENSG00000227372.12 | GFOD3P | -0.197992896639427 | 4.02974800194974 | -1.43111289814534 | 0.159710581675825 | 0.471368354263479 | -5.31363829715186 |
| ENSG00000124222.22 | STX16 | 0.280854956427421 | 5.14209321439581 | 1.43069625268101 | 0.159829351597882 | 0.471615759811971 | -5.38992135386155 |
| lnc-LRRC28-7 | lnc-LRRC28-7 | 0.224381517409919 | 1.98380786352964 | 1.43065808860236 | 0.159840234195613 | 0.471615759811971 | -5.09468772557974 |
| ENSG00000109332.19 | UBE2D3 | 0.132176589615711 | 7.5068816625394 | 1.43055348912563 | 0.159870064014378 | 0.471636165631067 | -5.40453312016909 |
| ENSG00000178982.9 | EIF3K | 0.186680697125064 | 6.29844675795951 | 1.43025962958088 | 0.159953890598992 | 0.471812632660278 | -5.41610196664408 |
| lnc-SLC35G2-1 | lnc-SLC35G2-1 | -0.446214611534281 | -0.68103675780653 | -1.43018310479718 | 0.159975725760874 | 0.471812632660278 | -4.65933129224253 |
| lnc-TBC1D3F-1 | lnc-TBC1D3F-1 | -0.218336380681171 | 0.913640442037963 | -1.43010005965368 | 0.159999424046973 | 0.471814920509865 | -4.95762981598005 |
| lnc-ESS2-5 | lnc-ESS2-5 | -0.20201639256389 | 1.15147712968028 | -1.42996447331542 | 0.160038121727176 | 0.471861432389408 | -4.98646551424724 |
| ENSG00000139323.13 | POC1B | -0.125911195975549 | 3.92594282111715 | -1.4297744008465 | 0.160092382623576 | 0.471921460940362 | -5.30838549226994 |
| ENSG00000082898.16 | XPO1 | 0.162862740187053 | 7.41381856515993 | 1.42972111200068 | 0.160107597829991 | 0.471921460940362 | -5.40740500483379 |
| ENSG00000211772.11 | TRBC2 | -0.26173061877118 | 5.44971448005963 | -1.42964402075342 | 0.160129611179837 | 0.471921460940362 | -5.40339486527596 |
| ENSG00000241978.9 | NA | -0.33222466179796 | 6.90667023026718 | -1.42966819367834 | 0.160150192132752 | 0.471921460940362 | -5.41283619788165 |
| ENSG00000123297.17 | TSFM | 0.176753079264559 | 2.9824807342414 | 1.429486331465 | 0.160174646622334 | 0.471925959265316 | -5.20193088830749 |
| ENSG00000131389.17 | SLC6A6 | -0.206110533350488 | 5.59697683646589 | -1.42915013739799 | 0.160270695434398 | 0.472089123541829 | -5.40753091613317 |
| ENSG00000151422.12 | FER | -0.213557295675164 | 4.78836303163575 | -1.42913193391187 | 0.160275897357543 | 0.472089123541829 | -5.37863686616459 |
| ENSG00000198947.15 | DMD | -1.05688253080819 | 5.99626522357819 | -1.42998829637989 | 0.160320991754239 | 0.472154381639363 | -5.41107713657489 |
| ENSG00000110844.13 | PRPF40B | -0.263759814977997 | 2.36986814823487 | -1.4287687126702 | 0.160379720984763 | 0.472212949273073 | -5.12813306190219 |
| lnc-MYLPF-3 | lnc-MYLPF-3 | -0.375242958795735 | 0.308744739896471 | -1.42874408569067 | 0.160386762297432 | 0.472212949273073 | -4.82566224867641 |
| ENSG00000280571.2 | | 0.237133866616649 | 1.5559999621376 | 1.42847153660607 | 0.160464705320943 | 0.472374861215813 | -5.03571343365523 |
| ENSG00000117593.10 | DARS2 | -0.152381184314583 | 3.61794731895965 | -1.42820939562407 | 0.160539699822539 | 0.472475520245357 | -5.27727199552835 |
| ENSG00000279162.1 | | 0.364964644547487 | 0.322815995913666 | 1.42819154269353 | 0.160544808268767 | 0.472475520245357 | -4.82136309092552 |
| ENSG00000171853.15 | TRAPPC12 | -0.175224971947403 | 5.10934977272282 | -1.42804257499757 | 0.16058743891725 | 0.472493829596756 | -5.39571011960232 |
| lnc-PFDN4-3 | lnc-PFDN4-3 | -1.86505891147775 | -2.45060757304449 | -1.4281279411526 | 0.160596940769514 | 0.472493829596756 | -4.43915796919125 |
| ENSG00000230149.2 | | -0.208438429585071 | 1.15571837990388 | -1.42785395248588 | 0.160641430443441 | 0.472557176335194 | -4.99145204209647 |
| ENSG00000285160.1 | THSD1P1 | 0.296090115797877 | -0.00186372612259226 | 1.42760239172155 | 0.160713459592283 | 0.472701505971414 | -4.78878923980801 |
| lnc-PAWR-7 | lnc-PAWR-7 | -0.290109726267417 | 2.37570312489474 | -1.42674172482616 | 0.160960084656002 | 0.473235768810479 | -5.15155808672569 |

| | | | | | | | |
|---|---|---|---|---|---|---|---|
| ENSG00000253981.5 | ALG1L13P | -0.623622527131761 | 0.115128623679075 | -1.42670645904503 | 0.16097019641612 | 0.473235768810479 | -4.76829271110033 |
| LINC01578 | LINC01578 | 0.220293383876217 | 7.35872781155878 | 1.4266438296782 | 0.160988155364194 | 0.473235768810479 | -5.41164368833514 |
| CCDC18-AS1 | CCDC18-AS1 | -0.212623880091694 | 7.6184721073159 | -1.42660135032491 | 0.161000337195328 | 0.473235768810479 | -5.40945736919286 |
| ENSG00000155085.15 | AK9 | -0.20570869048764 | 4.76886183273144 | -1.42656744359953 | 0.161010061166 | 0.473235768810479 | -5.37370552307115 |
| ENSG00000185710.9 | SMG1P4 | -0.688219017145495 | 2.03105806296399 | -1.42708937856397 | 0.161036906523475 | 0.473247094085191 | -5.08913030506413 |
| lnc-MUSTN1-3 | lnc-MUSTN1-3 | -0.250516987279239 | 0.385571615111251 | -1.42631066304426 | 0.16108371712326 | 0.47327085598431 | -4.86151555139412 |
| ENSG00000174231.16 | PRPF8 | 0.190578597651056 | 8.22999874937086 | 1.42622595286995 | 0.161108021507993 | 0.47327085598431 | -5.39792251707367 |
| ENSG00000154721.14 | JAM2 | -0.644798877862526 | 0.692954152449443 | -1.42638269310162 | 0.161113972107179 | 0.47327085598431 | -4.90059606924419 |
| GRIK1-AS1 | GRIK1-AS1 | -1.19202541778498 | -0.98968631353827 | -1.42621946665251 | 0.161200364268036 | 0.473429640641696 | -4.5789259337193 |
| ENSG00000272573.5 | MUSTN1 | -0.189136111612459 | 1.2741760453695 | -1.42571860323342 | 0.161253646362967 | 0.473429640641696 | -5.01092014085496 |
| LINC02449 | LINC02449 | 0.716173824490649 | -0.693564086648749 | 1.42571200403434 | 0.161255541212871 | 0.473429640641696 | -4.66627175733355 |
| ENSG00000131899.10 | LLGL1 | -0.262721645741077 | 3.02997031144781 | -1.42568077414603 | 0.161264508590416 | 0.473429640641696 | -5.21490807846619 |
| ENSG00000162736.16 | NCSTN | 0.135492289392631 | 5.89044191876301 | 1.42561626987978 | 0.161283031636767 | 0.473429640641696 | -5.41948390696182 |
| lnc-SLC25A39-2 | lnc-SLC25A39-2 | -0.275548163751008 | 2.04540938946592 | -1.42544994820852 | 0.16133080024781 | 0.473484958703947 | -5.11388251153489 |
| ENSG00000151846.8 | PABPC3 | 1.26967662464491 | -0.593433358217806 | 1.42554161795313 | 0.16135838150752 | 0.473484958703947 | -4.52017433538149 |
| ENSG00000100836.10 | PABPN1 | -0.156376809949257 | 6.02516491830971 | -1.42525143411741 | 0.161387829204215 | 0.473484958703947 | -5.42181992555776 |
| lnc-UNC80-1 | lnc-UNC80-1 | -0.860056503612972 | 1.23460211623393 | -1.42594288352992 | 0.161393891573964 | 0.473484958703947 | -5.01976740801677 |
| lnc-WDR24-1 | lnc-WDR24-1 | -0.219866260644195 | 1.70348127797839 | -1.42494707427954 | 0.161475296048279 | 0.47365626600566 | -5.07696509930638 |
| ENSG00000112893.9 | MAN2A1 | 0.201682710518195 | 7.587780681069 | 1.42474652152118 | 0.161532951100538 | 0.473757870120029 | -5.4118316028553 |
| ENSG00000266993.3 | | 0.231083807963424 | 1.60370096539083 | 1.42436979631098 | 0.161641295852304 | 0.474008091256301 | -5.04653180453083 |
| ENSG00000089091.16 | DZANK1 | -0.240344768627995 | 2.43170579750315 | -1.4238726953063 | 0.161784347169407 | 0.47436000253389 | -5.15031855673412 |
| lnc-ANKRD55-6 | lnc-ANKRD55-6 | 0.510329631964225 | 2.91492871771273 | 1.42420124627011 | 0.161809264758798 | 0.474365488752363 | -5.22288019659581 |
| ENSG00000064489.22 | BORCS8-MEF2B | -0.196658707792919 | 1.76045250741356 | -1.42367056502071 | 0.16184254272549 | 0.474375182951741 | -5.07296820778609 |
| ENSG00000242338.6 | BMS1P4 | 0.14644735260844 | 3.14104337706861 | 1.42361457629023 | 0.161858665397176 | 0.474375182951741 | -5.22771796609303 |
| LINC01569 | LINC01569 | -0.298896085381734 | -0.0448054557271712 | -1.42344483374023 | 0.161907552610673 | 0.474450904810006 | -4.75880007137186 |
| ENSG00000136877.14 | FPGS | -0.141878930138727 | 5.01561974181447 | -1.4230197158484 | 0.162030040618855 | 0.474733252969791 | -5.40092744762577 |
| ENSG00000162032.15 | SPSB3 | -0.161648351472854 | 5.0708969066842 | -1.42295035130774 | 0.162050033297661 | 0.474733252969791 | -5.40276993674825 |
| ENSG00000006837.11 | CDKL3 | -0.21534563612088 | 1.26107350717563 | -1.42284353582551 | 0.162080824089971 | 0.474749912806278 | -5.00393311927952 |
| lnc-BDH1-8 | lnc-BDH1-8 | 0.25582146544643 | 2.53603520433393 | 1.42276470072123 | 0.16210355214883 | 0.474749912806278 | -5.1540053644491 |
| ENSG00000164880.15 | INTS1 | -0.29239701247935 | 6.23426812287637 | -1.42262014699823 | 0.162145233271511 | 0.474749912806278 | -5.42586206134846 |
| ENSG00000262209.2 | PCDHGB3 | -1.25374507745797 | -1.06688162571978 | -1.42323060215149 | 0.162147980713605 | 0.474749912806278 | -4.67177298104752 |
| ENSG00000126012.11 | KDM5C | 0.212048209241924 | 7.17683191786607 | 1.42240176219918 | 0.162208218987621 | 0.474787606161024 | -5.42033278481 |
| ENSG00000164548.10 | TRA2A | 0.218182565406737 | 6.2307125486317 | 1.42235992433015 | 0.162220287889353 | 0.474787606161024 | -5.42674600765058 |
| ENSG00000006459.10 | KDM7A | 0.194779647002004 | 7.50772551382176 | 1.42232606556353 | 0.162230055586187 | 0.474787606161024 | -5.41756677373416 |
| ENSG00000259040.5 | BLOC1S5-TXNDC5 | -0.513676125954312 | 6.98885330352224 | -1.42265051453645 | 0.162333873781285 | 0.475008968140257 | -5.42120176751311 |
| ENSG00000154079.5 | SDHAF4 | -0.208575016957292 | 2.21935600693324 | -1.42190401718579 | 0.162351848169203 | 0.475008968140257 | -5.12697757214722 |
| lnc-PACRG-4 | lnc-PACRG-4 | 0.362228334544872 | -0.0575298037910264 | 1.42160119090547 | 0.16243928025676 | 0.475175775165012 | -4.76817499928266 |
| lnc-FNBP4-5 | lnc-FNBP4-5 | 0.231352837154288 | 1.35227909125844 | 1.4214984063821 | 0.162468964593795 | 0.475175775165012 | -5.01463356721584 |
| APTR | APTR | 0.198142064232137 | 2.783201688662 | 1.42141536546388 | 0.162492950040711 | 0.475175775165012 | -5.19213711880289 |
| lnc-SYCP1-4 | lnc-SYCP1-4 | -0.257570067145662 | 1.35897142515934 | -1.42138679142742 | 0.162501203973498 | 0.475175775165012 | -5.03346529900616 |
| ENSG00000265118.5 | | -0.363620726359677 | 5.44094044650437 | -1.42146496543692 | 0.162543290954509 | 0.475231329221115 | -5.41308629176643 |
| ENSG00000228205.1 | RPS3P3 | 0.947957684130593 | 0.590921526105071 | 1.4213860368226 | 0.162643964077147 | 0.475458132736816 | -4.77368125463205 |
| ENSG00000044459.14 | CNTLN | -2.09064339427831 | 1.00982243236418 | -1.42170074974532 | 0.162699644724884 | 0.47555336372796 | -4.82586003786162 |
| ENSG00000165819.11 | METTL3 | -0.147920900125011 | 6.32693984317453 | -1.42041544184039 | 0.162781984134596 | 0.475726477274229 | -5.42920214500564 |
| ENSG00000085662.13 | AKR1B1 | 0.218058014994436 | 3.83773834164707 | 1.41995034945846 | 0.162916558714893 | 0.476052176040409 | -5.30516347844835 |
| lnc-MOK-7 | lnc-MOK-7 | 0.28570072829936 | 4.34197050495671 | 1.41983222561687 | 0.162950751706455 | 0.476084502821003 | -5.33977188472173 |
| ENSG00000047849.21 | MAP4 | -0.159081916347443 | 6.39248014622228 | -1.41964813849259 | 0.163004050085178 | 0.476124385479288 | -5.43018513755862 |
| ENSG00000186468.12 | RPS23 | 0.198435132479 | 9.50896556498376 | 1.41962529004293 | 0.163010666300954 | 0.476124385479288 | -5.37336760994283 |
| ENSG00000116898.11 | MRPS15 | 0.151040164202689 | 4.04265225634821 | 1.41937965788771 | 0.163081807145513 | 0.47626352027518 | -5.32923339351204 |
| lnc-ZDHHC14-7 | lnc-ZDHHC14-7 | 0.427820485981579 | 1.04266840739627 | 1.41930104903671 | 0.16310457923625 | 0.47626352027518 | -4.96901243459533 |
| ENSG00000186001.13 | LRCH3 | -0.112030519136765 | 6.36007214932638 | -1.41911468154669 | 0.163158577701518 | 0.476353617706431 | -5.43102331586181 |
| RPARP-AS1 | RPARP-AS1 | -0.265915782442718 | 3.7467052459734 | -1.41901361910255 | 0.163187865559796 | 0.4763715553563 | -5.29438903915908 |
| lnc-ZNF266-2 | lnc-ZNF266-2 | 0.565772431100538 | 4.02531640566936 | 1.41930313983175 | 0.163299670048282 | 0.47663033304081 | -5.32940114301847 |
| ENSG00000077585.13 | GPR137B | -0.485359078068026 | 1.74865623150576 | -1.41846723319252 | 0.163365202626512 | 0.476754000519141 | -5.06043597654888 |
| lnc-UBAC2-6 | lnc-UBAC2-6 | -0.486247839011034 | 0.398466669005548 | -1.41820519938832 | 0.163422293059532 | 0.476791588937183 | -4.88182822136905 |
| ENSG00000249565.2 | SERBP1P5 | 0.97267340877109 | -0.617211730857118 | 1.41819789843395 | 0.163424411405132 | 0.476791588937183 | -4.54402539250157 |
| lnc-PLEKHA2-1 | lnc-PLEKHA2-1 | 0.273221248256938 | 0.564076381761391 | 1.41790252852533 | 0.163510129896437 | 0.476974065144326 | -4.90262409758475 |
| lnc-METTL16-2 | lnc-METTL16-2 | 0.238722853183854 | 1.19637543758048 | 1.4173186076947 | 0.163679691225848 | 0.477401032237727 | -4.98894322500976 |
| lnc-RASGRF1-4 | lnc-RASGRF1-4 | -0.36318454294218 | 4.18797183107178 | -1.41729314563961 | 0.163720393846662 | 0.477452092171415 | -5.35479196219716 |
| ENSG00000229808.1 | | 0.307873801137356 | 0.753867940634269 | 1.41666232944357 | 0.163870427763514 | 0.477821931528037 | -4.93236083599915 |
| ENSG00000243207.6 | PPAN-P2RY11 | -0.246081745767036 | 2.83834043281458 | -1.41644760862925 | 0.163932870498688 | 0.477888931250232 | -5.21324623675235 |
| ENSG00000134265.12 | NAPG | 0.164612541760165 | 5.19824023378696 | 1.4164236438657 | 0.163939840818048 | 0.477888931250232 | -5.41331979704083 |
| lnc-SHC4-3 | lnc-SHC4-3 | 0.234556831985588 | 3.73520660858797 | 1.41614761738911 | 0.16402014172507 | 0.477947045969321 | -5.29628650279328 |
| ENSG00000119878.5 | CRIPT | 0.197831946149717 | 3.3526792572949 | 1.41606430111206 | 0.164044385909125 | 0.477947045969321 | -5.26409066884262 |
| ENSG00000066136.20 | NFYC | -0.104139748295903 | 5.07488806272821 | -1.41595544955296 | 0.164076064811573 | 0.477947045969321 | -5.41034066426064 |
| ENSG00000129197.14 | RPAIN | -0.165166713302784 | 5.92633003212371 | -1.4159482964078 | 0.164078146747568 | 0.477947045969321 | -5.43269640995267 |
| ENSG00000084234.17 | APLP2 | -0.392615768888223 | 6.96477137603818 | -1.41624753764309 | 0.164093742883203 | 0.477947045969321 | -5.42974880231328 |
| lnc-RASL12-1 | lnc-RASL12-1 | -0.215161626967638 | -0.0087024963512538( | -1.41587630910846 | 0.164099099930292 | 0.477947045969321 | -4.79319235282741 |

| | | | | | | | |
|---|---|---|---|---|---|---|---|
| lnc-SSR3-10 | lnc-SSR3-10 | -0.339305820574822 | 0.609920554673093 | -1.41558112778215 | 0.164185039531782 | 0.478067196854798 | -4.91987150067226 |
| ENSG00000213240.8 | NA | 0.356145401602796 | 2.73746154546981 | 1.41557506388827 | 0.164186805352309 | 0.478067196854798 | -5.18018090440388 |
| lnc-ALG5-2 | lnc-ALG5-2 | 0.224361173052438 | 1.11951959219162 | 1.41534771135757 | 0.164253021649499 | 0.478192354259071 | -4.99807751784612 |
| ENSG00000134086.7 | VHL | 0.159512112683426 | 6.35112580737058 | 1.41516986724008 | 0.164304833169003 | 0.47827554534261 | -5.43651110439516 |
| lnc-PRKAG2-3 | lnc-PRKAG2-3 | 0.222988580327657 | 2.30063857955143 | 1.41506782622613 | 0.164334566643494 | 0.478294454924072 | -5.1369263553146 |
| ENSG00000147905.17 | ZCCHC7 | 0.173915966824115 | 7.42170787761686 | 1.41475856680217 | 0.164424706605957 | 0.478489146906605 | -5.42870678939408 |
| lnc-TUT1-2 | lnc-TUT1-2 | -0.257269797696013 | 8.40421198367449 | -1.41410102374705 | 0.164616489091597 | 0.478979529964991 | -5.40852242755466 |
| ENSG00000104522.15 | GFUS | -0.251423914739206 | 3.42126710544076 | -1.41402113684871 | 0.164639801206471 | 0.47897965063361 | -5.27480341193874 |
| lnc-VAMP4-4 | lnc-VAMP4-4 | 0.324844919203244 | 0.591288606318492 | 1.41389625660161 | 0.164676248167775 | 0.47901409401632 | -4.91516314721073 |
| lnc-ZCCHC7-2 | lnc-ZCCHC7-2 | 0.132537602910534 | 7.93808324979878 | 1.41378424922558 | 0.164708943465524 | 0.47901409401632 | -5.42118221637024 |
| ENSG00000136478.7 | TEX2 | -0.194063223876233 | 3.88760679203792 | -1.41371154719981 | 0.164730168119505 | 0.47901409401632 | -5.32714724394459 |
| ENSG00000132522.15 | GPS2 | -0.229751625280526 | 4.69803865465614 | -1.41366167278566 | 0.164744729706142 | 0.47901409401632 | -5.39542678881209 |
| lnc-AKR1B1-3 | lnc-AKR1B1-3 | -0.36794650421631 | 0.264518551935782 | -1.41317835308931 | 0.164885894169584 | 0.479356830465048 | -4.85969078717003 |
| ENSG00000144591.18 | GMPPA | -0.173234276445393 | 4.0244901855326 | -1.41301309639509 | 0.164934182759346 | 0.47942216999776 | -5.34128205231862 |
| ENSG00000163053.10 | SLC16A14 | -0.790229125303806 | -0.389795369332071 | -1.41305200102622 | 0.164954953501634 | 0.47942216999776 | -4.73819920576214 |
| lnc-GCM1-3 | lnc-GCM1-3 | -0.499781131378262 | 0.327206185942424 | -1.41276097720507 | 0.165007874117261 | 0.479497019517844 | -4.84077634855269 |
| ENSG00000166228.8 | PCBD1 | -0.273351490041259 | 2.24278852386669 | -1.41269453525435 | 0.165027298560191 | 0.479497019517844 | -5.14575674895958 |
| ENSG00000185414.19 | MRPL30 | 0.152537982810821 | 3.46839814699105 | 1.41247121006089 | 0.165092601211781 | 0.479619055856327 | -5.27614006374891 |
| ENSG00000114383.9 | TUSC2 | 0.193096261860978 | 3.44933089684582 | 1.41206341982524 | 0.16521189536512 | 0.479893284298152 | -5.28305718998623 |
| lnc-PYDC2-7 | lnc-PYDC2-7 | 0.319766363280845 | 2.68871540748133 | 1.41198916443978 | 0.165233625118836 | 0.479893284298152 | -5.18730441353399 |
| lnc-PRLHR-3 | lnc-PRLHR-3 | 0.155673850928811 | 1.73877358305741 | 1.41182809296801 | 0.165280767988538 | 0.479909258913902 | -5.08524624329485 |
| ENSG00000222724.1 | RNU2-63P | -0.651884677886869 | 3.02421692782983 | -1.4124939049518 | 0.165289930124277 | 0.479909258913902 | -5.24033261449995 |
| ENSG00000175906.4 | ARL4D | 0.808220381272722 | -0.75180952142626 | 1.41166559246382 | 0.165328339733922 | 0.479909258913902 | -4.63106284048622 |
| ENSG00000204316.12 | MRPL38 | -0.154422938660598 | 3.53238725635745 | -1.41165176354264 | 0.165332388619661 | 0.479909258913902 | -5.29122958546372 |
| ENSG00000117569.18 | PTBP2 | 0.234273686642595 | 5.425539481971 | 1.41149970721416 | 0.16537691336395 | 0.479945249543826 | -5.4297238264924 |
| lnc-TTLL1-3 | lnc-TTLL1-3 | 0.338491892080096 | 0.128351777735555 | 1.41145016305185 | 0.165391422776775 | 0.479945249543826 | -4.80425198650813 |
| lnc-ENDOU-6 | lnc-ENDOU-6 | -0.231068870007368 | 0.469717225157368 | -1.41124425049334 | 0.165451736582238 | 0.479952171040619 | -4.91433165434323 |
| lnc-FCRL2-1 | lnc-FCRL2-1 | -1.19811341168329 | 2.66356784309697 | -1.4121842301467 | 0.165464982256596 | 0.479952171040619 | -5.17386489580965 |
| ENSG00000180758.11 | GPR157 | -0.475268693779958 | 1.33610053630838 | -1.41124377886482 | 0.165466112158932 | 0.479952171040619 | -5.07076497106708 |
| ENSG00000100129.17 | EIF3L | 0.189692992490612 | 7.51433243573815 | 1.41110813592493 | 0.165491615281524 | 0.479952171040619 | -5.43176754299866 |
| ENSG00000102265.11 | TIMP1 | -0.750570121540646 | 0.704914849872052 | -1.41138651784835 | 0.165510397417593 | 0.479952171040619 | -4.88233219981655 |
| ENSG00000269404.6 | SPIB | -0.330178906164078 | 5.98274394565271 | -1.41098222511681 | 0.165557418240092 | 0.48000870005506 | -5.44025613655848 |
| ENSG00000280332.1 | | -0.319612103984692 | 1.23794379045582 | -1.41081838002769 | 0.165576532594419 | 0.48000870005506 | -5.01821623845481 |
| ENSG00000157881.13 | PANK4 | -0.211676275983125 | 3.58195700422747 | -1.41059165446759 | 0.165643001594903 | 0.48006570167195 | -5.29784457221036 |
| lnc-CYR61-3 | lnc-CYR61-3 | -0.339322300916724 | 2.1217536939445 | -1.41058036442715 | 0.16564631203282 | 0.48006570167195 | -5.13784582167055 |
| ENSG00000020426.10 | MNAT1 | -0.16252881652539 | 3.63329235442356 | -1.4105126600265 | 0.165666165232272 | 0.48006570167195 | -5.29526898671637 |
| ENSG00000106211.8 | HSPB1 | -0.389435250418708 | 2.19496641356988 | -1.41034865957099 | 0.165714263346736 | 0.480096602189961 | -5.15942224512308 |
| ENSG00000276234.4 | TADA2A | -0.130981107897137 | 4.27197752031983 | -1.41031724273763 | 0.165723478528867 | 0.480096602189961 | -5.36542964873282 |
| lnc-PUS10-5 | lnc-PUS10-5 | 0.322364290606831 | 3.72697595865707 | 1.41023559521231 | 0.1657474292391 | 0.480098414864677 | -5.2909087315264 |
| ENSG00000225921.6 | NOL7 | 0.122256619135818 | 4.86704893961864 | 1.4099810724066 | 0.165822108967126 | 0.480247146316357 | -5.40433687346894 |
| lnc-MAP3K3-1 | lnc-MAP3K3-1 | -0.229288798278275 | 2.06950572163477 | -1.40987281063117 | 0.165853882081076 | 0.480271589036969 | -5.11864476410082 |
| lnc-TUBGCP5-2 | lnc-TUBGCP5-2 | -0.414559302734591 | 0.155249206760158 | -1.40956456315536 | 0.165944373801453 | 0.480466035441737 | -4.85286982899031 |
| lnc-GTF2F2-4 | lnc-GTF2F2-4 | -0.350141182614061 | 2.08380274621413 | -1.40945622618237 | 0.165976187246148 | 0.48048975792442 | -5.13865026989864 |
| ENSG00000112812.15 | PRSS16 | -1.05851451834417 | 0.0269855235980139 | -1.41004798914062 | 0.16599925514262 | 0.48048975792442 | -4.82813576196887 |
| ENSG00000173480.10 | ZNF417 | 0.126218941616408 | 5.0086641318034 | 1.4092069739185 | 0.166049398884496 | 0.480543135369751 | -5.41095976978063 |
| ENSG00000204611.6 | ZNF616 | -0.243887282896434 | 2.46441153631367 | -1.40915594945183 | 0.166064389150514 | 0.480543135369751 | -5.16505339508642 |
| lnc-APBB3-3 | lnc-APBB3-3 | 0.295725467503535 | 1.35699000766002 | 1.40900712833634 | 0.166108116707799 | 0.480547100919347 | -5.02778094717568 |
| ENSG00000187118.13 | CMC1 | -0.298347565886096 | 2.32904383280347 | -1.40899237150028 | 0.166112453142941 | 0.480547100919347 | -5.15889544309854 |
| ENSG00000162373.12 | BEND5 | 0.168829784574526 | 2.08695906326621 | 1.40817507934647 | 0.166352759824669 | 0.481174656474305 | -5.13321594274571 |
| ENSG00000275131.3 | PDE4DIPP2 | -0.328639922671364 | 3.88588960655415 | -1.40796635537646 | 0.166414173864464 | 0.481212945359523 | -5.33283471825829 |
| ENSG00000250506.7 | CDK3 | -0.195925122229767 | 2.37725984009312 | -1.40781792095471 | 0.166457859308023 | 0.481212945359523 | -5.16625903244834 |
| ENSG00000237399.7 | PITRM1-AS1 | -0.340700755746225 | 0.839980955342701 | -1.40779283005927 | 0.166465244642385 | 0.481212945359523 | -4.96752709583482 |
| ENSG00000107362.13 | ABHD17B | 0.518721869998075 | 3.70980415322352 | 1.40832867426203 | 0.166468860396279 | 0.481212945359523 | -5.31787318036703 |
| lnc-PRKAA1-3 | lnc-PRKAA1-3 | -0.323480769952048 | 0.77254130869752 | -1.40773287558143 | 0.166482892868152 | 0.481212945359523 | -4.96271813942085 |
| lnc-TAL2-2 | lnc-TAL2-2 | 0.370336555962091 | 2.02089048361354 | 1.40761786537692 | 0.16651675139863 | 0.481243231401012 | -5.11302972301187 |
| ENSG00000095319.14 | NUP188 | 0.107965961480142 | 5.71629708681129 | 1.4073942201367 | 0.166582606984476 | 0.481358333510474 | -5.44128317023483 |
| ENSG00000138604.9 | GLCE | -1.08386356257248 | 2.11885135996373 | -1.40813644600456 | 0.166640017932013 | 0.481358333510474 | -5.10305345118546 |
| lnc-IFT80-4 | lnc-IFT80-4 | 0.19329853782606 | 1.9578311074166 | 1.40716984406969 | 0.166648698146943 | 0.481358333510474 | -5.11235364435176 |
| ENSG00000165512.4 | ZNF22 | 0.217673939331992 | 4.96617520018175 | 1.40697401641708 | 0.166706396902614 | 0.481358333510474 | -5.41824457638426 |
| lnc-OTOA-1 | lnc-OTOA-1 | 0.230578213579061 | 0.56995971502481 | 1.40696406110596 | 0.166709330555687 | 0.481358333510474 | -4.88899983521079 |
| ENSG00000172428.10 | COPS9 | 0.188818272975566 | 2.98396122350154 | 1.40691765223466 | 0.166723006954726 | 0.481358333510474 | -5.22792831621225 |
| lnc-GBP6-5 | lnc-GBP6-5 | 0.219901483854757 | 0.321722743551441 | 1.40689122804257 | 0.166730794386176 | 0.481358333510474 | -4.86462666691617 |
| ENSG00000143297.18 | FCRL5 | -0.562147403670954 | 8.40431189006181 | -1.40754008030959 | 0.166747183450913 | 0.481358333510474 | -5.41607409009105 |
| ENSG00000117450.13 | PRDX1 | -0.290423176325369 | 5.41074770108448 | -1.40653025946295 | 0.166837203213996 | 0.481358333510474 | -5.43575260149577 |
| lnc-B3GALNT2-3 | lnc-B3GALNT2-3 | 0.320034397149785 | 0.468025406988497 | 1.40640268172282 | 0.166874824106876 | 0.481358333510474 | -4.8849708929818 |
| ENSG00000185787.14 | MORF4L1 | 0.140134038353635 | 7.35059836534398 | 1.40634458771185 | 0.166891957406634 | 0.481358333510474 | -5.44030217958155 |
| lnc-MYLK4-29 | lnc-MYLK4-29 | 0.39086737022652 | 0.350562945453768 | 1.40628531412651 | 0.166909440001868 | 0.481358333510474 | -4.84332607781155 |
| lnc-TM2D3-1 | lnc-TM2D3-1 | -0.258767980597525 | 1.55308486698396 | -1.40627081174742 | 0.166913717659207 | 0.481358333510474 | -5.06779752849562 |

| | | | | | | | |
|---|---|---|---|---|---|---|---|
| ENSG00000127824.13 | TUBA4A | -0.324207492851981 | 5.45939723115259 | -1.4062779919978 | 0.166934323359014 | 0.481358333510474 | -5.43891906167312 |
| ENSG00000185418.15 | TARS3 | -0.250051149646943 | 4.98923239381341 | -1.40614066739988 | 0.166952109174031 | 0.481358333510474 | -5.41869327531607 |
| lnc-KHNYN-1 | lnc-KHNYN-1 | -0.325356214493369 | 0.444243998242267 | -1.40609719244233 | 0.166964935461026 | 0.481358333510474 | -4.88102653228888 |
| ENSG00000254328.1 | | 0.258844325025279 | 0.332399177106071 | 1.40595551226678 | 0.167006740262348 | 0.481358333510474 | -4.86967524486695 |
| lnc-MYOC-7 | lnc-MYOC-7 | -0.430428008063788 | 0.420243143735935 | -1.40594984108005 | 0.167008413798211 | 0.481358333510474 | -4.88739626899615 |
| lnc-IL17RA-10 | lnc-IL17RA-10 | 1.12332059354964 | -0.618627299884516 | 1.40616138936811 | 0.167012546743999 | 0.481358333510474 | -4.59476148562522 |
| lnc-TAF13-2 | lnc-TAF13-2 | -0.37942873623917 | -0.0068081290787981( | -1.40589600059021 | 0.167024302479318 | 0.481358333510474 | -4.80880244305885 |
| ENSG00000262621.5 | NA | -0.782975109429467 | 0.204587338123688 | -1.40569365308077 | 0.167158452099736 | 0.481664764667472 | -4.81660631247738 |
| ENSG00000104915.14 | STX10 | -0.195648842515464 | 3.91306889443243 | -1.40537730975359 | 0.167177431802187 | 0.481664764667472 | -5.33985672260964 |
| ENSG00000135631.16 | RAB11FIP5 | 0.705903892547506 | 0.0726711235557007 | 1.40538031081126 | 0.167231025989119 | 0.481751743587688 | -4.85672558544376 |
| ENSG00000169926.10 | KLF13 | 0.248852428154245 | 6.95503061425301 | 1.40431758384125 | 0.167490626531546 | 0.482270567141075 | -5.44797445785474 |
| lnc-MTSS1-3 | lnc-MTSS1-3 | -0.220197675608861 | 1.41867639632325 | -1.40431242455529 | 0.167492152438866 | 0.482270567141075 | -5.05090708287936 |
| ENSG00000166938.12 | DIS3L | -0.1261991997579 | 4.9718433393249 | -1.40422446640062 | 0.167518168553696 | 0.482270567141075 | -5.42030435470931 |
| lnc-KLRG2-1 | lnc-KLRG2-1 | 0.38992630897204 | 1.09246593081378 | 1.40420696757108 | 0.167523344702892 | 0.482270567141075 | -5.01943823826812 |
| lnc-TRAPPC6B-2 | lnc-TRAPPC6B-2 | -0.380244736016021 | -0.601007666590643 | -1.40419028865904 | 0.16752827843651 | 0.482270567141075 | -4.7105627916374 |
| lnc-MAP2K6-5 | lnc-MAP2K6-5 | -0.561617643995714 | 1.89098985363469 | -1.40437904326817 | 0.167566861547367 | 0.482308903939942 | -5.10791752199132 |
| ENSG00000258405.9 | ZNF578 | -1.13228137290716 | -0.111846080387668 | -1.40465420356495 | 0.167588460427463 | 0.482308903939942 | -4.78880119525516 |
| ENSG00000149346.14 | SLX4IP | -0.176914487657358 | 3.87062093013078 | -1.40383790822038 | 0.167632541346069 | 0.482368320778154 | -5.32863455414337 |
| lnc-PML-2 | lnc-PML-2 | -0.310418747281988 | 0.131996568353442 | -1.40364763547398 | 0.167688860542402 | 0.482445218923764 | -4.83260862720666 |
| ENSG00000165501.16 | LRR1 | 0.213443321395993 | 2.31202890383732 | 1.40358925716116 | 0.167706143001483 | 0.482445218923764 | -5.16432957639226 |
| lnc-NUDT3-1 | lnc-NUDT3-1 | -0.212846061896841 | 5.03401028567946 | -1.40336726496888 | 0.167771874765193 | 0.482510588151465 | -5.42363173218403 |
| ENSG00000260300.5 | NA | -0.217395429339959 | 0.912932409102792 | -1.40335417665423 | 0.167775750833213 | 0.482510588151465 | -4.9750603221614 |
| ENSG00000149269.9 | PAK1 | 0.750475556017265 | 4.50629963263859 | 1.40411355482898 | 0.167815387857358 | 0.482557156785134 | -5.36529101366507 |
| ENSG00000101040.19 | ZMYND8 | 0.184563708817913 | 6.68343799147357 | 1.40249563711276 | 0.168030157077861 | 0.482774987128453 | -5.45285295745485 |
| lnc-DYM-1 | lnc-DYM-1 | -0.201525587161371 | 2.64879861469231 | -1.40246196002728 | 0.168040142531524 | 0.482774987128453 | -5.20093225255707 |
| ENSG00000133874.1 | RNF122 | -0.320118654351882 | 2.38188536746115 | -1.40242930930024 | 0.168049824104378 | 0.482774987128453 | -5.17456262182029 |
| lnc-CYTIP-2 | lnc-CYTIP-2 | 0.564491329099516 | -0.911997074595371 | 1.40242439881591 | 0.168051280195592 | 0.482774987128453 | -4.60943376750475 |
| ENSG00000196267.12 | ZNF836 | -0.164834529414846 | 4.33287759388436 | -1.40238431537977 | 0.168063166383725 | 0.482774987128453 | -5.37586634403705 |
| FAM239A | FAM239A | -0.403751479583238 | 0.969949291988175 | -1.40230143022726 | 0.168087746901644 | 0.482774987128453 | -5.00711223171115 |
| ENSG00000204520.12 | MICA | -0.210868788083723 | 2.66550387472422 | -1.40229967146972 | 0.168088268511122 | 0.482774987128453 | -5.2031610433526 |
| lnc-QRICH1-2 | lnc-QRICH1-2 | 0.201576235477156 | 3.72086070244657 | 1.40229052992407 | 0.168090979716284 | 0.482774987128453 | -5.3177918867621 |
| ENSG00000132824.13 | SERINC3 | -0.111630759500977 | 6.00738238176061 | -1.402198843935 | 0.168118173883218 | 0.482774987128453 | -5.4522185079005 |
| lnc-EIF2S3B-8 | lnc-EIF2S3B-8 | -0.663627755513309 | 2.46098831775811 | -1.40262252351051 | 0.168182702833029 | 0.482774987128453 | -5.19010260835484 |
| ENSG00000067064.11 | IDI1 | 0.214784888083612 | 5.80448335128831 | 1.40194951089588 | 0.168192143640293 | 0.482774987128453 | -5.45087126147773 |
| ENSG00000144840.9 | RABL3 | 0.21074993566323 | 4.11508671934276 | 1.40185352675115 | 0.168220626048408 | 0.482774987128453 | -5.35023266958688 |
| ENSG00000005339.14 | CREBBP | 0.196055570293938 | 7.46085262603184 | 1.40178164794459 | 0.168241957875415 | 0.482774987128453 | -5.44554035089279 |
| ENSG00000154265.15 | ABCA5 | -0.292813935308761 | 5.7484990510791 | -1.4017211831756 | 0.168259903930794 | 0.482774987128453 | -5.44989799638954 |
| ENSG00000130165.10 | ELOF1 | -0.282491954732344 | 4.17929507524154 | -1.40151977439736 | 0.168319693163351 | 0.482774987128453 | -5.37949883734331 |
| ENSG00000117262.18 | GPR89A | 0.149612825184962 | 2.79260011188377 | 1.40144043356739 | 0.168343250430852 | 0.482774987128453 | -5.22098142855179 |
| lnc-DARS2-3 | lnc-DARS2-3 | -0.173891251429877 | 3.38413762950487 | -1.40143722721162 | 0.168344202491157 | 0.482774987128453 | -5.28415824696219 |
| ENSG00000154370.15 | TRIM11 | 0.143978267079555 | 4.8916225257906 | 1.40136418020974 | 0.16836589340494 | 0.482774987128453 | -5.41983102228758 |
| ENSG00000249209.2 | | 0.241530270642075 | 3.96815693667671 | 1.40134929085153 | 0.168370314986192 | 0.482774987128453 | -5.3416710744643 |
| lnc-SH2D3A-1 | lnc-SH2D3A-1 | -0.24858910234318 | 1.67159441788412 | -1.40126058669949 | 0.168396658664863 | 0.482774987128453 | -5.09648516265868 |
| ENSG00000165169.10 | DYNLT3 | 0.260697658457683 | 3.48822295796402 | 1.401229336561 | 0.168405940206707 | 0.482774987128453 | -5.30118005195424 |
| lnc-TAL2-4 | lnc-TAL2-4 | 0.497168953958716 | 0.444451068252948 | 1.40122525699058 | 0.168407151901195 | 0.482774987128453 | -4.88406303662728 |
| ENSG00000167562.12 | ZNF701 | -0.16900947595958 | 4.8542448732794 | -1.40088972212908 | 0.168506834061086 | 0.482895646470406 | -5.41357555139262 |
| ENSG00000164916.10 | FOXK1 | 0.2051519540424 | 5.71093363624086 | 1.40085897753211 | 0.168515970093 | 0.482895646470406 | -5.45060894658818 |
| lnc-TM9SF2-7 | lnc-TM9SF2-7 | 0.359600346231807 | 0.889890236992077 | 1.40084668048197 | 0.168519624379193 | 0.482895646470406 | -4.96622634600893 |
| lnc-SLC16A11-5 | lnc-SLC16A11-5 | -0.704206284676967 | -0.89236769682577 | -1.40052925032552 | 0.168613975686509 | 0.483098755784438 | -4.60290506316832 |
| ENSG00000196387.9 | ZNF140 | 0.164180469733776 | 4.22772966724819 | 1.40044362942718 | 0.168639432230764 | 0.483104444482368 | -5.36521867323487 |
| lnc-ST6GALNAC6-1 | lnc-ST6GALNAC6-1 | -0.16526441454115 | 3.44829814902454 | -1.40030789223014 | 0.168679795320548 | 0.483152828706211 | -5.29250855370195 |
| ENSG00000121864.9 | ZNF639 | 0.173454306617691 | 5.4610029138458 | 1.40016731889302 | 0.168721604407634 | 0.483176333426929 | -5.44768872663705 |
| ENSG00000186298.11 | PPP1CC | 0.134339002988023 | 7.89791587963513 | 1.40012245169898 | 0.168734950428643 | 0.483176333426929 | -5.43915771667982 |
| lnc-ANKRD30BL-3 | lnc-ANKRD30BL-3 | -1.48022134129984 | -0.844524803165246 | -1.40026365319859 | 0.168855582359307 | 0.483454507122217 | -4.58311865555425 |
| ENSG00000197043.13 | ANXA6 | -0.319577282672744 | 7.26712517295726 | -1.39967084099524 | 0.168891449630631 | 0.483489945444684 | -5.449154753233 |
| ENSG00000116698.21 | SMG7 | 0.229615990438996 | 5.26454953668283 | 1.39882643379964 | 0.169120813049258 | 0.484076550148649 | -5.43990504898713 |
| ENSG00000267120.3 | | -0.24431858768669 | 4.63322132599719 | -1.39875064969652 | 0.169143397399266 | 0.484076550148649 | -5.40409975416075 |
| lnc-TSTA3-5 | lnc-TSTA3-5 | -0.28449828275122 | 0.486906847018843 | -1.39843130554342 | 0.169238590588834 | 0.484244029184145 | -4.89039039444006 |
| ENSG00000213614.9 | HEXA | 0.13608344141057 | 6.37723343275274 | 1.39839649479702 | 0.169248969827078 | 0.484244029184145 | -5.45896522295102 |
| lnc-WNT5B-3 | lnc-WNT5B-3 | 0.267710784046055 | 1.90066686692222 | 1.39829635295711 | 0.169278831065174 | 0.484262151468309 | -5.11483710612776 |
| ENSG00000215114.8 | UBXN2B | -0.14385713434433 | 4.58463234206324 | -1.39813055109345 | 0.169328280424159 | 0.484320641057562 | -5.40723681739559 |
| ENSG00000100412.15 | ACO2 | 0.143553565252586 | 5.73789631644895 | 1.39807001756539 | 0.169346336962118 | 0.484320641057562 | -5.45418732931277 |
| ENSG00000123329.17 | ARHGAP9 | -0.236673217333295 | 6.02208532192938 | -1.39791999209978 | 0.169391094483902 | 0.484381341728559 | -5.45835179822658 |
| lnc-HMGN5-1 | lnc-HMGN5-1 | 0.206493242334424 | 0.792180495809246 | 1.39722645380021 | 0.16959811913184 | 0.484905971119691 | -4.95151342129144 |
| ENSG00000071994.10 | PDCD2 | -0.137667618423679 | 5.56449489177317 | -1.39707417712281 | 0.169643600774097 | 0.48496864371295 | -5.45307443008765 |
| lnc-PTPN12-1 | lnc-PTPN12-1 | -0.43297678657939 | 0.563569062853868 | -1.39653477481584 | 0.169804784384364 | 0.485272716863941 | -4.95054315641407 |
| lnc-STYX-4 | lnc-STYX-4 | -0.206012614147607 | 0.924376885351955 | -1.3964416144434 | 0.169832634501757 | 0.485272716863941 | -4.99154880119013 |
| ENSG00000132768.13 | DPH2 | -0.220734294124708 | 3.81492745613013 | -1.39635770490621 | 0.169857722134629 | 0.485272716863941 | -5.34420653114456 |

| | | | | | | | |
|---|---|---|---|---|---|---|---|
| lnc-BAZ2A-1 | lnc-BAZ2A-1 | 0.266653258369959 | 3.34808490688646 | 1.39634057256681 | 0.169862844788502 | 0.485272716863941 | -5.2855621440723 |
| ENSG00000085760.14 | MTIF2 | 0.134172212955696 | 5.2790992141473 | 1.39632383851764 | 0.16986784846741 | 0.485272716863941 | -5.44286195118388 |
| ENSG00000120697.8 | ALG5 | 0.119811167857248 | 3.35495202291206 | 1.3961998188357 | 0.169904935387688 | 0.485311307949595 | -5.2852840585644 |
| ENSG00000227063.5 | RPL41P1 | 0.364673103881115 | 1.4776350708818 | 1.39609972374588 | 0.169934872466959 | 0.485322254435401 | -5.08117129827794 |
| ENSG00000099624.7 | ATP5F1D | -0.318777885405836 | 2.50205786423413 | -1.39602934154656 | 0.169955925276702 | 0.485322254435401 | -5.20288626813449 |
| ENSG00000141736.13 | ERBB2 | -0.16563775592271 | 2.36787477168993 | -1.39587905128974 | 0.170000887059733 | 0.485348823016651 | -5.18446731671898 |
| lnc-AMDHD2-7 | lnc-AMDHD2-7 | 0.181673469344683 | 1.72753319435515 | 1.3957804861448 | 0.170030379444108 | 0.485348823016651 | -5.10506027359288 |
| ENSG00000229119.3 | RPLP0P9 | 0.969270779604178 | 1.66694082237034 | 1.39649346549055 | 0.170035969624111 | 0.485348823016651 | -4.99402355863701 |
| LINC00663 | LINC00663 | -0.198297920961688 | 2.72521410375355 | -1.39561765585481 | 0.170079109762904 | 0.485404647289221 | -5.21429132809591 |
| lnc-STARD10-2 | lnc-STARD10-2 | -0.450976300581574 | 0.369222923213787 | -1.39537695788953 | 0.170151163441454 | 0.485491833178942 | -4.88565459015117 |
| ENSG00000229106.1 | BTBD6P1 | -0.253842177922941 | 4.63526459060103 | -1.3953206041772 | 0.170168036521255 | 0.485491833178942 | -5.40589234147627 |
| ENSG00000138115.13 | CYP2C8 | 0.733598149883487 | 1.43656950310472 | 1.39581623372572 | 0.170180419588304 | 0.485491833178942 | -5.02657377769664 |
| ENSG00000202400.1 | SNORD82 | 0.254548572327827 | 0.39864244485276 | 1.39518624177683 | 0.170208271718733 | 0.485503999000372 | -4.90218678002381 |
| ENSG00000134313.15 | KIDINS220 | 0.136636575800041 | 6.37174031112661 | 1.39496487984099 | 0.170274575271792 | 0.485575332750865 | -5.46352827772455 |
| ENSG00000198265.11 | HELZ | 0.14016344295603 | 7.00592508379915 | 1.39494522916685 | 0.170280462119018 | 0.485575332750865 | -5.45997123554971 |
| ENSG00000187808.4 | SOWAHD | -0.466502003688201 | 1.18448458530059 | -1.39486459086966 | 0.170304620976472 | 0.485576951594226 | -5.05411106322398 |
| lnc-GTF2A2-1 | lnc-GTF2A2-1 | 0.169001055754848 | 2.72156822131845 | 1.39464770365679 | 0.170369612570662 | 0.485694977221874 | -5.22077390029119 |
| lnc-RHCE-2 | lnc-RHCE-2 | -0.885687894339206 | 0.388156726039186 | -1.39474799283757 | 0.170501729071284 | 0.485940997954409 | -4.89069743083476 |
| lnc-PTP4A2-2 | lnc-PTP4A2-2 | 0.170109521068108 | 4.69385885327167 | 1.39420234233713 | 0.170503128175035 | 0.485940997954409 | -5.41829800674161 |
| ENSG00000261796.1 | ISY1-RAB43 | -0.171182835417985 | 4.58596108109137 | -1.3940506620964 | 0.170548619172512 | 0.486003354344151 | -5.41227870531256 |
| ENSG00000159176.13 | CSRP1 | -0.213743341177767 | 4.58385573504666 | -1.39387700886486 | 0.170600711733919 | 0.486019649690178 | -5.41287073718905 |
| ENSG00000072803.17 | FBXW11 | 0.138740198317533 | 6.40593495067266 | 1.39387417182633 | 0.170601562892267 | 0.486019649690178 | -5.46496159268639 |
| ENSG00000139343.10 | SNRPF | 0.208964716842352 | 3.89214438806343 | 1.39355018580951 | 0.170698785712716 | 0.486229325536235 | -5.35167001158242 |
| ENSG00000204580.13 | DDR1 | -0.423862824971725 | 3.74877552888665 | -1.39366841325603 | 0.170750332502215 | 0.486301534869575 | -5.34952586202606 |
| ENSG00000142686.7 | C1orf216 | -0.308418578037398 | 2.2833304189589 | -1.39330833637991 | 0.170771388720657 | 0.486301534869575 | -5.178988645731 |
| lnc-TXNDC9-3 | lnc-TXNDC9-3 | 0.160887018137525 | 1.35695608626136 | 1.39292818970182 | 0.170885556865552 | 0.486542634129396 | -5.05834407520665 |
| lnc-TMEM39B-2 | lnc-TMEM39B-2 | 0.302735281445132 | -0.078080687350842 | 1.39286902751669 | 0.17090333016351 | 0.486542634129396 | -4.80041439442962 |
| ENSG00000136897.7 | MRPL50 | 0.188263408754699 | 4.24162396614958 | 1.39274649886016 | 0.170940144355942 | 0.486580139853181 | -5.38056841417426 |
| lnc-IYD-1 | lnc-IYD-1 | 0.359483655760974 | 4.51083300135908 | 1.39265347425911 | 0.171010475594329 | 0.486636972283401 | -5.40054043337539 |
| lnc-AK6-3 | lnc-AK6-3 | -0.302335094463496 | -0.296716802433857 | -1.39248196353993 | 0.171019645953631 | 0.486636972283401 | -4.74636234097864 |
| lnc-MTO1-1 | lnc-MTO1-1 | -0.354540345174317 | -0.25014380614378 | -1.39244406483384 | 0.171031038113886 | 0.486636972283401 | -4.76701642942125 |
| ENSG00000123552.17 | USP45 | -0.177364738643126 | 5.73886030032926 | -1.3922329517051 | 0.171094508428321 | 0.486702612233591 | -5.46227977201372 |
| ENSG00000130803.14 | ZNF317 | 0.15095793071624 | 6.23901910606437 | 1.39221003508806 | 0.17110139931605 | 0.486702612233591 | -5.46705235071053 |
| ENSG00000008382.15 | MPND | -0.1932300701909 | 2.20902575292011 | -1.39202023688561 | 0.171158478756859 | 0.486750956994226 | -5.16474631219378 |
| ENSG00000156265.15 | MAP3K7CL | 1.72933288180123 | 0.675928927549549 | 1.39293768889471 | 0.17116964327452 | 0.486750956994226 | -4.89229298919415 |
| ENSG00000197580.11 | BCO2 | 0.41144524322541 | -0.0733225592588487 | 1.39191764001171 | 0.171189339633737 | 0.486750956994226 | -4.76469717666318 |
| lnc-IRF8-1 | lnc-IRF8-1 | -0.474117779458506 | 2.02014676401984 | -1.3917503699453 | 0.171262296084687 | 0.486858693508437 | -5.11099109192578 |
| ENSG00000102871.15 | TRADD | -0.315098932442531 | 2.64004793242106 | -1.39163447535499 | 0.171274537224632 | 0.486858693508437 | -5.22503120702608 |
| UBAC2-AS1 | UBAC2-AS1 | -0.293854503127267 | 0.7960797623273 | -1.39142213412566 | 0.17133844727762 | 0.486896564376674 | -4.96797366981457 |
| lnc-PPM1K-1 | lnc-PPM1K-1 | -0.204467145082748 | 3.47131498892178 | -1.39138420345825 | 0.17134986552272 | 0.486896564376674 | -5.31789264624442 |
| ENSG00000241061.3 | RPL5P1 | 0.937327155918754 | -0.692564734586644 | 1.39135443922651 | 0.171358825843882 | 0.486896564376674 | -4.58800339934353 |
| lnc-PRAF2-1 | lnc-PRAF2-1 | 0.150271552523428 | 2.40891479505729 | 1.39117447414715 | 0.171413010855515 | 0.486983299163432 | -5.18713123474892 |
| ENSG00000212125.2 | TAS2R15P | -0.317412856940766 | 1.48016243722542 | -1.39108301117594 | 0.17144055418926 | 0.486994331614344 | -5.08925262157827 |
| ENSG00000173276.13 | ZBTB21 | 0.232992421113973 | 6.01525051233164 | 1.39093338050457 | 0.171485621652961 | 0.487046468303847 | -5.46727998984942 |
| IQCH-AS1 | IQCH-AS1 | -0.250937209219089 | 3.33914662567538 | -1.39086495657158 | 0.171506233409429 | 0.487046468303847 | -5.30321824366471 |
| lnc-HRH1-5 | lnc-HRH1-5 | -0.427001766890287 | -0.569218700402877 | -1.3907222503407 | 0.171549227852644 | 0.487101359758723 | -4.68558564422605 |
| lnc-HCK-1 | lnc-HCK-1 | 0.277607581695039 | 0.347061752400396 | 1.39033480422929 | 0.17166599950657 | 0.487365692116376 | -4.86775770662286 |
| lnc-C2orf66-2 | lnc-C2orf66-2 | 0.187304235046914 | 1.10288706412242 | 1.39020005902183 | 0.171706624546362 | 0.487413798515759 | -5.02563142507858 |
| ENSG00000121410.11 | A1BG | 0.300642768883287 | 1.21692013460982 | 1.38971370661118 | 0.171853319474304 | 0.487696495830178 | -5.0436008567409 |
| ENSG00000155304.5 | HSPA13 | 0.296918862326163 | 4.99891327163742 | 1.38962271995441 | 0.171880773896821 | 0.487696495830178 | -5.43954084394553 |
| ENSG00000272657.1 | NA | -0.376807571058299 | 0.0382737446851278 | -1.38959626094216 | 0.171888758308664 | 0.487696495830178 | -4.83908477820635 |
| ENSG00000172172.7 | MRPL13 | 0.202116695820937 | 3.36186589546973 | 1.38955570145501 | 0.171900998312574 | 0.487696495830178 | -5.2893501689097 |
| ENSG00000169696.15 | ASPSCR1 | -0.190478485395958 | 4.05991881046645 | -1.38947722048303 | 0.17192468414438 | 0.487696495830178 | -5.37527425532121 |
| lnc-CLEC16A-5 | lnc-CLEC16A-5 | 0.362535296129186 | 0.949592374608941 | 1.38932312669221 | 0.171971197546856 | 0.487761218011154 | -4.96666751617748 |
| lnc-SNX11-6 | lnc-SNX11-6 | 0.299647160085912 | 1.16606289351505 | 1.38878258483856 | 0.172134437876788 | 0.488156948858905 | -5.04204981416259 |
| ENSG00000203685.9 | STUM | -0.864771122042558 | 1.04971930516004 | -1.3886961750537 | 0.172349964813982 | 0.488681505404379 | -4.97881774946355 |
| ENSG00000161048.11 | NAPEPLD | -0.242691625725953 | 2.70828539629768 | -1.38801354001999 | 0.172366891572453 | 0.488681505404379 | -5.22364253521944 |
| ENSG00000159399.9 | HK2 | -0.514557424431324 | 2.16838768369895 | -1.38814797318998 | 0.172403495071736 | 0.488717964338457 | -5.16576546912248 |
| CCNT2-AS1 | CCNT2-AS1 | -0.215312964025602 | 2.49825376779396 | -1.38772170706582 | 0.172455165456327 | 0.488797117954775 | -5.20204086651827 |
| ENSG00000173272.15 | MZT2A | -0.195069558833021 | 3.07865782730856 | -1.38740648100839 | 0.172550554640072 | 0.488933312449177 | -5.2742111583418 |
| lnc-FCRL3-1 | lnc-FCRL3-1 | -1.24356031929366 | 1.18435822532292 | -1.38831633407823 | 0.172550725435108 | 0.488933312449177 | -5.02133625925407 |
| lnc-SNAPC4-1 | lnc-SNAPC4-1 | -0.223679359324525 | 1.78554245875566 | -1.38732617603941 | 0.172574861906719 | 0.488934395406193 | -5.12120273017882 |
| lnc-RSPH10B2-1 | lnc-RSPH10B2-1 | 0.207248965454287 | 0.745647046853389 | 1.38719186472009 | 0.172615522117859 | 0.48898228623065 | -4.97898826491812 |
| lnc-NMUR1-4 | lnc-NMUR1-4 | -0.999503102761253 | 0.0818316529644162 | -1.38749516454219 | 0.172699208628397 | 0.489152031264385 | -4.8319027340616 |
| ENSG00000279602.1 | | -0.969353600046841 | 4.38094142672913 | -1.38740878313518 | 0.172836337448267 | 0.489473078384382 | -5.43032321258184 |
| lnc-PTGR2-1 | lnc-PTGR2-1 | -0.388319551801502 | 1.28861087526313 | -1.38629927896313 | 0.172885924076697 | 0.489546151502359 | -5.05576421830537 |
| TRERNA1 | TRERNA1 | 0.5285752450894 | -0.124015398968366 | 1.38618613524724 | 0.172920223499867 | 0.489575923012071 | -4.75313025409849 |
| ENSG00000167632.15 | TRAPPC9 | 0.161635832982905 | 5.49808615844489 | 1.38585945626215 | 0.17301928552825 | 0.489748495438165 | -5.46182997166935 |

| | | | | | | | |
|---|---|---|---|---|---|---|---|
| ENSG00000165209.18 | STRBP | -0.150817069209127 | 7.82659342551032 | -1.38579942749583 | 0.173037493409078 | 0.489748495438165 | -5.45996728805745 |
| lnc-C19orf48-2 | lnc-C19orf48-2 | -0.371066720274008 | -0.0685453024098275 | -1.38574976443888 | 0.17305255829188 | 0.489748495438165 | -4.84016229454804 |
| lnc-ZNF562-2 | lnc-ZNF562-2 | 0.144743180592553 | 2.39417755180821 | 1.38556754999802 | 0.173107840251541 | 0.489797562912947 | -5.18924578228664 |
| lnc-C1orf159-1 | lnc-C1orf159-1 | -0.269989214208032 | 0.238611275137356 | -1.38553575239392 | 0.173117488713584 | 0.489797562912947 | -4.90410089449383 |
| ENSG00000278535.4 | DHRS11 | -0.342172229702076 | 0.823516135812814 | -1.38522487339071 | 0.173211841786176 | 0.489997160456962 | -5.01715042024059 |
| lnc-PIGN-7 | lnc-PIGN-7 | -0.376290941350931 | 0.0840238425927249 | -1.3850165341648 | 0.173275095882194 | 0.490108739665137 | -4.8464801648829 |
| lnc-RMDN3-5 | lnc-RMDN3-5 | 0.324080999473156 | 1.64564044199098 | 1.38437652384037 | 0.173469521928244 | 0.49059125719278 | -5.10201900507949 |
| ENSG00000198088.10 | NUP62CL | -1.2269611846504 | -1.27240677804672 | -1.38489365620758 | 0.173513394666656 | 0.490620568698666 | -4.75668157581076 |
| lnc-MAP3K9-3 | lnc-MAP3K9-3 | -0.461532446223967 | -0.34924661885932 | -1.38418313396125 | 0.173528304228903 | 0.490620568698666 | -4.73669739817631 |
| ENSG00000163516.13 | ANKZF1 | -0.220381837012718 | 6.32178206567313 | -1.38408608303148 | 0.173557809393175 | 0.490620568698666 | -5.47751095332729 |
| ENSG00000137880.5 | GCHFR | -0.36279707438957 | 2.67190986312784 | -1.38402878412096 | 0.173575231077281 | 0.490620568698666 | -5.23498148639415 |
| ENSG00000126391.13 | FRMD8 | -0.304167012554712 | 4.96337562259492 | -1.38330222311021 | 0.173798092099014 | 0.49113983364547 | -5.44878027391046 |
| ENSG00000142684.8 | ZNF593 | 0.27644293198636 | 1.44160494965556 | 1.38326803918997 | 0.173806663181927 | 0.49113983364547 | -5.09031644267892 |
| ENSG00000104356.10 | POP1 | -0.230046986093466 | 2.18849405886485 | -1.38315920652472 | 0.173839791519728 | 0.491166016314421 | -5.17679122085352 |
| ENSG00000119953.12 | SMNDC1 | 0.157247848812753 | 4.61266464220093 | 1.38285933298296 | 0.173931097402202 | 0.491296509394005 | -5.42659712266424 |
| ENSG00000117280.12 | RAB29 | 0.263326222615563 | 5.08956946042335 | 1.38278063429207 | 0.173955065825393 | 0.491296509394005 | -5.45111566038306 |
| ENSG00000215440.11 | NPEPL1 | -0.183507980920123 | 4.46760718579973 | -1.38276461063311 | 0.173959946293817 | 0.491296509394005 | -5.41924529045743 |
| lnc-STRN3-1 | lnc-STRN3-1 | -0.525883196435589 | 0.238953684196589 | -1.38258875509388 | 0.174013515140559 | 0.491296509394005 | -4.89622290762302 |
| ENSG00000113272.13 | THG1L | -0.166040185626986 | 3.41058353906281 | -1.38255682368685 | 0.174023243405804 | 0.491296509394005 | -5.32035497200579 |
| lnc-SAMD5-1 | lnc-SAMD5-1 | -0.417650533959451 | -0.224970676389804 | -1.38253730052574 | 0.174029191565452 | 0.491296509394005 | -4.81964313910617 |
| ENSG00000198898.13 | CAPZA2 | 0.14437565058206 | 5.67527668834261 | 1.38239392890588 | 0.174072877694998 | 0.491350181655823 | -5.47460033471523 |
| ENSG00000263883.1 | EEF1DP7 | -0.242600924039239 | 0.470932417719789 | -1.38231823034362 | 0.174095946888982 | 0.491350181655823 | -4.94465159888888 |
| ENSG00000165055.15 | METTL2B | -0.185117588495601 | 3.99202344976566 | -1.38202106182774 | 0.174186532081029 | 0.491538441160381 | -5.37564358478446 |
| lnc-PSPC1-2 | lnc-PSPC1-2 | -0.271526230036551 | 2.74532407631914 | -1.38192597838359 | 0.174215523843953 | 0.49155286186156 | -5.23378255890859 |
| ENSG00000162836.11 | ACP6 | -0.385776174410108 | 2.77746924779571 | -1.38169720130047 | 0.17428551451692 | 0.491682942064385 | -5.2313442539753 |
| ENSG00000125676.19 | THOC2 | 0.103303332068371 | 6.67346868474187 | 1.38157630156824 | 0.174322175421814 | 0.491687741573702 | -5.48023338071856 |
| ENSG00000107551.20 | RASSF4 | -0.639027036141227 | 1.02230541824496 | -1.38187496865969 | 0.174334991886745 | 0.491687741573702 | -5.03440913522606 |
| ENSG00000184990.12 | SIVA1 | 0.168535934158402 | 4.84833380764378 | 1.38142460901265 | 0.174368457414943 | 0.491714749824875 | -5.44302775675493 |
| lnc-ZNF621-1 | lnc-ZNF621-1 | 1.18382655453026 | -0.763999920840924 | 1.3813928914113 | 0.174472884020989 | 0.491941831753974 | -4.62016368681018 |
| ENSG00000241489.7 | | 0.195406763349341 | 4.65259741555231 | 1.38089013410072 | 0.17453160352484 | 0.491981642937997 | -5.42512805481075 |
| ENSG00000111405.8 | ENDOU | -0.57010765522913 | 2.78724859620772 | -1.38132784101346 | 0.174558042729421 | 0.491981642937997 | -5.24202549535421 |
| ENSG00000173559.12 | NABP1 | -0.347961695934647 | 5.27902095592745 | -1.38092895205322 | 0.17457385223264 | 0.491981642937997 | -5.46546639811639 |
| lnc-PTBP3-4 | lnc-PTBP3-4 | -0.256490618649889 | 2.0054148173415 | -1.38072310428433 | 0.174582612836765 | 0.491981642937997 | -5.16045199267477 |
| ENSG00000174903.15 | RAB1B | -0.206459652942888 | 5.05390593139588 | -1.38040175244136 | 0.174680783040434 | 0.492190904492986 | -5.45711264404202 |
| ENSG00000172500.12 | FIBP | -0.261232063080699 | 4.36774880504335 | -1.38025389829944 | 0.17472596553150 7 | 0.492233191915527 | -5.41277140570743 |
| lnc-MEAF6-2 | lnc-MEAF6-2 | 1.76232145220581 | -1.13408993324622 | 1.38066463222608 | 0.174743620139278 | 0.492233191915527 | -4.51587996871063 |
| ENSG00000279192.1 | PWAR5 | -0.287214368326602 | 4.33862523290236 | -1.37988998595447 | 0.174837211356343 | 0.492429436418667 | -5.40229288019053 |
| ENSG00000152520.13 | PAN3 | 0.211075557944595 | 7.23703997226393 | 1.37962564338098 | 0.174918053619829 | 0.492496035610423 | -5.47774624346197 |
| ENSG00000207468.1 | SNORA19 | 0.307586923496681 | 0.173733752269566 | 1.37960109164194 | 0.174925563593237 | 0.492496035610423 | -4.87951552633449 |
| lnc-TMEM201-2 | lnc-TMEM201-2 | 0.237423527431509 | 1.18515811580212 | 1.3795779601007 | 0.174932639379478 | 0.492496035610423 | -5.04160172980713 |
| ENSG00000279620.1 | | 0.471615564456258 | 1.37954972461508 | 1.37933550162074 | 0.175006819144814 | 0.492637494318614 | -5.05763385606036 |
| ENSG00000253731.2 | PCDHGA6 | -1.33370160499351 | -0.696354882471811 | -1.37961072481438 | 0.175147445745356 | 0.492936801244551 | -4.78477775201 |
| ENSG00000242259.8 | C22orf39 | 0.113014735765547 | 5.22915390038348 | 1.37883166849186 | 0.175161043788692 | 0.492936801244551 | -5.46671972730833 |
| ENSG00000206688.1 | SNORD116-18 | -0.388583346516364 | 2.70832115359102 | -1.37847211370854 | 0.175271168653931 | 0.493179284265737 | -5.22121957354031 |
| lnc-CD63-1 | lnc-CD63-1 | 0.179312883231422 | 2.3465762650065 | 1.37802266435794 | 0.175408901754699 | 0.493459847756192 | -5.19918589807518 |
| lnc-HMOX1-2 | lnc-HMOX1-2 | -0.821238434961846 | -0.666777306887197 | -1.37800675870345 | 0.175454341656078 | 0.493459847756192 | -4.75247886423845 |
| ENSG00000146826.16 | TRAPPC14 | 0.211120776938516 | 4.56053033539045 | 1.37757841076977 | 0.175545124780732 | 0.493459847756192 | -5.43398519756911 |
| lnc-WWP2-2 | lnc-WWP2-2 | -0.214061392699823 | 2.39480403140853 | -1.37748101030427 | 0.175575001944685 | 0.493459847756192 | -5.2112135405966 |
| ENSG00000187792.4 | ZNF70 | -0.24865225406657 | 3.16315730086676 | -1.37742842059369 | 0.175591135239734 | 0.493459847756192 | -5.29295007942851 |
| ENSG00000103061.12 | SLC7A6OS | -0.125800029115704 | 5.14855366396725 | -1.37723460520335 | 0.175650603162143 | 0.493459847756192 | -5.46470991670181 |
| ENSG00000141026.5 | MED9 | 0.140900044594397 | 3.99258930818457 | 1.377218749966 | 0.175655468675763 | 0.493459847756192 | -5.38228550366355 |
| ENSG00000196371.3 | FUT4 | 0.320757404155936 | 2.59131033633246 | 1.37721235948252 | 0.17565742975969 | 0.493459847756192 | -5.22668199632133 |
| lnc-PPP1R14D-1 | lnc-PPP1R14D-1 | -0.186736827002101 | 0.534265789483133 | -1.37716090064647 | 0.175673221841059 | 0.493459847756192 | -4.94623266636133 |
| ENSG00000144362.11 | PHOSPHO2 | -0.337248095035094 | 1.42674991548163 | -1.37710829203108 | 0.175689367909111 | 0.493459847756192 | -5.09787647243011 |
| RALY-AS1 | RALY-AS1 | -0.220629932323995 | 1.85429913408032 | -1.37687829104102 | 0.175759970788727 | 0.493459847756192 | -5.13666463942382 |
| ENSG00000198093.10 | ZNF649 | -0.119459510486836 | 4.14010802541526 | -1.37685573788835 | 0.175766895058134 | 0.493459847756192 | -5.39175838211906 |
| ENSG00000215375.6 | MYL5 | -0.321555080289605 | 3.1787001416082 | -1.37685249112418 | 0.175767891897086 | 0.493459847756192 | -5.29678654059354 |
| ENSG00000204147.10 | ASAH2B | -0.315503991718267 | 1.30806768991641 | -1.3768438729733 | 0.175770537909174 | 0.493459847756192 | -5.07048534660617 |
| lnc-TDRKH-1 | lnc-TDRKH-1 | 0.119232165432182 | 3.0993901001777 | 1.3768120962904 | 0.175780294500738 | 0.493459847756192 | -5.28535927246156 |
| ENSG00000075826.16 | SEC31B | -0.201069415041341 | 5.24918413919151 | -1.37675928404822 | 0.175796510695435 | 0.493459847756192 | -5.46835908888635 |
| ENSG00000178951.8 | ZBTB7A | -0.252332796422909 | 4.32509306099807 | -1.37663161174587 | 0.175835717721629 | 0.493459847756192 | -5.41997640461873 |
| ENSG00000104805.15 | NUCB1 | -0.192471976113131 | 5.56399136092102 | -1.37660994856519 | 0.175842370961418 | 0.493459847756192 | -5.47902785679509 |
| lnc-SLC15A2-1 | lnc-SLC15A2-1 | -0.559803381999736 | 0.252249730691508 | -1.37657672738601 | 0.175852574294717 | 0.493459847756192 | -4.87735069344535 |
| ENSG00000086102.18 | NFX1 | 0.170167228535676 | 7.27883086986983 | 1.37649078803453 | 0.175878971260108 | 0.493459847756192 | -5.48199063180995 |
| ENSG00000239014.1 | SNORD108 | -0.319980031696312 | 1.18081443543922 | -1.37646686864143 | 0.175886318839078 | 0.493459847756192 | -5.03755585297397 |
| ENSG00000014641.17 | MDH1 | 0.169154320662349 | 5.85524872354127 | 1.37642783599255 | 0.175898309429489 | 0.493459847756192 | -5.48502115256372 |
| ENSG00000278311.4 | GGNBP2 | 0.142336232798081 | 6.8316838583543 | 1.37607206931813 | 0.176007627869639 | 0.493699237454453 | -5.48643870187166 |
| ENSG00000181690.7 | PLAG1 | -0.441362137127574 | 0.797708690532352 | -1.37570516218656 | 0.176120424444094 | 0.4938920365289 | -5.02444089616086 |

| | | | | | | | |
|---|---|---|---|---|---|---|---|
| ENSG00000272578.5 | GUSBP11 | 0.157315236596657 | 5.30873172526641 | 1.37566840302905 | 0.176131728218295 | 0.4938920365289 | -5.47279562026958 |
| ENSG00000234776.4 | FREY1 | -0.245744005201467 | 0.425399290747677 | -1.37557238478722 | 0.176161257340691 | 0.4938920365289 | -4.93929979799909 |
| lnc-TBC1D8-4 | lnc-TBC1D8-4 | 0.352789824189274 | -0.442425542945607 | 1.37552113839537 | 0.176177019045439 | 0.4938920365289 | -4.72288132172961 |
| ENSG00000121594.11 | CD80 | -1.86493505231018 | -0.637204643685804 | -1.37618881811938 | 0.17619633796005 | 0.4938920365289 | -4.60563433651521 |
| lnc-AIG1-8 | lnc-AIG1-8 | -0.208908603073704 | 4.36038242785357 | -1.37503534726946 | 0.176326486503657 | 0.494155831328973 | -5.41606676312845 |
| lnc-TBC1D19-8 | lnc-TBC1D19-8 | -0.359120728532355 | 3.45125611686045 | -1.37501570969539 | 0.176350960874575 | 0.494155831328973 | -5.348829786725 |
| ENSG00000064490.13 | RFXANK | -0.204519891417939 | 4.0033604023443 | -1.37491844044519 | 0.176362470822652 | 0.494155831328973 | -5.38773960372327 |
| lnc-HNRNPA2B1-5 | lnc-HNRNPA2B1-5 | 0.21504527093517 | 4.94407230630846 | 1.37479978032238 | 0.176399000609516 | 0.494190911875006 | -5.46352824352875 |
| ENSG00000075303.12 | SLC25A40 | -0.161127279259452 | 3.81839854858903 | -1.37430115499984 | 0.176552567380713 | 0.494471193545345 | -5.37263354505843 |
| ENSG00000165915.13 | SLC39A13 | -0.176895869609139 | 3.99762742091386 | -1.37429816892466 | 0.176553487343722 | 0.494471193545345 | -5.39017626379193 |
| ENSG00000101935.9 | AMMECR1 | -0.164417575927847 | 4.26395033127061 | -1.37424095228901 | 0.176571115607943 | 0.494471193545345 | -5.41559851742631 |
| ENSG00000224331.2 | | 1.52933441806963 | -0.433519810886148 | 1.37453632694197 | 0.17666640305374 | 0.494659225148413 | -4.62158433632713 |
| lnc-RASGEF1B-1 | lnc-RASGEF1B-1 | 0.599328739071876 | -0.51462494238217 | 1.37386712256123 | 0.176686324796732 | 0.494659225148413 | -4.67796593599394 |
| ENSG00000133624.13 | ZNF767P | -0.214114425497492 | 4.42489173524392 | -1.37367476583508 | 0.176745629098669 | 0.494757960524671 | -5.41979762279429 |
| lnc-ZC3HC1-1 | lnc-ZC3HC1-1 | 0.199589023837792 | 1.17893273533529 | 1.37346710424796 | 0.176809669187695 | 0.494816655612768 | -5.0475474564265 |
| ENSG00000165650.11 | PDZD8 | 0.140277941082587 | 5.08087140717529 | 1.37338726287498 | 0.176834295976711 | 0.494816655612768 | -5.46595251096192 |
| ENSG00000176974.19 | SHMT1 | -0.302838591791406 | 2.66123494577166 | -1.3733729295368 | 0.176838717324371 | 0.494816655612768 | -5.24446607972683 |
| ENSG00000105447.12 | GRWD1 | 0.20912198381758 | 3.96246519327152 | 1.37323624232743 | 0.176880884968023 | 0.494858016099903 | -5.38315432468559 |
| ENSG00000205189.11 | ZBTB10 | 0.30978512152811 | 6.73616566029736 | 1.37323297417987 | 0.176901582979307 | 0.494858016099903 | -5.49084225243768 |
| ENSG00000135723.13 | FHOD1 | -0.203396840698399 | 2.78838126718863 | -1.37287020858764 | 0.176993843616747 | 0.49501849468028 | -5.25728332817124 |
| lnc-OVGP1-1 | lnc-OVGP1-1 | -0.223127900847967 | 3.39054544629407 | -1.3728274244121 | 0.177007050519694 | 0.49501849468028 | -5.31970962244719 |
| ENSG00000182712.15 | CMC4 | 0.135297659624351 | 2.12910817341949 | 1.37263237571557 | 0.177067269078329 | 0.495063237455174 | -5.17783782120181 |
| ENSG00000261236.7 | BOP1 | -0.342103251359447 | 3.22578675552008 | -1.37261979551989 | 0.177071153580381 | 0.495063237455174 | -5.31225918872971 |
| lnc-CTNNBIP1-2 | lnc-CTNNBIP1-2 | 0.186818365177015 | 1.14677675536069 | 1.37247259436398 | 0.177116611112519 | 0.495123075720355 | -5.05036926214387 |
| ENSG00000144218.18 | AFF3 | 0.196103967406351 | 8.3886189392386 | 1.37213391360255 | 0.177221234091593 | 0.495331607573193 | -5.46796778844839 |
| ENSG00000104980.7 | TIMM44 | 0.144671051016065 | 4.07821311500262 | 1.37207532477785 | 0.177239337792186 | 0.495331607573193 | -5.39284110095077 |
| lnc-APOL4-2 | lnc-APOL4-2 | 0.297434494372239 | 0.318956020877897 | 1.37193920747006 | 0.177281402974958 | 0.49535529094516 | -4.92939055711969 |
| lnc-MED20-3 | lnc-MED20-3 | -0.581812689505916 | -0.912119963647903 | -1.37185975173772 | 0.177305961235931 | 0.49535529094516 | -4.6408200095297 |
| ENSG00000207195.1 | | 0.630207738939735 | 0.0667028728639586 | 1.37174035406313 | 0.177344070817731 | 0.49535529094516 | -4.85052280290243 |
| ENSG00000159377.10 | PSMB4 | 0.131624184241698 | 6.65151622717709 | 1.37173644857736 | 0.177344077101243 | 0.49535529094516 | -5.49295788302262 |
| ENSG00000157870.15 | PRXL2B | -0.298103437338775 | 2.08777574769367 | -1.3715544725769 | 0.177400341647046 | 0.495445214670441 | -5.18431414765881 |
| LINC00926 | LINC00926 | -0.380549678190084 | 8.99294112402953 | -1.3715301070221 | 0.177486802314227 | 0.495525513768293 | -5.44864636888241 |
| ENSG00000163964.14 | PIGX | 0.185296861059059 | 2.56919680475329 | 1.37113970956337 | 0.177528632270624 | 0.495525513768293 | -5.22582272679266 |
| ENSG00000184047.17 | DIABLO | 0.173315824106589 | 4.7714358955292 | 1.37112647163358 | 0.177532728081355 | 0.495525513768293 | -5.4536205879107 |
| ENSG00000109920.12 | FNBP4 | 0.142894949981471 | 7.27436229904799 | 1.37109033034069 | 0.177543910555457 | 0.495525513768293 | -5.48828151488176 |
| ENSG00000279913.1 | | 0.456751544840747 | 0.353370071736833 | 1.3709009766084 | 0.177602507354213 | 0.495525513768293 | -4.88844898623044 |
| lnc-PPP1R12A-3 | lnc-PPP1R12A-3 | -0.284897241504616 | 0.802624267914231 | -1.37089441291869 | 0.177604538799972 | 0.495525513768293 | -5.03608516900646 |
| ENSG00000161533.11 | ACOX1 | -0.125376346253816 | 4.91836383540802 | -1.37083207687164 | 0.17762383254571 | 0.495525513768293 | -5.46197789240446 |
| ENSG00000123360.11 | PDE1B | 0.231599849707186 | 3.80960307869366 | 1.37083058350926 | 0.177624294778896 | 0.495525513768293 | -5.36279842136994 |
| ENSG00000122390.18 | NAA60 | 0.140066815316008 | 3.83166305202971 | 1.37053121579313 | 0.177716975341435 | 0.495525513768293 | -5.37215230447079 |
| ENSG00000235859.5 | | 0.192117928203429 | 4.64022328972897 | 1.37044069027813 | 0.177745008268182 | 0.495525513768293 | -5.44282117052972 |
| ENSG00000174125.7 | TLR1 | 0.255159853047269 | 4.52793720949959 | 1.37035861038248 | 0.177770428797093 | 0.495525513768293 | -5.43076013550317 |
| ENSG00000283726.1 | MIR5193 | 0.344916835136 | 0.644625466277799 | 1.37035720735093 | 0.177770863346914 | 0.495525513768293 | -4.96028491628965 |
| ENSG00000205302.6 | SNX2 | 0.213995695932383 | 6.86574544747001 | 1.37031576459784 | 0.177783699451098 | 0.495525513768293 | -5.49395460866717 |
| lnc-FOSB-1 | lnc-FOSB-1 | -1.27352320302719 | -2.1306562270509 | -1.37029786653483 | 0.177789243256481 | 0.495525513768293 | -4.49952632146849 |
| ENSG00000275888.1 | | -0.567186017487998 | -0.808969586691976 | -1.3701840898856 | 0.177824487929675 | 0.495525513768293 | -4.67712659136981 |
| ENSG00000272196.2 | NA | 1.18224293233236 | -0.364237671250219 | 1.37043836493904 | 0.177836373308642 | 0.495525513768293 | -4.62614369615265 |
| ENSG00000197355.10 | UAP1L1 | -0.246111812359904 | 4.13486053772166 | -1.37010716863551 | 0.177848318931459 | 0.495525513768293 | -5.4088788499643 |
| lnc-IRS1-6 | lnc-IRS1-6 | -0.288587030393725 | 2.67284008390174 | -1.37006160941694 | 0.177862434811259 | 0.495525513768293 | -5.24685688178045 |
| ENSG00000151092.16 | NGLY1 | 0.147903440147676 | 6.3198771997249 | 1.36992744368944 | 0.177904009182221 | 0.495574261875445 | -5.49649092862968 |
| ENSG00000137944.17 | KYAT3 | 0.139678901809156 | 4.19644300570247 | 1.36963138457014 | 0.177995776489985 | 0.49576279668381 | -5.41196136600974 |
| ENSG00000158528.11 | PPP1R9A | -2.875804697343 | 0.627413402864008 | -1.37029662665058 | 0.178074595439502 | 0.495915220935093 | -4.94021801989144 |
| ENSG00000164168.7 | TMEM184C | 0.136731993069445 | 4.36929733271889 | 1.36919052334681 | 0.178132494676312 | 0.496009353073934 | -5.42490050997755 |
| ENSG00000279133.1 | | 0.261828366301059 | 3.60475275566399 | 1.36887007907277 | 0.178231920407216 | 0.496219074495027 | -5.35214063322762 |
| ENSG00000135334.8 | AKIRIN2 | 0.254726912940485 | 4.58334575641237 | 1.36865422509612 | 0.178298918525612 | 0.496338469030656 | -5.44620051776966 |
| ENSG00000102710.19 | SUPT20H | -0.139998233077465 | 5.48356572569112 | -1.36845788750041 | 0.17835987588044 | 0.496441017612858 | -5.48846010472388 |
| lnc-HTR1D-1 | lnc-HTR1D-1 | -0.473903524122667 | 1.12624216329628 | -1.36824026586876 | 0.178427924982363 | 0.496563274731203 | -5.06754061054206 |
| lnc-TRIM17-1 | lnc-TRIM17-1 | 0.194494840026504 | 0.709457135990392 | 1.36800671914745 | 0.178500011937717 | 0.496696734583483 | -4.99061018275235 |
| ENSG00000258881.6 | | 0.175424125361104 | 1.63773398606657 | 1.36787203918091 | 0.17854186091848 | 0.496734290668022 | -5.12691924971615 |
| ENSG00000109118.13 | PHF12 | -0.167247269816635 | 5.60409749948336 | -1.36780795918833 | 0.178561775088797 | 0.496734290668022 | -5.49257716654409 |
| lnc-GRHL2-12 | lnc-GRHL2-12 | -0.377370800118565 | -0.369002353225011 | -1.36752206901686 | 0.178650642165327 | 0.49684833200228 | -4.77102295311507 |
| ENSG00000142634.12 | EFHD2 | 0.262863879752381 | 4.16373099989369 | 1.3675207661507 | 0.17865104723067 | 0.49684833200228 | -5.40683687709748 |
| ENSG00000107077.18 | KDM4C | 0.139476072474818 | 6.71829738364017 | 1.36736881957778 | 0.17869829276939 | 0.496883468022706 | -5.4986984653945 |
| ENSG00000196507.10 | TCEAL3 | -0.234264084503031 | 2.56279040268243 | -1.36731465571017 | 0.178715136549514 | 0.496883468022706 | -5.23821423636805 |
| lnc-ATG7-1 | lnc-ATG7-1 | -0.313625635201191 | 0.258937130239762 | -1.36717396149192 | 0.178758895093911 | 0.496883468022706 | -4.88609294330422 |
| ENSG00000062485.18 | CS | 0.170979055576343 | 6.1577708324002 | 1.36716962822177 | 0.178760242953318 | 0.496883468022706 | -5.4998294063055 |
| RAB30-AS1 | RAB30-AS1 | -0.285371693713954 | 5.35929163294323 | -1.36692678930136 | 0.178835790273358 | 0.497026339616057 | -5.48104459346 |
| lnc-WSB1-1 | lnc-WSB1-1 | 0.23446083431579 | 0.352922058220237 | 1.36678193074982 | 0.178880867556666 | 0.497056002998143 | -4.91757792680912 |

| | | | | | | | |
|---|---|---|---|---|---|---|---|
| ENSG00000124713.5 | GNMT | 0.283363346012745 | 0.833040127827538 | 1.36673728829569 | 0.178894761220922 | 0.497056002998143 | -5.02209908651368 |
| ENSG00000125245.12 | GPR18 | 0.334604704697819 | 4.43552229250827 | 1.36640112281468 | 0.179016477348282 | 0.497327055373152 | -5.41676672539077 |
| lnc-CDO1-1 | lnc-CDO1-1 | 0.121791310913635 | 4.08128603953977 | 1.36611901288448 | 0.179087266921832 | 0.49742969770282 | -5.40698886350488 |
| ENSG00000131469.14 | RPL27 | 0.202248857397569 | 7.86294500585418 | 1.36607249149914 | 0.179101758231337 | 0.49742969770282 | -5.48529553994034 |
| SNHG5 | SNHG5 | -0.163021512878856 | 5.9528572551031 | -1.36577025236056 | 0.179195927051523 | 0.497624091540947 | -5.50014590751095 |
| lnc-DCXR-2 | lnc-DCXR-2 | -0.247100852181741 | 1.28978145910526 | -1.36562643519301 | 0.179240749633284 | 0.497647122316262 | -5.09521379172116 |
| lnc-ZFAND4-1 | lnc-ZFAND4-1 | 0.156377552360609 | 2.73108826065901 | 1.36554536607815 | 0.179266019733061 | 0.497647122316262 | -5.2521918161975 |
| ENSG00000264668.1 | | -0.177874128319561 | 1.98137965886083 | -1.36551093452561 | 0.179276753241757 | 0.497647122316262 | -5.16607962009046 |
| ENSG00000174021.10 | GNG5 | -0.168082885205408 | 2.39369756237116 | -1.3653445103618 | 0.179328640400002 | 0.497706989554501 | -5.22706941266655 |
| ENSG00000279948.1 | NA | -0.301924941855347 | 0.380338495580868 | -1.36528665416524 | 0.179346681315928 | 0.497706989554501 | -4.92124306529172 |
| ENSG00000104450.12 | SPAG1 | 0.451006331835113 | 2.58550532001192 | 1.36494894868645 | 0.179519731034396 | 0.498120062534508 | -5.2740779759226 |
| ENSG00000086619.13 | ERO1B | 0.216740954849002 | 4.61641723705092 | 1.36453655742375 | 0.179580705548971 | 0.49822208684654 | -5.44940986460099 |
| ENSG00000002549.12 | LAP3 | 0.182302273873228 | 4.47386828727481 | 1.36436130725531 | 0.179635416072537 | 0.498306464313918 | -5.4423809553374 |
| ENSG00000100583.4 | SAMD15 | -0.397836919891328 | 0.0175827062722505 | -1.36428405216312 | 0.179659538049535 | 0.498306464313918 | -4.81975270088963 |
| lnc-PMM2-8 | lnc-PMM2-8 | -0.25028562694394 | 0.437857296430914 | -1.36418199521508 | 0.179691407931173 | 0.498327708090453 | -4.94427854855047 |
| ENSG00000279766.1 | | 0.444203518856619 | -0.281830096112029 | 1.36384059964927 | 0.179798049017333 | 0.498556276831978 | -4.75131977768295 |
| ENSG00000128228.4 | SDF2L1 | 0.411622312167529 | 0.532175153796091 | 1.36366518935127 | 0.179852860539083 | 0.498641086809797 | -4.96874599063589 |
| ENSG00000175334.7 | BANF1 | -0.326474443280992 | 3.2694748637621 | -1.36335773184378 | 0.179948964677052 | 0.498840342080507 | -5.32036993100691 |
| ENSG00000080546.13 | SESN1 | -0.57953660661026 | 5.76430964752906 | -1.36380280243827 | 0.180043547641358 | 0.499035327377063 | -5.49804290134835 |
| ENSG00000197857.13 | ZNF44 | -0.212669272002753 | 4.97526289174029 | -1.36241814280289 | 0.180242903494363 | 0.499503234240898 | -5.47406208868044 |
| ENSG00000140299.11 | BNIP2 | 0.131990972599732 | 7.04258521287954 | 1.36236066698225 | 0.180260896076441 | 0.499503234240898 | -5.50232288820438 |
| ENSG00000183309.11 | ZNF623 | -0.217815979119134 | 4.81068438312078 | -1.36181586551873 | 0.180431512683992 | 0.499908712555472 | -5.46671943456266 |
| lnc-SUMO4-3 | lnc-SUMO4-3 | 0.302634935972791 | 0.766641561736701 | 1.36156976955241 | 0.180508623786724 | 0.500045521793246 | -5.00174126283575 |
| ENSG00000112796.9 | ENPP5 | -1.25070173654561 | -1.37574806685181 | -1.36152538537819 | 0.180546552101738 | 0.500045521793246 | -4.54820818483574 |
| lnc-AP3S1-4 | lnc-AP3S1-4 | -0.308353844051539 | 0.330741371620046 | -1.36140437853387 | 0.180560461240349 | 0.500045521793246 | -4.94093800530241 |
| lnc-MCM7-2 | lnc-MCM7-2 | 0.164079916149944 | 1.36096461651363 | 1.36134821281913 | 0.180578067506641 | 0.500045521793246 | -5.09640462593652 |
| ENSG00000010671.15 | BTK | -0.232214132128286 | 7.11922650619037 | -1.36118950025416 | 0.180627826270725 | 0.500116027459018 | -5.50325426951652 |
| ENSG00000168818.9 | STX18 | 0.167194272097513 | 4.15287450996372 | 1.36087613032079 | 0.180726103397637 | 0.500295711289929 | -5.41990522427332 |
| lnc-HJURP-8 | lnc-HJURP-8 | -0.200267299405771 | 0.62409841385691 | -1.36082757234317 | 0.180741335526984 | 0.500295711289929 | -4.98438707039499 |
| ENSG00000132879.13 | FBXO44 | -0.216332934110354 | 4.32347549890704 | -1.36049336719042 | 0.180846198966968 | 0.500406318056171 | -5.42975849240773 |
| lnc-NR1D1-4 | lnc-NR1D1-4 | -0.143514737328014 | 0.878884958945253 | -1.36049264529358 | 0.180846425526926 | 0.500406318056171 | -5.02004405376515 |
| ENSG00000132341.11 | RAN | 0.185274074050202 | 6.96139955845258 | 1.36046778050827 | 0.180854229219252 | 0.500406318056171 | -5.50485468768485 |
| ENSG00000161914.9 | ZNF653 | -0.204496976151342 | 1.2321507015015 | -1.36004124496483 | 0.180988135652037 | 0.500612571285006 | -5.10193414694025 |
| ENSG00000243317.7 | STMP1 | -0.152144560666055 | 3.56176045195957 | -1.36001787849567 | 0.180995473515653 | 0.500612571285006 | -5.36015873372006 |
| lnc-ABCA10-4 | lnc-ABCA10-4 | -0.329795652953003 | 1.29346966912001 | -1.35999793338514 | 0.18100173713759 | 0.500612571285006 | -5.09689714052263 |
| lnc-AADACL4-2 | lnc-AADACL4-2 | 0.480850439744319 | -0.629828701713498 | 1.35989154368144 | 0.181035150892483 | 0.500637714741365 | -4.66833549274493 |
| ENSG00000105640.12 | RPL18A | -0.249463177310743 | 8.70826442298691 | -1.35980296626057 | 0.181062973971205 | 0.500647392967398 | -5.47094927492331 |
| ENSG00000028310.17 | BRD9 | 0.131273764495429 | 5.30399220179918 | 1.35964993878614 | 0.18111104921981 | 0.500713059246655 | -5.49259869242217 |
| ENSG00000085377.13 | PREP | -0.259881172397395 | 3.35253606316703 | -1.35946431120667 | 0.181169379317246 | 0.500771193699319 | -5.33157512322342 |
| ENSG00000201512.1 | SNORA71C | 0.272963177627533 | -0.192050961972452 | 1.3592529623152 | 0.181235809450248 | 0.500771193699319 | -4.81905897703228 |
| lnc-SALL3-7 | lnc-SALL3-7 | 0.433655119908705 | -0.172228620539222 | 1.35920857907164 | 0.181249762149254 | 0.500771193699319 | -4.77091086504128 |
| ENSG00000160606.10 | TLCD1 | -0.456687128140585 | 0.746559770241155 | -1.35920751736847 | 0.181250095925526 | 0.500771193699319 | -4.9957209907556 |
| ENSG00000131018.22 | SYNE1 | -0.170037789602159 | 7.28748003972737 | -1.3591959782273 | 0.181253723609772 | 0.500771193699319 | -5.50290778634643 |
| lnc-SH3YL1-1 | lnc-SH3YL1-1 | 0.279911862721815 | 3.32339400375157 | 1.35904016674812 | 0.181302713209537 | 0.500839316332291 | -5.31285516689793 |
| ENSG00000125703.14 | ATG4C | -0.235218890127838 | 4.22496828004631 | -1.35852001499962 | 0.181466330866855 | 0.500980097033572 | -5.41548217346922 |
| ENSG00000284029.1 | MIR6805 | -0.334695124440093 | -0.275744027781457 | -1.35833904809775 | 0.18152328197147 | 0.500980097033572 | -4.83094838679318 |
| ENSG00000175544.13 | CABP4 | -0.21132037971434 | 2.57163195112148 | -1.35831646441039 | 0.181530390124182 | 0.500980097033572 | -5.25926390620655 |
| ENSG00000103160.11 | HSDL1 | -0.179822898880183 | 3.70362200997623 | -1.35815309281678 | 0.181581817248989 | 0.500980097033572 | -5.38207143454718 |
| lnc-HIBADH-4 | lnc-HIBADH-4 | -0.32941585672364 | 1.48237041792724 | -1.35806845616008 | 0.181608464103368 | 0.500980097033572 | -5.12979507381595 |
| ENSG00000133740.10 | E2F5 | 0.159296761897928 | 5.73415877682401 | 1.35806377583027 | 0.181609937737747 | 0.500980097033572 | -5.50679316982111 |
| ENSG00000117395.12 | EBNA1BP2 | -0.194032920319065 | 3.81757736078225 | -1.35803569771742 | 0.181618778519692 | 0.500980097033572 | -5.38663079331419 |
| ENSG00000122335.15 | SERAC1 | 0.264234913687626 | 0.96890587032936 | 1.35803272718458 | 0.181619713852362 | 0.500980097033572 | -5.04172367264505 |
| ENSG00000188647.12 | PTAR1 | 0.155793542360963 | 7.11746858600983 | 1.3579929548054 | 0.181632237351243 | 0.500980097033572 | -5.50628824427059 |
| lnc-MRFAP1-2 | lnc-MRFAP1-2 | -0.248626321932882 | 1.78406326978448 | -1.35797084423687 | 0.181639199798543 | 0.500980097033572 | -5.17080679932232 |
| lnc-IRF1-7 | lnc-IRF1-7 | -0.343699464219579 | 1.32214639277397 | -1.35795586391675 | 0.181643917102212 | 0.500980097033572 | -5.098819068751 |
| ENSG00000180884.9 | ZNF792 | -0.221321086707663 | 3.06033959333356 | -1.3579426357939 | 0.181648082717128 | 0.500980097033572 | -5.30606947076392 |
| ENSG00000155893.12 | PXYLP1 | 0.401244519515659 | 0.799233646492266 | 1.3578688386772 | 0.181671323217098 | 0.500980097033572 | -5.01965999296452 |
| lnc-RNF149-3 | lnc-RNF149-3 | 0.220658889293632 | 1.38627267028876 | 1.35777215328219 | 0.181701775247872 | 0.500980097033572 | -5.11313202807193 |
| lnc-AGO3-2 | lnc-AGO3-2 | 0.170078835952607 | 2.27855612664184 | 1.35771820617838 | 0.181718768131597 | 0.500980097033572 | -5.20935619200033 |
| ENSG00000170379.20 | TCAF2 | -0.363031852623002 | 2.5248775326788 | -1.35749753584008 | 0.181788290143997 | 0.501064190512585 | -5.26414138685189 |
| ENSG00000090470.14 | PDCD7 | -0.133272449385957 | 3.84411815623694 | -1.35746685335472 | 0.181797958254287 | 0.501064190512585 | -5.38893633514606 |
| ENSG00000283236.1 | | -0.31696080577216 | 2.38670019964321 | -1.3572896843923 | 0.181853792262369 | 0.501116306870689 | -5.2253845471356 |
| lnc-ATOX1-4 | lnc-ATOX1-4 | -0.244872147882634 | 3.36893273102432 | -1.35711792948855 | 0.181907932629238 | 0.501116306870689 | -5.34287272722292 |
| ENSG00000112874.9 | NUDT12 | -1.88310808492681 | 0.082099519569501 | -1.35794763103644 | 0.181918970311131 | 0.501116306870689 | -4.74027310252479 |
| RUSC1-AS1 | RUSC1-AS1 | -0.367904415174687 | 0.0748685375919008 | -1.35702255138863 | 0.181938002941924 | 0.501116306870689 | -4.86642334236933 |
| lnc-SPAG9-2 | lnc-SPAG9-2 | 0.220593659826737 | 2.26823904283256 | 1.35702066429829 | 0.1819385979325 | 0.501116306870689 | -5.21017607478108 |
| ENSG00000167840.13 | ZNF232 | -0.293371103705069 | 1.87365273004515 | -1.35691925421975 | 0.18197057424471 | 0.501137319999848 | -5.17655620384993 |
| ENSG00000077063.10 | CTTNBP2 | -3.28940577729237 | -1.09292346376573 | -1.35758682474789 | 0.182043918686499 | 0.501272237902904 | -4.55231910866071 |

| | | | | | | | |
|---|---|---|---|---|---|---|---|
| ENSG00000221995.5 | NA | -0.163219388634796 | 3.34930105769148 | -1.35660505486975 | 0.18206967402061 | 0.501276096892218 | -5.34064432127434 |
| ENSG00000260342.2 | | 0.198065890096851 | 7.60039099764198 | 1.35589512931915 | 0.18229374012085 | 0.501678433748377 | -5.50366431907815 |
| ENSG00000205531.12 | NAP1L4 | 0.133675564675148 | 6.07483766048576 | 1.35580761161601 | 0.18232137703991 | 0.501678433748377 | -5.51396190643976 |
| ENSG00000106608.16 | URGCP | -0.109248720898831 | 4.3352725182818 | -1.35576881961118 | 0.182333628064864 | 0.501678433748377 | -5.44022695628985 |
| ENSG00000168172.8 | HOOK3 | 0.133062619304865 | 6.16890357863764 | 1.35570922841781 | 0.182352448978359 | 0.501678433748377 | -5.51444165807136 |
| ENSG00000123349.13 | PFDN5 | 0.179389587795479 | 7.73752988109031 | 1.35566484427526 | 0.182366467960448 | 0.501678433748377 | -5.50166870821661 |
| lnc-KLRG2-3 | lnc-KLRG2-3 | -0.392687949269085 | -0.138818788012598 | -1.35558882899883 | 0.182390479737312 | 0.501678433748377 | -4.85278657440877 |
| lnc-ZCCHC10-4 | lnc-ZCCHC10-4 | -0.287096722141309 | 1.42983181264651 | -1.35557371910913 | 0.182395252952691 | 0.501678433748377 | -5.12408216822348 |
| lnc-ZNF487-1 | lnc-ZNF487-1 | -0.248239004770725 | 0.189850514881666 | -1.35552452124126 | 0.182410795227754 | 0.501678433748377 | -4.90249103091141 |
| ENSG00000136159.3 | NUDT15 | 0.311862289886934 | 1.95821746547228 | 1.35541464650211 | 0.182445509822843 | 0.501706870899611 | -5.18497137150717 |
| ENSG00000159433.11 | STARD9 | -0.448510974648404 | 6.32265489406547 | -1.35581367268539 | 0.18248956356588 | 0.501760978743855 | -5.51358316228831 |
| ENSG00000160856.20 | FCRL3 | -1.09458103901069 | 6.28480114410146 | -1.3559717516222 | 0.182553149420505 | 0.501809831486368 | -5.50969803630247 |
| lnc-DNASE1L1-1 | lnc-DNASE1L1-1 | -0.267712312462483 | 1.03178201283811 | -1.35504860976978 | 0.182561194633535 | 0.501809831486368 | -5.06983221851033 |
| lnc-ZNF570-2 | lnc-ZNF570-2 | 0.27613598951474 | -0.380558297055011 | 1.35496157173885 | 0.182588711028386 | 0.501809831486368 | -4.75000688841761 |
| lnc-APOA5-1 | lnc-APOA5-1 | 0.218837109229252 | 2.16641573568124 | 1.35487177906644 | 0.182617101618519 | 0.501809831486368 | -5.20863200643179 |
| lnc-TENM1-9 | lnc-TENM1-9 | 0.210706737870198 | 1.43827547411371 | 1.35483342214251 | 0.182629230319408 | 0.501809831486368 | -5.12126229650457 |
| ENSG00000007516.13 | BAIAP3 | -0.604271998047883 | 5.65499544591284 | -1.35550906981964 | 0.182662095889998 | 0.501833144648135 | -5.50459638088958 |
| ZNF667-AS1 | ZNF667-AS1 | -1.7041247991854 | 0.229765666060361 | -1.35541157885951 | 0.182730026840339 | 0.501895850983383 | -4.9598271480205 |
| ENSG00000115839.17 | RAB3GAP1 | 0.0993848180148067 | 6.38359461448647 | 1.35445779860963 | 0.182748037530188 | 0.501895850983383 | -5.51657485460582 |
| ENSG00000279765.3 | NA | 0.178539506844442 | 7.08527149899793 | 1.35442608089474 | 0.182758072347105 | 0.501895850983383 | -5.5125105316265 |
| ENSG00000266338.6 | NBPF15 | -0.159001283157242 | 4.84800849916765 | -1.35402512774383 | 0.182884961952077 | 0.502121096665973 | -5.47501724863919 |
| lnc-MAVS-3 | lnc-MAVS-3 | -0.194604888529628 | 2.35191826799508 | -1.35401274381229 | 0.182888882169985 | 0.502121096665973 | -5.2408205890487 |
| ENSG00000232486.1 | DEPDC1P2 | 0.465337697993262 | 0.973966220085428 | 1.3538760554302 | 0.182932156093649 | 0.502172921962599 | -5.02268503369343 |
| lnc-SERTAD2-8 | lnc-SERTAD2-8 | -0.361050174284094 | 0.0833210969142889 | -1.35368821064454 | 0.182991638349699 | 0.502224174886983 | -4.90072768081474 |
| ENSG00000103404.14 | USP31 | -0.147135662304108 | 3.1762526163179 | -1.35357198458095 | 0.183028449511203 | 0.502224174886983 | -5.32547113774091 |
| TPT1-AS1 | TPT1-AS1 | -0.247177960724059 | 4.83579611427073 | -1.35356148899658 | 0.183031773956292 | 0.502224174886983 | -5.48163399690431 |
| ENSG00000285283.1 | | -0.398899405811716 | -0.328896545382031 | -1.35350891796494 | 0.183048426371381 | 0.502224174886983 | -4.80584563271327 |
| lnc-GABARAP-2 | lnc-GABARAP-2 | -0.287293374388021 | 0.0218476579304651 | -1.35336595102821 | 0.183093718499989 | 0.502281488455986 | -4.89569632487903 |
| ENSG00000182899.16 | RPL35A | 0.190601576803235 | 8.17805587098333 | 1.35324139916801 | 0.183133183715777 | 0.502309902054532 | -5.49563207302182 |
| ENSG00000260371.1 | | -0.185121962039838 | 2.0168419184589 | -1.35317923241448 | 0.183152884172339 | 0.502309902054532 | -5.19375084401134 |
| ENSG00000108963.17 | DPH1 | 0.111257036180058 | 5.09441038755684 | 1.35306135437419 | 0.183190243832933 | 0.502315122475364 | -5.4948580254325 |
| ENSG00000178307.9 | TMEM11 | 0.170105317717786 | 2.30945534893648 | 1.35300969615293 | 0.183206617966999 | 0.502315122475364 | -5.23461961365806 |
| lnc-PARL-1 | lnc-PARL-1 | 0.149904515818243 | 1.8702289938636 | 1.35294224162593 | 0.183228000755237 | 0.502315122475364 | -5.17258993908538 |
| ENSG00000172493.20 | AFF1 | 0.431877698860915 | 5.66806252661222 | 1.35320710497232 | 0.183302636553706 | 0.502417855497066 | -5.51115537455446 |
| lnc-USP53-4 | lnc-USP53-4 | 0.254848114237287 | 0.666986243767475 | 1.35263321753371 | 0.18332598345646 | 0.502417855497066 | -5.00939237769436 |
| ENSG00000173327.7 | MAP3K11 | -0.217070423375485 | 4.40000951206776 | -1.35259311661069 | 0.18333870245535 | 0.502417855497066 | -5.45413728906093 |
| lnc-GLRX3-2 | lnc-GLRX3-2 | -0.350755599240882 | 0.486530371798899 | -1.35175126673371 | 0.183605850016597 | 0.502885626616578 | -4.95807998926231 |
| ENSG00000066583.11 | ISOC1 | -0.197371874426954 | 2.55659139011608 | -1.35172677706241 | 0.183613625888383 | 0.502885626616578 | -5.26868598775636 |
| ENSG00000183978.7 | COA3 | -0.193980238430165 | 2.58038543773937 | -1.35167500379667 | 0.183630065579831 | 0.502885626616578 | -5.26481825115777 |
| ENSG00000168066.20 | SF1 | 0.211491164787997 | 8.19539198882696 | 1.35166599914117 | 0.183632924965331 | 0.502885626616578 | -5.49807547163597 |
| ENSG00000196378.11 | ZNF34 | -0.171068568460333 | 3.62302793322799 | -1.35166319625194 | 0.183633815016439 | 0.502885626616578 | -5.37059312420339 |
| lnc-VNN2-3 | lnc-VNN2-3 | 0.227016842004164 | 2.04002442414865 | 1.35159336564006 | 0.183655990636779 | 0.502885626616578 | -5.18725053716904 |
| lnc-LRRC38-8 | lnc-LRRC38-8 | -0.246395511093898 | 3.19263060649877 | -1.35143208405368 | 0.183707215555852 | 0.50290892110325 | -5.31981417206875 |
| ENSG00000207051.1 | SNORA27 | 0.22704279408768 | 1.15196750853113 | 1.3512981701022 | 0.183749756520428 | 0.50290892110325 | -5.08498096658153 |
| ENSG00000280077.1 | NA | -0.217842718449682 | 3.47143562468857 | -1.35121784162679 | 0.18377527840418 | 0.50290892110325 | -5.3718787445269 |
| ENSG00000237214.3 | RPS9P2 | 0.819745582424707 | -0.250130255435704 | 1.35113887753444 | 0.183802548018793 | 0.50290892110325 | -4.70604013332772 |
| lnc-LARP1B-1 | lnc-LARP1B-1 | -0.217797403257513 | 3.96811975985065 | -1.35103920964695 | 0.183832042925451 | 0.50290892110325 | -5.40925208237459 |
| ENSG00000279759.1 | | -0.266442845098945 | 1.58332118465934 | -1.35097674230676 | 0.183851896568007 | 0.50290892110325 | -5.13638000422379 |
| lnc-PAGR1-2 | lnc-PAGR1-2 | 0.535375281581107 | 0.759136207960235 | 1.35071306393827 | 0.183935935141507 | 0.50290892110325 | -4.98123328372024 |
| SREBF2-AS1 | SREBF2-AS1 | -0.373912424617012 | 2.56638761272877 | -1.3506098352415 | 0.183968541767791 | 0.50290892110325 | -5.26138506938777 |
| ENSG00000227081.5 | RPS27P3 | 0.321201113049195 | 5.0812157213913 | 1.35066189086035 | 0.183974324699207 | 0.50290892110325 | -5.48356956002327 |
| lnc-PIEZO2-14 | lnc-PIEZO2-14 | 0.207260681798208 | 0.797977312924685 | 1.3505584056677 | 0.183984896526279 | 0.50290892110325 | -5.02009954523685 |
| ENSG00000091140.13 | DLD | 0.168463330435518 | 4.96767084974235 | 1.35052900774078 | 0.183994245655866 | 0.50290892110325 | -5.49309190017681 |
| ENSG00000105639.18 | JAK3 | 0.233083669755269 | 5.41898065906185 | 1.35046714518881 | 0.184013920376348 | 0.50290892110325 | -5.50954511948214 |
| lnc-TTC36-2 | lnc-TTC36-2 | -0.164986843605984 | 3.17345270499295 | -1.35045282396324 | 0.184018475318339 | 0.50290892110325 | -5.33334591300818 |
| ENSG00000139190.16 | VAMP1 | -0.301458090458849 | 5.61313829255569 | -1.35039376315293 | 0.184051235800315 | 0.50290892110325 | -5.51484449104847 |
| ENSG00000011009.10 | LYPLA2 | -0.168830096734767 | 3.93959821590467 | -1.35033119174289 | 0.18405716457863 | 0.50290892110325 | -5.41652697026201 |
| lnc-VKORC1L1-5 | lnc-VKORC1L1-5 | -0.334513535952296 | 0.075008517061327 | -1.35023055198007 | 0.184089181189711 | 0.50290892110325 | -4.85794960733553 |
| lnc-LCAT-2 | lnc-LCAT-2 | -0.314023883230302 | -0.00607944080805511 | -1.35021281092319 | 0.18409482560954 | 0.50290892110325 | -4.87835186317963 |
| ENSG00000125454.11 | SLC25A19 | -0.292496103607147 | 3.56138488899958 | -1.3501830459593 | 0.184104295802992 | 0.50290892110325 | -5.35996831745671 |
| ENSG00000137161.16 | CNPY3 | -0.141489746188686 | 6.72319550163133 | -1.34979554585986 | 0.184227619168909 | 0.503154903060889 | -5.52068121071922 |
| lnc-UBP1-4 | lnc-UBP1-4 | 0.2200104171943 | 0.561533890864436 | 1.34974649306527 | 0.184243234920562 | 0.503154903060889 | -4.981620647129 |
| lnc-SHC4-2 | lnc-SHC4-2 | 0.220283387897234 | 3.52295881479127 | 1.34959567255636 | 0.184291254354413 | 0.50320404936489 | -5.35906077805938 |
| ENSG00000201217.1 | | -0.342709205845009 | 0.581594547975122 | -1.3495037478147 | 0.184320526790926 | 0.50320404936489 | -5.00231308582007 |
| lnc-TMEM167A-5 | lnc-TMEM167A-5 | -0.248912805885596 | 1.75279286161025 | -1.34938509890219 | 0.184358314518873 | 0.50320404936489 | -5.16919217694549 |
| ENSG00000165688.11 | PMPCA | 0.105336536678241 | 5.16261212089766 | 1.34938287974369 | 0.184359021340943 | 0.50320404936489 | -5.50077798272808 |
| ENSG00000197157.10 | SND1 | -0.13333773495626 | 6.84058684152708 | -1.34910197684286 | 0.184448508232519 | 0.503253213483422 | -5.5212112627886 |
| ENSG00000187626.8 | ZKSCAN4 | -0.263876897460476 | 2.04456559321272 | -1.34904690129577 | 0.184466057485151 | 0.503253213483422 | -5.21265108867038 |

| | | | | | | | |
|---|---|---|---|---|---|---|---|
| ENSG00000119669.4 | IRF2BPL | -0.327123083591324 | 3.79986663059972 | -1.34904837591221 | 0.184471659905664 | 0.503253213483422 | -5.41578672575187 |
| ENSG00000135679.23 | MDM2 | 0.208074780318499 | 6.80095939833707 | 1.34898415842482 | 0.184486051410596 | 0.503253213483422 | -5.52239665530169 |
| ENSG00000235453.10 | SMIM27 | -0.191739205041719 | 1.81222364853769 | -1.34894263822346 | 0.184499283337993 | 0.503253213483422 | -5.17109900123967 |
| ST3GAL5-AS1 | ST3GAL5-AS1 | -0.270028956603825 | 3.83436769899523 | -1.34871396601049 | 0.184572171119527 | 0.503385318424968 | -5.40245041042153 |
| ENSG00000164244.20 | PRRC1 | -0.158811972447437 | 5.23218869837644 | -1.34827562647422 | 0.184711950693893 | 0.503699798772178 | -5.50565194815963 |
| ENSG00000197037.10 | ZSCAN25 | 0.145035071518084 | 5.28540555352549 | 1.3480189819262 | 0.18479382824324 | 0.50385632093398 | -5.5068020868235 |
| ENSG00000146067.15 | FAM193B | -0.173982380505715 | 5.67084583771183 | -1.34773411702209 | 0.184884741523067 | 0.504037435068781 | -5.51903059617707 |
| ENSG00000160685.13 | ZBTB7B | -0.278455887854672 | 4.60096065265756 | -1.34752021377199 | 0.18495303025411 | 0.504111591250403 | -5.47362053658108 |
| ENSG00000153989.7 | NUS1 | 0.233605742758609 | 3.69578715513274 | 1.34747390355624 | 0.184967817360734 | 0.504111591250403 | -5.39190645910003 |
| ENSG00000110871.14 | COQ5 | 0.164750909666969 | 3.32744997874874 | 1.34741878714311 | 0.184985417515148 | 0.504111591250403 | -5.342386584999 |
| lnc-RNMT-2 | lnc-RNMT-2 | -0.459282503618204 | -0.432261045790855 | -1.347295310421 | 0.185024851613512 | 0.504152306163742 | -4.7801285840385 |
| lnc-TBCCD1-4 | lnc-TBCCD1-4 | -0.297162264129418 | -0.187650086219699 | -1.3468339897204 | 0.185172238112109 | 0.504487118075652 | -4.81314129618751 |
| lnc-IGFLR1-1 | lnc-IGFLR1-1 | -0.275921618247602 | 1.75154633900122 | -1.3467121131741 | 0.185211191239741 | 0.504526462319691 | -5.1747581338557 |
| lnc-OST4-6 | lnc-OST4-6 | -0.250089933120923 | -0.110996809100719 | -1.346425963486 | 0.185302672597101 | 0.504708867284124 | -4.8356579683684 |
| ENSG00000234648.1 | | 0.851624050763445 | -0.531686340333913 | 1.34595419121986 | 0.185453572602601 | 0.505053041132486 | -4.65347061213178 |
| lnc-PPIAL4D-19 | lnc-PPIAL4D-19 | -0.236485870471924 | -0.125766533088314 | -1.34575282885257 | 0.185518008561706 | 0.505142294218558 | -4.85387942751857 |
| ENSG00000010270.13 | STARD3NL | 0.181484989453601 | 3.35262538943012 | 1.34563652695016 | 0.185555232981678 | 0.505142294218558 | -5.35225394653383 |
| ENSG00000167074.14 | TEF | 0.26585120418177 | 2.06398803818198 | 1.34562172450148 | 0.185559971169728 | 0.505142294218558 | -5.19523491534672 |
| ENSG00000164659.14 | ELAPOR2 | -1.98509651374996 | -0.429768928621711 | -1.34582383046769 | 0.185777867288247 | 0.505668585346997 | -4.91544267687649 |
| lnc-RWDD2B-5 | lnc-RWDD2B-5 | 0.221552574950909 | 1.69100503529194 | 1.3445956012577 | 0.185888653842928 | 0.505869678475011 | -5.16298054138738 |
| ENSG00000243147.7 | MRPL33 | 0.128704741260269 | 3.94340603158749 | 1.34450956874761 | 0.185916231585613 | 0.505869678475011 | -5.41293439321188 |
| ENSG00000110107.8 | PRPF19 | -0.142083783006293 | 4.40436845148972 | -1.3444807248857 | 0.185925478193823 | 0.505869678475011 | -5.46323933086381 |
| lnc-LCT-1 | lnc-LCT-1 | 0.363915567155472 | -0.29498506783033 | 1.34421095100678 | 0.186011977880653 | 0.50597364889433 | -4.78610049912375 |
| lnc-VLDLR-4 | lnc-VLDLR-4 | -0.144939091190587 | 3.76481592797737 | -1.34414821457769 | 0.186032097961699 | 0.50597364889433 | -5.4042953008801 |
| ENSG00000183340.6 | JRKL | 0.237172291873228 | 2.50236643746781 | 1.34407662169517 | 0.186055060414302 | 0.50597364889433 | -5.25307726851724 |
| lnc-UBQLN2-4 | lnc-UBQLN2-4 | 0.288228066767963 | 0.287048295190012 | 1.34395230436947 | 0.186094938676171 | 0.50597364889433 | -4.9398507491028 |
| ENSG00000164331.9 | ANKRA2 | 0.112524670574502 | 5.09310066661984 | 1.34391555219204 | 0.186106729218649 | 0.50597364889433 | -5.50311909035999 |
| lnc-MOGAT2-5 | lnc-MOGAT2-5 | -0.419896921487936 | -0.448230996415986 | -1.34389276980056 | 0.186114038372117 | 0.50597364889433 | -4.78352692047874 |
| ENSG00000115073.7 | ACTR1B | -0.159908385113395 | 5.23852813552413 | -1.34376394379955 | 0.186155373066753 | 0.50597364889433 | -5.51360151309154 |
| ENSG00000130312.6 | MRPL34 | -0.353218127407371 | 1.8353207187805 | -1.34369443832243 | 0.186177677287608 | 0.50597364889433 | -5.19491147958959 |
| ENSG00000115977.18 | AAK1 | 0.375269389392019 | 1.44621971782564 | 1.34367183806331 | 0.186184930123192 | 0.50597364889433 | -5.12170166270251 |
| lnc-NSMCE1-1 | lnc-NSMCE1-1 | -0.451325591014534 | -0.19884859862507 | -1.34350304353758 | 0.186239106205922 | 0.506054062447062 | -4.82377640340278 |
| ENSG00000180938.5 | ZNF572 | -0.258323045955704 | 2.07959459465477 | -1.3433463065678 | 0.186289423112414 | 0.506123969894776 | -5.2137934933716 |
| ENSG00000147669.10 | POLR2K | 0.225286026810508 | 4.64256579943665 | 1.34303744576817 | 0.186388606428806 | 0.506318573358545 | -5.48052017508073 |
| lnc-TRIM7-3 | lnc-TRIM7-3 | -0.487184691102981 | -0.592635381936558 | -1.34277810250973 | 0.186471919568472 | 0.506318573358545 | -4.72584010420097 |
| ENSG00000004660.14 | CAMKK1 | -0.237170313798805 | 2.13068886637979 | -1.34274988663594 | 0.186480985540426 | 0.506318573358545 | -5.23119015774854 |
| lnc-VSX1-4 | lnc-VSX1-4 | -0.309368324945092 | 0.0182198090501578 | -1.34272404615575 | 0.186489288576462 | 0.506318573358545 | -4.91288005998567 |
| ENSG00000163320.10 | CGGBP1 | -0.133804765488691 | 7.16629965974524 | -1.34267575465455 | 0.18650480630935 | 0.506318573358545 | -5.52514858247622 |
| ENSG00000109133.12 | TMEM33 | 0.191826975518742 | 5.23778392390176 | 1.34266381158984 | 0.18650864418231 | 0.506318573358545 | -5.51478729635505 |
| lnc-ZFP36L2-3 | lnc-ZFP36L2-3 | 0.205610948354433 | 2.06677417402331 | 1.34258097822296 | 0.186535264133536 | 0.506324059298505 | -5.20059706989102 |
| lnc-RFT1-1 | lnc-RFT1-1 | -0.225580651642435 | 0.58779207211779 | -1.34236086816043 | 0.186606014477641 | 0.506408928531958 | -4.9883295330018 |
| lnc-KIAA1551-4 | lnc-KIAA1551-4 | -0.351565332527878 | 0.379988268803644 | -1.34233062434873 | 0.18661573740052 | 0.506408928531958 | -4.97116274319804 |
| PVT1 | PVT1 | 0.376092727705365 | 5.20026016850706 | 1.34255035296534 | 0.186641822558745 | 0.506412949344403 | -5.5074759000043 |
| ENSG00000101166.15 | PRELID3B | 0.208851924992579 | 5.77212651653936 | 1.34187142399251 | 0.186763410913834 | 0.506676062585929 | -5.52971793122419 |
| ENSG00000136448.11 | NMT1 | -0.189903687320513 | 5.8238341434244 | -1.34170764774842 | 0.186816101077671 | 0.506752215139919 | -5.52902009227412 |
| ENSG00000164347.17 | GFM2 | 0.106780333594958 | 4.51422185784087 | 1.34146870446723 | 0.186892994408915 | 0.506759051821586 | -5.4712179102189 |
| ENSG00000159256.12 | MORC3 | -0.188191233715278 | 7.97722229977136 | -1.3414614566814 | 0.186895327166587 | 0.506759051821586 | -5.5139938373962 |
| ENSG00000113328.18 | CCNG1 | 0.18329422582465 | 5.06367680141808 | 1.34145517747419 | 0.186897348197514 | 0.506759051821586 | -5.50877206531285 |
| ENSG00000115944.14 | COX7A2L | 0.203841912839524 | 5.02898288655247 | 1.34139380481766 | 0.186917102532647 | 0.506759051821586 | -5.50581751363543 |
| ENSG00000159140.19 | SON | 0.16793029764308 | 8.78598479645441 | 1.34131533884815 | 0.186942361107175 | 0.506760782124191 | -5.49625634520444 |
| ENSG00000108523.15 | RNF167 | -0.156774512900081 | 5.14785444557631 | -1.34118657870748 | 0.186983815275148 | 0.506806408981877 | -5.51271044940677 |
| ENSG00000123124.13 | WWP1 | 0.185763015902496 | 4.25133129393218 | 1.34078872994972 | 0.187111946613763 | 0.507065875118301 | -5.4502586843649 |
| lnc-ZNF14-4 | lnc-ZNF14-4 | -0.220895704633971 | 2.29018507536137 | -1.34073637077842 | 0.187128814429316 | 0.507065875118301 | -5.23549109533208 |
| ENSG00000166199.12 | ALKBH3 | -0.514016085311212 | 2.65865855509326 | -1.3409721502981 | 0.187167213060577 | 0.507103165252845 | -5.26588787035453 |
| lnc-SH3BGRL-1 | lnc-SH3BGRL-1 | -0.367482112335216 | 0.987941411473516 | -1.34031913994939 | 0.187263269355213 | 0.507147614385141 | -5.08330333528625 |
| ENSG00000065000.17 | AP3D1 | -0.135372844452326 | 6.14379300660533 | -1.34031582814205 | 0.187264336898745 | 0.507147614385141 | -5.53423961741766 |
| ENSG00000111684.10 | LPCAT3 | 0.247397983464706 | 3.30262423067256 | 1.3402810920713 | 0.187275534167307 | 0.507147614385141 | -5.35046088516728 |
| lnc-LIPJ-2 | lnc-LIPJ-2 | -0.319554905027576 | 0.163777628380526 | -1.34022132946994 | 0.187294800005372 | 0.507147614385141 | -4.95093891224787 |
| ENSG00000124571.17 | XPO5 | 0.107447184256364 | 4.77058938294323 | 1.34009934216545 | 0.187334130099875 | 0.507147614385141 | -5.48970641911599 |
| lnc-CRELD1-4 | lnc-CRELD1-4 | -0.273095076700209 | 0.422514891112775 | -1.34007076664666 | 0.187343344084977 | 0.507147614385141 | -4.9806362871546 |
| lnc-ARL14EPL-4 | lnc-ARL14EPL-4 | -0.190746897425444 | 3.2825560067675 | -1.33994056083194 | 0.187385332466505 | 0.507147614385141 | -5.35116417936073 |
| ENSG00000135077.8 | HAVCR2 | -0.368319435601894 | 0.850639916827157 | -1.3399184878969 | 0.18739245119375 | 0.507147614385141 | -5.05966713057215 |
| ENSG00000010017.12 | RANBP9 | 0.132567015243051 | 5.46937765042481 | 1.33987842871688 | 0.18740537118075 | 0.507147614385141 | -5.52371742401751 |
| ENSG00000161981.10 | SNRNP25 | 0.430076206891894 | 3.26521303939896 | 1.33998262923156 | 0.187438888913256 | 0.507171638030964 | -5.33184306673943 |
| ENSG00000186231.16 | KLHL32 | -0.386359632650625 | 2.37237185076926 | -1.33968504745908 | 0.187467750567517 | 0.507183058613461 | -5.24373370504923 |
| ENSG00000164091.11 | WDR82 | 0.146851471638869 | 7.08355798296132 | 1.33954988550068 | 0.187511359460079 | 0.507234368743172 | -5.53134576165604 |
| lnc-C10orf71-2 | lnc-C10orf71-2 | -0.256120840923805 | 2.41695881643532 | -1.3392672230399 | 0.187602583241874 | 0.507414450836728 | -5.24586679206346 |
| ENSG00000237188.4 | | -0.163849415773992 | 0.999449865606325 | -1.338897710173 | 0.187721887451194 | 0.507670028648974 | -5.09010032164193 |

| | | | | | | | |
|---|---|---|---|---|---|---|---|
| lnc-ULK2-6 | lnc-ULK2-6 | 0.162593443530503 | 1.66626488753791 | 1.33882179526561 | 0.18774640519244 | 0.507670028648974 | -5.16418948130416 |
| lnc-USP34-6 | lnc-USP34-6 | 0.152762764138645 | 3.73345663560357 | 1.33869112805096 | 0.18778861166016 | 0.507717456167224 | -5.4034817518295 |
| ENSG00000261459.1 | | -0.224016987571619 | 1.92192520550338 | -1.33859668028923 | 0.187819123494189 | 0.507733257010888 | -5.20679411186258 |
| lnc-HMGB1-4 | lnc-HMGB1-4 | -0.356650546757862 | 1.41257395404899 | -1.33814669242009 | 0.187964546415904 | 0.508059653168556 | -5.14796383371518 |
| lnc-MRPS27-10 | lnc-MRPS27-10 | 0.161969272626364 | 2.14010906031939 | 1.33801220332986 | 0.188008026054374 | 0.508106132024287 | -5.22249188857976 |
| lnc-MASTL-6 | lnc-MASTL-6 | 0.163802921319687 | 2.11522833642935 | 1.33790372295115 | 0.188043102808258 | 0.508106132024287 | -5.22532015797047 |
| ENSG00000105851.10 | PIK3CG | 0.279526347032693 | 4.82665915975794 | 1.33786446135013 | 0.188055799143032 | 0.508106132024287 | -5.49426432149715 |
| ENSG00000197599.12 | CCDC154 | -0.403480927295228 | 0.417268593425143 | -1.33771473191044 | 0.18810422434981 | 0.508131697398168 | -5.01739589641073 |
| ENSG00000172331.11 | BPGM | -0.220282150244474 | 3.47924780578449 | -1.33761804208216 | 0.188135500650374 | 0.508131697398168 | -5.38226396545574 |
| lnc-NTAN1-5 | lnc-NTAN1-5 | 0.908139875171102 | 0.840730308266808 | 1.33823397730546 | 0.18813932205565 | 0.508131697398168 | -5.05116943197871 |
| ENSG00000156675.15 | RAB11FIP1 | 0.205351556266249 | 7.52957584309312 | 1.33750655634623 | 0.188171567921387 | 0.508152110013894 | -5.52865484184571 |
| ENSG00000176731.11 | RBIS | 0.138005216362847 | 4.7825186886867 | 1.33709309866766 | 0.188305373672767 | 0.508358119526859 | -5.49646724706109 |
| ENSG00000107262.21 | BAG1 | 0.123986132270674 | 4.40708872778922 | 1.33707289152618 | 0.18831191509647 | 0.508358119526859 | -5.47528354142229 |
| ENSG00000269547.1 | | -0.234715794968227 | 1.53453227275446 | -1.33694233322508 | 0.188354183406338 | 0.508358119526859 | -5.16765989786784 |
| ENSG00000167114.12 | SLC27A4 | -0.196889928328416 | 2.47729454502524 | -1.33693670580722 | 0.188356005448368 | 0.508358119526859 | -5.27475963965031 |
| ENSG00000172638.12 | EFEMP2 | -0.365074698551097 | -0.0815309221726365 | -1.33677627320252 | 0.188407955886228 | 0.508358119526859 | -4.83291905418753 |
| LINC01003 | LINC01003 | -0.350231230274622 | 0.866781548113853 | -1.33676874640627 | 0.188410393442369 | 0.508358119526859 | -5.07502193146027 |
| ENSG00000169629.11 | RGPD8 | 0.365191889383933 | 2.61039629696676 | 1.33667487856582 | 0.188441981045548 | 0.508358119526859 | -5.30952010463001 |
| lnc-IL16-12 | lnc-IL16-12 | -0.412232593330179 | 1.13545230635033 | -1.33664637375713 | 0.188450027256191 | 0.508358119526859 | -5.09413433869221 |
| ENSG00000177058.11 | SLC38A9 | -0.154433529321971 | 3.97019489793572 | -1.33655622912824 | 0.188479227195306 | 0.508358119526859 | -5.42798836630585 |
| ENSG00000207370.1 | | 0.399675549860677 | 0.99161747927084 | 1.33641676027734 | 0.188524411205221 | 0.508358119526859 | -5.0722003812779 |
| ENSG00000144524.17 | COPS7B | 0.0876127631017001 | 4.41404149952365 | 1.33633816786364 | 0.188549876595291 | 0.508358119526859 | -5.4732729393519 |
| ENSG00000059377.16 | TBXAS1 | 1.04180913342875 | -0.827286037065956 | 1.33647518573245 | 0.188575086675386 | 0.508358119526859 | -4.68907585888439 |
| lnc-TRIM59-3 | lnc-TRIM59-3 | 0.16477424537401 | 1.66051934617323 | 1.33624390359286 | 0.18858042342108 | 0.508358119526859 | -5.16827632582365 |
| ENSG00000119636.15 | BBOF1 | 0.223274258747258 | 0.534391155569598 | 1.33620316592874 | 0.188593625842059 | 0.508358119526859 | -4.96913480475917 |
| lnc-SBDS-29 | lnc-SBDS-29 | 0.229869383541285 | 0.0875636838419834 | 1.33605259057219 | 0.188642431017542 | 0.508423092527703 | -4.89636445473859 |
| ENSG00000171130.18 | ATP6V0E2 | -0.300746653684517 | 3.23642332988444 | -1.3358573586654 | 0.188705724839497 | 0.508527092743045 | -5.36350801836611 |
| ENSG00000197930.12 | ERO1A | -0.186517623088973 | 4.88619192334006 | -1.33568369986917 | 0.18876203832684 | 0.508544551981025 | -5.50556737144604 |
| lnc-HECTD2-3 | lnc-HECTD2-3 | -0.234799277771009 | 3.89295264961709 | -1.33558113346566 | 0.188795304240994 | 0.508544551981025 | -5.42772129233818 |
| RNASEH1-AS1 | RNASEH1-AS1 | -0.268326779723938 | 2.05861769271791 | -1.33552273680425 | 0.188814246344704 | 0.508544551981025 | -5.21759199015159 |
| ENSG00000214530.9 | STARD10 | 0.33598912035101 | 1.46101847839898 | 1.33545733997975 | 0.18883546081134 | 0.508544551981025 | -5.13472629638668 |
| lnc-PDCL-1 | lnc-PDCL-1 | 0.217035134297814 | 2.04611117540926 | 1.3354383298167 | 0.188841627973144 | 0.508544551981025 | -5.21402881770667 |
| ENSG00000137309.19 | HMGA1 | -0.308152121241196 | 3.87871962979741 | -1.33525911965362 | 0.188899773806518 | 0.508544551981025 | -5.43548702841757 |
| ENSG00000276087.2 | | 0.262496634152989 | 2.98472521097007 | 1.33513040158602 | 0.18894154560563 | 0.508544551981025 | -5.32070168783282 |
| ENSG00000086544.2 | ITPKC | -0.261237245135701 | 2.21871050545725 | -1.33501611388183 | 0.188978640350244 | 0.508544551981025 | -5.24002346480698 |
| ENSG00000137996.12 | RTCA | 0.101807508539635 | 4.70680686122232 | 1.33493287081521 | 0.189005662332192 | 0.508544551981025 | -5.49692860339488 |
| lnc-SLC9A9-3 | lnc-SLC9A9-3 | -0.283975792553367 | 1.55600610109918 | -1.33485697697777 | 0.189030301211873 | 0.508544551981025 | -5.16439011102301 |
| lnc-ADRB1-4 | lnc-ADRB1-4 | -0.157019370623014 | 3.65495634197144 | -1.33478587053975 | 0.189053388090784 | 0.508544551981025 | -5.40321630494044 |
| ENSG00000136273.12 | HUS1 | 0.170344905293146 | 4.82227511183475 | 1.33471195469684 | 0.189077389410778 | 0.508544551981025 | -5.49652095118296 |
| ENSG00000153214.10 | TMEM87B | -0.123280369886926 | 5.54890639263443 | -1.33465596293063 | 0.189095572126427 | 0.508544551981025 | -5.53500572318333 |
| ENSG00000101161.7 | PRPF6 | -0.107228041779464 | 6.04732773833265 | -1.33457919312011 | 0.18912050445526 | 0.508544551981025 | -5.54068405147557 |
| ENSG00000280119.1 | | -0.295126287695595 | 2.62338533345449 | -1.33443580852638 | 0.189167077812205 | 0.508544551981025 | -5.29152860069546 |
| lnc-BARD1-1 | lnc-BARD1-1 | -0.592786437642825 | 0.68904150959607 | -1.33440390859455 | 0.189204787063052 | 0.508544551981025 | -4.98426052692551 |
| ENSG00000275052.4 | PPP4R3B | 0.138306965401359 | 7.22282443790949 | 1.33430937811768 | 0.189208151471835 | 0.508544551981025 | -5.53561187693399 |
| ENSG00000135972.8 | MRPS9 | 0.105672271979499 | 4.13932703151106 | 1.33429449082929 | 0.189212988378243 | 0.508544551981025 | -5.44284121503565 |
| ENSG00000041988.15 | THAP3 | -0.229855845980041 | 1.9604645652389 | -1.3342690771804 | 0.189221245535938 | 0.508544551981025 | -5.21810955050283 |
| ENSG00000019582.14 | CD74 | -0.234025567220353 | 11.6873922992306 | -1.334197019675 | 0.189244659261267 | 0.508544551981025 | -5.41040690357223 |
| ENSG00000279348.1 | | -0.251119413469863 | 1.2941963913695 | -1.33409028663858 | 0.189279344209204 | 0.508544551981025 | -5.1300600909799 |
| lnc-XRCC5-3 | lnc-XRCC5-3 | -0.251119413469863 | 1.2941963913695 | -1.33409028663858 | 0.189279344209204 | 0.508544551981025 | -5.1300600909799 |
| ENSG00000200530.1 | SNORD35B | 0.177204342169575 | 1.81358784353653 | 1.33408683772961 | 0.189280465079271 | 0.508544551981025 | -5.19515227384794 |
| ENSG00000159335.15 | PTMS | 0.949448018254843 | 1.08003538169621 | 1.33470124884325 | 0.189329068537407 | 0.508608746763958 | -5.13794670206703 |
| ENSG00000013441.15 | CLK1 | 0.214075461302989 | 8.02173706557865 | 1.33378322604825 | 0.189379156499804 | 0.50864389800023 | -5.52368913728082 |
| LINC00920 | LINC00920 | -1.01978912941409 | -1.90953450098003 | -1.33374502377582 | 0.189391577237223 | 0.50864389800023 | -4.56663316621179 |
| lnc-DGCR2-2 | lnc-DGCR2-2 | -0.297341526328623 | -0.312771774662801 | -1.33366364204239 | 0.189418039022669 | 0.508648597156373 | -4.81527863752162 |
| ENSG00000176248.8 | ANAPC2 | 0.167593457177317 | 5.2146851145469 | 1.33331526277114 | 0.189531348642084 | 0.508882811980874 | -5.52312722184194 |
| ENSG00000001084.12 | GCLC | -0.212867577780366 | 6.01991686039729 | -1.33317057580103 | 0.189578422972134 | 0.508882811980874 | -5.54193628297462 |
| ENSG00000285558.1 | | 0.178622814527341 | 1.70747866153048 | 1.33316748172387 | 0.189579429736644 | 0.508882811980874 | -5.17962152429675 |
| lnc-FNDC10-1 | lnc-FNDC10-1 | -0.210297188298841 | 3.02368974116753 | -1.33279063894987 | 0.189702079016524 | 0.509044721122193 | -5.329316977088 |
| lnc-DHX40-1 | lnc-DHX40-1 | 0.34911423814137 | 0.959809956161183 | 1.33278420684287 | 0.189704172970129 | 0.509044721122193 | -5.07778768086428 |
| lnc-AP1S3-2 | lnc-AP1S3-2 | 0.217041220574353 | 3.81082174087515 | 1.33272758573378 | 0.189722606567405 | 0.509044721122193 | -5.4237225519686 |
| ENSG00000183665.5 | TRMT12 | 0.187796392824318 | 3.98414116072457 | 1.33267823997364 | 0.189738672712892 | 0.509044721122193 | -5.4291204948186 |
| ENSG00000079974.17 | RABL2B | -0.135891431415165 | 3.74872522267025 | -1.33254446374294 | 0.189782233214382 | 0.509095230842125 | -5.411953783124 |
| ENSG00000121774.17 | KHDRBS1 | 0.154995681694811 | 5.8465330152533 | 1.33240586637106 | 0.189827371633403 | 0.509141786748171 | -5.54157939037384 |
| lnc-STON2-5 | lnc-STON2-5 | -0.257049610024962 | 3.17345344027254 | -1.33232180730224 | 0.189854751997342 | 0.509141786748171 | -5.35659052736549 |
| lnc-MC5R-2 | lnc-MC5R-2 | 0.163260300044987 | 1.99988687050347 | 1.33223117923724 | 0.189884275440341 | 0.509141786748171 | -5.21022879061391 |
| ENSG00000158796.16 | DEDD | 0.123818500355167 | 4.34208973873858 | 1.33206339292856 | 0.189938943575959 | 0.509141786748171 | -5.46960091735917 |
| ENSG00000143119.13 | CD53 | 0.146731304101333 | 7.8377481419589 | 1.33189735321652 | 0.189993054454673 | 0.509141786748171 | -5.53018540876054 |
| ENSG00000100632.10 | ERH | 0.20932954563997 | 5.15243144055241 | 1.33185624860472 | 0.19000645190111 | 0.509141786748171 | -5.52496558972197 |

| | | | | | | | |
|---|---|---|---|---|---|---|---|
| ENSG00000181924.6 | COA4 | -0.156251576188678 | 3.60239006035093 | -1.33182548216588 | 0.190016480242991 | 0.509141786748171 | -5.40131351638965 |
| ENSG00000112697.15 | TMEM30A | 0.124906527284168 | 6.11322777129269 | 1.33182225643957 | 0.190017531694083 | 0.509141786748171 | -5.54504964321947 |
| ENSG00000120329.6 | SLC25A2 | -0.374156180407172 | -0.50882712026226 | -1.33178569416861 | 0.190029449766729 | 0.509141786748171 | -4.75786767024472 |
| ENSG00000262227.1 | NA | -0.212384792894424 | 0.327290112404455 | -1.33168863018813 | 0.190061092131364 | 0.509141786748171 | -4.97642371022977 |
| lnc-RNF19B-1 | lnc-RNF19B-1 | -0.139376651098909 | 3.17824925192046 | -1.33154623053628 | 0.190107520967543 | 0.509141786748171 | -5.35252189492444 |
| ENSG00000109083.13 | IFT20 | 0.170738140781236 | 2.77867536211227 | 1.33154083052382 | 0.190109281790215 | 0.509141786748171 | -5.30914837186799 |
| ENSG00000105355.8 | PLIN3 | -0.182323811760548 | 3.74061251791201 | -1.33150324530612 | 0.190121537829414 | 0.509141786748171 | -5.41478141342362 |
| ENSG00000157212.18 | PAXIP1 | -0.166053125307628 | 4.28893636452177 | -1.33142856167186 | 0.190145892956964 | 0.509141786748171 | -5.46853538227051 |
| lnc-COX10-1 | lnc-COX10-1 | -0.301493123444911 | 0.191149109068406 | -1.33114491563677 | 0.190238414628483 | 0.509308254495624 | -4.94503423876349 |
| ENSG00000269307.1 | | -0.168631192232211 | 3.70786326473332 | -1.33108626241755 | 0.190257550834031 | 0.509308254495624 | -5.41141694708247 |
| ENSG00000198793.12 | MTOR | 0.0832468918002602 | 6.51686607622148 | 1.33094959130929 | 0.190302146860213 | 0.509330129189602 | -5.54644787800181 |
| ENSG00000114107.8 | CEP70 | 0.207873602250147 | 3.67875334626487 | 1.3308290977536 | 0.19034147073411 | 0.509330129189602 | -5.41006833412651 |
| ENSG00000152332.15 | UHMK1 | 0.112587246277351 | 7.38476607571231 | 1.33077205418576 | 0.190360089443268 | 0.509330129189602 | -5.53830423410054 |
| ENSG00000148672.8 | GLUD1 | 0.147490881438268 | 6.07795467188942 | 1.33075059608561 | 0.190367093609499 | 0.509330129189602 | -5.54631157773109 |
| ENSG00000212283.1 | SNORD89 | -0.295886144317948 | 1.14277354651156 | -1.33068211390284 | 0.190389448281825 | 0.509330129189602 | -5.11531050433509 |
| ENSG00000015479.18 | MATR3 | 0.127018512560165 | 8.23602760443916 | 1.33057059917906 | 0.190425854371299 | 0.509333748632855 | -5.52265072544931 |
| lnc-ERMARD-2 | lnc-ERMARD-2 | -0.320129844881839 | 1.44301647514203 | -1.33052638019818 | 0.190440291963196 | 0.509333748632855 | -5.15500658279697 |
| lnc-ZNF704-7 | lnc-ZNF704-7 | -0.259077216734925 | 2.02846460366827 | -1.33038747615561 | 0.190485649859579 | 0.509338368052433 | -5.22761664029595 |
| ENSG00000241258.6 | CRCP | 0.145530096077682 | 4.9193897030394 | 1.33035710546415 | 0.190495568239753 | 0.509338368052433 | -5.51225231174854 |
| lnc-CALML6-1 | lnc-CALML6-1 | -0.335065517788313 | 0.34023354174271 | -1.33029376238696 | 0.190516255921667 | 0.509338368052433 | -4.97140793625251 |
| ENSG00000072135.12 | PTPN18 | 0.287954336234353 | 5.35584331430392 | 1.32993019463789 | 0.19063682917064 | 0.509576779348489 | -5.53257197663146 |
| ENSG00000180357.9 | ZNF609 | -0.171808153768507 | 6.17701116183678 | -1.32986924260068 | 0.1906549471779 | 0.509576779348489 | -5.54729247206248 |
| ENSG00000169189.16 | NSMCE1 | -0.290585077065399 | 4.81287350606848 | -1.32971487408321 | 0.190705398685831 | 0.509645445488243 | -5.50576066147966 |
| ENSG00000261341.6 | SMIM47 | -0.280316414609472 | 0.0138259001023066 | -1.32946456745458 | 0.190787226808456 | 0.509790295739912 | -4.89568135106199 |
| lnc-UACA-5 | lnc-UACA-5 | 0.471514178864028 | -0.334278658161111 | 1.32933468301711 | 0.190829698074785 | 0.509790295739912 | -4.80778408845553 |
| ENSG00000238917.1 | SNORD10 | 0.430946530439568 | 3.01549889029789 | 1.32952083543477 | 0.190833903156781 | 0.509790295739912 | -5.32812860621417 |
| lnc-NPIPB12-2 | lnc-NPIPB12-2 | 0.298755347088419 | 1.06877495870384 | 1.3291075528718 | 0.190903985300599 | 0.509909798895645 | -5.11021889513545 |
| ENSG00000225573.4 | RPL35P5 | 1.15484076888488 | 1.17469575292171 | 1.32973333769763 | 0.19092818442835 | 0.509909798895645 | -4.93239825783534 |
| ENSG00000185324.21 | CDK10 | -0.172302828381475 | 5.26850367590796 | -1.32872120766813 | 0.191030397466834 | 0.510116589395414 | -5.53204208012924 |
| ENSG00000124279.11 | FASTKD3 | 0.174719246655453 | 2.4986869107148 | 1.32860798466897 | 0.191067456121985 | 0.510149364296123 | -5.27845180603647 |
| ENSG00000166669.13 | ATF7IP2 | 0.229515137953627 | 4.57915959566335 | 1.32849808281431 | 0.191103432984049 | 0.510162795959147 | -5.48374206030159 |
| ENSG00000186648.14 | CARMIL3 | 1.408966854245 | -0.9121883299287 | 1.32889850178787 | 0.191122057991594 | 0.510162795959147 | -4.63647706133151 |
| ENSG00000185800.11 | DMWD | -0.177431621259319 | 3.71156526931789 | -1.32803208641108 | 0.191256036338867 | 0.510454226654941 | -5.42171450933319 |
| SPINT1-AS1 | SPINT1-AS1 | -0.3864850879216 | 0.947398974555666 | -1.32790571756855 | 0.191297435296784 | 0.510484990549164 | -5.09322640400273 |
| ENSG00000157227.12 | MMP14 | -0.612734440663437 | 2.64640085056002 | -1.32836422651717 | 0.191327716476186 | 0.510484990549164 | -5.28670004425654 |
| lnc-KMT5B-1 | lnc-KMT5B-1 | -0.25359273039055 | 1.73983836196232 | -1.32770644928742 | 0.19136273028593 | 0.510484990549164 | -5.18167107685805 |
| ENSG00000205707.10 | ETFRF1 | -0.221716062383761 | 3.30981701389528 | -1.327694128431 | 0.191366768064779 | 0.510484990549164 | -5.38112746533076 |
| ENSG00000180329.13 | CCDC43 | 0.158009360122881 | 4.14180134027963 | 1.32755237411001 | 0.191413228320169 | 0.510542759947394 | -5.46132944147538 |
| ENSG00000198805.11 | PNP | 0.309983611191678 | 3.33962098823818 | 1.32737122214532 | 0.191472613738075 | 0.510634984266753 | -5.36794836723775 |
| lnc-AZIN2-3 | lnc-AZIN2-3 | -0.363549104079178 | 0.625158861558243 | -1.32722016254169 | 0.191522144979874 | 0.510700908164368 | -5.02030683222895 |
| lnc-ZNF709-1 | lnc-ZNF709-1 | 0.325453497039874 | -0.123178356025303 | 1.32696792280624 | 0.191604874138658 | 0.51082525462933 | -4.86901587159276 |
| ENSG00000203995.9 | ZYG11A | -1.3325157347644 | -2.08806658164766 | -1.32734251372556 | 0.191639087180991 | 0.51082525462933 | -4.6751625063555 |
| lnc-THTPA-2 | lnc-THTPA-2 | -0.445151473357971 | -0.214234387787997 | -1.32684747745611 | 0.191644387200887 | 0.51082525462933 | -4.85714360587771 |
| ENSG00000285725.1 | | -0.158351540076948 | 3.27384535985322 | -1.32677536127542 | 0.191668048462436 | 0.51082525462933 | -5.36929521562983 |
| ENSG00000054277.13 | OPN3 | -0.368292651605656 | 3.87647071681056 | -1.32666668769122 | 0.191754362008326 | 0.510989129106339 | -5.44069860273947 |
| LINC02033 | LINC02033 | -0.452393197024568 | -0.332781935929813 | -1.32622671971703 | 0.191848130186801 | 0.511114586088673 | -4.83149668887882 |
| ENSG00000112699.10 | GMDS | 0.200470558148276 | 3.69478714981635 | 1.32616253704515 | 0.191869205410886 | 0.511114586088673 | -5.40686974188652 |
| ENSG00000276547.1 | PCDHGB5 | -1.06318208928577 | -2.04766927891066 | -1.32618420979195 | 0.191875936778272 | 0.511114586088673 | -4.61531180093546 |
| KANSL1-AS1 | KANSL1-AS1 | 0.458259007743078 | 0.996107034901165 | 1.32593147288746 | 0.191945092965051 | 0.51118699127895 | -5.07982545597291 |
| lnc-ALB-6 | lnc-ALB-6 | -0.388581188319177 | 0.301026443466251 | -1.32590804376819 | 0.191952788980955 | 0.51118699127895 | -4.92470612003389 |
| ENSG00000265681.7 | RPL17 | 0.191200218414971 | 7.96285638266775 | 1.32569999274959 | 0.192021140055072 | 0.511248847742923 | -5.53534817538457 |
| ENSG00000168476.11 | REEP4 | -0.213049136504965 | 1.63610545155527 | -1.32568613580552 | 0.192025693139996 | 0.511248847742923 | -5.19238465481083 |
| ENSG00000142279.12 | WTIP | -0.306113009183349 | -0.0991639738014064 | -1.32554537775672 | 0.192071947776357 | 0.5112497826326 | -4.88074073124484 |
| ENSG00000229132.2 | EIF4A1P10 | 0.815198640296276 | -0.976735048697545 | 1.32545940036044 | 0.192100205063784 | 0.5112497826326 | -4.63227976998131 |
| ENSG00000229994.1 | RPL5P4 | 0.388721901921741 | 0.517382490672473 | 1.32545832159379 | 0.192100559630789 | 0.5112497826326 | -4.95909453315956 |
| lnc-USP33-2 | lnc-USP33-2 | 0.148478596151829 | 1.8675284749211 | 1.3252767389345 | 0.192160248981383 | 0.511342521626866 | -5.20611121255204 |
| ENSG00000131446.16 | MGAT1 | -0.194744875517224 | 5.51010029309253 | -1.32516650158763 | 0.192196492785925 | 0.511372855611775 | -5.54420509155403 |
| ENSG00000215472.10 | RPL17-C18orf32 | 0.193578923727842 | 7.97215147638294 | 1.3248938957584 | 0.19228614240464 | 0.511404559359015 | -5.53620647069301 |
| lnc-HSDL1-4 | lnc-HSDL1-4 | 0.253189664017227 | 0.641585162239137 | 1.32488391211369 | 0.192289426247305 | 0.511404559359015 | -5.01649921771558 |
| ENSG00000066322.14 | ELOVL1 | -0.169532619690485 | 4.34358910631268 | -1.32487067701544 | 0.192293779631127 | 0.511404559359015 | -5.48448367084331 |
| lnc-SLC25A11-1 | lnc-SLC25A11-1 | -0.238282290906572 | 1.34949563067548 | -1.32482807736104 | 0.192307792325646 | 0.511404559359015 | -5.14086014955818 |
| ENSG00000152953.12 | STK32B | -1.64310619309953 | 0.833725241254292 | -1.32545805808662 | 0.192381041517611 | 0.511428978931949 | -5.14614465292044 |
| ENSG00000119514.6 | GALNT12 | -0.325758590283243 | 1.58653156490471 | -1.32454262438026 | 0.192401709046989 | 0.511428978931949 | -5.16463620382498 |
| ENSG00000117174.10 | ZNHIT6 | -0.137722709353311 | 4.46607598173121 | -1.32445701552549 | 0.192429881971793 | 0.511428978931949 | -5.491905225684 |
| lnc-RAB1A-1 | lnc-RAB1A-1 | 0.209707743551148 | 3.80699395077756 | 1.32443403251778 | 0.192437445959434 | 0.511428978931949 | -5.41636364945381 |
| ENSG00000181638.17 | ZFP41 | -0.159096689786756 | 2.5274633097143 | -1.32442259340013 | 0.192441210796674 | 0.511428978931949 | -5.27410234841462 |
| ENSG00000136541.14 | ERMN | 0.342327884736264 | 4.81003340780204 | 1.32436799040437 | 0.192511398490494 | 0.511537048126215 | -5.51209172102666 |
| ENSG00000124160.11 | NCOA5 | -0.136993027789974 | 5.00857320999892 | -1.32414806574651 | 0.192531580033707 | 0.511537048126215 | -5.52662182108367 |

| | | | | | | | |
|---|---|---|---|---|---|---|---|
| HCG23 | HCG23 | 0.351838006983675 | 4.44092788033993 | 1.32420940251457 | 0.192560218997328 | 0.511547107333765 | -5.47902072153872 |
| lnc-VAMP2-2 | lnc-VAMP2-2 | 0.356827572794666 | 1.84440491353408 | 1.32385622222001 | 0.192627684741102 | 0.51161318502647 | -5.20009181312103 |
| ENSG00000196735.11 | HLA-DQA1 | -0.245503863758197 | 8.6896409588261 | -1.32383460545023 | 0.19263480464243 | 0.51161318502647 | -5.5227294842107 |
| ENSG00000198160.14 | MIER1 | 0.140744361742891 | 6.48944208172217 | 1.32370815863832 | 0.192676456358641 | 0.511657786250794 | -5.55560127888407 |
| ENSG00000204261.8 | PSMB8-AS1 | -0.224779175994874 | 4.94183567511678 | -1.32353848689594 | 0.192732357182089 | 0.511682479362458 | -5.52004636494487 |
| ENSG00000105193.8 | RPS16 | 0.251992682784199 | 6.76366217309036 | 1.32341386578263 | 0.192773423281724 | 0.511682479362458 | -5.55457765727809 |
| lnc-C2orf42-7 | lnc-C2orf42-7 | 0.370861563284724 | -0.157942513608676 | 1.32339163764837 | 0.192780748765754 | 0.511682479362458 | -4.87304168904223 |
| ENSG00000197712.11 | FAM114A1 | -0.477756453881765 | 1.69377560134375 | -1.32337391216497 | 0.192827575135742 | 0.511682479362458 | -5.19221147031108 |
| lnc-ZNF782-3 | lnc-ZNF782-3 | 0.18774757146604 | 3.30031983782928 | 1.32321160097492 | 0.192840089297186 | 0.511682479362458 | -5.37257227778039 |
| ENSG00000101160.13 | CTSZ | -0.201092528848988 | 6.84927826041904 | -1.32316935004971 | 0.192854017319781 | 0.511682479362458 | -5.55423913581723 |
| ENSG00000142856.16 | ITGB3BP | -0.196902776338968 | 3.21263741133769 | -1.32309550958722 | 0.192878360675028 | 0.511682479362458 | -5.36986675886857 |
| ENSG00000184227.7 | ACOT1 | -0.638595356310277 | -0.702046552767217 | -1.32298222077069 | 0.192915713712188 | 0.511682479362458 | -4.74457663083234 |
| ENSG00000197409.7 | H3C4 | 1.51561002373351 | 2.51039260021052 | 1.32378386522933 | 0.192931725309341 | 0.511682479362458 | -5.23713015747014 |
| lnc-SELENOF-2 | lnc-SELENOF-2 | -0.265565840565685 | 1.26944277433232 | -1.32291346957586 | 0.192938384705063 | 0.511682479362458 | -5.13880226254317 |
| ENSG00000183751.14 | TBL3 | -0.188535542963722 | 3.69962663624969 | -1.32278697495509 | 0.192980102129136 | 0.511682479362458 | -5.42951885130991 |
| ENSG00000169641.13 | LUZP1 | -0.367537112153363 | 5.12951985709778 | -1.32284310674779 | 0.193050998182509 | 0.511682479362458 | -5.53180305358715 |
| SP2-AS1 | SP2-AS1 | -0.221695989666423 | 0.937960883738904 | -1.32251301320124 | 0.193070477156212 | 0.511682479362458 | -5.08274192938478 |
| lnc-MTFMT-2 | lnc-MTFMT-2 | 0.146272506266229 | 2.38248603487426 | 1.32249851808182 | 0.193075259730668 | 0.511682479362458 | -5.26510297769865 |
| LINC00571 | LINC00571 | -0.203411011093017 | 1.62410687052708 | -1.32249256378505 | 0.193077224339948 | 0.511682479362458 | -5.18622744081573 |
| lnc-WNT9A-3 | lnc-WNT9A-3 | -0.218571193808521 | 0.788781741777754 | -1.3224735238252 | 0.193083506641802 | 0.511682479362458 | -5.0693238355146 |
| ENSG00000107164.15 | FUBP3 | 0.125560875420194 | 5.04191358088222 | 1.32223439249446 | 0.193162422109485 | 0.511809158220847 | -5.53178542656175 |
| ENSG00000153574.8 | RPIA | 0.205109753155064 | 3.2246068253253 | 1.32216539473689 | 0.193185196544187 | 0.511809158220847 | -5.36896672543642 |
| ENSG00000147099.20 | HDAC8 | -0.192223635270519 | 4.61296727591489 | -1.32206185755459 | 0.193219375411735 | 0.511809158220847 | -5.50279826283395 |
| ENSG00000124104.18 | SNX21 | 0.480162552795843 | -0.0403200202362975 | 1.32201435365534 | 0.193235058558805 | 0.511809158220847 | -4.94520253419557 |
| ENSG00000161920.9 | MED11 | -0.311209232669827 | 2.33303551938713 | -1.32190906399901 | 0.193269822802473 | 0.511809158220847 | -5.27127483606112 |
| ENSG00000113593.11 | PPWD1 | 0.14135169696737 | 6.21444713964581 | 1.32177183581451 | 0.193315139560539 | 0.511809158220847 | -5.55807029513823 |
| ENSG00000116151.13 | MORN1 | -0.258760025026083 | 2.72547724804664 | -1.32159987801616 | 0.193371936460874 | 0.511809158220847 | -5.3209124623835 |
| LINC01725 | LINC01725 | -0.634474434550582 | 4.11075883618789 | -1.32231608439949 | 0.193373087186276 | 0.511809158220847 | -5.45906264730065 |
| lnc-USP16-9 | lnc-USP16-9 | -0.329066538264374 | 0.176231967536764 | -1.32159621200356 | 0.193373147466786 | 0.511809158220847 | -4.91990139449391 |
| ENSG00000159110.19 | IFNAR2 | 0.201411483849023 | 4.7746421458598 | 1.32139762788885 | 0.193438755025321 | 0.511809158220847 | -5.51549885790072 |
| ENSG00000196943.13 | NOP9 | 0.101403515921792 | 5.4216971407748 | 1.32137538974603 | 0.193446103042816 | 0.511809158220847 | -5.545631894476 |
| ENSG00000212719.11 | LINC02693 | -0.904741747109948 | -1.40677522226302 | -1.32135367555148 | 0.193453278140037 | 0.511809158220847 | -4.61791463204363 |
| ENSG00000173473.10 | SMARCC1 | -0.125639805164313 | 7.20113583419624 | -1.32126112007147 | 0.193483863840919 | 0.511809158220847 | -5.55279559282718 |
| ENSG00000127445.13 | PIN1 | -0.134173398714015 | 4.50976256629906 | -1.32124319192174 | 0.193489788766978 | 0.511809158220847 | -5.50040053866291 |
| MIR22HG | MIR22HG | 0.360486667225437 | 3.00084810817381 | 1.32120831981775 | 0.193504293313639 | 0.511809158220847 | -5.34020011776328 |
| ENSG00000213672.7 | NCKIPSD | -0.228999572093302 | 2.96148514654872 | -1.32098336057942 | 0.19357567377214 | 0.511877741964559 | -5.33680207824199 |
| lnc-CLK3-2 | lnc-CLK3-2 | -0.176407098892953 | 0.146812120854846 | -1.32097039164545 | 0.193579961300691 | 0.511877741964559 | -4.94370924090016 |
| ENSG00000124782.19 | RREB1 | 0.126697816159545 | 6.40834163550416 | 1.32079264372912 | 0.193638732025715 | 0.51195980327298 | -5.55956760165262 |
| ENSG00000198205.6 | ZXDA | 0.160581703685786 | 4.70370056959359 | 1.32071874623377 | 0.193663169550141 | 0.51195980327298 | -5.51240021798934 |
| ENSG00000146243.13 | IRAK1BP1 | -0.250992451128001 | 2.11773948361843 | -1.32065088262979 | 0.193685613763139 | 0.51195980327298 | -5.25654020953967 |
| ENSG00000110092.3 | CCND1 | -0.550465624204958 | 1.87991383013172 | -1.32079774440009 | 0.193749093474386 | 0.512061837568476 | -5.21687005058333 |
| lnc-FAM102B-3 | lnc-FAM102B-3 | -0.30292591863749 | -0.413695532516367 | -1.320380348289 | 0.193775105981267 | 0.512064835847017 | -4.80429654713974 |
| ENSG00000144231.10 | POLR2D | 0.189308954602804 | 3.52365950411774 | 1.32025933407663 | 0.193815147415908 | 0.512104901060543 | -5.40139450127928 |
| CD27-AS1 | CD27-AS1 | -0.217979761055285 | 1.12526881965434 | -1.32013452844663 | 0.193856449951898 | 0.51214828768578 | -5.12863115439011 |
| ENSG00000089692.8 | LAG3 | 1.15180274766585 | 0.874264695984779 | 1.32065983615487 | 0.193950616015659 | 0.51233130511426 | -5.10833306840897 |
| ENSG00000100979.14 | PLTP | -0.786893785476652 | -0.271867925381669 | -1.31987564551344 | 0.193992791443452 | 0.512376957048707 | -4.83531145673979 |
| ENSG00000280670.2 | CCDC163 | 0.82539751307381 | 0.220139309855723 | 1.31982384135672 | 0.194023946829638 | 0.512393494687507 | -4.82228240903891 |
| ENSG00000207445.1 | SNORD15B | 0.350723401031718 | 3.08616123942454 | 1.31944504177946 | 0.194084745744059 | 0.51243659860795 | -5.32867713002255 |
| ENSG00000272899.3 | SPMIP1 | -0.322865616465985 | 0.346189624030363 | -1.31942899668807 | 0.194090060863216 | 0.51243659860795 | -4.99446629622544 |
| ENSG00000086589.11 | RBM22 | 0.103704496991835 | 5.95048316759796 | 1.31922857036474 | 0.194156463687451 | 0.512546170588534 | -5.55970205647153 |
| ENSG00000133119.12 | RFC3 | 0.213480045862309 | 3.33771967466202 | 1.3191364184399 | 0.194187000142225 | 0.512558318054766 | -5.38129609993831 |
| ENSG00000146215.13 | CRIP3 | -0.671510300117864 | 0.919135034944749 | -1.31929344026592 | 0.194238272951963 | 0.512558318054766 | -5.03778546243243 |
| ENSG00000157045.8 | NTAN1 | 0.15702015914338 | 3.5990240882014 | 1.31896275003811 | 0.194244558696919 | 0.512558318054766 | -5.40793179617625 |
| ENSG00000101019.21 | UQCC1 | -0.172580143583708 | 4.58451965726511 | -1.31890427222551 | 0.194263942793713 | 0.512558318054766 | -5.50757199472553 |
| lnc-ZC2HC1B-2 | lnc-ZC2HC1B-2 | 0.225056262109831 | 1.99488463894733 | 1.31883901650628 | 0.194285575351663 | 0.512558318054766 | -5.24650825957426 |
| ENSG00000185347.17 | TEDC1 | -0.30819536743194 | 2.44615471945681 | -1.31857253975533 | 0.19437393256936 | 0.512587238908263 | -5.29034450645228 |
| lnc-SIRT7-2 | lnc-SIRT7-2 | -0.204394728956106 | 0.904385410141313 | -1.31856570049745 | 0.194376200702999 | 0.512587238908263 | -5.0927905341129 |
| ENSG00000198513.11 | ATL1 | -0.293796469083841 | 2.08331304362187 | -1.31848264942222 | 0.194403744907071 | 0.512587238908263 | -5.23325207653863 |
| ENSG00000112078.13 | KCTD20 | 0.158113116966168 | 6.07975789123044 | 1.31830384211784 | 0.194463057094749 | 0.512587238908263 | -5.56216774841877 |
| ENSG00000231222.1 | ARF4P4 | -1.10033261892927 | 0.135354100683748 | -1.31884332842859 | 0.194487362536368 | 0.512587238908263 | -4.92245493511448 |
| ENSG00000178764.7 | ZHX2 | -0.201685664090315 | 6.22223265235502 | -1.31805385847902 | 0.194546002240006 | 0.512587238908263 | -5.56245484854299 |
| ENSG00000259431.5 | THTPA | -0.186317277825447 | 3.93754335608054 | -1.31804540775476 | 0.19454880667937 | 0.512587238908263 | -5.44549687938711 |
| ENSG00000232388.4 | SMIM26 | 0.228684542796919 | 3.2922960800113 | 1.31798893504079 | 0.194567548382503 | 0.512587238908263 | -5.37767773880106 |
| ENSG00000268434.5 | | -0.203143656234808 | 1.10374266834451 | -1.31796922770217 | 0.194574089015984 | 0.512587238908263 | -5.12415343629533 |
| lnc-SRM-1 | lnc-SRM-1 | 0.112758160362139 | 3.05790787972796 | 1.31790742319469 | 0.194594602287269 | 0.512587238908263 | -5.35646310274047 |
| lnc-OST4-5 | lnc-OST4-5 | -0.29949308117161 | -0.350956963762221 | -1.31784245163519 | 0.194616168492864 | 0.512587238908263 | -4.82223791101701 |
| lnc-APOL5-4 | lnc-APOL5-4 | 0.510123299942135 | 1.05703387397642 | 1.31792157249462 | 0.194621048297541 | 0.512587238908263 | -5.12239804745088 |
| lnc-SLC16A3-2 | lnc-SLC16A3-2 | -0.364300080494777 | -0.18930647146376 | -1.3177862730994 | 0.194634817467466 | 0.512587238908263 | -4.86622967630661 |

| | | | | | | | |
|---|---|---|---|---|---|---|---|
| ENSG00000184208.10 | C22orf46P | -0.256491843938196 | 2.82249308036619 | -1.3177550411027 | 0.1946451857993 | 0.512587238908263 | -5.3174048209921 |
| ENSG00000106628.10 | POLD2 | -0.223834253960654 | 3.90411022341171 | -1.31752580937573 | 0.194721298503568 | 0.512630693992376 | -5.45439778775363 |
| ENSG00000275074.1 | NUDT18 | -0.200966096010401 | 2.39003099667699 | -1.31751588508239 | 0.194724594215955 | 0.512630693992376 | -5.27257026414441 |
| ENSG00000174175.16 | SELP | -0.598059193515837 | 3.67418427656203 | -1.31802965007742 | 0.194751706748761 | 0.512630693992376 | -5.39571112526771 |
| ENSG00000131196.17 | NFATC1 | -0.179759492217487 | 5.47544268563986 | -1.31740533486552 | 0.194761309188183 | 0.512630693992376 | -5.55313884422498 |
| lnc-ZNF33B-5 | lnc-ZNF33B-5 | -0.300381509557178 | 0.202804418487701 | -1.3172144102165 | 0.194824729798094 | 0.512719044083911 | -4.93615473924232 |
| ENSG00000198919.12 | DZIP3 | -0.15345028322662 | 4.81308011438662 | -1.31715431489738 | 0.194844695273981 | 0.512719044083911 | -5.52234680497576 |
| ZSCAN16-AS1 | ZSCAN16-AS1 | -0.205661306195653 | 3.08344649973652 | -1.31684140788056 | 0.194948677529043 | 0.51292709057654З | -5.35171041756581 |
| ENSG00000168802.12 | CHTF8 | -0.181739813536784 | 4.8444084585761 | -1.31675164946515 | 0.194978512961725 | 0.512940022020858 | -5.52568337126109 |
| lnc-DUSP11-2 | lnc-DUSP11-2 | 0.26563427525269 | 1.37712672145316 | 1.31661233054096 | 0.195024829030328 | 0.512996301077475 | -5.14798708564095 |
| lnc-IGIP-1 | lnc-IGIP-1 | -0.286455855917207 | 0.598615144529779 | -1.31603636279822 | 0.195216396090775 | 0.513352325781942 | -5.04737134746563 |
| ENSG00000204463.12 | BAG6 | -0.151110089034611 | 6.60108871258424 | -1.31599532280088 | 0.195230051459016 | 0.513352325781942 | -5.56473911083201 |
| ENSG00000089057.14 | SLC23A2 | -0.171326489335247 | 7.46288593473469 | -1.31591518568759 | 0.19525671782537 | 0.513352325781942 | -5.55444623929032 |
| THUMPD3-AS1 | THUMPD3-AS1 | -0.149137619509228 | 6.41724176516761 | -1.31590550107499 | 0.195259940657184 | 0.513352325781942 | -5.56539562855091 |
| ENSG00000198042.10 | MAK16 | 0.162583872392854 | 5.03770890666617 | 1.31582933298828 | 0.195285289174505 | 0.513353398094361 | -5.5381734855947 |
| ENSG00000122224.17 | LY9 | 0.489862408666389 | 6.64496217757921 | 1.31617587874674 | 0.195367499465737 | 0.513503925616557 | -5.5650588327931 |
| ENSG00000201772.1 | SNORA5C | 0.285000212483266 | 2.65034433232942 | 1.31514011722962 | 0.195514771780636 | 0.513825401884681 | -5.31819383981929 |
| lnc-IGF2BP2-1 | lnc-IGF2BP2-1 | 0.204888662657087 | 5.05811953047495 | 1.31478848320825 | 0.195631931363254 | 0.514067667975214 | -5.54063421188619 |
| ENSG00000196154.11 | S100A4 | -0.484205344312479 | 4.80163480136048 | -1.31513974337478 | 0.195699447525371 | 0.514143631442453 | -5.52653491431245 |
| ENSG00000090061.17 | CCNK | 0.186107502665958 | 5.58337292088239 | 1.31455186920718 | 0.195710797859963 | 0.514143631442453 | -5.55904888467271 |
| ENSG00000168214.20 | RBPJ | -0.322457848933355 | 6.25100313571309 | -1.31462244293337 | 0.195736385038729 | 0.514145228847902 | -5.56670466998413 |
| lnc-C14orf177-5 | lnc-C14orf177-5 | 0.825852094181711 | -0.517478339199843 | 1.31391958978502 | 0.195921663146174 | 0.514538546374348 | -4.68915340318184 |
| ENSG00000267680.5 | ZNF224 | -0.10603995628456 | 5.57438666024226 | -1.31387626442342 | 0.195936118470687 | 0.514538546374348 | -5.55927224710196 |
| lnc-MEGF11-2 | lnc-MEGF11-2 | 0.340834351317981 | 1.9883814253782 | 1.31361438407661 | 0.196023510978348 | 0.514702376127992 | -5.22779775541896 |
| ENSG00000178404.9 | CEP295NL | -0.322273016367048 | 1.55789732869483 | -1.31352435216701 | 0.196053562501776 | 0.514715622063017 | -5.1775728863632 |
| ENSG00000105514.7 | RAB3D | -0.280924160269664 | 2.07710665398739 | -1.31338222429739 | 0.196101010119241 | 0.514774530198231 | -5.24294311239404 |
| ENSG00000164542.12 | MATCAP2 | 1.40765137604694 | -1.82500696799962 | 1.3129942333715 | 0.196257051424009 | 0.514992210750366 | -4.55193880398559 |
| ENSG00000198746.12 | GPATCH3 | 0.136104046016668 | 3.76864023601498 | 1.31289068708635 | 0.19626517086389 | 0.514992210750366 | -5.4378367045221 |
| ENSG00000200418.1 | SNORA63B | 0.231393921880266 | 3.27100450650892 | 1.3128508958277 | 0.196278464677813 | 0.514992210750366 | -5.39658691107657 |
| lnc-GPR152-1 | lnc-GPR152-1 | -0.130055902486324 | 2.45801559144942 | -1.31283428128767 | 0.196284015611749 | 0.514992210750366 | -5.29848681153357 |
| lnc-ATP5F1B-2 | lnc-ATP5F1B-2 | 0.338656305788862 | -0.386283646025206 | 1.31256894403429 | 0.196372681184976 | 0.515090925084594 | -4.80012228778709 |
| ENSG00000228502.1 | EEF1A1P11 | 0.320615448126978 | -0.235855488614522 | 1.31250383957819 | 0.196394441248435 | 0.515090925084594 | -4.88022113080407 |
| lnc-TAPBPL-3 | lnc-TAPBPL-3 | -0.303510501832334 | -0.121202927456755 | -1.31249703769702 | 0.196396714767716 | 0.515090925084594 | -4.87957639029522 |
| ENSG00000150093.18 | ITGB1 | -0.423994697416808 | 7.73135154702893 | -1.31273588360452 | 0.196473021376529 | 0.515225404381845 | -5.5565379557076 |
| lnc-VSIG10-1 | lnc-VSIG10-1 | -0.282759720850101 | 0.511232997333568 | -1.31211227484785 | 0.196525353722468 | 0.515241888967121 | -5.01278087091559 |
| ENSG00000102057.9 | KCND1 | -0.219170794092469 | 0.582299082853069 | -1.31210025820988 | 0.196529372311173 | 0.515241888967121 | -5.03446494514109 |
| ENSG00000227986.1 | TRIM60P18 | -0.183616686897804 | 1.3152820633384 | -1.31201181530811 | 0.196558951190841 | 0.515253806974156 | -5.15856490048353 |
| lnc-FUCA1-1 | lnc-FUCA1-1 | -0.28408707980641 | 1.20890764635449 | -1.31188285405263 | 0.196602087096191 | 0.515255115794193 | -5.13777844941425 |
| ENSG00000205784.2 | ARRDC5 | -0.335961780681375 | 3.40011423195465 | -1.31181001524254 | 0.19662645392756 | 0.515255115794193 | -5.38978403989224 |
| lnc-BACE2-1 | lnc-BACE2-1 | -0.625301344992377 | 0.615388099301003 | -1.31189374799499 | 0.196657832891678 | 0.515255115794193 | -5.01430023684144 |
| ENSG00000213397.10 | HAUS7 | 0.166456302896887 | 2.50961903196249 | 1.31171099583747 | 0.19665958264972 | 0.515255115794193 | -5.30258774636699 |
| lnc-LRRFIP2-2 | lnc-LRRFIP2-2 | 0.162257392281824 | 0.809136336367315 | 1.31139797379693 | 0.196764337644153 | 0.515420212511594 | -5.08608879806025 |
| ENSG00000129933.20 | MAU2 | -0.124449691154967 | 6.11367788375854 | -1.31137305731888 | 0.196772677934028 | 0.515420212511594 | -5.57062854605915 |
| ENSG00000106290.14 | TAF6 | -0.181862159638146 | 3.82891021128686 | -1.31114990655171 | 0.196847385117454 | 0.515526798964559 | -5.45046516854114 |
| ENSG00000196678.13 | ERI2 | -0.267483390815703 | 0.369967231818509 | -1.31103313379715 | 0.196886487274146 | 0.515526798964559 | -4.99218431189934 |
| ENSG00000198556.13 | ZNF789 | -0.141434962967274 | 3.94821383470729 | -1.31102709873642 | 0.19688850831562 | 0.515526798964559 | -5.45982851133642 |
| lnc-MAGT1-1 | lnc-MAGT1-1 | 0.175296033294381 | 2.22079356084302 | 1.31050150978501 | 0.197064579661416 | 0.515766768113684 | -5.25575309601695 |
| lnc-NDUFV3-1 | lnc-NDUFV3-1 | -0.355145804485706 | -0.452036672511001 | -1.31050112026926 | 0.197064710192741 | 0.515766768113684 | -4.79560264459026 |
| lnc-ERMN-5 | lnc-ERMN-5 | 0.389780648574432 | 1.85695684313416 | 1.31049904140819 | 0.197065406844719 | 0.515766768113684 | -5.21479920602248 |
| ENSG00000100814.17 | CCNB1IP1 | 0.149052690451303 | 4.03155691913664 | 1.31045433715365 | 0.197080388243411 | 0.515766768113684 | -5.46594958358391 |
| ENSG00000166145.14 | SPINT1 | -0.311609991623154 | 2.99012808730925 | -1.31029949400982 | 0.197132286330028 | 0.515837000957407 | -5.35744414974831 |
| ENSG00000207181.1 | SNORA14B | 0.279788654397374 | -0.261018349314641 | 1.31019690039168 | 0.197166677881903 | 0.515861412335477 | -4.84709798664799 |
| ENSG00000144233.9 | AMMECR1L | 0.138331658590039 | 3.67934727726426 | 1.31007624772792 | 0.197207129027631 | 0.515901669646127 | -5.43636675974152 |
| lnc-CCDC22-1 | lnc-CCDC22-1 | -0.234166601976306 | 2.88800754624073 | -1.3098818793074 | 0.197272308041008 | 0.516006597586488 | -5.3461243156856 |
| ENSG00000119689.14 | DLST | 0.165954437048393 | 5.5420508554896 | 1.30965519449607 | 0.197348344576835 | 0.516131289671975 | -5.56475872385846 |
| lnc-CXorf56-3 | lnc-CXorf56-3 | -0.190687566654171 | 0.208940761177473 | -1.30959025934208 | 0.197370129767678 | 0.516131289671975 | -4.95894413754431 |
| lnc-ASB5-2 | lnc-ASB5-2 | -0.270655761442478 | 2.35271075018789 | -1.30920335534564 | 0.197499970548293 | 0.51628423544907 | -5.29105658843164 |
| lnc-RRP8-6 | lnc-RRP8-6 | -0.339405673443942 | 0.847504880575582 | -1.30903886374365 | 0.197555191751029 | 0.51628423544907 | -5.10500189343083 |
| ENSG00000179295.17 | PTPN11 | 0.130218632856759 | 6.28020450639156 | 1.30902390425867 | 0.197560214355289 | 0.51628423544907 | -5.57439769284423 |
| ENSG00000088035.16 | ALG6 | -0.140218952973054 | 4.34982934723343 | -1.30892509923385 | 0.197593390289845 | 0.51628423544907 | -5.50508532048955 |
| ENSG00000188010.13 | MORN2 | 0.349664161768058 | -0.534211754311515 | 1.3088059364423 | 0.197633407403052 | 0.51628423544907 | -4.79807904145049 |
| ENSG00000188191.14 | PRKAR1B | -0.329044104015952 | 1.857644603632 | -1.30879572138787 | 0.19763683809679 | 0.51628423544907 | -5.23535031943818 |
| ENSG00000163754.17 | GYG1 | 0.168965407981998 | 3.44711358202508 | 1.30874568195454 | 0.197653644334252 | 0.51628423544907 | -5.41301455795105 |
| ENSG00000066557.5 | LRRC40 | 0.148013225832344 | 4.45519131728626 | 1.30873319532182 | 0.197657838261892 | 0.51628423544907 | -5.51247511612811 |
| ENSG00000175463.11 | TBC1D10C | -0.24552262261584 | 6.4637004664507 | -1.30853962319009 | 0.197722862615303 | 0.51628423544907 | -5.57439389408498 |
| lnc-GEMIN5-4 | lnc-GEMIN5-4 | -0.285811728107569 | 0.160665657502198 | -1.30852923064345 | 0.197726354116658 | 0.51628423544907 | -4.93468899565179 |
| ENSG00000207137.1 | SNORD116-13 | -0.366324389468173 | 1.94892719497197 | -1.30850345808646 | 0.19773501291952 | 0.51628423544907 | -5.22034218935087 |
| ENSG00000125846.15 | ZNF133 | -0.187832031688753 | 2.98438888666297 | -1.30845536420906 | 0.197751171782022 | 0.51628423544907 | -5.34706836058075 |

| | | | | | | | |
|---|---|---|---|---|---|---|---|
| lnc-URGCP-MRPS24-2 | lnc-URGCP-MRPS2 | -0.236666136406416 | 2.46993582899099 | -1.30844487476655 | 0.197754696219233 | 0.51628423544907 | -5.30064762805532 |
| lnc-SLC4A5-4 | lnc-SLC4A5-4 | -0.196626573470213 | 2.48577986278895 | -1.30830265371352 | 0.197802486980655 | 0.516324190618691 | -5.31088452524502 |
| ENSG00000099795.6 | NDUFB7 | -0.291366165710575 | 3.18442081949906 | -1.30821132814185 | 0.197833179867705 | 0.516324190618691 | -5.390905052688 |
| lnc-CCDC136-1 | lnc-CCDC136-1 | 0.220601035586745 | 0.182106958333402 | 1.3081754008468 | 0.197845255376263 | 0.516324190618691 | -4.95300104107122 |
| lnc-USP50-4 | lnc-USP50-4 | 0.131479033426726 | 2.62732352781947 | 1.3080837614017 | 0.197876058801731 | 0.51633911236258 | -5.31598335953427 |
| ENSG00000138376.10 | BARD1 | 0.216432305136634 | 4.16997080969622 | 1.30730420592567 | 0.198138243323264 | 0.516957721171598 | -5.48674155374018 |
| lnc-RBM45-1 | lnc-RBM45-1 | 0.226581574280784 | -0.231636901414788 | 1.30688712179444 | 0.198278627433931 | 0.517252271964352 | -4.881371972937 |
| lnc-ADGRD2-3 | lnc-ADGRD2-3 | -0.334455470331689 | -0.450841970533377 | -1.30674298312906 | 0.198327159804788 | 0.517252271964352 | -4.83428457740626 |
| ENSG00000249741.2 | | 1.31048830052421 | -1.45856766806508 | 1.30697094453415 | 0.198339605461738 | 0.517252271964352 | -4.62574340014172 |
| lnc-COX18-4 | lnc-COX18-4 | -0.159267022103498 | 1.30756478791489 | -1.30667023401825 | 0.198351658291533 | 0.517252271964352 | -5.15861436321106 |
| ENSG00000198056.13 | PRIM1 | -0.255354149045605 | 2.50567798125186 | -1.30615538576156 | 0.198525100527556 | 0.517604056732227 | -5.30783637261292 |
| lnc-KBTBD6-1 | lnc-KBTBD6-1 | -0.362897314097521 | 0.836990987368729 | -1.30611752269633 | 0.19853786038014 | 0.517604056732227 | -5.10445752203476 |
| ENSG00000136436.14 | CALCOCO2 | 0.167765666852888 | 5.75991429137365 | 1.30604589883863 | 0.19856199931777 | 0.517604056732227 | -5.57221976206932 |
| LINC00562 | LINC00562 | 0.33530016471925 | 1.34167765789521 | 1.30551787273226 | 0.198740025212651 | 0.517950981160502 | -5.16954773071222 |
| lnc-TMTC4-1 | lnc-TMTC4-1 | -0.202827781696695 | 1.8281079868992 | -1.30550189616506 | 0.198745413652317 | 0.517950981160502 | -5.22083511714884 |
| lnc-C8orf76-1 | lnc-C8orf76-1 | 0.214593027680648 | 1.31575538833805 | 1.30487763073026 | 0.19895604714863 | 0.518434272497817 | -5.16584376919966 |
| lnc-NMRAL1-5 | lnc-NMRAL1-5 | -0.295924212827863 | -0.256806949798933 | -1.30473147291787 | 0.199005386680486 | 0.518463880881397 | -4.83731345207577 |
| ENSG00000079257.7 | LXN | -0.300667724490949 | 1.38795732337746 | -1.30469474206179 | 0.199017787632702 | 0.518463880881397 | -5.18128798626917 |
| lnc-FAM124B-1 | lnc-FAM124B-1 | 0.186779530400173 | 2.77409385178164 | 1.3046062071335 | 0.199047680911224 | 0.518476134674226 | -5.33998328189954 |
| ENSG00000170270.4 | GON7 | -0.231030267679846 | 1.54723177465239 | -1.30435719745794 | 0.199131775729046 | 0.518629550782102 | -5.19369475117516 |
| ENSG00000112977.15 | DAP | 0.188127293501403 | 3.67454268479231 | 1.30398648230707 | 0.199257022370469 | 0.518890092541924 | -5.4353238551935 |
| lnc-HES5-4 | lnc-HES5-4 | -0.216311457821435 | 0.219063386014958 | -1.30376865568608 | 0.199330643202266 | 0.518977706748911 | -4.96665097681476 |
| ENSG00000171466.9 | ZNF562 | 0.137860336673369 | 5.17626407988354 | 1.30368396085901 | 0.199359273834815 | 0.518977706748911 | -5.55915124372781 |
| ENSG00000087448.10 | KLHL42 | -0.29218498196964 | 3.65930079046385 | -1.30355020700316 | 0.199404494946022 | 0.518977706748911 | -5.44551225440875 |
| ENSG00000179988.14 | PSTK | 0.231193275220103 | 2.69119523954816 | 1.30352471697904 | 0.199413113800125 | 0.518977706748911 | -5.32782590360886 |
| ENSG00000117533.14 | VAMP4 | -0.176705979117415 | 4.63988463858342 | -1.30351400386818 | 0.199416736271542 | 0.518977706748911 | -5.53716421340495 |
| ENSG00000102805.14 | CLN5 | -0.170392889619369 | 4.7739839014419 | -1.30317594540713 | 0.199531071044548 | 0.519209612554984 | -5.53815435489204 |
| lnc-EBNA1BP2-1 | lnc-EBNA1BP2-1 | -0.19728698784841 | 1.98006650962032 | -1.30298159100039 | 0.199596826107998 | 0.519315064060285 | -5.25382630900248 |
| ENSG00000229417.1 | NPM1P25 | -0.210044870002741 | 2.16800085733546 | -1.3028191020547 | 0.199651812851626 | 0.519326837346773 | -5.2757240256625 |
| lnc-BLNK-2 | lnc-BLNK-2 | -0.210044870002741 | 2.16800085733546 | -1.3028191020547 | 0.199651812851626 | 0.519326837346773 | -5.2757240256625 |
| lnc-ELOVL3-3 | lnc-ELOVL3-3 | 0.382313048989997 | -0.520374255117252 | 1.30268622957274 | 0.19969678581119 | 0.519338787227913 | -4.78058078186618 |
| ENSG00000123411.14 | IKZF4 | 0.277662997587593 | 0.953372443454922 | 1.30236957715139 | 0.199803993117785 | 0.519338787227913 | -5.12567419323908 |
| ENSG00000116668.12 | SWT1 | 0.163030565062605 | 3.19587051704967 | 1.30234381506108 | 0.199812717164559 | 0.519338787227913 | -5.38236259965737 |
| lnc-ZZEF1-4 | lnc-ZZEF1-4 | -0.187784208083833 | 0.416930943965918 | -1.30232893468542 | 0.199817756370356 | 0.519338787227913 | -5.02897727239772 |
| ENSG00000161217.11 | PCYT1A | -0.104582317675191 | 5.18397226856624 | -1.30232586368581 | 0.199818796369443 | 0.519338787227913 | -5.56152469624294 |
| ENSG00000241081.1 | RPL22P2 | 0.629277597284501 | -0.769909630360532 | 1.30229874128605 | 0.199827981592625 | 0.519338787227913 | -4.65790177945107 |
| ENSG00000175564.12 | UCP3 | -0.168256998680544 | 2.00119521010653 | -1.30228384344361 | 0.199833027005056 | 0.519338787227913 | -5.24584744413092 |
| ENSG00000279329.1 | | -0.248729678661018 | 0.433616428465765 | -1.30212692255924 | 0.199886176833308 | 0.519411334144677 | -5.01403641817175 |
| ENSG00000177854.7 | TMEM187 | -0.407371678559345 | -0.401737250226103 | -1.30197758941264 | 0.199936766587149 | 0.519437683785329 | -4.83798323612904 |
| ENSG00000144566.10 | RAB5A | 0.137370924086392 | 5.2373345363306 | 1.30194800666163 | 0.199946789517133 | 0.519437683785329 | -5.56444354270933 |
| ENSG00000100298.15 | APOBEC3H | 0.436994941217808 | 0.543911632385312 | 1.30172836377286 | 0.200021218594456 | 0.519473142026424 | -5.00146193805552 |
| ENSG00000171365.16 | CLCN5 | 0.293630481708292 | 2.39207086117401 | 1.30171248088028 | 0.200026601547825 | 0.519473142026424 | -5.2833073831331 |
| ENSG00000166260.11 | COX11 | -0.253196666198413 | 4.1178218660923 | -1.30168430118391 | 0.200036152344237 | 0.519473142026424 | -5.49325226522206 |
| ENSG00000205832.7 | C16orf96 | -0.315758748876568 | -0.0724403908834031 | -1.30121035123042 | 0.200196837388064 | 0.519713814264637 | -4.88744643792348 |
| ENSG00000173465.7 | ZNRD2 | -0.215736894371659 | 3.01616494458033 | -1.30118955547598 | 0.200203890085603 | 0.519713814264637 | -5.3815328890093 |
| ENSG00000272968.5 | RBAK-RBAKDN | 0.135884940672989 | 2.24501243939505 | 1.30118752648428 | 0.200204578210383 | 0.519713814264637 | -5.28261795299614 |
| lnc-GTF2F2-5 | lnc-GTF2F2-5 | -0.388665090823084 | -0.263849763515095 | -1.30100859807047 | 0.200265268130362 | 0.519734298366289 | -4.86952364877357 |
| LINC01128 | LINC01128 | 0.1441477391395 | 3.28819745965605 | 1.30100751363413 | 0.200265635997824 | 0.519734298366289 | -5.39844163323508 |
| ENSG00000278615.4 | C11orf98 | -0.164909991939065 | 1.63324835676248 | -1.30076704852675 | 0.200347220289671 | 0.519734298366289 | -5.21631808462763 |
| ENSG00000155115.6 | GTF3C6 | 0.212128199834697 | 3.39412162382856 | 1.30075532137048 | 0.200351199687087 | 0.519734298366289 | -5.41389106699792 |
| ENSG00000111875.7 | ASF1A | -0.236977027977575 | 4.26413434073983 | -1.30067572357764 | 0.20037821133113 | 0.519734298366289 | -5.50556988770499 |
| ENSG00000120217.13 | CD274 | 0.740019254696695 | 0.675846954696858 | 1.30098257284545 | 0.200410179147264 | 0.519734298366289 | -5.028490699914 |
| lnc-LEFTY1-2 | lnc-LEFTY1-2 | -0.134168763747947 | 1.66682434406094 | -1.30057768423877 | 0.200411484928143 | 0.519734298366289 | -5.21339570495927 |
| lnc-BBS10-5 | lnc-BBS10-5 | 0.173203548279124 | 0.907818581320049 | 1.30052154382755 | 0.20043054031762 | 0.519734298366289 | -5.10450917345019 |
| ENSG00000132581.9 | SDF2 | 0.159697237673924 | 3.90285094361236 | 1.3004054634328 | 0.200469945105266 | 0.519734298366289 | -5.46313953728869 |
| ENSG00000114861.20 | FOXP1 | -0.127346948870041 | 9.26503805278403 | -1.30035358341115 | 0.200487558251907 | 0.519734298366289 | -5.53442794576585 |
| ENSG00000279589.1 | | -0.285407143266833 | -0.125072594271879 | -1.3002739634677 | 0.2005145913125 | 0.519734298366289 | -4.88969179655195 |
| ENSG00000008869.11 | HEATR5B | 0.216143893260065 | 7.10662645575077 | 1.30027135497322 | 0.200515477011451 | 0.519734298366289 | -5.58141249176065 |
| ENSG00000154978.12 | VOPP1 | -0.189323905670353 | 8.39588391839505 | -1.30016807606659 | 0.200550547132822 | 0.519759747120987 | -5.55837458080629 |
| ENSG00000141040.14 | ZNF287 | -0.214476421392327 | 2.52804028571576 | -1.29962927274017 | 0.200733582215967 | 0.520168616738164 | -5.31406206760095 |
| ENSG00000204304.11 | PBX2 | -0.159071406951192 | 5.37273803751026 | -1.29949198181117 | 0.200780241036058 | 0.52022403087175 | -5.57265163702261 |
| lnc-EVPL-1 | lnc-EVPL-1 | -0.327804215527865 | 0.1007601120083 | -1.29928729916538 | 0.20084981839255 | 0.520248578137966 | -4.96213758130388 |
| ENSG00000161960.14 | EIF4A1 | 0.247124707402589 | 5.47184751985125 | 1.29927329964829 | 0.200854577885591 | 0.520248578137966 | -5.57441287357678 |
| lnc-RBBP7-3 | lnc-RBBP7-3 | -0.326993675931448 | 1.38500599969545 | -1.29924105095921 | 0.200865541974562 | 0.520248578137966 | -5.1828819655734 |
| ENSG00000138463.8 | SLC49A4 | 0.189206770635474 | 3.53100666835321 | 1.29915846561748 | 0.200893621861138 | 0.520255840307977 | -5.42178913272564 |
| ENSG00000171503.11 | ETFDH | 0.211832076590649 | 3.48076233596576 | 1.29857946328385 | 0.201090572056373 | 0.520700370441102 | -5.41859844421801 |
| ENSG00000162144.9 | CYB561A3 | -0.285562353048291 | 7.85808777709226 | -1.29806248591677 | 0.201273666611174 | 0.521086634377214 | -5.57299865169704 |
| ENSG00000111913.18 | RIPOR2 | 0.219747561993902 | 8.96611658038557 | 1.29791966080452 | 0.201315184564329 | 0.521086634377214 | -5.54926433658788 |

| | | | | | | | |
|---|---|---|---|---|---|---|---|
| ENSG00000264772.6 | | 0.157892424312682 | 6.29314470541024 | 1.29791816682151 | 0.20131569336674 | 0.521086634377214 | -5.58824962708572 |
| ENSG00000143164.15 | DCAF6 | -0.102770106251226 | 5.36576847645126 | -1.29775525228144 | 0.201371182630941 | 0.521164724266649 | -5.57374816965702 |
| lnc-MYOM1-4 | lnc-MYOM1-4 | -0.203814187212239 | 2.71796880981532 | -1.29764675804849 | 0.201408142555268 | 0.521194845136418 | -5.33812647602414 |
| ENSG00000236104.2 | ZBTB22 | -0.21980783307062 | 3.88557852300552 | -1.29754019306743 | 0.201444450243137 | 0.521223270817661 | -5.47574869063575 |
| ENSG00000099783.11 | HNRNPM | 0.13431878727029 | 6.75633829381831 | 1.2974477052993 | 0.201475965699287 | 0.521239291350984 | -5.58729405077951 |
| ENSG00000104894.11 | CD37 | -0.196793881890084 | 8.8028517014116 | -1.2973155744572 | 0.201520996099333 | 0.52129026802973 | -5.55191519640802 |
| lnc-A1BG-2 | lnc-A1BG-2 | -0.256117166636407 | 1.01589783215413 | -1.29705896519807 | 0.201608470654329 | 0.521410944835508 | -5.123456591701 |
| ENSG00000165923.15 | AGBL2 | -0.196714949662688 | 3.10537699512645 | -1.29698851191355 | 0.201632492222815 | 0.521410944835508 | -5.38172014474178 |
| ENSG00000130810.19 | PPAN | -0.201940387754998 | 2.85005438806778 | -1.29695580732995 | 0.201643643826976 | 0.521410944835508 | -5.36295069319747 |
| ENSG00000030582.17 | GRN | -0.363160815618153 | 6.48671764197148 | -1.29704575601872 | 0.201717330778978 | 0.521502902464119 | -5.58793949237884 |
| lnc-ZNF726-3 | lnc-ZNF726-3 | -0.818206750592378 | 0.216758875359808 | -1.29700905312745 | 0.201729879484007 | 0.521502902464119 | -4.90930748212044 |
| lnc-MED8-4 | lnc-MED8-4 | 0.365124489585818 | 0.674884435073634 | 1.29649980922277 | 0.201799178514973 | 0.521616537909542 | -5.04231015808789 |
| lnc-ARVCF-3 | lnc-ARVCF-3 | -0.200305430220242 | -0.156293150051783 | -1.29609702571297 | 0.201936637759821 | 0.521906305249925 | -4.88716606368256 |
| ENSG00000160256.12 | SLX9 | -0.274430876345907 | 1.78562196576142 | -1.29598351487961 | 0.201975388737231 | 0.521940919821521 | -5.23794373164406 |
| ENSG00000153790.11 | SPMIP4 | 0.214983416525383 | 2.71771985668851 | 1.29551209137435 | 0.20213638605983 | 0.522229906617742 | -5.33261350394167 |
| ENSG00000163703.17 | CRELD1 | -0.242999381572384 | 3.47453997909452 | -1.29547307465962 | 0.202149715119677 | 0.522229906617742 | -5.42561348679948 |
| ENSG00000120656.11 | TAF12 | -0.184545425436893 | 3.21827410239812 | -1.2954332126983 | 0.202163333621443 | 0.522229906617742 | -5.39728172161063 |
| ENSG00000279700.1 | | 0.520928049282979 | -0.227796362624664 | 1.29534680244782 | 0.202192857329833 | 0.522240630244693 | -4.84586929618542 |
| CAPN10-DT | CAPN10-DT | -0.279520360411594 | 1.92863898916386 | -1.29524126141518 | 0.202228921848949 | 0.522268243214167 | -5.26072828041018 |
| ENSG00000180229.12 | HERC2P3 | -0.708999056574343 | 1.82278788681891 | -1.29501053690046 | 0.202496408248316 | 0.522880177804174 | -5.21205574031095 |
| lnc-KLRB1-1 | lnc-KLRB1-1 | -0.393903309064759 | -0.320555249375567 | -1.294332804005 | 0.202539552401481 | 0.522880177804174 | -4.87753930825124 |
| ENSG00000200959.1 | SNORA74A | 0.603776216684465 | 2.04222858515069 | 1.2947741137325 | 0.202542081214239 | 0.522880177804174 | -5.23817919771287 |
| lnc-CC2D2B-2 | lnc-CC2D2B-2 | 0.310962545961021 | 0.214280616534301 | 1.29417328625125 | 0.202594133745976 | 0.522925609009311 | -4.96129718627566 |
| ENSG00000284012.1 | MIR6840 | 0.317021938260172 | -0.53190867981028 | 1.29412548798525 | 0.202610490786049 | 0.522925609009311 | -4.80634443584368 |
| ENSG00000133256.12 | PDE6B | -0.211421564954063 | 3.02920030803957 | -1.29377445644475 | 0.202730647786125 | 0.52305925854586 | -5.38451918524257 |
| ENSG00000147133.15 | TAF1 | 0.153764291279407 | 6.81100362593516 | 1.29370730671014 | 0.202753639054624 | 0.52305925854586 | -5.59181945623311 |
| lnc-AP3D1-1 | lnc-AP3D1-1 | -0.165063596785226 | 2.81545979898698 | -1.29362803726918 | 0.20278078249755 | 0.52305925854586 | -5.3581677232303 |
| ENSG00000213619.9 | NDUFS3 | 0.117519672146444 | 4.63099472392939 | 1.29360519573799 | 0.202788604403655 | 0.52305925854586 | -5.54213822943017 |
| ENSG00000277791.4 | PSMB3 | 0.195073897101437 | 4.81615140832727 | 1.29360306215799 | 0.202789335043286 | 0.52305925854586 | -5.55294648148274 |
| lnc-ZFYVE21-1 | lnc-ZFYVE21-1 | 0.227718627759365 | 1.62905540189946 | 1.29350371253933 | 0.202823359293996 | 0.523068096226392 | -5.21623092469011 |
| lnc-ARSI-2 | lnc-ARSI-2 | 0.747209314055423 | -0.0420718781578468 | 1.29350089382145 | 0.202843586653018 | 0.523068096226392 | -4.82812896408873 |
| ENSG00000182372.9 | CLN8 | 0.445885663578395 | 3.88000474118819 | 1.29374309243987 | 0.202882071110642 | 0.523101800033864 | -5.47595280719093 |
| lnc-PABPC1-3 | lnc-PABPC1-3 | 0.315509492925585 | 4.75927892550711 | 1.29318353322385 | 0.202959971679122 | 0.523237111356633 | -5.54846353563421 |
| lnc-ZNF649-3 | lnc-ZNF649-3 | -0.298422107316144 | 2.95972160839439 | -1.29293333299857 | 0.203018780433069 | 0.523244675324276 | -5.36996647529454 |
| ENSG00000203867.7 | RBM20 | -2.77437650126686 | -0.917574104405348 | -1.29369864174571 | 0.203022516687699 | 0.523244675324276 | -4.62738704179319 |
| ENSG00000152102.17 | FAM168B | 0.110612850362987 | 6.25914725669035 | 1.29281727744008 | 0.20305856030213 | 0.523244675324276 | -5.59456103818404 |
| KLF7-IT1 | KLF7-IT1 | -0.695124493336892 | 0.611069744789421 | -1.29313296371367 | 0.203064590511117 | 0.523244675324276 | -5.05169405785086 |
| SNHG11 | SNHG11 | -0.15419745927248 | 2.756042129245 | -1.29254172218234 | 0.20315303475157 | 0.523407048978792 | -5.3534930426377 |
| ENSG00000109805.9 | NCAPG | 0.604193809698915 | -0.453616556259023 | 1.29228570454682 | 0.20324084040568 | 0.523461248083024 | -4.83523090012258 |
| USP30-AS1 | USP30-AS1 | -0.202172912120246 | 2.08741243043599 | -1.29225912635299 | 0.203249957494065 | 0.523461248083024 | -5.27575017672249 |
| ENSG00000130958.12 | SLC35D2 | 0.233042910427861 | 3.76640226974201 | 1.29225550086598 | 0.20325120116476 | 0.523461248083024 | -5.45827925635577 |
| lnc-KIAA1614-3 | lnc-KIAA1614-3 | 0.274745922690058 | -0.325190919757552 | 1.2921838000246 | 0.203275798276617 | 0.523461248083024 | -4.84998394034639 |
| ENSG00000157933.9 | SKI | 0.21674162323905 | 4.78962182919329 | 1.29205218603608 | 0.203320954550462 | 0.523482463235897 | -5.54732519979585 |
| LINC00265 | LINC00265 | 0.187082312924326 | 2.6232641924724 | 1.2919802397605 | 0.20334564224933 | 0.523482463235897 | -5.32785423243721 |
| ENSG00000207165.1 | SNORA70 | 0.29777511233678 | 2.70945780523353 | 1.291937424346 | 0.203360335035429 | 0.523482463235897 | -5.33693836164657 |
| ENSG00000153395.9 | LPCAT1 | 0.281373810358588 | 5.13220451019327 | 1.29167133447381 | 0.20345187783299 | 0.52365261991202 | -5.56552854008434 |
| ENSG00000102753.9 | KPNA3 | 0.254197473949577 | 4.28079869044007 | 1.29152636122125 | 0.203501438578467 | 0.523704911485445 | -5.51960518625134 |
| ENSG00000186376.14 | ZNF75D | 0.126187377928709 | 4.29278064745235 | 1.29141033429455 | 0.203541279904209 | 0.523704911485445 | -5.51625657810351 |
| ENSG00000165699.14 | TSC1 | 0.128258596181169 | 6.7418399036852 | 1.2913892365255 | 0.203548525087867 | 0.523704911485445 | -5.59484770664782 |
| ENSG00000072518.20 | MARK2 | -0.178970204125315 | 4.79462947041382 | -1.29107968137688 | 0.203654851758147 | 0.523912987593794 | -5.55641409946374 |
| ENSG00000149532.15 | CPSF7 | -0.145470438857703 | 6.18854794002404 | -1.29067332210325 | 0.203794492463532 | 0.524143445536548 | -5.59666233513285 |
| ENSG00000173085.14 | COQ2 | 0.297257819732413 | 2.11818062032844 | 1.29067078409936 | 0.203795364846183 | 0.524143445536548 | -5.26170743826061 |
| ENSG00000103148.15 | NPRL3 | -0.204922615029268 | 3.22784545311121 | -1.29056880949162 | 0.203830418688578 | 0.524168104431161 | -5.40596482685915 |
| ENSG00000258186.2 | SLC7A5P2 | -0.301496783708747 | 2.69346338804459 | -1.29049315869082 | 0.20385642663994 | 0.52416949775514 | -5.36952787558896 |
| lnc-ITGAE-2 | lnc-ITGAE-2 | 0.143843801257404 | 2.35517123874783 | 1.29034414751967 | 0.203907662431482 | 0.524220573689245 | -5.31290827458096 |
| ENSG00000164305.18 | CASP3 | 0.31281055102752 | 5.3374830761867 | 1.2903415288454 | 0.20394484551037 | 0.524220573689245 | -5.579625585647 |
| ENSG00000070831.15 | CDC42 | 0.146867800928851 | 7.24299166532262 | 1.29021319608619 | 0.203952696599304 | 0.524220573689245 | -5.59101828612336 |
| ENSG00000104472.9 | CHRAC1 | 0.223520365422809 | 4.45731843928155 | 1.28995223755817 | 0.204042462527234 | 0.52425473608942 | -5.53225682178351 |
| ENSG00000268521.1 | VN1R83P | -0.227173888864221 | 0.220891419044495 | -1.2898809850222 | 0.20406697753389 | 0.52425473608942 | -4.99211430613146 |
| lnc-ZNF708-4 | lnc-ZNF708-4 | -0.227173888864221 | 0.220891419044495 | -1.2898809850222 | 0.20406697753389 | 0.52425473608942 | -4.99211430613146 |
| ENSG00000126091.20 | ST3GAL3 | -0.206737469193213 | 3.29989777613889 | -1.28987839440302 | 0.204067868899015 | 0.52425473608942 | -5.4139070767918 |
| lnc-SLC38A7-4 | lnc-SLC38A7-4 | -0.11561859928535 | 2.30927235599926 | -1.28964373055518 | 0.204148622835511 | 0.524390995782429 | -5.30171430447867 |
| lnc-ASXL1-1 | lnc-ASXL1-1 | -0.355633369992116 | -0.189012692631432 | -1.28948072988359 | 0.204204729777625 | 0.524390995782429 | -4.86872331256519 |
| ENSG00000109111.14 | SUPT6H | 0.143884450434521 | 6.55946910376584 | 1.28948060855331 | 0.204204771545406 | 0.524390995782429 | -5.5984277876891 |
| lnc-RPL3-2 | lnc-RPL3-2 | 0.170128762271762 | 3.28535739282617 | 1.28941978821253 | 0.204225709673579 | 0.524390995782429 | -5.41547002359873 |
| ENSG00000284525.1 | MIR4687 | 0.311262881454197 | -0.0673880043652587 | 1.28935419445367 | 0.204248292920747 | 0.524390995782429 | -4.91905501281803 |
| lnc-PSPC1-1 | lnc-PSPC1-1 | 0.188009962264297 | 2.40161671895177 | 1.28914806742608 | 0.204319272552735 | 0.524507805806054 | -5.31144776342428 |
| ENSG00000285756.1 | | -0.390501717659587 | 1.42789838395522 | -1.28902393998624 | 0.20436202467375 | 0.524552132916797 | -5.20579001334782 |

| | | | | | | | |
|---|---|---|---|---|---|---|---|
| ENSG00000271254.6 | | -0.335388182234484 | 3.02164415680395 | -1.28891252848874 | 0.204400402893133 | 0.524585223534832 | -5.3775926566911 |
| ENSG00000172007.5 | RAB33B | 0.155779758790767 | 4.35576939683736 | 1.28875687275344 | 0.20445403113566 | 0.524654024391759 | -5.52990399892695 |
| ENSG00000198807.12 | PAX9 | -1.56698458328749 | -1.07278520868651 | -1.28933974154634 | 0.204478189946591 | 0.524654024391759 | -4.75835491357069 |
| lnc-HP1BP3-2 | lnc-HP1BP3-2 | -0.207680582948561 | 2.03034929664102 | -1.28848437244381 | 0.204547941486259 | 0.524767578164237 | -5.27537995922101 |
| ENSG00000197763.15 | TXNRD3 | 1.75531317841691 | -1.86912792411124 | 1.28853044017115 | 0.204580616538369 | 0.524785995788789 | -4.53804774640604 |
| ENSG00000111676.14 | ATN1 | -0.329402443687324 | 3.73012761749223 | -1.28804713612236 | 0.204713117992168 | 0.525060449549257 | -5.4725301075435 |
| lnc-C1QTNF3-AMACR-3 | lnc-C1QTNF3-AMA | -0.334111853121928 | 1.57549510941214 | -1.28775856111232 | 0.204798232543024 | 0.525187752598117 | -5.20487494236348 |
| lnc-CA5B-1 | lnc-CA5B-1 | 0.203637741087976 | 0.898469959879613 | 1.28771348962466 | 0.204813782738429 | 0.525187752598117 | -5.12459383855117 |
| lnc-SLC4A5-5 | lnc-SLC4A5-5 | -0.202335147098924 | 2.2844573203576 | -1.28747621130822 | 0.204895661220511 | 0.525332261447655 | -5.30477709651306 |
| ENSG00000174776.11 | WDR49 | 0.588918497441146 | 0.112088015128798 | 1.28723167324641 | 0.204980070635907 | 0.525378468793451 | -4.94217342321213 |
| ENSG00000170275.14 | CRTAP | -0.164660733803124 | 4.92520725731207 | -1.28710268840553 | 0.205024604046196 | 0.525378468793451 | -5.56667446739581 |
| lnc-DRD5-4 | lnc-DRD5-4 | 0.605142809464342 | 2.04633456919236 | 1.2875059807341 | 0.205038150692246 | 0.525378468793451 | -5.23931631064313 |
| ENSG00000169519.20 | METTL15 | -0.130249920929765 | 4.0386598083565 | -1.28702977929337 | 0.205049779927373 | 0.525378468793451 | -5.50120759480306 |
| lnc-NCOA3-14 | lnc-NCOA3-14 | 0.303247528078135 | 4.81508711389401 | 1.28704070359177 | 0.205064530563551 | 0.525378468793451 | -5.55741460704697 |
| lnc-PRAG1-3 | lnc-PRAG1-3 | -0.669137012051441 | -0.367649034235522 | -1.28703493197559 | 0.205074856069843 | 0.525378468793451 | -4.92842836740567 |
| ENSG00000259577.1 | CERNA1 | -0.24596206934983 | 1.96273626548029 | -1.28690649005575 | 0.20509235761334 | 0.525378468793451 | -5.26097454402202 |
| ENSG00000102189.16 | EEA1 | -0.187287115032554 | 5.1483482068793 | -1.2867933768941 | 0.205131426866524 | 0.525413160676166 | -5.57907918889853 |
| ENSG00000139880.19 | CDH24 | -0.434211362583768 | 1.46408560019553 | -1.2867252316139 | 0.205162049506562 | 0.525426211894186 | -5.20947069046484 |
| ENSG00000198818.9 | SFT2D1 | -0.166864254877201 | 3.90081924310064 | -1.28652571560057 | 0.20522389931467 | 0.525519223636955 | -5.49893779355152 |
| lnc-EPHX3-7 | lnc-EPHX3-7 | -0.38396165594396 | 0.289392856163258 | -1.28641316677484 | 0.205262792399861 | 0.525553434005018 | -4.96391319794863 |
| ENSG00000112996.10 | MRPS30 | -0.114995902792466 | 4.68508029711983 | -1.2861242459338 | 0.205362659118017 | 0.525652226702551 | -5.55671706622642 |
| ENSG00000234936.1 | NA | -0.283973394919849 | 1.99943379732165 | -1.28610032642123 | 0.205370928638197 | 0.525652226702551 | -5.28045596708379 |
| ENSG00000135968.20 | GCC2 | 0.189434041871579 | 6.75880638449087 | 1.28607989659324 | 0.205377991893403 | 0.525652226702551 | -5.60154323990024 |
| ENSG00000106144.19 | CASP2 | 0.135670578578197 | 4.70735418249612 | 1.28597213401272 | 0.205415251943994 | 0.52568222438931 | -5.55239144578304 |
| lnc-CLEC4C-1 | lnc-CLEC4C-1 | 1.63626199827131 | 1.93149918699347 | 1.28661288082935 | 0.205469515671164 | 0.52569668104274 | -5.14152908589343 |
| ENSG00000126456.15 | IRF3 | -0.155974107495782 | 5.7231209885758 | -1.28580808993569 | 0.205471981683372 | 0.52569668104274 | -5.59800889630811 |
| ENSG00000268750.6 | | 0.126674464334132 | 2.26629618356218 | 1.28562038178735 | 0.205536909398326 | 0.52578106020156 | -5.3019297979058 |
| lnc-UCP3-7 | lnc-UCP3-7 | 0.15850161876943 | 0.878040259534668 | 1.28549575792655 | 0.205580024965096 | 0.52578106020156 | -5.10638945786345 |
| ENSG00000134419.15 | RPS15A | 0.179630144455654 | 8.24500637295613 | 1.28528000270554 | 0.205654684936744 | 0.52578106020156 | -5.5801534251783 |
| lnc-MCM4-7 | lnc-MCM4-7 | -0.363213292716948 | 0.208207075976831 | -1.28524271235906 | 0.205667590961918 | 0.52578106020156 | -4.96521455489369 |
| lnc-GLDN-5 | lnc-GLDN-5 | 0.262406682117733 | 0.899374835942822 | 1.28516227424547 | 0.205695432314974 | 0.52578106020156 | -5.11281145393375 |
| ENSG00000125834.12 | STK35 | 0.0861637338953489 | 4.57106511597956 | 1.28516026009761 | 0.205696129491059 | 0.52578106020156 | -5.54726525568628 |
| ENSG00000269713.7 | NBPF9 | 0.155691326241953 | 4.2427520679987 | 1.28512433508912 | 0.20570856485333 | 0.52578106020156 | -5.5190524116255 |
| ENSG00000142208.16 | AKT1 | -0.134290094895137 | 5.67723474406033 | -1.28512216139434 | 0.205709317291122 | 0.52578106020156 | -5.59887144714449 |
| ENSG00000168118.11 | RAB4A | 0.211782635767479 | 3.93204881594429 | 1.28487354085316 | 0.205795392482429 | 0.525877189245181 | -5.48777764820531 |
| lnc-EMC3-3 | lnc-EMC3-3 | 0.167645745620906 | 2.18637423584308 | 1.28486593677861 | 0.205798025524459 | 0.525877189245181 | -5.30324993629127 |
| ENSG00000204922.4 | UQCC3 | -0.247577951377608 | 1.26089843325999 | -1.28471670169804 | 0.205849705876489 | 0.525943954326684 | -5.19097123518207 |
| lnc-EPHX3-2 | lnc-EPHX3-2 | -0.294827557016886 | 0.0744342723151996 | -1.28455585803714 | 0.205905417230932 | 0.526021000727848 | -4.96102184101273 |
| ENSG00000076924.11 | XAB2 | 0.147573787761092 | 5.74408283251309 | 1.28442411435117 | 0.205951057688532 | 0.526072303351087 | -5.59995286284277 |
| ENSG00000230225.1 | MTND5P14 | -0.420734291256662 | -0.0293207357844746 | -1.28423876589443 | 0.206015281531952 | 0.526171055934007 | -4.91631606993908 |
| ENSG00000198416.9 | ZNF658B | -0.431279317149571 | -0.219518667404407 | -1.28401650574959 | 0.206092315258539 | 0.526212363459654 | -4.83860523777401 |
| ENSG00000068724.15 | TTC7A | 0.179892447045471 | 6.45303663081028 | 1.28392285433738 | 0.206124780634411 | 0.526212363459654 | -5.60537329406411 |
| ENSG00000278195.1 | SSTR3 | -0.488418932700759 | 2.61537480710176 | -1.28415097727348 | 0.206159307874417 | 0.526212363459654 | -5.34636827676218 |
| lnc-RBMX-1 | lnc-RBMX-1 | -0.287198238350755 | 0.86690906168347 | -1.28378105205869 | 0.206173945402445 | 0.526212363459654 | -5.13744721142215 |
| ENSG00000221887.5 | HMSD | -0.277564109773471 | 0.89182774680244 | -1.28377397385627 | 0.206176399742059 | 0.526212363459654 | -5.12328702648923 |
| ENSG00000090905.18 | TNRC6A | 0.151027460565515 | 6.97424081123503 | 1.28374961206707 | 0.206184847267265 | 0.526212363459654 | -5.6024825758154 |
| lnc-AKR7A3-1 | lnc-AKR7A3-1 | -0.187721442475954 | 2.16351803397986 | -1.28361285006064 | 0.206232274745592 | 0.526268151635076 | -5.29185550357931 |
| lnc-TAP1-1 | lnc-TAP1-1 | -0.35616701740749 | 0.472860579979303 | -1.28349955184378 | 0.206271571467461 | 0.526303180304295 | -5.08052905293364 |
| ENSG00000196405.12 | EVL | -0.169174803393646 | 7.94346821551103 | -1.28332453477855 | 0.206332286023302 | 0.526341883883473 | -5.58861002852621 |
| ENSG00000185000.11 | DGAT1 | 0.193507793146324 | 3.55494809210422 | 1.28330840062691 | 0.206337883741716 | 0.526341883883473 | -5.4475981424581 |
| ENSG00000142731.10 | PLK4 | 0.24648149672878 | 3.04668889605982 | 1.28315476599134 | 0.20639119275736 | 0.526363105263641 | -5.40366443361699 |
| lnc-FLCN-3 | lnc-FLCN-3 | 0.170686147141146 | 2.10873605440517 | 1.28301096069914 | 0.206441100515384 | 0.526363105263641 | -5.29145192115231 |
| lnc-SOCS2-4 | lnc-SOCS2-4 | 0.212459129800511 | 1.53804320905613 | 1.28300770934911 | 0.20644222900428 | 0.526363105263641 | -5.23199044468229 |
| lnc-TMEM101-1 | lnc-TMEM101-1 | 0.21199223817939 | 0.450150526788918 | 1.28298747013456 | 0.20644925379946 | 0.526363105263641 | -5.05005495279795 |
| lnc-PRSS2-6 | lnc-PRSS2-6 | -0.236296991562853 | 5.5109880274156 | -1.28283310387744 | 0.206502838439536 | 0.526363105263641 | -5.59502709842554 |
| ENSG00000175445.15 | LPL | -1.62580697962468 | 2.61609303959856 | -1.28357459104919 | 0.206520886599024 | 0.526363105263641 | -5.3028964914114 |
| ENSG00000090060.17 | PAPOLA | 0.105423555095748 | 7.34066079902517 | 1.28270109018735 | 0.20654867220342 | 0.526363105263641 | -5.59861046936529 |
| ENSG00000162104.9 | ADCY9 | 1.22338503599474 | -0.973606048122649 | 1.28300327310901 | 0.206631672650641 | 0.526363105263641 | -4.81662359446282 |
| ENSG00000101361.16 | NOP56 | 0.164326664954742 | 5.86758190866699 | 1.28241886987007 | 0.206646681781099 | 0.526363105263641 | -5.60516403669189 |
| ZNF674-AS1 | ZNF674-AS1 | 0.175623370544186 | 1.99101020334553 | 1.28239534094789 | 0.206654854492784 | 0.526363105263641 | -5.26934070850056 |
| ENSG00000038382.19 | TRIO | 0.889197332975582 | 6.50062105435258 | 1.28318776045294 | 0.206655035786918 | 0.526363105263641 | -5.60509575577708 |
| ENSG00000172432.18 | GTPBP2 | -0.128228061614641 | 5.52111356932242 | -1.28233848730234 | 0.206674603465941 | 0.526363105263641 | -5.59748000701229 |
| lnc-PALLD-8 | lnc-PALLD-8 | -0.250669739456132 | 1.67713181643419 | -1.28232684364658 | 0.206678648240818 | 0.526363105263641 | -5.25138180124474 |
| lnc-ERP44-3 | lnc-ERP44-3 | -0.329703719766971 | 1.44942591325725 | -1.28212720314515 | 0.206748008646039 | 0.526422713173985 | -5.20434055587982 |
| lnc-EIF2AK4-4 | lnc-EIF2AK4-4 | -0.27080903085627 | 1.38823352687611 | -1.28209530700277 | 0.206759091832878 | 0.526422713173985 | -5.20772993259321 |
| ENSG00000147813.15 | NAPRT | -0.340741278313778 | 3.01865110900028 | -1.28175848908064 | 0.206877844898605 | 0.526422713173985 | -5.40246836414696 |
| lnc-ST8SIA1-6 | lnc-ST8SIA1-6 | 0.166355890690923 | 2.14803946227801 | 1.28174694734528 | 0.206880168000024 | 0.526422713173985 | -5.2975048775647 |
| ENSG00000119927.13 | GPAM | 0.139554823195948 | 3.83118088131996 | 1.28160716692311 | 0.206928765177394 | 0.526422713173985 | -5.48386419366961 |

| | | | | | | | |
|---|---|---|---|---|---|---|---|
| lnc-TACSTD2-4 | lnc-TACSTD2-4 | 0.20477879640375 | 2.05376328974738 | 1.28155286109486 | 0.206947647889307 | 0.526422713173985 | -5.27364937633024 |
| ENSG00000142733.15 | MAP3K6 | 0.45459787260797 | 0.962351091486483 | 1.28153994945197 | 0.206952137594913 | 0.526422713173985 | -5.1255596970459 |
| ENSG00000158710.14 | TAGLN2 | 0.21789350141239 | 5.99615040313347 | 1.28153111639744 | 0.20695520911411 | 0.526422713173985 | -5.60759593384681 |
| ENSG00000164983.7 | TMEM65 | -0.339968607636772 | 4.15807686729741 | -1.28156911607349 | 0.206989512432908 | 0.526422713173985 | -5.53226791093514 |
| lnc-LRRC31-1 | lnc-LRRC31-1 | -0.481728863572972 | 0.140699970924326 | -1.28141973255092 | 0.206993943573657 | 0.526422713173985 | -4.99292962089144 |
| ENSG00000142657.20 | PGD | -0.26556507935691 | 4.16453563777916 | -1.28138135554397 | 0.207007290690306 | 0.526422713173985 | -5.52924124280337 |
| lnc-SERPINB9-3 | lnc-SERPINB9-3 | 0.314943030299105 | 3.79366741875364 | 1.28137655356352 | 0.20700896081379 | 0.526422713173985 | -5.46347707421423 |
| ENSG00000268614.1 | | -0.202796054387778 | 1.15584242173192 | -1.28123765859535 | 0.207057272712804 | 0.526480524304837 | -5.17485418108876 |
| ENSG00000240972.1 | MIF | -0.359816712942762 | 2.97413924198446 | -1.28109599344245 | 0.20712523894244 | 0.526588289668013 | -5.40652780268531 |
| lnc-ANKRD35-2 | lnc-ANKRD35-2 | -0.259930360771922 | -0.343193517006436 | -1.28089100034596 | 0.207177887972927 | 0.526657091286497 | -4.88505337486703 |
| lnc-RAP2C-3 | lnc-RAP2C-3 | -0.183271480737802 | 1.8124445922196 | -1.28076478607371 | 0.207221815707472 | 0.526694340056781 | -5.24832586200693 |
| ENSG00000148175.12 | STOM | 0.657519207999588 | 1.25829599731152 | 1.28109601087571 | 0.207260486341997 | 0.526694340056781 | -5.18288041571327 |
| ENSG00000204386.10 | NEU1 | -0.257860037578626 | 4.15087769106166 | -1.28059750468693 | 0.207280047264106 | 0.526694340056781 | -5.51831888275473 |
| lnc-ISG20-1 | lnc-ISG20-1 | 0.349006568759768 | -0.0128746813914271 | 1.28055485385874 | 0.207294896215322 | 0.526694340056781 | -4.9015949618881 |
| lnc-TNFRSF13C-1 | lnc-TNFRSF13C-1 | 0.232535158349721 | 4.37124752401151 | 1.28014559679297 | 0.207437420323377 | 0.526976701476223 | -5.53161655427548 |
| ENSG00000238390.1 | | -0.293729417938395 | -0.148923569879494 | -1.28008872360637 | 0.207457232282069 | 0.526976701476223 | -4.89274872024795 |
| lnc-MTPAP-1 | lnc-MTPAP-1 | -0.224607388347587 | 0.219813488722007 | -1.27992217695587 | 0.207515257523784 | 0.526998682854264 | -5.01523938086756 |
| lnc-CCNJ-1 | lnc-CCNJ-1 | -0.188271728195555 | 3.55783756708354 | -1.2798885023602 | 0.20752699131319 | 0.526998682854264 | -5.45984021006483 |
| ENSG00000053372.4 | MRTO4 | -0.209811525119162 | 3.66130491447756 | -1.27984343239809 | 0.207542696556219 | 0.526998682854264 | -5.47129732514284 |
| ENSG00000196236.12 | XPNPEP3 | -0.088927131460804 | 4.48630192742021 | -1.27961754544688 | 0.207621423397085 | 0.52713355837945 | -5.54454693131281 |
| ENSG00000100360.14 | IFT27 | -0.171188511082605 | 3.95503739572681 | -1.27938692260266 | 0.207701823947312 | 0.527178559251002 | -5.49917282801521 |
| lnc-PRPH-3 | lnc-PRPH-3 | -0.372059913940465 | -0.580763965751002 | -1.27937442594442 | 0.207706181244274 | 0.527178559251002 | -4.81914381568249 |
| ENSG00000196470.11 | SIAH1 | 0.206462593883057 | 5.93625534204576 | 1.27934631016881 | 0.207715984818813 | 0.527178559251002 | -5.60907909898433 |
| ENSG00000188846.13 | RPL14 | 0.167107871551279 | 8.19498423006204 | 1.27925965454725 | 0.207746202598625 | 0.527190246342928 | -5.58725236780489 |
| lnc-PPP1R27-2 | lnc-PPP1R27-2 | -0.263935823349844 | 1.53751329673333 | -1.27890618678284 | 0.207869494944796 | 0.527438093497131 | -5.24001960841478 |
| ENSG00000142252.10 | GEMIN7 | -0.19843601200788 | 3.70404695442949 | -1.27865289309289 | 0.207957879576091 | 0.527490624953976 | -5.47270945151061 |
| ENSG00000255561.6 | FDXACB1 | 0.279226637281403 | 1.58706116841527 | 1.27861559020491 | 0.207970898478373 | 0.527490624953976 | -5.23058572102545 |
| lnc-ARFIP1-2 | lnc-ARFIP1-2 | 0.888296360193471 | -0.251230256500327 | 1.2787027301817 | 0.207983768363445 | 0.527490624953976 | -4.77619380625528 |
| ENSG00000136152.14 | COG3 | 0.146842552666417 | 5.59464397607908 | 1.27855310313309 | 0.207992708163361 | 0.527490624953976 | -5.60623738186279 |
| ENSG00000203710.11 | CR1 | 1.38414130259218 | 0.917560624957112 | 1.27915016723561 | 0.208048504132643 | 0.527567125854651 | -5.06511757717663 |
| ENSG00000165525.17 | NEMF | 0.137295224956707 | 6.97667969256354 | 1.27823762036346 | 0.208102846443006 | 0.527570333614244 | -5.60921533374965 |
| ENSG00000127952.16 | STYXL1 | -0.182259434364469 | 2.94485957346769 | -1.27817672506961 | 0.208124110657624 | 0.527570333614244 | -5.38361957480775 |
| ENSG00000063854.12 | HAGH | -0.127965087191797 | 3.64594661061975 | -1.27813961065841 | 0.208137071552971 | 0.527570333614244 | -5.4741989214319 |
| LINC00996 | LINC00996 | -1.03464614192469 | 1.03101999121298 | -1.27878439120511 | 0.208156082966201 | 0.527570333614244 | -5.10386071747625 |
| ENSG00000165113.12 | GKAP1 | 0.206211816133261 | 2.54885624019016 | 1.27799628584249 | 0.208187128364346 | 0.527570333614244 | -5.34209656029612 |
| ENSG00000139187.9 | KLRG1 | -0.300697020486817 | -0.0105478117172731 | -1.27794925144073 | 0.20820355730207 | 0.527570333614244 | -4.95783843495729 |
| ENSG00000162676.11 | GFI1 | -0.556139759579844 | 0.699725705047255 | -1.27779932868392 | 0.208320067504798 | 0.52769407631365 | -5.14065639280104 |
| FOXD2-AS1 | FOXD2-AS1 | -0.619508366780998 | -0.856713355212928 | -1.2775617368797 | 0.208338951713245 | 0.52769407631365 | -4.80015259945668 |
| ENSG00000163946.13 | TASOR | -0.107808274909012 | 7.24065565368118 | -1.27755292387696 | 0.208342031672679 | 0.52769407631365 | -5.60635597341735 |
| ENSG00000055147.18 | FAM114A2 | -0.138198490508057 | 3.93481702790817 | -1.2774698896061 | 0.208371052085261 | 0.52769407631365 | -5.49914070200163 |
| ENSG00000166851.14 | PLK1 | 0.345114284403464 | 0.16366656388143 | 1.27744263347418 | 0.208380578743494 | 0.52769407631365 | -4.97815571001756 |
| ENSG00000136197.12 | C7orf25 | -0.198069514000754 | 1.17993464240648 | -1.27734885696111 | 0.208413358334877 | 0.52771216073401 | -5.1835015241964 |
| lnc-MYOCOS-2 | lnc-MYOCOS-2 | -0.540468431350921 | -0.67673821558569 | -1.27726459765422 | 0.208442814486946 | 0.527721826640197 | -4.81031183907239 |
| ENSG00000165506.14 | DNAAF2 | -0.214261375861138 | 3.04431179947713 | -1.27707789297338 | 0.208508095579453 | 0.527822178245219 | -5.41187245944581 |
| ENSG00000196821.9 | ILRUN | -0.155679319019709 | 7.48816161226419 | -1.27658467826652 | 0.208680621388782 | 0.528132161954163 | -5.60438015887238 |
| lnc-SCGN-6 | lnc-SCGN-6 | -0.594145465513456 | 0.313458696435792 | -1.27662877603799 | 0.208689897707245 | 0.528132161954163 | -5.01106729050248 |
| ENSG00000223612.3 | | 0.484326802698133 | 0.846627980060975 | 1.27650779159617 | 0.208707525887148 | 0.528132161954163 | -5.03683457166516 |
| ENSG00000271810.5 | | -0.336225734244514 | 2.64201617865406 | -1.27641515866837 | 0.208739943837646 | 0.528149264168441 | -5.34709870811804 |
| ENSG00000149476.15 | TKFC | -0.13004946254603 | 4.31784885095689 | -1.27618860513776 | 0.208819244753753 | 0.528269879269475 | -5.53621370004108 |
| ENSG00000241217.3 | RN7SL809P | 0.328727014546609 | -0.38382903904987 | 1.27613233345611 | 0.20883894513024 | 0.528269879269475 | -4.84038594985018 |
| ENSG00000095321.16 | CRAT | -0.462978286650723 | 2.11080529529553 | -1.27622597115451 | 0.208871441374423 | 0.528287156280383 | -5.30059112491251 |
| ENSG00000143933.16 | CALM2 | 0.153203756788481 | 7.56748160175319 | 1.27589097150627 | 0.208923460313499 | 0.52834603412689 | -5.60342185375593 |
| ENSG00000129317.14 | PUS7L | -0.167156224021499 | 5.7839915874675 | -1.27581441680555 | 0.208950272030349 | 0.52834603412689 | -5.61110414440476 |
| lnc-ZNF891-1 | lnc-ZNF891-1 | 0.202607989157833 | 0.954000273178222 | 1.27567229156624 | 0.20900005533144 | 0.52834603412689 | -5.14717925595562 |
| lnc-SLC17A6-3 | lnc-SLC17A6-3 | -0.265067596653527 | 0.0893444752257919 | -1.27559677455905 | 0.209026510874711 | 0.52834603412689 | -4.97501392835527 |
| ENSG00000215030.5 | RPL13P12 | 0.654230687176642 | 4.35991279700506 | 1.27618352542726 | 0.209061675290111 | 0.52834603412689 | -5.48830939380693 |
| lnc-STK38-1 | lnc-STK38-1 | 0.28618413001547 | 1.29466894622511 | 1.2754835877128 | 0.209066167832939 | 0.52834603412689 | -5.17190043006549 |
| ENSG00000186566.12 | GPATCH8 | 0.12452627603051 | 6.65946873979502 | 1.275460083816 | 0.209074403535127 | 0.52834603412689 | -5.61511796156114 |
| ENSG00000145919.10 | BOD1 | 0.138265872786071 | 4.18282598450162 | 1.2750715709498 | 0.209210572662553 | 0.528494121494384 | -5.52845646717792 |
| lnc-RNASET2-1 | lnc-RNASET2-1 | -0.203968445550996 | 4.18331670146192 | -1.27507126205462 | 0.209210680953094 | 0.528494121494384 | -5.51881814302902 |
| ENSG00000185658.13 | BRWD1 | 0.135866838886963 | 8.01704104126666 | 1.27496256476848 | 0.209248789978116 | 0.528494121494384 | -5.59756384023333 |
| ENSG00000174684.6 | B4GAT1 | -0.342667936884745 | 0.403406224286495 | -1.27490098706798 | 0.209270381292312 | 0.528494121494384 | -5.03842920921452 |
| ENSG00000169612.3 | RAMAC | 0.168181387523445 | 3.77585580823363 | 1.27489149082838 | 0.209273711157657 | 0.528494121494384 | -5.48594363336728 |
| ENSG00000135677.10 | GNS | 0.198501635330796 | 5.64939888498127 | 1.27484522410404 | 0.209289935196561 | 0.528494121494384 | -5.60968768598933 |
| ENSG00000114520.10 | SNX4 | 0.141809520520743 | 4.73463407435493 | 1.27478020209275 | 0.209312737619502 | 0.528494121494384 | -5.57353557643846 |
| ENSG00000163513.17 | TGFBR2 | 0.136336986898089 | 7.54448455095609 | 1.27447389105645 | 0.209420182214525 | 0.528671572826645 | -5.60634435796205 |
| ENSG00000196743.8 | GM2A | -0.294904299256195 | 6.78425443022199 | -1.27448937520384 | 0.209440759079242 | 0.528671572826645 | -5.61431571455211 |
| ENSG00000205710.3 | C17orf107 | 0.585361027145322 | 1.90744487131852 | 1.27475553417566 | 0.209460072700835 | 0.528671572826645 | -5.24428189088823 |

| | | | | | | | |
|---|---|---|---|---|---|---|---|
| lnc-ACTR1B-1 | lnc-ACTR1B-1 | -0.141297116101267 | 2.20944516228301 | -1.27407423996725 | 0.209560429846414 | 0.528860020540553 | -5.30908007503392 |
| lnc-CYTH1-1 | lnc-CYTH1-1 | 0.202340554847922 | 0.125813170886192 | 1.27398794415415 | 0.209590722464571 | 0.528870427702226 | -4.97190575708739 |
| ENSG00000184545.10 | DUSP8 | 0.822121075188789 | -0.0241765095532371 | 1.2742899304396 | 0.209615942729182 | 0.528870427702226 | -4.95967267600846 |
| ENSG00000119673.14 | ACOT2 | 0.207835957350795 | 1.9328293484352 | 1.27341422869139 | 0.209792198611927 | 0.52914324221216 | -5.27602896995525 |
| ENSG00000126353.3 | CCR7 | -0.233838351447718 | 7.92732033268488 | -1.27341417628536 | 0.209792217022392 | 0.52914324221216 | -5.60065544162917 |
| ENSG00000198931.10 | APRT | -0.238699231292889 | 3.76781114031154 | -1.27332636316379 | 0.209823067854805 | 0.52914324221216 | -5.48976011003169 |
| ENSG00000089351.14 | GRAMD1A | -0.143296445402612 | 3.83484735989722 | -1.27325109621295 | 0.209849513626845 | 0.52914324221216 | -5.50047173272628 |
| ENSG00000105393.15 | BABAM1 | -0.134692306479003 | 4.49918948504153 | -1.27324228212116 | 0.20985261070679 | 0.52914324221216 | -5.55816681474627 |
| ENSG00000169914.5 | OTUD3 | 0.115415072370319 | 4.43295014133185 | 1.27311582851302 | 0.209897047523994 | 0.529179095169636 | -5.54978633760339 |
| lnc-TMEM178A-1 | lnc-TMEM178A-1 | 1.24255099762882 | 0.585683544001612 | 1.27371446026132 | 0.209927110092381 | 0.529179095169636 | -4.96143751523562 |
| ENSG00000131848.9 | ZSCAN5A | -0.180754772433257 | 1.32174320296935 | -1.27288497951849 | 0.209978187914363 | 0.529179095169636 | -5.20422449233148 |
| ENSG00000180720.7 | CHRM4 | -0.259145510779238 | 0.0780130247927837 | -1.27285052893025 | 0.209990298859818 | 0.529179095169636 | -4.98837208401511 |
| ENSG00000204427.11 | ABHD16A | 0.140232295871967 | 3.7902754399423 | 1.27281765150692 | 0.210001857254697 | 0.529179095169636 | -5.48819764529661 |
| ENSG00000066777.8 | ARFGEF1 | 0.121705319116876 | 7.0357374632155 | 1.27270456125847 | 0.21004161893128 | 0.529179095169636 | -5.61540428245813 |
| ENSG00000103811.16 | CTSH | -0.331614611477606 | 7.3545707079358 | -1.2728415881131 | 0.210060532068161 | 0.529179095169636 | -5.60907524890405 |
| ENSG00000186517.13 | ARHGAP30 | -0.21383248361956 | 6.84614380768286 | -1.27260267922046 | 0.21007744471752 | 0.529179095169636 | -5.61680794826296 |
| lnc-CYS1-1 | lnc-CYS1-1 | 0.481754678999032 | 1.89867013912811 | 1.27269403139986 | 0.210110487014342 | 0.529179095169636 | -5.26865224599466 |
| ENSG00000213865.7 | NA | -0.171131641320566 | 2.73259135402563 | -1.2724705176116 | 0.210123924832213 | 0.529179095169636 | -5.37617844010964 |
| lnc-NAPRT-3 | lnc-NAPRT-3 | 0.163939110369215 | 4.18084306671955 | 1.27229105773721 | 0.210187051683812 | 0.529273315978456 | -5.52172999325285 |
| lnc-RBM48-2 | lnc-RBM48-2 | -0.26422620307593 | 0.673001636830698 | -1.27216631433563 | 0.210230939828138 | 0.529275261411265 | -5.1172772415261 |
| ENSG00000285901.1 | | 0.280902368179135 | 3.49705960374104 | 1.27200776374397 | 0.210286732179468 | 0.529275261411265 | -5.44597156193428 |
| ENSG00000007255.10 | TRAPPC6A | -0.221064066031321 | 3.19099040136492 | -1.27200105230901 | 0.210289094110396 | 0.529275261411265 | -5.42515498579186 |
| ENSG00000116213.15 | WRAP73 | 0.117925386529087 | 4.26919953410714 | 1.27199654307818 | 0.210290681038786 | 0.529275261411265 | -5.53002838639397 |
| ENSG00000170832.12 | USP32 | 0.169765069134225 | 5.13765085246867 | 1.27157578515485 | 0.210438797380473 | 0.529478839226448 | -5.59683249477328 |
| ENSG00000267809.1 | NDUFV2P1 | 0.354945728010739 | 0.431042737568901 | 1.27149567534701 | 0.210467006708153 | 0.529478839226448 | -5.00845321872306 |
| ENSG00000115267.6 | IFIH1 | -0.22702387476594 | 4.71632970282261 | -1.27147366798028 | 0.210474756730007 | 0.529478839226448 | -5.5790593358012 |
| ENSG00000259924.1 | | 0.345025931412148 | 3.10564280049188 | 1.27143995149951 | 0.210487783851385 | 0.529478839226448 | -5.39577633496504 |
| ENSG00000101189.6 | MRGBP | 0.224704789616445 | 4.21916707946764 | 1.27134666590732 | 0.2105194854244 | 0.529478839226448 | -5.53254420412433 |
| ENSG00000151413.16 | NUBPL | 0.195231538413557 | 3.9150590221073 | 1.27132842368404 | 0.210525910714145 | 0.529478839226448 | -5.50004524867654 |
| ENSG00000215424.9 | MCM3AP-AS1 | -0.31317950494694 | 0.0803662436814797 | -1.27118050681262 | 0.210578015533131 | 0.529544679227692 | -4.97622563170414 |
| ENSG00000155097.11 | ATP6V1C1 | 0.152705964957047 | 5.39865398936054 | 1.27110804604614 | 0.210603543903118 | 0.529544679227692 | -5.6066014192065 |
| ENSG00000169683.7 | LRRC45 | -0.245483582354938 | 2.3014124960025 | -1.27098897209016 | 0.210645499419725 | 0.529565533034083 | -5.33610216150103 |
| ENSG00000150967.17 | ABCB9 | -0.221578885973392 | 0.174813422626825 | -1.27090647630205 | 0.210674570349516 | 0.529565533034083 | -5.00493962600802 |
| ENSG00000240399.1 | | 0.234162191866836 | -0.0234393132656201 | 1.27076689856622 | 0.210723763396905 | 0.529565533034083 | -4.95232817857801 |
| lnc-COL2A1-3 | lnc-COL2A1-3 | 0.234162191866836 | -0.0234393132656201 | 1.27076689856622 | 0.210723763396905 | 0.529565533034083 | -4.95232817857801 |
| ENSG00000204308.7 | RNF5 | -0.231197080538775 | 3.44625132398278 | -1.27068264185125 | 0.210753463161408 | 0.529565533034083 | -5.46154617398625 |
| lnc-MYO18B-4 | lnc-MYO18B-4 | -0.331904337054585 | 0.146623504491388 | -1.27064450708819 | 0.21076690636686 | 0.529565533034083 | -4.96708287367061 |
| lnc-CCDC141-2 | lnc-CCDC141-2 | -0.613059542081185 | 1.24350515401196 | -1.27091269313522 | 0.210806322719204 | 0.529565533034083 | -5.19477623060605 |
| lnc-NAXD-4 | lnc-NAXD-4 | 0.243099272337219 | 1.93824923251464 | 1.27050053780083 | 0.210817663979074 | 0.529565533034083 | -5.28057268791968 |
| ENSG00000184271.17 | POU6F1 | 0.268212640581631 | 3.28904498881772 | 1.27035228756046 | 0.210869940446368 | 0.529568846090411 | -5.42843676140569 |
| ENSG00000225193.5 | RPS12P26 | 0.399062332126484 | 2.30995827548384 | 1.27038721650549 | 0.210870439807463 | 0.529568846090411 | -5.30176549064876 |
| lnc-ZNF135-2 | lnc-ZNF135-2 | -0.841160627531603 | 1.85592929464771 | -1.27085556772086 | 0.210932446908007 | 0.529604057718644 | -5.26392254612964 |
| lnc-NUP107-2 | lnc-NUP107-2 | 0.316308727334802 | 1.4804341531659 | 1.27016521290613 | 0.210935921157141 | 0.529604057718644 | -5.22187177174631 |
| ENSG00000259529.1 | | 0.149282361207416 | 5.84515318332746 | 1.26985323679566 | 0.211045988658075 | 0.529678569692552 | -5.61972852961277 |
| lnc-DLG2-3 | lnc-DLG2-3 | 0.175673500948469 | 2.44766253330623 | 1.2697765515967 | 0.211073050353524 | 0.529678569692552 | -5.33303070933977 |
| ENSG00000239195.1 | SNORD5 | 0.321911551984131 | 0.865627206482786 | 1.26973498262892 | 0.211087720851221 | 0.529678569692552 | -5.12187328726033 |
| ENSG00000173786.16 | CNP | -0.313385766872182 | 4.5885507912406 | -1.26963182784523 | 0.211150761998639 | 0.529678569692552 | -5.56798008020863 |
| lnc-UGDH-3 | lnc-UGDH-3 | -0.345248553117103 | 4.00807852352087 | -1.26963698961213 | 0.211171691805502 | 0.529678569692552 | -5.5302789512796 |
| lnc-ESRRB-3 | lnc-ESRRB-3 | 0.175668667837471 | 2.930804038406 | 1.2693106041 | 0.211237535986608 | 0.529678569692552 | -5.40130829055287 |
| lnc-BLNK-1 | lnc-BLNK-1 | -0.210464415760007 | 3.35363753728477 | -1.2693095707553 | 0.211237900877602 | 0.529678569692552 | -5.45089956115547 |
| ENSG00000188092.14 | GPR89B | 0.123322820925 | 2.86840282319976 | 1.26929837497706 | 0.211241854321105 | 0.529678569692552 | -5.39356230296845 |
| ENSG00000162695.11 | SLC30A7 | 0.133466617408071 | 5.20934723534254 | 1.26924639248359 | 0.211260211059941 | 0.529678569692552 | -5.60428796212535 |
| lnc-LIMA1-3 | lnc-LIMA1-3 | 0.109483324845251 | 2.7112924589305 | 1.26921625700661 | 0.211270853441138 | 0.529678569692552 | -5.36923671448937 |
| lnc-HEMGN-3 | lnc-HEMGN-3 | 0.199592349193999 | 2.2449891284664 | 1.26918718224687 | 0.211281121609201 | 0.529678569692552 | -5.31264035098784 |
| ENSG00000159792.9 | PSKH1 | -0.178381580103057 | 3.10013345240274 | -1.26915773107974 | 0.211291523091884 | 0.529678569692552 | -5.42476722709084 |
| lnc-GXYLT1-4 | lnc-GXYLT1-4 | 0.201645173982752 | 1.34927969511478 | 1.26913333995966 | 0.211300137771245 | 0.529678569692552 | -5.21089116010858 |
| ENSG00000178691.10 | SUZ12 | 0.125637449671091 | 5.31095002959994 | 1.26881866056282 | 0.211411302708937 | 0.529892698935466 | -5.60908742082908 |
| ENSG00000255823.4 | NA | 0.775598504368517 | 2.7408082303947 | 1.26933950178536 | 0.211478541042572 | 0.529970535712655 | -5.38092188863053 |
| ENSG00000126602.10 | TRAP1 | -0.190467834943006 | 4.58438007487552 | -1.2685850614721 | 0.211493853196509 | 0.529970535712655 | -5.56785923657225 |
| ENSG00000100461.17 | RBM23 | 0.107669262506237 | 6.42755769200164 | 1.26844312252996 | 0.21154402411831 | 0.530031728353887 | -5.62448626812581 |
| ENSG00000203812.2 | NA | 1.18811897705343 | -1.63028113984472 | 1.26831127836516 | 0.211590634836997 | 0.530066423128463 | -4.5763330862694 |
| ENSG00000122432.17 | SPATA1 | -0.354714268296857 | -0.416331693629065 | -1.26823005844966 | 0.211619352248405 | 0.530066423128463 | -4.83982022841791 |
| ENSG00000100316.15 | RPL3 | 0.146713068201882 | 9.49355279253098 | 1.26816819835347 | 0.211641226449854 | 0.530066423128463 | -5.56553566922763 |
| ENSG00000200983.1 | SNORA3A | 0.186281238263191 | 1.87616798842028 | 1.26807592962622 | 0.21167385651981 | 0.530066423128463 | -5.27871221233728 |
| lnc-WDPCP-6 | lnc-WDPCP-6 | 0.356703824884393 | 0.330414751473903 | 1.26803979993903 | 0.211686634509837 | 0.530066423128463 | -5.01799712699841 |
| ENSG00000073331.17 | ALPK1 | -0.256679688426491 | 4.68682962960045 | -1.26786871898082 | 0.211747148569031 | 0.530141016063556 | -5.57685554837782 |
| ENSG00000136451.8 | VEZF1 | 0.194507501390182 | 5.52583156785238 | 1.26777089550474 | 0.211781756108036 | 0.530141016063556 | -5.61686389730549 |
| ENSG00000262660.1 | | 0.266493011354366 | 0.318831120537018 | 1.26773715792661 | 0.211793692614809 | 0.530141016063556 | -5.02896984562994 |

| | | | | | | | |
|---|---|---|---|---|---|---|---|
| ENSG00000144848.10 | ATG3 | 0.122700552008749 | 4.3023555466786 | 1.26763870644835 | 0.211828528068853 | 0.530142472134851 | -5.55295435820073 |
| ENSG00000130529.15 | TRPM4 | -0.984169234748988 | -0.747954543708674 | -1.26800805589015 | 0.211845786975133 | 0.530142472134851 | -4.87121583829152 |
| ENSG00000204681.10 | GABBR1 | -0.21393524040483 | 6.04749283410705 | -1.26750139083941 | 0.211877122124427 | 0.530145808903425 | -5.62364209686617 |
| ENSG00000185278.15 | ZBTB37 | -0.13295623085129 | 6.84364746021951 | -1.26744061265038 | 0.21189863333083 | 0.530145808903425 | -5.62320601293117 |
| ENSG00000198342.9 | ZNF442 | -0.418318791196285 | 3.08577730303984 | -1.26756713542478 | 0.211929497508791 | 0.530154856525404 | -5.42151293701892 |
| ENSG00000004897.11 | CDC27 | 0.103198834696894 | 5.48364290552041 | 1.26721870535991 | 0.211977186789605 | 0.530154856525404 | -5.61470352350842 |
| lnc-SMIM10-2 | lnc-SMIM10-2 | -1.12000161296846 | -1.99724148150233 | -1.26716867597437 | 0.211994899813724 | 0.530154856525404 | -4.61225298197823 |
| ENSG00000082146.12 | STRADB | -0.249102720581249 | 3.34715193537818 | -1.26700112010214 | 0.212054231439111 | 0.530154856525404 | -5.44679086199992 |
| lnc-MEF2C-9 | lnc-MEF2C-9 | 0.173499714838709 | 1.45772656041436 | 1.2668717109045 | 0.212100063809554 | 0.530154856525404 | -5.23363835417754 |
| ENSG00000110400.10 | NECTIN1 | -0.772608093231578 | -1.12033776523058 | -1.26675951913705 | 0.21214155433679 | 0.530154856525404 | -4.7885429889573 |
| lnc-SETD7-3 | lnc-SETD7-3 | -0.244707051568991 | 5.11657314307348 | -1.26674965836569 | 0.2121432974912 | 0.530154856525404 | -5.60120294013598 |
| lnc-DDX41-2 | lnc-DDX41-2 | -0.172597086888306 | 2.44430566107705 | -1.26668733177393 | 0.212165377477722 | 0.530154856525404 | -5.35149614731621 |
| ENSG00000091106.18 | NLRC4 | 0.588330449848678 | -0.447497579070869 | 1.26668626165329 | 0.21216575659656 | 0.530154856525404 | -4.82721040225824 |
| ENSG00000125734.15 | GPR108 | -0.137416037861382 | 4.8641553432143 | -1.26667641409277 | 0.212169245381868 | 0.530154856525404 | -5.58942365377384 |
| lnc-C2orf68-2 | lnc-C2orf68-2 | 0.997711681342432 | 0.373131681285552 | 1.26718552023405 | 0.212185575866311 | 0.530154856525404 | -5.00903307824649 |
| ENSG00000129167.9 | TPH1 | -0.230161832054792 | 1.5872262786219 | -1.26621524550962 | 0.212332675867512 | 0.530458000653114 | -5.25204538813881 |
| ENSG00000140455.16 | USP3 | 0.15828268085551 | 6.89525884239017 | 1.26613104579367 | 0.212362525012291 | 0.530468185962133 | -5.62478328246938 |
| lnc-MYBPC2-1 | lnc-MYBPC2-1 | -0.169590374768589 | 1.09200423641291 | -1.26600087647725 | 0.212408676746178 | 0.530500201814591 | -5.17247932195948 |
| ENSG00000163050.16 | COQ8A | -0.165470754007616 | 5.17579095394226 | -1.26587742613919 | 0.212452453187163 | 0.530500201814591 | -5.60779086833422 |
| lnc-ZFP36L2-2 | lnc-ZFP36L2-2 | 0.197052262502696 | 0.32225570876859 | 1.26587683436817 | 0.212452663049977 | 0.530500201814591 | -5.0198009344648 |
| lnc-A2M-3 | lnc-A2M-3 | 0.204482998946733 | 1.63391044848565 | 1.2657001611103 | 0.212515324534033 | 0.530539804451926 | -5.2560368805329 |
| lnc-RCOR1-4 | lnc-RCOR1-4 | 0.350641977462494 | 2.36162979129822 | 1.2656867710149 | 0.212520074221743 | 0.530539804451926 | -5.33535207979594 |
| ENSG00000276314.1 | SNORD107 | -0.296779436433062 | 0.190238099671961 | -1.26549065108375 | 0.212589650294486 | 0.530607444586361 | -5.00056992271995 |
| ENSG00000233913.7 | RPL10P9 | 1.16121371625511 | 2.22065517491738 | 1.26623495216918 | 0.212598726886446 | 0.530607444586361 | -5.29942594108069 |
| lnc-RALGAPA1-4 | lnc-RALGAPA1-4 | -0.357139502087668 | 4.33243640607847 | -1.26545856024488 | 0.212674622920378 | 0.530708106025942 | -5.5634369385906 |
| lnc-FLNA-2 | lnc-FLNA-2 | -0.345516349382413 | 0.33562755344121 | -1.26518641320113 | 0.212697616313675 | 0.530708106025942 | -5.01744337608019 |
| ENSG00000169756.16 | LIMS1 | 0.212626326919919 | 5.08704800920781 | 1.26507124259331 | 0.212738498029598 | 0.530708106025942 | -5.60279977482748 |
| lnc-LAMP3-2 | lnc-LAMP3-2 | 0.183278524595121 | 0.821825619809744 | 1.26506083073563 | 0.212742194180637 | 0.530708106025942 | -5.14209977789855 |
| ENSG00000232593.7 | KANTR | -0.191471356300266 | 3.07320368845371 | -1.26493603988602 | 0.212786497968866 | 0.530737039052481 | -5.41253575843413 |
| lnc-OSBPL10-6 | lnc-OSBPL10-6 | 0.316063551455559 | 4.83510796587024 | 1.26494585141772 | 0.212820803776082 | 0.530737039052481 | -5.5880459670505 |
| OLMALINC | OLMALINC | -0.387952821516561 | 2.05935930545751 | -1.26485123918879 | 0.212831148051265 | 0.530737039052481 | -5.33924240485005 |
| ENSG00000279616.1 | | -0.302172537642092 | 0.148084996168634 | -1.26456824633906 | 0.212917113759112 | 0.530887092974415 | -4.99221745437362 |
| ENSG00000165233.17 | CARD19 | 0.203855145003063 | 2.86873186563072 | 1.26447472928877 | 0.212950334348673 | 0.53090561190634 | -5.40672723920446 |
| lnc-ADGRG2-3 | lnc-ADGRG2-3 | 0.140882278737666 | 1.16782105943978 | 1.26414275257156 | 0.213068295608474 | 0.531098279747943 | -5.18244767071188 |
| ENSG00000156471.12 | PTDSS1 | 0.145852628762996 | 6.25276298146559 | 1.26409510346634 | 0.213085230769315 | 0.531098279747943 | -5.62966441322813 |
| MIR4458HG | MIR4458HG | -1.16581528127358 | -0.748938846022337 | -1.26456462138292 | 0.213111302767493 | 0.531098279747943 | -4.88981135660423 |
| ENSG00000280987.4 | | 0.113714651655385 | 7.99894049547405 | 1.26396683000378 | 0.21313082595919 | 0.531098279747943 | -5.61051390734819 |
| ENSG00000154217.14 | PITPNC1 | -0.6553929658169 | 3.75337101939181 | -1.26451683691842 | 0.21317970386104 | 0.531155773462266 | -5.50895762432462 |
| ENSG00000105887.10 | MTPN | 0.161523715844287 | 7.84467793744109 | 1.26358414275115 | 0.213266896635347 | 0.531276673750574 | -5.61490684878748 |
| ENSG00000176024.17 | ZNF613 | 0.148147605317728 | 2.50924407453088 | 1.26354772166302 | 0.213279850128795 | 0.531276673750574 | -5.3567388287418 |
| ENSG00000244617.2 | ASPRV1 | 0.311296435703989 | -0.00678843993667059 | 1.26276938577209 | 0.213556812999836 | 0.531826032187579 | -4.9590192184457 |
| ENSG00000238090.1 | | 0.353164347121444 | 0.242002156697148 | 1.26275569854718 | 0.213561685862501 | 0.531826032187579 | -4.99777575084259 |
| lnc-PNMA1-3 | lnc-PNMA1-3 | -0.164942458272557 | 2.58702974904491 | -1.26271014781007 | 0.213577903224143 | 0.531826032187579 | -5.35578640952803 |
| ENSG00000102393.10 | GLA | 0.2350765830807 | 2.6971570653024 | 1.26238002317502 | 0.213695464513747 | 0.53205440257487 | -5.37228353966246 |
| ENSG00000163219.11 | ARHGAP25 | 0.194943437869989 | 8.18078257123403 | 1.2622237163456 | 0.213751144088665 | 0.532122289834525 | -5.61066980060932 |
| ENSG00000213516.9 | RBMXL1 | 0.138983238714382 | 5.10372887148035 | 1.26202560999854 | 0.213821729049853 | 0.532122289834525 | -5.60684323514181 |
| ENSG00000231043.3 | | 0.729075481445114 | -0.095997434200408 | 1.26196630097676 | 0.21384286414211 | 0.532122289834525 | -4.80762081318777 |
| ENSG00000263400.7 | TMEM220-AS1 | 0.486489621001857 | 0.189589574946678 | 1.26195341153708 | 0.213847457570435 | 0.532122289834525 | -5.02631888709061 |
| lnc-PROKR2-4 | lnc-PROKR2-4 | 0.263928841697539 | 3.00656463829945 | 1.26194068374073 | 0.213851993466025 | 0.532122289834525 | -5.41881423824514 |
| ENSG00000179598.5 | PLD6 | -0.396929228274116 | 1.82111634398455 | -1.26114343600402 | 0.2141383532044 | 0.5326085631847 | -5.2868929483867 |
| lnc-FUBP1-6 | lnc-FUBP1-6 | 0.205814639620937 | 1.41437374298814 | 1.26112084783815 | 0.214144315679162 | 0.5326085631847 | -5.22664228675206 |
| ENSG00000204301.6 | NOTCH4 | -0.314275370314364 | 0.13304480788988 | -1.26109697147 | 0.214152833551618 | 0.5326085631847 | -5.00550201356416 |
| lnc-STK3-2 | lnc-STK3-2 | -0.195788177984079 | 2.44883436798232 | -1.2610640042745 | 0.214164594984306 | 0.5326085631847 | -5.34711075589182 |
| lnc-VN1R4-2 | lnc-VN1R4-2 | -1.29731911537252 | -0.709479359319507 | -1.26150988786671 | 0.214176800149626 | 0.5326085631847 | -4.76710473027375 |
| ENSG00000087884.14 | AAMDC | 0.274451619142595 | -0.241918064342339 | 1.26088325804353 | 0.214229086890203 | 0.532674232358185 | -4.90952427433392 |
| ENSG00000255769.7 | GOLGA2P10 | 0.591970837845434 | 0.111913230546064 | 1.26079871097285 | 0.214269679622306 | 0.532710812376606 | -4.98464224482542 |
| ENSG00000160408.14 | ST6GALNAC6 | -0.154410011334067 | 5.3286282196412 | -1.26044918761536 | 0.21438402642179 | 0.532827518086562 | -5.61847462408887 |
| ENSG00000138111.14 | MFSD13A | 0.679625109527239 | -0.442326702879039 | 1.26043213230209 | 0.214390115948962 | 0.532827518086562 | -4.80837565883928 |
| lnc-ARPC1B-2 | lnc-ARPC1B-2 | -0.33601356062603 | 0.723526462440726 | -1.26038159880185 | 0.214408159478908 | 0.532827518086562 | -5.11031887479817 |
| ENSG00000201457.1 | SNORA55 | -0.240510941150927 | -0.037420420419249 | -1.26034796701694 | 0.214420168698003 | 0.532827518086562 | -4.96290114967303 |
| lnc-DIRAS1-1 | lnc-DIRAS1-1 | 0.571139465585629 | 0.356911109758659 | 1.26017518804404 | 0.214489345904528 | 0.5328809097117 | -5.00662163240823 |
| ENSG00000268107.5 | | -0.156578543741667 | 0.492632069818628 | -1.26014282024851 | 0.214493433268301 | 0.5328809097117 | -5.07205789832401 |
| ENSG00000197798.8 | FAM118B | -0.206922909413641 | 1.9932438416529 | -1.25969772961677 | 0.214652453847092 | 0.533133319114225 | -5.30605690614119 |
| ENSG00000148337.20 | CIZ1 | -0.173568645462107 | 5.54860501620271 | -1.25969296816915 | 0.214654155477783 | 0.533133319114225 | -5.62660019268324 |
| lnc-SPAG9-3 | lnc-SPAG9-3 | 0.175458469344903 | 0.902634871657338 | 1.2596409753643 | 0.214672737152927 | 0.533133319114225 | -5.14520022989395 |
| ENSG00000220472.1 | RPS3P4 | 1.02506731220596 | -0.163583188930536 | 1.25968324124944 | 0.214731003167651 | 0.533151580407379 | -4.7507722877374 |
| ENSG00000232629.8 | HLA-DQB2 | 1.24799324630764 | 3.53697587892694 | 1.26023731218483 | 0.21473189532999 | 0.533151580407379 | -5.42688547623591 |
| lnc-RBBP6-1 | lnc-RBBP6-1 | -0.77679863802188 | 0.0252347492498376 | -1.25956447337768 | 0.214796847582263 | 0.533248524157004 | -4.95608958403696 |

| | | | | | | | |
|---|---|---|---|---|---|---|---|
| ENSG00000183444.10 | OR7E38P | -0.163459918848076 | 2.25717364376906 | -1.25903288825235 | 0.214890150172801 | 0.533400942822305 | -5.33690145025782 |
| LINC02289 | LINC02289 | -0.376658507779031 | -0.343783652989431 | -1.25882019255005 | 0.214966235321973 | 0.533400942822305 | -4.85353482895162 |
| ENSG00000184897.5 | H1-10 | -0.39210293412118 | 5.84849736090945 | -1.25920493873934 | 0.214975153237555 | 0.533400942822305 | -5.63182693544209 |
| ENSG00000168071.21 | CCDC88B | -0.242013331937393 | 6.49431541839249 | -1.25878890085503 | 0.214977430630888 | 0.533400942822305 | -5.63550440298445 |
| lnc-BIRC2-9 | lnc-BIRC2-9 | -0.514238938404533 | -0.65989104875002 | -1.25875987341862 | 0.214987816239316 | 0.533400942822305 | -4.79582541892336 |
| lnc-MCPH1-1 | lnc-MCPH1-1 | -0.181709463587675 | 1.24250095408715 | -1.2586713168588 | 0.2150195028445 | 0.533415261787193 | -5.2022176366715 |
| lnc-ZCCHC10-3 | lnc-ZCCHC10-3 | -0.26703407141475 | 1.03036183768765 | -1.25857310998202 | 0.215054646525175 | 0.533438152497913 | -5.18425407754261 |
| lnc-ZNF8-1 | lnc-ZNF8-1 | -0.20962631336844 | 1.06186165150467 | -1.25835727486705 | 0.215131898952722 | 0.533548330148119 | -5.1848052507368 |
| ENSG00000125633.10 | CCDC93 | 0.0960756995159988 | 6.35834382289874 | 1.25817835585617 | 0.215195953941584 | 0.533548330148119 | -5.63692803208746 |
| ENSG00000050426.15 | LETMD1 | 0.134596210666547 | 5.27424894946017 | 1.25817040085382 | 0.215198802250997 | 0.533548330148119 | -5.61836340009793 |
| ENSG00000153283.12 | CD96 | 2.39550128432233 | -2.11194846980458 | 1.25855883093997 | 0.215214188404061 | 0.533548330148119 | -4.55042290787151 |
| lnc-ZNF121-1 | lnc-ZNF121-1 | 0.196200675552246 | 4.01240269140207 | 1.25808697879886 | 0.215228673430493 | 0.533548330148119 | -5.53817121614735 |
| lnc-ZKSCAN3-2 | lnc-ZKSCAN3-2 | -0.559785175009637 | -0.427017728032302 | -1.25786261448525 | 0.215309027567242 | 0.533683250772705 | -4.89012670579764 |
| ENSG00000118762.7 | PKD2 | -0.779257412383917 | -0.106435740304932 | -1.25801300691466 | 0.215363837609259 | 0.533734662520505 | -4.97488236077813 |
| ENSG00000228158.2 | TLE1P1 | -0.433551279170967 | 0.483213205767403 | -1.25765994589797 | 0.215381630806749 | 0.533734662520505 | -5.0526613245221 |
| ENSG00000100034.13 | PPM1F | -0.219676950289719 | 4.27536142860425 | -1.25729585528293 | 0.215512107150344 | 0.533993704282597 | -5.56386608134165 |
| ENSG00000143799.12 | PARP1 | -0.210509274480805 | 7.21795993844815 | -1.25720554582687 | 0.215544479777803 | 0.534009632554948 | -5.63142749209387 |
| ENSG00000197603.13 | CPLANE1 | -0.1921951886916 | 5.5694804741515 | -1.25712776148423 | 0.215572365519092 | 0.534014442108493 | -5.62936788597062 |
| ENSG00000198231.12 | DDX42 | 0.1273731135559 | 6.56142165211922 | 1.25669390666771 | 0.21572795214172 | 0.534335552218173 | -5.63832083914585 |
| ENSG00000139211.6 | AMIGO2 | 0.275635523289798 | 3.9043412694219 | 1.25660121158863 | 0.215761204795998 | 0.534353613080979 | -5.51903806599052 |
| SNHG4 | SNHG4 | -0.294312508577161 | 1.15508222964576 | -1.2561617603517 | 0.215918901864397 | 0.53465731122851 | -5.20796318723445 |
| ENSG00000213390.10 | ARHGAP19 | -0.12108519930233 | 4.48377832094789 | -1.25611473130175 | 0.215935783328115 | 0.53465731122851 | -5.57595431511864 |
| ENSG00000148358.19 | GPR107 | 0.123568859770323 | 5.25884465183018 | 1.25600210217274 | 0.215976216480002 | 0.53467820470108 | -5.62028762904245 |
| ENSG00000230715.3 | | -0.263156518732962 | 1.86249603478477 | -1.25592071261803 | 0.216005438334034 | 0.53467820470108 | -5.29873299868346 |
| ENSG00000249915.7 | PDCD6 | -0.120035697573978 | 5.00486572570809 | -1.25587416550676 | 0.21602215179003 | 0.53467820470108 | -5.61118292719165 |
| ENSG00000145390.11 | USP53 | -0.362574299158788 | 4.36115055963707 | -1.25584353258247 | 0.216113350488667 | 0.534819093154732 | -5.56491682829763 |
| ENSG00000167524.14 | RSKR | -0.183971764838949 | 2.722877307143 | -1.25557097389238 | 0.216131040997962 | 0.534819093154732 | -5.38992010022548 |
| lnc-TET1-2 | lnc-TET1-2 | -0.252352261854111 | 0.261562565076535 | -1.25538219325652 | 0.216198860950082 | 0.534889396095329 | -5.03757699329481 |
| ENSG00000067225.17 | PKM | -0.230192835238431 | 7.18293063213102 | -1.25534722337429 | 0.216211425716035 | 0.534889396095329 | -5.63381299283261 |
| ENSG00000168827.14 | GFM1 | 0.11394803993429 | 5.32789253471124 | 1.25519830589697 | 0.216264938232585 | 0.534927321684458 | -5.62535842915838 |
| lnc-C19orf47-3 | lnc-C19orf47-3 | 0.231446925881789 | 1.59438687776492 | 1.25515992022414 | 0.216278733472189 | 0.534927321684458 | -5.24139020560267 |
| ENSG00000023839.11 | ABCC2 | 0.224402297250739 | 2.77005874138273 | 1.25501688441789 | 0.216330144189077 | 0.534950762557358 | -5.39024945583221 |
| lnc-FAM111B-1 | lnc-FAM111B-1 | -0.211900801934304 | 2.53683008950501 | -1.25498893547971 | 0.21634019081414 | 0.534950762557358 | -5.36579582199457 |
| ENSG00000104218.14 | CSPP1 | 0.13120502163434 | 4.31401253389004 | 1.25484197890494 | 0.216393022071253 | 0.534989117986301 | -5.55929926439067 |
| ENSG00000067057.16 | PFKP | -0.265545112813331 | 6.30354284307361 | -1.25476481572714 | 0.216420974402639 | 0.534989117986301 | -5.64053237770781 |
| ENSG00000131116.11 | ZNF428 | -0.215185282196799 | 2.33868109843431 | -1.2547289088861 | 0.216433677572362 | 0.534989117986301 | -5.35683617244779 |
| lnc-GPR6-1 | lnc-GPR6-1 | 0.24823044321875 | 0.877742204762123 | 1.25446353243083 | 0.216529118798162 | 0.535078451426878 | -5.15773565357488 |
| lnc-CSRNP1-2 | lnc-CSRNP1-2 | 0.193561019595805 | 0.449001347937211 | 1.2544170304305 | 0.216545846223814 | 0.535078451426878 | -5.07409066347434 |
| ENSG00000231989.4 | PPP1R2B | 0.287954703296579 | 0.160899612754706 | 1.25437835292636 | 0.216559759801431 | 0.535078451426878 | -5.00078935539413 |
| lnc-CD63-7 | lnc-CD63-7 | 0.284081390748896 | 0.485517507432525 | 1.25433931809724 | 0.216573802596187 | 0.535078451426878 | -5.06209688203917 |
| lnc-EZH2-5 | lnc-EZH2-5 | -0.432686494919665 | 0.242617580581318 | -1.25401451619682 | 0.216690676522676 | 0.535223477564807 | -5.01308698327764 |
| ENSG00000113971.19 | NPHP3 | -0.16672787064214 | 5.01538108132271 | -1.25400990470588 | 0.216692336220304 | 0.535223477564807 | -5.6117727690318 |
| lnc-CLEC2D-8 | lnc-CLEC2D-8 | 0.259505089424475 | 5.22818934287883 | 1.25391678062473 | 0.216725854049345 | 0.535223477564807 | -5.61983714084406 |
| ENSG00000184900.15 | SUMO3 | 0.183732769381434 | 5.26666015095113 | 1.25379222430333 | 0.216770691215433 | 0.535223477564807 | -5.62487603926296 |
| ENSG00000101856.9 | PGRMC1 | 0.180298826628723 | 3.19909480956732 | 1.25365601936378 | 0.216819729504847 | 0.535223477564807 | -5.45058061667935 |
| lnc-SMIM14-11 | lnc-SMIM14-11 | 0.129567706267931 | 2.61395850698858 | 1.25362960313128 | 0.216829241180434 | 0.535223477564807 | -5.38116970784992 |
| lnc-MYO18B-5 | lnc-MYO18B-5 | -0.387017448260082 | 0.128266290619754 | -1.25360812511814 | 0.21683697498376 | 0.535223477564807 | -5.00268334150202 |
| lnc-TTPAL-1 | lnc-TTPAL-1 | 0.160761339338327 | 2.4229550933476 | 1.25359825914795 | 0.216840527591358 | 0.535223477564807 | -5.35152364029303 |
| ENSG00000183605.16 | SFXN4 | -0.17522437735617 | 2.76967759592173 | -1.2529029545812 | 0.217091006967979 | 0.535776364314102 | -5.41247767875777 |
| ENSG00000099194.5 | SCD | -0.38977154974911 | 2.68635774485343 | -1.25293257845011 | 0.21713164449717 | 0.535776364314102 | -5.43418361509884 |
| ENSG00000129810.14 | SGO1 | 0.433383502252431 | 0.105400645863131 | 1.25267315735477 | 0.217173837437744 | 0.535776364314102 | -4.98816099734815 |
| ENSG00000152767.16 | FARP1 | -2.35906105966178 | 0.956854559646587 | -1.25339232702738 | 0.217185940930655 | 0.535776364314102 | -5.01203314238425 |
| ENSG00000196233.13 | LCOR | 0.106913139572962 | 7.14187561219827 | 1.25261535939456 | 0.217194674432147 | 0.535776364314102 | -5.63824323302985 |
| lnc-ESRRB-2 | lnc-ESRRB-2 | -0.29107863063236 | 0.696459997505202 | -1.25252991070811 | 0.217225482638492 | 0.535788149688206 | -5.14616058004796 |
| ENSG00000163558.12 | PRKCI | -0.198197098768345 | 3.38021357510328 | -1.25219951892157 | 0.217344634794093 | 0.535956775715995 | -5.46913865545735 |
| ENSG00000036448.9 | MYOM2 | -1.16441157765213 | -1.88872545639676 | -1.25213197755654 | 0.217390387617503 | 0.535956775715995 | -4.63947340586152 |
| lnc-RPS10-2 | lnc-RPS10-2 | -0.348699483368999 | -0.518545434937809 | -1.25206772727117 | 0.217392177578666 | 0.535956775715995 | -4.85085211690929 |
| ENSG00000173950.15 | XXYLT1 | -0.343524128209044 | 1.69543271942806 | -1.25205157697889 | 0.217398004200206 | 0.535956775715995 | -5.29280037460323 |
| lnc-POLR1E-2 | lnc-POLR1E-2 | 0.212463738789358 | 4.53975938741889 | 1.25172140996835 | 0.217517145693873 | 0.536186276624788 | -5.57889976670297 |
| ENSG00000253558.1 | | 0.442092905215008 | 2.10952871351061 | 1.25162891689922 | 0.217601019293837 | 0.536190465713097 | -5.32105892356452 |
| ENSG00000161904.11 | LEMD2 | 0.106130080328963 | 5.80528779575343 | 1.25144009932498 | 0.217618695616016 | 0.536190465713097 | -5.64146637140567 |
| ENSG00000172869.14 | DMXL1 | 0.172731606185533 | 8.04410797182656 | 1.25132487909282 | 0.217660298989823 | 0.536190465713097 | -5.62601299535499 |
| lnc-MRPL14-1 | lnc-MRPL14-1 | -0.264763779471154 | 0.242106919217964 | -1.25130561703116 | 0.217667254654656 | 0.536190465713097 | -5.0223641491993 |
| ENSG00000100916.13 | BRMS1L | 0.257699477223953 | 1.15076953171005 | 1.25118527614421 | 0.217710714335345 | 0.536190465713097 | -5.20887254046474 |
| lnc-ZNF736-1 | lnc-ZNF736-1 | -0.295559133116023 | 0.31967682448694 | -1.25112164412585 | 0.217733696893349 | 0.536190465713097 | -5.0456595944409 |
| ENSG00000100503.23 | NIN | -0.164588307470618 | 7.74680909780336 | -1.25110167794819 | 0.217740908632628 | 0.536190465713097 | -5.6312645936041 |
| lnc-TBC1D8-5 | lnc-TBC1D8-5 | 0.182876967599161 | 2.38464568857166 | 1.25103076108499 | 0.21776652508462 | 0.536190465713097 | -5.3598067959493 |
| ENSG00000284491.2 | THSD8 | -0.263024548676385 | 0.723478722523573 | -1.25093702288984 | 0.217800388456767 | 0.536190465713097 | -5.12639922326165 |

| | | | | | | | |
|---|---|---|---|---|---|---|---|
| ENSG00000261221.3 | ZNF865 | -0.333299100346284 | 1.41792274118605 | -1.25093005295002 | 0.2178029065375 | 0.536190465713097 | -5.25256602219143 |
| lnc-IRS1-3 | lnc-IRS1-3 | -1.33349092711865 | -0.111328025509853 | -1.25159794236589 | 0.217822143709166 | 0.536190465713097 | -5.0495514087177 |
| ENSG00000275143.1 | SCARNA16 | 0.311996023566228 | 0.779809575846113 | 1.25080167809762 | 0.217849289327839 | 0.536190465713097 | -5.10535416430306 |
| ENSG00000137100.15 | DCTN3 | 0.114041460172934 | 3.16675738729475 | 1.2507744697612 | 0.21785912084678 | 0.536190465713097 | -5.44933726554551 |
| ENSG00000136261.14 | BZW2 | -0.306329611362697 | 3.57111729661262 | -1.25064046394022 | 0.217907547618633 | 0.536190465713097 | -5.50089996777346 |
| ENSG00000186416.14 | NKRF | -0.125545972588233 | 3.5202126256886 | -1.25063479198228 | 0.217909597516886 | 0.536190465713097 | -5.4886949977177 |
| ENSG00000132376.19 | INPP5K | 0.123571350287525 | 4.64651497002205 | 1.25049515697891 | 0.217960067430222 | 0.536250546009594 | -5.59977081162524 |
| ENSG00000261487.1 | | -0.194979891003891 | 2.69872852966471 | -1.25021010895663 | 0.218063122682936 | 0.536412523949552 | -5.39713312850333 |
| ENSG00000163568.14 | AIM2 | -0.373607118276924 | 5.81165325209352 | -1.25052879550294 | 0.218078025574982 | 0.536412523949552 | -5.6406219484527 |
| ENSG00000125629.14 | INSIG2 | 0.164822583157385 | 3.66063546419323 | 1.24981921812127 | 0.218204502932798 | 0.536659491440529 | -5.50639565813265 |
| lnc-NAPRT-1 | lnc-NAPRT-1 | -0.368417252753198 | 0.581047640845539 | -1.24944199436454 | 0.218341004644058 | 0.536926572470892 | -5.11891758791171 |
| ENSG00000074935.13 | TUBE1 | -0.23742357612307 | 3.81894697612399 | -1.2493749571122 | 0.218365269307382 | 0.536926572470892 | -5.52389989187562 |
| lnc-PPARA-10 | lnc-PPARA-10 | -0.344358709632697 | 0.150538252081687 | -1.24861309763977 | 0.218641171358286 | 0.537540758488723 | -4.99313065801058 |
| ENSG00000226608.3 | FTLP3 | 0.702934856599869 | 0.659559824855746 | 1.24870230703001 | 0.218673225048304 | 0.537555355448376 | -4.98018550762522 |
| ENSG00000285943.1 | | 0.157521556929327 | 4.45012951971639 | 1.24830604836968 | 0.218752440383758 | 0.537620496043062 | -5.57777812149457 |
| ENSG00000113580.14 | NR3C1 | 0.115213059657049 | 7.10737851788067 | 1.24823649144288 | 0.218777652389404 | 0.537620496043062 | -5.64423929883909 |
| lnc-SLC25A19-4 | lnc-SLC25A19-4 | -0.17969494334612 | 5.57678681333349 | -1.24823530450078 | 0.218778082634052 | 0.537620496043062 | -5.63988185230098 |
| ENSG00000258900.1 | HNRNPCP1 | -0.264905338417217 | 0.614397174407487 | -1.24807330540421 | 0.218836810227645 | 0.537700616559105 | -5.11022490379885 |
| ENSG00000230928.1 | | -0.329265181621795 | 0.620441998156465 | -1.24792429912374 | 0.218890838043072 | 0.53770761467206 | -5.12060810398316 |
| ENSG00000135698.9 | MPHOSPH6 | -0.157392742219073 | 3.20763981665821 | -1.24792135373577 | 0.21889190610393 | 0.53770761467206 | -5.45816799233061 |
| ENSG00000197024.8 | ZNF398 | -0.133106255764711 | 4.80378364897128 | -1.24777407417068 | 0.218945317776205 | 0.537774639115468 | -5.60786120651165 |
| ENSG00000124120.10 | TTPAL | 0.123093351943783 | 4.53987484287002 | 1.2476949235857 | 0.218974026136493 | 0.537780978399646 | -5.58259747396237 |
| ENSG00000168763.15 | CNNM3 | 0.209119357231435 | 4.47300392592033 | 1.24740895210648 | 0.219077772922512 | 0.537971581968988 | -5.57675298423779 |
| ENSG00000047230.14 | CTPS2 | -0.297019706023399 | 4.85959699758727 | -1.24720469382723 | 0.219173230701207 | 0.538141787847185 | -5.61188532116086 |
| ENSG00000084070.11 | SMAP2 | -0.247246648655052 | 8.89438807538874 | -1.24701882163315 | 0.219219366197505 | 0.538146935849774 | -5.60847739778958 |
| ENSG00000165417.11 | GTF2A1 | -0.17278952552831 | 5.15611344019394 | -1.2469867454947 | 0.219231010881971 | 0.538146935849774 | -5.62851510331152 |
| lnc-STX3-2 | lnc-STX3-2 | -0.257350216603644 | 0.492403870042422 | -1.2469240767426 | 0.219253763010067 | 0.538146935849774 | -5.09465057822484 |
| ENSG00000197860.9 | SGTB | -0.324682124336392 | 4.90415740450893 | -1.24688137529828 | 0.219327011611501 | 0.538262534875346 | -5.61438459493163 |
| ENSG00000108953.16 | YWHAE | 0.145365522462595 | 5.71146308145697 | 1.24655381065311 | 0.219388225348343 | 0.538300317325132 | -5.64623460103955 |
| ENSG00000107938.17 | EDRF1 | 0.0890731897477943 | 5.78020681758498 | 1.24653595210305 | 0.219394712240224 | 0.538300317325132 | -5.64661953665941 |
| ENSG00000131495.8 | NDUFA2 | 0.172092860916799 | 2.8795439993058 | 1.24610978138006 | 0.219549555657719 | 0.538554464284439 | -5.41874539648981 |
| ENSG00000184863.10 | RBM33 | 0.169129002390749 | 7.42683955323171 | 1.24610683964965 | 0.21955062477825 | 0.538554464284439 | -5.64302122433651 |
| ENSG00000000938.12 | FGR | 0.385620739603066 | 7.20393944034809 | 1.24640181038988 | 0.219587484283488 | 0.538580694590923 | -5.64727326892618 |
| ENSG00000090487.10 | SPG21 | 0.128787153146804 | 4.66743686222026 | 1.24566775114469 | 0.219710247226473 | 0.53881758830722 | -5.59901799312798 |
| ENSG00000163565.18 | IFI16 | -0.164594534216084 | 7.48389200366794 | -1.24552177746438 | 0.219763332338427 | 0.538883567967816 | -5.64194975282342 |
| ENSG00000129515.18 | SNX6 | 0.130539268890515 | 5.47380340824303 | 1.24519068162526 | 0.219883774714226 | 0.53907321511108 | -5.64184635326094 |
| ENSG00000112195.8 | TREML2 | 0.57613783014359 | 4.49340834045867 | 1.24576306736903 | 0.21989305320277 | 0.53907321511108 | -5.55296137676679 |
| ENSG00000107560.11 | RAB11FIP2 | 0.130730421700578 | 5.59934745660469 | 1.24487536728039 | 0.21999852186704 | 0.539267545026709 | -5.64595823656048 |
| ENSG00000137871.20 | ZNF280D | -0.154817707496702 | 5.40403848567422 | -1.24457466072654 | 0.220107994491217 | 0.539394952534474 | -5.6371098978998 |
| ENSG00000239672.7 | NME1 | -0.324559192175381 | 2.41551196556321 | -1.2445546274714 | 0.220115289062329 | 0.539394952534474 | -5.38497186142934 |
| lnc-MDGA1-1 | lnc-MDGA1-1 | -0.30381051157614 | 0.0896529969364158 | -1.24451665468648 | 0.220129116323645 | 0.539394952534474 | -5.01938989141898 |
| lnc-SPRYD3-1 | lnc-SPRYD3-1 | -0.421033946713634 | 1.9393587525653 | -1.24440401734746 | 0.220194438771347 | 0.539446589696297 | -5.30587294896547 |
| lnc-TCP11-2 | lnc-TCP11-2 | -0.28950178837199 | 1.73311524343658 | -1.2443148641789 | 0.220202606356133 | 0.539446589696297 | -5.28749644553699 |
| lnc-SEC24A-2 | lnc-SEC24A-2 | -0.188334299834454 | 1.57021603617801 | -1.24400985250329 | 0.220313723073211 | 0.539608519987978 | -5.26267088755629 |
| ENSG00000011600.11 | TYROBP | -1.01566688019613 | 0.934536436321557 | -1.24457753359034 | 0.220343750830979 | 0.539608519987978 | -5.12860741031249 |
| lnc-JAK1-2 | lnc-JAK1-2 | -0.202845218268217 | 5.54564858266309 | -1.24391029075572 | 0.220350002747723 | 0.539608519987978 | -5.64405917406009 |
| ENSG00000163002.12 | NUP35 | 0.140056730589743 | 3.22864922206008 | 1.2438456183411 | 0.220373571346205 | 0.539608519987978 | -5.46159004525195 |
| lnc-DDHD1-2 | lnc-DDHD1-2 | 0.155839416581521 | 3.87779000747001 | 1.24368829262789 | 0.220430913447868 | 0.53962783602982 | -5.52784215049988 |
| ENSG00000051341.13 | POLQ | 0.423323526166195 | 0.641008973571781 | 1.2436801152316 | 0.220433894249562 | 0.53962783602982 | -5.10954732152941 |
| ENSG00000100221.10 | JOSD1 | 0.204529486357385 | 5.58850343308791 | 1.24360800146428 | 0.220460182255134 | 0.539628009437202 | -5.64489265983926 |
| ENSG00000146143.17 | PRIM2 | -0.130517022595575 | 3.51684894373059 | -1.24315284430498 | 0.220626156784081 | 0.539827397081098 | -5.4932912487084 |
| ENSG00000153786.12 | ZDHHC7 | -0.185262698889541 | 4.31446037203857 | -1.24314406210401 | 0.220629360154301 | 0.539827397081098 | -5.57661213108086 |
| ENSG00000168566.12 | SNRNP48 | -0.134044696201246 | 4.3983964523252 | -1.24312108830458 | 0.220637740174285 | 0.539827397081098 | -5.5899066610474 |
| ENSG00000154258.16 | ABCA9 | -0.540687513949157 | 6.85957769031511 | -1.24372492895442 | 0.220646547716235 | 0.539827397081098 | -5.65126477107767 |
| ENSG00000134255.13 | CEPT1 | -0.153470587937223 | 6.10902848477878 | -1.24297361757392 | 0.220691537848471 | 0.539873297306285 | -5.65443493440829 |
| ENSG00000285508.1 | | -0.190341356923078 | 0.291558436845628 | -1.24264040731507 | 0.220813129613654 | 0.540106553397249 | -5.05472857855107 |
| ENSG00000073111.13 | MCM2 | 0.200458279822203 | 4.9646202700978 | 1.24237345504237 | 0.220910579058209 | 0.540193822347673 | -5.62203806727784 |
| lnc-HLA-DRB1-3 | lnc-HLA-DRB1-3 | -0.433962502045292 | 5.77643832740393 | -1.24275054102523 | 0.220949006490531 | 0.540193822347673 | -5.65039428440752 |
| ENSG00000160753.15 | RUSC1 | -0.192275442459049 | 1.64990977111183 | -1.24225584057198 | 0.2209535236988 | 0.540193822347673 | -5.28701875424911 |
| FAR1-IT1 | FAR1-IT1 | 0.265158218757089 | 0.789201356218065 | 1.24225512050384 | 0.22095378663679 | 0.540193822347673 | -5.151089735978 |
| ENSG00000117091.9 | CD48 | 0.262491229930735 | 6.70769454821023 | 1.24208292451398 | 0.221016878622693 | 0.540272171931054 | -5.6550871718308 |
| ENSG00000130783.13 | CCDC62 | 0.27933574581819 | 1.7104210677719 | 1.2420155163384 | 0.221041292697403 | 0.540272171931054 | -5.28022328036024 |
| lnc-RNF166-5 | lnc-RNF166-5 | 0.402084564664505 | 0.0272293586774956 | 1.24188377987483 | 0.221089415253596 | 0.540272171931054 | -5.00766666863291 |
| lnc-IQCG-5 | lnc-IQCG-5 | -0.159198440941263 | 0.699533124514235 | -1.24187991391169 | 0.22109082758467 | 0.540272171931054 | -5.12345946425968 |
| ENSG00000284163.1 | MIR3620 | 0.333983661521898 | -0.25743753559451 | 1.24177943242252 | 0.221127538284315 | 0.54029506849123 | -4.87460075908685 |
| lnc-GPR27-10 | lnc-GPR27-10 | -0.834248976682012 | 2.36440237351887 | -1.24238881603878 | 0.221152696498985 | 0.54029506849123 | -5.36441891024762 |
| ENSG00000204789.4 | ZNF204P | -1.14874663730047 | -0.307895265790994 | -1.24193992382047 | 0.221248573565579 | 0.540374808134931 | -4.86953791790468 |
| ENSG00000176714.9 | CCDC121 | -0.362414473192739 | 0.200358860837462 | -1.2413769701681 | 0.221274622374199 | 0.540374808134931 | -5.05339043647355 |

| | | | | | | | |
|---|---|---|---|---|---|---|---|
| lnc-NCOR1-1 | lnc-NCOR1-1 | -0.454704841155299 | 0.312516151993705 | -1.24129236920123 | 0.221305549926861 | 0.540374808134931 | -5.08172720953208 |
| lnc-FBXL20-2 | lnc-FBXL20-2 | 0.143671341538781 | 1.20522226829039 | 1.24127114385962 | 0.221313309772678 | 0.540374808134931 | -5.2190644966129 |
| lnc-DDX6-1 | lnc-DDX6-1 | 0.16819251010904 | 4.60547060616312 | 1.24126213705817 | 0.221316602661297 | 0.540374808134931 | -5.60254732740314 |
| lnc-CAGE1-4 | lnc-CAGE1-4 | 0.171906624811685 | 2.59572656923016 | 1.24113585588038 | 0.221362774917052 | 0.540423436854191 | -5.38807333209929 |
| ENSG00000149262.16 | INTS4 | 0.107780014098512 | 5.65013090013155 | 1.24083889942614 | 0.221471379437345 | 0.540624454810114 | -5.65014456975461 |
| ENSG00000109113.19 | RAB34 | 1.36957638880064 | -0.310055222951574 | 1.24124638486443 | 0.221518204854427 | 0.540674636608405 | -4.82316252912503 |
| ENSG00000100324.13 | TAB1 | -0.14277024570613 | 3.32774741139538 | -1.24034707874483 | 0.221651337720444 | 0.540878532344236 | -5.47989765918197 |
| lnc-MUC4-3 | lnc-MUC4-3 | -0.245767568189816 | 1.67579139420707 | -1.24028365458013 | 0.221674552659056 | 0.540878532344236 | -5.30572459674019 |
| ENSG00000111602.11 | TIMELESS | -0.233736945303061 | 4.99756968294373 | -1.2402003493006 | 0.221705047357562 | 0.540878532344236 | -5.62723208376998 |
| ENSG00000261611.6 | | 0.152019889398774 | 3.8735336955496 | 1.24017309324173 | 0.221715025375668 | 0.540878532344236 | -5.52603107139072 |
| ENSG00000148331.11 | ASB6 | 0.145882323908519 | 3.85294657366917 | 1.24010507850918 | 0.221739925961609 | 0.540878532344236 | -5.54214532974865 |
| ENSG00000076248.10 | UNG | -0.274837514530924 | 2.89602688813545 | -1.24005186297206 | 0.221759409923384 | 0.540878532344236 | -5.43936748583287 |
| ENSG00000284594.1 | MIR7847 | -0.282817373301754 | 3.83330609603451 | -1.23979884029201 | 0.221852067240916 | 0.541040414694287 | -5.53313526040675 |
| ENSG00000119285.10 | HEATR1 | 0.1095438530605 | 6.50271483859562 | 1.23962378648105 | 0.221916189037696 | 0.541132676100331 | -5.65888499700899 |
| lnc-ABHD17B-5 | lnc-ABHD17B-5 | -0.36622211093544 | 3.85208381473283 | -1.23973107878341 | 0.221948678840726 | 0.541147791587143 | -5.54439598154 |
| lnc-CREB3L4-1 | lnc-CREB3L4-1 | -0.337754472058464 | -0.469455978856511 | -1.2393542673927 | 0.22201494011744 | 0.541245234210265 | -4.89006143958401 |
| ENSG00000255072.1 | PIGY | 0.195419562241872 | 1.71427342160485 | 1.2387110526021 | 0.222250744015468 | 0.541755928951964 | -5.2967107101905 |
| lnc-FAM120AOS-3 | lnc-FAM120AOS-3 | 0.113585643589623 | 3.72401448762816 | 1.23859060159144 | 0.222294922248501 | 0.541775889123689 | -5.52303270813522 |
| ENSG00000198874.12 | TYW1 | 0.0921515688746375 | 4.73321733876605 | 1.23843519362984 | 0.222351931383457 | 0.541775889123689 | -5.61113800847196 |
| ENSG00000140451.12 | PIF1 | 0.536169280327916 | -0.691309792020182 | 1.23843409987082 | 0.222352332651353 | 0.541775889123689 | -4.84721445283897 |
| ENSG00000149084.12 | HSD17B12 | -0.183052046171561 | 4.35311239831885 | -1.23840170233623 | 0.222364218593836 | 0.541775889123689 | -5.5920366904276 |
| ENSG00000125257.14 | ABCC4 | -0.236533006500842 | 4.02216267680105 | -1.23811349748094 | 0.222469975350141 | 0.541938241096876 | -5.55199152082345 |
| ENSG00000281100.1 | | -0.307881051702895 | 0.540051211715433 | -1.23807661388527 | 0.22248351247479 | 0.541938241096876 | -5.12528181908113 |
| ENSG00000196505.10 | GDAP2 | 0.112542432493932 | 5.5971198003594 | 1.23781572499989 | 0.222579282110611 | 0.542043840986828 | -5.65260936589064 |
| ENSG00000153037.14 | VIII.19 | 0.12620938755118 | 4.13805163341981 | 1.23781503949951 | 0.222579533791025 | 0.542043840986828 | -5.56665794268261 |
| lnc-IGF2BP3-2 | lnc-IGF2BP3-2 | 0.22158494594446 | 2.37589889829143 | 1.23765090499479 | 0.222639801595946 | 0.542126468265628 | -5.38587373087348 |
| lnc-SEC11A-2 | lnc-SEC11A-2 | -0.322923560339588 | 0.910458760252067 | -1.23719787763835 | 0.222806209429431 | 0.542458357508902 | -5.17315426340981 |
| lnc-HMOX1-1 | lnc-HMOX1-1 | -0.182326352390179 | 1.06363744744123 | -1.23708645141384 | 0.222847153081366 | 0.542458357508902 | -5.20688672914249 |
| lnc-PPP1R10-1 | lnc-PPP1R10-1 | -0.232441636768982 | 4.19030930456941 | -1.23704801538005 | 0.222861277725041 | 0.542458357508902 | -5.57906309815581 |
| ENSG00000122965.10 | RBM19 | -0.117649934176736 | 6.4302196726491 | -1.23699293473725 | 0.222881520159976 | 0.542458357508902 | -5.66197395082399 |
| ENSG00000102081.14 | FMR1 | 0.176498740090509 | 5.9573819118051 | 1.23677662362086 | 0.222961028852131 | 0.542525849104035 | -5.66131069258262 |
| ENSG00000180354.15 | MTURN | 0.285138873884328 | 4.99407622728023 | 1.23674600996236 | 0.222986339077279 | 0.542525849104035 | -5.62969696456499 |
| ENSG00000103671.9 | TRIP4 | 0.175799872366201 | 3.58577817496773 | 1.23670237631104 | 0.222988324511497 | 0.542525849104035 | -5.50371181771402 |
| ENSG00000132254.12 | ARFIP2 | 0.116583465217633 | 4.57253059401673 | 1.23630107314776 | 0.223135899111878 | 0.542814964078697 | -5.6050716688403 |
| ENSG00000164086.9 | DUSP7 | -0.370623824191227 | 0.249218116599877 | -1.23620334982782 | 0.223171846867993 | 0.542814964078697 | -5.04939134354748 |
| ENSG00000283930.1 | PLD5P1 | 0.248151208879901 | 2.96192317740027 | 1.2361641373789 | 0.223186272214838 | 0.542814964078697 | -5.43625455398287 |
| ENSG00000138092.10 | CENPO | 0.165459745609009 | 2.91932915549804 | 1.23597770938564 | 0.223254865038753 | 0.54284515048409 | -5.4342344244214 |
| lnc-TMTC3-22 | lnc-TMTC3-22 | -0.539182069915771 | -0.761086147666222 | -1.23591680643407 | 0.223277276567674 | 0.54284515048409 | -4.82494392034489 |
| ENSG00000015153.14 | YAF2 | 0.180029274491658 | 4.9807358624207 | 1.23591537263694 | 0.223277804207273 | 0.54284515048409 | -5.63621047960299 |
| ENSG00000181754.6 | AMIGO1 | -0.602085512095035 | 1.93646992716604 | -1.23614144136385 | 0.223359770909256 | 0.542958591360791 | -5.32237041714672 |
| lnc-SLC7A7-1 | lnc-SLC7A7-1 | -0.352414358609589 | -0.375853759450766 | -1.23564526268916 | 0.223377221573297 | 0.542958591360791 | -4.86579167398262 |
| ENSG00000083642.18 | PDS5B | 0.161062170540599 | 5.93943643901447 | 1.2354022379586 | 0.223466697906848 | 0.543111942734285 | -5.6620205243688 |
| lnc-MYL7-1 | lnc-MYL7-1 | -0.237840792545093 | 0.523473473110878 | -1.23526749237567 | 0.2235163196915 | 0.543168407108636 | -5.11930796638828 |
| ENSG00000197937.12 | ZNF347 | -0.174626881848689 | 4.86364540600939 | -1.23514738403166 | 0.223560558005365 | 0.543200323518513 | -5.62112142701399 |
| ENSG00000103429.10 | BFAR | -0.140189637209029 | 5.07392515013089 | -1.23508853705434 | 0.223582234895246 | 0.543200323518513 | -5.63669943597767 |
| lnc-RHOB-1 | lnc-RHOB-1 | -0.771301761877294 | -0.735401201445346 | -1.23499372845641 | 0.223663795667666 | 0.543334345123047 | -4.89164633822948 |
| ENSG00000147124.12 | ZNF41 | -0.153936751882804 | 4.20681599763102 | -1.23457217078392 | 0.223772510493205 | 0.543475401565387 | -5.5762700528636 |
| lnc-RIT1-2 | lnc-RIT1-2 | -0.166637843550686 | 2.26662490525976 | -1.23456631279477 | 0.223774669788984 | 0.543475401565387 | -5.36532480599254 |
| lnc-COQ8A-4 | lnc-COQ8A-4 | -0.339663299569306 | -0.262025053039161 | -1.23440001871378 | 0.223835973387007 | 0.543560151041148 | -4.96865756751543 |
| ENSG00000035115.21 | SH3YL1 | 0.235028969149251 | 3.68322204014614 | 1.23419637814552 | 0.223911061545434 | 0.54367835080685 | -5.50995274343012 |
| lnc-BOLA2B-1 | lnc-BOLA2B-1 | -0.333135721109065 | 1.06139232480309 | -1.23409842341256 | 0.223947186927319 | 0.5437019283469 | -5.21445159257113 |
| lnc-TMEM155-3 | lnc-TMEM155-3 | -0.267217353976047 | 2.67264097351755 | -1.23390790338525 | 0.224017462443319 | 0.543776842017833 | -5.42150133303011 |
| lnc-SLC19A1-8 | lnc-SLC19A1-8 | 0.303289579891308 | 0.137171977741034 | 1.23384864826592 | 0.224039322708033 | 0.543776842017833 | -4.9831953425974 |
| ENSG00000123562.16 | MORF4L2 | 0.165898577169415 | 5.24204872770253 | 1.23377599781441 | 0.224066126903658 | 0.543776842017833 | -5.64732947148201 |
| ENSG00000228696.8 | ARL17B | -0.446034144110903 | 1.32425927053854 | -1.23376423565087 | 0.224083718310997 | 0.543776842017833 | -5.23208083241178 |
| lnc-NOL11-3 | lnc-NOL11-3 | -0.621088438578393 | -0.90688061030067 | -1.23328645097758 | 0.224246805843173 | 0.544012328573523 | -4.81904229166978 |
| ENSG00000176014.12 | TUBB6 | 1.01629165233658 | 0.292912341989417 | 1.23389885368916 | 0.224267374465524 | 0.544012328573523 | -5.17854398540007 |
| ENSG00000280417.1 | | -0.366788148697056 | 0.329171270566042 | -1.23322781082979 | 0.224268455619766 | 0.544012328573523 | -5.07749435872596 |
| ENSG00000182185.18 | RAD51B | -0.178092111459936 | 4.14958319435934 | -1.2331391353877 | 0.22430119728082 | 0.544012328573523 | -5.57258067907188 |
| ENSG00000182606.14 | TRAK1 | 0.14584337616471 | 7.46060800588005 | 1.23295482300284 | 0.224369262319823 | 0.544012328573523 | -5.65917689958079 |
| ENSG00000221946.7 | FXYD7 | -0.472166326123621 | 0.811763726497195 | -1.23291331435589 | 0.22438459322871 | 0.544012328573523 | -5.13746375597618 |
| lnc-CAMSAP2-1 | lnc-CAMSAP2-1 | -0.352587909177645 | -0.600563874074062 | -1.23286682441412 | 0.224401764862139 | 0.544012328573523 | -4.86446214665354 |
| ENSG00000105135.15 | ILVBL | -0.128661129678504 | 3.70952568254779 | -1.2328344516586 | 0.22441372271313 | 0.544012328573523 | -5.53072496967763 |
| ENSG00000251703.1 | RNU6-998P | -0.224907789492062 | 0.341160956001231 | -1.23282116411872 | 0.224418631002176 | 0.544012328573523 | -5.07978895299069 |
| ENSG00000238795.1 | SCARNA12 | 0.377271302921605 | 5.73387074034978 | 1.23295276865545 | 0.22450571357682 | 0.544159338501142 | -5.65785544639669 |
| ENSG00000167193.7 | CRK | 0.180159021037391 | 4.2628615920143 | 1.2324202553785 | 0.224566760189283 | 0.544243214997763 | -5.58853896771267 |
| ENSG00000261326.3 | LINC01355 | -0.413602145317689 | 0.572141358034928 | -1.23224121897913 | 0.224632934561341 | 0.544324739741277 | -5.12907370885779 |
| lnc-ZNF25-2 | lnc-ZNF25-2 | -0.389047566856761 | -0.39209502701651 | -1.23218615520745 | 0.224653289807227 | 0.544324739741277 | -4.89880543173587 |

| | | | | | | | |
|---|---|---|---|---|---|---|---|
| lnc-TMEM39B-1 | lnc-TMEM39B-1 | 0.284361608817577 | -0.443019646763998 | 1.23194402896768 | 0.224742812025299 | 0.544474727914333 | -4.89188654309566 |
| ENSG00000123131.12 | PRDX4 | -0.230323129384606 | 2.37317177579629 | -1.23187565193279 | 0.224768098094937 | 0.544474727914333 | -5.3816616215908 |
| lnc-CNPY2-4 | lnc-CNPY2-4 | 0.156579419118104 | 2.59013087820648 | 1.23174190984725 | 0.224817562476897 | 0.544492409163706 | -5.40637834994886 |
| ENSG00000174945.13 | AMZ1 | 0.632725254230111 | 0.427860022061948 | 1.23194328261249 | 0.224828304206498 | 0.544492409163706 | -5.10629210646597 |
| lnc-BRD1-23 | lnc-BRD1-23 | -0.547805211611341 | 1.37610472845771 | -1.23145328732381 | 0.225019066202126 | 0.544890287016279 | -5.23997863164443 |
| ENSG00000103942.12 | HOMER2 | 0.560522946376792 | 4.35843382965285 | 1.23138943730065 | 0.225159711951843 | 0.54516226687641 | -5.57597978731305 |
| lnc-RTP2-1 | lnc-RTP2-1 | 0.330415792584165 | 0.788070939540977 | 1.23075085085091 | 0.225184355681059 | 0.54516226687641 | -5.16668919375251 |
| ENSG00000072163.19 | LIMS2 | -0.385272702682709 | 0.138839859863895 | -1.23044231714177 | 0.225298635011584 | 0.545318827876624 | -5.05831771190047 |
| ENSG00000213015.8 | ZNF580 | -0.246457387304189 | 3.39432619526841 | -1.23043320173533 | 0.225302011964379 | 0.545318827876624 | -5.49454320525585 |
| ENSG00000173366.11 | | -0.307323600851555 | 3.29369836912284 | -1.23032642514661 | 0.22534157190183 | 0.545350449671414 | -5.48806297480343 |
| ENSG00000115956.9 | PLEK | -0.436746307699658 | 5.66653256327892 | -1.23053105066111 | 0.225442838737275 | 0.545451808190608 | -5.66160942082877 |
| ENSG00000112029.9 | FBXO5 | 0.228701029734473 | 2.26937062774778 | 1.23004956893286 | 0.22544416901071 | 0.545451808190608 | -5.37327867978155 |
| ENSG00000265612.1 | MIR4539 | -0.581603443248851 | -1.33427257862076 | -1.22998010793892 | 0.225469915218362 | 0.545451808190608 | -4.83604197916531 |
| ENSG00000131373.14 | HACL1 | 0.0967042512929215 | 4.32670515587385 | 1.22992739739433 | 0.225489454204629 | 0.545451808190608 | -5.59269000987125 |
| ENSG00000228486.10 | C2orf92 | -0.23269798523199 | 2.88608647737964 | -1.2297688957116 | 0.225548215877786 | 0.545509436194171 | -5.42653671963839 |
| ENSG00000236609.3 | ZNF853 | -0.313038265353555 | 2.73179754711568 | -1.22972016711654 | 0.225566283410999 | 0.545509436194171 | -5.43853730728372 |
| STX18-AS1 | STX18-AS1 | -0.209966704154971 | 4.32262649653604 | -1.22950934791884 | 0.225644463043881 | 0.545616575910106 | -5.59591960431279 |
| ENSG00000143801.16 | PSEN2 | -0.282608571786246 | 2.58008455196357 | -1.22943335109642 | 0.225672650416142 | 0.545616575910106 | -5.42878162173636 |
| ENSG00000162623.15 | TYW3 | 0.184223199259595 | 4.38286879110353 | 1.22936415520725 | 0.225698317572899 | 0.545616575910106 | -5.59972201299484 |
| ENSG00000252021.1 | | -0.305613331979836 | -0.267376796843902 | -1.22930866488571 | 0.225718902418252 | 0.545616575910106 | -4.9229230852566 |
| ENSG00000269837.1 | IPO5P1 | -0.216468161842604 | 3.05100614832229 | -1.22924337072023 | 0.225743125902466 | 0.545616575910106 | -5.45492202952447 |
| lnc-XPNPEP1-1 | lnc-XPNPEP1-1 | -1.21140083312081 | -1.45953065910351 | -1.22955787619893 | 0.225776337066933 | 0.545632775137806 | -4.78000356307 |
| ENSG00000103550.13 | KNOP1 | -0.155979728033281 | 3.52843243493106 | -1.22847568716985 | 0.226028072981061 | 0.546177016455645 | -5.51882512864185 |
| ENSG00000078401.6 | EDN1 | -0.301965631608972 | 1.21443681809407 | -1.22837968119382 | 0.226063726959425 | 0.546199048245784 | -5.24222801233467 |
| ENSG00000028528.14 | SNX1 | -0.107601536777401 | 6.18905928068604 | -1.22813114919711 | 0.226156044212901 | 0.546357964557997 | -5.6723359092547 |
| lnc-PRORY-6 | lnc-PRORY-6 | -2.0962248740552 | -1.62297119810748 | -1.2284851187243 | 0.226184360676418 | 0.546362245722651 | -4.5912608749108 |
| ENSG00000154889.16 | MPPE1 | -0.181700584490945 | 4.82570307213807 | -1.22766828337586 | 0.22632805005638 | 0.546585104146738 | -5.63040146150909 |
| ENSG00000155906.17 | RMND1 | -0.147417368927679 | 2.69830358993915 | -1.22759290872985 | 0.226356069223088 | 0.546585104146738 | -5.42186469669228 |
| ENSG00000237238.3 | BMS1P10 | 1.2620090731164 | -1.6873413192629 | 1.22759869374303 | 0.226385642495495 | 0.546585104146738 | -4.63807692119457 |
| ENSG00000032389.12 | EIPR1 | 0.11975807976943 | 3.74192896088651 | 1.22750676027127 | 0.22638809649904 | 0.546585104146738 | -5.53995102268738 |
| ENSG00000132694.18 | ARHGEF11 | -1.14402966712201 | -1.07586923011563 | -1.22785226583144 | 0.226409396004231 | 0.546585104146738 | -4.81640605115768 |
| ENSG00000131401.11 | NAPSB | -0.246913297131394 | 7.09856369185718 | -1.22733793760904 | 0.226450869141943 | 0.546612419405477 | -5.66843637310705 |
| ENSG00000132603.14 | NIP7 | 0.180948241725888 | 4.0273111818864 | 1.22727392681563 | 0.226474673376439 | 0.546612419405477 | -5.56748754063084 |
| GMDS-DT | GMDS-DT | -0.188462552544284 | 5.47670694572174 | -1.22720480551739 | 0.226500380173411 | 0.546612419405477 | -5.66118810483714 |
| MALINC1 | MALINC1 | -0.260399989252362 | 1.31766658771534 | -1.22701786106902 | 0.226569917198227 | 0.546716131968464 | -5.27331281102891 |
| ENSG00000135482.6 | ZC3H10 | 0.147968013138716 | 3.90574989701625 | 1.22693135771089 | 0.226602098864174 | 0.54672969182057 | -5.55865160476377 |
| ENSG00000153208.16 | MERTK | -1.03481214677652 | -0.236860232859277 | -1.22724386489466 | 0.226652290601839 | 0.5467866968422 | -4.91972430540549 |
| ENSG00000197044.10 | ZNF441 | -0.232010739450375 | 5.78800853294693 | -1.22664473227507 | 0.226708755664106 | 0.546858820793893 | -5.67012419558748 |
| lnc-NOP53-2 | lnc-NOP53-2 | -0.166667627315713 | 2.47171292014944 | -1.22656749411685 | 0.226737503260679 | 0.546864076589472 | -5.39812596863831 |
| ENSG00000163634.11 | THOC7 | -0.164065760500593 | 3.8642187518566 | -1.22601623843663 | 0.226942755628304 | 0.547294989935252 | -5.5486912212976 |
| lnc-TGFBRAP1-13 | lnc-TGFBRAP1-13 | 0.124761373349343 | 2.40765309176156 | 1.22587989854491 | 0.22699354104462 | 0.547311171734617 | -5.39310402571577 |
| ENSG00000223224.1 | SNORD71 | 0.283990611319903 | 0.762349846982041 | 1.2258279742741 | 0.227012884587553 | 0.547311171734617 | -5.13817050459251 |
| ENSG00000172845.14 | SP3 | 0.108236325711403 | 7.66675870585644 | 1.22578408182229 | 0.227029236956234 | 0.547311171734617 | -5.66207844262239 |
| ENSG00000198315.10 | ZKSCAN8 | 0.148206505056621 | 6.59363247206137 | 1.22534172128802 | 0.227194089307484 | 0.547644447852904 | -5.67548572953813 |
| ENSG00000198816.6 | ZNF358 | -0.721097521003233 | -0.327434311702428 | -1.22543347916865 | 0.227244789990873 | 0.547702519014301 | -5.01067956999098 |
| lnc-CDRT1-2 | lnc-CDRT1-2 | -0.405456510550537 | -0.0308975792792328 | -1.22510295589524 | 0.227283105563683 | 0.547730729635558 | -5.06295232477788 |
| ENSG00000119782.13 | FKBP1B | 0.378553772853726 | -0.0837290057754379 | 1.22495170665502 | 0.227339507462138 | 0.547801666139592 | -4.9572045009143 |
| ENSG00000198954.7 | KIFBP | -0.203863654283893 | 2.81745042404818 | -1.2248812909096 | 0.227365769510302 | 0.547801666139592 | -5.4468182470891 |
| ENSG00000117697.14 | NSL1 | -0.122576503949873 | 4.33857410464474 | -1.22480481923638 | 0.227394292696035 | 0.547806264813026 | -5.60272669274139 |
| ENSG00000131748.15 | STARD3 | 0.14666002364724 | 4.48522010495819 | 1.22465678065007 | 0.227449517145916 | 0.547853352820049 | -5.61260867844686 |
| ENSG00000207217.1 | SNORA80D | 0.438302878133082 | -0.402088045726229 | 1.22455345662101 | 0.22748806710206 | 0.547853352820049 | -4.89295585122674 |
| lnc-SORCS1-5 | lnc-SORCS1-5 | -1.22204567567545 | -1.11719770588078 | -1.22499455260068 | 0.227505834162398 | 0.547853352820049 | -4.84820752721629 |
| lnc-TCFL5-7 | lnc-TCFL5-7 | 0.183713919486798 | 1.42312023230577 | 1.22432249492988 | 0.227574255812879 | 0.547853352820049 | -5.2615583519608 |
| ENSG00000135766.8 | EGLN1 | -0.15054767980214 | 4.88550820639776 | -1.22426027028898 | 0.227597480501333 | 0.547853352820049 | -5.64692753813536 |
| ENSG00000132334.16 | PTPRE | 0.206619201400759 | 5.61926816894213 | 1.22425682081036 | 0.227598768033884 | 0.547853352820049 | -5.66755833861397 |
| ENSG00000102967.11 | DHODH | -0.223213158323314 | 2.86218317373126 | -1.22425310109881 | 0.227600156438043 | 0.547853352820049 | -5.4465426772146 |
| ENSG00000165895.18 | ARHGAP42 | 0.334799698654842 | 4.72488481623231 | 1.2238623991222 | 0.22781500698508 | 0.548306394851953 | -5.62860831113578 |
| ENSG00000103966.10 | EHD4 | -0.239327190806732 | 4.36975276918788 | -1.22320687566379 | 0.227990914394795 | 0.548665613350646 | -5.60661978809597 |
| LINC01134 | LINC01134 | -0.254367979685265 | -0.266973547720925 | -1.22312711137134 | 0.228020726100434 | 0.54867320614043 | -4.91111761065951 |
| ENSG00000283239.1 | | 0.600655907786805 | 2.00599162514605 | 1.22349194712439 | 0.228050473771603 | 0.548680643090697 | -5.34435121777273 |
| ENSG00000129625.12 | REEP5 | -0.186565094160809 | 5.24862658673158 | -1.22288038766033 | 0.228112956678546 | 0.548766828811889 | -5.66095894904877 |
| ENSG00000214021.15 | TTLL3 | 0.21801539630227 | 5.49584780066659 | 1.22265272435902 | 0.228198086480731 | 0.548868752039487 | -5.66546885068021 |
| ENSG00000105835.11 | NAMPT | 0.249465828663448 | 5.13489281549236 | 1.22262446082819 | 0.228208656656169 | 0.548868752039487 | -5.65314514358422 |
| lnc-LINS1-2 | lnc-LINS1-2 | -0.482784992205751 | -0.475126580839895 | -1.22191019266236 | 0.228475903122481 | 0.549380925239875 | -4.90466798310922 |
| ENSG00000175606.10 | TMEM70 | 0.194813707964345 | 2.81637410199829 | 1.22183638276605 | 0.2285035325606 | 0.549380925239875 | -5.44520367972323 |
| ENSG00000283765.1 | | 0.128506747764772 | 3.03710715139749 | 1.22179238676607 | 0.228520002862392 | 0.549380925239875 | -5.46482385783324 |
| ENSG00000165392.9 | WRN | 0.179724398479006 | 4.92394550322087 | 1.221688101186 | 0.228559046612494 | 0.549380925239875 | -5.64650595950086 |
| LINC01011 | LINC01011 | -0.3126905689946 | 0.209903047575873 | -1.22163772012793 | 0.228577910670329 | 0.549380925239875 | -5.0743113274675 |

| | | | | | | | |
|---|---|---|---|---|---|---|---|
| ENSG00000083123.14 | BCKDHB | -0.131350772695134 | 3.30291075657497 | -1.22162745580442 | 0.228581754056954 | 0.549380925239875 | -5.49042184610106 |
| ENSG00000235316.1 | DUSP8P5 | -0.198644965094787 | 2.23001613936221 | -1.22152787399956 | 0.228619044071167 | 0.549406396277505 | -5.37355522624024 |
| lnc-COL28A1-2 | lnc-COL28A1-2 | 0.285316385805218 | 1.43001956433267 | 1.22130945284106 | 0.228700851108988 | 0.549538830069612 | -5.27771193544487 |
| ENSG00000059769.19 | DNAJC25 | 0.162414660950819 | 1.62155102613704 | 1.22115870936073 | 0.228757322867268 | 0.549610362621335 | -5.3066074706195 |
| ENSG00000008311.14 | AASS | -1.48377025934214 | -1.87056269043176 | -1.22097430823384 | 0.22892777576977 | 0.549955696623385 | -4.64430724719274 |
| ENSG00000021762.19 | OSBPL5 | -1.68654255326521 | -0.273159671006638 | -1.22129105521271 | 0.228974240716824 | 0.550003127164708 | -5.01285155582888 |
| ENSG00000149679.11 | CABLES2 | 0.177653293685694 | 2.16219775001249 | 1.22027643638679 | 0.229088047532045 | 0.550097057278395 | -5.36511624938922 |
| lnc-MPV17-1 | lnc-MPV17-1 | 0.162694087587751 | 1.94544323755343 | 1.22025140884746 | 0.229097434374655 | 0.550097057278395 | -5.33817213298481 |
| ENSG00000176022.5 | B3GALT6 | -0.282971187482616 | 2.37253963364143 | -1.22016996054524 | 0.229127984381993 | 0.550097057278395 | -5.40391130148387 |
| lnc-NGFR-3 | lnc-NGFR-3 | -0.219612904086766 | 2.33414496889902 | -1.22016433648238 | 0.229130093992419 | 0.550097057278395 | -5.38258860524758 |
| ENSG00000173418.11 | NAA20 | 0.123251647931957 | 3.03478481055194 | 1.22005879352242 | 0.229169686270729 | 0.550097057278395 | -5.46980874998308 |
| ENSG00000125875.13 | TBC1D20 | -0.125995777979686 | 5.09524365712894 | -1.22004809481115 | 0.22917369995444 | 0.550097057278395 | -5.65861408331491 |
| ENSG00000090581.9 | GNPTG | -0.113253956575952 | 4.35733646994505 | -1.2197203685488 | 0.22929667346815 | 0.550274336455964 | -5.61230036197572 |
| SPAG5-AS1 | SPAG5-AS1 | -0.245008842095604 | 0.572622698958412 | -1.21963228290806 | 0.229329734345122 | 0.550274336455964 | -5.11558531701227 |
| lnc-EML1-5 | lnc-EML1-5 | -0.261393489883782 | 1.45685191743018 | -1.21962057665172 | 0.229334128277995 | 0.550274336455964 | -5.26847275071819 |
| lnc-FAM217A-1 | lnc-FAM217A-1 | -0.245783103690621 | 3.61634675875837 | -1.21956632300072 | 0.229354493147535 | 0.550274336455964 | -5.53193996960491 |
| lnc-PFDN4-13 | lnc-PFDN4-13 | -1.56908482366548 | -2.65171367920919 | -1.21949906179247 | 0.229397448884767 | 0.550313250628806 | -4.57876168519449 |
| lnc-RPL13-1 | lnc-RPL13-1 | -0.229470799059526 | 0.063578210097618 | -1.21912607462545 | 0.22951979583826 | 0.550374866309413 | -5.00027108169211 |
| lnc-TMPPE-1 | lnc-TMPPE-1 | 0.233395751910991 | 0.463432130590691 | 1.21901597194713 | 0.229561150467885 | 0.550374866309413 | -5.10417875028384 |
| lnc-RPL26L1-1 | lnc-RPL26L1-1 | -0.325468422012305 | -0.408207110059375 | -1.21900268112323 | 0.229566142880345 | 0.550374866309413 | -4.91762071623641 |
| ENSG00000159128.14 | IFNGR2 | 0.181047241710176 | 6.78711840696555 | 1.21890308168565 | 0.229603557815127 | 0.550374866309413 | -5.68226335803318 |
| ENSG00000124429.17 | POF1B | 2.26099096447834 | -0.226483004116426 | 1.21955946551525 | 0.229612979860314 | 0.550374866309413 | -4.77757647512686 |
| ENSG00000123607.14 | TTC21B | 0.14519551094849 | 5.51964622393316 | 1.21880785060849 | 0.229639335958633 | 0.550374866309413 | -5.67224108058355 |
| lnc-UBE2U-2 | lnc-UBE2U-2 | -0.530438185180057 | 3.9550763807504 | -1.21926483885526 | 0.229659434726781 | 0.550374866309413 | -5.57202214557442 |
| ENSG00000200502.1 | | 0.247907922881411 | 1.98792675257605 | 1.21871888398088 | 0.22967276427051 | 0.550374866309413 | -5.33966489349192 |
| ENSG00000153391.15 | INO80C | 0.177689704181977 | 2.27650336554539 | 1.21863561648397 | 0.229704054439162 | 0.550374866309413 | -5.37749413359113 |
| ENSG00000259378.1 | DCAF13P3 | 0.257190707093985 | 0.104692143750628 | 1.21860936154117 | 0.229713921143933 | 0.550374866309413 | -5.02360903616477 |
| ENSG00000167720.12 | SRR | -0.22807139006051 | 1.30587733408919 | -1.21856949554984 | 0.22972890352811 | 0.550374866309413 | -5.25763946317747 |
| lnc-NPIPA8-9 | lnc-NPIPA8-9 | 0.290082467464708 | 0.770968820991437 | 1.21848646761076 | 0.229760109289189 | 0.550374866309413 | -5.16895940644878 |
| lnc-AKAP1-5 | lnc-AKAP1-5 | 0.518771614716379 | 0.554306640712194 | 1.21846252010605 | 0.229770744118777 | 0.550374866309413 | -5.08148789337514 |
| ENSG00000175970.10 | UNC119B | 0.196408842575703 | 3.58725318827903 | 1.21827652212591 | 0.22983903049689 | 0.550474373367172 | -5.53098602161527 |
| lnc-SETMAR-2 | lnc-SETMAR-2 | 0.241358437449968 | 0.991115427612965 | 1.21807952979789 | 0.229913100602644 | 0.550587707935336 | -5.21181024596816 |
| ENSG00000100393.12 | EP300 | 0.130802907368208 | 7.87447324860641 | 1.21793246731126 | 0.229968408297791 | 0.550615360260641 | -5.66857938647374 |
| lnc-EVI5-2 | lnc-EVI5-2 | -0.387135131736782 | 0.956676919770602 | -1.21790656814208 | 0.229978149548692 | 0.550615360260641 | -5.18665817199613 |
| lnc-CD46-6 | lnc-CD46-6 | -0.576576230918786 | 0.479238211431474 | -1.21784648984434 | 0.230048642556207 | 0.550720075568087 | -5.11285410686723 |
| lnc-RSBN1L-3 | lnc-RSBN1L-3 | -0.357608333906214 | 0.188283712249617 | -1.21757597583822 | 0.230102519331229 | 0.550784993068344 | -5.05303676586515 |
| ENSG00000169203.16 | NPIPB12 | -0.262805190436098 | 1.06022315876001 | -1.21729485244624 | 0.230208317726557 | 0.550974163228573 | -5.24650830570407 |
| ENSG00000126903.15 | SLC10A3 | -0.241519428889982 | 3.51439998955532 | -1.21718844870759 | 0.230248371210748 | 0.551005955659902 | -5.53319459799216 |
| lnc-ITFG2-3 | lnc-ITFG2-3 | 0.213591030802401 | 0.945653592707721 | 1.21700717920191 | 0.230316618179191 | 0.551105202508985 | -5.19137884144283 |
| lnc-VIPR1-5 | lnc-VIPR1-5 | 0.467132717475068 | -0.288803512662471 | 1.21688296850328 | 0.230363391420348 | 0.551153049587937 | -4.86931779078053 |
| lnc-CRIPT-5 | lnc-CRIPT-5 | 0.206094279548292 | 3.55449048038333 | 1.21680440356162 | 0.230392979736458 | 0.551159774746108 | -5.53458387164806 |
| ENSG00000140471.16 | LINS1 | -0.122437658861839 | 4.13562630656599 | -1.21668597685611 | 0.230437585665231 | 0.551202420191452 | -5.59092175188059 |
| ENSG00000119411.10 | BSPRY | -0.400816296892641 | 2.44764419607952 | -1.21666070250542 | 0.230489709467933 | 0.551224870788913 | -5.40828470179777 |
| lnc-INO80D-9 | lnc-INO80D-9 | -0.20296787960655 | -0.303351675824749 | -1.21651888427649 | 0.230500532618188 | 0.551224870788913 | -4.95355549679382 |
| ENSG00000284448.1 | MIR6501 | 0.174676574004876 | 3.00490347140383 | 1.2163975248465 | 0.230546258958479 | 0.551270172878994 | -5.46520935735897 |
| ENSG00000111775.2 | COX6A1 | 0.19224055580396 | 4.26076867921974 | 1.21631721606888 | 0.230576521724434 | 0.551278493048441 | -5.60625603527551 |
| ENSG00000284987.1 | | -0.217534095458092 | 2.74901215953764 | -1.21619623987992 | 0.230622114715675 | 0.551308960231139 | -5.45080482202652 |
| TCL6 | TCL6 | -0.531296375093456 | 5.0332755053076 | -1.21672624934277 | 0.230642834210287 | 0.551308960231139 | -5.65680289892498 |
| lnc-PPP1R27-1 | lnc-PPP1R27-1 | -0.216890019461639 | 3.00003873563962 | -1.21590434353715 | 0.230732150688379 | 0.551436157822433 | -5.46684384636346 |
| ENSG00000100024.14 | UPB1 | -0.470779239150702 | 0.843246731438965 | -1.21591051638416 | 0.230749629661595 | 0.551436157822433 | -5.20984201297309 |
| lnc-RASA3-7 | lnc-RASA3-7 | -0.917090547029187 | 0.989717821142451 | -1.21628251582301 | 0.230800846116195 | 0.551450304459066 | -5.1577659355114 |
| ENSG00000110931.18 | CAMKK2 | -0.206268069609399 | 3.96422669655333 | -1.2157001922229 | 0.230809132435255 | 0.551450304459066 | -5.58124847165759 |
| lnc-HEATR1-4 | lnc-HEATR1-4 | -0.187329539256717 | 0.693902364046072 | -1.21549543663826 | 0.230886361026043 | 0.551514575818244 | -5.17646053292836 |
| ENSG00000172531.14 | PPP1CA | -0.204215124427298 | 5.93249042483408 | -1.21548679063785 | 0.230889622495545 | 0.551514575818244 | -5.68583278071026 |
| lnc-HNRNPLL-4 | lnc-HNRNPLL-4 | 0.569963558565804 | -0.950995585066652 | 1.21540860616298 | 0.230919117010781 | 0.551521024069727 | -4.80825567698214 |
| ENSG00000167395.10 | ZNF646 | -0.15315531177787 | 5.28796089648471 | -1.21508459436084 | 0.231041377621696 | 0.551705615702185 | -5.67103307145304 |
| LINC00909 | LINC00909 | -0.179759518832892 | 4.55046121287309 | -1.2150617151991 | 0.231050012503174 | 0.551705615702185 | -5.62939589033515 |
| ENSG00000069998.12 | HDHD5 | -0.193487826613814 | 2.58130968394798 | -1.21476867009317 | 0.231160632386111 | 0.551867852967157 | -5.42836801586551 |
| ENSG00000101079.20 | NDRG3 | -0.171410135820182 | 4.50003499603903 | -1.21469896824087 | 0.231186949465843 | 0.551867852967157 | -5.62429273781083 |
| lnc-LIPG-13 | lnc-LIPG-13 | -0.187404794103208 | 4.14450126690598 | -1.21466866486968 | 0.231198391689054 | 0.551867852967157 | -5.59515163674784 |
| ENSG00000114126.17 | TFDP2 | -0.206921833225411 | 5.15034761557867 | -1.21454377586372 | 0.231245552815488 | 0.551916420872123 | -5.66691385557716 |
| ENSG00000114670.13 | NEK11 | -0.275923739129384 | 1.78179860911702 | -1.21432954563004 | 0.231326467844126 | 0.551979295476228 | -5.33101862863304 |
| ENSG00000188459.4 | WASF4P | 0.95106244511992 | -1.03553412004189 | 1.21430155323858 | 0.23133704214388 | 0.551979295476228 | -4.6691070867953 |
| ENSG00000174151.14 | CYB561D1 | -0.194595242112561 | 3.82269462581826 | -1.21426103668101 | 0.231352348154954 | 0.551979295476228 | -5.56998948743921 |
| ENSG00000180370.10 | PAK2 | 0.100749500946125 | 7.13024396394956 | 1.21417632175961 | 0.231384353466572 | 0.551991672160692 | -5.6841495081227 |
| ENSG00000163322.13 | ABRAXAS1 | -0.181835671661173 | 4.14424266346891 | -1.21405166463127 | 0.231431454872676 | 0.5520400551216 | -5.5923529897547 |
| ENSG00000050748.17 | MAPK9 | 0.111964148605072 | 4.89811200803787 | 1.2138853693611 | 0.231494300132661 | 0.552125976782221 | -5.65529339438636 |
| ENSG00000197928.10 | ZNF677 | -1.41405743858208 | 1.00810847569786 | -1.2143513473935 | 0.231583650596693 | 0.552217102670354 | -5.18460696795058 |

| | | | | | | | |
|---|---|---|---|---|---|---|---|
| ENSG00000156642.16 | NPTN | -0.116047638053802 | 4.49113443567215 | -1.21364234534924 | 0.231586164808361 | 0.552217102670354 | -5.62496550326281 |
| lnc-RSC1A1-1 | lnc-RSC1A1-1 | -0.125747666741157 | 3.02898198256754 | -1.21331707118814 | 0.231709162534259 | 0.552401397503803 | -5.47690826797175 |
| lnc-ZC2HC1A-1 | lnc-ZC2HC1A-1 | 0.398757636236985 | 1.00944995867428 | 1.21327152880944 | 0.231726387560027 | 0.552401397503803 | -5.21293424757766 |
| ENSG00000165943.4 | MOAP1 | 0.140216331630521 | 5.30746629681212 | 1.21318772982657 | 0.231758084448186 | 0.552401397503803 | -5.67226552793912 |
| lnc-ZNF484-1 | lnc-ZNF484-1 | 0.186818188922395 | 2.25891288068386 | 1.21315410344802 | 0.231770804491223 | 0.552401397503803 | -5.38773236386226 |
| ITFG2-AS1 | ITFG2-AS1 | -0.198113689104712 | 2.83856792799471 | -1.21302695655948 | 0.231818905714132 | 0.552452070894291 | -5.44503529930699 |
| ENSG00000167797.7 | CDK2AP2 | -0.29410118119676 | 2.8923784083847 | -1.21292688354156 | 0.231856769722513 | 0.552478338874563 | -5.4742978430497 |
| lnc-TMEM63A-1 | lnc-TMEM63A-1 | -0.237295775806965 | 2.76350768934688 | -1.21281650911302 | 0.231898536685067 | 0.552513899825064 | -5.43469227572856 |
| ENSG00000134001.12 | EIF2S1 | 0.123648177523883 | 4.95953595867083 | 1.21265015495382 | 0.23196149747765 | 0.552599942428528 | -5.6576472860894 |
| ENSG00000164074.15 | ABHD18 | -0.0930992427647475 | 4.51870634577129 | -1.21249540388188 | 0.232020078080789 | 0.552675531435815 | -5.63143081441573 |
| lnc-LRCH3-1 | lnc-LRCH3-1 | 0.4139283353999 | 1.08641545455313 | 1.21216690433596 | 0.232144466748883 | 0.55290784067256 | -5.22458992552097 |
| ENSG00000254701.3 | | -0.224824122648801 | 1.45533757118053 | -1.21186515050012 | 0.232258771154007 | 0.553077801908543 | -5.28204919068043 |
| lnc-NTHL1-4 | lnc-NTHL1-4 | 0.236611464182515 | 0.0465005141296317 | 1.21183665319515 | 0.232269568075472 | 0.553077801908543 | -5.02273152228442 |
| ENSG00000143401.14 | ANP32E | -0.189251998687354 | 6.25643150145577 | -1.21154470372299 | 0.23238020173274 | 0.553277234501444 | -5.69193312159354 |
| ENSG00000132773.11 | TOE1 | 0.134293838342883 | 5.09431172213389 | 1.2114101963291 | 0.232431186065015 | 0.553324438284088 | -5.66657077850064 |
| lnc-OR2A4-2 | lnc-OR2A4-2 | 0.180800168994423 | 2.24258975529672 | 1.21135001536297 | 0.232454000010675 | 0.553324438284088 | -5.39059811831622 |
| ENSG00000186469.8 | GNG2 | -0.166096683444719 | 6.47787428108884 | -1.21114009753019 | 0.232533590444427 | 0.553324438284088 | -5.69216984157961 |
| lnc-TSPAN9-1 | lnc-TSPAN9-1 | -1.11773122780102 | -2.12946873870997 | -1.21113874120658 | 0.232534104760192 | 0.553324438284088 | -4.64538874243292 |
| lnc-CLK3-1 | lnc-CLK3-1 | -0.259720178090804 | 1.1618311772929 | -1.21113785551372 | 0.232534440613971 | 0.553324438284088 | -5.24075025129342 |
| ENSG00000188921.13 | HACD4 | -0.292910112694472 | 5.27580114709213 | -1.21100876928323 | 0.232618189873648 | 0.553459739009283 | -5.67699472156716 |
| lnc-MED21-1 | lnc-MED21-1 | 0.302401721894576 | -0.195397980561198 | 1.21081504923303 | 0.232656872148859 | 0.553487794664814 | -4.95783398819647 |
| ENSG00000047188.15 | YTHDC2 | 0.147469685787469 | 6.56762267444418 | 1.21053745601146 | 0.232762193485703 | 0.55361110319222 | -5.69302177146447 |
| lnc-BRF1-9 | lnc-BRF1-9 | -0.34945560807713 | 6.15042987333801 | -1.21082695791095 | 0.232762497547756 | 0.55361110319222 | -5.69171207282496 |
| lnc-PPEF2-2 | lnc-PPEF2-2 | -0.172445877159557 | 2.28066279743382 | -1.21032804499662 | 0.232841669062756 | 0.553728099970836 | -5.3920297244171 |
| ENSG00000169246.16 | NPIPB3 | -0.322816157659005 | 2.14080224649471 | -1.21017312208929 | 0.232900478164934 | 0.553728099970836 | -5.38695402405203 |
| ENSG00000283426.1 | SMBD1P | 0.433658108747665 | 0.767673172230863 | 1.2101444932254 | 0.232911346943747 | 0.553728099970836 | -5.1975340086221 |
| lnc-USP14-3 | lnc-USP14-3 | -0.300083650107039 | -0.339317985403807 | -1.2100766558581 | 0.232937102486318 | 0.553728099970836 | -4.95200221458515 |
| ENSG00000231563.1 | | -0.251725858463696 | 0.466970800839954 | -1.21005269745798 | 0.232946199176382 | 0.553728099970836 | -5.11625487643502 |
| lnc-SOX15-2 | lnc-SOX15-2 | -0.199202925270437 | 0.994285490560778 | -1.20958322283185 | 0.233124505187825 | 0.554041667969365 | -5.24084451274028 |
| ENSG00000157036.12 | EXOG | 0.151436022998778 | 4.34568244302976 | 1.20938048341847 | 0.233201536404199 | 0.554041667969365 | -5.61722616163318 |
| ENSG00000135476.11 | ESPL1 | 0.493049261637484 | -0.273769916071364 | 1.20935935639985 | 0.23320956472945 | 0.554041667969365 | -4.92519521101906 |
| ENSG00000110958.15 | PTGES3 | 0.142694964451938 | 6.21383643402516 | 1.20929161530941 | 0.233235307899098 | 0.554041667969365 | -5.69473920330766 |
| ENSG00000141560.14 | FN3KRP | -0.135332642779854 | 3.59133747452856 | -1.20928748920519 | 0.233236875980856 | 0.554041667969365 | -5.54316842001379 |
| lnc-SPOCK2-2 | lnc-SPOCK2-2 | -0.257464921848312 | 0.63401052630583 | -1.20926801701246 | 0.233244276283823 | 0.554041667969365 | -5.16104092982273 |
| ENSG00000175741.5 | RWDD4P2 | 0.346512141654145 | -0.201785703626724 | 1.20920945045911 | 0.233266535229195 | 0.554041667969365 | -4.91051031483731 |
| ENSG00000267264.1 | VN1R95P | -0.273848459097603 | -0.454356913868729 | -1.20859298353987 | 0.233500925760051 | 0.554534389629529 | -4.91248039742244 |
| ENSG00000204252.13 | HLA-DOA | -0.245402227881502 | 7.85698502926096 | -1.20842814666577 | 0.233563628651664 | 0.55461930878371 | -5.68119343364324 |
| ENSG00000232859.9 | LYRM9 | -0.175652970183413 | 2.32999529792216 | -1.20793981063785 | 0.233749461104465 | 0.554884119240388 | -5.39028710399494 |
| ENSG00000035664.11 | DAPK2 | 0.382189314668004 | 2.43585808270952 | 1.20798829979486 | 0.233754587253948 | 0.554884119240388 | -5.40545684835411 |
| lnc-TPGS2-3 | lnc-TPGS2-3 | -0.140040379470781 | 1.81169737759465 | -1.20778962219996 | 0.233806635955016 | 0.554884119240388 | -5.33576587664263 |
| ENSG00000269711.1 | | -0.306315381372017 | 2.04379642781013 | -1.20772241435462 | 0.233832224461126 | 0.554884119240388 | -5.38626229140888 |
| lnc-ARHGEF3-3 | lnc-ARHGEF3-3 | 0.156893707370658 | 3.3454679076355 | 1.20763228101828 | 0.233866544774083 | 0.554884119240388 | -5.51878018266985 |
| ENSG00000099860.8 | GADD45B | 0.560512128071226 | 3.48195607267126 | 1.2081174015216 | 0.233874271934347 | 0.554884119240388 | -5.54036792035567 |
| ENSG00000263712.1 | MIR4639 | 0.303218809713609 | -0.264293523712552 | 1.20751596621758 | 0.233910839737386 | 0.554884119240388 | -4.96396850767957 |
| ENSG00000278099.1 | RNVU1-2A | -1.27638142529103 | -2.45970719392754 | -1.20750856648304 | 0.233913657910107 | 0.554884119240388 | -4.59028432683043 |
| ENSG00000100401.19 | RANGAP1 | -0.114022557262742 | 4.83634686519693 | -1.20743542730136 | 0.233941514147916 | 0.554884119240388 | -5.65814136490071 |
| lnc-HES5-2 | lnc-HES5-2 | -0.174267811098574 | 0.176145852146254 | -1.20742698301899 | 0.233944730445906 | 0.554884119240388 | -5.0691162015957 |
| ENSG00000241170.3 | RPL31P46 | -0.214125088680523 | 0.984362604330274 | -1.20709648749281 | 0.23407063662763 | 0.555071893682279 | -5.23155952171602 |
| ENSG00000201302.1 | SNORA65 | 0.308400561835457 | -0.317307872556257 | 1.2070157236065 | 0.234101412144741 | 0.555071893682279 | -4.93761398890996 |
| ENSG00000140326.13 | CDAN1 | -0.141835568029141 | 4.14764721554078 | -1.20700683200885 | 0.234104800517702 | 0.555071893682279 | -5.6042863823589 |
| lnc-PLA2G10-7 | lnc-PLA2G10-7 | -0.157114199152967 | 2.28317721701782 | -1.20653777138034 | 0.234283599213031 | 0.555400182969721 | -5.38747344193767 |
| ENSG00000225215.1 | SMARCE1P1 | 0.44381979283204 | 1.00248459575807 | 1.20645726148737 | 0.234314298417044 | 0.555400182969721 | -5.18192203206875 |
| ENSG00000159753.13 | CARMIL2 | -0.157995784246694 | 5.76442943144007 | -1.20643127286785 | 0.234324208760096 | 0.555400182969721 | -5.69422608933562 |
| lnc-XKR6-3 | lnc-XKR6-3 | -0.24684839320583 | 5.09424569918749 | -1.20618285817274 | 0.23441895323633 | 0.555560773110349 | -5.67155696725591 |
| ENSG00000106780.8 | MEGF9 | 0.737900591181105 | 3.25436881121671 | 1.2065435145245 | 0.234524096436545 | 0.555711709019551 | -5.49108294019903 |
| ENSG00000134030.13 | CTIF | 0.499312359677292 | 0.915097673797034 | 1.20594179211196 | 0.234536637820184 | 0.555711709019551 | -5.18732939971532 |
| lnc-COL24A1-1 | lnc-COL24A1-1 | 0.222496742626941 | 1.58245636944728 | 1.20551395631605 | 0.234674209717541 | 0.55597367157184 | -5.31981199326028 |
| lnc-RPS16-1 | lnc-RPS16-1 | 0.243830825121085 | 6.32308476462844 | 1.20513361259741 | 0.234819441768377 | 0.556100493120707 | -5.69972931631606 |
| lnc-OPRD1-3 | lnc-OPRD1-3 | -0.449048756037142 | -0.100383196124721 | -1.2051113051233 | 0.234827961794204 | 0.556100493120707 | -4.97675432025311 |
| ENSG00000285509.1 | TBCEL-TECTA | 0.153139374522851 | 2.66537980728523 | 1.20509919848822 | 0.234832585848008 | 0.556100493120707 | -5.4329852095053 |
| lnc-NR1D1-33 | lnc-NR1D1-33 | -0.125460988971833 | 1.34075084261597 | -1.20509075601892 | 0.234835810436049 | 0.556100493120707 | -5.28398785812742 |
| ENSG00000167107.12 | ACSF2 | -0.241103441967488 | 4.70660890577348 | -1.20498497979942 | 0.234876214250757 | 0.55613218887879 | -5.65043606979114 |
| ENSG00000196826.7 | | 0.181654449596342 | 2.32289771546542 | 1.20490269617372 | 0.234907648019328 | 0.556139210890401 | -5.40028220801195 |
| ENSG00000137965.10 | IFI44 | 1.06672886191322 | 1.91044935804327 | 1.2054493356896 | 0.234962886586656 | 0.556139210890401 | -5.38598248630594 |
| lnc-EOMES-5 | lnc-EOMES-5 | 0.141722288603603 | 1.0168691161554 | 1.20462480424623 | 0.235013830343646 | 0.556139210890401 | -5.23981158753071 |
| lnc-B4GALT7-1 | lnc-B4GALT7-1 | -0.165032435573108 | 1.81424292380233 | -1.20458612787382 | 0.235028611349581 | 0.556139210890401 | -5.34315056923612 |
| ENSG00000200792.1 | SNORA80A | 0.565815827817676 | -0.760759789187126 | 1.20455057040192 | 0.235042201002087 | 0.556139210890401 | -4.83254701293609 |
| ENSG00000126266.3 | FFAR1 | 0.463357721977782 | 0.773363904264168 | 1.20447084005086 | 0.2350822741998 | 0.556139210890401 | -5.20021469725865 |

| | | | | | | | |
|---|---|---|---|---|---|---|---|
| ENSG00000181135.15 | ZNF707 | 0.171115672731586 | 3.01529348585971 | 1.2043772296633 | 0.235108458067663 | 0.556139210890401 | -5.47869394011789 |
| ENSG00000197020.10 | ZNF100 | -0.201933165788491 | 4.65726943893634 | -1.20436113447044 | 0.235114610924015 | 0.556139210890401 | -5.6437414010639 |
| lnc-FAM229A-2 | lnc-FAM229A-2 | 0.233900577334809 | 3.3011562483443 | 1.2043408797775 | 0.2351223540382 | 0.556139210890401 | -5.50994999852873 |
| ENSG00000130764.9 | LRRC47 | -0.126082301148116 | 4.52428878757074 | -1.20421472520422 | 0.235170585552178 | 0.5561545502165 | -5.64501682829311 |
| lnc-CNPY1-4 | lnc-CNPY1-4 | 0.247217615880991 | -0.121709067187536 | 1.20409765831918 | 0.235215349141986 | 0.5561545502165 | -4.98787737348719 |
| ENSG00000101752.11 | MIB1 | -0.111048530609196 | 6.17304487575123 | -1.20384276397932 | 0.235312836274064 | 0.5561545502165 | -5.70088761243707 |
| ENSG00000248283.2 | CCNL2P1 | -0.36044738307953 | -0.256456144930332 | -1.20368953846974 | 0.235371453299383 | 0.5561545502165 | -5.00766041838946 |
| lnc-BEND4-6 | lnc-BEND4-6 | -0.36044738307953 | -0.256456144930332 | -1.20368953846974 | 0.235371453299383 | 0.5561545502165 | -5.00766041838946 |
| ENSG00000249784.1 | SCARNA22 | 0.689139479313749 | -0.264255155674707 | 1.20371648115746 | 0.235372232241397 | 0.5561545502165 | -4.90979067949242 |
| ENSG00000266946.1 | MRPL37P1 | 0.240988626835658 | -0.339884488635771 | 1.20363862935562 | 0.235390931150993 | 0.5561545502165 | -4.93373596552657 |
| lnc-ZNF532-3 | lnc-ZNF532-3 | 0.240988626835658 | -0.339884488635771 | 1.20363862935562 | 0.235390931150993 | 0.5561545502165 | -4.93373596552657 |
| ENSG00000132003.9 | ZSWIM4 | -0.226424855998183 | 1.31112333008439 | -1.2035909369439 | 0.235409179362478 | 0.5561545502165 | -5.29976564008831 |
| ENSG00000249129.1 | SUDS3P1 | 0.317123117302758 | -0.365719467913503 | 1.2035459626122 | 0.235426388524028 | 0.5561545502165 | -4.90454185856884 |
| ENSG00000210049.1 | MT-TF | 0.291996270942116 | 4.53695505716377 | 1.20353211540246 | 0.23545249434451 | 0.5561545502165 | -5.6401401548025 |
| lnc-PAPSS2-10 | lnc-PAPSS2-10 | -0.247946459357112 | 2.00045356865458 | -1.20347621138865 | 0.235453080240324 | 0.5561545502165 | -5.37751667910199 |
| ENSG00000179271.2 | GADD45GIP1 | -0.242105771090984 | 2.73104057259401 | -1.20339403237129 | 0.235484530549043 | 0.556165013458515 | -5.46128709261599 |
| ENSG00000111450.13 | STX2 | 0.275515524443669 | 3.61063694252311 | 1.20307805752141 | 0.235605484334062 | 0.556325837349782 | -5.55775052016718 |
| lnc-MAP1LC3C-4 | lnc-MAP1LC3C-4 | -1.16944635404875 | -2.3667421096493 | -1.20307493083119 | 0.235606681444787 | 0.556325837349782 | -4.6138039569466 |
| lnc-FAM184A-2 | lnc-FAM184A-2 | 0.626289371839836 | 1.54927548182126 | 1.20321458853142 | 0.235709673820946 | 0.556477524924432 | -5.2890456088581 |
| ENSG00000160446.18 | ZDHHC12 | -0.187577495758863 | 2.40118559925995 | -1.20270259214028 | 0.23574927001616 | 0.556477524924432 | -5.43230239060309 |
| lnc-SALL2-2 | lnc-SALL2-2 | 0.164894955298244 | 2.69563571613419 | 1.20269538872597 | 0.235752029215423 | 0.556477524924432 | -5.45083646769977 |
| ENSG00000103037.11 | SETD6 | -0.133906894554499 | 3.9222920403295 | -1.20252305634349 | 0.23581804654818 | 0.556501729764954 | -5.6007259594255 |
| ENSG00000172216.5 | CEBPB | -0.283045158969973 | 0.903255269058684 | -1.20246243188193 | 0.23584127387285 | 0.556501729764954 | -5.23449569653822 |
| lnc-EMC7-2 | lnc-EMC7-2 | -0.253861457983674 | 1.20665877643756 | -1.20245689719072 | 0.235843394487669 | 0.556501729764954 | -5.28331922752235 |
| ENSG00000168899.4 | VAMP5 | 0.346067756104499 | 2.52208477097549 | 1.20223273669246 | 0.235930981596284 | 0.556571262589459 | -5.42826490201897 |
| lnc-HEATR4-2 | lnc-HEATR4-2 | -0.527561893579216 | -0.101735547899925 | -1.2022065979304 | 0.235939311155432 | 0.556571262589459 | -4.98631348743559 |
| ENSG00000173041.11 | ZNF680 | -0.17541827850722 | 4.72830531752937 | -1.20216431699742 | 0.235955516366302 | 0.556571262589459 | -5.65584227708364 |
| ENSG00000172775.16 | PSME3IP1 | -0.0868733896676179 | 6.32445932194092 | -1.20209777053108 | 0.235981023593169 | 0.556571262589459 | -5.70308409570176 |
| ENSG00000181885.18 | CLDN7 | -0.906249135861295 | -1.22854029731435 | -1.20178178159237 | 0.236128997653103 | 0.556820885605144 | -4.78628365355072 |
| DDX11-AS1 | DDX11-AS1 | -0.322801807055992 | 1.01146136126561 | -1.20168061177546 | 0.23614096635317 | 0.556820885605144 | -5.23266030894705 |
| lnc-CHST6-1 | lnc-CHST6-1 | 0.184991788117131 | 1.10259480328971 | 1.20141314326506 | 0.236243558171792 | 0.55691696585811 | -5.25298785550511 |
| ENSG00000267796.7 | LIN37 | -0.183662529666368 | 1.62557395325851 | -1.20141258295188 | 0.236243773123119 | 0.55691696585811 | -5.33210858804174 |
| lnc-MBD3L2B-5 | lnc-MBD3L2B-5 | -1.07896061484759 | 0.238936656873331 | -1.20170848848619 | 0.236316527860832 | 0.55691696585811 | -4.95998826386855 |
| lnc-TCL1A-8 | lnc-TCL1A-8 | 0.201386611012412 | 1.15721006103289 | 1.20112179139407 | 0.236355347976216 | 0.55691696585811 | -5.2664713096348 |
| lnc-ZNF837-1 | lnc-ZNF837-1 | -0.194450197438562 | 1.83693464630811 | -1.20110549415944 | 0.236361602262248 | 0.55691696585811 | -5.35465104767742 |
| lnc-HLA-DMA-2 | lnc-HLA-DMA-2 | -0.218261084726858 | 6.25413282441727 | -1.20109646052162 | 0.236365069096159 | 0.55691696585811 | -5.70383756623139 |
| lnc-IPO5-7 | lnc-IPO5-7 | -2.21365202195011 | -0.849810459780446 | -1.20174057606973 | 0.236371112750155 | 0.55691696585811 | -4.74754833653561 |
| ENSG00000103353.15 | UBFD1 | 0.118259077044714 | 5.02377070908222 | 1.20088167343227 | 0.236447508780456 | 0.557033200552607 | -5.67904693192903 |
| ENSG00000272325.1 | NUDT3 | -0.176060075368795 | 7.01757885669198 | -1.20067882633933 | 0.236525384979316 | 0.557152895288311 | -5.70089332227963 |
| ENSG00000123870.10 | ZNF137P | -0.244001912218794 | 2.2017420757507 | -1.20060588087607 | 0.236553394483987 | 0.557155111415941 | -5.3947854380337 |
| ENSG00000162852.13 | CNST | 0.145539995444302 | 6.03523704552672 | 1.20031223413323 | 0.236666173146537 | 0.557356961312949 | -5.70412077743433 |
| ENSG00000109685.17 | NSD2 | 0.174352931002742 | 5.7906036278453 | 1.20012161196301 | 0.236739405015778 | 0.5574656416245 | -5.70240590621577 |
| ENSG00000010803.16 | SCMH1 | -0.198028214665962 | 4.74484746078975 | -1.19996329520927 | 0.236800238628682 | 0.557527959139648 | -5.66207933312832 |
| ENSG00000115540.14 | MOB4 | 0.156872211184177 | 4.77235154032198 | 1.19983471671513 | 0.236849653660216 | 0.557527959139648 | -5.66554104646111 |
| lnc-JHY-5 | lnc-JHY-5 | -0.272059357047926 | 0.226584154276704 | -1.19970729304269 | 0.236898632319291 | 0.557527959139648 | -5.11513255938418 |
| ENSG00000185359.12 | HGS | -0.143698674142112 | 5.56573223258969 | -1.19968967666743 | 0.236905404222372 | 0.557527959139648 | -5.69806274064923 |
| ENSG00000137727.12 | ARHGAP20 | 0.719477440424561 | 3.51707726071978 | 1.20026058053626 | 0.236927773069036 | 0.557527959139648 | -5.51136880992748 |
| lnc-WDR34-2 | lnc-WDR34-2 | -0.285641196037115 | 0.164560356264624 | -1.19962988348248 | 0.236928390350994 | 0.557527959139648 | -5.06771318237979 |
| ENSG00000115274.14 | INO80B | -0.13232439363561 | 3.22177361605731 | -1.19940850595055 | 0.237013508113079 | 0.557664499027379 | -5.51361558909607 |
| ENSG00000141279.15 | NPEPPS | 0.132031403281199 | 5.97035600872217 | 1.19925297144631 | 0.237073323179842 | 0.557740277091379 | -5.70524387978105 |
| ENSG00000177432.7 | NAP1L5 | 0.305509108760571 | 0.702651389220075 | 1.19918385102182 | 0.237099908883524 | 0.557740277091379 | -5.19457886714933 |
| ENSG00000100162.14 | CENPM | 0.484029448604522 | 0.621508014469354 | 1.19893961831173 | 0.237202197645903 | 0.557873258047719 | -5.16514492191391 |
| lnc-RNF227-2 | lnc-RNF227-2 | -0.451139073482373 | 1.39064725758074 | -1.19896255326966 | 0.237210647351389 | 0.557873258047719 | -5.27774392181753 |
| ENSG00000023171.17 | GRAMD1B | -0.627922916586617 | 4.10241116727238 | -1.1990737238061 | 0.237383852060927 | 0.5581703780806 | -5.60462188552608 |
| lnc-ZNF572-3 | lnc-ZNF572-3 | -0.407482186512075 | -0.256219402988778 | -1.1984268386961 | 0.237391220410901 | 0.5581703780806 | -5.0138078431273 |
| ENSG00000267697.1 | NA | 0.208099966809518 | 3.49016699142752 | 1.19821395766879 | 0.237473187892713 | 0.558299329114301 | -5.53742096382036 |
| ENSG00000170542.5 | SERPINB9 | 0.248949452829929 | 5.55243084005504 | 1.19808487523457 | 0.237522899737075 | 0.558352426369143 | -5.6949805715147 |
| ENSG00000112305.14 | SMAP1 | 0.0926215506483752 | 4.76840299011349 | 1.19761095146666 | 0.237705481137755 | 0.558653116133009 | -5.66687896909172 |
| lnc-FAM122C-3 | lnc-FAM122C-3 | -0.276003998881542 | 0.146202229369833 | -1.19760262819946 | 0.237708688633308 | 0.558653116133009 | -5.06990769919089 |
| ENSG00000176020.8 | AMIGO3 | -0.228917415945228 | 1.65419725313876 | -1.19754152278689 | 0.237732237487685 | 0.558653116133009 | -5.33781034715657 |
| ENSG00000198910.13 | L1CAM | -0.623692537076703 | 2.10243817089223 | -1.19789538630009 | 0.23778170336952 | 0.558705570828178 | -5.41198606113678 |
| ENSG00000141577.13 | CEP131 | -0.2287923991451 | 3.29215672744598 | -1.19717560265593 | 0.237873291793614 | 0.558856975800476 | -5.52103782649237 |
| ENSG00000258052.1 | | 0.200347096323187 | 0.480707426013313 | 1.19676614474777 | 0.238031201523174 | 0.559020267193171 | -5.13649013167491 |
| ENSG00000123374.10 | CDK2 | 0.23464744928927 | 3.42925345372514 | 1.19673283066977 | 0.238044052654585 | 0.559020267193171 | -5.55441286321666 |
| ENSG00000235109.7 | ZSCAN31 | -0.424893224699714 | 1.15791007980394 | -1.19675738199013 | 0.238044135379686 | 0.559020267193171 | -5.27460198438912 |
| ENSG00000167702.12 | KIFC2 | -0.241217642703629 | 5.1723449411915 | -1.19671369958275 | 0.238051432830401 | 0.559020267193171 | -5.68883671198606 |
| lnc-NIT1-1 | lnc-NIT1-1 | -0.345947482761627 | 0.172774357715547 | -1.19660765871857 | 0.238092343117433 | 0.559052555143295 | -5.04561489108165 |
| ENSG00000187862.11 | TTC24 | 0.330739216823655 | 3.21974917364063 | 1.19637297795466 | 0.238205213193704 | 0.559253781586176 | -5.53646946050048 |

| | | | | | | | |
|---|---|---|---|---|---|---|---|
| ENSG00000100906.10 | NFKBIA | -0.304348443287028 | 6.31435784676578 | -1.19637568348055 | 0.238242688022358 | 0.559277970753216 | -5.70915560141814 |
| lnc-ASAP1-6 | lnc-ASAP1-6 | 0.171964526132814 | 3.19929727733995 | 1.19562706670609 | 0.238470896520794 | 0.559703849614335 | -5.50848555371957 |
| lnc-SF3B1-4 | lnc-SF3B1-4 | -0.379563603228497 | 2.75486258494109 | -1.19570960342551 | 0.238478490070336 | 0.559703849614335 | -5.45924343028281 |
| ENSG00000164543.6 | STK17A | 0.255693792307405 | 5.8311007531416 | 1.19549702011788 | 0.238522814815361 | 0.559744053967002 | -5.70652966805851 |
| ENSG00000175707.8 | KDF1 | -0.555720279389379 | -0.505918891714008 | -1.1953933042175 | 0.238561204445068 | 0.559757009312175 | -4.92626903340063 |
| ENSG00000171421.12 | MRPL36 | 0.235344584526833 | 1.44676032056561 | 1.19516218811907 | 0.238650514576535 | 0.559757009312175 | -5.30889567278478 |
| ENSG00000179262.9 | RAD23A | -0.169446924795562 | 5.29286306364636 | -1.19508573835844 | 0.238680062414321 | 0.559757009312175 | -5.69711924639487 |
| ENSG00000213231.12 | TCL1B | -0.461094161451735 | 0.801827460177819 | -1.19506484763598 | 0.238694495117483 | 0.559757009312175 | -5.19838846403824 |
| ENSG00000109684.14 | CLNK | -0.527112170196506 | 7.3830461946179 | -1.19556798770202 | 0.238716378956877 | 0.559757009312175 | -5.70119938996753 |
| lnc-EDC3-2 | lnc-EDC3-2 | -0.252995446880591 | 2.11921973716037 | -1.19496117166065 | 0.23872821317856 | 0.559757009312175 | -5.38403378757917 |
| ENSG00000167785.8 | ZNF558 | -0.461896275701749 | 4.67198183340009 | -1.1953572204931 | 0.238760667586822 | 0.559757009312175 | -5.65790662237145 |
| ENSG00000143515.16 | ATP8B2 | 0.137149815050582 | 5.45637177913328 | 1.19486629969382 | 0.238764890323643 | 0.559757009312175 | -5.70037058735071 |
| lnc-SERPINC1-6 | lnc-SERPINC1-6 | 0.156858793981995 | 2.28787835102259 | 1.194845087421 | 0.238773091471646 | 0.559757009312175 | -5.40751245223136 |
| ENSG00000115904.12 | SOS1 | 0.148012415593515 | 6.02541892632136 | 1.19469096463493 | 0.238832685033663 | 0.559806387957512 | -5.71038433566268 |
| ENSG00000167900.11 | TK1 | 0.450952152163044 | -0.121622193645043 | 1.1946499395896 | 0.238848549727585 | 0.559806387957512 | -4.99975922099696 |
| lnc-TBCA-5 | lnc-TBCA-5 | 0.391528145321092 | 0.746375195849501 | 1.19448454283434 | 0.238912517708151 | 0.559892559716141 | -5.19548058263546 |
| lnc-SOX30-3 | lnc-SOX30-3 | -0.309896534233135 | 0.860535311952496 | -1.19428059758268 | 0.238991411734658 | 0.56001368712824 | -5.22368805963043 |
| ENSG00000196922.10 | ZNF252P | -0.185432288650618 | 4.86797100678077 | -1.19401682542534 | 0.239093477381366 | 0.560189077397912 | -5.67347244641342 |
| ENSG00000237541.3 | HLA-DQA2 | 1.78769330331566 | 3.80207627111026 | 1.19456757114876 | 0.239142645148573 | 0.560240503652751 | -5.48844996279985 |
| ENSG00000210082.2 | MT-RNR2 | -0.252787674193222 | 17.8248781823665 | -1.19327398664902 | 0.239381087160363 | 0.56073528132716 | -5.34172153796325 |
| ENSG00000166347.18 | CYB5A | -0.601952851534888 | 2.15465123558356 | -1.19362718841215 | 0.239422827288147 | 0.560753503813483 | -5.39982701175894 |
| lnc-ZSWIM2-2 | lnc-ZSWIM2-2 | -0.326943239358807 | 0.319447908385608 | -1.19307837575074 | 0.23945686513126 | 0.560753503813483 | -5.13389810387314 |
| ENSG00000197451.11 | HNRNPAB | 0.147698277922213 | 4.79021235193748 | 1.19304293351123 | 0.239470597022811 | 0.560753503813483 | -5.67370314490985 |
| lnc-N4BP2-1 | lnc-N4BP2-1 | -0.207089582832425 | 1.51090583485302 | -1.19286523291273 | 0.239539454765891 | 0.560850938169303 | -5.32917538594572 |
| ENSG00000004455.16 | AK2 | -0.129838891305387 | 5.95008956686354 | -1.19276142400951 | 0.239579686681661 | 0.560866197699882 | -5.71207392962316 |
| lnc-PGPEP1-6 | lnc-PGPEP1-6 | -0.356019705693427 | -0.236554553294935 | -1.19269330090105 | 0.239606090982383 | 0.560866197699882 | -4.95592879637833 |
| ENSG00000055609.17 | KMT2C | 0.163666991986455 | 8.31907056200365 | 1.19257900797666 | 0.239650395330202 | 0.560866197699882 | -5.68890681460152 |
| ENSG00000138686.9 | BBS7 | -0.23375868181225 | 1.96637118509576 | -1.19256721233161 | 0.239654968117887 | 0.560866197699882 | -5.3789133454972 |
| ENSG00000181513.14 | ACBD4 | -0.180720755309949 | 3.19484208313182 | -1.19232043785997 | 0.239750649157744 | 0.561026331470425 | -5.51681985150084 |
| lnc-POLE4-5 | lnc-POLE4-5 | 0.160043837200104 | 1.92029629646009 | 1.19212794466095 | 0.239825303261412 | 0.561137230536505 | -5.37072106715259 |
| lnc-CEBPD-4 | lnc-CEBPD-4 | 0.133489863783971 | 2.02919808512781 | 1.19203973190989 | 0.239859520237944 | 0.561153501370493 | -5.38465146759489 |
| ENSG00000241127.7 | YAE1 | -0.166137434839317 | 2.12691850850745 | -1.19189024851466 | 0.239917511713038 | 0.561225382837762 | -5.40130223649501 |
| lnc-CAMK2N2-4 | lnc-CAMK2N2-4 | -0.259519006029425 | 0.0192632229883562 | -1.19179055373379 | 0.239956193586468 | 0.561252083248893 | -5.06843791586107 |
| ENSG00000170962.12 | PDGFD | -0.705822810644588 | 4.72177337315196 | -1.19230983060397 | 0.240006482524852 | 0.561301725056192 | -5.65617018822775 |
| ENSG00000013573.16 | DDX11 | -0.256652122355456 | 5.18649373686492 | -1.19155574774111 | 0.240047517690169 | 0.561301725056192 | -5.69747035096534 |
| ENSG00000207642.1 | MIR571 | 0.277599010306753 | -0.0126064543752506 | 1.19142085535792 | 0.240099677405417 | 0.561301725056192 | -5.02575972924804 |
| lnc-ERAP2-4 | lnc-ERAP2-4 | -0.16395269414021 | 3.81271392526397 | -1.19140241498547 | 0.240106835946744 | 0.561301725056192 | -5.5900826641635 |
| ENSG00000126247.10 | CAPNS1 | -0.229639702852353 | 6.17619056447088 | -1.19138455813043 | 0.240113768115424 | 0.561301725056192 | -5.71479753681067 |
| ENSG00000142546.13 | NOSIP | 0.37740288848947 | 6.12213657688568 | 1.19145379176487 | 0.240233387039257 | 0.561313838974314 | -5.71452367236064 |
| ENSG00000213862.4 | RPL7AP62 | 0.694933661516013 | -0.302430346576584 | 1.19103111413026 | 0.240251007858953 | 0.561313838974314 | -4.86946066249623 |
| lnc-SLC35F5-10 | lnc-SLC35F5-10 | -0.122385685791476 | 2.21001052410143 | -1.19102317637446 | 0.240254090689094 | 0.561313838974314 | -5.41631887010018 |
| ENSG00000204673.10 | AKT1S1 | -0.122350765548932 | 3.67457772968374 | -1.19089798255501 | 0.240302716723631 | 0.561313838974314 | -5.57309758852117 |
| ENSG00000187051.8 | RPS19BP1 | 0.153107317123609 | 3.3319750394807 | 1.19089217353585 | 0.240304973156046 | 0.561313838974314 | -5.53734066174693 |
| lnc-OR4F16-16 | lnc-OR4F16-16 | 0.225877274245916 | -0.00413557886461499 | 1.1908862862041 | 0.240307260023665 | 0.561313838974314 | -5.00601237379847 |
| ENSG00000124575.6 | H1-3 | 1.47048885513651 | 3.89228310833048 | 1.19147909136766 | 0.240338802687347 | 0.561313838974314 | -5.54927087252077 |
| lnc-RBM25-1 | lnc-RBM25-1 | 0.312286031735704 | 1.14671779785317 | 1.19080058166589 | 0.240340552783283 | 0.561313838974314 | -5.26025098637252 |
| ENSG00000231991.4 | ANXA2P2 | 0.99710497389447 | -0.187871601880596 | 1.19097214811497 | 0.240364386956207 | 0.561313838974314 | -4.82464216383327 |
| ENSG00000100060.17 | MFNG | -0.18431053972452 | 4.51712760818895 | -1.19059971496405 | 0.240418594590744 | 0.561376736524252 | -5.65537829543019 |
| lnc-B4GALNT3-2 | lnc-B4GALNT3-2 | -0.261443236484308 | 3.37755976187773 | -1.19036933433356 | 0.240508126071063 | 0.561475101393651 | -5.54796820886612 |
| ENSG00000173660.11 | UQCRH | 0.188925647920152 | 5.5285333082944 | 1.19028526787059 | 0.240540802387122 | 0.561475101393651 | -5.70777475382084 |
| ENSG00000142330.19 | CAPN10 | -0.132418162051851 | 4.399596371693 | -1.19028075483825 | 0.240542556677317 | 0.561475101393651 | -5.6525411623233 |
| ENSG00000043093.13 | DCUN1D1 | 0.112464169880971 | 5.59422498246034 | 1.19014992162008 | 0.240593417767474 | 0.561495065205998 | -5.71023958321926 |
| ENSG00000105619.13 | TFPT | -0.205199268382531 | 1.56489105548843 | -1.19011841124914 | 0.24060566851853 | 0.561495065205998 | -5.32900883833084 |
| ENSG00000197119.12 | SLC25A29 | -0.257923568839492 | 3.69577861223081 | -1.18980898350202 | 0.24072599347797 | 0.561564930472925 | -5.57213940992075 |
| ENSG00000168303.7 | MPLKIP | -0.174344637774463 | 3.3522331251255 | -1.18976026013955 | 0.240744944183441 | 0.561564930472925 | -5.54634234884327 |
| lnc-PFKFB3-453 | lnc-PFKFB3-453 | 0.305864181251992 | 0.00045928691048016 | 1.18975057414299 | 0.240748711632493 | 0.561564930472925 | -5.01247885188731 |
| ENSG00000182511.11 | FES | 0.724703118962901 | 0.382068766605225 | 1.19005387200898 | 0.240754834014839 | 0.561564930472925 | -5.09157836682238 |
| lnc-KCNC3-1 | lnc-KCNC3-1 | -0.247332227602766 | 4.27435036798456 | -1.18966345653431 | 0.240782598685012 | 0.561564930472925 | -5.638387027824 |
| lnc-ACSL4-1 | lnc-ACSL4-1 | 0.206377778161676 | 0.416372343616368 | 1.18962051315492 | 0.240799304103096 | 0.561564930472925 | -5.12813753401659 |
| lnc-ROPN1B-2 | lnc-ROPN1B-2 | -0.220542041631092 | 0.287385218373373 | -1.18948635246144 | 0.240851499444502 | 0.561622067344686 | -5.12064699297565 |
| lnc-PYURF-3 | lnc-PYURF-3 | 0.341100912743292 | 2.83549764887522 | 1.18946406047039 | 0.240878375869353 | 0.561622067344686 | -5.49673361713183 |
| lnc-DHRS7-1 | lnc-DHRS7-1 | -0.267076365205855 | 0.51015296504965 | -1.18929604346063 | 0.240925553490778 | 0.561668441216524 | -5.15881680142483 |
| ENSG00000144746.6 | ARL6IP5 | -0.200271373362197 | 6.07052480861809 | -1.1891364966009 | 0.240987650005256 | 0.561678786998193 | -5.71724357443611 |
| ENSG00000169951.9 | ZNF764 | -0.181112104006685 | 2.50926432536877 | -1.18910768706762 | 0.2409988640779 | 0.561678786998193 | -5.45403900947532 |
| ENSG00000196793.13 | ZNF239 | -0.282910024252827 | 1.507308925929 | -1.18907430953223 | 0.241011856715155 | 0.561678786998193 | -5.31136391412734 |
| lnc-ADAT1-3 | lnc-ADAT1-3 | 0.116691351628601 | 3.00852654053585 | 1.18896015332815 | 0.241056297354864 | 0.561718755627211 | -5.49626406838267 |
| ENSG00000180787.5 | ZFP3 | -0.183579197661499 | 4.50809031252043 | -1.18858120723815 | 0.241203862679379 | 0.56197892800552 | -5.65911220167065 |
| ENSG00000196458.10 | ZNF605 | -0.148051712911276 | 4.78143883434134 | -1.18844768090704 | 0.241255874832542 | 0.56197892800552 | -5.67785435545322 |

| | | | | | | | |
|---|---|---|---|---|---|---|---|
| ENSG00000176783.14 | RUFY1 | -0.110617093972687 | 5.20188691327552 | -1.18841951584694 | 0.241266846937041 | 0.56197892800552 | -5.70071966879088 |
| ENSG00000026950.16 | BTN3A1 | 0.180662516384074 | 6.08611990931747 | 1.18839304306556 | 0.241277160121692 | 0.56197892800552 | -5.71698980777606 |
| ENSG00000213246.6 | SUPT4H1 | 0.161178520264019 | 3.90137091489292 | 1.18831963311606 | 0.241305760625201 | 0.561981949643417 | -5.60353742863561 |
| ENSG00000108423.14 | TUBD1 | 0.171749395727635 | 3.17157209561008 | 1.18814507720482 | 0.241373777503833 | 0.562053242519622 | -5.50948711935123 |
| ENSG00000114450.9 | GNB4 | 1.23688112794392 | -0.174703644774527 | 1.18867717948839 | 0.241390985953139 | 0.562053242519622 | -4.99023884363421 |
| ENSG00000068796.16 | KIF2A | 0.0924691572199432 | 5.94096863869852 | 1.18751590583293 | 0.241619054316962 | 0.562520641896394 | -5.71785490020648 |
| ENSG00000130332.14 | LSM7 | 0.206773815077513 | 4.08417684674665 | 1.18727717116826 | 0.241712170411383 | 0.56267378461632 | -5.62139913226655 |
| ENSG00000104907.12 | TRMT1 | -0.150434037972112 | 4.28065387216768 | -1.18699132830264 | 0.241823694993836 | 0.562796788248669 | -5.64622916543078 |
| ENSG00000138802.11 | SEC24B | 0.123718322197147 | 6.12321447091161 | 1.18675398212967 | 0.241916326575652 | 0.562796788248669 | -5.72031773258714 |
| ENSG00000273784.4 | | 0.296710501275687 | -0.372031817913416 | 1.18662155318786 | 0.241968022244917 | 0.562796788248669 | -4.9693168292121 |
| ENSG00000236576.1 | | -0.370816392001285 | -0.70805558259739 | -1.18659234650523 | 0.241979424604884 | 0.562796788248669 | -4.87097357504834 |
| lnc-GSPT2-2 | lnc-GSPT2-2 | -0.370816392001285 | -0.70805558259739 | -1.18659234650523 | 0.241979424604884 | 0.562796788248669 | -4.87097357504834 |
| ENSG00000163874.10 | ZC3H12A | -0.264259535313226 | 4.9387859521767 | -1.18660658824199 | 0.241980232357619 | 0.562796788248669 | -5.69362538522768 |
| ENSG00000113552.15 | GNPDA1 | -0.212745630130088 | 3.12231262823902 | -1.18658410512675 | 0.241982642129921 | 0.562796788248669 | -5.50977877755998 |
| lnc-CDHR4-3 | lnc-CDHR4-3 | 0.251951824473593 | 2.02140896877999 | 1.18646919289021 | 0.242027508381739 | 0.562796788248669 | -5.39025994333743 |
| lnc-CLK4-2 | lnc-CLK4-2 | 0.10055256585438 | 3.17540722347905 | 1.18631615004411 | 0.242087271732587 | 0.562796788248669 | -5.52652616453333 |
| ENSG00000114573.9 | ATP6V1A | -0.212947732154377 | 4.95983712821956 | -1.18628849355024 | 0.242098072761941 | 0.562796788248669 | -5.69130707689223 |
| LINC00963 | LINC00963 | -0.727279730015036 | 1.18035393669992 | -1.18673808663159 | 0.242105376888832 | 0.562796788248669 | -5.25284029109618 |
| ENSG00000114853.13 | ZBTB47 | 0.816690825653673 | -1.51708949447167 | 1.18624905251418 | 0.242113476760274 | 0.562796788248669 | -4.7596330551127 |
| ENSG00000010318.20 | PHF7 | 0.128300961654738 | 3.47940715792532 | 1.18623115654996 | 0.242120466401495 | 0.562796788248669 | -5.54757086413448 |
| ENSG00000165458.13 | INPPL1 | -0.130132863366859 | 5.74777124234749 | -1.18563689621663 | 0.242352650637052 | 0.563272878055832 | -5.71772484173048 |
| lnc-BCL6-10 | lnc-BCL6-10 | -0.788816994661117 | -0.94004137324226 | -1.18547700681934 | 0.242416749859954 | 0.563358243464765 | -4.81705022455126 |
| ENSG00000264057.1 | | -0.335469337733095 | 1.78706504970873 | -1.18515221121824 | 0.242542142596217 | 0.563547956180796 | -5.36062310846573 |
| ENSG00000166582.9 | CENPV | -0.578472273665466 | -0.553054538178328 | -1.18512408253472 | 0.242553143069799 | 0.563547956180796 | -4.9141269242365 |
| ENSG00000285782.1 | NA | -0.568469736703164 | 3.09121135737828 | -1.18534457194032 | 0.242651271382122 | 0.563580145802924 | -5.50762521790047 |
| lnc-B3GNT5-3 | lnc-B3GNT5-3 | 0.304353436419089 | 1.05844095288043 | 1.18485273247575 | 0.24265928042283 | 0.563580145802924 | -5.25841965800913 |
| ENSG00000104613.11 | INTS10 | -0.158410381007638 | 5.3419394093204 | -1.18479658836753 | 0.242681245159866 | 0.563580145802924 | -5.70751488707142 |
| lnc-ANKRD30BL-4 | lnc-ANKRD30BL-4 | -1.04981128838124 | -1.64782095670578 | -1.18471411845121 | 0.242713511722918 | 0.563580145802924 | -4.67882942247985 |
| lnc-BTN2A2-1 | lnc-BTN2A2-1 | 0.211651832582355 | 1.96330653447104 | 1.18464191451689 | 0.242741764259237 | 0.563580145802924 | -5.37022592192278 |
| lnc-MNS1-11 | lnc-MNS1-11 | 0.332146251294629 | -0.0229804250921669 | 1.18458622085311 | 0.242763558159873 | 0.563580145802924 | -5.04221157928885 |
| ENSG00000163026.11 | WDCP | 0.2065405056185 | 3.33501975728959 | 1.1845674069565 | 0.242770920687212 | 0.563580145802924 | -5.54436825822787 |
| ENSG00000069966.18 | GNB5 | 0.099087506419159 | 5.67509926977529 | 1.18452876163417 | 0.242786044446122 | 0.563580145802924 | -5.71744021452395 |
| ENSG00000232587.1 | EEF1A1P3 | 0.363102642550035 | 3.51850794955829 | 1.18445682268194 | 0.242874689117975 | 0.563722341690942 | -5.55107314065314 |
| ENSG00000110367.12 | DDX6 | 0.173433540095564 | 8.61575485374562 | 1.18380263751493 | 0.243070339011514 | 0.564112841117826 | -5.69180028124871 |
| lnc-ZNF683-3 | lnc-ZNF683-3 | -0.306491692995211 | 0.312160223813554 | -1.18340317526715 | 0.243226841104216 | 0.564412409295162 | -5.12487430453832 |
| ENSG00000272655.2 | POLR2J4 | -0.142519584788162 | 2.36087484968036 | -1.18324311275515 | 0.243289571281196 | 0.56443812236512 | -5.43824045365622 |
| GHRLOS | GHRLOS | 0.361092441073352 | 0.159756650072953 | 1.18318975064151 | 0.243310487069903 | 0.56443812236512 | -5.04796567967657 |
| lnc-FAM104B-1 | lnc-FAM104B-1 | -0.267601709290366 | -0.311008306202779 | -1.18316499821274 | 0.243320189464398 | 0.56443812236512 | -4.95656344355327 |
| lnc-NBPF1-2 | lnc-NBPF1-2 | -0.246309058196351 | 1.97129720691886 | -1.18294183575786 | 0.243407676832537 | 0.56457744109129 | -5.38632414075665 |
| ENSG00000147687.18 | TATDN1 | -0.170224739112279 | 3.34214280302829 | -1.18286526698555 | 0.243437699702119 | 0.564583456109151 | -5.54567597333746 |
| lnc-SPCS2-3 | lnc-SPCS2-3 | -0.202767567516965 | 1.09373079808689 | -1.18257576111251 | 0.243551240270642 | 0.564704795208524 | -5.27483102414672 |
| ENSG00000148400.10 | NOTCH1 | 0.219508079310585 | 5.2761558909044 | 1.18257447434753 | 0.243551745009707 | 0.564704795208524 | -5.70929816621185 |
| ENSG00000275454.1 | | -0.238455571586177 | 0.223885608879221 | -1.18251969769466 | 0.243573232091762 | 0.564704795208524 | -5.11742182669316 |
| ENSG00000078900.14 | TP73 | 0.974884042268475 | 1.81931838968798 | 1.18307530290851 | 0.243603790221943 | 0.564704795208524 | -5.29292010482475 |
| ENSG00000196504.16 | PRPF40A | 0.0972399373225379 | 7.20047080742529 | 1.18238214315633 | 0.243627196300427 | 0.564704795208524 | -5.71943803727342 |
| ENSG00000125356.6 | NDUFA1 | 0.186818427901481 | 4.60358632775874 | 1.18218131859458 | 0.243705997677639 | 0.564823843058084 | -5.66814540414126 |
| lnc-KCND1-2 | lnc-KCND1-2 | 0.177466530522741 | 1.00297789070019 | 1.18181555140942 | 0.243849568387284 | 0.564842922420962 | -5.24705191913279 |
| ENSG00000256436.1 | TAS2R31 | -0.259980381822343 | 2.33851568960943 | -1.18179254937129 | 0.243858599188838 | 0.564842922420962 | -5.44068513777979 |
| lnc-CER1-111 | lnc-CER1-111 | -0.237497952433082 | 3.68585614293474 | -1.18178768056157 | 0.243860510757356 | 0.564842922420962 | -5.59263924277099 |
| lnc-C3orf33-2 | lnc-C3orf33-2 | -0.132950536456991 | 0.952260503990303 | -1.18173200147765 | 0.243882371984988 | 0.564842922420962 | -5.2532082823657 |
| ENSG00000187676.7 | B3GLCT | 0.1654291141896 | 3.50295186946855 | 1.18172701203453 | 0.243884331055054 | 0.564842922420962 | -5.56667658353126 |
| ENSG00000096746.17 | HNRNPH3 | 0.229732348930558 | 6.48379150366991 | 1.18172525607152 | 0.243885020524415 | 0.564842922420962 | -5.72642648022436 |
| ENSG00000175854.11 | SWI5 | 0.181529047186188 | 1.87119377996867 | 1.18167099805271 | 0.243906325340209 | 0.564842922420962 | -5.36053445826128 |
| ENSG00000092140.15 | G2E3 | 0.161310205064271 | 5.25910918428355 | 1.18144241897057 | 0.243996093485829 | 0.564930848573782 | -5.70921287588452 |
| ENSG00000200769.1 | | -0.358813494659495 | -0.725909724373363 | -1.1814345457072 | 0.243999185921436 | 0.564930848573782 | -4.93290483479523 |
| ENSG00000240342.3 | RPS2P5 | -0.251902730907967 | 5.28582421039007 | -1.18117809292205 | 0.244099930225962 | 0.565083416901019 | -5.70548942227169 |
| ENSG00000099942.12 | CRKL | 0.163427399642605 | 5.93013337295932 | 1.18111408908939 | 0.244125078056235 | 0.565083416901019 | -5.72568384210432 |
| lnc-NXPH2-3 | lnc-NXPH2-3 | -0.274693609374711 | 3.80097229024981 | -1.1810009025351 | 0.244169554946457 | 0.565083416901019 | -5.59308308076637 |
| lnc-GRAMD2A-6 | lnc-GRAMD2A-6 | -0.128950318001192 | 1.80717492344669 | -1.18098730825992 | 0.244174897241299 | 0.565083416901019 | -5.37490781925805 |
| ENSG00000187109.13 | NAP1L1 | 0.121436564720815 | 7.79067173904366 | 1.18072109790593 | 0.244279530041092 | 0.565199176714772 | -5.71171623476043 |
| lnc-DMXL1-8 | lnc-DMXL1-8 | -0.367223133642177 | 4.2793039571528 | -1.18092586561123 | 0.244302948231461 | 0.565199176714772 | -5.64202110474062 |
| ENSG00000214626.2 | POLR3DP1 | -0.383465778560372 | 1.04081273350923 | -1.18065046851726 | 0.244307296080811 | 0.565199176714772 | -5.24607156839759 |
| lnc-TKT-1 | lnc-TKT-1 | 0.183809191545273 | 2.74878708983726 | 1.18046817030967 | 0.244378972310588 | 0.565301459549206 | -5.48013434034283 |
| ENSG00000143756.11 | FBXO28 | 0.139155373893793 | 5.50326808739291 | 1.18035967764475 | 0.244421636844354 | 0.565328551211069 | -5.71781115112513 |
| ENSG00000269897.5 | COMMD3-BMI1 | -0.192405176658192 | 4.64172430799177 | -1.18029870755054 | 0.24444561559435 | 0.565328551211069 | -5.67739229183422 |
| lnc-C1QTNF8-4 | lnc-C1QTNF8-4 | 0.213588483051651 | 1.69719669825213 | 1.18020062975708 | 0.244484191911066 | 0.565352711936249 | -5.35088264215816 |
| ENSG00000133250.13 | ZNF414 | -0.161731804705445 | 2.02183191961005 | -1.18013248735941 | 0.244510996533862 | 0.565352711936249 | -5.40132204225801 |
| lnc-STK25-1 | lnc-STK25-1 | -0.243030078517351 | 0.935234533771193 | -1.1799549467319 | 0.244580844286126 | 0.56545069279359 | -5.24478473026023 |

| | | | | | | | |
|---|---|---|---|---|---|---|---|
| ENSG00000188428.19 | BLOC1S5 | -0.134418211950954 | 3.56890646171922 | -1.17980719981578 | 0.24463898170025 | 0.565521581349533 | -5.57728615572716 |
| ENSG00000072062.13 | PRKACA | -0.18155737077794 | 2.78375444830979 | -1.17960585734855 | 0.244718224772874 | 0.565641237563173 | -5.48207255518517 |
| ENSG00000252139.1 | | 0.226997367587732 | 1.55575902323427 | 1.1794603031539 | 0.244775522673726 | 0.565710148573243 | -5.33065593122818 |
| lnc-NT5DC3-5 | lnc-NT5DC3-5 | -0.571293352982784 | 2.05141518925666 | -1.17956754062762 | 0.244902861009717 | 0.565940899086449 | -5.42648822752241 |
| ENSG00000105404.10 | RABAC1 | -0.214045593789072 | 2.78675824758588 | -1.17874087929463 | 0.245058869506624 | 0.566207825106641 | -5.49680637862449 |
| ENSG00000269403.1 | | -0.272492363426813 | 1.51218451589526 | -1.17870403666555 | 0.245073386477922 | 0.566207825106641 | -5.32464861133624 |
| ENSG00000108055.9 | SMC3 | 0.11483734477133 | 6.22851582525139 | 1.17824160839735 | 0.245255648601432 | 0.566565321567148 | -5.73046417630301 |
| ENSG00000139618.14 | BRCA2 | 0.211968897897052 | 4.92512252155572 | 1.17779803320187 | 0.245430572463214 | 0.56690578756709 | -5.69533181578419 |
| ENSG00000172725.13 | CORO1B | -0.298835124438378 | 5.81743673118814 | -1.17783623796533 | 0.245475458421331 | 0.566945843883108 | -5.72770140410266 |
| ENSG00000115204.14 | MPV17 | -0.131021831968223 | 4.68499265155729 | -1.1775059920668 | 0.245545788336143 | 0.56704464953695 | -5.67966181036009 |
| ENSG00000146918.19 | NCAPG2 | 0.153455955697727 | 2.98252179929309 | 1.17711205364641 | 0.245701266884746 | 0.56727334883343 | -5.51194765007351 |
| lnc-ZNF131-10 | lnc-ZNF131-10 | 0.243951886875206 | 0.284202651251279 | 1.17704668426342 | 0.245727073610191 | 0.56727334883343 | -5.12090570428134 |
| ENSG00000187688.14 | TRPV2 | -0.183603209170573 | 4.57676377976662 | -1.17704559862056 | 0.245727502220224 | 0.56727334883343 | -5.67732003636013 |
| ENSG00000277925.1 | | -0.438373552629336 | 3.07291825499881 | -1.17721425519826 | 0.245791901371036 | 0.567358383528095 | -5.54001229563011 |
| lnc-NXPE3-1 | lnc-NXPE3-1 | -0.363096909885494 | 0.227447325054666 | -1.17664300252648 | 0.245886483933911 | 0.567472933277139 | -5.1002605246101 |
| lnc-GTF3C1-2 | lnc-GTF3C1-2 | 0.188621429638503 | -0.117562911933306 | 1.17656027848725 | 0.245919160188458 | 0.567472933277139 | -5.02658203493482 |
| ENSG00000224094.1 | RPS24P8 | 0.778391914168771 | -0.68270443425287 | 1.17654742727462 | 0.245924236740615 | 0.567472933277139 | -4.78222975193835 |
| lnc-RRP8-5 | lnc-RRP8-5 | 0.178157109086314 | 0.415743085813844 | 1.1764556737122 | 0.24596048391598 | 0.567492953754356 | -5.18171654898983 |
| ENSG00000120963.11 | ZNF706 | 0.169997082379277 | 5.83177086622602 | 1.17630238260119 | 0.246021050106355 | 0.567529171345378 | -5.72974411281086 |
| lnc-OCM-3 | lnc-OCM-3 | -0.181137432702902 | 0.556366895632942 | -1.17622350320727 | 0.24605222002266 | 0.567529171345378 | -5.18454567918056 |
| ENSG00000129691.15 | ASH2L | 0.0938446683086562 | 5.27811084735863 | 1.17620660130196 | 0.246058899338588 | 0.567529171345378 | -5.71349211763193 |
| ENSG00000109654.14 | TRIM2 | -0.526564019414189 | 2.55008702579492 | -1.17646159531483 | 0.246123008390738 | 0.567613432123985 | -5.48886690904338 |
| ENSG00000250251.6 | PKD1P6 | -0.249102905000996 | 2.89148839380562 | -1.1758704696737 | 0.246191759591959 | 0.567624346590219 | -5.51715232502971 |
| ENSG00000185697.16 | MYBL1 | -0.819803921286692 | 1.6108490799639 | -1.17642089475014 | 0.246194165635979 | 0.567624346590219 | -5.31277710923537 |
| ENSG00000090971.4 | NAT14 | -0.247936914734054 | 0.99597331321806 | -1.17582313593583 | 0.246210473029076 | 0.567624346590219 | -5.26989092027655 |
| ENSG00000137210.13 | TMEM14B | 0.121333608828963 | 4.05931732687305 | 1.17570030229917 | 0.246259040237296 | 0.567640209058344 | -5.63044432209469 |
| ENSG00000170043.11 | TRAPPC1 | -0.23064726100824 | 3.41365013142343 | -1.17566623918727 | 0.246272509687169 | 0.567640209058344 | -5.56479355712806 |
| ENSG00000148730.6 | EIF4EBP2 | 0.114484406641542 | 6.12314484897866 | 1.17534674288045 | 0.246398872955502 | 0.56786787616651 | -5.73306637835972 |
| ENSG00000122678.16 | POLM | -0.155432596247133 | 5.83983006820133 | -1.17522365228239 | 0.246447568796423 | 0.567916514614507 | -5.73097271268038 |
| lnc-MICALL2-5 | lnc-MICALL2-5 | -0.194233668330059 | 1.64655809660979 | -1.175050400714 | 0.246516120638757 | 0.568010893440897 | -5.35765803123385 |
| ENSG00000127720.7 | METTL25 | 0.160907736873354 | 3.68724122621099 | 1.17485819014277 | 0.246592190332372 | 0.568024526924497 | -5.59194222892835 |
| lnc-ARFGAP3-1 | lnc-ARFGAP3-1 | 0.157638013423513 | 2.17223642485636 | 1.17484035132885 | 0.246599251125663 | 0.568024526924497 | -5.42426556137477 |
| ENSG00000128159.11 | TUBGCP6 | -0.13299863270616 | 7.1341971153761 | -1.17482582670852 | 0.246605000234037 | 0.568024526924497 | -5.7291483406482 |
| lnc-ZNF652-2 | lnc-ZNF652-2 | -0.203654974756451 | 2.05158750820422 | -1.17475654462945 | 0.246632424676881 | 0.568024526924497 | -5.4137863085012 |
| ENSG00000066455.12 | GOLGA5 | -0.141975259805609 | 4.83133103291512 | -1.17464414151054 | 0.24667692275033 | 0.568063448307231 | -5.69147864720851 |
| ENSG00000169131.12 | ZNF354A | 0.141428350979853 | 3.62404402345783 | 1.17457172115032 | 0.246705595560706 | 0.568065921627252 | -5.58162796590031 |
| ENSG00000145029.13 | NICN1 | -0.194094345865092 | 2.83191933120638 | -1.17444272934877 | 0.246756672235559 | 0.568119975908782 | -5.48672430759042 |
| lnc-MACROD1-1 | lnc-MACROD1-1 | -0.236108045598597 | 1.28351062994248 | -1.17416679472442 | 0.246865959380228 | 0.568162301474732 | -5.30374841117082 |
| ENSG00000010310.8 | GIPR | 1.25265179227301 | -1.75063401828021 | 1.17413051441136 | 0.246888532147328 | 0.568162301474732 | -4.64234240768474 |
| lnc-CABLES2-7 | lnc-CABLES2-7 | -0.239992680991966 | -0.0723233465925862 | -1.17409474916185 | 0.246894499650261 | 0.568162301474732 | -5.03420392747188 |
| ENSG00000269335.5 | IKBKG | 0.153491372297122 | 2.69310608497536 | 1.17406627043395 | 0.246905781929253 | 0.568162301474732 | -5.47549468780361 |
| ENSG00000103266.10 | STUB1 | -0.149547567924848 | 4.1331484370253 | -1.17404786618638 | 0.246913073249355 | 0.568162301474732 | -5.64287575786925 |
| ENSG00000125388.19 | GRK4 | 0.332746483616279 | -0.201688125720472 | 1.17379011419422 | 0.247015204799263 | 0.568333776032108 | -5.00031201070356 |
| ENSG00000153975.9 | ZUP1 | 0.20762295992292 | 4.15880446353003 | 1.17349840760962 | 0.247130827461987 | 0.568411698584527 | -5.64170739748282 |
| lnc-PRAG1-7 | lnc-PRAG1-7 | -0.474334769183482 | 0.59155857989763 | -1.1734730397983 | 0.247141082612286 | 0.568411698584527 | -5.16493728136963 |
| lnc-ME2-6 | lnc-ME2-6 | -0.2605539701039 | 0.0450102566726968 | -1.17347214115374 | 0.247141240526181 | 0.568411698584527 | -5.09578128996933 |
| ENSG00000213985.4 | ACTR2P2 | 0.94141226116984 | -1.44736114975968 | 1.17342599696865 | 0.247159534680635 | 0.568411698584527 | -4.66252965150955 |
| lnc-TNFRSF14-2 | lnc-TNFRSF14-2 | -0.306158829848367 | 0.539346341283367 | -1.17329950725577 | 0.247209687364331 | 0.568422207297279 | -5.1647787704631 |
| ENSG00000018189.12 | RUFY3 | 0.210525061372141 | 5.12186718883831 | 1.17327517379995 | 0.247219336332179 | 0.568422207297279 | -5.71477446382393 |
| ENSG00000143889.15 | HNRNPLL | 0.37060379583103 | 2.78312939284145 | 1.17328317451833 | 0.247258446305479 | 0.568448631777691 | -5.4961473610343 |
| ENSG00000171150.8 | SOCS5 | 0.171204404664113 | 4.45035908974934 | 1.17297927616362 | 0.2473366907508 | 0.568461174762372 | -5.66871373568115 |
| ENSG00000101391.20 | CDK5RAP1 | 0.10465603827986 | 5.18772009884914 | 1.17294021243248 | 0.247352186631461 | 0.568461174762372 | -5.71530361323411 |
| ENSG00000129347.19 | KRI1 | -0.193840488484353 | 4.85516164175258 | -1.17289666886337 | 0.247369460413062 | 0.568461174762372 | -5.69809002525088 |
| ENSG00000132549.18 | VPS13B | 0.16213374692083 | 7.93041901014913 | 1.17288428286505 | 0.247374374111965 | 0.568461174762372 | -5.71955094911267 |
| ENSG00000157540.20 | DYRK1A | 0.110829639670771 | 7.944470109676 | 1.17274042987069 | 0.247431447792585 | 0.568528855761865 | -5.71833679352839 |
| lnc-C19orf48-3 | lnc-C19orf48-3 | -0.239521909728642 | 0.00354279941808845 | -1.17265869683473 | 0.247463879630555 | 0.568539907850845 | -5.07286798199276 |
| ENSG00000259151.2 | CAP2P1 | -0.318022280570065 | 1.18971028228091 | -1.17247226318711 | 0.247537868421093 | 0.568646422512429 | -5.28154446618142 |
| ENSG00000170160.17 | CCDC144A | -1.60846813757249 | -0.223508283593459 | -1.17293742868279 | 0.247611893023746 | 0.56875133898392 | -5.13173962601981 |
| lnc-EML5-3 | lnc-EML5-3 | 0.152238636200586 | 1.12149865143347 | 1.17219580955948 | 0.247647612416412 | 0.56875133898392 | -5.27711602963335 |
| ENSG00000101181.17 | MTG2 | -0.152896351287791 | 3.45117711193789 | -1.17214840052442 | 0.247666435957483 | 0.56875133898392 | -5.5704991737018 |
| ENSG00000139433.9 | GLTP | 0.148637594714016 | 4.38654381720909 | 1.17194940798922 | 0.24774545634864 | 0.568834817512348 | -5.66782812212226 |
| lnc-OTP-4 | lnc-OTP-4 | 0.16657957350253 | 1.22699483302666 | 1.17184241598143 | 0.247787950673592 | 0.568834817512348 | -5.29837213155328 |
| lnc-NCS1-2 | lnc-NCS1-2 | 0.281135741912612 | 0.459269149290046 | 1.17180195370395 | 0.247804022569979 | 0.568834817512348 | -5.14778414122663 |
| ENSG00000171219.8 | CDC42BPG | -0.197086923112765 | 2.81507122267133 | -1.17176716007171 | 0.247817843445484 | 0.568834817512348 | -5.49136048089042 |
| lnc-BRAT1-3 | lnc-BRAT1-3 | 0.308888092098781 | 0.654645084124091 | 1.17170894796589 | 0.247840967956602 | 0.568834817512348 | -5.22533639723632 |
| lnc-GOLGA8O-3 | lnc-GOLGA8O-3 | 0.235049757147235 | 1.01084728165827 | 1.17146466347074 | 0.247938025984212 | 0.5689870067403 | -5.28121767726931 |
| lnc-HOMEZ-1 | lnc-HOMEZ-1 | -0.135266460855902 | 3.77456050356513 | -1.17140291546285 | 0.247962563788587 | 0.5689870067403 | -5.59588247405104 |
| lnc-SPACA5-1 | lnc-SPACA5-1 | -0.186098135623002 | 1.75913933729718 | -1.17117889315723 | 0.248051601929465 | 0.569088988713045 | -5.38359594179218 |

| | | | | | | | |
|---|---|---|---|---|---|---|---|
| lnc-BHLHE41-4 | lnc-BHLHE41-4 | 0.218021470897245 | 2.03414177267085 | 1.17111083812793 | 0.248078655132955 | 0.569088988713045 | -5.40033626966497 |
| ENSG00000178125.14 | PPP1R42 | 0.149581540069261 | 1.80434953816197 | 1.17101287495465 | 0.24811760116168 | 0.569088988713045 | -5.37525238618508 |
| lnc-PPP1R42-1 | lnc-PPP1R42-1 | 0.149581540069261 | 1.80434953816197 | 1.17101287495465 | 0.24811760116168 | 0.569088988713045 | -5.37525238618508 |
| ENSG00000022556.15 | NLRP2 | 3.38399302019322 | 0.126904556310059 | 1.17141453726537 | 0.248216284929873 | 0.569157280644306 | -4.78690458325184 |
| MIR34AHG | MIR34AHG | 0.662499844100295 | -0.466960143995245 | 1.17079006652068 | 0.248217387003953 | 0.569157280644306 | -4.9514020099547 |
| ENSG00000109390.11 | NDUFC1 | 0.130308003457365 | 2.99084491349534 | 1.17072938127537 | 0.24823033126094 | 0.569157280644306 | -5.52311316627931 |
| ENSG00000201675.1 | SNORD32A | 0.218391353289255 | 1.77778379974873 | 1.17054444281947 | 0.248303891277336 | 0.569258194435653 | -5.38117329842166 |
| ENSG00000234009.1 | RPL5P34 | 0.437842434975128 | 1.60324113199674 | 1.17053980113579 | 0.248332851718185 | 0.569258194435653 | -5.31735991487076 |
| lnc-FAM169A-4 | lnc-FAM169A-4 | 0.133469711208333 | 0.949092792220884 | 1.17041015772782 | 0.24835731360988 | 0.569258194435653 | -5.25734969854839 |
| ENSG00000281910.1 | SNORA50A | 0.181056439982383 | 0.289656425659748 | 1.17029978854996 | 0.248401227762737 | 0.56929545385151 | -5.13317510533081 |
| ENSG00000126934.13 | MAP2K2 | -0.190433064049905 | 5.05358583778313 | -1.17019437619077 | 0.248443174927222 | 0.569328197453463 | -5.71479166887231 |
| ENSG00000204422.7 | | 0.137729445603716 | 3.47628459369134 | 1.1701018516315 | 0.248479997826 | 0.569349192391468 | -5.57196369460932 |
| ENSG00000172840.6 | PDP2 | -0.172489347851608 | 2.03267834979812 | -1.169293441875 | 0.248801896697949 | 0.570023312526034 | -5.40124882447537 |
| ENSG00000163545.8 | NUAK2 | 0.196344024167802 | 6.19052667368207 | 1.16905182994672 | 0.248898162200688 | 0.570180397615667 | -5.7409465447514 |
| ENSG00000116791.13 | CRYZ | -0.29038967222263 | 3.31147790400943 | -1.16892474196366 | 0.248948808721353 | 0.570232954586203 | -5.56484995781335 |
| ENSG00000111843.13 | TMEM14C | 0.240132099545878 | 3.37671088926882 | 1.16880088191267 | 0.248998176041016 | 0.570282570095942 | -5.56459324194229 |
| ENSG00000171206.14 | TRIM8 | -0.122840952225072 | 5.39924677761044 | -1.16864323864961 | 0.249061018700944 | 0.57029231874211 | -5.72909284741722 |
| ENSG00000166579.15 | NDEL1 | 0.186347006547679 | 5.7095903867033 | 1.16859440657292 | 0.249080487371589 | 0.57029231874211 | -5.73594805524639 |
| ENSG00000081386.12 | ZNF510 | 0.125935023747431 | 5.31872544874502 | 1.16854379433751 | 0.249100666929515 | 0.57029231874211 | -5.72384761110932 |
| ENSG00000112742.9 | TTK | 0.37933957986948 | 0.1711399472173 | 1.16848546851268 | 0.24912392343348 | 0.57029231874211 | -5.09627074680837 |
| ENSG00000148187.17 | MRRF | 0.127706850336912 | 4.88587349758928 | 1.16836439782673 | 0.249172203480345 | 0.57029231874211 | -5.70079571561278 |
| ENSG00000155755.18 | TMEM237 | -0.360194433157263 | 1.02974702225864 | -1.16836359222044 | 0.249172524759239 | 0.57029231874211 | -5.27455099942736 |
| ENSG00000100281.13 | HMGXB4 | 0.115502139526904 | 4.19600077720777 | 1.16830377427126 | 0.2491963812256 | 0.57029231874211 | -5.65197839343101 |
| lnc-C6orf62-1 | lnc-C6orf62-1 | -0.268108869375975 | 0.880089233383308 | -1.16816980800712 | 0.249249815362087 | 0.570341726767734 | -5.26028004981695 |
| lnc-BACE1-4 | lnc-BACE1-4 | 0.264458223182845 | 1.64187024184711 | 1.1679822543171 | 0.249324637453197 | 0.570341726767734 | -5.36262449384894 |
| ENSG00000138382.14 | METTL5 | -0.132912679138605 | 3.29081089188441 | -1.16798187601925 | 0.249324788386603 | 0.570341726767734 | -5.56153629527061 |
| ENSG00000100027.14 | YPEL1 | -0.19559845300211 | 4.46110958324803 | -1.16797180107356 | 0.249328808116216 | 0.570341726767734 | -5.66340862105866 |
| ENSG00000212607.1 | SNORA3B | 0.164026699087637 | 2.1031335709854 | 1.16787097140121 | 0.249369040000861 | 0.570355576829395 | -5.42463751126448 |
| lnc-RARA-1 | lnc-RARA-1 | -0.33245882384249 | 1.06027442781622 | -1.16781773709938 | 0.249390282828769 | 0.570355576829395 | -5.2698287740926 |
| ENSG00000116521.10 | SCAMP3 | -0.106721382299649 | 3.61545735435171 | -1.16770343202085 | 0.249435900008098 | 0.570355840071139 | -5.59427657657518 |
| lnc-CTSK-1 | lnc-CTSK-1 | -0.145134597402861 | 2.00855086179976 | -1.16762588662717 | 0.249466850471082 | 0.570355840071139 | -5.40573854690585 |
| ENSG00000263776.1 | SNORA4 | -0.103613630713719 | 3.4384016097702 | -1.16759502236699 | 0.249479170004944 | 0.570355840071139 | -5.58081638718346 |
| ENSG00000177380.13 | PPFIA3 | 0.345070467923138 | -0.339429689745272 | 1.16750553258065 | 0.249514892528973 | 0.570355840071139 | -4.92836910920517 |
| lnc-ADIPOQ-3 | lnc-ADIPOQ-3 | -0.177945545865632 | 2.30565010339322 | -1.16745499011682 | 0.249535069700324 | 0.570355840071139 | -5.44566402009475 |
| ENSG00000135900.3 | MRPL44 | 0.176171390897694 | 4.65210196828711 | 1.16740083900712 | 0.249556688797587 | 0.570355840071139 | -5.69559232667779 |
| ENSG00000250539.1 | KRT8P33 | 0.243357131153567 | 2.31038413868058 | 1.16733151300937 | 0.249584368241789 | 0.570355840071139 | -5.43642455630685 |
| ENSG00000171302.16 | CANT1 | 0.121900392147777 | 3.64812961153651 | 1.16718290715426 | 0.249643708843021 | 0.570428114910734 | -5.5993111278609 |
| ENSG00000104884.15 | ERCC2 | -0.139768493037494 | 2.93448837956545 | -1.16698764260988 | 0.249721696495369 | 0.570542976908 | -5.51891005904592 |
| ENSG00000213020.9 | ZNF611 | 0.137470154928224 | 5.06636270504258 | 1.1667119180078 | 0.249831849497869 | 0.570731293919493 | -5.71248528050177 |
| lnc-AHSP-5 | lnc-AHSP-5 | -0.293832687651437 | 0.643535983401501 | -1.16629603549962 | 0.249998062596384 | 0.571047622064296 | -5.2057849922883 |
| lnc-GADL1-11 | lnc-GADL1-11 | -0.256332997875845 | -0.204922116242492 | -1.16602018727294 | 0.250108353206465 | 0.571154684034973 | -5.00693846989615 |
| lnc-TOB1-4 | lnc-TOB1-4 | 0.150279634581944 | 1.30822069077586 | 1.16597583801731 | 0.250126088365588 | 0.571154684034973 | -5.32624475127543 |
| ENSG00000144118.13 | RALB | 0.23568860235718 | 4.27681142558268 | 1.16597060859805 | 0.250128179657545 | 0.571154684034973 | -5.65523024696079 |
| ENSG00000150457.8 | LATS2 | -0.254664990916546 | 2.830884427047 | -1.16571495599706 | 0.250230432862193 | 0.571296428943926 | -5.50809438815724 |
| ENSG00000159873.9 | CCDC117 | -0.217828397886072 | 5.09744238469934 | -1.16560094333737 | 0.250276044177873 | 0.571296428943926 | -5.72056150632914 |
| ENSG00000176732.6 | PFN4 | 0.399632738273183 | 0.503619611163141 | 1.16559888878084 | 0.250276866168294 | 0.571296428943926 | -5.13672185285102 |
| ENSG00000100721.10 | TCL1A | -0.358656287550957 | 7.17067213828102 | -1.16582844616615 | 0.250317560132958 | 0.571296428943926 | -5.73903800998694 |
| ENSG00000265727.2 | RN7SL648P | -0.331807665222571 | 0.801222654871205 | -1.16546850752914 | 0.250329033311241 | 0.571296428943926 | -5.2548565103108 |
| lnc-ACTRT3-2 | lnc-ACTRT3-2 | -0.427304685836368 | 3.10367959913306 | -1.16552858561612 | 0.250431532181836 | 0.571466987461057 | -5.55668606863666 |
| ENSG00000221116.1 | SNORD110 | 0.290845600604021 | -0.31747309305009 | 1.16502956473391 | 0.250504717521164 | 0.571570624181146 | -5.01053344993777 |
| ENSG00000285635.1 | | 0.105301625162794 | 2.35791285030722 | 1.16477417343495 | 0.250606977366401 | 0.57168459790679 | -5.45413444657486 |
| ENSG00000204560.9 | DHX16 | -0.114718954069189 | 5.20500902563991 | -1.16469795204506 | 0.250637502606756 | 0.57168459790679 | -5.7251162494089 |
| ENSG00000111266.8 | DUSP16 | 0.721128769991452 | 0.456368063543482 | 1.16507481608282 | 0.250637993077339 | 0.57168459790679 | -5.18206107366592 |
| ENSG00000284154.1 | MIR3605 | 0.189423387853272 | 0.153613825719985 | 1.16444944117449 | 0.25073704522908 | 0.571752136165551 | -5.09178175142303 |
| ENSG00000117859.18 | OSBPL9 | 0.0889757446672332 | 5.79486266954637 | 1.16438104206153 | 0.250764447947071 | 0.571752136165551 | -5.74278464032662 |
| ENSG00000085982.13 | USP40 | -0.139142416994532 | 4.71178447289522 | -1.164353833833 | 0.25077534897409 | 0.571752136165551 | -5.69892251357767 |
| ENSG00000197362.14 | ZNF786 | -0.229503335552659 | 2.65853770357878 | -1.16434543330408 | 0.250778714730729 | 0.571752136165551 | -5.48139766283714 |
| ENSG00000135637.13 | CCDC142 | -0.17092830811766 | 3.31133286068348 | -1.16415488930294 | 0.250855066877152 | 0.571822233782899 | -5.56987830956265 |
| ENSG00000273599.1 | | -0.326516792353525 | -0.459949651385506 | -1.16413004562532 | 0.250865023128775 | 0.571822233782899 | -4.97475462226743 |
| ENSG00000142409.5 | ZNF787 | -0.213970856453052 | 2.55705125282901 | -1.16399020920008 | 0.250921068736567 | 0.571886652397825 | -5.49192587621664 |
| ENSG00000114268.11 | PFKFB4 | -0.41071143670583 | -0.773760690855753 | -1.16367371654897 | 0.251047950528641 | 0.57211248513408 | -4.89632162700623 |
| ENSG00000006712.14 | PAF1 | -0.167932005483207 | 5.03965643765058 | -1.16348454164838 | 0.251123812816403 | 0.572143861333029 | -5.71673033269683 |
| ENSG00000006530.16 | AGK | -0.176294776778247 | 4.06067978654396 | -1.16338167791395 | 0.25116506984034 | 0.572143861333029 | -5.64048413480249 |
| ENSG00000068976.13 | PYGM | -0.249032402684154 | 3.24420883389426 | -1.16337821780415 | 0.251166457721032 | 0.572143861333029 | -5.546906549616 |
| lnc-RPL26L1-2 | lnc-RPL26L1-2 | 0.166404358064761 | 3.01855684502738 | 1.16336214102885 | 0.251172906330722 | 0.572143861333029 | -5.53177156718639 |
| lnc-CCDC144NL-8 | lnc-CCDC144NL-8 | 0.111512768537404 | 3.94091826648433 | 1.1628017982382 | 0.251397742080437 | 0.572592644156427 | -5.63749105767957 |
| ENSG00000063438.16 | AHRR | -0.205744491080849 | 1.22840662618841 | -1.16269295982574 | 0.251441430003694 | 0.572628784439702 | -5.31508179652687 |
| ENSG00000167065.13 | DUSP18 | -0.167685047999156 | 2.69430482428741 | -1.16225617268554 | 0.251616812274814 | 0.572964802196316 | -5.49624311955644 |

| | | | | | | | |
|---|---|---|---|---|---|---|---|
| lnc-CBWD5-1 | lnc-CBWD5-1 | -0.69407857914714 | 1.28179468380963 | -1.16230150352957 | 0.251773870723441 | 0.573197104734016 | -5.29066743452521 |
| ENSG00000165195.15 | PIGA | 0.176175025869357 | 4.58123563264557 | 1.16180390164379 | 0.251798504863625 | 0.573197104734016 | -5.69215109723673 |
| ENSG00000139651.10 | ZNF740 | -0.144668828221605 | 6.22220712888819 | -1.16179427831686 | 0.251802371909099 | 0.573197104734016 | -5.74886935867165 |
| lnc-TMED4-2 | lnc-TMED4-2 | 0.115580347820094 | 2.01955113698225 | 1.16168956925539 | 0.251844451055265 | 0.573209409519528 | -5.42112193390367 |
| ENSG00000123384.13 | LRP1 | 0.659303325379673 | -0.520438851411511 | 1.16158047589635 | 0.251888297508566 | 0.573209409519528 | -4.8803346062647 |
| ENSG00000153310.19 | CYRIB | 0.164436980220933 | 5.59589240136013 | 1.16157294780379 | 0.251891323378717 | 0.573209409519528 | -5.74066175885389 |
| ENSG00000164329.13 | TENT2 | 0.122326612931041 | 6.81928416714264 | 1.16145519668186 | 0.251938656128824 | 0.573229071410574 | -5.74727417163224 |
| ENSG00000170854.17 | RIOX2 | -0.168654899700059 | 4.51601193911951 | -1.16141289237173 | 0.251955662879632 | 0.573229071410574 | -5.69131417151069 |
| ENSG00000104946.12 | TBC1D17 | -0.166838081499243 | 4.71010721050429 | -1.16110097828789 | 0.252081081013335 | 0.573368271020533 | -5.69961997869169 |
| lnc-LGALS3BP-4 | lnc-LGALS3BP-4 | 0.142195847060025 | 1.43304682156989 | 1.16091532872089 | 0.252155750599946 | 0.573368271020533 | -5.34838367427609 |
| ENSG00000131013.3 | PPIL4 | 0.167559602757304 | 5.63593816575222 | 1.160861798461 | 0.252177283819876 | 0.573368271020533 | -5.74188914373299 |
| ENSG00000112367.10 | FIG4 | -0.193116091164439 | 5.96107692139517 | -1.16080489392241 | 0.25220017584153 | 0.573368271020533 | -5.74720756081932 |
| ENSG00000198246.8 | SLC29A3 | -0.15641136448893 | 2.49541331851702 | -1.16079146903568 | 0.252205576733107 | 0.573368271020533 | -5.47227665705535 |
| ENSG00000163738.18 | MTHFD2L | 0.160689039634685 | 3.74079226822146 | 1.1607371493786 | 0.252227430624934 | 0.573368271020533 | -5.61159839279272 |
| ENSG00000128654.13 | MTX2 | -0.155923053695894 | 2.97489679365914 | -1.16069668491891 | 0.252243711180335 | 0.573368271020533 | -5.52899971084673 |
| ENSG00000174885.12 | NLRP6 | -0.618181172030197 | 1.38242163433646 | -1.16113262527846 | 0.252246465441494 | 0.573368271020533 | -5.39760062537139 |
| ENSG00000185219.16 | ZNF445 | -0.135310645530367 | 4.81873053592607 | -1.16063742844911 | 0.252267553921292 | 0.573368271020533 | -5.70709077245391 |
| lnc-ATP9A-2 | lnc-ATP9A-2 | -0.225997291099148 | 0.0839331623643699 | -1.16050413816716 | 0.252321191236924 | 0.573426860906438 | -5.06866531076537 |
| ENSG00000163637.12 | PRICKLE2 | -1.16423561155621 | -0.601156797977972 | -1.16088714936889 | 0.252393213989289 | 0.573527216111894 | -5.0621012843864 |
| ENSG00000100097.11 | LGALS1 | 0.510855209045706 | 4.07112781889038 | 1.16070144893912 | 0.25243190683215 | 0.573549398892816 | -5.65487844260643 |
| ENSG00000285744.1 | MITA1 | 0.342332041853823 | 2.81303303850884 | 1.16020684963922 | 0.252458706406691 | 0.573549398892816 | -5.50370027556797 |
| ENSG00000106804.7 | C5 | -0.155540401098655 | 3.83937191710216 | -1.1600083262475 | 0.252520783076815 | 0.573627113792085 | -5.61972595101793 |
| ENSG00000198331.10 | HYLS1 | 0.315046872922172 | 1.21483849036383 | 1.15974250642633 | 0.252627837230998 | 0.573786361961709 | -5.32657371245417 |
| lnc-DUOXA1-1 | lnc-DUOXA1-1 | -0.477833808554051 | -0.530676790670195 | -1.15951406168698 | 0.252719865407968 | 0.573786361961709 | -4.90924676495522 |
| ENSG00000174255.6 | ZNF80 | -0.6611106529126 | -0.682847139422574 | -1.15947363727961 | 0.252736152760138 | 0.573786361961709 | -4.86652578674229 |
| ENSG00000186654.20 | PRR5 | 0.90650264684395 | -0.0265314758518821 | 1.15977795670094 | 0.252763797882448 | 0.573786361961709 | -5.01003440212645 |
| lnc-GLTP-3 | lnc-GLTP-3 | -0.17456849010688 | 2.58178473785179 | -1.15939391790543 | 0.252768274622533 | 0.573786361961709 | -5.48599835574259 |
| ENSG00000104679.10 | R3HCC1 | -0.13763684735293 | 3.74875828251623 | -1.15934204904061 | 0.252789176074124 | 0.573786361961709 | -5.62074832689094 |
| ENSG00000072506.12 | HSD17B10 | 0.173907863039088 | 3.96409534407142 | 1.15933032392168 | 0.252793901085363 | 0.573786361961709 | -5.63715797513076 |
| ENSG00000135441.7 | BLOC1S1 | 0.17528896822233 | 3.35568257727153 | 1.15928069679218 | 0.252813900628224 | 0.573786361961709 | -5.57193805904168 |
| ENSG00000145781.8 | COMMD10 | -0.118208911371274 | 4.40620116437263 | -1.15920818868319 | 0.252843123170559 | 0.573789416121237 | -5.68126350686717 |
| lnc-HNRNPH1-5 | lnc-HNRNPH1-5 | 0.4388818437868 | -0.103381922522158 | 1.15907231407152 | 0.252897890526617 | 0.573850433327016 | -5.1094905000981 |
| ENSG00000116977.18 | LGALS8 | 0.182187989911121 | 4.93582574814522 | 1.15865797152305 | 0.253064953568513 | 0.574166219058719 | -5.71671701512268 |
| ENSG00000177548.12 | RABEP2 | -0.140116009210849 | 5.71524934255444 | -1.15834846114174 | 0.253189800184097 | 0.574368163031992 | -5.74742169877977 |
| lnc-C7orf49-3 | lnc-C7orf49-3 | -0.297691633788538 | 2.62085861066258 | -1.15820664734149 | 0.253247018186215 | 0.574393649369486 | -5.49188278373739 |
| ENSG00000285446.1 | | 0.416812470966132 | 2.03596909151937 | 1.1582401496227 | 0.253272011893255 | 0.574393649369486 | -5.38961428198526 |
| ENSG00000130734.9 | ATG4D | 0.144095916976888 | 3.77320295017742 | 1.15813279616453 | 0.253276818815404 | 0.574393649369486 | -5.62915122517583 |
| ENSG00000070540.12 | WIPI1 | 0.180578996229349 | 1.94656536549536 | 1.1579265237743 | 0.253360067812569 | 0.574468893459111 | -5.43141438602878 |
| ENSG00000153914.15 | SREK1 | 0.176577207944676 | 7.77081551988214 | 1.157912279956 | 0.253365817170568 | 0.574468893459111 | -5.73713674886524 |
| lnc-TMEM161B-10 | lnc-TMEM161B-10 | -0.234923493929962 | 3.9656244622499 | -1.15783688590284 | 0.253396250704222 | 0.574474614852406 | -5.62761404680684 |
| ENSG00000153406.13 | NMRAL1 | -0.162201145934303 | 4.02506393538933 | -1.15769103817769 | 0.253455131021508 | 0.574518705378601 | -5.64497367088555 |
| ENSG00000283885.1 | MIR3714 | -0.448647080458364 | -0.456212658859543 | -1.1576275295235 | 0.253480773241033 | 0.574518705378601 | -4.92196590992208 |
| lnc-EIF4H-1 | lnc-EIF4H-1 | -0.393383952355301 | -0.163857374765502 | -1.1575813110426 | 0.253499435565683 | 0.574518705378601 | -5.07490122638306 |
| ENSG00000173511.9 | VEGFB | -0.198414341667756 | 3.13355666635944 | -1.15738334218941 | 0.253579383609713 | 0.574636623674857 | -5.55231251369872 |
| ENSG00000102554.13 | KLF5 | -0.463549510222258 | -0.211206044800951 | -1.15722712649216 | 0.253642482827144 | 0.574716339061108 | -5.00781279743717 |
| ENSG00000160124.9 | MIX23 | 0.115437224736469 | 2.94851406014802 | 1.15711475001006 | 0.25368788134835 | 0.574755934154441 | -5.52505945122753 |
| lnc-CHIC2-2 | lnc-CHIC2-2 | 0.191451911213104 | 2.5959962157111 | 1.15696131850744 | 0.253749874980716 | 0.574833114321822 | -5.47890604919308 |
| ENSG00000196814.14 | MVB12B | -0.749066580622829 | 0.621250622782113 | -1.15723973829664 | 0.253826264596944 | 0.574942885902817 | -5.30940254984377 |
| ENSG00000184785.5 | SMIM10 | -1.01875082349453 | -1.84252099172498 | -1.15671056370231 | 0.25386384339669 | 0.574964732464136 | -4.72664663207451 |
| ENSG00000179088.14 | C12orf42 | -0.577050041154873 | 2.65374330951838 | -1.15667846008385 | 0.254048431666006 | 0.575183459664066 | -5.52318036059854 |
| ENSG00000084764.11 | MAPRE3 | 0.502650976362666 | -0.590966517735075 | 1.15619020416718 | 0.254061607264108 | 0.575183459664066 | -4.92932978943982 |
| ENSG00000279366.1 | | 0.301685067392741 | 0.358166430986442 | 1.15613947881777 | 0.254082123270365 | 0.575183459664066 | -5.14283911075862 |
| ENSG00000184260.5 | H2AC20 | -0.250783691973892 | 6.96982265339048 | -1.15606807354921 | 0.254116946162669 | 0.575183459664066 | -5.75095321339792 |
| ENSG00000272916.5 | | 0.0979513883082177 | 5.1328420699368 | 1.15595697541622 | 0.254155947145899 | 0.575183459664066 | -5.73135065677154 |
| ENSG00000115459.17 | ELMOD3 | 0.123082835889574 | 4.53200099565809 | 1.15594096415471 | 0.254162424548286 | 0.575183459664066 | -5.69184516537144 |
| ENSG00000258539.1 | | -0.174233079498206 | 5.60122755534723 | -1.15588591955804 | 0.254184693907403 | 0.575183459664066 | -5.74767079194665 |
| ENSG00000116478.11 | HDAC1 | -0.143827964554633 | 6.50500083368692 | -1.15586609882438 | 0.25419271311392 | 0.575183459664066 | -5.7551536613955 |
| lnc-ZSCAN23-1 | lnc-ZSCAN23-1 | 1.37465272164478 | -1.5469202178824 | 1.15603529346046 | 0.254250316789291 | 0.575183459664066 | -4.72850956587796 |
| ENSG00000247596.8 | TWF2 | -0.27254713783992 | 3.75828688609751 | -1.15572275706941 | 0.254250712717767 | 0.575183459664066 | -5.62585803491384 |
| lnc-MRGPRE-2 | lnc-MRGPRE-2 | 0.13953257806679 | 1.17741924472073 | 1.15568051094217 | 0.25426780836046 | 0.575183459664066 | -5.31767194405025 |
| lnc-PEX16-3 | lnc-PEX16-3 | -0.189652422984452 | 2.10711702593104 | -1.15529894831482 | 0.254422251953111 | 0.575407749994753 | -5.43406945729925 |
| FUT8-AS1 | FUT8-AS1 | 0.636821859197244 | -0.795629539089242 | 1.15526088476152 | 0.254437662493464 | 0.575407749994753 | -4.91113212986985 |
| lnc-SPPL2A-1 | lnc-SPPL2A-1 | -0.183123869500675 | 2.26730711597038 | -1.15512045086412 | 0.254494524859073 | 0.575407749994753 | -5.44548345843123 |
| lnc-AGBL1-7 | lnc-AGBL1-7 | 0.0867941135821808 | 3.83935594709801 | 1.15509753243714 | 0.254503805510113 | 0.575407749994753 | -5.63402262833027 |
| lnc-FCGR1B-12 | lnc-FCGR1B-12 | 0.237823445403821 | 0.132940806493529 | 1.15492132452802 | 0.254575167770636 | 0.575407749994753 | -5.12173473060063 |
| ENSG00000278705.1 | H4C2 | 0.323213054264329 | 4.69100446556914 | 1.15508884450266 | 0.254577739774401 | 0.575407749994753 | -5.70133896215626 |
| ENSG00000174177.12 | CTU2 | -0.236177418535024 | 1.9953759553584 | -1.15489949175362 | 0.254584010807129 | 0.575407749994753 | -5.43205959029607 |
| ENSG00000259158.4 | ADAM20P1 | -0.388089633804129 | 0.94782499635999 | -1.15488321677526 | 0.254590602885984 | 0.575407749994753 | -5.27466700528735 |

| | | | | | | | |
|---|---|---|---|---|---|---|---|
| lnc-CTSZ-2 | lnc-CTSZ-2 | -0.199954271989536 | 1.7609885645011 | -1.15478386794506 | 0.25463084617862 | 0.575435518982711 | -5.38956818693685 |
| lnc-CTAGE5-5 | lnc-CTAGE5-5 | -0.378398886728517 | 0.22954187983498 | -1.15439066685339 | 0.254790165346211 | 0.575546124759545 | -5.15440110883076 |
| ENSG00000117335.19 | CD46 | -0.140636869903357 | 7.04397660464647 | -1.15437518591384 | 0.254796439458731 | 0.575546124759545 | -5.75166064035215 |
| ENSG00000164054.15 | SHISA5 | -0.192976739008784 | 5.94432067214823 | -1.15434826277807 | 0.254807351127609 | 0.575546124759545 | -5.75513760147851 |
| ENSG00000128833.12 | MYO5C | 0.646671924174295 | 2.96875478327398 | 1.1548750904632 | 0.254810642913644 | 0.575546124759545 | -5.50953307863081 |
| ENSG00000174106.2 | LEMD3 | 0.124306712357846 | 6.29585885053086 | 1.15431804027934 | 0.254819600395168 | 0.575546124759545 | -5.75751487914089 |
| lnc-NOX3-3 | lnc-NOX3-3 | -0.605901771572013 | 1.49626974609354 | -1.15436982690341 | 0.254948287811937 | 0.575773601934727 | -5.32674600851744 |
| lnc-TBC1D9B-4 | lnc-TBC1D9B-4 | 0.227352457429693 | 2.15482459236148 | 1.15363383505919 | 0.255097024270326 | 0.575970914212887 | -5.43993460238534 |
| TMEM44-AS1 | TMEM44-AS1 | -0.246757559150041 | 2.4289582290366 | -1.15362730875579 | 0.255099671529477 | 0.575970914212887 | -5.46267065615923 |
| ENSG00000162746.14 | FCRLB | -0.30897726688676 | 1.25049299570802 | -1.15357816880005 | 0.255119604765043 | 0.575970914212887 | -5.3300368696983 |
| lnc-DAD1-2 | lnc-DAD1-2 | -0.509115507793196 | 5.66098136535739 | -1.15397214877084 | 0.255176433128086 | 0.576036030168392 | -5.74867787664465 |
| ENSG00000270136.6 | MICOS10-NBL1 | 0.143543198750382 | 1.26144143455254 | 1.15318726726153 | 0.255278210836588 | 0.576100832842047 | -5.3283904258641 |
| ENSG00000244462.7 | RBM12 | -0.169790918521096 | 6.93597909585856 | -1.15304076394373 | 0.255337672007086 | 0.576100832842047 | -5.75538914506549 |
| ENSG00000119650.12 | IFT43 | -0.1303950337627 | 2.67803719441336 | -1.15301735939083 | 0.255347172115592 | 0.576100832842047 | -5.49945668256413 |
| lnc-FAM72D-8 | lnc-FAM72D-8 | 0.984093313063239 | 0.397201853350933 | 1.15346769720581 | 0.255356523937916 | 0.576100832842047 | -5.1053924296486 |
| ENSG00000213221.4 | DNLZ | -0.210554089591472 | 1.74612523596835 | -1.15297710209196 | 0.255363513493284 | 0.576100832842047 | -5.40642024907133 |
| ENSG00000052749.13 | RRP12 | 0.225527507336391 | 4.59558682905005 | 1.15289766562645 | 0.255395760817942 | 0.576100832842047 | -5.69722006195102 |
| ENSG00000103319.11 | EEF2K | -0.195632939049712 | 4.7399853477434 | -1.15269726227314 | 0.255477127815691 | 0.576100832842047 | -5.7134504798838 |
| ENSG00000149177.12 | PTPRJ | 0.19534499597393 | 7.52046113681253 | 1.15268087521665 | 0.255483782050426 | 0.576100832842047 | -5.74973779151416 |
| ENSG00000171858.17 | RPS21 | 0.201499471963486 | 7.14873681994244 | 1.15263505655407 | 0.255502388136589 | 0.576100832842047 | -5.75333300076715 |
| ENSG00000100994.11 | PYGB | -0.153892308588929 | 4.69049580197719 | -1.15263040946994 | 0.255504275283411 | 0.576100832842047 | -5.71017089687423 |
| ENSG00000004399.12 | PLXND1 | -0.448651231500376 | 4.30782363406651 | -1.15299001748188 | 0.255532035104264 | 0.576100832842047 | -5.67966158517752 |
| ENSG00000070915.9 | SLC12A3 | 0.550494883108965 | 0.924880509576926 | 1.15270396052857 | 0.255549720608396 | 0.576100832842047 | -5.24023234911054 |
| ENSG00000155666.11 | KDM8 | 0.142654414722611 | 3.5936920184479 | 1.15247104298884 | 0.255568998915646 | 0.576100832842047 | -5.60773985307701 |
| ENSG00000279176.1 | | 0.417037942654151 | 1.5970491919576 | 1.15209017027592 | 0.255747624383327 | 0.576309949387958 | -5.35240378677596 |
| ENSG00000160058.18 | BSDC1 | 0.132171132074074 | 6.22206893985943 | 1.15202724907286 | 0.255749299464593 | 0.576309949387958 | -5.7599348199907 |
| ENSG00000129355.6 | CDKN2D | 0.233618959602406 | 3.50097444773024 | 1.15201674150437 | 0.255753569492785 | 0.576309949387958 | -5.5935265778926 |
| ENSG00000144115.16 | THNSL2 | 0.470705960552856 | 3.2590975838264 | 1.15233946099047 | 0.255773764157752 | 0.576309949387958 | -5.55867989060543 |
| ENSG00000270672.1 | NA | 1.4054994247528 | -1.31611965671828 | 1.15195125522383 | 0.255818519736389 | 0.576327175341314 | -4.6481077887191 |
| lnc-TMEM272-2 | lnc-TMEM272-2 | 0.209057837888822 | 0.0657745562349241 | 1.15181045571593 | 0.255837409565835 | 0.576327175341314 | -5.10876862018775 |
| lnc-DNAJC24-6 | lnc-DNAJC24-6 | -0.338810383045851 | 0.471796518764509 | -1.15162099631584 | 0.255914428364426 | 0.576396807813293 | -5.1917497341531 |
| ENSG00000237506.3 | RPSAP15 | 0.526855108697599 | -0.754912718830572 | 1.15159664905356 | 0.255924327192619 | 0.576396807813293 | -4.83429956464868 |
| ENSG00000260428.2 | SCX | -0.441215163252696 | -0.80747786522982 | -1.15144840296032 | 0.255984605319343 | 0.576469489200004 | -4.95973815992319 |
| ENSG00000177932.6 | ZNF354C | -0.631387993415976 | 2.85127017355245 | -1.15164656399345 | 0.256106646104236 | 0.57662392923692 | -5.50130378251646 |
| lnc-KIAA1328-3 | lnc-KIAA1328-3 | -0.210261656233159 | -0.0633134095432718 | -1.151142022854 | 0.256109214452894 | 0.57662392923692 | -5.07143966113667 |
| ENSG00000198612.10 | COPS8 | 0.100464561299843 | 5.30875826357207 | 1.15088442116666 | 0.256214018474851 | 0.576796799985548 | -5.74604869656097 |
| ENSG00000167595.15 | PROSER3 | -0.160376318658335 | 3.40157699720983 | -1.15076950550752 | 0.256260781313651 | 0.576813689397705 | -5.5916699917116 |
| ENSG00000116062.14 | MSH6 | 0.22897866524031 | 4.78672876076668 | 1.15072825646386 | 0.256277568359423 | 0.576813689397705 | -5.72031823653549 |
| ENSG00000144579.7 | CTDSP1 | -0.163954244452572 | 5.5737826751476 | -1.15038277122773 | 0.256418200892615 | 0.577066316909367 | -5.75598561983978 |
| lnc-C21orf2-1 | lnc-C21orf2-1 | -0.215173713068665 | 1.27123582484009 | -1.15031478304415 | 0.256445882561822 | 0.577066316909367 | -5.3403804004943 |
| ENSG00000107581.12 | EIF3A | 0.104899486094093 | 7.80303762487737 | 1.15007555767776 | 0.256543301240415 | 0.577222427790933 | -5.7470472593281 |
| ENSG00000141994.15 | DUS3L | 0.164967927277758 | 4.01511113544738 | 1.14987024428994 | 0.256626931313371 | 0.577347483574502 | -5.66227721832739 |
| ENSG00000103044.10 | HAS3 | 0.277606386741895 | 0.140077774827349 | 1.14962259071509 | 0.256727833848355 | 0.577511366568382 | -5.09551799963284 |
| lnc-SLC35A3-1 | lnc-SLC35A3-1 | -0.299422710596422 | -0.464965242400621 | -1.14953584576311 | 0.256763183452498 | 0.5775277680038 | -4.97891238875887 |
| lnc-PBX2-1 | lnc-PBX2-1 | -0.156578379074899 | 1.65980024726206 | -1.149434929003 | 0.256804312641234 | 0.577557164236725 | -5.39443391521553 |
| ENSG00000167723.14 | TRPV3 | -0.799050353707862 | -0.711211336083304 | -1.14945252473023 | 0.256838128887606 | 0.577570108914409 | -4.90372706558803 |
| lnc-CAND1-3 | lnc-CAND1-3 | -0.17313984324642 | 1.33074274928079 | -1.14925882807637 | 0.256876094900004 | 0.577592381617519 | -5.35027152944694 |
| ENSG00000136492.8 | BRIP1 | 0.294912609702681 | 1.81401088668341 | 1.14900846158093 | 0.256978174118659 | 0.577758793870493 | -5.3974663382223 |
| ENSG00000206952.3 | SNORA50A | 0.177649591701286 | 0.288646062175929 | 1.14864896680607 | 0.257124798015397 | 0.578025307727274 | -5.15532558012761 |
| ENSG00000120265.17 | PCMT1 | -0.137500202639854 | 4.29260248468351 | -1.14855666416258 | 0.257162454362448 | 0.578033835491644 | -5.68836828701905 |
| ENSG00000206620.1 | SNORD45C | 0.281145765553999 | 0.289889816733825 | 1.14850199942865 | 0.257184757587935 | 0.578033835491644 | -5.18603543424352 |
| lnc-HIST1H2AI-3 | lnc-HIST1H2AI-3 | 0.248017342797484 | 2.60561338540899 | 1.14826901954563 | 0.257279829027869 | 0.578184378303378 | -5.50129778570041 |
| lnc-GLG1-1 | lnc-GLG1-1 | -0.192858741877704 | 2.75429581902068 | -1.14809353491854 | 0.257351455215259 | 0.578255234709122 | -5.51490466139059 |
| ENSG00000229809.8 | ZNF688 | -0.144559244288627 | 2.44522251503697 | -1.14803641406638 | 0.2573747728774 | 0.578255234709122 | -5.4828594506253 |
| ENSG00000150637.8 | CD226 | -0.33649268998499 | 0.0398255974309378 | -1.14798529913712 | 0.257395640111013 | 0.578255234709122 | -5.08838514461995 |
| ENSG00000125457.14 | MIF4GD | -0.256490564439122 | 4.10241989004105 | -1.14770905769962 | 0.257508434357718 | 0.578445089506001 | -5.66183271567455 |
| ENSG00000082701.15 | GSK3B | 0.0872868456138693 | 5.43742302381568 | 1.14763098866944 | 0.257540317753689 | 0.578445089506001 | -5.75303512160132 |
| DNAJC3-DT | DNAJC3-DT | -0.277407632151581 | 1.1941959618064 | -1.14757188322531 | 0.257564458306491 | 0.578445089506001 | -5.31191312273948 |
| lnc-TK1-6 | lnc-TK1-6 | 0.203196907300723 | 3.57405660202511 | 1.14736871312043 | 0.257647451870162 | 0.578568350626613 | -5.60811964999788 |
| ENSG00000169955.7 | ZNF747 | -0.174218705112318 | 3.54852657985692 | -1.14728677847716 | 0.257680927032927 | 0.578580399380248 | -5.61001981078747 |
| NFYC-AS1 | NFYC-AS1 | -0.209967780609047 | 3.55188602840113 | -1.14715550244966 | 0.257734567591265 | 0.578589382253917 | -5.6129544644855 |
| ENSG00000183684.7 | ALYREF | 0.164819463332274 | 4.11690487656043 | 1.14713939978817 | 0.25774114783492 | 0.578589382253917 | -5.6704697474659 |
| ENSG00000058799.14 | YIPF1 | 0.102085285208109 | 3.45080358179384 | 1.14701636808285 | 0.257791427892124 | 0.578639145071274 | -5.59301034193078 |
| ENSG00000039523.19 | RIPOR1 | -0.22370993225288 | 6.00425042004376 | -1.14657928533119 | 0.257970109924497 | 0.578974093287726 | -5.76459378619875 |
| lnc-FNTB-2 | lnc-FNTB-2 | -0.195383454429863 | 2.79905642480435 | -1.14650267781601 | 0.258001436691539 | 0.578974093287726 | -5.52789553703569 |
| ENSG00000198734.10 | F5 | -0.685547692084872 | 1.56549098923121 | -1.14688423486048 | 0.258025038027902 | 0.578974093287726 | -5.32916571328064 |
| ENSG00000165487.13 | MICU2 | -0.103183716047083 | 4.83960285666823 | -1.14626810218862 | 0.25809737763879 | 0.57902084386309 | -5.7262588848742 |
| ENSG00000165775.17 | FUNDC2 | 0.133564051527178 | 4.73007602473401 | 1.14625647233798 | 0.258102134890145 | 0.57902084386309 | -5.72195178849496 |

| | | | | | | | |
|---|---|---|---|---|---|---|---|
| DNAJC27-AS1 | DNAJC27-AS1 | -0.362298402697739 | 1.46387313488602 | -1.14604438045217 | 0.258188903232379 | 0.579152375243249 | -5.36513270959947 |
| ENSG00000156869.13 | FRRS1 | 1.22676411080393 | -0.335477320470023 | 1.14627449675145 | 0.258249545445397 | 0.579160117926357 | -4.83624214725135 |
| ENSG00000162377.5 | COA7 | 0.25182158639321 | 1.80835474520674 | 1.14580673811251 | 0.258286149330714 | 0.579160117926357 | -5.42083088925838 |
| ENSG00000105176.17 | URI1 | -0.12301923580062 | 5.88776176905727 | -1.14571785549033 | 0.25832252792296 | 0.579160117926357 | -5.76403270512801 |
| ENSG00000164556.7 | CFAP144P1 | -0.280580617754399 | 1.13054729760412 | -1.14567517170488 | 0.258339999192257 | 0.579160117926357 | -5.29694174415131 |
| ENSG00000135253.13 | KCP | 1.07085213316448 | -1.32442084029889 | 1.14561157992122 | 0.258386053691455 | 0.579160117926357 | -4.72285182715363 |
| ENSG00000266405.3 | CBX3P2 | -0.200575375426847 | 1.41240950989233 | -1.14555468925334 | 0.258389319482959 | 0.579160117926357 | -5.35751391188132 |
| lnc-METTL4-5 | lnc-METTL4-5 | -0.200575375426847 | 1.41240950989233 | -1.14555468925334 | 0.258389319482959 | 0.579160117926357 | -5.35751391188132 |
| lnc-CLASP1-3 | lnc-CLASP1-3 | 0.143775813429603 | 1.79241657257592 | 1.14527465091124 | 0.258503981149028 | 0.579354033535546 | -5.40454192389971 |
| lnc-NOP14-3 | lnc-NOP14-3 | 0.214298756022401 | 0.378872137283581 | 1.14520305027894 | 0.258533303881563 | 0.579356667805576 | -5.1676207150054 |
| lnc-MOCS2-8 | lnc-MOCS2-8 | -0.394900923777323 | -0.344611538085298 | -1.14500284737493 | 0.2586153059984 | 0.579477339795893 | -4.9883584480475 |
| ENSG00000170801.9 | HTRA3 | -1.46478259190092 | 0.104202464621442 | -1.14477511987454 | 0.258951446737014 | 0.580114443677987 | -5.01140053084581 |
| ENSG00000254413.8 | CHKB-CPT1B | -0.141373843420536 | 5.91273160142379 | -1.14410627368193 | 0.258982767130754 | 0.580114443677987 | -5.76672923854896 |
| ENSG00000225259.4 | ST13P6 | 0.883800760686154 | -0.649862198504705 | 1.14410414021308 | 0.258994063772738 | 0.580114443677987 | -4.79155344802549 |
| ENSG00000089159.16 | PXN | -0.52739906881047 | 3.12418896557779 | -1.144471379671 | 0.25901237610653 | 0.580114443677987 | -5.55350642753966 |
| ENSG00000117408.10 | IPO13 | -0.153324235898617 | 3.58395471707437 | -1.14382714662676 | 0.25909724391296 | 0.580168614056772 | -5.61878347730195 |
| lnc-ARSG-2 | lnc-ARSG-2 | -0.331469178185233 | 1.57515940567005 | -1.14380944063987 | 0.259104506792308 | 0.580168614056772 | -5.40289139909446 |
| ENSG00000177951.17 | BET1L | 0.144964232036743 | 4.75858820351077 | 1.14372712984249 | 0.259138272057637 | 0.580168614056772 | -5.72188535773737 |
| ENSG00000155090.14 | KLF10 | 0.483792544215794 | 4.5738802442963 | 1.14412045117499 | 0.259164863404459 | 0.580168614056772 | -5.71456736453219 |
| lnc-MTRNR2L8-3 | lnc-MTRNR2L8-3 | 0.260100165314903 | 1.30064236857707 | 1.14363152117174 | 0.259177496295585 | 0.580168614056772 | -5.35089000453389 |
| ENSG00000187713.6 | TMEM203 | 0.154571953526275 | 3.42070506472505 | 1.1432721994042 | 0.259324949066567 | 0.58043556183527 | -5.59790185693859 |
| ENSG00000213762.11 | ZNF134 | 0.121250650723444 | 5.94620634358899 | 1.14309839525884 | 0.259396293644178 | 0.580532120482562 | -5.767687007293 |
| ENSG00000252759.1 | | 0.206197086559347 | 2.5939976132618 | 1.1429990527811 | 0.259437078896666 | 0.580560273421405 | -5.4972900759408 |
| ENSG00000150672.17 | DLG2 | -0.690549709487995 | 4.41609318925878 | -1.14325152543136 | 0.259576957028409 | 0.580810143115093 | -5.72084293659218 |
| lnc-UBE2O-2 | lnc-UBE2O-2 | 0.282688920130153 | 0.455433829918848 | 1.14243613447768 | 0.259668272996555 | 0.580889507977348 | -5.226109179094 |
| ENSG00000274641.1 | H2BC17 | 0.548228361107422 | 3.31327125958653 | 1.14289177617155 | 0.259668870567924 | 0.580889507977348 | -5.59350981893804 |
| lnc-GALNT1-1 | lnc-GALNT1-1 | -0.261757452467115 | 1.32152231313034 | -1.14226922719489 | 0.25973685125752 | 0.580978440494841 | -5.34580232939147 |
| ENSG00000253910.2 | PCDHGB2 | -0.973062985558956 | -0.729535645788056 | -1.14236213117245 | 0.259877698814175 | 0.581203408239703 | -5.06025887762793 |
| lnc-TTC27-9 | lnc-TTC27-9 | 0.217554254770622 | 4.1865109131278 | 1.14183642773108 | 0.259914738828332 | 0.581203408239703 | -5.68035946502238 |
| lnc-TMCC1-7 | lnc-TMCC1-7 | -0.396374772454878 | 0.853303573556541 | -1.14175617962263 | 0.259947731693253 | 0.581203408239703 | -5.26829141904685 |
| ENSG00000001460.17 | STPG1 | -0.238182455263821 | 0.97543246403461 | -1.14174974975701 | 0.25995037537068 | 0.581203408239703 | -5.30176116716593 |
| ENSG00000111816.7 | FRK | 1.10101229577224 | 3.05245267450901 | 1.14223359875281 | 0.260004646577289 | 0.581261609699178 | -5.51097747143352 |
| lnc-PSMC3IP-3 | lnc-PSMC3IP-3 | -0.160289900634994 | 2.72416956212423 | -1.14153972012524 | 0.260036740898073 | 0.581270225663014 | -5.52569642499877 |
| lnc-RC3H2-1 | lnc-RC3H2-1 | 0.144352204983879 | 4.17076776128596 | 1.14145667067181 | 0.260070897038075 | 0.581283448119742 | -5.68280708915648 |
| ENSG00000058262.9 | SEC61A1 | 0.101322904424901 | 6.1163560027369 | 1.14135102787997 | 0.260114349893711 | 0.581317444501874 | -5.77134988260911 |
| ENSG00000054983.16 | GALC | 0.115675036759005 | 3.26275634345686 | 1.14120881965524 | 0.260172851012348 | 0.581385060513208 | -5.57825117323646 |
| ENSG00000182324.6 | KCNJ14 | 0.296370825075887 | 0.779722961663808 | 1.14102535262875 | 0.260248338961373 | 0.581490616678457 | -5.25917226217919 |
| ENSG00000068079.7 | IFI35 | -0.205368528627591 | 3.26510606750673 | -1.14075644477709 | 0.26035901010494 | 0.581674753607944 | -5.58826082540999 |
| ENSG00000124155.17 | PIGT | 0.122330082286439 | 5.48591656143194 | 1.14059366621175 | 0.260426019289233 | 0.581681240279118 | -5.76153512376951 |
| lnc-EPG5-1 | lnc-EPG5-1 | -0.258654350300373 | 0.664612112533993 | -1.14055680485084 | 0.260441195298657 | 0.581681240279118 | -5.23764261251344 |
| ENSG00000140406.3 | TLNRD1 | -0.25451406419556 | 3.34733979091376 | -1.14054344838595 | 0.260446694379456 | 0.581681240279118 | -5.61101186793846 |
| ENSG00000198700.9 | IPO9 | 0.0616193587401037 | 6.05666450875002 | 1.14018328776465 | 0.260595009922931 | 0.581949342437201 | -5.7721276129451 |
| ENSG00000207034.1 | | 0.332267694676134 | 1.4206898125622 | 1.13995102290007 | 0.260690689585323 | 0.582099855037395 | -5.35326305868117 |
| ENSG00000138778.11 | CENPE | 0.369046084430028 | 1.50153606720956 | 1.13971997914359 | 0.260785891187102 | 0.582191068763847 | -5.35753009444381 |
| ENSG00000106771.12 | TMEM245 | 0.101076411485668 | 5.60044727070845 | 1.13971459671615 | 0.260788109313641 | 0.582191068763847 | -5.76515320594441 |
| ENSG00000183208.12 | GDPGP1 | -0.131422418116299 | 1.66748403256746 | -1.13943536575227 | 0.260903200392964 | 0.582384836101115 | -5.396943697859 |
| lnc-HJURP-11 | lnc-HJURP-11 | -0.185461322192528 | 0.874615649790685 | -1.13929926621766 | 0.260959309943978 | 0.582446917867804 | -5.2778085358709 |
| ENSG00000131242.17 | RAB11FIP4 | 0.140702079139161 | 6.16894056546766 | 1.13911737434972 | 0.261034311723908 | 0.582523954953831 | -5.7739119198519 |
| ENSG00000257365.7 | FNTB | -0.152606868999876 | 2.00786243791415 | -1.13907829445429 | 0.261050428047133 | 0.582523954953831 | -5.43414601032219 |
| lnc-C1R-2 | lnc-C1R-2 | -0.18387844478302 | 0.48746080801271 | -1.13883826735742 | 0.261149429464733 | 0.582607428148112 | -5.20244274267068 |
| ENSG00000253305.2 | PCDHGB6 | -0.938263495398783 | -1.02079501522481 | -1.1391155610288 | 0.261159468079142 | 0.582607428148112 | -4.93855622102858 |
| MALAT1 | MALAT1 | 0.298198493693183 | 13.5124425852164 | 1.13878173354155 | 0.261172751276423 | 0.582607428148112 | -5.56635725178323 |
| ENSG00000055483.19 | USP36 | 0.164373445677998 | 5.4182810426535 | 1.13847355279415 | 0.261299910849422 | 0.582827922086438 | -5.76148423104609 |
| lnc-FAM192A-4 | lnc-FAM192A-4 | 0.346113291509145 | 2.75964685858022 | 1.13816515520075 | 0.261449681209595 | 0.583051791915229 | -5.51722630354085 |
| ENSG00000066855.15 | MTFR1 | 0.147319981823085 | 2.00902230218817 | 1.1380602203808 | 0.261470527016437 | 0.583051791915229 | -5.44702508470154 |
| ENSG00000166340.16 | TPP1 | -0.169611545725997 | 5.31426767163425 | -1.13802453932524 | 0.261485259251299 | 0.583051791915229 | -5.75510237650752 |
| ENSG00000125618.16 | PAX8 | -1.35100244804925 | -1.93735050844263 | -1.13777063020912 | 0.261658145954218 | 0.583323244290174 | -4.69153706987732 |
| lnc-NPIPB4-6 | lnc-NPIPB4-6 | -0.989226109077256 | -2.4792810011771 | -1.13759252481097 | 0.261663679470008 | 0.583323244290174 | -4.67291688729019 |
| ENSG00000183828.14 | NUDT14 | -0.439205754142962 | 0.646777236348158 | -1.13754102728101 | 0.261702613859537 | 0.583346859548501 | -5.29536682751747 |
| ENSG00000140941.12 | MAP1LC3B | 0.181653252356797 | 5.29368865989141 | 1.13720599596624 | 0.261823388136161 | 0.58355287471647 | -5.75959077337243 |
| lnc-PPP1R3G-4 | lnc-PPP1R3G-4 | -0.33565351195294 | 0.393985937030969 | -1.13710267050911 | 0.261866092678368 | 0.58357506242746 | -5.18380574974575 |
| ENSG00000058091.16 | CDK14 | 0.129955419673044 | 8.89564743589963 | 1.13700491711135 | 0.261906498869608 | 0.58357506242746 | -5.73590355764059 |
| lnc-DAPK2-3 | lnc-DAPK2-3 | 0.264781862448497 | 5.27925523746799 | 1.13702426689045 | 0.261918399975945 | 0.58357506242746 | -5.75611056455894 |
| ENSG00000154640.14 | BTG3 | 0.870217684828378 | 2.82858679724198 | 1.13714145504378 | 0.262092508012667 | 0.583899782706433 | -5.49166488862247 |
| ENSG00000117360.12 | PRPF3 | 0.148205952034982 | 6.38317095474904 | 1.13636637968095 | 0.262170546922114 | 0.584010429361242 | -5.77737786070451 |
| ENSG00000182362.13 | YBEY | -0.191605187451625 | 3.11469614799257 | -1.13628401075157 | 0.262204621991905 | 0.584023128931867 | -5.55948041677487 |
| ENSG00000165476.13 | REEP3 | -0.20196675165354 | 3.58086025478552 | -1.13612209479982 | 0.26227161397645 | 0.58405359244571 | -5.63360242673927 |
| lnc-DYNC1H1-4 | lnc-DYNC1H1-4 | 0.246688226514063 | 0.529805621973811 | 1.13608759470353 | 0.262285889815725 | 0.58405359244571 | -5.20276309426722 |

| | | | | | | | |
|---|---|---|---|---|---|---|---|
| ENSG00000227268.3 | KLLN | -0.283667725645654 | -0.304281368804776 | -1.13604521827593 | 0.262303425572956 | 0.58405359244571 | -5.03830171413754 |
| ENSG00000119655.10 | NPC2 | 0.228136668030524 | 3.62090670426955 | 1.1357346094359 | 0.26243198398202 | 0.584225477494763 | -5.62064379939652 |
| ENSG00000230581.2 | ACTG1P14 | 0.232727260006127 | 0.924325414040836 | 1.13572155431429 | 0.262437388374706 | 0.584225477494763 | -5.29738890353324 |
| lnc-STK10-3 | lnc-STK10-3 | 0.134507565886669 | 2.14106640507512 | 1.13557671425949 | 0.262497352753326 | 0.584240078996362 | -5.45386301029895 |
| ENSG00000241553.12 | ARPC4 | 0.122610933593899 | 5.82190167225339 | 1.13556858971614 | 0.262500716637922 | 0.584240078996362 | -5.7747815571327 |
| lnc-TGFBRAP1-1 | lnc-TGFBRAP1-1 | 0.118274140776947 | 3.45381118998799 | 1.13540209007857 | 0.262569660910573 | 0.584330341715032 | -5.60865402381779 |
| ENSG00000143207.20 | COP1 | 0.122784387962006 | 5.53116812783073 | 1.13518463398337 | 0.262659724764123 | 0.584411231018611 | -5.76904079390483 |
| ENSG00000143774.16 | GUK1 | 0.122667291158772 | 5.30925978186686 | 1.13517722327279 | 0.262662794449457 | 0.584411231018611 | -5.76293913913197 |
| ENSG00000260404.3 | | 0.291982721775798 | 2.79532861623963 | 1.13510013004252 | 0.26269472974523 | 0.584419111797025 | -5.51760780244579 |
| lnc-ZNF852-4 | lnc-ZNF852-4 | -0.148076862448963 | 2.42390818188186 | -1.13485718665703 | 0.262795385391249 | 0.584510081985514 | -5.49060856086312 |
| ENSG00000115825.9 | PRKD3 | 0.19194671756948 | 6.86381013768496 | 1.13482244770446 | 0.262809780593241 | 0.584510081985514 | -5.77605753408603 |
| NDUFA6-DT | NDUFA6-DT | -0.157375499769717 | 2.29773758371915 | -1.13473208381797 | 0.262847228415792 | 0.584510081985514 | -5.472067164253 |
| lnc-SERTAD2-3 | lnc-SERTAD2-3 | -0.184831835702457 | 4.89805301439635 | -1.13470298426885 | 0.262859288412021 | 0.584510081985514 | -5.7437955553323 |
| lnc-ZSCAN29-1 | lnc-ZSCAN29-1 | -0.102553998620777 | 2.18085334003692 | -1.13464383609596 | 0.262883802957579 | 0.584510081985514 | -5.46707956381522 |
| lnc-GOLGA8A-6 | lnc-GOLGA8A-6 | -0.367013293639064 | 3.19915156064395 | -1.13475770771132 | 0.262906006851377 | 0.584510081985514 | -5.59037572693564 |
| lnc-MALL-5 | lnc-MALL-5 | -1.23234975680039 | -2.20214375794595 | -1.13450596177696 | 0.262963369471367 | 0.584574471738756 | -4.68010904685667 |
| lnc-ITGB7-2 | lnc-ITGB7-2 | -0.262984619152053 | -0.0927925623365849 | -1.13422327144622 | 0.263058157268241 | 0.584722035750777 | -5.05781236925543 |
| ENSG00000108064.10 | TFAM | 0.131893327004616 | 5.56109602379048 | 1.13403631746957 | 0.26313568967869 | 0.584818952068543 | -5.77182400534715 |
| lnc-KIAA1324L-3 | lnc-KIAA1324L-3 | 0.137206205353305 | 4.80961078699443 | 1.1339811199488 | 0.263158583979927 | 0.584818952068543 | -5.74050011352658 |
| ENSG00000235505.7 | CASP4LP | 0.743734627003491 | -0.842482011562769 | 1.13379571798662 | 0.263236384040119 | 0.584928694018976 | -4.87057976193506 |
| ENSG00000160752.14 | FDPS | -0.129783885726252 | 4.94024550968295 | -1.13356094022719 | 0.263332908787866 | 0.58502943351201 | -5.74920621204835 |
| lnc-PDSS1-7 | lnc-PDSS1-7 | -0.284398061355448 | 2.48774825630319 | -1.13354321696654 | 0.263340263654921 | 0.58502943351201 | -5.49481447955043 |
| lnc-KMT2C-1 | lnc-KMT2C-1 | -0.269237694198475 | 1.46172740121855 | -1.1333310917331 | 0.263428303595702 | 0.58502943351201 | -5.37144519008525 |
| lnc-UNC5CL-4 | lnc-UNC5CL-4 | -0.223014884234975 | 1.28695846958809 | -1.13327139681163 | 0.263453083022493 | 0.58502943351201 | -5.34072275807543 |
| ENSG00000074800.15 | ENO1 | -0.232589620870044 | 7.17931012844866 | -1.13326420651234 | 0.263456067835933 | 0.58502943351201 | -5.77369027147004 |
| lnc-CD244-2 | lnc-CD244-2 | 0.362929148046435 | 3.4326693866937 | 1.13334322188842 | 0.263485438021823 | 0.58502943351201 | -5.58109942034205 |
| lnc-MUC20-6 | lnc-MUC20-6 | -0.270642270172713 | 1.26154548227844 | -1.1331433211336 | 0.263506252995715 | 0.58502943351201 | -5.35229092136553 |
| lnc-CASP10-1 | lnc-CASP10-1 | -0.206656681247056 | 2.83250397593734 | -1.13313645528096 | 0.263509103536406 | 0.58502943351201 | -5.54509161737152 |
| ENSG00000185862.6 | EVI2B | -0.23224040648402 | 7.04915675905067 | -1.1328664738013 | 0.263621210971182 | 0.58519392440913 | -5.77725899105691 |
| ENSG00000102699.5 | PARP4 | 0.156022604387531 | 7.55962685361799 | 1.13282109988189 | 0.263640055436324 | 0.58519392440913 | -5.77116608509088 |
| ENSG00000167555.13 | ZNF528 | -0.237554592694758 | 5.46145855755612 | -1.13271804677672 | 0.263682858502173 | 0.585217504455232 | -5.7679236723882 |
| PARTICL | PARTICL | -0.212213545877276 | 1.24826945607396 | -1.1326137332629 | 0.263726190134091 | 0.585217504455232 | -5.35974769238591 |
| lnc-ZSWIM7-5 | lnc-ZSWIM7-5 | 0.192713200399194 | 2.34644942707492 | 1.13259017980702 | 0.263735974898058 | 0.585217504455232 | -5.48342479698534 |
| ENSG00000204584.1 | | -0.240015271730436 | 1.72374236022406 | -1.13237878685537 | 0.263823805048359 | 0.585318641460744 | -5.41149152751401 |
| ENSG00000170430.9 | MGMT | -0.109953501511248 | 3.32471129108117 | -1.13219674256883 | 0.263899458073087 | 0.585318641460744 | -5.5968329199925 |
| ENSG00000113312.10 | TTC1 | -0.119372733057079 | 4.61700841814845 | -1.1321792057067 | 0.263906746769882 | 0.585318641460744 | -5.72805330628276 |
| lnc-ANXA6-5 | lnc-ANXA6-5 | -0.150221964647048 | 3.32447199561911 | -1.13216879944696 | 0.263911071903054 | 0.585318641460744 | -5.6018706737446 |
| ENSG00000260286.3 | ARMH2 | -0.337622313883701 | -0.43251832171751 | -1.13213832413024 | 0.263923738589961 | 0.585318641460744 | -4.9831626565101 |
| ENSG00000175928.5 | LRRN1 | 0.242823618773136 | 4.46610134014489 | 1.13157531045888 | 0.264157826346533 | 0.585774675683116 | -5.70702660890044 |
| ENSG00000171877.20 | FRMD5 | 1.43702616847132 | -1.39306293089767 | 1.13184423726959 | 0.264226979247779 | 0.585849733198518 | -4.839468968567 |
| lnc-GOLGB1-3 | lnc-GOLGB1-3 | 0.15400233452116 | 1.75808047047758 | 1.13129805762409 | 0.264273155979837 | 0.585849733198518 | -5.41915105302958 |
| lnc-ATPAF1-2 | lnc-ATPAF1-2 | 0.11433105612156 | 2.81506080816072 | 1.1312886683967 | 0.264277062272178 | 0.585849733198518 | -5.53842815239875 |
| ENSG00000081148.11 | IMPG2 | -0.228866080966076 | 1.73189306706516 | -1.13115398687042 | 0.264333099685411 | 0.585910854062547 | -5.40935181546009 |
| ENSG00000224557.7 | HLA-DPB2 | -0.496221930908299 | 1.46429824016739 | -1.13125539317272 | 0.264377698665164 | 0.585946610490478 | -5.35543260831455 |
| ENSG00000125731.12 | SH2D3A | -0.190032758524518 | 3.76200449068175 | -1.13024305623889 | 0.264712336517938 | 0.586625110093531 | -5.65426499783726 |
| lnc-CHP1-6 | lnc-CHP1-6 | -0.330011819146551 | 0.0992573586035236 | -1.13009579754442 | 0.264773679410534 | 0.586634518045694 | -5.10757561311109 |
| ENSG00000088682.13 | COQ9 | -0.14191815994873 | 3.56837058545484 | -1.12997762797747 | 0.264822912112658 | 0.586634518045694 | -5.63087847638703 |
| ENSG00000184575.11 | XPOT | -0.106385955363161 | 5.68847065470561 | -1.12994104466717 | 0.264838155053414 | 0.586634518045694 | -5.77859802930896 |
| lnc-NACA-2 | lnc-NACA-2 | -0.237639189550684 | 0.855651606001437 | -1.12990728897388 | 0.264852220383951 | 0.586634518045694 | -5.30383656842572 |
| lnc-TARSL2-1 | lnc-TARSL2-1 | 0.179691000717368 | 2.04736485106686 | 1.12989081144053 | 0.2648590864402 | 0.586634518045694 | -5.46035371186021 |
| ENSG00000164190.17 | NIPBL | 0.180165141930159 | 7.38782135274597 | 1.12980215304804 | 0.264896031863115 | 0.586653219694264 | -5.77616978907785 |
| lnc-FAM126A-5 | lnc-FAM126A-5 | -0.307180886547482 | 2.07803837206838 | -1.12966107046051 | 0.264954830900008 | 0.586690880598962 | -5.47161772267317 |
| lnc-COIL-7 | lnc-COIL-7 | 0.206880537725895 | 1.72727906428878 | 1.12955476541424 | 0.264999141841694 | 0.586690880598962 | -5.41444127499722 |
| lnc-ANKRD60-3 | lnc-ANKRD60-3 | -0.258457586845657 | -0.136959003539246 | -1.12953615683392 | 0.265006898966631 | 0.586690880598962 | -5.09928880080459 |
| ENSG00000269881.1 | NA | -0.16222887976021 | 2.43710113315449 | -1.12942601299305 | 0.265052816562575 | 0.586690880598962 | -5.51190078229097 |
| ENSG00000258311.5 | | 0.179040174019278 | 3.51845954364651 | 1.12941944347582 | 0.265055555491899 | 0.586690880598962 | -5.61891986644336 |
| ENSG00000127220.5 | ABHD8 | -0.293833329230883 | 0.194119287215891 | -1.12929752848179 | 0.265106387328712 | 0.586740297891062 | -5.15725696373476 |
| lnc-NFRKB-4 | lnc-NFRKB-4 | -0.145883784807654 | 0.479728558055715 | -1.12919682184532 | 0.265148381694002 | 0.586770147339816 | -5.22442348193209 |
| lnc-TAOK3-1 | lnc-TAOK3-1 | -0.31224009136642 | 2.18115657062691 | -1.12900431648008 | 0.265228669060295 | 0.586826707676157 | -5.48543205248564 |
| ENSG00000141252.19 | VPS53 | 0.165322721438835 | 4.95241389649228 | 1.12899659731973 | 0.265231888818191 | 0.586826707676157 | -5.75276298788033 |
| ENSG00000130717.12 | UCK1 | 0.109681849071182 | 3.64294199340148 | 1.12888314568921 | 0.265279214122528 | 0.586826707676157 | -5.64360699790186 |
| lnc-RPL18-1 | lnc-RPL18-1 | -0.213860113007818 | 6.91310417364805 | -1.12885605425118 | 0.265290515957676 | 0.586826707676157 | -5.78162904956256 |
| ENSG00000262919.7 | CCNQ | -0.250872019767752 | 3.3069546502225 | -1.12876141411381 | 0.265330000028701 | 0.586826707676157 | -5.61337670250267 |
| ENSG00000125375.14 | DMAC2L | 0.0904450365597587 | 5.44684721831157 | 1.1287254591419 | 0.26534500161988 | 0.586826707676157 | -5.77372252807946 |
| ENSG00000141526.16 | SLC16A3 | -0.292358685577435 | 3.99467184294812 | -1.12867921162848 | 0.265388705998339 | 0.586859769486489 | -5.68773979834428 |
| lnc-PROSER1-3 | lnc-PROSER1-3 | -1.30504529656591 | -1.11215284834568 | -1.12888418344167 | 0.265416974889459 | 0.586859769486489 | -4.78192471369607 |
| ENSG00000167461.11 | RAB8A | -0.142677417984152 | 5.99847645916907 | -1.12835389982323 | 0.265500063762337 | 0.586954308214677 | -5.7847061508676 |
| lnc-FSD2-31 | lnc-FSD2-31 | 0.493128781052059 | 1.58201013882412 | 1.1285423105596 | 0.26551676450405 | 0.586954308214677 | -5.38551228642547 |

| | | | | | | | |
|---|---|---|---|---|---|---|---|
| ENSG00000145675.14 | PIK3R1 | -0.136464414223962 | 5.58783591227632 | -1.12797660810058 | 0.265657584318733 | 0.587202540596271 | -5.77818634782515 |
| ENSG00000168878.16 | SFTPB | 0.966123076262525 | 2.64379871088322 | 1.12836353237913 | 0.265746632820893 | 0.587216822234982 | -5.51734990334013 |
| lnc-ENTPD6-2 | lnc-ENTPD6-2 | -0.185201737998336 | 0.50336948023112 | -1.12774140063025 | 0.265755817942487 | 0.587216822234982 | -5.21770613791447 |
| lnc-PDE6D-3 | lnc-PDE6D-3 | -0.276577979280246 | 0.998958425306326 | -1.1277334580321 | 0.26575913559442 | 0.587216822234982 | -5.31357471609841 |
| ENSG00000134882.15 | UBAC2 | 0.0893197680386617 | 5.13427981788843 | 1.12757229418045 | 0.265826460696727 | 0.587216822234982 | -5.76248355477883 |
| ENSG00000235183.3 | SRP14P3 | 0.228512123823989 | 0.904365779517399 | 1.12757177175328 | 0.265826678956893 | 0.587216822234982 | -5.29379920391836 |
| ENSG00000124215.16 | CDH26 | -0.276827824513993 | 1.1465414858661 | -1.12749189514826 | 0.265860051391407 | 0.587216822234982 | -5.32087831248167 |
| ENSG00000071073.12 | MGAT4A | 0.905396450349556 | -0.400924509992321 | 1.12780882816456 | 0.265863750007666 | 0.587216822234982 | -4.9914345086474 |
| ENSG00000141971.12 | MVB12A | -0.159568129390118 | 2.67269590245645 | -1.12729052989909 | 0.265944195021499 | 0.587256687575574 | -5.52916413369606 |
| ENSG00000174799.10 | CEP135 | 0.136899528296841 | 4.38132900436366 | 1.12724220409514 | 0.265964391541922 | 0.587256687575574 | -5.71627066946343 |
| lnc-TSPAN33-1 | lnc-TSPAN33-1 | -0.25301391403348 | 1.93716226806481 | -1.12723502348616 | 0.265967392585119 | 0.587256687575574 | -5.46148278457296 |
| ENSG00000220867.1 | HSPE1P26 | 0.457013095896251 | 0.0627104691512286 | 1.12696484788437 | 0.266080326546646 | 0.587380025880482 | -5.0936015430349 |
| lnc-SYNJ2-2 | lnc-SYNJ2-2 | 0.457013095896251 | 0.0627104691512286 | 1.12696484788437 | 0.266080326546646 | 0.587380025880482 | -5.0936015430349 |
| ENSG00000187796.14 | CARD9 | -0.260253435131102 | 1.72207320181442 | -1.12684903503588 | 0.266128747022889 | 0.587423914206126 | -5.43037235176716 |
| lnc-UAP1L1-4 | lnc-UAP1L1-4 | -0.195190028069356 | 1.68594941680779 | -1.12676741229307 | 0.26616287664747 | 0.587436252416349 | -5.42393211705633 |
| ENSG00000101940.17 | WDR13 | -0.141156712992076 | 4.91153250081371 | -1.12659535961256 | 0.266234828744239 | 0.58753205532783 | -5.75456186695801 |
| ENSG00000172671.19 | ZFAND4 | 0.143700903887925 | 3.57852300876139 | 1.12619439345794 | 0.266402565809469 | 0.587820107687327 | -5.63683291207372 |
| ENSG00000273899.4 | NOL12 | 0.125450227242718 | 3.33399718347742 | 1.12609530759499 | 0.266444028221733 | 0.587820107687327 | -5.60515218170584 |
| ENSG00000169379.15 | ARL13B | -0.139620596814961 | 2.45862739862005 | -1.12607856970409 | 0.266451032634833 | 0.587820107687327 | -5.49999190057655 |
| ENSG00000251349.3 | MSANTD3-TMEFF[illegible] | -0.192593163953656 | -0.0915526417631549 | -1.12562334608488 | 0.266641583311844 | 0.58817744178627 | -5.10372748369175 |
| PRKAR2A-AS1 | PRKAR2A-AS1 | -0.170563370702019 | 1.10684028911313 | -1.12547369703038 | 0.266704245656218 | 0.588203985433088 | -5.3222246116494 |
| ENSG00000196459.13 | TRAPPC2 | -0.143690418307921 | 4.74678613514929 | -1.12543689259005 | 0.266719658336552 | 0.588203985433088 | -5.74982306555319 |
| SMC5-AS1 | SMC5-AS1 | -0.269618497243327 | 1.34093743506664 | -1.12531378363563 | 0.266771217568313 | 0.588203985433088 | -5.35314002209542 |
| ENSG00000206838.1 | SNORA5A | 0.281745650718445 | 1.34083747975497 | 1.12531074314004 | 0.26677249104738 | 0.588203985433088 | -5.37741687663753 |
| ENSG00000224728.1 | IMPDH1P8 | -0.269359065063721 | 0.407485278118151 | -1.1252534168623 | 0.266796502356474 | 0.588203985433088 | -5.18921305490182 |
| ENSG00000278129.1 | ZNF8 | 0.119964385498438 | 4.29678126115912 | 1.12503420898785 | 0.266888332529535 | 0.588316599644303 | -5.71187740913065 |
| lnc-NSD3-5 | lnc-NSD3-5 | -0.413256135715709 | 0.742240025376362 | -1.12497495854176 | 0.266913157485803 | 0.588316599644303 | -5.26665413826944 |
| ENSG00000179841.8 | AKAP5 | -0.594004400823317 | 2.33982385869087 | -1.12534883063285 | 0.266933329644762 | 0.588316599644303 | -5.47225869953512 |
| ENSG00000156384.14 | SFR1 | 0.178523129666288 | 2.44984632616695 | 1.12482533546871 | 0.266975854397964 | 0.588347324526048 | -5.51401072315001 |
| lnc-PRKAR2B-1 | lnc-PRKAR2B-1 | 0.280312454684398 | 2.26802251212882 | 1.12470469072253 | 0.267026416091259 | 0.588395752318423 | -5.48021848835965 |
| lnc-ISCA1-2 | lnc-ISCA1-2 | 0.150108034461664 | 2.68771109435812 | 1.12460558717594 | 0.267067955062471 | 0.588404194359987 | -5.53313034724623 |
| ENSG00000274769.1 | | -0.203658097590051 | 2.14967998964113 | -1.12453771264067 | 0.267096407137084 | 0.588404194359987 | -5.46614609312615 |
| lnc-PRR14L-1 | lnc-PRR14L-1 | -0.172460397652033 | 0.663897643250379 | -1.12443618077219 | 0.267138971933989 | 0.588404194359987 | -5.26637054551477 |
| ADNP-AS1 | ADNP-AS1 | -0.202591172974132 | 0.421714415015902 | -1.12442276896269 | 0.267144594874123 | 0.588404194359987 | -5.18131881747864 |
| ENSG00000186106.11 | ANKRD46 | -0.163697306975883 | 3.28187212166368 | -1.1242409871425 | 0.267220815744088 | 0.588453554885146 | -5.60305068334936 |
| ENSG00000196419.12 | XRCC6 | 0.117358179770512 | 7.28326748582099 | 1.12411535033333 | 0.267273504136593 | 0.588453554885146 | -5.78393541890868 |
| lnc-PGAM1-4 | lnc-PGAM1-4 | 0.267300256395632 | 0.0935032357490269 | 1.12394538412466 | 0.267344794751173 | 0.588453554885146 | -5.12016574231271 |
| lnc-SSH3-5 | lnc-SSH3-5 | -0.413336457631563 | -0.0097353255621678[illegible] | -1.1239427326709 | 0.267345906983993 | 0.588453554885146 | -5.10338197429583 |
| ENSG00000137478.14 | FCHSD2 | 0.120036943577261 | 7.16748793024039 | 1.1238044701229 | 0.267403909974967 | 0.588453554885146 | -5.78638714563129 |
| ENSG00000158604.14 | TMED4 | 0.0870854624866011 | 6.10568129430395 | 1.12378775853176 | 0.267410921318179 | 0.588453554885146 | -5.79066210516668 |
| ENSG00000248710.1 | TRIM59-IFT80 | 0.118602395910359 | 3.9255878937111 | 1.12378759799771 | 0.267410988670828 | 0.588453554885146 | -5.6708896157039 |
| lnc-NUDT21-4 | lnc-NUDT21-4 | 0.164649873694906 | 1.71557245772938 | 1.12377611065665 | 0.267415808258203 | 0.588453554885146 | -5.41728421565687 |
| ENSG00000124802.11 | EEF1E1 | 0.148030397308017 | 2.20998194633269 | 1.12369014744505 | 0.2674518766362 | 0.588453554885146 | -5.49033487926466 |
| lnc-NRP1-6 | lnc-NRP1-6 | -0.352508572071164 | 3.54748079938628 | -1.1238005591912 | 0.267472132169418 | 0.588453554885146 | -5.62155017826157 |
| ENSG00000130695.15 | CEP85 | -0.153151271688597 | 3.58141289223015 | -1.12361958058955 | 0.267481487611399 | 0.588453554885146 | -5.63764036465202 |
| ENSG00000014919.12 | COX15 | -0.170423487954896 | 4.35312202541553 | -1.12342822742573 | 0.267561794188176 | 0.588567319629713 | -5.71687544067504 |
| ENSG00000267216.1 | ZNF8-ERVK3-1 | 0.124938928200673 | 2.57313682838122 | 1.12288377119509 | 0.267790383990457 | 0.589007210266679 | -5.52495545985288 |
| ENSG00000185480.11 | PARPBP | 0.197889195577864 | 2.10240911215659 | 1.12280319002901 | 0.267824227776104 | 0.589018707160545 | -5.47716082739836 |
| ENSG00000113360.16 | DROSHA | -0.135078654885321 | 5.09002618350103 | -1.12262266902755 | 0.267900056963499 | 0.589098645908045 | -5.76575472858689 |
| ENSG00000205269.5 | TMEM170B | 0.30644149222918 | 2.80027722650611 | 1.12258420003617 | 0.267917816855911 | 0.589098645908045 | -5.56357391667707 |
| ENSG00000211768.1 | TRBJ2-4 | -0.386337733318297 | -0.32736268324948 | -1.12243436057764 | 0.267979173606704 | 0.589156437527702 | -4.99975489046907 |
| lnc-ATXN3-3 | lnc-ATXN3-3 | -0.162607879991035 | 1.55048858542512 | -1.12238159204578 | 0.268001346964576 | 0.589156437527702 | -5.39757588426711 |
| lnc-PRDM8-4 | lnc-PRDM8-4 | -0.356652273363669 | 2.37772648871865 | -1.12229007843288 | 0.268072571302158 | 0.589250078504092 | -5.5252510495566 |
| ENSG00000172270.19 | BSG | -0.179123783646833 | 5.55474908134814 | -1.12199036574089 | 0.268165781132564 | 0.589349348430504 | -5.78372945702466 |
| lnc-OSBPL7-3 | lnc-OSBPL7-3 | -0.256486685222574 | 0.418969763443221 | -1.12189679123032 | 0.268205121553727 | 0.589349348430504 | -5.18718617127332 |
| ENSG00000131115.15 | ZNF227 | 0.10985893289413 | 5.01321022507361 | 1.12177139682218 | 0.268257846074314 | 0.589349348430504 | -5.76099660071764 |
| ENSG00000284276.1 | MIR4800 | 0.311031660066089 | -0.370689741810648 | 1.12176621802274 | 0.268260023759922 | 0.589349348430504 | -5.01781389007206 |
| ENSG00000164104.11 | HMGB2 | 0.239474768511203 | 6.61704047880784 | 1.12176794068582 | 0.268260897121187 | 0.589349348430504 | -5.79232635394845 |
| ENSG00000115295.19 | CLIP4 | -1.35423160564902 | 2.36443413350375 | -1.12222634299482 | 0.268316025159046 | 0.589407550250655 | -5.46969058259001 |
| ENSG00000117318.8 | ID3 | -0.372905693028678 | 4.21917970768711 | -1.12154236516418 | 0.268481945594597 | 0.589709090403755 | -5.70942181430808 |
| lnc-TSPAN3-2 | lnc-TSPAN3-2 | -0.173801539319336 | 0.382338544245903 | -1.1209209367154 | 0.268615633182951 | 0.589897358704131 | -5.21704596816836 |
| lnc-IGLL1-1 | lnc-IGLL1-1 | 0.121780016534283 | 4.8153251916854 | 1.12089873246146 | 0.268624979018308 | 0.589897358704131 | -5.75386995119886 |
| ENSG00000175792.11 | RUVBL1 | -0.225866157757867 | 3.78970941850889 | -1.12052054603976 | 0.268784194300163 | 0.590184027232798 | -5.66351589179691 |
| ENSG00000176401.5 | EID2B | -0.211544334245014 | 1.01117101812872 | -1.12018771061141 | 0.268924372462238 | 0.590371139302833 | -5.32651394623322 |
| ENSG00000180488.15 | MIGA1 | 0.0913397922125831 | 4.72169714802272 | 1.12018200805333 | 0.268926774624854 | 0.590371139302833 | -5.74620522914071 |
| ENSG00000144136.10 | SLC20A1 | 0.200248194821892 | 5.70627615798167 | 1.12010742329551 | 0.26895819433585 | 0.590377147703401 | -5.7894959116263 |
| ENSG00000173193.14 | PARP14 | 0.176567791745221 | 7.49669507817372 | 1.11999432272223 | 0.269005844268682 | 0.59040761030096 | -5.78363658728008 |
| lnc-NGLY1-2 | lnc-NGLY1-2 | 0.135283231007129 | 2.93794604642506 | 1.11988017774315 | 0.269053940300362 | 0.59040761030096 | -5.56151847421929 |

| | | | | | | | |
|---|---|---|---|---|---|---|---|
| ENSG00000266910.2 | | -0.258205592391058 | 0.0794558985012184 | -1.11987024725708 | 0.269058124890589 | 0.59040761030096 | -5.11271551343251 |
| ENSG00000128203.6 | ASPHD2 | 0.230027135977947 | 2.29195953506423 | 1.11964644317288 | 0.269152445575151 | 0.590509073015415 | -5.49348650849312 |
| lnc-TESC-1 | lnc-TESC-1 | -0.410806490444956 | 0.761151925561728 | -1.11962438908513 | 0.269161741390012 | 0.590509073015415 | -5.29455298901859 |
| ENSG00000100592.15 | DAAM1 | 0.113621899868428 | 3.72426058832928 | 1.11910578039372 | 0.269380401132293 | 0.590862938996702 | -5.66048168336319 |
| ENSG00000182566.13 | CLEC4G | -0.965874495903985 | -2.10443379503644 | -1.11904388359279 | 0.26940650696267 | 0.590862938996702 | -4.70139994167315 |
| ENSG00000126768.12 | TIMM17B | -0.127814041080758 | 4.22512156270882 | -1.11903760014973 | 0.269409157192054 | 0.590862938996702 | -5.71332109902071 |
| ENSG00000261448.1 | | -0.322456680917978 | 1.37553510570862 | -1.11879831728819 | 0.269510095657811 | 0.590938460103775 | -5.37619211538973 |
| ENSG00000099849.14 | RASSF7 | -0.229044813772525 | 2.8318265472226 | -1.11878632418052 | 0.26951515550657 | 0.590938460103775 | -5.56815224775885 |
| ENSG00000158079.15 | PTPDC1 | -0.182443356714822 | 3.21441902132567 | -1.11875179935433 | 0.269529721782745 | 0.590938460103775 | -5.59667611671346 |
| ENSG00000168591.15 | TMUB2 | -0.148980799827842 | 5.3783347033446 | -1.11866875644734 | 0.269564760482311 | 0.590952334115177 | -5.78132217682403 |
| lnc-SMUG1-9 | lnc-SMUG1-9 | 0.411432370666994 | -0.642671587324331 | 1.11837623841495 | 0.269688209824765 | 0.59116000243058 | -4.94057408855427 |
| ENSG00000023228.13 | NDUFS1 | 0.108174694910232 | 6.00239214270654 | 1.11824172481373 | 0.269744991140487 | 0.591221504913708 | -5.79566388624061 |
| ENSG00000138085.16 | ATRAID | 0.135568915310318 | 5.29224682298528 | 1.11805760013888 | 0.269822728211607 | 0.591282221910817 | -5.77839381634469 |
| ENSG00000158195.10 | WASF2 | -0.145006584532023 | 7.65282594224664 | -1.11804003089544 | 0.269830146742861 | 0.591282221910817 | -5.78368689895981 |
| ENSG00000160445.10 | ZER1 | -0.106825492527226 | 5.13301750320022 | -1.11749541717617 | 0.270060179211787 | 0.591723298777541 | -5.77362466298147 |
| ENSG00000137501.17 | SYTL2 | 0.830247409902972 | 3.47626136864174 | 1.11776705222768 | 0.270194017541953 | 0.591942112748068 | -5.61752994301449 |
| ENSG00000121316.10 | PLBD1 | 0.745921690827887 | 2.74883300946787 | 1.11766338515791 | 0.270217562618707 | 0.591942112748068 | -5.51621965834544 |
| lnc-SUMF1-2 | lnc-SUMF1-2 | -0.216700075829604 | 0.295836042658961 | -1.11703321247526 | 0.270255513215368 | 0.59196224630328 | -5.17597897040501 |
| lnc-MYO1C-1 | lnc-MYO1C-1 | 0.153947231048536 | 1.37666155691032 | 1.11688666355209 | 0.270317467729734 | 0.592034947678348 | -5.3915358774604 |
| lnc-ISY1-1 | lnc-ISY1-1 | -0.250911827444894 | 0.067627228796513 | -1.11670753209961 | 0.270393210417548 | 0.592137828494988 | -5.12133749999949 |
| ENSG00000183155.4 | RABIF | -0.149389884820065 | 2.76594526723185 | -1.11657601347589 | 0.270448830422791 | 0.592151565843914 | -5.55699354642051 |
| ENSG00000122862.4 | SRGN | -0.27980226010699 | 6.29989785258216 | -1.11660133541305 | 0.27049117601098 | 0.592151565843914 | -5.79827843504746 |
| ENSG00000100580.7 | TMED8 | -0.175176541428333 | 7.59938036323657 | -1.11643168685363 | 0.270509876353878 | 0.592151565843914 | -5.78746895400761 |
| LINC01215 | LINC01215 | -0.152329168244601 | 5.15266995757511 | -1.11642061629757 | 0.270514559278547 | 0.592151565843914 | -5.77770695642697 |
| OSGEPL1-AS1 | OSGEPL1-AS1 | -0.225850907817165 | 0.633138782096373 | -1.11616869000889 | 0.270621141463844 | 0.592321879492801 | -5.23118127785173 |
| ENSG00000140876.11 | NUDT7 | -1.13042464642192 | -1.5769206510432 | -1.11615797179695 | 0.270665474907019 | 0.592323727089123 | -4.7384007031809 |
| ENSG00000100297.15 | MCM5 | -0.133327388594669 | 7.2762195039934 | -1.11603068376975 | 0.270679540251678 | 0.592323727089123 | -5.7916982139445 |
| lnc-PGPEP1-5 | lnc-PGPEP1-5 | -0.335825845932446 | 1.15784087305451 | -1.11586365078785 | 0.270750233974833 | 0.592371992142905 | -5.35323699441312 |
| lnc-KCNS3-12 | lnc-KCNS3-12 | 0.276137091088427 | -0.300454694917492 | 1.11572862639283 | 0.270807390213554 | 0.592371992142905 | -5.07919288526384 |
| lnc-USP32-3 | lnc-USP32-3 | 0.118995936918766 | 2.23009552813309 | 1.11571442573154 | 0.270813401894485 | 0.592371992142905 | -5.49664089521613 |
| ENSG00000008952.16 | SEC62 | -0.145993261563041 | 7.25365262903655 | -1.11564390119272 | 0.270843259021812 | 0.592371992142905 | -5.79270991324316 |
| ENSG00000168936.10 | TMEM129 | -0.216112101343731 | 3.76423718354724 | -1.11562330776155 | 0.270851977856203 | 0.592371992142905 | -5.67625869027146 |
| ENSG00000197958.12 | RPL12 | -0.199544054152385 | 9.04527026780244 | -1.11554688488169 | 0.270884335469947 | 0.592371992142905 | -5.75196708682968 |
| ENSG00000040633.12 | PHF23 | -0.194110555479587 | 3.22716741838779 | -1.11547440204625 | 0.270915027395089 | 0.592371992142905 | -5.61719865442698 |
| lnc-CREG1-3 | lnc-CREG1-3 | -0.334196511041924 | 4.40564973472082 | -1.11561254231091 | 0.270950017288296 | 0.592371992142905 | -5.73585457191586 |
| ENSG00000285505.1 | | -0.216301739477164 | 2.31847590532231 | -1.1153394678755 | 0.270972170108707 | 0.592371992142905 | -5.51219691158268 |
| ENSG00000070814.19 | TCOF1 | -0.125091668737125 | 5.57433507194944 | -1.11523694616618 | 0.271015592320795 | 0.592371992142905 | -5.79282819200418 |
| ENSG00000222009.8 | BTBD19 | -0.37692202655354 | 1.02380062274782 | -1.11516464688511 | 0.271046217044936 | 0.592371992142905 | -5.3213176390034 |
| ENSG00000119616.11 | FCF1 | -0.104120591347617 | 4.33826149915674 | -1.11507653898536 | 0.271083541345888 | 0.592371992142905 | -5.72454271478473 |
| ENSG00000106701.11 | FSD1L | -0.30142740833797 | 2.77549717854594 | -1.11504366903044 | 0.27109905620138 | 0.592371992142905 | -5.57813272831636 |
| ENSG00000100522.9 | GNPNAT1 | -0.2586681155765 | 1.82099340227024 | -1.11502703986611 | 0.271104511780895 | 0.592371992142905 | -5.4589831105346 |
| ENSG00000169504.14 | CLIC4 | -0.238825678580368 | 6.24111620193014 | -1.11496224063857 | 0.271133874621752 | 0.592373266249817 | -5.80008348825009 |
| lnc-ACTA1-3 | lnc-ACTA1-3 | 0.217707617168424 | 1.59043653558628 | 1.11486080155983 | 0.27117494751044 | 0.592400121482422 | -5.4107129129256 |
| ENSG00000183495.13 | EP400 | 0.183174956623782 | 6.34151349361686 | 1.11448489753961 | 0.271334267179215 | 0.592641050070912 | -5.80118839687292 |
| ENSG00000154174.7 | TOMM70 | 0.0995073305240295 | 5.54985362492853 | 1.11446472310333 | 0.271342819602015 | 0.592641050070912 | -5.79247837064383 |
| lnc-GLDC-6 | lnc-GLDC-6 | -0.213592050334778 | 1.21194307010997 | -1.1143575242514 | 0.271388266949627 | 0.592677421604686 | -5.36947487635596 |
| lnc-ADAMTSL4-7 | lnc-ADAMTSL4-7 | -0.221417270336786 | 1.70590597914972 | -1.11428096375701 | 0.271420728357952 | 0.592685428791823 | -5.42992051172285 |
| ENSG00000059378.12 | PARP12 | 0.128292403963111 | 5.78842749656058 | 1.11416850541882 | 0.271468415332791 | 0.592726677924561 | -5.79805909462535 |
| lnc-CDC42SE2-1 | lnc-CDC42SE2-1 | -0.247679206584089 | 2.52855418894845 | -1.11405051314482 | 0.271518455313004 | 0.592773055335975 | -5.52754610353046 |
| lnc-IL1RN-5 | lnc-IL1RN-5 | -0.158059431628424 | 3.25764882542122 | -1.11394856104795 | 0.271561697996291 | 0.592804584776504 | -5.62070809641887 |
| ENSG00000142867.13 | BCL10 | 0.123616251355146 | 5.71068330418758 | 1.11384550771939 | 0.271605412727467 | 0.592831274618815 | -5.79714677478686 |
| ENSG00000158156.7 | XKR8 | 0.205547997956994 | 1.92838640180285 | 1.11377339807903 | 0.27163600426121 | 0.592831274618815 | -5.46530585663844 |
| ENSG00000132275.10 | RRP8 | 0.0831115137752052 | 5.02604072742889 | 1.1137151070792 | 0.271660735210721 | 0.592831274618815 | -5.77092662082489 |
| ZNF793-AS1 | ZNF793-AS1 | -0.260432039237424 | 0.623801354174665 | -1.11364817922333 | 0.271689132462677 | 0.592831274618815 | -5.25992593642758 |
| ENSG00000105568.17 | PPP2R1A | 0.0744264264736136 | 6.62896378221171 | 1.11342982630074 | 0.271781793470463 | 0.592928026202871 | -5.80125903144941 |
| ENSG00000284461.2 | | 0.137774986099466 | 3.87335125974363 | 1.11334136697333 | 0.271819338756626 | 0.592928026202871 | -5.67307481219205 |
| ENSG00000162688.16 | AGL | -0.0910899335868764 | 5.61599767001693 | -1.11334006147309 | 0.271819892884919 | 0.592928026202871 | -5.79396438317169 |
| ENSG00000136040.8 | PLXNC1 | 0.75616266158159 | 1.85621744815997 | 1.11358796502412 | 0.271942311022402 | 0.593132201735095 | -5.4724716205062 |
| lnc-XKRX-1 | lnc-XKRX-1 | 0.139212929335865 | 1.86785648490669 | 1.11290892503102 | 0.272002935439325 | 0.593175079673603 | -5.44313648051131 |
| ENSG00000121542.11 | SEC22A | 0.104465680058599 | 3.36027486952209 | 1.11285153295975 | 0.272027308308523 | 0.593175079673603 | -5.61927748027714 |
| LINC01033 | LINC01033 | -0.378175462855639 | 0.985274276923495 | -1.11280180900085 | 0.272048425988379 | 0.593175079673603 | -5.31559645033863 |
| SNHG15 | SNHG15 | 0.248117086443985 | 4.14906189427827 | 1.11222114432432 | 0.272295119310401 | 0.593650083758711 | -5.71123446125523 |
| ENSG00000177542.10 | SLC25A22 | -0.190748560242117 | 2.17154425358312 | -1.11205463234444 | 0.272365890575922 | 0.593698266948268 | -5.49815275775399 |
| lnc-CDC6-3 | lnc-CDC6-3 | -0.243167031108986 | 0.00410096990148478 | -1.1120334177882 | 0.272374908166569 | 0.593698266948268 | -5.14969691801291 |
| ENSG00000108557.18 | RAI1 | -0.233787344452552 | 3.91399499996413 | -1.11184734298477 | 0.272454011341818 | 0.593799934581127 | -5.69377532106004 |
| ENSG00000148908.14 | RGS10 | -0.190308098142729 | 4.03529722966134 | -1.11171143125809 | 0.272511799732126 | 0.593799934581127 | -5.69902540710295 |
| ENSG00000241112.1 | RPL29P14 | 0.199144545109218 | 1.39541612493961 | 1.1117081220167 | 0.272513206898804 | 0.593799934581127 | -5.39754948722725 |
| ENSG00000140987.19 | ZSCAN32 | 0.167943011025943 | 3.84318488591687 | 1.11165229390376 | 0.272536947091673 | 0.593799934581127 | -5.67629913287553 |

| | | | | | | | |
|---|---|---|---|---|---|---|---|
| lnc-ENTHD1-1 | lnc-ENTHD1-1 | -0.388516965376283 | -0.13549466817179 | -1.11152096613544 | 0.272592798328179 | 0.593843138073623 | -5.10795862173397 |
| ENSG00000090565.15 | RAB11FIP3 | -0.139049297543599 | 3.8047276798674 | -1.11146999275916 | 0.272614478533628 | 0.593843138073623 | -5.67837293806043 |
| ENSG00000282978.1 | BTF3P16 | 0.418222041527509 | 0.653203961519073 | 1.11133829409973 | 0.272670498803622 | 0.593902315013223 | -5.19189171942095 |
| lnc-SYNJ2-1 | lnc-SYNJ2-1 | -0.21300188738635 | 2.46960895537094 | -1.11114535432119 | 0.272752583754019 | 0.59401824477928 | -5.54663879618548 |
| lnc-ZNF836-1 | lnc-ZNF836-1 | -0.146108655660152 | 0.739050244057793 | -1.11099177531531 | 0.272817935437151 | 0.594091711699989 | -5.27712035451025 |
| ENSG00000128791.11 | TWSG1 | -0.296629697889679 | 2.67327445965635 | -1.11086836115993 | 0.272870459264755 | 0.594091711699989 | -5.5639234090156 |
| DARS-AS1 | DARS-AS1 | -0.246575626448811 | 0.108896190870633 | -1.11086261044772 | 0.27287290688489 | 0.594091711699989 | -5.16304445335048 |
| lnc-C17orf50-1 | lnc-C17orf50-1 | 0.304586122246305 | -0.506258639107278 | 1.11067021290917 | 0.272954804156736 | 0.594141049599223 | -4.96552715957882 |
| lnc-TRIML1-9 | lnc-TRIML1-9 | 0.350677590152314 | -0.555452765702408 | 1.11063075877743 | 0.272971600627343 | 0.594141049599223 | -4.9994289153868 |
| ENSG00000136826.14 | KLF4 | -1.21157723074536 | 0.561898035311836 | -1.11116235326871 | 0.272982165187769 | 0.594141049599223 | -5.15173065270745 |
| lnc-FAM124B-2 | lnc-FAM124B-2 | 0.353324056535304 | 1.30608607050038 | 1.11049461169426 | 0.273029566984486 | 0.594158770466844 | -5.35873316735881 |
| ENSG00000101911.12 | PRPS2 | -0.161930847810447 | 4.44658730007807 | -1.11045122765835 | 0.273048040122717 | 0.594158770466844 | -5.73880654558828 |
| ENSG00000113658.17 | SMAD5 | -0.155767584824392 | 5.72219867354086 | -1.11037661232297 | 0.273079813774087 | 0.594165095868079 | -5.80172866072738 |
| MINCR | MINCR | -0.20229400550683 | 1.60426042277006 | -1.11028826071791 | 0.273117440164601 | 0.594184152933937 | -5.42686519865287 |
| ENSG00000110619.17 | CARS1 | 0.112792363452683 | 5.09382673061655 | 1.11017496420375 | 0.273165695242446 | 0.594226326905016 | -5.77903143455103 |
| lnc-ZBTB10-1 | lnc-ZBTB10-1 | 0.299436865891674 | 1.20548234526883 | 1.11000776398266 | 0.273236919934392 | 0.594318453239945 | -5.35813829062858 |
| ENSG00000125818.17 | PSMF1 | -0.0982899924482847 | 5.53583880155068 | -1.10989725910737 | 0.273284000508831 | 0.594323710313645 | -5.79689771190806 |
| ENSG00000130175.9 | PRKCSH | -0.118684183373672 | 6.88711500087309 | -1.10986654831716 | 0.273297085853309 | 0.594323710313645 | -5.80363124973856 |
| lnc-FAM174A-6 | lnc-FAM174A-6 | -0.360875902350958 | 3.07152253284864 | -1.10983591441754 | 0.273383305847242 | 0.594448403153791 | -5.62327228910634 |
| lnc-MUC17-4 | lnc-MUC17-4 | -0.121015546519604 | 1.7488583853975 | -1.10954122166127 | 0.273435729242035 | 0.594499482077976 | -5.43953552376579 |
| lnc-SPOCK2-1 | lnc-SPOCK2-1 | -0.279968540932688 | 3.0144200219956 | -1.10947357837726 | 0.273464562809808 | 0.594499482077976 | -5.58256486116657 |
| lnc-LRRN1-1 | lnc-LRRN1-1 | 0.260811905729025 | 1.07524335242791 | 1.10939110462485 | 0.27349972091244 | 0.594513122350908 | -5.33257066137435 |
| lnc-MAPK6-3 | lnc-MAPK6-3 | -0.15893993842189 | 3.53920853312297 | -1.1090664958029 | 0.273638130934596 | 0.594597830563569 | -5.65223134322841 |
| ENSG00000187555.14 | USP7 | 0.111331788914199 | 8.0357976159937 | 1.10905097893353 | 0.273644748417231 | 0.594597830563569 | -5.78652218941131 |
| ENSG00000137500.9 | CCDC90B | 0.139425882417429 | 4.21148607363481 | 1.10896599681656 | 0.273680992768192 | 0.594597830563569 | -5.71247842925285 |
| ENSG00000275318.1 | | 0.222086565787895 | 0.715899784065744 | 1.10896586186693 | 0.27368105032608 | 0.594597830563569 | -5.27603670220192 |
| lnc-SLC38A1-5 | lnc-SLC38A1-5 | 0.198448307419685 | 3.71109940821827 | 1.10892185131098 | 0.273699821896223 | 0.594597830563569 | -5.68040591949009 |
| ENSG00000108798.8 | ABI3 | 1.39986227030984 | -1.58083956852041 | 1.10903616076897 | 0.27371763936264 | 0.594597830563569 | -4.69304980756533 |
| lnc-MVB12B-7 | lnc-MVB12B-7 | -0.295307591662451 | 2.3861069787452 | -1.10877545544166 | 0.273762269844858 | 0.594597830563569 | -5.54577944634668 |
| ENSG00000163704.11 | PRRT3 | -0.289451428422508 | -0.191277007557228 | -1.10875782164031 | 0.273769792559439 | 0.594597830563569 | -5.06246809030626 |
| ENSG00000181090.20 | EHMT1 | 0.162200713204495 | 7.44271051008184 | 1.10837028634583 | 0.273935155023028 | 0.594856766918033 | -5.79859684973321 |
| ENSG00000185973.10 | TMLHE | 0.155049958264225 | 3.52087726360593 | 1.10834296679587 | 0.273946815022885 | 0.594856766918033 | -5.64281697928127 |
| ENSG00000137601.16 | NEK1 | 0.123990068838726 | 4.68169197306884 | 1.10817202037986 | 0.274019783027667 | 0.594939432045832 | -5.75424569136398 |
| ENSG00000183196.9 | CHST6 | 0.988252155343315 | -1.11555247908729 | 1.10825207603322 | 0.274042693253002 | 0.594939432045832 | -4.82391815616832 |
| ENSG00000197142.10 | ACSL5 | -0.119536796131353 | 5.54124190067528 | -1.10790435569111 | 0.274134062584087 | 0.595075027438121 | -5.79876154983414 |
| ENSG00000112667.12 | DNPH1 | -0.265705460480802 | 2.52604516484218 | -1.10772845713336 | 0.274209180910174 | 0.595133079662014 | -5.54185256791627 |
| ENSG00000120437.8 | ACAT2 | -0.157212769149447 | 3.84247462076514 | -1.10770632583876 | 0.274218633217439 | 0.595133079662014 | -5.69478134529344 |
| ENSG00000178605.13 | GTPBP6 | -0.188011699434276 | 4.52907295548633 | -1.10744399580021 | 0.274330692309419 | 0.595266031697525 | -5.74957165614571 |
| ENSG00000128923.10 | MINDY2 | -0.123592534796301 | 4.44846651005819 | -1.10741355692179 | 0.274343696932374 | 0.595266031697525 | -5.74214333416899 |
| lnc-C11orf54-1 | lnc-C11orf54-1 | 0.215433646874591 | 5.5265216718548 | 1.10731303475213 | 0.274386646846703 | 0.595266031697525 | -5.79934759093158 |
| ENSG00000100888.13 | CHD8 | 0.0820365593552603 | 7.32506090248732 | 1.10729214154941 | 0.274395574442312 | 0.595266031697525 | -5.80119197510638 |
| ENSG00000141741.11 | MIEN1 | 0.160360807558031 | 3.56195700108074 | 1.1071748007351 | 0.274445717597157 | 0.595269067392347 | -5.65947917404483 |
| ENSG00000168806.7 | LCMT2 | -0.237281060932489 | 2.74082628827342 | -1.1071535142427 | 0.27445481463117 | 0.595269067392347 | -5.56408875781719 |
| lnc-EIF2AK2-1 | lnc-EIF2AK2-1 | 0.179322659578896 | 2.25355678216546 | 1.10700104747666 | 0.274519979340017 | 0.595346791821412 | -5.49293121845359 |
| ENSG00000010818.9 | HIVEP2 | -0.178446727397835 | 7.8751516096264 | -1.10693432847873 | 0.274548498662432 | 0.595346791821412 | -5.79192169729928 |
| ENSG00000151746.13 | BICD1 | -0.190721155294198 | 5.32990838972208 | -1.10686158231888 | 0.274579596709092 | 0.59535150522103 | -5.79544619231912 |
| ENSG00000111679.16 | PTPN6 | -0.307810935032093 | 7.57694806824195 | -1.10690120875634 | 0.274648421518233 | 0.595438009280575 | -5.79663942474773 |
| ENSG00000122644.12 | ARL4A | -0.726361079309622 | 3.49291239651468 | -1.10716485842473 | 0.274686961500154 | 0.595458844503178 | -5.65938483801278 |
| ENSG00000160293.16 | VAV2 | 0.200245465014047 | 5.61578966693858 | 1.10637299413006 | 0.274788526425466 | 0.595527835488578 | -5.79869564055233 |
| ENSG00000280138.1 | | 1.10191199499261 | 0.499564261853276 | 1.10683509272418 | 0.274817352453072 | 0.595527835488578 | -5.18691727393967 |
| ENSG00000197054.11 | ZNF763 | -0.191519491456678 | 2.2015407327278 | -1.10629308041732 | 0.274822709761936 | 0.595527835488578 | -5.49229890750599 |
| lnc-C12orf49-3 | lnc-C12orf49-3 | -0.239673088319632 | -0.0439993535807151 | -1.10626547390852 | 0.274834519229753 | 0.595527835488578 | -5.14113441740451 |
| ENSG00000250853.1 | RNF138P1 | 0.285471439750548 | -0.229243752598447 | 1.1061235946006 | 0.274895217792169 | 0.595575251371534 | -5.06421097676967 |
| lnc-TM6SF2-2 | lnc-TM6SF2-2 | 0.183059903176471 | 0.697788999546854 | 1.10599884861556 | 0.274948594235866 | 0.595575251371534 | -5.2800285595544 |
| lnc-SHARPIN-4 | lnc-SHARPIN-4 | -0.139588899865003 | 2.98303940227377 | -1.1059502306738 | 0.274969398916384 | 0.595575251371534 | -5.59371512573068 |
| ENSG00000135473.14 | PAN2 | 0.127225087421708 | 6.72547765568133 | 1.10589262542452 | 0.274994050902929 | 0.595575251371534 | -5.80934119453502 |
| ENSG00000148943.11 | LIN7C | 0.150024136968297 | 5.32806394266633 | 1.10587620554032 | 0.275001078027258 | 0.595575251371534 | -5.79582149164268 |
| ENSG00000163382.11 | NAXE | -0.166659090895401 | 3.70197504021691 | -1.10565003326733 | 0.275097884604782 | 0.595681347340063 | -5.67923099190819 |
| lnc-GPR149-5 | lnc-GPR149-5 | 0.185066710166655 | 1.94806853111044 | 1.10560901983963 | 0.275115441810385 | 0.595681347340063 | -5.46722161421318 |
| ENSG00000100300.17 | TSPO | -0.385920194804031 | 3.51454504742519 | -1.10582908575934 | 0.275136888168001 | 0.595681347340063 | -5.65383237071852 |
| ENSG00000185909.14 | KLHDC8B | -0.48580478251964 | 1.52749938341004 | -1.10575195599188 | 0.275180565087807 | 0.595713248969535 | -5.47498715940898 |
| ENSG00000162819.11 | BROX | -0.124708951907227 | 5.99947315157522 | -1.10536029116313 | 0.275221935626937 | 0.595731091969652 | -5.80947017998544 |
| ENSG00000162341.16 | TPCN2 | 0.182172445602101 | 3.23433011981705 | 1.10530247689304 | 0.27524669312692 | 0.595731091969652 | -5.61300633340297 |
| ENSG00000250410.1 | LRP2BP-AS1 | -0.427800357473449 | 0.262849332054624 | -1.10523288759519 | 0.275276495072566 | 0.595732951117509 | -5.1746683089717 |
| lnc-MRPS23-1 | lnc-MRPS23-1 | 0.140731218994131 | 1.12569706606111 | 1.10503492640975 | 0.275361285357424 | 0.595851071835885 | -5.36785587664404 |
| lnc-CLVS1-1 | lnc-CLVS1-1 | -0.362636814605102 | 2.61589409996073 | -1.1050567051414 | 0.275388973734381 | 0.595851071835885 | -5.5256374027275 |
| lnc-TRMT11-1 | lnc-TRMT11-1 | -0.232768144725696 | 3.13891078893312 | -1.10458304040357 | 0.275554905296467 | 0.596147426499598 | -5.60827724401383 |
| ENSG00000003402.19 | CFLAR | -0.198719938724944 | 8.43581216507718 | -1.10444585610368 | 0.275613703788441 | 0.596201563699124 | -5.78140138270236 |

| | | | | | | | |
|---|---|---|---|---|---|---|---|
| ENSG00000117362.12 | APH1A | 0.131163445517779 | 6.21231694041275 | 1.10435223621645 | 0.275653835248967 | 0.596201563699124 | -5.81188924622334 |
| ENSG00000283189.2 | | -0.180667488306362 | 3.62424991916157 | -1.10428908796892 | 0.27568090695092 | 0.596201563699124 | -5.65358604991826 |
| ENSG00000147364.16 | FBXO25 | -0.111350782960614 | 4.68205834157086 | -1.10425436892349 | 0.275695791832496 | 0.596201563699124 | -5.76411290362055 |
| ENSG00000175550.7 | DRAP1 | -0.219236342348614 | 4.68127111666212 | -1.10418156091825 | 0.275727008202097 | 0.596206430278786 | -5.76749233468573 |
| ENSG00000197226.12 | TBC1D9B | -0.119074613631301 | 6.58140038079424 | -1.10391532937412 | 0.275841176007634 | 0.59634327019686 | -5.81166310438428 |
| lnc-SPACA5B-3 | lnc-SPACA5B-3 | -0.228877751199037 | 0.475987930119355 | -1.10381601452392 | 0.275883773646244 | 0.59634327019686 | -5.20785729407307 |
| ENSG00000276057.1 | | 0.164673707940878 | 1.96324569424982 | 1.10377201067192 | 0.27590264904821 | 0.59634327019686 | -5.4618647699538 |
| lnc-DUSP4-6 | lnc-DUSP4-6 | -0.842880309032308 | 0.0889882678211003 | -1.10408054412553 | 0.275944233625423 | 0.59634327019686 | -5.17084846347087 |
| ENSG00000181751.9 | MACIR | -0.267798761470191 | 2.24538703019148 | -1.10365708311727 | 0.275951951393851 | 0.59634327019686 | -5.50907278856821 |
| lnc-CHMP3-1 | lnc-CHMP3-1 | -0.138505867117655 | 4.02777928324779 | -1.10362870051992 | 0.275964128097221 | 0.59634327019686 | -5.70798050285108 |
| ENSG00000137822.12 | TUBGCP4 | 0.16664144491664 | 3.91014608000981 | 1.10328966701526 | 0.276109609612627 | 0.596457594340052 | -5.68944862634563 |
| ENSG00000179454.13 | KLHL28 | 0.136287395624437 | 5.19958856692534 | 1.10328110552671 | 0.276113284104954 | 0.596457594340052 | -5.79283457841143 |
| ENSG00000133056.13 | PIK3C2B | 0.489914655806941 | 3.18999157227558 | 1.10368535368606 | 0.276113679092411 | 0.596457594340052 | -5.63830293990032 |
| ENSG00000234585.6 | CCT6P3 | -0.196205811921918 | 2.59818611638173 | -1.10323529634981 | 0.276132945463069 | 0.596457594340052 | -5.54140988274872 |
| ENSG00000189343.7 | RPS2P46 | 0.33362320184085 | 2.30553566378677 | 1.10304883112892 | 0.276214561461678 | 0.596531863266515 | -5.48943699671588 |
| lnc-GFAP-3 | lnc-GFAP-3 | 0.107476489868638 | 1.78572630529186 | 1.10302016772436 | 0.27622529217282 | 0.596531863266515 | -5.45188557575834 |
| lnc-KDM7A-1 | lnc-KDM7A-1 | 0.1445058029692 | 2.34668142418286 | 1.1024407953819 | 0.276474103601027 | 0.596975248980658 | -5.52433299352122 |
| ENSG00000187736.12 | NHEJ1 | -0.213095982011144 | 2.02444232679512 | -1.10240702897585 | 0.276488609460847 | 0.596975248980658 | -5.49400579187413 |
| lnc-SLC9B1-1 | lnc-SLC9B1-1 | -0.243424027817167 | 0.341946674233553 | -1.10233799253233 | 0.276518268804477 | 0.596976665282415 | -5.21964477338777 |
| ENSG00000177425.10 | PAWR | -0.131228663124277 | 5.92553259157222 | -1.1020608110856 | 0.276637373736515 | 0.597171165560429 | -5.81297399234891 |
| ENSG00000198346.10 | ZNF813 | -0.123961908592987 | 4.31294212633796 | -1.1017608432934 | 0.276766310764039 | 0.597224791456318 | -5.72667043764363 |
| ENSG00000092929.11 | UNC13D | -0.170566470168967 | 5.24473917518297 | -1.10164653755885 | 0.27681545467068 | 0.597224791456318 | -5.79834234709276 |
| ENSG00000162729.13 | IGSF8 | -0.190283321502208 | 3.77724399350116 | -1.10164372526251 | 0.276816663849749 | 0.597224791456318 | -5.68970898469711 |
| ENSG00000127124.15 | HIVEP3 | -0.34874386088113 | 4.0476598572361 | -1.10184412359402 | 0.276826320861542 | 0.597224791456318 | -5.70749851236868 |
| ENSG00000148429.14 | USP6NL | -0.294422506169106 | 6.31646136113699 | -1.10177109542242 | 0.276834606198071 | 0.597224791456318 | -5.81409600971632 |
| ENSG00000187024.14 | PTRH1 | -0.220782444365423 | 0.310077616327538 | -1.10159803744047 | 0.276836308375102 | 0.597224791456318 | -5.24171555035602 |
| ENSG00000124762.13 | CDKN1A | 0.532850179819857 | 2.87110552439352 | 1.10181025340794 | 0.276918666677987 | 0.597272026415345 | -5.58174050070307 |
| ENSG00000135297.15 | MTO1 | 0.0803012839570192 | 5.66624069046619 | 1.10138634834123 | 0.276927341791886 | 0.597272026415345 | -5.80748764464721 |
| ENSG00000023041.11 | ZDHHC6 | 0.106272928835969 | 6.02989499365549 | 1.10134469463502 | 0.276945256770542 | 0.597272026415345 | -5.81393521676195 |
| ENSG00000251593.1 | MSNP1 | 0.702960024112432 | -0.336113427179325 | 1.1011798310482 | 0.277016367670791 | 0.597357576770423 | -4.91456358576765 |
| lnc-SNX31-1 | lnc-SNX31-1 | -0.182808010564928 | 0.530559299609551 | -1.10111753997974 | 0.27704296885053 | 0.597357576770423 | -5.27275871953354 |
| ENSG00000229186.4 | ADAM1A | -0.219985579219021 | 1.56661082779102 | -1.10096522355128 | 0.277108502394989 | 0.597436294626172 | -5.43109674841649 |
| lnc-MANBA-2 | lnc-MANBA-2 | 0.11064612862059 | 6.24658036529824 | 1.10070533888189 | 0.27722034200311 | 0.597610617029239 | -5.81594759805073 |
| lnc-PATL2-3 | lnc-PATL2-3 | -0.109024885460612 | 3.005237467791 | -1.10058346218743 | 0.277272801781275 | 0.597610617029239 | -5.59502509427151 |
| ENSG00000180953.11 | ST20 | -0.215561756665955 | 1.24283934644592 | -1.10057496201489 | 0.277276460799166 | 0.597610617029239 | -5.40152924710834 |
| lnc-NOX3-6 | lnc-NOX3-6 | -0.624672225987655 | 1.47391236328445 | -1.10082966369438 | 0.277330955008626 | 0.59766548496938 | -5.39123655521114 |
| ENSG00000171241.8 | SHCBP1 | 0.369076741231636 | -0.0610030281729802 | 1.10021215293006 | 0.277432668790575 | 0.597801845454627 | -5.11942187523577 |
| ENSG00000202566.4 | MIR421 | -0.314784393276913 | -0.185183771805179 | -1.1001257814969 | 0.277469865285911 | 0.597801845454627 | -5.08309649233209 |
| lnc-CALHM3-2 | lnc-CALHM3-2 | -0.555895787394425 | 1.7822413678531 | -1.10045356802538 | 0.277481359931667 | 0.597801845454627 | -5.45494580049224 |
| ENSG00000060069.16 | CTDP1 | 0.159359340776371 | 4.17354712556928 | 1.09980372714631 | 0.277608591380956 | 0.597956230392291 | -5.73410869570154 |
| ENSG00000112118.18 | MCM3 | -0.240272226796967 | 5.17190721142387 | -1.09978780967061 | 0.277617019752406 | 0.597956230392291 | -5.79445240679423 |
| lnc-PEAK1-1 | lnc-PEAK1-1 | -0.14489752233145 | 1.91519583842726 | -1.0997304244223 | 0.277640173631595 | 0.597956230392291 | -5.47221582132113 |
| lnc-H3F3C-10 | lnc-H3F3C-10 | -0.414853629429124 | 0.603015131008435 | -1.09944206045658 | 0.277767254312796 | 0.597997036825363 | -5.31177030080183 |
| ENSG00000130714.15 | POMT1 | -0.133377941756464 | 3.89338547510274 | -1.09935162023886 | 0.277803420616832 | 0.597997036825363 | -5.70635385022259 |
| ENSG00000166598.14 | HSP90B1 | 0.165168317706232 | 7.59174555000725 | 1.09922504754185 | 0.277857982645767 | 0.597997036825363 | -5.80423582216189 |
| ENSG00000103275.19 | UBE2I | -0.0913650040733183 | 6.38153779448654 | -1.09921890134594 | 0.277860632295126 | 0.597997036825363 | -5.81731011634563 |
| ENSG00000181619.11 | GPR135 | -0.27254668866231 | 1.33983485395423 | -1.09910679913166 | 0.277908963128888 | 0.597997036825363 | -5.38866652396377 |
| ENSG00000165282.13 | PIGO | -0.0986445262820244 | 4.89391738126783 | -1.09908987663359 | 0.277916259471479 | 0.597997036825363 | -5.78070748003028 |
| ENSG00000132781.17 | MUTYH | 0.121535169228422 | 4.25334300886233 | 1.09904023689636 | 0.277937663032172 | 0.597997036825363 | -5.73029100874822 |
| ENSG00000129696.12 | TTI2 | 0.131630551254037 | 2.92991574432453 | 1.09900194396263 | 0.277954174895324 | 0.597997036825363 | -5.58532536606513 |
| ENSG00000248672.5 | LY75-CD302 | -0.107574953419807 | 5.66813403461428 | -1.0989918000826 | 0.277958549039048 | 0.597997036825363 | -5.8112827770802 |
| ENSG00000132623.15 | ANKEF1 | 1.18971263582919 | 0.718228000763196 | 1.09947992325857 | 0.277973233224086 | 0.597997036825363 | -5.10918904103841 |
| ENSG00000108961.13 | RANGRF | -0.146066378814885 | 3.65823063802305 | -1.09892660304682 | 0.277986663819038 | 0.597997036825363 | -5.67887411391428 |
| lnc-DOCK7-8 | lnc-DOCK7-8 | -0.306417535471923 | 0.0575726975350432 | -1.09887769377628 | 0.278007756176548 | 0.597997036825363 | -5.14499340345011 |
| lnc-UBE2D2-4 | lnc-UBE2D2-4 | 0.1602119338893 | 4.79278879443086 | 1.09867234407248 | 0.278096326534536 | 0.598125045878824 | -5.77675269221114 |
| ENSG00000243660.9 | ZNF487 | -0.16430619778631 | 3.53320272033781 | -1.09855934746371 | 0.278145072123216 | 0.598167382667657 | -5.65453636379613 |
| ENSG00000155749.12 | FLACC1 | -0.342491154260436 | 2.0439627150052 | -1.09832335035581 | 0.278248467051217 | 0.5983272249598 | -5.47135083365902 |
| ENSG00000164430.15 | CGAS | 0.221429844265489 | 2.04536466140668 | 1.09810063046005 | 0.278343019830458 | 0.598377727157235 | -5.48959528828892 |
| ENSG00000283580.3 | | 0.108679332966304 | 1.90716010974268 | 1.09799686473969 | 0.278387811066141 | 0.598377727157235 | -5.47051857034898 |
| ENSG00000169031.19 | COL4A3 | -1.57118770679032 | 3.99572384246949 | -1.09855314529689 | 0.278392616847697 | 0.598377727157235 | -5.75728969498682 |
| ENSG00000226721.2 | EEF1DP2 | -0.265151983102669 | 0.483978457360198 | -1.09790089618446 | 0.278429241119556 | 0.598377727157235 | -5.23544040883313 |
| lnc-HLA-A-4 | lnc-HLA-A-4 | -0.297763846292206 | 0.695560910387043 | -1.09769779339697 | 0.278516935827147 | 0.598377727157235 | -5.31885366612903 |
| lnc-CHST12-1 | lnc-CHST12-1 | 0.209889391693147 | 0.992963894543521 | 1.09764016379125 | 0.278541822395069 | 0.598377727157235 | -5.35124676639689 |
| ENSG00000125520.13 | SLC2A4RG | -0.206172146860063 | 2.48669474225383 | -1.09753720488966 | 0.278586287708361 | 0.598377727157235 | -5.55400482386391 |
| lnc-CPXM2-4 | lnc-CPXM2-4 | 0.266683113660314 | 3.67223172325782 | 1.09752625271588 | 0.278591017966085 | 0.598377727157235 | -5.65713020929254 |
| ENSG00000206602.1 | SNORD58A | 0.175314534970285 | 1.23954053989908 | 1.09751689930182 | 0.278595057762267 | 0.598377727157235 | -5.37769365662192 |
| lnc-SUPT16H-2 | lnc-SUPT16H-2 | -0.130519459963768 | 2.00289826374392 | -1.09749316314831 | 0.278605309736011 | 0.598377727157235 | -5.49367281587957 |
| ENSG00000188981.10 | MSANTD1 | -0.248981345425289 | 0.848367632844323 | -1.09748481607382 | 0.278608915016371 | 0.598377727157235 | -5.32894024178105 |

| | | | | | | | |
|---|---|---|---|---|---|---|---|
| ENSG00000130304.16 | SLC27A1 | -0.229687267789266 | 1.61692766209142 | -1.09741436729795 | 0.278639344661996 | 0.598377727157235 | -5.43174801399949 |
| ENSG00000065665.20 | SEC61A2 | 0.200077569295542 | 3.5328389869983 | 1.09738997390854 | 0.278649881688874 | 0.598377727157235 | -5.64967656190378 |
| ENSG00000228446.2 | ANKRD49P1 | 0.349147882350294 | 1.29908957691896 | 1.0971831625501 | 0.278739227698723 | 0.598507148312416 | -5.37124980738851 |
| lnc-LY6E-7 | lnc-LY6E-7 | -1.51793906814981 | -2.13333521221941 | -1.097255045004 | 0.278792921065892 | 0.598522483980615 | -4.68426772652622 |
| ENSG00000234465.10 | PINLYP | 0.994661692511519 | -1.95096147758836 | 1.09686396609186 | 0.27887716558693 | 0.598522483980615 | -4.65911793824707 |
| ENSG00000140807.6 | NKD1 | -0.306634263211958 | 0.134214365239913 | -1.09678144949375 | 0.27891283221592 | 0.598522483980615 | -5.143665058719 |
| ENSG00000284820.1 | | 0.160952626128612 | 3.32790098845359 | 1.09675128676231 | 0.278925870430614 | 0.598522483980615 | -5.64436361857912 |
| lnc-GRIK2-2 | lnc-GRIK2-2 | -0.194912197936044 | 1.02485311669512 | -1.09674444876596 | 0.278928826299027 | 0.598522483980615 | -5.36686688867835 |
| LINC00106 | LINC00106 | 0.298674307818167 | 0.894511087443752 | 1.09673859692993 | 0.278931355896324 | 0.598522483980615 | -5.31810417757006 |
| ENSG00000168615.11 | ADAM9 | 0.297570681407624 | 1.82341819962361 | 1.09669565449569 | 0.278949919293885 | 0.598522483980615 | -5.47677758477371 |
| lnc-ALKAL2-5 | lnc-ALKAL2-5 | 0.143277982160938 | 1.21925168674748 | 1.0965947056744 | 0.278993561446779 | 0.59855372892006 | -5.39024773385801 |
| lnc-SLC1A7-5 | lnc-SLC1A7-5 | 0.247555203739467 | 0.927279883938287 | 1.09649071855117 | 0.279038522145465 | 0.598587795864523 | -5.3196312467684 |
| ENSG00000135423.12 | GLS2 | -0.333551260021298 | 0.928420021870026 | -1.09633967323385 | 0.279103838388337 | 0.598665517460101 | -5.30200114860852 |
| ENSG00000241494.1 | | 0.644554731681838 | -1.02181572469006 | 1.09615766365549 | 0.279182558752669 | 0.598711564919186 | -4.82368128211415 |
| lnc-PREPL-3 | lnc-PREPL-3 | -0.345753416376486 | -0.304930190619055 | -1.09614623051012 | 0.279187504186682 | 0.598711564919186 | -5.0362089537145 |
| ENSG00000114784.3 | EIF1B | 0.191268489704867 | 4.66965452150861 | 1.09608828595118 | 0.279212569191044 | 0.598711564919186 | -5.76843863569046 |
| ENSG00000267552.6 | | -0.210284266903297 | 0.173917048907959 | -1.09566814614544 | 0.279394355935005 | 0.599038961271897 | -5.17025407361573 |
| lnc-TTC33-4 | lnc-TTC33-4 | 0.101504054032937 | 1.94814280495924 | 1.09553683909145 | 0.279451187163577 | 0.599098404894064 | -5.47597614727148 |
| ENSG00000133265.10 | HSPBP1 | -0.22466438873743 | 1.68235301695672 | -1.09516393027014 | 0.279612630839725 | 0.599282850313448 | -5.46176737442402 |
| ENSG00000247796.2 | MOCS2-DT | -0.262961435374003 | -0.230098021812439 | -1.09516325705454 | 0.279612922354786 | 0.599282850313448 | -5.04335005665257 |
| ENSG00000204475.9 | NCR3 | -0.331155648815821 | 2.9681494353832 | -1.09521446957868 | 0.279624568547362 | 0.599282850313448 | -5.60417287802457 |
| ENSG00000138378.18 | STAT4 | 1.44231579108504 | -0.227596206416308 | 1.09549589451927 | 0.279687681753741 | 0.599355705730062 | -4.94783401693024 |
| ENSG00000166135.13 | HIF1AN | 0.0936504038222647 | 6.08208216837925 | 1.0946157806379 | 0.279850060810988 | 0.599641245229291 | -5.8215494069774 |
| lnc-LAMTOR1-3 | lnc-LAMTOR1-3 | -0.102577632880123 | 3.80820662331681 | -1.09451818370392 | 0.279892349614142 | 0.599669431883823 | -5.69214000604774 |
| lnc-HIST1H3D-1 | lnc-HIST1H3D-1 | 0.928842248382143 | -1.46230664787936 | 1.09444423478256 | 0.279924394715554 | 0.599675667821634 | -4.76079774341787 |
| ENSG00000131471.6 | AOC3 | 0.292808107477066 | -0.41273679623154 | 1.09428735168516 | 0.279992387146855 | 0.599723435650375 | -5.00734079110102 |
| ENSG00000112624.12 | BICRAL | -0.119900781224551 | 5.94566470368337 | -1.09425833050695 | 0.280004966069091 | 0.599723435650375 | -5.82041537465052 |
| lnc-RRM2B-6 | lnc-RRM2B-6 | 0.288517705454655 | 0.0801964174769193 | 1.09418496366835 | 0.280036767921427 | 0.599729143078424 | -5.16850850554729 |
| ENSG00000262633.2 | | -0.0893664658268629 | 3.40451931048018 | -1.0940762386899 | 0.280083900913537 | 0.599730039584935 | -5.64919234428272 |
| ENSG00000279207.1 | | -0.203253241264354 | 2.35937350256975 | -1.09404957455608 | 0.280095460842928 | 0.599730039584935 | -5.54004970176044 |
| lnc-IRF2BP2-8 | lnc-IRF2BP2-8 | 0.204554565972073 | 0.889363138881938 | 1.0939802899474 | 0.280125499959831 | 0.599731970633783 | -5.34276285356585 |
| ENSG00000100372.14 | SLC25A17 | -0.158215852730367 | 3.12061510957797 | -1.0938845922979 | 0.280166994473526 | 0.599758424050815 | -5.6084132913614 |
| lnc-LCOR-1 | lnc-LCOR-1 | 0.117908544684632 | 2.5742211575565 | 1.09364673697089 | 0.280270147307032 | 0.599890252498396 | -5.5542244221346 |
| lnc-EFHD1-1 | lnc-EFHD1-1 | 0.121263662742804 | 1.36251390993795 | 1.09360819242906 | 0.28028686577647 | 0.599890252498396 | -5.40460224982764 |
| ENSG00000200090.1 | | 0.258704358739595 | 0.870019037518186 | 1.09346255988657 | 0.280350039372609 | 0.59990301352827 | -5.30598481953703 |
| lnc-FGD5-2 | lnc-FGD5-2 | 0.106476597970514 | 4.2708878799153 | 1.09346007071655 | 0.280351119230843 | 0.59990301352827 | -5.74324071587189 |
| ENSG00000169220.17 | RGS14 | -0.232756163978888 | 4.5629437208439 | -1.09330264863076 | 0.280419418434935 | 0.599986786865517 | -5.76468842162857 |
| lnc-MORN1-1 | lnc-MORN1-1 | -0.175928013496985 | 0.376848030506689 | -1.09291611053642 | 0.280587171644792 | 0.600283312957568 | -5.23280659530364 |
| ENSG00000282988.2 | | 1.292493958394 | 3.33634088101445 | 1.09338328407219 | 0.280628281493037 | 0.600308866968528 | -5.60612495942537 |
| ENSG00000005187.11 | ACSM3 | 0.321733665636254 | 6.89655072468598 | 1.09293007778823 | 0.280693328067644 | 0.600385614841144 | -5.8217830121603 |
| lnc-CDC42SE2-3 | lnc-CDC42SE2-3 | -0.343472522746179 | 1.31704315697278 | -1.09258280712583 | 0.2807318782557 | 0.600405678526295 | -5.4128247890891 |
| ENSG00000139842.14 | CUL4A | -0.0816271755269389 | 5.79856250404583 | -1.09215818117726 | 0.280916309358803 | 0.600737703431921 | -5.82106448699937 |
| ENSG00000225423.1 | TNPO1P1 | 0.295039089547017 | 0.744024704163577 | 1.09190578035099 | 0.281025976940303 | 0.600871010694348 | -5.29301618137829 |
| ENSG00000153044.9 | CENPH | 0.214634111427363 | 2.38235114289805 | 1.09188034196857 | 0.281037031528664 | 0.600871010694348 | -5.53902572815858 |
| ENSG00000140307.10 | GTF2A2 | 0.112063359067771 | 3.34875229921124 | 1.0917179444637 | 0.281107610735679 | 0.600959488135904 | -5.64870284762527 |
| lnc-TAPBPL-2 | lnc-TAPBPL-2 | 0.16739896220294 | -0.217161520832194 | 1.09107760925368 | 0.281386026774568 | 0.601335522004332 | -5.08088923733866 |
| ENSG00000273703.1 | H2BC14 | 0.590860014265094 | 2.40583472569105 | 1.09148850267401 | 0.281392488929939 | 0.601335522004332 | -5.55584700323765 |
| lnc-COMMD9-2 | lnc-COMMD9-2 | 0.154083843404001 | 1.10519530663931 | 1.0910576168115 | 0.28139472255371 | 0.601335522004332 | -5.36779806628647 |
| ENSG00000243302.3 | | -0.205610056346061 | 3.85358290667679 | -1.09104464058466 | 0.281400366707755 | 0.601335522004332 | -5.7026661528644 |
| lnc-PAICS-3 | lnc-PAICS-3 | -0.208125308417582 | 2.31462991240319 | -1.09063019969353 | 0.281580674228307 | 0.601658265415323 | -5.52652969187271 |
| ENSG00000119711.12 | ALDH6A1 | -0.160189925918799 | 4.13460128372327 | -1.0905245413515 | 0.281626655150906 | 0.601658265415323 | -5.72960510004461 |
| ENSG00000214113.10 | LYRM4 | 0.188364898239284 | 2.39757982372667 | 1.09049596996877 | 0.281639089893438 | 0.601658265415323 | -5.5488461514539 |
| ENSG00000077254.14 | USP33 | 0.0921729342642204 | 6.10344359322659 | 1.09030219142944 | 0.281723435727819 | 0.601775993937911 | -5.82652964365857 |
| ENSG00000141337.12 | ARSG | -0.441228546239308 | 2.7893168685254 | -1.09047433830828 | 0.281784669333211 | 0.601844334220762 | -5.59789063996227 |
| ENSG00000239291.1 | | 0.231180158371186 | 1.26608023427032 | 1.08988964262252 | 0.281903064632459 | 0.602001092588009 | -5.38417093627534 |
| lnc-FEM1B-4 | lnc-FEM1B-4 | 0.271356294367392 | 1.52992232796984 | 1.08980642356278 | 0.281939308999972 | 0.602001092588009 | -5.4156693477301 |
| ENSG00000166734.19 | GOLM2 | -0.115317058536845 | 6.48970135408835 | -1.08979150603456 | 0.281945806371686 | 0.602001092588009 | -5.82698954093564 |
| ENSG00000105364.13 | MRPL4 | -0.171025705138807 | 3.36800749631221 | -1.08950997064385 | 0.282068449632952 | 0.602200487376315 | -5.65631086270191 |
| ENSG00000239775.1 | | -0.16643238649928 | 3.46780187857214 | -1.0894344178268 | 0.282101368545456 | 0.602208304166265 | -5.64948280020316 |
| ENSG00000262302.1 | | -0.147114690554414 | 1.50392480646426 | -1.08909884398058 | 0.282247613119366 | 0.602313277701142 | -5.42647271936716 |
| lnc-ATP5PF-3 | lnc-ATP5PF-3 | -0.238843777783582 | 1.25219502333244 | -1.0889516700985 | 0.282311768943266 | 0.602313277701142 | -5.39448466234599 |
| ENSG00000131931.8 | THAP1 | -0.142396597486778 | 2.83524023150578 | -1.08895046588049 | 0.282312293926456 | 0.602313277701142 | -5.60155665557422 |
| ENSG00000007923.15 | DNAJC11 | -0.0899540724251864 | 4.75895387796588 | -1.08894201922713 | 0.28231597629475 | 0.602313277701142 | -5.78504512628954 |
| ENSG00000155629.14 | PIK3AP1 | -0.1919332609204 | 7.80917281086978 | -1.08892081739165 | 0.282325219508411 | 0.602313277701142 | -5.81237077336621 |
| ENSG00000170242.17 | USP47 | 0.104926543816739 | 7.16792562089921 | 1.08891875464673 | 0.28232611880001 | 0.602313277701142 | -5.82242021027579 |
| ENSG00000167216.16 | KATNAL2 | 0.333073427671356 | 0.637547969424906 | 1.08882227654217 | 0.282368182451761 | 0.602340584454825 | -5.28731327485683 |
| lnc-PHB2-1 | lnc-PHB2-1 | 0.26145386840104 | 6.33106975658707 | 1.08856398404994 | 0.282516692926282 | 0.602594931658204 | -5.82862024384081 |
| ENSG00000277958.1 | | 0.380357278841698 | -0.54565000515283 | 1.0883115692688 | 0.282590919875351 | 0.602669967858958 | -4.9588503328376 |

| | | | | | | | |
|---|---|---|---|---|---|---|---|
| ENSG00000085514.15 | PILRA | 0.259938874432977 | 2.6185020587607 | 1.08826684314561 | 0.282610432384719 | 0.602669967858958 | -5.57124078256002 |
| lnc-NAA10-2 | lnc-NAA10-2 | -0.180282269014324 | 3.24498876012639 | -1.08817028684709 | 0.282652559888241 | 0.602697362496599 | -5.63786033192634 |
| lnc-COX11-2 | lnc-COX11-2 | -0.285782359088344 | -0.421828126280243 | -1.0877883626021 | 0.282819236599352 | 0.602989783575055 | -5.02608884478126 |
| lnc-PDE6B-2 | lnc-PDE6B-2 | -0.286316145982907 | -0.144877610257045 | -1.08772180477777 | 0.282848290356953 | 0.602989783575055 | -5.11167005580665 |
| lnc-NOP16-1 | lnc-NOP16-1 | 0.171779395987981 | 0.235378002170184 | 1.08733762403041 | 0.283016033508519 | 0.603241874740293 | -5.21546911758356 |
| lnc-CDC27-7 | lnc-CDC27-7 | 0.103375010609203 | 3.54151707003947 | 1.08725442831433 | 0.283052368081857 | 0.603241874740293 | -5.66922999981953 |
| ENSG00000275714.1 | H3C1 | 0.273011824072687 | 3.21692927594961 | 1.08724963019107 | 0.283054463695368 | 0.603241874740293 | -5.64087715964509 |
| ENSG00000165630.13 | PRPF18 | 0.0957615862272787 | 4.32875912854631 | 1.08691259110887 | 0.283201695086758 | 0.603424785665701 | -5.74662912605629 |
| ENSG00000261408.7 | TEN1-CDK3 | -0.132485513775389 | 3.28925125325515 | -1.08679272441503 | 0.283254070332717 | 0.603424785665701 | -5.64750675976964 |
| ENSG00000124383.8 | MPHOSPH10 | 0.124165460219034 | 4.88454066705849 | 1.086765383107 | 0.283266017953599 | 0.603424785665701 | -5.7955234628255 |
| lnc-ESAM-65 | lnc-ESAM-65 | -0.178393910753553 | 3.21944634262615 | -1.08673118451733 | 0.283280962571111 | 0.603424785665701 | -5.6502643450019 |
| ENSG00000110696.9 | C11orf58 | 0.0995064250602677 | 7.14040764578363 | 1.08671766004876 | 0.283286872850783 | 0.603424785665701 | -5.82566710955702 |
| ENSG00000227500.9 | SCAMP4 | -0.164851639990092 | 3.97699008054643 | -1.08635748248138 | 0.283444304591531 | 0.60359743990493 | -5.71602270015464 |
| lnc-GTF2E2-2 | lnc-GTF2E2-2 | 0.20803902820593 | 2.25131154138492 | 1.08633641484627 | 0.283453515044628 | 0.60359743990493 | -5.51948373950174 |
| LINC01624 | LINC01624 | -0.514579188867731 | -0.333847246733512 | -1.08633095251072 | 0.283455903129819 | 0.60359743990493 | -5.08997610955501 |
| ENSG00000278236.1 | | -0.214018850320145 | 1.3037072801389 | -1.08618457549632 | 0.283519903132794 | 0.603671269906104 | -5.40831336910107 |
| lnc-RAB2B-1 | lnc-RAB2B-1 | 0.183866116263765 | -0.243912052338442 | 1.08559691520209 | 0.283776946180103 | 0.604156069841235 | -5.06528977767885 |
| ENSG00000279520.1 | | 0.25219998060134 | 3.24699544511405 | 1.08551947075206 | 0.28381083262248 | 0.604165722191386 | -5.63372949168539 |
| lnc-NFKBID-1 | lnc-NFKBID-1 | -0.262583469143384 | 3.00601232389392 | -1.08544589115024 | 0.283843030597804 | 0.604171778572347 | -5.60467610556743 |
| ENSG00000143674.10 | MAP3K21 | -0.527003017405181 | 0.771814543492394 | -1.08533311987207 | 0.283957696432206 | 0.604316151375421 | -5.28576836084271 |
| lnc-DDX3X-7 | lnc-DDX3X-7 | 0.283457854275623 | 1.22585785718801 | 1.08515675948809 | 0.2839695776176 | 0.604316151375421 | -5.38792681828727 |
| ENSG00000166508.17 | MCM7 | 0.229612326149306 | 4.99338298387815 | 1.08499919926117 | 0.284038555157238 | 0.604352032399974 | -5.80270726286005 |
| lnc-SND1-1 | lnc-SND1-1 | -0.304026111908798 | -0.407695244228686 | -1.08498411004157 | 0.284045161610909 | 0.604352032399974 | -5.04981946609234 |
| ENSG00000148288.12 | GBGT1 | -0.365167939991872 | 1.52635992731089 | -1.08468244416183 | 0.284187753028386 | 0.604573970751136 | -5.44421680798495 |
| ENSG00000048140.17 | TSPAN17 | 0.218391286033314 | 3.23330100838535 | 1.08458901362747 | 0.284218183457934 | 0.604573970751136 | -5.64565151705165 |
| ENSG00000130741.10 | EIF2S3 | 0.16693429290712 | 7.61265412972575 | 1.08453144969585 | 0.284243398197668 | 0.604573970751136 | -5.81942178311943 |
| ENSG00000213085.9 | CFAP45 | 0.24900475961614 | 0.881512296655443 | 1.08443150682181 | 0.284287179917621 | 0.604573970751136 | -5.33986490612834 |
| lnc-TUBA1C-1 | lnc-TUBA1C-1 | -0.186932564734049 | 2.55163211628326 | -1.08441060943422 | 0.284296334980335 | 0.604573970751136 | -5.55773994929205 |
| ENSG00000213639.9 | PPP1CB | 0.181848043836804 | 6.3377001717164 | 1.08425174059882 | 0.284365941544453 | 0.604637297136271 | -5.83333771298996 |
| lnc-IL16-11 | lnc-IL16-11 | -0.329949046428436 | -0.59547163476164 | -1.08420855503978 | 0.284384864867912 | 0.604637297136271 | -4.99193369953182 |
| ENSG00000116586.11 | LAMTOR2 | -0.220912828706728 | 2.81174410140097 | -1.08409274265513 | 0.284435616633267 | 0.604644112051801 | -5.5898317104118 |
| lnc-A4GNT-1 | lnc-A4GNT-1 | 0.134237962578098 | 0.409305732284143 | 1.08406717467934 | 0.284446821988903 | 0.604644112051801 | -5.24977063186313 |
| ENSG00000134297.6 | PLEKHA8P1 | 0.541026678310474 | 0.79631323035515 | 1.08406112692648 | 0.28450539495724 | 0.604706169393316 | -5.2358515670222 |
| ENSG00000159840.15 | ZYX | -0.276109764463566 | 2.18582574700885 | -1.08383101834198 | 0.284550333884943 | 0.604718046064977 | -5.52030530450183 |
| lnc-CFAP36-3 | lnc-CFAP36-3 | -0.175131808881678 | 3.81012024174773 | -1.08378674716274 | 0.284569741737617 | 0.604718046064977 | -5.70135876494509 |
| ENSG00000122481.16 | RWDD3 | -0.0943306165852444 | 3.26563572151803 | -1.08358420231534 | 0.284658546334671 | 0.604798757486578 | -5.64738970041671 |
| ENSG00000047578.12 | KATNIP | -0.095700730763876 | 4.78604970767478 | -1.08346290998884 | 0.284711735536126 | 0.604798757486578 | -5.79398776959508 |
| ENSG00000105755.7 | ETHE1 | -0.229480063688112 | 2.04870593398568 | -1.08343759794052 | 0.284722836272526 | 0.604798757486578 | -5.49634843507315 |
| lnc-WIZ-1 | lnc-WIZ-1 | -0.145996230170376 | 0.446783921884936 | -1.08343207878722 | 0.284725256767475 | 0.604798757486578 | -5.2595570650707 |
| lnc-PKP2-4 | lnc-PKP2-4 | -0.213804918981296 | 0.222997942398221 | -1.08333504951124 | 0.284767812546385 | 0.604813480048917 | -5.21965737570074 |
| ENSG00000088986.10 | DYNLL1 | -0.208801156172839 | 3.96539159606101 | -1.0832822854168 | 0.284790956062484 | 0.604813480048917 | -5.72768653916653 |
| ENSG00000179958.8 | DCTPP1 | -0.28008741894253 | 2.45204111619731 | -1.08315153231221 | 0.284848312987178 | 0.604813992148617 | -5.55699799693681 |
| ENSG00000157450.15 | RNF111 | -0.0932247561107449 | 5.46451951026271 | -1.083147765452 | 0.284849965500107 | 0.604813992148617 | -5.82265518033688 |
| lnc-TFCP2-4 | lnc-TFCP2-4 | 0.272631787387814 | -0.232592953247946 | 1.08297058986806 | 0.284927699608876 | 0.60491643415392 | -5.0705718584831 |
| ENSG00000168826.15 | ZBTB49 | 0.171475574312631 | 2.92388740003527 | 1.08290383622568 | 0.284956990990449 | 0.60491643415392 | -5.6079845715531 |
| ENSG00000214562.14 | NUTM2D | -0.180180249385736 | -0.0327524049598098 | -1.0827805641173 | 0.285011088119777 | 0.604938065634097 | -5.16699757956376 |
| lnc-EXTL3-4 | lnc-EXTL3-4 | 0.191203552892798 | 2.03774732656401 | 1.08270092795185 | 0.285046039735471 | 0.604938065634097 | -5.493848387862 |
| ENSG00000010932.16 | FMO1 | -0.797601298967689 | -0.554602351521432 | -1.08275818860234 | 0.285058535641994 | 0.604938065634097 | -4.92506823627174 |
| ENSG00000127616.18 | SMARCA4 | -0.145895775234591 | 5.30096954024499 | -1.08238423626267 | 0.28518506265023 | 0.604938065634097 | -5.82035015889748 |
| ENSG00000182287.14 | AP1S2 | -0.254853351390199 | 4.07833722410836 | -1.08237900792751 | 0.285188906629989 | 0.604938065634097 | -5.74389927056641 |
| lnc-RHEX-8 | lnc-RHEX-8 | 0.978934860481532 | 0.126579168038997 | 1.08273716751026 | 0.285209315322366 | 0.604938065634097 | -5.12367498544418 |
| lnc-PSMB7-2 | lnc-PSMB7-2 | -0.27328195388996 | 0.549667934946295 | -1.08224573123174 | 0.285245879209111 | 0.604938065634097 | -5.29194166620113 |
| ENSG00000070501.11 | POLB | 0.127332840619045 | 3.4311216956113 | 1.08224086376392 | 0.285248016644256 | 0.604938065634097 | -5.66725422872221 |
| lnc-CARD10-2 | lnc-CARD10-2 | -0.170441545806184 | 1.29044654781894 | -1.08219958650727 | 0.285266143041122 | 0.604938065634097 | -5.40674160466465 |
| ENSG00000111077.17 | TNS2 | -0.385614487653217 | -0.300886517784307 | -1.08216051377162 | 0.285283302092939 | 0.604938065634097 | -5.06583479010622 |
| ENSG00000198369.9 | SPRED2 | -0.698160347925432 | 0.16082925167835 | -1.08246684241373 | 0.28529047287131 | 0.604938065634097 | -5.23195110478647 |
| lnc-PPIAL4G-11 | lnc-PPIAL4G-11 | -0.228086045210675 | 0.998015347193154 | -1.08186272381238 | 0.28541410227891 | 0.605102067829436 | -5.36617924417391 |
| ENSG00000276903.1 | H2AC16 | 0.817185948164539 | 2.20840090636584 | 1.08236312986073 | 0.285426613057183 | 0.605102067829436 | -5.55336858314396 |
| ENSG00000137959.15 | IFI44L | 1.19717217271654 | 1.92925223597674 | 1.08225026546662 | 0.28548551385652 | 0.605102249442154 | -5.52145229250405 |
| lnc-MAP1LC3C-3 | lnc-MAP1LC3C-3 | -1.19252469442917 | -1.5850442466787 | -1.08191663540064 | 0.285496609956712 | 0.605102249442154 | -4.80645948256836 |
| lnc-ZNF79-1 | lnc-ZNF79-1 | -0.178213712109583 | 0.553150748842079 | -1.08163332049054 | 0.285514893192547 | 0.605102249442154 | -5.28196208463842 |
| ENSG00000113532.12 | ST8SIA4 | -0.211219003973992 | 4.2808904281633 | -1.08150776658671 | 0.285570067253175 | 0.605156871643376 | -5.7622584023202 |
| ENSG00000105063.18 | PPP6R1 | -0.126148931066616 | 5.98732730853491 | -1.08141501253051 | 0.285610832371546 | 0.605180951482761 | -5.83471511928954 |
| ENSG00000124370.10 | MCEE | -0.114983919418457 | 2.66649388216827 | -1.08126610795618 | 0.285676283985991 | 0.605257329210874 | -5.57199905872841 |
| ENSG00000178229.7 | ZNF543 | 0.150499385701304 | 4.57319451352778 | 1.08118000170996 | 0.285714137129108 | 0.605275224838249 | -5.77166739758758 |
| ENSG00000275111.4 | ZNF2 | -0.284430380636226 | 0.638842682064024 | -1.08105689807484 | 0.285768260781282 | 0.605327581729044 | -5.28446210459485 |
| lnc-VPS36-3 | lnc-VPS36-3 | 0.279195952962 | 0.716793012508504 | 1.08093158865376 | 0.285823361597212 | 0.605350081911748 | -5.30058689494978 |
| lnc-NOC2L-5 | lnc-NOC2L-5 | -0.291763618126742 | -0.043252832742914 | -1.08089897585072 | 0.285837703255127 | 0.605350081911748 | -5.154612171537 |

| | | | | | | | |
|---|---|---|---|---|---|---|---|
| ENSG00000147883.10 | CDKN2B | -0.420071749102739 | -0.7101435198523 | -1.08079036875254 | 0.285885467449824 | 0.605380205823318 | -4.98831926613577 |
| lnc-IQGAP3-2 | lnc-IQGAP3-2 | 0.171272579833304 | 4.09645582110084 | 1.08073288447034 | 0.285910750658722 | 0.605380205823318 | -5.73968231045837 |
| lnc-RTP5-8 | lnc-RTP5-8 | -0.237069810703825 | 0.925830625195151 | -1.08029805852173 | 0.286102050061645 | 0.60563392083636 | -5.34289598294202 |
| ENSG00000135317.12 | SNX14 | -0.0858279543399814 | 5.79278438870411 | -1.08029266847921 | 0.286104421944489 | 0.60563392083636 | -5.83421098145747 |
| ENSG00000187033.9 | SAMD7 | 0.642145689305971 | 0.351445295871761 | 1.08047526641205 | 0.286152672244196 | 0.60563392083636 | -5.25487353217543 |
| ENSG00000285385.1 | NA | 0.154792965632457 | 2.1787531828459 | 1.08004861234746 | 0.286211833017581 | 0.60563392083636 | -5.51137121319849 |
| lnc-TTC39A-3 | lnc-TTC39A-3 | 0.14906626362112 | 2.56163782499031 | 1.08002098571041 | 0.286223993500762 | 0.60563392083636 | -5.56048056587614 |
| lnc-CASKIN2-1 | lnc-CASKIN2-1 | -0.262881456658695 | 0.971464589133121 | -1.08002056137052 | 0.28622418028629 | 0.60563392083636 | -5.34753262998599 |
| ENSG00000017483.14 | SLC38A5 | -0.199798414136307 | 3.03139330683521 | -1.07998876985846 | 0.286238174486205 | 0.60563392083636 | -5.62259830453966 |
| ENSG00000280287.1 | | 0.364205320579436 | 1.73776600357203 | 1.07990922876421 | 0.286285096125853 | 0.60563392083636 | -5.45727195964869 |
| lnc-CNST-1 | lnc-CNST-1 | -0.277040335100089 | 0.316207460263537 | -1.07980230424873 | 0.286320263799638 | 0.60563392083636 | -5.24113329302102 |
| ENSG00000119048.7 | UBE2B | 0.169801820379762 | 4.55449570794207 | 1.07972921822029 | 0.286352443564702 | 0.60563392083636 | -5.78214443693766 |
| LINC01888 | LINC01888 | -0.30209048184072 | -0.143471313136931 | -1.0797251388412 | 0.28635423978912 | 0.60563392083636 | -5.1165000514501 |
| ENSG00000135002.11 | RFK | 0.250559894554352 | 2.05278048635236 | 1.07953809041484 | 0.286436609061382 | 0.605684221763112 | -5.52582962681346 |
| lnc-TCL1A-6 | lnc-TCL1A-6 | 0.164642603083124 | 0.938859172161925 | 1.07945882175671 | 0.286471521072819 | 0.605684221763112 | -5.35842914543391 |
| lnc-CPM-2 | lnc-CPM-2 | -0.261671373777962 | 0.00911114042337913 | -1.07940884795359 | 0.286493532385664 | 0.605684221763112 | -5.1619098572952 |
| ENSG00000162148.10 | SAXO4 | -0.160336951665788 | 1.09932036907901 | -1.07937980646587 | 0.286506324456503 | 0.605684221763112 | -5.37407564028327 |
| lnc-PRND-2 | lnc-PRND-2 | 0.268961743289371 | 1.63518132783434 | 1.07933705743504 | 0.28652515509437 | 0.605684221763112 | -5.45559728056704 |
| lnc-MED6-2 | lnc-MED6-2 | -0.401525947338245 | 0.00263562073224488 | -1.07919704229116 | 0.286586836809608 | 0.605752399060604 | -5.16361758680017 |
| ENSG00000081307.12 | UBA5 | 0.109085937478221 | 4.93004033466555 | 1.07885481334652 | 0.286737640076379 | 0.606008917311028 | -5.80751628758433 |
| lnc-BTG4-3 | lnc-BTG4-3 | 0.221690989275223 | 3.36292720342009 | 1.07872599234866 | 0.286794419452189 | 0.606066687431664 | -5.65633188516924 |
| ENSG00000132906.17 | CASP9 | -0.176397336726035 | 3.47786285876292 | -1.07838652757225 | 0.286944080213188 | 0.606248384145548 | -5.67187127182849 |
| ENSG00000111653.19 | ING4 | -0.183375362530637 | 4.33429390830981 | -1.07834139563337 | 0.286963981765508 | 0.606248384145548 | -5.76336618625741 |
| ENSG00000134744.13 | TUT4 | 0.115415287918407 | 7.66528058895974 | 1.07833055790649 | 0.286968760954675 | 0.606248384145548 | -5.82715752598464 |
| ENSG00000141642.8 | ELAC1 | -0.152792935994165 | 2.40920743017608 | -1.07824220070737 | 0.287007726530547 | 0.60626847651665 | -5.54952950483615 |
| ENSG00000035928.15 | RFC1 | 0.112787571319012 | 6.32335685650555 | 1.07803223353723 | 0.287100337011708 | 0.606401871391687 | -5.8398567167319 |
| ENSG00000145337.4 | PYURF | 0.146590548215675 | 3.90402706900704 | 1.07783662128352 | 0.287186634745829 | 0.606412877344465 | -5.72355437782549 |
| HCG11 | HCG11 | -0.380089481295057 | 3.4294119999756 | -1.07799173844552 | 0.287236598651122 | 0.606412877344465 | -5.68752558208377 |
| TRG-AS1 | TRG-AS1 | -0.592516038176865 | 1.41639855685822 | -1.0779861338479 | 0.287261801653269 | 0.606412877344465 | -5.36618027783562 |
| ENSG00000182552.14 | RWDD4 | -0.134618451030611 | 3.42669124852096 | -1.07765944723473 | 0.28726481379305 | 0.606412877344465 | -5.67073314065494 |
| lnc-ERGIC1-2 | lnc-ERGIC1-2 | -0.248630663067834 | 0.0276339484086834 | -1.07765929146023 | 0.287264882535967 | 0.606412877344465 | -5.16593039368871 |
| ENSG00000172667.10 | ZMAT3 | 0.178862089128003 | 4.617112653703 | 1.07761239370437 | 0.287285578927918 | 0.606412877344465 | -5.78689624157188 |
| lnc-ESX1-1 | lnc-ESX1-1 | -0.124048219379846 | 1.87718621913758 | -1.07755302446689 | 0.287311780588992 | 0.606412877344465 | -5.48456198942937 |
| lnc-FBXL16-3 | lnc-FBXL16-3 | -0.504065955453112 | -0.291551142326239 | -1.07747053832402 | 0.287348187297736 | 0.60642753400485 | -5.05726996795513 |
| ENSG00000141540.10 | TTYH2 | 0.363832296144171 | 1.05066566874644 | 1.07732578170897 | 0.287412085967006 | 0.606500201502858 | -5.35692197976829 |
| ENSG00000229917.2 | RPL7P46 | 0.388696149014277 | -0.335808818013969 | 1.07705208288598 | 0.287532929622601 | 0.606693007731625 | -4.98499173576058 |
| ENSG00000158769.17 | F11R | 0.198172728132321 | 5.63383279551589 | 1.07694229862792 | 0.287581411603786 | 0.606708206250906 | -5.83229032923785 |
| lnc-SLC25A47-5 | lnc-SLC25A47-5 | 0.105407728096222 | 4.69155539107481 | 1.07690228139887 | 0.287599085089156 | 0.606708206250906 | -5.79175250837865 |
| lnc-ZMYND19-1 | lnc-ZMYND19-1 | 0.155439108915929 | 2.63136818786772 | 1.0766271940871 | 0.287720597087691 | 0.606841794075101 | -5.58220936002655 |
| lnc-TNFAIP6-5 | lnc-TNFAIP6-5 | -0.211516887035125 | 1.67623554986748 | -1.0766109783649 | 0.287727761041193 | 0.606841794075101 | -5.48143450871789 |
| ENSG00000177963.14 | RIC8A | -0.105751052823861 | 5.54254825552818 | -1.0764637946561 | 0.287792791117674 | 0.606841794075101 | -5.83406312826027 |
| ENSG00000266714.8 | MYO15B | -0.402729187737271 | 6.11663655768531 | -1.07681809942794 | 0.287806342178946 | 0.606841794075101 | -5.83949401317991 |
| ENSG00000149792.8 | MRPL49 | -0.131043295434933 | 4.72110396374276 | -1.07642524917413 | 0.287809823337761 | 0.606841794075101 | -5.79374573554284 |
| ENSG00000166289.5 | PLEKHF1 | -0.416948976845468 | 0.15331699300739 | -1.07629791610479 | 0.287866093428965 | 0.606850948051045 | -5.20687689371059 |
| ENSG00000150990.7 | DHX37 | 0.104488639083009 | 5.00058610216788 | 1.0761680477956 | 0.287923491789949 | 0.606850948051045 | -5.8134287997625 |
| lnc-TPCN1-2 | lnc-TPCN1-2 | -0.243835554182613 | -0.230807786489811 | -1.07614557169944 | 0.287933426441957 | 0.606850948051045 | -5.06861600497247 |
| lnc-SH3D19-8 | lnc-SH3D19-8 | 0.136722021938473 | 2.25719158513568 | 1.07611002745321 | 0.287949137831468 | 0.606850948051045 | -5.53001323803753 |
| ENSG00000138385.15 | SSB | 0.106156793404853 | 5.64928408794058 | 1.07607083461026 | 0.287966462680343 | 0.606850948051045 | -5.83559382930777 |
| ENSG00000171169.8 | NAIF1 | 0.114792104631891 | 3.22106155240732 | 1.07601518459687 | 0.287991063526338 | 0.606850948051045 | -5.64280927213841 |
| ENSG00000241360.1 | PDXP | 0.349487397894479 | 0.969578947230162 | 1.0755436375931 | 0.288199576213211 | 0.607228158173459 | -5.36354081080148 |
| lnc-CPXM2-1 | lnc-CPXM2-1 | 0.350749801826442 | 1.21631330654826 | 1.07540908193478 | 0.288259094513156 | 0.607291396352538 | -5.37767366986735 |
| ENSG00000127561.14 | SYNGR3 | -0.468775568498433 | 1.42684676265991 | -1.07541932007273 | 0.288337801514108 | 0.607395043349185 | -5.41885526344521 |
| lnc-CA4-3 | lnc-CA4-3 | -0.32023566472415 | 2.54656694451074 | -1.07484401303787 | 0.288525448038354 | 0.607728131086107 | -5.58307971596011 |
| lnc-ARF6-2 | lnc-ARF6-2 | -0.31910288242794 | 0.662215387567387 | -1.0746718958145 | 0.288585327985197 | 0.607792060362152 | -5.26820255592112 |
| lnc-ARMCX5-3 | lnc-ARMCX5-3 | 0.416575815253844 | 1.83479362708902 | 1.07469890142845 | 0.288655209780984 | 0.607877039382237 | -5.51950248058912 |
| lnc-ZNF624-3 | lnc-ZNF624-3 | -0.194065734504562 | -0.0983841394821759 | -1.07431302023202 | 0.288744237598241 | 0.607981173704829 | -5.12390865731288 |
| lnc-USP33-1 | lnc-USP33-1 | 0.0928848349969096 | 2.31249023323121 | 1.07426899788692 | 0.288763734836438 | 0.607981173704829 | -5.54416944023645 |
| ENSG00000233680.4 | HNRNPA1P27 | -0.427125413931179 | 1.60792108585046 | -1.0740039086661 | 0.28893838498082 | 0.608250579861785 | -5.4455875109872 |
| ENSG00000104728.15 | ARHGEF10 | 0.77743721434159 | 0.98276777722777 | 1.07430649267897 | 0.288950792881919 | 0.608250579861785 | -5.37795680096053 |
| ENSG00000075089.9 | ACTR6 | 0.110595523530842 | 4.22777850056128 | 1.07348688822409 | 0.289110279792054 | 0.608410664085596 | -5.75842292642655 |
| lnc-ASXL1-3 | lnc-ASXL1-3 | 0.180294212973026 | 1.91616946747333 | 1.07346008228137 | 0.28912216237756 | 0.608410664085596 | -5.4948196308033 |
| ENSG00000150347.15 | ARID5B | 0.206619464095397 | 8.20244925925003 | 1.07344973850863 | 0.289126747674468 | 0.608410664085596 | -5.82178698550898 |
| ENSG00000151893.14 | CACUL1 | 0.0820072740489435 | 5.63791268352119 | 1.07340839173709 | 0.289145076815104 | 0.608410664085596 | -5.83848219253788 |
| ENSG00000163659.12 | TIPARP | 0.226090734877461 | 5.90752276485338 | 1.07308060474245 | 0.289290414404611 | 0.608546155366367 | -5.84225058670505 |
| ENSG00000179335.18 | CLK3 | 0.125499677582154 | 6.56285561004734 | 1.07301801195013 | 0.289318173238302 | 0.608546155366367 | -5.84466553686866 |
| LINC02482 | LINC02482 | -0.336408995696123 | -0.544901076442992 | -1.0729973963582 | 0.28932731630775 | 0.608546155366367 | -5.04045720654811 |
| ENSG00000267419.2 | ZNF56P | -0.147617516686579 | 0.829253382373863 | -1.07299646250845 | 0.289327730477349 | 0.608546155366367 | -5.34539314513923 |
| ENSG00000143878.9 | RHOB | -0.365359674474147 | 6.28956321650723 | -1.07324219173474 | 0.289368014699376 | 0.608568697921452 | -5.84326603297339 |

| | | | | | | | |
|---|---|---|---|---|---|---|---|
| ENSG00000171295.12 | ZNF440 | 0.127206347865613 | 4.20771660928955 | 1.07280830547619 | 0.28941118801574 | 0.608597311292192 | -5.75496445899108 |
| ENSG00000181827.14 | RFX7 | 0.151514728258499 | 6.641695798507 | 1.07266739519886 | 0.289473700128956 | 0.60865531177969 | -5.84490201826384 |
| ENSG00000115604.10 | IL18R1 | 1.01309296825699 | -0.559650616900323 | 1.0727895685278 | 0.289504923017253 | 0.60865531177969 | -4.86609000674735 |
| ENSG00000161551.14 | ZNF577 | -0.266849957725961 | 4.51311324089393 | -1.07259823556213 | 0.289527481787638 | 0.60865531177969 | -5.7785108168916 |
| lnc-PSEN2-4 | lnc-PSEN2-4 | -0.568606249018804 | -0.955427580729715 | -1.07224682189106 | 0.289660335323067 | 0.608766147504103 | -4.88514709243716 |
| ENSG00000179163.11 | FUCA1 | -0.137682740093726 | 2.97026928924195 | -1.0721831976815 | 0.289688576747409 | 0.608766147504103 | -5.63027031247179 |
| lnc-GNL2-1 | lnc-GNL2-1 | -0.208547132873259 | 1.13498143012005 | -1.07202260028518 | 0.289759871032784 | 0.608766147504103 | -5.39230492268256 |
| ENSG00000106829.18 | TLE4 | -0.299077485998475 | 7.51309679410008 | -1.07218649185678 | 0.289772312773786 | 0.608766147504103 | -5.83427898081934 |
| ENSG00000057252.12 | SOAT1 | 0.150953754550386 | 4.8818938790644 | 1.07199280615706 | 0.289773098937912 | 0.608766147504103 | -5.81472069418708 |
| ENSG00000127334.10 | DYRK2 | 0.186975841622779 | 5.89385374258592 | 1.07198282978948 | 0.289777528308923 | 0.608766147504103 | -5.84437529128158 |
| ENSG00000104064.17 | GABPB1 | 0.150866660757955 | 5.70265800026233 | 1.07196096152769 | 0.289787237683778 | 0.608766147504103 | -5.84129471633349 |
| LINC01284 | LINC01284 | -1.66499250832366 | 1.79984846673687 | -1.07209621150274 | 0.289966757816495 | 0.609065496487508 | -5.33739512457921 |
| ENSG00000122176.11 | FMOD | -0.651573459905125 | 7.49117413715179 | -1.0719750369088 | 0.290020530185247 | 0.609065496487508 | -5.83626418825028 |
| ENSG00000169564.6 | PCBP1 | 0.193467538115579 | 5.2019629860187 | 1.07133983757423 | 0.290063107651364 | 0.609065496487508 | -5.82361968131707 |
| lnc-NXN-6 | lnc-NXN-6 | -0.267432468561761 | 0.799989558203708 | -1.0713244067682 | 0.290069963518021 | 0.609065496487508 | -5.3558034523457 |
| ENSG00000105058.11 | FAM32A | 0.160389687852962 | 5.11749409121653 | 1.07130702014592 | 0.290077688484042 | 0.609065496487508 | -5.82052821400965 |
| ENSG00000040487.12 | SLC66A1 | -0.125822607132294 | 2.70267243534139 | -1.07104212198571 | 0.29019540184054 | 0.60924915929228 | -5.59902231006151 |
| ENSG00000168397.16 | ATG4B | -0.121518495000643 | 5.01127276370643 | -1.07097696645648 | 0.290224360241987 | 0.60924915929228 | -5.8169112261409 |
| ENSG00000241399.6 | CD302 | 0.233147458946741 | 1.39790614367128 | 1.07079654319339 | 0.290304559944764 | 0.60928869264335 | -5.41703165228807 |
| lnc-MISP3-1 | lnc-MISP3-1 | -0.180536089158993 | 0.0612924130063863 | -1.07073510721502 | 0.290331872293824 | 0.60928869264335 | -5.18540672949856 |
| ENSG00000104889.6 | RNASEH2A | -0.228442918471638 | 0.842657520648588 | -1.07073050390341 | 0.290333918841908 | 0.60928869264335 | -5.3394379544735 |
| ENSG00000276302.1 | | -0.134142811140716 | 1.15455063306747 | -1.07066824566033 | 0.290361598704689 | 0.60928869264335 | -5.40404095974556 |
| LINC01126 | LINC01126 | -0.24933734648729 | 2.74707591068256 | -1.07055482484457 | 0.290412030049037 | 0.609299138411248 | -5.60379996432954 |
| ENSG00000137955.15 | RABGGTB | 0.143223459229242 | 7.05443497129114 | 1.07052390150855 | 0.290425780836261 | 0.609299138411248 | -5.84284723245723 |
| lnc-ZNF461-2 | lnc-ZNF461-2 | 0.126401208805344 | 1.49049435731127 | 1.07036809621368 | 0.290495070219539 | 0.609322667706218 | -5.44335947759321 |
| ENSG00000177666.16 | PNPLA2 | 0.191922805327703 | 5.5757555823822 | 1.07036555012932 | 0.290496202604145 | 0.609322667706218 | -5.83893487002202 |
| ENSG00000182685.7 | BRICD5 | -0.242863284672398 | 1.08793018138686 | -1.06969472939705 | 0.290794660867628 | 0.609812386644842 | -5.39592975167591 |
| ENSG00000164163.10 | ABCE1 | 0.129910510045586 | 5.97313443099177 | 1.06966419377641 | 0.290808251706946 | 0.609812386644842 | -5.84731961974254 |
| ENSG00000167085.11 | PHB1 | -0.167714099781499 | 3.64500848925974 | -1.06958678441112 | 0.290842707164591 | 0.609812386644842 | -5.70859720997544 |
| ENSG00000167705.11 | RILP | -0.272423609781025 | -0.225975946815594 | -1.0695744773635 | 0.290848185380771 | 0.609812386644842 | -5.13655699699203 |
| ENSG00000152348.15 | ATG10 | -0.173706039811418 | 2.40709222228756 | -1.06949742972355 | 0.290882483105446 | 0.60982217862695 | -5.56681138305714 |
| lnc-SLC2A6-2 | lnc-SLC2A6-2 | 0.273624856493095 | 0.701683224711464 | 1.06911991721874 | 0.291050573355471 | 0.610029250991872 | -5.32180318851812 |
| lnc-GOLGA3-1 | lnc-GOLGA3-1 | 0.129001944943359 | 3.49532856058992 | 1.06901901876209 | 0.291095510597349 | 0.610029250991872 | -5.68597383185234 |
| ENSG00000049449.9 | RCN1 | -0.327388888238671 | 1.14602228230647 | -1.06897685961771 | 0.291114288486822 | 0.610029250991872 | -5.40517911815858 |
| ENSG00000247077.6 | PGAM5 | 0.188297668495896 | 4.26909622633245 | 1.06897113624141 | 0.291116837771698 | 0.610029250991872 | -5.76764603216284 |
| ENSG00000184220.11 | CMSS1 | -0.12435264888351 | 3.29036911647072 | -1.06894283608293 | 0.291129443350977 | 0.610029250991872 | -5.66476885708195 |
| ENSG00000069329.17 | VPS35 | -0.105584513329116 | 6.47437388224026 | -1.06884097998257 | 0.291174815682611 | 0.610046273000885 | -5.84887520035958 |
| ENSG00000179698.13 | WDR97 | -0.198623326550459 | 2.44847477008751 | -1.06860104952466 | 0.291281713417273 | 0.610046273000885 | -5.57892345341729 |
| lnc-ZNF273-7 | lnc-ZNF273-7 | -0.331724151413306 | -0.580796835328646 | -1.06853806330012 | 0.291309780595991 | 0.610046273000885 | -5.03498254648459 |
| ENSG00000279811.1 | | -0.369094894153022 | -0.653909082527911 | -1.06852787790334 | 0.291314319469272 | 0.610046273000885 | -5.02484565957304 |
| lnc-SRPRA-5 | lnc-SRPRA-5 | 0.179248105804501 | 0.614520115659221 | 1.0685188326342 | 0.291318350313794 | 0.610046273000885 | -5.32433798967694 |
| ENSG00000116237.15 | ICMT | -0.199494452135825 | 3.34627247144958 | -1.06848217661647 | 0.291334685737548 | 0.610046273000885 | -5.68109793802718 |
| ENSG00000255150.2 | EID3 | 0.268049561561563 | 3.43172520968237 | 1.06841428188205 | 0.291364944098239 | 0.610046273000885 | -5.6752807232253 |
| ENSG00000166484.19 | MAPK7 | -0.142910304725961 | 2.58795242751571 | -1.06839245119474 | 0.291374673753665 | 0.610046273000885 | -5.58720432458521 |
| ENSG00000213468.6 | FIRRE | 0.625807174551467 | 0.243303950931384 | 1.06850887875705 | 0.291416547750831 | 0.610071887953147 | -5.20873324440513 |
| ENSG00000099800.7 | TIMM13 | -0.271439008061181 | 2.39930885974677 | -1.06821757280635 | 0.291452622956864 | 0.610085359332974 | -5.56092594988338 |
| ENSG00000243753.5 | HLA-L | 0.241334389664607 | 3.7553234554769 | 1.06806427330497 | 0.291520965660998 | 0.610166365283743 | -5.69348626701151 |
| ENSG00000252835.1 | SCARNA21 | -0.381353745599985 | 7.69960514861239 | -1.06831404703496 | 0.29156744990997 | 0.610201608693127 | -5.83848062731674 |
| ENSG00000111300.9 | NAA25 | 0.117233263141578 | 5.28414057421496 | 1.0677303578402 | 0.291669867677517 | 0.610214687373615 | -5.83450987814259 |
| ENSG00000215908.10 | CROCCP2 | -0.169574177435491 | 5.25999705984334 | -1.06769334182234 | 0.291686377388613 | 0.610214687373615 | -5.8333199179927 |
| lnc-GALNT15-3 | lnc-GALNT15-3 | -0.216640547719764 | 0.193106574332484 | -1.06758739638875 | 0.291733634270069 | 0.610214687373615 | -5.23205624183715 |
| ENSG00000007376.7 | RPUSD1 | -0.270052945890509 | 2.7022023048811 | -1.06754184225607 | 0.291753955294944 | 0.610214687373615 | -5.60987615900953 |
| ENSG00000138835.22 | RGS3 | -0.273909396940615 | 1.38194054309753 | -1.06752749818653 | 0.291760354176156 | 0.610214687373615 | -5.44735889369171 |
| ENSG00000100578.14 | KIAA0586 | -0.0909040231237377 | 5.75092865366252 | -1.06749515157502 | 0.291774784339503 | 0.610214687373615 | -5.84474708023153 |
| lnc-RTL8C-1 | lnc-RTL8C-1 | -0.267887620458081 | 0.620394601787857 | -1.06747753776877 | 0.291782642252454 | 0.610214687373615 | -5.31000585056753 |
| ENSG00000261717.5 | | -0.170105137473488 | 1.76411031625489 | -1.06729524669904 | 0.291863975025598 | 0.610214687373615 | -5.47859727313248 |
| lnc-ERICH1-19 | lnc-ERICH1-19 | -0.207798346324826 | 2.15114205857348 | -1.06726591502408 | 0.291877063406983 | 0.610214687373615 | -5.53921967176962 |
| lnc-ARC-1 | lnc-ARC-1 | -0.187411085218291 | 0.459614381510066 | -1.06725936757743 | 0.291879985064697 | 0.610214687373615 | -5.27465869265323 |
| lnc-DGKQ-2 | lnc-DGKQ-2 | 0.174769883569408 | 1.78186389427648 | 1.06708180596724 | 0.291959225886596 | 0.610214687373615 | -5.48960140473276 |
| ENSG00000152556.16 | PFKM | 0.156552270529334 | 4.59430082284906 | 1.06707601864391 | 0.291961808860354 | 0.610214687373615 | -5.79009828832586 |
| ENSG00000275924.1 | MIR6807 | 0.181144270976316 | -0.104753073182131 | 1.06699921875304 | 0.291996087369573 | 0.610214687373615 | -5.12594249998956 |
| ENSG00000261884.2 | | 0.120155775341756 | 4.89693750056025 | 1.06695723751047 | 0.292014826267513 | 0.610214687373615 | -5.81593365082367 |
| ENSG00000077150.18 | NFKB2 | -0.253572853116114 | 5.635276745158 | -1.06693797499675 | 0.292051376449095 | 0.610214687373615 | -5.84537554542816 |
| ENSG00000005812.10 | FBXL3 | 0.111248402746643 | 6.23316754321854 | 1.06683661045741 | 0.292068674447515 | 0.610214687373615 | -5.85148337229355 |
| ENSG00000147535.16 | PLPP5 | -0.376934436750386 | 5.99818581634096 | -1.06714569944866 | 0.292088382186056 | 0.610214687373615 | -5.84685115604893 |
| ENSG00000067704.9 | IARS2 | -0.143996063505729 | 5.18706225112418 | -1.06675001106593 | 0.292107336864997 | 0.610214687373615 | -5.83082391642333 |
| ENSG00000073605.18 | GSDMB | -0.253538441041697 | 4.69349823065803 | -1.06671028615117 | 0.292140694053682 | 0.610222438167946 | -5.79645293981711 |
| ENSG00000102001.12 | CACNA1F | -0.253419837289271 | 2.95433650731267 | -1.06649683216347 | 0.29222038935546 | 0.610287419041272 | -5.61968072113667 |

| | | | | | | | |
|---|---|---|---|---|---|---|---|
| ENSG00000198862.13 | LTN1 | 0.134917216853377 | 6.04171518070943 | 1.06646134965379 | 0.292236235864173 | 0.610287419041272 | -5.85093851243406 |
| ENSG00000245958.6 | SEPTIN7P14 | -0.126670527897864 | 4.05003388688024 | -1.06640645391504 | 0.292260753509684 | 0.610287419041272 | -5.74522214747066 |
| lnc-ARPC5L-1 | lnc-ARPC5L-1 | -0.305128233146147 | -0.285738604081691 | -1.06623960965097 | 0.292335278614115 | 0.610381115816021 | -5.07974948358421 |
| ENSG00000183864.4 | TOB2 | -0.185766398241865 | 4.05011245377717 | -1.06611714329146 | 0.292389989644687 | 0.610382506182818 | -5.75680578628522 |
| lnc-SLC51A-14 | lnc-SLC51A-14 | -0.40546432255872 | 0.104178825599398 | -1.06610536051526 | 0.292395253897031 | 0.610382506182818 | -5.15147151189029 |
| lnc-RIC8A-2 | lnc-RIC8A-2 | -0.20969584676986 | 0.886316948561313 | -1.06582573115443 | 0.292520204710459 | 0.61057653294649 | -5.38377786410281 |
| ENSG00000279799.1 | | -0.207231918396424 | 0.520004853383636 | -1.06576459510912 | 0.292547527955997 | 0.61057653294649 | -5.3192786422121 |
| lnc-HBEGF-2 | lnc-HBEGF-2 | -0.179501778373483 | 1.26040656053383 | -1.06536702403562 | 0.29272525582793 | 0.610871356891902 | -5.41876851016096 |
| ENSG00000182308.6 | DCAF4L1 | -0.222442670722247 | -0.204685338358253 | -1.06526051152814 | 0.292772883311164 | 0.610871356891902 | -5.12484663923655 |
| lnc-BANP-1 | lnc-BANP-1 | 0.432017030810924 | -0.56916069558063 | 1.06516249645768 | 0.292816715892302 | 0.610871356891902 | -4.97430750700839 |
| lnc-NAGA-1 | lnc-NAGA-1 | 0.165405240833305 | 1.03143423572456 | 1.06509694109791 | 0.292846034956948 | 0.610871356891902 | -5.3876602756217 |
| ENSG00000170364.12 | SETMAR | 0.133704883691135 | 1.75345539611344 | 1.06508769707094 | 0.292850169431814 | 0.610871356891902 | -5.4853938759887 |
| lnc-ATRIP-24 | lnc-ATRIP-24 | -0.431524518854323 | -0.313357881299127 | -1.06505038375459 | 0.292866858563343 | 0.610871356891902 | -5.08970675120692 |
| ENSG00000230359.5 | TPI1P2 | -0.24510581273572 | 0.0201581442189819 | -1.06490916939773 | 0.292930025512271 | 0.610941201248259 | -5.17277018467049 |
| ENSG00000106609.16 | TMEM248 | -0.0978137930460781 | 7.18248088204187 | -1.06469362709934 | 0.293026458539377 | 0.611080404874975 | -5.84786080711313 |
| ENSG00000006194.9 | ZNF263 | 0.0899875615281243 | 5.68184411386338 | 1.0644694778251 | 0.293126765707722 | 0.611214977904235 | -5.84704952783408 |
| lnc-RPP40-5 | lnc-RPP40-5 | 0.239014119676897 | 1.01259136332321 | 1.06441671745331 | 0.293150379530224 | 0.611214977904235 | -5.39890505837076 |
| lnc-MBD3L2B-3 | lnc-MBD3L2B-3 | -1.03908650446496 | 1.26903560374262 | -1.06441189070321 | 0.293381252358999 | 0.611634388464022 | -5.31708510380125 |
| ENSG00000269313.5 | MAGIX | -0.246442195849416 | -0.135725492493665 | -1.06371705280144 | 0.293463651645067 | 0.611727161275846 | -5.11761742529672 |
| ENSG00000213965.3 | NUDT19 | -0.197148204315598 | 2.50213932707656 | -1.06366896230367 | 0.293485192518048 | 0.611727161275846 | -5.57986402364276 |
| ENSG00000128534.7 | LSM8 | -0.117579666021466 | 5.83704936896057 | -1.0633631987317 | 0.293622176961891 | 0.611950715715532 | -5.85145987967637 |
| ENSG00000178397.12 | FAM220A | -0.196459359224618 | 2.65718618310511 | -1.06279325763559 | 0.293877633527778 | 0.612421112777387 | -5.59803814175709 |
| lnc-FTHL17-1 | lnc-FTHL17-1 | -0.656076440708162 | 1.61483776901943 | -1.06306908173222 | 0.293938498887566 | 0.612485940737273 | -5.47508960611139 |
| ENSG00000143178.12 | TBX19 | 0.20148773380182 | 1.89674578117792 | 1.06244089795472 | 0.294035643877041 | 0.612626344561306 | -5.50033820567088 |
| ENSG00000110717.12 | NDUFS8 | -0.163474109351391 | 3.69000359009417 | -1.06181278232813 | 0.294317459096896 | 0.613087027251746 | -5.72567371484745 |
| ENSG00000178718.6 | RPP25 | 0.69064859147258 | -0.189608400203801 | 1.06204585748313 | 0.294326546211912 | 0.613087027251746 | -5.17044039352321 |
| lnc-MANEAL-3 | lnc-MANEAL-3 | 0.186363286245029 | 1.18175847354403 | 1.06174894569618 | 0.294346111028604 | 0.613087027251746 | -5.40944821868871 |
| ENSG00000129103.17 | SUMF2 | -0.115815008288573 | 6.45483722558249 | -1.06126598952476 | 0.29456294001985 | 0.613395479343308 | -5.85691590425619 |
| lnc-CDC26-2 | lnc-CDC26-2 | -0.205886768867293 | 0.51939714177184 | -1.06123226637209 | 0.294578084575442 | 0.613395479343308 | -5.29910775312559 |
| ENSG00000183726.10 | TMEM50A | 0.112694146437744 | 5.32155872674459 | 1.06116458477345 | 0.294608480984023 | 0.613395479343308 | -5.84369900880032 |
| lnc-EPHX2-5 | lnc-EPHX2-5 | -0.196437832108122 | -0.154418722778682 | -1.06109228192255 | 0.29464095524305 | 0.613395479343308 | -5.12187911725298 |
| lnc-C3orf62-2 | lnc-C3orf62-2 | 0.147587866096926 | 1.74908968579441 | 1.06104213365034 | 0.294663480405734 | 0.613395479343308 | -5.484678296521 |
| ENSG00000133858.15 | ZFC3H1 | 0.150203519795411 | 6.59883495754377 | 1.06096128932847 | 0.294699795868097 | 0.613395479343308 | -5.85716687948872 |
| ENSG00000005379.16 | TSPOAP1 | -0.29084358550869 | 3.60525803635694 | -1.06097158443667 | 0.29472803230242 | 0.613395479343308 | -5.72955118431884 |
| lnc-PPIG-1 | lnc-PPIG-1 | -0.243084142602103 | 0.0511598939744664 | -1.06088824776543 | 0.2947326089834 | 0.613395479343308 | -5.19905910301489 |
| ENSG00000102096.9 | PIM2 | 0.155347879794987 | 7.72852863331516 | 1.0607675230317 | 0.294786848794888 | 0.613414715789122 | -5.84282612544497 |
| lnc-ARPC3-3 | lnc-ARPC3-3 | 0.155948687925652 | 1.11879985915673 | 1.06073501358049 | 0.294801455987271 | 0.613414715789122 | -5.39517149068075 |
| ENSG00000036257.12 | CUL3 | 0.180492244723048 | 6.62666229428678 | 1.06057087150441 | 0.294875216206311 | 0.613488995520856 | -5.8570716132446 |
| ENSG00000004468.12 | CD38 | 1.00607283416158 | 2.34162275763061 | 1.0609920526051 | 0.294923258305244 | 0.613488995520856 | -5.59635638003317 |
| lnc-CCDC146-1 | lnc-CCDC146-1 | 0.202787509999566 | 0.178294281122599 | 1.0604306512904 | 0.294938236844341 | 0.613488995520856 | -5.17112765123832 |
| ENSG00000266302.6 | | -1.37159347791749 | -0.382715851116284 | -1.06088601027785 | 0.294970873946315 | 0.613488995520856 | -5.22851304150563 |
| ENSG00000163564.14 | PYHIN1 | -0.418130479288152 | 5.47332719475688 | -1.06070421393112 | 0.294986182221732 | 0.613488995520856 | -5.84375353269289 |
| lnc-NUP35-4 | lnc-NUP35-4 | -0.259233933619687 | 1.11252452265302 | -1.06012240288675 | 0.295076809001126 | 0.613566034676413 | -5.4160733292957 |
| ENSG00000137449.15 | CPEB2 | -0.116962535792199 | 4.24546120773055 | -1.06010898063295 | 0.29508284396161 | 0.613566034676413 | -5.78181061647532 |
| ENSG00000182004.12 | SNRPE | 0.141638258005561 | 4.29103846890088 | 1.05997782157217 | 0.295141820661778 | 0.61359075260737 | -5.78044919692097 |
| lnc-SYNE3-5 | lnc-SYNE3-5 | -0.330955164452919 | 5.36473432968003 | -1.06021120082974 | 0.295161041882795 | 0.61359075260737 | -5.8410304130601 |
| ENSG00000163815.5 | CLEC3B | 0.179709668052403 | 0.244814552450148 | 1.05988366652902 | 0.295184163256968 | 0.61359075260737 | -5.21964122658445 |
| lnc-IQUB-3 | lnc-IQUB-3 | 0.180148387557409 | 1.9674838817881 | 1.05981044169617 | 0.295217096211049 | 0.61359724238231 | -5.51281810078305 |
| lnc-GATD1-1 | lnc-GATD1-1 | -0.140379565684685 | 1.71224890723092 | -1.05963794515239 | 0.295294686806646 | 0.613696540644305 | -5.49959388318699 |
| lnc-SOX7-7 | lnc-SOX7-7 | 0.280507920448332 | -0.167284066327003 | 1.05947385057648 | 0.295368511243786 | 0.613787992623003 | -5.08967248977553 |
| ENSG00000187231.13 | SESTD1 | -0.618536904339948 | 5.982562868803 | -1.05984607007322 | 0.295428929274661 | 0.61379754042294 | -5.85262131783896 |
| ENSG00000106462.10 | EZH2 | -0.283804440978159 | 3.83068404149972 | -1.059220881477 | 0.295508737954141 | 0.61379754042294 | -5.73472757949128 |
| lnc-HEMGN-2 | lnc-HEMGN-2 | 0.132884567513277 | 2.62909170835728 | 1.0591593306073 | 0.295510046273805 | 0.61379754042294 | -5.59598413990071 |
| ENSG00000163001.11 | CFAP36 | 0.173413607817586 | 2.90012499553749 | 1.05914272006861 | 0.295517522378659 | 0.61379754042294 | -5.63085799508072 |
| ENSG00000113851.14 | CRBN | 0.098091958768344 | 5.83495893435537 | 1.05913230764458 | 0.295522208890411 | 0.61379754042294 | -5.855793825432 |
| ENSG00000275996.1 | SNORD27 | 0.223521345344761 | 2.1852467838627 | 1.05883854107343 | 0.295654451069426 | 0.614010247841993 | -5.54574037198576 |
| ENSG00000170222.11 | ADPRM | 0.165548341020239 | 3.75015428124322 | 1.05854967078114 | 0.295784529142131 | 0.614218418415303 | -5.72861384865793 |
| ENSG00000156374.15 | PCGF6 | 0.169868905180193 | 1.46744044353424 | 1.05833731264897 | 0.295880179149091 | 0.61435506178006 | -5.46178830770156 |
| ENSG00000157191.19 | NECAP2 | 0.151916051360017 | 5.41462487301351 | 1.05799937466599 | 0.296032436837364 | 0.61460920389585 | -5.84885730198419 |
| ENSG00000159674.11 | SPON2 | 0.618777932264838 | 1.46600338648293 | 1.05826674829509 | 0.296079347687542 | 0.614644600449085 | -5.46494475443381 |
| ENSG00000078699.21 | CBFA2T2 | -0.13093495302607 | 4.71966888216906 | -1.05767952515297 | 0.296176594796743 | 0.61471565330046 | -5.81375230785129 |
| ENSG00000206834.1 | SNORA1 | 0.243816535696867 | 0.483427012445767 | 1.05765001103555 | 0.296189899425434 | 0.61471565330046 | -5.26377765128577 |
| ENSG00000185085.2 | INTS5 | -0.179896748207392 | 3.17065861218062 | -1.05761700122542 | 0.29620478036335 | 0.61471565330046 | -5.67012626317428 |
| ENSG00000114302.15 | PRKAR2A | 0.0949277082875835 | 5.13737315507855 | 1.05755432760588 | 0.296233035276066 | 0.61471565330046 | -5.83769209796302 |
| ENSG00000168734.13 | PKIG | -0.272049010648233 | 5.31515506511744 | -1.05743984804077 | 0.296338803141641 | 0.614873143655685 | -5.84339732590973 |
| lnc-HBS1L-2 | lnc-HBS1L-2 | -0.261594049008637 | 1.290825212438 | -1.05722169656533 | 0.296383025341847 | 0.614902914082374 | -5.42763219681561 |
| ENSG00000185187.12 | SIGIRR | -0.224457032280458 | 2.80234847230791 | -1.05712359723473 | 0.296427270360963 | 0.614932725845565 | -5.63241762170732 |
| ENSG00000105221.16 | AKT2 | 0.115125957601103 | 6.98649332424476 | 1.05679567608933 | 0.296575203438041 | 0.615135003673297 | -5.85838126370761 |

| | | | | | | | |
|---|---|---|---|---|---|---|---|
| lnc-SRY-9 | lnc-SRY-9 | -1.09674558389642 | -2.11844236082552 | -1.05677496324883 | 0.296584549213127 | 0.615135003673297 | -4.69999876759579 |
| ENSG00000107175.11 | CREB3 | -0.152028096106395 | 3.83485872610522 | -1.05650266450309 | 0.296707431238389 | 0.615236456466563 | -5.73957099866387 |
| ENSG00000203799.12 | CCDC162P | 0.449799043754243 | 0.510980454479312 | 1.05647834421125 | 0.296722251330078 | 0.615236456466563 | -5.25630446833102 |
| ENSG00000198792.12 | TMEM184B | 0.16656684954964 | 2.16476999635667 | 1.05640520014954 | 0.296751423180295 | 0.615236456466563 | -5.54377538151984 |
| ENSG00000204001.9 | LCN8 | 1.05459438201184 | -0.510452656274948 | 1.056756112823 | 0.296810221546857 | 0.615236456466563 | -5.24602472160864 |
| lnc-BAHD1-3 | lnc-BAHD1-3 | -0.206758394663919 | 0.464936975302993 | -1.05623551015097 | 0.296828025988815 | 0.615236456466563 | -5.3178828042611 |
| lnc-CCNL2-4 | lnc-CCNL2-4 | 0.282307150275793 | -0.0657284273517163 | 1.05621331664246 | 0.296838045771571 | 0.615236456466563 | -5.17834973122092 |
| ENSG00000159579.13 | RSPRY1 | 0.0814945074036721 | 4.67869430035774 | 1.05620301268205 | 0.296842697817103 | 0.615236456466563 | -5.8104010036975 |
| TMEM254-AS1 | TMEM254-AS1 | 0.367219578398016 | 0.772399743990118 | 1.05612426456665 | 0.296878252786895 | 0.61524819544026 | -5.32442669085806 |
| VASH1-AS1 | VASH1-AS1 | -0.185383237587035 | 2.54044179443325 | -1.05573177496377 | 0.297055506801239 | 0.615540782633774 | -5.58750948474473 |
| lnc-ANAPC10-4 | lnc-ANAPC10-4 | 0.122488234066128 | 0.933607746167087 | 1.05567922056792 | 0.297079246693092 | 0.615540782633774 | -5.36598204234894 |
| ENSG00000262246.5 | CORO7 | 0.168629653735514 | 4.90164216797157 | 1.05549558769128 | 0.29716220772116 | 0.615580217462541 | -5.82556963593234 |
| lnc-TRAF5-8 | lnc-TRAF5-8 | 0.206176690427585 | 2.6975294328221 | 1.05545201985237 | 0.297181893003367 | 0.615580217462541 | -5.60845485091906 |
| lnc-SLC10A7-8 | lnc-SLC10A7-8 | 0.187491530258981 | 1.16924554914659 | 1.05543481771058 | 0.297189665705793 | 0.615580217462541 | -5.41107220752973 |
| ENSG00000162971.10 | TYW5 | 0.131673207717876 | 5.62421201278229 | 1.05535535534095 | 0.297225572210096 | 0.615580217462541 | -5.85685798002694 |
| lnc-PRND-3 | lnc-PRND-3 | 0.0978762884073165 | 3.65074782848435 | 1.05530613402969 | 0.297247815253374 | 0.615580217462541 | -5.71587086391163 |
| lnc-MST1-1 | lnc-MST1-1 | 0.33162038347699 | -0.500172419640016 | 1.05495443770875 | 0.297406779874973 | 0.615826504471399 | -5.02212150990879 |
| ENSG00000113638.13 | TTC33 | -0.143865050087199 | 2.98851228899402 | -1.05482925469724 | 0.297463376041603 | 0.615826504471399 | -5.64624031672659 |
| ENSG00000105963.14 | ADAP1 | 0.743037857298777 | -0.654809141826412 | 1.054874696899 | 0.29748713678205 | 0.615826504471399 | -4.99056129359382 |
| lnc-NUMA1-5 | lnc-NUMA1-5 | -0.166918025727729 | -0.189248851160634 | -1.05469167442703 | 0.29752558569915 | 0.615826504471399 | -5.12479019048745 |
| ENSG00000280365.1 | | -0.326095756474153 | 1.65784865307044 | -1.05465045728232 | 0.297544224604362 | 0.615826504471399 | -5.47620418936494 |
| ENSG00000102543.14 | CDADC1 | -0.180493251530202 | 2.43302651492721 | -1.05462820242702 | 0.297554288862565 | 0.615826504471399 | -5.59057051803161 |
| lnc-ZNF737-3 | lnc-ZNF737-3 | -0.216775150173268 | 1.48489270503 | -1.05457980755869 | 0.297576175167494 | 0.615826504471399 | -5.46310982071708 |
| ENSG00000214843.3 | ZFYVE9P2 | -0.920894903453517 | -0.889451733588239 | -1.05454546908921 | 0.297658591620506 | 0.615935135349841 | -4.90176538816214 |
| ENSG00000188933.15 | USP32P1 | -1.6729242393711 | -1.05400085104063 | -1.05480384599468 | 0.297710858146433 | 0.615944709393253 | -5.03657360150708 |
| ENSG00000187866.8 | PABIR1 | 0.118627423412877 | 3.39154667028565 | 1.05425506217807 | 0.297723068248237 | 0.615944709393253 | -5.69733014031586 |
| lnc-NCMAP-1 | lnc-NCMAP-1 | -0.168575380356396 | 0.802795889828957 | -1.05408635536419 | 0.297799399724377 | 0.616040707992649 | -5.35132260185638 |
| ENSG00000187531.13 | SIRT7 | 0.112777716875268 | 4.47568677667181 | 1.05401004680732 | 0.29783393001377 | 0.616050224246149 | -5.79856107513902 |
| ENSG00000135919.12 | SERPINE2 | -0.829139178553241 | -1.60108303280518 | -1.05372420960684 | 0.297969771571022 | 0.616238405286507 | -4.85131453309276 |
| ENSG00000204104.11 | TRAF3IP1 | -0.120846558031779 | 2.43716376628356 | -1.05362361118585 | 0.298008838050399 | 0.616238405286507 | -5.57539676918531 |
| lnc-SNIP1-1 | lnc-SNIP1-1 | 0.160815960421638 | 1.4063751058149 | 1.05361060733184 | 0.298014725075154 | 0.616238405286507 | -5.4495472114317 |
| ENSG00000138326.19 | RPS24 | 0.174687378580394 | 9.14233416281173 | 1.05325822446904 | 0.298174284306572 | 0.616506408225231 | -5.81841725167 |
| ENSG00000173153.13 | ESRRA | -0.159993527547198 | 3.03989714529757 | -1.05311483327483 | 0.298239228861987 | 0.61655720179209 | -5.65857693935682 |
| ENSG00000171155.7 | C1GALT1C1 | -0.147182937826388 | 2.75715872110484 | -1.05307171454151 | 0.298258760056544 | 0.61655720179209 | -5.61438275080352 |
| lnc-SMDT1-3 | lnc-SMDT1-3 | -0.213817815832382 | 0.0741667206985158 | -1.05279812552661 | 0.298382706396015 | 0.616709449174838 | -5.20221117376853 |
| ZNF710-AS1 | ZNF710-AS1 | -0.143787163915161 | 2.04721654005181 | -1.05271570608241 | 0.298420052553124 | 0.616709449174838 | -5.53090693164797 |
| lnc-CAPN9-4 | lnc-CAPN9-4 | 0.217492170430029 | 1.2943851685765 | 1.05265306902821 | 0.298448437013371 | 0.616709449174838 | -5.4313570211443 |
| lnc-TM2D2-7 | lnc-TM2D2-7 | -0.227053477418005 | 1.09828975672892 | -1.05264463776945 | 0.298452257845336 | 0.616709449174838 | -5.40396804150602 |
| ENSG00000085449.14 | WDFY1 | 0.177498634582317 | 6.99151528917861 | 1.0523583323278 | 0.298582024264983 | 0.616872132076223 | -5.86158575902102 |
| LINC01277 | LINC01277 | -0.350493384710331 | 0.14149991360593 | -1.05233869310117 | 0.298590927069689 | 0.616872132076223 | -5.2321640083536 |
| ENSG00000124181.14 | PLCG1 | -0.151396330577783 | 6.44636452967083 | -1.05215113389325 | 0.29867596018376 | 0.616985878007058 | -5.86635016892807 |
| lnc-NNT-3 | lnc-NNT-3 | -0.170509349348927 | 2.22203856875592 | -1.05155171602556 | 0.298947828578833 | 0.617479562561345 | -5.56020771025184 |
| ENSG00000026036.22 | RTEL1-TNFRSF6B | -0.137134217714053 | 2.99948546494756 | -1.05149194721718 | 0.29897494633855 | 0.617479562561345 | -5.65135476997088 |
| ENSG00000259084.6 | | -0.456522395069182 | 3.15200663649267 | -1.05152190454752 | 0.299116238392262 | 0.617631977678797 | -5.66376623752615 |
| ENSG00000183172.8 | SMDT1 | 0.164819061961151 | 3.31896524908253 | 1.05093202340649 | 0.299229072445159 | 0.617631977678797 | -5.67782305137577 |
| ENSG00000133313.14 | CNDP2 | 0.14514817924207 | 6.64187305547229 | 1.05092207929655 | 0.299233587011249 | 0.617631977678797 | -5.86749866013685 |
| ENSG00000176399.3 | DMRTA1 | -1.01783401056935 | -0.331657849604466 | -1.05125393798857 | 0.299238439938462 | 0.617631977678797 | -5.01435391909251 |
| ENSG00000155849.15 | ELMO1 | 0.118880400360352 | 6.37561475417697 | 1.05089826753966 | 0.299244397596922 | 0.617631977678797 | -5.86787413435595 |
| lnc-RFNG-2 | lnc-RFNG-2 | -0.168864659291676 | 2.33590648095439 | -1.05088644499665 | 0.299249765155564 | 0.617631977678797 | -5.57659748427256 |
| ENSG00000164494.11 | PDSS2 | -0.125895644050775 | 3.53378674775139 | -1.05080892083339 | 0.299284963587971 | 0.617631977678797 | -5.70245877303634 |
| lnc-ATAD2B-1 | lnc-ATAD2B-1 | 0.279796854014199 | 0.238552969932292 | 1.05080047423692 | 0.299288798783866 | 0.617631977678797 | -5.20721362495253 |
| lnc-RARRES2-2 | lnc-RARRES2-2 | -0.255492545359983 | 1.11989332958014 | -1.05066184278343 | 0.299351749555887 | 0.617696514109912 | -5.38806546536745 |
| lnc-VCAM1-3 | lnc-VCAM1-3 | 0.255152401025626 | 1.07409858272324 | 1.05057481310896 | 0.299391273293285 | 0.617696514109912 | -5.40265820191651 |
| ENSG00000165046.12 | LETM2 | -0.259658749128065 | 1.5589182904876 | -1.0504720641315 | 0.299437940450443 | 0.617696514109912 | -5.47632927586611 |
| ENSG00000163463.11 | KRTCAP2 | -0.105306743268447 | 3.48013193226318 | -1.05039046809528 | 0.299475003815727 | 0.617696514109912 | -5.70509565722547 |
| ENSG00000106328.9 | FSCN3 | -0.125877757450993 | 2.52706089223962 | -1.05028797377922 | 0.299521564296807 | 0.617696514109912 | -5.60182939957487 |
| ENSG00000171105.13 | INSR | -1.30183111465054 | -0.418026044710504 | -1.05056289489186 | 0.299622679639264 | 0.617696514109912 | -5.19747701110991 |
| lnc-PGAP1-4 | lnc-PGAP1-4 | -0.195022677655942 | 4.10239474566145 | -1.04997078255093 | 0.299665687641282 | 0.617696514109912 | -5.76774922969408 |
| ENSG00000138669.9 | PRKG2 | -1.0777723181592 | -1.05802580394758 | -1.05015811478114 | 0.299693921324508 | 0.617696514109912 | -4.87940744541358 |
| ENSG00000130517.13 | PGPEP1 | -0.138164684443788 | 4.68656231939315 | -1.04981671369257 | 0.29973570975642 | 0.617696514109912 | -5.82036981198008 |
| ENSG00000074054.18 | CLASP1 | 0.0822274750796641 | 6.63958612848782 | 1.04975938084376 | 0.299761769609504 | 0.617696514109912 | -5.86838951004146 |
| ENSG00000166140.17 | ZFYVE19 | -0.145291812788733 | 3.91844292824478 | -1.0496855816588 | 0.29979531631131 | 0.617696514109912 | -5.74873324629557 |
| lnc-RSRP1-1 | lnc-RSRP1-1 | 0.185003674183433 | 3.14196201716814 | 1.04966697428189 | 0.299803775027071 | 0.617696514109912 | -5.65700103852903 |
| lnc-ITGB3BP-5 | lnc-ITGB3BP-5 | -0.147912047930172 | 2.91935240033643 | -1.04966598514753 | 0.299804224681649 | 0.617696514109912 | -5.64238426836378 |
| ENSG00000105701.15 | FKBP8 | -0.149800631944114 | 5.8575002481673 | -1.04965942858564 | 0.299807205267355 | 0.617696514109912 | -5.86652342198709 |
| COX10-AS1 | COX10-AS1 | -0.175510708719345 | 3.54820925811488 | -1.04965001466006 | 0.299811484834464 | 0.617696514109912 | -5.71731614663157 |
| ENSG00000172830.12 | SSH3 | 1.00380518753181 | 0.952649942249631 | 1.05011361146219 | 0.299825809337353 | 0.617696514109912 | -5.31665750537837 |
| ENSG00000214338.10 | MTCL3 | -0.959390230082711 | -1.59935280155887 | -1.04968454049507 | 0.299848339368153 | 0.617696514109912 | -4.85254083757233 |

| | | | | | | | |
|---|---|---|---|---|---|---|---|
| ENSG00000139637.13 | MYG1 | -0.167796319711404 | 3.31843461555404 | -1.0495347133366 | 0.299863904196608 | 0.617696514109912 | -5.69614236518771 |
| lnc-PDGFA-17 | lnc-PDGFA-17 | 0.265412794970416 | 1.50797998875602 | 1.04947674183079 | 0.299890262133817 | 0.617696514109912 | -5.47495316115337 |
| lnc-SLC38A1-4 | lnc-SLC38A1-4 | 0.205515828748793 | 3.32757980126136 | 1.04939918658773 | 0.299925526721821 | 0.617707336053156 | -5.69959598214963 |
| lnc-FRAT1-7 | lnc-FRAT1-7 | 0.129341351321939 | 1.28795871859783 | 1.04922289866245 | 0.300005695991351 | 0.617779577342454 | -5.43955153063613 |
| ENSG00000128191.15 | DGCR8 | -0.0890152563451068 | 4.59294273427654 | -1.04919006020017 | 0.300020631351852 | 0.617779577342454 | -5.81390922469549 |
| ENSG00000049245.12 | VAMP3 | 0.103377197392692 | 4.90393279279414 | 1.04906809019551 | 0.300076109393928 | 0.617832005567193 | -5.83459945299374 |
| ENSG00000130309.10 | COLGALT1 | -0.108534048553096 | 6.16227488287198 | -1.04785599936568 | 0.300627813709395 | 0.618868065372592 | -5.87032161188929 |
| ENSG00000133818.13 | RRAS2 | -0.324835476174766 | 5.94635917056878 | -1.04810500950509 | 0.300639449334914 | 0.618868065372592 | -5.86818442084107 |
| lnc-OR13A1-4 | lnc-OR13A1-4 | 0.10133002782054 | 4.37271847656549 | 1.04761129702223 | 0.300739279126354 | 0.61897366564734 | -5.79982082858477 |
| ENSG00000120049.19 | KCNIP2 | -0.246304866403156 | 0.436789406197366 | -1.04751182918136 | 0.300784596301324 | 0.61897366564734 | -5.27886373327989 |
| ENSG00000086300.15 | SNX10 | 0.14851641261081 | 5.68193190857105 | 1.04748896678738 | 0.300795012988124 | 0.61897366564734 | -5.86638035209663 |
| ENSG00000166313.18 | APBB1 | 0.156205646247699 | 4.62674191197974 | 1.04745379827454 | 0.300811037140861 | 0.61897366564734 | -5.82072356115326 |
| lnc-GRK7-1 | lnc-GRK7-1 | 0.202607862822446 | 0.47385700469318 | 1.04736302450137 | 0.300852399951483 | 0.618996896061712 | -5.32608529785368 |
| ENSG00000146085.7 | MMUT | 0.139187550743495 | 3.59396711312918 | 1.04725688678219 | 0.300900768615114 | 0.619034534773103 | -5.716398479229 |
| ENSG00000156313.13 | RPGR | 0.209437101157046 | 3.71464051376226 | 1.04692804614463 | 0.301050660635364 | 0.61928100618206 | -5.73370795835246 |
| ENSG00000102221.13 | JADE3 | 0.227637571756002 | 6.02861130985056 | 1.04685206564135 | 0.301091319908667 | 0.619302751841719 | -5.87068535991445 |
| lnc-RCL1-1 | lnc-RCL1-1 | 0.142995741827461 | 1.06121560497714 | 1.04675509227478 | 0.301129517071319 | 0.61931942944434 | -5.40299284233383 |
| lnc-NCBP2-AS2-1 | lnc-NCBP2-AS2-1 | 0.228004761239455 | 2.51770618529641 | 1.04662571464908 | 0.301188514706939 | 0.619378878830345 | -5.58832952322657 |
| lnc-TCIRG1-1 | lnc-TCIRG1-1 | -0.210702338823824 | 3.06802728651002 | -1.04643452752987 | 0.301275712753273 | 0.619395029473509 | -5.65996938932657 |
| ENSG00000062725.9 | APPBP2 | 0.110613121616455 | 5.52818172823917 | 1.0463674371298 | 0.301306315969384 | 0.619395029473509 | -5.86406803184179 |
| ENSG00000078967.12 | UBE2D4 | 0.124646141860969 | 3.49509858314416 | 1.04634495199513 | 0.301316573019782 | 0.619395029473509 | -5.70372345536742 |
| ENSG00000124602.9 | UNC5CL | -0.154246835627854 | 2.73891021982826 | -1.04620599349926 | 0.301379967099543 | 0.619395029473509 | -5.6180067122731 |
| lnc-ADAT1-1 | lnc-ADAT1-1 | -0.183784257921115 | 1.27025652564069 | -1.04618603294345 | 0.301389074034893 | 0.619395029473509 | -5.43021984233334 |
| ENSG00000257800.1 | FNBP1P1 | -0.245540807556172 | 1.94735433689597 | -1.04612665927794 | 0.30141616418856 | 0.619395029473509 | -5.51092060447259 |
| ENSG00000198353.7 | HOXC4 | 1.33711717204917 | -0.435892305151993 | 1.04641116511872 | 0.301451992202299 | 0.619395029473509 | -4.88689287944343 |
| ENSG00000206341.7 | HLA-H | 0.369659819794224 | 5.17294499500087 | 1.0462941141837 | 0.301489171309713 | 0.619395029473509 | -5.84596202120514 |
| lnc-TNPO3-12 | lnc-TNPO3-12 | 1.17422170102508 | -1.84409061328311 | 1.04597003681382 | 0.301493647426553 | 0.619395029473509 | -4.70877836394517 |
| ENSG00000143387.12 | CTSK | 0.223733381556913 | 3.58314811533352 | 1.04594886877659 | 0.301497293897638 | 0.619395029473509 | -5.71349067885673 |
| lnc-PDGFD-1 | lnc-PDGFD-1 | -0.635640897800646 | 0.838500145099907 | -1.04610995616783 | 0.301567684251253 | 0.6194778088766 | -5.33393160461822 |
| lnc-CAPZA2-2 | lnc-CAPZA2-2 | -0.387148107948627 | 1.02282929565824 | -1.04572285275384 | 0.301613616835745 | 0.619510335827776 | -5.41337781475839 |
| ENSG00000181991.15 | MRPS11 | -0.158548382154787 | 3.18659533809194 | -1.04553245112886 | 0.301687373341281 | 0.619538415811868 | -5.67374081580125 |
| ENSG00000213967.10 | ZNF726 | -0.186240384214333 | 2.78651285128203 | -1.04553220252911 | 0.301687486842654 | 0.619538415811868 | -5.62594678084707 |
| ENSG00000147894.15 | C9orf72 | 0.1705630146252 | 5.58389322013325 | 1.04534917508506 | 0.301771058336224 | 0.619648213660663 | -5.86541872303959 |
| lnc-BCL7C-1 | lnc-BCL7C-1 | -0.189231553025894 | 1.88202200281447 | -1.04524204696089 | 0.30181998112147 | 0.619686850017278 | -5.53115511602276 |
| lnc-ISYNA1-2 | lnc-ISYNA1-2 | 0.211850317440919 | 0.505954411033276 | 1.04502335120772 | 0.301919871076408 | 0.619825242337124 | -5.32077379003587 |
| ENSG00000181023.7 | OR56B1 | 0.309497549719688 | 0.977568143458557 | 1.0449626242413 | 0.301947612345836 | 0.619825242337124 | -5.38655491304783 |
| lnc-SPATA7-4 | lnc-SPATA7-4 | -0.239056418903489 | 0.277278317744305 | -1.04460338800777 | 0.302111754438086 | 0.620064710327012 | -5.2439585802289 |
| ENSG00000285404.1 | | -0.165870854779389 | 4.64689926173641 | -1.04457545696259 | 0.302124519257135 | 0.620064710327012 | -5.81706846684063 |
| ENSG00000170191.4 | NANP | 0.181050655027657 | 1.77752577556257 | 1.04436265227939 | 0.302221785725076 | 0.620090566351446 | -5.50734415528729 |
| ENSG00000167118.10 | URM1 | 0.108370755528561 | 4.37843368283544 | 1.04435709130694 | 0.302224327763599 | 0.620090566351446 | -5.80609629948184 |
| lnc-CASD1-1 | lnc-CASD1-1 | -0.242894079103685 | 1.11454988429326 | -1.04435015928791 | 0.302227496557242 | 0.620090566351446 | -5.4296492478294 |
| ENSG00000120875.8 | DUSP4 | -0.715152506961422 | 2.37348609824039 | -1.04468903565045 | 0.302295801003953 | 0.620168889869867 | -5.59986301456544 |
| ENSG00000164402.13 | SEPTIN8 | -0.414869343602255 | 2.21250116200749 | -1.04429939182428 | 0.302348212782796 | 0.620179126344061 | -5.5792384038125 |
| ENSG00000126945.8 | HNRNPH2 | 0.146367111680378 | 5.81870560619592 | 1.04398848175618 | 0.302392859866243 | 0.620179126344061 | -5.87123026581449 |
| ENSG00000198771.10 | RCSD1 | -0.174915148881341 | 8.37715214134124 | -1.04392247080769 | 0.30242304758949 | 0.620179126344061 | -5.84671320293811 |
| ENSG00000128596.16 | CCDC136 | 0.497029859500528 | 0.151176651384582 | 1.04391453708897 | 0.302445115984733 | 0.620179126344061 | -5.22337062255805 |
| ENSG00000121895.7 | TMEM156 | 0.257631956754095 | 3.63794096697025 | 1.04386038120456 | 0.302451443921765 | 0.620179126344061 | -5.70324135213492 |
| ENSG00000107960.10 | STN1 | -0.119470276512644 | 3.28681516067482 | -1.04371867949307 | 0.302516257285152 | 0.620250236447881 | -5.69472157954803 |
| lnc-KMT2C-3 | lnc-KMT2C-3 | -0.359282101640282 | -0.550787548752755 | -1.04348084897311 | 0.302625060790665 | 0.62033992476207 | -5.03478542872065 |
| lnc-MANSC1-3 | lnc-MANSC1-3 | 0.768508717621829 | -1.9279185054849 | 1.04344642592008 | 0.302640810997277 | 0.62033992476207 | -4.78833109294763 |
| ENSG00000008086.11 | CDKL5 | 0.399298711884851 | 2.09904849390955 | 1.04357091142103 | 0.302650416579736 | 0.62033992476207 | -5.5376271054378 |
| ENSG00000139974.15 | SLC38A6 | -0.165772643541817 | 2.80588278060619 | -1.04329911624237 | 0.302708218655694 | 0.620384324536598 | -5.63174877994695 |
| ENSG00000108384.14 | RAD51C | 0.122031405808622 | 3.21771584257084 | 1.04324636541074 | 0.302732359502774 | 0.620384324536598 | -5.68051665747352 |
| lnc-EXOC7-2 | lnc-EXOC7-2 | 0.18613441801549 | 0.482254595649348 | 1.04308542905676 | 0.302806018476116 | 0.620473497092473 | -5.2864588743921 |
| ENSG00000044115.20 | CTNNA1 | 0.120845555046225 | 6.03827888905276 | 1.0429103590037 | 0.302886160317937 | 0.620575934483785 | -5.87504825421687 |
| lnc-FDFT1-1 | lnc-FDFT1-1 | -0.25885471476004 | 0.670036406490578 | -1.04280119588376 | 0.302936139326382 | 0.620594942025534 | -5.32894358588493 |
| lnc-NME5-3 | lnc-NME5-3 | 0.133139061522243 | 2.42921626571717 | 1.04270944877958 | 0.302978149015236 | 0.620594942025534 | -5.58052414337374 |
| ENSG00000167685.14 | ZNF444 | -0.111299822247079 | 3.18962426640638 | -1.0426925439579 | 0.302985889929291 | 0.620594942025534 | -5.67898683683758 |
| lnc-HMGB2-15 | lnc-HMGB2-15 | 0.306039765271977 | -0.438692160578316 | 1.04256882054663 | 0.303042548460755 | 0.620618716749966 | -5.06496516693818 |
| ENSG00000142459.8 | EVI5L | -0.187269083460594 | 2.25446006100263 | -1.04253551635666 | 0.30305780119778 | 0.620618716749966 | -5.57383461996511 |
| lnc-MFSD9-7 | lnc-MFSD9-7 | -0.168588523507792 | -0.06728605589585 | -1.04242225176004 | 0.303109678359141 | 0.620663202314585 | -5.17286913435547 |
| lnc-STX17-4 | lnc-STX17-4 | -0.349315091449778 | 0.215198870444453 | -1.04207210279119 | 0.303270091357771 | 0.620883014305755 | -5.22031134619343 |
| ENSG00000166263.13 | STXBP4 | -0.140348255201264 | 3.18347397436179 | -1.04205624779975 | 0.30327735635379 | 0.620883014305755 | -5.67474045680845 |
| ENSG00000100425.18 | BRD1 | 0.115810483190756 | 5.67643998599017 | 1.04177827549957 | 0.303404746912373 | 0.621071587753518 | -5.87164484196075 |
| ENSG00000087157.18 | PGS1 | 0.082562224968961 | 4.65588309063995 | 1.04166875821227 | 0.303454947163336 | 0.621071587753518 | -5.82897895895929 |
| ENSG00000276418.5 | | 0.205581426971391 | 3.26930299790824 | 1.04165775911666 | 0.303459989216204 | 0.621071587753518 | -5.67634035517378 |
| ENSG00000215284.2 | CHMP5P1 | 0.341593864681319 | 1.55173479181315 | 1.04144754319663 | 0.303556364564002 | 0.621158192420308 | -5.45916941585805 |
| lnc-TET3-3 | lnc-TET3-3 | -0.244825477503276 | 1.94807726980216 | -1.04143381022128 | 0.303562661300874 | 0.621158192420308 | -5.51565665479578 |

| | | | | | | | |
|---|---|---|---|---|---|---|---|
| ENSG00000169813.16 | HNRNPF | -0.151467997177406 | 6.72372949657511 | -1.04128131327083 | 0.303632589055241 | 0.621239126620573 | -5.87595588629454 |
| ENSG00000213799.12 | ZNF845 | 0.134255584713391 | 5.27571030987089 | 1.04121592037152 | 0.30366257848368 | 0.621239126620573 | -5.85623406008151 |
| ENSG00000128245.14 | YWHAH | -0.227135803532923 | 5.47332791168257 | -1.04110767428052 | 0.303718212312021 | 0.621288869394638 | -5.87059844689294 |
| ENSG00000182544.8 | MFSD5 | 0.181974221722305 | 3.37637739494672 | 1.04103128937573 | 0.303747261950068 | 0.621288869394638 | -5.70756234497458 |
| lnc-MBTPS1-1 | lnc-MBTPS1-1 | -0.173110790436936 | 0.869351287499672 | -1.0408437028095 | 0.303833317667431 | 0.621337155726687 | -5.38691357765938 |
| ENSG00000127946.16 | HIP1 | 0.536390955249522 | 2.65558460465892 | 1.04116183719499 | 0.303860184888024 | 0.621337155726687 | -5.63671145694181 |
| lnc-TXNDC9-2 | lnc-TXNDC9-2 | -0.116348536546605 | 2.01830283735897 | -1.04072485711034 | 0.303887847051567 | 0.621337155726687 | -5.54347762671051 |
| lnc-ZNF43-2 | lnc-ZNF43-2 | -0.185886575350337 | 0.714881182532299 | -1.04071664176194 | 0.30389161670799 | 0.621337155726687 | -5.33125456405547 |
| ENSG00000089820.15 | ARHGAP4 | -0.184945487357986 | 6.87475610265622 | -1.04058717626763 | 0.303951026888419 | 0.621374524294269 | -5.87495669007519 |
| ENSG00000129235.10 | TXNDC17 | 0.177360951330543 | 2.46065734696213 | 1.04054524391577 | 0.30397027086037 | 0.621374524294269 | -5.60407648609396 |
| ENSG00000159388.5 | BTG2 | 0.195443823723584 | 8.59115288783922 | 1.04044817723513 | 0.304014820789683 | 0.621403878482031 | -5.84462639135629 |
| lnc-GPR183-47 | lnc-GPR183-47 | 0.306343254781736 | 2.88479515692172 | 1.04031675147934 | 0.304097830144757 | 0.621511829795407 | -5.65611059236625 |
| ENSG00000185585.19 | OLFML2A | -0.461356068715783 | 0.734439556110743 | -1.03969355306873 | 0.304395807425745 | 0.622005030277615 | -5.32886981927041 |
| lnc-F11R-1 | lnc-F11R-1 | -0.222446663996523 | 2.62969124824838 | -1.03952885508772 | 0.304436977523355 | 0.622005030277615 | -5.61095276668783 |
| ENSG00000117322.17 | CR2 | 0.348215825674709 | 5.01834742582959 | 1.03978923427273 | 0.304452192690628 | 0.622005030277615 | -5.84779350958099 |
| ENSG00000267283.1 | | -0.256072163511603 | 0.412690808541629 | -1.03944367041778 | 0.304476115089354 | 0.622005030277615 | -5.27497496576295 |
| ENSG00000119917.13 | IFIT3 | 0.710926356615422 | -1.44661811177977 | 1.03941292081739 | 0.304490243651424 | 0.622005030277615 | -4.86133702236488 |
| lnc-LRCH4-2 | lnc-LRCH4-2 | 0.122549625358941 | 3.23450525648701 | 1.03911388640261 | 0.304627664919839 | 0.622177700465083 | -5.68178045348677 |
| ENSG00000175106.16 | TVP23C | -0.306009335514023 | 2.95334654311273 | -1.03915748371928 | 0.304635226414376 | 0.622177700465083 | -5.67557360129911 |
| ENSG00000153147.5 | SMARCA5 | 0.0960434975810118 | 7.60901282902812 | 1.03895253556815 | 0.304701831403793 | 0.622251988866383 | -5.86813059593332 |
| lnc-NXF1-1 | lnc-NXF1-1 | 0.350977938398519 | 0.112583697466318 | 1.03831553577519 | 0.30499475579202 | 0.622674504847997 | -5.19769618649547 |
| ENSG00000180211.5 | RPL23P6 | 0.713432666451579 | -0.942571133738059 | 1.03830525940842 | 0.304999482965433 | 0.622674504847997 | -4.84012758354732 |
| lnc-MOCS1-3 | lnc-MOCS1-3 | 0.713432666451579 | -0.942571133738059 | 1.03830525940842 | 0.304999482965433 | 0.622674504847997 | -4.84012758354732 |
| ENSG00000157800.17 | SLC37A3 | 0.490184652916195 | 1.74619748913579 | 1.03802344273163 | 0.305253033985602 | 0.622999376449578 | -5.49050647823189 |
| lnc-SPON2-1 | lnc-SPON2-1 | 0.688290555939479 | 1.35263339585599 | 1.03813131566682 | 0.305267967322001 | 0.622999376449578 | -5.45732222787693 |
| ENSG00000233369.7 | GTF2IP4 | 0.132309845206771 | 3.894324112719 | 1.03747359508345 | 0.305382219098702 | 0.622999376449578 | -5.76331106447693 |
| ENSG00000228232.1 | GAPDHP1 | 0.758265420797637 | -0.471138131903472 | 1.03750778368814 | 0.305384828085987 | 0.622999376449578 | -4.93331795836798 |
| ENSG00000131408.13 | NR1H2 | 0.154868902683814 | 5.27105808754162 | 1.03746296501302 | 0.305387113245503 | 0.622999376449578 | -5.8636176754582 |
| ENSG00000101224.17 | CDC25B | 0.174666617896185 | 6.66838205500477 | 1.03746002122508 | 0.305388468592273 | 0.622999376449578 | -5.88118689496949 |
| ENSG00000142168.14 | SOD1 | -0.175142540679709 | 5.90998291864506 | -1.03745262674496 | 0.305391873096465 | 0.622999376449578 | -5.87829454572287 |
| ENSG00000080802.18 | CNOT4 | -0.141317305371697 | 5.34493917457739 | -1.03730582957887 | 0.305459465606403 | 0.622999376449578 | -5.86564084756171 |
| ENSG00000051108.14 | HERPUD1 | -0.297972911187281 | 6.74587705125645 | -1.03746319494479 | 0.30548296222707 | 0.622999376449578 | -5.87952586364464 |
| lnc-RHOV-1 | lnc-RHOV-1 | 0.276222782982145 | 1.35751527025787 | 1.03721705496504 | 0.305500346714971 | 0.622999376449578 | -5.44939204124741 |
| lnc-RELN-2 | lnc-RELN-2 | -0.161927269190525 | 0.274852460431513 | -1.03713418569361 | 0.305538511778068 | 0.622999376449578 | -5.25412430039765 |
| ENSG00000108479.11 | GALK1 | -0.201140368382374 | 1.20831652698073 | -1.03709792300531 | 0.305555213422921 | 0.622999376449578 | -5.45327708938944 |
| ENSG00000140988.15 | RPS2 | -0.186545625463374 | 9.1828131749604 | -1.03697665984054 | 0.305611068616224 | 0.622999376449578 | -5.82996486768572 |
| lnc-SLC22A5-6 | lnc-SLC22A5-6 | 0.138765889150648 | 4.43300348600062 | 1.03697621584109 | 0.305611273140302 | 0.622999376449578 | -5.80618768601904 |
| ENSG00000107263.18 | RAPGEF1 | 0.153715547170294 | 7.65329820928 | 1.03691779097687 | 0.305638186812856 | 0.622999376449578 | -5.8697358334917 |
| ENSG00000124374.8 | PAIP2B | -0.316630877178071 | 4.57369491373926 | -1.03707641405611 | 0.305659588329341 | 0.622999376449578 | -5.81697518559263 |
| ENSG00000164114.18 | MAP9 | -0.259912424751121 | 2.99948795740482 | -1.03680714930319 | 0.305689158849985 | 0.622999376449578 | -5.67378926061713 |
| lnc-CCDC117-2 | lnc-CCDC117-2 | -0.233651345212227 | 0.227614429187774 | -1.03677617386125 | 0.305703430119066 | 0.622999376449578 | -5.24641767854319 |
| lnc-EML4-5 | lnc-EML4-5 | 0.33001004479735 | 0.0694607558302341 | 1.03629383267046 | 0.305925717489858 | 0.623390658656939 | -5.20106424072271 |
| ENSG00000274267.1 | NA | 0.75198302418129 | -0.502555288218195 | 1.03631848316782 | 0.30597415500577 | 0.623427641307045 | -5.01824051041872 |
| ENSG00000161654.9 | LSM12 | 0.121383572966187 | 3.29922477408286 | 1.03610939119398 | 0.306010746833817 | 0.623440483230768 | -5.69547093733028 |
| ENSG00000177112.7 | IRAG1-AS1 | -0.43203124441913 | 0.00562573963511248 | -1.03597369633774 | 0.306073313862327 | 0.623449389170988 | -5.11406359438534 |
| ENSG00000139631.18 | CSAD | 0.119818043262661 | 3.99283779191935 | 1.03596852752933 | 0.30607569730228 | 0.623449389170988 | -5.77114865301468 |
| lnc-ITGA11-4 | lnc-ITGA11-4 | -0.158471914594345 | 2.83796017305945 | -1.03573562175515 | 0.306183107990718 | 0.62360646267296 | -5.64456762571982 |
| lnc-OR8D4-4 | lnc-OR8D4-4 | -0.55049681911547 | 1.22558919912922 | -1.03582667710731 | 0.306277091804398 | 0.623728202463668 | -5.43586539883068 |
| ENSG00000275713.2 | H2BC9 | 2.42858516085969 | -0.269945692275304 | 1.03593123104181 | 0.306324898951689 | 0.623728202463668 | -4.83970176796443 |
| lnc-CCDC167-5 | lnc-CCDC167-5 | -0.594810823311845 | -0.699836720129312 | -1.03540898247926 | 0.306333789957986 | 0.623728202463668 | -4.98382867946458 |
| ENSG00000271819.1 | RNU6-94P | 0.274207414591557 | -0.488468048821075 | 1.03533283331156 | 0.306368925645423 | 0.623738041202744 | -5.04976904010559 |
| ENSG00000234825.3 | XRCC6P2 | 0.63518360253891 | -0.983925660927803 | 1.03517521616586 | 0.306441659938173 | 0.623764087333788 | -4.84280120469673 |
| lnc-ATL3-2 | lnc-ATL3-2 | -0.201226839028001 | 3.9725641123356 | -1.03517376711431 | 0.306442328675085 | 0.623764087333788 | -5.7781048848782 |
| DIP2A-IT1 | DIP2A-IT1 | 0.230728793395991 | 0.623464573076579 | 1.03499663179157 | 0.30652408412417 | 0.623862436235035 | -5.32384503529483 |
| ENSG00000170525.20 | PFKFB3 | 0.164902277895779 | 2.84415102686096 | 1.03491626747968 | 0.306561180590707 | 0.623862436235035 | -5.64209573080203 |
| ENSG00000253919.1 | THAP12P7 | 0.288308233526429 | 1.29763901230137 | 1.03487209033702 | 0.306581574236864 | 0.623862436235035 | -5.42442517762431 |
| lnc-PIK3R3-3 | lnc-PIK3R3-3 | -0.103511084975341 | 1.31788395189252 | -1.0344886408696 | 0.306758626437184 | 0.624144685743354 | -5.45787666166829 |
| lnc-IGLL5-1 | lnc-IGLL5-1 | 1.42434758472286 | 3.82487601780137 | 1.03486365725944 | 0.306817241980425 | 0.624144685743354 | -5.70525481857921 |
| lnc-PAPD7-7 | lnc-PAPD7-7 | -0.264496163409083 | -0.113181672523466 | -1.03431956200144 | 0.306836718419145 | 0.624144685743354 | -5.18528848723582 |
| ENSG00000196620.9 | UGT2B15 | 0.974701649823871 | -2.33208190438881 | 1.03430905452009 | 0.306841571930088 | 0.624144685743354 | -4.76081741623724 |
| ENSG00000130725.7 | UBE2M | -0.134133400628841 | 3.23138097641139 | -1.03422867780651 | 0.306878700481843 | 0.624158526879227 | -5.69336389870001 |
| ENSG00000151014.5 | NOCT | -0.242557729820072 | 1.25270998846027 | -1.03403996798493 | 0.306965883640983 | 0.624274160950731 | -5.44715358194248 |
| ENSG00000198721.12 | ECI2 | -0.111937354808274 | 4.19928977164835 | -1.03391664211067 | 0.307022868864305 | 0.624316545876827 | -5.79416770512093 |
| ENSG00000258366.8 | RTEL1 | -0.128713211178234 | 2.86373521403523 | -1.03365879312187 | 0.307142036665105 | 0.624316545876827 | -5.6523948480716 |
| ENSG00000106615.9 | RHEB | 0.131538062930616 | 3.38450411404276 | 1.03363350371914 | 0.307153726153929 | 0.624316545876827 | -5.70602944698193 |
| ENSG00000137288.9 | UQCC2 | -0.202383637526101 | 2.70738832306267 | -1.0336112799828 | 0.30716399885549 | 0.624316545876827 | -5.63655601938213 |
| lnc-LSM10-2 | lnc-LSM10-2 | -0.194349043594955 | 0.492451733565664 | -1.03357884543532 | 0.307178991826004 | 0.624316545876827 | -5.30493985781391 |
| ENSG00000100284.20 | TOM1 | 0.149359999973009 | 4.60312138916788 | 1.03356018168332 | 0.307187619430925 | 0.624316545876827 | -5.83096102522828 |

| | | | | | | | |
|---|---|---|---|---|---|---|---|
| ENSG00000261150.2 | EPPK1 | -1.0002570206918 | -0.0361256540150945 | -1.0339296037456 | 0.307199046622966 | 0.624316545876827 | -5.14135758357114 |
| ENSG00000186665.9 | C17orf58 | -0.252627865413884 | 0.466215393674444 | -1.0333522228457 | 0.307283762826364 | 0.624427059952123 | -5.34553832378991 |
| ENSG00000196116.7 | TDRD7 | 0.0932497517428423 | 4.61873428731482 | 1.03287510273077 | 0.307504422655244 | 0.624727885289625 | -5.83356712470511 |
| lnc-DNAH7-2 | lnc-DNAH7-2 | -1.14318764484056 | -1.07161816884339 | -1.03321904200045 | 0.307508982792161 | 0.624727885289625 | -4.993009531664 |
| ENSG00000106263.17 | EIF3B | 0.173652095407388 | 6.18179200345628 | 1.03283525717522 | 0.307522855451318 | 0.624727885289625 | -5.88599110472211 |
| ENSG00000151490.13 | PTPRO | 0.290068633711631 | 4.50706377701699 | 1.03260167098046 | 0.307685737933285 | 0.624928381759214 | -5.79751676527219 |
| ENSG00000143127.12 | ITGA10 | -0.292727221591543 | 0.575526757102593 | -1.03246900886032 | 0.307692319625217 | 0.624928381759214 | -5.31328067074602 |
| ENSG00000123130.16 | ACOT9 | -0.140962809798123 | 4.16339646695264 | -1.03242511406587 | 0.307712634170414 | 0.624928381759214 | -5.80445807235477 |
| lnc-RAB1A-9 | lnc-RAB1A-9 | -0.298421725579256 | 0.337923474320685 | -1.03211276922563 | 0.307857214150087 | 0.625160323485719 | -5.26888191016311 |
| ENSG00000100485.11 | SOS2 | -0.0876213800488088 | 6.60625570021798 | -1.03176144144079 | 0.308019894403129 | 0.625425743700852 | -5.8862067117782 |
| lnc-ZRANB1-3 | lnc-ZRANB1-3 | 0.246425999797087 | 1.05222241150484 | 1.03169926674051 | 0.30804869016369 | 0.625425743700852 | -5.40784013044426 |
| lnc-POLE2-3 | lnc-POLE2-3 | -0.15799707544562 | 2.64026045971253 | -1.03152680618619 | 0.308128573669718 | 0.625434445737916 | -5.6235851803779 |
| ENSG00000278985.1 | | 0.19679943447457 | -0.0885438765003693 | 1.03152556729016 | 0.308129147576102 | 0.625434445737916 | -5.15889814099415 |
| lnc-INPP5B-1 | lnc-INPP5B-1 | 0.107550726961412 | 2.02440590585996 | 1.03145563913091 | 0.308161542294202 | 0.625434445737916 | -5.54870966938692 |
| ENSG00000125351.11 | UPF3B | 0.136675318935814 | 3.85744522444111 | 1.03142762613073 | 0.308174520170721 | 0.625434445737916 | -5.75769008212337 |
| lnc-STAM2-2 | lnc-STAM2-2 | 0.134934334730985 | 2.11903543825287 | 1.03134041698046 | 0.308214924866088 | 0.625437654406718 | -5.56736053145115 |
| lnc-STAM2-1 | lnc-STAM2-1 | 0.112125257859625 | 3.27236756591245 | 1.03129304661828 | 0.308236873453906 | 0.625437654406718 | -5.69607269491771 |
| lnc-FNBP1L-2 | lnc-FNBP1L-2 | -0.252941993327886 | -0.018368097171422 | -1.03114572319273 | 0.308305141146085 | 0.625479108031636 | -5.20901289073347 |
| lnc-P2RX7-1 | lnc-P2RX7-1 | -0.293419855047106 | 0.147400077592044 | -1.03110329355922 | 0.308324804387923 | 0.625479108031636 | -5.24149707893068 |
| ENSG00000284292.1 | | -0.1855618139454 | 5.63350622532449 | -1.0310289852817 | 0.308359243277044 | 0.625479108031636 | -5.88088972746458 |
| ENSG00000176681.14 | LRRC37A | 0.216009939973781 | 2.71482921761603 | 1.03098667023946 | 0.308378855769569 | 0.625479108031636 | -5.61660709088479 |
| ENSG00000210174.1 | MT-TR | 0.574828749227944 | -0.376013557224294 | 1.03077398194582 | 0.308477447072008 | 0.625617429607168 | -4.99020349757025 |
| ENSG00000196301.3 | HLA-DRB9 | -0.387590894118339 | -0.183399619743536 | -1.03053295346545 | 0.308589201512481 | 0.625711046957564 | -5.14464793928339 |
| lnc-APH1A-1 | lnc-APH1A-1 | 0.150016821773227 | 3.49682300803561 | 1.03048568393029 | 0.30861112159491 | 0.625711046957564 | -5.72546529936575 |
| ENSG00000136237.18 | RAPGEF5 | 1.4644896735308 | 1.46079423740378 | 1.03094035586549 | 0.308631255434807 | 0.625711046957564 | -5.41026943908983 |
| ENSG00000121775.17 | TMEM39B | -0.142979149429625 | 2.64662915102211 | -1.03041218882991 | 0.308645205254829 | 0.625711046957564 | -5.6276326533571 |
| ENSG00000125863.19 | MKKS | -0.169499277940229 | 1.6396894370738 | -1.03033333248158 | 0.308681778082395 | 0.625717125521223 | -5.51151528105072 |
| lnc-PAPSS2-6 | lnc-PAPSS2-6 | -0.367449198081719 | 0.178027254948567 | -1.03022005880785 | 0.308734318532292 | 0.625717125521223 | -5.28037083122676 |
| ENSG00000218739.9 | CEBPZOS | -0.171507580855972 | 4.34159320222644 | -1.03020769016655 | 0.3087400559288 | 0.625717125521223 | -5.8116673552353 |
| ENSG00000185015.7 | CA13 | -0.188795355154322 | 3.25055209487975 | -1.03010329597207 | 0.308788483791298 | 0.625717125521223 | -5.692195787663 |
| ENSG00000263813.1 | MIR3679 | 0.403518787346852 | 1.7376515041723 | 1.03013485223713 | 0.308825702900911 | 0.625717125521223 | -5.4844228772555 |
| ENSG00000072415.8 | PALS1 | 0.150948832363445 | 3.82796870564256 | 1.03001251267706 | 0.308830601864165 | 0.625717125521223 | -5.75441575843821 |
| ENSG00000094914.12 | AAAS | -0.170900155680316 | 4.23792843794854 | -1.02964533464701 | 0.309000990849866 | 0.625918855170832 | -5.80904769416601 |
| lnc-TIGD7-1 | lnc-TIGD7-1 | -0.200046128802354 | 0.469817993975098 | -1.02964278321304 | 0.309002175068039 | 0.625918855170832 | -5.29857340551118 |
| lnc-CNMD-2 | lnc-CNMD-2 | 0.286936188925517 | 0.29880912695409 | 1.02959542834358 | 0.30902415483963 | 0.625918855170832 | -5.31135258066639 |
| ENSG00000164663.14 | USP49 | 0.207191101456717 | 2.06132445523062 | 1.02953585765252 | 0.309051806107038 | 0.625918855170832 | -5.55398308139823 |
| lnc-SLMAP-2 | lnc-SLMAP-2 | -0.542806726633038 | -0.916453628525301 | -1.02937066147099 | 0.309128495020942 | 0.626012575070449 | -5.04551673341168 |
| ENSG00000224003.1 | YES1P1 | -0.284603871912647 | 1.97181974448771 | -1.02918280395453 | 0.309215719817899 | 0.626093611725382 | -5.5286987224201 |
| lnc-MAML2-6 | lnc-MAML2-6 | 0.186236808514081 | 0.908229198500982 | 1.02910024685131 | 0.309254057529781 | 0.626093611725382 | -5.40785060393067 |
| ENSG00000178922.16 | HYI | 0.271039079291682 | 3.13170505121827 | 1.02908795610761 | 0.309259765360344 | 0.626093611725382 | -5.66907620446923 |
| ENSG00000234289.5 | H2BC12L | 1.10177153294886 | -1.27845422201256 | 1.02904202246692 | 0.309299286822494 | 0.626099960600756 | -4.7519997506499 |
| ENSG00000152894.14 | PTPRK | 2.10133886216548 | 0.0179244846251433 | 1.02944684446002 | 0.30932373800462 | 0.626099960600756 | -5.03559959190123 |
| lnc-GLRX3-10 | lnc-GLRX3-10 | -0.294026471775575 | 0.0620288183394945 | -1.02865928132464 | 0.309458887359532 | 0.626311925125 | -5.18910726032138 |
| LINC01311 | LINC01311 | -0.195241708994482 | 1.63864571212398 | -1.02855763980418 | 0.309506113316593 | 0.626345917844403 | -5.51497093999534 |
| lnc-POU4F1-2 | lnc-POU4F1-2 | -0.274297167867224 | 0.387952106425571 | -1.02845052923254 | 0.309555885708753 | 0.626385056581131 | -5.27071830386841 |
| ENSG00000134644.15 | PUM1 | 0.0833029453996019 | 7.00621251994393 | 1.02834885004192 | 0.309603139300413 | 0.626395073330728 | -5.88683012470944 |
| ENSG00000063322.13 | MED29 | -0.120227131518896 | 5.0886185035043 | -1.02829110797621 | 0.309629976093189 | 0.626395073330728 | -5.86480183927438 |
| lnc-RTRAF-2 | lnc-RTRAF-2 | -0.15137901859236 | 4.07854911259496 | -1.02824343589756 | 0.309652133855131 | 0.626395073330728 | -5.78849770620537 |
| ENSG00000278845.4 | MRPL45 | -0.112681968518396 | 3.39111589374072 | -1.02801829397456 | 0.309756793439521 | 0.626545212201813 | -5.71702560911628 |
| ENSG00000233954.6 | UQCRHL | 0.235382636241781 | 1.66243655117658 | 1.02776031098606 | 0.309876749283618 | 0.626726258278934 | -5.49162974883525 |
| lnc-TAF1D-1 | lnc-TAF1D-1 | -0.221774888106105 | 1.46141446268016 | -1.02749569571238 | 0.309999821984101 | 0.626885961963212 | -5.48617755132418 |
| lnc-RUBCN-2 | lnc-RUBCN-2 | -0.192913501773265 | 3.41489686909818 | -1.02740301533042 | 0.310042935581004 | 0.626885961963212 | -5.72325513313604 |
| LINC01560 | LINC01560 | -0.232110706451598 | -0.00540996184047875 | -1.02739410191071 | 0.310047082193339 | 0.626885961963212 | -5.23241277224849 |
| ENSG00000051180.16 | RAD51 | 0.250428571453442 | 0.69693094461584 | 1.0272327482309 | 0.31012215210231 | 0.626976157226388 | -5.36745290736512 |
| ENSG00000086758.15 | HUWE1 | -0.0719332196550813 | 8.39144185873789 | -1.02689213913251 | 0.31028066152398 | 0.62723500846082 | -5.86402299589091 |
| ENSG00000163191.5 | S100A11 | -0.416757805074395 | 1.26546916938198 | -1.02688592607443 | 0.310336718271352 | 0.62726862622118 | -5.4605077725877 |
| lnc-TTLL7-17 | lnc-TTLL7-17 | -0.398148041568609 | 3.70644492034612 | -1.02702683962999 | 0.310358241725526 | 0.62726862622118 | -5.74627231038756 |
| ENSG00000056050.6 | HPF1 | -0.197965549006024 | 1.65377376653917 | -1.026617372052 | 0.310408570371834 | 0.627308748548205 | -5.51380990036957 |
| ENSG00000172006.11 | ZNF554 | -0.142820000118144 | 2.1555197398414 | -1.02647824718394 | 0.310473349158542 | 0.627378062608509 | -5.57196032960423 |
| ENSG00000186364.11 | NUDT17 | -0.157647882239969 | 2.83963071107065 | -1.02640582747805 | 0.310507072600644 | 0.627384615229122 | -5.65352721885349 |
| lnc-PMVK-1 | lnc-PMVK-1 | 0.18393591263822 | 3.77637791108166 | 1.02611785109702 | 0.310641198371782 | 0.627594011198113 | -5.7608652607944 |
| LINC01684 | LINC01684 | -1.42121485697876 | -0.228337078144263 | -1.02637860661661 | 0.310749697718151 | 0.627700925049643 | -5.24420274120858 |
| lnc-ME1-2 | lnc-ME1-2 | 0.16509920740858 | -0.0463335196051612 | 1.02587334276643 | 0.310755109860364 | 0.627700925049643 | -5.18027309339764 |
| ENSG00000197894.10 | ADH5 | 0.109654580909414 | 5.23179561056696 | 1.02566181793964 | 0.310853678027693 | 0.627838411818664 | -5.87474038435841 |
| ENSG00000101132.9 | PFDN4 | -0.117894896437794 | 2.85764563245971 | -1.02544586292796 | 0.310954332648093 | 0.627980085252719 | -5.66471346357281 |
| lnc-INTS7-3 | lnc-INTS7-3 | 0.159872374134449 | 0.889639586978348 | 1.02525257273547 | 0.311044442290678 | 0.628043990030348 | -5.38876067499161 |
| ENSG00000274072.1 | | 0.324417097986672 | 1.00705431742424 | 1.02524708338796 | 0.31104700162121 | 0.628043990030348 | -5.41376864409226 |
| lnc-RAD50-4 | lnc-RAD50-4 | -0.289939250650768 | -0.335850290933435 | -1.02503692945832 | 0.311144993733225 | 0.628158035945176 | -5.12467828929777 |

| | | | | | | | |
|---|---|---|---|---|---|---|---|
| ENSG00000217128.11 | FNIP1 | 0.127825898630898 | 6.07794442278545 | 1.02499505680548 | 0.311164520939466 | 0.628158035945176 | -5.89323406898627 |
| lnc-TM2D3-3 | lnc-TM2D3-3 | -0.204308465884194 | 0.737992567857167 | -1.02480319991687 | 0.311254003621782 | 0.628276750022352 | -5.37192203058548 |
| ENSG00000080007.7 | DDX43 | 2.97725367105082 | -2.18800716118741 | 1.02519052417061 | 0.311284375296504 | 0.628276750022352 | -4.68370136002743 |
| ENSG00000198832.10 | SELENOM | 0.444012697790769 | 1.99631000782317 | 1.02490628098741 | 0.311316241449055 | 0.628279458549456 | -5.54606688584022 |
| lnc-SYNDIG1L-2 | lnc-SYNDIG1L-2 | -0.10613814740145 | 2.90665645099702 | -1.02438139734233 | 0.311450795521467 | 0.628471239883697 | -5.66283756687118 |
| ENSG00000168283.13 | BMI1 | -0.175690128923694 | 5.00953459898382 | -1.02429872026017 | 0.311489378453832 | 0.628471239883697 | -5.86927083888913 |
| lnc-SPINK8-2 | lnc-SPINK8-2 | -0.155804713113196 | 0.566964149637101 | -1.02421320319019 | 0.311529290157922 | 0.628471239883697 | -5.33145003994697 |
| ENSG00000004866.20 | ST7 | -0.298174478522999 | 3.34289561249152 | -1.02428186049903 | 0.311533404291569 | 0.628471239883697 | -5.72276622353329 |
| ENSG00000283977.1 | | 0.159046738005776 | 2.98248827942874 | 1.02407460194051 | 0.311593984215842 | 0.628481354899398 | -5.66844864437948 |
| ENSG00000161021.12 | MAML1 | 0.0907527581277368 | 5.30820757009346 | 1.02406281509805 | 0.311599486311439 | 0.628481354899398 | -5.87808637040255 |
| ENSG00000118412.12 | CASP8AP2 | -0.152158577040586 | 6.10584933945106 | -1.02390800124947 | 0.311671759541512 | 0.628565532690863 | -5.89334178496101 |
| ENSG00000148834.12 | GSTO1 | 0.181425060265211 | 3.02279944972374 | 1.02354526146575 | 0.311841145648432 | 0.628739860594438 | -5.67929914303455 |
| ENSG00000207650.1 | MIR570 | 0.373826469588451 | 0.0772519692399721 | 1.0234902433662 | 0.311866842560439 | 0.628739860594438 | -5.23057874906643 |
| lnc-VN1R4-3 | lnc-VN1R4-3 | -1.02490163297477 | -0.305733566579105 | -1.02387013406577 | 0.311867861460199 | 0.628739860594438 | -5.12029647326281 |
| ENSG00000106268.15 | NUDT1 | 0.329382927601885 | 2.65091815195066 | 1.02351878898609 | 0.311880385593413 | 0.628739860594438 | -5.60850880017102 |
| lnc-EPHB3-5 | lnc-EPHB3-5 | 0.123739943212932 | 0.898530217971738 | 1.02338653619172 | 0.311915284262252 | 0.628748633431586 | -5.40793875132973 |
| ENSG00000279696.1 | | -0.22080423226582 | 1.45140846567109 | -1.02323453181045 | 0.311986294907549 | 0.628830190764011 | -5.48905104509674 |
| ENSG00000146376.10 | ARHGAP18 | -0.268768289910974 | 2.62604439656275 | -1.02293659030577 | 0.312125513815715 | 0.629049197186808 | -5.62001159990305 |
| lnc-DCAF4L1-2 | lnc-DCAF4L1-2 | 0.161159574672662 | 2.03507580942398 | 1.02278971555473 | 0.31219415947246 | 0.629125943256476 | -5.56515449760474 |
| ENSG00000131236.16 | CAP1 | -0.197830764686621 | 7.26488202171091 | -1.02251395069774 | 0.3123230730334 | 0.629324112799459 | -5.88906715954968 |
| lnc-FAM177B-1 | lnc-FAM177B-1 | -0.204626291827352 | 3.88652111249047 | -1.02241395258069 | 0.312369828764977 | 0.629356713534604 | -5.78505280764209 |
| lnc-BTBD2-2 | lnc-BTBD2-2 | -0.191287573920204 | 1.33266563635919 | -1.02227312884308 | 0.312435681271625 | 0.629427779936758 | -5.47305332814539 |
| lnc-LUZP1-1 | lnc-LUZP1-1 | -0.377335821242639 | -0.573566012354182 | -1.02212582483404 | 0.312504574241088 | 0.629504957095743 | -5.07247515237503 |
| ENSG00000181788.3 | SIAH2 | 0.175597183578643 | 4.8812123465895 | 1.02205089310676 | 0.312539623218945 | 0.629513950945841 | -5.85044926450343 |
| ENSG00000262526.2 | | -0.115996180024541 | 4.64731357366773 | -1.0218076485279 | 0.312653418256357 | 0.629681536983423 | -5.84803819648196 |
| lnc-ATP8A2-2 | lnc-ATP8A2-2 | 0.231786812654995 | 0.58505670093484 | 1.02157315871896 | 0.312763144375623 | 0.629840896260977 | -5.35339760960937 |
| ENSG00000170145.4 | SIK2 | 0.197081887706231 | 4.12236317162246 | 1.021367870803 | 0.312859227450335 | 0.629903755148591 | -5.7929459056113 |
| ENSG00000053918.16 | KCNQ1 | 2.22163694413689 | -0.343333723517441 | 1.02184515095618 | 0.312864837727265 | 0.629903755148591 | -4.88066390632513 |
| RC3H1-IT1 | RC3H1-IT1 | 0.199630823829533 | 2.95049879799291 | 1.02131031857792 | 0.31288616783944 | 0.629903755148591 | -5.67694370064402 |
| lnc-RGS11-1 | lnc-RGS11-1 | -0.177995843207971 | 3.19306631684588 | -1.02123367286029 | 0.312922048412616 | 0.629914378726345 | -5.70375627845158 |
| lnc-TRAF6-2 | lnc-TRAF6-2 | 0.1142204096621 | 3.78042156698879 | 1.0211539933419 | 0.312959352189923 | 0.629927864866536 | -5.76777296354014 |
| lnc-ARMCX3-1 | lnc-ARMCX3-1 | 0.23006988980713 | 1.16524357652884 | 1.02082207354245 | 0.313114780613561 | 0.630179087647299 | -5.44611266146362 |
| ENSG00000101844.17 | ATG4A | 0.10550672715993 | 3.19978301439126 | 1.02052100509911 | 0.313255807753726 | 0.630313995515133 | -5.69830021368893 |
| ENSG00000169727.12 | GPS1 | -0.14287408985587 | 4.92416706871242 | -1.02047831112739 | 0.313275810062671 | 0.630313995515133 | -5.86592842649485 |
| ENSG00000279330.1 | | 0.239273383218753 | 2.52100697920855 | 1.02040381561095 | 0.313310713616762 | 0.630313995515133 | -5.60906106660549 |
| lnc-PDS5B-2 | lnc-PDS5B-2 | -0.263655257774176 | 1.74463867040626 | -1.02035106829314 | 0.31333542904074 | 0.630313995515133 | -5.53760564857457 |
| lnc-TUBE1-1 | lnc-TUBE1-1 | -0.234441230008414 | 0.251805119228293 | -1.02023355423165 | 0.313390496521484 | 0.630313995515133 | -5.26815584850736 |
| ENSG00000117713.19 | ARID1A | -0.122219995043307 | 7.16271789931616 | -1.02018982950837 | 0.31341098775629 | 0.630313995515133 | -5.89291207206686 |
| ENSG00000130177.15 | CDC16 | 0.132001568847373 | 5.93800744953544 | 1.02015530208768 | 0.313427169395566 | 0.630313995515133 | -5.89640088346014 |
| lnc-MON2-2 | lnc-MON2-2 | -0.170282754642464 | 5.73635575063942 | -1.02007706625355 | 0.313463837550078 | 0.630313995515133 | -5.89510563057778 |
| lnc-NKAIN1-1 | lnc-NKAIN1-1 | 0.131566238731301 | 1.79855285408916 | 1.01999139402642 | 0.313503994402564 | 0.630313995515133 | -5.54018146652024 |
| ENSG00000124523.15 | SIRT5 | -0.161444291827149 | 3.87006934099541 | -1.0198753087759 | 0.313558412242191 | 0.630313995515133 | -5.77406310533838 |
| ENSG00000112419.14 | PHACTR2 | 0.225391720842487 | 5.28103351259047 | 1.01986800516324 | 0.313569077793899 | 0.630313995515133 | -5.88563223434846 |
| ENSG00000176533.12 | GNG7 | 0.477423708846298 | 4.14865352113507 | 1.02025849924443 | 0.313572800162058 | 0.630313995515133 | -5.82041960498294 |
| ENSG00000164109.13 | MAD2L1 | 0.239715825358274 | 1.54549679751734 | 1.01978538527617 | 0.313600570536942 | 0.630313995515133 | -5.50457781954117 |
| ENSG00000197892.12 | KIF13B | -0.101006480740353 | 5.68865384958604 | -1.01976413374755 | 0.313610534327867 | 0.630313995515133 | -5.89406094542377 |
| ENSG00000197102.11 | DYNC1H1 | 0.104198649982234 | 8.58882238346102 | 1.0195228066282 | 0.313723695815899 | 0.630479870238102 | -5.86621594655882 |
| ENSG00000168280.16 | KIF5C | -0.960259357263635 | 0.619841403621485 | -1.0198400177837 | 0.313784892415054 | 0.630541290694041 | -5.31126323331498 |
| ENSG00000131844.15 | MCCC2 | -0.0709261608350644 | 5.25563363368848 | -1.0193123795959 | 0.313822390548211 | 0.630543360927323 | -5.88289312978468 |
| ENSG00000183604.14 | SMG1P5 | 0.293728443552762 | 3.51087231644946 | 1.01925220623798 | 0.313889209586193 | 0.630543360927323 | -5.74364137128643 |
| ENSG00000185624.14 | P4HB | -0.16183293393245 | 6.16937128895798 | -1.019152306687 | 0.313897482296309 | 0.630543360927323 | -5.89911291604329 |
| ENSG00000250644.3 | | -0.168673393382142 | 2.92668249119595 | -1.01912890904796 | 0.313908459380182 | 0.630543360927323 | -5.6713888524524 |
| ENSG00000205639.10 | MFSD2B | 0.283714186768729 | -0.300797611354217 | 1.01899752656911 | 0.31397010278509 | 0.630584282388174 | -5.10502133423529 |
| ENSG00000196172.9 | ZNF681 | -0.409145627745785 | 3.11732783137055 | -1.01921995701478 | 0.313990103978837 | 0.630584282388174 | -5.66471451910406 |
| lnc-ARID4A-4 | lnc-ARID4A-4 | 0.204090784902907 | 0.361507231484103 | 1.01853676204777 | 0.314186354270571 | 0.630916851702553 | -5.30372904143251 |
| ENSG00000176410.7 | DNAJC30 | -0.145035549917268 | 2.51090786370364 | -1.01791924034265 | 0.314476335955561 | 0.631437559552562 | -5.62457145233562 |
| ENSG00000198837.9 | DENND4B | -0.0995872729032055 | 6.49584740538797 | -1.01783158221988 | 0.314517514052234 | 0.631458641410177 | -5.90072783277471 |
| lnc-COL9A2-1 | lnc-COL9A2-1 | -0.168486166793381 | 2.61108105773043 | -1.01767482911904 | 0.31459115921355 | 0.631513151683906 | -5.63647873694262 |
| lnc-WDR34-1 | lnc-WDR34-1 | -0.252404573929592 | 0.43624570873612 | -1.01760345179077 | 0.314624697335386 | 0.631513151683906 | -5.27598290399307 |
| ENSG00000105854.12 | PON2 | 0.986748809436572 | 1.56889853253326 | 1.01796654806401 | 0.314682235235616 | 0.631513151683906 | -5.485442059994 |
| ENSG00000170906.15 | NDUFA3 | -0.154691308209411 | 3.42295598853742 | -1.01733336495116 | 0.31475162528415 | 0.631513151683906 | -5.73137426131002 |
| ITPKB-AS1 | ITPKB-AS1 | 0.486957265297348 | 2.16162718122667 | 1.01762645156219 | 0.314754325788834 | 0.631513151683906 | -5.5707765187768 |
| lnc-SRSF7-4 | lnc-SRSF7-4 | 0.163151053828884 | 0.200114673471377 | 1.01731764727563 | 0.314759012916048 | 0.631513151683906 | -5.25352789304621 |
| ENSG00000183735.9 | TBK1 | 0.116351653340316 | 5.22607904045673 | 1.01725609459177 | 0.314787945084081 | 0.631513151683906 | -5.88476775620861 |
| lnc-ALDH8A1-2 | lnc-ALDH8A1-2 | 0.165196087470866 | 1.93154312130444 | 1.0172266528372 | 0.314801784499858 | 0.631513151683906 | -5.56016089381277 |
| ENSG00000144642.21 | RBMS3 | -2.34186286069322 | 0.169399277636688 | -1.01764121143959 | 0.31483500478048 | 0.631513151683906 | -5.06524181814174 |
| ENSG00000119632.3 | IFI27L2 | -0.14283200849018 | 2.63757245218794 | -1.01712094387034 | 0.314851477558191 | 0.631513151683906 | -5.64779679229898 |
| ENSG00000187953.10 | PMS2CL | 0.14957117325414 | 3.07104150806047 | 1.01624784385356 | 0.315262120034043 | 0.632243282992352 | -5.6894075543414 |

| | | | | | | | |
|---|---|---|---|---|---|---|---|
| lnc-FUS-1 | lnc-FUS-1 | -0.240253200598449 | 1.1308225109815 | -1.01616925785542 | 0.315299099015121 | 0.632243282992352 | -5.44387749430206 |
| lnc-C7orf77-2 | lnc-C7orf77-2 | -0.137896793624549 | 0.705307460213656 | -1.01615109279362 | 0.315307647083345 | 0.632243282992352 | -5.3532987193977 |
| ENSG00000162616.8 | DNAJB4 | 0.217360118617575 | 3.19322790459718 | 1.0160745516183 | 0.315343667365316 | 0.63225391636278 | -5.68972004685685 |
| ENSG00000055955.15 | ITIH4 | -0.176175986262477 | 3.5967036022239 | -1.015678024754 | 0.315530317749636 | 0.632566526759595 | -5.75161461759821 |
| ENSG00000179151.12 | EDC3 | -0.140308399578899 | 3.53365846413341 | -1.01554448485016 | 0.315593193641919 | 0.632630960725712 | -5.74078612224431 |
| ENSG00000189077.10 | TMEM120A | -0.112511891075991 | 1.83255701667528 | -1.01543737629227 | 0.315643630764778 | 0.632670450095573 | -5.5442258679656 |
| lnc-PADI2-1 | lnc-PADI2-1 | 0.147749370085281 | 2.90315518123944 | 1.01534407974928 | 0.315687568315338 | 0.632670945808677 | -5.6740487250948 |
| lnc-PEAK1-3 | lnc-PEAK1-3 | 0.237308174823407 | 1.32300561013308 | 1.01526610509739 | 0.315724293287057 | 0.632670945808677 | -5.48297654011081 |
| lnc-SRPK2-7 | lnc-SRPK2-7 | -0.220649057363795 | 1.45553997580034 | -1.01524105802925 | 0.315736090722768 | 0.632670945808677 | -5.48923500283503 |
| ENSG00000120333.4 | MRPS14 | -0.0981669725948806 | 4.03339967277196 | -1.01509638482824 | 0.315804239207107 | 0.632745902423819 | -5.79501846879557 |
| ENSG00000270532.1 | PEBP1P2 | 0.888948629503463 | -1.69603370716461 | 1.0148494622727 | 0.31592057547544 | 0.632917384174711 | -4.7699140712206 |
| ENSG00000137460.8 | FHDC1 | 0.493139933756939 | 3.25559789576514 | 1.01504136798048 | 0.315991008067434 | 0.632996877766617 | -5.67527832559821 |
| ENSG00000141068.14 | KSR1 | 0.2426055875029 | 2.84467857681818 | 1.0144083174342 | 0.316128491081189 | 0.633210659003904 | -5.66780201396879 |
| ENSG00000164897.12 | TMUB1 | -0.184963542583834 | 2.97709988437829 | -1.01424020442073 | 0.316207748758769 | 0.633253127450845 | -5.69388575807945 |
| lnc-CASP10-2 | lnc-CASP10-2 | -0.130622439814288 | 2.40438997132489 | -1.01423283166502 | 0.316211224988572 | 0.633253127450845 | -5.61459449137354 |
| ENSG00000203965.12 | EFCAB7 | -0.128512594040827 | 3.05102411980409 | -1.01376726186857 | 0.316430792213291 | 0.633599458010753 | -5.69803605457028 |
| ENSG00000226396.1 | | 0.262490630242917 | 1.2998630621082 | 1.01373559912101 | 0.316445728433685 | 0.633599458010753 | -5.4631627519652 |
| ENSG00000163082.9 | SGPP2 | -0.46220699991736 | 3.48842619418155 | -1.01361921549722 | 0.316661264134168 | 0.633940227764598 | -5.73277478646614 |
| ENSG00000214160.9 | ALG3 | -0.155559917578757 | 2.41136973012827 | -1.01324435991907 | 0.316677521346528 | 0.633940227764598 | -5.62262019178052 |
| ENSG00000184277.12 | TM2D3 | 0.147592081385917 | 5.14196101144588 | 1.01299246697485 | 0.316796422633041 | 0.634116577560623 | -5.88447460782325 |
| ENSG00000114353.16 | GNAI2 | -0.149317997330879 | 7.17564239397318 | -1.01292509917973 | 0.316828227462609 | 0.634118573109965 | -5.89964137920271 |
| ENSG00000132434.9 | LANCL2 | 0.145146903171098 | 4.49143599034571 | 1.0127577341533 | 0.316907251105812 | 0.634215065582005 | -5.84348516571651 |
| lnc-ACER2-2 | lnc-ACER2-2 | -0.27376802661195 | 0.788207506041709 | -1.01268434478792 | 0.31694190710189 | 0.634222756550476 | -5.39149798304589 |
| ENSG00000107201.9 | RIGI | -0.151158735707045 | 4.88194583324359 | -1.01246836928261 | 0.317043910195782 | 0.634365199140642 | -5.86528815815908 |
| ENSG00000285851.1 | | -0.103248247350961 | 3.28246939529692 | -1.01223570862573 | 0.317153818466148 | 0.634424098976616 | -5.72301577155442 |
| lnc-REELD1-2 | lnc-REELD1-2 | 0.16812744325952 | 2.80932224332566 | 1.01221120683742 | 0.317165394551036 | 0.634424098976616 | -5.67494905459979 |
| lnc-FLNB-3 | lnc-FLNB-3 | -0.224917987155094 | 0.073375078724819 | -1.01209283030038 | 0.317221326626732 | 0.634424098976616 | -5.23358953693732 |
| ENSG00000149212.11 | SESN3 | 0.387187589324614 | 7.35458572250802 | 1.01237188260438 | 0.317249923656622 | 0.634424098976616 | -5.90031240388391 |
| ENSG00000177156.10 | TALDO1 | -0.186500253902273 | 4.39811355530873 | -1.01202328554286 | 0.317254189155579 | 0.634424098976616 | -5.83777192654915 |
| ENSG00000198744.5 | MTCO3P12 | -0.464143326237848 | 1.45298891921325 | -1.01224747820998 | 0.317258283572186 | 0.634424098976616 | -5.51058100778837 |
| ENSG00000275023.4 | MLLT6 | -0.10663072116256 | 7.32551143970881 | -1.0118381824667 | 0.317341668626473 | 0.634529198109452 | -5.89830348780045 |
| ENSG00000151929.9 | BAG3 | 0.213118052302561 | 1.9993398030542 | 1.01175543971062 | 0.317380778040933 | 0.634545755649979 | -5.5773849924357 |
| ENSG00000123992.19 | DNPEP | 0.141155228471144 | 4.60867528171622 | 1.01165600107333 | 0.317427783305185 | 0.634578094771817 | -5.85364667676884 |
| ENSG00000136631.14 | VPS45 | -0.115513624970578 | 4.82773895724777 | -1.01126083832228 | 0.317614625913752 | 0.634602551878862 | -5.86879188853594 |
| lnc-ABLIM2-4 | lnc-ABLIM2-4 | -0.167316713541648 | 0.0491499090171931 | -1.01126069051756 | 0.317614695813403 | 0.634602551878862 | -5.23051738226193 |
| lnc-CALR3-6 | lnc-CALR3-6 | -0.236582192207717 | 0.368447612164903 | -1.01119614407282 | 0.317645222046518 | 0.634602551878862 | -5.29865477070625 |
| ENSG00000106003.12 | LFNG | 0.200346148304509 | 5.6681165329535 | 1.01114908545466 | 0.317667478940355 | 0.634602551878862 | -5.90031237992693 |
| ENSG00000105726.16 | ATP13A1 | -0.122336614295587 | 6.23905996068731 | -1.0111160893307 | 0.317683085453372 | 0.634602551878862 | -5.90746555074613 |
| ENSG00000182858.13 | ALG12 | 0.120151153113941 | 3.76994437718477 | 1.01110972465271 | 0.317686095879854 | 0.634602551878862 | -5.77336399925676 |
| lnc-FGF9-3 | lnc-FGF9-3 | 1.57069487686544 | -2.71620505919993 | 1.01107793240446 | 0.317751541996711 | 0.634602551878862 | -4.68020009899266 |
| ENSG00000132182.11 | NUP210 | -0.103099127259781 | 7.86914081158305 | -1.01093739271838 | 0.317767614452322 | 0.634602551878862 | -5.8905909981912 |
| ENSG00000179344.16 | HLA-DQB1 | -0.394978460630128 | 8.7826193968788 | -1.01118679208005 | 0.317809897506435 | 0.634602551878862 | -5.86708263983983 |
| ENSG00000227615.1 | | 0.370023728640075 | 2.18410086330266 | 1.01093923811932 | 0.317819102678874 | 0.634602551878862 | -5.56700841370388 |
| ENSG00000138629.15 | UBL7 | -0.194988728354711 | 4.50935437514538 | -1.01080694750885 | 0.317829328690336 | 0.634602551878862 | -5.84588047152294 |
| ENSG00000255837.1 | TAS2R20 | -0.239011647642523 | 1.41243741200113 | -1.01079203437515 | 0.317836384681656 | 0.634602551878862 | -5.50649212227939 |
| ENSG00000103051.18 | COG4 | -0.151942282298116 | 5.53999794990893 | -1.01078264903979 | 0.317840825308225 | 0.634602551878862 | -5.89736132782917 |
| ENSG00000271698.1 | | -0.263399141245005 | -0.378069591530879 | -1.0103654453244 | 0.318038265789345 | 0.634910733775532 | -5.15083144907665 |
| lnc-GFPT2-1 | lnc-GFPT2-1 | 0.125434034364714 | 0.592745090781625 | 1.01032614043041 | 0.31805687100809 | 0.634910733775532 | -5.35525882509272 |
| ENSG00000154237.12 | LRRK1 | -0.122869087811947 | 6.17455546393715 | -1.01022131573857 | 0.318106494046259 | 0.634948212466462 | -5.90795020867849 |
| ENSG00000026297.15 | RNASET2 | 0.140772443994039 | 6.17492705578072 | 1.0101404051375 | 0.318144799967266 | 0.634963096841486 | -5.90815731166485 |
| lnc-RABEPK-2 | lnc-RABEPK-2 | 0.17399999949445 | -0.267052437359313 | 1.01007353721913 | 0.318176459950817 | 0.634964715451586 | -5.14494357120358 |
| ENSG00000173269.13 | MMRN2 | -0.228947129163105 | 0.499270349726712 | -1.00998066141956 | 0.318220437443774 | 0.634979615994114 | -5.33561407126692 |
| WEE2-AS1 | WEE2-AS1 | -0.320685397181651 | 0.821156195329632 | -1.00992247554902 | 0.318247991056719 | 0.634979615994114 | -5.38065746616732 |
| lnc-TSC22D3-2 | lnc-TSC22D3-2 | -0.496867251055945 | 0.0077598380324062 | -1.0098783573614 | 0.318282458059093 | 0.634979615994114 | -5.21485768909037 |
| ENSG00000279691.1 | | -0.233244493860651 | 0.447003135594103 | -1.0096792233469 | 0.318363199383828 | 0.634979615994114 | -5.34134012197504 |
| ENSG00000182326.14 | C1S | -0.22440837430713 | 0.935272477384849 | -1.00962132762161 | 0.318390623938783 | 0.634979615994114 | -5.4168438125104 |
| ENSG00000162511.7 | LAPTM5 | -0.172728630702764 | 10.1102239469164 | -1.0096116300319 | 0.318395217734812 | 0.634979615994114 | -5.82995219847698 |
| ENSG00000273542.1 | H4C12 | 0.32798370984351 | 0.743841415678897 | 1.00950518820361 | 0.318445642708186 | 0.634979615994114 | -5.3651851695644 |
| lnc-FBXL2-3 | lnc-FBXL2-3 | 0.224176456581793 | -0.0778501558838792 | 1.00942752661339 | 0.318482436959566 | 0.634979615994114 | -5.20025360060699 |
| lnc-LRRTM2-4 | lnc-LRRTM2-4 | 0.135248791128917 | 0.761770619435569 | 1.00935330298687 | 0.318517605078538 | 0.634979615994114 | -5.37594254128765 |
| ENSG00000280383.1 | | 0.204406273283452 | 3.03448137100915 | 1.00935129516575 | 0.318518556446864 | 0.634979615994114 | -5.68825736008906 |
| ENSG00000168301.12 | KCTD6 | 0.139231514462227 | 1.83960289024683 | 1.00934134043165 | 0.318523273339126 | 0.634979615994114 | -5.55597881407335 |
| ENSG00000105329.9 | TGFB1 | -0.223495105776147 | 4.05324570334083 | -1.00896176207123 | 0.318703165844296 | 0.635276705653025 | -5.80796228119837 |
| lnc-SFI1-3 | lnc-SFI1-3 | -0.103959154227283 | 2.69147955651434 | -1.00842153470647 | 0.318959313057855 | 0.635659010251207 | -5.65109157387414 |
| lnc-KATNAL1-5 | lnc-KATNAL1-5 | 0.122128610161965 | 4.32775221209326 | 1.00841559429199 | 0.318962130463435 | 0.635659010251207 | -5.84006463286741 |
| ENSG00000169231.13 | THBS3 | -0.137016574877606 | 3.42583959240203 | -1.00836187919476 | 0.318987607097348 | 0.635659010251207 | -5.73812120786337 |
| ENSG00000132646.10 | PCNA | 0.220493140214915 | 3.82146270334175 | 1.00805649432375 | 0.319132474873386 | 0.635841146553003 | -5.77646135628852 |
| ENSG00000034713.7 | GABARAPL2 | 0.193141584474135 | 5.54147655177539 | 1.00798303482238 | 0.319167329076084 | 0.635841146553003 | -5.89656965772298 |

| | | | | | | | |
|---|---|---|---|---|---|---|---|
| ENSG00000144935.14 | TRPC1 | -1.11205475711259 | -1.40477824479367 | -1.00824047684149 | 0.319171681785241 | 0.635841146553003 | -4.90995057618335 |
| ENSG00000140265.12 | ZSCAN29 | 0.0816458430121401 | 5.31030541951504 | 1.00783548800007 | 0.319237343157041 | 0.635910406871323 | -5.89313057621576 |
| ENSG00000237984.3 | PTENP1 | 0.445688635254454 | -0.385224651182202 | 1.00763025788943 | 0.319334746483056 | 0.635933208260332 | -5.06482564230791 |
| ENSG00000101004.14 | NINL | -0.996202627522158 | 2.20414033159405 | -1.00809696740992 | 0.319339227955751 | 0.635933208260332 | -5.5830817949245 |
| lnc-GABRB2-6 | lnc-GABRB2-6 | 0.144306719198241 | 1.77513218934821 | 1.0076160761726 | 0.319341477946791 | 0.635933208260332 | -5.54685643110394 |
| ENSG00000146006.7 | LRRTM2 | -0.347522908639816 | -0.368973108076868 | -1.00719516622372 | 0.319541309952912 | 0.636269592992241 | -5.10829398104572 |
| ENSG00000113732.8 | ATP6V0E1 | -0.149573053497676 | 5.50389004229547 | -1.00705567128586 | 0.31960755552244 | 0.636339941508839 | -5.90045672495195 |
| ENSG00000116688.16 | MFN2 | 0.0723507684850846 | 5.84669435477932 | 1.00664070751902 | 0.319804675104299 | 0.636670821904613 | -5.90916102651905 |
| lnc-PRSS12-3 | lnc-PRSS12-3 | 0.188698036007247 | -0.265861352584754 | 1.00647775590205 | 0.319882104266661 | 0.636725624655211 | -5.13020018907813 |
| ENSG00000181524.6 | RPL24P4 | 0.527510568330252 | 0.721608939287147 | 1.00644218028756 | 0.319952094817111 | 0.636725624655211 | -5.27250102879777 |
| lnc-FA2H-1 | lnc-FA2H-1 | -0.127926745674829 | 1.17695369848974 | -1.00631755386914 | 0.319958239295213 | 0.636725624655211 | -5.45235750153584 |
| lnc-SIX5-1 | lnc-SIX5-1 | -0.207348125736929 | 0.216410809608345 | -1.00625976516949 | 0.319985706030169 | 0.636725624655211 | -5.31024265036698 |
| ENSG00000221926.12 | TRIM16 | -0.175080076141877 | 1.41512625550323 | -1.00623772632195 | 0.319996181425768 | 0.636725624655211 | -5.49180591897233 |
| ENSG00000276259.1 | | -0.227200644225208 | -0.231765298151462 | -1.0061922238155 | 0.320017810185986 | 0.636725624655211 | -5.17625235763118 |
| ENSG00000164818.15 | DNAAF5 | -0.158451091860675 | 2.77546562167509 | -1.00573356228192 | 0.320235881599484 | 0.63709792682797 | -5.68174341945341 |
| lnc-HERC6-2 | lnc-HERC6-2 | -0.234927935130089 | 1.26884000239426 | -1.00557902152465 | 0.320309380924119 | 0.637182563792513 | -5.47535195798496 |
| lnc-UCN-2 | lnc-UCN-2 | -0.141276352873308 | 1.97650670185065 | -1.00515545034155 | 0.320510889212398 | 0.637465736444117 | -5.57094110729258 |
| lnc-INAFM2-1 | lnc-INAFM2-1 | 0.219494937154513 | 1.53787748285853 | 1.00510963665902 | 0.320532689596509 | 0.637465736444117 | -5.52783450464458 |
| lnc-IFITM3-1 | lnc-IFITM3-1 | -0.272019948137143 | 1.55367298956207 | -1.00508451887867 | 0.32054464228716 | 0.637465736444117 | -5.55717377363063 |
| ENSG00000168852.12 | TPTE2P5 | -0.158967187306437 | 1.49198897542059 | -1.00486408294329 | 0.320649553134141 | 0.637612767091105 | -5.50641541275879 |
| lnc-VPS54-8 | lnc-VPS54-8 | 0.18850183131475 | 0.919569687675366 | 1.00475068837996 | 0.320703529427829 | 0.637654550418279 | -5.40520008667358 |
| lnc-ZRANB1-4 | lnc-ZRANB1-4 | 0.197763321381099 | 4.07662671395117 | 1.00468977935125 | 0.320732524922531 | 0.637654550418279 | -5.80964360920767 |
| lnc-DEDD2-1 | lnc-DEDD2-1 | 0.158846704730109 | 1.48714336501963 | 1.00460751677516 | 0.320771688502356 | 0.637670819436353 | -5.5107661398105 |
| lnc-KLHL11-1 | lnc-KLHL11-1 | 0.179676614826681 | 3.66593400480219 | 1.00448200461165 | 0.32083144858759 | 0.637708766907452 | -5.77692246484884 |
| ENSG00000127995.16 | CASD1 | -0.142016695709088 | 5.90653625327309 | -1.00443728641894 | 0.320852742073243 | 0.637708766907452 | -5.91228313150996 |
| lnc-USP45-3 | lnc-USP45-3 | 0.143352235861717 | 2.56100611938267 | 1.00420602808773 | 0.320962875735798 | 0.63786606848218 | -5.65142932102187 |
| ENSG00000143811.18 | PYCR2 | -0.0965089763599547 | 4.43377233561556 | -1.00413988665721 | 0.320994379403618 | 0.637867089328507 | -5.84638417235746 |
| ENSG00000168924.14 | LETM1 | 0.172709864151168 | 5.10252789393176 | 1.00390930595895 | 0.321104223107879 | 0.638011207577218 | -5.89114193438146 |
| lnc-GAPT-6 | lnc-GAPT-6 | -0.446962540546799 | 0.868342004427014 | -1.00392551002218 | 0.321138873186349 | 0.638011207577218 | -5.38343101485616 |
| ENSG00000117143.13 | UAP1 | 0.128932389663549 | 3.88191300873952 | 1.00379246122511 | 0.321159895141989 | 0.638011207577218 | -5.78836125693939 |
| ENSG00000143353.11 | LYPLAL1 | -0.151709936642169 | 3.62634235424075 | -1.00345959444747 | 0.321318529187436 | 0.638231045769425 | -5.77290945348155 |
| ENSG00000153046.17 | CDYL | 0.0835463801562302 | 5.08234818112673 | 1.00343013355147 | 0.321332571894697 | 0.638231045769425 | -5.88726217336583 |
| ENSG00000073578.16 | SDHA | 0.0851652333494597 | 6.46200326251155 | 1.00318130575581 | 0.321451193666862 | 0.63835889327041 | -5.9153570229577 |
| OTUD6B-AS1 | OTUD6B-AS1 | 0.168032400949022 | 4.42965346476405 | 1.00312593002748 | 0.321477596542337 | 0.63835889327041 | -5.84023436461588 |
| ENSG00000101104.12 | PABPC1L | -0.149471949216067 | 5.03631961675281 | -1.00309995581722 | 0.321489981423557 | 0.63835889327041 | -5.88432579769272 |
| ENSG00000160190.13 | SLC37A1 | 0.168433634967749 | 3.87401712808429 | 1.00280174501326 | 0.32163219578871 | 0.638579674536415 | -5.79046226595001 |
| lnc-MAP3K8-21 | lnc-MAP3K8-21 | 0.321954401028869 | 1.44680670352527 | 1.00271782530124 | 0.321672224101311 | 0.638597549062237 | -5.5082557416559 |
| lnc-PDF-1 | lnc-PDF-1 | -0.149659824811056 | 1.96229825706115 | -1.0024514882038 | 0.321799284791929 | 0.638788183901271 | -5.57299303068633 |
| lnc-PAQR6-3 | lnc-PAQR6-3 | 0.121769243692216 | 0.838356164150625 | 1.00186427960155 | 0.322079542601126 | 0.639282856833074 | -5.41150431522875 |
| ENSG00000173517.10 | PEAK1 | -0.127883214686141 | 6.27477925606455 | -1.00150779436092 | 0.322249763219297 | 0.639447985083119 | -5.91689562661276 |
| lnc-ATP5F1E-1 | lnc-ATP5F1E-1 | -0.138180361111197 | 3.34685057791586 | -1.00145473992432 | 0.322275101747278 | 0.639447985083119 | -5.74094923430247 |
| ENSG00000169902.14 | TPST1 | 0.353643265322661 | -0.213139327474221 | 1.00138380820979 | 0.322308980474988 | 0.639447985083119 | -5.26304792383602 |
| ENSG00000096092.5 | TMEM14A | -0.200371953148307 | 1.83122612707238 | -1.00136660595574 | 0.322317197056262 | 0.639447985083119 | -5.5547695502523 |
| ENSG00000142687.17 | KIAA0319L | -0.0677159343111207 | 5.85045905657241 | -1.00136477731547 | 0.3223180705066 | 0.639447985083119 | -5.91423821080901 |
| lnc-C1GALT1-1 | lnc-C1GALT1-1 | -0.272159706139412 | -0.390504841575786 | -1.0012873452873 | 0.32235505739564 | 0.639459728833314 | -5.14416621989857 |
| ENSG00000223501.8 | VPS52 | 0.104819824450953 | 5.60068323041357 | 1.00122080730695 | 0.322386842825213 | 0.639461153114711 | -5.90900472965193 |
| lnc-EBPL-2 | lnc-EBPL-2 | 0.26903251641301 | 1.56807757516489 | 1.00111999758217 | 0.322435004023169 | 0.639495055676324 | -5.52621065543346 |
| ENSG00000120526.10 | NUDCD1 | -0.147834926140403 | 3.54862874219903 | -1.0010099803319 | 0.322487569603974 | 0.639515786966898 | -5.75947785232775 |
| ENSG00000167005.13 | NUDT21 | -0.111849617217791 | 6.2444897221642 | -1.00087798869548 | 0.322550642053278 | 0.639515786966898 | -5.91753654965848 |
| ENSG00000134294.13 | SLC38A2 | 0.185746217861405 | 7.05644498173076 | 1.00087331382057 | 0.322552876103746 | 0.639515786966898 | -5.91182687362998 |
| ENSG00000078061.12 | ARAF | 0.0846961422449555 | 5.19929010230628 | 1.00083803179454 | 0.322569737175841 | 0.639515786966898 | -5.89795358998202 |
| ENSG00000139278.9 | GLIPR1 | -0.415845930200704 | 4.94911889986032 | -1.00104986069646 | 0.322633201623322 | 0.639580004720489 | -5.86616060249316 |
| ENSG00000160803.7 | UBQLN4 | -0.131452530353729 | 4.03344123812233 | -1.00024913869727 | 0.322851253615756 | 0.639900646077225 | -5.81502302847308 |
| ENSG00000101421.3 | CHMP4B | 0.163139417430418 | 4.13575560852641 | 1.00023685970092 | 0.322857125273866 | 0.639900646077225 | -5.81919461205448 |
| LINC01435 | LINC01435 | -1.22131524167316 | -0.0884537091700837 | -1.00049704726738 | 0.322956994561129 | 0.640036086306467 | -5.32033706610032 |
| ENSG00000128944.13 | KNSTRN | -0.128579623876614 | 1.89697992961579 | -0.999906970981329 | 0.323014900810803 | 0.640036086306467 | -5.57049814820012 |
| ENSG00000143106.12 | PSMA5 | 0.091646631127337 | 4.5486328238634 | 0.999898930946868 | 0.323018746759538 | 0.640036086306467 | -5.86403898310267 |
| lnc-ADCY4-4 | lnc-ADCY4-4 | -0.162591562728569 | 0.588330353301249 | -0.999307436964953 | 0.323301772586463 | 0.640535218514502 | -5.35447123514141 |
| ENSG00000166848.5 | TERF2IP | 0.10827287977737 | 6.23096752035327 | 0.999127453659166 | 0.323387926575698 | 0.64054390473429 | -5.9191444432671 |
| ENSG00000268193.5 | | -0.171959345632843 | 3.30985836057215 | -0.99898794372989 | 0.323454717523414 | 0.64054390473429 | -5.73490937868289 |
| ENSG00000160072.19 | ATAD3B | -0.10490894879216 | 4.56645930105462 | -0.998953003499658 | 0.323471446758935 | 0.64054390473429 | -5.86360648267216 |
| lnc-ZMAT3-9 | lnc-ZMAT3-9 | -0.155983197550953 | 5.62817323607728 | -0.998809051098128 | 0.323540376723268 | 0.64054390473429 | -5.91297628582832 |
| lnc-CP-4 | lnc-CP-4 | 0.233384504071409 | 0.513794479606907 | 0.998665477611021 | 0.323609135121724 | 0.64054390473429 | -5.32361399411155 |
| ENSG00000243708.10 | PLA2G4B | -0.169462667308491 | 3.30088637506574 | -0.998647448312645 | 0.323617770182088 | 0.64054390473429 | -5.72898886719766 |
| ENSG00000162997.15 | PRORSD1P | -0.259970096070055 | 1.59570646309763 | -0.998420353941859 | 0.323726549436407 | 0.64054390473429 | -5.54529413069281 |
| lnc-MTHFS-5 | lnc-MTHFS-5 | -0.109855202575054 | 3.56551443560098 | -0.998396400215368 | 0.323738024820469 | 0.64054390473429 | -5.76801439217132 |
| ENSG00000131269.16 | ABCB7 | 0.101210999492541 | 4.95301314016407 | 0.998396225467674 | 0.323738108536923 | 0.64054390473429 | -5.88571624487945 |
| ENSG00000196455.7 | PIK3R4 | 0.116163331669675 | 4.89961427818091 | 0.998377213378069 | 0.323747216754303 | 0.64054390473429 | -5.88360666513892 |

| | | | | | | | |
|---|---|---|---|---|---|---|---|
| ENSG00000157911.10 | PEX10 | -0.128627986973773 | 3.3228592761236 | -0.998347044842053 | 0.323761670101428 | 0.64054390473429 | -5.74009377454958 |
| ENSG00000171045.14 | TSNARE1 | -0.188018592760463 | 3.3036164629052 | -0.998338907394794 | 0.323765568719478 | 0.64054390473429 | -5.74153961495916 |
| ENSG00000135341.17 | MAP3K7 | 0.0760421971153624 | 5.69691965998149 | 0.998299564849658 | 0.323784418020404 | 0.64054390473429 | -5.9148925157653 |
| ENSG00000210825.1 | SNORA40 | 0.225062514482306 | 1.10392907697596 | 0.998279486467331 | 0.323794038005475 | 0.64054390473429 | -5.43836488878611 |
| lnc-RPL14-1 | lnc-RPL14-1 | -0.172617590982738 | 0.680483354371933 | -0.998167297537238 | 0.323847793685281 | 0.64054390473429 | -5.39359406512567 |
| ENSG00000144736.13 | SHQ1 | -0.0971441974372946 | 3.4528609935402 | -0.998140170521902 | 0.323860792584145 | 0.64054390473429 | -5.7546112736057 |
| ENSG00000111678.10 | C12orf57 | -0.163396515510432 | 4.96942429147445 | -0.9981021370116 | 0.323879018320623 | 0.64054390473429 | -5.89572049493476 |
| lnc-KCNJ4-2 | lnc-KCNJ4-2 | 0.223024797865558 | 1.44123129358999 | 0.998073438700183 | 0.323892771068602 | 0.64054390473429 | -5.50172498339436 |
| ENSG00000152784.15 | PRDM8 | -0.329986650104145 | 4.454400421128 | -0.998321156288217 | 0.323897437714351 | 0.64054390473429 | -5.86872539204036 |
| ENSG00000197933.12 | ZNF823 | -0.288568663757471 | 1.47097814968161 | -0.997721096078381 | 0.324061652112889 | 0.640751152711936 | -5.52069062421695 |
| ENSG00000135040.15 | NAA35 | 0.122562932650094 | 4.45374345540665 | 0.997715166007031 | 0.324064494958522 | 0.640751152711936 | -5.85612423875294 |
| lnc-DKK4-1 | lnc-DKK4-1 | -0.18229750815939 | 1.49113581579395 | -0.99743791237943 | 0.32419742769685 | 0.640874436925265 | -5.51553796859533 |
| ENSG00000137269.14 | LRRC1 | -0.148941422705795 | 2.52143823388976 | -0.997424403745241 | 0.32420390552083 | 0.640874436925265 | -5.64625856082596 |
| ENSG00000280120.1 | | 0.175709447680489 | 1.433529884597 | 0.997371927214046 | 0.324229070531238 | 0.640874436925265 | -5.50830899111486 |
| ENSG00000164603.11 | BMT2 | 0.132002246797049 | 4.09070710474415 | 0.997325384171338 | 0.32425139125199 | 0.640874436925265 | -5.82455566905291 |
| ENSG00000130638.16 | ATXN10 | 0.09177153719037 | 5.01723036367075 | 0.997211012792757 | 0.324306244917134 | 0.640877404011411 | -5.89085542667898 |
| ENSG00000124380.10 | SNRNP27 | 0.0978640790548093 | 4.2477855956917 | 0.997143573007037 | 0.324338592650299 | 0.640877404011411 | -5.83931573763429 |
| ENSG00000101336.13 | HCK | 1.45483566512991 | -1.19422361945113 | 0.997429458202735 | 0.324363961456186 | 0.640877404011411 | -4.87683478893455 |
| ENSG00000143409.15 | MINDY1 | -0.228827286726488 | 3.12020001009121 | -0.997062593887171 | 0.324377437447936 | 0.640877404011411 | -5.71526885648886 |
| lnc-ERCC4-7 | lnc-ERCC4-7 | -0.156231187356697 | 0.885861100586919 | -0.9966688446526 | 0.324566359355935 | 0.641189113602382 | -5.41125045909886 |
| ENSG00000200397.1 | | 0.171360267425819 | 1.90652490003184 | 0.99650535616759 | 0.324644823346952 | 0.641275678357885 | -5.57564853568244 |
| lnc-NEK3-1 | lnc-NEK3-1 | 0.149147971675507 | 3.24850926747532 | 0.99644771773099 | 0.324672489149663 | 0.641275678357885 | -5.73928285051896 |
| ENSG00000132004.12 | FBXW9 | 0.169422965324072 | 0.492338458554251 | 0.996281366461231 | 0.324752344804623 | 0.641359639087631 | -5.36430447012319 |
| ENSG00000070785.16 | EIF2B3 | -0.103623760481956 | 2.72566357354169 | -0.996229351060964 | 0.324777317116571 | 0.641359639087631 | -5.66711767804408 |
| lnc-B3GALNT2-2 | lnc-B3GALNT2-2 | 0.13362123957981 | 2.40137108291022 | 0.995994076653875 | 0.324890287252478 | 0.641476352815875 | -5.62715992111002 |
| ENSG00000164692.17 | COL1A2 | 1.1678322724089 | -2.18738369188061 | 0.995845892403638 | 0.324961453482677 | 0.641476352815875 | -4.71353462106326 |
| lnc-RHNO1-2 | lnc-RHNO1-2 | -0.202014754823685 | 2.79013727853772 | -0.995813995358675 | 0.324976773572612 | 0.641476352815875 | -5.67370666970437 |
| ENSG00000101888.11 | NXT2 | 0.236599965104864 | 3.42104422628943 | 0.995751485759457 | 0.325006798225599 | 0.641476352815875 | -5.75471975557162 |
| ENSG00000128585.17 | MKLN1 | 0.139507264934059 | 6.72855488910901 | 0.995687222289002 | 0.325037667247892 | 0.641476352815875 | -5.92151153107605 |
| ENSG00000163728.10 | TTC14 | -0.116644327137641 | 7.01512464321894 | -0.995685692965564 | 0.325038401884001 | 0.641476352815875 | -5.9184083874297 |
| ENSG00000103064.14 | SLC7A6 | 0.15510974198634 | 6.19089104392619 | 0.995652020974846 | 0.325054577071835 | 0.641476352815875 | -5.92182565075152 |
| lnc-SHB-4 | lnc-SHB-4 | -0.174460396515825 | 2.12951286697714 | -0.995465034645207 | 0.325144410528548 | 0.641503127827211 | -5.59895293699374 |
| ENSG00000208018.1 | MIR645 | -0.263948149876796 | 1.19116700494685 | -0.995435485058019 | 0.325158608506286 | 0.641503127827211 | -5.46809465466568 |
| lnc-TBX6-1 | lnc-TBX6-1 | -0.172527164435159 | 3.87626618098556 | -0.995429166106993 | 0.325161644688398 | 0.641503127827211 | -5.79452071069054 |
| lnc-TSC22D2-1 | lnc-TSC22D2-1 | -0.225262496409848 | -0.325155226145403 | -0.995283604039435 | 0.325231590848708 | 0.64153992429826 | -5.1481586179579 |
| lnc-MRPL57-5 | lnc-MRPL57-5 | -0.26323351443701 | 0.799660843134372 | -0.995204286880586 | 0.325269708964541 | 0.64153992429826 | -5.44175857453392 |
| lnc-ZNF567-5 | lnc-ZNF567-5 | -0.224635088688831 | 3.7753337439362 | -0.995195772036137 | 0.325273801194026 | 0.64153992429826 | -5.77570703672912 |
| ENSG00000224083.1 | MTCO1P11 | -0.319875269539402 | 0.496314863045243 | -0.995006481386318 | 0.325364783130863 | 0.641601409338538 | -5.32855618318244 |
| lnc-ZCCHC8-1 | lnc-ZCCHC8-1 | 0.123580498328963 | 4.97249595421634 | 0.994944445215445 | 0.325394604344623 | 0.641601409338538 | -5.89082513833566 |
| lnc-BICD1-1 | lnc-BICD1-1 | -0.228601137313626 | 1.1333328775807 | -0.994834606306971 | 0.325447409178711 | 0.641601409338538 | -5.48761341128557 |
| ENSG00000132881.11 | CPLANE2 | -0.172754487581333 | 0.693564289356736 | -0.994828733495013 | 0.325450232683786 | 0.641601409338538 | -5.39695162245435 |
| ENSG00000141378.14 | PTRH2 | 0.143397610879433 | 3.38207333452317 | 0.994761567685678 | 0.325482525543885 | 0.641601409338538 | -5.74251279779297 |
| ENSG00000127125.8 | PPCS | -0.090052181098863 | 5.06976868947803 | -0.994703872418919 | 0.32551026675531 | 0.641601409338538 | -5.89675608515854 |
| lnc-POPDC2-4 | lnc-POPDC2-4 | -0.115361252626239 | 1.80934648882284 | -0.994636669393489 | 0.325542581523575 | 0.641601409338538 | -5.56192638457344 |
| ENSG00000101109.11 | STK4 | 0.0960882155413803 | 8.79804266693449 | 0.994612203097658 | 0.325554346748855 | 0.641601409338538 | -5.88669589304358 |
| ENSG00000059122.16 | FLYWCH1 | 0.187881017402596 | 4.09045100938493 | 0.994494684989047 | 0.325610862237926 | 0.641616683574731 | -5.81415629751902 |
| lnc-NMRAL1-4 | lnc-NMRAL1-4 | -0.186242114390664 | 0.871897514153876 | -0.994466450646439 | 0.325624441365284 | 0.641616683574731 | -5.40775669696558 |
| ENSG00000179222.17 | MAGED1 | -0.132995948141714 | 5.2297202752408 | -0.994153712135876 | 0.325774876469084 | 0.641851659075635 | -5.90396215371185 |
| lnc-TLCD2-3 | lnc-TLCD2-3 | 0.146911809551585 | 3.37052406201615 | 0.993885245816858 | 0.325904052857715 | 0.642044709032383 | -5.74172830438422 |
| lnc-FAM126B-3 | lnc-FAM126B-3 | 0.145802403633467 | 1.06716273590492 | 0.99380542356582 | 0.325942467119667 | 0.642058933938569 | -5.45847040471553 |
| ENSG00000197771.12 | MCMBP | 0.0963309816875952 | 5.12823456098097 | 0.99330000903813 | 0.326185767373586 | 0.642458710778507 | -5.90441504348008 |
| ENSG00000132382.14 | MYBBP1A | -0.101967170318992 | 5.11409564521927 | -0.99320397024295 | 0.326232013070171 | 0.642458710778507 | -5.90445034543472 |
| lnc-MYO1A-2 | lnc-MYO1A-2 | -0.172775211341035 | 0.583183504275754 | -0.993189349438932 | 0.326239053833598 | 0.642458710778507 | -5.34992670459325 |
| lnc-NUP62CL-3 | lnc-NUP62CL-3 | -0.136316926832118 | 0.54230872030897 | -0.993020825310424 | 0.326320215339361 | 0.642494990912692 | -5.34446406454393 |
| ENSG00000184988.8 | TMEM106A | -0.0984018148510062 | 4.23920484597215 | -0.993014389293424 | 0.326323315205419 | 0.642494990912692 | -5.83864089706612 |
| ENSG00000117682.16 | DHDDS | -0.0811464199775722 | 4.6129326855846 | -0.992896093780403 | 0.32638029500563 | 0.642494990912692 | -5.87226664955586 |
| ENSG00000144560.14 | VGLL4 | -0.19858779560083 | 3.7824980206672 | -0.992891856426629 | 0.326382336150372 | 0.642494990912692 | -5.80217227498958 |
| ENSG00000171817.16 | ZNF540 | -0.186847793076996 | 3.32986906198973 | -0.992654443094121 | 0.326496712517209 | 0.642658681623957 | -5.74258920198021 |
| ENSG00000133116.7 | KL | -0.843342183802347 | -2.08170274863481 | -0.992482878019513 | 0.326579382579268 | 0.642685133312154 | -4.83031516678345 |
| lnc-KLC2-2 | lnc-KLC2-2 | -0.300511801256595 | 1.43612888777768 | -0.992420112874414 | 0.326609630006297 | 0.642685133312154 | -5.50777054058341 |
| lnc-MTF1-2 | lnc-MTF1-2 | -0.155322112308391 | 1.33407817012596 | -0.992367537371663 | 0.326634968348643 | 0.642685133312154 | -5.4888222809188 |
| FMR1-IT1 | FMR1-IT1 | -0.182909780422657 | 0.0896036049643552 | -0.992319648864696 | 0.326658048981833 | 0.642685133312154 | -5.29286270407453 |
| lnc-ZKSCAN5-1 | lnc-ZKSCAN5-1 | -0.150435181688987 | 0.559022803004254 | -0.9923025892042 | 0.32666627142359 | 0.642685133312154 | -5.35211760141071 |
| ENSG00000132780.16 | NASP | 0.0987739594723305 | 5.93963437884453 | 0.991799257892157 | 0.32690893049348 | 0.643092264562597 | -5.92456443674979 |
| ENSG00000147010.17 | SH3KBP1 | 0.151830356673563 | 5.65270313687311 | 0.991702839215665 | 0.326955428354272 | 0.643092264562597 | -5.91957076540843 |
| ENSG00000198198.16 | SZT2 | 0.112588193171418 | 7.16303232238963 | 0.991678967635584 | 0.326966941099333 | 0.643092264562597 | -5.92153917772357 |
| ENSG00000160948.13 | VPS28 | -0.130474666463773 | 5.51346360318785 | -0.991216606810287 | 0.327189981474085 | 0.643469461941629 | -5.91743944329133 |
| ENSG00000101445.9 | PPP1R16B | -0.176889993922061 | 7.60638400775338 | -0.99114133955301 | 0.327226299674665 | 0.643479404433327 | -5.91353420353772 |

| | | | | | | | |
|---|---|---|---|---|---|---|---|
| lnc-NMRAL1-2 | lnc-NMRAL1-2 | 0.158798002913687 | 3.29474515202869 | 0.991020569793585 | 0.32728457955371 | 0.643503671104476 | -5.73409220505568 |
| ENSG00000103707.9 | MTFMT | -0.12714483683597 | 2.26728551360752 | -0.990951628746503 | 0.32731785157294 | 0.643503671104476 | -5.63156484343973 |
| ENSG00000284195.1 | MIR6852 | 0.17021903584086 | 2.12209786430093 | 0.990921420109467 | 0.327332431446527 | 0.643503671104476 | -5.60094236580267 |
| lnc-NMB-1 | lnc-NMB-1 | 0.142723127181816 | 3.54475494942765 | 0.990818034306834 | 0.327382332795839 | 0.643519806329475 | -5.76383330471397 |
| lnc-U2AF2-1 | lnc-U2AF2-1 | -0.147143102227949 | 2.77392715044006 | -0.99077487049091 | 0.327403168239919 | 0.643519806329475 | -5.66870477751511 |
| ENSG00000230076.1 | RPL10P6 | 1.06049680124589 | 0.11650483166571 | 0.991147340079981 | 0.327436977433494 | 0.643524807267603 | -5.37020660104817 |
| lnc-MMP24OS-73 | lnc-MMP24OS-73 | -0.143967852987013 | 0.437402511112457 | -0.990555345763808 | 0.327509147995324 | 0.643605193162856 | -5.31812498802122 |
| lnc-XXYLT1-6 | lnc-XXYLT1-6 | -0.284509864689482 | 0.770023518369436 | -0.990245879287095 | 0.327658588012305 | 0.643801662148836 | -5.3901690412508 |
| lnc-ATP6V1C1-8 | lnc-ATP6V1C1-8 | -0.256344761449561 | -0.183289428453562 | -0.990151540708488 | 0.327704152815554 | 0.643801662148836 | -5.16070197710642 |
| ENSG00000267587.1 | | 0.207969276662088 | -0.116492939559104 | 0.990089256854251 | 0.327734237768814 | 0.643801662148836 | -5.18887409237659 |
| lnc-ST8SIA5-3 | lnc-ST8SIA5-3 | 0.207969276662088 | -0.116492939559104 | 0.990089256854251 | 0.327734237768814 | 0.643801662148836 | -5.18887409237659 |
| ENSG00000085465.12 | OVGP1 | -0.205370521748873 | 3.69406289615959 | -0.989981158104718 | 0.327786457084859 | 0.643842794749275 | -5.78327938953441 |
| ENSG00000106591.3 | MRPL32 | 0.12036761982928 | 5.02100796833777 | 0.989816769256949 | 0.327865879202718 | 0.643883547420245 | -5.90283809646404 |
| ENSG00000273734.1 | | -0.15994819520913 | 4.55378047940858 | -0.989808718274198 | 0.327869769251887 | 0.643883547420245 | -5.87182467073104 |
| lnc-WDR49-4 | lnc-WDR49-4 | -0.154817462396095 | 1.96539059780487 | -0.989710289019225 | 0.327917330505019 | 0.643915513621904 | -5.58897277804354 |
| lnc-PGAM2-2 | lnc-PGAM2-2 | -0.126864526184173 | 2.71089188780861 | -0.989276225290956 | 0.328127126441212 | 0.644117489280858 | -5.68055558571486 |
| ENSG00000177888.7 | ZBTB41 | 0.109965704367569 | 4.9486158043057 | 0.989250671848012 | 0.328139479991593 | 0.644117489280858 | -5.89438534247519 |
| ENSG00000079616.12 | KIF22 | 0.114574713745615 | 5.2322745101453 | 0.98914328123403 | 0.328191400300128 | 0.644117489280858 | -5.91085580179808 |
| ENSG00000185304.14 | RGPD2 | 1.94615190438929 | 0.486544868272293 | 0.989592972493603 | 0.328195859355583 | 0.644117489280858 | -5.10237773732889 |
| ENSG00000120662.15 | MTRF1 | 0.107895386272166 | 3.94676250726815 | 0.98907870442714 | 0.32822262400703 | 0.644117489280858 | -5.81325976544358 |
| ENSG00000175806.14 | MSRA | -0.136526099745187 | 2.39135849806599 | -0.98905556391555 | 0.328233813224121 | 0.644117489280858 | -5.63434365497119 |
| ENSG00000113721.13 | PDGFRB | 0.375117619842641 | 1.35300330672368 | 0.989088985290074 | 0.328239243310835 | 0.644117489280858 | -5.48467307562123 |
| ENSG00000183258.11 | DDX41 | -0.136190206196277 | 5.1996071036301 | -0.98897475601305 | 0.328272888577473 | 0.644122103488096 | -5.90942937648343 |
| ENSG00000100325.14 | ASCC2 | 0.0855206375154181 | 4.82977692371607 | 0.988851226646217 | 0.328332628299712 | 0.64417791328691 | -5.88593133591356 |
| STAM-AS1 | STAM-AS1 | -0.242333253897106 | -0.0502927102872443 | -0.988683980942085 | 0.32841352120308 | 0.64423582337605 | -5.23581330799325 |
| lnc-TBC1D31-2 | lnc-TBC1D31-2 | 0.246332228148344 | 3.85057819035959 | 0.988610998770056 | 0.328450185929023 | 0.64423582337605 | -5.778976669035 |
| lnc-ADAM8-2 | lnc-ADAM8-2 | 0.16292494034197 | 1.67215925574439 | 0.988596078674338 | 0.32845604287559 | 0.64423582337605 | -5.55295990194832 |
| lnc-CORO6-1 | lnc-CORO6-1 | 0.183051765771163 | 2.61841408974874 | 0.987952286683476 | 0.328767582263979 | 0.64475725949055 | -5.65866807250399 |
| ENSG00000112144.15 | CILK1 | -0.129453280451364 | 4.56727590895735 | -0.987915540130958 | 0.3287853703857 | 0.64475725949055 | -5.87076384896288 |
| ENSG00000256269.8 | HMBS | -0.230878888796925 | 0.925872966349341 | -0.98785254735895 | 0.328815865173799 | 0.64475725949055 | -5.45901189455096 |
| lnc-CAMKK2-2 | lnc-CAMKK2-2 | 0.146586375991899 | 2.75197979549562 | 0.987682904779267 | 0.328897998554041 | 0.644856877904155 | -5.67876250550758 |
| ENSG00000204120.14 | GIGYF2 | -0.0717206025417374 | 6.66041586628273 | -0.987389572006883 | 0.329040049707402 | 0.645073944483043 | -5.92968114647742 |
| lnc-SH3BP2-2 | lnc-SH3BP2-2 | 0.170348965702645 | 1.82157272154783 | 0.987166811735511 | 0.32914795248797 | 0.645223676266444 | -5.57233870114248 |
| ENSG00000137764.19 | MAP2K5 | -0.132657888996749 | 3.82432206765415 | -0.987102477228914 | 0.329179119879217 | 0.645223676266444 | -5.79869010540335 |
| lnc-COBLL1-1 | lnc-COBLL1-1 | 0.951692641050265 | -1.11552589156147 | 0.987153307679952 | 0.329214156014674 | 0.64523090585127 | -4.89143045536866 |
| ENSG00000141753.6 | IGFBP4 | -0.357074947116414 | 4.55295062687712 | -0.987248201131418 | 0.329246267766079 | 0.645232403068572 | -5.86942970776234 |
| lnc-F11R-2 | lnc-F11R-2 | -0.249936374194716 | 1.58787144929792 | -0.986788990839543 | 0.329331019327177 | 0.64531115028137 | -5.52290707402739 |
| ENSG00000176058.12 | TPRN | -0.165040043513992 | 1.79507676995406 | -0.986751571734357 | 0.329349153850546 | 0.64531115028137 | -5.57940611296377 |
| FLVCR1-DT | FLVCR1-DT | 0.269494960097496 | 1.21784402684728 | 0.986454520110199 | 0.329493138563509 | 0.64553181715712 | -5.4757063474474 |
| ENSG00000114166.7 | KAT2B | 0.0775329789821102 | 6.38147916090316 | 0.986287212144737 | 0.3295742534575 | 0.645629281328231 | -5.93185114866558 |
| ENSG00000163344.5 | PMVK | -0.234741335747636 | 1.68451842631519 | -0.986217500103623 | 0.329608055471797 | 0.645634050797107 | -5.55281748993371 |
| lnc-MOV10-2 | lnc-MOV10-2 | 0.133163523302648 | 2.75226821614215 | 0.985992820927961 | 0.329717013864142 | 0.645786021159543 | -5.68549987529539 |
| ENSG00000134871.18 | COL4A2 | 0.740771864161951 | -1.34764419062073 | 0.985911897354424 | 0.329756263749539 | 0.645801444030138 | -4.86669039320804 |
| lnc-SORBS3-1 | lnc-SORBS3-1 | -0.250209870371693 | 1.55936036000586 | -0.9856670516386 | 0.329875038912547 | 0.645972593086953 | -5.54135269059855 |
| ENSG00000136935.13 | GOLGA1 | 0.105109969190547 | 4.77092106388366 | 0.985543669026254 | 0.329934902943196 | 0.645973343873136 | -5.88359332705571 |
| ENSG00000142507.9 | PSMB6 | 0.141370487280247 | 4.33030070188612 | 0.985536894744012 | 0.329938189969309 | 0.645973343873136 | -5.86043037885002 |
| lnc-PAQR6-2 | lnc-PAQR6-2 | 0.106299598037777 | 1.50921073969141 | 0.985469652045103 | 0.329970818750673 | 0.645975781086656 | -5.53486335684801 |
| ENSG00000109689.15 | STIM2 | -0.152745666330228 | 7.46820161759399 | -0.985247935084078 | 0.330078419800496 | 0.646124975249986 | -5.92131405615044 |
| ENSG00000113569.15 | NUP155 | 0.0875793635234086 | 5.70813889784385 | 0.985144022337739 | 0.330128857599863 | 0.646162255228031 | -5.92783715575972 |
| ENSG00000285396.1 | NA | 0.262240142617498 | 0.38376084683501 | 0.984962814127884 | 0.330216825902125 | 0.646218812413054 | -5.33963383083094 |
| lnc-MIXL1-7 | lnc-MIXL1-7 | 0.18181363072907 | 2.20798265276837 | 0.984937974473614 | 0.330228885643355 | 0.646218812413054 | -5.61096620630013 |
| lnc-ZNF439-1 | lnc-ZNF439-1 | -0.168809347950664 | 3.17443490724827 | -0.984831574120454 | 0.330280546737456 | 0.646218812413054 | -5.72611913247354 |
| ENSG00000051620.10 | HEBP2 | 0.101346467042706 | 5.62463536728384 | 0.984825829548093 | 0.330283336082463 | 0.646218812413054 | -5.92543445975114 |
| lnc-KLF9-5 | lnc-KLF9-5 | -0.300440835536561 | -0.318147013214017 | -0.984752678285892 | 0.330318856923287 | 0.646226882620664 | -5.14458970571963 |
| ENSG00000178075.19 | GRAMD1C | -0.606821718850853 | 1.49105718667838 | -0.984762068431269 | 0.330482155280742 | 0.646462720767056 | -5.5447749725891 |
| ENSG00000103226.17 | NOMO3 | 0.16233542249528 | 1.53189527190674 | 0.984364984428942 | 0.330507156324755 | 0.646462720767056 | -5.51225080862652 |
| ENSG00000214176.9 | PLEKHM1P1 | 0.117186720607344 | 5.69193199514009 | 0.984255237079438 | 0.330560472682401 | 0.646462720767056 | -5.9272657905868 |
| ENSG00000166181.12 | API5 | 0.0757002912553117 | 6.44150825295718 | 0.984245844407029 | 0.330565036004143 | 0.646462720767056 | -5.93371976004385 |
| ENSG00000104979.8 | C19orf53 | -0.131011840895582 | 4.13449991166214 | -0.984095174480709 | 0.330638243023284 | 0.6465444571678 | -5.84231153000681 |
| ENSG00000134278.15 | SPIRE1 | -0.388891751390585 | 0.695695600129972 | -0.98398224644933 | 0.330715804501174 | 0.646596836225938 | -5.45553912455095 |
| lnc-AGBL3-1 | lnc-AGBL3-1 | 0.26143599492943 | 2.42067542460552 | 0.983902757303729 | 0.330731749857629 | 0.646596836225938 | -5.63282657667718 |
| lnc-PLCG2-8 | lnc-PLCG2-8 | 0.125759031973683 | 4.71969821805827 | 0.983846130227156 | 0.330759271662192 | 0.646596836225938 | -5.88760365410767 |
| ENSG00000103274.10 | NUBP1 | -0.111925431999696 | 4.42387512470697 | -0.983720495346929 | 0.330820338012071 | 0.646645991647385 | -5.8674958749301 |
| ENSG00000144031.11 | ANKRD53 | -0.625792946637948 | 0.349014392015731 | -0.983944582601671 | 0.330847249576768 | 0.646645991647385 | -5.35665797394651 |
| ENSG00000276291.5 | FRG1HP | -0.132694340551161 | 3.89268387273512 | -0.983590678035514 | 0.330883445217709 | 0.64665533164794 | -5.81711411689223 |
| ENSG00000175595.14 | ERCC4 | -0.0978440874032951 | 3.58989871565127 | -0.983489968292865 | 0.330932408114053 | 0.646689618932076 | -5.77657515384952 |
| lnc-TESK2-1 | lnc-TESK2-1 | 0.675889940651905 | -0.795430568592244 | 0.983267331431362 | 0.331040666560284 | 0.646839760756629 | -4.93049426667535 |
| ENSG00000082781.11 | ITGB5 | 1.5748598024115 | -0.113731538492066 | 0.983516027807516 | 0.331131659657851 | 0.646905155600633 | -5.0594106700779 |

| | | | | | | | |
|---|---|---|---|---|---|---|---|
| ENSG00000270149.5 | | 0.171285019571269 | 4.51038097967238 | 0.983051627933607 | 0.331145576241968 | 0.646905155600633 | -5.86660055001291 |
| LINC00921 | LINC00921 | 0.203424871859022 | 2.19287507759575 | 0.982887285753104 | 0.331225520746942 | 0.646905155600633 | -5.60658906069865 |
| ENSG00000102898.11 | NUTF2 | -0.183738794499086 | 4.22654053523299 | -0.982858740826561 | 0.331239407787507 | 0.646905155600633 | -5.85528767866016 |
| lnc-GPCPD1-8 | lnc-GPCPD1-8 | -0.244229196883944 | 1.91971079293894 | -0.982858172345815 | 0.331239684356032 | 0.646905155600633 | -5.61135605718454 |
| ENSG00000215154.6 | | -0.250133773205675 | 1.91026889009824 | -0.982810846724159 | 0.331262709033216 | 0.646905155600633 | -5.57891433322212 |
| lnc-VCL-4 | lnc-VCL-4 | -0.356498562153112 | 0.0760882522914341 | -0.982595317955611 | 0.331367580805799 | 0.64697547434679 | -5.26415501908376 |
| lnc-ADH4-4 | lnc-ADH4-4 | 0.154270370873165 | 1.87259793118007 | 0.982593700293076 | 0.331368368010466 | 0.64697547434679 | -5.58265739450357 |
| lnc-PKM-2 | lnc-PKM-2 | 0.0985671424512795 | 1.74057168360596 | 0.982523411487367 | 0.331402573928525 | 0.64697547434679 | -5.56417897686288 |
| ENSG00000255439.6 | | -0.15641583711929 | 1.93117163632312 | -0.982478466767639 | 0.331424447432957 | 0.64697547434679 | -5.59017512254821 |
| ENSG00000167202.11 | TBC1D2B | 0.324879351738104 | 3.40329064632144 | 0.982423003470597 | 0.331517420395454 | 0.647095596396361 | -5.7365413476687 |
| ENSG00000279059.1 | | -0.223860754931052 | -0.0531815496665541 | -0.981960391289385 | 0.331676651988456 | 0.647212259032375 | -5.20668421749991 |
| ENSG00000131981.15 | LGALS3 | -0.612952024599888 | -0.895854113843785 | -0.981977678461763 | 0.331679102217267 | 0.647212259032375 | -5.05685514575026 |
| ENSG00000048471.13 | SNX29 | -0.134281009979605 | 7.37178502895885 | -0.981912563644772 | 0.331699941461617 | 0.647212259032375 | -5.92727485655265 |
| ENSG00000135387.20 | CAPRIN1 | 0.0786216465688376 | 7.12343700029641 | 0.981906355425941 | 0.331702964608294 | 0.647212259032375 | -5.93058364579023 |
| lnc-TRIM16-1 | lnc-TRIM16-1 | -0.191510721094301 | -0.138135936321238 | -0.980998221031094 | 0.332145387305588 | 0.648014076484447 | -5.19175839007108 |
| lnc-KRT222-1 | lnc-KRT222-1 | 0.118933894203112 | 3.28201763140155 | 0.980867857889939 | 0.332208929695667 | 0.648028509737363 | -5.74396417386782 |
| lnc-DNPH1-1 | lnc-DNPH1-1 | -0.234535794937582 | 1.48462397656873 | -0.980853861158451 | 0.332215752550583 | 0.648028509737363 | -5.53204857085089 |
| ENSG00000283773.1 | | -0.169354585872536 | 0.825809640313205 | -0.980696300032614 | 0.332292563856911 | 0.648116918588723 | -5.43279499243784 |
| ENSG00000105220.15 | GPI | -0.0998064944127917 | 5.79855568702123 | -0.980560168013476 | 0.332358938005139 | 0.648184955558061 | -5.93323433993327 |
| ENSG00000180185.11 | FAHD1 | -0.149834524874397 | 2.26221489102413 | -0.979791041226103 | 0.332734109217121 | 0.648855155851823 | -5.63171030571981 |
| ENSG00000101558.13 | VAPA | 0.113831489176659 | 6.64876053285737 | 0.979545978452839 | 0.332853707492438 | 0.649026890992501 | -5.93730881238983 |
| lnc-VPS26C-2 | lnc-VPS26C-2 | -0.259519836683722 | 0.440463442679786 | -0.979243744344738 | 0.333001246741031 | 0.649180016352935 | -5.35767966849241 |
| lnc-RNMT-4 | lnc-RNMT-4 | -0.315144426169361 | 2.54372278977437 | -0.979258534478844 | 0.33302428803648 | 0.649180016352935 | -5.66095500326516 |
| ENSG00000114209.14 | PDCD10 | -0.124389617693015 | 4.51494638776729 | -0.979191291349395 | 0.333026856759007 | 0.649180016352935 | -5.88255499341719 |
| ENSG00000173113.6 | TRMT112 | -0.156656387024761 | 4.94269004219058 | -0.979103414554676 | 0.333069765291692 | 0.649202176041562 | -5.90533581888352 |
| lnc-GGACT-3 | lnc-GGACT-3 | 0.228342900895945 | -0.107641440465655 | 0.978952561648077 | 0.333143432454169 | 0.649243707839092 | -5.22288079615786 |
| ENSG00000229807.11 | XIST | 3.00199619950174 | -0.161099599997594 | 0.97938020006146 | 0.333154158392678 | 0.649243707839092 | -5.20473315665801 |
| lnc-IFT20-5 | lnc-IFT20-5 | 0.186097089517435 | -0.163147823397536 | 0.978788065196409 | 0.333223774686351 | 0.649275700858413 | -5.20544780494203 |
| ENSG00000112297.14 | CRYBG1 | 0.228687970957905 | 7.31178192514099 | 0.978811558205673 | 0.333233663890991 | 0.649275700858413 | -5.9316173501842 |
| ENSG00000159788.19 | RGS12 | 0.417499327942545 | 1.48558784326662 | 0.978732623675696 | 0.333325940554113 | 0.649344570367688 | -5.52817974506775 |
| lnc-SLC22A1-2 | lnc-SLC22A1-2 | -0.186487600822781 | 0.504887480616262 | -0.978474810031202 | 0.333376808441972 | 0.649344570367688 | -5.36620149479249 |
| lnc-E2F2-1 | lnc-E2F2-1 | -0.328853318345408 | 2.9197245531972 | -0.978567578018062 | 0.3333906219469 | 0.649344570367688 | -5.71037571012842 |
| ENSG00000183426.16 | NPIPA1 | -0.15464552439196 | 2.9429643615267 | -0.978369863590967 | 0.33342808816324 | 0.649344570367688 | -5.70880186892268 |
| ENSG00000151229.12 | SLC2A13 | -0.145505050425218 | 2.94142627330213 | -0.978322702392693 | 0.333451134140162 | 0.649344570367688 | -5.72224023827678 |
| ENSG00000160613.12 | PCSK7 | 0.225553804243284 | 5.95140605218351 | 0.978348190107334 | 0.333458296107781 | 0.649344570367688 | -5.93712609119702 |
| lnc-NDRG1-5 | lnc-NDRG1-5 | -0.168535957193015 | 4.23846293924482 | -0.977928830795099 | 0.333643646551425 | 0.64964404285006 | -5.83841238213216 |
| ENSG00000065308.4 | TRAM2 | -0.134887652263064 | 5.96779242852889 | -0.977840053476305 | 0.333687048446923 | 0.649667094039255 | -5.9380929037918 |
| ENSG00000261056.2 | | -0.207684052344519 | 1.09049980680108 | -0.9777258366223 | 0.33374289289617 | 0.649714363423992 | -5.46974229311177 |
| XIST | XIST | 2.9967018756016 | -0.134540296687186 | 0.977925457520226 | 0.333864495028914 | 0.649878599457212 | -5.20302835930653 |
| lnc-NFS1-2 | lnc-NFS1-2 | -0.118677532943533 | 2.72685362985929 | -0.977424197928302 | 0.333890404181121 | 0.649878599457212 | -5.68922290822261 |
| ENSG00000233830.2 | EIF4HP1 | 0.545746487152688 | 0.545154092111487 | 0.977338411741251 | 0.333980946356771 | 0.649912552709245 | -5.23586028553081 |
| ENSG00000127580.16 | WDR24 | -0.125518334940288 | 3.2535818528056 | -0.977197206433885 | 0.334001439231727 | 0.649912552709245 | -5.75259343726626 |
| ENSG00000088179.8 | PTPN4 | 0.124621343140668 | 5.22386799008902 | 0.977194886671011 | 0.334002574093106 | 0.649912552709245 | -5.91998354852607 |
| lnc-PIP4K2A-1 | lnc-PIP4K2A-1 | -0.19832757655384 | 1.22555135443622 | -0.976961452581672 | 0.334116786576778 | 0.650073335675378 | -5.52386174440683 |
| lnc-CRABP2-1 | lnc-CRABP2-1 | 0.243541848478897 | -0.299266191425973 | 0.97660658080082 | 0.334290464931594 | 0.65026411267112 | -5.1498415525561 |
| lnc-SEPSECS-2 | lnc-SEPSECS-2 | -0.214479815359182 | 4.65643455426969 | -0.976580433070737 | 0.33430326431705 | 0.65026411267112 | -5.89206674908125 |
| ENSG00000133111.3 | RFXAP | -0.181712465701015 | 1.32288777288036 | -0.97656745612737 | 0.334309616687839 | 0.65026411267112 | -5.51725443247026 |
| ENSG00000109775.10 | UFSP2 | -0.149755217490675 | 3.62548795416211 | -0.976458752888382 | 0.334362831403471 | 0.650306166377931 | -5.78551333750839 |
| lnc-ITGA2B-3 | lnc-ITGA2B-3 | 0.240512653120292 | 0.919198895362211 | 0.976213487619408 | 0.334482919634231 | 0.650478262928134 | -5.45785004106838 |
| ENSG00000143493.12 | INTS7 | -0.148909739456138 | 3.8850828796489 | -0.976040081316453 | 0.334567841224864 | 0.650515199971965 | -5.81507291054923 |
| lnc-ZKSCAN8-1 | lnc-ZKSCAN8-1 | -0.200286505001067 | 0.936889622329961 | -0.976029802816556 | 0.33457287532618 | 0.650515199971965 | -5.45586146046183 |
| RAP2C-AS1 | RAP2C-AS1 | -0.169751159799935 | 2.8456200698401 | -0.975847285864035 | 0.334662275081272 | 0.650515199971965 | -5.70932766985711 |
| lnc-CMBL-1 | lnc-CMBL-1 | -0.21475823114116 | 0.118011970163864 | -0.975829964307434 | 0.334670760288088 | 0.650515199971965 | -5.26578546711756 |
| ENSG00000135976.18 | ANKRD36 | -0.155565107130529 | 6.94107110551781 | -0.975814486955879 | 0.334678342206728 | 0.650515199971965 | -5.93833006064884 |
| lnc-TTC14-1 | lnc-TTC14-1 | -0.205962044899382 | 0.0484876708874798 | -0.975774899151333 | 0.334697735678873 | 0.650515199971965 | -5.26874244991853 |
| ENSG00000145332.13 | KLHL8 | 0.120011010118104 | 4.16864164603087 | 0.975694650156711 | 0.334737050759717 | 0.650515199971965 | -5.84913089717818 |
| ENSG00000149231.13 | CCDC82 | 0.170531394301763 | 5.29067608252652 | 0.975658527044626 | 0.334754748972601 | 0.650515199971965 | -5.92536079221601 |
| ENSG00000197520.10 | FAM177B | -0.504962143639556 | 0.422356112667811 | -0.9754792826872 | 0.334882953444368 | 0.65070290104271 | -5.28238750925037 |
| ENSG00000207523.1 | SNORA66 | 0.214011653560738 | -0.233862160523911 | 0.975163253786707 | 0.334997466885778 | 0.650863966481968 | -5.21818029948994 |
| ENSG00000205572.9 | SERF1B | -0.263577759236134 | -0.676552906006563 | -0.974961737077541 | 0.335096257511944 | 0.650994456665251 | -5.12149212546495 |
| lnc-CREBBP-2 | lnc-CREBBP-2 | -0.163598486781024 | 0.833613482333906 | -0.974638671432579 | 0.335254676281422 | 0.651173901991511 | -5.43676970226494 |
| lnc-ZNF596-6 | lnc-ZNF596-6 | -0.155428398512526 | 1.54795786001903 | -0.974600611610541 | 0.33527334261977 | 0.651173901991511 | -5.55667978422359 |
| ENSG00000176927.15 | EFCAB5 | -0.285755137133495 | 0.399772463095395 | -0.974553509651289 | 0.335296444610034 | 0.651173901991511 | -5.31830011973342 |
| ENSG00000198624.12 | CCDC69 | -0.133506472463107 | 7.5669357799817 | -0.97451532833135 | 0.335315172093865 | 0.651173901991511 | -5.93127074104427 |
| ENSG00000165792.17 | METTL17 | -0.0836700465123554 | 5.35130528379953 | -0.974383800135303 | 0.335379690452458 | 0.651194984077452 | -5.92889071007899 |
| ENSG00000189091.12 | SF3B3 | -0.0697450843263168 | 7.35372758115825 | -0.974315703370629 | 0.335413097124419 | 0.651194984077452 | -5.93491097525869 |
| ENSG00000100605.16 | ITPK1 | -0.146881809653722 | 4.81320161517197 | -0.974284126105297 | 0.33542858894159 | 0.651194984077452 | -5.90493040439735 |
| ENSG00000100028.11 | SNRPD3 | -0.137335584802556 | 5.45094602826823 | -0.97422612171736 | 0.335457047157451 | 0.651194984077452 | -5.93062270883378 |

| | | | | | | | |
|---|---|---|---|---|---|---|---|
| ENSG00000205534.6 | SMG1P2 | 0.165045743587962 | 2.57802727927001 | 0.974146411888019 | 0.3354961571621 | 0.651194984077452 | -5.67978205687937 |
| lnc-LIPI-11 | lnc-LIPI-11 | -0.854045477667323 | 0.191592479203214 | -0.974458010057441 | 0.335516922691775 | 0.651194984077452 | -5.27557594211676 |
| lnc-DCT-12 | lnc-DCT-12 | -0.150494905392108 | -0.189596696254985 | -0.974026996438368 | 0.335554754606717 | 0.651194984077452 | -5.20996296221444 |
| ENSG00000144455.13 | SUMF1 | -0.0792136938414547 | 3.97790164546595 | -0.973977329668582 | 0.335579128217924 | 0.651194984077452 | -5.82763174092579 |
| ENSG00000120868.13 | APAF1 | 0.128758540619374 | 4.89450376706549 | 0.973707019602426 | 0.335711801640571 | 0.651388832860279 | -5.90621247002552 |
| ENSG00000285589.1 | | -0.0725374427346237 | 4.89308323702413 | -0.973622272042738 | 0.335753404586217 | 0.651388832860279 | -5.90822480070607 |
| ENSG00000175203.15 | DCTN2 | 0.095751865582586 | 4.8110995867501 | 0.973529989781515 | 0.335798710258566 | 0.651388832860279 | -5.90478462272635 |
| ENSG00000224051.6 | CPTP | -0.189261892092405 | 2.04592640008443 | -0.973515933109319 | 0.335805611693165 | 0.651388832860279 | -5.62037633023554 |
| lnc-CFL2-5 | lnc-CFL2-5 | 0.118539639964014 | 2.109634493794 | 0.973417181775247 | 0.335854098513669 | 0.651421495449194 | -5.60990309073341 |
| ENSG00000280434.1 | | -0.207778907660358 | 2.22214714295014 | -0.973032679813764 | 0.336042933107472 | 0.651660434913015 | -5.63634806399414 |
| lnc-VIPR1-6 | lnc-VIPR1-6 | 0.27018752998766 | 4.11306987117052 | 0.972982632689303 | 0.336106995470475 | 0.651660434913015 | -5.83054377287312 |
| ENSG00000269242.1 | | 0.185466835650571 | 3.3673455731324 | 0.97289627282074 | 0.336109941598672 | 0.651660434913015 | -5.75462715550317 |
| ENSG00000260005.6 | | -0.555666166121485 | 1.1616544241278 | -0.973138982698617 | 0.336125492144595 | 0.651660434913015 | -5.46326199497481 |
| lnc-SLC9B1-3 | lnc-SLC9B1-3 | 0.167006722537272 | 3.0901613715966 | 0.972844067142681 | 0.336135589441358 | 0.651660434913015 | -5.72852756183953 |
| lnc-BAZ2B-2 | lnc-BAZ2B-2 | -0.265416202817634 | 0.575616380153935 | -0.972740464069162 | 0.33618649189238 | 0.651686402838524 | -5.417242991609 |
| ENSG00000071537.13 | SEL1L | 0.096608268424486 | 6.28802660809734 | 0.972687926228988 | 0.336212306842651 | 0.651686402838524 | -5.94479757229184 |
| ENSG00000168385.17 | SEPTIN2 | 0.093480666188726 | 7.58282135013706 | 0.972452445393473 | 0.336328028741918 | 0.651849323502345 | -5.93355611165987 |
| ENSG00000152193.7 | OBI1 | 0.155549013359101 | 4.35477924421475 | 0.97238404844422 | 0.336361645897741 | 0.651853098344148 | -5.86358574082414 |
| ENSG00000085433.15 | WDR47 | 0.11661199552773 | 3.19026356697355 | 0.972240587812606 | 0.336432164186319 | 0.651928378407739 | -5.74409627034931 |
| lnc-RET-4 | lnc-RET-4 | 0.313241379866066 | -0.624247788414664 | 0.972125619685473 | 0.336488684050596 | 0.651976521115825 | -5.09798114094062 |
| ENSG00000126756.11 | UXT | 0.118445377497545 | 4.81440018942891 | 0.97199716152317 | 0.336551843278054 | 0.652037517902127 | -5.90588373743547 |
| lnc-ZNF614-1 | lnc-ZNF614-1 | 0.178441710316825 | 0.388614937202086 | 0.971912439753725 | 0.336593502885647 | 0.652056853637202 | -5.33777936088612 |
| ENSG00000213123.10 | DYNLT2B | -0.186768723882068 | 0.128467768527618 | -0.971841343212304 | 0.336628465311545 | 0.652063213015955 | -5.29109879371139 |
| ENSG00000284828.1 | NA | -0.218405863371132 | 1.21441487009373 | -0.971594639655542 | 0.336749802979641 | 0.652134433076428 | -5.50103359332495 |
| ENSG00000261349.1 | | -1.00531893874177 | -1.92518524058816 | -0.971614067181433 | 0.336757428889848 | 0.652134433076428 | -4.80319238152321 |
| ENSG00000073849.14 | ST6GAL1 | -0.120983582900172 | 8.76106017452691 | -0.971573336034813 | 0.336760282231422 | 0.652134433076428 | -5.90818655576261 |
| ENSG00000170088.13 | TMEM192 | 0.135610105129874 | 4.01670446106821 | 0.971292790267396 | 0.336898302966132 | 0.652340335837431 | -5.83727815301675 |
| ENSG00000007047.14 | MARK4 | -0.123118241375718 | 2.94017122783746 | -0.971226022345491 | 0.336931156479943 | 0.652342582431254 | -5.71986793551109 |
| ENSG00000204209.12 | DAXX | -0.176324169277408 | 4.70908493430599 | -0.971160071104046 | 0.33696361023607 | 0.652344054692346 | -5.90016626898401 |
| ENSG00000281469.1 | | -0.207963810496174 | 0.349861330343066 | -0.971064643341874 | 0.337010572692975 | 0.652373612126352 | -5.33940038988054 |
| lnc-MYO19-1 | lnc-MYO19-1 | -0.129192522367765 | 1.460300493157 | -0.97089946480241 | 0.337091871601566 | 0.652469625086989 | -5.54839628169709 |
| ENSG00000107223.12 | EDF1 | -0.13602696809188 | 6.11144249123638 | -0.9706848219535 | 0.337197535760254 | 0.652528785317058 | -5.94611200477824 |
| ENSG00000115310.17 | RTN4 | 0.132828683650989 | 5.44611804843468 | 0.970628210004429 | 0.337225408305977 | 0.652528785317058 | -5.93590839466298 |
| lnc-ITK-2 | lnc-ITK-2 | 0.21059422470815 | 2.29497051554382 | 0.970578065419182 | 0.337250097965663 | 0.652528785317058 | -5.64031879386793 |
| lnc-CCDC117-1 | lnc-CCDC117-1 | -0.223690352864653 | 2.73612049679448 | -0.970568954109692 | 0.33725458422489 | 0.652528785317058 | -5.7028842377339 |
| lnc-CCDC171-5 | lnc-CCDC171-5 | -0.231812134344991 | -0.136092246371463 | -0.970466411816492 | 0.3373050771062 | 0.652528785317058 | -5.22771849436791 |
| lnc-SLC38A1-1 | lnc-SLC38A1-1 | 0.128736307443338 | 2.87253678644143 | 0.970451033381228 | 0.337312650040009 | 0.652528785317058 | -5.71226096770639 |
| ENSG00000111897.6 | SERINC1 | 0.160261458192438 | 7.62561606599766 | 0.970379641682122 | 0.337347807544396 | 0.652535468723457 | -5.93499440994286 |
| lnc-CHCHD1-2 | lnc-CHCHD1-2 | 0.173979584526826 | 0.351299478044148 | 0.970242237244914 | 0.337415480498345 | 0.652541605062663 | -5.33263508815562 |
| ENSG00000278771.1 | RN7SL3 | 0.373068000284198 | 6.41172986048822 | 0.970523077338328 | 0.33743386210841 | 0.652541605062663 | -5.94658665227399 |
| ENSG00000170476.15 | MZB1 | 0.260658403979599 | 5.27467457124594 | 0.9703041023773 | 0.33744608872793 | 0.652541605062663 | -5.93255387629088 |
| ENSG00000254772.9 | EEF1G | -0.158362120532356 | 9.10644537281854 | -0.970065154455535 | 0.337502708731814 | 0.652566434567642 | -5.89866705853465 |
| ENSG00000189050.15 | RNFT1 | -0.106203307207274 | 4.02984507671398 | -0.96998972388313 | 0.337539869218317 | 0.652566434567642 | -5.84347895339458 |
| ENSG00000149357.9 | LAMTOR1 | -0.107239148082229 | 4.805724390186 | -0.969960958321097 | 0.337554041142772 | 0.652566434567642 | -5.9077786046669 |
| ENSG00000198087.7 | CD2AP | 0.109394248177854 | 6.27420578458875 | 0.969797168995854 | 0.337634742747228 | 0.652661148569327 | -5.9475708065875 |
| ENSG00000198740.8 | ZNF652 | 0.104768888149895 | 6.34351673749362 | 0.969589689672261 | 0.337736989539048 | 0.652797488560637 | -5.94776331263685 |
| ENSG00000062822.13 | POLD1 | -0.207964148557925 | 3.25975154774887 | -0.969469990193621 | 0.337795987378452 | 0.652850216733398 | -5.76527069447696 |
| ENSG00000188313.12 | PLSCR1 | 0.26063618679877 | 1.79167089272187 | 0.969315570512837 | 0.337872108343444 | 0.652893183510402 | -5.58143979966352 |
| LINC02551 | LINC02551 | -0.125428572840587 | 4.70044537899532 | -0.969296197247979 | 0.337881659172754 | 0.652893183510402 | -5.89640820799629 |
| ENSG00000258634.3 | | -0.117582563553625 | 3.37834681355075 | -0.96915770636384 | 0.337949939049429 | 0.652959435786468 | -5.77037072386122 |
| ENSG00000204138.12 | PHACTR4 | 0.0975863925930609 | 5.14377824322466 | 0.96909797287508 | 0.337979392161931 | 0.652959435786468 | -5.92518804150334 |
| lnc-RALGDS-2 | lnc-RALGDS-2 | -0.176887985133778 | 1.44238628033955 | -0.969015144382733 | 0.338020235677276 | 0.652977054054001 | -5.54223485380395 |
| ENSG00000266728.5 | | 0.190878296809963 | 2.87892703003859 | 0.968891902079683 | 0.338081013702482 | 0.653033174271601 | -5.70769603384828 |
| ENSG00000153815.16 | CMIP | -0.190636189519066 | 4.57272671449565 | -0.968778832517948 | 0.338136781343826 | 0.653079606868723 | -5.89012282599932 |
| lnc-DAZAP2-3 | lnc-DAZAP2-3 | -0.29960751070949 | 0.685887832191018 | -0.968590027377145 | 0.338229916570594 | 0.653195219994537 | -5.40793134091713 |
| ENSG00000173542.8 | MOB1B | 0.114482606836916 | 4.59165341931826 | 0.968528825651168 | 0.338260110281386 | 0.653195219994537 | -5.89657239479806 |
| lnc-PTK2-5 | lnc-PTK2-5 | 0.366527158175581 | 1.16789154585812 | 0.96832219424835 | 0.338376551069714 | 0.653358775860029 | -5.47227597497802 |
| ENSG00000155542.11 | SETD9 | 0.164110840047681 | 1.10497538544242 | 0.967966763957629 | 0.338537485756851 | 0.653601034152551 | -5.48620304854149 |
| ENSG00000148219.16 | ASTN2 | 0.22295191992377 | 1.38189675565386 | 0.967909961021281 | 0.338565526217485 | 0.653601034152551 | -5.4913413735103 |
| ENSG00000136891.13 | TEX10 | 0.0962880140718143 | 4.7518784825815 | 0.967661118987494 | 0.338688383931386 | 0.653776892662452 | -5.90647494626746 |
| ENSG00000280143.1 | | -0.311420569657913 | 1.42645733530963 | -0.967478650733376 | 0.338778490565606 | 0.653889504061503 | -5.51473662961511 |
| lnc-PRLHR-1 | lnc-PRLHR-1 | -0.133052046523314 | 1.20251249093619 | -0.967374524353431 | 0.338829917486234 | 0.653927444127441 | -5.50424637565352 |
| ENSG00000166398.12 | GARRE1 | -0.0945510128194131 | 5.86868967045366 | -0.967305016687151 | 0.338864249476898 | 0.653932387678885 | -5.94663901433948 |
| ENSG00000165288.10 | BRWD3 | 0.0939481405123903 | 6.53849563716591 | 0.967059404688534 | 0.338985583372863 | 0.654105208827564 | -5.95001248216056 |
| lnc-FAM53B-1 | lnc-FAM53B-1 | 0.101094806377882 | 2.04672033432636 | 0.966958959722477 | 0.339035212143922 | 0.654139648627516 | -5.61252927129879 |
| ENSG00000130254.11 | SAFB2 | 0.135080459396308 | 6.10579223725646 | 0.966690427743265 | 0.339167914595192 | 0.654295938620491 | -5.94961790233362 |
| lnc-CASTOR3-4 | lnc-CASTOR3-4 | -0.140297387430318 | 1.0185981721839 | -0.966629944361011 | 0.33919780888044 | 0.654295938620491 | -5.47344890542582 |
| ENSG00000282246.1 | NA | 0.0823154240058659 | 3.75201994028963 | 0.966602082269895 | 0.339211580479972 | 0.654295938620491 | -5.80702700049587 |

| | | | | | | | |
|---|---|---|---|---|---|---|---|
| lnc-PEX16-4 | lnc-PEX16-4 | -0.130203973329679 | 1.55188273254813 | -0.966441071167716 | 0.339291171890644 | 0.654388136340435 | -5.55023719452483 |
| ENSG00000075790.10 | BCAP29 | 0.0963810194739754 | 4.74624881550048 | 0.966221295216318 | 0.339399832110705 | 0.654536376307939 | -5.90583783008007 |
| ENSG00000203814.6 | H2BC18 | -0.219084662912032 | 5.79514349482669 | -0.966066138948197 | 0.339492861531346 | 0.65465444715193 | -5.9487229047659 |
| lnc-EPB41-3 | lnc-EPB41-3 | -0.143459321946007 | 3.7255140584579 | -0.965393124501306 | 0.339809498614657 | 0.655123534297464 | -5.81229847532305 |
| ENSG00000078804.12 | TP53INP2 | 0.23790812363548 | 2.92810936709384 | 0.965340165775081 | 0.339835706570936 | 0.655123534297464 | -5.71071113686795 |
| LOXL1-AS1 | LOXL1-AS1 | -0.717494836570983 | -1.70642590450131 | -0.965199299082376 | 0.339905424525302 | 0.655123534297464 | -4.89616926498919 |
| lnc-HOMER2-3 | lnc-HOMER2-3 | -0.0938289063759719 | 2.51172180761874 | -0.965142525799281 | 0.339933525527985 | 0.655123534297464 | -5.67462327570578 |
| lnc-TEFM-7 | lnc-TEFM-7 | 0.116136506492786 | 2.64005373018425 | 0.965124977563066 | 0.33994221166908 | 0.655123534297464 | -5.68564855066349 |
| ENSG00000203667.9 | COX20 | -0.104884523175695 | 4.77877413626916 | -0.965095062644438 | 0.339957019491462 | 0.655123534297464 | -5.91190520473271 |
| ENSG00000159055.3 | MIS18A | -0.171257463567747 | 1.05311038947877 | -0.965021115763444 | 0.339993624880886 | 0.655123534297464 | -5.48585601501707 |
| ENSG00000243696.4 | | -0.156765935411256 | 3.7502347370659 | -0.964999409101053 | 0.340004370669641 | 0.655123534297464 | -5.81599150745501 |
| ENSG00000140391.14 | TSPAN3 | 0.210003211562209 | 5.81750029897004 | 0.964973850425302 | 0.340022577704893 | 0.655123534297464 | -5.94886318778275 |
| lnc-RCL1-2 | lnc-RCL1-2 | -0.208165900922123 | 1.52393551527768 | -0.964897044062952 | 0.340055049074779 | 0.655124773035023 | -5.55384144813109 |
| ENSG00000239732.3 | TLR9 | -0.278272760671031 | 2.23401081451991 | -0.964694984795643 | 0.340155098349489 | 0.655256189923025 | -5.64350947223198 |
| ENSG00000104756.15 | KCTD9 | 0.140128300412142 | 4.05126906205938 | 0.964626951158277 | 0.340188789476051 | 0.655259765467486 | -5.84501201716451 |
| ENSG00000165804.15 | ZNF219 | -0.332561646701123 | 1.13773958037762 | -0.964314888704225 | 0.340346237104586 | 0.65544191204105 | -5.51877774406997 |
| lnc-MAP6D1-2 | lnc-MAP6D1-2 | 0.124472524399158 | 1.29535757673787 | 0.964292361641232 | 0.340354514708791 | 0.65544191204105 | -5.52111249878779 |
| ENSG00000122359.17 | ANXA11 | -0.0933723666029156 | 7.61937943026678 | -0.964188044702226 | 0.340406194791543 | 0.65544191204105 | -5.94062722123467 |
| lnc-STOML3-5 | lnc-STOML3-5 | 0.160107860292753 | 0.937010930240095 | 0.964173205764239 | 0.340413546633696 | 0.65544191204105 | -5.46749190199419 |
| lnc-TEC-2 | lnc-TEC-2 | -0.237404000581681 | 0.00837148361333548 | -0.964114620949245 | 0.340442573076367 | 0.65544191204105 | -5.27335775865108 |
| lnc-HOXB1-1 | lnc-HOXB1-1 | -0.465665219559922 | 0.129236697601041 | -0.963803639120198 | 0.340602226809863 | 0.655628582098691 | -5.24699755750824 |
| ENSG00000122435.9 | TRMT13 | -0.113793688274514 | 4.34719279534448 | -0.96379040863129 | 0.340603237058801 | 0.655628582098691 | -5.88004033938579 |
| lnc-PLPP4-8 | lnc-PLPP4-8 | -0.18101874183429 | 2.23272845860502 | -0.963675044518288 | 0.34066041807903 | 0.655677331711303 | -5.64644212416129 |
| lnc-ERLEC1-7 | lnc-ERLEC1-7 | -0.231677702485926 | 0.87549110558712 | -0.963476622028014 | 0.340758782453322 | 0.655788876954501 | -5.45859164732064 |
| ENSG00000261794.1 | GOLGA8H | -0.475692113588328 | -0.529504952088857 | -0.963429604364743 | 0.340782093373432 | 0.655788876954501 | -5.19190949639051 |
| ENSG00000130940.14 | CASZ1 | 0.16589639316316 | 3.74027704623729 | 0.963251609092399 | 0.340870351355264 | 0.655897395713321 | -5.81793362346903 |
| lnc-RNF145-6 | lnc-RNF145-6 | -0.272728897903487 | -0.39113888491529 | -0.96303733848699 | 0.340976616369928 | 0.656040539796432 | -5.15675966717938 |
| ENSG00000160050.14 | CCDC28B | 0.295328118958387 | -0.0928781976365666 | 0.962901802126049 | 0.341043845393922 | 0.656108558719796 | -5.22887928458171 |
| ENSG00000197879.16 | MYO1C | -0.197861703538283 | 4.89438737269496 | -0.962814004188686 | 0.341087399797874 | 0.65613102336819 | -5.92139257533816 |
| ENSG00000260261.2 | | -0.187783086273656 | 1.79637402789813 | -0.962692039765137 | 0.341147909478112 | 0.65618145619457 | -5.59417842607835 |
| lnc-WDR77-4 | lnc-WDR77-4 | -0.242755071191128 | 0.736868711247763 | -0.96263265032584 | 0.341177376679507 | 0.65618145619457 | -5.41826234353998 |
| ENSG00000003393.14 | ALS2 | 0.0844349179785517 | 4.14261859055978 | 0.962509638542409 | 0.341238416686243 | 0.656210318652218 | -5.85544626454773 |
| lnc-IDI2-2 | lnc-IDI2-2 | 0.267389651898778 | -0.164707923499784 | 0.96243184938687 | 0.341277020388299 | 0.656210318652218 | -5.19702749182508 |
| ENSG00000080298.15 | RFX3 | 0.165849279896937 | 5.34908558132401 | 0.962361346315875 | 0.341312010793909 | 0.656210318652218 | -5.93705851971092 |
| ENSG00000116030.16 | SUMO1 | 0.110927546893075 | 5.16895174177438 | 0.962305809535587 | 0.341339575160965 | 0.656210318652218 | -5.93248865417122 |
| ENSG00000149929.15 | HIRIP3 | -0.164574818900001 | 1.91136080995521 | -0.962228179409141 | 0.34137810751074 | 0.656210318652218 | -5.60163990095976 |
| lnc-PLAC8L1-3 | lnc-PLAC8L1-3 | -0.115667508319687 | 2.59482563447182 | -0.96211081475806 | 0.341436367892314 | 0.656210318652218 | -5.68327205393648 |
| lnc-KDM5A-1 | lnc-KDM5A-1 | -0.190778652248442 | 2.90698038879166 | -0.962077856300266 | 0.341452729815688 | 0.656210318652218 | -5.72792128590844 |
| ENSG00000220515.2 | PGAM1P10 | -0.343014311128112 | -0.741538770250938 | -0.962069076228853 | 0.341457088689089 | 0.656210318652218 | -5.0697576016673 |
| lnc-POLDIP3-1 | lnc-POLDIP3-1 | 0.135414878531131 | 1.35060935801772 | 0.961966624903615 | 0.341507953456684 | 0.656210318652218 | -5.53511929232258 |
| ENSG00000102977.14 | ACD | -0.187680686546512 | 2.95496718582583 | -0.961960095503775 | 0.341511195326364 | 0.656210318652218 | -5.73880153042588 |
| ENSG00000279838.1 | NA | 0.187537256733753 | 0.287985669914539 | 0.961737909112642 | 0.34162152379717 | 0.656299778263689 | -5.32871667723404 |
| lnc-IVNS1ABP-3 | lnc-IVNS1ABP-3 | 0.187537256733753 | 0.287985669914539 | 0.961737909112642 | 0.34162152379717 | 0.656299778263689 | -5.32871667723404 |
| ENSG00000184281.14 | TSSC4 | 0.143949487894451 | 4.16011370098142 | 0.961614572403117 | 0.341682777844065 | 0.656350135924908 | -5.86702648111683 |
| lnc-KBTBD4-4 | lnc-KBTBD4-4 | 0.090574428680819 | 4.68454204936356 | 0.961508431782508 | 0.341735497437505 | 0.656350135924908 | -5.90414365320306 |
| lnc-IFNAR1-1 | lnc-IFNAR1-1 | -0.229221176229341 | 0.754395631829935 | -0.96149252200832 | 0.341743400218979 | 0.656350135924908 | -5.43671614931328 |
| ENSG00000153064.11 | BANK1 | 0.338122394413208 | 6.70223942753501 | 0.961636490558043 | 0.341814309144206 | 0.656425072319015 | -5.95400517852633 |
| lnc-ISM2-1 | lnc-ISM2-1 | -0.210537229507311 | 1.05898489908964 | -0.961169234770451 | 0.341904011281518 | 0.65647525880588 | -5.50288709193835 |
| ENSG00000198440.9 | ZNF583 | -0.174237678269753 | 2.82634281512803 | -0.96116879377742 | 0.341904230403684 | 0.65647525880588 | -5.71066279723277 |
| lnc-GTPBP4-7 | lnc-GTPBP4-7 | 0.192833797711945 | 0.291426598182089 | 0.960940381759329 | 0.342017737073649 | 0.656543841134014 | -5.33310802541311 |
| lnc-GOLGA8O-5 | lnc-GOLGA8O-5 | -0.185743704725605 | 0.320738735798946 | -0.960889577489012 | 0.34204298705246 | 0.656543841134014 | -5.3495666802274 |
| ENSG00000164975.15 | SNAPC3 | -0.165605548842029 | 5.087121044024 | -0.960863185427356 | 0.342056104527694 | 0.656543841134014 | -5.92737794028663 |
| ENSG00000158427.14 | TMSB15B | -0.189040426951433 | -0.349863378379736 | -0.960840180361239 | 0.342067538858338 | 0.656543841134014 | -5.17634283978924 |
| ENSG00000157601.13 | MX1 | 0.263048860911716 | 6.36892678822218 | 0.960823190263892 | 0.342150607627937 | 0.656642047254623 | -5.95593897067133 |
| ENSG00000164949.7 | GEM | -0.392199355188692 | 0.913096298816371 | -0.960525212515067 | 0.342249114826424 | 0.656690534956247 | -5.45413210218226 |
| ENSG00000189136.9 | UBE2Q2P1 | 0.244298266323489 | 2.34079833474451 | 0.960447505062658 | 0.342262751400768 | 0.656690534956247 | -5.66082812317183 |
| ENSG00000122707.11 | RECK | -0.296340843946372 | 1.31355358645167 | -0.960429339068141 | 0.342271784133485 | 0.656690534956247 | -5.53534040103155 |
| ENSG00000167967.15 | E4F1 | -0.139543302601411 | 4.36490310247461 | -0.960348304538811 | 0.342312079101925 | 0.656690534956247 | -5.88780314270541 |
| ENSG00000166272.17 | WBP1L | -0.194842625407468 | 4.83029488115294 | -0.960196257500235 | 0.342387693998776 | 0.656690534956247 | -5.91750212683342 |
| ENSG00000100442.10 | FKBP3 | 0.0721011578982758 | 4.48970634517887 | 0.960139462882415 | 0.342415941510874 | 0.656690534956247 | -5.88966132370586 |
| lnc-C21orf58-2 | lnc-C21orf58-2 | 0.134459990863141 | 1.78874571000115 | 0.96013197514904 | 0.342419665743668 | 0.656690534956247 | -5.58016107866542 |
| ENSG00000128915.11 | ICE2 | -0.0810437035568468 | 5.49391071673055 | -0.960108968792575 | 0.342431108763805 | 0.656690534956247 | -5.9455889213975 |
| ENSG00000057704.12 | TMCC3 | 0.190640559201761 | 5.13638047091662 | 0.959999322745481 | 0.342485648558924 | 0.656733939285293 | -5.93148672762002 |
| ENSG00000075239.13 | ACAT1 | 0.185803496639221 | 3.43969031370117 | 0.959686150578785 | 0.342641457356377 | 0.656971505986614 | -5.78142507962674 |
| ENSG00000173258.12 | ZNF483 | -0.446333912707926 | 3.59278521551992 | -0.959893334352818 | 0.342693304338182 | 0.656977708209621 | -5.78276120992956 |
| ENSG00000137760.14 | ALKBH8 | 0.181588059335603 | 3.96717568023698 | 0.959551367054914 | 0.342708529031079 | 0.656977708209621 | -5.83047433879044 |
| ENSG00000167972.13 | ABCA3 | -0.40148401367838 | 1.66200585313739 | -0.959592610207444 | 0.342776802054949 | 0.657047393993017 | -5.60854680673162 |
| lnc-LSS-1 | lnc-LSS-1 | 0.115092862304049 | 1.826982194224 | 0.959271964687222 | 0.342847594364528 | 0.657049981031847 | -5.59018990846483 |

| | | | | | | | |
|---|---|---|---|---|---|---|---|
| ENSG00000087460.24 | GNAS | 0.109289198201504 | 8.51897622476078 | 0.959228201397159 | 0.342869379798108 | 0.657049981031847 | -5.92584997537567 |
| lnc-APOLD1-1 | lnc-APOLD1-1 | 0.17123743815474 | 3.21499084448242 | 0.959219085914714 | 0.342873917614685 | 0.657049981031847 | -5.75997342778535 |
| lnc-COIL-2 | lnc-COIL-2 | 0.183897189532903 | 0.670296165059519 | 0.958986682936947 | 0.342989624542519 | 0.657169098153601 | -5.39306736980687 |
| ENSG00000225830.12 | ERCC6 | 0.0831582240371311 | 5.64363764963003 | 0.95896120071 8382 | 0.343002312996686 | 0.657169098153601 | -5.94997828035126 |
| lnc-SGSH-2 | lnc-SGSH-2 | -0.188429860251728 | 1.13929742303651 | -0.95890186231966 | 0.343031860786197 | 0.657169098153601 | -5.48908155333783 |
| KCNJ2-AS1 | KCNJ2-AS1 | -0.778391609033069 | -1.97093442865614 | -0.958751068579452 | 0.343106956702685 | 0.657196093597388 | -4.84964211226786 |
| ENSG00000167548.15 | KMT2D | 0.183489780340761 | 6.80475932469404 | 0.958745339295371 | 0.34310981012474 | 0.657196093597388 | -5.95635206267639 |
| ENSG00000151005.4 | TKTL2 | 0.280553154755888 | 0.715067080612639 | 0.958675564424316 | 0.343144562176278 | 0.65720150025815 | -5.43462778159984 |
| ENSG00000160789.19 | LMNA | -0.580227492747527 | 1.26466598180335 | -0.958890218879382 | 0.34319691166051 | 0.657232492038792 | -5.5700154846575 |
| lnc-RLN1-2 | lnc-RLN1-2 | -0.253063404359801 | 0.235009115169068 | -0.958469965851866 | 0.343246976082318 | 0.657232492038792 | -5.3363045000364 |
| ENSG00000155158.20 | TTC39B | 0.765210365796436 | 1.52763288089946 | 0.958796281132222 | 0.343290884604635 | 0.657232492038792 | -5.58760362851318 |
| ENSG00000172469.15 | MANEA | -0.16773388346259 | 2.66266403084631 | -0.958378673211636 | 0.343292457763757 | 0.657232492038792 | -5.70947030828991 |
| lnc-EFCAB11-9 | lnc-EFCAB11-9 | -0.263210673639435 | 1.22947796912012 | -0.958320565365166 | 0.343321408970812 | 0.657232492038792 | -5.50331968959608 |
| ENSG00000100380.13 | ST13 | -0.139853171185785 | 6.60759204923863 | -0.958258509572993 | 0.343352328955601 | 0.657232492038792 | -5.95727779437526 |
| ENSG00000197063.10 | MAFG | 0.182357125946423 | 3.38232356295838 | 0.958126455304617 | 0.343418132586457 | 0.657250605393611 | -5.78686489137742 |
| ENSG00000167302.10 | TEPSIN | 0.137080239720923 | 5.56212494484725 | 0.958111360039723 | 0.343425655201297 | 0.657250605393611 | -5.9503620732237 |
| ENSG00000147419.17 | CCDC25 | 0.112844450341968 | 4.46379707219427 | 0.957935218288007 | 0.34351344221174 | 0.657313295464022 | -5.89341464785416 |
| lnc-COQ3-1 | lnc-COQ3-1 | -0.137914712938041 | 4.47468390076278 | -0.957917484275395 | 0.343522281460743 | 0.657313295464022 | -5.89472811634967 |
| ENSG00000143110.11 | C1orf162 | -0.350053266408927 | 4.08930102837804 | -0.957984695911042 | 0.343608914722783 | 0.65741794866509 | -5.83463346526067 |
| lnc-DDX5-4 | lnc-DDX5-4 | 0.142969092788584 | 1.49127606766236 | 0.957381893087912 | 0.343789309629765 | 0.657538021307337 | -5.55150036956518 |
| ENSG00000213047.12 | DENND1B | 0.260693402998259 | 3.88904935060287 | 0.957455340609221 | 0.343790574421752 | 0.657538021307337 | -5.86110584534802 |
| ENSG00000233892.1 | PAIP1P1 | 0.190857798824281 | 1.28620484362325 | 0.957352842124375 | 0.343803797409089 | 0.657538021307337 | -5.49773353599134 |
| lnc-ANAPC7-4 | lnc-ANAPC7-4 | 0.312474910960089 | -0.389420980543251 | 0.957326197634801 | 0.343817085429499 | 0.657538021307337 | -5.15542152948184 |
| lnc-ZNF800-2 | lnc-ZNF800-2 | -0.174197013487241 | 4.9017521352851 | -0.957297493850191 | 0.343831400832282 | 0.657538021307337 | -5.92177477363031 |
| ENSG00000100784.11 | RPS6KA5 | -0.134791711046854 | 5.03204897884533 | -0.956969483784771 | 0.343995016869256 | 0.657773667748763 | -5.93078573755881 |
| ENSG00000145723.16 | GIN1 | -0.151062442328466 | 2.45438243176801 | -0.956922341505273 | 0.344018536331702 | 0.657773667748763 | -5.67372353814311 |
| ENSG00000187187.13 | ZNF546 | -0.167744450096239 | 3.49338566715849 | -0.956368589549292 | 0.344294884871599 | 0.658240907979949 | -5.78230581666369 |
| ENSG00000071553.16 | ATP6AP1 | -0.09703419298065 | 4.5938196844123 | -0.956105919159844 | 0.3444260211679 | 0.658383641693213 | -5.90457950274236 |
| lnc-EXOSC1-5 | lnc-EXOSC1-5 | 0.219748116347341 | -0.223520736293888 | 0.956090909348287 | 0.34443351570483 | 0.658383641693213 | -5.1904664930744 |
| ENSG00000165669.13 | FAM204A | 0.0795787236073453 | 5.31151020377064 | 0.955962967127461 | 0.344497402806407 | 0.658444613424113 | -5.94501300343358 |
| ENSG00000167775.10 | CD320 | -0.236304982749652 | 2.11272559427464 | -0.955767083749754 | 0.34459523104593 | 0.658558637467758 | -5.64387952757233 |
| lnc-IER3IP1-1 | lnc-IER3IP1-1 | -0.113431705703335 | 0.925232291881209 | -0.955715390918075 | 0.344621050583745 | 0.658558637467758 | -5.46612248848911 |
| ENSG00000119686.9 | FLVCR2 | -0.183921225539039 | 0.610980218151359 | -0.955513281530075 | 0.344722012457656 | 0.658643370738649 | -5.41270479340143 |
| lnc-GLT8D1-2 | lnc-GLT8D1-2 | 0.135091431693652 | 0.243334124069633 | 0.955498514549876 | 0.34472938993186 | 0.658643370738649 | -5.32139565922034 |
| ENSG00000157426.13 | AASDH | -0.10482021391373 | 4.02453063408924 | -0.95534243028613 | 0.34480737485454 | 0.658683364819953 | -5.85100846679796 |
| lnc-SNAPC5-3 | lnc-SNAPC5-3 | 0.151023705234449 | 7.12511904205446 | 0.955236497340972 | 0.344860309137917 | 0.658683364819953 | -5.95622908180808 |
| lnc-CIR1-3 | lnc-CIR1-3 | 0.250751767930823 | -0.25466482173671 | 0.955202600869264 | 0.344877248207397 | 0.658683364819953 | -5.18745917048126 |
| lnc-SH3RF1-2 | lnc-SH3RF1-2 | -0.149837673227616 | 1.35091717658504 | -0.955200440401526 | 0.344878327875657 | 0.658683364819953 | -5.52628753226188 |
| ENSG00000055070.16 | SZRD1 | 0.117406816867781 | 6.43396767485041 | 0.955071249020719 | 0.344942893802205 | 0.658745554196585 | -5.96123630012452 |
| ENSG00000160551.11 | TAOK1 | -0.088640617701425 | 7.06271758357289 | -0.954804222626085 | 0.345076370783183 | 0.65893094094509 | -5.95717276863712 |
| ENSG00000197782.14 | ZNF780A | -0.134407327628787 | 4.39481495537975 | -0.954748966331775 | 0.345103995694897 | 0.65893094094509 | -5.89344528546136 |
| ENSG00000161558.10 | TMEM143 | -0.153045712962638 | 2.03235200306623 | -0.954622236784878 | 0.345167358563457 | 0.658935457519149 | -5.61621974054631 |
| ENSG00000167113.10 | COQ4 | -0.126335127994948 | 4.38289044209334 | -0.954616177456904 | 0.345170388328789 | 0.658935457519149 | -5.89319663140242 |
| ENSG00000020129.15 | NCDN | -0.163369634980308 | 3.18461180049342 | -0.954500315198101 | 0.345228324774299 | 0.658984940075062 | -5.76273561074643 |
| ENSG00000164332.7 | UBLCP1 | 0.116588819952953 | 4.35976671187057 | 0.954365910217225 | 0.345295541475936 | 0.659041730829073 | -5.8858667894061 |
| ENSG00000119771.14 | KLHL29 | -0.402128762643898 | 3.05909050789563 | -0.95450583872702 | 0.345356748488906 | 0.659041730829073 | -5.74033591482126 |
| ENSG00000112651.11 | MRPL2 | 0.100680322821576 | 2.65986021250811 | 0.954111159507572 | 0.345422967475111 | 0.659041730829073 | -5.6955651154247 |
| ENSG00000133063.15 | CHIT1 | -0.169322852807079 | 2.91551121807362 | -0.954041982968767 | 0.345457574855595 | 0.659041730829073 | -5.72399000386061 |
| ENSG00000073861.2 | TBX21 | 0.222399858842532 | 2.85536366310053 | 0.954014926343665 | 0.345471111265623 | 0.659041730829073 | -5.72546630096875 |
| ENSG00000164051.13 | CCDC51 | -0.165406431239877 | 1.41682307387179 | -0.954007890843472 | 0.345474631178657 | 0.659041730829073 | -5.53919627870588 |
| ENSG00000092529.23 | CAPN3 | -0.126259306392257 | 5.26893261400562 | -0.953909480832715 | 0.34552386891997 | 0.659041730829073 | -5.94171937345996 |
| ENSG00000168040.4 | FADD | -0.170643969483242 | 2.27331424194596 | -0.95389543645109 | 0.345530896160314 | 0.659041730829073 | -5.65273456079463 |
| ENSG00000110422.11 | HIPK3 | -0.103547579298048 | 7.31413123427917 | -0.953864761653376 | 0.345546244916064 | 0.659041730829073 | -5.95463229884866 |
| lnc-FAM120AOS-4 | lnc-FAM120AOS-4 | 0.116267688333361 | 2.05638685486823 | 0.953546560130928 | 0.345705490034448 | 0.659273395736612 | -5.62487774178981 |
| lnc-NBPF3-3 | lnc-NBPF3-3 | -0.19654997609448 | 0.128884367070327 | -0.953494056013434 | 0.345731770567021 | 0.659273395736612 | -5.28024399318826 |
| lnc-KCNK7-2 | lnc-KCNK7-2 | -0.202029448393822 | 0.654260154780082 | -0.953378297162614 | 0.345789717428214 | 0.659322342203897 | -5.44456745579823 |
| lnc-IL22-1 | lnc-IL22-1 | -0.124716209307816 | 1.75174103537743 | -0.953314804448632 | 0.34582150349479 | 0.659322342203897 | -5.59940524211762 |
| ENSG00000285721.1 | NA | 0.102484213879164 | 7.2418161608718 | 0.953198051845793 | 0.345879957847958 | 0.659372712085258 | -5.95619626734275 |
| lnc-P2RX5-3 | lnc-P2RX5-3 | 0.206967546981196 | 1.53140530520864 | 0.953002685661149 | 0.345977786128745 | 0.659498126679752 | -5.55859037382052 |
| ENSG00000149187.18 | CELF1 | 0.0765806014492618 | 6.96202922930391 | 0.952699527448022 | 0.346129626670888 | 0.659726465947485 | -5.9600450292024 |
| ENSG00000175489.9 | LRRC25 | -0.58885538808977 | 1.02035144594915 | -0.952532783513102 | 0.346357569471214 | 0.660099801150556 | -5.46150744812246 |
| ENSG00000169188.4 | APEX2 | -0.110078491778633 | 3.37424222662638 | -0.952089160127449 | 0.346435470029446 | 0.660140390280108 | -5.78428400296517 |
| lnc-SLC35E3-8 | lnc-SLC35E3-8 | 0.162342234618907 | 3.45785807541772 | 0.95206088111711 | 0.346449644417811 | 0.660140390280108 | -5.79071950549791 |
| ENSG00000116497.17 | S100PBP | -0.155919304518912 | 5.41272603332928 | -0.952010130802835 | 0.346475083136375 | 0.660140390280108 | -5.9486923545044 |
| lnc-ANXA2R-3 | lnc-ANXA2R-3 | -0.276919293407039 | 0.882721494729707 | -0.951903917567702 | 0.346528326765525 | 0.660180724714439 | -5.45485292089968 |
| ENSG00000213740.2 | SERBP1P1 | 0.710061693931164 | -1.15805331243219 | 0.951798498258201 | 0.346581177742483 | 0.66022030369954 | -4.86178083767713 |
| lnc-RPLP1-12 | lnc-RPLP1-12 | 0.403526791329798 | 1.2736500167368 | 0.951679753481589 | 0.346693405791942 | 0.660312847241379 | -5.51023119143886 |
| lnc-NAP1L4-1 | lnc-NAP1L4-1 | 0.0770801454066499 | 4.89405999893461 | 0.951573653576042 | 0.346693919260437 | 0.660312847241379 | -5.92822966584708 |

| | | | | | | | |
|---|---|---|---|---|---|---|---|
| ENSG00000169217.8 | CD2BP2 | -0.0878654835568922 | 5.82969689313365 | -0.95128654914336 | 0.346837914179701 | 0.660428991641794 | -5.96161698313919 |
| ENSG00000282100.1 | HSP90AB4P | 0.282237641316846 | 0.204787605839243 | 0.951247346989189 | 0.346857578763547 | 0.660428991641794 | -5.27352193148641 |
| lnc-TKTL2-1 | lnc-TKTL2-1 | 0.142642856197399 | 2.8259156820998 | 0.951206866002725 | 0.346877885606509 | 0.660428991641794 | -5.70731917284559 |
| lnc-ZNF782-1 | lnc-ZNF782-1 | -0.116703077790379 | 2.85589941930997 | -0.951109643914832 | 0.346926659201941 | 0.660428991641794 | -5.72262491180896 |
| ENSG00000163138.18 | PACRGL | -0.123047244313414 | 3.19161621080302 | -0.951047250943533 | 0.346957962387413 | 0.660428991641794 | -5.7702136559357 |
| ENSG00000096968.13 | JAK2 | -0.112631858510659 | 6.92843299140861 | -0.951029551399751 | 0.346966842766707 | 0.660428991641794 | -5.96261666929181 |
| ENSG00000254101.6 | LINC02055 | 1.32351444681489 | -0.950931274747901 | 0.951305128596482 | 0.346986545521583 | 0.660428991641794 | -4.94711722553083 |
| ENSG00000064393.15 | HIPK2 | 0.190028084883139 | 5.38503472914786 | 0.950913160272973 | 0.34702524333425 | 0.660428991641794 | -5.95097706753204 |
| lnc-COMMD10-14 | lnc-COMMD10-14 | -0.251150261593218 | -0.380338241713128 | -0.950788133156665 | 0.347087984324572 | 0.660428991641794 | -5.1503865015547 |
| ENSG00000238862.1 | SNORD19B | 0.181605843225396 | 0.63171694344342 | 0.950785802153502 | 0.347089154137378 | 0.660428991641794 | -5.39833331286958 |
| ENSG00000108651.9 | UTP6 | -0.0937636278393272 | 5.18650114541281 | -0.950687151362348 | 0.347138664370963 | 0.660428991641794 | -5.94263980326109 |
| ENSG00000118707.9 | TGIF2 | -0.173700049897726 | 4.57650771290728 | -0.950665066252773 | 0.347149748943289 | 0.660428991641794 | -5.90745213338155 |
| ENSG00000284378.1 | MIR1244-1 | 0.247916692191492 | 4.22878487349064 | 0.950640174932684 | 0.347194269647793 | 0.660428991641794 | -5.89033303983661 |
| lnc-TNXB-1 | lnc-TNXB-1 | -0.147219662678701 | 2.27125856557881 | -0.950452138311618 | 0.347256629980054 | 0.660428991641794 | -5.65764596757755 |
| ENSG00000204439.3 | C6orf47 | -0.141157464220034 | 4.72400117451561 | -0.950438316778339 | 0.347263568568552 | 0.660428991641794 | -5.92106433617183 |
| lnc-TSSK4-3 | lnc-TSSK4-3 | -0.152003359717897 | 1.78704889856418 | -0.950314093479311 | 0.347325934371733 | 0.660428991641794 | -5.59393968165692 |
| lnc-DLX2-8 | lnc-DLX2-8 | -0.14366380278707 | 0.871821482343488 | -0.95026542470337 | 0.347350370345072 | 0.660428991641794 | -5.46412608860167 |
| lnc-ZNF24-1 | lnc-ZNF24-1 | -0.142554289962823 | 2.40170111493172 | -0.950202206481703 | 0.347382113101246 | 0.660428991641794 | -5.66958440610047 |
| ENSG00000162572.20 | SCNN1D | -0.242539604840963 | 1.6951232671547 | -0.950139608943185 | 0.347413546086762 | 0.660428991641794 | -5.58813971158722 |
| ENSG00000213714.1 | FAM209B | -0.147690104067491 | 0.132587336603308 | -0.950128574354391 | 0.347419087234839 | 0.660428991641794 | -5.28554165290013 |
| LINC00641 | LINC00641 | -0.158401558064154 | 6.03184990385457 | -0.95010941353806 | 0.34742870920164 | 0.660428991641794 | -5.96467759876498 |
| lnc-SSR3-6 | lnc-SSR3-6 | 0.185583624579969 | 0.123705071708095 | 0.950000084107065 | 0.347483614404704 | 0.660472364511222 | -5.28076580704062 |
| lnc-CEND1-3 | lnc-CEND1-3 | -0.159419336353549 | 1.43851373923395 | -0.9497775905613 | 0.347595368239567 | 0.660603163797799 | -5.56241201459202 |
| lnc-XPO5-2 | lnc-XPO5-2 | -0.0944575625113999 | 1.68369280002956 | -0.949735287387714 | 0.347616618913373 | 0.660603163797799 | -5.58007741797837 |
| ENSG00000184787.18 | UBE2G2 | 0.0831823975264783 | 6.13911788469958 | 0.949622420913958 | 0.347673320705445 | 0.660649922459396 | -5.96598356103914 |
| ENSG00000146859.6 | TMEM140 | 0.215295503820506 | 4.65897837428826 | 0.949454046727513 | 0.347759282338861 | 0.660752266997209 | -5.90659643533019 |
| ENSG00000111057.10 | KRT18 | -0.689751074406953 | 0.0845416513727121 | -0.949547368775701 | 0.347861409700025 | 0.660885305137125 | -5.36784720235236 |
| ENSG00000114270.17 | COL7A1 | -0.269946313192492 | -0.0267400510946413 | -0.948979476956583 | 0.34799643869115 | 0.661080821188735 | -5.2670683639126 |
| ENSG00000167930.15 | FAM234A | -0.136153394508284 | 3.54929184672172 | -0.948764260225149 | 0.348104642191306 | 0.6612253460893 | -5.81706332328523 |
| ENSG00000203778.7 | FAM229B | 0.346539757546284 | -0.403846588137288 | 0.948661755888373 | 0.348156185599041 | 0.66126222830904 | -5.11492831425342 |
| ENSG00000196187.11 | TMEM63A | -0.176940664559421 | 6.68726628729436 | -0.948145837260041 | 0.348415687039007 | 0.661694047455609 | -5.96667595148425 |
| ENSG00000079246.15 | XRCC5 | 0.104931212319295 | 7.77049478095171 | 0.947922392959248 | 0.348528116579583 | 0.661846500927905 | -5.95172787436004 |
| ENSG00000105136.20 | ZNF419 | -0.0989306488650268 | 3.93930011082007 | -0.947822900966614 | 0.348578185233239 | 0.661880515374148 | -5.85201031956289 |
| lnc-BRD1-7 | lnc-BRD1-7 | -0.61106615197906 | -0.647548968754815 | -0.947476904737674 | 0.348760061893679 | 0.661881809295995 | -5.05481413197818 |
| ENSG00000105085.10 | MED26 | 0.151294365537732 | 3.63004642193501 | 0.947460415283076 | 0.348760643666846 | 0.661881809295995 | -5.81757711683443 |
| ENSG00000239521.8 | CASTOR3P | 0.161335918645306 | 4.48257609062211 | 0.947416062983097 | 0.348782972865419 | 0.661881809295995 | -5.90627267934336 |
| ENSG00000124208.16 | PEDS1-UBE2V1 | -0.0996538859834004 | 4.5033227429247 | -0.947307956024623 | 0.348837403334561 | 0.661881809295995 | -5.90550396928513 |
| ENSG00000187987.9 | ZSCAN23 | 0.929486431961682 | -2.20366655932914 | 0.947273849478036 | 0.348854576703506 | 0.661881809295995 | -4.7926886428661 |
| ENSG00000197563.10 | PIGN | -0.0971248313122706 | 4.99639453622706 | -0.947260061735312 | 0.348861519281998 | 0.661881809295995 | -5.93865570714757 |
| lnc-ENSA-4 | lnc-ENSA-4 | 0.172377568273356 | 4.35219747578324 | 0.947143340665533 | 0.348920295791934 | 0.661881809295995 | -5.89345401435737 |
| ENSG00000112303.13 | VNN2 | -0.18605256584287 | 4.47438490724014 | -0.947141706434039 | 0.348921118778056 | 0.661881809295995 | -5.89932262211836 |
| lnc-PLEKHD1-3 | lnc-PLEKHD1-3 | 0.154019988449183 | 0.268213971759741 | 0.947087132653729 | 0.348948602437706 | 0.661881809295995 | -5.3332519507341 |
| lnc-SLC5A10-3 | lnc-SLC5A10-3 | 0.927783674730213 | 1.05066748761507 | 0.947468177408122 | 0.348951662698961 | 0.661881809295995 | -5.30721667071842 |
| ENSG00000151093.7 | OXSM | 0.114989073218144 | 2.95761926834841 | 0.946997136084218 | 0.348993928321503 | 0.661881809295995 | -5.73270711361587 |
| ENSG00000185684.14 | EP400P1 | -0.112583886416616 | 4.8456106866734 | -0.94695583469334 | 0.349014730655616 | 0.661881809295995 | -5.92804915475016 |
| ENSG00000271383.6 | NBPF19 | -0.24966296949476 | 3.67899161740678 | -0.946971043823395 | 0.349021476093604 | 0.661881809295995 | -5.81938469782501 |
| ENSG00000197696.9 | NMB | 0.354010107156689 | 1.10962325289762 | 0.946943380221007 | 0.349029060783109 | 0.661881809295995 | -5.4794849626459 |
| ENSG00000170296.9 | GABARAP | -0.108160757119585 | 4.72737249137964 | -0.946818250685083 | 0.349084033684906 | 0.661925072808514 | -5.9250170969924 |
| lnc-SPTBN4-1 | lnc-SPTBN4-1 | -0.185202395616852 | 0.364227474613164 | -0.946683901510586 | 0.349151716015607 | 0.661992425455899 | -5.3476305384684 |
| ENSG00000182093.15 | GET1 | -0.150900461059067 | 2.78402927763912 | -0.946563734825566 | 0.349212260812985 | 0.66203565889423 | -5.72068214275295 |
| ENSG00000277634.1 | | 0.223053320444508 | 0.405077300842064 | 0.94651097238498 | 0.349238846828623 | 0.66203565889423 | -5.36726148912316 |
| lnc-THAP4-4 | lnc-THAP4-4 | 0.12645643464137 | 2.12011317209407 | 0.946367833493447 | 0.349310978565395 | 0.662041846233682 | -5.64337959823577 |
| ENSG00000188026.12 | RILPL1 | -0.387086927251801 | 0.742185027672758 | -0.94641823214657 | 0.349319553579649 | 0.662041846233682 | -5.49249630234139 |
| lnc-C2orf74-5 | lnc-C2orf74-5 | -0.171717416920389 | 2.93587139952256 | -0.946260926757144 | 0.349364858284091 | 0.662041846233682 | -5.74185569935453 |
| ENSG00000163956.11 | LRPAP1 | 0.122705050893457 | 4.9493368952534 | 0.94624919996556 | 0.349370768779588 | 0.662041846233682 | -5.93545566001528 |
| ENSG00000160633.12 | SAFB | 0.125130935216887 | 6.07453692402838 | 0.946087893078984 | 0.349452076770786 | 0.662124320871088 | -5.96864004166428 |
| ENSG00000123965.13 | PMS2P5 | -0.481873762120372 | -0.348584450200638 | -0.946035221349215 | 0.349478629060074 | 0.662124320871088 | -5.16177887639163 |
| lnc-ZFP69B-1 | lnc-ZFP69B-1 | -0.152682163573093 | 0.930161798092508 | -0.945775146733271 | 0.349609754438632 | 0.662298887573556 | -5.47626907625393 |
| ENSG00000196792.11 | STRN3 | -0.0717196045160087 | 4.92527637378732 | -0.945724840371718 | 0.349635121817726 | 0.662298887573556 | -5.93437808407892 |
| ENSG00000086730.16 | LAT2 | -0.290884849904115 | 4.34040880100723 | -0.94511665948701 | 0.350029193563903 | 0.662984346289299 | -5.90099819045914 |
| ENSG00000145362.17 | ANK2 | 0.722718315104151 | 0.80208656933186 | 0.945165041656875 | 0.350097083449833 | 0.663051920192115 | -5.45643551341343 |
| lnc-SAMD8-1 | lnc-SAMD8-1 | 0.287809914305676 | 0.736944273720862 | 0.944498082452799 | 0.350254098629151 | 0.663145765918021 | -5.41949053358203 |
| ENSG00000103194.15 | USP10 | 0.0686787813031877 | 6.12764990929245 | 0.944484293940948 | 0.350261059892267 | 0.663145765918021 | -5.970672583102 |
| lnc-CWC15-1 | lnc-CWC15-1 | 0.357108505553409 | 6.35314864132452 | 0.944692410948914 | 0.350305580832314 | 0.663145765918021 | -5.97050427970499 |
| ENSG00000214941.7 | ZSWIM7 | 0.11935955338881 | 3.09689316467529 | 0.944244352372977 | 0.350382211256665 | 0.663145765918021 | -5.75806746186313 |
| ENSG00000165934.12 | CPSF2 | -0.116522219550823 | 5.84459883050467 | -0.944222628302625 | 0.350393181539041 | 0.663145765918021 | -5.96740062561281 |
| ENSG00000256361.1 | | 0.199005596513007 | 0.831001320698852 | 0.944158424145728 | 0.350425604854034 | 0.663145765918021 | -5.45000713822842 |
| lnc-UEVLD-4 | lnc-UEVLD-4 | 0.199005596513007 | 0.831001320698852 | 0.944158424145728 | 0.350425604854034 | 0.663145765918021 | -5.45000713822842 |

| | | | | | | | |
|---|---|---|---|---|---|---|---|
| ENSG00000158006.13 | PAFAH2 | -0.123965146182602 | 2.5417571576869 | -0.944149535238107 | 0.35043009393754 | 0.663145765918021 | -5.69046717394066 |
| lnc-MAP2K2-2 | lnc-MAP2K2-2 | 0.16255892377697 | 1.51678115003916 | 0.944136656641939 | 0.350436597963869 | 0.663145765918021 | -5.56638010395142 |
| ENSG00000054392.12 | HHAT | 1.49875154942632 | -1.45076942434608 | 0.944325151869589 | 0.3504909314068 | 0.663187611798692 | -4.85590075855574 |
| ENSG00000137414.5 | FAM8A1 | 0.100896238244146 | 3.93195143790773 | 0.943663552398839 | 0.350675582874394 | 0.663475087414646 | -5.85624557059388 |
| ENSG00000255036.6 | SUGT1P4-STRA6LP | 0.150599447089427 | 3.81676992934302 | 0.943600722576745 | 0.350707328915675 | 0.663475087414646 | -5.83294625517763 |
| lnc-DAPK3-1 | lnc-DAPK3-1 | -0.170055441659589 | 6.31260813591795 | -0.943412907255865 | 0.350802237664594 | 0.663593645607052 | -5.97184176233459 |
| ENSG00000180628.14 | PCGF5 | -0.135272668749938 | 7.21720777625166 | -0.943254409014703 | 0.350882344726845 | 0.663684185031488 | -5.965447311037 |
| lnc-ZNF385C-6 | lnc-ZNF385C-6 | 0.177642033690836 | 1.36590510404 | 0.942997702870871 | 0.351012112806876 | 0.663868631618481 | -5.5558784310489 |
| ENSG00000050130.17 | JKAMP | -0.130773237685192 | 3.95925146364188 | -0.942755236358409 | 0.351134711505039 | 0.664039486117991 | -5.86290785902484 |
| ENSG00000096717.11 | SIRT1 | 0.105983396302913 | 5.69655114229112 | 0.942485479593395 | 0.351271142035044 | 0.664236464539027 | -5.96716424106973 |
| lnc-AMDHD2-6 | lnc-AMDHD2-6 | 0.1182411865923 | 0.968955635568728 | 0.942395041827111 | 0.351316889070808 | 0.664243068667722 | -5.49652229347648 |
| ENSG00000111912.19 | NCOA7 | 0.169998107477793 | 5.04161053522431 | 0.942350982593334 | 0.351339177407836 | 0.664243068667722 | -5.93995445213025 |
| ENSG00000163875.15 | MEAF6 | -0.109313758130773 | 5.27973468243287 | -0.94210827833335 | 0.351461971353285 | 0.664414195110641 | -5.95541812138707 |
| ENSG00000105520.10 | PLPPR2 | -0.198771278733431 | 1.55334510676493 | -0.941640795687935 | 0.351698569153722 | 0.664783172975694 | -5.5725141073743 |
| lnc-ZNF514-11 | lnc-ZNF514-11 | -0.18126102550479 | 0.0464038011281725 | -0.941595007371199 | 0.351721748710359 | 0.664783172975694 | -5.26484833447997 |
| ENSG00000183077.15 | AFMID | -0.298840305796148 | 1.61754035809392 | -0.941363305809498 | 0.351839059049384 | 0.664869252842559 | -5.58059420287764 |
| lnc-TRMT1-1 | lnc-TRMT1-1 | -0.257173907945036 | 0.0371518729680012 | -0.941251180220414 | 0.351895837369167 | 0.664869252842559 | -5.29456123073832 |
| ENSG00000130827.6 | PLXNA3 | -0.16347187099771 | 4.93061480809933 | -0.941246308178466 | 0.351898304617102 | 0.664869252842559 | -5.93942852226236 |
| lnc-SYNDIG1L-1 | lnc-SYNDIG1L-1 | -0.116713175138473 | 0.807277807412274 | -0.941231063126253 | 0.351906024928432 | 0.664869252842559 | -5.46772025660988 |
| ENSG00000251357.4 | | -0.174339088114163 | 3.10107565614247 | -0.941186089344256 | 0.351928800938623 | 0.664869252842559 | -5.76947171279062 |
| lnc-ANAPC11-7 | lnc-ANAPC11-7 | -0.278902216525706 | 1.6789965531139 | -0.941069867965618 | 0.351987663247237 | 0.664875213369702 | -5.60297812312784 |
| ENSG00000111231.8 | GPN3 | 0.18449652106112 | 2.06033147534083 | 0.941026243803119 | 0.352009759118061 | 0.664875213369702 | -5.64170626858351 |
| ENSG00000113161.15 | HMGCR | -0.15846877171273 | 4.17590214728643 | -0.940988529344378 | 0.352028862425449 | 0.664875213369702 | -5.89499539122486 |
| ENSG00000151247.12 | EIF4E | 0.0933710617452057 | 5.176820684443 | 0.940854143126036 | 0.352096937899669 | 0.664942772069812 | -5.95523936133425 |
| ENSG00000157741.14 | UBN2 | -0.137232356946677 | 5.81485145349561 | -0.940691416515306 | 0.352179381202654 | 0.665037449843507 | -5.97029938531153 |
| lnc-MMD-9 | lnc-MMD-9 | -0.215108791927121 | -0.0592422252085093 | -0.940613476133516 | 0.352218873156875 | 0.665044742108331 | -5.23999884503856 |
| PAN3-AS1 | PAN3-AS1 | -0.22658941084163 | 1.72395232213531 | -0.940514986978657 | 0.352268781211113 | 0.665044742108331 | -5.59495120755675 |
| ENSG00000145020.15 | AMT | -0.176651079222785 | 2.36992557165893 | -0.940492505425106 | 0.352280174085757 | 0.665044742108331 | -5.66782948658794 |
| lnc-ANXA6-3 | lnc-ANXA6-3 | -0.125140848129237 | 1.95224987643759 | -0.940371793411864 | 0.352341350893831 | 0.665099231974295 | -5.63513320640291 |
| lnc-TRMT11-2 | lnc-TRMT11-2 | -0.250956857787673 | 0.338621341634499 | -0.940221877600139 | 0.352417337871327 | 0.665158133248081 | -5.35813354162627 |
| lnc-QRSL1-2 | lnc-QRSL1-2 | -0.445828495962457 | 0.00934305666949485 | -0.940186085579524 | 0.352437186085778 | 0.665158133248081 | -5.27072038209899 |
| lnc-TRMT61B-1 | lnc-TRMT61B-1 | -0.225133255797693 | 0.751179839677429 | -0.939948620636084 | 0.352555869708684 | 0.665316068104087 | -5.45004020104892 |
| lnc-ZNF316-1 | lnc-ZNF316-1 | 0.183993363764009 | 1.43486976046238 | 0.939890152317965 | 0.352585515759577 | 0.665316068104087 | -5.5590517296706 |
| ENSG00000147872.9 | PLIN2 | 0.349391371874138 | 3.72112414036302 | 0.940042775509753 | 0.352625929174698 | 0.665331381946357 | -5.82802599000208 |
| lnc-EGR1-3 | lnc-EGR1-3 | 0.141677838667625 | 0.254757743233809 | 0.939736367081752 | 0.352663499547741 | 0.665341229256752 | -5.33322271949895 |
| ENSG00000156983.15 | BRPF1 | 0.112695945625639 | 4.79761918630431 | 0.939663757572932 | 0.352700323424337 | 0.665349716528561 | -5.93384507823332 |
| lnc-NRDE2-4 | lnc-NRDE2-4 | -0.303218341339269 | 1.04834250779058 | -0.939560458759855 | 0.352752715708003 | 0.665387568495036 | -5.53005260123604 |
| ENSG00000120334.15 | CENPL | 0.120149063132433 | 2.59359196644798 | 0.939427062706133 | 0.3528203806012 | 0.665438188984777 | -5.7057451080525 |
| ENSG00000283399.1 | | 0.367528753456072 | 4.83715826810491 | 0.939673347505283 | 0.352844210978956 | 0.665438188984777 | -5.92784700064403 |
| ENSG00000114779.19 | ABHD14B | -0.171323732437925 | 5.10411219304481 | -0.939260224866718 | 0.352905020743905 | 0.665491895737224 | -5.95057251596199 |
| lnc-CDRT4-4 | lnc-CDRT4-4 | -0.304849040123311 | 0.235282366566923 | -0.939108328136372 | 0.352982092533017 | 0.665576256010176 | -5.40552047440566 |
| ENSG00000215790.7 | SLC35E2A | 0.206226667678286 | 4.9414952923107 | 0.939029980271635 | 0.353022011412478 | 0.665586720747028 | -5.93798611823608 |
| ENSG00000021300.13 | PLEKHB1 | 0.189634091030041 | 2.31825394625379 | 0.938876855763273 | 0.353099561883875 | 0.665586720747028 | -5.65065917684185 |
| ENSG00000169994.18 | MYO7B | -0.462622860683358 | -0.107644304761386 | -0.938900241423428 | 0.353114616585507 | 0.665586720747028 | -5.32355235775887 |
| ENSG00000110171.19 | TRIM3 | -0.151621632277192 | 2.55479169477536 | -0.938745965166418 | 0.353165998565638 | 0.665586720747028 | -5.70199618997227 |
| ENSG00000151657.11 | KIN | 0.0844094309059695 | 4.29974677819765 | 0.938716290872814 | 0.353181061607929 | 0.665586720747028 | -5.89478316366634 |
| ENSG00000166847.9 | DCTN5 | 0.0935102472261562 | 6.17718459574192 | 0.938715106672903 | 0.353181662731334 | 0.665586720747028 | -5.97619180162905 |
| lnc-DPH5-5 | lnc-DPH5-5 | 0.87784255377144 | -0.280612585755767 | 0.938848089070142 | 0.353253079071127 | 0.665660361303763 | -5.13168183761236 |
| ENSG00000125954.12 | CHURC1-FNTB | 0.0987467137846347 | 3.10855585520033 | 0.938507508611831 | 0.353287053986973 | 0.665663441249897 | -5.76246277990406 |
| ENSG00000120949.14 | TNFRSF8 | -0.304955374770027 | 0.245438146087523 | -0.938257284028236 | 0.353414112807055 | 0.665819677243017 | -5.32752041310452 |
| ENSG00000256514.1 | | 0.186991264723258 | 3.62978680435667 | 0.938216806869579 | 0.353434669074343 | 0.665819677243017 | -5.82380154225708 |
| ENSG00000165178.9 | NCF1C | -0.176082129283746 | 4.95811273246892 | -0.938134925577217 | 0.353476254765364 | 0.665837078048457 | -5.94670533601555 |
| ENSG00000118680.13 | MYL12B | -0.167686850043724 | 6.47740614963607 | -0.937920490392977 | 0.353585176815612 | 0.665981304391997 | -5.97679075844759 |
| ENSG00000131143.8 | COX4I1 | 0.119513633276003 | 7.39554148659918 | 0.937745384618721 | 0.353674137842449 | 0.666032326994681 | -5.96793478735882 |
| lnc-TTC21B-2 | lnc-TTC21B-2 | -0.189939467651 | 1.70440485834422 | -0.937739785522774 | 0.353676982658109 | 0.666032326994681 | -5.58830212316401 |
| ENSG00000072134.15 | EPN2 | -0.242448160520936 | 2.26229177145897 | -0.937480473270494 | 0.3538087516876 | 0.666210230358184 | -5.67512738977325 |
| ENSG00000259384.6 | GH1 | -0.26250887387829 | -0.514630834867069 | -0.937389935651381 | 0.353854765773721 | 0.666210230358184 | -5.15713982826084 |
| ENSG00000254999.3 | BRK1 | 0.145031216694607 | 5.13924491525362 | 0.937304334402246 | 0.353898274642034 | 0.666210230358184 | -5.95259815092415 |
| lnc-FNBP4-3 | lnc-FNBP4-3 | 0.150231032118284 | 1.20295217384475 | 0.937257379870205 | 0.353922141885154 | 0.666210230358184 | -5.52664502271129 |
| ENSG00000279819.1 | | 0.25727063437881 | 1.14906157309121 | 0.937222274601688 | 0.353939986771551 | 0.666210230358184 | -5.50886399737248 |
| ENSG00000229358.3 | DPY19L1P1 | -0.452639875759649 | 3.62697115472219 | -0.937480274702851 | 0.353977172342622 | 0.666210230358184 | -5.83406890174638 |
| lnc-ZDHHC7-2 | lnc-ZDHHC7-2 | -0.154062597284267 | 1.28617636489493 | -0.93710811283179 | 0.353998022126389 | 0.666210230358184 | -5.54296073804636 |
| ENSG00000109917.10 | ZPR1 | 0.135003147019224 | 5.90917681587373 | 0.936979402080686 | 0.354063461078529 | 0.666272464744867 | -5.97594157442972 |
| ENSG00000031698.12 | SARS1 | -0.0839735784363533 | 6.03989748838781 | -0.936767443312734 | 0.354171242127299 | 0.666393219839748 | -5.97719776409508 |
| lnc-PAPSS2-7 | lnc-PAPSS2-7 | -0.29204085887028 | 0.332499807058762 | -0.936725872748521 | 0.354192383279738 | 0.666393219839748 | -5.37836112307866 |
| ENSG00000162777.16 | DENND2D | -0.179056530025034 | 5.86679621405154 | -0.936372205887029 | 0.354372277711904 | 0.666661652018616 | -5.97564661672415 |
| lnc-ZNF320-1 | lnc-ZNF320-1 | -0.10865651809017 | 2.21545924284831 | -0.936261141046349 | 0.354428783732064 | 0.666661652018616 | -5.66298261669354 |
| ENSG00000170260.8 | ZNF212 | 0.158160649261583 | 3.32316652552096 | 0.93625438025292 | 0.354432223584498 | 0.666661652018616 | -5.78827056514515 |

| | | | | | | | |
|---|---|---|---|---|---|---|---|
| ENSG00000133627.17 | ACTR3B | 0.131165959876892 | 1.57789934446513 | 0.935982204521621 | 0.354570723149937 | 0.666861220266369 | -5.57051170711976 |
| ENSG00000231887.7 | PRH1 | -0.16127226380274 | 2.97862254234889 | -0.935878780236275 | 0.354623360995023 | 0.66689208633158 | -5.75547912936655 |
| ENSG00000177054.13 | ZDHHC13 | -0.104294163802932 | 4.31205783527657 | -0.935822641556343 | 0.354651934945609 | 0.66689208633158 | -5.90055512550693 |
| ENSG00000253953.2 | PCDHGB4 | -0.856908366929931 | -2.00556264827183 | -0.9357497858189 | 0.354764479286423 | 0.667042776756413 | -4.99010051791265 |
| SCAMP1-AS1 | SCAMP1-AS1 | -0.150992230815739 | 3.36680713913562 | -0.935449651392115 | 0.354841820933782 | 0.667127256145418 | -5.79271943462441 |
| ENSG00000198178.10 | CLEC4C | 1.27555154134007 | 2.06920064722035 | 0.935769326662816 | 0.354888527996345 | 0.667154130217258 | -5.55173313538177 |
| PAX8-AS1 | PAX8-AS1 | -1.64057731643144 | 1.56835858602377 | -0.935479661845104 | 0.355035927342579 | 0.667278128159781 | -5.52859831911209 |
| ENSG00000107957.16 | SH3PXD2A | -0.488507880576626 | 5.44073897431535 | -0.935415896868319 | 0.355059484144057 | 0.667278128159781 | -5.96677949241334 |
| lnc-XXYLT1-4 | lnc-XXYLT1-4 | -0.245464605102536 | -0.151269419067286 | -0.935009489617494 | 0.355065988879562 | 0.667278128159781 | -5.2346399989435 |
| lnc-ENOPH1-3 | lnc-ENOPH1-3 | -0.182292407751802 | 2.45231817538245 | -0.934939028556124 | 0.355101882259307 | 0.667278128159781 | -5.70656800578513 |
| lnc-SAA2-3 | lnc-SAA2-3 | 0.13641973124288 | 1.81838187542996 | 0.934903706788961 | 0.355119876318089 | 0.667278128159781 | -5.60597317273805 |
| ENSG00000197858.10 | GPAA1 | -0.174773547967757 | 4.34371794071815 | -0.934846537829169 | 0.355149001311296 | 0.667278128159781 | -5.90475495245794 |
| lnc-CTAGE1-9 | lnc-CTAGE1-9 | -0.183733802554788 | 1.15893684891082 | -0.934688521125112 | 0.355229511779059 | 0.667368477633112 | -5.54084772065879 |
| ENSG00000140632.16 | GLYR1 | -0.0709262748133348 | 5.54250460701562 | -0.934496571608207 | 0.355327327263817 | 0.667491318524335 | -5.97103312049825 |
| ENSG00000229222.1 | KRT18P4 | -0.237873940818769 | -0.39537955199338 | -0.934189560919806 | 0.355483813313348 | 0.667538313645505 | -5.18110700102534 |
| ENSG00000119397.16 | CNTRL | 0.099264645897378 | 8.06970239700719 | 0.934175407116865 | 0.355491028716851 | 0.667538313645505 | -5.96064828807043 |
| ENSG00000234353.2 | | 0.144180017497274 | 1.93314345466728 | 0.934170609210038 | 0.355493474641895 | 0.667538313645505 | -5.62757971085381 |
| lnc-TAF6-2 | lnc-TAF6-2 | 0.210129488794782 | 1.90746237233848 | 0.934135779175347 | 0.355511230975792 | 0.667538313645505 | -5.63256910415474 |
| lnc-CEP97-4 | lnc-CEP97-4 | 0.129130720892731 | 0.161574251054061 | 0.934074582436926 | 0.355542430445181 | 0.667538313645505 | -5.32425177026191 |
| ENSG00000285897.1 | | 0.105229015370701 | 4.10951690921231 | 0.934019154275825 | 0.355570690508269 | 0.667538313645505 | -5.88345438026047 |
| lnc-GALNS-1 | lnc-GALNS-1 | -0.192557204728371 | 3.38076204262251 | -0.933920396513687 | 0.355621045816684 | 0.667538313645505 | -5.80490718588626 |
| lnc-STN1-1 | lnc-STN1-1 | -0.185324213276971 | 0.85331616656949 | -0.933903917153973 | 0.35562944888304 | 0.667538313645505 | -5.4776770543472 |
| ENSG00000241547.1 | ACTG1P20 | 0.166019991977137 | 0.929171473961787 | 0.93379452925635 | 0.355685230658688 | 0.667538313645505 | -5.48619149028317 |
| ENSG00000005206.16 | SPPL2B | 0.16903887469949 | 4.56574226911946 | 0.933713720164557 | 0.35572644249493 | 0.667538313645505 | -5.92632656199662 |
| lnc-PRRT3-3 | lnc-PRRT3-3 | -0.234909992059843 | -0.0766191934424859 | -0.933702866810209 | 0.355731977835699 | 0.667538313645505 | -5.25495505729322 |
| lnc-HBEGF-4 | lnc-HBEGF-4 | 0.113771934309757 | 3.51537575861323 | 0.933658779620387 | 0.355754463409211 | 0.667538313645505 | -5.81508716776503 |
| lnc-HIST1H2AI-1 | lnc-HIST1H2AI-1 | -0.253153086923037 | 3.92259035559909 | -0.93367531911398 | 0.355773954267657 | 0.667538313645505 | -5.85839196700364 |
| ENSG00000228327.3 | | -0.156671869807486 | 1.66376236908715 | -0.933533168701404 | 0.35581853322137 | 0.667561103754941 | -5.58579926747642 |
| lnc-DNAJB4-1 | lnc-DNAJB4-1 | -0.18640386736559 | 2.79538287310848 | -0.933352418588773 | 0.355910740864451 | 0.667673239082482 | -5.72900755900593 |
| lnc-PDCD2L-3 | lnc-PDCD2L-3 | -0.197222958411656 | 1.56478366932785 | -0.93294288447409 | 0.356119717827443 | 0.668004388229496 | -5.5862662867964 |
| lnc-PLAC8L1-2 | lnc-PLAC8L1-2 | -0.11667504202231 | 2.91494327869502 | -0.932795160936348 | 0.356195117813601 | 0.668027032839782 | -5.74757609879612 |
| lnc-IGFBP4-4 | lnc-IGFBP4-4 | -0.432807226497172 | -0.568202672631113 | -0.932792060451137 | 0.356196700452636 | 0.668027032839782 | -5.11115700771875 |
| lnc-ZNF19-2 | lnc-ZNF19-2 | 0.136786234918956 | 0.710984427863697 | 0.932696093497401 | 0.356245688941644 | 0.668035543068633 | -5.43270631635597 |
| ENSG00000102780.16 | DGKH | -0.847128593693122 | 2.74321484051511 | -0.933011062784106 | 0.356293728603154 | 0.668035543068633 | -5.74921048384847 |
| ENSG00000254806.5 | SYS1-DBNDD2 | 0.140349441988935 | 0.384718550395244 | 0.9325924417491 | 0.356298605247411 | 0.668035543068633 | -5.37717871472238 |
| lnc-SFT2D3-2 | lnc-SFT2D3-2 | -0.181542631041647 | 1.52428057266837 | -0.932390818573071 | 0.356401552637849 | 0.668140195423972 | -5.57509427042748 |
| lnc-HIST1H4D-1 | lnc-HIST1H4D-1 | -0.196763779063039 | 0.253651980213435 | -0.932355979042347 | 0.356419343426868 | 0.668140195423972 | -5.3214221817249 |
| ENSG00000167037.18 | SGSM1 | 1.09156317358853 | 4.48376090115132 | 0.93268241077179 | 0.35646140292814 | 0.668158187457418 | -5.90418996360676 |
| lnc-MTERF4-5 | lnc-MTERF4-5 | -0.19017290492678 | 1.24184298727462 | -0.932126470301492 | 0.356536556437562 | 0.668219144498154 | -5.54427693982215 |
| lnc-UEVLD-2 | lnc-UEVLD-2 | 0.240941557716444 | 0.912887173464291 | 0.932082818821296 | 0.356558852646516 | 0.668219144498154 | -5.50393216758918 |
| ENSG00000119335.16 | SET | 0.117875175664137 | 8.22160945295104 | 0.931941147831534 | 0.356631221318072 | 0.668293921011459 | -5.95718676347429 |
| ENSG00000169764.15 | UGP2 | 0.111441244498593 | 5.15279334042819 | 0.931679830177569 | 0.356764733292934 | 0.668322133711233 | -5.95972365705535 |
| ENSG00000103126.14 | AXIN1 | 0.141909796858629 | 5.34394812600831 | 0.931646467393859 | 0.356781781297445 | 0.668322133711233 | -5.96779429652112 |
| lnc-TNFRSF17-3 | lnc-TNFRSF17-3 | -0.331660298244864 | -0.59770910385089 | -0.931571598130182 | 0.356820040577716 | 0.668322133711233 | -5.11913839712843 |
| ENSG00000100142.14 | POLR2F | -0.110809789199851 | 2.78945920358942 | -0.931535264382756 | 0.35683860861462 | 0.668322133711233 | -5.73735688451281 |
| lnc-NDST2-5 | lnc-NDST2-5 | -0.151733690376878 | 1.15530275496358 | -0.931522055140836 | 0.3568453592359 | 0.668322133711233 | -5.52315120471845 |
| ENSG00000005700.14 | IBTK | 0.113759504549241 | 6.15868585087849 | 0.931489447241226 | 0.356862023953987 | 0.668322133711233 | -5.98269310205053 |
| ENSG00000166845.14 | C18orf54 | -0.276845420898952 | 0.402135864886913 | -0.931342081026821 | 0.356937343828311 | 0.668322133711233 | -5.39774731343109 |
| ENSG00000273727.1 | | -0.47026630866386 | 1.18046807610038 | -0.931484274283105 | 0.356986867680902 | 0.668322133711233 | -5.57920535358487 |
| ENSG00000169136.10 | ATF5 | 0.327495644559161 | 2.93644945441189 | 0.931357989689275 | 0.35700506505448 | 0.668322133711233 | -5.76084597500353 |
| ENSG00000099954.18 | CECR2 | -0.664909504490729 | 1.03230015412609 | -0.931580483863977 | 0.357007006993838 | 0.668322133711233 | -5.59003248834507 |
| ENSG00000115234.10 | SNX17 | 0.124895987633555 | 5.62769160325461 | 0.931196601820066 | 0.357011709411717 | 0.668322133711233 | -5.9763175071276 |
| lnc-CDC42BPA-5 | lnc-CDC42BPA-5 | -0.286041753562452 | 3.58296440665463 | -0.931245181515502 | 0.357035912271716 | 0.668322133711233 | -5.8083375213695 |
| ENSG00000284755.1 | NA | 0.275369793131609 | 0.250140623713772 | 0.931070239291411 | 0.357076311195519 | 0.668336974932925 | -5.35206048422844 |
| ENSG00000170632.13 | ARMC10 | -0.122501606404474 | 3.38340351668717 | -0.930986567684348 | 0.35711909179912 | 0.668356270822085 | -5.80961722785753 |
| ENSG00000093167.17 | LRRFIP2 | 0.105297980008154 | 5.15692281774038 | 0.930649930116553 | 0.3572912455951 | 0.668617665977266 | -5.96166894461886 |
| ENSG00000113269.13 | RNF130 | 0.15230950864715 | 4.78456486435995 | 0.930568374265646 | 0.357332960756167 | 0.668634939204016 | -5.94089242952306 |
| ENSG00000118961.14 | LDAH | -0.116861517967994 | 4.96981611619925 | -0.930492552282952 | 0.35737174593855 | 0.668646727265991 | -5.94939747942879 |
| ENSG00000153113.23 | CAST | -0.0754394728278767 | 7.50824382856313 | -0.930165477515857 | 0.357539085796257 | 0.668899018627919 | -5.97298597423606 |
| ENSG00000256537.4 | SMIM10L1 | -0.120373546411814 | 4.00647422070523 | -0.929961046741449 | 0.357643703773924 | 0.669033932089309 | -5.87718499504715 |
| ENSG00000166164.15 | BRD7 | 0.0810863647745066 | 6.00391914926883 | 0.929786907939198 | 0.357732835493749 | 0.669092152006618 | -5.98307754345243 |
| ENSG00000067445.20 | TRO | -0.452204918471984 | 1.75644070324544 | -0.929753184006625 | 0.357866541343639 | 0.669092152006618 | -5.59114434117847 |
| ENSG00000285064.1 | | -0.181974361809702 | 1.53083574198038 | -0.929405864645785 | 0.35792792032007 | 0.669092152006618 | -5.59177266139058 |
| ENSG00000177885.14 | GRB2 | -0.119164540241622 | 7.76364370898906 | -0.929352233353629 | 0.357955383791894 | 0.669092152006618 | -5.97049823345867 |
| lnc-CASP3-5 | lnc-CASP3-5 | 0.26288059895259 | 1.31088094453174 | 0.929345098473528 | 0.357959037519433 | 0.669092152006618 | -5.5391981725055 |
| ENSG00000165272.15 | AQP3 | 0.37319580336357 | 1.20912845225021 | 0.929414808416683 | 0.357962919315802 | 0.669092152006618 | -5.54101420171443 |
| lnc-SLC35B1-2 | lnc-SLC35B1-2 | 0.0974543912539851 | 1.39316376941052 | 0.929304940628052 | 0.357979602554635 | 0.669092152006618 | -5.56131053808365 |
| ENSG00000164080.13 | RAD54L2 | 0.0830972027280064 | 5.82109237443061 | 0.929295196838036 | 0.35798459251471 | 0.669092152006618 | -5.98135162519606 |

| | | | | | | | |
|---|---|---|---|---|---|---|---|
| ENSG00000284048.2 | | -0.140995340952907 | 1.98781117462436 | -0.929280435401403 | 0.357992152183039 | 0.669092152006618 | -5.62873190960606 |
| ENSG00000011523.13 | CEP68 | 0.188335840663914 | 6.10989372086789 | 0.92926531358308 | 0.357999896518918 | 0.669092152006618 | -5.98415596908348 |
| ENSG00000256618.2 | NA | -1.01395217181226 | 0.270544087661617 | -0.929476695701349 | 0.358082702896785 | 0.669143703617666 | -5.28192160712338 |
| ENSG00000041353.9 | RAB27B | -0.801707098087618 | -1.22302301376194 | -0.92913265238747 | 0.358092498437963 | 0.669143703617666 | -4.93801031912256 |
| ENSG00000130338.12 | TULP4 | 0.171021364067952 | 6.21693458046463 | 0.928620390893217 | 0.35833028234432 | 0.669527251406422 | -5.98558832329345 |
| ENSG00000163468.14 | CCT3 | -0.180211227196669 | 6.2588752876962 | -0.928484559218035 | 0.358399892525621 | 0.669596531528988 | -5.98543117595637 |
| lnc-RAB3B-3 | lnc-RAB3B-3 | 0.147260645202492 | 1.67554185954763 | 0.928267740552597 | 0.358511024671476 | 0.669729482041453 | -5.59553206300012 |
| ENSG00000252906.1 | SCARNA3 | -0.195349596197753 | 0.0100222103915293 | -0.928178064207876 | 0.358556995561358 | 0.669729482041453 | -5.27021064260423 |
| lnc-FA2H-4 | lnc-FA2H-4 | -0.132358424410239 | 0.482731399844172 | -0.928155295865402 | 0.358568667934549 | 0.669729482041453 | -5.38991367889948 |
| ENSG00000221390.1 | MIR1295A | -0.440919154121417 | 0.773855835438061 | -0.92808111071215 | 0.358656130001162 | 0.669824375784518 | -5.42676493969496 |
| lnc-EXOSC1-1 | lnc-EXOSC1-1 | -0.189963420517594 | 3.16123207051624 | -0.927894599948559 | 0.35870233342253 | 0.669824375784518 | -5.78235746365885 |
| ENSG00000183576.12 | SETD3 | 0.109008069435363 | 4.41303909070397 | 0.92786580131049 | 0.35871710121327 | 0.669824375784518 | -5.91556050647177 |
| ENSG00000198856.12 | OSTC | 0.174114909455584 | 3.82463757712967 | 0.92760479514435 | 0.358850961860937 | 0.669966416045391 | -5.8545195676202 |
| ENSG00000130347.12 | RTN4IP1 | -0.142225588729131 | 3.08931188594105 | -0.927585379285971 | 0.358860920853775 | 0.669966416045391 | -5.77380695927936 |
| ENSG00000249485.1 | RBBP4P1 | 0.400797620431099 | -0.278538731298024 | 0.927489611292848 | 0.358910045845013 | 0.669966416045391 | -5.11402329857649 |
| ENSG00000275291.1 | RNVU1-26 | -1.36107233069204 | -1.77697024231243 | -0.927702241711788 | 0.358923367328987 | 0.669966416045391 | -4.83339547698658 |
| lnc-PCYT2-1 | lnc-PCYT2-1 | 0.146987245392448 | 2.90249353263437 | 0.927386399456832 | 0.35896299411894 | 0.669979625312852 | -5.75018546043401 |
| lnc-LAP3-1 | lnc-LAP3-1 | 0.113899357137127 | 3.95841750619537 | 0.927223395996739 | 0.359046626201038 | 0.670006138053679 | -5.87401201949061 |
| ENSG00000173898.12 | SPTBN2 | -0.337804127037254 | 4.3188859120127 | -0.927357941568083 | 0.359099932204958 | 0.670006138053679 | -5.88558231447462 |
| ENSG00000199545.1 | RNA5SP195 | 0.29690802436505 | 0.697599766571052 | 0.92706023338124 | 0.359130352641534 | 0.670006138053679 | -5.42884555499298 |
| ENSG00000160972.9 | PPP1R16A | -0.171102090313711 | 1.78177414922609 | -0.927016231080823 | 0.359152934474577 | 0.670006138053679 | -5.62849971579848 |
| ENSG00000117676.13 | RPS6KA1 | -0.168929136072344 | 5.50668799304833 | -0.927009214521588 | 0.359156535434681 | 0.670006138053679 | -5.97800860239057 |
| ENSG00000088899.15 | LZTS3 | 0.359429305778254 | 2.18846326666219 | 0.927118187905132 | 0.359172507762925 | 0.670006138053679 | -5.66314391141498 |
| lnc-PLEKHA3-20 | lnc-PLEKHA3-20 | 0.134785153095746 | 3.40844523896545 | 0.926820550934742 | 0.359253368062224 | 0.670067228220248 | -5.81591607642333 |
| ENSG00000104866.10 | PPP1R37 | -0.129690138777856 | 3.66248594578631 | -0.926787437669287 | 0.359270365381075 | 0.670067228220248 | -5.84381342110059 |
| ENSG00000178773.14 | CPNE7 | 0.647397065021769 | -1.02323764771438 | 0.926647041630842 | 0.359403828394438 | 0.670255413594521 | -5.15456011562134 |
| lnc-FAM69A-2 | lnc-FAM69A-2 | 0.106599252921824 | 1.03133448240927 | 0.92641467918104 | 0.359461741565755 | 0.670285843643743 | -5.51737854946792 |
| lnc-TAOK1-5 | lnc-TAOK1-5 | -0.251612426649723 | -0.244427924756133 | -0.926368848974154 | 0.359485275614987 | 0.670285843643743 | -5.21919866930549 |
| ENSG00000013364.18 | MVP | -0.123790494948102 | 6.75992419598522 | -0.926254714311882 | 0.359543888713565 | 0.670334407734721 | -5.9856349877444 |
| lnc-NUDT5-5 | lnc-NUDT5-5 | 0.253668903443395 | -0.0365194946754329 | 0.92600031249099 | 0.359674557453611 | 0.670517291555808 | -5.25539258079676 |
| ENSG00000198791.11 | CNOT7 | -0.121876647912013 | 6.51112674829424 | -0.925913245428097 | 0.359719284917436 | 0.670539942171307 | -5.98754837158251 |
| lnc-PKDREJ-3 | lnc-PKDREJ-3 | -0.228605297549442 | -0.414983689666191 | -0.92580578651092 | 0.359774492925607 | 0.670582123325887 | -5.18128906484932 |
| lnc-DEGS2-2 | lnc-DEGS2-2 | -0.212464268391105 | 1.56166994181211 | -0.925677310638309 | 0.359840505821069 | 0.670644434200929 | -5.59659100110777 |
| lnc-CORIN-1 | lnc-CORIN-1 | 0.131430124650894 | 2.44339646821032 | 0.925357375292421 | 0.360004927794418 | 0.670890124834088 | -5.69979448673799 |
| ENSG00000138594.13 | TMOD3 | -0.087257467733437 | 5.78171552777815 | -0.925252571417085 | 0.360058799485626 | 0.670929772751461 | -5.98473672626603 |
| lnc-INPP5K-1 | lnc-INPP5K-1 | -0.196835767737876 | 1.21709192195498 | -0.925060655860621 | 0.360157462245466 | 0.670997137951764 | -5.54936824535888 |
| ENSG00000105497.7 | ZNF175 | -0.225789790452623 | 2.9141746781438 | -0.925055427012673 | 0.360160150614152 | 0.670997137951764 | -5.75820115718878 |
| ENSG00000258111.1 | | 0.34166143042096 | -0.148481940564082 | 0.924816963183101 | 0.360282768674055 | 0.67099776421044 | -5.21815012469535 |
| lnc-TXNRD1-5 | lnc-TXNRD1-5 | 0.34166143042096 | -0.148481940564082 | 0.924816963183101 | 0.360282768674055 | 0.67099776421044 | -5.21815012469535 |
| lnc-NDUFB1-1 | lnc-NDUFB1-1 | -0.119963455398472 | 1.22790006247617 | -0.924809416939488 | 0.360286649394151 | 0.67099776421044 | -5.54944002318179 |
| ENSG00000232381.1 | OR52U1P | 0.300534519240771 | 0.403639134590099 | 0.924701783924467 | 0.360342003547307 | 0.67099776421044 | -5.37370664424463 |
| lnc-CACNB2-1 | lnc-CACNB2-1 | 0.198557665526653 | 3.28288040998572 | 0.924678755848894 | 0.360353847282147 | 0.67099776421044 | -5.81164354461849 |
| ENSG00000267127.7 | | 0.196351658459755 | 1.42789788412342 | 0.924671313460737 | 0.360357675084325 | 0.67099776421044 | -5.56864536807041 |
| BACH1-IT3 | BACH1-IT3 | -0.524331747565153 | 0.598641345859729 | -0.924813577158327 | 0.360389144809774 | 0.67099776421044 | -5.42073375533624 |
| ENSG00000248121.8 | SMURF2P1 | 0.144766804264846 | 0.81309525906354 | 0.924493224543453 | 0.360449278451453 | 0.67099776421044 | -5.49239570826325 |
| lnc-OTP-2 | lnc-OTP-2 | -0.211586718772215 | 1.64678899413616 | -0.924427013459992 | 0.360483339220382 | 0.67099776421044 | -5.60370671659611 |
| lnc-SLC25A41-4 | lnc-SLC25A41-4 | -0.155187899493225 | 1.13331500593404 | -0.924420902717616 | 0.36048648285668 | 0.67099776421044 | -5.54345413666022 |
| ENSG00000174586.10 | ZNF497 | -0.236404074661362 | -0.371298559329561 | -0.92426297000486 | 0.360567736614179 | 0.671088319263011 | -5.21122850265611 |
| ENSG00000006007.11 | GDE1 | 0.120652315093817 | 4.2868359493911 | 0.923965282236329 | 0.360720924385069 | 0.671312729350622 | -5.90982364772327 |
| ENSG00000145414.8 | NAF1 | 0.159558754420968 | 3.63841195814452 | 0.923843903454124 | 0.360783397081266 | 0.671367449846989 | -5.84593561791647 |
| lnc-DERL2-1 | lnc-DERL2-1 | -0.138864727115953 | 3.87938155666763 | -0.923757617466667 | 0.360827812069165 | 0.671367449846989 | -5.86034194119441 |
| ENSG00000100365.14 | NCF4 | -0.166096502822064 | 5.36537739061676 | -0.923715389065973 | 0.360849550078695 | 0.671367449846989 | -5.97484373789265 |
| ENSG00000184743.12 | ATL3 | -0.12868522092752 | 5.85731667833821 | -0.923643350428072 | 0.360886635535337 | 0.671367449846989 | -5.98675889875759 |
| lnc-GPR31-51 | lnc-GPR31-51 | 0.488727555015578 | -0.693795152955541 | 0.923485731048327 | 0.360967786552779 | 0.671367449846989 | -5.01619737179914 |
| ENSG00000123444.13 | KBTBD4 | 0.100065341861986 | 5.12862786226742 | 0.923477210709141 | 0.360972173623848 | 0.671367449846989 | -5.96572888068928 |
| lnc-ROS1-3 | lnc-ROS1-3 | -0.145158156283278 | 1.82687052547928 | -0.923464631385443 | 0.360978650704787 | 0.671367449846989 | -5.64060372428345 |
| lnc-ALYREF-1 | lnc-ALYREF-1 | -0.138483868185599 | 2.59927206996273 | -0.923299618432597 | 0.361063622696669 | 0.671464812609824 | -5.72538090933307 |
| lnc-CMAS-6 | lnc-CMAS-6 | -0.180521997992589 | 3.98209748926341 | -0.923134882538592 | 0.36114846497787 | 0.671561916581398 | -5.89035266016329 |
| ENSG00000119326.14 | CTNNAL1 | 0.400604987645397 | -0.0956279778670093 | 0.92251123222843 | 0.36146977464988 | 0.672056815978111 | -5.28338123067204 |
| ENSG00000198168.8 | SVIP | -0.201481223284981 | 4.90996948343058 | -0.92234404704371 | 0.361556099505874 | 0.672056815978111 | -5.9583235895464 |
| ENSG00000027847.13 | B4GALT7 | -0.105884806498777 | 3.73898273961677 | -0.922309632834126 | 0.361573679969063 | 0.672056815978111 | -5.85464139369792 |
| lnc-FGD4-9 | lnc-FGD4-9 | -0.214049194071963 | 1.25833413261179 | -0.922275509166647 | 0.361591269393708 | 0.672056815978111 | -5.56199169872328 |
| ENSG00000110090.12 | CPT1A | -0.299800876276447 | 5.053852268971 | -0.922486533592317 | 0.361597984237419 | 0.672056815978111 | -5.96783777785615 |
| ENSG00000168159.11 | RNF187 | -0.123798178818393 | 4.5637557532354 | -0.922238173091384 | 0.361610515326122 | 0.672056815978111 | -5.93415034666665 |
| ENSG00000242852.6 | ZNF709 | 0.138838290040422 | 2.23058818728555 | 0.922139909717276 | 0.361661171130281 | 0.672090274952561 | -5.67569007848959 |
| ENSG00000004779.9 | NDUFAB1 | 0.138340089644783 | 3.14382872127165 | 0.922036640167329 | 0.361714412634887 | 0.672128532568734 | -5.78695709041473 |
| lnc-TMEM116-5 | lnc-TMEM116-5 | 0.153240783865122 | 0.613430442497341 | 0.921900694640536 | 0.361784508276923 | 0.672198098380926 | -5.45346555062329 |
| lnc-VPS33A-2 | lnc-VPS33A-2 | 0.196411509031851 | 2.1792421711692 | 0.921732595477301 | 0.361871195027528 | 0.672298475246107 | -5.67072144393997 |

| | | | | | | | |
|---|---|---|---|---|---|---|---|
| ENSG00000132185.16 | FCRLA | -0.172433523517936 | 7.8296656672024 | -0.921569532442507 | 0.361955297585678 | 0.672339339619754 | -5.97587184790445 |
| ENSG00000122484.8 | RPAP2 | -0.106626379494217 | 5.46302146871335 | -0.9215353716185 | 0.36197291822369 | 0.672339339619754 | -5.97869099657872 |
| ENSG00000070669.16 | ASNS | -0.12509581436155 | 4.79560427521004 | -0.92143846543049 | 0.362022906840507 | 0.672339339619754 | -5.95484133236126 |
| ENSG00000100068.12 | LRP5L | 0.212751464393776 | 4.05084442460809 | 0.921385266098476 | 0.362050351378763 | 0.672339339619754 | -5.88614012117311 |
| ENSG00000188342.11 | GTF2F2 | -0.0991879982502804 | 4.07713308556198 | -0.921334762945754 | 0.362076406257737 | 0.672339339619754 | -5.89957215333055 |
| ENSG00000111325.16 | OGFOD2 | -0.135806638216661 | 2.66138622832795 | -0.9212463076093 | 0.3621220438271 | 0.672339339619754 | -5.73187188616915 |
| ENSG00000082212.12 | ME2 | -0.155651631413889 | 5.51458876125264 | -0.921144083120227 | 0.362174790089823 | 0.672339339619754 | -5.98344687866402 |
| ENSG00000090104.11 | RGS1 | -0.870237271789304 | 4.42573776134937 | -0.921479240233133 | 0.362207953927613 | 0.672339339619754 | -5.91350385856099 |
| lnc-ZNF585B-2 | lnc-ZNF585B-2 | -0.135325100757633 | 0.0763703429950767 | -0.921074323451674 | 0.36221078786622 | 0.672339339619754 | -5.29523930162312 |
| ENSG00000136518.16 | ACTL6A | -0.131086728651943 | 3.98294769481856 | -0.921056784929478 | 0.362219838558201 | 0.672339339619754 | -5.88747973918047 |
| ENSG00000099917.17 | MED15 | 0.144486486870487 | 5.49047217670961 | 0.920924402039752 | 0.362288159010698 | 0.672405516403714 | -5.98045265004776 |
| ENSG00000064666.14 | CNN2 | 0.187560022473086 | 6.69835223755407 | 0.920828838245607 | 0.362337482978481 | 0.672435256764603 | -5.99196246972493 |
| ENSG00000115526.10 | CHST10 | 0.203341858182453 | 2.91137468697473 | 0.920766768377354 | 0.362369521840012 | 0.672435256764603 | -5.7506002115801 |
| lnc-OGFOD1-2 | lnc-OGFOD1-2 | -0.827386819133314 | -1.67500991922847 | -0.920580912247664 | 0.362465466958635 | 0.672552664419867 | -4.83698119406709 |
| ENSG00000133195.11 | SLC39A11 | -0.118686056805957 | 3.28795712673994 | -0.920446162956597 | 0.362535039328964 | 0.672621120831806 | -5.80022413841453 |
| lnc-CD226-3 | lnc-CD226-3 | 0.174042794564849 | -0.0334876065952433 | 0.92026345545785 | 0.362629386820283 | 0.67273552671671 | -5.26169354666068 |
| ENSG00000153982.10 | GDPD1 | 0.180168178389707 | 0.504176468363765 | 0.92008081279154 | 0.362723716745479 | 0.672798553091741 | -5.41039308329176 |
| ENSG00000159885.13 | ZNF222 | -0.146493155946368 | 1.51032063314216 | -0.920071098085395 | 0.36272873457023 | 0.672798553091741 | -5.58231816035422 |
| ENSG00000135945.9 | REV1 | 0.0830092677890963 | 5.4259871766272 | 0.919560196755367 | 0.362992687947222 | 0.672980819389453 | -5.98064557908672 |
| ENSG00000114030.12 | KPNA1 | 0.0958877270206067 | 5.51024375341779 | 0.919518877184228 | 0.36301404083753 | 0.672980819389453 | -5.98428628949058 |
| ENSG00000101596.15 | SMCHD1 | 0.132937759037979 | 10.0388759438206 | 0.919507966433445 | 0.363019679368445 | 0.672980819389453 | -5.91813807533863 |
| ENSG00000128272.14 | ATF4 | -0.12449538672422 | 7.75998657148948 | -0.91940949121964 | 0.363070572617876 | 0.672980819389453 | -5.97688854909107 |
| ENSG00000198142.4 | SOWAHC | -1.65593500238952 | -0.5321340577644 | -0.91977529819716 | 0.363087218887529 | 0.672980819389453 | -5.10182951851654 |
| ENSG00000134864.10 | GGACT | -0.146499637688332 | 1.5876584855501 | -0.91922088049228 | 0.363168061963911 | 0.672980819389453 | -5.59210425875657 |
| ENSG00000156110.13 | ADK | -0.176136837079643 | 5.14986982405426 | -0.919187778867021 | 0.363185173322806 | 0.672980819389453 | -5.96751011630688 |
| ENSG00000137494.13 | ANKRD42 | 0.21876003614579 | 3.49532724592943 | 0.919108058323033 | 0.363226385735071 | 0.672980819389453 | -5.81721002279846 |
| ENSG00000167264.17 | DUS2 | -0.18511569546978 | 5.69297993791195 | -0.919087690593003 | 0.363236915543247 | 0.672980819389453 | -5.98695710842801 |
| ENSG00000139174.11 | PRICKLE1 | 0.352277805445495 | 3.45618309730129 | 0.919296512848803 | 0.3632494603062 | 0.672980819389453 | -5.83903137102222 |
| lnc-ELK1-2 | lnc-ELK1-2 | 0.111771291962988 | 3.99736584768762 | 0.919011288208206 | 0.363276416182721 | 0.672980819389453 | -5.88635927002409 |
| lnc-MARCH8-3 | lnc-MARCH8-3 | 0.153292429027822 | 1.06706467366655 | 0.918946888034058 | 0.363309713746007 | 0.672980819389453 | -5.5177017852433 |
| ENSG00000089195.14 | TRMT6 | 0.113462157549214 | 4.38427530815978 | 0.918888597247687 | 0.363339854207379 | 0.672980819389453 | -5.92346039811805 |
| lnc-NR2C2AP-1 | lnc-NR2C2AP-1 | 0.157314781785095 | 0.334583574456208 | 0.918813895875594 | 0.363378482471503 | 0.672980819389453 | -5.38328006792913 |
| ENSG00000136286.15 | MYO1G | 0.23208764192891 | 5.78196126717059 | 0.918898059567882 | 0.363380666071867 | 0.672980819389453 | -5.98988060632482 |
| ENSG00000103316.10 | CRYM | 0.580440965848068 | 1.62600027105876 | 0.919066491621522 | 0.363390551596252 | 0.672980819389453 | -5.48397253881616 |
| TMPO-AS1 | TMPO-AS1 | -0.217836579127776 | 1.73768088818263 | -0.918762903932232 | 0.36340485205957 | 0.672980819389453 | -5.63078814581042 |
| ENSG00000196417.12 | ZNF765 | 0.0919343297787041 | 5.14401061202682 | 0.918742260613329 | 0.363415527742009 | 0.672980819389453 | -5.96967647190124 |
| ENSG00000136824.18 | SMC2 | 0.120777453790557 | 4.09528626183491 | 0.918639990864982 | 0.363468419490167 | 0.673018215038332 | -5.8904895312614 |
| ENSG00000285794.1 | | -0.173548476393886 | 1.13942460262666 | -0.918127066415878 | 0.36373376838431 | 0.67344896596692 | -5.53152991127781 |
| ENSG00000143324.13 | XPR1 | 0.0929372605194363 | 4.80924362420682 | 0.918023205865325 | 0.363787513367167 | 0.673487892393992 | -5.9544876491984 |
| ENSG00000125779.22 | PANK2 | -0.103222581969749 | 5.7372050698466 | -0.917901743152198 | 0.363850373510383 | 0.673543685400144 | -5.99066763008318 |
| lnc-DHRS7B-1 | lnc-DHRS7B-1 | -0.16749856365463 | 2.77922201339781 | -0.917790170848948 | 0.363908121305141 | 0.67359000546976 | -5.74156081664898 |
| lnc-ATE1-5 | lnc-ATE1-5 | 0.116842077520085 | 2.55016679499086 | 0.917539847071135 | 0.364037705935455 | 0.673653698941998 | -5.72125188652755 |
| lnc-RNF19A-8 | lnc-RNF19A-8 | 0.226969804205085 | 4.0163062199418 | 0.917537766150446 | 0.364048911790722 | 0.673653698941998 | -5.88836127799842 |
| ENSG00000146282.17 | RARS2 | 0.105843944463648 | 5.00675137459388 | 0.917416619672395 | 0.364101507800105 | 0.673653698941998 | -5.96712722547465 |
| lnc-LYST-4 | lnc-LYST-4 | -0.214919744483256 | 2.76605699628427 | -0.917415309259488 | 0.364102186314638 | 0.673653698941998 | -5.76972403429038 |
| ENSG00000168090.9 | COPS6 | 0.109851225401536 | 3.80718461442446 | 0.917407605890279 | 0.364106175034239 | 0.673653698941998 | -5.86381844684199 |
| ENSG00000122566.21 | HNRNPA2B1 | 0.121382159728682 | 9.11452156974942 | 0.917304668226635 | 0.364159477735907 | 0.673691760762015 | -5.94666154792607 |
| lnc-KLHDC4-3 | lnc-KLHDC4-3 | -0.140755387140012 | -0.19628745661731 | -0.917039666219815 | 0.364296723083886 | 0.673885094925464 | -5.22666222932087 |
| ENSG00000228590.2 | MIR4432HG | 0.281392547675933 | 2.98293711364528 | 0.916894633636155 | 0.364395490243899 | 0.673994884872757 | -5.7463967379383 |
| ENSG00000137275.13 | RIPK1 | 0.0983608036047183 | 4.92134460148667 | 0.916757413906521 | 0.364442939237625 | 0.673994884872757 | -5.95996456351095 |
| ENSG00000130396.20 | AFDN | -0.350002870952854 | 5.2786205637773 | -0.917015303383181 | 0.364454310286828 | 0.673994884872757 | -5.99250317057365 |
| ENSG00000101945.16 | SUV39H1 | 0.122776343215868 | 2.9777970954522 | 0.916291479632172 | 0.364684391893788 | 0.674346223870121 | -5.77778371649884 |
| ENSG00000090554.12 | FLT3LG | 0.196412404624113 | 0.191261660073061 | 0.916242428022643 | 0.36470981703941 | 0.674346223870121 | -5.35761103801045 |
| ENSG00000090372.14 | STRN4 | 0.110247730221761 | 4.87679013396651 | 0.915977754844202 | 0.364847026091941 | 0.674518840972672 | -5.96162261668373 |
| ENSG00000167703.14 | SLC43A2 | 0.120918289492534 | 5.66931962187878 | 0.915855873153553 | 0.364910221930002 | 0.674518840972672 | -5.99141037321447 |
| ENSG00000010295.19 | IFFO1 | -0.146445942276487 | 4.29859494666896 | -0.915848478369067 | 0.364914056364608 | 0.674518840972672 | -5.91681379505853 |
| ENSG00000176809.10 | LRRC37A3 | -0.203608892415292 | 3.09044272591411 | -0.915756247506718 | 0.364961883235701 | 0.674518840972672 | -5.78758764114844 |
| lnc-RRM2B-5 | lnc-RRM2B-5 | 0.34145565896737 | -0.595353879553498 | 0.915746327152037 | 0.364967027737096 | 0.674518840972672 | -5.0918282618985 |
| ENSG00000075785.12 | RAB7A | -0.13044463634039 | 5.07683297153697 | -0.915610229629022 | 0.365037609979812 | 0.674544737233852 | -5.97257638576494 |
| ENSG00000138078.15 | PREPL | 0.138605286438531 | 5.69335788739288 | 0.915537789317864 | 0.365075182225807 | 0.674544737233852 | -5.99108871756786 |
| ENSG00000067066.16 | SP100 | 0.120453375460679 | 8.65230959741595 | 0.915529743567208 | 0.365079355428514 | 0.674544737233852 | -5.96170035073745 |
| lnc-ZNF202-1 | lnc-ZNF202-1 | 0.128561256222371 | 0.912364183420136 | 0.915374112457341 | 0.365160084877023 | 0.674633338750898 | -5.50753696069177 |
| ENSG00000159069.13 | FBXW5 | -0.17640223465999 | 4.6526443965017 | -0.915211027870726 | 0.365244693003129 | 0.674729089578478 | -5.95260378100444 |
| ENSG00000092094.10 | OSGEP | -0.0984497583416051 | 5.52442375526023 | -0.915132872432738 | 0.365285244462129 | 0.674734196452502 | -5.98854673473076 |
| lnc-VGLL4-2 | lnc-VGLL4-2 | -0.205753298657638 | -0.413720170063364 | -0.915004605929809 | 0.365351802683904 | 0.674734196452502 | -5.2087808774252 |
| ENSG00000151503.12 | NCAPD3 | 0.125190614074233 | 4.33047957549736 | 0.914992976218381 | 0.365357837795376 | 0.674734196452502 | -5.91316523095982 |
| A1BG-AS1 | A1BG-AS1 | -0.165381133745803 | 2.03765877455639 | -0.914953002872776 | 0.365378582017179 | 0.674734196452502 | -5.66778241142895 |
| ENSG00000135486.17 | HNRNPA1 | 0.10803598445742 | 9.01985589301794 | 0.914873278980339 | 0.365419957113132 | 0.674750065242631 | -5.95120056991454 |

| | | | | | | | |
|---|---|---|---|---|---|---|---|
| ENSG00000196663.15 | TECPR2 | -0.0972335350616245 | 3.18585743752649 | -0.914725057152693 | 0.365496889310506 | 0.674774511745654 | -5.79899989985648 |
| ENSG00000134717.17 | BTF3L4 | 0.102840976036021 | 5.21498902903774 | 0.914711930917556 | 0.365503702780119 | 0.674774511745654 | -5.97998404562355 |
| ENSG00000185803.9 | SLC52A2 | -0.202539742916018 | 2.99690306910737 | -0.914609287907353 | 0.365556984790732 | 0.674774511745654 | -5.78273058811 |
| ENSG00000139793.18 | MBNL2 | -0.0880376928122546 | 5.62843045322211 | -0.914541128368973 | 0.36559236919723 | 0.674774511745654 | -5.9914389383227 |
| ENSG00000271581.1 | | -0.224234687226529 | -0.237681165192746 | -0.914438358186204 | 0.365645725591963 | 0.674774511745654 | -5.25501986913715 |
| ENSG00000142937.11 | RPS8 | 0.150879691230454 | 9.92651479172638 | 0.914417035418495 | 0.365656796612546 | 0.674774511745654 | -5.9252184712857 |
| ENSG00000197448.13 | GSTK1 | -0.0962212711049597 | 5.95110961590513 | -0.914374095090403 | 0.36567909237093 | 0.674774511745654 | -5.99610255116521 |
| ENSG00000108788.11 | MLX | -0.106951046960954 | 3.97648471729003 | -0.914342570699035 | 0.365695461231248 | 0.674774511745654 | -5.8877560353045 |
| ENSG00000167792.12 | NDUFV1 | -0.135886387298748 | 6.0068448312189 | -0.914206774166438 | 0.365765978220045 | 0.674818037513794 | -5.99715989897461 |
| ENSG00000100883.11 | VIII.54 | 0.0844371190062825 | 5.27245866630122 | 0.914143847028169 | 0.365798658266124 | 0.674818037513794 | -5.9793154552619 |
| LINC01869 | LINC01869 | -0.27966454380776 | -0.359002711423248 | -0.914107749404838 | 0.365817405751296 | 0.674818037513794 | -5.22665147593681 |
| lnc-RNF13-4 | lnc-RNF13-4 | -0.215410320701623 | 0.987319739387015 | -0.913951374283791 | 0.365898627141275 | 0.674859479289936 | -5.52961878340777 |
| ENSG00000185236.11 | RAB11B | -0.130172107928796 | 4.10112234043747 | -0.913927400255346 | 0.365911080305194 | 0.674859479289936 | -5.90268251214279 |
| ENSG00000113119.12 | TMCO6 | -0.0895789883621307 | 3.29061339571894 | -0.913875129604478 | 0.3659382329279 | 0.674859479289936 | -5.80903706954539 |
| lnc-LRP2BP-1 | lnc-LRP2BP-1 | -0.247737355456002 | 1.74698548978667 | -0.913598898850124 | 0.366081745959966 | 0.675017571530025 | -5.61458806553386 |
| ENSG00000100601.9 | ALKBH1 | 0.162112480502432 | 2.85975742935836 | 0.913583885107868 | 0.366089547247227 | 0.675017571530025 | -5.7591196146699 |
| lnc-GJC1-2 | lnc-GJC1-2 | 0.141283260587258 | 1.22846381076957 | 0.913262659896487 | 0.366256484741419 | 0.675262575600081 | -5.5502963166503 |
| ENSG00000148090.11 | AUH | 0.106865963359486 | 3.89542546536444 | 0.913104182662931 | 0.366338861869606 | 0.675262575600081 | -5.87787438281671 |
| lnc-CCER2-4 | lnc-CCER2-4 | 0.139637551103127 | 0.571541491591031 | 0.913100150293661 | 0.366340958068125 | 0.675262575600081 | -5.44374221988942 |
| lnc-RNF121-1 | lnc-RNF121-1 | 0.119978503912656 | 1.45754926558862 | 0.91307573532144 | 0.366353650183458 | 0.675262575600081 | -5.57826329940566 |
| ENSG00000181523.12 | SGSH | -0.143872519369047 | 4.41802999517099 | -0.912895927431691 | 0.366447132004688 | 0.675374401688081 | -5.92574295465261 |
| ENSG00000265185.5 | SNORD3B-1 | 0.73934314456649 | 1.37369653282669 | 0.91312030347133 | 0.36650948940464 | 0.675428849531356 | -5.45647964392507 |
| lnc-KCNE4-1 | lnc-KCNE4-1 | -0.255666234974274 | 0.975127260388058 | -0.91258138591845 | 0.366610698655798 | 0.675554880074511 | -5.55115824635852 |
| lnc-TMED10-1 | lnc-TMED10-1 | 0.126988074895773 | 1.3282681431689 | 0.912412585031916 | 0.366698497271401 | 0.675641847153749 | -5.5648167415315 |
| ENSG00000149577.15 | SIDT2 | -0.300654082263447 | 6.81731988012164 | -0.912597143009743 | 0.366723544497968 | 0.675641847153749 | -5.99812274014243 |
| lnc-MAP4K2-3 | lnc-MAP4K2-3 | 0.157249392539731 | 4.7228266541134 | 0.912264876842457 | 0.366775336053334 | 0.675676787074714 | -5.94888304411135 |
| ENSG00000230092.7 | | 0.133134547796308 | 0.841622336185787 | 0.911952821465541 | 0.366937703511829 | 0.675915406424196 | -5.47330578807732 |
| ENSG00000266086.2 | | 0.122498353143489 | 5.61977762480024 | 0.911372930956607 | 0.367239553127202 | 0.676281395422901 | -5.99446052302384 |
| ENSG00000135686.12 | KLHL36 | -0.102646646155981 | 5.8844889612383 | -0.911349139671399 | 0.367251940593466 | 0.676281395422901 | -5.99862528257439 |
| lnc-C12orf76-1 | lnc-C12orf76-1 | 0.138834658049746 | 3.19378644457376 | 0.911234192583804 | 0.367311794171296 | 0.676281395422901 | -5.79204137363448 |
| ENSG00000187189.10 | TSPYL4 | -0.106527289128167 | 4.44461349936374 | -0.911220233109606 | 0.367319063374543 | 0.676281395422901 | -5.93441123663711 |
| lnc-SKIL-4 | lnc-SKIL-4 | -0.295340281984509 | 1.3891382474825 | -0.91120197303637 | 0.367328728324417 | 0.676281395422901 | -5.60361305856254 |
| ENSG00000182180.13 | MRPS16 | 0.0985549525631635 | 5.36839044070722 | 0.911156850161694 | 0.367352070336235 | 0.676281395422901 | -5.98699676294518 |
| lnc-AFG1L-8 | lnc-AFG1L-8 | 0.253877281512035 | 3.23581120052105 | 0.911105315708349 | 0.367394974034329 | 0.676281395422901 | -5.81336514987757 |
| ENSG00000129003.17 | VPS13C | 0.107897514566645 | 9.18448488891568 | 0.910955227442009 | 0.367457079021205 | 0.676281395422901 | -5.95293561987904 |
| ENSG00000178105.10 | DDX10 | 0.178355145048336 | 4.4966144813315 | 0.910947365430577 | 0.367461174088336 | 0.676281395422901 | -5.9265039319609 |
| ENSG00000197121.14 | PGAP1 | 0.570018133959865 | 0.247951471038417 | 0.911158942312081 | 0.36746495294222 | 0.676281395422901 | -5.39004102751782 |
| ENSG00000115539.13 | PDCL3 | 0.16052242513122 | 2.4707935305662 | 0.910677425316256 | 0.367601795011878 | 0.676472753395572 | -5.70807062912632 |
| lnc-ANXA2R-1 | lnc-ANXA2R-1 | 0.125001462567071 | 5.16548643815926 | 0.91055694460912 | 0.367664568684729 | 0.676527786271926 | -5.97746200069204 |
| lnc-HIP1-2 | lnc-HIP1-2 | -0.121614321947303 | 1.106734198737 | -0.910431793545329 | 0.367729783059712 | 0.676553239524354 | -5.53083598059984 |
| ENSG00000147443.12 | DOK2 | -0.744024176004774 | 1.08188146455642 | -0.910731140812591 | 0.367756089568944 | 0.676553239524354 | -5.47094974812446 |
| ENSG00000133703.11 | KRAS | 0.187561780533018 | 6.0342615241759 | 0.910341170576677 | 0.367777010010105 | 0.676553239524354 | -6.00184793586698 |
| ENSG00000163807.5 | KIAA1143 | -0.115536052596137 | 5.38829678700756 | -0.910256939663185 | 0.36782090933383 | 0.676573527865794 | -5.98933350551059 |
| lnc-ANKRD36B-3 | lnc-ANKRD36B-3 | 0.358892235819508 | 0.928521176068566 | 0.910174187149785 | 0.367877175617494 | 0.676616558460805 | -5.48485712240022 |
| ENSG00000150455.13 | TIRAP | -0.0950854408029231 | 3.04243860298247 | -0.909859294242338 | 0.368028199128348 | 0.676833847628556 | -5.79485156856308 |
| lnc-EGR2-5 | lnc-EGR2-5 | 0.319635102603658 | 1.69936080706046 | 0.90974765176824 | 0.368106206184546 | 0.676916826757483 | -5.62214338324491 |
| ENSG00000144426.18 | NBEAL1 | 0.13756287281468 | 4.49361086038628 | 0.909615014008582 | 0.368155578059131 | 0.676947138037439 | -5.93391381780604 |
| ENSG00000100345.21 | MYH9 | 0.117024478592313 | 9.06629592275388 | 0.909338236672455 | 0.368299936813453 | 0.677087898248033 | -5.95854385945039 |
| ENSG00000143373.17 | ZNF687 | 0.0922382751510371 | 5.6579730777563 | 0.909204942100273 | 0.368369472278819 | 0.677087898248033 | -5.99744490702011 |
| ENSG00000117020.16 | AKT3 | 0.899851610289158 | 4.50065429254982 | 0.909529560417797 | 0.36840337966048 | 0.677087898248033 | -5.88189357760047 |
| ENSG00000171608.15 | PIK3CD | 0.132905737283473 | 6.7571818552235 | 0.909050142526032 | 0.36845023682709 | 0.677087898248033 | -6.00179156396512 |
| MAST4-IT1 | MAST4-IT1 | -0.726900625868643 | -2.27841763715921 | -0.908978913823194 | 0.368487403256402 | 0.677087898248033 | -4.89175508401269 |
| ENSG00000172403.10 | SYNPO2 | 0.517110724300155 | -0.580338117953917 | 0.90897546477743 | 0.36849045004926 | 0.677087898248033 | -5.15119472974117 |
| ENSG00000160781.16 | PAQR6 | -0.162183194783658 | 1.27386523942284 | -0.908970489262027 | 0.368491799268481 | 0.677087898248033 | -5.57887016610553 |
| lnc-TSR3-3 | lnc-TSR3-3 | -0.166719295736107 | 0.749525175870394 | -0.908963791991266 | 0.368495293988946 | 0.677087898248033 | -5.48206390866036 |
| ENSG00000123200.16 | ZC3H13 | -0.165560859475163 | 5.98975761071738 | -0.908761240806037 | 0.368600997847159 | 0.67722166729341 | -6.00231147520687 |
| ENSG00000197323.11 | TRIM33 | 0.0673713081893732 | 6.91423644904734 | 0.908637096488 | 0.368665793762117 | 0.677280259996934 | -6.00029967597075 |
| ENSG00000111224.13 | PARP11 | 0.14785365774876 | 4.24727959810206 | 0.908443292332458 | 0.368766962620576 | 0.677405657440367 | -5.91701482018284 |
| lnc-KLRG1-6 | lnc-KLRG1-6 | 0.528288787032824 | -0.495264054971094 | 0.908314808771997 | 0.368834042946134 | 0.677468419236147 | -5.12718825675132 |
| ENSG00000266066.1 | POLRMTP1 | -0.54050557388309 | -0.85185714315992 | -0.908226421373781 | 0.368880193922911 | 0.677492730571542 | -5.02569396591641 |
| lnc-MPLKIP-5 | lnc-MPLKIP-5 | -0.272337772960164 | 0.438883814613422 | -0.908094541495172 | 0.368949061166665 | 0.677529445238946 | -5.39608983750273 |
| ENSG00000152942.18 | RAD17 | -0.107547105889496 | 5.09429517346471 | -0.908060899242284 | 0.368966630367465 | 0.677529445238946 | -5.97710568742145 |
| ENSG00000163635.18 | ATXN7 | 0.0840905772575624 | 5.83206828166441 | 0.907989008197253 | 0.369004176276661 | 0.677529445238946 | -6.00057783324587 |
| ENSG00000204435.13 | CSNK2B | -0.0988521232965391 | 4.87316114481249 | -0.907883944456714 | 0.369059051426395 | 0.677529445238946 | -5.96784813330178 |
| lnc-HAND1-9 | lnc-HAND1-9 | -0.107956154606466 | 0.702235029691166 | -0.907813444580003 | 0.369095876698891 | 0.677529445238946 | -5.47002285715183 |
| ENSG00000131067.16 | GGT7 | 0.109598953697633 | 4.11543577617795 | 0.907809981030402 | 0.369097685928402 | 0.677529445238946 | -5.90511756518949 |
| ENSG00000150768.15 | DLAT | 0.175610907059484 | 4.09160166228667 | 0.907347885368185 | 0.369339118732809 | 0.677871277230622 | -5.90062121797624 |
| lnc-RPP25L-1 | lnc-RPP25L-1 | 0.0984427889489126 | 1.46187193957201 | 0.907324971067662 | 0.369351093496559 | 0.677871277230622 | -5.5924685064852 |

| | | | | | | | |
|---|---|---|---|---|---|---|---|
| lnc-NFKBIA-6 | lnc-NFKBIA-6 | 0.228447808267806 | -0.553183270383769 | 0.907217047074167 | 0.369407496760861 | 0.677871277230622 | -5.1945129856261 |
| ENSG00000196683.10 | TOMM7 | 0.157630781284558 | 6.43250420632953 | 0.907043919000751 | 0.369497988580375 | 0.677871277230622 | -6.00501655571707 |
| lnc-ABCC6-9 | lnc-ABCC6-9 | 0.298857032194604 | -0.384040673879107 | 0.907037289420571 | 0.369501454060431 | 0.677871277230622 | -5.18669392549889 |
| lnc-NAA25-2 | lnc-NAA25-2 | 0.204096236093491 | 1.33826297249466 | 0.907022714937683 | 0.369509072652575 | 0.677871277230622 | -5.56582060213046 |
| ENSG00000169057.21 | MECP2 | 0.101601871244415 | 6.45712650916899 | 0.906998842221236 | 0.369521551975211 | 0.677871277230622 | -6.00498462403012 |
| lnc-USP34-3 | lnc-USP34-3 | 0.115912763079486 | 4.01031401487769 | 0.906934123188476 | 0.369555384835153 | 0.677871277230622 | -5.89561396730329 |
| lnc-CELF6-1 | lnc-CELF6-1 | 0.116131766642837 | 3.19930321601379 | 0.906886450594387 | 0.369580307685082 | 0.677871277230622 | -5.80679924233229 |
| lnc-GLYATL2-2 | lnc-GLYATL2-2 | 0.212628858450977 | 3.86964096192349 | 0.906608264602633 | 0.369725762668245 | 0.678044645560536 | -5.86754971241764 |
| ENSG00000168026.18 | TTC21A | 0.153737754188844 | 5.18746256562883 | 0.906529288314546 | 0.369767063678202 | 0.678044645560536 | -5.97988276320379 |
| ENSG00000166477.12 | LEO1 | -0.107597314169054 | 3.51486966480506 | -0.90648591054307 | 0.369789749544864 | 0.678044645560536 | -5.83869434080313 |
| lnc-PLEKHD1-4 | lnc-PLEKHD1-4 | 0.247976897930534 | 0.595674496786658 | 0.906453697073331 | 0.369806597243482 | 0.678044645560536 | -5.44311785403519 |
| ENSG00000145041.15 | DCAF1 | 0.101070739536741 | 5.67224025469032 | 0.906344744411602 | 0.369863583328816 | 0.678088726788725 | -5.99917433360904 |
| lnc-DNAH1-1 | lnc-DNAH1-1 | 0.217201550825348 | 0.251562028960478 | 0.906179416212035 | 0.369950066602087 | 0.678142081231143 | -5.33364398120611 |
| lnc-RNF170-2 | lnc-RNF170-2 | -0.148371336926568 | 0.619154665965165 | -0.906163144721052 | 0.369958578931376 | 0.678142081231143 | -5.47007248100474 |
| lnc-ESAM-2 | lnc-ESAM-2 | -0.176107948224141 | 0.203025795646095 | -0.906050793202294 | 0.370017358368945 | 0.678189428967765 | -5.37803201353274 |
| lnc-GGCT-1 | lnc-GGCT-1 | -0.126125661954491 | 7.37827259770692 | -0.905883311770033 | 0.370104991519939 | 0.678289648335403 | -5.99704702661528 |
| lnc-TMEM88B-6 | lnc-TMEM88B-6 | -0.182915358107651 | 1.79584254388859 | -0.90573035399758 | 0.370185036961868 | 0.678375945137654 | -5.64411333518152 |
| ENSG00000258964.1 | | 0.249286500960184 | 2.02824928964428 | 0.905650055908703 | 0.370227062795085 | 0.678392560625945 | -5.67581367435278 |
| ENSG00000197279.3 | ZNF165 | 0.217131006677864 | 1.96847576907921 | 0.90549341285253 | 0.370309054338286 | 0.678449989433671 | -5.64194276353645 |
| lnc-SERP1-1 | lnc-SERP1-1 | -0.218417776443564 | 0.154199445499596 | -0.905464236427662 | 0.37032432742007 | 0.678449989433671 | -5.29915613990797 |
| lnc-RFLNB-3 | lnc-RFLNB-3 | 0.2054120336802 | 0.999050610153989 | 0.905390438087973 | 0.370362960694998 | 0.678460379137162 | -5.51966717091755 |
| ENSG00000167550.10 | RHEBL1 | 0.190478265630933 | 1.27895155367709 | 0.905173831753178 | 0.370476368623295 | 0.678571642144629 | -5.57887819319104 |
| ENSG00000138303.17 | ASCC1 | -0.168327967276524 | 4.41689692446779 | -0.90511682297061 | 0.370506220252418 | 0.678571642144629 | -5.93635577640035 |
| lnc-TMPPE-3 | lnc-TMPPE-3 | -0.20218803007565 | -0.0112928062729041 | -0.905065653854494 | 0.37053301536192 | 0.678571642144629 | -5.30155808443583 |
| lnc-RNF38-1 | lnc-RNF38-1 | 0.174813741481354 | 4.06499169216088 | 0.905016244999057 | 0.370558889879659 | 0.678571642144629 | -5.89768204256672 |
| ENSG00000276368.1 | H2AC14 | 0.521683419757981 | 2.40949764910994 | 0.905287325232036 | 0.370588535653101 | 0.678571642144629 | -5.73521531191443 |
| ENSG00000171097.13 | KYAT1 | 0.178971680630305 | 3.00325412037328 | 0.904773593639348 | 0.370685978841636 | 0.678689690668688 | -5.7767666489435 |
| lnc-FNDC1-8 | lnc-FNDC1-8 | -0.337855376823086 | -0.506423200700911 | -0.904668724016644 | 0.370740913113753 | 0.678729895456762 | -5.17403782073849 |
| LINC00623 | LINC00623 | -0.143070964414182 | 0.728917853173562 | -0.904549951150488 | 0.37080313669791 | 0.67877485794977 | -5.48057894291776 |
| lnc-ERMP1-4 | lnc-ERMP1-4 | -0.140735477008219 | -0.039385474182952 | -0.90443833785822 | 0.370861615591599 | 0.67877485794977 | -5.27514738210677 |
| lnc-RECQL4-3 | lnc-RECQL4-3 | -0.215909188295557 | 0.706772283825063 | -0.90443210017642 | 0.370864883950112 | 0.67877485794977 | -5.44593402949926 |
| lnc-MRM3-3 | lnc-MRM3-3 | 0.205052249856509 | -0.298207475480214 | 0.904307439568325 | 0.370930206259254 | 0.67877485794977 | -5.25428159444085 |
| ENSG00000158467.16 | AHCYL2 | 0.131767277713528 | 3.30842006473255 | 0.904223681748673 | 0.370974099611508 | 0.67877485794977 | -5.81886337089266 |
| BIN3-IT1 | BIN3-IT1 | 0.19981685677192 | 1.44030666216702 | 0.904152104017938 | 0.371011612623952 | 0.67877485794977 | -5.58353195734493 |
| ENSG00000159314.11 | ARHGAP27 | 0.140616817900099 | 5.64797146077982 | 0.904149281017665 | 0.371013092173811 | 0.67877485794977 | -6.00118155682205 |
| ENSG00000106415.12 | GLCCI1 | -0.208718175538782 | 4.86056416362044 | -0.904143610610669 | 0.371029292464308 | 0.67877485794977 | -5.97922464243875 |
| lnc-MTRF1-2 | lnc-MTRF1-2 | 0.130944056267414 | 1.22048230964567 | 0.903986451407939 | 0.371098438479205 | 0.678841020193519 | -5.57330281800717 |
| lnc-DNAL4-4 | lnc-DNAL4-4 | 0.120049145648016 | 4.12115128304401 | 0.903922941113107 | 0.371131730494937 | 0.678841170635387 | -5.90857454400111 |
| lnc-SLC25A51-6 | lnc-SLC25A51-6 | -0.0930699163858362 | 1.52645463478678 | -0.903860465385909 | 0.371164482064356 | 0.678841170635387 | -5.60211394790476 |
| ENSG00000161040.16 | FBXL13 | 0.173924205928687 | 1.17590125447338 | 0.903793371140771 | 0.371199656861799 | 0.678845183224026 | -5.55774347825167 |
| ENSG00000145088.8 | EAF2 | 0.299136061919096 | 4.17465609018716 | 0.903753566295242 | 0.371313751539851 | 0.678947174032979 | -5.89561639780401 |
| lnc-MORC2-3 | lnc-MORC2-3 | -0.402914463122534 | -0.0089730057363242 | -0.903522474008665 | 0.371341699048989 | 0.678947174032979 | -5.28383538661869 |
| MAN1B1-DT | MAN1B1-DT | -0.226109598922002 | 0.0192035670430772 | -0.903324655654668 | 0.371445445172953 | 0.678947174032979 | -5.30734593966695 |
| ENSG00000189306.10 | RRP7A | 0.112360917244532 | 5.54494085995191 | 0.90330669013535 | 0.371454868138012 | 0.678947174032979 | -5.99839616116702 |
| ENSG00000120798.16 | NR2C1 | 0.0859243975015165 | 4.7324359959558 | 0.903264478376622 | 0.371477008927676 | 0.678947174032979 | -5.96480606425851 |
| ENSG00000161921.14 | CXCL16 | 0.202490366820589 | 2.44769973271406 | 0.903262987257875 | 0.371477791060474 | 0.678947174032979 | -5.71381235780369 |
| ENSG00000196110.7 | ZNF699 | 0.18760244610877 | 4.64498095914431 | 0.903246713317928 | 0.371486327258388 | 0.678947174032979 | -5.94976251874655 |
| lnc-ACOT12-9 | lnc-ACOT12-9 | -0.507626138903388 | -0.0659781467509545 | -0.903116630886455 | 0.371611214860652 | 0.679078476381823 | -5.33388729886249 |
| lnc-KYNU-5 | lnc-KYNU-5 | -0.218340637345782 | 0.736570478610597 | -0.902937812646582 | 0.37164837935855 | 0.679078476381823 | -5.49575396076541 |
| ENSG00000175701.10 | MTLN | 0.573257923984066 | 0.303907621881906 | 0.903092086023065 | 0.371657146015704 | 0.679078476381823 | -5.31364701445957 |
| ENSG00000183010.16 | PYCR1 | -0.416571785856257 | -0.19593819304722 | -0.902832781954626 | 0.371703489756711 | 0.679102869666464 | -5.24742747535625 |
| ENSG00000169020.9 | ATP5ME | 0.148736227798692 | 4.02365466129493 | 0.902537685286588 | 0.371858357297695 | 0.679272644471338 | -5.90203175771056 |
| lnc-ARMCX3-3 | lnc-ARMCX3-3 | 0.254300945325159 | 1.15778411167587 | 0.902506015522469 | 0.371874980138479 | 0.679272644471338 | -5.56543898203677 |
| lnc-ANKRD20A2-22 | lnc-ANKRD20A2-2 | -0.127946218038672 | 4.39636916233139 | -0.902422649076238 | 0.371918739839466 | 0.679272644471338 | -5.94084575491681 |
| ENSG00000103199.13 | ZNF500 | -0.104940968508059 | 3.67954128117554 | -0.902404205111113 | 0.371928421667977 | 0.679272644471338 | -5.8669829212667 |
| ENSG00000165660.7 | ABRAXAS2 | 0.0892643110003952 | 4.32706845164919 | 0.902135441635105 | 0.372069522592655 | 0.679470054433912 | -5.92942835761248 |
| ENSG00000141627.13 | DYM | 0.0629009170204338 | 5.59332340641854 | 0.902051902035299 | 0.372113387914079 | 0.679489874328911 | -6.00155643210647 |
| ENSG00000085415.15 | SEH1L | 0.113555986168511 | 4.81846935110248 | 0.901912471809159 | 0.372186607913206 | 0.679563288448285 | -5.96674552655291 |
| ENSG00000162384.13 | CZIB | -0.0978848678668522 | 3.5343031805637 | -0.901836666580243 | 0.372226419939927 | 0.679573427958179 | -5.84673539159151 |
| lnc-MFN1-1 | lnc-MFN1-1 | -0.258912310208815 | 0.734310860825138 | -0.901776170641905 | 0.372258193666058 | 0.679573427958179 | -5.47489510583836 |
| ENSG00000196950.13 | SLC39A10 | -0.106848967084671 | 6.25553106776465 | -0.901674085128351 | 0.372311815050377 | 0.67961104018995 | -6.00948334143645 |
| ENSG00000175414.6 | ARL10 | -0.160988676731074 | 3.966636123706 | -0.901309521247729 | 0.372503346148275 | 0.679869512466955 | -5.89109382707834 |
| lnc-ZNF813-3 | lnc-ZNF813-3 | 0.34720325666963 | 0.994325858617452 | 0.901243668124534 | 0.372546410161232 | 0.679869512466955 | -5.49537931849129 |
| lnc-COMMD1-3 | lnc-COMMD1-3 | -0.199588189987319 | 0.315972401346947 | -0.901215968855272 | 0.372552506005674 | 0.679869512466955 | -5.37959399391629 |
| ENSG00000100941.8 | PNN | -0.105060375848494 | 7.6820643931129 | -0.901016583749313 | 0.372657292666995 | 0.679966640721382 | -5.99539775344577 |
| ENSG00000169567.11 | HINT1 | 0.110372255018629 | 6.35742761435809 | 0.900961348159006 | 0.372686325025983 | 0.679966640721382 | -6.01036904880173 |
| lnc-AFG3L2-6 | lnc-AFG3L2-6 | -0.151503981587068 | 2.48827099320109 | -0.900926131384617 | 0.372704836059789 | 0.679966640721382 | -5.72491953426567 |
| ENSG00000010626.14 | LRRC23 | 0.145178868046375 | 2.43212849989919 | 0.900757632773831 | 0.372793412342211 | 0.680067961201784 | -5.71488997694567 |

| | | | | | | | |
|---|---|---|---|---|---|---|---|
| lnc-UBASH3A-6 | lnc-UBASH3A-6 | -0.553758017325761 | 0.582790962631527 | -0.900727879652806 | 0.372940346831503 | 0.680200137088607 | -5.48832323157092 |
| lnc-CORO1C-1 | lnc-CORO1C-1 | 0.128485343855447 | 0.465158912869777 | 0.900348843240302 | 0.373008360779465 | 0.680200137088607 | -5.4054800843638 |
| ENSG00000104689.9 | TNFRSF10A | 0.117575160337382 | 5.90765015642004 | 0.900322001141417 | 0.373022477588743 | 0.680200137088607 | -6.0080937687298 |
| ENSG00000107882.11 | SUFU | 0.155027109246161 | 2.58885217830217 | 0.900217716628465 | 0.373077326200478 | 0.680200137088607 | -5.72490877190768 |
| ENSG00000113595.14 | TRIM23 | 0.115739764890189 | 4.83787579127801 | 0.90015274425596 | 0.373111501141961 | 0.680200137088607 | -5.97062931941536 |
| lnc-CYGB-4 | lnc-CYGB-4 | 0.256710434601835 | 2.62856536599413 | 0.900141741881744 | 0.373125829648668 | 0.680200137088607 | -5.76613421792403 |
| ENSG00000133983.14 | COX16 | 0.106067912831705 | 2.9945913161081 | 0.900073567067086 | 0.373153150415629 | 0.680200137088607 | -5.78868798256803 |
| ENSG00000232176.1 | | 0.563347195047609 | 0.902596944821446 | 0.900178851023406 | 0.373204462919987 | 0.680200137088607 | -5.37118656014161 |
| UBL7-AS1 | UBL7-AS1 | -0.11145703559592 | 2.17212388091217 | -0.899972127034477 | 0.373206514884301 | 0.680200137088607 | -5.68629474930291 |
| lnc-ZNF654-3 | lnc-ZNF654-3 | -0.127866021397877 | 4.34533690219067 | -0.899960249172693 | 0.373212763780634 | 0.680200137088607 | -5.94335535563836 |
| lnc-KLF6-18 | lnc-KLF6-18 | -0.349036342146563 | 1.43948105260148 | -0.899985468280354 | 0.373242420904208 | 0.680200137088607 | -5.60535046014788 |
| ENSG00000119041.10 | GTF3C3 | 0.105158570620223 | 5.27384053977941 | 0.89981238055529 | 0.373290562494746 | 0.680200137088607 | -5.99320210319166 |
| lnc-BAIAP2L2-3 | lnc-BAIAP2L2-3 | 0.109136453501928 | 0.895129726956042 | 0.899771662970455 | 0.373311987228214 | 0.680200137088607 | -5.49600512914583 |
| lnc-ZNF700-1 | lnc-ZNF700-1 | -0.165896135509175 | 0.834897391955314 | -0.899725206788651 | 0.373336432453141 | 0.680200137088607 | -5.5056065270353 |
| lnc-TLR6-2 | lnc-TLR6-2 | 0.277932440429054 | 1.93315833470597 | 0.89962297928232 | 0.373390228141393 | 0.680200137088607 | -5.63209973300644 |
| ENSG00000219626.8 | FAM228B | 0.122386334165396 | 4.3679211153827 | 0.899614644456332 | 0.373394614437359 | 0.680200137088607 | -5.93439719903882 |
| ENSG00000119705.9 | SLIRP | 0.105844972578375 | 3.2256774576561 | 0.899450073556782 | 0.373481228490567 | 0.680295353324406 | -5.81682836253257 |
| lnc-NGLY1-3 | lnc-NGLY1-3 | 0.110040999643001 | 2.95530869241975 | 0.899366184919144 | 0.373525384234218 | 0.680295353324406 | -5.78533204948209 |
| lnc-CRYGN-3 | lnc-CRYGN-3 | 0.143207408861386 | 1.4154193782858 | 0.899237742449292 | 0.373592997880877 | 0.680295353324406 | -5.58633590170114 |
| ENSG00000119900.8 | OGFRL1 | 0.214423286314918 | 5.33032915662719 | 0.899255648031452 | 0.373607049892894 | 0.680295353324406 | -5.99940155908972 |
| ENSG00000204920.10 | ZNF155 | -0.225800752302868 | 3.32485193379685 | -0.899201382106735 | 0.373612139823111 | 0.680295353324406 | -5.82422960739635 |
| ENSG00000180287.16 | PLD5 | -1.15900778985589 | -0.0743590996577608 | -0.899372090506755 | 0.373717343634711 | 0.680426721276714 | -5.31044927065351 |
| ENSG00000155957.17 | TMBIM4 | 0.0733108628067241 | 5.17603291348777 | 0.898867534246006 | 0.37378792377862 | 0.680495032295714 | -5.99016993096819 |
| ENSG00000148120.16 | AOPEP | 0.131093816873471 | 3.84214425585652 | 0.898803754256007 | 0.37382151246688 | 0.680495992845652 | -5.87513811637591 |
| lnc-ATAD5-9 | lnc-ATAD5-9 | 0.104700579766781 | 3.62522460572925 | 0.898628952242332 | 0.373913579001786 | 0.680602456169384 | -5.86268767415191 |
| ENSG00000104731.13 | KLHDC4 | 0.112411788358704 | 4.14619733713995 | 0.898567157493809 | 0.373946129175512 | 0.680602456169384 | -5.91085430679811 |
| ENSG00000167778.8 | SPRYD3 | -0.10349579906634 | 3.95524549650925 | -0.898260337195169 | 0.374107772609998 | 0.680795063968932 | -5.90471922646512 |
| ENSG00000139946.9 | PELI2 | -0.908220162163045 | 2.76983261271103 | -0.898537166847795 | 0.374162526822523 | 0.680795063968932 | -5.82132438858205 |
| lnc-SP7-2 | lnc-SP7-2 | -0.152809614759216 | 2.09065032288401 | -0.898115611583805 | 0.374184034549653 | 0.680795063968932 | -5.69365143663555 |
| ENSG00000196586.13 | MYO6 | 0.833276889979253 | -0.714178964529212 | 0.898334056650081 | 0.374192852861137 | 0.680795063968932 | -5.1377556714773 |
| ENSG00000110066.14 | KMT5B | -0.0740816974806968 | 5.58285674913035 | -0.898052426637775 | 0.374217332446412 | 0.680795063968932 | -6.0045536260278 |
| lnc-SGSM2-2 | lnc-SGSM2-2 | -0.280276748053488 | -0.581418412962351 | -0.897764718414592 | 0.374368976081837 | 0.681010749862346 | -5.14746172687048 |
| ENSG00000112208.11 | BAG2 | -0.119023880609356 | 3.13064006224849 | -0.897675146013238 | 0.374416195432799 | 0.681011612980056 | -5.8118547063167 |
| ENSG00000200742.1 | | 0.225903264237339 | -0.27452427640452 | 0.897638295511787 | 0.374435622800141 | 0.681011612980056 | -5.26631707145294 |
| ENSG00000277149.4 | TYW1B | -0.0991128368486982 | 2.87015495048017 | -0.89695709994389 | 0.374794861343928 | 0.681533203546783 | -5.77047310031604 |
| lnc-TTN-1 | lnc-TTN-1 | -0.461358861504373 | 3.70258478695314 | -0.897242644223532 | 0.374814200183062 | 0.681533203546783 | -5.85904804091786 |
| lnc-NPIPB5-3 | lnc-NPIPB5-3 | 0.301394451277297 | -0.101487313234722 | 0.896861674643455 | 0.374845202898349 | 0.681533203546783 | -5.27892177194228 |
| ENSG00000106610.15 | STAG3L4 | 0.104310458559722 | 3.93148131135742 | 0.896664911812931 | 0.374949018662797 | 0.681533203546783 | -5.88987799748426 |
| lnc-MYL7-3 | lnc-MYL7-3 | -0.16827503025172 | 1.21712335910691 | -0.896627050435043 | 0.374968997147805 | 0.681533203546783 | -5.58081930281499 |
| ENSG00000057935.13 | MTA3 | 0.142185658965442 | 4.60981593569056 | 0.896579152418612 | 0.374994272687548 | 0.681533203546783 | -5.95821438247903 |
| lnc-CARHSP1-2 | lnc-CARHSP1-2 | 0.106918747396621 | 4.61026849402586 | 0.896474831960867 | 0.375049325836244 | 0.681533203546783 | -5.95684365319635 |
| ENSG00000168612.4 | ZSWIM1 | -0.141200475039359 | 3.45054903202085 | -0.89632288864309 | 0.375129520301499 | 0.681533203546783 | -5.84621345135605 |
| ENSG00000223959.8 | AFG3L1P | 0.107986682783447 | 4.69324064299691 | 0.896286627242265 | 0.375148660399915 | 0.681533203546783 | -5.9664717857146 |
| ENSG00000115241.10 | PPM1G | 0.137829599301676 | 5.22123048106908 | 0.89627884689452 | 0.375152767234476 | 0.681533203546783 | -5.99534547432547 |
| lnc-OR4F16-4 | lnc-OR4F16-4 | -0.196646685889343 | 0.618186547394052 | -0.896271403691952 | 0.375156696134834 | 0.681533203546783 | -5.44016361853154 |
| lnc-GSG1L-7 | lnc-GSG1L-7 | -0.179212473646727 | 1.47666804254509 | -0.896246022196434 | 0.375170093975276 | 0.681533203546783 | -5.6031232257412 |
| lnc-POU4F3-2 | lnc-POU4F3-2 | 0.168044701499721 | -0.10598735842584 | 0.896165695249142 | 0.375212497262138 | 0.681533203546783 | -5.29811744107924 |
| lnc-DUSP14-9 | lnc-DUSP14-9 | -0.158091691158062 | -0.0201854012413383 | -0.896057462791911 | 0.37526963626248 | 0.681533203546783 | -5.33406417792898 |
| lnc-NDUFB3-2 | lnc-NDUFB3-2 | -0.187277803674524 | 3.02552205177131 | -0.896031684906066 | 0.375283245964302 | 0.681533203546783 | -5.80489072665629 |
| ENSG00000104960.15 | PTOV1 | -0.105660623935484 | 4.81082425906946 | -0.895962048851781 | 0.375320012619168 | 0.681533203546783 | -5.97211642495567 |
| lnc-GOLGA8N-1 | lnc-GOLGA8N-1 | -0.205714397765331 | -0.298355610552362 | -0.895890221759981 | 0.375357938518999 | 0.681533203546783 | -5.2368328337802 |
| ENSG00000163832.15 | ELP6 | -0.122901174625428 | 3.07096020348965 | -0.895737483773628 | 0.375438594867524 | 0.681533203546783 | -5.79306525064993 |
| ENSG00000146476.10 | ARMT1 | -0.124566627034994 | 2.96860022535432 | -0.895718661592879 | 0.375448535063776 | 0.681533203546783 | -5.80283162959815 |
| ENSG00000065183.15 | WDR3 | -0.0987341725125983 | 4.39649934328265 | -0.895639462013434 | 0.375490363058728 | 0.681533203546783 | -5.94453014248335 |
| ENSG00000092108.20 | SCFD1 | 0.0826625734038569 | 5.90497016415106 | 0.895625318173357 | 0.375497833216011 | 0.681533203546783 | -6.01255826161904 |
| ENSG00000125870.10 | SNRPB2 | 0.10913783751448 | 4.70477250742279 | 0.895540457417091 | 0.375542654949483 | 0.681533203546783 | -5.96455755597003 |
| ENSG00000196876.15 | SCN8A | 0.771580299085647 | -1.19634856507555 | 0.895596375211258 | 0.375563407407698 | 0.681533203546783 | -5.01467799964321 |
| lnc-MFSD14C-4 | lnc-MFSD14C-4 | -0.125972629530188 | 1.70967040990746 | -0.895326749778697 | 0.37565554615749 | 0.681533203546783 | -5.62983154345044 |
| ENSG00000084072.16 | PPIE | -0.198785087452127 | 5.10991685762285 | -0.89526438227293 | 0.375693627914902 | 0.681533203546783 | -5.99350779440851 |
| ENSG00000124549.14 | BTN2A3P | 0.203603818227978 | 1.03649899961153 | 0.895226394100329 | 0.375708566602355 | 0.681533203546783 | -5.51083178888542 |
| lnc-AK6-6 | lnc-AK6-6 | 0.163204284316143 | 0.965737978048599 | 0.895146507483438 | 0.375750776144889 | 0.681533203546783 | -5.50318871328862 |
| ENSG00000143603.18 | KCNN3 | -0.483405043581576 | 0.916636441821771 | -0.895215869263961 | 0.375807835911385 | 0.681533203546783 | -5.46296576493708 |
| ENSG00000115966.16 | ATF2 | 0.0606291209384056 | 5.69667014628485 | 0.895013891420635 | 0.375820852941464 | 0.681533203546783 | -6.01015993551047 |
| ENSG00000205476.8 | CCDC85C | 0.34423690926028 | 1.37661110156274 | 0.895039641736819 | 0.375842244722891 | 0.681533203546783 | -5.59741684692169 |
| ENSG00000198876.12 | DCAF12 | -0.368489559972212 | 2.05738429336336 | -0.895133681343045 | 0.375852341644848 | 0.681533203546783 | -5.70674899244433 |
| lnc-SELENOI-1 | lnc-SELENOI-1 | 0.259269506886544 | 0.952200181429516 | 0.894890611885167 | 0.375886003637775 | 0.681533203546783 | -5.51287928974425 |
| lnc-FOXO6-2 | lnc-FOXO6-2 | -0.338158671727322 | 1.92497991539005 | -0.894978724154437 | 0.375892689729978 | 0.681533203546783 | -5.67166759801788 |
| lnc-NOX1-2 | lnc-NOX1-2 | -0.468238956063716 | 0.668439678438411 | -0.895013352630592 | 0.37590085694222 | 0.681533203546783 | -5.45719678028403 |

| | | | | | | | |
|---|---|---|---|---|---|---|---|
| lnc-NENF-2 | lnc-NENF-2 | 0.266069655574768 | 0.0266360447311574 | 0.89484416916339 | 0.375910549532626 | 0.681533203546783 | -5.29806195248847 |
| ENSG00000163794.6 | UCN | -0.199489669265303 | -0.0295476982098634 | -0.894834687584747 | 0.375915560859995 | 0.681533203546783 | -5.31897062123449 |
| ENSG00000104885.17 | DOT1L | 0.156956027057809 | 3.63645122124378 | 0.894743311937175 | 0.375963858096071 | 0.681533203546783 | -5.86651928172342 |
| ENSG00000170899.10 | GSTA4 | -0.417139782221208 | -0.272647285773056 | -0.89464555864108 | 0.37601553067478 | 0.681533203546783 | -5.23897653672011 |
| ENSG00000182774.12 | RPS17 | -0.154841203585311 | 8.11395888350629 | -0.894606370799534 | 0.376036246717376 | 0.681533203546783 | -5.99411582757163 |
| ENSG00000230562.3 | FAM133DP | 0.289508386803909 | 1.79687119253683 | 0.894587817147179 | 0.376047591295033 | 0.681533203546783 | -5.63261220058808 |
| lnc-MAFA-3 | lnc-MAFA-3 | -0.164906949457712 | 0.26700833838107 | -0.894503608568936 | 0.376090573836625 | 0.681533203546783 | -5.39732721363524 |
| ENSG00000196436.8 | NPIPB15 | -1.29144334573265 | -1.11344279791958 | -0.89476830286125 | 0.376113086483404 | 0.681533203546783 | -4.99223995327037 |
| ENSG00000162620.15 | LRRIQ3 | -0.253273638615093 | -0.0356513561099859 | -0.8943064832014 | 0.376194801784548 | 0.681608497945741 | -5.29194997187006 |
| ENSG00000196150.13 | ZNF250 | 0.103699653790721 | 4.29211136617738 | 0.894257188106516 | 0.376220868928803 | 0.681608497945741 | -5.93011693427254 |
| lnc-IGFBP4-3 | lnc-IGFBP4-3 | -0.3885167187578 | -0.0504155861684001 | -0.894141155977556 | 0.376282231035468 | 0.681659669195832 | -5.27373206789154 |
| lnc-ABT1-1 | lnc-ABT1-1 | -0.270340020035206 | 0.105253556769889 | -0.893968723172986 | 0.376373431732867 | 0.681664743205053 | -5.36280584174709 |
| ENSG00000169018.5 | FEM1B | 0.101390576280859 | 6.37729996695217 | 0.893961789716857 | 0.376377099172902 | 0.681664743205053 | -6.0164716331384 |
| ENSG00000272256.1 | | -0.340701483150812 | 0.783188175493927 | -0.89388589236238 | 0.376423366757239 | 0.681664743205053 | -5.50405435504192 |
| lnc-PAOX-2 | lnc-PAOX-2 | -0.213727368539332 | -0.29887572334113 | -0.893825717266565 | 0.376449079090173 | 0.681664743205053 | -5.24009959643851 |
| ENSG00000103091.14 | WDR59 | -0.0867588527010259 | 4.76482422580631 | -0.893779321205604 | 0.376473623795881 | 0.681664743205053 | -5.96991456943942 |
| ENSG00000260836.2 | | -0.162934501858557 | 8.03712639157924 | -0.893758172840356 | 0.376484812162566 | 0.681664743205053 | -5.9965616067466 |
| ENSG00000157823.16 | AP3S2 | -0.082617729168633 | 4.19065896869219 | -0.893682390855858 | 0.37652490573568 | 0.681664743205053 | -5.92699218695207 |
| lnc-SLC38A2-4 | lnc-SLC38A2-4 | 0.26290001533001 | -0.00564719415844115 | 0.893515042305561 | 0.376613453610583 | 0.681664743205053 | -5.31368362177272 |
| ENSG00000106400.11 | ZNHIT1 | 0.113010287091112 | 4.31407277875405 | 0.893483216390981 | 0.376630294922027 | 0.681664743205053 | -5.93416391549364 |
| ENSG00000267314.1 | | -0.145737118586351 | 1.36680046193583 | -0.89347768165662 | 0.376633223785169 | 0.681664743205053 | -5.59611365488842 |
| ENSG00000107521.18 | HPS1 | -0.101785992669621 | 6.21398344331969 | -0.893369921486927 | 0.376690251063667 | 0.681664743205053 | -6.0167744395697 |
| ENSG00000138867.16 | GUCD1 | 0.203073424677017 | 6.23190483083131 | 0.893391537879358 | 0.376694541052751 | 0.681664743205053 | -6.01701675238826 |
| ENSG00000133884.9 | DPF2 | -0.0890383629970817 | 5.20522971379043 | -0.893322092856616 | 0.376715564007068 | 0.681664743205053 | -5.99762593553191 |
| ENSG00000110934.10 | BIN2 | -0.233556690580513 | 6.21040750714818 | -0.893321475857562 | 0.376769036350515 | 0.681701571308251 | -6.0157780449153 |
| ENSG00000121741.16 | ZMYM2 | 0.106804514965165 | 7.51827011213815 | 0.893019294404523 | 0.376875843028644 | 0.681768874744338 | -6.00668946870193 |
| ENSG00000259330.2 | INAFM2 | 0.199100029648277 | 1.6857661483647 | 0.893013039963012 | 0.376879154124726 | 0.681768874744338 | -5.6487599709866 |
| ENSG00000134308.13 | YWHAQ | 0.129246698267074 | 6.71223305217156 | 0.892895097193925 | 0.376941596406318 | 0.681768874744338 | -6.01586431467393 |
| ENSG00000085491.16 | SLC25A24 | -0.12575813407186 | 5.03840813898583 | -0.892840187494048 | 0.376970669428875 | 0.681768874744338 | -5.99288085380673 |
| ENSG00000198932.12 | GPRASP1 | 0.382212341779829 | 5.97051164044953 | 0.893157404227413 | 0.376971848781291 | 0.681768874744338 | -6.01685165147884 |
| ENSG00000131437.15 | KIF3A | -0.111678093637393 | 3.61162834225687 | -0.892759722405938 | 0.377013275842127 | 0.681783891816771 | -5.86934042689876 |
| ENSG00000070214.15 | SLC44A1 | 0.246705481824216 | 3.88799536998404 | 0.892735445949505 | 0.377057597307169 | 0.681804139978342 | -5.8932500822152 |
| ENSG00000137221.14 | TJAP1 | -0.0795119529413952 | 4.19306215566338 | -0.892395867128878 | 0.377205976277154 | 0.682012527205962 | -5.92373556058108 |
| ENSG00000129559.12 | NEDD8 | -0.14811493529188 | 3.66288910788941 | -0.892240101666537 | 0.377288490033437 | 0.682068474686955 | -5.87188104863522 |
| ENSG00000257246.1 | PHB1P19 | -0.268913613290313 | -0.266308867913345 | -0.8921818878119 | 0.377319330660589 | 0.682068474686955 | -5.24432054583093 |
| ENSG00000284503.1 | MIR3652 | -0.174382508619966 | 2.28894392218753 | -0.892149797912968 | 0.377336331985636 | 0.682068474686955 | -5.72322785468911 |
| ENSG00000182810.6 | DDX28 | 0.115787696922131 | 3.67370722007087 | 0.892019748168305 | 0.377405237738694 | 0.682073820711843 | -5.87996699984976 |
| ENSG00000136068.14 | FLNB | 0.356902660467345 | 6.63931655186402 | 0.892297688219087 | 0.377405564985045 | 0.682073820711843 | -6.01429652906122 |
| ENSG00000135446.16 | CDK4 | -0.139245225454083 | 3.97751056234618 | -0.891783013527032 | 0.377530690195662 | 0.682211851442396 | -5.91284770247168 |
| ENSG00000137337.14 | MDC1 | -0.112405781050713 | 5.41656561867487 | -0.891749920777089 | 0.377548229110447 | 0.682211851442396 | -6.00403560547355 |
| ENSG00000241357.1 | | -0.2164306357153 | 0.417965992658723 | -0.891282778291263 | 0.377795866683999 | 0.682568545985297 | -5.40912075196409 |
| ENSG00000134602.15 | STK26 | 0.112460250708683 | 5.51040704950752 | 0.891252438442609 | 0.377811953768182 | 0.682568545985297 | -6.01136825637255 |
| ENSG00000113048.16 | MRPS27 | -0.0816184902942076 | 5.59231209173993 | -0.890960456670602 | 0.377966793473408 | 0.682746837488325 | -6.01036444480398 |
| ENSG00000182405.5 | PGBD4 | -0.125092035535553 | 3.64502235115083 | -0.890890066346402 | 0.378004127944894 | 0.682746837488325 | -5.87147401100018 |
| lnc-SERGEF-1 | lnc-SERGEF-1 | -0.0857890887417672 | 1.8907870087489 | -0.890874880646773 | 0.378012182628428 | 0.682746837488325 | -5.66134534341997 |
| lnc-SEC14L1-1 | lnc-SEC14L1-1 | -0.219008765985198 | 0.792118969602429 | -0.890706365477893 | 0.378101572514009 | 0.682746837488325 | -5.48565556420601 |
| ENSG00000162600.11 | OMA1 | 0.13886701435617 | 5.66339684078336 | 0.890701831718788 | 0.378103977660059 | 0.682746837488325 | -6.01131518838157 |
| ENSG00000161970.14 | RPL26 | 0.131399714251178 | 5.65631750175398 | 0.890691114146041 | 0.378109663340107 | 0.682746837488325 | -6.01298504213197 |
| ENSG00000175216.14 | CKAP5 | 0.0791643341194082 | 5.78730463709412 | 0.890513151652398 | 0.378204080541759 | 0.682848071678712 | -6.01503912037185 |
| ENSG00000156500.15 | PABIR3 | 0.083938969595752 | 3.08491910251949 | 0.890456787304702 | 0.378233987528485 | 0.682848071678712 | -5.80214959834116 |
| lnc-ERVW-1-2 | lnc-ERVW-1-2 | -0.104460407006772 | 1.84772465642684 | -0.890397864733824 | 0.378265253523814 | 0.682848071678712 | -5.66038879336734 |
| ENSG00000240694.8 | PNMA2 | 0.845446824544414 | 0.196973550665909 | 0.890656958783419 | 0.378318737575861 | 0.68288472994159 | -5.48333098676912 |
| ENSG00000100290.2 | BIK | -0.708486503005781 | -0.398019716999464 | -0.890380036354399 | 0.378380394451518 | 0.682936132847737 | -5.1911412230984 |
| lnc-GLT8D2-1 | lnc-GLT8D2-1 | 0.116859658423643 | 0.968826349082267 | 0.890106341691086 | 0.378419968208632 | 0.68294767256802 | -5.53767174929712 |
| ENSG00000163249.11 | CCNYL1 | 0.114095349736362 | 4.05177782420209 | 0.889787850739662 | 0.378589041192655 | 0.683164430945521 | -5.91090703569686 |
| ENSG00000140153.17 | WDR20 | 0.0806121586705482 | 4.93030626459464 | 0.889755052611975 | 0.378606455025777 | 0.683164430945521 | -5.98361561312287 |
| ENSG00000266865.6 | | -0.180503364779103 | 0.608032093887374 | -0.889610365114387 | 0.378683281479529 | 0.683243161175767 | -5.44702114340978 |
| ENSG00000107672.14 | NSMCE4A | 0.102936095879595 | 5.36774341473938 | 0.88929185287049 | 0.378852440756079 | 0.683482224355458 | -6.00685588919094 |
| lnc-GNG13-6 | lnc-GNG13-6 | 0.239922061239033 | -0.213994989942865 | 0.889235841224328 | 0.378882193066889 | 0.683482224355458 | -5.31700525777403 |
| ENSG00000197385.5 | ZNF860 | -0.208438099888051 | 5.03307826749639 | -0.888674533724816 | 0.379196071097223 | 0.683972643361888 | -5.98940464848008 |
| lnc-SERPINB9-5 | lnc-SERPINB9-5 | 0.137696513380323 | 2.28927885178557 | 0.888548660754102 | 0.379247331332445 | 0.683972643361888 | -5.71056848038951 |
| ENSG00000006576.16 | PHTF2 | 0.109183840083831 | 6.66659199613293 | 0.888536599165926 | 0.379253742345102 | 0.683972643361888 | -6.02067691204634 |
| lnc-C12orf73-2 | lnc-C12orf73-2 | -0.164018173967165 | 0.999427480618045 | -0.888400254672618 | 0.379326217393263 | 0.683985615845032 | -5.54293050180978 |
| ENSG00000100101.15 | | 0.102243424889592 | 2.73963292745395 | 0.888393578760557 | 0.379329766256339 | 0.683985615845032 | -5.76551250961936 |
| ENSG00000180425.11 | C11orf71 | 0.187298130235217 | 2.58772733978097 | 0.888335526573633 | 0.37936062724029 | 0.683985615845032 | -5.73831624284523 |
| ENSG00000244165.1 | P2RY11 | -0.158789719389744 | 2.30977755102398 | -0.8880995131101114 | 0.379486110259766 | 0.684151931985351 | -5.72788258065404 |
| lnc-CARNMT1-2 | lnc-CARNMT1-2 | 0.266598664344396 | 0.400483271658591 | 0.887934900063229 | 0.379573646940469 | 0.684209133502096 | -5.3869551442298 |
| ENSG00000070367.15 | EXOC5 | 0.0878311665630616 | 5.59791357246593 | 0.887914827618619 | 0.379584321792763 | 0.684209133502096 | -6.01508987644965 |

| | | | | | | | |
|---|---|---|---|---|---|---|---|
| LINC02021 | LINC02021 | 0.307587963449618 | 0.873651479604864 | 0.887357773067277 | 0.379880648731716 | 0.684683309355946 | -5.49398463181333 |
| EXOC3-AS1 | EXOC3-AS1 | -0.163997873351879 | 0.179735433336504 | -0.88710539561305 | 0.380014950184347 | 0.684865398795589 | -5.36966642854456 |
| lnc-USP53-6 | lnc-USP53-6 | 0.181338948479464 | 1.35542466739225 | 0.886871835913204 | 0.380139264782896 | 0.68502945955405 | -5.59790914442242 |
| ENSG00000166925.8 | TSC22D4 | -0.16444461706633 | 3.95831261555965 | -0.886347643566891 | 0.380418365874787 | 0.685447469444663 | -5.91153619004107 |
| ENSG00000133247.13 | KMT5C | 0.190757347780744 | 1.05443404679356 | 0.886311092957014 | 0.380437831751243 | 0.685447469444663 | -5.54826085261527 |
| ENSG00000279415.1 | | 0.291870714374101 | -0.381397071203185 | 0.886107508766412 | 0.380546266837076 | 0.685566962309618 | -5.20420679622795 |
| ENSG00000157224.15 | CLDN12 | 0.131024456100469 | 1.69403682665597 | 0.886061514134216 | 0.38057076768934 | 0.685566962309618 | -5.63542522399048 |
| ENSG00000143367.15 | TUFT1 | -0.507587715850422 | -0.843995167825542 | -0.885978890435992 | 0.380627541183812 | 0.685580297426692 | -5.2022986361814 |
| ENSG00000212309.1 | SNORD70B | 0.20747003571885 | 0.59774109058835 | 0.885885073584498 | 0.380664765005181 | 0.685580297426692 | -5.46249142922542 |
| lnc-GLRA4-2 | lnc-GLRA4-2 | 0.124414933249658 | 3.00999443506811 | 0.885764763865418 | 0.38072886752957 | 0.685580297426692 | -5.79730945469875 |
| ENSG00000101928.12 | MOSPD1 | 0.10921372959533 | 2.02258406318929 | 0.885742952716637 | 0.380740489517836 | 0.685580297426692 | -5.68824765813334 |
| lnc-TLE3-6 | lnc-TLE3-6 | -0.230632648003609 | -0.516842776881979 | -0.885735030910912 | 0.380744710677963 | 0.685580297426692 | -5.15935422320437 |
| GNAS-AS1 | GNAS-AS1 | -0.24858322919021 | 0.197502017724708 | -0.885596154899566 | 0.380818716050321 | 0.685614233915404 | -5.37381035268321 |
| lnc-PKP2-3 | lnc-PKP2-3 | -0.198186360531122 | -0.273419505701732 | -0.885574648910542 | 0.38083017715371 | 0.685614233915404 | -5.28456637032322 |
| lnc-CCDC89-4 | lnc-CCDC89-4 | 0.666539500741972 | -0.9033976692604 | 0.885517886174223 | 0.380879655750007 | 0.685643340414763 | -5.04071101418356 |
| ENSG00000253861.1 | SLC2A3P1 | 0.0939987005913365 | 3.74475495206941 | 0.885401338451948 | 0.380922546831823 | 0.685660584297281 | -5.88880071247118 |
| lnc-CLEC18B-7 | lnc-CLEC18B-7 | -0.0974768619404163 | 3.29459600228505 | -0.885316491786249 | 0.380967772950896 | 0.685682027863613 | -5.83523752111294 |
| ENSG00000107872.12 | FBXL15 | -0.19106080473702 | 1.97843801672756 | -0.884994325070042 | 0.381139529712677 | 0.685800449756303 | -5.69459905253248 |
| ENSG00000168679.17 | SLC16A4 | -0.873286463655994 | 1.49248136275161 | -0.885269804539429 | 0.381184421545204 | 0.685800449756303 | -5.54657848331427 |
| ENSG00000139832.4 | RAB20 | 0.664415466573799 | 0.270821787062245 | 0.885088244313205 | 0.381207616819051 | 0.685800449756303 | -5.2930671890671 |
| lnc-SLCO4C1-5 | lnc-SLCO4C1-5 | -0.28876165479566 | 0.356430612610398 | -0.884858710798227 | 0.381211844479218 | 0.685800449756303 | -5.45334650260486 |
| lnc-EDN1-2 | lnc-EDN1-2 | 0.244362004474623 | 0.339172635543588 | 0.88477494747901 | 0.381256514672825 | 0.685800449756303 | -5.39881951042461 |
| ENSG00000199477.1 | SNORA31 | 0.15514371348561 | 0.275036035563915 | 0.884756702778507 | 0.381266244842018 | 0.685800449756303 | -5.40956798044249 |
| ENSG00000197808.11 | ZNF461 | 0.108018652644888 | 3.99483286779198 | 0.884755661633191 | 0.381266800105008 | 0.685800449756303 | -5.91119763600657 |
| ENSG00000276966.2 | H4C5 | 1.06815636544191 | 4.23669627717327 | 0.885006682416187 | 0.381330273429993 | 0.685854685250746 | -5.90234325065642 |
| ENSG00000166949.15 | SMAD3 | 0.752444302157486 | 1.48280585707115 | 0.884873762555836 | 0.381393471718116 | 0.685908416633813 | -5.61256124457373 |
| ENSG00000070087.14 | PFN2 | -0.269624953936796 | 3.0080050548494 | -0.884476144530832 | 0.381451497732289 | 0.685952837482414 | -5.82257231513276 |
| ENSG00000148935.11 | GAS2 | -0.18879226715213 | -0.342833949352719 | -0.88426203559207 | 0.381530118342793 | 0.685989501384217 | -5.23025380066704 |
| lnc-CCNL2-2 | lnc-CCNL2-2 | 0.120147688624384 | 1.25099502117198 | 0.884246247700078 | 0.381538542090391 | 0.685989501384217 | -5.56879074878613 |
| ENSG00000251451.1 | GTF2IP9 | 0.171864439360864 | -0.324160247836322 | 0.884158712152959 | 0.381585249479162 | 0.686004108030723 | -5.21573095520836 |
| ENSG00000183624.13 | HMCES | 0.318869781174977 | 3.81336365819279 | 0.884303044252396 | 0.381613323468483 | 0.686004108030723 | -5.87704404495074 |
| lnc-DDR2-3 | lnc-DDR2-3 | -0.229786451254122 | 1.4428949663849 | -0.88403751415811 | 0.381649924540528 | 0.686008074441374 | -5.60487242406627 |
| ENSG00000144036.15 | EXOC6B | 0.165525840372116 | 4.05929919972078 | 0.883946176413075 | 0.381698669835882 | 0.686008074441374 | -5.92820494075853 |
| lnc-MMGT1-3 | lnc-MMGT1-3 | 0.184258473829706 | -0.0574043559211368 | 0.883914611226614 | 0.38171551652223 | 0.686008074441374 | -5.28434053163257 |
| ENSG00000230795.3 | HLA-K | -0.29298886108582 | 2.09094442090511 | -0.883830861920551 | 0.381777763221754 | 0.686060040194986 | -5.68274940579868 |
| lnc-FAM83F-4 | lnc-FAM83F-4 | -0.274544558802047 | -0.58163005880848 | -0.883667753734591 | 0.381847283368182 | 0.686066987940297 | -5.14692723702789 |
| ENSG00000176340.3 | COX8A | 0.129229090571182 | 4.04158835147707 | 0.883665862546003 | 0.381848292952633 | 0.686066987940297 | -5.92419179806927 |
| ENSG00000204983.13 | PRSS1 | -0.67865440909111 | -1.04828598773364 | -0.883532766573843 | 0.38192144526044 | 0.686125129401977 | -4.97835447710072 |
| lnc-POPDC2-3 | lnc-POPDC2-3 | -0.139067685218855 | 2.75145209220493 | -0.883445728122707 | 0.381965820207244 | 0.686125129401977 | -5.77571185101536 |
| lnc-MGAT4A-1 | lnc-MGAT4A-1 | -0.0884291396162449 | 2.74404771344393 | -0.883417941601347 | 0.381980656746696 | 0.686125129401977 | -5.77091212579599 |
| ENSG00000285258.1 | NA | 0.0775138159987927 | 5.57840719622552 | 0.883241677838972 | 0.382074780841328 | 0.686234311872344 | -6.01709297616112 |
| lnc-THOC5-3 | lnc-THOC5-3 | 0.112893218629477 | 1.55079227978615 | 0.882958862824516 | 0.382225833624246 | 0.686445714465391 | -5.6221194324652 |
| ENSG00000104331.8 | BPNT2 | 0.146771266093038 | 5.45727258350497 | 0.882813496373544 | 0.382303489258966 | 0.686525276515207 | -6.01540428594204 |
| lnc-FAM13C-5 | lnc-FAM13C-5 | -0.222358840220903 | -0.310069940956282 | -0.882713872840095 | 0.3823567145362 | 0.686560957454296 | -5.26302552204641 |
| lnc-SLX1B-3 | lnc-SLX1B-3 | -0.200029047871535 | 2.79643224296272 | -0.882572726910044 | 0.382432131793798 | 0.686636476684556 | -5.77060257496851 |
| ENSG00000059145.18 | UNKL | -0.146485670632274 | 4.96360069551409 | -0.882431703189684 | 0.382507493183335 | 0.686711882441569 | -5.99596404644729 |
| ENSG00000076685.18 | NT5C2 | 0.133409396820472 | 6.65490987481573 | 0.882082908903672 | 0.382693925168285 | 0.686971212176971 | -6.0259429611808 |
| ENSG00000176293.19 | ZNF135 | -0.989467150643905 | 1.81767258131777 | -0.882384742888576 | 0.382729287276734 | 0.686971212176971 | -5.70656094250492 |
| ENSG00000207133.1 | SNORD116-7 | -0.280367946085417 | 0.519425742959078 | -0.881950640616615 | 0.382764638192794 | 0.686971212176971 | -5.42706845953452 |
| lnc-HIST4H4-5 | lnc-HIST4H4-5 | -0.129442352515814 | 3.1636736510546 | -0.881911722924173 | 0.38278544587561 | 0.686971212176971 | -5.83548367244199 |
| ENSG00000138468.15 | SENP7 | 0.0806416363675169 | 6.37421919119817 | 0.881661895729165 | 0.382919035253016 | 0.687083204191773 | -6.02720851753596 |
| ENSG00000198668.10 | CALM1 | 0.101125929019714 | 7.90862516210359 | 0.881611398871014 | 0.382946040886429 | 0.687083204191773 | -6.00858834625646 |
| ENSG00000075188.8 | NUP37 | 0.103691362833741 | 3.631632203134 | 0.881607750718803 | 0.382947991958802 | 0.687083204191773 | -5.87890481476882 |
| ENSG00000168970.22 | JMJD7-PLA2G4B | -0.152617578477424 | 4.35524397777005 | -0.881313801824082 | 0.383105219825144 | 0.687305390016642 | -5.94821125161175 |
| lnc-SRSF7-5 | lnc-SRSF7-5 | 0.259027368934215 | 0.446400550455796 | 0.881140009462751 | 0.383198197429557 | 0.687339053227799 | -5.45050764519382 |
| lnc-CEP170-10 | lnc-CEP170-10 | 0.847432770985836 | 0.893077141872787 | 0.881436446658888 | 0.383220095546721 | 0.687339053227799 | -5.41211140378105 |
| ENSG00000233836.7 | | -0.499676451219376 | 3.01623904763887 | -0.881406200825414 | 0.383224164351271 | 0.687339053227799 | -5.78612387684866 |
| lnc-ZNF506-8 | lnc-ZNF506-8 | -0.299201664509673 | 0.12030270302764 | -0.880973970437642 | 0.383287040426766 | 0.68735653477417 | -5.33544681716449 |
| ENSG00000169740.13 | ZNF32 | 0.0911731826376307 | 3.84041021243219 | 0.880861990357854 | 0.383346965318857 | 0.68735653477417 | -5.89509649222746 |
| ENSG00000140030.5 | GPR65 | 0.281970142003989 | 2.44814273797759 | 0.880889648190394 | 0.383353155863676 | 0.68735653477417 | -5.73551534844263 |
| ENSG00000224861.1 | YBX1P1 | 0.318833172206416 | 0.459594812640376 | 0.880804682395746 | 0.383377635334831 | 0.68735653477417 | -5.37677407638888 |
| ENSG00000156860.15 | FBRS | -0.131202188864797 | 5.91617014625818 | -0.880761245197879 | 0.383400883046312 | 0.68735653477417 | -6.02582481442644 |
| ENSG00000169964.7 | TMEM42 | -0.103038409556394 | 3.14955807866895 | -0.880595875238201 | 0.383489397698545 | 0.687455345134048 | -5.8186652708626 |
| ENSG00000226318.1 | RPS3AP38 | -0.279577564346041 | -0.768447099081664 | -0.880443967507732 | 0.383570718079932 | 0.687481373235739 | -5.1069312801876 |
| lnc-DNAJC12-2 | lnc-DNAJC12-2 | -0.279577564346041 | -0.768447099081664 | -0.880443967507732 | 0.383570718079932 | 0.687481373235739 | -5.1069312801876 |
| ENSG00000023191.16 | RNH1 | -0.154577024573478 | 5.82393609821074 | -0.880290882945707 | 0.383652679512497 | 0.687568402473289 | -6.02496893609854 |
| ENSG00000285565.1 | NA | 0.150907534536857 | 4.48841044654928 | 0.880125611174971 | 0.383741178437588 | 0.687637637625832 | -5.96226648886574 |
| ENSG00000105655.18 | ISYNA1 | 0.191870431690534 | 1.7442961888386 | 0.880042036605175 | 0.383785935468939 | 0.687637637625832 | -5.6699550702511 |

| | | | | | | | |
|---|---|---|---|---|---|---|---|
| ENSG00000167799.9 | NUDT8 | -0.35435953622928 | -0.141926872954226 | -0.87992076248139 | 0.38385088778154 | 0.687637637625832 | -5.34726839172342 |
| ENSG00000172765.17 | TMCC1 | -0.0979668521775592 | 4.49191154525612 | -0.87987669978192 | 0.38387448872314 | 0.687637637625832 | -5.96450616970112 |
| ENSG00000241549.8 | GUSBP2 | -0.298926598384066 | 1.55958185953738 | -0.87986493958183 | 0.383885824166323 | 0.687637637625832 | -5.61421137943335 |
| ENSG00000188002.10 | PDCD6P1 | 0.0843787346122129 | 2.91923896401449 | 0.879714623408167 | 0.383961308273223 | 0.687637637625832 | -5.79116829862722 |
| ENSG00000233010.1 | RPEP4 | 0.216881469708907 | -0.370291449398889 | 0.879612022714025 | 0.384016274882088 | 0.687637637625832 | -5.19837615244474 |
| ENSG00000214212.8 | C19orf38 | 0.241610371240862 | 0.654172920665549 | 0.879575960175052 | 0.384035595969685 | 0.687637637625832 | -5.47529199818376 |
| lnc-SRPK2-6 | lnc-SRPK2-6 | -0.197448696384495 | 0.0789800951364027 | -0.879567397996698 | 0.384040183385502 | 0.687637637625832 | -5.34183706534312 |
| ENSG00000162694.13 | EXTL2 | 0.174940059142006 | 1.31529848012967 | 0.879427946762828 | 0.384114902998786 | 0.687637637625832 | -5.60633471809458 |
| ENSG00000160087.20 | UBE2J2 | 0.083274547850043 | 3.7121649246002 | 0.879384103106938 | 0.384138396849347 | 0.687637637625832 | -5.88699085664687 |
| ENSG00000168291.12 | PDHB | -0.0902793588066748 | 5.05196953412705 | -0.879263199282509 | 0.384203188508868 | 0.687637637625832 | -6.00356197876197 |
| ENSG00000160783.19 | PMF1 | -0.104300265213546 | 4.44539324296968 | -0.879224251545997 | 0.384224061849936 | 0.687637637625832 | -5.96360245530752 |
| ENSG00000053900.10 | ANAPC4 | -0.101792406647808 | 5.72159439089611 | -0.879186539867336 | 0.384244273432661 | 0.687637637625832 | -6.02526553840168 |
| lnc-SYNCRIP-4 | lnc-SYNCRIP-4 | 0.130836655878732 | 1.49374271641352 | 0.879087225461406 | 0.38429750423409 | 0.687637637625832 | -5.62887220901356 |
| lnc-LAMTOR1-2 | lnc-LAMTOR1-2 | -0.0976149384209326 | 2.6985325203281 | -0.878992483800658 | 0.384348288477565 | 0.687637637625832 | -5.76538770281102 |
| ENSG00000248909.1 | HMGB1P21 | -0.395136455673661 | 0.960230465990594 | -0.879089415820665 | 0.384349145666463 | 0.687637637625832 | -5.53565057549143 |
| ENSG00000104897.9 | SF3A2 | -0.141980216393652 | 3.81624375775008 | -0.87888561529786 | 0.384405578158369 | 0.687637637625832 | -5.90645106834174 |
| lnc-RCC1L-5 | lnc-RCC1L-5 | 0.141664956057113 | -0.0396207582983122 | 0.878760874664774 | 0.384472455508313 | 0.687637637625832 | -5.28882141818286 |
| ENSG00000164466.12 | SFXN1 | -0.110555787241166 | 4.74415738279488 | -0.878735813400296 | 0.38448589252412 | 0.687637637625832 | -5.98763590384774 |
| lnc-POLR1E-4 | lnc-POLR1E-4 | 0.228543024373507 | 0.289217698351644 | 0.878706666906101 | 0.38450152027828 | 0.687637637625832 | -5.3728108386941 |
| ENSG00000269858.5 | EGLN2 | 0.0959627368956699 | 5.30192295585994 | 0.878630788309653 | 0.384542206717245 | 0.687637637625832 | -6.01102117877249 |
| lnc-SMARCC1-2 | lnc-SMARCC1-2 | -0.12125842089821 | 0.877665957262864 | -0.878592092130097 | 0.384562956829779 | 0.687637637625832 | -5.52110683723293 |
| ENSG00000172915.18 | NBEA | 0.258140624995124 | 4.55971936396023 | 0.878706727744655 | 0.384570684587246 | 0.687637637625832 | -5.96711929696285 |
| lnc-DNAAF5-1 | lnc-DNAAF5-1 | -0.122691228484328 | 2.73031574760652 | -0.878533202753565 | 0.384594536532612 | 0.687637637625832 | -5.77287580776415 |
| ENSG00000103197.17 | TSC2 | -0.106867501629787 | 5.49881739545442 | -0.878524109414146 | 0.384599413024681 | 0.687637637625832 | -6.01910398020324 |
| ENSG00000115289.13 | PCGF1 | 0.120743140670471 | 2.70018351158241 | 0.878453397728368 | 0.384637334970695 | 0.687637637625832 | -5.78077514861804 |
| ENSG00000141504.11 | SAT2 | 0.118545509567137 | 3.42565860351061 | 0.878391497622026 | 0.3846705333016 | 0.687637637625832 | -5.85745987646028 |
| ENSG00000102781.13 | KATNAL1 | -0.222951571034925 | 3.26246817497988 | -0.878374245499525 | 0.384679786301661 | 0.687637637625832 | -5.84350886280638 |
| FTX | FTX | -0.210264121754079 | 9.14336138487541 | -0.878377371306024 | 0.384693552798983 | 0.687637637625832 | -5.97967652417077 |
| ENSG00000105122.12 | RASAL3 | -0.108302250631512 | 4.59331829438909 | -0.877982430873064 | 0.384889970035708 | 0.687875550830252 | -5.97656008880187 |
| ENSG00000115762.16 | PLEKHB2 | 0.105356653741246 | 5.20895922656953 | 0.877975868985281 | 0.384893490691931 | 0.687875550830252 | -6.01012804788976 |
| lnc-ERGIC2-7 | lnc-ERGIC2-7 | 0.216190871192655 | 0.853910889720789 | 0.877903502510784 | 0.38493231890137 | 0.687880206832972 | -5.53144681526534 |
| ENSG00000133067.17 | LGR6 | -0.396781599527865 | 1.01670202002654 | -0.877925715867665 | 0.38496293555405 | 0.687880206832972 | -5.49176564227961 |
| ENSG00000121897.14 | LIAS | -0.0766208318693908 | 3.54164231350971 | -0.877553111035049 | 0.385120356411924 | 0.688069679515756 | -5.86881130328719 |
| ENSG00000163697.16 | APBB2 | 0.478152729414836 | 6.05334773736954 | 0.877820148345023 | 0.385165053599055 | 0.688069679515756 | -6.02956995678544 |
| ENSG00000064419.13 | TNPO3 | 0.0736351033590283 | 5.97690556581087 | 0.87745639330324 | 0.385172270200389 | 0.688069679515756 | -6.02912394939684 |
| lnc-HACD2-5 | lnc-HACD2-5 | -0.866885039631131 | -1.64195312039843 | -0.877527328648574 | 0.385202687512027 | 0.688069679515756 | -4.97980711991241 |
| ENSG00000211829.9 | TRDC | -1.10391082788852 | -1.40590412892608 | -0.87738655464234 | 0.385305608700403 | 0.688193799364839 | -4.89356422381597 |
| lnc-TMEM138-10 | lnc-TMEM138-10 | -0.163192878346608 | 0.512627807179627 | -0.877097067383268 | 0.385365179211549 | 0.688240475810092 | -5.45504717604099 |
| lnc-C2orf42-10 | lnc-C2orf42-10 | -0.247361413946274 | 0.920583722977191 | -0.876943552454338 | 0.38544761442875 | 0.688296766015818 | -5.53812719891286 |
| lnc-ATOX1-1 | lnc-ATOX1-1 | -0.221255117708611 | 0.440097154471298 | -0.876913827077355 | 0.385463577800337 | 0.688296766015818 | -5.44787134213846 |
| TMC3-AS1 | TMC3-AS1 | -0.274795876748516 | 0.234393702640177 | -0.876829130032591 | 0.38550906481276 | 0.688303562554411 | -5.37380839601841 |
| lnc-PMM2-1 | lnc-PMM2-1 | -0.171610503536167 | -0.249587405029906 | -0.876782210224046 | 0.385534264815129 | 0.688303562554411 | -5.26268735268899 |
| ENSG00000272182.1 | | -0.164375281097905 | 1.55058558497837 | -0.876582330713242 | 0.385641629107364 | 0.688359991189654 | -5.62103976083198 |
| ENSG00000233382.6 | NKAPP1 | -0.111104578438795 | 2.0660398341862 | -0.876573771168544 | 0.385646227246857 | 0.688359991189654 | -5.69771800021377 |
| lnc-UCP2-2 | lnc-UCP2-2 | -0.147718125792668 | 0.736223983648195 | -0.87653659001631 | 0.38566620115625 | 0.688359991189654 | -5.5024190200613 |
| ENSG00000214753.3 | HNRNPUL2 | -0.0977049170338118 | 4.93583334423993 | -0.876464265394222 | 0.385705056196095 | 0.688369649846894 | -5.99757630983376 |
| ENSG00000090316.15 | MAEA | -0.122457253607696 | 4.8858217925433 | -0.876247282007926 | 0.385821641288834 | 0.688518019997232 | -5.99900575712958 |
| LINC00174 | LINC00174 | -0.0917954627152406 | 3.91538388364174 | -0.875966787501588 | 0.38597238391021 | 0.688727314956555 | -5.91183558653781 |
| ENSG00000007341.18 | ST7L | -0.109504406163039 | 3.35730125456633 | -0.875869259112262 | 0.386024806087058 | 0.688728877659885 | -5.85510860472437 |
| LINC02427 | LINC02427 | -0.257563769963017 | 1.03590018987435 | -0.87584065512621 | 0.38604018177924 | 0.688728877659885 | -5.56966044048324 |
| ENSG00000157107.13 | FCHO2 | 0.188749376806222 | 4.52983598922322 | 0.875666285757158 | 0.38613392009008 | 0.688817831983023 | -5.97202288385908 |
| lnc-INTS4-2 | lnc-INTS4-2 | 0.254096629389959 | -0.377122225988999 | 0.875623408568415 | 0.38615697241666 | 0.688817831983023 | -5.16656739068753 |
| lnc-TRMU-3 | lnc-TRMU-3 | -0.150318376647119 | 0.425173851142434 | -0.875459102868303 | 0.386245317156358 | 0.688829740883932 | -5.44271936531607 |
| lnc-UCP3-5 | lnc-UCP3-5 | -0.12172660583669 | 0.772706060392472 | -0.875425747203917 | 0.386263253562986 | 0.688829740883932 | -5.5027907351198 |
| lnc-ADGRV1-17 | lnc-ADGRV1-17 | -0.201882679052983 | 0.475630318440556 | -0.875424272615604 | 0.386264046508397 | 0.688829740883932 | -5.46461992219673 |
| ENSG00000256574.7 | OR13A1 | -0.332588224540111 | 1.78509181318823 | -0.875364333874799 | 0.386343060542564 | 0.688908833591695 | -5.66884675352458 |
| ENSG00000168255.20 | POLR2J3 | 0.210150526969383 | 2.72939156733945 | 0.875217330783678 | 0.386375337656441 | 0.688908833591695 | -5.7726863743555 |
| ENSG00000172794.19 | RAB37 | 0.254858872522876 | 3.84732153805221 | 0.87514677685165 | 0.386459661572987 | 0.688999498844243 | -5.9113393646427 |
| ENSG00000161692.17 | DBF4B | -0.12822412778182 | 2.33918866655349 | -0.874902797803792 | 0.386544529009009 | 0.689091117321362 | -5.72781619937682 |
| ENSG00000136933.16 | RABEPK | -0.106338922641156 | 2.61524467640214 | -0.874790833988098 | 0.386604767090917 | 0.689098463833596 | -5.76119334238476 |
| lnc-ANKRD20A4-5 | lnc-ANKRD20A4-5 | -0.518348608220737 | 1.02515545218517 | -0.874993776545924 | 0.386615608043063 | 0.689098463833596 | -5.51001398120837 |
| ENSG00000168887.10 | C2orf68 | -0.112668190224439 | 4.9264171962737 | -0.874673040682514 | 0.3866681479227 | 0.689113631751138 | -6.00062662829886 |
| ENSG00000168067.11 | MAP4K2 | -0.0985393180252627 | 6.1612924328975 | -0.874630429189536 | 0.386691077429221 | 0.689113631751138 | -6.03278249096814 |
| LINC01136 | LINC01136 | -0.199565500354276 | 2.77203154334644 | -0.874560978173559 | 0.386728451289642 | 0.689120570763977 | -5.78345473316749 |
| ENSG00000226572.1 | SNORD57 | 0.197406227063317 | -0.201184989450495 | 0.874461669624236 | 0.386781896429351 | 0.689151774551527 | -5.27638284621173 |
| lnc-C16orf45-4 | lnc-C16orf45-4 | 0.149713213746044 | 1.31304300697317 | 0.874404016720665 | 0.386812925783112 | 0.689151774551527 | -5.59632537729129 |
| ENSG00000264364.2 | DYNLL2 | 0.111692444070767 | 5.07734957175207 | 0.874316903471703 | 0.386859813965139 | 0.689175657853943 | -6.00961076405752 |
| ENSG00000091542.8 | ALKBH5 | 0.0881938751186837 | 6.17551419675189 | 0.874120544655163 | 0.386965516085419 | 0.689304302690279 | -6.03341854180887 |

| | | | | | | | |
|---|---|---|---|---|---|---|---|
| lnc-ZNF599-5 | lnc-ZNF599-5 | 0.171994611238839 | 1.77126636434809 | 0.87394633792702 | 0.387059308764085 | 0.689411712728553 | -5.66088595640232 |
| ENSG00000181007.8 | ZFP82 | 0.091124416916017 | 4.15642866433728 | 0.873861086766021 | 0.387105213112377 | 0.689417902197403 | -5.93240196997382 |
| ENSG00000213923.11 | CSNK1E | -0.1620996050449 | 6.01294779540752 | -0.873807079448336 | 0.38713429567701 | 0.689417902197403 | -6.03326608658864 |
| ENSG00000067601.7 | PMS2P4 | 0.129370941808151 | 0.701513953027089 | 0.873746730996596 | 0.387166794530744 | 0.689417902197403 | -5.47844546415549 |
| lnc-NANOS1-3 | lnc-NANOS1-3 | -0.214034491098795 | -0.0950286594613115 | -0.873675781558057 | 0.387205004432512 | 0.689417902197403 | -5.28460603658947 |
| ENSG00000100599.15 | RIN3 | -0.152805599780602 | 5.40826899434267 | -0.873628895293084 | 0.387230256391399 | 0.689417902197403 | -6.01902725834027 |
| lnc-LGALSL-2 | lnc-LGALSL-2 | -0.326780131193326 | 5.93941849632143 | -0.873713401426054 | 0.387322335221089 | 0.68952219562841 | -6.03071170625579 |
| ENSG00000172345.13 | STARD5 | 0.194017512038806 | 3.57584535942869 | 0.873339928100273 | 0.387385911017495 | 0.689523000445865 | -5.87285499832274 |
| ENSG00000196866.2 | H2AC7 | 1.00602735082693 | 2.09725322014278 | 0.873670931412472 | 0.38740219489981 | 0.689523000445865 | -5.67372735259632 |
| ENSG00000066427.22 | ATXN3 | -0.0840614012368777 | 4.86630600022567 | -0.873246099889934 | 0.387436460873555 | 0.689523000445865 | -5.99713346233335 |
| lnc-PIGU-1 | lnc-PIGU-1 | -0.147628346216841 | -0.0313980566536143 | -0.873208375824948 | 0.387456785850307 | 0.689523000445865 | -5.31525011812627 |
| ENSG00000081189.15 | MEF2C | -0.157904090132887 | 7.48608809644176 | -0.87305319916016 | 0.387540399034532 | 0.689612175441149 | -6.02548796462892 |
| lnc-TMUB2-1 | lnc-TMUB2-1 | -0.20134151749975 | 2.60247582370071 | -0.872765594312682 | 0.387695398054527 | 0.689828352191937 | -5.76399116737861 |
| RHOXF1-AS1 | RHOXF1-AS1 | -0.43949124451994 | 0.740231466077796 | -0.872540872750777 | 0.387867163470636 | 0.690074321524427 | -5.43426449462321 |
| ENSG00000263956.6 | NBPF11 | -0.122650454729827 | 5.14920377219981 | -0.872150242437613 | 0.388027161346676 | 0.690299313915179 | -6.00970221029315 |
| ENSG00000188706.12 | ZDHHC9 | -0.148547937832213 | 2.00730882778508 | -0.87200452464997 | 0.388105750448512 | 0.690312073454197 | -5.69640234758847 |
| lnc-TACC3-1 | lnc-TACC3-1 | -0.203582423136371 | 1.22053265875542 | -0.871962992075383 | 0.388128151802255 | 0.690312073454197 | -5.60643258439947 |
| ENSG00000070614.14 | NDST1 | -0.180357867723093 | 3.84189630518291 | -0.871870693075948 | 0.388177937876658 | 0.690312073454197 | -5.90638229801808 |
| lnc-FOS-1 | lnc-FOS-1 | -0.697021237056308 | -0.655702651291087 | -0.871975793295371 | 0.388213659322896 | 0.690312073454197 | -5.17290874008492 |
| ENSG00000250011.1 | HMGB1P3 | -0.175998422179433 | 1.86352230764597 | -0.871712025204904 | 0.388263532731918 | 0.690312073454197 | -5.6831322969914 |
| ENSG00000101158.13 | NELFCD | -0.0558326429111435 | 5.66874952164334 | -0.871696211867396 | 0.388272064035961 | 0.690312073454197 | -6.029756267675 |
| ENSG00000166435.15 | XRRA1 | 0.294955296974809 | 4.06497504071849 | 0.87184321081121 | 0.388290808176591 | 0.690312073454197 | -5.92878262227886 |
| ENSG00000159884.11 | CCDC107 | -0.157426894096336 | 2.41367038965764 | -0.871639543834447 | 0.388302637441223 | 0.690312073454197 | -5.74875107113171 |
| lnc-IL17RD-2 | lnc-IL17RD-2 | 0.144627376049936 | 2.81282909142163 | 0.87156284717031 | 0.388344019065468 | 0.69032601644568 | -5.77669846043418 |
| ENSG00000120899.17 | PTK2B | -0.120048654321078 | 7.59988240349814 | -0.871312039380136 | 0.388479361630618 | 0.690506968950174 | -6.02382266013177 |
| lnc-PPP2CB-1 | lnc-PPP2CB-1 | 0.144349973400853 | -0.0692237873681541 | 0.871017336717704 | 0.38863842904677 | 0.690730056564172 | -5.31518661517584 |
| ENSG00000259164.1 | | 0.185578097918286 | 0.120727131222058 | 0.870953963948098 | 0.388672640219633 | 0.690731216857254 | -5.37569715274811 |
| ENSG00000276664.1 | | 0.12670506063547 | 1.94636518961372 | 0.870873972714621 | 0.388715825419545 | 0.690748323803031 | -5.6840759682463 |
| ENSG00000169255.14 | B3GALNT1 | -1.05568834077355 | -1.43418557706799 | -0.870861475426752 | 0.388833743842902 | 0.690898217327214 | -4.96003338190855 |
| lnc-MFNG-1 | lnc-MFNG-1 | -0.187845317110069 | 0.716377376989328 | -0.870478976220935 | 0.388929118171796 | 0.691008031016839 | -5.48771614370498 |
| ENSG00000162585.16 | FAAP20 | -0.131428428110535 | 3.18933822775887 | -0.870042378093613 | 0.389164961065323 | 0.69136737386566 | -5.84040799327953 |
| ENSG00000164465.18 | DCBLD1 | 0.958452430383408 | -1.19239469511485 | 0.869981897840028 | 0.389291566764686 | 0.691499168216715 | -4.96843068923588 |
| ENSG00000280649.2 | | -0.204296193758434 | 0.6480929833081 | -0.869732217518787 | 0.389332559284574 | 0.691499168216715 | -5.50363114328948 |
| ENSG00000138614.14 | INTS14 | 0.086736926607224 | 4.5814199017338 | 0.86967213708009 | 0.389365029578864 | 0.691499168216715 | -5.9798441838734 |
| ENSG00000131238.17 | PPT1 | -0.115432059633567 | 4.94299044573146 | -0.869648564953028 | 0.389377769530807 | 0.691499168216715 | -6.0040855433494 |
| ENSG00000134028.14 | ADAMDEC1 | -0.611894624173156 | -0.458137411635832 | -0.869721911142786 | 0.389407125239272 | 0.691499168216715 | -5.21766006676768 |
| ENSG00000141503.15 | MINK1 | -0.104484642579607 | 5.76892102629389 | -0.869345317014961 | 0.389541688393729 | 0.691578210280217 | -6.03316004818616 |
| ENSG00000115561.15 | CHMP3 | 0.0731584511755496 | 5.41450176106712 | 0.869289467869845 | 0.38957188205528 | 0.691578210280217 | -6.02450961063625 |
| ENSG00000167470.12 | MIDN | 0.191670496034737 | 2.8966507222988 | 0.869281603363461 | 0.38957613395308 | 0.691578210280217 | -5.81059434110532 |
| ENSG00000130779.20 | CLIP1 | 0.166088482183121 | 6.69890967365345 | 0.869263291056827 | 0.38958603450416 | 0.691578210280217 | -6.03656988185232 |
| ENSG00000232653.8 | GOLGA8N | -0.242264747442239 | 1.26370238894751 | -0.869153078492914 | 0.389645624289239 | 0.691624343286082 | -5.60311629262972 |
| ENSG00000172466.15 | ZNF24 | 0.0822214511024226 | 7.20836333890838 | 0.869030110621735 | 0.389712117396124 | 0.691682720735054 | -6.03154707541934 |
| ENSG00000136485.14 | DCAF7 | -0.0997681000098565 | 5.56781290452569 | -0.868783083478143 | 0.389845715366746 | 0.691832681863566 | -6.02991169354372 |
| ENSG00000279738.1 | | 0.10605114176129 | 6.09832554181211 | 0.8687098638673 | 0.389885319759828 | 0.691832681863566 | -6.03761876597035 |
| ENSG00000167987.10 | VPS37C | -0.111636391393983 | 2.22861425515605 | -0.868687448036602 | 0.389897444960276 | 0.691832681863566 | -5.72283995861575 |
| ENSG00000130758.7 | MAP3K10 | -0.143243386491112 | 2.96011435369848 | -0.868271629793402 | 0.390122412963175 | 0.692130736229924 | -5.81188232665466 |
| ENSG00000134070.4 | IRAK2 | -0.266584782437525 | 3.25665630995063 | -0.868251581005883 | 0.390174543462572 | 0.692130736229924 | -5.84448856129321 |
| lnc-RNF4-1 | lnc-RNF4-1 | 0.190627215996673 | 0.7493969003149 | 0.868116146883452 | 0.390206554067918 | 0.692130736229924 | -5.50983450604773 |
| ENSG00000165275.9 | TRMT10B | -0.14507384747807 | 2.91251651684867 | -0.868061160797848 | 0.390236313062622 | 0.692130736229924 | -5.79484435057369 |
| ENSG00000102043.15 | MTMR8 | -0.124479128008455 | 2.93437325279446 | -0.868060413804745 | 0.39023671735226 | 0.692130736229924 | -5.80148122155729 |
| ENSG00000176018.12 | LYSMD3 | 0.128573511862037 | 4.85730889302853 | 0.868004134043799 | 0.390267177995652 | 0.692130736229924 | -5.99904457076593 |
| ENSG00000206799.1 | SNORA32 | 0.298859747653618 | 2.99675102667723 | 0.86790722826634 | 0.390384585593938 | 0.692279307769451 | -5.81707607778986 |
| ENSG00000203761.5 | MSTO2P | -0.124416173689525 | 4.16139787171918 | -0.867713942595047 | 0.390424263861849 | 0.692290026108583 | -5.94384728763514 |
| lnc-TBCA-9 | lnc-TBCA-9 | 0.235550788818021 | 4.23002361897763 | 0.867484686458225 | 0.390583252882833 | 0.692512283063774 | -5.94543651202575 |
| ENSG00000113068.9 | PFDN1 | 0.098694258288885 | 3.6004723587392 | 0.867335907102848 | 0.39062896096693 | 0.692512847052741 | -5.88309563027352 |
| lnc-AIRE-3 | lnc-AIRE-3 | -0.177467088743842 | 0.37801089078634 | -0.867259859835161 | 0.390670146885739 | 0.692512847052741 | -5.39082558017688 |
| ENSG00000186470.13 | BTN3A2 | -0.204305626593275 | 5.31534569885877 | -0.867267402808434 | 0.390684505658866 | 0.692512847052741 | -6.02324272039562 |
| lnc-TCP1-2 | lnc-TCP1-2 | 0.118948432785617 | 1.97694606011197 | 0.867084737324624 | 0.390765000675996 | 0.69259588469078 | -5.68791320165372 |
| ENSG00000182871.14 | COL18A1 | -1.14911924236248 | 1.39346480251731 | -0.867136791541514 | 0.390929788347972 | 0.692828296329112 | -5.59258942967935 |
| lnc-CLTRN-3 | lnc-CLTRN-3 | -0.190520560855869 | 1.73654839411335 | -0.86665381081248 | 0.390998470461939 | 0.692881397706751 | -5.67084620158848 |
| ENSG00000089847.12 | ANKRD24 | 0.355707837511953 | -0.281393487760256 | 0.866601022376413 | 0.391027076507877 | 0.692881397706751 | -5.18238837234279 |
| ENSG00000279519.1 | | -0.156738624178729 | 1.5236007986618 | -0.866429577593507 | 0.39111999149002 | 0.692928407905531 | -5.64206271035568 |
| ENSG00000141401.11 | IMPA2 | -0.33691696922475 | 0.212622927587243 | -0.866368777516499 | 0.391152945597333 | 0.692928407905531 | -5.39038004350057 |
| ENSG00000202078.1 | | 0.260138166213017 | -0.205970303829985 | 0.866365721686711 | 0.391154601926559 | 0.692928407905531 | -5.25124958843709 |
| NALT1 | NALT1 | 0.216004350321013 | 0.63227418591899 | 0.866232212010665 | 0.391226971512217 | 0.692996966836141 | -5.49640020689839 |
| LINC01224 | LINC01224 | -0.185079956429549 | 5.66446918308364 | -0.866123380077061 | 0.391286118404254 | 0.693025606188993 | -6.03072339511915 |
| ENSG00000276058.1 | STMN1P1 | 0.1789469336824 | 0.601174472473231 | 0.866066301763747 | 0.391316915719621 | 0.693025606188993 | -5.42437547837088 |
| lnc-EZR-4 | lnc-EZR-4 | -0.329188507772095 | -0.167035216912929 | -0.865996090086059 | 0.391354983188717 | 0.693025606188993 | -5.31587869876288 |

| | | | | | | | |
|---|---|---|---|---|---|---|---|
| ENSG00000117133.10 | RPF1 | 0.0848535846447342 | 5.0896753663406 | 0.865953973944881 | 0.391377818896218 | 0.693025606188993 | -6.01549624317987 |
| ENSG00000128039.10 | SRD5A3 | -0.227250167083813 | 1.11545714142747 | -0.865803809787075 | 0.391459245917444 | 0.69311016419086 | -5.59894416348263 |
| lnc-MSMP-3 | lnc-MSMP-3 | -0.105724309423996 | 2.28280809776282 | -0.865707856585998 | 0.391511282440578 | 0.693142673875316 | -5.74276118661181 |
| ENSG00000279265.1 | | -0.208395290997979 | 0.463931532978327 | -0.865551014058826 | 0.391596349306238 | 0.693156671746349 | -5.43750551286942 |
| ENSG00000145833.15 | DDX46 | 0.068303006318061 | 6.37134785052558 | 0.865511558253122 | 0.391617750826572 | 0.693156671746349 | -6.04100714714768 |
| ENSG00000074966.10 | TXK | -0.176122906667156 | 1.68776533897507 | -0.865507010880693 | 0.391620217448297 | 0.693156671746349 | -5.66226615546126 |
| ENSG00000185246.17 | PRPF39 | 0.0767213692221285 | 5.91270747308358 | 0.865293396479837 | 0.391736098864069 | 0.693302160182212 | -6.0391117355901 |
| ENSG00000125844.15 | RRBP1 | -0.280133503458561 | 6.44825233507937 | -0.865365342521062 | 0.39180751212686 | 0.693326585991036 | -6.04007952318481 |
| lnc-ANKS4B-5 | lnc-ANKS4B-5 | 0.218973940849951 | 2.67738160100295 | 0.865143792043473 | 0.391817269020441 | 0.693326585991036 | -5.74709590367392 |
| ENSG00000108854.15 | SMURF2 | 0.0896728051891227 | 5.16752424676814 | 0.86507142758804 | 0.391856535245545 | 0.693336462216028 | -6.02034627659059 |
| lnc-LSM5-2 | lnc-LSM5-2 | -0.344766996160181 | 0.448090615045276 | -0.864784371442835 | 0.392012321310382 | 0.69355248491762 | -5.45173250377505 |
| lnc-TIMM10B-1 | lnc-TIMM10B-1 | -0.211149880889228 | 0.717058626244873 | -0.864716757787111 | 0.3920490210659 | 0.693557799793675 | -5.50638740239219 |
| lnc-GHRL-1 | lnc-GHRL-1 | 0.0905225435096333 | 1.67569598022603 | 0.864559064715177 | 0.392134623067229 | 0.693649617273357 | -5.65701670114924 |
| lnc-C1QBP-4 | lnc-C1QBP-4 | -0.226588338652284 | -0.257112941236663 | -0.86439608692665 | 0.392223106156789 | 0.69372327721799 | -5.29076736042407 |
| lnc-ARF6-3 | lnc-ARF6-3 | 0.136813229029555 | 4.61140411135818 | 0.864358209870856 | 0.392243671975075 | 0.69372327721799 | -5.98912703676825 |
| lnc-KLF6-8 | lnc-KLF6-8 | -0.262115911808549 | 5.59959341971251 | -0.864456575916939 | 0.392286560779425 | 0.693739520622296 | -6.03578621498206 |
| lnc-SYVN1-3 | lnc-SYVN1-3 | 0.378126092089294 | 0.191721259812068 | 0.864204003638759 | 0.392327407157035 | 0.69375214961454 | -5.33497163953178 |
| ENSG00000239857.6 | GET4 | 0.16893601372231 | 3.00961898234216 | 0.86404355634371 | 0.392414543200402 | 0.693846623373754 | -5.8256383119048 |
| ENSG00000070269.13 | TMEM260 | -0.0904069140829966 | 4.65727876319341 | -0.863979623522527 | 0.392449267352695 | 0.693848416931844 | -5.99146578054859 |
| lnc-TNFRSF8-2 | lnc-TNFRSF8-2 | -0.220012919313919 | 1.25907177022322 | -0.863843762581843 | 0.392523064589848 | 0.693919285274657 | -5.59221235402955 |
| lnc-MECR-3 | lnc-MECR-3 | -0.128350187794576 | -0.0410546245469133 | -0.863748262701838 | 0.392574943633981 | 0.69395139684114 | -5.30683399837356 |
| LINC-PINT | LINC-PINT | 0.225450780041871 | 9.91192548172573 | 0.8632980937527 | 0.39285033284552 | 0.694343244003425 | -5.97247435778219 |
| ENSG00000254870.5 | ATP6V1G2-DDX39 | 0.0881646813560657 | 5.58750817726477 | 0.863086801809809 | 0.392934391655391 | 0.694343244003425 | -6.03458733027733 |
| ENSG00000198646.13 | NCOA6 | -0.101119748709973 | 5.19600633577337 | -0.863062679986602 | 0.392947503727464 | 0.694343244003425 | -6.02073333449068 |
| ENSG00000144749.13 | LRIG1 | 0.256972252299684 | 4.41102488907798 | 0.863150615769639 | 0.392967983597147 | 0.694343244003425 | -5.96882873650461 |
| ENSG00000158373.8 | H2BC5 | -0.15849374475453 | 6.77820050171128 | -0.863002090044481 | 0.392980440247013 | 0.694343244003425 | -6.04035005151825 |
| ENSG00000260272.1 | | -0.145811011060335 | 2.93172793827016 | -0.862931862993797 | 0.393018617638938 | 0.694343244003425 | -5.81023997853816 |
| ENSG00000277157.1 | H4C4 | 1.07771072747632 | 1.40601700965252 | 0.863201481429871 | 0.39306407967582 | 0.694343244003425 | -5.5560164842177 |
| ENSG00000197601.12 | FAR1 | 0.135239419512533 | 5.48877727420955 | 0.86281835605863 | 0.39308032810913 | 0.694343244003425 | -6.03481956996436 |
| ENSG00000196664.4 | TLR7 | 0.25543620493384 | 5.26977840378311 | 0.862930816426525 | 0.393100219713934 | 0.694343244003425 | -6.02103659385188 |
| ENSG00000138073.13 | PREB | -0.138681744665214 | 3.95822362838252 | -0.86258965942312 | 0.393204682385618 | 0.694468163509797 | -5.93142146595632 |
| ENSG00000280069.1 | | 0.151507205240915 | 2.66708896399619 | 0.862404996885062 | 0.393305111037996 | 0.6944807904164 | -5.78135390757593 |
| ENSG00000108551.4 | RASD1 | 1.19294957939342 | -0.563028962021548 | 0.862582011718604 | 0.393373710925984 | 0.6944807904164 | -5.07308939502804 |
| ENSG00000249850.1 | KRT18P31 | 0.204833783277477 | 3.14891964457834 | 0.862255498656904 | 0.393386427379219 | 0.6944807904164 | -5.83079120301732 |
| ENSG00000138640.14 | FAM13A | -0.456752286416869 | -0.191746445825363 | -0.862283932897615 | 0.39338978143511 | 0.6944807904164 | -5.3127823014107 |
| ENSG00000173226.16 | IQCB1 | 0.0778185720429372 | 5.29236057133953 | 0.862248567825864 | 0.393390197511254 | 0.6944807904164 | -6.02417065543971 |
| ENSG00000182318.5 | ZSCAN22 | -0.149717789710629 | 1.92972910561162 | -0.862176904594994 | 0.393429181155483 | 0.6944807904164 | -5.68755725599641 |
| ENSG00000173933.20 | RBM4 | 0.0902722409776748 | 6.30143082260341 | 0.862142283744613 | 0.393448015208941 | 0.6944807904164 | -6.0438234112487 |
| ENSG00000155660.10 | PDIA4 | -0.146616705218372 | 4.99606805080164 | -0.862003768681898 | 0.393523374304089 | 0.694508713214774 | -6.01072435576173 |
| ENSG00000166405.14 | RIC3 | -0.179339685859455 | 6.02452174889214 | -0.861989168264234 | 0.393531318186072 | 0.694508713214774 | -6.04362933030565 |
| lnc-PIEZO1-1 | lnc-PIEZO1-1 | -0.179214631724002 | 0.555606584087424 | -0.861855293261074 | 0.393604162381967 | 0.694577715561388 | -5.48818010302774 |
| lnc-BRD3-4 | lnc-BRD3-4 | 0.174750428703806 | 0.612147769826101 | 0.861756439308055 | 0.393657956317099 | 0.694602884167278 | -5.46523584161626 |
| ENSG00000174529.7 | TMEM81 | 0.141057896207737 | 0.149280063655707 | 0.861576653707676 | 0.393755803117603 | 0.694602884167278 | -5.37176367347095 |
| ENSG00000210196.2 | MT-TP | 0.378829202064566 | 0.260662133636502 | 0.861466570952223 | 0.393815722257644 | 0.694602884167278 | -5.37527949713737 |
| ENSG00000166801.15 | FAM111A | -0.105500547086211 | 6.38968943329125 | -0.861452547463151 | 0.393823355792896 | 0.694602884167278 | -6.04426350248083 |
| ENSG00000107614.21 | TRDMT1 | -0.112271949489733 | 3.82537180642003 | -0.861443572137686 | 0.39382824146334 | 0.694602884167278 | -5.915112302572 |
| lnc-KIAA0355-5 | lnc-KIAA0355-5 | 0.141963182941718 | 3.29084378391234 | 0.861428201702281 | 0.393836608365959 | 0.694602884167278 | -5.85748462339575 |
| ENSG00000154611.14 | PSMA8 | 0.928178990203741 | -3.34979645771734 | 0.861395058963741 | 0.393854650008079 | 0.694602884167278 | -4.78402319597898 |
| lnc-PLLP-3 | lnc-PLLP-3 | -0.0976376029970991 | 2.92525134296031 | -0.861289411862712 | 0.393912163716638 | 0.694619356289735 | -5.81820236987326 |
| ENSG00000078124.11 | ACER3 | 0.135217798405911 | 3.15554123417496 | 0.861253923583452 | 0.393931484524611 | 0.694619356289735 | -5.84249026619225 |
| lnc-HAUS2-1 | lnc-HAUS2-1 | -0.178985308259077 | 1.03958023153024 | -0.861177321618558 | 0.39397319079359 | 0.694633389249996 | -5.54840379746826 |
| lnc-PACRG-3 | lnc-PACRG-3 | 0.226973745723069 | -0.247661864636924 | 0.86102838003045 | 0.394054290603933 | 0.694716870535396 | -5.27390410822546 |
| ENSG00000111011.17 | RSRC2 | 0.0967735326664902 | 7.71341732787226 | 0.860900252126249 | 0.394124065579742 | 0.694780373572099 | -6.03035834610665 |
| ENSG00000032219.18 | ARID4A | 0.0977195354224794 | 6.36170937373991 | 0.860631422440849 | 0.394270488089002 | 0.69494234392444 | -6.0451390212942 |
| ENSG00000108344.14 | PSMD3 | 0.0744937726975579 | 5.67446330993142 | 0.860607588236244 | 0.394283471425429 | 0.69494234392444 | -6.03914667597932 |
| lnc-SHOC2-1 | lnc-SHOC2-1 | 0.161060932150047 | 3.21838420717164 | 0.860340560330616 | 0.394428949309218 | 0.695139229697032 | -5.84798390247907 |
| ENSG00000158864.12 | NDUFS2 | -0.0885262281595862 | 4.97098049813278 | -0.859960793930175 | 0.394635905498977 | 0.695444421479919 | -6.01244226040328 |
| lnc-ZC3H15-2 | lnc-ZC3H15-2 | -0.149206634785596 | 2.57490124675996 | -0.859747283340914 | 0.394752289360164 | 0.69554603727487 | -5.77540312671687 |
| lnc-ERAP2-3 | lnc-ERAP2-3 | -0.190251410060531 | 0.648293388569175 | -0.859731024698322 | 0.394761152769034 | 0.69554603727487 | -5.48966334446592 |
| ENSG00000111752.10 | PHC1 | 0.133759244832544 | 4.47553024266616 | 0.859588718242389 | 0.394838736539556 | 0.695623188752347 | -5.98496631902404 |
| ENSG00000146909.7 | NOM1 | 0.067938801402005 | 4.76771373024462 | 0.859367966084061 | 0.394959106858949 | 0.695709360560867 | -6.00278487196393 |
| lnc-LRRC8C-4 | lnc-LRRC8C-4 | 0.165323346978256 | 2.44437934338309 | 0.859345160546096 | 0.394971543425654 | 0.695709360560867 | -5.74848715291062 |
| ENSG00000196642.18 | RABL6 | -0.10048388911337 | 4.86533226413175 | -0.859169648960909 | 0.395067263538049 | 0.695709360560867 | -6.01257890357889 |
| ENSG00000149657.19 | LSM14B | -0.0599597908791578 | 4.89288522890473 | -0.859135898999833 | 0.395085671680512 | 0.695709360560867 | -6.01194808925515 |
| lnc-CHD9-1 | lnc-CHD9-1 | 0.173810772308727 | 0.900523675873531 | 0.859131330830727 | 0.395088163324875 | 0.695709360560867 | -5.55729444195517 |
| lnc-SINHCAF-10 | lnc-SINHCAF-10 | 0.324871364579464 | 0.195337049007832 | 0.859094020144539 | 0.395108514288535 | 0.695709360560867 | -5.44236104914022 |
| lnc-INTS4-1 | lnc-INTS4-1 | -0.197987829887357 | 0.791387634298538 | -0.85899414961204 | 0.395162991498057 | 0.695709360560867 | -5.51667625500458 |
| lnc-ANK2-8 | lnc-ANK2-8 | 0.112241799632751 | 1.18495454019206 | 0.858985121222144 | 0.39516791652069 | 0.695709360560867 | -5.58610733190613 |

| | | | | | | | |
|---|---|---|---|---|---|---|---|
| ENSG00000115541.10 | HSPE1 | 0.135171142749049 | 3.4819357405317 | 0.858941248589956 | 0.395191849763283 | 0.695709360560867 | -5.8859367148899 |
| ENSG00000134108.12 | ARL8B | 0.0970319181827728 | 5.34572440369786 | 0.858814780014346 | 0.39526084552956 | 0.695727286378875 | -6.03272164302474 |
| ENSG00000146830.10 | GIGYF1 | -0.101373103794091 | 6.31889565165379 | -0.858752700810503 | 0.395294716007887 | 0.695727286378875 | -6.04653670644658 |
| ENSG00000243789.10 | JMJD7 | -0.144668534927859 | 2.75967728320296 | -0.858736719789975 | 0.395303435563375 | 0.695727286378875 | -5.79289208014086 |
| ENSG00000176896.8 | TCEANC | 0.12191186626416 | 3.24070483318187 | 0.858534612429319 | 0.395413719657192 | 0.695817016747171 | -5.85201614478617 |
| ENSG00000214717.11 | ZBED1 | -0.134781980595976 | 5.44677146663263 | -0.858519384144771 | 0.395422030068098 | 0.695817016747171 | -6.03445393067475 |
| ENSG00000228784.8 | LINC00954 | -0.302761538309642 | 2.4678346912073 | -0.858479935662602 | 0.395488786794526 | 0.695847340055357 | -5.74598439543066 |
| lnc-RPL30-3 | lnc-RPL30-3 | 0.128775906954093 | 8.39221178402498 | 0.858351064471055 | 0.395513893109451 | 0.695847340055357 | -6.01976569286818 |
| ENSG00000142102.15 | PGGHG | -0.215897342887732 | 8.06443474640455 | -0.858361974998138 | 0.39554239523161 | 0.695847340055357 | -6.02470108422346 |
| MBNL1-AS1 | MBNL1-AS1 | -0.173582737909628 | 0.23926987047845 | -0.858227629202587 | 0.395581268286071 | 0.695847340055357 | -5.401191136852 |
| lnc-MYLK4-32 | lnc-MYLK4-32 | 0.286257892676899 | -0.469686387438348 | 0.858129673768833 | 0.3956347408112 | 0.695847340055357 | -5.23866654000788 |
| ENSG00000072571.19 | HMMR | -0.249859079790916 | 0.0537482138677561 | -0.858116186444825 | 0.395642103710239 | 0.695847340055357 | -5.35324470667616 |
| ENSG00000125772.12 | GPCPD1 | 0.119909521642652 | 7.72371915567292 | 0.857730018903192 | 0.395852953774836 | 0.696140306270209 | -6.03254196468873 |
| ENSG00000135437.9 | RDH5 | 0.172866422113923 | 2.41176279927109 | 0.857687232924961 | 0.39587631953033 | 0.696140306270209 | -5.74749169455831 |
| ENSG00000083223.17 | TUT7 | 0.0941326766595578 | 7.19541491128298 | 0.857519165126818 | 0.395968111012246 | 0.696242237225786 | -6.04186210064106 |
| lnc-TIAM1-2 | lnc-TIAM1-2 | 0.14805804153322 | 2.86014476696199 | 0.857423430297035 | 0.396020403254477 | 0.696274704039199 | -5.79485745955563 |
| ENSG00000133739.15 | LRRCC1 | 0.282317216775374 | 2.04121146523656 | 0.85732418612816 | 0.396093174539825 | 0.696343168052034 | -5.72019517422615 |
| ENSG00000130826.17 | DKC1 | 0.084223589956868 | 5.30059133573454 | 0.857124987096307 | 0.396183446502995 | 0.696413542223059 | -6.03284756913844 |
| ENSG00000123739.10 | PLA2G12A | -0.1546617738448 | 2.07832457550312 | -0.857093092303885 | 0.396200873508819 | 0.696413542223059 | -5.71187328506483 |
| ENSG00000108405.3 | P2RX1 | 0.446079585325899 | 5.30350457442032 | 0.857215368645803 | 0.396315917357563 | 0.696556274184162 | -6.02477717488198 |
| ENSG00000211454.13 | AKR7L | 0.182265485695293 | -0.0516024318546147 | 0.856815440490183 | 0.396352600020028 | 0.696561267606919 | -5.31926248914184 |
| lnc-BTK-1 | lnc-BTK-1 | -0.148350768861322 | 4.65949660519984 | -0.856577929936611 | 0.396482419632423 | 0.696729927712299 | -5.98963633444337 |
| ENSG00000263843.1 | MIF4GD-DT | -0.269275647188088 | 0.58667971481792 | -0.856446537830703 | 0.396554247952902 | 0.696796660885656 | -5.48539257394211 |
| ENSG00000140612.13 | SEC11A | 0.10712649824552 | 5.19650270069115 | 0.856209924051257 | 0.396683618474891 | 0.696964483061774 | -6.02527966678544 |
| lnc-CCDC146-2 | lnc-CCDC146-2 | 0.245078653134098 | -0.540562791188558 | 0.855959117592361 | 0.396820777754787 | 0.697120636332801 | -5.14292922219211 |
| lnc-MED16-2 | lnc-MED16-2 | 0.117527049076341 | 2.57794164377769 | 0.855805614310268 | 0.39690473917212 | 0.697120636332801 | -5.76977222262637 |
| ENSG00000266038.1 | MIR4659A | 0.333456633443501 | 0.283239679479156 | 0.855804505788402 | 0.39690534553872 | 0.697120636332801 | -5.40149208103202 |
| ENSG00000163961.4 | RNF168 | 0.0695739523964782 | 5.18997119407335 | 0.85576387954406 | 0.396927568682037 | 0.697120636332801 | -6.02667009131145 |
| ENSG00000155530.3 | LRGUK | -0.281399571697686 | 0.158452464295595 | -0.855737793922054 | 0.396941838304447 | 0.697120636332801 | -5.39184528913718 |
| ENSG00000156482.10 | RPL30 | 0.125471593750099 | 8.4549238529773 | 0.855589368797067 | 0.397023037426804 | 0.697203752184618 | -6.02049832604935 |
| ENSG00000124444.15 | ZNF576 | 0.125805895064496 | 1.6161860167741 | 0.855496272188335 | 0.397073973202441 | 0.697233713566443 | -5.65192143830746 |
| ENSG00000158517.13 | NCF1 | -0.174409094245301 | 5.44836071257752 | -0.855195436190756 | 0.397238596953981 | 0.69739407542983 | -6.03826782301765 |
| ENSG00000147485.12 | PXDNL | -0.442874823855702 | 0.677318541597806 | -0.85536619329689 | 0.397246470827771 | 0.69739407542983 | -5.57251602593393 |
| lnc-IFRD2-3 | lnc-IFRD2-3 | -0.141406008351258 | 0.681825122695051 | -0.85514363990497 | 0.397266945266227 | 0.69739407542983 | -5.51503012098726 |
| ENSG00000048740.18 | CELF2 | -0.0819673495405908 | 7.72528687255736 | -0.854687309079477 | 0.397516751531191 | 0.697653392847009 | -6.03534887853203 |
| lnc-ZNF512B-1 | lnc-ZNF512B-1 | -0.163761763344985 | 1.60369468607828 | -0.854673850858563 | 0.397524120367634 | 0.697653392847009 | -5.66648399236963 |
| ENSG00000240729.1 | RPL21P60 | 0.217274177452531 | 0.116699174734481 | 0.854626143142612 | 0.397550242661545 | 0.697653392847009 | -5.36516805232288 |
| lnc-PRELID1-2 | lnc-PRELID1-2 | 0.217274177452531 | 0.116699174734481 | 0.854626143142612 | 0.397550242661545 | 0.697653392847009 | -5.36516805232288 |
| lnc-EXTL3-2 | lnc-EXTL3-2 | 0.430049835841427 | 0.494123286229113 | 0.854508478591276 | 0.39768278142255 | 0.697826486787753 | -5.5017510727928 |
| ENSG00000136710.9 | CCDC115 | -0.0769408787878243 | 4.98786987476033 | -0.853999129095615 | 0.397893662935615 | 0.698051328485353 | -6.02039276267353 |
| ENSG00000110077.14 | MS4A6A | 1.43758030860475 | -0.880487196567748 | 0.854245722539987 | 0.397909489172589 | 0.698051328485353 | -4.9253963114741 |
| ENSG00000099203.6 | TMED1 | 0.119651435242761 | 2.58900124577069 | 0.853964458137592 | 0.397912657878767 | 0.698051328485353 | -5.78661722923567 |
| lnc-ZNF726-5 | lnc-ZNF726-5 | -0.509222236500995 | 1.00901699294965 | -0.85409629762553 | 0.39795770273397 | 0.698070853534456 | -5.52644835727298 |
| lnc-RTN4-1 | lnc-RTN4-1 | 0.268221969492934 | 0.0274921254493824 | 0.853448570914267 | 0.398195360376555 | 0.698369392151701 | -5.33963145253163 |
| ENSG00000274602.4 | PI4KAP1 | 0.333577834030797 | 2.25744075425763 | 0.853386349923361 | 0.398295189953538 | 0.698369392151701 | -5.71860889137976 |
| lnc-ILKAP-2 | lnc-ILKAP-2 | 0.161490085613157 | 2.29093172373574 | 0.853053017409155 | 0.398412205714912 | 0.698369392151701 | -5.73379430082919 |
| ENSG00000229127.1 | KANSL1L-AS1 | -0.153330445956301 | 2.83513377859795 | -0.853044565703014 | 0.398416839805926 | 0.698369392151701 | -5.8107229774486 |
| ENSG00000121067.17 | SPOP | 0.0672567586890773 | 5.20855498259218 | 0.853040422943035 | 0.398419111303568 | 0.698369392151701 | -6.02948225960768 |
| lnc-PREX1-1 | lnc-PREX1-1 | 0.19153919696129 | 1.13937347464041 | 0.852971015092849 | 0.398457169203587 | 0.698369392151701 | -5.57211788262213 |
| ENSG00000100227.17 | POLDIP3 | 0.0631479793029177 | 5.67721258680006 | 0.8529175809019 | 0.398486469928316 | 0.698369392151701 | -6.04570080988455 |
| ENSG00000069275.12 | NUCKS1 | -0.120540725754147 | 7.37577545612987 | -0.85290715998743 | 0.398492184410156 | 0.698369392151701 | -6.04177363329255 |
| ENSG00000091129.19 | NRCAM | -1.17591065746634 | -1.30164830112402 | -0.853196430407449 | 0.398495261403919 | 0.698369392151701 | -5.0794973076549 |
| ENSG00000136167.13 | LCP1 | 0.192500719579536 | 8.35816986472256 | 0.852907257344713 | 0.398500096842653 | 0.698369392151701 | -6.02383205506242 |
| lnc-SYT2-2 | lnc-SYT2-2 | -0.150071712382218 | 0.930970125162085 | -0.852890868988727 | 0.398501117952763 | 0.698369392151701 | -5.5721337117183 |
| ENSG00000125898.12 | FAM110A | -0.239004345676386 | 0.283017515662051 | -0.852776188671131 | 0.398564009068488 | 0.698420142913049 | -5.44716330965982 |
| lnc-EXOSC6-1 | lnc-EXOSC6-1 | 0.153213384789406 | 0.447176993470168 | 0.852711507924569 | 0.398599482962183 | 0.698422844795319 | -5.456785020882 |
| ENSG00000122034.14 | GTF3A | 0.0952602626115677 | 6.47545739032403 | 0.852593398989197 | 0.398664264410043 | 0.698476894309833 | -6.05169930202139 |
| lnc-EIF3E-5 | lnc-EIF3E-5 | 0.186748351575361 | 2.78029126375398 | 0.852509701696501 | 0.398710175426103 | 0.698497875631583 | -5.80396962414507 |
| ENSG00000100889.11 | PCK2 | -0.152064656332938 | 3.96197574633461 | -0.852319427312619 | 0.398814560127891 | 0.698621284349991 | -5.9361415148339 |
| ENSG00000135678.11 | CPM | -0.956922710701567 | -0.182140865584695 | -0.852501920090352 | 0.39886916912409 | 0.698625363368354 | -5.19579100086615 |
| ENSG00000059573.8 | ALDH18A1 | -0.127376561072823 | 5.24349575476881 | -0.852191460538544 | 0.398884772399791 | 0.698625363368354 | -6.03305863044122 |
| ENSG00000147394.18 | ZNF185 | 0.392523235115742 | 0.1958071189931 | 0.85211203762258 | 0.39892835370706 | 0.698640984539856 | -5.31687210635329 |
| ENSG00000183765.21 | CHEK2 | -0.236419935837613 | 2.94453865103949 | -0.852056006731591 | 0.398961576654592 | 0.698640984539856 | -5.81151828071113 |
| lnc-RNMT-6 | lnc-RNMT-6 | -0.201987503146318 | 0.854257423683915 | -0.851757085591855 | 0.399123160890399 | 0.698717439639169 | -5.54603586689477 |
| ENSG00000123983.13 | ACSL3 | -0.149631255222778 | 5.14242965802069 | -0.851749662434818 | 0.399127235550687 | 0.698717439639169 | -6.03386430351334 |
| lnc-NDRG1-7 | lnc-NDRG1-7 | -0.119892918490265 | 3.33219835362858 | -0.851749159722912 | 0.399127511496243 | 0.698717439639169 | -5.85435297190703 |
| ENSG00000280968.1 | | 0.162590070908327 | -0.237862304172505 | 0.851724546880145 | 0.399141021973344 | 0.698717439639169 | -5.29313989142905 |
| ENSG00000100056.11 | ESS2 | 0.112458225088649 | 3.80038247694943 | 0.851500249947241 | 0.399264156138071 | 0.698817750188976 | -5.92532093344358 |

| | | | | | | | |
|---|---|---|---|---|---|---|---|
| ENSG00000129534.13 | MIS18BP1 | 0.124689606705485 | 6.19021583829359 | 0.851481111151356 | 0.399274664019054 | 0.698817750188976 | -6.0523871243319 |
| FOXP1-IT1 | FOXP1-IT1 | -0.543224477395209 | 3.13532133306736 | -0.85176400625047 | 0.399300177815311 | 0.698817750188976 | -5.84209282792972 |
| ENSG00000204822.6 | MRPL53 | 0.117478499933519 | 2.16969904915027 | 0.851369012201556 | 0.39933621379799 | 0.698821398452987 | -5.7279210631542 |
| lnc-METTL2B-4 | lnc-METTL2B-4 | 0.287660009164291 | -0.334962297192323 | 0.851097464979535 | 0.399485335788257 | 0.69900953624792 | -5.23288878500437 |
| ENSG00000217094.2 | PPIAP31 | 0.452860801517923 | -0.809925767319663 | 0.851049563559374 | 0.399511644775812 | 0.69900953624792 | -5.07076500033173 |
| lnc-ILK-9 | lnc-ILK-9 | -0.315239217092478 | -0.58554740663862 | -0.850970592192023 | 0.399555020725812 | 0.699026008635732 | -5.2113606366749 |
| lnc-TMEM161B-8 | lnc-TMEM161B-8 | -0.192302389810623 | 4.01724177718441 | -0.850695770816666 | 0.399705992458645 | 0.69912229367661 | -5.93305397035149 |
| lnc-APBB2-3 | lnc-APBB2-3 | 0.186346402702235 | 0.305273234638349 | 0.850539308693577 | 0.399791960005057 | 0.69912229367661 | -5.40553070663169 |
| ENSG00000048828.16 | FAM120A | -0.0747319294028754 | 7.04996222853653 | -0.850499577503091 | 0.399813791996856 | 0.69912229367661 | -6.0485683216999 |
| ENSG00000135439.11 | AGAP2 | -0.150082004936158 | 3.89290514748222 | -0.850497086118278 | 0.399815161018902 | 0.69912229367661 | -5.92653251374834 |
| lnc-FBXO30-3 | lnc-FBXO30-3 | 0.154964610715484 | 0.11589796365632 | 0.850476397296024 | 0.399826529690095 | 0.69912229367661 | -5.37157761509402 |
| ENSG00000065413.19 | ANKRD44 | -0.0908639795184172 | 8.71633368588648 | -0.85037705165164 | 0.399881123710498 | 0.69912229367661 | -6.01759341427159 |
| ENSG00000280496.1 | SNORA17B | 0.252344779970799 | 1.51404952344726 | 0.850372435221072 | 0.399883660718725 | 0.69912229367661 | -5.64166150913772 |
| ENSG00000169857.7 | AVEN | 0.0813314868063872 | 2.31048037100202 | 0.850337716647708 | 0.399902740998939 | 0.69912229367661 | -5.74837093261564 |
| ENSG00000109466.13 | KLHL2 | 0.124574444654144 | 4.89606678907144 | 0.850314045686421 | 0.399915750169966 | 0.69912229367661 | -6.01854477651516 |
| LINC01507 | LINC01507 | -0.940477344885965 | -0.863992484965825 | -0.850376422742181 | 0.40000476326494 | 0.699181518958076 | -5.0741121720501 |
| lnc-MYO5C-2 | lnc-MYO5C-2 | 0.11114907842012 | 2.46242507032938 | 0.850128801725804 | 0.400017566304732 | 0.699181518958076 | -5.75550892298252 |
| ENSG00000147684.8 | NDUFB9 | 0.156921352268371 | 4.32669001882497 | 0.849965910751667 | 0.400107109842342 | 0.699278648372669 | -5.96990391692318 |
| ENSG00000237004.4 | ZNRF2P1 | -0.309990126802812 | -0.0992387870162882 | -0.849756681965551 | 0.400222144256317 | 0.699420308645591 | -5.3283833271496 |
| ENSG00000031691.6 | CENPQ | 0.132748094460907 | 1.66343324133839 | 0.849659208383436 | 0.400275742459999 | 0.699454589273636 | -5.65800802079714 |
| ENSG00000152433.14 | ZNF547 | -0.0959408265070516 | 1.56129747539779 | -0.849466000394855 | 0.400381995744065 | 0.699477422846184 | -5.64858605753026 |
| ENSG00000116747.12 | RO60 | 0.0956340583313757 | 6.45771312085751 | 0.849440926884873 | 0.40039578601956 | 0.699477422846184 | -6.0542337993912 |
| lnc-BACE1-2 | lnc-BACE1-2 | 0.204954499110227 | 3.81080891292732 | 0.849359587821621 | 0.400440524036448 | 0.699477422846184 | -5.91557456812535 |
| ENSG00000107651.12 | SEC23IP | -0.0563235035763912 | 5.3088066546071 | -0.849249747178748 | 0.400500943395813 | 0.699477422846184 | -6.03724149075776 |
| ENSG00000119203.13 | CPSF3 | 0.0903981168216008 | 4.71774507481104 | 0.84920654456868 | 0.400524709141255 | 0.699477422846184 | -6.0077359129459 |
| ENSG00000229117.8 | RPL41 | 0.135222234803935 | 7.89763472078578 | 0.849159800484866 | 0.400550424038966 | 0.699477422846184 | -6.036631511681 |
| lnc-SEPT7-3 | lnc-SEPT7-3 | -0.136560521557072 | 2.05468796652972 | -0.849137780537995 | 0.400562538028034 | 0.699477422846184 | -5.71226599072492 |
| lnc-WNT5B-7 | lnc-WNT5B-7 | 0.259414768669931 | 0.506979928641549 | 0.849132609239901 | 0.400565382983875 | 0.699477422846184 | -5.4782936300757 |
| ENSG00000171824.13 | EXOSC10 | 0.0683086817410707 | 5.92918652654813 | 0.849071987360553 | 0.400598734652117 | 0.699477422846184 | -6.0529790671618 |
| ENSG00000197150.12 | ABCB8 | -0.0896889804333911 | 4.21158747399479 | -0.849017628774524 | 0.400628641982882 | 0.699477422846184 | -5.96519812883975 |
| lnc-CDC27-2 | lnc-CDC27-2 | -0.205012244337614 | 0.64021527154756 | -0.848692873137467 | 0.400807346915693 | 0.699658417376657 | -5.50213206396644 |
| ENSG00000225450.1 | | -0.17001670190046 | 0.427766574574176 | -0.848661947047898 | 0.400824367357574 | 0.699658417376657 | -5.46182039076408 |
| ENSG00000259371.2 | | -0.13813671817053 | 2.64561102677075 | -0.848564863621794 | 0.400877800985044 | 0.699658417376657 | -5.78866894719513 |
| lnc-CRTAP-1 | lnc-CRTAP-1 | -0.132002905534788 | 1.27306205188347 | -0.848539145750445 | 0.400891956554934 | 0.699658417376657 | -5.61675992909058 |
| ENSG00000085382.11 | HACE1 | -0.102475630112669 | 4.10028706496607 | -0.848497911992583 | 0.400914652991231 | 0.699658417376657 | -5.9576330151499 |
| lnc-POU5F1-8 | lnc-POU5F1-8 | 0.562337270843212 | 1.11419884142854 | 0.848700769704115 | 0.400936259678262 | 0.699658417376657 | -5.53350030364506 |
| ENSG00000244115.1 | DNAJC25-GNG10 | 0.148119212085343 | 1.11112054746559 | 0.848341934264509 | 0.401000515567568 | 0.699711225154904 | -5.59688750032374 |
| ENSG00000187147.17 | RNF220 | -0.0839382004239013 | 6.06005458586846 | -0.847745694249477 | 0.401328839011098 | 0.700224759969943 | -6.05514634301475 |
| lnc-MAP3K2-4 | lnc-MAP3K2-4 | 0.176559137619199 | 1.99268763099811 | 0.847500694683432 | 0.401463798049485 | 0.700400860760514 | -5.70295957375313 |
| ENSG00000042493.15 | CAPG | -0.250410372881402 | 5.11575153364815 | -0.84717700611525 | 0.40171982655968 | 0.700788133746748 | -6.02972139549311 |
| lnc-ITGB3BP-12 | lnc-ITGB3BP-12 | 0.13179235216983 | 1.31284115958648 | 0.84677932885857 | 0.401861329230679 | 0.700975571148734 | -5.63087141351449 |
| ENSG00000115419.12 | GLS | -0.079032681722726 | 7.37755785968702 | -0.846547973161411 | 0.40198887680108 | 0.701124793657938 | -6.0469691313988 |
| ENSG00000168268.10 | NT5DC2 | 0.332409254677585 | 0.341907528835083 | 0.846500588353849 | 0.402015003388767 | 0.701124793657938 | -5.41444564121403 |
| lnc-TGM1-1 | lnc-TGM1-1 | -0.119469560681319 | 1.41202270147322 | -0.846403356344526 | 0.402068617558845 | 0.701158888012684 | -5.63559908595033 |
| lnc-SORT1-4 | lnc-SORT1-4 | 0.125487343014596 | 1.59881351234671 | 0.846237944393217 | 0.402159836662442 | 0.701179186101174 | -5.67593573641024 |
| ENSG00000166997.7 | CNPY4 | -0.145837720022052 | 2.19216138121806 | -0.84618508764601 | 0.402188988082941 | 0.701179186101174 | -5.74250596628713 |
| ENSG00000186716.20 | BCR | 0.280519192355015 | 4.17387880020215 | 0.84631881200906 | 0.402210124051104 | 0.701179186101174 | -5.96994591262043 |
| lnc-CAB39L-4 | lnc-CAB39L-4 | -0.209327718034624 | 1.20578492681937 | -0.84607556128935 | 0.402249397961547 | 0.701179186101174 | -5.60259893165529 |
| ENSG00000188827.10 | SLX4 | -0.109185599937307 | 4.17485817030402 | -0.846073405842448 | 0.402250586866961 | 0.701179186101174 | -5.96473392989893 |
| lnc-FCHSD1-1 | lnc-FCHSD1-1 | 0.0995407499244054 | 2.83948010086427 | 0.845933949966394 | 0.402327512836962 | 0.701253890822524 | -5.81805809240774 |
| lnc-MX1-4 | lnc-MX1-4 | 0.226661709847279 | 2.88439904378929 | 0.845763960054786 | 0.402421294197145 | 0.701309613340395 | -5.81973326666989 |
| ENSG00000221184.1 | | 0.162449010863121 | 0.754979125888392 | 0.84575248227768 | 0.402427626835904 | 0.701309613340395 | -5.52727316456179 |
| ENSG00000187608.9 | ISG15 | 0.320979809886403 | 1.96147181729686 | 0.845508443944645 | 0.402612883429496 | 0.701573059569024 | -5.71564866417643 |
| ENSG00000091164.12 | TXNL1 | 0.0812904020984595 | 4.80392290773194 | 0.845152544263235 | 0.402758717028743 | 0.701649111842979 | -6.01939610358658 |
| lnc-NR1D1-2 | lnc-NR1D1-2 | 0.150290461284936 | 2.08766009753767 | 0.845142686175322 | 0.402764158862797 | 0.701649111842979 | -5.72958161312343 |
| ENSG00000147601.13 | TERF1 | 0.0726185796596877 | 4.861215404425 | 0.845132012520857 | 0.402770050955072 | 0.701649111842979 | -6.02204390657391 |
| lnc-RTN4RL1-1 | lnc-RTN4RL1-1 | 0.159556711719845 | -0.20229794305622 | 0.845064896428191 | 0.402807101741886 | 0.701649111842979 | -5.29938532017002 |
| lnc-GDI1-1 | lnc-GDI1-1 | -0.125395010780175 | 0.972149417380057 | -0.84502890467149 | 0.402826971512825 | 0.701649111842979 | -5.57017192861182 |
| ENSG00000278986.1 | | 0.0970404517024703 | 1.67776268962162 | 0.844845897021921 | 0.402928012948618 | 0.701752689260146 | -5.67771710397962 |
| lnc-EXOC2-3 | lnc-EXOC2-3 | -0.299848072489698 | 1.43645371182901 | -0.844824495920259 | 0.402954624416541 | 0.701752689260146 | -5.6462287053626 |
| lnc-ENO1-5 | lnc-ENO1-5 | -0.168380927264185 | 1.51243682959819 | -0.844671058131447 | 0.403024558972249 | 0.701760816365651 | -5.63517689726951 |
| lnc-FAM98B-1 | lnc-FAM98B-1 | 0.190694102644641 | 0.716271802018322 | 0.844665769666478 | 0.403027479485904 | 0.701760816365651 | -5.50776298242072 |
| lnc-TBC1D19-2 | lnc-TBC1D19-2 | -0.233756691802677 | 2.33707767317228 | -0.844544154359976 | 0.403094644216579 | 0.701818394680244 | -5.76527693320787 |
| LINC01962 | LINC01962 | -0.43805352676558 | -1.29228681919516 | -0.844390665486264 | 0.403179421739699 | 0.701881894948572 | -4.96941566609063 |
| ENSG00000197296.5 | FITM2 | -0.140850034838502 | 1.9321225580681 | -0.844347795891213 | 0.403203102162375 | 0.701881894948572 | -5.69970239132494 |
| ENSG00000152683.14 | SLC30A6 | 0.0994958335134601 | 4.78959876047845 | 0.844292919296608 | 0.403233416303107 | 0.701881894948572 | -6.01480739701079 |
| ENSG00000115042.9 | FAHD2A | -0.0826837828025524 | 3.64049321513786 | -0.844142962883743 | 0.403316260282895 | 0.701966733079894 | -5.90521186824817 |
| ENSG00000066933.15 | MYO9A | -0.0583483367594377 | 6.5262927342227 | -0.843969868247473 | 0.403411900186838 | 0.702073826122066 | -6.05858050668462 |

| | | | | | | | |
|---|---|---|---|---|---|---|---|
| ENSG00000149196.15 | HIKESHI | 0.156342227124905 | 3.59748240643876 | 0.843569702098489 | 0.403633057660229 | 0.70239932582182 | -5.89072834233027 |
| ENSG00000158201.9 | ABHD3 | -0.0927818701859442 | 4.41533295662758 | -0.843399557790425 | 0.403727113089608 | 0.70241403896438 | -5.99426253079079 |
| ENSG00000105676.13 | ARMC6 | -0.15985007108193 | 3.20551083655158 | -0.843366610271782 | 0.40374532798169 | 0.70241403896438 | -5.86858258721711 |
| ENSG00000104671.7 | DCTN6 | 0.114649363367743 | 3.84204288617103 | 0.843337777463884 | 0.403761268491919 | 0.70241403896438 | -5.93463150638438 |
| ENSG00000166188.2 | ZNF319 | -0.114382825234919 | 3.16178252169287 | -0.843271095975712 | 0.403798135523556 | 0.70241403896438 | -5.86172729655601 |
| ENSG00000158716.8 | DUSP23 | -0.188901441406932 | 1.1775913141895 | -0.843197037934187 | 0.403839083368912 | 0.70241403896438 | -5.60895847751163 |
| lnc-APOL2-2 | lnc-APOL2-2 | 0.122405928310645 | 5.26837947739137 | 0.843089525146388 | 0.403898533445994 | 0.70241403896438 | -6.03744651535623 |
| ENSG00000130699.18 | TAF4 | -0.125887562119635 | 2.95412630591048 | -0.843054487696951 | 0.403917908862869 | 0.70241403896438 | -5.8291939211028 |
| lnc-EIF1AD-1 | lnc-EIF1AD-1 | 0.154262382932407 | 0.772716709274259 | 0.843028749476244 | 0.40393214225101 | 0.70241403896438 | -5.52398778829841 |
| ENSG00000175182.14 | FAM131A | -0.115944017871177 | 2.32822715844937 | -0.84299890659097 | 0.403948645932146 | 0.70241403896438 | -5.76418937556697 |
| ENSG00000161513.11 | FDXR | 0.228881564892571 | 0.718332129361694 | 0.84285948830067 | 0.404025752426422 | 0.702450973623433 | -5.49353105109552 |
| ENSG00000168038.10 | ULK4 | 0.123100517384498 | 4.58384277887937 | 0.842837087928269 | 0.404038141997173 | 0.702450973623433 | -5.99909531129617 |
| lnc-GATAD1-4 | lnc-GATAD1-4 | -0.198418711817636 | 0.651404422573579 | -0.842509125242686 | 0.404219564019079 | 0.70270703430783 | -5.52309432358386 |
| lnc-ATAD2-2 | lnc-ATAD2-2 | -0.227112511578512 | 0.775225462344981 | -0.842348152563358 | 0.404308629167688 | 0.702802509429822 | -5.50595705550217 |
| ENSG00000100109.16 | TFIP11 | 0.124824231978105 | 5.58293764276839 | 0.841970569703936 | 0.404517591044242 | 0.703106365180175 | -6.05069159404337 |
| INE1 | INE1 | 0.174732087287552 | 1.32379023704367 | 0.841759976188382 | 0.404634166749248 | 0.703249603495717 | -5.62043300520033 |
| ENSG00000119402.16 | FBXW2 | 0.0663682817011741 | 6.19018817373575 | 0.841506898752808 | 0.404774287289832 | 0.703364736651313 | -6.06090832268408 |
| lnc-STON1-1 | lnc-STON1-1 | -0.195734337936521 | 0.942726749691726 | -0.841494548870718 | 0.404781125776795 | 0.703364736651313 | -5.57011836944653 |
| ENSG00000125651.13 | GTF2F1 | -0.0931571809254734 | 5.1350425778284 | -0.841455175980674 | 0.404802928162632 | 0.703364736651313 | -6.03855673047467 |
| lnc-C19orf44-5 | lnc-C19orf44-5 | 0.129999102609359 | 2.64625491101063 | 0.841362206413547 | 0.404854412115079 | 0.703371076985848 | -5.78653031241872 |
| lnc-COPZ2-1 | lnc-COPZ2-1 | -0.214881857460993 | -0.348662805746925 | -0.841272523730147 | 0.404904079725055 | 0.703371076985848 | -5.27177178808061 |
| ENSG00000101849.16 | TBL1X | -0.13957269228866 | 4.79302161453148 | -0.841263469221606 | 0.404909094456848 | 0.703371076985848 | -6.02008390143052 |
| ENSG00000111669.14 | TPI1 | -0.183223409357797 | 6.08396614226606 | -0.841122149422026 | 0.404992995513019 | 0.703451548857606 | -6.06070752243834 |
| ENSG00000254469.7 | XNDC1N | -0.109595058615733 | 2.78811839887442 | -0.841056427916765 | 0.40502377231266 | 0.703451548857606 | -5.80405049803234 |
| ENSG00000138002.14 | IFT172 | 0.270953559066161 | 3.56709446186056 | 0.840926418656879 | 0.405150875881514 | 0.703520468841893 | -5.88138765244099 |
| lnc-PHF2-1 | lnc-PHF2-1 | -0.150501813764892 | 0.640998161834843 | -0.840812405021327 | 0.405158959670533 | 0.703520468841893 | -5.54216728422878 |
| ENSG00000106348.17 | IMPDH1 | -0.157729850234164 | 3.7513032781789 | -0.840799710408414 | 0.405165993181353 | 0.703520468841893 | -5.93650068687078 |
| lnc-RETSAT-1 | lnc-RETSAT-1 | -0.249856568417242 | 2.2529562129669 | -0.840690174907135 | 0.405227879882436 | 0.703568574577831 | -5.75201530315093 |
| lnc-ZNF91-4 | lnc-ZNF91-4 | -0.153282631479088 | 4.68344176562908 | -0.840523634698071 | 0.405318972940041 | 0.703622985931183 | -6.00580590607708 |
| ENSG00000137133.10 | HINT2 | -0.128788340279678 | 2.08767993236011 | -0.840506260802746 | 0.40532860140162 | 0.703622985931183 | -5.73154363095159 |
| lnc-PDE4D-7 | lnc-PDE4D-7 | 0.368737659181568 | -0.969546068582916 | 0.84039886128482 | 0.405388124421933 | 0.703622985931183 | -5.07059690628239 |
| lnc-DGAT1-2 | lnc-DGAT1-2 | -0.227964223194242 | 0.0744999255479057 | -0.840360201658363 | 0.405409551705527 | 0.703622985931183 | -5.37365986261428 |
| ENSG00000198146.4 | ZNF770 | 0.0924754121361087 | 6.25033063637699 | 0.840298295802849 | 0.405443864782934 | 0.703622985931183 | -6.06199575130629 |
| ENSG00000136003.15 | ISCU | 0.101395867453236 | 7.7363525851966 | 0.840261381057353 | 0.405464326683659 | 0.703622985931183 | -6.04749087962295 |
| LINC01679 | LINC01679 | 0.764926915575729 | -0.437348040686568 | 0.840362014861612 | 0.405540351405188 | 0.703695586998228 | -5.27986193673346 |
| lnc-SLC38A9-3 | lnc-SLC38A9-3 | -0.105286681901586 | 0.708899491489704 | -0.8399229480717 | 0.405651950396369 | 0.70382989926734 | -5.51861308812428 |
| ENSG00000102225.15 | CDK16 | 0.123209237596706 | 4.82195986654898 | 0.839777178758644 | 0.405732779928877 | 0.703910806580923 | -6.02590666173624 |
| ENSG00000087266.15 | SH3BP2 | -0.20955641926652 | 6.22880732027858 | -0.839713035480866 | 0.40580735290428 | 0.703925877579732 | -6.06191833399089 |
| lnc-ST8SIA1-4 | lnc-ST8SIA1-4 | 0.158249538795914 | 0.794707167183324 | 0.839637888410753 | 0.405810026163667 | 0.703925877579732 | -5.53071641506688 |
| ENSG00000005175.9 | RPAP3 | -0.123299638409648 | 4.6872410594324 | -0.839546566043071 | 0.405860675737286 | 0.703925877579732 | -6.0117721099603 |
| lnc-EIF3B-2 | lnc-EIF3B-2 | 0.175517157478819 | 2.05174530226874 | 0.839476703468905 | 0.405899425834862 | 0.703925877579732 | -5.738788001919 |
| lnc-MED29-3 | lnc-MED29-3 | 0.230339308066777 | 1.54607593783259 | 0.839453198363993 | 0.405912463731907 | 0.703925877579732 | -5.63950474838334 |
| ENSG00000133943.20 | DGLUCY | -0.0900503584995187 | 5.37569139351611 | -0.839295399724443 | 0.405999998761202 | 0.704018363479513 | -6.04786750092038 |
| ENSG00000266019.1 | MIR3609 | 0.216454960385559 | 7.3551848615428 | 0.839231499565587 | 0.406079554491086 | 0.704092713825032 | -6.05350338968249 |
| lnc-MED6-1 | lnc-MED6-1 | 0.105910500365398 | 2.35845267036893 | 0.839094804907567 | 0.406111290799727 | 0.704092713825032 | -5.75853228517384 |
| ENSG00000141258.12 | SGSM2 | -0.0939639217137103 | 6.09014880033524 | -0.838947069204587 | 0.40619326812748 | 0.704125795136444 | -6.06255653054125 |
| lnc-SPDYA-1 | lnc-SPDYA-1 | 0.228318113639135 | 0.108740804066716 | 0.838937118880282 | 0.406198789848425 | 0.704125795136444 | -5.36325670799096 |
| ENSG00000221883.3 | ARIH2OS | -0.131492843877596 | 1.25227420639046 | -0.838636059188931 | 0.406365878468284 | 0.70430343120958 | -5.61749754345255 |
| ENSG00000198466.11 | ZNF587 | 0.0823050131844045 | 5.72401077529443 | 0.838616322931202 | 0.406376833607309 | 0.70430343120958 | -6.05770413019584 |
| ENSG00000226210.3 | WASH8P | -0.244148138221018 | 3.75021555071098 | -0.838646369967745 | 0.40640391840238 | 0.70430343120958 | -5.93723102129591 |
| ENSG00000226117.1 | | 0.357801532046976 | -0.597406642048999 | 0.838350378516416 | 0.406524470989526 | 0.704432629269864 | -5.14965955570489 |
| ENSG00000161265.14 | U2AF1L4 | -0.146963826083716 | 3.64379083605794 | -0.838309950038185 | 0.406546917509436 | 0.704432629269864 | -5.91382253953704 |
| ENSG00000165632.7 | TAF3 | -0.104724005606887 | 4.16898614559491 | -0.838243850889896 | 0.406583618434602 | 0.704436920727224 | -5.96558049158781 |
| lnc-DDX52-3 | lnc-DDX52-3 | -0.165825395178465 | 1.13733450679625 | -0.838139810286028 | 0.406641390140938 | 0.704461963943788 | -5.60467138166207 |
| lnc-AGAP2-1 | lnc-AGAP2-1 | -0.1593731788942 | 0.159665100782132 | -0.838080616126097 | 0.406674261760802 | 0.704461963943788 | -5.38622566631519 |
| lnc-SEMA4B-2 | lnc-SEMA4B-2 | -0.173928899228678 | 1.52475206804828 | -0.837970101043181 | 0.406735637239924 | 0.704461963943788 | -5.65311354635036 |
| ENSG00000267519.6 | MIR23AHG | -0.413119506428842 | 2.28865220674208 | -0.838189498903949 | 0.406739682121706 | 0.704461963943788 | -5.76033028514199 |
| ENSG00000197774.12 | EME2 | -0.128485216777003 | 4.75417774053853 | -0.837822516873731 | 0.406817608272305 | 0.704461963943788 | -6.01727551662615 |
| ENSG00000134809.8 | TIMM10 | 0.106947927914399 | 2.29816166055208 | 0.837791061794864 | 0.406835080336085 | 0.704461963943788 | -5.75871065758205 |
| lnc-SMARCA5-4 | lnc-SMARCA5-4 | -0.270102986417339 | 0.416492736248009 | -0.837773333616248 | 0.406844927848931 | 0.704461963943788 | -5.51178618599857 |
| lnc-DHX57-4 | lnc-DHX57-4 | -0.182857011049334 | 2.03994591153284 | -0.837700623454709 | 0.406885317873064 | 0.704461963943788 | -5.72775226356068 |
| ENSG00000176946.11 | THAP4 | -0.107420802302367 | 3.7210620500843 | -0.837663210298208 | 0.40690610160461 | 0.704461963943788 | -5.93122691267204 |
| ENSG00000130413.15 | STK33 | 0.847701950461692 | -0.835300849945144 | 0.837751639098806 | 0.406990792956595 | 0.704549326444542 | -5.21863690701625 |
| ENSG00000089123.15 | TASP1 | 0.0754828112353776 | 3.45830264417165 | 0.837159678880191 | 0.407185886849973 | 0.70482777807022 | -5.89135320455225 |
| ENSG00000197959.13 | DNM3 | 0.157959139114279 | 1.25550363330446 | 0.837088769082917 | 0.407225297140953 | 0.704836721413744 | -5.59818392904254 |
| C1RL-AS1 | C1RL-AS1 | -0.164437561144058 | 3.91971623011026 | -0.837007844543651 | 0.407270276309533 | 0.70485530120904 | -5.94956665596774 |
| ENSG00000106617.13 | PRKAG2 | -0.0930481938945955 | 3.63476368719216 | -0.836902247672875 | 0.407328973372683 | 0.704861484597063 | -5.91153201917769 |
| lnc-ELOVL5-2 | lnc-ELOVL5-2 | 0.27621300281321 | 0.181698883561911 | 0.836830271111726 | 0.407368985248457 | 0.704861484597063 | -5.37680639750902 |

| | | | | | | | |
|---|---|---|---|---|---|---|---|
| lnc-UNC93B1-6 | lnc-UNC93B1-6 | -0.135806230642721 | 1.51659257423481 | -0.836816603036549 | 0.407376583625645 | 0.704861484597063 | -5.66324371763609 |
| lnc-ZNF143-3 | lnc-ZNF143-3 | -0.289327081017716 | -0.270649950760427 | -0.836736718862512 | 0.407420994707832 | 0.704878815895849 | -5.25197820443922 |
| ENSG00000213977.7 | TAX1BP3 | 0.197180548062467 | 0.831177945300412 | 0.836627638303391 | 0.4074816421563 | 0.704878815895849 | -5.55121943461549 |
| ENSG00000271852.1 | SNORD11B | -0.217624900797178 | 0.529186558024775 | -0.836476147444115 | 0.407565878454683 | 0.704878815895849 | -5.4813859843656 |
| lnc-ZNF136-1 | lnc-ZNF136-1 | -0.268597480260398 | 0.727991987665636 | -0.836453037264084 | 0.407578729785701 | 0.704878815895849 | -5.53153942010616 |
| lnc-ESAM-1 | lnc-ESAM-1 | -0.139931274220725 | 0.133614728735292 | -0.836433689635775 | 0.407589488993173 | 0.704878815895849 | -5.39221544080816 |
| lnc-PIDD1-2 | lnc-PIDD1-2 | -0.16312302528247 | 0.633099158037905 | -0.836389396022094 | 0.407614121312139 | 0.704878815895849 | -5.51124107351571 |
| ENSG00000125166.12 | GOT2 | 0.0897078496529122 | 4.11758484668694 | 0.836323025186375 | 0.407651032813539 | 0.704878815895849 | -5.96720735722031 |
| ENSG00000012817.15 | KDM5D | -1.9659871512371 | 3.06742245939498 | -0.836639440009774 | 0.407660565730175 | 0.704878815895849 | -5.65400137469429 |
| ENSG00000100084.14 | HIRA | 0.0949620471193224 | 4.47962066184471 | 0.836159907558513 | 0.407741757878262 | 0.70492051159645 | -6.0035586483959 |
| ENSG00000231234.2 | SKP1P1 | 0.221024923805751 | -0.479908017812918 | 0.836129919596379 | 0.407758438362388 | 0.70492051159645 | -5.1852991692501 |
| ENSG00000110848.8 | CD69 | -0.310509564505774 | 7.26213320579631 | -0.836285387461891 | 0.407803057940486 | 0.70492051159645 | -6.05731223677916 |
| lnc-KCNK9-4 | lnc-KCNK9-4 | -0.260345403699373 | 0.273560350519344 | -0.836016247823227 | 0.407821670897854 | 0.70492051159645 | -5.39855358659719 |
| ENSG00000126709.14 | IFI6 | 0.221711314673045 | 1.74043329024779 | 0.83570844449088 | 0.407992923950461 | 0.705076772458618 | -5.68193084526312 |
| ENSG00000233261.3 | FAM238A | -0.225992365189657 | 0.441303371038245 | -0.835686789077435 | 0.408004974079409 | 0.705076772458618 | -5.45488890889861 |
| ENSG00000131473.16 | ACLY | -0.0922173588678943 | 6.2119431905011 | -0.83566906052822 | 0.408014839272924 | 0.705076772458618 | -6.06551344223192 |
| lnc-FOXD4L5-27 | lnc-FOXD4L5-27 | -0.444801867600989 | 0.568396379642967 | -0.83570807441726 | 0.408050450610861 | 0.705079115591282 | -5.44358136427161 |
| lnc-C19orf57-6 | lnc-C19orf57-6 | -0.415642377316902 | 2.28292069465435 | -0.835723717190445 | 0.408111441604949 | 0.705125308700971 | -5.76236404901686 |
| ENSG00000104447.12 | TRPS1 | 0.739698862879605 | 5.243073776579 | 0.835589093662281 | 0.408244565007081 | 0.705296112266305 | -6.04396165708889 |
| SNHG19 | SNHG19 | -0.189761702186405 | 2.46208236183792 | -0.835127222246196 | 0.408316420518086 | 0.705361047714962 | -5.78199065145506 |
| ENSG00000119844.15 | AFTPH | 0.102929679871028 | 7.31053427472691 | 0.83495046285809 | 0.408414832590219 | 0.705456801890331 | -6.05887459611841 |
| ENSG00000107371.12 | EXOSC3 | -0.0812604402675543 | 4.17264989499514 | -0.834886445131214 | 0.4084504785324 | 0.705456801890331 | -5.97420496535933 |
| ENSG00000070444.14 | MNT | 0.102650002269004 | 3.58705134614129 | 0.8348244723352 | 0.408484987656879 | 0.705456801890331 | -5.9057567438055 |
| ENSG00000104835.14 | SARS2 | -0.136192944529892 | 2.50806109201739 | -0.83478145013464 | 0.408508945330168 | 0.705456801890331 | -5.79255407221031 |
| lnc-SLFNL1-2 | lnc-SLFNL1-2 | -0.140325667760554 | 0.918255976499876 | -0.834520196465531 | 0.408654447662731 | 0.705600935071467 | -5.57020806458105 |
| lnc-SLC35E3-1 | lnc-SLC35E3-1 | -0.259069540797362 | -0.243211623203998 | -0.834444615254019 | 0.40869654773854 | 0.705600935071467 | -5.24811352820487 |
| ENSG00000275835.4 | TUBGCP5 | 0.130231246222349 | 4.81766972595644 | 0.834398008544207 | 0.408722509835642 | 0.705600935071467 | -6.02197107512942 |
| ENSG00000156232.7 | WHAMM | 0.133509470396221 | 5.93016313844029 | 0.834385403513014 | 0.408729531596808 | 0.705600935071467 | -6.06460631632901 |
| ENSG00000130119.15 | GNL3L | 0.0688360478035186 | 5.16872996180551 | 0.834293283566637 | 0.408780850210126 | 0.705630345511156 | -6.04198269954862 |
| lnc-APOL4-3 | lnc-APOL4-3 | 0.175739210907183 | 1.58346727442505 | 0.834197637387145 | 0.408834137436961 | 0.705663148919494 | -5.68317625859661 |
| ENSG00000189403.14 | HMGB1 | 0.080384690157878 | 8.27355476774121 | 0.834094955118441 | 0.408891349436512 | 0.705702720564458 | -6.03940414126204 |
| ENSG00000145740.18 | SLC30A5 | -0.090980205840542 | 4.42157592004207 | -0.833997298886495 | 0.408945765632067 | 0.705737460719781 | -6.00018310297396 |
| lnc-AJUBA-2 | lnc-AJUBA-2 | 0.170538248572102 | 1.41995778746184 | 0.833695107344656 | 0.409114181665651 | 0.705940001189161 | -5.63096439733309 |
| ENSG00000241973.10 | PI4KA | 0.0696290854376 | 6.86234430259833 | 0.833603638946922 | 0.40916516684789 | 0.705940001189161 | -6.06536795420279 |
| lnc-RINT1-3 | lnc-RINT1-3 | -0.209051525417642 | -0.553724870228217 | -0.833602106297705 | 0.409166021191599 | 0.705940001189161 | -5.20748848824822 |
| lnc-HIPK2-1 | lnc-HIPK2-1 | 0.179410680389983 | 2.73115534987772 | 0.833451782846137 | 0.40924982124172 | 0.70602540194605 | -5.81227385592991 |
| lnc-SCAF8-2 | lnc-SCAF8-2 | 0.158588523509397 | 2.07793269048241 | 0.833348031918707 | 0.409307664911151 | 0.706066012979066 | -5.7351603896276 |
| ENSG00000116584.17 | ARHGEF2 | -0.143874148532002 | 6.83106904302454 | -0.833248551764474 | 0.409363132250553 | 0.706102518320047 | -6.06539243518061 |
| lnc-TM9SF1-4 | lnc-TM9SF1-4 | -0.0839752075642273 | 2.73023774184399 | -0.833120085866706 | 0.40943476807714 | 0.706166903915851 | -5.80909420522453 |
| ENSG00000135250.16 | SRPK2 | 0.0997015437068827 | 5.69795654736039 | 0.832925862691203 | 0.409543086483169 | 0.706294541188359 | -6.06018287230221 |
| ENSG00000179950.13 | PUF60 | 0.111621739469027 | 5.59500714434703 | 0.832802620853449 | 0.409611827698062 | 0.706344262357725 | -6.06067829978077 |
| lnc-GPR75-ASB3-4 | lnc-GPR75-ASB3-4 | -0.163779615190037 | 0.83693986049793 | -0.832527329222386 | 0.409765404148406 | 0.706344262357725 | -5.57612486420533 |
| ENSG00000204219.10 | TCEA3 | 0.897749334328158 | -2.1415555912003 | 0.832548590671264 | 0.409766275513882 | 0.706344262357725 | -4.85766252060649 |
| lnc-KYNU-11 | lnc-KYNU-11 | 0.225265271593484 | 0.512300798410112 | 0.832428843814395 | 0.409820354637287 | 0.706344262357725 | -5.45929162837791 |
| ENSG00000204604.10 | ZNF468 | 0.0762244253745106 | 5.67861022533422 | 0.832291594555827 | 0.409896941202574 | 0.706344262357725 | -6.06155272646677 |
| ENSG00000047621.11 | FERRY3 | 0.119494386730831 | 3.98509308125496 | 0.832271287177409 | 0.409908273686886 | 0.706344262357725 | -5.95001966329994 |
| ENSG00000096070.19 | BRPF3 | -0.107683147670383 | 5.04356215887368 | -0.832246997367288 | 0.409921828811104 | 0.706344262357725 | -6.03803952041863 |
| ENSG00000132849.20 | PATJ | -0.120134955219693 | 6.20671752429958 | -0.832156121297702 | 0.409972545381527 | 0.706344262357725 | -6.06839141732483 |
| RB1-DT | RB1-DT | -0.205044612101243 | 1.72517923949102 | -0.832079307827057 | 0.410015416838984 | 0.706344262357725 | -5.69316943351651 |
| ENSG00000163346.16 | PBXIP1 | 0.153612060658556 | 7.14996889839081 | 0.832012621898363 | 0.410052638108042 | 0.706344262357725 | -6.06287230805454 |
| ENSG00000113460.12 | BRIX1 | 0.0973707015152572 | 4.71152251472191 | 0.831979361838808 | 0.410071203246015 | 0.706344262357725 | -6.02474895181666 |
| NPTN-IT1 | NPTN-IT1 | -0.141845114063395 | 1.995844927912 | -0.831971365779018 | 0.410075666572934 | 0.706344262357725 | -5.72163554826711 |
| lnc-MED26-1 | lnc-MED26-1 | -0.0946800357374551 | 3.25167398737146 | -0.831960891581018 | 0.410081513219015 | 0.706344262357725 | -5.87501693762911 |
| lnc-ECD-2 | lnc-ECD-2 | 0.107749826906367 | 2.09522729911068 | 0.831837329660916 | 0.410150488751313 | 0.706344262357725 | -5.72953782548438 |
| lnc-SUSD2-1 | lnc-SUSD2-1 | 0.14114565761506 | 1.93592590483416 | 0.831780367705068 | 0.410182288826483 | 0.706344262357725 | -5.71924592527105 |
| lnc-CCDC89-8 | lnc-CCDC89-8 | 0.738698719775606 | -1.0911358585581 | 0.83180517920665 | 0.410209851323855 | 0.706344262357725 | -5.02087131366691 |
| LINC02240 | LINC02240 | -0.774928132833087 | -0.807765390187545 | -0.831767610855725 | 0.410231995491861 | 0.706344262357725 | -4.99719604944729 |
| ENSG00000006715.15 | VPS41 | -0.0883516614812319 | 6.59332806892731 | -0.831656607184275 | 0.410251385677586 | 0.706344262357725 | -6.06854412244493 |
| lnc-H2AFJ-6 | lnc-H2AFJ-6 | -0.249704432302235 | 1.01624503465621 | -0.831602024138858 | 0.41028186226453 | 0.706344262357725 | -5.5704367505714 |
| lnc-SEMA5B-2 | lnc-SEMA5B-2 | 0.278114256719127 | -0.585971876176716 | 0.831501212913609 | 0.410338154140242 | 0.706344262357725 | -5.21123616291826 |
| ENSG00000184436.11 | THAP7 | -0.148286923615737 | 4.34935080571055 | -0.831500414922547 | 0.410338599748605 | 0.706344262357725 | -5.99709985858952 |
| lnc-TRMT2A-1 | lnc-TRMT2A-1 | -0.16210941134991 | 0.0910642570487155 | -0.831451272041452 | 0.410366042332512 | 0.706344262357725 | -5.39277843303 |
| ENSG00000144567.10 | RETREG2 | -0.113543043355598 | 6.23739643877928 | -0.831414553129458 | 0.410386547806087 | 0.706344262357725 | -6.06909687330649 |
| lnc-TATDN1-3 | lnc-TATDN1-3 | -0.465572378170859 | 3.61001836856917 | -0.831701623939399 | 0.41039552220454 | 0.706344262357725 | -5.93107728799661 |
| ENSG00000114739.13 | ACVR2B | -0.23836059212332 | 3.06563650834364 | -0.831341375263051 | 0.410441947075844 | 0.706365100055359 | -5.85391823559013 |
| lnc-GRHL2-19 | lnc-GRHL2-19 | 0.517792990142552 | 0.774769234626676 | 0.831425518289514 | 0.410483520740257 | 0.706377586104566 | -5.45930561440431 |
| ENSG00000093183.13 | SEC22C | 0.100422192528861 | 5.38536064579136 | 0.831118066479502 | 0.410552142235943 | 0.70642909865974 | -6.05669544131426 |
| lnc-TRIM73-1 | lnc-TRIM73-1 | -0.122908277544239 | 2.79186632229917 | -0.831026162762899 | 0.410603480863825 | 0.70642909865974 | -5.81474006908957 |

| | | | | | | | |
|---|---|---|---|---|---|---|---|
| lnc-ST8SIA1-1 | lnc-ST8SIA1-1 | -0.231486658338939 | -0.169496690566972 | -0.831003004089794 | 0.410616418226941 | 0.70642909865974 | -5.32586320055952 |
| ENSG00000135775.13 | COG2 | 0.0772558592098589 | 4.93288543370903 | 0.83093721910974 | 0.410653169721687 | 0.706433279764436 | -6.03486051431762 |
| ENSG00000176407.17 | KCMF1 | 0.0758675808297049 | 5.61864190911218 | 0.830809663614088 | 0.4107244357522 | 0.706493250650268 | -6.06297484741713 |
| ENSG00000118579.12 | MED28 | 0.0885175164330345 | 6.3924080227591 | 0.830751956933853 | 0.410756679324285 | 0.706493250650268 | -6.06983767347002 |
| ENSG00000184678.10 | H2BC21 | -0.170447561105092 | 6.42001286960233 | -0.83066624628614 | 0.410804572969455 | 0.706516588020579 | -6.06945211535087 |
| ENSG00000029993.14 | HMGB3 | -0.324317504888178 | 2.55535166887477 | -0.830653906991888 | 0.410896449915464 | 0.70661555924455 | -5.80293866745647 |
| ENSG00000188629.12 | ZNF177 | -0.226342841345595 | 2.03336724177614 | -0.830325316575404 | 0.410995112665183 | 0.70671043295214 | -5.72798258223096 |
| ENSG00000102384.13 | CENPI | 0.218630383243444 | 0.0853137118600193 | 0.830280277579319 | 0.41102028824127 | 0.70671043295214 | -5.3655864961803 |
| lnc-GRAMD2A-3 | lnc-GRAMD2A-3 | -0.087939480694875 | 1.67374527443477 | -0.830218455783326 | 0.41105484649095 | 0.706710817350753 | -5.68760837711126 |
| ENSG00000105875.13 | WDR91 | -0.200791660468211 | 6.46980440665993 | -0.829815419893423 | 0.411308795252476 | 0.707088359866509 | -6.07003643724576 |
| lnc-TBL1Y-6 | lnc-TBL1Y-6 | -0.907311168863694 | -1.99374528484882 | -0.829700702812506 | 0.411344339407637 | 0.70709040738495 | -4.82328880614931 |
| ENSG00000123178.14 | SPRYD7 | -0.148870558996926 | 1.5088458669925 | -0.829473048292878 | 0.411471668281664 | 0.70717635844392 | -5.6851575401741 |
| ENSG00000251634.2 | NAIPP4 | -0.199300116276783 | 1.27946808441488 | -0.829391930950302 | 0.411517043670132 | 0.70717635844392 | -5.64146881752514 |
| lnc-ZSWIM7-3 | lnc-ZSWIM7-3 | -0.127694471640512 | 2.67061714339458 | -0.829352306031282 | 0.411539210160288 | 0.70717635844392 | -5.8088040573619 |
| ENSG00000124532.14 | MRS2 | -0.116522724072766 | 3.52480053627332 | -0.829333510643927 | 0.411549724703793 | 0.70717635844392 | -5.91203807963353 |
| lnc-SYNE3-6 | lnc-SYNE3-6 | -0.254123453718338 | 5.34596186020615 | -0.829368822446892 | 0.411620851735258 | 0.70717635844392 | -6.0518346428593 |
| ENSG00000109971.13 | HSPA8 | -0.148853169995181 | 8.46956995344541 | -0.829192846211323 | 0.411628420650397 | 0.70717635844392 | -6.0401390163204 |
| ENSG00000223508.5 | RPL23AP53 | 0.145738943693238 | 0.0249603470921255 | 0.829181369356944 | 0.411634841884886 | 0.70717635844392 | -5.35740520859923 |
| ENSG00000186230.6 | ZNF749 | 0.124869545833905 | 2.27228331076182 | 0.829033925286861 | 0.411717341474715 | 0.70724456116916 | -5.75748367727368 |
| ENSG00000132485.13 | ZRANB2 | -0.105911362268617 | 6.46410192984814 | -0.828973910045039 | 0.411750924797041 | 0.70724456116916 | -6.07079726378032 |
| ENSG00000134824.13 | FADS2 | -0.562111298264525 | 0.408341439843127 | -0.829160121346359 | 0.411777623332421 | 0.70724456116916 | -5.54361049463551 |
| ENSG00000168002.11 | POLR2G | 0.0862412472035965 | 4.90741848227048 | 0.828615359410144 | 0.411951597581822 | 0.707484333168682 | -6.03580905299554 |
| lnc-ADAMTS12-6 | lnc-ADAMTS12-6 | -0.235142028790271 | -0.518773100866223 | -0.828487785156139 | 0.412023012530691 | 0.707515297332793 | -5.21956228430083 |
| lnc-SMAD7-3 | lnc-SMAD7-3 | 0.116305730281036 | 0.797342746371222 | 0.828459394674436 | 0.412038906307744 | 0.707515297332793 | -5.5445405472866 |
| lnc-ABCB1-1 | lnc-ABCB1-1 | -0.140186910537096 | 0.542707410272473 | -0.828364165588977 | 0.412092220933979 | 0.707515297332793 | -5.51790873257498 |
| ENSG00000278463.1 | H2AC4 | -0.401482841320722 | 1.5534444107318 | -0.828520981501097 | 0.412107122369084 | 0.707515297332793 | -5.67503088252064 |
| lnc-QRICH2-1 | lnc-QRICH2-1 | -0.138918842235317 | 0.795837064711736 | -0.827967916948658 | 0.41231410881344 | 0.707729179141517 | -5.54010333931252 |
| ENSG00000186871.6 | ERCC6L | 0.508692857200285 | -0.383943263306278 | 0.828007627733662 | 0.412335487255328 | 0.707729179141517 | -5.29377874045418 |
| ENSG00000269469.1 | | 0.162875318159137 | 1.3086927712184 | 0.827913351951859 | 0.412344669392582 | 0.707729179141517 | -5.64979178184425 |
| ENSG00000044446.11 | PHKA2 | 0.074592703538447 | 5.24659528472833 | 0.827869485533161 | 0.412369238956625 | 0.707729179141517 | -6.05134308531456 |
| ENSG00000205593.11 | DENND6B | -0.252896325912606 | 2.02684347811815 | -0.827728428232622 | 0.41245295849529 | 0.707756079508036 | -5.7505579359611 |
| lnc-TBC1D20-2 | lnc-TBC1D20-2 | -0.0975170243184649 | 2.68526328915942 | -0.827718730217747 | 0.412453683802113 | 0.707756079508036 | -5.81676673074735 |
| ENSG00000185947.14 | ZNF267 | 0.103139200293771 | 5.91453787010761 | 0.827615929794796 | 0.412511273038437 | 0.707784369994994 | -6.06960690498136 |
| ENSG00000168228.14 | ZCCHC4 | 0.113334346683519 | 3.99692009795362 | 0.827566538299832 | 0.412538944120388 | 0.707784369994994 | -5.9661981887399 |
| lnc-POLR1A-5 | lnc-POLR1A-5 | 0.23712983153261 | 0.295349920191435 | 0.82735606084769 | 0.41265676747 4122 | 0.707927048350016 | -5.41888768780134 |
| lnc-EPS15L1-3 | lnc-EPS15L1-3 | -0.344002559883144 | 0.134104119545471 | -0.827281120524509 | 0.4126988688359 | 0.707927048350016 | -5.40843973967549 |
| lnc-FASTK-2 | lnc-FASTK-2 | -0.226168853553754 | 0.068267348439181 | -0.827212006256348 | 0.412737600525482 | 0.707927048350016 | -5.41265181450071 |
| ENSG00000231841.1 | FAM192BP | 0.390653221389039 | -0.363868103785669 | 0.827172607404548 | 0.412759680670871 | 0.707927048350016 | -5.16294436881657 |
| lnc-PNN-4 | lnc-PNN-4 | -0.263735468246772 | -0.418158449313822 | -0.827007227220473 | 0.412852371951805 | 0.708027026485919 | -5.26855274682839 |
| lnc-GMNN-6 | lnc-GMNN-6 | -0.216190735146817 | -0.0300041787086903 | -0.826763839594441 | 0.412988807597463 | 0.708148223504186 | -5.33828788329716 |
| ENSG00000116701.14 | NCF2 | 0.238641531534022 | 4.58308931968162 | 0.82679975619575 | 0.413022878950454 | 0.708148223504186 | -5.99809782266355 |
| ENSG00000229180.7 | RABGEF1P1 | -0.0799053360800137 | 4.30318485317515 | -0.826654006766096 | 0.41305038557263 | 0.708148223504186 | -5.99232314475434 |
| lnc-PBLD-1 | lnc-PBLD-1 | 0.100459294416792 | 2.41003662633714 | 0.826604868177353 | 0.413077937035447 | 0.708148223504186 | -5.78149067785121 |
| ENSG00000187446.11 | CHP1 | 0.13271412098045 | 4.59511374588836 | 0.826536516906747 | 0.41311626271097 | 0.708148223504186 | -6.01424561029265 |
| ENSG00000164611.12 | PTTG1 | 0.195373353197058 | 1.65967543745779 | 0.826499142226057 | 0.413137220228219 | 0.708148223504186 | -5.68590158690031 |
| ENSG00000160439.15 | RDH13 | -0.169128715182187 | 2.70471623254936 | -0.826414429193825 | 0.413184724716074 | 0.708148223504186 | -5.79667496737265 |
| lnc-DAZL-5 | lnc-DAZL-5 | 0.0781631781993593 | 2.26012415072989 | 0.826330472678465 | 0.4132318082778 | 0.708148223504186 | -5.75830988831804 |
| lnc-TRAF3IP2-7 | lnc-TRAF3IP2-7 | -0.225286211051026 | -0.33053396114591 | -0.826328913128635 | 0.413232682918368 | 0.708148223504186 | -5.31400711881144 |
| ENSG00000108622.10 | ICAM2 | -0.158636975952702 | 3.71641756422186 | -0.826236411520444 | 0.413284562521602 | 0.70817816769748 | -5.92158574814367 |
| lnc-MTCL1-4 | lnc-MTCL1-4 | 0.127108116319795 | 1.59855531501585 | 0.826009534417629 | 0.413411823644323 | 0.708337265135361 | -5.67763017941355 |
| ENSG00000285171.1 | | -0.0771835143224449 | 5.51514971597071 | -0.825915261470048 | 0.413464710798483 | 0.708355409309951 | -6.06314974656543 |
| ENSG00000234284.6 | ZNF879 | -0.23878018912797 | 3.32825868116178 | -0.825862763375934 | 0.413515588225492 | 0.708355409309951 | -5.88200854717683 |
| ENSG00000131791.7 | PRKAB2 | -0.0787518909817688 | 4.33359567536349 | -0.825806632176302 | 0.413525657011532 | 0.708355409309951 | -6.00574289815379 |
| lnc-TMPRSS2-83 | lnc-TMPRSS2-83 | 0.409176557545537 | 0.0754357054532653 | 0.825575300416994 | 0.413656869650855 | 0.70842098618686 | -5.32420088232429 |
| ENSG00000258790.1 | | 0.0838145234475585 | 4.75342701381944 | 0.825562313566573 | 0.413662751524322 | 0.70842098618686 | -6.02921900645729 |
| lnc-GOLPH3-4 | lnc-GOLPH3-4 | -0.114034212287487 | 1.00897614449886 | -0.825554399209353 | 0.413667192973807 | 0.70842098618686 | -5.5838320877388 |
| ENSG00000181610.12 | MRPS23 | -0.0970934060931259 | 3.55888238748624 | -0.825451305746594 | 0.413725050552115 | 0.708461124418817 | -5.92303559496118 |
| lnc-PPP1R9B-2 | lnc-PPP1R9B-2 | -0.427444869655055 | -0.854995042192323 | -0.825338763785146 | 0.413788216430702 | 0.708483827794491 | -5.0553603653538 |
| ENSG00000120860.10 | WASHC3 | -0.0714607188001128 | 4.52649059039523 | -0.825305031296603 | 0.413807150451605 | 0.708483827794491 | -6.01304046053229 |
| ENSG00000108733.9 | PEX12 | -0.160975674986502 | 1.70415521686434 | -0.825097593312571 | 0.413923596911867 | 0.708578977177144 | -5.69447971864862 |
| ENSG00000084754.11 | HADHA | 0.126065431827977 | 7.00050954822069 | 0.82496275050921 | 0.413999302434435 | 0.708578977177144 | -6.07062584693836 |
| ENSG00000136810.12 | TXN | 0.183188386480045 | 2.90430748023942 | 0.824856231194179 | 0.414059112146447 | 0.708578977177144 | -5.84425116740961 |
| FIRRE | FIRRE | 0.445318501354504 | 0.233253213286733 | 0.824933225417935 | 0.414069949696617 | 0.708578977177144 | -5.43479628412904 |
| ENSG00000163029.15 | SMC6 | 0.171182541987263 | 6.9063882342336 | 0.824782662157571 | 0.414100423641362 | 0.708578977177144 | -6.07289443612538 |
| lnc-NKX3-2-3 | lnc-NKX3-2-3 | 0.222323143340175 | 1.85242212942971 | 0.824751379212175 | 0.414117990835336 | 0.708578977177144 | -5.72606557132333 |
| lnc-TM9SF2-2 | lnc-TM9SF2-2 | -0.219441020113544 | 0.40831477455347 | -0.824729959517764 | 0.414130019503365 | 0.708578977177144 | -5.47375139418075 |
| lnc-PSME2-2 | lnc-PSME2-2 | -0.245784742240572 | -0.0526038397484066 | -0.824671543112475 | 0.414162825520002 | 0.708578977177144 | -5.37871668018267 |
| lnc-AGO2-6 | lnc-AGO2-6 | 0.117799859664296 | 1.6777153567518 | 0.824569967195792 | 0.414219873239096 | 0.708578977177144 | -5.66910676996251 |

| | | | | | | | |
|---|---|---|---|---|---|---|---|
| lnc-SLIT3-2 | lnc-SLIT3-2 | 0.0804650586421936 | 2.84311911421865 | 0.824551785765339 | 0.414230084918839 | 0.708578977177144 | -5.83640584442603 |
| ENSG00000120686.11 | UFM1 | 0.0784376530930073 | 6.34174038127817 | 0.824490534743607 | 0.414264487957398 | 0.708578977177144 | -6.07494965937949 |
| ENSG00000250317.8 | SMIM20 | 0.131503191974909 | 2.70121834575434 | 0.824424110096612 | 0.414301798859343 | 0.708578977177144 | -5.8157919233188 |
| lnc-NSF-2 | lnc-NSF-2 | 0.240548836103081 | 1.89434781668783 | 0.824401559005996 | 0.414314466335328 | 0.708578977177144 | -5.71458898268592 |
| ENSG00000105204.13 | DYRK1B | 0.158301359186447 | 3.44480018713112 | 0.824347771467697 | 0.414344681013657 | 0.708578977177144 | -5.91115028571918 |
| ENSG00000269237.1 | | -0.116057920478377 | 1.22401570350543 | -0.823977804755428 | 0.41455254314024 | 0.708845134964311 | -5.61833472074193 |
| ENSG00000266820.1 | KPNA2P3 | -0.139407921878379 | 2.99058102585394 | -0.823948172111133 | 0.414569194709245 | 0.708845134964311 | -5.84932706064294 |
| lnc-SH2B2-2 | lnc-SH2B2-2 | 0.133273643992782 | 1.42834153375487 | 0.823795370695568 | 0.414655065411257 | 0.708896292781189 | -5.66460204414992 |
| lnc-ERLEC1-6 | lnc-ERLEC1-6 | -0.20095654643565 | 1.68463345733164 | -0.823772362924098 | 0.414667996166036 | 0.708896292781189 | -5.68489883629689 |
| ENSG00000140104.13 | CLBA1 | -0.253614552081319 | -0.303879461509039 | -0.823673173420945 | 0.414723745162804 | 0.70893271710705 | -5.30480414129063 |
| lnc-CEP170-4 | lnc-CEP170-4 | 0.74103402380628 | 2.26867139448578 | 0.823795023786278 | 0.414837563563104 | 0.709068391531255 | -5.70308416792617 |
| ENSG00000155621.14 | C9orf85 | -0.0951269735002316 | 2.19747518860735 | -0.823171019345096 | 0.415006048931486 | 0.709297476140229 | -5.75636884442733 |
| ENSG00000106733.20 | NMRK1 | 0.199344527888583 | 2.86075527665143 | 0.822883874843905 | 0.415167530268914 | 0.709514552933 | -5.82305079651505 |
| ENSG00000178222.12 | RNF212 | 0.90651962268578 | -2.93311109927589 | 0.822585315334227 | 0.41533547182869 | 0.709742633179737 | -4.78251178324056 |
| lnc-NANOS3-2 | lnc-NANOS3-2 | -0.54333710287792 | -0.895712017122876 | -0.822482828285179 | 0.415400780713693 | 0.709771009824031 | -5.13013859582963 |
| ENSG00000204599.14 | TRIM39 | 0.0920783573118182 | 4.6811992731525 | 0.822371563313806 | 0.415455734182599 | 0.709771009824031 | -6.02393386670205 |
| lnc-ANXA6-6 | lnc-ANXA6-6 | -0.123752573610528 | 3.50046215832412 | -0.822318831093553 | 0.415485405954103 | 0.709771009824031 | -5.9140157483131 |
| ENSG00000144909.7 | OSBPL11 | 0.10882471762956 | 5.16810280380255 | 0.822224203371543 | 0.415538655059282 | 0.709771009824031 | -6.05160089391977 |
| ENSG00000182325.10 | FBXL6 | -0.138807428132329 | 2.56652586712906 | -0.82220240813001 | 0.415550920313609 | 0.709771009824031 | -5.81018008941891 |
| ENSG00000147439.12 | BIN3 | 0.0914693031456082 | 4.54590044450446 | 0.822186470453475 | 0.415559889368201 | 0.709771009824031 | -6.01443772403513 |
| ENSG00000213297.8 | ZNF625-ZNF20 | -0.108698333848 | 1.91241618606327 | -0.822126816552062 | 0.415593461128078 | 0.709771009824031 | -5.72099020195766 |
| ENSG00000242779.6 | ZNF702P | -0.587429561639208 | -1.04646352953534 | -0.821997853793978 | 0.415670269711756 | 0.709843288930315 | -5.06696618520295 |
| lnc-NOL10-4 | lnc-NOL10-4 | 0.125952056525623 | 2.18760642003473 | 0.821835017707375 | 0.415757702298993 | 0.709849448941572 | -5.75479953127344 |
| lnc-TM9SF2-6 | lnc-TM9SF2-6 | 0.2001491801821 | 0.596283584756412 | 0.82180508941769 | 0.415774549905676 | 0.709849448941572 | -5.4938075544628 |
| lnc-ROCK1-1 | lnc-ROCK1-1 | -0.178376062528294 | 0.649030658917093 | -0.821800135961456 | 0.415777338407404 | 0.709849448941572 | -5.5547681854793 |
| ENSG00000134779.14 | TPGS2 | -0.087497612656616 | 4.24955244786277 | -0.821454514444923 | 0.415971931035458 | 0.710031529193146 | -5.98746223147621 |
| lnc-PLEKHM1-6 | lnc-PLEKHM1-6 | 0.148139147255617 | 2.89896951199875 | 0.821407557698984 | 0.41599837302725 | 0.710031529193146 | -5.83809422358637 |
| lnc-MYLK4-31 | lnc-MYLK4-31 | 0.23556351666725 | 0.144711213001226 | 0.821387933155583 | 0.416009424182992 | 0.710031529193146 | -5.38771054665797 |
| ENSG00000008838.19 | MED24 | -0.0668071025070907 | 4.93030757410251 | -0.821365651897134 | 0.41602197163044 | 0.710031529193146 | -6.04301510130683 |
| ENSG00000265527.1 | MIR5690 | -0.515032130313642 | 0.115442404182591 | -0.821342191777042 | 0.416126946243127 | 0.710151806195364 | -5.41761083605898 |
| ENSG00000090863.11 | GLG1 | 0.0963348019267389 | 7.24802569653995 | 0.820861763641908 | 0.416305792576686 | 0.710398120428282 | -6.07188778622972 |
| ENSG00000243244.6 | STON1 | 0.192851126457169 | 0.964309829524274 | 0.820674925050347 | 0.416411061663656 | 0.71045022423517 | -5.60692926652246 |
| lnc-HNRNPU-12 | lnc-HNRNPU-12 | -0.207505035980465 | 0.476789695036098 | -0.820600558149033 | 0.416452966182631 | 0.71045022423517 | -5.49062767956784 |
| ENSG00000237788.1 | GLYR1P1 | 0.101155087846249 | 2.3375098740822 | 0.820541147937814 | 0.416486444705554 | 0.71045022423517 | -5.76973918471866 |
| ENSG00000187325.4 | TAF9B | -0.136302048113677 | 3.02825279265243 | -0.820519264364392 | 0.416498776833652 | 0.71045022423517 | -5.86931589217595 |
| ENSG00000119138.4 | KLF9 | 0.177026848421261 | 6.07556334941164 | 0.820423860352322 | 0.416556164187357 | 0.71045022423517 | -6.07701963933385 |
| ENSG00000198563.13 | DDX39B | 0.0991919776057326 | 7.45827499129533 | 0.820416532478022 | 0.416556672681747 | 0.71045022423517 | -6.06820522431283 |
| lnc-DUOX1-2 | lnc-DUOX1-2 | -0.324025420683205 | -0.168218426176018 | -0.820367055378223 | 0.416584557878559 | 0.71045022423517 | -5.3329370189779 |
| lnc-FBXO15-6 | lnc-FBXO15-6 | -0.367315982688302 | -0.306828798447548 | -0.820317555244314 | 0.416612457198586 | 0.71045022423517 | -5.26664998609687 |
| lnc-WEE1-3 | lnc-WEE1-3 | -0.198462185314447 | 2.62032518761297 | -0.820087321687407 | 0.41674223669101 | 0.710565787705047 | -5.79979719491121 |
| ENSG00000270804.1 | UBE2CP5 | 0.143350631218168 | 1.19969985039007 | 0.820024606401443 | 0.416777592715164 | 0.710565787705047 | -5.62140689363722 |
| ENSG00000197724.10 | PHF2 | 0.0990371508294041 | 4.81936356510155 | 0.820013613392567 | 0.416783790262014 | 0.710565787705047 | -6.03803333464527 |
| ENSG00000132604.10 | TERF2 | -0.123839985846988 | 4.18326217056723 | -0.819786043157282 | 0.41691210055513 | 0.7107256721655 | -5.99019911458572 |
| ENSG00000143630.9 | HCN3 | -0.112321182548565 | 1.42248214571618 | -0.819700755854408 | 0.416960194072171 | 0.7107487929265 | -5.67053790973506 |
| lnc-LAMA4-3 | lnc-LAMA4-3 | -0.180158171226686 | 0.146676932423228 | -0.819595143503665 | 0.417019753580513 | 0.710785150467997 | -5.41774226592315 |
| ENSG00000151687.14 | ANKAR | -0.109145462443496 | 4.78377615677274 | -0.819540469979488 | 0.417050588456591 | 0.710785150467997 | -6.03087833868525 |
| lnc-SLC9A3-1 | lnc-SLC9A3-1 | -0.199525890169999 | 2.23204656374742 | -0.819429020074115 | 0.417113448493324 | 0.710833425263958 | -5.77118307306885 |
| lnc-CEP83-5 | lnc-CEP83-5 | -0.188471466980527 | 0.251081750971583 | -0.819353719075082 | 0.417155923078652 | 0.710846954609478 | -5.43445925391011 |
| lnc-RRN3-3 | lnc-RRN3-3 | 0.150549719253683 | 0.750346438286125 | 0.819265837939518 | 0.417205497005766 | 0.710853055665803 | -5.55266296208667 |
| ENSG00000175220.11 | ARHGAP1 | -0.0741646116241679 | 5.33415994815256 | -0.819224928545585 | 0.417228575304794 | 0.710853055665803 | -6.06268456864996 |
| lnc-KAT7-2 | lnc-KAT7-2 | -0.217791300509337 | 0.926975818796668 | -0.819110430375599 | 0.417293171531778 | 0.710904266647789 | -5.58480057394907 |
| lnc-HNRNPD-1 | lnc-HNRNPD-1 | -0.345837838667314 | -0.759078895037705 | -0.818927264471378 | 0.41739652060534 | 0.710977941581098 | -5.2522516527693 |
| ENSG00000259399.1 | TGIF2-RAB5IF | 0.126830253160922 | 2.94471565357289 | 0.818911348161217 | 0.417405501922265 | 0.710977941581098 | -5.85405934868576 |
| ENSG00000166762.17 | CATSPER2 | -0.171722733597595 | 2.81388432805314 | -0.818838689841846 | 0.417446503337443 | 0.71098894316877 | -5.82472467202642 |
| ENSG00000051382.8 | PIK3CB | 0.164574873755232 | 3.84647351592975 | 0.818762141606167 | 0.417489702508834 | 0.711003685813282 | -5.94789436069756 |
| lnc-HLA-DPB1-13 | lnc-HLA-DPB1-13 | 0.311985633176056 | -0.452343643701989 | 0.818632050314526 | 0.417563124384627 | 0.711069892380969 | -5.16525207195204 |
| ENSG00000186767.6 | SPIN4 | -0.188945536109901 | 3.86457238809709 | -0.818540872275386 | 0.41761458881057 | 0.711078085444752 | -5.95116492079151 |
| lnc-GRAMD2A-2 | lnc-GRAMD2A-2 | -0.0943904999832326 | 1.48602375815488 | -0.818484621310608 | 0.417646340969924 | 0.711078085444752 | -5.66337064712121 |
| ENSG00000111737.11 | RAB35 | -0.0834768911884591 | 3.92453801101551 | -0.81839238714661 | 0.417698407867114 | 0.711078085444752 | -5.96782819507742 |
| ENSG00000013016.15 | EHD3 | 0.201543697158546 | 5.7563558297707 | 0.818378523764395 | 0.417737939308815 | 0.711078085444752 | -6.07380805871966 |
| ENSG00000200469.1 | RNU6-112P | 0.196607088722668 | 0.0365916350112426 | 0.818317526960512 | 0.417740669931397 | 0.711078085444752 | -5.37500293865889 |
| lnc-CLDN10-7 | lnc-CLDN10-7 | 0.223166449203002 | 0.297863919297444 | 0.818113944616115 | 0.417855614848031 | 0.711146641737604 | -5.40542463575146 |
| ENSG00000127585.11 | FBXL16 | -0.327175183307677 | -0.600195236447678 | -0.81807130281842 | 0.417879693337756 | 0.711146641737604 | -5.25366962080614 |
| lnc-ADSS-1 | lnc-ADSS-1 | 0.104483616857699 | 1.28110467046632 | 0.818062621291467 | 0.417884595628252 | 0.711146641737604 | -5.65329378828869 |
| ENSG00000116095.10 | PLEKHA3 | 0.106131261002361 | 5.24948156433257 | 0.817822936579883 | 0.418019954807582 | 0.711257279906098 | -6.06304701517356 |
| HDAC2-AS2 | HDAC2-AS2 | -0.228203062658437 | -0.544756252357664 | -0.817787969672333 | 0.418039704200805 | 0.711257279906098 | -5.18465567662034 |
| ENSG00000230291.5 | | 0.589570435815055 | -0.778578447981722 | 0.817763941623001 | 0.418053275630568 | 0.711257279906098 | -5.04627888101897 |
| ENSG00000172831.11 | CES2 | -0.0909532591043449 | 4.45702478204194 | -0.817594549393571 | 0.418148958718136 | 0.711361270955897 | -6.01566264421272 |

| | | | | | | | |
|---|---|---|---|---|---|---|---|
| ENSG00000171159.4 | BBLN | -0.161715211973911 | 3.31302231500887 | -0.817531494832835 | 0.418184579199838 | 0.711363073857047 | -5.90815655191065 |
| lnc-SLC46A2-4 | lnc-SLC46A2-4 | 0.100552682582577 | 1.48718029263037 | 0.817451589076406 | 0.418229721836825 | 0.711374059095135 | -5.67228062186051 |
| ENSG00000034677.12 | RNF19A | 0.215395464140195 | 7.36317865996746 | 0.817483001250759 | 0.418260159508785 | 0.711374059095135 | -6.0715450150481 |
| ENSG00000151743.10 | AMN1 | -0.129988756382782 | 4.60061020441 | -0.817326484738328 | 0.418300405312529 | 0.711383726559348 | -6.01991489131082 |
| ENSG00000267793.1 | | -0.936390903911319 | -1.93606824390459 | -0.817083944764414 | 0.41844637849186 | 0.711514399710695 | -4.83265143936022 |
| lnc-EIF1AY-4 | lnc-EIF1AY-4 | -0.936390903911319 | -1.93606824390459 | -0.817083944764414 | 0.41844637849186 | 0.711514399710695 | -4.83265143936022 |
| ENSG00000178896.8 | EXOSC4 | -0.185015379213998 | 0.654477877709928 | -0.816776095138555 | 0.41861145984807 | 0.711636451502181 | -5.56026211413531 |
| ENSG00000221968.8 | FADS3 | -0.272772001707094 | 4.55245688730778 | -0.816945879686946 | 0.418616575306844 | 0.711636451502181 | -6.04010619373787 |
| ENSG00000258727.1 | AP1G2-AS1 | -0.131954051371185 | 3.94996688843209 | -0.816709084975314 | 0.418649340492584 | 0.711636451502181 | -5.9573103428464 |
| ENSG00000135114.12 | OASL | 0.397369574920984 | -0.65433537369562 | 0.816678895091361 | 0.418666407426175 | 0.711636451502181 | -5.20983504176222 |
| ENSG00000175279.21 | CENPS | 0.180811115591363 | -0.328193209366088 | 0.816635344953035 | 0.418691027920683 | 0.711636451502181 | -5.26551123357936 |
| ENSG00000214787.9 | MS4A4E | 1.48500442248719 | -0.059233034791119 | 0.816787955494718 | 0.41877831953319 | 0.71168733243722 | -5.10368794594136 |
| ENSG00000129055.12 | ANAPC13 | 0.0895414107743706 | 4.08792611924574 | 0.816428192058343 | 0.418808151149685 | 0.71168733243722 | -5.9785571223854 |
| ENSG00000128016.5 | ZFP36 | 0.12914213832387 | 6.96565694405977 | 0.81629908005396 | 0.418881160541523 | 0.71168733243722 | -6.07860287085238 |
| ENSG00000169981.10 | ZNF35 | -0.169202748669839 | 3.77812733063479 | -0.81629813566847 | 0.418881694594875 | 0.71168733243722 | -5.94832317146279 |
| ENSG00000254979.5 | | -0.176780701797174 | 1.00344762618875 | -0.816276648083704 | 0.418893846012579 | 0.71168733243722 | -5.59409581594964 |
| ENSG00000184752.12 | NDUFA12 | 0.133893154353387 | 3.49723032780345 | 0.816007969835927 | 0.419045804051535 | 0.711886743545126 | -5.91588302460162 |
| lnc-NR5A1-6 | lnc-NR5A1-6 | 0.11210711354591 | 2.25904063637524 | 0.815883895780654 | 0.419115988725269 | 0.711947214321383 | -5.7714803392198 |
| ENSG00000175518.6 | UBQLNL | -0.185748307883344 | 0.04097532392216 | -0.815780312623086 | 0.419174587846052 | 0.711987996503984 | -5.41079307493368 |
| ENSG00000091039.16 | OSBPL8 | 0.099318733879703 | 6.62901223550243 | 0.815675580140351 | 0.419233842237337 | 0.712023886922956 | -6.08091067119416 |
| ENSG00000104299.14 | INTS9 | -0.0988288313120167 | 3.89994195062704 | -0.815620682713275 | 0.419264903537202 | 0.712023886922956 | -5.96231806436603 |
| lnc-HEY1-2 | lnc-HEY1-2 | 0.250453040079048 | 0.52407281482713 | 0.815506615528997 | 0.41932944792915 | 0.712037609352562 | -5.47163066947841 |
| ENSG00000120071.14 | KANSL1 | 0.111213244575921 | 7.28869735617658 | 0.815425266115311 | 0.419375482817483 | 0.712037609352562 | -6.07515505701852 |
| lnc-SLC30A9-1 | lnc-SLC30A9-1 | -0.151387174444852 | 0.447291550731225 | -0.815385098352121 | 0.41939821452183 | 0.712037609352562 | -5.48150793681256 |
| ENSG00000155729.12 | KCTD18 | -0.0620246455746747 | 4.74338506050608 | -0.815361874452984 | 0.419411357712212 | 0.712037609352562 | -6.03529715198803 |
| ENSG00000160014.16 | CALM3 | -0.15435785632098 | 5.41086678387504 | -0.815274010539685 | 0.419461085143389 | 0.712063300247949 | -6.06932637754118 |
| lnc-LSM3-5 | lnc-LSM3-5 | -0.175905042951207 | 0.0911047767405945 | -0.815184523270314 | 0.419511735018669 | 0.712090552687552 | -5.39562122972818 |
| ENSG00000162066.15 | AMDHD2 | 0.138880519672832 | 4.41106009131041 | 0.815114983253984 | 0.419551097301871 | 0.712097600453415 | -6.01416665003697 |
| ENSG00000166197.16 | NOLC1 | 0.116464410033461 | 5.72681761422534 | 0.815023385431683 | 0.419602948570296 | 0.712097600453415 | -6.07756200972241 |
| ENSG00000078142.12 | PIK3C3 | 0.0651465244026017 | 6.10113017912896 | 0.814993836661296 | 0.419619676232788 | 0.712097600453415 | -6.08162367926558 |
| ENSG00000100413.16 | POLR3H | -0.113799288092686 | 3.69608226709678 | -0.8143100558464 | 0.420006880335907 | 0.712695928932727 | -5.94432432906715 |
| ENSG00000112584.13 | FAM120B | 0.102325115685099 | 5.33298834816223 | 0.814094685079763 | 0.420128883207151 | 0.712820148229781 | -6.06599086876102 |
| lnc-ZNF2-1 | lnc-ZNF2-1 | -0.189746662124194 | 0.104022844637253 | -0.813998613202782 | 0.420183312797035 | 0.712820148229781 | -5.39781417741153 |
| lnc-LRRC47-78 | lnc-LRRC47-78 | -0.11589044791458 | 2.84802326511071 | -0.813931174197261 | 0.420221522975992 | 0.712820148229781 | -5.84298596817856 |
| ENSG00000163808.16 | KIF15 | 0.266524358025494 | 1.75752505390916 | 0.813867717243638 | 0.420258631174065 | 0.712820148229781 | -5.68329473886285 |
| lnc-ITGB1-6 | lnc-ITGB1-6 | -0.57937312426532 | -1.24690348816038 | -0.81381880469821 | 0.420285194960638 | 0.712820148229781 | -5.05302229175148 |
| lnc-FARSB-1 | lnc-FARSB-1 | -0.157476644016732 | 2.12034295567973 | -0.8138140761889 | 0.420287874406871 | 0.712820148229781 | -5.76859794696474 |
| ENSG00000145246.13 | ATP10D | -0.212955272764753 | 5.36520074959116 | -0.813459745605983 | 0.420531472337208 | 0.713050758970827 | -6.06478600677995 |
| lnc-NENF-1 | lnc-NENF-1 | 0.200485359906416 | -0.194156842867655 | 0.813289705925841 | 0.420585077325262 | 0.713050758970827 | -5.31305407704518 |
| lnc-ZNF587B-2 | lnc-ZNF587B-2 | 0.113769763065296 | 1.68899002384818 | 0.813253294726398 | 0.420605719232837 | 0.713050758970827 | -5.70357079642239 |
| ENSG00000181220.16 | ZNF746 | 0.0766179363258006 | 4.40875523673968 | 0.813236606023692 | 0.420615180446986 | 0.713050758970827 | -6.0127674843764 |
| ENSG00000038210.12 | PI4K2B | 0.102192336412668 | 4.2973783874397 | 0.813215485769477 | 0.420627154196398 | 0.713050758970827 | -6.01079038833143 |
| ENSG00000147118.11 | ZNF182 | -0.0859725769919182 | 3.75752503799888 | -0.813167882694194 | 0.420654142666391 | 0.713050758970827 | -5.94330878419373 |
| ENSG00000184481.16 | FOXO4 | -0.124858489867936 | 3.7702455401444 | -0.813146361570616 | 0.420666344370731 | 0.713050758970827 | -5.95093047323933 |
| ENSG00000134490.13 | TMEM241 | -0.133919534663676 | 1.55238292081858 | -0.812981691702938 | 0.420759713396067 | 0.713150294864233 | -5.68237921177327 |
| ENSG00000213149.3 | CNN2P9 | 0.181007899338585 | 0.00759469816091575 | 0.81280761953164 | 0.420858427294917 | 0.713258617941209 | -5.38046084820997 |
| ENSG00000143476.17 | DTL | 0.311332769978799 | 0.371179062477433 | 0.812729865564752 | 0.420902525032834 | 0.713258617941209 | -5.45637576293765 |
| lnc-TRAPPC9-1 | lnc-TRAPPC9-1 | 0.168779163752138 | 0.201424866764536 | 0.812626662025129 | 0.420961060695083 | 0.713258617941209 | -5.42987391913646 |
| ENSG00000285994.1 | | 0.244802600059267 | 1.76411935059987 | 0.812624591047787 | 0.420962235376272 | 0.713258617941209 | -5.69696177386136 |
| lnc-RRAS-1 | lnc-RRAS-1 | 0.243441803602527 | 4.69671113555309 | 0.812607866479632 | 0.421043471741398 | 0.71333754044392 | -6.03251569226255 |
| ENSG00000233822.4 | H2BC15 | 0.335934649329677 | 4.11146022108803 | 0.812220238701258 | 0.421318495761921 | 0.713656258256829 | -6.00228031234635 |
| MHENCR | MHENCR | 0.106321293739317 | 3.93001963735622 | 0.811956596767125 | 0.421341233058893 | 0.713656258256829 | -5.95012357082656 |
| lnc-CRYBA1-1 | lnc-CRYBA1-1 | 0.206538864569952 | -0.157251524050951 | 0.811831853779326 | 0.421412031036474 | 0.713656258256829 | -5.33160292948685 |
| ENSG00000197081.13 | IGF2R | 0.0931497020336529 | 6.70799499506745 | 0.81181992743798 | 0.421418800219073 | 0.713656258256829 | -6.08393825356422 |
| lnc-GREB1L-2 | lnc-GREB1L-2 | -0.205414791330094 | -0.283862681208399 | -0.8117944383235 | 0.421433267615763 | 0.713656258256829 | -5.32032647038802 |
| ENSG00000119777.19 | TMEM214 | 0.115128787381912 | 4.53154221029605 | 0.811783236333963 | 0.421439625861719 | 0.713656258256829 | -6.01911157416587 |
| ENSG00000168653.10 | NDUFS5 | 0.124026078860434 | 4.30665455451607 | 0.811541746004176 | 0.421576709871039 | 0.713829665948963 | -6.00802840705487 |
| lnc-MRPL40-1 | lnc-MRPL40-1 | -0.143546526517793 | 0.640480848226884 | -0.811474037693107 | 0.42161514992215 | 0.71383603116292 | -5.53065899906581 |
| ENSG00000169733.11 | RFNG | -0.163209185564267 | 2.43144120556467 | -0.811323669433663 | 0.421700526133461 | 0.713921856341917 | -5.79415670016264 |
| ENSG00000203875.11 | SNHG5 | 0.130491182396945 | 4.92737189501809 | 0.81115071537614 | 0.421798739109993 | 0.713941841437494 | -6.04869922312863 |
| ENSG00000153832.11 | FBXO36 | 0.171178473390219 | 0.68712096006677 | 0.811129715943171 | 0.421810664706783 | 0.713941841437494 | -5.5481200584905 |
| lnc-PDLIM1-2 | lnc-PDLIM1-2 | 0.301650299528276 | 0.0793283179615057 | 0.811034918586954 | 0.421864502758599 | 0.713941841437494 | -5.35608067057037 |
| ENSG00000131966.13 | ACTR10 | -0.0810509271868334 | 4.45857015822261 | -0.810989265923736 | 0.421890431662622 | 0.713941841437494 | -6.02234635779459 |
| ENSG00000198885.9 | ITPRIPL1 | -0.180075608718039 | 2.31151958328823 | -0.810971298075964 | 0.42190063695366 | 0.713941841437494 | -5.76754944858973 |
| lnc-C3orf38-2 | lnc-C3orf38-2 | -0.126156662850805 | -0.185952092437813 | -0.810936420577124 | 0.421920446934153 | 0.713941841437494 | -5.33211259489284 |
| ENSG00000111886.10 | GABRR2 | 0.272014834212039 | 0.728701468917675 | 0.810805742881234 | 0.421994675243113 | 0.713982770106259 | -5.49634499102831 |
| ENSG00000180694.13 | TMEM64 | -0.213943543727301 | 4.32251657394484 | -0.810813095326865 | 0.422020614700652 | 0.713982770106259 | -6.01020101691806 |
| ENSG00000176595.3 | KBTBD11 | 0.964243015555504 | -0.715543864334747 | 0.810950175422734 | 0.422048698629104 | 0.713982770106259 | -5.15217569216581 |

| | | | | | | | |
|---|---|---|---|---|---|---|---|
| ENSG00000197635.9 | DPP4 | -0.793057378215397 | -0.587045196928372 | -0.810537782246206 | 0.42227959813912 | 0.714194090182262 | -5.28464212434638 |
| ENSG00000165138.17 | ANKS6 | -0.12803399256876 | 3.91582018966345 | -0.810294785053142 | 0.422284988592961 | 0.714194090182262 | -5.9653673265358 |
| ENSG00000186522.14 | SEPTIN10 | -1.77146680657256 | 0.554487906427651 | -0.810609266814792 | 0.422285317833195 | 0.714194090182262 | -5.39687993919269 |
| ENSG00000143537.13 | ADAM15 | 0.198563634382478 | 2.41325197665291 | 0.810174071957132 | 0.42235359242612 | 0.714194090182262 | -5.79296766500275 |
| lnc-FAM208B-5 | lnc-FAM208B-5 | -0.139275730978364 | 2.38100873064947 | -0.810163363773356 | 0.422359678442473 | 0.714194090182262 | -5.79234335021353 |
| ENSG00000278970.1 | HEIH | -0.106221250939332 | 3.24807067491476 | -0.810124436253685 | 0.422381803419748 | 0.714194090182262 | -5.89848873474598 |
| ENSG00000144199.11 | FAHD2B | -0.228387368489629 | 1.40162572271727 | -0.809860872510355 | 0.422531621904558 | 0.714388727916997 | -5.65369316823443 |
| ENSG00000123136.14 | DDX39A | 0.109792804885398 | 5.83449168531454 | 0.80969750092787 | 0.422624503984233 | 0.714487077249073 | -6.08387005962662 |
| lnc-NREP-1 | lnc-NREP-1 | 0.279496691437541 | 0.743123610035846 | 0.80958037762789 | 0.422691100027573 | 0.714499924237414 | -5.57545838231099 |
| ENSG00000275307.1 | | 0.108155445958453 | 2.89077938445877 | 0.809562036561179 | 0.422701529293057 | 0.714499924237414 | -5.83719862871063 |
| ENSG00000174744.13 | BRMS1 | 0.0885130067344146 | 4.89797894300684 | 0.809443767773083 | 0.422768784123966 | 0.714508098476768 | -6.05143997924474 |
| lnc-TIGD5-1 | lnc-TIGD5-1 | 0.191964990750258 | 0.25764944864725 | 0.809431444563872 | 0.422775792224096 | 0.714508098476768 | -5.46103179075981 |
| ENSG00000105373.18 | NOP53 | -0.103492970332944 | 7.42795021041318 | -0.809314074358903 | 0.422842543153463 | 0.714562238565496 | -6.07747739135149 |
| ENSG00000014123.9 | UFL1 | 0.0906486522928258 | 6.1717905169287 | 0.809187900079235 | 0.42291430827594 | 0.714589267561996 | -6.0870602540542 |
| ENSG00000101017.13 | CD40 | -0.152241585464965 | 5.36610300149272 | -0.809151162840767 | 0.422935204988033 | 0.714589267561996 | -6.07373503489259 |
| ENSG00000150753.11 | CCT5 | -0.156778757033995 | 5.49052499999055 | -0.809048932490293 | 0.422993358495473 | 0.714589267561996 | -6.0777299343531 |
| ENSG00000277468.1 | | -0.334798990727206 | 0.488731564547729 | -0.809048906246447 | 0.422997407373821 | 0.714589267561996 | -5.49191065232325 |
| lnc-CCDC198-3 | lnc-CCDC198-3 | 0.109739213799106 | 1.04597492684586 | 0.808911137263316 | 0.42307175067584 | 0.714656203870399 | -5.61046957659852 |
| ENSG00000185896.10 | LAMP1 | 0.0710544724317526 | 6.06452284552001 | 0.808666959534838 | 0.423210685862327 | 0.714780096610381 | -6.08658134117842 |
| ENSG00000177697.18 | CD151 | -0.238102616349589 | 2.03398974143717 | -0.808566264114599 | 0.423267988810429 | 0.714780096610381 | -5.74070338609829 |
| lnc-ANKRD42-1 | lnc-ANKRD42-1 | -0.149077824003277 | 1.9895011712276 | -0.808447702902056 | 0.423335464710249 | 0.714780096610381 | -5.74609765498945 |
| lnc-FAM3B-7 | lnc-FAM3B-7 | 0.310560294970317 | 0.286817148474088 | 0.80820510980157 | 0.4234735503147 | 0.714780096610381 | -5.3967548402067 |
| ENSG00000106524.8 | ANKMY2 | -0.118609120351728 | 3.87368342161821 | -0.808200753859635 | 0.423476029995239 | 0.714780096610381 | -5.96666225746944 |
| ENSG00000111196.9 | MAGOHB | -0.118735869463178 | 4.32767748780254 | -0.808195885063479 | 0.42347880163559 | 0.714780096610381 | -6.01267784105143 |
| ENSG00000151135.9 | TMEM263 | -0.130668471459904 | 5.13382638727444 | -0.80818970157131 | 0.423482321703595 | 0.714780096610381 | -6.0676870518283 |
| ENSG00000279328.1 | | -0.308560127030111 | -0.508855270301432 | -0.808162937340306 | 0.423497557944715 | 0.714780096610381 | -5.22009574739336 |
| ENSG00000185596.16 | WASH3P | 0.123004966504893 | 3.95890053768319 | 0.808139789431047 | 0.423510735767657 | 0.714780096610381 | -5.98227358981458 |
| lnc-EMP3-1 | lnc-EMP3-1 | -0.201108369174521 | 0.141099284060104 | -0.808121307931858 | 0.423521257238559 | 0.714780096610381 | -5.3856940289529 |
| ENSG00000110057.7 | UNC93B1 | 0.118330622428445 | 5.66310747901051 | 0.808111065514129 | 0.423527088289375 | 0.714780096610381 | -6.08192681054668 |
| ENSG00000132357.13 | CARD6 | 0.951450541807225 | -0.409461337548264 | 0.807815805378451 | 0.423821295221786 | 0.714990254350355 | -5.10415199321638 |
| ENSG00000181904.8 | C5orf24 | -0.0837748134188567 | 5.04029902800297 | -0.80758609433232 | 0.423826021704179 | 0.714990254350355 | -6.06313534828771 |
| lnc-NXPE3-2 | lnc-NXPE3-2 | -0.238753749596388 | -0.489816075285543 | -0.807480875788353 | 0.423885951487419 | 0.714990254350355 | -5.25614882946528 |
| ENSG00000111364.15 | DDX55 | 0.0791863135545938 | 4.64735590619121 | 0.807452192885613 | 0.423902289423328 | 0.714990254350355 | -6.03710442335068 |
| lnc-MUC20-67 | lnc-MUC20-67 | 0.235214556436991 | 5.24848835879839 | 0.807556071329 | 0.423916648212803 | 0.714990254350355 | -6.07096069005029 |
| ENSG00000197837.3 | H4C16 | 0.121506780813752 | 4.68298765445116 | 0.807343822426352 | 0.423964021263795 | 0.714990254350355 | -6.04231762007978 |
| ENSG00000126107.14 | HECTD3 | -0.101008665262831 | 4.804368886623 | -0.807323009925428 | 0.423975877460943 | 0.714990254350355 | -6.04717838435501 |
| ENSG00000173588.14 | CEP83 | 0.124498042112874 | 3.44007059119715 | 0.807315104109667 | 0.42398038119675 | 0.714990254350355 | -5.91577658748753 |
| lnc-FAM49B-4 | lnc-FAM49B-4 | -0.170498487397782 | 0.039906515506993 | -0.807297563862095 | 0.423990373519037 | 0.714990254350355 | -5.36996959313662 |
| ENSG00000122986.13 | HVCN1 | -0.16026263281783 | 6.2035680781073 | -0.807282453134775 | 0.42399898190742 | 0.714990254350355 | -6.08815044283734 |
| lnc-H3F3C-5 | lnc-H3F3C-5 | -0.224498659759811 | -0.47135954631371 | -0.807086611710822 | 0.424110559842108 | 0.715119820859352 | -5.25608277292026 |
| lnc-MYOCOS-1 | lnc-MYOCOS-1 | 0.158822841230008 | 1.68877215937604 | 0.807015453618249 | 0.424151105582157 | 0.715129604046325 | -5.70196359290169 |
| ENSG00000107518.17 | ATRNL1 | 0.441913149111158 | -0.926529089807598 | 0.806935026352694 | 0.424196935704389 | 0.715143400241407 | -5.19914787659917 |
| ENSG00000100387.8 | RBX1 | 0.0953277480525561 | 4.55927149389258 | 0.806804696586109 | 0.42427120829242 | 0.715143400241407 | -6.03083914560021 |
| lnc-EIF4E3-6 | lnc-EIF4E3-6 | -0.219461194059721 | 1.9706811012714 | -0.806769399251598 | 0.424291324962956 | 0.715143400241407 | -5.73640518272933 |
| lnc-CARS-2 | lnc-CARS-2 | -0.79257723568687 | -2.54689545258506 | -0.806757220982038 | 0.424298265741051 | 0.715143400241407 | -4.84865899654684 |
| ENSG00000284252.1 | MIR4444-1 | 0.14164911717456 | 0.885027632002189 | 0.806561997086981 | 0.42440953935056 | 0.715272377667452 | -5.57961092415668 |
| lnc-HSD17B12-1 | lnc-HSD17B12-1 | -0.159027598573358 | 0.448287879181554 | -0.806404094852713 | 0.424499553302005 | 0.715365507255213 | -5.51827898515667 |
| lnc-NLRP3-1 | lnc-NLRP3-1 | -0.153789603606865 | 1.79955121857169 | -0.806204622699597 | 0.424613281173936 | 0.715498580958094 | -5.72527171942708 |
| ENSG00000012211.12 | PRICKLE3 | -0.164764334591238 | 1.14170052832853 | -0.806101189684986 | 0.424672260156977 | 0.715539385298874 | -5.62956079863271 |
| ENSG00000140044.12 | JDP2 | -0.185345520855177 | 3.15288327104663 | -0.805870547629172 | 0.424803793405807 | 0.715622368772641 | -5.8685145623266 |
| ENSG00000124942.13 | AHNAK | -0.108910558782954 | 11.8288223244501 | -0.805859396093315 | 0.424810153658798 | 0.715622368772641 | -5.94701033681006 |
| lnc-RDH13-1 | lnc-RDH13-1 | -0.149352038020785 | 1.47599518278099 | -0.805831939461887 | 0.424825813731375 | 0.715622368772641 | -5.65630027370779 |
| ENSG00000101442.9 | ACTR5 | 0.0852145704524268 | 3.87011794067253 | 0.805760914836219 | 0.424866324720071 | 0.715632042693389 | -5.96805838203854 |
| ENSG00000132530.16 | XAF1 | -0.184674762702221 | 5.34435766355473 | -0.805594585020127 | 0.424973584038716 | 0.715754134708198 | -6.07474870813135 |
| lnc-CYP2C8-2 | lnc-CYP2C8-2 | 0.362162284404753 | -0.256882760194804 | 0.80521375449686 | 0.425178492103576 | 0.71602855781309 | -5.27438321583857 |
| lnc-USP25-1 | lnc-USP25-1 | -0.22381182397266 | 2.49085771053778 | -0.805093225513769 | 0.425247275262942 | 0.71602855781309 | -5.8089990275592 |
| ENSG00000278144.1 | | -0.253676447367502 | 1.85650519713428 | -0.80502768252692 | 0.425289016918895 | 0.71602855781309 | -5.73780094698694 |
| lnc-HSF5-2 | lnc-HSF5-2 | -0.295257439861781 | -0.44397218920182 | -0.804916622297662 | 0.425348070869866 | 0.71602855781309 | -5.22858680072749 |
| ENSG00000255374.3 | TAS2R43 | -0.388800560912907 | 1.07930744624115 | -0.804961759515769 | 0.425390690557232 | 0.71602855781309 | -5.59145990522922 |
| lnc-IMMT-2 | lnc-IMMT-2 | 0.110273917826571 | -0.0545496204348621 | 0.80471171492853 | 0.425465349305586 | 0.71602855781309 | -5.36782442477819 |
| lnc-MYO5B-2 | lnc-MYO5B-2 | -0.14572101012479 | 2.33222433032311 | -0.804697043612125 | 0.425473414707664 | 0.71602855781309 | -5.79383666907917 |
| lnc-TSEN2-3 | lnc-TSEN2-3 | 0.153768970114431 | 1.3845164348467 | 0.804686794006297 | 0.425479266116746 | 0.71602855781309 | -5.67940515184165 |
| lnc-PEX19-6 | lnc-PEX19-6 | -0.0693312947372431 | 3.69031228811169 | -0.804657226131688 | 0.425496146426809 | 0.71602855781309 | -5.94637089354302 |
| ENSG00000268361.1 | | -0.0949976576778201 | 0.229790918543559 | -0.804639824883377 | 0.425506080994885 | 0.71602855781309 | -5.4156929067478 |
| ENSG00000062598.17 | ELMO2 | 0.0855792057185818 | 4.52270016047602 | 0.804592036816574 | 0.425533364460274 | 0.71602855781309 | -6.02969463184683 |
| lnc-RGS9-11 | lnc-RGS9-11 | -0.251463621516643 | 0.0812022142457503 | -0.804555948056082 | 0.425553969184644 | 0.71602855781309 | -5.44241806462278 |
| lnc-PIGBOS1-3 | lnc-PIGBOS1-3 | -0.142870988366855 | 0.294934370435492 | -0.804434027755673 | 0.425623583522646 | 0.716073758105326 | -5.44192491102833 |
| ENSG00000136193.16 | SCRN1 | 0.355801076156845 | 6.5456466450783 | 0.804657644736 | 0.425650412010818 | 0.716073758105326 | -6.08869738260443 |

| | | | | | | | |
|---|---|---|---|---|---|---|---|
| LINC01588 | LINC01588 | 0.180084798273991 | 4.26156365988647 | 0.804156724057313 | 0.425781944669634 | 0.716224410151005 | -5.99202236520594 |
| ENSG00000168894.9 | RNF181 | 0.10302469445473 | 3.92259110400362 | 0.804058629380037 | 0.425837972591279 | 0.716224410151005 | -5.9762135030062 |
| lnc-HECTD1-1 | lnc-HECTD1-1 | 0.117375207140059 | 0.597139157885227 | 0.803960363079406 | 0.425894103004831 | 0.716224410151005 | -5.52141902565867 |
| lnc-ZFP2-3 | lnc-ZFP2-3 | 0.14358139083366 | -0.306675197633758 | 0.80395451357179 | 0.425897444426401 | 0.716224410151005 | -5.28805793328228 |
| lnc-CLN5-4 | lnc-CLN5-4 | -0.213436240846113 | 0.303533555491679 | -0.803923436340709 | 0.425915196977749 | 0.716224410151005 | -5.45740273560377 |
| ENSG00000117597.17 | UTP25 | -0.0899386569409151 | 5.30466976496319 | -0.803855962180587 | 0.425953742443063 | 0.716224410151005 | -6.07589619856091 |
| ENSG00000270179.1 | | 0.269271382520531 | 0.0713128851685449 | 0.803758800519055 | 0.42600925097119 | 0.716224410151005 | -5.329293096583 |
| ENSG00000148399.12 | DPH7 | 0.11429222900189 | 4.35111712195595 | 0.803701446924508 | 0.426042019173051 | 0.716224410151005 | -6.01806276562509 |
| lnc-GPR156-1 | lnc-GPR156-1 | 0.116967358703443 | 2.98174089406962 | 0.803630707359295 | 0.426082437366024 | 0.716224410151005 | -5.86707207638069 |
| lnc-COLEC12-5 | lnc-COLEC12-5 | -0.14080985540507 | 1.12351979444812 | -0.803621091187277 | 0.426087931900066 | 0.716224410151005 | -5.64051036440231 |
| lnc-NEDD8-MDP1-1 | lnc-NEDD8-MDP1- | -0.0746852509388525 | 4.82687847509471 | -0.803403728226408 | 0.426212141200948 | 0.71637469396857 | -6.04888983772241 |
| lnc-TNFRSF11A-5 | lnc-TNFRSF11A-5 | -0.180966432479329 | 1.49006807887958 | -0.803273456266226 | 0.426286593938586 | 0.716441329553189 | -5.6882214036464 |
| ENSG00000226918.1 | | -2.00865348020216 | 1.13953595774007 | -0.803411380281887 | 0.426384997867099 | 0.716548204694521 | -5.26286721307645 |
| ENSG00000170315.13 | UBB | 0.113741731558957 | 7.87987264981183 | 0.802945998778167 | 0.426473776394791 | 0.716638888034451 | -6.07301572968092 |
| ENSG00000176978.13 | DPP7 | -0.130832819652619 | 5.84637700898048 | -0.80284250337179 | 0.426532947142088 | 0.716655999305594 | -6.08945353594433 |
| ENSG00000180902.17 | D2HGDH | -0.1120228423921 | 4.00231504759747 | -0.802806390159063 | 0.426553595078115 | 0.716655999305594 | -5.97948519069716 |
| lnc-FAM124B-4 | lnc-FAM124B-4 | 0.13569935635025 | 3.1214862968807 | 0.802686533177442 | 0.426622128318965 | 0.716696460567422 | -5.88465351717076 |
| ENSG00000213137.3 | ARF1P2 | 0.306208474156357 | -0.31925109842566 | 0.802600443696178 | 0.426671357682044 | 0.716696460567422 | -5.2453889883879 |
| ENSG00000235613.2 | NSRP1P1 | -0.204056915544456 | 1.04399937204906 | -0.802521610786417 | 0.426716440459389 | 0.716696460567422 | -5.596612728217 |
| lnc-C7orf77-4 | lnc-C7orf77-4 | -0.118416365690596 | 0.528567031380769 | -0.802376822855877 | 0.426799248928719 | 0.716696460567422 | -5.49629095504707 |
| ENSG00000184445.11 | KNTC1 | -0.11198102583112 | 7.08933405977961 | -0.802283172814523 | 0.42685281530373 | 0.716696460567422 | -6.08753579157137 |
| lnc-TNNT1-1 | lnc-TNNT1-1 | 0.132801172162474 | 1.40230154017003 | 0.802276551099677 | 0.426856602975805 | 0.716696460567422 | -5.67795217674581 |
| lnc-EAF2-3 | lnc-EAF2-3 | -0.266104415443417 | -0.0341984460553694 | -0.8022717084518 | 0.426859373020666 | 0.716696460567422 | -5.36114101270676 |
| MIR646HG | MIR646HG | -1.42046895204479 | -2.25905747147517 | -0.802396170215394 | 0.426887238767862 | 0.716696460567422 | -4.81104184623243 |
| lnc-EPC2-1 | lnc-EPC2-1 | -0.225898687428177 | -0.513017161832997 | -0.802116419759454 | 0.426948205517623 | 0.716696460567422 | -5.26431613079359 |
| ENSG00000141232.4 | TOB1 | 0.155142003237237 | 2.38024590747764 | 0.802112221861645 | 0.426950607069149 | 0.716696460567422 | -5.80513000820209 |
| ENSG00000197822.10 | OCLN | 0.663001308741539 | -1.75894440327787 | 0.802094587160302 | 0.426960695694395 | 0.716696460567422 | -4.93083653461004 |
| lnc-ZNF736-3 | lnc-ZNF736-3 | 0.131800178305733 | 1.54594257933447 | 0.801958255437284 | 0.427038694480145 | 0.71676893488419 | -5.67502374247609 |
| ENSG00000205981.7 | DNAJC19 | -0.0770973156284667 | 3.59292497389846 | -0.801715328352506 | 0.427177700448016 | 0.71694378737129 | -5.9450963403696 |
| PITPNA-AS1 | PITPNA-AS1 | -0.129569366356182 | 1.41029291725977 | -0.801534903732564 | 0.427280959377892 | 0.717011952370425 | -5.67278306654947 |
| ENSG00000180979.9 | LRRC57 | 0.0725232647844578 | 4.37997684600191 | 0.801522627736321 | 0.4272879856083 | 0.717011952370425 | -6.01990365045658 |
| ENSG00000158089.14 | GALNT14 | 0.313192488312908 | 0.971467790621717 | 0.801320574847748 | 0.427420408519797 | 0.717155816237711 | -5.60746010116155 |
| ENSG00000002822.15 | MAD1L1 | -0.188560433778476 | 4.89809413711767 | -0.801262928597942 | 0.427450583390873 | 0.717155816237711 | -6.06301424595184 |
| ENSG00000116830.11 | TTF2 | -0.0880730552843376 | 4.71362394970564 | -0.801190259765364 | 0.427478244639774 | 0.717155816237711 | -6.04618146717924 |
| ZNF236-DT | ZNF236-DT | -0.139358245713787 | 2.85977615069951 | -0.800909037115567 | 0.427639266207234 | 0.717224748300988 | -5.8550572550499 |
| lnc-RNFT1-2 | lnc-RNFT1-2 | 0.137701179738827 | 1.44502707906316 | 0.800804072480747 | 0.427699375895979 | 0.717224748300988 | -5.68108830566807 |
| ENSG00000256372.1 | | -0.224498659759811 | -0.478252561008829 | -0.800798129423095 | 0.427702779436068 | 0.717224748300988 | -5.26039375617964 |
| ENSG00000076928.17 | ARHGEF1 | 0.109671365844538 | 7.13315798289896 | 0.8007701936749 | 0.42771877822742 | 0.717224748300988 | -6.08912384473173 |
| ENSG00000139641.12 | ESYT1 | -0.135701895574079 | 7.72157082190098 | -0.800754330305779 | 0.427727863332951 | 0.717224748300988 | -6.07978515608819 |
| ENSG00000249624.9 | | 0.108730462868663 | 4.65171631196622 | 0.80075338237084 | 0.42772840622818 | 0.717224748300988 | -6.03732819565047 |
| ENSG00000110925.6 | CSRNP2 | 0.0761802061857431 | 3.69625970549892 | 0.80049489487865 | 0.427876461034114 | 0.717359528727594 | -5.9452486121295 |
| ENSG00000185721.12 | DRG1 | 0.0696368533385288 | 4.64033148308811 | 0.800491355061966 | 0.427878488762021 | 0.717359528727594 | -6.03859194953933 |
| ENSG00000164211.12 | STARD4 | 0.306726269626622 | 3.72972565446702 | 0.800486179856233 | 0.427990642082765 | 0.717448708270317 | -5.97357577239798 |
| lnc-CALM1-7 | lnc-CALM1-7 | -0.217614919918182 | 0.680893452005444 | -0.800204109891343 | 0.428043051916522 | 0.717448708270317 | -5.56373810956793 |
| ENSG00000181274.6 | FRAT2 | 0.155747265725207 | 2.51020527362241 | 0.800173621035725 | 0.428060521263339 | 0.717448708270317 | -5.80831997269084 |
| ENSG00000182952.4 | HMGN4 | 0.0958282813898084 | 5.55649453432147 | 0.800133278418155 | 0.428083637228249 | 0.717448708270317 | -6.08510465412766 |
| ENSG00000257489.6 | | -0.0964374813100131 | 2.29204209108511 | -0.8000943162371 | 0.428105962929409 | 0.717448708270317 | -5.7927284264898 |
| ENSG00000171033.12 | PKIA | 0.291713548098869 | 3.67995490019107 | 0.80010179330062 | 0.428200413587513 | 0.717548572243896 | -5.96316573505231 |
| lnc-TMEM170A-3 | lnc-TMEM170A-3 | -0.126838321575079 | 0.716105501367458 | -0.799839649902435 | 0.428251906450853 | 0.717565186097847 | -5.55073659605949 |
| ENSG00000105612.8 | DNASE2 | 0.184538838137008 | 3.05384947585173 | 0.799780682419469 | 0.428285703657335 | 0.717565186097847 | -5.87521784003582 |
| ENSG00000100359.20 | SGSM3 | -0.101809067516892 | 4.61437425448236 | -0.79972971996981 | 0.428314914074632 | 0.717565186097847 | -6.04069818694203 |
| ENSG00000125743.10 | SNRPD2 | 0.1415186466611 | 5.73461863786024 | 0.799654483925238 | 0.428358039711268 | 0.717579029167171 | -6.08945812390599 |
| lnc-DRD5-9 | lnc-DRD5-9 | -0.667408349924886 | 1.19778776599517 | -0.799759738702089 | 0.428467153146842 | 0.717693192803327 | -5.67876464507299 |
| ENSG00000271793.1 | | -0.070082289228985 | 3.63578435363267 | -0.799413961007434 | 0.428495926072977 | 0.717693192803327 | -5.94962927361368 |
| ENSG00000254665.1 | | -0.220482666605427 | 0.562563323979063 | -0.799186117075772 | 0.428626568517934 | 0.717810012971812 | -5.52422923329105 |
| lnc-RPS6KA3-3 | lnc-RPS6KA3-3 | 0.236346150003092 | 1.50507772809001 | 0.799156593202811 | 0.428643498839318 | 0.717810012971812 | -5.68306726683899 |
| ENSG00000153443.12 | UBALD1 | 0.150693107300133 | 2.44841571127988 | 0.799109866476033 | 0.428670294876817 | 0.717810012971812 | -5.81975477774204 |
| ENSG00000174912.7 | METTL15P1 | 0.202311710285285 | -0.30702151030614 | 0.799015719088502 | 0.428724287961536 | 0.717842025470779 | -5.27350706739135 |
| ENSG00000168237.17 | GLYCTK | 0.12555534890002 | 3.7427447567949 | 0.798872620069626 | 0.428806362420863 | 0.717921047479147 | -5.95165831687056 |
| ENSG00000111371.15 | SLC38A1 | 0.0957852356975142 | 9.25480445261703 | 0.79874358277119 | 0.428880379891616 | 0.71793468755336 | -6.04241894784834 |
| ENSG00000278272.1 | NA | -0.498072605145354 | 1.14195020186266 | -0.798953905910424 | 0.428884269453244 | 0.71793468755336 | -5.61238477483403 |
| lnc-KIAA1551-14 | lnc-KIAA1551-14 | -0.20844047310528 | 0.546608293436614 | -0.798465803643265 | 0.429039743713955 | 0.718109599438511 | -5.54920407010199 |
| ENSG00000275994.1 | SNORA24 | -0.130783092211853 | 1.30846327952898 | -0.798401559933936 | 0.429076605853613 | 0.718109599438511 | -5.66100568980365 |
| ENSG00000183579.15 | ZNRF3 | 0.21015956106909 | 1.02615205528627 | 0.798372248737443 | 0.429093424840609 | 0.718109599438511 | -5.62328654221813 |
| ENSG00000127483.17 | HP1BP3 | 0.066267139606742 | 7.69376426950793 | 0.798253009066687 | 0.429161849560027 | 0.71816571947106 | -6.08214762678924 |
| lnc-KDM5D-4 | lnc-KDM5D-4 | -0.927542126641443 | -1.95083466805018 | -0.798160704549042 | 0.429223507631556 | 0.718177681408557 | -4.83371613837768 |
| lnc-LRIF1-1 | lnc-LRIF1-1 | 0.218156677042198 | 0.800641805457477 | 0.798061482838217 | 0.429271769082514 | 0.718177681408557 | -5.56703189775627 |
| lnc-TCF21-3 | lnc-TCF21-3 | -0.137399250853667 | 0.327706942282696 | -0.798058165415184 | 0.429273673146261 | 0.718177681408557 | -5.49375444609836 |

| | | | | | | | |
|---|---|---|---|---|---|---|---|
| ENSG00000154328.15 | NEIL2 | 0.147029164229052 | 2.74501138625237 | 0.797779045350718 | 0.429433894736561 | 0.718263610802677 | -5.84174190712478 |
| ENSG00000148158.16 | SNX30 | 0.1060059517569 | 5.00126955923809 | 0.797775046138813 | 0.429436190641287 | 0.718263610802677 | -6.0635777247008 |
| ENSG00000065534.18 | MYLK | -1.28286565522374 | 2.58017326801796 | -0.797998362615073 | 0.429483876697333 | 0.718263610802677 | -5.87074200873893 |
| ENSG00000153006.15 | SREK1IP1 | 0.0910669986138108 | 4.13219675100609 | 0.797655097985372 | 0.429505055024847 | 0.718263610802677 | -6.00334749817112 |
| ENSG00000138834.12 | MAPK8IP3 | 0.14528569404869 | 7.46766499725908 | 0.79763043387339 | 0.429519215956979 | 0.718263610802677 | -6.08638248346979 |
| ENSG00000196712.17 | NF1 | 0.129910194315159 | 5.77966840617448 | 0.797512070731098 | 0.429587178218955 | 0.718263610802677 | -6.09284933608257 |
| ENSG00000196126.11 | HLA-DRB1 | -0.170054623610206 | 9.27514365628108 | -0.797508713752788 | 0.42958910583766 | 0.718263610802677 | -6.0441367036052 |
| lnc-PERP-2 | lnc-PERP-2 | -0.147090354166411 | 2.25699684130871 | -0.797482421930684 | 0.429604203108962 | 0.718263610802677 | -5.78576920975518 |
| ENSG00000095906.16 | NUBP2 | -0.131214476684392 | 3.61785844269277 | -0.797297170251408 | 0.429710587244338 | 0.718372883259031 | -5.95094686133277 |
| lnc-EPGN-4 | lnc-EPGN-4 | 0.178520271868963 | 0.897655410964551 | 0.797247066172304 | 0.429739363142809 | 0.718372883259031 | -5.60773531748181 |
| ENSG00000159403.16 | C1R | 0.187341062443586 | 0.845230013046362 | 0.797137623876227 | 0.429802222343964 | 0.718400734042466 | -5.57350949388291 |
| ENSG00000105518.13 | TMEM205 | -0.128618613865345 | 3.03038854172853 | -0.797091322804643 | 0.42982881746009 | 0.718400734042466 | -5.87239946095857 |
| ENSG00000129493.14 | HEATR5A | -0.100166382700022 | 4.1030179793937 | -0.797035763627912 | 0.429860731694457 | 0.718400734042466 | -5.99406405514896 |
| ENSG00000120742.10 | SERP1 | 0.102247809758357 | 6.11264429407838 | 0.796911557363255 | 0.429932083259659 | 0.718431461032449 | -6.09661226167875 |
| lnc-MSS51-1 | lnc-MSS51-1 | 0.178132827120227 | -0.0840478870876395 | 0.796882240509311 | 0.429948925666701 | 0.718431461032449 | -5.37144587517709 |
| lnc-LAMTOR5-1 | lnc-LAMTOR5-1 | -0.752651168389645 | -0.299376851781507 | -0.797009610358967 | 0.429985066866276 | 0.71843352798998 | -5.20119117307807 |
| ENSG00000100811.12 | YY1 | 0.0682839231742669 | 6.48305420749584 | 0.796706048595742 | 0.430050155524906 | 0.718470468538033 | -6.09681410819252 |
| LINC00476 | LINC00476 | 0.0999423439055818 | 4.85952996775472 | 0.796540659311098 | 0.430145191846829 | 0.718470468538033 | -6.06055079320772 |
| ENSG00000125740.13 | FOSB | -0.745013655493371 | 4.92467558754472 | -0.796835212042268 | 0.430151531915318 | 0.718470468538033 | -6.05516247802471 |
| ENSG00000134250.19 | NOTCH2 | 0.248376753749248 | 7.36557923217536 | 0.7966189604145 | 0.430193753368275 | 0.718470468538033 | -6.08899572505401 |
| ENSG00000159063.12 | ALG8 | 0.0841590409393009 | 3.46444193037103 | 0.796429508946734 | 0.43020906838633 | 0.718470468538033 | -5.92371276574639 |
| ENSG00000156531.16 | PHF6 | -0.0819014298612482 | 5.91153607486213 | -0.79641638310714 | 0.430216611996855 | 0.718470468538033 | -6.09472375096921 |
| ENSG00000113108.19 | APBB3 | 0.190486660835895 | 4.05159476671278 | 0.795788526100133 | 0.430578666784937 | 0.719016769303452 | -5.98846474393365 |
| ENSG00000018510.14 | AGPS | -0.0907008228476052 | 4.98590768710358 | -0.795509659593588 | 0.430737911633096 | 0.71920898849529 | -6.06924836100539 |
| lnc-CDK2AP1-2 | lnc-CDK2AP1-2 | 0.12403013737949 | 0.340283304110795 | 0.79545907853374 | 0.430767003211985 | 0.71920898849529 | -5.48084297933895 |
| lnc-TMED4-1 | lnc-TMED4-1 | 0.0979934974747021 | 1.97682330741011 | 0.795332464300733 | 0.430839830269445 | 0.71920898849529 | -5.75210613124807 |
| lnc-SEMA3B-1 | lnc-SEMA3B-1 | -0.188242370744683 | -0.189699810124068 | -0.795265139783633 | 0.43087855757316 | 0.71920898849529 | -5.39215415485496 |
| ENSG00000140157.14 | NIPA2 | 0.0976328435754157 | 5.23202023691493 | 0.795247911131611 | 0.43088846840346 | 0.71920898849529 | -6.07865235157635 |
| ENSG00000282278.1 | | 0.0771866012652307 | 4.378364233639 | 0.795221907373293 | 0.430903427397557 | 0.71920898849529 | -6.02176034104827 |
| lnc-COA6-1 | lnc-COA6-1 | 1.21979365825282 | -0.653501001267133 | 0.795386582478885 | 0.430938588254673 | 0.719209353932208 | -4.9912507255492 |
| ENSG00000100258.17 | LMF2 | -0.141091258783006 | 5.10957886381638 | -0.795065737448119 | 0.430993272704457 | 0.719242300314241 | -6.07652725115849 |
| ENSG00000234335.1 | RPS4XP11 | 0.687659080797335 | -0.499586298510766 | 0.795075460744898 | 0.43102918216356 | 0.719243912158294 | -5.06989162676791 |
| ENSG00000080200.9 | CRYBG3 | 0.811395267001778 | 2.87909377069519 | 0.795226698714817 | 0.431075856916232 | 0.719263485968451 | -5.87565444392325 |
| ENSG00000283971.1 | MIR4442 | -0.200517536564294 | -0.438916561479346 | -0.794829213087337 | 0.431129367728096 | 0.719294461858426 | -5.31439126591033 |
| ENSG00000173875.13 | ZNF791 | 0.0862694230603331 | 6.23420337547811 | 0.7947256656722 | 0.431188956553993 | 0.719335572438875 | -6.09843763401902 |
| ENSG00000175354.19 | PTPN2 | 0.101060674050986 | 6.50115529113878 | 0.794536510585192 | 0.431297823122519 | 0.719441507473865 | -6.09865621579214 |
| ENSG00000177683.13 | THAP5 | -0.125651058533375 | 4.9493523921021 | -0.794493876249508 | 0.431322363223412 | 0.719441507473865 | -6.06447588083905 |
| ENSG00000168014.16 | C2CD3 | 0.0755552765570524 | 5.64309948253548 | 0.794142151685335 | 0.431524846030949 | 0.719678260170282 | -6.0909005426424 |
| ENSG00000199574.1 | SNORD18C | 0.212050121265357 | 0.354785897305097 | 0.79412585097082 | 0.431534231502775 | 0.719678260170282 | -5.50716764999371 |
| ENSG00000066230.11 | SLC9A3 | -0.190668887088282 | 2.23296506601246 | -0.793973766048094 | 0.431621803431417 | 0.719713821618717 | -5.79093501971335 |
| ENSG00000141522.11 | ARHGDIA | -0.121993339143414 | 6.00006303791353 | -0.793967367975487 | 0.43162548773558 | 0.719713821618717 | -6.09787510303225 |
| ENSG00000214954.8 | LRRC69 | 0.124496289846581 | 1.75172409867339 | 0.793685276710628 | 0.431787947608809 | 0.719853841739705 | -5.71940691810493 |
| lnc-MOCS2-9 | lnc-MOCS2-9 | -0.167436400407826 | 2.2086369967395 | -0.793644437552782 | 0.431811470428944 | 0.719853841739705 | -5.79178385814792 |
| ENSG00000106638.16 | TBL2 | -0.0991009493676884 | 3.78079355175758 | -0.793624148744342 | 0.431823156802833 | 0.719853841739705 | -5.9679076532099 |
| ENSG00000063241.7 | ISOC2 | -0.22276823401978 | 1.7539676420795 | -0.793578670089176 | 0.431849353243059 | 0.719853841739705 | -5.7329155932155 |
| ENSG00000179057.13 | IGSF22 | -0.550311894683885 | -0.345347878162859 | -0.793702313859028 | 0.431885547269589 | 0.719855876544655 | -5.43343397130418 |
| ENSG00000168564.5 | CDKN2AIP | 0.102489490975193 | 4.76772031889093 | 0.793380529612408 | 0.431963496486561 | 0.719927501877156 | -6.05575131199461 |
| ENSG00000104290.10 | FZD3 | 0.38071368450936 | 4.0512037909285 | 0.793549029649529 | 0.432014722417471 | 0.719954581128556 | -6.0039434427585 |
| lnc-CARD8-3 | lnc-CARD8-3 | -0.144087967303438 | 2.12656900840613 | -0.793104615786411 | 0.432122472952692 | 0.720022678747803 | -5.77857357520815 |
| ENSG00000110906.12 | KCTD10 | 0.0701926961213326 | 4.87428827211066 | 0.793099279775339 | 0.432125547809921 | 0.720022678747803 | -6.06361408852231 |
| ENSG00000167110.17 | GOLGA2 | 0.0658054563391222 | 4.84112032804847 | 0.792947972898333 | 0.43221274331721 | 0.720103493547106 | -6.05903563664176 |
| ENSG00000232366.1 | VDAC1P9 | -0.490699643909185 | -0.751736276642418 | -0.792849433202213 | 0.432269535694866 | 0.720103493547106 | -5.11638497404711 |
| lnc-CD69-1 | lnc-CD69-1 | -0.276702020587834 | 5.95989446691079 | -0.79297576475111 | 0.432310717412911 | 0.720103493547106 | -6.09719575373898 |
| ENSG00000105438.8 | KDELR1 | -0.0965263543214432 | 3.46740895498595 | -0.792772305402825 | 0.43231399066033 | 0.720103493547106 | -5.92599793193526 |
| ENSG00000100207.18 | TCF20 | 0.0697943048305953 | 6.73331977172582 | 0.792621552007087 | 0.432400889906029 | 0.720189959292426 | -6.09869754042285 |
| ENSG00000095002.14 | MSH2 | -0.125011120533817 | 4.40358941170361 | -0.792409862036464 | 0.432522932689579 | 0.720313933979863 | -6.03040893980201 |
| ENSG00000149308.16 | NPAT | 0.138257326869716 | 5.71073735488264 | 0.792351159897415 | 0.43255677910486 | 0.720313933979863 | -6.09224474509582 |
| ENSG00000132153.14 | DHX30 | -0.0781414878457838 | 5.7066716870786 | -0.792213903598184 | 0.432635924389084 | 0.720313933979863 | -6.09523017226903 |
| ENSG00000284946.1 | | 0.120772258452718 | 4.95808577096258 | 0.79218405776609 2 | 0.432653135362934 | 0.720313933979863 | -6.06421293282922 |
| ENSG00000023909.9 | GCLM | -0.16939569315997 | 0.921173094585733 | -0.792130320259415 | 0.432684124805734 | 0.720313933979863 | -5.60437152005434 |
| ENSG00000196535.16 | MYO18A | -0.0880114130273926 | 6.27206679877219 | -0.792128286676047 | 0.432685297562407 | 0.720313933979863 | -6.10040451028731 |
| ENSG00000115271.10 | GCA | -0.145891178132146 | 3.55128692689991 | -0.792028874349363 | 0.432742630443916 | 0.720351117058401 | -5.95749515562807 |
| ENSG00000144713.12 | RPL32 | 0.111958592031497 | 9.2700613326369 | 0.791907301147088 | 0.432812750091129 | 0.720409577480649 | -6.04858265012788 |
| ENSG00000272296.1 | SNORD96A | 0.115027673709477 | 1.1351082646928 | 0.791828276627166 | 0.432858332631564 | 0.72042719014761 | -5.63815329901028 |
| ENSG00000260144.1 | | 0.22905707584977 | -0.505676634052675 | 0.791675836381247 | 0.432946270602889 | 0.720507341994068 | -5.24628255827336 |
| lnc-CUL2-3 | lnc-CUL2-3 | -0.109546527109907 | 5.10841194666964 | -0.791572730555845 | 0.433005755171588 | 0.720507341994068 | -6.07653695531501 |
| ENSG00000147454.13 | SLC25A37 | 0.127713547203158 | 5.22073834557547 | 0.791519127953639 | 0.433036681912585 | 0.720507341994068 | -6.07974792117948 |
| ENSG00000235333.3 | PVRIG2P | -0.138374888566422 | 3.34923780841705 | -0.79150209322303 | 0.433046510606258 | 0.720507341994068 | -5.93704163877542 |

| | | | | | | | |
|---|---|---|---|---|---|---|---|
| ENSG00000264781.1 | MIR4537 | 0.621981054990645 | -1.27574385281835 | 0.79145502575127 | 0.433082270879578 | 0.72050859857051 | -5.04276197910246 |
| lnc-PON3-2 | lnc-PON3-2 | 0.631836656601129 | -0.365041844981873 | 0.791284157440248 | 0.433268294146168 | 0.720759823681449 | -5.27311940302193 |
| ENSG00000169919.16 | GUSB | -0.0897912729769024 | 4.37277246274298 | -0.790981642147796 | 0.433346864732802 | 0.720832270631587 | -6.02491570844763 |
| ENSG00000279649.1 | | -0.139216637153776 | 0.653232831407058 | -0.790748778104575 | 0.433481291777754 | 0.720954604691598 | -5.5487024535914 |
| ENSG00000128581.15 | IFT22 | 0.161997337422563 | 1.66546267097268 | 0.790732893672455 | 0.433490462404082 | 0.720954604691598 | -5.70488045631558 |
| ENSG00000155016.17 | CYP2U1 | 0.119280218239553 | 3.75848512105574 | 0.790440060503051 | 0.433659545829188 | 0.721177541553056 | -5.97490622582582 |
| ENSG00000172349.17 | IL16 | -0.161082218247292 | 7.74604345932758 | -0.790313767269396 | 0.433732480398315 | 0.721240559382657 | -6.08678413713979 |
| ENSG00000117707.15 | PROX1 | 1.99746306895619 | -0.646722689790005 | 0.790519508613265 | 0.433787677035855 | 0.721243475696639 | -5.00478556990327 |
| ENSG00000079691.17 | CARMIL1 | -0.211921074273048 | 6.50194498524603 | -0.790158082425682 | 0.433875136621921 | 0.721243475696639 | -6.10123271931773 |
| SNHG8 | SNHG8 | -0.115640524572428 | 4.91541597887818 | -0.790041983378899 | 0.433889460988168 | 0.721243475696639 | -6.06182297320179 |
| ENSG00000143149.12 | ALDH9A1 | 0.113152648450718 | 4.59830366942447 | 0.789974667690249 | 0.433928347342042 | 0.721243475696639 | -6.04400785729293 |
| lnc-FUBP1-56 | lnc-FUBP1-56 | 0.0875805520685287 | 1.96133732998435 | 0.789972061160925 | 0.433929853101658 | 0.721243475696639 | -5.76183593584513 |
| lnc-PITPNA-2 | lnc-PITPNA-2 | 0.110971825236496 | 1.52200615839926 | 0.78990734829414 | 0.433967237923751 | 0.721243475696639 | -5.70679215954456 |
| lnc-MBOAT4-3 | lnc-MBOAT4-3 | -0.161106446576512 | -0.174559264521268 | -0.789886089982331 | 0.433979519336485 | 0.721243475696639 | -5.35608847224096 |
| lnc-METTL16-1 | lnc-METTL16-1 | -0.117982215764882 | 2.2451660040542 | -0.789782182001785 | 0.434039552345731 | 0.721285007745211 | -5.79743561410208 |
| ENSG00000152454.3 | ZNF256 | 0.0840486248008574 | 4.68102036181091 | 0.789543612461219 | 0.434177405100009 | 0.721381992591903 | -6.04816986087856 |
| ENSG00000131626.18 | PPFIA1 | 0.0719901802411558 | 5.25054040011808 | 0.789340674807149 | 0.434294689282862 | 0.721381992591903 | -6.08252969627201 |
| ENSG00000077463.14 | SIRT6 | -0.108477324807265 | 2.16755092165583 | -0.789309033262635 | 0.434312977654846 | 0.721381992591903 | -5.78136628881035 |
| ENSG00000197483.9 | ZNF628 | 0.201313235276069 | 0.96533948292833 | 0.789213200312589 | 0.43436837057604 | 0.721381992591903 | -5.61990840405248 |
| lnc-TCL1B-5 | lnc-TCL1B-5 | -0.605069809714877 | -0.992910919663086 | -0.789324519671695 | 0.434378755378546 | 0.721381992591903 | -5.24662929267016 |
| lnc-CLDN34-2 | lnc-CLDN34-2 | -0.181254042782485 | 0.145388414171059 | -0.789169868853033 | 0.434393418214589 | 0.721381992591903 | -5.45204008353899 |
| lnc-HACD2-6 | lnc-HACD2-6 | -0.75642674712047 | -1.76668780053396 | -0.789235447029516 | 0.43439666880905 | 0.721381992591903 | -5.00876130474938 |
| lnc-RC3H1-2 | lnc-RC3H1-2 | -0.100180630923467 | 4.55660320075229 | -0.789154361410886 | 0.434402382461508 | 0.721381992591903 | -6.04711611561774 |
| ENSG00000170604.4 | IRF2BP1 | -0.195900671004241 | 2.28893897223761 | -0.789135404229819 | 0.434413341018153 | 0.721381992591903 | -5.81242039748898 |
| lnc-TBL1Y-11 | lnc-TBL1Y-11 | -1.00773662102484 | -1.73718802707591 | -0.789056801236832 | 0.434499675518476 | 0.721404073542705 | -4.85921334785702 |
| ENSG00000135838.13 | NPL | 0.251562882472033 | -0.0676122499420054 | 0.78898042801874 | 0.434502934146619 | 0.721404073542705 | -5.30776674936395 |
| lnc-CORO6-4 | lnc-CORO6-4 | 0.122511074604778 | -0.164040341743141 | 0.788930076964679 | 0.434532044918193 | 0.721404073542705 | -5.32875468767532 |
| lnc-ETV5-3 | lnc-ETV5-3 | 0.36262265748994 | 1.12907524589349 | 0.788950867987328 | 0.434566832233202 | 0.721404073542705 | -5.58605219616753 |
| ENSG00000165526.8 | RPUSD4 | 0.0920414934701856 | 4.54784267819056 | 0.788605839272711 | 0.43471953287542 | 0.721557920519544 | -6.04975429753777 |
| ENSG00000202031.1 | SNORD38A | 0.165805882577539 | 0.461598340763198 | 0.788588396856696 | 0.434729620189616 | 0.721557920519544 | -5.51632778271605 |
| ENSG00000198815.8 | FOXJ3 | 0.0670309169628558 | 6.19747592756807 | 0.788466553267301 | 0.434800088790964 | 0.721599410237228 | -6.10325537213391 |
| ENSG00000183161.5 | FANCF | -0.145058317282605 | 3.51464783102936 | -0.788423944545459 | 0.434824733283406 | 0.721599410237228 | -5.95307006797872 |
| lnc-ZDHHC11-3 | lnc-ZDHHC11-3 | 0.086571990205317 | 2.37460152941556 | 0.787999137766257 | 0.435070483282751 | 0.72193222462844 | -5.80770279350151 |
| lnc-CRIPT-1 | lnc-CRIPT-1 | 0.177408491185426 | 3.06092871184357 | 0.787956021644197 | 0.435095430525958 | 0.72193222462844 | -5.87311862833272 |
| lnc-OGDH-2 | lnc-OGDH-2 | -0.165611963938203 | -0.0566861972800866 | -0.787695607012338 | 0.435246126148852 | 0.722075504953976 | -5.35181781765933 |
| lnc-TOP3A-2 | lnc-TOP3A-2 | -0.266339395548526 | -0.627461045466986 | -0.787685552125691 | 0.435251945292911 | 0.722075504953976 | -5.22336408879138 |
| ENSG00000187240.14 | DYNC2H1 | -0.303664647685395 | 2.58186346313169 | -0.787684135769676 | 0.435325775087037 | 0.722139783092882 | -5.83228453096213 |
| ENSG00000278963.1 | | 0.15085215532608 | 3.53475908718508 | 0.787415880807773 | 0.435408031651751 | 0.722161138606829 | -5.94957221246546 |
| ENSG00000256060.2 | TRAPPC2B | -0.115761923774011 | 2.25086362747543 | -0.787346524496134 | 0.435448180645485 | 0.722161138606829 | -5.78663195119904 |
| ENSG00000182087.13 | TMEM259 | -0.128080563302936 | 6.54152628211349 | -0.787265757448704 | 0.435494937875531 | 0.722161138606829 | -6.10341699540411 |
| lnc-UBQLN2-6 | lnc-UBQLN2-6 | 0.183230107016082 | 0.564544472303981 | 0.787246269431846 | 0.435506220223895 | 0.722161138606829 | -5.52922501210591 |
| ENSG00000104432.13 | IL7 | 0.206826439011624 | 2.55477533195297 | 0.787232701388977 | 0.435514075379029 | 0.722161138606829 | -5.81129865230605 |
| ENSG00000242612.6 | DECR2 | 0.192300080469843 | 0.513047440865213 | 0.787018545383095 | 0.435638071210963 | 0.722308556447176 | -5.52865880243316 |
| LINC00589 | LINC00589 | -0.300804018939878 | -0.212146526870952 | -0.786933444269079 | 0.435687350413365 | 0.722332076806951 | -5.35867899227794 |
| ENSG00000197153.4 | H3C12 | 0.462763000494488 | 3.85953157495503 | 0.786990463616503 | 0.435820101657753 | 0.722493971683435 | -5.98710485105076 |
| lnc-CTDSPL2-2 | lnc-CTDSPL2-2 | 0.199859803160263 | 1.00554948358887 | 0.786601494968025 | 0.435879602947127 | 0.722534417931924 | -5.60758578262521 |
| ENSG00000054148.17 | PHPT1 | 0.16937290737572 | 2.24042025609543 | 0.786492999403974 | 0.435942450474876 | 0.722559233483156 | -5.79783166931189 |
| ENSG00000123179.13 | EBPL | -0.187474664324858 | 0.888645294409738 | -0.786454449007308 | 0.43596478261805 | 0.722559233483156 | -5.61059522264457 |
| lnc-TSC22D2-3 | lnc-TSC22D2-3 | 0.236088823174329 | -0.325900311145794 | 0.786333843615167 | 0.436034653415059 | 0.722599889265896 | -5.32345534299291 |
| lnc-C12orf74-3 | lnc-C12orf74-3 | -0.130588028359028 | 3.95160629847736 | -0.786290913252838 | 0.436059526044391 | 0.722599889265896 | -5.9908693345098 |
| lnc-CCR7-1 | lnc-CCR7-1 | 0.270470257746782 | -0.538535415249916 | 0.786009770666387 | 0.436222432988705 | 0.722727732161616 | -5.21218173774594 |
| ENSG00000131148.8 | EMC8 | 0.0999142599173708 | 2.97594316622821 | 0.785983510603643 | 0.436237651131619 | 0.722727732161616 | -5.88214059164554 |
| ENSG00000149548.14 | CCDC15 | -0.225776503191716 | 0.336974403734722 | -0.785975984740441 | 0.43624201255288 | 0.722727732161616 | -5.4804700254689 |
| ENSG00000221994.10 | ZNF630 | -0.11933971577691 | 2.19886457361566 | -0.785844183074473 | 0.436318399051826 | 0.722796105246175 | -5.77920414187272 |
| ENSG00000225973.3 | PIGBOS1 | 0.117692284279557 | 0.951332833203311 | 0.785783430856198 | 0.436353611076386 | 0.722796264025854 | -5.61973647223138 |
| lnc-PDCL2-2 | lnc-PDCL2-2 | 0.121490502361573 | 0.811967814754729 | 0.785556357907022 | 0.436485237718157 | 0.722956115551044 | -5.57443381041853 |
| ENSG00000196363.9 | WDR5 | -0.108156319434773 | 3.62433095864511 | -0.78542206173338 | 0.436563095895353 | 0.722986312173414 | -5.96037354561008 |
| lnc-MEF2C-8 | lnc-MEF2C-8 | 0.163275564753681 | 1.6159992122351 | 0.785335884514013 | 0.436613061483828 | 0.722986312173414 | -5.71375662723798 |
| ENSG00000204356.13 | NELFE | 0.0779399661943403 | 4.81414536503105 | 0.785252517448386 | 0.436661400990651 | 0.722986312173414 | -6.06758592528217 |
| ENSG00000163848.19 | ZNF148 | -0.0694006175879256 | 6.8194471015599 | -0.785206575946605 | 0.43668804104663 | 0.722986312173414 | -6.10349203095811 |
| lnc-COL4A3-4 | lnc-COL4A3-4 | -0.100994298213698 | 1.71531731577272 | -0.785145698316888 | 0.436723343594876 | 0.722986312173414 | -5.73408890326303 |
| ENSG00000131080.14 | EDA2R | 0.640145004556634 | 0.0893771426961355 | 0.785310459338044 | 0.436749920909987 | 0.722986312173414 | -5.39542323964552 |
| lnc-ZNF692-1 | lnc-ZNF692-1 | -0.133794328347182 | 0.546869162327927 | -0.785079661524783 | 0.436761639833366 | 0.722986312173414 | -5.53288312479139 |
| ENSG00000165675.16 | ENOX2 | 0.0884724527788165 | 3.65713519950121 | 0.785008973266024 | 0.436802635782987 | 0.722986312173414 | -5.95547462100336 |
| ENSG00000259581.3 | TYRO3P | 0.206193551469496 | 1.29027079708311 | 0.784979727533754 | 0.436819597638273 | 0.722986312173414 | -5.65477367220677 |
| ENSG00000102401.19 | ARMCX3 | 0.125149772607864 | 4.38187865086313 | 0.784844716751549 | 0.43689790589462 | 0.723057778968318 | -6.03323231875891 |
| ENSG00000156973.13 | PDE6D | -0.141654105233298 | 3.07991637076812 | -0.784564513909511 | 0.437060454417232 | 0.723245129888675 | -5.8858378585454 |
| ENSG00000130818.11 | ZNF426 | 0.101662726150039 | 5.6193431936402 | 0.784528436123478 | 0.437081386128612 | 0.723245129888675 | -6.09694977589 |

| | | | | | | | |
|---|---|---|---|---|---|---|---|
| lnc-ESRP2-3 | lnc-ESRP2-3 | -0.137658473986683 | 0.614950177383595 | -0.784219050424276 | 0.437260910934714 | 0.723444377513689 | -5.54557395184143 |
| ENSG00000147912.12 | FBXO10 | 0.17047377044707 | 4.221285776669 | 0.784069387075136 | 0.437347770682686 | 0.723444377513689 | -6.01071696753695 |
| ENSG00000197885.10 | NKIRAS1 | 0.135218462590274 | 0.933679305661568 | 0.784057570809344 | 0.437354628897704 | 0.723444377513689 | -5.61234494412491 |
| ENSG00000103549.21 | RNF40 | -0.0800488145176255 | 5.87749017757562 | -0.783934099683289 | 0.43742629594189 | 0.723444377513689 | -6.10434840921853 |
| lnc-UQCRFS1-16 | lnc-UQCRFS1-16 | 1.02231680913707 | -2.72598985682985 | 0.783938840265832 | 0.437453866812924 | 0.723444377513689 | -4.83549556187468 |
| ENSG00000148773.13 | MKI67 | 0.863065557281594 | -1.02521825420223 | 0.784045945586759 | 0.437455557808812 | 0.723444377513689 | -5.05001359575415 |
| ENSG00000170100.13 | ZNF778 | -0.123813592502886 | 3.22228488125456 | -0.783868414664642 | 0.437464424721056 | 0.723444377513689 | -5.90590345054435 |
| ENSG00000161714.11 | PLCD3 | -0.487192160552446 | -0.723136259196972 | -0.783847581754975 | 0.437487724108015 | 0.723444377513689 | -5.21383152762336 |
| lnc-STX5-3 | lnc-STX5-3 | 0.0926754721770466 | 2.70090456146222 | 0.783708258957377 | 0.437557400061235 | 0.723444377513689 | -5.85070263530723 |
| lnc-AKT3-7 | lnc-AKT3-7 | 0.089519983219589 | 2.0984568036414 | 0.783673826718567 | 0.437577390578457 | 0.723444377513689 | -5.76978958906653 |
| ENSG00000004777.18 | ARHGAP33 | -0.161230914608102 | 1.73027386263528 | -0.783654825445435 | 0.437588422486782 | 0.723444377513689 | -5.72763504362187 |
| lnc-TFDP2-1 | lnc-TFDP2-1 | -0.16319435427743 | 1.57244500415425 | -0.783548434573906 | 0.437650194796198 | 0.723488391252923 | -5.71113338787235 |
| ENSG00000162924.14 | REL | 0.109538396321422 | 7.59454724413159 | 0.783277335321186 | 0.437807623018612 | 0.72369051594861 | -6.09674370738702 |
| ENSG00000124217.4 | MOCS3 | 0.133428920348071 | 2.50225010361178 | 0.783143008711716 | 0.437885639418028 | 0.7237072695564 | -5.83450241221098 |
| ENSG00000122490.18 | SLC66A2 | -0.161477419989969 | 3.41744495274415 | -0.78313880811466 | 0.437888079242842 | 0.7237072695564 | -5.94398608052851 |
| lnc-TMEM161B-1 | lnc-TMEM161B-1 | -0.159837116482387 | 0.698013471697261 | -0.782990727870623 | 0.437974093584342 | 0.723741210367196 | -5.55883691201746 |
| ENSG00000172382.9 | PRSS27 | -0.193707879444008 | -0.527685055819275 | -0.782969696593891 | 0.43798631069191 | 0.723741210367196 | -5.28369949860319 |
| ENSG00000043514.16 | TRIT1 | 0.0891788096450861 | 3.6470903891723 | 0.782921856564592 | 0.438014101809462 | 0.723741210367196 | -5.96011266900957 |
| ENSG00000104205.13 | SGK3 | 0.119679483751372 | 4.26871578443985 | 0.782775464389422 | 0.438099150132487 | 0.723804261591142 | -6.01651556507683 |
| ENSG00000251287.8 | ALG1L2 | -0.281504570987551 | -0.432396707990245 | -0.782735118628348 | 0.438122591224021 | 0.723804261591142 | -5.26019520636314 |
| lnc-SUMO2-1 | lnc-SUMO2-1 | -0.111536357763755 | 0.909017273533676 | -0.78255570780573 | 0.438226838866273 | 0.723918380769222 | -5.61920741166185 |
| ENSG00000154479.12 | CFAP210 | -0.626825272684021 | -1.16832051938295 | -0.782083181450828 | 0.438515036373077 | 0.724282218469427 | -5.05311700100644 |
| lnc-USP10-3 | lnc-USP10-3 | -0.107904801468701 | 3.93405313857032 | -0.782055670560916 | 0.438517466188894 | 0.724282218469427 | -5.99226924930905 |
| ENSG00000148296.6 | SURF6 | -0.0828233399452767 | 4.16822771205405 | -0.781703694568792 | 0.438722107415465 | 0.724462130331431 | -6.01560627435476 |
| lnc-CHRNE-3 | lnc-CHRNE-3 | -0.171489485351472 | 0.441334640880843 | -0.78156602376038 | 0.438802165615618 | 0.724462130331431 | -5.49934818237239 |
| lnc-PPP1R18-1 | lnc-PPP1R18-1 | -0.177386073993802 | 2.50476359027579 | -0.781543645431699 | 0.438815179861773 | 0.724462130331431 | -5.82973603592086 |
| lnc-FOXD4L5-26 | lnc-FOXD4L5-26 | -0.481162988415456 | -0.415174361982186 | -0.781533788500645 | 0.438834127227228 | 0.724462130331431 | -5.23054308952106 |
| lnc-TCOF1-2 | lnc-TCOF1-2 | -0.224309912415127 | -0.123672116487191 | -0.781492158829136 | 0.438845123064614 | 0.724462130331431 | -5.33652659806232 |
| ENSG00000146729.9 | NIPSNAP2 | 0.0590660443240223 | 5.25096228067835 | 0.781473640155402 | 0.438855893317104 | 0.724462130331431 | -6.09080491399645 |
| ENSG00000099308.10 | MAST3 | -0.174597370099422 | 4.98410016699731 | -0.781449445056288 | 0.438872773798893 | 0.724462130331431 | -6.07290741042729 |
| lnc-FCHO2-1 | lnc-FCHO2-1 | 0.11790600399227 | 3.59618937610796 | 0.781255492760577 | 0.438982777227983 | 0.724532625995476 | -5.95669512344087 |
| ENSG00000183878.15 | UTY | -1.84567531689444 | 3.1725796872778 | -0.781507272225168 | 0.439008086284251 | 0.724532625995476 | -5.70280568167642 |
| lnc-RAB6D-1 | lnc-RAB6D-1 | -0.180867938603775 | 0.222294540982303 | -0.781189645599072 | 0.43902108105794 | 0.724532625995476 | -5.47735074156161 |
| lnc-ATP6V1G2-DDX39B-[illegible] | lnc-ATP6V1G2-DD[illegible] | -0.225096602543584 | 0.0256010662268797 | -0.780965393099167 | 0.439151545478018 | 0.724689830880626 | -5.42777185245866 |
| ENSG00000130487.7 | KLHDC7B | -0.25175270917828 | 0.507338022490176 | -0.780880717254018 | 0.43920081357825 | 0.7247130307745 | -5.60759895059872 |
| ENSG00000185101.12 | ANO9 | -0.13286136955463 | 5.14320817664782 | -0.780270363627534 | 0.439556042470003 | 0.725194213335737 | -6.08644662044342 |
| ENSG00000110060.8 | PUS3 | 0.154433058826549 | 1.93259845650578 | 0.780258597693456 | 0.439562891977684 | 0.725194213335737 | -5.76007369096714 |
| ENSG00000267740.5 | | -0.130394746217028 | 2.79632350345896 | -0.779970013727176 | 0.439730910240325 | 0.725413266448394 | -5.86647529598228 |
| ENSG00000037241.7 | RPL26L1 | 0.0947833660441764 | 1.90758221141114 | 0.779862789887386 | 0.439793347422943 | 0.725458123898576 | -5.74965698053453 |
| lnc-NR2C2-6 | lnc-NR2C2-6 | 0.233383168519647 | -0.0439016469583442 | 0.779793238494888 | 0.43983385049646 | 0.725466795798432 | -5.37967376091198 |
| lnc-C1orf35-2 | lnc-C1orf35-2 | -0.107903433287734 | 2.32786822889018 | -0.779621910065835 | 0.439933632619732 | 0.725539844401061 | -5.80513111464484 |
| ENSG00000091317.7 | CMTM6 | 0.129122144646983 | 5.62453241372483 | 0.779596148946043 | 0.439948637126252 | 0.725539844401061 | -6.1037468200188 |
| ENSG00000123975.4 | CKS2 | 0.248332403330093 | 0.771363597369241 | 0.77949706640413 | 0.440006350364811 | 0.725576887482688 | -5.60060628648134 |
| lnc-DHX15-1 | lnc-DHX15-1 | 0.123997657539686 | 4.17987960991069 | 0.779357906721963 | 0.440087415185521 | 0.725652428644042 | -6.02624514920338 |
| lnc-MYOM2-4 | lnc-MYOM2-4 | -1.54929637198203 | 1.37963154688147 | -0.779258595013863 | 0.440316462254037 | 0.725971943493659 | -5.72831987022807 |
| ENSG00000188243.12 | COMMD6 | -0.127765590463824 | 4.35238244013227 | -0.778862080599108 | 0.44037632136401 | 0.726012480787778 | -6.03364212552292 |
| ENSG00000175602.3 | CCDC85B | -0.275470784801423 | 0.878673237883921 | -0.778767205608273 | 0.440431615617908 | 0.726045486485937 | -5.62734301416949 |
| lnc-IL7-2 | lnc-IL7-2 | 0.242439167064109 | -0.0616603365083572 | 0.778635637843812 | 0.440508301672257 | 0.726113748163696 | -5.34488278710148 |
| lnc-MTHFD1-4 | lnc-MTHFD1-4 | -0.267123354631459 | -0.387201074625501 | -0.778512816494858 | 0.440579896914904 | 0.72614267112288 | -5.27995009420099 |
| ENSG00000182154.7 | MRPL41 | 0.146041409788888 | 2.52549775475 | 0.778484497328041 | 0.440596405754927 | 0.72614267112288 | -5.84307683213405 |
| ENSG00000144827.8 | ABHD10 | -0.146609734605977 | 3.90740000215124 | -0.778199261030827 | 0.44076270656402 | 0.726312844080447 | -5.98722349899086 |
| ENSG00000231500.6 | RPS18 | 0.136613358403896 | 9.42432887662353 | 0.778186350864413 | 0.440770234436616 | 0.726312844080447 | -6.05496993665617 |
| ENSG00000105829.12 | BET1 | -0.109943304753679 | 3.39081220326464 | -0.777958468908745 | 0.440903124165786 | 0.726473663521003 | -5.93646606245218 |
| lnc-CYSTM1-1 | lnc-CYSTM1-1 | -0.267178925700098 | 0.751887258853376 | -0.777759014173164 | 0.441019456028752 | 0.726607177094357 | -5.57177541892814 |
| lnc-IBTK-10 | lnc-IBTK-10 | -0.424733004795133 | -0.637133226786187 | -0.777637275639456 | 0.441090468911324 | 0.72666600941266 | -5.19792667138562 |
| ENSG00000274466.1 | MIR1273H | 0.202741832192507 | 0.350069895788015 | 0.777455178528787 | 0.441196703002277 | 0.726782852172539 | -5.46266479220204 |
| lnc-COL23A1-1 | lnc-COL23A1-1 | -0.128142410877936 | 2.27459453492305 | -0.777246747267169 | 0.441318318869956 | 0.726874981795779 | -5.79787797014838 |
| lnc-TCEA1-1 | lnc-TCEA1-1 | 0.12282432814013 | 2.80744660765018 | 0.77723828072448 | 0.441323259364614 | 0.726874981795779 | -5.86865981792109 |
| ENSG00000179388.8 | EGR3 | -0.675515500508395 | -0.0440919307446716 | -0.777244108861499 | 0.441426091267382 | 0.726943026259031 | -5.27888447794737 |
| ENSG00000255389.1 | | -0.22089738495058 | 0.755466204740723 | -0.777013890479008 | 0.441454210106622 | 0.726943026259031 | -5.56658640489817 |
| lnc-SLC38A9-4 | lnc-SLC38A9-4 | 0.124254785802921 | 0.0441328505286411 | 0.776974811089204 | 0.441477018601628 | 0.726943026259031 | -5.40185329381874 |
| lnc-KAT7-1 | lnc-KAT7-1 | -0.155131294067078 | 1.01129009197673 | -0.77687569096465 | 0.441534872712769 | 0.726943026259031 | -5.62182160446363 |
| ENSG00000125967.16 | NECAB3 | -0.136764701059005 | 1.25968478400121 | -0.776815091010493 | 0.441570245711174 | 0.726943026259031 | -5.67378669564268 |
| lnc-PRR18-1 | lnc-PRR18-1 | -0.104772325790098 | 2.97074157832101 | -0.776551949865362 | 0.441723864185435 | 0.726943026259031 | -5.89937150047397 |
| ENSG00000202252.1 | SNORD14C | 0.301574052781354 | -0.457715013399106 | 0.776485341983396 | 0.441762754050053 | 0.726943026259031 | -5.28068569485726 |
| lnc-ST7L-3 | lnc-ST7L-3 | -0.100965949098166 | 2.4985206488401 | -0.776445869280996 | 0.441785801650639 | 0.726943026259031 | -5.83219639583216 |
| ENSG00000003056.7 | M6PR | -0.103308163806079 | 5.82074971385328 | -0.776388324167592 | 0.44181940277555 | 0.726943026259031 | -6.10919694819239 |
| ENSG00000151466.11 | SCLT1 | 0.0684822113430652 | 4.60144520450768 | 0.776346730558272 | 0.441843690612745 | 0.726943026259031 | -6.0533226446384 |

| | | | | | | | |
|---|---|---|---|---|---|---|---|
| ENSG00000149179.13 | CSTPP1 | -0.112445094149789 | 3.11747156856629 | -0.77632001928904 | 0.44185928859352 | 0.726943026259031 | -5.90780049470962 |
| ENSG00000274471.2 | | -0.348655140237826 | 0.759718253935187 | -0.776412277192388 | 0.441859925500985 | 0.726943026259031 | -5.64190308467618 |
| lnc-FNDC1-3 | lnc-FNDC1-3 | -0.138481279052401 | 1.37581413631086 | -0.776142521522192 | 0.44196294627473 | 0.726943026259031 | -5.69673717901925 |
| ENSG00000119929.12 | CUTC | -0.0635451204365527 | 4.5302735557048 | -0.776110793169579 | 0.441981476969776 | 0.726943026259031 | -6.05033749826505 |
| ENSG00000160932.10 | LY6E | 0.187070131754774 | 4.60849234682329 | 0.776104797098246 | 0.441998423573002 | 0.726943026259031 | -6.05658074782065 |
| ENSG00000182600.9 | SNORC | -0.102698085551599 | 1.48846111410856 | -0.776068892021264 | 0.442005949711903 | 0.726943026259031 | -5.70720558755704 |
| ENSG00000096433.10 | ITPR3 | 0.194638943782937 | 6.6650596368062 | 0.776089276506814 | 0.442030506948164 | 0.726943026259031 | -6.11198759051354 |
| ENSG00000096063.15 | SRPK1 | -0.0671484459805205 | 5.12947173376865 | -0.776025494901367 | 0.442031297036362 | 0.726943026259031 | -6.08979553591299 |
| lnc-MAPK15-2 | lnc-MAPK15-2 | -0.172805946485017 | 0.667469783478092 | -0.775909543960599 | 0.442099025713303 | 0.726943026259031 | -5.55966081915379 |
| lnc-SLC41A1-3 | lnc-SLC41A1-3 | 0.16594553233949 | 1.13925704766529 | 0.775837595835115 | 0.442141054780372 | 0.726943026259031 | -5.64507735700336 |
| ENSG00000163006.11 | CCDC138 | -0.108474322111969 | 2.53244565940796 | -0.775760325939964 | 0.442186195238303 | 0.726943026259031 | -5.82952748692328 |
| lnc-CAST-4 | lnc-CAST-4 | -0.16270240582831 | 3.45678958014693 | -0.775744664986289 | 0.442195344575675 | 0.726943026259031 | -5.94278884583714 |
| lnc-NBPF15-4 | lnc-NBPF15-4 | -0.104003906091179 | 2.27858057471768 | -0.775723720718149 | 0.442207580670941 | 0.726943026259031 | -5.80280117437028 |
| lnc-CBX5-3 | lnc-CBX5-3 | -0.141692407031492 | 0.388899313469926 | -0.775715820718693 | 0.442212196073912 | 0.726943026259031 | -5.4920546714507 |
| ENSG00000280198.1 | | -0.112245603563194 | 1.25200464048722 | -0.775308116908049 | 0.442450426911491 | 0.727223704304686 | -5.6693318826508 |
| ENSG00000130997.16 | POLN | 0.126901022832279 | 1.05092957095913 | 0.775264140128616 | 0.44247612811054 | 0.727223704304686 | -5.65063010967266 |
| ENSG00000108582.11 | CPD | 0.150365353315196 | 3.83255841232044 | 0.775242233445536 | 0.442488931288534 | 0.727223704304686 | -5.99555646825432 |
| lnc-GTF2H5-24 | lnc-GTF2H5-24 | 0.183569449715836 | 2.95368315700145 | 0.775006448386054 | 0.442626747789348 | 0.727392123732387 | -5.88271152998087 |
| lnc-ZNF274-1 | lnc-ZNF274-1 | -0.102165350659915 | 1.5404530125671 | -0.774585817764127 | 0.442872669747 | 0.727592964930868 | -5.70857556328088 |
| ENSG00000163528.12 | CHCHD4 | -0.138337658548625 | 1.1651058719788 | -0.774561624025112 | 0.442886817091749 | 0.727592964930868 | -5.65447072765714 |
| ENSG00000160117.14 | ANKLE1 | -0.150465886303979 | 0.566816664620793 | -0.774520478992283 | 0.442910877359166 | 0.727592964930868 | -5.52162615686758 |
| ENSG00000165716.10 | DIPK1B | 0.625359908140801 | -1.15708923026787 | 0.774652924608392 | 0.442919250282025 | 0.727592964930868 | -5.24793552936028 |
| lnc-EFCAB11-11 | lnc-EFCAB11-11 | 0.0680558072980944 | 2.78604463239232 | 0.774434328505075 | 0.442961257848603 | 0.727592964930868 | -5.8575602001663 |
| lnc-GDF11-1 | lnc-GDF11-1 | -0.233516198229436 | 0.617210396926085 | -0.774379145848772 | 0.442993530246462 | 0.727592964930868 | -5.56167575849254 |
| lnc-PRSS37-3 | lnc-PRSS37-3 | 0.127122706674204 | 3.98499935572808 | 0.774374227724329 | 0.442996406574048 | 0.727592964930868 | -6.00128391734881 |
| lnc-CCDC149-1 | lnc-CCDC149-1 | 0.125577776004864 | 3.63363445416848 | 0.774179404477771 | 0.443110356358027 | 0.727710797066762 | -5.97147942057364 |
| ENSG00000145241.10 | CENPC | 0.0943487730490338 | 6.57620380283994 | 0.774130677684391 | 0.443138858788067 | 0.727710797066762 | -6.11392741064246 |
| lnc-SFRP4-3 | lnc-SFRP4-3 | -0.477609315937843 | 1.28265707872943 | -0.774256893942048 | 0.443185190484719 | 0.72772882135995 | -5.630844268503 |
| ENSG00000258704.6 | SRP54-AS1 | -0.125460018469904 | 0.808809895309767 | -0.773957515919966 | 0.443240157448715 | 0.727761021120525 | -5.58743744179403 |
| lnc-CCK-4 | lnc-CCK-4 | 0.147646261033886 | 1.04558777480267 | 0.773769620917696 | 0.443350090472562 | 0.727870103134514 | -5.62561223649587 |
| lnc-SNAI3-1 | lnc-SNAI3-1 | 0.158578698614973 | -0.105821100821557 | 0.77372308712534 9 | 0.443377318811666 | 0.727870103134514 | -5.39211698993181 |
| ENSG00000242616.3 | GNG10 | 0.133628745827956 | 1.0802533938703 | 0.773562110663845 | 0.443471518673842 | 0.727966685450489 | -5.64806602222612 |
| ENSG00000177663.13 | IL17RA | -0.104086448652937 | 4.89836894028171 | -0.773074805734294 | 0.44375675090143 | 0.728376810510697 | -6.07979829991999 |
| lnc-NOTCH2-2 | lnc-NOTCH2-2 | 0.327281947646031 | 1.0602837664998 | 0.772972575185166 | 0.443843003813776 | 0.72842229922852 | -5.59726100000335 |
| ENSG00000170921.15 | TANC2 | 1.16492531811048 | -0.249888285751272 | 0.773185152976891 | 0.443855243498231 | 0.72842229922852 | -5.25166947026066 |
| ENSG00000057657.16 | PRDM1 | 0.417638957135124 | 1.7560370615963 | 0.772931629994234 | 0.443952674096651 | 0.728514229495795 | -5.7109649202361 |
| ENSG00000197181.11 | PIWIL2 | 0.185858879024215 | 0.0448089128760556 | 0.772690027837698 | 0.443982048039414 | 0.728514229495795 | -5.38332636871157 |
| lnc-CRIPT-4 | lnc-CRIPT-4 | 0.181710559624416 | 1.19733627845361 | 0.772625505678911 | 0.444019834007082 | 0.728518154114609 | -5.66352211411083 |
| lnc-TOR1AIP2-8 | lnc-TOR1AIP2-8 | -0.0691286591320734 | 2.32179342116746 | -0.772512695367552 | 0.444085903425867 | 0.728568480010729 | -5.80997385449946 |
| ENSG00000198909.7 | MAP3K3 | 0.0648980199277869 | 5.86453792312622 | 0.77224194852842 | 0.44424449496838 | 0.728770577822123 | -6.11364627598908 |
| ENSG00000133193.12 | VCF1 | -0.118239877083447 | 4.51393421915597 | -0.772099317733591 | 0.44432805759494 | 0.728822674066168 | -6.05297430631543 |
| ENSG00000149823.8 | VPS51 | -0.110000653822858 | 6.19042462204803 | -0.772066863686649 | 0.444347069759333 | 0.728822674066168 | -6.11577836632703 |
| NUP50-DT | NUP50-DT | -0.106359334677657 | 2.98792887028471 | -0.771906574622115 | 0.444440988603735 | 0.728918634934715 | -5.88665053768324 |
| ENSG00000280202.1 | | -0.122340015435377 | 1.55217193144447 | -0.771545609063209 | 0.444652533619154 | 0.72919926757668 | -5.73078865138175 |
| ENSG00000163938.16 | GNL3 | 0.105225117047045 | 5.94750368206509 | 0.771493713154147 | 0.444682952272384 | 0.72919926757668 | -6.1141568372481 |
| ENSG00000214837.8 | LINC01347 | -0.145466798339379 | 0.54503959588309 | -0.771410610424747 | 0.444731665283335 | 0.729211411855956 | -5.51951466458804 |
| ENSG00000185917.13 | SETD4 | -0.0886308028325695 | 3.67711655615498 | -0.771360204220991 | 0.444761213838589 | 0.729211411855956 | -5.97028147184457 |
| lnc-PSMB2-2 | lnc-PSMB2-2 | -0.130086193834194 | -0.0911352600405737 | -0.771218314570506 | 0.444844397013162 | 0.729242230478116 | -5.38737844388723 |
| lnc-RPL13-5 | lnc-RPL13-5 | 0.13998548227462 | 3.48031017062917 | 0.771207274872925 | 0.444850869449702 | 0.729242230478116 | -5.94230608171462 |
| H1FX-AS1 | H1FX-AS1 | -0.323566719708988 | 0.692162495168103 | -0.770974337048893 | 0.444995800229696 | 0.729376594481528 | -5.53240433054186 |
| ENSG00000134825.15 | TMEM258 | -0.110729149053641 | 3.57935325169741 | -0.77080128634055 | 0.445088934079718 | 0.729376594481528 | -5.96395797870014 |
| ENSG00000158417.10 | EIF5B | 0.0994548937670441 | 5.48889692339948 | 0.770742685050129 | 0.44512330307152 | 0.729376594481528 | -6.10583216651986 |
| lnc-TMF1-1 | lnc-TMF1-1 | 0.173745259088813 | 0.229316330846061 | 0.77073083465367 | 0.44513025338504 | 0.729376594481528 | -5.48023456393696 |
| LINC00158 | LINC00158 | -0.362262689325983 | 1.80960294175859 | -0.770875173220507 | 0.445137469939342 | 0.729376594481528 | -5.73078283994657 |
| lnc-BAZ1A-4 | lnc-BAZ1A-4 | -0.132176933674165 | 0.422582978001663 | -0.770658574296374 | 0.445172635812693 | 0.729376594481528 | -5.50007336627042 |
| ENSG00000174749.5 | FAM241A | -0.185333770762455 | 2.45383603175547 | -0.770625778276996 | 0.445191872248421 | 0.729376594481528 | -5.83160920704397 |
| ENSG00000162783.10 | IER5 | 0.15414358313889 | 4.90346527072099 | 0.77058409720852 | 0.445216320899087 | 0.729376594481528 | -6.08170847747857 |
| lnc-DCAF16-2 | lnc-DCAF16-2 | 0.124085975352056 | 1.18986210170989 | 0.770273285929714 | 0.445398656870405 | 0.729597312369766 | -5.66285225019207 |
| lnc-GOLGA4-3 | lnc-GOLGA4-3 | 0.158229559222361 | 2.00935099856074 | 0.770233599739706 | 0.445421941759631 | 0.729597312369766 | -5.76170772947008 |
| lnc-CDK14-1 | lnc-CDK14-1 | -0.199298824571914 | -0.181370037334515 | -0.77016693925312 | 0.445461054766379 | 0.729603317439038 | -5.37937911074138 |
| lnc-INPP5F-3 | lnc-INPP5F-3 | -0.108835925877009 | 0.126841104481835 | -0.770014407870673 | 0.445550560103452 | 0.729675048850923 | -5.43271216959177 |
| ENSG00000278677.1 | H2AC17 | 0.414864387817156 | 3.4884929230311 | 0.770229311200888 | 0.445575751299969 | 0.729675048850923 | -5.96017505057915 |
| ENSG00000136238.17 | RAC1 | 0.0789652983000639 | 5.53925726123686 | 0.769815159679475 | 0.445667494825582 | 0.729755671126648 | -6.1095485585422 |
| ENSG00000197183.14 | NOL4L | 0.255478904003141 | 3.54921044373891 | 0.769867548529239 | 0.445695891839532 | 0.729755671126648 | -5.93788021154836 |
| ENSG00000116819.7 | TFAP2E | -0.110426696963345 | 1.02939835929805 | -0.769470302392751 | 0.445869927362104 | 0.729982557659416 | -5.62807504940811 |
| lnc-NOD1-4 | lnc-NOD1-4 | -0.18599209613074 | 0.768721279457224 | -0.769299083074065 | 0.445970453900189 | 0.730089068052779 | -5.58871737622532 |
| ENSG00000151881.14 | TMEM267 | 0.0838514341941581 | 3.67319894803697 | 0.769021810594392 | 0.44613327487718 | 0.730230109572546 | -5.97487546904647 |
| ENSG00000285978.1 | | 0.112650620068962 | 0.312662539142692 | 0.768963468957454 | 0.446167538933929 | 0.730230109572546 | -5.47470053277462 |

| | | | | | | | |
|---|---|---|---|---|---|---|---|
| ENSG00000269743.2 | SLC25A53 | -0.135759920680659 | 3.08845861053785 | -0.768901204628633 | 0.44620410850032 | 0.730230109572546 | -5.89338827514476 |
| lnc-MCEE-1 | lnc-MCEE-1 | -0.18017718977784 | 1.9120087825617 | -0.768808361578315 | 0.446258641080096 | 0.730230109572546 | -5.77160344623572 |
| ENSG00000138459.8 | SLC35A5 | -0.0776379158210272 | 4.55704218752214 | -0.768737046983318 | 0.446300531308513 | 0.730230109572546 | -6.05725055545198 |
| lnc-ATXN7-16 | lnc-ATXN7-16 | -0.0950170228555525 | 2.84303208882105 | -0.768730297195081 | 0.446304496257566 | 0.730230109572546 | -5.87603240569712 |
| ENSG00000131653.12 | TRAF7 | -0.128964148768732 | 4.50552147924188 | -0.768729525376652 | 0.44630494963915 | 0.730230109572546 | -6.05803920087209 |
| ENSG00000112576.12 | CCND3 | -0.159189391554277 | 6.68104077259149 | -0.768553993477686 | 0.446408067641173 | 0.730286052275484 | -6.11737148822346 |
| ENSG00000020633.18 | RUNX3 | 0.142508491146117 | 4.92177771039966 | 0.768506306055759 | 0.446436084529 | 0.730286052275484 | -6.08760613163956 |
| ENSG00000132680.10 | KHDC4 | -0.0903663795237213 | 5.90973805851996 | -0.768485584115431 | 0.446448259221206 | 0.730286052275484 | -6.11580807132111 |
| ENSG00000212232.1 | SNORD17 | 0.267569412378372 | 6.56575235341503 | 0.768417702074813 | 0.446598117839643 | 0.730286052275484 | -6.11868523507444 |
| lnc-RRBP1-3 | lnc-RRBP1-3 | 0.267569412378372 | 6.56575235341503 | 0.768417702074813 | 0.446598117839643 | 0.730286052275484 | -6.11868523507444 |
| lnc-IGDCC4-2 | lnc-IGDCC4-2 | 0.0737761118614992 | 1.94801871403817 | 0.768092121726364 | 0.446679465998647 | 0.730286052275484 | -5.7617911756405 |
| lnc-CYB5R4-5 | lnc-CYB5R4-5 | -0.194050599561116 | 0.967528923174204 | -0.768076635851085 | 0.446688567267833 | 0.730286052275484 | -5.61207695166091 |
| lnc-LYAR-3 | lnc-LYAR-3 | 0.120522661862381 | 1.06214185933283 | 0.768001831081996 | 0.446732532633668 | 0.730286052275484 | -5.64222240218205 |
| ENSG00000174032.16 | SLC25A30 | -0.121313166126063 | 4.0241610836849 | -0.767980214907831 | 0.446745237686312 | 0.730286052275484 | -6.02076301330879 |
| ENSG00000213801.4 | ZNF321P | -0.0868154733498745 | 2.54873738176958 | -0.767974885620437 | 0.44674837004376 | 0.730286052275484 | -5.84201514686667 |
| ENSG00000172456.17 | FGGY | 0.087731221678534 | 3.8023783242336 | 0.767940230808572 | 0.44676873917241 | 0.730286052275484 | -5.97808561532266 |
| lnc-HOOK1-6 | lnc-HOOK1-6 | -0.210680221930186 | -0.606991017980863 | -0.767913115259598 | 0.446784677314925 | 0.730286052275484 | -5.24396244329136 |
| lnc-SIAH1-4 | lnc-SIAH1-4 | -0.0793267413152381 | 1.66941771056247 | -0.767875883067344 | 0.446806562425488 | 0.730286052275484 | -5.73535423445117 |
| ENSG00000074842.7 | MYDGF | -0.105959043596329 | 2.96386213531664 | -0.767798966501936 | 0.446851776040723 | 0.730286052275484 | -5.89630527417689 |
| lnc-CPEB3-1 | lnc-CPEB3-1 | 0.0923084319249515 | 0.57962750574986 | 0.767765683004335 | 0.446871341806817 | 0.730286052275484 | -5.5640491326176 |
| ENSG00000198855.6 | FICD | 0.130835707254686 | 2.52850064003198 | 0.767567473664076 | 0.446987869989874 | 0.730418492219878 | -5.83264336681115 |
| ENSG00000155034.18 | FBXL18 | 0.154145428772053 | 1.52017775246351 | 0.767286399744275 | 0.447153145327524 | 0.730630562060525 | -5.7270533887359 |
| ENSG00000176155.18 | CCDC57 | -0.143987748372225 | 3.79970508427066 | -0.767149116648006 | 0.447233882761344 | 0.730704477605711 | -5.99617696907014 |
| ENSG00000222932.1 | RNU6-172P | 0.29643712844728 | -0.652879201194246 | 0.766956483335465 | 0.44734718663332 | 0.730831585242768 | -5.25152486259038 |
| LACTB2-AS1 | LACTB2-AS1 | 0.244036698341099 | 0.672230334213208 | 0.766865219465022 | 0.447400872500929 | 0.730861282435447 | -5.52912181059053 |
| ENSG00000274618.1 | H4C6 | 0.927087396991456 | 1.62155883495269 | 0.766991510907681 | 0.447494695961759 | 0.730890048475781 | -5.66094307337241 |
| lnc-VWCE-1 | lnc-VWCE-1 | 0.105223621062402 | 2.84344763402262 | 0.76668093754014 | 0.4475092876958 | 0.730890048475781 | -5.8791433032351 |
| lnc-SERGEF-3 | lnc-SERGEF-3 | -0.0978734697888586 | 1.49058572862176 | -0.766654215263146 | 0.447525010005357 | 0.730890048475781 | -5.71262886331214 |
| SNHG6 | SNHG6 | 0.12198153384291 | 5.53373905388896 | 0.766515670840941 | 0.447606529166578 | 0.730965184848911 | -6.11159455602398 |
| lnc-CERS3-2 | lnc-CERS3-2 | -0.126149292706371 | 0.0999537714581675 | -0.766307960740751 | 0.447728761599211 | 0.731067443589716 | -5.43400855862045 |
| ENSG00000135913.10 | USP37 | -0.101260048364536 | 4.98565746470822 | -0.766288553415654 | 0.447740183349947 | 0.731067443589716 | -6.09217110647902 |
| ENSG00000102547.18 | CAB39L | 0.195399721078644 | 2.55759969047385 | 0.76621343452201 | 0.447784394525167 | 0.731081636591696 | -5.84426631423311 |
| ENSG00000162063.12 | CCNF | -0.177224188093473 | 1.12847523018955 | -0.766126521216852 | 0.447835550499349 | 0.731086801133788 | -5.67156346328298 |
| lnc-RPS10-1 | lnc-RPS10-1 | 0.153460075702069 | 1.96160634121705 | 0.766087369626149 | 0.447858595709854 | 0.731086801133788 | -5.75974466422545 |
| ENSG00000280214.1 | | -0.173124301440113 | 1.4129324977651 | -0.765882435275122 | 0.447979234513014 | 0.731192272826552 | -5.70812352758111 |
| lnc-TRAF3IP2-6 | lnc-TRAF3IP2-6 | -0.102536579372489 | 0.87312662684824 | -0.765856921436842 | 0.44799425509477 | 0.731192272826552 | -5.61118679355113 |
| lnc-SIN3A-1 | lnc-SIN3A-1 | -0.117474888237486 | 1.73654025357059 | -0.765788304804204 | 0.448034652750197 | 0.731200226574477 | -5.74463620900498 |
| lnc-DLG2-1 | lnc-DLG2-1 | 0.238247705525475 | -0.469728862906091 | 0.765562397877928 | 0.448167669321519 | 0.73135932273407 | -5.27242161478492 |
| lnc-AGAP9-39 | lnc-AGAP9-39 | -0.135060950254873 | 0.186543755779394 | -0.765483942329193 | 0.448213870276525 | 0.731376731560307 | -5.4635089047156 |
| ENSG00000127663.14 | KDM4B | 0.107143480143248 | 5.88338727785032 | 0.765270719858645 | 0.448339447017155 | 0.731523649461284 | -6.11783947598502 |
| ENSG00000132613.14 | MTSS2 | 0.212998070537497 | 2.10619577806071 | 0.765187606051155 | 0.448388402251477 | 0.731545536108287 | -5.79505117310035 |
| lnc-PRMT9-2 | lnc-PRMT9-2 | 0.65121036967871 | -1.08644486548417 | 0.76514060227808 | 0.448505443347911 | 0.731678492544191 | -5.2308221834355 |
| lnc-CXXC5-1 | lnc-CXXC5-1 | 0.241733797574547 | -0.0407209053900226 | 0.764889722904915 | 0.448563885553467 | 0.731698625359227 | -5.36374519487538 |
| ENSG00000102038.15 | SMARCA1 | -0.560309249900834 | -1.53749551305248 | -0.764771881098723 | 0.448654478368379 | 0.731698625359227 | -5.14947956203382 |
| lnc-LEFTY1-3 | lnc-LEFTY1-3 | -0.154356126914739 | 2.83909730543178 | -0.764699205974427 | 0.448676140463432 | 0.731698625359227 | -5.86788615340237 |
| ENSG00000226015.2 | CCT8P1 | 0.194888271342054 | 0.371774334018588 | 0.764659633986564 | 0.448699458834792 | 0.731698625359227 | -5.47525965732296 |
| lnc-ECHDC3-9 | lnc-ECHDC3-9 | 0.185924373404247 | 3.38788947009349 | 0.764629283766759 | 0.448717343627466 | 0.731698625359227 | -5.92617968974076 |
| ENSG00000273398.6 | | 0.0948343081384375 | 3.30379833555922 | 0.764605979749057 | 0.448731076515062 | 0.731698625359227 | -5.92573883935663 |
| lnc-PPT1-4 | lnc-PPT1-4 | -0.229228538227831 | -0.18144866837585 | -0.764466141559754 | 0.448813487328814 | 0.731736282523816 | -5.3512996235294 |
| ENSG00000125686.11 | MED1 | 0.0734349243056744 | 6.10225173330247 | 0.764446146613089 | 0.448825271673866 | 0.731736282523816 | -6.12106748533426 |
| ENSG00000264522.5 | OTUD7B | -0.38729196713148 | 1.05753684383677 | -0.764388222543197 | 0.448951112550816 | 0.73186223320046 | -5.6714023141032 |
| lnc-TMEM41B-5 | lnc-TMEM41B-5 | -0.144426869568226 | 1.88418508221078 | -0.764166822647672 | 0.448989914783472 | 0.73186223320046 | -5.77538792741459 |
| ENSG00000108587.15 | GOSR1 | 0.0687081561557532 | 5.59426055340815 | 0.764134115223965 | 0.449009195979955 | 0.73186223320046 | -6.11372441995463 |
| lnc-DOLK-1 | lnc-DOLK-1 | -0.0984786342199802 | 3.00325395092989 | -0.764068967105655 | 0.449047602584472 | 0.731866878137317 | -5.89951724665046 |
| ENSG00000067048.16 | DDX3Y | -1.79829888056478 | 3.58904288541867 | -0.764170291768722 | 0.449155274927713 | 0.731971463082326 | -5.78975219482039 |
| ENSG00000123570.3 | RAB9B | -0.181983553133886 | 0.704596817623556 | -0.763839498351561 | 0.449182896088659 | 0.731971463082326 | -5.59984954142377 |
| ENSG00000161912.18 | ADCY10P1 | 0.191704436656699 | 3.01635900290933 | 0.763657651816214 | 0.449290128803846 | 0.732057926056162 | -5.89088366375408 |
| ENSG00000115392.11 | FANCL | 0.126342490751651 | 4.20325781223064 | 0.7636288955659 | 0.449307087395788 | 0.732057926056162 | -6.03658365732678 |
| lnc-TXNDC9-1 | lnc-TXNDC9-1 | 0.105196013781604 | 1.41398379328273 | 0.763360045465526 | 0.44946565613027 | 0.732199886416924 | -5.70125292443981 |
| ENSG00000106355.9 | LSM5 | 0.111883197068014 | 4.10558547141495 | 0.763353020440439 | 0.449469799955339 | 0.732199886416924 | -6.02281014513107 |
| lnc-PAPSS2-9 | lnc-PAPSS2-9 | -0.149600894346918 | 1.86854825027325 | -0.763300236808262 | 0.449500935955121 | 0.732199886416924 | -5.76830957877431 |
| ENSG00000278259.4 | MYO19 | -0.0810026863092759 | 5.08965265968213 | -0.763013675156136 | 0.449669994996549 | 0.732376094077443 | -6.09876888487558 |
| ENSG00000164088.17 | PPM1M | -0.161317487885099 | 5.41511596748815 | -0.762996254115522 | 0.449680273864389 | 0.732376094077443 | -6.10313185641104 |
| lnc-EPB42-1 | lnc-EPB42-1 | -0.182165557678731 | 1.04098775141977 | -0.762918056133134 | 0.449726414410719 | 0.732386729305145 | -5.63867819809328 |
| lnc-SHC3-7 | lnc-SHC3-7 | 0.15695416913436 | 1.12262692296719 | 0.762864582263476 | 0.4497579681493 | 0.732386729305145 | -5.64846281394198 |
| lnc-LIPI-7 | lnc-LIPI-7 | -0.823786667972105 | 1.10950386537422 | -0.762972995986162 | 0.449861101609005 | 0.732496721336773 | -5.64295144010735 |
| ENSG00000136856.17 | SLC2A8 | 0.114231037296493 | 2.23132876416562 | 0.762501046646967 | 0.449972516864407 | 0.73255921918403 | -5.81442536332601 |
| ENSG00000131669.9 | NINJ1 | 0.372377569437825 | -0.140915578288323 | 0.762494542733575 | 0.449976355841969 | 0.73255921918403 | -5.33349539795254 |

| | | | | | | | |
|---|---|---|---|---|---|---|---|
| ENSG00000183291.16 | SELENOF | 0.109651022440802 | 6.22979910144717 | 0.762443887632695 | 0.450006256005581 | 0.73255921918403 | -6.12323485770228 |
| ENSG00000136270.13 | TBRG4 | 0.104835842847926 | 5.54819599297911 | 0.762314934791397 | 0.450082378203659 | 0.732567975087178 | -6.11544068807653 |
| ENSG00000227827.3 | PKD1P2 | -0.465310816025429 | -0.386981884283458 | -0.762390287628345 | 0.450082908415403 | 0.732567975087178 | -5.36306327636177 |
| LY86-AS1 | LY86-AS1 | 0.51459085483982 | -0.166810393645724 | 0.762370704315693 | 0.450118407468665 | 0.732567975087178 | -5.35673322467864 |
| lnc-RAP1GAP2-4 | lnc-RAP1GAP2-4 | 0.175228690185805 | 0.612105622934102 | 0.762087926472359 | 0.4502164019249 | 0.732669528844102 | -5.55815745743049 |
| ENSG00000115085.13 | ZAP70 | -0.613074170270638 | 5.183876448212 | -0.762290894257228 | 0.450263503803075 | 0.732688251939181 | -6.09251833806674 |
| lnc-BRI3-2 | lnc-BRI3-2 | 0.119435676351977 | 2.62841931486209 | 0.761929893465966 | 0.450309717049831 | 0.732705526168906 | -5.85445271076856 |
| ENSG00000278048.1 | | -0.324027470936786 | -0.120491582130172 | -0.761851574849796 | 0.450355966729747 | 0.732722856943987 | -5.46511169608426 |
| ENSG00000143452.15 | HORMAD1 | -0.206880050372649 | 0.137436439650445 | -0.761617881746032 | 0.450493986639784 | 0.732889482058701 | -5.46223180938422 |
| ENSG00000130684.13 | ZNF337 | -0.0733325337972932 | 5.21547831159466 | -0.761494646104883 | 0.45056678000214 | 0.73291260366522 | -6.09998391254085 |
| ENSG00000126746.17 | ZNF384 | 0.0746755736120264 | 4.92682026819667 | 0.761451319204596 | 0.450592374162249 | 0.73291260366522 | -6.09061960057453 |
| ENSG00000189362.11 | NEMP2 | 0.0937529454723189 | 3.96879369443653 | 0.761374514350928 | 0.450637746586659 | 0.73291260366522 | -6.00653678198564 |
| lnc-ECHDC3-6 | lnc-ECHDC3-6 | -0.185527985016698 | 3.07704916550796 | -0.761352707022066 | 0.450650629742362 | 0.73291260366522 | -5.90023735505738 |
| lnc-ZNF724-6 | lnc-ZNF724-6 | -0.157406713028909 | 0.434037258133918 | -0.761066950386036 | 0.450819466672416 | 0.733083571181485 | -5.52394463402143 |
| ENSG00000240370.6 | RPL13P5 | 0.14811012417802 | 0.0504086748854972 | 0.761054226103299 | 0.450826985571357 | 0.733083571181485 | -5.4335164736479 |
| ENSG00000184009.11 | ACTG1 | -0.197480109284015 | 8.11807073570229 | -0.760874472609416 | 0.450971705035924 | 0.733211288477318 | -6.09988779946118 |
| ENSG00000121653.11 | MAPK8IP1 | -0.23411008393367 | -0.504762867059274 | -0.760800766300776 | 0.450976772682449 | 0.733211288477318 | -5.31372684836925 |
| ENSG00000267787.6 | NA | -0.107113913420472 | 4.14447459211832 | -0.760338183981919 | 0.451250220098114 | 0.733522742546315 | -6.02308305179043 |
| lnc-MCTP2-1 | lnc-MCTP2-1 | 0.105815936897582 | 4.77410312820311 | 0.760303976416706 | 0.451270445157094 | 0.733522742546315 | -6.07626386292186 |
| ENSG00000087299.11 | L2HGDH | -0.0854132001705561 | 1.82788319354885 | -0.760145019523166 | 0.451364434627395 | 0.733522742546315 | -5.7636931539122 |
| lnc-CCDC154-2 | lnc-CCDC154-2 | -0.301185258630991 | -0.36799563874013 | -0.760090546561464 | 0.451396646531671 | 0.733522742546315 | -5.34706151612674 |
| ENSG00000175893.11 | ZDHHC21 | -0.118174671415286 | 6.19862973366813 | -0.760086491113165 | 0.451399044724143 | 0.733522742546315 | -6.12470291054478 |
| ENSG00000252874.2 | | 0.188604895876516 | -0.138186742773486 | 0.759982341343278 | 0.45146063632821 | 0.733522742546315 | -5.36617268043983 |
| ENSG00000240065.7 | PSMB9 | 0.137233089734879 | 4.93378851853867 | 0.759942772167994 | 0.451484037852652 | 0.733522742546315 | -6.09116893735614 |
| ENSG00000169962.4 | TAS1R3 | -0.292939004267123 | 0.17285462519495 | -0.759924145718748 | 0.451495053928614 | 0.733522742546315 | -5.47832522933343 |
| lnc-MSANTD3-1 | lnc-MSANTD3-1 | 0.149378661797611 | 0.108789703716541 | 0.759888591747677 | 0.451516081736062 | 0.733522742546315 | -5.39582010842861 |
| ENSG00000173744.17 | AGFG1 | 0.0975509410260278 | 6.62922853136986 | 0.759849995765345 | 0.451538909340114 | 0.733522742546315 | -6.12389600511267 |
| ENSG00000087338.4 | GMCL1 | -0.0917626633728523 | 3.43109726445679 | -0.759658741296736 | 0.451652036821519 | 0.733522742546315 | -5.95659637329393 |
| ENSG00000164828.17 | SUN1 | -0.123894272940114 | 6.87170093787715 | -0.759642822102482 | 0.451661453811799 | 0.733522742546315 | -6.1225131195311 |
| lnc-DYNLL2-4 | lnc-DYNLL2-4 | 0.261060615123244 | 2.97300008996377 | 0.759509287746705 | 0.4517933238261 | 0.733522742546315 | -5.89754780215671 |
| ENSG00000174109.4 | UQCC4 | 0.128849715960577 | 1.73566756886091 | 0.759399196136755 | 0.451805584959023 | 0.733522742546315 | -5.75379436121242 |
| ENSG00000155307.18 | SAMSN1 | -0.597099038559834 | 3.88140986766973 | -0.759666245771215 | 0.45181387197928 | 0.733522742546315 | -5.95590858655522 |
| lnc-TINF2-1 | lnc-TINF2-1 | 0.0862880547714564 | 4.69157846641862 | 0.759331774136202 | 0.451845477132983 | 0.733522742546315 | -6.0714995199 |
| ENSG00000175048.16 | ZDHHC14 | 0.730567993935037 | -1.39119806095011 | 0.75934062167745 | 0.451851128210575 | 0.733522742546315 | -4.94875989600148 |
| lnc-SPINT2-4 | lnc-SPINT2-4 | 0.197983012321607 | -0.356278141310948 | 0.759249564431651 | 0.451894121669691 | 0.733522742546315 | -5.28695282778759 |
| lnc-BCL2L10-4 | lnc-BCL2L10-4 | 0.0900775145447541 | 1.90433084572405 | 0.759247594297784 | 0.451895287460794 | 0.733522742546315 | -5.76905555720098 |
| ENSG00000081913.13 | PHLPP1 | -0.680740999775108 | -1.65202187442035 | -0.759327926224285 | 0.451910283433946 | 0.733522742546315 | -5.12594711708191 |
| LINC01685 | LINC01685 | 0.331903373663966 | -0.693653187895009 | 0.75921137066925 | 0.451916722452015 | 0.733522742546315 | -5.26109981805105 |
| ENSG00000135272.10 | MDFIC | -0.143851282927439 | 6.1202064865871 | -0.759109223029162 | 0.451977170562738 | 0.733563010699639 | -6.12448198003672 |
| ENSG00000230148.8 | HOXB-AS1 | 0.772239092565752 | -2.62919407350389 | 0.75900451037011 | 0.452039141493099 | 0.733605743858114 | -4.85924013653071 |
| ENSG00000129636.12 | ITFG1 | 0.0705150840501414 | 4.97335849377479 | 0.758472785029916 | 0.452353903348546 | 0.734058687529416 | -6.08963583819739 |
| lnc-TRAF6-17 | lnc-TRAF6-17 | 0.0960282891941338 | 1.99052140713568 | 0.758359546237647 | 0.452420953113112 | 0.73407739119145 | -5.78147261297452 |
| ENSG00000206503.12 | HLA-A | 0.10029806685438 | 9.69447111744049 | 0.758182482397094 | 0.452525805949851 | 0.73407739119145 | -6.06185958460184 |
| lnc-NEFL-6 | lnc-NEFL-6 | -0.216406869167079 | 4.70091384303136 | -0.758224104308161 | 0.452553927856948 | 0.73407739119145 | -6.07362316727935 |
| lnc-RNF219-7 | lnc-RNF219-7 | -0.159587642599638 | 2.9189609938008 | -0.758119980706287 | 0.452562821303508 | 0.73407739119145 | -5.89956439634943 |
| ENSG00000274428.1 | RNVU1-25 | -1.27671270614967 | -1.87750765366052 | -0.758294537463927 | 0.452570354900611 | 0.73407739119145 | -4.89777372341961 |
| lnc-WRNIP1-34 | lnc-WRNIP1-34 | -0.211095730288436 | 1.83486026065676 | -0.75809196304068 | 0.452579414770418 | 0.73407739119145 | -5.76100216665523 |
| ENSG00000117500.12 | TMED5 | 0.0955121847807101 | 5.97782053978489 | 0.757798175987962 | 0.452753431631638 | 0.734253954489816 | -6.12565170518211 |
| lnc-EXOC7-3 | lnc-EXOC7-3 | -0.0807642105000896 | 2.16162473364312 | -0.757787735164626 | 0.452759616692646 | 0.734253954489816 | -5.79887852870427 |
| ENSG00000116863.10 | ADPRS | 0.101699329935181 | 3.69658908158322 | 0.757718607827489 | 0.452800568426146 | 0.734261910689992 | -5.97988726334205 |
| ENSG00000121858.10 | TNFSF10 | -0.228779754916285 | 2.0727862846354 | -0.757539932657698 | 0.452906554510233 | 0.734261910689992 | -5.79031902964919 |
| ENSG00000174989.12 | FBXW8 | -0.0901894271420467 | 4.72240402533033 | -0.757496232375901 | 0.452932320584948 | 0.734261910689992 | -6.07942609526148 |
| ENSG00000222078.1 | RN7SKP110 | 0.20318197191306 | 1.51701001897972 | 0.757478013747995 | 0.452943115681533 | 0.734261910689992 | -5.70027611785686 |
| lnc-MDN1-1 | lnc-MDN1-1 | 0.20318197191306 | 1.51701001897972 | 0.757478013747995 | 0.452943115681533 | 0.734261910689992 | -5.70027611785686 |
| ENSG00000273841.4 | TAF9 | 0.0860146542250693 | 5.17613725605574 | 0.757418193600214 | 0.452978562014357 | 0.734261910689992 | -6.1047792708247 |
| ENSG00000164050.12 | PLXNB1 | 0.322065547770983 | -0.532259015425662 | 0.757296988858522 | 0.453050386662501 | 0.734320506234684 | -5.36442552671554 |
| ENSG00000206621.1 | SNORD116-14 | -0.200968948748125 | 1.05805671150442 | -0.757048853166098 | 0.45319745001777 | 0.734501032576044 | -5.63109362422123 |
| ENSG00000065911.11 | MTHFD2 | -0.182069224218663 | 3.92923631847448 | -0.756957422824975 | 0.453252662078375 | 0.734532678022139 | -6.00813555662666 |
| ENSG00000119242.8 | CCDC92 | -0.131971848106095 | 3.30394558887945 | -0.756866265482518 | 0.453305682650079 | 0.734560767279686 | -5.93681654905924 |
| lnc-TIAF1-1 | lnc-TIAF1-1 | -0.110804746225312 | 1.3194146987522 | -0.756643576356224 | 0.453437706690857 | 0.734678170307639 | -5.69664369819192 |
| lnc-INTS13-3 | lnc-INTS13-3 | 0.227115019665714 | 0.452949376381969 | 0.756598279605032 | 0.453464564187775 | 0.734678170307639 | -5.50252989588327 |
| ENSG00000248871.1 | TNFSF12-TNFSF13 | -0.149959383185249 | 1.60180674441392 | -0.756563454037865 | 0.45348521370833 | 0.734678170307639 | -5.72508465730656 |
| ENSG00000100596.6 | SPTLC2 | -0.13320351370447 | 5.65087756296468 | -0.756239788934971 | 0.453677154430613 | 0.73492516464158 | -6.1228490935392 |
| ENSG00000130449.5 | ZSWIM6 | -0.10326046198719 | 4.16835477463955 | -0.756182801923076 | 0.453710953930135 | 0.73492516464158 | -6.0271373684601 |
| lnc-SOX30-5 | lnc-SOX30-5 | 0.223942497235039 | 1.70682690682298 | 0.756010442473357 | 0.453813190793127 | 0.73492516464158 | -5.74069151768036 |
| ENSG00000275146.1 | | 0.155093310034609 | 4.9893996682145 | 0.755995045739904 | 0.453822324186 | 0.73492516464158 | -6.09484774698246 |
| ENSG00000139629.15 | GALNT6 | -0.376116261492756 | -0.156463790583874 | -0.755913070546038 | 0.4538791165478 | 0.73492516464158 | -5.39524776206991 |
| lnc-TRMO-1 | lnc-TRMO-1 | 0.129091100943529 | 3.51980094916567 | 0.755893082943245 | 0.453882811561861 | 0.73492516464158 | -5.96134241962652 |

| | | | | | | | |
|---|---|---|---|---|---|---|---|
| ENSG00000103202.12 | NME4 | -0.330659899393028 | 1.24958168450024 | -0.755993573498507 | 0.453887610791613 | 0.73492516464158 | -5.717543804169 |
| lnc-SLC4A5-3 | lnc-SLC4A5-3 | -0.0980996134324021 | 2.26174632509288 | -0.755816548228085 | 0.453928217334021 | 0.734933099770797 | -5.81077804941673 |
| lnc-PLK1-1 | lnc-PLK1-1 | -0.0965329188297246 | 2.45140670085068 | -0.755662208662443 | 0.454019790480523 | 0.734979406884445 | -5.83814963354074 |
| lnc-PIAS3-1 | lnc-PIAS3-1 | 0.185097783543444 | 0.865853045764933 | 0.755647977185589 | 0.454028234880033 | 0.734979406884445 | -5.59453825176184 |
| ENSG00000103005.11 | USB1 | 0.0889765292430131 | 3.39091937632805 | 0.75557414009202 | 0.454072048384187 | 0.734992526886735 | -5.94170108755017 |
| ENSG00000247626.4 | MARS2 | 0.107812460230801 | 3.2095407193413 | 0.755396480047402 | 0.454177478536838 | 0.735105373965851 | -5.93514394752232 |
| ENSG00000142552.7 | RCN3 | -0.260363499699029 | 1.67689191807422 | -0.755263573847293 | 0.454264513963921 | 0.735188433001996 | -5.71578743896731 |
| ENSG00000186073.12 | CDIN1 | -0.113321392219803 | 2.60486656066628 | -0.755079879463957 | 0.45436539665648 | 0.735293887835547 | -5.85631007518976 |
| RFX3-AS1 | RFX3-AS1 | -0.132889346043751 | 6.02458661946824 | -0.754943958939153 | 0.454446086139403 | 0.735366650236425 | -6.12755919565873 |
| ENSG00000087263.16 | OGFOD1 | -0.0748547208153667 | 4.51440984179879 | -0.754847869969493 | 0.454503134588308 | 0.735401149220278 | -6.06838170737527 |
| ENSG00000279089.1 | | 0.130097065851552 | 3.1420917866734 | 0.754719057802436 | 0.454579617501726 | 0.735444451047096 | -5.91926154502375 |
| lnc-LHFPL2-3 | lnc-LHFPL2-3 | -0.319210558027383 | 1.12244918049127 | -0.754746673266912 | 0.454601357949385 | 0.735444451047096 | -5.65147992741862 |
| ENSG00000239704.10 | CDRT4 | -0.280342509495345 | 1.22016484645004 | -0.754649587395609 | 0.454638197301287 | 0.735446244502703 | -5.69783983999806 |
| ENSG00000186017.14 | ZNF566 | 0.101967002528303 | 3.44154920949723 | 0.754310576926892 | 0.454822204873794 | 0.735654299313378 | -5.9450165846621 |
| lnc-SPINT2-9 | lnc-SPINT2-9 | -0.24701063139534 | -0.317083355683306 | -0.754283489111958 | 0.454838294372154 | 0.735654299313378 | -5.33040045272459 |
| ENSG00000176485.11 | PLAAT3 | 0.714879040298292 | -1.47880360454853 | 0.754158807438027 | 0.454952197509675 | 0.73578070883489 | -5.03992143670587 |
| lnc-LRFN1-3 | lnc-LRFN1-3 | 0.10398621165169 | 4.16481662809227 | 0.753924093713581 | 0.455051797858183 | 0.735843559567051 | -6.03154744035914 |
| lnc-RIMKLB-6 | lnc-RIMKLB-6 | 1.16281023696605 | -2.67757932789583 | 0.753951052770794 | 0.455064501285242 | 0.735843559567051 | -4.78412322578064 |
| ENSG00000162664.16 | ZNF326 | 0.104732722761047 | 6.00732339278472 | 0.753845811905568 | 0.455098309929969 | 0.735843559567051 | -6.12869560649326 |
| lnc-CDK12-2 | lnc-CDK12-2 | 0.138668050153411 | -0.114908891548746 | 0.753721407861073 | 0.45517223178347 | 0.735905274275325 | -5.36755174536208 |
| ENSG00000089177.18 | KIF16B | -0.0874451655618635 | 4.2092898919358 | -0.753232886323019 | 0.45546258281461 | 0.736316866326823 | -6.03183188710764 |
| lnc-GJA9-1 | lnc-GJA9-1 | -0.136420835522726 | 1.11627775737861 | -0.752832939939469 | 0.455700369873719 | 0.736613015723991 | -5.66785890142026 |
| ENSG00000065135.10 | GNAI3 | 0.0975483226302032 | 5.97856581147115 | 0.752804390787384 | 0.455717346462095 | 0.736613015723991 | -6.12912764348113 |
| ENSG00000097007.17 | ABL1 | 0.0663382264811299 | 4.65262313385372 | 0.752564956671677 | 0.455859739081644 | 0.736730943387882 | -6.07904373929914 |
| ENSG00000151689.12 | INPP1 | 0.142097041684168 | 1.19867632893319 | 0.752561339151028 | 0.455861890637325 | 0.736730943387882 | -5.6846730363224 |
| ENSG00000105409.18 | ATP1A3 | -0.316478831374668 | -0.315697789129505 | -0.752345778300816 | 0.455990108234614 | 0.736880301310596 | -5.32085698966331 |
| ZBED3-AS1 | ZBED3-AS1 | -0.257785067587501 | 2.54207975622946 | -0.752280349739774 | 0.45606799098698 | 0.736915299405804 | -5.85436859519252 |
| lnc-ERCC6-1 | lnc-ERCC6-1 | -0.115838030735755 | 3.59136741429078 | -0.752189008220261 | 0.456083369728928 | 0.736915299405804 | -5.97246719532348 |
| ENSG00000135905.19 | DOCK10 | 0.123874016635165 | 8.67713803822818 | 0.752054282808726 | 0.45616352586969 | 0.736986958632326 | -6.0941701591693 |
| ENSG00000221978.11 | CCNL2 | 0.0814045483221735 | 6.90310832312088 | 0.751779095320994 | 0.456327276724982 | 0.737193653310596 | -6.12843374561044 |
| ENSG00000005075.15 | POLR2J | -0.126393063818974 | 3.29263397885627 | -0.751599272866579 | 0.456434298918067 | 0.737271098543879 | -5.94501687470607 |
| ENSG00000278861.1 | | -0.118288590096786 | 0.283273808241313 | -0.751571207439047 | 0.456451003509623 | 0.737271098543879 | -5.49431901615295 |
| ENSG00000165752.16 | STK32C | -0.151527080666992 | 1.24078797532172 | -0.751469393114236 | 0.456511606561413 | 0.737271098543879 | -5.68768813300992 |
| ENSG00000161847.13 | RAVER1 | -0.12441569953637 | 3.52544108505001 | -0.751427870107619 | 0.45653632369064 | 0.737271098543879 | -5.97195561092681 |
| lnc-MFAP2-53 | lnc-MFAP2-53 | -0.16624040176821 | 2.08092171960748 | -0.751397650378323 | 0.456554312881518 | 0.737271098543879 | -5.77941081148586 |
| lnc-OR4F3-5 | lnc-OR4F3-5 | -0.184114457930243 | 0.368612205271322 | -0.751300067715443 | 0.456612404675502 | 0.737307062480259 | -5.49483749824975 |
| ENSG00000137825.10 | ITPKA | 0.453735001404592 | -0.926776890446364 | 0.751035084362661 | 0.456770173234461 | 0.737503959498346 | -5.18068335869402 |
| ENSG00000173681.16 | BCLAF3 | -0.0829782944127175 | 3.9362788707663 | -0.750826328331628 | 0.456894486920169 | 0.737591478721425 | -6.01559685790084 |
| lnc-HEBP2-2 | lnc-HEBP2-2 | -0.102776840665648 | 4.29483901208155 | -0.750823707256375 | 0.456896047888946 | 0.737591478721425 | -6.04614255417819 |
| lnc-ELOA3D-2 | lnc-ELOA3D-2 | 0.130169743712377 | 0.0872390452938899 | 0.750612760928693 | 0.457021686135147 | 0.737672684849307 | -5.45114169970252 |
| ENSG00000002016.17 | RAD52 | -0.15524652424036 | 2.73780393704098 | -0.750575182465159 | 0.457044069730007 | 0.737672684849307 | -5.87666750731235 |
| ENSG00000196873.15 | ZNG1C | -0.126810955486834 | 2.23274671774032 | -0.750558734315782 | 0.457053867263431 | 0.737672684849307 | -5.81987683384045 |
| ENSG00000126461.14 | SCAF1 | -0.0978079932102385 | 3.85814438331983 | -0.750356257913628 | 0.457174484720304 | 0.737718415497432 | -6.0092758180464 |
| ENSG00000196588.15 | MRTFA | -0.0869776779229277 | 4.76239029515507 | -0.750200512022381 | 0.457267276886198 | 0.737718415497432 | -6.08970439693495 |
| lnc-KL-5 | lnc-KL-5 | 0.15794226959709 | 1.87179342319588 | 0.75018366732273 | 0.457277313480924 | 0.737718415497432 | -5.75977801378579 |
| lnc-CCDC146-96 | lnc-CCDC146-96 | -0.266159467391387 | -0.280833445034302 | -0.750175583417737 | 0.45728213016785 | 0.737718415497432 | -5.35852596616019 |
| ENSG00000120314.18 | WDR55 | -0.056622199772776 | 4.99661276869855 | -0.750173111895659 | 0.457283602797192 | 0.737718415497432 | -6.10408054393595 |
| ENSG00000255104.8 | ZNF286A-TBC1D26 | 0.121592013115853 | 1.65673236899891 | 0.75015021066374 | 0.457297248376414 | 0.737718415497432 | -5.73145630921561 |
| ENSG00000055044.10 | NOP58 | 0.112005756655098 | 7.3564356949628 | 0.750037888851376 | 0.457364178169523 | 0.737768564205587 | -6.12407517093753 |
| lnc-FHDC1-4 | lnc-FHDC1-4 | -0.0918395231477697 | 0.776685625155303 | -0.749753275140918 | 0.457533797950554 | 0.737984340037321 | -5.6064836878233 |
| lnc-MARCH6-3 | lnc-MARCH6-3 | -0.254121224503057 | -0.488749270520542 | -0.749632392261941 | 0.4576058509673 | 0.738042722963481 | -5.29827039539224 |
| ENSG00000188938.16 | FAM120AOS | 0.0808582852954078 | 5.46705347505619 | 0.749497326569955 | 0.457686365543086 | 0.738114742769986 | -6.12078415598757 |
| ENSG00000164219.9 | PGGT1B | 0.0794987689704945 | 5.76828774640886 | 0.749335844296349 | 0.457782638225118 | 0.738207611062225 | -6.12868353696434 |
| lnc-PTBP3-6 | lnc-PTBP3-6 | 0.200106870304228 | -0.512142483019774 | 0.749256465944837 | 0.457829966413732 | 0.738207611062225 | -5.31637538388301 |
| lnc-RBP4-1 | lnc-RBP4-1 | -0.112316128386489 | 2.33236509477164 | -0.749185643945821 | 0.457872195401843 | 0.738207611062225 | -5.82890807647846 |
| lnc-PTPN4-1 | lnc-PTPN4-1 | -0.178621968312975 | 0.97608972636996 | -0.749160127711352 | 0.457887410505341 | 0.738207611062225 | -5.64777793076806 |
| ENSG00000189164.14 | ZNF527 | 0.0869051112620616 | 3.43351456839305 | 0.749094056175564 | 0.457926809739433 | 0.738213308651844 | -5.95122532602123 |
| ENSG00000070413.19 | DGCR2 | -0.0834678090817444 | 5.63537921174698 | -0.748792164006074 | 0.458106856654702 | 0.738445722493831 | -6.126101632016 |
| lnc-DCANP1-4 | lnc-DCANP1-4 | 0.125938456057227 | 3.22560957165965 | 0.748694056088469 | 0.458165376558309 | 0.738446169576812 | -5.92804753497909 |
| ENSG00000182518.13 | VCF2 | -0.167141309848824 | 0.275116241145042 | -0.748671406984774 | 0.458178887026452 | 0.738446169576812 | -5.47466430107606 |
| ENSG00000170734.11 | POLH | -0.183630357017077 | 3.35403433352999 | -0.748490422567374 | 0.458286854764469 | 0.738562349797765 | -5.94408339771017 |
| lnc-CHKA-1 | lnc-CHKA-1 | -0.0747667968574248 | 2.38914211726873 | -0.748295826685403 | 0.458402959018198 | 0.738691623383041 | -5.83338940834572 |
| ENSG00000078589.12 | P2RY10 | 0.0944351699151813 | 7.04495072957789 | 0.748187323910968 | 0.458467703846677 | 0.738738120265864 | -6.13060060390983 |
| ENSG00000115947.13 | ORC4 | -0.0959771965840543 | 4.59744261932499 | -0.74810723312559 | 0.458515498327411 | 0.738757299575193 | -6.07954281253658 |
| ENSG00000132763.14 | MMACHC | -0.25628621243179 | 0.633648511848132 | -0.748034592548412 | 0.458558849369965 | 0.738769317202724 | -5.53900271953327 |
| ENSG00000163600.12 | ICOS | -0.919071380236023 | -1.11074972960695 | -0.748084172507662 | 0.458644772947763 | 0.738849914814417 | -5.1166970116544 |
| lnc-SYN1-3 | lnc-SYN1-3 | -0.108443837789099 | 1.15917023387052 | -0.747815018505598 | 0.458689903045793 | 0.738864789043008 | -5.66651526736793 |
| ENSG00000184083.11 | FAM120C | -0.0876757918197733 | 3.14522364809335 | -0.747700196581573 | 0.458758443643527 | 0.738917367987692 | -5.93207237310164 |

| | | | | | | | |
|---|---|---|---|---|---|---|---|
| lnc-MAP1LC3C-1 | lnc-MAP1LC3C-1 | -0.733107287656448 | -1.95319543685872 | -0.747539597076657 | 0.458867198939459 | 0.739034707024326 | -4.95214680811908 |
| lnc-SLC35G5-2 | lnc-SLC35G5-2 | -0.151619401259302 | 4.74527963883159 | -0.747345660429828 | 0.458970114259225 | 0.739056806016661 | -6.08703131062771 |
| ENSG00000103495.13 | MAZ | -0.140749002523057 | 4.77083630837944 | -0.747344890955335 | 0.45897057372433 | 0.739056806016661 | -6.09370481989112 |
| ENSG00000183837.9 | PNMA3 | -0.25301969370667 | 1.07665992187987 | -0.747314637492101 | 0.4589886387461 | 0.739056806016661 | -5.61996473516147 |
| lnc-PDIA4-4 | lnc-PDIA4-4 | -0.204219551330311 | 0.0353013356137163 | -0.747092607278965 | 0.459121230594401 | 0.739205502934337 | -5.42895153469428 |
| ENSG00000143322.19 | ABL2 | 0.0951886120826693 | 4.84241360484373 | 0.746981807313371 | 0.459187406352531 | 0.739205502934337 | -6.09294500771732 |
| lnc-RSF1-3 | lnc-RSF1-3 | 0.115716530423746 | 1.22407473661731 | 0.746979596805441 | 0.459188726644242 | 0.739205502934337 | -5.66530789903653 |
| ENSG00000183323.12 | CCDC125 | -0.110691947318949 | 3.21108423455818 | -0.746857817756901 | 0.45926146621158 | 0.739264781343 | -5.92912689136123 |
| lnc-INTS14-1 | lnc-INTS14-1 | 0.1317055214225 | 2.850952169531 | 0.746704002082271 | 0.45935335105624 | 0.739354865873521 | -5.88664575187286 |
| ENSG00000119522.16 | DENND1A | 0.13546378152332 | 4.51064468582737 | 0.74662275325714 | 0.459401890957408 | 0.739375175664737 | -6.0690246034313 |
| lnc-IL31RA-1 | lnc-IL31RA-1 | -0.265774039916586 | -0.0513564599333394 | -0.746497346044255 | 0.459476817926019 | 0.739437947097049 | -5.4334966383773 |
| ENSG00000175105.6 | ZNF654 | 0.0839322089204311 | 4.43945018370893 | 0.746408852578058 | 0.459529694326462 | 0.739465225417995 | -6.06825894423701 |
| lnc-PLEKHM1-8 | lnc-PLEKHM1-8 | -0.189263346925217 | 1.23732577441594 | -0.746090312424031 | 0.459720056873715 | 0.739621772312236 | -5.69842778187553 |
| ENSG00000198952.8 | SMG5 | 0.0671981317251822 | 5.912831471238 | 0.746081870780349 | 0.459725102299996 | 0.739621772312236 | -6.13337304338764 |
| ENSG00000196357.11 | ZNF565 | 0.0954705482768626 | 1.0377357127237 | 0.745999643211768 | 0.459774249993017 | 0.739621772312236 | -5.63790610357832 |
| ENSG00000186111.9 | PIP5K1C | 0.160953489333433 | 3.89520397293374 | 0.745997348104314 | 0.459775621830121 | 0.739621772312236 | -6.00448804759485 |
| lnc-PDK3-1 | lnc-PDK3-1 | 0.0881846590198974 | 4.19381778850383 | 0.745945444869784 | 0.45980664618896 | 0.739621772312236 | -6.04531814252368 |
| ENSG00000116874.11 | WARS2 | 0.094854231532902 | 3.16665170068625 | 0.745838382870302 | 0.459870644683142 | 0.739666912519583 | -5.92897036706812 |
| lnc-WBP4-2 | lnc-WBP4-2 | -0.183486716169021 | 1.51234840814893 | -0.745750280605262 | 0.459923313481947 | 0.739693824144313 | -5.70879259248679 |
| ENSG00000185619.18 | PCGF3 | -0.11162770404687 | 6.06733460095033 | -0.745681846215007 | 0.459964226955489 | 0.739701826972797 | -6.13491355959792 |
| ENSG00000171428.13 | NAT1 | -0.146311647715716 | 1.60169604454488 | -0.745419099406584 | 0.460121329628593 | 0.739803072511091 | -5.73252961731171 |
| lnc-RGS17-2 | lnc-RGS17-2 | 0.0989679479027221 | 0.468508061643006 | 0.745398339277282 | 0.460133743937485 | 0.739803072511091 | -5.55814379212292 |
| ENSG00000224430.2 | MKRN5P | -0.280838442764603 | 1.18507803842951 | -0.745361669924824 | 0.460171108010292 | 0.739803072511091 | -5.68188023174421 |
| ENSG00000152495.10 | CAMK4 | -1.83559278866731 | -2.15713876855397 | -0.745485483305774 | 0.460219700543605 | 0.739803072511091 | -4.82794945503136 |
| ENSG00000108256.8 | NUFIP2 | 0.0703953757224589 | 7.59724578597718 | 0.74522010045388 | 0.460240336607397 | 0.739803072511091 | -6.12366101305528 |
| lnc-TPD52-3 | lnc-TPD52-3 | 0.207131319349584 | 3.3790580616393 | 0.74523619174535 | 0.46024283843485 | 0.739803072511091 | -5.9400016232485 |
| lnc-EMC9-2 | lnc-EMC9-2 | 0.130281484505375 | 2.75182594847301 | 0.744644585551074 | 0.460584611081558 | 0.740294631414314 | -5.88042414650043 |
| ENSG00000130921.7 | MTRFR | -0.0846520271297075 | 4.14961640688773 | -0.744552756710185 | 0.460639557121325 | 0.740325135022116 | -6.03680995327251 |
| ENSG00000123415.15 | SMUG1 | -0.125494308200277 | 2.79350467938314 | -0.744453536348332 | 0.460698930164352 | 0.740362748248974 | -5.8854091365323 |
| ENSG00000105778.18 | AVL9 | 0.0702009446803746 | 6.30729172050328 | 0.743843555988581 | 0.461064037168947 | 0.740841599522931 | -6.13704832577889 |
| lnc-TEX29-7 | lnc-TEX29-7 | 0.194686835088151 | 0.945062364164747 | 0.743835456221435 | 0.461068886454295 | 0.740841599522931 | -5.64928979289134 |
| lnc-EIF4E3-8 | lnc-EIF4E3-8 | 0.221808455562982 | 1.25112060130763 | 0.743537903434253 | 0.461247050105916 | 0.74107002047694 | -5.67246489506485 |
| ENSG00000125868.15 | DSTN | 0.164614985055119 | 1.95875694090145 | 0.743405838071375 | 0.461326138744811 | 0.741139237729039 | -5.78732042282517 |
| ENSG00000008988.9 | RPS20 | 0.132277311799683 | 9.11339935104874 | 0.743277117431151 | 0.461402034020077 | 0.741203314308533 | -6.09030249321261 |
| ENSG00000124209.3 | RAB22A | 0.070895229168622 | 5.08873366746587 | 0.743154478550782 | 0.461476689537971 | 0.741205208524602 | -6.11167706061147 |
| ENSG00000244405.7 | ETV5 | -0.298816161660168 | 0.786262248903613 | -0.743055791132061 | 0.461554802077334 | 0.741205208524602 | -5.64305163486891 |
| lnc-ZNF669-2 | lnc-ZNF669-2 | 0.14269500436352 | 0.619301294734046 | 0.742942276795106 | 0.461603809064626 | 0.741205208524602 | -5.56899064892356 |
| ENSG00000237550.5 | | 0.232332645482229 | 5.56911910572154 | 0.743060406301246 | 0.461619438610658 | 0.741205208524602 | -6.124740153428 |
| ENSG00000119509.12 | INVS | 0.116596027873155 | 4.70779718077231 | 0.742894375139317 | 0.46163250736793 | 0.741205208524602 | -6.08371234713739 |
| lnc-FBXL20-3 | lnc-FBXL20-3 | -0.108151600301372 | 1.33317778189934 | -0.742876400335207 | 0.461643276497429 | 0.741205208524602 | -5.70698990191923 |
| ENSG00000264653.1 | MIR5194 | 0.177028494921707 | 0.426173034168893 | 0.742856353757386 | 0.461655287046854 | 0.741205208524602 | -5.50702415666439 |
| ENSG00000033100.16 | CHPF2 | -0.08690493952568 | 5.43421722867218 | -0.742785392630356 | 0.461697803592096 | 0.74121565332939 | -6.12544747923519 |
| ENSG00000145907.14 | G3BP1 | -0.0867069546545354 | 7.37546854577024 | -0.742690217698139 | 0.46175483145739 | 0.741249391349825 | -6.12837309271066 |
| ENSG00000137413.15 | TAF8 | 0.0689207985692573 | 5.39221001115223 | 0.742399352657054 | 0.461929140118645 | 0.741471378855344 | -6.12371641671279 |
| lnc-RAP2C-5 | lnc-RAP2C-5 | 0.08534295351176 | 2.68779835266142 | 0.742223085517007 | 0.462034791430382 | 0.741583134124419 | -5.87733101207743 |
| ENSG00000224769.1 | MUC20P1 | -0.427319663143505 | -0.372907035747166 | -0.742034090791364 | 0.462163846055164 | 0.741732432230288 | -5.33004156140005 |
| ENSG00000137171.14 | KLC4 | -0.0708116916622918 | 3.84085648513483 | -0.741786253326429 | 0.462296680881351 | 0.741806160585558 | -6.01475912904747 |
| ENSG00000084090.13 | STARD7 | -0.133221447728959 | 7.87116265918258 | -0.741762427641585 | 0.462310967312473 | 0.741806160585558 | -6.12098374189295 |
| ENSG00000101049.15 | SGK2 | 0.21313090134789 | 0.0554765000563499 | 0.741750858575408 | 0.462317904483872 | 0.741806160585558 | -5.44296088586917 |
| lnc-ARMCX1-2 | lnc-ARMCX1-2 | 0.114802539284076 | 1.8522775155124 | 0.74157412279635 | 0.462423888224428 | 0.741831138235186 | -5.77065886892417 |
| lnc-MAP7D2-1 | lnc-MAP7D2-1 | -0.153279635314556 | -0.19694312285265 | -0.741560319409849 | 0.462432166338728 | 0.741831138235186 | -5.3832533626636 |
| lnc-HOMER3-3 | lnc-HOMER3-3 | 0.0869218704654351 | 2.94084545353989 | 0.741543831910969 | 0.462442054270879 | 0.741831138235186 | -5.90752337823092 |
| ENSG00000251682.1 | | 0.211741121062965 | 1.34651671400852 | 0.741484504615028 | 0.46247763522489 | 0.741831138235186 | -5.68562395946171 |
| ENSG00000071127.16 | WDR1 | 0.0751920212188768 | 6.71152639027296 | 0.741280008008792 | 0.462600292146191 | 0.741957301457733 | -6.13729848240525 |
| lnc-CLEC3A-2 | lnc-CLEC3A-2 | 0.219833868942381 | 0.51855155238594 | 0.741156210840154 | 0.46267455473176 | 0.741957301457733 | -5.55896252212001 |
| lnc-ZNF736-4 | lnc-ZNF736-4 | 0.107735926228894 | 3.72773322626567 | 0.741155865001013 | 0.462674762200986 | 0.741957301457733 | -6.00136127145402 |
| lnc-FAM96B-4 | lnc-FAM96B-4 | -0.198853962553538 | -0.193265156495551 | -0.741096086928754 | 0.462710623930166 | 0.741957301457733 | -5.36847237544979 |
| ENSG00000164118.12 | CEP44 | -0.103730579560457 | 4.51513179307776 | -0.74105291522304 5 | 0.46273652425851 | 0.741957301457733 | -6.07356858195855 |
| ENSG00000197647.11 | ZNF433 | -0.284426603974184 | 2.62589586666274 | -0.741049993792014 | 0.462805016201198 | 0.742006598796769 | -5.86250294869569 |
| lnc-HNRNPF-1 | lnc-HNRNPF-1 | -0.333221205109368 | 0.401513849819858 | -0.740885592573055 | 0.462852244328743 | 0.742006598796769 | -5.52759184992832 |
| ENSG00000197530.12 | MIB2 | -0.147690012963401 | 3.73881001674721 | -0.740802572611771 | 0.462886730701692 | 0.742006598796769 | -6.00506107646739 |
| lnc-STYXL1-2 | lnc-STYXL1-2 | -0.0927409306992404 | 0.681137555151647 | -0.740739543079623 | 0.46292455307788 | 0.742006598796769 | -5.5934002278528 |
| lnc-WIPF1-3 | lnc-WIPF1-3 | 0.266856130458858 | -1.03304319556391 | 0.740650551834996 | 0.462977957438003 | 0.742006598796769 | -5.20865416183896 |
| lnc-GJA10-23 | lnc-GJA10-23 | -0.242533033340563 | 1.58870142144954 | -0.740644297080188 | 0.462983566455177 | 0.742006598796769 | -5.74334810637338 |
| ENSG00000272693.4 | NUPR2P1 | -0.293810483478097 | -0.0706683607023354 | -0.74038863352184 | 0.463135157359588 | 0.742061718639962 | -5.38196760082263 |
| lnc-GPR68-2 | lnc-GPR68-2 | 0.0936047666512001 | 1.59486978078182 | 0.740255617113641 | 0.463215003851697 | 0.742061718639962 | -5.74530135311597 |
| lnc-CCDC13-5 | lnc-CCDC13-5 | 0.106303295174976 | 1.54625046112528 | 0.740068220676069 | 0.463327506824766 | 0.742061718639962 | -5.73797877818325 |
| ENSG00000256745.1 | ARPC3P3 | 0.621390944092073 | -1.54156274436323 | 0.739951360623386 | 0.463397671441435 | 0.742061718639962 | -4.92896896060319 |

| | | | | | | | |
|---|---|---|---|---|---|---|---|
| lnc-GPR83-5 | lnc-GPR83-5 | 0.621390944092073 | -1.54156274436323 | 0.739951360623386 | 0.463397671441435 | 0.742061718639962 | -4.92896896060319 |
| lnc-ICAM3-2 | lnc-ICAM3-2 | -0.090805191876806 | 2.57970731645205 | -0.739852789033185 | 0.463456860144142 | 0.742061718639962 | -5.85900605316952 |
| ENSG00000134186.11 | PRPF38B | -0.0698414827566909 | 6.62826407588715 | -0.739814868678015 | 0.463479631118499 | 0.742061718639962 | -6.13874431352876 |
| lnc-CLEC16A-1 | lnc-CLEC16A-1 | 0.150967834486727 | 2.79645644433716 | 0.739782165605528 | 0.463499269660404 | 0.742061718639962 | -5.87614860648339 |
| ENSG00000111361.12 | EIF2B1 | 0.0806504791894476 | 5.35615910952443 | 0.739702678256408 | 0.463547004658333 | 0.742061718639962 | -6.12448389862297 |
| lnc-SLC7A6-3 | lnc-SLC7A6-3 | -0.0994033134621392 | 0.243840509613053 | -0.73969732597699 | 0.463550218995599 | 0.742061718639962 | -5.48554843539179 |
| lnc-RDH13-4 | lnc-RDH13-4 | -0.0869313006723229 | 1.68980922017252 | -0.739630201653624 | 0.463590531924261 | 0.742061718639962 | -5.76297337026782 |
| ENSG00000139055.6 | ERP27 | -0.15722846979279 | 2.17927145621473 | -0.739593358097334 | 0.463612659959102 | 0.742061718639962 | -5.81622259117376 |
| LINC01473 | LINC01473 | -0.166820263201614 | 7.48367171567602 | -0.739538990308556 | 0.463653904712853 | 0.742061718639962 | -6.12824502280896 |
| ENSG00000226501.2 | USF1P1 | 0.164205084649239 | 0.106245787283948 | 0.739501529911177 | 0.463667814108467 | 0.742061718639962 | -5.45151764487161 |
| lnc-PLA2G15-3 | lnc-PLA2G15-3 | 0.324344636366377 | -0.650693131182299 | 0.739458557863226 | 0.463693625422294 | 0.742061718639962 | -5.16112901413652 |
| ENSG00000186577.13 | SMIM29 | -0.149244871047748 | 2.5773901163617 | -0.739440387538983 | 0.463704539741667 | 0.742061718639962 | -5.88496016227747 |
| ENSG00000147642.16 | SYBU | -0.297833247363886 | 0.884322216957958 | -0.739454271791617 | 0.463710848300723 | 0.742061718639962 | -5.62185561069614 |
| lnc-A1BG-1 | lnc-A1BG-1 | -0.11234566298831 | 0.658189214686162 | -0.739421191256 | 0.46371607048187 | 0.742061718639962 | -5.58070878389097 |
| RNF139-AS1 | RNF139-AS1 | -0.130942562973982 | 2.36466279512552 | -0.739395485476491 | 0.46373151157731 | 0.742061718639962 | -5.84606938778233 |
| ENSG00000136811.16 | ODF2 | -0.091770398011227 | 5.16562279888628 | -0.739383015164807 | 0.463739002422671 | 0.742061718639962 | -6.11757681315723 |
| ENSG00000234545.7 | FAM133B | 0.0762092696013346 | 4.47109899467806 | 0.739271275935335 | 0.463806126657373 | 0.742111435400242 | -6.07581206156259 |
| lnc-BLVRB-1 | lnc-BLVRB-1 | 0.171210341978033 | 1.02530236736787 | 0.739208658729479 | 0.463843744652367 | 0.742113936741521 | -5.66281066816872 |
| lnc-SUN2-2 | lnc-SUN2-2 | 0.16362268154088 | 0.948065376597229 | 0.738705056766424 | 0.464146353209341 | 0.74244863369042 | -5.63571978601677 |
| ENSG00000128908.16 | INO80 | 0.0772909324070836 | 6.49278123043854 | 0.738652707617498 | 0.464177815732712 | 0.74244863369042 | -6.14074931027102 |
| lnc-NOP14-8 | lnc-NOP14-8 | 0.100985393443518 | 3.46004026474128 | 0.7386275088881 | 0.464192960937089 | 0.74244863369042 | -5.96901846875945 |
| ENSG00000181804.14 | SLC9A9 | 0.210288663858873 | 3.90630615678048 | 0.738619239238168 | 0.464231798747553 | 0.74244863369042 | -6.01493490416466 |
| lnc-TXNRD2-2 | lnc-TXNRD2-2 | -0.114504817358225 | 0.437690156072312 | -0.738525444230388 | 0.464254307820708 | 0.74244863369042 | -5.54477340230408 |
| ENSG00000188186.10 | LAMTOR4 | 0.0895010688025524 | 4.68442935097435 | 0.73850039236112 | 0.464269366187116 | 0.74244863369042 | -6.09157594105031 |
| lnc-UGT3A2-3 | lnc-UGT3A2-3 | -0.115833679128019 | 2.00004770405502 | -0.738377295479624 | 0.464343362278852 | 0.742453551782707 | -5.80195121038908 |
| ENSG00000268049.1 | | -0.16246612273862 | 0.652215512004511 | -0.738375263199176 | 0.464344583981868 | 0.742453551782707 | -5.6019140407403 |
| ENSG00000259080.1 | | 0.0809382705896415 | 4.03230642474486 | 0.738210296337264 | 0.464443759796714 | 0.742554443676849 | -6.03087178631208 |
| ENSG00000181744.8 | DIPK2A | -0.103163928113387 | 4.27589597574894 | -0.737929499308118 | 0.464612599189369 | 0.742687958458807 | -6.06515317422456 |
| ENSG00000170653.18 | ATF7 | -0.107472087826026 | 4.02362600046298 | -0.737829776709442 | 0.464672569525569 | 0.742687958458807 | -6.03240346910002 |
| lnc-CABLES2-5 | lnc-CABLES2-5 | 0.245676340679071 | 0.143681457494391 | 0.73772732782034 | 0.464734184019257 | 0.742687958458807 | -5.42583111516746 |
| ENSG00000104687.13 | GSR | 0.128740341536997 | 4.81739603770936 | 0.737674918937126 | 0.464765705427129 | 0.742687958458807 | -6.10018997034101 |
| ENSG00000183066.14 | WBP2NL | 0.471271859705675 | 1.29873894427567 | 0.73785583683755 | 0.464776086601892 | 0.742687958458807 | -5.70237815661196 |
| lnc-TPD52-4 | lnc-TPD52-4 | 0.106055540481514 | 1.65446742622174 | 0.737617747763838 | 0.464800092526613 | 0.742687958458807 | -5.74206572231411 |
| ENSG00000226124.7 | FTCDNL1 | -0.220059131703015 | 0.0558796284924948 | -0.737605823420455 | 0.46480726491995 | 0.742687958458807 | -5.40995166490027 |
| ENSG00000107742.12 | SPOCK2 | -0.157805038078969 | 8.69613399019147 | -0.7374846403711 | 0.46488015913199 | 0.742687958458807 | -6.10634837087249 |
| lnc-SYNJ2BP-2 | lnc-SYNJ2BP-2 | -0.160518037353638 | 1.16117136271281 | -0.737464680455902 | 0.464892166082054 | 0.742687958458807 | -5.68444173256643 |
| lnc-PAM16-4 | lnc-PAM16-4 | 0.144504598512285 | 1.5070626292232 | 0.73737628578269 2 | 0.464945342324076 | 0.742687958458807 | -5.73364485918202 |
| ENSG00000132535.19 | DLG4 | -0.13933881988961 | 0.78535203025732 | -0.73731702589972 | 0.464980993694655 | 0.742687958458807 | -5.62480618803623 |
| lnc-GTF3A-4 | lnc-GTF3A-4 | -0.16157996235873 | 0.3368892550991 | -0.737312940359556 | 0.464983451656715 | 0.742687958458807 | -5.52071390599767 |
| ENSG00000143393.16 | PI4KB | 0.0620009530022374 | 6.05155426855639 | 0.737238170055668 | 0.465028436643297 | 0.742687958458807 | -6.14082271250701 |
| ENSG00000136628.17 | EPRS1 | -0.101848771110491 | 6.66097864039393 | -0.737218900184532 | 0.465040030621371 | 0.742687958458807 | -6.14083457294003 |
| lnc-KIAA1586-1 | lnc-KIAA1586-1 | -0.15761383084785 | 0.789051351789249 | -0.737171570576051 | 0.46506850782566 | 0.742687958458807 | -5.62582529891743 |
| ENSG00000133606.10 | MKRN1 | 0.0729285835533418 | 6.84292723550022 | 0.737005405904399 | 0.465168493474352 | 0.742790000091744 | -6.13965138612119 |
| lnc-LYPLAL1-1 | lnc-LYPLAL1-1 | 0.179243203412174 | 0.864307131101058 | 0.736409227289055 | 0.465527331656674 | 0.743305334534894 | -5.62198811211153 |
| ENSG00000121281.12 | ADCY7 | -0.0697194966097454 | 6.68903889902963 | -0.736249560955902 | 0.465623461398891 | 0.743401156218847 | -6.14164008743002 |
| lnc-MAP4K5-1 | lnc-MAP4K5-1 | 0.121371065390474 | 2.08031817615173 | 0.736180237471386 | 0.465665202299073 | 0.743407739353204 | -5.80553083996714 |
| lnc-LHX1-2 | lnc-LHX1-2 | 0.20809592990612 | -0.0822274527955098 | 0.736122747251936 | 0.465699819813449 | 0.743407739353204 | -5.36868779106218 |
| ENSG00000154874.15 | CCDC144BP | -0.722389061599226 | 1.90896215511579 | -0.736131455051484 | 0.465854937263329 | 0.743597686986514 | -5.91543541459576 |
| ENSG00000153560.11 | UBP1 | 0.059429380998135 | 5.30694838868404 | 0.735793798843901 | 0.465897923303109 | 0.743608634874992 | -6.12591092838722 |
| ENSG00000253106.1 | | 0.159152211666369 | 1.50887629490772 | 0.735649383065449 | 0.465984910517942 | 0.743646612420778 | -5.73709001024286 |
| ENSG00000230006.7 | ANKRD36BP2 | -1.19628705273962 | 0.327485572325659 | -0.735899820565132 | 0.465994370952797 | 0.743646612420778 | -5.38417934990308 |
| ENSG00000257529.5 | RPL36A-HNRNPH2 | 0.132033775178741 | 6.80608246940151 | 0.735574357318993 | 0.466030105116631 | 0.743646612420778 | -6.14154719385851 |
| ENSG00000281207.1 | SLFNL1-AS1 | 0.163185011852462 | 1.14615337723222 | 0.735476424201538 | 0.466089102630279 | 0.743683100731709 | -5.68281188099776 |
| ENSG00000082068.8 | WDR70 | -0.0624499414509927 | 5.22013481770609 | -0.735326262018124 | 0.466179572663289 | 0.743769796459847 | -6.12149984048796 |
| ENSG00000097033.14 | SH3GLB1 | 0.0620274310679116 | 6.43977155130503 | 0.735000177033303 | 0.466376067809229 | 0.744025624222397 | -6.14332958641186 |
| ENSG00000115446.11 | UNC50 | 0.0717452493266633 | 4.592455518634 | 0.734735474460392 | 0.466535609609669 | 0.744222463969296 | -6.08741855122243 |
| lnc-SESN3-8 | lnc-SESN3-8 | -0.302622052969431 | -0.0330588451714209 | -0.734665488549418 | 0.466577796830778 | 0.744232082468065 | -5.38591128178399 |
| lnc-AKAP11-1 | lnc-AKAP11-1 | -0.593413685031849 | -0.140241309039993 | -0.734305729550226 | 0.466907756597158 | 0.744700686093707 | -5.44427624942489 |
| ENSG00000139626.15 | ITGB7 | -0.169669740134965 | 6.81061764074597 | -0.733885640580357 | 0.467063196356472 | 0.744890885681486 | -6.14263413314364 |
| lnc-CSPG5-2 | lnc-CSPG5-2 | -0.144140277722805 | 0.738293313522915 | -0.733781701985563 | 0.467110728377991 | 0.744908973596048 | -5.60634735434509 |
| ENSG00000152234.15 | ATP5F1A | 0.106177525960378 | 7.4195938777946 | 0.733300305829595 | 0.467401161769052 | 0.745314387410319 | -6.13467919726683 |
| ENSG00000278419.1 | | -0.147812559675281 | 2.29455755030998 | -0.73317203676713 | 0.467478565872293 | 0.745380069823332 | -5.84372739967662 |
| ENSG00000251474.6 | RPL32P3 | -0.106217917143342 | 3.8983050564522 | -0.732999165483877 | 0.467582896873894 | 0.7454886728393 | -6.02214551510036 |
| ENSG00000103479.15 | RBL2 | 0.105070990839617 | 7.63630849092531 | 0.73243957142632 | 0.467920713823881 | 0.745914726777884 | -6.13312223827355 |
| lnc-COL4A3BP-3 | lnc-COL4A3BP-3 | -0.0841542447280155 | 1.29424215245676 | -0.732436441481128 | 0.467922603709665 | 0.745914726777884 | -5.70826318080169 |
| ENSG00000208797.1 | SNORD73A | 0.126457060432168 | 1.38734859885041 | 0.732280022357214 | 0.46801705639772 | 0.746007517372746 | -5.71660312854355 |
| lnc-CUL2-1 | lnc-CUL2-1 | -0.227066925062868 | -0.509996835633321 | -0.732173699867396 | 0.468081264796765 | 0.746052088687611 | -5.24884735770345 |
| ENSG00000151292.17 | CSNK1G3 | -0.101478487654135 | 6.42049770744994 | -0.732047860731646 | 0.468157265872443 | 0.746115447421795 | -6.14523875014745 |

| | | | | | | | |
|---|---|---|---|---|---|---|---|
| ENSG00000102879.15 | CORO1A | -0.142373599015683 | 7.43798955291637 | -0.731987182076941 | 0.468193915530371 | 0.746116085735647 | -6.13529917855249 |
| ENSG00000196705.8 | ZNF431 | 0.068204948423876 | 5.89619383273888 | 0.731704779962 | 0.468364506894752 | 0.746330157576424 | -6.14310092677597 |
| ENSG00000136643.11 | RPS6KC1 | -0.0686728506082854 | 4.46800495869689 | -0.731481930459808 | 0.468499149316003 | 0.746409334359622 | -6.0772408348526 |
| ENSG00000198598.6 | MMP17 | -0.374512914222827 | 2.81027181882817 | -0.73164767873971 | 0.468513709466633 | 0.746409334359622 | -5.88427249841414 |
| ENSG00000173163.10 | COMMD1 | 0.0822676159262388 | 3.45915833590005 | 0.731442483370125 | 0.468522984984032 | 0.746409334359622 | -5.97131460439271 |
| ENSG00000243943.9 | ZNF512 | -0.0904381304823509 | 5.34886990119103 | -0.731225624373239 | 0.468654033151281 | 0.746531971902136 | -6.13057779084084 |
| lnc-KHDRBS1-2 | lnc-KHDRBS1-2 | 0.152889548157981 | -0.0453105782777937 | 0.73116992285425 | 0.468687697040924 | 0.746531971902136 | -5.41092596049596 |
| ENSG00000139168.7 | ZCRB1 | -0.0943435521855269 | 4.23418214426163 | -0.731111469117803 | 0.468723025756922 | 0.746531971902136 | -6.0632134798475 |
| ENSG00000130755.12 | GMFG | 0.0864700225574788 | 5.34166109233513 | 0.731018321958166 | 0.468779325902883 | 0.746531971902136 | -6.12903076843706 |
| ENSG00000198604.10 | BAZ1A | 0.0702109935986002 | 6.28195443259634 | 0.730909004903337 | 0.468845404423822 | 0.746531971902136 | -6.14632539949461 |
| ENSG00000060749.15 | QSER1 | -0.0969505047469434 | 5.08234037985349 | -0.730855949309791 | 0.468877476687868 | 0.746531971902136 | -6.1171247936589 |
| ENSG00000131725.13 | WDR44 | -0.0533732968081493 | 5.08677920519988 | -0.730760464972909 | 0.468935200420473 | 0.746531971902136 | -6.12084360283275 |
| lnc-ATXN7-17 | lnc-ATXN7-17 | -0.126181156603852 | 4.04424241751042 | -0.730714063009808 | 0.468963253554292 | 0.746531971902136 | -6.03652206006729 |
| ENSG00000129158.10 | SERGEF | 0.0649129904092511 | 4.18791734483438 | 0.73067079771418 | 0.468989411226091 | 0.746531971902136 | -6.04899306141718 |
| ENSG00000140319.10 | VIII.14 | -0.102679738870765 | 6.12242955190326 | -0.730644134619096 | 0.469005531825556 | 0.746531971902136 | -6.14599232452462 |
| lnc-TPRKB-1 | lnc-TPRKB-1 | -0.109368714797427 | 0.982996808114324 | -0.730640818956468 | 0.469007536508837 | 0.746531971902136 | -5.67068538962903 |
| ENSG00000168556.6 | ING2 | -0.181695512441384 | 1.18723643200448 | -0.730595070422067 | 0.469035197038256 | 0.746531971902136 | -5.68168553290288 |
| lnc-CCDC89-5 | lnc-CCDC89-5 | 0.103298692515502 | 4.77519272381223 | 0.730496763829253 | 0.46909463844075 | 0.746568850462071 | -6.10174884047207 |
| ENSG00000168807.16 | SNTB2 | -0.10378947259122 | 4.33226812794655 | -0.73043513871792 | 0.469131902465963 | 0.746570430528601 | -6.07642444034432 |
| ENSG00000256100.1 | | 0.0844265886629705 | 2.68159239659282 | 0.730360814629711 | 0.469176847678146 | 0.746584233147219 | -5.88436145670704 |
| ENSG00000131002.11 | TXLNGY | -1.74495080255311 | 2.58768437076462 | -0.730467580281706 | 0.469271222482722 | 0.746621362438011 | -5.63263373744761 |
| lnc-SPHAR-2 | lnc-SPHAR-2 | 0.185048288808934 | -0.0509975889008449 | 0.73020227439233 | 0.469272728264128 | 0.746621362438011 | -5.44094813293896 |
| ENSG00000196313.11 | POM121 | 0.0846785993050867 | 4.85512939032748 | 0.730109830277766 | 0.469328640990788 | 0.746652606083892 | -6.10615693733343 |
| ENSG00000104324.15 | CPQ | 0.135230162472832 | 3.49487241562959 | 0.729942613797444 | 0.469429787778558 | 0.746755801982075 | -5.96455373630348 |
| ENSG00000163257.10 | DCAF16 | 0.0846099805224192 | 6.02020893157127 | 0.729838435637401 | 0.469492809924253 | 0.746766381612306 | -6.14640351876212 |
| ENSG00000105576.15 | TNPO2 | -0.0740168770175345 | 5.07456196482104 | -0.729803529186852 | 0.469513927520146 | 0.746766381612306 | -6.12153285225526 |
| lnc-ZMYM2-12 | lnc-ZMYM2-12 | -0.135049866095104 | 2.34545864823247 | -0.729751705573663 | 0.469545280611498 | 0.746766381612306 | -5.84320303423259 |
| lnc-STRAP-5 | lnc-STRAP-5 | 0.219645686503111 | 1.80368012597995 | 0.729634948608066 | 0.469615922529404 | 0.746770098169807 | -5.73592793206773 |
| lnc-ZFAND3-1 | lnc-ZFAND3-1 | -0.18971886252548 | 0.211123795311348 | -0.729627913324302 | 0.469620179308652 | 0.746770098169807 | -5.48195241259568 |
| lnc-IMMT-1 | lnc-IMMT-1 | 0.104371518532755 | -0.156132370786419 | 0.729349609739979 | 0.469788587807213 | 0.746980185618838 | -5.39342373921142 |
| lnc-SLC15A3-6 | lnc-SLC15A3-6 | -0.110160768214323 | 1.30259414173088 | -0.729278200496472 | 0.469831804908063 | 0.746984647823212 | -5.70065229101465 |
| ENSG00000068489.12 | PRR11 | 0.258513434990119 | -0.011584138440962 | 0.729191046316052 | 0.469884553977655 | 0.746984647823212 | -5.41642952403837 |
| ENSG00000132635.16 | PCED1A | -0.102404891177483 | 5.17944260162858 | -0.729165083742334 | 0.469900268183207 | 0.746984647823212 | -6.12596985944028 |
| lnc-ZNF500-1 | lnc-ZNF500-1 | -0.132847455145385 | -0.382951642241732 | -0.728765496773773 | 0.470142161632209 | 0.747311461338772 | -5.34543830080714 |
| lnc-OVOL2-1 | lnc-OVOL2-1 | 0.0971477775605565 | 3.50691008734692 | 0.728427220004137 | 0.470346996085363 | 0.747457289706404 | -5.97853411068268 |
| lnc-TMSB4Y-1 | lnc-TMSB4Y-1 | -0.799730997774079 | -2.01813915196776 | -0.72841185807031 | 0.470356299305048 | 0.747457289706404 | -4.86726336798567 |
| ENSG00000111652.9 | COPS7A | -0.0809397956616187 | 3.69968729375857 | -0.728392523175018 | 0.470368008708201 | 0.747457289706404 | -6.00944283420096 |
| ENSG00000166275.15 | BORCS7 | 0.113841444891388 | 3.63289859991979 | 0.728374108236649 | 0.470379161131373 | 0.747457289706404 | -5.99413421992743 |
| ENSG00000158691.14 | ZSCAN12 | 0.115287156828372 | 4.13605645728727 | 0.728199684962311 | 0.470484802552883 | 0.747479258469926 | -6.0475016792888 |
| ENSG00000198171.12 | DDRGK1 | -0.0996343418491552 | 3.40055573066664 | -0.728153058677087 | 0.470513044585782 | 0.747479258469926 | -5.96708328646256 |
| lnc-FTSJ3-1 | lnc-FTSJ3-1 | 0.12381788374741 | -0.165596654965534 | 0.72812830159755 | 0.470528040604047 | 0.747479258469926 | -5.39194840536564 |
| ENSG00000242330.3 | RN7SL683P | 0.341988661091043 | -0.653902605777408 | 0.72805273876568 | 0.470573812700978 | 0.747479258469926 | -5.184803138475 |
| ENSG00000197461.13 | PDGFA | 0.469008845147706 | -0.44982485772119 | 0.728100551524339 | 0.470587539461896 | 0.747479258469926 | -5.34331382679676 |
| ENSG00000100014.19 | SPECC1L | 0.0686451684992599 | 5.61186223773086 | 0.727991551931087 | 0.470610878419655 | 0.747479258469926 | -6.13963173049941 |
| ENSG00000213066.12 | CEP43 | 0.10030509806432 | 3.81518030758553 | 0.727814230772741 | 0.470718305336697 | 0.747580921403785 | -6.02236370527156 |
| lnc-GLOD4-2 | lnc-GLOD4-2 | 0.113211681077722 | 2.35115462992635 | 0.727766002953884 | 0.470747525740391 | 0.747580921403785 | -5.85182727516654 |
| ENSG00000114686.8 | MRPL3 | 0.0688219949110573 | 4.94231209924815 | 0.727604267864343 | 0.470845525810428 | 0.747678865742635 | -6.11655993144051 |
| lnc-RNF14-1 | lnc-RNF14-1 | -0.183737672500783 | -0.0226464936710112 | -0.727491846553419 | 0.470913652073696 | 0.747710170735419 | -5.39158132357075 |
| lnc-CLDN10-10 | lnc-CLDN10-10 | 0.172753575556335 | -0.133888733450212 | 0.727358761645292 | 0.470994307564764 | 0.747710170735419 | -5.41273409694474 |
| ENSG00000139977.13 | NAA30 | 0.0815609324036819 | 4.98539623111231 | 0.727347236621138 | 0.471001292623561 | 0.747710170735419 | -6.12232632881983 |
| lnc-TBCCD1-5 | lnc-TBCCD1-5 | 0.117078714019258 | 0.465133342207174 | 0.727331968692249 | 0.471010546264171 | 0.747710170735419 | -5.56117587868403 |
| lnc-OPN3-1 | lnc-OPN3-1 | -0.121779348571536 | 0.289078988480092 | -0.727222795177173 | 0.47107671756343 | 0.747757544351668 | -5.51759103932901 |
| ENSG00000129521.13 | EGLN3 | -0.802223094648186 | -2.94911681826046 | -0.727023565896201 | 0.471197486368374 | 0.747891568624324 | -4.79835416320858 |
| lnc-ZFHX4-6 | lnc-ZFHX4-6 | -0.152951673182539 | 3.05971519095422 | -0.726816273677087 | 0.47132316153885 | 0.748024903424393 | -5.91925618738076 |
| ENSG00000112473.17 | SLC39A7 | 0.100979971205263 | 4.48627794065575 | 0.726765122971939 | 0.471354175650507 | 0.748024903424393 | -6.08277797379858 |
| ENSG00000082512.14 | TRAF5 | 0.100161202498564 | 7.56600523337527 | 0.726414413419 | 0.471566852111201 | 0.748168043407389 | -6.13768274579534 |
| lnc-WDR73-3 | lnc-WDR73-3 | 0.190230196297155 | 1.86006944877233 | 0.726367993217368 | 0.471595006246218 | 0.748168043407389 | -5.79127430363007 |
| ENSG00000112242.14 | E2F3 | -0.120488399626209 | 2.76094520761241 | -0.72632623948785 | 0.471620330957349 | 0.748168043407389 | -5.89722983669354 |
| ENSG00000115365.11 | LANCL1 | 0.0927486409921139 | 4.59600707035215 | 0.726271945059294 | 0.471653263086374 | 0.748168043407389 | -6.09584101954397 |
| ENSG00000119862.12 | LGALSL | 0.188056716738539 | 1.53317701871561 | 0.726243245364156 | 0.471670671334085 | 0.748168043407389 | -5.75388655594301 |
| lnc-PMM1-1 | lnc-PMM1-1 | -0.108253702751295 | 1.70086799899042 | -0.726227567368994 | 0.471680181221508 | 0.748168043407389 | -5.76434047517222 |
| ENSG00000236287.7 | ZBED5 | -0.0768764071306451 | 6.9008787702443 | -0.726196849043969 | 0.471698814521581 | 0.748168043407389 | -6.14641840376078 |
| ENSG00000109046.14 | WSB1 | 0.0830697677843411 | 6.51818728081687 | 0.726088878468181 | 0.471764311275084 | 0.748170237781014 | -6.14938149241396 |
| ENSG00000163319.10 | MRPS18C | 0.105794648871157 | 4.04157760609 | 0.726074727468125 | 0.471772895892716 | 0.748170237781014 | -6.04291230918208 |
| ENSG00000135596.17 | MICAL1 | -0.114636567425515 | 7.34984008574862 | -0.726012530680522 | 0.47181062825079 | 0.748172263653011 | -6.14185191542462 |
| lnc-NPFF-3 | lnc-NPFF-3 | -0.122351425226695 | -0.0673213503451207 | -0.725952791293294 | 0.471846871422034 | 0.748172263653011 | -5.40036989494109 |
| ENSG00000089289.15 | IGBP1 | 0.107859672260704 | 5.44017898838229 | 0.725887725091699 | 0.471886348115768 | 0.748177222559456 | -6.13791597791136 |
| ENSG00000197549.9 | PRAMENP | 0.629554264181645 | -0.968270684437936 | 0.725674773178456 | 0.472074973861938 | 0.748341505121046 | -5.16881630250726 |

| | | | | | | | |
|---|---|---|---|---|---|---|---|
| ENSG00000055211.13 | GINM1 | 0.0883206821497851 | 5.08921479917434 | 0.72557657256396 | 0.472075155336314 | 0.748341505121046 | -6.12734316881088 |
| ENSG00000078902.15 | TOLLIP | 0.112678848296038 | 3.70769259769527 | 0.725487349965618 | 0.47212930352253 | 0.748341505121046 | -6.00871502713371 |
| ENSG00000268790.5 | | 0.0707233867434018 | 2.72702369285924 | 0.725430876455251 | 0.472163578488863 | 0.748341505121046 | -5.88774326978185 |
| lnc-BEND4-3 | lnc-BEND4-3 | -0.080017048533751 | 2.98665022777758 | -0.725417413183495 | 0.472171749842444 | 0.748341505121046 | -5.92234090895448 |
| ENSG00000161929.14 | SCIMP | -0.140611271019646 | 6.25366638806532 | -0.725293669845019 | 0.472246857993305 | 0.748378939918891 | -6.14991576443049 |
| ENSG00000205629.11 | LCMT1 | -0.0732576156732434 | 3.40166249126032 | -0.725258695068546 | 0.472268087771776 | 0.748378939918891 | -5.97384856365718 |
| ENSG00000135801.9 | TAF5L | 0.0676011492987056 | 4.90530846783214 | 0.724857789766894 | 0.472511477197848 | 0.748706985001025 | -6.1161298968245 |
| lnc-RAD54L2-1 | lnc-RAD54L2-1 | 0.125265849740591 | 1.21018908684867 | 0.724680560093957 | 0.472619096049557 | 0.748807907402762 | -5.70426331770776 |
| lnc-CD40LG-2 | lnc-CD40LG-2 | -0.180835862603536 | 0.588752800231205 | -0.724573058926499 | 0.472684380597337 | 0.748807907402762 | -5.55913219027286 |
| ENSG00000134245.17 | WNT2B | -0.585051628212903 | 0.209939800527752 | -0.72475513834493 | 0.472690556783345 | 0.748807907402762 | -5.49920679239072 |
| ENSG00000178234.12 | GALNT11 | -0.395374697424199 | 2.70424869160231 | -0.724733333893231 | 0.472720689345163 | 0.748807907402762 | -5.93360687914105 |
| lnc-NIPA1-3 | lnc-NIPA1-3 | 0.0721631807821113 | 2.51445502778875 | 0.724138523210487 | 0.472948322845811 | 0.749110837178555 | -5.86659962243335 |
| lnc-KIAA1551-6 | lnc-KIAA1551-6 | -0.170404353255865 | 2.74565383756422 | -0.723857654847579 | 0.473118970373385 | 0.74932346623541 | -5.89704104603663 |
| ENSG00000115677.16 | HDLBP | -0.0760972700531317 | 6.61278782329817 | -0.723700171600337 | 0.47321466800861 | 0.749417366440042 | -6.150833763768 |
| lnc-ARMCX4-3 | lnc-ARMCX4-3 | -0.203610299324687 | 1.652272852219 | -0.723417080342265 | 0.473386721425712 | 0.749567120467639 | -5.74789205115279 |
| ENSG00000211766.1 | TRBJ2-2P | 0.174369097455112 | 0.25397912247533 | 0.723411125428077 | 0.473390341006424 | 0.749567120467639 | -5.47682843044365 |
| ENSG00000186141.8 | POLR3C | 0.0592267295290059 | 4.98984111915174 | 0.72330744184128 | 0.473453365618792 | 0.749567120467639 | -6.11969434777136 |
| ENSG00000119487.16 | MAPKAP1 | -0.0643528034534357 | 5.89826224561108 | -0.723304923559427 | 0.473454896429081 | 0.749567120467639 | -6.14939868170803 |
| ENSG00000176994.10 | SMCR8 | -0.0640578525789461 | 6.56105101622573 | -0.723210963940401 | 0.473512014509612 | 0.749577667338266 | -6.15134454052463 |
| ENSG00000167642.12 | SPINT2 | -0.122839146201049 | 4.93601795253499 | -0.723076204900543 | 0.473593941420844 | 0.749577667338266 | -6.12184599931791 |
| ENSG00000146278.10 | PNRC1 | 0.0870339731208567 | 8.47635763262765 | 0.723060713930925 | 0.473603359690701 | 0.749577667338266 | -6.12063615017724 |
| lnc-URGCP-MRPS24-5 | lnc-URGCP-MRPS2 | -0.0682844025589118 | 2.61157190881226 | -0.723049925969932 | 0.473609918667082 | 0.749577667338266 | -5.87862282714944 |
| lnc-GTF2F2-14 | lnc-GTF2F2-14 | -0.121706697723993 | 1.31762831780063 | -0.722964988067898 | 0.473661561900425 | 0.749577667338266 | -5.73044984966531 |
| lnc-SMYD1-2 | lnc-SMYD1-2 | -0.241958511730344 | 1.44652875291221 | -0.722940821505628 | 0.473680062142002 | 0.749577667338266 | -5.74214752409567 |
| lnc-RPS7-3 | lnc-RPS7-3 | -0.149554471832938 | 2.11843600415845 | -0.722522918274112 | 0.473930397252192 | 0.749769787548907 | -5.82095123833795 |
| CTBP1-DT | CTBP1-DT | -0.124253698421907 | 3.96334082717436 | -0.722453236132493 | 0.473972780889165 | 0.749769787548907 | -6.04776974392173 |
| lnc-UPRT-2 | lnc-UPRT-2 | -0.16008900325255 | 0.734807304779009 | -0.722429133476602 | 0.473987441649976 | 0.749769787548907 | -5.60419959582813 |
| lnc-CETP-1 | lnc-CETP-1 | -0.222350878838395 | 3.47332868903536 | -0.722484770988201 | 0.473990955432676 | 0.749769787548907 | -5.97912834228377 |
| ENSG00000134256.12 | CD101 | -0.699814507035135 | 0.0253241442464485 | -0.722538342743825 | 0.474031988310363 | 0.749769787548907 | -5.31066947784841 |
| ENSG00000197375.12 | SLC22A5 | -0.185962985916483 | 2.24956737515812 | -0.722324813624514 | 0.474050898562791 | 0.749769787548907 | -5.83000801800822 |
| ENSG00000149311.18 | ATM | 0.156997536628001 | 9.12404656349485 | 0.722228783728371 | 0.474109317037145 | 0.749769787548907 | -6.10896332772832 |
| ENSG00000114374.12 | USP9Y | -1.76304402818427 | 2.78154913186853 | -0.722458838260214 | 0.474126297801063 | 0.749769787548907 | -5.68953563995153 |
| ENSG00000102030.15 | NAA10 | 0.1140531857293 | 3.40054988037057 | 0.722194543393127 | 0.474130147667761 | 0.749769787548907 | -5.97413914398962 |
| ENSG00000149932.16 | TMEM219 | 0.0841760670223755 | 4.72092438808222 | 0.722136048638582 | 0.474165735049513 | 0.749769787548907 | -6.10402915198104 |
| lnc-NAA15-2 | lnc-NAA15-2 | 0.255147885579216 | -0.597956853662802 | 0.722023329139236 | 0.474234316286075 | 0.749779387595607 | -5.25490100103815 |
| ENSG00000206875.1 | RNU6-761P | -0.25968172051336 | -0.149425087444044 | -0.722006328285121 | 0.474244660501735 | 0.749779387595607 | -5.38129806240361 |
| ENSG00000117602.11 | RCAN3 | -0.0967316494367676 | 5.15961400940721 | -0.721757562366213 | 0.474396037480888 | 0.749961109022206 | -6.13014330779976 |
| ENSG00000132424.15 | PNISR | -0.103582375352483 | 8.6507987749244 | -0.721564874821833 | 0.474513309001063 | 0.749989037999877 | -6.11691948915255 |
| lnc-ZNF335-1 | lnc-ZNF335-1 | -0.166908369271796 | 0.712384528547423 | -0.721559796948263 | 0.474516399667464 | 0.749989037999877 | -5.60575917429705 |
| ENSG00000228314.1 | CYP4F29P | -0.793417575896407 | 1.68159084672905 | -0.721764262156528 | 0.474548710115556 | 0.749989037999877 | -5.76637786986698 |
| lnc-MED20-5 | lnc-MED20-5 | -0.154946831712286 | 4.41032389394266 | -0.721489062768154 | 0.474559453476675 | 0.749989037999877 | -6.0791192614161 |
| lnc-DYNC1H1-7 | lnc-DYNC1H1-7 | -0.244348890914225 | -0.524456194439259 | -0.721172991469218 | 0.474751863897458 | 0.750165967058003 | -5.23924223172343 |
| lnc-PACRG-2 | lnc-PACRG-2 | 0.202244790368722 | -0.4276979451179 | 0.7211453058364 | 0.474768719815933 | 0.750165967058003 | -5.32378639784775 |
| ENSG00000170482.16 | SLC23A1 | -0.210901567746588 | 1.21628630143413 | -0.721125556678581 | 0.474780743956887 | 0.750165967058003 | -5.72067421056985 |
| ENSG00000120832.9 | MTERF2 | -0.200349756834926 | 2.38362334863974 | -0.720988126234932 | 0.474864422346393 | 0.750191619067468 | -5.86174917587423 |
| lnc-MFSD6-1 | lnc-MFSD6-1 | 0.188900926117396 | -0.332180574416764 | 0.720946833176729 | 0.474889566423936 | 0.750191619067468 | -5.31695980281914 |
| lnc-PPP1R18-3 | lnc-PPP1R18-3 | -0.0860023960021297 | 0.883266610296334 | -0.720919319226735 | 0.474906320577618 | 0.750191619067468 | -5.64304380637923 |
| lnc-TUBGCP6-5 | lnc-TUBGCP6-5 | 0.144653212788212 | 2.16546827426591 | 0.720834324709427 | 0.474958078671257 | 0.750215803337463 | -5.82243111092231 |
| ENSG00000054611.13 | TBC1D22A | -0.0745904732948866 | 6.06098820647431 | -0.720556879478755 | 0.475127053488207 | 0.750373432150065 | -6.15210035329212 |
| lnc-PIGP-6 | lnc-PIGP-6 | 0.134623155421986 | 0.422326163086769 | 0.720432704773805 | 0.475202691726345 | 0.750373432150065 | -5.53255255259006 |
| lnc-KATNBL1-1 | lnc-KATNBL1-1 | -0.0962354774220624 | 2.4227770833156 | -0.720358125603218 | 0.475248123245365 | 0.750373432150065 | -5.86681216791172 |
| ENSG00000232119.7 | MCTS1 | -0.0979152991226342 | 3.51534852346237 | -0.720312886370241 | 0.475275682907807 | 0.750373432150065 | -5.99087550820037 |
| ENSG00000206634.1 | SNORA22 | 0.172585503802238 | -0.0695532429708215 | 0.720296452521588 | 0.475285694604833 | 0.750373432150065 | -5.40008257249417 |
| lnc-KLHL22-5 | lnc-KLHL22-5 | -0.10168854017289 | 0.98633732648689 | -0.720296160481336 | 0.475285872520329 | 0.750373432150065 | -5.67781307209909 |
| lnc-DTNB-1 | lnc-DTNB-1 | -0.170704165531577 | 1.26882024177353 | -0.720200350055561 | 0.475344243775214 | 0.750373432150065 | -5.69456637684183 |
| lnc-AKAP12-1 | lnc-AKAP12-1 | -0.105185885032056 | 5.50556528420953 | -0.720178075816604 | 0.475357814648914 | 0.750373432150065 | -6.14435972008794 |
| ENSG00000053770.11 | AP5M1 | 0.0671983679142354 | 6.20248221021336 | 0.72013185409321 | 0.47538597654554 | 0.750373432150065 | -6.15398661300259 |
| lnc-SP110-8 | lnc-SP110-8 | 0.339500172134952 | -0.50053824672925 | 0.719998469702325 | 0.475467250085204 | 0.750442369333251 | -5.26345841550887 |
| ENSG00000128595.16 | CALU | 0.10058474356302 | 4.79318444962056 | 0.719831052229802 | 0.475569271772294 | 0.750442369333251 | -6.10975948603872 |
| ENSG00000144357.16 | UBR3 | 0.0930719260664983 | 5.89986768764643 | 0.719830242791104 | 0.475569765062253 | 0.750442369333251 | -6.15166063006938 |
| lnc-SPTBN5-4 | lnc-SPTBN5-4 | -0.172563970400042 | 1.02346431051874 | -0.719770699410951 | 0.475606052920443 | 0.750442369333251 | -5.67689384250643 |
| ENSG00000136938.8 | ANP32B | 0.139120692722575 | 3.98588099663195 | 0.719761028054049 | 0.475611947138525 | 0.750442369333251 | -6.04643918046149 |
| ENSG00000150054.18 | MPP7 | 0.540909103813588 | 2.66172006579345 | 0.719769102421211 | 0.475757218938995 | 0.750557494521753 | -5.90363705349763 |
| ENSG00000158882.14 | TOMM40L | -0.113708692384303 | 2.44910961094435 | -0.719489997074921 | 0.47577714411077 | 0.750557494521753 | -5.8663325016294 |
| ENSG00000078618.21 | NRDC | 0.0602859732178382 | 6.0771940354925 | 0.719366851204852 | 0.475852213914291 | 0.750557494521753 | -6.15351144798449 |
| ENSG00000163655.15 | GMPS | -0.0508576982299571 | 5.1540021378824 | -0.719342328401624 | 0.47586716383426 | 0.750557494521753 | -6.13239836160848 |
| ENSG00000204394.12 | VARS1 | -0.133219414420259 | 4.45789765458886 | -0.71925357362744 | 0.475921273938446 | 0.750557494521753 | -6.08968435593334 |
| ENSG00000105722.9 | ERF | -0.183671007490509 | 2.93349639446945 | -0.719224864107109 | 0.475938777689808 | 0.750557494521753 | -5.93176633955995 |

| | | | | | | | |
|---|---|---|---|---|---|---|---|
| ENSG00000235655.3 | H3P6 | 0.16147989976684 | 3.60493400206863 | 0.719222588493574 | 0.475940165111884 | 0.750557494521753 | -6.01748450312234 |
| lnc-YIPF1-1 | lnc-YIPF1-1 | 0.113563471098357 | 0.86359010746889 | 0.719095232585869 | 0.476017816577985 | 0.750577497695864 | -5.6531299334019 |
| ENSG00000214706.10 | IFRD2 | -0.119579362957171 | 3.19091494662718 | -0.719082170464654 | 0.476025781223427 | 0.750577497695864 | -5.95838407188965 |
| lnc-KDSR-2 | lnc-KDSR-2 | -0.192321198483383 | 2.40728964778147 | -0.718897991126184 | 0.476138092868529 | 0.750671319043269 | -5.86901194450551 |
| ENSG00000109854.13 | HTATIP2 | 0.0925888666713099 | 3.35047631049458 | 0.718860987743656 | 0.476160659161555 | 0.750671319043269 | -5.9699501434055 |
| ENSG00000284570.1 | MIR7705 | 0.33382018308498 | 1.17215011891186 | 0.71892048446628 | 0.47619469527027 | 0.750671319043269 | -5.72098073881388 |
| ENSG00000130227.16 | XPO7 | -0.0773393766900821 | 5.6226857245858 | -0.71863122957069 | 0.476300789403777 | 0.750773712483753 | -6.14765449956825 |
| ENSG00000198824.6 | CHAMP1 | -0.0947252402851069 | 4.45146494999084 | -0.718570745296183 | 0.476337682846512 | 0.750773712483753 | -6.08672775743148 |
| ENSG00000280028.1 | | 0.275682873995393 | 1.33001745736983 | 0.718527902033292 | 0.476387267135003 | 0.750773712483753 | -5.71914576496207 |
| LINC00938 | LINC00938 | -0.0887748488955156 | 4.86084788220056 | -0.718459486845125 | 0.4764055511223 | 0.750773712483753 | -6.12140789045786 |
| lnc-KMT2E-1 | lnc-KMT2E-1 | -0.17243409918371 | 5.15241276511211 | -0.71825993960642 | 0.476545354924523 | 0.750914128437864 | -6.13577837189101 |
| ENSG00000131149.18 | GSE1 | 0.18744420145371 | 5.44452427092634 | 0.718221479312218 | 0.476588414083952 | 0.750914128437864 | -6.14200965096746 |
| ENSG00000060339.13 | CCAR1 | 0.0664166954310454 | 6.81815171632233 | 0.718134052789184 | 0.476604099203821 | 0.750914128437864 | -6.15332494036867 |
| ENSG00000276710.4 | CSPG4P10 | 0.42141022250065 | 0.254014527604988 | 0.717892979683252 | 0.476824350154748 | 0.751064255391157 | -5.52345641875769 |
| ENSG00000196597.11 | ZNF782 | -0.0960163044492219 | 4.20188703359668 | -0.717732108705385 | 0.476849391008426 | 0.751064255391157 | -6.05882207417354 |
| ENSG00000119004.15 | CYP20A1 | 0.0613505687063247 | 4.95799114683282 | 0.717719214591348 | 0.476857261000542 | 0.751064255391157 | -6.12076436355497 |
| lnc-NDRG1-3 | lnc-NDRG1-3 | -0.129072387565017 | 1.46224246734579 | -0.717648291141973 | 0.476900550827502 | 0.751064255391157 | -5.71336933039703 |
| ENSG00000136158.11 | SPRY2 | -0.669566848548681 | 1.03590792002153 | -0.717848598530497 | 0.476928023026953 | 0.751064255391157 | -5.6511493403174 |
| ENSG00000136816.15 | TOR1B | -0.108748830343239 | 3.07835049000081 | -0.717503316807538 | 0.476989046319249 | 0.751064255391157 | -5.94207231399359 |
| ENSG00000185305.10 | ARL15 | 0.134276106957234 | 3.76889676184934 | 0.717451860253872 | 0.477020458759101 | 0.751064255391157 | -6.01311655350365 |
| ENSG00000186174.12 | BCL9L | 0.227303900231425 | 2.39399134371773 | 0.717468444420546 | 0.477021388302967 | 0.751064255391157 | -5.83639149897678 |
| lnc-TESC-3 | lnc-TESC-3 | -0.210456758046519 | 1.38605870567862 | -0.717368740750706 | 0.477071202807796 | 0.751064255391157 | -5.73281491113325 |
| lnc-FKBP7-2 | lnc-FKBP7-2 | 0.0787464248079779 | 2.45888664601288 | 0.71736414711431 | 0.477074007289498 | 0.751064255391157 | -5.87220576391503 |
| ENSG00000186300.11 | ZNF555 | 0.0992162108477648 | 3.19148057226432 | 0.71732031233258 | 0.477100769530164 | 0.751064255391157 | -5.94322223541606 |
| lnc-KLF10-2 | lnc-KLF10-2 | 0.428808075452207 | -0.895413443398054 | 0.717245077065079 | 0.477146704543721 | 0.751079123556394 | -5.23797109254399 |
| lnc-TMCC3-6 | lnc-TMCC3-6 | 0.142707251463167 | -0.176772922536025 | 0.716961091729285 | 0.477320114835526 | 0.751240368847247 | -5.39090954278719 |
| lnc-MAP1LC3B2-28 | lnc-MAP1LC3B2-28 | 0.392295640346731 | 0.400879906881364 | 0.717067116513004 | 0.47732213689862 | 0.751240368847247 | -5.57009916441695 |
| ENSG00000179967.11 | PPP1R14BP3 | -0.180810213187103 | -0.0163948881648498 | -0.716758219215807 | 0.477444016979076 | 0.75137473824301 | -5.42102024909374 |
| ENSG00000198908.11 | GPRASP3 | -0.18594300825233 | 0.671290988450722 | -0.716575736847144 | 0.477555481633903 | 0.751492697130781 | -5.60632633946645 |
| lnc-ECHDC3-8 | lnc-ECHDC3-8 | -0.149604453894785 | 2.53470730443288 | -0.716393249458355 | 0.47766696409698 | 0.751610666004751 | -5.86970676926212 |
| ENSG00000204574.12 | ABCF1 | 0.0837718904451883 | 5.46010571679667 | 0.716235936037148 | 0.477763079487245 | 0.751704438548078 | -6.14517861638324 |
| ENSG00000010361.13 | FUZ | -0.096637951510421 | 2.45316797127882 | -0.716036503978104 | 0.477884944276618 | 0.751784819072135 | -5.86889004566681 |
| ENSG00000148291.9 | SURF2 | -0.127693034852977 | 1.66114937520794 | -0.716032786108032 | 0.47788721628236 | 0.751784819072135 | -5.77293301719077 |
| lnc-CAPN2-3 | lnc-CAPN2-3 | 0.163131359685346 | -0.0871977441130768 | 0.71563010971078 | 0.478133329696094 | 0.752000198810345 | -5.46406764861042 |
| lnc-TRIM59-1 | lnc-TRIM59-1 | 0.0957826704563938 | 3.31124507694863 | 0.715532853495794 | 0.478192782871113 | 0.752000198810345 | -5.96401300409495 |
| ENSG00000147155.10 | EBP | -0.0980277536200793 | 3.30684042099805 | -0.715516043378924 | 0.478203059397612 | 0.752000198810345 | -5.97797330886058 |
| lnc-GTF2H2-7 | lnc-GTF2H2-7 | -0.128707545377697 | 2.96848319097441 | -0.71543208021767 | 0.478254390457283 | 0.752000198810345 | -5.9311829094056 |
| lnc-LEFTY1-5 | lnc-LEFTY1-5 | -0.133778418071242 | 2.1708921771478 | -0.715397043610661 | 0.478275811084705 | 0.752000198810345 | -5.82825219284314 |
| lnc-ACBD6-7 | lnc-ACBD6-7 | 0.105560428422055 | 0.552591498658825 | 0.715310529927236 | 0.478328706014483 | 0.752000198810345 | -5.56822816498732 |
| ENSG00000238961.1 | SNORA47 | -0.371971499401885 | 0.146548094617442 | -0.715374605772915 | 0.478331356713969 | 0.752000198810345 | -5.5310261170861 |
| lnc-AKAP13-6 | lnc-AKAP13-6 | 0.150912480114113 | 2.71768827554034 | 0.715234719893196 | 0.478375059397227 | 0.752000198810345 | -5.89528800248305 |
| ENSG00000175309.14 | PHYKPL | -0.101270018995021 | 6.33323796522009 | -0.715051113667813 | 0.478487334352332 | 0.752000198810345 | -6.15753052002424 |
| lnc-SMAP2-1 | lnc-SMAP2-1 | -0.251225953845311 | -0.659275183819832 | -0.715019935598628 | 0.47850640117972 | 0.752000198810345 | -5.25749355608608 |
| ENSG00000105369.9 | CD79A | -0.135354941557887 | 8.14957708929331 | -0.714943895573316 | 0.478552904965281 | 0.752000198810345 | -6.1333726050332 |
| ENSG00000107789.15 | MINPP1 | -0.156195689709052 | 1.76243867701048 | -0.714943329911047 | 0.478553250916841 | 0.752000198810345 | -5.79376315690974 |
| ENSG00000204469.12 | PRRC2A | -0.084410358158495 | 7.22217257297696 | -0.714916176185066 | 0.478569857942867 | 0.752000198810345 | -6.15128715583069 |
| ENSG00000089157.15 | RPLP0 | -0.146607852480269 | 9.24591879892154 | -0.714889843026805 | 0.478585963426924 | 0.752000198810345 | -6.10404108405065 |
| lnc-SLC35E1-3 | lnc-SLC35E1-3 | 0.20661377037422 | -0.0893639735795828 | 0.714804125644988 | 0.478638390706379 | 0.752000198810345 | -5.41926329146455 |
| ENSG00000155850.7 | SLC26A2 | -0.113043210421957 | 4.41827353128201 | -0.714738474354414 | 0.478678547169566 | 0.752000198810345 | -6.08784100313353 |
| lnc-CCK-6 | lnc-CCK-6 | 0.120342122765813 | 0.773805946231598 | 0.714718889130063 | 0.478690527095075 | 0.752000198810345 | -5.61405215418716 |
| lnc-UHRF2-9 | lnc-UHRF2-9 | -0.235197054164657 | 0.968749613872787 | -0.714691523853689 | 0.478707266220943 | 0.752000198810345 | -5.66960229643743 |
| lnc-PJA2-1 | lnc-PJA2-1 | 0.152217127806501 | 3.53333733382854 | 0.7146734990431 | 0.478718292037698 | 0.752000198810345 | -5.98842608559268 |
| lnc-UPF2-5 | lnc-UPF2-5 | -0.220442605520161 | 1.62186925217743 | -0.714576341541163 | 0.478777725971812 | 0.752036167099955 | -5.74873259972116 |
| LINC01410 | LINC01410 | 0.468121001538041 | 3.73909293223526 | 0.714692834544096 | 0.47885543122703 | 0.752100827237386 | -5.98281180707684 |
| ENSG00000235449.2 | COX7CP2 | 0.201026525513311 | -0.325317306420921 | 0.714206189443728 | 0.479004196494254 | 0.75227707740281 | -5.33885306312293 |
| ENSG00000110880.10 | CORO1C | 0.132164719825944 | 5.08139385329376 | 0.71403667903777 | 0.479107928430988 | 0.75238258113184 | -6.13397950780389 |
| ENSG00000141552.17 | ANAPC11 | 0.167003878178484 | 2.16243013738682 | 0.713881081183823 | 0.479203157756528 | 0.752465450624407 | -5.82682372923128 |
| ENSG00000135127.11 | BICDL1 | -0.168484323127015 | 3.62730275309082 | -0.71383099491636 | 0.479233813935546 | 0.752465450624407 | -6.00460653057818 |
| lnc-FAM46C-5 | lnc-FAM46C-5 | 0.259797380265164 | 0.665180837907857 | 0.713644347665874 | 0.479348064423482 | 0.752587430208126 | -5.62080268423456 |
| lnc-UCP3-4 | lnc-UCP3-4 | 0.124058673900755 | 4.10800654513451 | 0.713390206885642 | 0.479503653789206 | 0.752660863819883 | -6.04757723784428 |
| ENSG00000049541.10 | RFC2 | -0.109178856785557 | 3.11206846965561 | -0.713378005673587 | 0.479511124299369 | 0.752660863819883 | -5.95153107294749 |
| ENSG00000135617.3 | PRADC1 | 0.141673295738328 | 0.953844940654726 | 0.713310568524052 | 0.479552415638521 | 0.752660863819883 | -5.65831132954795 |
| E2F3-IT1 | E2F3-IT1 | -0.175158622118413 | -0.230822843171646 | -0.713306836723454 | 0.479554700655032 | 0.752660863819883 | -5.35964156142616 |
| ENSG00000157514.16 | TSC22D3 | 0.144198856970972 | 9.87509803541745 | 0.713211780069565 | 0.479612906809984 | 0.752660863819883 | -6.08697040152785 |
| lnc-GABARAPL2-3 | lnc-GABARAPL2-3 | -0.172078322692791 | 0.364389143950317 | -0.713192417439834 | 0.479624763641605 | 0.752660863819883 | -5.5224200315574 |
| lnc-KHDRBS3-11 | lnc-KHDRBS3-11 | 0.839083202807945 | -1.30303569357043 | 0.71325392355535 | 0.479673637843264 | 0.752660863819883 | -5.05464240701601 |
| lnc-ORC1-4 | lnc-ORC1-4 | 0.0875809251660939 | 2.21208735412228 | 0.713039824069354 | 0.479718210974586 | 0.752660863819883 | -5.83458702316898 |
| lnc-SUSD1-3 | lnc-SUSD1-3 | 0.187667650376172 | 0.610432761102284 | 0.712994547792242 | 0.479745939893839 | 0.752660863819883 | -5.57778767980066 |

| | | | | | | | |
|---|---|---|---|---|---|---|---|
| lnc-ZNF860-1 | lnc-ZNF860-1 | -0.14885312655653 | 2.38869644942744 | -0.712970761545191 | 0.479760507861675 | 0.752660863819883 | -5.85667643700818 |
| lnc-PDLIM1-3 | lnc-PDLIM1-3 | 0.230621438879657 | -0.531882065123963 | 0.712895338689817 | 0.47980670249811 | 0.752675966581708 | -5.28407374537057 |
| ENSG00000136828.18 | RALGPS1 | 0.132085198141586 | 5.73475123049022 | 0.712816560528722 | 0.479854954858863 | 0.752694294761467 | -6.15483988548673 |
| ENSG00000285932.1 | | -0.106029957884185 | 2.51053454192452 | -0.712721573269434 | 0.479913139088428 | 0.752719467228362 | -5.8735760105022 |
| lnc-GRK4-2 | lnc-GRK4-2 | 0.129445300004941 | 3.84687281360716 | 0.712670961830263 | 0.479944142637372 | 0.752719467228362 | -6.03697840594827 |
| ENSG00000259706.1 | HSP90B2P | 0.148204561340008 | -0.0994238416909404 | 0.712539086390675 | 0.480024932190546 | 0.752788813659277 | -5.40040063778316 |
| lnc-ATF2-1 | lnc-ATF2-1 | -0.12916173728987 | 2.62152843796243 | -0.712462084338185 | 0.480072108751275 | 0.752805440684709 | -5.89495289249545 |
| ENSG00000141698.16 | NT5C3B | -0.112131755808394 | 3.66676520335598 | -0.712398738503765 | 0.480110920570362 | 0.752808949348652 | -6.00803137759279 |
| lnc-VEZF1-1 | lnc-VEZF1-1 | 0.151651769186877 | 5.25075148582425 | 0.712327073524213 | 0.480154831642704 | 0.752820452445291 | -6.14145980771933 |
| lnc-ARF6-5 | lnc-ARF6-5 | -0.18210126278621 | 0.422988552826166 | -0.71225318240359 | 0.480200109106009 | 0.752834095950109 | -5.55315628893503 |
| lnc-ANGPTL1-5 | lnc-ANGPTL1-5 | 0.104304133948309 | 2.23414117392089 | 0.711931108676865 | 0.480397490889886 | 0.753086180882446 | -5.83788838366671 |
| CYP1B1-AS1 | CYP1B1-AS1 | 0.189703860408165 | -0.387839199970407 | 0.711422254488729 | 0.480709433746019 | 0.753491218059641 | -5.31062693749582 |
| ENSG00000165609.12 | NUDT5 | -0.0706432974244136 | 4.81495324844656 | -0.711390211201101 | 0.480729081064918 | 0.753491218059641 | -6.11758736276621 |
| lnc-RABGAP1-1 | lnc-RABGAP1-1 | -0.099792410533198 | 2.39007678634552 | -0.711299816118139 | 0.480784509186954 | 0.753520715190365 | -5.8655094405489 |
| FAM99B | FAM99B | 0.142016434253794 | 0.165133046167238 | 0.711029586017957 | 0.480950229374424 | 0.753723052475542 | -5.47741117833989 |
| ENSG00000077514.8 | POLD3 | 0.0788251891045282 | 3.88087783787169 | 0.710892333398549 | 0.4810344126803 | 0.753795354089074 | -6.03723152721211 |
| ENSG00000125611.15 | CHCHD5 | 0.104965350607463 | 1.6387633331514 | 0.710828473628433 | 0.481073583628918 | 0.753795354089074 | -5.78437708961606 |
| ENSG00000214654.8 | B3GALT9 | -0.125930164370429 | 1.62219571146866 | -0.71077525027411 | 0.481106231679938 | 0.753795354089074 | -5.76041969085198 |
| lnc-NAV1-9 | lnc-NAV1-9 | -0.0932102299213975 | 2.69287252596935 | -0.710574266396134 | 0.481229529649396 | 0.753856407817478 | -5.89624793801175 |
| ENSG00000121060.17 | TRIM25 | 0.113589662328065 | 4.76903336898921 | 0.710495144698242 | 0.481278073477457 | 0.753856407817478 | -6.1153745747968 |
| ENSG00000143258.15 | USP21 | -0.0549134044041981 | 4.495885025341 | -0.710481122404101 | 0.481286676915426 | 0.753856407817478 | -6.09847882555379 |
| lnc-ZFYVE21-3 | lnc-ZFYVE21-3 | 0.103006363756495 | 2.83655395072869 | 0.710423748774788 | 0.481321879651819 | 0.753856407817478 | -5.91511678083146 |
| ENSG00000254634.4 | SMG1P6 | 0.14850869062727 | 1.73849936545747 | 0.710413244517582 | 0.481328324905859 | 0.753856407817478 | -5.76420787515269 |
| ENSG00000162645.12 | GBP2 | -0.159895756948196 | 6.42522017562168 | -0.710350370373991 | 0.481377357077713 | 0.753875838144303 | -6.16071074844613 |
| ENSG00000159199.13 | ATP5MC1 | -0.105240374755874 | 3.68570267431456 | -0.710268709495018 | 0.48141701435925 | 0.753880584795834 | -6.02101063398062 |
| ENSG00000156017.12 | CARNMT1 | -0.114196384863017 | 3.30878099216588 | -0.710084219937124 | 0.481530234060383 | 0.753970460649813 | -5.98304862203341 |
| ENSG00000135148.11 | TRAFD1 | 0.0895115142090388 | 4.8843786240258 | 0.710055811804689 | 0.48154766922715 | 0.753970460649813 | -6.12394181401289 |
| lnc-SEC24C-3 | lnc-SEC24C-3 | 0.0983759304513132 | 1.17256714772491 | 0.709884161587642 | 0.481653025155666 | 0.753990574396175 | -5.69376109610391 |
| lnc-TRPM7-1 | lnc-TRPM7-1 | 0.220525894575742 | -0.352593822746961 | 0.709876855113835 | 0.481657510030839 | 0.753990574396175 | -5.37130352139538 |
| ENSG00000122122.9 | SASH3 | -0.157723496272356 | 6.52410735645348 | -0.709832180077199 | 0.4816925239106 | 0.753990574396175 | -6.16055176479196 |
| ENSG00000182568.16 | SATB1 | 0.368996726592951 | 4.17237657475297 | 0.709986069237527 | 0.481731372020089 | 0.753990574396175 | -6.04920864767977 |
| lnc-GAN-1 | lnc-GAN-1 | -0.134526841322652 | 2.71030328862721 | -0.709704547367744 | 0.481763283155575 | 0.753990574396175 | -5.90034689523198 |
| ENSG00000181381.13 | DDX60L | -0.0977427034203835 | 5.09366031117566 | -0.709641443061076 | 0.481802023745712 | 0.753990574396175 | -6.13632889943423 |
| ENSG00000086712.12 | TXLNG | 0.117445435121548 | 4.5360358124589 | 0.709617150864139 | 0.481816937522853 | 0.753990574396175 | -6.10479313919455 |
| lnc-SFXN5-1 | lnc-SFXN5-1 | -0.13880217683385 | -0.169842696146979 | -0.709215117099886 | 0.482063797011792 | 0.754319532757618 | -5.40343576348898 |
| lnc-GTF3C2-1 | lnc-GTF3C2-1 | -0.10360401593881 | 1.42492660547296 | -0.70906324213954 | 0.482157070833964 | 0.754408132951386 | -5.73724952414045 |
| ENSG00000163923.9 | RPL39L | 0.24567166750759 | 0.191407348079439 | 0.708894957829786 | 0.482260434305453 | 0.754512505268253 | -5.4819434359523 |
| ENSG00000275464.4 | | -0.312697112487804 | 2.69612429355086 | -0.708794709690769 | 0.482422072727907 | 0.754708027890743 | -5.93231537162022 |
| ENSG00000105793.15 | GTPBP10 | -0.0809906017591393 | 4.26250959092126 | -0.708536675558795 | 0.482480539741002 | 0.75474001295327 | -6.07905102719987 |
| lnc-KDM5D-1 | lnc-KDM5D-1 | -1.03597215967577 | -1.08077851401311 | -0.708587946561981 | 0.48255956968295 | 0.75474001295327 | -5.04345381814289 |
| ENSG00000255669.2 | | -0.194360966227011 | 0.411022101009211 | -0.708402470082645 | 0.482563001452335 | 0.75474001295327 | -5.52293816741941 |
| ENSG00000161618.9 | ALDH16A1 | 0.141509032898265 | 3.93677780251813 | 0.708359850221548 | 0.482589190617402 | 0.75474001295327 | -6.04779202117959 |
| ENSG00000265145.1 | SNORD53 | 0.205543536099175 | -0.180522130534252 | 0.708285824339293 | 0.482634680138533 | 0.75474156548104 | -5.36029197002557 |
| lnc-C1QBP-2 | lnc-C1QBP-2 | 0.0965549062108764 | 3.69476011265067 | 0.708238895222202 | 0.482663519721466 | 0.75474156548104 | -6.01795725529233 |
| ENSG00000147471.11 | PLPBP | 0.10834663936484 | 3.706507046788 | 0.708148740542094 | 0.482718925644577 | 0.754770863456572 | -6.02246281170477 |
| lnc-KIAA1671-2 | lnc-KIAA1671-2 | 0.618095154689625 | 0.672246576535325 | 0.70821910603155 | 0.482816550725181 | 0.754807445657509 | -5.59983642056051 |
| lnc-ZBTB20-1 | lnc-ZBTB20-1 | 0.177413409337047 | 5.98857920755122 | 0.707858740143763 | 0.482929910472176 | 0.754807445657509 | -6.15986726432944 |
| lnc-NUP54-2 | lnc-NUP54-2 | 0.10377673082935 | 0.321671115785606 | 0.70769537925482 | 0.482997599923168 | 0.754807445657509 | -5.52677296264758 |
| ENSG00000164941.13 | INTS8 | -0.0882252736275779 | 5.8145702307569 | -0.707679738986901 | 0.48300721537206 | 0.754807445657509 | -6.15885679890826 |
| ENSG00000235106.9 | BRD3OS | -0.139670521068742 | 2.51807999260673 | -0.707660414716137 | 0.48301909585035 | 0.754807445657509 | -5.88312419444685 |
| SLC25A25-AS1 | SLC25A25-AS1 | -0.122059883146692 | 4.0027417858068 | -0.707629828523314 | 0.483037900445667 | 0.754807445657509 | -6.05758293847897 |
| ENSG00000126368.5 | NR1D1 | -0.211117878652863 | 1.04887007651423 | -0.707612349278407 | 0.48304864698688 | 0.754807445657509 | -5.66826449926682 |
| lnc-HSPA13-6 | lnc-HSPA13-6 | -0.500156507174019 | 1.61216100043761 | -0.707759719883436 | 0.483088067278289 | 0.754807445657509 | -5.70076290540057 |
| LINC01481 | LINC01481 | -0.131638266826361 | 5.4161698104199 | -0.707514733533196 | 0.48310866530277 | 0.754807445657509 | -6.1487922527048 |
| lnc-NXF5-2 | lnc-NXF5-2 | -0.187672207724372 | 1.88159546095836 | -0.707514130553096 | 0.483109036053638 | 0.754807445657509 | -5.81187754818995 |
| ENSG00000123064.12 | DDX54 | 0.0924257892379741 | 5.76932284001064 | 0.707018876592689 | 0.483413604006422 | 0.755225974289502 | -6.15995855477635 |
| ENSG00000140382.14 | HMG20A | -0.0655103380795108 | 5.12023810528569 | -0.706545848580724 | 0.483704604345266 | 0.755530351345933 | -6.13827646930937 |
| lnc-SYAP1-2 | lnc-SYAP1-2 | 0.120536789680502 | 2.78719166199299 | 0.706524893639616 | 0.483717497813816 | 0.755530351345933 | -5.92222657517575 |
| ENSG00000229816.1 | DDX50P1 | 0.149117537227738 | 1.90442446996791 | 0.706523179228329 | 0.483718552690896 | 0.755530351345933 | -5.79541126488569 |
| ENSG00000213753.10 | CENPBD2P | -0.1367466738369 | 1.48992802713728 | -0.706190816287781 | 0.483923079986516 | 0.755669901246667 | -5.76393952452771 |
| lnc-ZPR1-1 | lnc-ZPR1-1 | -0.108144302024979 | 1.94067583090726 | -0.706178093191305 | 0.483930910406308 | 0.755669901246667 | -5.82440835954137 |
| ENSG00000125148.6 | MT2A | -0.249171260998193 | -0.710629005885465 | -0.706165396129887 | 0.483938724873801 | 0.755669901246667 | -5.29204166032058 |
| ENSG00000275379.1 | H3C11 | 0.43567646957769 | 2.09130222694111 | 0.70633944389914 | 0.483967443965558 | 0.755669901246667 | -5.8699516095017 |
| ENSG00000072694.20 | FCGR2B | 0.103073626965853 | 6.489846448153 | 0.706079707470401 | 0.483991464224593 | 0.755669901246667 | -6.16384470185532 |
| lnc-HSPA12A-1 | lnc-HSPA12A-1 | -1.07484315268938 | 1.45282778884889 | -0.706231160811644 | 0.484051098441052 | 0.755705685619855 | -5.83358339051719 |
| lnc-PCNX4-3 | lnc-PCNX4-3 | -0.159296321773762 | 1.88955808650826 | -0.705753107499647 | 0.484192508409842 | 0.75576197556304 | -5.82484837888368 |
| ENSG00000280381.1 | | -0.28787243452442 | 3.21310764177441 | -0.705849221093542 | 0.484222875277549 | 0.75576197556304 | -5.94978834432426 |
| ENSG00000143507.17 | DUSP10 | 0.316264227357737 | 4.38851291755257 | 0.705859930320317 | 0.484242862980158 | 0.75576197556304 | -6.0643689859565 |

| | | | | | | | |
|---|---|---|---|---|---|---|---|
| ENSG00000117475.13 | BLZF1 | -0.0784749917302925 | 4.10969811036305 | -0.7056641357142 | 0.484247284637257 | 0.75576197556304 | -6.06211870981642 |
| lnc-TTF1-1 | lnc-TTF1-1 | 0.08870755801754 | 3.40880399810212 | 0.705626035144554 | 0.484270742637173 | 0.75576197556304 | -5.98342400677571 |
| lnc-SLC26A3-2 | lnc-SLC26A3-2 | 0.18216653671166 | 1.60475159019722 | 0.705508609088067 | 0.484343044268408 | 0.755817504183218 | -5.76824527809112 |
| lnc-LYSMD4-1 | lnc-LYSMD4-1 | 0.177975763584375 | -0.393121691827037 | 0.705254152728756 | 0.484499739060707 | 0.755971787238898 | -5.33433523968648 |
| ENSG00000166444.18 | DENND2B | -0.259215009326743 | 0.0468758166789204 | -0.70522877563986 | 0.484515367888818 | 0.755971787238898 | -5.4383667446931 |
| LINC00954 | LINC00954 | -0.267315456042204 | 3.42634377017761 | -0.705249544128597 | 0.484584080760159 | 0.756021688341268 | -5.97143025944201 |
| lnc-JMJD8-2 | lnc-JMJD8-2 | -0.150559533840998 | 0.462530982450731 | -0.704918566259855 | 0.484706437458483 | 0.756109229464503 | -5.59243511592404 |
| ENSG00000144559.10 | TAMM41 | 0.0812729638287966 | 3.80476361429427 | 0.704906530992243 | 0.484713851282292 | 0.756109229464503 | -6.03104378365768 |
| lnc-NUP160-2 | lnc-NUP160-2 | 0.10311616001516 | 3.49490819015762 | 0.704833884643446 | 0.484758603381753 | 0.756109229464503 | -5.98996774127514 |
| lnc-LRRC31-3 | lnc-LRRC31-3 | -0.114486692872058 | 4.09635336397307 | -0.704787578827411 | 0.484787130216346 | 0.756109229464503 | -6.05788033108739 |
| ENSG00000130706.12 | ADRM1 | -0.129600103990722 | 4.4520880103501 | -0.704578867063088 | 0.484915719404402 | 0.756252481626065 | -6.10586567634011 |
| ENSG00000144048.10 | DUSP11 | 0.0717327388883646 | 5.16292459803739 | 0.704281831794868 | 0.485098758484344 | 0.756475926491471 | -6.14297166744587 |
| ENSG00000108599.14 | AKAP10 | 0.0684376520401608 | 5.21115681956698 | 0.704227084431693 | 0.485132499134597 | 0.756475926491471 | -6.14417234683498 |
| lnc-CRACR2A-6 | lnc-CRACR2A-6 | -0.173798926622562 | 0.780629411261416 | -0.704119023242468 | 0.485199100812739 | 0.756522467391001 | -5.62797020533104 |
| lnc-EHD3-1 | lnc-EHD3-1 | 0.173348822678985 | 3.1215505237393 | 0.703719136345953 | 0.485445608898008 | 0.756639457414812 | -5.9415086253998 |
| ENSG00000198589.11 | LRBA | 0.071424670350086 | 7.97299334398843 | 0.703704828731709 | 0.485454430049157 | 0.756639457414812 | -6.14695207866415 |
| lnc-RSBN1-1 | lnc-RSBN1-1 | -0.142389636906608 | 2.76971819422266 | -0.703645000150472 | 0.485491317462895 | 0.756639457414812 | -5.92422087420913 |
| ENSG00000124702.17 | KLHDC3 | -0.111091714131528 | 4.7015355012338 | -0.703626396288847 | 0.48550278802573 | 0.756639457414812 | -6.11840948451529 |
| lnc-TPP2-3 | lnc-TPP2-3 | -0.156036455418158 | 0.0750718823347808 | -0.70356550261373 | 0.485540334235518 | 0.756639457414812 | -5.48656064108816 |
| ENSG00000178802.17 | MPI | -0.0991393031173793 | 4.56470367630292 | -0.703494741923561 | 0.485583966356696 | 0.756639457414812 | -6.09950245975647 |
| ENSG00000189367.14 | KIAA0408 | -0.619082461610957 | -0.701308850415489 | -0.703496100112867 | 0.485678783979914 | 0.756639457414812 | -5.32241966740074 |
| lnc-GPC2-2 | lnc-GPC2-2 | 0.128302246540632 | 1.22691214101351 | 0.703269750800848 | 0.485722713917513 | 0.756639457414812 | -5.72452429060302 |
| lnc-MGAT1-1 | lnc-MGAT1-1 | -0.156201283132043 | 0.124367214183883 | -0.703263080245601 | 0.485726827855647 | 0.756639457414812 | -5.48250516899959 |
| ENSG00000137270.10 | GCM1 | -0.250285566825859 | 1.07343362284001 | -0.703258047504534 | 0.48573090626855 | 0.756639457414812 | -5.68702509775087 |
| ENSG00000167766.18 | ZNF83 | -0.0874995028114851 | 7.06611965666016 | -0.70312790574218 | 0.48581019836765 | 0.756639457414812 | -6.16189325631616 |
| lnc-RXRA-1 | lnc-RXRA-1 | -0.335894532456697 | -0.491472251435156 | -0.703122791269604 | 0.485813352937969 | 0.756639457414812 | -5.36379171675841 |
| lnc-VCP-2 | lnc-VCP-2 | 0.0912137972588077 | -0.00909532440381015 | 0.703101950556337 | 0.485826207460399 | 0.756639457414812 | -5.42522378463862 |
| ENSG00000196562.14 | SULF2 | 0.575921642324404 | 1.70300440568497 | 0.703300908973956 | 0.485849714828416 | 0.756639457414812 | -5.81985560128032 |
| ENSG00000007168.12 | PAFAH1B1 | 0.0833096768506856 | 7.02644716733942 | 0.702992275616053 | 0.485893857948574 | 0.756639457414812 | -6.16130677580504 |
| ENSG00000071794.15 | HLTF | 0.146545600667002 | 3.95059861250891 | 0.702964961478156 | 0.485910706873081 | 0.756639457414812 | -6.0370962917753 |
| ENSG00000137265.14 | IRF4 | -0.201939912578163 | 6.47252590823867 | -0.703045900584997 | 0.48592357529509 | 0.756639457414812 | -6.16523900523974 |
| ENSG00000149600.11 | COMMD7 | -0.0941836777081861 | 3.11173961624336 | -0.702924250701745 | 0.485935820218938 | 0.756639457414812 | -5.95539277986833 |
| ENSG00000273802.2 | H2BC8 | 0.940885103051223 | 3.85011118573891 | 0.702916420046158 | 0.48609266668033 | 0.756747458795315 | -6.01604330654051 |
| ENSG00000179134.15 | SAMD4B | 0.113435144567796 | 4.6339931704571 | 0.702645123655122 | 0.486108025514386 | 0.756747458795315 | -6.1083150159245 |
| ENSG00000132964.11 | CDK8 | 0.067401067129178 | 3.73961831860462 | 0.702539617712079 | 0.48617312549917 | 0.756747458795315 | -6.03176832576298 |
| ENSG00000204516.9 | MICB | 0.105690672519 | 4.28073484153281 | 0.702390646567579 | 0.486265052987836 | 0.756747458795315 | -6.08637991910914 |
| ENSG00000108106.13 | UBE2S | 0.133449141180372 | 2.55421313469493 | 0.702287877651387 | 0.486328475562327 | 0.756747458795315 | -5.89491556846782 |
| ENSG00000091483.6 | FH | 0.119130440881571 | 3.6857047793673 | 0.702273941281677 | 0.486337076578369 | 0.756747458795315 | -6.02392710945678 |
| lnc-STEAP1B-5 | lnc-STEAP1B-5 | 0.753770317299578 | -1.55931302820864 | 0.70231390128896 | 0.486348110734925 | 0.756747458795315 | -4.98695437322081 |
| lnc-PTH-3 | lnc-PTH-3 | -0.106366888753186 | 0.475947428612941 | -0.702128792370112 | 0.486426662223796 | 0.756747458795315 | -5.56374339444154 |
| lnc-SERF1A-5 | lnc-SERF1A-5 | -0.172637533353642 | 1.4998111474129 | -0.70206246208262 | 0.486467604232625 | 0.756747458795315 | -5.7533031367133 |
| lnc-RDH5-2 | lnc-RDH5-2 | 0.160949510141606 | 0.188358002983056 | 0.702053699085697 | 0.486473013289033 | 0.756747458795315 | -5.48668283071441 |
| ENSG00000203791.14 | EEF1AKMT2 | 0.0748065904954363 | 3.47236371605437 | 0.702039750538695 | 0.486481623252818 | 0.756747458795315 | -5.99467666465331 |
| lnc-ADI1-4 | lnc-ADI1-4 | 0.0988538355781037 | 1.37287561142553 | 0.702022660528937 | 0.486492172451035 | 0.756747458795315 | -5.74034869847715 |
| ENSG00000166855.9 | CLPX | 0.0619821455693383 | 5.13315357371182 | 0.701987274697932 | 0.486514015569188 | 0.756747458795315 | -6.14306160669572 |
| ENSG00000134013.15 | LOXL2 | -0.182435572297874 | 1.98265118708298 | -0.701920445655486 | 0.486555269585786 | 0.756747458795315 | -5.82210889692731 |
| ENSG00000110063.9 | DCPS | -0.0888172420088562 | 3.26347363256947 | -0.701909965704152 | 0.486561739107714 | 0.756747458795315 | -5.97914095586705 |
| ENSG00000065882.15 | TBC1D1 | 0.120836404396165 | 7.81123757082663 | 0.701802662564829 | 0.486627982644981 | 0.756747458795315 | -6.15214847404408 |
| lnc-LGALS2-5 | lnc-LGALS2-5 | -0.132502509542649 | 3.25168812369197 | -0.701799074276399 | 0.486630197960199 | 0.756747458795315 | -5.97319303766236 |
| lnc-PPIA-1 | lnc-PPIA-1 | -0.180303474061935 | 0.657021992220168 | -0.70169298904613 | 0.486695694750796 | 0.756792134551306 | -5.59695143105151 |
| ENSG00000108439.10 | PNPO | 0.124557508071311 | 2.2501367750652 | 0.701404383955401 | 0.486873903871964 | 0.75701205343682 | -5.84993695709199 |
| lnc-RFX3-4 | lnc-RFX3-4 | -0.2022760036577 | -0.32522373084806 | -0.701195530570132 | 0.48700289033229 | 0.757077191048369 | -5.34685282261467 |
| lnc-FUBP1-2 | lnc-FUBP1-2 | 0.135179583429517 | 5.03580296996882 | 0.701120716680053 | 0.487049099542746 | 0.757077191048369 | -6.14448038232062 |
| lnc-ZNF33A-8 | lnc-ZNF33A-8 | 0.349809295706103 | 1.76228476266607 | 0.701158595177638 | 0.487115940988692 | 0.757077191048369 | -5.77683976393157 |
| lnc-ITGA9-5 | lnc-ITGA9-5 | 0.152087353444565 | 1.70535948525492 | 0.700977263490641 | 0.487137711053651 | 0.757077191048369 | -5.79318484766879 |
| ENSG00000263142.5 | LRRC37A17P | -0.289396348463866 | 0.642199877902905 | -0.700986868043524 | 0.487143733791648 | 0.757077191048369 | -5.61321151639358 |
| ENSG00000112685.13 | EXOC2 | 0.045462075016883 | 5.51580416292688 | 0.700888468187198 | 0.48719256472496 | 0.757077191048369 | -6.15706779502382 |
| LGALSL-DT | LGALSL-DT | -0.166879054979557 | 1.79603762142825 | -0.700734626973648 | 0.487287608956921 | 0.757077191048369 | -5.80534702628333 |
| lnc-SMCO4-5 | lnc-SMCO4-5 | 0.168423452431826 | -0.238640731348636 | 0.700665634953022 | 0.487330236101564 | 0.757077191048369 | -5.36770881664097 |
| lnc-GBA2-1 | lnc-GBA2-1 | 0.0995855879605401 | 2.33036427364971 | 0.70062908751023 | 0.487352818008895 | 0.757077191048369 | -5.85310297105045 |
| ENSG00000199753.1 | SNORD104 | -0.16370914278786 | 0.110724305245299 | -0.700575623548408 | 0.487385853336674 | 0.757077191048369 | -5.48452217630647 |
| lnc-DDX5-2 | lnc-DDX5-2 | -0.0712645904129507 | 2.6669893563375 | -0.700565594909275 | 0.487392050162077 | 0.757077191048369 | -5.90425374171882 |
| lnc-NPR2-1 | lnc-NPR2-1 | -0.12948310345258 | 2.18377701195581 | -0.700553130737467 | 0.487399751995916 | 0.757077191048369 | -5.84065010582744 |
| lnc-ESCO1-5 | lnc-ESCO1-5 | 0.0756856752746544 | 3.01475351251463 | 0.700504138076152 | 0.487430026093678 | 0.757077191048369 | -5.94209889940623 |
| ENSG00000035687.9 | ADSS2 | 0.0637724103680908 | 5.18205401915439 | 0.700502981651529 | 0.487430740697322 | 0.757077191048369 | -6.14843382345656 |
| ENSG00000165995.19 | CACNB2 | 0.713653806837432 | 2.95851254575733 | 0.700549230287901 | 0.487553583926066 | 0.757137805248308 | -5.83110880106975 |
| lnc-MYLK4-33 | lnc-MYLK4-33 | 0.252762312937921 | -0.248851522374802 | 0.700241108040156 | 0.487592578552365 | 0.757137805248308 | -5.38215058824284 |
| lnc-TMEM248-4 | lnc-TMEM248-4 | 0.174324143394459 | 2.37467013323431 | 0.700160418555693 | 0.487642450695005 | 0.757137805248308 | -5.85139186314086 |

| | | | | | | | |
|---|---|---|---|---|---|---|---|
| lnc-TSPAN3-5 | lnc-TSPAN3-5 | -0.102288218794467 | 2.16782032966501 | -0.70014403301819 | 0.487652578532241 | 0.757137805248308 | -5.84473445740857 |
| ENSG00000205544.3 | TMEM256 | -0.115610040535609 | 1.85886440088274 | -0.700142236161504 | 0.487653689169549 | 0.757137805248308 | -5.81009055393301 |
| ENSG00000100083.18 | GGA1 | -0.0922548133530219 | 6.65092115310872 | -0.699874613517171 | 0.487819122539936 | 0.757320092523581 | -6.16720816390284 |
| ENSG00000197608.11 | ZNF841 | 0.0847311777874644 | 5.26095711736347 | 0.699820530104402 | 0.487852558502082 | 0.757320092523581 | -6.14639317295773 |
| ENSG00000102755.11 | FLT1 | -0.186859989932747 | 4.18069153837564 | -0.699816790868431 | 0.487881476308735 | 0.757320092523581 | -6.08599286765892 |
| lnc-AGO2-2 | lnc-AGO2-2 | -0.134542687449263 | 3.23356593728809 | -0.699600043352712 | 0.487988883171347 | 0.757429694767802 | -5.95914037217801 |
| ENSG00000147457.13 | CHMP7 | -0.10833537339129 | 7.53202833873088 | -0.69943510906227 | 0.488090874243761 | 0.75749103009151 | -6.15681808468483 |
| lnc-EXOC3-3 | lnc-EXOC3-3 | -0.103121178846919 | 0.918648013393725 | -0.699403030298553 | 0.488110712299407 | 0.75749103009151 | -5.66446503778521 |
| ENSG00000160683.4 | CXCR5 | 0.187215800596653 | 6.56681761532639 | 0.699326242726183 | 0.488203759221065 | 0.75749103009151 | -6.16811220541809 |
| ENSG00000123545.5 | NDUFAF4 | 0.109157786394615 | 2.36053724874454 | 0.69919774769758 | 0.488237673230234 | 0.75749103009151 | -5.87195023523531 |
| ENSG00000101639.18 | CEP192 | 0.082932310296866 | 6.45601328474289 | 0.699149743800268 | 0.488267364814438 | 0.75749103009151 | -6.16878249818945 |
| lnc-COQ5-2 | lnc-COQ5-2 | 0.111381275484912 | 1.65129184840752 | 0.699147428164628 | 0.488268797117194 | 0.75749103009151 | -5.77616217902626 |
| ENSG00000122376.11 | SHLD2 | 0.0702008028463057 | 5.30154395182438 | 0.69911959660113 | 0.48828601210971 | 0.75749103009151 | -6.1519037044235 |
| lnc-SLC25A52-1 | lnc-SLC25A52-1 | -0.0786025377192066 | 1.65472663502442 | -0.6988660704474 | 0.488442844304041 | 0.757677222421439 | -5.77233277566366 |
| ENSG00000081019.13 | RSBN1 | 0.0634570443890592 | 5.61549446156221 | 0.698755780698772 | 0.488511078718639 | 0.757708871383966 | -6.16178511298434 |
| lnc-MYLK4-36 | lnc-MYLK4-36 | 0.133292246463039 | 0.277681590763829 | 0.698607892161385 | 0.488602583223797 | 0.757708871383966 | -5.49371029275823 |
| ENSG00000181915.4 | ADO | 0.116497879881418 | 4.16950175305553 | 0.698558540291604 | 0.488633121312577 | 0.757708871383966 | -6.07634266762747 |
| ENSG00000147679.11 | UTP23 | 0.108113465624624 | 4.38170243612574 | 0.698520815765286 | 0.488656465319503 | 0.757708871383966 | -6.09197717337418 |
| lnc-UBAP1L-2 | lnc-UBAP1L-2 | 0.0853396074660395 | 0.739377083487631 | 0.698476196748587 | 0.488684076453399 | 0.757708871383966 | -5.62323492159453 |
| lnc-ATOH7-4 | lnc-ATOH7-4 | -0.0775820939913257 | 1.67436634474167 | -0.698476124364586 | 0.488684121246764 | 0.757708871383966 | -5.77928112320126 |
| lnc-TSPAN8-3 | lnc-TSPAN8-3 | 0.129792198932292 | 1.76586429304461 | 0.698391899027998 | 0.488736243934092 | 0.757732608383204 | -5.77945349472846 |
| ENSG00000103024.7 | NME3 | -0.153314918884403 | 2.85281916877575 | -0.698182006714317 | 0.488866148887599 | 0.757876925702602 | -5.94416250985818 |
| lnc-ZNF253-2 | lnc-ZNF253-2 | -0.108096251341906 | 0.437515389308947 | -0.698024706997255 | 0.488963516260583 | 0.757970782888355 | -5.5668375286788 |
| lnc-C2CD6-2 | lnc-C2CD6-2 | 0.213126507869855 | 1.9953948576114 | 0.697905688500928 | 0.489037195028244 | 0.758027907618522 | -5.80217743848319 |
| lnc-SLC4A11-1 | lnc-SLC4A11-1 | -0.083434101733205 | 2.1312054078386 | -0.697575999061275 | 0.489241322631802 | 0.758287209618251 | -5.84359072922958 |
| ENSG00000160294.10 | MCM3AP | 0.0549550344528784 | 6.74732344660872 | 0.69746117852 2122 | 0.48931242505886 | 0.758299912497135 | -6.16862814372761 |
| lnc-ZNF506-5 | lnc-ZNF506-5 | -0.613515854957087 | 0.794864040957675 | -0.697450614577943 | 0.489463210249774 | 0.758299912497135 | -5.68638583387257 |
| ENSG00000132716.18 | DCAF8 | -0.0523718491713927 | 7.02532320790868 | -0.697207442035658 | 0.489469571390601 | 0.758299912497135 | -6.16555666955973 |
| ENSG00000118418.14 | HMGN3 | -0.0985445169199046 | 5.56517211737958 | -0.697181642421336 | 0.489485551412512 | 0.758299912497135 | -6.16150806679128 |
| ENSG00000171940.13 | ZNF217 | 0.0838276150191399 | 6.79496419368858 | 0.697175258228757 | 0.489489505762151 | 0.758299912497135 | -6.16846586138735 |
| ENSG00000185869.14 | ZNF829 | 0.125514632030031 | 3.77400729260088 | 0.697147394247171 | 0.489506764836855 | 0.758299912497135 | -6.0301322687715 |
| ENSG00000119965.12 | C10orf88 | 0.0817339134587987 | 3.05388348604777 | 0.697058711749173 | 0.489561697430614 | 0.758299912497135 | -5.94745492674725 |
| lnc-GEMIN4-3 | lnc-GEMIN4-3 | 0.103692901756387 | 0.492940650286008 | 0.697053881423961 | 0.489564689577323 | 0.758299912497135 | -5.57754733899523 |
| NORAD | NORAD | -0.100146397286839 | 6.26025167371173 | -0.696986726295206 | 0.489606289902392 | 0.758299912497135 | -6.17009056155018 |
| ENSG00000149136.8 | SSRP1 | -0.0778147656382355 | 5.82583645634652 | -0.696967937595294 | 0.489617929217652 | 0.758299912497135 | -6.16674262976574 |
| lnc-REST-6 | lnc-REST-6 | -0.0832572271575698 | 0.844578755351918 | -0.696706948814811 | 0.489779623768653 | 0.758493265821246 | -5.64709119717154 |
| BRE-AS1 | BRE-AS1 | 0.277187720268349 | -0.470234304564245 | 0.696543517876764 | 0.48988089190111 | 0.758591227999311 | -5.28720492848561 |
| lnc-USP6-4 | lnc-USP6-4 | 0.129305213671296 | 0.886495539302694 | 0.696485909501668 | 0.489916591060333 | 0.758591227999311 | -5.66019952924983 |
| ENSG00000262814.7 | MRPL12 | 0.158550791515145 | -0.0487560299028303 | 0.69638161409006 | 0.489981225255275 | 0.758634237959179 | -5.44405887341245 |
| ENSG00000012232.8 | EXTL3 | -0.119103742233076 | 4.1293126751194 | -0.696168007830949 | 0.490113616651311 | 0.758772813993743 | -6.08174004516037 |
| lnc-RSPH10B-2 | lnc-RSPH10B-2 | 0.08847687519454 | 1.76800887712059 | 0.696118255082866 | 0.490144455854871 | 0.758772813993743 | -5.79262709461053 |
| ENSG00000161547.16 | SRSF2 | -0.104201331280764 | 6.03560334631798 | -0.695910081228248 | 0.490273503985175 | 0.758915509703456 | -6.17044555895072 |
| ENSG00000183458.13 | PKD1P3 | -0.140103157864901 | 2.76900569449181 | -0.695785372991118 | 0.490350820377603 | 0.758972404884449 | -5.92587978954853 |
| ENSG00000123146.19 | ADGRE5 | -0.14707916938961 | 5.63118634393201 | -0.695731847936433 | 0.490384006828853 | 0.758972404884449 | -6.16554508777845 |
| ENSG00000188848.15 | BEND4 | -0.0521065348069163 | 5.69509338545092 | -0.69561865960113 | 0.490454189655442 | 0.758975240753974 | -6.16483627178714 |
| ENSG00000111674.8 | ENO2 | 0.209520582793246 | 3.6476184782733 | 0.695675261118698 | 0.490459586905145 | 0.758975240753974 | -6.04036408495357 |
| ENSG00000169714.16 | CNBP | 0.0827814607674471 | 7.87021340857182 | 0.695271933944254 | 0.490669212924588 | 0.759242550716193 | -6.15457403738503 |
| ENSG00000182551.13 | ADI1 | -0.125966612306194 | 4.83402124246166 | -0.695198327328064 | 0.490714867106398 | 0.759256115887469 | -6.13216075289685 |
| ENSG00000250850.2 | | -0.212237521477346 | 2.73738164767367 | -0.695044872168556 | 0.490824664254456 | 0.759368916442384 | -5.91236551741922 |
| ENSG00000280537.2 | | -0.126604495634224 | 2.20161225063361 | -0.694431148414059 | 0.491190847192521 | 0.75970902303884 | -5.86251054322255 |
| ENSG00000029364.11 | SLC39A9 | 0.0734282102980215 | 5.68454660561357 | 0.69441059968644 | 0.491203599750691 | 0.75970902303884 | -6.16588294136776 |
| ENSG00000228716.6 | DHFR | 0.121142653764272 | 3.36759514774087 | 0.694381134087562 | 0.491221886448267 | 0.75970902303884 | -5.99437767981211 |
| ENSG00000122483.17 | CCDC18 | -0.0748608324879427 | 4.46435191466975 | -0.694379449513958 | 0.491222931925922 | 0.75970902303884 | -6.10187888967683 |
| ENSG00000138443.15 | ABI2 | -0.10448487843902 | 5.17118735925957 | -0.694323267505977 | 0.491257800232117 | 0.75970902303884 | -6.15180783238373 |
| ERICH6-AS1 | ERICH6-AS1 | 0.243898116660584 | 0.31421715809842 | 0.694310132665675 | 0.491265952322157 | 0.75970902303884 | -5.53160099368729 |
| ENSG00000226976.3 | COX6A1P2 | 0.208690512566371 | -0.381434208545754 | 0.694239458150096 | 0.49130981749037 | 0.759719778634535 | -5.30544016410324 |
| ENSG00000176101.11 | SSNA1 | -0.135729173132119 | 3.08433053055917 | -0.694084441209263 | 0.491406038642591 | 0.759772127061786 | -5.97207160279939 |
| lnc-PPP1R2C-9 | lnc-PPP1R2C-9 | 0.0957947360527616 | 2.10515048166763 | 0.694021902632583 | 0.491444860165689 | 0.759772127061786 | -5.82565575735689 |
| ENSG00000162129.13 | CLPB | 0.0812054495564052 | 4.24210668460719 | 0.694006520652445 | 0.491454408963738 | 0.759772127061786 | -6.08178986222866 |
| lnc-ZNF154-2 | lnc-ZNF154-2 | -0.0802020293250783 | 0.541090187594203 | -0.693693262845854 | 0.491648895025684 | 0.760015712077631 | -5.58703978048864 |
| ENSG00000141458.12 | NPC1 | 0.0798299714824928 | 4.25121455710402 | 0.693292509523813 | 0.491897765032308 | 0.760246650840061 | -6.08623870313029 |
| lnc-DGAT2-4 | lnc-DGAT2-4 | -0.183844517969166 | 3.30341386323483 | -0.693233030062948 | 0.491937345164314 | 0.760246650840061 | -5.97469750595346 |
| ENSG00000153234.13 | NR4A2 | 0.342654146286407 | 3.77406543983034 | 0.69342566767873 | 0.491937594590558 | 0.760246650840061 | -6.02020809509446 |
| ENSG00000188725.7 | SMIM15 | 0.100809067933596 | 4.62997886803669 | 0.693214801339384 | 0.491946030342456 | 0.760246650840061 | -6.12202271838394 |
| LINC00852 | LINC00852 | 0.0961753742660747 | 1.84403910654907 | 0.693021499447525 | 0.492066103444246 | 0.760375120660129 | -5.79609363709041 |
| lnc-MFSD1-2 | lnc-MFSD1-2 | -0.155644193001395 | 0.885588340817481 | -0.692824009442165 | 0.492188794885231 | 0.760507616929863 | -5.6521247868506 |
| ENSG00000165591.6 | FAAH2 | -0.445084994020014 | -0.786512408143545 | -0.692707360776593 | 0.492274579347107 | 0.760583070612634 | -5.26413831300132 |
| ENSG00000122257.19 | RBBP6 | 0.0623797981036276 | 7.28499455269534 | 0.692601308861696 | 0.492327168878518 | 0.76060722937976 | -6.16601037576872 |

| | | | | | | | |
|---|---|---|---|---|---|---|---|
| ENSG00000280392.1 | | 0.179875424895137 | 1.99593175171602 | 0.692446245861805 | 0.49242352932656 | 0.760670985800533 | -5.81375119040021 |
| lnc-ZC2HC1A-4 | lnc-ZC2HC1A-4 | -0.395049554131983 | -0.482336152199814 | -0.692422561014599 | 0.492442349793135 | 0.760670985800533 | -5.3133530635872 |
| ENSG00000180573.9 | H2AC6 | -0.15898667234658 | 6.46103877321528 | -0.692311502591742 | 0.492520570234288 | 0.760678253933783 | -6.17286732632184 |
| lnc-AMFR-5 | lnc-AMFR-5 | 0.088078082056259 | 1.4361236920349 | 0.692289464461562 | 0.492520968283318 | 0.760678253933783 | -5.75322988301715 |
| ENSG00000148950.10 | IMMP1L | 0.0957469038213601 | 2.33127793224602 | 0.692125790563088 | 0.492622702318469 | 0.760778292453561 | -5.87105709583499 |
| ENSG00000175003.13 | SLC22A1 | -0.228124216653567 | -0.607982111816135 | -0.691805358295972 | 0.49282190570684 | 0.761028830757288 | -5.2815915888505 |
| LINC00115 | LINC00115 | 0.0844225207849403 | 1.73505685549534 | 0.691586489596506 | 0.492957995683061 | 0.761175534564247 | -5.78630020140034 |
| lnc-SCGN-2 | lnc-SCGN-2 | -0.240787098348343 | 1.47507911798695 | -0.691503289979385 | 0.493019653221108 | 0.761175534564247 | -5.76102975752457 |
| ENSG00000121892.14 | PDS5A | 0.0468623573213875 | 7.55767993033817 | 0.691436441033824 | 0.493051306157418 | 0.761175534564247 | -6.16236374832462 |
| ENSG00000196693.14 | ZNF33B | -0.1246816383573 | 4.83210807328552 | -0.691396525445687 | 0.493076130054379 | 0.761175534564247 | -6.13354030758059 |
| ENSG00000013810.18 | TACC3 | 0.159010766448902 | 4.43823030827119 | 0.691323689412464 | 0.493122384655164 | 0.761175534564247 | -6.09892085114828 |
| ENSG00000082293.12 | COL19A1 | -0.207457692164352 | 6.71869728352397 | -0.691378666090645 | 0.493160566145904 | 0.761175534564247 | -6.17160544880551 |
| ENSG00000284981.1 | UPK3BL2 | 0.315267563751054 | 0.0960606366057663 | 0.691236316301521 | 0.493175772673993 | 0.761175534564247 | -5.4546392322988 |
| lnc-ZNF510-1 | lnc-ZNF510-1 | -0.179962825778591 | -0.244617403316608 | -0.691151658627759 | 0.493228430311338 | 0.76119972865699 | -5.38945493075639 |
| lnc-UBE2W-1 | lnc-UBE2W-1 | -0.0744057161575259 | 0.959615260519496 | -0.69109017149906 | 0.493266677670309 | 0.761201681398109 | -5.6826905569106 |
| lnc-GP5-4 | lnc-GP5-4 | 0.136821498705805 | 2.17103814425894 | 0.691020118147754 | 0.493310255553993 | 0.761211859214921 | -5.84698658670685 |
| ENSG00000111331.12 | OAS3 | -0.258057528364724 | 4.43422853356755 | -0.691116211347091 | 0.493349757119121 | 0.761215745935748 | -6.1219191354795 |
| lnc-GLRA1-1 | lnc-GLRA1-1 | -0.136534150524114 | 1.01476250985894 | -0.690832796764862 | 0.493426792507773 | 0.761277540678172 | -5.68623462416938 |
| ENSG00000214135.8 | SDHAP4 | 0.0943857211411832 | 3.66649238054853 | 0.690480578687945 | 0.493645956833884 | 0.761558591628829 | -6.03046579535775 |
| ENSG00000077147.15 | TM9SF3 | 0.0786015995215212 | 6.52560564296592 | 0.690269029168055 | 0.493777617433941 | 0.761666189111802 | -6.17425670022161 |
| ENSG00000239884.3 | RN7SL608P | 0.150573355743127 | 0.312770206482265 | 0.690249598577151 | 0.493789711291254 | 0.761666189111802 | -5.54209973919488 |
| ENSG00000077097.15 | TOP2B | -0.0507868719607103 | 7.64609026295421 | -0.690026404025764 | 0.493928642312026 | 0.761823397879987 | -6.16130239431275 |
| ENSG00000100077.14 | GRK3 | -0.0860297026496442 | 6.66838662831955 | -0.689962080129016 | 0.493968685771771 | 0.761824887006369 | -6.17432284087171 |
| lnc-AZIN1-1 | lnc-AZIN1-1 | 0.102009547102919 | 3.58808796898337 | 0.689905945807156 | 0.494003632457562 | 0.761824887006369 | -6.01011895992442 |
| ENSG00000171823.6 | FBXL14 | 0.114475413597491 | 3.02005782390624 | 0.689844334785484 | 0.494041990265476 | 0.76182696176463 | -5.96247461648857 |
| ENSG00000177971.8 | IMP3 | 0.107921255682525 | 4.1841199824494 | 0.689681416262982 | 0.494143428077958 | 0.76192630010702 | -6.08623780787311 |
| ENSG00000269476.1 | | 0.0676447392728263 | 1.60451096509509 | 0.689611785286601 | 0.494186785869264 | 0.761936075921128 | -5.77120429103095 |
| lnc-ARF6-1 | lnc-ARF6-1 | 0.249901271856312 | 1.36515511021901 | 0.689536826865107 | 0.494243561055945 | 0.761966535631377 | -5.728431838122 |
| ENSG00000111801.15 | BTN3A3 | 0.142136839578378 | 5.03812283670014 | 0.689329994561841 | 0.494362272736734 | 0.762092470022483 | -6.1395361544639 |
| ENSG00000172262.11 | ZNF131 | 0.0506283566857251 | 5.53463394234703 | 0.689252329025071 | 0.494410645482863 | 0.762109961504813 | -6.16576831465414 |
| lnc-SEMA4D-5 | lnc-SEMA4D-5 | 0.274227741719078 | -0.0937522747366053 | 0.688952502180082 | 0.494597412461674 | 0.762284190312507 | -5.42674266298453 |
| ENSG00000148362.10 | PAXX | -0.220794841005662 | 2.13515668311445 | -0.688975183742117 | 0.494597743848007 | 0.762284190312507 | -5.874226608675 |
| ENSG00000104081.13 | BMF | -0.233424566557343 | 8.16078033421512 | -0.688876373809237 | 0.494733907582997 | 0.762403439766346 | -6.1510292978048 |
| ENSG00000163683.11 | SMIM14 | -0.180940982448623 | 6.12692570254085 | -0.688773120637512 | 0.494749198122678 | 0.762403439766346 | -6.17534895077651 |
| lnc-OR2A4-3 | lnc-OR2A4-3 | 0.0788120432192736 | 2.12265920949921 | 0.688489022301 | 0.49488619840211 | 0.76255746531746 | -5.84882735545283 |
| ENSG00000062582.13 | MRPS24 | 0.0987298227579609 | 2.41532658303392 | 0.68834796008178 | 0.4949741102426 | 0.762592394765228 | -5.88297186369088 |
| lnc-TMEM120B-1 | lnc-TMEM120B-1 | -0.160611768293466 | 1.07279899281566 | -0.688333750634147 | 0.494982966237353 | 0.762592394765228 | -5.69429923596085 |
| ENSG00000103043.14 | VAC14 | 0.067299868129246 | 5.09101643699286 | 0.688133732790902 | 0.495107636049438 | 0.762727376154898 | -6.14920079851783 |
| ENSG00000275227.1 | | 0.121685096796691 | 1.28433535310934 | 0.688072362407351 | 0.495145891290364 | 0.762729223202332 | -5.73144060931874 |
| lnc-TSPOAP1-1 | lnc-TSPOAP1-1 | -0.207647768322977 | 1.72312498520451 | -0.687882481943011 | 0.495264263672392 | 0.762836188983187 | -5.82790584822763 |
| ENSG00000125912.10 | NCLN | -0.0867215033365776 | 4.71649109252386 | -0.687781368317311 | 0.495327304772934 | 0.762836188983187 | -6.13138920190129 |
| lnc-SNX5-1 | lnc-SNX5-1 | -0.342512965561758 | 2.28647221125523 | -0.687890666885592 | 0.495357143205763 | 0.762836188983187 | -5.84790659822962 |
| lnc-SH2D3A-2 | lnc-SH2D3A-2 | -0.171850942545582 | -0.141065722145866 | -0.687723193633059 | 0.495363576832788 | 0.762836188983187 | -5.41396918855098 |
| ENSG00000165617.14 | DACT1 | 0.283353250194315 | 2.7880748415578 | 0.687735804650995 | 0.495436236667407 | 0.762860985590234 | -5.91149624562291 |
| lnc-RPS5-3 | lnc-RPS5-3 | -0.162935920838382 | 1.31065743797182 | -0.687553233429571 | 0.495469555855975 | 0.762860985590234 | -5.72739529644831 |
| ENSG00000173013.5 | CFAP184 | -0.136733098917561 | 0.861437585653613 | -0.687519058775505 | 0.495490867043474 | 0.762860985590234 | -5.66834600635315 |
| ENSG00000168803.15 | ADAL | -0.0916732088660455 | 2.60965418028259 | -0.687435788641766 | 0.495542796112131 | 0.76288387227943 | -5.89193968807962 |
| ENSG00000198276.15 | UCKL1 | -0.0898142814756717 | 5.01955270070364 | -0.687203011483065 | 0.49568797698002 | 0.762976048278299 | -6.14883727529099 |
| ENSG00000218730.1 | | -0.283193457850685 | 1.40630037877664 | -0.687248826103915 | 0.495695407218502 | 0.762976048278299 | -5.73634683755571 |
| lnc-ACMSD-6 | lnc-ACMSD-6 | 0.359082061206193 | 3.72677078105216 | 0.68736299394339 | 0.495713875218661 | 0.762976048278299 | -6.00204726125253 |
| lnc-MDN1-3 | lnc-MDN1-3 | -0.283193457850685 | 1.40677093246286 | -0.687119517606479 | 0.495776068907429 | 0.763014717087006 | -5.7364308663018 |
| ENSG00000163935.13 | SFMBT1 | 0.115012834052873 | 7.71557944492088 | 0.686971593245846 | 0.495832333607228 | 0.763044255898136 | -6.16329103162476 |
| lnc-RNF39-13 | lnc-RNF39-13 | -0.214487679152999 | -0.0279925244081102 | -0.686679950436241 | 0.496014290873282 | 0.763247258533318 | -5.48244657302399 |
| ENSG00000103342.12 | GSPT1 | -0.0658990299437052 | 6.33748711175741 | -0.686615480601822 | 0.49605451887338 | 0.763247258533318 | -6.17720447037031 |
| lnc-MN1-7 | lnc-MN1-7 | 0.108631604077181 | 0.445579537179918 | 0.686581872100575 | 0.496075490679625 | 0.763247258533318 | -5.55488664019209 |
| ENSG00000002330.13 | BAD | -0.111971233116813 | 2.26749974405651 | -0.686334135373643 | 0.496230094258892 | 0.763421768148401 | -5.86794217998935 |
| ENSG00000159842.15 | ABR | -0.110815233374187 | 7.45532764580611 | -0.686281262586303 | 0.496263093709644 | 0.763421768148401 | -6.16558340067432 |
| ENSG00000108797.11 | CNTNAP1 | 0.420796782116821 | 3.20115562871187 | 0.686197342520798 | 0.4964512680218 | 0.763637772642183 | -5.92183832124113 |
| ENSG00000120942.13 | UBIAD1 | 0.0736966123187184 | 4.50206994438239 | 0.685937447244805 | 0.4964777084729 | 0.763637772642183 | -6.11216081295006 |
| ENSG00000162639.15 | HENMT1 | 0.129645837837273 | 2.57677576466813 | 0.685711598472072 | 0.496618714597901 | 0.763797579210088 | -5.88874275589747 |
| ENSG00000165861.13 | ZFYVE1 | 0.0833942118038129 | 3.9075057605585 | 0.68551546539904 | 0.496741185998539 | 0.763807077641168 | -6.05373266110652 |
| ENSG00000185115.5 | NSMCE3 | -0.0889175045616106 | 5.4563245401286 | -0.685454726662322 | 0.49677911647673 | 0.763807077641168 | -6.16513373746916 |
| lnc-KIF3B-1 | lnc-KIF3B-1 | -0.108091098667134 | 0.775800919705977 | -0.685401498673675 | 0.496812357913844 | 0.763807077641168 | -5.63949552442554 |
| ENSG00000010322.15 | NISCH | -0.0692882777217019 | 6.5410393016628 | -0.685393336692415 | 0.496817455265301 | 0.763807077641168 | -6.1776268500025 |
| ENSG00000244754.8 | N4BP2L2 | -0.0613422162347685 | 7.74122363064477 | -0.685379171502028 | 0.496826301832186 | 0.763807077641168 | -6.16298484102118 |
| ENSG00000151006.7 | PRSS53 | -0.138318625096312 | 1.1057515550852 | -0.685264656218614 | 0.496897823097386 | 0.763807077641168 | -5.70199352907476 |
| ENSG00000157077.14 | ZFYVE9 | 0.428987842573885 | 1.31209175411815 | 0.685422961793008 | 0.496910928413471 | 0.763807077641168 | -5.76414164055386 |
| lnc-ADAMTS12-2 | lnc-ADAMTS12-2 | -0.21431050420699 | 0.498288230298355 | -0.685226332740463 | 0.496921759543938 | 0.763807077641168 | -5.56310259504349 |

| | | | | | | | |
|---|---|---|---|---|---|---|---|
| lnc-HIST1H4D-2 | lnc-HIST1H4D-2 | -0.314725095047208 | 0.469444095473899 | -0.684789008803223 | 0.497221001046943 | 0.764209965990833 | -5.59785108337336 |
| lnc-AGO3-3 | lnc-AGO3-3 | -0.0636038319732357 | 3.6181760397537 | -0.684667662956315 | 0.497270771134754 | 0.764229394629034 | -6.02217216014574 |
| lnc-RUNX1-5 | lnc-RUNX1-5 | 0.101409223727563 | 0.0567303224224396 | 0.684607503143519 | 0.49730836217154 | 0.764230104418157 | -5.48146552604519 |
| lnc-KCNK9-2 | lnc-KCNK9-2 | -0.158281060274749 | -0.275045089599345 | -0.684447807432762 | 0.497408156110765 | 0.764326396209621 | -5.37822501241532 |
| ENSG00000175536.6 | LIPT2 | 0.168438120856532 | 0.816967329845589 | 0.684253278767766 | 0.497529732101747 | 0.764422013313408 | -5.66256746489211 |
| ENSG00000168060.15 | NAALADL1 | -0.189499540755833 | 2.67682715951933 | -0.684230127644672 | 0.497544658813571 | 0.764422013313408 | -5.92421150271791 |
| ENSG00000184857.7 | TMEM186 | 0.1135996739618 | 1.18005693530792 | 0.684141260078131 | 0.4975997486688 | 0.764449591494993 | -5.71183946639792 |
| ENSG00000165025.14 | SYK | 0.0708904015073948 | 8.91600229848948 | 0.684073249424869 | 0.497642260943978 | 0.764457844392112 | -6.13753119727424 |
| ENSG00000146426.18 | TIAM2 | -0.134926999684325 | 4.12368544622461 | -0.68397180004652 | 0.497705678905144 | 0.764498208127207 | -6.08148667757609 |
| lnc-NOX3-4 | lnc-NOX3-4 | -0.347780749283637 | 5.61124750737409 | -0.68403643529843 | 0.497800571249326 | 0.764507188087091 | -6.16146274324061 |
| MIR4432HG | MIR4432HG | 0.223066813516311 | 2.44228669056205 | 0.683836247156862 | 0.497804758666656 | 0.764507188087091 | -5.85977113826392 |
| lnc-DDX23-1 | lnc-DDX23-1 | -0.0883438243083399 | 2.9368532837953 | -0.68378421604075 | 0.497822953016144 | 0.764507188087091 | -5.9445724885797 |
| ENSG00000156876.9 | SASS6 | -0.0777289449378702 | 3.32233254225796 | -0.683670593360604 | 0.497893995257083 | 0.76455924383591 | -5.99618210663458 |
| ENSG00000110777.11 | POU2AF1 | 0.135078580290918 | 6.27608122710476 | 0.683563608658477 | 0.497960892229264 | 0.764604926874669 | -6.17940292983099 |
| lnc-NFYA-1 | lnc-NFYA-1 | 0.174293337976888 | 2.25674402261082 | 0.683467451101585 | 0.498021023270965 | 0.764618509236477 | -5.85041841256663 |
| ENSG00000166321.13 | NUDT13 | -0.167848931065076 | 0.529515627502759 | -0.683430655451511 | 0.498044034073432 | 0.764618509236477 | -5.5790839568045 |
| ENSG00000109445.10 | ZNF330 | 0.099059783731744 | 4.46390246546228 | 0.683330913504043 | 0.498106412388766 | 0.76465724091572 | -6.11247398556665 |
| ENSG00000181004.9 | BBS12 | 0.127168243930645 | 1.20449328604284 | 0.682904841717797 | 0.498372924870867 | 0.765009315580361 | -5.71135457888254 |
| ENSG00000011304.19 | PTBP1 | -0.066155080300728 | 6.53170447179292 | -0.682640756851988 | 0.498538152211389 | 0.765205875239897 | -6.17943296743447 |
| lnc-MNS1-5 | lnc-MNS1-5 | -0.122506166345283 | 1.24825567112025 | -0.682560051973882 | 0.498588652043437 | 0.765226323541128 | -5.71061346738167 |
| lnc-PSMC6-1 | lnc-PSMC6-1 | 0.130451513897649 | 1.16124390965962 | 0.682333405647777 | 0.498730487545171 | 0.765370263742655 | -5.72466698107307 |
| ENSG00000198879.11 | SFMBT2 | -0.57478513658582 | 2.26341249948801 | -0.682516989539903 | 0.498756806470399 | 0.765370263742655 | -5.81131604230044 |
| ENSG00000177830.17 | CHID1 | -0.0571484015430577 | 3.71941216639477 | -0.682012240316652 | 0.498931511169063 | 0.765581280333444 | -6.03873388356318 |
| ENSG00000101084.17 | RAB5IF | 0.109360099306554 | 2.7859733865236 | 0.681583830026212 | 0.499199730898715 | 0.765829739192258 | -5.94057903524328 |
| ENSG00000168765.16 | GSTM4 | 0.163496472667397 | 2.75647842189075 | 0.681581540730529 | 0.499201164397295 | 0.765829739192258 | -5.9097998385515 |
| ENSG00000099725.14 | PRKY | -1.30931293098256 | 1.14229531788741 | -0.681809666219235 | 0.499205053235317 | 0.765829739192258 | -5.44227175593578 |
| ENSG00000280238.1 | | -0.141766052773555 | -0.14204359440313 | -0.681414926496778 | 0.499305500061696 | 0.765926748231472 | -5.43793438769509 |
| LINC01522 | LINC01522 | -0.747385976682672 | -2.48964497523805 | -0.681224989520955 | 0.499424455313292 | 0.765978051078572 | -4.86544979103589 |
| lnc-NIN-3 | lnc-NIN-3 | 0.148465536858915 | -0.279405011140577 | 0.681053295151958 | 0.499531998850175 | 0.765978051078572 | -5.39188951850116 |
| ENSG00000229676.2 | ZNF492 | -0.769025617125463 | -2.27152680759938 | -0.681079620763325 | 0.499532407396384 | 0.765978051078572 | -4.9268340093924 |
| ENSG00000109063.14 | MYH3 | -0.111076365036852 | 2.95411539883154 | -0.681043630556792 | 0.499538052803772 | 0.765978051078572 | -5.9554071192205 |
| ENSG00000278715.1 | SNORD116-20 | -0.209556982650657 | 1.58183586552381 | -0.681015650969595 | 0.499555579592848 | 0.765978051078572 | -5.76337200270931 |
| ENSG00000165527.6 | ARF6 | 0.103750189164685 | 6.37070713775164 | 0.681005036206918 | 0.499562228911177 | 0.765978051078572 | -6.18112542535305 |
| lnc-FAM167A-6 | lnc-FAM167A-6 | -0.192984156850649 | 1.3947288693055 | -0.680854969680089 | 0.499656239056454 | 0.766020512347117 | -5.75384821090447 |
| ENSG00000140995.16 | DEF8 | 0.170592410430871 | 6.21881845794779 | 0.680890449243397 | 0.499665303530718 | 0.766020512347117 | -6.18060939886407 |
| ENSG00000220804.8 | LINC01881 | 0.124291595026741 | 1.69690868817243 | 0.680782614034894 | 0.499701570193108 | 0.766020512347117 | -5.78056800447 |
| ENSG00000111725.10 | PRKAB1 | -0.0552691808582679 | 5.02628407757037 | -0.680371925473858 | 0.499958911252747 | 0.766298764717237 | -6.15381366633824 |
| ENSG00000185130.5 | H2BC13 | 0.373772661524014 | 1.86153212279109 | 0.680528037773149 | 0.499967603757134 | 0.766298764717237 | -5.86219283987509 |
| ADAMTSL4-AS1 | ADAMTSL4-AS1 | -0.175969242363933 | 1.66531116433435 | -0.680314703413893 | 0.499994772878224 | 0.766298764717237 | -5.79934600602254 |
| ENSG00000114857.17 | NKTR | 0.093681671823834 | 8.45072103023717 | 0.679913019710385 | 0.50024655177497 | 0.766488991413956 | -6.15017065451693 |
| lnc-PLCG2-1 | lnc-PLCG2-1 | -0.218079667348778 | 2.39075564892753 | -0.67987905193824 | 0.500281551431613 | 0.766488991413956 | -5.86944180342212 |
| lnc-RBM28-1 | lnc-RBM28-1 | -0.159211474241962 | -0.241491544999698 | -0.679836818452657 | 0.500294323255559 | 0.766488991413956 | -5.39711869055753 |
| SSSCA1-AS1 | SSSCA1-AS1 | -0.152994522069778 | 0.961406063211928 | -0.679833672119489 | 0.500296295783353 | 0.766488991413956 | -5.67907577964125 |
| ENSG00000132670.20 | PTPRA | -0.0552016036660889 | 5.96176364857578 | -0.679819650008051 | 0.500305086704878 | 0.766488991413956 | -6.17975956762061 |
| lnc-MS4A6A-1 | lnc-MS4A6A-1 | 0.845950567659202 | -2.24454426705056 | 0.679529394784763 | 0.500490618372687 | 0.766660090913572 | -4.84463658703846 |
| ENSG00000120725.12 | SIL1 | -0.0707949214553625 | 3.00628430987287 | -0.679522719979112 | 0.500491261798847 | 0.766660090913572 | -5.95204119640432 |
| ENSG00000203880.11 | PCMTD2 | 0.106206750159661 | 6.51066201410763 | 0.679396241257552 | 0.500570575495555 | 0.766672894732307 | -6.18170949108508 |
| ENSG00000065600.12 | PACC1 | -0.101690388781015 | 3.57188263641259 | -0.679390595390473 | 0.500574116130179 | 0.766672894732307 | -6.01994230157463 |
| ENSG00000058729.10 | RIOK2 | 0.0830714499273659 | 4.47770202136658 | 0.679270956220077 | 0.500649147447134 | 0.766730759070265 | -6.11492811647965 |
| ENSG00000138964.16 | PARVG | -0.102166343265134 | 7.21135183537101 | -0.679159424822378 | 0.500719099561642 | 0.766780836714327 | -6.17730482919894 |
| lnc-ECH1-5 | lnc-ECH1-5 | -0.077422490218535 | 3.3670242333444 | -0.678936297738938 | 0.500859060215256 | 0.766883866574653 | -6.0040751983669 |
| ENSG00000133812.15 | SBF2 | 0.156101720674037 | 4.29421760047929 | 0.678934862886263 | 0.500860895805425 | 0.766883866574653 | -6.10748234544463 |
| lnc-ZNF518B-2 | lnc-ZNF518B-2 | -0.185528730033927 | 2.7343292507587 | -0.678784558781703 | 0.500954253591395 | 0.766940944778819 | -5.93282475425067 |
| ENSG00000284129.1 | MIR6789 | -0.163962584420119 | -0.0530371440864124 | -0.678755163213027 | 0.500972696038052 | 0.766940944778819 | -5.47106320256691 |
| ENSG00000144821.9 | MYH15 | 0.415141830079451 | 1.06956460395095 | 0.678750659220519 | 0.50107872610179 | 0.766977091583979 | -5.70308038184835 |
| ENSG00000164535.14 | DAGLB | 0.11009466635251 | 3.19690954540188 | 0.678548932384884 | 0.501102093398602 | 0.766977091583979 | -5.98581668120326 |
| ENSG00000145782.12 | ATG12 | 0.0591660016411621 | 5.81650358811356 | 0.678481142791412 | 0.50114463126927 | 0.766977091583979 | -6.17978999334654 |
| ENSG00000066336.11 | SPI1 | -0.148509693433798 | 3.3157849290595 | -0.67846541563017 | 0.501154500323054 | 0.766977091583979 | -6.00375648241331 |
| ENSG00000140497.16 | SCAMP2 | -0.0670191248854701 | 5.26859061627992 | -0.678374388544123 | 0.501211623419574 | 0.766977091583979 | -6.16464602721761 |
| lnc-XRN1-3 | lnc-XRN1-3 | 0.0735330492522934 | 1.89885036351491 | 0.678361226616473 | 0.50121988334529 | 0.766977091583979 | -5.81973940276736 |
| ENSG00000143633.12 | C1orf131 | 0.0629631383142395 | 3.64982451489093 | 0.678136339811717 | 0.501361025297259 | 0.76713603803847 | -6.03029688067551 |
| ENSG00000135916.15 | ITM2C | 0.566429289725031 | 0.421089136124816 | 0.678249104485157 | 0.501424216565582 | 0.767142750068387 | -5.67121294174327 |
| SGO1-AS1 | SGO1-AS1 | 0.165389517770978 | 1.67578802499399 | 0.678010595931748 | 0.501439953331394 | 0.767142750068387 | -5.78686698115158 |
| ENSG00000004059.10 | ARF5 | -0.0985007359401342 | 4.81327555294131 | -0.677875164307499 | 0.501524969876458 | 0.767149855727269 | -6.14357984559471 |
| ENSG00000048649.13 | RSF1 | 0.0572705346887368 | 6.87886634216774 | 0.677849658261735 | 0.501540982057377 | 0.767149855727269 | -6.18108624519766 |
| ENSG00000231875.1 | | -0.178179779070025 | 0.236263591041435 | -0.677776646434676 | 0.501586818959015 | 0.767149855727269 | -5.52482371351523 |
| lnc-RHOXF1-3 | lnc-RHOXF1-3 | -0.107258062788523 | 0.988658476558504 | -0.677747345456555 | 0.501605214789683 | 0.767149855727269 | -5.69366879050527 |
| lnc-PHB-3 | lnc-PHB-3 | -0.174236520244129 | 0.0860085474222945 | -0.677586455786955 | 0.501706231631397 | 0.767149855727269 | -5.47029950858108 |

| | | | | | | | |
|---|---|---|---|---|---|---|---|
| lnc-ZNF814-1 | lnc-ZNF814-1 | 0.0597276841476226 | 3.11575050150098 | 0.677502894393932 | 0.501758701227282 | 0.767149855727269 | -5.97087189050837 |
| ENSG00000175691.8 | ZNF77 | -0.110266365459698 | 1.9795119659281 | -0.677493387156305 | 0.501764671171344 | 0.767149855727269 | -5.82583735213656 |
| ENSG00000122971.8 | ACADS | 0.135793767606448 | 1.89897295926704 | 0.677433948358434 | 0.501801995860613 | 0.767149855727269 | -5.82861880502821 |
| ENSG00000158987.20 | RAPGEF6 | -0.0470364563448147 | 5.6977450766096 | -0.677396913781413 | 0.501825252552688 | 0.767149855727269 | -6.17709570007953 |
| ENSG00000215845.10 | TSTD1 | -0.108770555285135 | 5.21058069344389 | -0.677375642403544 | 0.501838610657939 | 0.767149855727269 | -6.15632922259029 |
| ENSG00000125753.13 | VASP | -0.13028849081768 | 5.19854428162969 | -0.677350214301033 | 0.501854579379472 | 0.767149855727269 | -6.16375110104964 |
| ENSG00000147955.16 | SIGMAR1 | -0.147351422333215 | 2.75215669371713 | -0.677251739948101 | 0.501916423409589 | 0.767184576510386 | -5.93466721286205 |
| lnc-PIK3R4-3 | lnc-PIK3R4-3 | -0.148869854268379 | 0.891916211860006 | -0.677190618952479 | 0.5019548108216 | 0.767184576510386 | -5.67106576803026 |
| ENSG00000239969.5 | | -0.0866244357108612 | 2.48371138090227 | -0.67713600751089 | 0.501989111229747 | 0.767184576510386 | -5.88837870436718 |
| ENSG00000186866.16 | POFUT2 | -0.0645236926017319 | 4.26384860596593 | -0.677062586262868 | 0.502035227743874 | 0.767198091369231 | -6.09034791102392 |
| ENSG00000140983.13 | RHOT2 | -0.0841247237668599 | 5.67513673242392 | -0.676931776971134 | 0.50211739593157 | 0.767229964331513 | -6.17815939842849 |
| lnc-PSMC3IP-2 | lnc-PSMC3IP-2 | 0.114006694144722 | 2.51264219915322 | 0.676910702695373 | 0.502130634480387 | 0.767229964331513 | -5.89694996655918 |
| lnc-RBAK-RBAKDN-2 | lnc-RBAK-RBAKDN | 0.072000726294124 | 3.08141775270487 | 0.676811775190208 | 0.502192781834041 | 0.767267965297659 | -5.9708529856786 |
| lnc-MED4-5 | lnc-MED4-5 | 0.218330718966939 | -0.58070905128283 | 0.676619776767211 | 0.50231340939302 | 0.767395302125475 | -5.31161475936524 |
| ENSG00000138623.10 | SEMA7A | 0.272910571695837 | 2.91407981668103 | 0.676635522816448 | 0.502381026499481 | 0.767430422171042 | -5.92866442164186 |
| ENSG00000204619.7 | PPP1R11 | -0.0828643939950133 | 4.94709450852272 | -0.676448937591536 | 0.502420756496092 | 0.767430422171042 | -6.14927048545947 |
| ENSG00000178809.11 | TRIM73 | 0.129275327193391 | 0.324976994681882 | 0.676405182805344 | 0.502448251915511 | 0.767430422171042 | -5.54613850371504 |
| ENSG00000135709.12 | KIAA0513 | 0.291836259572142 | 0.670794674547429 | 0.676291453300155 | 0.502533513694751 | 0.76750369610292 | -5.60639523407526 |
| lnc-GPR155-3 | lnc-GPR155-3 | 0.109364088630823 | 1.95410539196939 | 0.676156772175974 | 0.502604368272819 | 0.767554957127128 | -5.82374486727763 |
| ENSG00000092010.14 | PSME1 | 0.104849440473781 | 6.06536293579729 | 0.675935026999712 | 0.502743748869461 | 0.767710852658218 | -6.18378387969085 |
| ENSG00000163527.9 | STT3B | 0.0842563662020321 | 6.91375187306093 | 0.675614394968122 | 0.502945323399883 | 0.767961690767046 | -6.18092620094464 |
| lnc-FER-1 | lnc-FER-1 | 0.17102589483149 | 0.117809330886596 | 0.675544931713894 | 0.502988999297141 | 0.767971409578892 | -5.54235721687745 |
| ENSG00000134291.11 | TMEM106C | -0.140099471726475 | 3.08565093820325 | -0.675396230134815 | 0.503082504267703 | 0.768055180315247 | -5.96508859848716 |
| ENSG00000165572.7 | KBTBD6 | -0.166902074465233 | 1.07319393768977 | -0.675338996812622 | 0.503118495661006 | 0.768055180315247 | -5.71700074632013 |
| ENSG00000117984.13 | CTSD | -0.112003344482539 | 3.70186528729166 | -0.675244072067993 | 0.503178192555178 | 0.768089345695878 | -6.04379848471966 |
| lnc-COLQ-2 | lnc-COLQ-2 | -0.0778394040940108 | 0.820275391126478 | -0.674990100080553 | 0.503337931171851 | 0.768216535716306 | -5.6520275164617 |
| ENSG00000167377.18 | ZNF23 | 0.0776342645417403 | 4.05413417405052 | 0.674870822522527 | 0.503412961741786 | 0.768216535716306 | -6.06993256608689 |
| ENSG00000108349.16 | CASC3 | 0.0647057678276139 | 6.31139964138771 | 0.674866814399218 | 0.503415483125115 | 0.768216535716306 | -6.18519964164148 |
| lnc-TRMT1L-1 | lnc-TRMT1L-1 | -0.0937402150937439 | 1.30203878219137 | -0.674860689723762 | 0.503419335977654 | 0.768216535716306 | -5.72939367092401 |
| lnc-PRDM14-4 | lnc-PRDM14-4 | -0.213567785885813 | -0.520326727838855 | -0.674762182382471 | 0.503481306253817 | 0.768216535716306 | -5.37925951071134 |
| ENSG00000177570.14 | SAMD12 | -1.35447184037311 | 0.469226039170851 | -0.674986118076086 | 0.503485452403098 | 0.768216535716306 | -5.23146727092348 |
| lnc-TEAD2-2 | lnc-TEAD2-2 | 0.133219497601587 | 2.9055441313101 | 0.674641165299562 | 0.503557442960429 | 0.768269427652103 | -5.93569243427195 |
| lnc-DNAH10OS-6 | lnc-DNAH10OS-6 | -0.17369759788815 | -0.255939226176844 | -0.674473239094569 | 0.503663102507675 | 0.768318246683734 | -5.42881783957793 |
| ENSG00000182796.14 | TMEM198B | -0.112480045776137 | 2.82774966861025 | -0.674471659092346 | 0.503664096706195 | 0.768318246683734 | -5.93706883999412 |
| ENSG00000085832.16 | EPS15 | 0.0750205216809343 | 7.22465108723143 | 0.673974794511678 | 0.50397679633549 | 0.768738283605557 | -6.17968774946028 |
| ENSG00000147164.11 | SNX12 | -0.0626945612446423 | 4.0522699333742 | -0.673827747491768 | 0.504069360099013 | 0.768822500105075 | -6.08057557995004 |
| ENSG00000146066.2 | HIGD2A | 0.110855800929872 | 4.53835977021498 | 0.67358895735147 | 0.504219694488556 | 0.768976535413716 | -6.12912053219269 |
| ENSG00000166750.9 | SLFN5 | -0.255435235508759 | 4.63568566663993 | -0.673706220696449 | 0.504245071101342 | 0.768976535413716 | -6.13550839346021 |
| ENSG00000179195.15 | ZNF664 | 0.0808789199867812 | 5.4498807842466 | 0.673489151981511 | 0.504282535912177 | 0.768976695560849 | -6.17472041058778 |
| ENSG00000212124.2 | TAS2R19 | -0.171284177188781 | 2.1962515554211 | -0.673406630929521 | 0.504334497673772 | 0.768998960339228 | -5.87836038445585 |
| ENSG00000107745.18 | MICU1 | -0.0571113805744217 | 4.70111795164823 | -0.673257809715704 | 0.504428214637746 | 0.769084884584351 | -6.13494656440412 |
| ENSG00000204388.6 | HSPA1B | 0.227850401273541 | 1.76808568070422 | 0.673143362537702 | 0.504511697092778 | 0.769155193042045 | -5.81537149872393 |
| ENSG00000034152.18 | MAP2K3 | 0.117073162326833 | 4.12251525170226 | 0.672962869274389 | 0.504613975146618 | 0.769254143863341 | -6.08946221546994 |
| lnc-PCSK7-6 | lnc-PCSK7-6 | 0.233729529671296 | 1.55909514350551 | 0.672798545652027 | 0.504726512166862 | 0.769276636210436 | -5.7723283453563 |
| ENSG00000185442.12 | FAM174B | -0.434766081967005 | 1.71510902602471 | -0.67297612153539 | 0.504728252246426 | 0.769276636210436 | -5.8108507515719 |
| ENSG00000118496.4 | FBXO30 | 0.109645069815017 | 4.41405602310428 | 0.67271070485325 | 0.504772823912847 | 0.769276636210436 | -6.11499151351914 |
| ENSG00000089053.12 | ANAPC5 | 0.0903747751632693 | 6.8736983668312 | 0.672702128307514 | 0.504778227112576 | 0.769276636210436 | -6.18395813135611 |
| lnc-ZNF345-29 | lnc-ZNF345-29 | -0.0994752068497622 | 4.49815459333812 | -0.672641149048221 | 0.504816644784901 | 0.769278226075933 | -6.12384380302792 |
| ENSG00000134470.20 | IL15RA | 0.117153117138171 | 2.05573842650656 | 0.672516807089445 | 0.504894986670553 | 0.769313572530427 | -5.85967774470666 |
| ENSG00000177732.8 | SOX12 | -0.120367673983009 | 3.84455128304563 | -0.672485691406914 | 0.504914592202961 | 0.769313572530427 | -6.06389854863547 |
| ENSG00000177076.5 | ACER2 | -0.119834706940389 | 1.37484970266392 | -0.672364583506965 | 0.504990904459137 | 0.769340051836662 | -5.74563429121265 |
| ENSG00000139971.15 | ARMH4 | 0.0561671580276095 | 3.13674953132536 | 0.672336446971197 | 0.505008634695051 | 0.769340051836662 | -5.97325102628545 |
| STXBP5-AS1 | STXBP5-AS1 | 0.199976009288638 | -0.260993886715589 | 0.672280162441653 | 0.505044103406548 | 0.769340051836662 | -5.38314533574041 |
| lnc-LRRC41-3 | lnc-LRRC41-3 | 0.235337068877638 | 0.298386566918135 | 0.672090547971393 | 0.50516360238747 | 0.769465139342951 | -5.53689990834609 |
| lnc-FGA-3 | lnc-FGA-3 | 0.0870562892593489 | 2.17713037900057 | 0.672017585619794 | 0.505209588887775 | 0.769478242420977 | -5.85523067564875 |
| ENSG00000188493.14 | ACTMAP | 0.128418733899412 | 0.495151741305881 | 0.671609820691463 | 0.505466635830569 | 0.769812783226089 | -5.5797954468383 |
| ENSG00000198429.9 | ZNF69 | 0.181785004713893 | 2.11891125703182 | 0.67136688888649 | 0.505619809129657 | 0.769836028197381 | -5.82142024211244 |
| ENSG00000140396.12 | NCOA2 | 0.0657751663241557 | 7.22841507852323 | 0.67130414729696 | 0.505659373052179 | 0.769836028197381 | -6.18024909971225 |
| ENSG00000251467.1 | CTDNEP1P2 | -0.320368151475978 | -0.43385843781397 | -0.67114588576196 | 0.505759177943485 | 0.769836028197381 | -5.35092968338155 |
| lnc-RANBP2-2 | lnc-RANBP2-2 | 0.134285807955808 | 2.54235228046322 | 0.671099655870453 | 0.505788334048875 | 0.769836028197381 | -5.90771514007428 |
| ENSG00000145741.15 | BTF3 | 0.0924832822727568 | 7.34911634827706 | 0.671087974430022 | 0.50579570140364 | 0.769836028197381 | -6.17945534301433 |
| ENSG00000139714.12 | MORN3 | 0.244132213848098 | 0.036611455438538 | 0.671063117243691 | 0.505811378749702 | 0.769836028197381 | -5.47663730346919 |
| ENSG00000080189.14 | SLC35C2 | -0.0450857526535513 | 5.26506434456974 | -0.671007040321297 | 0.505846747253849 | 0.769836028197381 | -6.16808378819521 |
| ENSG00000051128.18 | HOMER3 | 0.915652067210982 | -0.229391192765539 | 0.671174638116846 | 0.505863287314661 | 0.769836028197381 | -5.18645814474171 |
| lnc-PDCL2-4 | lnc-PDCL2-4 | -0.0840877786348493 | 2.39339418361202 | -0.670934977495147 | 0.505892200271512 | 0.769836028197381 | -5.89500752353163 |
| ENSG00000141568.20 | FOXK2 | -0.10735770099763 | 4.4080604008223 | -0.670909284905444 | 0.505908406193268 | 0.769836028197381 | -6.12102650583511 |
| ENSG00000185515.14 | BRCC3 | -0.0705866780236986 | 3.60182238887669 | -0.670879313288132 | 0.505927311522647 | 0.769836028197381 | -6.03401642108039 |
| ENSG00000137507.11 | LRRC32 | -0.789263575509446 | -0.402959695205907 | -0.671067685407284 | 0.50593071726308 | 0.769836028197381 | -5.29358558074582 |

| | | | | | | | |
|---|---|---|---|---|---|---|---|
| lnc-ACER2-1 | lnc-ACER2-1 | -0.161006996878946 | 0.864521389419753 | -0.670578260580866 | 0.506117229222516 | 0.770062901322224 | -5.6691990992598 |
| ENSG00000172167.7 | MTBP | 0.0739387354908132 | 3.58124879350622 | 0.670399030419854 | 0.506230314179086 | 0.77012716501269 | -6.03144308728249 |
| ENSG00000178607.15 | ERN1 | 0.112712149103736 | 3.82703955914352 | 0.67039271780555 | 0.506234297362955 | 0.77012716501269 | -6.06264984033332 |
| ENSG00000143379.12 | SETDB1 | 0.041818908126011 | 6.00503484049579 | 0.670266716848287 | 0.506313806026092 | 0.770191195730919 | -6.18663143544747 |
| ENSG00000241015.2 | TPM3P9 | -0.107528869405833 | 3.65939365408038 | -0.670207095577201 | 0.506351430393028 | 0.770191508408194 | -6.04632240273493 |
| ENSG00000008083.13 | JARID2 | -0.0913707848094782 | 5.97676715437031 | -0.669884371864815 | 0.506555113587961 | 0.770444388013079 | -6.18735241687829 |
| ENSG00000142528.15 | ZNF473 | 0.0817249260382793 | 2.81624375653673 | 0.669787088559441 | 0.506616521527565 | 0.770454991955273 | -5.94016997970865 |
| lnc-ULK2-5 | lnc-ULK2-5 | -0.0817245257338221 | 0.463018151071234 | -0.66975472889532 | 0.506636948749678 | 0.770454991955273 | -5.60384593280415 |
| lnc-NTN1-2 | lnc-NTN1-2 | -0.476267868480773 | 0.401893633888102 | -0.669625213113213 | 0.50679803825898 | 0.770643027591946 | -5.44362216605778 |
| lnc-RPIA-1 | lnc-RPIA-1 | -1.13020696580319 | -0.678523594303728 | -0.669536147076721 | 0.506912491411072 | 0.77071793382557 | -5.17439388895955 |
| lnc-POLE2-4 | lnc-POLE2-4 | 0.0786549545361162 | 2.72503365801244 | 0.669302943565553 | 0.506922187636913 | 0.77071793382557 | -5.92770048849212 |
| lnc-SIPA1L1-5 | lnc-SIPA1L1-5 | -0.137498937609404 | 0.247404375417243 | -0.669133248765508 | 0.507029348607823 | 0.770823922179986 | -5.5332327052466 |
| ENSG00000011007.12 | ELOA | -0.0898478741178241 | 5.75857343317755 | -0.668944481059461 | 0.507148568454135 | 0.770948226328764 | -6.1843093399964 |
| ENSG00000163141.19 | BNIPL | 0.14809976115177 | -0.105130328948287 | 0.668880822382161 | 0.50718877674483 | 0.770952410585544 | -5.42081459309542 |
| ENSG00000277075.2 | H2AC8 | 0.854244160497584 | 3.43062040659267 | 0.669034599213043 | 0.50723517938462 | 0.770964071113081 | -5.99448097844188 |
| lnc-LRRC8C-5 | lnc-LRRC8C-5 | 0.16133365273384 | -0.423208558001402 | 0.668725723689903 | 0.50728674792003 | 0.770964071113081 | -5.32738918866148 |
| lnc-MSX1-5 | lnc-MSX1-5 | -0.217870229272344 | -0.436749973495159 | -0.66869078834862 | 0.507308816943865 | 0.770964071113081 | -5.34661081519158 |
| ENSG00000104881.15 | PPP1R13L | -0.204641688414237 | -0.355875605107309 | -0.668571952316734 | 0.507383890828729 | 0.770969209119003 | -5.37809818205997 |
| ENSG00000204217.13 | BMPR2 | 0.104678971460455 | 7.67959000856208 | 0.668566855175877 | 0.507387111049216 | 0.770969209119003 | -6.17541919119781 |
| lnc-YIPF1-2 | lnc-YIPF1-2 | -0.0803720036730016 | 2.07136839558203 | -0.668277162561794 | 0.507570148407304 | 0.771190401171295 | -5.85781573324572 |
| ENSG00000204130.13 | RUFY2 | 0.0713426831219452 | 4.73878449597027 | 0.668169099180123 | 0.507638435612478 | 0.771237224698231 | -6.14322437542803 |
| lnc-PGM2-1 | lnc-PGM2-1 | -0.106753833060658 | -0.0373911419242077 | -0.66789864754255 | 0.507809360783482 | 0.771323482842956 | -5.45098843778871 |
| ENSG00000258064.1 | | 0.141086045114719 | 3.27920670062407 | 0.667795583808424 | 0.507874505193551 | 0.771323482842956 | -5.97815589161343 |
| ENSG00000126883.16 | NUP214 | 0.0989405532207259 | 6.36051237102758 | 0.667788297834508 | 0.507879110675699 | 0.771323482842956 | -6.18991349192958 |
| lnc-CCDC68-5 | lnc-CCDC68-5 | -0.188775732148771 | 3.81964858631706 | -0.667767247228454 | 0.507916106536519 | 0.771323482842956 | -6.04714510093702 |
| GATA2-AS1 | GATA2-AS1 | 0.353516007172633 | -0.213928090972304 | 0.667711121071719 | 0.507931573459508 | 0.771323482842956 | -5.38384675788126 |
| ENSG00000009844.15 | VTA1 | 0.0547655482967873 | 4.29816181008264 | 0.667682237762045 | 0.507946154079912 | 0.771323482842956 | -6.1120467610667 |
| ENSG00000198001.13 | IRAK4 | 0.094266225172583 | 5.92123294283448 | 0.667664244561139 | 0.507957528539876 | 0.771323482842956 | -6.18702624262233 |
| BHLHE40-AS1 | BHLHE40-AS1 | -0.193297058918946 | -0.478146956714222 | -0.667426231725468 | 0.508108002145405 | 0.771495058140961 | -5.30128604480837 |
| ENSG00000215559.8 | ANKRD20A11P | -0.791979590597869 | 0.457820934950997 | -0.667538018753041 | 0.5081804812717 | 0.771548192521606 | -5.56750419324747 |
| ENSG00000221923.8 | ZNF880 | 0.0979041624264276 | 4.92855302586847 | 0.667116024879404 | 0.508304153722267 | 0.771679037916021 | -6.15028482153906 |
| ENSG00000272679.1 | | -0.0902300713477957 | 1.93645208434874 | -0.666975943482209 | 0.508392744184578 | 0.771756608418848 | -5.83789340437559 |
| ENSG00000166946.13 | CCNDBP1 | 0.0680691332607521 | 6.80590592603165 | 0.666652012220556 | 0.508597637331593 | 0.772010705692933 | -6.188359154689 |
| lnc-RBMX-4 | lnc-RBMX-4 | 0.0671211777087866 | 5.83887910863403 | 0.666433747142885 | 0.508735719733562 | 0.77216335958085 | -6.18771520760952 |
| lnc-SUGCT-5 | lnc-SUGCT-5 | -0.122924322643184 | 2.27177388912228 | -0.666363252598731 | 0.508780321485357 | 0.772174115700716 | -5.88607780523532 |
| RBFADN | RBFADN | -0.192294963379566 | 0.0889201337892791 | -0.666273143766576 | 0.50883733624672 | 0.772198190454119 | -5.46281508067956 |
| ENSG00000134970.13 | TMED7 | 0.147024410051247 | 5.08078579547668 | 0.666132902045098 | 0.508929396848174 | 0.772198190454119 | -6.16954872080198 |
| ENSG00000049247.13 | UTS2 | 1.20406929153378 | -0.712863339132815 | 0.666289198592254 | 0.508948384197763 | 0.772198190454119 | -5.00697956350887 |
| ENSG00000111785.19 | RIC8B | 0.42150380848909 | 2.61575826169561 | 0.666284794343825 | 0.508955248748457 | 0.772198190454119 | -5.8651981714291 |
| ENSG00000185829.17 | ARL17A | 0.166058370305669 | 2.77895998114479 | 0.666041744783322 | 0.508983765723706 | 0.772198190454119 | -5.92567436549587 |
| ENSG00000173889.15 | PHC3 | 0.0638856308786249 | 8.38574495452868 | 0.665784809110997 | 0.5091463816253 | 0.772227050745535 | -6.16357265301344 |
| ENSG00000157954.14 | WIPI2 | 0.09379279701984 | 5.79212302071187 | 0.665776068530317 | 0.509151914079206 | 0.772227050745535 | -6.18758729583555 |
| ENSG00000189007.15 | ADAT2 | -0.084519855657919 | 4.13358937551009 | -0.665743430358058 | 0.509172573084564 | 0.772227050745535 | -6.10284867092356 |
| lnc-BRD2-5 | lnc-BRD2-5 | 0.166210188240239 | 3.39818780755946 | 0.66574088039873 | 0.509174187153147 | 0.772227050745535 | -5.98678167704576 |
| ENSG00000158985.13 | CDC42SE2 | 0.0702464080336753 | 8.10193734479701 | 0.665715303157126 | 0.509190377142224 | 0.772227050745535 | -6.16791889815003 |
| lnc-TSR3-4 | lnc-TSR3-4 | -0.201332214610037 | -0.0352836750667499 | -0.665438506020013 | 0.509365603210545 | 0.772344302142655 | -5.4805151641184 |
| ENSG00000079462.7 | PAFAH1B3 | -0.16041404315405 | 2.01293457523621 | -0.66541054670347 | 0.509383304637875 | 0.772344302142655 | -5.85002987696491 |
| ENSG00000167280.16 | ENGASE | -0.100838194002625 | 5.00004260421544 | -0.665381949565716 | 0.509401410223677 | 0.772344302142655 | -6.15952981239713 |
| ENSG00000265257.5 | | 0.0732949622353569 | 5.14418644660199 | 0.66529535107171 | 0.509456240093233 | 0.772344302142655 | -6.16518409668556 |
| ENSG00000084073.8 | ZMPSTE24 | 0.0798817424474683 | 4.25330247863606 | 0.665195031327412 | 0.509519761578402 | 0.772344302142655 | -6.1087528501607 |
| ENSG00000182022.17 | CHST15 | 0.155559401911802 | 6.74667732885357 | 0.665209128731739 | 0.509525354584234 | 0.772344302142655 | -6.19085486975651 |
| ENSG00000089022.13 | MAPKAPK5 | 0.0548023909130251 | 5.68349746431773 | 0.665178303146727 | 0.509530354117238 | 0.772344302142655 | -6.18619691129559 |
| ENSG00000093000.18 | NUP50 | 0.0671615257116533 | 5.65458430745135 | 0.665069958854302 | 0.509598962264472 | 0.772378877531145 | -6.18575033987526 |
| ENSG00000110002.15 | VWA5A | -0.194970062834271 | 3.12881867467431 | -0.665041524785195 | 0.509628214339527 | 0.772378877531145 | -5.94704067360472 |
| ENSG00000140463.13 | BBS4 | -0.06756127728399 | 4.08723048387893 | -0.664877654020122 | 0.509720750078922 | 0.772392364934946 | -6.0892625447281 |
| ENSG00000169814.13 | BTD | -0.0659377485004837 | 3.93987992627992 | -0.664852938186733 | 0.509736403909754 | 0.772392364934946 | -6.06599089006676 |
| ENSG00000277072.4 | STAG3L2 | -0.0850895100340483 | 3.45319019595636 | -0.664805395132347 | 0.509766516147048 | 0.772392364934946 | -6.0194765489605 |
| lnc-SMIM14-9 | lnc-SMIM14-9 | 0.066483607247966 | 2.65199538743851 | 0.664733444000545 | 0.509812089507021 | 0.772392364934946 | -5.92605321131592 |
| ENTPD1-AS1 | ENTPD1-AS1 | -0.0842479390375406 | 4.58768834200086 | -0.664659302703631 | 0.509859052412678 | 0.772392364934946 | -6.12911739378432 |
| ENSG00000144043.11 | TEX261 | 0.0896795511344184 | 4.77731141731598 | 0.664646634720088 | 0.509867076856918 | 0.772392364934946 | -6.14717933799939 |
| ENSG00000118873.15 | RAB3GAP2 | -0.126512730470829 | 6.91964732081706 | -0.664561184871386 | 0.509921206244785 | 0.772392364934946 | -6.18878139460714 |
| ENSG00000241343.9 | RPL36A | 0.112580037795857 | 6.86201464649588 | 0.664509509465075 | 0.50995394225621 | 0.772392364934946 | -6.19003804405806 |
| lnc-SHB-3 | lnc-SHB-3 | -0.136247616229576 | -0.280425829675915 | -0.664476514024362 | 0.509974845234704 | 0.772392364934946 | -5.41835802953011 |
| ENSG00000140990.14 | NDUFB10 | -0.123279528675672 | 3.79179642209412 | -0.664337563834873 | 0.510062876818417 | 0.772468853914611 | -6.05454098160446 |
| ENSG00000140280.13 | LYSMD2 | -0.166105672759031 | 2.30148847404571 | -0.664177917880514 | 0.51016403032073 | 0.772565202773071 | -5.89279575025036 |
| ENSG00000144867.11 | SRPRB | -0.0784205951354275 | 3.90140950159447 | -0.663935698787865 | 0.510317523856654 | 0.772740792580115 | -6.07938619450229 |
| ENSG00000114391.12 | RPL24 | 0.0906996388638239 | 7.94403236792249 | 0.663583065113006 | 0.510541031497471 | 0.772888731762773 | -6.17384577705188 |
| lnc-SLC30A4-4 | lnc-SLC30A4-4 | -0.223803827571471 | 0.270635049984937 | -0.663571586568187 | 0.510548307763757 | 0.772888731762773 | -5.53438159144714 |

| | | | | | | | |
|---|---|---|---|---|---|---|---|
| lnc-MPHOSPH8-7 | lnc-MPHOSPH8-7 | 0.176244230302012 | 0.703332096097919 | 0.663545546449279 | 0.510564814841233 | 0.772888731762773 | -5.63519222863637 |
| ENSG00000117009.11 | KMO | -0.162015801010169 | 5.00922401035641 | -0.663570450059713 | 0.510565422230891 | 0.772888731762773 | -6.15635698064036 |
| lnc-LEKR1-4 | lnc-LEKR1-4 | 0.118163285646228 | 2.05590201792717 | 0.663431238979626 | 0.510637278844493 | 0.772930252514892 | -5.84606107465846 |
| lnc-NFKB2-1 | lnc-NFKB2-1 | -0.177668533377526 | 0.363536951959745 | -0.663382852951629 | 0.510667954328816 | 0.772930252514892 | -5.58287374604124 |
| lnc-TOP3A-4 | lnc-TOP3A-4 | -0.0739719832445869 | 2.59370132150187 | -0.663301869106372 | 0.510719298214011 | 0.772951126113161 | -5.92340593878722 |
| ENSG00000184110.14 | EIF3C | -0.123540646073414 | 1.44464096840108 | -0.663138185242535 | 0.510823082571669 | 0.7730513571482 | -5.76905732030306 |
| ENSG00000182158.14 | CREB3L2 | -0.176406672019186 | 7.9243583889747 | -0.663098016513372 | 0.510886422437728 | 0.773090371492115 | -6.17372332900982 |
| ENSG00000249863.2 | | -0.106611590939199 | -0.0337933145012534 | -0.662882894081239 | 0.510984973654895 | 0.773120333969386 | -5.45533178118742 |
| lnc-CDH24-2 | lnc-CDH24-2 | -0.0790351907090443 | 3.59267912029922 | -0.662768029282325 | 0.511057823408473 | 0.773120333969386 | -6.03726862495365 |
| lnc-CDC42SE1-1 | lnc-CDC42SE1-1 | 0.0807187320684868 | 5.72054354145948 | 0.662690284722507 | 0.511107133887539 | 0.773120333969386 | -6.18691095798486 |
| lnc-RAB6B-1 | lnc-RAB6B-1 | 0.0797715493417879 | 1.67804418871349 | 0.66267161506898 | 0.511118975736758 | 0.773120333969386 | -5.80291519735537 |
| lnc-GNRH2-1 | lnc-GNRH2-1 | -0.200336549497882 | -0.633460463013229 | -0.662507094434279 | 0.511223334842298 | 0.773120333969386 | -5.27709728103111 |
| ENSG00000150527.17 | MIA2 | 0.0651586750053701 | 5.57378377510182 | 0.662438557798636 | 0.511266812558273 | 0.773120333969386 | -6.1849798155149 |
| ENSG00000124535.15 | WRNIP1 | -0.0767593916450418 | 4.59359293999569 | -0.662400121026089 | 0.511291196642073 | 0.773120333969386 | -6.14103053081084 |
| ENSG00000120685.19 | PROSER1 | 0.0821185651679963 | 4.69714144350384 | 0.662352825506485 | 0.511321201530921 | 0.773120333969386 | -6.14359300839924 |
| ENSG00000125877.12 | ITPA | -0.10322314931783 | 4.08668688062367 | -0.662332495548653 | 0.511334099411536 | 0.773120333969386 | -6.0921924798056 |
| ENSG00000156639.11 | ZFAND3 | 0.0666129031678544 | 4.5155699070877 | 0.662256488942867 | 0.511382321637067 | 0.773120333969386 | -6.13001548658415 |
| ENSG00000119421.6 | NDUFA8 | -0.15635466099405 | 1.48650659177056 | -0.662247784999178 | 0.511387843992118 | 0.773120333969386 | -5.77436892601097 |
| lnc-TSPYL1-2 | lnc-TSPYL1-2 | -0.0992097672212315 | 1.65099321784179 | -0.662221743731565 | 0.51140436647974 | 0.773120333969386 | -5.80728144068291 |
| ENSG00000140443.14 | IGF1R | -0.703642384977183 | 1.52262651780877 | -0.662381963052606 | 0.511444583659129 | 0.773120333969386 | -5.9113700803165 |
| lnc-PSMB1-2 | lnc-PSMB1-2 | -0.134857883374817 | 0.21161582044629 | -0.662137702981517 | 0.511457690056254 | 0.773120333969386 | -5.54919600721276 |
| ENSG00000175087.9 | PDIK1L | 0.11211684683495 | 4.19634962318049 | 0.662118870847938 | 0.511469639394701 | 0.773120333969386 | -6.10649755373872 |
| ENSG00000023902.13 | PLEKHO1 | -0.123005056611022 | 6.10405337941399 | -0.661935462390644 | 0.511586023359953 | 0.773239471201198 | -6.19330962002751 |
| ENSG00000200534.1 | SNORA33 | -0.117840805804667 | 1.82763344676563 | -0.661814054547089 | 0.511663071990688 | 0.773299141697946 | -5.83522886128799 |
| ENSG00000115694.14 | STK25 | 0.0876703565913166 | 5.36943081576347 | 0.661664101184305 | 0.511758245020872 | 0.773386193631762 | -6.17921035225018 |
| lnc-DGKG-1 | lnc-DGKG-1 | 0.241690399352373 | 0.35844289219714 | 0.661600805810168 | 0.511798420466282 | 0.773390124694037 | -5.53820573939586 |
| ENSG00000240429.1 | LRRFIP1P1 | -0.161389343590015 | 2.14141824868687 | -0.661522337602398 | 0.51184822892412 | 0.773408610772658 | -5.88720483585725 |
| ENSG00000283900.1 | TPTEP2-CSNK1E | -0.128344718448667 | 4.5537696373272 | -0.661359308812529 | 0.511951721408021 | 0.773503962856913 | -6.14235807931869 |
| ENSG00000279432.1 | | 0.14882001229629 | 2.91030053293603 | 0.661261612695363 | 0.512013745404184 | 0.773503962856913 | -5.94569884969183 |
| lnc-RAP1GAP2-5 | lnc-RAP1GAP2-5 | 0.142787296693027 | 0.447176550239272 | 0.661196504221055 | 0.512055082850438 | 0.773503962856913 | -5.58796745644817 |
| lnc-SLC35F5-16 | lnc-SLC35F5-16 | -0.271010498681392 | 0.0461106599336706 | -0.661186156624344 | 0.512061652717694 | 0.773503962856913 | -5.44824753643756 |
| lnc-CCDC92-6 | lnc-CCDC92-6 | -0.144108160126058 | -0.15860459187208 | -0.661120085505995 | 0.512103603481639 | 0.773510565088616 | -5.42507937006152 |
| ENSG00000279789.1 | | -0.0927354435849939 | 1.82162531933768 | -0.661019024091957 | 0.512167774346774 | 0.773550726400033 | -5.82381946938179 |
| lnc-RIT1-1 | lnc-RIT1-1 | -0.0936695816860449 | 2.55989502720621 | -0.660936736442285 | 0.51222002766143 | 0.773572883509812 | -5.90959942323428 |
| lnc-TBC1D7-8 | lnc-TBC1D7-8 | 0.117233848762973 | 2.30359869832611 | 0.660796888976063 | 0.512308838523828 | 0.773572896440271 | -5.87585077665652 |
| ENSG00000123268.8 | ATF1 | -0.0982136804341014 | 3.65867350860441 | -0.660643761554915 | 0.512406092427846 | 0.773572896440271 | -6.05079108440912 |
| ENSG00000167460.15 | TPM4 | -0.0863933293682475 | 6.8473008793643 | -0.660641596598651 | 0.512407467500908 | 0.773572896440271 | -6.19195308766354 |
| lnc-SLC26A3-1 | lnc-SLC26A3-1 | -0.14187535547418 | 0.280110828499456 | -0.660600471582387 | 0.512433588454135 | 0.773572896440271 | -5.5730176610561 |
| lnc-STX3-1 | lnc-STX3-1 | -0.273187296939392 | -0.408781675156691 | -0.660552359805548 | 0.512464148028701 | 0.773572896440271 | -5.35764115859847 |
| ENSG00000113300.11 | CNOT6 | 0.0681520308477603 | 4.6913701342636 | 0.66046046827592 | 0.512522518302889 | 0.773572896440271 | -6.14452121566814 |
| lnc-EIF1AX-2 | lnc-EIF1AX-2 | 0.112039131296393 | 0.793836345009963 | 0.660438406308373 | 0.512536532784948 | 0.773572896440271 | -5.65569930626095 |
| ENSG00000136717.14 | BIN1 | 0.172232090178872 | 5.21642167235675 | 0.660409730084942 | 0.512588015243513 | 0.773572896440271 | -6.17219362182601 |
| ENSG00000006534.15 | ALDH3B1 | 0.324964084879013 | -0.313916924179899 | 0.660337171173699 | 0.512600843300232 | 0.773572896440271 | -5.37371330825944 |
| ENSG00000107290.13 | SETX | 0.0778680030507726 | 8.91324064436542 | 0.66032561197908 | 0.512608186657992 | 0.773572896440271 | -6.15309702696945 |
| ENSG00000276293.4 | PIP4K2B | -0.078695711106735 | 4.83476112000447 | -0.660233664229007 | 0.512666601506382 | 0.773572896440271 | -6.15653642954146 |
| lnc-NAA25-5 | lnc-NAA25-5 | -0.120763689552227 | 0.777800905862133 | -0.660200510026525 | 0.512687665412207 | 0.773572896440271 | -5.66200313264939 |
| ENSG00000260345.1 | | -0.0845915914433208 | 0.862647436682578 | -0.660118463748988 | 0.512739793996742 | 0.773572896440271 | -5.68448837355538 |
| ENSG00000009335.17 | UBE3C | -0.0543143171383687 | 6.00625613247823 | -0.660108382163409 | 0.51274619958872 | 0.773572896440271 | -6.19330332275411 |
| ENSG00000258677.2 | | 0.0602151223443874 | 2.7203880660338 | 0.660019312477764 | 0.512802794156677 | 0.773601576673034 | -5.93451083795137 |
| lnc-YTHDF3-4 | lnc-YTHDF3-4 | -0.163563305172497 | 0.036560543616036 | -0.659840479120771 | 0.5129164344269 | 0.773716304126705 | -5.5210870980939 |
| lnc-AKTIP-1 | lnc-AKTIP-1 | -0.0968396022089291 | 0.993845354657796 | -0.659630763063538 | 0.513049716523738 | 0.773819908122905 | -5.7212938487163 |
| lnc-CTSK-2 | lnc-CTSK-2 | -0.1015782555051 | 2.93108088845981 | -0.659614101454259 | 0.513060306376781 | 0.773819908122905 | -5.9523062194053 |
| ENSG00000106868.16 | SUSD1 | -0.169314971142785 | 3.58497657244829 | -0.659502933319101 | 0.513137934653318 | 0.773880283482507 | -6.05072319157138 |
| ENSG00000127527.13 | EPS15L1 | 0.0916530555858506 | 5.14948129296838 | 0.659200556838831 | 0.513323186516976 | 0.774006641048541 | -6.16834556347948 |
| ENSG00000099204.19 | ABLIM1 | 0.128147195839992 | 6.85245314885586 | 0.659183989662288 | 0.513333719376618 | 0.774006641048541 | -6.19394285601273 |
| ENSG00000205220.11 | PSMB10 | 0.0825606155484193 | 4.69022178089389 | 0.659101157078271 | 0.513386383325614 | 0.774006641048541 | -6.14814728241855 |
| HCG26 | HCG26 | -0.140329560933595 | 2.6689876148443 | -0.659087319724602 | 0.513395181230678 | 0.774006641048541 | -5.92381792341003 |
| ENSG00000028203.17 | VEZT | -0.0704496325634896 | 5.91692840263165 | -0.659064422584226 | 0.513409739602377 | 0.774006641048541 | -6.19387394497372 |
| ENSG00000172339.9 | ALG14 | 0.0892732753742154 | 1.38795032954729 | 0.658754359034857 | 0.513606904929639 | 0.774231666411934 | -5.76163841370023 |
| ENSG00000035720.7 | STAP1 | -0.0767326307614491 | 6.43109660195851 | -0.658711388778927 | 0.513634232369186 | 0.774231666411934 | -6.19570881580608 |
| ENSG00000066294.14 | CD84 | -0.148983416949911 | 7.20290555652444 | -0.658588000862703 | 0.513722278764032 | 0.774242658315092 | -6.18948449538651 |
| ENSG00000258469.1 | CHMP4BP1 | -0.168597189796899 | 0.450514626861787 | -0.658525177851063 | 0.513752664454882 | 0.774242658315092 | -5.58308285142177 |
| ENSG00000166337.9 | TAF10 | -0.0847156787979424 | 5.45953382118047 | -0.658522494131463 | 0.513754371436017 | 0.774242658315092 | -6.18510794717265 |
| ENSG00000260899.1 | | -0.0806866578492786 | 1.20719085749287 | -0.658414145198327 | 0.513823289379365 | 0.774289828339957 | -5.74375380278558 |
| lnc-PLEKHG5-3 | lnc-PLEKHG5-3 | -0.0729149572127181 | 1.71773261927751 | -0.658257840450127 | 0.51392271954829 | 0.774335852023818 | -5.82023067182451 |
| RAB11B-AS1 | RAB11B-AS1 | 0.121486547317122 | 2.33612992230097 | 0.658247856022155 | 0.513929071296574 | 0.774335852023818 | -5.87833706371775 |
| ENSG00000254463.1 | PPIAP41 | -0.15818641454494 | 1.0427997152175 | -0.658114564997157 | 0.514013870494945 | 0.774406931371501 | -5.71017569873238 |
| lnc-RNF111-2 | lnc-RNF111-2 | -0.151980778999107 | 0.35094332624287 | -0.657968320346867 | 0.514106919407609 | 0.774480591423044 | -5.58544643680849 |

| | | | | | | | |
|---|---|---|---|---|---|---|---|
| lnc-TTLL7-16 | lnc-TTLL7-16 | -0.25199199598468 | 4.76254159072305 | -0.6580730691494 | 0.514138016868507 | 0.774480591423044 | -6.1406738677547 |
| lnc-MASTL-4 | lnc-MASTL-4 | -0.192819333804308 | 0.184124212124787 | -0.657806353225835 | 0.514209982411008 | 0.774532313791861 | -5.57591105418124 |
| lnc-ATAD2-1 | lnc-ATAD2-1 | -0.124577574397081 | 2.45945795159692 | -0.657688201424843 | 0.514285171845679 | 0.774588884245544 | -5.89190818101024 |
| lnc-NT5C-3 | lnc-NT5C-3 | -0.223019099515635 | 0.158234502198627 | -0.657502652844443 | 0.514403263186725 | 0.774710058255094 | -5.5147824033362 |
| lnc-ZNF213-3 | lnc-ZNF213-3 | 0.305905525277501 | -0.645061733060478 | 0.6573884245654 | 0.514475970386759 | 0.774762869364257 | -5.35475722890045 |
| lnc-CTDSP2-2 | lnc-CTDSP2-2 | -0.217778181862964 | 1.36796519795857 | -0.657195739747391 | 0.514598628328682 | 0.774890889376638 | -5.76535968339092 |
| lnc-STYX-5 | lnc-STYX-5 | -0.144773735335304 | 2.31082777147257 | -0.656931958741687 | 0.514766569700874 | 0.77504857997467 | -5.87062261134095 |
| lnc-PDGFB-2 | lnc-PDGFB-2 | 0.100894924475948 | 6.49219001745275 | 0.656912972295141 | 0.514778658933766 | 0.77504857997467 | -6.19669435396882 |
| lnc-KMT2E-2 | lnc-KMT2E-2 | -0.169192814405354 | 1.6521076941055 | -0.656626770765992 | 0.514960910432475 | 0.775221627841528 | -5.81953726435619 |
| ENSG00000174928.15 | C3orf33 | 0.140715058072774 | 0.349601845622108 | 0.656614191233406 | 0.514968921803295 | 0.775221627841528 | -5.58232393717273 |
| lnc-STC1-1 | lnc-STC1-1 | 0.226188767928874 | 0.148103312046851 | 0.656546932615821 | 0.515011757102082 | 0.775229413224525 | -5.5188974005242 |
| lnc-ZNF821-2 | lnc-ZNF821-2 | -0.0985916007354193 | 0.661271198318261 | -0.65617669840449 | 0.515247584142791 | 0.775470472581294 | -5.63315415584774 |
| ENSG00000165533.18 | TTC8 | -0.157961294559207 | 1.97833748862716 | -0.656153968318814 | 0.515262064353135 | 0.775470472581294 | -5.85235203340838 |
| ENSG00000227394.1 | RPL11P1 | -0.176384105435544 | 1.36680982853861 | -0.656080275669088 | 0.51530901179046 | 0.775470472581294 | -5.74852426688378 |
| ENSG00000257411.1 | | 0.106145069820801 | 3.37540967154251 | 0.656058943472065 | 0.515322602340132 | 0.775470472581294 | -6.01416889177013 |
| ENSG00000178537.9 | SLC25A20 | -0.254438619161021 | 1.86954350998561 | -0.655631873535346 | 0.515634541513263 | 0.775883161632246 | -5.83761573584224 |
| ENSG00000117868.15 | ESYT2 | -0.0869648367754774 | 6.18464160862064 | -0.655427428886609 | 0.515725021857196 | 0.775897177888586 | -6.19758994995199 |
| ENSG00000207008.1 | SNORA54 | 0.335749634644029 | 4.65975165362926 | 0.65561944634414 | 0.51573141955442 | 0.775897177888586 | -6.1301500366277 |
| ENSG00000259075.6 | POC1B-GALNT4 | 0.0821509042071198 | 2.71406717973306 | 0.655377344269754 | 0.5157569444625 | 0.775897177888586 | -5.94504581244285 |
| lnc-BAIAP2L2-2 | lnc-BAIAP2L2-2 | 0.095824900242457 | 0.534523761002604 | 0.655093100224978 | 0.515938134182435 | 0.776071012381089 | -5.60162535056474 |
| ENSG00000187605.15 | TET3 | -0.0824374000012122 | 5.73703639819067 | -0.655077773268607 | 0.515947905233582 | 0.776071012381089 | -6.19268053600668 |
| ENSG00000271913.6 | TAGAP-AS1 | -0.128062641484505 | -0.288851117262942 | -0.654870208891798 | 0.51608023888934 | 0.776213339938567 | -5.37307249461708 |
| lnc-MCL1-7 | lnc-MCL1-7 | -0.18146407272757 | 0.1924773882796 | -0.654803279227396 | 0.516122914101947 | 0.776220805272568 | -5.5324593522119 |
| lnc-GTF2H5-2 | lnc-GTF2H5-2 | 0.174153304977529 | 1.99718475631263 | 0.654600445280378 | 0.516252255202884 | 0.776358600777451 | -5.85379294046566 |
| ENSG00000213557.4 | RPL31P43 | -0.52996748504459 | 0.512947602005762 | -0.654671588556632 | 0.516303766363254 | 0.776379341251816 | -5.46118397190893 |
| lnc-RDH11-3 | lnc-RDH11-3 | -0.108663332237772 | 0.905893321701117 | -0.654202873245331 | 0.516505825403076 | 0.776609183992882 | -5.69400067172209 |
| lnc-KDM5C-2 | lnc-KDM5C-2 | -0.367591832121778 | 0.331552143903635 | -0.654265619115099 | 0.516532076415613 | 0.776609183992882 | -5.60373251019616 |
| lnc-PLPP5-1 | lnc-PLPP5-1 | 0.0811039354550265 | 3.67517761425355 | 0.653707162156891 | 0.516822082006271 | 0.776988453285741 | -6.047378863838 |
| lnc-CCDC88C-2 | lnc-CCDC88C-2 | -0.0622217602776258 | 4.65063750593612 | -0.653580381337466 | 0.516902983041591 | 0.777053323104372 | -6.14346008254102 |
| ENSG00000058866.14 | DGKG | 0.294579230098685 | 5.19652044366846 | 0.653583587561966 | 0.517019454553735 | 0.777102423109879 | -6.16185396376689 |
| ENSG00000172613.7 | RAD9A | -0.13153587514542 | 3.45672265292601 | -0.653311388059506 | 0.517074654804828 | 0.777102423109879 | -6.02556246474741 |
| lnc-P4HTM-2 | lnc-P4HTM-2 | 0.064147472185641 | 2.50570175233309 | 0.653298849919739 | 0.517082657403258 | 0.777102423109879 | -5.91601934039965 |
| lnc-ZNF382-5 | lnc-ZNF382-5 | -0.13059773132397 | 0.946639334792532 | -0.653292573974293 | 0.517086663115819 | 0.777102423109879 | -5.68943436711474 |
| ENSG00000114767.6 | RRP9 | -0.0932501537195494 | 3.06826288529754 | -0.653143821153707 | 0.517181611617672 | 0.777138092311308 | -5.98485433727896 |
| lnc-AFF3-1 | lnc-AFF3-1 | 0.093347650337595 | 4.17295992392597 | 0.65307861376595 | 0.51722323625493 | 0.777138092311308 | -6.09314491506232 |
| ENSG00000148019.13 | CEP78 | 0.0927832620212241 | 4.24941779657247 | 0.65303192218432 | 0.517253042572603 | 0.777138092311308 | -6.11133180217795 |
| ENSG00000145216.15 | FIP1L1 | -0.0623070552797342 | 5.71188244311637 | -0.653018794885131 | 0.517261422759798 | 0.777138092311308 | -6.19441987668054 |
| ENSG00000186088.15 | GSAP | -0.0645443982232772 | 7.82524589158521 | -0.652949565255709 | 0.517305618679651 | 0.777147766534067 | -6.18263700222107 |
| ENSG00000279276.1 | | -0.225820273516602 | 2.06017773035674 | -0.652811698903265 | 0.517412284175042 | 0.777217234481269 | -5.84691631683884 |
| ENSG00000088970.15 | KIZ | 0.0590806269929764 | 5.18452421479204 | 0.652758851463774 | 0.517427380075637 | 0.777217234481269 | -6.17609954346681 |
| lnc-HDHD2-7 | lnc-HDHD2-7 | 0.161557654556245 | -0.364427356327384 | 0.652663208638317 | 0.517488449109423 | 0.777252243725866 | -5.37107387879841 |
| lnc-LRMP-6 | lnc-LRMP-6 | -0.142629097410718 | 1.02521012718584 | -0.652507663193622 | 0.517587774898226 | 0.777344704175862 | -5.71645893669434 |
| lnc-KIF5B-3 | lnc-KIF5B-3 | 0.0754041623239488 | 2.67653784843265 | 0.652351512442587 | 0.517687497490257 | 0.777389149999949 | -5.93696248204717 |
| lnc-SUPT16H-3 | lnc-SUPT16H-3 | 0.0571237976161161 | 5.78773293879116 | 0.652328854568656 | 0.517701968349089 | 0.777389149999949 | -6.1955483611535 |
| ENSG00000137145.20 | DENND4C | -0.0885960848239176 | 6.45292336445381 | -0.652251871461949 | 0.517751136614474 | 0.777389149999949 | -6.19974299469007 |
| ENSG00000006125.17 | AP2B1 | -0.0647676400038186 | 6.72852123444356 | -0.652156023785083 | 0.517812356970507 | 0.777389149999949 | -6.19856812728647 |
| ENSG00000128872.9 | TMOD2 | 0.282638049491226 | 3.30798530496789 | 0.652260466527295 | 0.517842386711967 | 0.777389149999949 | -6.02546756875298 |
| lnc-GLRX-4 | lnc-GLRX-4 | -0.300665264858964 | -1.05864134472727 | -0.652106516943172 | 0.517843979772108 | 0.777389149999949 | -5.19451926069575 |
| lnc-ENSA-1 | lnc-ENSA-1 | -0.0838688203733251 | 3.4990318848906 | -0.651753255031307 | 0.518069658031925 | 0.77763370266762 | -6.04311063451601 |
| ENSG00000239203.1 | | -0.124686835122188 | 0.718036322545449 | -0.651733241677032 | 0.518082444963371 | 0.77763370266762 | -5.66956256345427 |
| ENSG00000061794.12 | MRPS35 | 0.0815109402845522 | 4.10445977230998 | 0.651325468233828 | 0.518343016355894 | 0.777968084122311 | -6.10536766849537 |
| lnc-DISP1-1 | lnc-DISP1-1 | -0.0735080110963398 | 2.05547328337084 | -0.650862334956438 | 0.5186390483206 | 0.778190152003204 | -5.86142987676313 |
| lnc-PUF60-1 | lnc-PUF60-1 | -0.175872350643701 | -0.0828321044740015 | -0.650833446981077 | 0.518657516335293 | 0.778190152003204 | -5.47499962171349 |
| lnc-RAD1-1 | lnc-RAD1-1 | -0.118896027640543 | 1.9523765515896 | -0.650814487044457 | 0.518669637569442 | 0.778190152003204 | -5.85397364827467 |
| lnc-HSPA5-3 | lnc-HSPA5-3 | -0.0692072559420316 | 4.07440639321729 | -0.650801346040744 | 0.518678038803707 | 0.778190152003204 | -6.10168926750621 |
| ENSG00000143951.15 | WDPCP | 0.0969748913482862 | 3.72532199338899 | 0.650798258998744 | 0.518680012404992 | 0.778190152003204 | -6.06014513922343 |
| ENSG00000204498.10 | NFKBIL1 | -0.145331907475672 | 1.33502299127376 | -0.650689874744513 | 0.518749306948582 | 0.778237389571622 | -5.76526365842082 |
| lnc-RBM43-3 | lnc-RBM43-3 | -0.111516076340022 | 0.847665808659851 | -0.650505476309938 | 0.518867211887597 | 0.778338354538369 | -5.68599169526244 |
| ENSG00000145916.18 | RMND5B | -0.0526763966249835 | 4.69506446320892 | -0.650466341958459 | 0.518892236358913 | 0.778338354538369 | -6.15190237305777 |
| lnc-ALKAL1-3 | lnc-ALKAL1-3 | 0.102660573795868 | 1.59978338564251 | 0.650327561153516 | 0.518980984985003 | 0.778391198391123 | -5.81729641869039 |
| ENSG00000179387.9 | ELMOD2 | 0.0741328964583694 | 4.33418144052601 | 0.650269109388813 | 0.519018366602715 | 0.778391198391123 | -6.12789793232041 |
| lnc-FAM13A-2 | lnc-FAM13A-2 | -1.15697362154463 | -1.55026967652863 | -0.650387397460252 | 0.519040917160675 | 0.778391198391123 | -4.95275249251905 |
| ENSG00000106605.10 | BLVRA | -0.308645127738888 | 0.943080566477654 | -0.650152813211096 | 0.519141643638886 | 0.778485534825818 | -5.69067819918429 |
| ENSG00000143970.16 | ASXL2 | -0.101667901201123 | 6.13907908213409 | -0.650007008121714 | 0.51918600575375 | 0.778495341766551 | -6.20070515595155 |
| ENSG00000158555.14 | GDPD5 | 0.397226020516435 | -0.979484714738437 | 0.649817611842599 | 0.519307161038327 | 0.77856233634901 | -5.18056642651007 |
| ENSG00000166166.12 | TRMT61A | -0.102091921031557 | 3.38428035978512 | -0.649772875220943 | 0.519335780903836 | 0.77856233634901 | -6.0293459940909 |
| lnc-TESC-4 | lnc-TESC-4 | -0.177919645957799 | 0.332801722585127 | -0.649759775348501 | 0.519344161593155 | 0.77856233634901 | -5.55399134979534 |
| lnc-RAB1A-6 | lnc-RAB1A-6 | 0.202314473024582 | -0.707626187485071 | 0.649568920671948 | 0.519466269740524 | 0.778668880544244 | -5.30495883670712 |

| | | | | | | | |
|---|---|---|---|---|---|---|---|
| ENSG00000119599.16 | DCAF4 | 0.102558165628832 | 4.3242297133362 | 0.649489519635663 | 0.519517074762336 | 0.778668880544244 | -6.11888471966351 |
| ENSG00000229944.3 | EIF4EP2 | 0.2670681316432 | -0.653368365778193 | 0.649422590246466 | 0.519559901821065 | 0.778668880544244 | -5.21141407404111 |
| ENSG00000166889.13 | PATL1 | 0.0822250494869561 | 5.20545184737159 | 0.649355295480675 | 0.5196029645806 | 0.778668880544244 | -6.1792441070297 |
| ENSG00000248905.8 | FMN1 | -0.471573642056203 | -1.19741317838263 | -0.649343743178347 | 0.519622719818842 | 0.778668880544244 | -5.19820233232974 |
| lnc-CWC15-6 | lnc-CWC15-6 | 0.284658523971847 | 2.22411465466353 | 0.649342276624717 | 0.519682669394305 | 0.778668880544244 | -5.86076786299362 |
| ENSG00000082805.19 | ERC1 | 0.135721696918127 | 5.79412220175367 | 0.649191243717648 | 0.519707951340333 | 0.778668880544244 | -6.19682077509889 |
| lnc-GK5-7 | lnc-GK5-7 | -0.0893629375939722 | 1.25417025618287 | -0.649151658462062 | 0.519733286064409 | 0.778668880544244 | -5.75395421748624 |
| ENSG00000160688.18 | FLAD1 | -0.066557293211047 | 4.68787847987069 | -0.649116624356142 | 0.519755708584626 | 0.778668880544244 | -6.1544315786753 |
| ENSG00000213638.5 | ADAT3 | -0.148567117236354 | 0.140392296543422 | -0.649016481931565 | 0.519819804565439 | 0.778697693511174 | -5.52666977548367 |
| ENSG00000185651.14 | UBE2L3 | -0.0896672681902917 | 5.11898430376109 | -0.648867160167991 | 0.519915385536173 | 0.778697693511174 | -6.17764802362923 |
| ENSG00000109084.13 | TMEM97 | -0.162181052983033 | 1.50589428019285 | -0.648832433370839 | 0.519937615529902 | 0.778697693511174 | -5.80021721575114 |
| lnc-KLF10-6 | lnc-KLF10-6 | 0.10408018633638 | 3.31946130020087 | 0.64880711428967 | 0.519953823593242 | 0.778697693511174 | -6.00552749639724 |
| ENSG00000169446.5 | MMGT1 | 0.0800653358556467 | 4.93599552596882 | 0.648791059212946 | 0.519964101424359 | 0.778697693511174 | -6.16805847047372 |
| lnc-SCAI-2 | lnc-SCAI-2 | -0.167551092560148 | -0.564239060221578 | -0.648716244171379 | 0.520011996513719 | 0.778712762767688 | -5.36010670090351 |
| ENSG00000111144.9 | LTA4H | 0.119874926676613 | 7.47063075729494 | 0.648555614684058 | 0.520114836249495 | 0.778809160304467 | -6.19396917681046 |
| ENSG00000256594.8 | | 0.118801172077945 | 7.14298385384639 | 0.648449757462063 | 0.520182615101535 | 0.778809160304467 | -6.19692811731874 |
| lnc-MYO1C-2 | lnc-MYO1C-2 | 0.0746366293155494 | 1.22448047825557 | 0.648438408839744 | 0.520189881740554 | 0.778809160304467 | -5.76264004748842 |
| ENSG00000114744.8 | COMMD2 | 0.0602610607355194 | 4.75499311140688 | 0.648309500823202 | 0.520272426666897 | 0.778876089758145 | -6.16011537809156 |
| ENSG00000145687.16 | SSBP2 | -0.377582732505223 | 3.62995035431685 | -0.64826045373396 | 0.520435865453027 | 0.779064103172339 | -6.06939197880925 |
| lnc-C1QTNF1-8 | lnc-C1QTNF1-8 | -0.165527175382429 | 1.01670208362804 | -0.647898020841277 | 0.520535960429728 | 0.779089726196526 | -5.70458275285171 |
| lnc-SLC45A1-1 | lnc-SLC45A1-1 | -0.177420563217907 | 0.391322248842277 | -0.647842815116893 | 0.520571322543723 | 0.779089726196526 | -5.56517215406307 |
| ENSG00000134697.12 | GNL2 | 0.0677453488337418 | 4.91096566608517 | 0.647827416603193 | 0.520581186314645 | 0.779089726196526 | -6.16594150617493 |
| ENSG00000162604.12 | TM2D1 | -0.0556162702099909 | 4.59485263371387 | -0.647791198468008 | 0.520604386829278 | 0.779089726196526 | -6.14620509238341 |
| ENSG00000189319.13 | FAM53B | -0.100284779427819 | 7.84942736314185 | -0.647710813993409 | 0.520655881268627 | 0.779110142068496 | -6.18526328162461 |
| lnc-RSPO2-1 | lnc-RSPO2-1 | 0.114522197115922 | 4.49945777808494 | 0.647622219515783 | 0.520712638198636 | 0.779138429197625 | -6.13888199818764 |
| lnc-PPARA-4 | lnc-PPARA-4 | 0.126515055436162 | -0.33369222372002 | 0.647481698928684 | 0.520802667702245 | 0.779216494098664 | -5.42531486887438 |
| lnc-CEP72-2 | lnc-CEP72-2 | 0.159651026184689 | 0.0857804865081342 | 0.64739195268391 | 0.520860171165233 | 0.779245886254832 | -5.49844193593751 |
| ENSG00000232303.2 | DFFBP1 | 0.0921363179500728 | 0.672554537273559 | 0.647249444711425 | 0.520951487814355 | 0.779295632204028 | -5.66081186964118 |
| ENSG00000164040.16 | PGRMC2 | 0.0925005228622142 | 2.67558231070591 | 0.647191283102686 | 0.520988759222632 | 0.779295632204028 | -5.94517493409051 |
| ENSG00000158050.4 | DUSP2 | -0.233685118085771 | 4.27727976246006 | -0.647290702059332 | 0.521007005526873 | 0.779295632204028 | -6.11316838489554 |
| ENSG00000110801.13 | PSMD9 | 0.0962328778029278 | 3.29811346671107 | 0.646973536028517 | 0.52112830960074 | 0.779420432823139 | -6.0170040084803 |
| lnc-PAPSS2-2 | lnc-PAPSS2-2 | -0.220232769700729 | 1.07007731287953 | -0.646842316747058 | 0.521213299583449 | 0.779452848789089 | -5.76501678950908 |
| ENSG00000146535.13 | GNA12 | 0.135734887201798 | 5.83183663990618 | 0.646748915359194 | 0.521272286029079 | 0.779452848789089 | -6.20063948009244 |
| ENSG00000083635.7 | NUFIP1 | 0.0890688434915305 | 2.41382787051761 | 0.646691346662932 | 0.521309189587395 | 0.779452848789089 | -5.91894785443447 |
| ENSG00000177189.13 | RPS6KA3 | -0.0765800357736248 | 6.66024851360707 | -0.646670875900864 | 0.521322312401224 | 0.779452848789089 | -6.20247972273805 |
| CDKN2B-AS1 | CDKN2B-AS1 | 0.124073662841359 | 3.60111959533075 | 0.646581024396456 | 0.521379913930186 | 0.779452848789089 | -6.05026811237515 |
| RABGAP1L-IT1 | RABGAP1L-IT1 | -0.211343738127256 | 0.831692550309504 | -0.646573882674271 | 0.521384492453328 | 0.779452848789089 | -5.68524241929926 |
| ENSG00000275043.1 | SNORD25 | 0.131971585669981 | 2.02271615888561 | 0.646526196125964 | 0.521415064615312 | 0.779452848789089 | -5.86378484023757 |
| ENSG00000078043.15 | PIAS2 | -0.0724041198253751 | 5.5167389235438 | -0.646404558093577 | 0.521493051889942 | 0.779512816779279 | -6.19533673175901 |
| ENSG00000075240.16 | GRAMD4 | 0.237208538411941 | 3.17267877654113 | 0.646248087082075 | 0.521651571877044 | 0.779693145301372 | -5.97656142838729 |
| ENSG00000111237.18 | VPS29 | -0.0577501033015224 | 5.65295441110688 | -0.645989681783022 | 0.521759093344587 | 0.779762252491406 | -6.19840359081244 |
| ENSG00000196456.11 | ZNF775 | 0.167169205661145 | 0.237168440460293 | 0.645955984066103 | 0.521780705343801 | 0.779762252491406 | -5.53543031159655 |
| lnc-ABCA3-1 | lnc-ABCA3-1 | 0.0894275447850941 | 2.15220046988389 | 0.645908033442168 | 0.521811459253589 | 0.779762252491406 | -5.86889794220816 |
| ENSG00000251728.1 | | 0.156671905063097 | 0.870550307033855 | 0.645817589718971 | 0.521869469429131 | 0.779792325899079 | -5.68000175835173 |
| lnc-TMCC3-3 | lnc-TMCC3-3 | -0.262267607402165 | -0.71062258864125 | -0.645710239448574 | 0.52193832784145 | 0.779806843763919 | -5.27580578401779 |
| lnc-CRISPLD2-2 | lnc-CRISPLD2-2 | 0.103119706030057 | 4.15486952454848 | 0.645684315219503 | 0.521954957320872 | 0.779806843763919 | -6.10356876599956 |
| ENSG00000213741.10 | RPS29 | 0.114030205333291 | 8.08741998007646 | 0.645524481874548 | 0.522057491011358 | 0.779877724394585 | -6.18274318244347 |
| ENSG00000124641.15 | MED20 | -0.0966919977486609 | 2.5339066815597 | -0.645492234231874 | 0.522078179295194 | 0.779877724394585 | -5.92248210090038 |
| ENSG00000134684.10 | YARS1 | 0.0804305327036507 | 5.15772839733788 | 0.645358803334867 | 0.522163785734982 | 0.779948998678021 | -6.18026439341024 |
| ENSG00000128626.11 | MRPS12 | -0.130654348399263 | 2.33217594865954 | -0.64519153747908 | 0.522271110522778 | 0.780052700666885 | -5.91187182811063 |
| ENSG00000279554.1 | | 0.148553267003972 | 2.59304831155172 | 0.645056105446578 | 0.522358017996084 | 0.780125894965418 | -5.91736564632954 |
| lnc-SERAC1-12 | lnc-SERAC1-12 | -0.252482720324985 | -0.226454184561797 | -0.64465075009174 | 0.522618182721724 | 0.780403130269769 | -5.43842989498829 |
| ENSG00000152684.10 | PELO | 0.146459002251533 | 1.42308573588617 | 0.644609440587002 | 0.522644699812428 | 0.780403130269769 | -5.79187253143488 |
| lnc-ACAA1-1 | lnc-ACAA1-1 | -0.0921327241031892 | 0.804185774298271 | -0.644589664649062 | 0.522657394489082 | 0.780403130269769 | -5.68598830081542 |
| lnc-ORC1-2 | lnc-ORC1-2 | 0.079299112488334 | 3.51861739963522 | 0.644495167550682 | 0.522718056840195 | 0.780437092988665 | -6.03470522290909 |
| lnc-SAR1B-3 | lnc-SAR1B-3 | -0.126813406876225 | -0.022654848863394 | -0.64435931315516 | 0.522805275043483 | 0.780482240698449 | -5.49709239060463 |
| lnc-CD83-5 | lnc-CD83-5 | -0.298720245027983 | 0.173004692126541 | -0.644263339165788 | 0.522876029601228 | 0.780482240698449 | -5.53758335213353 |
| lnc-LHX4-1 | lnc-LHX4-1 | 0.126495563775692 | 0.125323348831314 | 0.644191684991546 | 0.522912902597465 | 0.780482240698449 | -5.51020227392563 |
| ENSG00000158623.14 | COPG2 | 0.0961267559253578 | 3.60710746257367 | 0.644162238611225 | 0.52293181019482 | 0.780482240698449 | -6.04595616269224 |
| ENSG00000146263.11 | MMS22L | -0.0973944984768185 | 4.43066376449379 | -0.644152770778582 | 0.522937889592008 | 0.780482240698449 | -6.13698191211677 |
| ENSG00000138606.19 | SHF | -0.153202774891922 | 1.75738682999539 | -0.643900253308287 | 0.523100047643299 | 0.780667653758848 | -5.82143364658222 |
| lnc-LRRD1-1 | lnc-LRRD1-1 | 0.121261545311983 | 2.28860653577186 | 0.643785287768432 | 0.523173883421372 | 0.780721238487791 | -5.89533790310817 |
| ENSG00000104872.10 | PIH1D1 | -0.0819516116811672 | 5.01228208259568 | -0.643710654422586 | 0.523221819107914 | 0.780736168094693 | -6.17574538892221 |
| ENSG00000073350.13 | LLGL2 | -0.134255022578003 | 3.15760532427755 | -0.643179894590644 | 0.523562784143966 | 0.781183320854253 | -5.99927022509122 |
| ENSG00000162378.12 | ZYG11B | 0.0663400889999215 | 5.14404836101647 | 0.643102230788118 | 0.523612685989032 | 0.781183320854253 | -6.18353254096181 |
| lnc-ADH5-1 | lnc-ADH5-1 | 0.14373346992802 | 0.99337880747687 | 0.643066969298959 | 0.523635343624648 | 0.781183320854253 | -5.72659775438327 |
| lnc-ITGB6-2 | lnc-ITGB6-2 | -0.239846365663201 | -0.497694220768703 | -0.64279404589366 | 0.523810731003845 | 0.781340997568425 | -5.39123963607728 |
| lnc-SHOC2-3 | lnc-SHOC2-3 | -0.166090279432078 | 2.8135331880139 | -0.642617285691828 | 0.523924338156252 | 0.781340997568425 | -5.9604544962148 |

| | | | | | | | |
|---|---|---|---|---|---|---|---|
| ENSG00000141867.17 | BRD4 | -0.108230132053134 | 5.93557351730508 | -0.642612651885081 | 0.523927316568187 | 0.781340997568425 | -6.20373195724813 |
| ENSG00000070081.16 | NUCB2 | 0.297084168556809 | -0.0429193261253832 | 0.642553821650082 | 0.523965130897842 | 0.781340997568425 | -5.47300515742621 |
| ENSG00000138107.12 | ACTR1A | 0.0656373317761272 | 5.17661278036659 | 0.642501465736231 | 0.523998784944551 | 0.781340997568425 | -6.18559232017213 |
| ENSG00000121413.12 | ZSCAN18 | -0.109793344505948 | 5.9472702826583 | -0.642491960717843 | 0.524004894832688 | 0.781340997568425 | -6.20403136982127 |
| ENSG00000198853.11 | RUSC2 | -0.623646135574854 | -1.55207329645067 | -0.64250076092477 | 0.524049891496263 | 0.781340997568425 | -5.13301969359282 |
| lnc-ARID2-2 | lnc-ARID2-2 | -0.190338388080741 | 0.153804936809864 | -0.642222180889496 | 0.524178326845607 | 0.781340997568425 | -5.54390293520652 |
| ENSG00000133961.20 | NUMB | 0.0835482021391964 | 5.51720418862164 | 0.642180826216862 | 0.524204915010682 | 0.781340997568425 | -6.19469230194293 |
| ENSG00000171604.11 | CXXC5 | 0.101768798506959 | 6.85910580705936 | 0.642163769217393 | 0.524215881678078 | 0.781340997568425 | -6.20335260251929 |
| lnc-HACD1-5 | lnc-HACD1-5 | 0.0828810990672582 | 1.30712744341863 | 0.642103120954422 | 0.524254876001248 | 0.781340997568425 | -5.76007937113129 |
| ENSG00000214309.4 | MBLAC1 | -0.128468742844079 | -0.0790243542283464 | -0.642097991675232 | 0.524258173986095 | 0.781340997568425 | -5.45846314891616 |
| ENSG00000116580.18 | GON4L | -0.0476394349182216 | 6.68908144682907 | -0.642076158853991 | 0.524272212010299 | 0.781340997568425 | -6.2053681352887 |
| lnc-SLC25A51-7 | lnc-SLC25A51-7 | -0.143506153005554 | 0.118749283237254 | -0.642055627491629 | 0.524285413407052 | 0.781340997568425 | -5.50180790085766 |
| ENSG00000103769.9 | RAB11A | 0.106251859738738 | 5.3905291327491 | 0.642016700412215 | 0.524310443492936 | 0.781340997568425 | -6.19608189690023 |
| lnc-ARL4D-3 | lnc-ARL4D-3 | -0.132774444252377 | 0.939525229541408 | -0.641784423053019 | 0.52445981085883 | 0.781471119382634 | -5.70657057343717 |
| ENSG00000198039.11 | ZNF273 | 0.0793943479113324 | 4.35470848031511 | 0.641762835553264 | 0.524473693978123 | 0.781471119382634 | -6.1263043957771 |
| ENSG00000249709.7 | ZNF564 | 0.0753942403585111 | 4.11533841772313 | 0.641621840521497 | 0.52456437397277 | 0.78154965685715 | -6.10054192077764 |
| ENSG00000135048.13 | CEMIP2 | -0.221143844819771 | 7.31207637440497 | -0.64161439133014 | 0.524655097999175 | 0.781628248560873 | -6.1985278609284 |
| lnc-AKIRIN2-2 | lnc-AKIRIN2-2 | -0.0667574929345647 | 1.81294646659252 | -0.641390564999062 | 0.524713135258352 | 0.781658135848784 | -5.84404002860314 |
| ENSG00000198502.5 | HLA-DRB5 | 0.24628591702111 | 7.1646096854679 | 0.641459778761925 | 0.524766021245948 | 0.781680345585856 | -6.20227489341889 |
| ENSG00000167978.16 | SRRM2 | 0.132734706187509 | 8.91360225275511 | 0.641047476759144 | 0.524933858046002 | 0.78187376801222 | -6.16703829299696 |
| ENSG00000118308.15 | IRAG2 | -0.143073206432316 | 8.80983231509745 | -0.640981437928625 | 0.5249773997653 | 0.781882041922517 | -6.16716575670992 |
| ENSG00000185482.7 | STAC3 | -0.149937641684581 | 1.05916292989362 | -0.640895501310352 | 0.52503164583819 | 0.781894171047569 | -5.72530439830187 |
| ENSG00000264343.5 | NOTCH2NLA | 0.163950454089977 | 2.81022692592333 | 0.640849078379144 | 0.525061518351042 | 0.781894171047569 | -5.9473063588386 |
| ENSG00000151240.16 | DIP2C | 0.386986410996379 | 2.98509532266061 | 0.640729206224627 | 0.525269272219722 | 0.782009344712561 | -6.04783535109773 |
| lnc-DRGX-4 | lnc-DRGX-4 | -0.0925056492523382 | 0.838061582472037 | -0.640475530060759 | 0.525301924305264 | 0.782009344712561 | -5.67847882259663 |
| ENSG00000154122.13 | ANKH | 0.282052084817297 | 3.36023260487869 | 0.640518774959586 | 0.525365357400155 | 0.782009344712561 | -6.00240265041247 |
| ENSG00000146776.14 | ATXN7L1 | -0.203953748763408 | 2.90448764867931 | -0.640378718185461 | 0.52539004545433 | 0.782009344712561 | -5.96768505847399 |
| ENSG00000129824.15 | RPS4Y1 | -1.49947524364652 | 2.74386504649129 | -0.6405465891467 | 0.525392619331714 | 0.782009344712561 | -5.72219786300728 |
| lnc-EPCAM-6 | lnc-EPCAM-6 | 0.13486545067549 | 1.03485077910797 | 0.64031674845859 | 0.525404129665674 | 0.782009344712561 | -5.72562027716238 |
| ENSG00000272047.2 | GTF2H5 | 0.101199176711738 | 2.96684128119111 | 0.640315689780667 | 0.525404811156294 | 0.782009344712561 | -5.97286181064891 |
| lnc-LPCAT1-4 | lnc-LPCAT1-4 | 0.0750865411022001 | 4.900369071292 | 0.640026437655516 | 0.525591025658226 | 0.782169567348012 | -6.16829030675896 |
| ENSG00000121310.16 | ECHDC2 | 0.138672484549056 | 4.50516768564445 | 0.64001785980663 | 0.525596548432584 | 0.782169567348012 | -6.14000188703273 |
| ENSG00000159131.16 | GART | -0.071214347158886 | 5.33682721369595 | -0.639954619170297 | 0.525637266321956 | 0.782169567348012 | -6.1932551974339 |
| lnc-FBXL12-2 | lnc-FBXL12-2 | -0.230536970156867 | -0.454100455175796 | -0.639891519894881 | 0.525677894860208 | 0.782169567348012 | -5.33377337301133 |
| ENSG00000151882.11 | CCL28 | -0.662474317472356 | -0.218852013435842 | -0.640047839017622 | 0.525702462939921 | 0.782169567348012 | -5.48037845968673 |
| ENSG00000168661.14 | ZNF30 | -0.148032692722151 | 1.03159847572278 | -0.639590958670382 | 0.525871443889431 | 0.782261746417902 | -5.71162391709227 |
| lnc-CROT-4 | lnc-CROT-4 | -0.070551776520146 | 3.06127583102939 | -0.639574776520164 | 0.525881865596862 | 0.782261746417902 | -5.9895199314443 |
| ENSG00000075142.13 | SRI | -0.0930105259970606 | 4.68480163301019 | -0.639524014854068 | 0.525914558082519 | 0.782261746417902 | -6.16171920777358 |
| ENSG00000205581.10 | HMGN1 | -0.0831860964985855 | 6.33806331574439 | -0.639521095684658 | 0.52591643817378 | 0.782261746417902 | -6.20794319192553 |
| lnc-PRKD2-1 | lnc-PRKD2-1 | 0.0794039633226804 | 2.26145832734286 | 0.639397755722536 | 0.5259958785361 | 0.782323373647557 | -5.89530932729384 |
| MIR3936HG | MIR3936HG | -0.19953433806199 | 2.78270955572678 | -0.63888426151753 | 0.526342615793822 | 0.782777578004412 | -5.93980093813327 |
| lnc-ABCA9-1 | lnc-ABCA9-1 | -0.264078472655349 | 4.50978716282966 | -0.638954943799838 | 0.526377323867224 | 0.782777578004412 | -6.13019765018994 |
| lnc-NFKBIA-8 | lnc-NFKBIA-8 | 0.100405930922608 | 1.43583562038926 | 0.638241159947057 | 0.526741123553705 | 0.783233104670542 | -5.78076929472789 |
| ENSG00000105245.9 | NUMBL | -0.116066112785203 | 1.66813574711865 | -0.638212268402158 | 0.526759746779104 | 0.783233104670542 | -5.81346674525619 |
| ENSG00000258461.5 | | -0.0813750828837421 | 5.48898314268793 | -0.638101740355108 | 0.526830995359994 | 0.783282460090636 | -6.19588512910989 |
| ADIRF-AS1 | ADIRF-AS1 | -0.12209475626808 | 2.80331689928679 | -0.63794789556174 | 0.526930175259166 | 0.783373333142609 | -5.96048755394425 |
| lnc-TXNDC2-3 | lnc-TXNDC2-3 | 0.14254096217638 | 0.780411892084708 | 0.637876973204883 | 0.526975900456381 | 0.783384729098201 | -5.67557335588277 |
| ENSG00000278828.1 | H3C10 | 0.352714172307133 | 4.41024254076193 | 0.63784155226416 | 0.527128728993105 | 0.783520975851105 | -6.15799898585617 |
| lnc-IL27-1 | lnc-IL27-1 | -0.103559160052784 | 1.35073665096811 | -0.637522024314455 | 0.52720477529295 | 0.783520975851105 | -5.77388230164192 |
| ENSG00000205356.9 | TECPR1 | 0.104905443679644 | 4.95083551162471 | 0.637518865343305 | 0.527206812467082 | 0.783520975851105 | -6.17063089364876 |
| lnc-CRYBA4-52 | lnc-CRYBA4-52 | -0.0971653893229041 | 6.44842570321729 | -0.63749869816898 | 0.527219818079862 | 0.783520975851105 | -6.20921642654107 |
| ENSG00000151366.12 | NDUFC2 | -0.121389797095588 | 3.0377644681458 | -0.637229120370318 | 0.527393682433236 | 0.78372277564965 | -5.99797925934387 |
| ENSG00000148660.20 | CAMK2G | -0.0759602019236398 | 4.36000984120334 | -0.637121636772307 | 0.527463012484965 | 0.783769216429254 | -6.13309296809232 |
| ENSG00000004478.7 | FKBP4 | -0.105588718429117 | 4.51382202246845 | -0.636948323121287 | 0.527574814977074 | 0.783872849024301 | -6.14799531811752 |
| lnc-RTL9-2 | lnc-RTL9-2 | 0.0915921532329418 | 1.72609619415808 | 0.636874442414909 | 0.527622478311031 | 0.783872849024301 | -5.83456411030981 |
| lnc-FAM169A-5 | lnc-FAM169A-5 | 0.0755349799384656 | 0.620182142650996 | 0.636751178745521 | 0.527702005587788 | 0.783872849024301 | -5.63342554711745 |
| lnc-SIVA1-1 | lnc-SIVA1-1 | 0.216126093547201 | 0.164499849443358 | 0.636720514104547 | 0.527721790789526 | 0.783872849024301 | -5.53480168742124 |
| lnc-MINDY1-2 | lnc-MINDY1-2 | -0.171913675350296 | 1.16939627522994 | -0.636709115450532 | 0.527729145440487 | 0.783872849024301 | -5.75151849882978 |
| lnc-STIM2-12 | lnc-STIM2-12 | -0.891090714937624 | 0.638974955929187 | -0.636854692639153 | 0.527770734961306 | 0.783872849024301 | -5.55531729246501 |
| ENSG00000065526.10 | SPEN | 0.0906299012443696 | 7.73899553737488 | 0.636600328216772 | 0.527799339971228 | 0.783872849024301 | -6.19664872149348 |
| lnc-SLC38A2-5 | lnc-SLC38A2-5 | 0.120776166884801 | 3.37346065151692 | 0.636405341084588 | 0.527925166978197 | 0.784003153817621 | -6.03624782326118 |
| ENSG00000198286.9 | CARD11 | 0.104116042731501 | 7.60133505370819 | 0.63618250366666 | 0.528068985417852 | 0.784106794836241 | -6.19742761979745 |
| ENSG00000119638.12 | NEK9 | -0.05496054113976 | 6.07207740922209 | -0.636179156837418 | 0.528071145606567 | 0.784106794836241 | -6.20895709088018 |
| ENSG00000108639.7 | SYNGR2 | -0.110020297331094 | 6.56593810538528 | -0.635518884450462 | 0.5284974052256 | 0.784683119942186 | -6.20954672580483 |
| ENSG00000084112.14 | SSH1 | 0.0786301901083331 | 4.61662436381946 | 0.635416694270843 | 0.528563393524051 | 0.784700566438825 | -6.15279868173906 |
| lnc-NINJ2-4 | lnc-NINJ2-4 | -0.155002083205479 | 0.421764069218559 | -0.635382611378628 | 0.528585403180989 | 0.784700566438825 | -5.59177163495878 |
| ENSG00000214174.8 | AMZ2P1 | -0.164155883527118 | 1.08560239568653 | -0.635247663782797 | 0.528672552827194 | 0.784741648061916 | -5.7267471155097 |
| lnc-HSD17B10-6 | lnc-HSD17B10-6 | -0.0564046286517472 | 2.16089719849217 | -0.635221689815094 | 0.528689327778973 | 0.784741648061916 | -5.8876835990042 |

| | | | | | | | |
|---|---|---|---|---|---|---|---|
| lnc-PRCD-5 | lnc-PRCD-5 | 0.313779659213348 | -1.09466418692549 | 0.635030978715798 | 0.528812504674957 | 0.784867881722284 | -5.40872865830627 |
| ENSG00000278879.1 | | 0.257188081260344 | -0.0546234792610559 | 0.634790994123086 | 0.52896752793382 | 0.785041360405351 | -5.46846073901368 |
| ENSG00000175137.10 | SH3BP5L | -0.0836690955074459 | 4.04583231615731 | -0.634678878235661 | 0.529039959833169 | 0.785058045646988 | -6.107297924196 |
| ENSG00000280355.1 | | -0.164188162934764 | 0.519889756729742 | -0.634655517198845 | 0.529055052770216 | 0.785058045646988 | -5.66871793739201 |
| ENSG00000111788.10 | DDX12B | 0.258792018122668 | 3.12049130354937 | 0.634692418960268 | 0.529107255688405 | 0.785078910311018 | -5.98470752539652 |
| ENSG00000277972.1 | CISD3 | -0.12079075684137 | 1.55009976032858 | -0.634500320124804 | 0.529155327182384 | 0.785093642283212 | -5.81545048748939 |
| lnc-DENND4A-7 | lnc-DENND4A-7 | 0.107231495255782 | 1.19395680646602 | 0.63433056132229 | 0.529265021541252 | 0.785197870448619 | -5.75878185246562 |
| ENSG00000198551.9 | ZNF627 | 0.103237351236855 | 4.27413794730761 | 0.634273535531934 | 0.529301873025049 | 0.785197870448619 | -6.12738209227601 |
| lnc-TRMT9B-2 | lnc-TRMT9B-2 | -0.210636465541812 | -0.279365848798998 | -0.634003057837061 | 0.529476680843786 | 0.785400585313321 | -5.41127593380665 |
| ENSG00000174444.14 | RPL4 | 0.0969546428420873 | 9.89539563460564 | 0.633913220433563 | 0.529534748846641 | 0.785430117137018 | -6.13908832630485 |
| lnc-AGO1-1 | lnc-AGO1-1 | -0.0825066211280174 | 2.15899573030723 | -0.633634042218048 | 0.529715222049052 | 0.78564118860323 | -5.88535145397567 |
| lnc-BICD1-6 | lnc-BICD1-6 | 0.078877978149956 | 4.09995650262396 | 0.633497931648613 | 0.529803221708218 | 0.785713641987859 | -6.10990401970378 |
| ENSG00000136770.10 | DNAJC1 | -0.108020023762696 | 2.96610267749424 | -0.633391101765395 | 0.529872295894755 | 0.785713641987859 | -6.00215287657871 |
| ENSG00000283782.2 | | 0.0674509413635631 | 4.58542062670939 | 0.633381364198615 | 0.529878592257371 | 0.785713641987859 | -6.14967883535897 |
| ENSG00000270316.1 | BORCS7-ASMT | 0.105982258478716 | 1.80243660128321 | 0.633210188742906 | 0.52998928164963 | 0.78582116295882 | -5.82119997505847 |
| ENSG00000145050.16 | MANF | 0.0980775073656934 | 2.96964486549469 | 0.632993446599121 | 0.530129453871413 | 0.785972379819585 | -5.98290586764708 |
| lnc-TONSL-1 | lnc-TONSL-1 | 0.11619546540021 | 0.244663162760453 | 0.63286691461974 | 0.530211294082252 | 0.786037097817271 | -5.55104708971334 |
| lnc-MYL7-2 | lnc-MYL7-2 | -0.116222853868909 | 1.09463886203748 | -0.632792247354197 | 0.530259591590073 | 0.786052083089555 | -5.75535745765333 |
| ENSG00000163932.13 | PRKCD | -0.11087969935794 | 3.48361156354531 | -0.632582278168694 | 0.530395419711398 | 0.786196811459002 | -6.03662418149181 |
| lnc-MRPL39-2 | lnc-MRPL39-2 | -0.180568684063924 | -0.555309590180005 | -0.632382363879039 | 0.530524760367309 | 0.786246211378888 | -5.33100367621683 |
| lnc-ZNF131-3 | lnc-ZNF131-3 | 0.128041130443396 | 2.44693048994497 | 0.632236749281 | 0.530618980616732 | 0.786246211378888 | -5.91380232001917 |
| ENSG00000241468.7 | ATP5MF | -0.113835128532503 | 3.02063732175944 | -0.632226946895606 | 0.530625323588 | 0.786246211378888 | -5.99388570580925 |
| ENSG00000179021.9 | C3orf38 | 0.0698628006223778 | 5.89570902633238 | 0.632198106474695 | 0.530643986007374 | 0.786246211378888 | -6.20980991948432 |
| lnc-TFPI2-2 | lnc-TFPI2-2 | 0.102549061006634 | 0.955049602164684 | 0.632168692092046 | 0.530663020188472 | 0.786246211378888 | -5.70015370932814 |
| lnc-EBLN1-6 | lnc-EBLN1-6 | 0.137177492852787 | 1.92681685899069 | 0.632143049274168 | 0.530679614065631 | 0.786246211378888 | -5.85810225237551 |
| lnc-ELF2-5 | lnc-ELF2-5 | 0.175903683973239 | 1.8167829516027 | 0.632117514491029 | 0.530696138302817 | 0.786246211378888 | -5.84469569262355 |
| ENSG00000240344.8 | PPIL3 | -0.0470233503434936 | 3.9464571656573 | -0.631586106222583 | 0.531040088096303 | 0.786699160305614 | -6.10108845726535 |
| lnc-TSKS-1 | lnc-TSKS-1 | -0.13583230393077 | 0.145491160282511 | -0.631416613912069 | 0.531149815280713 | 0.786805084413308 | -5.55008111805056 |
| ENSG00000239127.1 | SNORD125 | 0.119844664616928 | -0.332211209485304 | 0.631167096515385 | 0.531311371365468 | 0.786987763155976 | -5.41082854249451 |
| ENSG00000284649.1 | | -0.12728824755396 | 1.23104252871717 | -0.630899810378161 | 0.531484460875151 | 0.787187497891144 | -5.7504209983285 |
| lnc-GNPNAT1-2 | lnc-GNPNAT1-2 | 0.111747968458804 | 0.829792999322019 | 0.630709931466632 | 0.531607440894888 | 0.78729040205347 | -5.67428405481756 |
| lnc-UTY-1 | lnc-UTY-1 | -1.21638437049805 | -0.556968364422919 | -0.630872572958872 | 0.531630437649574 | 0.78729040205347 | -5.09499484289732 |
| ENSG00000122417.15 | ODF2L | -0.102133083761035 | 4.93139476111462 | -0.63055553526516 | 0.531707450623762 | 0.787347802603518 | -6.1812860244873 |
| lnc-BRD2-3 | lnc-BRD2-3 | 0.153496736894227 | 3.5626005063283 | 0.630258954800953 | 0.531899587529062 | 0.78757565715493 | -6.04200006159647 |
| ENSG00000180182.10 | MED14 | 0.0809263334283177 | 5.61501894390111 | 0.63019312708974 | 0.531942238341387 | 0.787582153055731 | -6.20735712969995 |
| ENSG00000206754.1 | SNORD101 | 0.128292363291148 | 1.14752957004005 | 0.630004764217153 | 0.532064291529603 | 0.787674738590045 | -5.74904868315644 |
| ENSG00000180817.11 | PPA1 | 0.124521335643542 | 3.54853392769809 | 0.629936205157758 | 0.532108719279466 | 0.787674738590045 | -6.05705862592426 |
| lnc-ULK4-15 | lnc-ULK4-15 | -0.191909289372379 | -0.26384704612631 | -0.629919451427533 | 0.532119576354009 | 0.787674738590045 | -5.39105505404968 |
| lnc-NOX3-2 | lnc-NOX3-2 | -0.26586397784303 | 0.785424273646601 | -0.629838641012153 | 0.532180569298541 | 0.787708374649207 | -5.62446292674813 |
| ENSG00000104361.9 | NIPAL2 | -0.111840606701869 | 5.85983147741274 | -0.629557538491688 | 0.532354138498728 | 0.787908623910212 | -6.2120015747268 |
| lnc-CHRDL2-5 | lnc-CHRDL2-5 | 0.253647349564159 | -0.0604277246508968 | 0.629157979395718 | 0.532613162734379 | 0.788160668602653 | -5.46996196206692 |
| ENSG00000285382.1 | | -0.0822866442523573 | 1.12607995296496 | -0.62910890418129 | 0.532644981530999 | 0.788160668602653 | -5.74709919418227 |
| lnc-PTCD1-1 | lnc-PTCD1-1 | -0.127912987238175 | 2.07899489885744 | -0.629105526460141 | 0.53264717157381 | 0.788160668602653 | -5.87849638939669 |
| lnc-PPARA-11 | lnc-PPARA-11 | -0.161865785230335 | 1.05664474502469 | -0.628980645218776 | 0.53272814523677 | 0.788160668602653 | -5.74942770096855 |
| ENSG00000196290.14 | NIF3L1 | -0.0682653470056787 | 3.54274268666216 | -0.628959011640561 | 0.532742173219766 | 0.788160668602653 | -6.04931299441383 |
| lnc-RPS14-2 | lnc-RPS14-2 | 0.112168540789612 | 7.4259524106825 | 0.628940488201659 | 0.532754184631429 | 0.788160668602653 | -6.20393212538761 |
| ENSG00000156928.4 | MALSU1 | 0.0841025678843357 | 3.01657501403912 | 0.628824028120538 | 0.532829705715725 | 0.788215741896419 | -5.99992679876978 |
| lnc-RFNG-1 | lnc-RFNG-1 | -0.132149979309715 | 1.65699547504885 | -0.628745204767745 | 0.532880823623967 | 0.788234710215746 | -5.82078707259799 |
| ENSG00000270106.6 | TSNAX-DISC1 | 0.062560827433662 | 3.16495882792903 | 0.628643140788779 | 0.532947017179574 | 0.788275974030409 | -6.00701983585818 |
| ENSG00000233006.7 | MIR3936HG | -0.224895195858489 | 0.277859573267832 | -0.62808785339526 | 0.533307224063513 | 0.788752072494021 | -5.54256734613861 |
| lnc-RAD23B-18 | lnc-RAD23B-18 | -0.473041254672548 | 2.20164269261101 | -0.627937385404826 | 0.533527310654543 | 0.788957465127891 | -5.82197750451992 |
| lnc-MUSTN1-5 | lnc-MUSTN1-5 | -0.136785488793659 | -0.132978556914149 | -0.627725722520505 | 0.533542201764488 | 0.788957465127891 | -5.45707334832394 |
| ENSG00000176087.14 | SLC35A4 | -0.0742124770097572 | 4.44560920892018 | -0.627696617010892 | 0.533561089956772 | 0.788957465127891 | -6.14530348581908 |
| ENSG00000075151.20 | EIF4G3 | -0.0667195260877911 | 6.20688143981381 | -0.627542272364335 | 0.533661258674544 | 0.789048896436903 | -6.2152221954073 |
| ENSG00000102796.10 | DHRS12 | 0.112534070705707 | 2.62320219706433 | 0.627210144693212 | 0.533876840838323 | 0.7892504536173 | -5.94782523596957 |
| ENSG00000135390.18 | ATP5MC2 | 0.0971204277497923 | 5.77997460730538 | 0.627132329247326 | 0.533927356983891 | 0.7892504536173 | -6.2123252468585 |
| ENSG00000207392.1 | SNORA20 | -0.141134361136934 | -0.537994463936766 | -0.627131756103503 | 0.533927729066043 | 0.7892504536173 | -5.37900841490603 |
| ENSG00000125648.14 | SLC25A23 | 0.198616073555927 | 2.54337512885475 | 0.627120001503867 | 0.533950957908026 | 0.7892504536173 | -5.93539693240301 |
| lnc-UBTD2-7 | lnc-UBTD2-7 | -0.0742513126854383 | 2.524054676738 | -0.626895046409465 | 0.53408141143017 | 0.789365440433446 | -5.93415303741602 |
| ENSG00000167315.17 | ACAA2 | -0.118424567575906 | 3.82659681127544 | -0.626858023291072 | 0.53410545056195 | 0.789365440433446 | -6.08987745012399 |
| ENSG00000174775.16 | HRAS | 0.124765079340539 | 2.16326081527471 | 0.626772155450173 | 0.534161206777664 | 0.789391163060358 | -5.8908661124771 |
| SLC25A5-AS1 | SLC25A5-AS1 | -0.0908328485035603 | 1.10141370475612 | -0.626640201348685 | 0.534246893929103 | 0.789428324023101 | -5.74069175241242 |
| lnc-ASB11-2 | lnc-ASB11-2 | 0.0997614243236742 | 1.33625980862816 | 0.626615308389447 | 0.534263059497731 | 0.789428324023101 | -5.78177000621979 |
| lnc-CACNA2D4-3 | lnc-CACNA2D4-3 | 0.108384481556644 | 0.641340717857174 | 0.626291017204716 | 0.534473678610857 | 0.789682845944101 | -5.67704693543508 |
| ENSG00000175155.8 | YPEL2 | 0.135595701598417 | 7.05906541806512 | 0.62619085521384 | 0.534541648818775 | 0.789715223017525 | -6.21251202742527 |
| ENSG00000198695.2 | MT-ND6 | -0.267896390374016 | 2.34107708976902 | -0.626250297573993 | 0.534572326789252 | 0.789715223017525 | -5.91569902958372 |
| lnc-WARS-2 | lnc-WARS-2 | -0.148693471975783 | 1.11889408842335 | -0.626032064715719 | 0.534641893011346 | 0.789744391816159 | -5.7209504281054 |
| ENSG00000181722.16 | ZBTB20 | 0.14076967119339 | 7.36857041537447 | 0.625960024444212 | 0.534696164127409 | 0.789744391816159 | -6.20923230235175 |

| | | | | | | | |
|---|---|---|---|---|---|---|---|
| ENSG00000204856.11 | FAM216A | -0.116932072523888 | 2.6771261265402 | -0.625931575246071 | 0.5347071779794 | 0.789744391816159 | -5.94890984613043 |
| KLHL7-DT | KLHL7-DT | -0.180711127271033 | -0.303839486961677 | -0.625397334431777 | 0.535054327990176 | 0.790176368054558 | -5.37011098397899 |
| ENSG00000108591.9 | DRG2 | 0.0686520500785424 | 5.4232495277065 | 0.625338225307371 | 0.535092744365574 | 0.790176368054558 | -6.2053978562748 |
| ENSG00000166825.13 | ANPEP | -0.878963582501681 | -1.67949024988207 | -0.62541155340162 | 0.535114822635791 | 0.790176368054558 | -4.97483465427286 |
| ENSG00000253203.6 | GUSBP3 | 0.112649227279584 | 0.256373902792939 | 0.625048145763211 | 0.535281294588073 | 0.790249625442176 | -5.53544912913471 |
| ENSG00000008277.14 | ADAM22 | -0.497686466164112 | -0.916235988405494 | -0.625095617312735 | 0.535294119969045 | 0.790249625442176 | -5.26538811795538 |
| lnc-FANCI-5 | lnc-FANCI-5 | 0.100690061091391 | 1.54188630337226 | 0.625010227793587 | 0.535305943637481 | 0.790249625442176 | -5.81337295237353 |
| lnc-SCGN-4 | lnc-SCGN-4 | -0.312649844402028 | -0.33849781434473 | -0.62498078290086 | 0.535325085066172 | 0.790249625442176 | -5.375996831224 |
| ENSG00000157216.15 | SSBP3 | 0.294764526803369 | 1.50480430469927 | 0.624949913610218 | 0.535410971619927 | 0.790249625442176 | -5.79329012561483 |
| ENSG00000100731.15 | PCNX1 | -0.0775961063445427 | 7.19204467118106 | -0.624778559483998 | 0.535456555380733 | 0.790249625442176 | -6.21085389848596 |
| lnc-PAIP2-2 | lnc-PAIP2-2 | 0.167358057086657 | 2.24099844102861 | 0.624768381808622 | 0.535463172577989 | 0.790249625442176 | -5.90240066167388 |
| lnc-ZNF675-4 | lnc-ZNF675-4 | -0.284355858080458 | 0.463174911814599 | -0.624765513872344 | 0.535471579752321 | 0.790249625442176 | -5.55558563411537 |
| ENSG00000153107.12 | ANAPC1 | 0.0409845827649322 | 5.78546741484088 | 0.624521908661413 | 0.535623434522165 | 0.790406577127904 | -6.21351507695978 |
| ENSG00000151151.5 | IPMK | -0.0987751625052575 | 3.58961789924825 | -0.624473779859063 | 0.535654731786286 | 0.790406577127904 | -6.07042795581819 |
| ENSG00000092470.11 | WDR76 | 0.0906345490082319 | 4.03847570740662 | 0.624353424934055 | 0.535733000529071 | 0.790427615689606 | -6.10665873120093 |
| LINC00278 | LINC00278 | -1.14818047514433 | -0.262862110979222 | -0.624529504861932 | 0.535745793445353 | 0.790427615689606 | -5.15357134107278 |
| ENSG00000129680.15 | MAP7D3 | -0.0898981051762 | 3.26651886927571 | -0.624114962874819 | 0.535888093858008 | 0.790549531106102 | -6.02161615492177 |
| ENSG00000171435.13 | KSR2 | -0.382792242187103 | 4.74861098968528 | -0.624261654169567 | 0.535919949040211 | 0.790549531106102 | -6.16609831594848 |
| lnc-ECE1-2 | lnc-ECE1-2 | -0.574110866818972 | -0.134126581839986 | -0.624164057781505 | 0.535943650442333 | 0.790549531106102 | -5.35298805688977 |
| ENSG00000205022.9 | PABPN1L | -0.177202334162804 | -0.283347443087009 | -0.62388048890951 | 0.536040616199175 | 0.790558189495676 | -5.41703734281109 |
| lnc-THYN1-4 | lnc-THYN1-4 | 0.13309917706345 | -0.0752740376442857 | 0.623791787679626 | 0.536098321124691 | 0.790558189495676 | -5.46332521055236 |
| lnc-C4orf46-2 | lnc-C4orf46-2 | 0.144487980727794 | 1.0422588771206 | 0.623786510658376 | 0.536101754213182 | 0.790558189495676 | -5.75390161613739 |
| lnc-HSFY1-2 | lnc-HSFY1-2 | -1.4224349551422 | 1.01929633576255 | -0.623984644573162 | 0.536105188895784 | 0.790558189495676 | -5.39357666917542 |
| ENSG00000221874.4 | ZNF816-ZNF321P | -0.0587474587202035 | 1.96615070979198 | -0.623725323412571 | 0.53614156183113 | 0.790558189495676 | -5.86163715831015 |
| lnc-UNC93B1-7 | lnc-UNC93B1-7 | 0.119426407334124 | 0.938889051302777 | 0.623528168316178 | 0.536269838826731 | 0.790675080212076 | -5.70859438784984 |
| TTTY10 | TTTY10 | -0.861249537492429 | -1.50679583664865 | -0.623580821327926 | 0.536297662869397 | 0.790675080212076 | -4.96327423078186 |
| ENSG00000196715.6 | VKORC1L1 | -0.101341455816585 | 2.78994280516097 | -0.623357059714429 | 0.536381181895609 | 0.790688901364033 | -5.96931926973481 |
| ENSG00000279845.1 | | 0.146939554014069 | 0.7506934501794 | 0.623339632739322 | 0.536392522579411 | 0.790688901364033 | -5.71053515134404 |
| lnc-ACO1-1 | lnc-ACO1-1 | -0.151567348998463 | 0.0301670933327469 | -0.623293903894239 | 0.536422281428721 | 0.790688901364033 | -5.49364646498198 |
| ENSG00000031003.10 | FAM13B | 0.0504017831170005 | 6.5055400363269 | 0.623229591943634 | 0.536464135016169 | 0.790693969999126 | -6.21806513917493 |
| ENSG00000139722.6 | VPS37B | 0.637002056894153 | -0.351135439092336 | 0.623252112435319 | 0.536547432107157 | 0.790760117074438 | -5.36712479516629 |
| ENSG00000260729.1 | | 0.0698444635816289 | 4.20074752127989 | 0.622919325291331 | 0.536666077417599 | 0.790878346923924 | -6.12478085484961 |
| ENSG00000078177.13 | N4BP2 | 0.0722731931977014 | 6.34141651415965 | 0.622669342417277 | 0.536828811882157 | 0.791061528849545 | -6.21861216216791 |
| lnc-RAB20-3 | lnc-RAB20-3 | -0.0622965547273489 | 2.8208197773919 | -0.622509370203115 | 0.536932964469359 | 0.791158365500238 | -5.96778237444428 |
| ENSG00000137806.8 | NDUFAF1 | -0.122033941061269 | 2.1773603616542 | -0.622329244149375 | 0.53705025114325 | 0.791274539676558 | -5.88455190828818 |
| lnc-ZMAT5-4 | lnc-ZMAT5-4 | 0.0536721990613043 | 1.67435650311055 | 0.622137676959215 | 0.537175002193925 | 0.791347735753252 | -5.82878338259027 |
| ENSG00000121903.14 | ZSCAN20 | -0.163724643662512 | 0.895285144775678 | -0.622113595066808 | 0.537190685703249 | 0.791347735753252 | -5.69762969139706 |
| lnc-N4BP2L2-2 | lnc-N4BP2L2-2 | 0.0577137419193372 | 6.14704759571464 | 0.622010873453724 | 0.537257586584657 | 0.791347735753252 | -6.21840763774288 |
| ENSG00000168209.4 | DDIT4 | 0.345774047679508 | 3.52915210691947 | 0.622151158595167 | 0.537287366035632 | 0.791347735753252 | -6.06699360570278 |
| ENSG00000221500.1 | SNORD100 | 0.11015337598215 | 1.21665380431359 | 0.621935377096371 | 0.537306758873331 | 0.791347735753252 | -5.77726932910811 |
| ENSG00000121716.20 | PILRB | 0.090365783053473 | 5.89402134836442 | 0.621839929907487 | 0.537368928889876 | 0.791347735753252 | -6.21729185016791 |
| lnc-HIP1-1 | lnc-HIP1-1 | -0.0718712724896742 | 2.67472178546274 | -0.62180796472908 | 0.537389750409604 | 0.791347735753252 | -5.96061046314249 |
| ENSG00000101412.12 | E2F1 | 0.280173895916452 | -0.144840101605139 | 0.621780710275638 | 0.537407503782683 | 0.791347735753252 | -5.4656557028735 |
| ENSG00000213204.8 | | -0.0772889908695409 | 2.43764062901964 | -0.621711617765375 | 0.537452511560259 | 0.791357396483641 | -5.92696278334495 |
| ENSG00000048991.16 | R3HDM1 | -0.059742234561412 | 5.15215689996902 | -0.621420307240621 | 0.53764229666273 | 0.791550577877106 | -6.1941970368074 |
| lnc-TBK1-1 | lnc-TBK1-1 | 0.202093504211334 | -0.0669858505760327 | 0.621365829371607 | 0.537677792170078 | 0.791550577877106 | -5.43903580749377 |
| ENSG00000184584.12 | STING1 | 0.41536070127439 | -1.04254045931789 | 0.62133315687803 | 0.537699080788889 | 0.791550577877106 | -5.20662641506302 |
| ENSG00000197019.4 | SERTAD1 | 0.159445128545656 | 2.22105173648179 | 0.62111861730113 | 0.537838880557574 | 0.791699755311202 | -5.89681097709844 |
| ENSG00000185379.20 | RAD51D | -0.0596249543209684 | 4.39882740792372 | -0.620908980272981 | 0.537975503947663 | 0.791792510130369 | -6.14261630867334 |
| ENSG00000123472.12 | ATPAF1 | -0.0815293469568621 | 3.92113407554188 | -0.620903877349385 | 0.537978829819424 | 0.791792510130369 | -6.09783502221988 |
| ENSG00000139597.17 | N4BP2L1 | 0.0442343908249105 | 5.3484422099537 | 0.620803306632224 | 0.538044379779587 | 0.791832365866097 | -6.20384910589111 |
| ENSG00000205268.10 | PDE7A | -0.115503413740832 | 6.69204208927469 | -0.620725484181474 | 0.538095105728696 | 0.7918504011735 | -6.21856788126491 |
| ENSG00000132141.13 | CCT6B | 0.133322941456252 | 1.76304771233736 | 0.620637901405541 | 0.538152196590469 | 0.791877799715586 | -5.84180322226586 |
| ENSG00000166797.10 | CIAO2A | -0.0972159229183482 | 4.98127407654144 | -0.620523217012529 | 0.538226958410057 | 0.791931194864123 | -6.18490890829516 |
| lnc-MALT1-1 | lnc-MALT1-1 | 0.133085755238645 | 1.38093288129345 | 0.620429076627544 | 0.538288331791851 | 0.791964884436859 | -5.78311759353526 |
| ENSG00000204387.13 | SNHG32 | 0.104389820461435 | 4.9926361624576 | 0.620147827001925 | 0.538471709834338 | 0.792139204733672 | -6.19019518114287 |
| ENSG00000198839.9 | ZNF277 | 0.0757000856488128 | 4.29200433098703 | 0.620034601097623 | 0.538545543607921 | 0.792139204733672 | -6.13765143960938 |
| ENSG00000163535.17 | SGO2 | 0.124299011356466 | 1.44721660924565 | 0.619998658120583 | 0.538568982856556 | 0.792139204733672 | -5.81019931634434 |
| ENSG00000272391.5 | POM121C | 0.0797217041055749 | 4.74665662296081 | 0.619965015073546 | 0.538590922747582 | 0.792139204733672 | -6.17227291421474 |
| ENSG00000213613.2 | RPL11P3 | -0.1342536799422 | 1.44089542153834 | -0.619952260433349 | 0.538599240647512 | 0.792139204733672 | -5.79099095255277 |
| ENSG00000147400.8 | CETN2 | 0.0891822853661477 | 2.44714111892133 | 0.61975416372176 | 0.538728437381644 | 0.792205996551917 | -5.92275545105874 |
| ENSG00000138160.5 | KIF11 | -0.225636322301646 | 2.38393676205829 | -0.619760803378255 | 0.538759106408368 | 0.792205996551917 | -5.90572542059696 |
| ENSG00000231466.2 | PHF10P2 | 0.150182448433367 | 0.335488172749155 | 0.619682432212092 | 0.538775223937594 | 0.792205996551917 | -5.56089519353594 |
| ENSG00000228451.4 | SDAD1P1 | -0.103808199957137 | 2.06198051824613 | -0.619623413420267 | 0.538813720262196 | 0.792205996551917 | -5.86002599542173 |
| ENSG00000142751.14 | GPN2 | -0.04986086580438 | 4.91764301302269 | -0.61956662481307 | 0.538850763247201 | 0.792205996551917 | -6.1876171084938 |
| ENSG00000212296.1 | SNORD72 | 0.176055279306818 | -0.322437776284451 | 0.619528573673971 | 0.538875584595219 | 0.792205996551917 | -5.43947596091668 |
| ENSG00000197982.13 | C1orf122 | 0.11516907267216 | 1.52101105878907 | 0.619322549435693 | 0.539009987706708 | 0.792330610278075 | -5.81034877576691 |
| lnc-SUMO3-1 | lnc-SUMO3-1 | 0.074389930081401 | 1.74843293298154 | 0.619229691882218 | 0.539070570461539 | 0.792330610278075 | -5.84679913763295 |

| | | | | | | | |
|---|---|---|---|---|---|---|---|
| ENSG00000196544.7 | BORCS6 | -0.110410179998593 | 2.27003591981123 | -0.61914160825884 | 0.539128041842391 | 0.792330610278075 | -5.92558358937305 |
| lnc-HSFY2-13 | lnc-HSFY2-13 | -1.33178538104544 | 0.100740870424264 | -0.619257976559948 | 0.539183276110783 | 0.792330610278075 | -5.21874596952654 |
| lnc-RWDD2B-4 | lnc-RWDD2B-4 | 0.101155466167527 | 3.22210955835565 | 0.61904894047784 | 0.539188507652351 | 0.792330610278075 | -6.01547209872858 |
| ENSG00000156858.11 | PRR14 | -0.0902097914485549 | 3.57374461555308 | -0.619016022467163 | 0.539209987530731 | 0.792330610278075 | -6.07051874701659 |
| lnc-SLC10A7-7 | lnc-SLC10A7-7 | 0.126215412265265 | 0.518080046922018 | 0.61898564433644 | 0.539229810463745 | 0.792330610278075 | -5.61133111562096 |
| lnc-TMEM44-3 | lnc-TMEM44-3 | 0.0848430336428418 | 1.06045273993101 | 0.618825227928049 | 0.539334494791263 | 0.792427861109898 | -5.74800067837841 |
| ENSG00000197442.9 | MAP3K5 | 0.059036819694442 | 6.94766507669776 | 0.618642098822793 | 0.539454013826977 | 0.792502012441803 | -6.21838272061556 |
| ENSG00000111670.15 | GNPTAB | -0.120688791421191 | 7.36536662032883 | -0.618615138250564 | 0.539471610778753 | 0.792502012441803 | -6.21004107642147 |
| ENSG00000126070.19 | AGO3 | 0.0549417477657759 | 6.0672605698687 | 0.618570921751379 | 0.539500471181779 | 0.792502012441803 | -6.22006501133309 |
| ENSG00000177954.12 | RPS27 | 0.0783143193400839 | 10.3865153422093 | 0.618504870288699 | 0.539543584902153 | 0.792508785273748 | -6.13159549786482 |
| LINC02550 | LINC02550 | 0.302770580717412 | -0.289321901008764 | 0.618432730089165 | 0.539590674960419 | 0.792521397367655 | -5.38133582352105 |
| ENSG00000092931.11 | MFSD11 | -0.0577362048992387 | 3.81024788313373 | -0.618297867503695 | 0.539678713244243 | 0.792587368776262 | -6.08797615061408 |
| ENSG00000227671.4 | ZNF731P | 0.0709660949701316 | 5.80167649306691 | 0.618190524155952 | 0.539748792294934 | 0.792587368776262 | -6.2179073257743 |
| ENSG00000134184.12 | GSTM1 | 1.17890923051223 | -0.442027951914329 | 0.61837122242368 | 0.539751112478106 | 0.792587368776262 | -5.06993045224768 |
| ENSG00000281842.2 | MIR1291 | 0.238621474215738 | 1.4731938708591 | 0.618112081235109 | 0.539822729724525 | 0.79263358703583 | -5.79121335244064 |
| lnc-LRRC31-4 | lnc-LRRC31-4 | 0.134218417184021 | 2.21843957445297 | 0.618020801399716 | 0.539859605337186 | 0.79263358703583 | -5.89211793196085 |
| ENSG00000170310.14 | STX8 | 0.0661025188132562 | 3.45077405598122 | 0.617907697994543 | 0.539933457800621 | 0.792657436779716 | -6.04497231100025 |
| lnc-NUP107-26 | lnc-NUP107-26 | -0.169920023077789 | -0.386521950225665 | -0.617877970131638 | 0.539952869896925 | 0.792657436779716 | -5.37960921279726 |
| ENSG00000196961.12 | AP2A1 | -0.079384746014278 | 4.6943819498501 | -0.61749823716411 | 0.540200864809218 | 0.792953447270094 | -6.17111312087065 |
| lnc-DDR2-5 | lnc-DDR2-5 | -0.175171897989318 | -0.653543464215153 | -0.617432086595052 | 0.540244072266766 | 0.792953447270094 | -5.34139523326921 |
| ENSG00000123609.10 | NMI | 0.0790690719616789 | 4.14530602773593 | 0.617392263569959 | 0.540270084269339 | 0.792953447270094 | -6.12144019953421 |
| lnc-GALNT3-1 | lnc-GALNT3-1 | -0.217848861674684 | -0.0370546770987187 | -0.616925555060922 | 0.540574981959312 | 0.793112588183737 | -5.47753010081077 |
| ENSG00000197299.11 | BLM | -0.0640028812588662 | 4.48113937817319 | -0.616911790391078 | 0.540583975682649 | 0.793112588183737 | -6.15284863492437 |
| lnc-MTA2-2 | lnc-MTA2-2 | 0.091883063045775 | 1.2467021576551 | 0.616899239596346 | 0.540592176337388 | 0.793112588183737 | -5.76882622972503 |
| ENSG00000260290.2 | | 0.357748019180563 | -0.661525798763173 | 0.616853899722713 | 0.540621801822679 | 0.793112588183737 | -5.19219096714229 |
| ENSG00000074416.14 | MGLL | 0.350930096199373 | -0.174671161058917 | 0.616802202585809 | 0.540664593679331 | 0.793112588183737 | -5.38558089236615 |
| ENSG00000198040.10 | ZNF84 | 0.0570800243919631 | 6.11593395668113 | 0.616718753381437 | 0.540710112656363 | 0.793112588183737 | -6.22161391176006 |
| ENSG00000067955.13 | CBFB | 0.0784556723635008 | 5.54409148857937 | 0.616708121143774 | 0.540717060566192 | 0.793112588183737 | -6.21576587547105 |
| lnc-TMC7-5 | lnc-TMC7-5 | -0.0936463354561182 | 0.860237870026453 | -0.616666850371053 | 0.540744030454568 | 0.793112588183737 | -5.68872000887084 |
| lnc-GATA2-2 | lnc-GATA2-2 | 0.0779342185753986 | 3.30292280493269 | 0.616566736160845 | 0.540809456621867 | 0.793112588183737 | -6.031510778717 |
| ENSG00000184117.11 | NIPSNAP1 | -0.0939642468793567 | 4.29127466509026 | -0.616421632726265 | 0.540904291207622 | 0.793112588183737 | -6.14529332927044 |
| ENSG00000187801.14 | ZFP69B | -0.0882215875488663 | 1.7864776160951 | -0.616379846848744 | 0.540931602607761 | 0.793112588183737 | -5.84344839226136 |
| ENSG00000230397.1 | SPTLC1P1 | 0.215134120100473 | 0.68533740849299 | 0.616273255569928 | 0.541001274283603 | 0.793112588183737 | -5.64625060876614 |
| ENSG00000145375.7 | AFG2A | 0.050568705943439 | 5.0853229190426 | 0.61623817454031 | 0.541024205450191 | 0.793112588183737 | -6.19534883931909 |
| lnc-PLXND1-1 | lnc-PLXND1-1 | -0.228539390290717 | -0.382768302778243 | -0.616233045007997 | 0.541027558477161 | 0.793112588183737 | -5.42126081877561 |
| ENSG00000284353.1 | MIR142 | -0.247048845079455 | -0.77515559909663 | -0.616166369576565 | 0.541071143257185 | 0.793112588183737 | -5.3725626919467 |
| ENSG00000164209.16 | SLC25A46 | -0.0768634491412108 | 5.53267317661725 | -0.616133561735578 | 0.541092589942465 | 0.793112588183737 | -6.21471098631483 |
| lnc-GPALPP1-5 | lnc-GPALPP1-5 | 0.125410366352091 | 1.33472864103094 | 0.615941460103273 | 0.541218176774851 | 0.793112588183737 | -5.78673860478282 |
| ENSG00000145293.15 | ENOPH1 | 0.0763579218159958 | 4.13986529433102 | 0.615893569978964 | 0.54124948738866 | 0.793112588183737 | -6.12838032796586 |
| lnc-SDC3-1 | lnc-SDC3-1 | -0.0690528971464888 | 2.69869946041608 | -0.615870196353851 | 0.541264769429649 | 0.793112588183737 | -5.96477648544322 |
| lnc-PMM2-2 | lnc-PMM2-2 | -0.124913484617044 | 0.279948303222793 | -0.615825902451186 | 0.541293730086876 | 0.793112588183737 | -5.58512396812604 |
| ENSG00000149100.12 | EIF3M | 0.0473608321018464 | 6.40561920301197 | 0.615816746339906 | 0.541299716721869 | 0.793112588183737 | -6.22272126378162 |
| ENSG00000170142.11 | UBE2E1 | 0.0522117607972108 | 4.90482470952905 | 0.61577140046305 | 0.54132936619057 | 0.793112588183737 | -6.19297507229581 |
| ENSG00000136715.18 | SAP130 | 0.0745913679304238 | 4.43337015925066 | 0.615757086047631 | 0.541338725866693 | 0.793112588183737 | -6.15092193377429 |
| ENSG00000169398.19 | PTK2 | -0.971235095359682 | 0.0799058913186165 | -0.615950495378337 | 0.541342613180505 | 0.793112588183737 | -5.49622323701271 |
| ENSG00000168575.9 | SLC20A2 | -0.07502330875338 | 3.43214136366498 | -0.615687641774419 | 0.541384134141537 | 0.793112588183737 | -6.04859724129544 |
| lnc-CRY1-1 | lnc-CRY1-1 | 0.179617280084287 | -0.347212181829754 | 0.615683576037109 | 0.541386792710014 | 0.793112588183737 | -5.40026015384719 |
| ENSG00000006831.9 | ADIPOR2 | 0.0828054181320734 | 4.41673308040071 | 0.615634493301685 | 0.541418888236393 | 0.793112588183737 | -6.15530303820804 |
| lnc-RCSD1-4 | lnc-RCSD1-4 | 0.0970294009510488 | 2.98822125312302 | 0.615528571974065 | 0.541488154247932 | 0.793127945359516 | -5.99426546224271 |
| ENSG00000103426.12 | CORO7-PAM16 | 0.10240051432215 | 4.75137331379709 | 0.615500613490259 | 0.541506438135222 | 0.793127945359516 | -6.17599681431848 |
| ENSG00000221491.2 | SNORA2C | 0.224629626219359 | 1.6103929272343 | 0.615420623508121 | 0.541574193980953 | 0.793141784162149 | -5.81396941192511 |
| ENSG00000197548.12 | ATG7 | 0.0618055628308989 | 4.62933160046899 | 0.615368325586334 | 0.541592954204878 | 0.793141784162149 | -6.16813846754638 |
| lnc-COMMD5-1 | lnc-COMMD5-1 | 0.0761759668517923 | 1.75574410339734 | 0.615131696834543 | 0.541747726836766 | 0.793287016417675 | -5.83085218767299 |
| lnc-SLC11A2-7 | lnc-SLC11A2-7 | -0.132147486433442 | 0.554294787065849 | -0.615098858851953 | 0.541769207102135 | 0.793287016417675 | -5.66523995967722 |
| ENSG00000196141.13 | SPATS2L | -0.185662322966718 | 4.60894859870773 | -0.614950490441767 | 0.541919993653757 | 0.793369453431789 | -6.17430333997707 |
| ENSG00000067167.7 | TRAM1 | 0.0914458250346897 | 6.72987558096727 | 0.614773495917031 | 0.541982060105133 | 0.793369453431789 | -6.22123411789498 |
| lnc-SLC38A6-1 | lnc-SLC38A6-1 | -0.158433442128597 | 0.170453340720948 | -0.614757081379618 | 0.541992799669068 | 0.793369453431789 | -5.56917719626268 |
| ENSG00000163867.16 | ZMYM6 | -0.0723434301776509 | 5.89423180381294 | -0.614739467440226 | 0.542004324089817 | 0.793369453431789 | -6.22078516198076 |
| ENSG00000185477.4 | GPRIN3 | -1.4446401040636 | 0.418623890658739 | -0.614917048342621 | 0.542018231266473 | 0.793369453431789 | -5.33575812954498 |
| lnc-SPAG16-7 | lnc-SPAG16-7 | -0.353481193166389 | 0.59233947425512 | -0.614488988558638 | 0.542244410268484 | 0.793644079965599 | -5.69188434751393 |
| ENSG00000119408.16 | NEK6 | -0.112572811366955 | 4.38331422951565 | -0.614232311129926 | 0.542336199818764 | 0.7937219852723 | -6.14814035212824 |
| ENSG00000257093.6 | DENND11 | 0.053086563481731 | 7.51719686504454 | 0.614105876558454 | 0.54241895310523 | 0.793786655653391 | -6.21266354153409 |
| LIX1L-AS1 | LIX1L-AS1 | -0.153429411289712 | -0.0699669973787986 | -0.613989395055291 | 0.542495197731475 | 0.793791826956923 | -5.51191294797441 |
| ENSG00000101003.9 | GINS1 | -0.195653697222937 | 0.0132512615036816 | -0.613894952090351 | 0.542557020780051 | 0.793791826956923 | -5.51977414027284 |
| lnc-CCDC92-7 | lnc-CCDC92-7 | -0.136703935227434 | 1.69382309842473 | -0.613826035134335 | 0.542602136613476 | 0.793791826956923 | -5.83898429037874 |
| ENSG00000189180.15 | ZNF33A | 0.0711804479114866 | 6.91140949858919 | 0.613667217299659 | 0.542706112579242 | 0.793791826956923 | -6.221470149186 |
| ENSG00000138381.9 | ASNSD1 | 0.0721166430678313 | 5.09844420612888 | 0.613632278955636 | 0.54272898763732 | 0.793791826956923 | -6.19791967503314 |
| ENSG00000152133.14 | GPATCH11 | -0.155226778983336 | 4.75856087583416 | -0.613654687949131 | 0.542733983463414 | 0.793791826956923 | -6.17889658312721 |

| | | | | | | | |
|---|---|---|---|---|---|---|---|
| lnc-INVS-1 | lnc-INVS-1 | -0.122193835198783 | 1.88819812699532 | -0.613623115192835 | 0.542734987477607 | 0.793791826956923 | -5.85505872826508 |
| ENSG00000270170.1 | NCBP2AS2 | 0.0823524175886168 | 3.01784813675967 | 0.613595998099368 | 0.542752742200286 | 0.793791826956923 | -6.00744848513757 |
| ENSG00000163785.12 | RYK | 0.082949243725162 | 5.57967439693515 | 0.613549105869765 | 0.542783445258946 | 0.793791826956923 | -6.21595115411609 |
| lnc-HLX-3 | lnc-HLX-3 | 0.22582601100126 | 3.49959748319989 | 0.613613436488846 | 0.542808140913312 | 0.793791826956923 | -6.04681071906386 |
| ENSG00000238650.1 | SNORD54 | 0.130392786677014 | 0.215660230910104 | 0.613363300758269 | 0.542905111437655 | 0.793858970512519 | -5.5696421939795 |
| lnc-RBL2-4 | lnc-RBL2-4 | -0.0777827545782455 | 1.93188550823107 | -0.613323473844746 | 0.542931192144232 | 0.793858970512519 | -5.86857681003684 |
| ENSG00000139921.12 | TMX1 | 0.0849067895750553 | 5.21584214588871 | 0.613219248428548 | 0.542999447353686 | 0.793902374263455 | -6.20553710021199 |
| lnc-MRPS14-1 | lnc-MRPS14-1 | -0.113727683724426 | 4.07352612054605 | -0.613111443766177 | 0.543070051198865 | 0.793911472717408 | -6.12215740340788 |
| lnc-KYNU-10 | lnc-KYNU-10 | -0.185752914333899 | 0.639592045730479 | -0.613054551776713 | 0.543107313028666 | 0.793911472717408 | -5.65859040471473 |
| ENSG00000146414.15 | SHPRH | 0.0588048255729641 | 6.54165427259201 | 0.613033068470027 | 0.543121384022437 | 0.793911472717408 | -6.2240357606358 |
| lnc-TRIM52-2 | lnc-TRIM52-2 | 0.102407244539233 | 6.56630680185082 | 0.612849162149979 | 0.543241845457818 | 0.793976517953623 | -6.22383782545251 |
| ENSG00000123575.8 | FAM199X | 0.0680252667454884 | 5.63716972701806 | 0.612800702625331 | 0.543273589476387 | 0.793976517953623 | -6.21917918520001 |
| ENSG00000174306.21 | ZHX3 | 0.123567921507088 | 3.31847019991043 | 0.612788466350464 | 0.543281605152739 | 0.793976517953623 | -6.01799474842182 |
| ENSG00000130589.16 | HELZ2 | -0.106925822631409 | 5.63633726763979 | -0.612501303356504 | 0.543469735924233 | 0.794094935703116 | -6.2185130025062 |
| lnc-TTC7B-8 | lnc-TTC7B-8 | 0.133838553635098 | -0.326300975021603 | 0.612487970143266 | 0.543478471805916 | 0.794094935703116 | -5.39733296823838 |
| ENSG00000005007.12 | UPF1 | 0.0618526033429134 | 5.88796069044579 | 0.612436573047491 | 0.543512147707574 | 0.794094935703116 | -6.22201862166466 |
| ENSG00000171606.17 | ZNF274 | 0.0809373432231606 | 6.7769471105354 | 0.612393346384445 | 0.543540471090967 | 0.794094935703116 | -6.22344030640556 |
| ENSG00000113924.11 | HGD | 0.158813080337324 | 0.1220915403044 | 0.612337485382117 | 0.543577073992463 | 0.794094935703116 | -5.48783152297297 |
| ENSG00000145703.15 | IQGAP2 | 0.646628370992656 | 1.4018321561288 | 0.612416712758252 | 0.543654633178848 | 0.794094935703116 | -5.78041295135186 |
| ENSG00000159259.7 | CHAF1B | -0.128326719521001 | 1.63979590934464 | -0.612174191239898 | 0.543684079703834 | 0.794094935703116 | -5.82314111792212 |
| ENSG00000109065.11 | NAT9 | -0.0806137908022819 | 4.25265154540794 | -0.612119550602984 | 0.54371988782219 | 0.794094935703116 | -6.14205034528588 |
| ENSG00000132305.20 | IMMT | -0.0801606745239347 | 5.32179811926081 | -0.612082629623383 | 0.543744084253127 | 0.794094935703116 | -6.21007902894825 |
| ENSG00000102763.17 | VWA8 | -0.0998542045833055 | 5.42165034767485 | -0.612075991995655 | 0.543748434329287 | 0.794094935703116 | -6.21258586083758 |
| ENSG00000064601.18 | CTSA | -0.0918360566966507 | 4.57460739104283 | -0.611955696701674 | 0.543827274893388 | 0.794153728210756 | -6.16991833345334 |
| ENSG00000130748.6 | TMEM160 | 0.162633774479012 | -0.277976167195628 | 0.611852747897069 | 0.543894751384742 | 0.794195918540873 | -5.44757120731114 |
| lnc-TTC3-8 | lnc-TTC3-8 | 0.1309379324425 | 1.58175152824965 | 0.611734614343169 | 0.543972185841365 | 0.794225949309812 | -5.81118081921495 |
| ENSG00000233264.2 | | -0.143553114938682 | 2.42403998524742 | -0.611703639003817 | 0.543992490568417 | 0.794225949309812 | -5.93927518906717 |
| lnc-ZNF860-2 | lnc-ZNF860-2 | -0.107016616909385 | 4.91105187142713 | -0.611507724104519 | 0.544120924306624 | 0.79435711646239 | -6.18202728649915 |
| ENSG00000157693.14 | TMEM268 | -0.0879115549585014 | 6.29388182489619 | -0.611355801519918 | 0.544220529242953 | 0.794384629955594 | -6.22530316668085 |
| ENSG00000131724.10 | IL13RA1 | 0.425240584797633 | 2.16513145293841 | 0.611520503995496 | 0.544231280477896 | 0.794384629955594 | -5.89313040025802 |
| ENSG00000136169.16 | SETDB2 | -0.112938651599412 | 5.53582339934573 | -0.61130032227171 | 0.544256905417595 | 0.794384629955594 | -6.21692759066387 |
| lnc-XRCC2-10 | lnc-XRCC2-10 | 0.170554091836057 | 0.284862416023396 | 0.611243524487052 | 0.54429414741601 | 0.794384629955594 | -5.60745318407378 |
| ENSG00000070961.15 | ATP2B1 | 0.0913104551806692 | 7.92880506338951 | 0.611117380396126 | 0.544376864102248 | 0.794449021115753 | -6.20771534732326 |
| ENSG00000151883.17 | PARP8 | 0.0617710796931528 | 6.0847203438242 | 0.610941123741896 | 0.544492452023385 | 0.794561371144795 | -6.22518590767601 |
| lnc-SERF1A-2 | lnc-SERF1A-2 | -0.344954801367282 | -0.100921968602897 | -0.610851734734584 | 0.544578569970208 | 0.794573178015779 | -5.47994969232667 |
| ENSG00000117616.17 | RSRP1 | -0.0800377088334754 | 6.49077636817263 | -0.610764869478481 | 0.544608051005822 | 0.794573178015779 | -6.22546471985775 |
| ENSG00000273088.1 | | -0.0712445430103692 | 2.96273574030118 | -0.610752212011339 | 0.544616353080047 | 0.794573178015779 | -5.99653653651366 |
| lnc-TAOK1-1 | lnc-TAOK1-1 | -0.127485317096981 | 1.30790862727431 | -0.610689412736061 | 0.544657544293741 | 0.794576953306263 | -5.7977924731768 |
| lnc-HAND1-8 | lnc-HAND1-8 | -0.075560721752854 | 1.7401756220778 | -0.610444406652319 | 0.544818263669314 | 0.794709405926148 | -5.83532398301542 |
| ENSG00000205464.12 | ATP6AP1L | -0.15357907966711 | 1.45319577438821 | -0.610402568325422 | 0.544845711261987 | 0.794709405926148 | -5.80292296882331 |
| ENSG00000241859.6 | ANOS2P | -0.981315936013399 | -1.10272229854454 | -0.610439624323186 | 0.544912603912321 | 0.794709405926148 | -5.02746386663782 |
| lnc-CNR2-1 | lnc-CNR2-1 | 0.0565280814255065 | 5.16346909275954 | 0.610259184210911 | 0.544939782291856 | 0.794709405926148 | -6.20311535782027 |
| ENSG00000271141.1 | | -0.184507057789102 | 0.55250015071359 | -0.61025673939799 | 0.54494138634998 | 0.794709405926148 | -5.65141875913298 |
| lnc-LYSMD3-1 | lnc-LYSMD3-1 | 0.181108028139905 | 5.65563021767837 | 0.61006827914475 | 0.545125569332732 | 0.794921685694341 | -6.22062377037573 |
| ENSG00000005882.11 | PDK2 | -0.09991758697551 | 3.9622026790984 | -0.609881564100278 | 0.545187570148628 | 0.794955777583538 | -6.11053799697822 |
| ENSG00000169598.15 | DFFB | -0.0992123784191219 | 2.93113850303625 | -0.609786948819878 | 0.545249664150424 | 0.794990000510603 | -5.98619976465263 |
| lnc-LILRA4-1 | lnc-LILRA4-1 | 0.830207607168474 | 0.350092275364777 | 0.60986870932656 | 0.545319857503227 | 0.795008080839477 | -5.44186448765903 |
| ENSG00000110455.13 | ACCS | 0.114845259916756 | 4.61855864974523 | 0.609583542153656 | 0.545383167938679 | 0.795008080839477 | -6.17237583900762 |
| lnc-ORMDL3-1 | lnc-ORMDL3-1 | -0.156897322504423 | 0.884792544085944 | -0.609454612869419 | 0.545467797992398 | 0.795008080839477 | -5.68660648376089 |
| ENSG00000011275.18 | RNF216 | 0.0673117246942025 | 6.72235570606162 | 0.609393926397574 | 0.545507635338355 | 0.795008080839477 | -6.22525201317419 |
| lnc-RHOV-3 | lnc-RHOV-3 | 0.150585236467335 | -0.104709162869161 | 0.609372196773016 | 0.545521900010568 | 0.795008080839477 | -5.44641380891017 |
| ENSG00000156599.10 | ZDHHC5 | 0.0567489149795026 | 5.91198569034968 | 0.609365216193679 | 0.545526482536127 | 0.795008080839477 | -6.22430550576414 |
| lnc-AMZ2-5 | lnc-AMZ2-5 | 0.0856750676887792 | 3.99149660482706 | 0.60935614703798 | 0.545532436174356 | 0.795008080839477 | -6.11416767224383 |
| ENSG00000177733.6 | HNRNPA0 | 0.0704281068374453 | 5.44627502070454 | 0.609216482332187 | 0.545624126227004 | 0.795085408150826 | -6.21613478701726 |
| ENSG00000063177.12 | RPL18 | -0.106242721410283 | 7.88485510356524 | -0.609132753014707 | 0.545679098423392 | 0.795109222911559 | -6.20750538351314 |
| lnc-SUMO4-2 | lnc-SUMO4-2 | 0.152153978016534 | 1.79076144584179 | 0.608599141333795 | 0.546029506173133 | 0.795530628216397 | -5.83945657064449 |
| ENSG00000103264.17 | FBXO31 | -0.0647919438105609 | 4.622912872869 | -0.608392598233452 | 0.546165168210353 | 0.795530628216397 | -6.17607675563896 |
| ENSG00000188322.4 | SBK1 | 0.241597808278586 | 0.349330004273826 | 0.608365382480444 | 0.546183045403659 | 0.795530628216397 | -5.55974511039744 |
| ENSG00000283498.1 | MIR1244-2 | 0.282604675928577 | 0.379540111896016 | 0.608407188206938 | 0.546188802761063 | 0.795530628216397 | -5.68378274805778 |
| ENSG00000143553.10 | SNAPIN | 0.0897668806871597 | 2.83450127596509 | 0.608349824376843 | 0.546193265179593 | 0.795530628216397 | -5.98190926778886 |
| lnc-CISD2-1 | lnc-CISD2-1 | -0.105080932388487 | 1.87507487482532 | -0.60833019141143 | 0.546206161782669 | 0.795530628216397 | -5.85782528942548 |
| ENSG00000227097.5 | RPS28P7 | 0.199689186960362 | 1.97142933353336 | 0.608143873244021 | 0.546338132764259 | 0.795530628216397 | -5.87394071846621 |
| lnc-ALG5-4 | lnc-ALG5-4 | 0.0843190416710589 | 2.61601796488696 | 0.608101602273461 | 0.546356330091493 | 0.795530628216397 | -5.95166824232823 |
| ENSG00000278206.1 | | 0.0852137858080638 | 0.933693851326434 | 0.608092000902667 | 0.546362638035944 | 0.795530628216397 | -5.72803397018458 |
| ENSG00000274997.1 | H2AC12 | -0.211697125614748 | 3.72471570188674 | -0.608154459086787 | 0.54638772895427 | 0.795530628216397 | -6.10092019386787 |
| lnc-TCF4-8 | lnc-TCF4-8 | -0.229116133266438 | 1.88455020047828 | -0.608030034724157 | 0.546436646794564 | 0.795530628216397 | -5.87503869850693 |
| ENSG00000006282.20 | SPATA20 | -0.108544883259747 | 5.36868299149711 | -0.607933593315238 | 0.546466714638594 | 0.795530628216397 | -6.21608877604748 |
| ENSG00000223865.10 | HLA-DPB1 | -0.161357418270111 | 8.75943693902236 | -0.607883083048099 | 0.546530317901562 | 0.795530628216397 | -6.19154625330849 |

| | | | | | | | |
|---|---|---|---|---|---|---|---|
| lnc-SBDS-28 | lnc-SBDS-28 | 0.115025428993124 | 0.0112433002693276 | 0.607818987539677 | 0.546542019012765 | 0.795530628216397 | -5.47621666536805 |
| ENSG00000169991.10 | IFFO2 | -0.15590815855934 | 2.68059216956269 | -0.607809801899166 | 0.546548054880633 | 0.795530628216397 | -5.97515643107843 |
| ENSG00000102316.16 | MAGED2 | -0.0812135556976177 | 6.27483423649123 | -0.607643673194631 | 0.546657223655076 | 0.795633265060984 | -6.22758436379801 |
| ENSG00000162999.12 | DUSP19 | -0.183357825331971 | 0.088914060901292 | -0.607563904040266 | 0.546709646630745 | 0.795653302453554 | -5.47554312132869 |
| lnc-NUMB-1 | lnc-NUMB-1 | 0.10721985048601 | 1.20967516656857 | 0.607438835002016 | 0.546791845134595 | 0.795675996014785 | -5.7583314210819 |
| ENSG00000165868.14 | HSPA12A | -0.792295510660237 | 0.244328740222072 | -0.607560157517118 | 0.546840385848032 | 0.795675996014785 | -5.70153825715342 |
| lnc-MAMDC2-1 | lnc-MAMDC2-1 | 0.134540377389086 | 1.90474954933164 | 0.607363727532757 | 0.546841210689654 | 0.795675996014785 | -5.87356466314719 |
| ENSG00000212304.1 | SNORD12 | 0.136307019064054 | 1.06484843969267 | 0.607257649912283 | 0.546910935752178 | 0.795719528250277 | -5.75112203942426 |
| ENSG00000116171.17 | SCP2 | 0.0749927704802653 | 5.27402559423756 | 0.607200584212453 | 0.546948447052661 | 0.795719528250277 | -6.20992775122355 |
| lnc-NLN-3 | lnc-NLN-3 | 0.0565022421367867 | 3.15495382680009 | 0.60711263164862 | 0.547006263962935 | 0.795747397777164 | -6.01805131579343 |
| lnc-H2AFV-3 | lnc-H2AFV-3 | -0.0893799131707276 | 2.80389107333839 | -0.606960080634422 | 0.547106553042891 | 0.795837044613556 | -5.98114828748225 |
| ENSG00000100299.17 | ARSA | 0.0849415922548662 | 4.22365338051196 | 0.606819635726663 | 0.547198891737584 | 0.795915114736393 | -6.14232674650466 |
| ENSG00000111247.14 | RAD51AP1 | 0.148755485045085 | 1.66575022267965 | 0.606518734924143 | 0.547396752686741 | 0.79610695385099 | -5.83466123543774 |
| lnc-SYF2-7 | lnc-SYF2-7 | 0.120781202195855 | 1.53029937286174 | 0.606501421198184 | 0.547408138650977 | 0.79610695385099 | -5.82702643388437 |
| ENSG00000268797.1 | | -0.100860388772672 | 0.734266382442931 | -0.606252378001165 | 0.54757192946881 | 0.796288135204875 | -5.68757146341017 |
| LIMD1-AS1 | LIMD1-AS1 | -0.142332244512964 | 0.242721948640986 | -0.606194355296342 | 0.547610093466696 | 0.796288135204875 | -5.57526018342731 |
| ENSG00000182389.19 | CACNB4 | 0.188121355302576 | 0.104719788844577 | 0.606061389838274 | 0.54769755563449 | 0.796359055356365 | -5.54651075533823 |
| ENSG00000146112.11 | PPP1R18 | -0.146318058581867 | 3.90919436724383 | -0.605924895885446 | 0.547788330119463 | 0.796434781298927 | -6.11252814365319 |
| ENSG00000108468.14 | CBX1 | -0.0998370032612028 | 4.13738713172443 | -0.605820635682632 | 0.547855937371611 | 0.796476815858163 | -6.13170832092243 |
| ENSG00000264943.1 | SH3GL1P2 | 0.172701802677109 | -0.176404532302969 | 0.60561918527295 | 0.547988480890931 | 0.796516166291575 | -5.48531774957745 |
| ENSG00000025770.18 | NCAPH2 | 0.0753159344608534 | 4.95597905536771 | 0.605598569474895 | 0.548002045901359 | 0.796516166291575 | -6.19736887693144 |
| ENSG00000172062.16 | SMN1 | 0.0941536264193914 | 2.08712781702169 | 0.605564140770753 | 0.548024700063403 | 0.796516166291575 | -5.90022002546857 |
| ENSG00000244486.8 | SCARF2 | -0.154344885978496 | -0.129216225540857 | -0.605544239123551 | 0.54803779560906 | 0.796516166291575 | -5.48659879208235 |
| NARF-IT1 | NARF-IT1 | 0.07273682821262 | 1.00550985798126 | 0.605391647264774 | 0.548138208380599 | 0.796605856322663 | -5.74410611637788 |
| ENSG00000100714.15 | MTHFD1 | -0.0954332066392043 | 4.13675450575491 | -0.605306549725142 | 0.548194210737388 | 0.796630996865832 | -6.13057037546769 |
| lnc-MCM6-1 | lnc-MCM6-1 | 0.0823646007007679 | 2.61985631624327 | 0.605240992112593 | 0.548237355941177 | 0.796637451276897 | -5.95596919196248 |
| ENSG00000264773.1 | MIR4420 | 0.174155725464793 | 2.81146136370812 | 0.604992285469387 | 0.548404145691871 | 0.796773987791101 | -5.95635144196814 |
| ENSG00000269590.1 | | 0.0989267638033002 | 2.69103410444325 | 0.604980606919981 | 0.548408739495532 | 0.796773987791101 | -5.96381027768049 |
| lnc-TNRC6A-3 | lnc-TNRC6A-3 | -0.127985030144545 | 2.71928219076431 | -0.604881965573993 | 0.548473671635835 | 0.796812082388509 | -5.98179067998155 |
| ENSG00000012779.10 | ALOX5 | 0.0740907205267618 | 8.12902506637736 | 0.604774966417576 | 0.548544109875262 | 0.79685817019991 | -6.20814981515236 |
| ENSG00000177455.12 | CD19 | -0.0954590816781868 | 7.29781330746498 | -0.604676407208861 | 0.548608996135099 | 0.796896186834774 | -6.22093028867962 |
| lnc-PTGER3-7 | lnc-PTGER3-7 | -0.107212527049763 | 5.29320609324722 | -0.604609509347795 | 0.548653040445281 | 0.796903925727558 | -6.21601209817089 |
| lnc-TMEM150A-2 | lnc-TMEM150A-2 | -0.0913897225420771 | 0.69211299696344 | -0.604533787936753 | 0.548702896198275 | 0.796904396661649 | -5.69836190356257 |
| lnc-PLSCR3-1 | lnc-PLSCR3-1 | 0.132238315256719 | -0.200183249706407 | 0.604491412038801 | 0.548730797934487 | 0.796904396661649 | -5.43489309335448 |
| ENSG00000200840.1 | RNU6-82P | 0.204613622731517 | 0.378433161114987 | 0.604297640937412 | 0.548858392675957 | 0.796979787594534 | -5.60977631293228 |
| ENSG00000138031.14 | ADCY3 | 0.141978784048698 | 3.75913353216764 | 0.604248919566785 | 0.548890477196935 | 0.796979787594534 | -6.07900599866797 |
| lnc-SAG-6 | lnc-SAG-6 | -0.144222936868656 | 2.14298703415324 | -0.604236173021481 | 0.548898871347234 | 0.796979787594534 | -5.90221972220636 |
| ENSG00000118482.11 | PHF3 | 0.060850196736153 | 8.41715969557978 | 0.604068014022254 | 0.549009617448781 | 0.797051073953891 | -6.20147931870575 |
| ENSG00000120688.8 | WBP4 | -0.0623618316199805 | 3.52992386579121 | -0.604044027424859 | 0.549025415465105 | 0.797051073953891 | -6.07161364328893 |
| ENSG00000148153.13 | INIP | -0.0954383566039643 | 3.31937883173931 | -0.603880306174476 | 0.549133251509641 | 0.797107069331165 | -6.04920905080458 |
| ENSG00000176371.13 | ZSCAN2 | -0.106890912387488 | 3.17954354806458 | -0.603867875942569 | 0.549141439203027 | 0.797107069331165 | -6.03249011857221 |
| lnc-CASTOR3-2 | lnc-CASTOR3-2 | -0.159954254191032 | -0.487832480117988 | -0.603562042780366 | 0.549342908680756 | 0.79730507069039 | -5.36489853527484 |
| ENSG00000198498.9 | TMA16 | -0.102064028797644 | 3.50794512084693 | -0.603530904351995 | 0.549363423429369 | 0.79730507069039 | -6.0708565213122 |
| ENSG00000107758.15 | PPP3CB | 0.0454983844287495 | 4.8508209255569 | 0.603365413930968 | 0.54947245908043 | 0.79730507069039 | -6.19128546007667 |
| ENSG00000269514.2 | | -0.169526327717164 | 0.0298020604127294 | -0.603291867157488 | 0.549520919935151 | 0.79730507069039 | -5.50268035147025 |
| lnc-DNAJB6-3 | lnc-DNAJB6-3 | -0.119532275057656 | 1.89237727228031 | -0.603283700067624 | 0.549526301463631 | 0.79730507069039 | -5.87037516855102 |
| ENSG00000087074.7 | PPP1R15A | -0.192625020228802 | 7.02179701644397 | -0.60325461536752 | 0.549616671081733 | 0.79730507069039 | -6.22626054548196 |
| ENSG00000136144.11 | RCBTB1 | -0.131715485666132 | 5.14485633109925 | -0.602951902555381 | 0.54974576969079 | 0.79730507069039 | -6.20746610053368 |
| lnc-KIF5B-6 | lnc-KIF5B-6 | -0.0892764938029287 | 1.45750441803371 | -0.602904733983753 | 0.549776042615708 | 0.79730507069039 | -5.81568659065056 |
| lnc-LTBP3-11 | lnc-LTBP3-11 | -0.254870126195816 | -0.415133954454065 | -0.602711946392231 | 0.549903113138504 | 0.79730507069039 | -5.37639194128082 |
| ENSG00000139625.12 | MAP3K12 | -0.130654825628032 | 2.29143092146441 | -0.602705319472373 | 0.549907481352906 | 0.79730507069039 | -5.90900189655429 |
| ENSG00000125812.15 | GZF1 | 0.107605031125834 | 4.09647049247879 | 0.602697325914253 | 0.549912750427014 | 0.79730507069039 | -6.12754561629101 |
| ENSG00000204348.9 | DXO | -0.0879598232069747 | 3.80248218596577 | -0.602669005113542 | 0.549931418716247 | 0.79730507069039 | -6.10406995753955 |
| lnc-C13orf42-2 | lnc-C13orf42-2 | 0.132119695825272 | 0.27844838565594 | 0.602640933781334 | 0.549949922881844 | 0.79730507069039 | -5.62945309207811 |
| ENSG00000081177.18 | EXD2 | 0.148819463964323 | 2.408875822621 | 0.602620430183186 | 0.549963438721007 | 0.79730507069039 | -5.92507899700658 |
| ENSG00000197969.12 | VPS13A | 0.139632697992932 | 6.03219339716662 | 0.602622001469719 | 0.549974458907614 | 0.79730507069039 | -6.22978510467641 |
| lnc-C16orf74-1 | lnc-C16orf74-1 | -0.188659750912039 | 1.13769013806063 | -0.6024999322763 | 0.55004287358585 | 0.79730507069039 | -5.76779314858731 |
| ENSG00000119383.19 | PTPA | -0.0813995913356873 | 4.38214535792036 | -0.602458796197726 | 0.550069992731815 | 0.79730507069039 | -6.15981846711569 |
| lnc-SPATA45-3 | lnc-SPATA45-3 | 0.0857870987702096 | 1.53041223019385 | 0.602416877039624 | 0.550097628828188 | 0.79730507069039 | -5.82301043606413 |
| lnc-ORMDL2-1 | lnc-ORMDL2-1 | -0.101962952585938 | 2.5818187495806 | -0.602406643705278 | 0.550104375479021 | 0.79730507069039 | -5.95283129900716 |
| ENSG00000054967.12 | RELT | 0.105771161751729 | 5.06831402794575 | 0.602397239468195 | 0.550110575558356 | 0.79730507069039 | -6.19980303567392 |
| lnc-ENO1-4 | lnc-ENO1-4 | 0.111515254313755 | 2.12952824513303 | 0.602383573033384 | 0.550119585707384 | 0.79730507069039 | -5.89085242108756 |
| MIR9-3HG | MIR9-3HG | -0.879206033541836 | 1.29124552185466 | -0.602560429415541 | 0.550130040030361 | 0.79730507069039 | -5.6350487482445 |
| ENSG00000258732.1 | | -1.19890405771246 | -0.920418602842846 | -0.602379013860646 | 0.550233403162471 | 0.79739872824707 | -4.99449867745329 |
| ENSG00000022277.12 | RTF2 | 0.0550767878943699 | 5.50540506924775 | 0.602138515384923 | 0.550281162634239 | 0.797408236175956 | -6.21994249175069 |
| lnc-KCNE5-2 | lnc-KCNE5-2 | 0.150433785905937 | 0.00294462966089413 | 0.601995944544788 | 0.550375176785695 | 0.797408236175956 | -5.52574395347159 |
| lnc-UCP3-6 | lnc-UCP3-6 | 0.0845402733166966 | 0.322421245032932 | 0.601992989097056 | 0.550377125755508 | 0.797408236175956 | -5.57484731545778 |
| ENSG00000281896.1 | | 0.120886931836601 | 2.54866584395326 | 0.601893472664675 | 0.550442753914533 | 0.797408236175956 | -5.94394771241115 |

| | | | | | | | |
|---|---|---|---|---|---|---|---|
| ENSG00000099219.13 | ERMP1 | -0.0686036129008132 | 4.00413417638561 | -0.601844405958174 | 0.55047511343226 | 0.797408236175956 | -6.11502056789262 |
| lnc-BCL2L15-3 | lnc-BCL2L15-3 | 0.124824734196031 | 3.74040509735265 | 0.6018423310793 | 0.550476481837299 | 0.797408236175956 | -6.08093105919533 |
| lnc-CCDC85A-7 | lnc-CCDC85A-7 | 0.188165558636464 | -0.344050510523511 | 0.601784974292121 | 0.550514309944813 | 0.797408236175956 | -5.41174467849623 |
| ENSG00000162076.12 | FLYWCH2 | 0.105378183534955 | 1.33061112162703 | 0.601722542907255 | 0.550555486371273 | 0.797408236175956 | -5.78762927678988 |
| ENSG00000244560.7 | | -0.070378340269293 | 3.28240256284269 | -0.601663891781942 | 0.550594170969451 | 0.797408236175956 | -6.04191249786383 |
| ENSG00000011451.19 | WIZ | -0.0886142688992142 | 3.25647140442218 | -0.601613551535312 | 0.550627375055573 | 0.797408236175956 | -6.03800712665078 |
| ENSG00000183648.9 | NDUFB1 | -0.0703039291403176 | 3.57186777358064 | -0.601499571774458 | 0.550702559106103 | 0.797434861123271 | -6.06944050003949 |
| lnc-COIL-8 | lnc-COIL-8 | 0.242693382308026 | -0.82123502431547 | 0.601468213218109 | 0.550723244952986 | 0.797434861123271 | -5.28873816243768 |
| lnc-HNRNPD-2 | lnc-HNRNPD-2 | -0.0705626047031647 | 4.9684586152537 | -0.601331643292788 | 0.550813338684471 | 0.797509211461907 | -6.20273911182727 |
| ENSG00000255330.9 | | -0.566168839571593 | -0.305048057754217 | -0.601267265433139 | 0.550959055171312 | 0.797594150643452 | -5.49131935768322 |
| ENSG00000144635.8 | DYNC1LI1 | -0.0447863547681866 | 4.81974475247897 | -0.601110490247596 | 0.550959246928561 | 0.797594150643452 | -6.19357500852169 |
| lnc-SCGB2B2-8 | lnc-SCGB2B2-8 | -0.110227363213525 | -0.101179141294876 | -0.601066528101677 | 0.550988253801644 | 0.797594150643452 | -5.47101877636246 |
| ENSG00000167861.15 | HID1 | -0.312926801381843 | 0.100360597421161 | -0.601002258732462 | 0.551075984980222 | 0.797665049145423 | -5.64200386136128 |
| lnc-XKR5-10 | lnc-XKR5-10 | -0.188804792338513 | 2.87570450755667 | -0.600892659137408 | 0.551121686192484 | 0.797675104908227 | -5.97578975785123 |
| lnc-MSTN-2 | lnc-MSTN-2 | 0.16781671727982 | 2.32356750711025 | 0.600666527110051 | 0.551252216098372 | 0.797751091295262 | -5.91019632688179 |
| ENSG00000120451.10 | SNX19 | -0.075584451062074 | 6.50390172327144 | -0.600647302753462 | 0.551264903951739 | 0.797751091295262 | -6.23161181364985 |
| ENSG00000070610.14 | GBA2 | -0.0580215161552404 | 5.71924708723652 | -0.60057518859574 | 0.55131249978816 | 0.797751091295262 | -6.22784956570186 |
| ENSG00000176124.12 | DLEU1 | -0.410852915939756 | 2.20515022052792 | -0.600726068634872 | 0.551329217007973 | 0.797751091295262 | -5.92439929890726 |
| ENSG00000170385.9 | SLC30A1 | -0.104880110928768 | 3.61152043850788 | -0.600332043067835 | 0.551472992914782 | 0.797855828155927 | -6.09953247985113 |
| lnc-CD47-5 | lnc-CD47-5 | 0.107196392824851 | 5.40369890196972 | 0.600314931292979 | 0.551484288783766 | 0.797855828155927 | -6.21936780478334 |
| ENSG00000182831.11 | HAPSTR1 | 0.0636869652504437 | 8.15002159315464 | 0.600264031373176 | 0.551517889666352 | 0.797855828155927 | -6.2079443492889 |
| ENSG00000197256.10 | KANK2 | 0.629447021561405 | 1.363963565499 | 0.600329995912261 | 0.551600844991698 | 0.797869788309846 | -5.78786985118112 |
| ENSG00000164951.15 | PDP1 | 0.0951132230232776 | 6.33608449967766 | 0.600078815030949 | 0.551640166483118 | 0.797869788309846 | -6.23226962333629 |
| lnc-MFSD14B-7 | lnc-MFSD14B-7 | 0.17172702195484 | -0.107964242423644 | 0.600051631882801 | 0.551658113516278 | 0.797869788309846 | -5.45016661547971 |
| lnc-NFAM1-3 | lnc-NFAM1-3 | 0.112321316082991 | 3.61321301579846 | 0.600014554061384 | 0.551682593753376 | 0.797869788309846 | -6.0715113879054 |
| lnc-RFC4-1 | lnc-RFC4-1 | -0.164840429759956 | -0.521513252070988 | -0.599888182318981 | 0.551766033504116 | 0.797934396656729 | -5.41359784787619 |
| ENSG00000197124.11 | ZNF682 | -0.213610347406185 | 3.13847409777612 | -0.599801395060174 | 0.551874607574987 | 0.797995124788513 | -6.01909362220641 |
| lnc-APIP-1 | lnc-APIP-1 | -0.310248602193261 | -0.270543907202333 | -0.599707164350467 | 0.551885565824442 | 0.797995124788513 | -5.40468847102164 |
| ENSG00000170234.12 | PWWP2A | -0.0575055591171285 | 4.70008244187636 | -0.599301055256599 | 0.552153781385116 | 0.798326867255538 | -6.18386864211371 |
| ENSG00000104691.14 | UBXN8 | -0.108440971179385 | 1.24196649638429 | -0.599024965997461 | 0.552336162925346 | 0.798534469445277 | -5.78227439524971 |
| lnc-GPR3-2 | lnc-GPR3-2 | 0.442637465426302 | -1.32121022013946 | 0.598938047700775 | 0.552394929325195 | 0.798546945983297 | -5.16217874438792 |
| SOS1-IT1 | SOS1-IT1 | -0.0990027002119352 | 0.930018863081922 | -0.598894458129055 | 0.552422385651079 | 0.798546945983297 | -5.72768852736074 |
| ENSG00000006652.13 | IFRD1 | 0.117587088277034 | 5.54695637494359 | 0.598820182062459 | 0.552471460726938 | 0.798561803099899 | -6.22569655358215 |
| MIF-AS1 | MIF-AS1 | -0.695316246618853 | 1.09924783998129 | -0.598674988653421 | 0.552688605637215 | 0.798669059022235 | -5.55050829765382 |
| ENSG00000283321.1 | | 0.731557183426323 | -0.954945754784352 | 0.598592299291404 | 0.552711183811294 | 0.798669059022235 | -5.2069234660685 |
| ENSG00000126861.4 | OMG | 0.160116142130765 | -0.280289316458927 | 0.598447897914381 | 0.55271746664475 | 0.798669059022235 | -5.46268605718643 |
| ENSG00000111885.6 | MAN1A1 | 0.192680204436457 | 6.89596185698049 | 0.598446376150742 | 0.552789464155936 | 0.798669059022235 | -6.23006196585323 |
| lnc-SHISA5-1 | lnc-SHISA5-1 | -0.0907606512852817 | 3.90683130216666 | -0.598311179105187 | 0.552807824605577 | 0.798669059022235 | -6.1111079746068 |
| lnc-BCL7B-2 | lnc-BCL7B-2 | -0.105054258081766 | -0.0169583440761422 | -0.598297469826258 | 0.552816885532019 | 0.798669059022235 | -5.51560084949197 |
| ENSG00000049192.14 | ADAMTS6 | -0.128581056301416 | 6.01498785558424 | -0.598299062447697 | 0.552817280468823 | 0.798669059022235 | -6.23217205589424 |
| ENSG00000284976.1 | | 0.104303791454638 | 2.94473601903155 | 0.598179850507004 | 0.552894627220761 | 0.798724741162614 | -6.00627742992308 |
| ENSG00000113761.11 | ZNF346 | 0.0535076445917095 | 4.159103154239 | 0.598031814984101 | 0.552992480700522 | 0.798810037915562 | -6.13693923736536 |
| ENSG00000145214.13 | DGKQ | -0.156339529747957 | 4.36511225300357 | -0.598003900476912 | 0.553032210432491 | 0.798811367532067 | -6.16682329890685 |
| ENSG00000141446.10 | ESCO1 | 0.0576369020902749 | 5.66012549915704 | 0.597740071623407 | 0.553185352741175 | 0.798946231078941 | -6.22689753967585 |
| ENSG00000152457.17 | DCLRE1C | -0.0772984942716217 | 5.59982157757994 | -0.597700768257986 | 0.553211338873896 | 0.798946231078941 | -6.22425041409819 |
| ENSG00000135953.10 | MFSD9 | -0.0647199897107815 | 3.29603021333971 | -0.597654355201347 | 0.553242026505764 | 0.798946231078941 | -6.04550362915683 |
| lnc-PTGER1-1 | lnc-PTGER1-1 | 0.0862763171749812 | 3.06636050907609 | 0.597409241210487 | 0.553404106608644 | 0.799081299657942 | -6.02092304687885 |
| lnc-KMT5A-20 | lnc-KMT5A-20 | -0.0781273891753482 | 1.10167788241858 | -0.597395488060793 | 0.553413201507262 | 0.799081299657942 | -5.76470021555943 |
| ENSG00000157617.16 | C2CD2 | -0.100708667678384 | 3.76767528244093 | -0.59733004035536 | 0.553456482832025 | 0.799087737523259 | -6.09858376881263 |
| ENSG00000107874.10 | CUEDC2 | -0.0957207516212009 | 3.36576710507645 | -0.597075849433939 | 0.553624598516774 | 0.799122325446749 | -6.04690377208667 |
| lnc-CHRNA10-1 | lnc-CHRNA10-1 | -0.0763131750206205 | 1.26287924657037 | -0.597011508195012 | 0.553667156353155 | 0.799122325446749 | -5.78479987678027 |
| ENSG00000177600.8 | RPLP2 | 0.0907333915905451 | 8.53133114076757 | 0.596938408252926 | 0.553715509555229 | 0.799122325446749 | -6.20094731250713 |
| ENSG00000108179.13 | PPIF | 0.108861030834101 | 3.85528897644811 | 0.596924641257057 | 0.55372461620885 | 0.799122325446749 | -6.11544262512951 |
| lnc-SNX2-1 | lnc-SNX2-1 | -0.132996800551534 | 2.02983982103139 | -0.596907859896771 | 0.553735716920375 | 0.799122325446749 | -5.88549027528951 |
| ENSG00000131398.13 | KCNC3 | 0.202157022280432 | 2.33652091590949 | 0.596932557680512 | 0.553738280626588 | 0.799122325446749 | -5.91191547994271 |
| lnc-TSTD1-2 | lnc-TSTD1-2 | -0.184030804378152 | -0.370758873250681 | -0.596867484736977 | 0.553762425170145 | 0.799122325446749 | -5.37886538032898 |
| ENSG00000135457.9 | TFCP2 | 0.0844440659619767 | 3.91302019721232 | 0.596798673021417 | 0.553807945764527 | 0.799122325446749 | -6.10801446540556 |
| ENSG00000149054.15 | ZNF215 | -0.524876384350698 | -1.59984688662683 | -0.596870252985228 | 0.553829858256711 | 0.799122325446749 | -5.31215806433702 |
| ENSG00000136895.18 | GARNL3 | 0.146986404293128 | 3.34796023113045 | 0.596498041930563 | 0.554006842668228 | 0.799161184653164 | -6.05172558360429 |
| ENSG00000100219.16 | XBP1 | -0.178699380975954 | 5.61606306236273 | -0.59654567456033 | 0.554036223925134 | 0.799161184653164 | -6.22881204647528 |
| ENSG00000129351.17 | ILF3 | 0.0536166607310289 | 8.0903456151168 | 0.596403045473332 | 0.554069699655174 | 0.799161184653164 | -6.21251694230367 |
| ENSG00000128050.8 | PAICS | -0.105372271305808 | 4.36866472810197 | -0.5963853234268 | 0.554081426328865 | 0.799161184653164 | -6.16705888981019 |
| lnc-STAU2-3 | lnc-STAU2-3 | -0.121629423355865 | 1.2649605273401 | -0.59637915845559 | 0.554085505718243 | 0.799161184653164 | -5.79366395374728 |
| ENSG00000204568.11 | MRPS18B | -0.114104262576292 | 3.86169198229749 | -0.596372748578648 | 0.554089747178997 | 0.799161184653164 | -6.11496770869759 |
| lnc-HEXA-5 | lnc-HEXA-5 | 0.080641692842454 | 1.91191287371667 | 0.596206178447923 | 0.554199973578989 | 0.799264157523566 | -5.87483343740362 |
| lnc-ABHD17B-2 | lnc-ABHD17B-2 | -0.202755945659569 | 3.35044252920647 | -0.596074266613138 | 0.554335364993995 | 0.799366692118236 | -6.0523970654956 |
| lnc-MGARP-1 | lnc-MGARP-1 | 0.0873732501150044 | 1.6092984854507 | 0.595976027414953 | 0.554352292395808 | 0.799366692118236 | -5.83794370035953 |
| lnc-EIF1AX-4 | lnc-EIF1AX-4 | 0.211140203420545 | 0.105166954040116 | 0.595922715181961 | 0.554387578583677 | 0.799366692118236 | -5.48863811822731 |

| | | | | | | | |
|---|---|---|---|---|---|---|---|
| lnc-SESN3-1 | lnc-SESN3-1 | -0.170294580462838 | -0.232707932548708 | -0.595591903949524 | 0.554606560657124 | 0.799626424629139 | -5.44328127234106 |
| ENSG00000174373.16 | RALGAPA1 | 0.0661413953272607 | 6.74444657719833 | 0.595331170854979 | 0.55477918503852 | 0.799813931697455 | -6.23355780721535 |
| ENSG00000170509.11 | HSD17B13 | -0.127374073148252 | 0.90233100123979 | -0.595278095448335 | 0.554814328172582 | 0.799813931697455 | -5.74959042991443 |
| ENSG00000136319.11 | TTC5 | -0.0573261923436066 | 4.40952063960935 | -0.595097934478514 | 0.554933627630728 | 0.799929887073554 | -6.16310700666206 |
| ENSG00000095787.21 | WAC | 0.0602773608756051 | 7.84735790715858 | 0.594840152661331 | 0.555104348842345 | 0.80005568961922 | -6.21787053292309 |
| ENSG00000213906.9 | LTB4R2 | -0.0689871045605392 | 2.5326807874838 | -0.594776313125922 | 0.555146631969086 | 0.80005568961922 | -5.95430553222694 |
| ENSG00000112290.12 | WASF1 | 0.152413447677609 | 1.93292269282762 | 0.594743607455124 | 0.555168294694535 | 0.80005568961922 | -5.8847549477947 |
| lnc-CBR3-3 | lnc-CBR3-3 | -0.190404403245683 | -0.234248346863983 | -0.594729588870662 | 0.555177580088236 | 0.80005568961922 | -5.42544375950953 |
| ENSG00000105011.8 | ASF1B | 0.188850444661402 | 0.138234925217606 | 0.594672719578915 | 0.555215249017195 | 0.80005568961922 | -5.53406408590514 |
| lnc-TMEM63A-2 | lnc-TMEM63A-2 | -0.223897893761111 | -0.698709713860194 | -0.594330474956114 | 0.555441971398303 | 0.80019378699126 | -5.27102896656872 |
| ENSG00000115866.10 | DARS1 | 0.0627700265519691 | 5.40598651915165 | 0.594326064485112 | 0.555444893451734 | 0.80019378699126 | -6.2227767091416 |
| lnc-PUS7-3 | lnc-PUS7-3 | -0.204278267301746 | -0.760591359596781 | -0.594320994657418 | 0.555448252355979 | 0.80019378699126 | -5.32359598211763 |
| lnc-ZNF507-7 | lnc-ZNF507-7 | 0.119640093054016 | 1.56079338496712 | 0.594293316168715 | 0.555466590318764 | 0.80019378699126 | -5.83252634383977 |
| ENSG00000127774.6 | EMC6 | -0.108897040091674 | 1.87725874455239 | -0.594174248930678 | 0.555545480002653 | 0.800251425214823 | -5.87741078407787 |
| ENSG00000164649.19 | CDCA7L | 0.121043926882886 | 6.47108473693651 | 0.593830864334109 | 0.55577302605592 | 0.800474971960826 | -6.23577844125782 |
| ENSG00000174915.11 | PTDSS2 | 0.0908430144736361 | 2.99713759978344 | 0.593822680934285 | 0.555778449414184 | 0.800474971960826 | -6.00954419579884 |
| ENSG00000175756.13 | AURKAIP1 | 0.0743077114115924 | 3.07773867677296 | 0.593519819192104 | 0.555979182817877 | 0.80070805485169 | -6.01962744232287 |
| ENSG00000172053.17 | QARS1 | -0.0673504881211366 | 6.31545544667342 | -0.593312187760659 | 0.556116819793537 | 0.800807435428035 | -6.23612976833566 |
| lnc-EOMES-6 | lnc-EOMES-6 | 0.0783013862937676 | 0.174757315751383 | 0.593229987583228 | 0.556171314295257 | 0.800807435428035 | -5.56816417941522 |
| ENSG00000285920.1 | | -0.0774600692629357 | 3.08592641551019 | -0.59317462132523 | 0.556208020805086 | 0.800807435428035 | -6.02188654991435 |
| ENSG00000255835.1 | | -0.0737647677752014 | 3.05744717375891 | -0.593158446288344 | 0.556218744700122 | 0.800807435428035 | -6.01755985615885 |
| lnc-ZNF491-2 | lnc-ZNF491-2 | -0.0933831846917674 | 0.968912429736853 | -0.593122284603397 | 0.556242719929778 | 0.800807435428035 | -5.74206207210026 |
| lnc-IGFBP4-6 | lnc-IGFBP4-6 | -0.322864343540479 | -0.734461813342709 | -0.592960272626541 | 0.556350140424272 | 0.800854402672332 | -5.25777452419751 |
| ENSG00000163354.14 | DCST2 | -0.141760286325955 | 0.190364045770514 | -0.592860064074939 | 0.55641658799167 | 0.800854402672332 | -5.5849243001808 |
| ENSG00000161328.10 | LRRC56 | 0.159315492610914 | 2.6533771287395 | 0.592845964281518 | 0.55642593778455 | 0.800854402672332 | -5.96322649613785 |
| ENSG00000100100.12 | PIK3IP1 | 0.146778296204265 | 7.56608230603047 | 0.592871438820206 | 0.556430977632854 | 0.800854402672332 | -6.22581476562378 |
| lnc-GSAP-5 | lnc-GSAP-5 | -0.120160220169543 | 1.18130673339399 | -0.592732288124845 | 0.556501321114835 | 0.800899642882579 | -5.78480444536425 |
| ENSG00000078246.16 | TULP3 | -0.0708158748644832 | 3.30186170017478 | -0.592259142616634 | 0.556815138730801 | 0.801295252778863 | -6.0470134203035 |
| ENSG00000076242.14 | MLH1 | 0.0520643071847884 | 5.40220437651338 | 0.592137813198077 | 0.556895625841902 | 0.801345359997583 | -6.2229566427693 |
| ENSG00000117528.12 | ABCD3 | 0.0630326508218881 | 3.81902323066547 | 0.592089280863441 | 0.55692782270638 | 0.801345359997583 | -6.10539004189963 |
| lnc-NOX3-9 | lnc-NOX3-9 | -0.324906916110334 | 6.46590362099099 | -0.592161344904659 | 0.556999688767874 | 0.801392743877335 | -6.23612621799787 |
| lnc-PCBP1-12 | lnc-PCBP1-12 | -0.199080073304563 | 0.713465731782747 | -0.591896042999473 | 0.55705602806495 | 0.801402733279855 | -5.6892766301039 |
| lnc-NOS1-2 | lnc-NOS1-2 | -0.353812239773108 | 2.62471257904467 | -0.591988643792865 | 0.55710374851306 | 0.801402733279855 | -5.95957535374702 |
| ENSG00000164440.14 | TXLNB | -0.119839915264679 | 4.10585087959403 | -0.591794449211696 | 0.557123437326991 | 0.801402733279855 | -6.13495990095459 |
| lnc-CYP2C18-1 | lnc-CYP2C18-1 | -0.131168816512512 | 0.30088477528406 | -0.591688658073432 | 0.55719363597767 | 0.801447701560335 | -5.59940742393782 |
| ENSG00000279977.1 | | 0.129406306716203 | 1.36323606634371 | 0.591569717053079 | 0.557272565676639 | 0.80145008986256 | -5.78956147311204 |
| lnc-EFEMP1-8 | lnc-EFEMP1-8 | -0.10558288415425 | 0.730035789584946 | -0.591568804366683 | 0.55727317136049 | 0.80145008986256 | -5.70184575159053 |
| ENSG00000251992.1 | SCARNA17 | -0.0898944891512548 | 3.60896634106883 | -0.591485918192531 | 0.557328178294484 | 0.801473198758846 | -6.08508662214978 |
| lnc-SLTM-2 | lnc-SLTM-2 | 0.0975485739580191 | 1.83338292103862 | 0.59130463957613 | 0.557448492349628 | 0.801590213639261 | -5.87523807160789 |
| ENSG00000188886.3 | ASTL | -0.199601284638509 | -0.46916749104295 | -0.591075018620802 | 0.557600909860716 | 0.80168466781927 | -5.38435203053916 |
| ENSG00000167130.17 | DOLPP1 | -0.120507944622655 | 1.10678665428896 | -0.591066696059444 | 0.557606434594541 | 0.80168466781927 | -5.77885313539379 |
| lnc-ABR-2 | lnc-ABR-2 | -0.141334595502969 | 1.29963499880545 | -0.591029653004766 | 0.557631025077418 | 0.80168466781927 | -5.81004334913899 |
| ENSG00000110274.15 | CEP164 | -0.0871293789797853 | 5.19024597018846 | -0.590919563562747 | 0.557704109547765 | 0.80173373982551 | -6.21584224941576 |
| lnc-TARBP1-1 | lnc-TARBP1-1 | -0.0832541601286416 | 2.0501570118292 | -0.590794492947296 | 0.55778714534378 | 0.801797109617363 | -5.9007543715205 |
| ENSG00000227376.1 | FTH1P16 | 0.145226332962586 | -0.0641466301526714 | 0.590690809421952 | 0.55785598673121 | 0.801840068078805 | -5.4665500411772 |
| lnc-CHML-1 | lnc-CHML-1 | -0.150293974438013 | 2.89251766283727 | -0.590479689188408 | 0.557996174688363 | 0.801938930612789 | -6.00768327134826 |
| ENSG00000196411.9 | EPHB4 | -0.162448618791776 | 0.868734795803527 | -0.590446211627047 | 0.55801840607268 | 0.801938930612789 | -5.7237910887047 |
| ENSG00000087269.15 | NOP14 | 0.0846353054391527 | 5.06786659454184 | 0.590411208474336 | 0.558041651030801 | 0.801938930612789 | -6.21066802860175 |
| lnc-SMCO1-2 | lnc-SMCO1-2 | -0.0791541795442861 | 2.68119921867645 | -0.590303306153094 | 0.55811331007185 | 0.801985916029663 | -5.98016049299265 |
| lnc-ACTR3C-1 | lnc-ACTR3C-1 | 0.121432743004204 | 2.92975913160411 | 0.590032449339305 | 0.558293209281141 | 0.802093901595382 | -5.99480894221005 |
| ENSG00000186715.11 | MST1L | -0.29854966981829 | -0.538447662402241 | -0.589987986793142 | 0.558322743455707 | 0.802093901595382 | -5.32877508743833 |
| ENSG00000146386.7 | ABRACL | -0.191820532401823 | 2.63500075239932 | -0.5899953792024 | 0.558341042212159 | 0.802093901595382 | -5.97588419731625 |
| ENSG00000068438.14 | FTSJ1 | 0.0591743227744113 | 4.3669347579511 | 0.589955484162339 | 0.558344333773436 | 0.802093901595382 | -6.16594618448702 |
| lnc-RAB4A-1 | lnc-RAB4A-1 | -0.118182965046248 | 0.425553136614637 | -0.589891506407492 | 0.558386833101333 | 0.802098973112201 | -5.62571149608521 |
| ENSG00000164715.5 | LMTK2 | 0.0817304779718953 | 4.89478804499768 | 0.58966969225715 | 0.558534193055109 | 0.802175725903985 | -6.20273658341854 |
| lnc-C11orf24-4 | lnc-C11orf24-4 | 0.236895383441705 | -0.146074159524585 | 0.589606045372795 | 0.558576479822705 | 0.802175725903985 | -5.44193865840957 |
| lnc-ROGDI-4 | lnc-ROGDI-4 | -0.0819034276774981 | 1.80195293489872 | -0.58958173191839 | 0.55859263402214 | 0.802175725903985 | -5.86052346696742 |
| ENSG00000108932.11 | SLC16A6 | -0.36151936387675 | 0.83081416294124 | -0.589703724198207 | 0.558596156021077 | 0.802175725903985 | -5.73656218561975 |
| ENSG00000138346.14 | DNA2 | 0.0705326801934691 | 3.21462560753799 | 0.58949883693249 | 0.558647712378222 | 0.802193795443069 | -6.03388959997067 |
| lnc-FAM122C-1 | lnc-FAM122C-1 | -0.118769096895917 | 0.4990073607294 | -0.589413793498393 | 0.558704221078848 | 0.802218973314679 | -5.65024171107476 |
| ENSG00000125995.15 | ROMO1 | 0.0948459570958886 | 3.15144454194975 | 0.589342682990081 | 0.55875147399395 | 0.80223085862287 | -6.03422872673857 |
| ENSG00000005448.16 | WDR54 | -0.0867409691930918 | 2.64390072016902 | -0.589098624341695 | 0.558913666195547 | 0.802370172607447 | -5.97890218363886 |
| lnc-C1QBP-3 | lnc-C1QBP-3 | -0.072939256382512 | 3.40639956250567 | -0.589059274644644 | 0.558939818741299 | 0.802370172607447 | -6.05945602488139 |
| lnc-SEMA4D-9 | lnc-SEMA4D-9 | -0.170622737547663 | 0.0320638056480015 | -0.589020706701628 | 0.5589654523159 | 0.802370172607447 | -5.54656533247549 |
| ENSG00000186642.15 | PDE2A | -0.143480155983337 | -0.238241931858008 | -0.588705942211122 | 0.55917467766612 | 0.802614532106118 | -5.44770506640185 |
| ENSG00000285053.1 | | -0.0546966025392148 | 3.59405295759761 | -0.588586357513254 | 0.559254176463601 | 0.80267266677012 | -6.077313625793 |
| lnc-ACACB-1 | lnc-ACACB-1 | 0.485675900931402 | -1.29911489599168 | 0.58827600662972 | 0.559460521353345 | 0.802796166079591 | -5.07997131935104 |
| lnc-TK1-3 | lnc-TK1-3 | 0.10135355721079 | 3.36497995379987 | 0.588270434534183 | 0.559464226456167 | 0.802796166079591 | -6.05386363866929 |

| | | | | | | | |
|---|---|---|---|---|---|---|---|
| ENSG00000231105.1 | ECE1-AS1 | -0.118283021007819 | 1.16031777982475 | -0.588239390995047 | 0.559484868737218 | 0.802796166079591 | -5.75610718811703 |
| ENSG00000151470.12 | C4orf33 | 0.11637394815655 | 1.98175468228827 | 0.588222297920903 | 0.559496234874009 | 0.802796166079591 | -5.87528859833181 |
| ERI3-IT1 | ERI3-IT1 | 0.0655483317299741 | 1.59156302000391 | 0.587976295426845 | 0.559659828480446 | 0.8029749232966 | -5.83196644071425 |
| ENSG00000156873.15 | PHKG2 | -0.0588443935786973 | 4.05223927205155 | -0.587884768254266 | 0.559720700907707 | 0.803006286107432 | -6.13348597488742 |
| lnc-PABPC4-1 | lnc-PABPC4-1 | -0.316565470924564 | 3.32821974804478 | -0.587848131013734 | 0.559849814740892 | 0.803030686433576 | -6.03727404502979 |
| lnc-FBXL3-7 | lnc-FBXL3-7 | 0.0699249494933509 | 3.35527615833962 | 0.587652187557534 | 0.559875399457977 | 0.803030686433576 | -6.05098429775914 |
| ENSG00000153898.12 | MCOLN2 | 0.133665183646656 | 6.35682568708628 | 0.587658537653297 | 0.559880911383156 | 0.803030686433576 | -6.23955865825803 |
| ENSG00000058056.8 | USP13 | -0.0617694084635341 | 4.22925594917256 | -0.58761534224539 | 0.559899908691298 | 0.803030686433576 | -6.15252893962797 |
| ENSG00000103351.12 | CLUAP1 | 0.122214117915087 | 1.45449681537282 | 0.587565927228653 | 0.559932780046382 | 0.803030686433576 | -5.81686875568947 |
| ENSG00000118689.14 | FOXO3 | 0.157462163529309 | 4.90107142589259 | 0.587486969422979 | 0.5600170883435 | 0.803045729079045 | -6.20747895600193 |
| ENSG00000257594.3 | GALNT4 | 0.0749770315805032 | 2.71296238103638 | 0.587399339306225 | 0.560043603110046 | 0.803045729079045 | -5.98651303420078 |
| lnc-SYNM-7 | lnc-SYNM-7 | 0.150565626873304 | 0.324529689007562 | 0.587317987764207 | 0.560097726443804 | 0.803045729079045 | -5.60305077551339 |
| ENSG00000157625.15 | TAB3 | -0.0534125889160206 | 5.73552413470861 | -0.587291707214074 | 0.560115211504295 | 0.803045729079045 | -6.23508209553739 |
| ENSG00000111424.10 | VDR | -0.32228449790529 | -0.460577973706322 | -0.587256939196801 | 0.560138343894857 | 0.803045729079045 | -5.32887079447135 |
| ENSG00000108298.11 | RPL19 | 0.076632492607061 | 9.92449998784207 | 0.587067310114792 | 0.56026451926423 | 0.803121923971383 | -6.16550432532978 |
| ENSG00000167615.16 | LENG8 | -0.091479830107728 | 8.01768670648096 | -0.587003774685948 | 0.560306797648032 | 0.803121923971383 | -6.21924258133028 |
| ENSG00000166436.16 | TRIM66 | -0.11568034327619 | 4.60458700133852 | -0.586896241036828 | 0.560378357417639 | 0.803121923971383 | -6.17790254178665 |
| ENSG00000074603.18 | DPP8 | 0.0419006236107053 | 6.14627681426633 | 0.586889782182859 | 0.560382655698263 | 0.803121923971383 | -6.2396122888936 |
| lnc-PQLC1-104 | lnc-PQLC1-104 | -0.107133395875481 | 0.753757097781156 | -0.586883878309843 | 0.560386584660983 | 0.803121923971383 | -5.6884732426363 |
| lnc-RPL9-2 | lnc-RPL9-2 | 0.0749827875780809 | 2.24378064710814 | 0.586811743927398 | 0.560434590417191 | 0.803134802935114 | -5.91832503729578 |
| ENSG00000265491.4 | RNF115 | 0.0424071590153527 | 6.10898650419743 | 0.586708410740667 | 0.560503362712735 | 0.803177437671695 | -6.23961399405505 |
| ENSG00000185088.13 | RPS27L | 0.098602932624984 | 4.95963184762931 | 0.586603433801899 | 0.560573233324754 | 0.803202695425877 | -6.20519386778938 |
| lnc-MTFR2-2 | lnc-MTFR2-2 | 0.175032780794741 | -0.190057293281546 | 0.586460721027478 | 0.560668227182588 | 0.803202695425877 | -5.45726949420471 |
| lnc-CAMSAP1-3 | lnc-CAMSAP1-3 | -0.107564855140011 | -0.107513251689892 | -0.586445666762227 | 0.560678248219006 | 0.803202695425877 | -5.47490179639353 |
| lnc-DZIP1-8 | lnc-DZIP1-8 | -0.534681360333449 | -1.66757872170266 | -0.58642377958497 | 0.560693606514227 | 0.803202695425877 | -5.03045126235028 |
| lnc-SCAP-2 | lnc-SCAP-2 | -0.108250318362573 | -0.159022096822878 | -0.586388801293591 | 0.560716102151019 | 0.803202695425877 | -5.46210568885009 |
| lnc-SVIP-1 | lnc-SVIP-1 | -0.107851443117405 | 3.10578999286279 | -0.586288243213161 | 0.560783044307917 | 0.803242686220588 | -6.02431699895752 |
| ENSG00000099365.10 | STX1B | -0.353099558309894 | 0.965972927018925 | -0.58605246050584 | 0.56101576945419 | 0.803520115696583 | -5.69205780848351 |
| lnc-DPP7-2 | lnc-DPP7-2 | -0.109736505307988 | 1.51501169890992 | -0.585746451898102 | 0.561143787196058 | 0.80359784348506 | -5.82788758728925 |
| ENSG00000134575.9 | ACP2 | -0.0709275538453249 | 3.00404343566535 | -0.585739941941655 | 0.561148122451089 | 0.80359784348506 | -6.01990273388963 |
| ENSG00000094841.13 | UPRT | -0.0562888960832488 | 3.88911456353602 | -0.585528026632566 | 0.561289254973868 | 0.803712823992217 | -6.11902965089405 |
| ENSG00000108262.15 | GIT1 | -0.0860475276728998 | 4.79523821999399 | -0.585447446125784 | 0.561342925089352 | 0.803712823992217 | -6.19715819154561 |
| ENSG00000130158.13 | DOCK6 | -0.132086992979196 | 3.01446522363623 | -0.58540060568585 | 0.561374124037216 | 0.803712823992217 | -6.01815944796323 |
| lnc-SLC25A17-1 | lnc-SLC25A17-1 | -0.0868009880121638 | 4.86171250942737 | -0.585340757102196 | 0.561413988565562 | 0.803712823992217 | -6.20351652814297 |
| ENSG00000214745.2 | GOT2P6 | -0.190795214725604 | 0.829950155640649 | -0.585326253053868 | 0.561423649777005 | 0.803712823992217 | -5.70438367681101 |
| KDM5C-IT1 | KDM5C-IT1 | 0.148149341087205 | 0.722354216136422 | 0.585213464577554 | 0.561498781527085 | 0.803764477374781 | -5.69251803604818 |
| lnc-VTI1B-1 | lnc-VTI1B-1 | 0.102822107313498 | 0.593643441172429 | 0.585015629530735 | 0.561630577482552 | 0.80386911284438 | -5.68944184496434 |
| ENSG00000188820.12 | CALHM6 | 0.190400725127711 | 1.85616497646309 | 0.584987911852133 | 0.561649988428073 | 0.80386911284438 | -5.84388830990566 |
| ENSG00000134014.16 | ELP3 | -0.102707329462007 | 4.76929356001144 | -0.58483678165532 | 0.561749737683003 | 0.803909850877116 | -6.19554383761337 |
| ENSG00000129187.14 | DCTD | -0.0664360250489908 | 5.2052099356262 | -0.584826534525055 | 0.561756565377523 | 0.803909850877116 | -6.21932544057715 |
| lnc-MGMT-2 | lnc-MGMT-2 | -0.138284854927588 | 1.20929298054699 | -0.584687517555202 | 0.56184919691838 | 0.803969982528278 | -5.76157601775313 |
| ENSG00000198917.12 | SPOUT1 | 0.0539453706818493 | 4.76316024291122 | 0.584457163892166 | 0.562002705906057 | 0.803969982528278 | -6.19891306879962 |
| ASH1L-AS1 | ASH1L-AS1 | -0.0782783947036503 | 3.63141543681606 | -0.584417062993488 | 0.562029431517232 | 0.803969982528278 | -6.09544028511832 |
| ENSG00000167522.15 | ANKRD11 | -0.105096437715599 | 6.99089293631562 | -0.584365155879606 | 0.562064026433032 | 0.803969982528278 | -6.23814469938329 |
| ENSG00000150459.12 | SAP18 | 0.0540294276849327 | 5.91400835635664 | 0.584179397994087 | 0.562187838585611 | 0.803969982528278 | -6.23954631563682 |
| lnc-APOA1-4 | lnc-APOA1-4 | -0.127359045635109 | 3.41826505032756 | -0.584179070723381 | 0.562188056731543 | 0.803969982528278 | -6.06447472223118 |
| ENSG00000101363.12 | MANBAL | 0.0593187297065159 | 2.50849855658026 | 0.584162490928365 | 0.5621991082339 | 0.803969982528278 | -5.94727439119705 |
| ENSG00000177565.16 | TBL1XR1 | -0.0589446972483119 | 6.83609298611476 | -0.584134658984508 | 0.562217660263289 | 0.803969982528278 | -6.23936466764447 |
| ENSG00000124614.15 | RPS10 | 0.0895051759816559 | 7.73753170382792 | 0.584116486226579 | 0.562229773902948 | 0.803969982528278 | -6.22685099168097 |
| ENSG00000280433.1 | | 0.0914164453498825 | 2.61353407501286 | 0.584092143259258 | 0.56224600069986 | 0.803969982528278 | -5.98072921888327 |
| ENSG00000147251.15 | DOCK11 | 0.0439021439462755 | 7.89314234590247 | 0.584091411281021 | 0.562246488633412 | 0.803969982528278 | -6.22435132364502 |
| lnc-NPHS2-7 | lnc-NPHS2-7 | 0.0655288423288041 | 2.82273685392917 | 0.583970893293934 | 0.562326828294904 | 0.803969982528278 | -5.99313734805459 |
| lnc-OGFRL1-4 | lnc-OGFRL1-4 | 0.157601350535865 | 2.21730017552951 | 0.583932024578559 | 0.562352740169824 | 0.803969982528278 | -5.93915184856293 |
| ENSG00000106546.13 | AHR | 0.724461449636142 | 0.0485264509960751 | 0.584084848546636 | 0.562368640369354 | 0.803969982528278 | -5.51466128747498 |
| lnc-NSMF-2 | lnc-NSMF-2 | -0.18216887195868 | 0.66623623741805 | -0.583829057649002 | 0.562421386081065 | 0.803969982528278 | -5.66926621806039 |
| lnc-GLE1-1 | lnc-GLE1-1 | 0.0509418608945467 | 2.28302403767097 | 0.583789404364068 | 0.562447823219337 | 0.803969982528278 | -5.92852188854245 |
| ENSG00000109458.8 | GAB1 | -0.257899989068691 | 7.61337586174284 | -0.58390082362333 | 0.562477469506129 | 0.803969982528278 | -6.22833091446447 |
| ENSG00000278864.1 | | 0.148793912047648 | 1.27089877282336 | 0.583708653319909 | 0.562501662458812 | 0.803969982528278 | -5.79484275259371 |
| ENSG00000175634.14 | RPS6KB2 | -0.0557215989252535 | 4.88215975780663 | -0.583531784372238 | 0.562619595504896 | 0.804082706171175 | -6.20522699773809 |
| ENSG00000146463.11 | ZMYM4 | 0.0415815025586133 | 6.11154885057247 | 0.58335746831292 | 0.56273583842419 | 0.804114850025196 | -6.2411603294245 |
| ENSG00000104412.7 | EMC2 | -0.0770596457535015 | 4.47658535924196 | -0.583346142462125 | 0.562743391499674 | 0.804114850025196 | -6.17491162221856 |
| lnc-C3-2 | lnc-C3-2 | -0.0694647181237891 | 1.63335879236331 | -0.583316986146858 | 0.562762835733428 | 0.804114850025196 | -5.84793457702562 |
| ENSG00000277301.1 | | -0.330615362332692 | -0.635873944014897 | -0.583263727716665 | 0.562798354441188 | 0.804114850025196 | -5.28538704779036 |
| ENSG00000181396.12 | OGFOD3 | -0.0951907697411176 | 3.43818519859783 | -0.583168838312973 | 0.56286164013783 | 0.804138269849579 | -6.07586065892727 |
| ENSG00000125962.14 | ARMCX5 | 0.0540668987411701 | 4.45446029442334 | 0.583102766021298 | 0.562905708602024 | 0.804138269849579 | -6.17103056055624 |
| ENSG00000170606.14 | HSPA4 | -0.0760789071081316 | 7.45737420232278 | -0.583010296338255 | 0.562967386320111 | 0.804138269849579 | -6.23260044788449 |
| lnc-TAF5L-3 | lnc-TAF5L-3 | -0.0800569071056213 | 0.634199086741164 | -0.583004851370442 | 0.562971018244786 | 0.804138269849579 | -5.68740564000977 |
| ENSG00000101695.8 | RNF125 | -0.201763938047901 | 3.65304898501991 | -0.582963728228903 | 0.563051475095723 | 0.804143213194884 | -6.07370896403249 |

| | | | | | | | |
|---|---|---|---|---|---|---|---|
| ENSG00000207092.1 | | 0.227000164186801 | -0.413206076294385 | 0.582863582762834 | 0.563065251900427 | 0.804143213194884 | -5.35488384825093 |
| ENSG00000278527.1 | | 0.128831022052962 | 2.21709423475244 | 0.582802623151588 | 0.563105917628695 | 0.804143213194884 | -5.92943874903486 |
| ENSG00000284969.1 | | 0.187757485121107 | 2.15471036324861 | 0.582777331740843 | 0.563131299182086 | 0.804143213194884 | -5.90439102178925 |
| ENSG00000147853.16 | AK3 | 0.0646377715406388 | 5.03287037696628 | 0.582706834335356 | 0.563169820638597 | 0.804143213194884 | -6.21626032079331 |
| ENSG00000177646.18 | ACAD9 | -0.0499134702377591 | 4.78171854103342 | -0.582557685724882 | 0.563269328453462 | 0.804179022477461 | -6.19583136030845 |
| ENSG00000198692.9 | EIF1AY | -1.25373772743839 | 1.06471195576383 | -0.58273528152338 | 0.563273039258493 | 0.804179022477461 | -5.45990244881083 |
| ENSG00000213420.7 | GPC2 | 0.189846065219403 | -0.721819926291086 | 0.582330765795466 | 0.563420739974506 | 0.804334102572844 | -5.39529115403726 |
| ENSG00000133641.17 | RLIG1 | 0.0511141127937829 | 3.34724613286158 | 0.581940736294501 | 0.563681033251441 | 0.804649886081865 | -6.06415434620018 |
| lnc-U2AF1L4-2 | lnc-U2AF1L4-2 | -0.126994604647575 | 2.3062605781922 | -0.581809050531863 | 0.563768929688508 | 0.804719547834852 | -5.92942633813137 |
| ENSG00000128563.13 | PRKRIP1 | -0.0689693214207887 | 4.99017037933739 | -0.581705068915026 | 0.563838339241568 | 0.804762813716746 | -6.21004225661333 |
| ENSG00000102575.11 | ACP5 | -0.221581580626502 | 4.98701990036302 | -0.58164681859057 | 0.563963439156595 | 0.804885554580515 | -6.21305816741614 |
| lnc-PDHX-1 | lnc-PDHX-1 | 0.166784785246689 | 1.32543305056109 | 0.581266043031766 | 0.564131443713663 | 0.805021727318498 | -5.80346971273014 |
| ENSG00000092208.17 | GEMIN2 | 0.065090667880745 | 2.87832608344689 | 0.581235733271639 | 0.564151682058284 | 0.805021727318498 | -6.00745577582989 |
| ENSG00000124074.11 | ENKD1 | -0.121954826854404 | 0.500522794525235 | -0.581199037352778 | 0.564176185034705 | 0.805021727318498 | -5.68266301240011 |
| ENSG00000184005.10 | ST6GALNAC3 | 0.845450399873448 | -1.57133020951385 | 0.581175316036099 | 0.564267958315449 | 0.805096865798342 | -5.0329245477268 |
| DGCR11 | DGCR11 | 0.0928933326567227 | 2.35390154294655 | 0.580771508155391 | 0.564461698329876 | 0.805317469794402 | -5.93219339324493 |
| ENSG00000137747.15 | TMPRSS13 | -0.162505412414706 | 2.01757557067394 | -0.580671703035203 | 0.564528360726431 | 0.805356754146946 | -5.89587002069576 |
| ENSG00000086065.13 | CHMP5 | 0.0584981384145706 | 4.11218918002522 | 0.580415017095147 | 0.564699825860715 | 0.80554553438846 | -6.14297479707627 |
| lnc-RIPPLY3-5 | lnc-RIPPLY3-5 | 0.143871602367255 | 0.463669435017087 | 0.580306592690409 | 0.564772260703581 | 0.805593031327915 | -5.62892157897498 |
| lnc-SYNRG-2 | lnc-SYNRG-2 | -0.166492846291257 | -0.513989573509864 | -0.580204151085299 | 0.5648407028835 | 0.80563343219455 | -5.32478425270198 |
| lnc-DRD5-23 | lnc-DRD5-23 | 0.56795778715132 | 0.531359897397033 | 0.580287903968222 | 0.564901179778214 | 0.80563343219455 | -5.65341617607755 |
| ENSG00000157978.11 | LDLRAP1 | -0.135374583415381 | 4.43552629143567 | -0.580075707021166 | 0.564929645083861 | 0.80563343219455 | -6.17286191569686 |
| lnc-CYFIP1-1 | lnc-CYFIP1-1 | 0.105380526914125 | 0.482285796281107 | 0.580023752275145 | 0.564961239032781 | 0.80563343219455 | -5.63240355378556 |
| ENSG00000175809.5 | CBLL2 | 0.376768590748682 | 0.367549149944349 | 0.580064527848372 | 0.564996287894298 | 0.80563343219455 | -5.54427435126653 |
| lnc-CES5A-3 | lnc-CES5A-3 | -0.534541422778055 | -1.17090745303946 | -0.57989641026077 | 0.565074409468599 | 0.805650881676248 | -5.13202158925362 |
| lnc-ROPN1L-2 | lnc-ROPN1L-2 | -0.105019359684665 | 3.57179900054618 | -0.579835840844143 | 0.565086808471068 | 0.805650881676248 | -6.09694865778397 |
| ENSG00000106399.11 | RPA3 | 0.115249226652971 | 2.8174827304159 | 0.579749715805021 | 0.565144365083392 | 0.805677134403066 | -6.00052066616166 |
| lnc-HTRA4-2 | lnc-HTRA4-2 | 0.128997556999047 | 4.84222720765092 | 0.579478446153835 | 0.565326605398129 | 0.805796897362748 | -6.2061717221088 |
| ENSG00000279561.2 | | -0.410252313145023 | -0.677469938815775 | -0.579497992240135 | 0.565330866789155 | 0.805796897362748 | -5.28638521554287 |
| ENSG00000169045.17 | HNRNPH1 | 0.0609207192024819 | 9.91331212641853 | 0.579439769917736 | 0.56535152336449 | 0.805796897362748 | -6.16752661530839 |
| LINC00886 | LINC00886 | -0.154920593973572 | 2.54772190337026 | -0.579315796743894 | 0.56543439379401 | 0.805796897362748 | -5.96085549760058 |
| ENSG00000169718.17 | DUS1L | -0.0803303008194032 | 5.1525274755563 | -0.579294835571638 | 0.565448405981931 | 0.805796897362748 | -6.22312580845403 |
| ENSG00000231307.1 | RPS3P2 | 0.165115028920458 | 0.495396041743138 | 0.579183970838655 | 0.565522520048046 | 0.805796897362748 | -5.63989052263381 |
| ENSG00000230373.8 | GOLGA6L5P | -0.149280139983974 | -0.621469060584273 | -0.579181809815483 | 0.565523964759398 | 0.805796897362748 | -5.4366019632556 |
| ENSG00000222255.1 | RNU6-101P | -0.210788386712164 | -0.311596166886539 | -0.579155487162915 | 0.565541562420554 | 0.805796897362748 | -5.50304753543707 |
| ENSG00000277161.1 | PIGW | -0.111827724705911 | 2.42645266691593 | -0.579084729403247 | 0.565588867935186 | 0.805808518634313 | -5.95334800934975 |
| ENSG00000228663.1 | PSMD10P1 | -0.14322555790187 | 2.64382139862604 | -0.578943013387786 | 0.565683618926452 | 0.805887730368436 | -5.96177683740549 |
| ENSG00000134453.15 | RBM17 | 0.0477558379457834 | 6.53463595181388 | 0.578831971264365 | 0.565757866943021 | 0.805908058343683 | -6.24420220986996 |
| ENSG00000082458.11 | DLG3 | 0.130873280367284 | 2.48048120898943 | 0.578804560176717 | 0.565776196038781 | 0.805908058343683 | -5.95119950087245 |
| ENSG00000172661.18 | WASHC2C | 0.0407587162533565 | 6.09090087442121 | 0.578401920486113 | 0.566045464963011 | 0.806235817959564 | -6.24429834160938 |
| ENSG00000234773.7 | | -0.192927536106324 | 0.0250871807539583 | -0.578187360556363 | 0.566188979865805 | 0.806384429323129 | -5.5098096275224 |
| lnc-C12orf45-3 | lnc-C12orf45-3 | -0.109582626270854 | 2.65746540549659 | -0.577921271118021 | 0.566366986956526 | 0.806558109290599 | -5.98926231067958 |
| ENSG00000215018.9 | COL28A1 | -0.179344233493794 | -0.417451804524406 | -0.577870509282065 | 0.566400948496308 | 0.806558109290599 | -5.49054961786527 |
| ENSG00000285799.1 | | -0.169611744673782 | 2.94272168532624 | -0.577850548906718 | 0.566428483204373 | 0.806558109290599 | -6.03625872218304 |
| ENSG00000155393.12 | HEATR3 | -0.117709304776906 | 3.03149415181936 | -0.57767744626467 | 0.566530124013806 | 0.806629659648979 | -6.02005262256759 |
| ENSG00000050030.14 | NEXMIF | -0.804973817013404 | -0.9016437007169 | -0.577803233959811 | 0.566557109728674 | 0.806629659648979 | -5.29416708620993 |
| ENSG00000196367.13 | TRRAP | -0.116202809277304 | 7.81512623344994 | -0.577335881646566 | 0.56675869552496 | 0.806825476158989 | -6.22925578113065 |
| lnc-ARHGAP17-7 | lnc-ARHGAP17-7 | 0.153419652313645 | -0.0418759277764922 | 0.577314442714956 | 0.566773043763437 | 0.806825476158989 | -5.48375475262716 |
| lnc-ROPN1B-16 | lnc-ROPN1B-16 | -0.161804689105588 | 0.120560754760142 | -0.577179960995296 | 0.566863051232734 | 0.806897799690412 | -5.55815232273651 |
| ENSG00000139083.10 | ETV6 | 0.0897748336735216 | 6.55414880994446 | 0.57712122971551 | 0.566902361804037 | 0.806897954015109 | -6.24506844961904 |
| LINC00957 | LINC00957 | -0.181908196768449 | -0.202042704254787 | -0.577048169018575 | 0.566951265363997 | 0.806911761512041 | -5.42598124247231 |
| ENSG00000200087.1 | SNORA73B | 0.185947583737923 | 5.80233005792891 | 0.576773354374074 | 0.567202813807381 | 0.807136913027796 | -6.23817473339725 |
| ENSG00000108799.12 | EZH1 | 0.0517360491064727 | 7.26269651492277 | 0.576652835005545 | 0.567215920569899 | 0.807136913027796 | -6.23917260564312 |
| lnc-TLCD2-4 | lnc-TLCD2-4 | -0.105334520080808 | 0.558134199001159 | -0.576625279527361 | 0.567234369789542 | 0.807136913027796 | -5.66006920973445 |
| lnc-PHACTR4-3 | lnc-PHACTR4-3 | 0.186497975047814 | 5.79425708969204 | 0.576659144165384 | 0.567279633390642 | 0.807136913027796 | -6.23802997739038 |
| ENSG00000103111.14 | MON1B | 0.0618675827062605 | 5.10789022058135 | 0.576486055357346 | 0.567327589101682 | 0.807136913027796 | -6.21907179892343 |
| lnc-YDJC-1 | lnc-YDJC-1 | -0.0835995623626309 | 1.57257413606737 | -0.576460436130884 | 0.56734474360813 | 0.807136913027796 | -5.84070361303481 |
| ENSG00000104343.19 | UBE2W | 0.0871426362391414 | 4.69227976924296 | 0.576184990662939 | 0.567529196774462 | 0.807343524584197 | -6.20104227455407 |
| ENSG00000200320.1 | SNORA63 | 0.184024653712378 | 4.63233724070936 | 0.576136282200657 | 0.567620943860227 | 0.80741823686766 | -6.18107565957468 |
| ENSG00000173020.10 | GRK2 | -0.0712142819597776 | 7.29883473672343 | -0.575951482448712 | 0.567685589777692 | 0.807437162245733 | -6.23861500755652 |
| ENSG00000012963.14 | UBR7 | -0.0598710620250147 | 4.09013468136819 | -0.575910999969448 | 0.567712705243174 | 0.807437162245733 | -6.13960392098301 |
| ENSG00000154655.15 | L3MBTL4 | -0.70007480292311 | 1.8593215332647 | -0.575973278117743 | 0.567791541337165 | 0.80748049844424 | -5.91689834823593 |
| ENSG00000236047.1 | RPL13AP12 | 0.140980605019579 | 0.487156131409472 | 0.575689817127653 | 0.567860866490812 | 0.80748049844424 | -5.62776358831719 |
| lnc-FAM117B-7 | lnc-FAM117B-7 | 0.140980605019579 | 0.487156131409472 | 0.575689817127653 | 0.567860866490812 | 0.80748049844424 | -5.62776358831719 |
| lnc-CSRNP1-3 | lnc-CSRNP1-3 | 0.07928349545906 | 0.994005682803601 | 0.575588434409087 | 0.567928784990083 | 0.807488452015295 | -5.75424213746801 |
| lnc-ISCU-1 | lnc-ISCU-1 | -0.0845960721880843 | 6.56610676309118 | -0.57556434809719 | 0.567944921528709 | 0.807488452015295 | -6.2461357693076 |
| DLEU1 | DLEU1 | -0.395508278731015 | 2.24045712674569 | -0.575626931724782 | 0.568013657994646 | 0.807530399399351 | -5.94253609458252 |
| lnc-EML3-1 | lnc-EML3-1 | 0.0828748244985822 | 2.83230424497995 | 0.57519517064482 | 0.568192279069185 | 0.807647912199485 | -5.99591479554703 |

| | | | | | | | |
|---|---|---|---|---|---|---|---|
| ENSG00000164845.16 | FAM86FP | -0.148445247409377 | -0.496448186102145 | -0.575189721427605 | 0.568195930571978 | 0.807647912199485 | -5.36269741708867 |
| lnc-MASTL-3 | lnc-MASTL-3 | -0.13564294642031 | 0.325256335777859 | -0.575162708825025 | 0.568214031801037 | 0.807647912199485 | -5.63996687542858 |
| ENSG00000253816.3 | GUSBP14 | -0.122286285128866 | 2.38369968453815 | -0.57510036575281 | 0.568255809178049 | 0.807651520529746 | -5.93918789969845 |
| ENSG00000188599.17 | NPIPP1 | -0.118826994381164 | 1.83137537119834 | -0.574976722420267 | 0.568338669622737 | 0.807713514937843 | -5.88517945079992 |
| ENSG00000074964.16 | ARHGEF10L | -0.231080593281678 | 3.46307972952451 | -0.575003093555334 | 0.568395338087196 | 0.807734015816725 | -6.07926474330281 |
| ENSG00000186105.7 | LRRC70 | 0.128338982871011 | 0.667409117151784 | 0.574838092741394 | 0.568431580386982 | 0.807734015816725 | -5.69129308538676 |
| ENSG00000184368.15 | MAP7D2 | 0.467911422833806 | -1.86974380556837 | 0.574770333314282 | 0.568476996055024 | 0.807742786814894 | -5.0927727604972 |
| ENSG00000142327.12 | RNPEPL1 | -0.0850039206161082 | 5.11416046539151 | -0.574611966527588 | 0.568583148193801 | 0.80777476673881 | -6.22207380466795 |
| ENSG00000255863.2 | NA | 0.126257292930018 | 0.264902652947643 | 0.574605836921624 | 0.568587257022627 | 0.80777476673881 | -5.60306292365648 |
| ENSG00000233757.6 | ZNF892 | 0.13616921505809 | 0.851607363556151 | 0.574535180823128 | 0.568634620641191 | 0.80777476673881 | -5.69751626873345 |
| ENSG00000025708.13 | TYMP | -0.16078875093593 | 3.29345227563371 | -0.574520931661217 | 0.568658428082582 | 0.80777476673881 | -6.08575274164653 |
| lnc-TMC8-4 | lnc-TMC8-4 | 0.12474127418826 | 2.0986329910371 | 0.574444028175029 | 0.56869572680426 | 0.80777476673881 | -5.91111492129943 |
| lnc-CCDC17-5 | lnc-CCDC17-5 | -0.125765222433194 | 0.40321753158673 | -0.574373984759433 | 0.5687426841416 | 0.807785720927861 | -5.66227076659115 |
| ENSG00000234420.7 | ZNF37BP | -0.0516206343990531 | 5.89025849601058 | -0.574182046492519 | 0.568871370022693 | 0.80791274471656 | -6.24390312786219 |
| lnc-C19orf12-12 | lnc-C19orf12-12 | -0.221216457845749 | -0.850451035800999 | -0.574100661996048 | 0.568925938976302 | 0.807923635133113 | -5.29560949639837 |
| ENSG00000197841.14 | ZNF181 | -0.0908829082009932 | 4.35318160124564 | -0.57405353049526 | 0.568957542207379 | 0.807923635133113 | -6.17029214105503 |
| ENSG00000241069.1 | | 0.105879721631462 | -0.177850586886544 | 0.573951188519044 | 0.569026168879623 | 0.807947609257865 | -5.48287338794117 |
| ENSG00000168906.12 | MAT2A | 0.0740748871815367 | 6.74839890728882 | 0.573888336632901 | 0.569068317015122 | 0.807947609257865 | -6.24552119204653 |
| lnc-SIRT3-1 | lnc-SIRT3-1 | 0.0991120670605095 | 1.62459275555385 | 0.573756700522288 | 0.569156596483625 | 0.807947609257865 | -5.83953805153976 |
| lnc-KCNG4-4 | lnc-KCNG4-4 | -0.0915509323512329 | 0.3682381547384 | -0.573573566571786 | 0.569279423347184 | 0.807947609257865 | -5.62505092548101 |
| lnc-CSE1L-4 | lnc-CSE1L-4 | -0.127573231293367 | 0.21934555511222 | -0.573549923377263 | 0.569295281653899 | 0.807947609257865 | -5.59193802364274 |
| ENSG00000135469.13 | COQ10A | 0.0872233799331039 | 3.64924421254993 | 0.573535579993619 | 0.56930490236288 | 0.807947609257865 | -6.09642178290682 |
| ENSG00000128973.12 | CLN6 | -0.0680930285871764 | 2.72541479155003 | -0.573512626970408 | 0.569320298085453 | 0.807947609257865 | -5.99207267103851 |
| LINC01089 | LINC01089 | -0.095677214444482 | 5.32966577935492 | -0.573494434005639 | 0.569332501148745 | 0.807947609257865 | -6.23110454936022 |
| ENSG00000117748.9 | RPA2 | -0.0895167445193854 | 5.20077683479249 | -0.573404829419925 | 0.569392605962486 | 0.807947609257865 | -6.22415527660251 |
| ENSG00000116857.16 | TMEM9 | -0.0754078281024131 | 3.19743611313558 | -0.573329391361425 | 0.569443210608863 | 0.807947609257865 | -6.05181739227185 |
| ENSG00000056097.15 | ZFR | 0.0441447628924605 | 7.35341446512591 | 0.573188302306939 | 0.569537860603398 | 0.807947609257865 | -6.2393233634582 |
| ENSG00000177728.16 | TMEM94 | -0.0759226545683971 | 5.09422702626803 | -0.573154609882076 | 0.569560464412739 | 0.807947609257865 | -6.22214343939311 |
| ENSG00000027001.9 | MIPEP | -0.0996951347154953 | 3.5509370824222 | -0.573130817765229 | 0.569576426501359 | 0.807947609257865 | -6.08543887635737 |
| ENSG00000198720.12 | ANKRD13B | -0.250620379239399 | 0.288496811803909 | -0.573130548674231 | 0.569577374986579 | 0.807947609257865 | -5.58496738227805 |
| lnc-LGALS3BP-1 | lnc-LGALS3BP-1 | 0.098949945680562 | 0.518120561564077 | 0.573127446115531 | 0.569578688552993 | 0.807947609257865 | -5.68384908443544 |
| lnc-RBM4-1 | lnc-RBM4-1 | -0.070206044062898 | 4.20629829157021 | -0.57309199108035 | 0.569602475729528 | 0.807947609257865 | -6.15828158518854 |
| lnc-CASP9-4 | lnc-CASP9-4 | -0.117344133641042 | 0.244471361219804 | -0.572934494070631 | 0.5697081481547 | 0.807948187506633 | -5.5885007099846 |
| lnc-C16orf95-7 | lnc-C16orf95-7 | -0.0816638428791434 | 1.62882084927533 | -0.572907976830363 | 0.569725940817153 | 0.807948187506633 | -5.8597880229396 |
| lnc-TXLNB-3 | lnc-TXLNB-3 | -0.137124159477741 | 0.397908855396092 | -0.572812838815704 | 0.569789777228644 | 0.807948187506633 | -5.60865412700041 |
| LINC02001 | LINC02001 | 0.0630267365810733 | 2.75613312351057 | 0.57280801143437 | 0.569793018536499 | 0.807948187506633 | -5.98751912347147 |
| ENSG00000151461.19 | UPF2 | 0.0635000833692292 | 6.31150171447579 | 0.572798875110033 | 0.569799149290496 | 0.807948187506633 | -6.24799560164217 |
| ENSG00000173039.18 | RELA | 0.0666529952567059 | 5.54073868766061 | 0.572694575362374 | 0.569869139931204 | 0.807991768767926 | -6.23851348502778 |
| ENSG00000166783.21 | MARF1 | -0.0667741347412081 | 7.72992499074713 | -0.572610984533629 | 0.569925236858726 | 0.808015646112629 | -6.2342514085545 |
| ENSG00000180233.10 | ZNRF2 | 0.0845361328488314 | 4.73640076085409 | 0.572428158723401 | 0.570047938845721 | 0.808125220976705 | -6.20663596848 |
| lnc-VPREB1-7 | lnc-VPREB1-7 | -0.111862200546279 | 4.07733296316787 | -0.572378829627528 | 0.57008104788329 | 0.808125220976705 | -6.14948083878352 |
| ENSG00000188687.17 | SLC4A5 | 0.0904007702369802 | 2.71199265828792 | 0.57223933246901 | 0.570174681672522 | 0.808159635946482 | -5.98248329496296 |
| ENSG00000140830.8 | TXNL4B | -0.0592083828548926 | 3.22940526333629 | -0.572185774645872 | 0.570210632966084 | 0.808159635946482 | -6.0507485623334 |
| lnc-ZNF652-4 | lnc-ZNF652-4 | -0.125343788384082 | 1.07445014622467 | -0.572066702889856 | 0.570290565236343 | 0.808159635946482 | -5.77973607786632 |
| lnc-ZNF611-1 | lnc-ZNF611-1 | -0.101436119734022 | 2.60776410453607 | -0.572050719516596 | 0.57030129521536 | 0.808159635946482 | -5.96671076819533 |
| lnc-RNF223-1 | lnc-RNF223-1 | -0.178983256326542 | -0.270850974565391 | -0.572010654210107 | 0.570328192347214 | 0.808159635946482 | -5.49264368257853 |
| ENSG00000063169.10 | BICRA | 0.112705453504154 | 1.67676770377527 | 0.571957138343774 | 0.570364120250191 | 0.808159635946482 | -5.85729322156572 |
| ENSG00000103544.14 | VPS35L | -0.048343917243562 | 4.96707088189496 | -0.571933232776833 | 0.570380169625154 | 0.808159635946482 | -6.21392731145759 |
| lnc-TBC1D7-2 | lnc-TBC1D7-2 | 0.113310216499788 | 3.39014845354891 | 0.571828923994487 | 0.570450201559868 | 0.808203228159882 | -6.07552500781325 |
| ENSG00000023318.7 | ERP44 | -0.0554481518841934 | 5.18174799259131 | -0.571688826481991 | 0.570544268387422 | 0.80820354551534 | -6.22647394914961 |
| ENSG00000068366.19 | ACSL4 | 0.0589471606129273 | 4.79558304635897 | 0.571633518710016 | 0.570581406247622 | 0.80820354551534 | -6.2078319682442 |
| lnc-ARHGEF4-1 | lnc-ARHGEF4-1 | 0.0865038217163758 | 0.149919454702506 | 0.571551818476478 | 0.570636268203317 | 0.80820354551534 | -5.56752295911905 |
| ENSG00000174326.11 | SLC16A11 | -0.294255022936437 | -1.06713369523881 | -0.571532920340134 | 0.570648958730017 | 0.80820354551534 | -5.34229724703784 |
| ENSG00000231925.11 | TAPBP | -0.0813795739352283 | 7.46650701638779 | -0.571495442762986 | 0.570674126183964 | 0.80820354551534 | -6.23791028808284 |
| lnc-BBS9-1 | lnc-BBS9-1 | -0.137560548497981 | 0.341081247615798 | -0.571477733008974 | 0.570686019070104 | 0.80820354551534 | -5.62782063448719 |
| ENSG00000261609.6 | GAN | -0.0951837622715077 | 4.61841396400583 | -0.57139001865556 | 0.570744924955256 | 0.808231358125492 | -6.1899092797126 |
| lnc-PAQR9-1 | lnc-PAQR9-1 | -0.25880809066769 | -0.752005501130625 | -0.571284111669935 | 0.570816052370762 | 0.808276472616083 | -5.28317311989769 |
| ENSG00000188783.5 | PRELP | 0.559340755641373 | -1.7369790913966 | 0.571225043320818 | 0.570861317465399 | 0.808284962329938 | -5.01255027505957 |
| ENSG00000163902.11 | RPN1 | 0.0693395603378635 | 6.07480075969816 | 0.571073726075248 | 0.570957360866475 | 0.808365343149997 | -6.24819710613086 |
| HOTAIRM1 | HOTAIRM1 | 0.38106580607798 | -0.297056159725241 | 0.57099729567062 | 0.571027463851486 | 0.808408988820079 | -5.33967216756715 |
| ENSG00000174123.10 | TLR10 | -0.238632545352059 | 4.13904300063803 | -0.570912308329747 | 0.571150159960267 | 0.808487316026615 | -6.15831488648284 |
| lnc-POLE3-3 | lnc-POLE3-3 | -0.101466820922914 | 0.988087863299193 | -0.570770064718741 | 0.571161349771317 | 0.808487316026615 | -5.76222938231149 |
| ENSG00000164587.12 | RPS14 | 0.0914773828142437 | 8.41264515624071 | 0.570650679446221 | 0.571241558390014 | 0.808506860798345 | -6.21871348078522 |
| ENSG00000198912.10 | C1orf174 | -0.0665178372266864 | 4.08793286190687 | -0.570587653887231 | 0.571283904151642 | 0.808506860798345 | -6.14802192776768 |
| ENSG00000135211.5 | TMEM60 | 0.11328482363718 | 2.18322400340559 | 0.570574118899611 | 0.571292998272901 | 0.808506860798345 | -5.92291299559583 |
| lnc-PMM1-2 | lnc-PMM1-2 | -0.0928178335160581 | 1.75813669944883 | -0.57035752277665 | 0.571438538321384 | 0.808622516456322 | -5.87109566205428 |
| ENSG00000201208.1 | | -0.0955049806282732 | 0.160921926209584 | -0.570335566428084 | 0.571453292735445 | 0.808622516456322 | -5.60765337661791 |
| ENSG00000277577.1 | | 1.05162274335119 | -2.18153674526763 | 0.570411210512467 | 0.57150360542367 | 0.808629821406179 | -5.04018523761435 |

| | | | | | | | |
|---|---|---|---|---|---|---|---|
| lnc-CNPY3-1 | lnc-CNPY3-1 | -0.0745523351420032 | 1.99717281562171 | -0.570210963937565 | 0.571537027732453 | 0.808629821406179 | -5.91069733761659 |
| lnc-MAP3K14-3 | lnc-MAP3K14-3 | -0.0679867342174578 | 2.39373489420083 | -0.570028511004033 | 0.571659650116112 | 0.808746135140424 | -5.95068486381307 |
| ENSG00000128284.19 | APOL3 | 0.113076538263171 | 4.33778164319006 | 0.569971718314784 | 0.571697821808928 | 0.808746135140424 | -6.17887425354036 |
| ENSG00000163110.14 | PDLIM5 | -0.0723146474554106 | 4.0264132525275 | -0.569897146337252 | 0.571747945282635 | 0.808750940025076 | -6.14243304169904 |
| ENSG00000200795.1 | RNU4-1 | -0.211659542734967 | 1.88572465729898 | -0.569834812147771 | 0.571816723394723 | 0.808750940025076 | -5.88227507680591 |
| ENSG00000114405.10 | CEP15 | 0.32661308287225 | 0.995625206761858 | 0.569844662947655 | 0.571840950553805 | 0.808750940025076 | -5.63984725442533 |
| lnc-MAP3K3-2 | lnc-MAP3K3-2 | -0.102822856056394 | 2.53636874504707 | -0.569609577978939 | 0.571941254301351 | 0.808750940025076 | -5.97052952321978 |
| ENSG00000133422.12 | MORC2 | 0.0703672565069358 | 4.33667677206484 | 0.569593009126583 | 0.571952393184897 | 0.808750940025076 | -6.17392633854409 |
| lnc-ZBTB7C-4 | lnc-ZBTB7C-4 | 0.168337643955168 | 0.112954540922342 | 0.569584383447821 | 0.571958192085717 | 0.808750940025076 | -5.54354689307687 |
| lnc-MNT-4 | lnc-MNT-4 | 0.10036984537126 | 1.88997994369945 | 0.569557502907478 | 0.571976263612935 | 0.808750940025076 | -5.88757556984219 |
| ENSG00000177082.12 | WDR73 | -0.0512276180155142 | 4.20226905701884 | -0.56933477045969 | 0.572126015294217 | 0.808870078521388 | -6.1607336631354 |
| ENSG00000124635.8 | H2BC11 | -0.149959004082977 | 4.2408875174249 | -0.569343094567556 | 0.572139118359466 | 0.808870078521388 | -6.16884927440938 |
| ENSG00000163872.15 | YEATS2 | -0.0563091658426949 | 5.24624603810323 | -0.569182036573937 | 0.572228715371649 | 0.808895521720721 | -6.23138913801586 |
| ENSG00000146731.10 | CCT6A | -0.0783218222615316 | 5.71378190659535 | -0.569034095746566 | 0.57232820118671 | 0.808895521720721 | -6.24527922765078 |
| ENSG00000152213.3 | ARL11 | -0.213560961941039 | 2.32643643813714 | -0.569088043146019 | 0.572332606722863 | 0.808895521720721 | -5.94646126908175 |
| ENSG00000122507.20 | BBS9 | -0.0604690654854734 | 4.94805027756127 | -0.568985005864082 | 0.572361214555491 | 0.808895521720721 | -6.21607082759786 |
| lnc-SLC38A11-2 | lnc-SLC38A11-2 | 0.348783559997127 | 1.2655896146073 | 0.569077611758907 | 0.572382991203318 | 0.808895521720721 | -5.79487403763952 |
| ENSG00000138036.18 | DYNC2LI1 | 0.109148619726916 | 2.41094734484997 | 0.568937876475986 | 0.572392910356231 | 0.808895521720721 | -5.9550629775338 |
| lnc-SPICE1-1 | lnc-SPICE1-1 | -0.109569575646678 | 0.220967910080723 | -0.568836595513364 | 0.57246102749093 | 0.808936243913621 | -5.58566527208698 |
| lnc-H3F3A-7 | lnc-H3F3A-7 | -0.217498756301135 | 2.92344214628648 | -0.568801766591441 | 0.572542875134137 | 0.808996361562844 | -6.01549315293854 |
| lnc-TMEM65-4 | lnc-TMEM65-4 | -0.138659171629755 | 1.5571126705856 | -0.568559803217174 | 0.572647206186332 | 0.809088237570928 | -5.85861524694493 |
| lnc-PCED1B-8 | lnc-PCED1B-8 | -0.520770078535004 | -0.883386660630955 | -0.568482729503008 | 0.572772417152034 | 0.809209599989039 | -5.35234470238691 |
| ENSG00000159423.16 | ALDH4A1 | -0.168745514060411 | 0.0675839261694751 | -0.568278735260832 | 0.572836291275452 | 0.809222494246963 | -5.52753439221399 |
| lnc-ELL2-4 | lnc-ELL2-4 | 0.240917989690078 | -0.437268841015995 | 0.568158164377259 | 0.57291741329247 | 0.809222494246963 | -5.35494162742365 |
| ENSG00000248751.6 | | -0.0965623107020632 | 2.08223971698251 | -0.568157757829665 | 0.572917686833737 | 0.809222494246963 | -5.9128584532435 |
| ENSG00000239961.2 | LILRA4 | 0.808650160096102 | 2.28503776597022 | 0.568302776494785 | 0.572938804288053 | 0.809222494246963 | -5.85225122957864 |
| ENSG00000073282.12 | TP63 | -0.6856941677261 | -0.141981231026219 | -0.568134037275787 | 0.573047695262641 | 0.809320756936389 | -5.51480999419066 |
| ENSG00000140464.19 | PML | -0.077036597956929 | 4.84098694298696 | -0.567899989203562 | 0.573091136659334 | 0.80932323626281 | -6.21263317114806 |
| ENSG00000177556.11 | ATOX1 | -0.0867000514136409 | 5.36111673147979 | -0.567773224592705 | 0.573176444708722 | 0.80932323626281 | -6.23722688751187 |
| ENSG00000125817.7 | CENPB | -0.10661276344083 | 4.64100659138805 | -0.567664539565956 | 0.573249590811024 | 0.80932323626281 | -6.20164951074356 |
| ENSG00000185825.16 | BCAP31 | 0.0729234702438912 | 5.14120670958455 | 0.567550010666644 | 0.573326674858401 | 0.80932323626281 | -6.22657555757087 |
| ENSG00000275066.4 | SYNRG | 0.0663592717399517 | 6.45048300537346 | 0.567481320590017 | 0.573372909379322 | 0.80932323626281 | -6.25102467929969 |
| lnc-PQBP1-2 | lnc-PQBP1-2 | 0.168481918862284 | -0.58652568655841 | 0.567469053543255 | 0.573381166383725 | 0.80932323626281 | -5.36783790090383 |
| lnc-NQO2-5 | lnc-NQO2-5 | -0.0724577794303527 | 3.19227269382988 | -0.567458650031181 | 0.573388169080296 | 0.80932323626281 | -6.06241539383259 |
| ENSG00000185880.12 | TRIM69 | 0.115265161301806 | 3.25128547923302 | 0.567448640509999 | 0.573394906617732 | 0.80932323626281 | -6.05312096265336 |
| ENSG00000279077.1 | | -0.132021358944663 | 1.64600272109806 | -0.567436125579521 | 0.573403330633074 | 0.80932323626281 | -5.84339669162132 |
| ENSG00000239827.8 | SUGT1P3 | 0.0935767322051801 | 1.52731613262192 | 0.567335729485523 | 0.573470911172883 | 0.809363121549057 | -5.84179069880914 |
| lnc-LTBP2-4 | lnc-LTBP2-4 | 0.0619075968172824 | 2.09608110904396 | 0.56717192175651 | 0.573581184960668 | 0.809374444921887 | -5.91825192995515 |
| ENSG00000125386.15 | FAM193A | -0.0819840947178321 | 5.52625712528678 | -0.567001249245616 | 0.573696091124972 | 0.809374444921887 | -6.24185272811403 |
| ENSG00000172757.12 | CFL1 | -0.0929402551985161 | 7.32007960721561 | -0.566939085489614 | 0.573737946006828 | 0.809374444921887 | -6.24267859600945 |
| ENSG00000196924.15 | FLNA | 0.119869450662707 | 9.13372830854852 | 0.56686588009784 | 0.573787237147854 | 0.809374444921887 | -6.20488991275342 |
| ENSG00000188986.6 | NELFB | -0.0663216364875927 | 4.39118064088179 | -0.56684532406159 | 0.573801078448498 | 0.809374444921887 | -6.17921841736961 |
| ENSG00000259232.2 | | -0.131927065674622 | 0.294583561077964 | -0.566839658701839 | 0.573804893217972 | 0.809374444921887 | -5.59640814404915 |
| ENSG00000132010.15 | ZNF20 | -0.0807595799731431 | 1.54852096696669 | -0.566824217811889 | 0.573815290402997 | 0.809374444921887 | -5.84611920970952 |
| ENSG00000149970.15 | CNKSR2 | -0.0876306329843644 | 4.80373443601527 | -0.566816117756191 | 0.573820744644875 | 0.809374444921887 | -6.21048641330244 |
| ENSG00000132965.9 | ALOX5AP | -0.0979483389589699 | 4.33163452260618 | -0.566784536431118 | 0.573842010443536 | 0.809374444921887 | -6.163086069583 |
| lnc-INTS12-4 | lnc-INTS12-4 | -0.100501412434056 | -0.123997404011557 | -0.566609364398922 | 0.573959972399694 | 0.809374444921887 | -5.49572930909575 |
| TTTY15 | TTTY15 | -1.36764422452821 | 1.5211182226637 | -0.566783631346624 | 0.573960939376262 | 0.809374444921887 | -5.53052849576072 |
| ENSG00000198176.12 | TFDP1 | -0.165703694420886 | 4.64912414557066 | -0.566656769389839 | 0.573970530943752 | 0.809374444921887 | -6.19778167372881 |
| lnc-LRGUK-4 | lnc-LRGUK-4 | 0.158853915046438 | 0.701653868688206 | 0.566564588859371 | 0.573990126440499 | 0.809374444921887 | -5.7097351983506 |
| ENSG00000160818.16 | GPATCH4 | -0.094846928352248 | 4.04560436642 | -0.566479656355641 | 0.574047326282294 | 0.809379929221837 | -6.15311404704793 |
| ENSG00000197747.8 | S100A10 | -0.199024368143457 | 2.52564527714165 | -0.56649110108691 | 0.574072661259758 | 0.809379929221837 | -5.97985659069486 |
| ENSG00000135624.15 | CCT7 | -0.0945152330315642 | 6.16544770316415 | -0.56626347793492 | 0.57419292946387 | 0.809494045695537 | -6.25119751554455 |
| lnc-CENPC-1 | lnc-CENPC-1 | -0.146791109917654 | 0.551729019986266 | -0.566191885060143 | 0.574241153579941 | 0.809506586133548 | -5.64594733468505 |
| ENSG00000268870.1 | | 0.126504984456526 | -0.397279351204959 | 0.566068929987172 | 0.574323979300754 | 0.809517904461741 | -5.4278199037661 |
| lnc-KIAA1551-3 | lnc-KIAA1551-3 | -0.0868569888134698 | 2.68274751702274 | -0.566063196948589 | 0.574327841366895 | 0.809517904461741 | -5.99463379250523 |
| ENSG00000108592.16 | FTSJ3 | 0.0658777348973203 | 4.89139949545471 | 0.565820289967441 | 0.57449148755104 | 0.809687057291005 | -6.21480342320914 |
| lnc-NKX6-3-3 | lnc-NKX6-3-3 | 0.183971322814061 | 0.0323790596229512 | 0.565714458802055 | 0.574562793045622 | 0.809687057291005 | -5.50385726962866 |
| ENSG00000269755.1 | | 0.0887908545333252 | 0.284913144974901 | 0.565662110322842 | 0.574598065300813 | 0.809687057291005 | -5.59674478269414 |
| ENSG00000285668.1 | | 0.68168915911844 | -2.63010097326311 | 0.5656651116023 | 0.574605200692304 | 0.809687057291005 | -4.91921868159562 |
| ENSG00000163611.11 | SPICE1 | 0.0699897295546714 | 4.25209992110416 | 0.565526766402344 | 0.574689264564235 | 0.809750077527769 | -6.16419772653554 |
| ENSG00000159348.12 | CYB5R1 | 0.0826361844724777 | 4.01763168635788 | 0.565396158806643 | 0.574777279057197 | 0.809763076574813 | -6.13983370785267 |
| ENSG00000171552.12 | BCL2L1 | -0.0746757290538453 | 3.02965086850763 | -0.565321721822562 | 0.574827443974464 | 0.809763076574813 | -6.03686288544329 |
| lnc-TMEM150A-4 | lnc-TMEM150A-4 | -0.117598661215611 | 1.44742478996551 | -0.565229206627423 | 0.574889795232567 | 0.809763076574813 | -5.8133780731015 |
| ENSG00000279416.1 | | 0.151475305566506 | 1.1268355982096 | 0.565205766408531 | 0.574905593456794 | 0.809763076574813 | -5.76725085468257 |
| lnc-IDI1-2 | lnc-IDI1-2 | -0.0860686827347688 | 3.912032971227 | -0.565180523010173 | 0.574922607224598 | 0.809763076574813 | -6.1391785028191 |
| lnc-GDAP2-2 | lnc-GDAP2-2 | 0.0804573264955302 | 3.44969048299693 | 0.565135472038717 | 0.574952971687603 | 0.809763076574813 | -6.07001645958269 |
| lnc-LRRC63-4 | lnc-LRRC63-4 | -0.215092535194545 | -0.015603412126901 | -0.565104452161068 | 0.574973879616231 | 0.809763076574813 | -5.6152682366313 |

| | | | | | | | |
|---|---|---|---|---|---|---|---|
| ENSG00000082996.19 | RNF13 | 0.0510094821020903 | 6.10147203605695 | 0.564939429619342 | 0.57508511389355 | 0.809835669583685 | -6.25146633019071 |
| lnc-AURKB-4 | lnc-AURKB-4 | 0.0913934515727113 | 3.7398837805068 | 0.564863441259041 | 0.575136337774144 | 0.809835669583685 | -6.10824771972623 |
| lnc-OVOL2-2 | lnc-OVOL2-2 | -0.0835693099042076 | 1.78563567725413 | -0.564852877398673 | 0.575143459066915 | 0.809835669583685 | -5.88338426672305 |
| ENSG00000055130.15 | CUL1 | 0.0574941207726107 | 5.95290301386977 | 0.564707373021318 | 0.575241550638652 | 0.809865150530679 | -6.25017507921649 |
| ENSG00000279608.1 | | -0.13194242380306 | 3.08402145309001 | -0.564705091195178 | 0.575243088993759 | 0.809865150530679 | -6.03708299352746 |
| ENSG00000163406.10 | SLC15A2 | 0.177507556359554 | 4.38132638657265 | 0.564661123631399 | 0.575321390200328 | 0.809919990048105 | -6.16694518809391 |
| ENSG00000277611.1 | | 0.116925436029615 | -0.124577506406206 | 0.564414930869478 | 0.575438724870871 | 0.810029768432303 | -5.50362431771922 |
| lnc-APCDD1L-3 | lnc-APCDD1L-3 | -0.153083846435271 | -0.381831630055126 | -0.56424196473789 | 0.575555359956478 | 0.810135308522165 | -5.42989306448627 |
| lnc-LRGUK-5 | lnc-LRGUK-5 | -0.161412015557719 | 0.720325764082413 | -0.564187011550939 | 0.575592418589522 | 0.810135308522165 | -5.71984044516594 |
| lnc-CLEC16A-2 | lnc-CLEC16A-2 | -0.204707717462511 | -0.390136997278668 | -0.563841620572356 | 0.575825365696729 | 0.810407760829796 | -5.39968919221534 |
| lnc-ZNF514-1 | lnc-ZNF514-1 | -0.104056447951188 | 0.143030871706831 | -0.563739242249343 | 0.575894423068733 | 0.810417550165591 | -5.56334405168891 |
| ENSG00000197978.9 | GOLGA6L9 | 0.206615691749181 | 1.14954565148198 | 0.563723172615611 | 0.575911067690429 | 0.810417550165591 | -5.7821412777791 |
| lnc-ZNF726-4 | lnc-ZNF726-4 | -0.353483566109982 | 1.0725113411488 | -0.563605741069238 | 0.576066846127208 | 0.810578860950143 | -5.73902083219659 |
| lnc-DCAF8-1 | lnc-DCAF8-1 | -0.13063827301998 | 0.271879221157769 | -0.563427892964079 | 0.57610446275274 | 0.810578860950143 | -5.62746940127619 |
| ENSG00000145819.16 | ARHGAP26 | 0.155951851700603 | 2.56689400701072 | 0.563192607409858 | 0.576263213835028 | 0.810702578809186 | -5.97356615303764 |
| ENSG00000161813.21 | LARP4 | 0.0591397927549755 | 4.94671967246427 | 0.563146280256882 | 0.576294474055411 | 0.810702578809186 | -6.22156351771149 |
| lnc-GOLGB1-1 | lnc-GOLGB1-1 | -0.104177253195913 | 3.5927628798054 | -0.563107814568515 | 0.576320430216872 | 0.810702578809186 | -6.09035032851934 |
| ENSG00000226259.10 | GTF2H2B | 0.274384686140127 | 1.39982758230832 | 0.563150193053726 | 0.576349941005433 | 0.810702578809186 | -5.79759050379832 |
| ENSG00000134042.12 | MRO | -0.574963429012446 | -0.738007451322908 | -0.563057219106445 | 0.576437595273949 | 0.810770467645462 | -5.33676035145656 |
| lnc-RBL1-3 | lnc-RBL1-3 | -0.253722918073954 | -0.726949637991742 | -0.562785817666711 | 0.576537732066949 | 0.810822977679671 | -5.26635405896116 |
| ENSG00000100644.16 | HIF1A | 0.218028610547487 | 6.04169537800737 | 0.562788855543664 | 0.576620216459365 | 0.810822977679671 | -6.25289692397734 |
| lnc-CERK-1 | lnc-CERK-1 | 0.113583175006911 | 1.51489252698585 | 0.562657702271235 | 0.576624202760647 | 0.810822977679671 | -5.83140287824624 |
| ENSG00000162613.16 | FUBP1 | 0.0465435010374033 | 7.66663909322522 | 0.562599747638313 | 0.576663320962073 | 0.810822977679671 | -6.23940260658988 |
| lnc-NBDY-7 | lnc-NBDY-7 | -0.131256031807148 | 0.00210983295751536 | -0.562569252900701 | 0.576683904811597 | 0.810822977679671 | -5.542032389427 |
| ENSG00000136450.12 | SRSF1 | 0.0826540632545099 | 6.60369090494302 | 0.562485975204229 | 0.576740118815716 | 0.810822977679671 | -6.25290682296205 |
| ENSG00000049246.14 | PER3 | 0.314569162786092 | 2.75251772580405 | 0.562587792592478 | 0.576763853038964 | 0.810822977679671 | -5.96100696557981 |
| ENSG00000198743.6 | SLC5A3 | -0.136658792129587 | 4.9795716862153 | -0.562432333239335 | 0.576790071378601 | 0.810822977679671 | -6.22440519771618 |
| ENSG00000238829.1 | RNU7-45P | -0.156660961373385 | 0.247963250010387 | -0.562279076893474 | 0.576879790628073 | 0.810893719217212 | -5.61229375831045 |
| lnc-ZNF131-2 | lnc-ZNF131-2 | -0.0575860845314109 | 5.57893260149812 | -0.562164216031016 | 0.576957337415724 | 0.810928812775941 | -6.24450589178535 |
| ENSG00000182872.15 | RBM10 | -0.0601952121313847 | 5.47708486436964 | -0.562125388295365 | 0.576983552596981 | 0.810928812775941 | -6.24347767860832 |
| lnc-ZMAT3-8 | lnc-ZMAT3-8 | -0.0764561695983614 | 1.71998938125014 | -0.561856908410721 | 0.577164837091103 | 0.811128215338398 | -5.86284878207892 |
| ENSG00000164291.16 | ARSK | -0.110928051533948 | 1.10707866288561 | -0.561728802368149 | 0.577251347364498 | 0.811166279092175 | -5.79009657579536 |
| ENSG00000207460.1 | SNORD116-19 | -0.171182434555003 | 1.71018269070103 | -0.561556799616468 | 0.577367511122921 | 0.811166279092175 | -5.85149292148892 |
| ENSG00000162510.5 | MATN1 | 0.155058734493466 | 1.63546323894848 | 0.561487932746993 | 0.577414024239999 | 0.811166279092175 | -5.84753908016549 |
| ENSG00000213719.8 | CLIC1 | -0.128079189967151 | 6.48142087754633 | -0.561488786374608 | 0.577422681739115 | 0.811166279092175 | -6.25385889097998 |
| ENSG00000186153.16 | WWOX | -0.0783255389600117 | 2.93421244727581 | -0.56144746074193 | 0.57744136013947 | 0.811166279092175 | -6.0210825441216 |
| lnc-CDK6-1 | lnc-CDK6-1 | 0.075144634231822 | 2.45451096922486 | 0.561418957069664 | 0.577460612678286 | 0.811166279092175 | -5.95881909711455 |
| lnc-HEATR1-1 | lnc-HEATR1-1 | -0.0837241355209114 | 0.448911005544295 | -0.561408333503761 | 0.577467788346579 | 0.811166279092175 | -5.64470630047821 |
| lnc-PTP4A2-3 | lnc-PTP4A2-3 | 0.072656035015501 | 4.59233823660687 | 0.561284236185494 | 0.577551612875209 | 0.811228664379039 | -6.20018614549034 |
| lnc-SYNRG-5 | lnc-SYNRG-5 | -0.160263688014414 | -0.376294969143137 | -0.561203163391457 | 0.577606378654388 | 0.811250228034341 | -5.36373231633637 |
| ENSG00000153130.17 | SCOC | 0.115086596042859 | 3.48272875099269 | 0.560801014164377 | 0.577878073367712 | 0.811576445423554 | -6.09367980452173 |
| ENSG00000180855.15 | ZNF443 | 0.0942722530962066 | 1.84376747558147 | 0.560594615577351 | 0.578017541816533 | 0.811711222945087 | -5.88471758440583 |
| ENSG00000255710.2 | ISG20L2P2 | 0.165664974512239 | 1.93156415988341 | 0.560542273984568 | 0.578052912885832 | 0.811711222945087 | -5.8828711827834 |
| ENSG00000174720.15 | LARP7 | -0.06065019487363 | 5.58793410595899 | -0.560202816923324 | 0.578282334614681 | 0.811890778671576 | -6.24674530910593 |
| ENSG00000155111.14 | CDK19 | -0.0735507722559454 | 5.27502529258401 | -0.560158412788196 | 0.578312348385911 | 0.811890778671576 | -6.23560051674938 |
| ENSG00000086967.9 | MYBPC2 | -0.167025506551245 | 1.24124630626486 | -0.560084611073092 | 0.578362234332271 | 0.811890778671576 | -5.80551642996695 |
| ENSG00000127837.9 | AAMP | -0.0816862293943519 | 4.66283147712466 | -0.560069647894525 | 0.578372348882805 | 0.811890778671576 | -6.20699792522275 |
| ENSG00000165410.14 | CFL2 | 0.0608506362688388 | 4.2673515090206 | 0.560061279678691 | 0.578378005522097 | 0.811890778671576 | -6.1767301944074 |
| ENSG00000165898.13 | ISCA2 | -0.0728736414387572 | 3.58646839932873 | -0.559754987099037 | 0.578585067788522 | 0.812121643000533 | -6.09760331362837 |
| ENSG00000280067.1 | | -0.147723705198016 | 1.49543361128973 | -0.55970127634947 | 0.578621381460565 | 0.812121643000533 | -5.8350918867781 |
| ENSG00000161405.16 | IKZF3 | 0.0865610985563929 | 9.09730428710177 | 0.559231089725157 | 0.578939320533927 | 0.812490682758495 | -6.20853010444197 |
| ENSG00000267645.5 | | -0.202137640217254 | 1.64481555329914 | -0.559171015508512 | 0.578997464665066 | 0.812490682758495 | -5.87013174546851 |
| ENSG00000128335.13 | APOL2 | -0.0747167312600441 | 3.98498123764786 | -0.559137320898225 | 0.579002736952903 | 0.812490682758495 | -6.14632356082436 |
| lnc-PAWR-14 | lnc-PAWR-14 | 0.133964914308468 | 1.87430532105274 | 0.559034778754435 | 0.579072090685874 | 0.812532609079511 | -5.90590835576642 |
| lnc-COL4A3-2 | lnc-COL4A3-2 | -0.167011466788004 | 2.7415216843771 | -0.558844335897442 | 0.579210089870153 | 0.812670843885301 | -5.99968797227961 |
| lnc-ARMC4-3 | lnc-ARMC4-3 | 0.413829544475586 | -1.56293345661151 | 0.558501123559301 | 0.579433090220942 | 0.812928314090222 | -5.13353667376799 |
| ENSG00000089639.10 | GMIP | -0.0727193403162429 | 5.78758418524828 | -0.558137865162812 | 0.579678884684257 | 0.813202353260005 | -6.25173462658896 |
| lnc-MLXIPL-2 | lnc-MLXIPL-2 | -0.0864065745699785 | 0.326198812703608 | -0.558079552902046 | 0.579718345697958 | 0.813202353260005 | -5.62312975682311 |
| ENSG00000198752.10 | CDC42BPB | 0.334006341097811 | 2.70489728390152 | 0.55819433840997 | 0.57974745286714 | 0.813202353260005 | -6.07488293148416 |
| lnc-ZNF316-2 | lnc-ZNF316-2 | 0.0934657358121189 | 2.95252424526753 | 0.557950672227829 | 0.579805566325403 | 0.813202353260005 | -6.02462584860427 |
| lnc-BCL2L13-3 | lnc-BCL2L13-3 | -0.118145050336431 | 1.33554369942334 | -0.557914983165766 | 0.57982972019973 | 0.813202353260005 | -5.803532490555 |
| lnc-METTL8-4 | lnc-METTL8-4 | -0.124568422038281 | 3.08188882028669 | -0.557859584001007 | 0.579867214572908 | 0.813202353260005 | -6.02828778946368 |
| ENSG00000138430.15 | OLA1 | -0.060033001992772 | 4.89508856248988 | -0.557803790567739 | 0.579904976978592 | 0.813202353260005 | -6.21913161865681 |
| ENSG00000169508.6 | GPR183 | -0.51876089385158 | 3.92651342187261 | -0.557818858695612 | 0.580010929813561 | 0.813279385631696 | -6.11273989056125 |
| ENSG00000213366.12 | GSTM2 | 0.214906485768445 | 2.5686060871762 | 0.55766019558462 | 0.580043203976493 | 0.813279385631696 | -5.93570516177922 |
| ENSG00000234473.1 | ZNF101P2 | -0.159565049357753 | -0.0819448992646791 | -0.557516292650848 | 0.580099581763447 | 0.813279385631696 | -5.50381209293413 |
| ENSG00000229391.7 | HLA-DRB6 | -0.400657075074895 | 5.21222281740734 | -0.557660677114872 | 0.580117958501898 | 0.813279385631696 | -6.22738701048521 |
| ENSG00000215305.9 | VPS16 | -0.0504571614260234 | 4.85673601441474 | -0.557390433426216 | 0.580184784730041 | 0.813292472195273 | -6.21694171221228 |

| | | | | | | | |
|---|---|---|---|---|---|---|---|
| LINC00865 | LINC00865 | 0.460706712611899 | 1.02983303022426 | 0.557512191636417 | 0.580206318890123 | 0.813292472195273 | -5.70205186793287 |
| lnc-STAR-1 | lnc-STAR-1 | 0.114342820701429 | -0.150808857409743 | 0.557108479956447 | 0.580375680919025 | 0.813303218593534 | -5.46629977734004 |
| ENSG00000179041.3 | RRS1 | -0.144089726055785 | 1.27674074109855 | -0.557037469409412 | 0.580423763311117 | 0.813303218593534 | -5.81760558705377 |
| lnc-SLC7A5-3 | lnc-SLC7A5-3 | 0.222695645149864 | -0.655418144468302 | 0.557015375830932 | 0.580438723625317 | 0.813303218593534 | -5.33349082914678 |
| ENSG00000238083.7 | LRRC37A2 | 0.104082074578276 | 4.70159904975827 | 0.556994282341341 | 0.580453006919847 | 0.813303218593534 | -6.20129642934125 |
| ENSG00000149480.6 | MTA2 | 0.072145391621123 | 6.12405437692526 | 0.55698981048933 | 0.58045603502201 | 0.813303218593534 | -6.25643922296733 |
| ENSG00000176209.11 | SMIM19 | 0.0532219496617037 | 3.63654885239537 | 0.556961977811921 | 0.580474882014468 | 0.813303218593534 | -6.1013745869451 |
| ENSG00000173275.12 | ZNF449 | -0.0588153042344386 | 2.50077599243783 | -0.556888080150822 | 0.580524923508242 | 0.813303218593534 | -5.97566508163335 |
| ENSG00000118363.11 | SPCS2 | 0.0669370962520954 | 4.54650119404722 | 0.556880447589146 | 0.58053009219143 | 0.813303218593534 | -6.19461240451529 |
| lnc-GTF2IRD2-5 | lnc-GTF2IRD2-5 | -0.0611375954950493 | 2.3912658226693 | -0.556812463639058 | 0.580576131127261 | 0.813312360102935 | -5.95408249872132 |
| ENSG00000162368.13 | CMPK1 | 0.0788092216044056 | 6.33596715589509 | 0.556668012551472 | 0.580673959717728 | 0.81339404606438 | -6.25707669482942 |
| lnc-SPIRE1-3 | lnc-SPIRE1-3 | 0.0813855315961121 | 2.60358511124569 | 0.5565401688536 | 0.580760547725977 | 0.813438776512412 | -5.97898953072504 |
| lnc-PMM2-6 | lnc-PMM2-6 | 0.0619486569469707 | 7.2594390262369 | 0.556504167819271 | 0.580784932207472 | 0.813438776512412 | -6.2493784886905 |
| ENSG00000176422.13 | SPRYD4 | -0.0545282385622388 | 5.01312288110356 | -0.556293265131473 | 0.580927792316194 | 0.813576099043292 | -6.22634918412969 |
| lnc-DYNLRB1-2 | lnc-DYNLRB1-2 | 0.136710168597161 | 0.316605246968075 | 0.556196680992692 | 0.580993221637976 | 0.813576099043292 | -5.60339123672444 |
| ENSG00000141682.11 | PMAIP1 | 0.16960019532373 | 7.80566402514577 | 0.556268639505861 | 0.581001558559848 | 0.813576099043292 | -6.24156140622285 |
| ENSG00000198478.7 | SH3BGRL2 | -1.73233865999123 | 0.557584134816959 | -0.556215386646382 | 0.581096314052159 | 0.813653430757523 | -5.28729703976487 |
| ENSG00000154620.5 | TMSB4Y | -0.941101417895112 | -0.750144345436916 | -0.556055404828483 | 0.581191002224441 | 0.81373065769578 | -5.12280371375702 |
| ENSG00000100439.10 | ABHD4 | 0.090063886577334 | 3.48045950154052 | 0.555735938159343 | 0.58130539337437 | 0.813835458565138 | -6.07987198648657 |
| lnc-FNDC9-2 | lnc-FNDC9-2 | -0.128904551814775 | 1.46635301430005 | -0.555637592708181 | 0.581372036880919 | 0.813847246058147 | -5.81967295327638 |
| ENSG00000159761.14 | C16orf86 | -0.116634525941107 | 1.63512362848876 | -0.555606817329899 | 0.581392892485984 | 0.813847246058147 | -5.85929607324427 |
| ENSG00000165832.5 | TRUB1 | -0.076255542743457 | 3.89655289214615 | -0.555425949988001 | 0.581515468500087 | 0.813869337753468 | -6.14875138041239 |
| lnc-EFCAB12-2 | lnc-EFCAB12-2 | -0.0811856202411141 | 4.42373787917448 | -0.555386735520719 | 0.58154204627587 | 0.813869337753468 | -6.18707923038238 |
| ENSG00000091879.13 | ANGPT2 | 1.15287038301258 | -1.7441893769365 | 0.555425664781253 | 0.581599646359063 | 0.813869337753468 | -4.92406454256048 |
| ENSG00000167112.9 | TRUB2 | -0.0597079087822108 | 4.04728227310382 | -0.555297573145486 | 0.58160247865262 | 0.813869337753468 | -6.15502626918824 |
| ENSG00000107625.12 | DDX50 | 0.0425480732041859 | 5.30652805413079 | 0.555291819410825 | 0.581606378517017 | 0.813869337753468 | -6.23971468667541 |
| ENSG00000236397.3 | DDX11L2 | -0.246823176477707 | -0.243248894892052 | -0.555183928149561 | 0.581679509240561 | 0.813916338456728 | -5.42273287317368 |
| ENSG00000196787.3 | H2AC11 | -0.184898677528752 | 4.86056490146393 | -0.554823163844452 | 0.581989276139476 | 0.814181338345336 | -6.22560578785262 |
| lnc-MTURN-6 | lnc-MTURN-6 | 0.195366490913888 | -0.00328499802006218 | 0.554654044533535 | 0.582038738914512 | 0.814181338345336 | -5.52546822573351 |
| ENSG00000132432.13 | SEC61G | 0.0885172807281078 | 3.50816941205163 | 0.554642883916138 | 0.582046306305436 | 0.814181338345336 | -6.10028536357085 |
| ENSG00000128463.12 | EMC4 | 0.0476427857366541 | 4.60665270889987 | 0.554623651762637 | 0.582059346665588 | 0.814181338345336 | -6.2013335050398 |
| ENSG00000280071.3 | | -0.153303197340232 | 1.57888834788059 | -0.554523194809577 | 0.582127463791936 | 0.814181338345336 | -5.86220483325219 |
| ENSG00000272602.5 | ZNF595 | 0.775051783727092 | 3.58067379150809 | 0.55462767836404 | 0.582171998097387 | 0.814181338345336 | -6.15895630397945 |
| ENSG00000264536.1 | MIR5706 | -0.221331177646362 | -0.637216719142627 | -0.554455044396075 | 0.582173676928224 | 0.814181338345336 | -5.32527079764811 |
| lnc-XPC-7 | lnc-XPC-7 | 0.185297597261714 | -0.139318874066174 | 0.55443783924527 | 0.582185344107596 | 0.814181338345336 | -5.45006506969478 |
| ENSG00000140006.11 | WDR89 | 0.0780854193395522 | 4.10459763605262 | 0.554351658410467 | 0.582243786870052 | 0.814207749517378 | -6.15394248202174 |
| ENSG00000122783.16 | CYREN | -0.0686979372400387 | 4.24014261484512 | -0.554196101143216 | 0.582349283822363 | 0.814299953051338 | -6.17075674382659 |
| lnc-DICER1-1 | lnc-DICER1-1 | 0.0862830463957511 | 1.34694829717097 | 0.553936825027201 | 0.582525142113161 | 0.814454930377362 | -5.82762481838818 |
| ENSG00000143420.18 | ENSA | 0.0474716491471536 | 6.3252386252622 | 0.553904052322536 | 0.582547372566666 | 0.814454930377362 | -6.25858340591447 |
| ENSG00000217512.1 | | -0.147777448365196 | -0.140669383565238 | -0.553799355386814 | 0.582618393571213 | 0.814454930377362 | -5.5040607063158 |
| lnc-HMGN3-6 | lnc-HMGN3-6 | -0.147777448365196 | -0.140669383565238 | -0.553799355386814 | 0.582618393571213 | 0.814454930377362 | -5.5040607063158 |
| ENSG00000205413.7 | SAMD9 | 0.0833463668028687 | 6.7357814837649 | 0.553595000534547 | 0.582757029424358 | 0.814593408260887 | -6.25757319347162 |
| ENSG00000233937.6 | NA | -0.104619286699899 | 1.52896896787952 | -0.553467296410587 | 0.582843672945437 | 0.814621387326024 | -5.85057682725105 |
| ENSG00000185753.12 | CXorf38 | -0.0801924158495098 | 5.02541216614064 | -0.553448833040362 | 0.582856200318241 | 0.814621387326024 | -6.23330248014002 |
| ENSG00000140365.15 | COMMD4 | 0.0931251610794352 | 3.54024047677359 | 0.553275607713108 | 0.582973739813818 | 0.814688102866444 | -6.09838312346207 |
| ENSG00000111540.15 | RAB5B | -0.0740489807182794 | 6.12673820230407 | -0.553175940468475 | 0.583041372743906 | 0.814688102866444 | -6.25841800598916 |
| lnc-CFAP53-1 | lnc-CFAP53-1 | 0.0512611322041747 | 1.95886363066456 | 0.553175866253477 | 0.583041423106676 | 0.814688102866444 | -5.90391002770307 |
| TSPOAP1-AS1 | TSPOAP1-AS1 | -0.116617046241113 | 3.93388051132367 | -0.553119162607442 | 0.583079903175683 | 0.814688102866444 | -6.135264875144 |
| ENSG00000095139.13 | ARCN1 | 0.0745102122189882 | 6.45070362238829 | 0.553086840600739 | 0.5831018379939 | 0.814688102866444 | -6.25898971717266 |
| lnc-CISD3-1 | lnc-CISD3-1 | -0.0662103738165762 | 3.13587018925466 | -0.55289768133214 | 0.583230215926903 | 0.814787000876172 | -6.05577650353288 |
| ENSG00000250959.2 | GLUD1P3 | -0.0699470142449417 | 1.35183637025477 | -0.552750128779795 | 0.58333036583645 | 0.814787000876172 | -5.81986058989929 |
| ENSG00000248643.5 | RBM14-RBM4 | 0.0567237370418692 | 5.36085605392864 | 0.552722673772464 | 0.583349001580933 | 0.814787000876172 | -6.24271382741553 |
| ENSG00000100239.15 | PPP6R2 | -0.0473166589206695 | 5.83647150615613 | -0.552653990938462 | 0.583395622959967 | 0.814787000876172 | -6.25550330845619 |
| lnc-SLC15A3-2 | lnc-SLC15A3-2 | -0.233839716814953 | 0.0883100838451344 | -0.552643616414481 | 0.583402665263331 | 0.814787000876172 | -5.52023838465371 |
| lnc-AKIRIN2-4 | lnc-AKIRIN2-4 | -0.531632263122344 | 1.1481251789241 | -0.552655339367478 | 0.583509614093047 | 0.814787000876172 | -5.84854769040309 |
| lnc-MBOAT4-5 | lnc-MBOAT4-5 | 0.0715961258679059 | 5.36851176531542 | 0.552473169591543 | 0.583518371697756 | 0.814787000876172 | -6.24140990126651 |
| ENSG00000145979.17 | TBC1D7 | -0.0779328331071723 | 2.96151123960832 | -0.552429131682558 | 0.583548268272947 | 0.814787000876172 | -6.01945208988632 |
| ENSG00000155329.11 | ZCCHC10 | -0.10949617033759 | 4.68943380447543 | -0.552426055903468 | 0.58355035639382 | 0.814787000876172 | -6.21177509912661 |
| lnc-LRRC8D-9 | lnc-LRRC8D-9 | 0.0654645747936318 | 3.13374805573329 | 0.552382763473474 | 0.58357974764743 | 0.814787000876172 | -6.04998922005969 |
| ENSG00000131153.8 | GINS2 | 0.173696566188537 | 1.26638939434485 | 0.552341057295709 | 0.583608062669067 | 0.814787000876172 | -5.80561981141027 |
| ENSG00000138685.15 | FGF2 | 0.342477581969644 | 1.85820734424282 | 0.55225233035003 | 0.583773487027766 | 0.814888950924511 | -5.97921165528231 |
| ENSG00000067191.15 | CACNB1 | 0.104826597395772 | 0.574065781735357 | 0.552074614115543 | 0.583788971043411 | 0.814888950924511 | -5.68939062383346 |
| ENSG00000104856.13 | RELB | -0.100709294858265 | 5.08845966141113 | -0.552004838876608 | 0.583836351185655 | 0.814888950924511 | -6.23594063898738 |
| ENSG00000067082.14 | KLF6 | -0.177520411132007 | 7.69717421698325 | -0.552094879560985 | 0.583839448053431 | 0.814888950924511 | -6.24236089781624 |
| ENSG00000149115.13 | TNKS1BP1 | -0.0847975211671441 | 2.40441717701299 | -0.551923715984443 | 0.583891439164015 | 0.814906257954497 | -5.97024222904089 |
| ENSG00000205560.12 | CPT1B | -0.127309409717291 | 4.54858373063685 | -0.551840771349908 | 0.583951401351026 | 0.814934686691177 | -6.20511813040773 |
| ENSG00000214050.7 | FBXO16 | 0.092289163750789 | 4.90987652980048 | 0.551706291046298 | 0.584039097892224 | 0.815001813689197 | -6.22584611522978 |
| ENSG00000175727.13 | MLXIP | 0.125605845133165 | 4.85032800974418 | 0.55154087618992 | 0.584156266103019 | 0.815035708786638 | -6.22369769335382 |

| | | | | | | | |
|---|---|---|---|---|---|---|---|
| ENSG00000187510.9 | PLEKHG7 | 0.183367346742448 | 2.1751175543624 | 0.551530982879945 | 0.584167396715441 | 0.815035708786638 | -5.90482050290743 |
| ENSG00000178057.14 | NDUFAF3 | 0.061625815798755 | 3.94907688741854 | 0.551450375392042 | 0.584212919694214 | 0.815035708786638 | -6.14204564271357 |
| lnc-AP3B1-3 | lnc-AP3B1-3 | 0.164117197354397 | 3.54009186319917 | 0.551391127675331 | 0.584275402665635 | 0.815035708786638 | -6.08573682311864 |
| lnc-TLR1-1 | lnc-TLR1-1 | 0.0951163873079683 | 1.76619672521415 | 0.551352519803333 | 0.584279391286897 | 0.815035708786638 | -5.87519145811959 |
| ENSG00000198901.13 | PRC1 | 0.12293196201651 | 3.964722286174 | 0.551320750223128 | 0.584300972591733 | 0.815035708786638 | -6.13582045810659 |
| lnc-LIPI-4 | lnc-LIPI-4 | -0.469223331164796 | -1.43543549996519 | -0.551203802925421 | 0.584394005501177 | 0.815058254256647 | -5.15360624108172 |
| ENSG00000158747.14 | NBL1 | 0.118711367831742 | -0.243998499664929 | 0.551120154957598 | 0.584437247271283 | 0.815058254256647 | -5.50407716734992 |
| ENSG00000183484.11 | GPR132 | 0.151029102979682 | 3.68908388934191 | 0.551080854027983 | 0.584475099637745 | 0.815058254256647 | -6.10459500268973 |
| ENSG00000099385.11 | BCL7C | -0.0805927015835421 | 3.20420532369883 | -0.551063807217543 | 0.584475529943551 | 0.815058254256647 | -6.06286471414804 |
| lnc-CWF19L2-1 | lnc-CWF19L2-1 | 0.125573592406925 | 3.58407541376282 | 0.550972967001729 | 0.584537249358318 | 0.815083506353547 | -6.096824458158 |
| ENSG00000163492.15 | CCDC141 | -0.240881536207849 | 8.12646742159278 | -0.551063613471702 | 0.584572837817182 | 0.815083506353547 | -6.23921348552424 |
| lnc-C11orf58-8 | lnc-C11orf58-8 | 0.127776074085763 | 1.18332151057979 | 0.550860297322962 | 0.584613804689988 | 0.815085412310102 | -5.81847666861961 |
| ENSG00000185010.14 | F8 | 0.195566804045214 | 1.05730615835261 | 0.550646639051272 | 0.584758991701167 | 0.815232614886557 | -5.75806452786982 |
| lnc-TYR-2 | lnc-TYR-2 | 0.186969990262713 | -0.696321994965905 | 0.550453674461089 | 0.584890131673563 | 0.815360174991186 | -5.2422335945281 |
| lnc-RNF13-3 | lnc-RNF13-3 | -0.149796507484754 | -0.238875258646006 | -0.55039543263217 | 0.584929715975312 | 0.815360174991186 | -5.50916590266665 |
| lnc-CD164-2 | lnc-CD164-2 | -0.154818192541494 | 0.165499349511273 | -0.550265247096554 | 0.585018201779975 | 0.815428296013898 | -5.56196046373305 |
| ENSG00000112306.7 | RPS12 | 0.105234581559897 | 9.08436871543555 | 0.550174218098633 | 0.585080077105392 | 0.81545931927548 | -6.21370752220948 |
| ENSG00000145868.16 | FBXO38 | 0.0529510262506632 | 5.98511879698444 | 0.550050875273492 | 0.585163922198872 | 0.815520956775636 | -6.25866037186681 |
| lnc-MTMR7-3 | lnc-MTMR7-3 | -0.084974798977643 | 3.18663378892041 | -0.549909697593933 | 0.585259898036351 | 0.815561320474004 | -6.04840476319445 |
| lnc-GRHPR-5 | lnc-GRHPR-5 | -0.17129739256204 | 0.661498166187728 | -0.549891704605587 | 0.585272130628262 | 0.815561320474004 | -5.66918808644679 |
| ENSG00000257218.5 | GATC | 0.0604335573908354 | 4.68073910362309 | 0.54980971322174 | 0.585327874295748 | 0.815583782604209 | -6.20959123969582 |
| ENSG00000168502.17 | MTCL1 | -0.278345695433369 | 3.34601294655948 | -0.549863862180505 | 0.585383011307684 | 0.815605396449338 | -6.07221810570699 |
| ENSG00000258610.1 | | 0.728327398259008 | -0.513413270860819 | 0.549532575917552 | 0.585621177241737 | 0.81588200156807 | -5.33704431099375 |
| lnc-MYEF2-6 | lnc-MYEF2-6 | -0.327452620848838 | -0.74336134300193 | -0.549308754271694 | 0.585671448265463 | 0.815896813512556 | -5.35245475917837 |
| ENSG00000112249.13 | ASCC3 | 0.0689463906761159 | 6.8242211574576 | 0.549064210406038 | 0.585834838129098 | 0.816069198241149 | -6.2592978078583 |
| ENSG00000114395.10 | CYB561D2 | 0.0825027701212746 | 2.86449166344101 | 0.548905471018331 | 0.585942812826723 | 0.816164371415878 | -6.01205204619317 |
| ENSG00000120699.12 | EXOSC8 | 0.055121717180648 | 4.74035685911314 | 0.548844550991148 | 0.585984253232084 | 0.816166861840302 | -6.21521843280822 |
| ENSG00000146701.11 | MDH2 | -0.0869941681729509 | 5.12917707599795 | -0.548756247816796 | 0.586044323324074 | 0.816168735998754 | -6.2369690978338 |
| ENSG00000134330.18 | IAH1 | -0.0568474264810126 | 4.59544013843008 | -0.548725995245032 | 0.586064903952853 | 0.816168735998754 | -6.20631529159329 |
| lnc-REXO4-2 | lnc-REXO4-2 | -0.0844145540156664 | 0.870680969792536 | -0.548647521753055 | 0.586118290580052 | 0.816177188137473 | -5.74745873821696 |
| ENSG00000172530.20 | BANP | 0.0782322499477361 | 4.58330626571868 | 0.548600503156148 | 0.58615027911617 | 0.816177188137473 | -6.20041003361132 |
| ENSG00000137312.14 | FLOT1 | -0.0801982578327909 | 4.94882655076295 | -0.548480729968974 | 0.58623176915043 | 0.816201162610123 | -6.22962917577386 |
| lnc-CD164-1 | lnc-CD164-1 | -0.0733369715137329 | 5.96770572758267 | -0.548452377717867 | 0.586251059956348 | 0.816201162610123 | -6.26066083539142 |
| ENSG00000146094.14 | DOK3 | -0.174772056018532 | 3.76652304097527 | -0.548463777331304 | 0.586286459174594 | 0.816201162610123 | -6.13816146321572 |
| ENSG00000114999.7 | TTL | -0.0763762623854367 | 5.06046681643833 | -0.548317626483636 | 0.586342748551334 | 0.816224319858793 | -6.23155646849674 |
| ENSG00000142784.15 | WDTC1 | 0.0681404780405845 | 5.36643486786986 | 0.54802965431218 | 0.586538716134764 | 0.816407102263596 | -6.246911073518 |
| ENSG00000275981.1 | | 0.190417457046546 | 0.0922893483141041 | 0.548008107231806 | 0.58655338037575 | 0.816407102263596 | -5.58738953709957 |
| ENSG00000160193.11 | WDR4 | -0.0744110081841507 | 3.1064215675159 | -0.547856404000754 | 0.586656629633949 | 0.816446276211074 | -6.04964768039268 |
| ENSG00000143157.11 | POGK | 0.0318650862897059 | 6.52231041136218 | 0.54785019264713 | 0.586660857268706 | 0.816446276211074 | -6.2614867146561 |
| ENSG00000125910.5 | S1PR4 | -0.133765688220342 | 4.19159388368869 | -0.547582958814422 | 0.586849151810339 | 0.816510765293971 | -6.17505591735701 |
| ENSG00000146410.11 | MTFR2 | 0.121204770573846 | 0.711055541730379 | 0.547440304787775 | 0.5869398716306 | 0.816510765293971 | -5.70967039791077 |
| ENSG00000152795.17 | HNRNPDL | -0.0569515082013918 | 8.1486864196231 | -0.547439017688072 | 0.586940747871223 | 0.816510765293971 | -6.23805102765673 |
| SNHG20 | SNHG20 | -0.102461393235287 | 2.61530342366276 | -0.547357944353913 | 0.586995942800008 | 0.816510765293971 | -5.98653234070805 |
| ENSG00000109771.15 | LRP2BP | -0.152979055687058 | 0.12071872609932 | -0.547298433319748 | 0.58703645964538 | 0.816510765293971 | -5.54725571838 |
| lnc-UCK1-5 | lnc-UCK1-5 | 0.0846173322855877 | 2.5219738093272 | 0.547262399047261 | 0.587060993479578 | 0.816510765293971 | -5.97007326601818 |
| ENSG00000102174.8 | PHEX | 0.408623820043406 | 5.69772208716656 | 0.547353387543965 | 0.58711267617437 | 0.816510765293971 | -6.24547459553397 |
| ENSG00000061938.17 | TNK2 | -0.0830181175571028 | 5.82238735133066 | -0.547100967254716 | 0.58717090990225 | 0.816510765293971 | -6.25899554077333 |
| ENSG00000256683.6 | ZNF350 | 0.111780812262596 | 5.21862615327455 | 0.547081272558087 | 0.587184320393398 | 0.816510765293971 | -6.23878886759844 |
| ENSG00000167397.14 | VKORC1 | -0.104374534376648 | 2.03529125228725 | -0.547062967929253 | 0.587196784492789 | 0.816510765293971 | -5.91986962010649 |
| lnc-ZNF785-3 | lnc-ZNF785-3 | -0.100168285411116 | 0.525747715199861 | -0.546962400789655 | 0.587265265545204 | 0.816510765293971 | -5.68387648228553 |
| ENSG00000104518.10 | GSDMD | -0.0847205588425277 | 5.37905938899344 | -0.546947658981614 | 0.587275304280441 | 0.816510765293971 | -6.2479836391401 |
| ENSG00000130939.18 | UBE4B | -0.0416572988161166 | 5.72747966101412 | -0.546887558327701 | 0.587316231899448 | 0.816510765293971 | -6.25717865515516 |
| ENSG00000108506.12 | INTS2 | -0.04603958800584 | 4.7400328195307 | -0.546868775040653 | 0.587329023308866 | 0.816510765293971 | -6.21515080766505 |
| ENSG00000155099.7 | PIP4P2 | 0.102429044527205 | 2.27490502837445 | 0.546851656851109 | 0.587340680903999 | 0.816510765293971 | -5.95858132071512 |
| lnc-MOB3C-3 | lnc-MOB3C-3 | -0.0834585681442365 | -0.0675668712717033 | -0.546849862209684 | 0.587341903072562 | 0.816510765293971 | -5.51057725018081 |
| ENSG00000107669.17 | ATE1 | 0.0562437706241745 | 4.80881937955321 | 0.546535470632204 | 0.587556025657338 | 0.816531157981477 | -6.22117184502179 |
| ENSG00000112514.15 | CUTA | -0.0744953336375884 | 5.0663846726955 | -0.546490386710631 | 0.587586734017654 | 0.816531157981477 | -6.23519345761087 |
| ENSG00000132005.8 | RFX1 | -0.092917644933364 | 2.02660355312045 | -0.546482828312633 | 0.587591882403032 | 0.816531157981477 | -5.92467955230624 |
| ENSG00000188529.14 | SRSF10 | 0.0504502561471671 | 8.10944146353569 | 0.546473272339348 | 0.587598391463151 | 0.816531157981477 | -6.23925502944115 |
| ENSG00000260916.7 | CCPG1 | -0.0733336597315057 | 5.60743989245301 | -0.546455197174469 | 0.587610703473049 | 0.816531157981477 | -6.25408652869421 |
| ENSG00000204923.3 | FBXO48 | -0.083398167582394 | 1.59901309637572 | -0.546353658122515 | 0.58767986972114 | 0.816531157981477 | -5.85910243375044 |
| lnc-TEX29-3 | lnc-TEX29-3 | 0.105082670515043 | 1.05678813453298 | 0.546333104558734 | 0.587693870847399 | 0.816531157981477 | -5.78754519738983 |
| ENSG00000137812.19 | KNL1 | 0.172457603587547 | 5.56729533822383 | 0.546347051213156 | 0.587746974995764 | 0.816531157981477 | -6.25586124391625 |
| ENSG00000160813.6 | PPP1R35 | -0.0957798988155223 | 3.06071322482415 | -0.546130735371755 | 0.587831733649382 | 0.816531157981477 | -6.0532836317107 |
| lnc-EIF4E3-5 | lnc-EIF4E3-5 | 0.186764732587293 | 0.594194449772431 | 0.546011028516693 | 0.587913290520997 | 0.816531157981477 | -5.6573690631727 |
| ENSG00000135316.17 | SYNCRIP | 0.0700624214337138 | 6.86149416181802 | 0.546004369986182 | 0.587917827169803 | 0.816531157981477 | -6.26000058781035 |
| lnc-HNRNPH1-1 | lnc-HNRNPH1-1 | 0.13134965676863 | 0.148726397714806 | 0.545991373511282 | 0.587926682091142 | 0.816531157981477 | -5.56206243488857 |
| ENSG00000136738.14 | STAM | 0.103320821335164 | 6.59412577931111 | 0.545982045262368 | 0.587933037770098 | 0.816531157981477 | -6.26261105755296 |

| | | | | | | | |
|---|---|---|---|---|---|---|---|
| lnc-LRP1-1 | lnc-LRP1-1 | 0.179480649502155 | -0.551444495016312 | 0.54597344350629 | 0.587938898491932 | 0.816531157981477 | -5.36042985090709 |
| ENSG00000239789.5 | MRPS17 | 0.0790826656976549 | 2.21979958346419 | 0.545949199062164 | 0.587955417355835 | 0.816531157981477 | -5.94231770420902 |
| ENSG00000114867.20 | EIF4G1 | 0.0576865121458878 | 7.01834784564535 | 0.54588694791026 | 0.587997832969843 | 0.816531157981477 | -6.25852950841084 |
| lnc-CYB5R4-2 | lnc-CYB5R4-2 | -0.213022243060126 | 0.0841024465809343 | -0.54583832312442 | 0.588030965105156 | 0.816531157981477 | -5.52005677806345 |
| lnc-PLPP2-5 | lnc-PLPP2-5 | -0.118848716919587 | -0.194174824747511 | -0.545600617352217 | 0.588192946801852 | 0.816695000597989 | -5.45944264494892 |
| lnc-ARHGAP22-1 | lnc-ARHGAP22-1 | 0.313428381535757 | -0.742665772665021 | 0.545548720901705 | 0.588228313844686 | 0.816695000597989 | -5.34260261510503 |
| ENSG00000180385.8 | EMC3-AS1 | 0.0545575303910506 | 3.22426691101 | 0.545484799009102 | 0.588271877535188 | 0.816700394935031 | -6.05634545905268 |
| ENSG00000048462.10 | TNFRSF17 | -0.186453028378306 | 1.66982330659937 | -0.545427178369517 | 0.588316735204044 | 0.81670758485903 | -5.85227006920282 |
| lnc-UBD-1 | lnc-UBD-1 | 0.0869034495593806 | 1.47045321450669 | 0.545332898843698 | 0.588375405879398 | 0.81673394788344 | -5.82765264651636 |
| ENSG00000149418.10 | ST14 | 0.0950972370112462 | 6.38022568603117 | 0.545239844771878 | 0.588438831669647 | 0.816766907563311 | -6.26331304825759 |
| ENSG00000168792.4 | ABHD15 | -0.103755136327994 | 4.85369432302796 | -0.544994472945241 | 0.588606093145529 | 0.816912986672556 | -6.22539115457184 |
| ENSG00000151498.11 | ACAD8 | 0.0741816329373801 | 4.30416083401934 | 0.544927483109471 | 0.588651761747692 | 0.816912986672556 | -6.17553787865587 |
| ENSG00000111605.17 | CPSF6 | -0.0509621809675824 | 7.22658119233536 | -0.544858924375948 | 0.588698501660178 | 0.816912986672556 | -6.25567004515236 |
| ENSG00000285554.1 | LINC02988 | 0.146994876876051 | -0.240865656219874 | 0.544852576964217 | 0.588702829097509 | 0.816912986672556 | -5.45569152901859 |
| lnc-ERCC6L-2 | lnc-ERCC6L-2 | -0.129135708484269 | 6.84402814547257 | -0.544770005912797 | 0.588772527512544 | 0.816930956871558 | -6.26079486661291 |
| ENSG00000279801.1 | | -0.156833198342666 | 1.12527910998379 | -0.544717155235293 | 0.588795158392342 | 0.816930956871558 | -5.7911450752972 |
| ENSG00000243970.3 | PPIEL | -0.302291146167927 | 2.62309798191186 | -0.544786429868435 | 0.588836827651003 | 0.816933703393138 | -5.97603521424542 |
| lnc-TENM1-11 | lnc-TENM1-11 | -0.0756379700822263 | 1.8239111903785 | -0.544524877216137 | 0.588926263678031 | 0.817002715190734 | -5.89200578434243 |
| ENSG00000198131.13 | ZNF544 | -0.0584370673574854 | 5.16347921851843 | -0.544424411601997 | 0.588994771978001 | 0.817042687129209 | -6.23967175337967 |
| ENSG00000130305.16 | NSUN5 | 0.0780086860518576 | 3.92288604155973 | 0.544136443229411 | 0.589191161001257 | 0.817260035507034 | -6.1507868386067 |
| ENSG00000188227.12 | ZNF793 | -0.109452031963523 | 4.19233321104432 | -0.543884617010006 | 0.589362927393946 | 0.817423936596941 | -6.17061681093491 |
| ENSG00000078487.17 | ZCWPW1 | -0.0745159065314556 | 3.50480783984983 | -0.543846761465304 | 0.589388750086732 | 0.817423936596941 | -6.09580382887191 |
| ENSG00000060237.16 | WNK1 | -0.0656700014447695 | 7.69719432408092 | -0.543711491186812 | 0.589481027453624 | 0.817444487308283 | -6.24999290348554 |
| ENSG00000236946.2 | HNRNPA1P70 | -0.472105324889173 | -0.303540399925724 | -0.543834215882089 | 0.589482996896314 | 0.817444487308283 | -5.55627735066239 |
| ENSG00000198707.14 | CEP290 | 0.0631317889198808 | 5.56946343433824 | 0.543585152349701 | 0.589567218287028 | 0.817506201428315 | -6.25421118699583 |
| lnc-ENO1-2 | lnc-ENO1-2 | -0.152393364156338 | -0.308001792664738 | -0.543379596626872 | 0.589707465292381 | 0.817594181591022 | -5.43428979705238 |
| ENSG00000236809.2 | SNX25P1 | 0.214561899101477 | 0.248867306843987 | 0.54337571874952 | 0.589710111252019 | 0.817594181591022 | -5.57591157752447 |
| ENSG00000256981.1 | | 0.117906696902782 | 1.5961430252305 | 0.543071818105183 | 0.589917486860898 | 0.817826606860501 | -5.86982299814978 |
| GASAL1 | GASAL1 | 0.203273141997634 | 1.56906911122984 | 0.542770838350926 | 0.590140418871061 | 0.818043969575415 | -5.8504204376225 |
| ENSG00000197498.12 | RPF2 | -0.0826677872474844 | 2.61618445661986 | -0.542692968782335 | 0.590176054640529 | 0.818043969575415 | -5.98608708870579 |
| ENSG00000196628.16 | TCF4 | -0.119944486495445 | 9.85856936459714 | -0.542667401175185 | 0.59019350668974 | 0.818043969575415 | -6.19430844261425 |
| ENSG00000167491.17 | GATAD2A | -0.0632470733829624 | 5.16141285077411 | -0.542374464980746 | 0.590393477922852 | 0.818266039733759 | -6.24429337299541 |
| lnc-CXCR4-3 | lnc-CXCR4-3 | -0.164718652646448 | -0.767319765287944 | -0.542227018513342 | 0.590494143638934 | 0.818350455057917 | -5.39008300684499 |
| ENSG00000118246.13 | FASTKD2 | 0.0483807480066654 | 5.03698767826139 | 0.541875572447678 | 0.590734118420234 | 0.818589034166395 | -6.23746618938185 |
| ENSG00000148356.13 | LRSAM1 | -0.065909476932835 | 4.22042503869681 | -0.5418584157027 | 0.590745834596882 | 0.818589034166395 | -6.17663885612664 |
| ENSG00000271699.5 | SNX29P2 | 0.180946752425178 | 1.36166420166029 | 0.541493354314643 | 0.590995157756719 | 0.818842090677457 | -5.81743811127624 |
| ENSG00000074201.8 | CLNS1A | -0.0636100496480116 | 5.61879686343689 | -0.54139574492029 | 0.591061829782199 | 0.818842090677457 | -6.25723225354899 |
| lnc-FAM46C-3 | lnc-FAM46C-3 | 0.14847184995944 | -0.110255292192565 | 0.541366303850235 | 0.591081940186667 | 0.818842090677457 | -5.52574089711801 |
| ENSG00000171700.13 | RGS19 | -0.140753123723511 | 3.08597195227931 | -0.541309933931892 | 0.591120445873513 | 0.818842090677457 | -6.05971463648575 |
| ENSG00000140009.18 | ESR2 | 0.281219337425176 | 1.25959027500963 | 0.54139173356594 | 0.591127368477692 | 0.818842090677457 | -5.80860167959936 |
| lnc-C2orf78-2 | lnc-C2orf78-2 | -0.111148201025437 | 0.0550511623742886 | -0.541197313347648 | 0.59119737936387 | 0.8188839609318 | -5.52657680837546 |
| lnc-EXOC4-1 | lnc-EXOC4-1 | 0.14993501373736 | -0.546093809844291 | 0.540971691567444 | 0.591351520688242 | 0.819042349123618 | -5.35218938388257 |
| lnc-CPNE4-1 | lnc-CPNE4-1 | 0.0633244770041843 | 1.52892053960526 | 0.540716622324641 | 0.59152580309829 | 0.819228610225549 | -5.85737122809707 |
| lnc-PCP2-1 | lnc-PCP2-1 | -0.186554587342668 | -0.169124827546605 | -0.540563623078712 | 0.591630355367417 | 0.819313411080956 | -5.42513159999896 |
| ENSG00000164902.13 | PHAX | -0.0630003977102414 | 4.37145044919085 | -0.540508466451082 | 0.591668048889218 | 0.819313411080956 | -6.19171039411208 |
| ENSG00000270096.1 | | -0.161059765855665 | 0.250605652927788 | -0.540452276462307 | 0.59170644977498 | 0.819313411080956 | -5.63446541736143 |
| lnc-RBBP6-4 | lnc-RBBP6-4 | -0.39768620618794 | 1.4795923874241 | -0.540411116487957 | 0.591832782802339 | 0.819384272073354 | -5.79716797500283 |
| lnc-HSPD1-1 | lnc-HSPD1-1 | 0.0439085984882919 | 5.78426231394955 | 0.540260906637195 | 0.59183724301193 | 0.819384272073354 | -6.26216264769222 |
| lnc-MYL12B-3 | lnc-MYL12B-3 | -0.0927924486036713 | 3.42251275719418 | -0.540192949842939 | 0.591883691934068 | 0.819393464560056 | -6.08844721405084 |
| lnc-GAPVD1-6 | lnc-GAPVD1-6 | 0.145995350283263 | 1.19764171835417 | 0.540099526334043 | 0.591947550354962 | 0.819408064395501 | -5.82745709444066 |
| ENSG00000285329.1 | | -0.0833479135989768 | 1.15107598766938 | -0.54005380437149 | 0.591978804199919 | 0.819408064395501 | -5.78906429167375 |
| ENSG00000103245.13 | CIAO3 | -0.0825834093948004 | 3.95294075632769 | -0.540002802877701 | 0.592013667863067 | 0.819408064395501 | -6.15698709486363 |
| ENSG00000144802.11 | NFKBIZ | 0.11262362318412 | 6.8390627451353 | 0.539888437981191 | 0.59209184910641 | 0.819461170666839 | -6.26398498975881 |
| ENSG00000141959.16 | PFKL | -0.0705581822041488 | 6.93681248431262 | -0.539829172586421 | 0.592132365586303 | 0.819462144884212 | -6.26329149598024 |
| ENSG00000277194.1 | SNORD22 | 0.0961494319437199 | 2.68450801310835 | 0.539692235488205 | 0.592225986963334 | 0.81948800912283 | -6.00744034405652 |
| lnc-YME1L1-1 | lnc-YME1L1-1 | 0.0730911907182765 | 0.747980031564985 | 0.539685367898616 | 0.592230682393339 | 0.81948800912283 | -5.69816669231899 |
| lnc-TMEM138-9 | lnc-TMEM138-9 | -0.0711170866438496 | -0.030497183210538 | -0.539561696201292 | 0.592315240822975 | 0.819501029878432 | -5.51664487528135 |
| lnc-MYH11-1 | lnc-MYH11-1 | -0.0791880599793875 | 3.12084678583277 | -0.539544701091631 | 0.592326861389842 | 0.819501029878432 | -6.04424078581578 |
| ENSG00000119392.14 | GLE1 | 0.0465126762843235 | 4.66782099121515 | 0.539496915731407 | 0.592359535662018 | 0.819501029878432 | -6.21465928362243 |
| ENSG00000170873.18 | MTSS1 | -0.307598937163805 | 6.8242013283192 | -0.539528050477825 | 0.59244566916194 | 0.81956510574368 | -6.26263299531257 |
| ENSG00000204444.10 | APOM | -0.108283847526198 | 2.07447940761637 | -0.539174214296834 | 0.592580212152775 | 0.819673528746032 | -5.92636190001515 |
| ENSG00000164308.16 | ERAP2 | -0.155453809287152 | 6.49992653689299 | -0.539207724638071 | 0.592603691457499 | 0.819673528746032 | -6.26578076286304 |
| ENSG00000100650.15 | SRSF5 | -0.0677801781057976 | 8.85557624990383 | -0.53890446554695 | 0.592764707377479 | 0.819800818082516 | -6.22567530050617 |
| ENSG00000083535.15 | PIBF1 | 0.0527087514009832 | 4.86056696227371 | 0.538888867469039 | 0.592775376549681 | 0.819800818082516 | -6.2244734913376 |
| ENSG00000006025.11 | OSBPL7 | -0.0600799822713629 | 4.43831369811583 | -0.538563914836116 | 0.592997666692085 | 0.819977669385173 | -6.19658517468476 |
| ENSG00000183255.11 | PTTG1IP | 0.066348916742391 | 4.82237937315968 | 0.538537762394818 | 0.593015558495966 | 0.819977669385173 | -6.22430099916108 |
| lnc-EMC7-4 | lnc-EMC7-4 | -0.0806327048309932 | 0.0883537004225531 | -0.53852722772929 | 0.593022765703138 | 0.819977669385173 | -5.56320049520934 |
| lnc-GSR-3 | lnc-GSR-3 | 0.100946601555385 | 1.80520463998485 | 0.53834064964072 | 0.593150418497063 | 0.820099084028014 | -5.88329475381526 |

| | | | | | | | |
|---|---|---|---|---|---|---|---|
| lnc-SLC6A3-4 | lnc-SLC6A3-4 | 0.139890405385348 | 0.387655495372986 | 0.538112000579325 | 0.593306873138703 | 0.820256983776578 | -5.65497467159799 |
| lnc-NFE2L3-4 | lnc-NFE2L3-4 | 0.0948325148002658 | 4.1053122892722 | 0.538057271888023 | 0.593344324512922 | 0.820256983776578 | -6.1779801898239 |
| ENSG00000261879.6 | ZNF594-DT | -0.0969920368827025 | 1.02460655803559 | -0.537882327249568 | 0.593464048345306 | 0.820367394700566 | -5.77946881778771 |
| ENSG00000283704.1 | | -0.113850367537607 | 0.261432274365801 | -0.537751202079574 | 0.593553791716733 | 0.820434823086432 | -5.63456666494665 |
| ENSG00000144468.16 | RHBDD1 | 0.0798504767035917 | 6.52967689166135 | 0.537694579827436 | 0.593592546538559 | 0.820434823086432 | -6.26702380120834 |
| lnc-GOLGA8R-1 | lnc-GOLGA8R-1 | -0.107966979766369 | -0.241445558470634 | -0.537515477763236 | 0.593715139962492 | 0.820549165772375 | -5.44453905815398 |
| ENSG00000139173.9 | TMEM117 | 0.0824574606410238 | 2.91665722980549 | 0.5373914400443 | 0.593800049470801 | 0.820611415218041 | -6.02793501930265 |
| ENSG00000258644.5 | SYNJ2BP-COX16 | 0.0520288424094644 | 3.12868877179154 | 0.536873832937656 | 0.594154437373614 | 0.821046041252843 | -6.05507009231801 |
| ENSG00000127399.14 | LRRC61 | -0.113486495139321 | 2.00668144650772 | -0.536608029447198 | 0.594336462860295 | 0.821242441934307 | -5.91858806332527 |
| lnc-CYFIP1-2 | lnc-CYFIP1-2 | 0.100972742777708 | 0.77988741231892 | 0.536360393577252 | 0.594506070690092 | 0.821419577040775 | -5.72388731684521 |
| lnc-SFXN4-1 | lnc-SFXN4-1 | -0.101039551145487 | 2.34560439035889 | -0.536255303243072 | 0.594578054841127 | 0.821419577040775 | -5.96355662496385 |
| ENSG00000108825.17 | PTGES3L-AARSD1 | -0.0505760778037954 | 2.17174700379947 | -0.536246070451103 | 0.594584379261065 | 0.821419577040775 | -5.94200123001602 |
| lnc-GDF5-2 | lnc-GDF5-2 | -0.10156511616624 | 1.55439370959223 | -0.536065603843746 | 0.594708004447038 | 0.821535225203583 | -5.8583555799583 |
| ENSG00000226767.1 | | -0.412687045227607 | -0.486399899812007 | -0.535976942282166 | 0.594814639285382 | 0.82158791839518 | -5.37979177760358 |
| ENSG00000148842.17 | CNNM2 | -0.0717710056057808 | 3.86055885937096 | -0.535810331661693 | 0.594882894520236 | 0.82158791839518 | -6.13879461467366 |
| ENSG00000154262.12 | ABCA6 | -0.215134132191645 | 8.43100497563645 | -0.535906540683208 | 0.594897556761094 | 0.82158791839518 | -6.24049970513728 |
| lnc-CSRNP3-9 | lnc-CSRNP3-9 | 0.240655703728277 | 3.07331412192176 | 0.535880647655041 | 0.594905812357856 | 0.82158791839518 | -6.01366284099638 |
| ENSG00000176953.12 | NFATC2IP | 0.0406673511308178 | 5.73033661663415 | 0.535697546138541 | 0.594960172999579 | 0.82160786587389 | -6.26273878994774 |
| ENSG00000143156.13 | NME7 | -0.068368134791716 | 2.03142367160626 | -0.535626937787161 | 0.595008554905206 | 0.821619554918413 | -5.9133629201815 |
| ENSG00000166377.20 | ATP9B | 0.0708039712141995 | 5.96292822985041 | 0.535529496874219 | 0.595075325941132 | 0.821621698186087 | -6.26596521728545 |
| lnc-ZNF263-3 | lnc-ZNF263-3 | -0.0628151796202916 | 2.30826126607414 | -0.535395864286394 | 0.595166902952722 | 0.821621698186087 | -5.95447518665658 |
| ENSG00000082258.12 | CCNT2 | 0.0496652729761218 | 6.87736300821749 | 0.535389572046013 | 0.595171215123321 | 0.821621698186087 | -6.26544586309432 |
| ENSG00000224892.6 | RPS4XP16 | 0.121806484784543 | 1.63149750356156 | 0.535364798273397 | 0.595188193120127 | 0.821621698186087 | -5.86821856239795 |
| LINC01816 | LINC01816 | -0.136151215099665 | -0.193491274324892 | -0.535333424615306 | 0.595209694488303 | 0.821621698186087 | -5.46845603647749 |
| ENSG00000082213.17 | C5orf22 | 0.0691687772216456 | 4.25753069564206 | 0.535038634054652 | 0.595411741788358 | 0.821845485597491 | -6.18499540715852 |
| ENSG00000175764.14 | TTLL11 | 0.110683543010115 | 0.113890060039535 | 0.534565942560342 | 0.595735788718786 | 0.822143256370077 | -5.58094553706863 |
| ENSG00000243927.5 | MRPS6 | -0.113073472841295 | 2.94573530525196 | -0.534395140459281 | 0.595852900150255 | 0.822143256370077 | -6.03740900522646 |
| ENSG00000101096.19 | NFATC2 | -0.0848426087387191 | 5.33009677795172 | -0.53438284771705 | 0.595861329158122 | 0.822143256370077 | -6.24928743476203 |
| ENSG00000279598.1 | | -0.138020950523234 | 2.02773629424436 | -0.534377415836091 | 0.595865053761382 | 0.822143256370077 | -5.92423698881253 |
| ENSG00000090975.12 | PITPNM2 | -0.151853092762973 | 6.95533622931815 | -0.534243085364371 | 0.59600126098052 | 0.822143256370077 | -6.26598086217767 |
| lnc-WDR5-8 | lnc-WDR5-8 | -0.119862479224892 | -0.422292893867893 | -0.534097269882644 | 0.596057162807319 | 0.822143256370077 | -5.42562425385172 |
| lnc-SPINT2-5 | lnc-SPINT2-5 | -0.149000069851055 | 1.96639772127736 | -0.534050541123343 | 0.596089209724669 | 0.822143256370077 | -5.90215198454534 |
| lnc-PDZD9-4 | lnc-PDZD9-4 | -0.238430376680721 | 0.640504327425814 | -0.534047434959339 | 0.596118789970688 | 0.822143256370077 | -5.75563930573619 |
| ENSG00000158828.7 | PINK1 | -0.0701652438111204 | 3.63891770943592 | -0.533989685124386 | 0.596130946423028 | 0.822143256370077 | -6.115461930409 |
| LINC02391 | LINC02391 | 0.1371475110638 | 0.0702005320482226 | 0.533986595200862 | 0.596133065613363 | 0.822143256370077 | -5.56593680218943 |
| ENSG00000168904.14 | LRRC28 | -0.0743283584251673 | 3.10239073092916 | -0.533971451603568 | 0.596143451735739 | 0.822143256370077 | -6.05323187571587 |
| lnc-BFSP1-4 | lnc-BFSP1-4 | -0.171480197463802 | 3.32656160583826 | -0.534010917615614 | 0.596149086721963 | 0.822143256370077 | -6.091540609318 |
| ENSG00000279549.1 | | -0.114487026514776 | 0.0182514732753999 | -0.533912333397705 | 0.59618399833018 | 0.822143256370077 | -5.54311157658507 |
| lnc-NUP43-1 | lnc-NUP43-1 | -0.0744889348612594 | 1.65092167621427 | -0.533908436954747 | 0.596186670775872 | 0.822143256370077 | -5.87470070515191 |
| ENSG00000136444.9 | RSAD1 | 0.0582039288461926 | 4.43250118480962 | 0.533811246776435 | 0.596253332241748 | 0.822167819182222 | -6.19990650046968 |
| ENSG00000162086.14 | ZNF75A | -0.0584054315314537 | 5.95634559813807 | -0.533765995309427 | 0.596284370827975 | 0.822167819182222 | -6.26719894343308 |
| LINC02288 | LINC02288 | -0.160296296968596 | -0.159525455162578 | -0.53369257783879 | 0.5963347304742 | 0.822182179472869 | -5.50132162779067 |
| ENSG00000279319.1 | | -0.124726038478698 | 2.99355005175302 | -0.533589938030209 | 0.596405138121061 | 0.822215415072946 | -6.03421814494499 |
| lnc-SLC30A5-5 | lnc-SLC30A5-5 | 0.137416024886955 | 0.433832634687943 | 0.533536214063342 | 0.596441992618196 | 0.822215415072946 | -5.66534918759898 |
| lnc-FAM167A-2 | lnc-FAM167A-2 | 0.149683706226703 | 1.53207114771563 | 0.533463279823743 | 0.596492027029684 | 0.822215415072946 | -5.87782694784316 |
| ENSG00000257267.3 | ZNF271P | 0.0447976452905117 | 4.44264963446027 | 0.533424514236591 | 0.596518621837647 | 0.822215415072946 | -6.1961858538296 |
| ENSG00000105186.15 | ANKRD27 | 0.049879784793462 | 4.37788371448518 | 0.533312310225322 | 0.596595601615647 | 0.822219457268155 | -6.1938994075683 |
| ENSG00000282907.1 | | -0.105860797559928 | 0.625689277215044 | -0.533200411578287 | 0.596672376557021 | 0.822219457268155 | -5.71629637468553 |
| ENSG00000267041.5 | ZNF850 | 0.104051144917846 | 2.57562820550818 | 0.533189034484053 | 0.596680182772336 | 0.822219457268155 | -5.99418181177954 |
| lnc-SNRPD2-2 | lnc-SNRPD2-2 | -0.0938064034591362 | 1.27358806868159 | -0.533187347050279 | 0.596681340582735 | 0.822219457268155 | -5.80801846943019 |
| ENSG00000126062.3 | TMEM115 | -0.0792372846358145 | 3.62374813704722 | -0.532802756711444 | 0.596945249789009 | 0.8224788171253 | -6.12100229317462 |
| ENSG00000111335.12 | OAS2 | -0.137541811850125 | 5.32775627839917 | -0.53283368988421 | 0.596949475248256 | 0.8224788171253 | -6.25287104388073 |
| ENSG00000147383.10 | NSDHL | 0.0656201495085715 | 1.90889784415725 | 0.532701794336133 | 0.597014540155143 | 0.822513405623381 | -5.91239154703004 |
| ENSG00000119723.16 | COQ6 | 0.0784232757532068 | 4.09020618879993 | 0.532543975748466 | 0.597122858467204 | 0.822517570375499 | -6.16352999408958 |
| lnc-CCDC184-3 | lnc-CCDC184-3 | -0.153098372708642 | 0.114714895533271 | -0.532464829813997 | 0.597177183526013 | 0.822517570375499 | -5.55724689223788 |
| ENSG00000132386.10 | SERPINF1 | 0.21950550278253 | 4.7507661127211 | 0.532579684761684 | 0.597177654329816 | 0.822517570375499 | -6.21081670049178 |
| ENSG00000180574.3 | EIF2S3B | 0.32052365481371 | 1.06693112626022 | 0.532515971225814 | 0.597214834441231 | 0.822517570375499 | -5.76469394838558 |
| ENSG00000066926.10 | FECH | 0.0845635125627964 | 3.14540120042102 | 0.53237582864463 | 0.597238275910167 | 0.822517570375499 | -6.06867220454261 |
| ENSG00000077420.15 | APBB1IP | -0.10530614768553 | 5.25603520382584 | -0.532291861732836 | 0.597295915366073 | 0.822517570375499 | -6.24681395963412 |
| ENSG00000284630.1 | | -0.112610799354547 | -0.330684794964614 | -0.532286988482905 | 0.597299260710348 | 0.822517570375499 | -5.46296208152728 |
| ENSG00000143363.16 | PRUNE1 | -0.0652802668168517 | 3.82131746457577 | -0.532177707618522 | 0.597374281161109 | 0.822517570375499 | -6.14003668474459 |
| lnc-PLA2G15-1 | lnc-PLA2G15-1 | -0.0713883964177704 | 2.88032382932125 | -0.532173438102274 | 0.59737721224036 | 0.822517570375499 | -6.02269049380202 |
| lnc-ZSCAN25-2 | lnc-ZSCAN25-2 | -0.0943858346102704 | 1.53111068777889 | -0.532097782575095 | 0.59742915188127 | 0.822522236325645 | -5.85118881359995 |
| lnc-ERRFI1-3 | lnc-ERRFI1-3 | 0.0812144859593462 | 1.82600685546391 | 0.531979416214462 | 0.597510417972 | 0.822522236325645 | -5.88857084014459 |
| lnc-AK7-6 | lnc-AK7-6 | -0.105668478488323 | 2.04798103227953 | -0.531894547362921 | 0.597568689079245 | 0.822522236325645 | -5.93984588430796 |
| ENSG00000124224.16 | PPP4R1L | -0.0920198164108665 | 4.48217399986377 | -0.531859336279146 | 0.597592865858201 | 0.822522236325645 | -6.20858120214521 |
| ENSG00000182199.10 | SHMT2 | -0.154144717882162 | 6.14271441824683 | -0.53190942533851 | 0.59760528895128 | 0.822522236325645 | -6.2696753315425 |
| ENSG00000134007.3 | ADAM20 | -0.22984020099007 | 0.503988252445324 | -0.531829927907912 | 0.597620368471556 | 0.822522236325645 | -5.65896019891567 |

| | | | | | | | |
|---|---|---|---|---|---|---|---|
| ENSG00000187514.16 | PTMA | -0.097846582932787 | 8.18587711773879 | -0.531646117848113 | 0.597739276606407 | 0.822525080896205 | -6.24417733735467 |
| ENSG00000182359.14 | KBTBD3 | 0.103281753209635 | 1.73261406704363 | 0.531629049315274 | 0.597750997789317 | 0.822525080896205 | -5.87947672606702 |
| ENSG00000163482.11 | STK36 | -0.0915597059880957 | 3.49070444999749 | -0.531607946946223 | 0.597765489209381 | 0.822525080896205 | -6.10735828238134 |
| lnc-MAG-1 | lnc-MAG-1 | 0.184165612656349 | 2.2495032163003 | 0.531613526614029 | 0.597782280771818 | 0.822525080896205 | -5.94887540025732 |
| ENSG00000181929.12 | PRKAG1 | 0.0619435056041775 | 4.56647192161704 | 0.531335130679078 | 0.597952852492137 | 0.822628293689282 | -6.20915271803653 |
| ENSG00000161542.16 | PRPSAP1 | -0.0830383636709349 | 3.48727991444246 | -0.531321823843183 | 0.597961991995496 | 0.822628293689282 | -6.09877071389266 |
| ENSG00000100379.17 | KCTD17 | -0.117469264345292 | 2.32180884338159 | -0.53129969421816 | 0.597977191380218 | 0.822628293689282 | -5.97604797638643 |
| ENSG00000164978.17 | NUDT2 | -0.103673920527702 | 2.42265008989832 | -0.53097773611691 | 0.598198343785315 | 0.82282414685094 | -5.97304947949018 |
| ENSG00000102738.7 | MRPS31 | -0.0551962427778108 | 3.47925160863326 | -0.5309760371124 | 0.598199510930256 | 0.82282414685094 | -6.09887997423995 |
| ENSG00000160214.12 | RRP1 | -0.07791311036156 | 3.21327172565654 | -0.53083081269858 | 0.598299277976812 | 0.822866477880977 | -6.06547124854111 |
| ENSG00000122406.13 | RPL5 | 0.077664859521775 | 9.42017591541438 | 0.530814853996544 | 0.598310241848453 | 0.822866477880977 | -6.21287945645912 |
| ENSG00000198755.10 | RPL10A | 0.0810925293378695 | 8.79524038459004 | 0.530411562289274 | 0.598587340743123 | 0.823120975953071 | -6.23059885636641 |
| ENSG00000119969.14 | HELLS | -0.0915789535487423 | 4.43799181144575 | -0.530390824360399 | 0.598601591257879 | 0.823120975953071 | -6.19827019670225 |
| lnc-ZNF33A-7 | lnc-ZNF33A-7 | 0.188606761433899 | 0.00218709639318341 | 0.530339958307032 | 0.598636545637563 | 0.823120975953071 | -5.53140986167425 |
| ENSG00000179104.8 | TMTC2 | -0.543957503634982 | 1.13216082356719 | -0.530472219867815 | 0.598655249964411 | 0.823120975953071 | -5.76890810430822 |
| lnc-APLN-1 | lnc-APLN-1 | 0.138862309945453 | -0.219511418692445 | 0.530111757922066 | 0.598793373276413 | 0.8232195641886 | -5.43448925600471 |
| lnc-ATP6V1G1-2 | lnc-ATP6V1G1-2 | -0.0921666255804592 | 2.64451936957504 | -0.530092013381251 | 0.598806943352851 | 0.8232195641886 | -6.0014464953646 |
| lnc-GPR155-4 | lnc-GPR155-4 | -0.0833455303045483 | 3.14507268749239 | -0.529928311535049 | 0.598919458321227 | 0.82331925536805 | -6.05909032459879 |
| ENSG00000149554.12 | CHEK1 | -0.127184987323941 | 2.10201321808727 | -0.529804362574355 | 0.599004657080954 | 0.823381384846886 | -5.94458038291655 |
| ENSG00000188312.13 | CENPP | 0.0934429754374833 | 1.97444574823302 | 0.529681913111442 | 0.59908883071768 | 0.823395913392197 | -5.91466452790894 |
| ENSG00000270277.1 | | 0.116125179163403 | 1.63833481406831 | 0.529672598603343 | 0.599095233880461 | 0.823395913392197 | -5.86574512596504 |
| lnc-NINL-2 | lnc-NINL-2 | 0.087195642800407 | 0.208205411553861 | 0.529535754348272 | 0.599189309758397 | 0.823455535437253 | -5.59378058402862 |
| ENSG00000010404.17 | IDS | 0.0823234662989236 | 6.24482395222982 | 0.529493110363745 | 0.599218627497409 | 0.823455535437253 | -6.27149421505727 |
| ENSG00000149636.15 | DSN1 | 0.0586045644750652 | 3.46418730911747 | 0.529434403057809 | 0.59925898987097 | 0.823456024335015 | -6.09086909361551 |
| lnc-C5orf63-4 | lnc-C5orf63-4 | -0.102896680578759 | 1.61741764268055 | -0.529238072899119 | 0.59939397980197 | 0.823500945886017 | -5.86442374863566 |
| ENSG00000156011.16 | PSD3 | -0.543223024652585 | 3.36156777546856 | -0.529351859791161 | 0.599425065507621 | 0.823500945886017 | -6.07145790291803 |
| ENSG00000134262.12 | AP4B1 | 0.0597930032736569 | 5.07991021986982 | 0.529144759881495 | 0.599458143655776 | 0.823500945886017 | -6.24199769206407 |
| ENSG00000135336.14 | ORC3 | 0.0522710293815667 | 4.74025606091837 | 0.529101030969747 | 0.599488213618369 | 0.823500945886017 | -6.22618521603569 |
| ENSG00000160323.18 | ADAMTS13 | 0.121633729337052 | 1.31343699427088 | 0.529062208018394 | 0.599514910615228 | 0.823500945886017 | -5.82365282601475 |
| lnc-TAOK1-4 | lnc-TAOK1-4 | -0.0599424283074663 | 4.81337895209454 | -0.529037744238059 | 0.599531733668657 | 0.823500945886017 | -6.22957859342437 |
| ENSG00000263020.6 | | -0.0583263061207715 | 4.78249634526225 | -0.528871909553267 | 0.599645779352293 | 0.823602634219155 | -6.22823401269575 |
| ENSG00000085978.21 | ATG16L1 | 0.0479129314356879 | 5.23912970281987 | 0.528696564104932 | 0.59976637671813 | 0.823632738957448 | -6.25335125393993 |
| ENSG00000196218.12 | RYR1 | 0.311434838853266 | 1.05362555542103 | 0.528785068894788 | 0.599777670580477 | 0.823632738957448 | -5.7810313510519 |
| lnc-FAP-3 | lnc-FAP-3 | -0.121272240983604 | 1.46293144119196 | -0.528665499096994 | 0.599787743488957 | 0.823632738957448 | -5.86884172736764 |
| ENSG00000132465.10 | JCHAIN | 0.445618231058158 | 3.18016010879391 | 0.528672569714348 | 0.599892041827363 | 0.823659794711657 | -6.07924855714634 |
| ENSG00000213145.9 | CRIP1 | -0.192793428290225 | 4.0448129322796 | -0.528597362165917 | 0.599898841150519 | 0.823659794711657 | -6.16537943477616 |
| ENSG00000157538.13 | VPS26C | -0.0512229536822728 | 5.43122543509394 | -0.52846232718322 | 0.599927495618576 | 0.823659794711657 | -6.25821851243543 |
| lnc-STAU1-5 | lnc-STAU1-5 | -0.202511475037804 | 0.612957723129343 | -0.528288294469831 | 0.600047216435171 | 0.823769216026488 | -5.68754668191662 |
| ENSG00000197128.11 | ZNF772 | -0.0778965087744972 | 4.23763350304417 | -0.528222087664477 | 0.600092764448299 | 0.823776802096794 | -6.18298715793418 |
| lnc-IPP-2 | lnc-IPP-2 | -0.0527657881547889 | 3.17526811872532 | -0.528129094149259 | 0.60015674353515 | 0.823807655268021 | -6.06941731040002 |
| lnc-PPP1CC-2 | lnc-PPP1CC-2 | 0.111376242751375 | 3.2998378383329 | 0.528028559197258 | 0.600225914688285 | 0.823807655268021 | -6.06754688738652 |
| lnc-LIPI-10 | lnc-LIPI-10 | -0.540790929184804 | 0.626332010638091 | -0.52816724429193 | 0.600235310987442 | 0.823807655268021 | -5.67963020819037 |
| ENSG00000130723.19 | NA | 0.0634014489476948 | 7.84928306768118 | 0.527853331700877 | 0.600346485545135 | 0.823825877664696 | -6.25569654694842 |
| lnc-GRK3-5 | lnc-GRK3-5 | -0.125002039282445 | 1.40565147614101 | -0.527845321383773 | 0.600351997569893 | 0.823825877664696 | -5.84038895739364 |
| lnc-RORA-5 | lnc-RORA-5 | -0.0720733125321846 | 2.29215023392585 | -0.527821104460853 | 0.600368661758269 | 0.823825877664696 | -5.95834900514951 |
| ENSG00000170027.6 | YWHAG | 0.076829519353292 | 5.53979616214041 | 0.527723580680045 | 0.600435772173289 | 0.823863042373362 | -6.26461281712454 |
| ENSG00000184216.13 | IRAK1 | 0.0953716267578759 | 5.52959306760152 | 0.527656100774419 | 0.600482210128575 | 0.823871839159876 | -6.26260349888911 |
| ENSG00000235119.1 | | 0.093359943925237 | 0.405178057512189 | 0.527560875411531 | 0.60054774466258 | 0.823906833859221 | -5.63039560391285 |
| ENSG00000152147.10 | GEMIN6 | -0.113414640790635 | 2.53844444630986 | -0.527493164134391 | 0.600594345912529 | 0.823915850567687 | -5.991314818941 |
| ENSG00000243175.1 | RPSAP36 | -0.229748006775708 | 2.87220727417045 | -0.527267850854818 | 0.600816442213478 | 0.824034082095866 | -6.02034630090049 |
| ENSG00000125352.5 | RNF113A | 0.0679180848923324 | 3.69217784927622 | 0.527127971655917 | 0.600845713223226 | 0.824034082095866 | -6.12429171131998 |
| lnc-ACTR1B-5 | lnc-ACTR1B-5 | 0.0833316873256198 | 2.77068640741536 | 0.527091788423201 | 0.600870621350013 | 0.824034082095866 | -6.01405078241181 |
| ENSG00000270393.1 | | 0.213452460118998 | -0.574823048628165 | 0.527009544888007 | 0.600927238663156 | 0.824034082095866 | -5.3404244521934 |
| ENSG00000214198.8 | TTC41P | 0.105350916798163 | 1.34841623635063 | 0.526998892224634 | 0.600934572251022 | 0.824034082095866 | -5.83209950798588 |
| ENSG00000278610.1 | NA | -0.0796061059156415 | 0.299276149638148 | -0.526987938711748 | 0.600942112995662 | 0.824034082095866 | -5.61584673182964 |
| lnc-BDH1-5 | lnc-BDH1-5 | 0.0680154400288426 | 4.05927027277805 | 0.52695824060541 | 0.600962558333868 | 0.824034082095866 | -6.17113805690373 |
| ENSG00000285330.1 | | -0.103352577300213 | -0.0582111320353499 | -0.526856787590237 | 0.601032405012596 | 0.824034082095866 | -5.51139355405409 |
| ENSG00000113141.17 | IK | -0.0534549609040207 | 6.64928982504076 | -0.526844531486322 | 0.601040843147512 | 0.824034082095866 | -6.27221563635837 |
| ENSG00000144815.15 | NXPE3 | 0.0569600660313947 | 4.99565360333476 | 0.526603296442751 | 0.601206940960163 | 0.824206904607902 | -6.24294977059999 |
| lnc-MEST-6 | lnc-MEST-6 | 0.133909042597331 | 3.00838508712313 | 0.526422835493962 | 0.601331207953407 | 0.824278329403714 | -6.05531517953982 |
| ENSG00000171161.12 | ZNF672 | 0.0829328732628851 | 3.96979623871664 | 0.52641132590985 | 0.601339133961335 | 0.824278329403714 | -6.15226290664939 |
| ENSG00000198189.10 | HSD17B11 | 0.0708225433768872 | 5.56519510795866 | 0.526155045544526 | 0.60151563258754 | 0.824465356965395 | -6.26472804781191 |
| ENSG00000268758.7 | ADGRE4P | -0.506370392347424 | 3.36300921704787 | -0.526151875397556 | 0.601626377521132 | 0.824469995264274 | -6.01933964468675 |
| lnc-RGPD3-7 | lnc-RGPD3-7 | -0.0824341578439005 | 0.400671200389043 | -0.52598721339376 | 0.601631230617528 | 0.824469995264274 | -5.62864701961601 |
| ENSG00000116288.12 | PARK7 | 0.0804396853681464 | 5.04068684057914 | 0.525975666425956 | 0.601639184223359 | 0.824469995264274 | -6.24334582550717 |
| ENSG00000069974.15 | RAB27A | -0.0837001634550734 | 2.48624022335984 | -0.525702922360632 | 0.601827065902492 | 0.824672558249849 | -5.98892946391897 |
| ENSG00000101337.15 | TM9SF4 | -0.0389425252901177 | 5.5122346957782 | -0.525551282652576 | 0.601931535839192 | 0.824729655419233 | -6.26259679190788 |
| ENSG00000141127.14 | PRPSAP2 | 0.0535605626210973 | 4.78654103781156 | 0.525476674700929 | 0.601982938993096 | 0.824729655419233 | -6.2314331153219 |

| | | | | | | | |
|---|---|---|---|---|---|---|---|
| lnc-KIF16B-41 | lnc-KIF16B-41 | 0.31022437748674 | 1.64175397159812 | 0.525582324433892 | 0.601988939562676 | 0.824729655419233 | -5.88193095637989 |
| lnc-ACTR2-4 | lnc-ACTR2-4 | 0.153864494318147 | 0.494422954577769 | 0.525241135899178 | 0.602145233229261 | 0.824824191148188 | -5.66923642222784 |
| ENSG00000064545.14 | TMEM161A | -0.107309063491157 | 1.73879734432674 | -0.525224340822721 | 0.602156806387491 | 0.824824191148188 | -5.90484543056269 |
| lnc-STYK1-1 | lnc-STYK1-1 | -0.187260032459028 | 5.35565360370068 | -0.525297281670484 | 0.602178162579984 | 0.824824191148188 | -6.25962166227995 |
| lnc-FBXO25-3 | lnc-FBXO25-3 | -0.154780724367588 | -0.55595997806962 | -0.525105899640035 | 0.602238424834134 | 0.824851843116914 | -5.44531071132281 |
| lnc-NDUFC2-1 | lnc-NDUFC2-1 | 0.070218540700244 | 1.30468535083 | 0.524989391253456 | 0.602318716417371 | 0.824856670877896 | -5.82696898356023 |
| lnc-FAM19A4-3 | lnc-FAM19A4-3 | -0.108507795971494 | 0.0429227350462789 | -0.524984483091665 | 0.602322098979487 | 0.824856670877896 | -5.56797704245603 |
| ENSG00000141424.12 | SLC39A6 | 0.0709599098845386 | 4.70257179947087 | 0.524783239937925 | 0.602460797529628 | 0.824960262699749 | -6.2292283059844 |
| ENSG00000099992.15 | TBC1D10A | -0.0782912003439282 | 3.6659368762004 | -0.524680265033217 | 0.602531774500684 | 0.824960262699749 | -6.12696333566416 |
| ENSG00000213413.2 | PVRIG | 0.154095958017329 | 2.82896926932397 | 0.52464680148226 1 | 0.602565107009221 | 0.824960262699749 | -6.04462657608383 |
| ENSG00000182890.4 | GLUD2 | 0.249253698980514 | -0.123398790317493 | 0.524549310231906 | 0.602622042661203 | 0.824960262699749 | -5.42739867505105 |
| ENSG00000103227.18 | LMF1 | -0.0735671707598278 | 3.52572772459891 | -0.524407965108687 | 0.602719480027564 | 0.824960262699749 | -6.11616478571843 |
| ENSG00000063761.15 | ADCK1 | 0.0858586756608428 | 2.28541282370434 | 0.52436851674132 | 0.60274667537928 | 0.824960262699749 | -5.95560596061227 |
| ARF4-AS1 | ARF4-AS1 | -0.113086192383129 | 2.16011498783112 | -0.524333473041023 | 0.602770834672745 | 0.824960262699749 | -5.9473076074201 |
| ENSG00000163376.11 | KBTBD8 | 0.105425786103099 | 6.24542879752216 | 0.524283141206466 | 0.602805534468419 | 0.824960262699749 | -6.27383338432787 |
| lnc-ZFYVE1-1 | lnc-ZFYVE1-1 | -0.136874581848465 | -0.364647643784475 | -0.524266404317865 | 0.602817073427886 | 0.824960262699749 | -5.46104095240121 |
| ENSG00000153487.12 | ING1 | -0.098155776002983 | 3.97941409498357 | -0.524258271540298 | 0.602822680468423 | 0.824960262699749 | -6.16663075295114 |
| ENSG00000175826.11 | CTDNEP1 | -0.0578047872543315 | 3.4922337183093 | -0.524227736614708 | 0.602843732603313 | 0.824960262699749 | -6.10634729640396 |
| lnc-LILRA4-2 | lnc-LILRA4-2 | -0.734921172578164 | -1.64785602046507 | -0.524273914207239 | 0.602878699486451 | 0.824960262699749 | -5.07035354948405 |
| lnc-SASH3-1 | lnc-SASH3-1 | -0.100026185934528 | 2.72697351677147 | -0.523892659918901 | 0.60307477188179 | 0.825173704024655 | -6.01943442626282 |
| lnc-ITGB4-2 | lnc-ITGB4-2 | 0.0624895365475278 | 1.75120423477384 | 0.523657443337303 | 0.603236981092623 | 0.825340785816902 | -5.89450370454973 |
| ENSG00000100911.15 | PSME2 | 0.0790986457784344 | 4.74902688678447 | 0.523578401447246 | 0.603291494237607 | 0.825360506872227 | -6.23026558722329 |
| ENSG00000105397.13 | TYK2 | -0.0643060829291748 | 6.72980015931208 | -0.523345369547648 | 0.60345222365727 | 0.825525529678159 | -6.27339278539577 |
| lnc-TWF2-1 | lnc-TWF2-1 | -0.0835997778766516 | 0.964133294883817 | -0.523047213566087 | 0.603657900272181 | 0.825593490131447 | -5.78652788165882 |
| ENSG00000285437.1 | POLR2J3 | 0.108547788909196 | 3.24119672888406 | 0.52303324085017 | 0.603667539855865 | 0.825593490131447 | -6.07245484663797 |
| ENSG00000197705.9 | KLHL14 | 0.174765882772694 | 6.72641598563463 | 0.523117013214686 | 0.603676014394375 | 0.825593490131447 | -6.27375501092531 |
| lnc-HOXB2-1 | lnc-HOXB2-1 | 0.498661466550981 | -2.61331948871808 | 0.522916009570286 | 0.60374841891839 | 0.825593490131447 | -4.95945730008818 |
| ENSG00000101546.12 | RBFA | 0.0923589366560824 | 3.30137308975392 | 0.522874846366208 | 0.603776819031281 | 0.825593490131447 | -6.08732684731577 |
| ENSG00000115514.11 | TXNDC9 | -0.0794596898973197 | 3.38352316026298 | -0.522812040307254 | 0.603820152598523 | 0.825593490131447 | -6.09823826864837 |
| lnc-ZC3H12B-7 | lnc-ZC3H12B-7 | 0.121945098521929 | 0.821871828827692 | 0.522788987323246 | 0.603836058561289 | 0.825593490131447 | -5.73574734851909 |
| ENSG00000119688.20 | ABCD4 | -0.0514410348738776 | 4.89802843957867 | -0.522785297683831 | 0.603838604335162 | 0.825593490131447 | -6.23814782989533 |
| ENSG00000089009.15 | RPL6 | 0.0834110894194424 | 8.33876772158412 | 0.522750091275631 | 0.603862896270171 | 0.825593490131447 | -6.24795347434898 |
| lnc-GSDMB-1 | lnc-GSDMB-1 | -0.139872854181579 | 2.53086165313719 | -0.522583252815763 | 0.603978018683528 | 0.825662208335248 | -5.9809422429703 |
| lnc-NUAK1-7 | lnc-NUAK1-7 | 0.178276645454035 | 3.47557519345698 | 0.522599245406831 | 0.604006552089788 | 0.825662208335248 | -6.07814474200396 |
| lnc-AMN1-2 | lnc-AMN1-2 | -0.234014552170206 | 0.917555174322484 | -0.522547014545989 | 0.604033500126908 | 0.825662208335248 | -5.78048317535341 |
| lnc-DPYSL2-1 | lnc-DPYSL2-1 | -0.22254171596297 | -0.0999261826660233 | -0.52227814208387 | 0.604188578579935 | 0.825779270236441 | -5.60206293234904 |
| lnc-XPO1-4 | lnc-XPO1-4 | 0.196074636704323 | -0.441600651092 | 0.522217506346051 | 0.604230427960302 | 0.825779270236441 | -5.4632345387928 |
| ENSG00000153094.22 | BCL2L11 | 0.178021864869159 | 6.00496725204281 | 0.522280665054365 | 0.604253831390523 | 0.825779270236441 | -6.27437988873941 |
| ENSG00000214331.8 | PDPR2P | 0.0470007986956222 | 4.30381695975834 | 0.52214623782638 | 0.604279617563099 | 0.825779270236441 | -6.19040898164958 |
| lnc-ACBD4-1 | lnc-ACBD4-1 | -0.0934345644392478 | 0.277674890363476 | -0.521694981756092 | 0.604591118153362 | 0.826150101902055 | -5.62981780597896 |
| ENSG00000279513.1 | | 0.143057588643344 | -0.108234127964315 | 0.521575879245829 | 0.604673346670216 | 0.826207613815259 | -5.53635556650374 |
| ENSG00000143624.13 | INTS3 | 0.0649390646116785 | 6.06832906615433 | 0.521424541320942 | 0.604777838050782 | 0.826253112964827 | -6.27473076984423 |
| ENSG00000005844.17 | ITGAL | 0.167243370097096 | 5.5253646567822 | 0.521498477018651 | 0.604786930958951 | 0.826253112964827 | -6.26365468407428 |
| ENSG00000157653.11 | C9orf43 | 0.11915595983709 | 0.372804274067908 | 0.521312895397646 | 0.604854929435457 | 0.826291166958918 | -5.64995813112672 |
| lnc-HSPA13-4 | lnc-HSPA13-4 | -0.347061222522604 | 0.859141226801318 | -0.521140198821705 | 0.605041010780587 | 0.826490517978287 | -5.64742564110716 |
| ENSG00000163617.10 | CCDC191 | -0.217861394705656 | 6.51307684396472 | -0.521028881542597 | 0.605138378522333 | 0.826543329744371 | -6.27509846000709 |
| ENSG00000205236.6 | | -0.179589265265933 | 1.14192502984395 | -0.520873241433164 | 0.605159985388314 | 0.826543329744371 | -5.82911583013883 |
| lnc-ARHGEF2-3 | lnc-ARHGEF2-3 | 0.0719173643615783 | 4.55944730093534 | 0.520621794494136 | 0.605332235360455 | 0.826707625437912 | -6.21374778339305 |
| ENSG00000266315.1 | MIR4668 | -0.102242532832191 | 0.54943260668399 | -0.520565035578303 | 0.605371443431442 | 0.826707625437912 | -5.68433991294805 |
| lnc-GRHPR-8 | lnc-GRHPR-8 | -0.108788159122559 | 1.52824174923906 | -0.520522583322513 | 0.605400769480913 | 0.826707625437912 | -5.86929337356008 |
| lnc-CABLES2-2 | lnc-CABLES2-2 | 0.160772692879397 | -0.342249225896102 | 0.520314382786709 | 0.6055446041057 | 0.826832753200637 | -5.40025060132544 |
| ENSG00000133275.15 | CSNK1G2 | -0.082138347555202 | 5.62795105194273 | -0.520218916100951 | 0.605610562237294 | 0.826832753200637 | -6.27040434193648 |
| ENSG00000152413.14 | HOMER1 | 0.111067221267563 | 2.30167935093345 | 0.520013853948365 | 0.605752251368289 | 0.826832753200637 | -5.94916189104651 |
| lnc-CCDC173-3 | lnc-CCDC173-3 | -0.0779396468482563 | 0.716015043888498 | -0.519886200891227 | 0.60584046190358 | 0.826832753200637 | -5.73934492041978 |
| ENSG00000174718.11 | RESF1 | 0.0714367426338711 | 9.74493309779782 | 0.51984646142802 | 0.60586792379556 | 0.826832753200637 | -6.2097091712898 |
| lnc-DDX31-2 | lnc-DDX31-2 | -0.0666560829502196 | 1.45497540363985 | -0.519782785431631 | 0.605911928193195 | 0.826832753200637 | -5.85691682134173 |
| lnc-CDK19-4 | lnc-CDK19-4 | -0.0760725733517975 | 1.02568084421751 | -0.51975821415544 | 0.605928908994302 | 0.826832753200637 | -5.78666739192808 |
| ENSG00000171174.14 | RBKS | -0.130704946887653 | 0.245909569052777 | -0.519687125667322 | 0.605978038310559 | 0.826832753200637 | -5.60310665998769 |
| lnc-CCDC146-4 | lnc-CCDC146-4 | -0.130737419391448 | 0.653262625615642 | -0.519682831936815 | 0.605981005770584 | 0.826832753200637 | -5.72488211863361 |
| lnc-GGCT-4 | lnc-GGCT-4 | 0.0888454001037933 | 2.35229348331736 | 0.519634958697696 | 0.606014092123254 | 0.826832753200637 | -5.96614446360918 |
| ENSG00000124171.8 | PARD6B | 0.0731692596996281 | 1.88300994619848 | 0.519587806906298 | 0.606046680684273 | 0.826832753200637 | -5.92029979801748 |
| ENSG00000250462.8 | LRRC37BP1 | -0.0695763057850791 | 3.78260052609782 | -0.519574896548624 | 0.606055603710177 | 0.826832753200637 | -6.15354814264288 |
| ENSG00000210077.1 | MT-TV | 0.175524072588864 | 2.22474407181903 | 0.51954985233843 | 0.606088563996951 | 0.826832753200637 | -5.95076475257809 |
| ENSG00000104365.14 | IKBKB | -0.065941233494673 | 6.59637893416688 | -0.519486886397847 | 0.606116433767343 | 0.826832753200637 | -6.27639553549918 |
| ENSG00000284184.1 | MIR6506 | -0.0819147441376271 | 1.65199387093657 | -0.519436403631386 | 0.606151327264288 | 0.826832753200637 | -5.86727273761608 |
| ENSG00000131781.12 | FMO5 | 0.0917618323905489 | 2.97781553104837 | 0.51943291863103 | 0.606153736117673 | 0.826832753200637 | -6.03464077647709 |
| ENSG00000159588.14 | CCDC17 | -0.0917173371985274 | 2.61984542652536 | -0.519401718218179 | 0.606175302229879 | 0.826832753200637 | -6.0042640558049 |
| ENSG00000129128.12 | SPCS3 | 0.0910478110896582 | 6.88949912181962 | 0.519239534807036 | 0.606287411140384 | 0.826930871612386 | -6.27316914737303 |

| | | | | | | | |
|---|---|---|---|---|---|---|---|
| ENSG00000147606.8 | SLC26A7 | -0.166809256567332 | -0.0479787323041685 | -0.519120231495554 | 0.606369885410507 | 0.826988560257386 | -5.536320703702 |
| ENSG00000180900.18 | SCRIB | -0.0983189162568596 | 4.1633338333228 | -0.518955061044832 | 0.606484076167126 | 0.827089494450934 | -6.1849129640743 |
| ENSG00000266094.7 | RASSF5 | -0.107347843998486 | 6.54706445277908 | -0.518834595216164 | 0.606567366617057 | 0.827097538605053 | -6.27661267179704 |
| ENSG00000184203.7 | PPP1R2 | 0.0667606934990133 | 6.8681621883845 | 0.518774519324939 | 0.60660890525451 | 0.827097538605053 | -6.27385243201007 |
| ENSG00000170322.14 | NFRKB | -0.0416440709399373 | 5.08944555345839 | -0.518772176310519 | 0.606610525325845 | 0.827097538605053 | -6.25197648345458 |
| ENSG00000072042.12 | RDH11 | -0.075214783199165 | 4.1732768477562 | -0.518561301269421 | 0.606756342535411 | 0.827241557819855 | -6.18446341786369 |
| ENSG00000239282.7 | CASTOR1 | -0.116465879837051 | 0.0861106812436454 | -0.518490727401525 | 0.606805147023272 | 0.82725330117764 | -5.57957409133512 |
| ENSG00000266208.1 | GJD3-AS1 | -0.170883470373911 | 0.255665289172307 | -0.518354559810274 | 0.606899317174628 | 0.827326885582182 | -5.60999600818771 |
| ENSG00000213096.10 | ZNF254 | 0.0700914283307501 | 5.83878602439071 | 0.518294199262353 | 0.606941063205231 | 0.827329000261806 | -6.27350051551455 |
| ENSG00000121964.14 | GTDC1 | -0.11902855643417 | 3.43719356120826 | -0.51798562664591 | 0.60715449623165 | 0.827565127868092 | -6.11836848016222 |
| ENSG00000122550.17 | KLHL7 | 0.0502668873156025 | 3.62393154512873 | 0.517924008518866 | 0.60719712031871 | 0.827568423223415 | -6.13216849146224 |
| ENSG00000167985.6 | SDHAF2 | -0.0607118362740366 | 3.71920501893754 | -0.517683037514692 | 0.60736382430842 | 0.827740819422645 | -6.12844864105238 |
| ENSG00000186272.12 | ZNF17 | -0.0521813096676662 | 3.35862349356665 | -0.517604771514717 | 0.607417973359348 | 0.827742560704423 | -6.08793644189317 |
| ENSG00000283856.1 | MIR7703 | 0.0966081423263416 | 1.21381191890636 | 0.517564940423563 | 0.60744553172231 | 0.827742560704423 | -5.81184538052452 |
| ENSG00000006625.17 | GGCT | -0.123301309983245 | 1.89620030739293 | -0.517174697474931 | 0.607715563843965 | 0.828055703071649 | -5.92687542368823 |
| lnc-PACSIN3-3 | lnc-PACSIN3-3 | 0.0461721661539705 | 2.47178842558224 | 0.517029957428299 | 0.607815732117998 | 0.828115529912348 | -5.9864370340307 |
| lnc-PDPK1-1 | lnc-PDPK1-1 | 0.21697549620289 | -0.324914821884261 | 0.516994740313427 | 0.607840105500714 | 0.828115529912348 | -5.40039874494408 |
| lnc-ANKRD42-2 | lnc-ANKRD42-2 | 0.13613735591174 | 1.39716980675343 | 0.516936852348212 | 0.607880170113476 | 0.828115529912348 | -5.82317829319821 |
| lnc-DNAH9-4 | lnc-DNAH9-4 | -0.0896027244871991 | 0.32254076174906 | -0.516827474940729 | 0.607955874211161 | 0.82816384903298 | -5.65634382622868 |
| ENSG00000166432.14 | ZMAT1 | -0.122751003135464 | 5.66295364163153 | -0.516599274554837 | 0.60812629003506 | 0.828297109404896 | -6.27230664122431 |
| lnc-ZNF18-1 | lnc-ZNF18-1 | 0.0804899959835578 | 0.0618677641184114 | 0.516569876938987 | 0.608134184391332 | 0.828297109404896 | -5.55556171509843 |
| ENSG00000272822.1 | | 0.0881669261787428 | 2.55972044426003 | 0.516336742393679 | 0.608295581693447 | 0.828377431157084 | -5.99783449523521 |
| ENSG00000269891.2 | ARHGAP19-SLIT1 | -0.0539301645102808 | 2.33610258254583 | -0.516296223161561 | 0.608323634870869 | 0.828377431157084 | -5.96809813838429 |
| ENSG00000140488.15 | CELF6 | 0.0927144053392712 | 1.48096793468181 | 0.516269197435877 | 0.608342346255182 | 0.828377431157084 | -5.85879817543097 |
| ENSG00000186501.14 | TMEM222 | -0.0536773654480838 | 4.20762646559449 | -0.516252164376741 | 0.608354139307704 | 0.828377431157084 | -6.1945554032984 |
| ENSG00000167613.15 | LAIR1 | -0.836701845431577 | -0.127438141420176 | -0.516260617990947 | 0.608454500253363 | 0.828411191534101 | -5.37900496124453 |
| ENSG00000136026.13 | CKAP4 | 0.164874206929853 | 5.6084128355991 | 0.516185943201563 | 0.608459427372712 | 0.828411191534101 | -6.26585061450578 |
| TUG1 | TUG1 | 0.0517086029068414 | 7.89202831440957 | 0.51600912371451 | 0.608522423053165 | 0.828427219291414 | -6.26152430715336 |
| ENSG00000168405.17 | CMAHP | 0.0880204378401963 | 2.65224716087896 | 0.515966850188731 | 0.608551695850273 | 0.828427219291414 | -5.99886649539342 |
| ENSG00000257341.5 | | -0.185460036230296 | 3.98180463116747 | -0.515878527089428 | 0.608672255575306 | 0.828536541003011 | -6.16550687615807 |
| ENSG00000140564.11 | FURIN | 0.0988244541083221 | 3.91841632049987 | 0.515699899394989 | 0.608736564046336 | 0.828569283015854 | -6.16440457809658 |
| ENSG00000114904.12 | NEK4 | -0.0592731346345854 | 4.17044985393288 | -0.515557490835174 | 0.608835195109122 | 0.828636608514641 | -6.18530336631926 |
| lnc-ZNF783-3 | lnc-ZNF783-3 | -0.0596136886891627 | 2.70679691978292 | -0.515512230494231 | 0.608866543612468 | 0.828636608514641 | -6.01790697401099 |
| lnc-ERLEC1-4 | lnc-ERLEC1-4 | -0.119518531297488 | 1.32877378873053 | -0.515416474025832 | 0.608932869514783 | 0.828672082857704 | -5.83031093621601 |
| ENSG00000141933.9 | TPGS1 | -0.104734457305382 | 1.555216628059 | -0.515172669944981 | 0.609101755911699 | 0.828769140508201 | -5.88193911653252 |
| lnc-KCTD1-3 | lnc-KCTD1-3 | 0.10815376961889 | 0.444612211333813 | 0.515148268469733 | 0.609118660332935 | 0.828769140508201 | -5.69204666994341 |
| ENSG00000081760.16 | AACS | 0.126901027666336 | 3.54573718258927 | 0.515139139520691 | 0.609124984579899 | 0.828769140508201 | -6.12177580989479 |
| ENSG00000250479.8 | CHCHD10 | -0.0902336650043805 | 3.10014460200455 | -0.515040545536148 | 0.609193289299499 | 0.828793383033964 | -6.06440702106902 |
| ENSG00000112972.14 | HMGCS1 | 0.085232758894997 | 4.74523227468842 | 0.514839461185413 | 0.609332609040149 | 0.828793383033964 | -6.23840446667432 |
| lnc-BRD3OS-1 | lnc-BRD3OS-1 | -0.105257787873795 | 0.122999824529051 | -0.514787070442005 | 0.60936890997188 | 0.828793383033964 | -5.58652935321107 |
| lnc-ZFY-3 | lnc-ZFY-3 | -0.552637258045231 | -1.9451595372674 | -0.514802409491121 | 0.60936926380039 | 0.828793383033964 | -5.014123241887 |
| ENSG00000169891.17 | REPS2 | -0.235678812769116 | 1.66347833740293 | -0.514726093677396 | 0.609465662805645 | 0.828793383033964 | -5.89439127279172 |
| lnc-AKAP1-7 | lnc-AKAP1-7 | 0.175529836106166 | 5.09312330455126 | 0.514532774011377 | 0.60960993444474 | 0.828793383033964 | -6.2541138730644 |
| lnc-L1TD1-3 | lnc-L1TD1-3 | -0.618131681128379 | -2.56304773785602 | -0.514357179005794 | 0.609669717832497 | 0.828793383033964 | -4.91661623570825 |
| lnc-FARP2-2 | lnc-FARP2-2 | 0.133331648643909 | -0.208141690408774 | 0.51432048661491 | 0.60969224435387 | 0.828793383033964 | -5.45083592575198 |
| lnc-LSM14A-2 | lnc-LSM14A-2 | -0.0526065038702423 | 2.3646568500814 | -0.51430943544416 | 0.609699903577892 | 0.828793383033964 | -5.97309704984364 |
| ENSG00000114812.12 | VIPR1 | 0.532761826581193 | 3.72093278340586 | 0.514442885888347 | 0.609713199924708 | 0.828793383033964 | -6.17807924902541 |
| ENSG00000175482.8 | POLD4 | 0.095738628935524 | 4.31553834697279 | 0.514172257843188 | 0.609794980808733 | 0.828793383033964 | -6.19852088555689 |
| ENSG00000162521.18 | RBBP4 | 0.0416062805738822 | 7.18483602104043 | 0.51409564544792 | 0.609848083513572 | 0.828793383033964 | -6.27362949073968 |
| lnc-MCOLN2-1 | lnc-MCOLN2-1 | 0.114744167274419 | 3.0927533642824 | 0.513901986936865 | 0.609982324420978 | 0.828793383033964 | -6.06130739689265 |
| TNKS2-AS1 | TNKS2-AS1 | -0.0940622263823779 | 0.678229749842899 | -0.513879207451995 | 0.609998115680116 | 0.828793383033964 | -5.71907906108005 |
| ENSG00000055118.14 | KCNH2 | 0.246152763029396 | 3.49839439726994 | 0.513949585346676 | 0.610027654052001 | 0.828793383033964 | -6.10466898493739 |
| lnc-STARD10-3 | lnc-STARD10-3 | -0.151925221477462 | 1.08410126155291 | -0.513826955198559 | 0.610034338845839 | 0.828793383033964 | -5.79753485040207 |
| ENSG00000197816.13 | CCDC180 | 0.0822803845320652 | 3.93248891590463 | 0.513772945844475 | 0.610071781137499 | 0.828793383033964 | -6.15057555194313 |
| ENSG00000100150.17 | DEPDC5 | -0.0996282405434794 | 6.60093269043577 | -0.513723928343259 | 0.610105763713649 | 0.828793383033964 | -6.27942475448991 |
| ENSG00000188917.14 | TRMT2B | 0.0386788867022361 | 5.5840485901927 | 0.513617268458516 | 0.610179711283322 | 0.828793383033964 | -6.27101042737366 |
| ENSG00000005471.17 | ABCB4 | -0.0687272471122817 | 6.32768235763699 | -0.513597733879575 | 0.61019325510314 | 0.828793383033964 | -6.27971969076272 |
| ENSG00000214087.8 | ARL16 | 0.0830569876194227 | 3.189632235537 | 0.513595332320983 | 0.61019492017426 | 0.828793383033964 | -6.07987137678911 |
| ENSG00000088448.14 | ANKRD10 | -0.0633737133058005 | 6.71677085523673 | -0.513570854326969 | 0.610211891605736 | 0.828793383033964 | -6.27785415836651 |
| lnc-TESK2-2 | lnc-TESK2-2 | 0.0661905489984154 | 0.877620212625294 | 0.51355068897192 | 0.610225873099975 | 0.828793383033964 | -5.74883968603324 |
| ENSG00000139517.8 | LNX2 | -0.0528050750959181 | 4.0366951439156 | -0.513518747450136 | 0.610248019810502 | 0.828793383033964 | -6.17552937004076 |
| lnc-CAMK2B-1 | lnc-CAMK2B-1 | 0.0738951803141176 | 0.457130538890664 | 0.513446606826069 | 0.610298039999285 | 0.828793383033964 | -5.66810226284739 |
| ENSG00000125249.6 | RAP2A | 0.0966956926334741 | 3.72639292895475 | 0.513445399695009 | 0.610298877004521 | 0.828793383033964 | -6.14274720196351 |
| lnc-MIOS-1 | lnc-MIOS-1 | 0.0990765179680815 | 1.46835761813424 | 0.513326609176102 | 0.610381247018236 | 0.828793383033964 | -5.85694107234284 |
| ENSG00000065559.14 | MAP2K4 | 0.0619088844163036 | 4.91585350519037 | 0.513247045086146 | 0.610436420062813 | 0.828793383033964 | -6.24719658200656 |
| ENSG00000231389.7 | HLA-DPA1 | -0.114548874983237 | 9.50000527852052 | -0.513211062465665 | 0.610461453535537 | 0.828793383033964 | -6.22421412163503 |
| ENSG00000148335.14 | NTMT1 | 0.0836797826079114 | 2.70039697334996 | 0.513191321916844 | 0.610475062187481 | 0.828793383033964 | -6.01262411780936 |
| ENSG00000186918.13 | ZNF395 | 0.0894848841009992 | 6.78826622510108 | 0.513182737707755 | 0.610481015144752 | 0.828793383033964 | -6.2778594025754 |

| | | | | | | | |
|---|---|---|---|---|---|---|---|
| ENSG00000120733.13 | KDM3B | 0.0377641010869629 | 7.2081201005054 | 0.513152427735842 | 0.610502034646669 | 0.828793383033964 | -6.27364024012191 |
| ENSG00000204745.4 | | 0.173317949012561 | 0.17881848261417 | 0.513139359364404 | 0.610511097464981 | 0.828793383033964 | -5.51661446169383 |
| lnc-ABHD17A-1 | lnc-ABHD17A-1 | -0.11568542391966 | 1.49546154718876 | -0.51311981607634 | 0.610524650705129 | 0.828793383033964 | -5.85770735253444 |
| PSMA3-AS1 | PSMA3-AS1 | 0.0410080773283248 | 7.28641989170766 | 0.513068454732197 | 0.610560270377233 | 0.828793383033964 | -6.27295395716448 |
| ENSG00000197785.13 | ATAD3A | -0.100216441037052 | 2.55980263830643 | -0.513022162650052 | 0.610592375277026 | 0.828793383033964 | -6.00198787130793 |
| lnc-RCAN1-6 | lnc-RCAN1-6 | -0.146189303879718 | 2.15436318620581 | -0.512958614956987 | 0.610636448705931 | 0.828798550854875 | -5.94215608633424 |
| ENSG00000234219.1 | CDCA4P4 | -0.101998667466819 | 0.00824361240978376 | -0.512808441100194 | 0.61074060741773 | 0.828846609660853 | -5.5795936778234 |
| ENSG00000075391.16 | RASAL2 | 0.486611406462654 | -0.824069871965492 | 0.512732884571265 | 0.6108470594509 | 0.828846609660853 | -5.27196633330443 |
| ENSG00000013392.7 | RWDD2A | -0.088865082778397 | 1.90560861659493 | -0.512587244896529 | 0.610894041202995 | 0.828846609660853 | -5.9181934661549 |
| lnc-EPHX3-3 | lnc-EPHX3-3 | -0.107481852001142 | 1.39078802453431 | -0.512577787285553 | 0.610900601913816 | 0.828846609660853 | -5.83520923409698 |
| lnc-ZSWIM7-1 | lnc-ZSWIM7-1 | 0.123529402134917 | 1.64916527127484 | 0.512569193298249 | 0.610906563560051 | 0.828846609660853 | -5.87661977104872 |
| lnc-FBXO43-6 | lnc-FBXO43-6 | 0.133811690666961 | 4.66009134519327 | 0.5125875299794 | 0.610913468161337 | 0.828846609660853 | -6.22573722589253 |
| LINC01801 | LINC01801 | -0.155259226689684 | 1.40096138823405 | -0.512392237866505 | 0.611029323423811 | 0.82892364664726 | -5.86001308628346 |
| ENSG00000259024.6 | TVP23C-CDRT4 | -0.184011084619412 | 2.00078686868561 | -0.512396363136106 | 0.611050793887109 | 0.82892364664726 | -5.94914391276965 |
| ENSG00000146802.12 | TMEM168 | 0.0750984359840681 | 4.93283187239859 | 0.512294764862126 | 0.611096948522873 | 0.828931625902616 | -6.24579506901606 |
| lnc-GPAT2-3 | lnc-GPAT2-3 | 0.101538172606992 | 1.75250732529041 | 0.512197399788813 | 0.611164502170693 | 0.828968629204572 | -5.90203729771459 |
| ENSG00000087088.19 | BAX | 0.0853143545541357 | 4.51006342489999 | 0.512062399864128 | 0.611258173228747 | 0.828994525548825 | -6.21329581817549 |
| ENSG00000176619.12 | LMNB2 | -0.126098328708774 | 3.18603402888836 | -0.512053792352419 | 0.611264145860881 | 0.828994525548825 | -6.08233987337757 |
| HLA-F-AS1 | HLA-F-AS1 | -0.0992913552803611 | 1.8236754372101 | -0.51191451486733 | 0.611360792284248 | 0.82899605502797 | -5.90660906845233 |
| lnc-GOLGA8H-8 | lnc-GOLGA8H-8 | 0.0802502284501707 | 3.09322527361397 | 0.511889934559668 | 0.611377849601026 | 0.82899605502797 | -6.07610114958403 |
| lnc-RAB28-5 | lnc-RAB28-5 | -0.120594975777022 | 0.762341685387622 | -0.511878044170278 | 0.611386100924286 | 0.82899605502797 | -5.7296633459356 |
| ENSG00000122008.15 | POLK | -0.060357220417167 | 5.93079024517165 | -0.511645311339032 | 0.611547615913763 | 0.829124965281903 | -6.2788250123305 |
| ENSG00000178425.13 | NT5DC1 | -0.0646280450232704 | 5.65382900626144 | -0.511624965775069 | 0.611561736525767 | 0.829124965281903 | -6.27340225131347 |
| lnc-CHRDL2-3 | lnc-CHRDL2-3 | 0.180413576782485 | 1.75677526484755 | 0.511520212847228 | 0.611644412761614 | 0.829182437454542 | -5.88064836018212 |
| lnc-MEST-7 | lnc-MEST-7 | 0.359133594400791 | -1.45912921370057 | 0.511357376257623 | 0.611747467963289 | 0.82922172178966 | -5.15869648171786 |
| ENSG00000243508.1 | DNAJB6P7 | 0.173418853072495 | -0.592846995263445 | 0.511289980325513 | 0.611794250939988 | 0.82922172178966 | -5.33908068698258 |
| lnc-SNX4-2 | lnc-SNX4-2 | 0.173418853072495 | -0.592846995263445 | 0.511289980325513 | 0.611794250939988 | 0.82922172178966 | -5.33908068698258 |
| ENSG00000101365.20 | IDH3B | 0.0549374209052376 | 5.54368593708848 | 0.511140868707969 | 0.61189776279716 | 0.829269780752077 | -6.27304654734047 |
| ENSG00000102878.16 | HSF4 | 0.198703038275381 | -0.188014165233088 | 0.51112282854858 | 0.611910286647356 | 0.829269780752077 | -5.510599971034 |
| ENSG00000110448.10 | CD5 | 0.156552063150864 | 6.15180710590212 | 0.511106622180974 | 0.611975598530626 | 0.829303689812093 | -6.27896257000597 |
| ENSG00000119986.6 | AVPI1 | 0.12790438391786 | 0.815108618692918 | 0.510757379326217 | 0.612164014313847 | 0.829454053907226 | -5.73773985523041 |
| lnc-RTP5-1 | lnc-RTP5-1 | 0.128693674868462 | 0.540027067155657 | 0.510752857457311 | 0.612167154103127 | 0.829454053907226 | -5.72356793522365 |
| lnc-SMCR8-1 | lnc-SMCR8-1 | 0.0447939294851381 | 4.30142467885602 | 0.510574760151383 | 0.612290822999982 | 0.829567009597724 | -6.19732320274673 |
| lnc-HSD11B1L-1 | lnc-HSD11B1L-1 | -0.0812795088523561 | 4.6154549975426 | -0.510309436127182 | 0.612475082429411 | 0.829762036572406 | -6.22384225849962 |
| ENSG00000048392.11 | RRM2B | 0.0630451479017955 | 5.93543823708487 | 0.510216034799251 | 0.612539952839005 | 0.829795304020353 | -6.27981668981421 |
| ENSG00000127022.14 | CANX | 0.0639423575930813 | 7.68084098346049 | 0.510142487957436 | 0.612591035843681 | 0.829809890804243 | -6.26715731814372 |
| ENSG00000173821.19 | RNF213 | 0.0598669773467275 | 8.81116991069333 | 0.509999325872522 | 0.612690476696662 | 0.829881906456215 | -6.24193195416579 |
| ENSG00000173846.12 | PLK3 | 0.138046799115734 | 4.5794178594078 | 0.5099846992863 | 0.612724837604582 | 0.829881906456215 | -6.21579070367084 |
| ENSG00000130340.15 | SNX9 | 0.0999512297977683 | 6.51343012336338 | 0.509820893301562 | 0.61281442688471 | 0.829948634594551 | -6.28117373805038 |
| ENSG00000142583.17 | SLC2A5 | 0.51521802924396 | -1.0830349898306 | 0.509819594466886 | 0.612868985417095 | 0.829967914128565 | -5.22588213593816 |
| ENSG00000110583.12 | NAA40 | -0.0820558737865564 | 5.22432770957704 | -0.509631072380092 | 0.612946300711045 | 0.8300180070747 | -6.26057899416412 |
| lnc-PGAP1-3 | lnc-PGAP1-3 | -0.135830343102522 | 0.742469086510484 | -0.509547883261433 | 0.613004098575352 | 0.830041665744127 | -5.70551186772956 |
| lnc-MCM10-4 | lnc-MCM10-4 | -0.191395563371589 | -0.0873655671641889 | -0.509486085897026 | 0.613047035559962 | 0.830045200166471 | -5.51362756576006 |
| ENSG00000068097.14 | HEATR6 | -0.0652865210526103 | 5.69923619519677 | -0.509393547040583 | 0.613111334392355 | 0.83007765544944 | -6.27665138506643 |
| ENSG00000277846.1 | SNORD30 | 0.116838864842221 | 1.1213354455823 | 0.509316305164587 | 0.613165006775207 | 0.830095720498164 | -5.81319763347452 |
| ENSG00000173209.23 | AHSA2P | 0.0610785196947656 | 7.70215994332949 | 0.509088508032794 | 0.61332330667237 | 0.830141431153076 | -6.26834634457724 |
| ENSG00000117054.13 | ACADM | -0.0706314171462381 | 5.83398764684522 | -0.509082885714882 | 0.613327213947874 | 0.830141431153076 | -6.2786487549815 |
| ENSG00000005100.12 | DHX33 | 0.0624364160612658 | 2.83913220337126 | 0.509061489396541 | 0.613342083597146 | 0.830141431153076 | -6.04409771299828 |
| lnc-PTHLH-1 | lnc-PTHLH-1 | -0.113637862119161 | 2.04607179187452 | -0.509035569179432 | 0.613360097409318 | 0.830141431153076 | -5.92244745632949 |
| ENSG00000119912.16 | IDE | 0.0397755492808072 | 5.08583243921125 | 0.508865735208422 | 0.613478133161062 | 0.830246591811042 | -6.25685140601332 |
| ENSG00000130702.15 | LAMA5 | 0.150631921020932 | 5.07263099694375 | 0.508788689391181 | 0.613574819837354 | 0.83025884217824 | -6.24970597077474 |
| lnc-SF3B2-2 | lnc-SF3B2-2 | -0.154750651122335 | 2.11291365823153 | -0.50869896912525 | 0.613594046808849 | 0.83025884217824 | -5.93160562733612 |
| ENSG00000166557.12 | TMED3 | -0.090996742850364 | 4.09885229272319 | -0.508678612907805 | 0.613608196434698 | 0.83025884217824 | -6.1792703547649 |
| ENSG00000207241.1 | SNORD45A | 0.0919785622081828 | 1.65982867120709 | 0.50849814844175 | 0.613733643972108 | 0.830373995522113 | -5.89627014023799 |
| ENSG00000090615.14 | GOLGA3 | 0.0587219815093105 | 6.63536567613152 | 0.508398106863452 | 0.613803191626291 | 0.830413506851844 | -6.28185078519767 |
| lnc-NLN-2 | lnc-NLN-2 | 0.124917532701908 | 2.28657499525628 | 0.508180621013107 | 0.6139543974846 | 0.8305340622102 | -5.95795071127913 |
| ENSG00000124356.15 | STAMBP | -0.0426640085012799 | 5.07889526935144 | -0.508153863701982 | 0.613973001534781 | 0.8305340622102 | -6.25507328648908 |
| lnc-NDRG1-1 | lnc-NDRG1-1 | -0.0694322755937256 | 1.72069550464257 | -0.507948821585853 | 0.614115573497296 | 0.83058511084589 | -5.89230969299755 |
| ENSG00000090989.17 | EXOC1 | 0.0493622577807864 | 5.49221579628869 | 0.507935151290192 | 0.614125079403076 | 0.83058511084589 | -6.2719041351638 |
| ENSG00000105323.16 | HNRNPUL1 | 0.0574656663755997 | 7.10271816741105 | 0.507898234683347 | 0.614150750418672 | 0.83058511084589 | -6.27786673293275 |
| ENSG00000130787.13 | HIP1R | 0.250543766919003 | 4.44682251353012 | 0.507952871677227 | 0.61420852531838 | 0.83058511084589 | -6.23344129375225 |
| ENSG00000048162.20 | NOP16 | 0.0892506332972025 | 2.7094713754771 | 0.507809431696489 | 0.614212504114332 | 0.83058511084589 | -6.01752331952085 |
| ENSG00000140829.11 | DHX38 | 0.0425752531011643 | 6.2355137378261 | 0.507345670775146 | 0.614535049998391 | 0.830919787896039 | -6.28289760671054 |
| ENSG00000171055.14 | FEZ2 | -0.053797884495407 | 4.3797390058388 | -0.50733749900996 | 0.61454073415641 | 0.830919787896039 | -6.21335742034518 |
| ENSG00000136936.10 | XPA | -0.0465040141637925 | 4.24566857188688 | -0.506993421277265 | 0.614780091250655 | 0.831017101227935 | -6.19835503163513 |
| ENSG00000198833.6 | UBE2J1 | 0.107462362473654 | 6.94610332138529 | 0.506977396804037 | 0.614791319432436 | 0.831017101227935 | -6.2811224003303 |
| ENSG00000088832.16 | FKBP1A | -0.0845643771535722 | 3.67798036614164 | -0.506968653655272 | 0.614797322444478 | 0.831017101227935 | -6.14520884594023 |
| ENSG00000068885.14 | IFT80 | 0.0588638318586669 | 4.624165719659 | 0.506952521656057 | 0.614808545828674 | 0.831017101227935 | -6.22627268283339 |

| | | | | | | | |
|---|---|---|---|---|---|---|---|
| lnc-ZNF236-1 | lnc-ZNF236-1 | 0.303083994425104 | 2.96311427419872 | 0.5070812395106 | 0.614814575984987 | 0.831017101227935 | -6.05855475301888 |
| ENSG00000143226.13 | FCGR2A | 0.325653152077092 | 1.45615405045775 | 0.506988137061924 | 0.614872034538032 | 0.831040192192285 | -5.90626758131699 |
| ENSG00000154016.13 | GRAP | -0.111005828538416 | 4.37748078964718 | -0.506753650249946 | 0.614946912680207 | 0.831086822304445 | -6.21390299006186 |
| ENSG00000123338.12 | NCKAP1L | 0.0789780882273394 | 8.24565838683484 | 0.506681371941854 | 0.614997204581068 | 0.831100220726946 | -6.25846642882117 |
| ENSG00000279908.1 | | 0.131906872066924 | 0.37990273957742 | 0.506512892519422 | 0.615114441362235 | 0.831183004132036 | -5.66854522763449 |
| ENSG00000169228.13 | RAB24 | -0.0806687331608829 | 3.11789299004482 | -0.506477275797998 | 0.615139226640592 | 0.831183004132036 | -6.077161628473 |
| lnc-CDK2AP1-1 | lnc-CDK2AP1-1 | -0.121510918402306 | 1.11704719968465 | -0.506374317216199 | 0.615210876924886 | 0.831225251394574 | -5.83697148564622 |
| lnc-MOAP1-2 | lnc-MOAP1-2 | -0.135400545479074 | -0.285234338577196 | -0.506031060726381 | 0.615449781261978 | 0.831347506928233 | -5.44107270854381 |
| ENSG00000169100.13 | SLC25A6 | -0.103430927995992 | 6.74906230142179 | -0.50599445467308 | 0.615475261334483 | 0.831347506928233 | -6.28136176533551 |
| ENSG00000115307.16 | AUP1 | 0.0507349814573717 | 6.70081172163171 | 0.505977949652256 | 0.615486750004763 | 0.831347506928233 | -6.28201301780891 |
| lnc-ZNRF2-1 | lnc-ZNRF2-1 | -0.157967215023244 | -0.453765997666035 | -0.505970922460553 | 0.61549164146042 | 0.831347506928233 | -5.42027708125019 |
| ENSG00000204536.13 | CCHCR1 | 0.0782759333979515 | 3.14875800688619 | 0.505954157258788 | 0.615503311377319 | 0.831347506928233 | -6.06732190366442 |
| lnc-IFT57-4 | lnc-IFT57-4 | -0.0472589512691224 | 6.80153367854486 | -0.505829084734991 | 0.615590375002362 | 0.831410543876222 | -6.28132540981264 |
| lnc-MYO15B-3 | lnc-MYO15B-3 | -0.0968826743994504 | 2.06011247885609 | -0.505674145452768 | 0.615698236802267 | 0.831501660527595 | -5.93312814211005 |
| lnc-WDR77-3 | lnc-WDR77-3 | -0.105722957404937 | 0.907081498156044 | -0.505568394585029 | 0.615771860764725 | 0.831532319265205 | -5.76726910943288 |
| lnc-GUF1-4 | lnc-GUF1-4 | -0.139051523734904 | 0.288595725239884 | -0.505525483758826 | 0.615801736508746 | 0.831532319265205 | -5.66437443814914 |
| lnc-DR1-2 | lnc-DR1-2 | -0.0547121484200392 | 3.5559365214817 | -0.50532143590248 | 0.615943809481767 | 0.831669603159486 | -6.1277222226579 |
| ENSG00000271723.5 | MROH7-TTC4 | 0.0506125484334876 | 2.60916356635887 | 0.505157567514207 | 0.61605791737564 | 0.831684327937139 | -6.01169432384709 |
| ENSG00000143222.11 | UFC1 | 0.0639232820048647 | 5.35375563133513 | 0.505149361335561 | 0.616063631907793 | 0.831684327937139 | -6.26789287612035 |
| ENSG00000100030.14 | MAPK1 | 0.0319879607706142 | 7.17034232672923 | 0.505119817144831 | 0.616084205779376 | 0.831684327937139 | -6.27787852127863 |
| ENSG00000270011.7 | ZNF559-ZNF177 | -0.114418654258241 | 3.25228136959259 | -0.505061260302337 | 0.616124984294587 | 0.831684327937139 | -6.08207082417136 |
| CDC37L1-DT | CDC37L1-DT | -0.0827267479472088 | 1.09422050932154 | -0.504851057883409 | 0.616271377679853 | 0.831684327937139 | -5.79900793661368 |
| lnc-GTPBP1-4 | lnc-GTPBP1-4 | -0.0813560751477425 | 1.7001368055216 | -0.504739466866647 | 0.616349100557763 | 0.831684327937139 | -5.8936446303909 |
| ENSG00000095951.16 | HIVEP1 | 0.0706337805231669 | 8.24381528618757 | 0.504715958529488 | 0.616365474626737 | 0.831684327937139 | -6.25815015557265 |
| ENSG00000165060.12 | FXN | 0.0705883109428589 | 2.84270966998216 | 0.504675357369461 | 0.616393754684714 | 0.831684327937139 | -6.04082967825977 |
| ENSG00000160223.17 | ICOSLG | 0.397347523198188 | 1.78241270064314 | 0.50481685065835 | 0.61639431625825 | 0.831684327937139 | -5.88739241997949 |
| ENSG00000132792.18 | CTNNBL1 | 0.0528470698985242 | 4.79872902199669 | 0.50467361595038 | 0.616394967654193 | 0.831684327937139 | -6.23819417675925 |
| ENSG00000139436.20 | GIT2 | -0.0674034453496466 | 6.55886272241651 | -0.504582289711964 | 0.616458581627327 | 0.831684327937139 | -6.28388964256361 |
| ENSG00000009954.10 | BAZ1B | 0.0497185180132984 | 6.64174219956929 | 0.504544952959915 | 0.61648458967966 | 0.831684327937139 | -6.28354991910656 |
| ENSG00000276610.1 | SNORD64 | 0.126850570228955 | 0.997532225979071 | 0.504320084851484 | 0.616641238953567 | 0.831684327937139 | -5.75604751124322 |
| ENSG00000007312.12 | CD79B | -0.132196785284289 | 6.68944791073319 | -0.504347993460581 | 0.616649672825401 | 0.831684327937139 | -6.28299399387235 |
| ENSG00000134333.13 | LDHA | -0.10101254913313 | 5.21112102966125 | -0.504293134338411 | 0.616660014631132 | 0.831684327937139 | -6.26454849659178 |
| lnc-CREB3L4-12 | lnc-CREB3L4-12 | -0.106386619118695 | 0.810266427910118 | -0.504190749929489 | 0.616731345385613 | 0.831684327937139 | -5.75085188882265 |
| ENSG00000114541.14 | FRMD4B | -0.755401872215718 | -0.638246044605046 | -0.504312932356517 | 0.616745608175337 | 0.831684327937139 | -5.38400006017305 |
| ENSG00000010219.13 | DYRK4 | 0.119844660878966 | 3.81812600815548 | 0.504122978434772 | 0.616778563540143 | 0.831684327937139 | -6.13205606234064 |
| lnc-BRF1-31 | lnc-BRF1-31 | -0.171271586077053 | 7.74560030829399 | -0.504107222926892 | 0.616853180983602 | 0.831684327937139 | -6.27102532550184 |
| lnc-TM9SF2-5 | lnc-TM9SF2-5 | 0.107833053573061 | 1.21893431792182 | 0.503894632604107 | 0.616937670071957 | 0.831684327937139 | -5.8091873404378 |
| ENSG00000069509.5 | FUNDC1 | 0.0913650539608644 | 2.08566316253127 | 0.503881252039397 | 0.616946993943471 | 0.831684327937139 | -5.94696698206382 |
| ENSG00000075945.12 | KIFAP3 | 0.0600824211955966 | 3.83154962479234 | 0.503868267583116 | 0.616956041859076 | 0.831684327937139 | -6.16013032323994 |
| ENSG00000171867.16 | PRNP | 0.0889162595441659 | 4.44397789222933 | 0.503834885484842 | 0.616979303672365 | 0.831684327937139 | -6.21293093445824 |
| ENSG00000167658.15 | EEF2 | -0.081899144309407 | 10.4244466705269 | -0.503735947368426 | 0.617048249549478 | 0.831684327937139 | -6.19120971003828 |
| ENSG00000164879.6 | CA3 | -0.149117454406444 | 1.17982465504002 | -0.503727938563097 | 0.6170538307073 | 0.831684327937139 | -5.80362681877156 |
| lnc-ARSF-6 | lnc-ARSF-6 | 0.193237583440556 | -0.630317139720261 | 0.503676275313671 | 0.617089834224167 | 0.831684327937139 | -5.44788527436614 |
| ENSG00000169026.12 | SLC49A3 | 0.310599984658832 | 1.31082753355611 | 0.503707349369766 | 0.617141185304053 | 0.831684327937139 | -5.81337961203894 |
| lnc-DNAJC7-1 | lnc-DNAJC7-1 | 0.0919025147749994 | 1.50602308162648 | 0.503584073239163 | 0.617154091144743 | 0.831684327937139 | -5.88048152701983 |
| lnc-HTR4-1 | lnc-HTR4-1 | -0.126912126206415 | 0.0154747829063726 | -0.503532823318532 | 0.617189809246839 | 0.831684327937139 | -5.54822463923486 |
| ENSG00000139133.6 | ALG10 | 0.0472620249610824 | 3.83607125760979 | 0.503524414334518 | 0.617195669890838 | 0.831684327937139 | -6.157314456662 |
| lnc-PRDX3-4 | lnc-PRDX3-4 | -0.0755770463834043 | 0.576718040229142 | -0.503507709639308 | 0.617207312308206 | 0.831684327937139 | -5.72771127226545 |
| lnc-PDGFB-5 | lnc-PDGFB-5 | -0.0786273412813081 | 4.48668312663523 | -0.503370469708301 | 0.617302966107264 | 0.83172626377583 | -6.22215844925832 |
| lnc-POLDIP2-6 | lnc-POLDIP2-6 | 0.0654492749754287 | 1.7486198957882 | 0.503347106620614 | 0.617319250434988 | 0.83172626377583 | -5.91087188536192 |
| lnc-SLC25A16-1 | lnc-SLC25A16-1 | 0.0618563707364745 | 0.879738306471628 | 0.503104882274565 | 0.617488094935221 | 0.831840908509536 | -5.77082082651933 |
| ENSG00000077713.18 | SLC25A43 | 0.0885657756291005 | 2.95244043388167 | 0.502980544373404 | 0.6175747738385 | 0.831840908509536 | -6.05869757626517 |
| ENSG00000114993.16 | RTKN | -0.293520509213325 | -0.253034172260801 | -0.502981637194391 | 0.61759602996648 | 0.831840908509536 | -5.50788206149293 |
| lnc-EIF5A2-1 | lnc-EIF5A2-1 | 0.0969159576717162 | 3.23703668953196 | 0.502892379506605 | 0.617636239002827 | 0.831840908509536 | -6.09256169350809 |
| lnc-ZFP57-15 | lnc-ZFP57-15 | 0.162181341049597 | 0.801826935583384 | 0.502887730675239 | 0.617639480066753 | 0.831840908509536 | -5.7406940261418 |
| ENSG00000086475.14 | SEPHS1 | 0.0467645126960221 | 4.14755819711881 | 0.502877195082978 | 0.617646825280631 | 0.831840908509536 | -6.19447897839765 |
| ENSG00000164031.16 | DNAJB14 | 0.0685979553458793 | 6.39245580611439 | 0.502614912450714 | 0.617829696488049 | 0.832004319751781 | -6.28525862442505 |
| ENSG00000213339.8 | QTRT1 | -0.0774316122725089 | 4.29811307218758 | -0.502587224274775 | 0.617849002934751 | 0.832004319751781 | -6.20796887504503 |
| lnc-TIMMDC1-2 | lnc-TIMMDC1-2 | -0.0871219912605442 | -0.152306211428017 | -0.502444209377186 | 0.617948728897588 | 0.832084174205093 | -5.51854077846788 |
| lnc-LYSMD3-2 | lnc-LYSMD3-2 | 0.141691816422999 | 5.55983118620215 | 0.502423436719973 | 0.618001990332522 | 0.832101456598044 | -6.27735878101677 |
| lnc-B3GLCT-6 | lnc-B3GLCT-6 | -0.246429051332658 | 0.373109027887113 | -0.50226333772723 | 0.618091684443884 | 0.83211853217628 | -5.63248591401515 |
| ENSG00000130038.9 | CRACR2A | -0.0497173857757307 | 4.39849291784706 | -0.50223370790788 | 0.618095527301324 | 0.83211853217628 | -6.21369725199354 |
| lnc-HMGB2-1 | lnc-HMGB2-1 | -0.227339367770753 | -0.333180774069251 | -0.501848806645821 | 0.618363988545883 | 0.832385379290905 | -5.46864425194054 |
| lnc-FBXO31-1 | lnc-FBXO31-1 | 0.101960800800641 | -0.159846988078969 | 0.501833563377446 | 0.618374621519566 | 0.832385379290905 | -5.50352230693534 |
| ENSG00000062282.14 | DGAT2 | -0.203020612731214 | 1.32427647184319 | -0.501783728856219 | 0.618432334851477 | 0.83240862858017 | -5.83585152528206 |
| ENSG00000077152.10 | UBE2T | 0.190737546659066 | -0.074454467988139 | 0.501575467312404 | 0.618554669531108 | 0.832518849340142 | -5.54366188541675 |
| lnc-LRRD1-4 | lnc-LRRD1-4 | 0.0608289758568843 | 1.99005390441938 | 0.50121845255638 | 0.618808372798786 | 0.832805854408095 | -5.93545844225428 |
| ENSG00000274544.1 | SNORD28 | 0.132454514617081 | 1.34740164204332 | 0.501138892446213 | 0.618859278476018 | 0.832813091038863 | -5.86502717802368 |

| | | | | | | | |
|---|---|---|---|---|---|---|---|
| lnc-AFAP1L2-1 | lnc-AFAP1L2-1 | 0.121808046550203 | 1.09517749086968 | 0.501043672211919 | 0.618925724969157 | 0.832813091038863 | -5.793343167007 |
| ENSG00000136425.13 | CIB2 | 0.126240585047039 | -0.0185685508324551 | 0.501021827804509 | 0.618940968867623 | 0.832813091038863 | -5.59558209672471 |
| ENSG00000165424.6 | ZCCHC24 | -0.135150877209801 | 0.197422415957181 | -0.500972210045821 | 0.618975594748701 | 0.832813091038863 | -5.66866753628092 |
| lnc-B2M-1 | lnc-B2M-1 | 0.106539870618528 | 2.81081609520448 | 0.500904936979588 | 0.619022542830426 | 0.832821818359282 | -6.03786257050329 |
| ENSG00000164082.14 | GRM2 | -0.108854427735494 | 0.240338266524122 | -0.500545787488346 | 0.619273210931442 | 0.833065939716519 | -5.61119771228081 |
| ENSG00000143499.13 | SMYD2 | -0.0674182205208176 | 4.70937065832589 | -0.500528982409986 | 0.619284941145725 | 0.833065939716519 | -6.23947364652474 |
| ENSG00000198130.15 | HIBCH | 0.11777816685094 | 4.66684050698911 | 0.500457292113466 | 0.61934004295259 | 0.833079533472302 | -6.2290738089259 |
| ENSG00000161202.18 | DVL3 | -0.0526089077466665 | 5.04216864568947 | -0.500398540954284 | 0.619375994788255 | 0.833079533472302 | -6.25790059290511 |
| ENSG00000108556.8 | CHRNE | -0.283937930106581 | 1.78695506281657 | -0.500223386665232 | 0.6195706967219 | 0.833159004858003 | -5.89402737409562 |
| ENSG00000167182.14 | SP2 | 0.0928009603685531 | 3.77148284047135 | 0.500108863567507 | 0.619578223459056 | 0.833159004858003 | -6.1414868357858 |
| ENSG00000120129.5 | DUSP1 | 0.172327064838582 | 7.41099193953943 | 0.500198830462035 | 0.619579627984447 | 0.833159004858003 | -6.27802094766095 |
| ENSG00000100395.14 | L3MBTL2 | -0.0347613880491249 | 4.84957939146023 | -0.500081981136162 | 0.61959699204982 | 0.833159004858003 | -6.24594875932503 |
| ENSG00000169258.6 | GPRIN1 | 0.165140464204655 | 0.471112652793492 | 0.499877137821655 | 0.61974001657546 | 0.833209534157401 | -5.67999474708 |
| ENSG00000005020.12 | SKAP2 | 0.0826081546085664 | 5.54874404548131 | 0.499782203579426 | 0.61980630607236 | 0.833209534157401 | -6.27807152111414 |
| lnc-EXOC2-37 | lnc-EXOC2-37 | 0.2102326805944 | 0.331912128908822 | 0.499620161496732 | 0.619919462187315 | 0.833209534157401 | -5.59109536082539 |
| ENSG00000159685.10 | CHCHD6 | 0.0833516230282128 | 1.27952939315677 | 0.499555998038866 | 0.619964270949404 | 0.833209534157401 | -5.82770119145656 |
| lnc-DHX36-3 | lnc-DHX36-3 | -0.157500095540985 | -0.122681057597926 | -0.499543275100624 | 0.619973156228858 | 0.833209534157401 | -5.49164348799168 |
| lnc-CCDC117-3 | lnc-CCDC117-3 | -0.112340267792542 | -0.239349639934086 | -0.499541610431921 | 0.619974318782698 | 0.833209534157401 | -5.48991267476689 |
| ENSG00000158019.20 | BABAM2 | 0.0468241806540597 | 4.2396755910228 | 0.499535143238891 | 0.619978835282358 | 0.833209534157401 | -6.1962914549223 |
| ENSG00000112679.14 | DUSP22 | -0.146296033571068 | 5.7626402855521 | -0.499579129295559 | 0.619993605885055 | 0.833209534157401 | -6.28210899723164 |
| ENSG00000117543.20 | DPH5 | 0.0757724743994559 | 3.52137339144627 | 0.499506422040676 | 0.619998893511867 | 0.833209534157401 | -6.11538231145341 |
| ENSG00000164815.10 | ORC5 | -0.0541413834750197 | 3.17904481454948 | -0.499193529366494 | 0.620217429584817 | 0.833437695497109 | -6.0870000175903 |
| ENSG00000188283.11 | ZNF383 | 0.0439702436962026 | 4.76356235629278 | 0.499047158360271 | 0.620319672555864 | 0.833437695497109 | -6.24006766498373 |
| ENSG00000139131.12 | YARS2 | 0.0475242681108674 | 3.52403650827317 | 0.498991351150278 | 0.620358656969757 | 0.833437695497109 | -6.12676072047747 |
| lnc-ITGB2-1 | lnc-ITGB2-1 | 0.0780561925778414 | 3.03628230812092 | 0.498985611435159 | 0.62036266653996 | 0.833437695497109 | -6.06077994956868 |
| ENSG00000111641.11 | NOP2 | 0.061760884518594 | 3.29605989070281 | 0.498973498523765 | 0.620371128247156 | 0.833437695497109 | -6.09867768211482 |
| ENSG00000107242.17 | PIP5K1B | 0.183659145572376 | 3.22586787800193 | 0.498731861878879 | 0.620585410931346 | 0.833671159979108 | -6.06333478242357 |
| ENSG00000083093.9 | PALB2 | 0.0539999711052577 | 3.91424603505123 | 0.498355963845242 | 0.620802587909427 | 0.83389335400797 | -6.1627085059444 |
| lnc-RWDD1-1 | lnc-RWDD1-1 | -0.130320361179361 | 0.241517789707911 | -0.498314103658041 | 0.620831839713265 | 0.83389335400797 | -5.64840944439416 |
| lnc-NUDT14-3 | lnc-NUDT14-3 | 0.11528896014555 | -0.0475747669389729 | 0.498084614899238 | 0.620992216969091 | 0.833996464521033 | -5.55083525564061 |
| ENSG00000238317.2 | SNORD11 | 0.12091808169332 | 2.31238953790173 | 0.49806009539117 | 0.62100935342759 | 0.833996464521033 | -5.96561388171051 |
| ENSG00000203666.12 | EFCAB2 | 0.104142200034395 | 1.80292774993562 | 0.498010171902556 | 0.621044245152464 | 0.833996464521033 | -5.92458952790505 |
| lnc-FZD1-5 | lnc-FZD1-5 | 0.127900116781936 | 2.68170435783156 | 0.497972349333191 | 0.621070680084458 | 0.833996464521033 | -6.01431502104398 |
| ENSG00000107902.13 | LHPP | -0.131491229677017 | 4.4779560679168 | -0.497940966945785 | 0.621111742060176 | 0.833997193999908 | -6.22371513590151 |
| ENSG00000111639.7 | MRPL51 | -0.101328587053679 | 3.2462681838975 | -0.497446422087854 | 0.621438313393253 | 0.834376924611074 | -6.08959604990306 |
| ENSG00000152217.18 | SETBP1 | -0.06723456824027 | 7.38523546653802 | -0.49739306383214 | 0.621475617315861 | 0.834376924611074 | -6.28052661919301 |
| lnc-PARP1-5 | lnc-PARP1-5 | 0.170529622591777 | 0.437811386959554 | 0.497305773627938 | 0.62153664597738 | 0.834382631770272 | -5.61266722084483 |
| lnc-ST8SIA1-7 | lnc-ST8SIA1-7 | -0.169721332894741 | -0.296825046811244 | -0.497271022087622 | 0.621560943153747 | 0.834382631770272 | -5.54176802164847 |
| ENSG00000182484.15 | WASH6P | -0.0907350409704129 | 2.59468285512397 | -0.496925459183456 | 0.621802573037143 | 0.834580717989265 | -6.01673499644242 |
| ENSG00000003400.14 | CASP10 | 0.287506293349362 | 1.48842113732874 | 0.497005153080236 | 0.621818088673043 | 0.834580717989265 | -5.84478213791377 |
| ENSG00000274211.4 | SOCS7 | 0.0673986902537429 | 4.40356183539704 | 0.496886030294778 | 0.621830145803982 | 0.834580717989265 | -6.21478850138974 |
| lnc-SLC1A3-1 | lnc-SLC1A3-1 | 0.132207289986448 | 3.16356148049709 | 0.496778606499767 | 0.621906352079156 | 0.834628574352564 | -6.08133388290092 |
| ENSG00000205746.9 | PKD1P4 | -0.164947163287424 | 0.762851480959838 | -0.496524705108116 | 0.622082847432651 | 0.834811008521727 | -5.7831001821344 |
| ENSG00000132466.18 | ANKRD17 | -0.0472450811093044 | 7.16078828290816 | -0.49642537416504 | 0.622152325049809 | 0.83484981462287 | -6.28264547890525 |
| lnc-SLC6A4-4 | lnc-SLC6A4-4 | 0.124112573783436 | -0.344883834504737 | 0.495887276072366 | 0.622528761539172 | 0.83525225247099 | -5.46302831651741 |
| ENSG00000166348.18 | USP54 | -0.0880405187683873 | 2.59208687077313 | -0.495880658123543 | 0.622533391884441 | 0.83525225247099 | -6.01603931946253 |
| ENSG00000165629.19 | ATP5F1C | 0.0577405021468662 | 5.60630147971452 | 0.495359316168982 | 0.622898205032557 | 0.835687247698156 | -6.28022405559591 |
| ENSG00000254692.1 | | -0.0443075632180738 | 4.28093095286008 | -0.495266841273105 | 0.622962925085925 | 0.835703467771612 | -6.201317799172 |
| ENSG00000130544.11 | ZNF557 | 0.0488688843761489 | 4.7545490194447 | 0.495226015964712 | 0.622991498298965 | 0.835703467771612 | -6.24095184961638 |
| lnc-SHCBP1-2 | lnc-SHCBP1-2 | -0.128609543539579 | -0.203740941698 | -0.495166647904516 | 0.623033050442143 | 0.83570474273383 | -5.5413883214326 |
| ENSG00000207166.1 | SNORA68 | -0.138565509744309 | 1.65859005252588 | -0.495068406285047 | 0.623101813206642 | 0.835742514090467 | -5.88779856916122 |
| lnc-SSBP2-5 | lnc-SSBP2-5 | -0.287697481280285 | 1.30162170484266 | -0.495074543811276 | 0.623169090806677 | 0.835745331045199 | -5.86269428359566 |
| ENSG00000007202.14 | BLTP2 | 0.0322973208273378 | 7.5594456691959 | 0.494949390965459 | 0.623185120773537 | 0.835745331045199 | -6.27731753313496 |
| ENSG00000177685.16 | CRACR2B | 0.200381587331826 | 2.79055038568859 | 0.49475890725736 | 0.623371628376978 | 0.835887124876195 | -6.02406239949073 |
| lnc-MEF2A-9 | lnc-MEF2A-9 | 0.0906103031266603 | 1.1698201243056 | 0.494646274044714 | 0.623397317133347 | 0.835887124876195 | -5.81882101875321 |
| ENSG00000160570.13 | DEDD2 | 0.0677329966032252 | 4.38160190844224 | 0.494574672719111 | 0.623447446218368 | 0.835887124876195 | -6.21734095997316 |
| ENSG00000183354.11 | BRD10 | -0.0371165687406023 | 6.71371701262766 | -0.494566320918328 | 0.623453293548042 | 0.835887124876195 | -6.2883350765928 |
| lnc-DACT3-1 | lnc-DACT3-1 | 0.0796265079679692 | 1.34161744564234 | 0.494374162508545 | 0.623587835821286 | 0.836013054432979 | -5.84470318345222 |
| ENSG00000166685.11 | COG1 | 0.0601798142075864 | 5.02861991183278 | 0.494256916599556 | 0.623669933497998 | 0.836034738424511 | -6.25868046764409 |
| ENSG00000278987.1 | | -0.1053361074057 | 0.0655062880380872 | -0.494235049854633 | 0.623685245518553 | 0.836034738424511 | -5.58715550021892 |
| ENSG00000256500.5 | | 0.0416201722303073 | 4.90683013330094 | 0.494008951870681 | 0.623843578771161 | 0.836062197578369 | -6.25133383774727 |
| ENSG00000116157.5 | GPX7 | 0.0681507945957476 | 3.62450654185318 | 0.493993044397308 | 0.623854719229829 | 0.836062197578369 | -6.13225129095836 |
| ENSG00000211459.2 | MT-RNR1 | 0.14079529658189 | 14.5968271687888 | 0.493993917767013 | 0.623856700786095 | 0.836062197578369 | -6.02647964819195 |
| ENSG00000107833.10 | NPM3 | -0.13072125532594 | 1.95352450239788 | -0.493973786203117 | 0.623868206413359 | 0.836062197578369 | -5.94229204125973 |
| ENSG00000156804.7 | FBXO32 | -0.0853582008867884 | 3.88043981245068 | -0.49385836462603 | 0.623949042896483 | 0.836063087300032 | -6.14864850463705 |
| ENSG00000172315.5 | TP53RK | 0.0817170047386528 | 3.54251023204519 | 0.49385684310651 | 0.623950108536938 | 0.836063087300032 | -6.13278000636403 |
| ENSG00000126767.17 | ELK1 | 0.0751689068454953 | 3.80255774527 | 0.493564815090936 | 0.624154653937788 | 0.836265292636499 | -6.16352605750267 |
| ENSG00000085224.22 | ATRX | 0.0405601703402851 | 7.83511716335754 | 0.493525389219682 | 0.624182271329472 | 0.836265292636499 | -6.27381547628646 |

| | | | | | | | |
|---|---|---|---|---|---|---|---|
| ENSG00000279145.2 | | -0.113110930229536 | 1.55600065165928 | -0.493326562269335 | 0.624321555766618 | 0.836397460449381 | -5.8930913338854 |
| ENSG00000183569.17 | SERHL2 | 0.356850273164939 | -1.40894252122515 | 0.493257305837281 | 0.624370075307777 | 0.836408022132116 | -5.20235685497249 |
| ENSG00000181192.11 | DHTKD1 | -0.0577330479791227 | 5.98090907362995 | -0.493046686361973 | 0.624517641069239 | 0.836551256418596 | -6.28797823721994 |
| lnc-NPY1R-5 | lnc-NPY1R-5 | 0.0882991563541672 | 0.502223619949695 | 0.49291767234448 | 0.624608039519299 | 0.836617900528779 | -5.69084690986649 |
| lnc-LRRC37A2-1 | lnc-LRRC37A2-1 | -0.110846036410705 | 0.479437989693161 | -0.492652832148857 | 0.624793627943554 | 0.836812027847616 | -5.64360958603457 |
| lnc-POLR2B-3 | lnc-POLR2B-3 | -0.114766674028509 | -0.339360879828074 | -0.492536262035731 | 0.624875322991445 | 0.836866990248742 | -5.47176760143202 |
| ENSG00000226048.1 | YBX1P8 | -0.17796183588737 | 0.289622390563728 | -0.492452647953654 | 0.624933924626066 | 0.836891019556169 | -5.61713120838701 |
| ENSG00000148985.19 | PGAP2 | -0.0501180975645359 | 3.11042844226438 | -0.492371067147451 | 0.624991103589088 | 0.836913140665812 | -6.07288920656062 |
| ENSG00000269693.1 | | 0.0434656243431428 | 4.37808225873323 | 0.49205197370234 | 0.625214774628869 | 0.837128501387436 | -6.2104812816022 |
| ENSG00000177465.4 | ACOT4 | -0.277087134020523 | -1.15217607262704 | -0.492025586289051 | 0.625233272692467 | 0.837128501387436 | -5.26742130278378 |
| lnc-RPL7-3 | lnc-RPL7-3 | -0.137535388304889 | -0.644344162169758 | -0.491913548778427 | 0.625311815776617 | 0.837179205420197 | -5.40129179727246 |
| ENSG00000209582.1 | SNORA48 | 0.141845902557975 | 3.71104083881333 | 0.491524519187472 | 0.625603278757045 | 0.83746118966296 | -6.1399057059598 |
| lnc-SAMD11-7 | lnc-SAMD11-7 | -0.116168406841862 | 1.4196964155057 | -0.491429907729315 | 0.625650919373989 | 0.83746118966296 | -5.85409780591594 |
| ENSG00000273217.1 | | 0.0364424350361094 | 5.02008914043181 | 0.491412190358991 | 0.625663343417735 | 0.83746118966296 | -6.25742308914002 |
| lnc-AMMECR1L-3 | lnc-AMMECR1L-3 | 0.0978842314352208 | 1.3149161209214 | 0.491381042624617 | 0.625685185572416 | 0.83746118966296 | -5.85195941030214 |
| lnc-PPIAL4C-4 | lnc-PPIAL4C-4 | -0.0830739348188845 | 0.336206073611281 | -0.490815609409415 | 0.626081751225604 | 0.837937491740466 | -5.65787403732599 |
| ENSG00000165490.12 | DDIAS | 0.135733768593736 | 1.74259158478481 | 0.490360583556509 | 0.62640096421244 | 0.838310211078326 | -5.90941446189744 |
| ENSG00000126814.6 | TRMT5 | -0.0607949883036469 | 2.56916177124318 | -0.490077684252901 | 0.626599462304853 | 0.838521340135284 | -6.02179928148038 |
| lnc-ETV6-5 | lnc-ETV6-5 | 0.150865558639689 | -0.163393295830507 | 0.489972979665346 | 0.626672936046354 | 0.838528800344438 | -5.53813913732099 |
| ENSG00000114480.12 | GBE1 | -0.0508475966521962 | 3.61414120102903 | -0.489953629598532 | 0.626686514876281 | 0.838528800344438 | -6.13850948039856 |
| ENSG00000064607.16 | SUGP2 | 0.0494769386403275 | 6.74214557388809 | 0.489844405020723 | 0.626763165232303 | 0.838576848022393 | -6.29019946807909 |
| lnc-PLPP4-6 | lnc-PLPP4-6 | 0.138396547108457 | 1.45811291482588 | 0.489665181593862 | 0.626888947611356 | 0.838649876097161 | -5.85586965504911 |
| ENSG00000166896.8 | ATP23 | 0.0998148068373437 | 0.768843065037488 | 0.489650521376783 | 0.626899236925177 | 0.838649876097161 | -5.77539372363583 |
| ENSG00000165168.7 | CYBB | 0.765459853579208 | 1.40400909329847 | 0.489659554045646 | 0.626992855940669 | 0.83872060530492 | -5.79540811375277 |
| lnc-SFMBT1-5 | lnc-SFMBT1-5 | 0.101305481686872 | 2.42971290751078 | 0.489088360230767 | 0.627293847971824 | 0.839068707616588 | -5.9898291413428 |
| ENSG00000183773.15 | AIFM3 | -0.216691882259162 | -0.838692566531548 | -0.488930550151023 | 0.627404643184899 | 0.839162373817322 | -5.33124205980722 |
| ENSG00000077044.10 | DGKD | 0.0812198799236254 | 7.52293000217728 | 0.48882587529206 | 0.627478138051725 | 0.839206141359237 | -6.28147520728869 |
| ENSG00000143437.20 | ARNT | -0.0590639023169679 | 6.03940876898072 | -0.488552507160486 | 0.627670094782906 | 0.83940832700387 | -6.29074133347001 |
| lnc-SERF1A-4 | lnc-SERF1A-4 | 0.174792553874897 | -0.356696724995752 | 0.488263256289413 | 0.627873232649211 | 0.839576172475656 | -5.5557241893788 |
| lnc-TRIM41-4 | lnc-TRIM41-4 | -0.117074692334068 | -0.305847266323868 | -0.488257634146715 | 0.627877181310682 | 0.839576172475656 | -5.44621583891165 |
| ENSG00000224442.1 | RPL4P7 | -0.129777877706267 | 0.745198868717535 | -0.487871864910031 | 0.628148149216909 | 0.839876433707834 | -5.71613771017674 |
| lnc-RAPSN-5 | lnc-RAPSN-5 | 0.0482570879728552 | 2.69946805680483 | 0.487821769552366 | 0.628183340471851 | 0.839876433707834 | -6.02855293652921 |
| lnc-FUBP1-57 | lnc-FUBP1-57 | -0.0658882857995119 | 0.647669083008765 | -0.487567363159633 | 0.628362070745049 | 0.840060827627003 | -5.73759768236137 |
| ENSG00000281398.4 | SNHG4 | -0.107542839768219 | 3.69615900384211 | -0.487232984339903 | 0.628597019044123 | 0.840320350911553 | -6.15468405242902 |
| lnc-C16orf45-1 | lnc-C16orf45-1 | 0.105156256859919 | 0.222906916461209 | 0.487135080116703 | 0.628665817949735 | 0.840357743269216 | -5.63601925668416 |
| ENSG00000248971.2 | KRT8P46 | -0.0615428718433466 | 2.06181999096678 | -0.486822768313263 | 0.628885306900899 | 0.84059654990202 | -5.94809421519793 |
| lnc-SAT2-2 | lnc-SAT2-2 | 0.0730235904909684 | 1.03607055876638 | 0.486651998815068 | 0.629005336001349 | 0.840702391617907 | -5.81154477113447 |
| ENSG00000279235.1 | | 0.134886370421146 | 0.464930324465617 | 0.486221857220788 | 0.629307715480206 | 0.840997319031884 | -5.68083048052261 |
| lnc-MORF4L1-3 | lnc-MORF4L1-3 | 0.134886370421146 | 0.464930324465617 | 0.486221857220788 | 0.629307715480206 | 0.840997319031884 | -5.68083048052261 |
| lnc-SCAF4-2 | lnc-SCAF4-2 | 0.094535897736282 | 1.5709295974757 | 0.486128323879546 | 0.629373475754031 | 0.841027510663841 | -5.88812247319174 |
| lnc-PRSS12-4 | lnc-PRSS12-4 | 0.0714914828237379 | -0.0167787458614523 | 0.485966801914798 | 0.629487043800899 | 0.841027510663841 | -5.54449644253424 |
| lnc-L3HYPDH-1 | lnc-L3HYPDH-1 | -0.084821348742262 | 2.5708564414248 | -0.485965555357033 | 0.629487920306027 | 0.841027510663841 | -6.02635838534052 |
| lnc-KLHL22-4 | lnc-KLHL22-4 | 0.0469670129840956 | 1.86596360479861 | 0.485957266300897 | 0.629493748690042 | 0.841027510663841 | -5.92290004064744 |
| ENSG00000184967.6 | NOC4L | -0.091712425282465 | 2.33312961146102 | -0.485786732717512 | 0.629613663328085 | 0.841101878454335 | -6.01041487576792 |
| lnc-SYCP3-4 | lnc-SYCP3-4 | -0.0948656292080448 | 2.94904605380038 | -0.485749746231593 | 0.62963967256664 | 0.841101878454335 | -6.07888855622972 |
| ENSG00000167378.8 | IRGQ | 0.0557158426177629 | 3.29867818960849 | 0.485660087011824 | 0.629702723732163 | 0.841101878454335 | -6.10539011884581 |
| ENSG00000207313.1 | SNORA2B | -0.0949789141470521 | 0.176812345688536 | -0.485645663200291 | 0.629712867268197 | 0.841101878454335 | -5.62680920111977 |
| lnc-BCKDK-1 | lnc-BCKDK-1 | 0.0821230122314025 | 1.89381577801821 | 0.485450269676802 | 0.629850284761273 | 0.841197891984568 | -5.92877538042629 |
| ENSG00000166343.9 | MSS51 | -0.0881761336584876 | 2.20195368169143 | -0.485330049237691 | 0.629934840684094 | 0.841197891984568 | -5.96949386677462 |
| lnc-CAMTA1-2 | lnc-CAMTA1-2 | 0.0906356845148981 | 0.487273971606124 | 0.485076425518261 | 0.630113241029582 | 0.841197891984568 | -5.70065109484132 |
| ENSG00000070061.14 | ELP1 | 0.0483477269590313 | 5.62002995681093 | 0.48506765689441 | 0.630119409328361 | 0.841197891984568 | -6.28697758363933 |
| lnc-KLF11-1 | lnc-KLF11-1 | 0.393782871186753 | 1.36479809554378 | 0.485021479045895 | 0.630246916081719 | 0.841197891984568 | -5.88755556263176 |
| lnc-KHDRBS3-4 | lnc-KHDRBS3-4 | -0.173842238696344 | -0.0308931134243736 | -0.484864186604003 | 0.630262548242692 | 0.841197891984568 | -5.5553923338299 |
| lnc-CCDC167-4 | lnc-CCDC167-4 | 0.373705909021532 | 0.470261664724746 | 0.484924817076715 | 0.630291228236099 | 0.841197891984568 | -5.65576718385034 |
| ENSG00000215529.12 | EFCAB8 | -0.117114193764013 | -0.244369351773383 | -0.48478128908291 | 0.630320869778332 | 0.841197891984568 | -5.46973178366475 |
| lnc-CELF6-3 | lnc-CELF6-3 | 0.0476435208524055 | 2.51879244804214 | 0.484777195579484 | 0.630323749774346 | 0.841197891984568 | -6.01092483315695 |
| ENSG00000156261.12 | CCT8 | 0.0694611515871637 | 5.9628268351145 | 0.484696707676679 | 0.6303803784497 | 0.841197891984568 | -6.29223454458739 |
| ENSG00000135535.16 | CD164 | -0.0549299430752264 | 6.54652093650855 | -0.48469466267667 | 0.630381817274611 | 0.841197891984568 | -6.29311236148199 |
| ENSG00000198026.7 | ZNF335 | 0.0800497069522859 | 6.03038554060052 | 0.484679001467043 | 0.630392836265741 | 0.841197891984568 | -6.29253998544507 |
| ENSG00000202538.1 | RNU4-2 | 0.170963347645137 | 3.27818755185555 | 0.484677269436501 | 0.630434856754223 | 0.841197891984568 | -6.09713561880211 |
| ENSG00000196747.4 | H2AC13 | 0.311867815280971 | 3.29696874438895 | 0.484723104164046 | 0.630456784277667 | 0.841197891984568 | -6.14970358184498 |
| ENSG00000100376.11 | FAM118A | 0.156529476164696 | 4.03911760859954 | 0.484576238588973 | 0.630502343647986 | 0.841197891984568 | -6.15573352369121 |
| lnc-RWDD2B-1 | lnc-RWDD2B-1 | -0.0876745448060869 | 0.528955265660534 | -0.484483117839066 | 0.630530664226386 | 0.841197891984568 | -5.7239533143199 |
| ZBED5-AS1 | ZBED5-AS1 | -0.0664445440297978 | 2.94055396130836 | -0.484459656923045 | 0.630547172727673 | 0.841197891984568 | -6.05983315406756 |
| ENSG00000103540.16 | CCP110 | 0.0553821702045258 | 3.96064301620949 | 0.484451285392812 | 0.630553063482418 | 0.841197891984568 | -6.1782795509262 |
| lnc-DICER1-5 | lnc-DICER1-5 | 0.089429792587062 | 1.37548646928207 | 0.484439646737963 | 0.630561253239562 | 0.841197891984568 | -5.8616848454361 |
| ENSG00000162650.16 | ATXN7L2 | -0.074912927685884 | 3.41331942541808 | -0.48423835398265 | 0.630702904086863 | 0.841306544140268 | -6.1164830280666 |
| ENSG00000202503.1 | SNORD34 | 0.0728227925063523 | 1.26757147746151 | 0.484183327897431 | 0.630741628699846 | 0.841306544140268 | -5.85091185762333 |

| | | | | | | | |
|---|---|---|---|---|---|---|---|
| ENSG00000139645.9 | ANKRD52 | 0.0578001049192371 | 5.53712050286932 | 0.484105805229264 | 0.63079618706736 | 0.841306544140268 | -6.28530810370151 |
| VAV3-AS1 | VAV3-AS1 | -0.172868282957407 | 3.10505488955185 | -0.484005412504367 | 0.630907983321246 | 0.841306544140268 | -6.08993448099694 |
| ENSG00000063127.15 | SLC6A16 | -0.0920689092443645 | 4.4400682445996 | -0.483936493647796 | 0.630915351248381 | 0.841306544140268 | -6.21780910689902 |
| ENSG00000149483.11 | TMEM138 | 0.0540842307221343 | 5.06793367177402 | 0.483926972982963 | 0.63092205233874 | 0.841306544140268 | -6.26586811249871 |
| lnc-PHF11-2 | lnc-PHF11-2 | -0.0923521400521518 | 1.02098209988815 | -0.483917363767205 | 0.63092881578726 | 0.841306544140268 | -5.80729179684536 |
| ENSG00000265408.1 | | 0.068865994068811 | 0.296563242208135 | 0.483732072144833 | 0.631059239594287 | 0.841393196029746 | -5.63513726698609 |
| lnc-DDX53-3 | lnc-DDX53-3 | 0.386023065944851 | -0.401251629213679 | 0.483667380656706 | 0.631145482448462 | 0.841393196029746 | -5.36997043070503 |
| lnc-ANKS1B-2 | lnc-ANKS1B-2 | 0.0632488325603837 | 0.905274341507092 | 0.483490211321403 | 0.631229499460326 | 0.841393196029746 | -5.77909242722122 |
| ENSG00000176108.9 | CHMP6 | -0.0530315495054414 | 3.1865503142848 | -0.483488741163411 | 0.631230534451814 | 0.841393196029746 | -6.09088421855107 |
| ENSG00000176105.13 | YES1 | -0.69701428799374 | -0.677146411532419 | -0.483571122706755 | 0.631263060020516 | 0.841393196029746 | -5.29331770392664 |
| lnc-ZBTB49-4 | lnc-ZBTB49-4 | 0.168864774819753 | -0.556067840088351 | 0.483347167406816 | 0.631330205909803 | 0.841393196029746 | -5.39657619154693 |
| SNHG7 | SNHG7 | -0.104214006349872 | 5.118027264725 | -0.483344398125999 | 0.631332570495261 | 0.841393196029746 | -6.27856217783768 |
| lnc-RAD23B-17 | lnc-RAD23B-17 | -0.352494721122001 | 1.59510548948935 | -0.48345069686132 | 0.631340907389948 | 0.841393196029746 | -5.79759775594413 |
| ENSG00000185420.18 | SMYD3 | 0.072858870894005 | 3.13798422132328 | 0.483184922795478 | 0.631444438702945 | 0.841393196029746 | -6.07392993351495 |
| ENSG00000156931.15 | VPS8 | 0.0453991047721161 | 6.16513496852524 | 0.483096375905684 | 0.631506786428164 | 0.841393196029746 | -6.29447925238866 |
| ENSG00000197653.15 | DNAH10 | -0.188816112418743 | 0.72515612700304 | -0.483087280560325 | 0.631521876069022 | 0.841393196029746 | -5.79211569093526 |
| ENSG00000157600.11 | TMEM164 | 0.0608562150602183 | 3.85599220487276 | 0.483019805623452 | 0.631560703361364 | 0.841393196029746 | -6.1702384486072 |
| ENSG00000198464.13 | ZNF480 | -0.0552818880393353 | 5.47453609276979 | -0.483013400937073 | 0.631565213310683 | 0.841393196029746 | -6.28245836573522 |
| ENSG00000252361.1 | RNU6-118P | -0.152854302695213 | 0.94093227370401 | -0.483012152607215 | 0.631566092341232 | 0.841393196029746 | -5.79009447091708 |
| lnc-MROH6-1 | lnc-MROH6-1 | -0.127823523473682 | 0.973459307124183 | -0.482948890919637 | 0.631610639731888 | 0.841398084111155 | -5.80651606563444 |
| lnc-TERF2IP-5 | lnc-TERF2IP-5 | 0.104909990383096 | 3.57344257875266 | 0.482850902472838 | 0.631679643623569 | 0.841435549100694 | -6.12316746285616 |
| lnc-THAP5-1 | lnc-THAP5-1 | 0.121300203603647 | 1.54090884634838 | 0.482734719804686 | 0.631761464271699 | 0.841452544042364 | -5.88311925812181 |
| ENSG00000139197.10 | PEX5 | 0.0623504761956593 | 4.07122159924495 | 0.482716687333538 | 0.631774163903741 | 0.841452544042364 | -6.19286663189379 |
| ENSG00000112308.12 | C6orf62 | -0.0404774381669055 | 7.30680138617024 | -0.482293093980344 | 0.632072518091987 | 0.84179544742073 | -6.2866299825714 |
| ENSG00000167552.13 | TUBA1A | 0.0835628099141843 | 6.63613010879227 | 0.482114190399607 | 0.632198545869313 | 0.841908816616724 | -6.29416228718166 |
| ENSG00000100796.17 | PPP4R3A | -0.0427163715643024 | 7.32532799113974 | -0.482009591889334 | 0.632272234919286 | 0.841922248009899 | -6.28746795802852 |
| lnc-DDX3X-2 | lnc-DDX3X-2 | -0.064278486912754 | 0.980879673519276 | -0.481983752526512 | 0.632290439184619 | 0.841922248009899 | -5.8079864028438 |
| lnc-CRACR2A-5 | lnc-CRACR2A-5 | -0.0634531356927723 | 0.716891888696392 | -0.481837812598396 | 0.632393260649056 | 0.84200468878579 | -5.75457560008803 |
| ENSG00000278217.1 | | 0.163742917617622 | 8.05425726067355 | 0.481826783077986 | 0.632460856198941 | 0.842040220125666 | -6.27195742326698 |
| ENSG00000112130.16 | RNF8 | -0.045200135163141 | 3.20270908016898 | -0.48151229541019 | 0.632622629158061 | 0.842201123857472 | -6.09424043848185 |
| ENSG00000198618.5 | PPIAP22 | 0.12077216827816 | 3.97420627523557 | 0.481449144672605 | 0.632671737133999 | 0.84221202725579 | -6.17075899376825 |
| lnc-ASPHD1-1 | lnc-ASPHD1-1 | -0.127260273564802 | -0.115071471905276 | -0.481361756650902 | 0.632728715353552 | 0.842233405427288 | -5.53731908051762 |
| KDM7A-DT | KDM7A-DT | -0.117237782275359 | 1.97854350693287 | -0.481150313846169 | 0.632877734464774 | 0.842377289736708 | -5.93833591578082 |
| lnc-GMCL1-1 | lnc-GMCL1-1 | 0.0923226319916339 | 0.644857145114706 | 0.480602661268205 | 0.633263776781952 | 0.842836619301837 | -5.7444254096232 |
| ENSG00000255872.3 | | 0.0648038064736263 | 1.54462314782644 | 0.480477281346828 | 0.633352172106125 | 0.842899764545478 | -5.88572268346232 |
| ENSG00000225828.1 | FAM229A | -0.072534519640866 | 3.87268669403357 | -0.480395458053496 | 0.633409862069522 | 0.842922039889893 | -6.17110375452625 |
| ENSG00000162994.15 | CLHC1 | -0.0981731090337663 | 1.94153657953286 | -0.480159703128681 | 0.633576095284101 | 0.843076863418327 | -5.94541060912383 |
| ENSG00000115841.19 | RMDN2 | -0.0775715326622533 | 3.71865886630046 | -0.480074835338114 | 0.633635941123206 | 0.843076863418327 | -6.15381017552272 |
| ENSG00000234857.2 | HNRNPUL2-BSCL2 | -0.053650404671612 | 4.68490995319829 | -0.480056199108216 | 0.633649083082229 | 0.843076863418327 | -6.24437769018628 |
| ENSG00000259066.5 | | 0.058957852918156 | 1.70779386897279 | 0.479853381706645 | 0.633792114275456 | 0.843195907238149 | -5.90199857378056 |
| ENSG00000054654.16 | SYNE2 | 0.104658674437886 | 9.98510366481793 | 0.47981315222783 | 0.633820486653483 | 0.843195907238149 | -6.22162704911657 |
| ENSG00000150995.19 | ITPR1 | 0.0850262464008757 | 8.58194129122494 | 0.479517367194707 | 0.634029110121409 | 0.843418934507139 | -6.26234218245378 |
| ENSG00000267179.1 | | 0.0722967285728622 | 2.84391432324307 | 0.479361748360191 | 0.634138883486665 | 0.843435598609922 | -6.04083529710032 |
| ENSG00000285245.1 | | -0.0630156100860546 | 4.69818759382649 | -0.479287290036282 | 0.634191409260326 | 0.843435598609922 | -6.25332619267969 |
| lnc-POU2AF1-1 | lnc-POU2AF1-1 | -0.132315409611758 | 2.98983357967673 | -0.479219161172271 | 0.634245376145016 | 0.843435598609922 | -6.09369451930214 |
| ENSG00000228049.7 | POLR2J2 | -0.131857822759697 | 1.29823527695081 | -0.479182481814173 | 0.634265348256374 | 0.843435598609922 | -5.83651008578083 |
| ENSG00000128609.14 | NDUFA5 | 0.0624895843321633 | 5.03286859637698 | 0.479159414083408 | 0.634281622342716 | 0.843435598609922 | -6.26963311484436 |
| ENSG00000188306.6 | LRRIQ4 | 0.302716313664812 | -1.00073948690985 | 0.479046324999208 | 0.634361408371716 | 0.843435598609922 | -5.25792390146634 |
| lnc-ORC6-5 | lnc-ORC6-5 | -0.196995324692046 | 1.627262249842 | -0.479089363482946 | 0.634361522658182 | 0.843435598609922 | -5.89359136032082 |
| ENSG00000169499.14 | PLEKHA2 | 0.0930703569194282 | 8.48113567627474 | 0.47901568172958 | 0.634383028411454 | 0.843435598609922 | -6.26434395095328 |
| ENSG00000063245.14 | EPN1 | 0.0688137815087728 | 4.89757965436785 | 0.478976840687099 | 0.634410432768732 | 0.843435598609922 | -6.25412066253358 |
| ENSG00000164414.17 | SLC35A1 | -0.065949764248709 | 2.94129265653009 | -0.478908189153447 | 0.634458871233328 | 0.843445517444654 | -6.06729481360459 |
| lnc-ITGA11-3 | lnc-ITGA11-3 | 0.0492965275048052 | 3.68762883616051 | 0.478803105364919 | 0.634533018345189 | 0.843489609700274 | -6.15357149107115 |
| lnc-VTI1A-6 | lnc-VTI1A-6 | 0.101884997231257 | 1.30166468487882 | 0.478540618587816 | 0.634718245582837 | 0.843590303452885 | -5.84591979431712 |
| ENSG00000184432.9 | COPB2 | -0.0481453208936067 | 5.86927327826708 | -0.478533924457518 | 0.634722969693494 | 0.843590303452885 | -6.29379720301095 |
| ENSG00000163795.13 | ZNF513 | 0.0619672393148347 | 2.76606788825425 | 0.478521522675094 | 0.634731721788604 | 0.843590303452885 | -6.04779502053612 |
| lnc-BORCS5-1 | lnc-BORCS5-1 | -0.0892317989870781 | 1.62476362622744 | -0.478127252790738 | 0.635009990601365 | 0.843884793514381 | -5.88681835961422 |
| ENSG00000198252.11 | STYX | -0.0696566903990625 | 4.83822076381517 | -0.478091397400402 | 0.635035299360844 | 0.843884793514381 | -6.25931780232406 |
| ENSG00000121680.15 | PEX16 | 0.0479138988686421 | 4.41004387941682 | 0.47795499122946 | 0.635131586576196 | 0.84390633086065 | -6.23093795403638 |
| ENSG00000103381.11 | CPPED1 | -0.106886144577718 | 2.91931982035308 | -0.477952270956999 | 0.635133506843625 | 0.84390633086065 | -6.07448253032435 |
| ENSG00000092036.18 | HAUS4 | 0.0838238898521793 | 3.16067898514745 | 0.477780121346049 | 0.635255034138469 | 0.844013320918675 | -6.0745533388982 |
| ENSG00000176171.11 | BNIP3 | -0.0914947552060088 | 2.28288499639712 | -0.47769492981706 | 0.63531517800999 | 0.844038747110284 | -6.00025533187217 |
| ENSG00000284034.1 | MIR5196 | -0.157253620453248 | 1.44962081484574 | -0.477556525852574 | 0.635413516721104 | 0.84411490994388 | -5.88466461890734 |
| ENSG00000198185.11 | ZNF334 | 0.559027923743671 | -2.16217610770187 | 0.47726180478622 | 0.635628654770405 | 0.844346214981558 | -4.96103051245701 |
| lnc-SMARCD2-2 | lnc-SMARCD2-2 | -0.0826439684541336 | 1.12263443338353 | -0.476907496939027 | 0.635871211080464 | 0.844522960051654 | -5.82251762365819 |
| ENSG00000197062.11 | ZSCAN26 | -0.05082749342969 | 4.0664141901286 | -0.476886367407989 | 0.635886134289656 | 0.844522960051654 | -6.19260647178226 |
| ENSG00000181472.4 | ZBTB2 | 0.0687061286781905 | 4.8557020132018 | 0.476829566397033 | 0.635926252047264 | 0.844522960051654 | -6.25809140515731 |
| lnc-LCP1-2 | lnc-LCP1-2 | -0.0892175893824971 | 5.42874947599743 | -0.476783142814979 | 0.635959041193882 | 0.844522960051654 | -6.28812896365631 |

| | | | | | | | |
|---|---|---|---|---|---|---|---|
| ENSG00000265558.1 | MIR3918 | -0.149711313893936 | 0.555221038805431 | -0.476711348545051 | 0.636009751205503 | 0.844522960051654 | -5.70693632245997 |
| ENSG00000148516.21 | ZEB1 | 0.0507974293617073 | 6.48883029235605 | 0.47667602184029 | 0.63603470395145 | 0.844522960051654 | -6.2977184631313 |
| ENSG00000279901.1 | | 0.0856647701392096 | 0.862827489346141 | 0.476655895385656 | 0.636048920309029 | 0.844522960051654 | -5.77531977957473 |
| lnc-RPS7-4 | lnc-RPS7-4 | -0.135559845252473 | -0.0615590080729329 | -0.476554700089341 | 0.636120401893424 | 0.844563389806641 | -5.54852677235684 |
| ENSG00000160213.7 | CSTB | -0.139169321883824 | 2.61799413112018 | -0.476479353127625 | 0.636177066226442 | 0.844584143068812 | -6.01958107172842 |
| lnc-ETFRF1-7 | lnc-ETFRF1-7 | -0.198548255556093 | 0.99063970990832 | -0.476198257612582 | 0.636394687739782 | 0.844818565478744 | -5.83057737521276 |
| ENSG00000285399.2 | NA | -0.0599930011196856 | 2.97777331376969 | -0.47603619466424 | 0.636486714967166 | 0.844878291609035 | -6.06918289433454 |
| ENSG00000180644.7 | PRF1 | -0.531616484968954 | -1.08361411490037 | -0.476085034625467 | 0.636522548770568 | 0.844878291609035 | -5.35428219917944 |
| ENSG00000112877.7 | CEP72 | 0.0723535847639001 | 2.56223390320593 | 0.475875898730828 | 0.636599979274505 | 0.844878291609035 | -6.00605408386397 |
| lnc-KLHL1-9 | lnc-KLHL1-9 | -0.295977933411508 | 2.91035414610551 | -0.475996067667846 | 0.636603972509353 | 0.844878291609035 | -6.07088975903516 |
| ENSG00000173436.14 | MICOS10 | 0.0575089283402535 | 3.06685123940477 | 0.475812306244007 | 0.636644915845899 | 0.844878291609035 | -6.0840032489153 |
| ENSG00000199631.1 | SNORD33 | -0.0820914373167614 | 0.498802698190614 | -0.475434439643484 | 0.6369119575308 | 0.845126632839214 | -5.70935005371754 |
| FARP1-AS1 | FARP1-AS1 | -0.152877304092162 | -0.901870665664915 | -0.475422533133848 | 0.636920372760977 | 0.845126632839214 | -5.3519125275232 |
| ENSG00000274944.4 | | 0.102064709206497 | 0.101092237774908 | 0.475373218198164 | 0.636955227869345 | 0.845126632839214 | -5.60157059748673 |
| ENSG00000107771.16 | CCSER2 | 0.0627750819904325 | 5.96899620618316 | 0.475251607616001 | 0.637041184086925 | 0.845186199049967 | -6.29731217919016 |
| ENSG00000196083.9 | IL1RAP | -0.0622839620619269 | 3.56829322165421 | -0.475010000897173 | 0.637211970413121 | 0.845238386050261 | -6.13936775005597 |
| ENSG00000092820.17 | EZR | 0.0917190112558259 | 9.48455953028966 | 0.474985751559249 | 0.637229112823908 | 0.845238386050261 | -6.23709892825196 |
| lnc-DNAJB13-3 | lnc-DNAJB13-3 | 0.16617428172704 | -0.31880274390726 | 0.474953300984079 | 0.637252053191111 | 0.845238386050261 | -5.47883540467043 |
| lnc-MTMR7-2 | lnc-MTMR7-2 | 0.0604076013662804 | 2.512770821459 | 0.474911325190515 | 0.637281727779034 | 0.845238386050261 | -6.00470069918042 |
| ENSG00000188234.13 | AGAP4 | 0.0731335414552237 | 2.76575321848739 | 0.47490550367005 | 0.637285843322839 | 0.845238386050261 | -6.02848000161281 |
| lnc-ITGA7-2 | lnc-ITGA7-2 | -0.104519338263577 | 2.17448308236697 | -0.474420394089063 | 0.637628833893434 | 0.845638807218334 | -5.98042278942973 |
| ENSG00000103363.14 | ELOB | 0.077246888396048 | 4.38630854755535 | 0.47430905243425 | 0.637707567940586 | 0.845688735965535 | -6.22612740984644 |
| ENSG00000164506.14 | STXBP5 | -0.056518049795689 | 5.36826095478246 | -0.474243230737986 | 0.637754115032477 | 0.845695976659804 | -6.2854716181702 |
| lnc-TK1-4 | lnc-TK1-4 | 0.0988328321995362 | 3.38182987660419 | 0.474035371794405 | 0.637901116236163 | 0.845789258312647 | -6.12093876195371 |
| lnc-SDR39U1-1 | lnc-SDR39U1-1 | -0.0705356188728206 | 2.9291619755857 | -0.474027556480485 | 0.637906643640166 | 0.845789258312647 | -6.06586445931107 |
| lnc-AGO2-5 | lnc-AGO2-5 | -0.494818368855957 | -2.55356377904342 | -0.47396562664021 | 0.637950444440957 | 0.845792850108098 | -4.94980126013814 |
| ENSG00000146834.13 | MEPCE | -0.0739380398214529 | 5.58425860790602 | -0.47385228868858 | 0.638030607783103 | 0.845844647687724 | -6.29041323313451 |
| ENSG00000184634.15 | MED12 | 0.0574616710679887 | 5.86469994211625 | 0.473729049732961 | 0.638117779028305 | 0.845893489000999 | -6.29603161362115 |
| ENSG00000206527.9 | HACD2 | 0.0754520120801416 | 4.77766902283862 | 0.473684003969298 | 0.638149642773527 | 0.845893489000999 | -6.25009157622697 |
| ENSG00000178026.12 | LRRC75B | -0.367623328810317 | -1.7829566142818 | -0.473533937050001 | 0.638258169323427 | 0.845982864265832 | -5.18508222982409 |
| ENSG00000073670.13 | ADAM11 | -0.138549311770242 | -0.206477444847537 | -0.47342376222644 | 0.638333741927573 | 0.846000491623362 | -5.51518244435166 |
| ENSG00000235145.2 | RPSAP16 | 0.0876464582603771 | 0.933477447434129 | 0.473395590805816 | 0.63835367222477 | 0.846000491623362 | -5.78830533348039 |
| lnc-ZNF699-1 | lnc-ZNF699-1 | 0.108268488958828 | 0.636106765992911 | 0.473292908135922 | 0.638426318921934 | 0.846042294770163 | -5.70825850120743 |
| ENSG00000119977.20 | TCTN3 | -0.0557318562264615 | 3.73787192248752 | -0.473199810749962 | 0.638492187258402 | 0.846075110431963 | -6.16195808705104 |
| lnc-HECTD4-1 | lnc-HECTD4-1 | 0.080489633481152 | 8.22036696374256 | 0.47303303395618 | 0.638610192694938 | 0.846132044140645 | -6.27529543067124 |
| ENSG00000116903.7 | EXOC8 | 0.0530255932222339 | 5.93454889945447 | 0.473022892161598 | 0.638617368980466 | 0.846132044140645 | -6.29780428700474 |
| lnc-PCED1B-7 | lnc-PCED1B-7 | -0.142870925003545 | 3.02254570899599 | -0.472928726595555 | 0.638696620268821 | 0.846136131481065 | -6.07032136397909 |
| lnc-OXA1L-3 | lnc-OXA1L-3 | -0.0894145333795146 | 0.611014827557286 | -0.472848662537087 | 0.638740658505826 | 0.846136131481065 | -5.71290147531434 |
| ENSG00000273611.4 | ZNHIT3 | 0.0440432586437538 | 4.68166974690013 | 0.472844252388443 | 0.638743779378749 | 0.846136131481065 | -6.24713075508535 |
| ENSG00000127990.17 | SGCE | 0.184975264581892 | 5.5096016472799 | 0.472799921832695 | 0.638844332549348 | 0.846214872055038 | -6.29127834980132 |
| lnc-DYRK2-6 | lnc-DYRK2-6 | -0.112432449369405 | 1.33256288768443 | -0.472607964316717 | 0.638910999993364 | 0.846227214614746 | -5.87150665734565 |
| ENSG00000088727.12 | KIF9 | -0.171855718974667 | 1.02320794228044 | -0.472290887314265 | 0.639137115632397 | 0.846227214614746 | -5.80438778869417 |
| ENSG00000156162.16 | DPY19L4 | -0.102629568402878 | 3.56963807334299 | -0.47227515044321 | 0.639146563983059 | 0.846227214614746 | -6.15804854284167 |
| ENSG00000135452.9 | TSPAN31 | 0.0739633339660251 | 3.85654083775879 | 0.472247719926373 | 0.63916598084728 | 0.846227214614746 | -6.16859342309264 |
| ENSG00000117519.15 | CNN3 | 0.378795025010509 | 0.454750791461507 | 0.472353235061066 | 0.639174997752778 | 0.846227214614746 | -5.74285542605301 |
| ENSG00000114331.13 | ACAP2 | 0.0375463480717197 | 7.40457071444623 | 0.472186373771728 | 0.639209406036289 | 0.846227214614746 | -6.29154137313835 |
| lnc-PLEKHA5-3 | lnc-PLEKHA5-3 | 0.192823840796005 | -0.377309842251662 | 0.472163431249098 | 0.639225646721213 | 0.846227214614746 | -5.43689277666122 |
| ENSG00000118855.18 | MFSD1 | -0.0653550851162708 | 4.66934507158528 | -0.472139458631866 | 0.639242616786805 | 0.846227214614746 | -6.25027003713427 |
| lnc-CDS2-3 | lnc-CDS2-3 | -0.124195905541479 | 0.959939285858757 | -0.472079927098194 | 0.639284759632468 | 0.846227214614746 | -5.8008863686769 |
| ENSG00000171425.9 | ZNF581 | -0.065814006347694 | 3.88729251096351 | -0.472012417864324 | 0.63933255141464 | 0.846227214614746 | -6.1786945722461 |
| lnc-SEL1L2-2 | lnc-SEL1L2-2 | 0.0639904891410809 | 0.521264268501214 | 0.471997828416357 | 0.639342879919877 | 0.846227214614746 | -5.69466352640506 |
| ENSG00000264545.1 | | -0.070884374769614 | 1.5311347377565 | -0.471958467941669 | 0.639370745274344 | 0.846227214614746 | -5.89299522631502 |
| ENSG00000166974.12 | MAPRE2 | -0.0712657646586603 | 5.66476845654641 | -0.471907738918763 | 0.63940665980065 | 0.846227214614746 | -6.29406873165527 |
| ENSG00000126215.13 | XRCC3 | 0.095746692054461 | 2.79565980465575 | 0.471875857390574 | 0.639429231351268 | 0.846227214614746 | -6.04990054799553 |
| lnc-TMEM167A-4 | lnc-TMEM167A-4 | 0.0526035153375748 | 4.95901468812178 | 0.471724064617897 | 0.639536702655691 | 0.84631502833754 | -6.26718896384379 |
| lnc-LONRF1-3 | lnc-LONRF1-3 | -0.102168654280806 | 1.64200675871465 | -0.471652338938443 | 0.639587488118069 | 0.846327821789407 | -5.90777395364805 |
| lnc-CDKN1B-1 | lnc-CDKN1B-1 | -0.105132517812852 | 1.02867067525999 | -0.471330348852327 | 0.639815495176034 | 0.846536394670514 | -5.80213980608241 |
| lnc-CAPN12-2 | lnc-CAPN12-2 | 0.0794008247529741 | 3.20555507795856 | 0.471313585113385 | 0.639827366850759 | 0.846536394670514 | -6.08841919311346 |
| ENSG00000169692.12 | AGPAT2 | -0.087715236299053 | 2.34531813530417 | -0.471226245310928 | 0.63988922033276 | 0.846563814250495 | -6.00833958878307 |
| ENSG00000219545.11 | UMAD1 | -0.0519756734478349 | 4.00455099766628 | -0.471063446278304 | 0.640004520470987 | 0.846640389832784 | -6.19208853356613 |
| ENSG00000252391.1 | RNU6-638P | 0.125980254535369 | 2.36991560594074 | 0.4710283653388 | 0.640029367235005 | 0.846640389832784 | -5.98995648755169 |
| ENSG00000164889.13 | SLC4A2 | 0.0635461147687177 | 4.35241540743554 | 0.470930806113401 | 0.640098467669209 | 0.846677383203864 | -6.22899485537559 |
| ENSG00000063180.8 | CA11 | -0.111988654905251 | 2.19731213478797 | -0.470755604731562 | 0.640222569548922 | 0.846763589904261 | -5.97064353397855 |
| ENSG00000110917.7 | MLEC | -0.0593706791582136 | 5.56497484444492 | -0.47070933862863 | 0.64025534334942 | 0.846763589904261 | -6.29158639730157 |
| ENSG00000160352.15 | ZNF714 | 0.0700033534701054 | 4.02921655418171 | 0.470664568788561 | 0.640287057924983 | 0.846763589904261 | -6.19987690543653 |
| ENSG00000180096.11 | SEPTIN1 | 0.075927166982833 | 4.9275892536532 | 0.470461329021335 | 0.640431039810347 | 0.846849296634555 | -6.26703338246831 |
| ENSG00000185262.8 | UBALD2 | -0.104477828835483 | 4.78133789663639 | -0.470447264546322 | 0.640441615919918 | 0.846849296634555 | -6.26191343375793 |
| ENSG00000067646.11 | ZFY | -0.942975595802176 | 1.62633773703524 | -0.470533684181634 | 0.64047529518098 | 0.846849296634555 | -5.66377615961653 |

| | | | | | | | |
|---|---|---|---|---|---|---|---|
| ENSG00000227345.8 | PARG | -0.0362072713664547 | 4.9359926513415 | -0.470340023377119 | 0.640516983488698 | 0.846850017417001 | -6.26472611869283 |
| ENSG00000115840.13 | SLC25A12 | 0.0431588139902077 | 4.29898751708487 | 0.470212870135058 | 0.640607075483919 | 0.846914730212955 | -6.21838774786344 |
| ENSG00000168275.14 | COA6 | 0.069151311169247 | 1.69217784313887 | 0.470067474933082 | 0.640710099192186 | 0.846917986129241 | -5.9236993735187 |
| lnc-HIVEP1-1 | lnc-HIVEP1-1 | -0.0805500727031594 | 1.09781920764761 | -0.470000145354037 | 0.640757809817958 | 0.846917986129241 | -5.82372995092217 |
| ENSG00000232838.3 | PET117 | 0.0539060080750424 | 2.26939253843766 | 0.469998147264153 | 0.640759225714235 | 0.846917986129241 | -5.98872120390423 |
| ENSG00000071889.16 | FAM3A | -0.0622914867410951 | 3.53545137948441 | -0.469969323398032 | 0.640779651174581 | 0.846917986129241 | -6.13934023879259 |
| ENSG00000174695.9 | TMEM167A | 0.0523995685457725 | 4.95888210775528 | 0.469919059080443 | 0.640815270659126 | 0.846917986129241 | -6.2680239770722 |
| lnc-ZNF385C-7 | lnc-ZNF385C-7 | 0.0641843985370523 | 2.776194878448 | 0.469769924074043 | 0.640920959261214 | 0.847003281186104 | -6.04482875809928 |
| ENSG00000181544.14 | FANCB | 0.0912689610624664 | 1.86201308688982 | 0.469450168437489 | 0.641147588251332 | 0.847197010656207 | -5.92201678439124 |
| ENSG00000100418.7 | DESI1 | -0.0611956352222078 | 3.33052701948807 | -0.4693785885721 | 0.641198325715708 | 0.847197010656207 | -6.11367478747295 |
| ENSG00000231697.3 | NANOGP5 | -0.169850709773551 | 0.166481400500026 | -0.469345699032369 | 0.641221639164308 | 0.847197010656207 | -5.60231624661953 |
| ENSG00000167283.7 | ATP5MG | 0.0844754201562958 | 4.72639827476226 | 0.469330810303089 | 0.641232193024003 | 0.847197010656207 | -6.25693854079554 |
| ENSG00000271741.1 | | -0.0529176672950182 | 5.03129293916056 | -0.469095921968461 | 0.641398703302942 | 0.847362612970761 | -6.27137254314632 |
| ENSG00000186104.10 | CYP2R1 | 0.0716026376852175 | 3.52771570256772 | 0.468934794955004 | 0.641512935632825 | 0.847459133183392 | -6.12755937369585 |
| ENSG00000188856.6 | RPSAP47 | -0.595059002588301 | -0.345187784028015 | -0.468910773431967 | 0.641617365914984 | 0.847542693019389 | -5.44049636211378 |
| lnc-GNA13-2 | lnc-GNA13-2 | -0.116861442357834 | 0.196051062262278 | -0.468600631629807 | 0.641749871500997 | 0.847663325746328 | -5.61493048276562 |
| lnc-RXFP2-2 | lnc-RXFP2-2 | 0.202383777534124 | 2.55848519864533 | 0.46835353774129 | 0.641983017478451 | 0.847916866578475 | -6.04884965500381 |
| ENSG00000278540.4 | ACACA | -0.112004352870393 | 5.4973140547834 | -0.467815582539784 | 0.642317594981255 | 0.848270298871744 | -6.28466589714751 |
| ENSG00000151806.13 | GUF1 | 0.0522354792275941 | 5.48321215800705 | 0.467778390483751 | 0.642333035913861 | 0.848270298871744 | -6.29128458462977 |
| ENSG00000075131.9 | TIPIN | 0.0723798084698326 | 1.99946129123447 | 0.467671417942881 | 0.64240892167209 | 0.848316086147214 | -5.9550310682294 |
| lnc-PGRMC2-3 | lnc-PGRMC2-3 | -0.0731132321045975 | 0.700572874478749 | -0.467479214434039 | 0.642545279543912 | 0.848416721554504 | -5.73838968250911 |
| lnc-ZNF23-5 | lnc-ZNF23-5 | 0.0621569255627632 | 1.28571415699828 | 0.46744779798768 | 0.642567568979894 | 0.848416721554504 | -5.83795654171236 |
| ENSG00000033050.8 | ABCF2 | 0.0455878489726089 | 4.02343718569401 | 0.467355136063757 | 0.642633312985059 | 0.848449100780722 | -6.18406098629445 |
| lnc-C2orf42-4 | lnc-C2orf42-4 | -0.0517249487077995 | 2.44972532453699 | -0.467249692729414 | 0.642708128978665 | 0.8484934525539 | -6.00873669434059 |
| lnc-ZNF644-2 | lnc-ZNF644-2 | -0.158377446441389 | -0.447228840042536 | -0.467065079981912 | 0.642839127673764 | 0.84851185177375 | -5.43481723324188 |
| lnc-PLEKHG7-1 | lnc-PLEKHG7-1 | -0.151337304690557 | 0.325913832280037 | -0.467061438720228 | 0.642841711579225 | 0.84851185177375 | -5.61675230407161 |
| ENSG00000092203.13 | TOX4 | 0.0413394170263661 | 6.20138374020085 | 0.466971840676273 | 0.642905293389124 | 0.84851185177375 | -6.30229491455583 |
| ENSG00000184226.14 | PCDH9 | -0.258307556991127 | 8.48868836516178 | -0.46708312086629 | 0.64291735245956 | 0.84851185177375 | -6.26897897456122 |
| lnc-CTNND2-2 | lnc-CTNND2-2 | -0.0780118740707652 | 1.43486976386498 | -0.46693958269657 | 0.642928185408512 | 0.84851185177375 | -5.88823338688091 |
| lnc-WASL-1 | lnc-WASL-1 | -0.118493021944951 | 0.445657725279539 | -0.466833672901522 | 0.643003347236448 | 0.848556638851561 | -5.69649352749722 |
| lnc-ULK1-2 | lnc-ULK1-2 | -0.0960299968739682 | 0.159674936052887 | -0.466665650759428 | 0.643122596580635 | 0.848613619096269 | -5.61064502097741 |
| lnc-KLHL5-1 | lnc-KLHL5-1 | 0.104502777713135 | 0.584527179355541 | 0.466656653559516 | 0.643128982377821 | 0.848613619096269 | -5.69472992782569 |
| ENSG00000141429.13 | GALNT1 | 0.0525977949360881 | 5.68355702882782 | 0.466579693838563 | 0.643183605950828 | 0.848631292390121 | -6.29636317381201 |
| lnc-NME5-4 | lnc-NME5-4 | 0.0579450891748821 | 1.71367980020915 | 0.466258706020031 | 0.643411454476016 | 0.848871855982126 | -5.90491911351427 |
| LINC01422 | LINC01422 | 0.101922224745859 | 2.63683437154393 | 0.466206643465377 | 0.643448413595352 | 0.848871855982126 | -6.02320335523507 |
| ENSG00000241370.5 | RPP21 | -0.0461913173388981 | 2.45815815534554 | -0.465975699511856 | 0.643612371329392 | 0.849033739606029 | -6.01696897282732 |
| ENSG00000137103.19 | TMEM8B | 0.112287579006153 | 2.61405070173331 | 0.465755709292499 | 0.643768569214247 | 0.849168577395039 | -6.01610347699627 |
| ENSG00000125450.10 | NUP85 | 0.054757016218737 | 5.16854327220267 | 0.465715532849311 | 0.643797097131108 | 0.849168577395039 | -6.27886562668045 |
| ENSG00000057757.9 | PITHD1 | 0.0510383883247956 | 4.84033860098488 | 0.465624827771929 | 0.643861505700108 | 0.849199113919347 | -6.26422605631861 |
| lnc-NEMP2-38 | lnc-NEMP2-38 | -0.115325988678128 | 0.85517422659696 | -0.465507570725252 | 0.643944772603821 | 0.849209766107933 | -5.80106804037825 |
| lnc-MTA3-5 | lnc-MTA3-5 | -0.237017494329627 | 3.04454751312401 | -0.465547652079767 | 0.643988261047763 | 0.849209766107933 | -6.08288531142396 |
| ENSG00000147138.2 | GPR174 | -0.105080406888001 | 5.50426076515502 | -0.465446237277984 | 0.643993355641973 | 0.849209766107933 | -6.29027594368771 |
| lnc-TAS2R30-1 | lnc-TAS2R30-1 | 0.102560546624241 | 1.72435869630062 | 0.465336441739425 | 0.644066303506969 | 0.849251551895191 | -5.92588483899424 |
| ENSG00000164576.11 | SAP30L | 0.0655808541998608 | 4.29466147311309 | 0.465161405286937 | 0.644190619576821 | 0.849361060966608 | -6.22232792250258 |
| lnc-RIPPLY3-6 | lnc-RIPPLY3-6 | -0.109233555994011 | -0.19390524994398 | -0.465013093327584 | 0.644295963188068 | 0.849379815703198 | -5.5252999329968 |
| ENSG00000170325.14 | PRDM10 | 0.0435117804355953 | 3.81363448993029 | 0.464975670202409 | 0.644322545404053 | 0.849379815703198 | -6.17014624334755 |
| ENSG00000182179.12 | UBA7 | 0.0930088133466143 | 6.0645827233672 | 0.464967087071136 | 0.644328642199375 | 0.849379815703198 | -6.30265676603192 |
| lnc-GGA2-2 | lnc-GGA2-2 | -0.0775065044723994 | 0.698826796316121 | -0.464777697344623 | 0.644463176377652 | 0.849457067805199 | -5.75582995847147 |
| ENSG00000170791.17 | CHCHD7 | -0.0460801044910433 | 4.82704858524494 | -0.464768395475885 | 0.644469784329106 | 0.849457067805199 | -6.26778672109384 |
| lnc-PDLIM1-1 | lnc-PDLIM1-1 | 0.146927003584212 | 1.40966965120489 | 0.464327187409125 | 0.644783247389383 | 0.849815814882222 | -5.86202242530522 |
| lnc-HACL1-9 | lnc-HACL1-9 | 0.100651268217451 | 1.8877340193293 | 0.464086652068339 | 0.644954166974306 | 0.849986657627874 | -5.95355171069341 |
| lnc-ADCY5-1 | lnc-ADCY5-1 | 0.0821433044544442 | 2.33275653137873 | 0.463963389924966 | 0.645041762109757 | 0.850047672034389 | -5.98901263055206 |
| ENSG00000172578.11 | KLHL6 | -0.0655283025603612 | 5.68944775726987 | -0.463764473746758 | 0.645183130864769 | 0.850121448849739 | -6.29926666677526 |
| ENSG00000131044.17 | TTLL9 | 0.246604902387068 | -0.0326157799366321 | 0.46361004334469 | 0.645294428851863 | 0.850121448849739 | -5.47525476874833 |
| LIPE-AS1 | LIPE-AS1 | -0.0754221643150095 | 1.07090210913992 | -0.463526455554173 | 0.645352306684216 | 0.850121448849739 | -5.80799992045162 |
| ENSG00000112033.13 | PPARD | -0.0633927263631193 | 4.64383800888401 | -0.463506137623999 | 0.645366748908444 | 0.850121448849739 | -6.24422957173987 |
| ENSG00000145247.11 | OCIAD2 | 0.0437373319920464 | 3.24863931746132 | 0.46349922909842 | 0.645371659601402 | 0.850121448849739 | -6.10578343944397 |
| ENSG00000198783.5 | ZNF830 | -0.0531299599969222 | 4.51557623359389 | -0.463412928642539 | 0.645433004726131 | 0.850121448849739 | -6.24110189761265 |
| ENSG00000198865.9 | CCDC152 | 0.852691973342337 | -1.1473775457342 | 0.463451712573555 | 0.64547875485039 | 0.850121448849739 | -5.04127985337329 |
| ENSG00000284191.1 | MIR671 | 0.0802906481276812 | 0.172493728140552 | 0.463344006389936 | 0.645481998662558 | 0.850121448849739 | -5.62378389632113 |
| ENSG00000272921.1 | | 0.114489317297902 | 0.125196201265102 | 0.46334297744791 | 0.645482730106328 | 0.850121448849739 | -5.58320256316478 |
| ENSG00000108826.15 | MRPL27 | 0.0568532209159658 | 4.33872535546341 | 0.463303537084264 | 0.64551076733579 | 0.850121448849739 | -6.22461848492981 |
| ENSG00000169599.12 | NFU1 | -0.0488366401031879 | 3.52367517399186 | -0.463186663300607 | 0.645593853231057 | 0.850176473515986 | -6.1423786150725 |
| ENSG00000131779.10 | PEX11B | -0.0459991273503038 | 3.81340577868677 | -0.462957480801033 | 0.645756792999624 | 0.850299479329681 | -6.17709961708706 |
| lnc-CCT8-6 | lnc-CCT8-6 | 0.113160125709006 | 1.73308799346684 | 0.462939072965704 | 0.645769881012563 | 0.850299479329681 | -5.91447624614498 |
| ENSG00000108433.16 | GOSR2 | -0.0363788320533622 | 6.33210224451748 | -0.462798471903386 | 0.645869852435643 | 0.85030276167114 | -6.30435923462303 |
| ENSG00000187778.13 | MCRS1 | 0.0632105411757014 | 5.03835892183895 | 0.462796646607904 | 0.645871150317104 | 0.85030276167114 | -6.27587065005024 |
| ENSG00000164037.16 | SLC9B1 | 0.0701321459802935 | 1.38586133358975 | 0.462761267995447 | 0.645896306599052 | 0.85030276167114 | -5.85650803495039 |

| | | | | | | | |
|---|---|---|---|---|---|---|---|
| lnc-FLII-1 | lnc-FLII-1 | -0.099538064590468 | 1.08324012216798 | -0.462564978236037 | 0.64603588782416 | 0.850432123246655 | -5.83682424605525 |
| ENSG00000136870.10 | ZNF189 | 0.0646501834935389 | 4.75009660300071 | 0.462269575390464 | 0.64624597247877 | 0.85050821480144 | -6.25301662295975 |
| ENSG00000226278.1 | PSPHP1 | -0.531863763849119 | -2.52367612396694 | -0.462232039882907 | 0.64627266908102 | 0.85050821480144 | -4.93577329917755 |
| ENSG00000171792.10 | RHNO1 | -0.0547229044271775 | 4.54466842584437 | -0.462231242076857 | 0.646273236514481 | 0.85050821480144 | -6.23997038985341 |
| lnc-SLC44A2-1 | lnc-SLC44A2-1 | -0.135058771575461 | -0.323336082674996 | -0.462218630170564 | 0.646282206664839 | 0.85050821480144 | -5.48369218024692 |
| ENSG00000083845.8 | RPS5 | 0.079598952994445 | 8.28887776534023 | 0.462155954425414 | 0.646326785240581 | 0.85050821480144 | -6.27658560685491 |
| ENSG00000111412.5 | SPRING1 | 0.0836079114296604 | 5.64352925742535 | 0.462060558054579 | 0.646394639121343 | 0.85050821480144 | -6.29852507947457 |
| ENSG00000164970.14 | FAM219A | -0.0709518062904495 | 2.48328510405712 | -0.462032341203461 | 0.646414709890527 | 0.85050821480144 | -6.02107948424814 |
| ENSG00000164938.13 | TP53INP1 | 0.14897273455196 | 6.87641768105168 | 0.46209301155433 | 0.64642425848291 | 0.85050821480144 | -6.30060967507083 |
| ENSG00000206611.1 | SNORD24 | -0.0861723822553899 | -0.0971698874709013 | -0.46194506885567 | 0.646476788778552 | 0.850522962187852 | -5.58055736915341 |
| ENSG00000186818.12 | LILRB4 | -0.200176774546872 | 3.34695497920133 | -0.46174177745558 | 0.646681291160268 | 0.85056078775012 | -6.08251793444226 |
| lnc-KLF12-5 | lnc-KLF12-5 | 0.15039301463685 | 1.45007368689465 | 0.461613310625119 | 0.6467127995235 | 0.85056078775012 | -5.88002841278316 |
| lnc-CYB5R2-3 | lnc-CYB5R2-3 | 0.0937793518573176 | 3.92962824107497 | 0.461607874847575 | 0.646716666808467 | 0.85056078775012 | -6.17544767521199 |
| ENSG00000165312.6 | OTUD1 | 0.073192629259518 | 5.31306186195713 | 0.461583236974702 | 0.64673419555171 | 0.85056078775012 | -6.2884284204961 |
| ENSG00000175387.15 | SMAD2 | 0.0412875538783615 | 7.1203537049123 | 0.461583229206986 | 0.646734201078124 | 0.85056078775012 | -6.29972344047801 |
| SRRM2-AS1 | SRRM2-AS1 | -0.0691533978125765 | 1.97347251276609 | -0.461556130966359 | 0.646753480497358 | 0.85056078775012 | -5.94280725101534 |
| ENSG00000132359.14 | RAP1GAP2 | 0.123116838052874 | 3.80902137663987 | 0.461480483704826 | 0.646816399124426 | 0.850589186249557 | -6.17526450690348 |
| lnc-RCAN1-3 | lnc-RCAN1-3 | -0.101816652029446 | 2.20150166484356 | -0.461294501817648 | 0.646939632814645 | 0.850696892750517 | -5.96193778483162 |
| ENSG00000168488.18 | ATXN2L | -0.0701967900059115 | 5.9354207975602 | -0.461224199535851 | 0.64698965764289 | 0.850708325237231 | -6.30262228769529 |
| lnc-SNAPC1-4 | lnc-SNAPC1-4 | 0.164609875251666 | 1.51252410237935 | 0.461089532215587 | 0.64709457544586 | 0.850722520703031 | -5.89719317087737 |
| ENSG00000111269.2 | CREBL2 | -0.0792311551914295 | 4.88803771012251 | -0.461035956674568 | 0.647123613303346 | 0.850722520703031 | -6.26678344695196 |
| ENSG00000119899.12 | SLC17A5 | 0.0641947556396527 | 2.53511600982515 | 0.461034784205115 | 0.647124447682425 | 0.850722520703031 | -6.01988071517129 |
| ENSG00000117523.16 | PRRC2C | 0.0611357021716579 | 8.75476006597532 | 0.460960483753519 | 0.647177323977503 | 0.850737696987415 | -6.26843055662617 |
| ENSG00000230521.1 | | -0.13120616254818 | 1.93716882107723 | -0.460831826255892 | 0.647268888103315 | 0.85080372462038 | -5.97966637839392 |
| ENSG00000165802.22 | NSMF | -0.125338563647332 | 2.89119031759159 | -0.46074726588172 | 0.64732966932587 | 0.85082928376337 | -6.06899714677948 |
| ENSG00000155506.16 | LARP1 | -0.0520771719323285 | 6.42825348605844 | -0.460514425952585 | 0.64749480205082 | 0.850991684636813 | -6.3052863673674 |
| lnc-CCDC173-2 | lnc-CCDC173-2 | 0.0620446393641175 | 0.482542057424535 | 0.460408655346994 | 0.647570093178427 | 0.850991684636813 | -5.68031948900797 |
| ENSG00000214194.8 | SMIM30 | -0.119853984214969 | 2.25179487679582 | -0.460398586566644 | 0.647577260674654 | 0.850991684636813 | -5.99615753805446 |
| ENSG00000133114.17 | GPALPP1 | -0.0617011837564689 | 4.90427122056801 | -0.46032728128005 | 0.647628020558661 | 0.851004056892168 | -6.27363639163377 |
| lnc-WDR89-3 | lnc-WDR89-3 | -0.134606835385627 | 1.22358901034415 | -0.460131463919301 | 0.647767425213574 | 0.851048440546354 | -5.83860193647907 |
| ENSG00000105202.8 | FBL | 0.0656946099119589 | 4.99116399493972 | 0.460113214171484 | 0.647780418072909 | 0.851048440546354 | -6.27346930548519 |
| lnc-MSMB-3 | lnc-MSMB-3 | 0.0802011318186753 | 2.99148844273012 | 0.46001553849161 | 0.647849959901692 | 0.851048440546354 | -6.07726739117201 |
| lnc-TMEM88B-3 | lnc-TMEM88B-3 | -0.0765510147036426 | 3.0929964786288 | -0.460008239803087 | 0.647855156452466 | 0.851048440546354 | -6.08650046400611 |
| ENSG00000279281.1 | | 0.147741757497067 | 2.26420947456135 | 0.459950706940056 | 0.647904950778274 | 0.851048440546354 | -5.99060218153159 |
| lnc-UBE2B-1 | lnc-UBE2B-1 | 0.0835014012320511 | 0.69320904794787 | 0.459931380468902 | 0.647909880161365 | 0.851048440546354 | -5.7280847759361 |
| ENSG00000179922.5 | ZNF784 | 0.0915702316800397 | 1.52548137708795 | 0.459722130554556 | 0.648058875710235 | 0.851189830817675 | -5.89587884794015 |
| ENSG00000272941.1 | | 0.116599431009859 | 2.85751077113774 | 0.459604681551107 | 0.64814251119133 | 0.851204161969304 | -6.04355822972664 |
| lnc-ZNF784-1 | lnc-ZNF784-1 | 0.109245031505814 | -0.451175162738073 | 0.459523238491762 | 0.648200509525428 | 0.851204161969304 | -5.45534981300499 |
| ENSG00000154767.14 | XPC | 0.0620558161753095 | 6.21864847057626 | 0.45945241696161 | 0.648250945709407 | 0.851204161969304 | -6.3055146937719 |
| lnc-DCAF16-1 | lnc-DCAF16-1 | 0.0768599381589379 | 2.46238512600943 | 0.459426012592958 | 0.648269750243675 | 0.851204161969304 | -6.02081758548338 |
| ENSG00000181017.5 | OR56B2P | 0.158149342096602 | 0.742981125919071 | 0.459393912637688 | 0.648292611345901 | 0.851204161969304 | -5.74981470404137 |
| ENSG00000149089.12 | APIP | 0.0487974643011175 | 3.04321159370518 | 0.459347135438715 | 0.648325925973641 | 0.851204161969304 | -6.08620031434999 |
| ENSG00000145990.10 | GFOD1 | 0.0792609318863375 | 5.61454884349413 | 0.459251544842561 | 0.648394007672575 | 0.851204161969304 | -6.29947741002643 |
| ENSG00000198799.11 | LRIG2 | 0.0438878437016221 | 5.61072911829045 | 0.459242254568831 | 0.648400624569631 | 0.851204161969304 | -6.29758949610135 |
| ENSG00000115652.14 | UXS1 | -0.0685873966798183 | 4.00141581671276 | -0.459054265359327 | 0.648534524029095 | 0.851246237482646 | -6.18761696276202 |
| lnc-SMARCA5-7 | lnc-SMARCA5-7 | -0.232512292178181 | 1.97496569523517 | -0.459127508227938 | 0.648542064676544 | 0.851246237482646 | -5.93962101416858 |
| ENSG00000160714.9 | UBE2Q1 | 0.0449631416836445 | 5.02589549597917 | 0.459023068703522 | 0.648556745673753 | 0.851246237482646 | -6.27573020019511 |
| lnc-ACOT12-4 | lnc-ACOT12-4 | -0.257383189828831 | -0.0475095111498807 | -0.458922550697105 | 0.648653377367775 | 0.851318782526361 | -5.59936950437075 |
| NUTM2B-AS1 | NUTM2B-AS1 | -0.0882848906482279 | 4.62704041172628 | -0.458668089524524 | 0.648809623199126 | 0.851469553322342 | -6.25669897432338 |
| ENSG00000010539.11 | ZNF200 | 0.122814941624166 | 3.31677745473347 | 0.45855947484752 | 0.648889387976019 | 0.851519940880485 | -6.10519164163128 |
| ENSG00000173207.12 | CKS1B | -0.0934588063820654 | 1.99184750578652 | -0.458334312162101 | 0.6490474353819 | 0.851673043635449 | -5.96098176135218 |
| ENSG00000256667.6 | KLRA1P | -0.0940415282150138 | 1.90195533658736 | -0.458072143018537 | 0.649234253592588 | 0.851863877203814 | -5.95128646546173 |
| ENSG00000243678.11 | NME2 | 0.0984874508095999 | 4.25455196050989 | 0.457954528555187 | 0.649318071505692 | 0.851919547794599 | -6.22235735280708 |
| lnc-ACSBG1-5 | lnc-ACSBG1-5 | 0.0683609232860103 | 1.72449292709472 | 0.457601862325756 | 0.649569426553261 | 0.852183855526492 | -5.91386481293249 |
| ENSG00000139233.6 | LLPH | 0.0495714047559055 | 4.28417949554342 | 0.457555705547561 | 0.649602326833341 | 0.852183855526492 | -6.22706866853754 |
| lnc-TRIM52-3 | lnc-TRIM52-3 | 0.0717073627260225 | 6.66575609286988 | 0.457351474200521 | 0.649747910229276 | 0.852311075804644 | -6.30572673108638 |
| lnc-CCDC167-3 | lnc-CCDC167-3 | -0.335127560783903 | -0.903896728968537 | -0.45728387531244 | 0.649807757721631 | 0.852311075804644 | -5.2999567616602 |
| lnc-AFG3L2-7 | lnc-AFG3L2-7 | -0.0771311136869288 | 2.07930869429431 | -0.457173969716371 | 0.649874453013743 | 0.852311075804644 | -5.96241583061623 |
| ENSG00000280248.1 | | -0.0705954349203868 | 2.76183516229812 | -0.457119146446981 | 0.649913538586654 | 0.852311075804644 | -6.05884659663855 |
| ENSG00000198521.11 | ZNF43 | -0.0534917500546834 | 6.07985960860046 | -0.457068582656319 | 0.649949588301218 | 0.852311075804644 | -6.30624103460986 |
| lnc-POLE4-15 | lnc-POLE4-15 | -0.133809233087341 | 0.859595643158439 | -0.457056564273048 | 0.649958156994196 | 0.852311075804644 | -5.77917859533882 |
| ENSG00000033867.16 | SLC4A7 | -0.321434751035353 | 2.76248949778098 | -0.457137211370025 | 0.649989163722757 | 0.852311075804644 | -6.02478097963948 |
| ENSG00000147140.15 | NONO | 0.0359555800630957 | 7.65785241942535 | 0.456864600320861 | 0.650095027172313 | 0.852395588246128 | -6.29292067583994 |
| ENSG00000125970.11 | RALY | -0.0383382345117438 | 5.87914324614527 | -0.456801694181965 | 0.650139881870095 | 0.852400101186838 | -6.30451011457655 |
| lnc-TMEM201-1 | lnc-TMEM201-1 | -0.102268266959242 | 0.277509216567327 | -0.456607944721665 | 0.650278041555188 | 0.852526938173913 | -5.67929938318235 |
| ENSG00000159086.14 | PAXBP1 | -0.055273219761844 | 5.80856126269513 | -0.456472567300051 | 0.650374584448691 | 0.852599202070403 | -6.30336344161307 |
| ENSG00000115053.15 | NCL | -0.0519136198227041 | 8.24185284755501 | -0.456351357435225 | 0.650461029063738 | 0.852658219412744 | -6.28159652597439 |
| lnc-ZNF136-2 | lnc-ZNF136-2 | -0.149080109302438 | 0.514082612959874 | -0.456224418422639 | 0.650551564821257 | 0.852722591779655 | -5.73962768420848 |

| | | | | | | | |
|---|---|---|---|---|---|---|---|
| SNHG3 | SNHG3 | -0.109482047892142 | 6.37529256508308 | -0.456097648248452 | 0.650655471305627 | 0.852804480761826 | -6.3070591456461 |
| ENSG00000147804.9 | SLC39A4 | 0.122631943690632 | 0.435072890349472 | 0.455863898496335 | 0.650808724856503 | 0.852908998109224 | -5.70306456866981 |
| CACNA1C-AS2 | CACNA1C-AS2 | 0.175033443656804 | -0.347549199245601 | 0.455850771885588 | 0.650818088923066 | 0.852908998109224 | -5.42362661663514 |
| lnc-ARMC9-1 | lnc-ARMC9-1 | -0.0978053453321006 | 0.574061042134611 | -0.455698115105405 | 0.650926993139944 | 0.852995581199315 | -5.75353458401094 |
| ENSG00000160221.17 | GATD3 | 0.403602152279683 | -0.45149132516912 | 0.455703773937892 | 0.650967040250636 | 0.852995581199315 | -5.32664328875803 |
| ENSG00000163428.3 | LRRC58 | 0.0621180410111215 | 5.09764224620189 | 0.455488097415474 | 0.651076830815733 | 0.853085136461916 | -6.28371998801103 |
| ENSG00000177302.14 | TOP3A | 0.0602579294556493 | 4.53401904979779 | 0.455313715658298 | 0.651201255035738 | 0.853193853112705 | -6.24571697255244 |
| ENSG00000243364.7 | EFNA4 | -0.123834598410542 | 1.03153676956508 | -0.455213405137277 | 0.651272832783328 | 0.853233321696852 | -5.81953837850569 |
| lnc-CNN1-3 | lnc-CNN1-3 | -0.121569604513985 | 0.019367072222878 | -0.455066525398352 | 0.651377646548149 | 0.853263488186913 | -5.59583347337448 |
| ENSG00000141499.16 | WRAP53 | 0.11653264338491 | 2.60256214519058 | 0.455064953362933 | 0.651378768395324 | 0.853263488186913 | -6.04049891761001 |
| ENSG00000181704.11 | YIPF6 | -0.040007230087781 | 3.85542643739558 | -0.454910277243125 | 0.651489153472567 | 0.853277190139163 | -6.18249211720386 |
| ENSG00000004534.14 | RBM6 | -0.0648085679782753 | 7.2372052988398 | -0.454892147147367 | 0.651502092587895 | 0.853277190139163 | -6.30138288135818 |
| ENSG00000078269.14 | SYNJ2 | -0.224954907353432 | 1.9537394119642 | -0.454877318394053 | 0.651567109437748 | 0.853277190139163 | -5.91746081137572 |
| lnc-NRBP2-1 | lnc-NRBP2-1 | -0.110764909718213 | -0.262835545425779 | -0.454791131577802 | 0.651574187528718 | 0.853277190139163 | -5.46541259421568 |
| ENSG00000199673.1 | SNORD16 | 0.0700457581650435 | 0.681137852897903 | 0.45462135751251 | 0.651695363095162 | 0.853277190139163 | -5.76311763368461 |
| lnc-CSMD1-1 | lnc-CSMD1-1 | -0.544416010491832 | -2.14895943201072 | -0.454649460083231 | 0.651699046024586 | 0.853277190139163 | -5.02973996694837 |
| ENSG00000243417.1 | RPS3AP18 | 0.205509378424176 | -0.613076331673534 | 0.454556554067785 | 0.651741618809454 | 0.853277190139163 | -5.32713589924769 |
| ENSG00000155846.16 | PPARGC1B | -0.149398702457479 | 3.50810259831032 | -0.454518958092373 | 0.651802807154897 | 0.853277190139163 | -6.14983029657696 |
| MIR210HG | MIR210HG | 0.139723621972145 | -0.347325054037424 | 0.454397010183478 | 0.65185550472444 | 0.853277190139163 | -5.48276045042762 |
| ENSG00000164068.15 | RNF123 | -0.0424780944747134 | 4.61293627907582 | -0.45435081105278 | 0.65188848424498 | 0.853277190139163 | -6.25350033745702 |
| ENSG00000204843.12 | DCTN1 | -0.0450893188571859 | 5.87012277956912 | -0.454295380774678 | 0.651928054404804 | 0.853277190139163 | -6.30525002770399 |
| ENSG00000076650.6 | GPATCH1 | 0.0494142699992803 | 3.43686188502118 | 0.454156022949439 | 0.652027542628939 | 0.853277190139163 | -6.12835359904378 |
| lnc-ST8SIA5-4 | lnc-ST8SIA5-4 | 0.0584868499133333 | 1.32297765181566 | 0.45414959378141 | 0.652032132597817 | 0.853277190139163 | -5.86751213012696 |
| lnc-INTS13-1 | lnc-INTS13-1 | 0.0757237013725786 | 0.185352541162527 | 0.454145495731398 | 0.652035058321289 | 0.853277190139163 | -5.63007960152216 |
| ENSG00000146223.14 | RPL7L1 | 0.0417812913536437 | 5.74742340187153 | 0.454129260853763 | 0.652046648952397 | 0.853277190139163 | -6.30365371271495 |
| lnc-TIAL1-1 | lnc-TIAL1-1 | 0.185319448995175 | -0.306897754682865 | 0.454083460377326 | 0.652079347938829 | 0.853277190139163 | -5.50332907917029 |
| ENSG00000083844.10 | ZNF264 | -0.0388222426800748 | 5.74940586814468 | -0.454058576538591 | 0.652097113902627 | 0.853277190139163 | -6.30376185378493 |
| lnc-AGRP-1 | lnc-AGRP-1 | -0.0872298851559226 | 4.17050494191646 | -0.45400491547886 | 0.652135426229372 | 0.853277190139163 | -6.21840782193744 |
| ENSG00000259112.2 | NDUFC2-KCTD14 | -0.086311566594925 | 2.03788242045102 | -0.453920073177158 | 0.652196002932443 | 0.853302207497995 | -5.97167228876497 |
| ENSG00000180917.17 | CMTR2 | -0.0570267897351986 | 5.14848976320921 | -0.453830947402819 | 0.652259640558928 | 0.853331226188548 | -6.28606040040021 |
| ENSG00000160993.3 | ALKBH4 | 0.114312065093626 | 1.05440846127656 | 0.453692451911609 | 0.65235853437602 | 0.853406362848711 | -5.82022371996878 |
| ENSG00000120253.13 | NUP43 | -0.0616245666736939 | 4.88723071673588 | -0.453552415708255 | 0.652458534786682 | 0.853482937496935 | -6.27357965728289 |
| ENSG00000112659.13 | CUL9 | 0.0410555106067108 | 6.09796189153306 | 0.45343723046413 | 0.652540793877859 | 0.853483691307908 | -6.30780439502732 |
| ENSG00000274290.2 | H2BC6 | 0.206827100395314 | 3.13006360098554 | 0.453398193052636 | 0.652632249149048 | 0.853483691307908 | -6.08503395460371 |
| lnc-HLA-DRB1-8 | lnc-HLA-DRB1-8 | -0.142447180048221 | 0.934106086913049 | -0.453308652153252 | 0.652632622744429 | 0.853483691307908 | -5.79981650023518 |
| ENSG00000137364.4 | TPMT | 0.0751272158451983 | 2.64553701786369 | 0.453278319416191 | 0.652654286764166 | 0.853483691307908 | -6.03867466494708 |
| ENSG00000060762.18 | MPC1 | -0.0616144973331938 | 4.0434571671549 | -0.453238672243173 | 0.652682603727884 | 0.853483691307908 | -6.2088696708827 |
| lnc-IGSF9-1 | lnc-IGSF9-1 | 0.0748045771181456 | 3.90727422391772 | 0.453131917507219 | 0.652758853102361 | 0.853483691307908 | -6.19960909061179 |
| ENSG00000131788.15 | PIAS3 | -0.0620002326524332 | 3.76033619623355 | -0.453100968031453 | 0.652780959411254 | 0.853483691307908 | -6.17812274372053 |
| ENSG00000122674.11 | CCZ1 | 0.0727612658097745 | 3.40246732713911 | 0.453087142371795 | 0.652790834779206 | 0.853483691307908 | -6.137853455965 |
| lnc-PNPLA2-2 | lnc-PNPLA2-2 | 0.103228556673891 | -0.0377726460943099 | 0.452788936143758 | 0.653003852335793 | 0.853618661939874 | -5.5751706857393 |
| lnc-MAP3K8-25 | lnc-MAP3K8-25 | -0.101845796963022 | 1.91332883749128 | -0.452749692578451 | 0.653031887360825 | 0.853618661939874 | -5.95324481097743 |
| ENSG00000160007.18 | ARHGAP35 | -0.0478795034728332 | 5.60247434371771 | -0.452675471577597 | 0.653084911137875 | 0.853618661939874 | -6.30149458739421 |
| ENSG00000143228.12 | NUF2 | 0.120129651773101 | 0.330440807660574 | 0.452674770823876 | 0.653085411767757 | 0.853618661939874 | -5.6561097858905 |
| lnc-PTRH2-3 | lnc-PTRH2-3 | -0.123383511542785 | 0.584659718780549 | -0.452652352608951 | 0.653101427791339 | 0.853618661939874 | -5.73577649234856 |
| ENSG00000152056.16 | AP1S3 | -0.0599957214646825 | 4.99368943039121 | -0.452411432907243 | 0.653273556119243 | 0.853700090768508 | -6.27840816286516 |
| ENSG00000140400.16 | MAN2C1 | -0.069010129802181 | 5.67065408978452 | -0.452359432260535 | 0.653310711190125 | 0.853700090768508 | -6.30343125322411 |
| lnc-N4BP2L1-1 | lnc-N4BP2L1-1 | -0.117537498567842 | 2.01198607850899 | -0.451887585076281 | 0.653647892207826 | 0.853700090768508 | -5.96037718265556 |
| lnc-TESMIN-2 | lnc-TESMIN-2 | -0.166008957421054 | 0.82645658427184 | -0.451840482482327 | 0.653682257495011 | 0.853700090768508 | -5.79888218137849 |
| ENSG00000072401.14 | UBE2D1 | 0.0867688346975894 | 2.63453791539094 | 0.45183769610891 | 0.653683547042695 | 0.853700090768508 | -6.0523641320785 |
| ENSG00000065613.13 | SLK | 0.0433924500546228 | 5.87506467998315 | 0.45182919019382 | 0.653689626163784 | 0.853700090768508 | -6.30735951471901 |
| ENSG00000215032.2 | GNL3LP1 | 0.075442581751538 | 0.0373979117505022 | 0.451755226381237 | 0.653742488613166 | 0.853700090768508 | -5.59890750916847 |
| lnc-HS3ST5-10 | lnc-HS3ST5-10 | 0.0743415431087766 | 1.57869075144391 | 0.451735832624293 | 0.653756349763393 | 0.853700090768508 | -5.90294619830453 |
| lnc-ARL17B-2 | lnc-ARL17B-2 | -0.0438699363224218 | 3.36922622763474 | -0.451719408353126 | 0.653768088652708 | 0.853700090768508 | -6.13140938290358 |
| ENSG00000072786.12 | STK10 | -0.0412745534029666 | 7.31190018431876 | -0.45166129693542 | 0.653809623232306 | 0.853700090768508 | -6.30125097847522 |
| ENSG00000136560.13 | TANK | -0.0444640088412793 | 6.9417780998747 | -0.451634626728573 | 0.653828685879165 | 0.853700090768508 | -6.30642056085306 |
| ENSG00000151117.8 | TMEM86A | 0.223962573715429 | -0.623104676522918 | 0.451528263589697 | 0.653904711723671 | 0.853700090768508 | -5.49150908166515 |
| lnc-ZNF674-10 | lnc-ZNF674-10 | -0.16146795828174 | 0.304471351580115 | -0.451483099751316 | 0.653936994887268 | 0.853700090768508 | -5.63931234452038 |
| lnc-FAM49B-5 | lnc-FAM49B-5 | -0.100740337712558 | -0.276476938701008 | -0.451482600000207 | 0.653937352113686 | 0.853700090768508 | -5.48278711955319 |
| lnc-GRHPR-6 | lnc-GRHPR-6 | 0.13668222864335 | 1.53120325452986 | 0.451461189147047 | 0.653952656853958 | 0.853700090768508 | -5.88597393438202 |
| lnc-LTBP1-4 | lnc-LTBP1-4 | -0.105031142391013 | 0.156787500575976 | -0.451419666630956 | 0.653982338086904 | 0.853700090768508 | -5.64085050637502 |
| ENSG00000139163.15 | ETNK1 | -0.0584395371956576 | 5.47391885394108 | -0.451416683333859 | 0.653984470636916 | 0.853700090768508 | -6.29987872739978 |
| lnc-TFIP11-1 | lnc-TFIP11-1 | -0.0412638586470822 | 3.03519448792146 | -0.451393167082742 | 0.654001280859136 | 0.853700090768508 | -6.08573414179868 |
| ENSG00000262265.2 | | 0.0882504444398193 | 1.94497749345576 | 0.45133860630432 | 0.654040283473482 | 0.853700090768508 | -5.94734039947503 |
| lnc-HLA-DOA-2 | lnc-HLA-DOA-2 | 0.170759511533289 | 0.279056870326861 | 0.451183217404852 | 0.654151368138717 | 0.853700090768508 | -5.59219785714796 |
| ENSG00000111271.14 | ACAD10 | 0.0522508964460862 | 4.97672683738535 | 0.451178221152027 | 0.654154939999592 | 0.853700090768508 | -6.27332205674017 |
| ENSG00000242294.6 | STAG3L5P | 0.0562714752130979 | 4.45802706409762 | 0.451172589270176 | 0.654158966286547 | 0.853700090768508 | -6.24560899100926 |
| ENSG00000168958.19 | MFF | -0.0563833256730435 | 5.23872289731604 | -0.451094255399404 | 0.654214969010957 | 0.853700090768508 | -6.29220376487334 |

| | | | | | | | |
|---|---|---|---|---|---|---|---|
| ENSG00000108774.14 | RAB5C | -0.0525019215406335 | 4.61260830871645 | -0.45105808397193 | 0.654240829494536 | 0.853700090768508 | -6.25775196085155 |
| ENSG00000171659.14 | GPR34 | -0.616358641273855 | -1.35985125599338 | -0.451096975878707 | 0.654275174185087 | 0.853700090768508 | -5.15076338960716 |
| ENSG00000179091.4 | CYC1 | -0.0932743141962362 | 4.40925907443047 | -0.450999173907964 | 0.654282947711195 | 0.853700090768508 | -6.24331636447319 |
| ENSG00000164180.13 | TMEM161B | 0.0599418889004228 | 5.28509429567136 | 0.450998289226421 | 0.654283580229957 | 0.853700090768508 | -6.29317619685709 |
| lnc-FKBP3-4 | lnc-FKBP3-4 | 0.0735186812949515 | 2.04915632915113 | 0.45060475457307 | 0.654564970296904 | 0.85400560993261 | -5.96612755058329 |
| ENSG00000164221.12 | CCDC112 | -0.0880916210976204 | 2.01682817540059 | -0.450554769950371 | 0.654600714565748 | 0.85400560993261 | -5.9681275446355 |
| ENSG00000165555.9 | NOXRED1 | -0.0948124374154171 | 0.317890438928818 | -0.45047903143038 | 0.654654877145765 | 0.854022140720704 | -5.65443506821939 |
| ENSG00000244270.1 | RPL32P29 | 0.111292343189815 | 2.12890439191045 | 0.450320888707907 | 0.654767975176854 | 0.854083096046696 | -5.9726938043737 |
| ENSG00000112182.14 | BACH2 | 0.0976973627219895 | 7.63784362093945 | 0.450265450046668 | 0.654808591924445 | 0.854083096046696 | -6.29870403983574 |
| ENSG00000013375.15 | PGM3 | -0.0504858525165455 | 4.61494989244773 | -0.45023963724559 | 0.654826086513769 | 0.854083096046696 | -6.25576408286051 |
| ENSG00000151332.18 | MBIP | 0.0664300885454079 | 3.49825744769891 | 0.450173594827982 | 0.654873321882925 | 0.854090583216085 | -6.14798212773052 |
| lnc-MTA2-4 | lnc-MTA2-4 | -0.0657514565133277 | 1.4787165253341 | -0.449828670074762 | 0.655120044878224 | 0.854358226175535 | -5.89121846973885 |
| ENSG00000111445.13 | RFC5 | 0.05607081228941 | 4.0423341197049 | 0.449580608602249 | 0.655297506151115 | 0.854535515022073 | -6.20049049727973 |
| ENSG00000090020.10 | SLC9A1 | 0.0769361100773285 | 2.85423502248976 | 0.449220667322703 | 0.655555041229489 | 0.854801521559274 | -6.06299622326261 |
| lnc-EBLN1-8 | lnc-EBLN1-8 | -0.134391170565472 | -0.113564709463622 | -0.449179426664814 | 0.655584551296444 | 0.854801521559274 | -5.55308313713003 |
| ENSG00000105258.8 | POLR2I | -0.0697681155441194 | 2.60037566799528 | -0.449065485082129 | 0.655666085958624 | 0.854853680071348 | -6.03571255743008 |
| ENSG00000272779.1 | BMS1P20 | -0.0835572110231904 | 3.34708528066695 | -0.44878494570319 | 0.655866853287759 | 0.855026128737222 | -6.13119736287172 |
| lnc-SNW1-5 | lnc-SNW1-5 | 0.0922367147497932 | 0.841974892688183 | 0.448764573590874 | 0.655881433543798 | 0.855026128737222 | -5.77653177194392 |
| lnc-GTSF1L-10 | lnc-GTSF1L-10 | -0.158000270157864 | -0.481700447530777 | -0.448488648981692 | 0.656078925282377 | 0.855229418561568 | -5.44490056499864 |
| ENSG00000207500.1 | SNORD102 | -0.101248549366618 | 1.10981650800314 | -0.448339424728601 | 0.656185742212789 | 0.85528772297654 | -5.85391176020206 |
| ENSG00000204351.11 | SKIC2 | 0.0491087611656287 | 5.71588462894844 | 0.448253999462037 | 0.656246894160686 | 0.85528772297654 | -6.30489613838182 |
| ENSG00000197566.9 | ZNF624 | 0.0527797052204289 | 3.32898048105311 | 0.448252018664373 | 0.656248312149274 | 0.85528772297654 | -6.12542770409323 |
| lnc-EP400-1 | lnc-EP400-1 | 0.0615844076845244 | 0.263034289373626 | 0.448019968578211 | 0.65641443812954 | 0.855450068381685 | -5.6509574142394 |
| lnc-C3orf38-1 | lnc-C3orf38-1 | 0.108899761266485 | 1.17675012902621 | 0.447909746103412 | 0.656493353192099 | 0.85548570431102 | -5.86488467056861 |
| ENSG00000138363.14 | ATIC | 0.0944936179080025 | 3.95246488161479 | 0.447753670899818 | 0.65660510383804 | 0.85548570431102 | -6.19038350281061 |
| ENSG00000134597.15 | RBMX2 | -0.0563856734192401 | 3.61519030144224 | -0.447718186169702 | 0.656630512196858 | 0.85548570431102 | -6.15727403176583 |
| lnc-SERINC1-5 | lnc-SERINC1-5 | 0.0884191583511572 | 0.0206872773184326 | 0.447695033072417 | 0.656647090879251 | 0.85548570431102 | -5.57285022392815 |
| lnc-POC1B-GALNT4-3 | lnc-POC1B-GALNT4 | 0.078581057761852 | 3.2543282722126 | 0.447661099056285 | 0.656671389511637 | 0.85548570431102 | -6.11237129194059 |
| ENSG00000175866.15 | BAIAP2 | -0.143073721681653 | -0.599517530747701 | -0.447633490395144 | 0.656691159117433 | 0.85548570431102 | -5.38930049796364 |
| ENSG00000257727.5 | CNPY2 | -0.0568349649587243 | 3.43573922452649 | -0.447527090928614 | 0.656767350417271 | 0.855530812837079 | -6.1344677469191 |
| ENSG00000153339.13 | TRAPPC8 | 0.0296266079002443 | 6.14957525225196 | 0.447422002164251 | 0.656842606768017 | 0.855574697829775 | -6.31086832145155 |
| LINC01191 | LINC01191 | -0.145712604940432 | 1.69891425883604 | -0.447235616616279 | 0.656978406105768 | 0.85569743294786 | -5.92543210952435 |
| ENSG00000261832.6 | | -0.0636429272926606 | 2.09149288274988 | -0.447094180355215 | 0.657077390257619 | 0.855727224928675 | -5.97033490162596 |
| lnc-SLC25A29-5 | lnc-SLC25A29-5 | 0.218857186041923 | 0.796827442324497 | 0.447129106639912 | 0.657084428411685 | 0.855727224928675 | -5.77059666709026 |
| ENSG00000104419.14 | NDRG1 | -0.0443850007677372 | 4.96287115959209 | -0.446973829057863 | 0.657163593741831 | 0.855776176767564 | -6.27897648720526 |
| ENSG00000130724.8 | CHMP2A | 0.061428316514256 | 4.11961943806721 | 0.446837410104521 | 0.657261311646348 | 0.855845934478687 | -6.20830463627366 |
| lnc-TRA2A-6 | lnc-TRA2A-6 | -0.462280067984434 | -2.64381418422236 | -0.446622329042078 | 0.657415388126492 | 0.855845934478687 | -4.96807771873183 |
| ENSG00000105499.13 | PLA2G4C | -0.238880801586326 | 1.05383783086561 | -0.446635184171541 | 0.65745349790503 | 0.855845934478687 | -5.78745467780715 |
| lnc-FRRS1L-2 | lnc-FRRS1L-2 | 0.0943352066962662 | 1.50807101656649 | 0.446548543273122 | 0.657468249120594 | 0.855845934478687 | -5.91284934773808 |
| ENSG00000100417.11 | PMM1 | -0.0680296824522135 | 2.2851242729194 | -0.446528689357421 | 0.657482473002655 | 0.855845934478687 | -6.00269142142025 |
| ENSG00000279861.1 | | 0.0745557768953122 | 2.12850468982899 | 0.446458078812007 | 0.657533061347579 | 0.855845934478687 | -5.98834544517076 |
| ENSG00000131351.14 | HAUS8 | -0.069419652791376 | 3.7517131234702 | -0.446407017778865 | 0.657569644617367 | 0.855845934478687 | -6.17393702522159 |
| lnc-RGS9-3 | lnc-RGS9-3 | 0.249146498368596 | -0.695256010570869 | 0.44640077299811 | 0.657574118821317 | 0.855845934478687 | -5.31317225247598 |
| lnc-FBXW7-9 | lnc-FBXW7-9 | 0.111043055030244 | 3.31287878798107 | 0.446376675845864 | 0.6575913838498 | 0.855845934478687 | -6.10007366412701 |
| ENSG00000154473.17 | BUB3 | 0.0401825373754967 | 6.5928694228356 | 0.446240511131378 | 0.657688946138893 | 0.85588231781717 | -6.31091647141167 |
| ENSG00000279885.1 | | -0.41386606826969 | -0.480994822498504 | -0.446278041860695 | 0.657726849787885 | 0.85588231781717 | -5.5010936973907 |
| ENSG00000233719.4 | GOT2P3 | -0.304001827847673 | 3.35130259910392 | -0.446283702392703 | 0.657744463639683 | 0.85588231781717 | -6.11893624217205 |
| ENSG00000168710.17 | AHCYL1 | -0.0449368942689983 | 5.28846810050449 | -0.446082365858924 | 0.657802265122006 | 0.85588231781717 | -6.29357589579936 |
| ENSG00000021574.12 | SPAST | 0.0658831643689144 | 5.33436734806942 | 0.446025863643138 | 0.657842753755348 | 0.85588231781717 | -6.29685800585148 |
| lnc-GOLPH3-1 | lnc-GOLPH3-1 | 0.136032597626873 | -0.553254056717564 | 0.445989473288932 | 0.657868831083197 | 0.85588231781717 | -5.42914818110119 |
| ENSG00000272821.1 | | -0.0922974778007261 | 0.734058706377123 | -0.445876544207062 | 0.657949758808394 | 0.855922249729427 | -5.78072098722359 |
| ENSG00000134061.5 | CD180 | 0.149945445067067 | 4.74551041717818 | 0.445900772671023 | 0.65798269239026 | 0.855922249729427 | -6.25694612351503 |
| lnc-SLC2A9-4 | lnc-SLC2A9-4 | -0.0496636529509615 | 3.41315441657975 | -0.445433567114739 | 0.658267247034513 | 0.85623829282807 | -6.1389341781538 |
| ENSG00000121931.15 | LRIF1 | 0.0571519246068064 | 3.70716143331226 | 0.445156948013254 | 0.658465536407835 | 0.856350853091408 | -6.16870721502243 |
| lnc-NFIL3-3 | lnc-NFIL3-3 | 0.0924374690980461 | 2.84156803157597 | 0.445111006988476 | 0.658498470805363 | 0.856350853091408 | -6.06724772743801 |
| ENSG00000173064.12 | HECTD4 | 0.0708734141554137 | 6.96349235170735 | 0.445082400514961 | 0.658518978682583 | 0.856350853091408 | -6.30926082932372 |
| lnc-MTMR2-2 | lnc-MTMR2-2 | 0.0936243210622666 | 0.0968850764298447 | 0.445080695138022 | 0.658520201269533 | 0.856350853091408 | -5.627371337243 |
| FOXP4-AS1 | FOXP4-AS1 | -0.144657876436306 | -0.28727450136351 | -0.444916677775317 | 0.658637789948226 | 0.856415408979484 | -5.49016154057459 |
| ENSG00000162734.12 | PEA15 | -0.0915794046509539 | 4.71175427251065 | -0.444895380061247 | 0.65865305952219 | 0.856415408979484 | -6.27092059993345 |
| lnc-DEFB115-5 | lnc-DEFB115-5 | 0.211232351780592 | 1.18855267298683 | 0.444820546879162 | 0.658751972392229 | 0.856454836729234 | -5.88415671497614 |
| ENSG00000110911.15 | SLC11A2 | -0.0607692097321193 | 5.09288495720024 | -0.444737019184213 | 0.658766602297879 | 0.856454836729234 | -6.28689365612327 |
| ENSG00000166881.9 | NEMP1 | -0.0546092655591011 | 5.2626182253194 | -0.444608505201959 | 0.658858751210629 | 0.856520537874589 | -6.29338491869262 |
| lnc-AGBL1-3 | lnc-AGBL1-3 | -0.09215795771367 | 2.0249063906861 | -0.444465636896684 | 0.658961198959541 | 0.85657474261962 | -5.96094293414427 |
| lnc-SPATA9-6 | lnc-SPATA9-6 | 0.036096564731452 | 2.6118383367282 | 0.44437720857867 | 0.659024612300576 | 0.85657474261962 | -6.03721035009958 |
| lnc-HNRNPUL2-BSCL2-1 | lnc-HNRNPUL2-BS | 0.0621564210003074 | 2.3365598343685 | 0.44428910421268 | 0.659087795855323 | 0.85657474261962 | -5.99680079026654 |
| AKT3-IT1 | AKT3-IT1 | 0.335045383354246 | -0.665678535631321 | 0.444288708588637 | 0.659101540123348 | 0.85657474261962 | -5.2956442946569 |
| ENSG00000165322.17 | ARHGAP12 | 0.0461962482768632 | 6.62434159550431 | 0.443981490909865 | 0.659308418774817 | 0.85657474261962 | -6.3117530253281 |
| ENSG00000277224.2 | H2BC7 | 0.52456778192371 | 2.7654856893361 | 0.44410229406327 | 0.659311228517943 | 0.85657474261962 | -6.02179523222838 |

| | | | | | | | |
|---|---|---|---|---|---|---|---|
| lnc-ACTR1B-7 | lnc-ACTR1B-7 | 0.0522354251966908 | 2.95496286469997 | 0.443953426004467 | 0.659328548696016 | 0.85657474261962 | -6.07208863527855 |
| ENSG00000128011.4 | LRFN1 | -0.147981198754282 | -0.343048663213829 | -0.443902590309942 | 0.659365011914749 | 0.85657474261962 | -5.50390655170098 |
| lnc-ADAD1-3 | lnc-ADAD1-3 | -0.101593086976643 | 1.7059419346793 | -0.443866519401278 | 0.659390885216819 | 0.85657474261962 | -5.94154213608594 |
| ENSG00000248167.7 | TRIM39-RPP21 | 0.0406899380473642 | 3.44593741614499 | 0.443866029728768 | 0.659391236456917 | 0.85657474261962 | -6.1394287014017 |
| ENSG00000188612.11 | SUMO2 | -0.0631434777910478 | 6.04438949461275 | -0.443844262840588 | 0.659406849835778 | 0.85657474261962 | -6.31211059158957 |
| ENSG00000106266.10 | SNX8 | -0.0809749682530283 | 5.0073622775436 | -0.443799259905912 | 0.659439130907398 | 0.85657474261962 | -6.28606876384406 |
| ENSG00000155827.11 | RNF20 | 0.0437626222527641 | 5.52796517150294 | 0.443788799036358 | 0.659446634691675 | 0.85657474261962 | -6.30291835757648 |
| lnc-ECE1-1 | lnc-ECE1-1 | -0.108876336621801 | 1.38987485167738 | -0.443688838660609 | 0.659518339994481 | 0.85657474261962 | -5.87623742224197 |
| ENSG00000163602.10 | RYBP | 0.0922285919739103 | 4.51252408377737 | 0.443679998421189 | 0.659524681583625 | 0.85657474261962 | -6.24808096019829 |
| ENSG00000117586.10 | TNFSF4 | -0.0826342157648461 | 0.278335540007759 | -0.443506101237996 | 0.659649432733266 | 0.856666602154365 | -5.64719887777416 |
| ENSG00000119950.20 | MXI1 | 0.0700840779659471 | 5.32849922129871 | 0.443429523705294 | 0.659704371379865 | 0.856666602154365 | -6.29439641595454 |
| ENSG00000102893.15 | PHKB | -0.0621650251552435 | 7.10925894304974 | -0.443381999944434 | 0.659738467076135 | 0.856666602154365 | -6.30897520004153 |
| ENSG00000196365.11 | LONP1 | -0.0431840189304964 | 5.15500196019326 | -0.443335739191312 | 0.659771657336551 | 0.856666602154365 | -6.29074046985514 |
| ENSG00000249437.7 | NAIP | 0.0853975187325338 | 0.890600892742078 | 0.443291343857518 | 0.659803509884523 | 0.856666602154365 | -5.78260927151452 |
| lnc-NAT10-5 | lnc-NAT10-5 | 0.056123776861855 | 0.988425384144094 | 0.443180885959017 | 0.659882763451025 | 0.856680992789557 | -5.80196962570095 |
| lnc-GPAT2-1 | lnc-GPAT2-1 | 0.0768752979193312 | 2.68235935443647 | 0.443159486212818 | 0.659898118230744 | 0.856680992789557 | -6.04599570005583 |
| ENSG00000125821.12 | DTD1 | -0.0482558911927335 | 3.24779052752175 | -0.443048980362474 | 0.65997741092413 | 0.856680992789557 | -6.12183600731013 |
| ENSG00000183020.13 | AP2A2 | -0.0510759655978879 | 4.97966334488208 | -0.442995920292486 | 0.660015485208633 | 0.856680992789557 | -6.28343241185705 |
| ENSG00000280347.1 | | -0.0770771428469712 | 3.44385842309732 | -0.442951075512594 | 0.660047665163034 | 0.856680992789557 | -6.13332691301521 |
| ENSG00000186787.8 | SPIN2B | -0.0559129795143152 | 1.38015764470955 | -0.442846738012279 | 0.660122538738757 | 0.856680992789557 | -5.87552275104632 |
| lnc-ABCG1-2 | lnc-ABCG1-2 | -0.150615038478308 | -0.322180543339561 | -0.442817093038254 | 0.660143812895168 | 0.856680992789557 | -5.44710134043061 |
| ENSG00000113522.13 | RAD50 | 0.0441923661450101 | 4.74316711984397 | 0.442811871645084 | 0.660147559958955 | 0.856680992789557 | -6.26351269575146 |
| ENSG00000270231.3 | NBPF8 | 0.0612170909551146 | 3.9133820480389 | 0.442541582338179 | 0.660341541560127 | 0.856767280676143 | -6.18940981497667 |
| lnc-PARP16-2 | lnc-PARP16-2 | -0.104943030814411 | 0.910052234678916 | -0.442531317657692 | 0.660348908796154 | 0.856767280676143 | -5.82470209726744 |
| ENSG00000164024.11 | METAP1 | -0.0456062157464245 | 4.66725648549743 | -0.442482928487283 | 0.660383639458613 | 0.856767280676143 | -6.26573149458332 |
| lnc-ZDHHC19-1 | lnc-ZDHHC19-1 | 0.0703401797529022 | 3.19333966654216 | 0.442421405942513 | 0.660427797514134 | 0.856767280676143 | -6.1045754098761 |
| lnc-TRIM59-2 | lnc-TRIM59-2 | -0.100569883708753 | 8.02893297649971 | -0.442395206662304 | 0.660450999612192 | 0.856767280676143 | -6.29168180521236 |
| ENSG00000238741.1 | SCARNA7 | -0.100554282056482 | 8.02890642896368 | -0.44231703905779 | 0.660507106278296 | 0.856767280676143 | -6.2917164674348 |
| ENSG00000251595.7 | ABCA11P | -0.0565518722950387 | 3.24308445891471 | -0.44230855142254 | 0.660508802485997 | 0.856767280676143 | -6.11857597558115 |
| lnc-TMEM140-1 | lnc-TMEM140-1 | 0.0966137397633209 | 3.41800453566352 | 0.442255264565182 | 0.660547052278565 | 0.856767280676143 | -6.12363096156063 |
| ENSG00000231752.6 | EMBP1 | -0.336136157912034 | -0.802015763501722 | -0.4421790699279 | 0.660613611313266 | 0.85679158807991 | -5.28052226157302 |
| lnc-SIN3A-2 | lnc-SIN3A-2 | -0.0641079635638828 | 2.15189153572405 | -0.442113181821905 | 0.66064904505739 | 0.85679158807991 | -5.98594988995004 |
| ENSG00000151287.16 | TEX30 | 0.089851681320011 | 1.801041120342 | 0.44197299420717 | 0.660749683833544 | 0.856808680026917 | -5.94436465474361 |
| lnc-PAQR9-7 | lnc-PAQR9-7 | 0.125794471941109 | 0.179475517298043 | 0.441942984852373 | 0.660771227965582 | 0.856808680026917 | -5.62518927605376 |
| ENSG00000144589.21 | STK11IP | 0.0639401127324524 | 4.32747080988658 | 0.441920869290946 | 0.660787105220195 | 0.856808680026917 | -6.24052652883992 |
| ENSG00000091640.7 | SPAG7 | -0.0704977051388411 | 4.2567365180809 | -0.44185210971473 | 0.660836470262831 | 0.856818712908336 | -6.23363675095551 |
| lnc-LY6E-2 | lnc-LY6E-2 | -0.120056193669238 | 0.539399946561178 | -0.441724270272551 | 0.660928254987583 | 0.856846492125096 | -5.73201410365082 |
| lnc-DUSP27-1 | lnc-DUSP27-1 | -0.115744385863072 | -0.409282914047444 | -0.441706306053221 | 0.660941153158925 | 0.856846492125096 | -5.47196218131572 |
| ENSG00000112787.13 | FBRSL1 | 0.0937977880370369 | 4.22274790060664 | 0.441622596817443 | 0.66100125712152 | 0.856870441827198 | -6.22640878304369 |
| lnc-FNDC5-3 | lnc-FNDC5-3 | 0.06922257655487 | 1.75755369452314 | 0.441398526750206 | 0.661162152526635 | 0.857025038444218 | -5.92567214788254 |
| lnc-FKBP3-6 | lnc-FKBP3-6 | 0.0765520729057405 | 2.39874432942467 | 0.441115303241802 | 0.661365546759046 | 0.857128217595434 | -6.00391862279743 |
| ENSG00000114062.19 | UBE3A | -0.0471617253180909 | 7.01149533328161 | -0.441109235340083 | 0.661369904647523 | 0.857128217595434 | -6.30987166099898 |
| lnc-MAST4-6 | lnc-MAST4-6 | -0.139481499091153 | -0.336766919978367 | -0.441047174913884 | 0.661414476324698 | 0.857128217595434 | -5.44161784956159 |
| ENSG00000117016.9 | RIMS3 | -0.246964712046056 | 0.620037157431729 | -0.441070379795667 | 0.661447838549942 | 0.857128217595434 | -5.79106314444644 |
| ENSG00000173171.14 | MTX1 | -0.05292051902572 | 2.97280838363112 | -0.440997764098673 | 0.661449963964722 | 0.857128217595434 | -6.08498852041205 |
| lnc-SLC2A6-1 | lnc-SLC2A6-1 | 0.130862186221691 | -0.316430721454897 | 0.440905050714722 | 0.661516554330277 | 0.8571605437696 | -5.53066909447101 |
| ENSG00000279407.1 | | -0.0804762496058132 | 0.756491129961448 | -0.440740620742225 | 0.661634661165664 | 0.857247534879737 | -5.75466133344316 |
| RBM15-AS1 | RBM15-AS1 | -0.0815187451486767 | 2.12045639107605 | -0.440608499871135 | 0.661729567324207 | 0.857247534879737 | -5.9628961876998 |
| ENSG00000127947.15 | PTPN12 | 0.0759632836354465 | 6.42184944380162 | 0.440591528588713 | 0.661741758684395 | 0.857247534879737 | -6.31433275701824 |
| lnc-APOD-3 | lnc-APOD-3 | 0.0567971887499068 | 4.43742444336494 | 0.440515424390062 | 0.661796429454526 | 0.857247534879737 | -6.2463410281103 |
| ENSG00000278831.1 | | -0.211284327201727 | 1.27122851822972 | -0.440545500643227 | 0.661819591949648 | 0.857247534879737 | -5.87730070048101 |
| ENSG00000284214.1 | MIR29C | -0.163918914459667 | -0.0754830106429746 | -0.4404637106492 | 0.661833579982214 | 0.857247534879737 | -5.56745777196314 |
| ENSG00000063244.12 | U2AF2 | -0.0427993335384835 | 6.05380880136601 | -0.440317705784422 | 0.661938472769281 | 0.857329447836025 | -6.31338667610443 |
| lnc-ROGDI-3 | lnc-ROGDI-3 | -0.0437576298143072 | 1.98820719447034 | -0.440110575207956 | 0.662087291265973 | 0.857468238603632 | -5.96052934414744 |
| ENSG00000283764.1 | MIR6850 | 0.157883000754316 | -0.244200431796459 | 0.439740677496422 | 0.662353088541236 | 0.857676171175627 | -5.5858105147256 |
| lnc-PCGF2-4 | lnc-PCGF2-4 | -0.0565793165965589 | 3.68406891143152 | -0.439721037838626 | 0.66236720223549 | 0.857676171175627 | -6.17224151276815 |
| lnc-JSRP1-4 | lnc-JSRP1-4 | 0.0897661979499351 | 2.351519908132 | 0.439713175768564 | 0.66237285220851 | 0.857676171175627 | -6.0197833977025 |
| lnc-CROT-1 | lnc-CROT-1 | -0.189296346237847 | 0.916819956906219 | -0.439595530502104 | 0.66247069891853 | 0.857748908337429 | -5.76752287328595 |
| ARRDC3-AS1 | ARRDC3-AS1 | 0.067518184600059 | 1.69239295437945 | 0.439436450380528 | 0.662571729936869 | 0.857825759044569 | -5.92805549936284 |
| ENSG00000112763.16 | BTN2A1 | 0.0598470692765982 | 4.59471834734096 | 0.439250183967948 | 0.662705610248192 | 0.857895317438538 | -6.25268366161749 |
| ENSG00000145725.19 | PPIP5K2 | 0.0335237686233123 | 6.52698096179618 | 0.439194072870526 | 0.662745942687011 | 0.857895317438538 | -6.31455242952308 |
| lnc-RNF135-1 | lnc-RNF135-1 | -0.0566497776122534 | 2.34199477008655 | -0.43918773944179 | 0.662750495194588 | 0.857895317438538 | -6.01149197836155 |
| lnc-NOD1-18 | lnc-NOD1-18 | -0.10456246180141 | 0.240288617187122 | -0.43907649152971 | 0.662830462982491 | 0.857897581372072 | -5.64952352532782 |
| lnc-NOBOX-2 | lnc-NOBOX-2 | -0.0878962844272793 | 1.19101391550127 | -0.439063441386204 | 0.662839844013623 | 0.857897581372072 | -5.83710343845316 |
| ENSG00000285444.1 | | 0.0922841433062959 | 0.983238478267119 | 0.438922801944784 | 0.662940945455944 | 0.857897581372072 | -5.81559085690595 |
| ENSG00000254858.9 | MPV17L2 | -0.0615248190555349 | 1.76334903863918 | -0.438889147078154 | 0.66296513986505 | 0.857897581372072 | -5.94173824737281 |
| ENSG00000147533.16 | GOLGA7 | 0.0661782406330218 | 3.58516673295409 | 0.438841550282144 | 0.662999357717957 | 0.857897581372072 | -6.15912574716553 |
| lnc-FOXD4-5 | lnc-FOXD4-5 | -0.0745024437442531 | 0.347761160915486 | -0.438837424793181 | 0.663002323611017 | 0.857897581372072 | -5.6751131445589 |

| | | | | | | | |
|---|---|---|---|---|---|---|---|
| ENSG00000089916.17 | GPATCH2L | -0.0320944891272141 | 6.52088413503436 | -0.438663595190027 | 0.663127298014712 | 0.858005354226604 | -6.31477738562735 |
| ENSG00000101557.14 | USP14 | 0.0724817808567506 | 4.79060039711143 | 0.438499320461519 | 0.663245411887075 | 0.858104237404719 | -6.269652809621 |
| ENSG00000207750.1 | MIR553 | 0.0992754578180034 | -0.274493621489691 | 0.438297251056671 | 0.663390712101172 | 0.858183106370475 | -5.48398618686238 |
| ENSG00000023330.14 | ALAS1 | -0.0748822192197696 | 2.96398312076076 | -0.438255891286553 | 0.663420453912731 | 0.858183106370475 | -6.09182139657253 |
| lnc-DRD5-30 | lnc-DRD5-30 | -0.431016938077488 | -0.77203815490148 | -0.438273405513141 | 0.66347017395201 | 0.858183106370475 | -5.42730912937145 |
| lnc-ZNF33B-4 | lnc-ZNF33B-4 | -0.0412055611409148 | 5.61241456997055 | -0.438182617605288 | 0.663473146367069 | 0.858183106370475 | -6.30643743956017 |
| lnc-SCARF2-5 | lnc-SCARF2-5 | -0.0599125995688499 | 0.202101161545565 | -0.4380866367988 | 0.663542170541475 | 0.858196131692208 | -5.61939246684957 |
| ENSG00000139112.10 | GABARAPL1 | 0.27747872590787 | 2.76016808289491 | 0.43816520444539 | 0.663566605251008 | 0.858196131692208 | -6.01736405962599 |
| ENSG00000111348.8 | ARHGDIB | 0.0751356243704269 | 7.95643193405482 | 0.43780784085501 | 0.663742682186124 | 0.858369918788451 | -6.29744715266185 |
| ENSG00000109103.11 | UNC119 | -0.0723059524266727 | 4.19959644669858 | -0.437709687834193 | 0.663813280338922 | 0.858407284614942 | -6.22814103275987 |
| ENSG00000151651.15 | ADAM8 | 0.140642674619909 | 4.83240529567494 | 0.437649895788639 | 0.663902509691997 | 0.858468737089483 | -6.27052261052076 |
| ENSG00000155096.13 | AZIN1 | 0.0608075729016413 | 7.42287109931181 | 0.437497315776951 | 0.663966042964017 | 0.858470523300882 | -6.3068777290597 |
| lnc-COG5-4 | lnc-COG5-4 | -0.0510772649116795 | 2.77626238156255 | -0.437467756515667 | 0.663987306561242 | 0.858470523300882 | -6.05547618333908 |
| ENSG00000153485.5 | LYSET | 0.0848148900815642 | 1.38853610277335 | 0.437373023119565 | 0.66405545537455 | 0.858504706863536 | -5.88133927854752 |
| lnc-DNAJC5-2 | lnc-DNAJC5-2 | 0.0986318404796494 | -0.292337871384152 | 0.43729043976522 | 0.664114866101957 | 0.858527590062591 | -5.51499915792008 |
| ENSG00000100294.12 | MCAT | 0.091457935365135 | 0.973745598621208 | 0.437182489133144 | 0.6641925294261 | 0.858574064760247 | -5.82990951327128 |
| lnc-ARAF-1 | lnc-ARAF-1 | 0.0733193921613597 | 1.21637039095648 | 0.437074036682202 | 0.664270557536523 | 0.85860855582729 | -5.86570914349963 |
| ENSG00000136783.9 | NIPSNAP3A | -0.0675007978777695 | 3.46466385724069 | -0.437029445461337 | 0.664302640603876 | 0.85860855582729 | -6.15307808154032 |
| ENSG00000266931.2 | | 0.0607326422999583 | 1.20588027988841 | 0.43643991474227 | 0.664726863646424 | 0.859102915636967 | -5.86513574088141 |
| lnc-GLRX5-6 | lnc-GLRX5-6 | 0.171079857443543 | -0.395587414054047 | 0.43621030913008 | 0.664892116678648 | 0.859262537677943 | -5.43855920727993 |
| ENSG00000160999.10 | SH2B2 | 0.13667636083211 | 3.05679664059473 | 0.436153243266909 | 0.664948610078835 | 0.859263608553464 | -6.08058226746092 |
| ENSG00000266953.6 | | -0.0574734536966859 | 2.64875351554942 | -0.436093160741891 | 0.664976437868538 | 0.859263608553464 | -6.0465254395938 |
| ENSG00000052723.11 | SIKE1 | 0.0492412510957925 | 5.25894843725981 | 0.435573188766506 | 0.665350756811546 | 0.859654457554908 | -6.29961775327802 |
| ENSG00000267228.7 | | -0.0431447445814381 | 3.18123282090574 | -0.435509472218369 | 0.665396631209945 | 0.859654457554908 | -6.11042923385938 |
| lnc-TPST1-1 | lnc-TPST1-1 | 0.127955740721892 | -0.387533527334971 | 0.435444813421902 | 0.665443185330793 | 0.859654457554908 | -5.43798533811273 |
| lnc-DOLK-2 | lnc-DOLK-2 | 0.070214743707633 | 1.57678355066453 | 0.435440941028006 | 0.665445973484057 | 0.859654457554908 | -5.91677576546817 |
| ENSG00000224315.2 | RPL7P7 | -0.174798758822674 | -0.626029622768834 | -0.435361300544056 | 0.665503316304846 | 0.859674580111877 | -5.33622121057623 |
| ENSG00000087502.17 | ERGIC2 | 0.038993661753536 | 5.43277430160756 | 0.4353025660474 | 0.665545607673852 | 0.859675258409218 | -6.30443835885385 |
| ENSG00000179361.17 | ARID3B | -0.0572008038047898 | 4.35381624164528 | -0.435014908653765 | 0.665752749305314 | 0.859888857865918 | -6.23827160065689 |
| ENSG00000180104.15 | EXOC3 | 0.0351172082447553 | 5.37506144368852 | 0.4346761370096 | 0.66599673209841 | 0.860098475021307 | -6.30182451501998 |
| ENSG00000181038.13 | METTL23 | -0.0508583731334333 | 3.68797549160233 | -0.434639314191103 | 0.666023254039585 | 0.860098475021307 | -6.16880271206959 |
| lnc-DTYMK-6 | lnc-DTYMK-6 | -0.0690864975610521 | 0.513408214476039 | -0.434615506508238 | 0.666040401951349 | 0.860098475021307 | -5.72854765527943 |
| lnc-RAP1GAP2-3 | lnc-RAP1GAP2-3 | 0.102721798967021 | 1.23065831840466 | 0.434504317164554 | 0.666120490469795 | 0.86014630895197 | -5.84549979218193 |
| ENSG00000221944.7 | TIGD1 | -0.0456124483562377 | 4.49903556554508 | -0.434446353613341 | 0.666162242571522 | 0.86014630895197 | -6.25254106810908 |
| ENSG00000197363.9 | ZNF517 | -0.0608505404536896 | 2.76544188716832 | -0.434390034856328 | 0.666202810927091 | 0.86014630895197 | -6.0624937592029 |
| LINC01353 | LINC01353 | -0.0978141141644984 | 0.730415273642575 | -0.434207511518773 | 0.666334295772372 | 0.860228601717775 | -5.77177503336306 |
| ENSG00000209082.1 | MT-TL1 | 0.162118503308429 | 8.34436690489078 | 0.4342711359332 | 0.666350134858291 | 0.860228601717775 | -6.29380292426389 |
| ENSG00000163348.3 | PYGO2 | -0.0618606946299423 | 4.41001332511454 | -0.433932152280144 | 0.66653267716761 | 0.860410291260014 | -6.25133117424746 |
| lnc-STYX-2 | lnc-STYX-2 | -0.0837094820498602 | 0.0950572591484389 | -0.433754739160959 | 0.666660506493159 | 0.860521334826833 | -5.59283058412181 |
| ENSG00000196498.13 | NCOR2 | -0.0936500275239096 | 5.15287455373771 | -0.433376402062876 | 0.666933205332417 | 0.860789426121539 | -6.29458855043157 |
| ENSG00000272150.5 | NBPF25P | -0.107829359972831 | -0.189631162194438 | -0.433332730617139 | 0.66696461147296 | 0.860789426121539 | -5.52461952461158 |
| ENSG00000162804.13 | SNED1 | 0.403728657392854 | 4.24527264767031 | 0.433413181846121 | 0.666993662595949 | 0.860789426121539 | -6.24085123740108 |
| ENSG00000159479.16 | MED8 | 0.0493583189619518 | 4.21650348835104 | 0.433203546374836 | 0.667057714691427 | 0.860818115454147 | -6.22837515605942 |
| ENSG00000067365.14 | METTL22 | -0.0469436424796879 | 3.91709887098509 | -0.432982605319241 | 0.667216959446441 | 0.860858851012859 | -6.19254915165627 |
| ENSG00000137177.19 | KIF13A | 0.0987494582641926 | 4.8804511527157 | 0.43297731807411 | 0.66722506213403 | 0.860858851012859 | -6.28663806852984 |
| ENSG00000229623.1 | METTL21AP1 | 0.148296691004952 | -0.258226062297281 | 0.432951233799219 | 0.66723957193977 | 0.860858851012859 | -5.46267480093818 |
| lnc-ZBTB8OS-3 | lnc-ZBTB8OS-3 | -0.0465604322236133 | 2.4228093093829 | -0.432897266777326 | 0.667278471932332 | 0.860858851012859 | -6.01464699697294 |
| lnc-FYCO1-5 | lnc-FYCO1-5 | 0.0721189696443947 | -0.131088017207116 | 0.43284926633235 | 0.667313071932167 | 0.860858851012859 | -5.539675097351 |
| ENSG00000058272.18 | PPP1R12A | 0.0387634628944715 | 6.89267596559282 | 0.432774678599248 | 0.667366838211357 | 0.860858851012859 | -6.3145809626312 |
| ENSG00000211456.10 | SACM1L | 0.0411490234644312 | 5.05874563369798 | 0.43275357969862 | 0.667382047593266 | 0.860858851012859 | -6.28855605570047 |
| lnc-PLPP5-4 | lnc-PLPP5-4 | -0.041470693081024 | 3.92533902965538 | -0.432677576228774 | 0.667436836729869 | 0.860875574032516 | -6.18990216884539 |
| lnc-SFMBT2-3 | lnc-SFMBT2-3 | -0.34457402788118 | 0.186413750478776 | -0.432670241411261 | 0.667495795524521 | 0.860897672741477 | -5.52485880293527 |
| ENSG00000028137.18 | TNFRSF1B | -0.127895514249859 | 6.77443230511256 | -0.432502582236475 | 0.667603757976438 | 0.860940493828973 | -6.31559305566966 |
| ENSG00000106665.15 | CLIP2 | -0.263916104928002 | -0.283776529588469 | -0.432411591925835 | 0.667647841987578 | 0.860940493828973 | -5.50169666454885 |
| lnc-TBC1D3F-8 | lnc-TBC1D3F-8 | -0.0737197135500308 | 1.04749041171563 | -0.432375686668538 | 0.667654480031971 | 0.860940493828973 | -5.82039617473781 |
| ENSG00000166130.14 | IKBIP | 0.078334132105907 | 2.51502678630516 | 0.432257136008774 | 0.667739955486515 | 0.860996774026119 | -6.02927114737034 |
| ENSG00000285304.1 | | 0.0857965087507315 | 3.38287100562795 | 0.432114053006283 | 0.66784312479363 | 0.860999366244295 | -6.13266297751174 |
| ENSG00000149541.9 | B3GAT3 | -0.0429145308704406 | 4.25475157378965 | -0.432070607829475 | 0.667874452014933 | 0.860999366244295 | -6.23211809596125 |
| ENSG00000185986.11 | SDHAP3 | 0.0749343415852008 | 3.25768984184422 | 0.43207043866876 | 0.667874573993629 | 0.860999366244295 | -6.11710920906033 |
| lnc-SLC16A11-2 | lnc-SLC16A11-2 | -0.156049901064433 | -0.529239272129301 | -0.432022297210519 | 0.667909288287551 | 0.860999366244295 | -5.45788665773707 |
| lnc-PRSS27-4 | lnc-PRSS27-4 | 0.0470218955242627 | 5.16981769140678 | 0.431950369150861 | 0.667961156217822 | 0.861012304510986 | -6.28984128434337 |
| ENSG00000171843.15 | MLLT3 | -0.430714044614318 | 1.7833073006219 | -0.431886761671734 | 0.668093710205448 | 0.861068412424926 | -5.92367169933874 |
| ENSG00000007968.6 | E2F2 | 0.174351422612461 | 0.149553791621177 | 0.431706164754146 | 0.668137266318667 | 0.861068412424926 | -5.63493071444843 |
| lnc-XRCC1-2 | lnc-XRCC1-2 | 0.0568430956256483 | 1.84795267463536 | 0.431688145165678 | 0.668150262049564 | 0.861068412424926 | -5.95153989448834 |
| lnc-PHF20L1-6 | lnc-PHF20L1-6 | 0.114415585449176 | 2.89213789733603 | 0.431657976697255 | 0.66817201978579 | 0.861068412424926 | -6.08669747339814 |
| ENSG00000105556.11 | MIER2 | -0.0756739902999386 | 2.02200229264458 | -0.431240941884308 | 0.668472818142806 | 0.861402117062128 | -5.9944246929978 |
| ENSG00000225614.2 | ZNF469 | -0.31455902051855 | -0.126173118537585 | -0.43118219972683 | 0.668566210842382 | 0.861468531223785 | -5.57524971348883 |
| ENSG00000050393.11 | MCUR1 | 0.0556466777232854 | 2.79398225805948 | 0.430938972257381 | 0.66869065684016 | 0.861574947714618 | -6.06582416370652 |

| | | | | | | | |
|---|---|---|---|---|---|---|---|
| ENSG00000171813.13 | PWWP2B | 0.110324910438327 | 2.48607872618141 | 0.430682899463647 | 0.668875408533046 | 0.861737889081551 | -6.04098895895987 |
| ENSG00000111726.12 | CMAS | -0.0631702238465834 | 2.60832598307722 | -0.43064763496253 | 0.668900852832723 | 0.861737889081551 | -6.04682306513886 |
| lnc-CXorf38-2 | lnc-CXorf38-2 | -0.0717247320896239 | 1.44552710216 | -0.430367851093332 | 0.66910273848146 | 0.861944027172606 | -5.89927972489926 |
| ENSG00000285730.1 | | -0.0854485837993109 | 0.489762324349845 | -0.430236383883467 | 0.669197610773403 | 0.862012292543273 | -5.6987865717908 |
| ENSG00000259248.6 | USP3-AS1 | -0.105817877607665 | -0.0252640681680277 | -0.43008620151607 | 0.66930599540544 | 0.862034088246212 | -5.55372942289283 |
| ENSG00000143153.12 | ATP1B1 | -0.371155482315064 | 0.952924678228437 | -0.430089885305992 | 0.669382595160065 | 0.862034088246212 | -5.78744200201544 |
| ENSG00000091157.13 | WDR7 | -0.0296436427952935 | 5.46324055648064 | -0.429868823971099 | 0.66946288654663 | 0.862034088246212 | -6.30633044588684 |
| ENSG00000060971.17 | ACAA1 | -0.0552360042307742 | 3.87244707385412 | -0.42986856856473 | 0.669463070893671 | 0.862034088246212 | -6.20179649800581 |
| lnc-CCDC173-5 | lnc-CCDC173-5 | -0.0759925480891073 | 0.380642073639552 | -0.429865368821601 | 0.669465380403912 | 0.862034088246212 | -5.66496117947845 |
| ENSG00000008710.19 | PKD1 | -0.0893029461279351 | 4.98820830245596 | -0.429864736569424 | 0.669465836751201 | 0.862034088246212 | -6.28879788988327 |
| ENSG00000142188.16 | TMEM50B | 0.0618084588476222 | 4.7512379713004 | 0.429774996805931 | 0.66953061044421 | 0.862034088246212 | -6.27394201762137 |
| ENSG00000171603.17 | CLSTN1 | -0.0584799959459623 | 5.72138985186022 | -0.429748718827171 | 0.669549578234086 | 0.862034088246212 | -6.31342502252748 |
| lnc-RABGGTB-1 | lnc-RABGGTB-1 | 0.103992307181169 | 0.885155859729901 | 0.429554404176422 | 0.669689843873242 | 0.862160749089501 | -5.82528533778596 |
| lnc-GRAMD4-4 | lnc-GRAMD4-4 | -0.133768840498361 | 1.74034068807097 | -0.429477572234868 | 0.669745308147378 | 0.862178227381168 | -5.92216097946071 |
| ENSG00000122696.13 | SLC25A51 | 0.05230274334621 | 3.01122905046534 | 0.429190110306575 | 0.669952840819529 | 0.862391452319473 | -6.09216365959959 |
| lnc-SLC36A1-4 | lnc-SLC36A1-4 | -0.09399516685813 | 0.440370944196715 | -0.429080941503102 | 0.67003166189753 | 0.862438977977416 | -5.71504873227812 |
| ENSG00000080371.5 | RAB21 | 0.0508214964006521 | 6.15394226423351 | 0.428874189409094 | 0.670180949496344 | 0.862453815989753 | -6.31915495420456 |
| lnc-ZMYM3-1 | lnc-ZMYM3-1 | 0.0729868965068263 | 2.27028708618 | 0.428850712123933 | 0.670197902378567 | 0.862453815989753 | -5.9978016901472 |
| ENSG00000172977.12 | KAT5 | 0.0449667697527962 | 5.32316666338463 | 0.42883016338556 | 0.670212740708068 | 0.862453815989753 | -6.30164218079852 |
| ENSG00000168092.13 | PAFAH1B2 | 0.0627326093105083 | 5.99635280857602 | 0.428803104144155 | 0.670232280502175 | 0.862453815989753 | -6.3184435766263 |
| ENSG00000175548.8 | ALG10B | -0.069428385061383 | 4.31305574200025 | -0.428712700843529 | 0.670297563456355 | 0.862453815989753 | -6.24152613951246 |
| ENSG00000180596.7 | H2BC4 | -0.0713197239204071 | 6.95152439174301 | -0.428674136490551 | 0.670325412725182 | 0.862453815989753 | -6.31587102430444 |
| lnc-KIAA0355-3 | lnc-KIAA0355-3 | 0.100969117620883 | 0.108720258627695 | 0.42863743685459 | 0.670351915823302 | 0.862453815989753 | -5.62401419638357 |
| lnc-ZFAT-1 | lnc-ZFAT-1 | 0.0930149802910094 | 6.06355030444484 | 0.428603820702031 | 0.670378399748339 | 0.862453815989753 | -6.31893757720242 |
| lnc-MTX3-4 | lnc-MTX3-4 | 0.159529141384878 | 2.83092718006827 | 0.428395575335966 | 0.670560108612774 | 0.86258193799338 | -6.05376356617692 |
| lnc-TMEM132D-5 | lnc-TMEM132D-5 | -0.123470725732829 | 0.703956854726892 | -0.428278163966396 | 0.670611391638717 | 0.86258193799338 | -5.7781038070754 |
| lnc-NPIPB11-5 | lnc-NPIPB11-5 | -0.111112565961557 | 0.226704948212431 | -0.428254836139553 | 0.670628240982617 | 0.86258193799338 | -5.63230236143107 |
| ENSG00000104529.17 | EEF1D | 0.0664050327592474 | 8.19858682831716 | 0.42820398426399 | 0.670664971133905 | 0.86258193799338 | -6.29585806814956 |
| ENSG00000229241.1 | PNPT1P1 | -0.136125415539459 | -0.728547578769389 | -0.428073677759969 | 0.670759094836059 | 0.86258193799338 | -5.37084685780358 |
| ENSG00000273340.1 | MICE | -0.100926442151628 | 0.199396106964812 | -0.427967626148256 | 0.670835702562248 | 0.86258193799338 | -5.63259840959666 |
| ENSG00000276410.3 | H2BC3 | -0.201107415363262 | 3.21888158068141 | -0.42804837558877 | 0.670838051871952 | 0.86258193799338 | -6.11354790809652 |
| ENSG00000125733.17 | TRIP10 | 0.113725142618823 | 1.91022407730262 | 0.427952919629502 | 0.6708463262823 | 0.86258193799338 | -5.96039027489116 |
| ENSG00000184428.12 | TOP1MT | 0.0808044381340141 | 5.16479576212855 | 0.427940697481434 | 0.670855155390275 | 0.86258193799338 | -6.29244181636592 |
| ENSG00000104427.11 | ZC2HC1A | 0.213206620038666 | 3.09299359468023 | 0.427841741902792 | 0.670990219272339 | 0.862682457065805 | -6.09717698134614 |
| ENSG00000131374.14 | TBC1D5 | 0.0245093297491996 | 7.15511176273966 | 0.427716450290099 | 0.671017156761577 | 0.862682457065805 | -6.31417348542782 |
| ENSG00000207751.3 | | 0.0441415125389526 | 1.44212407868761 | 0.427525359072506 | 0.671155217986138 | 0.862774763301066 | -5.88476617585587 |
| ENSG00000163479.13 | SSR2 | 0.0530280950064981 | 5.97158182507838 | 0.427497564467701 | 0.671175300228726 | 0.862774763301066 | -6.31811805852047 |
| lnc-WNT5B-5 | lnc-WNT5B-5 | -0.1386409750015 | 0.617897610559691 | -0.427443026921412 | 0.671214705569643 | 0.862774763301066 | -5.73665226021748 |
| ENSG00000239900.12 | ADSL | 0.0513820247808781 | 4.05837053508331 | 0.427145544934333 | 0.671429663494949 | 0.86299717520398 | -6.21800656482118 |
| lnc-TRIB1-3 | lnc-TRIB1-3 | -0.118352366908011 | 0.368553851091513 | -0.427053428725313 | 0.671496231520997 | 0.863021463512018 | -5.69058014852221 |
| lnc-OR1A2-1 | lnc-OR1A2-1 | 0.121904914887031 | -0.541154415312914 | 0.427003355770868 | 0.671532417995846 | 0.863021463512018 | -5.43841806434997 |
| ENSG00000243609.1 | RPS2P44 | 0.114577869315764 | 0.197883331843435 | 0.426900779389487 | 0.671606549849543 | 0.863036217492446 | -5.65101485382216 |
| lnc-TRMT2B-1 | lnc-TRMT2B-1 | -0.0665501794704193 | 2.92813571715365 | -0.426790536516989 | 0.671686225957746 | 0.863036217492446 | -6.08423709442151 |
| ENSG00000172888.11 | ZNF621 | 0.0430073090852972 | 5.13889623916476 | 0.426733809594603 | 0.67172722583775 | 0.863036217492446 | -6.29617355779902 |
| ENSG00000089048.14 | ESF1 | 0.0445149715310897 | 4.24096088968739 | 0.426715390776763 | 0.671740538417211 | 0.863036217492446 | -6.22626469284022 |
| ENSG00000026652.13 | AGPAT4 | -0.308833937851454 | 2.68876531976841 | -0.426811158134719 | 0.671753546151993 | 0.863036217492446 | -6.09724197242958 |
| ENSG00000118432.12 | CNR1 | -0.426374606801033 | 3.35341813976905 | -0.426684168483051 | 0.671848616461705 | 0.863104485871382 | -6.17662184083917 |
| ENSG00000088854.12 | DNAAF9 | -0.0535565998899719 | 4.78942577823841 | -0.426307860064207 | 0.672035116897532 | 0.863209026812854 | -6.27916067300613 |
| ENSG00000222345.1 | SNORD19C | 0.119353557543833 | -0.279499290482172 | 0.426296331260203 | 0.672043451108271 | 0.863209026812854 | -5.50541707192748 |
| ENSG00000171766.15 | GATM | 0.15755502495321 | 3.92580184939182 | 0.426342391563448 | 0.672058412762548 | 0.863209026812854 | -6.18769367189085 |
| ENSG00000125459.15 | MSTO1 | 0.0460419788104907 | 3.63750361580058 | 0.426172066495805 | 0.6721332851693 | 0.863209026812854 | -6.17560006907091 |
| ENSG00000256274.1 | TAS2R64P | -0.14844564263235 | 2.34552640855195 | -0.426172939694414 | 0.672152386860789 | 0.863209026812854 | -6.01832342444918 |
| lnc-HSPA5-5 | lnc-HSPA5-5 | 0.0425957499336334 | 2.84234830561241 | 0.426092340926442 | 0.672190923301899 | 0.863209026812854 | -6.07546198824851 |
| ENSG00000263366.2 | ABHD17AP5 | -0.436201416525229 | -1.49449454459544 | -0.426086982357552 | 0.672223557828462 | 0.863209026812854 | -5.1684590722231 |
| ENSG00000281344.1 | HELLPAR | -0.0816268390914416 | 2.6724931532589 | -0.42596628125321 | 0.67228206305632 | 0.863230299681117 | -6.06568942097097 |
| lnc-TMCC1-4 | lnc-TMCC1-4 | 0.0532138394022565 | 2.32390756145269 | 0.425808066881607 | 0.672396457361803 | 0.863294528712287 | -6.00959712792732 |
| ENSG00000184939.15 | ZFP90 | -0.0444012942321175 | 5.51142646040834 | -0.425718181035813 | 0.672461451346817 | 0.863294528712287 | -6.30878234556642 |
| ENSG00000106526.10 | ACTR3C | 0.0839142820134047 | 2.75749553154909 | 0.42567922402633 | 0.67248962087722 | 0.863294528712287 | -6.0616272005235 |
| lnc-BAIAP2L1-2 | lnc-BAIAP2L1-2 | 0.104120927566395 | 0.101818176547067 | 0.425665073752862 | 0.672499852955002 | 0.863294528712287 | -5.61609687235222 |
| LINC00894 | LINC00894 | -0.0750994159726606 | 3.47382596705913 | -0.425495427780503 | 0.67262252902803 | 0.863316487969228 | -6.14265121950344 |
| ENSG00000132423.11 | COQ3 | -0.0781367872826082 | 0.507935326374956 | -0.425413908465095 | 0.67268148129214 | 0.863316487969228 | -5.73513538644424 |
| lnc-AVIL-2 | lnc-AVIL-2 | 0.0632183656890621 | 2.70652438252862 | 0.425412541401746 | 0.672682469928116 | 0.863316487969228 | -6.05944932825631 |
| ENSG00000132436.11 | FIGNL1 | -0.0576948494872629 | 3.91738499238636 | -0.42540941377934 | 0.672684731771388 | 0.863316487969228 | -6.20962823222085 |
| lnc-DALRD3-1 | lnc-DALRD3-1 | -0.0881326375520692 | 0.486167624969788 | -0.425335732728413 | 0.672738017540113 | 0.863331044019462 | -5.71208043452277 |
| ENSG00000138617.15 | PARP16 | -0.0492932926764116 | 3.16385157427607 | -0.425138523161259 | 0.672880646889494 | 0.863460246566488 | -6.11229873140121 |
| lnc-FZD5-1 | lnc-FZD5-1 | 0.0813322924514611 | 0.323417564923674 | 0.425062167294913 | 0.67293587358683 | 0.863477282341358 | -5.66704218143283 |
| lnc-HIST1H2AH-5 | lnc-HIST1H2AH-5 | -0.209157920964488 | -0.776942886106388 | -0.424768632913683 | 0.673148198243806 | 0.863695883583859 | -5.38273387973664 |
| lnc-PRR14L-4 | lnc-PRR14L-4 | -0.135049716705923 | 0.0456550265419605 | -0.424580154343873 | 0.673284546236558 | 0.863786292179275 | -5.61185442737468 |

| | | | | | | | |
|---|---|---|---|---|---|---|---|
| FGD5-AS1 | FGD5-AS1 | -0.0428870284592193 | 6.77381203238704 | -0.424515444546279 | 0.673331360759936 | 0.863786292179275 | -6.31923495328741 |
| ENSG00000223336.1 | RNU2-6P | 0.10993083982692 | 2.63863790211079 | 0.424497201479168 | 0.673344559005804 | 0.863786292179275 | -6.0536859624519 |
| ENSG00000178498.15 | DTX3 | 0.0978617937721262 | 4.25966661599501 | 0.424324524759269 | 0.673469753380407 | 0.863893053335966 | -6.23900754756214 |
| lnc-SLC10A1-1 | lnc-SLC10A1-1 | -0.0969437076807159 | 0.648041448437446 | -0.424264085540807 | 0.673513219720979 | 0.863894971117151 | -5.75718939493403 |
| lnc-MYC-8 | lnc-MYC-8 | 0.153816966903986 | 1.57093395708489 | 0.423729065320026 | 0.673910418296006 | 0.864281197900946 | -5.91318170626905 |
| ENSG00000197756.9 | RPL37A | 0.0633166153098372 | 9.11727021342401 | 0.423697038016225 | 0.673923552813876 | 0.864281197900946 | -6.27183476020825 |
| ENSG00000281593.1 | | 0.065409388580532 | 3.01195094841065 | 0.423673895585769 | 0.673940301526026 | 0.864281197900946 | -6.09318660275421 |
| lnc-PAWR-9 | lnc-PAWR-9 | 0.0502427613228091 | 2.31294897942232 | 0.423615249394856 | 0.673982745880232 | 0.864281780478027 | -6.01440570016806 |
| ENSG00000141219.15 | MTNAP1 | -0.0766529460908068 | 3.68773028443742 | -0.423378233235439 | 0.674154293933167 | 0.864345097968789 | -6.16851946707801 |
| lnc-ADAM10-6 | lnc-ADAM10-6 | 0.0442382582707852 | 3.6728179013924 | 0.423329419209087 | 0.674189626830777 | 0.864345097968789 | -6.17084358160334 |
| ENSG00000205208.4 | C4orf46 | 0.0972852117663827 | 1.16141035307708 | 0.423256784954874 | 0.674242202823776 | 0.864345097968789 | -5.87386157727363 |
| ENSG00000216866.5 | RPS2P55 | 0.171988507582535 | -0.834158882947474 | 0.423240224337155 | 0.674254190386434 | 0.864345097968789 | -5.32182600134225 |
| ENSG00000183092.16 | BEGAIN | 0.434056358427328 | -0.733677503564626 | 0.4232948413735 | 0.674266732863055 | 0.864345097968789 | -5.31667501888223 |
| lnc-PPP4R3B-4 | lnc-PPP4R3B-4 | -0.16612859901771 | -0.595503104765263 | -0.423198931545094 | 0.674284080944704 | 0.864345097968789 | -5.43487930744027 |
| ENSG00000177721.4 | ANXA2R | 0.0883138087104749 | 2.3883360224852 | 0.423049285126119 | 0.674392409758757 | 0.864430126420756 | -6.01438002335558 |
| lnc-HIST2H2AA4-1 | lnc-HIST2H2AA4-1 | -0.125317276268059 | 0.550250723142629 | -0.422923085032881 | 0.674483771246708 | 0.864493397071485 | -5.72043549664011 |
| ENSG00000254004.6 | ZNF260 | -0.0392957139241412 | 4.61011338722482 | -0.422863130523613 | 0.674527176547132 | 0.864495197688021 | -6.26257102509246 |
| ENSG00000067177.14 | PHKA1 | -0.106583444921682 | 0.341128217214341 | -0.422539019677689 | 0.674761842679693 | 0.864742108702829 | -5.67956475922603 |
| lnc-HP1BP3-1 | lnc-HP1BP3-1 | 0.0847918136645686 | -0.443699950233647 | 0.422468060930382 | 0.674813223347564 | 0.864754113819133 | -5.43967103603115 |
| ENSG00000264293.2 | RN7SL657P | 0.0580345164877525 | 0.929128459030225 | 0.422224947061031 | 0.67498927208058 | 0.864852760927439 | -5.79648867742752 |
| lnc-RANBP9-1 | lnc-RANBP9-1 | 0.0887235281639123 | -0.231568782748654 | 0.422185076496073 | 0.675018145751802 | 0.864852760927439 | -5.50721411783333 |
| ENSG00000197372.9 | ZNF675 | 0.0677943171549851 | 5.58764070802959 | 0.422151901681544 | 0.675042170837551 | 0.864852760927439 | -6.30866645219708 |
| lnc-GOLGA8F-6 | lnc-GOLGA8F-6 | -0.0665226291603277 | -0.112038700875748 | -0.422024773369375 | 0.675134239895763 | 0.864852760927439 | -5.54625991299413 |
| ENSG00000099977.14 | DDT | 0.0650455912745096 | 1.92462755208487 | 0.421842871160143 | 0.675265986144295 | 0.864852760927439 | -5.9524869144234 |
| ENSG00000263798.2 | | 0.0603153530130278 | 6.43043702603106 | 0.421820722010752 | 0.675282028804689 | 0.864852760927439 | -6.32240651066073 |
| lnc-ZNF275-1 | lnc-ZNF275-1 | -0.0909042198866487 | 0.298609033176406 | -0.42182047916234 | 0.675282204700986 | 0.864852760927439 | -5.6737636758194 |
| ENSG00000275126.1 | H4C13 | 0.24445317886107 | 0.83346687831174 | 0.421845861992708 | 0.675319617778205 | 0.864852760927439 | -5.85125041728911 |
| ENSG00000161671.16 | EMC10 | -0.0486002178867277 | 4.99704759017275 | -0.42174555331152 | 0.675336474739406 | 0.864852760927439 | -6.29200473147021 |
| ENSG00000152382.5 | TADA1 | 0.0360818652979713 | 3.91845559860217 | 0.42174144447694 | 0.675339450887108 | 0.864852760927439 | -6.21010349621706 |
| ENSG00000100726.14 | TELO2 | -0.068650376581469 | 3.57667304112862 | -0.421694067720105 | 0.67537376762181 | 0.864852760927439 | -6.1776626770329 |
| lnc-MRPS16-2 | lnc-MRPS16-2 | 0.0521812199566996 | 1.75830698714832 | 0.421636054284031 | 0.675415789847875 | 0.864852760927439 | -5.94866221638114 |
| TRIM52-AS1 | TRIM52-AS1 | 0.0578377978480265 | 3.90136524190222 | 0.421557980312545 | 0.675472344643477 | 0.864852760927439 | -6.19682689972608 |
| lnc-H2AFB3-2 | lnc-H2AFB3-2 | 0.067983794028404 | 0.734403638526358 | 0.421549548775565 | 0.675478452347544 | 0.864852760927439 | -5.78151590125234 |
| ENSG00000147044.21 | CASK | -0.0552817681707403 | 5.6046407627047 | -0.421274330026135 | 0.675677829649243 | 0.865028675750346 | -6.31310560962055 |
| lnc-AEN-1 | lnc-AEN-1 | 0.0953450346989091 | 3.29442876972717 | 0.421243865829058 | 0.67569990034077 | 0.865028675750346 | -6.11907112225241 |
| ENSG00000115685.14 | PPP1R7 | -0.0686235111586631 | 4.00604351309235 | -0.421081534014549 | 0.675817511307911 | 0.865104890257356 | -6.20633341981234 |
| ENSG00000007952.17 | NOX1 | -0.233297160672943 | -0.0826161326623573 | -0.420967474137761 | 0.675907735429326 | 0.865104890257356 | -5.51404423484077 |
| lnc-CCDC154-1 | lnc-CCDC154-1 | 0.0781951760561881 | 1.23963307768207 | 0.420931672261489 | 0.675926094869665 | 0.865104890257356 | -5.87306944972692 |
| ENSG00000264922.1 | MIR4540 | -0.156319516006994 | -0.391382365016843 | -0.420929658430133 | 0.67592755405523 | 0.865104890257356 | -5.41747067846317 |
| SNHG9 | SNHG9 | 0.0538149855392307 | 2.37846391220238 | 0.420763514977174 | 0.676047942920974 | 0.865205173732663 | -6.01227474548669 |
| lnc-TIMM10-1 | lnc-TIMM10-1 | 0.140519110352278 | 0.104372610233374 | 0.420656051036243 | 0.676125816715272 | 0.865250020021089 | -5.62725511311341 |
| lnc-ERV3-1-7 | lnc-ERV3-1-7 | 0.0627752080781996 | 1.16200932165906 | 0.420599139917773 | 0.676167058836866 | 0.865250020021089 | -5.85277984778785 |
| ENSG00000090612.20 | ZNF268 | -0.0600358964253491 | 5.71832844432776 | -0.420491679133232 | 0.676244935831955 | 0.865295879288147 | -6.31641638711327 |
| ENSG00000156603.17 | MED19 | -0.0666827309074947 | 2.38214528415439 | -0.420263842762347 | 0.676410061062285 | 0.865453365254244 | -6.02427712153905 |
| ENSG00000213139.7 | CRYGS | -0.050665492742155 | 2.25644438060862 | -0.41996884990964 | 0.676623882136402 | 0.865641892153026 | -6.00215292674661 |
| ENSG00000173334.3 | TRIB1 | -0.143238938757998 | 5.72286525892425 | -0.419932542652021 | 0.67670247682821 | 0.865641892153026 | -6.31763749443376 |
| lnc-WASHC4-2 | lnc-WASHC4-2 | 0.0806787529250801 | 1.42534040721346 | 0.419848737615492 | 0.676710951439113 | 0.865641892153026 | -5.87683625202268 |
| ENSG00000090266.12 | NDUFB2 | 0.068532484390881 | 3.51239321779251 | 0.419828486144816 | 0.676725632154416 | 0.865641892153026 | -6.15667704026447 |
| lnc-MID1-1 | lnc-MID1-1 | 0.120001647921601 | -0.539532262820706 | 0.419619779758374 | 0.676876935196727 | 0.865771897127279 | -5.4113803824429 |
| ENSG00000100744.14 | GSKIP | 0.0630713913758933 | 3.28448451715892 | 0.41957225563526 | 0.676911390004824 | 0.865771897127279 | -6.14407154537356 |
| ENSG00000161395.13 | PGAP3 | -0.0475841506257568 | 4.15831380882384 | -0.419487607251206 | 0.676972761495076 | 0.865796591888478 | -6.2361308399675 |
| lnc-KLHL18-8 | lnc-KLHL18-8 | 0.103776256114552 | 3.16276803338388 | 0.419400600214871 | 0.67703584536263 | 0.865823473445108 | -6.09674319197674 |
| ENSG00000095380.10 | NANS | 0.0707759934390508 | 2.32982412204177 | 0.419281413829388 | 0.677122264459884 | 0.86587945369912 | -6.01345592436599 |
| ENSG00000161526.14 | SAP30BP | 0.0398619916012253 | 5.94223667744596 | 0.419174683001414 | 0.677199656076881 | 0.86587945369912 | -6.32147484695419 |
| ENSG00000163251.3 | FZD5 | -0.216564001600744 | 0.00448718309719231 | -0.419173866713702 | 0.677205822554945 | 0.86587945369912 | -5.55214413799822 |
| ENSG00000173611.17 | SCAI | -0.0411998296379288 | 4.75979986653645 | -0.418996933146365 | 0.677328552151147 | 0.865913544609111 | -6.27620412329384 |
| ENSG00000198089.15 | SFI1 | -0.0608133404955611 | 5.9072318326475 | -0.418911821458315 | 0.677390274723755 | 0.865913544609111 | -6.32135737450504 |
| ENSG00000166261.10 | ZNF202 | 0.0546298715364273 | 4.10623264149799 | 0.418911319413035 | 0.67739063881115 | 0.865913544609111 | -6.2262660378446 |
| ENSG00000279500.1 | | -0.121074007426183 | 0.731747687346362 | -0.418889858363156 | 0.677406202615201 | 0.865913544609111 | -5.78581855781374 |
| ENSG00000226254.1 | PTP4A1P3 | 0.0995043614658906 | 1.6029997919236 | 0.418722539759824 | 0.677527548910695 | 0.865913544609111 | -5.92822918876953 |
| lnc-FZD1-3 | lnc-FZD1-3 | 0.0995043614658906 | 1.6029997919236 | 0.418722539759824 | 0.677527548910695 | 0.865913544609111 | -5.92822918876953 |
| ENSG00000006451.7 | RALA | -0.0698471843459392 | 3.15360710290676 | -0.41871556330118 | 0.677532608712105 | 0.865913544609111 | -6.12132958137368 |
| ENSG00000145451.12 | GLRA3 | 0.428338303043936 | -0.20463378137104 | 0.418677519384397 | 0.67763710171696 | 0.865913544609111 | -5.64669450938575 |
| ENSG00000240230.5 | COX19 | 0.0438909858370273 | 4.61922246098067 | 0.418555087272348 | 0.677649000980978 | 0.865913544609111 | -6.27076900929799 |
| ENSG00000164167.11 | LSM6 | 0.0402961769548771 | 4.0359808949954 | 0.418541561070958 | 0.677658811815191 | 0.865913544609111 | -6.22139250544981 |
| lnc-MYLK4-22 | lnc-MYLK4-22 | -0.158755194448377 | -0.802885946395569 | -0.418491327135248 | 0.677695248025466 | 0.865913544609111 | -5.36666019796319 |
| ENSG00000171970.12 | ZNF57 | -0.0969160375873535 | 1.51504427269837 | -0.418412220795363 | 0.677752627856606 | 0.865933106093887 | -5.90736324533791 |
| lnc-CXorf36-46 | lnc-CXorf36-46 | -0.107382719841144 | 4.23715567094043 | -0.41811445102695 | 0.6779782713843 | 0.866067849576596 | -6.23746093211632 |

| | | | | | | | |
|---|---|---|---|---|---|---|---|
| ENSG00000174501.14 | ANKRD36C | 0.0688695369714676 | 6.19469748198839 | 0.418100207892358 | 0.67797896547704 | 0.866067849576596 | -6.32369669204681 |
| ENSG00000160051.11 | IQCC | -0.117331255486168 | 0.0539029305937567 | -0.418092826919638 | 0.677984320081023 | 0.866067849576596 | -5.59227531973902 |
| lnc-C15orf40-1 | lnc-C15orf40-1 | -0.0815493999783647 | 0.0664692921664612 | -0.417984601221922 | 0.678062835481223 | 0.866076253017517 | -5.62795317162096 |
| ENSG00000153936.16 | HS2ST1 | -0.0615954344114087 | 5.35360217050123 | -0.417911427194302 | 0.678115923705341 | 0.866076253017517 | -6.30805189552026 |
| lnc-COX19-4 | lnc-COX19-4 | -0.151534394898758 | 1.41145555085721 | -0.417893750497338 | 0.678138655846159 | 0.866076253017517 | -5.91725052555394 |
| ENSG00000214413.7 | BBIP1 | -0.0692823899770101 | 4.268307074237 | -0.417797664247514 | 0.678198462735692 | 0.866076253017517 | -6.23557460548898 |
| lnc-IL23A-6 | lnc-IL23A-6 | -0.090710462807631 | 0.0938358028583131 | -0.417793774640685 | 0.678201284853342 | 0.866076253017517 | -5.61588715924986 |
| ENSG00000283023.1 | FRG1GP | 0.176459215697859 | -0.839771858708753 | 0.417708580067934 | 0.67826309924192 | 0.866101456150905 | -5.44458839540559 |
| lnc-ZNF816-2 | lnc-ZNF816-2 | -0.0966701639353212 | 0.138380884400874 | -0.417468678259889 | 0.67843717617405 | 0.8662010680805 | -5.63415274754041 |
| ENSG00000102172.15 | SMS | -0.0522305680259754 | 4.38761131241851 | -0.417457468473836 | 0.678445310624823 | 0.8662010680805 | -6.25083454876534 |
| lnc-SLC35A5-1 | lnc-SLC35A5-1 | -0.105111732175194 | 1.18773030369408 | -0.417427086747142 | 0.678467357508323 | 0.8662010680805 | -5.86443797397548 |
| ENSG00000105618.13 | PRPF31 | -0.0411077135750189 | 5.16197830722248 | -0.417347271873956 | 0.678525277536442 | 0.866216532461916 | -6.30099728841418 |
| ENSG00000140320.11 | BAHD1 | 0.0426279681026669 | 4.47122707792528 | 0.417294411368911 | 0.678563638411112 | 0.866216532461916 | -6.25812500273254 |
| ENSG00000164236.11 | ANKRD33B | 0.367920716235049 | -1.48243598909144 | 0.417246148390897 | 0.678618501607333 | 0.866223314523987 | -5.23366115568555 |
| ENSG00000085788.13 | DDHD2 | -0.0479305318203904 | 6.07248763424791 | -0.417171112108351 | 0.678653120051198 | 0.866223314523987 | -6.32331837008874 |
| ENSG00000089693.10 | MLF2 | 0.0579613830690251 | 4.82963273170941 | 0.417042991576338 | 0.678746105590255 | 0.866287423975169 | -6.28459211298857 |
| lnc-GPR83-1 | lnc-GPR83-1 | 0.103036134920781 | 0.998558548286216 | 0.41698592746918 | 0.678787522415174 | 0.866287423975169 | -5.82414309666534 |
| lnc-MTHFD1-3 | lnc-MTHFD1-3 | -0.123564664420517 | 2.10178029065317 | -0.416861418131334 | 0.678877894116326 | 0.866314540458954 | -5.99179832367875 |
| ENSG00000283551.1 | SNORD98 | -0.0919229502759227 | 0.126913996322769 | -0.416840679070447 | 0.678892947461639 | 0.866314540458954 | -5.64041521690789 |
| ENSG00000181027.10 | FKRP | -0.0548642979435483 | 3.46387398932627 | -0.416709078648393 | 0.678988472071929 | 0.866376875936164 | -6.15883202474621 |
| ENSG00000198900.5 | TOP1 | 0.0752554297280801 | 6.11298264849775 | 0.416596782012125 | 0.679069988893431 | 0.866376875936164 | -6.32337347848629 |
| ENSG00000182150.15 | ERCC6L2 | 0.0431106368937471 | 5.59367072596024 | 0.416543695998769 | 0.67910852571154 | 0.866376875936164 | -6.31599222640949 |
| ENSG00000105072.8 | C19orf44 | 0.105514206638452 | 1.24225612794894 | 0.416529289885962 | 0.679118983714864 | 0.866376875936164 | -5.84814248642013 |
| lnc-MKNK2-3 | lnc-MKNK2-3 | -0.0659515413180973 | 1.75610040499102 | -0.416459261423024 | 0.679169821233305 | 0.866376875936164 | -5.93967134669129 |
| ENSG00000180543.4 | TSPYL5 | -0.844117662616781 | -0.748570541687618 | -0.416501852834333 | 0.67921579810483 | 0.866376875936164 | -5.12242607108902 |
| ENSG00000188997.7 | KCTD21 | -0.127727151278746 | 0.420206216436979 | -0.416324590497318 | 0.679267590523787 | 0.866376875936164 | -5.68177635239332 |
| ENSG00000142166.12 | IFNAR1 | -0.0597914142300615 | 6.18221918112766 | -0.416309519814602 | 0.679278531985513 | 0.866376875936164 | -6.32420526576016 |
| ENSG00000260914.3 | | 0.0361825746225701 | 3.91616636976706 | 0.416158296671352 | 0.679388325326896 | 0.866440445306222 | -6.20315973982973 |
| ENSG00000213073.4 | CHP1P2 | 0.0768028132585921 | 1.36109707746159 | 0.416116772376582 | 0.679418474663735 | 0.866440445306222 | -5.88429171968074 |
| ENSG00000267855.5 | NDUFA7 | -0.0513125443519518 | 2.5046538950573 | -0.415901798429581 | 0.679574568211553 | 0.866440445306222 | -6.04139060795144 |
| ENSG00000152904.11 | GGPS1 | -0.0493747689340176 | 3.50289823589944 | -0.415874750395415 | 0.679594208918332 | 0.866440445306222 | -6.15095334118255 |
| lnc-IL2RG-1 | lnc-IL2RG-1 | 0.063030962794643 | 1.0329603677351 | 0.415862104878702 | 0.679603391433187 | 0.866440445306222 | -5.83348166862619 |
| ENSG00000167565.12 | SERTAD3 | 0.0971759659126121 | 3.05965423312805 | 0.415825998230567 | 0.679629610469267 | 0.866440445306222 | -6.10583448977732 |
| lnc-TBCA-10 | lnc-TBCA-10 | 0.0407067586216563 | 2.39464867206913 | 0.415820409963103 | 0.6796336684555 | 0.866440445306222 | -6.02172801559739 |
| lnc-CFAP44-2 | lnc-CFAP44-2 | -0.0803994919426749 | -0.360269069291397 | -0.415777080650318 | 0.679665132872481 | 0.866440445306222 | -5.48803130667924 |
| lnc-TATDN1-4 | lnc-TATDN1-4 | -0.231605508748804 | 2.86808176246227 | -0.415597710976453 | 0.679865838946318 | 0.866642631017035 | -6.08376205836795 |
| lnc-BCL2L15-2 | lnc-BCL2L15-2 | -0.0641848214654395 | 1.14718684803853 | -0.415405415096534 | 0.679935048720375 | 0.866677180320255 | -5.84396538141374 |
| ENSG00000135821.18 | GLUL | -0.620614696792018 | 5.03522847390239 | -0.415342626200945 | 0.680063618190536 | 0.86678328118344 | -6.26276988313758 |
| ENSG00000197134.11 | ZNF257 | 0.199742617612714 | 2.79286984322027 | 0.415251248991852 | 0.680102511349782 | 0.86678328118344 | -6.03454720272702 |
| ENSG00000166529.14 | ZSCAN21 | -0.0808044863313011 | 2.0390107297278 | -0.415101378434919 | 0.680155881817709 | 0.866797629586645 | -5.97926486769131 |
| ENSG00000213654.9 | GPSM3 | -0.0718434193403263 | 5.64572885112459 | -0.414869416220283 | 0.680324383681432 | 0.866883766722079 | -6.31881303159691 |
| ENSG00000114446.4 | IFT57 | 0.0675625518372946 | 7.21248943514851 | 0.414854225827539 | 0.68033541885494 | 0.866883766722079 | -6.31846371997335 |
| ENSG00000151665.12 | PIGF | -0.0660140413970955 | 2.93232080649275 | -0.414834400667098 | 0.680349821096462 | 0.866883766722079 | -6.09597566376157 |
| lnc-ZNF841-1 | lnc-ZNF841-1 | 0.0952521021491171 | -0.0283642967549678 | 0.414762581644735 | 0.680401995955348 | 0.86689658203336 | -5.58809738746696 |
| ENSG00000204531.17 | POU5F1 | 0.241637224465568 | -0.37731186974222 | 0.414486149424137 | 0.680604958183505 | 0.867078478152924 | -5.42612070693037 |
| ENSG00000177946.6 | CENPBD1P | 0.0823018712642531 | 0.872699229792874 | 0.414450116836487 | 0.680629012851227 | 0.867078478152924 | -5.78642228798576 |
| ENSG00000204165.5 | CXorf65 | -0.0603550589371391 | 1.15227787546957 | -0.414352273103831 | 0.680700105996937 | 0.867115378248233 | -5.85091776215384 |
| ENSG00000187097.12 | ENTPD5 | -0.0530673303537221 | 3.17913647578741 | -0.413938828030733 | 0.681000547153381 | 0.867444412529119 | -6.11438729178417 |
| LINC02145 | LINC02145 | -0.134394748222532 | -0.284320318456014 | -0.413836025543375 | 0.681075259528413 | 0.86748589522731 | -5.53079324566655 |
| lnc-GEMIN4-1 | lnc-GEMIN4-1 | -0.14201358052513 | -0.736017503739029 | -0.413720566404088 | 0.681159174070645 | 0.867487544958291 | -5.37720239661997 |
| HCG27 | HCG27 | -0.108317936004845 | 2.84941715541811 | -0.413617950242631 | 0.681233757895511 | 0.867487544958291 | -6.06198674625675 |
| lnc-FAM122A-1 | lnc-FAM122A-1 | 0.159871785433497 | -0.313465540051058 | 0.413591934499578 | 0.681252667258523 | 0.867487544958291 | -5.45806501798876 |
| ENSG00000145734.18 | BDP1 | 0.0347880746130551 | 7.69922774640314 | 0.413550809905409 | 0.681282558808613 | 0.867487544958291 | -6.31147494772549 |
| ENSG00000138434.16 | ITPRID2 | 0.0571257102131755 | 5.36683000951418 | 0.4135443092392 | 0.681287283887226 | 0.867487544958291 | -6.31125271971361 |
| ENSG00000164111.14 | ANXA5 | -0.0784605300646755 | 5.51777679618672 | -0.41329198732275 | 0.681470696769458 | 0.86763396948 | -6.31451279880452 |
| LINC00899 | LINC00899 | 0.290462482021627 | 0.637301480917893 | 0.413332591789739 | 0.68150374583788 | 0.86763396948 | -5.79921506740742 |
| lnc-PKD1L1-1 | lnc-PKD1L1-1 | -0.0683257974786273 | 3.0440195235308 | -0.413170477152798 | 0.681559029507893 | 0.86763396948 | -6.09594219376608 |
| ENSG00000212534.1 | SNORD70 | -0.086931405533283 | 1.22377879984254 | -0.41314861140923 | 0.68157492545581 | 0.86763396948 | -5.87267708964272 |
| ENSG00000267640.6 | | -0.390663217837949 | -2.26618470791634 | -0.413085770577918 | 0.681635498579739 | 0.86763396948 | -5.15741296196408 |
| ENSG00000159882.12 | ZNF230 | 0.0842204917352372 | 2.73340601772081 | 0.413022645258268 | 0.681666503119367 | 0.86763396948 | -6.05914822519644 |
| ENSG00000112339.14 | HBS1L | 0.0441845505756371 | 5.87187207472153 | 0.412950542621475 | 0.681718924077819 | 0.86763396948 | -6.32344312785769 |
| ENSG00000186951.16 | PPARA | 0.0591395823798782 | 3.93026622223921 | 0.412901184983471 | 0.681754809600972 | 0.86763396948 | -6.21152629266181 |
| ENSG00000171314.8 | PGAM1 | 0.0674461822918957 | 4.55253995931247 | 0.412864260806694 | 0.681781655850436 | 0.86763396948 | -6.26611614666818 |
| ENSG00000282034.1 | | 0.065830036980393 | 5.09731106057536 | 0.412733844132341 | 0.68187648049303 | 0.867700995238827 | -6.29632726036655 |
| NUTM2A-AS1 | NUTM2A-AS1 | -0.0769390903242639 | 5.09757907687756 | -0.412427202030858 | 0.682099457356851 | 0.867931078806631 | -6.30611941784846 |
| ENSG00000171456.18 | ASXL1 | 0.0699904385000698 | 7.64223017288138 | 0.412296222779729 | 0.682194708562864 | 0.867998620655835 | -6.31367376596976 |
| lnc-MED4-3 | lnc-MED4-3 | -0.127823202022562 | -0.548043449249302 | -0.412093808325218 | 0.682341919450011 | 0.86808443154784 | -5.46098328772782 |
| lnc-IYD-2 | lnc-IYD-2 | 0.143473942265457 | 3.16831311237884 | 0.4120023867351 | 0.682437921812429 | 0.86808443154784 | -6.10478064090118 |

| | | | | | | | |
|---|---|---|---|---|---|---|---|
| ENSG00000105321.13 | CCDC9 | 0.0669609356842122 | 3.5263543363637 | 0.411961084010845 | 0.682438453265738 | 0.86808443154784 | -6.15830541110757 |
| lnc-RAB3B-1 | lnc-RAB3B-1 | 0.0854344253161867 | 1.48214861137547 | 0.411908106101105 | 0.682476986968062 | 0.86808443154784 | -5.89075305836021 |
| ENSG00000111203.12 | ITFG2 | -0.0428865128031859 | 5.71125062821085 | -0.41186664029961 | 0.682507147888796 | 0.86808443154784 | -6.32083671189982 |
| lnc-TMTC3-5 | lnc-TMTC3-5 | 0.083380899123251 | 3.48410255966699 | 0.411855570619169 | 0.682515199715236 | 0.86808443154784 | -6.16323579170428 |
| lnc-PATL2-8 | lnc-PATL2-8 | -0.158277076259109 | -0.64265882935522 | -0.411603122742574 | 0.682698834548325 | 0.868169815432865 | -5.5064765687904 |
| ENSG00000179331.2 | RAB39A | -0.113907434406472 | 1.2960011713239 | -0.411588378323717 | 0.682709560488067 | 0.868169815432865 | -5.87149875184395 |
| ENSG00000156697.12 | UTP14A | -0.0452141279964664 | 4.17850573042226 | -0.411579115161775 | 0.682716299079681 | 0.868169815432865 | -6.23204173750722 |
| lnc-CHMP7-1 | lnc-CHMP7-1 | -0.0544077750649592 | 3.46104594398738 | -0.411489685548955 | 0.682781357006953 | 0.868169815432865 | -6.15577416902356 |
| ENSG00000141380.13 | SS18 | -0.0435378532444757 | 6.20516826222022 | -0.411473370515002 | 0.68279322607138 | 0.868169815432865 | -6.32649951484379 |
| ENSG00000150403.17 | TMCO3 | -0.0468193709971683 | 4.0604692180981 | -0.411318068296046 | 0.682906211333429 | 0.868259839883623 | -6.2238981633225 |
| lnc-PI4K2B-2 | lnc-PI4K2B-2 | 0.0791198213811601 | 1.18296818506638 | 0.410912416364114 | 0.683201365387724 | 0.868385321311715 | -5.87657527027707 |
| lnc-FRMD4A-5 | lnc-FRMD4A-5 | 0.0763645201005885 | 1.49706961568939 | 0.410890271231602 | 0.683217479723714 | 0.868385321311715 | -5.88791165615805 |
| ENSG00000125977.6 | EIF2S2 | 0.04440556057083 | 5.19176052717225 | 0.410788237405449 | 0.683291728550991 | 0.868385321311715 | -6.30518631097577 |
| lnc-FANCD2-1 | lnc-FANCD2-1 | -0.0456181622194451 | 1.20891497385904 | -0.410782252346267 | 0.68329608390739 | 0.868385321311715 | -5.86059986127669 |
| ENSG00000134627.11 | PIWIL4 | 0.10333840804757 | 1.20536989346139 | 0.410627567555929 | 0.683408652561158 | 0.868385321311715 | -5.85903496080426 |
| ENSG00000164953.15 | TMEM67 | -0.092485896537426 | 2.566410260052 | -0.410588656413169 | 0.683436970487323 | 0.868385321311715 | -6.05351533361689 |
| ENSG00000101347.9 | SAMHD1 | -0.166369902149913 | 3.32652639588627 | -0.410641611434938 | 0.683446452955949 | 0.868385321311715 | -6.11807409387434 |
| ENSG00000073910.21 | FRY | 0.193127341593886 | 7.36523758694027 | 0.410667374349219 | 0.683454938986266 | 0.868385321311715 | -6.31445740132821 |
| ENSG00000125652.7 | ALKBH7 | 0.0914808639753137 | 2.29995847570695 | 0.410484267774283 | 0.683512942515224 | 0.868385321311715 | -6.02722424420883 |
| ENSG00000167232.13 | ZNF91 | -0.0585528143292642 | 7.17337050900163 | -0.410441911352949 | 0.683543769644213 | 0.868385321311715 | -6.3225256680419 |
| lnc-NPFFR1-1 | lnc-NPFFR1-1 | 0.0875066554318567 | 1.14001473796833 | 0.410424611409666 | 0.683556360751366 | 0.868385321311715 | -5.86046064660316 |
| LINC01361 | LINC01361 | -0.18298186348019 | 0.522877745893505 | -0.410419953316017 | 0.683567422969501 | 0.868385321311715 | -5.70833142907516 |
| ENSG00000102786.14 | INTS6 | 0.07106680707835 | 6.12488779472574 | 0.410271185549414 | 0.683668029929046 | 0.868385321311715 | -6.32707083211798 |
| ENSG00000134057.14 | CCNB1 | 0.0922131751257087 | 0.982220994603588 | 0.410251566619382 | 0.68368230984949 | 0.868385321311715 | -5.83807368669089 |
| lnc-EIF2S3-1 | lnc-EIF2S3-1 | 0.0876221466689797 | 3.27089567079352 | 0.410213702520786 | 0.683709870109628 | 0.868385321311715 | -6.15019963471625 |
| ENSG00000136521.12 | NDUFB5 | -0.0638018613159792 | 4.39927719091826 | -0.410166467425704 | 0.683744251878158 | 0.868385321311715 | -6.25243960469136 |
| ENSG00000143337.18 | TOR1AIP1 | 0.034612338393873 | 5.96978777010196 | 0.410111637788302 | 0.683784162460048 | 0.868385321311715 | -6.32557118600007 |
| ENSG00000160888.6 | IER2 | -0.128423135630679 | 3.53623361446351 | -0.410032439297261 | 0.683867802930535 | 0.868385321311715 | -6.18229791484617 |
| ENSG00000240891.7 | PLCXD2 | 0.302982269268516 | 1.91914838728417 | 0.410102749872374 | 0.68386879671658 | 0.868385321311715 | -5.97377792852308 |
| ENSG00000279481.1 | | 0.179352734789198 | 0.398893226131684 | 0.409993850053071 | 0.683873940387076 | 0.868385321311715 | -5.68867061066959 |
| lnc-ZHX1-6 | lnc-ZHX1-6 | 0.0739364220933256 | 0.638479688501397 | 0.409943784314496 | 0.68390634896304 | 0.868385321311715 | -5.75456072964155 |
| ENSG00000147475.15 | ERLIN2 | 0.040003970205766 | 4.67305507289198 | 0.409907073641017 | 0.683933073108109 | 0.868385321311715 | -6.27556639061458 |
| ENSG00000148300.11 | REXO4 | -0.0413068559530121 | 4.23318605602115 | -0.409557810993471 | 0.684187345159869 | 0.868636446668491 | -6.23940846831856 |
| ENSG00000167751.12 | KLK2 | 0.178700399183548 | 2.4145320701932 | 0.409576632167219 | 0.684222110940938 | 0.868636446668491 | -6.05650340764521 |
| lnc-MTURN-3 | lnc-MTURN-3 | -0.0915527674761727 | 1.0203434148928 | -0.409461508566963 | 0.684257462288438 | 0.868636446668491 | -5.85097368167616 |
| ENSG00000176715.16 | ACSF3 | -0.0602835771983017 | 6.11086569823868 | -0.40934130557536 | 0.684344985207623 | 0.868662774334479 | -6.32684834544429 |
| ENSG00000140259.6 | MFAP1 | 0.0703996656867183 | 4.0977421664041 | 0.409317104151155 | 0.684362607423986 | 0.868662774334479 | -6.2141166927936 |
| ENSG00000283411.1 | | -0.0512244185725435 | 1.73364592858697 | -0.409137551385486 | 0.684493353952091 | 0.868734362751014 | -5.95037203659342 |
| lnc-USP50-5 | lnc-USP50-5 | -0.0383151218967077 | 3.23370962827655 | -0.409076858433674 | 0.684537551491217 | 0.868734362751014 | -6.12845339753611 |
| lnc-ERGIC2-8 | lnc-ERGIC2-8 | -0.0993264823465277 | 0.577624541744509 | -0.409065769887558 | 0.684545626461579 | 0.868734362751014 | -5.72293774317504 |
| lnc-ZNF713-3 | lnc-ZNF713-3 | -0.614902629981547 | -1.85977958742211 | -0.408893326620375 | 0.684718995070733 | 0.868823141879904 | -5.04132227475202 |
| lnc-PAPD5-2 | lnc-PAPD5-2 | -0.0784265882248487 | 0.893344506748288 | -0.408796783576256 | 0.684741520792597 | 0.868823141879904 | -5.81495412172806 |
| ENSG00000131508.15 | UBE2D2 | -0.0391062133345148 | 6.32466173367154 | -0.408795830651651 | 0.684742214816878 | 0.868823141879904 | -6.32771356705416 |
| lnc-ZSWIM7-4 | lnc-ZSWIM7-4 | 0.101071272857698 | 1.79690890207466 | 0.408639553465685 | 0.68485603674395 | 0.868913998909001 | -5.9447980571373 |
| ENSG00000158470.5 | B4GALT5 | 0.117021478883553 | 2.80767310143154 | 0.40854320324139 | 0.684931550388287 | 0.868956243937507 | -6.0922867534503 |
| lnc-ZSWIM7-6 | lnc-ZSWIM7-6 | 0.0600108944740319 | 2.24873729522715 | 0.408362966575543 | 0.685057502435762 | 0.869062469808031 | -6.01030910905515 |
| lnc-SLC15A1-1 | lnc-SLC15A1-1 | 0.100565058349833 | 2.30233710701048 | 0.40797764172746 | 0.685338211450749 | 0.86916458153804 | -6.00331952177719 |
| ENSG00000151164.18 | RAD9B | -0.0775596296696216 | 0.514569683062036 | -0.407969894650618 | 0.685343855655 | 0.86916458153804 | -5.72267831629555 |
| ENSG00000157315.4 | TMED6 | -0.0939224023469512 | -0.10784421663728 | -0.407922413359303 | 0.685378448984824 | 0.86916458153804 | -5.58522559874855 |
| ENSG00000106991.13 | ENG | 0.259663501916379 | -0.735943325536081 | 0.40790204623296 | 0.685404156783355 | 0.86916458153804 | -5.43195387360105 |
| lnc-MTPN-1 | lnc-MTPN-1 | 0.0935744796469584 | 2.82713796099934 | 0.407827369117723 | 0.685447697190781 | 0.86916458153804 | -6.08441089928631 |
| lnc-SENP6-3 | lnc-SENP6-3 | -0.0833502280468585 | 1.09775880767013 | -0.407737250960423 | 0.685513358832638 | 0.86916458153804 | -5.83572711012767 |
| lnc-CDCA5-2 | lnc-CDCA5-2 | -0.124145650358161 | 0.196288893647114 | -0.407718260716899 | 0.685527195765025 | 0.86916458153804 | -5.6818971383752 |
| lnc-DMGDH-2 | lnc-DMGDH-2 | -0.0718671232402373 | 0.232114153614556 | -0.407697670615314 | 0.685542198531412 | 0.86916458153804 | -5.65033523735993 |
| ENSG00000128891.15 | CCDC32 | 0.0901894657111457 | 6.05481341724206 | 0.407701725953096 | 0.685543255383177 | 0.86916458153804 | -6.32708126411845 |
| ENSG00000105248.15 | YJU2 | -0.0527053825066224 | 4.39990028565684 | -0.407672404397029 | 0.685560608677739 | 0.86916458153804 | -6.25409718346372 |
| ENSG00000184992.10 | BRI3BP | -0.113258354681424 | 4.46243889717338 | -0.407542516211113 | 0.685675329757057 | 0.86916458153804 | -6.25804320743952 |
| ENSG00000174886.12 | NDUFA11 | -0.0421554276477893 | 3.98558558780068 | -0.407478950248159 | 0.685701574809567 | 0.86916458153804 | -6.21808239306205 |
| lnc-GRTP1-3 | lnc-GRTP1-3 | 0.0452668630117978 | 1.66098056898675 | 0.40741967165841 | 0.685744772188852 | 0.86916458153804 | -5.92916806190736 |
| ENSG00000161980.5 | POLR3K | -0.0708592814772604 | 1.96669454126333 | -0.407392328168679 | 0.6857646982423 | 0.86916458153804 | -5.97307831818166 |
| ENSG00000152219.4 | ARL14EP | -0.0521268897332527 | 5.09834520238598 | -0.407383126565294 | 0.685771403788455 | 0.86916458153804 | -6.30337344597034 |
| ENSG00000011295.15 | TTC19 | -0.0374287992730942 | 4.41665962754424 | -0.40697154505614 | 0.686071364545218 | 0.869491219532925 | -6.2627563191873 |
| ENSG00000022567.9 | SLC45A4 | -0.0659510076439678 | 2.76355288584227 | -0.40686869367297 | 0.68614633067388 | 0.869532688354911 | -6.07139910371004 |
| ENSG00000182173.12 | TSEN54 | -0.0880469624712688 | 2.23679346432937 | -0.406790749027685 | 0.686203144955345 | 0.869551150194906 | -6.0109649976795 |
| lnc-COL4A1-6 | lnc-COL4A1-6 | 0.197240181695881 | 0.65985194539856 | 0.406767214685818 | 0.686250324389013 | 0.869557401311195 | -5.80384368459411 |
| ENSG00000137804.12 | NUSAP1 | 0.12932221375634 | 1.12856823344325 | 0.406569922988111 | 0.686364116244374 | 0.86956199264059 | -5.83923068720863 |
| LRRC75A-AS1 | LRRC75A-AS1 | -0.0744324586917573 | 6.80281809451109 | -0.406569458590513 | 0.686364454782843 | 0.86956199264059 | -6.32565264492871 |
| ENSG00000282980.1 | | 0.115641830128237 | -0.200237002789746 | 0.406525236897227 | 0.686396691993028 | 0.86956199264059 | -5.51464440897039 |

| | | | | | | | |
|---|---|---|---|---|---|---|---|
| ENSG00000131845.14 | ZNF304 | 0.0428706738269393 | 3.84184524801448 | 0.406489239384871 | 0.686422934286756 | 0.86956199264059 | -6.19614971494113 |
| ENSG00000102100.15 | SLC35A2 | 0.0628643171629464 | 3.75383247483098 | 0.406414736737875 | 0.686477248181955 | 0.869565218189426 | -6.19344781878928 |
| lnc-COL28A1-3 | lnc-COL28A1-3 | 0.0827711918401535 | 0.176765574683847 | 0.406295119061007 | 0.686564455328537 | 0.869565218189426 | -5.67117767982438 |
| ENSG00000100353.17 | EIF3D | 0.0487658788205758 | 7.48052990308544 | 0.406213347577473 | 0.686624073232894 | 0.869565218189426 | -6.31817452587796 |
| lnc-KIAA0100-1 | lnc-KIAA0100-1 | -0.0709158967875337 | 0.835627381504772 | -0.406149483041027 | 0.68667063695526 | 0.869565218189426 | -5.79506410059011 |
| ENSG00000126749.15 | EMG1 | 0.0454143236390028 | 5.31267743447385 | 0.406063237391944 | 0.686733520732362 | 0.869565218189426 | -6.31269789139201 |
| lnc-ABHD17B-3 | lnc-ABHD17B-3 | -0.14560528055471 | 3.09035507661676 | -0.406081863031023 | 0.68675335093982 | 0.869565218189426 | -6.11241873697852 |
| MIR762HG | MIR762HG | 0.0670469331288537 | 0.9041535299964 | 0.405994177784076 | 0.686783875372312 | 0.869565218189426 | -5.81260261411125 |
| ENSG00000139318.7 | DUSP6 | -0.217379983502221 | 1.86617122578622 | -0.406058812376684 | 0.68679501989781 | 0.869565218189426 | -6.01877596237197 |
| ENSG00000119596.17 | YLPM1 | -0.05365222553817 | 6.65228984100559 | -0.405964244759689 | 0.686805701409197 | 0.869565218189426 | -6.32820593193874 |
| lnc-LRGUK-6 | lnc-LRGUK-6 | -0.120145469797252 | 1.32100431360419 | -0.405782403985563 | 0.686938298681087 | 0.86967960399513 | -5.87920964738193 |
| LINC00944 | LINC00944 | 0.613063122589061 | -2.46878621456282 | 0.405713270889796 | 0.687010236424449 | 0.869691418326759 | -4.92027546232604 |
| lnc-DYNLT1-4 | lnc-DYNLT1-4 | 0.124250307905768 | 5.27213763497266 | 0.405708520467237 | 0.687032136325759 | 0.869691418326759 | -6.30948953954782 |
| ENSG00000047579.19 | DTNBP1 | -0.0611673817139167 | 3.94633976914287 | -0.405379910789999 | 0.687231829933318 | 0.869888270586675 | -6.21467195046168 |
| ENSG00000196914.8 | ARHGEF12 | -0.0591161807945726 | 6.06195855217428 | -0.405252105797835 | 0.687325046097493 | 0.869888270586675 | -6.32851756641691 |
| lnc-IYD-4 | lnc-IYD-4 | 0.129354323825982 | -0.317257034309003 | 0.405226175933072 | 0.687343958968313 | 0.869888270586675 | -5.49763163363052 |
| ENSG00000277462.1 | ZNF670 | -0.0782685931234774 | 1.27834978659039 | -0.405079766783976 | 0.687450751499803 | 0.869888270586675 | -5.87042131281243 |
| ENSG00000106624.10 | AEBP1 | 0.208025681719176 | 4.98563569475013 | 0.405125498580323 | 0.687491515825606 | 0.869888270586675 | -6.30086727810835 |
| MIR4453HG | MIR4453HG | 0.0877781238683988 | 3.05099386818905 | 0.404970868982846 | 0.687530187007098 | 0.869888270586675 | -6.11128534412607 |
| ENSG00000126705.14 | AHDC1 | -0.0857993937536534 | 3.48870356863033 | -0.404966179221514 | 0.687533608034002 | 0.869888270586675 | -6.1751409103948 |
| ENSG00000207067.1 | SNORA72 | -0.102162372864517 | 0.973684348524037 | -0.404801268369228 | 0.687653909297557 | 0.869888270586675 | -5.85300502671974 |
| ENSG00000149646.12 | CNBD2 | -0.0729791145269543 | 1.22527342015584 | -0.404760565678981 | 0.687683602873177 | 0.869888270586675 | -5.87116250408864 |
| ENSG00000184207.8 | PGP | -0.0733727135124336 | 2.610080253749 | -0.404747308367582 | 0.687693274503221 | 0.869888270586675 | -6.06089326172935 |
| ENSG00000142541.16 | RPL13A | -0.0641567368661811 | 10.5758206910099 | -0.404739064249875 | 0.687699288875807 | 0.869888270586675 | -6.22958332969325 |
| lnc-URGCP-MRPS24-3 | lnc-URGCP-MRPS2 | -0.0487191710428125 | 1.12248691642875 | -0.404727252951402 | 0.687707905667346 | 0.869888270586675 | -5.84686208313117 |
| ENSG00000076043.9 | REXO2 | 0.0886909712341031 | 2.87263880382127 | 0.404687295889496 | 0.687737056175337 | 0.869888270586675 | -6.07942838064941 |
| ENSG00000084207.16 | GSTP1 | 0.0860317393037899 | 5.71592854877956 | 0.40459611292375 | 0.687803641896834 | 0.869919034113466 | -6.32402734974329 |
| ENSG00000177599.12 | ZNF491 | -0.107894807531773 | 0.163473318762662 | -0.404388156166783 | 0.687955307578378 | 0.87005739452447 | -5.64042968608699 |
| lnc-MAFG-1 | lnc-MAFG-1 | 0.0645704650247434 | -0.0759187874335722 | 0.40368037216477 | 0.688471811738799 | 0.870442216529753 | -5.59249608356536 |
| ENSG00000125445.10 | MRPS7 | 0.0495128868827592 | 4.54956336568058 | 0.403629250553037 | 0.68850912347721 | 0.870442216529753 | -6.26545304157372 |
| lnc-THSD1-1 | lnc-THSD1-1 | -0.071189380010733 | 1.75532326665579 | -0.403585304565587 | 0.688541198625263 | 0.870442216529753 | -5.961543533969 |
| LINC02161 | LINC02161 | 0.730694503360265 | -2.84349149012302 | 0.403612936897599 | 0.688545943798107 | 0.870442216529753 | -4.854338001056 |
| ENSG00000126947.12 | ARMCX1 | 0.314960831454094 | -1.04156598862876 | 0.403457908908343 | 0.688646615456899 | 0.870442216529753 | -5.28508465163258 |
| ENSG00000167699.13 | GLOD4 | -0.0390083644876285 | 4.93773182393971 | -0.403436113035411 | 0.688650094359472 | 0.870442216529753 | -6.29726337246643 |
| ENSG00000075975.15 | MKRN2 | 0.0347650842373568 | 4.62286495579433 | 0.403429067034356 | 0.688655237440426 | 0.870442216529753 | -6.27579016780063 |
| ENSG00000080608.9 | PUM3 | -0.0354689987463921 | 4.87629176151962 | -0.403407945731973 | 0.688670654582216 | 0.870442216529753 | -6.29237442034026 |
| ENSG00000267554.1 | | 0.155574008450892 | -0.283246782191118 | 0.403346942387149 | 0.688715183706622 | 0.870442216529753 | -5.49352847357413 |
| lnc-SLFN13-2 | lnc-SLFN13-2 | 0.155574008450892 | -0.283246782191118 | 0.403346942387149 | 0.688715183706622 | 0.870442216529753 | -5.49352847357413 |
| lnc-RNF111-3 | lnc-RNF111-3 | 0.0615004291489695 | 0.938470530426184 | 0.403290647714683 | 0.688756276747425 | 0.870442216529753 | -5.83207458301287 |
| lnc-RAP2C-4 | lnc-RAP2C-4 | -0.05729573439256 | 5.1199055938676 | -0.403275876629817 | 0.688767059253757 | 0.870442216529753 | -6.30721781947736 |
| ENSG00000196123.12 | MATCAP1 | 0.0952130715609452 | 0.750453993088323 | 0.403130247515998 | 0.688873368210392 | 0.870523117502118 | -5.77859758893022 |
| ENSG00000141034.9 | GID4 | -0.060561236771103 | 2.56925392289186 | -0.403025198051557 | 0.688950058056671 | 0.870525410324409 | -6.0548534210873 |
| ENSG00000110315.6 | RNF141 | 0.0501639891759558 | 5.6576831431723 | 0.403011895771758 | 0.688959769430337 | 0.870525410324409 | -6.32181857655031 |
| ENSG00000170779.10 | CDCA4 | -0.108233895111816 | 1.10088082462807 | -0.402785192269618 | 0.689125283237299 | 0.870681094154645 | -5.88043730135487 |
| ENSG00000258892.1 | | -0.0538991257942892 | 0.717051988370125 | -0.402691327816942 | 0.689193817184002 | 0.870714236379715 | -5.76334663328767 |
| ENSG00000234769.7 | WASH4P | -0.0785376426828764 | 1.0031116656209 | -0.402467737785251 | 0.689357079265588 | 0.870821933562507 | -5.82661728610603 |
| ENSG00000100897.17 | DCAF11 | -0.0505461440731061 | 5.91519226990318 | -0.402312362299363 | 0.689470540933553 | 0.870821933562507 | -6.32772064886586 |
| ENSG00000160271.15 | RALGDS | -0.0535024842972138 | 5.11116401852125 | -0.402296921315754 | 0.689481816979671 | 0.870821933562507 | -6.30850309995644 |
| ENSG00000111640.14 | GAPDH | -0.0837464387834288 | 7.79770946356389 | -0.402292634923553 | 0.689484947204593 | 0.870821933562507 | -6.31304035007643 |
| ENSG00000152256.13 | PDK1 | 0.0474550627299477 | 4.9922909338724 | 0.402284892159955 | 0.689490601528843 | 0.870821933562507 | -6.30154016347034 |
| lnc-SMARCC2-1 | lnc-SMARCC2-1 | 0.053066242839531 | 2.10161168885769 | 0.402226633714906 | 0.689533146621873 | 0.87082223321377 | -5.9979572652644 |
| lnc-RPL32-1 | lnc-RPL32-1 | -0.090804509266317 | 0.155579466363508 | -0.402079418088477 | 0.689640660071831 | 0.87088467165924 | -5.66861141940194 |
| ENSG00000154127.9 | UBASH3B | -0.329974369756751 | 6.03554042155575 | -0.402108000008687 | 0.689699803856458 | 0.87088467165924 | -6.3288109761898 |
| ENSG00000103042.8 | SLC38A7 | -0.0495366598206846 | 2.50108676745032 | -0.40198102518184 | 0.689712521284783 | 0.87088467165924 | -6.04439663186536 |
| lnc-ZNF500-3 | lnc-ZNF500-3 | -0.0707120894398193 | 0.0394023377294394 | -0.401874111012089 | 0.68979060928375 | 0.87088467165924 | -5.60886769544596 |
| lnc-ST5-2 | lnc-ST5-2 | -0.0780051037948858 | 0.609680882584446 | -0.401869275816647 | 0.689794140895913 | 0.87088467165924 | -5.73385324396035 |
| ENSG00000189058.8 | APOD | -0.423521175952314 | 2.71311336189565 | -0.401714131691511 | 0.689987392084262 | 0.871031023000959 | -6.05466571911841 |
| lnc-TACSTD2-5 | lnc-TACSTD2-5 | -0.153207866507196 | 0.38694657410456 | -0.401533709018248 | 0.690039254962985 | 0.871031023000959 | -5.68956023596006 |
| ENSG00000276043.4 | UHRF1 | 0.203043073228094 | -0.438229100874663 | 0.401498210724308 | 0.690065186587874 | 0.871031023000959 | -5.42584316022234 |
| ENSG00000196776.15 | CD47 | 0.0417338310636071 | 7.54585496925804 | 0.401478846917253 | 0.690079332073878 | 0.871031023000959 | -6.31855892858682 |
| ENSG00000113583.7 | C5orf15 | -0.0529339692060296 | 4.12627393044666 | -0.4014141016302 | 0.690126630072466 | 0.871037308485502 | -6.23664620069545 |
| ENSG00000167969.12 | ECI1 | 0.0611829088940889 | 3.9775910116276 | 0.40126489774848 | 0.690235631890056 | 0.871115558046899 | -6.21458086428068 |
| ENSG00000145014.17 | TMEM44 | -0.0753958701183941 | 2.34595331969315 | -0.401121509699516 | 0.690340391185612 | 0.871115558046899 | -6.01726912509801 |
| ENSG00000172197.10 | MBOAT1 | -0.097559674552615 | 4.65323890433898 | -0.401098087479766 | 0.690365074857475 | 0.871115558046899 | -6.28283732900518 |
| lnc-LONP2-12 | lnc-LONP2-12 | -0.11590271943933 | 0.0228610987214179 | -0.401043205799972 | 0.690397602601394 | 0.871115558046899 | -5.57939928613563 |
| ENSG00000283496.1 | ZNF511-PRAP1 | 0.0569920060914484 | 1.61070969535426 | 0.401039598875472 | 0.690400237983728 | 0.871115558046899 | -5.92726058887804 |
| ENSG00000158062.20 | UBXN11 | 0.0465385589826806 | 4.12652919506611 | 0.40095065384465 | 0.690465226479732 | 0.871144155733255 | -6.23122717956899 |
| ENSG00000100055.20 | CYTH4 | -0.0690958420447814 | 6.25431727953205 | -0.400737464740827 | 0.690621004657996 | 0.871287290154798 | -6.33064678082045 |

| | | | | | | | |
|---|---|---|---|---|---|---|---|
| ENSG00000162482.4 | AKR7A3 | 0.0705492814432075 | 0.146848971471937 | 0.40067711728046 | 0.690665103266829 | 0.871289520749028 | -5.64174376197337 |
| ENSG00000173991.5 | TCAP | 0.0778084457102166 | 0.397272306366477 | 0.400473386559614 | 0.690813986516167 | 0.871423931143119 | -5.73456707531757 |
| ENSG00000200259.1 | SNORD35A | 0.0691669514782618 | 1.51207922222233 | 0.400242650730574 | 0.690982619637703 | 0.871563066155343 | -5.91788093860231 |
| ENSG00000178538.9 | CA8 | -0.451906467082829 | -0.619952553642472 | -0.400283615058701 | 0.691008972287278 | 0.871563066155343 | -5.31873125965822 |
| ENSG00000120647.9 | CCDC77 | -0.0597484697001542 | 3.54269447787787 | -0.399723205438722 | 0.691362313977915 | 0.871955300772514 | -6.15993532730767 |
| ENSG00000167863.11 | ATP5PD | 0.0775490135766637 | 4.3726141616707 | 0.399303809072699 | 0.691668935104014 | 0.872288566340661 | -6.2497946915437 |
| ENSG00000162825.16 | NBPF20 | -0.0558010085157852 | 2.66823086374837 | -0.399240573402375 | 0.691715171313464 | 0.872293430409572 | -6.05958052872795 |
| ENSG00000107890.16 | ANKRD26 | 0.0457015395838241 | 4.48705824796061 | 0.399132720157221 | 0.691794033431488 | 0.872322741332365 | -6.25573563072756 |
| ENSG00000087258.14 | GNAO1 | -0.412548532269546 | 2.12444286649097 | -0.399147515292716 | 0.691862576972246 | 0.872322741332365 | -5.92563277680029 |
| lnc-COL24A1-3 | lnc-COL24A1-3 | -0.0799161802291876 | 0.742310113485931 | -0.399034907834674 | 0.691865556636587 | 0.872322741332365 | -5.79066706125235 |
| lnc-CERK-2 | lnc-CERK-2 | 0.0963788199885833 | 2.16030457111999 | 0.39880370266089 | 0.692034631876647 | 0.872482471390239 | -5.98961473184711 |
| ENSG00000111845.4 | PAK1IP1 | -0.0688150644320578 | 2.41811830353867 | -0.398608901806164 | 0.692177097782117 | 0.872608636225229 | -6.03675130214601 |
| lnc-DENND4B-3 | lnc-DENND4B-3 | -0.0891326327216866 | 0.748329365367666 | -0.398388889907982 | 0.692338015129409 | 0.872729875581859 | -5.78475900330898 |
| lnc-TRPC5OS-5 | lnc-TRPC5OS-5 | -0.104054880620083 | 0.0771493282441996 | -0.39836147264823 | 0.692358069201582 | 0.872729875581859 | -5.6227484341871 |
| ENSG00000125124.11 | BBS2 | -0.052207825904568 | 5.61562770338281 | -0.39827473128366 | 0.692421516750073 | 0.872756404119214 | -6.32415897504503 |
| ENSG00000177700.5 | POLR2L | -0.0794669135573143 | 2.41614873236133 | -0.398121469713701 | 0.692533626408954 | 0.87284426136645 | -6.03877719149087 |
| ENSG00000156398.12 | SFXN2 | -0.0543942597473395 | 3.53588833871258 | -0.398032612605789 | 0.692598627894242 | 0.872872738057015 | -6.16658826850428 |
| lnc-TAS1R3-4 | lnc-TAS1R3-4 | 0.0995138715182417 | -0.130320577174097 | 0.397612813909792 | 0.692905754196688 | 0.873206339229194 | -5.55213987138605 |
| ENSG00000198218.10 | QRICH1 | -0.0448764175220262 | 6.95602758571427 | -0.397507368474928 | 0.692982906689848 | 0.873250102142595 | -6.32832146738582 |
| ENSG00000136231.13 | IGF2BP3 | -0.483989696471067 | 0.602628513964812 | -0.397537457956028 | 0.693039896040265 | 0.873268453027045 | -5.80571896978637 |
| ENSG00000240053.8 | LY6G5B | 0.0490362801534094 | 3.68979890251949 | 0.397356339866017 | 0.693093417300605 | 0.873282431947744 | -6.18506447589428 |
| lnc-HMGN1-1 | lnc-HMGN1-1 | -0.0500485507396853 | 5.40185721489671 | -0.39725875932112 | 0.693164822505784 | 0.873318941154224 | -6.31974173046674 |
| lnc-SVBP-1 | lnc-SVBP-1 | -0.098193610635208 | -0.171646171145609 | -0.397171687688874 | 0.693228540128159 | 0.87334576089227 | -5.48493335059584 |
| ENSG00000169087.10 | HSPBAP1 | 0.0587230913085044 | 4.47575676467265 | 0.397069868195521 | 0.69330305284864 | 0.87338617643574 | -6.26460763030048 |
| ENSG00000225138.7 | SLC9A3-AS1 | -0.120845596732473 | 2.1322689218035 | -0.397012876198251 | 0.693346496584752 | 0.873387450379679 | -6.00311671235421 |
| ENSG00000198034.10 | RPS4X | 0.0761787190574719 | 9.53463488153084 | 0.39688421379824 | 0.693438924859486 | 0.873450424722037 | -6.26980022578344 |
| ENSG00000156467.9 | UQCRB | 0.0677349131539415 | 6.05646889949861 | 0.396704265797307 | 0.693570630362099 | 0.87356286162912 | -6.33152832033874 |
| ENSG00000116918.13 | TSNAX | -0.0401227401435067 | 4.80685360666737 | -0.396565318988132 | 0.693672333296994 | 0.873570331578263 | -6.29126166823673 |
| ENSG00000182118.7 | FAM89A | 0.0823413682862962 | 0.0258300951272758 | 0.396513589077919 | 0.693710198769257 | 0.873570331578263 | -5.66910385853573 |
| ENSG00000100023.18 | PPIL2 | -0.0347959107215763 | 5.62504656379786 | -0.396475554293315 | 0.693738040132464 | 0.873570331578263 | -6.32465046667982 |
| ENSG00000274180.1 | NATD1 | 0.0842166137997298 | 3.56566596192525 | 0.396393303671533 | 0.693798248837233 | 0.873570331578263 | -6.16689104756194 |
| lnc-MPHOSPH9-1 | lnc-MPHOSPH9-1 | 0.0659494705090509 | -0.0550518699564085 | 0.39634322374303 | 0.693834909086273 | 0.873570331578263 | -5.58372026786023 |
| lnc-CLDN15-1 | lnc-CLDN15-1 | 0.053149208223797 | 1.9008731888438 | 0.396300915195841 | 0.693865880992227 | 0.873570331578263 | -5.97369341528601 |
| ENSG00000111780.8 | | 0.0650693012645722 | 3.48649357185595 | 0.396290301748736 | 0.693873650632707 | 0.873570331578263 | -6.16921812342036 |
| ENSG00000164077.14 | MON1A | -0.086832599448555 | 0.569338978187622 | -0.395801452319493 | 0.694231552026017 | 0.873961968365317 | -5.74999628982404 |
| CCDC183-AS1 | CCDC183-AS1 | -0.115466625275481 | 1.30307279118043 | -0.395697568617389 | 0.694307617515064 | 0.873961968365317 | -5.87459394287645 |
| ENSG00000168404.12 | MLKL | 0.115584613219827 | 2.52390791108356 | 0.395694193574745 | 0.694312107404638 | 0.873961968365317 | -6.04815757317211 |
| lnc-COMMD1-7 | lnc-COMMD1-7 | 0.216515800888597 | -1.05476876870307 | 0.395591862996163 | 0.694385020318655 | 0.873975302727261 | -5.28797151113223 |
| ENSG00000124608.4 | AARS2 | -0.0343891639071927 | 4.46397817076252 | -0.39556099657992 | 0.694407622834457 | 0.873975302727261 | -6.2643475538374 |
| ENSG00000169032.9 | MAP2K1 | 0.0532906280128875 | 4.77459570846894 | 0.395347853688406 | 0.694563708419562 | 0.874118300849883 | -6.28718649057187 |
| ENSG00000126561.16 | STAT5A | -0.0427758355247532 | 5.22813865596079 | -0.395075243643276 | 0.694763361641134 | 0.874289361679496 | -6.31534512776935 |
| ENSG00000181873.12 | IBA57 | 0.0669132917739601 | 3.36606188996412 | 0.39502317749886 | 0.694801496162178 | 0.874289361679496 | -6.14616428869895 |
| lnc-ZNF561-1 | lnc-ZNF561-1 | 0.0454237406530313 | 1.83761608787747 | 0.394988274919181 | 0.694827060115299 | 0.874289361679496 | -5.96555800519713 |
| ENSG00000256625.1 | | 0.0957870606490211 | 0.380185041823297 | 0.394776968837421 | 0.694981836256375 | 0.874376340314834 | -5.69379491886403 |
| lnc-LDLRAD2-3 | lnc-LDLRAD2-3 | -0.128954744094115 | 0.296135971527337 | -0.394774816886031 | 0.694983412571847 | 0.874376340314834 | -5.66436690747669 |
| ENSG00000260238.6 | PMF1-BGLAP | -0.0510002193375816 | 3.19119160194939 | -0.39471366028342 | 0.695028210668972 | 0.874376340314834 | -6.12643064926152 |
| lnc-CTNNBIP1-9 | lnc-CTNNBIP1-9 | -0.0642365982051867 | 2.03050655270401 | -0.394661926886157 | 0.695066106992728 | 0.874376340314834 | -5.98409641492214 |
| lnc-LDHAL6A-1 | lnc-LDHAL6A-1 | -0.0972558344480743 | 0.534034901094922 | -0.394598113370223 | 0.695112853466356 | 0.874381706465685 | -5.75181456871784 |
| ENSG00000018699.12 | TTC27 | 0.042708749502685 | 4.5611022284766 | 0.39426625562376 | 0.695355974597748 | 0.874571871906429 | -6.27220642772523 |
| ENSG00000274949.1 | NA | -0.0543603782215956 | 0.124803736517626 | -0.394192427904956 | 0.695410065674305 | 0.874571871906429 | -5.62404073470762 |
| lnc-MSTN-3 | lnc-MSTN-3 | 0.103515028979388 | 1.99332785386841 | 0.394117529046639 | 0.695464943179089 | 0.874571871906429 | -5.97122151201454 |
| lnc-ZNF169-7 | lnc-ZNF169-7 | 0.0485408821843247 | 6.45445322750901 | 0.394063911181724 | 0.695504229367158 | 0.874571871906429 | -6.33322019999778 |
| ENSG00000214465.3 | SMARCE1P6 | -0.118802084201808 | -0.133255434139227 | -0.394022900839917 | 0.695534278505414 | 0.874571871906429 | -5.54368368051857 |
| ENSG00000121210.15 | TMEM131L | 0.0323557598425275 | 8.06102783655254 | 0.394013206414752 | 0.695541381886585 | 0.874571871906429 | -6.31243335274806 |
| ENSG00000108239.8 | TBC1D12 | -0.0566540734923207 | 4.40747878971273 | -0.393915656877043 | 0.695612860748591 | 0.874571871906429 | -6.2695074155487 |
| ENSG00000254577.1 | | 0.0869064926908856 | 1.20302787446169 | 0.393818291695321 | 0.695684207317997 | 0.874571871906429 | -5.87684700883657 |
| ENSG00000169249.12 | ZRSR2 | -0.0706973075161317 | 4.2591179877589 | -0.393788808762065 | 0.695705812163707 | 0.874571871906429 | -6.25311181525891 |
| lnc-ALCAM-8 | lnc-ALCAM-8 | 0.356266547620144 | -1.50835692170535 | 0.393695223954677 | 0.695781806181714 | 0.874571871906429 | -5.14956370635212 |
| ENSG00000204946.9 | ZNF783 | -0.0326770764152409 | 4.77405904449522 | -0.393681593704116 | 0.695784380613516 | 0.874571871906429 | -6.2872957474654 |
| ENSG00000272104.1 | | 0.0564308273315681 | 1.05801416993336 | 0.393646327466905 | 0.695810224864287 | 0.874571871906429 | -5.84616852631751 |
| lnc-HECTD2-4 | lnc-HECTD2-4 | -0.0992254384303517 | 3.35534044196072 | -0.39363790027721 | 0.695816400638375 | 0.874571871906429 | -6.14201389167293 |
| ENSG00000084623.11 | EIF3I | 0.0546936164657641 | 5.75124817599824 | 0.393509016780444 | 0.695910854106705 | 0.874637180807126 | -6.32943503748794 |
| ENSG00000130383.7 | FUT5 | 0.101458998214201 | -0.00117522337477792 | 0.393429376320149 | 0.695969221800343 | 0.874657131048752 | -5.60166062958048 |
| LINC02390 | LINC02390 | 0.111669524243511 | 2.67773103518409 | 0.392999527148538 | 0.696284382405408 | 0.874976517847122 | -6.05133592436933 |
| ENSG00000156787.16 | TBC1D31 | 0.0675106784486323 | 2.86439809506703 | 0.392909168034282 | 0.696350523095042 | 0.874976517847122 | -6.07995375409907 |
| ENSG00000183598.3 | H3C13 | -0.102142442816583 | 2.67287945184699 | -0.392908669508471 | 0.696350888541695 | 0.874976517847122 | -6.09687474575032 |
| ENSG00000117481.10 | NSUN4 | 0.0389953002136059 | 4.77239543278363 | 0.392792797764654 | 0.696435830840087 | 0.875029831899738 | -6.28566460185167 |
| lnc-MYCL-3 | lnc-MYCL-3 | 0.0442375045403794 | 0.661951184645655 | 0.392732963965434 | 0.69647969484983 | 0.875031530189467 | -5.77634482640353 |

| | | | | | | | |
|---|---|---|---|---|---|---|---|
| ENSG00000255339.6 | | 0.0281397120947711 | 4.42951860261993 | 0.392570825589888 | 0.696598563374041 | 0.875127455439934 | -6.26118516625386 |
| ENSG00000178950.16 | GAK | 0.0631239492507154 | 6.00796244587185 | 0.392463796975575 | 0.696677033501388 | 0.875172620113461 | -6.33331343326302 |
| ENSG00000162980.16 | ARL5A | -0.0595886048419878 | 5.64065778178676 | -0.392211612830953 | 0.696861940542572 | 0.875335225644199 | -6.32879155556511 |
| ENSG00000157470.11 | FAM81A | -0.149729993532946 | 1.37493385354774 | -0.392173108845699 | 0.696900950830339 | 0.875335225644199 | -5.88968133600072 |
| ENSG00000134463.14 | ECHDC3 | -0.193817532495453 | -0.233371765747554 | -0.392092207051644 | 0.696954594109594 | 0.875335225644199 | -5.59829700647244 |
| ENSG00000147996.16 | ZNG1E | 0.0582477657160165 | 3.8406756434107 | 0.392055270967033 | 0.696976583252334 | 0.875335225644199 | -6.20158969257351 |
| lnc-KIF13B-4 | lnc-KIF13B-4 | -0.0622325280345669 | 0.0352865390373358 | -0.391955309426168 | 0.697049887029505 | 0.875354959881977 | -5.61861580557732 |
| ENSG00000285947.1 | | -0.103012337312387 | 5.58701929826684 | -0.391946128631374 | 0.6970773525446 | 0.875354959881977 | -6.32563186511583 |
| ENSG00000198553.8 | KCNRG | -0.142895525677124 | 0.430759333215758 | -0.391747406904504 | 0.697202355445677 | 0.875420714067204 | -5.72956952113869 |
| lnc-ITGAE-1 | lnc-ITGAE-1 | -0.0457742888313732 | 2.04893053546574 | -0.391727759613315 | 0.697216764734877 | 0.875420714067204 | -5.99646958003324 |
| ENSG00000169139.11 | UBE2V2 | -0.0661233600275883 | 4.03740389192447 | -0.391672478014375 | 0.697257308770235 | 0.875420714067204 | -6.2266124262046 |
| lnc-CNKSR3-6 | lnc-CNKSR3-6 | -0.117605123580126 | 0.19927986715383 | -0.391559619554013 | 0.697340082987164 | 0.875471236848112 | -5.63953793290049 |
| ENSG00000126267.9 | COX6B1 | 0.0613350327035596 | 4.91038419664929 | 0.391378543484843 | 0.697472898102768 | 0.875584573167191 | -6.29834050972 |
| ENSG00000128731.16 | HERC2 | -0.0747117530211767 | 6.22239602829229 | -0.391275471300961 | 0.697548503457294 | 0.875599199040458 | -6.33427380935596 |
| lnc-LONRF2-3 | lnc-LONRF2-3 | -0.112783553303528 | 2.01551546353614 | -0.391177619453358 | 0.697620282476034 | 0.875599199040458 | -5.9787241431029 |
| ENSG00000090273.13 | NUDC | -0.035412359719178 | 4.71661716835996 | -0.391121449143262 | 0.697661487355832 | 0.875599199040458 | -6.28735751427529 |
| ENSG00000252213.1 | SNORA74D | 0.131154311220203 | 0.903721470717812 | 0.391116222169112 | 0.69766532175658 | 0.875599199040458 | -5.78088751726403 |
| ENSG00000154814.13 | OXNAD1 | -0.0453516997582954 | 3.26231943466827 | -0.391072700889429 | 0.697697248382769 | 0.875599199040458 | -6.13458787988391 |
| ENSG00000053501.12 | USE1 | -0.0384939156585182 | 2.83103478051017 | -0.390603487469724 | 0.698041492229636 | 0.875950371133532 | -6.08955853675539 |
| lnc-LRPPRC-4 | lnc-LRPPRC-4 | 0.059584186262315 | 1.22295775037043 | 0.390575286544086 | 0.698062184215291 | 0.875950371133532 | -5.87743537193859 |
| ENSG00000175581.13 | MRPL48 | -0.064366436615845 | 3.04673773802956 | -0.390448954618177 | 0.698154881131782 | 0.87601328446327 | -6.11855981137563 |
| lnc-FRMD4A-4 | lnc-FRMD4A-4 | -0.0834273073413585 | 2.11133085335341 | -0.390368017796538 | 0.698214271530349 | 0.876016220962202 | -5.98274265693092 |
| ENSG00000172650.13 | AGAP5 | 0.0508055294679657 | 1.86623940890612 | 0.390329764746977 | 0.698242341792056 | 0.876016220962202 | -5.96600475358048 |
| lnc-AARS-1 | lnc-AARS-1 | -0.0841920280891325 | 0.443002115847311 | -0.390225088297741 | 0.698319156031633 | 0.876059193551478 | -5.70628533963302 |
| lnc-MED14-6 | lnc-MED14-6 | -0.0645991908130736 | 1.62914860888259 | -0.39015153198436 | 0.698373135438852 | 0.876068917460877 | -5.92577882501386 |
| ENSG00000148459.15 | PDSS1 | 0.0527692588231957 | 0.922448454013589 | 0.390094289474662 | 0.69841514403077 | 0.876068917460877 | -5.83430767527732 |
| ENSG00000185163.9 | DDX51 | 0.0388418323582966 | 4.6795095610532 | 0.389985058307649 | 0.698495308221616 | 0.876068917460877 | -6.2849737214724 |
| ENSG00000188167.8 | TMPPE | 0.0583715412132522 | 3.20098213826967 | 0.38998253778008 | 0.69849715806493 | 0.876068917460877 | -6.12435621590567 |
| ENSG00000248124.7 | RRN3P1 | -0.0634522773488344 | 4.34432210530185 | -0.389587497919754 | 0.698787105142941 | 0.87628617024983 | -6.25516930538361 |
| ENSG00000117480.15 | FAAH | 0.0672210140627316 | 1.81024448080075 | 0.389554703796562 | 0.698811177064843 | 0.87628617024983 | -5.94777641125654 |
| LINC01000 | LINC01000 | 0.0789153836721397 | 5.55348884412843 | 0.389521241168193 | 0.698835740013947 | 0.87628617024983 | -6.32552123249963 |
| lnc-DOCK4-8 | lnc-DOCK4-8 | -0.0817955058844126 | 2.00235896323913 | -0.38941318630617 | 0.698915058967982 | 0.87628617024983 | -5.9994842036233 |
| ENSG00000127418.14 | FGFRL1 | 0.143561524949359 | 1.80367601709414 | 0.389414114327636 | 0.698933640279086 | 0.87628617024983 | -5.99650717264152 |
| lnc-D2HGDH-1 | lnc-D2HGDH-1 | -0.0563855196253765 | 0.898033056698179 | -0.389357192163568 | 0.698956163472073 | 0.87628617024983 | -5.83276743441629 |
| lnc-RCSD1-5 | lnc-RCSD1-5 | 0.0387846434776504 | 4.43120869366162 | 0.389300252326485 | 0.698997963134309 | 0.87628617024983 | -6.26364891357383 |
| ENSG00000276612.3 | NA | -0.148290641636097 | 0.393049712864271 | -0.389261401482514 | 0.699026484168465 | 0.87628617024983 | -5.70664448840273 |
| ENSG00000188033.9 | ZNF490 | 0.0526743516930033 | 3.08323793940101 | 0.389224553571782 | 0.69905353522335 | 0.87628617024983 | -6.11351929132575 |
| BACE2-IT1 | BACE2-IT1 | 0.157916186004145 | -0.502308954725278 | 0.388981883406504 | 0.699231695883612 | 0.876456123035892 | -5.39126920319024 |
| ENSG00000092841.18 | MYL6 | 0.0518255006457808 | 7.63697264621605 | 0.388867923255982 | 0.699315367694362 | 0.876507624726164 | -6.32239705497186 |
| ENSG00000213920.8 | MDP1 | -0.0473793060840015 | 1.17461487733131 | -0.388807007444634 | 0.699360094846619 | 0.876510310675757 | -5.86129467020483 |
| ENSG00000197903.7 | H2BC12 | -0.0674598407179596 | 6.93347622826067 | -0.388709719828249 | 0.699431530068846 | 0.876546467572276 | -6.33158880060358 |
| lnc-SYPL1-3 | lnc-SYPL1-3 | -0.155520640199047 | -1.01334218386432 | -0.388589052506913 | 0.699520136097529 | 0.876604137673999 | -5.33348712448487 |
| ENSG00000015676.17 | NUDCD3 | -0.0259213141624253 | 5.93961348697457 | -0.388372551080515 | 0.699679123742602 | 0.876698705676988 | -6.3336119126402 |
| ENSG00000184384.13 | MAML2 | -0.143628357500469 | 6.11313191217333 | -0.388444423997208 | 0.699680786967686 | 0.876698705676988 | -6.33393772627263 |
| ENSG00000109016.17 | DHRS7B | -0.048920254585832 | 1.92360804929207 | -0.388163301584248 | 0.699832798896305 | 0.876801290440255 | -5.98037175025711 |
| ENSG00000263620.1 | | 0.0746114800052808 | 2.64422655147899 | 0.388135652256292 | 0.69985310582217 | 0.876801290440255 | -6.06643393170943 |
| ENSG00000204287.13 | HLA-DRA | 0.082875338560779 | 10.7966788492001 | 0.387954571950361 | 0.699986104940949 | 0.876801290440255 | -6.23470275956736 |
| ENSG00000125246.15 | CLYBL | -0.058712506301553 | 2.93404196684973 | -0.387887736933895 | 0.700035196057099 | 0.876801290440255 | -6.09851300734854 |
| ENSG00000279880.1 | | -0.0408459109288963 | 3.12272289540978 | -0.387871478201609 | 0.700047138486036 | 0.876801290440255 | -6.11566883151813 |
| lnc-CMTM7-2 | lnc-CMTM7-2 | -0.0810543150285937 | -0.292947892211553 | -0.387790262116577 | 0.700106794799126 | 0.876801290440255 | -5.53316611336086 |
| lnc-FKBP3-5 | lnc-FKBP3-5 | 0.0655263870150797 | 2.71048427221443 | 0.387789535334994 | 0.700107328656578 | 0.876801290440255 | -6.06461951442115 |
| lnc-DNAL4-3 | lnc-DNAL4-3 | -0.0564393895070574 | 3.33615091261913 | -0.38776125419822 | 0.700128102686373 | 0.876801290440255 | -6.14905278864923 |
| ENSG00000112425.14 | EPM2A | -0.0540655788412536 | 4.10914631908486 | -0.387736830614608 | 0.700146043320995 | 0.876801290440255 | -6.24141441207931 |
| ENSG00000168393.12 | DTYMK | -0.0660037393668327 | 1.83596002273431 | -0.387660542978245 | 0.700202082429991 | 0.876818121473293 | -5.96612603048666 |
| lnc-BPNT1-2 | lnc-BPNT1-2 | -0.0790914816412398 | 3.54523503829134 | -0.387558543440052 | 0.700277011540064 | 0.87685860375527 | -6.16902996815666 |
| lnc-CROCC-3 | lnc-CROCC-3 | -0.104590905206775 | -0.0153049890487451 | -0.387426532796403 | 0.700373991366649 | 0.876908400622957 | -5.6187448585395 |
| ENSG00000170515.13 | PA2G4 | 0.0533342753902444 | 5.28887757291628 | 0.387342068440141 | 0.700436044612955 | 0.876908400622957 | -6.31946780371458 |
| ENSG00000166900.16 | STX3 | -0.0606805465267701 | 2.45871133421186 | -0.387330435792726 | 0.700444590907158 | 0.876908400622957 | -6.03304173661108 |
| ENSG00000174748.20 | RPL15 | -0.0565218534734857 | 9.05292312534706 | -0.387266416588382 | 0.700491625353485 | 0.876913947491229 | -6.28815182792719 |
| ENSG00000102743.14 | SLC25A15 | -0.0854171066677685 | 0.490578958229782 | -0.387024074252625 | 0.700669683285825 | 0.877083506116411 | -5.75042559619395 |
| ENSG00000167711.13 | SERPINF2 | -0.144855700574468 | 1.90669086033068 | -0.386910303276974 | 0.700772228598045 | 0.87715237455857 | -5.98152016763053 |
| lnc-SLFN12L-3 | lnc-SLFN12L-3 | 0.142043540711436 | -0.742594132426215 | 0.386833209781854 | 0.700809930506519 | 0.87715237455857 | -5.36721333113137 |
| ENSG00000165795.23 | NDRG2 | -0.202204960093406 | 0.387651901411638 | -0.386503001818341 | 0.701076927584794 | 0.87738146669895 | -5.71439554086367 |
| lnc-TNFAIP8-8 | lnc-TNFAIP8-8 | -0.105825563312184 | 0.506051517413469 | -0.386468132052646 | 0.701078219054445 | 0.87738146669895 | -5.71590311176527 |
| lnc-PIGC-2 | lnc-PIGC-2 | -0.049428089945708 | 1.26149095901481 | -0.386248432909914 | 0.701239690307771 | 0.877530188813525 | -5.888342870852 |
| ENSG00000018869.16 | ZNF582 | -0.0698847315382267 | 2.06393846705654 | -0.386161302176481 | 0.701303732247073 | 0.877556977375008 | -5.98315760431243 |
| ENSG00000124459.11 | ZNF45 | -0.033778463131882 | 4.05976781899494 | -0.385951180523481 | 0.701458182752572 | 0.87769688605448 | -6.22862351210485 |
| ENSG00000163870.15 | TPRA1 | -0.0629686102935603 | 2.39874486430736 | -0.385788874230431 | 0.701577495190802 | 0.877706659389469 | -6.03918109386219 |

| | | | | | | | |
|---|---|---|---|---|---|---|---|
| ENSG00000238105.7 | GOLGA2P5 | 0.0451414083834338 | 6.32333047526405 | 0.385751010455058 | 0.701605330200839 | 0.877706659389469 | -6.33687014800292 |
| ENSG00000067829.18 | IDH3G | -0.0407679870604563 | 4.78201478421733 | -0.385713282582944 | 0.701633065715996 | 0.877706659389469 | -6.29414050386442 |
| ENSG00000006071.13 | ABCC8 | 0.290857969524525 | -0.475645906697917 | 0.385733021612052 | 0.701636562871998 | 0.877706659389469 | -5.35929154619665 |
| lnc-SLC16A4-3 | lnc-SLC16A4-3 | 0.0687221765974171 | 0.999340754456132 | 0.385520417496612 | 0.701774856250019 | 0.877768125650341 | -5.82769390292318 |
| lnc-KIAA1147-1 | lnc-KIAA1147-1 | -0.0675005765100902 | 2.54231702271506 | -0.385438323580227 | 0.701835213309956 | 0.877768125650341 | -6.0628723447153 |
| ENSG00000202363.1 | SNORA62 | -0.0609176538590253 | 1.11366710917932 | -0.385426529103049 | 0.701843885001308 | 0.877768125650341 | -5.86015403047817 |
| lnc-CAB39L-10 | lnc-CAB39L-10 | -0.100376333034874 | 0.770228672101288 | -0.385355590907032 | 0.701896041967227 | 0.877768125650341 | -5.81029083801585 |
| ENSG00000185621.11 | LMLN | -0.0785962074083102 | 2.4898114680012 | -0.385315385104068 | 0.701925603735335 | 0.877768125650341 | -6.0522233063684 |
| ENSG00000156504.16 | PABIR2 | 0.0487753266298308 | 4.24328086338295 | 0.385267579099866 | 0.701960754244657 | 0.877768125650341 | -6.24916318729731 |
| lnc-ANKRD46-5 | lnc-ANKRD46-5 | -0.168349603311107 | -0.307414019959252 | -0.38523567084254 | 0.701984215922376 | 0.877768125650341 | -5.53425437959196 |
| ENSG00000112701.17 | SENP6 | -0.0365523755943977 | 7.04011482611241 | -0.38507529794395 | 0.702102140237346 | 0.8778622495751 | -6.33233395294338 |
| ENSG00000160255.17 | ITGB2 | 0.2434169558622 | 3.40574175340057 | 0.384820495199329 | 0.702362481056616 | 0.877932210134966 | -6.14110944158901 |
| ENSG00000112139.15 | MDGA1 | -0.293806249224327 | 3.64383194835196 | -0.384791829666223 | 0.702386793948309 | 0.877932210134966 | -6.17366602816651 |
| ENSG00000143374.16 | TARS2 | -0.0274376837687331 | 4.53491693227515 | -0.384686037090824 | 0.70238839978571 | 0.877932210134966 | -6.2728894685296 |
| lnc-ZC3HC1-2 | lnc-ZC3HC1-2 | -0.0582630312586323 | 2.29994171263877 | -0.38467365186445 | 0.702397508507393 | 0.877932210134966 | -6.01338917337103 |
| lnc-FAAP100-1 | lnc-FAAP100-1 | -0.106828782578449 | 5.30660529032512 | -0.384697153514349 | 0.7024065641199 | 0.877932210134966 | -6.32188185549529 |
| ENSG00000205057.5 | CLLU1-AS1 | -0.262293123374659 | 2.02196383967361 | -0.384746374486328 | 0.702414013336376 | 0.877932210134966 | -6.01627870526286 |
| ENSG00000215695.1 | RSC1A1 | -0.0407360791437426 | 3.40041770148603 | -0.384552158314154 | 0.702486863353692 | 0.877969950112882 | -6.15668499395289 |
| lnc-CGAS-3 | lnc-CGAS-3 | -0.0625696378339651 | 9.23850376490516 | -0.384349667785133 | 0.70263579850011 | 0.878022764777291 | -6.28174437022241 |
| lnc-DDX18-1 | lnc-DDX18-1 | -0.0752520502363949 | 2.32620524554547 | -0.384337485560141 | 0.702644759105981 | 0.878022764777291 | -6.02459612846955 |
| ENSG00000171862.10 | PTEN | -0.0537876223568299 | 7.22007243345183 | -0.384299651018008 | 0.702672588483967 | 0.878022764777291 | -6.32959714722914 |
| lnc-PLA2G1B-2 | lnc-PLA2G1B-2 | -0.117749834695442 | 4.46145466862576 | -0.384304963073981 | 0.702699752425609 | 0.878022764777291 | -6.26809165907878 |
| ENSG00000176834.13 | VSIG10 | -0.183445832612362 | 3.51405840004594 | -0.38411505663718 | 0.702868196126416 | 0.878179924782689 | -6.19213367327254 |
| ENSG00000204172.12 | AGAP9 | 0.0738282455857897 | 2.54165433060113 | 0.383906283413244 | 0.702961956349138 | 0.87819054802681 | -6.03972069437391 |
| lnc-PDE7A-4 | lnc-PDE7A-4 | -0.0730363006891466 | 4.14915272695005 | -0.383906182916728 | 0.702962030281773 | 0.87819054802681 | -6.23834529703262 |
| lnc-TRPM7-3 | lnc-TRPM7-3 | -0.0878281884831189 | -0.061079430035253 | -0.383793905264858 | 0.703044631804392 | 0.878208021364618 | -5.58353392956988 |
| lnc-FAS-2 | lnc-FAS-2 | -0.133956110598005 | -0.518475074405896 | -0.383745414905579 | 0.703080306789493 | 0.878208021364618 | -5.4248660177216 |
| ENSG00000111667.13 | USP5 | -0.06641913925107 | 5.04279563776888 | -0.383682722805816 | 0.70312643118255 | 0.878208021364618 | -6.31090673057813 |
| ENSG00000011052.21 | NME1-NME2 | 0.082209369837833 | 4.3907023927825 | 0.383602090929456 | 0.703185756052431 | 0.878208021364618 | -6.26588466900023 |
| lnc-KLHL18-6 | lnc-KLHL18-6 | -0.0882924306666298 | 0.554475299761862 | -0.383597205670033 | 0.703189350440182 | 0.878208021364618 | -5.73152206903157 |
| lnc-XRRA1-1 | lnc-XRRA1-1 | 0.130669181097481 | 0.740441611768941 | 0.38341258074271 | 0.703325195462889 | 0.878261595427657 | -5.78638710165981 |
| lnc-RASA2-1 | lnc-RASA2-1 | 0.0523146791877878 | 3.59879153923206 | 0.383405103178236 | 0.703330697580582 | 0.878261595427657 | -6.19078110870235 |
| ENSG00000186448.14 | ZNF197 | -0.0384904236664951 | 4.39201958749272 | -0.383364933333116 | 0.703360255503547 | 0.878261595427657 | -6.26094600338485 |
| lnc-AIDA-1 | lnc-AIDA-1 | -0.0890211082301327 | 1.86893098700906 | -0.383218500916342 | 0.70346800787305 | 0.878342857509918 | -5.96589522799953 |
| ENSG00000279476.1 | | 0.123842684438854 | 1.06481765279097 | 0.383158196477119 | 0.70351238471894 | 0.878344984512309 | -5.84129481796959 |
| ENSG00000105866.14 | SP4 | -0.0554890085181494 | 6.84383786661771 | -0.383067983042112 | 0.703578772956762 | 0.878363461245333 | -6.33484142785744 |
| ENSG00000285779.1 | | -0.0492186443161247 | 3.73370517199257 | -0.38302210969312 | 0.703612532138806 | 0.878363461245333 | -6.19642959897485 |
| ENSG00000159352.15 | PSMD4 | -0.0564061273976861 | 5.68953725807941 | -0.382895904080665 | 0.703705412679467 | 0.878368822677697 | -6.33139094225468 |
| ENSG00000197713.14 | RPE | -0.0555056149739492 | 4.81789507650318 | -0.382875427981838 | 0.703720482418237 | 0.878368822677697 | -6.30016203344229 |
| ENSG00000133789.14 | SWAP70 | 0.0578392266047851 | 8.06900346670455 | 0.382842318251838 | 0.703744850351159 | 0.878368822677697 | -6.31840429925562 |
| ENSG00000087077.13 | TRIP6 | -0.110051341926588 | 1.6297546987191 | -0.382694189399913 | 0.703853873337779 | 0.878451629572612 | -5.94672974900589 |
| lnc-SHISA9-4 | lnc-SHISA9-4 | -0.101821803118523 | 1.07156942554232 | -0.382423342421174 | 0.704053233291537 | 0.878644872719301 | -5.84130544253826 |
| ENSG00000148688.13 | RPP30 | -0.0350331063535236 | 4.6136790922221 | -0.382367846911955 | 0.704094083983489 | 0.878644872719301 | -6.28486621272316 |
| lnc-CCDC80-3 | lnc-CCDC80-3 | -0.0485642010098553 | 0.97635053643184 | -0.382241685657943 | 0.704186955589899 | 0.878703848648576 | -5.86085364627673 |
| lnc-LDLRAD1-1 | lnc-LDLRAD1-1 | -0.0577586685397021 | 0.527115797809846 | -0.382187662439848 | 0.704226725322203 | 0.878703848648576 | -5.74922108163344 |
| TOB1-AS1 | TOB1-AS1 | -0.0801971917628264 | -0.0058086983946122 | -0.38207206301253 | 0.704311827810315 | 0.878748315328019 | -5.60665809091744 |
| ENSG00000165637.13 | VDAC2 | 0.0399871999443926 | 5.38691854486116 | 0.382023271583702 | 0.704347748446857 | 0.878748315328019 | -6.32501326003816 |
| ENSG00000188897.9 | | -0.0823755154260084 | 2.21842756730534 | -0.381692726362829 | 0.704591116439958 | 0.878998663536194 | -6.01632286725994 |
| lnc-TMEM200C-2 | lnc-TMEM200C-2 | -0.308254933368585 | -1.07666683311773 | -0.381667512956841 | 0.704636589505759 | 0.879002116472548 | -5.36843624876862 |
| lnc-FAM90A1-1 | lnc-FAM90A1-1 | -0.0830519873506979 | 1.35457811171548 | -0.381519425656578 | 0.704718723738202 | 0.879051299357821 | -5.87944313978888 |
| lnc-AGAP5-1 | lnc-AGAP5-1 | -0.0289717841406048 | 3.623835082144 | -0.381334890437532 | 0.704854612857734 | 0.879167524933386 | -6.18803310116409 |
| ENSG00000138760.9 | SCARB2 | 0.0394015861066963 | 4.89899234510873 | 0.38124431377541 | 0.704921315796998 | 0.879197445497765 | -6.30475226453856 |
| lnc-ATP6V1E2-8 | lnc-ATP6V1E2-8 | -0.0625774623571023 | 1.33049346101488 | -0.381086305408951 | 0.705037682768861 | 0.879250931564081 | -5.89742684470552 |
| ENSG00000119772.16 | DNMT3A | -0.0813509059359383 | 4.49966802220197 | -0.381035508471029 | 0.705075094242061 | 0.879250931564081 | -6.27685628403154 |
| lnc-SRPK2-5 | lnc-SRPK2-5 | -0.0509424617347149 | 2.67290259228421 | -0.380967946998083 | 0.705124853784181 | 0.879250931564081 | -6.06471221862692 |
| lnc-DNASE1L1-5 | lnc-DNASE1L1-5 | 0.0793447629805306 | 3.67474216740224 | 0.380927782406825 | 0.705154435932372 | 0.879250931564081 | -6.18158566615725 |
| ENSG00000116459.10 | ATP5PB | 0.0518596011216816 | 5.74712140955853 | 0.380896079150329 | 0.705177786438316 | 0.879250931564081 | -6.33372444725062 |
| lnc-GOLGA6B-6 | lnc-GOLGA6B-6 | -0.0806914112297669 | 0.0707523215367264 | -0.38072532344459 | 0.7053035586576 | 0.879315317926885 | -5.62964471333551 |
| ENSG00000181045.14 | SLC26A11 | -0.0842986914457149 | 2.49793416231481 | -0.380709971693671 | 0.705314866587615 | 0.879315317926885 | -6.0446152033436 |
| ENSG00000130766.4 | SESN2 | 0.11759533603831 | 0.333073605023939 | 0.38059119955149 | 0.705402355112766 | 0.879347701438359 | -5.69978925570325 |
| lnc-RNMT-7 | lnc-RNMT-7 | -0.252114870317259 | -1.33195289983822 | -0.380557530350499 | 0.705427156853775 | 0.879347701438359 | -5.24169261913814 |
| ENSG00000104964.14 | TLE5 | -0.0555737382524875 | 7.4762396729238 | -0.380456865644055 | 0.705501311422291 | 0.879347701438359 | -6.32756431809797 |
| lnc-P3H4-1 | lnc-P3H4-1 | 0.0665929918211781 | 0.285777216043899 | 0.3804427225614 | 0.705511730143956 | 0.879347701438359 | -5.66739215402806 |
| ENSG00000142798.19 | HSPG2 | 0.141439314505361 | 2.49699518956657 | 0.380084900068936 | 0.705799251424102 | 0.879623988717281 | -6.03179536315793 |
| lnc-SGSH-4 | lnc-SGSH-4 | 0.0672754701203249 | 1.56839600663412 | 0.379932723188913 | 0.705887467304369 | 0.879623988717281 | -5.92814276685058 |
| lnc-EIF4EBP3-1 | lnc-EIF4EBP3-1 | -0.116220794308508 | -0.577495862100261 | -0.379845950990171 | 0.705951403287799 | 0.879623988717281 | -5.43619814164404 |
| lnc-STK33-1 | lnc-STK33-1 | -0.0875649669373515 | 0.405300153887515 | -0.379818448484172 | 0.705971668288856 | 0.879623988717281 | -5.68665121704673 |
| lnc-NSD3-9 | lnc-NSD3-9 | -0.150018457367648 | 4.06829003075744 | -0.379867955560353 | 0.705984224698777 | 0.879623988717281 | -6.21632706765509 |

| | | | | | | | |
|---|---|---|---|---|---|---|---|
| lnc-ZNF33B-3 | lnc-ZNF33B-3 | 0.0602022043461148 | 1.90180454300071 | 0.379782520119912 | 0.705998142141325 | 0.879623988717281 | -5.97323500618351 |
| ENSG00000050438.16 | SLC4A8 | -0.193122099129731 | 1.65201433440649 | -0.379781317922105 | 0.706045431490875 | 0.879623988717281 | -5.94440667489397 |
| lnc-MTA3-6 | lnc-MTA3-6 | 0.104719893249807 | 0.448441426587369 | 0.37965254684461 | 0.706093916161488 | 0.879623988717281 | -5.69272760159622 |
| lnc-CLASP2-7 | lnc-CLASP2-7 | 0.0359738502075357 | 4.37750726659944 | 0.379619839613244 | 0.70611801805255 | 0.879623988717281 | -6.26424288015408 |
| ENSG00000283050.1 | GTF2IP12 | 0.102918562770077 | 1.24463043335576 | 0.379489403082185 | 0.706214139498144 | 0.879652726090876 | -5.86557796625408 |
| ENSG00000147123.10 | NDUFB11 | 0.0642385922668307 | 3.97069741074606 | 0.37947254804239 | 0.706226560688516 | 0.879652726090876 | -6.22490910048243 |
| lnc-TOR1AIP2-5 | lnc-TOR1AIP2-5 | -0.126064689268369 | 0.819048874166783 | -0.37924967907858 | 0.706390809855742 | 0.879787062727855 | -5.81434839469233 |
| lnc-ZNF408-3 | lnc-ZNF408-3 | 0.0395192913035305 | 2.05487542069548 | 0.379210209849105 | 0.706419899222818 | 0.879787062727855 | -6.00459811649087 |
| ENSG00000254536.1 | | 0.0487940840495873 | 2.95441434222213 | 0.37906244804206 | 0.706528805658128 | 0.879869458304771 | -6.10537917415914 |
| ENSG00000206990.1 | | 0.0810484514898698 | -0.179249980520227 | 0.378938618326167 | 0.706620077965711 | 0.879929884733996 | -5.5784026992415 |
| ENSG00000103174.12 | NAGPA | -0.0388177304631254 | 4.17094167811343 | -0.378706037907053 | 0.70679151996261 | 0.880032286262486 | -6.24805188015626 |
| ENSG00000255275.3 | NA | -0.0814601016736121 | 0.886173597929048 | -0.37870073194262 | 0.706795431327758 | 0.880032286262486 | -5.82779134538717 |
| ENSG00000178952.10 | TUFM | -0.0526121680549929 | 5.95767641375716 | -0.37861853201703 | 0.70685602716516 | 0.880032286262486 | -6.33794281501438 |
| ENSG00000057663.15 | ATG5 | 0.0379442972977442 | 4.76031522513261 | 0.378404348251916 | 0.707013927397887 | 0.880032286262486 | -6.29576028625388 |
| ENSG00000183530.13 | PRR14L | -0.0344535669639114 | 6.45546894902509 | -0.378394506336325 | 0.707021183353855 | 0.880032286262486 | -6.33952840932527 |
| lnc-ARHGEF4-2 | lnc-ARHGEF4-2 | 0.0574171810249684 | 1.09897608767322 | 0.378220739094176 | 0.707149297857624 | 0.880032286262486 | -5.84887319510984 |
| ENSG00000243716.10 | NPIPB5 | -0.0845370649670323 | 2.36738605590018 | -0.37817566454305 | 0.707182531667983 | 0.880032286262486 | -6.04638393630666 |
| ENSG00000112110.9 | MRPL18 | 0.0498118400661842 | 3.06530450150562 | 0.37816094858658 | 0.707193381980074 | 0.880032286262486 | -6.11969348311286 |
| ENSG00000187954.12 | ZFTRAF1 | 0.0451528463836963 | 4.52681628208169 | 0.378114155681958 | 0.707227883552785 | 0.880032286262486 | -6.27621084041448 |
| ENSG00000137992.14 | DBT | -0.0377335315335792 | 4.31263989942693 | -0.378091415110802 | 0.707244650966502 | 0.880032286262486 | -6.25529577024798 |
| ENSG00000146007.10 | ZMAT2 | -0.0521521788462467 | 5.65845664918691 | -0.37808600276155 | 0.707248641701439 | 0.880032286262486 | -6.33284600127251 |
| ENSG00000250264.1 | | -0.0484525598561429 | 5.95808366731853 | -0.378077208345457 | 0.707255126183144 | 0.880032286262486 | -6.33693784730261 |
| ENSG00000187837.3 | H1-2 | 0.0873007849235474 | 8.27167519511044 | 0.378024405582364 | 0.707299135158025 | 0.880032286262486 | -6.31357876693827 |
| ENSG00000196428.12 | TSC22D2 | 0.0456802655949993 | 5.01947110845339 | 0.378015150226991 | 0.707300884790376 | 0.880032286262486 | -6.31189055963835 |
| ENSG00000250091.3 | DNAH10OS | -0.116639641454553 | -0.146958143574457 | -0.377566898435204 | 0.707631436147194 | 0.880351349899037 | -5.5651135420449 |
| lnc-CRNKL1-6 | lnc-CRNKL1-6 | -0.0846344337663601 | 0.0275876363689334 | -0.377551204668899 | 0.707643010132003 | 0.880351349899037 | -5.60933075486006 |
| ENSG00000165959.11 | CLMN | 0.052012803831655 | 6.6217384640077 | 0.377493407543955 | 0.707685635496743 | 0.880351349899037 | -6.33903244203094 |
| lnc-COQ10B-3 | lnc-COQ10B-3 | -0.0964144442414605 | 0.682495538320136 | -0.377369308416879 | 0.70777716177622 | 0.880411997391668 | -5.80085035694222 |
| ENSG00000156508.17 | EEF1A1 | 0.0572585072959964 | 12.6004342488042 | 0.377260086005555 | 0.707857719715215 | 0.880458994736419 | -6.16112013836846 |
| ENSG00000261864.1 | | 0.0386278714204904 | 3.52709570524959 | 0.377122129921382 | 0.707959475263991 | 0.880532351160718 | -6.17097301164092 |
| ENSG00000232869.2 | TRBV29-1 | -0.258282528626055 | -0.548392136626897 | -0.377008851215331 | 0.708050100896534 | 0.88056953822313 | -5.36049067287295 |
| ENSG00000165507.8 | DEPP1 | -0.387052835000058 | -1.09385105156475 | -0.376916215160011 | 0.708164025016888 | 0.88056953822313 | -5.41911330434462 |
| ENSG00000166128.12 | RAB8B | 0.0615878368071767 | 6.64122818620313 | 0.376783517492929 | 0.708209256575528 | 0.88056953822313 | -6.33899996445326 |
| ENSG00000270647.5 | TAF15 | 0.0504767248144535 | 5.14098385990201 | 0.376716400249171 | 0.708258770235761 | 0.88056953822313 | -6.31402317693748 |
| ENSG00000111911.6 | HINT3 | 0.0505469683475974 | 4.04071118117551 | 0.376702730297201 | 0.708268854974447 | 0.88056953822313 | -6.23580990083793 |
| lnc-SERINC5-1 | lnc-SERINC5-1 | -0.166552625159548 | 0.801294479181013 | -0.376699397883906 | 0.708281742266248 | 0.88056953822313 | -5.76164143560587 |
| ENSG00000141425.17 | RPRD1A | -0.0388393473400024 | 5.15546383849713 | -0.376675635290505 | 0.708288843940249 | 0.88056953822313 | -6.31803781414557 |
| ENSG00000101997.12 | CCDC22 | -0.045024313695774 | 4.00747641389125 | -0.37654149463615 | 0.708387807414901 | 0.880601519687981 | -6.23242494942524 |
| ENSG00000005893.15 | LAMP2 | 0.0442610866349753 | 5.31991206468806 | 0.376486809096429 | 0.708428153627117 | 0.880601519687981 | -6.32549392169915 |
| ENSG00000110013.12 | SIAE | -0.0760460152233342 | 1.43627387733894 | -0.376460305694873 | 0.708447707764119 | 0.880601519687981 | -5.9132475583113 |
| ENSG00000178966.16 | RMI1 | 0.0661352500645269 | 1.19909870823322 | 0.376408812132161 | 0.708485700142494 | 0.880601519687981 | -5.88031701078807 |
| ENSG00000147592.8 | LACTB2 | -0.0718716668792745 | 1.56208777761734 | -0.376326902927842 | 0.708546134981759 | 0.880623458506706 | -5.93487034009698 |
| ENSG00000125901.5 | MRPS26 | -0.0824635180444986 | 2.39434848644649 | -0.376215793908677 | 0.70862811727611 | 0.880672173523377 | -6.04903077289241 |
| lnc-DAZL-1 | lnc-DAZL-1 | -0.357030407821841 | -0.449044988835734 | -0.376182963669171 | 0.708688942503924 | 0.880683249770654 | -5.30864275197421 |
| lnc-CAVIN2-2 | lnc-CAVIN2-2 | -0.163152608120749 | -0.233459218053227 | -0.376085305157316 | 0.708724403457653 | 0.880683249770654 | -5.51236662082606 |
| lnc-DRD5-10 | lnc-DRD5-10 | -0.381503346325463 | 0.094029418874976 | -0.376122257760403 | 0.708765390489767 | 0.880683249770654 | -5.66817145745629 |
| lnc-CHRNA10-2 | lnc-CHRNA10-2 | 0.0537458708773177 | 0.851810218989492 | 0.375872091379134 | 0.70888174187835 | 0.880760597929525 | -5.81593891738907 |
| ENSG00000136827.11 | TOR1A | -0.0335900800022868 | 4.2159620871667 | -0.375829435345095 | 0.708913220905526 | 0.880760597929525 | -6.25482797687912 |
| ENSG00000101298.14 | SNPH | -0.199103743335805 | 0.812466293856332 | -0.375753325329239 | 0.709007428126413 | 0.880824474476458 | -5.8152585830773 |
| ENSG00000254633.1 | STRA6LP | 0.072027011964179 | 2.03209253898784 | 0.375593747546399 | 0.70908716155918 | 0.880855151897018 | -5.98547050197598 |
| ENSG00000177150.12 | FAM210A | -0.0697947968408634 | 2.80139439176301 | -0.375552354155526 | 0.709117712040693 | 0.880855151897018 | -6.08990315713837 |
| ENSG00000273189.1 | | -0.0721908005764735 | 6.47179125104835 | -0.37531440203071 | 0.709293342516573 | 0.880970581452454 | -6.34018623572172 |
| lnc-SLC38A8-2 | lnc-SLC38A8-2 | 0.209667457250222 | 0.744125622159482 | 0.375361123758371 | 0.70929623844095 | 0.880970581452454 | -5.76058541794834 |
| ENSG00000124772.11 | CPNE5 | -0.16889464774161 | 5.10996199971187 | -0.375319940989113 | 0.709352151345457 | 0.880986866055847 | -6.31154438329854 |
| ENSG00000248593.3 | DSTNP2 | 0.073237750938351 | 0.00305086006068037 | 0.374916646588743 | 0.709586958211744 | 0.881225314400405 | -5.61678695854769 |
| lnc-POLR2F-2 | lnc-POLR2F-2 | 0.0613696263668218 | 0.710989315263783 | 0.374712274949715 | 0.709737838943396 | 0.881359513663465 | -5.78987486056137 |
| ENSG00000149782.11 | PLCB3 | 0.0707702100553846 | 1.74079454134246 | 0.374515868987236 | 0.709882849990788 | 0.88147239830394 | -5.95850609923342 |
| ENSG00000177272.8 | KCNA3 | 0.055858743799444 | 5.48169283690933 | 0.374473148726637 | 0.709914392784551 | 0.88147239830394 | -6.32840617796729 |
| ENSG00000204622.11 | HLA-J | 0.191822182523116 | 2.00371442824899 | 0.374394569831869 | 0.710018421808273 | 0.88149179722108 | -5.95102154040625 |
| ENSG00000142892.14 | PIGK | -0.0440626309801955 | 4.36533703748193 | -0.374291510440235 | 0.71004851239049 | 0.88149179722108 | -6.26194294648669 |
| ENSG00000184293.7 | CLECL1P | -0.0823795382532957 | 4.72610743956933 | -0.374277991794788 | 0.710058494773624 | 0.88149179722108 | -6.28073185103642 |
| ENSG00000168096.14 | ANKS3 | -0.0444360192215223 | 3.05666419032224 | -0.374049837287335 | 0.710226975468007 | 0.881647779751401 | -6.12051787428227 |
| ENSG00000271895.2 | NA | -0.0550496436575436 | 2.63825455588757 | -0.373962734278221 | 0.710291300553983 | 0.881674456597675 | -6.07023238171953 |
| lnc-XRCC2-13 | lnc-XRCC2-13 | 0.103081850108317 | 4.87979434650661 | 0.373899386687208 | 0.710358969502804 | 0.881705280665513 | -6.30246569818565 |
| ENSG00000116198.12 | CEP104 | 0.0485788691084563 | 4.62752668291051 | 0.373776691204917 | 0.710428699483206 | 0.881738659036591 | -6.28251846221566 |
| lnc-COIL-5 | lnc-COIL-5 | 0.0616843278028873 | 0.203234542501797 | 0.373579738437993 | 0.710574166201751 | 0.881866027309656 | -5.64046308063643 |
| lnc-ABR-5 | lnc-ABR-5 | -0.0639705498724741 | 3.78927051279184 | -0.373458056659724 | 0.710664044217334 | 0.881924395401265 | -6.21492535244049 |
| lnc-VEZF1-2 | lnc-VEZF1-2 | -0.0675940986175121 | 4.69408970632216 | -0.373340582855957 | 0.71075081804288 | 0.88197890442977 | -6.29112977935244 |

| | | | | | | | |
|---|---|---|---|---|---|---|---|
| ENSG00000237854.3 | LINC00674 | 0.0601098979853525 | 2.08948923226095 | 0.373239063441353 | 0.710825810041847 | 0.882018787562776 | -6.00752377461944 |
| ENSG00000256632.3 | | 0.0877628284654745 | 0.230164568576555 | 0.373082065257065 | 0.710941789701337 | 0.882109521817024 | -5.6655636482942 |
| ENSG00000208892.1 | SNORA49 | 0.124259803040161 | 2.97916469320787 | 0.373034051580119 | 0.710994775355263 | 0.882122089278926 | -6.08948596652767 |
| lnc-AKT3-2 | lnc-AKT3-2 | 0.0784178342097687 | 1.04669860555606 | 0.372889023662115 | 0.71108440529904 | 0.882180116585507 | -5.82834697923033 |
| ENSG00000101444.12 | AHCY | -0.0812783355500795 | 4.71658667063195 | -0.372634652188007 | 0.711272346281781 | 0.882310097210656 | -6.29442058190645 |
| ENSG00000009830.11 | POMT2 | -0.0443742754910779 | 3.14404315915628 | -0.372561500411784 | 0.711326397439937 | 0.882310097210656 | -6.13724245667077 |
| ENSG00000122970.15 | IFT81 | -0.0825932267738547 | 1.24245964278918 | -0.372554562785397 | 0.71133152366473 | 0.882310097210656 | -5.90339648124379 |
| ENSG00000163607.15 | GTPBP8 | 0.041224352007533 | 2.76127412196877 | 0.372464760293143 | 0.711397880109899 | 0.882310097210656 | -6.09119288438769 |
| ENSG00000163131.10 | CTSS | 0.0498726663519573 | 8.15090050323066 | 0.372457146108599 | 0.711403506452318 | 0.882310097210656 | -6.32057667398898 |
| ENSG00000110660.14 | SLC35F2 | 0.0903114333930208 | 4.014814700803 | 0.372293062489559 | 0.711526372249459 | 0.882342191413447 | -6.2284506215099 |
| ENSG00000105053.10 | VRK3 | 0.0263098963838857 | 5.08719844953165 | 0.372238697110355 | 0.711564931702493 | 0.882342191413447 | -6.31415351676474 |
| ENSG00000071051.13 | NCK2 | -0.0987810959676692 | 5.96231257662029 | -0.372235289577615 | 0.711588233046858 | 0.882342191413447 | -6.34007455047673 |
| ENSG00000175213.2 | ZNF408 | 0.0430125683707967 | 3.08449957764196 | 0.372190087692179 | 0.711600853979654 | 0.882342191413447 | -6.13038338413213 |
| lnc-ETV5-2 | lnc-ETV5-2 | 0.186835101010916 | 0.125631683352135 | 0.372049865382934 | 0.711707152268165 | 0.882366965916812 | -5.57082640718724 |
| lnc-ZNF683-1 | lnc-ZNF683-1 | -0.0910139838712316 | 1.05193236017827 | -0.372030685717932 | 0.711718656418589 | 0.882366965916812 | -5.84620906768535 |
| ENSG00000169905.12 | TOR1AIP2 | -0.0463881576743121 | 6.27262253287813 | -0.371989032367958 | 0.711749440563418 | 0.882366965916812 | -6.34180676705684 |
| ENSG00000113645.14 | WWC1 | -0.38794878667921 | 0.0807323899017034 | -0.371964978068203 | 0.711837767846379 | 0.882423318211396 | -5.75422886058247 |
| lnc-KIF13B-2 | lnc-KIF13B-2 | 0.126641482093056 | -0.236163637727854 | 0.371724872741062 | 0.7119446805628 | 0.882471572462523 | -5.55799546051209 |
| ENSG00000165416.14 | SUGT1 | 0.066043423636045 | 5.7644408672188 | 0.371662925923753 | 0.711990468197891 | 0.882471572462523 | -6.33877172258779 |
| ENSG00000196476.11 | C20orf96 | -0.0970716440892194 | 0.724246337420303 | -0.371596426036963 | 0.712039622405183 | 0.882471572462523 | -5.76236926181824 |
| ENSG00000081087.14 | OSTM1 | 0.0520700712936861 | 3.96482782468373 | 0.371584836525321 | 0.712048189061729 | 0.882471572462523 | -6.23036384866808 |
| ENSG00000258674.5 | | -0.0547635306151317 | 4.03071941461107 | -0.371399435071951 | 0.712185237965988 | 0.882504026824873 | -6.23175166387215 |
| ENSG00000202337.1 | RNU6-8 | 0.124413926172538 | -0.20678255845228 | 0.371329582208254 | 0.712236875756661 | 0.882504026824873 | -5.55188025527588 |
| FAM30A | FAM30A | 0.6628045698092 | 3.25449101969182 | 0.371400611683575 | 0.712257639670137 | 0.882504026824873 | -6.09963505629612 |
| ENSG00000267361.1 | SEC24AP1 | -0.157267243431903 | 3.53771292296599 | -0.371257910347007 | 0.712338266392516 | 0.882504026824873 | -6.16601589521799 |
| ENSG00000087087.19 | SRRT | 0.0432336122494927 | 5.95018715209037 | 0.371141391397785 | 0.712376000083345 | 0.882504026824873 | -6.34054567876334 |
| ENSG00000107020.9 | PLGRKT | 0.0582400640312226 | 2.43087758700466 | 0.371107561659683 | 0.712401010536232 | 0.882504026824873 | -6.04583920747175 |
| lnc-ZEB2-8 | lnc-ZEB2-8 | -0.0949850232571899 | 0.0348927014319548 | -0.371101193908054 | 0.712405718274838 | 0.882504026824873 | -5.66431019582876 |
| ENSG00000239556.4 | UPK3BL3P | 0.0466516965383397 | 1.80307047594874 | 0.371084148628648 | 0.712418320065897 | 0.882504026824873 | -5.95577325136966 |
| ENSG00000137656.11 | BUD13 | -0.0558636359857996 | 3.80229613913043 | -0.37093946076995 | 0.712525292907878 | 0.882504026824873 | -6.20890384259387 |
| lnc-CD74-3 | lnc-CD74-3 | 0.0744681854426162 | 0.696885548514536 | 0.370764331612655 | 0.712654779912219 | 0.882504026824873 | -5.78522279053195 |
| lnc-TRAF3IP2-5 | lnc-TRAF3IP2-5 | 0.0629066131137918 | 0.848810079431628 | 0.370742940653773 | 0.712670596548322 | 0.882504026824873 | -5.8083918510377 |
| ENSG00000060138.12 | YBX3 | -0.126028225595666 | 8.29651224492298 | -0.370761198874332 | 0.712700960544622 | 0.882504026824873 | -6.31487223122004 |
| ENSG00000142961.14 | MOB3C | -0.0392529746686634 | 4.03865512123442 | -0.370654781897668 | 0.71273578314701 | 0.882504026824873 | -6.23560410606427 |
| ENSG00000170017.12 | ALCAM | 0.0722535836282914 | 5.31456842026611 | 0.370631864235616 | 0.712752729345918 | 0.882504026824873 | -6.3248340431115 |
| ENSG00000179532.12 | DNHD1 | -0.0718672438358539 | 6.41015071475256 | -0.37062194763662 | 0.712760062105339 | 0.882504026824873 | -6.34240925978452 |
| ENSG00000134056.11 | KGD4 | 0.0641202345148848 | 2.93432858774358 | 0.370594558787838 | 0.712780314739885 | 0.882504026824873 | -6.11080865407396 |
| ENSG00000168028.13 | RPSA | 0.0597799006289481 | 8.97743153139564 | 0.370563532471567 | 0.712803257346524 | 0.882504026824873 | -6.29736013825839 |
| lnc-UBXN4-14 | lnc-UBXN4-14 | -0.124742784253878 | 0.740014361675444 | -0.37044148809553 | 0.712893506446704 | 0.882541502677677 | -5.79541188333173 |
| ENSG00000221420.2 | SNORA81 | 0.0488535469799252 | 3.50099935311308 | 0.370406633776839 | 0.712919281203991 | 0.882541502677677 | -6.18216383392913 |
| lnc-NOS1AP-2 | lnc-NOS1AP-2 | -0.0714194400522932 | 0.232186456986855 | -0.370210538741112 | 0.713064299746175 | 0.882667938517893 | -5.69177409759217 |
| lnc-DRD3-1 | lnc-DRD3-1 | -0.0557150624584841 | 0.511504424062445 | -0.370019188351184 | 0.713205819825898 | 0.882790028833752 | -5.73065816825903 |
| ENSG00000131941.7 | RHPN2 | 0.220631682904903 | 2.07036081451535 | 0.369877743140189 | 0.713364503811977 | 0.882845785242956 | -5.95569920025179 |
| ENSG00000285447.1 | NA | 0.353261226319374 | -1.43320709668879 | 0.369806857829962 | 0.713373535627953 | 0.882845785242956 | -5.14647069630966 |
| ENSG00000178904.18 | DPY19L3 | -0.0465223422715253 | 5.20648443745854 | -0.369719907758679 | 0.713427184075644 | 0.882845785242956 | -6.32512660513741 |
| ENSG00000176623.11 | RMDN1 | -0.0362816262082175 | 5.09502314461863 | -0.369574323971512 | 0.71353487482152 | 0.882845785242956 | -6.31722010825977 |
| lnc-EN1-7 | lnc-EN1-7 | 0.0454476487855278 | 1.19686711208856 | 0.369567315272374 | 0.713540059421871 | 0.882845785242956 | -5.87000576805932 |
| ENSG00000174514.12 | MFSD4A | 0.0927809093681788 | 0.559501241978765 | 0.369566890682664 | 0.71354037350741 | 0.882845785242956 | -5.71748415377023 |
| ENSG00000118564.14 | FBXL5 | 0.0316714683886374 | 5.97753603713155 | 0.369552377737537 | 0.713551109328175 | 0.882845785242956 | -6.34193615549882 |
| lnc-ALG12-5 | lnc-ALG12-5 | 0.0867750249247736 | 1.95920738023518 | 0.36939798263117 | 0.713665325357331 | 0.88293188055589 | -5.98561042243278 |
| ENSG00000069849.10 | ATP1B3 | 0.0440776706820092 | 4.506914608706 | 0.369235361698778 | 0.713785633747896 | 0.88293188055589 | -6.27609934883792 |
| lnc-EXT1-6 | lnc-EXT1-6 | -0.678678828891419 | -1.98295279582596 | -0.369244223459613 | 0.713815223307424 | 0.88293188055589 | -4.90326568166173 |
| ENSG00000116717.12 | GADD45A | 0.12445093706087 | 0.993287816327553 | 0.369189623510647 | 0.713819472588351 | 0.88293188055589 | -5.85124923524683 |
| ENSG00000155330.9 | C16orf87 | -0.0599674595127082 | 4.50639016424805 | -0.369165917857588 | 0.713837011155927 | 0.88293188055589 | -6.28148576357305 |
| ENSG00000168256.17 | NKIRAS2 | -0.0433225658921748 | 4.01837499445863 | -0.369110419603682 | 0.713878072011423 | 0.88293188055589 | -6.23502998313294 |
| ENSG00000148481.13 | MINDY3 | 0.0317028425359522 | 4.8831154030163 | 0.369049275279724 | 0.713923311153478 | 0.882934778193357 | -6.305079533948 |
| ENSG00000279611.1 | | -0.0867154938005024 | -0.216596212125734 | -0.368972766753049 | 0.713979919348301 | 0.882951735156578 | -5.58210180056121 |
| ENSG00000136104.20 | RNASEH2B | -0.160449924088336 | 7.03625678434337 | -0.368619826722771 | 0.71430273845186 | 0.883297883181414 | -6.33864247738823 |
| ENSG00000109323.9 | MANBA | 0.0415056795391537 | 5.59595601917778 | 0.368356902121435 | 0.714435653213992 | 0.883363574684855 | -6.3350940809743 |
| ENSG00000167394.12 | ZNF668 | -0.092346168329617 | 1.52798417667632 | -0.368348737110535 | 0.71444169595194 | 0.883363574684855 | -5.93589282980215 |
| LINC00294 | LINC00294 | -0.0524614178273444 | 3.41477307428726 | -0.368281160461899 | 0.714491708596821 | 0.883372347311892 | -6.1623099970645 |
| lnc-NOC2L-6 | lnc-NOC2L-6 | -0.0628341096044678 | 2.12804336648865 | -0.368156047638666 | 0.714584306407416 | 0.883420936493097 | -6.01074016322611 |
| lnc-ZNF460-2 | lnc-ZNF460-2 | 0.0387158155157064 | 5.02052738105383 | 0.36810192391819 | 0.714624365504545 | 0.883420936493097 | -6.31323203724967 |
| lnc-TMEM132D-4 | lnc-TMEM132D-4 | -0.0764430756860371 | 0.886811150180577 | -0.368023789940384 | 0.714682196974982 | 0.883420936493097 | -5.83246234332405 |
| lnc-GCN1-1 | lnc-GCN1-1 | -0.0754622223134289 | 6.44988289141939 | -0.367981282147224 | 0.714713660160838 | 0.883420936493097 | -6.34316667650196 |
| ENSG00000102524.11 | TNFSF13B | -0.0666398284673257 | 1.12775401243285 | -0.367899123523236 | 0.714774473304915 | 0.883420936493097 | -5.86023105764153 |
| lnc-ZFHX4-9 | lnc-ZFHX4-9 | -0.096110567918693 | 1.18527242095603 | -0.367880135677848 | 0.714788528220184 | 0.883420936493097 | -5.85088924104279 |
| lnc-KCTD7-1 | lnc-KCTD7-1 | 0.0660583056575659 | 2.11256861396124 | 0.367630930112015 | 0.714973000934227 | 0.883553431373726 | -6.00849910859633 |

| | | | | | | | |
|---|---|---|---|---|---|---|---|
| lnc-AGO2-8 | lnc-AGO2-8 | -0.112842065929901 | 1.04401532955161 | -0.367601596004665 | 0.714994716440944 | 0.883553431373726 | -5.84572347695918 |
| lnc-SLC7A9-4 | lnc-SLC7A9-4 | 0.0448740308593126 | -0.0244765801689248 | 0.367351831665603 | 0.715179622119024 | 0.883553431373726 | -5.57747799114672 |
| ENSG00000084731.14 | KIF3C | 0.0702773774724967 | 1.33052101314445 | 0.367293810542758 | 0.71522257882789 | 0.883553431373726 | -5.90449319197058 |
| lnc-NUP210L-2 | lnc-NUP210L-2 | 0.0609954167866977 | 4.27973691037082 | 0.36729134974202 | 0.715224400735046 | 0.883553431373726 | -6.25533747144667 |
| ENSG00000180398.12 | MCFD2 | 0.0513538423355495 | 5.41839163443297 | 0.36721073215611 | 0.715284088640949 | 0.883553431373726 | -6.33090093054807 |
| ENSG00000171490.12 | RSL1D1 | -0.0493913447593251 | 6.49648202804116 | -0.367146830960708 | 0.715331401293115 | 0.883553431373726 | -6.34344561557142 |
| lnc-PBDC1-3 | lnc-PBDC1-3 | 0.0556967978620535 | 2.71017286052928 | 0.367128858922813 | 0.715344708052279 | 0.883553431373726 | -6.08064120322814 |
| lnc-TGOLN2-1 | lnc-TGOLN2-1 | 0.0737080319774732 | 1.27850358887648 | 0.367092754825267 | 0.715371440323786 | 0.883553431373726 | -5.88029391737848 |
| ENSG00000266472.5 | MRPS21 | -0.0387655558467817 | 5.12543986791007 | -0.367077421913036 | 0.715382793260128 | 0.883553431373726 | -6.32040359116061 |
| lnc-SBDS-17 | lnc-SBDS-17 | 0.0526909306372154 | 0.830972475253688 | 0.366987450748745 | 0.715449411853298 | 0.883553431373726 | -5.80238626365499 |
| ENSG00000139725.7 | RHOF | -0.0794312716118863 | 4.18230759550398 | -0.366982502767934 | 0.715453075620255 | 0.883553431373726 | -6.25471507059854 |
| lnc-DCANP1-2 | lnc-DCANP1-2 | 0.0862335849075299 | 0.311554796324234 | 0.366973372403749 | 0.715459836280018 | 0.883553431373726 | -5.66154681951304 |
| ENSG00000186704.9 | DTX2P1 | -0.0706721299373068 | 2.28731874005446 | -0.366923586807479 | 0.715496700876319 | 0.883553431373726 | -6.02622677511417 |
| ENSG00000104524.13 | PYCR3 | -0.0716349542740611 | 2.97362927994424 | -0.366775628562041 | 0.715606263144865 | 0.883635713858705 | -6.10835732965375 |
| ENSG00000105341.18 | DMAC2 | -0.0565721832777757 | 3.49170263177341 | -0.366615012948485 | 0.715725204992146 | 0.883729567747651 | -6.17608274874698 |
| ENSG00000279665.1 | | -0.0839719832069025 | -0.0599712732272285 | -0.366261799406916 | 0.715986797916329 | 0.883906894432678 | -5.58979560165299 |
| ENSG00000135090.13 | TAOK3 | 0.0447748638668353 | 6.45299949796815 | 0.366229037440765 | 0.716011063454878 | 0.883906894432678 | -6.34409662395571 |
| ENSG00000284108.1 | MIR611 | -0.0818612432728741 | 0.444308845022539 | -0.366216234236951 | 0.716020546380266 | 0.883906894432678 | -5.73823072755057 |
| lnc-HEMGN-4 | lnc-HEMGN-4 | 0.104972648192143 | -0.221598258790783 | 0.366141936218122 | 0.716075577452489 | 0.883906894432678 | -5.56013834059423 |
| ENSG00000205084.10 | TMEM231 | -0.140044774541583 | 1.03937207104497 | -0.366135539940018 | 0.716085337334141 | 0.883906894432678 | -5.84264705862193 |
| ENSG00000167962.13 | ZNF598 | -0.0553843050287885 | 3.52822804599055 | -0.366057880560375 | 0.716137837656298 | 0.883906894432678 | -6.18832971703636 |
| ENSG00000166311.9 | SMPD1 | -0.055796915256193 | 3.13944341992985 | -0.366015236190642 | 0.716169425178728 | 0.883906894432678 | -6.12960223166812 |
| ENSG00000174136.11 | RGMB | 0.0630230108385597 | 3.78266977921624 | 0.365880187243975 | 0.716269461905502 | 0.883977355462342 | -6.20777783208494 |
| ENSG00000090924.14 | PLEKHG2 | 0.0670551508898365 | 4.54436276309206 | 0.365806379111792 | 0.716324136984424 | 0.883991828739757 | -6.2803429304079 |
| ENSG00000279433.1 | | 0.113167595933335 | 0.839374330149964 | 0.3655121567388 | 0.716542103970833 | 0.884171914279111 | -5.77068045278564 |
| lnc-DKK4-4 | lnc-DKK4-4 | -0.0460624691498805 | 1.82970665966433 | -0.365493429543261 | 0.716555978335998 | 0.884171914279111 | -5.9685963073171 |
| ENSG00000079432.7 | CIC | -0.0656906391360383 | 5.7722290453135 | -0.365300153623582 | 0.716699175787597 | 0.88427070279194 | -6.33907105182186 |
| ZBTB40-IT1 | ZBTB40-IT1 | 0.0603474148762397 | 0.873619020962703 | 0.365246204803159 | 0.716739148121442 | 0.88427070279194 | -5.81302034972324 |
| ENSG00000130640.13 | TUBGCP2 | 0.0399157471946721 | 5.51408079033682 | 0.365211418520338 | 0.716764922770282 | 0.88427070279194 | -6.33420183942823 |
| ENSG00000197943.9 | PLCG2 | 0.0503063042968126 | 8.3474045006112 | 0.365093150232064 | 0.71685255531608 | 0.884280285076456 | -6.31695631276783 |
| ENSG00000160695.14 | VPS11 | -0.052599216520191 | 5.82825772673864 | -0.365039464791153 | 0.716892335570078 | 0.884280285076456 | -6.34131300248988 |
| ENSG00000189283.9 | FHIT | 0.163920014748375 | 0.976608191827699 | 0.365047647096961 | 0.716901574943926 | 0.884280285076456 | -5.79016592115458 |
| lnc-TLR10-1 | lnc-TLR10-1 | -0.159431457234577 | 0.626171249903906 | -0.364910396461697 | 0.71699755712845 | 0.884345680631285 | -5.77170981266391 |
| ENSG00000242802.7 | AP5Z1 | 0.0638827289243578 | 4.97198102594209 | 0.364838480029044 | 0.717041269878956 | 0.884346603027057 | -6.31565918553176 |
| ENSG00000201403.1 | SNORD14B | 0.0525944326825254 | 1.20807072296344 | 0.36474444030516 | 0.717110959293816 | 0.884379561123104 | -5.88857817430747 |
| lnc-PUS7-2 | lnc-PUS7-2 | -0.0746480877736648 | 1.06789457819614 | -0.364548464489047 | 0.717256197671637 | 0.884505680706687 | -5.86405957746901 |
| lnc-ZNF611-5 | lnc-ZNF611-5 | -0.0396292400055684 | 1.41606072444339 | -0.364317715999429 | 0.717427219773933 | 0.884663579235973 | -5.91077492055637 |
| lnc-HNRNPU-1 | lnc-HNRNPU-1 | -0.0419898056922205 | 6.01527582114731 | -0.364002964860863 | 0.717660525132041 | 0.884848153545878 | -6.34388510548094 |
| ENSG00000086598.10 | TMED2 | 0.043505379646982 | 6.23580860231621 | 0.36399978704207 | 0.717662880789724 | 0.884848153545878 | -6.3449541507426 |
| ENSG00000158717.10 | RNF166 | -0.0677146814386536 | 3.14412452977203 | -0.36392287535739 | 0.717719894832133 | 0.884865444464199 | -6.13924023415765 |
| ENSG00000111644.7 | ACRBP | -0.172571317586009 | 1.19208267372707 | -0.363888887786942 | 0.717771117502211 | 0.88487559364806 | -5.83622068428194 |
| lnc-CRLF3-4 | lnc-CRLF3-4 | 0.0504652348130936 | 1.67206849381095 | 0.363705509498487 | 0.717881035280347 | 0.884958097213594 | -5.94888485586052 |
| ENSG00000256028.2 | | -0.0525696039760613 | 4.7492415959352 | -0.363524496711748 | 0.718015235922717 | 0.885042136837542 | -6.30060054177206 |
| lnc-CSGALNACT2-1 | lnc-CSGALNACT2-1 | -0.105719506973887 | 0.136482586748992 | -0.363497561804136 | 0.718035205900506 | 0.885042136837542 | -5.64328738225753 |
| ENSG00000178980.14 | SELENOW | -0.0458707045826237 | 3.52351884534575 | -0.363328171927626 | 0.718160798876938 | 0.885139845038359 | -6.18097679689366 |
| ENSG00000261732.1 | | -0.0562675347462118 | 4.52390217210514 | -0.363232297701165 | 0.718231887668246 | 0.885139845038359 | -6.28052397594457 |
| ENSG00000185201.16 | IFITM2 | -0.100497266942202 | 4.5173194811476 | -0.363242651036806 | 0.718243486947028 | 0.885139845038359 | -6.29551637772094 |
| ENSG00000203485.12 | INF2 | -0.0798166754991545 | 4.43133816922497 | -0.363113836554619 | 0.718319727690774 | 0.885180803152679 | -6.27437655338184 |
| lnc-NIF3L1-3 | lnc-NIF3L1-3 | -0.0753109048430267 | 2.93547367938313 | -0.362907187035483 | 0.718472969425808 | 0.885316638511219 | -6.11269559526545 |
| ENSG00000256188.3 | TAS2R30 | 0.092354752172422 | 3.1407011759394 | 0.362709155073794 | 0.718619831781977 | 0.885444597280361 | -6.14172012491383 |
| ENSG00000169925.16 | BRD3 | -0.0375408860364121 | 3.49319560520592 | -0.362530656093409 | 0.718752217520437 | 0.885535025984168 | -6.18365933757165 |
| ENSG00000196284.15 | SUPT3H | -0.045782699927821 | 3.24401673734998 | -0.362494184325323 | 0.718779268285357 | 0.885535025984168 | -6.1479235605568 |
| lnc-TSPAN8-2 | lnc-TSPAN8-2 | 0.0552725349224552 | 2.49667529115328 | 0.362430913349493 | 0.718826196634444 | 0.88553983752255 | -6.04911404333698 |
| ENSG00000167196.13 | FBXO22 | -0.0519609491325097 | 5.12837718212819 | -0.362361374277808 | 0.718877775323239 | 0.885550377011086 | -6.32108814088558 |
| ENSG00000148308.17 | GTF3C5 | 0.0306335462732266 | 4.92527838912965 | 0.362223535394941 | 0.718980017433022 | 0.885573656457146 | -6.31171394207961 |
| lnc-HMBOX1-1 | lnc-HMBOX1-1 | -0.0489328538413361 | 5.52614312673899 | -0.362215150301871 | 0.718986237251103 | 0.885573656457146 | -6.33489998296495 |
| lnc-SNX17-1 | lnc-SNX17-1 | -0.079735439978808 | 0.0818359058283259 | -0.362161887071393 | 0.719025746815907 | 0.885573656457146 | -5.64560368927893 |
| ENSG00000229474.6 | PATL2 | 0.082611561622346 | 6.24008703219919 | 0.361890459692986 | 0.719232716204101 | 0.885768736722146 | -6.34513220385046 |
| ENSG00000160741.16 | CRTC2 | 0.0493363459190785 | 5.04026101913042 | 0.361782426844486 | 0.71930724507846 | 0.885768736722146 | -6.31890843477107 |
| ENSG00000232479.2 | KRT8P52 | -0.0822782111413369 | 0.371273872172164 | -0.361774345248486 | 0.719313240747737 | 0.885768736722146 | -5.69439955388959 |
| ENSG00000127419.16 | TMEM175 | 0.110840170993576 | 4.4677854599055 | 0.361726668775015 | 0.71937705665334 | 0.88579432622013 | -6.271109070198 |
| lnc-C6orf120-3 | lnc-C6orf120-3 | -0.0637974900024665 | 0.945813318776082 | -0.361621133305428 | 0.719426910805534 | 0.885802722085919 | -5.85620676852363 |
| lnc-GPAT2-6 | lnc-GPAT2-6 | -0.0826213760023442 | 1.44834860674594 | -0.361460344883418 | 0.719546208868721 | 0.885896615453098 | -5.92832172294914 |
| ENSG00000187699.10 | C2orf88 | -0.21593478346508 | 2.22385888304528 | -0.361412439436086 | 0.719639681329413 | 0.885958703319774 | -6.00888504840879 |
| ENSG00000213754.2 | KPNA4P1 | -0.0920413218943347 | 0.567707657628484 | -0.361139339404225 | 0.719784402226706 | 0.886046736260516 | -5.73912389405872 |
| ENSG00000207088.1 | SNORA7B | 0.0977371989361557 | 0.888893998418435 | 0.361059750776793 | 0.719843463156409 | 0.886046736260516 | -5.81647304569855 |
| lnc-MYCL-2 | lnc-MYCL-2 | 0.052395164081155 | 0.411524444747554 | 0.360996332587656 | 0.719890525607481 | 0.886046736260516 | -5.72569391950545 |
| ENSG00000196584.2 | XRCC2 | -0.0876803637290371 | 0.710646036696799 | -0.360988633552399 | 0.719896239113674 | 0.886046736260516 | -5.80181657429374 |

| | | | | | | | |
|---|---|---|---|---|---|---|---|
| ENSG00000196967.10 | ZNF585A | -0.0439509452751734 | 4.20497998734816 | -0.360872209037249 | 0.719982640509632 | 0.886046736260516 | -6.24791822118158 |
| ENSG00000066739.11 | ATG2B | 0.0407607192443082 | 6.58589286556586 | 0.360867273198796 | 0.719986303594354 | 0.886046736260516 | -6.34524042710304 |
| ENSG00000152229.18 | PSTPIP2 | -0.239652230814531 | 0.129903663172708 | -0.360875510464953 | 0.720012520560336 | 0.886046736260516 | -5.58535144762769 |
| ENSG00000154359.12 | LONRF1 | -0.0670120923920026 | 5.48237102800859 | -0.360618932055769 | 0.7201706161622 | 0.88615592580812 | -6.33612239917728 |
| ENSG00000170581.13 | STAT2 | 0.0661441284836739 | 7.27819958243376 | 0.3605268381616 | 0.720238970222603 | 0.88615592580812 | -6.34002688935005 |
| ENSG00000174371.16 | EXO1 | 0.14184910558148 | 0.534943718804941 | 0.360479256139357 | 0.720274287525869 | 0.88615592580812 | -5.70668469036698 |
| lnc-SGTB-7 | lnc-SGTB-7 | 0.117335942581484 | 1.26558731696515 | 0.360443374557109 | 0.720300920697711 | 0.88615592580812 | -5.87756199555042 |
| ENSG00000265972.5 | TXNIP | -0.0873176960273483 | 11.4026518533868 | -0.360371759466307 | 0.720354132524147 | 0.88615592580812 | -6.21416617064177 |
| lnc-FGD6-2 | lnc-FGD6-2 | -0.0430522158729665 | 2.03154236191923 | -0.360352298168058 | 0.720368523896884 | 0.88615592580812 | -6.01002042956415 |
| lnc-ANGPTL1-3 | lnc-ANGPTL1-3 | -0.0876038754966554 | 0.967742987247547 | -0.360217434111885 | 0.720468633491363 | 0.88615592580812 | -5.82800954929322 |
| ENSG00000136243.16 | NUP42 | 0.0454773849881898 | 4.22665177794862 | 0.360182164088325 | 0.720494815250027 | 0.88615592580812 | -6.26357403676635 |
| ENSG00000257949.6 | TEN1 | -0.0544855447531134 | 2.26144682759295 | -0.360137259416145 | 0.720528149523531 | 0.88615592580812 | -6.03169209583506 |
| ENSG00000174238.14 | PITPNA | -0.05008440304203 | 4.59944732919725 | -0.360132372334016 | 0.720531777404883 | 0.88615592580812 | -6.29300319401353 |
| ENSG00000070190.12 | DAPP1 | 0.0819417013765591 | 5.73717610617416 | 0.359917830742518 | 0.720696372829973 | 0.886273901517972 | -6.34000155994185 |
| ENSG00000133226.16 | SRRM1 | 0.0454808700966947 | 6.86969839039835 | 0.359887155440916 | 0.720713820318117 | 0.886273901517972 | -6.34402797014868 |
| lnc-HMG20A-6 | lnc-HMG20A-6 | -0.043654633864822 | 3.74185245568258 | -0.359783555792579 | 0.720790735043213 | 0.88631553255239 | -6.20929715450295 |
| lnc-ZNF329-2 | lnc-ZNF329-2 | 0.168206264270767 | -0.634039563176864 | 0.359693887379206 | 0.720857309265842 | 0.886344444242463 | -5.37427120743166 |
| ENSG00000141556.20 | TBCD | -0.0521617790663216 | 4.27009230454208 | -0.359605407501341 | 0.720923003209238 | 0.886362105660803 | -6.25814200462387 |
| lnc-UCK1-4 | lnc-UCK1-4 | -0.122374962594784 | 0.389966278026513 | -0.359558544211037 | 0.720957798813252 | 0.886362105660803 | -5.68719262139749 |
| ENSG00000164117.13 | FBXO8 | 0.0528021638112341 | 3.8624568645393 | 0.359334322966441 | 0.72112428950419 | 0.886472262268743 | -6.21332038664912 |
| CDC42-IT1 | CDC42-IT1 | 0.0774310958471032 | 1.86075743276656 | 0.359321871582889 | 0.721133535414072 | 0.886472262268743 | -5.98629346685741 |
| ENSG00000197312.11 | DDI2 | -0.0297441728678942 | 5.86275493080546 | -0.359126003878693 | 0.721278984655735 | 0.886566583818417 | -6.34378463932047 |
| LINC01237 | LINC01237 | -0.0716472390088603 | 3.2077943872261 | -0.358969738097751 | 0.721395033424649 | 0.886566583818417 | -6.14301732500566 |
| ENSG00000122299.11 | ZC3H7A | 0.0577538318818105 | 6.37259222019463 | 0.358963736397519 | 0.721399490642484 | 0.886566583818417 | -6.34678590882609 |
| lnc-MMADHC-12 | lnc-MMADHC-12 | -0.0600052358897929 | 1.11177533248338 | -0.358925485691944 | 0.72142789811075 | 0.886566583818417 | -5.8799194919125 |
| ENSG00000146192.14 | FGD2 | -0.0794760048091874 | 6.56944264933907 | -0.358913845446989 | 0.721440026691103 | 0.886566583818417 | -6.34613440566343 |
| lnc-SNX20-1 | lnc-SNX20-1 | 0.0635636209668222 | 3.86276894399839 | 0.35887054505748 | 0.721468701304887 | 0.886566583818417 | -6.21338720959955 |
| ENSG00000072864.14 | NDE1 | -0.0367586781475584 | 5.08533213908513 | -0.358662358482514 | 0.721623324318232 | 0.886643241236203 | -6.32031701678446 |
| ENSG00000227161.1 | | 0.0828290672570761 | -0.195404515990202 | 0.35863097314112 | 0.721646635662182 | 0.886643241236203 | -5.54418822668756 |
| ENSG00000161011.19 | SQSTM1 | -0.0528412671741609 | 7.42475799518076 | -0.358561742823988 | 0.721698057165797 | 0.886643241236203 | -6.33658300220687 |
| ENSG00000175040.5 | CHST2 | -0.0957145335735907 | 6.00823049985213 | -0.358533023251581 | 0.721740260679334 | 0.886643241236203 | -6.34539745124333 |
| lnc-BORCS5-4 | lnc-BORCS5-4 | -0.0794957771588968 | -0.0703649717922277 | -0.358488400006135 | 0.721752534690971 | 0.886643241236203 | -5.56305872134859 |
| ENSG00000237757.2 | EEF1A1P30 | -0.0903595544794508 | 0.201878937088081 | -0.358410979811087 | 0.721810042393223 | 0.886643241236203 | -5.66995299401934 |
| ENSG00000104177.17 | MYEF2 | -0.246775707498839 | 4.01975149799378 | -0.358475457689999 | 0.721832618830834 | 0.886643241236203 | -6.25882916846636 |
| ENSG00000260007.3 | | -0.0511094586757253 | 2.03244093035199 | -0.358316439268463 | 0.721880269293847 | 0.886648859223968 | -6.0020306811186 |
| ENSG00000130811.11 | EIF3G | -0.050923036741465 | 6.29466315477267 | -0.358026844004011 | 0.722095402456332 | 0.886860174757366 | -6.34702469331109 |
| lnc-EXOC1-3 | lnc-EXOC1-3 | -0.0712157715678484 | -0.268037383512378 | -0.357874916261256 | 0.722208274924902 | 0.88694587844745 | -5.58439042859125 |
| ENSG00000236670.1 | KRT18P5 | -0.081754257350762 | 0.239152347368409 | -0.357770413654766 | 0.722285917225739 | 0.886974213261682 | -5.6539766054749 |
| ENSG00000267261.5 | | -0.0410529022627459 | 4.18021905617701 | -0.35772786281087 | 0.72231753207464 | 0.886974213261682 | -6.25720200153054 |
| lnc-HEYL-4 | lnc-HEYL-4 | -0.0518406139925278 | 2.87836141533067 | -0.357484791086459 | 0.722498141333981 | 0.887105389091396 | -6.09079218037784 |
| ENSG00000114120.12 | SLC25A36 | -0.0501511718855934 | 6.62627513101918 | -0.357468085740087 | 0.722510554473505 | 0.887105389091396 | -6.34582323095837 |
| ENSG00000089597.17 | GANAB | -0.0345174593651505 | 6.71022716875535 | -0.357408605546388 | 0.722554752671266 | 0.887106738695656 | -6.34606556037625 |
| ENSG00000104442.9 | ARMC1 | -0.0455664889195946 | 4.40156999014013 | -0.357332350725438 | 0.722611417064432 | 0.887123392427365 | -6.2783275379923 |
| ENSG00000112511.17 | PHF1 | 0.0576033947247785 | 6.28433171919093 | 0.35701421401427 | 0.722847839096259 | 0.887269835401283 | -6.3474553788936 |
| ENSG00000164597.13 | COG5 | -0.0296606022185459 | 5.65745173762412 | -0.357012654213959 | 0.722848998323184 | 0.887269835401283 | -6.34028805746265 |
| ENSG00000156970.12 | BUB1B | -0.122690349433623 | 1.57187096587119 | -0.356967568178962 | 0.722884206006298 | 0.887269835401283 | -5.90585638893313 |
| ENSG00000234882.1 | EIF3EP1 | -0.0553641335204962 | 1.04798029263489 | -0.356864782398346 | 0.72295889806764 | 0.887269835401283 | -5.84180128468103 |
| ENSG00000140386.12 | SCAPER | -0.0494457845104579 | 5.73760914628431 | -0.356818576936982 | 0.722993239621122 | 0.887269835401283 | -6.34271404042522 |
| lnc-POLRMT-4 | lnc-POLRMT-4 | 0.0631430971245872 | 1.90960765015216 | 0.356781312820267 | 0.723020936067967 | 0.887269835401283 | -5.99025134020299 |
| ENSG00000163563.7 | MNDA | -0.326589076311969 | 2.93189281722128 | -0.356860165883246 | 0.723032451498116 | 0.887269835401283 | -6.08904089431821 |
| ENSG00000168003.16 | SLC3A2 | 0.0587402434288427 | 5.65713419247518 | 0.356457949468832 | 0.723261290717212 | 0.88747026926151 | -6.34219075182405 |
| lnc-LDLRAP1-1 | lnc-LDLRAP1-1 | 0.124530172606891 | -0.534038142522847 | 0.356430066238554 | 0.723282017532032 | 0.88747026926151 | -5.4669011964974 |
| lnc-HIST1H3A-4 | lnc-HIST1H3A-4 | 0.0656897261656024 | 1.28695692097907 | 0.356346150710018 | 0.723344396853427 | 0.887493903220915 | -5.88866998938337 |
| lnc-KLHL5-2 | lnc-KLHL5-2 | 0.0904225244207841 | -0.0520021473318543 | 0.356082039352267 | 0.723540738697988 | 0.887681887382768 | -5.57534417988139 |
| ENSG00000160208.12 | RRP1B | -0.0425546792591494 | 4.98499740682847 | -0.355901882852251 | 0.72367467884862 | 0.887720658040192 | -6.31689449543175 |
| lnc-FARS2-2 | lnc-FARS2-2 | 0.303529323996343 | -1.2101790824417 | 0.355910259810436 | 0.723682017296137 | 0.887720658040192 | -5.25341556739643 |
| ENSG00000143569.18 | UBAP2L | 0.036208610352138 | 6.94777207523626 | 0.355865503428247 | 0.723701726760648 | 0.887720658040192 | -6.34462966121862 |
| lnc-SLC25A52-2 | lnc-SLC25A52-2 | -0.0397390311390922 | 2.35491953308058 | -0.355609684126026 | 0.723891937167475 | 0.887901063269063 | -6.03393596156327 |
| lnc-RNF139-4 | lnc-RNF139-4 | -0.0487095394528876 | 1.88802281101452 | -0.35538726961649 | 0.724057324318894 | 0.888051001457263 | -5.9783480333888 |
| lnc-YEATS2-1 | lnc-YEATS2-1 | 0.0452103721944801 | 1.91269078354857 | 0.355238952099189 | 0.724167620454584 | 0.888133357076607 | -5.9859843289425 |
| lnc-RAD51D-2 | lnc-RAD51D-2 | -0.0476956801439697 | -0.0255758926035409 | -0.355077191664112 | 0.724287920160378 | 0.888208699400507 | -5.61514843714179 |
| ENSG00000284031.1 | MIR4761 | -0.0752545980684289 | -0.0979316484106236 | -0.355040298455894 | 0.72431535827807 | 0.888208699400507 | -5.60615569136483 |
| lnc-URGCP-1 | lnc-URGCP-1 | -0.0452320469569422 | 0.217677401203214 | -0.354978493698407 | 0.724361324368105 | 0.888212149378572 | -5.65984697236898 |
| lnc-SOCS4-1 | lnc-SOCS4-1 | 0.0505081348665408 | 2.45594863199977 | 0.35481475936699 | 0.724483103576725 | 0.888308555484586 | -6.05253879826012 |
| ENSG00000013306.15 | SLC25A39 | -0.0502439747908383 | 4.65419528380948 | -0.354719105407735 | 0.724554250601536 | 0.888342872297559 | -6.29767668856527 |
| ENSG00000080345.17 | RIF1 | 0.0365676518018354 | 7.03623700392994 | 0.354659189699185 | 0.724598816916121 | 0.888344597568915 | -6.34380464975783 |
| lnc-DUSP2-2 | lnc-DUSP2-2 | -0.0666504638825485 | 4.34824318545034 | -0.354531644250837 | 0.724693690590101 | 0.888364813636577 | -6.26740628060133 |
| lnc-PSD-2 | lnc-PSD-2 | -0.1332318679172 | 1.05995865826235 | -0.35452835933044 | 0.72470162686546 | 0.888364813636577 | -5.86803562408333 |

| | | | | | | | |
|---|---|---|---|---|---|---|---|
| ENSG00000038532.15 | CLEC16A | 0.0364708941687617 | 5.44045976655114 | 0.35433976533372 | 0.724836426453703 | 0.888477141835193 | -6.33427183146906 |
| lnc-DMC1-3 | lnc-DMC1-3 | -0.0382254317852233 | 6.17372915163775 | -0.354266110751671 | 0.724891219629179 | 0.888491393653748 | -6.34775560657464 |
| lnc-FAM105A-9 | lnc-FAM105A-9 | 0.242750925809948 | -1.29630283989161 | 0.35413244606813 | 0.724990659293642 | 0.888550204561809 | -5.24412299769961 |
| ENSG00000185900.9 | POMK | -0.0529267509828646 | 3.45447212028589 | -0.35404078047679 | 0.725058856565873 | 0.888550204561809 | -6.17270164239733 |
| ENSG00000139737.21 | SLAIN1 | 0.0608113010791658 | 3.4250515834877 | 0.354027537934177 | 0.725068708925819 | 0.888550204561809 | -6.17846620825372 |
| lnc-SAMD7-1 | lnc-SAMD7-1 | 0.233554513977614 | 0.550307341338495 | 0.353938842071146 | 0.725183835847508 | 0.888623744617029 | -5.82858250429254 |
| ENSG00000102910.13 | LONP2 | -0.0293855512362544 | 6.94131901002382 | -0.353821048738585 | 0.725222341553633 | 0.888623744617029 | -6.34517001135435 |
| ENSG00000213445.9 | SIPA1 | -0.08948138347239 | 5.25805192346793 | -0.35376912291036 | 0.725275340511815 | 0.888623744617029 | -6.3333980795804 |
| ENSG00000127603.25 | MACF1 | 0.0760591884873953 | 8.12471078431952 | 0.353714783987806 | 0.725301409394456 | 0.888623744617029 | -6.32662656068446 |
| lnc-TTBK2-1 | lnc-TTBK2-1 | -0.0617569761887325 | -0.17676765095938 | -0.353444391673184 | 0.725502612423459 | 0.888770036449073 | -5.56725232021174 |
| ENSG00000150456.10 | EEF1AKMT1 | -0.0731510070574961 | 0.011196133770625 | -0.353323953662727 | 0.725592238494741 | 0.888770036449073 | -5.58617057236168 |
| ENSG00000170006.11 | TMEM154 | -0.0758737838826402 | 7.20265205827162 | -0.353266397022078 | 0.725635515971276 | 0.888770036449073 | -6.34380676931239 |
| lnc-C10orf55-1 | lnc-C10orf55-1 | 0.0424207289436742 | 2.07563138952087 | 0.353260911752787 | 0.725639153796736 | 0.888770036449073 | -6.0030853341657 |
| ENSG00000174842.16 | GLMN | 0.0570309819345556 | 3.17003088312429 | 0.353237849118032 | 0.725656317097619 | 0.888770036449073 | -6.1460434000459 |
| ENSG00000128266.8 | GNAZ | -0.172158676775639 | 2.24552863219135 | -0.3532720595590201 | 0.725679892754366 | 0.888770036449073 | -6.0748426542652 |
| ENSG00000166902.4 | MRPL16 | -0.0326153181241353 | 3.86835599067617 | -0.353118648709703 | 0.725745028771386 | 0.888796925519215 | -6.22267489757279 |
| ENSG00000188177.13 | ZC3H6 | -0.0494393517456038 | 5.17617187869138 | -0.353050993819726 | 0.725795380785246 | 0.888805706967082 | -6.32334714551903 |
| ENSG00000176182.5 | MYPOP | 0.0830777501873436 | -0.0680311310043192 | 0.352892361793281 | 0.725913447159912 | 0.88883168511076 | -5.57994626829021 |
| ENSG00000125695.12 | | -0.035321816324664 | 3.78112780016256 | -0.352837388004775 | 0.725954364531628 | 0.88883168511076 | -6.21184619893485 |
| lnc-MAT1A-1 | lnc-MAT1A-1 | -0.0619011741683966 | 1.24635376890043 | -0.352826263325681 | 0.725962644805906 | 0.88883168511076 | -5.89185389886632 |
| lnc-CLK1-4 | lnc-CLK1-4 | 0.03160302059272 | 2.1433058076224 | 0.352738801218582 | 0.726027745388412 | 0.88883168511076 | -6.01665158248224 |
| lnc-CCDC114-2 | lnc-CCDC114-2 | 0.0669528254669425 | 3.59258757649812 | 0.352732402169654 | 0.72603250846656 | 0.88883168511076 | -6.18379262088489 |
| ENSG00000176871.8 | WSB2 | 0.0899647238839041 | 5.72928656966139 | 0.352623852065245 | 0.72612930121664 | 0.888859839588373 | -6.34398812664781 |
| lnc-PTGR2-2 | lnc-PTGR2-2 | 0.0701922401414122 | -0.41652427613695 | 0.352551088529825 | 0.726167472305654 | 0.888859839588373 | -5.49154166024887 |
| lnc-ZNF366-11 | lnc-ZNF366-11 | -0.0482789949599391 | 2.78156302787152 | -0.352527463465387 | 0.726185058665796 | 0.888859839588373 | -6.09193140627797 |
| ENSG00000204954.9 | UQCC6 | -0.0558485831955331 | 0.569684735484416 | -0.352451287927715 | 0.726241764309601 | 0.88887638905777 | -5.76903236679624 |
| ENSG00000139370.11 | SLC15A4 | -0.0689931118023743 | 4.55118281884554 | -0.352232849243749 | 0.726404380325595 | 0.889000683924786 | -6.28969016443176 |
| ENSG00000163755.8 | HPS3 | -0.0467421745920227 | 5.59704453625532 | -0.352198840193702 | 0.726429699408782 | 0.889000683924786 | -6.34103423641153 |
| ENSG00000124126.13 | PREX1 | 0.0673275049347657 | 7.3176102334351 | 0.352080190834475 | 0.726518033996367 | 0.889055927095554 | -6.34172901881622 |
| ENSG00000183060.15 | LYSMD4 | 0.0723575896514439 | 3.28607633024944 | 0.35194290885684 | 0.726620245289791 | 0.88912814391533 | -6.16293963659899 |
| ENSG00000258924.4 | | 0.0323827498178506 | 1.9701066085894 | 0.351784903956475 | 0.726737891771502 | 0.889219238279207 | -5.99796626585527 |
| lnc-RAD51-2 | lnc-RAD51-2 | -0.100386232855605 | 0.335854057074028 | -0.351661885562389 | 0.726829492800002 | 0.889278455109221 | -5.75349259755275 |
| lnc-ERRFI1-4 | lnc-ERRFI1-4 | 0.065969587418984 | 1.04715959203196 | 0.351410547402496 | 0.727016654895473 | 0.889423615540681 | -5.84455722301525 |
| lnc-STK26-6 | lnc-STK26-6 | 0.0722887953775806 | 1.57157774194298 | 0.351386505108713 | 0.727034559173685 | 0.889423615540681 | -5.94199347370514 |
| ENSG00000221914.9 | PPP2R2A | 0.0429966061361053 | 5.14370227228815 | 0.35126260218418 | 0.727126832036501 | 0.889436742836438 | -6.32653463474515 |
| ENSG00000159202.17 | UBE2Z | 0.0356814224257915 | 6.19415020076377 | 0.351244228268391 | 0.727140515789801 | 0.889436742836438 | -6.34939594570524 |
| ENSG00000104738.17 | MCM4 | 0.0826851237487496 | 4.04739077552023 | 0.351198094851592 | 0.727174926320341 | 0.889436742836438 | -6.24584852790034 |
| ENSG00000284431.1 | | -0.0405207116693367 | 5.15505959645464 | -0.350943070249022 | 0.7273648124529 | 0.889616134928875 | -6.32753236260109 |
| lnc-CDKN2B-2 | lnc-CDKN2B-2 | 0.0982220485951788 | 0.889047969990902 | 0.350729946440742 | 0.727523557560377 | 0.889638312714106 | -5.84331467028252 |
| ENSG00000125354.22 | SEPTIN6 | -0.041677991426412 | 7.69327789332585 | -0.350718585678537 | 0.727532019955563 | 0.889638312714106 | -6.33602802467987 |
| lnc-PSORS1C2-1 | lnc-PSORS1C2-1 | 0.0560842052006419 | 0.856013938589957 | 0.350669704665897 | 0.72756843079651 | 0.889638312714106 | -5.81003069939417 |
| ENSG00000254995.4 | STX16-NPEPL1 | 0.0508557429638001 | 5.15585411532189 | 0.350632533624264 | 0.727596119456228 | 0.889638312714106 | -6.32534817897272 |
| ENSG00000092758.17 | COL9A3 | -0.116239894306251 | 5.33280370028824 | -0.350682723974258 | 0.727599055348066 | 0.889638312714106 | -6.33302697740063 |
| ENSG00000172301.10 | COPRS | -0.0655201860734187 | 2.35291531952998 | -0.350528158592964 | 0.727673870247877 | 0.889644839054721 | -6.05198609012268 |
| ENSG00000011376.10 | LARS2 | 0.0253002219581351 | 4.51245350340023 | 0.350287845266002 | 0.727852894896445 | 0.889644839054721 | -6.28307365043668 |
| ENSG00000157916.19 | RER1 | -0.0253323224562072 | 5.60841117529737 | -0.350281783279439 | 0.727857411053883 | 0.889644839054721 | -6.34163674836774 |
| lnc-FAM3C-2 | lnc-FAM3C-2 | -0.229085575644194 | -1.30503879700263 | -0.350261622883144 | 0.727872430544518 | 0.889644839054721 | -5.23736845035783 |
| ENSG00000164258.11 | NDUFS4 | 0.0416223371015878 | 3.587860408623 | 0.35021749960473 | 0.727905302754543 | 0.889644839054721 | -6.19095724352698 |
| lnc-STX17-2 | lnc-STX17-2 | 0.0676764594722725 | -0.194032568933263 | 0.35015931873467 | 0.727948648779946 | 0.889644839054721 | -5.54554312344686 |
| lnc-NAA25-4 | lnc-NAA25-4 | -0.102378473185963 | -0.296231919906564 | -0.350101239198722 | 0.727991920208116 | 0.889644839054721 | -5.52954808689435 |
| ENSG00000165548.10 | TMEM63C | 0.0800523449200923 | 1.13887241889299 | 0.350024964645255 | 0.728048748969385 | 0.889644839054721 | -5.88202666317287 |
| ENSG00000167333.12 | TRIM68 | 0.0418362218075803 | 3.18016487089351 | 0.349987152929267 | 0.728076921363357 | 0.889644839054721 | -6.13743525062996 |
| ENSG00000132612.15 | VPS4A | 0.0281875710563044 | 5.29409644672429 | 0.349985998840492 | 0.728077781246837 | 0.889644839054721 | -6.3329266512744 |
| ENSG00000130856.16 | ZNF236 | -0.0423265257557137 | 5.76299105927135 | -0.349930227165658 | 0.728119335787886 | 0.889644839054721 | -6.34489305005884 |
| ENSG00000110723.11 | EXPH5 | 0.217951151033963 | 2.11377652675046 | 0.350000286025082 | 0.728123060715922 | 0.889644839054721 | -5.97395499499658 |
| ENSG00000200120.1 | | -0.0798317831008356 | -0.326381940653802 | -0.349747794597619 | 0.728255269020787 | 0.889703596345205 | -5.56623096456163 |
| LINC00630 | LINC00630 | -0.0259598308506082 | 4.2141058990552 | -0.349744665673094 | 0.728257600506438 | 0.889703596345205 | -6.25700456902754 |
| ENSG00000092978.10 | GPATCH2 | 0.0373656902257666 | 4.57972649723038 | 0.349367523619813 | 0.72853864306173 | 0.889946976434583 | -6.29289664378098 |
| ENSG00000167770.11 | OTUB1 | 0.0326827921004265 | 4.84035661406413 | 0.349267056144377 | 0.728613516814815 | 0.889946976434583 | -6.31087464559658 |
| lnc-UBP1-3 | lnc-UBP1-3 | 0.057818974474478 | 0.116626079039581 | 0.349230737757912 | 0.728640583884914 | 0.889946976434583 | -5.64181584533456 |
| ENSG00000158941.16 | CCAR2 | -0.0308748008270268 | 5.9159120974686 | -0.34919760236635 | 0.728665279064264 | 0.889946976434583 | -6.34779795322304 |
| ENSG00000105669.13 | COPE | -0.0439366407563044 | 4.74712918177571 | -0.349055208278305 | 0.728771405998548 | 0.889946976434583 | -6.30586592413203 |
| lnc-CCSAP-1 | lnc-CCSAP-1 | -0.120539432568646 | 0.312253224913764 | -0.34896125904131 | 0.728841429722202 | 0.889946976434583 | -5.65286311627737 |
| ENSG00000165672.6 | PRDX3 | -0.0702541727129975 | 3.49760997119279 | -0.34894217021167 | 0.728855657595497 | 0.889946976434583 | -6.18405456644086 |
| lnc-SP140L-7 | lnc-SP140L-7 | 0.0533000239720325 | 2.40408158609687 | 0.348930550952485 | 0.728864318066283 | 0.889946976434583 | -6.05705333602233 |
| ENSG00000010292.12 | NCAPD2 | 0.0495762359445111 | 5.37456335220257 | 0.348910342197702 | 0.728879380844339 | 0.889946976434583 | -6.33637620637423 |
| lnc-SEMA4D-4 | lnc-SEMA4D-4 | -0.0853509991842654 | 1.0300252377374 | -0.348792552030531 | 0.728967178967259 | 0.889946976434583 | -5.86250070513248 |
| ENSG00000165502.6 | RPL36AL | -0.0629744155629623 | 6.50054684632967 | -0.348734010317729 | 0.729010816003757 | 0.889946976434583 | -6.35002848509774 |

| | | | | | | | |
|---|---|---|---|---|---|---|---|
| lnc-NT5C2-2 | lnc-NT5C2-2 | 0.0748427522650931 | 0.33597349603345 | 0.348727873563827 | 0.729015390397358 | 0.889946976434583 | -5.73212765379139 |
| ENSG00000074356.16 | NCBP3 | 0.0298722060179957 | 5.79840201068254 | 0.348723168479836 | 0.729018897617621 | 0.889946976434583 | -6.34731459119329 |
| lnc-NIF3L1-5 | lnc-NIF3L1-5 | 0.101253452977145 | -0.332784488189732 | 0.348413714210166 | 0.729249581054528 | 0.89010675843386 | -5.52274024591479 |
| ENSG00000120837.7 | NFYB | -0.0417417037355206 | 4.18657780051136 | -0.348385404540585 | 0.729270685834185 | 0.89010675843386 | -6.26272669777543 |
| ENSG00000227232.5 | WASH7P | 0.0723964104117407 | 1.47688588667537 | 0.348373553676309 | 0.729279520683507 | 0.89010675843386 | -5.94594410767626 |
| lnc-DNTTIP2-1 | lnc-DNTTIP2-1 | -0.0869252539487895 | 1.37262200083512 | -0.348052554742834 | 0.729518840357981 | 0.890346059358928 | -5.92075478919675 |
| ENSG00000188566.13 | NDOR1 | -0.0543562699005418 | 3.77766800808722 | -0.347761696357369 | 0.729735712416256 | 0.890473325985491 | -6.22057157591712 |
| ENSG00000196275.13 | GTF2IRD2 | -0.0511063109153735 | 2.13930215361853 | -0.347760473688305 | 0.729736624119402 | 0.890473325985491 | -5.99729209408186 |
| ENSG00000164713.9 | BRI3 | 0.0602122533257923 | 3.00456784216356 | 0.347738638529868 | 0.729752905928352 | 0.890473325985491 | -6.12991851140179 |
| lnc-RNF130-4 | lnc-RNF130-4 | 0.0379775951214025 | 2.61911504339699 | 0.347475658854092 | 0.729949011727997 | 0.890659820285574 | -6.07200954941715 |
| lnc-PLEKHG2-1 | lnc-PLEKHG2-1 | -0.0660170042333446 | -0.0072865513392303 | -0.34736566308923 | 0.730031041771706 | 0.890696324420811 | -5.61646608986909 |
| ENSG00000140263.13 | SORD | 0.0827868759211547 | 2.68979191836868 | 0.347306056924069 | 0.730075494786327 | 0.890696324420811 | -6.07915592114428 |
| ENSG00000105983.20 | LMBR1 | 0.0381450288519725 | 5.01567981567709 | 0.347261467509149 | 0.73010874924102 | 0.890696324420811 | -6.32017496485428 |
| ENSG00000204084.12 | INPP5B | -0.0511523125334037 | 5.5132143132065 | -0.347070825931201 | 0.730250934241072 | 0.890797906092995 | -6.33697867587801 |
| ENSG00000088247.17 | KHSRP | -0.0417320392340423 | 5.42438113157033 | -0.347000175095745 | 0.730303629744913 | 0.890797906092995 | -6.33839598569193 |
| lnc-ATPAF1-1 | lnc-ATPAF1-1 | 0.0559269797305435 | 0.249957884330264 | 0.346898634162734 | 0.730379367190097 | 0.890797906092995 | -5.6571836747266 |
| ENSG00000067560.10 | RHOA | 0.0468387689625622 | 7.92639734831645 | 0.346832664705089 | 0.730428574009499 | 0.890797906092995 | -6.33062493229604 |
| lnc-NPIPB4-3 | lnc-NPIPB4-3 | -0.0484255905324766 | 2.92179217821723 | -0.346825615774404 | 0.730433831896324 | 0.890797906092995 | -6.11170433095414 |
| lnc-ZFC3H1-10 | lnc-ZFC3H1-10 | 0.0594726494103601 | 2.14162070967258 | 0.346801079941364 | 0.730452133587775 | 0.890797906092995 | -6.00978943225704 |
| lnc-MLLT6-1 | lnc-MLLT6-1 | 0.0708272250741024 | 0.883279369333202 | 0.346743660111114 | 0.730494964628269 | 0.890797906092995 | -5.82636452647995 |
| lnc-PLCG1-2 | lnc-PLCG1-2 | -0.0900260296550801 | 2.57601984932247 | -0.346542139445308 | 0.730645291356886 | 0.890896048224286 | -6.06317328425606 |
| ENSG00000165097.14 | KDM1B | 0.055636273654553 | 4.16567339167062 | 0.346479399873104 | 0.730692094871762 | 0.890896048224286 | -6.25260582492284 |
| ENSG00000168813.16 | ZNF507 | 0.0414177284517878 | 5.53493583576362 | 0.346461705561193 | 0.730705294958091 | 0.890896048224286 | -6.34133461491402 |
| ENSG00000037042.8 | TUBG2 | 0.118364794734826 | 2.38888779592559 | 0.346191951869055 | 0.730918524195943 | 0.891092142035689 | -6.03341233392848 |
| ENSG00000136156.13 | ITM2B | -0.0767028965835683 | 6.23491212604597 | -0.346134554912954 | 0.730952714673788 | 0.891092142035689 | -6.35114842670018 |
| ENSG00000164776.9 | PHKG1 | 0.071267979118974 | 0.933295882281614 | 0.346052058552064 | 0.731010917711066 | 0.891110317989155 | -5.84384763370309 |
| ENSG00000163104.17 | SMARCAD1 | 0.0288927525757323 | 5.49550472095114 | 0.345923522778943 | 0.73110682269933 | 0.891174448159441 | -6.34111975408534 |
| ENSG00000074370.17 | ATP2A3 | 0.05211188474604 | 7.90630809734976 | 0.345863253144037 | 0.73115179345852 | 0.891176488704726 | -6.33171358700041 |
| lnc-TXNDC11-2 | lnc-TXNDC11-2 | 0.0670295369514389 | 2.06991287363417 | 0.345747842219286 | 0.731237911083819 | 0.891228678578769 | -6.00457109153618 |
| lnc-ZNF652-1 | lnc-ZNF652-1 | -0.0491153317411885 | 1.88053785818254 | -0.345665780957105 | 0.731299145904424 | 0.891250537044867 | -5.98458894468954 |
| ENSG00000102385.12 | DRP2 | -0.287269282848623 | 0.124163985152393 | -0.345680174902551 | 0.731345867601524 | 0.891254706503385 | -5.72692105641177 |
| ENSG00000196653.11 | ZNF502 | 0.324827944770702 | 1.36581519880297 | 0.345546354405877 | 0.731453334495684 | 0.891332898225366 | -5.86991287256734 |
| ENSG00000091732.16 | ZC3HC1 | -0.0417223440435386 | 3.31828034173129 | -0.345330056428737 | 0.731549684898398 | 0.891397535180188 | -6.15840257550439 |
| lnc-FST-1 | lnc-FST-1 | 0.0736142231957044 | 0.782529600965428 | 0.344879519051589 | 0.731885951128528 | 0.891754486004765 | -5.81422834863038 |
| lnc-ZNF468-3 | lnc-ZNF468-3 | -0.0566762849495665 | 0.492542545449226 | -0.344693684913041 | 0.732024667136589 | 0.891795126090554 | -5.73424424466742 |
| ENSG00000186496.11 | ZNF396 | -0.0860434714361158 | 1.00651282406781 | -0.344682236152008 | 0.732033213368001 | 0.891795126090554 | -5.83746105824829 |
| ENSG00000130511.15 | SSBP4 | -0.10578951641385 | 2.36740091718249 | -0.34466235049269 | 0.732053383346106 | 0.891795126090554 | -6.06593735493697 |
| lnc-RSL24D1-1 | lnc-RSL24D1-1 | -0.0751067855368093 | 2.06149307069241 | -0.344602664593671 | 0.732092612619739 | 0.891795126090554 | -6.02153473861214 |
| ENSG00000132467.3 | UTP3 | 0.0398822048308523 | 4.15496208641706 | 0.344323622774761 | 0.732300927258332 | 0.891996093369526 | -6.25571573017684 |
| ENSG00000124562.9 | SNRPC | -0.0450894890682729 | 4.33437871737416 | -0.344220911714454 | 0.732377609861279 | 0.892025097360529 | -6.27626674680215 |
| ENSG00000185905.3 | C16orf54 | 0.0991308432758622 | 4.30325275802132 | 0.344088191616371 | 0.732495592616987 | 0.892025097360529 | -6.26907502823955 |
| ENSG00000227262.3 | HCG4B | -0.106627702775455 | -0.287774324716028 | -0.344046383137539 | 0.732507916736734 | 0.892025097360529 | -5.56187979872556 |
| ENSG00000116337.15 | AMPD2 | -0.0591300602703515 | 3.77656203972152 | -0.343960399450537 | 0.732572116986012 | 0.892025097360529 | -6.21422121583207 |
| ENSG00000137076.20 | TLN1 | -0.0428815259215511 | 8.31587873956142 | -0.34392962387426 | 0.732595096230512 | 0.892025097360529 | -6.32504936968743 |
| lnc-GABPB2-1 | lnc-GABPB2-1 | -0.0502476119849087 | 1.41519638643051 | -0.343903173564251 | 0.732614846120505 | 0.892025097360529 | -5.9084837283519 |
| ENSG00000137274.12 | BPHL | -0.0716752554469564 | 2.61024965580284 | -0.343885417499245 | 0.732628104303539 | 0.892025097360529 | -6.07653167043856 |
| DANCR | DANCR | 0.0497021756992773 | 4.11949289946936 | 0.343812367020378 | 0.732682650851625 | 0.892038743787945 | -6.24886983487428 |
| lnc-MRPS28-2 | lnc-MRPS28-2 | -0.0802556215658228 | 1.13076997496627 | -0.343650041800207 | 0.732803863555793 | 0.89209060087947 | -5.86491915280646 |
| ENSG00000229992.1 | HMGB3P9 | 0.0944636431225653 | -0.347079270999238 | 0.343623346112319 | 0.732823798620727 | 0.89209060087947 | -5.49682550112703 |
| ENSG00000198355.4 | PIM3 | -0.109452746965927 | 3.85027128083367 | -0.343612808640295 | 0.732857158767362 | 0.89209060087947 | -6.22898500795636 |
| ENSG00000146833.15 | TRIM4 | 0.0253347862718609 | 4.85974384925414 | 0.343499941339976 | 0.732915953837372 | 0.89209060087947 | -6.31075743017319 |
| ENSG00000198373.12 | WWP2 | -0.0322692560410687 | 5.59134649513148 | -0.343403712009481 | 0.732987817964592 | 0.89209060087947 | -6.34440535125949 |
| ENSG00000167106.11 | EEIG1 | -0.0948595096190799 | 6.13240946172883 | -0.343341719397144 | 0.733058068589623 | 0.89209060087947 | -6.35147518339817 |
| ENSG00000111671.9 | SPSB2 | -0.0902350818307269 | 0.78227922403679 | -0.343214665279056 | 0.73312900524509 | 0.89209060087947 | -5.81225830833024 |
| lnc-PLEKHM1-2 | lnc-PLEKHM1-2 | -0.0672229111429795 | 0.880883545067439 | -0.343134736016249 | 0.733188702267979 | 0.89209060087947 | -5.8421042138022 |
| lnc-DCHS1-1 | lnc-DCHS1-1 | -0.0544662811265244 | 2.4575897510798 | -0.34303189745095 | 0.733265512092824 | 0.89209060087947 | -6.04620380514548 |
| lnc-FAM105A-10 | lnc-FAM105A-10 | 0.236394503244857 | -0.875266318492885 | 0.342994478705727 | 0.7332979426905 | 0.89209060087947 | -5.34224505403774 |
| lnc-HEATR4-10 | lnc-HEATR4-10 | 0.053674034299716 | 1.92984557832871 | 0.342888854065335 | 0.733372355379727 | 0.89209060087947 | -5.98150766535987 |
| ENSG00000072501.17 | SMC1A | 0.0362694178161416 | 6.46693531948069 | 0.342856157822912 | 0.733396777910281 | 0.89209060087947 | -6.35218873524078 |
| lnc-GOLGA8A-3 | lnc-GOLGA8A-3 | -0.0522783061715667 | 0.269567323830994 | -0.342840200167718 | 0.733408697617582 | 0.89209060087947 | -5.68378276822835 |
| ENSG00000276712.1 | MIR7111 | 0.0676616249375507 | 0.679982276959784 | 0.342811309616807 | 0.733430277831509 | 0.89209060087947 | -5.7787742753277 |
| lnc-PDAP1-1 | lnc-PDAP1-1 | 0.0581651041436113 | 1.67584553885704 | 0.342679738205557 | 0.733528559754694 | 0.89209060087947 | -5.95535330120971 |
| ENSG00000189159.15 | JPT1 | -0.0675612792508065 | 3.76752551882232 | -0.342617523142645 | 0.733575035073483 | 0.89209060087947 | -6.22759657283979 |
| ENSG00000124177.14 | CHD6 | -0.0327944239765534 | 7.01129970658079 | -0.342602061146754 | 0.733586585507017 | 0.89209060087947 | -6.34850691998251 |
| ENSG00000166483.10 | WEE1 | -0.103958062671803 | 5.66020825104798 | -0.342596746866164 | 0.733623850605059 | 0.89209060087947 | -6.34364021390329 |
| lnc-AAMP-1 | lnc-AAMP-1 | -0.0805392608389008 | 1.21397890849849 | -0.342546674026909 | 0.733627961353688 | 0.89209060087947 | -5.89306996345999 |
| ENSG00000088387.18 | DOCK9 | -0.0463868213947212 | 5.32234143722695 | -0.342542084697927 | 0.733631389756607 | 0.89209060087947 | -6.33618108169942 |
| ENSG00000012983.11 | MAP4K5 | -0.039314534335352 | 6.30167778711529 | -0.342483944761569 | 0.733674822971812 | 0.89209060087947 | -6.35245054884902 |

| | | | | | | | |
|---|---|---|---|---|---|---|---|
| lnc-MAGT1-3 | lnc-MAGT1-3 | -0.0544176683658737 | 2.35321659347069 | -0.342478689350899 | 0.733678749049588 | 0.89209060087947 | -6.03524145212785 |
| ENSG00000214655.10 | ZSWIM8 | 0.0436218736755736 | 6.01834695701538 | 0.34229437250677 | 0.733816448324669 | 0.892205325528186 | -6.35140620224703 |
| ENSG00000111666.10 | CHPT1 | -0.0791257661766425 | 6.06122703805289 | -0.341945481605391 | 0.734083471235134 | 0.892463343892701 | -6.35234995799587 |
| ENSG00000266173.6 | STRADA | 0.0443497722550719 | 5.08198337376897 | 0.34186396823312 | 0.734138028916367 | 0.892463343892701 | -6.32469151593911 |
| ENSG00000178996.13 | SNX18 | 0.0699943860534633 | 7.45886911532908 | 0.341824049933167 | 0.734167856697186 | 0.892463343892701 | -6.34362316870341 |
| lnc-PPP1R37-3 | lnc-PPP1R37-3 | -0.0571767836469956 | 0.469150027974745 | -0.341755966120898 | 0.734218731290901 | 0.892463343892701 | -5.73843045374467 |
| ENSG00000089063.14 | TMEM230 | -0.0353993189548678 | 5.36281791105367 | -0.341720199058953 | 0.734245458168343 | 0.892463343892701 | -6.33799707835117 |
| ENSG00000145349.16 | CAMK2D | -0.0507701452873528 | 7.90406588173875 | -0.34148930929434 | 0.73441799818259 | 0.892565032055439 | -6.33469871936702 |
| ENSG00000141506.13 | PIK3R5 | -0.315716928811177 | 2.30793464867861 | -0.341536575914821 | 0.734449510365819 | 0.892565032055439 | -5.95990706653482 |
| ENSG00000171530.13 | TBCA | -0.0516761547646247 | 4.29857707864687 | -0.341395345822757 | 0.734488219471063 | 0.892565032055439 | -6.27361231066224 |
| ENSG00000113838.12 | TBCCD1 | 0.0290328464413543 | 4.09172311857376 | 0.341376136231021 | 0.734502575569889 | 0.892565032055439 | -6.24598061638357 |
| ENSG00000139508.14 | SLC46A3 | 0.0445096273990577 | 3.9515951678932 | 0.341184001975976 | 0.734646170495893 | 0.892642116704173 | -6.233836497307 |
| lnc-SEL1L3-8 | lnc-SEL1L3-8 | 0.07739878415887 | 0.166228888770135 | 0.341175204971781 | 0.73465274532229 | 0.892642116704173 | -5.64311852787138 |
| ENSG00000119185.12 | ITGB1BP1 | 0.0486508073155996 | 3.79633782918355 | 0.341073752911314 | 0.734728571418813 | 0.892681552772176 | -6.21584452093156 |
| ENSG00000254732.1 | | 0.0518183819647202 | 2.77308470912948 | 0.340841297338192 | 0.734902320705501 | 0.892828539153948 | -6.08903669036958 |
| ENSG00000078319.9 | PMS2P1 | 0.0446056949124589 | 2.84230751731917 | 0.340795834403008 | 0.734936303691656 | 0.892828539153948 | -6.10527138848186 |
| DLEU2 | DLEU2 | -0.193954951863228 | 6.4520469388717 | -0.34080757740783 | 0.734991620703303 | 0.892843043492451 | -6.3523684507454 |
| lnc-MSMO1-1 | lnc-MSMO1-1 | 0.0863729908576223 | 1.16211193223701 | 0.340609766077771 | 0.735075393081814 | 0.892892110699497 | -5.87722671309936 |
| LINC00528 | LINC00528 | 0.207466386162912 | 1.27690052238091 | 0.340418837358764 | 0.735263196113548 | 0.89301743875547 | -5.84422149659373 |
| ENSG00000173917.10 | HOXB2 | 0.341613929076916 | -2.0314153987828 | 0.340369572880833 | 0.735270341352702 | 0.89301743875547 | -5.15864432788545 |
| ENSG00000172771.11 | EFCAB12 | -0.0947353683944727 | 2.2233186095224 | -0.340297646579856 | 0.735308728175082 | 0.89301743875547 | -6.01219182426975 |
| ENSG00000085871.8 | MGST2 | -0.251889462134145 | -0.222743127188563 | -0.340120338268784 | 0.735473136887718 | 0.893121814898881 | -5.50955334779184 |
| ENSG00000099381.17 | SETD1A | 0.0520908364834973 | 4.72157410949669 | 0.340066622890948 | 0.73548145374999 | 0.893121814898881 | -6.30360482605861 |
| ENSG00000204311.13 | PJVK | 0.0561446221373734 | 1.26726453867095 | 0.339828616198698 | 0.73565941467529 | 0.893285218114654 | -5.90210738754743 |
| lnc-DEF8-2 | lnc-DEF8-2 | 0.0405218258001696 | 4.06366040753264 | 0.339739543245123 | 0.735726019544593 | 0.893313394306651 | -6.25224458860461 |
| ENSG00000113648.16 | MACROH2A1 | 0.0467681488651337 | 6.25574118232645 | 0.339325247174795 | 0.736035839106496 | 0.893600200063894 | -6.35349209801103 |
| lnc-HLA-F-2 | lnc-HLA-F-2 | -0.107686188421783 | 0.814146735324997 | -0.33925605439277 | 0.736087587299698 | 0.893600200063894 | -5.81051139545471 |
| ENSG00000237765.6 | FAM200B | 0.0515690837774088 | 2.36807965157021 | 0.33924952046251 | 0.736092473987432 | 0.893600200063894 | -6.05722846676606 |
| IFNG-AS1 | IFNG-AS1 | 0.315776726486869 | 2.24494358776455 | 0.339138467651633 | 0.736241854694249 | 0.893728833166532 | -6.05320547740754 |
| lnc-CEP120-1 | lnc-CEP120-1 | -0.364222596517625 | -2.79442276956165 | -0.338823323450568 | 0.736411248081949 | 0.893830493131161 | -4.88753815644109 |
| ENSG00000079335.19 | CDC14A | 0.0455372988163094 | 3.70292618919351 | 0.338821713751188 | 0.73641245214586 | 0.893830493131161 | -6.22246338364594 |
| lnc-MOAP1-3 | lnc-MOAP1-3 | -0.0499557585656272 | 1.50079894156613 | -0.338706652954765 | 0.736498519985057 | 0.893882247588445 | -5.93698068441102 |
| lnc-KIAA1551-13 | lnc-KIAA1551-13 | 0.0522668321385305 | 3.39262743710937 | 0.338638374986219 | 0.736549594926852 | 0.893891527852559 | -6.16916489618161 |
| lnc-ZNF616-1 | lnc-ZNF616-1 | -0.0513610630236319 | 1.54465165734948 | -0.338512658729861 | 0.736643639402617 | 0.89395295264572 | -5.92910336672998 |
| ENSG00000126698.10 | DNAJC8 | -0.0437692327870076 | 5.72281810557218 | -0.33834495410203 | 0.736769100451192 | 0.894052493490561 | -6.34867689622711 |
| ENSG00000168594.15 | ADAM29 | -0.432682049011313 | -0.375840252184871 | -0.338240879193948 | 0.736911207241027 | 0.894131254983498 | -5.62560376189257 |
| ENSG00000100926.14 | TM9SF1 | 0.0360751530148025 | 2.24176274554323 | 0.338034212827349 | 0.737001587457929 | 0.894131254983498 | -6.03340208064088 |
| lnc-IER5-4 | lnc-IER5-4 | -0.0513806184630372 | 2.2044078062011 | -0.337987074066557 | 0.737036857398408 | 0.894131254983498 | -6.01092070912848 |
| ENSG00000168234.12 | TTC39C | -0.0999993359049332 | 3.3840211938877 | -0.337997730195338 | 0.737038822807062 | 0.894131254983498 | -6.15736485743277 |
| ENSG00000140374.15 | ETFA | 0.0366221657837151 | 4.67223849174741 | 0.337956499201556 | 0.737059734285872 | 0.894131254983498 | -6.30363441306165 |
| ENSG00000213901.10 | SLC23A3 | 0.0991506574130979 | -0.507810725323148 | 0.337909838128594 | 0.737094647746198 | 0.894131254983498 | -5.46327200732212 |
| ENSG00000258352.1 | IFITM3P6 | 0.102047649804459 | 0.358995714841926 | 0.337791985534297 | 0.737182831715261 | 0.894149732278818 | -5.68199238357333 |
| ENSG00000116661.10 | FBXO2 | 0.107502492362311 | 0.397231875218669 | 0.337714816865187 | 0.737240575614656 | 0.894149732278818 | -5.73099149378689 |
| ENSG00000137547.8 | MRPL15 | -0.062866652573526 | 2.68880329151182 | -0.337711478366087 | 0.737243073786955 | 0.894149732278818 | -6.09558192391336 |
| ENSG00000081791.8 | DELE1 | 0.0364420070053191 | 6.13601738427313 | 0.337657261053895 | 0.737283644572516 | 0.894149732278818 | -6.35334355539554 |
| ENSG00000124795.15 | DEK | 0.041870867930367 | 6.58292145600901 | 0.337498425077203 | 0.737402505812225 | 0.894192672505767 | -6.35313988961237 |
| ENSG00000146556.14 | WASH2P | 0.0500387747635331 | 2.70621199951286 | 0.337370699594181 | 0.737498090949889 | 0.894192672505767 | -6.09334556702288 |
| ENSG00000135956.8 | TMEM127 | 0.0421435542697139 | 6.21621966316832 | 0.337295108022195 | 0.737554662932409 | 0.894192672505767 | -6.35401674582312 |
| ENSG00000072736.18 | NFATC3 | -0.0377251193975923 | 6.96677530683058 | -0.337280865708799 | 0.737565321903307 | 0.894192672505767 | -6.35121559036127 |
| ENSG00000067606.16 | PRKCZ | 0.0372951636520753 | 4.17393458788132 | 0.337167630691728 | 0.737650069036917 | 0.894192672505767 | -6.25816694140992 |
| ENSG00000182809.10 | CRIP2 | -0.309278285678163 | 1.0944523027835 | -0.337251591183672 | 0.737650582644213 | 0.894192672505767 | -5.83683126059327 |
| ENSG00000159593.14 | NAE1 | 0.0358835954695564 | 5.22980025323619 | 0.337165624232794 | 0.737651570736836 | 0.894192672505767 | -6.33686374679812 |
| ENSG00000155744.9 | HYCC2 | 0.036514701346408 | 5.42293145553548 | 0.33714554649176 | 0.737666597636298 | 0.894192672505767 | -6.34312640536255 |
| ENSG00000169221.13 | TBC1D10B | -0.0387668294359846 | 3.75941731013903 | -0.33706223445681 | 0.737728952452169 | 0.894215595566986 | -6.21694140728987 |
| lnc-GRTP1-2 | lnc-GRTP1-2 | -0.0353563797145222 | 1.71648901775356 | -0.336766828679467 | 0.737950063037016 | 0.89443093554887 | -5.95631324118779 |
| lnc-MN1-8 | lnc-MN1-8 | 0.0917475722776386 | -0.109405692155432 | 0.336545150885095 | 0.738116003135324 | 0.894549167978616 | -5.58933744044326 |
| ENSG00000156587.15 | UBE2L6 | 0.0585308076804862 | 4.86281725059885 | 0.336520399607629 | 0.738134531844182 | 0.894549167978616 | -6.3183726527313 |
| ENSG00000132510.10 | KDM6B | 0.0934923791882416 | 3.91601168516108 | 0.336292466816121 | 0.738315910337023 | 0.894681210170757 | -6.22554998009943 |
| ENSG00000197818.11 | SLC9A8 | -0.0329140520757356 | 5.35996001221326 | -0.336219037146403 | 0.738360143229757 | 0.894681210170757 | -6.33912151612772 |
| ENSG00000110888.17 | CAPRIN2 | -0.0366566307548322 | 5.78209446980389 | -0.336200679719633 | 0.738373887050996 | 0.894681210170757 | -6.35007362918054 |
| ENSG00000176749.8 | CDK5R1 | 0.177391655917975 | 2.93046974788466 | 0.336202260554692 | 0.738427231977223 | 0.894688822502347 | -6.14323042312486 |
| ENSG00000167904.14 | TMEM68 | 0.0409771322398003 | 2.83385788740202 | 0.33607617396669 | 0.738467104187552 | 0.894688822502347 | -6.10845305708553 |
| ENSG00000119820.10 | YIPF4 | 0.0421025624423308 | 5.31741979688234 | 0.335912280060668 | 0.738589817203222 | 0.894738618080508 | -6.33824447363957 |
| ENSG00000078304.19 | PPP2R5C | 0.0209131417094417 | 7.73709053848257 | 0.335905165164601 | 0.738595144527324 | 0.894738618080508 | -6.34064298879623 |
| lnc-HIST1H2BK-1 | lnc-HIST1H2BK-1 | -0.157296026564098 | 3.08031424863858 | -0.335838943939748 | 0.738691127215245 | 0.89480222878245 | -6.1464456496879 |
| lnc-ADIPOQ-2 | lnc-ADIPOQ-2 | -0.0835692357197919 | 3.42816585860257 | -0.335636211037139 | 0.738796535186308 | 0.89487724849878 | -6.17153120430378 |
| ENSG00000197694.15 | SPTAN1 | 0.0407812839666688 | 8.05581373499688 | 0.335447087606865 | 0.738938160384147 | 0.894996125408197 | -6.33430975962184 |
| lnc-CCK-8 | lnc-CCK-8 | 0.0745647422089749 | 0.353785764126626 | 0.335265860729173 | 0.739073880849547 | 0.894997066151135 | -5.70572929862957 |

| | | | | | | | |
|---|---|---|---|---|---|---|---|
| ENSG00000172508.10 | CARNS1 | 0.106266368205903 | 4.22851042078409 | 0.335258715641746 | 0.739106281303201 | 0.894997066151135 | -6.2598540765204 |
| ENSG00000172171.10 | TEFM | 0.0476968517318155 | 2.89120265774193 | 0.335044071940788 | 0.739239989508024 | 0.894997066151135 | -6.10462910590988 |
| ENSG00000280137.1 | | 0.0601956480041228 | 2.03754930337636 | 0.335021389322394 | 0.739256978359871 | 0.894997066151135 | -6.00523935902848 |
| ENSG00000235750.9 | KIAA0040 | -0.0885681457795666 | 7.72064435856098 | -0.334881643958757 | 0.739379087334134 | 0.894997066151135 | -6.34088284479756 |
| lnc-TCTN3-2 | lnc-TCTN3-2 | -0.118116006400213 | -0.50480427646113 | -0.334858260633425 | 0.739379162541934 | 0.894997066151135 | -5.48827459363755 |
| ENSG00000161791.13 | FMNL3 | -0.0644718743861734 | 5.58836756382263 | -0.334849826261938 | 0.739385480112508 | 0.894997066151135 | -6.34642530253486 |
| ENSG00000091592.15 | NLRP1 | -0.0477791599439046 | 7.69710367824771 | -0.334829135337195 | 0.739400978247513 | 0.894997066151135 | -6.34166198677403 |
| ENSG00000261556.9 | SMG1P7 | 0.0476194777951711 | 3.37830553929218 | 0.334826468804524 | 0.739402975569954 | 0.894997066151135 | -6.17318809649209 |
| ENSG00000162576.16 | MXRA8 | 0.497939009755398 | -2.06357863760165 | 0.334807641214242 | 0.739436388727713 | 0.894997066151135 | -4.95348831460265 |
| lnc-GNA12-2 | lnc-GNA12-2 | -0.113118670462497 | 1.20770968189971 | -0.334773700920279 | 0.739442919836655 | 0.894997066151135 | -5.9140623298161 |
| lnc-U2AF2-3 | lnc-U2AF2-3 | 0.0703825616355558 | 0.997232175557053 | 0.334713937221383 | 0.739487267160789 | 0.894997066151135 | -5.81927947205799 |
| ENSG00000166454.9 | ATMIN | -0.0368292286746223 | 5.84821539191559 | -0.334691321857892 | 0.739504207551492 | 0.894997066151135 | -6.35235389570761 |
| ENSG00000002745.12 | WNT16 | 0.161145382518816 | 2.52363248123351 | 0.334628743556842 | 0.739597080031 | 0.89503764323219 | -6.08424797159129 |
| lnc-DYSF-9 | lnc-DYSF-9 | 0.0391249095062296 | 2.2885198740939 | 0.334526405336117 | 0.739627744801058 | 0.89503764323219 | -6.03391612717304 |
| ENSG00000104765.15 | BNIP3L | 0.0439462800679236 | 6.61107884700467 | 0.334472417696129 | 0.739668187891928 | 0.89503764323219 | -6.35413218390939 |
| lnc-MICU2-2 | lnc-MICU2-2 | 0.0600386911421049 | 4.1378094779394 | 0.334101656776644 | 0.739945951554012 | 0.895224937123598 | -6.26255530936346 |
| ENSG00000040275.16 | SPDL1 | -0.0537592644605214 | 2.10908133497467 | -0.334091852727553 | 0.739953296948947 | 0.895224937123598 | -6.01490640328756 |
| ENSG00000129925.10 | PGAP6 | -0.0579507825021479 | 3.51844322731308 | -0.334091648840894 | 0.73995344970528 | 0.895224937123598 | -6.20220077153896 |
| lnc-SEPSECS-3 | lnc-SEPSECS-3 | -0.0706581524147172 | 3.55898101252041 | -0.333979480596495 | 0.740037490211367 | 0.895234847347952 | -6.19748587985515 |
| lnc-FAM45A-4 | lnc-FAM45A-4 | -0.0517726653397912 | 0.330048664180449 | -0.333964614259159 | 0.740048628850285 | 0.895234847347952 | -5.70963954159266 |
| ENSG00000171763.18 | AFG2B | -0.0431068555199698 | 2.34310787960727 | -0.333905925217219 | 0.740092602307107 | 0.895235427438128 | -6.05534095006997 |
| ENSG00000258465.7 | | -0.0242002426424461 | 5.5282297377563 | -0.333711795893041 | 0.740238062259466 | 0.895310251572876 | -6.34527900771566 |
| ENSG00000269800.1 | PLEKHA3P1 | 0.0683741030218157 | 1.61231637987084 | 0.333707268584129 | 0.74024145466002 | 0.895310251572876 | -5.95053402015511 |
| ENSG00000240463.1 | RPS19P3 | 0.0735166964634611 | 0.310880742676628 | 0.333544341240672 | 0.740363542766675 | 0.895314569476829 | -5.67308102678594 |
| lnc-TBC1D3F-2 | lnc-TBC1D3F-2 | 0.0471773702003931 | 0.861286655654255 | 0.333533324537575 | 0.740371798276351 | 0.895314569476829 | -5.8285977302418 |
| ENSG00000214253.8 | FIS1 | 0.0488100519439776 | 4.50306219420228 | 0.333520187615241 | 0.740381642640354 | 0.895314569476829 | -6.28877246852588 |
| lnc-FA2H-2 | lnc-FA2H-2 | 0.0444790885412665 | 1.15211226716497 | 0.333385550723849 | 0.740482537497837 | 0.895314569476829 | -5.87721801456065 |
| lnc-RABAC1-1 | lnc-RABAC1-1 | -0.0885258438473662 | -0.412136680786826 | -0.33333451480096 | 0.740520784257879 | 0.895314569476829 | -5.52856565717161 |
| ENSG00000151148.13 | UBE3B | 0.0353697903635037 | 5.69426758556873 | 0.33330123584952 | 0.74054572414841 | 0.895314569476829 | -6.34848772895976 |
| ENSG00000134318.13 | ROCK2 | -0.0470054838145728 | 5.45037983535169 | -0.333296185400272 | 0.740549509077541 | 0.895314569476829 | -6.34433964019524 |
| ENSG00000204344.14 | STK19 | -0.035056188405894 | 3.17290188322442 | -0.333220534131912 | 0.740606204750797 | 0.895330524632072 | -6.14610121720226 |
| ENSG00000127526.14 | SLC35E1 | -0.035207494141047 | 4.86370889446875 | -0.333112312362124 | 0.740687312419164 | 0.895375988225974 | -6.31841553053535 |
| ENSG00000198561.13 | CTNND1 | 0.049411787383919 | 3.82947653623844 | 0.332999705589517 | 0.740771709631715 | 0.895420814971096 | -6.21900851834302 |
| ENSG00000168297.15 | PXK | -0.0666735378455297 | 5.88525710818946 | -0.332905885772701 | 0.740842028749053 | 0.895420814971096 | -6.35166201082387 |
| ENSG00000109180.14 | OCIAD1 | 0.0211170663553126 | 5.95725925485368 | 0.332888708580676 | 0.740854903510556 | 0.895420814971096 | -6.35401973693212 |
| ENSG00000114859.15 | CLCN2 | -0.060062652402094 | 0.979101681695246 | -0.332675482844805 | 0.741014728116803 | 0.895507700425368 | -5.85435872669046 |
| ENSG00000243679.1 | | -0.08347514858256 | 2.8426999866282 | -0.332537017397597 | 0.741118521907215 | 0.895507700425368 | -6.11856316126925 |
| ENSG00000186575.17 | NF2 | 0.0272103184560155 | 5.21229574263258 | 0.332357554415967 | 0.741253054735993 | 0.895507700425368 | -6.33579188182899 |
| ENSG00000106028.10 | SSBP1 | -0.030415930709971 | 6.39222876002671 | -0.332344525596879 | 0.741262821993402 | 0.895507700425368 | -6.35581985826663 |
| ENSG00000163681.14 | SLMAP | -0.0262364386389962 | 5.69029594723583 | -0.332287274864239 | 0.741305741409606 | 0.895507700425368 | -6.35060567567618 |
| ENSG00000137496.17 | IL18BP | -0.0453268992979372 | 3.80774803021633 | -0.332240231267971 | 0.741341009419503 | 0.895507700425368 | -6.22072409180098 |
| ENSG00000122687.17 | MRM2 | -0.0730056105344072 | 3.78122687033543 | -0.33220980703682 | 0.741363818393891 | 0.895507700425368 | -6.21708086968876 |
| lnc-ABHD15-2 | lnc-ABHD15-2 | -0.0677923617153544 | 1.79902320423941 | -0.332195247417148 | 0.741374733789708 | 0.895507700425368 | -5.97569426057126 |
| ENSG00000106537.7 | TSPAN13 | -0.308733667148388 | -0.187386924058341 | -0.332261931919933 | 0.741382830801161 | 0.895507700425368 | -5.69111223103442 |
| lnc-HMGCS1-1 | lnc-HMGCS1-1 | -0.0717042692027586 | -0.154926924855314 | -0.332167859197302 | 0.741395266976252 | 0.895507700425368 | -5.62173843325774 |
| lnc-RAB15-2 | lnc-RAB15-2 | -0.0580182757008432 | 3.44068295864277 | -0.332153971552129 | 0.741405678737451 | 0.895507700425368 | -6.17691267039526 |
| ENSG00000173456.4 | RNF26 | 0.0456555400746379 | 3.14584321049756 | 0.33206528573828 | 0.741472168884713 | 0.895507700425368 | -6.14692067255007 |
| ENSG00000169252.5 | ADRB2 | -0.0636181965816034 | 4.290361351877 | -0.332038323063047 | 0.741492383926033 | 0.895507700425368 | -6.26747955338428 |
| ENSG00000119328.11 | ABITRAM | 0.0556639777574177 | 1.89454805902369 | 0.331829741027679 | 0.74164877281288 | 0.895635732952432 | -5.99555647745169 |
| ENSG00000180479.13 | ZNF571 | -0.0418176391182326 | 3.79134848718035 | -0.331768593549848 | 0.741694621550236 | 0.895635732952432 | -6.21364788435643 |
| lnc-COG5-3 | lnc-COG5-3 | -0.0690963688603197 | 0.374436317725183 | -0.331666111136941 | 0.741771465593514 | 0.895635732952432 | -5.73534874948469 |
| lnc-RCAN1-4 | lnc-RCAN1-4 | -0.0893956042050618 | -0.486660884936895 | -0.331664798230981 | 0.741772450062679 | 0.895635732952432 | -5.42579935679279 |
| lnc-TP53BP1-4 | lnc-TP53BP1-4 | 0.0409302644355918 | 1.53218153764253 | 0.331549449414171 | 0.741858944906472 | 0.895647735317387 | -5.9455677960812 |
| lnc-RABGEF1-3 | lnc-RABGEF1-3 | -0.0528440224128118 | 0.940894765086629 | -0.331535482322175 | 0.74186941842424 | 0.895647735317387 | -5.8513321515834 |
| RASA2-IT1 | RASA2-IT1 | 0.0818871029475589 | 0.0790448314025651 | 0.331413906316168 | 0.741960586840996 | 0.895705264454441 | -5.6459340785764 |
| ENSG00000075275.16 | CELSR1 | 0.0577570532242486 | 7.60023349907830 | 0.331264402827415 | 0.742072702889722 | 0.895750408350322 | -6.34307884132419 |
| lnc-RAB40AL-1 | lnc-RAB40AL-1 | -0.0628895215349271 | 0.481972738853201 | -0.331209717222018 | 0.742113714278573 | 0.895750408350322 | -5.73589114844645 |
| lnc-APBB1IP-1 | lnc-APBB1IP-1 | -0.088037122513188 | 2.24301168579821 | -0.331189950119288 | 0.742128538774622 | 0.895750408350322 | -6.02796095268997 |
| ENSG00000173281.4 | PPP1R3B | 0.0417173364942958 | 3.10213757068197 | 0.33111109244207 | 0.742187679703397 | 0.8957692630078 | -6.1372315871872 |
| lnc-ZNF529-3 | lnc-ZNF529-3 | -0.0524566089330352 | 0.041346508350481 | -0.330913909710817 | 0.742335567826612 | 0.895895220907368 | -5.63145183606538 |
| ENSG00000250321.1 | | 0.118852519624842 | 1.61881272391303 | 0.33077554991327 | 0.742443728377068 | 0.895960059348538 | -5.90954058209347 |
| lnc-CAMKK2-4 | lnc-CAMKK2-4 | 0.0483524182703206 | 2.45796442921311 | 0.330726212401287 | 0.74247635098421 | 0.895960059348538 | -6.06794072212776 |
| lnc-ADAMTS8-5 | lnc-ADAMTS8-5 | -0.0429126721487845 | 3.3694048029796 | -0.33066494144621 | 0.742522309454006 | 0.895962990707691 | -6.16598169707888 |
| ENSG00000112149.9 | CD83 | 0.114420996332068 | 7.08429481798494 | 0.33031036902667 | 0.742829943131207 | 0.896193663846597 | -6.35311206343732 |
| ENSG00000131873.6 | CHSY1 | -0.0544704666581102 | 3.68727969442821 | -0.33024612912607 | 0.742836480052 | 0.896193663846597 | -6.22257754500866 |
| lnc-SLC16A11-1 | lnc-SLC16A11-1 | 0.0585281045516606 | 0.403336198456196 | 0.330235972653606 | 0.742844099445503 | 0.896193663846597 | -5.74515946892471 |
| ENSG00000153933.9 | DGKE | 0.0430450464581646 | 4.70077768685748 | 0.33007115966874 | 0.742967745920628 | 0.896290300919253 | -6.3099506502184 |
| lnc-SLC38A3-6 | lnc-SLC38A3-6 | 0.0772726714921562 | 0.562493017593977 | 0.329815344855231 | 0.74315967765192 | 0.896435800164953 | -5.7705855787728 |

| | | | | | | | |
|---|---|---|---|---|---|---|---|
| lnc-TFIP11-2 | lnc-TFIP11-2 | 0.033679039886342 | 1.4034287708554 | 0.329794310393865 | 0.743175460040555 | 0.896435800164953 | -5.92160675848821 |
| ENSG00000126001.15 | CEP250 | -0.0334545990000406 | 5.58459272617042 | -0.329630976223022 | 0.743298015275802 | 0.89649011309908 | -6.34758031880654 |
| lnc-RPIA-2 | lnc-RPIA-2 | -0.0774003514610566 | 2.5278288792607 | -0.329618206590575 | 0.74330759705334 | 0.89649011309908 | -6.07076524612239 |
| ENSG00000053371.12 | AKR7A2 | -0.0517782810677589 | 3.60536482730113 | -0.329345590464571 | 0.743512166173486 | 0.896610745228809 | -6.19983714351841 |
| lnc-ZNF527-5 | lnc-ZNF527-5 | -0.062370825035594 | 1.05344351791809 | -0.329340179180427 | 0.743516226950848 | 0.896610745228809 | -5.85603981953927 |
| ENSG00000102858.12 | MGRN1 | 0.0491095001464235 | 4.49721236896948 | 0.32931076659662 | 0.743538299096854 | 0.896610745228809 | -6.29528252793623 |
| ENSG00000037897.16 | METTL1 | -0.0692475217671168 | 0.640805503084387 | -0.329241940279138 | 0.743589949425329 | 0.896620499649769 | -5.80686986361265 |
| ENSG00000042753.11 | AP2S1 | 0.0862707572716297 | 3.23663435391327 | 0.329062221462278 | 0.743724873253431 | 0.896730658202529 | -6.14882488380561 |
| ENSG00000276180.1 | H4C9 | 0.103078001025164 | 1.12117073876508 | 0.328980704382867 | 0.743786003328936 | 0.896751833793316 | -5.914241014902 |
| ENSG00000137575.11 | SDCBP | 0.0665772966908786 | 5.43208598751902 | 0.328824284878687 | 0.743903402224081 | 0.896840843904308 | -6.34261791952116 |
| ENSG00000068831.18 | RASGRP2 | -0.0641797058538796 | 7.4491627467596 | -0.328618032051895 | 0.744058212381071 | 0.896928336432252 | -6.34829110566669 |
| ENSG00000272333.5 | KMT2B | -0.054047479817096 | 5.49657140272581 | -0.328611484462875 | 0.744063127074805 | 0.896928336432252 | -6.34571690546803 |
| ENSG00000145743.15 | FBXL17 | -0.0280601420425963 | 4.8515403142223 | -0.328433191464855 | 0.744196959947145 | 0.896973504520241 | -6.3164639006184 |
| ENSG00000166189.7 | HPS6 | 0.0488038942923161 | 3.7728342542989 | 0.32838164324096 | 0.744235655312768 | 0.896973504520241 | -6.22577671793924 |
| lnc-ABCA9-2 | lnc-ABCA9-2 | -0.136063556400117 | 3.55479333360955 | -0.328362444047283 | 0.744291496739989 | 0.896973504520241 | -6.18284101680285 |
| MID1IP1-AS1 | MID1IP1-AS1 | -0.145306926033258 | 0.324349823851187 | -0.328255546698884 | 0.744333913875217 | 0.896973504520241 | -5.71372005651196 |
| ENSG00000173120.14 | KDM2A | -0.0445287491769966 | 7.60159169746829 | -0.328199408977297 | 0.744372457267182 | 0.896973504520241 | -6.34410462872654 |
| lnc-WDR1-1 | lnc-WDR1-1 | 0.0588400745117039 | 3.95555510036622 | 0.328150773422763 | 0.744408969041467 | 0.896973504520241 | -6.24362851276105 |
| ENSG00000176986.15 | SEC24C | 0.0281397870496338 | 6.77422013195632 | 0.328113158580664 | 0.744437207733934 | 0.896973504520241 | -6.35559451268665 |
| lnc-ZNF600-1 | lnc-ZNF600-1 | -0.0556492450288084 | 1.70523730834851 | -0.328086605281145 | 0.744457142378779 | 0.896973504520241 | -5.97879556401802 |
| ENSG00000228014.1 | ZNF680P1 | 0.380101578889065 | -2.71713233841889 | 0.328017167290147 | 0.744509322183697 | 0.896973504520241 | -4.96299831866067 |
| ENSG00000187840.4 | EIF4EBP1 | -0.0717141918356572 | 2.09622113009736 | -0.327981060720168 | 0.744536380737906 | 0.896973504520241 | -6.0264922324774 |
| ENSG00000099341.11 | PSMD8 | -0.0427667612957929 | 5.32893697491892 | -0.327849420409654 | 0.744635214598436 | 0.897040069184105 | -6.34147812475985 |
| ENSG00000170638.9 | TRABD | 0.0630047777368201 | 4.88278419839685 | 0.327607252610725 | 0.744817042456402 | 0.897206600625043 | -6.31862335809141 |
| lnc-NUMA1-3 | lnc-NUMA1-3 | -0.0573464197710622 | 0.159732915996491 | -0.327530533050582 | 0.744874649196101 | 0.897223484574166 | -5.66665283148976 |
| lnc-RABL2A-7 | lnc-RABL2A-7 | 0.0644343760810547 | -0.296308713679435 | 0.327367585958611 | 0.744997006868835 | 0.897279524404151 | -5.5514908927078 |
| lnc-JMJD6-1 | lnc-JMJD6-1 | -0.0453702037841461 | 5.76902936419151 | -0.327299903851441 | 0.745047831613431 | 0.897279524404151 | -6.35462832911449 |
| ENSG00000144034.14 | TPRKB | 0.0315587086114051 | 2.80316215515063 | 0.327294415372183 | 0.745051953145379 | 0.897279524404151 | -6.10490331666994 |
| ENSG00000204632.11 | HLA-G | -0.108404577939537 | 3.3516026990423 | -0.327099904816737 | 0.745217422131116 | 0.897426292986471 | -6.15318343641529 |
| ENSG00000206530.10 | CFAP44 | 0.053874078313676 | 4.57775733106638 | 0.326856559812724 | 0.745380781808799 | 0.897529241417517 | -6.29390043498874 |
| ENSG00000173141.4 | MRPL57 | -0.0494392527399166 | 3.25557864459772 | -0.32684412554224 | 0.74539012062338 | 0.897529241417517 | -6.16192012633952 |
| lnc-NBPF14-3 | lnc-NBPF14-3 | 0.0533251242616481 | 0.614490603586142 | 0.32677102143758 | 0.74544502657242 | 0.897542847729753 | -5.79797101086639 |
| lnc-MTERF1-1 | lnc-MTERF1-1 | -0.0512786687233278 | 0.844761962985441 | -0.326644204794164 | 0.745540277316451 | 0.897605026205224 | -5.82539223735016 |
| ENSG00000256950.2 | | 0.0592909312344903 | 2.52556135685952 | 0.326306990253023 | 0.74579357546371 | 0.897857469952011 | -6.07632387465257 |
| ENSG00000166794.4 | PPIB | -0.0599526977360183 | 6.45399726774601 | -0.326239787269224 | 0.745844058275935 | 0.897865729997284 | -6.35776400086122 |
| ENSG00000163644.14 | PPM1K | -0.0333071028657024 | 7.47292392937028 | -0.32602024226704 | 0.746008988130391 | 0.897983025396442 | -6.34738328951949 |
| ENSG00000189420.8 | ZFP92 | -0.127902548224025 | 0.0476866993011842 | -0.325993939239598 | 0.746028748689655 | 0.897983025396442 | -5.68765796503916 |
| ENSG00000117298.15 | ECE1 | -0.0638198309184324 | 5.67925704691243 | -0.325879159049104 | 0.746114981120872 | 0.898034305386288 | -6.35114974188341 |
| ENSG00000163798.13 | SLC4A1AP | -0.0442710736710374 | 5.04154447182303 | -0.325648097998685 | 0.746288583397562 | 0.898190732784003 | -6.3316166576234 |
| ENSG00000206885.1 | SNORA75 | -0.0499545036035249 | 1.66625394783811 | -0.325491671589781 | 0.746406118266111 | 0.898279666273248 | -5.96314216437249 |
| ENSG00000174516.14 | PELI3 | -0.08199250395282 | 3.01407163104266 | -0.325014772019142 | 0.746764486305051 | 0.898657585171467 | -6.12383067411533 |
| ENSG00000105656.12 | ELL | 0.0552702717778919 | 3.99652282504375 | 0.324957586767258 | 0.746807462194916 | 0.898657585171467 | -6.26012993211763 |
| lnc-OR4F16-15 | lnc-OR4F16-15 | -0.0564191303723065 | 2.00921198217153 | -0.3245716052019 | 0.747097556578211 | 0.898831605200876 | -5.99527282351069 |
| ENSG00000279544.1 | | 0.0707629567073029 | 2.46572870837716 | 0.324541108428937 | 0.747120478795447 | 0.898831605200876 | -6.07070702833536 |
| lnc-CPLX1-2 | lnc-CPLX1-2 | -0.0547805678848712 | 3.11383723289347 | -0.324497348936283 | 0.747153370043915 | 0.898831605200876 | -6.14648218008634 |
| ZNF528-AS1 | ZNF528-AS1 | -0.0888502651658074 | 2.9438676472241 | -0.324495429431779 | 0.747154812825207 | 0.898831605200876 | -6.09786885350629 |
| lnc-EIF3H-3 | lnc-EIF3H-3 | 0.234637099180045 | -0.662826311231286 | 0.324473956159665 | 0.747193430098101 | 0.898831605200876 | -5.47327987565048 |
| ENSG00000185883.11 | ATP6V0C | -0.0579582839426731 | 2.91482262177406 | -0.324387278532764 | 0.747236105126603 | 0.898831605200876 | -6.1167253328158 |
| lnc-ASIP-1 | lnc-ASIP-1 | 0.0589020800028961 | -0.0981861424858746 | 0.324268820289892 | 0.747325148349021 | 0.898831605200876 | -5.57776958071527 |
| ENSG00000144791.9 | LIMD1 | -0.0463528640423574 | 6.78051556696304 | -0.324225865604264 | 0.74735743758228 | 0.898831605200876 | -6.35680732606685 |
| ENSG00000113441.15 | LNPEP | 0.0287634177103726 | 8.12327557359386 | 0.324195045379709 | 0.747380605568669 | 0.898831605200876 | -6.33652359095837 |
| lnc-PILRB-1 | lnc-PILRB-1 | 0.0413632727929416 | 4.6680922114745 | 0.324184192758959 | 0.747388763689112 | 0.898831605200876 | -6.31090448410346 |
| ENSG00000160305.17 | DIP2A | -0.0429860770085106 | 5.85526067663307 | -0.324031392035041 | 0.747503629996154 | 0.898917224597501 | -6.35405710333389 |
| lnc-ADCY5-2 | lnc-ADCY5-2 | 0.0548718401859092 | 1.71914217556595 | 0.323966158202697 | 0.747552670599554 | 0.898923679321763 | -5.95780576776862 |
| lnc-MAP3K12-3 | lnc-MAP3K12-3 | -0.0875871343628369 | -0.193684611194693 | -0.323738925456472 | 0.747723504805196 | 0.89907658017323 | -5.54682296180923 |
| ENSG00000143155.12 | TIPRL | -0.0387454275136037 | 4.41768092711622 | -0.323582100289703 | 0.747841413882709 | 0.899147111823399 | -6.29167221942646 |
| ENSG00000152620.12 | NADK2 | -0.0645199803930967 | 3.52953749416582 | -0.323544704409882 | 0.747869530895234 | 0.899147111823399 | -6.20753943364801 |
| ENSG00000147548.16 | NSD3 | -0.0325780270485225 | 7.84519724719223 | -0.323411085602006 | 0.747969998321753 | 0.899176653731741 | -6.343430236566 |
| lnc-ZNF765-5 | lnc-ZNF765-5 | 0.0556634250668158 | -0.20244370837724 | 0.323353251823678 | 0.748013484674075 | 0.899176653731741 | -5.55738272872883 |
| ENSG00000181031.15 | RPH3AL | 0.187546457337028 | -0.859298076056279 | 0.323338240523778 | 0.748025158715863 | 0.899176653731741 | -5.44020834063459 |
| lnc-MSMB-6 | lnc-MSMB-6 | -0.205809370848624 | 2.53978740133499 | -0.32317975968634 | 0.748201603603185 | 0.899265635970644 | -6.05134673603507 |
| LINC01765 | LINC01765 | 0.152719116584376 | -0.928222808941152 | 0.322995485655851 | 0.748282514523817 | 0.899265635970644 | -5.35253893128001 |
| lnc-BEND4-4 | lnc-BEND4-4 | 0.0626973025108912 | 1.24800341853843 | 0.322988822561301 | 0.748287525280134 | 0.899265635970644 | -5.90039511741697 |
| ENSG00000066654.13 | THUMPD1 | 0.0349317767630417 | 6.16713429357277 | 0.322985046711643 | 0.748290364785854 | 0.899265635970644 | -6.35860857738629 |
| ENSG00000152484.13 | USP12 | 0.030468756857732 | 5.55471812769356 | 0.322948788633647 | 0.748317631681737 | 0.899265635970644 | -6.35010988931819 |
| ENSG00000285025.1 | | 0.0461294905749573 | 0.199420753675858 | 0.322744396826758 | 0.74847134504745 | 0.899389488529413 | -5.68361773097279 |
| ENSG00000226450.2 | CYP2D8P | -0.064285365013502 | 0.620934806922296 | -0.322661037584554 | 0.748534038545142 | 0.899389488529413 | -5.79298699951941 |
| lnc-PRDM9-17 | lnc-PRDM9-17 | 0.0507624985651048 | 0.0979287438234928 | 0.322637445804433 | 0.748551781955855 | 0.899389488529413 | -5.63981009360991 |

| | | | | | | | |
|---|---|---|---|---|---|---|---|
| lnc-BID-4 | lnc-BID-4 | -0.074889013414404 | 0.436101961899105 | -0.322555981871494 | 0.748613052153512 | 0.899394646544655 | -5.73221411282063 |
| lnc-CCKAR-11 | lnc-CCKAR-11 | -0.118535292722286 | 3.62920524547148 | -0.322466296339359 | 0.748714124394869 | 0.899394646544655 | -6.19356891978134 |
| ENSG00000196865.4 | NHLRC2 | 0.0273984983584921 | 5.66526163558682 | 0.322327713955505 | 0.748784744497293 | 0.899394646544655 | -6.35305569477038 |
| ENSG00000258738.1 | BAZ1A-AS1 | 0.0816812579992054 | 1.20934876386392 | 0.322291123317648 | 0.748812267446418 | 0.899394646544655 | -5.90919450798037 |
| ENSG00000120254.15 | MTHFD1L | 0.0583618297967123 | 3.23685230069737 | 0.322285868446347 | 0.748816220111902 | 0.899394646544655 | -6.15254556815666 |
| lnc-JAKMIP3-5 | lnc-JAKMIP3-5 | -0.117001492581683 | -0.620328515280744 | -0.322283168646905 | 0.748818250878674 | 0.899394646544655 | -5.51522555552649 |
| ENSG00000163913.11 | IFT122 | 0.0401744568756572 | 3.91451269145517 | 0.322157059753392 | 0.748913110953693 | 0.899456095388019 | -6.2326752752097 |
| ENSG00000196372.12 | ASB13 | -0.281788138115437 | -2.22376060803816 | -0.321962125330574 | 0.749059749860638 | 0.899577236359573 | -5.02795159267444 |
| ENSG00000149531.15 | FRG1BP | 0.14094886954875 | 2.02651372343737 | 0.321931866256228 | 0.749117592104019 | 0.899577236359573 | -6.04314809135323 |
| ENSG00000264663.1 | KRT8P34 | -0.0935922516604712 | 1.76868079601277 | -0.321796799996891 | 0.749184122772909 | 0.899577236359573 | -5.97473002420498 |
| ENSG00000188761.12 | BCL2L15 | -0.0631951923159727 | 0.528758109489059 | -0.321747508474417 | 0.749221205694465 | 0.899577236359573 | -5.76870440545088 |
| lnc-FBXL6-1 | lnc-FBXL6-1 | 0.0542954056576298 | 0.847160324002481 | 0.321732494971722 | 0.749232500748781 | 0.899577236359573 | -5.83339723547243 |
| lnc-MCM7-3 | lnc-MCM7-3 | 0.0525231466803722 | -0.11566856026533 | 0.321596745449174 | 0.749334631226039 | 0.899647381855201 | -5.58516380135697 |
| lnc-WASHC2C-4 | lnc-WASHC2C-4 | -0.0637750747103456 | 3.5623352014313 | -0.321537258360437 | 0.749379387475729 | 0.899648639977423 | -6.19349131983488 |
| ENSG00000184677.17 | ZBTB40 | -0.0380193341974033 | 7.1806165143133 | -0.321313793597179 | 0.749547523275047 | 0.899798009539851 | -6.35394074334364 |
| ENSG00000072364.12 | AFF4 | 0.0525466349032975 | 7.99642762803242 | 0.32123028338897 | 0.749610359860864 | 0.899801821521705 | -6.33941249996177 |
| ENSG00000136720.6 | HS6ST1 | 0.0530837953813879 | 2.84085223618558 | 0.321193377066755 | 0.749638130275188 | 0.899801821521705 | -6.11377847709404 |
| lnc-PTPN2-4 | lnc-PTPN2-4 | -0.0448488460062628 | 3.5252873354075 | -0.321040625666342 | 0.749753072684338 | 0.89988731092552 | -6.1927241416599 |
| lnc-CALCOCO1-3 | lnc-CALCOCO1-3 | 0.054361495273362 | 4.4167320694605 | 0.320639645139932 | 0.75005482998207 | 0.900197000905012 | -6.29135123158148 |
| lnc-MSANTD4-2 | lnc-MSANTD4-2 | 0.0756358452808044 | -0.317304725982327 | 0.320377816787575 | 0.750251889901229 | 0.900381006928867 | -5.49229002953125 |
| ENSG00000243650.3 | RN7SL834P | 0.048918816670199 | 2.41871252261948 | 0.320296127875424 | 0.750313374896806 | 0.900382671815647 | -6.06846885609362 |
| ENSG00000213090.2 | SCYL2P1 | 0.104040632330602 | 2.10575207290319 | 0.320263825634567 | 0.750340765177758 | 0.900382671815647 | -5.99418447388886 |
| ENSG00000126790.11 | L3HYPDH | -0.0598639313698692 | 3.89464580272527 | -0.320177970150303 | 0.750402311870289 | 0.90040403318329 | -6.24657011274121 |
| ENSG00000123240.16 | OPTN | -0.0597046024332215 | 5.27448973365328 | -0.320102851134243 | 0.750458855520022 | 0.900416243384572 | -6.34304430328729 |
| ENSG00000144029.11 | MRPS5 | -0.033673744592325 | 4.37746015129568 | -0.320048218942226 | 0.750499979182127 | 0.900416243384572 | -6.28585163496314 |
| lnc-SLC47A2-2 | lnc-SLC47A2-2 | -0.14454451717153 | -0.407477767375655 | -0.319965525579347 | 0.750562226917238 | 0.90043844009078 | -5.51261595772819 |
| ENSG00000148925.10 | BTBD10 | -0.0316264703774065 | 4.40794537480911 | -0.319807633199383 | 0.750681085661683 | 0.900492530278188 | -6.28481046231506 |
| ENSG00000163170.11 | BOLA3 | 0.0496697210194594 | 0.767523806338815 | 0.319789398855346 | 0.750694812565876 | 0.900492530278188 | -5.82767640623834 |
| ENSG00000178028.13 | DMAP1 | -0.0313861007769622 | 5.68464514136894 | -0.319716479300239 | 0.750749707583788 | 0.900496936033006 | -6.35383730459255 |
| ENSG00000205045.8 | SLFN12L | 0.111931125078592 | 3.00829244959216 | 0.319696930913004 | 0.750785984514523 | 0.900496936033006 | -6.12269290839883 |
| lnc-TCTE1-1 | lnc-TCTE1-1 | 0.045953936721054 | 0.302347649055162 | 0.31949350175493 | 0.750917576784857 | 0.900574123805509 | -5.71464394039942 |
| lnc-MALT1-2 | lnc-MALT1-2 | -0.0637629585152963 | 1.51541198115696 | -0.31944417280652 | 0.750954715856546 | 0.900574123805509 | -5.94848369299698 |
| lnc-METTL2B-5 | lnc-METTL2B-5 | -0.109882892225904 | -0.491853904049668 | -0.319403881621252 | 0.750985050966816 | 0.900574123805509 | -5.44897796655275 |
| ENSG00000070019.4 | GUCY2C | -0.130157876561453 | -0.58815191328383 | -0.319350353524086 | 0.751025352724168 | 0.900574123805509 | -5.38452309693435 |
| lnc-KDM4C-5 | lnc-KDM4C-5 | -0.0688061240423314 | 0.374517032069546 | -0.318956708083081 | 0.751321753309677 | 0.900861030016038 | -5.73847145857288 |
| ENSG00000221829.9 | FANCG | -0.0330185859436867 | 2.97191574544598 | -0.318916341596156 | 0.751352149943415 | 0.900861030016038 | -6.12734455078487 |
| ENSG00000181163.13 | NPM1 | 0.0446585862711171 | 8.48900573382611 | 0.318642021754217 | 0.751558727913049 | 0.901056226504793 | -6.32852387553069 |
| ENSG00000152078.9 | TLCD4 | 0.134359783111624 | 3.68281320578942 | 0.318631401036687 | 0.751610550674391 | 0.901065872475449 | -6.21768354067765 |
| ENSG00000114023.15 | FAM162A | -0.0331378170832086 | 3.96301513787409 | -0.318457210292792 | 0.751697911532255 | 0.901118119573 | -6.23664368902104 |
| lnc-ACVR2B-1 | lnc-ACVR2B-1 | -0.117477918718067 | -0.145566726158201 | -0.318379193617275 | 0.751756669291381 | 0.901136073846864 | -5.57915645319216 |
| ENSG00000106723.16 | SPIN1 | 0.0354217019293488 | 4.91571039216762 | 0.317970728869106 | 0.752064326217729 | 0.901327384179571 | -6.32743775806938 |
| ENSG00000149091.15 | DGKZ | -0.0441475446758401 | 4.47177721093267 | -0.317906740622834 | 0.752112526063017 | 0.901327384179571 | -6.29591856694415 |
| ENSG00000221838.9 | AP4M1 | 0.0413364595233152 | 2.29469841474832 | 0.317805069917799 | 0.752189112691012 | 0.901327384179571 | -6.05070554223848 |
| lnc-YWHAZ-1 | lnc-YWHAZ-1 | -0.0630434715541819 | 2.92974836140605 | -0.317750925614531 | 0.75222989960652 | 0.901327384179571 | -6.11923304147255 |
| ENSG00000133131.14 | MORC4 | 0.0898569418945994 | 2.45109714042427 | 0.317693091502278 | 0.752273466847381 | 0.901327384179571 | -6.05509362251759 |
| lnc-ZNF397-5 | lnc-ZNF397-5 | 0.0744921532465095 | 1.39761949882842 | 0.317643465854235 | 0.75231085118963 | 0.901327384179571 | -5.92669078860246 |
| lnc-EIF4E3-4 | lnc-EIF4E3-4 | 0.0675391371315195 | 2.92896233882756 | 0.317640009347019 | 0.752313455092375 | 0.901327384179571 | -6.10764146981436 |
| lnc-INTS8-5 | lnc-INTS8-5 | -0.0966915210321408 | 2.56374511893411 | -0.317627001380282 | 0.752329115020775 | 0.901327384179571 | -6.09475160658847 |
| ENSG00000273025.1 | | 0.0284651445619968 | 3.23143032257469 | 0.317488940748468 | 0.752427262982902 | 0.901327384179571 | -6.15953678223425 |
| lnc-RFX7-1 | lnc-RFX7-1 | 0.0707503721911786 | 2.48229755533684 | 0.317352244148626 | 0.752530248497537 | 0.901327384179571 | -6.05829005489544 |
| ENSG00000285815.1 | GET1-SH3BGR | -0.0456966386927188 | 0.864376237522609 | -0.317324144524594 | 0.752551418969402 | 0.901327384179571 | -5.83047736194121 |
| lnc-DNASE1L1-4 | lnc-DNASE1L1-4 | 0.0653861676854222 | 4.68282797608641 | 0.317293109284419 | 0.752574801380958 | 0.901327384179571 | -6.30409202970065 |
| lnc-ZNF415-4 | lnc-ZNF415-4 | 0.0439590499755949 | 3.04680749751231 | 0.317287151187857 | 0.752579290326428 | 0.901327384179571 | -6.13110834847218 |
| ENSG00000106330.11 | MOSPD3 | -0.0721095212588239 | 1.01287479322565 | -0.317211389248444 | 0.752636371596103 | 0.901327384179571 | -5.88229364050296 |
| ENSG00000198382.8 | UVRAG | 0.0480161379192996 | 7.17314106820221 | 0.317182975085456 | 0.752657780022006 | 0.901327384179571 | -6.35550178293756 |
| ENSG00000149182.14 | ARFGAP2 | 0.0215796011775744 | 6.68152933588851 | 0.317182579476739 | 0.752658078091649 | 0.901327384179571 | -6.35947389279124 |
| ENSG00000213689.13 | TREX1 | -0.0638900206898616 | 2.11571736424042 | -0.317179106605193 | 0.752660694713038 | 0.901327384179571 | -6.03363303070456 |
| ENSG00000179397.17 | CATSPERE | -0.0453986160679445 | 1.11459511855606 | -0.316964337207151 | 0.752822517587912 | 0.901411050383085 | -5.87511451238496 |
| lnc-PKIA-3 | lnc-PKIA-3 | -0.202034012239197 | -0.431427344151466 | -0.316954909253053 | 0.752850981977731 | 0.901411050383085 | -5.57705511932261 |
| ENSG00000119922.9 | IFIT2 | 0.0879504635214672 | 2.35081667713111 | 0.316850569306093 | 0.752908243151821 | 0.901411050383085 | -6.0503588286488 |
| lnc-CERS4-6 | lnc-CERS4-6 | -0.106756607356933 | 0.27175167386471 | -0.316795071907128 | 0.752950062294646 | 0.901411050383085 | -5.75828328565262 |
| lnc-BBS10-1 | lnc-BBS10-1 | -0.0462613203646036 | 2.34587211193939 | -0.31672623051466 | 0.753001937625281 | 0.901411050383085 | -6.04902166018965 |
| lnc-NID1-3 | lnc-NID1-3 | -0.101448622409888 | -0.0975168831101632 | -0.31656917906428 | 0.753120287838777 | 0.901411050383085 | -5.58670300114855 |
| ENSG00000104660.18 | LEPROTL1 | -0.0402618456083434 | 4.3756030946418 | -0.316534474607703 | 0.753146441097155 | 0.901411050383085 | -6.28541445192315 |
| lnc-ZNF490-1 | lnc-ZNF490-1 | -0.0440859880863288 | -0.0320272159429928 | -0.316531900398445 | 0.753148381031448 | 0.901411050383085 | -5.6129741635247 |
| ENSG00000176393.10 | RNPEP | 0.0295716404292863 | 3.95231242451114 | 0.316493634348203 | 0.753177218668813 | 0.901411050383085 | -6.24134317721096 |
| ENSG00000204231.10 | RXRB | -0.023420645785996 | 4.81882184960765 | -0.316460281627153 | 0.753202353868286 | 0.901411050383085 | -6.31991728808839 |
| lnc-CNDP1-5 | lnc-CNDP1-5 | 0.0872501058245148 | 1.40635765717667 | 0.316447091314817 | 0.75321229439531 | 0.901411050383085 | -5.9333762455316 |

| | | | | | | | |
|---|---|---|---|---|---|---|---|
| ENSG00000198242.13 | RPL23A | 0.0480507238840815 | 9.24415627533003 | 0.316351283678735 | 0.753284498535879 | 0.901445048451395 | -6.30795054862799 |
| ENSG00000087086.14 | FTL | -0.0394851990144947 | 8.6060036691075 | -0.316272329270946 | 0.75334400314652 | 0.901463846099925 | -6.32795650352518 |
| lnc-ECHDC3-5 | lnc-ECHDC3-5 | -0.0579534153560912 | 2.98209700054575 | -0.316195261738493 | 0.753402087162067 | 0.901480941754263 | -6.12464316511048 |
| ENSG00000254681.6 | PKD1P5 | -0.127199690928892 | 1.18522945922703 | -0.31608385558045 | 0.753499797433362 | 0.901545447338888 | -5.91397116583516 |
| ENSG00000241839.9 | PLEKHO2 | 0.0523354387158657 | 3.46935432863458 | 0.315928163885518 | 0.75360340381504 | 0.901586016581866 | -6.19110670441219 |
| lnc-SAMD9L-1 | lnc-SAMD9L-1 | 0.0584573037504996 | 2.68117290958859 | 0.315904408425897 | 0.753621309590001 | 0.901586016581866 | -6.09015901170204 |
| ENSG00000248019.2 | FAM13A-AS1 | 0.0584557712472566 | 3.83093466225402 | 0.315656142589882 | 0.753808449189032 | 0.901723005443453 | -6.22622806989324 |
| lnc-NUP62-1 | lnc-NUP62-1 | -0.050886133325367 | 2.8441486903858 | -0.315636263268067 | 0.753823434614269 | 0.901723005443453 | -6.12063128836449 |
| TRAF3IP2-AS1 | TRAF3IP2-AS1 | 0.0681195187822468 | 4.54617116700592 | 0.315565092918546 | 0.753877085015932 | 0.901734776899286 | -6.29602224113719 |
| ENSG00000232112.3 | TMA7 | -0.0427066346383587 | 5.31346710163128 | -0.315478987229385 | 0.753941995781443 | 0.901760015060111 | -6.3436254623639 |
| ENSG00000104763.18 | ASAH1 | -0.0361043094083024 | 5.79324493771838 | -0.315283836427682 | 0.754089116858721 | 0.9018405198974 | -6.35746970430444 |
| lnc-HYAL3-2 | lnc-HYAL3-2 | -0.0599365472537866 | 0.550710294226432 | -0.315273467909243 | 0.754096933778077 | 0.9018405198974 | -5.78757157899637 |
| ENSG00000147654.14 | EBAG9 | -0.0376852639637173 | 4.42760343410312 | -0.315189445335097 | 0.754160280115245 | 0.901863876691384 | -6.28693075033875 |
| lnc-JMJD7-PLA2G4B-2 | lnc-JMJD7-PLA2G4 | 0.253357344987819 | 2.687606184321 | 0.31519133663364 | 0.754220111345743 | 0.901883027469321 | -6.11860589713001 |
| ENSG00000178127.12 | NDUFV2 | 0.0368425442068032 | 5.69319576634665 | 0.314851565414701 | 0.754415031989302 | 0.902063704161485 | -6.35501252896878 |
| ENSG00000274582.1 | SNORA16A | -0.0648278164408808 | 0.661911984876408 | -0.314781854473427 | 0.754467595506183 | 0.902074151513434 | -5.79748934500788 |
| ENSG00000131023.12 | LATS1 | 0.0269366499580288 | 5.97914070415011 | 0.314689768550739 | 0.754537032035521 | 0.902104770584754 | -6.36020713586313 |
| ENSG00000076944.15 | STXBP2 | -0.0352494375580923 | 4.4637663351763 | -0.314565271890885 | 0.754630910845013 | 0.902109608418882 | -6.29203692442847 |
| lnc-RGN-3 | lnc-RGN-3 | 0.127421887953788 | -0.495893977623018 | 0.314538956538106 | 0.754650754862024 | 0.902109608418882 | -5.45988531147989 |
| ENSG00000171298.12 | GAA | 0.0588306392235913 | 3.69870679483791 | 0.314487111602321 | 0.754689850849448 | 0.902109608418882 | -6.21032184559652 |
| ENSG00000100347.14 | SAMM50 | -0.0388442711748951 | 4.07905803069991 | -0.314451918595181 | 0.754716390077887 | 0.902109608418882 | -6.25787488748809 |
| ENSG00000183513.8 | COA5 | -0.0345513567170981 | 4.48335380733028 | -0.314318341364499 | 0.754817124086619 | 0.902177624126061 | -6.29618741325 |
| lnc-COG8-1 | lnc-COG8-1 | 0.0657698596608626 | -0.162333688700868 | 0.31417517613944 | 0.75492509344239 | 0.902254279312781 | -5.58428480492719 |
| ENSG00000222328.1 | RNU2-2P | -0.102255124087331 | 5.65388553468909 | -0.31382398115336 | 0.755224883012345 | 0.90256016762719 | -6.35358265704367 |
| ENSG00000105429.12 | MEGF8 | -0.0473328250184575 | 3.76454948574803 | -0.3136671097479 | 0.755308296343731 | 0.902587664010065 | -6.23066288649061 |
| ENSG00000102125.15 | TAFAZZIN | 0.0287546845507807 | 4.76321167437649 | 0.313588743916035 | 0.755367408373981 | 0.902587664010065 | -6.31446179725954 |
| ENSG00000115657.13 | ABCB6 | 0.0685377632989966 | 0.705903193801113 | 0.313572788038823 | 0.755379444213058 | 0.902587664010065 | -5.79868149709488 |
| ENSG00000131697.17 | NPHP4 | -0.0736159684405138 | 2.24978853525247 | -0.313364961335326 | 0.755536217691001 | 0.902623005726465 | -6.03581306065369 |
| lnc-VARS-1 | lnc-VARS-1 | 0.0715884822528567 | -0.335056507337341 | 0.313362021168534 | 0.755538435672202 | 0.902623005726465 | -5.53537252520278 |
| ENSG00000159658.10 | EFCAB14 | -0.0265592526047707 | 7.1440616033965 | -0.31330995301965 | 0.755577714801558 | 0.902623005726465 | -6.35624279072419 |
| lnc-GOLGA4-4 | lnc-GOLGA4-4 | -0.0860805571522214 | 1.12400524049195 | -0.31328486743944 | 0.755596639075476 | 0.902623005726465 | -5.89619529934728 |
| lnc-SLC25A19-3 | lnc-SLC25A19-3 | -0.0752849709632697 | 1.34758912339137 | -0.313242917397204 | 0.755628286045412 | 0.902623005726465 | -5.90938245190657 |
| ENSG00000130956.13 | HABP4 | 0.0506134189983669 | 2.93935119756403 | 0.313173862741881 | 0.75568038156862 | 0.902632851313075 | -6.1220056283998 |
| ENSG00000112782.16 | CLIC5 | 0.201555147462471 | 0.265639811714674 | 0.313133261415749 | 0.755744058369527 | 0.902656528371147 | -5.68210248796131 |
| ENSG00000127054.20 | INTS11 | -0.0389686514238683 | 5.170624643276 | -0.313022089412735 | 0.75579488493017 | 0.902664855316101 | -6.34217066438283 |
| ENSG00000270800.3 | RPS10-NUDT3 | 0.045524528425072 | 8.00678989326557 | 0.312897999349653 | 0.755888507149115 | 0.90270011645774 | -6.34202166678011 |
| ENSG00000196812.4 | ZSCAN16 | 0.0589025029886785 | 2.95231890457314 | 0.312865557653674 | 0.755912984044783 | 0.90270011645774 | -6.13191726789185 |
| ENSG00000115268.9 | RPS15 | -0.0562927890163261 | 7.79840675098873 | -0.312763121422831 | 0.755990272696906 | 0.90270011645774 | -6.34407639129752 |
| ENSG00000174243.9 | DDX23 | -0.0369770517169057 | 6.12265392650276 | -0.312677064034456 | 0.756055205384446 | 0.90270011645774 | -6.36157529453311 |
| ENSG00000143390.17 | RFX5 | -0.0330971405139815 | 6.85347759645446 | -0.312600682066494 | 0.75611283919344 | 0.90270011645774 | -6.35967387800969 |
| ENSG00000149503.12 | INCENP | 0.0454785758017859 | 3.18878254173975 | 0.312578119846441 | 0.756129863724921 | 0.90270011645774 | -6.15651153412365 |
| ENSG00000177034.15 | MTX3 | 0.0367944728605414 | 5.66604172112885 | 0.312534254482004 | 0.756162963091614 | 0.90270011645774 | -6.35613409931469 |
| ENSG00000137310.11 | TCF19 | -0.0690732219306671 | 3.55849871872399 | -0.312495281168587 | 0.756192371465656 | 0.90270011645774 | -6.19597550847524 |
| ENSG00000130844.17 | ZNF331 | 0.0829517168292959 | 4.95700993525563 | 0.312464506902115 | 0.756229972692645 | 0.90270011645774 | -6.32500977327664 |
| ENSG00000033800.13 | PIAS1 | 0.0287601503479085 | 6.99934995009763 | 0.312401716536617 | 0.756262974697434 | 0.90270011645774 | -6.35809946109618 |
| ENSG00000259630.2 | FABP5P9 | 0.100140173688458 | 2.02294624012144 | 0.312026189654242 | 0.756548570096529 | 0.902823327549636 | -5.99426226233385 |
| lnc-KPNA2-6 | lnc-KPNA2-6 | -0.0403609073194859 | 6.37032833016428 | -0.31196996358643 | 0.756588799813927 | 0.902823327549636 | -6.36233061559815 |
| LINC00426 | LINC00426 | -0.0965606941438395 | 3.14424282730592 | -0.311928577339859 | 0.756631318482773 | 0.902823327549636 | -6.14420100243062 |
| lnc-NPC1-7 | lnc-NPC1-7 | -0.063422242111563 | 0.294442710294582 | -0.311890063706722 | 0.75664910168383 | 0.902823327549636 | -5.69936578034201 |
| ENSG00000268400.5 | | 0.0303315390674537 | 3.87974144281701 | 0.311869691551112 | 0.756664477159987 | 0.902823327549636 | -6.23367382945074 |
| lnc-CCNL1-9 | lnc-CCNL1-9 | -0.0350355149812582 | 3.00064703602826 | -0.311723129312108 | 0.756775095012617 | 0.902823327549636 | -6.13337153764978 |
| ENSG00000115935.17 | WIPF1 | 0.0366502968780829 | 6.82379938930048 | 0.311648747593434 | 0.756831236590597 | 0.902823327549636 | -6.36052275456936 |
| ENSG00000178338.10 | ZNF354B | -0.0398100289340761 | 3.41726426345005 | -0.311599642385935 | 0.756868300778672 | 0.902823327549636 | -6.17930889958745 |
| ENSG00000129474.15 | AJUBA | -0.145987977762291 | -0.0644801255811586 | -0.311591037650291 | 0.756874795618731 | 0.902823327549636 | -5.57692478817048 |
| ENSG00000170089.15 | | 0.0567057012710708 | 1.57270387675406 | 0.311469428339964 | 0.756966588023964 | 0.902823327549636 | -5.95562792684136 |
| ENSG00000154310.16 | TNIK | -0.332917333770459 | -1.09307221195931 | -0.311495383472058 | 0.756993852561219 | 0.902823327549636 | -5.47597792766536 |
| lnc-PAPLN-2 | lnc-PAPLN-2 | 0.0344771547024947 | 3.16294178303344 | 0.311431793030144 | 0.75699499640076 | 0.902823327549636 | -6.15254923451549 |
| ENSG00000279236.1 | | -0.0622906483422989 | 1.75235869256735 | -0.311417346934365 | 0.757005900882029 | 0.902823327549636 | -5.96644999342546 |
| ENSG00000155463.12 | OXA1L | -0.0323946820504366 | 6.34625619040158 | -0.311403044503977 | 0.757016696968363 | 0.902823327549636 | -6.3625477238619 |
| ENSG00000101152.10 | DNAJC5 | -0.0460462949565876 | 4.9843178107906 | -0.311383032405662 | 0.757031803038734 | 0.902823327549636 | -6.32930338943959 |
| lnc-ESCO1-3 | lnc-ESCO1-3 | -0.0603001434802921 | 1.90352461326623 | -0.311257577238805 | 0.757126504672508 | 0.902823327549636 | -5.99306951707389 |
| ENSG00000108474.16 | PIGL | -0.0454453096158591 | 4.65669052193715 | -0.311235611394319 | 0.757143086294101 | 0.902823327549636 | -6.30804482072667 |
| ENSG00000186166.8 | CENATAC | -0.0575356898488915 | 5.79449451480721 | -0.311218869357295 | 0.757155724634981 | 0.902823327549636 | -6.35852063836118 |
| ENSG00000102981.9 | PARD6A | -0.0686429611812982 | 1.38347388073451 | -0.311132484112344 | 0.757220936775033 | 0.902828300063082 | -5.93252259743109 |
| lnc-ABCC8-1 | lnc-ABCC8-1 | 0.0849241713816852 | -0.361487690730022 | 0.311097137417771 | 0.757247620477532 | 0.902828300063082 | -5.52184807578399 |
| lnc-ATP6V1G3-1 | lnc-ATP6V1G3-1 | 0.227592434822852 | -1.0814044664497 | 0.310929052625978 | 0.757390168233763 | 0.902945950347845 | -5.41843120142644 |
| lnc-KLHL1-10 | lnc-KLHL1-10 | 0.169635250137295 | 0.561947833710011 | 0.310823643719902 | 0.757486586928348 | 0.902996677483756 | -5.82668310901335 |
| ENSG00000160767.20 | ENTREP3 | 0.081344102098993 | 0.168695196165555 | 0.310705667771184 | 0.75754316628988 | 0.902996677483756 | -5.67730715596511 |

| | | | | | | | |
|---|---|---|---|---|---|---|---|
| ENSG00000110497.14 | AMBRA1 | 0.0242983229846905 | 5.31619309053248 | 0.310677635485883 | 0.757564331088252 | 0.902996677483756 | -6.34580972285712 |
| ENSG00000113916.17 | BCL6 | 0.08050960537956 | 5.06664082291503 | 0.310388013678098 | 0.757796612618695 | 0.903221244862505 | -6.33534958548937 |
| ENSG00000279088.1 | | 0.0369023711710756 | 2.7309116225509 | 0.310305539029588 | 0.757845287353627 | 0.903226957127951 | -6.10580658503694 |
| ENSG00000105698.15 | USF2 | -0.0391175969042585 | 5.9278664009894 | -0.310220123932045 | 0.75790978580988 | 0.903240486472117 | -6.36079125544947 |
| ENSG00000006757.11 | PNPLA4 | -0.0605897036153091 | 1.50258788304479 | -0.310157549513854 | 0.757957037971422 | 0.903240486472117 | -5.95005648902304 |
| ENSG00000105127.8 | AKAP8 | 0.0306119278022683 | 6.04527323630824 | 0.310116167436883 | 0.757988287559291 | 0.903240486472117 | -6.36224548158642 |
| ENSG00000123080.10 | CDKN2C | 0.226249612086799 | 0.148120084806168 | 0.309850955570278 | 0.758227827400978 | 0.903473623445312 | -5.66821787279941 |
| ENSG00000278896.1 | | -0.0881549418740484 | 2.27769466107753 | -0.309619652593306 | 0.758363261614791 | 0.903558018582554 | -6.05568772318579 |
| lnc-CDC73-5 | lnc-CDC73-5 | 0.0607817780638804 | 3.92528578882765 | 0.309588948844506 | 0.75838645139349 | 0.903558018582554 | -6.23479745851078 |
| ENSG00000010165.19 | METTL13 | -0.0443508189493134 | 3.36423298342832 | -0.309467446005047 | 0.758478221677493 | 0.903608519601893 | -6.17707004137213 |
| ENSG00000179627.9 | ZBTB42 | -0.0811205842347564 | 1.21315830823911 | -0.309416581921012 | 0.75851664002728 | 0.903608519601893 | -5.91626892770176 |
| ENSG00000130560.8 | UBAC1 | -0.0453207065306662 | 3.25086679066501 | -0.309298817150681 | 0.758605591765149 | 0.903662185039763 | -6.16486135400293 |
| lnc-STYK1-2 | lnc-STYK1-2 | -0.109916256111929 | 4.68305999242777 | -0.309200291761793 | 0.758717088273956 | 0.903742698376321 | -6.3126456405803 |
| lnc-GSK3A-1 | lnc-GSK3A-1 | 0.0576490367707236 | 0.921780525123032 | 0.309072258326155 | 0.758776728658878 | 0.90376143776319 | -5.85107412174622 |
| lnc-TBC1D22A-1 | lnc-TBC1D22A-1 | -0.0978487886793916 | 0.832971011508963 | -0.308995094172396 | 0.758835019333306 | 0.903778567465422 | -5.80693811995088 |
| lnc-METTL16-5 | lnc-METTL16-5 | -0.0605019883035905 | -0.216930692345992 | -0.308766883870566 | 0.759007420281242 | 0.903861635185609 | -5.5841370905932 |
| ENSG00000124422.11 | USP22 | 0.0323738205972681 | 7.19279791695863 | 0.30874006548333 | 0.759027680985039 | 0.903861635185609 | -6.35628736894093 |
| ENSG00000244038.9 | DDOST | -0.0365325538274277 | 6.07093525724374 | -0.308728386685712 | 0.759036504114233 | 0.903861635185609 | -6.36255580611648 |
| lnc-ZNF652-3 | lnc-ZNF652-3 | -0.0583359553087193 | 1.90388350457395 | -0.308651206718762 | 0.759094813058371 | 0.903878776881896 | -6.00378208840051 |
| lnc-GOLGA8H-119 | lnc-GOLGA8H-119 | 0.0637832350204949 | 0.932186873232053 | 0.308547040818367 | 0.759173511942274 | 0.903920194152127 | -5.8492814155861 |
| lnc-BCAS2-4 | lnc-BCAS2-4 | 0.0306455420842404 | 2.82407817234561 | 0.308332017981499 | 0.759335973055905 | 0.904017605743978 | -6.11229607596298 |
| lnc-COPE-1 | lnc-COPE-1 | -0.0543378010880806 | 2.54007970369664 | -0.308322498215281 | 0.759343165996581 | 0.904017605743978 | -6.08425094364907 |
| ENSG00000172244.8 | C5orf34 | -0.0651487535785505 | 0.798844422229449 | -0.308127197813077 | 0.759490735744167 | 0.904125877654084 | -5.82408089537746 |
| lnc-SMIM14-2 | lnc-SMIM14-2 | 0.0572817226941882 | 1.26980520874087 | 0.308043233789552 | 0.75955418207668 | 0.904125877654084 | -5.9177349370699 |
| lnc-FAM83F-5 | lnc-FAM83F-5 | 0.101555737033305 | 0.1419426677842 | 0.308027742333083 | 0.759565888179295 | 0.904125877654084 | -5.63216004671443 |
| ENSG00000007545.15 | CRAMP1 | -0.0484855459155735 | 5.40051495826158 | -0.307877720308323 | 0.759679255113682 | 0.904208529925687 | -6.34929978981222 |
| ENSG00000171953.15 | ATPAF2 | 0.0340884950829312 | 3.92522669341112 | 0.307687583488527 | 0.759822943222738 | 0.904327260341946 | -6.24845692870651 |
| ENSG00000163378.13 | EOGT | 0.0608548961617297 | 3.98176491892908 | 0.307603829110041 | 0.75988623988575 | 0.904328397488514 | -6.25594563843898 |
| ENSG00000150477.14 | KIAA1328 | -0.0279683053577138 | 4.35211412994498 | -0.307570048124585 | 0.759911770051078 | 0.904328397488514 | -6.28187940054058 |
| ENSG00000182827.8 | ACBD3 | -0.034666941618104 | 6.31180285446333 | -0.307308747672039 | 0.760109258453436 | 0.904511121393714 | -6.36378803740055 |
| ENSG00000011143.16 | MKS1 | 0.0626078116650182 | 2.16022239971604 | 0.307202060413999 | 0.760189896327689 | 0.904539362854173 | -6.03333505678134 |
| lnc-ZNF516-14 | lnc-ZNF516-14 | 0.0797704838524611 | 2.27014321690627 | 0.307113546541283 | 0.760256800185122 | 0.904539362854173 | -6.04386149040965 |
| lnc-ZNF639-4 | lnc-ZNF639-4 | -0.0653735254500676 | 2.35184827990972 | -0.307102524102904 | 0.760265131705769 | 0.904539362854173 | -6.07421510879135 |
| ENSG00000181817.5 | LSM10 | -0.0709200560364227 | 3.15921150443708 | -0.307044785444514 | 0.760308775044456 | 0.904539362854173 | -6.14971397080871 |
| ENSG00000101447.14 | FAM83D | -0.109933760227578 | -0.0980917949007314 | -0.306969049841394 | 0.760366023062345 | 0.904555187417191 | -5.63939486339884 |
| ENSG00000170345.9 | FOS | -0.286598883905896 | 3.70412471384122 | -0.306858046716027 | 0.760509444074878 | 0.904651201145003 | -6.21310592019939 |
| ENSG00000138621.11 | PPCDC | -0.0617573313441917 | 2.41498984364552 | -0.306745997561806 | 0.760534634631159 | 0.904651201145003 | -6.05892820557144 |
| ENSG00000108848.15 | LUC7L3 | -0.033183595751483 | 7.99512107401347 | -0.306621917526788 | 0.760628435363025 | 0.904710493214513 | -6.34196666983163 |
| ENSG00000222365.1 | SNORD12B | 0.0568441016724366 | 2.37506974795441 | 0.306520389427148 | 0.760705190229091 | 0.904749504824071 | -6.05501316179054 |
| lnc-LMF2-1 | lnc-LMF2-1 | 0.0427544021366133 | 2.23776844782317 | 0.306291865547758 | 0.760877962356707 | 0.904797184934113 | -6.03871445109081 |
| lnc-UQCR11-1 | lnc-UQCR11-1 | -0.0718234450973415 | 0.0653644782710007 | -0.306213879571826 | 0.760936925337147 | 0.904797184934113 | -5.66827176211481 |
| ENSG00000160584.15 | SIK3 | -0.0529091033523903 | 7.10644159583253 | -0.306206811318258 | 0.760942269513906 | 0.904797184934113 | -6.35881640824743 |
| ENSG00000251149.1 | MTND5P5 | -0.111451061760984 | 0.745413769828253 | -0.30613165565658 | 0.760999094060453 | 0.904797184934113 | -5.81997016766843 |
| lnc-NFKB1-6 | lnc-NFKB1-6 | -0.111451061760984 | 0.745413769828253 | -0.30613165565658 | 0.760999094060453 | 0.904797184934113 | -5.81997016766843 |
| ENSG00000091527.15 | CDV3 | -0.0361828581833743 | 7.68642113971568 | -0.306118514578889 | 0.761009030052926 | 0.904797184934113 | -6.34881608240431 |
| ENSG00000168273.7 | UQCC5 | 0.0456309839622997 | 1.08970107582951 | 0.306048931127648 | 0.761061642905836 | 0.904798459493299 | -5.87460637087145 |
| lnc-RENBP-2 | lnc-RENBP-2 | -0.0592118534407517 | 0.957057802310683 | -0.306000822419725 | 0.761098019125831 | 0.904798459493299 | -5.8711331588507 |
| ENSG00000285547.1 | | 0.0315568140898115 | 1.88657421723791 | 0.305850208810712 | 0.761211905449544 | 0.904875255602806 | -5.99495155936864 |
| ENSG00000250896.1 | RNPS1P1 | -0.172205940547941 | 0.215872532581889 | -0.30581333996962 | 0.761255591650682 | 0.904875255602806 | -5.64597231697069 |
| ENSG00000203711.13 | LINC02901 | -0.093680505406076 | 0.111147499907989 | -0.30573059765786 | 0.761302353115558 | 0.904875255602806 | -5.69332285638037 |
| ENSG00000075426.11 | FOSL2 | -0.278073572411288 | 0.479037088672861 | -0.305751597414831 | 0.761338467498877 | 0.904875255602806 | -5.69569944842648 |
| ENSG00000154102.10 | C16orf74 | -0.0793314977429693 | 4.17841133756198 | -0.305533535629123 | 0.761458316363499 | 0.904946340719433 | -6.27277976832697 |
| GABPB1-IT1 | GABPB1-IT1 | -0.0483242385032874 | 6.0134525425948 | -0.305487475174891 | 0.761486208097002 | 0.904946340719433 | -6.36320121588763 |
| ENSG00000244879.7 | GABPB1-AS1 | -0.0696714943705565 | 6.89120013927913 | -0.305360348755383 | 0.761587020256648 | 0.905013892843519 | -6.36164460469219 |
| ENSG00000125952.18 | MAX | 0.0339527455522584 | 5.44390141019225 | 0.304984658018501 | 0.761866494579445 | 0.905166985073506 | -6.35188569979411 |
| GABPB1-AS1 | GABPB1-AS1 | -0.0688660428977936 | 6.94459026152931 | -0.304936092214333 | 0.761907243559642 | 0.905166985073506 | -6.3612562209333 |
| CD81-AS1 | CD81-AS1 | 0.0751589079294462 | 1.02247409502972 | 0.304903936741731 | 0.76192755057396 | 0.905166985073506 | -5.87165412561888 |
| ENSG00000255112.2 | CHMP1B | -0.0401710094794518 | 6.2464473693333 | -0.304897794668123 | 0.76193219638111 | 0.905166985073506 | -6.36440203383624 |
| ENSG00000189067.12 | LITAF | -0.0413014735176649 | 6.81431638047189 | -0.304893119600063 | 0.761935732564912 | 0.905166985073506 | -6.36278279204514 |
| lnc-EXOC8-4 | lnc-EXOC8-4 | -0.0451956942627965 | 0.608174217170945 | -0.304822308967604 | 0.761989293786034 | 0.905178370981586 | -5.78144897085039 |
| lnc-RABEP1-5 | lnc-RABEP1-5 | -0.0701819180428142 | 0.253654282262542 | -0.304763102294001 | 0.762034078669417 | 0.905179330635538 | -5.69792061384704 |
| lnc-WDR91-3 | lnc-WDR91-3 | -0.0508154409997297 | 2.44062509248763 | -0.304601390585187 | 0.762156404220772 | 0.905272391256054 | -6.06364653273102 |
| lnc-ZNF227-2 | lnc-ZNF227-2 | 0.0353753108830174 | 0.93955729826164 | 0.304533925724292 | 0.762207439304046 | 0.90528076879372 | -5.85352032148853 |
| lnc-PDE7B-8 | lnc-PDE7B-8 | -0.101998892853734 | 2.03631720886715 | -0.30444718494807 | 0.762279972381742 | 0.905314677256557 | -6.02679205091948 |
| MATN1-AS1 | MATN1-AS1 | 0.0777883997879255 | 0.386503878871159 | 0.304037572573213 | 0.762582948038442 | 0.905574747248171 | -5.70656580694021 |
| ENSG00000105738.10 | SIPA1L3 | 0.0543816844167262 | 7.68889433903622 | 0.304032289411748 | 0.76258694524863 | 0.905574747248171 | -6.35246053618003 |
| lnc-ESS2-2 | lnc-ESS2-2 | 0.0748145609245994 | 0.558574825034448 | 0.303955433045373 | 0.762645095083761 | 0.905591553715764 | -5.76521110186381 |
| lnc-YWHAZ-3 | lnc-YWHAZ-3 | -0.051228798733143 | 1.89978830199822 | -0.303881718132354 | 0.762700869386177 | 0.905605537616133 | -6.00037693904842 |

| | | | | | | | |
|---|---|---|---|---|---|---|---|
| ENSG00000220205.8 | VAMP2 | -0.0556372702322427 | 5.05492291814414 | -0.303740865514584 | 0.762807445080809 | 0.905631743542708 | -6.33862531427587 |
| ENSG00000108819.10 | PPP1R9B | -0.0576124156181139 | 6.79111381608252 | -0.303736249244578 | 0.762810938045957 | 0.905631743542708 | -6.36297518380562 |
| lnc-SELENOP-6 | lnc-SELENOP-6 | 0.10814700745404 | -0.336297459962424 | 0.303624793635897 | 0.762895274006457 | 0.905679629996246 | -5.59892264524758 |
| ENSG00000168350.7 | DEGS2 | 0.0737130791126579 | 3.11204591017637 | 0.303396660849062 | 0.763067906060171 | 0.905832326572265 | -6.15092701214407 |
| lnc-CALM3-1 | lnc-CALM3-1 | -0.0443391856578936 | 0.697187017758774 | -0.303252328471754 | 0.76317713118767 | 0.905909739978998 | -5.809345131148 |
| ENSG00000205133.11 | TRIQK | -0.463813710990626 | -1.0396123902669 | -0.303167845599115 | 0.763287115839659 | 0.905942579310744 | -5.2054409670726 |
| lnc-SHROOM2-1 | lnc-SHROOM2-1 | -0.0644083759715905 | -0.276075805380711 | -0.303099455696173 | 0.763292824680898 | 0.905942579310744 | -5.55880776185051 |
| lnc-EIF4E3-3 | lnc-EIF4E3-3 | -0.195588797312239 | 1.25729483961515 | -0.302795101185083 | 0.763566659876185 | 0.906172674769495 | -5.90480964965057 |
| lnc-PIGP-5 | lnc-PIGP-5 | 0.0646696000697481 | 0.122524274542066 | 0.302726974442811 | 0.763574739892247 | 0.906172674769495 | -5.64910103508668 |
| lnc-MRPL48-4 | lnc-MRPL48-4 | 0.0645740613049237 | -0.0281835942451456 | 0.302401226946445 | 0.763821310926338 | 0.90641303215894 | -5.57802993135923 |
| ENSG00000118513.18 | MYB | -0.29594189771359 | 1.22489450519047 | -0.302278614961969 | 0.763972684118318 | 0.906540398778239 | -5.93500753158419 |
| ENSG00000101265.15 | RASSF2 | 0.0713523153715867 | 6.76293454917274 | 0.301967196729533 | 0.764156738161585 | 0.906683323943713 | -6.36402524834265 |
| ENSG00000214711.9 | CAPN14 | 0.0516176816648737 | 2.60780079228721 | 0.301849887185971 | 0.764238698093373 | 0.906683323943713 | -6.08052429399031 |
| ENSG00000070047.11 | PHRF1 | 0.0570920858052695 | 5.06652709630981 | 0.301841067194795 | 0.76424537577103 | 0.906683323943713 | -6.33831273533241 |
| lnc-REEP1-2 | lnc-REEP1-2 | -0.046981544523867 | 1.53081707801465 | -0.301809424960855 | 0.764269332479397 | 0.906683323943713 | -5.95542908986516 |
| ENSG00000272434.1 | | -0.0671817029929602 | 0.523692996923154 | -0.301708800303864 | 0.764345518146498 | 0.90672144545037 | -5.75422487632375 |
| ENSG00000120616.15 | EPC1 | 0.029590485848014 | 6.98599784216957 | 0.301525467923403 | 0.764484330154989 | 0.906833850137168 | -6.36185489051539 |
| ENSG00000136514.2 | RTP4 | 0.0538933135615987 | 1.83139178183136 | 0.301417797381236 | 0.764565857673242 | 0.906878294732228 | -5.9884743897193 |
| lnc-GINS1-9 | lnc-GINS1-9 | -0.0808843753675309 | -0.506755737564764 | -0.301329702468807 | 0.764632564646822 | 0.90690515605195 | -5.46915434847244 |
| ENSG00000131828.13 | PDHA1 | -0.0328417285615794 | 4.72524401220257 | -0.300760642745932 | 0.765063509950504 | 0.907250959882054 | -6.31725864142877 |
| ENSG00000100348.9 | TXN2 | -0.0507350901262513 | 3.50259445403765 | -0.300709059016168 | 0.76510257770657 | 0.907250959882054 | -6.19838734572982 |
| ENSG00000100368.13 | CSF2RB | -0.0689136783792921 | 3.849965612468 | -0.300599897386897 | 0.765185255038985 | 0.907250959882054 | -6.24472461242134 |
| lnc-ITGA11-6 | lnc-ITGA11-6 | -0.0362530939281163 | 3.65631291703348 | -0.300575536756293 | 0.765203705782661 | 0.907250959882054 | -6.21183024677792 |
| lnc-PTGES3L-AARSD1-2 | lnc-PTGES3L-AARS | 0.0405260109222835 | 0.35699572724262 | 0.300565753611721 | 0.765211115576021 | 0.907250959882054 | -5.72287840053726 |
| LINC01754 | LINC01754 | -0.200038993420447 | 2.11547172612198 | -0.300598111987433 | 0.765240078004204 | 0.907250959882054 | -6.04511093240805 |
| ENSG00000120053.11 | GOT1 | 0.0358093185431856 | 2.54325930553578 | 0.300410533697591 | 0.76532868274534 | 0.907250959882054 | -6.07666477702984 |
| ENSG00000249140.1 | PRDX2P3 | 0.0557055404230842 | -0.278007262083294 | 0.300393926790258 | 0.765341261534426 | 0.907250959882054 | -5.54249068526963 |
| lnc-CANX-5 | lnc-CANX-5 | 0.0557055404230842 | -0.278007262083294 | 0.300393926790258 | 0.765341261534426 | 0.907250959882054 | -5.54249068526963 |
| lnc-TBC1D28-1 | lnc-TBC1D28-1 | 0.0869115243928204 | 0.350801606205633 | 0.300362128401812 | 0.765365347186476 | 0.907250959882054 | -5.68262402759409 |
| lnc-MAFA-5 | lnc-MAFA-5 | -0.0706708503797361 | 0.17575186585854 | -0.300304531729931 | 0.765408974316274 | 0.907250959882054 | -5.67820501608631 |
| ENSG00000157343.8 | ARMC12 | 0.0819559502948686 | -0.202462893304421 | 0.300093095206126 | 0.765569135466194 | 0.907290077093286 | -5.59675705376225 |
| ENSG00000064547.13 | LPAR2 | -0.0369109115677544 | 3.23648242589518 | -0.300063611434935 | 0.765591469966403 | 0.907290077093286 | -6.16690981605659 |
| ENSG00000164691.17 | TAGAP | -0.0608828457969128 | 8.66477774288081 | -0.299981588734089 | 0.765653604736001 | 0.907290077093286 | -6.32726270301783 |
| ENSG00000078808.16 | SDF4 | 0.0316269802665907 | 5.03745931673376 | 0.299929676142298 | 0.765692930960121 | 0.907290077093286 | -6.33579028964509 |
| ENSG00000267561.2 | | -0.0445518022982623 | 2.77117512503128 | -0.299909292805969 | 0.765708372464812 | 0.907290077093286 | -6.10096429810838 |
| lnc-SCML4-1 | lnc-SCML4-1 | -0.337809781360533 | -0.4935850087831 | -0.29986440870373 | 0.765781212737768 | 0.907290077093286 | -5.32768688258035 |
| lnc-GSTT2B-4 | lnc-GSTT2B-4 | -0.27521936521733 | 0.281996944594496 | -0.299829698728083 | 0.765807035410485 | 0.907290077093286 | -5.48674989991385 |
| ENSG00000165175.15 | MID1IP1 | 0.14354159759924 | 3.75973043519833 | 0.299814706189702 | 0.765826750034801 | 0.907290077093286 | -6.22886386501753 |
| lnc-ZC3HC1-4 | lnc-ZC3HC1-4 | -0.0447575312816695 | 3.5246399607591 | -0.299737271317909 | 0.76583869209633 | 0.907290077093286 | -6.19345537968462 |
| lnc-ANKRD12-2 | lnc-ANKRD12-2 | 0.0423714857830615 | 2.51523409262871 | 0.299553793500326 | 0.765977698350808 | 0.907402530368615 | -6.07624272310218 |
| ENSG00000101608.12 | MYL12A | -0.0539583965012476 | 6.72013637095343 | -0.299481031327978 | 0.766032826505861 | 0.907415611646532 | -6.36504020855434 |
| lnc-CSNK1A1-6 | lnc-CSNK1A1-6 | -0.099680459233233 | -0.138157132679933 | -0.299377331847131 | 0.766111396388491 | 0.90745645806896 | -5.61035175266173 |
| ENSG00000105173.13 | CCNE1 | 0.0771078426399088 | 2.32750795211949 | 0.299043312990493 | 0.766364489056446 | 0.907699397625159 | -6.0491602710907 |
| lnc-PSMB2-6 | lnc-PSMB2-6 | 0.0313000193961601 | 1.1725430422543 | 0.298990254740063 | 0.766404694715427 | 0.907699397625159 | -5.89561578040314 |
| ENSG00000282218.1 | | -0.0443807429220047 | 2.20204179505452 | -0.29880714016192 | 0.766543457446671 | 0.907811506595214 | -6.03421097734158 |
| ENSG00000077549.17 | CAPZB | 0.0473459899160718 | 4.74217510249792 | 0.298633769532839 | 0.766674843440083 | 0.907914866953986 | -6.3183384978705 |
| lnc-SPSB4-1 | lnc-SPSB4-1 | -0.0679087247191116 | 0.517378306319223 | -0.298574698284723 | 0.766719611171656 | 0.907915645883453 | -5.79117964761196 |
| ENSG00000163281.11 | GNPDA2 | 0.0367982662307291 | 4.24395582205993 | 0.298388099888378 | 0.76686103191477 | 0.908030870384958 | -6.28628232458136 |
| ENSG00000104886.11 | PLEKHJ1 | 0.0550772427886681 | 3.37309526552932 | 0.298228038728528 | 0.766982346805161 | 0.908091578799503 | -6.18977680320621 |
| lnc-BAMBI-3 | lnc-BAMBI-3 | 0.0844477039975757 | 0.984332603849033 | 0.298204036670904 | 0.767000539155097 | 0.908091578799503 | -5.86708645815487 |
| lnc-TACSTD2-3 | lnc-TACSTD2-3 | -0.0861797341002374 | -0.363513084043477 | -0.297876487509332 | 0.767248818198269 | 0.908320300430794 | -5.50844749720159 |
| ENSG00000115159.15 | GPD2 | -0.0724606326027519 | 5.69045337818633 | -0.297841246691578 | 0.767284426468317 | 0.908320300430794 | -6.36005482566339 |
| ENSG00000106244.12 | PDAP1 | 0.0402705153234573 | 4.35903980239618 | 0.297729055514126 | 0.767360578277398 | 0.908320300430794 | -6.29112383862098 |
| ENSG00000280135.1 | | -0.335651284262194 | -0.492220918619164 | -0.297725943060067 | 0.767401518242044 | 0.908320300430794 | -5.3284520554869 |
| ENSG00000143742.12 | VIII.09 | 0.0399417823451493 | 5.76906539514872 | 0.297658093747116 | 0.767414372272822 | 0.908320300430794 | -6.36373750249078 |
| ENSG00000239920.2 | | 0.0474902536603965 | 1.4322188147882 | 0.2975648041785 | 0.767485094073302 | 0.908351773406404 | -5.93439551068155 |
| lnc-ZNF35-1 | lnc-ZNF35-1 | -0.0668373241979399 | 1.86010172406314 | -0.297440944364686 | 0.767578993940424 | 0.908410673428918 | -5.99794760964407 |
| ENSG00000256229.7 | ZNF486 | -0.0787567802601652 | 3.81670563572278 | -0.297306970570194 | 0.767685561763517 | 0.908484558027854 | -6.22702499771979 |
| ENSG00000158042.8 | MRPL17 | -0.0433413336011199 | 2.71668818700061 | -0.297237769961067 | 0.767733030965101 | 0.908488500422828 | -6.11124060389903 |
| ENSG00000141644.17 | MBD1 | 0.0232287580643522 | 5.79986934409163 | 0.297117555102935 | 0.767824176541907 | 0.908511294297138 | -6.36312758126293 |
| ENSG00000101220.17 | ADISSP | -0.0678724521289396 | 1.3515319845066 | -0.297057085998485 | 0.767870024805668 | 0.908511294297138 | -5.92866050346391 |
| ENSG00000136709.11 | WDR33 | 0.0221533333869619 | 7.42488047690524 | 0.297037717923587 | 0.76788471004651 | 0.908511294297138 | -6.35769137052467 |
| ENSG00000125630.15 | POLR1B | -0.0335719108229187 | 4.19399556153335 | -0.29688020179759 | 0.768004144957809 | 0.908600374506644 | -6.27806625460768 |
| ENSG00000104154.6 | SLC30A4 | -0.101433528595634 | 2.76996391860332 | -0.296635605141305 | 0.768204403637274 | 0.908785058915225 | -6.10075373126455 |
| ENSG00000256525.6 | POLG2 | -0.0293068049897141 | 4.71666822402715 | -0.296347628358232 | 0.768408005286501 | 0.908924720466776 | -6.31911983916142 |
| ENSG00000187764.11 | SEMA4D | 0.048844065274869 | 6.39583682918146 | 0.2963439711522 | 0.768410778838963 | 0.908924720466776 | -6.36712351348606 |
| lnc-COL28A1-1 | lnc-COL28A1-1 | -0.0545071264954942 | 2.96098516561941 | -0.296194009464265 | 0.768524509450184 | 0.909007009424959 | -6.15889232350379 |
| ENSG00000148110.15 | MFSD14B | 0.0363739502839815 | 4.5121534033388 | 0.296117179632527 | 0.768582779023174 | 0.909023693652473 | -6.30599066014069 |

| | | | | | | | |
|---|---|---|---|---|---|---|---|
| lnc-TP53I13-6 | lnc-TP53I13-6 | 0.0728953282985297 | 0.778723658141318 | 0.295869017692217 | 0.768771000193965 | 0.909111737384619 | -5.79414498120674 |
| ENSG00000158850.14 | B4GALT3 | 0.050657968546612 | 5.45301111079096 | 0.295839915110431 | 0.768793074294303 | 0.909111737384619 | -6.354732366635 |
| ENSG00000162910.18 | MRPL55 | -0.0358909654938691 | 3.0573879090405 | -0.29580189919501 | 0.768821909386924 | 0.909111737384619 | -6.14440150892225 |
| ENSG00000070366.13 | SMG6 | 0.0374865310485344 | 5.03988071683227 | 0.295785125719561 | 0.768834632182526 | 0.909111737384619 | -6.33796780662872 |
| ENSG00000116128.10 | BCL9 | 0.0651568459774023 | 3.15928752318596 | 0.295602358302159 | 0.768973266693995 | 0.909111737384619 | -6.14812275328464 |
| ENSG00000153048.10 | CARHSP1 | -0.0346888476395546 | 3.67787131076878 | -0.295545699367541 | 0.769016245731499 | 0.909111737384619 | -6.22411518594518 |
| ENSG00000100105.17 | PATZ1 | -0.0267689730200527 | 5.58190181148527 | -0.295481133322668 | 0.769065223659844 | 0.909111737384619 | -6.35976750802925 |
| ENSG00000104325.6 | DECR1 | 0.0350749745493006 | 4.93809954481521 | 0.295466495936911 | 0.769076327288717 | 0.909111737384619 | -6.33042752581936 |
| lnc-CCDC167-2 | lnc-CCDC167-2 | -0.258343199464339 | 1.28948844254392 | -0.295523784082144 | 0.769087794995902 | 0.909111737384619 | -5.8861182757966 |
| lnc-SLC46A3-1 | lnc-SLC46A3-1 | 0.0471955149726995 | 2.57595629143562 | 0.295198658029748 | 0.76927951244094 | 0.909111737384619 | -6.08102974746784 |
| lnc-EVL-6 | lnc-EVL-6 | -0.0479246151485313 | 4.17032468329321 | -0.295192816947897 | 0.769283943740832 | 0.909111737384619 | -6.27279602350754 |
| lnc-DENND4C-1 | lnc-DENND4C-1 | -0.0805669484426654 | -0.280964354143675 | -0.295189616186412 | 0.769286371981648 | 0.909111737384619 | -5.57244002682859 |
| ENSG00000237190.3 | CDKN2AIPNL | 0.0491085430753135 | 2.68058079375869 | 0.29508643306936 | 0.769364652564421 | 0.909111737384619 | -6.10758724511083 |
| lnc-ZNF142-4 | lnc-ZNF142-4 | -0.0443524208782884 | 1.22732858822658 | -0.29507080739803 | 0.769376507299931 | 0.909111737384619 | -5.90787583914265 |
| ENSG00000184708.17 | EIF4ENIF1 | -0.0390063298191503 | 4.95607255131525 | -0.295067592021625 | 0.769378946718005 | 0.909111737384619 | -6.33222373364681 |
| lnc-C8orf59-8 | lnc-C8orf59-8 | 0.101212188132948 | 0.179231541727859 | 0.295041777315916 | 0.769398531714364 | 0.909111737384619 | -5.66197027137335 |
| lnc-PPP1R3G-9 | lnc-PPP1R3G-9 | 0.0623922207415551 | 0.644424624194413 | 0.294997210091711 | 0.769432344154079 | 0.909111737384619 | -5.78090987175282 |
| lnc-IGFBP3-4 | lnc-IGFBP3-4 | 0.0283034651228817 | 3.58770270109076 | 0.294961355988059 | 0.769459546418274 | 0.909111737384619 | -6.20852418525064 |
| ENSG00000197122.11 | SRC | -0.110304790883023 | 1.07265612530923 | -0.294917996710034 | 0.769496413968558 | 0.909111737384619 | -5.88638357498901 |
| lnc-GRAMD2A-5 | lnc-GRAMD2A-5 | 0.0309803337953927 | 0.808659028536616 | 0.294776681827872 | 0.769599662099505 | 0.90914842902741 | -5.82615002819603 |
| ENSG00000231181.1 | | -0.0754120101347497 | 0.145421392264138 | -0.294727319684281 | 0.769637115385343 | 0.90914842902741 | -5.65431839311798 |
| ENSG00000158423.16 | RIBC1 | -0.0673435539534982 | 0.372522097874334 | -0.294692481985438 | 0.769663548654547 | 0.90914842902741 | -5.73094320816382 |
| lnc-HMGN5-3 | lnc-HMGN5-3 | -0.0610873721028824 | 1.95134365074194 | -0.294638971891005 | 0.76970415023231 | 0.90914842902741 | -6.0223484156941 |
| ENSG00000101187.15 | SLCO4A1 | -0.119491894346753 | 3.62518232427897 | -0.294494995568502 | 0.769849905780792 | 0.909171684795661 | -6.208799033873 |
| lnc-SUGT1-3 | lnc-SUGT1-3 | -0.0532406255302789 | 2.55751614258708 | -0.294434795173625 | 0.769859078351215 | 0.909171684795661 | -6.09265244165858 |
| ENSG00000266265.3 | KLF14 | -0.148095867879443 | -0.102034459397172 | -0.294443351109671 | 0.769859257031401 | 0.909171684795661 | -5.65900011390883 |
| ENSG00000118058.21 | KMT2A | -0.0429910640854789 | 8.12431075697456 | -0.294380178143792 | 0.76990052305244 | 0.909171684795661 | -6.34441870765095 |
| ENSG00000159228.12 | CBR1 | -0.0888672315825826 | 0.762227340027294 | -0.294236275526817 | 0.770009723039622 | 0.909239339151958 | -5.85686143721983 |
| lnc-MTG2-2 | lnc-MTG2-2 | -0.0621140229180072 | -0.363127306622611 | -0.294182891418121 | 0.770050234581081 | 0.909239339151958 | -5.54360440486061 |
| ENSG00000183889.12 | NPIPA6 | -0.0619356419974706 | 1.33104295519353 | -0.2941300477247 | 0.770090336658217 | 0.909239339151958 | -5.92021543274219 |
| ENSG00000116120.9 | FARSB | -0.0382098398412953 | 4.17907953831515 | -0.293860548381807 | 0.770294864501509 | 0.909428656420475 | -6.28061952839393 |
| ENSG00000054282.15 | SDCCAG8 | 0.030406688636428 | 4.35552757869991 | 0.293475419499266 | 0.770587174372315 | 0.909721583602257 | -6.28218157128875 |
| ENSG00000285077.1 | ARHGAP11B | -0.0678198114429308 | 0.388675687171628 | -0.293250916111126 | 0.770757586297231 | 0.909767017515696 | -5.7256599500705 |
| lnc-NSMCE4A-6 | lnc-NSMCE4A-6 | -0.0417490706303141 | 0.239985769493161 | -0.293226550619077 | 0.770776081898745 | 0.909767017515696 | -5.70195560726546 |
| ENSG00000091127.13 | PUS7 | -0.0661530488122954 | 2.40144964963866 | -0.293196743357169 | 0.770798708476941 | 0.909767017515696 | -6.0680131769186 |
| ENSG00000131370.15 | SH3BP5 | -0.0376975350293547 | 8.01265348785981 | -0.293191802297267 | 0.77080245923608 | 0.909767017515696 | -6.34742232638386 |
| ENSG00000148303.16 | RPL7A | -0.0460162521438168 | 9.66675737436081 | -0.292906906465277 | 0.771018733078194 | 0.909970102233283 | -6.29985328073788 |
| lnc-FBXO42-1 | lnc-FBXO42-1 | 0.0696947270054261 | 0.161166306299109 | 0.292656780690286 | 0.771208627019545 | 0.910069709397125 | -5.65960457884147 |
| ENSG00000102595.19 | UGGT2 | -0.472377760610811 | 1.70704167052322 | -0.292699090880778 | 0.771233072161064 | 0.910069709397125 | -5.90661444814353 |
| ENSG00000134242.15 | PTPN22 | -0.0502161502387244 | 7.3822884630744 | -0.292584892884471 | 0.771263206424537 | 0.910069709397125 | -6.35973265376798 |
| lnc-THUMPD3-11 | lnc-THUMPD3-11 | -0.0776603957304167 | -0.149031866873716 | -0.292529526837252 | 0.771305242808105 | 0.910069709397125 | -5.61229126189895 |
| lnc-SOWAHB-6 | lnc-SOWAHB-6 | 0.0552809146441615 | 2.80840204701383 | 0.292466561289684 | 0.771353049917998 | 0.910069709397125 | -6.1293640791452 |
| ENSG00000100241.20 | SBF1 | 0.033592428609829 | 6.68555212715311 | 0.292433306740198 | 0.771378299071128 | 0.910069709397125 | -6.36713025208798 |
| ENSG00000151414.14 | NEK7 | -0.0303428552300602 | 5.35275301028292 | -0.292388087893913 | 0.771412632747978 | 0.910069709397125 | -6.35508982954401 |
| ENSG00000143418.19 | CERS2 | -0.025254772731599 | 5.64210126439031 | -0.292255287418755 | 0.771513467912422 | 0.910090703472939 | -6.36281055428151 |
| ENSG00000197136.4 | PCNX3 | 0.0345752922284257 | 5.60429535440547 | 0.292248186828187 | 0.771518859490808 | 0.910090703472939 | -6.3604033107103 |
| ENSG00000285762.1 | NA | 0.0460657850997177 | 1.91151062331212 | 0.292188860288518 | 0.771563907414246 | 0.910091685175211 | -5.99548682185632 |
| lnc-RNF219-6 | lnc-RNF219-6 | 0.0593819790238506 | 3.84713272356878 | 0.291948570546191 | 0.771746372764171 | 0.910190563920181 | -6.25300086456318 |
| ENSG00000144040.12 | SFXN5 | 0.0472655323123075 | 3.11693951097203 | 0.291891815308924 | 0.771789472078013 | 0.910190563920181 | -6.1517721778041 |
| ENSG00000188677.14 | PARVB | 0.0670578448863548 | 3.80645033261955 | 0.291830572589158 | 0.771835979955691 | 0.910190563920181 | -6.23434271162005 |
| ENSG00000112599.8 | GUCA1B | 0.0570563268590124 | 0.77384487621505 | 0.291819026553782 | 0.771844748139553 | 0.910190563920181 | -5.80812516166637 |
| lnc-EIF4ENIF1-1 | lnc-EIF4ENIF1-1 | -0.056030492593558 | -0.172418779013392 | -0.291750268864744 | 0.771896964095512 | 0.910190563920181 | -5.58583138191688 |
| lnc-HNRNPA2B1-7 | lnc-HNRNPA2B1-7 | -0.0748094848723335 | 3.90092245395947 | -0.291734932069256 | 0.771913058533289 | 0.910190563920181 | -6.25713885924401 |
| lnc-SMARCD3-1 | lnc-SMARCD3-1 | -0.0465488437868889 | -0.215483386128462 | -0.29159309222964 | 0.772016331175067 | 0.910260190443742 | -5.54057606939673 |
| ENSG00000203668.2 | CHML | -0.0490212578745322 | 5.74483647084743 | -0.291316405610182 | 0.772226473143899 | 0.910397054386838 | -6.36348254197269 |
| ENSG00000125843.10 | AP5S1 | 0.0317564682765528 | 2.98330733749191 | 0.291314626074423 | 0.772227824747558 | 0.910397054386838 | -6.13681089721545 |
| ENSG00000204257.14 | HLA-DMA | -0.0424610354103532 | 8.19880449349789 | -0.291265548409474 | 0.772265100791634 | 0.910397054386838 | -6.34571648721397 |
| lnc-UQCC3-1 | lnc-UQCC3-1 | -0.107977460676376 | 2.28446995327329 | -0.291089596686585 | 0.772414313840361 | 0.910520807615609 | -6.04215356468061 |
| lnc-FHL3-2 | lnc-FHL3-2 | 0.0320756248145577 | 2.14547644227402 | 0.290886487876191 | 0.772553027627663 | 0.910632170865891 | -6.0299134300996 |
| ENSG00000080603.16 | SRCAP | 0.0466165715095013 | 5.33092698419826 | 0.290785100529974 | 0.772630044936777 | 0.910657652308723 | -6.34988594123455 |
| lnc-TRIM10-1 | lnc-TRIM10-1 | 0.0292730956786214 | 2.50901965527878 | 0.290741545287367 | 0.772663131706726 | 0.910657652308723 | -6.08078670693286 |
| lnc-STRN-2 | lnc-STRN-2 | 0.0421342031913969 | 0.76381542635336 | 0.290638060601982 | 0.772741745635413 | 0.910677717255211 | -5.8300483590043 |
| lnc-ERO1B-2 | lnc-ERO1B-2 | 0.0321796086725483 | 1.40588293845258 | 0.290531189492843 | 0.772822934654549 | 0.910677717255211 | -5.93944317875181 |
| lnc-SERPINC1-5 | lnc-SERPINC1-5 | 0.0331784862281532 | 2.4536755196872 | 0.290448419782742 | 0.772885815827429 | 0.910677717255211 | -6.07079887406806 |
| ENSG00000149273.14 | RPS3 | -0.0425337644209556 | 9.25019997613031 | -0.290443693550104 | 0.772889406451451 | 0.910677717255211 | -6.31575929279211 |
| lnc-DHX57-7 | lnc-DHX57-7 | 0.0528572364511593 | 1.75924030276328 | 0.290427937121337 | 0.772901376997099 | 0.910677717255211 | -5.97642505893195 |
| ENSG00000167635.11 | ZNF146 | -0.0332654222570001 | 5.93895220498057 | -0.290263210576018 | 0.773026527146445 | 0.910711394182686 | -6.366367048919 |
| lnc-RNF38-4 | lnc-RNF38-4 | 0.0539555327995487 | 4.20825388914402 | 0.290246956419629 | 0.773038876489929 | 0.910711394182686 | -6.26666408232592 |

| | | | | | | | |
|---|---|---|---|---|---|---|---|
| ENSG00000110435.11 | PDHX | -0.0341498391873269 | 4.23008053672127 | -0.29019427760378 | 0.773078900433548 | 0.910711394182686 | -6.28070460747506 |
| ENSG00000140961.12 | OSGIN1 | -0.0768313166295819 | -0.289275506958893 | -0.290145425417941 | 0.773116017565446 | 0.910711394182686 | -5.54711162857492 |
| ENSG00000124831.18 | LRRFIP1 | 0.064379634626464 | 5.92583416695094 | 0.29006304138242 | 0.773181378619958 | 0.910711394182686 | -6.36700326277848 |
| ENSG00000133112.16 | TPT1 | 0.0504417199528753 | 9.9208236434555 | 0.289973456655691 | 0.773246681034704 | 0.910711394182686 | -6.29132896167911 |
| ENSG00000270617.1 | URGCP-MRPS24 | 0.0409639795289523 | 2.49610626392384 | 0.289969507618252 | 0.773249681628369 | 0.910711394182686 | -6.07974735891588 |
| ENSG00000127337.6 | YEATS4 | 0.0489982056482487 | 3.43165185325434 | 0.289924440242508 | 0.773283925381665 | 0.910711394182686 | -6.20428635286461 |
| lnc-CHD4-2 | lnc-CHD4-2 | 0.0408504511023298 | 0.823263701749141 | 0.289822792678328 | 0.773361162400056 | 0.910750246348572 | -5.84174264293262 |
| ENSG00000164442.9 | CITED2 | 0.0694011485747423 | 2.69292948897842 | 0.289684495871845 | 0.773466251119693 | 0.91077003048624 | -6.10738185200709 |
| lnc-GOLGA1-1 | lnc-GOLGA1-1 | 0.0417051821650636 | 0.49539218873776 | 0.289611975027178 | 0.773521359843081 | 0.91077003048624 | -5.75119312685083 |
| ENSG00000121390.18 | PSPC1 | 0.0238980703036384 | 6.02250791844242 | 0.289608479488288 | 0.773524016139389 | 0.91077003048624 | -6.36757701910448 |
| ENSG00000127980.15 | PEX1 | -0.0318400056441588 | 5.66356688916094 | -0.289567763740526 | 0.773554956661344 | 0.91077003048624 | -6.36336778281672 |
| lnc-GP5-2 | lnc-GP5-2 | -0.0485156252634898 | 0.0677593875398903 | -0.289313652010892 | 0.773748068485199 | 0.91094088050791 | -5.66878900674334 |
| lnc-PATL2-9 | lnc-PATL2-9 | -0.112981932005845 | -0.157843201303916 | -0.289260345385526 | 0.773788580615085 | 0.91094088050791 | -5.59040191246481 |
| ENSG00000106305.9 | AIMP2 | 0.052663144553835 | 2.34398723424092 | 0.289107742853722 | 0.77390455947517 | 0.911025310133109 | -6.07052745122514 |
| lnc-GDPD1-2 | lnc-GDPD1-2 | 0.108689294459809 | -0.302439780784197 | 0.288979226291967 | 0.774002236905049 | 0.911088187247891 | -5.52527080250551 |
| lnc-DYSF-2 | lnc-DYSF-2 | 0.0605981496156573 | 4.10347619107687 | 0.288735705940468 | 0.774187331711282 | 0.911253951427536 | -6.26066255250015 |
| ENSG00000278863.1 | | 0.108689294459809 | -0.30179165057622 | 0.288581091989447 | 0.774304857493697 | 0.911340169361435 | -5.525371231384 |
| ENSG00000145416.13 | MARCHF1 | 0.0393291548166044 | 8.39347692829051 | 0.288463980152384 | 0.774393880583011 | 0.911392832296879 | -6.34280779790833 |
| ENSG00000175066.15 | GK5 | -0.0365956721743808 | 3.98919372422333 | -0.28836176901795 | 0.774471579347042 | 0.911432162247008 | -6.2550350815555 |
| ITPKB-IT1 | ITPKB-IT1 | -0.0699733883678793 | 1.61379311864582 | -0.288129462253815 | 0.774648182784171 | 0.911536809450259 | -5.93955274096036 |
| lnc-NFAT5-2 | lnc-NFAT5-2 | -0.0635453909582061 | 0.241725193471644 | -0.288076886074887 | 0.774688153739448 | 0.911536809450259 | -5.7034257697582 |
| ENSG00000154518.9 | ATP5MC3 | 0.040124823484164 | 4.52782450008556 | 0.288070039382719 | 0.774693358971646 | 0.911536809450259 | -6.3105338825525 |
| lnc-SLC16A11-7 | lnc-SLC16A11-7 | -0.0384216372488662 | 3.20810418377337 | -0.287865308639673 | 0.774849011394419 | 0.911667840489958 | -6.16880895894068 |
| lnc-TSTD2-6 | lnc-TSTD2-6 | -0.0508061669902049 | 1.06226795809709 | -0.287529179667916 | 0.775104583411742 | 0.911908090650197 | -5.902171333579 |
| lnc-MTMR9-3 | lnc-MTMR9-3 | 0.0857873194573514 | -0.320062068380418 | 0.287433279695691 | 0.775177504543137 | 0.911908090650197 | -5.55867122214556 |
| HYI-AS1 | HYI-AS1 | -0.0773933547343347 | 0.780800247780877 | -0.287421952451946 | 0.775186117772263 | 0.911908090650197 | -5.80707616555473 |
| ENSG00000108578.14 | BLMH | -0.065230355724327 | 3.0319068381522 | -0.287363048967674 | 0.77523090840699 | 0.911908663146707 | -6.15106458335007 |
| lnc-MFSD14B-5 | lnc-MFSD14B-5 | 0.102976359321384 | -0.844148033656857 | 0.28729407011704 | 0.775283361407575 | 0.911918248348598 | -5.35578764395093 |
| ENSG00000128815.19 | WDFY4 | -0.0405935661451853 | 9.32408082909075 | -0.287143036759013 | 0.775398214126802 | 0.912001225221255 | -6.31615376805815 |
| ENSG00000136942.14 | RPL35 | 0.0456543770803783 | 8.29714680933523 | 0.286601269246246 | 0.775810240932535 | 0.912433700309375 | -6.34306390341082 |
| lnc-MYLK4-34 | lnc-MYLK4-34 | 0.0827033389751115 | 2.26416560466476 | 0.286310479799269 | 0.776031420039312 | 0.912641682017436 | -6.05508210405684 |
| lnc-MCM7-1 | lnc-MCM7-1 | -0.0462213502727795 | 0.491733241056179 | -0.286245009917492 | 0.776081220074259 | 0.912648103341625 | -5.7596925506528 |
| lnc-EFCAB11-3 | lnc-EFCAB11-3 | -0.0872334555961244 | 1.11604859556383 | -0.286008030438166 | 0.776261487822035 | 0.912758025254169 | -5.86898842412102 |
| lnc-CLHC1-4 | lnc-CLHC1-4 | -0.050748464197198 | 1.66529511889035 | -0.285998191167744 | 0.776268972719446 | 0.912758025254169 | -5.98789760873085 |
| NCK1-DT | NCK1-DT | -0.0527866045603802 | 1.26787013825567 | -0.285896846042563 | 0.776346068901571 | 0.912758025254169 | -5.9109915415312 |
| ENSG00000101199.12 | ARFGAP1 | 0.0309704693052035 | 5.43975315423954 | 0.285888951722069 | 0.776352074436415 | 0.912758025254169 | -6.35812061693332 |
| ENSG00000107099.15 | DOCK8 | 0.0324821074421404 | 8.39114684603497 | 0.285701559734622 | 0.776494635306326 | 0.912819461523162 | -6.34231593773926 |
| ENSG00000258555.6 | SPECC1L-ADORA2/ | -0.0215111679078985 | 5.42006774200037 | -0.285679213265374 | 0.776511636191294 | 0.912819461523162 | -6.35547382723291 |
| ENSG00000133302.12 | SLF1 | -0.0452843966641092 | 3.83845882679452 | -0.285627504640373 | 0.776550975823658 | 0.912819461523162 | -6.23825913048266 |
| lnc-SUDS3-1 | lnc-SUDS3-1 | -0.0633598915056996 | -0.142421312054481 | -0.285587091496233 | 0.776581722330666 | 0.912819461523162 | -5.61033441066811 |
| lnc-SLC30A5-6 | lnc-SLC30A5-6 | 0.0409891259719892 | 0.692276301322838 | 0.285520234304952 | 0.776632588386983 | 0.912827122360054 | -5.81424769865494 |
| lnc-NLRP2-2 | lnc-NLRP2-2 | 0.304453934736009 | -1.19130313768814 | 0.285323240223574 | 0.776809018692784 | 0.912982357777284 | -5.2391692760384 |
| ENSG00000160746.12 | ANO10 | -0.148049939113625 | 0.658121712510055 | -0.285115525503046 | 0.776961358206624 | 0.91310926321611 | -5.78803219419111 |
| ENSG00000143190.22 | POU2F1 | -0.0263836547949081 | 6.46757334993125 | -0.284704137339672 | 0.777253568338715 | 0.913284281118961 | -6.37025464093906 |
| lnc-SLFN5-1 | lnc-SLFN5-1 | 0.159681795631301 | -0.414146314156603 | 0.284694331138065 | 0.77726103091603 | 0.913284281118961 | -5.46327229602303 |
| ENSG00000155275.18 | TRMT44 | -0.0349456216428017 | 4.02879990184612 | -0.284638406420343 | 0.777303590363159 | 0.913284281118961 | -6.26818956874195 |
| lnc-SLC30A7-2 | lnc-SLC30A7-2 | -0.0629769316648714 | 3.12643688268152 | -0.284560965773126 | 0.777362524858863 | 0.913284281118961 | -6.15200758515729 |
| ENSG00000078403.16 | MLLT10 | -0.026806244288891 | 5.46557458048881 | -0.284400785642695 | 0.777484430622451 | 0.913284281118961 | -6.35841303748392 |
| ENSG00000279425.1 | | 0.0845587938291703 | 2.85285509905518 | 0.284329704105634 | 0.777542658957322 | 0.913284281118961 | -6.10740654150436 |
| lnc-KIF17-2 | lnc-KIF17-2 | -0.0434602949718075 | 3.13904383153412 | -0.284321780102255 | 0.777544560211541 | 0.913284281118961 | -6.16245461427163 |
| ENSG00000106261.16 | ZKSCAN1 | -0.0316588398158322 | 6.05016826123199 | -0.284308507084704 | 0.777554662183929 | 0.913284281118961 | -6.36943809903816 |
| ENSG00000134460.17 | IL2RA | -0.100270642420094 | 5.68298594810522 | -0.284316275690699 | 0.777584992687573 | 0.913284281118961 | -6.36668942206313 |
| ENSG00000147874.10 | HAUS6 | 0.0313242769847095 | 4.8276484977969 | 0.284259372517008 | 0.777592058393387 | 0.913284281118961 | -6.33002361483989 |
| ENSG00000106123.11 | EPHB6 | 0.0604966102422035 | 3.43861912898509 | 0.284251094060078 | 0.777598359160948 | 0.913284281118961 | -6.18575186215932 |
| ENSG00000188215.9 | DCUN1D3 | 0.0461326360805845 | 1.73242046299568 | 0.284150139242383 | 0.777675197505727 | 0.913307079711252 | -5.97778694912423 |
| ENSG00000137818.11 | RPLP1 | -0.0603206032140164 | 8.26714409173201 | -0.284108994067399 | 0.777706514409906 | 0.913307079711252 | -6.34533747482549 |
| lnc-SLC26A5-1 | lnc-SLC26A5-1 | 0.0435551475185492 | 1.43675859793163 | 0.283863576930747 | 0.777893316981716 | 0.913331434227484 | -5.94537472950672 |
| ENSG00000267534.3 | S1PR2 | -0.183886885050974 | -0.761206908054201 | -0.283869491431323 | 0.777896361804234 | 0.913331434227484 | -5.45622026529617 |
| ENSG00000182378.13 | PLCXD1 | -0.0569864551398016 | 4.20325756792316 | -0.283832646849054 | 0.777916860775196 | 0.913331434227484 | -6.27498237488521 |
| ENSG00000248919.7 | ATP5MF-PTCD1 | -0.0380229933802112 | 3.32385314195771 | -0.283776497645422 | 0.777959601753273 | 0.913331434227484 | -6.18113930214776 |
| ENSG00000144306.14 | SCRN3 | 0.0312470610030062 | 2.99987208097931 | 0.283708117569129 | 0.778011653855675 | 0.913331434227484 | -6.13951326391669 |
| ENSG00000151552.11 | QDPR | 0.0788360865432826 | -0.121237092574586 | 0.283637184359967 | 0.778065650537307 | 0.913331434227484 | -5.62425142921551 |
| ENSG00000185024.16 | BRF1 | -0.0335500229696111 | 4.28223657822662 | -0.283598400455907 | 0.778095174584442 | 0.913331434227484 | -6.2939532791013 |
| ENSG00000155657.26 | TTN | -0.137869511563959 | 11.5789870177499 | -0.283640704294434 | 0.778108369303441 | 0.913331434227484 | -6.23896804170754 |
| lnc-CAPN14-2 | lnc-CAPN14-2 | 0.0977632917944186 | -0.267225936005998 | 0.283418798681049 | 0.778231899834806 | 0.913331434227484 | -5.56359446604265 |
| lnc-ARHGEF39-1 | lnc-ARHGEF39-1 | 0.0693242353467038 | 10.5704188239736 | 0.283412567547208 | 0.77823934153195 | 0.913331434227484 | -6.27518016188556 |
| lnc-SNRPG-1 | lnc-SNRPG-1 | 0.0375251629868436 | 3.85280414795751 | 0.283387462399474 | 0.778255755900863 | 0.913331434227484 | -6.23972205781065 |
| ENSG00000277027.1 | RMRP | 0.0693179015270644 | 10.5704121604354 | 0.283385785674925 | 0.778259730112998 | 0.913331434227484 | -6.27518794638115 |

| | | | | | | | |
|---|---|---|---|---|---|---|---|
| ENSG00000165283.15 | STOML2 | 0.0318409846231145 | 4.26408481477015 | 0.283166225227449 | 0.778424188186063 | 0.913472352647305 | -6.28606635519637 |
| ENSG00000058453.16 | CROCC | -0.0516443505223183 | 4.03644698441896 | -0.28292332022673 | 0.778609129037887 | 0.91363728994851 | -6.26700937076993 |
| ENSG00000131375.9 | CAPN7 | 0.0352201422245049 | 5.83316449957833 | 0.282781341855249 | 0.778717233282798 | 0.913661156815521 | -6.36793615317077 |
| ENSG00000005884.17 | ITGA3 | 0.110446643352124 | 3.3668316580324 | 0.282818918248739 | 0.77871824680633 | 0.913661156815521 | -6.1524060641618 |
| lnc-NPBWR1-7 | lnc-NPBWR1-7 | -0.0490279602080084 | 1.52656494785381 | -0.282707258010894 | 0.778773643480648 | 0.913674071121876 | -5.95259299164752 |
| ENSG00000155008.13 | APOOL | -0.0429300254743921 | 3.3660430063804 | -0.282534491667518 | 0.778905198896986 | 0.913727219707637 | -6.1813292709934 |
| ENSG00000175283.7 | DOLK | -0.0408350123936234 | 2.78734443920285 | -0.282531168370505 | 0.778907729533605 | 0.913727219707637 | -6.11911389753551 |
| ENSG00000069020.18 | MAST4 | -0.230201731383228 | 3.53357036394605 | -0.282421386458298 | 0.779045774692354 | 0.913785171259386 | -6.28752930003767 |
| ILF3-DT | ILF3-DT | -0.0466612747700763 | 3.70232887925573 | -0.282296537546584 | 0.779086403223673 | 0.913785171259386 | -6.22169846149401 |
| ENSG00000150867.13 | PIP4K2A | 0.0526568694715754 | 5.34793618333191 | 0.282291399784545 | 0.779090315815525 | 0.913785171259386 | -6.3562954737118 |
| lnc-TCF7L2-2 | lnc-TCF7L2-2 | 0.0773041670516116 | 0.324174102801923 | 0.282140258613999 | 0.779205417883672 | 0.913794716070643 | -5.70607171603152 |
| ENSG00000154001.13 | PPP2R5E | 0.0261466158898616 | 5.51899981831785 | 0.282121901199021 | 0.779219398376523 | 0.913794716070643 | -6.36036305662923 |
| lnc-PTK2-4 | lnc-PTK2-4 | 0.123281362147965 | -0.145066763206664 | 0.282105826466746 | 0.779231640502936 | 0.913794716070643 | -5.55673642257883 |
| ENSG00000249773.3 | | -0.0359733764162884 | 1.6463151610333 | -0.281928146596512 | 0.779366960962197 | 0.913876995846292 | -5.96337579092245 |
| lnc-PAIP2-3 | lnc-PAIP2-3 | 0.0837795735373636 | 1.99803298098187 | 0.281897104590921 | 0.779390603171832 | 0.913876995846292 | -6.03741887456751 |
| ENSG00000265942.2 | RN7SL577P | -0.067758208589201 | 0.703671129507946 | -0.281622566975725 | 0.779599705684941 | 0.91402929048214 | -5.83137061440084 |
| ENSG00000135424.16 | ITGA7 | -0.0797262469526088 | -0.0871109509829481 | -0.281454727056521 | 0.779727549669726 | 0.91402929048214 | -5.60025233218739 |
| ENSG00000120519.14 | SLC10A7 | -0.0243770757821262 | 3.8489252839018 | -0.281398575932321 | 0.779770321465075 | 0.91402929048214 | -6.24310167281164 |
| lnc-SMKR1-3 | lnc-SMKR1-3 | -0.0602965779495258 | 0.607419972572479 | -0.281393274481441 | 0.779774359755999 | 0.91402929048214 | -5.78375006039199 |
| ENSG00000165389.6 | SPTSSA | -0.0598129284186009 | 1.44027398728479 | -0.281358572504299 | 0.779800793554645 | 0.91402929048214 | -5.96129188871547 |
| ENSG00000143126.7 | CELSR2 | 0.163767795568434 | -0.681036584126366 | 0.281341554955731 | 0.779813756556559 | 0.91402929048214 | -5.42646815193845 |
| ENSG00000125510.15 | OPRL1 | -0.0922178551585339 | 0.567432889247028 | -0.281318478370199 | 0.779831335089971 | 0.91402929048214 | -5.80255866185558 |
| lnc-MCL1-2 | lnc-MCL1-2 | -0.0770126691654007 | 2.27163663333641 | -0.281049182554819 | 0.78003647914465 | 0.914199555195267 | -6.0576727276812 |
| lnc-TPP1-2 | lnc-TPP1-2 | -0.050255603757583 | 1.57761049911882 | -0.280913074646353 | 0.780140169406459 | 0.914199555195267 | -5.94992284665742 |
| ENSG00000185670.7 | ZBTB3 | -0.045638749304258 | 3.19860398343998 | -0.280906163331927 | 0.780145434719253 | 0.914199555195267 | -6.16736311967393 |
| ENSG00000162757.4 | C1orf74 | -0.0773714655487623 | 0.244568501292612 | -0.28085258150055 | 0.780186255832819 | 0.914199555195267 | -5.71190582701786 |
| lnc-SLC16A3-3 | lnc-SLC16A3-3 | -0.0746019165303455 | -0.14985821379896 | -0.28083627648899 | 0.780198677868146 | 0.914199555195267 | -5.58868592453172 |
| lnc-FRMD4A-6 | lnc-FRMD4A-6 | 0.0693662359562088 | 1.13766091203399 | 0.280704966911806 | 0.780298718689187 | 0.91422479561451 | -5.88597881549017 |
| ENSG00000168884.14 | TNIP2 | -0.0512881474181044 | 3.86337495232257 | -0.28063424505548 | 0.780352601107362 | 0.91422479561451 | -6.25409401582261 |
| ENSG00000100124.14 | ANKRD54 | -0.0361866667704587 | 2.89738987566294 | -0.28063310710446 | 0.780353468112099 | 0.91422479561451 | -6.12578638158604 |
| lnc-UBR2-4 | lnc-UBR2-4 | -0.0554328049159385 | 3.91811213102927 | -0.28046346412416 | 0.780482722240804 | 0.914249953971447 | -6.25150004235792 |
| lnc-RBPMS2-3 | lnc-RBPMS2-3 | 0.039953972817633 | 1.95307578140785 | 0.280454245121975 | 0.780489746549345 | 0.914249953971447 | -6.0087307357411 |
| lnc-MRPS5-3 | lnc-MRPS5-3 | 0.0476873801470951 | 0.452826238501509 | 0.280389857675146 | 0.780538806312279 | 0.914249953971447 | -5.76427668177302 |
| ENSG00000132507.17 | EIF5A | 0.0320583463727626 | 6.18531528605887 | 0.28035216756522 | 0.780567524563804 | 0.914249953971447 | -6.37165407181426 |
| lnc-NUMA1-4 | lnc-NUMA1-4 | -0.0835481052458554 | -0.360829977612959 | -0.280313443329036 | 0.780597031098198 | 0.914249953971447 | -5.50679000662978 |
| ENSG00000179912.20 | R3HDM2 | 0.0358509409153873 | 5.26773189486772 | 0.280176345355496 | 0.780701497662392 | 0.914291208403074 | -6.35122505055981 |
| lnc-KCTD1-4 | lnc-KCTD1-4 | -0.0360720791883702 | 1.72282845466936 | -0.280121368444637 | 0.780743390381423 | 0.914291208403074 | -5.99162725327319 |
| ENSG00000075292.18 | ZNF638 | -0.0316174616846228 | 7.64239636095849 | -0.28009233568118 | 0.780765513778401 | 0.914291208403074 | -6.35936352597008 |
| ENSG00000168939.11 | SPRY3 | -0.0565195762263622 | 1.07484508576347 | -0.279994816629609 | 0.78083982608958 | 0.914326211195917 | -5.88114461353872 |
| lnc-MRPS9-1 | lnc-MRPS9-1 | -0.0724109350669844 | -0.219744928394274 | -0.27989531309856 | 0.780915652767711 | 0.914362983157051 | -5.5974389770583 |
| lnc-SCARB1-4 | lnc-SCARB1-4 | -0.0751236520530049 | 0.271923424947114 | -0.279705561014236 | 0.781060259346749 | 0.914443229109286 | -5.70576153864701 |
| ENSG00000083099.10 | LYRM2 | -0.0342953150756502 | 4.95627123189734 | -0.279647027445911 | 0.781104868287039 | 0.914443229109286 | -6.34085939439776 |
| lnc-ESCO1-6 | lnc-ESCO1-6 | 0.0318334162219882 | 2.78216605461885 | 0.279630494264576 | 0.781117468503108 | 0.914443229109286 | -6.11280224417986 |
| ENSG00000237285.1 | HNRNPA1P2 | -0.11491209117815 | 0.846113860728212 | -0.279297994863003 | 0.78137505834392 | 0.914613571283768 | -5.80682011891027 |
| ENSG00000159712.10 | ANKRD18CP | -0.0636518889285504 | 0.408174158981788 | -0.279237402848043 | 0.781417067616833 | 0.914613571283768 | -5.73237003824302 |
| lnc-TMX2-1 | lnc-TMX2-1 | 0.0329658460111601 | 1.80534314475886 | 0.279226695965744 | 0.781425228459741 | 0.914613571283768 | -5.99109581241717 |
| lnc-MTFR2-3 | lnc-MTFR2-3 | -0.0882074346149027 | -0.0918617622897618 | -0.279206376218446 | 0.781440716348265 | 0.914613571283768 | -5.66532753585002 |
| ENSG00000115317.11 | HTRA2 | -0.0311644073235025 | 4.07548894488682 | -0.279033318310363 | 0.781572626231881 | 0.91469235532415 | -6.27483258555988 |
| ENSG00000099901.16 | RANBP1 | -0.0377516612502715 | 4.36033743492338 | -0.278988935050991 | 0.781606457520659 | 0.91469235532415 | -6.29483094628022 |
| ENSG00000271870.1 | NA | -0.0508887721800519 | 0.988406328913247 | -0.278943164618074 | 0.781641346635402 | 0.91469235532415 | -5.87337990884857 |
| lnc-UHRF1-2 | lnc-UHRF1-2 | 0.0704527992010462 | 2.43360113187274 | 0.278693311912416 | 0.78183180816881 | 0.914694449600538 | -6.07357474543504 |
| lnc-FGL2-8 | lnc-FGL2-8 | -0.0275657419323196 | 3.09506643688557 | -0.278688876184896 | 0.781835189625323 | 0.914694449600538 | -6.15797400847938 |
| ENSG00000064932.15 | SBNO2 | 0.0407602689466549 | 4.68637846498794 | 0.278666934913421 | 0.781851916021079 | 0.914694449600538 | -6.32528020456259 |
| ENSG00000156795.7 | NTAQ1 | -0.0511851853551799 | 2.28084950776954 | -0.278652709124401 | 0.781862760761886 | 0.914694449600538 | -6.05752616532351 |
| ENSG00000134046.11 | MBD2 | 0.0273524382513184 | 6.11098293429648 | 0.27857879899711 | 0.781919105350407 | 0.914694449600538 | -6.37158132751673 |
| lnc-ARHGDIA-2 | lnc-ARHGDIA-2 | -0.0390475906379502 | 3.42308129917442 | -0.278495887188221 | 0.781982313684405 | 0.914694449600538 | -6.1921573226442 |
| ENSG00000223599.1 | | 0.0824720414540838 | 1.83865080479888 | 0.278452680417257 | 0.782015253226432 | 0.914694449600538 | -5.98631250570376 |
| ENSG00000171408.13 | PDE7B | -0.111886759472151 | 3.79279136795874 | -0.278488551562057 | 0.782023357307797 | 0.914694449600538 | -6.24052875988937 |
| lnc-SCML4-2 | lnc-SCML4-2 | 0.0457000636627414 | 1.68602070982678 | 0.278330975583859 | 0.782108039505445 | 0.914694449600538 | -5.96500588454743 |
| ENSG00000106077.18 | ABHD11 | -0.0400274957656199 | 2.10119621692509 | -0.278309054066778 | 0.782124752543746 | 0.914694449600538 | -6.04109537616092 |
| lnc-CLPS-1 | lnc-CLPS-1 | -0.0373632653555357 | 0.580602031177373 | -0.278299589020365 | 0.782131968759144 | 0.914694449600538 | -5.78877227414203 |
| ENSG00000205339.9 | IPO7 | -0.0251206117507792 | 6.40465200696605 | -0.278178034491043 | 0.782224644480353 | 0.914750858322772 | -6.3721655005047 |
| ENSG00000101811.13 | CSTF2 | -0.0295963584581953 | 3.00210165651785 | -0.278102235483218 | 0.78228243685366 | 0.914766469592028 | -6.15384989440414 |
| lnc-CCDC142-3 | lnc-CCDC142-3 | -0.0579694921450061 | -0.276920071901593 | -0.277885792549768 | 0.782447468961886 | 0.914897354734516 | -5.59325050424582 |
| ENSG00000090339.8 | ICAM1 | 0.0883327305274606 | 2.20497621436962 | 0.277845105754889 | 0.782483264477793 | 0.914897354734516 | -6.06085809498294 |
| ENSG00000163840.9 | DTX3L | 0.0532725494229354 | 6.10515130649784 | 0.277773403688815 | 0.782533166546404 | 0.91490373002128 | -6.37220930589007 |
| lnc-KLHDC9-5 | lnc-KLHDC9-5 | -0.0510553823048965 | 2.71391023312383 | -0.277371759892398 | 0.782839446125327 | 0.915209832988617 | -6.10874674056221 |
| ENSG00000272949.1 | NA | 0.0561618390584555 | 1.24414639854168 | 0.277303298857353 | 0.782891655606631 | 0.915218887223904 | -5.90891752671043 |

| | | | | | | | |
|---|---|---|---|---|---|---|---|
| ENSG00000081721.11 | DUSP12 | 0.0284636692034868 | 3.90357679398179 | 0.277210310322143 | 0.782962571774046 | 0.91524980774791 | -6.25973403116339 |
| lnc-COL8A1-5 | lnc-COL8A1-5 | 0.0699743325462181 | 0.0364754224813056 | 0.276999548311467 | 0.783123312846423 | 0.91531716346731 | -5.64630833215822 |
| ENSG00000157778.8 | PSMG3 | -0.0431469843769485 | 3.00711013812308 | -0.276999236985936 | 0.783123550290992 | 0.91531716346731 | -6.1536851621407 |
| ENSG00000220848.5 | RPS18P9 | -0.0387635987234085 | 2.5666500857858 | -0.276871523421017 | 0.783220957790946 | 0.91531716346731 | -6.09963485975945 |
| lnc-GDF7-1 | lnc-GDF7-1 | 0.0866850008887985 | 4.02056887023725 | 0.276882594539122 | 0.783231880989255 | 0.91531716346731 | -6.25524022412455 |
| ENSG00000128951.13 | DUT | 0.042218762090882 | 4.92670015045465 | 0.276806847878086 | 0.783270287354263 | 0.91531716346731 | -6.34047806321513 |
| ENSG00000170348.8 | TMED10 | 0.0282304663430963 | 6.58238074596106 | 0.27678492384366 | 0.783287009537638 | 0.91531716346731 | -6.37220569770218 |
| ENSG00000088992.17 | TESC | 0.0728743931234117 | 4.8926954347098 | 0.27661742417933 | 0.783427076226558 | 0.915391956755734 | -6.33646540561337 |
| ENSG00000006062.15 | MAP3K14 | 0.027969447020638 | 5.93981496977854 | 0.276509137883837 | 0.783497369427498 | 0.915391956755734 | -6.37052720668809 |
| ENSG00000206760.1 | SNORA6 | -0.0377450126175791 | 1.73079007287768 | -0.276485860803223 | 0.783515125121197 | 0.915391956755734 | -5.99359739413759 |
| ENSG00000234268.1 | RPS27P19 | 0.100957958733268 | 0.504930440589093 | 0.276467793272149 | 0.783528907065176 | 0.915391956755734 | -5.71947380689416 |
| ENSG00000117625.13 | RCOR3 | 0.027944856188256 | 6.3108318451129 | 0.276339232894507 | 0.783626975174356 | 0.915454567796911 | -6.37285779939974 |
| ALMS1-IT1 | ALMS1-IT1 | -0.0637894309602957 | 0.526123594989821 | -0.276259356807857 | 0.783687907846121 | 0.915473791554864 | -5.809139887475 |
| lnc-ENDOD1-1 | lnc-ENDOD1-1 | 0.0898892488611875 | 2.14852534165016 | 0.276198224593834 | 0.783739277015287 | 0.915481842052418 | -6.03847569006625 |
| lnc-TAF1B-2 | lnc-TAF1B-2 | -0.0474611292266369 | 1.45596086152238 | -0.276099480200906 | 0.783809872238873 | 0.915512348217723 | -5.94211302629096 |
| ENSG00000206941.1 | SNORD15A | 0.0501760460825366 | 0.191260765267255 | 0.275913059739855 | 0.783952092975803 | 0.915572793671045 | -5.68719413850312 |
| ENSG00000166986.14 | MARS1 | -0.0384301804096134 | 6.84623002310471 | -0.275910234506011 | 0.783954248412804 | 0.915572793671045 | -6.37114930129751 |
| ENSG00000207719.1 | MIR623 | 0.0769137594060373 | -0.167430254494517 | 0.275823782854815 | 0.784020205230096 | 0.915572793671045 | -5.5844019414478 |
| lnc-DEGS2-6 | lnc-DEGS2-6 | 0.077931521409638 | 0.497503686870913 | 0.275798427233925 | 0.7840395501747 | 0.915572793671045 | -5.76806530291976 |
| ENSG00000276788.1 | SNORD26 | -0.0506468226398904 | 2.49570654047601 | -0.27560044814751 | 0.784190602084058 | 0.915589425395023 | -6.08605359989928 |
| ENSG00000276965.1 | | -0.0359188658410985 | 3.31209415486798 | -0.275568752674628 | 0.784214785530051 | 0.915589425395023 | -6.17611859363461 |
| ENSG00000077684.15 | JADE1 | 0.0215234275886358 | 6.13405970932708 | 0.275544026082988 | 0.784233651913443 | 0.915589425395023 | -6.37261107514831 |
| ENSG00000153404.14 | PLEKHG4B | 0.294069486880979 | -2.64396661601972 | 0.275523081821273 | 0.784265125721844 | 0.915589425395023 | -5.16177282501471 |
| ENSG00000121691.5 | CAT | -0.0593542967423408 | 5.78493519674189 | -0.275488993548531 | 0.784276206536445 | 0.915589425395023 | -6.36869047151942 |
| ENSG00000203326.11 | ZNF525 | -0.0301337762604241 | 5.27452492194927 | -0.275366425324666 | 0.784369165107407 | 0.915646014371924 | -6.35171024699526 |
| ENSG00000099910.16 | KLHL22 | -0.0285049951475336 | 3.7182322774156 | -0.275233914433968 | 0.784470278174811 | 0.915675583770895 | -6.22191141280563 |
| ENSG00000144401.14 | METTL21A | 0.0312685713086398 | 3.68323146993342 | 0.275216627818729 | 0.784483469086915 | 0.915675583770895 | -6.22651090573946 |
| ENSG00000101310.16 | SEC23B | -0.0277660830460418 | 4.93528116781634 | -0.274979259444641 | 0.784664604456237 | 0.915797592586259 | -6.33943783418908 |
| ENSG00000180747.15 | SMG1P3 | 0.0457953792333319 | 5.33457105881166 | 0.274963038479931 | 0.784676983085617 | 0.915797592586259 | -6.36088014575512 |
| ENSG00000231587.1 | SNORD62B | -0.0687521498654409 | -0.295835820606091 | -0.274904125866519 | 0.784721941268834 | 0.915798135575007 | -5.55896838722325 |
| ENSG00000094916.15 | CBX5 | -0.0323404310720772 | 7.00993522405435 | -0.274708968579472 | 0.784870877626828 | 0.915900299644689 | -6.36900099859225 |
| lnc-KCNK13-3 | lnc-KCNK13-3 | -0.0567781517011648 | 0.926082224604204 | -0.274626190026538 | 0.784934053423813 | 0.915900299644689 | -5.85437436571345 |
| ENSG00000114745.13 | GORASP1 | -0.0355995718950091 | 5.05753191814455 | -0.274614498345231 | 0.784942976521028 | 0.915900299644689 | -6.34491917736101 |
| ENSG00000109066.13 | TMEM104 | 0.0545780001888967 | 2.26925554591304 | 0.274474236331791 | 0.785050026843834 | 0.915973283970672 | -6.05323632920811 |
| ENSG00000225697.12 | SLC26A6 | 0.0479832122444955 | 2.79227401002341 | 0.274405569090494 | 0.785102436370373 | 0.915982510362282 | -6.1160158684558 |
| ENSG00000164002.11 | EXO5 | -0.0322068448109136 | 2.77862916858228 | -0.274337397738027 | 0.785154468413237 | 0.91599129532107 | -6.1169017384402 |
| lnc-SERAC1-1 | lnc-SERAC1-1 | -0.175470114490835 | -1.1408985330833 | -0.274217518816643 | 0.785245968865143 | 0.916046122033056 | -5.30799841638126 |
| lnc-ZNF599-3 | lnc-ZNF599-3 | -0.0619469659127834 | 2.08908265449918 | -0.274022978913643 | 0.785394462635531 | 0.916155150596181 | -6.03554250722147 |
| ENSG00000110074.10 | FOXRED1 | -0.0368409031724997 | 3.28215380432536 | -0.273978453898395 | 0.785428450051995 | 0.916155150596181 | -6.18210890005557 |
| ENSG00000115091.11 | ACTR3 | -0.0321782396802277 | 7.69450359630723 | -0.273889719716026 | 0.785496185050896 | 0.916182239298611 | -6.35788749982338 |
| lnc-UROC1-1 | lnc-UROC1-1 | 0.0384549703950978 | 3.48515429274094 | 0.273715702452752 | 0.785629025542892 | 0.916285257975371 | -6.19518152739005 |
| ENSG00000012822.15 | CALCOCO1 | 0.0515118240220636 | 7.0117361495214 | 0.273295916172944 | 0.785949506649226 | 0.916607099289536 | -6.37051761822214 |
| ENSG00000180992.6 | MRPL14 | -0.0616439627662264 | 2.43360555605829 | -0.273126034630957 | 0.786079211479806 | 0.916643808743942 | -6.07228451801218 |
| ENSG00000105865.10 | DUS4L | -0.0334372413105219 | 3.23456691978719 | -0.273088780006825 | 0.786107656267488 | 0.916643808743942 | -6.17665902662153 |
| ENSG00000117215.14 | PLA2G2D | 0.204855593165265 | -0.734302770579835 | 0.273110692218386 | 0.786116917948922 | 0.916643808743942 | -5.52800587658646 |
| ENSG00000100890.15 | PRORP | -0.0348972881227034 | 4.34823128243767 | -0.272966863583438 | 0.786200744410535 | 0.916643808743942 | -6.29461062106885 |
| ENSG00000130816.15 | DNMT1 | -0.0306230577119511 | 5.53252323591349 | -0.272942625892325 | 0.786219251249775 | 0.916643808743942 | -6.36472711278142 |
| lnc-ARHGEF3-1 | lnc-ARHGEF3-1 | -0.0270767918248085 | 2.8081164160844 | -0.272826131112972 | 0.786308203305754 | 0.916643808743942 | -6.1140081301088 |
| ENSG00000157240.3 | FZD1 | -0.0692496221084549 | 0.658076838719845 | -0.272824507798548 | 0.786309442842273 | 0.916643808743942 | -5.82368160289712 |
| ENSG00000176390.11 | CRLF3 | 0.0267919949730597 | 5.93930433934755 | 0.272788084078823 | 0.786337255550298 | 0.916643808743942 | -6.37195424030577 |
| ENSG00000042832.11 | TG | -0.105987631940381 | 0.625473488921843 | -0.272653465212183 | 0.786440051324524 | 0.91671172139612 | -5.7746459151164 |
| lnc-NPIPB12-1 | lnc-NPIPB12-1 | -0.0545889289332404 | 2.35338311758031 | -0.272520837711229 | 0.786541330250008 | 0.916746052276982 | -6.07578755879626 |
| ENSG00000170113.15 | NIPA1 | 0.0294962108115246 | 5.32889487888647 | 0.272498247206997 | 0.786558581509571 | 0.916746052276982 | -6.3597265688767 |
| lnc-JUNB-1 | lnc-JUNB-1 | -0.0780943052198566 | 2.54250773278184 | -0.272112144589202 | 0.786854301699627 | 0.917038791228323 | -6.09696696519288 |
| ENSG00000177989.13 | CIMAP1B | -0.093075464430848 | 0.71887251689513 | -0.272049169003708 | 0.786901543004361 | 0.917041923885107 | -5.88818993598862 |
| lnc-CD2BP2-1 | lnc-CD2BP2-1 | 0.0464053680055166 | -0.110946254388012 | 0.271814480957556 | 0.787080791549232 | 0.917198886511796 | -5.58467085917606 |
| lnc-P4HA1-1 | lnc-P4HA1-1 | 0.0414002792939312 | 1.29006980770376 | 0.271750363540984 | 0.787129764774743 | 0.917204027873558 | -5.90935533792873 |
| ENSG00000171222.10 | SCAND1 | -0.0726549144173894 | 3.00995803092892 | -0.271522731936472 | 0.787304558462062 | 0.917355772817697 | -6.16482399308891 |
| ENSG00000158321.16 | AUTS2 | 0.0713328552675803 | 4.38270727802102 | 0.271304438295628 | 0.787478234953292 | 0.917506198915635 | -6.290422569624 |
| ENSG00000161179.13 | YDJC | -0.0502687197148908 | 2.84131978465031 | -0.271091222347401 | 0.787633270860578 | 0.917598137723558 | -6.12992107907156 |
| ENSG00000119979.17 | DENND10 | -0.0465778790748734 | 3.46544117846851 | -0.271074160824549 | 0.787646305072138 | 0.917598137723558 | -6.19293594768876 |
| ENSG00000250461.1 | | 0.090937872931319 | 0.250507885553489 | 0.270999477483262 | 0.787703360403363 | 0.917612670053904 | -5.66729425961449 |
| ENSG00000101350.7 | KIF3B | 0.0399455219971419 | 5.12156062574263 | 0.270647927335416 | 0.78797194765541 | 0.91787360490047 | -6.34852051151453 |
| lnc-CPLX1-10 | lnc-CPLX1-10 | -0.0874347242290923 | -0.521530285311762 | -0.270557601969289 | 0.788040961208076 | 0.917902048811376 | -5.52046258549204 |
| ENSG00000171570.11 | RAB4B-EGLN2 | -0.0336073023409167 | 3.75651579815008 | -0.270118173600318 | 0.788376733250619 | 0.918036583393245 | -6.23612268190073 |
| ENSG00000147789.15 | ZNF7 | -0.0220824570819662 | 4.32598996205705 | -0.270034529498023 | 0.788440651243437 | 0.918036583393245 | -6.28868947987888 |
| ENSG00000072210.18 | ALDH3A2 | -0.0654574976414377 | 4.95672605078505 | -0.269955083250303 | 0.788506415260654 | 0.918036583393245 | -6.33510296258986 |
| lnc-GDPD3-2 | lnc-GDPD3-2 | -0.0520810141907179 | 0.372796048166365 | -0.2697773414455 | 0.78863719489044 | 0.918036583393245 | -5.75755815367217 |

| | | | | | | | |
|---|---|---|---|---|---|---|---|
| ENSG00000242588.6 | | -0.0591761823137822 | 1.95539396419874 | -0.269742220335092 | 0.788664035598006 | 0.918036583393245 | -6.01742724121231 |
| ENSG00000111859.16 | NEDD9 | -0.0580750946146802 | 4.5907365271147 | -0.269622307252904 | 0.788755679076624 | 0.918036583393245 | -6.31360747014563 |
| ENSG00000111145.7 | ELK3 | -0.121477965223099 | 2.66561711881007 | -0.269624395343016 | 0.788788335214863 | 0.918036583393245 | -6.13332345119704 |
| lnc-CELA1-1 | lnc-CELA1-1 | -0.0561363600723024 | 1.71112241038338 | -0.26955293282648 | 0.788808699809903 | 0.918036583393245 | -5.97517843656447 |
| lnc-ITGA9-2 | lnc-ITGA9-2 | 0.0606547582848189 | 2.63921970531019 | 0.269490526061012 | 0.788856396240519 | 0.918036583393245 | -6.09304190987817 |
| ENSG00000180035.12 | ZNF48 | -0.0582239339326225 | 1.80985337806331 | -0.269471563516172 | 0.788870889153756 | 0.918036583393245 | -6.01842179165271 |
| lnc-OSBPL10-3 | lnc-OSBPL10-3 | 0.0836456690556434 | -0.33596414772556 | 0.269369560598253 | 0.788948850422593 | 0.918036583393245 | -5.59571818698832 |
| lnc-SRM-2 | lnc-SRM-2 | 0.0307106760271467 | 2.37496773088593 | 0.269366256237539 | 0.788951375996302 | 0.918036583393245 | -6.07524334494054 |
| lnc-CASP3-2 | lnc-CASP3-2 | 0.0870731696241034 | 0.53426366221584 | 0.269364821739064 | 0.788952472406262 | 0.918036583393245 | -5.74960921594917 |
| ENSG00000123358.19 | NR4A1 | -0.152256990848828 | 1.77961542464563 | -0.269401947506308 | 0.78896070791407 | 0.918036583393245 | -5.97077793200009 |
| ENSG00000278050.1 | | 0.0811685400608586 | 2.52931690718974 | 0.269354559702371 | 0.788962871595205 | 0.918036583393245 | -6.08207499887381 |
| ENSG00000007392.16 | LUC7L | 0.0382554021190797 | 6.41627892939381 | 0.26922575016947 | 0.789058769069919 | 0.918036583393245 | -6.37476076442679 |
| lnc-BORCS8-MEF2B-2 | lnc-BORCS8-MEF2 | -0.0664021824246567 | -0.178092600112055 | -0.269224401544964 | 0.789059799884881 | 0.918036583393245 | -5.63815858736162 |
| ENSG00000141968.7 | VAV1 | -0.0472793957291504 | 5.53313972884595 | -0.269192182315067 | 0.789084426619338 | 0.918036583393245 | -6.36303268970635 |
| ENSG00000184831.13 | APOO | -0.0401845097911233 | 0.563931971590646 | -0.269161810555799 | 0.789107641441398 | 0.918036583393245 | -5.78362285749512 |
| lnc-RFWD3-1 | lnc-RFWD3-1 | 0.075525976729202 | 1.01660709816077 | 0.269155287333267 | 0.789112627527891 | 0.918036583393245 | -5.88368001013286 |
| lnc-CD226-6 | lnc-CD226-6 | 0.0284419246599488 | 0.964129989926198 | 0.269067365354359 | 0.78917983239209 | 0.918036583393245 | -5.88464113240103 |
| lnc-NEIL2-2 | lnc-NEIL2-2 | 0.0419308964377111 | 0.729521742069896 | 0.269025787787374 | 0.789211613574936 | 0.918036583393245 | -5.83987224889383 |
| lnc-REL-6 | lnc-REL-6 | -0.0492397811991001 | 4.76231445885462 | -0.269019685475521 | 0.789216278108574 | 0.918036583393245 | -6.32824122736749 |
| ENSG00000283293.1 | RN7SK | 0.0582134473137685 | 13.4346893256488 | 0.268948700231636 | 0.789270538948358 | 0.918036583393245 | -6.16709353687332 |
| ENSG00000202198.1 | | 0.0582131864307769 | 13.4346890047572 | 0.268947436922464 | 0.789271504626481 | 0.918036583393245 | -6.16709387498872 |
| lnc-MTIF2-2 | lnc-MTIF2-2 | 0.0889754161200331 | 4.79176638737973 | 0.268803087057414 | 0.789410274054941 | 0.918101460837997 | -6.31983627601471 |
| ENSG00000131171.12 | SH3BGRL | -0.0338268644007904 | 6.45934809915134 | -0.268757768115263 | 0.789416491955054 | 0.918101460837997 | -6.37454143075363 |
| ENSG00000180376.16 | CCDC66 | 0.0380900573431537 | 4.96576182232512 | 0.26857811922759 | 0.789553826783981 | 0.918166769664724 | -6.34329661794044 |
| lnc-MDGA1-2 | lnc-MDGA1-2 | -0.0535711193862714 | 0.0576403564602137 | -0.268567607452348 | 0.789561862851169 | 0.918166769664724 | -5.6704079379351 |
| ENSG00000112578.9 | BYSL | 0.0955189385485608 | -0.11721525657896 | 0.268491471127756 | 0.789620068423435 | 0.918182581117427 | -5.61989708271839 |
| ENSG00000225447.1 | RPS15AP10 | 0.0557594221589862 | 0.0643140202177654 | 0.268341992066097 | 0.789734347430423 | 0.918240892284881 | -5.66047213162052 |
| ENSG00000122729.18 | ACO1 | -0.0293781630096284 | 3.38389328530981 | -0.268309173716572 | 0.78975943818293 | 0.918240892284881 | -6.19380036852868 |
| lnc-RNF121-2 | lnc-RNF121-2 | 0.0431081817749318 | 0.371116800590769 | 0.268142526612686 | 0.789886849085992 | 0.918337156277506 | -5.73359109284301 |
| ENSG00000140262.17 | TCF12 | 0.0191020929250652 | 6.37748188429658 | 0.268034380566724 | 0.789969535816301 | 0.918379780231346 | -6.37501302672868 |
| ENSG00000167986.13 | DDB1 | -0.0238441028975744 | 7.02420211188797 | -0.267977864798989 | 0.790012747839295 | 0.918379780231346 | -6.37053897196265 |
| ENSG00000153551.13 | CMTM7 | -0.185846518045921 | -0.407671032199676 | -0.267831274389499 | 0.79015013756283 | 0.918437281932316 | -5.59307357706103 |
| lnc-UBXN4-12 | lnc-UBXN4-12 | -0.0778896104962059 | 2.50612477667577 | -0.26779928857768 | 0.790151454523512 | 0.918437281932316 | -6.1039250222236 |
| ENSG00000204272.12 | NBDY | 0.0299408650713699 | 3.66076437811113 | 0.26773272078388 | 0.790200192983413 | 0.918442067433372 | -6.2223551897037 |
| ENSG00000100099.20 | HPS4 | -0.0220132705049139 | 6.27744783701272 | -0.267566911603361 | 0.790326983241915 | 0.918533978967375 | -6.37514194479452 |
| ENSG00000091490.10 | SEL1L3 | -0.0610413635490339 | 8.11546245536642 | -0.267516171383027 | 0.790368522639614 | 0.918533978967375 | -6.35201519862409 |
| lnc-UQCRHL-1 | lnc-UQCRHL-1 | -0.0423374841668585 | 2.15510715541883 | -0.267356708363212 | 0.790487728822873 | 0.918585345206213 | -6.04280477133386 |
| ENSG00000177125.5 | ZBTB34 | 0.0813024681191683 | 5.64254800478886 | 0.267331150854682 | 0.790533248298315 | 0.918585345206213 | -6.37179883983714 |
| ENSG00000174796.12 | THAP6 | -0.0330120387173132 | 4.47402935044949 | -0.26726441062677 | 0.790558313207754 | 0.918585345206213 | -6.3065398831069 |
| lnc-FNTA-1 | lnc-FNTA-1 | -0.0430968609276909 | 1.79663767342574 | -0.267221362122516 | 0.790591235015696 | 0.918585345206213 | -6.0015196322448 |
| lnc-PLA2G4C-3 | lnc-PLA2G4C-3 | 0.0385758525116844 | 1.42862631181398 | 0.266887761436 | 0.790846372853596 | 0.918797887473088 | -5.95380341992563 |
| ENSG00000080815.18 | PSEN1 | 0.027112461576972 | 5.73617148477138 | 0.266857444132354 | 0.790869560681437 | 0.918797887473088 | -6.36983968167328 |
| ENSG00000146733.13 | PSPH | -0.0648049461324092 | 1.25656702189201 | -0.266783865855065 | 0.790925836945738 | 0.918797887473088 | -5.90233464705216 |
| ENSG00000162702.7 | ZNF281 | 0.0415520875090003 | 5.18256253541213 | 0.266698046070036 | 0.790991477539838 | 0.918797887473088 | -6.35596471975342 |
| ENSG00000029534.20 | ANK1 | -0.0697217272685648 | 2.69612964872978 | -0.26669036135077 | 0.790997355391494 | 0.918797887473088 | -6.09751645282164 |
| ENSG00000133805.15 | AMPD3 | -0.078618212612052 | 4.06576727287193 | -0.26656955495395 | 0.791105633788751 | 0.91887180521832 | -6.28458319287093 |
| ENSG00000154511.11 | DIPK1A | -0.0327260867803396 | 5.51416397322776 | -0.266428946407372 | 0.791197312543872 | 0.918926435170439 | -6.36200626323564 |
| ENSG00000012174.11 | MBTPS2 | -0.0447447795128793 | 2.72472261290378 | -0.266182907888361 | 0.791385521218905 | 0.919093166125866 | -6.11361592367348 |
| lnc-EMILIN3-5 | lnc-EMILIN3-5 | 0.0506525055944781 | 0.0850842739963881 | 0.266112147066857 | 0.791439652476185 | 0.919099119358076 | -5.68175772437524 |
| ENSG00000188807.12 | TMEM201 | 0.0573502062568685 | 1.77441767294223 | 0.266059465632641 | 0.791479953875621 | 0.919099119358076 | -5.99604426488723 |
| ENSG00000273820.1 | USP27X | -0.0666291807250453 | 0.738108789538547 | -0.265929258503377 | 0.791579565058391 | 0.919123769667558 | -5.84935650191357 |
| lnc-NOP9-1 | lnc-NOP9-1 | -0.0366015915736733 | 3.33669716260495 | -0.265881955497383 | 0.791615753722222 | 0.919123769667558 | -6.18254534022141 |
| lnc-HIST2H2AA3-1 | lnc-HIST2H2AA3-1 | -0.0600779879378961 | 3.34981168761607 | -0.26585660903296 | 0.791635144957804 | 0.919123769667558 | -6.18743823665179 |
| ENSG00000274760.1 | | -0.0432323962198269 | 3.87875780066244 | -0.265784092467951 | 0.791690624272919 | 0.919136337041541 | -6.25255107494367 |
| ENSG00000065802.11 | ASB1 | -0.0323995288359358 | 4.83055305115403 | -0.265695807573653 | 0.791758168735626 | 0.919162909593084 | -6.33467129223426 |
| ENSG00000114054.13 | PCCB | 0.0355645534578888 | 3.91273550559202 | 0.265569692684998 | 0.79185465874533 | 0.919223080534382 | -6.25689669895316 |
| ENSG00000144785.8 | | 0.0336614887029305 | 3.59257965559755 | 0.265494402071608 | 0.791912264874456 | 0.919238108950537 | -6.21779888219199 |
| ENSG00000136878.12 | USP20 | -0.0222452364546584 | 4.82498641138083 | -0.264799237910439 | 0.792444202347042 | 0.919730377801752 | -6.33755531309227 |
| lnc-AACS-5 | lnc-AACS-5 | -0.0775447840976926 | 1.88629293024588 | -0.26474435356618 | 0.792486203943306 | 0.919730377801752 | -6.00897040953587 |
| ENSG00000172534.13 | HCFC1 | -0.0390784291982609 | 6.21531965743001 | -0.264683219236462 | 0.792532989229078 | 0.919730377801752 | -6.37576592619954 |
| lnc-PPP1R3G-3 | lnc-PPP1R3G-3 | -0.0835782722210268 | 2.17731497156121 | -0.264650061085117 | 0.79256087344179 | 0.919730377801752 | -6.03909515294342 |
| lnc-CDHR5-3 | lnc-CDHR5-3 | -0.0739564664237669 | 0.190180722868984 | -0.264435754935332 | 0.792722378254301 | 0.919730377801752 | -5.71405752329832 |
| ENSG00000164169.12 | PRMT9 | 0.0258686450228858 | 3.51220803995839 | 0.264409475529257 | 0.79274249111585 | 0.919730377801752 | -6.20822591132424 |
| ENSG00000127957.17 | PMS2P3 | -0.029660792800231 | 3.02102334237731 | -0.264383108021813 | 0.792762671549035 | 0.919730377801752 | -6.15411764163325 |
| lnc-TMCO1-4 | lnc-TMCO1-4 | 0.0484829396596109 | 2.12309889854444 | 0.264371649775785 | 0.792771441189108 | 0.919730377801752 | -6.04028511310528 |
| ENSG00000146757.13 | ZNF92 | 0.0541648795766205 | 7.55654156289277 | 0.264368360539216 | 0.792773958631849 | 0.919730377801752 | -6.36381156951817 |
| lnc-FUBP1-58 | lnc-FUBP1-58 | -0.0261161811392627 | 2.4475737932729 | -0.264356301393759 | 0.792783188211567 | 0.919730377801752 | -6.08038489873036 |
| ENSG00000172183.14 | ISG20 | -0.0426780925196635 | 6.52391085063263 | -0.264278088447225 | 0.792843049951235 | 0.919747984960056 | -6.3756769703668 |

| | | | | | | | |
|---|---|---|---|---|---|---|---|
| ENSG00000184786.5 | DYNLT2 | -0.0410099999082454 | 2.53846864122231 | -0.264047087705356 | 0.793019858064722 | 0.919901247663783 | -6.09862264708238 |
| lnc-CENPV-4 | lnc-CENPV-4 | 0.0385517964609118 | 0.25451220348456 | 0.263965004538109 | 0.793082687226885 | 0.919922285217863 | -5.72731672364379 |
| ENSG00000252558.1 | RNU6-914P | 0.0639638667161181 | 0.542638864745772 | 0.263867337420553 | 0.793157446670645 | 0.919957157941051 | -5.78075702535323 |
| ENSG00000165714.10 | BORCS5 | 0.0396632637603744 | 3.41001076351178 | 0.26370547952813 | 0.79328134537531 | 0.920049018530457 | -6.18693010777308 |
| ENSG00000136098.16 | NEK3 | -0.0550699887529928 | 4.73679276266076 | -0.263520666341593 | 0.793422822467398 | 0.920161255062342 | -6.33642628858095 |
| ENSG00000116754.13 | SRSF11 | 0.0314774754295453 | 7.64660089237699 | 0.263414893954082 | 0.793503795885111 | 0.920203314592554 | -6.36173502892424 |
| ENSG00000053524.12 | MCF2L2 | 0.148639112278188 | 1.13704782597275 | 0.263082354327619 | 0.793788668761099 | 0.920427970324144 | -5.86575863100682 |
| ENSG00000237296.9 | SMG1P1 | 0.0498499921906381 | 4.19774029413016 | 0.262968580148221 | 0.793845494123614 | 0.920427970324144 | -6.29382679631449 |
| ENSG00000169972.11 | PUSL1 | 0.0469244316687407 | 1.2601670596466 | 0.26294561806915 | 0.793863075020402 | 0.920427970324144 | -5.92018444064335 |
| lnc-DHRS7C-2 | lnc-DHRS7C-2 | 0.0363593327207303 | 1.99989336051729 | 0.262928226335726 | 0.79387639105886 | 0.920427970324144 | -6.0174186548243 |
| ENSG00000131480.8 | AOC2 | 0.03588454766945 | 1.44734463420752 | 0.262650752412064 | 0.794088848303029 | 0.920606327269352 | -5.94321090254659 |
| ENSG00000133393.12 | CEP20 | 0.036224960461845 | 4.1478280815303 | 0.262610489321465 | 0.794119678407781 | 0.920606327269352 | -6.27619361354915 |
| PDCD4-AS1 | PDCD4-AS1 | -0.0369420159867975 | 3.28317093995128 | -0.262540550848398 | 0.794173232228429 | 0.920616559977346 | -6.17431466880498 |
| ENSG00000105879.11 | CBLL1 | -0.0315863961265039 | 5.11323871762405 | -0.262443232088693 | 0.794247753562743 | 0.920651095994929 | -6.35423035901137 |
| lnc-STAG2-3 | lnc-STAG2-3 | 0.0295971298572164 | 1.44760879584242 | 0.262379091161259 | 0.794296870207906 | 0.92065618197372 | -5.94940250195364 |
| lnc-TGFBR2-1 | lnc-TGFBR2-1 | -0.0570681771858128 | 2.78719951518914 | -0.26216723931183 | 0.79445910417401 | 0.920752124246626 | -6.12829163293861 |
| lnc-NOX3-7 | lnc-NOX3-7 | -0.135147606805081 | 1.78238380284734 | -0.262184974069484 | 0.794476502539345 | 0.920752124246626 | -5.96364142096512 |
| lnc-MAP7-2 | lnc-MAP7-2 | -0.0292904719255715 | 2.67675500815876 | -0.262058407725211 | 0.794542449863624 | 0.920752124246626 | -6.10304589015449 |
| ENSG00000105643.9 | ARRDC2 | 0.042687357831027 | 5.59856142807244 | 0.262037346944044 | 0.794558578966553 | 0.920752124246626 | -6.3698169649448 |
| lnc-SYF2-3 | lnc-SYF2-3 | -0.0350699876716867 | 6.17409508126934 | -0.261900656700278 | 0.794663263477144 | 0.92082159275742 | -6.37633807611425 |
| ENSG00000274487.2 | NPEPPSP1 | -0.0447627549695988 | 0.607533276220148 | -0.261717831095901 | 0.794803286837055 | 0.920932000279616 | -5.81028338390799 |
| lnc-SLC16A6-1 | lnc-SLC16A6-1 | 0.0734157949475206 | 0.391582725266352 | 0.26150727172553 | 0.794964559551213 | 0.921038792333347 | -5.79282808069492 |
| LINC00342 | LINC00342 | -0.0496221166016033 | 5.85273475523451 | -0.261416415757889 | 0.795034151223561 | 0.921038792333347 | -6.37393801674886 |
| ENSG00000134851.12 | TMEM165 | -0.0291067777927703 | 5.06926791297968 | -0.261323436932316 | 0.795105370660391 | 0.921038792333347 | -6.34776127293998 |
| ENSG00000177096.8 | PHETA2 | -0.0650415446245363 | 0.780679224368013 | -0.261316672572248 | 0.795110552058718 | 0.921038792333347 | -5.82979820776547 |
| ENSG00000160298.17 | C21orf58 | 0.0614171661555578 | 1.57835833127925 | 0.261305394855489 | 0.795119190641366 | 0.921038792333347 | -5.95418228883087 |
| lnc-PEX7-4 | lnc-PEX7-4 | 0.0794771258212372 | 3.45551345137288 | 0.261166964466684 | 0.795234561920486 | 0.92107337814404 | -6.17432033489211 |
| lnc-RGS10-3 | lnc-RGS10-3 | 0.0825866401886059 | -0.274696761264514 | 0.261149578582022 | 0.795238546518862 | 0.92107337814404 | -5.57938738199099 |
| ENSG00000272031.2 | ANKRD34A | -0.0588384896877141 | -0.0418567953507695 | -0.260945589541659 | 0.795394810426602 | 0.921202531116968 | -5.62013068984318 |
| lnc-WSB2-1 | lnc-WSB2-1 | 0.0682872139155015 | 2.80011124551523 | 0.260767465162668 | 0.795531267903108 | 0.921256908335859 | -6.12121738209033 |
| ENSG00000167081.16 | PBX3 | -0.0328292321021069 | 5.47197948885288 | -0.260751488595166 | 0.795543507544255 | 0.921256908335859 | -6.3672635656059 |
| LINC02328 | LINC02328 | -0.0555977493562372 | 3.02186817432193 | -0.260709029307242 | 0.795576035838793 | 0.921256908335859 | -6.14724997992751 |
| ENSG00000130649.9 | CYP2E1 | -0.101144172196778 | 0.468062305772068 | -0.260575737070918 | 0.795678154146281 | 0.921323326214722 | -5.77429081369427 |
| ENSG00000178177.15 | LCORL | 0.0325067758234976 | 5.17944531074464 | 0.260417608112706 | 0.795799305149168 | 0.921411773521141 | -6.35804392862859 |
| ENSG00000144021.2 | CIAO1 | 0.0188555702232449 | 5.62674959746062 | 0.260326300047094 | 0.79586926343533 | 0.921440941010766 | -6.36880132473653 |
| ENSG00000227398.3 | KIF9-AS1 | -0.045434139437569 | -0.0058970708748905 | -0.260122493557805 | 0.796025421735493 | 0.921532309294615 | -5.6461080308105 |
| LINC02035 | LINC02035 | 0.0374128497440744 | 3.21408806102162 | 0.260106438880724 | 0.796037723327904 | 0.921532309294615 | -6.16945473087624 |
| lnc-MAG-2 | lnc-MAG-2 | 0.0895191368888948 | 2.04238276968171 | 0.260030943038068 | 0.796102446325442 | 0.921533684647744 | -6.02625192763046 |
| lnc-OR13C9-2 | lnc-OR13C9-2 | 0.129646666846867 | 0.40847373717555 | 0.259955972312093 | 0.796164538435444 | 0.921533684647744 | -5.73790740127749 |
| ENSG00000143891.16 | GALM | -0.0627771639435664 | 2.17243763902347 | -0.259929600109506 | 0.796173226161921 | 0.921533684647744 | -6.03030187069087 |
| lnc-SCD5-2 | lnc-SCD5-2 | 0.0276978291131202 | 2.98898700799563 | 0.259861588530313 | 0.796225341777724 | 0.921542184537274 | -6.14408452071101 |
| ENSG00000152240.12 | HAUS1 | 0.0331695410304413 | 3.48416475060886 | 0.25963056576371 | 0.796402375921661 | 0.921695254630056 | -6.20248990061347 |
| ENSG00000209042.1 | SNORD12C | -0.0602214515310915 | 1.93706648087617 | -0.25956149734085 | 0.796455305579535 | 0.921704686536802 | -6.01769226684075 |
| ENSG00000113318.9 | MSH3 | 0.0283935964573745 | 4.83584272628331 | 0.259488890291582 | 0.796510948061698 | 0.921717256645524 | -6.33398485982755 |
| ENSG00000169621.9 | APLF | 0.0376101971026399 | 3.90130141668819 | 0.259394433222205 | 0.796583336931218 | 0.921739430092711 | -6.25033142158447 |
| ENSG00000168101.14 | NUDT16L1 | 0.0314601223061142 | 3.37739080012307 | 0.259329871818122 | 0.796632815764214 | 0.921739430092711 | -6.20003839127934 |
| ENSG00000115084.13 | SLC35F5 | -0.029335477852821 | 5.47165132176207 | -0.259288589621416 | 0.796664454226757 | 0.921739430092711 | -6.36816908453565 |
| ENSG00000226498.2 | RPSAP21 | 0.0833151543398091 | -0.128893016484774 | 0.259178684808943 | 0.796748686394949 | 0.921785071781644 | -5.59547286451932 |
| ENSG00000176624.10 | MEX3C | 0.0371236830484873 | 7.86179216826748 | 0.258983129354546 | 0.796898568150097 | 0.921869609288516 | -6.35942968951449 |
| ENSG00000130349.9 | MTRES1 | -0.0344437669718509 | 2.66220844917647 | -0.258966475694783 | 0.796911332559421 | 0.921869609288516 | -6.10759359946004 |
| ENSG00000177426.20 | TGIF1 | 0.0675054589405919 | 5.61144204531027 | 0.258834533137837 | 0.797024729492706 | 0.921948971965178 | -6.36786832924471 |
| lnc-CCDC71L-1 | lnc-CCDC71L-1 | -0.0675176351372857 | 0.968716106633695 | -0.258700071379992 | 0.797115529180129 | 0.921977323421512 | -5.90851874815023 |
| lnc-STAT6-1 | lnc-STAT6-1 | -0.0600812239794969 | -0.298815906512225 | -0.258510093758986 | 0.797261154181739 | 0.921977323421512 | -5.5197447994465 |
| ENSG00000101126.16 | ADNP | 0.0226716095817983 | 6.32198319262181 | 0.258490101632535 | 0.797276479324748 | 0.921977323421512 | -6.37759316420903 |
| ENSG00000116221.15 | MRPL37 | -0.0276934070225883 | 4.33276919578394 | -0.258482742012072 | 0.797282120927897 | 0.921977323421512 | -6.29946492487998 |
| ENSG00000143294.14 | PRCC | -0.024877977820867 | 4.47202815718333 | -0.25846135630318 | 0.797298514455428 | 0.921977323421512 | -6.3108369067185 |
| lnc-RRM1-5 | lnc-RRM1-5 | -0.0669567775348509 | -0.12453063344675 | -0.258425089202629 | 0.797326315742777 | 0.921977323421512 | -5.62101353634036 |
| lnc-TRAPPC6B-5 | lnc-TRAPPC6B-5 | 0.0367328838586867 | 1.4024088244772 | 0.25837750688669 | 0.79736279134365 | 0.921977323421512 | -5.94403289279052 |
| INTS6-AS1 | INTS6-AS1 | -0.0555327781075126 | 3.28815326709273 | -0.258249078039585 | 0.797461244479448 | 0.922030294235672 | -6.18969284764106 |
| ENSG00000123395.14 | ATG101 | 0.0511870981530103 | 2.35231232996844 | 0.258200879665688 | 0.797498194057119 | 0.922030294235672 | -6.07296491410595 |
| lnc-MBOAT4-4 | lnc-MBOAT4-4 | 0.034256326711871 | 4.14271361798129 | 0.258132266809334 | 0.797550794485923 | 0.92203931717051 | -6.27648296893572 |
| lnc-MEF2A-6 | lnc-MEF2A-6 | -0.0746474018356071 | -0.0767401427733298 | -0.257677402392485 | 0.797899529633572 | 0.922353874835649 | -5.64567204863699 |
| lnc-DRD5-22 | lnc-DRD5-22 | -0.240460667772899 | -1.3939164529582 | -0.257656437959914 | 0.797950525532595 | 0.922353874835649 | -5.51944477968239 |
| ENSG00000204977.9 | TRIM13 | -0.11268353744442 | 4.81194962656929 | -0.257654511423712 | 0.797957316678735 | 0.922353874835649 | -6.33475919140156 |
| ENSG00000117139.17 | KDM5B | -0.0402903893488409 | 5.94361853249863 | -0.257263931123428 | 0.798216565756546 | 0.922574447182294 | -6.37658858641109 |
| ENSG00000100823.11 | APEX1 | -0.0511154532458677 | 5.51794433498772 | -0.257236258999447 | 0.798237785059003 | 0.922574447182294 | -6.36790119958557 |
| ENSG00000196923.13 | PDLIM7 | -0.0902109463272999 | 1.07331696978087 | -0.257094422101157 | 0.798346549644401 | 0.92260227305619 | -5.88563498624325 |
| lnc-USP36-4 | lnc-USP36-4 | -0.0340496280901741 | 4.90515848200832 | -0.257087956410494 | 0.798351507817315 | 0.92260227305619 | -6.33882582278078 |

| | | | | | | | |
|---|---|---|---|---|---|---|---|
| ENSG00000101966.12 | XIAP | 0.0217511779979027 | 6.01257251517641 | 0.256976878966637 | 0.798436688149184 | 0.922648908161613 | -6.37667593208498 |
| lnc-SLC9B1-2 | lnc-SLC9B1-2 | 0.0355043725556893 | 0.0364134206582897 | 0.256856821794154 | 0.79852875742818 | 0.922703498239726 | -5.6474515932029 |
| ENSG00000198642.6 | KLHL9 | 0.0281290857684429 | 5.21336349122677 | 0.256659105039385 | 0.798680388503905 | 0.92279914574535 | -6.3566911085488 |
| lnc-ZNF106-5 | lnc-ZNF106-5 | -0.0266629597101122 | 1.05268006186526 | -0.256603063292199 | 0.798723368944659 | 0.92279914574535 | -5.89829869766827 |
| lnc-GATM-1 | lnc-GATM-1 | 0.0859836997271031 | 0.627976490046237 | 0.256495525746053 | 0.798805845143041 | 0.92279914574535 | -5.7828348323749 |
| ENSG00000179886.5 | TIGD5 | 0.0706883003164291 | 2.6374270879436 | 0.256464322802836 | 0.798829776756025 | 0.92279914574535 | -6.11175035828364 |
| ENSG00000156709.13 | AIFM1 | 0.0446344331532758 | 3.31024789670851 | 0.256456602059436 | 0.79883569833798 | 0.92279914574535 | -6.18519640601824 |
| lnc-KYNU-9 | lnc-KYNU-9 | 0.0940166109821403 | -0.566042854119353 | 0.256065566754106 | 0.799135626522191 | 0.923093810017749 | -5.4955846671562 |
| ENSG00000088256.8 | GNA11 | 0.0896537555268372 | 0.697642984779563 | 0.255916162585369 | 0.799250229194142 | 0.923097331742225 | -5.84429161763911 |
| ENSG00000052841.14 | TTC17 | 0.0180661001409925 | 7.19671275504925 | 0.2559160532607 | 0.799250313054882 | 0.923097331742225 | -6.37212189856519 |
| ENSG00000174456.13 | C12orf76 | -0.0505517410645382 | 2.28541178747721 | -0.255886193437496 | 0.799273218010491 | 0.923097331742225 | -6.06069568757697 |
| lnc-CDC37-1 | lnc-CDC37-1 | 0.0286507174476975 | 2.7131552699863 | 0.255747649889002 | 0.799379494717874 | 0.923129018995556 | -6.10977562730545 |
| ENSG00000174080.10 | CTSF | -0.0758098871378032 | 3.9030358415003 | -0.255750716857162 | 0.79939035294062 | 0.923129018995556 | -6.25832325210638 |
| ENSG00000179630.10 | LACC1 | 0.0883784926734807 | 0.951352943057489 | 0.255573693026327 | 0.799512942415508 | 0.923181615040605 | -5.86080478470177 |
| ENSG00000172939.8 | OXSR1 | -0.028428217051796 | 5.19831716324902 | -0.255515168843368 | 0.799557839492303 | 0.923181615040605 | -6.35700150017967 |
| ENSG00000197386.12 | HTT | 0.0419245163014228 | 7.22154234079622 | 0.255450363050963 | 0.799607556333145 | 0.923181615040605 | -6.37141880091683 |
| lnc-STAM2-3 | lnc-STAM2-3 | 0.0617322689061472 | -0.0799746319952411 | 0.255397522462322 | 0.79964809449156 | 0.923181615040605 | -5.61464387688354 |
| ENSG00000175416.12 | CLTB | 0.0334280728490106 | 3.49068480482214 | 0.255343767902483 | 0.799689334402622 | 0.923181615040605 | -6.21581488438834 |
| lnc-LHPP-8 | lnc-LHPP-8 | 0.0679130603316436 | 2.57942049745384 | 0.255323337153045 | 0.799705008802118 | 0.923181615040605 | -6.0769242243361 |
| lnc-TMEM161B-9 | lnc-TMEM161B-9 | 0.0281918370225538 | 1.44658229778617 | 0.255162426923246 | 0.79982846148768 | 0.923272347193142 | -5.94525160976939 |
| lnc-ZNF615-1 | lnc-ZNF615-1 | 0.0556515588418414 | 2.33475429568979 | 0.255094737190438 | 0.799880395590404 | 0.923280517184684 | -6.05927481035216 |
| ENSG00000011485.14 | PPP5C | -0.0393504139452681 | 3.82822546447707 | -0.255029328102205 | 0.799930580767714 | 0.923286667635387 | -6.24700877706785 |
| lnc-PGGHG-1 | lnc-PGGHG-1 | -0.0958822664055816 | 4.20822386260215 | -0.254644586268736 | 0.800256455894792 | 0.923611003234412 | -6.28572222268839 |
| ENSG00000167528.12 | ZNF641 | -0.0281189724052622 | 4.93751772428735 | -0.254434476883368 | 0.800387020104486 | 0.923709898223192 | -6.3432383702313 |
| ENSG00000139990.17 | DCAF5 | -0.0279050120972153 | 5.91715257419925 | -0.254226139675804 | 0.80054689744753 | 0.923842609899221 | -6.37642365036481 |
| ENSG00000089220.4 | PEBP1 | -0.0571154263615368 | 7.69951376921872 | -0.254160619025567 | 0.800599582906321 | 0.923851612656882 | -6.36329061122543 |
| ENSG00000068971.13 | PPP2R5B | 0.0443897773697564 | 3.37154729708531 | 0.254083196188479 | 0.800656596853711 | 0.923865608983066 | -6.20061630386551 |
| lnc-RNF186-3 | lnc-RNF186-3 | -0.0294945659115372 | 0.356879580021997 | -0.253810237015596 | 0.800866085767142 | 0.924039462651372 | -5.72482088755399 |
| ENSG00000114770.16 | ABCC5 | -0.0474288355777412 | 4.23436405588889 | -0.253715705328244 | 0.800938639784495 | 0.924039462651372 | -6.27814499513389 |
| ENSG00000280173.1 | | -0.0510537732951352 | -0.223892955605612 | -0.253711399018805 | 0.8009419449625 | 0.924039462651372 | -5.61183296265551 |
| ENSG00000164096.12 | C4orf3 | -0.0293305081858337 | 5.73197017322821 | -0.253628347619568 | 0.801005689282432 | 0.924061209645797 | -6.3728215052519 |
| ENSG00000156738.17 | MS4A1 | 0.146720056291858 | 8.39923659400618 | 0.253504821326313 | 0.801147169073006 | 0.924128411182615 | -6.35444858886436 |
| ENSG00000181789.14 | COPG1 | 0.0283247167210137 | 6.07642147374909 | 0.253317170402418 | 0.801244538879926 | 0.924128411182615 | -6.37796058928602 |
| ENSG00000109881.16 | CCDC34 | -0.0711612077517707 | -0.330468997821524 | -0.253283689447196 | 0.801270238928469 | 0.924128411182615 | -5.52279983619228 |
| ENSG00000175567.8 | UCP2 | 0.0448430214041116 | 6.83427459836825 | 0.2531971299014 | 0.801336683242081 | 0.924128411182615 | -6.37733966433307 |
| ENSG00000272533.1 | SNORA28 | 0.0403976818086118 | 2.98957086467622 | 0.253178769579632 | 0.801350777072268 | 0.924128411182615 | -6.13562524556572 |
| ENSG00000174996.11 | KLC2 | -0.0337270032029298 | 3.68688424765352 | -0.253123500523081 | 0.801393203350368 | 0.924128411182615 | -6.24211916330514 |
| lnc-EIF6-1 | lnc-EIF6-1 | -0.0438485767147743 | 1.72092738138311 | -0.253102325497673 | 0.801409458131181 | 0.924128411182615 | -5.99625511234689 |
| ENSG00000096060.14 | FKBP5 | 0.107190145931916 | 6.0640634558677 | 0.253136891396143 | 0.80142312294659 | 0.924128411182615 | -6.37858304822659 |
| ENSG00000114491.13 | UMPS | 0.0307569026998502 | 4.09418286932894 | 0.252995455471983 | 0.801491497116962 | 0.924132756020204 | -6.27849733896921 |
| lnc-TM9SF2-8 | lnc-TM9SF2-8 | 0.0951882862633296 | 0.219396796970205 | 0.252788724729167 | 0.801650200812221 | 0.924132756020204 | -5.77285985049521 |
| ENSG00000237836.5 | PHKA2-AS1 | 0.0793655942287035 | -0.167274280537464 | 0.25264564327434 | 0.80176004698934 | 0.924132756020204 | -5.59238992926462 |
| ENSG00000268975.2 | MIA-RAB4B | -0.0362139225302434 | 2.88699571626509 | -0.25264298944527 | 0.801762084419448 | 0.924132756020204 | -6.14128747886989 |
| lnc-COPS2-5 | lnc-COPS2-5 | -0.0607126529308187 | 0.69549962529812 | -0.252631931960916 | 0.801770573621583 | 0.924132756020204 | -5.83655816398075 |
| lnc-EPM2AIP1-1 | lnc-EPM2AIP1-1 | 0.0646086672458753 | 0.439334474576079 | 0.252595302347988 | 0.801798695578854 | 0.924132756020204 | -5.76943726982802 |
| lnc-COG2-4 | lnc-COG2-4 | -0.0612494904650848 | 0.971469964053195 | -0.252585509961441 | 0.801806213614241 | 0.924132756020204 | -5.86623706426916 |
| ENSG00000162769.12 | FLVCR1 | 0.0261720409529005 | 3.83065707617104 | 0.252544938294359 | 0.801837362426644 | 0.924132756020204 | -6.25049203323531 |
| ENSG00000114554.11 | PLXNA1 | 0.0653484254918708 | 2.54712015211018 | 0.252482681521942 | 0.801885160567496 | 0.924132756020204 | -6.09004656076277 |
| lnc-GOLPH3L-1 | lnc-GOLPH3L-1 | -0.0342412658462097 | 2.90760824655727 | -0.25243714050656 | 0.801920125536904 | 0.924132756020204 | -6.13683893454064 |
| ENSG00000283385.1 | MIR4524A | 0.121424458101434 | 2.10998163100255 | 0.252474994253175 | 0.801920767394299 | 0.924132756020204 | -6.02767352159173 |
| lnc-ZNF142-1 | lnc-ZNF142-1 | -0.0517512006090505 | -0.0945291072012827 | -0.252375828153198 | 0.801967199895923 | 0.92413452443499 | -5.61531522586373 |
| ENSG00000136718.9 | IMP4 | -0.0268979200217059 | 4.95010535976479 | -0.25224595189681 | 0.802066918662531 | 0.924150071845038 | -6.34953590061746 |
| ENSG00000136842.13 | TMOD1 | -0.173135065300039 | 0.892557496678655 | -0.25226717135353 | 0.802091608962875 | 0.924150071845038 | -5.94820340786894 |
| ENSG00000162493.16 | PDPN | 0.298245563943461 | -2.18978597847394 | 0.252139551982745 | 0.802150997970863 | 0.924150071845038 | -4.99074297390589 |
| ENSG00000109107.13 | ALDOC | -0.0817353564877048 | 2.55335244816135 | -0.252133806099727 | 0.802160286818416 | 0.924150071845038 | -6.09449232666655 |
| ENSG00000110318.13 | CEP126 | 0.123195092141947 | 2.34368257155885 | 0.251839746392416 | 0.802412913726318 | 0.924347938956586 | -6.11899091076401 |
| ENSG00000175895.3 | PLEKHF2 | 0.0431920813499582 | 6.4433114304778 | 0.251783713368716 | 0.802421851687134 | 0.924347938956586 | -6.37919641515825 |
| PIK3IP1-AS1 | PIK3IP1-AS1 | 0.0618900436843894 | 4.21809646572353 | 0.251693939069037 | 0.802493309331885 | 0.924351517624005 | -6.28258990305942 |
| ENSG00000109171.14 | SLAIN2 | -0.0257745666704441 | 5.30829198997367 | -0.251662705614655 | 0.802514775297137 | 0.924351517624005 | -6.36376891188553 |
| ENSG00000167566.16 | NCKAP5L | 0.034887644146635 | 3.53150315245751 | 0.251476762485706 | 0.802657569359714 | 0.924425296732025 | -6.22231351400717 |
| ENSG00000151575.14 | TEX9 | -0.0973760078975991 | 0.130967769425681 | -0.25146232882704 | 0.802668653898594 | 0.924425296732025 | -5.61648961357914 |
| ENSG00000101624.10 | CEP76 | -0.032873784707391 | 2.34417014102258 | -0.251144376705735 | 0.802912840296828 | 0.924654786092408 | -6.07971936510849 |
| lnc-ZNF720-4 | lnc-ZNF720-4 | -0.0536152114151726 | 0.256722525934409 | -0.250988650903783 | 0.803032444577831 | 0.924735777967865 | -5.71682906401678 |
| ENSG00000113407.13 | TARS1 | 0.0286502990541262 | 4.88798130906603 | 0.250935818940749 | 0.803073022940076 | 0.924735777967865 | -6.34431985328674 |
| ENSG00000129535.12 | NRL | 0.0424890784156179 | 0.922300576278118 | 0.250875374601955 | 0.803119448773253 | 0.924737503585806 | -5.85234811678129 |
| ENSG00000116906.12 | GNPAT | 0.0226528490188462 | 5.20314735633548 | 0.250811120157568 | 0.80316880184525 | 0.924742599338859 | -6.35857503440911 |
| ENSG00000163389.11 | POGLUT1 | -0.0304248679795709 | 4.07825116503665 | -0.250649393729196 | 0.803293025582439 | 0.924787424245077 | -6.27535165330601 |
| ENSG00000139437.17 | TCHP | -0.0449870485670862 | 4.05562581605701 | -0.250643447466794 | 0.803297593066012 | 0.924787424245077 | -6.27111138368047 |

| | | | | | | | |
|---|---|---|---|---|---|---|---|
| ENSG00000161996.18 | WDR90 | 0.0589344153285358 | 2.34478552198253 | 0.250418050042073 | 0.803470731999678 | 0.924845285338436 | -6.07935298296059 |
| ENSG00000179933.5 | C14orf119 | -0.039497330884855 | 3.63719878590477 | -0.250407903628824 | 0.80347852619759 | 0.924845285338436 | -6.22966877553164 |
| ENSG00000100483.13 | VCPKMT | 0.0315731984185224 | 4.8188984142529 | 0.250402534839875 | 0.803482650363014 | 0.924845285338436 | -6.33792706464039 |
| ENSG00000125107.17 | CNOT1 | 0.0196932356746675 | 8.20354682875533 | 0.250163832406111 | 0.803666021151032 | 0.924969525675673 | -6.35556230413243 |
| LINC02367 | LINC02367 | 0.218712968928856 | -1.61959161772392 | 0.250150817363414 | 0.803680464324138 | 0.924969525675673 | -5.20614881172124 |
| ENSG00000070476.14 | ZXDC | -0.0221770748789227 | 5.75444852059236 | -0.249924001314051 | 0.803850270243291 | 0.925045546445652 | -6.37494241901951 |
| ENSG00000272808.4 | GCAWKR | 0.205834564362291 | -1.15734320063787 | 0.249894593439384 | 0.803877083069481 | 0.925045546445652 | -5.24559996304521 |
| ENSG00000162896.5 | PIGR | -0.0991065656315282 | 6.49677711851395 | -0.249929700113189 | 0.80388134330629 | 0.925045546445652 | -6.37921878270815 |
| ENSG00000136754.17 | ABI1 | -0.0270881277448209 | 7.53821084981796 | -0.24972575835058 | 0.804002577975784 | 0.925133333099036 | -6.36684829290729 |
| ENSG00000162300.12 | ZFPL1 | 0.0330133067056773 | 4.45120412634212 | 0.249493434748145 | 0.804181079268797 | 0.925287000647865 | -6.31259154347526 |
| lnc-ABCA10-3 | lnc-ABCA10-3 | -0.108081826830315 | 4.46324088270844 | -0.248954520297197 | 0.804630489785912 | 0.92574756116889 | -6.29649014218675 |
| ENSG00000074696.12 | HACD3 | -0.0320043217281465 | 3.91558032583929 | -0.248799821009525 | 0.804714066688609 | 0.92574756116889 | -6.26293906312669 |
| lnc-ACAP3-2 | lnc-ACAP3-2 | -0.0309916513808453 | 1.5442003253974 | -0.248766776178009 | 0.804739461381922 | 0.92574756116889 | -5.97315935664643 |
| ENSG00000148484.17 | RSU1 | -0.0325163093936922 | 4.87298517374615 | -0.248709630024573 | 0.804783378255224 | 0.92574756116889 | -6.33879470329192 |
| ENSG00000137154.12 | RPS6 | -0.0435617620844848 | 10.1980061381411 | -0.248640434740702 | 0.80483655575061 | 0.92574756116889 | -6.29639810497491 |
| ENSG00000111846.17 | GCNT2 | -0.0571149532048452 | 4.42548965816378 | -0.24862175357175 | 0.804851217369543 | 0.92574756116889 | -6.30744050992091 |
| ENSG00000187607.15 | ZNF286A | 0.0371342229715124 | 2.91065273869117 | 0.247984436729189 | 0.805340745361652 | 0.926258860179866 | -6.1327255161339 |
| lnc-ADTRP-5 | lnc-ADTRP-5 | 0.125410886567404 | -0.0566457524455917 | 0.247662075949789 | 0.805595019873404 | 0.926408023846213 | -5.67473187656174 |
| ENSG00000181019.12 | NQO1 | -0.067191781167577 | 0.830832544961572 | -0.247653155658021 | 0.805595394582218 | 0.926408023846213 | -5.84229911665979 |
| ENSG00000265623.1 | MIR3139 | 0.114206141342187 | 0.363343107173609 | 0.24759806696141 | 0.805640080154739 | 0.926408023846213 | -5.68801261157645 |
| ENSG00000269955.2 | FMC1-LUC7L2 | 0.0345837443567678 | 5.27634335442299 | 0.247581509813881 | 0.805650470137843 | 0.926408023846213 | -6.36063185567783 |
| ENSG00000198951.11 | NAGA | -0.0369093068690858 | 3.33999575527092 | -0.247442084245186 | 0.805757652149678 | 0.926465353436501 | -6.19783032251489 |
| lnc-TFCP2L1-3 | lnc-TFCP2L1-3 | 0.0301165982177836 | 1.93656579338297 | 0.247386499829751 | 0.80580038316707 | 0.926465353436501 | -6.01140614344798 |
| lnc-NOTCH2-1 | lnc-NOTCH2-1 | 0.0808797827319959 | 0.958188014147189 | 0.24730467297484 | 0.805863289387682 | 0.926465353436501 | -5.86608794051049 |
| ENSG00000142864.14 | SERBP1 | -0.0329445431693589 | 6.95998679294372 | -0.247282453193253 | 0.805880371565229 | 0.926465353436501 | -6.37644758807144 |
| ENSG00000115998.7 | C2orf42 | 0.0232113395952407 | 3.75069907219518 | 0.246922796724887 | 0.806156882474127 | 0.926684241336922 | -6.23736658241267 |
| ENSG00000271092.5 | TLCD4-RWDD3 | 0.0415518146680331 | 3.05570062997547 | 0.246917683823012 | 0.806160813553852 | 0.926684241336922 | -6.15699999490175 |
| lnc-P2RX5-2 | lnc-P2RX5-2 | -0.0756607748958853 | 3.88969846056525 | -0.246794114104786 | 0.806270832550443 | 0.926758951604722 | -6.25720888854323 |
| lnc-TBX15-2 | lnc-TBX15-2 | 0.0776371191449008 | 0.0278533661014622 | 0.246685063650079 | 0.806339670080201 | 0.926762695345503 | -5.61754028963571 |
| ENSG00000109184.14 | DCUN1D4 | 0.0305297935047317 | 4.41481054637722 | 0.246653238124622 | 0.806364140841598 | 0.926762695345503 | -6.3075466909261 |
| lnc-CTNNBIP1-1 | lnc-CTNNBIP1-1 | -0.0362510617385121 | 2.33300619381572 | -0.246442562862159 | 0.806526134761708 | 0.92689712070353 | -6.07060307370381 |
| ENSG00000177239.14 | MAN1B1 | -0.0278773686295004 | 5.27913255436316 | -0.246300698526042 | 0.806635222939033 | 0.926922132988044 | -6.36197823799894 |
| ENSG00000261040.7 | WFDC21P | 0.123148814523893 | 2.75678579193937 | 0.246322997488173 | 0.806651378520471 | 0.926922132988044 | -6.1050324855215 |
| lnc-BIRC6-3 | lnc-BIRC6-3 | 0.0561329546644164 | 0.0588112600479074 | 0.246171627166319 | 0.806734477198022 | 0.926922132988044 | -5.667544335677 |
| ENSG00000182117.5 | NOP10 | -0.0599145991430201 | 3.56309966279226 | -0.246132714879777 | 0.80676440089143 | 0.926922132988044 | -6.22548347635217 |
| ENSG00000198933.9 | TBKBP1 | -0.127035858733195 | 0.884409701974686 | -0.246112986853237 | 0.806805909736717 | 0.926922132988044 | -5.92095096045406 |
| lnc-FAM186B-3 | lnc-FAM186B-3 | 0.0383851128160515 | 2.25182757078025 | 0.246042882250048 | 0.806833483637206 | 0.926922132988044 | -6.06183298427501 |
| ENSG00000081665.13 | ZNF506 | -0.0320500107397867 | 6.50029806549451 | -0.246004328563968 | 0.806863132524257 | 0.926922132988044 | -6.38053610057047 |
| ENSG00000107738.19 | VSIR | -0.0603163347410138 | 3.74158251948131 | -0.24588532960051 | 0.806954647942416 | 0.926975528440604 | -6.22373313520555 |
| lnc-ERVMER34-1-4 | lnc-ERVMER34-1-4 | 0.0507952763165669 | 0.440540137019533 | 0.245760449421527 | 0.807050689216186 | 0.92703209365923 | -5.74614133520661 |
| ENSG00000146285.13 | SCML4 | -0.311697455103525 | 2.24759121595614 | -0.245765273304536 | 0.807093966786834 | 0.92703209365923 | -5.97422753930577 |
| lnc-WNT10B-1 | lnc-WNT10B-1 | -0.0418734731243746 | 2.02941267067172 | -0.245626577013141 | 0.807153649466247 | 0.927048912837663 | -6.04269371313952 |
| ENSG00000277599.1 | | -0.0811827568622592 | 3.0645666251933 | -0.245510223795137 | 0.807255651241927 | 0.927114332636569 | -6.15225310011803 |
| lnc-LPAR3-1 | lnc-LPAR3-1 | 0.0581174011058945 | 3.04402359891179 | 0.245338497557185 | 0.807375220915457 | 0.927168866756005 | -6.15201448854869 |
| ENSG00000102931.7 | ARL2BP | -0.0304643867967949 | 4.72612767040471 | -0.245206906539534 | 0.807476437251435 | 0.927168866756005 | -6.33714864464667 |
| SND1-IT1 | SND1-IT1 | -0.066734336263248 | 1.64636760656258 | -0.245188366278944 | 0.807490698198636 | 0.927168866756005 | -5.96908817281163 |
| ENSG00000143971.8 | ETAA1 | 0.0328658328581728 | 5.20934846136749 | 0.245097558591964 | 0.807560547344772 | 0.927168866756005 | -6.3628153787875 |
| ENSG00000225224.1 | RPS27AP12 | 0.064125599970548 | -0.482044612027677 | 0.245080840129188 | 0.807573407336232 | 0.927168866756005 | -5.4850412218487 |
| lnc-DNAJB6-6 | lnc-DNAJB6-6 | 0.064125599970548 | -0.482044612027677 | 0.245080840129188 | 0.807573407336232 | 0.927168866756005 | -5.4850412218487 |
| lnc-SH3GL1-3 | lnc-SH3GL1-3 | -0.0342709355202935 | 2.42235173501179 | -0.244934863129789 | 0.807685696440661 | 0.92720005274854 | -6.08131091460322 |
| lnc-BEND4-5 | lnc-BEND4-5 | 0.044317874033547 | 1.19006785703611 | 0.244928404826873 | 0.807690664421188 | 0.92720005274854 | -5.92201824329138 |
| ENSG00000115806.12 | GORASP2 | 0.0257114437575545 | 5.30959390030726 | 0.244722796838292 | 0.807848830367222 | 0.927288186835554 | -6.36547628705497 |
| lnc-CCBE1-3 | lnc-CCBE1-3 | 0.0641885482506073 | 1.75539297154156 | 0.244711473753582 | 0.807857540996704 | 0.927288186835554 | -5.99319878782994 |
| ENSG00000164073.10 | MFSD8 | 0.0244641301677282 | 4.83765658167249 | 0.244617850846881 | 0.807929564228369 | 0.927297531520193 | -6.34219257578269 |
| ENSG00000199691.1 | RN7SKP173 | -0.0500727272057049 | -0.307381290294491 | -0.244525205629414 | 0.808000836992405 | 0.927297531520193 | -5.53678460372072 |
| lnc-SLC32A1-2 | lnc-SLC32A1-2 | -0.0500727272057049 | -0.307381290294491 | -0.244525205629414 | 0.808000836992405 | 0.927297531520193 | -5.53678460372072 |
| ENSG00000184924.5 | PTRHD1 | 0.0534619611423284 | 3.00031760861879 | 0.244302302017878 | 0.808172325422153 | 0.927394456353066 | -6.151431794659 |
| ENSG00000168246.5 | UBTD2 | 0.0417935479126554 | 2.55953638996232 | 0.244298299085801 | 0.808175405120971 | 0.927394456353066 | -6.09870135179907 |
| ENSG00000141956.13 | PRDM15 | 0.055518349376432 | 4.81203176202399 | 0.244029073858581 | 0.80838395004072 | 0.927408916681076 | -6.33858563036001 |
| lnc-MRPS31-2 | lnc-MRPS31-2 | 0.0536210025403741 | 1.69965089841541 | 0.244004525263972 | 0.808401431567912 | 0.927408916681076 | -5.97968951404316 |
| ENSG00000051009.10 | FHIP1B | 0.0226698663017402 | 4.6393279097866 | 0.243976749079281 | 0.808422803125542 | 0.927408916681076 | -6.3271035586918 |
| ENSG00000143614.9 | GATAD2B | -0.0338415674990236 | 5.65692607734874 | -0.243966390118275 | 0.808430773557309 | 0.927408916681076 | -6.3738253001856 |
| ENSG00000135404.11 | CD63 | -0.0493430661790724 | 3.99485452410522 | -0.243934440506421 | 0.80845535648067 | 0.927408916681076 | -6.27554320750322 |
| lnc-SLCO4C1-4 | lnc-SLCO4C1-4 | 0.0617069797302792 | 1.06164215421481 | 0.24393055159013 | 0.808458348734807 | 0.927408916681076 | -5.89400118631472 |
| N4BP2L2-IT2 | N4BP2L2-IT2 | -0.0402137179772142 | 4.51777615043618 | -0.243653991713888 | 0.808671150012464 | 0.92760133084633 | -6.3138012483943 |
| ENSG00000100479.12 | POLE2 | -0.0603311532031832 | 0.57005515311386 | -0.24355671235897 | 0.808746005899724 | 0.927616547219289 | -5.80128392327082 |
| ENSG00000085231.13 | AK6 | -0.0269160748517292 | 3.40624820149893 | -0.243519618741264 | 0.808774549696223 | 0.927616547219289 | -6.20389322902967 |
| lnc-SETBP1-2 | lnc-SETBP1-2 | -0.0308285293236898 | 4.115709472189 | -0.243295634486916 | 0.808946912734692 | 0.927743304972137 | -6.26935260316102 |

| | | | | | | | |
|---|---|---|---|---|---|---|---|
| ENSG00000100354.20 | TNRC6B | -0.0219069112058587 | 8.53684421235766 | -0.243208572306161 | 0.809013912455431 | 0.927743304972137 | -6.34941715596957 |
| ENSG00000100554.11 | ATP6V1D | -0.0362570784468383 | 3.83234015427107 | -0.243200288199996 | 0.8090202876619 | 0.927743304972137 | -6.24646685707572 |
| lnc-ANPEP-1 | lnc-ANPEP-1 | 0.0321829110355079 | 1.12278805116681 | 0.242844102595189 | 0.809294410125413 | 0.928005952290327 | -5.89668547869154 |
| lnc-SOGA1-1 | lnc-SOGA1-1 | 0.0316300140458048 | 1.14513968591548 | 0.242666129372691 | 0.809431388410347 | 0.92811131790152 | -5.89225291679707 |
| lnc-ADAMTS4-1 | lnc-ADAMTS4-1 | -0.0389186059724445 | 2.94475475192536 | -0.242452866888392 | 0.809595535276185 | 0.928247822113955 | -6.14929055044577 |
| ENSG00000267169.1 | ADGRL1-AS1 | -0.050187771289792 | 1.03958962586134 | -0.242356270025427 | 0.809669888162215 | 0.928251429174115 | -5.88613155513962 |
| ENSG00000126214.21 | KLC1 | 0.0266925371306774 | 6.11830972395622 | 0.242331600453704 | 0.809688877199245 | 0.928251429174115 | -6.38107329676295 |
| ENSG00000118181.10 | RPS25 | -0.0452813174131776 | 8.73381833799552 | -0.242105602262463 | 0.80986284135483 | 0.92830608436981 | -6.34367209690549 |
| ENSG00000092531.9 | SNAP23 | -0.0271629483886056 | 5.95121534704612 | -0.241995138060237 | 0.809947875715979 | 0.92830608436981 | -6.37919880728639 |
| ENSG00000179632.9 | MAF1 | -0.0317346904510707 | 6.38497960431956 | -0.241945063525361 | 0.8099864234133 | 0.92830608436981 | -6.38162778299808 |
| ENSG00000149806.10 | FAU | 0.0371390773897309 | 7.68855058189346 | 0.2419333627363 | 0.809995430824387 | 0.92830608436981 | -6.36688416597694 |
| ENSG00000134444.14 | RELCH | 0.0185385589445595 | 6.53758361790432 | 0.241926851328004 | 0.81000044339773 | 0.92830608436981 | -6.38130370719809 |
| ENSG00000116251.10 | RPL22 | -0.0353014503425877 | 7.10111816714193 | -0.241918132624758 | 0.810007155190293 | 0.92830608436981 | -6.37652727890873 |
| KTN1-AS1 | KTN1-AS1 | -0.0455393072221483 | 0.770570653280858 | -0.241620692071819 | 0.810236138190722 | 0.928444709298007 | -5.83485335665437 |
| ENSG00000188976.10 | NOC2L | 0.0272341726625571 | 4.99160553712598 | 0.241573702338713 | 0.810272314521643 | 0.928444709298007 | -6.35075529653493 |
| TAF1A-AS1 | TAF1A-AS1 | -0.0606759969404119 | -0.366817063781865 | -0.241566335565843 | 0.810277986071237 | 0.928444709298007 | -5.51994515697342 |
| ENSG00000133657.15 | ATP13A3 | 0.026586654878443 | 6.132977887667 | 0.241499834288802 | 0.810329184708244 | 0.928444709298007 | -6.38158619298848 |
| ENSG00000168522.12 | FNTA | 0.0243987212505932 | 5.74133016129269 | 0.241468055457441 | 0.810353651194612 | 0.928444709298007 | -6.37693733637501 |
| lnc-KCNK5-1 | lnc-KCNK5-1 | 0.0332936968762442 | 0.752313045103014 | 0.241389217856999 | 0.810414349002645 | 0.928462570718103 | -5.83303191795132 |
| ENSG00000033030.13 | ZCCHC8 | -0.0214051245948559 | 5.49581724888396 | -0.241199716054044 | 0.810560253047335 | 0.928562544640242 | -6.37125576579157 |
| ENSG00000162244.11 | RPL29 | -0.0401381536502479 | 7.55174499492042 | -0.241078272016679 | 0.810653760630002 | 0.928562544640242 | -6.36922982441484 |
| lnc-KIAA0040-2 | lnc-KIAA0040-2 | -0.0641454301762376 | 4.29208593116465 | -0.24105949488348 | 0.810675633963068 | 0.928562544640242 | -6.29379701190886 |
| lnc-LYPD6-2 | lnc-LYPD6-2 | 0.130347847545861 | 3.57036033242323 | 0.241095561949878 | 0.810682064188172 | 0.928562544640242 | -6.22406191906757 |
| ENSG00000141524.15 | TMC6 | 0.04295321148944 | 6.52545229815124 | 0.240716337668167 | 0.810932453777268 | 0.928790022190077 | -6.38156472027423 |
| ENSG00000173530.5 | TNFRSF10D | -0.0781752038831467 | 3.57082082159504 | -0.240687137964432 | 0.810970911859304 | 0.928790022190077 | -6.2182948399522 |
| ENSG00000151611.14 | MMAA | 0.0360354542473535 | 3.11977641450986 | 0.240482785613349 | 0.811112304423694 | 0.928861921902536 | -6.15772412568514 |
| ENSG00000131759.17 | RARA | -0.0415758552391927 | 3.4234905615607 | -0.240442920091212 | 0.811143004564331 | 0.928861921902536 | -6.20904280429548 |
| ENSG00000022267.16 | FHL1 | -0.234817811102449 | 0.223467130352882 | -0.240428945981366 | 0.811195944698915 | 0.928861921902536 | -5.72966038198082 |
| lnc-PRLHR-2 | lnc-PRLHR-2 | 0.0420510530023246 | 2.27932963433518 | 0.240350469236358 | 0.811214201434192 | 0.928861921902536 | -6.06677149570141 |
| ENSG00000120708.16 | TGFBI | 0.224459319018477 | 6.47260758548661 | 0.24023304530608 | 0.811350508117997 | 0.928959354260347 | -6.37521966460549 |
| lnc-CCDC68-2 | lnc-CCDC68-2 | -0.0801746841759554 | 3.86446659204536 | -0.240147939995212 | 0.811389557930939 | 0.928959354260347 | -6.24363608966492 |
| ENSG00000138336.8 | TET1 | -0.0640993143369837 | 2.208361670027 | -0.239926437103896 | 0.81154077142767 | 0.929080799727223 | -6.04107499497833 |
| lnc-RIT1-3 | lnc-RIT1-3 | -0.0226951887467037 | 2.5119842642008 | -0.239837869288247 | 0.811608986564317 | 0.929107217489063 | -6.08986829774625 |
| ENSG00000134698.10 | AGO4 | 0.0284667947805701 | 5.5541052110869 | 0.239683029438843 | 0.81172824813265 | 0.929146112769117 | -6.37294440060187 |
| ENSG00000143498.17 | TAF1A | 0.033158575297513 | 2.91628312054928 | 0.239676540893072 | 0.811733245873501 | 0.929146112769117 | -6.13941161781773 |
| ENSG00000272807.1 | | -0.0669628669113831 | 1.14742871627648 | -0.239529121171823 | 0.811846796656135 | 0.929162252137833 | -5.93995067223041 |
| lnc-VPS13A-2 | lnc-VPS13A-2 | -0.0600083412189841 | -0.590930302440352 | -0.239501925530237 | 0.811867744683864 | 0.929162252137833 | -5.49463315740345 |
| ENSG00000104973.17 | MED25 | 0.0307043358695749 | 4.56540208020147 | 0.239482415981862 | 0.811882772418935 | 0.929162252137833 | -6.32290580221736 |
| ENSG00000186417.13 | GLDN | -0.213248516825953 | -0.666354435471098 | -0.239185005718424 | 0.812140575870876 | 0.929351002869087 | -5.49989669429836 |
| ENSG00000185963.13 | BICD2 | -0.046352925303661 | 7.05690018920616 | -0.238992155469117 | 0.812260431827279 | 0.929351002869087 | -6.37655412803393 |
| lnc-FKBP1A-1 | lnc-FKBP1A-1 | -0.0399898317378673 | 2.37156732523798 | -0.238930853862618 | 0.812307657105463 | 0.929351002869087 | -6.08367813844121 |
| ENSG00000118640.10 | VAMP8 | -0.0476625787937692 | 4.61589702248692 | -0.238918780642458 | 0.81231695810624 | 0.929351002869087 | -6.32836202930428 |
| PCED1B-AS1 | PCED1B-AS1 | 0.0633406638835954 | 4.7244799038913 | 0.238922870948342 | 0.812323549713054 | 0.929351002869087 | -6.32597678430744 |
| ENSG00000150977.10 | RILPL2 | 0.0428435180538731 | 4.76287724652849 | 0.238904482518832 | 0.812327973169779 | 0.929351002869087 | -6.33782833780263 |
| ENSG00000276805.2 | | 0.062441042889997 | 0.79824499224984 | 0.238858031496889 | 0.812363758617335 | 0.929351002869087 | -5.83090480067864 |
| lnc-PSAT1-5 | lnc-PSAT1-5 | -0.0738425782834649 | 1.72997808342994 | -0.238741805494529 | 0.812453299857292 | 0.929401782413307 | -6.00230572105347 |
| lnc-NSD3-14 | lnc-NSD3-14 | 0.159598063174727 | -1.00363607311106 | 0.238510029769303 | 0.812631868908245 | 0.929554393561099 | -5.31038299722216 |
| ENSG00000106459.14 | NRF1 | -0.029224344059771 | 4.1537855945816 | -0.238163732888101 | 0.812898688318655 | 0.929759971943752 | -6.28580036040902 |
| lnc-PEX3-1 | lnc-PEX3-1 | 0.0391456674029938 | 0.326156741242015 | 0.238159523901149 | 0.81290193145313 | 0.929759971943752 | -5.74016540794678 |
| lnc-DCTD-1 | lnc-DCTD-1 | -0.0631724871765161 | -0.332887161025426 | -0.23806538971674 | 0.812974465171139 | 0.92979126613422 | -5.53503467226678 |
| lnc-COMMD6-2 | lnc-COMMD6-2 | 0.0346466078058838 | 3.57667334255934 | 0.237708750965743 | 0.813249283007637 | 0.930053894440838 | -6.21612714747562 |
| lnc-SERF1B-1 | lnc-SERF1B-1 | 0.0563717706979849 | 0.852843344632602 | 0.237407208554888 | 0.813481663420298 | 0.930267963674649 | -5.82476891946815 |
| ENSG00000215021.8 | PHB2 | 0.0435856914462773 | 5.67300461151048 | 0.237255344936282 | 0.813598701923037 | 0.930305133355618 | -6.3773239886817 |
| ENSG00000148814.17 | LRRC27 | -0.0332754112048037 | 3.38931466417382 | -0.237247740953785 | 0.813604562286084 | 0.930305133355618 | -6.19635588306292 |
| ENSG00000268655.2 | SAXO3 | 0.132944281018573 | -1.00087332585044 | 0.237083121843043 | 0.813731436315063 | 0.930366221278387 | -5.3797713667286 |
| ENSG00000110987.8 | BCL7A | 0.257230737179706 | 2.84319950272625 | 0.237119839443349 | 0.813748388557295 | 0.930366221278387 | -6.20622452184801 |
| ENSG00000277502.1 | | -0.0458273937352306 | 0.231036565149257 | -0.236992903904729 | 0.813800970567545 | 0.930374659919561 | -5.70191867861885 |
| ENSG00000118596.11 | SLC16A7 | -0.0317307698564262 | 5.40833109878927 | -0.236878354060828 | 0.813889260495987 | 0.930423918288748 | -6.36953175624616 |
| lnc-BICRAL-3 | lnc-BICRAL-3 | 0.0282788729587073 | 3.6459789197284 | 0.236787604320039 | 0.813959208101333 | 0.930452203729297 | -6.23481699920909 |
| lnc-CT55-5 | lnc-CT55-5 | 0.037553488618118 | 0.794294178628896 | 0.236615311911311 | 0.814092010960111 | 0.930483357838564 | -5.84113733578846 |
| ENSG00000102882.11 | MAPK3 | -0.0307347116600765 | 4.35498899081465 | -0.236536447505217 | 0.814152801431478 | 0.930483357838564 | -6.30514629620693 |
| ENSG00000001036.13 | FUCA2 | 0.0338825626250325 | 3.1900851300148 | 0.236516460550719 | 0.814168208013058 | 0.930483357838564 | -6.18097729190043 |
| ENSG00000146540.14 | C7orf50 | 0.0597021548138197 | 3.26419852065592 | 0.236474596440868 | 0.814200513036969 | 0.930483357838564 | -6.20970053680913 |
| lnc-USP53-8 | lnc-USP53-8 | -0.0319596068289885 | 4.85004833120256 | -0.236459009264887 | 0.814212493709876 | 0.930483357838564 | -6.34062837935547 |
| ENSG00000159111.12 | MRPL10 | -0.03098272444955 | 4.08988894287724 | -0.236378885760742 | 0.814274257065649 | 0.930502278102502 | -6.2843697049103 |
| ENSG00000135503.12 | ACVR1B | 0.0505021708210949 | 2.65922665206342 | 0.236149740405441 | 0.814450900797715 | 0.930625693952848 | -6.12388433055418 |
| ENSG00000197948.10 | FCHSD1 | -0.0278093497814949 | 6.48963695372275 | -0.23612148397664 | 0.814472683810261 | 0.930625693952848 | -6.3828231284087 |
| ENSG00000039319.16 | ZFYVE16 | 0.0246810970372993 | 5.23991561733888 | 0.235987573629308 | 0.814575917940621 | 0.930691985510508 | -6.3644508635446 |

| | | | | | | | |
|---|---|---|---|---|---|---|---|
| ENSG00000186056.10 | MATN1-AS1 | 0.0520620175340412 | 4.04921989315005 | 0.235763576198393 | 0.814748609435831 | 0.930800921353545 | -6.26791159808087 |
| LINC02019 | LINC02019 | -0.0577207518612322 | -0.253704259249051 | -0.235680970171342 | 0.814812297154992 | 0.930800921353545 | -5.59939995040567 |
| ENSG00000107862.4 | GBF1 | 0.0168675652867174 | 6.24592632567034 | 0.235644631728352 | 0.814840313823936 | 0.930800921353545 | -6.38308853607442 |
| lnc-DDRGK1-1 | lnc-DDRGK1-1 | -0.0272791301015973 | 1.8556900587607 | -0.235534327581559 | 0.814925359001549 | 0.930800921353545 | -6.00843126282078 |
| lnc-SLC7A5-4 | lnc-SLC7A5-4 | 0.031430627935269 | 0.527065632016436 | 0.235525040999296 | 0.814932519116854 | 0.930800921353545 | -5.76825194392074 |
| ENSG00000284776.1 | | 0.0292115152654753 | 2.75677280445819 | 0.235511974575099 | 0.814942593582677 | 0.930800921353545 | -6.12624006826327 |
| lnc-RAD52-2 | lnc-RAD52-2 | -0.0617868759506137 | 0.570794872698242 | -0.235430319141794 | 0.815005552219117 | 0.93082117863312 | -5.80526396000714 |
| lnc-HPD-2 | lnc-HPD-2 | -0.189895907702236 | -1.70629947491139 | -0.235290267942719 | 0.81511779714328 | 0.930897720612558 | -5.26996286684977 |
| ENSG00000111832.12 | RWDD1 | -0.0334158320427118 | 4.56066657201385 | -0.235151104759331 | 0.815220843709423 | 0.930938099522797 | -6.32834708106575 |
| ENSG00000254786.1 | | 0.07429464400985 | 2.89095981703205 | 0.235075468651286 | 0.81528614572993 | 0.930938099522797 | -6.12580046406071 |
| ENSG00000167740.9 | CYB5D2 | 0.0284554524448266 | 2.55712248933371 | 0.235062924103762 | 0.815288839430498 | 0.930938099522797 | -6.09893817418358 |
| lnc-RET-1 | lnc-RET-1 | 0.0654682071940564 | 1.74166915601485 | 0.234897350987171 | 0.81541651603451 | 0.931025624100442 | -5.9883121748592 |
| lnc-POF1B-1 | lnc-POF1B-1 | -0.0330549541943149 | 0.449100192948359 | -0.234846205360334 | 0.815455956434085 | 0.931025624100442 | -5.74945532089021 |
| lnc-GNRH1-1 | lnc-GNRH1-1 | 0.0704241964847394 | -0.681400725162278 | 0.234758119668414 | 0.815523883902338 | 0.931051533771247 | -5.41532373534137 |
| ENSG00000180353.10 | HCLS1 | 0.0318055292203994 | 7.52177661821185 | 0.234469373595627 | 0.815746561114268 | 0.931235361572945 | -6.37283749469283 |
| ENSG00000136960.12 | ENPP2 | 0.143842627612748 | 3.86189407246787 | 0.234450476404136 | 0.815804108760361 | 0.931235361572945 | -6.24687917176234 |
| ENSG00000155926.13 | SLA | 0.060598575091662 | 5.66076141100839 | 0.234387129258383 | 0.815820630611833 | 0.931235361572945 | -6.37580458478658 |
| ENSG00000167984.17 | NLRC3 | 0.0367932367214729 | 5.93437521985662 | 0.23373699693712 | 0.816311429533507 | 0.931743922480351 | -6.38065647138106 |
| ENSG00000131652.13 | THOC6 | -0.0353908537038699 | 2.90532220458044 | -0.233595537884838 | 0.81642054565462 | 0.931816795564436 | -6.14225916919513 |
| RHOA-IT1 | RHOA-IT1 | 0.0555858385294439 | 1.6765925027639 | 0.233379768436962 | 0.816586989080496 | 0.931861414459728 | -5.98998233232912 |
| lnc-NUMB-2 | lnc-NUMB-2 | -0.0646530011355474 | 0.900510722483859 | -0.233372228390369 | 0.81659280558806 | 0.931861414459728 | -5.85666870777133 |
| ENSG00000237298.9 | TTN-AS1 | 0.0440739974410498 | 5.47301251153857 | 0.233368788726956 | 0.816595459000637 | 0.931861414459728 | -6.3724282868092 |
| lnc-AVEN-2 | lnc-AVEN-2 | 0.0394855901168016 | 0.831055202912917 | 0.233297018398923 | 0.816650824301366 | 0.93187293029133 | -5.85663952368906 |
| ENSG00000111731.12 | C2CD5 | 0.0267632461387253 | 5.18209061545108 | 0.233229443309553 | 0.816702954163558 | 0.931880753120933 | -6.36317123168267 |
| ENSG00000111344.11 | RASAL1 | -0.165176666535699 | 0.398327033116447 | -0.233121199813959 | 0.816819407996541 | 0.931913001382711 | -5.83429910002851 |
| ENSG00000174446.12 | SNAPC5 | 0.0267047491570063 | 4.20817300572864 | 0.233073186016831 | 0.816823499893238 | 0.931913001382711 | -6.2952313291539 |
| ENSG00000162517.12 | PEF1 | 0.0243126504614137 | 4.83469719459767 | 0.232968291663359 | 0.816904423858872 | 0.931913001382711 | -6.34103006300227 |
| ENSG00000127452.8 | FBXL12 | 0.0278458672465867 | 4.0467468036985 | 0.232958056668931 | 0.816912320067447 | 0.931913001382711 | -6.27902127594677 |
| ENSG00000284427.1 | MIR4647 | -0.0907055463567952 | -0.0988495839126342 | -0.232264940906891 | 0.817447097549098 | 0.932471381559138 | -5.63869721133135 |
| lnc-VCP-3 | lnc-VCP-3 | -0.0282151729602582 | 1.2781607265153 | -0.232145009534605 | 0.817539640233236 | 0.932525265443097 | -5.92660048423421 |
| lnc-EIF1AD-4 | lnc-EIF1AD-4 | 0.0438498374900744 | 0.327858932441254 | 0.232036592969654 | 0.817623300009515 | 0.932557789405229 | -5.73601784111371 |
| ENSG00000016864.18 | GLT8D1 | -0.0406609259324591 | 3.88909764278523 | -0.231941128226948 | 0.817696967277438 | 0.932557789405229 | -6.24966172644171 |
| ENSG00000116678.19 | LEPR | -0.0484065632698838 | 1.98405961116266 | -0.231742854910557 | 0.817849974153762 | 0.932557789405229 | -6.03145918839187 |
| ENSG00000176894.9 | PXMP2 | 0.0713574127811543 | -0.171237470381991 | 0.231664476710236 | 0.817910460339558 | 0.932557789405229 | -5.59554649899062 |
| lnc-FBXL2-7 | lnc-FBXL2-7 | 0.0546741016196861 | -0.174872903478751 | 0.231664047849326 | 0.817910791304069 | 0.932557789405229 | -5.60997321804023 |
| ENSG00000106560.10 | GIMAP2 | -0.0412013181627162 | 2.87232261141446 | -0.231603029661085 | 0.817957881167718 | 0.932557789405229 | -6.14266548021121 |
| ENSG00000285836.1 | | 0.0760760570163695 | -0.242148529707363 | 0.231398569724327 | 0.818115674988515 | 0.932557789405229 | -5.54786812910179 |
| lnc-SLC35F5-14 | lnc-SLC35F5-14 | -0.0453847733052504 | 1.68586307735501 | -0.231296023071027 | 0.818194819177515 | 0.932557789405229 | -5.99072978263293 |
| ENSG00000066651.19 | TRMT11 | -0.0377507985789922 | 5.13417734922363 | -0.231289356466388 | 0.818199964443731 | 0.932557789405229 | -6.36050643795854 |
| lnc-IDS-1 | lnc-IDS-1 | -0.0673291425652691 | -0.315463456339435 | -0.231233412992852 | 0.818243141774671 | 0.932557789405229 | -5.52305975591932 |
| ENSG00000076555.15 | ACACB | -0.0655682825208297 | 3.96686800651772 | -0.231225306168595 | 0.818259256480411 | 0.932557789405229 | -6.26860749040798 |
| lnc-CLEC3A-14 | lnc-CLEC3A-14 | -0.0426985604256184 | 0.103406349764388 | -0.231210906836747 | 0.818260512250606 | 0.932557789405229 | -5.69308667859161 |
| lnc-CSPP1-1 | lnc-CSPP1-1 | -0.0672815850147878 | -0.119615367158711 | -0.230981621342927 | 0.818437482347721 | 0.932557789405229 | -5.5857395595106 |
| ENSG00000267360.6 | | -0.0235194982519888 | 2.5599968836446 | -0.230965421055532 | 0.818449986625697 | 0.932557789405229 | -6.09019993293694 |
| lnc-CGREF1-2 | lnc-CGREF1-2 | -0.0404372574627798 | 4.48298888546838 | -0.230959719372754 | 0.818454387511249 | 0.932557789405229 | -6.31884603869028 |
| lnc-SYT2-3 | lnc-SYT2-3 | -0.0416826793876554 | 0.898590450920425 | -0.230886337961516 | 0.818511028018913 | 0.932557789405229 | -5.87179342148222 |
| ENSG00000112081.16 | SRSF3 | 0.0224920076533497 | 8.04114671653051 | 0.230857316848973 | 0.818533428656698 | 0.932557789405229 | -6.36230508326351 |
| ENSG00000037637.10 | FBXO42 | -0.0178594268014702 | 5.43012057056359 | -0.230833261124085 | 0.818551996757901 | 0.932557789405229 | -6.37133369629886 |
| ENSG00000064115.10 | TM7SF3 | 0.0281877916682887 | 3.86556211455593 | 0.230737307738729 | 0.818626062179046 | 0.932557789405229 | -6.2556278531448 |
| lnc-FGL2-6 | lnc-FGL2-6 | -0.0237896190633542 | 4.9211048304081 | -0.230714292968261 | 0.818643827290511 | 0.932557789405229 | -6.34784858648841 |
| lnc-GJC1-4 | lnc-GJC1-4 | -0.0827675106392327 | -0.0186464976087236 | -0.230697521945402 | 0.818656772911184 | 0.932557789405229 | -5.71914310113373 |
| ENSG00000279369.1 | | -0.0390473226739099 | 2.04809164070778 | -0.230664873271927 | 0.818681974701721 | 0.932557789405229 | -6.04409871060388 |
| ENSG00000152422.15 | XRCC4 | -0.0327446662040298 | 2.37255444782985 | -0.230596868636189 | 0.81873446868509 | 0.932557789405229 | -6.07053515546287 |
| ENSG00000118420.16 | UBE3D | 0.0350453873251722 | 2.3896209206051 | 0.230514296204021 | 0.818798208939869 | 0.932557789405229 | -6.08404905107655 |
| ENSG00000164284.14 | GRPEL2 | -0.0444251459856573 | 3.18237539528358 | -0.230512383649359 | 0.818799685315602 | 0.932557789405229 | -6.18584455139674 |
| ENSG00000177613.8 | CSTF2T | 0.0263869269283461 | 5.42985037300762 | 0.230484094297784 | 0.818821523050026 | 0.932557789405229 | -6.37146880448476 |
| lnc-PUF60-2 | lnc-PUF60-2 | 0.0463858244915759 | -0.188525867237997 | 0.23043032345107 | 0.818863031418232 | 0.932557789405229 | -5.6054889117534 |
| ENSG00000113649.11 | TCERG1 | -0.0233520168980776 | 6.8160744907905 | -0.230424628505053 | 0.818867427658693 | 0.932557789405229 | -6.3817198586417 |
| TMEM202-AS1 | TMEM202-AS1 | -0.0466967373895852 | 2.5618188164684 | -0.230361414927436 | 0.818916226074334 | 0.932557789405229 | -6.09239929319771 |
| lnc-FBXL3-4 | lnc-FBXL3-4 | 0.0274425785927215 | 3.04772254930227 | 0.230346914144755 | 0.818927420215253 | 0.932557789405229 | -6.15982635636312 |
| ENSG00000085117.11 | CD82 | 0.0933555007956377 | 5.05842046307168 | 0.230277579592554 | 0.819012555887485 | 0.932557789405229 | -6.34077506915421 |
| lnc-DNAH10OS-5 | lnc-DNAH10OS-5 | 0.0572867092303295 | -0.415764142539374 | 0.230211436691212 | 0.81903200632722 | 0.932557789405229 | -5.53209423995243 |
| ENSG00000158109.14 | TPRG1L | 0.032242819438971 | 4.08129659141256 | 0.230151133044427 | 0.819078560710255 | 0.932557789405229 | -6.27849479717741 |
| ENSG00000105723.11 | GSK3A | -0.0306707092258551 | 3.55631461346739 | -0.230068897777466 | 0.819142047356815 | 0.932557789405229 | -6.22609122746971 |
| lnc-WDFY2-4 | lnc-WDFY2-4 | -0.0605761203880784 | 2.34386976537807 | -0.230003138168099 | 0.819192815474815 | 0.932557789405229 | -6.08513332578589 |
| ENSG00000272556.2 | GTF2IP13 | 0.0508239675525085 | 0.520555695862678 | 0.229994852256584 | 0.819199212468345 | 0.932557789405229 | -5.768109186091 |
| ANKRD44-IT1 | ANKRD44-IT1 | -0.0681584637826633 | 4.50788215282324 | -0.229751467543938 | 0.81940271879752 | 0.9327378697605 | -6.31405124642543 |
| ENSG00000198301.11 | SDAD1 | 0.0168718795279803 | 5.47999871611363 | 0.229576466537781 | 0.819522236060883 | 0.932822329527244 | -6.37335656561408 |

| | | | | | | | |
|---|---|---|---|---|---|---|---|
| lnc-COMMD10-6 | lnc-COMMD10-6 | -0.0554746483728363 | 1.51651073963986 | -0.229368113476894 | 0.819683111307342 | 0.932902224649762 | -5.96271586934773 |
| ENSG00000157184.6 | CPT2 | -0.0271497234070668 | 3.57997544987962 | -0.229363298264971 | 0.819686829360568 | 0.932902224649762 | -6.22236937102687 |
| ENSG00000279570.1 | | -0.0471638612026148 | 3.19298737454951 | -0.229309462329771 | 0.819728398922198 | 0.932902224649762 | -6.16805839236928 |
| ENSG00000160209.18 | PDXK | -0.0590031178593939 | 4.60199389384213 | -0.22888441647673 | 0.820064017281503 | 0.933232579626537 | -6.3316279746794 |
| ENSG00000101882.9 | NKAP | -0.0186114730289493 | 4.01934971101124 | -0.228785856664745 | 0.820132729785552 | 0.933259176093322 | -6.27691059549113 |
| lnc-TP53BP2-8 | lnc-TP53BP2-8 | 0.0369343721072983 | 1.93902773751982 | 0.228711620405774 | 0.820190059399401 | 0.93326289477209 | -6.00757192187218 |
| ENSG00000085872.14 | CHERP | -0.0322194982060249 | 3.83824268868782 | -0.228651513477008 | 0.82023647824086 | 0.93326289477209 | -6.25441448669275 |
| ENSG00000279981.1 | | -0.0590081436302421 | 1.5015603809011 | -0.228547503314934 | 0.820316803819334 | 0.93326289477209 | -5.96596936700849 |
| ENSG00000166471.10 | TMEM41B | -0.0342983887696233 | 3.99275706071792 | -0.228546778661481 | 0.820317363465805 | 0.93326289477209 | -6.27778655642991 |
| ENSG00000178974.9 | FBXO34 | 0.0294051381440239 | 4.89860431201579 | 0.228275992900905 | 0.820526496671943 | 0.933378434927875 | -6.34662254169803 |
| ENSG00000092201.9 | SUPT16H | 0.0181103611981879 | 6.88171331861825 | 0.228248391416944 | 0.820547814587139 | 0.933378434927875 | -6.38239003794924 |
| ENSG00000111276.10 | CDKN1B | 0.0291719993875734 | 6.79904003790653 | 0.228239137483732 | 0.820554961862697 | 0.933378434927875 | -6.38259344300519 |
| ENSG00000196177.12 | ACADSB | 0.0182847637206374 | 4.41519573394194 | 0.228065098799224 | 0.820689383518898 | 0.933437499858112 | -6.30790406278727 |
| ENSG00000169403.11 | PTAFR | -0.125225906322467 | -0.162126391638492 | -0.228057254257793 | 0.820697587083139 | 0.933437499858112 | -5.54379346077778 |
| lnc-PTCH1-8 | lnc-PTCH1-8 | 0.0504318684914341 | 0.285101573455547 | 0.227952056476422 | 0.820776696548604 | 0.933475894853571 | -5.73159056606851 |
| lnc-ZNF582-5 | lnc-ZNF582-5 | 0.0408738148143005 | -0.276091093364425 | 0.227870215559972 | 0.820839911284994 | 0.933496209402676 | -5.5520647475025 |
| ENSG00000103253.18 | HAGHL | -0.0650523321156458 | 1.9421743855361 | -0.227534212335477 | 0.821099456138021 | 0.93373978484469 | -6.03633934008599 |
| lnc-IL15RA-4 | lnc-IL15RA-4 | 0.0927612200295471 | 0.257709230202315 | 0.227392250061104 | 0.821209120640539 | 0.933812901505122 | -5.70122246677884 |
| ENSG00000147586.9 | MRPS28 | 0.0214404894926519 | 2.76270145584711 | 0.227120310990467 | 0.82141920108244 | 0.933949541602358 | -6.13309959253249 |
| ENSG00000126458.3 | RRAS | 0.064901984961327 | -0.392040163332425 | 0.227119233368093 | 0.82142003360188 | 0.933949541602358 | -5.50497500883133 |
| ENSG00000186666.5 | BCDIN3D | 0.0332034672959916 | 2.89411045102102 | 0.226920350503432 | 0.82157368454454 | 0.934072644110709 | -6.13580397388742 |
| ENSG00000183336.8 | BOLA2 | 0.0405746431164808 | 0.260588714799037 | 0.226525180190251 | 0.821879002319978 | 0.934368158229887 | -5.73632540578256 |
| ENSG00000216977.1 | RPL21P65 | 0.0617720710661933 | 0.285144875957177 | 0.22614287574173 | 0.822174406221308 | 0.934545184361134 | -5.70068124228661 |
| lnc-ESS2-3 | lnc-ESS2-3 | 0.0463479288161965 | 0.834803656798602 | 0.226131444934145 | 0.822183239126441 | 0.934545184361134 | -5.8520126310952 |
| ENSG00000165730.15 | STOX1 | 0.237613177598231 | -1.1324402639068 | 0.22612138637941 | 0.822206653600739 | 0.934545184361134 | -5.21121969269708 |
| lnc-CARS2-2 | lnc-CARS2-2 | -0.0605603449992298 | 3.06387316483977 | -0.225888245623731 | 0.822371447124777 | 0.934545184361134 | -6.16520830159745 |
| ENSG00000114850.6 | SSR3 | -0.0272983861713954 | 5.68185605370173 | -0.22575850784028 | 0.822471430859528 | 0.934545184361134 | -6.37995888198164 |
| lnc-FAM3B-9 | lnc-FAM3B-9 | 0.0755781566869566 | 0.892535394618204 | 0.225753231009381 | 0.822475508773675 | 0.934545184361134 | -5.83027691648164 |
| ENSG00000180257.13 | ZNF816 | -0.0259189614644804 | 3.97339569724577 | -0.225726136627612 | 0.822496447283465 | 0.934545184361134 | -6.26231177817578 |
| lnc-TIAF1-3 | lnc-TIAF1-3 | 0.0281681804904548 | 1.8595973775658 | 0.225689294114758 | 0.822524919348762 | 0.934545184361134 | -6.01676765342948 |
| lnc-PGGT1B-1 | lnc-PGGT1B-1 | 0.0335021153575878 | 1.99884324530758 | 0.225687087333923 | 0.822526624766938 | 0.934545184361134 | -6.03810025982895 |
| ENSG00000279495.1 | | -0.0514807163967207 | 0.649804182665618 | -0.225587666079449 | 0.822603459221627 | 0.934545184361134 | -5.81630817209376 |
| MYCBP2-AS2 | MYCBP2-AS2 | -0.0538320299299851 | 0.0522250722964017 | -0.225581294817201 | 0.822608383102699 | 0.934545184361134 | -5.66887120439267 |
| lnc-CXCR4-4 | lnc-CXCR4-4 | -0.0759340410360019 | 1.5763238606394 | -0.225576237632771 | 0.822614579749326 | 0.934545184361134 | -5.98993609643588 |
| lnc-CER1-1 | lnc-CER1-1 | 0.0324138120135253 | 2.18752355806386 | 0.225559838991532 | 0.822624964789148 | 0.934545184361134 | -6.05749470011353 |
| lnc-IL10RA-2 | lnc-IL10RA-2 | -0.0609807168579757 | 1.70346271477644 | -0.22524696092803 | 0.822866775378474 | 0.934730666029663 | -5.99222108010742 |
| lnc-SLC25A51-1 | lnc-SLC25A51-1 | 0.0250694479833511 | 2.41621203592519 | 0.225231067970223 | 0.822879058857188 | 0.934730666029663 | -6.08051261733489 |
| ENSG00000204564.11 | C6orf136 | 0.0268642819857114 | 2.47508948859523 | 0.225168996163612 | 0.822927033851384 | 0.934733576389991 | -6.09788761850429 |
| ENSG00000285645.1 | | -0.0458437666338056 | 1.5221959798667 | -0.224971104382309 | 0.823079988011897 | 0.934831003221004 | -5.97092603196799 |
| ENSG00000142765.17 | SYTL1 | -0.0595497472626452 | 5.62587322553004 | -0.224955437253048 | 0.823103642392958 | 0.934831003221004 | -6.37668563490544 |
| MIR29B2CHG | MIR29B2CHG | 0.0634678669469873 | 7.81972368767808 | 0.224833257144599 | 0.8232022958717 | 0.934891461924918 | -6.3703071053832 |
| ENSG00000270574.1 | | -0.0555746930959309 | 1.31885431528403 | -0.224471740380164 | 0.823465986475349 | 0.935139332392944 | -5.9368994967565 |
| ENSG00000144381.16 | HSPD1 | -0.0321928772911724 | 5.95893911802659 | -0.224412567028778 | 0.823511729234944 | 0.935139684588043 | -6.38418028327965 |
| ENSG00000236017.8 | ASMTL-AS1 | 0.0645054249085435 | 1.67165979623035 | 0.224266915031901 | 0.823624325203297 | 0.935215947793869 | -5.98178194972932 |
| ENSG00000101843.18 | PSMD10 | 0.0350004325485194 | 3.5618162479323 | 0.224143830330741 | 0.823719478502828 | 0.935272397728581 | -6.22735983116835 |
| ENSG00000084444.13 | FAM234B | -0.0538447040766905 | 2.17736717761316 | -0.223985696951558 | 0.823841730882287 | 0.935342753665503 | -6.0466380598244 |
| lnc-UGT8-1 | lnc-UGT8-1 | 0.0571365729179946 | 0.737923036550517 | 0.223946120699654 | 0.823872327841207 | 0.935342753665503 | -5.82650043907377 |
| ENSG00000198467.14 | TPM2 | -0.119247294242755 | 1.0901805803091 | -0.223864307075033 | 0.823960701845893 | 0.935391491152943 | -5.9216331001532 |
| ENSG00000101353.14 | MROH8 | 0.05336554355347 | 0.528719262619086 | 0.223755455836353 | 0.824019737422303 | 0.9354069190033048 | -5.78621253866067 |
| lnc-SLC25A47-3 | lnc-SLC25A47-3 | 0.0389277837528571 | 0.723438112272473 | 0.223518310568325 | 0.824203091578084 | 0.935563460759483 | -5.83419310149237 |
| ENSG00000102897.9 | LYRM1 | 0.0251441202375573 | 3.94902610458788 | 0.223443448405103 | 0.824260974993998 | 0.935577569664799 | -6.26603041757144 |
| lnc-ADAR-2 | lnc-ADAR-2 | 0.0239868674274204 | 3.47573934091493 | 0.223242478565568 | 0.824416369745251 | 0.935675030484071 | -6.21536117283184 |
| ACAP2-IT1 | ACAP2-IT1 | -0.0635983911181592 | 1.27323149780539 | -0.223214819419357 | 0.824437757025658 | 0.935675030484071 | -5.94827825341654 |
| lnc-CAPS2-4 | lnc-CAPS2-4 | 0.0614678678606892 | 0.0845275547307021 | 0.223084165270389 | 0.824538786441649 | 0.935738095678914 | -5.70350853863963 |
| lnc-MAGOHB-1 | lnc-MAGOHB-1 | -0.0459264667600126 | 1.30402121875702 | -0.222978331697358 | 0.824620625346257 | 0.935779376530048 | -5.9467031328023 |
| lnc-KYNU-7 | lnc-KYNU-7 | 0.0500821198128534 | 1.21371817201458 | 0.222854881133611 | 0.824716089598733 | 0.935836114020107 | -5.92333661767711 |
| lnc-RUNDC3A-2 | lnc-RUNDC3A-2 | -0.0571423887991617 | -0.275257200173163 | -0.222732316328002 | 0.824810871551175 | 0.935861492926599 | -5.59213760738305 |
| LINC01732 | LINC01732 | -0.0891404771738369 | 0.311052532501653 | -0.222708369525639 | 0.824829390428093 | 0.935861492926599 | -5.70123245040733 |
| ENSG00000103257.8 | SLC7A5 | -0.0860144558489109 | 1.77751691303955 | -0.222587413266181 | 0.824934349290526 | 0.935928988614645 | -6.0179482989693 |
| ENSG00000169762.16 | TAPT1 | -0.0732814939353323 | 5.63500137924729 | -0.222537375444012 | 0.824986704660271 | 0.935936798876832 | -6.38256794290537 |
| ENSG00000167747.14 | C19orf48P | -0.0397991110192519 | 4.80818272148211 | -0.222442107509289 | 0.825035306703831 | 0.93594035041253 | -6.34853785181449 |
| lnc-BAG3-4 | lnc-BAG3-4 | 0.0397455075066464 | 2.39682008228836 | 0.222270815715212 | 0.825167783442072 | 0.936039045830156 | -6.07997268981179 |
| ENSG00000178988.10 | MRFAP1L1 | -0.0345886629748486 | 4.95954444340324 | -0.222191628924209 | 0.8252290280944 | 0.936056931845422 | -6.35418254829963 |
| ENSG00000117151.12 | CTBS | 0.0279672622891747 | 4.05093990560601 | 0.222055220077403 | 0.825334532015544 | 0.936125016392568 | -6.28404573276945 |
| ENSG00000079805.16 | DNM2 | -0.0257976286533799 | 7.12798572483883 | -0.221971060135232 | 0.825399626226318 | 0.936131258510549 | -6.38039257570221 |
| ENSG00000230701.2 | FBXW4P1 | 0.0809164880324397 | -0.696443346370139 | 0.221899341901148 | 0.825455098275103 | 0.936131258510549 | -5.45639507820753 |
| ENSG00000108465.14 | CDK5RAP3 | 0.0341553347487107 | 6.86538678109928 | 0.221756558369892 | 0.825565540031037 | 0.936131258510549 | -6.38415065975612 |
| ENSG00000167487.11 | KLHL26 | -0.0398368559892075 | 1.5513129948825 | -0.221752280747083 | 0.825568848788337 | 0.936131258510549 | -5.9789245178683 |

| | | | | | | | |
|---|---|---|---|---|---|---|---|
| ENSG00000241058.3 | NSUN6 | -0.0187103772863226 | 4.64258547666653 | -0.221706572671836 | 0.825604204356439 | 0.936131258510549 | -6.32932268348113 |
| lnc-RIPK4-3 | lnc-RIPK4-3 | -0.058749362128742 | 0.715413667683621 | -0.221695304850361 | 0.82561292016431 | 0.936131258510549 | -5.84453097121556 |
| ENSG00000121481.10 | RNF2 | 0.0294055134218718 | 4.12538736533079 | 0.221565896370423 | 0.825713020918365 | 0.936158515160945 | -6.29475276477757 |
| ENSG00000249459.8 | ZNF286B | 0.052348764590221 | 1.37690791661578 | 0.221546631319475 | 0.825727923176746 | 0.936158515160945 | -5.93831972775612 |
| lnc-SIDT2-2 | lnc-SIDT2-2 | 0.0342967363637028 | 4.16920174044237 | 0.221414385513613 | 0.825830222152434 | 0.936168800572356 | -6.29917298656439 |
| ENSG00000171960.10 | PPIH | -0.0189825460710464 | 3.92545855539022 | -0.22138055863922 | 0.825856389482526 | 0.936168800572356 | -6.26681800280211 |
| ENSG00000107554.16 | DNMBP | -0.0420651721786722 | 8.16028861215637 | -0.221331913426824 | 0.825894020130724 | 0.936168800572356 | -6.36419694406637 |
| ENSG00000214013.9 | GANC | -0.0220360546765317 | 6.8287374993281 | -0.221272236216803 | 0.825940185405928 | 0.936168800572356 | -6.38454611120257 |
| lnc-ELMSAN1-2 | lnc-ELMSAN1-2 | 0.0350550411659922 | 1.76407716376955 | 0.221201157135832 | 0.825995171787889 | 0.936168800572356 | -6.00178443938853 |
| lnc-PABPC4-4 | lnc-PABPC4-4 | 0.0297118266563956 | 3.27449481175545 | 0.221182130243488 | 0.826009891036028 | 0.936168800572356 | -6.19325597065488 |
| ENSG00000106299.7 | WASL | -0.0258841323565657 | 3.82922601928988 | -0.220833577741177 | 0.826279543262846 | 0.936422852052591 | -6.26031110828811 |
| ENSG00000175711.8 | B3GNTL1 | -0.0492688278109105 | 1.07589398688268 | -0.220749695500541 | 0.826344440623932 | 0.9364448395839 | -5.87119246093717 |
| ENSG00000258230.2 | | -0.0560849867485911 | 2.28864266515705 | -0.220405331721568 | 0.826610878190629 | 0.936607104613879 | -6.07713315259697 |
| ENSG00000240303.7 | ACAD11 | 0.0267135868123001 | 4.39306795824395 | 0.220362208489038 | 0.826644244507543 | 0.936607104613879 | -6.31060442745175 |
| ENSG00000049239.12 | H6PD | -0.0237250681760797 | 5.72380291634814 | -0.220305028123109 | 0.826688487938643 | 0.936607104613879 | -6.38104431045765 |
| ENSG00000145817.16 | YIPF5 | 0.0236987445820693 | 3.97707460284569 | 0.220182667137993 | 0.826783166928133 | 0.936607104613879 | -6.27389667295314 |
| ENSG00000155367.15 | PPM1J | 0.0714098122517102 | 1.6572158914462 | 0.220165027337196 | 0.826796816252988 | 0.936607104613879 | -5.96598568489698 |
| ENSG00000173406.15 | DAB1 | 0.0430214449942754 | 1.75620930953782 | 0.220114644622512 | 0.826835801687812 | 0.936607104613879 | -5.99102242103695 |
| ENSG00000062716.12 | VMP1 | 0.0548803879590274 | 6.48608684022646 | 0.220058776176273 | 0.826887764765022 | 0.936607104613879 | -6.38627186559845 |
| ENSG00000257122.5 | RRN3P3 | 0.0359940233954696 | 3.76216806873364 | 0.220012966127665 | 0.826914480420719 | 0.936607104613879 | -6.23264341180315 |
| ENSG00000158483.15 | FAM86C1P | 0.0471792845595975 | 0.738376249094756 | 0.219989082644165 | 0.826932961700969 | 0.936607104613879 | -5.8262229913276 |
| ENSG00000132950.18 | ZMYM5 | 0.018956270615031 | 4.81446837717415 | 0.219975870507763 | 0.826943185427928 | 0.936607104613879 | -6.34401147761339 |
| ENSG00000186810.7 | CXCR3 | -0.119178520152845 | 2.50428742957861 | -0.219958121010558 | 0.82698817078427 | 0.936607104613879 | -6.09562604933625 |
| lnc-SERPINB9-4 | lnc-SERPINB9-4 | 0.16083310775146 | -1.40579917696456 | 0.219835040294677 | 0.827057803897427 | 0.936634430680646 | -5.37058687332758 |
| ENSG00000124767.6 | GLO1 | 0.0570555278346578 | 7.31220069974066 | 0.219693607100788 | 0.827171677218053 | 0.936711852562675 | -6.37737671918449 |
| ENSG00000107404.19 | DVL1 | -0.0500737535052802 | 3.1459920000071 | -0.2194860357423 | 0.827322247513986 | 0.9368308202993 | -6.18700841973622 |
| lnc-RHOA-1 | lnc-RHOA-1 | 0.037730852562168 | 1.26453906944632 | 0.219331877623395 | 0.827441552485319 | 0.936876162034151 | -5.93358006099504 |
| ENSG00000163156.11 | SCNM1 | 0.0225243937905262 | 4.02526327179211 | 0.219316668674283 | 0.827453323112221 | 0.936876162034151 | -6.27741860043196 |
| ENSG00000265996.1 | MIR3671 | -0.0471493925161594 | 0.454422793090036 | -0.219112017353377 | 0.82761171233988 | 0.93691064614545 | -5.78315880779491 |
| ENSG00000232373.2 | MTCYBP3 | 0.07632849722272 | -0.196050565231031 | 0.219103095533263 | 0.827618617518921 | 0.93691064614545 | -5.56954485114807 |
| ENSG00000102241.11 | HTATSF1 | -0.0240656649007081 | 4.98153828484025 | -0.219100875658209 | 0.827620335627293 | 0.93691064614545 | -6.35343435379439 |
| ENSG00000241527.1 | CA15P1 | 0.0621506585978467 | 0.989731989688071 | 0.218961964463983 | 0.827727849902495 | 0.936980824646266 | -5.86443876252324 |
| ENSG00000164172.18 | MOCS2 | -0.0202752365586787 | 3.78305574045748 | -0.218338633062091 | 0.828210336127829 | 0.937461294404076 | -6.2525230769766 |
| ENSG00000182923.17 | CEP63 | -0.0238228270127792 | 4.36181900996477 | -0.218295937048557 | 0.828243387200025 | 0.937461294404076 | -6.30383760785506 |
| ENSG00000284284.1 | MIR4712 | -0.0583273750375728 | 1.47991943495989 | -0.218182520809732 | 0.828331184473894 | 0.937509115254931 | -5.96193497686669 |
| ENSG00000141979.4 | | -0.0282167421088291 | 2.52937808151802 | -0.218054326043765 | 0.828430424709596 | 0.937519811713658 | -6.10305939687801 |
| ENSG00000163818.16 | LZTFL1 | 0.0287493481033736 | 3.273603671783 | 0.21799543773793 | 0.828476013241954 | 0.937519811713658 | -6.19284274648627 |
| lnc-THSD1-3 | lnc-THSD1-3 | -0.0348170913266714 | 2.43916072712003 | -0.217909742068582 | 0.828542355832106 | 0.937519811713658 | -6.09863321626735 |
| CNNM3-DT | CNNM3-DT | 0.0521535142458647 | 2.42187783354269 | 0.217866655616609 | 0.828575712334895 | 0.937519811713658 | -6.06171976473384 |
| lnc-TRIM62-3 | lnc-TRIM62-3 | 0.0467679935152518 | -0.223825687022218 | 0.21783049338302 | 0.828603708520168 | 0.937519811713658 | -5.57197819463831 |
| ENSG00000268465.1 | | -0.0301417786360569 | 1.2488360045135 | -0.217817297213398 | 0.828613924826064 | 0.937519811713658 | -5.93167178435077 |
| ENSG00000179832.17 | MROH1 | -0.0380523752797206 | 5.0936180230369 | -0.217632803677161 | 0.828756760507573 | 0.937629879707985 | -6.36462983942364 |
| ENSG00000115233.11 | PSMD14 | 0.0195369030400396 | 4.54644568680082 | 0.217474737301326 | 0.828879140833051 | 0.937686852402336 | -6.32928254123113 |
| ENSG00000167553.15 | TUBA1C | 0.038423807149143 | 3.5720155533326 | 0.217450081528938 | 0.828898230552422 | 0.937686852402336 | -6.24088385398254 |
| ENSG00000123643.12 | SLC36A1 | -0.0243118014613479 | 3.39269700268649 | -0.217276426914152 | 0.829032685513823 | 0.937760558554262 | -6.20478318477282 |
| lnc-VDR-2 | lnc-VDR-2 | -0.0559516077035493 | 0.837762669540674 | -0.217224232477561 | 0.829073098934704 | 0.937760558554262 | -5.83550041091715 |
| ENSG00000130653.15 | PNPLA7 | 0.0452358266643555 | 4.85151187798992 | 0.217104130269141 | 0.829166094170799 | 0.937760558554262 | -6.34141532139885 |
| ENSG00000113369.8 | ARRDC3 | 0.0630972728581281 | 8.19338803971854 | 0.217103119338747 | 0.829183075267092 | 0.937760558554262 | -6.36068751339727 |
| ENSG00000153015.15 | CWC27 | 0.0239689567674652 | 4.28049395551188 | 0.217071726015857 | 0.829191185234784 | 0.937760558554262 | -6.29864902645868 |
| ENSG00000153767.9 | GTF2E1 | -0.0314739763129044 | 3.36236127144106 | -0.21696112263341 | 0.829276828330954 | 0.937805887453218 | -6.1970577664655 |
| lnc-PTRHD1-1 | lnc-PTRHD1-1 | 0.0617287498219402 | 0.796714789622856 | 0.216875684528488 | 0.829342986732079 | 0.937829177887396 | -5.82784178400492 |
| ENSG00000283791.1 | MIR612 | -0.0800137965873623 | 0.253339459472558 | -0.216811382372818 | 0.829392779484838 | 0.937833960343703 | -5.72436006045784 |
| ENSG00000238942.1 | SNORD2 | 0.0274814448181901 | 3.66893547641577 | 0.216478664857528 | 0.829650432584056 | 0.938073766967569 | -6.23824978601866 |
| ENSG00000145012.13 | LPP | 0.0421980098063885 | 7.5326863743605 | 0.216386152012095 | 0.829722076974153 | 0.938103241436913 | -6.37726408907767 |
| ENSG00000136240.9 | KDELR2 | 0.0218661909071267 | 4.23000255903511 | 0.21627234490388 | 0.829810214215 | 0.938151358848036 | -6.29938666213802 |
| SCOC-AS1 | SCOC-AS1 | 0.0535578305429064 | -0.257177625224286 | 0.215966000945881 | 0.830047471496175 | 0.938281586989764 | -5.53769727090349 |
| ENSG00000090674.15 | MCOLN1 | 0.0379791877660963 | 0.951211338868014 | 0.21595783107738 | 0.830053799116079 | 0.938281586989764 | -5.87266571942727 |
| lnc-ZC4H2-3 | lnc-ZC4H2-3 | 0.0286584520442661 | 1.76350782975109 | 0.215925493570455 | 0.830078844851812 | 0.938281586989764 | -6.00289752595842 |
| lnc-SLC24A1-5 | lnc-SLC24A1-5 | 0.0503157588044481 | 0.0354595261230007 | 0.215888180790541 | 0.830107744210446 | 0.938281586989764 | -5.72012166411172 |
| lnc-KLF7-1 | lnc-KLF7-1 | 0.0521449061539991 | 4.37389888135573 | 0.215445069447171 | 0.83045326137157 | 0.938619780737032 | -6.3090862637715 |
| ENSG00000180879.13 | SSR4 | 0.0340995926325958 | 5.13108235568915 | 0.215384145329327 | 0.830498151230764 | 0.938619780737032 | -6.36394515173331 |
| ENSG00000075568.16 | TMEM131 | 0.0224874976651801 | 7.12562578500274 | 0.21496890710173 | 0.830819811794828 | 0.938848868445701 | -6.38307686283889 |
| ENSG00000132801.6 | ZSWIM3 | -0.0329162211247337 | 1.02096477623804 | -0.214893716350457 | 0.830878060775959 | 0.938848868445701 | -5.87566194517656 |
| lnc-RIOK1-2 | lnc-RIOK1-2 | -0.0475078421256254 | 1.32944678581886 | -0.214884941955982 | 0.830884858211075 | 0.938848868445701 | -5.94007244835558 |
| ENSG00000100528.11 | CNIH1 | -0.025345846295318 | 4.40564124230284 | -0.214732549582658 | 0.831002917139074 | 0.938848868445701 | -6.31397258230412 |
| ENSG00000071909.18 | MYO3B | -0.384669749650721 | -1.03271455592045 | -0.214757183502596 | 0.831021529548429 | 0.938848868445701 | -5.25454150301136 |
| lnc-CDC42-2 | lnc-CDC42-2 | 0.0628588645164516 | -0.383670956589441 | 0.21466038481877 | 0.831058824823431 | 0.938848868445701 | -5.53101784308144 |
| ENSG00000277138.1 | MIR6509 | 0.0834168029278733 | -0.740658404755092 | 0.214654962166824 | 0.831063025911033 | 0.938848868445701 | -5.42297791763476 |

| | | | | | | | |
|---|---|---|---|---|---|---|---|
| ENSG00000179218.13 | CALR | 0.0374483616520118 | 7.7632319891936 | 0.214651442801193 | 0.831065752469547 | 0.938848868445701 | -6.37183783774548 |
| ENSG00000124334.17 | IL9R | -0.162920205568897 | -1.16086269734337 | -0.214478274618506 | 0.831199913674772 | 0.938948895404634 | -5.21165665257911 |
| ENSG00000184922.13 | FMNL1 | 0.0325355488124384 | 6.36903047907101 | 0.214344088432486 | 0.831303877271814 | 0.939014801113255 | -6.38797105151376 |
| ENSG00000263126.1 | | -0.0219048993627502 | 3.19889453659504 | -0.214104894064627 | 0.831489205768591 | 0.939052719939598 | -6.18786805903972 |
| ENSG00000148341.17 | SH3GLB2 | 0.0262156789178443 | 6.17068820870255 | 0.214067039190102 | 0.831518536725936 | 0.939052719939598 | -6.38784583575112 |
| ENSG00000127884.4 | ECHS1 | 0.0389717317948239 | 4.30660261908608 | 0.214019502693527 | 0.831555369604131 | 0.939052719939598 | -6.30484057902474 |
| ENSG00000103512.14 | NOMO1 | 0.0205243227344787 | 3.95820943311793 | 0.214009384510741 | 0.83156320956231 | 0.939052719939598 | -6.26659038978299 |
| ENSG00000201793.1 | RN7SKP9 | 0.0802036364798722 | 0.219213120637673 | 0.214006350755711 | 0.831565560236071 | 0.939052719939598 | -5.78233418484383 |
| ENSG00000239912.1 | RPL39P36 | 0.040842399114307 | -0.338321647599282 | 0.213878388167056 | 0.83166471214736 | 0.939113164918209 | -5.54374590952011 |
| ENSG00000100422.13 | CERK | 0.0547327846615264 | 5.26285234859829 | 0.213643988295743 | 0.831855011775259 | 0.939179224576681 | -6.36494293867134 |
| ENSG00000092330.16 | TINF2 | -0.0252946380313821 | 5.6674666067109 | -0.213536672944294 | 0.831929504046525 | 0.939179224576681 | -6.3808428638977 |
| ENSG00000160216.18 | AGPAT3 | 0.0308712185110132 | 5.17412822232462 | 0.213525604722386 | 0.831938081037631 | 0.939179224576681 | -6.36516093459495 |
| ENSG00000105327.17 | BBC3 | -0.0596896279338663 | 2.56884909577867 | -0.213451667580127 | 0.831995376970132 | 0.939179224576681 | -6.11427949310655 |
| ENSG00000254505.9 | CHMP4A | -0.0154261086540854 | 3.45421638969586 | -0.213429559353004 | 0.832012509425175 | 0.939179224576681 | -6.21179551712697 |
| lnc-LRIG2-5 | lnc-LRIG2-5 | -0.0408455220034678 | 0.823669445996211 | -0.213401453022768 | 0.832034290143943 | 0.939179224576681 | -5.84039976989901 |
| lnc-TXNDC2-4 | lnc-TXNDC2-4 | 0.0474585858032393 | 1.15792037765222 | 0.213390709507786 | 0.832042615758431 | 0.939179224576681 | -5.91701823747208 |
| ENSG00000196396.9 | PTPN1 | 0.0327070947679795 | 7.8215436058283 | 0.213323919315007 | 0.832094374805675 | 0.939186143706142 | -6.37193498712842 |
| ENSG00000197776.7 | KLHDC1 | 0.0297276230607411 | 3.42104438249465 | 0.213169295498761 | 0.832214203395776 | 0.939269888056986 | -6.20549309954462 |
| ENSG00000101745.16 | ANKRD12 | 0.02591926214415 | 8.23729672341534 | 0.21300630179608 | 0.832340522756506 | 0.939360948563283 | -6.36385028560981 |
| LINC02365 | LINC02365 | -0.0796389630184331 | 3.5646852682703 | -0.212890646318899 | 0.832452245473285 | 0.939376891377009 | -6.21357976642362 |
| ENSG00000003249.13 | DBNDD1 | 0.116909006152662 | 1.58497759645127 | 0.212894504054897 | 0.832455755488444 | 0.939376891377009 | -5.99748140107131 |
| lnc-GRM2-2 | lnc-GRM2-2 | -0.0280142168471694 | 0.597904390268353 | -0.212634432202212 | 0.832628736740939 | 0.939376891377009 | -5.80885865313648 |
| ENSG00000279957.1 | | -0.0481837575941491 | 0.858192064817618 | -0.21260863195643 | 0.832648733840408 | 0.939376891377009 | -5.88396394161047 |
| ENSG00000145908.12 | ZNF300 | 0.319677530255887 | -0.516385382081628 | 0.212526146575878 | 0.832753024856823 | 0.939376891377009 | -5.6306170382864 |
| ENSG00000175215.10 | CTDSP2 | 0.0264915218651112 | 6.74956505887376 | 0.21243125601466 | 0.832786216354827 | 0.939376891377009 | -6.38638656116627 |
| ENSG00000264678.1 | MIR3140 | 0.0815355656892606 | 0.266500001966782 | 0.212423092972166 | 0.832792543583963 | 0.939376891377009 | -5.70144086312184 |
| ENSG00000105197.10 | TIMM50 | 0.0208663104277047 | 4.09788076677336 | 0.212399333554166 | 0.832810959732484 | 0.939376891377009 | -6.28882412313088 |
| lnc-AFG1L-7 | lnc-AFG1L-7 | 0.0677316766435366 | 0.425483587563198 | 0.212287103534331 | 0.832897951561546 | 0.939376891377009 | -5.79608879920973 |
| ENSG00000242889.3 | RN7SL449P | -0.0785938816209671 | -0.0892012892765909 | -0.212243136594257 | 0.832932031831544 | 0.939376891377009 | -5.63579558007173 |
| ENSG00000100092.22 | SH3BP1 | -0.0406490750563919 | 3.76091196694104 | -0.212227792435468 | 0.832943925687223 | 0.939376891377009 | -6.2476801000674 |
| ENSG00000156136.9 | DCK | -0.0209313051115005 | 5.59249382028125 | -0.212163292031486 | 0.83299392289802 | 0.939376891377009 | -6.38180651744046 |
| ENSG00000285292.1 | ABCF2-H2BK1 | -0.0230189864155759 | 3.41350017450174 | -0.21206203175566 | 0.833072415776533 | 0.939376891377009 | -6.20031694594635 |
| lnc-MED7-1 | lnc-MED7-1 | -0.0229506499649599 | 2.8652055439828 | -0.211999203589498 | 0.833121118501339 | 0.939376891377009 | -6.138236644363 |
| lnc-UBB-1 | lnc-UBB-1 | 0.0398683953663037 | 1.47150901258749 | 0.211952660834929 | 0.833157197635522 | 0.939376891377009 | -5.94822563325576 |
| ENSG00000164252.12 | AGGF1 | -0.0161869721660723 | 4.63597350449957 | -0.211940696400078 | 0.833166472316214 | 0.939376891377009 | -6.3344023843279 |
| lnc-CARS2-4 | lnc-CARS2-4 | -0.0213933963083432 | 5.31492670270595 | -0.211929393056246 | 0.833175234549482 | 0.939376891377009 | -6.37311328657934 |
| lnc-PASK-2 | lnc-PASK-2 | 0.0234167282786535 | 3.33397785433949 | 0.211928222553415 | 0.833176141912194 | 0.939376891377009 | -6.19310629854756 |
| lnc-SLC38A9-1 | lnc-SLC38A9-1 | -0.0636036784659684 | -0.607488362039146 | -0.211763871854751 | 0.833303547301773 | 0.939390785819668 | -5.42534232672422 |
| ENSG00000161955.16 | TNFSF13 | 0.0535984305452812 | 1.09474590083639 | 0.211763184302929 | 0.833304080304517 | 0.939390785819668 | -5.88339150739783 |
| lnc-CTC1-2 | lnc-CTC1-2 | 0.04067283273596 | 2.18638334070266 | 0.211735704777993 | 0.833325382999626 | 0.939390785819668 | -6.05852999866858 |
| lnc-MATN2-2 | lnc-MATN2-2 | 0.0603450869851565 | -0.223656440570186 | 0.211607158519195 | 0.83342503639635 | 0.939421049518391 | -5.59340242057347 |
| lnc-STRBP-13 | lnc-STRBP-13 | 0.037599156318193 | 0.61344894012213 | 0.211578072912378 | 0.833447584924611 | 0.939421049518391 | -5.81102189899203 |
| ENSG00000106443.16 | PHF14 | -0.0331260811787111 | 5.92305054674528 | -0.211524456125273 | 0.833489151547629 | 0.939421049518391 | -6.3872312133374 |
| ENSG00000013563.13 | DNASE1L1 | 0.031956754245412 | 4.33226566394148 | 0.211459340791565 | 0.833539633111817 | 0.939426505412065 | -6.30151965996393 |
| ENSG00000133872.13 | SARAF | 0.0248895788884465 | 7.37160279071295 | 0.211287271434213 | 0.833673035683859 | 0.939476438175753 | -6.38019688559387 |
| lnc-KPNA2-3 | lnc-KPNA2-3 | 0.0453689347061642 | 0.51375225757176 | 0.211218164644097 | 0.833726614449388 | 0.939476438175753 | -5.78200876305246 |
| lnc-CD180-8 | lnc-CD180-8 | -0.0730890571304172 | -0.459588690572373 | -0.211174209294868 | 0.833760693620182 | 0.939476438175753 | -5.50026238868662 |
| ENSG00000164300.16 | SERINC5 | 0.0747943115734948 | 6.59451681467036 | 0.211201272072521 | 0.833766511011829 | 0.939476438175753 | -6.38853357707951 |
| ENSG00000103018.16 | CYB5B | -0.0199824692458916 | 4.9569044102727 | -0.210712218985949 | 0.834118900444815 | 0.93972068193265 | -6.35700910114148 |
| lnc-DCDC1-5 | lnc-DCDC1-5 | -0.0294153195066099 | 0.851573359983145 | -0.210599504047559 | 0.834206300029635 | 0.93972068193265 | -5.8687771763656 |
| lnc-CYP7A1-6 | lnc-CYP7A1-6 | -0.0260852694235258 | 2.42835640651498 | -0.210579997266571 | 0.834221425875931 | 0.93972068193265 | -6.08375182795928 |
| ENSG00000143669.13 | LYST | -0.0508653359871092 | 7.75754317142906 | -0.2105541633585 | 0.834246574943324 | 0.93972068193265 | -6.37192834840295 |
| ENSG00000076201.14 | PTPN23 | -0.0277837471352376 | 4.51314990971878 | -0.21053561273891 | 0.834255842531417 | 0.93972068193265 | -6.32266957704381 |
| ENSG00000163945.16 | UVSSA | 0.0350309643149167 | 4.81197303706437 | 0.210533857528182 | 0.834257203563878 | 0.93972068193265 | -6.34466126051713 |
| ENSG00000143013.12 | LMO4 | 0.0436496146791887 | 4.62483814971876 | 0.210434703795989 | 0.834334090578958 | 0.939755860040861 | -6.33013664809398 |
| lnc-DNAJC5-1 | lnc-DNAJC5-1 | -0.0344991778579389 | 2.49460240329026 | -0.210303507480667 | 0.834435826971711 | 0.939819021973118 | -6.09755174056051 |
| ENSG00000147202.17 | DIAPH2 | -0.0239315642595104 | 5.19307209873761 | -0.210198061804588 | 0.834517597070518 | 0.939856353162773 | -6.36662004236179 |
| ENSG00000104774.12 | MAN2B1 | -0.0369141300861209 | 5.72983025232881 | -0.210085269827414 | 0.83460506606648 | 0.939856353162773 | -6.38364679281187 |
| lnc-MMRN2-1 | lnc-MMRN2-1 | 0.0283569297289752 | 2.89002904373015 | 0.210077997190203 | 0.834610705993142 | 0.939856353162773 | -6.14673753647134 |
| lnc-SRD5A3-4 | lnc-SRD5A3-4 | 0.0518102617867864 | 1.29777511812567 | 0.209959017125007 | 0.834702976227424 | 0.939856353162773 | -5.94105583617228 |
| ENSG00000112062.10 | MAPK14 | -0.0233847703361096 | 5.48457210863712 | -0.209948229036843 | 0.834711342614586 | 0.939856353162773 | -6.37910285965945 |
| ENSG00000102032.12 | RENBP | -0.0481356615069082 | 2.37646716833519 | -0.209887087605139 | 0.834758759434924 | 0.939856353162773 | -6.10556178584747 |
| ENSG00000101057.15 | MYBL2 | 0.0891443935925977 | 3.82984641245775 | 0.209885605704826 | 0.834788604601944 | 0.939856353162773 | -6.29274753347393 |
| ENSG00000021355.12 | SERPINB1 | 0.0236487392841171 | 5.38289038634739 | 0.209547732893897 | 0.835021949421431 | 0.939951902806262 | -6.3752495124545 |
| lnc-NFKBIA-7 | lnc-NFKBIA-7 | -0.0247539541305932 | 1.83830287276783 | -0.20944783697256 | 0.835099428363071 | 0.939951902806262 | -6.0131851840025 |
| ENSG00000065978.18 | YBX1 | -0.0324386950054842 | 7.61470681456988 | -0.209370297784384 | 0.835159568640086 | 0.939951902806262 | -6.37602240872956 |
| lnc-GRAMD2A-4 | lnc-GRAMD2A-4 | -0.0210264102398419 | 1.75951345028794 | -0.209319347880747 | 0.83519908650885 | 0.939951902806262 | -6.00244500197457 |
| ENSG00000211767.1 | TRBJ2-3 | 0.0524511336244501 | -0.216688547911689 | 0.209311486918156 | 0.835205183683154 | 0.939951902806262 | -5.55464144648965 |

| | | | | | | | |
|---|---|---|---|---|---|---|---|
| lnc-NPIPB5-6 | lnc-NPIPB5-6 | -0.0326468155843112 | 1.80415611302493 | -0.209260579357967 | 0.835244669206217 | 0.939951902806262 | -6.02132249449677 |
| ENSG00000106049.8 | HIBADH | 0.030921862578202 | 2.86822150798783 | 0.20923122213935 | 0.83526743979437 | 0.939951902806262 | -6.15074266278703 |
| ENSG00000066923.17 | STAG3 | 0.0737555177273248 | 3.64673813004723 | 0.209225232896282 | 0.835293247304447 | 0.939951902806262 | -6.26197371291917 |
| lnc-SH2B2-5 | lnc-SH2B2-5 | -0.0539029198758453 | 0.134351449463277 | -0.209177612119915 | 0.835309022156878 | 0.939951902806262 | -5.70983742881398 |
| ENSG00000163900.10 | TMEM41A | 0.0264397491692092 | 4.10349050817093 | 0.209150390298964 | 0.835330136818353 | 0.939951902806262 | -6.29004096856232 |
| ENSG00000108187.15 | PBLD | -0.0359418173799217 | 1.64864616610986 | -0.209060564349901 | 0.835399811372626 | 0.939978916387841 | -5.98910541338945 |
| ENSG00000105171.9 | POP4 | 0.0238033312481027 | 4.63327945628578 | 0.208873403969765 | 0.835544988820742 | 0.940070437844914 | -6.33015486773864 |
| lnc-ZC3HC1-5 | lnc-ZC3HC1-5 | -0.0728125792220694 | 0.304619991931186 | -0.208837944562521 | 0.835572494795351 | 0.940070437844914 | -5.6979505221075 |
| ENSG00000131379.9 | C3orf20 | -0.0600004733700143 | 0.469832919655312 | -0.208609751615554 | 0.835749509768953 | 0.940218198490072 | -5.71077997380387 |
| ENSG00000251192.7 | ZNF674 | -0.0218275675107571 | 1.60528070562639 | -0.208427569271865 | 0.83589083935318 | 0.940279612739096 | -5.97605292344923 |
| ENSG00000127328.21 | RAB3IP | 0.0301261123232628 | 4.43725913143253 | 0.208421606791638 | 0.835895464893358 | 0.940279612739096 | -6.32207129846383 |
| ENSG00000280828.1 | | -0.0436793239032505 | 3.92020375692105 | -0.208326266613054 | 0.835969428171957 | 0.940311423578523 | -6.26259765571808 |
| lnc-LCMT2-1 | lnc-LCMT2-1 | -0.0243732650217794 | 2.61123190186012 | -0.208232796137219 | 0.836041942425371 | 0.940341601144339 | -6.10881700744091 |
| ENSG00000244378.1 | RPS2P45 | -0.0516924469960605 | -0.279256680063971 | -0.208136909359939 | 0.836116332741359 | 0.940372598554031 | -5.5611305966056 |
| lnc-ARSD-3 | lnc-ARSD-3 | -0.0341800401103622 | 2.75783843805281 | -0.208079496187052 | 0.836160875418349 | 0.940372598554031 | -6.12461847480752 |
| lnc-NPBWR1-8 | lnc-NPBWR1-8 | -0.0536960509785229 | 0.59184361840793 | -0.207990157420771 | 0.836230187907628 | 0.9403991672241 | -5.77780940851007 |
| ENSG00000205702.10 | CYP2D7 | -0.0451513370865361 | 0.73445963552975 | -0.207923354262579 | 0.836282017256293 | 0.940406073054321 | -5.86126184157888 |
| ENSG00000077782.20 | FGFR1 | 0.208348868595999 | 2.07310250538091 | 0.207910756286042 | 0.836331238350369 | 0.940410045286296 | -6.09642523017056 |
| ENSG00000213443.2 | NPM1P7 | 0.0384950903485679 | 5.1711319508953 | 0.20770527550615 | 0.836451219168543 | 0.940493578288328 | -6.36157653439097 |
| ENSG00000168286.2 | THAP11 | -0.0480616933973073 | 3.91711655460194 | -0.207279234382646 | 0.836781796539449 | 0.940813880929234 | -6.28042176358091 |
| ENSG00000200879.1 | SNORD14E | -0.0944009881936132 | -0.736356507619954 | -0.207123855485417 | 0.836902366857047 | 0.940831299044624 | -5.40518831384473 |
| ENSG00000123933.16 | MXD4 | 0.0467956040712707 | 5.28803502164459 | 0.207123332402327 | 0.8369048268647 | 0.940831299044624 | -6.37489731417883 |
| lnc-PLEK2-3 | lnc-PLEK2-3 | 0.0436308329026475 | -0.0299443739774895 | 0.207050166918027 | 0.836959548814649 | 0.940831299044624 | -5.63720611618606 |
| lnc-CBARP-6 | lnc-CBARP-6 | 0.0583313492122063 | 0.193064353697696 | 0.206969484741597 | 0.837022158800762 | 0.940831299044624 | -5.69817312819718 |
| ENSG00000134905.16 | CARS2 | 0.0419022396578095 | 6.28214035228096 | 0.206917745482487 | 0.837062309424236 | 0.940831299044624 | -6.38949587220919 |
| ENSG00000147403.16 | RPL10 | -0.0359790437161767 | 9.49288012633651 | -0.206841280833852 | 0.837121648212719 | 0.940831299044624 | -6.32739238737373 |
| lnc-MNX1-21 | lnc-MNX1-21 | -0.0257278849748208 | 1.21816480996068 | -0.206805592867202 | 0.837149343439983 | 0.940831299044624 | -5.91756775552201 |
| lnc-GPR89B-2 | lnc-GPR89B-2 | -0.103555500228118 | 0.322276578029805 | -0.206795012668383 | 0.837162961764675 | 0.940831299044624 | -5.69413325856495 |
| ENSG00000123636.17 | BAZ2B | -0.0565176214583891 | 5.90968830734623 | -0.206622014700761 | 0.837305349464411 | 0.940939943657239 | -6.38789708121181 |
| ENSG00000284671.1 | | 0.0399994453328465 | 3.46415725394942 | 0.206459615795238 | 0.837417845711586 | 0.941014987076572 | -6.21037092977738 |
| ENSG00000130159.13 | ECSIT | -0.0299694740352243 | 3.64969668683254 | -0.206199898802208 | 0.83761941705066 | 0.941190111428854 | -6.24708374487316 |
| lnc-COX17-2 | lnc-COX17-2 | 0.0188684377881079 | 2.69935519834289 | 0.206131279233542 | 0.837672675857866 | 0.941198574550056 | -6.12311381524621 |
| lnc-NDRG1-4 | lnc-NDRG1-4 | -0.0553585966972316 | 2.24098310335894 | -0.205922745413483 | 0.837834533285093 | 0.9412811692115 | -6.07186452942463 |
| ENSG00000167136.6 | ENDOG | 0.0546178553652593 | 0.596086296647244 | 0.205918733047386 | 0.837837647627843 | 0.9412811692115 | -5.80923318149824 |
| ENSG00000100813.14 | ACIN1 | 0.0181537808759771 | 7.50410549815934 | 0.205810626847448 | 0.83792155914668 | 0.941293876413201 | -6.37884111605049 |
| ENSG00000272645.3 | GTF2IP20 | -0.035422498703928 | 1.77922414577737 | -0.205742453848217 | 0.8379744756864 | 0.941293876413201 | -5.99550172965136 |
| ENSG00000174292.12 | TNK1 | -0.0419595647255637 | 1.8629467686675 | -0.205711060568959 | 0.837998843707858 | 0.941293876413201 | -6.02131484813316 |
| ENSG00000174564.12 | IL20RB | -0.0285230908636884 | 1.51352972225223 | -0.205642683806671 | 0.838051919523205 | 0.941293876413201 | -5.96584118303 |
| ENSG00000213988.10 | ZNF90 | -0.0647930544315942 | 2.31648197627126 | -0.20561286953615 | 0.838077616438241 | 0.941293876413201 | -6.06189379126196 |
| ENSG00000109536.11 | FRG1 | -0.0314853436995763 | 4.26127383070266 | -0.20535797377282 | 0.838272927033004 | 0.941461868124642 | -6.30435916046438 |
| lnc-OXLD1-3 | lnc-OXLD1-3 | 0.0372388532648277 | 0.363311416415852 | 0.205129965153379 | 0.838449929330575 | 0.941580791918667 | -5.78081849801741 |
| lnc-SLC35C2-3 | lnc-SLC35C2-3 | -0.0325970411677399 | 0.388091619538612 | -0.205093366847306 | 0.838478341256858 | 0.941580791918667 | -5.76250303009413 |
| ENSG00000132356.11 | PRKAA1 | 0.0170061368674386 | 5.87176067502459 | 0.205044789774749 | 0.8385160528528 | 0.941580791918667 | -6.38731339693842 |
| lnc-EPHX2-4 | lnc-EPHX2-4 | 0.0402279476455575 | 0.632633002678868 | 0.204903025323675 | 0.838626110336242 | 0.941631883392685 | -5.79438057665463 |
| ENSG00000198858.9 | R3HDM4 | 0.0293549491387193 | 5.55426408975633 | 0.204868327745131 | 0.838653047969427 | 0.941631883392685 | -6.38093299368873 |
| ENSG00000089127.12 | OAS1 | -0.0614684218882591 | 5.29042907978767 | -0.204657750394984 | 0.838834398838124 | 0.941709463831583 | -6.36765602415059 |
| lnc-MYC-1 | lnc-MYC-1 | -0.0757362917377787 | 1.07224974965819 | -0.204586372200595 | 0.838872365650093 | 0.941709463831583 | -5.8710276045505 |
| ENSG00000188321.13 | ZNF559 | -0.0410455195170343 | 5.24692505512627 | -0.204508640968208 | 0.838932304070391 | 0.941709463831583 | -6.3686067177331 |
| ENSG00000184719.11 | RNLS | 0.0312055512576808 | 1.32371290807663 | 0.204500132836283 | 0.838938909926992 | 0.941709463831583 | -5.948196285963 |
| lnc-COIL-6 | lnc-COIL-6 | 0.0359489234506758 | 2.13120060814558 | 0.204453490280327 | 0.838975124200039 | 0.941709463831583 | -6.05195859392345 |
| ENSG00000168918.13 | INPP5D | 0.0297468571831061 | 8.02890583201021 | 0.204367437751184 | 0.839041938141635 | 0.941709463831583 | -6.36885125650235 |
| ENSG00000189227.5 | C15orf61 | -0.0373914668770625 | 1.2342710381278 | -0.204364293034388 | 0.839044379823302 | 0.941709463831583 | -5.93881781092983 |
| lnc-KLRB1-12 | lnc-KLRB1-12 | -0.0736683769931217 | 0.597097330511299 | -0.204307909466385 | 0.83908815851291 | 0.941709463831583 | -5.84890768654036 |
| ENSG00000180867.11 | PDIA3P1 | 0.037342332568882 | 2.4736300529722 | 0.20416505157672 | 0.839199081988916 | 0.941782602070654 | -6.09036579929423 |
| lnc-WDR49-5 | lnc-WDR49-5 | -0.0315268001523072 | 1.38028783938367 | -0.203815703623155 | 0.839470350728415 | 0.94203566835912 | -5.96071952950434 |
| ENSG00000261740.6 | BOLA2-SMG1P6 | 0.0379735981474737 | 0.498508085708219 | 0.20372115626937 | 0.839543770160498 | 0.942049058339117 | -5.79368826755552 |
| ENSG00000178104.19 | PDE4DIP | 0.0641848254296145 | 5.78342739152399 | 0.203676574571592 | 0.839599800324368 | 0.942049058339117 | -6.38756608637208 |
| lnc-PGM1-1 | lnc-PGM1-1 | -0.0306762494216892 | 1.80345867209865 | -0.203623522558513 | 0.839619587777831 | 0.942049058339117 | -6.02282161917028 |
| ENSG00000147274.14 | RBMX | -0.0183375059714521 | 7.93807540659604 | -0.203491375748037 | 0.839722209053619 | 0.942111462547755 | -6.37116056087599 |
| lnc-LMF2-2 | lnc-LMF2-2 | 0.0439250016761314 | 0.65369938653512 | 0.203397295884127 | 0.839795270392559 | 0.942111462547755 | -5.83479238277452 |
| GAS5-AS1 | GAS5-AS1 | -0.029190987211108 | -0.107826496382353 | -0.203375083141767 | 0.839812520761124 | 0.942111462547755 | -5.62554102832594 |
| ENSG00000162591.15 | MEGF6 | 0.0525851080932416 | 5.57449934769268 | 0.203306851741517 | 0.839874677795377 | 0.942129843210107 | -6.3804902554328 |
| ENSG00000139726.10 | DENR | -0.030738080745901 | 5.0936107056056 | -0.203161378706571 | 0.839978487241768 | 0.942194942937184 | -6.36614326832547 |
| ENSG00000179978.11 | NAIPP2 | -0.0509285218796396 | 1.35229536118362 | -0.202987383923078 | 0.840113619977018 | 0.942198147422657 | -5.95349791533084 |
| lnc-RGPD3-5 | lnc-RGPD3-5 | -0.0295904492815595 | 1.87597134166101 | -0.202937900651647 | 0.840152051962176 | 0.942198147422657 | -6.00964032161048 |
| ENSG00000141699.10 | RETREG3 | -0.0316344353641762 | 6.15710721196721 | -0.202921708495503 | 0.840164627948511 | 0.942198147422657 | -6.38992370588653 |
| ENSG00000099875.14 | MKNK2 | -0.028753540748709 | 6.87989323229171 | -0.202908967155267 | 0.840174523814155 | 0.942198147422657 | -6.38679874726729 |
| ENSG00000172809.12 | RPL38 | 0.0302255022413718 | 7.79237020537275 | 0.202828284995553 | 0.840237188144198 | 0.942198147422657 | -6.3746071685246 |

| | | | | | | | |
|---|---|---|---|---|---|---|---|
| lnc-DOCK4-2 | lnc-DOCK4-2 | -0.0668655256340566 | 0.685709550547694 | -0.202709483457293 | 0.840329460993914 | 0.942198147422657 | -5.8471834506781 |
| lnc-CCDC17-1 | lnc-CCDC17-1 | -0.0330202463998721 | 3.42415330356052 | -0.202697154954777 | 0.84033903664032 | 0.942198147422657 | -6.21183361452017 |
| lnc-VCX-2 | lnc-VCX-2 | -0.0466362956463131 | 0.611212928267316 | -0.202652773978809 | 0.840373507901519 | 0.942198147422657 | -5.81460618144206 |
| ENSG00000179409.10 | GEMIN4 | 0.0315151239114448 | 3.68684148640249 | 0.202568734254799 | 0.840438783478682 | 0.942198147422657 | -6.25065832499319 |
| ENSG00000071859.14 | FAM50A | -0.0318919117735357 | 4.34853484972286 | -0.202568327288858 | 0.840439099581207 | 0.942198147422657 | -6.31550467423973 |
| ENSG00000120137.6 | PANK3 | 0.0261893883651805 | 6.47553699545001 | 0.202385215403505 | 0.840581330720257 | 0.942306275813684 | -6.38999750746025 |
| ENSG00000100934.14 | SEC23A | 0.0254565360771508 | 5.63772973426516 | 0.202190718599912 | 0.840732410934142 | 0.942378057204206 | -6.38412853567221 |
| ENSG00000010244.18 | ZNF207 | -0.0143786062807294 | 8.12574048491182 | -0.202184898848414 | 0.840736931664035 | 0.942378057204206 | -6.36633365711144 |
| ENSG00000127415.12 | IDUA | -0.062963514675296 | 2.60906218399854 | -0.202115324787926 | 0.840795519533559 | 0.942392407893664 | -6.12613565912679 |
| ENSG00000189308.10 | LIN54 | 0.0191146168743739 | 4.59423148567123 | 0.201806991659531 | 0.841030498159659 | 0.942604451054738 | -6.33180135611579 |
| ENSG00000082516.8 | GEMIN5 | -0.0202208105630728 | 4.30159279376692 | -0.201740163883983 | 0.841082413807646 | 0.942611310214678 | -6.30590092765816 |
| ENSG00000232818.2 | RPS2P32 | 0.0798533561661682 | -0.686452934889739 | 0.201674962030139 | 0.841133067036222 | 0.942616753895931 | -5.44173657468657 |
| ENSG00000151725.11 | CENPU | 0.0598854355748486 | 2.32104985719368 | 0.201606791843158 | 0.841186056285978 | 0.942624814706788 | -6.06255828012724 |
| ENSG00000106086.19 | PLEKHA8 | -0.0364342098655035 | 2.55885082017901 | -0.201473383295209 | 0.841289671395623 | 0.942689602391862 | -6.11721114172242 |
| ENSG00000136997.17 | MYC | -0.106236674938522 | 3.67493329539964 | -0.201446589286548 | 0.841343957925757 | 0.942699111920841 | -6.22562144706457 |
| ENSG00000065057.7 | NTHL1 | -0.0465137972629797 | 1.87970112940306 | -0.201341002248738 | 0.841392520350681 | 0.942702207086063 | -6.01916241189771 |
| lnc-ARL16-1 | lnc-ARL16-1 | -0.0345517080018221 | 2.37216733956208 | -0.200916352157893 | 0.841722456716361 | 0.9429637048614 | -6.08477664810136 |
| lnc-ZNF592-6 | lnc-ZNF592-6 | -0.0473374437796988 | 0.805597665370153 | -0.200877956345234 | 0.841752290167785 | 0.9429637048614 | -5.86176368630999 |
| ENSG00000186010.18 | NDUFA13 | -0.0266666017802528 | 4.19124236224555 | -0.20086371735179 | 0.841763353890267 | 0.9429637048614 | -6.2964361708289 |
| ENSG00000160321.14 | ZNF208 | 0.492750231652984 | -0.521818308793685 | 0.200769303186585 | 0.841874755994071 | 0.943037175653113 | -5.24413286214257 |
| ENSG00000130559.18 | CAMSAP1 | -0.0215196283226007 | 4.97091854600235 | -0.200304744829085 | 0.842197702050536 | 0.943176181970903 | -6.35886649848409 |
| ENSG00000175115.11 | PACS1 | 0.0318848850014479 | 6.71630884781067 | 0.200288701532112 | 0.842210169187969 | 0.943176181970903 | -6.38989780346799 |
| ENSG00000170291.14 | ELP5 | 0.0211731246537319 | 3.68268349812596 | 0.200230378048416 | 0.842255492317314 | 0.943176181970903 | -6.24751861509916 |
| ENSG00000168434.12 | COG7 | -0.018045531699868 | 3.68305399437615 | -0.200174554067926 | 0.842298873591796 | 0.943176181970903 | -6.24237647098374 |
| ENSG00000164253.13 | WDR41 | 0.0215303323569289 | 4.61109186216326 | 0.200168511351728 | 0.842303569466 | 0.943176181970903 | -6.33535697124918 |
| lnc-SPINK4-4 | lnc-SPINK4-4 | 0.0528763693977512 | -0.413166694838774 | 0.200141780823311 | 0.842324342180745 | 0.943176181970903 | -5.50786317052357 |
| ENSG00000102531.16 | FNDC3A | 0.0329422930696408 | 6.55555330616424 | 0.200112253084071 | 0.842347288784187 | 0.943176181970903 | -6.39033399831585 |
| ENSG00000284626.1 | | -0.0270190776246102 | 0.356501687867069 | -0.200088902334949 | 0.842365435221838 | 0.943176181970903 | -5.73095579530613 |
| ENSG00000254673.1 | | -0.0203455015471068 | 3.38888723935314 | -0.19998111934146 | 0.842449197142973 | 0.943211017381793 | -6.20755029047497 |
| lnc-PROC-1 | lnc-PROC-1 | -0.0461746718991627 | 3.38074091575967 | -0.199930936263845 | 0.842488196791734 | 0.943211017381793 | -6.20286134271117 |
| ENSG00000226243.1 | RPL37AP1 | 0.151852227901445 | -1.03319555503388 | 0.199809351293412 | 0.842582687895422 | 0.943265499018355 | -5.22120651419738 |
| ENSG00000171812.12 | COL8A2 | -0.109401432465892 | -0.430367087762454 | -0.199617762865284 | 0.842733454581221 | 0.943382971427926 | -5.54962782265688 |
| ENSG00000138646.8 | HERC5 | 0.0457141211568906 | 3.5203764354362 | 0.199509051870946 | 0.842816078966745 | 0.943385399006866 | -6.23576171175769 |
| lnc-POU4F3-5 | lnc-POU4F3-5 | -0.0338945648228217 | 0.20748703056038 | -0.199494627815021 | 0.842827289624314 | 0.943385399006866 | -5.73056956565832 |
| lnc-KLHDC3-1 | lnc-KLHDC3-1 | 0.0336213202562663 | 1.11564664136056 | 0.199314821802517 | 0.842967041131752 | 0.943431499032112 | -5.92050926247854 |
| ENSG00000261371.5 | PECAM1 | -0.0477371295713587 | 6.75513344595214 | -0.199213261488243 | 0.843051333153653 | 0.943431499032112 | -6.3902882289936 |
| ENSG00000160226.15 | CFAP410 | -0.035069224225186 | 4.55680143799495 | -0.199129324941618 | 0.843111221132382 | 0.943431499032112 | -6.33032265592496 |
| lnc-IL2RG-2 | lnc-IL2RG-2 | -0.0428088410817708 | 0.058676328896265 | -0.199077640205875 | 0.843151394778376 | 0.943431499032112 | -5.63515139318547 |
| ENSG00000164620.8 | RELL2 | -0.0359597182463158 | 3.22385346520984 | -0.198953043538751 | 0.843248243337795 | 0.943431499032112 | -6.19822910227284 |
| ENSG00000174547.13 | MRPL11 | -0.0294540858116403 | 3.83161488839064 | -0.198887385962443 | 0.843299279733101 | 0.943431499032112 | -6.26375121849935 |
| ENSG00000139547.7 | RDH16 | -0.0331403278310692 | 0.126911807098934 | -0.19884196077383 | 0.843334589657743 | 0.943431499032112 | -5.69494313194541 |
| lnc-GMDS-6 | lnc-GMDS-6 | 0.0730126513584533 | 1.249437511769 | 0.198842542632995 | 0.843337108809939 | 0.943431499032112 | -5.93210222208412 |
| ENSG00000141985.9 | SH3GL1 | -0.0301306689558909 | 3.76569547850474 | -0.198815927009541 | 0.843354826380182 | 0.943431499032112 | -6.26019491458142 |
| lnc-WHRN-2 | lnc-WHRN-2 | -0.0350559356789557 | 1.73658061466732 | -0.198793911131138 | 0.843371939978957 | 0.943431499032112 | -6.00674437214028 |
| ENSG00000065485.19 | PDIA5 | -0.0274304617292518 | 1.83314488474735 | -0.198792977270207 | 0.843372665898599 | 0.943431499032112 | -6.00950204712272 |
| ENSG00000179820.15 | MYADM | -0.19579677204401 | -1.97363453070563 | -0.198707264122593 | 0.84344286490942 | 0.943458751612989 | -5.13497508739141 |
| lnc-ZNF445-3 | lnc-ZNF445-3 | -0.0395918618330508 | -0.0413404663168315 | -0.198566991218727 | 0.843548336055034 | 0.943479767389755 | -5.68797668123817 |
| ENSG00000079739.16 | PGM1 | -0.0343747347006511 | 4.0734597937293 | -0.198526035048885 | 0.843580174178525 | 0.943479767389755 | -6.29079136279582 |
| ENSG00000181481.13 | RNF135 | 0.0660793194494369 | 3.21875930397735 | 0.19847630705605 | 0.843629962984119 | 0.943479767389755 | -6.17793430015698 |
| ENSG00000213585.10 | VDAC1 | -0.0358146971089415 | 5.16974706196582 | -0.19844005709105 | 0.843647011778737 | 0.943479767389755 | -6.37119032916786 |
| ENSG00000133316.15 | WDR74 | -0.0416322026058626 | 6.62676697831865 | -0.198383676119543 | 0.843690841887764 | 0.943479767389755 | -6.39049888543405 |
| lnc-PCDH12-4 | lnc-PCDH12-4 | 0.0452507930396475 | -0.00825304357228761 | 0.198048292271627 | 0.843951576921322 | 0.943720068874427 | -5.64990186056381 |
| ENSG00000132313.14 | MRPL35 | -0.0375066167039724 | 3.63828514369648 | -0.197912329286046 | 0.844057282686329 | 0.943786997475701 | -6.24209314237712 |
| ENSG00000057608.16 | GDI2 | -0.0364345274155689 | 7.70837703504587 | -0.197745013180523 | 0.844187368235823 | 0.943881177706445 | -6.37731402383758 |
| ENSG00000136536.14 | MARCHF7 | -0.0155305586756338 | 7.10034070647505 | -0.197578520101057 | 0.844316818264831 | 0.943927238923037 | -6.38603093304237 |
| ENSG00000112357.12 | PEX7 | -0.0237548935967156 | 2.21306771981 | -0.19755751013715 | 0.844333154030711 | 0.943927238923037 | -6.0698297733437 |
| ENSG00000137207.11 | YIPF3 | -0.0221245372942374 | 5.02140852051056 | -0.197515082109294 | 0.844366143081716 | 0.943927238923037 | -6.3625463004282 |
| ENSG00000174606.13 | ANGEL2 | -0.0182105965092996 | 5.96535414190199 | -0.197416872337358 | 0.844442505178902 | 0.943954927010285 | -6.38986827258887 |
| ENSG00000142599.17 | RERE | -0.0298655662301037 | 6.02414183158861 | -0.197365265042384 | 0.844482632559267 | 0.943954927010285 | -6.38962294053719 |
| lnc-ARHGAP10-5 | lnc-ARHGAP10-5 | -0.146920184460529 | -0.461596893636948 | -0.197004495998436 | 0.844789536503389 | 0.944208046038553 | -5.72024277283765 |
| lnc-PRRT4-2 | lnc-PRRT4-2 | -0.0378484220102422 | 1.16630013512421 | -0.196956061276521 | 0.844800824750813 | 0.944208046038553 | -5.93402825448145 |
| lnc-MRPS17-1 | lnc-MRPS17-1 | 0.030866584769637 | 1.05681275455128 | 0.196857003127519 | 0.844877855184825 | 0.944242867636926 | -5.900467723735 |
| lnc-MRPS7-4 | lnc-MRPS7-4 | -0.0399470623266921 | 0.719557610699817 | -0.196634788972809 | 0.845050660818362 | 0.944384718841587 | -5.81572995009397 |
| ENSG00000038358.14 | EDC4 | 0.0169408098264539 | 5.81324610455828 | 0.196366381670904 | 0.845259399013637 | 0.944566708827715 | -6.38783984068732 |
| lnc-PKN3-1 | lnc-PKN3-1 | 0.0786406414929879 | -0.477483961103732 | 0.196246224793876 | 0.845352847694938 | 0.944590937186922 | -5.46955107494158 |
| ENSG00000100031.18 | GGT1 | 0.0532157656263474 | 1.38230498722785 | 0.196177381914382 | 0.845406389352144 | 0.944590937186922 | -5.93452728310342 |
| ENSG00000140264.19 | SERF2 | -0.0207161652209251 | 7.62243716854468 | -0.196120222875728 | 0.845450844615444 | 0.944590937186922 | -6.3787211876257 |
| lnc-KCTD18-1 | lnc-KCTD18-1 | 0.20567558178109 | -1.01922180934994 | 0.196137719428345 | 0.845464647320345 | 0.944590937186922 | -5.4912662466971 |

| | | | | | | | |
|---|---|---|---|---|---|---|---|
| lnc-INPP5K-3 | lnc-INPP5K-3 | -0.0354923444283459 | 0.415163096821308 | -0.196021721501122 | 0.84552745494678 | 0.944609835278418 | -5.76422367110694 |
| ENSG00000284518.1 | MIR4658 | 0.0416997330427809 | 0.784119305037228 | 0.195900323907756 | 0.845621875099379 | 0.944664046413597 | -5.87507734580335 |
| ENSG00000168517.10 | HEXIM2 | -0.0393318103944977 | -0.0282640298809751 | -0.195700431065836 | 0.845777351986868 | 0.944786455874618 | -5.63627371053012 |
| ENSG00000100209.9 | HSCB | 0.0340316334191777 | 1.62072200533051 | 0.195475327104345 | 0.845952445558396 | 0.944904824090678 | -5.99607375656815 |
| ENSG00000171291.8 | ZNF439 | -0.0364907583265603 | 5.24830808783627 | -0.195394289975586 | 0.846015480942034 | 0.944904824090678 | -6.36778479497955 |
| ENSG00000266967.6 | AARSD1 | -0.0155843798629425 | 2.85274818632174 | -0.195387147360366 | 0.846021036931793 | 0.944904824090678 | -6.14639684903793 |
| ENSG00000181852.17 | RNF41 | 0.0471498392691201 | 7.60470074535348 | 0.195289645636586 | 0.846101868297314 | 0.944938829857522 | -6.3793850044633 |
| ENSG00000185049.14 | NELFA | 0.029265843715776 | 3.73069512919592 | 0.195213834233467 | 0.846155853561771 | 0.944938829857522 | -6.25937646920381 |
| ENSG00000241120.1 | HMGN1P8 | 0.0435197404418074 | -0.395991209554588 | 0.195108112773311 | 0.846238094349006 | 0.944938829857522 | -5.54031478142375 |
| lnc-WDR49-3 | lnc-WDR49-3 | 0.0435197404418074 | -0.395991209554588 | 0.195108112773311 | 0.846238094349006 | 0.944938829857522 | -5.54031478142375 |
| lnc-RWDD2B-6 | lnc-RWDD2B-6 | 0.0286083256300453 | 3.16622491562975 | 0.195050959318108 | 0.846282554778536 | 0.944938829857522 | -6.18371148589079 |
| ENSG00000244723.3 | ASLP1 | 0.0334189141997239 | 1.36444082877767 | 0.194990875958778 | 0.846329294965767 | 0.944938829857522 | -5.94426419253922 |
| ENSG00000137522.17 | RNF121 | -0.0190115282325212 | 3.78483267537377 | -0.19493489419665 | 0.84637284493287 | 0.944938829857522 | -6.25531574052537 |
| ENSG00000280153.1 | | 0.0408788960502351 | 4.32169289198097 | 0.194858757006141 | 0.846432075219282 | 0.944953702025193 | -6.29944630880125 |
| lnc-PTP4A2-5 | lnc-PTP4A2-5 | -0.048089477409622 | 0.811637083419671 | -0.19467539763442 | 0.846574721786263 | 0.945054033212514 | -5.87981150878745 |
| lnc-NCBP3-1 | lnc-NCBP3-1 | 0.0381167372162144 | -0.0626319640528495 | 0.19456161913692 | 0.84666323971699 | 0.945054033212514 | -5.62562774520757 |
| LINC00513 | LINC00513 | -0.0675131087908897 | 0.127170547787261 | -0.194447141188572 | 0.846752303832153 | 0.945054033212514 | -5.67614299395365 |
| ENSG00000157429.15 | ZNF19 | -0.0244871348222389 | 3.52464951293935 | -0.194359839079883 | 0.846820226442328 | 0.945054033212514 | -6.21525755647623 |
| lnc-SDF2-3 | lnc-SDF2-3 | 0.0374943992773485 | 0.036242973928644 | 0.194294660222107 | 0.846870937533998 | 0.945054033212514 | -5.65190355855034 |
| ENSG00000120278.16 | PLEKHG1 | 0.030331110102904 | 7.85711265805028 | 0.194272090797325 | 0.846888497371436 | 0.945054033212514 | -6.37403956710415 |
| lnc-CLEC2D-2 | lnc-CLEC2D-2 | -0.0613300812439698 | 0.746619545035363 | -0.194248979390622 | 0.846906478972214 | 0.945054033212514 | -5.83225562719087 |
| lnc-ZFP36L1-2 | lnc-ZFP36L1-2 | -0.0519510481038274 | 2.1907701769268 | -0.194223456656908 | 0.846926336777979 | 0.945054033212514 | -6.05205191879681 |
| ENSG00000196655.11 | TRAPPC4 | -0.0282230532433765 | 3.91639038240583 | -0.194212097804681 | 0.846935174495362 | 0.945054033212514 | -6.27954619565695 |
| ENSG00000167306.19 | MYO5B | -0.196810626674428 | 0.0879957172960087 | -0.193894324095906 | 0.847217687966173 | 0.945318028482015 | -5.77346489015069 |
| ENSG00000101190.12 | TCFL5 | -0.0465439284084335 | 4.44167019910361 | -0.193721928457551 | 0.847318580521521 | 0.945361914450696 | -6.31951720285083 |
| lnc-PRR4-1 | lnc-PRR4-1 | -0.0337176514384816 | 3.52358589256133 | -0.193680405547255 | 0.847348878185925 | 0.945361914450696 | -6.22777111849759 |
| lnc-MTIF2-3 | lnc-MTIF2-3 | -0.0385150602059005 | 0.236023395869742 | -0.193560830177442 | 0.847441924424566 | 0.945414478614138 | -5.724157933743 |
| lnc-LIN9-2 | lnc-LIN9-2 | -0.027801840577951 | 3.69266815570027 | -0.193480252730519 | 0.847504626107167 | 0.94542738683 | -6.23961349784118 |
| lnc-ZNF709-2 | lnc-ZNF709-2 | -0.039237494560828 | 0.0551032234036961 | -0.193376286855959 | 0.847585529071035 | 0.94542738683 | -5.66389629882178 |
| ENSG00000165264.10 | NDUFB6 | -0.0355838888533802 | 2.31065745894418 | -0.19334461996983 | 0.84761017157337 | 0.94542738683 | -6.08936675968088 |
| lnc-C16orf89-1 | lnc-C16orf89-1 | -0.0279198996311134 | -0.0094394304126396 | -0.19330985550049 | 0.847637224727242 | 0.94542738683 | -5.67289539962075 |
| ENSG00000078140.13 | UBE2K | 0.0187396172843197 | 5.34929766886911 | 0.193139692935538 | 0.847769645194582 | 0.945513672203296 | -6.37718940514441 |
| lnc-PLEKHO1-3 | lnc-PLEKHO1-3 | -0.0360529161151797 | 0.703689188446241 | -0.193092388294715 | 0.847806458445515 | 0.945513672203296 | -5.8338213812355 |
| ENSG00000127528.5 | KLF2 | 0.0583370714526428 | 5.83680766708401 | 0.193043412740338 | 0.847862776457059 | 0.945525249380486 | -6.38852574768768 |
| ENSG00000240445.3 | FOXO3B | 0.0441199204935663 | 0.848063495871616 | 0.192866496971052 | 0.847982255535897 | 0.945590048492014 | -5.84984710119116 |
| ENSG00000068383.18 | INPP5A | -0.0258156426505165 | 4.10312579524296 | -0.192767526823632 | 0.848059280280325 | 0.945590048492014 | -6.29901922312597 |
| lnc-FAM207A-12 | lnc-FAM207A-12 | -0.0615582682701632 | 0.171707662606711 | -0.192669006877458 | 0.848135956142588 | 0.945590048492014 | -5.69887179749119 |
| ENSG00000141542.10 | RAB40B | 0.0605546619844938 | 0.315059495478038 | 0.192628476365433 | 0.848167500562312 | 0.945590048492014 | -5.74526840161586 |
| ENSG00000144460.12 | NYAP2 | -0.186584421045095 | -1.05978446404562 | -0.192606132922766 | 0.848195649911803 | 0.945590048492014 | -5.24487991433158 |
| ENSG00000264538.6 | SUZ12P1 | 0.0199704771517384 | 3.5728473036115 | 0.192591184538512 | 0.848196524574068 | 0.945590048492014 | -6.23841258069943 |
| ENSG00000166323.12 | C11orf65 | 0.0526525573027288 | 0.218702897808247 | 0.192507306378693 | 0.84826180723637 | 0.945611610612338 | -5.65010895100166 |
| ENSG00000132205.10 | EMILIN2 | -0.0586357978202967 | 2.31778046338708 | -0.192329917062802 | 0.848402744271165 | 0.945717502590489 | -6.10774751520433 |
| ENSG00000107798.17 | LIPA | -0.0289849031743987 | 5.35613716804813 | -0.192022616524558 | 0.848639064126928 | 0.945929700905695 | -6.37605258312907 |
| lnc-RPS16-4 | lnc-RPS16-4 | -0.0434698414563522 | 0.689164093665291 | -0.191583050170106 | 0.848981230329194 | 0.94625985075355 | -5.82066167666131 |
| ENSG00000106948.16 | AKNA | 0.0250694027983087 | 7.78868228850184 | 0.1915060603297 | 0.849041163613296 | 0.946270860296115 | -6.37718529877749 |
| ENSG00000179364.13 | PACS2 | -0.0350823324898624 | 5.43608076117789 | -0.191452247803397 | 0.849083054890393 | 0.946270860296115 | -6.38084490534098 |
| ENSG00000215252.11 | GOLGA8B | 0.0332352004919709 | 6.5385945306696 | 0.19137168899276 | 0.849145768092701 | 0.946276845453375 | -6.39234839772032 |
| ENSG00000185432.11 | TMT1A | 0.0436879594838251 | 7.34469481165374 | 0.191285752933791 | 0.84921552310465 | 0.946276845453375 | -6.38507686864612 |
| ENSG00000198663.16 | C6orf89 | 0.0159111108296042 | 6.22739892134943 | 0.191268183185455 | 0.849226346461387 | 0.946276845453375 | -6.39240505792113 |
| ENSG00000139546.10 | TARBP2 | 0.0322783710109094 | 3.28522608610829 | 0.190941046875885 | 0.849481029948723 | 0.94650939422046 | -6.20534844020315 |
| ENSG00000267168.1 | | -0.0193488321973445 | 1.97309207310306 | -0.190778018251204 | 0.849607957731101 | 0.946569972741283 | -6.03407042585829 |
| ENSG00000197008.9 | ZNF138 | -0.0199892122426668 | 4.44030997132804 | -0.190748371251535 | 0.849631040172449 | 0.946569972741283 | -6.3163032428601 |
| ENSG00000135632.11 | SMYD5 | -0.0221306868607698 | 4.78086991840667 | -0.190693505280549 | 0.849673757850803 | 0.946569972741283 | -6.35307453715536 |
| lnc-SNRPC-2 | lnc-SNRPC-2 | 0.0496905149172426 | 0.311395724860241 | 0.190634947698084 | 0.849719350257041 | 0.946569972741283 | -5.72637353629245 |
| lnc-GUF1-5 | lnc-GUF1-5 | 0.0593135248075738 | -0.372132169304167 | 0.190210545394836 | 0.850049801614057 | 0.946874535578756 | -5.56792796311206 |
| ENSG00000255508.7 | | 0.0277979610823111 | 7.74445054500971 | 0.190105860689382 | 0.850131316214426 | 0.946874535578756 | -6.37767001709184 |
| lnc-DYNLL2-6 | lnc-DYNLL2-6 | -0.0708382934379917 | -0.0818227214061758 | -0.190096415291031 | 0.850138671122786 | 0.946874535578756 | -5.64142458682389 |
| ENSG00000109586.11 | GALNT7 | 0.02305399015445 | 3.98273076053519 | 0.189973398536281 | 0.850234462594097 | 0.946874535578756 | -6.28366686450274 |
| lnc-JPH3-8 | lnc-JPH3-8 | 0.0405803963317953 | 0.800897673356089 | 0.189945116788638 | 0.850256485529483 | 0.946874535578756 | -5.84543211350293 |
| lnc-ZNF540-1 | lnc-ZNF540-1 | -0.0519904004120298 | 0.366691752719358 | -0.189929344430551 | 0.850268767482978 | 0.946874535578756 | -5.73064382106289 |
| ENSG00000179476.7 | C14orf28 | 0.0334417619464871 | 2.85149219089376 | 0.189759298968768 | 0.850401184469792 | 0.946970762818748 | -6.15400212331312 |
| ENSG00000214435.7 | AS3MT | -0.0557853130597654 | -0.103530701707353 | -0.189620270763911 | 0.850509451093965 | 0.94704008828061 | -5.59746942160378 |
| lnc-SEC24D-1 | lnc-SEC24D-1 | 0.0479529105698671 | -0.245310613249559 | 0.189516542161744 | 0.850590230457774 | 0.947040915955974 | -5.58210524329801 |
| ENSG00000213471.10 | TTLL13 | 0.0393171375731437 | -0.252232527288745 | 0.18945903963868 | 0.850635011643626 | 0.947040915955974 | -5.51645233577001 |
| ENSG00000157483.8 | MYO1E | -0.0385276098101492 | 6.87380213520053 | -0.189438405038374 | 0.850651081353566 | 0.947040915955974 | -6.39137093591012 |
| ENSG00000064687.12 | ABCA7 | -0.0343747802684424 | 6.39790684440318 | -0.189382989729997 | 0.850694237725793 | 0.947040915955974 | -6.3929123550229 |
| ENSG00000135144.7 | DTX1 | -0.108772928500604 | 4.20762547128304 | -0.18935279685205 | 0.850752168667868 | 0.947054185380785 | -6.34515038077754 |
| lnc-NKRF-2 | lnc-NKRF-2 | -0.0439506966857617 | -0.417010216492779 | -0.189168715133504 | 0.850861115001347 | 0.947069857725833 | -5.57061703338011 |

| | | | | | | | |
|---|---|---|---|---|---|---|---|
| ENSG00000152642.10 | GPD1L | -0.033936333200717 | 2.86933134775394 | -0.18911540695698 | 0.850902632538789 | 0.947069857725833 | -6.15088332804215 |
| ENSG00000179918.18 | SEPHS2 | 0.0451050155197854 | 3.13979689598083 | 0.189010632451218 | 0.850984234391287 | 0.947069857725833 | -6.18238737602276 |
| ENSG00000221821.3 | C6orf226 | 0.0357613285809386 | 0.206710092982593 | 0.189006990369532 | 0.850987070994762 | 0.947069857725833 | -5.74236646220203 |
| ENSG00000175265.17 | GOLGA8A | -0.0448397130844551 | 5.73554932001658 | -0.188950949178729 | 0.851034904914235 | 0.947069857725833 | -6.38772551252893 |
| ENSG00000200913.1 | SNORD46 | -0.0396768707983987 | 0.631172748135422 | -0.188865593715147 | 0.851097198086014 | 0.947069857725833 | -5.84582099541154 |
| ENSG00000026508.18 | CD44 | 0.0348812135374246 | 7.79143551444307 | 0.188829990132832 | 0.851124928487494 | 0.947069857725833 | -6.37746265385506 |
| lnc-THUMPD3-9 | lnc-THUMPD3-9 | 0.0298807783192383 | 0.634547619673785 | 0.188705700753805 | 0.851221734689526 | 0.947069857725833 | -5.82581519406198 |
| ENSG00000148483.8 | TMEM236 | 0.0641098817452359 | -0.317598901103004 | 0.188700949787774 | 0.851225435156202 | 0.947069857725833 | -5.49506465607459 |
| lnc-NBN-1 | lnc-NBN-1 | 0.0263210585408302 | 5.29043539512432 | 0.188668744160292 | 0.851250519796984 | 0.947069857725833 | -6.37673375033136 |
| ENSG00000183785.14 | TUBA8 | 0.0341934639330569 | -0.0631583412888072 | 0.188640675971546 | 0.851272381955285 | 0.947069857725833 | -5.6530849518415 |
| ENSG00000120802.13 | TMPO | 0.0243361287144603 | 7.49744883310879 | 0.18849197056703 | 0.851388209754022 | 0.947118297088893 | -6.38129241270962 |
| ENSG00000141480.17 | ARRB2 | 0.0287763254275708 | 4.09986209062462 | 0.188466626487488 | 0.851407950786367 | 0.947118297088893 | -6.29161337807869 |
| lnc-ATP6V1D-8 | lnc-ATP6V1D-8 | -0.205272884719661 | 2.72232678696757 | -0.188182833382268 | 0.851664586166024 | 0.94712421378349 | -6.06219911691479 |
| SLC2A1-AS1 | SLC2A1-AS1 | 0.0672673427569916 | 1.87819594942654 | 0.188124268404843 | 0.851679115563796 | 0.94712421378349 | -5.99994971554198 |
| lnc-SLC15A1-3 | lnc-SLC15A1-3 | 0.0365209362950535 | -0.300535511374957 | 0.188100990849429 | 0.851692762738394 | 0.94712421378349 | -5.56203481385585 |
| ENSG00000279024.1 | | -0.0824013859207769 | -0.308841923411715 | -0.188100043299926 | 0.851693500858308 | 0.94712421378349 | -5.59112698456519 |
| lnc-UBXN4-13 | lnc-UBXN4-13 | -0.0824013859207769 | -0.308841923411715 | -0.188100043299926 | 0.851693500858308 | 0.94712421378349 | -5.59112698456519 |
| ENSG00000174891.12 | RSRC1 | 0.0140686830304775 | 4.89502122059287 | 0.188049598548255 | 0.85173279638813 | 0.94712421378349 | -6.35530800825729 |
| ENSG00000001631.15 | KRIT1 | -0.0186749404519615 | 5.12671404160311 | -0.187931483197725 | 0.851824807555718 | 0.94712421378349 | -6.36804874737934 |
| lnc-ZNF529-1 | lnc-ZNF529-1 | -0.0220243466846007 | 0.0247689450443765 | -0.187872693807847 | 0.851870604925394 | 0.94712421378349 | -5.66046828819188 |
| lnc-NPIPA7-12 | lnc-NPIPA7-12 | 0.0842365533230973 | 0.299894328365062 | 0.18781705051822 | 0.851913951936219 | 0.94712421378349 | -5.67580556137551 |
| ENSG00000092439.14 | TRPM7 | 0.0134408902391172 | 7.23561194560732 | 0.187803192905647 | 0.851924747309677 | 0.94712421378349 | -6.38611351832648 |
| lnc-PHF8-1 | lnc-PHF8-1 | -0.0350569458890401 | 0.512655489680312 | -0.187761919482289 | 0.851956900350147 | 0.94712421378349 | -5.8051515911557 |
| lnc-ERN1-4 | lnc-ERN1-4 | -0.0444788237206715 | 0.776159519708071 | -0.187750947174807 | 0.851965448098009 | 0.94712421378349 | -5.827762432467 |
| lnc-ATP6V0E1-2 | lnc-ATP6V0E1-2 | -0.044693837772348 | 0.363744706172097 | -0.187626604170785 | 0.852062316186449 | 0.947168048585635 | -5.73287721324377 |
| lnc-NAV1-8 | lnc-NAV1-8 | -0.0259418451174989 | 1.38509592071628 | -0.187582195591693 | 0.852096912775602 | 0.947168048585635 | -5.95799177770838 |
| lnc-GNRH1-3 | lnc-GNRH1-3 | 0.0304027824683606 | 1.09777248613716 | 0.187253196738275 | 0.852353229142247 | 0.947335424095437 | -5.90540084944315 |
| lnc-CLEC2D-1 | lnc-CLEC2D-1 | 0.0450579791650236 | 7.40572673571922 | 0.187247917749855 | 0.852362117290436 | 0.947335424095437 | -6.38448979712944 |
| ENSG00000142227.10 | EMP3 | 0.0412568664277879 | 5.55202217368858 | 0.187166985803434 | 0.85242167683941 | 0.947335424095437 | -6.38423516030803 |
| ENSG00000109906.13 | ZBTB16 | 0.167774195601176 | -0.186175365766959 | 0.187184731060014 | 0.85243158867374 | 0.947335424095437 | -5.57649009257165 |
| ENSG00000132109.9 | TRIM21 | -0.0227562789746552 | 4.37586930628944 | -0.186958417665065 | 0.85258289928673 | 0.947354003869829 | -6.31719837029629 |
| lnc-C2CD2-3 | lnc-C2CD2-3 | 0.0443697354744513 | 0.862364057957819 | 0.186940341609403 | 0.852596983239505 | 0.947354003869829 | -5.86328503105908 |
| lnc-ATF4-1 | lnc-ATF4-1 | 0.0343458122484637 | -0.184806550026221 | 0.186899758627157 | 0.852628603633961 | 0.947354003869829 | -5.59538656664041 |
| ENSG00000196497.16 | IPO4 | 0.0282263198818997 | 3.28895217864919 | 0.186842335615787 | 0.852673345424151 | 0.947354003869829 | -6.20720497812347 |
| ENSG00000088833.17 | NSFL1C | -0.0158869012361329 | 4.92164488503938 | -0.186835800536627 | 0.85267843733627 | 0.947354003869829 | -6.35880537191645 |
| ENSG00000272034.1 | SNORD14A | 0.0313265451186469 | 1.28532362545575 | 0.186573627345943 | 0.85288271903968 | 0.94746109952259 | -5.95210828917486 |
| ENSG00000244491.1 | | -0.0323837256409095 | 0.387153234571531 | -0.18652049645658 | 0.852924119135438 | 0.94746109952259 | -5.75113348515847 |
| ENSG00000188603.19 | CLN3 | -0.0297105824336275 | 1.3730020202202 | -0.18647643864611 | 0.852958449719815 | 0.94746109952259 | -5.94284411571178 |
| lnc-ARHGEF3-2 | lnc-ARHGEF3-2 | -0.0275674501938874 | 3.0384439382159 | -0.186427060685184 | 0.852996926197399 | 0.94746109952259 | -6.17363419439731 |
| ENSG00000271936.1 | | -0.035396061302811 | -0.19427509669549 | -0.18636624005541 | 0.853044319571641 | 0.94746109952259 | -5.6167406637562 |
| ENSG00000241945.7 | PWP2 | 0.186899859481551 | 0.421802213765066 | 0.186399174523491 | 0.853051017653046 | 0.94746109952259 | -5.68049404396256 |
| ENSG00000234127.8 | TRIM26 | -0.0184285872902574 | 5.70462682974917 | -0.186282707190851 | 0.853109411938907 | 0.947474830083555 | -6.3878441489171 |
| lnc-RMDN2-3 | lnc-RMDN2-3 | -0.0898556590360053 | 4.78584214203066 | -0.186199033574461 | 0.853206146256506 | 0.947531137822255 | -6.34225112562671 |
| ENSG00000173852.14 | DPY19L1 | -0.0481042498534592 | 2.59402056684158 | -0.18602964392154 | 0.853306615941836 | 0.947591587587311 | -6.13657025139246 |
| lnc-GBA2-2 | lnc-GBA2-2 | 0.0225389830001319 | 2.33848023178877 | 0.185822919574546 | 0.853467716571246 | 0.947707012445816 | -6.07987371172358 |
| ENSG00000154319.15 | FAM167A | -0.144840282636001 | 1.07747891903739 | -0.185776792723525 | 0.853534628000848 | 0.947707012445816 | -5.87214846857161 |
| ENSG00000142676.14 | RPL11 | 0.0324323517662943 | 9.59528601644621 | 0.185719023110951 | 0.853548685649348 | 0.947707012445816 | -6.33080379552969 |
| lnc-MED10-16 | lnc-MED10-16 | -0.0353812902605708 | 0.999443745225841 | -0.185526957102599 | 0.853698371636057 | 0.947759442150946 | -5.88827168261246 |
| ENSG00000141027.20 | NCOR1 | 0.0246453939596103 | 7.68133179462554 | 0.185504518728863 | 0.85371585926186 | 0.947759442150946 | -6.37937753328562 |
| ENSG00000277578.1 | | 0.0301328552985642 | -0.228436388187809 | 0.185452708969209 | 0.85375623812716 | 0.947759442150946 | -5.60712853640342 |
| ENSG00000102078.15 | SLC25A14 | -0.018631482515632 | 2.91078536144682 | -0.185396516661983 | 0.853800033058046 | 0.947759442150946 | -6.15357102271879 |
| lnc-PEX16-1 | lnc-PEX16-1 | 0.0241569427744891 | 0.720346320504453 | 0.185363026178404 | 0.853826134956274 | 0.947759442150946 | -5.8141459225206 |
| ENSG00000255438.2 | | 0.106248019530784 | 1.07379606593052 | 0.185201010719478 | 0.853974408026737 | 0.94782281128863 | -5.90066030241872 |
| lnc-FAM177B-3 | lnc-FAM177B-3 | -0.0243940480596991 | 1.83198774954969 | -0.185072792457644 | 0.854052344989215 | 0.94782281128863 | -6.01954777384103 |
| ENSG00000128309.16 | MPST | -0.0944879894181958 | 1.44650209709823 | -0.185086297175634 | 0.854064242255323 | 0.94782281128863 | -6.01578878390601 |
| ENSG00000144895.11 | EIF2A | 0.0193057419962157 | 5.97391289856494 | 0.185044281543671 | 0.854074567250001 | 0.94782281128863 | -6.39245387368558 |
| ENSG00000085063.15 | CD59 | 0.0614213040846652 | 4.5715219843123 | 0.185018740970924 | 0.854113467608294 | 0.94782281128863 | -6.33078787097442 |
| ENSG00000183431.11 | SF3A3 | -0.0187958549726224 | 5.55718919491281 | -0.184763672576374 | 0.854293288683202 | 0.947860634764042 | -6.38456739789772 |
| lnc-PROM1-4 | lnc-PROM1-4 | -0.0583474906136918 | 2.66301883292491 | -0.184749785900942 | 0.854310100620395 | 0.947860634764042 | -6.14570774631655 |
| ENSG00000162614.18 | NEXN | 0.077853380290906 | -0.287802850589883 | 0.184722866422413 | 0.854325096118711 | 0.947860634764042 | -5.59150435665504 |
| ENSG00000165891.15 | E2F7 | -0.0886972511395889 | 2.29182446409327 | -0.18465624626452 | 0.854398582697028 | 0.947860634764042 | -6.04984121574689 |
| lnc-RNF7-1 | lnc-RNF7-1 | 0.0472244484435246 | -0.22641376208221 | 0.184556043290965 | 0.854455133368617 | 0.947860634764042 | -5.65544806548915 |
| ENSG00000182541.17 | LIMK2 | 0.0171395739272651 | 3.34483981700019 | 0.184507290098681 | 0.854493136851722 | 0.947860634764042 | -6.20526479727988 |
| ENSG00000273136.6 | NBPF26 | -0.050168094634309 | 3.11512616230274 | -0.184492832513215 | 0.854505674680021 | 0.947860634764042 | -6.1829428831446 |
| lnc-DPM2-3 | lnc-DPM2-3 | -0.0577489747033746 | 3.02366037469778 | -0.184474006990588 | 0.854525212613074 | 0.947860634764042 | -6.15823579475696 |
| ENSG00000157796.17 | WDR19 | 0.0352918347161815 | 5.69685192182347 | 0.184418939827797 | 0.854562007448688 | 0.947860634764042 | -6.38807536390852 |
| lnc-GALNT4-1 | lnc-GALNT4-1 | 0.0176512346195272 | 2.04883246103909 | 0.184057371898573 | 0.854843867989989 | 0.948122175602864 | -6.04690732160549 |
| ENSG00000142444.6 | TIMM29 | -0.0250997663840251 | 2.65190963270558 | -0.183926400168013 | 0.854945971832592 | 0.948184327724028 | -6.13032781454469 |

| | | | | | | | |
|---|---|---|---|---|---|---|---|
| ENSG00000108559.11 | NUP88 | 0.0231449352713797 | 7.14105178740771 | 0.18375104289007 | 0.855082682003906 | 0.94828484474561 | -6.38843204083912 |
| lnc-SEC63-3 | lnc-SEC63-3 | 0.0341661539547151 | 2.07803932448824 | 0.183690291725854 | 0.85513004522978 | 0.94828484474561 | -6.05906390179448 |
| lnc-PGAP2-1 | lnc-PGAP2-1 | 0.0339358967177654 | 0.549068863404493 | 0.183632863308684 | 0.855174818450567 | 0.94828484474561 | -5.81333585008698 |
| lnc-TIMM13-2 | lnc-TIMM13-2 | -0.0497595520493073 | 0.457599798322903 | -0.183524343827617 | 0.85525942538616 | 0.948294325463328 | -5.79623161386271 |
| lnc-RNPEPL1-1 | lnc-RNPEPL1-1 | 0.0201813190988948 | 0.908744619927656 | 0.183486959395864 | 0.855288572461817 | 0.948294325463328 | -5.87282377387843 |
| ENSG00000126226.21 | PCID2 | 0.0151676612317484 | 5.04711920931451 | 0.183405470033916 | 0.855352107012873 | 0.948294325463328 | -6.36276263961352 |
| ENSG00000149930.17 | TAOK2 | -0.0235691971899454 | 4.79044906800747 | -0.183385528158622 | 0.855367655179136 | 0.948294325463328 | -6.35074344010414 |
| ENSG00000132541.10 | RIDA | -0.0488616952716008 | 1.31435030704056 | -0.183253705977563 | 0.855470434999077 | 0.948357190907847 | -5.9450403342801 |
| ENSG00000166479.9 | TMX3 | -0.0186105531638514 | 5.45388975895465 | -0.18313180209738 | 0.855565483925317 | 0.948411479730439 | -6.38390473437118 |
| ENSG00000198863.7 | RUNDC1 | -0.0252888742923449 | 4.33064411873943 | -0.182786540854686 | 0.85583469722249 | 0.948654338221881 | -6.31477810437482 |
| ENSG00000065970.8 | FOXJ2 | 0.022796259135017 | 4.21621002538974 | 0.182732616195281 | 0.855876745896144 | 0.948654338221881 | -6.3001746074038 |
| ENSG00000196465.10 | MYL6B | -0.0286541929107802 | 5.56556102849034 | -0.182550225422231 | 0.856018971354658 | 0.948760889957079 | -6.38550895102684 |
| lnc-C12orf76-2 | lnc-C12orf76-2 | -0.0264351382477428 | 3.17054859495603 | -0.182024281220299 | 0.856429121480526 | 0.949119337033492 | -6.17899525045305 |
| ENSG00000208028.1 | MIR616 | 0.0598152602508143 | 0.457333632158051 | 0.181999770097895 | 0.856448237111959 | 0.949119337033492 | -5.78008816042774 |
| lnc-MAFA-6 | lnc-MAFA-6 | 0.0364119296463934 | 2.34267678631085 | 0.18194940639688 | 0.856487514817762 | 0.949119337033492 | -6.07371926856309 |
| ENSG00000047644.18 | WWC3 | 0.0357983939594471 | 6.02915127569823 | 0.181801173952819 | 0.856603120656408 | 0.949119337033492 | -6.39375789375348 |
| lnc-INTS8-1 | lnc-INTS8-1 | -0.0607370614749516 | 0.392494123366042 | -0.181796907066348 | 0.856606448429806 | 0.949119337033492 | -5.77492120664578 |
| ENSG00000182903.15 | ZNF721 | -0.0258722081282263 | 6.55459378486965 | -0.181773187476332 | 0.856624947545397 | 0.949119337033492 | -6.39399212136018 |
| ENSG00000205903.6 | ZNF316 | 0.0340939041968102 | 4.5370652392873 | 0.181657180854965 | 0.856715423298327 | 0.949119337033492 | -6.33683116097753 |
| lnc-RWDD2B-3 | lnc-RWDD2B-3 | 0.0397416344610306 | 2.05399730481791 | 0.181573664887365 | 0.856780560184652 | 0.949119337033492 | -6.0515323174558 |
| ENSG00000075624.14 | ACTB | 0.0323048273613367 | 9.80735983384166 | 0.181544526094931 | 0.856803286737673 | 0.949119337033492 | -6.32151768257518 |
| ENSG00000009780.15 | FAM76A | 0.0184713501033409 | 3.93780095204188 | 0.181522752157488 | 0.856820269215032 | 0.949119337033492 | -6.28055875969235 |
| ENSG00000128218.7 | VPREB3 | -0.0570149919987877 | 3.73706475738894 | -0.181498870544461 | 0.856849609908971 | 0.949119337033492 | -6.24383983057637 |
| lnc-LSM12-3 | lnc-LSM12-3 | -0.0581107667300331 | -0.0716222105553126 | -0.181310182377317 | 0.856986065593804 | 0.949218526338975 | -5.58534292186638 |
| lnc-GHRL-2 | lnc-GHRL-2 | -0.0244484851918643 | 1.22320237776952 | -0.181252073230686 | 0.857031389665429 | 0.949218526338975 | -5.93626099004268 |
| lnc-NPHS2-6 | lnc-NPHS2-6 | 0.0289889571102262 | 3.2207125642453 | 0.181157464851302 | 0.857105183515541 | 0.949249179031498 | -6.1913753575659 |
| ENSG00000274810.4 | NPHP3-ACAD11 | -0.021073344327073 | 5.23554836822205 | -0.180869457143016 | 0.8573298354001 | 0.949417975225913 | -6.37297452699244 |
| ENSG00000269378.1 | ITGB1P1 | 0.0725760252598171 | 0.701685214356084 | 0.180812771052304 | 0.857374053116897 | 0.949417975225913 | -5.76029059932008 |
| ENSG00000182197.11 | EXT1 | 0.0425278529033091 | 2.85662087778639 | 0.180784669789145 | 0.857395973546095 | 0.949417975225913 | -6.14362511120874 |
| lnc-CXXC1-3 | lnc-CXXC1-3 | -0.0231415671048889 | 1.76971379959103 | -0.180673197363934 | 0.857482928887405 | 0.949463183887754 | -6.02135205381385 |
| lnc-MAP3K7CL-1 | lnc-MAP3K7CL-1 | -0.110372278778242 | -1.19082621655128 | -0.180466415831266 | 0.857644235931145 | 0.949590710498696 | -5.33757440590892 |
| SLC16A1-AS1 | SLC16A1-AS1 | -0.0160798446417614 | 4.37919009229656 | -0.180381161955374 | 0.857710742940861 | 0.949598409523162 | -6.31682094837774 |
| ENSG00000105607.12 | GCDH | 0.0221373935549765 | 3.20900456408667 | 0.180339223623715 | 0.857743459644106 | 0.949598409523162 | -6.18689713122146 |
| ENSG00000170889.13 | RPS9 | -0.0269748896131752 | 8.03430652434954 | -0.180239463038087 | 0.857821285346574 | 0.94960021261962 | -6.3722449315696 |
| ENSG00000139719.10 | VPS33A | 0.014489343757746 | 4.47870325246899 | 0.180218859740626 | 0.857837358667309 | 0.94960021261962 | -6.32800211817401 |
| ENSG00000226479.3 | TMEM185B | 0.0322378986589831 | 3.72801087171276 | 0.1800325673768 | 0.857982694334918 | 0.949710018687584 | -6.25650095517992 |
| lnc-WNK1-3 | lnc-WNK1-3 | 0.0328195689272912 | -0.252544295263207 | 0.179752970753339 | 0.858200830514819 | 0.949900392261052 | -5.58892460667567 |
| lnc-USF2-2 | lnc-USF2-2 | -0.0799342223581878 | 1.79655516944903 | -0.179667986164309 | 0.858282435287921 | 0.949939633571612 | -5.99221644040231 |
| ENSG00000159596.6 | TMEM69 | -0.0228970301419267 | 3.04284262533185 | -0.17948007817281 | 0.858413747135797 | 0.950033883067863 | -6.1712070381194 |
| ENSG00000008128.22 | CDK11A | 0.0295501808023806 | 5.02765352231211 | 0.179393000536235 | 0.858481689213497 | 0.950057992853455 | -6.36525783486348 |
| lnc-RAB1A-2 | lnc-RAB1A-2 | -0.0349654275891913 | 2.13851933368021 | -0.179176328086364 | 0.858650751984623 | 0.950194001510726 | -6.04626042231136 |
| ENSG00000142534.6 | RPS11 | 0.0295474610533244 | 10.0188573016362 | 0.179087666303405 | 0.858719933951181 | 0.950219472099197 | -6.31490750661807 |
| ENSG00000101986.11 | ABCD1 | 0.0434655012150649 | 3.09463094959511 | 0.178998010975571 | 0.858789892314036 | 0.950237999890829 | -6.17592603347854 |
| ENSG00000004142.11 | POLDIP2 | -0.0215023467848425 | 4.65142136087804 | -0.178933932742496 | 0.858839893484908 | 0.950237999890829 | -6.34212015980412 |
| ENSG00000214114.8 | MYCBP | -0.0336661239925422 | 1.6396755092019 | -0.178888717282029 | 0.858875176114706 | 0.950237999890829 | -5.99888781217772 |
| ENSG00000174469.21 | CNTNAP2 | 0.334691413183037 | 1.9919050911855 | 0.178839855594225 | 0.85894706078624 | 0.950266452682783 | -5.66399817368931 |
| ENSG00000174851.15 | YIF1A | -0.0319928681775514 | 3.18905856669908 | -0.178634450288852 | 0.859073591740406 | 0.950355355196859 | -6.19949763145332 |
| ENSG00000145685.13 | LHFPL2 | -0.0695736330541195 | 4.57287110102526 | -0.178558751651515 | 0.859156494029334 | 0.950384577643491 | -6.33104544098935 |
| ENSG00000129595.12 | EPB41L4A | 0.130206009625375 | 1.31769740278645 | 0.178355850495961 | 0.85931776676459 | 0.950384577643491 | -5.85487457425865 |
| ENSG00000125691.12 | RPL23 | 0.0299563578244584 | 8.87588364414706 | 0.178265877977049 | 0.859361221153327 | 0.950384577643491 | -6.35246689640792 |
| lnc-CD53-4 | lnc-CD53-4 | -0.0628110121662444 | 0.218384056094072 | -0.178256551648676 | 0.85936849955907 | 0.950384577643491 | -5.68558276739255 |
| ENSG00000130522.5 | JUND | -0.0453193097985647 | 6.58560400869992 | -0.178266189584845 | 0.859368752416926 | 0.950384577643491 | -6.39367768386338 |
| lnc-ARNTL-4 | lnc-ARNTL-4 | 0.0365939127715934 | -0.091557657674835 | 0.178245599193792 | 0.859377047033992 | 0.950384577643491 | -5.65686317948129 |
| lnc-PAWR-10 | lnc-PAWR-10 | -0.0220904513501958 | 4.1220212769474 | -0.178095522961882 | 0.859494170698606 | 0.950416075676115 | -6.29558415393096 |
| lnc-TMEM123-2 | lnc-TMEM123-2 | 0.0566535873006888 | 4.35919849494269 | 0.178064013093591 | 0.85953353458347 | 0.950416075676115 | -6.30792178158572 |
| ENSG00000130520.10 | LSM4 | -0.030829042387996 | 3.84236976807759 | -0.178031607233919 | 0.859544053282363 | 0.950416075676115 | -6.26048178954416 |
| ENSG00000188994.12 | ZNF292 | 0.0262285448558259 | 7.68976734720083 | 0.177696215075394 | 0.859805817262641 | 0.950654444387459 | -6.38123029225923 |
| lnc-BRCA2-2 | lnc-BRCA2-2 | 0.0757653403822829 | 1.11066773662663 | 0.177595082853561 | 0.859895410502161 | 0.95070243510587 | -5.95769277861296 |
| ENSG00000213625.8 | LEPROT | 0.0233190285509384 | 4.84712691895778 | 0.177393224871835 | 0.860042306163742 | 0.950813770961881 | -6.35562450595235 |
| ENSG00000146072.6 | TNFRSF21 | -0.151079755840482 | -2.04966858491991 | -0.177298253836604 | 0.860116435315943 | 0.950844652672434 | -5.12653018950628 |
| ENSG00000274523.4 | RCC1L | -0.0133938625016645 | 4.67387852063444 | -0.17715660941223 | 0.860226997513256 | 0.950861270431523 | -6.3438255632162 |
| ENSG00000283803.1 | MIR3198-2 | 0.0338182142731306 | 0.297106894529056 | 0.177054827346966 | 0.860306446436331 | 0.950861270431523 | -5.7480739515604 |
| lnc-IER5-3 | lnc-IER5-3 | -0.06173757843876 | -0.543549463831785 | -0.177035418324753 | 0.860321596874006 | 0.950861270431523 | -5.50706131468379 |
| lnc-CCKAR-1 | lnc-CCKAR-1 | -0.0609418867322235 | -0.183364254362893 | -0.176994039432415 | 0.860353896892653 | 0.950861270431523 | -5.65066414367583 |
| ENSG00000105677.11 | TMEM147 | 0.0278977158717928 | 3.34160052767763 | 0.176945063134025 | 0.86039212769583 | 0.950861270431523 | -6.21304828277813 |
| ENSG00000197744.5 | PTMAP2 | 0.070355676595558 | 1.50430827156913 | 0.176878832362966 | 0.860448846723614 | 0.950861270431523 | -5.9158336978803 |
| ENSG00000134802.17 | SLC43A3 | -0.0375767246069598 | 2.04585997516411 | -0.176864722087481 | 0.860454842494173 | 0.950861270431523 | -6.04501559181538 |
| ENSG00000114942.13 | EEF1B2 | -0.0311167882828691 | 8.45737653928821 | -0.176703983732212 | 0.860580318745066 | 0.950948875327733 | -6.36391624701242 |

| | | | | | | | |
|---|---|---|---|---|---|---|---|
| ENSG00000168282.5 | MGAT2 | 0.0256822830183685 | 4.78581949451664 | 0.176626381058291 | 0.86064089857568 | 0.950964763548595 | -6.35338285075964 |
| lnc-PAIP2-6 | lnc-PAIP2-6 | 0.0448448613690265 | 0.818471114709894 | 0.17648993194508 | 0.860747418409981 | 0.951031408724711 | -5.86352485738066 |
| ENSG00000135124.14 | P2RX4 | -0.020629918772304 | 4.42614066754732 | -0.176273722084941 | 0.860916209330537 | 0.951045519902744 | -6.32298539391024 |
| ENSG00000162881.6 | OXER1 | 0.0910599089961001 | -0.397730864771959 | 0.176243033809411 | 0.860941580583042 | 0.951045519902744 | -5.59825982284057 |
| ENSG00000118971.7 | CCND2 | -0.04490656315983 | 6.3992661574056 | -0.176197404001292 | 0.860983612918939 | 0.951045519902744 | -6.39533076553416 |
| ENSG00000135931.17 | ARMC9 | -0.0731192238723362 | -0.421026537814192 | -0.176162471058326 | 0.861003063452781 | 0.951045519902744 | -5.4938942446222 |
| ENSG00000114902.13 | SPCS1 | 0.0204704162831337 | 4.66086551325095 | 0.176072729811416 | 0.861073126072487 | 0.951045519902744 | -6.34296402479753 |
| ENSG00000107736.20 | CDH23 | -0.054520374214733 | 0.339907480270469 | -0.176064281080285 | 0.861079722208499 | 0.951045519902744 | -5.78473921812198 |
| lnc-TFCP2-2 | lnc-TFCP2-2 | -0.0505333182135494 | -0.176861426990469 | -0.17602549803497 | 0.861110001232147 | 0.951045519902744 | -5.61689582963149 |
| ENSG00000285839.1 | | 0.0178291293661594 | 3.88724864538156 | 0.175951628148484 | 0.861167674132824 | 0.951045519902744 | -6.27187069999488 |
| lnc-ZNF679-1 | lnc-ZNF679-1 | 0.0363422576498986 | 3.46114897929706 | 0.175940914826358 | 0.861176038475798 | 0.951045519902744 | -6.22619822031595 |
| lnc-GPR88-1 | lnc-GPR88-1 | -0.0452856632328253 | 0.155741038339398 | -0.175777949252065 | 0.861303274568865 | 0.951135001901977 | -5.71289514917429 |
| ENSG00000199627.1 | RNU6-1010P | 0.0719498832699356 | 0.0558173499006905 | 0.175694811965397 | 0.861368185809422 | 0.951152672286098 | -5.61267709105852 |
| ENSG00000100216.5 | TOMM22 | 0.0295575740266107 | 4.71316901463552 | 0.175584921373032 | 0.861453986761915 | 0.951152672286098 | -6.35123706140009 |
| ENSG00000103994.17 | ZNF106 | 0.021336216599968 | 6.78583781003461 | 0.17557989947687 | 0.861457907824779 | 0.951152672286098 | -6.39372653457244 |
| ENSG00000144659.12 | SLC25A38 | -0.0161977551014703 | 4.97530313260423 | -0.175500539052029 | 0.861519872382017 | 0.951170065613853 | -6.36614206922207 |
| lnc-SPEN-3 | lnc-SPEN-3 | -0.0413059361336339 | 1.47573889464602 | -0.175318487336296 | 0.861662021552718 | 0.951267789365631 | -5.987252851047 |
| lnc-INCA1-1 | lnc-INCA1-1 | 0.029780670287575 | 0.6159111447349 | 0.175154984097221 | 0.861789691703201 | 0.951267789365631 | -5.79927087296529 |
| ENSG00000104142.10 | VPS18 | -0.019050706229045 | 4.93981405937263 | -0.175154879898195 | 0.861789773067341 | 0.951267789365631 | -6.36182579903378 |
| ENSG00000132017.10 | DCAF15 | -0.0189115733676468 | 4.50063224968078 | -0.175150426943525 | 0.861793250172517 | 0.951267789365631 | -6.33308871452192 |
| lnc-OGDH-5 | lnc-OGDH-5 | -0.0313068034229608 | -0.0239295703605609 | -0.174906715448907 | 0.861983557370899 | 0.951290841426575 | -5.65031719140434 |
| ENSG00000146054.17 | TRIM7 | -0.0576414984737441 | 0.388371324277507 | -0.174875666920881 | 0.862007802857013 | 0.951290841426575 | -5.73163743261389 |
| lnc-CAVIN1-1 | lnc-CAVIN1-1 | 0.0304749810260816 | 2.29636030037854 | 0.174870071807298 | 0.862012172039769 | 0.951290841426575 | -6.08976521109564 |
| lnc-CARHSP1-4 | lnc-CARHSP1-4 | -0.0249506077693806 | 3.54540951007826 | -0.174738967448988 | 0.862114551713401 | 0.951290841426575 | -6.23528498025777 |
| lnc-PATL2-1 | lnc-PATL2-1 | 0.0369171518596092 | 3.6177556814322 | 0.174711449653076 | 0.862136040723703 | 0.951290841426575 | -6.22650029833827 |
| lnc-RANBP3L-4 | lnc-RANBP3L-4 | -0.0530083371225252 | 2.87134855131587 | -0.174671020860486 | 0.862171473072703 | 0.951290841426575 | -6.15200680691682 |
| ENSG00000090530.9 | P3H2 | -0.195588125385424 | -1.9969206111135 | -0.174656866521659 | 0.862181791419795 | 0.951290841426575 | -5.05758314294477 |
| lnc-NAT10-1 | lnc-NAT10-1 | 0.0218235636567436 | 1.3647167943077 | 0.17465019911345 | 0.862183872458473 | 0.951290841426575 | -5.96532386151996 |
| ENSG00000022976.15 | ZNF839 | -0.0134850405194351 | 4.29816406943848 | -0.174562960919608 | 0.862251999360105 | 0.951315014088177 | -6.31046941701646 |
| ENSG00000100462.15 | PRMT5 | 0.0243846985099321 | 3.49621751391834 | 0.174362689644944 | 0.86240840117157 | 0.951366543247883 | -6.22151343073123 |
| ENSG00000264324.1 | | 0.0163802062508673 | 3.0912840129688 | 0.174344137015921 | 0.862422890126263 | 0.951366543247883 | -6.17405492525703 |
| ENSG00000136848.17 | DAB2IP | -0.120454727658446 | -0.140867895841211 | -0.174355932633552 | 0.862437367267272 | 0.951366543247883 | -5.83708818457839 |
| ENSG00000124570.19 | SERPINB6 | -0.139337814927535 | 2.76474377472473 | -0.174185184188249 | 0.862579881459188 | 0.951401084265515 | -6.22834418353434 |
| lnc-ZNF490-3 | lnc-ZNF490-3 | -0.0252504421048663 | 0.0725517167181443 | -0.174118919359613 | 0.862598781063889 | 0.951401084265515 | -5.6694162171005 |
| lnc-FAXDC2-2 | lnc-FAXDC2-2 | 0.0216655708994418 | 0.319707766417855 | 0.17406022641253 | 0.862644620358799 | 0.951401084265515 | -5.7253331739369 |
| ENSG00000151726.13 | ACSL1 | 0.0283600471680892 | 5.15755343721463 | 0.174036278216303 | 0.862663324079774 | 0.951401084265515 | -6.37304141885556 |
| lnc-EEF1B2-2 | lnc-EEF1B2-2 | 0.0264983238775187 | 2.08873440365624 | 0.173989583959177 | 0.862699792874293 | 0.951401084265515 | -6.06071697631902 |
| ENSG00000100296.13 | THOC5 | 0.0123824791361198 | 5.01115840457427 | 0.17375656566927 | 0.862881787579494 | 0.951512224105258 | -6.36375641471419 |
| lnc-ERMN-2 | lnc-ERMN-2 | 0.0698441914977599 | 1.74916831373014 | 0.173724494747228 | 0.86291742436363 | 0.951512224105258 | -6.01741259817503 |
| ENSG00000103852.12 | TTC23 | 0.0659184951198084 | -0.190599204515212 | 0.173619970105756 | 0.862988476569311 | 0.951512224105258 | -5.55738809726895 |
| lnc-PPP1R1C-2 | lnc-PPP1R1C-2 | -0.0369174490736082 | 0.106340238379722 | -0.173565222378286 | 0.863031238419248 | 0.951512224105258 | -5.70217483939805 |
| lnc-PKIA-1 | lnc-PKIA-1 | 0.0705540888515231 | -0.0091766005904385 | 0.173528909955197 | 0.863059601220954 | 0.951512224105258 | -5.72521946030816 |
| ENSG00000163154.5 | TNFAIP8L2 | -0.0621603517454736 | -0.103411177858993 | -0.173505432177554 | 0.863077939272466 | 0.951512224105258 | -5.62170453860818 |
| ENSG00000272305.5 | | 0.0295513473143432 | 3.928989213413 | 0.173386411198714 | 0.863170905500631 | 0.951521970355024 | -6.26891558490209 |
| ENSG00000204852.15 | TCTN1 | -0.0334840236468201 | 5.13050157570555 | -0.173354055530514 | 0.86319617856462 | 0.951521970355024 | -6.36693721767988 |
| ENSG00000052802.12 | MSMO1 | -0.0363207239247245 | 3.29992536507091 | -0.173261077339107 | 0.863268804784964 | 0.951521970355024 | -6.22506370427065 |
| ENSG00000279203.1 | | -0.0391881012077716 | -0.0779525020075295 | -0.173257378712555 | 0.863271693844907 | 0.951521970355024 | -5.64524910277749 |
| ENSG00000144182.16 | LIPT1 | 0.0195869678580176 | 2.23808350483273 | 0.173121656588341 | 0.86337771000782 | 0.95156064642583 | -6.0731530167516 |
| ENSG00000071246.10 | VASH1 | 0.261552386071545 | 2.36004812838814 | 0.173135886702158 | 0.863399243679192 | 0.95156064642583 | -6.07225408078079 |
| ENSG00000186301.8 | MST1P2 | -0.0566141002392692 | 0.774349596591451 | -0.172995607382912 | 0.863476172678447 | 0.951563441178438 | -5.8636494316254 |
| ENSG00000125447.17 | GGA3 | 0.0257233320855637 | 6.16416904487011 | 0.17297247762773 | 0.863494240602967 | 0.951563441178438 | -6.39529699640034 |
| ENSG00000253043.1 | RNU7-181P | -0.0534804459128065 | -0.614897795675748 | -0.172884292450881 | 0.86356312757535 | 0.951588407028397 | -5.4632586988839 |
| ENSG00000205571.13 | SMN2 | 0.0291290733400633 | 1.85706962720723 | 0.172780054313326 | 0.863644555907417 | 0.95162718919927 | -6.02627646789116 |
| ENSG00000115145.9 | STAM2 | 0.0136815326454576 | 5.17759239591012 | 0.172658024286625 | 0.863739884742249 | 0.951681282996077 | -6.37373208931833 |
| ENSG00000283228.1 | | 0.0254509429203118 | 3.82007663983191 | 0.172579178830049 | 0.86380147924516 | 0.951698204009375 | -6.26644261713181 |
| ENSG00000283845.1 | MIR4721 | 0.0284472363846746 | 1.89715279217858 | 0.172244035344465 | 0.864063304735051 | 0.951850175467218 | -6.04066610049112 |
| ENSG00000115286.19 | NDUFS7 | -0.0253647249031852 | 3.67396255355715 | -0.172238316937829 | 0.864067772283033 | 0.951850175467218 | -6.2524946300598 |
| ENSG00000138592.13 | USP8 | 0.0198774396022907 | 7.36337182754446 | 0.172158002525578 | 0.86413051899413 | 0.951850175467218 | -6.38749843035154 |
| ENSG00000205771.6 | CATSPER2P1 | 0.038180095490746 | 0.382347577372063 | 0.172094389751056 | 0.864180217956676 | 0.951850175467218 | -5.74583513226125 |
| ENSG00000140395.8 | SKIC8 | 0.0151438512286042 | 4.26678865844742 | 0.172057329955116 | 0.864209172042458 | 0.951850175467218 | -6.30801046034469 |
| ENSG00000154781.15 | CCDC174 | 0.0200855835453195 | 5.15457849969582 | 0.172023410345481 | 0.864235672928285 | 0.951850175467218 | -6.36958507245328 |
| ENSG00000127989.13 | MTERF1 | 0.0241069601291954 | 3.93150997100565 | 0.171988271780264 | 0.864263126332742 | 0.951850175467218 | -6.27097794973451 |
| ENSG00000198018.6 | ENTPD7 | 0.0274642756273056 | 2.29560422971698 | 0.171857480037948 | 0.864365314084992 | 0.951911784901626 | -6.07434948334018 |
| ENSG00000126016.15 | AMOT | 0.100977204797595 | 1.94901644177757 | 0.171736568687993 | 0.864485524260211 | 0.95199323448975 | -6.05840844033099 |
| ENSG00000249212.1 | ATP1B1P1 | 0.0389171256554841 | 0.473826799608525 | 0.171635954862989 | 0.864538397340422 | 0.952000526024925 | -5.7580837968188 |
| ENSG00000146963.17 | LUC7L2 | 0.024835903543627 | 6.4466448474408 | 0.171446210835891 | 0.864686654528701 | 0.952081368017807 | -6.39608761510329 |
| lnc-ZCCHC13-17 | lnc-ZCCHC13-17 | -0.0570602295519445 | 1.95337125312338 | -0.171427624960231 | 0.864704323651794 | 0.952081368017807 | -6.05405069691717 |
| lnc-RCAN1-5 | lnc-RCAN1-5 | -0.0293098219806017 | 1.0184067525293 | -0.171304604362942 | 0.864797302478175 | 0.952132809676826 | -5.88482169330601 |

| | | | | | | | |
|---|---|---|---|---|---|---|---|
| lnc-PTTG2-2 | lnc-PTTG2-2 | 0.0455341471673854 | 0.651791417885068 | 0.17119767234029 | 0.864880858451544 | 0.952173871924911 | -5.8076086820253 |
| lnc-CDNF-3 | lnc-CDNF-3 | -0.0278431367114038 | 1.31962553187372 | -0.170998554440583 | 0.865036452026307 | 0.952294233944348 | -5.94148304857594 |
| ENSG00000126254.11 | RBM42 | 0.0229738067732841 | 4.06883425210105 | 0.170675016857507 | 0.865289280474105 | 0.952487093142561 | -6.29273283423614 |
| lnc-VWA7-2 | lnc-VWA7-2 | -0.0405308419801683 | 0.778690307403766 | -0.170570564947919 | 0.865370907470587 | 0.952487093142561 | -5.85812964260174 |
| ENSG00000145191.13 | EIF2B5 | 0.0126686662936697 | 6.2266112614499 | 0.17056679963152 | 0.86537385001488 | 0.952487093142561 | -6.39621548994677 |
| ENSG00000171720.9 | HDAC3 | 0.0177035365747624 | 5.07702668365347 | 0.170532621792465 | 0.865400559626357 | 0.952487093142561 | -6.36945842356315 |
| lnc-NAA35-4 | lnc-NAA35-4 | -0.0407665900439254 | -0.514754909849473 | -0.17047829359039 | 0.865443016856249 | 0.952487093142561 | -5.51549831609174 |
| lnc-AGAP4-5 | lnc-AGAP4-5 | 0.0427075429587978 | -0.436837664388322 | 0.170380633222457 | 0.865519338977624 | 0.952520160090704 | -5.52991387085151 |
| ENSG00000009413.15 | REV3L | 0.0193796474723129 | 6.5631809238892 | 0.170060749442642 | 0.865769339024494 | 0.952744349077265 | -6.39595301128838 |
| ENSG00000157017.15 | GHRL | -0.0378420783663906 | -0.388027299483057 | -0.16979391862374 | 0.865977887021676 | 0.95290088272472 | -5.54479774947108 |
| ENSG00000122025.14 | FLT3 | 0.164315432746635 | 3.0139296890265 | 0.169801232490606 | 0.86600417394202 | 0.95290088272472 | -6.14535734365737 |
| lnc-FAM3B-3 | lnc-FAM3B-3 | -0.0393374757583032 | -0.33025667904923 | -0.169573709403235 | 0.866150004075283 | 0.952909896514607 | -5.56442696109683 |
| lnc-IL4R-2 | lnc-IL4R-2 | 0.0422539817983261 | 0.192270186901965 | 0.169511248317011 | 0.866198825290521 | 0.952909896514607 | -5.68704426794503 |
| ENSG00000075336.11 | TIMM21 | 0.0196492856333244 | 3.31946747806923 | 0.169456503785112 | 0.866241615480278 | 0.952909896514607 | -6.20907681978203 |
| ENSG00000213071.11 | LPAL2 | -0.104235257990389 | -0.549719253284936 | -0.169439782642092 | 0.866260491861951 | 0.952909896514607 | -5.54762362100344 |
| lnc-TIA1-1 | lnc-TIA1-1 | 0.0299625780222499 | -0.320652229476958 | 0.169372256259907 | 0.866307467007114 | 0.952909896514607 | -5.54658044681993 |
| ENSG00000179051.13 | RCC2 | -0.0172877002266398 | 5.91688916892996 | -0.169342231313034 | 0.866330936042709 | 0.952909896514607 | -6.39451171963742 |
| ENSG00000104133.14 | SPG11 | 0.0138235254786059 | 7.83536544527936 | 0.169335193058826 | 0.866336437520179 | 0.952909896514607 | -6.38005116396431 |
| ENSG00000168300.13 | PCMTD1 | -0.0162622786913449 | 6.92942744562864 | -0.168978604107058 | 0.866615175366571 | 0.95311586235295 | -6.3930380093282 |
| ENSG00000140105.17 | WARS1 | 0.0516503657720673 | 6.00695253039234 | 0.168998138928777 | 0.866616302965334 | 0.95311586235295 | -6.39551572076406 |
| lnc-DIP2B-1 | lnc-DIP2B-1 | -0.0432219762812973 | 0.46714584205861 | -0.168826896955797 | 0.866733766766282 | 0.953194118473519 | -5.8049163742976 |
| ENSG00000160211.17 | G6PD | -0.023584870337376 | 3.99605976716989 | -0.168683026708585 | 0.866846234829206 | 0.953261590461795 | -6.2819835516393 |
| lnc-UAP1-4 | lnc-UAP1-4 | 0.0223542584990488 | 5.54777828917523 | 0.1686101138899 | 0.8669032342229 | 0.953261590461795 | -6.38857202273912 |
| ENSG00000135899.17 | SP110 | 0.0216930393370571 | 6.90020839222636 | 0.168515113032686 | 0.866977501954721 | 0.953261590461795 | -6.39416306450224 |
| ENSG00000140743.7 | CDR2 | 0.0285750886001503 | 4.41608962450386 | 0.168431863607467 | 0.867042583903984 | 0.953261590461795 | -6.31378211762757 |
| ENSG00000166439.5 | RNF169 | 0.0148253296145988 | 6.24076625258422 | 0.168340527309973 | 0.867113989005687 | 0.953261590461795 | -6.39660593629014 |
| ENSG00000143183.16 | TMCO1 | 0.0143199697193106 | 5.02743684722008 | 0.168311142368951 | 0.86713696187157 | 0.953261590461795 | -6.36711381166521 |
| ENSG00000138942.15 | RNF185 | 0.0173049351928638 | 4.34172887030458 | 0.168289654087284 | 0.867153761278252 | 0.953261590461795 | -6.31704188945125 |
| ENSG00000136141.14 | LRCH1 | -0.0479323120010733 | 4.97163748790483 | -0.168278993971059 | 0.867173956708675 | 0.953261590461795 | -6.37587519160927 |
| lnc-MPHOSPH8-3 | lnc-MPHOSPH8-3 | 0.0351339498039966 | 0.957652720326557 | 0.168070634848331 | 0.867324992717669 | 0.953261590461795 | -5.87983514599539 |
| lnc-FAAP20-2 | lnc-FAAP20-2 | 0.0344243946536811 | 0.186621488840587 | 0.168013213242371 | 0.867369886572654 | 0.953261590461795 | -5.70770120252145 |
| lnc-IMPA2-1 | lnc-IMPA2-1 | 0.0315667096487395 | 1.57590948331023 | 0.167922592915977 | 0.867440737038647 | 0.953261590461795 | -5.98842994881478 |
| lnc-CHST2-2 | lnc-CHST2-2 | 0.043527192384146 | 2.64177416851081 | 0.167914696293732 | 0.867446910974698 | 0.953261590461795 | -6.1223583509226 |
| ENSG00000198554.11 | WDHD1 | 0.0193610294790894 | 3.0522774896584 | 0.167865115907561 | 0.86748567535204 | 0.953261590461795 | -6.1704510610081 |
| lnc-HLA-B-2 | lnc-HLA-B-2 | -0.0358306671122239 | 3.91857349610442 | -0.167807488096596 | 0.867530732015422 | 0.953261590461795 | -6.27671572119513 |
| ENSG00000197283.16 | SYNGAP1 | 0.0182500081923882 | 3.48970706103408 | 0.167740418173861 | 0.867583171613953 | 0.953261590461795 | -6.22574009701508 |
| lnc-PCED1B-5 | lnc-PCED1B-5 | 0.0434054365355809 | 4.61869492749673 | 0.16769670108402 | 0.867623168112653 | 0.953261590461795 | -6.34891679969013 |
| ENSG00000206786.1 | RNU6-701P | 0.0540576065976944 | -0.61142219658282 | 0.167596818123997 | 0.867695449438812 | 0.953261590461795 | -5.44451564513986 |
| ENSG00000178917.16 | ZNF852 | 0.0244249329524591 | 3.28632165084437 | 0.167533541920043 | 0.867744924634336 | 0.953261590461795 | -6.19962724499015 |
| ENSG00000163125.15 | RPRD2 | -0.0186198707191201 | 6.56299780587874 | -0.16752849437602 | 0.867748871294753 | 0.953261590461795 | -6.39653849098851 |
| lnc-CSNK1D-4 | lnc-CSNK1D-4 | 0.0329725956430376 | -0.119460167096021 | 0.167519374597996 | 0.867756002032093 | 0.953261590461795 | -5.64544650284878 |
| lnc-EIF5B-2 | lnc-EIF5B-2 | 0.0292135722052624 | 1.25169378952136 | 0.167447286935182 | 0.867812367632606 | 0.953261590461795 | -5.94370663615516 |
| ENSG00000253030.1 | MIR2116 | 0.0407071136293216 | 0.130669920094798 | 0.167392883348308 | 0.867854906451914 | 0.953261590461795 | -5.68749249750777 |
| ENSG00000243725.6 | TTC4 | 0.0159565524689205 | 2.86866464319009 | 0.167375585253778 | 0.867868432123235 | 0.953261590461795 | -6.1544241830427 |
| ENSG00000080823.22 | MOK | 0.153187568320168 | -0.780807539136309 | 0.1673420085177 | 0.867906631941605 | 0.953261590461795 | -5.3301864194051 |
| ENSG00000154114.12 | TBCEL | -0.0208794160488583 | 4.20177845366408 | -0.167162377755742 | 0.86803514594343 | 0.9533518707088 | -6.30276063426518 |
| ENSG00000167881.14 | VIII.68 | -0.0220449375304854 | 4.22707182426541 | -0.167004626899656 | 0.868158500338815 | 0.953405649247316 | -6.30408690036999 |
| ENSG00000256185.1 | | 0.0311590347562354 | 0.109014986619148 | 0.166955769744018 | 0.868196705210845 | 0.953405649247316 | -5.72195623939269 |
| lnc-STRN3-8 | lnc-STRN3-8 | 0.0122046124727122 | 2.92523585594902 | 0.166922051695858 | 0.868223071927887 | 0.953405649247316 | -6.15595214236379 |
| ENSG00000166295.8 | ANAPC16 | 0.0221467848076173 | 6.45578696088075 | 0.166700896557313 | 0.868396013782705 | 0.953446970037222 | -6.39687906269257 |
| ENSG00000101473.16 | ACOT8 | 0.0152276160734206 | 3.91198823993272 | 0.16669049694892 | 0.868404146368262 | 0.953446970037222 | -6.27238889610135 |
| ENSG00000134884.14 | ARGLU1 | -0.0169498559954686 | 8.42481725065837 | -0.166657213723643 | 0.86843017424113 | 0.953446970037222 | -6.36613654727541 |
| lnc-DNAJB12-3 | lnc-DNAJB12-3 | 0.0247129740010148 | 1.1041266036124 | 0.166636917871751 | 0.86844604590804 | 0.953446970037222 | -5.90654623593363 |
| lnc-RANBP9-8 | lnc-RANBP9-8 | -0.0245129539435642 | 1.38931010373848 | -0.166553200251915 | 0.868511514948737 | 0.953446970037222 | -5.96435263022042 |
| ENSG00000147536.11 | GINS4 | 0.03806307050837 | 2.33187407915312 | 0.166477312300603 | 0.868570861821351 | 0.953446970037222 | -6.07651009050643 |
| lnc-PAWR-11 | lnc-PAWR-11 | 0.0207918627150776 | 2.58425706393627 | 0.166459290836913 | 0.868584955311079 | 0.953446970037222 | -6.11545835078783 |
| ENSG00000250021.7 | ARPIN-AP3S2 | -0.018022033748029 | 2.74950777027817 | -0.166325913338471 | 0.86868926307761 | 0.953464080229332 | -6.13692465524935 |
| ENSG00000167384.10 | ZNF180 | 0.019870061957965 | 3.88848481849006 | 0.166320894137967 | 0.868693188385594 | 0.953464080229332 | -6.27150252199199 |
| ENSG00000105219.9 | CCNP | -0.036137379486237 | 0.185416429805946 | -0.166229263902861 | 0.868764849170921 | 0.953491889222836 | -5.7004118954661 |
| ENSG00000162298.18 | SYVN1 | 0.0178516299052305 | 6.48946246205458 | 0.166037752895947 | 0.868914626800834 | 0.953565616957116 | -6.3969175999143 |
| ENSG00000175556.16 | LONRF3 | 0.0429374585746682 | 3.73509665939276 | 0.166026871354891 | 0.868924681127516 | 0.953565616957116 | -6.24839639162945 |
| ENSG00000177688.6 | SUMO4 | 0.0334664680899637 | 1.65270767405826 | 0.165908481704909 | 0.869015730441883 | 0.953599995613878 | -6.00271750050756 |
| ENSG00000258466.5 | | -0.0190405337502058 | 2.55747143918382 | -0.1658232400805 | 0.869082399553609 | 0.953599995613878 | -6.11314171446506 |
| lnc-COP1-6 | lnc-COP1-6 | -0.0221415998341532 | 1.37365496391595 | -0.165807133751309 | 0.869094996731319 | 0.953599995613878 | -5.96118823042593 |
| ENSG00000156853.12 | ZNF689 | -0.0249002095335737 | 3.44518860802961 | -0.165484915167765 | 0.869347019466846 | 0.953809143119497 | -6.22412959137012 |
| ENSG00000274847.1 | MAFIP | -0.0568791590746614 | 0.12022307224577 | -0.165385692326245 | 0.869424629218676 | 0.953809143119497 | -5.6644552725696 |
| ENSG00000277400.1 | | -0.0568791590746614 | 0.12022307224577 | -0.165385692326245 | 0.869424629218676 | 0.953809143119497 | -5.6644552725696 |
| lnc-HORMAD2-3 | lnc-HORMAD2-3 | 0.0237047393714841 | 0.81846804017151 | 0.165319861505498 | 0.869476121243254 | 0.953814795264611 | -5.86547969032856 |

| | | | | | | | |
|---|---|---|---|---|---|---|---|
| lnc-ZBTB25-2 | lnc-ZBTB25-2 | -0.04017019124909 | 0.207996656718333 | -0.164730486990321 | 0.869937147950575 | 0.954269682171535 | -5.72561270393549 |
| ENSG00000129480.12 | DTD2 | 0.0273152314270231 | 2.87287034154205 | 0.164614531323982 | 0.870027857401858 | 0.954284595973163 | -6.15362852790662 |
| AFF1-AS1 | AFF1-AS1 | -0.0661650258514512 | 0.770449706255778 | -0.164599217838685 | 0.87004346927233 | 0.954284595973163 | -5.88929869226826 |
| ENSG00000138814.16 | PPP3CA | 0.0168612226670799 | 5.8448127062906 | 0.164476699892963 | 0.870135682064498 | 0.954322658694659 | -6.39425594300895 |
| LINC02416 | LINC02416 | -0.163156995958676 | -0.360337178256877 | -0.164412341159596 | 0.870211129807653 | 0.954322658694659 | -5.59018042200002 |
| ENSG00000182054.9 | IDH2 | 0.0324913201131176 | 4.41917497754171 | 0.164372414217193 | 0.870217265755238 | 0.954322658694659 | -6.32487479428596 |
| ITCH-IT1 | ITCH-IT1 | 0.0406765989397893 | -0.264375641650582 | 0.164162594040279 | 0.87038141443836 | 0.954347227742605 | -5.57143563175443 |
| ENSG00000132704.15 | FCRL2 | -0.103899464882813 | 7.12310788605483 | -0.164198312868755 | 0.870384390348875 | 0.954347227742605 | -6.39411520173434 |
| lnc-DDX52-2 | lnc-DDX52-2 | 0.0262486138931886 | 2.15259806941983 | 0.164148330908795 | 0.870392573128392 | 0.954347227742605 | -6.05416022878986 |
| ENSG00000177409.11 | SAMD9L | 0.0287867609236274 | 6.79139668702526 | 0.164106713182682 | 0.870425132702324 | 0.954347227742605 | -6.39572607057676 |
| ENSG00000196092.12 | PAX5 | 0.0261548997661212 | 8.7173473697072 | 0.164026505077579 | 0.870487884050676 | 0.954365192149518 | -6.36122661373472 |
| lnc-WDR41-4 | lnc-WDR41-4 | -0.071016654789483 | -0.444782203715582 | -0.163477860282958 | 0.870917142604495 | 0.954784955857701 | -5.54106610768407 |
| ENSG00000111834.12 | RSPH4A | 0.0449025909351885 | 0.413246092046386 | 0.163302632967435 | 0.871054248398171 | 0.954852832278537 | -5.77604475016673 |
| lnc-MBD5-1 | lnc-MBD5-1 | -0.0159865506205712 | 3.03892185959246 | -0.163278752007992 | 0.871072934254051 | 0.954852832278537 | -6.17321079897336 |
| ENSG00000206418.4 | RAB12 | -0.0198709413844043 | 4.72880971897309 | -0.163089208679396 | 0.87122124665468 | 0.954852832278537 | -6.34489008300682 |
| ENSG00000216490.3 | IFI30 | 0.0576793164762308 | 3.01936474159714 | 0.163016763336457 | 0.871289415452995 | 0.954852832278537 | -6.16773590854219 |
| lnc-USP28-3 | lnc-USP28-3 | 0.0609242662454438 | -0.591409448592501 | 0.163000700436654 | 0.871290503507801 | 0.954852832278537 | -5.45672359231869 |
| ENSG00000067113.16 | PLPP1 | 0.0205740724869581 | 1.96691755342814 | 0.162991519634398 | 0.871297687456242 | 0.954852832278537 | -6.05306629489564 |
| lnc-PRRT4-3 | lnc-PRRT4-3 | 0.0179011725483388 | 0.58704432078752 | 0.162952338099358 | 0.871328347005857 | 0.954852832278537 | -5.80376894024893 |
| lnc-DEPDC4-2 | lnc-DEPDC4-2 | 0.0252861101281199 | 2.02612173341904 | 0.162897347702099 | 0.871371377327238 | 0.954852832278537 | -6.04923731420285 |
| ENSG00000180098.9 | TRNAU1AP | 0.0154803943516599 | 4.34303674931486 | 0.162801834242754 | 0.871446118129043 | 0.954852832278537 | -6.31933951784153 |
| ENSG00000144741.17 | SLC25A26 | -0.0160132740238978 | 4.58495020898284 | -0.162751101123011 | 0.87148581808601 | 0.954852832278537 | -6.34169930709616 |
| ENSG00000110200.8 | ANAPC15 | 0.0207315907999237 | 3.58711061012371 | 0.162694517404068 | 0.871530096681233 | 0.954852832278537 | -6.24393512536011 |
| ENSG00000122694.15 | GLIPR2 | -0.0546708492346242 | 2.28282896319481 | -0.162637467579904 | 0.871579648004024 | 0.954852832278537 | -6.08408741610589 |
| ENSG00000077157.21 | PPP1R12B | 0.0186473305966626 | 5.1692938187491 | 0.162588529337576 | 0.871613036902573 | 0.954852832278537 | -6.37240825783401 |
| lnc-DHX9-1 | lnc-DHX9-1 | -0.0547821965864078 | -0.262701927684209 | -0.162568741534708 | 0.871628521871517 | 0.954852832278537 | -5.58508152953286 |
| ENSG00000173915.15 | ATP5MK | 0.0243616387750469 | 3.46649630968523 | 0.162342721737941 | 0.871805397548822 | 0.954980272631377 | -6.22303180745018 |
| lnc-DNPEP-4 | lnc-DNPEP-4 | -0.0445162116182766 | 0.181509181416045 | -0.162286406157908 | 0.871849469307544 | 0.954980272631377 | -5.73478593906497 |
| lnc-PDE7A-1 | lnc-PDE7A-1 | 0.0494005746200817 | -0.294588918577793 | 0.162184387823656 | 0.871929308430079 | 0.954980272631377 | -5.57152339289683 |
| ENSG00000244257.5 | PKD1P1 | -0.0359537017826639 | 1.28688178191966 | -0.16218294085946 | 0.871930440827939 | 0.954980272631377 | -5.94436884156664 |
| ENSG00000102572.14 | STK24 | 0.0137308184420295 | 6.90482524758095 | 0.162079648103915 | 0.872011278703337 | 0.955017992420495 | -6.39499046879181 |
| ENSG00000166452.11 | AKIP1 | 0.0405541574008186 | 1.86944936767439 | 0.162010612477714 | 0.872065307401158 | 0.955026348616017 | -6.02959465031877 |
| lnc-COX5B-2 | lnc-COX5B-2 | -0.0378155358952266 | 0.926275324306375 | -0.161939532971593 | 0.872120936327194 | 0.955036456289776 | -5.87224551217397 |
| ENSG00000099290.16 | WASHC2A | 0.0113740436239942 | 5.60935631046933 | 0.161700882233185 | 0.872307716250893 | 0.955190175219031 | -6.38956231209692 |
| lnc-C10orf105-4 | lnc-C10orf105-4 | -0.0557324918432147 | 1.10578829302034 | -0.161564141496898 | 0.872414739667441 | 0.955237796117754 | -5.9033885680619 |
| ENSG00000203737.3 | GPR52 | 0.0589713600431896 | 2.67845739020981 | 0.161540153039724 | 0.872444023185039 | 0.955237796117754 | -6.11860976397308 |
| lnc-MED9-4 | lnc-MED9-4 | -0.0386903450021185 | 0.356778089265367 | -0.161225181374349 | 0.872680045360393 | 0.955349330168285 | -5.75667347831109 |
| ENSG00000067533.5 | RRP15 | 0.0185826389703347 | 4.62394951892453 | 0.161190950158163 | 0.872706839120419 | 0.955349330168285 | -6.34569595732609 |
| ENSG00000135940.6 | COX5B | 0.0222920999343472 | 4.48861451869559 | 0.161172095619851 | 0.872721597175849 | 0.955349330168285 | -6.33719808726324 |
| lnc-THUMPD3-6 | lnc-THUMPD3-6 | 0.0390656290491371 | 1.49770801703505 | 0.161159382552477 | 0.872731548129732 | 0.955349330168285 | -5.96642567735965 |
| LINC00863 | LINC00863 | -0.0257183757339987 | 1.08809193849712 | -0.161014067561447 | 0.872845292638579 | 0.95542303012018 | -5.92132838191329 |
| ENSG00000176261.15 | ZBTB8OS | -0.011903465000989 | 4.06142107300022 | -0.160892634900712 | 0.872940345473031 | 0.955431751376878 | -6.29083112556678 |
| ENSG00000154723.12 | ATP5PF | -0.0263533363599331 | 3.38414643508676 | -0.160882680078427 | 0.872948137810791 | 0.955431751376878 | -6.21276575859224 |
| lnc-DNAJB6-5 | lnc-DNAJB6-5 | -0.03898720549316 | 0.582366229201823 | -0.160825987160516 | 0.872992515578144 | 0.955431751376878 | -5.81743294321447 |
| ENSG00000134955.11 | SLC37A2 | 0.0884556126994271 | 0.525509952005778 | 0.160385997160992 | 0.873351785210624 | 0.955774127764264 | -5.81857686632865 |
| ENSG00000109320.11 | NFKB1 | 0.0185450047655689 | 5.79642411436562 | 0.160201882372107 | 0.873481076183749 | 0.955864798292844 | -6.39288116120286 |
| ENSG00000127951.6 | FGL2 | -0.14505750933249 | 1.6680705982205 | -0.159953274528613 | 0.873704639521953 | 0.956058494164004 | -5.950883617757 |
| ENSG00000232882.1 | PHKA1P1 | 0.0422720001786322 | 0.754672400141624 | 0.1598532227501 | 0.873754035059405 | 0.956058494164004 | -5.83008601413235 |
| ENSG00000214736.7 | TOMM6 | 0.0206681470299115 | 3.82056511514188 | 0.159771529129734 | 0.873817993666569 | 0.956058494164004 | -6.26218151793389 |
| lnc-ZNF142-3 | lnc-ZNF142-3 | 0.0269410900981829 | 0.77905867562944 | 0.159547396360303 | 0.873993473438907 | 0.956058494164004 | -5.85877762081603 |
| ENSG00000263647.1 | BPTFP1 | -0.0284105691998264 | 2.15119820765442 | -0.15944995972443 | 0.874069761290588 | 0.956058494164004 | -6.06195687232011 |
| ENSG00000263503.1 | MAPK8IP1P2 | 0.163847286363022 | -2.74682882054289 | 0.159432500564477 | 0.874083431038884 | 0.956058494164004 | -5.01814731316487 |
| ENSG00000137513.9 | NARS2 | 0.0165869579456257 | 3.19641636000548 | 0.159370225635413 | 0.874132189864744 | 0.956058494164004 | -6.18955691269319 |
| lnc-RPS14-4 | lnc-RPS14-4 | 0.0223801267178585 | 5.52363535251038 | 0.159316208645763 | 0.874174483448985 | 0.956058494164004 | -6.38607071514349 |
| ENSG00000250486.4 | FAM218A | -0.170975060333645 | -2.32916911146835 | -0.159210366100208 | 0.874257378841029 | 0.956058494164004 | -5.02845381072639 |
| lnc-PTAFR-1 | lnc-PTAFR-1 | 0.0225971386957444 | 3.06371261452194 | 0.159133314050929 | 0.874317686861294 | 0.956058494164004 | -6.16976617592007 |
| ENSG00000008130.15 | NADK | -0.0296669925110278 | 5.45126618146435 | -0.15908461779686 | 0.874355815933606 | 0.956058494164004 | -6.38426578892972 |
| lnc-TMA16-1 | lnc-TMA16-1 | -0.0510648560421138 | -0.516966501534249 | -0.159016095642628 | 0.874409469158918 | 0.956058494164004 | -5.56453693879058 |
| ENSG00000100764.13 | PSMC1 | 0.0123913867117702 | 5.31630698807385 | 0.159001220495598 | 0.874421116560229 | 0.956058494164004 | -6.38278427089285 |
| ENSG00000069702.10 | TGFBR3 | -0.0832631576609399 | 5.68991607984379 | -0.1589939450822 | 0.874455458567772 | 0.956058494164004 | -6.39033559890996 |
| lnc-KDM6B-1 | lnc-KDM6B-1 | 0.0415313626650418 | 0.634375498948215 | 0.15892605090747 | 0.874479975593221 | 0.956058494164004 | -5.81870722161896 |
| ENSG00000065060.16 | BLTP3A | -0.0116540125071234 | 5.18044905521638 | -0.158916946373476 | 0.874487104643034 | 0.956058494164004 | -6.37727404820968 |
| ENSG00000228474.5 | OST4 | -0.0267395488625928 | 4.62839785618576 | -0.158901100130102 | 0.874499512624324 | 0.956058494164004 | -6.34417937592513 |
| lnc-FAM186B-4 | lnc-FAM186B-4 | 0.0371218079347495 | 0.354642956199594 | 0.15887157491418 | 0.874522631648116 | 0.956058494164004 | -5.74770144280731 |
| ENSG00000285953.1 | | 0.0137276313539525 | 4.49402282888081 | 0.158848617876206 | 0.874540607691778 | 0.956058494164004 | -6.32943960763505 |
| ENSG00000064652.10 | SNX24 | -0.0250241200768037 | 0.856673908876642 | -0.158664337069704 | 0.874684907474014 | 0.956142360831342 | -5.87542325420239 |
| ENSG00000141013.16 | GAS8 | 0.05251303249453 | 2.05871022693558 | 0.15857208451851 | 0.874760051755431 | 0.956142360831342 | -6.06011605763662 |
| ENSG00000139684.13 | ESD | 0.0214151135830258 | 4.93629435492747 | 0.158460820496339 | 0.874844274702464 | 0.956142360831342 | -6.36994869157848 |

| | | | | | | | |
|---|---|---|---|---|---|---|---|
| lnc-CEP41-3 | lnc-CEP41-3 | 0.0291707805657013 | 0.154483833247159 | 0.158398591547549 | 0.874893005223991 | 0.956142360831342 | -5.67820388208645 |
| lnc-SCCPDH-3 | lnc-SCCPDH-3 | 0.0247790352200012 | 2.20935259449659 | 0.158397554609434 | 0.874893817238257 | 0.956142360831342 | -6.0803353796032 |
| ENSG00000126953.6 | TIMM8A | 0.0298498762328727 | 1.70463697984292 | 0.158393733912791 | 0.874896809182864 | 0.956142360831342 | -5.99811108092687 |
| lnc-CTNNA1-4 | lnc-CTNNA1-4 | 0.0323224445970878 | 2.55626741201781 | 0.158300840137562 | 0.874969553824211 | 0.956142360831342 | -6.11404427833783 |
| ENSG00000185507.20 | IRF7 | -0.0342665579824719 | 3.62353743815703 | -0.158276074813121 | 0.87498894760818 | 0.956142360831342 | -6.24688414570281 |
| ENSG00000126088.13 | UROD | -0.0182175631826862 | 4.3319377841918 | -0.157945996362209 | 0.875247440241924 | 0.956374054387616 | -6.32198033584062 |
| lnc-STXBP5-4 | lnc-STXBP5-4 | 0.0432773981364919 | -0.179513163016716 | 0.157829024949707 | 0.875339046770244 | 0.956378560136754 | -5.61449751551624 |
| ENSG00000267699.2 | | -0.0189805590549827 | 4.13889819604036 | -0.157692009854353 | 0.87544635277157 | 0.956378560136754 | -6.30042331964178 |
| lnc-CHP2-1 | lnc-CHP2-1 | -0.0593385663975452 | 4.56954514850366 | -0.157704434772195 | 0.875456215246042 | 0.956378560136754 | -6.32621729009853 |
| ENSG00000135597.18 | REPS1 | 0.0140903205429357 | 5.23607887563157 | 0.157652159829189 | 0.875477562528815 | 0.956378560136754 | -6.38091691586322 |
| lnc-ATP10A-69 | lnc-ATP10A-69 | 0.0479174075493739 | 0.742325874999076 | 0.157601952942473 | 0.875516883862197 | 0.956378560136754 | -5.8470455167769 |
| ENSG00000183793.13 | NPIPA5 | -0.0789072219724428 | -0.0301931480113073 | -0.157517298881914 | 0.875586698739512 | 0.956378560136754 | -5.69369305180423 |
| ENSG00000058804.11 | NDC1 | -0.0188706079906167 | 3.93065203313174 | -0.157481131742804 | 0.875611510644887 | 0.956378560136754 | -6.28437396710664 |
| JPX | JPX | 0.0297245634876342 | 6.79621472957462 | 0.157466104981198 | 0.87562327968601 | 0.956378560136754 | -6.3962225031715 |
| ENSG00000107829.13 | FBXW4 | 0.0128602230355055 | 5.230036694542 | 0.157405900018855 | 0.875670432824457 | 0.956379312258612 | -6.37755486513889 |
| ENSG00000151623.14 | NR3C2 | 0.153229446051243 | 2.92532872030822 | 0.157364740239408 | 0.875732272632938 | 0.956396103762072 | -6.2435788935456 |
| ENSG00000188352.12 | FOCAD | 0.0458883020850296 | 4.50563503656228 | 0.157264359900472 | 0.875791091470852 | 0.956409594426175 | -6.3291361362551 |
| ENSG00000274322.1 | | -0.0305572771817677 | 0.0577723322205931 | -0.157022833103376 | 0.875970465426026 | 0.956506259353695 | -5.69511188804086 |
| lnc-ARL14EP-4 | lnc-ARL14EP-4 | 0.0238452407980829 | 2.61584694204565 | 0.156899113859137 | 0.876067370997577 | 0.956506259353695 | -6.12763140495921 |
| lnc-RIPOR2-4 | lnc-RIPOR2-4 | -0.0314104344141768 | 0.579252103408834 | -0.156849585357507 | 0.876106165726137 | 0.956506259353695 | -5.78983178175661 |
| ENSG00000013503.9 | POLR3B | 0.0184441810304579 | 3.64230876917717 | 0.156827582534397 | 0.876123400215787 | 0.956506259353695 | -6.24693960894303 |
| lnc-GOLGA8A-1 | lnc-GOLGA8A-1 | 0.0681742947477175 | 1.83685384365923 | 0.156778651662084 | 0.876172944488794 | 0.956506259353695 | -5.98037687353804 |
| lnc-RRP1-3 | lnc-RRP1-3 | 0.0336428824089902 | -0.184193750743147 | 0.156672902036967 | 0.876244560904543 | 0.956506259353695 | -5.59196271977659 |
| ENSG00000243554.1 | CCZ1P1 | -0.0214229827181516 | 2.19093686013893 | -0.156620955902649 | 0.87628525080244 | 0.956506259353695 | -6.07770684718797 |
| lnc-CLLU1OS-2 | lnc-CLLU1OS-2 | 0.0569528907222178 | -0.166426658920772 | 0.156617520397628 | 0.876287941877928 | 0.956506259353695 | -5.60097348650334 |
| lnc-OAZ1-1 | lnc-OAZ1-1 | -0.0311856046113006 | 0.524508987111624 | -0.156581350667233 | 0.876316274185133 | 0.956506259353695 | -5.8075481671683 |
| lnc-RASA1-9 | lnc-RASA1-9 | 0.0532226143395018 | 0.675626045457013 | 0.156491680899312 | 0.876386514603989 | 0.956506259353695 | -5.83560295023351 |
| ENSG00000116266.10 | STXBP3 | 0.0162405347229955 | 5.17004727743417 | 0.156474894321179 | 0.876399664033893 | 0.956506259353695 | -6.37650561182303 |
| ENSG00000160075.11 | SSU72 | 0.0128523451779925 | 5.56224166770274 | 0.156390146508378 | 0.876466050075724 | 0.956506259353695 | -6.38894899561476 |
| lnc-EGR3-3 | lnc-EGR3-3 | 0.0354395286527018 | 1.24617518829608 | 0.156322474549138 | 0.876519060627177 | 0.956506259353695 | -5.94271890383524 |
| ENSG00000132405.18 | TBC1D14 | -0.0238531120267451 | 6.23885181541687 | -0.156308257040791 | 0.876530197926901 | 0.956506259353695 | -6.39865153324882 |
| ENSG00000006744.18 | ELAC2 | -0.0148535678950115 | 5.64501412351223 | -0.15619515735078 | 0.876618795586511 | 0.956552227617937 | -6.39307485904145 |
| ENSG00000105583.10 | WDR83OS | 0.0236114071747013 | 3.43446032516694 | 0.15589709635271 | 0.876852292017301 | 0.956727015344595 | -6.22768655695004 |
| lnc-RAB15-1 | lnc-RAB15-1 | -0.0353432055328004 | 0.542091341854446 | -0.155780599865606 | 0.876943556602866 | 0.956727015344595 | -5.80186512525036 |
| ENSG00000196547.14 | MAN2A2 | -0.0272024715658166 | 5.84810079100334 | -0.155767670565723 | 0.876953685658222 | 0.956727015344595 | -6.39499202674089 |
| ENSG00000104870.12 | FCGRT | -0.0318104752395971 | 5.39632499666726 | -0.155668740110714 | 0.877031190317644 | 0.956727015344595 | -6.38027284809145 |
| ENSG00000185513.16 | L3MBTL1 | 0.0203557117761237 | 5.03774261202657 | 0.155624355424201 | 0.877065962817878 | 0.956727015344595 | -6.37266557475375 |
| ENSG00000184381.18 | PLA2G6 | 0.0183354343433164 | 4.5326167429794 | 0.155536555674896 | 0.877134748905217 | 0.956727015344595 | -6.33452501876798 |
| ENSG00000197050.10 | ZNF420 | 0.0241159446719506 | 3.10966919612089 | 0.155496685581725 | 0.877165985162611 | 0.956727015344595 | -6.18228956966064 |
| ENSG00000053254.15 | FOXN3 | 0.0159912297514673 | 7.23269921640892 | 0.155380461510647 | 0.877257042137239 | 0.956727015344595 | -6.39239004066714 |
| lnc-ACAP3-1 | lnc-ACAP3-1 | 0.0339505983351202 | 0.209923289138711 | 0.155374776698737 | 0.877261496006265 | 0.956727015344595 | -5.72818507029479 |
| ENSG00000104969.9 | SGTA | 0.0271364530229334 | 4.36798391309959 | 0.155327773191529 | 0.877298321910915 | 0.956727015344595 | -6.32569370648235 |
| ENSG00000274266.1 | SNORA73A | -0.0434225884130957 | 6.03028076222609 | -0.155290775379109 | 0.877338126763779 | 0.956727015344595 | -6.39458677725816 |
| ENSG00000068323.16 | TFE3 | -0.0179358134933487 | 4.33730249975841 | -0.155212084874149 | 0.877388961572848 | 0.956727015344595 | -6.32328240532698 |
| ENSG00000168264.10 | IRF2BP2 | -0.0315707965569674 | 5.69768571209149 | -0.155195853506887 | 0.8774016786828 | 0.956727015344595 | -6.39434397924535 |
| ENSG00000189369.8 | GSPT2 | -0.084994576688147 | 2.78610252497434 | -0.155193163631958 | 0.877429717177278 | 0.956727015344595 | -6.21454083172758 |
| ENSG00000128185.9 | DGCR6L | 0.0226714310435834 | 2.80910303374279 | 0.155065458779194 | 0.877503842804782 | 0.956757156290434 | -6.14537707450037 |
| ENSG00000122565.18 | CBX3 | 0.0170329826396696 | 6.47546463494495 | 0.15496232611621 | 0.877584648626556 | 0.956794577185254 | -6.39845112084124 |
| ENSG00000131475.6 | VPS25 | -0.015539542910108 | 3.84181985819472 | -0.154864656243028 | 0.877661175497375 | 0.956827329198224 | -6.27180830172673 |
| ENSG00000197779.13 | ZNF81 | 0.014073750668735 | 4.64756193428234 | 0.154760108096648 | 0.87774309298517 | 0.956838320114307 | -6.34141795080008 |
| ENSG00000162882.14 | HAAO | 0.0927394876487216 | 0.501440585181965 | 0.154752208816762 | 0.877767202087998 | 0.956838320114307 | -5.84420326817474 |
| ENSG00000167526.13 | RPL13 | -0.0232414039211744 | 9.53249833030467 | -0.1546215733351 | 0.877851642379666 | 0.956838320114307 | -6.33602309941198 |
| lnc-SLC3A2-7 | lnc-SLC3A2-7 | 0.0582685223796328 | -0.440100116454213 | 0.154614475030181 | 0.877857204345264 | 0.956838320114307 | -5.47881979016145 |
| LINC01504 | LINC01504 | -0.115059412537247 | -0.847973160822518 | -0.15449524921648 | 0.877950673574137 | 0.956874645652642 | -5.27535146539076 |
| lnc-EPB41L1-2 | lnc-EPB41L1-2 | -0.0226784425603959 | 2.46585077594221 | -0.15445328484185 | 0.877983508613701 | 0.956874645652642 | -6.10183624051402 |
| ENSG00000172785.18 | ZNG1A | -0.0237672412975338 | 3.79558769958092 | -0.154272885095466 | 0.878124868786311 | 0.956978036439278 | -6.25771694874144 |
| ENSG00000132475.10 | H3-3B | 0.0276376767177346 | 10.0250019183962 | 0.15385913858791 | 0.878449093291146 | 0.957231261386871 | -6.31936133577962 |
| ENSG00000138796.16 | HADH | 0.0192865449141086 | 4.08682757148424 | 0.153720900282646 | 0.878557425792797 | 0.957231261386871 | -6.29131104411263 |
| ENSG00000275700.4 | AATF | -0.016016634644294 | 5.41053231936 | -0.153697679905369 | 0.878575623017487 | 0.957231261386871 | -6.38518360866823 |
| ENSG00000273167.1 | | -0.0628366682247528 | 1.98807224227287 | -0.153708083207437 | 0.878578354980172 | 0.957231261386871 | -6.03447011932758 |
| ENSG00000197956.9 | S100A6 | -0.0261243340759867 | 6.09674444623784 | -0.153639683943647 | 0.878621073281783 | 0.957231261386871 | -6.3983081586366 |
| ENSG00000118985.15 | ELL2 | 0.0343534946000581 | 3.67573156194063 | 0.153620300261827 | 0.878636263974258 | 0.957231261386871 | -6.25971963180047 |
| lnc-MFSD14B-16 | lnc-MFSD14B-16 | 0.0383185583368808 | 1.34424831370004 | 0.153423350953657 | 0.878790612732006 | 0.957348744673594 | -5.93431135329146 |
| ENSG00000130830.14 | MPP1 | -0.084167745970863 | -0.27977910460255 | -0.152901063114579 | 0.879202750424748 | 0.957609866510066 | -5.59678088476337 |
| lnc-KIF14-1 | lnc-KIF14-1 | -0.0317545333764457 | 1.46117285040929 | -0.152889602627576 | 0.879208934138814 | 0.957609866510066 | -5.99826663985038 |
| lnc-ELMOD2-4 | lnc-ELMOD2-4 | 0.0334823363831711 | 1.68124088027495 | 0.152874093735174 | 0.879221089644229 | 0.957609866510066 | -5.99106895251687 |
| lnc-SYNE3-1 | lnc-SYNE3-1 | -0.0320804430581436 | 2.83681668061992 | -0.15286821024048 | 0.879225700996824 | 0.957609866510066 | -6.14560128696851 |
| lnc-SEZ6L2-1 | lnc-SEZ6L2-1 | -0.0233045978346762 | 0.85922033346704 | -0.152820712306492 | 0.879262928978951 | 0.957609866510066 | -5.87158975998225 |

| | | | | | | | |
|---|---|---|---|---|---|---|---|
| ENSG00000141543.10 | EIF4A3 | -0.0210804698240858 | 4.36645681524669 | -0.152553775919854 | 0.879472153808873 | 0.957787055124234 | -6.32184774665322 |
| ENSG00000126005.16 | MMP24OS | 0.0131920123746987 | 3.46194577216058 | 0.152456390154803 | 0.879548486975123 | 0.957819507296384 | -6.21892176147068 |
| lnc-ANKRD29-1 | lnc-ANKRD29-1 | 0.0276866820527604 | -0.253433231309505 | 0.15230740598994 | 0.879665266377954 | 0.957895999251795 | -5.58679120099837 |
| ENSG00000135912.10 | TTLL4 | -0.0210162171980435 | 3.39229784202052 | -0.152107305286158 | 0.879822117115285 | 0.957966982843697 | -6.2227450197592 |
| ENSG00000145996.11 | CDKAL1 | -0.012316512144694 | 5.33454563632467 | -0.152098935599859 | 0.879828677875662 | 0.957966982843697 | -6.38236872942826 |
| ENSG00000148606.12 | POLR3A | 0.0136505426509244 | 4.52449482397439 | 0.152046121226688 | 0.879870077766122 | 0.957966982843697 | -6.33383413655998 |
| ENSG00000285952.1 | | 0.0449125344562497 | 0.285042872335243 | 0.151760041673827 | 0.880094334411298 | 0.957989883317373 | -5.74142385170024 |
| lnc-RRN3-2 | lnc-RRN3-2 | 0.0204232812196448 | 1.41182632057182 | 0.151759986824913 | 0.880094377408103 | 0.957989883317373 | -5.96717524635277 |
| ENSG00000122545.19 | SEPTIN7 | 0.0163750948075698 | 8.469646907226 | 0.151754065921492 | 0.880099018886498 | 0.957989883317373 | -6.3674196133397 |
| ENSG00000129245.11 | FXR2 | -0.0219460592359964 | 3.60592263025899 | -0.151731639170975 | 0.880116599566271 | 0.957989883317373 | -6.24758480056439 |
| ENSG00000023516.8 | AKAP11 | 0.0143505341630199 | 7.29522033760892 | 0.151670970968915 | 0.880164158623471 | 0.957989883317373 | -6.39199369593588 |
| ENSG00000158290.16 | CUL4B | 0.0144052133004759 | 6.73170488170624 | 0.151594640595604 | 0.880223996216279 | 0.957989883317373 | -6.39785500381158 |
| ENSG00000108010.11 | GLRX3 | 0.0176941020689876 | 4.82398925242202 | 0.151538991747543 | 0.880267621408248 | 0.957989883317373 | -6.36169678049127 |
| lnc-THAP12-1 | lnc-THAP12-1 | -0.0237566030416634 | 2.33952361488735 | -0.151422494640075 | 0.880358949021633 | 0.957989883317373 | -6.07329608209026 |
| ENSG00000162735.18 | PEX19 | -0.0154896949507712 | 5.25788942239552 | -0.151415931796544 | 0.880364093995304 | 0.957989883317373 | -6.38065805012914 |
| ENSG00000274258.1 | MIR6728 | 0.0476596163653168 | -0.0004539764218889 | 0.151372615256522 | 0.88039805234244 | 0.957989883317373 | -5.61204209771576 |
| ENSG00000159216.18 | RUNX1 | -0.0175679629315866 | 4.9830007591297 | -0.151366199804121 | 0.8804030818069 | 0.957989883317373 | -6.36515997005857 |
| lnc-STK4-1 | lnc-STK4-1 | -0.0140709705758999 | 5.3914511152062 | -0.151235445163829 | 0.8805055894334 | 0.95801706314387 | -6.38637156987267 |
| lnc-CENPP-1 | lnc-CENPP-1 | -0.0374467113201003 | -0.0903423791604978 | -0.151166440241256 | 0.880559688011722 | 0.95801706314387 | -5.60989592408204 |
| ENSG00000110514.19 | MADD | -0.0125894724364475 | 6.3598119460865 | -0.15115623007639 | 0.880567692640474 | 0.95801706314387 | -6.39945385125133 |
| ENSG00000165516.10 | KLHDC2 | 0.0126593818795389 | 5.6580657553261 | 0.151046513311476 | 0.880653709871733 | 0.958027453210744 | -6.39347288886355 |
| lnc-LIPH-2 | lnc-LIPH-2 | 0.0254622469018884 | 0.930101425573037 | 0.150899398076395 | 0.880769049549182 | 0.958027453210744 | -5.88881978238684 |
| ENSG00000197965.11 | MPZL1 | -0.0347719745905293 | 2.97382052600381 | -0.150831026267541 | 0.880822654558612 | 0.958027453210744 | -6.1698402599131 |
| ENSG00000152455.15 | SUV39H2 | -0.0327770891314851 | 0.609857797981023 | -0.150800104521826 | 0.880846898076475 | 0.958027453210744 | -5.85676004478222 |
| lnc-CSAD-1 | lnc-CSAD-1 | -0.0389226860714076 | 4.07189872912238 | -0.150773692443417 | 0.880869841540565 | 0.958027453210744 | -6.28023264988565 |
| ENSG00000119280.16 | C1orf198 | 0.0317128337872554 | 2.28472391364329 | 0.150695379414455 | 0.880929006363459 | 0.958027453210744 | -6.09541937164713 |
| ENSG00000173137.11 | ADCK5 | 0.0350699917243151 | 2.06774287795309 | 0.150676615190336 | 0.880943718336601 | 0.958027453210744 | -6.05671832379485 |
| ENSG00000175305.17 | CCNE2 | -0.0823039325730535 | -0.704405154706897 | -0.150658259355089 | 0.880958110156162 | 0.958027453210744 | -5.44832623192271 |
| lnc-DKKL1-1 | lnc-DKKL1-1 | 0.0600810382675761 | 0.206782974366356 | 0.150604138453813 | 0.881000543670359 | 0.958027453210744 | -5.66989919425152 |
| ENSG00000199933.1 | RNY1P16 | 0.0415194324065253 | 1.51774977992722 | 0.150380980369517 | 0.88117551459109 | 0.958027453210744 | -6.00398355230776 |
| lnc-CYP7A1-7 | lnc-CYP7A1-7 | -0.0200319612399285 | 1.6174651537435 | -0.150365210196397 | 0.881187879692961 | 0.958027453210744 | -5.98620307702725 |
| lnc-AVEN-1 | lnc-AVEN-1 | -0.0289323021867865 | 0.177838661188925 | -0.150338074299904 | 0.881209156519103 | 0.958027453210744 | -5.70017281394414 |
| ENSG00000167965.17 | MLST8 | -0.0190437977391471 | 3.83228683570903 | -0.150263877823795 | 0.881267333252749 | 0.958027453210744 | -6.27547220560069 |
| ENSG00000164674.15 | SYTL3 | -0.0428808077908691 | 1.75499289964335 | -0.150190515604302 | 0.881324856506294 | 0.958027453210744 | -6.03534029000405 |
| ENSG00000258289.8 | CHURC1 | -0.0185868802300497 | 5.28731981893976 | -0.150060250861918 | 0.881426998576949 | 0.958027453210744 | -6.38087261413423 |
| ENSG00000172361.5 | CFAP53 | -0.0299835904859884 | -0.151724446668633 | -0.150040350990658 | 0.881442602472926 | 0.958027453210744 | -5.60837427233114 |
| lnc-TIMM10B-2 | lnc-TIMM10B-2 | 0.0314927697587799 | 1.40553631546204 | 0.149834331848837 | 0.88160414909455 | 0.958027453210744 | -5.96100135989121 |
| ENSG00000252782.1 | RNU6-341P | 0.0345410105210821 | 0.0862330842536117 | 0.149809384660246 | 0.881623711380692 | 0.958027453210744 | -5.68710538106553 |
| LINC02576 | LINC02576 | -0.0446552384831111 | 2.41729207139535 | -0.14979841128878 | 0.881633039055993 | 0.958027453210744 | -6.09344363228394 |
| ENSG00000094631.19 | HDAC6 | -0.0178720611291618 | 6.0773501585472 | -0.14968516106688 | 0.881721122166734 | 0.958027453210744 | -6.39903057657884 |
| ENSG00000196296.13 | ATP2A1 | -0.0428825165334342 | 0.419238282156857 | -0.14967795406042 | 0.881726773647355 | 0.958027453210744 | -5.77978632360618 |
| ENSG00000167258.13 | CDK12 | 0.0143063975471369 | 7.25001726365901 | 0.149441636340576 | 0.881912089132955 | 0.958027453210744 | -6.39272656784285 |
| lnc-SCYL3-3 | lnc-SCYL3-3 | -0.0254108735081732 | 0.227481648170817 | -0.149424088481785 | 0.881925850069334 | 0.958027453210744 | -5.72472992303128 |
| ENSG00000107815.9 | TWNK | 0.0329498927526815 | 2.32896224534517 | 0.149383206742491 | 0.881957909452324 | 0.958027453210744 | -6.09632865152614 |
| lnc-ZNF124-1 | lnc-ZNF124-1 | 0.0519502955939126 | 2.10785074785813 | 0.149345784022178 | 0.881993740737211 | 0.958027453210744 | -6.09515868013403 |
| ENSG00000160712.12 | IL6R | -0.0586647632035451 | 3.84148261696779 | -0.149337993267685 | 0.882010704042468 | 0.958027453210744 | -6.24788486357542 |
| lnc-DTWD2-15 | lnc-DTWD2-15 | 0.0485257995264674 | 0.190869168610685 | 0.149309048626081 | 0.882016064618751 | 0.958027453210744 | -5.68566521653097 |
| ENSG00000167716.18 | WDR81 | -0.0243906224694932 | 5.89771333020073 | -0.149142295588331 | 0.882146835591892 | 0.958027453210744 | -6.39800548588009 |
| ENSG00000120159.12 | CAAP1 | -0.0148071792165569 | 3.73261030939927 | -0.149028931230366 | 0.882235740024712 | 0.958027453210744 | -6.26263598632774 |
| ENSG00000124541.6 | RRP36 | 0.0202930710869897 | 3.65688887216645 | 0.148872044464755 | 0.882358778839343 | 0.958027453210744 | -6.24906062872893 |
| lnc-POLR3GL-3 | lnc-POLR3GL-3 | 0.0314337356288595 | 0.73647475126226 | 0.148870070940024 | 0.882360326599481 | 0.958027453210744 | -5.83504291721265 |
| ENSG00000196204.11 | RNF216P1 | 0.0189917748062727 | 3.58326556333547 | 0.148763183262943 | 0.882444155220554 | 0.958027453210744 | -6.24333499202499 |
| ENSG00000259522.3 | | 0.0214549961622912 | 3.39468409963487 | 0.148694028841972 | 0.882498391570069 | 0.958027453210744 | -6.2259891845425 |
| lnc-RMDN3-6 | lnc-RMDN3-6 | 0.0385045560062895 | 2.72898095733339 | 0.148653028812732 | 0.882530547295128 | 0.958027453210744 | -6.1326902837668 |
| lnc-PRDM14-1 | lnc-PRDM14-1 | -0.0236090200131798 | 1.64078042842477 | -0.148649953677694 | 0.882532959086779 | 0.958027453210744 | -6.00487695932966 |
| ENSG00000166123.13 | GPT2 | -0.0554821987986953 | 6.41196486031915 | -0.148641444724872 | 0.882559378561461 | 0.958027453210744 | -6.39967984566614 |
| ENSG00000171723.15 | GPHN | 0.147128335295903 | 0.698211794409195 | 0.148604531468857 | 0.882594230884968 | 0.958027453210744 | -5.69356143660409 |
| lnc-MYO6-1 | lnc-MYO6-1 | -0.0455292525568032 | 2.5744473996506 | -0.148562033492729 | 0.882604562488492 | 0.958027453210744 | -6.12837982823324 |
| ENSG00000166326.6 | TRIM44 | -0.0101003836286679 | 7.24473151421601 | -0.148481019532988 | 0.882665453849279 | 0.958027453210744 | -6.39286455165261 |
| ENSG00000187627.15 | RGPD1 | 0.158910489293521 | -1.34013839599825 | 0.148487288558924 | 0.882672154438229 | 0.958027453210744 | -5.23718465723992 |
| ENSG00000140718.20 | FTO | -0.0136998245215532 | 5.45518589251993 | -0.148455530053569 | 0.882685445499945 | 0.958027453210744 | -6.38628923112906 |
| ENSG00000254341.2 | SNORD87 | -0.0338408746440044 | 1.07960895333793 | -0.148450180030183 | 0.882689641585997 | 0.958027453210744 | -5.94325225519517 |
| ENSG00000136161.12 | RCBTB2 | 0.0523758474206548 | 1.99647260826793 | 0.148399839007495 | 0.882733835230465 | 0.958027453210744 | -6.04461801402758 |
| lnc-EFCAB12-1 | lnc-EFCAB12-1 | -0.0283978270453877 | 5.11138623634963 | -0.148386697977652 | 0.882739431564101 | 0.958027453210744 | -6.37122533939125 |
| ENSG00000099331.13 | MYO9B | 0.0176328697717496 | 7.63958969502153 | 0.148385519000473 | 0.882740356259094 | 0.958027453210744 | -6.38676691106977 |
| lnc-MRM2-2 | lnc-MRM2-2 | -0.0297885030872524 | 0.0177650170243115 | -0.148361984450252 | 0.88275881490438 | 0.958027453210744 | -5.65679814731823 |
| ENSG00000218713.1 | | 0.0471958132942657 | 1.06072542712723 | 0.148354304038558 | 0.882764838827915 | 0.958027453210744 | -5.87157290614551 |
| ENSG00000127084.18 | FGD3 | -0.0167420905583393 | 6.75701806701878 | -0.148232860501616 | 0.882860090720106 | 0.958029839334955 | -6.39836766350376 |

| | | | | | | | |
|---|---|---|---|---|---|---|---|
| ENSG00000124299.13 | PEPD | 0.0160942396358994 | 4.08322789248096 | 0.148198553536509 | 0.882886999043281 | 0.958029839334955 | -6.29548515119654 |
| lnc-NHLRC3-1 | lnc-NHLRC3-1 | 0.0392276952385559 | -0.0948295420022269 | 0.148089118730001 | 0.882972834068669 | 0.958029839334955 | -5.64537343221503 |
| lnc-LRRC56-1 | lnc-LRRC56-1 | -0.0207420198400552 | 4.01692067405278 | -0.14806140757513 | 0.882994569497024 | 0.958029839334955 | -6.29235387830552 |
| ENSG00000134452.19 | FBH1 | -0.0191714122189763 | 7.12382021800155 | -0.14802877505585 | 0.883020165147209 | 0.958029839334955 | -6.39528809605259 |
| lnc-DNAJC28-6 | lnc-DNAJC28-6 | 0.021489089668746 | 0.971601875491778 | 0.147995447891818 | 0.883046305779661 | 0.958029839334955 | -5.89334347725715 |
| ENSG00000259132.1 | | 0.0282430070222439 | 1.45714655059378 | 0.147908606469477 | 0.883114421697021 | 0.958052741847483 | -5.9521466527549 |
| ENSG00000146221.9 | TCTE1 | 0.0354026495005961 | 0.504367498418126 | 0.147849852156762 | 0.883160507399998 | 0.958052741847483 | -5.78925064479159 |
| lnc-HAX1-3 | lnc-HAX1-3 | 0.0292378721019571 | -0.0408330697736363 | 0.147713803623588 | 0.883267222709115 | 0.958118010277824 | -5.65088813731434 |
| lnc-PMS2-4 | lnc-PMS2-4 | 0.0186130845175336 | 2.58908654910049 | 0.147634677610345 | 0.883329289488254 | 0.95813484219734 | -6.12310817271877 |
| ENSG00000168961.16 | LGALS9 | 0.0281769717815452 | 5.28669759605958 | 0.147484765374559 | 0.88344688331302 | 0.958192921893028 | -6.38341911774439 |
| lnc-ST8SIA1-5 | lnc-ST8SIA1-5 | 0.0196822516252275 | 1.74119808987127 | 0.147383775777637 | 0.883526102849102 | 0.958192921893028 | -6.01587998046026 |
| ENSG00000162191.13 | UBXN1 | -0.0185838343144464 | 5.8216745227671 | -0.147343751745098 | 0.883557499340401 | 0.958192921893028 | -6.39684569769627 |
| ENSG00000087365.15 | SF3B2 | 0.0149570876717648 | 6.96905083381527 | 0.147266186668462 | 0.883618345104299 | 0.958192921893028 | -6.39676340935414 |
| lnc-GPR68-3 | lnc-GPR68-3 | -0.0209896451266094 | 2.01594186344082 | -0.147241263232831 | 0.883637896392775 | 0.958192921893028 | -6.04472472033826 |
| lnc-ADAP2-5 | lnc-ADAP2-5 | -0.062340886753934 | -0.884641691892643 | -0.147164348307208 | 0.883698233075636 | 0.958192921893028 | -5.38539902465273 |
| ENSG00000241057.2 | RASA4EP | 0.0207599316764686 | 1.08080576224089 | 0.147151001582054 | 0.883708703118853 | 0.958192921893028 | -5.91523213823217 |
| ENSG00000186395.7 | KRT10 | -0.0267315999364548 | 1.92290875837727 | -0.146848929446477 | 0.883945673860026 | 0.958324974537566 | -6.0392744619589 |
| lnc-CENPK-3 | lnc-CENPK-3 | 0.0395130904882856 | 0.00316225667325097 | 0.146822772177907 | 0.883966194323547 | 0.958324974537566 | -5.65933140450832 |
| ENSG00000068784.12 | SRBD1 | 0.0179470283513248 | 3.815845809431 | 0.146817707061687 | 0.883970167933257 | 0.958324974537566 | -6.26327779708557 |
| ENSG00000168016.14 | TRANK1 | 0.032575301943865 | 6.21118491354809 | 0.146702070438171 | 0.884062246597422 | 0.958374320414743 | -6.40003451159243 |
| ENSG00000127914.16 | AKAP9 | 0.0138792404156352 | 8.2269760827565 | 0.146148180776286 | 0.884495441315556 | 0.958792926078273 | -6.37510259606446 |
| ENSG00000132002.7 | DNAJB1 | 0.0270353167177939 | 4.47120253748737 | 0.146039735803638 | 0.884580526212138 | 0.958792926078273 | -6.3313992442479 |
| ENSG00000205250.8 | E2F4 | 0.0170180808715603 | 5.35825809036884 | 0.146030032935287 | 0.884588139057785 | 0.958792926078273 | -6.38705690738608 |
| ENSG00000207496.1 | SNORA7A | 0.0298503464285322 | 2.77913802165078 | 0.145272015864707 | 0.885182911357731 | 0.959376787124445 | -6.14362171717115 |
| ENSG00000120509.10 | PDZD11 | -0.0204153308689445 | 2.48219716542473 | -0.145166304582926 | 0.885265862238603 | 0.959376787124445 | -6.10925821435457 |
| lnc-SAMD12-4 | lnc-SAMD12-4 | 0.0620747680628631 | 0.213046208722433 | 0.145092457854787 | 0.885323810006571 | 0.959376787124445 | -5.69360037176172 |
| ENSG00000121057.12 | AKAP1 | -0.0195667902642072 | 3.75808596941427 | -0.145089497607926 | 0.885326132935163 | 0.959376787124445 | -6.26459572739977 |
| ENSG00000178163.7 | ZNF518B | 0.0387094831935348 | 6.02797040487612 | 0.144894877434123 | 0.885486911996942 | 0.959376787124445 | -6.39914357059963 |
| ENSG00000106460.18 | TMEM106B | -0.0221701624349793 | 5.47375730113032 | -0.144869207468812 | 0.885498999172674 | 0.959376787124445 | -6.39152316084073 |
| ENSG00000092098.16 | RNF31 | -0.0157690630463385 | 4.53002151280642 | -0.144826175598255 | 0.885532767832485 | 0.959376787124445 | -6.34076057407684 |
| lnc-SRP54-1 | lnc-SRP54-1 | 0.0381351318322813 | 0.96781727898281 | 0.144802679073918 | 0.885551206492888 | 0.959376787124445 | -5.91160796702314 |
| ENSG00000120992.17 | LYPLA1 | -0.0273127372529203 | 4.92796134967296 | -0.144726260160827 | 0.885611175907653 | 0.959376787124445 | -6.37320286023802 |
| ENSG00000184898.6 | RBM43 | 0.0196224386598667 | 2.86367178531187 | 0.144715686680637 | 0.885619473454393 | 0.959376787124445 | -6.15823788493308 |
| lnc-TYW1-7 | lnc-TYW1-7 | -0.0265495394430909 | 0.341678871490047 | -0.144680602837988 | 0.885647005620035 | 0.959376787124445 | -5.7348605934487 |
| ENSG00000083812.11 | ZNF324 | -0.0176852662569917 | 3.45785310470957 | -0.144578030142053 | 0.885727500714274 | 0.959376787124445 | -6.226682124801 |
| ENSG00000096996.15 | IL12RB1 | -0.0284059938070516 | 3.04451107745766 | -0.144571347831609 | 0.885732744776069 | 0.959376787124445 | -6.16697000306264 |
| ENSG00000186130.4 | ZBTB6 | -0.0226519605019675 | 4.54402279301599 | -0.144500489130111 | 0.885788352719297 | 0.95938653252059 | -6.3403752219395 |
| ENSG00000164039.14 | BDH2 | -0.0335054226920977 | 2.72547688118961 | -0.144401186125811 | 0.885866283942382 | 0.9594204536904 | -6.12183434043 |
| SNHG1 | SNHG1 | -0.0235020352328137 | 7.15586843101751 | -0.144233999292648 | 0.885997491763415 | 0.959512068450298 | -6.39355720265454 |
| lnc-GPAT2-5 | lnc-GPAT2-5 | -0.027757240647582 | 0.995256734577675 | -0.143994513493211 | 0.886185445275991 | 0.959604338656067 | -5.90296024018737 |
| ENSG00000234709.2 | UPF3AP3 | 0.0336962054969069 | -0.104509867596816 | 0.143944265877771 | 0.886224881507601 | 0.959604338656067 | -5.62728483957195 |
| ENSG00000172264.17 | MACROD2 | -0.0713917666456088 | 1.4866273130617 | -0.143960593363957 | 0.886227294596073 | 0.959604338656067 | -5.98492271519885 |
| ENSG00000103335.22 | PIEZO1 | -0.0553021090738728 | 5.49096272435454 | -0.143912169413408 | 0.886269177372192 | 0.959604338656067 | -6.39143071991386 |
| lnc-MAP3K1-6 | lnc-MAP3K1-6 | 0.0547299660867619 | 1.78642404247856 | 0.143596475922948 | 0.886504191719025 | 0.959747706722989 | -6.01533414806368 |
| ENSG00000185864.16 | NPIPB4 | -0.0365536189896307 | 2.76741469235026 | -0.143536406256618 | 0.886544995973523 | 0.959747706722989 | -6.15420616197026 |
| ENSG00000122884.12 | P4HA1 | -0.0129034814425397 | 4.67170207984519 | -0.143467917996084 | 0.886598751847344 | 0.959747706722989 | -6.34711267719657 |
| lnc-CSRNP1-4 | lnc-CSRNP1-4 | 0.0209952077268167 | 1.5791021182759 | 0.143460736580996 | 0.886604388512436 | 0.959747706722989 | -5.98614102393197 |
| lnc-NAT1-6 | lnc-NAT1-6 | 0.0273125549311457 | 1.26754810323161 | 0.143422080283966 | 0.886634729793404 | 0.959747706722989 | -5.95224154545235 |
| lnc-PSMB2-3 | lnc-PSMB2-3 | -0.015543607732821 | 0.496904029213261 | -0.143266770395039 | 0.886756634072324 | 0.959829185901906 | -5.7985936195512 |
| ENSG00000141646.13 | SMAD4 | -0.0175656425270851 | 7.02363679531279 | -0.143129956499063 | 0.88686402296117 | 0.959894945817414 | -6.39592084352243 |
| LINC01918 | LINC01918 | -0.0593924892736855 | 3.08858919325016 | -0.142923931409734 | 0.887041562145592 | 0.960016726157991 | -6.17052115616453 |
| ENSG00000198920.9 | KIAA0753 | 0.0176872422745076 | 3.62146605978285 | 0.142801239265693 | 0.887122050706287 | 0.960016726157991 | -6.25001044709876 |
| ENSG00000170396.7 | ZNF804A | -0.182087150924096 | 2.48448205122115 | -0.142784333150363 | 0.887162125917236 | 0.960016726157991 | -6.08340111152001 |
| lnc-SEM1-1 | lnc-SEM1-1 | -0.0332233425068159 | 0.703021548406071 | -0.142748941758481 | 0.887163102961237 | 0.960016726157991 | -5.84209150843808 |
| ENSG00000136490.8 | LIMD2 | -0.0193071873236939 | 7.78760245980121 | -0.142626642781948 | 0.887259105869021 | 0.96007013858175 | -6.38341627307074 |
| lnc-PRDX2-1 | lnc-PRDX2-1 | 0.0696201885295904 | 0.756401901815148 | 0.142573683700872 | 0.887313488653144 | 0.960078512166728 | -5.92381210391848 |
| lnc-SLC25A10-4 | lnc-SLC25A10-4 | 0.0340477373313922 | -0.244174116291758 | 0.142380531634994 | 0.887452304710867 | 0.960178237377196 | -5.57674215486075 |
| lnc-RNFT2-6 | lnc-RNFT2-6 | -0.0339319051545843 | 0.0326324901012842 | -0.142144076170052 | 0.887637930303855 | 0.960285523911917 | -5.670180788134 |
| ENSG00000108091.10 | CCDC6 | 0.0250036362798607 | 6.54758206626676 | 0.142114642439749 | 0.887661037231504 | 0.960285523911917 | -6.40030993608723 |
| ENSG00000103168.16 | TAF1C | -0.0199147213648052 | 5.22650800947513 | -0.142016608022834 | 0.887737999789184 | 0.960285523911917 | -6.38063061675377 |
| ENSG00000167578.17 | RAB4B | -0.0195006053960234 | 3.22949091927024 | -0.141993010393764 | 0.887756525425339 | 0.960285523911917 | -6.20005415104122 |
| lnc-TASP1-3 | lnc-TASP1-3 | -0.0281475885966223 | 1.53569329096778 | -0.141957075584737 | 0.887784736652571 | 0.960285523911917 | -5.98780874154209 |
| ENSG00000171574.17 | ZNF584 | 0.022848779319568 | 1.30026844892604 | 0.141848348564432 | 0.887870095528115 | 0.960324201488785 | -5.94949307905435 |
| lnc-LIN9-1 | lnc-LIN9-1 | 0.0312173868698693 | 2.67797432146402 | 0.141738016445773 | 0.887956715903182 | 0.960324201488785 | -6.13031150782477 |
| ENSG00000197586.12 | ENTPD6 | -0.0233725185547602 | 4.98140085768364 | -0.141651513292348 | 0.888024629423048 | 0.960324201488785 | -6.36490172889173 |
| lnc-CHD2-16 | lnc-CHD2-16 | 0.0316027879126792 | 0.969139077745658 | 0.14163349087088 | 0.888038778911264 | 0.960324201488785 | -5.8989802949874 |
| lnc-SIPA1L3-6 | lnc-SIPA1L3-6 | -0.142129221524828 | -1.69845752549346 | -0.141570378933534 | 0.888104691598963 | 0.960324201488785 | -5.40151294675259 |
| ENSG00000167996.15 | FTH1 | -0.0221177529735455 | 6.5879470753325 | -0.14153564342213 | 0.888115600050175 | 0.960324201488785 | -6.40032195746019 |

| | | | | | | | |
|---|---|---|---|---|---|---|---|
| ENSG00000013523.9 | ANGEL1 | 0.0246128218413055 | 4.85723103167771 | 0.141363355260392 | 0.88825086807998 | 0.960324201488785 | -6.35598676742432 |
| ENSG00000166886.12 | NAB2 | 0.0458573385224297 | 1.81598883429622 | 0.141320316753683 | 0.8882853404774 | 0.960324201488785 | -6.01509363194131 |
| ENSG00000145912.8 | NHP2 | 0.0253439984689448 | 3.91535977037221 | 0.141264547316131 | 0.888328446352739 | 0.960324201488785 | -6.28031455935521 |
| ENSG00000217624.2 | YWHAZP10 | -0.0412202011062697 | 0.0921620114074879 | -0.141205505050492 | 0.888374803447533 | 0.960324201488785 | -5.62862385738814 |
| lnc-AGO2-1 | lnc-AGO2-1 | -0.0239880262594134 | 3.52594968988547 | -0.141170381216727 | 0.888402381148848 | 0.960324201488785 | -6.23991656545507 |
| ENSG00000173548.8 | SNX33 | 0.0293287662888873 | 1.5832446660232 | 0.141153492095859 | 0.888415641799247 | 0.960324201488785 | -5.98697239586093 |
| lnc-GDF5-1 | lnc-GDF5-1 | 0.0190639658932132 | 1.68245874203661 | 0.14113899484805 | 0.888427024474054 | 0.960324201488785 | -5.99412562800525 |
| ENSG00000117862.12 | TXNDC12 | -0.0145301844749008 | 3.67183034811384 | -0.141060598594303 | 0.888488578574391 | 0.960340304195593 | -6.25466869197458 |
| ENSG00000214765.8 | SEPTIN7P2 | -0.0138730388258921 | 4.35351458885789 | -0.140935946875494 | 0.888586452343151 | 0.960395659975797 | -6.32311748540276 |
| lnc-URGCP-MRPS24-4 | lnc-URGCP-MRPS2 | 0.0216353548693844 | 1.18607204784708 | 0.140846493873478 | 0.888656689943307 | 0.960421141984935 | -5.92306887164755 |
| ENSG00000103490.13 | PYCARD | 0.0423218057969925 | 2.16488341117247 | 0.140327639520031 | 0.889064268694257 | 0.960811185683812 | -6.06335299193586 |
| lnc-EXOC8-1 | lnc-EXOC8-1 | -0.0241469012249469 | 1.65707745404723 | -0.140202756347128 | 0.889162172859716 | 0.960866540871084 | -6.01235311486841 |
| ENSG00000153029.14 | MR1 | 0.0187669562770978 | 4.99232846936951 | 0.13997163942225 | 0.889343664523246 | 0.960965043732044 | -6.36261441693137 |
| ENSG00000084093.16 | REST | -0.012790191834851 | 6.75264969439012 | -0.139949472940055 | 0.88936107175002 | 0.960965043732044 | -6.39940217697152 |
| ENSG00000269026.2 | | 0.0145762992766398 | 2.82711173666275 | 0.139908322759557 | 0.889393386925883 | 0.960965043732044 | -6.15082126489705 |
| ENSG00000121988.17 | ZRANB3 | -0.0197608641903946 | 3.16406019280799 | -0.139839338197122 | 0.889447560827671 | 0.960973132356094 | -6.18760267901039 |
| ENSG00000082641.15 | NFE2L1 | 0.0207560575864312 | 5.76990593872222 | 0.139655464881879 | 0.889591960019223 | 0.96102553512486 | -6.39838650291068 |
| ENSG00000209480.1 | SNORD83B | -0.0258957420960019 | -0.33136087100104 | -0.139613377760552 | 0.889625012365191 | 0.96102553512486 | -5.60763424581865 |
| ENSG00000143641.9 | GALNT2 | 0.0177416069803087 | 5.13114728430693 | 0.139576637405946 | 0.889653865888776 | 0.96102553512486 | -6.3780638640649 |
| ENSG00000155876.5 | RRAGA | 0.0175834549262106 | 4.21066805366408 | 0.139502946297056 | 0.889711738626649 | 0.96102553512486 | -6.30900212840627 |
| ENSG00000175197.12 | DDIT3 | 0.0330342554812431 | 4.62961659537068 | 0.139482575465034 | 0.889729514518745 | 0.96102553512486 | -6.35175980762591 |
| ENSG00000169118.17 | CSNK1G1 | 0.0122481189985075 | 5.21125792243549 | 0.139216865730797 | 0.889936415534302 | 0.961154032042432 | -6.37969630025076 |
| ENSG00000163393.12 | SLC22A15 | -0.0309894547363951 | 4.48830251262045 | -0.139201701466645 | 0.889948325232188 | 0.961154032042432 | -6.33093273830938 |
| ENSG00000278979.1 | | 0.0410762123480533 | 0.383975666278525 | 0.139136152935193 | 0.889999805982247 | 0.961154032042432 | -5.73363337317435 |
| lnc-YARS2-4 | lnc-YARS2-4 | 0.0351572024462767 | -0.286300569052724 | 0.139084510186219 | 0.890040365695321 | 0.961154032042432 | -5.58916603188133 |
| ENSG00000144792.9 | ZNF660 | 0.0417950405316736 | 2.33648288007597 | 0.138977671714462 | 0.890125563876947 | 0.961154032042432 | -6.10311036161259 |
| lnc-MASTL-7 | lnc-MASTL-7 | 0.0184112520101383 | 1.5409924442452 | 0.138972093661638 | 0.890128657571435 | 0.961154032042432 | -5.99199038998637 |
| lnc-FBXO30-2 | lnc-FBXO30-2 | -0.0339791050974831 | 1.35738095785503 | -0.138881205175193 | 0.890200042370074 | 0.96118068887915 | -5.98948267539719 |
| ENSG00000174028.6 | FAM3C2P | -0.0427406617165937 | 1.67090851520607 | -0.138809096600257 | 0.890256677879282 | 0.961191418421593 | -5.99133915627165 |
| ENSG00000279254.1 | | 0.0644628196140128 | -0.799167776164272 | 0.138589325379144 | 0.890429294144426 | 0.961327362254116 | -5.37851052278789 |
| ENSG00000108448.21 | TRIM16L | -0.0361729670483991 | -0.323918715597261 | -0.138273446749062 | 0.890677406063752 | 0.961544794346491 | -5.54243653420001 |
| ENSG00000256751.5 | PLBD1-AS1 | 0.0850059834428202 | -0.457585176288684 | 0.138099383808159 | 0.890817167331708 | 0.961645238117614 | -5.51602684184315 |
| ENSG00000169019.10 | COMMD8 | 0.0333030362238378 | 2.38875575002015 | 0.137801366506097 | 0.891048229695845 | 0.961844226548646 | -6.10496358690354 |
| ENSG00000125149.11 | PHAF1 | -0.0161850379876158 | 3.38052547769593 | -0.137620109301551 | 0.89119061555856 | 0.961947477059198 | -6.21773831039128 |
| ENSG00000198538.10 | ZNF28 | -0.0226019056159847 | 4.5549616135361 | -0.137496742754939 | 0.891287527727997 | 0.962001635197974 | -6.33306751671519 |
| lnc-ATP6V1E2-5 | lnc-ATP6V1E2-5 | 0.0288870770342253 | 0.876098216313379 | 0.137393969279588 | 0.891368264030975 | 0.962026421196589 | -5.85454335456703 |
| ENSG00000186281.12 | GPAT2 | -0.419486162327225 | -2.54984547874569 | -0.137372915005275 | 0.89140722988014 | 0.962026421196589 | -4.87605607805126 |
| ENSG00000131558.14 | EXOC4 | -0.0158908338272106 | 6.1561900142066 | -0.137289023019462 | 0.891450708423579 | 0.962026421196589 | -6.40067777783803 |
| lnc-NEDD9-2 | lnc-NEDD9-2 | -0.0289380657708138 | 2.00078807779722 | -0.137191827618047 | 0.89152706492817 | 0.962057791008569 | -6.03827478711967 |
| lnc-BST1-2 | lnc-BST1-2 | 0.0219980531445042 | 0.974804464536014 | 0.137089973816965 | 0.891607082179565 | 0.962057791008569 | -5.88775694941544 |
| ENSG00000205937.11 | RNPS1 | 0.0115032154221728 | 6.409264646987 | 0.137073533279852 | 0.89161999811881 | 0.962057791008569 | -6.40141802985989 |
| lnc-PJA2-3 | lnc-PJA2-3 | 0.0349549363679735 | 0.564221213061491 | 0.136727326306071 | 0.891891990561899 | 0.962199678658386 | -5.8196675501405 |
| ENSG00000226435.10 | ANKRD18DP | 0.100894459037723 | 0.399525833297476 | 0.136713321757497 | 0.891921125346391 | 0.962199678658386 | -5.75052754212038 |
| ENSG00000164823.10 | OSGIN2 | -0.0184324296793319 | 4.16234268090524 | -0.136643341543646 | 0.89195797394815 | 0.962199678658386 | -6.3099489214044 |
| RSF1-IT2 | RSF1-IT2 | 0.0292617390228221 | 0.473180639960525 | 0.136616966671976 | 0.891978695763883 | 0.962199678658386 | -5.78076059504682 |
| ENSG00000130489.14 | NA | -0.0330872583612413 | 0.496157756684481 | -0.136608644961485 | 0.891985233857098 | 0.962199678658386 | -5.83979092317792 |
| ENSG00000147316.12 | MCPH1 | -0.0138312203729063 | 6.00523004543499 | -0.136500361595329 | 0.892070309216285 | 0.962206186242617 | -6.39969383650356 |
| ENSG00000257704.3 | INAFM1 | -0.0259918411511391 | 0.977736152771617 | -0.136481966606936 | 0.89208476179701 | 0.962206186242617 | -5.89730818121702 |
| ENSG00000154305.16 | MIA3 | 0.0116728295298101 | 6.83889241439567 | 0.136353338713883 | 0.892185823240755 | 0.962264766284032 | -6.39917356949853 |
| ENSG00000112031.15 | MTRF1L | -0.0154115997473383 | 3.81682457466985 | -0.136197216676197 | 0.892308488951188 | 0.962346640192942 | -6.28106050672702 |
| ENSG00000239040.1 | | -0.0322067041515922 | -0.393014706287471 | -0.136039303243359 | 0.892432564882585 | 0.962399835780596 | -5.55692566347558 |
| ENSG00000257950.3 | P2RX5-TAX1BP3 | 0.0387802286396104 | 3.94616990871031 | 0.135904883604905 | 0.892543946921263 | 0.962399835780596 | -6.28216258895253 |
| ENSG00000143384.12 | MCL1 | 0.0197634558800619 | 8.26998093076421 | 0.135749740942023 | 0.892660087203069 | 0.962399835780596 | -6.37468319082852 |
| lnc-CEP76-1 | lnc-CEP76-1 | -0.0190657972369606 | 0.763123445186082 | -0.135731242734765 | 0.892674622400214 | 0.962399835780596 | -5.86709221089817 |
| ENSG00000056277.15 | ZNF280C | -0.0115955614297949 | 3.70421156551828 | -0.135710422952366 | 0.892690981848651 | 0.962399835780596 | -6.2538196878696 |
| lnc-DDX41-4 | lnc-DDX41-4 | -0.0195932113216566 | 2.23561640798384 | -0.135698366177605 | 0.892700455657331 | 0.962399835780596 | -6.07419589593125 |
| ENSG00000177873.12 | ZNF619 | 0.0128617511334976 | 3.9262816363861 | 0.135577257586019 | 0.892795619600594 | 0.962399835780596 | -6.2779124255387 |
| ENSG00000217930.7 | PAM16 | -0.0218525657077104 | 1.19890753061215 | -0.135575189232784 | 0.892797244872008 | 0.962399835780596 | -5.92531109133466 |
| lnc-RSPH6A-2 | lnc-RSPH6A-2 | -0.0214332901348948 | 1.80159316968865 | -0.135539861671121 | 0.892825004649219 | 0.962399835780596 | -6.02139330976851 |
| ENSG00000141582.14 | CBX4 | -0.0240455880368904 | 5.07402990352631 | -0.135539379809365 | 0.89282538328866 | 0.962399835780596 | -6.37747690295165 |
| ENSG00000001629.9 | ANKIB1 | 0.00898348325710252 | 6.46086015953946 | 0.1349074219877 | 0.89332198761565 | 0.962834835838944 | -6.40160390409627 |
| ENSG00000106976.20 | DNM1 | -0.0693159255559261 | -0.221190277685723 | -0.134906779713505 | 0.893322492348847 | 0.962834835838944 | -5.57106873657764 |
| NOP14-AS1 | NOP14-AS1 | -0.0241568513190185 | 3.87235298002284 | -0.134805756570157 | 0.893401882260543 | 0.96284362321084 | -6.27817636828142 |
| ENSG00000176476.8 | SGF29 | 0.0266607390126481 | 2.9052597114026 | 0.134777354486846 | 0.893424202482817 | 0.96284362321084 | -6.15918223511261 |
| LNCOC1 | LNCOC1 | 0.0526897057422733 | 0.448033743769285 | 0.134668037928797 | 0.893510111428658 | 0.962873473494387 | -5.78736535906882 |
| lnc-ONECUT2-6 | lnc-ONECUT2-6 | -0.0208152550047656 | 1.43613676594379 | -0.134623057541955 | 0.893545460681936 | 0.962873473494387 | -5.9589779411315 |
| lnc-MED6-3 | lnc-MED6-3 | -0.0185538329944035 | 2.91205381652503 | -0.134526797546864 | 0.893621110365697 | 0.96290458143949 | -6.15795244647696 |
| ENSG00000236804.1 | RPS3AP12 | 0.0353775373408186 | -0.410388990283057 | 0.134313867456042 | 0.893788453369021 | 0.963034483363585 | -5.5490127106934 |

| | | | | | | | |
|---|---|---|---|---|---|---|---|
| lnc-PMM2-4 | lnc-PMM2-4 | -0.0357460608414779 | 1.16023018564501 | -0.134140818927636 | 0.893924456826358 | 0.963130605886565 | -5.93678126050291 |
| lnc-NPIPB4-5 | lnc-NPIPB4-5 | 0.0309633614663603 | 3.35919357495428 | 0.134053618288905 | 0.893992991359972 | 0.96315402989596 | -6.224846034963 |
| ANP32A-IT1 | ANP32A-IT1 | -0.0318268714694825 | 3.01158562282069 | -0.133968426909206 | 0.894059947526616 | 0.96317282392466 | -6.16066169639977 |
| ENSG00000146416.18 | AIG1 | -0.0254415603187907 | 2.72837255137228 | -0.133848442374474 | 0.894154250700053 | 0.96317282392466 | -6.14209105939782 |
| lnc-TMLHE-2 | lnc-TMLHE-2 | 0.0405662376672135 | -0.0041310157932461 | 0.133779105816286 | 0.894208747242108 | 0.96317282392466 | -5.74548894005439 |
| ENSG00000175198.16 | PCCA | -0.0178663720932617 | 3.91167354130135 | -0.13377634705532 | 0.894210915559642 | 0.96317282392466 | -6.27940423259219 |
| ENSG00000280063.1 | | 0.0334276491209082 | 1.18853179384682 | 0.133733733644547 | 0.894244408745093 | 0.96317282392466 | -5.9081147833583 |
| ENSG00000106603.18 | COA1 | -0.0164782206712628 | 4.41801339999961 | -0.133585055362575 | 0.894361268070143 | 0.963201859980254 | -6.32695155899947 |
| ENSG00000118515.11 | SGK1 | -0.125886261306809 | -0.418196847177524 | -0.133601423827515 | 0.894364958846748 | 0.963201859980254 | -5.49107345006773 |
| ENSG00000086232.12 | EIF2AK1 | -0.0163952954906295 | 5.96703792146393 | -0.133304897568843 | 0.89458147514651 | 0.963384633649381 | -6.40039670140545 |
| ENSG00000273682.1 | BLVRBP1 | 0.0223677915672368 | 0.213054600385366 | 0.13297670840923 | 0.894839446099756 | 0.963612028830767 | -5.71925709378091 |
| lnc-IFT27-3 | lnc-IFT27-3 | -0.0216667071442742 | 0.0542249267337237 | -0.132840702838926 | 0.894946355770657 | 0.963654714740831 | -5.67869001984898 |
| lnc-ZNF460-1 | lnc-ZNF460-1 | 0.0258704445941773 | 3.51453029008474 | 0.132807161934524 | 0.894972721517064 | 0.963654714740831 | -6.24107246936338 |
| lnc-MAP3K7CL-5 | lnc-MAP3K7CL-5 | 0.0537216575332117 | -0.706170628063606 | 0.132723843616428 | 0.895038216682547 | 0.963674824186685 | -5.47915385396239 |
| lnc-TBX15-5 | lnc-TBX15-5 | -0.0280623368061514 | 0.588010070930914 | -0.132467863944439 | 0.895239442744458 | 0.963841063396233 | -5.81250564814048 |
| ENSG00000118162.13 | KPTN | 0.0232198393600168 | 0.553225661639431 | 0.132330282038886 | 0.895347599003134 | 0.963907088774596 | -5.79574624673633 |
| ENSG00000015133.18 | CCDC88C | -0.0160994975949338 | 7.34061971450286 | -0.132121283832055 | 0.89551190109122 | 0.963934522999733 | -6.39384225687418 |
| ENSG00000158473.6 | CD1D | 0.0626944377418597 | 2.09865462909715 | 0.131909200165328 | 0.895694023346761 | 0.963934522999733 | -6.07928801572946 |
| lnc-FAM20B-2 | lnc-FAM20B-2 | -0.0404181078327217 | 3.87914925500588 | -0.131884222192114 | 0.895706133672906 | 0.963934522999733 | -6.28038617656056 |
| lnc-ANXA7-2 | lnc-ANXA7-2 | 0.0316456751336938 | 0.789138650535362 | 0.131865159111145 | 0.895713257599771 | 0.963934522999733 | -5.8671436322091 |
| lnc-ZNF45-1 | lnc-ZNF45-1 | -0.0155855053142752 | 0.830522323379063 | -0.13186182671022 | 0.895715877465454 | 0.963934522999733 | -5.86914808955187 |
| lnc-FOXO1-2 | lnc-FOXO1-2 | -0.027050643016838 | 2.1721579052134 | -0.131861635744418 | 0.89571602759891 | 0.963934522999733 | -6.07608265975944 |
| ENSG00000099622.13 | CIRBP | -0.0158787768521513 | 8.01463561510264 | -0.13184617616284 | 0.895728181622553 | 0.963934522999733 | -6.38117966258018 |
| ENSG00000180901.10 | KCTD2 | -0.0177424874609029 | 4.74911353577372 | -0.131748444859701 | 0.89580501666685 | 0.963934522999733 | -6.35662852792808 |
| ENSG00000160131.13 | VMA21 | 0.0209662207723476 | 5.61743594834472 | 0.131674464117745 | 0.895863180010535 | 0.963934522999733 | -6.3961625999184 |
| ENSG00000163013.11 | FBXO41 | 0.0195004347073882 | 4.62233547516757 | 0.131551967153847 | 0.895959487875338 | 0.963934522999733 | -6.34747144167775 |
| lnc-ESRRB-1 | lnc-ESRRB-1 | -0.0157080259870666 | 1.13828877765664 | -0.131508755109253 | 0.895993461828937 | 0.963934522999733 | -5.93248610997161 |
| lnc-MYLPF-1 | lnc-MYLPF-1 | -0.0253216731113874 | 0.281035171459713 | -0.131494987688148 | 0.896004286022128 | 0.963934522999733 | -5.71945062711758 |
| ENSG00000177105.9 | RHOG | 0.0187863174670251 | 3.42348839261531 | 0.13148339151323 | 0.896013403158212 | 0.963934522999733 | -6.23192624149472 |
| ENSG00000164053.20 | ATRIP | -0.016219616687815 | 3.80371135597524 | -0.131433373989793 | 0.896052728061521 | 0.963934522999733 | -6.27493526019956 |
| ENSG00000138398.15 | PPIG | 0.0136338830718188 | 6.27528130779757 | 0.131404339560469 | 0.896075555705043 | 0.963934522999733 | -6.40222812965529 |
| ENSG00000158792.15 | SPATA2L | 0.0336805091904817 | 1.32094384887484 | 0.13115289081183 | 0.896273255149981 | 0.96409680746026 | -5.96169106686114 |
| ENSG00000141376.22 | BCAS3 | 0.0165566109814055 | 5.28413444956253 | 0.13107136136149 | 0.896337358421174 | 0.964115376692256 | -6.38386766408687 |
| ENSG00000119718.10 | EIF2B2 | 0.0133657966455116 | 4.25793954822363 | 0.130901201684214 | 0.896471150271015 | 0.964165023775226 | -6.31492002896487 |
| ENSG00000157764.13 | BRAF | 0.0186282822692485 | 6.55987008071163 | 0.130695529317461 | 0.896632868836341 | 0.964165023775226 | -6.40190225190979 |
| ENSG00000140750.16 | ARHGAP17 | 0.0180865677615403 | 6.66965193924015 | 0.130621730590917 | 0.896690897279556 | 0.964165023775226 | -6.40161161330852 |
| ENSG00000137267.5 | TUBB2A | -0.0437691271809517 | 0.909773389486321 | -0.130605103828165 | 0.896703971095786 | 0.964165023775226 | -5.92012268707898 |
| ENSG00000179715.12 | PCED1B | 0.0295926250647116 | 3.97545313744845 | 0.130569769971793 | 0.896731754612964 | 0.964165023775226 | -6.27523491541602 |
| ENSG00000169435.13 | RASSF6 | 0.187033041609617 | -1.587743945149 | 0.130588353609981 | 0.896737910386101 | 0.964165023775226 | -5.31869718114159 |
| ENSG00000279233.1 | | 0.0412279074396991 | 0.27939485255428 | 0.130561870130024 | 0.896737966389596 | 0.964165023775226 | -5.75332319615953 |
| lnc-DTX1-1 | lnc-DTX1-1 | 0.0701887125603091 | 0.901194550666202 | 0.130556198588041 | 0.896758257550062 | 0.964165023775226 | -5.97975949967328 |
| ENSG00000185522.8 | LMNTD2 | -0.0339020867084729 | 1.85794942952489 | -0.130387162126175 | 0.896875343866031 | 0.964216176347 | -6.02672295639658 |
| lnc-CSPG4-1 | lnc-CSPG4-1 | 0.0552369001356614 | -0.423956081739293 | 0.130356411171071 | 0.896899524478436 | 0.964216176347 | -5.48803727614725 |
| ENSG00000155189.11 | AGPAT5 | -0.0287296268126482 | 6.69560295474571 | -0.130098993968837 | 0.897103132080144 | 0.964384695649742 | -6.40095129703313 |
| lnc-TMEM167B-1 | lnc-TMEM167B-1 | -0.0416652236892725 | 6.49549709317607 | -0.129959491897049 | 0.897225277594397 | 0.964437389215513 | -6.40141413452655 |
| lnc-HIBCH-1 | lnc-HIBCH-1 | 0.0219541372189233 | 0.498905565272473 | 0.129897898978384 | 0.897260081310712 | 0.964437389215513 | -5.79797464673228 |
| lnc-HLA-A-8 | lnc-HLA-A-8 | -0.0429198911332698 | 2.39934915679977 | -0.129838336959699 | 0.897309475338806 | 0.964437389215513 | -6.07046673940874 |
| ENSG00000141696.12 | P3H4 | -0.0617399786391564 | -0.912069324262101 | -0.129796814302116 | 0.897339573476475 | 0.964437389215513 | -5.46844461079818 |
| ENSG00000242574.8 | HLA-DMB | 0.0215206194394995 | 8.1955285243411 | 0.129657663477433 | 0.897449002296547 | 0.964504637336561 | -6.38037710546106 |
| ENSG00000179943.7 | FIZ1 | -0.0189174328411865 | 1.88362823538085 | -0.129335466301216 | 0.897702387321823 | 0.96454283201719 | -6.03615230594909 |
| lnc-KCNA3-3 | lnc-KCNA3-3 | -0.0228703519672759 | 5.48302253168588 | -0.129206570461818 | 0.897803757703704 | 0.96454283201719 | -6.39041357453887 |
| ENSG00000137166.14 | FOXP4 | 0.0265111964433791 | 3.1635918249428 | 0.129147843090143 | 0.897849944531482 | 0.96454283201719 | -6.19294466393654 |
| lnc-NPTX1-2 | lnc-NPTX1-2 | -0.101100067135739 | -1.46193468803241 | -0.129083830842456 | 0.897900288119995 | 0.96454283201719 | -5.21954963125088 |
| ENSG00000139266.5 | MARCHF9 | -0.0240466160672073 | 4.59145097098425 | -0.129016144410656 | 0.897953521799924 | 0.96454283201719 | -6.34564430566288 |
| lnc-BAG6-1 | lnc-BAG6-1 | 0.0179172703000747 | 1.87993416900327 | 0.12897509385622 | 0.897985807262487 | 0.96454283201719 | -6.0376920247189 |
| lnc-SRGAP3-5 | lnc-SRGAP3-5 | 0.0179770888084156 | 1.21274298083872 | 0.128972742092922 | 0.897987656883872 | 0.96454283201719 | -5.93276678319623 |
| ENSG00000283941.1 | MIR600 | -0.030425504308502 | 0.516391652647469 | -0.128948646120479 | 0.898006607984833 | 0.96454283201719 | -5.79727525880799 |
| lnc-TRMO-2 | lnc-TRMO-2 | 0.0277289714007751 | 3.7376980247309 | 0.12887802131113 | 0.898062153626636 | 0.96454283201719 | -6.25821882692175 |
| ENSG00000184060.10 | ADAP2 | 0.0767438724501119 | 0.41649766923155 | 0.128878803497214 | 0.898071512088958 | 0.96454283201719 | -5.71564367703535 |
| ENSG00000206149.10 | HERC2P9 | -0.0178775844096393 | 3.38641782275457 | -0.12885806783133 | 0.898077846914469 | 0.96454283201719 | -6.21497602907458 |
| ENSG00000250569.1 | NTAN1P2 | -0.0260009711005376 | 0.474078775414075 | -0.128793012032122 | 0.898129013182852 | 0.96454283201719 | -5.77371044647152 |
| ENSG00000180448.10 | ARHGAP45 | -0.0232276406155885 | 7.94158909607084 | -0.128738538394569 | 0.898171856937654 | 0.96454283201719 | -6.38427560762167 |
| lnc-SAMD5-5 | lnc-SAMD5-5 | -0.0330764238370351 | -0.378109424133325 | -0.128737136457459 | 0.898172959571396 | 0.96454283201719 | -5.52108272615612 |
| ENSG00000110446.10 | SLC15A3 | -0.0228753478483048 | 4.37731428121586 | -0.128718701721717 | 0.898187458644196 | 0.96454283201719 | -6.32485586936752 |
| ENSG00000109929.9 | SC5D | 0.0243614163342372 | 4.28824255001071 | 0.128535083391341 | 0.898331877907005 | 0.964647591974118 | -6.32665275036601 |
| ENSG00000124103.8 | FAM209A | 0.0137567541641848 | 3.13242694809484 | 0.128311278850787 | 0.898507909093575 | 0.964786284776099 | -6.18961186481142 |
| ENSG00000165434.7 | PGM2L1 | -0.0231766284448631 | 3.84704509090797 | -0.128155805678457 | 0.898630197981972 | 0.964832420256554 | -6.27891952960144 |
| lnc-THOC2-5 | lnc-THOC2-5 | -0.0306895254977947 | -0.0848720451978299 | -0.128137463193657 | 0.898644625600661 | 0.964832420256554 | -5.64230077709446 |

| | | | | | | | |
|---|---|---|---|---|---|---|---|
| lnc-SFTA3-3 | lnc-SFTA3-3 | -0.0180246359214089 | 2.29802862192395 | -0.12799301967686 | 0.898758241516694 | 0.964858746430624 | -6.09504404295887 |
| lnc-ADPRM-6 | lnc-ADPRM-6 | 0.0268558263756946 | 1.43775155006226 | 0.1279870988256 | 0.89876289876667 | 0.964858746430624 | -5.97558658050175 |
| ENSG00000198730.8 | CTR9 | 0.0120026775453503 | 5.79688101862334 | 0.127876558157943 | 0.898849849008055 | 0.964879636414912 | -6.39905283016568 |
| lnc-UNCX-3 | lnc-UNCX-3 | -0.0265296558526662 | 1.42285761556086 | -0.127843169157296 | 0.898876112726811 | 0.964879636414912 | -5.98276551005799 |
| ENSG00000068400.13 | GRIPAP1 | 0.0118980789636209 | 5.59924039688144 | 0.127612907871379 | 0.899057238898923 | 0.964979905812422 | -6.39407436581316 |
| lnc-EIF2AK1-1 | lnc-EIF2AK1-1 | 0.0361764369539584 | 0.627901860886683 | 0.127605218110095 | 0.899063287847486 | 0.964979905812422 | -5.84529422600951 |
| ENSG00000134574.11 | DDB2 | -0.020701949556531 | 5.67990031641202 | -0.127488163038313 | 0.89915536689213 | 0.965028413637538 | -6.39555017019802 |
| lnc-SRPRA-6 | lnc-SRPRA-6 | -0.0166579769249824 | 1.51958954730586 | -0.127240707389829 | 0.899350027583546 | 0.965187007547428 | -5.99498190853209 |
| ENSG00000168538.15 | TRAPPC11 | 0.0163087246192664 | 5.69540954263612 | 0.127068396485569 | 0.899485579455786 | 0.965282152342984 | -6.39637955521359 |
| lnc-RAB9A-1 | lnc-RAB9A-1 | -0.0223614245179352 | 0.756613540616841 | -0.126865780252906 | 0.89964497551451 | 0.965318098817743 | -5.85129425808474 |
| ENSG00000188735.12 | TMEM120B | -0.0191607748891962 | 3.28417666962113 | -0.126858237859191 | 0.899650909117272 | 0.965318098817743 | -6.19709490809013 |
| ENSG00000196912.12 | ANKRD36B | 0.0198030456599589 | 5.57377729802228 | 0.126787633138483 | 0.899706454147928 | 0.965318098817743 | -6.39323873068593 |
| ENSG00000173540.12 | GMPPB | 0.020217074707495 | 4.85977712348692 | 0.126787357584483 | 0.899706670928415 | 0.965318098817743 | -6.36451318285159 |
| ENSG00000228300.13 | FAM174C | 0.0228119413149925 | 2.20203030569744 | 0.126718853472355 | 0.899760563889657 | 0.965325602634392 | -6.08220250900561 |
| ENSG00000105281.12 | SLC1A5 | -0.0244603992718009 | 3.3640478330979 | -0.126639983998538 | 0.899822611991155 | 0.965341854613465 | -6.21035093403799 |
| ENSG00000172175.13 | MALT1 | 0.0171058066781118 | 6.61147223381067 | 0.126350452315817 | 0.900050397456229 | 0.965535900914241 | -6.40232039672473 |
| lnc-FRY-1 | lnc-FRY-1 | 0.0487230879934579 | 0.821790853063585 | 0.126200215796768 | 0.900170499058571 | 0.965567486159573 | -5.90367701628166 |
| ENSG00000145736.14 | GTF2H2 | -0.0286850239317893 | 2.32960449881803 | -0.12619377831936 | 0.900173662338631 | 0.965567486159573 | -6.10986267678938 |
| ENSG00000154589.6 | LY96 | -0.0356125163579717 | 1.5964215734987 | -0.125911890242763 | 0.900395446966947 | 0.965721986431277 | -5.99027362077287 |
| ENSG00000166167.17 | BTRC | 0.0119548748114919 | 4.5325830977073 | 0.125829848968303 | 0.900459997126529 | 0.965721986431277 | -6.33747921900318 |
| lnc-PPIL2-7 | lnc-PPIL2-7 | -0.0659216653037468 | -0.640818245733816 | -0.125741528981198 | 0.900529488145761 | 0.965721986431277 | -5.37881920576995 |
| lnc-ZBTB41-4 | lnc-ZBTB41-4 | 0.0427162193313975 | 0.108463207874849 | 0.125645678899475 | 0.900604904809632 | 0.965721986431277 | -5.7750988765016 |
| ENSG00000168778.11 | TCTN2 | -0.0262834606392723 | 2.91763592130292 | -0.125535080298175 | 0.900691927043411 | 0.965721986431277 | -6.16263763157454 |
| lnc-RXFP4-3 | lnc-RXFP4-3 | 0.0830484781930486 | 0.0670303746201738 | 0.125524185338886 | 0.900712185043424 | 0.965721986431277 | -5.66408174648294 |
| ENSG00000140521.13 | POLG | -0.0113390455124645 | 6.44809993070486 | -0.12548823704086 | 0.900728785069326 | 0.965721986431277 | -6.40287919396201 |
| ENSG00000198133.8 | TMEM229B | -0.0655190062824216 | 0.266936664811427 | -0.125463149269338 | 0.900753955470485 | 0.965721986431277 | -5.72113329100031 |
| ENSG00000114388.12 | NPRL2 | 0.0113074948412208 | 4.06730865413052 | 0.125415103632791 | 0.900786329611793 | 0.965721986431277 | -6.29836226819182 |
| lnc-UPK2-1 | lnc-UPK2-1 | -0.037476568806039 | 3.04166237068905 | -0.125418485735713 | 0.900786883228591 | 0.965721986431277 | -6.18557502366378 |
| lnc-FNDC5-2 | lnc-FNDC5-2 | 0.0260686461621822 | 0.995776096615681 | 0.12530299302319 | 0.900874544157966 | 0.965765663666845 | -5.89171875228718 |
| lnc-CCDC122-2 | lnc-CCDC122-2 | 0.0532224489471476 | -0.181601587835552 | 0.125094241042078 | 0.901038804614732 | 0.965891449163326 | -5.60205342279641 |
| ENSG00000173457.10 | PPP1R14B | -0.0255407638320144 | 2.60746220929767 | -0.124910339885997 | 0.901183514364807 | 0.965904439974758 | -6.14277891361425 |
| ENSG00000204160.11 | ZDHHC18 | -0.0210180019788915 | 4.38157436812532 | -0.124781601754792 | 0.901284818975561 | 0.965904439974758 | -6.33238375341257 |
| lnc-RBL2-7 | lnc-RBL2-7 | -0.0323959671796867 | 0.722246584467044 | -0.124763255615473 | 0.901299255770167 | 0.965904439974758 | -5.81239931714679 |
| ENSG00000071655.17 | MBD3 | -0.0194864129471939 | 3.92382926899986 | -0.124728401904464 | 0.90132668266183 | 0.965904439974758 | -6.2898202719118 |
| lnc-TYMP-1 | lnc-TYMP-1 | 0.0368025890691792 | 0.0727753163825883 | 0.124709710793885 | 0.901341391018606 | 0.965904439974758 | -5.75862999540382 |
| lnc-LY6G5B-1 | lnc-LY6G5B-1 | -0.0148010534491227 | 1.23212660882677 | -0.124709486647081 | 0.901341567403798 | 0.965904439974758 | -5.94078372302569 |
| ENSG00000127666.9 | TICAM1 | 0.0263809403904238 | 2.32574036463999 | 0.124661391803114 | 0.901379414227039 | 0.965904439974758 | -6.09238925212333 |
| lnc-SLC44A3-1 | lnc-SLC44A3-1 | -0.0854153652374087 | -1.13790928325234 | -0.124592057177822 | 0.90143505669828 | 0.965913778542386 | -5.37192727096488 |
| lnc-AP3M1-4 | lnc-AP3M1-4 | -0.0158783596695695 | 0.438083719718602 | -0.124425140766963 | 0.901565328402649 | 0.96598115497536 | -5.78676386871497 |
| ENSG00000163534.14 | FCRL1 | 0.0630141592796676 | 7.30103452018666 | 0.124367340528651 | 0.901632723976969 | 0.96598115497536 | -6.39719601567851 |
| lnc-IL5RA-8 | lnc-IL5RA-8 | 0.0138187622975718 | 2.62798488491365 | 0.124259990922904 | 0.901695293893015 | 0.96598115497536 | -6.1274100319171 |
| ENSG00000125089.16 | SH3TC1 | 0.0358914611410226 | 5.71527020871448 | 0.124176646279149 | 0.90177048767831 | 0.96598115497536 | -6.39806334672843 |
| ENSG00000213347.10 | MXD3 | -0.0147946212579924 | 3.60662619640778 | -0.124128355558039 | 0.901798886952026 | 0.96598115497536 | -6.25270097512924 |
| ENSG00000201564.1 | RN7SKP50 | -0.041796544655352 | -0.274732342028162 | -0.124000180666416 | 0.901899758387257 | 0.96598115497536 | -5.56873361130572 |
| lnc-WEE1-2 | lnc-WEE1-2 | -0.041796544655352 | -0.274732342028162 | -0.124000180666416 | 0.901899758387257 | 0.96598115497536 | -5.56873361130572 |
| lnc-ICA1L-4 | lnc-ICA1L-4 | -0.0285247067558793 | -0.0847176625412748 | -0.12392873586128 | 0.901955984932227 | 0.96598115497536 | -5.6310955501844 |
| ENSG00000217527.1 | RPS16P5 | 0.0341801904521435 | 2.4336560858463 | 0.123917825984553 | 0.90196457097115 | 0.96598115497536 | -6.09640576602306 |
| ENSG00000285854.1 | | 0.0296908947154702 | 4.76898924290192 | 0.12391705293191 | 0.90196724566251 | 0.96598115497536 | -6.36190342220519 |
| ENSG00000148334.14 | PTGES2 | 0.0157975847228764 | 4.2678042998912 | 0.123743863762134 | 0.902101480300915 | 0.965982617788842 | -6.320588651864 |
| lnc-NUGGC-1 | lnc-NUGGC-1 | 0.0258980728971713 | 0.702298237793313 | 0.123738026175445 | 0.90210607456955 | 0.965982617788842 | -5.85035760447028 |
| ENSG00000118600.11 | RXYLT1 | -0.0132533459675259 | 3.36240191039994 | -0.123733794699786 | 0.90210940480674 | 0.965982617788842 | -6.2191338542827 |
| lnc-ADSL-2 | lnc-ADSL-2 | 0.0387298379133769 | 0.037317044178392 | 0.123555433776222 | 0.902249779250754 | 0.966033100093845 | -5.65201823231765 |
| ENSG00000178882.14 | RFLNA | 0.0193036366655719 | 0.181227491516125 | 0.123554624499632 | 0.902250416178598 | 0.966033100093845 | -5.74240178475455 |
| lnc-MIER3-6 | lnc-MIER3-6 | -0.043212997293757 | 0.530524111252541 | -0.1234927221881 | 0.902299135568961 | 0.966035012089255 | -5.81115036254204 |
| lnc-CATSPERB-1 | lnc-CATSPERB-1 | 0.024107945118796 | 2.70523840007107 | 0.123411584339481 | 0.902362994609807 | 0.966053132115555 | -6.14220234913294 |
| ENSG00000203705.10 | TATDN3 | -0.0147587789998864 | 4.99254384103673 | -0.123311724921276 | 0.90244158924403 | 0.96608702509023 | -6.37156411316956 |
| ENSG00000117620.14 | SLC35A3 | -0.0174755928960399 | 4.99678771424704 | -0.123209029287824 | 0.902522417162193 | 0.966123305203319 | -6.37548088278153 |
| lnc-TERF1-1 | lnc-TERF1-1 | 0.0268070578204433 | 1.75029935606378 | 0.123095024341977 | 0.902612147445603 | 0.966169110763579 | -6.01845139723479 |
| ENSG00000116984.12 | MTR | -0.0100269208471673 | 6.57239264310907 | -0.122946238925689 | 0.902729254456953 | 0.96622523289402 | -6.40277893090382 |
| lnc-LTBP3-10 | lnc-LTBP3-10 | -0.0424806750234716 | 0.114057237388484 | -0.122791391588201 | 0.902851135052981 | 0.96622523289402 | -5.69792677285104 |
| lnc-DND1-2 | lnc-DND1-2 | 0.0120129610220174 | 2.18335017698388 | 0.122758240428636 | 0.902877228693888 | 0.96622523289402 | -6.08128982463273 |
| ENSG00000140403.12 | DNAJA4 | -0.0883075143596888 | 1.77265231246479 | -0.12277749122959 | 0.902883195446263 | 0.96622523289402 | -6.03452554983258 |
| lnc-GOLGA6L9-2 | lnc-GOLGA6L9-2 | 0.0530222577861111 | -0.0895490641613243 | 0.122730209653105 | 0.902899292109196 | 0.96622523289402 | -5.59304551762776 |
| ENSG00000267748.4 | | -0.0254890660790354 | 2.64649018771027 | -0.122546100369199 | 0.903044209029924 | 0.966330073016008 | -6.14048868098473 |
| ENSG00000102974.15 | CTCF | -0.0106716784909127 | 6.245481049433 | -0.122369639466502 | 0.903183108862594 | 0.966355187644166 | -6.40332685513371 |
| lnc-RD3L-2 | lnc-RD3L-2 | 0.0211047395832993 | 2.89057880865886 | 0.122354179302 | 0.903195278357602 | 0.966355187644166 | -6.15694725176611 |
| lnc-HTRA4-1 | lnc-HTRA4-1 | -0.0255313190077349 | 4.70629659095304 | -0.12233734891937 | 0.903208526448937 | 0.966355187644166 | -6.35920974230757 |
| ENSG00000145864.12 | GABRB2 | -0.181024749980795 | 3.78343595999796 | -0.122291054509502 | 0.903267865032073 | 0.966368442536263 | -6.25203570327171 |

| | | | | | | | |
|---|---|---|---|---|---|---|---|
| lnc-CHD7-2 | lnc-CHD7-2 | -0.0375717403958682 | 1.1745586666213 | -0.12207028446969 | 0.903418750846175 | 0.966392655600601 | -5.94226320506285 |
| ENSG00000042088.13 | TDP1 | -0.0144655368108833 | 5.13562223046981 | -0.122056611716686 | 0.90342951377883 | 0.966392655600601 | -6.37714600210391 |
| ENSG00000279196.1 | | -0.0805582214201194 | -0.886835222478977 | -0.12205427894049 | 0.903431350097982 | 0.966392655600601 | -5.35676213882474 |
| ENSG00000279377.1 | | 0.269215382205195 | -1.35062066309107 | 0.121783819181039 | 0.903665200652102 | 0.966582241571143 | -5.08909582389843 |
| ENSG00000109919.9 | MTCH2 | 0.0123146668693472 | 4.3997497504876 | 0.121577444208981 | 0.903806716974567 | 0.966582241571143 | -6.33298819051142 |
| lnc-UCHL5-10 | lnc-UCHL5-10 | -0.0678418360189546 | -0.183074294920543 | -0.121565785813073 | 0.903819480344453 | 0.966582241571143 | -5.62356686431118 |
| ENSG00000184840.11 | TMED9 | 0.0151173488418088 | 4.52341125171194 | 0.121512423491623 | 0.903857903361519 | 0.966582241571143 | -6.34198661195353 |
| ENSG00000169371.13 | SNUPN | 0.0155626558251356 | 2.69479711160059 | 0.121476069391503 | 0.903886522651483 | 0.966582241571143 | -6.13043963160283 |
| lnc-C1orf198-4 | lnc-C1orf198-4 | 0.0251419848046248 | 0.749628092700811 | 0.121369217596556 | 0.903970641078367 | 0.966582241571143 | -5.8395218817109 |
| ENSG00000234426.2 | | 0.0765559142358649 | 0.581372769052329 | 0.121382513417555 | 0.903975193592637 | 0.966582241571143 | -5.84562696795386 |
| lnc-AMPH-12 | lnc-AMPH-12 | 0.0197040814620082 | 1.80668556531056 | 0.121308369282451 | 0.904018544043099 | 0.966582241571143 | -6.01488394619121 |
| lnc-MRTO4-1 | lnc-MRTO4-1 | 0.0198380249872203 | 3.810050832963 | 0.121292259739728 | 0.904031226375459 | 0.966582241571143 | -6.2658453181665 |
| ENSG00000079263.18 | SP140 | 0.0250602578796135 | 7.67275591174213 | 0.121036711710876 | 0.904232411454887 | 0.966676690753442 | -6.39053246732393 |
| ENSG00000183826.17 | BTBD9 | -0.01655652492353 | 5.87492707749942 | -0.12098581846527 | 0.904272478897795 | 0.966676690753442 | -6.40089395238055 |
| lnc-GCN1-3 | lnc-GCN1-3 | 0.0140724235769349 | 2.34498079802906 | 0.120943906093184 | 0.90430547603135 | 0.966676690753442 | -6.0993931645264 |
| ENSG00000132361.16 | CLUH | -0.0205791387486771 | 4.59506198336509 | -0.120816928122807 | 0.904405445386758 | 0.966676690753442 | -6.35163230595795 |
| ENSG00000268533.1 | | 0.015056302763076 | 0.622029444863042 | 0.120787121057559 | 0.904428912622646 | 0.966676690753442 | -5.82018279212717 |
| ENSG00000188130.13 | MAPK12 | 0.0249174982246507 | 3.05568371122374 | 0.120760376138338 | 0.904449969090844 | 0.966676690753442 | -6.18373589117835 |
| ENSG00000182196.13 | ARL6IP4 | -0.0184892052108063 | 4.46912042730787 | -0.120741965524971 | 0.904464463940245 | 0.966676690753442 | -6.33839052218101 |
| ENSG00000150961.14 | SEC24D | -0.0128697826321158 | 4.38523125404513 | -0.120702822282058 | 0.904495281893822 | 0.966676690753442 | -6.327242481686 |
| lnc-KCTD1-5 | lnc-KCTD1-5 | -0.0163097629322655 | 2.26053893038196 | -0.120587568863278 | 0.904586023184069 | 0.966722537689033 | -6.08703207632499 |
| ENSG00000229018.5 | PMS2P7 | -0.0237371782197136 | -0.0872939266775147 | -0.120515709356655 | 0.904642600244387 | 0.966722537689033 | -5.62801845170765 |
| ENSG00000117479.13 | SLC19A2 | 0.0481238332199236 | 1.84128096573187 | 0.120477857356633 | 0.90467908084177 | 0.966722537689033 | -6.01741532634672 |
| lnc-RNF41-1 | lnc-RNF41-1 | 0.0304229996544412 | 1.30002529205909 | 0.120333737097142 | 0.904785874489671 | 0.966765883365871 | -5.95197412300873 |
| lnc-AFF4-2 | lnc-AFF4-2 | 0.0213645543185557 | 3.10136040142464 | 0.120298545165038 | 0.904813582916005 | 0.966765883365871 | -6.18309760936491 |
| lnc-FANCL-1 | lnc-FANCL-1 | -0.03166482293788 | 1.4769503504947 | -0.119871027513669 | 0.905150199225857 | 0.966919510741399 | -5.97875031165195 |
| lnc-DIS3-6 | lnc-DIS3-6 | -0.0400930566474442 | -0.151881359258922 | -0.119828599647887 | 0.905183606790691 | 0.966919510741399 | -5.63988463417546 |
| lnc-C11orf94-4 | lnc-C11orf94-4 | 0.0148321144470867 | 2.18473024227303 | 0.119795377001835 | 0.905209766316486 | 0.966919510741399 | -6.08540489362547 |
| ENSG00000130204.12 | TOMM40 | 0.0151913500911456 | 3.80309906902322 | 0.119751305943022 | 0.905244468051533 | 0.966919510741399 | -6.27280752080808 |
| lnc-SAAL1-2 | lnc-SAAL1-2 | 0.0180188161652621 | 0.116768162432532 | 0.119748930131461 | 0.905246338780764 | 0.966919510741399 | -5.70181406945218 |
| ENSG00000273748.1 | | -0.0815527246749266 | 2.38091210839173 | -0.119728986273904 | 0.905283585301001 | 0.966919510741399 | -6.19326504096497 |
| ENSG00000089248.6 | ERP29 | -0.0223435596586405 | 6.93580770569809 | -0.119698304581056 | 0.90528620179262 | 0.966919510741399 | -6.40074897550213 |
| ENSG00000160957.12 | RECQL4 | 0.0328219090439925 | 1.33070121326257 | 0.119460986925809 | 0.90547307113336 | 0.966993954625105 | -5.93496373593644 |
| ENSG00000133597.10 | ADCK2 | -0.0189987694360783 | 3.05922528975028 | -0.119441288946617 | 0.905488582015751 | 0.966993954625105 | -6.18097649552362 |
| lnc-HEYL-5 | lnc-HEYL-5 | -0.0152190577880575 | 2.48019812978387 | -0.119304643142972 | 0.905596182760523 | 0.966993954625105 | -6.10667309331268 |
| ENSG00000114346.13 | ECT2 | -0.027360010475086 | 1.19267785436412 | -0.119295403012779 | 0.905603458898686 | 0.966993954625105 | -5.95294843217308 |
| lnc-CCDC32-3 | lnc-CCDC32-3 | -0.0282653224035241 | 2.20635945437061 | -0.119273734513976 | 0.905620521784726 | 0.966993954625105 | -6.08340183106388 |
| lnc-YWHAZ-2 | lnc-YWHAZ-2 | 0.0239136727285012 | 1.99585349085155 | 0.119196605291594 | 0.905681257651873 | 0.966993954625105 | -6.0522280986836 |
| ENSG00000102445.18 | RUBCNL | 0.0229291676952581 | 9.35620330945527 | 0.119192155036352 | 0.905684762049399 | 0.966993954625105 | -6.34448011908255 |
| ENSG00000137802.13 | MAPKBP1 | 0.0134913121738807 | 4.61188801090284 | 0.119080935164709 | 0.905772343855396 | 0.967037302431434 | -6.34578706873759 |
| lnc-PAFAH1B1-1 | lnc-PAFAH1B1-1 | -0.0235309305706075 | 0.209185150077273 | -0.118926626348755 | 0.905893858681617 | 0.96704100271534 | -5.73210965008692 |
| lnc-NEMF-2 | lnc-NEMF-2 | 0.0551298258761981 | 2.50825871182495 | 0.118923174712635 | 0.90591042984096 | 0.96704100271534 | -6.10147034167401 |
| ENSG00000185272.13 | RBM11 | -0.13026781736973 | -0.738714870645095 | -0.118857778403187 | 0.905961920996566 | 0.96704100271534 | -5.34413153704658 |
| ENSG00000234518.2 | PTGES3P1 | -0.028853923343332 | 1.22514424139946 | -0.118837887333024 | 0.905963739754162 | 0.96704100271534 | -5.9321605143418 |
| ENSG00000185504.16 | FAAP100 | -0.0190644615023743 | 4.36335079703731 | -0.118515780634197 | 0.906217401835936 | 0.967261119575678 | -6.33025004594223 |
| ENSG00000047056.15 | WDR37 | -0.0109402482365306 | 4.47364442704631 | -0.118456685768285 | 0.906263940680025 | 0.967261119575678 | -6.33489060401355 |
| lnc-ZNF391-5 | lnc-ZNF391-5 | -0.0307287359031667 | 1.49065384379803 | -0.118336338438962 | 0.906358718566493 | 0.967312117818838 | -5.96777205045758 |
| ENSG00000206561.12 | COLQ | -0.0200145473036349 | 0.619623397606934 | -0.118232524318873 | 0.906440477060654 | 0.967349216536498 | -5.83056523727097 |
| ENSG00000259956.1 | RBM15B | -0.0149796131097739 | 5.8616297344248 | -0.118131162151115 | 0.906520305517941 | 0.967384251787421 | -6.40091292911529 |
| ENSG00000279457.4 | WASH9P | 0.0171582775066814 | 3.17693843912523 | 0.117909739583831 | 0.906694691751453 | 0.967510020903431 | -6.21667063378368 |
| ENSG00000173674.10 | EIF1AX | 0.0206674637166664 | 5.80401892138974 | 0.117810714459253 | 0.906772682683603 | 0.967510020903431 | -6.4020329106145 |
| lnc-LYRM7-4 | lnc-LYRM7-4 | -0.0391824113677205 | 0.332431794300652 | -0.117796517270364 | 0.906783864285827 | 0.967510020903431 | -5.78351999751985 |
| SNHG16 | SNHG16 | -0.0209090816641035 | 5.16546082262475 | -0.117742785852634 | 0.906826182931011 | 0.967510020903431 | -6.38472345664651 |
| ENSG00000096080.11 | MRPS18A | -0.0115891807779507 | 4.01006817036994 | -0.117171069732832 | 0.907276481152203 | 0.967898294865156 | -6.29635279327704 |
| ENSG00000148803.11 | FUOM | -0.0213629952557275 | 2.24429306427189 | -0.117161330834783 | 0.90728415202489 | 0.967898294865156 | -6.08742352509049 |
| ENSG00000116001.15 | TIA1 | -0.0135590344959535 | 6.0879477739119 | -0.117047084341049 | 0.907374139291918 | 0.967944125889896 | -6.40350224521348 |
| ENSG00000154930.14 | ACSS1 | -0.0168757648791491 | 5.40326811579887 | -0.116849828427451 | 0.907529512564375 | 0.968031166587689 | -6.38990294152528 |
| ENSG00000151090.18 | THRB | -0.164546873818654 | -1.24034319189978 | -0.116848940451992 | 0.907549794570404 | 0.968031166587689 | -5.3817349436238 |
| ENSG00000114841.17 | DNAH1 | -0.0173388917351902 | 6.39272176084891 | -0.116445684744983 | 0.907847857305429 | 0.96826775516754 | -6.40407477202693 |
| lnc-ZNF266-1 | lnc-ZNF266-1 | 0.113785102964181 | 0.74054043522222 | 0.116316954486699 | 0.907970177253851 | 0.96826775516754 | -5.8285537538878 |
| lnc-AMPH-13 | lnc-AMPH-13 | -0.021118979339299 | 3.23770926443861 | -0.116218364540804 | 0.908026924598625 | 0.96826775516754 | -6.19720895377456 |
| ENSG00000213190.3 | MLLT11 | 0.0262409216305812 | 2.13483283146832 | 0.116160038281672 | 0.90807287081293 | 0.96826775516754 | -6.04883698772232 |
| ENSG00000111321.10 | LTBR | -0.0933321222641393 | -0.930267970883582 | -0.116101058925214 | 0.908131122462923 | 0.96826775516754 | -5.52449475129269 |
| lnc-TRAF3IP2-3 | lnc-TRAF3IP2-3 | -0.0163846717956213 | 1.510457439834 | -0.116041558196717 | 0.908166203878411 | 0.96826775516754 | -5.9876526980555 |
| ENSG00000156304.14 | SCAF4 | -0.0155455089247072 | 5.93026422159949 | -0.116037022048926 | 0.908169777269432 | 0.96826775516754 | -6.40158182008504 |
| ENSG00000131188.11 | PRR7 | -0.0177022155993196 | 2.06086371319275 | -0.115956170647451 | 0.908233469007354 | 0.96826775516754 | -6.06129490805522 |
| ENSG00000125991.19 | ERGIC3 | 0.0129563872229985 | 6.06360712656739 | 0.11593129686273 | 0.908253063776111 | 0.96826775516754 | -6.40357951724152 |
| lnc-TRAK1-1 | lnc-TRAK1-1 | -0.0233854247776113 | 0.691367552944799 | -0.115876492622961 | 0.908296237000118 | 0.96826775516754 | -5.84582919440786 |

| | | | | | | | |
|---|---|---|---|---|---|---|---|
| ENSG00000198586.13 | TLK1 | 0.0157566354312655 | 7.22713245255645 | 0.115760701897675 | 0.908387454568147 | 0.96826775516754 | -6.39782785155951 |
| ENSG00000116871.15 | MAP7D1 | 0.0197925204265665 | 4.15603463478512 | 0.115740858974869 | 0.908403086542987 | 0.96826775516754 | -6.31609694279277 |
| lnc-MELTF-3 | lnc-MELTF-3 | -0.0159189437468783 | 1.47830155063708 | -0.115700431124867 | 0.908434935146921 | 0.96826775516754 | -5.97644281769188 |
| lnc-RBL2-6 | lnc-RBL2-6 | 0.0293639324876399 | 0.660041115397884 | 0.115670061834033 | 0.908458859831733 | 0.96826775516754 | -5.81903105619231 |
| ENSG00000147180.16 | ZNF711 | 0.172771615630453 | -0.59943705099603 | 0.115615580442807 | 0.908522572804047 | 0.96826775516754 | -5.49708181068733 |
| ENSG00000101605.12 | MYOM1 | -0.0267680284217207 | 2.89600472788447 | -0.115559059016015 | 0.9085463076968 | 0.96826775516754 | -6.15523412176269 |
| ENSG00000168781.22 | PPIP5K1 | 0.0123718932490826 | 3.44389151319178 | 0.115527313137151 | 0.908571317266961 | 0.96826775516754 | -6.2226521102004 |
| ENSG00000119401.10 | TRIM32 | -0.0172166417686248 | 3.37667068190318 | -0.115459217154275 | 0.908624963956587 | 0.968274793327384 | -6.21635706708483 |
| ENSG00000163866.8 | SMIM12 | 0.0105739724660037 | 4.73157633915824 | 0.115357722673268 | 0.908704923114685 | 0.968293187362905 | -6.35823423999029 |
| ENSG00000198546.14 | ZNF511 | -0.0151399641528278 | 3.13799533148202 | -0.11501027956887 | 0.908978652221899 | 0.968293187362905 | -6.19190779764674 |
| ENSG00000152465.17 | NMT2 | -0.0379879070348816 | 2.92363656968376 | -0.114961395814975 | 0.909023067509925 | 0.968293187362905 | -6.185945882321 |
| ENSG00000122705.16 | CLTA | -0.0131356457062664 | 4.43686343598927 | -0.114889481270679 | 0.909073824414174 | 0.968293187362905 | -6.33557823744415 |
| ENSG00000152061.23 | RABGAP1L | 0.0241458351063404 | 8.14842294901303 | 0.114866671161202 | 0.909091795747274 | 0.968293187362905 | -6.38190745724058 |
| lnc-PPA1-1 | lnc-PPA1-1 | 0.0258132906661625 | 0.337330080089651 | 0.114835383400589 | 0.909116446420523 | 0.968293187362905 | -5.76956157565492 |
| lnc-ASXL2-6 | lnc-ASXL2-6 | 0.0195226161232926 | 0.898062174354796 | 0.114790786488497 | 0.909151583125334 | 0.968293187362905 | -5.88178263512917 |
| ENSG00000079313.14 | REXO1 | -0.0161344851621292 | 4.42107446614391 | -0.114759946632209 | 0.909175881124385 | 0.968293187362905 | -6.33485421442206 |
| ENSG00000198356.11 | GET3 | -0.0183904469537453 | 3.86732509020884 | -0.11471161143404 | 0.909213963469757 | 0.968293187362905 | -6.28365934283614 |
| ENSG00000177917.10 | ARL6IP6 | -0.0148191301642333 | 3.57108746234665 | -0.114701167761691 | 0.909222191860754 | 0.968293187362905 | -6.25170939703599 |
| ENSG00000254462.1 | NA | 0.0123865957166169 | 1.78900776574706 | 0.114696896593028 | 0.909225557044469 | 0.968293187362905 | -6.02567421515506 |
| lnc-OTP-5 | lnc-OTP-5 | 0.0236865836163267 | 1.01634530600586 | 0.114648596963752 | 0.909263611647339 | 0.968293187362905 | -5.91333119093196 |
| lnc-PPM1F-2 | lnc-PPM1F-2 | 0.0168521881952304 | 0.501990383348834 | 0.114600483593883 | 0.909301519713397 | 0.968293187362905 | -5.78894533199315 |
| lnc-SMNDC1-2 | lnc-SMNDC1-2 | -0.0235142819982075 | 2.84161974127056 | -0.114515419134692 | 0.90936854171421 | 0.968293187362905 | -6.14532940581141 |
| lnc-PPARA-2 | lnc-PPARA-2 | 0.0530297218122046 | -0.4977298230932 | 0.114471196855484 | 0.909403384562959 | 0.968293187362905 | -5.61101037030388 |
| lnc-TEX2-2 | lnc-TEX2-2 | -0.0260219964733444 | 3.97314498028835 | -0.11439727220575 | 0.909461630389639 | 0.968293187362905 | -6.27315203481974 |
| lnc-DMTN-5 | lnc-DMTN-5 | -0.0156307255497206 | 2.1481717609216 | -0.114360247804724 | 0.909490802402732 | 0.968293187362905 | -6.06795440655804 |
| ENSG00000170445.13 | HARS1 | 0.00686819354911172 | 5.19043758698115 | 0.114327098324485 | 0.90951692142195 | 0.968293187362905 | -6.38400064404616 |
| ENSG00000129255.15 | MPDU1 | 0.0157835096988874 | 4.1104754386737 | 0.114302823000051 | 0.909536048412496 | 0.968293187362905 | -6.30770602266008 |
| ENSG00000128607.13 | KLHDC10 | 0.014941106367603 | 5.00347572881379 | 0.113942703426686 | 0.9098197998564 | 0.968545174059699 | -6.37274621534178 |
| ENSG00000102103.15 | PQBP1 | -0.0178860624804438 | 4.1902866534956 | -0.113857922138467 | 0.909886603882484 | 0.968566196096047 | -6.31387147163964 |
| ENSG00000162733.17 | DDR2 | 0.0431856101860726 | -0.0371924506100692 | 0.113674947679842 | 0.910030782171805 | 0.96862686981731 | -5.62097381599337 |
| ENSG00000034533.11 | ASTE1 | 0.0132623107341665 | 4.27461143557932 | 0.113666141991141 | 0.910037720863195 | 0.96862686981731 | -6.31877875818362 |
| ENSG00000111328.6 | CDK2AP1 | -0.0290481914115892 | 2.25289812412655 | -0.113386600123209 | 0.910257997472686 | 0.968766935334866 | -6.1051555516648 |
| lnc-SRBD1-5 | lnc-SRBD1-5 | -0.0172108602758811 | 0.0116768771764601 | -0.113379684533477 | 0.910263446989083 | 0.968766935334866 | -5.67033497313797 |
| ENSG00000095059.15 | DHPS | -0.0103059860631456 | 4.30933688189732 | -0.113275538318258 | 0.910345515212449 | 0.968804184872439 | -6.32266962976878 |
| lnc-SLC12A8-4 | lnc-SLC12A8-4 | 0.0299098469197698 | 2.83002354097043 | 0.112809362632835 | 0.910712878283509 | 0.969145030178341 | -6.16505775197041 |
| lnc-LUC7L3-6 | lnc-LUC7L3-6 | -0.0222883088187134 | 3.5432933351081 | -0.112673349027956 | 0.910820065585339 | 0.969208985676629 | -6.25526378561219 |
| lnc-PTGES2-2 | lnc-PTGES2-2 | 0.0134813278413026 | 2.08038359121118 | 0.112568675121029 | 0.910902556372552 | 0.969246656214652 | -6.05174869823144 |
| ENSG00000159445.12 | THEM4 | 0.0259802219092204 | 2.67438151989315 | 0.112495644971455 | 0.910960110125592 | 0.969257789956839 | -6.13046825646845 |
| ENSG00000188315.7 | C3orf62 | 0.0129918068290844 | 3.27489459613553 | 0.112401143057267 | 0.911034586109865 | 0.969286926801373 | -6.21338260554842 |
| ENSG00000166441.12 | RPL27A | -0.0172614383675267 | 9.12420978018542 | -0.112177908752112 | 0.911210517977572 | 0.969397252653705 | -6.35477230277393 |
| lnc-C13orf42-4 | lnc-C13orf42-4 | 0.0265752767186882 | 0.182532295379183 | 0.112069034234064 | 0.911296324065978 | 0.969397252653705 | -5.71681988756692 |
| ENSG00000163466.15 | ARPC2 | 0.0141518134518259 | 7.69594118385966 | 0.11201826093518 | 0.91133633983952 | 0.969397252653705 | -6.38986499904756 |
| lnc-TTLL7-3 | lnc-TTLL7-3 | -0.0420614767682355 | -0.505736156029962 | -0.112015594721644 | 0.911338441159037 | 0.969397252653705 | -5.63751328343445 |
| lnc-MTX3-5 | lnc-MTX3-5 | -0.0420926624130371 | 3.42987273635962 | -0.111963221072979 | 0.911390858565166 | 0.969397252653705 | -6.21064255452689 |
| ENSG00000115919.14 | KYNU | -0.0216400704533983 | 4.94840490331759 | -0.111911015574769 | 0.911420863496794 | 0.969397252653705 | -6.36552396147403 |
| lnc-DPP8-2 | lnc-DPP8-2 | 0.0147478874941729 | 0.884665002924471 | 0.111671423513447 | 0.911609697766161 | 0.969547998197742 | -5.87699375819909 |
| ENSG00000175931.12 | UBE2O | -0.021067211659159 | 4.8710770240591 | -0.111527566875587 | 0.911723080731557 | 0.969618485620421 | -6.3649343559394 |
| ENSG00000085511.19 | MAP3K4 | 0.0102786857962469 | 5.74094702748251 | 0.111118931308883 | 0.912045163692028 | 0.969875901943906 | -6.39967858860227 |
| ENSG00000077454.15 | LRCH4 | 0.0138892647485465 | 5.4781171543129 | 0.111100911181038 | 0.912059367343256 | 0.969875901943906 | -6.3943332599782 |
| lnc-BRIP1-1 | lnc-BRIP1-1 | 0.0149592525322127 | 0.184612939302156 | 0.111027777843857 | 0.912117012094979 | 0.96988198392127 | -5.6981822316097 |
| ENSG00000281039.1 | | 0.0185646540469697 | 1.71223103146714 | 0.110974092666766 | 0.91215932783112 | 0.96988198392127 | -6.01577023610473 |
| ENSG00000159733.13 | ZFYVE28 | -0.114365798852741 | 0.0160902716305659 | -0.110890359857935 | 0.912242221531922 | 0.969920018895168 | -5.56950515569641 |
| ENSG00000177192.13 | PUS1 | 0.0133457853369432 | 3.45025372396627 | 0.110753865855503 | 0.912332917730557 | 0.96996634533306 | -6.23462735510029 |
| ENSG00000127838.13 | PNKD | -0.0234474346016502 | 2.50577919130294 | -0.110554423330003 | 0.912490128522007 | 0.970083379751484 | -6.12024682549779 |
| ENSG00000278249.1 | SCARNA2 | -0.0361431050953716 | 6.45849613421306 | -0.11043431313634 | 0.912596983218685 | 0.970104565817278 | -6.40396148844193 |
| ENSG00000259318.1 | HMGN1P1 | 0.0249277138379284 | 0.653906228416988 | 0.110313951540425 | 0.912679685392237 | 0.970104565817278 | -5.81220359049257 |
| lnc-TBPL2-2 | lnc-TBPL2-2 | 0.0249277138379284 | 0.653906228416988 | 0.110313951540425 | 0.912679685392237 | 0.970104565817278 | -5.81220359049257 |
| ENSG00000122512.15 | PMS2 | 0.0131362137549029 | 4.10718403220199 | 0.110227460856866 | 0.912747864724174 | 0.970104565817278 | -6.30738127743915 |
| ENSG00000162437.14 | RAVER2 | 0.0415756474675733 | 2.17448059132552 | 0.110206097890262 | 0.912771702103909 | 0.970104565817278 | -6.09121144754906 |
| ENSG00000168175.14 | MAPK1IP1L | -0.0141813290889692 | 6.67798033187821 | -0.110169103981384 | 0.912793866965387 | 0.970104565817278 | -6.40346739416004 |
| ENSG00000143376.13 | SNX27 | 0.0121416802358647 | 4.44637854096219 | 0.110065166563426 | 0.912875800719805 | 0.970104565817278 | -6.33066864779572 |
| ENSG00000183091.19 | NEB | 0.0272559477474568 | 4.39220361957616 | 0.110017445340407 | 0.912915042606129 | 0.970104565817278 | -6.32522047635986 |
| ENSG00000277967.1 | | -0.0191243002973498 | 2.24965959818825 | -0.109945134976234 | 0.912970422683772 | 0.970104565817278 | -6.08230391711106 |
| lnc-METTL2B-3 | lnc-METTL2B-3 | -0.0211544129258083 | 4.67009364021761 | -0.109931247637203 | 0.912981370278701 | 0.970104565817278 | -6.35010815970164 |
| ENSG00000184076.13 | UQCR10 | 0.0180148914579141 | 3.87300041852117 | 0.109677417592664 | 0.91318147127095 | 0.970267098036855 | -6.27498546985104 |
| ENSG00000100997.18 | ABHD12 | -0.013820592399342 | 2.77034710865344 | -0.109313840113404 | 0.913468099018958 | 0.970479350502037 | -6.15075291170848 |
| ENSG00000103047.7 | TANGO6 | -0.0103961587925427 | 4.91913016867675 | -0.109091810221049 | 0.913643142900647 | 0.970479350502037 | -6.36636181373338 |
| lnc-EDN2-3 | lnc-EDN2-3 | -0.0440102055404306 | 0.0580294729859246 | -0.109031269093212 | 0.913690873055916 | 0.970479350502037 | -5.68923002917356 |

| | | | | | | | |
|---|---|---|---|---|---|---|---|
| ENSG00000274070.1 | CASTOR2 | -0.0257818001162657 | 1.72864096862102 | -0.10898230457279 | 0.913729476539575 | 0.970479350502037 | -6.01715161660944 |
| ENSG00000108561.8 | C1QBP | -0.0235092749949901 | 4.40091378396581 | -0.108969001277197 | 0.913739964855049 | 0.970479350502037 | -6.33412721340004 |
| ENSG00000236184.1 | TCEA1P4 | 0.0276382962603995 | 0.362346865596177 | 0.108967063691907 | 0.913741492448186 | 0.970479350502037 | -5.73147944755497 |
| ENSG00000168216.11 | LMBRD1 | -0.010900017082573 | 6.37691602390129 | -0.108812290671512 | 0.913863516634486 | 0.970479350502037 | -6.4048797979842 |
| ENSG00000141101.12 | NOB1 | 0.015080864011435 | 3.36066185152496 | 0.108789360779814 | 0.913881594908469 | 0.970479350502037 | -6.22371519460802 |
| ENSG00000280607.1 | | 0.0193979932793906 | 1.4568461762596 | 0.108785083919272 | 0.913884966855218 | 0.970479350502037 | -5.98472041863652 |
| ENSG00000012048.21 | BRCA1 | 0.0174716250953417 | 2.83821802581609 | 0.108775197725597 | 0.913892761298027 | 0.970479350502037 | -6.15046559104417 |
| ENSG00000171928.13 | TVP23B | 0.0153818927422655 | 3.89893088103161 | 0.108766167348939 | 0.913899881007676 | 0.970479350502037 | -6.28400514070263 |
| lnc-HELZ-4 | lnc-HELZ-4 | -0.0119051467380658 | 4.33450342509464 | -0.108544055304077 | 0.914075000347222 | 0.970488247126849 | -6.32755602630881 |
| ENSG00000151773.12 | CCDC122 | 0.039853091217469 | 1.06929626984014 | 0.108515373544256 | 0.914098285631412 | 0.970488247126849 | -5.90797584049145 |
| ENSG00000104643.9 | MTMR9 | 0.0136668116994985 | 5.20252942176953 | 0.10845611576816 | 0.914144335527719 | 0.970488247126849 | -6.38283557505566 |
| ENSG00000143702.15 | CEP170 | -0.00828603085571089 | 6.43766643160517 | -0.108343500989836 | 0.914233126681634 | 0.970488247126849 | -6.40495850932791 |
| ENSG00000169682.17 | SPNS1 | -0.0159568355855209 | 2.62744712749727 | -0.108312741119649 | 0.914257379502804 | 0.970488247126849 | -6.14038483738609 |
| ENSG00000230424.1 | EMC1-AS1 | -0.0211757115862948 | -0.193446563633716 | -0.108297985416555 | 0.914269013762925 | 0.970488247126849 | -5.59950568815844 |
| lnc-FAM161B-3 | lnc-FAM161B-3 | -0.0185138228082525 | 0.448604180677147 | -0.108233941410018 | 0.914319510027266 | 0.970488247126849 | -5.77859790957953 |
| ENSG00000283818.1 | MIR7844 | 0.0264639816002709 | 0.407168804659382 | 0.10821308175697 | 0.914335957147993 | 0.970488247126849 | -5.78710680075058 |
| ENSG00000113163.16 | CERT1 | 0.00906810868099406 | 5.99870266924784 | 0.108176122193124 | 0.914365098589272 | 0.970488247126849 | -6.40359267607282 |
| ENSG00000167657.13 | DAPK3 | 0.0164157305498942 | 2.7832739811691 | 0.108142119599052 | 0.914391908659172 | 0.970488247126849 | -6.15108152387707 |
| ENSG00000060491.16 | OGFR | 0.0179210306581412 | 5.74717724440695 | 0.108097729860812 | 0.914426908846043 | 0.970488247126849 | -6.4007693840031 |
| ENSG00000127554.12 | GFER | 0.0152773761302689 | 3.2934321406495 | 0.108001322542183 | 0.914502924172389 | 0.970511140125893 | -6.21831752734399 |
| lnc-DEFB115-4 | lnc-DEFB115-4 | 0.0727315438346099 | -1.07277041906184 | 0.107955193991162 | 0.914542781610155 | 0.970511140125893 | -5.47209245661152 |
| ENSG00000073584.19 | SMARCE1 | -0.00967416951233524 | 6.86515241707836 | -0.107531297962008 | 0.914873541148059 | 0.970775407203333 | -6.40226130207825 |
| lnc-CNDP1-8 | lnc-CNDP1-8 | 0.0243283565802543 | 1.86156128437627 | 0.107515324604791 | 0.914886136565625 | 0.970775407203333 | -6.03879834711478 |
| lnc-EMCN-2 | lnc-EMCN-2 | 0.0298169022555936 | 1.13090260099358 | 0.107309467299092 | 0.915048462513118 | 0.970788806794867 | -5.93493161639669 |
| lnc-MFSD5-3 | lnc-MFSD5-3 | 0.013666107935447 | 2.05515423601089 | 0.107229923180522 | 0.915111186917157 | 0.970788806794867 | -6.06819119357306 |
| ENSG00000177943.13 | MAMDC4 | -0.0295627301993719 | 1.5111872042712 | -0.1072231665467 | 0.915116514876589 | 0.970788806794867 | -6.00838684033818 |
| ENSG00000213024.11 | NUP62 | 0.0146088023962037 | 5.1774049031576 | 0.107139750575552 | 0.915182293061401 | 0.970788806794867 | -6.38491810954325 |
| ENSG00000128059.8 | PPAT | -0.014625110868641 | 3.04631957980188 | -0.107121215959412 | 0.915196908730364 | 0.970788806794867 | -6.19835734708826 |
| lnc-CXorf38-4 | lnc-CXorf38-4 | 0.0204944610660818 | 0.304420938572053 | 0.107087801381283 | 0.915223258226852 | 0.970788806794867 | -5.7438805587685 |
| ENSG00000132463.13 | GRSF1 | 0.0117452873693589 | 5.23965089558961 | 0.107050171637778 | 0.915252931761857 | 0.970788806794867 | -6.38746771178566 |
| lnc-OR2AG2-1 | lnc-OR2AG2-1 | 0.0221367768755427 | 1.5473633841919 | 0.106967061437029 | 0.915318470070859 | 0.970788806794867 | -5.98854973129345 |
| NEAT1 | NEAT1 | 0.0396519338967334 | 8.51485221158394 | 0.106978314075886 | 0.915323246051012 | 0.970788806794867 | -6.37279332125685 |
| ENSG00000097021.19 | ACOT7 | 0.0239131496100444 | 0.984215125217106 | 0.106861242240013 | 0.915401916902564 | 0.970822220507289 | -5.89672180139186 |
| lnc-KIF5B-4 | lnc-KIF5B-4 | -0.0133983003326659 | 3.28211844856373 | -0.106676735001906 | 0.915547417806117 | 0.970926503204869 | -6.21129253396329 |
| ENSG00000143869.6 | GDF7 | 0.0333534655084851 | 7.48645020085839 | 0.106593818007783 | 0.915623473587212 | 0.970927251985357 | -6.39515966434183 |
| ENSG00000013561.17 | RNF14 | -0.00867298863303212 | 4.49267912007799 | -0.106492911667066 | 0.915692382303333 | 0.970927251985357 | -6.34099743687155 |
| lnc-MYLK3-6 | lnc-MYLK3-6 | 0.0228965764437921 | 2.30162901235378 | 0.106438384323588 | 0.915735383544894 | 0.970927251985357 | -6.08624080195665 |
| lnc-GLUD2-1 | lnc-GLUD2-1 | -0.115046987143179 | -2.8364978662908 | -0.106436576599678 | 0.915736809152783 | 0.970927251985357 | -4.92706197322922 |
| ENSG00000227540.1 | NA | 0.0202716734690475 | -0.429845396463481 | 0.106171938842343 | 0.915945510903166 | 0.971018087205843 | -5.52550073151921 |
| ENSG00000116514.16 | RNF19B | 0.0142615882849304 | 4.32273918247398 | 0.10606510615264 | 0.916029764260262 | 0.971018087205843 | -6.32658376440921 |
| ENSG00000127586.16 | CHTF18 | 0.0239420588114561 | 3.20669849770568 | 0.106054562458957 | 0.91603807957322 | 0.971018087205843 | -6.20621189855598 |
| lnc-LSG1-4 | lnc-LSG1-4 | -0.0218476083080686 | 0.671937233958877 | -0.105915759021657 | 0.916147548171701 | 0.971018087205843 | -5.82202994648404 |
| lnc-CHN1-5 | lnc-CHN1-5 | -0.0216466293609368 | 2.57815151752319 | -0.105907088469129 | 0.916154386336808 | 0.971018087205843 | -6.14535677672418 |
| ENSG00000088038.18 | CNOT3 | -0.0164395347236663 | 4.16107311896223 | -0.105902985705889 | 0.916157622046616 | 0.971018087205843 | -6.30324994736101 |
| ENSG00000230069.3 | LRRC37A15P | 0.0147775499574787 | 1.38024033504562 | 0.105854915357089 | 0.916195533602701 | 0.971018087205843 | -5.96071103567738 |
| ENSG00000284906.1 | | -0.0248004752381924 | 0.785185467185195 | -0.105792890063157 | 0.916244451272091 | 0.971018087205843 | -5.86136205556641 |
| TP53TG1 | TP53TG1 | 0.019645940248737 | 1.99240558327218 | 0.105756568357884 | 0.916273097369779 | 0.971018087205843 | -6.04323090261297 |
| ENSG00000253276.2 | CCDC71L | 0.0242225127844078 | 3.38575487817989 | 0.105651182989651 | 0.916356213035001 | 0.971018087205843 | -6.23760777616768 |
| ENSG00000095261.13 | PSMD5 | -0.021417495324171 | 3.99170167792128 | -0.105623506350927 | 0.916378041290924 | 0.971018087205843 | -6.27988571303225 |
| ENSG00000100445.17 | SDR39U1 | 0.0120407778158391 | 4.27806046219298 | 0.105581170029461 | 0.916411431601261 | 0.971018087205843 | -6.32087366298855 |
| ENSG00000124783.13 | SSR1 | -0.0126369066547651 | 6.69407922525947 | -0.105550316609161 | 0.916435765537614 | 0.971018087205843 | -6.40442452707452 |
| lnc-TTC3-6 | lnc-TTC3-6 | 0.0278333873718093 | 1.60949329930273 | 0.105438022395603 | 0.916524332099677 | 0.971033058672994 | -5.98585022216537 |
| lnc-PCED1B-1 | lnc-PCED1B-1 | 0.0318271612516647 | 2.24660298730823 | 0.105404026201117 | 0.916551371445935 | 0.971033058672994 | -6.07718224399712 |
| lnc-KRBA2-1 | lnc-KRBA2-1 | -0.0192722170119417 | 8.15226998160029 | -0.105352955941005 | 0.916591424814536 | 0.971033058672994 | -6.38129108482016 |
| ENSG00000189114.6 | BLOC1S3 | 0.0164241498352844 | 2.14578722554503 | 0.105098477418594 | 0.916792138096198 | 0.971195706558623 | -6.06657060237281 |
| lnc-KLF12-4 | lnc-KLF12-4 | 0.0273809619762684 | 2.02680085824491 | 0.105019556734624 | 0.916854385838991 | 0.971211662998505 | -6.06054335089438 |
| ENSG00000136603.13 | SKIL | 0.0154583938535238 | 5.25541095730902 | 0.104919244001896 | 0.916933507065554 | 0.971245490733342 | -6.38599883432652 |
| lnc-MPP4-3 | lnc-MPP4-3 | 0.0214573532824913 | 1.1504811011221 | 0.104482077924028 | 0.917278329823976 | 0.971560740120768 | -5.93340675087903 |
| ENSG00000154945.6 | ANKRD40 | -0.013339650656552 | 4.67828564570489 | -0.104253360768519 | 0.91745874109945 | 0.971623509506984 | -6.35602829095391 |
| ENSG00000125850.10 | OVOL2 | 0.0154091428445831 | 1.36779920679308 | 0.104246308915969 | 0.91746430364556 | 0.971623509506984 | -5.96840899117125 |
| ENSG00000154277.12 | UCHL1 | -0.132000785003358 | -1.89574827112497 | -0.104202481863087 | 0.9175036705325 | 0.971623509506984 | -5.05460555411202 |
| lnc-CYP2C18-2 | lnc-CYP2C18-2 | -0.017116218126542 | 0.0283910662334468 | -0.104105023142401 | 0.917575751643078 | 0.971623509506984 | -5.70510196977234 |
| ENSG00000170571.11 | EMB | 0.0176515266444985 | 6.27815578076989 | 0.104080899593983 | 0.917594780769593 | 0.971623509506984 | -6.40546375440563 |
| ENSG00000267059.2 | | -0.0193968809372986 | 3.13640503579392 | -0.103949173105814 | 0.917698690060272 | 0.971623509506984 | -6.19891974594819 |
| ENSG00000060982.14 | BCAT1 | 0.0734047084525109 | 5.02973455693356 | 0.103903529377441 | 0.917754111084052 | 0.971623509506984 | -6.39134228957825 |
| ENSG00000171223.5 | JUNB | -0.030631131366083 | 6.38765722099459 | -0.103882796569038 | 0.917760164215089 | 0.971623509506984 | -6.40527950036713 |
| lnc-MSH6-2 | lnc-MSH6-2 | -0.0218951726368188 | 1.77254051695966 | -0.103868359693348 | 0.917762438509196 | 0.971623509506984 | -6.0079748779962 |
| ENSG00000187164.19 | SHTN1 | -0.127972130340989 | -1.86673366948675 | -0.103394120926346 | 0.918142721977039 | 0.971936705049078 | -5.09850203860546 |

| | | | | | | | |
|---|---|---|---|---|---|---|---|
| lnc-PLPP5-3 | lnc-PLPP5-3 | 0.0135304892479808 | 1.94661737952917 | 0.103357072406883 | 0.918165772493815 | 0.971936705049078 | -6.03439092475543 |
| ENSG00000129071.9 | MBD4 | 0.0173348767133019 | 6.24829431706547 | 0.103313769493436 | 0.918199933424638 | 0.971936705049078 | -6.40531518819059 |
| lnc-AFG1L-5 | lnc-AFG1L-5 | 0.0543320805170174 | 0.636265258535283 | 0.102993990131502 | 0.918459897688918 | 0.97211125728985 | -5.81669701747866 |
| lnc-RBP2-2 | lnc-RBP2-2 | -0.0153924229869435 | 1.337456083116 | -0.102981194647852 | 0.918462301263406 | 0.97211125728985 | -5.96282192947672 |
| lnc-ACSBG1-6 | lnc-ACSBG1-6 | -0.0172320244855146 | -0.0221977127701043 | -0.10291961232135 | 0.918510884486014 | 0.97211125728985 | -5.64947599688275 |
| lnc-ZEB1-12 | lnc-ZEB1-12 | -0.0239627565847349 | 1.13472682758771 | -0.102800272339627 | 0.918605034487381 | 0.97211125728985 | -5.92153119103471 |
| ENSG00000137720.7 | CFAP68 | 0.0270053462964059 | 1.62746038669451 | 0.102674690093682 | 0.918704110427953 | 0.97211125728985 | -5.99589467671541 |
| ENSG00000102226.9 | USP11 | -0.0129091508730657 | 6.18541843890134 | -0.102663990514988 | 0.918712551735933 | 0.97211125728985 | -6.40525739008487 |
| ENSG00000279744.1 | | -0.0219732188035698 | -0.151621369074007 | -0.102625820378399 | 0.91874266569788 | 0.97211125728985 | -5.62270933658589 |
| lnc-HEXDC-1 | lnc-HEXDC-1 | -0.0219732188035698 | -0.151621369074007 | -0.102625820378399 | 0.91874266569788 | 0.97211125728985 | -5.62270933658589 |
| lnc-HSP90AA1-16 | lnc-HSP90AA1-16 | 0.032237397519614 | 0.217433946755117 | 0.1024189518093 | 0.91890587476349 | 0.972203578673577 | -5.7219633234575 |
| lnc-CD82-3 | lnc-CD82-3 | -0.0493756073618605 | 0.359692494397776 | -0.102380135106455 | 0.918938362051264 | 0.972203578673577 | -5.72042917719122 |
| lnc-UNC93B1-5 | lnc-UNC93B1-5 | -0.0201111975464097 | 1.82450665763641 | -0.102335622266417 | 0.918971618650008 | 0.972203578673577 | -6.02351146880125 |
| ENSG00000046647.13 | GEMIN8 | 0.0127257885605556 | 3.21524795848024 | 0.102196767812116 | 0.919081170885027 | 0.972269504051319 | -6.20847988782539 |
| lnc-KMT5B-2 | lnc-KMT5B-2 | 0.0176600870777681 | 1.70243695555007 | 0.102110429104784 | 0.919149290484057 | 0.972291594512968 | -6.00295067477875 |
| lnc-ALG13-3 | lnc-ALG13-3 | 0.0267371689223972 | 1.65548826288581 | 0.101934376464716 | 0.919288194574374 | 0.972299347141792 | -6.00827383423252 |
| lnc-ABCB1-3 | lnc-ABCB1-3 | -0.0120233954858421 | 1.00007358728781 | -0.101930299471502 | 0.919291411318847 | 0.972299347141792 | -5.90260600526088 |
| ENSG00000041802.10 | LSG1 | 0.0104188197002228 | 5.71403918657243 | 0.101921526336667 | 0.919298333320041 | 0.972299347141792 | -6.40026726715972 |
| ENSG00000124593.16 | | 0.0124328640465241 | 3.90802934604292 | 0.101582347543782 | 0.919565950153509 | 0.972393812333712 | -6.27965619688104 |
| lnc-ZFP3-3 | lnc-ZFP3-3 | 0.0252812180356528 | 0.605870353653199 | 0.101512480056604 | 0.919621077755629 | 0.972393812333712 | -5.80060667951096 |
| ENSG00000223547.9 | ZNF844 | 0.0179400075118901 | 6.50516028788292 | 0.101461853367353 | 0.9196610240248 | 0.972393812333712 | -6.40559682197039 |
| ENSG00000135315.11 | CEP162 | -0.0121186354424414 | 3.89672439213634 | -0.101416593596726 | 0.919696735780919 | 0.972393812333712 | -6.27456305504786 |
| lnc-LZTFL1-1 | lnc-LZTFL1-1 | -0.0180048068687801 | -0.0444577036415172 | -0.101331367069063 | 0.919763983350527 | 0.972393812333712 | -5.65276684254411 |
| ENSG00000163584.17 | RPL22L1 | 0.0183219669483212 | 3.69072196182338 | 0.101296945873886 | 0.919791143392633 | 0.972393812333712 | -6.26506397145466 |
| ENSG00000104408.9 | EIF3E | 0.0153879259347312 | 7.57541168431363 | 0.1012805412418 | 0.91980408750075 | 0.972393812333712 | -6.393479014305 |
| ENSG00000150593.17 | PDCD4 | -0.0173838104513369 | 8.24410842732283 | -0.101136700682976 | 0.919917586126608 | 0.972393812333712 | -6.37958446090962 |
| ENSG00000025796.13 | SEC63 | -0.0158176869267006 | 4.88945723893088 | -0.101108315542246 | 0.919939983864885 | 0.972393812333712 | -6.36490893460411 |
| ENSG00000130726.11 | TRIM28 | -0.0123800493462101 | 6.26681934349951 | -0.101008059251629 | 0.920019093181942 | 0.972393812333712 | -6.40572968997062 |
| ENSG00000129007.14 | CALML4 | -0.0187854738723965 | 1.02947519685301 | -0.100932168347813 | 0.920078977027561 | 0.972393812333712 | -5.91136026320341 |
| lnc-KCNK7-3 | lnc-KCNK7-3 | -0.0184692924040855 | 0.985790080725604 | -0.100911479212243 | 0.920095302451538 | 0.972393812333712 | -5.90688706491025 |
| lnc-ANKRD26-5 | lnc-ANKRD26-5 | -0.026928662647324 | -0.155138266070419 | -0.100793696560434 | 0.920188243283421 | 0.972393812333712 | -5.58174299101224 |
| lnc-FBXO11-6 | lnc-FBXO11-6 | -0.0193426645510393 | 0.81243927146756 | -0.100762438479958 | 0.920212908837249 | 0.972393812333712 | -5.87457136104326 |
| ENSG00000184979.9 | USP18 | 0.0552335410018309 | -0.0686512645850043 | 0.100712459362384 | 0.92025784368108 | 0.972393812333712 | -5.72272551833232 |
| ENSG00000280213.1 | UCKL1-AS1 | 0.0270280033772776 | 0.616622838534767 | 0.100700491175392 | 0.920261791298788 | 0.972393812333712 | -5.81192984251757 |
| LINC01948 | LINC01948 | -0.0167443361147104 | 2.19081219808329 | -0.100687381297087 | 0.920272136310652 | 0.972393812333712 | -6.06499453931827 |
| ENSG00000114013.15 | CD86 | -0.0590264652546146 | 3.58597338695767 | -0.100705355693649 | 0.920275234038926 | 0.972393812333712 | -6.24657647206055 |
| ENSG00000250565.6 | ATP6V1E2 | 0.018304559927288 | 2.07909651763994 | 0.100615860396422 | 0.920328573736982 | 0.972393812333712 | -6.04722247404968 |
| ENSG00000153814.12 | JAZF1 | -0.0157934924476999 | 5.21476275815678 | -0.100610884168446 | 0.920332500513679 | 0.972393812333712 | -6.38708030734827 |
| ENSG00000086200.16 | IPO11 | 0.0122864246460948 | 4.11893603523915 | 0.100438336004376 | 0.920468660734092 | 0.972410405064235 | -6.30488777531869 |
| ENSG00000130024.14 | PHF10 | -0.0143155552730407 | 6.21207370626212 | -0.100423458180064 | 0.920480401150863 | 0.972410405064235 | -6.4056637975222 |
| ENSG00000123416.15 | TUBA1B | 0.0193229273471855 | 7.05930687489805 | 0.100356372291894 | 0.920533340316322 | 0.972410405064235 | -6.39990770135141 |
| ENSG00000183891.5 | TTC32 | 0.0154462469398933 | 4.6208052877864 | 0.100351508610698 | 0.920537178383939 | 0.972410405064235 | -6.35371529946781 |
| ENSG00000136014.11 | USP44 | -0.054350149472886 | 2.87111541207846 | -0.100282610239784 | 0.920607095840144 | 0.972434355623405 | -6.16836405046541 |
| ENSG00000111880.15 | RNGTT | -0.0227613114582521 | 5.79167161969794 | -0.0999572326279412 | 0.920849792076541 | 0.972620524324445 | -6.40204904713492 |
| lnc-ADAT2-2 | lnc-ADAT2-2 | -0.0176862850102042 | 2.03522644209047 | -0.0999198124554038 | 0.920877849586298 | 0.972620524324445 | -6.0574637333178 |
| ENSG00000180776.15 | ZDHHC20 | 0.0130964452990607 | 6.53550166211715 | 0.0996866600426237 | 0.921061847088628 | 0.972764944257029 | -6.40536907426873 |
| ENSG00000143919.14 | CAMKMT | 0.0119028022703285 | 2.98605317354579 | 0.0995449894318874 | 0.921173651794885 | 0.972806243293197 | -6.18274467061457 |
| ENSG00000204178.10 | MACO1 | 0.00784101186614006 | 4.62231365181449 | 0.0994268665818139 | 0.921266874131178 | 0.972806243293197 | -6.35496074421802 |
| lnc-DMKN-1 | lnc-DMKN-1 | 0.0780066942154323 | 0.121081521364255 | 0.0994195037627555 | 0.921288971905452 | 0.972806243293197 | -5.87226807921158 |
| lnc-XXYLT1-5 | lnc-XXYLT1-5 | -0.0199059097768353 | 2.087353383638 | -0.0993975619294441 | 0.9212900014825 | 0.972806243293197 | -6.06288268437563 |
| lnc-ENKD1-1 | lnc-ENKD1-1 | 0.0239343925440288 | 0.0111448677482082 | 0.0991063063687738 | 0.921519865304428 | 0.972964662690003 | -5.69899203199015 |
| ENSG00000285596.1 | NA | 0.0113383427420857 | 1.53357241757617 | 0.0988725198632866 | 0.921704378487835 | 0.972964662690003 | -5.98757716465518 |
| ENSG00000174007.7 | CEP19 | 0.0276271297445775 | -0.10506250848617 | 0.0988696211396989 | 0.921706666297925 | 0.972964662690003 | -5.61790163539496 |
| ENSG00000137338.5 | PGBD1 | -0.0139320144259685 | 2.44044303141227 | -0.098795429347345 | 0.921765222206318 | 0.972964662690003 | -6.10685166787767 |
| lnc-SLC25A12-8 | lnc-SLC25A12-8 | -0.0132285655908629 | 0.290639771924832 | -0.0987653730277009 | 0.921788944296318 | 0.972964662690003 | -5.74902512529264 |
| ENSG00000146350.13 | TBC1D32 | 0.0182887649883328 | 4.03276909224647 | 0.0987333163022949 | 0.921814245295278 | 0.972964662690003 | -6.29069416131955 |
| lnc-ROPN1L-9 | lnc-ROPN1L-9 | -0.0343041714546665 | 0.248885963522194 | -0.0986750297144881 | 0.921860248609982 | 0.972964662690003 | -5.70511784418327 |
| ENSG00000122643.19 | NT5C3A | -0.0149786297190673 | 5.15409459301561 | -0.0986408848176671 | 0.921887197961863 | 0.972964662690003 | -6.38191146762692 |
| ENSG00000069424.14 | KCNAB2 | 0.0113314056646283 | 5.2396835436111 | 0.0985881219668932 | 0.921928841984852 | 0.972964662690003 | -6.38751210706481 |
| ENSG00000265818.1 | EEF1E1-BLOC1S5 | -0.0102824028189517 | 2.27991611938362 | -0.0985685326649894 | 0.921944303247928 | 0.972964662690003 | -6.09218034657436 |
| ENSG00000116273.5 | PHF13 | -0.0189384321978197 | 3.16073006184841 | -0.098548638816347 | 0.921960004912103 | 0.972964662690003 | -6.19364785138209 |
| LINC02397 | LINC02397 | 0.0375263168482913 | 7.26145421426888 | 0.0984334869238206 | 0.922063890273454 | 0.972988669827598 | -6.39583271759497 |
| lnc-AZIN2-5 | lnc-AZIN2-5 | 0.0297288146189801 | -0.495053167725981 | 0.0983684034143491 | 0.922102261161641 | 0.972988669827598 | -5.49666384380202 |
| lnc-B4GALT1-2 | lnc-B4GALT1-2 | -0.0171376233289706 | 0.297963983741323 | -0.0983047150444326 | 0.9221525297538 | 0.972988669827598 | -5.77732187613424 |
| lnc-HNRNPA3-1 | lnc-HNRNPA3-1 | 0.037313859361441 | 0.493507290909236 | 0.0982802505364239 | 0.922171839430442 | 0.972988669827598 | -5.82104711607124 |
| ENSG00000171310.10 | CHST11 | -0.0172905211075411 | 5.03975458895797 | -0.0980811288992438 | 0.922329006606207 | 0.973104615458279 | -6.37972570010796 |
| ENSG00000142694.6 | EVA1B | -0.0472699087124698 | -0.782767916073954 | -0.0979446910680412 | 0.922436699120582 | 0.973134653806255 | -5.5504158098649 |
| ENSG00000268173.3 | | -0.0281703810109906 | 3.43021646484954 | -0.0979284563351354 | 0.922452034708926 | 0.973134653806255 | -6.22627528736409 |

| | | | | | | | |
|---|---|---|---|---|---|---|---|
| ENSG00000102119.10 | EMD | -0.0151609757484774 | 4.10049773508809 | -0.0978588276973165 | 0.922504473177344 | 0.973140096935695 | -6.30632270291919 |
| lnc-NSUN6-2 | lnc-NSUN6-2 | 0.0106902486263727 | 2.28104115483731 | 0.0977829012193678 | 0.92256440428731 | 0.973153443009568 | -6.08635008699396 |
| ENSG00000178385.14 | PLEKHM3 | 0.00937672030567782 | 5.00129998106967 | 0.0977012182615385 | 0.922628879672721 | 0.973157149671562 | -6.37215320644489 |
| lnc-HJURP-10 | lnc-HJURP-10 | 0.0137502246984036 | 1.02577385915342 | 0.0976586538219475 | 0.922662477570837 | 0.973157149671562 | -5.90032866047202 |
| ENSG00000223396.4 | RPS10P7 | 0.014574432257674 | 0.73540785703583 | 0.0975073212478424 | 0.922781931878183 | 0.973215782839776 | -5.84863133303147 |
| ENSG00000188659.9 | SAXO2 | 0.0321420571113424 | 1.35457983405763 | 0.0974684268844196 | 0.922812633420002 | 0.973215782839776 | -5.95655753770843 |
| ENSG00000205765.8 | RIMOC1 | 0.0112055987039685 | 4.53386079545663 | 0.0973082095048546 | 0.92293910340098 | 0.973271750137571 | -6.34243870497621 |
| ENSG00000187961.13 | KLHL17 | -0.0232799012312668 | 1.38725161835645 | -0.0972813917411134 | 0.922960272600459 | 0.973271750137571 | -5.97492389552783 |
| lnc-FZD5-6 | lnc-FZD5-6 | -0.023307633205554 | 0.0485600275844441 | -0.0969724690480397 | 0.923204131684202 | 0.973453737078112 | -5.66914636314039 |
| lnc-ARHGEF10L-1 | lnc-ARHGEF10L-1 | 0.0377061985735654 | 1.43031479191285 | 0.0969197790295849 | 0.923249590896787 | 0.973453737078112 | -5.98282229438803 |
| lnc-LGR6-5 | lnc-LGR6-5 | -0.022635674021702 | 1.84171449002274 | -0.0968830308955367 | 0.923274734258316 | 0.973453737078112 | -6.02347425270197 |
| ENSG00000125247.15 | TMTC4 | 0.016248274064631 | 3.66604564586856 | 0.0967779181673948 | 0.923357711159535 | 0.973484499545887 | -6.2615294183632 |
| ENSG00000065357.19 | DGKA | 0.0183797185024082 | 7.75166705912286 | 0.0967262457649377 | 0.923398502119899 | 0.973484499545887 | -6.39279185506185 |
| lnc-CMTR1-10 | lnc-CMTR1-10 | 0.029328780368771 | 2.07409097432655 | 0.0966433940625273 | 0.923464202029284 | 0.973503901171255 | -6.07553718639907 |
| ENSG00000095585.16 | BLNK | -0.0133113615025833 | 6.48569257088563 | -0.0964632659511621 | 0.923606105513131 | 0.973603629508182 | -6.40594091340967 |
| ENSG00000123353.9 | ORMDL2 | 0.0138400291174956 | 3.12736128250659 | 0.0963385554124422 | 0.923704557277483 | 0.973648638227805 | -6.19901721013433 |
| ENSG00000185009.12 | AP3M1 | 0.00798722494601378 | 5.81598171124658 | 0.0962893404685501 | 0.923743409965145 | 0.973648638227805 | -6.4028390242065 |
| ENSG00000252498.1 | RNU6-1016P | -0.0185966875196408 | 1.82480253899032 | -0.096172772799308 | 0.923835434945132 | 0.973666477336593 | -6.03726752358789 |
| ENSG00000139579.12 | NABP2 | 0.0133905029797425 | 3.05441928200504 | 0.096082017334324 | 0.923907083071187 | 0.973666477336593 | -6.19067022429165 |
| ENSG00000259494.1 | MRPL46 | -0.0113722445178497 | 3.06970826476676 | -0.0960381993837745 | 0.923941675972213 | 0.973666477336593 | -6.19117822986295 |
| ENSG00000115504.14 | EHBP1 | 0.0143977295749096 | 4.41333143200346 | 0.0960282226440838 | 0.923949552316879 | 0.973666477336593 | -6.32971292049154 |
| ENSG00000178585.14 | CTNNBIP1 | -0.0108295067900328 | 2.93762937736875 | -0.0957118383031947 | 0.92419933251661 | 0.973810355300714 | -6.16713140349983 |
| ENSG00000242485.5 | MRPL20 | 0.00844526569775894 | 4.08464214671342 | 0.0956514994121909 | 0.924246969951833 | 0.973810355300714 | -6.30173452744009 |
| lnc-MNX1-22 | lnc-MNX1-22 | -0.0193021259682549 | 1.92842608204258 | -0.095633773308974 | 0.924260964728772 | 0.973810355300714 | -6.04127317055889 |
| ENSG00000174276.6 | ZNHIT2 | -0.0248851515388734 | 0.33916588026228 | -0.0955416732944778 | 0.924333678176736 | 0.973810355300714 | -5.79771865547242 |
| ENSG00000077235.17 | GTF3C1 | 0.00672810481553299 | 6.10647344132853 | 0.0955378955359803 | 0.924336660751009 | 0.973810355300714 | -6.40552837152244 |
| ENSG00000119661.14 | DNAL1 | 0.0137513102050688 | 2.59821650687272 | 0.0954848202310213 | 0.924378564297537 | 0.973810355300714 | -6.1307901999172 |
| ENSG00000198894.7 | CIPC | 0.0153785153467521 | 4.73006944034054 | 0.0953645476200387 | 0.924473521672286 | 0.973810355300714 | -6.3643916100682 |
| ENSG00000072274.12 | TFRC | 0.0137754555642617 | 7.15525801033352 | 0.0953613421859034 | 0.924476052435047 | 0.973810355300714 | -6.40097057224706 |
| ENSG00000242372.7 | EIF6 | -0.0152706491909447 | 3.98081078757827 | -0.0953159557157809 | 0.924511886164857 | 0.973810355300714 | -6.29362508667211 |
| ENSG00000196338.12 | NLGN3 | -0.025404439942898 | 0.767434874374628 | -0.0950532893510422 | 0.924719270806531 | 0.973978955634573 | -5.84341541180093 |
| ENSG00000079134.11 | THOC1 | 0.0127468417661376 | 5.78523173814847 | 0.0949368516756969 | 0.924811204279071 | 0.974025943697289 | -6.40218983780875 |
| ENSG00000186350.10 | RXRA | 0.0425187455578886 | 5.26273492121943 | 0.0948917464917868 | 0.924862274930337 | 0.974029891777074 | -6.3956683663318 |
| ENSG00000166927.12 | MS4A7 | -0.0727743175409926 | -0.489437491044021 | -0.0946810819450419 | 0.92501851101955 | 0.974043004397371 | -5.44085098020423 |
| ENSG00000273184.1 | | -0.0132053482015488 | 1.63197606274337 | -0.0946437736379884 | 0.925042608910741 | 0.974043004397371 | -5.99991505566219 |
| ENSG00000100151.15 | PICK1 | -0.013191960703391 | 3.43164845804876 | -0.0946190971712089 | 0.925062092924964 | 0.974043004397371 | -6.22239464863903 |
| ENSG00000070759.16 | TESK2 | -0.0236170053155744 | 2.920641761797 | -0.0946166610724218 | 0.925064016419366 | 0.974043004397371 | -6.14728672309569 |
| lnc-ELF1-2 | lnc-ELF1-2 | -0.0141447720751973 | 0.830630755208611 | -0.0944770145812447 | 0.925174279226537 | 0.974050763003323 | -5.87275965391972 |
| lnc-HIP1-4 | lnc-HIP1-4 | -0.017140339315824 | 0.196734358828182 | -0.0944056979363584 | 0.925230590370546 | 0.974050763003323 | -5.72839362496986 |
| lnc-ISM2-3 | lnc-ISM2-3 | -0.0159920353998544 | 0.984028734386827 | -0.0944043249812215 | 0.925231674450512 | 0.974050763003323 | -5.91780487702747 |
| ENSG00000185885.15 | IFITM1 | -0.0494082441177763 | 2.35233709826894 | -0.0940644078852292 | 0.925514036278858 | 0.974050763003323 | -6.17326604255556 |
| ENSG00000171497.4 | PPID | 0.0136686838146542 | 3.54272010904782 | 0.0940440382210442 | 0.925516160420019 | 0.974050763003323 | -6.23855248096886 |
| ENSG00000010256.10 | UQCRC1 | -0.0120955117801511 | 5.03857907665212 | -0.0940354297192725 | 0.925522957898497 | 0.974050763003323 | -6.37839309993511 |
| ZNF346-IT1 | ZNF346-IT1 | -0.0169191721491381 | 0.447782467204337 | -0.0939329886736072 | 0.925603848225411 | 0.974050763003323 | -5.78937608205863 |
| lnc-HSPA13-5 | lnc-HSPA13-5 | -0.0790057401222075 | -1.58793582757822 | -0.0939026578131462 | 0.925627798477602 | 0.974050763003323 | -5.11969468948742 |
| lnc-TGS1-1 | lnc-TGS1-1 | 0.0219263286302924 | 4.37348873464419 | 0.0937376806957833 | 0.925758454147244 | 0.974050763003323 | -6.32758419552526 |
| ENSG00000137504.13 | CREBZF | -0.0160635015998372 | 6.34595343369434 | -0.0936839441964445 | 0.925800504059825 | 0.974050763003323 | -6.40642245270749 |
| ENSG00000175787.16 | ZNF169 | 0.0094657619449281 | 3.57267892705747 | 0.0936387893181428 | 0.925836160725214 | 0.974050763003323 | -6.24163142066408 |
| RUNX1-IT1 | RUNX1-IT1 | 0.0231872464795191 | 2.13746441407167 | 0.0936322895438513 | 0.925841293301347 | 0.974050763003323 | -6.05979362775922 |
| lnc-NPIPA7-3 | lnc-NPIPA7-3 | -0.0224627366877843 | 1.48885174969261 | -0.0936241458380418 | 0.925847724019941 | 0.974050763003323 | -5.96269241944738 |
| ENSG00000054116.11 | TRAPPC3 | 0.00855697913968179 | 4.3304801961036 | 0.0935940505841892 | 0.925871488933307 | 0.974050763003323 | -6.32597415116015 |
| lnc-FRAT2-2 | lnc-FRAT2-2 | 0.019671906862906 | -0.183132751088893 | 0.0935689348434054 | 0.925891321794159 | 0.974050763003323 | -5.61031465476604 |
| lnc-KAT2B-3 | lnc-KAT2B-3 | -0.0330040489653419 | -0.524717718938843 | -0.0935664025128864 | 0.925893321473394 | 0.974050763003323 | -5.48375412026936 |
| lnc-RPS16-5 | lnc-RPS16-5 | -0.0126409732522217 | 2.07478923031409 | -0.0935130619747175 | 0.925935442456806 | 0.974050763003323 | -6.05068321523961 |
| ENSG00000242114.5 | MTFP1 | 0.0331462666219465 | 0.0701152483829222 | 0.0932111073966468 | 0.92617388849119 | 0.974050763003323 | -5.72327087985111 |
| lnc-SSMEM1-3 | lnc-SSMEM1-3 | -0.0150830235368482 | 1.04689229291784 | -0.0932032108375292 | 0.926180124300142 | 0.974050763003323 | -5.91821947009932 |
| ENSG00000117419.15 | ERI3 | 0.00927804876592431 | 3.62841886463381 | 0.0931644898510826 | 0.926210701822204 | 0.974050763003323 | -6.25217349278994 |
| ENSG00000083454.21 | P2RX5 | -0.0284311150394849 | 4.74802026991202 | -0.0931652081988076 | 0.926217779322373 | 0.974050763003323 | -6.35972717061382 |
| lnc-FBXO21-2 | lnc-FBXO21-2 | 0.0256275589853805 | 1.49620327783467 | 0.0931109373568289 | 0.926252991803092 | 0.974050763003323 | -5.99481449602307 |
| ENSG00000176273.14 | SLC35G1 | -0.0580143133371287 | -1.07227072010611 | -0.0930323950938644 | 0.926315016396851 | 0.974050763003323 | -5.29433755906909 |
| ENSG00000157992.12 | KRTCAP3 | 0.0216849753436771 | 0.718286948364892 | 0.0929670182380853 | 0.926366644661033 | 0.974050763003323 | -5.87030248366642 |
| lnc-PUM3-2 | lnc-PUM3-2 | -0.0179657893595433 | 2.71993075697958 | -0.0929528782873662 | 0.926377811057219 | 0.974050763003323 | -6.14532076555398 |
| lnc-TET3-2 | lnc-TET3-2 | 0.0183241990978966 | 1.12693170745997 | 0.0929083231944937 | 0.926412996555223 | 0.974050763003323 | -5.93406713677245 |
| ENSG00000153317.14 | ASAP1 | 0.0142375822682519 | 3.82291556436243 | 0.0929054984102354 | 0.926415227314006 | 0.974050763003323 | -6.26897468790253 |
| ENSG00000204003.8 | | -0.0766389221535406 | 0.753878950906007 | -0.0928980116640136 | 0.926437808460786 | 0.974050763003323 | -6.0011852679653 |
| ENSG00000165948.10 | IFI27L1 | 0.0119045451748603 | 1.04312536704905 | 0.0928423741413579 | 0.926465077301026 | 0.974050763003323 | -5.91703665587064 |
| lnc-TMC6-12 | lnc-TMC6-12 | -0.0676688473759178 | 0.570425817559186 | -0.0928035377629621 | 0.926509075568982 | 0.974050763003323 | -5.84943516838233 |
| ENSG00000262554.1 | NA | -0.012673437508415 | 1.74414982972395 | -0.0926953017195118 | 0.926581223303751 | 0.974050763003323 | -6.01051107362696 |

| | | | | | | | |
|---|---|---|---|---|---|---|---|
| ENSG00000183479.12 | TREX2 | 0.0111225058633089 | 2.28437762646187 | 0.0926896037313408 | 0.926585723150418 | 0.974050763003323 | -6.09330232130523 |
| ENSG00000278588.1 | H2BC10 | -0.102718726844848 | 1.85336737594468 | -0.09254705257019 | 0.926715575090254 | 0.974137515044313 | -6.02349627389088 |
| ENSG00000138698.14 | RAP1GDS1 | 0.0102036431535977 | 5.57207460947241 | 0.0922100411360566 | 0.926964454671146 | 0.974349370365448 | -6.39805746445758 |
| ENSG00000109079.9 | TNFAIP1 | 0.0124872781729723 | 3.05091503877694 | 0.0919591414262094 | 0.927162607961666 | 0.974462643573057 | -6.18901887216667 |
| ENSG00000125945.14 | ZNF436 | 0.0124655715176538 | 2.88160625801723 | 0.0918279019960729 | 0.927266258909453 | 0.974462643573057 | -6.15998248281638 |
| lnc-NPM2-3 | lnc-NPM2-3 | -0.0177916477174738 | 2.29600469685483 | -0.0918069193543136 | 0.927282830806057 | 0.974462643573057 | -6.09835867630984 |
| lnc-SLC22A31-7 | lnc-SLC22A31-7 | 0.0157087271767544 | 2.52883008442429 | 0.0917789775324012 | 0.927304899049567 | 0.974462643573057 | -6.11200978164238 |
| lnc-PLXNA4-1 | lnc-PLXNA4-1 | -0.0234065126639518 | 0.826146781186815 | -0.091773867981258 | 0.927308934541392 | 0.974462643573057 | -5.87340371759861 |
| lnc-KRR1-1 | lnc-KRR1-1 | -0.0156685337733111 | 1.98155402318273 | -0.0916734174669336 | 0.927388270128135 | 0.974466250535652 | -6.04329767687812 |
| lnc-TNS2-1 | lnc-TNS2-1 | 0.0201984851518909 | -0.179010931465084 | 0.0916496351233504 | 0.927407053478254 | 0.974466250535652 | -5.63415544886172 |
| lnc-SCARF2-3 | lnc-SCARF2-3 | 0.0130314356547634 | 0.745098136543652 | 0.0915064093416233 | 0.927520174418372 | 0.974535362432534 | -5.82810277646952 |
| ENSG00000049860.13 | HEXB | -0.0128713478507567 | 3.93391453333305 | -0.0911893448510131 | 0.927770600042215 | 0.974748724448415 | -6.28657270945455 |
| ENSG00000161016.17 | RPL8 | 0.0145919666091068 | 8.66214261220175 | 0.0911023681016317 | 0.927839297794042 | 0.974771144676132 | -6.36994447885798 |
| ENSG00000134996.11 | OSTF1 | 0.0109229787086429 | 5.01392299677648 | 0.0910360822106958 | 0.927891653443713 | 0.974776394775274 | -6.37469613037373 |
| ENSG00000125871.13 | MGME1 | -0.0145831265643128 | 4.43083279187355 | -0.0908716102947711 | 0.928021562334035 | 0.974807776860589 | -6.33568466741969 |
| lnc-SP110-1 | lnc-SP110-1 | -0.0800501499209019 | -1.0915554112452 | -0.0908427597707035 | 0.928047067638438 | 0.974807776860589 | -5.2581429972887 |
| ENSG00000167815.11 | PRDX2 | 0.04624002220881 | 2.11628909361639 | 0.0908335570748997 | 0.928063605662931 | 0.974807776860589 | -6.11726809283298 |
| lnc-STRN3-11 | lnc-STRN3-11 | -0.0194836248676106 | 0.760533579811364 | -0.0907343965578843 | 0.928129942748163 | 0.974827708913886 | -5.8668365663893 |
| ENSG00000136840.18 | ST6GALNAC4 | 0.0194549951929917 | 5.7667300542765 | 0.0906540374921276 | 0.928193536146305 | 0.974844757347658 | -6.40279399951051 |
| lnc-CCDC78-1 | lnc-CCDC78-1 | -0.0228711824748788 | -0.177194435933251 | -0.0904491322374427 | 0.928355267646135 | 0.974926049645957 | -5.6327797280762 |
| ENSG00000147854.16 | UHRF2 | -0.00887564112895234 | 7.58264417456969 | -0.0904359636729042 | 0.92836566939031 | 0.974926049645957 | -6.39487523660262 |
| lnc-FAM86B2-7 | lnc-FAM86B2-7 | 0.0184111047918285 | 2.42963108124299 | 0.0902960297499689 | 0.928476202863472 | 0.97499238220187 | -6.10495349259147 |
| lnc-RAB11FIP2-3 | lnc-RAB11FIP2-3 | -0.0135482932747112 | 0.434818649654663 | -0.090203897586764 | 0.928548978620648 | 0.975019060654464 | -5.79691616036822 |
| ENSG00000207721.1 | MIR186 | -0.0414381402651177 | 0.513935516238942 | -0.0898808044478838 | 0.928808304394821 | 0.97524161247557 | -5.85373291600597 |
| lnc-TEX44-2 | lnc-TEX44-2 | -0.0707640796784873 | 0.920977971281731 | -0.0898236015613497 | 0.928863970663468 | 0.975250311577543 | -5.85648114172342 |
| GAS5 | GAS5 | -0.0112978779868564 | 7.10480383636351 | -0.0896337365755569 | 0.928999366020847 | 0.975283938483723 | -6.40118036317213 |
| ENSG00000167671.11 | UBXN6 | 0.011604994784931 | 4.92737276325931 | 0.0895740886790106 | 0.929046485059127 | 0.975283938483723 | -6.37328337126743 |
| ENSG00000115170.13 | ACVR1 | 0.0155623324651946 | 2.16273758369424 | 0.0895496859149809 | 0.929065762171466 | 0.975283938483723 | -6.07739905968345 |
| lnc-JSRP1-3 | lnc-JSRP1-3 | 0.0179454602030678 | 1.89888900245725 | 0.0894710290616308 | 0.929127897926049 | 0.975283938483723 | -6.04349186553165 |
| ENSG00000085563.14 | ABCB1 | -0.0459159882944209 | 3.19891570886671 | -0.0894813080355388 | 0.929132913070365 | 0.975283938483723 | -6.19160184114208 |
| lnc-ATP10A-3 | lnc-ATP10A-3 | -0.0109483110204334 | 3.02104793637377 | -0.0893159855739671 | 0.929250377363358 | 0.975357497056094 | -6.17982500749001 |
| lnc-SLC3A1-3 | lnc-SLC3A1-3 | -0.028492044412189 | -0.181547561314473 | -0.0892108099089941 | 0.929333463790785 | 0.97536228814463 | -5.66299377512968 |
| ENSG00000172057.9 | ORMDL3 | -0.0232151076151697 | 5.53029887503039 | -0.0891952869993277 | 0.929349715546445 | 0.97536228814463 | -6.39457245779071 |
| lnc-ZNF250-3 | lnc-ZNF250-3 | 0.0121681280614454 | 0.838275609305496 | 0.0891268577725431 | 0.929399784677057 | 0.975365103146274 | -5.88345069449178 |
| lnc-GNB5-1 | lnc-GNB5-1 | -0.0180154966441272 | 0.0848711843138602 | -0.0888524913437224 | 0.92961653344153 | 0.975406437706669 | -5.6823057552484 |
| lnc-FCRL1-1 | lnc-FCRL1-1 | 0.0438718173723336 | 6.57673497275697 | 0.0888208349283045 | 0.92965676747482 | 0.975406437706669 | -6.40681994271554 |
| ENSG00000084092.6 | NOA1 | 0.0125567630994867 | 4.00572027840289 | 0.0887837159242634 | 0.929670866692111 | 0.975406437706669 | -6.29684031850347 |
| lnc-XRN1-1 | lnc-XRN1-1 | -0.0215564189751721 | -0.276273672591554 | -0.0887362347349282 | 0.929708377491815 | 0.975406437706669 | -5.6447845894025 |
| ENSG00000051596.9 | THOC3 | 0.0177942271633458 | 1.81240301485862 | 0.0886953161668138 | 0.929740703859265 | 0.975406437706669 | -6.03152071910134 |
| lnc-DLST-2 | lnc-DLST-2 | -0.020020872989376 | 3.37935978469956 | -0.0885989676276236 | 0.92981682132435 | 0.975406437706669 | -6.22702693874975 |
| ENSG00000170390.15 | DCLK2 | 0.0297045136887064 | 5.61788735377267 | 0.0885909706460618 | 0.929833296982035 | 0.975406437706669 | -6.40304485517084 |
| ENSG00000157873.17 | TNFRSF14 | -0.0134709863603852 | 5.99027165195742 | -0.0885413801058842 | 0.929862317050659 | 0.975406437706669 | -6.40552023880366 |
| ENSG00000071626.16 | DAZAP1 | 0.0107639284347529 | 5.62399258667629 | 0.0885107091255547 | 0.929886548065723 | 0.975406437706669 | -6.40019974624677 |
| ENSG00000283549.1 | | -0.0122095029541329 | 2.1613658577677 | -0.0883801969681013 | 0.929989657424981 | 0.975406437706669 | -6.07722902206437 |
| lnc-NPC1L1-3 | lnc-NPC1L1-3 | 0.00960775111633416 | 0.967222277031721 | 0.0883663681926895 | 0.930000582732716 | 0.975406437706669 | -5.89132302047432 |
| ENSG00000254093.8 | PINX1 | -0.0103126785292895 | 2.24307550500951 | -0.0883571841611177 | 0.930007838507185 | 0.975406437706669 | -6.08559145533735 |
| ENSG00000162929.13 | SANBR | 0.0176785352892672 | 2.26648131179318 | 0.0881777557614786 | 0.9301495957611 | 0.975473042741849 | -6.07708202209275 |
| ENSG00000115993.12 | TRAK2 | 0.0226780087975769 | 6.33682850135257 | 0.0881618469498116 | 0.930166127867379 | 0.975473042741849 | -6.40673732805276 |
| ENSG00000173545.4 | ZNF622 | 0.00932831870254702 | 4.4543727491754 | 0.0880276111880112 | 0.930268219117227 | 0.975522107390369 | -6.33765806349754 |
| ENSG00000214046.8 | SMIM7 | -0.00831357230503293 | 5.37501061257732 | -0.0879776361927529 | 0.930307702762744 | 0.975522107390369 | -6.39186627768317 |
| lnc-TP53I13-3 | lnc-TP53I13-3 | -0.00974761843021366 | 2.87489550933383 | -0.0876800125307901 | 0.930542849375796 | 0.975688521905352 | -6.15880030405907 |
| LINC02453 | LINC02453 | -0.0169889651079992 | -0.0859760061406178 | -0.087656774665519 | 0.93056120942156 | 0.975688521905352 | -5.65892033532131 |
| ENSG00000254206.5 | NPIPB11 | 0.0234433456835907 | 0.493656603434445 | 0.087533650284496 | 0.930658489618251 | 0.975724732516427 | -5.76438371691222 |
| ENSG00000261067.7 | | 0.0163767700427355 | 2.55980437601738 | 0.0874930677054839 | 0.930690554024924 | 0.975724732516427 | -6.1370868007179 |
| ENSG00000260413.1 | | 0.0205822544167903 | 0.371857238881091 | 0.0871220996741985 | 0.930983662274514 | 0.975883833716663 | -5.77005525977851 |
| lnc-CDH17-5 | lnc-CDH17-5 | -0.0158972307548584 | 0.794007486688497 | -0.0870949870444707 | 0.931005084812256 | 0.975883833716663 | -5.86090168149328 |
| ENSG00000102908.20 | NFAT5 | -0.0135426170860091 | 6.99808185173112 | -0.0870791168967646 | 0.931017624336931 | 0.975883833716663 | -6.4034400744688 |
| lnc-ATP6V1C1-9 | lnc-ATP6V1C1-9 | -0.0122453015984392 | 3.49570922189865 | -0.0870086640338499 | 0.93107329167075 | 0.975883833716663 | -6.23948036952473 |
| ENSG00000147437.9 | GNRH1 | 0.0167622404472088 | 1.0825566755568 | 0.0870009679868457 | 0.931079372614673 | 0.975883833716663 | -5.90906202919445 |
| lnc-CALR3-5 | lnc-CALR3-5 | 0.017987901809975 | 0.268810902899078 | 0.0869091843832369 | 0.931151894705188 | 0.975889738987473 | -5.7244949165841 |
| ENSG00000122140.10 | MRPS2 | -0.0117278600165394 | 3.01657326288607 | -0.0868738277922228 | 0.931179831594712 | 0.975889738987473 | -6.18798679003998 |
| lnc-P2RY10-1 | lnc-P2RY10-1 | -0.0208727844493461 | 1.05195890511028 | -0.0866471125521031 | 0.931358971952005 | 0.976027784720132 | -5.90718722624816 |
| ENSG00000125741.4 | OPA3 | -0.0134746377891455 | 4.14125777346789 | -0.0863906896665312 | 0.931561590316589 | 0.976190419177597 | -6.31624942272722 |
| lnc-PSMB9-6 | lnc-PSMB9-6 | 0.0164921933747335 | 3.4375471883911 | 0.0862926441659216 | 0.93163906440948 | 0.97622190412566 | -6.23140209222864 |
| lnc-ZNF529-2 | lnc-ZNF529-2 | -0.0115148916885781 | 1.86983702987786 | -0.0862106665207169 | 0.931703842437506 | 0.976240082920545 | -6.03897688001911 |
| lnc-HMGN2-6 | lnc-HMGN2-6 | 0.0286977690261442 | -0.462928240644645 | 0.0859772159079036 | 0.931888315663058 | 0.976383670210879 | -5.51794662055277 |
| ENSG00000067369.13 | TP53BP1 | -0.00641937918016566 | 6.70386509809103 | -0.0858359016442305 | 0.931999984369638 | 0.976450966012433 | -6.40594001993611 |
| lnc-PPP1R13L-3 | lnc-PPP1R13L-3 | 0.0106565598276587 | 1.67134857994917 | 0.0855957497955257 | 0.932189759235696 | 0.976504857372687 | -6.01805458967365 |

| | | | | | | | |
|---|---|---|---|---|---|---|---|
| lnc-AKIRIN1-16 | lnc-AKIRIN1-16 | -0.0189639267845719 | 0.422063037178304 | -0.0855554155071661 | 0.932221632936928 | 0.976504857372687 | -5.78879043181454 |
| LINC01176 | LINC01176 | 0.0193582692531526 | 3.47500424509136 | 0.0855429513530998 | 0.932231482612162 | 0.976504857372687 | -6.23394102821877 |
| lnc-STRN-1 | lnc-STRN-1 | -0.0147182001691415 | 2.72943656399197 | -0.0855306652023094 | 0.932241191632576 | 0.976504857372687 | -6.15386862453999 |
| ENSG00000156831.7 | NSMCE2 | -0.0087153658000045 | 3.71991484640893 | -0.085103056693988 | 0.932579111914104 | 0.976706819845528 | -6.25636923924703 |
| lnc-WDR73-7 | lnc-WDR73-7 | -0.0085629279761514 | 3.20868992136886 | -0.0850820481833304 | 0.932595714345075 | 0.976706819845528 | -6.20346889384652 |
| lnc-DDIT3-1 | lnc-DDIT3-1 | -0.0108050074013293 | 2.37725225690139 | -0.0850625061763789 | 0.932611157866964 | 0.976706819845528 | -6.10987579753426 |
| ENSG00000158869.10 | FCER1G | 0.0680474412288581 | -1.27393251970867 | 0.0850464990810009 | 0.932623807862994 | 0.976706819845528 | -5.22177940182971 |
| ENSG00000081059.19 | TCF7 | 0.0635671183242295 | 3.65881635020151 | 0.0849077130908779 | 0.932749326013885 | 0.976788571729524 | -6.24151336683793 |
| lnc-CYB561A3-1 | lnc-CYB561A3-1 | 0.00875402038144335 | 2.66697392966668 | 0.0847677737515734 | 0.93284408014869 | 0.976793323322455 | -6.13266456209693 |
| ENSG00000142453.11 | CARM1 | 0.0118431756737358 | 4.02350540684181 | 0.0847618318400089 | 0.932848776006875 | 0.976793323322455 | -6.29880036894575 |
| ENSG00000142655.12 | PEX14 | 0.00781653413569439 | 3.33382192537396 | 0.0846896768064112 | 0.932905799903777 | 0.976803341103847 | -6.2189937232173 |
| ENSG00000135506.15 | OS9 | -0.0101728743237022 | 6.92936328786282 | -0.0844539609890852 | 0.933092087859122 | 0.976908800082209 | -6.40428058737843 |
| ENSG00000127540.11 | UQCR11 | 0.0111898625465305 | 3.63041185964752 | 0.0844421232583264 | 0.93310144340554 | 0.976908800082209 | -6.25472996339202 |
| lnc-IFI27L2-3 | lnc-IFI27L2-3 | 0.0120902993513559 | 4.124902825275 | 0.0842804630234213 | 0.933229207011041 | 0.976992867499529 | -6.30675555739702 |
| ENSG00000119729.11 | RHOQ | 0.0138152120928678 | 5.58519985441804 | 0.0841744073809 | 0.933313026063713 | 0.977030923378568 | -6.3966405237157 |
| ENSG00000127948.15 | POR | -0.0110408999075364 | 4.2221517284675 | -0.0840919856259977 | 0.933378167053279 | 0.977043498084974 | -6.32212009326745 |
| lnc-DYNC1H1-3 | lnc-DYNC1H1-3 | 0.0188720829804496 | 2.91448896202298 | 0.0840264039334946 | 0.933429999043022 | 0.977043498084974 | -6.15938402995281 |
| ENSG00000176148.15 | TCP11L1 | 0.0149874980869189 | 4.24037952623002 | 0.0839790266315168 | 0.933467443513629 | 0.977043498084974 | -6.31465492781923 |
| lnc-APPBP2-2 | lnc-APPBP2-2 | 0.0162138353868962 | -0.200234600199391 | 0.0838686045423852 | 0.93355471578384 | 0.977054065668186 | -5.5910574837249 |
| lnc-DRD5-8 | lnc-DRD5-8 | 0.0759859658517097 | -1.11008263063646 | 0.083843711264288 | 0.933583143130315 | 0.977054065668186 | -5.44867482454901 |
| CBR3-AS1 | CBR3-AS1 | 0.021073969622411 | -0.26757927862313 | 0.0837860711353638 | 0.933619946730889 | 0.977054065668186 | -5.6309469489237 |
| lnc-ZDHHC21-2 | lnc-ZDHHC21-2 | -0.0187795707576224 | 2.10761180281547 | -0.0835892449910465 | 0.933775511731315 | 0.977167184806837 | -6.05657223618154 |
| ENSG00000281991.1 | TMEM265 | -0.0150579322539436 | 0.131479530003801 | -0.083146961558548 | 0.934125087757254 | 0.977448921980805 | -5.6930004006994 |
| ENSG00000243646.9 | IL10RB | -0.0110978844446579 | 3.9408284603467 | -0.0831284561286064 | 0.934139714535511 | 0.977448921980805 | -6.28162713596418 |
| ENSG00000163820.14 | FYCO1 | -0.00917898415538065 | 4.97765690762134 | -0.0830046977591235 | 0.934237534320453 | 0.977461235433567 | -6.37499726399214 |
| ENSG00000143740.14 | SNAP47 | -0.0160173094836082 | 4.42990101772954 | -0.0829934054401155 | 0.934246459927346 | 0.977461235433567 | -6.3398023070758 |
| ENSG00000110536.13 | PTPMT1 | 0.00879344259079629 | 3.41206166788799 | 0.0829088899556195 | 0.934313262420143 | 0.977481441516408 | -6.23109310299913 |
| ENSG00000107679.14 | PLEKHA1 | 0.0242236755359393 | 4.20079507812339 | 0.0828058766511836 | 0.934399259684291 | 0.977521726154092 | -6.3190338373804 |
| ENSG00000256591.5 | | -0.00928745839473555 | 3.5425822646425 | -0.082676593567824 | 0.934496875947369 | 0.97753015609324 | -6.24060333286941 |
| ENSG00000285629.1 | | -0.0142852316184028 | 6.21468922502034 | -0.0826284182518162 | 0.934534955509494 | 0.97753015609324 | -6.40706430483516 |
| lnc-KLHL31-2 | lnc-KLHL31-2 | 0.023369834634606 | -0.181258359176202 | 0.082477051533909 | 0.934654602403876 | 0.97753015609324 | -5.60657780781236 |
| ENSG00000227165.8 | WDR11-DT | -0.023811650437916 | 1.17153856189756 | -0.0823687231178055 | 0.934740230875015 | 0.97753015609324 | -5.9095167216929 |
| lnc-ZNF80-4 | lnc-ZNF80-4 | -0.0148438490048619 | 4.24152794129635 | -0.0823613137374512 | 0.934746087667081 | 0.97753015609324 | -6.31579476051031 |
| lnc-RAB11A-2 | lnc-RAB11A-2 | -0.0120512322800053 | 1.11055339694703 | -0.0823412191815902 | 0.934761971555627 | 0.97753015609324 | -5.94647046066516 |
| lnc-CEMP1-1 | lnc-CEMP1-1 | 0.071105873161327 | -1.76623286360637 | 0.0823124426612364 | 0.934784718213295 | 0.97753015609324 | -5.09753098770681 |
| ENSG00000006432.15 | MAP3K9 | 0.00928935915099238 | 5.40330628694326 | 0.0823092338550658 | 0.934787254646225 | 0.97753015609324 | -6.39227350735808 |
| lnc-NUAK1-5 | lnc-NUAK1-5 | -0.024851073657576 | 3.49020407321087 | -0.082245076373603 | 0.934841169198987 | 0.977536871856469 | -6.2213889288023 |
| ENSG00000198937.8 | CCDC167 | -0.02539938600518 | 2.10274775232405 | -0.0821459177308031 | 0.934916550761165 | 0.977566033239373 | -6.05745099885013 |
| ENSG00000269179.1 | | 0.0143953478403452 | 0.743334104543267 | 0.0820826144341796 | 0.934966389966307 | 0.977568485455476 | -5.86907354587077 |
| UBR5-AS1 | UBR5-AS1 | 0.0158606879109966 | 5.10290442232395 | 0.0818279229766307 | 0.935167719392533 | 0.977729322306929 | -6.37646921235261 |
| ENSG00000136379.11 | ABHD17C | 0.0273687975888309 | -0.505459156423359 | 0.0815521466107459 | 0.935385720935004 | 0.977807480976945 | -5.57742028065321 |
| ENSG00000129353.14 | SLC44A2 | 0.00752589553464755 | 7.19500583874104 | 0.0815029942724028 | 0.935424576443755 | 0.977807480976945 | -6.40143707937691 |
| ENSG00000151835.15 | SACS | -0.0149208884486169 | 6.33225749850449 | -0.0814982329963911 | 0.935428340297602 | 0.977807480976945 | -6.40740170434188 |
| lnc-DGKH-1 | lnc-DGKH-1 | -0.0202693592976874 | -0.214356839024161 | -0.0814929734067766 | 0.93543249807691 | 0.977807480976945 | -5.60270326655203 |
| ENSG00000196081.9 | ZNF724 | -0.0106267017103185 | 2.97953992139453 | -0.0813342727946781 | 0.935557953992276 | 0.977817677328195 | -6.1681733786367 |
| ENSG00000086015.20 | MAST2 | -0.0159987311819987 | 3.34697675741511 | -0.0812992993006113 | 0.935585601441376 | 0.977817677328195 | -6.23090028631097 |
| lnc-ADAM10-7 | lnc-ADAM10-7 | -0.0086471164830213 | 3.08454614985844 | -0.0810169067933795 | 0.935808842946143 | 0.977817677328195 | -6.18671588568402 |
| ENSG00000067836.12 | ROGDI | 0.0222581611951236 | 1.45360816427285 | 0.0810002990261869 | 0.93582197215053 | 0.977817677328195 | -5.98200886267979 |
| ENSG00000139117.13 | CPNE8 | 0.101719555009489 | -1.11634878982411 | 0.0809827782250781 | 0.935847191166203 | 0.977817677328195 | -5.37222039197129 |
| lnc-UBLCP1-9 | lnc-UBLCP1-9 | 0.0175352503780919 | 0.310740507106735 | 0.0809281346783243 | 0.935879021598946 | 0.977817677328195 | -5.76532790467183 |
| ENSG00000235217.6 | NA | 0.0191302976024037 | 0.504819211345569 | 0.0808050588722123 | 0.935976319832981 | 0.977817677328195 | -5.82127875615967 |
| ENSG00000012124.16 | CD22 | 0.0271342757398161 | 6.7832251098494 | 0.0807742119378594 | 0.93600998146105 | 0.977817677328195 | -6.40552961445584 |
| lnc-TUBB2A-7 | lnc-TUBB2A-7 | -0.0204122366912348 | -0.242819760131087 | -0.0807543512571749 | 0.936016407296252 | 0.977817677328195 | -5.61162165652917 |
| ENSG00000143862.7 | ARL8A | 0.0113390332515992 | 4.47882957121679 | 0.0807529049859963 | 0.936017550664317 | 0.977817677328195 | -6.34227752314665 |
| ENSG00000165879.8 | FRAT1 | -0.016545132875045 | 2.08969920707736 | -0.0806831770858473 | 0.936072675104254 | 0.977817677328195 | -6.06725844188267 |
| ENSG00000178567.7 | EPM2AIP1 | -0.00799945139144143 | 6.27744805784045 | -0.0806358445709525 | 0.936110094716483 | 0.977817677328195 | -6.40757768417571 |
| lnc-RPS6KA2-2 | lnc-RPS6KA2-2 | -0.018107910044137 | 3.38143315063831 | -0.0806047176689246 | 0.936134702755528 | 0.977817677328195 | -6.21331018195183 |
| ENSG00000141002.19 | TCF25 | 0.00917108035452063 | 6.08197609080935 | 0.0805267645022135 | 0.936196330578207 | 0.977817677328195 | -6.40666695053947 |
| ENSG00000171488.14 | LRRC8C | 0.0208981954803411 | 2.87935995506853 | 0.0805172031810786 | 0.936203889546671 | 0.977817677328195 | -6.17340584675257 |
| lnc-SRRD-1 | lnc-SRRD-1 | 0.0134002110293244 | 0.771581904932536 | 0.0804583923992154 | 0.936250384177613 | 0.977817677328195 | -5.8437856530068 |
| ENSG00000228623.6 | ZNF883 | -0.0778487772938896 | -1.06651957644501 | -0.0804432297791033 | 0.936268488635952 | 0.977817677328195 | -5.30915782338666 |
| ENSG00000196312.13 | MFSD14CP | -0.00901631413983406 | 4.67909655463009 | -0.080396393104485 | 0.936299399827005 | 0.977817677328195 | -6.35545120573364 |
| ENSG00000011478.11 | QPCTL | -0.0155033186751705 | 2.34436203874131 | -0.0803388811232443 | 0.936344868101769 | 0.977817677328195 | -6.0943716856729 |
| ENSG00000215126.10 | ZNG1F | -0.011770456112332 | 1.90731740276137 | -0.0802215332675264 | 0.936437642559816 | 0.977864948344006 | -6.04434211432211 |
| ENSG00000141664.9 | ZCCHC2 | -0.00608073505487461 | 5.73870433908782 | -0.0800762635574851 | 0.936552493093786 | 0.97789030828049 | -6.40207289278258 |
| ENSG00000167670.15 | CHAF1A | 0.0122739712623376 | 3.29283641062201 | 0.0800706295976688 | 0.936556947341656 | 0.97789030828049 | -6.21581993890929 |
| lnc-NIN-4 | lnc-NIN-4 | 0.0155985878257765 | 0.364472835535467 | 0.0799415629215245 | 0.936658988917588 | 0.977899278261608 | -5.78981632478791 |
| ENSG00000134109.10 | EDEM1 | -0.0106831151663469 | 7.5244094064639 | -0.0799395779427197 | 0.93666055827273 | 0.977899278261608 | -6.3962424409559 |

| | | | | | | | |
|---|---|---|---|---|---|---|---|
| lnc-HEATR6-1 | lnc-HEATR6-1 | 0.0385860641862195 | 3.04269598711384 | 0.0798113261606075 | 0.936772479015659 | 0.977956559937707 | -6.16754827502154 |
| lnc-AKAP8-1 | lnc-AKAP8-1 | -0.0179595256412522 | -0.159947598619675 | -0.0797499907941649 | 0.936810449997171 | 0.977956559937707 | -5.62869999831133 |
| ENSG00000175467.14 | SART1 | 0.0102677680796948 | 5.19910785090285 | 0.0795744176006643 | 0.936949264052442 | 0.978051866416037 | -6.38596231254714 |
| ENSG00000156172.5 | CFAP418 | 0.0169831410194545 | 1.92834018420541 | 0.0794280313460287 | 0.937065003452397 | 0.978077144773367 | -6.04056496652031 |
| ENSG00000207118.1 | SNORD14D | 0.0319365155886809 | -0.408382596426072 | 0.0793674190836116 | 0.937112926571542 | 0.978077144773367 | -5.54784957787288 |
| lnc-DDX52-4 | lnc-DDX52-4 | -0.0137312027024982 | 0.161268796578153 | -0.0793518291701983 | 0.937125252783445 | 0.978077144773367 | -5.70987095392507 |
| ENSG00000132256.18 | TRIM5 | 0.0132607823980115 | 3.7033500885035 | 0.0792581286874315 | 0.937199337680456 | 0.978077144773367 | -6.25995393455386 |
| lnc-PTGR2-3 | lnc-PTGR2-3 | -0.0169675521741265 | 0.909987847133231 | -0.0792432855884777 | 0.937211073525002 | 0.978077144773367 | -5.88718629979681 |
| ENSG00000162512.15 | SDC3 | 0.0749137634450212 | 0.190291448925079 | 0.079038310442897 | 0.937386678070943 | 0.978108724163 | -5.77136617351854 |
| ENSG00000162889.10 | MAPKAPK2 | 0.00915106068335803 | 6.69590088074254 | 0.0789610208623166 | 0.937434251637619 | 0.978108724163 | -6.40662529885766 |
| ENSG00000107341.4 | UBE2R2 | -0.00708375597795552 | 5.33962147334507 | -0.078940784060384 | 0.937450252455658 | 0.978108724163 | -6.39186665327827 |
| lnc-PANK3-13 | lnc-PANK3-13 | -0.0114153430143693 | 4.96039608973433 | -0.0789182749795202 | 0.937468049948078 | 0.978108724163 | -6.37915857672428 |
| ENSG00000205730.6 | ITPRIPL2 | 0.0892224524218829 | 1.08671591678819 | 0.0788766515444706 | 0.937515667159837 | 0.978108724163 | -5.85618454577045 |
| ENSG00000121579.12 | NAA50 | -0.00884293211504605 | 6.42865336563981 | -0.0788261919829417 | 0.937540858519204 | 0.978108724163 | -6.40740519009502 |
| ENSG00000205138.3 | SDHAF1 | -0.0139243032356022 | 2.1828953294266 | -0.0787843088687581 | 0.937573975015109 | 0.978108724163 | -6.08694730929566 |
| lnc-TBC1D31-3 | lnc-TBC1D31-3 | -0.0214246655909212 | 1.7641454270897 | -0.0784922381059489 | 0.937804915108482 | 0.97830006424803 | -6.01008372650808 |
| ENSG00000149485.18 | FADS1 | -0.0863154182214103 | -1.46222153176546 | -0.0783580106423014 | 0.93792157394923 | 0.978335583066108 | -5.45021118624374 |
| lnc-LSM3-2 | lnc-LSM3-2 | 0.0121651816222943 | 1.18169757532615 | 0.078328953873015 | 0.937934026169339 | 0.978335583066108 | -5.93181637566604 |
| lnc-AGAP4-2 | lnc-AGAP4-2 | 0.0137501445185456 | 0.962199264080258 | 0.0780903179506549 | 0.938122721873232 | 0.978482820731567 | -5.89006229466979 |
| lnc-NPIPB4-1 | lnc-NPIPB4-1 | -0.019974762876794 | 0.174864101881779 | -0.0779222566748162 | 0.938255614507089 | 0.978571842584212 | -5.73303143191017 |
| lnc-SLC22A1-3 | lnc-SLC22A1-3 | 0.0157517758103902 | 0.250924728154275 | 0.0777333030300163 | 0.938405029682164 | 0.978678087046412 | -5.73377659086576 |
| lnc-SIGIRR-1 | lnc-SIGIRR-1 | 0.0222731366558868 | 0.380576108213096 | 0.0774587480268339 | 0.938622138168999 | 0.978840781116768 | -5.7578832237617 |
| BISPR | BISPR | -0.0115459224772922 | 3.37531042001512 | -0.0774157494186794 | 0.93865614039172 | 0.978840781116768 | -6.22261926942061 |
| lnc-SCO1-1 | lnc-SCO1-1 | 0.00945091301593415 | 3.05376162741664 | 0.0772501777623547 | 0.938787071393117 | 0.978927721287022 | -6.19169749194136 |
| ENSG00000170860.3 | LSM3 | -0.0108807741413468 | 4.08864207873193 | -0.0771497412865787 | 0.938866495534632 | 0.978954037098924 | -6.29936266838588 |
| ENSG00000147416.10 | ATP6V1B2 | 0.00908493692151513 | 5.17307635877652 | 0.0770979766683614 | 0.938907430713203 | 0.978954037098924 | -6.38483632255659 |
| lnc-UBXN4-11 | lnc-UBXN4-11 | -0.0205264160485726 | -0.543732260227066 | -0.0769153816628955 | 0.939051827184778 | 0.979054997413853 | -5.50017850662662 |
| ENSG00000248993.1 | | 0.0120272245460677 | 6.56638981642354 | 0.0767700720500557 | 0.939166739781403 | 0.97905817087438 | -6.40772671896173 |
| ENSG00000255529.8 | POLR2M | -0.00747730324896612 | 5.1713036785349 | -0.0766731519458354 | 0.939243386099722 | 0.97905817087438 | -6.38800781373188 |
| ENSG00000078369.17 | GNB1 | 0.00692176802678546 | 7.14889538241389 | 0.0766327392604679 | 0.93927534541575 | 0.97905817087438 | -6.40190887931175 |
| lnc-PHOSPHO2-2 | lnc-PHOSPHO2-2 | 0.0119257137715882 | 1.24558120111714 | 0.0766201682142435 | 0.939285286919767 | 0.97905817087438 | -5.96000124722393 |
| ENSG00000131165.14 | CHMP1A | 0.00780488073366481 | 4.87194952225736 | 0.0766107909216971 | 0.93929270272829 | 0.97905817087438 | -6.37165270113239 |
| lnc-PI16-2 | lnc-PI16-2 | 0.0159340228629701 | 1.89406006308614 | 0.0763208739004951 | 0.939521979387902 | 0.97924756439848 | -6.0407022987021 |
| ENSG00000073169.13 | SELENOO | -0.0116721763261913 | 3.57283620050731 | -0.0761223500364687 | 0.939678982094403 | 0.979283235608144 | -6.25002344351444 |
| ENSG00000237973.1 | MTCO1P12 | 0.0280507823549843 | 2.75023790526694 | 0.0760552591176828 | 0.939737293193563 | 0.979283235608144 | -6.136940397141 |
| ENSG00000167272.10 | POP5 | 0.0119152531998847 | 3.17638443551796 | 0.0760174869411814 | 0.939761914111127 | 0.979283235608144 | -6.2006306099855 |
| ENSG00000131943.17 | C19orf12 | 0.0126140193164734 | 3.93412190467789 | 0.0759948830305754 | 0.939779790726205 | 0.979283235608144 | -6.28748549349705 |
| lnc-ZP3-4 | lnc-ZP3-4 | -0.0133914669357059 | -0.258836143904553 | -0.0759768025749637 | 0.939794089927011 | 0.979283235608144 | -5.56811193515891 |
| lnc-ARMC1-5 | lnc-ARMC1-5 | -0.0231608087866917 | 4.22940980274007 | -0.0759223025303101 | 0.939842427833584 | 0.979284028151193 | -6.31861614044883 |
| ENSG00000110046.12 | ATG2A | -0.0124518690475517 | 4.34673705375324 | -0.075521498289467 | 0.940154180797247 | 0.979533591367654 | -6.33123734781126 |
| lnc-APOF-1 | lnc-APOF-1 | 0.0143952661065505 | 4.01340481778109 | 0.0754924959988113 | 0.940177118547314 | 0.979533591367654 | -6.28506197305189 |
| ENSG00000173273.15 | TNKS | 0.00940868052973798 | 5.93698145500024 | 0.0751664661871595 | 0.940434977240362 | 0.979728234042735 | -6.40585007963546 |
| ENSG00000184357.4 | H1-5 | -0.0657207676227598 | 2.85587355325987 | -0.0750888234707579 | 0.940510382284323 | 0.979728234042735 | -6.21410538380734 |
| ENSG00000197927.12 | NBEAP2 | -0.0702427232116576 | -0.249295217787771 | -0.0750472273736345 | 0.940539176132307 | 0.979728234042735 | -5.55194785547606 |
| lnc-SP110-2 | lnc-SP110-2 | -0.0117096765758302 | 3.39575824397175 | -0.0750155544350363 | 0.940554336330197 | 0.979728234042735 | -6.22333764773541 |
| ENSG00000155636.14 | RBM45 | 0.0083305981410291 | 3.6541536500322 | 0.0747732350839862 | 0.940745994389838 | 0.979847097462446 | -6.26260987665996 |
| ENSG00000118454.12 | ANKRD13C | 0.0061323514340339 | 5.08884150962932 | 0.0747204565144487 | 0.940787739106027 | 0.979847097462446 | -6.38192942814191 |
| lnc-LONRF2-2 | lnc-LONRF2-2 | -0.0206337076438616 | 1.71420371238205 | -0.0746907173472278 | 0.940811261096552 | 0.979847097462446 | -6.02318469111556 |
| ENSG00000153187.19 | HNRNPU | 0.0068559559897956 | 9.03750594404273 | 0.0745122370308097 | 0.940952430002445 | 0.979871303319427 | -6.36040058677082 |
| lnc-RAB1A-8 | lnc-RAB1A-8 | 0.0176361681426469 | 2.1372931915117 | 0.0744490914065227 | 0.941002375458826 | 0.979871303319427 | -6.05817795484296 |
| ENSG00000122952.16 | ZWINT | 0.0289058821359061 | 0.315867448641895 | 0.0743971098832126 | 0.94104349077476 | 0.979871303319427 | -5.7363361989305 |
| ENSG00000049759.17 | NEDD4L | 0.0511793288758815 | 1.91141235960819 | 0.0743727776129235 | 0.941075470575086 | 0.979871303319427 | -6.04544527327495 |
| ENSG00000005436.13 | GCFC2 | -0.0104856432556542 | 4.23758675825187 | -0.0742941736124676 | 0.94112490974784 | 0.979871303319427 | -6.31945448480245 |
| lnc-RAPSN-2 | lnc-RAPSN-2 | -0.00609930806122971 | 4.75807215600329 | -0.0742787300648378 | 0.941137125106339 | 0.979871303319427 | -6.36529153649428 |
| lnc-FNIP1-6 | lnc-FNIP1-6 | 0.00740385227309748 | 1.78055004554517 | 0.0742400200039083 | 0.941167743605163 | 0.979871303319427 | -6.02282795731274 |
| ENSG00000162946.22 | DISC1 | 0.043191758745351 | 1.07262058481461 | 0.0740347824464292 | 0.94133975027712 | 0.979871645139407 | -5.95305052202136 |
| EPB41L4A-AS1 | EPB41L4A-AS1 | 0.00784958004367464 | 4.70113359687094 | 0.0740199317853924 | 0.941341828504431 | 0.979871645139407 | -6.3590005578635 |
| lnc-BIRC6-1 | lnc-BIRC6-1 | -0.0106360948819368 | 2.6996662657296 | -0.0739808028454435 | 0.941372778932167 | 0.979871645139407 | -6.14391623449975 |
| lnc-TRIM3-1 | lnc-TRIM3-1 | -0.0105427221822008 | 2.35216925230001 | -0.0739615911032965 | 0.941387975176969 | 0.979871645139407 | -6.09964708831418 |
| ENSG00000279355.1 | NA | -0.0391617400634233 | -0.509055096199936 | -0.0738644042941732 | 0.941465285223975 | 0.979871645139407 | -5.58074022589298 |
| lnc-WDR41-1 | lnc-WDR41-1 | -0.0119913006225322 | 0.499082855800807 | -0.0738114128478259 | 0.941506765036755 | 0.979871645139407 | -5.80201682104938 |
| lnc-ZNF587-1 | lnc-ZNF587-1 | 0.0123604683933152 | 1.47368626051967 | 0.0736804204896771 | 0.941610380097876 | 0.979871645139407 | -5.98002014657737 |
| ENSG00000095383.19 | TBC1D2 | 0.0122036576305302 | 1.31990504870763 | 0.0736361484942494 | 0.941645399513519 | 0.979871645139407 | -5.93838258395071 |
| ENSG00000166501.13 | PRKCB | 0.00729772280462093 | 9.90608046342286 | 0.0736138817080466 | 0.941663012719988 | 0.979871645139407 | -6.33421506252905 |
| lnc-NDUFV3-2 | lnc-NDUFV3-2 | 0.0178780575857852 | 0.0105469430491258 | 0.0735925517700399 | 0.941679884899728 | 0.979871645139407 | -5.66008046315051 |
| lnc-SPOPL-18 | lnc-SPOPL-18 | -0.0223269883384684 | -0.0734627453873989 | -0.0735775688997251 | 0.941691736505981 | 0.979871645139407 | -5.62176860487767 |
| ENSG00000187922.13 | LCN10 | -0.0612873381744554 | 0.826791817527695 | -0.0734313203178209 | 0.94182057636854 | 0.979956168405301 | -6.01551645622647 |
| ENSG00000132846.5 | ZBED3 | -0.0248256101236175 | 0.525904952060973 | -0.073101694604 | 0.942068164965441 | 0.980025810699925 | -5.82589048619453 |

| | | | | | | | |
|---|---|---|---|---|---|---|---|
| lnc-EGLN2-2 | lnc-EGLN2-2 | 0.0148187879909631 | 3.33456690466632 | 0.0730854191693319 | 0.942081039478474 | 0.980025810699925 | -6.20486983156571 |
| ENSG00000100568.10 | VTI1B | -0.0073869548753853 | 5.21986442434427 | -0.0730597047228074 | 0.942101380655439 | 0.980025810699925 | -6.38825443889152 |
| ENSG00000185238.12 | PRMT3 | 0.00952961274612418 | 3.89818307832702 | 0.0730339932844272 | 0.942121719491941 | 0.980025810699925 | -6.28542214188701 |
| lnc-RNF139-1 | lnc-RNF139-1 | -0.0178743595763616 | 0.44997563722819 | -0.0729842077356821 | 0.942161102080256 | 0.980025810699925 | -5.7963228865653 |
| ENSG00000132024.17 | CC2D1A | 0.00921863950720914 | 3.82074039522847 | 0.0729372494582847 | 0.942198248305117 | 0.980025810699925 | -6.27001572466145 |
| lnc-SLC6A18-2 | lnc-SLC6A18-2 | 0.0150147418079269 | 2.6042281293092 | 0.0729087383367264 | 0.94222080201821 | 0.980025810699925 | -6.10226506357147 |
| ENSG00000150907.7 | FOXO1 | -0.0110474007117087 | 7.29495020654076 | -0.0725957675294804 | 0.942468380619725 | 0.980233788696099 | -6.40067924130763 |
| ENSG00000158481.12 | CD1C | 0.0658602516084272 | 0.541522926363928 | 0.0724901665520531 | 0.942564328468432 | 0.980237758583965 | -5.79039388886686 |
| lnc-RBM28-3 | lnc-RBM28-3 | -0.0086562607475375 | 2.78608368592726 | -0.0724705397629423 | 0.94256744487654 | 0.980237758583965 | -6.14933366842155 |
| lnc-TXNDC12-1 | lnc-TXNDC12-1 | 0.00981071233769581 | 1.82258234603895 | 0.0723887644077582 | 0.94263213561734 | 0.980240927271593 | -6.0262632055522 |
| ENSG00000179981.10 | TSHZ1 | -0.0123205376308975 | 3.1444135417231 | -0.07229106112477 | 0.942709427116197 | 0.980240927271593 | -6.20084679548552 |
| ENSG00000148600.14 | CDHR1 | 0.0642798536622466 | 0.686241951997242 | 0.072301731678209 | 0.942713363228936 | 0.980240927271593 | -5.83875419092801 |
| ENSG00000177576.11 | C18orf32 | -0.0067420319348965 | 4.05282187398368 | -0.0721562948027203 | 0.942816039506969 | 0.980298168376033 | -6.30349967012336 |
| lnc-IL11RA-2 | lnc-IL11RA-2 | 0.0163438075102773 | 2.06951749563436 | 0.0719145750375141 | 0.943007264434145 | 0.980447467992525 | -6.04333566038087 |
| lnc-ARID2-12 | lnc-ARID2-12 | -0.0171083729833228 | 3.12550867031133 | -0.0717553180856094 | 0.943133254745866 | 0.980527083274935 | -6.20602465240049 |
| ENSG00000140526.17 | ABHD2 | 0.0066622762522579 | 5.70344959497416 | 0.0715819595574852 | 0.943270402678601 | 0.980527083274935 | -6.4027700919977 |
| ENSG00000140650.11 | PMM2 | -0.0076307075102128 | 4.93999184207454 | -0.0715295018531853 | 0.943311903522662 | 0.980527083274935 | -6.37398876319207 |
| ENSG00000206737.1 | RNVU1-18 | -0.0744485705746367 | -1.95950740081948 | -0.0714546892114018 | 0.943371747638485 | 0.980527083274935 | -5.08519460488345 |
| ENSG00000069493.14 | CLEC2D | -0.013668563434365 | 8.24376315656677 | -0.0714237338364616 | 0.943395580218813 | 0.980527083274935 | -6.38435925407232 |
| lnc-PIGBOS1-1 | lnc-PIGBOS1-1 | 0.0105380902124327 | 0.454812328948964 | 0.0712532260631419 | 0.943530476112691 | 0.980527083274935 | -5.78436859952868 |
| ENSG00000101052.12 | IFT52 | -0.0094042193363881 | 3.46813479167824 | -0.0712295654572649 | 0.943549195148948 | 0.980527083274935 | -6.24025614763487 |
| lnc-FRAT1-6 | lnc-FRAT1-6 | -0.0121231392008635 | 0.405892286399709 | -0.0712105171765988 | 0.943564265177726 | 0.980527083274935 | -5.76132396097736 |
| lnc-MOCOS-2 | lnc-MOCOS-2 | -0.0094172991538563 | 2.35723326187872 | -0.0712005506080746 | 0.943572150227087 | 0.980527083274935 | -6.09972935206369 |
| ENSG00000149499.11 | EML3 | -0.0091580178059969 | 4.97298328614784 | -0.0711862003905296 | 0.943583503409844 | 0.980527083274935 | -6.3772013249094 |
| ENSG00000131504.16 | DIAPH1 | -0.0101336149533671 | 6.41831522344744 | -0.0710698252764189 | 0.943675574085656 | 0.980527083274935 | -6.4083015290379 |
| ENSG00000145911.5 | N4BP3 | 0.0245466198723236 | 3.16387287288979 | 0.0710663354576801 | 0.943683502533213 | 0.980527083274935 | -6.21326910733299 |
| ENSG00000279792.1 | | 0.0116702992075073 | 2.35701700275175 | 0.071034995770759 | 0.943703129751565 | 0.980527083274935 | -6.10028211284985 |
| lnc-NSFL1C-1 | lnc-NSFL1C-1 | -0.0075367278420386 | 2.49591728719351 | -0.0708547943262198 | 0.9438456988031 | 0.980578500624182 | -6.1168381162992 |
| ENSG00000120694.19 | HSPH1 | 0.00986755427103247 | 5.3959800168649 | 0.0708333231711444 | 0.94386268615091 | 0.980578500624182 | -6.39333907951907 |
| ENSG00000113368.11 | LMNB1 | 0.0119519833614028 | 3.70790567502121 | 0.0707918017609649 | 0.943895536747166 | 0.980578500624182 | -6.26496774911296 |
| lnc-CAB39-1 | lnc-CAB39-1 | 0.0137441352754146 | 1.28788413409439 | 0.0706926339658607 | 0.943973995975373 | 0.980594607379769 | -5.97849454375497 |
| lnc-C5orf60-3 | lnc-C5orf60-3 | -0.0080686191177230 | 6.69171912975269 | -0.0706496182480998 | 0.944008029175123 | 0.980594607379769 | -6.40669826813733 |
| ENSG00000135930.13 | EIF4E2 | -0.0075456096159998 | 5.63811683476099 | -0.0705320531583607 | 0.944101044928676 | 0.980594607379769 | -6.40016440840548 |
| lnc-NAA25-3 | lnc-NAA25-3 | 0.0137488572773837 | 0.405700496489682 | 0.0705313453503046 | 0.944101604938293 | 0.980594607379769 | -5.77047478896464 |
| ENSG00000101901.11 | ALG13 | -0.0099080433473854 | 6.06488050557991 | -0.0701329991749584 | 0.944416776394859 | 0.980872464856967 | -6.40731966479321 |
| lnc-HIVEP1-5 | lnc-HIVEP1-5 | 0.0175913407986819 | 0.695781199137714 | 0.070063006846187 | 0.944472155251038 | 0.980880464983809 | -5.85074443022891 |
| lnc-TARS-3 | lnc-TARS-3 | 0.0133738385024949 | 0.541580914756317 | 0.0697804258988676 | 0.944695739857029 | 0.981051259222272 | -5.82437325487895 |
| ENSG00000163577.7 | EIF5A2 | 0.0625312275322824 | 0.370846170959545 | 0.0697470695455451 | 0.944731915126301 | 0.981051259222272 | -5.65477938868805 |
| ENSG00000124164.15 | VAPB | -0.0085688300842932 | 4.40551498825577 | -0.0696657580646105 | 0.94478646899763 | 0.981058414457583 | -6.3351067478874 |
| ENSG00000147065.16 | MSN | -0.0098189077046004 | 8.94865477085124 | -0.0694704288215774 | 0.944941021930801 | 0.981089404897835 | -6.36434920811983 |
| ENSG00000120696.8 | KBTBD7 | 0.0179376466050321 | 1.73450987858488 | 0.0694637341968434 | 0.944946319045101 | 0.981089404897835 | -6.02715259758684 |
| ENSG00000139291.13 | TMEM19 | -0.012936176660407 | 5.72538269107622 | -0.0694473174923272 | 0.944959308754778 | 0.981089404897835 | -6.40045246451464 |
| ENSG00000131732.11 | ZCCHC9 | -0.0072557586421848 | 3.67775518729996 | -0.0692608229801763 | 0.945106873512418 | 0.981193119010698 | -6.25870297019049 |
| lnc-ANKRD37-2 | lnc-ANKRD37-2 | -0.0249133551889216 | 1.49632748720768 | -0.0691178807905167 | 0.945220875241859 | 0.981261979881132 | -5.9604138034093 |
| ENSG00000102760.12 | RGCC | 0.0343715659241299 | 4.26594904328729 | 0.0690032417147626 | 0.945322203445709 | 0.981305299977381 | -6.32260760405801 |
| lnc-SLC25A52-3 | lnc-SLC25A52-3 | 0.0085575512181657 | 1.62875879576086 | 0.0688413051175273 | 0.94543882633584 | 0.981305299977381 | -6.00322693864405 |
| ENSG00000122741.15 | DCAF10 | -0.0081052938210168 | 4.40525450736384 | -0.068836328675467 | 0.945442764115113 | 0.981305299977381 | -6.33491840549973 |
| lnc-LRRC7-5 | lnc-LRRC7-5 | 0.0138866432971345 | 3.57874165166626 | 0.0688230057840205 | 0.945453306313532 | 0.981305299977381 | -6.26088331650403 |
| ENSG00000116127.17 | ALMS1 | 0.00980475804207881 | 6.80080463413421 | 0.0686746797605226 | 0.945570675074864 | 0.981377632365164 | -6.40634437793511 |
| ENSG00000214756.7 | CSKMT | -0.022348699283893 | 2.66315839936759 | -0.0685499723212005 | 0.945671513406079 | 0.981399543912421 | -6.12897531983779 |
| ENSG00000221909.2 | FAM200A | -0.0132332845600267 | 1.62365805655813 | -0.0685274882062897 | 0.945687147352619 | 0.981399543912421 | -6.00540755115201 |
| lnc-ZBTB34-1 | lnc-ZBTB34-1 | 0.0218412980027428 | -0.431580870498197 | 0.0683899947777213 | 0.945795946615997 | 0.981414341638783 | -5.56704316447538 |
| ENSG00000138101.18 | DTNB | 0.0111065471231928 | 4.09868451675558 | 0.0683889564639457 | 0.94579676824306 | 0.981414341638783 | -6.30258592724748 |
| ENSG00000228144.2 | | 0.00608921978357202 | 4.23238395138284 | 0.0682551877690207 | 0.945902621117944 | 0.981456609333513 | -6.31868576090382 |
| ENSG00000159592.10 | GPBP1L1 | -0.0055799608292082 | 7.09318601733668 | -0.068180163021108 | 0.945961989591181 | 0.981456609333513 | -6.40310053903841 |
| ENSG00000086189.9 | DIMT1 | -0.0093590430237040 | 3.32319458402121 | -0.0681474507354682 | 0.945987875526288 | 0.981456609333513 | -6.2230298557062 |
| ENSG00000099995.18 | SF3A1 | 0.00644007384237089 | 6.42226506493014 | 0.068096450213782 | 0.946028233453671 | 0.981456609333513 | -6.40851428251792 |
| ENSG00000138399.17 | FASTKD1 | -0.0084340071266062 | 3.6385868980274 | -0.0679880538932049 | 0.946114010519886 | 0.981496128145295 | -6.2486443628503 |
| ENSG00000174227.15 | PIGG | 0.00973817147252376 | 5.82617516993427 | 0.0677329658156101 | 0.946315871462259 | 0.981656062005226 | -6.40515903530384 |
| ENSG00000228808.1 | HMGB3P4 | -0.018531567987951 | -0.280184917574045 | -0.0676090027773821 | 0.946413969440481 | 0.981658875881547 | -5.5850757290288 |
| lnc-TM9SF2-4 | lnc-TM9SF2-4 | -0.018531567987951 | -0.280184917574045 | -0.0676090027773821 | 0.946413969440481 | 0.981658875881547 | -5.5850757290288 |
| ENSG00000220323.4 | H2BC19P | 0.0174986144489204 | 3.60264067050006 | 0.0674882625691332 | 0.946510046317115 | 0.981706766461286 | -6.24662108439286 |
| ENSG00000174353.17 | STAG3L3 | 0.00818744205129729 | 3.47368828056306 | 0.067430118634589 | 0.946555530641339 | 0.981706766461286 | -6.24384552562758 |
| ENSG00000042317.16 | SPATA7 | -0.0195111073878915 | 0.560330124329517 | -0.0672274669147735 | 0.94671590258929 | 0.981756498119457 | -5.8349417923112 |
| lnc-MRNIP-2 | lnc-MRNIP-2 | -0.0088519134720439 | 1.6964120924788 | -0.067154109869104 | 0.946773955509322 | 0.981756498119457 | -6.01530358274019 |
| lnc-KMT2E-3 | lnc-KMT2E-3 | -0.0220111284053269 | 0.500667897523177 | -0.0670904488714511 | 0.94682433545816 | 0.981756498119457 | -5.83023196566223 |
| ENSG00000134982.16 | APC | 0.00497306357565108 | 6.58052286915951 | 0.0670829477869778 | 0.946830271670117 | 0.981756498119457 | -6.40821926211913 |
| lnc-GRM1-12 | lnc-GRM1-12 | -0.0183918839111715 | -0.57117025676236 | -0.0670439281353213 | 0.946861151110737 | 0.981756498119457 | -5.55990791662302 |

| | | | | | | | |
|---|---|---|---|---|---|---|---|
| ENSG00000150627.15 | WDR17 | -0.0747733473604764 | 2.36597747283265 | -0.0668721411983411 | 0.947009558526097 | 0.981756498119457 | -6.10774544804147 |
| ENSG00000148218.15 | ALAD | -0.00998323362304143 | 3.7706450549611 | -0.0667273147705254 | 0.947111716205593 | 0.981756498119457 | -6.26682964617711 |
| LINC00525 | LINC00525 | 0.0588979126627742 | 2.95051550392068 | 0.0667413901804606 | 0.947113009852159 | 0.981756498119457 | -6.15342453568738 |
| ENSG00000108771.12 | DHX58 | 0.0094251112269208 | 3.32452750844756 | 0.066713915006325 | 0.947122320783455 | 0.981756498119457 | -6.22163268638631 |
| lnc-MAP3K11-1 | lnc-MAP3K11-1 | -0.00938663914092963 | 2.99343690107023 | -0.0666925076475993 | 0.947139262594427 | 0.981756498119457 | -6.17973805069461 |
| ENSG00000169490.16 | TM2D2 | 0.0102666478018842 | 2.70676521376818 | 0.0666569809028284 | 0.947167378560132 | 0.981756498119457 | -6.14490425444386 |
| lnc-WNT5B-6 | lnc-WNT5B-6 | 0.0219924618546039 | 1.28061653584035 | 0.0666462754129934 | 0.947175850928251 | 0.981756498119457 | -5.95365834656436 |
| ENSG00000224877.3 | NDUFAF8 | 0.00864027109693355 | 2.71058606740657 | 0.0664823874181556 | 0.947305553324429 | 0.981831451597633 | -6.14427458740794 |
| lnc-DNAJB6-4 | lnc-DNAJB6-4 | -0.0164890269462107 | 0.104813349978569 | -0.06643435550771 | 0.94734356647374 | 0.981831451597633 | -5.6836084015662 |
| ENSG00000280190.1 | | 0.0208782145966454 | -0.206361475924829 | 0.0662983902817625 | 0.947451171994774 | 0.981867605179998 | -5.58827816248483 |
| ENSG00000279202.1 | | 0.0185428723375125 | 2.58899594531347 | 0.0662697282535137 | 0.947473855807469 | 0.981867605179998 | -6.11947209543422 |
| ENSG00000156411.9 | ATP5MJ | 0.00872038984273971 | 3.58262393600758 | 0.0660487867283435 | 0.947648715678188 | 0.981942761470861 | -6.24775796294303 |
| ENSG00000254877.1 | | -0.0221501816225973 | 0.994444462057355 | -0.0660236310878435 | 0.947668624786108 | 0.981942761470861 | -5.86795668005243 |
| lnc-GLYATL2-4 | lnc-GLYATL2-4 | -0.0221501816225974 | 0.994981189882381 | -0.0659972561381704 | 0.947689498936921 | 0.981942761470861 | -5.86795832268858 |
| ENSG00000181404.16 | WASHC1 | 0.0186714382645516 | 3.5003498032034 | 0.0657597116760723 | 0.947878914938485 | 0.982089585532006 | -6.22528237116614 |
| ENSG00000134107.4 | BHLHE40 | 0.0214450969582192 | 4.1307271256782 | 0.0655945143520026 | 0.948013996297861 | 0.982180101968031 | -6.30984457873059 |
| ENSG00000114626.17 | ABTB1 | 0.00935663200673886 | 4.73071892178077 | 0.0653495156507972 | 0.948202157336162 | 0.982241997198176 | -6.36120034605488 |
| CASC11 | CASC11 | 0.0202706327503653 | -0.480394173917244 | 0.0653474122297239 | 0.948203822138674 | 0.982241997198176 | -5.51182096119367 |
| lnc-SLC13A4-2 | lnc-SLC13A4-2 | 0.00987456926595964 | 2.37885412403192 | 0.0652997238287976 | 0.948241566317837 | 0.982241997198176 | -6.09728950217159 |
| ENSG00000163918.10 | RFC4 | -0.00859237260605994 | 3.04317186574538 | -0.0652705931675523 | 0.94826462256838 | 0.982241997198176 | -6.18199721829745 |
| ENSG00000130255.12 | RPL36 | -0.00946450688856526 | 7.09146022579657 | -0.0651000895576001 | 0.948399573165956 | 0.982327260550813 | -6.40400936201731 |
| lnc-SCAPER-7 | lnc-SCAPER-7 | -0.00884452379959644 | 2.03810205390433 | -0.0650459969013235 | 0.948442386869082 | 0.982327260550813 | -6.07064886347964 |
| lnc-TWSG1-2 | lnc-TWSG1-2 | -0.0167484566860638 | 2.55788046868133 | -0.0649670406291914 | 0.948504880109311 | 0.98234255546392 | -6.13337908463699 |
| ENSG00000273066.5 | | -0.0109475646218817 | 1.31062118586061 | -0.0649048587089965 | 0.948554096818845 | 0.982344099362128 | -5.95953092902934 |
| lnc-PRMT5-1 | lnc-PRMT5-1 | 0.00774858605968154 | 1.90724734440048 | 0.0646547887190206 | 0.948752028139366 | 0.982499647574591 | -6.03329339927082 |
| ENSG00000152558.14 | TMEM123 | 0.0210434366435708 | 6.95953727273217 | 0.0644962124309021 | 0.948884584803598 | 0.982567835631321 | -6.40556115521463 |
| ENSG00000035141.7 | FAM136A | -0.00965249379830713 | 4.81099012441903 | -0.0644159989703417 | 0.948941034164347 | 0.982567835631321 | -6.37240925631218 |
| ENSG00000121274.12 | TENT4B | 0.00746901775757269 | 4.33977323819095 | 0.0643906675080079 | 0.948961084609387 | 0.982567835631321 | -6.33219595520923 |
| ENSG00000095777.15 | MYO3A | -0.0803882381081498 | 2.37115814466428 | -0.0642921481221314 | 0.949051040099039 | 0.982588271142632 | -6.15831239442392 |
| ENSG00000267368.1 | UPK3BL1 | -0.0227445540716769 | 2.10365429835938 | -0.0642377901068831 | 0.949084792392818 | 0.982588271142632 | -6.08162694350729 |
| ENSG00000254884.1 | PRR13P2 | -0.0166923243626083 | -0.480727907290331 | -0.0641514555784444 | 0.949150428096351 | 0.982588271142632 | -5.56580422276979 |
| lnc-RWDD3-3 | lnc-RWDD3-3 | 0.0353407531392724 | -0.872127387196322 | 0.063862340552442 | 0.949379275363459 | 0.982588271142632 | -5.42097590710244 |
| ENSG00000167088.10 | SNRPD1 | -0.00786720681564734 | 4.37262737547634 | -0.0638421875820364 | 0.949395227487921 | 0.982588271142632 | -6.33556895048607 |
| lnc-PAPPA-6 | lnc-PAPPA-6 | -0.0134385066951765 | 0.00402121372808036 | -0.0637472000101541 | 0.949470415375163 | 0.982588271142632 | -5.66967185506678 |
| lnc-ANKFY1-5 | lnc-ANKFY1-5 | 0.0128242887920829 | 1.94728824263658 | 0.063683339643609 | 0.949520964629428 | 0.982588271142632 | -6.06599459545701 |
| ENSG00000014164.6 | ZC3H3 | 0.0089535126811735 | 3.50750611067851 | 0.0635417132555041 | 0.949633071036125 | 0.982588271142632 | -6.23523789302076 |
| ENSG00000129250.11 | KIF1C | 0.012002829224997 | 4.44105361620628 | 0.063503010272146 | 0.949663707119592 | 0.982588271142632 | -6.33248176103564 |
| ENSG00000167977.8 | KCTD5 | -0.00876972313750901 | 3.61647160481923 | -0.0633380887585108 | 0.949794254747426 | 0.982588271142632 | -6.25261461964315 |
| ENSG00000248592.7 | STIMATE-MUSTN1 | -0.00591164874929628 | 4.27739491379924 | -0.0632819623400129 | 0.949838683295026 | 0.982588271142632 | -6.32291860510409 |
| ENSG00000134852.14 | CLOCK | -0.00580934810542288 | 5.07983250281461 | -0.0632318835045446 | 0.949878324833488 | 0.982588271142632 | -6.38211599003138 |
| ENSG00000280037.1 | | -0.0123650512811198 | 0.267375385068519 | -0.0632024458600013 | 0.949901627222815 | 0.982588271142632 | -5.75729255991271 |
| lnc-DUSP11-1 | lnc-DUSP11-1 | -0.0123650512811198 | 0.267375385068519 | -0.0632024458600013 | 0.949901627222815 | 0.982588271142632 | -5.75729255991271 |
| ENSG00000112167.9 | SAYSD1 | 0.00733747123264417 | 3.14612560939677 | 0.0631529178830273 | 0.949940832912236 | 0.982588271142632 | -6.19473641186465 |
| lnc-ALK-1 | lnc-ALK-1 | 0.0248011890401385 | 0.882676344459131 | 0.063138287792986 | 0.949952964714211 | 0.982588271142632 | -5.85840169211058 |
| ENSG00000255526.6 | NEDD8-MDP1 | -0.0100058757569981 | 3.3780072858401 | -0.0631268686742633 | 0.949961453170633 | 0.982588271142632 | -6.222327583868 |
| lnc-STRA8-1 | lnc-STRA8-1 | 0.0304704069144108 | -0.930626544486324 | 0.0631032119131538 | 0.949980179624654 | 0.982588271142632 | -5.38280487136516 |
| lnc-FOXH1-1 | lnc-FOXH1-1 | -0.015082050392059 | -0.451263705791564 | -0.0629643896392179 | 0.950090070515843 | 0.982588271142632 | -5.54041791937602 |
| ENSG00000031081.10 | ARHGAP31 | 0.0182054082931806 | 4.05843443503274 | 0.0629310194012637 | 0.950119276090193 | 0.982588271142632 | -6.29292022738804 |
| ENSG00000105671.11 | DDX49 | -0.00827084015099677 | 3.66787506019401 | -0.0629194477363545 | 0.950125646471152 | 0.982588271142632 | -6.26131013909444 |
| lnc-TRANK1-1 | lnc-TRANK1-1 | 0.0139560831770133 | -0.140736663378292 | 0.0629104259443224 | 0.950132788123095 | 0.982588271142632 | -5.64622375322186 |
| ENSG00000143458.11 | GABPB2 | 0.00766694948841984 | 5.36467267875279 | 0.0628929850692026 | 0.95014659433425 | 0.982588271142632 | -6.38897585866507 |
| ENSG00000138801.8 | PAPSS1 | 0.012126396683227 | 5.34963827639373 | 0.0628038006140861 | 0.950217193064779 | 0.982588271142632 | -6.39161008847156 |
| lnc-PLEKHO1-1 | lnc-PLEKHO1-1 | -0.015644287379416 | -0.0707430356261791 | -0.0627797526362171 | 0.950236229598527 | 0.982588271142632 | -5.66427373168071 |
| lnc-RSRC2-1 | lnc-RSRC2-1 | -0.00766530646217012 | 2.03168809857708 | -0.0627636652819019 | 0.950248964467958 | 0.982588271142632 | -6.06025752718619 |
| lnc-TTC27-10 | lnc-TTC27-10 | -0.0126552567853002 | 2.46224256211769 | -0.0627323961508126 | 0.950273717382542 | 0.982588271142632 | -6.12183470690558 |
| lnc-VPS54-7 | lnc-VPS54-7 | -0.0118107681959696 | 0.139140863405593 | -0.0626771117580426 | 0.950317481107535 | 0.982588271142632 | -5.71567836740362 |
| ENSG00000132406.11 | TMEM128 | 0.011861034841119 | 3.07543953319794 | 0.0625280413990342 | 0.950435487593768 | 0.982640643585423 | -6.20176784514117 |
| lnc-TEKT2-1 | lnc-TEKT2-1 | -0.00844183039197788 | 0.756394276493872 | -0.0624925108651846 | 0.950463614300257 | 0.982640643585423 | -5.86166344650623 |
| lnc-ICE2-1 | lnc-ICE2-1 | 0.0142287760038622 | 3.47944123093571 | 0.0622208579261764 | 0.950678662502953 | 0.982813606745267 | -6.24893023186795 |
| ENSG00000103356.15 | EARS2 | -0.00850180399990652 | 3.2813264614277 | -0.0620477462508615 | 0.950815704576711 | 0.982905913681003 | -6.2141792405647 |
| ENSG00000279632.1 | | -0.0115975648113503 | 0.773096628086259 | -0.0619209816048863 | 0.950916057458053 | 0.982960285790433 | -5.87248826829783 |
| lnc-HIST1H2BI-2 | lnc-HIST1H2BI-2 | 0.0164696614530584 | 1.60112214300477 | 0.0618103563404816 | 0.951003634301422 | 0.98300144653373 | -6.00726308854656 |
| ENSG00000263809.1 | | -0.0114243195941039 | 8.07894175207341 | -0.0617002292338335 | 0.951090817385351 | 0.983035086727487 | -6.38669819208852 |
| lnc-RGPD8-6 | lnc-RGPD8-6 | -0.00751080040955165 | 0.921125162539859 | -0.061562021635916 | 0.951200231492717 | 0.983035086727487 | -5.89075130220972 |
| ENSG00000206612.1 | SNORA2A | 0.0132660821995125 | -0.449829281532771 | 0.061559557974493 | 0.951202181895697 | 0.983035086727487 | -5.58563309820415 |
| ENSG00000184307.14 | ZDHHC23 | -0.0133395383832529 | 5.27730406025861 | -0.0613884673602257 | 0.951337702916929 | 0.983035086727487 | -6.3902835053416 |
| ENSG00000230189.7 | RABGEF1P2 | 0.00905641681742464 | 0.666759548892736 | 0.0613829421173706 | 0.951342003882722 | 0.983035086727487 | -5.81849151413654 |
| ENSG00000265421.1 | | 0.0249708584671839 | -0.542851492591962 | 0.0613219469916306 | 0.951390292431166 | 0.983035086727487 | -5.48484651973455 |

| | | | | | | | |
|---|---|---|---|---|---|---|---|
| ENSG00000115355.16 | CCDC88A | -0.0175469988375969 | 9.19093080842931 | -0.0613236859538598 | 0.951392793364776 | 0.983035086727487 | -6.36119450611 |
| lnc-ERVW-1-3 | lnc-ERVW-1-3 | 0.00885679654213867 | 1.53415414724766 | 0.0612655324011512 | 0.951434954831466 | 0.983035086727487 | -5.9963202304838 |
| ENSG00000203546.7 | | -0.00799162140187049 | 2.9396120638252 | -0.0612262986904662 | 0.951466015536365 | 0.983035086727487 | -6.16726549741671 |
| ENSG00000172878.13 | METAP1D | -0.00952575080088325 | 2.40279805518356 | -0.0609674715668404 | 0.951670926765818 | 0.983057957870886 | -6.09708244359061 |
| lnc-TRIM44-2 | lnc-TRIM44-2 | 0.00807386661610243 | 2.92648491272661 | 0.0609492776017795 | 0.951685330897659 | 0.983057957870886 | -6.17538637507228 |
| ENSG00000266997.2 | | 0.0143713389433071 | 1.43519198898777 | 0.0609444927678551 | 0.951689119045424 | 0.983057957870886 | -5.98424845621995 |
| ENSG00000167483.17 | NIBAN3 | -0.0099996017362578 | 8.01286988047062 | -0.0608983661625483 | 0.951725637486174 | 0.983057957870886 | -6.38860608787723 |
| ENSG00000165983.14 | PTER | 0.00753044690422788 | 3.88846774819437 | 0.0608967012877267 | 0.951726955569798 | 0.983057957870886 | -6.2863497978995 |
| ENSG00000215193.12 | PEX26 | 0.00458212208951725 | 5.64391922893001 | 0.0608081303000404 | 0.951797077536365 | 0.983076551197663 | -6.40167267145676 |
| lnc-TPD52-1 | lnc-TPD52-1 | 0.0245025026607595 | -0.442402460648274 | 0.0607533096115023 | 0.95184047946452 | 0.983076551197663 | -5.49214682147944 |
| ENSG00000162004.16 | CCDC78 | -0.0154448479201317 | 1.84286032444812 | -0.0606162693365144 | 0.951948975869601 | 0.983139275982137 | -6.04198316428451 |
| ENSG00000099251.14 | HSD17B7P2 | 0.0162874942038331 | 2.19456718935563 | 0.0604400522808421 | 0.952088490350228 | 0.983187446173158 | -6.08744659519948 |
| lnc-LDLRAD1-9 | lnc-LDLRAD1-9 | -0.0097562447755607 | 0.573936669659965 | -0.0604366907156913 | 0.952091151781732 | 0.983187446173158 | -5.82345641893905 |
| ENSG00000112658.7 | SRF | 0.00856912635816087 | 4.44856591872532 | 0.060181893527014 | 0.95229288241647 | 0.983346430481021 | -6.34310547680053 |
| ENSG00000158636.16 | EMSY | -0.0063094711243652 | 5.40918903177607 | -0.0597816529283009 | 0.952609771381835 | 0.983594999253136 | -6.3942261337422 |
| ENSG00000128524.4 | ATP6V1F | 0.00923412596159068 | 4.18396316049859 | 0.0597090166928255 | 0.952667281677307 | 0.983594999253136 | -6.31304705187561 |
| ENSG00000151690.14 | MFSD6 | -0.00611621670422302 | 5.67067071318205 | -0.059675892021015 | 0.952693508474554 | 0.983594999253136 | -6.40300535802557 |
| ENSG00000152939.15 | MARVELD2 | 0.0230245756990681 | -0.0781848381575845 | 0.0595924142389508 | 0.952759603096438 | 0.983594999253136 | -5.70406062406773 |
| ENSG00000165801.9 | ARHGEF40 | 0.0324821446165234 | -0.716549207210338 | 0.0595760807987023 | 0.952772535349999 | 0.983594999253136 | -5.49889351127172 |
| lnc-KLHL42-6 | lnc-KLHL42-6 | -0.0229081299417386 | 0.0077360644344624 | -0.0594810252045777 | 0.952847797340098 | 0.983602431270618 | -5.70927658020392 |
| lnc-PPIL1-1 | lnc-PPIL1-1 | 0.0102039978719961 | 1.47137198088362 | 0.0593726988942514 | 0.952933567191277 | 0.983602431270618 | -5.98644973729646 |
| lnc-CHURC1-1 | lnc-CHURC1-1 | 0.0153806643896829 | -0.388046359720706 | 0.0593537803923789 | 0.952948546408805 | 0.983602431270618 | -5.51815470620053 |
| ENSG00000119707.13 | RBM25 | 0.00690471921313307 | 7.89305249188253 | 0.0593088243554724 | 0.952984141593181 | 0.983602431270618 | -6.39000623636088 |
| ENSG00000110719.9 | TCIRG1 | -0.0142815432864731 | 6.14007135909399 | -0.0591804213636878 | 0.953087466039618 | 0.983602431270618 | -6.40877941957307 |
| ENSG00000146083.11 | RNF44 | 0.00753826331102569 | 4.98707218108666 | 0.0589661384049676 | 0.953255475849898 | 0.983602431270618 | -6.37595407656037 |
| ENSG00000113719.15 | ERGIC1 | 0.0065589217243005 | 7.26209767912467 | 0.0589545895831975 | 0.953264620153957 | 0.983602431270618 | -6.40117989211748 |
| ENSG00000204628.11 | RACK1 | -0.00941111401703054 | 9.19154898162539 | -0.0588581697965921 | 0.953340965146898 | 0.983602431270618 | -6.35611077240711 |
| ENSG00000196976.7 | LAGE3 | 0.0177636065992508 | 0.880595484735418 | 0.0588262662772171 | 0.953366226386987 | 0.983602431270618 | -5.86080101403353 |
| ENSG00000065154.11 | OAT | -0.0111425566546504 | 3.25409490510554 | -0.0588012151015246 | 0.953386061968173 | 0.983602431270618 | -6.20288012727706 |
| lnc-CWC25-1 | lnc-CWC25-1 | -0.0118870830819521 | 0.382677715980125 | -0.058788705618189 | 0.953395967018371 | 0.983602431270618 | -5.78758168743904 |
| ENSG00000249319.2 | | -0.00534795853669223 | 3.05488650516587 | -0.0587085918459446 | 0.953459401544375 | 0.983602431270618 | -6.19108983515138 |
| ENSG00000196369.11 | SRGAP2B | 0.0118056006112475 | 1.22315573140206 | 0.0586426433267666 | 0.953511620174158 | 0.983602431270618 | -5.94351332469378 |
| ENSG00000111716.12 | LDHB | 0.0132342254807947 | 4.95647208729183 | 0.0586080509992014 | 0.953539469454013 | 0.983602431270618 | -6.37979938013585 |
| ENSG00000241878.11 | PISD | 0.00559600073167221 | 4.91975393118208 | 0.0585626946038878 | 0.953574924568107 | 0.983602431270618 | -6.37553508748912 |
| lnc-RNF152-8 | lnc-RNF152-8 | -0.00855279734444094 | -0.0324565844563424 | -0.0585344130360436 | 0.953597318338219 | 0.983602431270618 | -5.6623375999295 |
| lnc-INTS8-4 | lnc-INTS8-4 | 0.0191145964408829 | -0.0738575574728329 | 0.0584876877224703 | 0.953634316226277 | 0.983602431270618 | -5.65703013121567 |
| ENSG00000176386.8 | CDC26 | 0.0088812462281358 | 2.65333881536077 | 0.0584768396167816 | 0.953642905953364 | 0.983602431270618 | -6.13600980390884 |
| ENSG00000143870.12 | PDIA6 | 0.0131064194859718 | 5.92976214513485 | 0.0584211769174133 | 0.953687689883288 | 0.983602431270618 | -6.40743452620044 |
| lnc-DNAJC8-1 | lnc-DNAJC8-1 | 0.0106969846788658 | 3.31269211646934 | 0.05834657125446 | 0.953746055243614 | 0.983613167156339 | -6.20764590164633 |
| ENSG00000198890.8 | PRMT6 | -0.00840692328036674 | 3.92866578713721 | -0.0582504355479115 | 0.953822178077796 | 0.983613167156339 | -6.29411181823476 |
| ENSG00000119471.14 | HSDL2 | 0.011439372058283 | 4.21376711877401 | 0.0581192029322449 | 0.953926092291276 | 0.983613167156339 | -6.30926962077191 |
| lnc-ETFA-4 | lnc-ETFA-4 | 0.00967910493566659 | 1.62031860639828 | 0.0581163009981243 | 0.953928390145389 | 0.983613167156339 | -6.01139042265838 |
| ENSG00000284159.1 | MIR3940 | -0.01105798596271 | 1.16233892029908 | -0.0581053803610875 | 0.953937037495741 | 0.983613167156339 | -5.93514957842104 |
| ENSG00000133878.8 | DUSP26 | 0.0518173696928265 | -0.0580521116939143 | 0.0579384879305113 | 0.95407797762164 | 0.983698549743143 | -5.69825090673741 |
| ENSG00000147889.17 | CDKN2A | -0.0162491772516742 | 1.79539121952312 | -0.0577789666041158 | 0.954195506151674 | 0.983698549743143 | -6.03700064718288 |
| lnc-NIM1K-7 | lnc-NIM1K-7 | 0.00787916552264578 | 1.27545953810432 | 0.0577734218069334 | 0.954199896807205 | 0.983698549743143 | -5.97029381849498 |
| ENSG00000196247.11 | ZNF107 | -0.00966683893022324 | 7.8907010349924 | -0.0577593858183027 | 0.95421101123119 | 0.983698549743143 | -6.39318287115738 |
| lnc-WNT5B-2 | lnc-WNT5B-2 | 0.0156882374774695 | 0.513414320416133 | 0.0575011248576939 | 0.954415517301978 | 0.98380720446199 | -5.81478106225139 |
| lnc-SLC25A3-7 | lnc-SLC25A3-7 | 0.00984736399666386 | 1.59469008122197 | 0.0574872482437019 | 0.954426505701282 | 0.98380720446199 | -6.00373830107303 |
| ENSG00000197620.10 | EOLA1 | -0.00830587297905794 | 2.46073131718046 | -0.0574383441715045 | 0.954465231176063 | 0.98380720446199 | -6.11941853427367 |
| lnc-RP9-6 | lnc-RP9-6 | 0.0114273653710943 | 1.50078726658967 | 0.0572339381666707 | 0.954627094557017 | 0.98380720446199 | -5.97318974057852 |
| ENSG00000101138.11 | CSTF1 | 0.00575877065644992 | 4.27825767329121 | 0.0571510293219365 | 0.954692748298149 | 0.98380720446199 | -6.32563464936376 |
| ENSG00000267157.1 | | -0.0140435147140798 | -0.26501464430378 | -0.0570623819145308 | 0.954762946636532 | 0.98380720446199 | -5.65025800724231 |
| ENSG00000109736.14 | MFSD10 | -0.0138845754384311 | 4.23276642814898 | -0.0570176724607722 | 0.954799064288815 | 0.98380720446199 | -6.3308843829756 |
| ENSG00000168916.15 | ZNF608 | 0.0826605399294932 | 0.267608146677883 | 0.0570028396811941 | 0.954820298438134 | 0.98380720446199 | -5.56093815981226 |
| ENSG00000182095.14 | TNRC18 | 0.0115598164038264 | 4.70476636297632 | 0.0569754782293946 | 0.954831764504385 | 0.98380720446199 | -6.35877997789674 |
| lnc-SYNCRIP-7 | lnc-SYNCRIP-7 | -0.00816663571043108 | 2.15441282181543 | -0.0569658286454388 | 0.954839405902591 | 0.98380720446199 | -6.08170541855102 |
| lnc-CXorf65-2 | lnc-CXorf65-2 | -0.0068146045689006 | 2.80745217293393 | -0.0569623296423065 | 0.954842176725316 | 0.98380720446199 | -6.16084672812335 |
| ENSG00000137824.15 | RMDN3 | -0.00702591302919053 | 4.14300543826568 | -0.0568587274838531 | 0.954924218409698 | 0.983813044749279 | -6.31154003511824 |
| lnc-MPEG1-1 | lnc-MPEG1-1 | -0.0337704146878512 | 0.242257193233197 | -0.0568384002940405 | 0.954943439782629 | 0.983813044749279 | -5.65663751096576 |
| lnc-FBLN1-3 | lnc-FBLN1-3 | -0.0121390438046267 | 1.86973378234597 | -0.0567412278524016 | 0.955017265985471 | 0.983839859148095 | -6.03828278610345 |
| ENSG00000261582.1 | | 0.00755522393233132 | 2.91936869827878 | 0.0566454016548874 | 0.955093150909538 | 0.983868791610581 | -6.16924962796651 |
| ENSG00000254612.2 | DNAJB6P1 | -0.0156565740371423 | -0.530492164308918 | -0.0564368963999842 | 0.955258268020871 | 0.983925531034007 | -5.51345173063759 |
| lnc-ALOX15-1 | lnc-ALOX15-1 | 0.00641931454029376 | 0.627722709175842 | 0.0564034251880492 | 0.955284774347977 | 0.983925531034007 | -5.82968122935957 |
| lnc-UBR5-5 | lnc-UBR5-5 | 0.00553435507869438 | 2.10360694325569 | 0.0562949408767112 | 0.955370684963476 | 0.983925531034007 | -6.07153325242854 |
| ENSG00000165029.15 | ABCA1 | -0.0229998552662094 | 6.13703043291535 | -0.056304294535416 | 0.955371463687863 | 0.983925531034007 | -6.40848373603173 |
| ENSG00000188707.5 | ZBED10P | -0.0109836688466625 | 3.88831601837856 | -0.0562210931755604 | 0.955429166551381 | 0.983925531034007 | -6.2857823937282 |
| lnc-SLC2A9-5 | lnc-SLC2A9-5 | 0.00696876406039681 | 3.8880860012877 | 0.0561920864450123 | 0.955452137675153 | 0.983925531034007 | -6.28580431655853 |

| | | | | | | | |
|---|---|---|---|---|---|---|---|
| lnc-COX14-2 | lnc-COX14-2 | 0.00713761124895895 | 1.48202948940141 | 0.0561358475319189 | 0.955496674724839 | 0.983925531034007 | -5.98713818303203 |
| ENSG00000242193.11 | CRYZL2P | -0.0710785660627495 | -2.07035062239118 | -0.0560565377195993 | 0.955562859631564 | 0.983925531034007 | -5.08704950157224 |
| lnc-ANKRD26-1 | lnc-ANKRD26-1 | -0.0113880956185256 | 0.619239776815021 | -0.0559755379206851 | 0.955623628841022 | 0.983925531034007 | -5.80169639151097 |
| ENSG00000128699.13 | ORMDL1 | -0.00712434709377607 | 6.61927852060259 | -0.0559128637119513 | 0.955673262793515 | 0.983925531034007 | -6.40849933287838 |
| ENSG00000135318.11 | NT5E | 0.0904213054057812 | -0.306564711353808 | 0.0559244914502552 | 0.955674061916721 | 0.983925531034007 | -5.42838578040842 |
| lnc-NUTM2E-1 | lnc-NUTM2E-1 | -0.00983383635395163 | 1.53013979095834 | -0.0558060231940224 | 0.955757874039701 | 0.983962602979001 | -5.99609815329612 |
| ENSG00000023697.12 | DERA | -0.00836086950383086 | 3.88802753770211 | -0.0557157219454404 | 0.955829387584769 | 0.983987010335966 | -6.28587214963584 |
| lnc-CRHR2-3 | lnc-CRHR2-3 | 0.0105601681681038 | 2.37439441967914 | 0.0556343082440745 | 0.955893863004535 | 0.984004170153655 | -6.11227744473915 |
| ENSG00000035681.8 | NSMAF | 0.00623487142511259 | 5.47138715544434 | 0.0554971960725852 | 0.956002449392873 | 0.984064128490472 | -6.39610453006999 |
| LINC01415 | LINC01415 | -0.0206920831665505 | 0.974780001139091 | -0.0554407179028144 | 0.956047727611328 | 0.984064128490472 | -5.916977202878 |
| ENSG00000207405.1 | SNORA64 | -0.00850727132027274 | 0.811623632278477 | -0.0553068498852213 | 0.956153196018422 | 0.984106550003859 | -5.87806360625763 |
| lnc-NAT9-3 | lnc-NAT9-3 | -0.00813343184794466 | 0.103940772361938 | -0.055267241340958 | 0.956184564617752 | 0.984106550003859 | -5.71153325314939 |
| ENSG00000118200.14 | CAMSAP2 | 0.0428738740522818 | -1.57689252591196 | 0.0549639567914903 | 0.956424794037753 | 0.984288045095697 | -5.24988252710675 |
| lnc-NHLRC1-2 | lnc-NHLRC1-2 | 0.0124006309560444 | -0.135986555341097 | 0.0548554917065517 | 0.956510660263359 | 0.984288045095697 | -5.66410389106621 |
| lnc-LMBR1-1 | lnc-LMBR1-1 | -0.0121278200514854 | 0.248380901244996 | -0.0548372910764359 | 0.956525074891901 | 0.984288045095697 | -5.71900928955362 |
| ENSG00000141385.9 | AFG3L2 | 0.00643184394093055 | 5.24043752989469 | 0.0548030518271688 | 0.956552191908333 | 0.984288045095697 | -6.38948931160443 |
| lnc-MED14-10 | lnc-MED14-10 | 0.024370756596462 | -0.909108293359305 | 0.054567303250702 | 0.956738902963733 | 0.98441643239045 | -5.40869804299344 |
| ENSG00000110344.9 | UBE4A | 0.00775197002373229 | 6.64335737945069 | 0.0545146950416369 | 0.956780568595671 | 0.98441643239045 | -6.40835420670909 |
| lnc-GRIFIN-4 | lnc-GRIFIN-4 | -0.0166119180076333 | 0.461796376407126 | -0.0544083921685281 | 0.95686476070356 | 0.98441643239045 | -5.76648863827277 |
| lnc-SLC25A41-2 | lnc-SLC25A41-2 | 0.00520796793121914 | 2.02419066983063 | 0.0543960146243167 | 0.956874563778045 | 0.98441643239045 | -6.05704800705766 |
| ENSG00000259562.2 | | -0.00903308201450789 | 0.0259825341316521 | -0.0543435768779963 | 0.956916094799987 | 0.98441643239045 | -5.66748268093147 |
| ENSG00000104998.3 | IL27RA | 0.0203026156746269 | 4.26172370023657 | 0.054263460950708 | 0.956985788604637 | 0.984438926825391 | -6.30749627259434 |
| ENSG00000029725.16 | RABEP1 | 0.00676007377209054 | 7.06766126195878 | 0.0540683355116591 | 0.957134089661522 | 0.984535298377543 | -6.40454790531653 |
| ENSG00000101266.18 | CSNK2A1 | -0.00467192679984674 | 6.87722780867381 | -0.0538960712923875 | 0.957270526928905 | 0.984535298377543 | -6.40630692607905 |
| ENSG00000105732.12 | ZNF574 | -0.00955584672617338 | 2.92513742887841 | -0.0537744367189707 | 0.957366865120601 | 0.984535298377543 | -6.18025661567111 |
| lnc-DENND2A-1 | lnc-DENND2A-1 | 0.00615395533668917 | 4.32186549379375 | 0.0537668899138467 | 0.957372842435476 | 0.984535298377543 | -6.32711226700007 |
| ENSG00000137168.7 | PPIL1 | 0.00971328148711548 | 2.59401548480004 | 0.0537263060512399 | 0.957404986215906 | 0.984535298377543 | -6.13506645809309 |
| ENSG00000107185.9 | RGP1 | 0.00516737781474941 | 5.75667521278131 | 0.0536789060932719 | 0.957442528663293 | 0.984535298377543 | -6.40502826329673 |
| lnc-RBM23-2 | lnc-RBM23-2 | 0.00720626362952603 | 1.2046084445167 | 0.053664534332911 | 0.957453911626403 | 0.984535298377543 | -5.93724326170654 |
| ENSG00000106554.12 | CHCHD3 | -0.00458230244185476 | 4.83273693359151 | -0.0536225978398939 | 0.957487126923554 | 0.984535298377543 | -6.3671173389132 |
| ENSG00000156976.16 | EIF4A2 | 0.00461884079136254 | 9.42184635219793 | 0.053593763734556 | 0.957509964675784 | 0.984535298377543 | -6.34969851165891 |
| ENSG00000115568.15 | ZNF142 | 0.00684361949900492 | 4.84589742817448 | 0.0534216011753887 | 0.95764632498515 | 0.984626320354131 | -6.37089282855481 |
| lnc-REELD1-3 | lnc-REELD1-3 | 0.00738826221868494 | 1.28829845515876 | 0.0533586431147552 | 0.957696190855437 | 0.98462840641246 | -5.95848850207029 |
| lnc-TRPC4AP-6 | lnc-TRPC4AP-6 | 0.00616566916705308 | 1.12197017176377 | 0.0532602947379268 | 0.95777408793404 | 0.984659310321479 | -5.92812300586747 |
| ENSG00000213588.5 | ZBTB9 | -0.00831012653910501 | 1.77873949262104 | -0.0531873518497894 | 0.957831862801354 | 0.984669525124376 | -6.0262325838507 |
| lnc-OSBPL9-8 | lnc-OSBPL9-8 | -0.0161125983674041 | -0.320504153951602 | -0.0530763684317786 | 0.957919768346428 | 0.98471071227461 | -5.54290913559947 |
| lnc-STX6-2 | lnc-STX6-2 | 0.00832444896285423 | 1.18481418135147 | 0.05287512747833 | 0.958079164641429 | 0.984825381832527 | -5.94699723269811 |
| ENSG00000267757.4 | EML2-AS1 | 0.016309010223068 | -0.298767325683501 | 0.0527874655388397 | 0.95814859930388 | 0.984847571509201 | -5.61576007614375 |
| ENSG00000133106.14 | EPSTI1 | 0.00764460878400985 | 5.31436828736086 | 0.0526044821861263 | 0.958293536550723 | 0.984947361571134 | -6.3921623470161 |
| ENSG00000108669.16 | CYTH1 | 0.00553367699938079 | 8.55998562551964 | 0.0524426431091731 | 0.958421727053631 | 0.984960047490399 | -6.37607939395104 |
| lnc-FJX1-1 | lnc-FJX1-1 | -0.0159773237956654 | 1.00966962030601 | -0.0523570167354163 | 0.9584895509751 | 0.984960047490399 | -5.88912266674077 |
| ENSG00000129932.8 | DOHH | -0.0126036515891536 | 0.715760602732015 | -0.0523367298063293 | 0.958505620124487 | 0.984960047490399 | -5.86384572683219 |
| lnc-FGFR2-12 | lnc-FGFR2-12 | -0.0174399973200736 | 3.3444530326409 | -0.0523118865019322 | 0.95852834098908 | 0.984960047490399 | -6.20117960181231 |
| ENSG00000185722.17 | ANKFY1 | 0.00572122395917596 | 5.32768518878232 | 0.0522868311661235 | 0.95854514459895 | 0.984960047490399 | -6.39329661349816 |
| SNHG21 | SNHG21 | -0.00765579952141282 | 2.49984786216147 | -0.0520399673882655 | 0.958740685768461 | 0.985111797882001 | -6.12733826072277 |
| lnc-MAP3K7-3 | lnc-MAP3K7-3 | -0.0146464199844445 | 4.28157192141268 | -0.0519735446758688 | 0.95879557586375 | 0.985119020516326 | -6.31241017927056 |
| ENSG00000089006.16 | SNX5 | 0.0060280775075201 | 6.24079159353188 | 0.0516548692702832 | 0.959045727695682 | 0.985326855004504 | -6.40937253646449 |
| ENSG00000127804.12 | METTL16 | -0.00684936261968883 | 5.52894217122243 | -0.0515243790900742 | 0.959149092312604 | 0.98538386658699 | -6.40079349043054 |
| ENSG00000177595.17 | PIDD1 | -0.00960377094465859 | 3.36882113006133 | -0.0511906422240684 | 0.959413457071264 | 0.985582497980444 | -6.22849195924209 |
| ENSG00000239306.4 | RBM14 | -0.00777380072902112 | 5.73013470127174 | -0.0511594042472515 | 0.959438202013028 | 0.985582497980444 | -6.40330216254751 |
| ENSG00000274386.5 | TMEM269 | -0.0104835411623236 | 0.633398857651261 | -0.0509639489041419 | 0.959593031494603 | 0.985692352892176 | -5.79723363319953 |
| ENSG00000254838.5 | GVINP1 | -0.0105426062023382 | 6.33310928105587 | -0.0508833562798814 | 0.959656873208707 | 0.985708739021648 | -6.40954566492489 |
| ENSG00000219085.1 | NPM1P37 | 0.0121815066271371 | 2.35493074913808 | 0.0506796571577015 | 0.959818235336203 | 0.98575218181758 | -6.09407370674768 |
| ENSG00000116455.13 | WDR77 | -0.00425922098536446 | 4.88325915329041 | -0.050676768844718 | 0.959820523352092 | 0.98575218181758 | -6.37148469703626 |
| lnc-PLPP5-2 | lnc-PLPP5-2 | 0.00677893174118127 | 1.90154373288204 | 0.0506472868803866 | 0.959843877905373 | 0.98575218181758 | -6.03164341075775 |
| ENSG00000284194.1 | SCO2 | -0.0129922083525639 | 0.552767972843428 | -0.0505881373971646 | 0.959890734109205 | 0.98575218181758 | -5.86076658581008 |
| ENSG00000266421.3 | MIR4451 | 0.0145143353909658 | 2.68978211979452 | 0.05044846625218 | 0.960001840909744 | 0.985817096958953 | -6.15780263285864 |
| lnc-FBXW11-4 | lnc-FBXW11-4 | -0.00612056702515273 | 3.09721385393788 | -0.050138868350295 | 0.960246634401956 | 0.986019280413842 | -6.18820750814294 |
| ENSG00000283709.1 | FAM238C | -0.0176498687545729 | -0.304196226614554 | -0.0500759197946058 | 0.960296501413151 | 0.98602129544981 | -5.56094262452165 |
| ENSG00000230202.1 | RPL29P4 | 0.0314977935483451 | 3.00238924520881 | 0.0497828009798238 | 0.960537219234457 | 0.986219263904965 | -6.16321838583652 |
| lnc-H6PD-2 | lnc-H6PD-2 | -0.00553872193031202 | 3.13188378134046 | -0.0496300254486372 | 0.960649737596228 | 0.986244054058796 | -6.19534057927267 |
| lnc-CCL25-1 | lnc-CCL25-1 | 0.00846648666216598 | -0.349618121864799 | 0.049620612284056 | 0.960657194763873 | 0.986244054058796 | -5.53388122321747 |
| ENSG00000198934.4 | MAGEE1 | -0.00947897436128486 | 1.28935376726098 | -0.049503055245687 | 0.960750324482346 | 0.986290470265343 | -5.96630092890596 |
| ENSG00000223496.2 | EXOSC6 | -0.00676809368316356 | 5.3682343463062 | -0.049303912543758 | 0.960908088345182 | 0.986403230881696 | -6.39634096391083 |
| ENSG00000078668.13 | VDAC3 | 0.0079929045551772 | 4.30098428047702 | 0.0491155698955992 | 0.961057297700786 | 0.986496897896699 | -6.32236354207182 |
| lnc-NEDD4L-2 | lnc-NEDD4L-2 | 0.0157040565600115 | -0.316815407395769 | 0.0490189262609805 | 0.961133861548328 | 0.986496897896699 | -5.53918613711129 |
| lnc-LRRC31-2 | lnc-LRRC31-2 | 0.00875088366482402 | 3.16592774590759 | 0.0490072426778217 | 0.961143117641763 | 0.986496897896699 | -6.20503573333556 |
| ENSG00000177370.4 | TIMM22 | -0.00734835488253482 | 2.87613346743389 | -0.0489085408566744 | 0.961221312472401 | 0.986527961835923 | -6.1695630111264 |

| | | | | | | | |
|---|---|---|---|---|---|---|---|
| ENSG00000201544.1 | SNORA16B | -0.0121544027698519 | -0.313616609011488 | -0.0488037133243795 | 0.961304360716015 | 0.986531604270006 | -5.57284405688728 |
| ENSG00000088451.10 | TGDS | 0.00520992546080024 | 2.87617806117144 | 0.0486528012934608 | 0.961423919565267 | 0.986531604270006 | -6.16616418722411 |
| lnc-TXNDC16-3 | lnc-TXNDC16-3 | 0.00771835465888582 | 0.927114122616289 | 0.0486468573004506 | 0.961428628664588 | 0.986531604270006 | -5.89310781534771 |
| lnc-AMOTL2-5 | lnc-AMOTL2-5 | 0.00845161778107051 | 1.78170328807442 | 0.0485803636491968 | 0.961481308028757 | 0.986531604270006 | -6.03526557515247 |
| lnc-CASTOR3-3 | lnc-CASTOR3-3 | 0.00552826840519938 | 3.72834597839269 | 0.0485751614548813 | 0.961485429456478 | 0.986531604270006 | -6.26758901893831 |
| lnc-ZDHHC6-1 | lnc-ZDHHC6-1 | 0.0115508313699172 | 0.647791921532674 | 0.0484434905060708 | 0.961589745848795 | 0.986531604270006 | -5.83367518751759 |
| ENSG00000175662.17 | TOM1L2 | -0.00848857613946041 | 4.48085089750402 | -0.0484272315983592 | 0.961602627025969 | 0.986531604270006 | -6.34088029150877 |
| ENSG00000092199.17 | HNRNPC | 0.00444590319503746 | 8.66888032272952 | 0.0484056112677166 | 0.961619755825513 | 0.986531604270006 | -6.37330158801931 |
| ENSG00000255468.7 | B4GAT1-DT | 0.0137912116354416 | -0.215936576042935 | 0.0483595774144821 | 0.961656226403993 | 0.986531604270006 | -5.53045959440679 |
| ENSG00000222489.1 | SNORA79B | 0.0142568634332187 | 2.23810240357204 | 0.0482641904870825 | 0.961731797496751 | 0.986559959525324 | -6.06119360449409 |
| ENSG00000152763.16 | DNAI4 | -0.0111400164949727 | 0.237750015168195 | -0.0480441550403579 | 0.961906123780649 | 0.986567005483094 | -5.71530015579862 |
| lnc-SLC26A3-4 | lnc-SLC26A3-4 | 0.00444987692252874 | 1.81631842774267 | 0.0479816557836271 | 0.961955640070411 | 0.986567005483094 | -6.02854667198765 |
| ENSG00000135932.10 | CAB39 | -0.00605442466345263 | 6.33212936005994 | -0.0479092137651613 | 0.962013033902743 | 0.986567005483094 | -6.40961169824764 |
| lnc-EFHD1-2 | lnc-EFHD1-2 | -0.00846223238794463 | 0.763456150976775 | -0.0478645181447238 | 0.962048445123581 | 0.986567005483094 | -5.85762184698933 |
| ENSG00000111802.13 | TDP2 | -0.00521734527252438 | 4.67799404877115 | -0.0478486762643988 | 0.962060996267276 | 0.986567005483094 | -6.35930026340646 |
| ENSG00000243056.1 | EIF4EBP3 | 0.00581084458982128 | 1.02255999403822 | 0.0478399166611533 | 0.96206793629605 | 0.986567005483094 | -5.90016031772981 |
| ENSG00000253626.3 | EIF5AL1 | -0.037545480146449 | -1.26274589125366 | -0.0478212838256867 | 0.962082698665686 | 0.986567005483094 | -5.17833612275407 |
| lnc-KDM7A-3 | lnc-KDM7A-3 | -0.00550346172933835 | 2.51085008191745 | -0.0477359629372816 | 0.962150296627218 | 0.986567005483094 | -6.12743789246372 |
| ENSG00000201944.1 | | 0.0148585318423268 | 0.0979208157167662 | 0.0477110350239982 | 0.96217004654655 | 0.986567005483094 | -5.75347152186436 |
| ENSG00000100201.20 | DDX17 | 0.0043164944826799 | 10.3799518770994 | 0.0476341976406492 | 0.962230923519073 | 0.986580278893802 | -6.31968317523974 |
| ENSG00000261716.2 | H2BC20P | 0.0112973780292407 | 4.22693068374981 | 0.0475635841885733 | 0.962287084068706 | 0.986588715450596 | -6.31784236241758 |
| lnc-SORBS3-2 | lnc-SORBS3-2 | -0.0121841536751337 | -0.251245335323064 | -0.0474885105191722 | 0.962346349612331 | 0.986600334416028 | -5.54333850029612 |
| ENSG00000076554.15 | TPD52 | 0.0132100248891005 | 4.39342125705378 | 0.0474254775519417 | 0.962398394493117 | 0.986604549947795 | -6.3306669235808 |
| ENSG00000185122.10 | HSF1 | -0.00586804836711075 | 5.19622414184695 | -0.0472213141662389 | 0.962558048079789 | 0.986717721601212 | -6.38836394481431 |
| ENSG00000115129.13 | TP53I3 | -0.0190911863093963 | 2.73319326613384 | -0.0470697916955784 | 0.962683002062674 | 0.986717721601212 | -6.16724699163392 |
| ENSG00000172954.13 | LCLAT1 | -0.00785381204801006 | 4.26333655135792 | -0.0470290764276911 | 0.962710358866417 | 0.986717721601212 | -6.31689296855824 |
| LINC02520 | LINC02520 | 0.0514760527168173 | -0.239823932331571 | 0.0469676558018222 | 0.962765415831978 | 0.986717721601212 | -5.46652935693886 |
| lnc-PAIP2-5 | lnc-PAIP2-5 | 0.0163239202362653 | 1.1178809433351 | 0.0469091479785857 | 0.962805435330576 | 0.986717721601212 | -5.93995522912535 |
| ENSG00000164008.15 | C1orf50 | 0.00421344267281754 | 4.18353469277298 | 0.0468971515179532 | 0.962814884355311 | 0.986717721601212 | -6.31354152605866 |
| ENSG00000065029.14 | ZNF76 | -0.00634628849416163 | 4.84002276749967 | -0.0468599509722678 | 0.962844358857326 | 0.986717721601212 | -6.36688418750789 |
| ENSG00000108861.8 | DUSP3 | 0.00785660019519696 | 3.33384497435031 | 0.0467421683044635 | 0.962937680025348 | 0.986764227221036 | -6.22949916289037 |
| lnc-PTS-1 | lnc-PTS-1 | -0.0131968432960141 | 3.34810525471386 | -0.0466124124424446 | 0.963041207407138 | 0.986821186442033 | -6.20428701284547 |
| ENSG00000104825.16 | NFKBIB | -0.00737964366540613 | 3.28252739283274 | -0.0463120949893043 | 0.963278438726425 | 0.987015138605436 | -6.21846467251609 |
| lnc-FJX1-2 | lnc-FJX1-2 | -0.010778187625375 | 2.8187145670175 | -0.0460923411764284 | 0.963452558292086 | 0.987128327891 | -6.1540638738499 |
| ENSG00000132718.8 | SYT11 | -0.0260006817535503 | 3.90300108634693 | -0.0460615575817987 | 0.963484822782403 | 0.987128327891 | -6.26220214878004 |
| ENSG00000107779.13 | BMPR1A | -0.0174707490731514 | 3.305967505951 | -0.0459881291804951 | 0.963539874934181 | 0.987135595329762 | -6.22577086548555 |
| ENSG00000100029.17 | PES1 | 0.00627835191843573 | 4.9988669574082 | 0.0458896878765391 | 0.963613130050585 | 0.987161509846266 | -6.38020186819564 |
| lnc-C8orf86-7 | lnc-C8orf86-7 | 0.0076529236981287 | -0.076705807899072 | 0.045781915079532 | 0.963698524138835 | 0.987199856783439 | -5.65457788745932 |
| lnc-SLC35E1-1 | lnc-SLC35E1-1 | -0.00586353252513727 | 1.74509835670736 | -0.0456774945116774 | 0.963781262489925 | 0.98723547953768 | -6.02366460312662 |
| lnc-ATP6V1G2-DDX39B- | lnc-ATP6V1G2-DD | 0.00533251870615237 | 3.52488063207635 | 0.0455925434322172 | 0.963848574356733 | 0.987255297635464 | -6.2453566372087 |
| lnc-FGL1-2 | lnc-FGL1-2 | 0.00580847426963871 | 3.29509297895771 | 0.0454286015666875 | 0.963978476144117 | 0.987337379240389 | -6.21050914813946 |
| lnc-LPP-2 | lnc-LPP-2 | -0.00894681592371516 | 4.44750564286343 | -0.0453431064629462 | 0.964046219860413 | 0.987337379240389 | -6.3367022831182 |
| ENSG00000125885.13 | MCM8 | 0.00489918789761834 | 4.57606806741439 | 0.0453097940780639 | 0.964072615652399 | 0.987337379240389 | -6.35054371330307 |
| lnc-PDZD9-3 | lnc-PDZD9-3 | -0.0087803069248642 | 1.96733559606407 | -0.0452166079335452 | 0.964146453935275 | 0.987363871901576 | -6.05480446247642 |
| ENSG00000146842.16 | TMEM209 | 0.00541755272786926 | 4.94699023427575 | 0.0451155717104758 | 0.964226512776471 | 0.987396731964085 | -6.37611248488813 |
| lnc-PLXND1-3 | lnc-PLXND1-3 | -0.00956354362607392 | 2.72454489478906 | -0.0449230657119507 | 0.964379051253018 | 0.987503806374851 | -6.15275193720367 |
| ENSG00000243414.5 | TICAM2 | 0.0101508922302562 | 0.428271059092016 | 0.0448200605910209 | 0.964460671312423 | 0.987538254781793 | -5.80765091896953 |
| ENSG00000030419.16 | IKZF2 | 0.0220988751196582 | 3.68810884045683 | 0.0446873368557464 | 0.964572839326372 | 0.98756142351622 | -6.26577911247637 |
| lnc-VCP-4 | lnc-VCP-4 | -0.006509828872656 | -0.157804234390858 | -0.0446704047323729 | 0.964579257560613 | 0.98756142351622 | -5.61943356813762 |
| ENSG00000113504.20 | SLC12A7 | -0.0509386731099875 | -0.279404343073337 | -0.0445360137993309 | 0.964693711943829 | 0.987603359003937 | -5.69229667204783 |
| lnc-MTA3-2 | lnc-MTA3-2 | -0.0139980569612602 | 0.0232753676462246 | -0.0444498605700704 | 0.964754016679406 | 0.987603359003937 | -5.64714739767784 |
| ENSG00000249240.2 | | -0.00819236721616101 | 0.902529196298796 | -0.0444296294660708 | 0.96477004788723 | 0.987603359003937 | -5.89733646649854 |
| ENSG00000285000.1 | | -0.00347653716330321 | 3.64431040727142 | -0.0443765060447847 | 0.964812143169177 | 0.987603359003937 | -6.25617304541974 |
| ENSG00000213533.11 | STIMATE | 0.00407434283529184 | 4.26166305327267 | 0.0443016047666518 | 0.964871495513401 | 0.98761499786923 | -6.32175866548545 |
| ENSG00000180228.12 | PRKRA | -0.00369363521538713 | 4.53518279308039 | -0.0442063169926374 | 0.964947002563781 | 0.987629206762841 | -6.35290096495314 |
| ENSG00000074657.13 | ZNF532 | -0.0240345237327899 | 4.68441502784585 | -0.0441270816658624 | 0.965017650915686 | 0.987629206762841 | -6.37218696132653 |
| lnc-WWP2-5 | lnc-WWP2-5 | -0.0148623783520778 | 0.735705259011842 | -0.044066085800231 | 0.96505812385101 | 0.987629206762841 | -5.85583743626982 |
| ENSG00000100344.10 | PNPLA3 | 0.0102065785220434 | 0.062631365036293 | 0.0440418758524909 | 0.965077308245689 | 0.987629206762841 | -5.67803584048676 |
| ENSG00000201592.1 | | 0.0147535379093493 | 0.798376136032701 | 0.043930690214129 | 0.965165413998606 | 0.98767026530443 | -5.85363235422226 |
| ENSG00000117691.9 | NENF | 0.00596406882564837 | 3.51862885353877 | 0.0437493223676395 | 0.96530913450016 | 0.987751327777685 | -6.24038745078986 |
| ENSG00000174130.12 | TLR6 | 0.0132883426126069 | 4.41237500495649 | 0.0437134717707649 | 0.965340606790843 | 0.987751327777685 | -6.3288943713519 |
| ENSG00000101493.10 | ZNF516 | -0.0516505083134358 | 0.438846483736699 | -0.0433765418210773 | 0.965612609117666 | 0.987945451908692 | -5.76504132190125 |
| lnc-C1QTNF12-1 | lnc-C1QTNF12-1 | 0.0040550030041917 | 2.06231747719222 | 0.0433204821038493 | 0.965648963082872 | 0.987945451908692 | -6.06464756968522 |
| ENSG00000011638.10 | LDAF1 | 0.0112747492504774 | 2.78717328396869 | 0.0432884826214091 | 0.965674320893503 | 0.987945451908692 | -6.13948973191809 |
| ENSG00000036549.12 | ZZZ3 | -0.00282987988653005 | 5.75814585256374 | -0.042859581620447 | 0.966014204549093 | 0.98821611357752 | -6.40506623609324 |
| ENSG00000106100.10 | NOD1 | -0.00699467530495923 | 4.15196096949002 | -0.042833461608694 | 0.966034903624024 | 0.98821611357752 | -6.31267099808906 |
| ENSG00000285133.1 | | -0.0109508380757486 | 1.63572787287855 | -0.0427454624949181 | 0.966104639611179 | 0.988238336006207 | -6.01923967782372 |
| ENSG00000103021.9 | CFAP263 | -0.0138773972980729 | 0.641150500354795 | -0.0426337334254196 | 0.966193181082291 | 0.988279791592546 | -5.868106274321 |

| | | | | | | | |
|---|---|---|---|---|---|---|---|
| ENSG00000184270.4 | H2AC21 | -0.0085260064343444 | 4.48908348878897 | -0.042512872773087 | 0.966288959506875 | 0.98830720217626 | -6.3505483110557 |
| ENSG00000018610.14 | STEEP1 | -0.0043654661019720 | 4.07311294768456 | -0.0424787378647977 | 0.966316010484246 | 0.98830720217626 | -6.30823963976475 |
| ENSG00000197045.12 | GMFB | 0.00556329613602724 | 6.56759302376824 | 0.0423799463904252 | 0.966394300240681 | 0.988338163661629 | -6.40918057829105 |
| ENSG00000132661.3 | NXT1 | -0.0075710844619911 | 2.73542630340181 | -0.0423150809604875 | 0.966445704643466 | 0.988341627598572 | -6.1569528613207 |
| lnc-ZMAT3-4 | lnc-ZMAT3-4 | -0.0080426430416027 | 1.71105434008828 | -0.0421772723925988 | 0.966554915331038 | 0.988385402797803 | -6.03145885707465 |
| ENSG00000139613.11 | SMARCC2 | 0.00536496135746223 | 6.83417675806404 | 0.0420686382998452 | 0.966641006256058 | 0.988385402797803 | -6.40793616755548 |
| ENSG00000058600.15 | POLR3E | -0.0031316605770497 | 6.02704288926522 | -0.0419732375761588 | 0.966716610262332 | 0.988385402797803 | -6.40886503110835 |
| ENSG00000136807.13 | CDK9 | -0.0044770624677799 | 5.23749204032219 | -0.0419602768649057 | 0.966726881504931 | 0.988385402797803 | -6.39250504118866 |
| ENSG00000158526.7 | TSR2 | 0.00495630454309026 | 4.70811208734608 | 0.0419580988697678 | 0.966728607546294 | 0.988385402797803 | -6.36124264089256 |
| ENSG00000138780.14 | GSTCD | 0.00492536421873835 | 3.36004159568905 | 0.0417671414818099 | 0.966879940168301 | 0.98843831563374 | -6.23093496169932 |
| ENSG00000185127.6 | C6orf120 | -0.0068555563205192 | 4.47129235757837 | -0.0417173537997449 | 0.966919396820726 | 0.98843831563374 | -6.34193547210775 |
| ENSG00000116473.14 | RAP1A | -0.0065862021897729 | 5.04564053665573 | -0.0416879720053413 | 0.966942681881629 | 0.98843831563374 | -6.38246949812653 |
| lnc-APOL4-1 | lnc-APOL4-1 | 0.00819628100383689 | -0.0316010802483358 | 0.041650410650682 | 0.966972449283438 | 0.98843831563374 | -5.67046704461382 |
| lnc-SOX15-1 | lnc-SOX15-1 | -0.0080814362176843 | -0.149875326967229 | -0.0415581999525007 | 0.967045526533119 | 0.988442201347361 | -5.63097201336643 |
| ENSG00000004766.16 | VPS50 | 0.00377639588211753 | 5.1303415815203 | 0.0415244227547276 | 0.967072295133516 | 0.988442201347361 | -6.38673519300417 |
| ENSG00000146676.8 | PURB | 0.00501313010959059 | 5.48837991880904 | 0.0414271192175914 | 0.967149408894459 | 0.988471934220897 | -6.3985971902483 |
| lnc-PTBP3-8 | lnc-PTBP3-8 | 0.00775526615949353 | -0.0590102131443606 | 0.0412599503865168 | 0.967281892151424 | 0.98855825154681 | -5.63065055876765 |
| lnc-ABHD17B-6 | lnc-ABHD17B-6 | -0.0168193523145681 | 2.95231748268658 | -0.0410678547410921 | 0.967438407347076 | 0.988669119627867 | -6.17122098008753 |
| lnc-TBC1D12-1 | lnc-TBC1D12-1 | -0.0097738093339242 | -0.268214481552936 | -0.0409938643657608 | 0.967492770367559 | 0.988675587949334 | -5.57334920708518 |
| ENSG00000142002.17 | DPP9 | 0.00395546794595614 | 5.28068101798955 | 0.040901841026866 | 0.967565701156961 | 0.988701028988419 | -6.39213991546504 |
| ENSG00000139182.14 | CLSTN3 | -0.0069552298735819 | 4.07998840193146 | -0.0407168089236905 | 0.967712344559697 | 0.98880178654052 | -6.30745660795507 |
| ENSG00000178852.15 | EFCAB13 | -0.0075137088283607 | 6.15624550314017 | -0.0405511870580812 | 0.967843605735496 | 0.988843700213672 | -6.40953612816125 |
| ENSG00000185404.16 | SP140L | 0.00578719522363258 | 6.77904071386839 | 0.0405031428839232 | 0.967881682608718 | 0.988843700213672 | -6.40831455767646 |
| ENSG00000204256.12 | BRD2 | 0.00419554387764443 | 8.06464476001521 | 0.0403896298122017 | 0.967971646426027 | 0.988843700213672 | -6.38761448042766 |
| ENSG00000135045.6 | C9orf40 | 0.00832772118160685 | 1.05398651250349 | 0.0403771180809842 | 0.967981562517793 | 0.988843700213672 | -5.91150609493406 |
| ENSG00000083814.13 | ZNF671 | 0.00357389882628034 | 4.47455612667319 | 0.0403619635835681 | 0.967993573123708 | 0.988843700213672 | -6.33990066672928 |
| lnc-EFNB2-6 | lnc-EFNB2-6 | -0.0048169396835646 | 4.47725324365851 | -0.0402796361963345 | 0.968058821331334 | 0.988861276400142 | -6.33817771775521 |
| lnc-SPPL2A-2 | lnc-SPPL2A-2 | -0.0077151984940440 | 1.0849289223882 | -0.0402008762704325 | 0.968121242370491 | 0.988865224765107 | -5.93386833574023 |
| ENSG00000148824.18 | MTG1 | 0.00550832535024526 | 3.52134055664308 | 0.0401535229447821 | 0.968158772261887 | 0.988865224765107 | -6.24609307785214 |
| ENSG00000136888.6 | ATP6V1G1 | 0.00633132449803409 | 6.36065410925583 | 0.0400039403008847 | 0.968277324507194 | 0.988887861736419 | -6.40999080405955 |
| ENSG00000184178.15 | SCFD2 | -0.0039120641802261 | 4.04394510975275 | -0.0399124093420764 | 0.968349868045582 | 0.988887861736419 | -6.29949884043631 |
| ENSG00000150625.16 | GPM6A | 0.0568271221098424 | 5.19276506281916 | 0.0399084952846411 | 0.968360394037358 | 0.988887861736419 | -6.3914378340147 |
| lnc-MNX1-23 | lnc-MNX1-23 | -0.0065499708351353 | 0.527214069426524 | -0.0398707294928969 | 0.96838290180824 | 0.988887861736419 | -5.80941335590628 |
| ENSG00000089060.11 | SLC8B1 | 0.00750635812628397 | 3.61143588449205 | 0.0397899608756168 | 0.968446915907373 | 0.988887861736419 | -6.258237158515 |
| ENSG00000172354.9 | GNB2 | 0.0071055763169127 | 4.71171287856337 | 0.0397618460299576 | 0.968469198701975 | 0.988887861736419 | -6.36178248295001 |
| ENSG00000235683.1 | | 0.0104751733038458 | 0.0822584192851231 | 0.0396775695521949 | 0.968535993297229 | 0.9889070067978 | -5.66271522750372 |
| lnc-ZNF732-9 | lnc-ZNF732-9 | 0.0313913253144565 | -0.740075883832034 | 0.0395336929476813 | 0.968653733640075 | 0.988978164658415 | -5.51568638526464 |
| ENSG00000136295.14 | TTYH3 | -0.0127677118076828 | 3.06489492362402 | -0.039435933390613 | 0.968729484623245 | 0.989006447200052 | -6.20917674364457 |
| ENSG00000100416.13 | TRMU | 0.00380105789306739 | 4.63338097642935 | 0.039350722199805 | 0.96879504326073 | 0.989008889779092 | -6.35522667523182 |
| lnc-BRD8-1 | lnc-BRD8-1 | -0.0048816960225157 | 1.06625368374089 | -0.0393091698660078 | 0.968827976709704 | 0.989008889779092 | -5.92537149694459 |
| ENSG00000125755.18 | SYMPK | -0.0050299635179582 | 6.11520348322622 | -0.0391465805435851 | 0.968956841893225 | 0.98905345992002 | -6.40951744609561 |
| ENSG00000123700.4 | KCNJ2 | -0.0337090035653562 | -1.97623378270937 | -0.0391328358348604 | 0.968967741305311 | 0.98905345992002 | -5.17800654513305 |
| ENSG00000259429.5 | UBE2Q2P2 | 0.015923179924497 | 0.0780922909227325 | 0.0389587429971797 | 0.969105719380508 | 0.989138879270857 | -5.65986536791502 |
| ENSG00000112592.13 | TBP | -0.0031595585039347 | 4.02886224413438 | -0.0389059808088518 | 0.969147538174968 | 0.989138879270857 | -6.30510634425103 |
| ENSG00000136866.13 | ZFP37 | 0.0433385277856655 | 1.02882682143891 | 0.0387535724718641 | 0.969275544730571 | 0.98917577688729 | -5.93497468280584 |
| ENSG00000099956.19 | SMARCB1 | -0.0046590617630499 | 6.74901691770054 | -0.0387289402656385 | 0.96928785941942 | 0.98917577688729 | -6.40872942975374 |
| lnc-FERMT2-4 | lnc-FERMT2-4 | 0.00714363808137727 | 0.187821582556284 | 0.0386311419175021 | 0.969365374210496 | 0.98917577688729 | -5.71915391157121 |
| ENSG00000161203.13 | AP2M1 | 0.00540428475762944 | 5.98542000135626 | 0.0385945785455996 | 0.969394354348918 | 0.98917577688729 | -6.40831308098274 |
| lnc-RNF219-3 | lnc-RNF219-3 | 0.0078589319839506 | 3.28868319649543 | 0.0384210235871111 | 0.969531914625973 | 0.98917577688729 | -6.22908318854655 |
| ENSG00000188690.13 | UROS | 0.00365621095247359 | 4.32817095642023 | 0.0383792194515428 | 0.969565048872893 | 0.98917577688729 | -6.32452354975164 |
| ENSG00000116017.10 | ARID3A | -0.0080318565310770 | 4.29657481188436 | -0.0383432151013426 | 0.969593586213439 | 0.98917577688729 | -6.33668379255104 |
| ENSG00000178623.11 | GPR35 | -0.0072771186000695 | 0.649169195892323 | -0.0382394293279001 | 0.969675847886356 | 0.98917577688729 | -5.84512290310409 |
| ENSG00000240682.9 | ISY1 | 0.00328113271424524 | 4.27350657754606 | 0.0382279758491335 | 0.969684926052075 | 0.98917577688729 | -6.32445120204559 |
| ENSG00000170584.10 | NUDCD2 | 0.00633340132228044 | 3.05437321969093 | 0.0382092212244585 | 0.969699791203079 | 0.98917577688729 | -6.19594792801937 |
| SEMA6A-AS1 | SEMA6A-AS1 | -0.0056893414246994 | 1.42396038427969 | -0.0381934054784824 | 0.9697123269704 | 0.98917577688729 | -5.97126250297674 |
| ENSG00000104814.12 | MAP4K1 | 0.00651493244023005 | 5.65100833940506 | 0.0380450717083076 | 0.969829898635403 | 0.989246682373383 | -6.40275905914055 |
| lnc-B9D2-2 | lnc-B9D2-2 | 0.00694494044198546 | 2.42825892198111 | 0.0377476882352632 | 0.970065611467663 | 0.989438081310154 | -6.10601044644587 |
| ENSG00000185798.7 | WDR53 | -0.0038286763452526 | 2.99580299883264 | -0.0376165746050852 | 0.970169535942658 | 0.989441853325019 | -6.17653368716276 |
| ENSG00000198015.12 | MRPL42 | 0.00368287609142381 | 5.37020244597659 | 0.0375959492487538 | 0.970185884247503 | 0.989441853325019 | -6.39686346161871 |
| lnc-GPAT2-4 | lnc-GPAT2-4 | -0.0076800483346311 | 2.55833648632778 | -0.037561080788205 | 0.970213522113339 | 0.989441853325019 | -6.13708434613163 |
| ENSG00000083457.11 | ITGAE | 0.00655483440283477 | 2.70181794098265 | 0.0374746716900967 | 0.970282012909536 | 0.989462676957837 | -6.14587010168612 |
| ENSG00000143079.14 | CTTNBP2NL | -0.0326689357434225 | -0.120396290802802 | -0.0370619916472595 | 0.970614086058141 | 0.989743303355559 | -5.641222020632 |
| ENSG00000105486.13 | LIG1 | 0.0060989783989287 | 5.20463842681605 | 0.0369500692416693 | 0.970697835543398 | 0.989743303355559 | -6.39114480995837 |
| ENSG00000134954.14 | ETS1 | 0.00568746585851088 | 9.02034136827911 | 0.0368566615229916 | 0.970771875417565 | 0.989743303355559 | -6.36464012332262 |
| ENSG00000125434.10 | SLC25A35 | -0.0087621605518502 | 1.53872259496826 | -0.036828910387153 | 0.970793872478759 | 0.989743303355559 | -5.97090836005627 |
| ENSG00000251079.6 | BMS1P2 | 0.00685323037401941 | 2.16133200731766 | 0.0368241740592382 | 0.970797626752436 | 0.989743303355559 | -6.05970742749753 |
| lnc-PPT1-1 | lnc-PPT1-1 | -0.0164524173268649 | -0.296469802987341 | -0.0367531440398879 | 0.970853929127867 | 0.989751680200043 | -5.54978535692161 |
| ENSG00000214140.10 | PRCD | 0.0193397832136802 | -0.0745532449180853 | 0.0365304820913713 | 0.971033765531951 | 0.98988598860602 | -5.86743556569891 |

| | | | | | | | |
|---|---|---|---|---|---|---|---|
| ENSG00000129116.18 | PALLD | 0.0329511252099712 | -2.11847056483806 | 0.036424372519353 | 0.971114577227398 | 0.989886919360703 | -5.14921407481935 |
| lnc-U2AF2-2 | lnc-U2AF2-2 | 0.0081254867805714 | 0.449658874991628 | 0.036356724979402 | 0.97116815585381 | 0.989886919360703 | -5.74070462287094 |
| ENSG00000115325.13 | DOK1 | 0.00620605146131181 | 2.53921740336475 | 0.0363431000505457 | 0.971178955913619 | 0.989886919360703 | -6.14073311479145 |
| ENSG00000183814.15 | LIN9 | 0.0051560055198272 | 2.37842714491127 | 0.0362461676995491 | 0.97125579134889 | 0.989916214574608 | -6.10792099472767 |
| ENSG00000161249.20 | DMKN | 0.0297533330515274 | 0.537335620147507 | 0.0360910219314715 | 0.971385177607458 | 0.989989046288037 | -5.94707561224836 |
| lnc-SATB1-8 | lnc-SATB1-8 | -0.00358764030344405 | 3.7967467675342 | -0.0360300314828293 | 0.971427117190584 | 0.989989046288037 | -6.27196882901746 |
| lnc-CNNM1-1 | lnc-CNNM1-1 | -0.00906673777576972 | -0.123651464120577 | -0.0359739871333307 | 0.971471542385744 | 0.989989046288037 | -5.62764666541629 |
| lnc-C18orf25-1 | lnc-C18orf25-1 | -0.0100049912244214 | 0.997254711617563 | -0.0357841717792674 | 0.971622005765985 | 0.990048336294975 | -5.90553327806457 |
| ENSG00000162714.12 | ZNF496 | 0.00392484726691134 | 5.29106548890256 | 0.0357278880099785 | 0.971666621147258 | 0.990048336294975 | -6.39180748048641 |
| lnc-CECR2-1 | lnc-CECR2-1 | -0.00953115810215149 | -0.459557572818417 | -0.0357185487419723 | 0.971674024267346 | 0.990048336294975 | -5.56049430789615 |
| lnc-CAPN1-1 | lnc-CAPN1-1 | 0.00610236312326151 | -0.130426073991623 | 0.0352230462331121 | 0.972066806471694 | 0.990359674460655 | -5.65429344112057 |
| ENSG00000117632.22 | STMN1 | 0.00771854834598162 | 4.87523020011093 | 0.0352116880518297 | 0.972075815144955 | 0.990359674460655 | -6.37323702365182 |
| ENSG00000179583.19 | CIITA | 0.00605073338212314 | 8.6899285607059 | 0.0350939612604939 | 0.972169132599204 | 0.990394644673965 | -6.3758591286922 |
| ENSG00000276023.4 | DUSP14 | 0.00977412498435148 | 1.21166249262795 | 0.0350189501984051 | 0.972228594359137 | 0.990394644673965 | -5.96731399086426 |
| ENSG00000146576.12 | INTS15 | 0.00521460971909082 | 3.85480504889557 | 0.0349472793737411 | 0.972285408440349 | 0.990394644673965 | -6.28849258084588 |
| lnc-NYAP2-10 | lnc-NYAP2-10 | 0.0242581783414937 | -0.375974755559063 | 0.0349277911926156 | 0.97230260821682 | 0.990394644673965 | -5.51367237854361 |
| lnc-E2F6-4 | lnc-E2F6-4 | 0.00654309197127056 | 0.79708011286067 | 0.034604403714868 | 0.972557210943956 | 0.990603859184758 | -5.88028202628245 |
| ENSG00000280387.1 | | -0.00532334070316953 | 0.776856511341802 | -0.0345450580274486 | 0.972604255466393 | 0.990603859184758 | -5.85911444763052 |
| ENSG00000172046.18 | USP19 | -0.00290952668959932 | 5.81551242865691 | -0.0344140903404319 | 0.972708076540399 | 0.990660580872392 | -6.4065796520695 |
| ENSG00000148180.19 | GSN | -0.00902149770445915 | 2.5039452339727 | -0.0342329740357435 | 0.972851652356701 | 0.990740408099505 | -6.10935134292161 |
| ATP6V0E2-AS1 | ATP6V0E2-AS1 | -0.00958099977910876 | 1.37558360360651 | -0.0341310990238866 | 0.972932411847592 | 0.990740408099505 | -5.97304807141061 |
| lnc-AKR7A2-2 | lnc-AKR7A2-2 | 0.00806421099079868 | 0.975106808115286 | 0.0340820167649819 | 0.972971320983656 | 0.990740408099505 | -5.89574037751725 |
| ENSG00000013725.14 | CD6 | 0.0105079374484269 | 4.85629681719235 | 0.0339539391796146 | 0.973075721759892 | 0.990740408099505 | -6.35572365693019 |
| ENSG00000111877.17 | MCM9 | -0.0044378165320591 | 5.6710831790225 | -0.0338936281376766 | 0.973120663520349 | 0.990740408099505 | -6.40357602026648 |
| ENSG00000111481.9 | COPZ1 | 0.00517891681758265 | 5.54464777058118 | 0.0338585224886346 | 0.973148493156933 | 0.990740408099505 | -6.40016181672659 |
| ENSG00000137776.16 | SLTM | 0.00317575220135335 | 7.24937085168777 | 0.033857642702178 | 0.973149190598892 | 0.990740408099505 | -6.40317262531611 |
| ENSG00000164967.9 | RPP25L | -0.00501007342348991 | 1.96790479419802 | -0.033753245584326 | 0.973231950526026 | 0.990740408099505 | -6.05403321906936 |
| ENSG00000134287.9 | ARF3 | -0.00518741148376911 | 5.1012034174061 | -0.0337239369348198 | 0.97325518476231 | 0.990740408099505 | -6.38700262708866 |
| ENSG00000121807.5 | CCR2 | 0.0312502458505457 | -0.304588983531398 | 0.0336514414338624 | 0.973316509122343 | 0.990740408099505 | -5.47304051016668 |
| ENSG00000182718.16 | ANXA2 | 0.00799517244183079 | 6.08437293399898 | 0.0336282678060584 | 0.973331847444119 | 0.990740408099505 | -6.41009077984613 |
| ENSG00000125944.19 | HNRNPR | 0.00378034497070612 | 6.90661277699346 | 0.033586592247713 | 0.97336406415917 | 0.990740408099505 | -6.40707002768421 |
| ENSG00000149218.4 | ENDOD1 | -0.00782709063420217 | 5.20891895115485 | -0.0335171175250862 | 0.973419631929388 | 0.990747974286831 | -6.38864891861175 |
| ENSG00000136122.15 | BORA | -0.00375093259246739 | 3.31051182299456 | -0.0332292289364568 | 0.973647364756644 | 0.990930760917029 | -6.21986779328673 |
| ENSG00000136732.15 | GYPC | -0.0123888017607241 | 3.75981611811277 | -0.0331122927042442 | 0.973743859097551 | 0.990945625292115 | -6.30156974023394 |
| ENSG00000159692.15 | CTBP1 | -0.00545313216574961 | 6.31447100179466 | -0.0330893467223004 | 0.973758257647491 | 0.990945625292115 | -6.41015966914619 |
| lnc-MAGOH-3 | lnc-MAGOH-3 | 0.00352969308737139 | 2.41622710490867 | 0.0329199568396511 | 0.973892543712559 | 0.991033283593 | -6.10532769972169 |
| ENSG00000158552.12 | ZFAND2B | 0.00483253496795122 | 4.36070614723812 | 0.0328541863888439 | 0.973944684312253 | 0.991037346114253 | -6.33742641471064 |
| ENSG00000187486.5 | KCNJ11 | 0.00901767597288261 | 2.3672726623559 | 0.0327752729929795 | 0.974007244348585 | 0.991052010006769 | -6.09897135095659 |
| ENSG00000144535.19 | DIS3L2 | -0.00296023919893162 | 5.2018506496344 | -0.0326992212553423 | 0.974067535909789 | 0.991064364391062 | -6.38775045118133 |
| ENSG00000126249.7 | PDCD2L | 0.00775002596814123 | 0.812315246921715 | 0.0325884396501198 | 0.974155360554089 | 0.991065481421349 | -5.88130346185245 |
| ENSG00000131061.13 | ZNF341 | 0.00762332995934489 | 2.26910912014365 | 0.0325763647793044 | 0.97416493320297 | 0.991065481421349 | -6.08559301710897 |
| ENSG00000270956.1 | | -0.0077201001963646 | 0.73510440660638 | -0.0322360999870535 | 0.974434688010382 | 0.991290919948485 | -5.84816266472928 |
| ENSG00000160867.14 | FGFR4 | -0.0088811512009252 | -0.498103719484525 | -0.0320888250437329 | 0.974551445444723 | 0.991311888187055 | -5.53666296580108 |
| ENSG00000197345.12 | MRPL21 | 0.0031359960340307 | 3.23965037404094 | 0.0320576229435226 | 0.974576182088817 | 0.991311888187055 | -6.21343779478343 |
| ENSG00000106246.17 | PTCD1 | 0.00392208698731356 | 3.47001178916581 | 0.0320278512564873 | 0.974599784742421 | 0.991311888187055 | -6.23805553113701 |
| ENSG00000147799.11 | ARHGAP39 | 0.037062309845679 | -1.98323348846418 | 0.0319086061782033 | 0.974695291020727 | 0.991355700915409 | -5.07482884527622 |
| ENSG00000177606.6 | JUN | -0.00914999244347481 | 7.28476701419127 | -0.0318527410916067 | 0.974741011675635 | 0.991355700915409 | -6.40142250645147 |
| ENSG00000169679.14 | BUB1 | 0.0148752737755316 | -0.0446604704865423 | 0.0317912626817128 | 0.974787350280667 | 0.991355700915409 | -5.65514175254186 |
| ENSG00000142632.16 | ARHGEF19 | -0.00527461956840503 | 2.03144968246044 | -0.0314867639455983 | 0.975028756577689 | 0.991516748821335 | -6.06118323094173 |
| lnc-PRSS55-1 | lnc-PRSS55-1 | -0.00678128097925246 | 1.8946314960488 | -0.0314699964630197 | 0.975042049890329 | 0.991516748821335 | -6.04276919457814 |
| ENSG00000124406.16 | ATP8A1 | 0.00562705670346553 | 6.61160344013725 | 0.0312935370580868 | 0.975181947893168 | 0.991610020427086 | -6.40926443456343 |
| ENSG00000163812.13 | ZDHHC3 | -0.0030126520240108 | 4.88581455837306 | -0.0310180935881942 | 0.975400322594853 | 0.991694266694331 | -6.37323277714552 |
| lnc-NAA35-1 | lnc-NAA35-1 | -0.00638912227147928 | 0.0572194199242526 | -0.0309578778067424 | 0.975448062603143 | 0.991694266694331 | -5.68500394347746 |
| LLPH-DT | LLPH-DT | 0.00813626729077799 | -0.631991767268895 | 0.0309396185514444 | 0.975462538842802 | 0.991694266694331 | -5.49879287835909 |
| ENSG00000226942.2 | IL9RP3 | 0.00914898031270006 | 2.35025127245572 | 0.0308303703140438 | 0.975549152827373 | 0.991694266694331 | -6.07463272188279 |
| ENSG00000119537.17 | KDSR | 0.00404111112609806 | 3.06363572646764 | 0.0308005918342986 | 0.975572761798182 | 0.991694266694331 | -6.19561044941841 |
| lnc-TMEM50B-30 | lnc-TMEM50B-30 | -0.00543093973113029 | 1.9094893440736 | -0.030764570941634 | 0.975601319907683 | 0.991694266694331 | -6.05700476868877 |
| ENSG00000164885.12 | CDK5 | 0.006283636031267 | 1.26235216700077 | 0.0307636371238959 | 0.975602060258151 | 0.991694266694331 | -5.94437892335641 |
| ENSG00000120306.10 | CYSTM1 | -0.010344510508471 | 1.56562792815712 | -0.0306019357268435 | 0.975730657117836 | 0.991760444327916 | -6.02166388923867 |
| ENSG00000172336.4 | POP7 | 0.00623659884219219 | 1.74333576118062 | 0.030559971775295 | 0.975763530976273 | 0.991760444327916 | -6.04102961398479 |
| ENSG00000115425.13 | PECR | -0.0100093579588403 | 0.680244381798995 | -0.0304175998472245 | 0.975876407428341 | 0.991784020663543 | -5.85150318326475 |
| ENSG00000139718.10 | SETD1B | -0.00582309404461245 | 5.00809865513074 | -0.0303493350269968 | 0.975930529867353 | 0.991784020663543 | -6.37774172329258 |
| ENSG00000103249.17 | CLCN7 | -0.00610712732519225 | 5.25368138337079 | -0.030306537804155 | 0.975964460874159 | 0.991784020663543 | -6.39410999509537 |
| lnc-HLA-DMB-1 | lnc-HLA-DMB-1 | -0.0051564206239183 | 7.08682898432037 | -0.030287612458854 | 0.975979465507461 | 0.991784020663543 | -6.40675431035596 |
| ENSG00000201785.1 | SNORD117 | 0.00680312269342161 | -0.0185996993006695 | 0.0301899904215572 | 0.976056863594119 | 0.991813705734486 | -5.67807740442653 |
| ENSG00000117640.17 | MTFR1L | -0.00384952944637185 | 4.28972553758901 | -0.0300502560636918 | 0.97616765017347 | 0.991824641047819 | -6.32343115191769 |
| ENSG00000069399.14 | BCL3 | 0.00587602858895808 | 3.44094074082303 | 0.0300280482219072 | 0.976185257417664 | 0.991824641047819 | -6.23562391086522 |
| ENSG00000011114.14 | BTBD7 | -0.00274046564540686 | 4.7718545083673 | -0.0299940848128842 | 0.976212184957866 | 0.991824641047819 | -6.36729809270837 |

| | | | | | | | |
|---|---|---|---|---|---|---|---|
| LINC02363 | LINC02363 | -0.0288560127794852 | 3.77037044928646 | -0.0298497014456352 | 0.976332208923905 | 0.991897623705861 | -6.30709190414014 |
| ENSG00000161800.12 | RACGAP1 | 0.00663419610077343 | 2.45528041429448 | 0.0297138953952138 | 0.976434331329745 | 0.99195241309709 | -6.11293401890327 |
| ENSG00000083444.16 | PLOD1 | -0.0064734631081897 | 1.87579094511197 | -0.0295765507656835 | 0.976543224825535 | 0.991955457030039 | -6.02654495725941 |
| ENSG00000110721.11 | CHKA | 0.00474928863572854 | 3.77411544446165 | 0.0295060945522335 | 0.976599086110816 | 0.991955457030039 | -6.27622569334108 |
| lnc-AMN1-1 | lnc-AMN1-1 | 0.00436419819724643 | 1.50743019017547 | 0.0294974092220498 | 0.976605972292511 | 0.991955457030039 | -5.97622153246009 |
| ENSG00000116922.14 | AIRIM | -0.0039937014909416 | 2.92485860783623 | -0.029382985845807 | 0.9766966932674 | 0.991955457030039 | -6.16750770559498 |
| lnc-DNMT3A-3 | lnc-DNMT3A-3 | -0.0055837560792564 | -0.0085915656784144 | -0.0292374480242038 | 0.976812083890247 | 0.991955457030039 | -5.67707169583927 |
| ENSG00000085365.17 | SCAMP1 | -0.0042788638622379 | 4.59307461139053 | -0.0291123478615289 | 0.976911270782001 | 0.991955457030039 | -6.35716277009092 |
| ENSG00000056972.18 | TRAF3IP2 | -0.0050624937320038 | 3.97701116411985 | -0.0290963418177352 | 0.976923961357624 | 0.991955457030039 | -6.29709479720763 |
| ENSG00000141384.12 | TAF4B | 0.00753764656323508 | 4.22376390929261 | 0.0290484081092742 | 0.976962675664543 | 0.991955457030039 | -6.32539489496486 |
| ENSG00000129911.8 | KLF16 | -0.0068434360706817 | 2.0865603623433 | -0.0289971177199234 | 0.977002632457843 | 0.991955457030039 | -6.07017558423819 |
| ENSG00000240563.1 | L1TD1 | 0.0528229792670633 | 0.180083212624324 | 0.0289293573866028 | 0.977061736734714 | 0.991955457030039 | -5.67342456129024 |
| ENSG00000135773.12 | CAPN9 | 0.0281846100166927 | -1.95890830594437 | 0.0288930884169876 | 0.977085113675416 | 0.991955457030039 | -5.08200838071792 |
| ENSG00000275221.1 | H2AC15 | 0.0102214029297563 | 3.42669020409817 | 0.0288848770579463 | 0.977094389734768 | 0.991955457030039 | -6.26616834788463 |
| ENSG00000100038.19 | TOP3B | -0.0027074259669257 | 4.88766784753606 | -0.0288046401543453 | 0.977155241427458 | 0.991955457030039 | -6.36937797057651 |
| ENSG00000160404.17 | TOR2A | -0.0055631760737822 | 1.82088597455774 | -0.0287346447163122 | 0.977210738664405 | 0.991955457030039 | -6.05125490253595 |
| lnc-ZSCAN21-2 | lnc-ZSCAN21-2 | 0.00474614175063781 | 2.13601112100505 | 0.0287011002967503 | 0.977237335046808 | 0.991955457030039 | -6.07299756971494 |
| lnc-CLNS1A-3 | lnc-CLNS1A-3 | -0.0037287670341128 | 3.35409094065223 | -0.0287008994503956 | 0.977237494292046 | 0.991955457030039 | -6.21884384555457 |
| ENSG00000277147.5 | LINC00869 | 0.00488113692923076 | 2.58022147777794 | 0.0285957614636182 | 0.977320855275647 | 0.991955457030039 | -6.11930840936249 |
| ENSG00000186591.12 | UBE2H | -0.0034919429699868 | 5.76235379824619 | -0.0285682713450505 | 0.977342651467919 | 0.991955457030039 | -6.4045960557224 |
| lnc-HEATR4-6 | lnc-HEATR4-6 | 0.00481481592830058 | 2.48718292516535 | 0.028537377944272 | 0.977367146054087 | 0.991955457030039 | -6.11063109735398 |
| lnc-YAF2-1 | lnc-YAF2-1 | 0.00413699514466481 | 1.42710713195 | 0.0284818128926529 | 0.977411202215874 | 0.991955457030039 | -5.98237862017528 |
| lnc-MIEN1-1 | lnc-MIEN1-1 | -0.0046760347499104 | 7.68058779250186 | -0.028382776313961 | 0.977489726058355 | 0.991955457030039 | -6.39684881273691 |
| lnc-HLCS-5 | lnc-HLCS-5 | -0.0046982150301714 | 0.0859925216740939 | -0.0283728801561646 | 0.977497572508395 | 0.991955457030039 | -5.66835852918538 |
| ENSG00000154229.11 | PRKCA | 0.0360772931978568 | -0.472640684807466 | 0.0282283516341551 | 0.977617192296222 | 0.991991542607311 | -5.58074086282587 |
| ENSG00000106344.8 | RBM28 | 0.00217177866779217 | 4.88979351011235 | 0.0282064628513265 | 0.977629521536671 | 0.991991542607311 | -6.37144209453136 |
| ENSG00000274998.1 | SNORA17A | 0.00724466218054837 | 0.099391387921485 | 0.0281223344683151 | 0.977696225528693 | 0.99201032288559 | -5.75176047159839 |
| ENSG00000076604.14 | TRAF4 | -0.0068861036907730 | 5.60891884975615 | -0.0277791955259463 | 0.977969231599209 | 0.992238413467073 | -6.40362494201262 |
| ENSG00000283208.1 | | 0.00303849089443942 | 2.18794551867159 | 0.0277052588624303 | 0.97802692019768 | 0.992248033242746 | -6.07198740536874 |
| DLGAP4-AS1 | DLGAP4-AS1 | 0.00433334050160106 | 2.19115181408083 | 0.0274213453172888 | 0.978252034371589 | 0.9923857111748 | -6.0690917215282 |
| ANKRD10-IT1 | ANKRD10-IT1 | 0.0059161483591601 | 3.75244310536858 | 0.0272683270723733 | 0.978373362810938 | 0.9923857111748 | -6.2852105508205 |
| ENSG00000185271.8 | KLHL33 | -0.018491397536895 | -0.980581840931444 | -0.0272632097494381 | 0.978377420354392 | 0.9923857111748 | -5.31535777117427 |
| lnc-ACTR3-2 | lnc-ACTR3-2 | -0.0068205544977890 | 1.53366023027607 | -0.0272385498042983 | 0.97839697332023 | 0.9923857111748 | -5.99998203210463 |
| ENSG00000104388.14 | RAB2A | 0.00311029089762016 | 5.1692760297724 | 0.0272300735744111 | 0.978403694158782 | 0.9923857111748 | -6.39072941588195 |
| ENSG00000088812.17 | ATRN | -0.0024816377111971 | 5.0731534774651 | -0.0268679920608975 | 0.978690791599394 | 0.992627996624142 | -6.38233866023747 |
| lnc-MYLK4-30 | lnc-MYLK4-30 | 0.0083457865015976 | 0.149761348571546 | 0.0266362505639947 | 0.978874542860582 | 0.992718895922431 | -5.71777799574579 |
| ENSG00000136682.14 | ZNG1B | -0.0025122122528458 | 4.33038711533662 | -0.0264494503698333 | 0.979022660343502 | 0.992718895922431 | -6.33202868384114 |
| ENSG00000200084.1 | SNORD68 | 0.00525090495727842 | 0.682937214581918 | 0.0263801490237082 | 0.979077610910512 | 0.992718895922431 | -5.85963393943938 |
| lnc-ZNF644-1 | lnc-ZNF644-1 | -0.0102292668027555 | 0.514076095829947 | -0.0263438621884401 | 0.979106383584175 | 0.992718895922431 | -5.80418206551443 |
| lnc-METTL26-7 | lnc-METTL26-7 | 0.00499299517865093 | 1.29683507435529 | 0.0262888158634929 | 0.979150031146221 | 0.992718895922431 | -5.96568177698039 |
| lnc-REL-1 | lnc-REL-1 | 0.00374547886452081 | 4.74773166493464 | 0.026229596247578 | 0.979196987883765 | 0.992718895922431 | -6.35767888779846 |
| ENSG00000258830.1 | | -0.0037108629414559 | 4.02294109859119 | -0.0262158059183553 | 0.979207922596781 | 0.992718895922431 | -6.29469044632299 |
| ENSG00000135164.18 | DMTF1 | -0.0026303956868591 | 7.77013656940337 | -0.0262037025793434 | 0.979217519654434 | 0.992718895922431 | -6.39459912353889 |
| ENSG00000007541.16 | PIGQ | -0.0039641650222201 | 3.53952935142874 | -0.026199004590834 | 0.979221244814631 | 0.992718895922431 | -6.25172749414793 |
| ENSG00000182240.15 | BACE2 | 0.0120388452196076 | 5.18043085397119 | 0.0261520832277741 | 0.979262714998253 | 0.992718895922431 | -6.38100174869954 |
| ZFAS1 | ZFAS1 | 0.00326032828821265 | 6.10947401449176 | 0.0259354343763996 | 0.979430237405784 | 0.992839821547563 | -6.40976998722124 |
| lnc-OSTC-2 | lnc-OSTC-2 | 0.00503497202312267 | 0.182835619768283 | 0.0257557009761179 | 0.97957275415554 | 0.992863655973034 | -5.72477404448235 |
| ENSG00000162139.9 | NEU3 | 0.00255351007646265 | 4.73752959245657 | 0.0257010235064469 | 0.979616109922278 | 0.992863655973034 | -6.36001901518036 |
| ENSG00000167774.2 | | 0.00369509558105596 | 2.2320702633478 | 0.025699973107858 | 0.979616942822372 | 0.992863655973034 | -6.09304339011834 |
| lnc-CD72-2 | lnc-CD72-2 | 0.011807173695054 | 3.21854047098279 | 0.0256667031673159 | 0.979646698205138 | 0.992863655973034 | -6.20250504777188 |
| ENSG00000172465.13 | TCEAL1 | 0.00484047311972313 | 2.32584260529928 | 0.0255632219238328 | 0.979725378116595 | 0.99289450801447 | -6.10317715251142 |
| lnc-HPGD-5 | lnc-HPGD-5 | -0.0055542362671994 | 1.46769435400896 | -0.0254571752337798 | 0.979809466899147 | 0.992930838323329 | -5.98349804326038 |
| ENSG00000126858.17 | RHOT1 | -0.0028472787797113 | 4.52157554267359 | -0.0253650069942744 | 0.979882551076159 | 0.992956013627441 | -6.34893745033557 |
| ENSG00000134215.15 | VAV3 | -0.0038758628120634 | 6.23601519780765 | -0.025283810642241 | 0.97994693531473 | 0.992972370874961 | -6.41052300470252 |
| lnc-RFTN1-2 | lnc-RFTN1-2 | 0.0168337989982858 | 0.373675855030365 | 0.0251458091943994 | 0.980058855033553 | 0.99303689146183 | -5.68117909292575 |
| ENSG00000143643.12 | TTC13 | -0.0039633659699234 | 4.57647975819088 | -0.0249920844818048 | 0.980178259218092 | 0.993108988899138 | -6.3479276898207 |
| lnc-SCPEP1-3 | lnc-SCPEP1-3 | 0.0100755274693032 | -0.289712118474681 | 0.0248859314899328 | 0.980262433533026 | 0.993145386365932 | -5.65096849390101 |
| ENSG00000163512.13 | AZI2 | 0.00229557084386522 | 3.65097249042552 | 0.0246671757114757 | 0.980435897262531 | 0.993174799432119 | -6.26667481982807 |
| ENSG00000131797.12 | CLUHP3 | 0.00221875976713903 | 5.6753649826523 | 0.0246667492229678 | 0.980436235450134 | 0.993174799432119 | -6.40381796907602 |
| lnc-SCCPDH-2 | lnc-SCCPDH-2 | -0.0064792494833133 | -0.0590868115853332 | -0.0246501580286445 | 0.980449391578256 | 0.993174799432119 | -5.65261935684691 |
| ENSG00000147174.11 | GCNA | 0.0047502720594288 | 2.40563076863976 | 0.0246059158552665 | 0.98048447381143 | 0.993174799432119 | -6.11824908486997 |
| lnc-ZNF124-2 | lnc-ZNF124-2 | -0.0025772395806997 | 5.44399039519908 | -0.0244997491073977 | 0.980568659857638 | 0.993176352355564 | -6.39980927745189 |
| lnc-ZBTB41-1 | lnc-ZBTB41-1 | 0.00583521690531095 | 0.00228567601860915 | 0.0244817455965199 | 0.980582935954359 | 0.993176352355564 | -5.67043560433254 |
| lnc-ASB3-2 | lnc-ASB3-2 | 0.00512866624333408 | 1.65507399009902 | 0.0244039712667842 | 0.980644608102225 | 0.993176352355564 | -6.01797312220242 |
| ENSG00000173349.5 | SFT2D3 | 0.00362602351570418 | 3.34032042355651 | 0.0243605798291992 | 0.980679015948816 | 0.993176352355564 | -6.23215458956627 |
| ENSG00000117505.12 | DR1 | -0.0025854644553415 | 6.21816345723571 | -0.0242889280819206 | 0.980735833284218 | 0.993185026149523 | -6.41051668687902 |
| lnc-WDR77-1 | lnc-WDR77-1 | -0.0021042319611557 | 3.26158691515511 | -0.024065294584804 | 0.980913167493577 | 0.993229730682636 | -6.21173558009548 |
| ENSG00000123908.11 | AGO2 | -0.0038596578351581 | 5.03839967749622 | -0.0240612135125646 | 0.980916403662416 | 0.993229730682636 | -6.38248761475474 |

| | | | | | | | |
|---|---|---|---|---|---|---|---|
| ENSG00000130595.18 | TNNT3 | 0.00558977559196516 | 1.45703258791843 | 0.0240220984740071 | 0.980947420740473 | 0.993229730682636 | -5.97316677542664 |
| lnc-M6PR-4 | lnc-M6PR-4 | 0.00313648267109001 | 1.67256815285646 | 0.0239898449100166 | 0.980972996892936 | 0.993229730682636 | -6.01597349606433 |
| ENSG00000185608.8 | MRPL40 | -0.00314804416799674 | 3.63130469461355 | -0.0239218700437839 | 0.981026899081269 | 0.993235448292659 | -6.25696435932502 |
| ENSG00000213152.5 | RPL7AP60 | -0.00696679900975311 | -0.214708961540539 | -0.0234798185526304 | 0.981377435861262 | 0.99348602239493 | -5.57715473474167 |
| ENSG00000034693.14 | PEX3 | -0.00208101918183967 | 3.73515573982236 | -0.0234367257432265 | 0.981411607679497 | 0.99348602239493 | -6.27157990521318 |
| ENSG00000076770.14 | MBNL3 | 0.00284199482618952 | 6.48995306773296 | 0.0233862393997855 | 0.981451642481392 | 0.99348602239493 | -6.41026180163066 |
| lnc-SYTL3-3 | lnc-SYTL3-3 | -0.00708171057126794 | 2.6043664314873 | -0.0233665813952216 | 0.981467462438833 | 0.99348602239493 | -6.11411982845363 |
| ENSG00000166839.16 | ANKDD1A | 0.00874865403519198 | 1.15476344697731 | 0.0230358648607498 | 0.981729947874445 | 0.993638572034538 | -5.94875767072811 |
| ENSG00000254470.2 | AP5B1 | -0.00304937675877959 | 4.50206810506635 | -0.0229724807069523 | 0.981779747820983 | 0.993638572034538 | -6.34819893326508 |
| ENSG00000244687.11 | UBE2V1 | 0.00169565956342588 | 4.91238982416347 | 0.0229141479855191 | 0.981826005179388 | 0.993638572034538 | -6.37400188592824 |
| ENSG00000118242.15 | MREG | -0.00913415216595348 | -0.186608817974415 | -0.0228260291428862 | 0.98189588280181 | 0.993638572034538 | -5.64746142783896 |
| lnc-SIPA1L1-2 | lnc-SIPA1L1-2 | -0.00735006647234382 | 5.05730372324143 | -0.0225873771083834 | 0.982087447792616 | 0.993638572034538 | -6.37356538070921 |
| ENSG00000111046.3 | MYF6 | -0.0515342626323401 | -2.64437836769358 | -0.0225770791052384 | 0.982095774246514 | 0.993638572034538 | -4.87610341335138 |
| ENSG00000251807.1 | RNU6-202P | -0.0074995982898084 | -0.592340562072718 | -0.0225725897373051 | 0.982096859267798 | 0.993638572034538 | -5.50862364433796 |
| ENSG00000263482.3 | ANTXRLP1 | 0.0137215853950559 | 1.53350292222468 | 0.0225740622217173 | 0.982098659179497 | 0.993638572034538 | -5.95443813342896 |
| ENSG00000227124.9 | ZNF717 | 0.00332706837234653 | 4.28157370677179 | 0.0225630768481378 | 0.982104402975021 | 0.993638572034538 | -6.3056558124801 |
| ENSG00000162227.7 | TAF6L | 0.00311033955504492 | 2.79786748911004 | 0.0225535627709738 | 0.982111947625981 | 0.993638572034538 | -6.16380854952144 |
| CYTOR | CYTOR | -0.0116301728739177 | 1.56403612516341 | -0.022509919590443 | 0.982149188555734 | 0.993638572034538 | -6.0157781615543 |
| lnc-CD83-3 | lnc-CD83-3 | -0.00559505262577825 | -0.355480176512147 | -0.0224031518073987 | 0.982231223546444 | 0.993672725560624 | -5.59373747390983 |
| lnc-PROM1-3 | lnc-PROM1-3 | 0.00808638259705285 | 0.43851936167201 | 0.0223334792619452 | 0.982286474026771 | 0.993679780552073 | -5.83390645269471 |
| ENSG00000127511.9 | SIN3B | -0.0030125424849361 | 5.43057145690894 | -0.022245629455225 | 0.982356139240273 | 0.993683600539255 | -6.39712111288384 |
| ENSG00000240695.1 | DNAJC8P2 | -0.00593089230815158 | 0.807869176827642 | -0.0222069603414177 | 0.982386804031157 | 0.993683600539255 | -5.86059772532716 |
| lnc-PPP2R2A-2 | lnc-PPP2R2A-2 | -0.00991835819434439 | -1.1674591783365 | -0.021990035333769 | 0.982558827090716 | 0.993756903461892 | -5.3971373524144 |
| lnc-ICAM2-1 | lnc-ICAM2-1 | 0.00361840506095068 | 1.00134438197991 | 0.0219584855827771 | 0.982583846335714 | 0.993756903461892 | -5.92132584238433 |
| ENSG00000275895.6 | | 0.0120520345149178 | 2.73224481600265 | 0.0217991954536345 | 0.982713604134082 | 0.993756903461892 | -6.17686551680243 |
| ENSG00000092964.17 | DPYSL2 | 0.00784872301245709 | 4.50617885387706 | 0.0217046047042831 | 0.982787691959278 | 0.993756903461892 | -6.35320058096938 |
| ENSG00000111358.13 | GTF2H3 | 0.00399989628389896 | 4.00990455183945 | 0.0216447694700106 | 0.98283262708235 | 0.993756903461892 | -6.30152315794059 |
| ENSG00000121064.12 | SCPEP1 | -0.00589077096655366 | 3.02785174307455 | -0.0215984186181694 | 0.982869409753856 | 0.993756903461892 | -6.17268370601776 |
| lnc-METTL6-5 | lnc-METTL6-5 | -0.00504577258831462 | 3.72755479103587 | -0.0215617911036377 | 0.982898430240472 | 0.993756903461892 | -6.25764755711885 |
| lnc-FCHO2-3 | lnc-FCHO2-3 | 0.00387520848615219 | 3.00278403927962 | 0.0215387243281269 | 0.982916722580035 | 0.993756903461892 | -6.18582246170735 |
| lnc-S1PR2-3 | lnc-S1PR2-3 | 0.00313953336144057 | 0.692296294701086 | 0.0215072536171063 | 0.982941679396302 | 0.993756903461892 | -5.85401610173602 |
| lnc-GATM-5 | lnc-GATM-5 | 0.00813734522683728 | -0.105220565057168 | 0.0214741788883294 | 0.982967908244929 | 0.993756903461892 | -5.61155581607982 |
| ENSG00000270629.5 | NBPF14 | 0.00390612333030571 | 5.25735399243027 | 0.0214339599311442 | 0.982999802619534 | 0.993756903461892 | -6.39256430554797 |
| ENSG00000180881.19 | CAPS2 | 0.00665686405082555 | 1.12102952814838 | 0.0213788904582741 | 0.983043473772526 | 0.993756903461892 | -5.95148024116877 |
| lnc-LGALS3BP-3 | lnc-LGALS3BP-3 | 0.00365110036860655 | 0.164061216097341 | 0.0213241050549071 | 0.983086919705148 | 0.993756903461892 | -5.73733321971944 |
| ENSG00000100429.17 | HDAC10 | -0.00405605349460725 | 3.03532044994516 | -0.0211992685662257 | 0.983185917758685 | 0.99376881343782 | -6.18669447817858 |
| ENSG00000170619.9 | COMMD5 | 0.00270955760397085 | 3.48232907859505 | 0.0211608566798127 | 0.983216379274959 | 0.99376881343782 | -6.23684705384299 |
| lnc-CALCA-2 | lnc-CALCA-2 | -0.00370812039568247 | 0.695998526407021 | -0.0211266008696272 | 0.983243544947831 | 0.99376881343782 | -5.84106308635989 |
| ENSG00000198860.12 | TSEN15 | -0.00280443811662145 | 3.86825373143683 | -0.0210603499723187 | 0.983296083543435 | 0.99377311634953 | -6.29036714444433 |
| lnc-EPSTI1-4 | lnc-EPSTI1-4 | 0.00332897265473031 | 2.68595834242815 | 0.020976581814294 | 0.98336251386622 | 0.993791457893082 | -6.13818043441356 |
| ENSG00000147421.17 | HMBOX1 | 0.00262296202259993 | 5.66445411671764 | 0.0208365686698301 | 0.983473548189096 | 0.993854872465444 | -6.40326558858525 |
| ENSG00000183486.12 | MX2 | -0.00525405615746521 | 5.68001169200621 | -0.0206898144032589 | 0.983590797509452 | 0.993903825856634 | -6.40380420543436 |
| lnc-MRPL20-1 | lnc-MRPL20-1 | -0.0023147454968564 | 5.08158331454087 | -0.0206537040149767 | 0.983618565452054 | 0.993903825856634 | -6.3839153215043 |
| ENSG00000154144.12 | TBRG1 | 0.00262266986056581 | 6.23116947759183 | 0.0203870666312295 | 0.983830018053915 | 0.994068688901071 | -6.41057584708953 |
| ENSG00000169330.8 | MINAR1 | 0.00622729607296951 | 0.178112622166893 | 0.0202539829390634 | 0.983935558433113 | 0.994094995585634 | -5.72189346898346 |
| ENSG00000284261.1 | MIR4657 | 0.00338696018343485 | -0.1157663466959 | 0.0202324340053096 | 0.983952647576452 | 0.994094995585634 | -5.6273991839145 |
| lnc-CXCL16-4 | lnc-CXCL16-4 | -0.00411141884807964 | 0.826294420143048 | -0.0200536904681805 | 0.984094398452866 | 0.994189408233795 | -5.87401002739695 |
| lnc-CCDC12-2 | lnc-CCDC12-2 | -0.00292485067934933 | 2.76361238272858 | -0.0199139206677991 | 0.984205241937467 | 0.994231497347848 | -6.13986429049122 |
| ENSG00000143549.19 | TPM3 | 0.00291304379182744 | 7.76404703655519 | 0.0198793383616792 | 0.984232667247716 | 0.994231497347848 | -6.3950439308739 |
| ENSG00000101194.17 | SLC17A9 | 0.00431957286717077 | 3.45378536683771 | 0.0198140895039013 | 0.98428441255291 | 0.994234973920427 | -6.2533624540976 |
| ENSG00000102890.14 | ELMO3 | 0.00371696953230916 | 1.10712194076022 | 0.0196778692328934 | 0.984392441629888 | 0.994295300130925 | -5.94487211225703 |
| ENSG00000214517.9 | PPME1 | 0.00181067438255145 | 3.5113446618247 | 0.0195994756222502 | 0.98445461158196 | 0.994309302232273 | -6.24881609785433 |
| ENSG00000198482.11 | ZNF808 | 0.00185501318029591 | 5.23701422165117 | 0.0193114431056052 | 0.984683036235293 | 0.994456032098135 | -6.38689770602024 |
| ENSG00000135823.13 | STX6 | -0.00195643784154331 | 4.49399346942344 | -0.0192944457376542 | 0.984696516066272 | 0.994456032098135 | -6.3461826200371 |
| lnc-CASC1-4 | lnc-CASC1-4 | -0.0114913789952302 | -0.755717007471636 | -0.0192314684489973 | 0.984746615409797 | 0.99445783460798 | -5.48858089347319 |
| ENSG00000163481.7 | RNF25 | 0.00328954879737653 | 3.50487015046805 | 0.0191356491347974 | 0.984822450568559 | 0.994480926394599 | -6.24809881702389 |
| lnc-UBE2E3-2 | lnc-UBE2E3-2 | -0.00361928511421561 | 1.10142712013184 | -0.0190805932988456 | 0.984866112987782 | 0.994480926394599 | -5.92496326270731 |
| ENSG00000068120.14 | COASY | -0.00214924347495504 | 4.24937162405507 | -0.0187436106228116 | 0.985133360495067 | 0.994529622070184 | -6.32208601430428 |
| ENSG00000107331.16 | ABCA2 | 0.0119206769038145 | 0.993256758926394 | 0.0186259386927085 | 0.985229130184875 | 0.994529622070184 | -5.89842631884733 |
| lnc-SMIM30-1 | lnc-SMIM30-1 | -0.00488067957265303 | 0.484221625383381 | -0.018601682097652 | 0.985245918861144 | 0.994529622070184 | -5.79366355433277 |
| lnc-NCAPD3-1 | lnc-NCAPD3-1 | -0.00420270201394687 | -0.0387184462084577 | -0.0184922797251345 | 0.985332682122873 | 0.994529622070184 | -5.63784177573603 |
| ENSG00000160410.14 | SHKBP1 | 0.00214757151459022 | 5.33172137129276 | 0.0184893241173984 | 0.985335026116288 | 0.994529622070184 | -6.39420638732533 |
| ENSG00000198055.10 | GRK6 | -0.002463143851128 | 6.49459444922325 | -0.0184041955338706 | 0.985402538798315 | 0.994529622070184 | -6.41040577729131 |
| ENSG00000105483.17 | CARD8 | -0.00338841327830153 | 6.09380080552133 | -0.0183782906860477 | 0.985423083103324 | 0.994529622070184 | -6.40993127818786 |
| lnc-AURKB-5 | lnc-AURKB-5 | 0.00290313966320355 | 3.71219582975116 | 0.0183691140511724 | 0.985430360800894 | 0.994529622070184 | -6.26828018418986 |
| ENSG00000169442.8 | CD52 | 0.00390657591327326 | 8.12740728431853 | 0.0183169771351205 | 0.985471708956395 | 0.994529622070184 | -6.38942834249468 |
| ENSG00000185950.8 | IRS2 | 0.0131745268475072 | 4.38132918732922 | 0.0182184063956922 | 0.985553269348317 | 0.994529622070184 | -6.3708109017019 |
| ENSG00000109927.10 | TECTA | -0.00400343405220896 | 0.175657701783888 | -0.0181972893198588 | 0.985566629767462 | 0.994529622070184 | -5.73156845078479 |

| | | | | | | | |
|---|---|---|---|---|---|---|---|
| ENSG00000182628.12 | SKA2 | -0.00334732960132822 | 2.97045841849237 | -0.0181143540542857 | 0.985632403359551 | 0.994529622070184 | -6.18126776757425 |
| lnc-HSFX3-1 | lnc-HSFX3-1 | 0.00506544646322837 | -0.255393199173442 | 0.0181056950536438 | 0.985639270571664 | 0.994529622070184 | -5.60547159374245 |
| ENSG00000154822.17 | PLCL2 | 0.00250327814424622 | 7.72101685989738 | 0.0180908697855369 | 0.98565102808123 | 0.994529622070184 | -6.39742085463668 |
| lnc-EVPL-2 | lnc-EVPL-2 | 0.00275821626956096 | 2.11223124562177 | 0.0180658354672883 | 0.98567088211306 | 0.994529622070184 | -6.06965964154945 |
| ENSG00000148343.18 | MIGA2 | -0.00227597176546546 | 4.74855111685391 | -0.0180449749276644 | 0.98568742604245 | 0.994529622070184 | -6.36506844363983 |
| lnc-MRC2-2 | lnc-MRC2-2 | 0.00536620930850551 | 0.594333234669509 | 0.0175902365736928 | 0.986048068294849 | 0.994823983863997 | -5.81382624712408 |
| ENSG00000144712.12 | CAND2 | -0.00628077941926741 | 1.47431951225226 | -0.0175556255919696 | 0.986075835339516 | 0.994823983863997 | -6.00067884714224 |
| ENSG00000176438.12 | SYNE3 | 0.00257139852161981 | 7.38230202394006 | 0.017218635431824 | 0.986342778582865 | 0.995044523431412 | -6.40306498139942 |
| ENSG00000238578.1 | SNORD4A | -0.0028369798341612 | 1.22349439541534 | -0.0169406126699729 | 0.986563274778577 | 0.99513955525104 | -5.96254567942785 |
| ENSG00000200623.1 | SNORD18A | 0.00280279990852594 | 0.307063657809846 | 0.0169308952652464 | 0.986570981543784 | 0.99513955525104 | -5.76964051147373 |
| lnc-EMCN-4 | lnc-EMCN-4 | -0.00391913987674597 | 2.77093284253797 | -0.0169169740917064 | 0.986582022273367 | 0.99513955525104 | -6.16711177423268 |
| ENSG00000100983.10 | GSS | 0.00244553419184257 | 3.56063253041196 | 0.0167467448445625 | 0.986717029431114 | 0.995226962159094 | -6.25454387939553 |
| ENSG00000196591.11 | HDAC2 | -0.00217669849026603 | 4.761325654554 | -0.016448350524172 | 0.986953684008068 | 0.995350728089294 | -6.36853708815892 |
| ENSG00000109762.15 | SNX25 | 0.00397111232558522 | 5.75935586474025 | 0.0163554130313257 | 0.987027861597667 | 0.995350728089294 | -6.40598519347064 |
| ENSG00000260314.2 | MRC1 | 0.0165669357848711 | 1.43087378186946 | 0.0163418966648534 | 0.987041028754134 | 0.995350728089294 | -5.78907263527772 |
| lnc-HERC3-14 | lnc-HERC3-14 | -0.00319347442940687 | 2.05607146500908 | -0.0162506139600637 | 0.987110508235465 | 0.995350728089294 | -6.06387868341486 |
| ENSG00000152082.13 | MZT2B | 0.00395420123734569 | 3.28834404066777 | 0.0161042104159686 | 0.987226620743916 | 0.995350728089294 | -6.23218889862668 |
| ENSG00000005302.18 | MSL3 | -0.0017545981747622 | 6.39842533143569 | -0.016086808133991 | 0.987240422495512 | 0.995350728089294 | -6.41067444875449 |
| ENSG00000233927.4 | RPS28 | 0.0025951252950307 | 6.84113222908783 | 0.0160210789652091 | 0.987292552341559 | 0.995350728089294 | -6.40813618943563 |
| ENSG00000135926.14 | TMBIM1 | -0.00340671788964375 | 4.16916443561299 | -0.0159440958014979 | 0.987353607809649 | 0.995350728089294 | -6.31268593530195 |
| ENSG00000167525.13 | PROCA1 | 0.0034461062257003 | 1.19906661603274 | 0.0159286534251245 | 0.987365855191861 | 0.995350728089294 | -5.9263878001303 |
| ENSG00000198324.13 | PHETA1 | -0.00644471578679934 | 0.378541494470303 | -0.0157508758573104 | 0.987506851187999 | 0.995350728089294 | -5.72874363535921 |
| ENSG00000204161.13 | TMEM273 | -0.0212223190153158 | -2.7022769598809 | -0.0157392770813233 | 0.987516050230935 | 0.995350728089294 | -4.89808597567875 |
| ENSG00000282995.1 | FRG1EP | 0.00994912609878418 | -1.09445209744819 | 0.015675131792273 | 0.987566926421632 | 0.995350728089294 | -5.40687774963617 |
| ENSG00000242265.5 | PEG10 | -0.00557624436106519 | 5.94794243349816 | -0.0156087640527907 | 0.987621524601011 | 0.995350728089294 | -6.40777395963759 |
| ENSG00000119321.8 | FKBP15 | -0.00276939091528838 | 5.20817379696146 | -0.015530857712555 | 0.987681348888986 | 0.995350728089294 | -6.38919754627065 |
| ENSG00000204406.13 | MBD5 | 0.00167134754111053 | 5.86203752910707 | 0.0155255443661162 | 0.987685562943274 | 0.995350728089294 | -6.4077761227682 |
| lnc-ASB1-2 | lnc-ASB1-2 | 0.00236743242280002 | 1.0197603588983 | 0.0154942509320959 | 0.987710382004042 | 0.995350728089294 | -5.91440042403628 |
| ENSG00000026751.16 | SLAMF7 | -0.0115884428076846 | 0.924629157047771 | -0.0154384625561933 | 0.987757063462103 | 0.995350728089294 | -5.88584994200342 |
| ENSG00000084774.13 | CAD | -0.00190274381006805 | 5.34520604734712 | -0.0153546541960275 | 0.987821097387886 | 0.995350728089294 | -6.3942616616507 |
| ENSG00000256019.1 | TAS2R63P | 0.00384010967169646 | 1.12909781445454 | 0.0153038916672178 | 0.987861357650722 | 0.995350728089294 | -5.95806602614465 |
| lnc-CFAP43-4 | lnc-CFAP43-4 | 0.00304053788163761 | 1.70741429309403 | 0.0152772124140814 | 0.98788251724266 | 0.995350728089294 | -6.00712178756677 |
| lnc-ZNF432-2 | lnc-ZNF432-2 | 0.00195710313958575 | 1.14216072836341 | 0.0151760726045959 | 0.987962732352941 | 0.995350728089294 | -5.92927109621782 |
| ENSG00000266412.5 | NCOA4 | -0.00216442166429864 | 6.36465461716887 | -0.0151253685293846 | 0.988002946367652 | 0.995350728089294 | -6.41070624819032 |
| lnc-HHIPL1-2 | lnc-HHIPL1-2 | 0.00414634913675749 | 0.195065250658358 | 0.01509525174869 | 0.988026832365275 | 0.995350728089294 | -5.72843172640351 |
| ENSG00000143321.18 | HDGF | -0.00249903549076258 | 5.54244584312432 | -0.0150909094664 | 0.9880302762849 | 0.995350728089294 | -6.4026939815721 |
| ENSG00000070761.7 | CFAP20 | 0.00326604660120333 | 3.52125052069591 | 0.0150676991870614 | 0.988048684658235 | 0.995350728089294 | -6.24810969384134 |
| ENSG00000184162.14 | NR2C2AP | 0.00194312549494491 | 2.21331569778652 | 0.0147233640625047 | 0.988321782076411 | 0.995577117431545 | -6.08808892422403 |
| ENSG00000109534.16 | GAR1 | -0.00227627996496249 | 1.83486262687604 | -0.0145603363134815 | 0.988451082357092 | 0.995658638942744 | -6.04362571662125 |
| ENSG00000137185.12 | ZSCAN9 | 0.00153627187598548 | 4.37211534461379 | 0.0142773394768177 | 0.988675533052822 | 0.99571146385744 | -6.33472448862485 |
| ENSG00000226761.3 | TAS2R46 | 0.00401073852574063 | 1.70619031609372 | 0.0142743361895476 | 0.988677915027628 | 0.99571146385744 | -6.03877664568017 |
| ENSG00000111906.17 | HDDC2 | -0.00222074358810431 | 3.39228082325953 | -0.0142443811669604 | 0.988701673036724 | 0.99571146385744 | -6.2260673677981 |
| ENSG00000078674.17 | PCM1 | -0.00172228898833815 | 8.46856160697132 | -0.0142105586105191 | 0.988728498487275 | 0.99571146385744 | -6.38047111175192 |
| lnc-NFE2L2-1 | lnc-NFE2L2-1 | -0.00361810681783004 | 1.82579630340539 | -0.0141892459967102 | 0.988745402020222 | 0.99571146385744 | -6.04352273031001 |
| ENSG00000257906.2 | LINC02156 | -0.0120772042720776 | -1.48112828128262 | -0.0140821471318425 | 0.988830925209558 | 0.995748871506278 | -5.30691689245966 |
| lnc-RAPH1-6 | lnc-RAPH1-6 | 0.00346463025923224 | 0.14169422608827 | 0.0135591915035525 | 0.989245115318313 | 0.99584909006094 | -5.70357451929867 |
| lnc-KCNIP3-4 | lnc-KCNIP3-4 | 0.00215416109987093 | 0.325141223134304 | 0.0135416565561042 | 0.98925902282391 | 0.99584909006094 | -5.75156637203233 |
| lnc-FAM25A-4 | lnc-FAM25A-4 | -0.00275546129192517 | 0.301443922629232 | -0.0134938128742357 | 0.989296969135216 | 0.99584909006094 | -5.7632946529301 |
| ENSG00000171634.17 | BPTF | 0.00121139013281457 | 8.07479234878339 | 0.0134851251974381 | 0.989303859604432 | 0.99584909006094 | -6.38961975520044 |
| ENSG00000251247.10 | ZNF345 | -0.00126481322326244 | 3.75394460980045 | -0.0134784705382127 | 0.989309137624145 | 0.99584909006094 | -6.26766131603867 |
| ENSG00000136147.17 | PHF11 | 0.00253195106556614 | 5.5455653932418 | 0.0134472237725813 | 0.989333920424053 | 0.99584909006094 | -6.40191115382528 |
| ENSG00000173145.11 | NOC3L | -0.00133125385363186 | 5.81667957352292 | -0.0133714228389448 | 0.989394040590591 | 0.99584909006094 | -6.40703759327399 |
| ENSG00000046651.14 | OFD1 | 0.0017633692150687 | 6.88162988641052 | 0.0133356151363536 | 0.989422440856731 | 0.99584909006094 | -6.40746132905447 |
| ENSG00000258984.5 | UBE2F-SCLY | 0.00187071231780234 | 2.85091269004396 | 0.0133264194348618 | 0.989429734272144 | 0.99584909006094 | -6.15990581865161 |
| ENSG00000100342.20 | APOL1 | -0.00450278069935761 | 1.92729852386813 | -0.0133224680417369 | 0.989433336817005 | 0.99584909006094 | -6.11109089217732 |
| ENSG00000265808.3 | SEC22B | -0.00103984059844375 | 5.69610196157136 | -0.013284917589233 | 0.989462650771816 | 0.99584909006094 | -6.40436079657727 |
| ENSG00000160691.18 | SHC1 | -0.0015296441212147 | 4.54740812564778 | -0.0131421308866862 | 0.989575899815326 | 0.995914372476964 | -6.3503964208394 |
| ENSG00000129472.14 | RAB2B | -0.00121723372794591 | 4.77930434198255 | -0.013048382149437 | 0.989650255280825 | 0.99594050772235 | -6.36660774624343 |
| ENSG00000117411.16 | B4GALT2 | 0.00532912568294108 | 1.73127485761177 | 0.0127658567580136 | 0.989875261190659 | 0.996059368734462 | -6.03425487865089 |
| ENSG00000135740.16 | SLC9A5 | -0.00260760706842511 | 1.70516188968713 | -0.0127562001910695 | 0.989881995802001 | 0.996059368734462 | -6.01964607844659 |
| ENSG00000165724.5 | ZMYND19 | 0.00190292434561723 | 2.67310437923758 | 0.0127164256280497 | 0.989913542575531 | 0.996059368734462 | -6.14378157943864 |
| lnc-KIAA0391-3 | lnc-KIAA0391-3 | 0.00241993698340615 | 1.70338756946593 | 0.0125424573587195 | 0.990051523857647 | 0.996127640476165 | -6.02338782171715 |
| ENSG00000063046.17 | EIF4B | 0.00176352257790668 | 9.48576616171813 | 0.0125088435475069 | 0.990078184368721 | 0.996127640476165 | -6.34839413442495 |
| ENSG00000196975.15 | ANXA4 | 0.00913824928920315 | 2.83807376895803 | 0.0123957033451237 | 0.990170135573712 | 0.996156801899485 | -6.15127279423141 |
| ENSG00000172340.14 | SUCLG2 | 0.00105326091202418 | 5.48673335962221 | 0.0123017461800299 | 0.990242442152301 | 0.996156801899485 | -6.39897175603002 |
| ENSG00000110104.12 | CCDC86 | -0.00197162206044865 | 2.40172844777631 | -0.0122892418410129 | 0.990252359892404 | 0.996156801899485 | -6.1075836583396 |
| ENSG00000145365.10 | TIFA | 0.00266366006480645 | 5.03484019517588 | 0.012168738651713 | 0.990347950833164 | 0.996204274850895 | -6.38486936149779 |
| ENSG00000259784.1 | | -0.00219947353098449 | 1.59126678153508 | -0.0120332585766989 | 0.990455391795275 | 0.99622735836348 | -6.01266608569608 |

| | | | | | | | |
|---|---|---|---|---|---|---|---|
| lnc-CELA2B-3 | lnc-CELA2B-3 | -0.00289270544615261 | -0.124906554945001 | -0.0119692036498883 | 0.990506196755049 | 0.99622735836348 | -5.62282335537657 |
| lnc-TXNDC11-1 | lnc-TXNDC11-1 | 0.002081613747559 | 2.29041097207318 | 0.0119567174781913 | 0.990516100126737 | 0.99622735836348 | -6.09433407321176 |
| ENSG00000100242.15 | SUN2 | 0.00153377529823473 | 6.52783944269624 | 0.0118678679294331 | 0.990586570938083 | 0.996249554852856 | -6.41047561059487 |
| GPRC5D-AS1 | GPRC5D-AS1 | 0.0222492912441754 | -2.72886493711187 | 0.0117260950873143 | 0.990699742639821 | 0.996314692077756 | -4.87618538264664 |
| lnc-DGKI-1 | lnc-DGKI-1 | -0.00497409459040954 | 3.78899322166832 | -0.011578557963317 | 0.99081755881918 | 0.996384493510611 | -6.2709468334661 |
| ENSG00000184205.14 | TSPYL2 | -0.00254846803832276 | 6.68947374026497 | -0.0115048627686642 | 0.990874593993302 | 0.996393168604433 | -6.40987017961338 |
| lnc-HUS1B-3 | lnc-HUS1B-3 | -0.00174217159696921 | 0.525654671564298 | -0.0112434288629482 | 0.991081845433402 | 0.996459722845338 | -5.82630335412401 |
| ENSG00000129484.13 | PARP2 | -0.0015537144271951 | 3.19273891369427 | -0.0112223489575735 | 0.991098565045496 | 0.996459722845338 | -6.20811987964618 |
| lnc-GRHPR-7 | lnc-GRHPR-7 | -0.00276214585403523 | 1.43795527222963 | -0.0111809932300055 | 0.991131366519616 | 0.996459722845338 | -5.98232159365607 |
| lnc-HLA-DRB1-7 | lnc-HLA-DRB1-7 | -0.00397761895265769 | -0.3441144533302 | -0.0111481867231308 | 0.991157387153426 | 0.996459722845338 | -5.57884462082664 |
| ENSG00000170264.12 | FAM161A | -0.00267365244224393 | 1.46260031003723 | -0.0111160979364793 | 0.99118283853352 | 0.996459722845338 | -5.99231063247317 |
| ENSG00000161956.12 | SENP3 | 0.0010669950520521 | 4.09519480864359 | 0.0110512709249845 | 0.991234256427147 | 0.996462744812697 | -6.31225359182989 |
| ENSG00000169062.14 | UPF3A | 0.00164664420605387 | 5.88073645164084 | 0.010904439108077 | 0.991350717013312 | 0.996531149066469 | -6.40708942531174 |
| lnc-ZNF557-1 | lnc-ZNF557-1 | 0.0017619775599973 | 0.667509075088961 | 0.0107416271225305 | 0.991479852594545 | 0.996588372472879 | -5.86188816935354 |
| lnc-SOCS5-3 | lnc-SOCS5-3 | -0.00541658460516409 | -0.149079219304378 | -0.0105788719780075 | 0.991608944830476 | 0.996588372472879 | -5.58433935640416 |
| ENSG00000174206.12 | KICS2 | -0.00125520545205848 | 3.56615082830488 | -0.0105645315352166 | 0.991620317586959 | 0.996588372472879 | -6.24669996124711 |
| ENSG00000116786.12 | PLEKHM2 | 0.00129056362046587 | 5.92779799229649 | 0.0105600721713451 | 0.991623854575881 | 0.996588372472879 | -6.40911224425368 |
| ENSG00000140199.11 | SLC12A6 | 0.00155530247521884 | 7.75146736460487 | 0.0105274450462616 | 0.991649733110675 | 0.996588372472879 | -6.39695067876597 |
| lnc-COL24A1-2 | lnc-COL24A1-2 | 0.00156021709376719 | 0.831525261170732 | 0.0100656338063517 | 0.992016024336529 | 0.996819466309223 | -5.87658370736248 |
| lnc-MEMO1-1 | lnc-MEMO1-1 | -0.00168412225782388 | 0.999617152778311 | -0.0099213864940568 | 0.992130436215167 | 0.996819466309223 | -5.91443766346725 |
| ENSG00000070423.17 | RNF126 | 0.00173003285198181 | 4.13098671217745 | 0.0098400940561344 | 0.992194914582681 | 0.996819466309223 | -6.31612612703572 |
| lnc-MEMO1-4 | lnc-MEMO1-4 | -0.00195290936828174 | 3.40233908279642 | -0.0098372407956035 | 0.992197177691822 | 0.996819466309223 | -6.23836725797877 |
| ENSG00000198517.9 | MAFK | 0.0017261320866222 | 4.71696184593755 | 0.0098316050503214 | 0.992201647773354 | 0.996819466309223 | -6.36947047525292 |
| ENSG00000280378.1 | | 0.00157722933632454 | 0.593012838754508 | 0.0098050361476417 | 0.992222721326979 | 0.996819466309223 | -5.82779295863366 |
| lnc-DAPK3-2 | lnc-DAPK3-2 | 0.00141333304467928 | 6.9393268839556 | 0.0097772670838425 | 0.992244746813756 | 0.996819466309223 | -6.4072559587903 |
| ENSG00000185298.12 | CCDC137 | 0.00098008475019195 | 3.96211243406152 | 0.0097111713986533 | 0.992297171711243 | 0.996819466309223 | -6.29732172966881 |
| lnc-UGDH-4 | lnc-UGDH-4 | -0.00286514728605436 | 3.84646512547638 | -0.0096886121266868 | 0.9923155450943 | 0.996819466309223 | -6.28938419806009 |
| ENSG00000283201.1 | | 0.00131432772518636 | 2.71723695735581 | 0.0095331260850013 | 0.992438391498456 | 0.996864128597109 | -6.139013157972 |
| lnc-RAB6B-2 | lnc-RAB6B-2 | 0.00112933981996921 | 1.75806459387807 | 0.0095098309743621 | 0.992456868445414 | 0.996864128597109 | -6.03517294976411 |
| ENSG00000127249.14 | ATP13A4 | -0.00244084142538603 | 1.28781612459369 | -0.0093173555810514 | 0.992609534015476 | 0.996928092087925 | -5.9285785043254 |
| ENSG00000136573.13 | BLK | -0.00179865938545731 | 7.88833792501037 | -0.0091771548628649 | 0.992720737095584 | 0.996928092087925 | -6.39259730174917 |
| ENSG00000136100.13 | VPS36 | 0.00143892273876123 | 5.80216667129411 | 0.0090221379940371 | 0.992843692080362 | 0.996928092087925 | -6.40830812318384 |
| lnc-AP3M1-1 | lnc-AP3M1-1 | 0.00182931129066133 | 1.57056854680982 | 0.0089842815508265 | 0.992873718764536 | 0.996928092087925 | -5.99762284438674 |
| lnc-KLHL9-1 | lnc-KLHL9-1 | 0.00117604206484545 | 1.88398939369506 | 0.0089810967520307 | 0.99287624485937 | 0.996928092087925 | -6.04950157867066 |
| ENSG00000264058.2 | | 0.00104720456742857 | 5.72980319883495 | 0.0089649251537976 | 0.992889071727572 | 0.996928092087925 | -6.40553905389704 |
| lnc-SCN4A-2 | lnc-SCN4A-2 | -0.00236588527429671 | 3.84866310932307 | -0.0089612232555425 | 0.992892166014175 | 0.996928092087925 | -6.2803158350945 |
| lnc-ANKRD60-4 | lnc-ANKRD60-4 | 0.00165853933664262 | 1.78866654483548 | 0.0089410290713267 | 0.992908025448305 | 0.996928092087925 | -6.0367869180497 |
| ENSG00000182512.4 | GLRX5 | -0.00177378318409538 | 2.42752810861433 | -0.0088200918693982 | 0.99300394960106 | 0.996975771652047 | -6.11202962067864 |
| ENSG00000269533.5 | | -0.00093819479832379 | 1.3848108030031 | -0.0085160626250445 | 0.993245097922401 | 0.997169244490136 | -5.97268627881566 |
| ENSG00000090621.13 | PABPC4 | 0.00085549604763635 | 5.55663963782017 | 0.0083182671298111 | 0.993401984660379 | 0.997278108094649 | -6.40078037740175 |
| lnc-NLN-1 | lnc-NLN-1 | -0.00208101156623314 | 0.397609529211141 | -0.0081475501883041 | 0.99353739353863 | 0.997365400468475 | -5.76697016963529 |
| lnc-GP5-3 | lnc-GP5-3 | -0.00134179950960189 | 1.12576585010549 | -0.0079542515572966 | 0.993690713942873 | 0.997420239416357 | -5.94333161575028 |
| ENSG00000182141.10 | ZNF708 | 0.00099229833920322 | 5.47346110389808 | 0.0079068031426145 | 0.993728349063488 | 0.997420239416357 | -6.3984442649683 |
| ENSG00000130772.13 | MED18 | -0.00108036617595264 | 2.54528287417668 | -0.0078953955299732 | 0.99373739735273 | 0.997420239416357 | -6.13411773765901 |
| ENSG00000140350.15 | ANP32A | -0.00058080457981457 | 5.92223911467192 | -0.0076668867969018 | 0.993918646064621 | 0.997553515308567 | -6.40853869876936 |
| ENSG00000184402.14 | SS18L1 | -0.00103367848737779 | 3.10305105966255 | -0.0074876167681640 | 0.994060839815605 | 0.997605131846617 | -6.19849826101108 |
| lnc-ADAP2-1 | lnc-ADAP2-1 | 0.0026629596682588 | 1.36981660406516 | 0.0074799250583891 | 0.99406700938513 | 0.997605131846617 | -5.9581645913941 |
| ENSG00000065615.13 | CYB5R4 | -0.00106945430968786 | 5.84854282768235 | -0.0073862276752075 | 0.99414125990086 | 0.997628551175403 | -6.40674546896416 |
| lnc-CCDC17-2 | lnc-CCDC17-2 | 0.00131050677142341 | 1.01769446787987 | 0.0073282046531437 | 0.994187282792104 | 0.997628551175403 | -5.92028376664368 |
| ENSG00000137101.12 | CD72 | -0.00344217403954605 | 4.78780417628588 | -0.0072239183343036 | 0.994271199767708 | 0.997635187276745 | -6.36320981099467 |
| ENSG00000212498.1 | SNORD86 | -0.00120760403548636 | 0.613485841931164 | -0.0071976535916219 | 0.994290833784927 | 0.997635187276745 | -5.84972426158918 |
| ENSG00000114933.15 | INO80D | 0.00086317281625714 | 7.0914953733842 | 0.0071168849644118 | 0.994354898205118 | 0.997650834499437 | -6.40569785917055 |
| ENSG00000132286.11 | TIMM10B | -0.00096775434997997 | 5.3333388641697 | -0.0067693622834132 | 0.994630548220804 | 0.997867805672155 | -6.39492713232622 |
| lnc-RXFP2-3 | lnc-RXFP2-3 | 0.00295517157914854 | -0.575701807009626 | 0.0067139673681847 | 0.994674486736782 | 0.997867805672155 | -5.58080462024744 |
| ENSG00000102870.5 | ZNF629 | -0.0036262134092917 | 2.58291544627326 | -0.0066621652892718 | 0.994716593148777 | 0.997867805672155 | -6.16499608934806 |
| lnc-ABHD14B-1 | lnc-ABHD14B-1 | 0.00112343511487545 | 6.06510479750069 | 0.0065064461252543 | 0.994839089996589 | 0.997916511516277 | -6.41002628415681 |
| lnc-ZNF80-3 | lnc-ZNF80-3 | 0.00201405597477256 | 0.947113898909751 | 0.0064311378524429 | 0.994898823635292 | 0.997916511516277 | -5.9134729216039 |
| ENSG00000159023.21 | EPB41 | 0.00095151813451366 | 8.60831676963079 | 0.0062448881832979 | 0.995046554814776 | 0.997916511516277 | -6.37758811908776 |
| lnc-SGSH-5 | lnc-SGSH-5 | 0.00107889655328952 | 1.77999351915237 | 0.0061888407288079 | 0.99509101107101 | 0.997916511516277 | -6.02918631328519 |
| ENSG00000166913.12 | YWHAB | 0.00061134352727432 | 6.9665239684382 | 0.0060680770389594 | 0.995186799623489 | 0.997916511516277 | -6.40699186121905 |
| lnc-ERGIC2-2 | lnc-ERGIC2-2 | -0.00128388397073602 | 0.0231314203299609 | -0.0060585539422297 | 0.995194353251764 | 0.997916511516277 | -5.65022565407772 |
| ENSG00000277209.1 | RPPH1 | 0.00187517071169552 | 10.2919508438771 | 0.0059999108955573 | 0.995241328338428 | 0.997916511516277 | -6.32431220554502 |
| ENSG00000140691.16 | ARMC5 | 0.00095565262069330 | 2.94049443830173 | 0.0059623996139044 | 0.995270621960423 | 0.997916511516277 | -6.18016493232232 |
| lnc-CCNB1IP1-1 | lnc-CCNB1IP1-1 | 0.00185323176294361 | 10.2919591571849 | 0.0059297496117994 | 0.995296974245188 | 0.997916511516277 | -6.32431230897863 |
| lnc-POU5F1-9 | lnc-POU5F1-9 | -0.00092266139134622 | 0.972883743843509 | -0.0058347275662146 | 0.995371890296622 | 0.997916511516277 | -5.89240882780484 |
| ENSG00000102034.16 | ELF4 | -0.00102327340275554 | 4.72707910645365 | -0.0058287240454533 | 0.995376652237972 | 0.997916511516277 | -6.36082056634338 |
| ENSG00000151465.13 | CDC123 | 0.00059678802769003 | 5.24535527749322 | 0.0057648220671863 | 0.995427338751441 | 0.997916511516277 | -6.39185006826537 |
| ENSG00000119333.11 | DYNC2I2 | 0.00123624248568355 | 3.93241957826025 | 0.005725433379276 | 0.995458581538029 | 0.997916511516277 | -6.29472661783269 |

| | | | | | | | |
|---|---|---|---|---|---|---|---|
| ENSG00000100263.13 | RHBDD3 | 0.00092375529693256 | 1.9136167098276 | 0.0057200291236636 | 0.995462868150149 | 0.997916511516277 | -6.05432031759482 |
| lnc-NETO2-3 | lnc-NETO2-3 | 0.00142706513799795 | 1.61381741436805 | 0.0056828181901924 | 0.995492383567201 | 0.997916511516277 | -5.99849954324546 |
| ENSG00000232334.1 | | -0.0020911815240001 | -0.371844643033585 | -0.0054195261417963 | 0.995701224910097 | 0.997983660124334 | -5.52786909537016 |
| lnc-CHST15-2 | lnc-CHST15-2 | -0.0020911815240001 | -0.371844643033585 | -0.0054195261417963 | 0.995701224910097 | 0.997983660124334 | -5.52786909537016 |
| ENSG00000163312.10 | HELQ | -0.00063498325117628 | 5.13312553702133 | -0.0054149854929607 | 0.995704826522534 | 0.997983660124334 | -6.38161500967256 |
| ENSG00000133619.17 | KRBA1 | 0.00078787743904085 | 2.85986153688609 | 0.0053082176923428 | 0.995789514057371 | 0.998019942924617 | -6.1604323170943 |
| ENSG00000100804.18 | PSMB5 | 0.00069329048570443 | 3.17729134104568 | 0.0051936560158931 | 0.99588038369915 | 0.998032270922005 | -6.20973212162683 |
| ENSG00000259753.1 | | -0.00094833585692023 | 3.01006935255918 | -0.0051585145345904 | 0.995908257724685 | 0.998032270922005 | -6.1815082289196 |
| lnc-SIRT3-2 | lnc-SIRT3-2 | 0.00067044608721911 | 2.68891133816741 | 0.0050673403310438 | 0.995980576614725 | 0.998032270922005 | -6.14857057854706 |
| ENSG00000142230.11 | SAE1 | 0.00061955338183050 | 4.95896558698617 | 0.0050481891472479 | 0.99599576723553 | 0.998032270922005 | -6.37857769389736 |
| lnc-RBMS3-5 | lnc-RBMS3-5 | 0.0012987987034474 | 1.33561191977151 | 0.0047361918878907 | 0.996243242069755 | 0.998144187732627 | -5.93189930697082 |
| ENSG00000156966.6 | B3GNT7 | 0.003563140135037 | -0.735933063668183 | 0.0047011755071210 | 0.996271501465204 | 0.998144187732627 | -5.59066398988965 |
| lnc-ETFBKMT-8 | lnc-ETFBKMT-8 | 0.00087405874359092 | 2.85729482498306 | 0.0046460277834906 | 0.996314759905923 | 0.998144187732627 | -6.15150132290596 |
| ENSG00000285972.1 | CERNA2 | 0.00424805784143278 | 0.917085261351088 | 0.0045134498836088 | 0.996420726695513 | 0.998144187732627 | -5.90225102212947 |
| ENSG00000102390.10 | PBDC1 | 0.00046341497513458 | 3.20986093421511 | 0.0045046240259841 | 0.996426920908105 | 0.998144187732627 | -6.20648598310497 |
| lnc-MSL2-6 | lnc-MSL2-6 | -0.0011902092900631 | 0.806307549646033 | -0.0044512947761944 | 0.996469221515319 | 0.998144187732627 | -5.86015915638933 |
| lnc-COP1-4 | lnc-COP1-4 | 0.00135655052342276 | -0.23579099597206 | 0.0043947429580319 | 0.996514078265702 | 0.998144187732627 | -5.6057432103 |
| ENSG00000088298.12 | EDEM2 | 0.00037339845003714 | 4.45252334055746 | 0.0043633562743058 | 0.996538974103926 | 0.998144187732627 | -6.34235921504598 |
| ENSG00000182749.5 | PAQR7 | -0.00080297143193552 | 3.03783768897595 | -0.0043196905760448 | 0.996573609631251 | 0.998144187732627 | -6.1862228108955 |
| ENSG00000215630.6 | GUSBP9 | -0.00097041820945920 | 2.4223970077196 | -0.0042489440492039 | 0.996629725621139 | 0.998144187732627 | -6.11142233893647 |
| lnc-MMGT1-4 | lnc-MMGT1-4 | 0.00084600424344161 | -0.16157458678124 | 0.0042348745719812 | 0.996640885501674 | 0.998144187732627 | -5.61940476182946 |
| ENSG00000181826.9 | RELL1 | 0.00234200298354004 | -0.756538005051723 | 0.0041635241184379 | 0.996697480545812 | 0.998152300981581 | -5.4725584573176 |
| lnc-SMIM7-1 | lnc-SMIM7-1 | -0.00052146429886612 | 0.575936438401284 | -0.0040889049471364 | 0.996756668349066 | 0.998163009858364 | -5.82824682166811 |
| ENSG00000182957.15 | SPATA13 | -0.0015643904305262 | 4.20422360096001 | -0.0040256100019220 | 0.996807368360009 | 0.998165218338801 | -6.31207652222314 |
| ENSG00000249590.7 | | 0.00135605888084672 | -0.0623635184846735 | 0.0039536336250117 | 0.996863965384926 | 0.998173331363978 | -5.68954081465124 |
| ENSG00000285827.1 | | -0.00056084705674575 | 5.94694091304796 | -0.0038403824435281 | 0.996953796119273 | 0.998214719342462 | -6.40907175892215 |
| ENSG00000254272.1 | BTF3P3 | 0.00092393948397384 | -0.25816821948261 | 0.0035736160871591 | 0.997165395106914 | 0.998348786895071 | -5.5407181320311 |
| ENSG00000119013.8 | NDUFB3 | -0.00070625402031376 | 2.82241585752559 | -0.0035492762835733 | 0.997184701441761 | 0.998348786895071 | -6.16270076108325 |
| ENSG00000178971.14 | CTC1 | 0.00054066881765013 | 7.05357849219707 | 0.0034291230687925 | 0.997280007009562 | 0.998364294714472 | -6.4064683320377 |
| lnc-HACL1-10 | lnc-HACL1-10 | 0.00127250686127367 | -0.673509122402158 | 0.0034074478467356 | 0.997297199806843 | 0.998364294714472 | -5.5048637340902 |
| lnc-MGARP-3 | lnc-MGARP-3 | 0.00049658363981253 | 2.10541277819831 | 0.0032203142466352 | 0.997445634327844 | 0.998464326980353 | -6.07830834161487 |
| lnc-MOCS2-7 | lnc-MOCS2-7 | 0.00131500102259818 | -0.327068466498586 | 0.0030691533603972 | 0.997565535330586 | 0.998535788246338 | -5.55301212179433 |
| ENSG00000174607.10 | UGT8 | -0.00040904648492201 | 4.5103584546124 | -0.0026656950240165 | 0.99788555925534 | 0.998772849029475 | -6.34660520148147 |
| ENSG00000164733.20 | CTSB | -0.00057685156514875 | 5.25990606243901 | -0.0025721592499514 | 0.99795977237844 | 0.998772849029475 | -6.39055059137905 |
| lnc-MRPL3-1 | lnc-MRPL3-1 | 0.00045492659092172 | 2.44993777225745 | 0.00255388002731 | 0.997974251189392 | 0.998772849029475 | -6.11435749967565 |
| ENSG00000107537.13 | PHYH | 0.00058189476670156 | 2.27589148177053 | 0.0025258780648318 | 0.99799646241992 | 0.998772849029475 | -6.09746395122676 |
| ENSG00000168137.15 | SETD5 | -0.00020278960540309 | 6.56003670486482 | -0.0022932912746694 | 0.998180950936078 | 0.998908912539736 | -6.41035404926831 |
| ENSG00000112335.14 | SNX3 | -0.00037955120531219 | 6.1459316020275 | -0.0021235868310433 | 0.998315561056822 | 0.998995050475598 | -6.4104623734567 |
| ENSG00000156042.17 | CFAP70 | 0.00032625153719401 | 3.05501371470734 | 0.0015978837136844 | 0.998732550789579 | 0.999363738108989 | -6.17341890737499 |
| ENSG00000105443.14 | CYTH2 | -0.00017382831250678 | 4.3883800878435 | -0.0015356079199272 | 0.998781948214536 | 0.999364583155889 | -6.34020106584609 |
| ENSG00000166619.13 | BLCAP | -0.00012420629705101 | 5.12892517911194 | -0.0011796690950305 | 0.999064280493925 | 0.99959848752705 | -6.38621387288101 |
| ENSG00000261072.2 | | -0.00026143522069310 | 1.28689011552848 | -0.0010054377512111 | 0.999202481627249 | 0.999673746472267 | -5.94979503229616 |
| lnc-OAZ3-2 | lnc-OAZ3-2 | 0.00053470705153152 | -0.250223688127708 | 0.0009623979965296 | 0.999236635083342 | 0.999673746472267 | -5.58059976071299 |
| ENSG00000135870.11 | RC3H1 | 7.11709939453797e-05 | 6.80127807077421 | 0.0007617745023230 | 0.999395756519595 | 0.999784342961984 | -6.40882998318896 |
| lnc-CEP76-3 | lnc-CEP76-3 | 9.55505162374415e-05 | 3.13673257104411 | 0.0006697329963210 | 0.99946876430689 | 0.999795581584837 | -6.19198092081381 |
| ENSG00000085185.15 | BCORL1 | -0.00016195879321134 | 1.84144568201431 | -0.0006251364970850 | 0.999504138476956 | 0.999795581584837 | -6.04304822839975 |
| lnc-FAM153C-5 | lnc-FAM153C-5 | -0.00013483708737710 | -0.382948370052591 | -0.0004813556850186 | 0.99961818615504 | 0.99980468863765 | -5.55645914843027 |
| lnc-EMILIN3-2 | lnc-EMILIN3-2 | -7.66406708211376e-0 | 0.66931517459838 | -0.0004751785101284 | 0.999623085922065 | 0.99980468863765 | -5.8481872657378 |
| ENSG00000251922.1 | | -0.00022013497352146 | -0.832659096717495 | -0.0004184899393050 | 0.999668051589878 | 0.99980468863765 | -5.42186586227294 |
| LINC01238 | LINC01238 | 0.00010693831560182 | 2.86854659336617 | 0.0003687121073026 | 0.999707540050113 | 0.99980468863765 | -6.16032683218901 |
| ENSG00000279410.1 | | -7.00960701831566e-0 | 1.17135098104403 | -0.0002105757281844 | 0.99983297022658 | 0.999881548254479 | -5.91838644175653 |
| ENSG00000159082.17 | SYNJ1 | -7.01415555859342e-0 | 2.87422599238907 | -4.44566401792254e- | 0.999964736759294 | 0.999964736759294 | -6.15787567518936 |

**Table S10.** Results of the pathway analysis in Group 1 and Group 2 of the *TP53* non-exp cases.

| ID | Description | GeneRatio | BgRatio | RichFactor | FoldEnrichment | zScore | pvalue | p.adjust | qvalue | geneID | Count |
|---|---|---|---|---|---|---|---|---|---|---|---|
| GO:0016887 | ATP hydrolysis activity | 103/1878 | 417/18639 | 0.247002398081535 | 2.45147907233319 | 10.0340964082769 | 2.73029195798152E-18 | 3.09888137230903E-15 | 2.41990087223204E-15 | RTEL1/CHD | 103 |
| GO:0140104 | molecular carrier activity | 66/1878 | 249/18639 | 0.265060240963855 | 2.6307017206205 | 8.67123233959424 | 1.01916181940979E-13 | 5.78374332515058E-11 | 4.51649606285814E-11 | TNPO2/RTE | 66 |
| GO:0140597 | protein carrier chaperone | 54/1878 | 188/18639 | 0.287234042553192 | 2.85077493032425 | 8.53729056813797 | 5.53489549152154E-13 | 2.09403546095898E-10 | 1.63522175574075E-10 | RTEL1/CHD | 54 |
| GO:0140097 | catalytic activity, acting on DNA | 78/1878 | 336/18639 | 0.232142857142857 | 2.3039993153811 | 8.07389964180474 | 1.19341953117978E-12 | 2.46952954738412E-10 | 1.92844319860647E-10 | RTEL1/CHD | 78 |
| GO:0061775 | cohesin loader activity | 47/1878 | 153/18639 | 0.30718954248366 | 3.04883167324438 | 8.51781111094955 | 1.2298335349311E-12 | 2.46952954738412E-10 | 1.92844319860647E-10 | RTEL1/CHD | 47 |
| GO:0140713 | histone chaperone activity | 49/1878 | 164/18639 | 0.298780487804878 | 2.9653724772072 | 8.46198117958283 | 1.30547817482861E-12 | 2.46952954738412E-10 | 1.92844319860647E-10 | RTEL1/CHD | 49 |
| GO:0008094 | ATP-dependent activity, acting on DNA | 57/1878 | 210/18639 | 0.271428571428571 | 2.69390689183021 | 8.26315276722564 | 1.69447541819635E-12 | 2.65675193334854E-10 | 2.07464421783397E-10 | RTEL1/CHD | 57 |
| GO:0003689 | DNA clamp loader activity | 48/1878 | 160/18639 | 0.3 | 2.97747603833866 | 8.40871190089324 | 1.87260048165536E-12 | 2.65675193334854E-10 | 2.07464421783397E-10 | RTEL1/CHD | 48 |
| GO:0000510 | H3-H4 histone complex chaperone activity | 46/1878 | 151/18639 | 0.304635761589404 | 3.023485601845 | 8.35682120592067 | 2.96213614589076E-12 | 2.80168710465501E-10 | 2.18782336389476E-10 | RTEL1/CHD | 46 |
| GO:0140584 | chromatin extrusion motor activity | 46/1878 | 151/18639 | 0.304635761589404 | 3.023485601845 | 8.35682120592067 | 2.96213614589076E-12 | 2.80168710465501E-10 | 2.18782336389476E-10 | RTEL1/CHD | 46 |
| GO:0140665 | ATP-dependent H3-H4 histone complex chaperone activity | 46/1878 | 151/18639 | 0.304635761589404 | 3.023485601845 | 8.35682120592067 | 2.96213614589076E-12 | 2.80168710465501E-10 | 2.18782336389476E-10 | RTEL1/CHD | 46 |
| GO:0140849 | ATP-dependent H2AZ histone chaperone activity | 46/1878 | 151/18639 | 0.304635761589404 | 3.023485601845 | 8.35682120592067 | 2.96213614589076E-12 | 2.80168710465501E-10 | 2.18782336389476E-10 | RTEL1/CHD | 46 |
| GO:0015616 | DNA translocase activity | 46/1878 | 152/18639 | 0.302631578947368 | 3.00359424920128 | 8.30225052803368 | 3.83840928914395E-12 | 3.11185324512741E-10 | 2.43003054245053E-10 | RTEL1/CHD | 46 |
| GO:0140674 | ATP-dependent histone chaperone activity | 46/1878 | 152/18639 | 0.302631578947368 | 3.00359424920128 | 8.30225052803368 | 3.83840928914395E-12 | 3.11185324512741E-10 | 2.43003054245053E-10 | RTEL1/CHD | 46 |
| GO:0004386 | helicase activity | 46/1878 | 155/18639 | 0.296774193548387 | 2.94546016695867 | 8.14118637119232 | 8.21286744187951E-12 | 6.21440303102217E-10 | 4.85279606039477E-10 | RTEL1/CHD | 46 |
| GO:0140658 | ATP-dependent chromatin remodeler activity | 46/1878 | 158/18639 | 0.291139240506329 | 2.88953370809237 | 7.98395238909089 | 1.71509722298817E-11 | 1.21664709255724E-09 | 9.50073593260554E-10 | RTEL1/CHD | 46 |
| GO:0003725 | double-stranded RNA binding | 27/1878 | 75/18639 | 0.36 | 3.57297124600639 | 7.47355559030473 | 1.52164688639417E-09 | 1.01592306826905E-07 | 7.93329212596836E-08 | OAS1/SLC3 | 27 |
| GO:0051020 | GTPase binding | 67/1878 | 315/18639 | 0.212698412698413 | 2.11101475734064 | 6.6567530750697 | 2.39393259801923E-09 | 1.50950749930657E-07 | 1.17876681142234E-07 | TNPO2/DVL | 67 |
| GO:0140098 | catalytic activity, acting on RNA | 77/1878 | 387/18639 | 0.198966408268734 | 1.97472571018154 | 6.48604800400844 | 3.83116131643124E-09 | 2.28861478639445E-07 | 1.78716777198621E-07 | THG1L/DDX | 77 |
| GO:0003712 | transcription coregulator activity | 92/1878 | 495/18639 | 0.185858585858586 | 1.84463161971149 | 6.37530403919271 | 4.44821714650894E-09 | 2.52436323064382E-07 | 1.97126254597923E-07 | TCF25/TDR | 92 |
| GO:0030695 | GTPase regulator activity | 91/1878 | 489/18639 | 0.186094069529652 | 1.84696877633823 | 6.35303136968507 | 5.06306535761089E-09 | 2.7364662766135E-07 | 2.13689274742274E-07 | SBF1/GMIP | 91 |
| GO:0060589 | nucleoside-triphosphatase regulator activity | 91/1878 | 490/18639 | 0.185714285714286 | 1.84319945230488 | 6.33139618337285 | 5.61803790860035E-09 | 2.89839683011882E-07 | 2.26334350193373E-07 | SBF1/GMIP | 91 |
| GO:0003735 | structural constituent of ribosome | 43/1878 | 175/18639 | 0.245714285714286 | 2.43869465997262 | 6.40062295694849 | 2.32383949568895E-08 | 1.14676427287259E-06 | 8.95502450970297E-07 | RPS18/RPS | 43 |
| GO:0044389 | ubiquitin-like protein ligase binding | 65/1878 | 322/18639 | 0.201863354037267 | 2.00347766554879 | 6.08003330973965 | 3.50981674572747E-08 | 1.65985083600028E-06 | 1.29616916662392E-06 | ACTG1/PRK | 65 |
| GO:0031267 | small GTPase binding | 59/1878 | 283/18639 | 0.208480565371025 | 2.06915295950507 | 6.06654635362998 | 4.53905686965169E-08 | 2.06073181882187E-06 | 1.60921510915652E-06 | TNPO2/DVL | 59 |
| GO:0031625 | ubiquitin protein ligase binding | 61/1878 | 302/18639 | 0.201986754966887 | 2.00470240991896 | 5.89215628403973 | 8.97825155053849E-08 | 3.91935211917738E-06 | 3.06060235042648E-06 | ACTG1/PRK | 61 |
| GO:0043021 | ribonucleoprotein complex binding | 42/1878 | 177/18639 | 0.23728813559322 | 2.35506579303623 | 6.06324513092187 | 9.72076956666724E-08 | 4.08632350302493E-06 | 3.19098946398979E-06 | SNRPN/DDX | 42 |
| GO:0003724 | RNA helicase activity | 24/1878 | 77/18639 | 0.311688311688312 | 3.09348159827393 | 6.16168668057352 | 2.85126214252477E-07 | 0.0000115577947563058 | 9.02542377445811E-06 | DDX5/DDX | 24 |
| GO:0008186 | ATP-dependent activity, acting on RNA | 24/1878 | 79/18639 | 0.30379746835443 | 3.01516560844421 | 6.00803960744864 | 4.8472524174396E-07 | 0.0000189711430820481 | 0.0000148144701832455 | DDX5/DDX | 24 |
| GO:0106310 | protein serine kinase activity | 67/1878 | 361/18639 | 0.185595567867036 | 1.84202118715319 | 5.40766486221983 | 5.86341404242123E-07 | 0.000022183249793827 | 0.0000173227881533989 | MINK1/PRK | 67 |
| GO:0045296 | cadherin binding | 63/1878 | 334/18639 | 0.188622754491018 | 1.87206577260814 | 5.3831327868489 | 7.10279675224369E-07 | 0.0000260054010122471 | 0.0000203074868094709 | PSEN1/DDX | 63 |
| GO:0004674 | protein serine/threonine kinase activity | 76/1878 | 433/18639 | 0.175519630484988 | 1.7420183134237 | 5.22936297474627 | 1.02767030459667E-06 | 0.0000364501811161631 | 0.0000284637630417893 | MINK1/PRK | 76 |
| GO:0003743 | translation initiation factor activity | 18/1878 | 51/18639 | 0.352941176470588 | 3.50291298628077 | 5.99117482128111 | 1.13337049752356E-06 | 0.0000389810762027044 | 0.0000304401262811751 | EIF2B2/EIF | 18 |
| GO:0140297 | DNA-binding transcription factor binding | 83/1878 | 488/18639 | 0.170081967213115 | 1.6880499397685 | 5.155574635872 | 1.24900161516804E-06 | 0.0000403570503560089 | 0.0000315146175745509 | PTMA/DDX | 83 |
| GO:0008135 | translation factor activity, RNA binding | 24/1878 | 83/18639 | 0.289156626506024 | 2.86985642249509 | 5.71482070162222 | 1.31439147320701E-06 | 0.0000403570503560089 | 0.0000315146175745509 | EIF2B2/EIF | 24 |
| GO:0061749 | forked DNA-dependent helicase activity | 17/1878 | 47/18639 | 0.361702127659574 | 3.58986472707498 | 5.95059563006987 | 1.51196587825912E-06 | 0.0000403570503560089 | 0.0000315146175745509 | RTEL1/CHD | 17 |
| GO:1990518 | single-stranded 3'-5' DNA helicase activity | 17/1878 | 47/18639 | 0.361702127659574 | 3.58986472707498 | 5.95059563006987 | 1.51196587825912E-06 | 0.0000403570503560089 | 0.0000315146175745509 | RTEL1/CHD | 17 |
| GO:0180051 | translation factor activity | 24/1878 | 84/18639 | 0.285714285714286 | 2.83569146508444 | 5.64425123930116 | 0.0000016659208647931 | 0.0000403570503560089 | 0.0000315146175745509 | EIF2B2/EIF | 24 |
| GO:0004694 | eukaryotic translation initiation factor 2alpha kinase activity | 49/1878 | 243/18639 | 0.201646090534979 | 2.0013213426419 | 5.25911168742817 | 1.69718061260334E-06 | 0.0000403570503560089 | 0.0000315146175745509 | MINK1/TNK | 49 |
| GO:0042162 | telomeric DNA binding | 15/1878 | 38/18639 | 0.394736842105263 | 3.91773162939297 | 6.02651830309269 | 1.72789510892121E-06 | 0.0000403570503560089 | 0.0000315146175745509 | TERF2/BLM | 15 |
| GO:0009378 | four-way junction helicase activity | 17/1878 | 48/18639 | 0.354166666666667 | 3.51507587859425 | 5.84006687015686 | 2.12230307850468E-06 | 0.0000403570503560089 | 0.0000315146175745509 | RTEL1/CHD | 17 |
| GO:0036121 | double-stranded DNA helicase activity | 17/1878 | 48/18639 | 0.354166666666667 | 3.51507587859425 | 5.84006687015686 | 2.12230307850468E-06 | 0.0000403570503560089 | 0.0000315146175745509 | RTEL1/CHD | 17 |
| GO:0043138 | 3'-5' DNA helicase activity | 17/1878 | 48/18639 | 0.354166666666667 | 3.51507587859425 | 5.84006687015686 | 2.12230307850468E-06 | 0.0000403570503560089 | 0.0000315146175745509 | RTEL1/CHD | 17 |
| GO:0003678 | DNA helicase activity | 22/1878 | 74/18639 | 0.297297297297297 | 2.95065192988516 | 5.62789063640363 | 2.15575904111868E-06 | 0.0000403570503560089 | 0.0000315146175745509 | RTEL1/CHD | 22 |
| GO:0004676 | 3-phosphoinositide-dependent protein kinase activity | 49/1878 | 245/18639 | 0.2 | 1.98498402555911 | 5.19483338517764 | 2.16896922618197E-06 | 0.0000403570503560089 | 0.0000315146175745509 | MINK1/TNK | 49 |
| GO:0004711 | ribosomal protein S6 kinase activity | 49/1878 | 245/18639 | 0.2 | 1.98498402555911 | 5.19483338517764 | 2.16896922618197E-06 | 0.0000403570503560089 | 0.0000315146175745509 | MINK1/TNK | 49 |
| GO:0035175 | histone H3S10 kinase activity | 49/1878 | 245/18639 | 0.2 | 1.98498402555911 | 5.19483338517764 | 2.16896922618197E-06 | 0.0000403570503560089 | 0.0000315146175745509 | MINK1/TNK | 49 |
| GO:0035979 | histone H2AXS139 kinase activity | 49/1878 | 245/18639 | 0.2 | 1.98498402555911 | 5.19483338517764 | 2.16896922618197E-06 | 0.0000403570503560089 | 0.0000315146175745509 | MINK1/TNK | 49 |
| GO:0044022 | histone H3S28 kinase activity | 49/1878 | 245/18639 | 0.2 | 1.98498402555911 | 5.19483338517764 | 2.16896922618197E-06 | 0.0000403570503560089 | 0.0000315146175745509 | MINK1/TNK | 49 |
| GO:0044023 | histone H4S1 kinase activity | 49/1878 | 245/18639 | 0.2 | 1.98498402555911 | 5.19483338517764 | 2.16896922618197E-06 | 0.0000403570503560089 | 0.0000315146175745509 | MINK1/TNK | 49 |
| GO:0044024 | histone H2AS1 kinase activity | 49/1878 | 245/18639 | 0.2 | 1.98498402555911 | 5.19483338517764 | 2.16896922618197E-06 | 0.0000403570503560089 | 0.0000315146175745509 | MINK1/TNK | 49 |

| | | | | | | | | | | | |
|---|---|---|---|---|---|---|---|---|---|---|---|
| GO:0044025 | histone H2BS14 kinase activity | 49/1878 | 245/18639 | 0.2 | 1.98498402555911 | 5.19483338517764 | 2.16896922618197E-06 | 0.0000403570503560089 | 0.0000315146175745509 | MINK1/TNK | 49 |
| GO:0072354 | histone H3T3 kinase activity | 49/1878 | 245/18639 | 0.2 | 1.98498402555911 | 5.19483338517764 | 2.16896922618197E-06 | 0.0000403570503560089 | 0.0000315146175745509 | MINK1/TNK | 49 |
| GO:0072371 | histone H2AS121 kinase activity | 49/1878 | 245/18639 | 0.2 | 1.98498402555911 | 5.19483338517764 | 2.16896922618197E-06 | 0.0000403570503560089 | 0.0000315146175745509 | MINK1/TNK | 49 |
| GO:0072518 | Rho-dependent protein serine/threonine kinase activity | 49/1878 | 245/18639 | 0.2 | 1.98498402555911 | 5.19483338517764 | 2.16896922618197E-06 | 0.0000403570503560089 | 0.0000315146175745509 | MINK1/TNK | 49 |
| GO:0140823 | histone H2BS36 kinase activity | 49/1878 | 245/18639 | 0.2 | 1.98498402555911 | 5.19483338517764 | 2.16896922618197E-06 | 0.0000403570503560089 | 0.0000315146175745509 | MINK1/TNK | 49 |
| GO:0140855 | histone H3S57 kinase activity | 49/1878 | 245/18639 | 0.2 | 1.98498402555911 | 5.19483338517764 | 2.16896922618197E-06 | 0.0000403570503560089 | 0.0000315146175745509 | MINK1/TNK | 49 |
| GO:0140995 | histone H2A kinase activity | 49/1878 | 245/18639 | 0.2 | 1.98498402555911 | 5.19483338517764 | 2.16896922618197E-06 | 0.0000403570503560089 | 0.0000315146175745509 | MINK1/TNK | 49 |
| GO:0140997 | histone H4 kinase activity | 49/1878 | 245/18639 | 0.2 | 1.98498402555911 | 5.19483338517764 | 2.16896922618197E-06 | 0.0000403570503560089 | 0.0000315146175745509 | MINK1/TNK | 49 |
| GO:0140998 | histone H2B kinase activity | 49/1878 | 245/18639 | 0.2 | 1.98498402555911 | 5.19483338517764 | 2.16896922618197E-06 | 0.0000403570503560089 | 0.0000315146175745509 | MINK1/TNK | 49 |
| GO:1990244 | histone H2AT120 kinase activity | 49/1878 | 245/18639 | 0.2 | 1.98498402555911 | 5.19483338517764 | 2.16896922618197E-06 | 0.0000403570503560089 | 0.0000315146175745509 | MINK1/TNK | 49 |
| GO:0004677 | DNA-dependent protein kinase activity | 49/1878 | 246/18639 | 0.199186991869919 | 1.9769149848048 | 5.16292152580801 | 2.44825525202911E-06 | 0.0000441074557310006 | 0.0000344432902624646 | MINK1/TNK | 49 |
| GO:0140857 | histone H3T45 kinase activity | 49/1878 | 246/18639 | 0.199186991869919 | 1.9769149848048 | 5.16292152580801 | 2.44825525202911E-06 | 0.0000441074557310006 | 0.0000344432902624646 | MINK1/TNK | 49 |
| GO:0035402 | histone H3T11 kinase activity | 49/1878 | 247/18639 | 0.198380566801619 | 1.96891128041288 | 5.13115925825162 | 2.76074212206271E-06 | 0.0000482068047467873 | 0.0000376444512838348 | MINK1/TNK | 49 |
| GO:0035403 | histone H3T6 kinase activity | 49/1878 | 247/18639 | 0.198380566801619 | 1.96891128041288 | 5.13115925825162 | 2.76074212206271E-06 | 0.0000482068047467873 | 0.0000376444512838348 | MINK1/TNK | 49 |
| GO:0004679 | AMP-activated protein kinase activity | 49/1878 | 248/18639 | 0.19758064516129 | 1.96097212202412 | 5.09954515548234 | 3.11003175394502E-06 | 0.000053483121829206 | 0.0000417647007467577 | MINK1/TNK | 49 |
| GO:0005085 | guanyl-nucleotide exchange factor activity | 46/1878 | 230/18639 | 0.2 | 1.98498402555911 | 5.03124528772745 | 4.38747275680812E-06 | 0.0000743250981936898 | 0.0000580400952275324 | SBF1/SH2D | 46 |
| GO:0042393 | histone binding | 48/1878 | 247/18639 | 0.194331983805668 | 1.92872941754731 | 4.91836419452958 | 6.27119835680593E-06 | 0.000104673678455511 | 0.0000817391488611546 | CHD3/TDRD | 48 |
| GO:0048027 | mRNA 5'-UTR binding | 11/1878 | 24/18639 | 0.458333333333333 | 4.54892172523962 | 5.82328446713011 | 7.58740762259085E-06 | 0.000124807357270154 | 0.0000974614373489168 | DDX3X/LAR | 11 |
| GO:0017116 | single-stranded DNA helicase activity | 18/1878 | 58/18639 | 0.310344827586207 | 3.08014762586758 | 5.31093635802747 | 9.18573403485294E-06 | 0.000148940116136544 | 0.000116306587328514 | RTEL1/CHD | 18 |
| GO:0019787 | ubiquitin-like protein transferase activity | 78/1878 | 479/18639 | 0.162839248434238 | 1.61616653438006 | 4.57304448626298 | 0.0000133790264149464 | 0.000213875985647382 | 0.00016701468111764 | TRIM26/TR | 78 |
| GO:0140101 | catalytic activity, acting on a tRNA | 31/1878 | 139/18639 | 0.223021582733813 | 2.21347139540764 | 4.80671711658882 | 0.0000169822251663195 | 0.000267705910607953 | 0.00020905019868481 | THG1L/RAR | 31 |
| GO:0140030 | modification-dependent protein binding | 36/1878 | 173/18639 | 0.208092485549133 | 2.06530129826959 | 4.71201405144079 | 0.0000189555250423346 | 0.000294719464699312 | 0.000230144947161438 | TDRD3/OPT | 36 |
| GO:0141003 | histone H2AX kinase activity | 61/1878 | 354/18639 | 0.172316384180791 | 1.71022634970488 | 4.51594260936024 | 0.0000207744725824392 | 0.00031863549163606 | 0.000248820852267621 | MINK1/TNK | 61 |
| GO:0140996 | histone H3 kinase activity | 61/1878 | 357/18639 | 0.170868347338936 | 1.69585469970736 | 4.44363424615903 | 0.0000269553688630276 | 0.000407924582127152 | 0.000318546253791849 | MINK1/TNK | 61 |
| GO:0035173 | histone kinase activity | 61/1878 | 360/18639 | 0.169444444444444 | 1.6817225772098 | 4.37200042743123 | 0.0000347962436228563 | 0.000519654427788709 | 0.000405795528122507 | MINK1/TNK | 61 |
| GO:0003713 | transcription coactivator activity | 47/1878 | 256/18639 | 0.18359375 | 1.82215330471246 | 4.43365688404267 | 0.0000359557470950102 | 0.00052999705133554 | 0.000413872030813378 | TDRD3/PM | 47 |
| GO:0141052 | histone H3 demethylase activity | 11/1878 | 28/18639 | 0.392857142857143 | 3.8990757644911 | 5.13867570847467 | 0.000044583784029017 | 0.00064875121631967 | 0.000506606560761569 | KDM7A/KD | 11 |
| GO:0016755 | aminoacyltransferase activity | 78/1878 | 496/18639 | 0.157258064516129 | 1.5607737297743 | 4.23712652705694 | 0.0000461398857907763 | 0.000662895827500393 | 0.000517652016466804 | TRIM26/TR | 78 |
| GO:0003697 | single-stranded DNA binding | 27/1878 | 121/18639 | 0.223140495867769 | 2.21465159876429 | 4.48688414705444 | 0.0000573536355918504 | 0.000813704704959377 | 0.000635417910109711 | TERF2/BLM | 27 |
| GO:0017124 | SH3 domain binding | 26/1878 | 115/18639 | 0.226086956521739 | 2.24389498541464 | 4.47881099481851 | 0.0000617767955826971 | 0.000865637814646435 | 0.000675972214173242 | ABI1/GRB2/ | 26 |
| GO:0035091 | phosphatidylinositol binding | 50/1878 | 284/18639 | 0.176056338028169 | 1.74734509292175 | 4.24815786891961 | 0.0000637594930701575 | 0.000876217483144206 | 0.000684233824073658 | KRIT1/WDR | 50 |
| GO:0140328 | floppase activity | 8/1878 | 16/18639 | 0.5 | 4.96246006389776 | 5.30759226475331 | 0.0000640758159479904 | 0.000876217483144206 | 0.000684233824073658 | ATP8B2/AB | 8 |
| GO:0004842 | ubiquitin-protein transferase activity | 71/1878 | 449/18639 | 0.158129175946548 | 1.56941944114361 | 4.08824200734255 | 0.0000834371927801985 | 0.00112739540244673 | 0.000880377397505353 | TRIM26/TR | 71 |
| GO:0042605 | peptide antigen binding | 14/1878 | 45/18639 | 0.311111111111111 | 3.0877529286475 | 4.69349575896824 | 0.000086834555787445 | 0.00115949671551471 | 0.000905445151368776 | HLA-C/HLA- | 14 |
| GO:0061659 | ubiquitin-like protein ligase activity | 61/1878 | 372/18639 | 0.163978494623656 | 1.62747346181593 | 4.09195362196366 | 0.0000919642612707136 | 0.00121371437839837 | 0.000947783451529264 | TRIM26/TR | 61 |
| GO:0032452 | histone demethylase activity | 11/1878 | 30/18639 | 0.366666666666667 | 3.63913738019169 | 4.84237693204699 | 0.0000937787225114018 | 0.00122343505805105 | 0.000955374281362376 | KDM7A/KD | 11 |
| GO:0003684 | damaged DNA binding | 19/1878 | 74/18639 | 0.256756756756757 | 2.54829030308264 | 4.4670237501912 | 0.0000987924551999228 | 0.00127419814377173 | 0.000995014919597308 | BLM/REV1/ | 19 |
| GO:0016874 | ligase activity | 33/1878 | 167/18639 | 0.197604790419162 | 1.96121176177995 | 4.17655145828186 | 0.000119530196044772 | 0.00152434575854849 | 0.00119035393340861 | ACSL1/DIP2 | 33 |
| GO:0004812 | aminoacyl-tRNA ligase activity | 13/1878 | 41/18639 | 0.317073170731707 | 3.14692589417907 | 4.60652407628035 | 0.000124345454059985 | 0.00155090209184707 | 0.00121109164046856 | RARS2/MA | 13 |
| GO:0016875 | ligase activity, forming carbon-oxygen bonds | 13/1878 | 41/18639 | 0.317073170731707 | 3.14692589417907 | 4.60652407628035 | 0.000124345454059985 | 0.00155090209184707 | 0.00121109164046856 | RARS2/MA | 13 |
| GO:0140457 | protein demethylase activity | 11/1878 | 31/18639 | 0.354838709677419 | 3.52174585179841 | 4.7035936777617 | 0.00013219315193352 | 0.00163086116787549 | 0.00127353128064101 | KDM7A/KD | 11 |
| GO:0019887 | protein kinase regulator activity | 45/1878 | 255/18639 | 0.176470588235294 | 1.75145649314039 | 4.04437493882641 | 0.000136756680082913 | 0.00166901969778608 | 0.00130332908466115 | PRKAG1/AC | 45 |
| GO:0032356 | oxidized DNA binding | 6/1878 | 10/18639 | 0.6 | 5.95495207667732 | 5.24616773002823 | 0.000152327328852583 | 0.00181991071839665 | 0.00142115912347784 | BLM/RPS3/ | 6 |
| GO:0098505 | G-rich strand telomeric DNA binding | 6/1878 | 10/18639 | 0.6 | 5.95495207667732 | 5.24616773002823 | 0.000152327328852583 | 0.00181991071839665 | 0.00142115912347784 | TERF2/HNR | 6 |
| GO:0003727 | single-stranded RNA binding | 21/1878 | 89/18639 | 0.235955056179775 | 2.34183508633378 | 4.24736588640621 | 0.000161313853945692 | 0.00190720025237876 | 0.00148932308138457 | MSI2/HNRN | 21 |
| GO:0019207 | kinase regulator activity | 49/1878 | 287/18639 | 0.170731707317073 | 1.69449855840411 | 3.96888103444381 | 0.000163065664944457 | 0.00190803638878308 | 0.00148997601609585 | PRKAG1/AC | 49 |
| GO:0061629 | RNA polymerase II-specific DNA-binding transcription factor bind | 57/1878 | 349/18639 | 0.163323782234957 | 1.62097549365142 | 3.91992544251758 | 0.000171302887886788 | 0.00197883460143874 | 0.00154526198415156 | DDX5/MED | 57 |
| GO:0036002 | pre-mRNA binding | 16/1878 | 59/18639 | 0.271186440677966 | 2.69150376346997 | 4.35585690277668 | 0.000172603194310516 | 0.00197883460143874 | 0.00154526198415156 | U2AF1L4/D | 16 |
| GO:0023023 | MHC protein complex binding | 12/1878 | 37/18639 | 0.324324324324324 | 3.21889301442017 | 4.52225335407422 | 0.000177021050001221 | 0.00199301751090668 | 0.00155633734679659 | CD74/HLA- | 12 |
| GO:0023026 | MHC class II protein complex binding | 10/1878 | 27/18639 | 0.37037037037037 | 3.67589634362797 | 4.65749214266567 | 0.000177352219032224 | 0.00199301751090668 | 0.00155633734679659 | CD74/HLA- | 10 |
| GO:0140142 | nucleocytoplasmic carrier activity | 11/1878 | 32/18639 | 0.34375 | 3.41169129392971 | 4.57041893773707 | 0.000183210571096875 | 0.00203866664897013 | 0.0015919845290358 | TNPO2/IPO | 11 |
| GO:0016922 | nuclear receptor binding | 28/1878 | 137/18639 | 0.204379562043796 | 2.0284508290385 | 4.04419383206888 | 0.000210866701698585 | 0.00232362821774654 | 0.00181450958436595 | DDX5/MED | 28 |
| GO:0061630 | ubiquitin protein ligase activity | 57/1878 | 353/18639 | 0.161473087818697 | 1.60260749938908 | 3.82613255318437 | 0.000232217716278807 | 0.00253429911515814 | 0.00197902142820603 | TRIM26/TR | 57 |
| GO:0034511 | U3 snoRNA binding | 6/1878 | 11/18639 | 0.545454545454545 | 5.41359279697938 | 4.9012062059875 | 0.000306488653916483 | 0.00331299640185912 | 0.00258710222153061 | XRCC5/ISG | 6 |
| GO:1990841 | promoter-specific chromatin binding | 17/1878 | 68/18639 | 0.25 | 2.48123003194888 | 4.09597035537772 | 0.0003187601409527 | 0.00341313924510674 | 0.00266530326397392 | DDX5/DNM | 17 |

| | | | | | | | | | | |
|---|---|---|---|---|---|---|---|---|---|---|
| GO:0051539 | 4 iron, 4 sulfur cluster binding | 13/1878 | 45/18639 | 0.288888888888889 | 2.86719914802982 | 4.19766686094624 | 0.000351154858178811 | 0.00372486695357898 | 0.00290872986312404 | RTEL1/NDU | 13 |
| GO:0043024 | ribosomal small subunit binding | 8/1878 | 20/18639 | 0.4 | 3.96996805111821 | 4.44821821325238 | 0.000433591595087236 | 0.00455672648540753 | 0.0035583247861935 | DDX3X/PTC | 8 |
| GO:0008139 | nuclear localization sequence binding | 9/1878 | 25/18639 | 0.36 | 3.57297124600639 | 4.30906025480019 | 0.000478608845030553 | 0.00493837308281525 | 0.00385635069393039 | TNPO2/IPO | 9 |
| GO:0017056 | structural constituent of nuclear pore | 9/1878 | 25/18639 | 0.36 | 3.57297124600639 | 4.30906025480019 | 0.000478608845030553 | 0.00493837308281525 | 0.00385635069393039 | NUP88/TPR | 9 |
| GO:0016877 | ligase activity, forming carbon-sulfur bonds | 12/1878 | 41/18639 | 0.292682926829268 | 2.90485467154991 | 4.08712705459496 | 0.000515700902640581 | 0.00526902299405315 | 0.00411455354601692 | ACSL1/DIP2 | 12 |
| GO:0015923 | mannosidase activity | 7/1878 | 16/18639 | 0.4375 | 4.34215255591054 | 4.4767096115354 | 0.000524581143901326 | 0.00526902299405315 | 0.00411455354601692 | MAN2A2/M | 7 |
| GO:0034450 | ubiquitin-ubiquitin ligase activity | 7/1878 | 16/18639 | 0.4375 | 4.34215255591054 | 4.4767096115354 | 0.000524581143901326 | 0.00526902299405315 | 0.00411455354601692 | ITCH/AMFR | 7 |
| GO:0015399 | primary active transmembrane transporter activity | 31/1878 | 166/18639 | 0.186746987951807 | 1.85344893952808 | 3.69709284057914 | 0.00053091003615025 | 0.00528581483360117 | 0.00412766620903518 | NDUFV1/TC | 31 |
| GO:0043047 | single-stranded telomeric DNA binding | 6/1878 | 12/18639 | 0.5 | 4.96246006389776 | 4.59601617576661 | 0.000560715255363029 | 0.00553401578119163 | 0.00432148508023497 | TERF2/HNR | 6 |
| GO:0019843 | rRNA binding | 16/1878 | 65/18639 | 0.246153846153846 | 2.44305726222659 | 3.90108135236351 | 0.000570117444117619 | 0.00557830430235773 | 0.00435606976358471 | RPS18/RPS1 | 16 |
| GO:0003714 | transcription corepressor activity | 36/1878 | 204/18639 | 0.176470588235294 | 1.75145649314039 | 3.61239172530996 | 0.000607845815352993 | 0.00589662393526194 | 0.00460464396335781 | TCF25/COP | 36 |
| GO:0042626 | ATPase-coupled transmembrane transporter activity | 21/1878 | 98/18639 | 0.214285714285714 | 2.12676859881333 | 3.74350727042686 | 0.000650737523950807 | 0.00625921262444208 | 0.00488778764644584 | TCIRG1/AB | 21 |
| GO:0043539 | protein serine/threonine kinase activator activity | 17/1878 | 73/18639 | 0.232876712328767 | 2.31128276948663 | 3.75747496907451 | 0.000771549507974344 | 0.00735889656765445 | 0.00574652530485977 | ACSL1/DDX | 17 |
| GO:0008173 | RNA methyltransferase activity | 16/1878 | 67/18639 | 0.238805970149254 | 2.37013017977207 | 3.76068866484983 | 0.000814888997305701 | 0.00770749176618309 | 0.00601874154150351 | METTL5/NS | 16 |
| GO:0031490 | chromatin DNA binding | 25/1878 | 128/18639 | 0.1953125 | 1.93846096246006 | 3.56618394807371 | 0.000904184285152361 | 0.0084813980466771 | 0.00662308106218606 | ZNF276/HN | 25 |
| GO:0140326 | ATPase-coupled intramembrane lipid transporter activity | 9/1878 | 27/18639 | 0.333333333333333 | 3.30830670926518 | 4.0176893947577 | 0.00091187635917838 | 0.00848343989891361 | 0.00662467553432438 | ATP8B2/AB | 9 |
| GO:0005096 | GTPase activator activity | 44/1878 | 271/18639 | 0.162361623616236 | 1.61142614621034 | 3.39386998552373 | 0.0010111560315333 | 0.00933058614463656 | 0.00728620777536191 | GMIP/DNM | 44 |
| GO:0019903 | protein phosphatase binding | 27/1878 | 144/18639 | 0.1875 | 1.86092252396166 | 3.47148771802571 | 0.00110339780368097 | 0.0100996492514347 | 0.00788676528607282 | CDC27/PPP | 27 |
| GO:0008525 | phosphatidylcholine transporter activity | 7/1878 | 18/18639 | 0.388888888888889 | 3.85969116080937 | 4.06304254659677 | 0.00121765593908032 | 0.0110563159268493 | 0.00863382147962635 | ATP8B2/AB | 7 |
| GO:0008757 | S-adenosylmethionine-dependent methyltransferase activity | 30/1878 | 167/18639 | 0.179640718562874 | 1.78291978343632 | 3.40185679498297 | 0.00124119296797497 | 0.0111519661695982 | 0.00870851427294383 | METTL5/KN | 30 |
| GO:0140034 | methylation-dependent protein binding | 17/1878 | 76/18639 | 0.223684210526316 | 2.22004792332268 | 3.56744337547931 | 0.00124784114849249 | 0.0111519661695982 | 0.00870851427294383 | TDRD3/GLY | 17 |
| GO:0005048 | signal sequence binding | 13/1878 | 51/18639 | 0.254901960784314 | 2.52988160120278 | 3.66204825293879 | 0.00129190577764493 | 0.0114555707627109 | 0.00894559757217955 | TNPO2/IPO | 13 |
| GO:0019902 | phosphatase binding | 33/1878 | 190/18639 | 0.173684210526316 | 1.72380191693291 | 3.35666189636091 | 0.00132335555989416 | 0.0116434772130222 | 0.0090923327738138 | CDC27/PPP | 33 |
| GO:0030983 | mismatched DNA binding | 6/1878 | 14/18639 | 0.428571428571429 | 4.25353719762665 | 4.07632815083501 | 0.00152573743861526 | 0.0131190302486994 | 0.010244584715423 | PMS2/MSH | 6 |
| GO:0032454 | histone H3K9 demethylase activity | 6/1878 | 14/18639 | 0.428571428571429 | 4.25353719762665 | 4.07632815083501 | 0.00152573743861526 | 0.0131190302486994 | 0.010244584715423 | KDM7A/KD | 6 |
| GO:0098847 | sequence-specific single stranded DNA binding | 6/1878 | 14/18639 | 0.428571428571429 | 4.25353719762665 | 4.07632815083501 | 0.00152573743861526 | 0.0131190302486994 | 0.010244584715423 | TERF2/HNR | 6 |
| GO:0032451 | demethylase activity | 11/1878 | 40/18639 | 0.275 | 2.72935303514377 | 3.66493556526775 | 0.0015422324765469 | 0.0131611568487273 | 0.0102774811654332 | KDM7A/KD | 11 |
| GO:0043022 | ribosome binding | 22/1878 | 112/18639 | 0.196428571428571 | 1.94953788224555 | 3.37377597303739 | 0.00165277470332519 | 0.0136656233968285 | 0.0106714165547225 | DDX3X/MA | 22 |
| GO:0005522 | profilin binding | 5/1878 | 10/18639 | 0.5 | 4.96246006389776 | 4.19534433372075 | 0.00168562755555594 | 0.0136656233968285 | 0.0106714165547225 | ACTG1/RH | 5 |
| GO:0008641 | ubiquitin-like modifier activating enzyme activity | 5/1878 | 10/18639 | 0.5 | 4.96246006389776 | 4.19534433372075 | 0.00168562755555594 | 0.0136656233968285 | 0.0106714165547225 | UBA6/ATG | 5 |
| GO:0015924 | mannosyl-oligosaccharide mannosidase activity | 5/1878 | 10/18639 | 0.5 | 4.96246006389776 | 4.19534433372075 | 0.00168562755555594 | 0.0136656233968285 | 0.0106714165547225 | MAN2A2/M | 5 |
| GO:0016427 | tRNA (cytidine) methyltransferase activity | 5/1878 | 10/18639 | 0.5 | 4.96246006389776 | 4.19534433372075 | 0.00168562755555594 | 0.0136656233968285 | 0.0106714165547225 | NSUN6/NS | 5 |
| GO:0032395 | MHC class II receptor activity | 5/1878 | 10/18639 | 0.5 | 4.96246006389776 | 4.19534433372075 | 0.00168562755555594 | 0.0136656233968285 | 0.0106714165547225 | HLA-DOB/H | 5 |
| GO:0051880 | G-quadruplex DNA binding | 5/1878 | 10/18639 | 0.5 | 4.96246006389776 | 4.19534433372075 | 0.00168562755555594 | 0.0136656233968285 | 0.0106714165547225 | BLM/XRN1/ | 5 |
| GO:0003756 | protein disulfide isomerase activity | 7/1878 | 19/18639 | 0.368421052631579 | 3.65654952076677 | 3.87795140676693 | 0.00176156143000888 | 0.0140800860778879 | 0.0109950683770754 | PDIA3/P4H | 7 |
| GO:0016864 | intramolecular oxidoreductase activity, transposing S-S bonds | 7/1878 | 19/18639 | 0.368421052631579 | 3.65654952076677 | 3.87795140676693 | 0.00176156143000888 | 0.0140800860778879 | 0.0109950683770754 | PDIA3/P4H | 7 |
| GO:0043495 | protein-membrane adaptor activity | 12/1878 | 47/18639 | 0.25531914893617 | 2.53402216028822 | 3.52464183476914 | 0.00193154209514116 | 0.0153307711747218 | 0.0119717220766202 | RB1CC1/SU | 12 |
| GO:0042277 | peptide binding | 40/1878 | 249/18639 | 0.160642570281124 | 1.59436467916394 | 3.16052541204028 | 0.00203851073097122 | 0.016067428330919 | 0.012546973943551 | TNPO2/IPO | 40 |
| GO:0016779 | nucleotidyltransferase activity | 25/1878 | 136/18639 | 0.183823529411765 | 1.82443384702124 | 3.22999371871867 | 0.00219377919405748 | 0.017171995760381 | 0.0134095250918069 | THG1L/TER | 25 |
| GO:0004559 | alpha-mannosidase activity | 6/1878 | 15/18639 | 0.4 | 3.96996805111821 | 3.85175282855886 | 0.00232748902241094 | 0.0180938358934001 | 0.0141293854136266 | MAN2A2/M | 6 |
| GO:0000339 | RNA cap binding | 7/1878 | 20/18639 | 0.35 | 3.47372204472843 | 3.70497435140489 | 0.00247662211035131 | 0.018993014157086 | 0.0148315491957027 | EIF3D/LARF | 7 |
| GO:0061608 | nuclear import signal receptor activity | 7/1878 | 20/18639 | 0.35 | 3.47372204472843 | 3.70497435140489 | 0.00247662211035131 | 0.018993014157086 | 0.0148315491957027 | TNPO2/IPO | 7 |
| GO:0005548 | phospholipid transporter activity | 15/1878 | 68/18639 | 0.220588235294118 | 2.18932061642548 | 3.28876803929563 | 0.00272418001015308 | 0.0207513041041862 | 0.0162045889689077 | ATP8B2/AB | 15 |
| GO:0060590 | ATPase regulator activity | 12/1878 | 49/18639 | 0.244897959183673 | 2.43059268435809 | 3.35638544892401 | 0.0028235889678811 | 0.0213087603975054 | 0.0166399037836304 | DNAJC7/DN | 12 |
| GO:0032404 | mismatch repair complex binding | 5/1878 | 11/18639 | 0.454545454545455 | 4.51132733081615 | 3.89925851719438 | 0.00283490997358882 | 0.0213087603975054 | 0.0166399037836304 | PMS2/WRN | 5 |
| GO:0050681 | nuclear androgen receptor binding | 8/1878 | 26/18639 | 0.307692307692308 | 3.05382157778324 | 3.50783116523208 | 0.00310405776353972 | 0.0231783260632735 | 0.0180998382056817 | DDX5/SNW | 8 |
| GO:1901981 | phosphatidylinositol phosphate binding | 31/1878 | 185/18639 | 0.167567567567568 | 1.66309472411709 | 3.03398663376313 | 0.00321926674750659 | 0.0238814886171241 | 0.0186489343061613 | KRIT1/WDR | 31 |
| GO:0031369 | translation initiation factor binding | 9/1878 | 32/18639 | 0.28125 | 2.79138378594249 | 3.39486832358458 | 0.00343771962907027 | 0.025172979219321 | 0.0196574528195394 | EIF2B5/DD | 9 |
| GO:0051059 | NF-kappaB binding | 9/1878 | 32/18639 | 0.28125 | 2.79138378594249 | 3.39486832358458 | 0.00343771962907027 | 0.025172979219321 | 0.0196574528195394 | NPM1/HIF1 | 9 |
| GO:0019209 | kinase activator activity | 26/1878 | 148/18639 | 0.175675675675676 | 1.74356704947759 | 3.03996286990003 | 0.00346826662739952 | 0.0252338629621696 | 0.0197049966280054 | ACSL1/DDX | 26 |
| GO:0008017 | microtubule binding | 42/1878 | 276/18639 | 0.152173913043478 | 1.51031393249062 | 2.8590229007272 | 0.00442652202634079 | 0.0320006528655847 | 0.0249891488178273 | DNM1L/CC | 42 |
| GO:1990247 | N6-methyladenosine-containing RNA reader activity | 5/1878 | 12/18639 | 0.416666666666667 | 4.13538338658147 | 3.63669854827088 | 0.00445933477232152 | 0.0320338289024362 | 0.0250150558180861 | HNRNPC/H | 5 |
| GO:0070034 | telomerase RNA binding | 7/1878 | 22/18639 | 0.318181818181818 | 3.1579291315713 | 3.38993330351064 | 0.00455262013487845 | 0.032295149081794 | 0.0252191194313661 | HNRNPC/X | 7 |
| GO:0140517 | protein-RNA adaptor activity | 7/1878 | 22/18639 | 0.318181818181818 | 3.1579291315713 | 3.38993330351064 | 0.00455262013487845 | 0.032295149081794 | 0.0252191194313661 | HNRNPC/H | 7 |
| GO:0120013 | lipid transfer activity | 14/1878 | 65/18639 | 0.215384615384615 | 2.13767510444827 | 3.07552808276187 | 0.00467064811820193 | 0.0329266187214857 | 0.0257122308958877 | ESYT1/ABC | 14 |

| | | | | | | | | | | | |
|---|---|---|---|---|---|---|---|---|---|---|---|
| GO:0008656 | cysteine-type endopeptidase activator activity involved in apopt | 6/1878 | 17/18639 | 0.352941176470588 | 3.50291298628077 | 3.45584723065309 | 0.00482451639014183 | 0.0338013956963641 | 0.0263953398343042 | CTSH/ATP2 | 6 |
| GO:0008175 | tRNA methyltransferase activity | 9/1878 | 34/18639 | 0.264705882352941 | 2.62718473971058 | 3.178768996821 | 0.00534847119584256 | 0.0370153341907397 | 0.0289050882342711 | NSUN6/NSU | 9 |
| GO:0052866 | phosphatidylinositol phosphate phosphatase activity | 9/1878 | 34/18639 | 0.264705882352941 | 2.62718473971058 | 3.178768996821 | 0.00534847119584256 | 0.0370153341907397 | 0.0289050882342711 | INPP5K/INP | 9 |
| GO:0003950 | NAD+ poly-ADP-ribosyltransferase activity | 7/1878 | 23/18639 | 0.304347826086957 | 3.02062786498125 | 3.24567144917198 | 0.00598377810785271 | 0.0406619629565951 | 0.0317527222902417 | PARP10/PA | 7 |
| GO:0044013 | histone H2B acetyltransferase activity | 7/1878 | 23/18639 | 0.304347826086957 | 3.02062786498125 | 3.24567144917198 | 0.00598377810785271 | 0.0406619629565951 | 0.0317527222902417 | KAT6B/CRE | 7 |
| GO:1990404 | NAD+-protein mono-ADP-ribosyltransferase activity | 7/1878 | 23/18639 | 0.304347826086957 | 3.02062786498125 | 3.24567144917198 | 0.00598377810785271 | 0.0406619629565951 | 0.0317527222902417 | PARP10/PA | 7 |
| GO:0140938 | histone H3 methyltransferase activity | 11/1878 | 47/18639 | 0.234042553191489 | 2.32285364693087 | 3.03945107570899 | 0.00603068732623338 | 0.0406619629565951 | 0.0317527222902417 | KMT2A/PRI | 11 |
| GO:0030295 | protein kinase activator activity | 24/1878 | 139/18639 | 0.172661870503597 | 1.71365527386398 | 2.82688082571038 | 0.00605451254595998 | 0.0406619629565951 | 0.0317527222902417 | ACSL1/DDX | 24 |
| GO:0031491 | nucleosome binding | 15/1878 | 74/18639 | 0.202702702702703 | 2.01180813401261 | 2.9192012352413 | 0.00631407033859415 | 0.0421557049076727 | 0.0329191778643732 | HNRNPC/G | 15 |
| GO:0004693 | cyclin-dependent protein serine/threonine kinase activity | 8/1878 | 29/18639 | 0.275862068965517 | 2.73790900077118 | 3.13509228624755 | 0.00649231313962972 | 0.0425268529470118 | 0.0332090055009357 | CDKL1/CDK | 8 |
| GO:0097472 | cyclin-dependent protein kinase activity | 8/1878 | 29/18639 | 0.275862068965517 | 2.73790900077118 | 3.13509228624755 | 0.00649231313962972 | 0.0425268529470118 | 0.0332090055009357 | CDKL1/CDK | 8 |
| GO:0106222 | lncRNA binding | 8/1878 | 29/18639 | 0.275862068965517 | 2.73790900077118 | 3.13509228624755 | 0.00649231313962972 | 0.0425268529470118 | 0.0332090055009357 | PCBP2/CSD | 8 |
| GO:0017069 | snRNA binding | 12/1878 | 54/18639 | 0.222222222222222 | 2.20553780617678 | 2.96957238835655 | 0.00655700375834984 | 0.0425268529470118 | 0.0332090055009357 | RNPC3/EFT | 12 |
| GO:0019829 | ATPase-coupled monoatomic cation transmembrane transporte | 12/1878 | 54/18639 | 0.222222222222222 | 2.20553780617678 | 2.96957238835655 | 0.00655700375834984 | 0.0425268529470118 | 0.0332090055009357 | TCIRG1/ATI | 12 |
| GO:0070628 | proteasome binding | 6/1878 | 18/18639 | 0.333333333333333 | 3.30830670926518 | 3.27963680094907 | 0.00662794661567259 | 0.042648816979974 | 0.0333042466006382 | BAG6/PSMI | 6 |
| GO:0140806 | NAD+-protein-aspartate ADP-ribosyltransferase activity | 5/1878 | 13/18639 | 0.384615384615385 | 3.81727697222905 | 3.40125275783308 | 0.00665096088586379 | 0.042648816979974 | 0.0333042466006382 | PARP10/PA | 5 |
| GO:0022853 | active monoatomic ion transmembrane transporter activity | 21/1878 | 118/18639 | 0.177966101694915 | 1.76629934477717 | 2.7951474746702 | 0.00694678027074703 | 0.0442954809398757 | 0.0345901182020639 | NDUFV1/TC | 21 |
| GO:0140693 | molecular condensate scaffold activity | 11/1878 | 48/18639 | 0.229166666666667 | 2.27446086261981 | 2.95932887132517 | 0.00712697437194063 | 0.0451905916880034 | 0.0352891056817054 | TARDBP/HN | 11 |
| GO:0044183 | protein folding chaperone | 15/1878 | 75/18639 | 0.2 | 1.98498402555911 | 2.86102414941633 | 0.00717542987173602 | 0.0452450716912243 | 0.0353316488421154 | CD74/TRAP | 15 |
| GO:0035198 | miRNA binding | 10/1878 | 42/18639 | 0.238095238095238 | 2.3630762209037 | 2.96019844265662 | 0.00763075767956136 | 0.0478503313055367 | 0.0373660829670873 | MATR3/HN | 10 |
| GO:0008168 | methyltransferase activity | 32/1878 | 204/18639 | 0.156862745098039 | 1.55685021612479 | 2.67688300164897 | 0.00767398245066198 | 0.0478569784697876 | 0.037371273704207 | METTL5/KN | 32 |

**Table S11.** Significantly different skipped exon (SE) events between TP53non-exp Group 1 and Group 2, sorted by p-value.

| ID | GeneID | geneSymbol | chr | strand | exonStart_0base | exonEnd | upstreamES | upstreamEE | downstreamES | downstreamEE | ID...12 | IJC_SAMPLE_1 | SJC_SAMPLE_1 | IJC_SAMPLE_2 | SJC_SAMPLE_2 | IncFormLen | SkipFormLen | PValue | FDR | IncLevel1 | IncLevel2 | IncLevelDifference |
|---|---|---|---|---|---|---|---|---|---|---|---|---|---|---|---|---|---|---|---|---|---|---|
| 93491 | ENSG00000179344 | HLA-DQB1 | chr6 | - | 32664797 | 32665067 | 32661966 | 32662248 | 32666498 | 32666638 | 93491 | 1803,1325,219,174,669,32 | 6,0,10,18,45,41,3,2,5,0,55, | 1856,2115,15,9,12,4,74,11 | 7,0,0,1,0,0,1,1,2,0,7,16,1,0 | 298 | 149 | <1E-15 | <1E-11 | 0.993,1.0,0.916,0.8 | 0.993,1.0,1.0,0.81 | -0,057 |
| 134101 | ENSG00000100201 | DDX17 | chr22 | - | 38490082 | 38490436 | 38488025 | 38488115 | 38492055 | 38492115 | 134101 | 17,9,31,5,11,8,16,24,3,3,4, | 223,109,242,214,130,97,12 | 37,64,23,43,9,1,10,19,15,2 | 123,52,139,40,64,84,60,14 | 298 | 149 | <1E-15 | <1E-11 | 0.037,0.04,0.06,0.0 | 0.131,0.381,0.076, | -0,086 |
| 211173 | ENSG00000197622 | CDC42SE1 | chr1 | - | 151054982 | 151055126 | 151054236 | 151054321 | 151055676 | 151055993 | 211173 | 540,301,317,269,152,200,2 | 1,0,3,3,0,3,3,1,2,10,2,0 | 270,246,262,130,197,152,1 | 14,15,2,0,7,7,3,8,5,6,4,0,7, | 293 | 149 | <1E-15 | <1E-11 | 0.996,1.0,0.982,0.9 | 0.907,0.893,0.985, | 0,054 |
| 221201 | ENSG00000234745 | HLA-B | chr6 | - | 31355158 | 31355223 | 31354632 | 31354665 | 31355316 | 31355592 | 221201 | 1278,1189,2614,2250,928, | 18,0,57,44,1,0,0,2,1,0,2,0 | 1560,1895,1075,1595,938, | 22,24,1,12,3,3,34,103,61,4 | 214 | 149 | <1E-15 | <1E-11 | 0.98,1.0,0.97,0.97 | 0.98,0.982,0.999,0 | 0,036 |
| 1617 | ENSG00000183765 | CHEK2 | chr22 | - | 28696900 | 28696987 | 28695709 | 28695873 | 28699837 | 28699937 | 1617 | 24,7,29,21,6,4,11,7,5,5,8,6 | 9,0,3,3,0,1,3,2,12,2,0,2 | 17,6,8,1,1,3,4,7,8,20,2,28,1 | 0,0,0,0,0,0,0,0,0,2,0,1,3,1,0 | 236 | 149 | 1.22124532709E-15 | 5.25699705989815E-11 | 0.627,1.0,0.859,0.8 | 1.0,1.0,1.0,1.0,1.0, | -0,214 |
| 82780 | ENSG00000197622 | CDC42SE1 | chr1 | - | 151055015 | 151055126 | 151054230 | 151054321 | 151055676 | 151055993 | 82780 | 829,467,598,537,441,452,5 | 1,0,3,3,0,3,3,1,2,10,2,0 | 514,498,485,311,377,292,3 | 14,15,2,0,7,7,3,8,5,6,4,0,7, | 260 | 149 | 2.44249065418E-15 | 8.76166176649692E-11 | 0.998,1.0,0.991,0.9 | 0.955,0.95,0.993,1 | 0,022 |
| 223900 | ENSG00000173200 | PARP15 | chr3 | + | 122620418 | 122620594 | 122619780 | 122619843 | 122621443 | 122621611 | 223900 | 0,6,11,5,14,21,3,3,3,2,0,4 | 335,269,265,198,186,171,1 | 38,39,0,4,29,13,2,2,19,25,1 | 187,244,191,208,238,155,8 | 298 | 149 | 3.44169137634E-15 | 1.05822668088719E-10 | 0.0,0.011,0.02,0.0 | 0.092,0.074,0.0,0.0 | -0,028 |
| 221184 | ENSG00000234745 | HLA-B | chr6 | - | 31355106 | 31355209 | 31354632 | 31354665 | 31355316 | 31356442 | 221184 | 919,915,2495,2204,913,11 | 18,0,57,44,1,0,0,2,1,0,2,0 | 1650,1709,793,1067,895,5 | 22,24,1,12,3,3,34,103,61,4 | 252 | 149 | 5.21804821574E-15 | 1.32752451007377E-10 | 0.968,1.0,0.963,0.9 | 0.978,0.977,0.998, | 0,026 |
| 221186 | ENSG00000234745 | HLA-B | chr6 | - | 31355106 | 31355221 | 31354632 | 31354665 | 31355316 | 31356442 | 221186 | 927,923,2563,2258,1020,1 | 18,0,57,44,1,0,0,2,1,0,2,0 | 1696,1762,818,1106,905,5 | 22,24,1,12,3,3,34,103,61,4 | 264 | 149 | 5.55111512313E-15 | 1.32752451007377E-10 | 0.967,1.0,0.962,0.9 | 0.978,0.976,0.998, | 0,026 |
| 221188 | ENSG00000234745 | HLA-B | chr6 | - | 31355106 | 31355251 | 31354632 | 31354665 | 31355316 | 31356442 | 221188 | 932,926,2588,2282,1035,1 | 18,0,57,44,1,0,0,2,1,0,2,0 | 1717,1772,821,1106,912,5 | 22,24,1,12,3,3,34,103,61,4 | 294 | 149 | 1.68753899743E-14 | 3.41888619978277E-10 | 0.963,1.0,0.958,0.9 | 0.975,0.974,0.998, | 0,027 |
| 221195 | ENSG00000234745 | HLA-B | chr6 | - | 31355124 | 31355223 | 31354632 | 31354665 | 31355316 | 31355592 | 221195 | 2472,2368,4595,3964,1508 | 18,0,57,44,1,0,0,2,1,0,2,0 | 3091,3653,1610,2579,1826 | 22,24,1,12,3,3,34,103,61,4 | 248 | 149 | 1.79856129989E-14 | 3.41888619978277E-10 | 0.988,1.0,0.98,0.98 | 0.988,0.989,0.999, | 0,02 |
| 221190 | ENSG00000234745 | HLA-B | chr6 | - | 31355106 | 31355257 | 31354632 | 31354665 | 31355316 | 31356442 | 221190 | 933,926,2598,2289,1035,1 | 18,0,57,44,1,0,0,2,1,0,2,0 | 1717,1776,821,1106,911,5 | 22,24,1,12,3,3,34,103,61,4 | 298 | 149 | 1.94289029309E-14 | 3.41888619978277E-10 | 0.963,1.0,0.958,0.9 | 0.975,0.974,0.998, | 0,028 |
| 221192 | ENSG00000234745 | HLA-B | chr6 | - | 31355106 | 31355260 | 31354632 | 31354665 | 31355316 | 31356442 | 221192 | 932,926,2587,2279,1035,1 | 18,0,57,44,1,0,0,2,1,0,2,0 | 1717,1772,821,1106,912,5 | 22,24,1,12,3,3,34,103,61,4 | 298 | 149 | 2.0650148258E-14 | 3.41888619978277E-10 | 0.963,1.0,0.958,0.9 | 0.975,0.974,0.998, | 0,028 |
| 211186 | ENSG00000197622 | CDC42SE1 | chr1 | - | 151055015 | 151055129 | 151054230 | 151054321 | 151055676 | 151055993 | 211186 | 250,150,268,227,229,200,2 | 1,0,3,3,0,3,3,1,2,10,2,0 | 228,226,195,169,166,129,1 | 14,15,2,0,7,7,3,8,5,6,4,0,7, | 263 | 149 | 3.16413562018E-14 | 4.86442909762115E-10 | 0.993,1.0,0.981,0.9 | 0.902,0.895,0.982, | 0,039 |
| 247073 | ENSG00000183648 | NDUFB1 | chr14 | - | 92121195 | 92121413 | 92117497 | 92117642 | 92121641 | 92121917 | 247073 | 0,0,28,13,0,0,0,5,2,4,3,1 | 21,26,98,81,26,12,15,19,22 | 0,0,0,0,0,1,0,0,0,0,0,0,0,2,1 | 28,24,24,9,13,12,5,21,16,1 | 298 | 149 | 4.2965631053E-14 | 6.16502382477883E-10 | 0.0,0.0,0.125,0.07 | 0.0,0.0,0.0,0.0,0.0, | 0,041 |
| 211190 | ENSG00000197622 | CDC42SE1 | chr1 | - | 151055015 | 151055224 | 151054230 | 151054321 | 151055676 | 151055993 | 211190 | 264,148,282,245,266,209,2 | 1,0,3,3,0,3,3,1,2,10,2,0 | 238,246,217,195,182,148,1 | 14,15,2,0,7,7,3,8,5,6,4,0,7, | 298 | 149 | 5.46229728116E-14 | 7.34784816325842E-10 | 0.992,1.0,0.979,0.9 | 0.895,0.891,0.982, | 0,04 |
| 211191 | ENSG00000197622 | CDC42SE1 | chr1 | - | 151055015 | 151055285 | 151054230 | 151054321 | 151055676 | 151055993 | 211191 | 259,147,276,243,270,206,2 | 1,0,3,3,0,3,3,1,2,10,2,0 | 236,236,213,192,176,140,1 | 14,15,2,0,7,7,3,8,5,6,4,0,7, | 298 | 149 | 6.29496454962E-14 | 7.35728057424857E-10 | 0.992,1.0,0.979,0.9 | 0.894,0.887,0.982, | 0,04 |
| 211189 | ENSG00000197622 | CDC42SE1 | chr1 | - | 151055015 | 151055166 | 151054230 | 151054321 | 151055676 | 151055993 | 211189 | 259,147,274,243,263,206,2 | 1,0,3,3,0,3,3,1,2,10,2,0 | 236,235,212,192,176,140,1 | 14,15,2,0,7,7,3,8,5,6,4,0,7, | 298 | 149 | 6.45039577307E-14 | 7.35728057424857E-10 | 0.992,1.0,0.979,0.9 | 0.894,0.887,0.981, | 0,041 |
| 211188 | ENSG00000197622 | CDC42SE1 | chr1 | - | 151055015 | 151055153 | 151054230 | 151054321 | 151055676 | 151055993 | 211188 | 260,149,275,243,252,201,2 | 1,0,3,3,0,3,3,1,2,10,2,0 | 238,236,211,190,176,140,1 | 14,15,2,0,7,7,3,8,5,6,4,0,7, | 287 | 149 | 6.49480469406E-14 | 7.35728057424857E-10 | 0.993,1.0,0.979,0.9 | 0.898,0.891,0.982, | 0,039 |
| 232413 | ENSG00000164754 | RAD21 | chr8 | - | 116861153 | 116861297 | 116858351 | 116858458 | 116861840 | 116861940 | 232413 | 16,0,14,10,8,9,7,15,0,1,3,2 | 150,96,331,232,120,111,10 | 21,12,31,41,11,34,4,13,10, | 173,185,122,92,91,72,89,1 | 293 | 149 | 6.98330282489E-14 | 7.5151162515195E-10 | 0.051,0.0,0.021,0.0 | 0.058,0.032,0.114, | -0,046 |
| 211181 | ENSG00000197622 | CDC42SE1 | chr1 | - | 151055015 | 151055125 | 151054230 | 151054321 | 151055676 | 151055993 | 211181 | 233,132,251,217,209,183,2 | 1,0,3,3,0,3,3,1,2,10,2,0 | 216,216,181,155,156,127,1 | 14,15,2,0,7,7,3,8,5,6,4,0,7, | 259 | 149 | 8.06021915878E-14 | 8.26099537982561E-10 | 0.993,1.0,0.98,0.97 | 0.899,0.892,0.981, | 0,04 |
| 179953 | ENSG00000204252 | HLA-DOA | chr6 | - | 33007077 | 33007215 | 33006800 | 33006841 | 33007310 | 33007592 | 179953 | 99,46,375,357,89,74,161,1 | 0,0,0,0,0,0,0,0,0,0,0,3,7 | 136,177,73,27,53,56,37,12 | 12,11,0,0,0,0,3,16,0,8,4,14 | 287 | 149 | 1.03805852802E-13 | 1.01555625020124E-09 | 1.0,1.0,1.0,1.0,1.0, | 0.855,0.893,1.0,1.0 | 0,042 |
| 40110 | ENSG00000076108 | BAZ2A | chr12 | - | 56605991 | 56606063 | 56605214 | 56605327 | 56606246 | 56606312 | 40110 | 102,165,83,49,19,35,24,41 | 2,0,2,0,0,3,0,5,0,0,1,0 | 54,74,61,125,49,34,34,30,2 | 1,9,3,26,3,1,0,6,7,1,2,10,4, | 221 | 149 | 1.32338584535E-13 | 1.21568057010154E-09 | 0.972,1.0,0.965,1.0 | 0.973,0.847,0.932, | 0,11 |
| 179954 | ENSG00000204252 | HLA-DOA | chr6 | - | 33007079 | 33007188 | 33004181 | 33006841 | 33007310 | 33007592 | 179954 | 132,72,352,355,60,53,125, | 0,0,0,0,0,0,0,0,0,0,0,3,7 | 150,150,111,54,46,52,40,1 | 12,11,0,0,0,0,3,16,0,8,4,14 | 258 | 149 | 1.35558231307E-13 | 1.21568057010154E-09 | 1.0,1.0,1.0,1.0,1.0, | 0.878,0.887,1.0,1.0 | 0,035 |
| 132397 | ENSG00000213923 | CSNK1E | chr22 | - | 38295386 | 38295447 | 38294341 | 38294534 | 38298785 | 38298934 | 132397 | 4,2,11,16,3,7,7,12,7,4,3,0 | 37,33,73,45,23,28,19,52,32 | 1,0,0,0,3,5,0,0,3,0,3,0,5,0,0 | 32,29,37,38,37,23,18,52,18 | 210 | 149 | 1.49658063719E-13 | 1.2637012163196E-09 | 0.071,0.041,0.097, | 0.022,0.0,0.0,0.0,0 | 0,083 |
| 179955 | ENSG00000204252 | HLA-DOA | chr6 | - | 33007079 | 33007207 | 33004181 | 33006841 | 33007310 | 33007592 | 179955 | 145,84,388,389,69,64,145, | 0,0,0,0,0,0,0,0,0,0,0,3,7 | 167,156,123,58,52,57,46,1 | 12,11,0,0,0,0,3,16,0,8,4,14 | 277 | 149 | 1.52655665886E-13 | 1.2637012163196E-09 | 1.0,1.0,1.0,1.0,1.0, | 0.882,0.884,1.0,1.0 | 0,034 |
| 29271 | ENSG00000113240 | CLK4 | chr5 | - | 178622392 | 178622459 | 178619837 | 178620673 | 178623255 | 178623416 | 29271 | 10,1,44,24,12,11,13,27,1,1 | 5,0,13,9,1,0,2,3,0,0,0,2 | 30,45,15,13,15,9,14,26,9,1 | 1,4,0,1,0,0,2,0,0,1,0,0,1,0,0 | 216 | 149 | 1.71862524212E-13 | 1.3700052943953E-09 | 0.58,1.0,0.7,0.648, | 0.954,0.886,1.0,0.9 | -0,099 |
| 179956 | ENSG00000204252 | HLA-DOA | chr6 | - | 33007079 | 33007213 | 33004181 | 33006841 | 33007310 | 33007592 | 179956 | 154,84,427,430,71,67,149, | 0,0,0,0,0,0,0,0,0,0,0,3,7 | 177,174,127,62,58,63,49,1 | 12,11,0,0,0,0,3,16,0,8,4,14 | 283 | 149 | 1.85962356625E-13 | 1.42945942781269E-09 | 1.0,1.0,1.0,1.0,1.0, | 0.886,0.893,1.0,1.0 | 0,033 |
| 24701 | ENSG00000136098 | NEK3 | chr13 | - | 52141019 | 52141069 | 52136799 | 52136902 | 52143915 | 52143987 | 24701 | 8,3,12,8,4,5,4,4,2,3,2,1 | 3,1,0,0,0,0,0,0,0,2,0,0 | 10,10,4,5,6,1,3,7,5,3,11,10 | 1,2,3,1,4,2,0,5,5,5,6,13,3,2 | 199 | 149 | 3.06865644006E-13 | 2.27748273879501E-09 | 0.666,0.692,1.0,1.0 | 0.882,0.789,0.5,0.7 | 0,273 |
| 12618 | ENSG00000011260 | UTP18 | chr17 | + | 51280004 | 51280105 | 51275867 | 51275991 | 51280388 | 51280479 | 12618 | 21,15,48,38,15,25,15,30,15 | 2,1,0,0,0,3,0,0,0,0,0,0 | 36,31,18,28,22,15,17,21,23 | 3,4,0,2,4,2,0,3,1,1,6,13,1,3 | 250 | 149 | 4.0079051189E-13 | 2.87541808881989E-09 | 0.862,0.899,1.0,1.0 | 0.877,0.822,1.0,0.8 | 0,117 |
| 205915 | ENSG00000128815 | WDFY4 | chr10 | + | 48824680 | 48824925 | 48822379 | 48822537 | 48826670 | 48826909 | 205915 | 0,0,2,0,19,22,0,1,0,0,9,18 | 97,60,97,107,74,48,37,52,8 | 1,1,0,2,0,0,0,0,0,0,0,0,1,0,2 | 65,92,63,46,53,31,34,47,30 | 298 | 149 | 4.39426273147E-13 | 3.05090826437748E-09 | 0.0,0.0,0.01,0.0,0.1 | 0.008,0.005,0.0,0.0 | 0,046 |
| 185503 | ENSG00000134352 | IL6ST | chr5 | - | 55948858 | 55948982 | 55947564 | 55947589 | 55951463 | 55951506 | 185503 | 1,1,0,0,1,0,1,0,0,1,0,0 | 7,7,12,5,6,9,18,59,13,5,2,1 | 5,3,7,6,2,1,3,8,3,8,7,0,0,0,1 | 9,25,10,13,17,4,5,13,3,2,26 | 273 | 149 | 4.93494134446E-13 | 3.31922612659209E-09 | 0.072,0.072,0.0,0.0 | 0.233,0.061,0.276, | -0,137 |
| 93277 | ENSG00000291107 | | chr12 | + | 9653682 | 9653808 | 9648046 | 9648933 | 9655872 | 9655979 | 93277 | 179,107,690,303,134,194,1 | 8,6,0,0,12,7,12,14,2,3,1,0 | 657,412,110,123,170,141,1 | 0,0,4,4,5,9,0,0,0,0,0,0,0,0,0 | 275 | 149 | 6.99884594724E-13 | 4.56475336991034E-09 | 0.924,0.906,1.0,1.0 | 1.0,1.0,0.937,0.94 | -0,046 |
| 233141 | ENSG00000065548 | ZC3H15 | chr2 | + | 186489161 | 186489245 | 186486266 | 186486457 | 186495232 | 186495334 | 233141 | 3,5,0,5,1,4,2,3,0,0,1,0 | 156,121,118,94,56,66,86,6 | 3,12,10,6,9,3,0,7,5,5,3,26,5 | 71,111,81,101,72,53,37,69 | 233 | 149 | 8.09352584952E-13 | 5.12346371211188E-09 | 0.012,0.026,0.0,0.0 | 0.026,0.065,0.073, | -0,035 |
| 40109 | ENSG00000076108 | BAZ2A | chr12 | - | 56605829 | 56606063 | 56605072 | 56605327 | 56606246 | 56606312 | 40109 | 157,237,235,154,72,104,55 | 2,0,2,0,0,3,0,5,0,0,1,0 | 122,250,116,236,122,80,12 | 1,9,3,26,3,1,0,6,7,1,2,10,4, | 298 | 149 | 9.35806987457E-13 | 5.75470496335307E-09 | 0.975,1.0,0.983,1.0 | 0.984,0.933,0.951, | 0,061 |
| 201480 | ENSG00000131127 | ZNF141 | chr4 | + | 342291 | 342426 | 337833 | 337986 | 343781 | 343908 | 201480 | 1,0,26,13,0,1,1,4,2,0,0,1 | 8,9,98,59,25,22,11,32,20,8 | 0,3,0,0,1,0,0,0,0,0,0,1,3,2,0 | 31,34,9,20,26,17,18,19,13, | 284 | 149 | 1.24955601422E-12 | 7.47064418046069E-09 | 0.062,0.0,0.122,0.1 | 0.0,0.044,0.0,0.0,0 | 0,03 |
| 221287 | ENSG00000173041 | ZNF680 | chr7 | - | 64543706 | 64544201 | 64519877 | 64522500 | 64544305 | 64544432 | 221287 | 7,8,24,12,6,4,7,6,5,5,6,2 | 2,4,5,6,0,1,1,5,0,4,0,0 | 17,6,6,6,1,5,3,9,1,5,12,3,9, | 0,0,0,0,1,0,0,0,1,0,0,3,1,0,0 | 298 | 149 | 2.02571293073E-12 | 1.17836816160527E-08 | 0.636,0.5,0.706,0.5 | 1.0,1.0,1.0,1.0,0.33 | -0,151 |
| 191064 | ENSG00000144559 | TAMM41 | chr3 | - | 11844890 | 11844999 | 11844028 | 11844211 | 11846501 | 11846919 | 191064 | 3,3,10,3,3,7,6,5,0,0,2,2 | 5,5,12,13,4,5,2,0,4,2,3,2 | 0,3,0,0,0,0,0,2,1,1,0,2,0,0,3 | 4,9,5,4,11,10,1,3,7,3,7,8,5, | 258 | 149 | 2.58559840205E-12 | 1.4644761307148E-08 | 0.257,0.257,0.325, | 0.0,0.161,0.0,0.0,0 | 0,258 |
| 168250 | ENSG00000138639 | ARHGAP24 | chr4 | + | 85664004 | 85664157 | 85604152 | 85604303 | 85721884 | 85722386 | 168250 | 0,0,13,8,0,0,5,19,2,4,0,0 | 3,3,70,41,8,36,37,108,47,2 | 0,0,0,2,0,0,0,0,0,0,2,0,0,0,0 | 85,24,38,48,50,70,38,54,59 | 298 | 149 | 3.16147108492E-12 | 1.72447425145462E-08 | 0.0,0.0,0.085,0.08 | 0.0,0.0,0.0,0.02,0.0 | 0,034 |
| 221254 | ENSG00000118873 | RAB3GAP2 | chr1 | - | 220206533 | 220206677 | 220205906 | 220206006 | 220210387 | 220211002 | 221254 | 6,0,4,2,0,1,30,28,0,3,2,2 | 26,23,128,101,34,43,38,34 | 0,0,2,4,2,0,0,1,1,2,0,0,2,3,1 | 45,27,73,63,34,25,14,53,31 | 293 | 149 | 3.20488080519E-12 | 1.72447425145462E-08 | 0.105,0.0,0.016,0.0 | 0.0,0.0,0.014,0.03 | 0,08 |
| 135175 | ENSG00000093000 | NUP50 | chr22 | + | 45175893 | 45176080 | 45171599 | 45171683 | 45178799 | 45178900 | 135175 | 26,11,35,23,33,14,8,26,22, | 1,0,3,1,1,0,2,0,0,0,0,0 | 25,10,9,4,7,21,8,18,3,11,12 | 4,3,4,1,0,2,0,4,3,0,3,6,2,1,5 | 298 | 149 | 3.56215057451E-12 | 1.86996397634722E-08 | 0.929,1.0,0.854,0.9 | 0.758,0.625,0.529, | 0,199 |
| 205916 | ENSG00000128815 | WDFY4 | chr10 | + | 48824680 | 48824999 | 48822379 | 48822537 | 48826670 | 48826909 | 205916 | 2,0,2,0,19,20,0,1,0,0,6,18 | 97,60,97,107,74,48,37,52,8 | 1,1,0,2,0,0,0,0,0,0,0,0,1,0,2 | 65,92,63,46,53,31,34,47,30 | 298 | 149 | 3.71369601737E-12 | 1.90310120836801E-08 | 0.01,0.0,0.01,0.0,0 | 0.008,0.005,0.0,0.0 | 0,043 |
| 76044 | ENSG00000135541 | AHI1 | chr6 | - | 135433026 | 135433256 | 135431207 | 135431314 | 135438374 | 135438498 | 76044 | 49,24,108,48,40,41,23,50,1 | 6,3,3,2,5,5,2,5,2,0,4,0 | 20,17,25,66,25,39,23,51,11 | 1,0,2,1,1,0,1,1,0,0,2,0,0,1,2 | 298 | 149 | 7.00872693216E-12 | 3.50812861938542E-08 | 0.803,0.8,0.947,0.9 | 0.909,1.0,0.862,0.9 | -0,105 |
| 12177 | ENSG00000170037 | CNTROB | chr17 | + | 7949081 | 7949157 | 7948486 | 7948619 | 7949384 | 7949578 | 12177 | 3,3,6,8,7,9,1,4,9,6,3,4 | 6,0,3,0,2,2,1,4,0,2,4,2 | 5,3,8,1,5,1,1,9,8,4,8,4,6,7,4 | 0,2,0,0,0,2,0,3,0,0,0,1,2,0,0 | 225 | 149 | 7.65831842386E-12 | 3.74615348337684E-08 | 0.249,1.0,0.57,1.0, | 1.0,0.498,1.0,1.0,1 | -0,227 |
| 50618 | ENSG00000152223 | EPG5 | chr18 | - | 45887750 | 45887907 | 45884616 | 45884811 | 45889797 | 45889940 | 50618 | 8,11,41,24,6,13,23,10,12,1 | 0,3,0,0,0,1,0,1,0,0,2,0 | 21,19,8,7,5,9,10,11,15,13,2 | 1,8,0,2,1,0,3,2,5,0,4,3,2,2,2 | 298 | 149 | 9.66760005383E-12 | 4.62392717152419E-08 | 1.0,0.647,1.0,1.0,1 | 0.913,0.543,1.0,0.6 | 0,129 |
| 94281 | ENSG00000173041 | ZNF680 | chr7 | - | 64543706 | 64543802 | 64519877 | 64522500 | 64544305 | 64544432 | 94281 | 8,11,36,33,16,6,6,14,7,6,6, | 2,4,5,6,0,1,1,5,0,4,0,0 | 12,4,8,8,4,10,8,10,5,23,18, | 0,0,0,0,1,0,0,0,1,0,0,3,1,0,0 | 245 | 149 | 1.26890720153E-11 | 5.93713404114138E-08 | 0.709,0.626,0.814, | 1.0,1.0,1.0,1.0,0.70 | -0,149 |
| 128796 | ENSG00000149100 | EIF3M | chr11 | + | 32602078 | 32602114 | 32601761 | 32601822 | 32602278 | 32602377 | 128796 | 2,4,6,2,1,9,0,1,2,0,2,2 | 160,139,342,280,109,98,10 | 12,12,36,16,3,1,8,6,7,2,11, | 169,238,172,102,198,82,63 | 185 | 149 | 1.49193990495E-11 | 6.83216420600625E-08 | 0.01,0.023,0.014,0 | 0.054,0.039,0.144, | -0,028 |
| 204585 | ENSG00000135541 | AHI1 | chr6 | - | 135433026 | 135433170 | 135431207 | 135431314 | 135438374 | 135438498 | 204585 | 33,17,56,34,18,21,17,23,7, | 6,3,3,2,5,5,2,5,2,0,4,0 | 7,6,17,36,17,26,14,19,6,10 | 1,0,2,1,1,0,1,1,0,0,2,0,0,1,2 | 293 | 149 | 1.87135862362E-11 | 8.39113308167409E-08 | 0.737,0.742,0.905, | 0.781,1.0,0.812,0.9 | -0,135 |
| 73760 | ENSG00000138398 | PPIG | chr2 | + | 169603641 | 169603694 | 169584341 | 169584490 | 169604025 | 169604102 | 73760 | 34,29,32,39,16,17,20,26,20 | 3,0,1,0,0,1,0,0,0,0,0,0 | 49,49,11,20,26,6,16,15,9,2 | 0,4,4,3,5,0,0,5,4,0,4,3,0,4,4 | 202 | 149 | 2.08216777153E-11 | 9.14585819661578E-08 | 0.893,1.0,0.959,1.0 | 1.0,0.9,0.67,0.831, | 0,115 |
| 1749 | ENSG00000284554 | | chr22 | + | 39025069 | 39025349 | 39022821 | 39023014 | 39029362 | 39029519 | 1749 | 9,4,12,23,13,2,7,12,7,7,3,4 | 5,1,6,8,2,0,3,4,0,4,0,0 | 8,7,2,6,7,11,2,8,3,4,5,21,6, | 1,2,0,2,0,0,0,2,0,0,0,1,1,0,0 | 298 | 149 | 2.87826429357E-11 | 1.23898340433873E-07 | 0.474,0.667,0.5,0.5 | 0.8,0.636,1.0,0.6,1 | -0,119 |
| 146276 | ENSG00000131148 | EMC8 | chr16 | - | 85780378 | 85780469 | 85778623 | 85779867 | 85781210 | 85781760 | 146276 | 15,8,27,12,12,5,7,5,9,8,2,8 | 0,4,5,5,3,3,2,0,4,0,0,1 | 10,20,8,6,7,6,3,11,11,7,2,2 | 0,1,0,0,0,0,0,0,2,0,3,1,4,1,0 | 240 | 149 | 3.58799656652E-11 | 1.51421193923268E-07 | 1.0,0.554,0.77,0.59 | 1.0,0.925,1.0,1.0,1 | -0,137 |
| 13035 | ENSG00000074370 | ATP2A3 | chr17 | - | 3928226 | 3928327 | 3923869 | 3925441 | 3928662 | 3928780 | 13035 | 2,2,22,31,1,2,4,0,4,5,0,0 | 80,60,111,70,80,62,49,34,6 | 2,1,1,0,6,8,0,0,1,1,0,0,6,1,0 | 101,125,32,24,62,56,16,71 | 250 | 149 | 4.23263646354E-11 | 1.73933563786614E-07 | 0.015,0.019,0.106, | 0.012,0.005,0.018, | 0,029 |
| 220337 | ENSG00000291107 | | chr12 | + | 9653682 | 9653780 | 9648046 | 9648933 | 9655872 | 9655979 | 220337 | 27,9,154,71,46,87,52,134,1 | 8,6,0,0,12,7,12,14,2,3,1,0 | 144,80,35,33,31,40,54,107 | 0,0,4,4,5,9,0,0,0,0,0,0,0,0,0 | 247 | 149 | 4.28306279332E-11 | 1.73933563786614E-07 | 0.671,0.475,1.0,1.0 | 1.0,1.0,0.841,0.83 | -0,137 |
| 31261 | ENSG00000163297 | ANTXR2 | chr4 | - | 79907576 | 79907664 | 79905590 | 79907467 | 79977620 | 79977701 | 31261 | 0,1,0,0,2,0,2,1,0,0,1,0 | 19,17,21,40,13,14,8,21,12, | 12,5,0,8,0,0,1,5,1,3,11,4,0, | 23,13,16,19,16,13,30,13,6, | 237 | 149 | 4.88075135863E-11 | 1.94534999198017E-07 | 0.0,0.036,0.0,0.0,0 | 0.247,0.195,0.0,0.2 | -0,063 |
| 76792 | ENSG00000116984 | MTR | chr1 | + | 236852520 | 236852637 | 236850343 | 236850523 | 236852947 | 236853088 | 76792 | 31,11,80,55,30,47,34,29,11 | 0,1,1,0,0,2,1,0,0,0,1,0 | 57,49,36,20,31,10,21,22,17 | 3,5,2,2,7,4,0,5,0,2,2,0,1,3,0 | 266 | 149 | 5.36207744872E-11 | 2.09833689339174E-07 | 1.0,0.86,0.978,1.0, | 0.914,0.846,0.91,0 | 0,081 |
| 246634 | ENSG00000054654 | SYNE2 | chr14 | + | 64082423 | 64082719 | 64081442 | 64081580 | 64087670 | 64087924 | 246634 | 2,3,15,18,5,3,1,13,8,0,0,8 | 34,67,92,80,51,100,74,121 | 0,1,4,7,0,0,2,1,9,5,1,0,2,3,2 | 66,86,38,60,97,89,27,62,90 | 298 | 149 | 5.53183054919E-11 | 2.12610968023699E-07 | 0.029,0.022,0.075, | 0.0,0.006,0.05,0.05 | 0,029 |
| 9907 | ENSG00000168961 | LGALS9 | chr17 | + | 27645313 | 27645349 | 27642237 | 27642348 | 27645860 | 27645899 | 9907 | 85,31,48,65,69,23,22,11,18 | 3,0,4,0,1,1,2,1,6,1,0,0 | 36,17,31,19,10,14,11,45,6, | 6,3,3,3,0,0,0,7,0,3,1,10,10, | 185 | 149 | 5.93749494016E-11 | 2.24198767274663E-07 | 0.958,1.0,0.906,1.0 | 0.829,0.82,0.893,0 | 0,106 |
| 145064 | ENSG00000154945 | ANKRD40 | chr17 | - | 50701872 | 50702008 | 50700567 | 50700716 | 50707520 | 50707914 | 145064 | 0,0,1,1,0,4,0,1,1,0,0,0 | 18,29,31,15,12,6,15,8,9,13 | 10,10,1,4,2,1,2,3,2,5,3,0,2, | 18,21,11,3,5,8,13,23,2,11,1 | 285 | 149 | 6.14496231677E-11 | 2.28032135241504E-07 | 0.0,0.0,0.017,0.03 | 0.225,0.199,0.045, | -0,101 |
| 132060 | ENSG00000272917 | | chr12 | + | 9653682 | 9653808 | 9642871 | 9643103 | 9655872 | 9655979 | 132060 | 179,114,625,275,119,93,13 | 19,15,0,0,16,3,19,7,14,4,0, | 664,440,73,85,179,122,127 | 0,0,7,16,29,12,0,0,0,0,0,0,0 | 275 | 149 | 6.71380728789E-11 | 2.44918551928789E-07 | 0.836,0.805,1.0,1.0 | 1.0,1.0,0.85,0.742, | -0,065 |
| 131254 | ENSG00000111229 | ARPC3 | chr12 | - | 110436556 | 110436683 | 110436109 | 110436204 | 110437083 | 110437152 | 131254 | 653,416,1198,1051,471,36 | 82,40,42,42,27,8,19,7,10,8 | 663,799,636,256,578,464,1 | 30,34,18,10,3,14,3,7,6,8,7, | 276 | 149 | 6.98704427649E-11 | 2.5063808777887E-07 | 0.811,0.849,0.939, | 0.923,0.927,0.95,0 | -0,038 |
| 130793 | ENSG00000123349 | PFDN5 | chr12 | + | 53295838 | 53296275 | 53295478 | 53295639 | 53297849 | 53297924 | 130793 | 1400,890,1219,1007,340,3 | 62,26,120,60,24,30,21,42,2 | 726,606,613,389,460,252,2 | 50,56,40,22,61,26,27,59,28 | 298 | 149 | 8.88222828621E-11 | 3.13398504306437E-07 | 0.919,0.945,0.836, | 0.879,0.844,0.885, | 0,06 |
| 94550 | ENSG00000231925 | TAPBP | chr6 | - | 33304127 | 33304217 | 33302853 | 33303989 | 33304988 | 33305387 | 94550 | 71,77,96,58,74,65,81,89,49 | 1,0,0,3,0,0,2,0,0,0,0,0 | 56,81,52,58,39,37,32,75,27 | 2,6,0,3,4,0,7,8,1,1,2,10,0,0 | 239 | 149 | 9.32283139576E-11 | 3.23639084538842E-07 | 0.978,1.0,1.0,0.92 | 0.946,0.894,1.0,0.9 | 0,065 |
| 23390 | ENSG00000070367 | EXOC5 | chr14 | - | 57244164 | 57244359 | 57239527 | 57239659 | 57246710 | 57246827 | 23390 | 49,35,82,59,45,25,31,34,44 | 0,0,3,0,2,2,1,1,0,0,0,0 | 60,48,33,12,27,19,14,29,19 | 0,5,5,3,2,1,1,1,12,2,1,1,3,1 | 298 | 149 | 9.82974812658E-11 | 3.35820082386022E-07 | 1.0,1.0,0.932,1.0,0 | 1.0,0.828,0.767,0.0 | 0,117 |
| 172651 | ENSG00000168216 | LMBRD1 | chr6 | - | 69738401 | 69738832 | 69737941 | 69738015 | 69741788 | 69741877 | 172651 | 11,4,16,13,5,4,4,8,0,0,1,8 | 31,37,138,108,72,60,44,65 | 1,7,1,0,2,0,1,0,0,0,0,2,5,5,4 | 94,100,38,43,69,47,56,58,4 | 298 | 149 | 1.00286889904E-10 | 3.37263243764497E-07 | 0.151,0.051,0.055, | 0.005,0.034,0.013, | 0,031 |
| 117118 | ENSG00000080608 | PUM3 | chr9 | - | 2824716 | 2824815 | 2823780 | 2823834 | 2827072 | 2827151 | 117118 | 18,9,41,16,18,24,15,31,22, | 2,2,8,2,2,2,2,6,0,0,0,2 | 29,15,21,7,5,3,10,20,5,11,2 | 1,0,2,1,0,0,0,0,0,0,0,0,0,1,0 | 248 | 149 | 1.27836075059E-10 | 4.23296711861902E-07 | 0.844,0.73,0.755,0 | 0.946,1.0,0.863,0.8 | -0,101 |
| 93468 | ENSG00000196735 | HLA-DQA1 | chr6 | + | 32642609 | 32642784 | 32641971 | 32642253 | 32642951 | 32643272 | 93468 | 505,335,722,980,566,191,5 | 0,2,0,0,0,1,0,1,10,4,0,4 | 438,302,134,102,63,38,109 | 1,3,0,0,0,0,0,0,0,0,3,61,0,0 | 298 | 149 | 1.31309962903E-10 | 4.28211736751145E-07 | 1.0,0.988,1.0,1.0,1 | 0.995,0.981,1.0,1.0 | 0,004 |
| 226178 | ENSG00000106443 | PHF14 | chr7 | + | 11061790 | 11061851 | 11051611 | 11051780 | 11061963 | 11063072 | 226178 | 15,3,22,6,9,15,3,7,7,12,1,8 | 0,0,2,0,0,0,1,4,0,0,0,0 | 9,6,6,1,19,7,5,4,3,8,12,7,4, | 3,3,2,4,1,2,0,0,0,0,4,5,2,3,0 | 210 | 149 | 1.47945877771E-10 | 4.75261779381046E-07 | 1.0,1.0,0.886,1.0,1 | 0.68,0.587,0.68,0.1 | 0,194 |
| 221179 | ENSG00000234745 | HLA-B | chr6 | - | 31354778 | 31355223 | 31354632 | 31354665 | 31355316 | 31355592 | 221179 | 3789,3687,6091,5420,1942 | 18,0,57,44,1,0,0,2,1,0,2,0 | 4100,4897,2232,3825,2537 | 22,24,1,12,3,3,34,103,61,4 | 298 | 149 | 1.62715951824E-10 | 5.15022309221049E-07 | 0.991,1.0,0.982,0.9 | 0.989,0.99,0.999,0 | 0,017 |
| 234648 | ENSG00000101199 | ARFGAP1 | chr20 | + | 63276919 | 63277305 | 63276479 | 63276651 | 63278116 | 63278203 | 234648 | 15,20,36,20,14,11,14,16,15 | 3,4,4,0,0,0,1,1,9,4,2,1 | 13,17,8,14,12,4,10,24,9,15 | 2,4,0,0,0,0,0,3,0,0,0,1,0,1,0 | 298 | 149 | 1.67239888604E-10 | 5.21669687885906E-07 | 0.714,0.714,0.818, | 0.765,0.68,1.0,1.0, | -0,142 |
| 119771 | ENSG00000166266 | CUL5 | chr11 | + | 108104189 | 108104902 | 108098405 | 108098529 | 108106206 | 108106852 | 119771 | 19,19,23,17,4,6,2,9,10,6,3, | 2,0,5,1,0,0,0,0,0,0,0,0 | 31,30,5,3,7,12,4,5,8,21,9,2 | 6,1,4,1,6,2,2,0,2,0,4,0,2,1,3 | 298 | 149 | 1.87334814328E-10 | 5.76003706037568E-07 | 0.826,1.0,0.697,0.8 | 0.721,0.938,0.385, | 0,238 |
| 232414 | ENSG00000164754 | RAD21 | chr8 | - | 116861153 | 116861318 | 116858351 | 116858458 | 116861840 | 116861940 | 232414 | 8,0,8,4,6,1,3,5,0,2,2,1 | 150,96,331,232,120,111,10 | 16,8,16,14,1,18,2,8,2,6,8,1 | 173,185,122,92,91,72,89,1 | 298 | 149 | 2.23975282765E-10 | 6.78963719504137E-07 | 0.026,0.0,0.012,0.0 | 0.044,0.021,0.062, | -0,024 |
| 239515 | ENSG00000108091 | CCDC6 | chr10 | - | 59794574 | 59794597 | 59791751 | 59793111 | 59804419 | 59804520 | 239515 | 6,2,10,8,5,7,14,18,7,2,2,2 | 3,0,4,0,4,4,2,1,7,1,1,3 | 11,9,6,2,5,6,2,11,5,4,26,9,6 | 0,0,3,1,0,0,0,1,1,0,3,1,0,4,0 | 172 | 149 | 2.56321186498E-10 | 7.66225906821542E-07 | 0.634,1.0,0.684,1.0 | 1.0,1.0,0.634,0.63 | -0,21 |
| 48513 | ENSG00000138074 | SLC5A6 | chr2 | - | 27206482 | 27206534 | 27206025 | 27206093 | 27206876 | 27206942 | 48513 | 12,7,11,8,12,8,7,8,3,5,5,9 | 0,0,3,0,0,0,2,2,0,0,0,0 | 25,19,4,1,10,6,9,13,2,2,14, | 4,6,2,0,3,1,4,2,3,0,0,6,3,0,0 | 201 | 149 | 2.89629653594E-10 | 8.53935342091647E-07 | 1.0,1.0,0.731,1.0,1 | 0.822,0.701,0.597, | 0,181 |
| 87646 | ENSG00000268362 | | chr19 | + | 24048936 | 24049014 | 24046183 | 24046279 | 24049404 | 24049548 | 87646 | 1,0,5,2,2,1,3,0,3,1,0,1 | 21,6,28,19,8,18,13,31,11,3 | 12,14,5,3,0,1,2,1,1,13,2,3,7 | 4,6,10,15,10,5,3,8,11,8,8,1 | 227 | 149 | 3.35145022845E-10 | 9.74778356918273E-07 | 0.03,0.0,0.105,0.0 | 0.663,0.605,0.247, | -0,193 |
| 201021 | ENSG00000120029 | ARMH3 | chr10 | - | 102001694 | 102002072 | 101995296 | 101995355 | 102006539 | 102006633 | 201021 | 12,2,17,15,10,7,6,21,6,3,7, | 2,0,2,0,3,2,0,0,0,0,0,0 | 18,12,4,5,6,7,5,11,6,12,7,6 | 3,11,2,1,2,0,2,2,1,1,1,0,1,1 | 298 | 149 | 3.44672179686E-10 | 9.88662245569262E-07 | 0.75,1.0,0.81,1.0,0 | 0.75,0.353,0.5,0.7 | 0,18 |

| | | | | | | | | | | | | | | | | | | | | | | |
|---|---|---|---|---|---|---|---|---|---|---|---|---|---|---|---|---|---|---|---|---|---|---|
| 92144 | ENSG00000137200 | CMTR1 | chr6 | + | 37461548 | 37461645 | 37459565 | 37459684 | 37471020 | 37471077 | 92144 | 39,19,41,27,13,10,10,17,18 | 1,1,0,1,0,1,0,1,0,0,0,3 | 22,23,10,15,25,26,10,20,10 | 7,7,0,1,5,1,0,5,1,0,1,0,1,2,2 | 246 | 149 | 3.49105522268E-10 | 9.88662245569262E-07 | 0.959,0.92,1.0,0.9 | 0.656,0.666,1.0,0. | 0,093 |
| 213603 | ENSG00000272752 | STAG3L5P-PVR | chr7 | + | 100351257 | 100351410 | 100349798 | 100349887 | 100352162 | 100352211 | 213603 | 21,10,28,24,15,13,10,13,12 | 3,0,0,1,0,0,0,1,0,0,0,0 | 6,13,3,9,13,19,6,14,3,9,11, | 3,5,1,0,0,0,1,4,0,1,0,3,0,1, | 298 | 149 | 4.1020720154E-10 | 1.14661436616436E-06 | 0.778,1.0,1.0,0.92 | 0.5,0.565,0.6,1.0,1 | 0,194 |
| 12179 | ENSG00000170037 | CNTROB | chr17 | + | 7949084 | 7949157 | 7948486 | 7948619 | 7949384 | 7949576 | 12179 | 12,2,10,11,13,7,2,7,18,21,4 | 6,0,3,0,2,2,1,4,0,2,4,2 | 7,3,9,3,5,2,3,13,8,15,6,7,4, | 0,2,0,0,0,2,0,3,0,0,0,1,2,0, | 222 | 149 | 4.29683177927E-10 | 0.0000011727005357357 | 0.573,1.0,0.691,1. | 1.0,0.502,1.0,1.0,1 | -0,13 |
| 248903 | ENSG00000119777 | TMEM214 | chr2 | + | 27039715 | 27039837 | 27039046 | 27039164 | 27040029 | 27040198 | 248903 | 3,2,9,4,8,5,5,5,3,2,2,7 | 0,2,3,0,2,0,0,1,0,0,0,0 | 5,8,5,1,0,3,4,3,3,4,3,4,11,1 | 1,6,0,3,2,1,4,0,1,0,2,2,2,0,1 | 271 | 149 | 4.30436797316E-10 | 0.0000011727005357357 | 1.0,0.355,0.623,1. | 0.733,0.423,1.0,0. | 0,255 |
| 219223 | ENSG00000137200 | CMTR1 | chr6 | + | 37461548 | 37461710 | 37459565 | 37459684 | 37471020 | 37471077 | 219223 | 39,18,39,25,11,10,9,19,18, | 1,1,0,1,0,1,0,1,0,0,0,3 | 20,21,11,15,25,26,13,13,10 | 7,7,0,1,5,1,0,5,1,0,1,0,1,2,2 | 298 | 149 | 4.57780369167E-10 | 1.23160658295228E-06 | 0.951,0.9,1.0,0.92 | 0.588,0.6,1.0,0.88 | 0,127 |
| 183755 | ENSG00000171163 | ZNF692 | chr1 | - | 248856287 | 248856384 | 248855557 | 248855946 | 248856513 | 248856562 | 183755 | 9,6,7,2,3,14,2,9,12,0,3,11 | 1,1,5,1,3,2,1,5,0,3,0,2 | 4,3,3,2,1,5,4,6,6,3,16,12,1, | 2,0,0,0,0,0,4,0,0,2,3,0,0, | 246 | 149 | 4.99015051503E-10 | 1.32596924135855E-06 | 0.845,0.784,0.459, | 0.548,1.0,1.0,1.0,1 | -0,272 |
| 198378 | ENSG00000069248 | NUP133 | chr1 | - | 229477596 | 229477755 | 229475637 | 229475732 | 229484053 | 229484145 | 198378 | 23,24,86,45,41,15,26,34,26 | 2,2,1,0,0,0,0,4,1,0,0,0 | 50,68,16,20,36,16,28,28,45 | 4,1,3,6,2,3,1,3,2,2,4,5,1,1,4 | 298 | 149 | 5.68326830042E-10 | 1.44684844156149E-06 | 0.852,0.857,0.977, | 0.862,0.971,0.727, | 0,114 |
| 198380 | ENSG00000069248 | NUP133 | chr1 | - | 229477596 | 229477759 | 229475637 | 229475732 | 229484053 | 229484145 | 198380 | 23,24,86,45,41,16,26,34,25 | 2,2,1,0,0,0,0,4,1,0,0,0 | 50,68,16,20,36,16,28,28,45 | 4,1,3,6,2,3,1,3,2,2,4,5,1,1,4 | 298 | 149 | 5.68506353105E-10 | 1.44684844156149E-06 | 0.852,0.857,0.977, | 0.862,0.971,0.727, | 0,114 |
| 198382 | ENSG00000069248 | NUP133 | chr1 | - | 229477596 | 229477897 | 229475637 | 229475732 | 229484053 | 229484145 | 198382 | 23,24,86,45,42,15,26,34,25 | 2,2,1,0,0,0,0,4,1,0,0,0 | 50,68,16,20,36,16,28,28,45 | 4,1,3,6,2,3,1,3,2,2,4,5,1,1,4 | 298 | 149 | 5.69024272146E-10 | 1.44684844156149E-06 | 0.852,0.857,0.977, | 0.862,0.971,0.727, | 0,114 |
| 213343 | ENSG00000135426 | TESPA1 | chr12 | - | 54983993 | 54984075 | 54974419 | 54974607 | 54984584 | 54984762 | 213343 | 1,1,15,4,4,3,0,6,1,1,2,1 | 41,14,38,23,21,11,9,10,10, | 2,2,1,1,0,2,0,0,0,0,0,0,2,0 | 18,23,14,14,13,2,13,14,8,2 | 231 | 149 | 5.71395930571E-10 | 1.44684844156149E-06 | 0.015,0.044,0.203, | 0.067,0.053,0.044, | 0,058 |
| 31897 | ENSG00000247774 | PCED1B-AS1 | chr12 | - | 47209280 | 47209482 | 47206510 | 47206588 | 47210639 | 47210761 | 31897 | 24,9,14,7,21,5,10,9,5,4,7,7 | 0,0,0,1,2,0,2,0,1,1,0,0 | 14,11,12,5,8,8,6,6,5,15,7,7 | 1,4,0,1,0,3,2,2,1,1,0,13,2,1 | 298 | 149 | 5.95870242037E-10 | 1.49127614027751E-06 | 1.0,1.0,1.0,0.778,0 | 0.875,0.579,1.0,0. | 0,1 |
| 241503 | ENSG00000266714 | MYO15B | chr17 | + | 75591912 | 75592080 | 75591615 | 75591712 | 75592237 | 75592301 | 241503 | 22,12,26,23,14,23,24,20,3, | 2,5,7,2,0,0,3,8,0,0,0,0 | 23,28,1,5,15,11,3,1,10,8,8, | 0,0,0,0,0,0,0,1,0,0,0,0,0,0, | 298 | 149 | 8.55328141647E-10 | 2.11601300292903E-06 | 0.846,0.545,0.65,0 | 1.0,1.0,1.0,1.0,1.0, | -0,092 |
| 205908 | ENSG00000128815 | WDFY4 | chr10 | + | 48820233 | 48820425 | 48817244 | 48817409 | 48821061 | 48821176 | 205908 | 80,70,95,67,91,81,50,64,94 | 5,0,3,0,14,18,1,1,2,2,8,9 | 79,75,22,50,20,31,66,84,40 | 0,4,0,2,2,2,0,0,3,0,2,1,0,2,1 | 298 | 149 | 8.91197782238E-10 | 2.17969761214622E-06 | 0.889,1.0,0.941,1. | 1.0,0.904,1.0,0.92 | -0,059 |
| 183910 | ENSG00000138095 | LRPPRC | chr2 | - | 43945382 | 43945417 | 43943686 | 43943894 | 43946112 | 43946243 | 183910 | 6,4,8,11,0,6,6,7,4,1,4,4 | 1,0,0,1,1,2,0,1,0,0,0,0 | 9,20,6,10,4,1,9,9,5,6,4,8,7, | 5,3,0,4,1,0,2,3,4,0,5,2,0,3,1 | 184 | 149 | 9.24211374098E-10 | 2.23504425009536E-06 | 0.829,1.0,1.0,0.89 | 0.593,0.844,1.0,0. | 0,091 |
| 241500 | ENSG00000266714 | MYO15B | chr17 | + | 75591791 | 75592080 | 75591615 | 75591712 | 75592237 | 75592301 | 241500 | 21,12,26,25,14,23,24,18,3, | 2,5,7,2,0,0,3,8,0,0,0,0 | 23,28,1,5,15,11,3,1,10,8,8, | 0,0,0,0,0,0,0,1,0,0,0,0,0,0, | 298 | 149 | 9.75057701247E-10 | 2.33180715663437E-06 | 0.84,0.545,0.65,0. | 1.0,1.0,1.0,1.0,1.0, | -0,094 |
| 154307 | ENSG00000075413 | MARK3 | chr14 | + | 103448920 | 103448967 | 103428386 | 103428440 | 103451917 | 103451983 | 154307 | 26,8,45,26,7,4,11,18,4,4,5, | 2,0,2,3,0,0,0,0,0,0,0,0 | 16,17,12,9,10,9,4,19,7,17,1 | 2,3,1,4,0,0,0,2,4,3,3,2,1,1,2 | 196 | 149 | 1.02166441973E-09 | 2.41641598596602E-06 | 0.908,1.0,0.945,0. | 0.859,0.812,0.901, | 0,156 |
| 219221 | ENSG00000137200 | CMTR1 | chr6 | + | 37461548 | 37461609 | 37459565 | 37459684 | 37471020 | 37471077 | 219221 | 33,17,25,16,7,5,9,11,17,4,4 | 1,1,0,1,0,1,0,1,0,0,0,3 | 14,16,7,14,23,17,5,6,7,7,17 | 7,7,0,1,5,1,0,5,1,0,1,0,1,2,2 | 210 | 149 | 0.0000000010388220 | 2.43029026081044E-06 | 0.959,0.923,1.0,0. | 0.587,0.619,1.0,0. | 0,135 |
| 33230 | ENSG00000132376 | INPP5K | chr17 | - | 1513452 | 1513561 | 1509177 | 1509353 | 1513871 | 1513979 | 33230 | 2,2,26,15,1,3,12,5,10,2,1,2 | 0,0,0,1,0,2,0,0,0,0,0,1 | 7,13,7,5,12,6,2,7,6,2,7,16,1 | 3,2,5,4,3,1,0,2,0,2,3,2,0,0, | 258 | 149 | 1.05905706427E-09 | 2.44442152405314E-06 | 1.0,1.0,1.0,0.897,1 | 0.574,0.79,0.447,0 | 0,164 |
| 203554 | ENSG00000275052 | PPP4R3B | chr2 | - | 55568193 | 55568363 | 55564901 | 55565211 | 55573618 | 55573738 | 203554 | 85,61,158,66,36,46,32,40,4 | 13,4,7,7,4,9,0,2,1,1,0,1 | 132,171,35,54,30,40,26,65 | 0,2,2,0,3,0,3,1,4,0,0,2,1,1,2 | 298 | 149 | 1.06757680474E-09 | 2.44442152405314E-06 | 0.766,0.884,0.919, | 1.0,0.977,0.897,1. | -0,05 |
| 150041 | ENSG00000159322 | ADPGK | chr15 | - | 72756250 | 72756447 | 72753068 | 72753141 | 72760406 | 72760527 | 150041 | 34,7,63,45,16,20,13,14,21, | 0,0,2,3,2,0,0,0,4,0,0,0 | 35,21,23,3,9,17,6,39,8,12,3 | 5,5,1,0,4,1,3,6,2,1,4,2,6,2,2 | 298 | 149 | 1.18529386306E-09 | 2.68538929937123E-06 | 1.0,1.0,0.94,0.882, | 0.778,0.677,0.92,1 | 0,174 |
| 170548 | ENSG00000076108 | BAZ2A | chr12 | - | 56605829 | 56606089 | 56604584 | 56605327 | 56606246 | 56606412 | 170548 | 18,22,106,69,38,41,13,68,2 | 2,0,2,0,0,3,0,5,0,0,1,0 | 53,119,28,53,47,38,68,62,2 | 1,9,3,26,3,1,0,6,7,1,2,10,4, | 298 | 149 | 1.25266308437E-09 | 2.80845758658375E-06 | 0.818,1.0,0.964,1. | 0.964,0.869,0.824, | 0,115 |
| 224033 | ENSG00000154978 | VOPP1 | chr7 | - | 55492748 | 55493209 | 55492385 | 55492418 | 55497612 | 55497690 | 224033 | 5,4,10,2,2,7,7,14,12,0,1,2 | 279,248,213,208,203,291,2 | 4,8,14,62,2,5,2,5,6,6,9,0,3, | 156,208,299,279,146,68,79 | 298 | 149 | 1.27945087858E-09 | 0.0000028389432169861 | 0.009,0.008,0.023, | 0.013,0.019,0.023, | -0,013 |
| 186149 | ENSG00000173276 | ZBTB21 | chr21 | - | 42002349 | 42002639 | 41988636 | 41994108 | 42002896 | 42002961 | 186149 | 20,10,17,8,4,3,11,11,5,5,2, | 0,0,1,1,0,0,2,1,2,0,0,0 | 10,12,7,9,19,6,12,10,4,6,20 | 4,0,4,0,4,3,0,0,0,0,3,8,1,0,5 | 298 | 149 | 1.34415778508E-09 | 0.0000029520859616383 | 1.0,1.0,0.895,0.8,1 | 0.556,1.0,0.467,1. | 0,111 |
| 56396 | ENSG00000173276 | ZBTB21 | chr21 | - | 42002349 | 42002563 | 41993947 | 41994108 | 42002896 | 42002961 | 56396 | 20,12,18,12,6,4,11,14,5,5,2 | 0,0,1,1,0,0,2,1,2,0,0,0 | 10,15,12,11,24,6,17,12,4,7 | 4,0,4,0,4,3,0,0,0,0,3,8,1,0,5 | 298 | 149 | 1.37717859339E-09 | 2.98714895494396E-06 | 1.0,1.0,0.9,0.857,1 | 0.556,1.0,0.6,1.0,0 | 0,106 |
| 163253 | ENSG00000113391 | FAM172A | chr5 | - | 94055659 | 94055801 | 94053125 | 94053227 | 94074645 | 94074771 | 163253 | 13,5,9,3,2,5,0,0,3,2,1,0 | 42,26,40,10,28,32,19,15,22 | 3,1,0,1,0,2,0,1,0,0,2,0,0,0,1 | 27,25,18,17,33,7,13,14,18, | 291 | 149 | 1.39626776807E-09 | 2.98714895494396E-06 | 0.137,0.09,0.103,0 | 0.054,0.02,0.0,0.0 | 0,047 |
| 186150 | ENSG00000173276 | ZBTB21 | chr21 | - | 42002349 | 42002781 | 41988636 | 41994108 | 42002896 | 42002961 | 186150 | 20,10,17,8,4,3,11,11,5,5,2, | 0,0,1,1,0,0,2,1,2,0,0,0 | 10,12,7,9,19,6,12,10,4,6,20 | 4,0,4,0,4,3,0,0,0,0,3,8,1,0,5 | 298 | 149 | 1.40546985161E-09 | 2.98714895494396E-06 | 1.0,1.0,0.895,0.8,1 | 0.556,1.0,0.467,1. | 0,111 |
| 201903 | ENSG00000134759 | ELP2 | chr18 | + | 36142168 | 36142347 | 36141136 | 36141201 | 36142825 | 36142857 | 201903 | 19,8,36,20,15,18,15,19,3,2 | 4,0,0,0,1,3,0,0,0,0,0,0 | 41,21,14,11,10,11,20,29,6, | 1,2,0,7,1,3,0,4,1,0,5,3,5,0,0 | 298 | 149 | 1.41563805123E-09 | 2.98714895494396E-06 | 0.704,1.0,1.0,1.0,0 | 0.953,0.84,1.0,0.4 | 0,117 |
| 56395 | ENSG00000173276 | ZBTB21 | chr21 | - | 42002349 | 42002503 | 41988636 | 41994108 | 42002896 | 42002961 | 56395 | 20,10,18,10,4,3,11,11,5,5,2 | 0,0,1,1,0,0,2,1,2,0,0,0 | 10,12,7,9,19,6,10,10,4,6,20 | 4,0,4,0,4,3,0,0,0,0,3,8,1,0,5 | 298 | 149 | 1.52053269975E-09 | 3.17733760679507E-06 | 1.0,1.0,0.9,0.833,1 | 0.556,1.0,0.467,1. | 0,119 |
| 234649 | ENSG00000101199 | ARFGAP1 | chr20 | + | 63277204 | 63277297 | 63276479 | 63276651 | 63278116 | 63278553 | 234649 | 16,3,24,15,6,3,5,4,4,2,2,8 | 3,4,4,0,0,0,1,1,9,4,2,1 | 10,8,5,3,7,3,3,11,4,1,11,16 | 2,4,0,0,0,0,0,3,0,0,0,1,0,1,0 | 242 | 149 | 2.05851435897E-09 | 4.26015484611031E-06 | 0.767,0.316,0.787, | 0.755,0.552,1.0,1. | -0,233 |
| 9636 | ENSG00000091592 | NLRP1 | chr17 | - | 5533896 | 5533988 | 5533291 | 5533384 | 5536850 | 5536940 | 9636 | 270,87,172,141,140,129,12 | 4,2,7,1,0,3,2,1,2,0,2,7 | 120,143,101,112,102,39,58 | 2,2,4,15,1,0,10,7,1,3,9,7,3, | 241 | 149 | 2.14089401762E-09 | 4.38844533625115E-06 | 0.977,0.964,0.938, | 0.974,0.978,0.94,0 | 0,068 |
| 249830 | ENSG00000077147 | TM9SF3 | chr10 | - | 96560415 | 96560703 | 96559658 | 96559736 | 96561977 | 96562138 | 249830 | 4,0,4,0,0,1,1,3,0,0,0,0 | 67,88,101,72,41,48,29,50,9 | 4,2,5,33,3,4,2,2,0,0,4,1,0,3 | 69,62,56,42,21,12,22,36,25 | 298 | 149 | 2.24638074808E-09 | 4.56123372443402E-06 | 0.029,0.0,0.019,0. | 0.028,0.016,0.043, | -0,04 |
| 47638 | ENSG00000137818 | RPLP1 | chr15 | + | 69453646 | 69453721 | 69452817 | 69453020 | 69455169 | 69455287 | 47638 | 2123,1573,2521,2810,1074 | 26,16,43,55,17,17,14,10,23 | 1828,2119,656,709,693,48 | 8,13,2,8,7,5,12,22,6,8,18,2 | 224 | 149 | 2.44333131505E-09 | 4.88032149408873E-06 | 0.982,0.985,0.975, | 0.993,0.991,0.995, | -0,007 |
| 46926 | ENSG00000100814 | CCNB1IP1 | chr14 | - | 20315484 | 20315618 | 20313467 | 20313801 | 20316226 | 20316408 | 46926 | 1,3,9,6,0,3,2,6,2,1,0,1 | 18,15,30,32,14,8,17,13,13, | 1,0,2,0,0,2,1,1,0,0,0,0,1,0,1 | 28,30,17,11,7,4,11,18,8,12 | 283 | 149 | 2.44887921053E-09 | 4.88032149408873E-06 | 0.028,0.095,0.136, | 0.018,0.0,0.058,0. | 0,065 |
| 31906 | ENSG00000247774 | PCED1B-AS1 | chr12 | - | 47210639 | 47210761 | 47205897 | 47206588 | 47216164 | 47216225 | 31906 | 10,9,31,42,29,19,11,5,9,11 | 0,1,1,1,1,2,0,0,0,0,2,3 | 27,19,9,5,16,21,12,10,18,3 | 2,6,2,2,0,3,3,1,0,4,2,33,4,3 | 271 | 149 | 2.53430976205E-09 | 5.00423875592462E-06 | 1.0,0.832,0.945,0. | 0.881,0.635,0.712, | 0,174 |
| 198376 | ENSG00000069248 | NUP133 | chr1 | - | 229477596 | 229477680 | 229475637 | 229475732 | 229484053 | 229484145 | 198376 | 12,10,46,24,19,7,10,10,7,6 | 2,2,1,0,0,0,0,4,1,0,0,0 | 27,33,7,5,15,11,15,17,19,3 | 4,1,3,6,2,3,1,3,2,2,4,5,1,1,4 | 233 | 149 | 2.80407341791E-09 | 5.48657750736534E-06 | 0.793,0.762,0.967, | 0.812,0.955,0.599, | 0,161 |
| 247220 | ENSG00000110330 | BIRC2 | chr11 | + | 102350826 | 102350943 | 102350000 | 102350749 | 102362895 | 102362974 | 247220 | 15,11,39,14,7,6,5,7,5,2,3,1 | 5,0,2,0,0,0,2,0,0,0,0,0 | 24,24,5,3,17,6,15,7,7,13,16 | 2,0,2,0,3,3,4,1,0,0,2,1,2,3,6 | 266 | 149 | 3.07234770958E-09 | 5.93634658619728E-06 | 0.627,1.0,0.916,1. | 0.87,1.0,0.583,1.0, | 0,136 |
| 108513 | ENSG00000101199 | ARFGAP1 | chr20 | + | 63277204 | 63277305 | 63276479 | 63276651 | 63278116 | 63278203 | 108513 | 33,28,61,40,27,14,22,19,19 | 3,4,4,0,0,0,1,1,9,4,2,1 | 27,29,16,19,18,8,16,39,15, | 2,4,0,0,0,0,0,3,0,0,0,1,0,1,0 | 250 | 149 | 3.08910341751E-09 | 5.93634658619728E-06 | 0.868,0.807,0.901, | 0.889,0.812,1.0,1. | -0,104 |
| 189265 | ENSG00000118308 | IRAG2 | chr12 | + | 25076530 | 25076640 | 25075739 | 25075802 | 25079243 | 25079290 | 189265 | 62,21,82,31,12,25,30,43,14 | 0,0,10,18,1,2,0,0,5,0,0,0 | 46,76,46,44,18,3,12,13,23, | 0,0,0,0,1,0,0,0,1,0,0,0,0,0,1 | 259 | 149 | 3.94929189351E-09 | 7.52221277461992E-06 | 1.0,1.0,0.825,0.49 | 1.0,1.0,1.0,1.0,0.9 | -0,096 |
| 112972 | ENSG00000154589 | LY96 | chr8 | + | 74004795 | 74004885 | 73991391 | 73991554 | 74010000 | 74010129 | 112972 | 13,21,40,69,20,8,11,14,8,1 | 0,0,1,2,2,0,0,1,0,0,0,4 | 7,17,14,5,28,8,5,5,19,32,4, | 0,1,1,1,2,0,0,0,7,2,1,22,4,3 | 239 | 149 | 0.0000000040608678 | 7.66688290596167E-06 | 1.0,1.0,0.961,0.95 | 1.0,0.914,0.897,0. | 0,09 |
| 3248 | ENSG00000181896 | ZNF101 | chr19 | + | 19678725 | 19678786 | 19677863 | 19677990 | 19679180 | 19679523 | 3248 | 151,192,591,346,102,182,1 | 4,1,11,3,1,11,5,10,3,6,4,5 | 394,417,174,206,205,156,9 | 13,13,5,33,13,4,6,8,1,6,19, | 210 | 149 | 0.0000000043406853 | 8.12391349704529E-06 | 0.964,0.993,0.974, | 0.956,0.958,0.961, | 0,034 |
| 50194 | ENSG00000204252 | HLA-DOA | chr6 | - | 33007079 | 33007215 | 33004181 | 33006841 | 33007310 | 33007592 | 50194 | 245,127,797,779,167,142,3 | 0,0,0,0,0,0,0,0,0,0,3,7 | 312,345,195,89,112,118,88 | 12,11,0,0,0,0,3,16,0,8,4,14 | 285 | 149 | 4.57989923763E-09 | 8.49772666219261E-06 | 1.0,1.0,1.0,1.0,1.0, | 0.931,0.943,1.0,1. | 0,021 |
| 50382 | ENSG00000266094 | RASSF5 | chr1 | + | 206585179 | 206585295 | 206583268 | 206583379 | 206586825 | 206589267 | 50382 | 92,104,92,105,32,24,14,19 | 0,0,19,20,0,0,0,0,0,0,0,0 | 65,103,20,10,39,39,7,18,25 | 0,3,0,0,0,0,0,0,1,0,1,2,0,4,0 | 265 | 149 | 4.65962601837E-09 | 8.57176040649396E-06 | 1.0,1.0,0.731,0.74 | 1.0,0.951,1.0,1.0,1 | -0,025 |
| 64647 | ENSG00000058673 | ZC3H11A | chr1 | + | 203796292 | 203796496 | 203795698 | 203795775 | 203801574 | 203801628 | 64647 | 23,10,28,6,36,20,10,17,12, | 0,0,6,2,0,0,1,0,0,0,0,0 | 18,19,23,17,10,8,13,14,26, | 8,2,2,0,0,1,0,5,0,2,3,6,2,4,7 | 298 | 149 | 4.89024887251E-09 | 8.91977250067118E-06 | 1.0,1.0,0.7,0.6,1.0, | 0.529,0.826,0.852, | 0,108 |
| 75219 | ENSG00000162613 | FUBP1 | chr1 | - | 77949154 | 77949300 | 77944054 | 77948774 | 77955254 | 77955282 | 75219 | 154,154,303,194,135,134,6 | 1,0,0,5,2,1,0,2,0,1,0,1 | 200,245,142,181,92,91,75, | 8,10,9,16,2,0,0,1,1,0,1,6,1, | 295 | 149 | 5.28014931778E-09 | 0.0000095500152757572 | 0.987,1.0,1.0,0.95 | 0.927,0.925,0.889, | 0,029 |
| 189263 | ENSG00000118308 | IRAG2 | chr12 | + | 25076522 | 25076640 | 25075739 | 25075802 | 25079243 | 25079290 | 189263 | 66,29,104,40,13,33,35,46,1 | 0,0,10,18,1,2,0,0,5,0,0,0 | 65,100,63,50,18,4,12,19,27 | 0,0,0,0,1,0,0,0,1,0,0,0,0,0,1 | 267 | 149 | 5.48638723341E-09 | 9.80567498567912E-06 | 1.0,1.0,0.853,0.55 | 1.0,1.0,1.0,1.0,0.9 | -0,088 |
| 201006 | ENSG00000120733 | KDM3B | chr5 | + | 138428689 | 138428801 | 138427966 | 138428086 | 138429825 | 138429965 | 201006 | 13,10,9,6,2,2,1,7,5,0,2,0 | 73,57,98,65,45,34,49,44,10 | 11,12,17,26,3,4,7,9,7,7,16, | 45,42,42,51,46,20,28,46,20 | 261 | 149 | 5.51261980508E-09 | 9.80567498567912E-06 | 0.092,0.091,0.05,0 | 0.122,0.14,0.188,0 | -0,067 |
| 155093 | ENSG00000129460 | NGDN | chr14 | + | 23471067 | 23471143 | 23470905 | 23470977 | 23475170 | 23478139 | 155093 | 3,0,37,7,23,28,1,1,3,0,1,2 | 36,27,80,50,16,28,50,68,20 | 0,2,1,2,0,0,0,2,6,8,1,2,2,2,6 | 91,95,46,28,73,25,56,79,14 | 225 | 149 | 5.71653424597E-09 | 0.000010085044117167 | 0.052,0.0,0.234,0. | 0.0,0.014,0.014,0. | 0,078 |
| 91124 | ENSG00000231789 | PIK3CD-AS2 | chr1 | - | 9680711 | 9681135 | 9672404 | 9672719 | 9687334 | 9687583 | 91124 | 10,16,21,24,5,3,2,15,20,11 | 0,2,3,3,0,1,2,2,10,1,0,2 | 4,3,7,5,10,8,4,6,2,11,10,2,3 | 2,0,0,0,0,0,1,0,0,2,2,0,0,0,1 | 298 | 149 | 5.89279836039E-09 | 0.0000103114868610171 | 1.0,0.8,0.778,0.8,1 | 0.5,1.0,1.0,1.0,1.0, | -0,112 |
| 102688 | ENSG00000162600 | OMA1 | chr1 | - | 58506059 | 58506209 | 58480718 | 58481174 | 58527260 | 58527335 | 102688 | 108,49,93,74,71,78,48,69,6 | 4,4,8,8,0,4,0,6,2,0,2,0 | 157,81,68,51,80,71,41,75,3 | 0,0,0,2,0,0,5,1,0,0,2,3,1,1,0 | 298 | 149 | 6.03436889346E-09 | 0.0000104740584782927 | 0.931,0.86,0.853,0 | 1.0,1.0,1.0,0.927,1 | -0,051 |
| 23436 | ENSG00000125746 | EML2 | chr19 | - | 45618821 | 45618943 | 45617629 | 45617697 | 45619059 | 45619191 | 23436 | 4,0,6,5,8,3,5,14,4,6,2,2 | 17,24,34,25,27,34,19,32,20 | 3,2,0,0,3,0,0,2,4,0,5,2,2,0,0 | 32,15,17,8,13,14,14,28,20, | 271 | 149 | 6.32258878586E-09 | 0.0000108865368557555 | 0.115,0.0,0.088,0. | 0.049,0.068,0.0,0. | 0,072 |
| 16043 | ENSG00000090857 | PDPR | chr16 | + | 70130422 | 70130544 | 70128783 | 70129122 | 70131301 | 70131419 | 16043 | 28,11,60,27,11,24,16,33,15 | 9,3,8,5,0,2,5,4,0,2,0,0 | 26,36,6,9,25,14,21,26,22,2 | 4,3,0,0,1,1,0,1,0,0,0,2,0,2,0 | 271 | 149 | 6.42698383402E-09 | 0.0000109784615680949 | 0.631,0.668,0.805, | 0.781,0.868,1.0,1. | -0,105 |
| 69559 | ENSG00000243943 | ZNF512 | chr2 | + | 27599969 | 27600053 | 27599582 | 27599678 | 27600690 | 27600728 | 69559 | 55,31,65,45,47,55,35,52,45 | 1,0,2,0,0,0,1,6,0,2,0,0 | 105,64,27,10,22,42,24,52,2 | 10,7,3,2,11,0,0,5,2,0,1,0,1, | 233 | 149 | 7.08776570768E-09 | 0.0000120118653624384 | 0.972,1.0,0.954,1. | 0.87,0.854,0.852,0 | 0,083 |
| 71419 | ENSG00000176444 | CLK2 | chr1 | - | 155265859 | 155265954 | 155264644 | 155264774 | 155268009 | 155268126 | 71419 | 18,29,36,30,17,25,6,14,20, | 0,1,2,0,0,1,0,4,1,1,1,0 | 10,21,17,23,18,14,9,8,8,21 | 3,6,2,0,0,1,5,1,2,3,4,5,2,0,2 | 244 | 149 | 7.14810921565E-09 | 0.0000120194898015122 | 1.0,0.947,0.917,1. | 0.671,0.681,0.838, | 0,16 |
| 108715 | ENSG00000155085 | AK9 | chr6 | - | 109564079 | 109564280 | 109563596 | 109563712 | 109564755 | 109564845 | 108715 | 38,8,31,21,12,6,22,22,10,8 | 1,4,5,0,2,7,0,1,2,3,1,0 | 29,16,13,14,6,12,9,13,11,2 | 0,0,2,4,0,0,0,1,0,0,3,4,0,0,0 | 298 | 149 | 7.46651229733E-09 | 0.0000124575574284235 | 0.95,0.5,0.756,1.0, | 1.0,1.0,0.765,0.63 | -0,154 |
| 107141 | ENSG00000215301 | DDX3X | chrX | + | 41347314 | 41347451 | 41346228 | 41346622 | 41347639 | 41347694 | 107141 | 251,213,374,268,106,138,1 | 3,1,7,4,3,0,2,0,0,1,0,2 | 229,278,187,176,192,127,9 | 8,5,1,31,0,6,5,8,1,6,10,0,3, | 286 | 149 | 7.86368503647E-09 | 0.0000130192984160344 | 0.978,0.991,0.965, | 0.937,0.967,0.99,0 | 0,038 |
| 59563 | ENSG00000158941 | CCAR2 | chr8 | + | 22613016 | 22613136 | 22607968 | 22608065 | 22614091 | 22614314 | 59563 | 39,18,70,75,67,42,31,35,66 | 1,1,0,1,3,5,2,3,1,0,0,4 | 51,35,24,25,24,22,10,27,5, | 7,8,3,3,1,2,5,6,1,1,8,6,1,5,2 | 269 | 149 | 8.38963976069E-09 | 0.0000137840500407105 | 0.956,0.909,1.0,0. | 0.801,0.708,0.816, | 0,139 |
| 107134 | ENSG00000215301 | DDX3X | chrX | + | 41346858 | 41347012 | 41346228 | 41346622 | 41347639 | 41347694 | 107134 | 298,265,589,392,142,180,1 | 3,1,7,4,3,0,2,0,0,1,0,2 | 405,479,236,336,238,182,1 | 8,5,1,31,0,6,5,8,1,6,10,0,3, | 298 | 149 | 0.0000000100648357 | 0.0000164110959220874 | 0.98,0.993,0.977,0 | 0.962,0.98,0.992,0 | 0,027 |
| 107137 | ENSG00000215301 | DDX3X | chrX | + | 41347311 | 41347451 | 41346493 | 41346622 | 41347639 | 41347915 | 107137 | 274,223,395,286,109,142,1 | 3,1,7,4,3,0,2,0,0,1,0,2 | 249,306,209,213,206,134,1 | 8,5,1,31,0,6,5,8,1,6,10,0,3, | 289 | 149 | 0.0000000103437555 | 0.0000167390740155402 | 0.979,0.991,0.967, | 0.941,0.969,0.991, | 0,034 |
| 192926 | ENSG00000106479 | ZNF862 | chr7 | + | 149852860 | 149853068 | 149850193 | 149850338 | 149859421 | 149859526 | 192926 | 2,0,8,3,4,0,2,7,13,3,1,0 | 6,7,4,2,5,11,6,10,7,7,7,3 | 0,0,0,1,2,2,0,0,0,2,2,1,0,1,1 | 4,4,3,1,4,4,8,5,3,3,0,4,5,13 | 298 | 149 | 0.0000000104489292 | 0.0000167830858007767 | 0.143,0.0,0.5,0.42 | 0.0,0.0,0.0,0.333,0 | 0,069 |
| 203557 | ENSG00000275052 | PPP4R3B | chr2 | - | 55569655 | 55569840 | 55568193 | 55568363 | 55577314 | 55577356 | 203557 | 8,13,5,7,1,2,4,13,6,9,7,8 | 7,2,6,2,5,2,0,0,0,2,1,2 | 17,36,11,42,12,11,7,12,9,1 | 2,0,0,4,0,1,0,0,1,3,0,3,0,1,0 | 298 | 149 | 0.0000000107350247 | 0.0000171148897310304 | 0.364,0.765,0.294, | 0.81,1.0,1.0,0.84,1 | -0,247 |
| 200291 | ENSG00000176444 | CLK2 | chr1 | - | 155265859 | 155266033 | 155264644 | 155264774 | 155268009 | 155268126 | 200291 | 16,29,37,29,17,25,6,15,20, | 0,1,2,0,0,1,0,4,1,1,1,0 | 13,21,16,24,16,14,9,8,8,21 | 3,6,2,0,0,1,5,1,2,3,4,5,2,0,2 | 298 | 149 | 0.0000000108681521 | 0.0000171997298285885 | 1.0,0.935,0.902,1. | 0.684,0.636,0.8,1. | 0,177 |
| 113569 | ENSG00000134014 | ELP3 | chr8 | + | 28183193 | 28183264 | 28161996 | 28162078 | 28189648 | 28190087 | 113569 | 9,1,6,4,4,0,0,0,6,7,0,2 | 23,4,21,22,28,20,20,12,25, | 2,0,0,0,3,2,0,1,1,0,0,0,0,0,0 | 30,8,21,8,1,13,13,10,14,28 | 220 | 149 | 0.0000000113151104 | 0.0000177763688131852 | 0.209,0.145,0.162, | 0.043,0.0,0.0,0.0,0 | 0,031 |
| 154163 | ENSG00000006432 | MAP3K9 | chr14 | - | 70761001 | 70761168 | 70749932 | 70750081 | 70800666 | 70801080 | 154163 | 14,21,27,9,27,17,13,7,16,7 | 0,0,0,0,3,2,0,0,0,0,0,0 | 14,25,18,12,17,6,13,8,15,1 | 3,4,4,4,1,2,0,4,1,0,0,0,1,1,1 | 298 | 149 | 0.0000000115804656 | 0.0000180519724635965 | 1.0,1.0,1.0,1.0,0.8 | 0.7,0.758,0.692,0. | 0,154 |
| 194053 | ENSG00000164241 | C5orf63 | chr5 | - | 127056396 | 127056544 | 127052612 | 127052669 | 127058881 | 127059002 | 194053 | 2,0,7,8,1,3,9,5,3,0,0,4 | 7,4,22,20,10,10,6,15,4,4,3, | 3,0,0,2,0,0,0,0,3,5,2,0,1,0 | 8,11,7,2,7,9,1,5,5,4,10,10,9 | 297 | 149 | 0.0000000116582842 | 0.0000180519724635965 | 0.125,0.0,0.138,0. | 0.158,0.0,0.0,0.33 | 0,069 |
| 24992 | ENSG00000167895 | TMC8 | chr17 | + | 78137234 | 78137323 | 78134869 | 78135009 | 78138004 | 78138188 | 24992 | 23,5,33,46,12,16,16,8,7,5,6 | 0,3,1,0,0,1,0,0,0,3,0,0 | 18,29,11,28,10,17,7,17,29, | 3,0,1,8,0,0,2,10,2,1,2,5,3,2 | 238 | 149 | 0.0000000121981177 | 0.0000187529505634929 | 1.0,0.511,0.954,1. | 0.79,1.0,0.873,0.6 | 0,067 |
| 39791 | ENSG00000114416 | FXR1 | chr3 | + | 180914651 | 180914995 | 180912481 | 180912736 | 180933333 | 180933386 | 39791 | 2,4,7,3,2,1,0,2,0,2,1,1 | 114,103,74,65,44,25,23,35 | 10,7,4,6,1,1,0,2,3,6,3,5,5,1 | 38,54,58,33,38,26,19,16,13 | 298 | 149 | 0.0000000126554656 | 0.00001925243613629 | 0.009,0.019,0.045, | 0.116,0.061,0.033, | -0,04 |
| 189735 | ENSG00000144744 | UBA3 | chr3 | - | 69057259 | 69057309 | 69056778 | 69056815 | 69061813 | 69061927 | 189735 | 4,1,17,14,6,9,5,6,1,2,3,5 | 0,0,6,0,1,0,0,3,0,0,1,0 | 9,6,1,4,6,2,1,4,1,2,2,9,4,5,8 | 1,3,3,2,2,4,3,2,0,3,1,3,3,4,0 | 199 | 149 | 0.0000000127019152 | 0.00001925243613629 | 1.0,1.0,0.68,1.0,0. | 0.871,0.6,0.2,0.6,0 | 0,296 |
| 165501 | ENSG00000109684 | CLNK | chr4 | - | 10531925 | 10532283 | 10528075 | 10528094 | 10540493 | 10540604 | 165501 | 7,8,35,10,3,13,14,21,2,1,4, | 0,1,2,1,0,1,2,2,0,0,1,0 | 8,10,7,8,11,12,9,11,11,5,20 | 0,4,2,4,6,0,4,1,1,0,1,0,1,0,1 | 298 | 149 | 0.0000000130793919 | 0.0000196859482423292 | 1.0,0.8,0.897,0.83 | 1.0,0.556,0.636,0. | 0,119 |
| 162529 | ENSG00000132952 | USPL1 | chr13 | + | 30643164 | 30643302 | 30642627 | 30642757 | 30646931 | 30647057 | 162529 | 0,0,4,1,0,0,1,2,2,3,0,0 | 27,21,62,42,28,19,20,27,14 | 5,4,3,8,5,2,10,5,1,7,5,2,2,3 | 30,52,18,39,22,21,30,35,9, | 287 | 149 | 0.0000000140674415 | 0.000021026038272027 | 0.0,0.0,0.032,0.01 | 0.08,0.038,0.08,0. | -0,048 |
| 65087 | ENSG00000103657 | HERC1 | chr15 | - | 63780461 | 63780552 | 63775187 | 63775649 | 63833821 | 63833915 | 65087 | 0,0,1,0,2,4,0,1,0,0,0,2 | 9,9,13,9,5,11,7,7,8,2,5,5 | 0,4,8,5,8,3,0,2,2,3,7,0,0,2,0 | 7,9,6,15,9,3,2,2,2,7,16,16,1 | 240 | 149 | 0.0000000142822281 | 0.0000211998500203972 | 0.0,0.0,0.046,0.0,0 | 0.0,0.216,0.453,0. | -0,131 |
| 197932 | ENSG00000135905 | DOCK10 | chr2 | - | 224946846 | 224947189 | 224931548 | 224931668 | 225042251 | 225042468 | 197932 | 200,144,266,143,138,146,1 | 2,4,8,5,9,10,12,10,5,6,5,35 | 281,273,151,188,181,103,8 | 2,6,4,6,4,3,2,7,2,1,3,11,2,1 | 298 | 149 | 0.0000000144294628 | 0.0000212716966984905 | 0.98,0.947,0.943,0 | 0.986,0.958,0.95,0 | -0,077 |
| 153502 | ENSG00000162086 | ZNF75A | chr16 | + | 3316844 | 3317022 | 3313048 | 3313175 | 3317189 | 3317252 | 153502 | 33,23,61,40,31,36,23,33,25 | 2,2,0,2,0,1,0,0,0,3,0,1 | 26,39,26,29,22,8,21,11,7,1 | 1,0,0,3,0,0,4,4,0,3,8,1,0,3,7 | 298 | 149 | 0.0000000151213891 | 0.0000221400796337655 | 0.892,0.852,1.0,0. | 0.929,1.0,1.0,0.82 | 0,092 |
| 141013 | ENSG00000130813 | SHFL | chr19 | + | 10089213 | 10089378 | 10087250 | 10087300 | 10089656 | 10089695 | 141013 | 4,2,9,6,8,1,8,5,3,4,4,8 | 22,16,40,31,21,24,16,22,14 | 5,4,0,3,0,0,0,1,3,3,2,0,0,2,5 | 17,18,22,22,24,8,11,15,7,2 | 298 | 149 | 0.0000000154544116 | 0.0000224747869698526 | 0.083,0.059,0.101, | 0.128,0.1,0.0,0.06 | 0,083 |
| 33226 | ENSG00000132376 | INPP5K | chr17 | - | 1509682 | 1509799 | 1508976 | 1509353 | 1513871 | 1513979 | 33226 | 17,6,23,20,20,21,17,17,12, | 0,0,0,1,0,2,0,0,0,0,0,1 | 12,19,16,9,12,6,7,10,7,14,1 | 3,2,5,4,3,1,0,2,0,2,3,2,0,0, | 266 | 149 | 0.0000000157564209 | 0.0000226168904310892 | 1.0,1.0,1.0,0.918,1 | 0.691,0.842,0.642, | 0,152 |
| 31887 | ENSG00000247774 | PCED1B-AS1 | chr12 | - | 47209280 | 47209458 | 47205897 | 47206588 | 47216164 | 47216251 | 31887 | 23,15,28,14,23,9,8,10,6,9,7 | 0,1,1,1,1,2,0,0,0,0,2,3 | 23,16,19,10,19,8,2,9,7,16,8 | 2,6,2,2,0,3,3,1,0,4,2,33,4,3 | 298 | 149 | 0.0000000157622905 | 0.0000226168904310892 | 1.0,0.882,0.933,0. | 0.852,0.571,0.826, | 0,176 |
| 153501 | ENSG00000162086 | ZNF75A | chr16 | + | 3316633 | 3317022 | 3313048 | 3313175 | 3317189 | 3317252 | 153501 | 33,23,61,40,32,34,23,33,25 | 2,2,0,2,0,1,0,0,0,3,0,1 | 26,39,26,29,22,8,21,11,7,1 | 1,0,0,3,0,0,4,4,0,3,8,1,0,3,7 | 298 | 149 | 0.0000000162517455 | 0.0000231647644818883 | 0.892,0.852,1.0,0. | 0.929,1.0,1.0,0.82 | 0,088 |
| 162696 | ENSG00000163297 | ANTXR2 | chr4 | - | 79984118 | 79984209 | 79983877 | 79983970 | 79984818 | 79984863 | 162696 | 10,0,0,1,0,0,1,1,0,1,1,2 | 9,15,12,20,15,19,19,15,18, | 1,9,3,13,10,5,2,0,3,8,10,4,3 | 21,19,4,14,4,7,17,7,3,6,16, | 240 | 149 | 0.0000000163738869 | 0.0000231853161458678 | 0.408,0.0,0.0,0.03, | 0.029,0.227,0.318, | -0,145 |
| 153500 | ENSG00000162086 | ZNF75A | chr16 | + | 3316550 | 3317022 | 3313048 | 3313175 | 3317189 | 3317252 | 153500 | 33,23,65,40,33,38,24,35,31 | 2,2,0,2,0,1,0,0,0,3,0,1 | 31,39,28,30,22,8,21,11,7,1 | 1,0,0,3,0,0,4,4,0,3,8,1,0,3,7 | 298 | 149 | 0.0000000169499101 | 0.00002370331764474 | 0.892,0.852,1.0,0. | 0.939,1.0,1.0,0.83 | 0,085 |
| 31898 | ENSG00000247774 | PCED1B-AS1 | chr12 | - | 47209280 | 47209482 | 47206510 | 47206588 | 47216164 | 47216225 | 31898 | 22,9,13,7,19,5,7,9,5,4,7,6 | 0,1,1,1,1,2,0,0,0,0,2,3 | 10,7,12,4,7,5,2,5,5,11,7,69 | 2,6,2,2,0,3,3,1,0,4,2,33,4,3 | 298 | 149 | 0.0000000169599682 | 0.00002370331764474 | 1.0,0.818,0.867,0. | 0.714,0.368,0.75,0 | 0,228 |
| 60087 | ENSG00000177200 | CHD9 | chr16 | + | 53321162 | 53321217 | 53318211 | 53318340 | 53321525 | 53321630 | 60087 | 0,1,4,0,0,0,0,0,0,0,0,0 | 1,9,17,15,7,11,9,4,6,1,4,11 | 2,7,1,3,1,0,0,2,2,0,1,2,4,4,2 | 26,21,6,3,13,3,4,3,17,11,6, | 204 | 149 | 0.0000000175262653 | 0.0000241980921082474 | 0.0,0.075,0.147,0. | 0.053,0.196,0.109, | -0,116 |

| | | | | | | | | | | | | | | | | | | | | | | |
|---|---|---|---|---|---|---|---|---|---|---|---|---|---|---|---|---|---|---|---|---|---|---|
| 247071 | ENSG00000183648 | NDUFB1 | chr14 | - | 92121195 | 92121399 | 92117497 | 92117642 | 92121641 | 92121917 | 247071 | 0,0,21,11,0,0,0,3,2,1,4,1 | 21,26,98,81,26,12,15,19,22 | 0,0,0,0,0,1,0,0,0,0,0,0,0,2,2 | 28,24,24,9,13,12,5,21,16,1 | 298 | 149 | 0.0000000175388413 | 0.0000241980921082474 | 0.0,0.0,0.097,0.064 | 0.0,0.0,0.0,0.0,0.0,0.0, | 0,029 |
| 175029 | ENSG00000101596 | SMCHD1 | chr18 | + | 2673016 | 2673120 | 2666869 | 2667031 | 2673280 | 2673363 | 175029 | 52,48,156,66,26,69,45,144 | 514,412,843,542,254,319,3 | 76,127,61,139,230,72,37,3 | 231,322,330,168,299,197,1 | 253 | 149 | 0.0000000185654416 | 0.0000254513284531445 | 0.056,0.064,0.098, | 0.162,0.188,0.098, | -0,06 |
| 199254 | ENSG00000076321 | KLHL20 | chr1 | + | 173756797 | 173756878 | 173755922 | 173756038 | 173756975 | 173757159 | 199254 | 1,0,0,0,0,1,1,1,0,0,0,0 | 4,7,13,12,9,11,3,4,13,5,3,1 | 5,2,4,1,1,4,0,2,0,0,3,8,2,0,4 | 18,10,12,10,13,1,8,8,3,8,11 | 230 | 149 | 0.0000000187776099 | 0.0000255792642964591 | 0.139,0.0,0.0,0.0,0.0 | 0.153,0.115,0.178, | -0,122 |
| 251989 | ENSG00000221817 | PPP3CB-AS1 | chr10 | + | 73504804 | 73504983 | 73504183 | 73504272 | 73505277 | 73505557 | 251989 | 10,2,16,7,3,2,6,13,6,7,2,6 | 0,3,2,6,4,2,3,0,0,1,4,0 | 9,2,12,3,5,8,1,8,3,5,5,4,9,5 | 0,0,0,0,0,1,1,6,1,0,0,0,0,2,1 | 298 | 149 | 0.0000000195463435 | 0.0000264589878443654 | 1.0,0.25,0.8,0.368, | 1.0,1.0,1.0,1.0,1.0, | -0,193 |
| 207727 | ENSG00000143924 | EML4 | chr2 | + | 42304483 | 42304547 | 42303314 | 42303446 | 42315961 | 42316050 | 207727 | 34,25,35,23,12,16,9,12,16, | 2,0,2,0,1,1,0,0,0,0,0,2 | 43,34,13,37,23,18,13,24,28 | 1,6,1,7,1,2,0,1,3,3,0,8,1,5,0 | 213 | 149 | 0.0000000197400235 | 0.0000265541562433755 | 0.922,1.0,0.924,1.0 | 0.968,0.799,0.901, | 0,047 |
| 207726 | ENSG00000143924 | EML4 | chr2 | + | 42304464 | 42304551 | 42303314 | 42303446 | 42315961 | 42316050 | 207726 | 19,20,35,13,26,32,7,22,4,1 | 2,0,2,0,1,1,0,0,0,0,0,2 | 32,39,9,28,27,25,18,38,20, | 1,6,1,7,1,2,0,1,3,3,0,8,1,5,0 | 236 | 149 | 0.0000000230563632 | 0.0000308226342949085 | 0.857,1.0,0.917,1.0 | 0.953,0.804,0.85,0 | 0,045 |
| 124745 | ENSG00000119777 | TMEM214 | chr2 | + | 27039740 | 27039837 | 27039046 | 27039164 | 27040029 | 27040198 | 124745 | 11,4,20,9,25,13,16,10,13,2 | 0,2,3,0,2,0,0,1,0,0,0,0 | 16,20,5,3,6,4,22,21,6,16,15 | 1,6,0,3,2,1,4,0,1,0,2,2,2,0,1 | 246 | 149 | 0.0000000247266779 | 0.0000327523217851984 | 1.0,0.548,0.802,1.0 | 0.906,0.669,1.0,0. | 0,122 |
| 72109 | ENSG00000120029 | ARMH3 | chr10 | - | 102001970 | 102002072 | 101995296 | 101995355 | 102006539 | 102006633 | 72109 | 26,8,36,27,25,18,7,35,5,3,1 | 2,0,2,0,3,2,0,0,0,0,0,0 | 32,21,13,20,25,13,8,20,14, | 3,11,2,1,2,0,2,2,1,1,1,0,1,1 | 251 | 149 | 0.0000000248041799 | 0.0000327523217851984 | 0.885,1.0,0.914,1.0 | 0.864,0.531,0.794, | 0,096 |
| 33293 | ENSG00000128739 | SNRPN | chr15 | + | 24967028 | 24967240 | 24962113 | 24962209 | 24967931 | 24968082 | 33293 | 12,4,110,124,5,8,2,13,30,3 | 126,81,198,89,72,86,83,10 | 6,7,11,0,22,20,4,5,5,3,4,3,2 | 161,170,110,16,59,53,58,1 | 298 | 149 | 0.0000000249873532 | 0.0000327930063268033 | 0.045,0.024,0.217, | 0.018,0.02,0.048,0 | 0,064 |
| 146748 | ENSG00000132600 | PRMT7 | chr16 | + | 68347210 | 68347334 | 68346120 | 68346280 | 68347630 | 68347678 | 146748 | 6,3,18,11,2,7,9,4,3,3,2,5 | 2,0,3,0,0,0,0,0,0,0,0,0 | 8,5,7,2,7,5,0,8,3,5,5,7,10,6 | 1,4,1,1,2,1,7,1,1,0,2,0,0,1,0 | 273 | 149 | 0.0000000253364149 | 0.0000330495874518009 | 0.621,1.0,0.766,1.0 | 0.814,0.406,0.793, | 0,252 |
| 240748 | ENSG00000021776 | AQR | chr15 | - | 34927034 | 34927136 | 34920331 | 34920434 | 34930257 | 34930371 | 240748 | 14,10,36,21,7,14,12,18,8,3 | 2,0,0,1,0,0,1,1,0,1,0,0 | 29,21,7,6,19,10,7,17,16,14 | 0,8,0,4,7,1,2,1,1,0,2,1,0,1,0 | 251 | 149 | 0.0000000276691122 | 0.0000358750041746846 | 0.806,1.0,1.0,0.92 | 1.0,0.609,1.0,0.47 | 0,113 |
| 252331 | ENSG00000151490 | PTPRO | chr12 | + | 15570418 | 15570590 | 15569416 | 15569498 | 15578852 | 15578943 | 252331 | 1,2,3,1,1,0,2,0,0,0,0,0 | 23,7,36,31,5,5,10,3,4,5,3,4 | 4,1,8,4,2,2,3,2,1,0,3,0,7,6,2 | 11,9,7,3,11,4,7,12,7,24,5,1 | 298 | 149 | 0.0000000280151717 | 0.0000361061881936615 | 0.021,0.125,0.04,0 | 0.154,0.053,0.364, | -0,098 |
| 80238 | ENSG00000124151 | NCOA3 | chr20 | + | 47639007 | 47639202 | 47637647 | 47637783 | 47639576 | 47639822 | 80238 | 364,239,900,579,318,247,2 | 2,1,7,5,2,2,4,8,2,0,1,3 | 470,623,345,395,423,287,3 | 9,12,8,18,12,1,3,10,6,1,10, | 298 | 149 | 0.0000000288731535 | 0.0000369904625501608 | 0.989,0.992,0.985, | 0.963,0.963,0.956, | 0,026 |
| 241506 | ENSG00000266714 | MYO15B | chr17 | + | 75592005 | 75592080 | 75591615 | 75591712 | 75592237 | 75592301 | 241506 | 17,6,17,14,10,20,19,11,3,7 | 2,5,7,2,0,0,3,8,0,0,0,0 | 17,21,1,3,8,10,2,0,7,7,6,16 | 0,0,0,0,0,0,0,1,0,0,0,0,0,0,0 | 224 | 149 | 0.0000000297433694 | 0.0000378798529388803 | 0.85,0.444,0.618,0 | 1.0,1.0,1.0,1.0,1.0, | -0,085 |
| 155898 | ENSG00000196199 | MPHOSPH8 | chr13 | + | 19661962 | 19662077 | 19661697 | 19661838 | 19663039 | 19663126 | 155898 | 15,0,9,2,4,3,2,3,1,1,2,3 | 26,18,91,52,45,23,25,30,48 | 3,0,2,2,1,2,0,0,0,0,1,1,2,2,0 | 67,48,18,24,13,7,28,62,22, | 264 | 149 | 0.0000000332410200 | 0.0000420852822697941 | 0.246,0.0,0.053,0.0 | 0.025,0.0,0.059,0.0 | 0,033 |
| 174869 | ENSG00000127483 | HP1BP3 | chr1 | - | 20775723 | 20776005 | 20773586 | 20773610 | 20776596 | 20776750 | 174869 | 5,5,23,19,1,7,8,8,10,1,0,6 | 173,118,231,180,122,133,1 | 3,1,5,2,2,1,2,3,0,1,4,6,8,8,5 | 163,118,115,70,105,81,96, | 298 | 149 | 0.0000000347858272 | 0.0000435783736834381 | 0.014,0.021,0.047, | 0.009,0.004,0.021, | 0,014 |
| 254154 | ENSG00000123349 | PFDN5 | chr12 | + | 53295838 | 53295948 | 53295290 | 53295639 | 53297849 | 53299100 | 254154 | 1189,772,818,686,175,230 | 62,26,120,60,24,30,21,42,2 | 483,371,485,288,321,190,1 | 50,56,40,22,61,26,27,59,28 | 259 | 149 | 0.0000000350050983 | 0.0000435783736834381 | 0.917,0.945,0.797, | 0.847,0.792,0.875, | 0,067 |
| 105952 | ENSG00000196776 | CD47 | chr3 | - | 108050577 | 108050602 | 108047078 | 108047292 | 108057476 | 108057569 | 105952 | 0,0,1,3,0,0,1,3,2,0,0,0 | 14,2,42,24,15,8,8,6,8,2,1,4 | 2,2,0,0,1,0,3,1,1,2,7,3,2,2,3 | 10,2,14,1,7,5,4,4,8,12,8,6,7 | 174 | 149 | 0.0000000350277545 | 0.0000435783736834381 | 0.0,0.0,0.02,0.097, | 0.146,0.461,0.0,0.0 | -0,186 |
| 45570 | ENSG00000100578 | KIAA0586 | chr14 | + | 58492143 | 58492275 | 58490163 | 58490240 | 58498782 | 58498960 | 45570 | 47,13,35,39,17,24,25,24,48 | 1,1,0,1,0,1,0,0,1,0,0,1 | 34,26,27,15,21,15,18,23,16 | 1,7,3,0,2,0,0,1,3,3,3,5,0,0,4 | 281 | 149 | 0.0000000356514924 | 0.0000440994618741589 | 0.961,0.873,1.0,0.9 | 0.947,0.663,0.827, | 0,091 |
| 70573 | ENSG00000144591 | GMPPA | chr2 | + | 219505292 | 219505362 | 219504082 | 219504213 | 219505457 | 219505555 | 70573 | 9,10,11,7,7,4,4,7,12,5,2,9 | 4,1,3,0,0,3,4,3,2,0,0,1 | 11,8,4,2,4,3,11,4,2,3,2,7,9, | 0,0,0,0,0,0,0,0,0,0,2,0,0,0 | 219 | 149 | 0.0000000360776671 | 0.000044371613636267 | 0.605,0.872,0.714, | 1.0,1.0,1.0,1.0,1.0, | -0,186 |
| 82231 | ENSG00000004897 | CDC27 | chr17 | - | 47141852 | 47142022 | 47138738 | 47138891 | 47142228 | 47142381 | 82231 | 48,51,88,37,32,38,25,40,18 | 0,0,2,10,0,1,0,0,0,0,0,0 | 56,52,29,34,20,17,22,48,18 | 3,4,3,3,3,5,0,0,7,0,3,5,6,4,2 | 298 | 149 | 0.0000000363468337 | 0.0000444486669208791 | 1.0,1.0,0.957,0.64 | 0.903,0.867,0.829, | 0,119 |
| 116358 | ENSG00000266714 | MYO15B | chr17 | + | 75591976 | 75592080 | 75591600 | 75591712 | 75592237 | 75592301 | 116358 | 31,25,37,34,29,58,20,23,11 | 2,5,7,2,0,0,3,8,0,0,0,0 | 48,54,1,12,38,22,3,2,20,25 | 0,0,0,0,0,0,1,0,0,0,0,0,0,0 | 253 | 149 | 0.0000000392311235 | 0.000047579734323884 | 0.901,0.746,0.757, | 1.0,1.0,1.0,1.0,1.0, | -0,069 |
| 76497 | ENSG00000168256 | NKIRAS2 | chr17 | + | 42022564 | 42022640 | 42022398 | 42022472 | 42023653 | 42024265 | 76497 | 5,6,18,12,4,2,5,4,4,1,1,6 | 2,3,6,6,0,2,2,0,0,0,3,4 | 7,6,3,2,2,3,4,8,1,10,9,11,5, | 2,2,0,0,1,0,0,2,0,0,2,1,0,2,0 | 225 | 149 | 0.0000000393493163 | 0.000047579734323884 | 0.623,0.57,0.665,0 | 0.699,0.665,1.0,1.0 | -0,165 |
| 194264 | ENSG00000103657 | HERC1 | chr15 | - | 63780461 | 63780555 | 63775187 | 63775649 | 63833821 | 63833917 | 194264 | 0,0,1,0,4,2,0,1,0,0,0,2 | 9,9,13,9,5,11,7,7,8,2,5,5 | 0,3,6,6,6,3,0,2,2,3,7,0,0,2,0 | 7,9,6,15,9,3,2,2,2,7,16,16,1 | 243 | 149 | 0.0000000410690385 | 0.0000493817331067249 | 0.0,0.0,0.045,0.0,0 | 0.0,0.17,0.38,0.19 | -0,116 |
| 110512 | ENSG00000101190 | TCFL5 | chr20 | - | 62859363 | 62859526 | 62857394 | 62857638 | 62860124 | 62860308 | 110512 | 48,35,119,60,31,24,6,30,35 | 0,0,8,0,0,1,0,2,0,0,0,0 | 60,82,21,7,9,17,19,28,7,15 | 5,17,6,1,4,2,0,2,1,0,0,1,0,0 | 298 | 149 | 0.0000000419679141 | 0.0000501822007172541 | 1.0,1.0,0.881,1.0,1 | 0.857,0.707,0.636, | 0,121 |
| 136111 | ENSG00000105486 | LIG1 | chr19 | - | 48151467 | 48151894 | 48151231 | 48151339 | 48153871 | 48154207 | 136111 | 2,6,4,1,0,0,4,2,7,1,5,4 | 26,18,20,5,27,19,12,15,26, | 0,0,1,0,0,0,0,1,1,0,3,0,0,4,0 | 14,11,19,13,6,14,4,16,10,1 | 298 | 149 | 0.0000000424439079 | 0.0000503112462748388 | 0.037,0.143,0.091, | 0.0,0.0,0.026,0.0,0 | 0,05 |
| 12339 | ENSG00000181610 | MRPS23 | chr17 | - | 57841182 | 57841260 | 57840799 | 57841052 | 57849239 | 57849410 | 12339 | 31,25,94,71,26,32,27,31,28 | 0,0,4,2,0,0,2,1,0,0,0,2 | 57,51,42,21,21,19,21,38,11 | 5,4,4,7,6,1,0,3,1,4,1,3,1,0,0 | 227 | 149 | 0.0000000425433456 | 0.0000503112462748388 | 1.0,1.0,0.939,0.95 | 0.882,0.893,0.873, | 0,071 |
| 189198 | ENSG00000118308 | IRAG2 | chr12 | + | 25062166 | 25062243 | 25061591 | 25061653 | 25062819 | 25062900 | 189198 | 1,0,26,26,4,14,0,1,6,3,3,6 | 208,78,469,405,181,94,186 | 17,38,43,14,10,1,2,11,10,1 | 169,249,150,55,67,53,65,1 | 226 | 149 | 0.0000000443876048 | 0.0000522054020312679 | 0.003,0.0,0.035,0.0 | 0.062,0.091,0.159, | -0,03 |
| 23312 | ENSG00000090061 | CCNK | chr14 | + | 99491945 | 99492320 | 99481424 | 99481479 | 99492625 | 99492874 | 23312 | 1,3,2,5,0,4,0,2,1,2,2,0 | 28,17,27,23,14,14,11,24,14 | 5,4,2,15,5,4,8,5,7,1,7,1,1,3 | 20,27,13,21,17,10,13,5,17, | 298 | 149 | 0.0000000472245250 | 0.000055155440284988 | 0.018,0.081,0.036, | 0.111,0.069,0.071, | -0,096 |
| 144683 | ENSG00000181610 | MRPS23 | chr17 | - | 57841182 | 57841256 | 57840799 | 57841052 | 57849239 | 57849410 | 144683 | 16,18,38,24,10,10,4,10,12, | 0,0,4,2,0,0,2,1,0,0,0,2 | 9,17,20,10,8,7,4,10,6,7,11, | 5,4,4,7,6,1,0,3,1,4,1,3,1,0,0 | 223 | 149 | 0.0000000474083958 | 0.000055155440284988 | 1.0,1.0,0.864,0.88 | 0.546,0.74,0.77,0.4 | 0,112 |
| 212632 | ENSG00000117133 | RPF1 | chr1 | + | 84488570 | 84488699 | 84482914 | 84482995 | 84489632 | 84489728 | 212632 | 0,0,3,1,0,0,34,36,1,0,3,0 | 22,19,100,58,68,65,19,40,1 | 2,0,0,3,0,0,0,1,0,0,2,1,1,1,0 | 71,68,39,19,30,26,29,58,25 | 278 | 149 | 0.0000000492125763 | 0.0000569466237625121 | 0.0,0.0,0.016,0.00 | 0.015,0.0,0.0,0.0,0.07 | 0,073 |
| 145443 | ENSG00000074370 | ATP2A3 | chr17 | - | 3928212 | 3928327 | 3925395 | 3925441 | 3928662 | 3928780 | 145443 | 0,2,19,26,1,2,4,0,1,3,0,0 | 80,60,111,70,80,62,49,34,6 | 2,1,1,0,5,7,0,0,1,1,0,0,5,0,0 | 101,125,32,24,62,56,16,71 | 264 | 149 | 0.0000000494890693 | 0.0000569603309926335 | 0.0,0.018,0.088,0. | 0.011,0.004,0.017, | 0,021 |
| 165578 | ENSG00000047365 | ARAP2 | chr4 | - | 36080215 | 36080446 | 36073688 | 36073823 | 36082250 | 36082286 | 165578 | 13,27,23,16,6,12,3,10,12,5 | 0,0,1,0,0,0,0,2,1,0,0,0 | 19,14,10,12,5,13,4,13,8,15 | 2,5,0,5,1,0,6,2,1,0,1,2,0,0,1 | 298 | 149 | 0.0000000509402028 | 0.0000580101100322756 | 1.0,1.0,0.92,1.0,1.0 | 0.826,0.583,1.0,0. | 0,17 |
| 165581 | ENSG00000047365 | ARAP2 | chr4 | - | 36080215 | 36080626 | 36073688 | 36073823 | 36082250 | 36082286 | 165581 | 13,27,23,16,6,12,3,10,12,5 | 0,0,1,0,0,0,0,2,1,0,0,0 | 18,14,10,12,5,13,4,13,8,15 | 2,5,0,5,1,0,6,2,1,0,1,2,0,0,1 | 298 | 149 | 0.0000000509402028 | 0.0000580101100322756 | 1.0,1.0,0.92,1.0,1.0 | 0.818,0.583,1.0,0. | 0,17 |
| 172121 | ENSG00000145012 | LPP | chr3 | + | 188341655 | 188341719 | 188154156 | 188154252 | 188406111 | 188406313 | 172121 | 57,31,8,0,4,2,2,8,0,8,3,3 | 0,0,0,3,2,0,1,2,3,7,2,2 | 17,13,17,18,13,5,9,7,4,3,4, | 0,0,0,0,0,0,1,1,1,0,0,1,0,0 | 213 | 149 | 0.0000000523851676 | 0.0000593416422004723 | 1.0,1.0,1.0,0.0,0.5 | 1.0,1.0,1.0,1.0,1.0, | -0,325 |
| 12361 | ENSG00000266967 | AARSD1 | chr17 | - | 42954875 | 42954967 | 42951799 | 42951894 | 42955157 | 42955224 | 12361 | 29,11,20,8,11,7,4,13,18,8,3 | 11,2,8,6,0,2,0,4,7,0,0,1 | 14,9,15,8,11,11,2,20,3,17,1 | 1,2,0,3,1,0,0,2,1,2,0,2,0,0,2 | 241 | 149 | 0.0000000527205241 | 0.0000594088541220441 | 0.62,0.773,0.607,0 | 0.896,0.736,1.0,0. | -0,092 |
| 152957 | ENSG00000257621 | PSMA3-AS1 | chr14 | - | 58293765 | 58295811 | 58291634 | 58291722 | 58297954 | 58298120 | 152957 | 16,4,69,30,7,11,14,45,27,1 | 0,0,21,20,0,0,0,0,0,0,0,3 | 31,26,17,11,26,13,28,28,10 | 0,0,0,0,8,0,0,0,0,0,0,0,0,0,0 | 298 | 149 | 0.0000000549295958 | 0.000061366267053381 | 1.0,1.0,0.622,0.42 | 1.0,1.0,1.0,1.0,0.6 | -0,072 |
| 189410 | ENSG00000177200 | CHD9 | chr16 | + | 53208264 | 53208709 | 53157163 | 53157541 | 53209481 | 53209813 | 189410 | 2,1,4,4,0,2,1,5,5,0,2,2 | 63,57,85,61,38,40,26,38,39 | 9,15,6,13,6,1,2,8,21,17,3,0 | 69,78,42,38,79,54,24,27,43 | 298 | 149 | 0.0000000550278052 | 0.000061366267053381 | 0.016,0.009,0.023, | 0.061,0.088,0.067, | -0,039 |
| 244244 | ENSG00000166266 | CUL5 | chr11 | + | 108067022 | 108067143 | 108054874 | 108054955 | 108070095 | 108070189 | 244244 | 1,0,5,1,0,1,0,4,0,0,0,0 | 13,14,55,37,16,15,10,9,3,4 | 8,5,4,4,2,1,0,0,3,0,2,6,3,7,0 | 25,33,8,12,25,8,7,5,13,10,2 | 270 | 149 | 0.0000000578395482 | 0.0000641694010926728 | 0.041,0.0,0.048,0.0 | 0.15,0.077,0.216,0 | -0,054 |
| 43805 | ENSG00000114867 | EIF4G1 | chr3 | + | 184317716 | 184317816 | 184314550 | 184314674 | 184319688 | 184319801 | 43805 | 43,33,39,55,18,26,27,34,37 | 1,0,0,2,2,0,0,4,0,2,0,2 | 30,56,24,41,37,19,27,24,12 | 6,1,1,10,0,1,3,2,1,1,5,4,2,2 | 249 | 149 | 0.0000000584501876 | 0.0000645143197265891 | 0.963,1.0,1.0,0.94 | 0.749,0.971,0.935, | 0,077 |
| 233318 | ENSG00000215301 | DDX3X | chrX | + | 41346858 | 41346952 | 41346228 | 41346622 | 41347639 | 41351668 | 233318 | 99,89,352,188,56,77,56,91 | 3,1,7,4,3,0,2,0,0,1,0,2 | 166,241,93,126,94,84,38,5 | 8,5,1,31,0,6,5,8,1,6,10,0,3, | 243 | 149 | 0.0000000594000031 | 0.0000651232871642908 | 0.953,0.982,0.969, | 0.927,0.967,0.983, | 0,052 |
| 98738 | ENSG00000125812 | GZF1 | chr20 | + | 23369583 | 23369741 | 23368761 | 23368929 | 23370090 | 23373045 | 98738 | 12,21,28,18,15,7,9,16,18,5 | 4,1,4,8,0,2,1,3,0,1,0,0 | 10,15,21,6,12,12,9,19,6,11 | 0,0,0,0,2,2,0,3,0,0,2,0,0,3,0 | 298 | 149 | 0.0000000597359383 | 0.0000651232871642908 | 0.6,0.913,0.778,0. | 1.0,1.0,1.0,1.0,0.7 | -0,077 |
| 235436 | ENSG00000156171 | DRAM2 | chr1 | - | 111119614 | 111119728 | 111118864 | 111118897 | 111119876 | 111119959 | 235436 | 19,1,22,9,12,7,5,14,6,2,2,1 | 70,41,198,95,101,55,57,10 | 12,1,8,6,0,0,3,8,0,0,3,4,6,0 | 97,77,75,37,50,48,39,61,54 | 263 | 149 | 0.0000000599177051 | 0.0000651232871642908 | 0.133,0.014,0.059, | 0.065,0.007,0.057, | 0,024 |
| 21679 | ENSG00000162086 | ZNF75A | chr16 | + | 3316911 | 3317022 | 3313048 | 3313175 | 3317189 | 3317252 | 21679 | 40,22,54,45,26,30,19,38,22 | 2,2,0,2,0,1,0,0,0,3,0,1 | 35,42,29,20,24,9,23,8,3,25 | 1,0,0,3,0,0,4,4,0,3,8,1,0,3,7 | 260 | 149 | 0.0000000602122098 | 0.0000651232871642908 | 0.92,0.863,1.0,0.9 | 0.953,1.0,1.0,0.79 | 0,077 |
| 241265 | ENSG00000107077 | KDM4C | chr9 | + | 6986343 | 6986433 | 6984165 | 6984404 | 6990415 | 6990524 | 241265 | 27,13,42,35,9,23,11,25,8,1 | 1,0,0,5,0,1,0,0,0,0,0,1 | 42,45,23,31,30,26,16,19,24 | 1,5,0,3,7,0,2,0,1,0,2,2,2,4,0 | 239 | 149 | 0.0000000627564744 | 0.0000675356937741155 | 0.944,1.0,1.0,0.81 | 0.963,0.849,1.0,0. | 0,075 |
| 82232 | ENSG00000004897 | CDC27 | chr17 | - | 47141852 | 47142025 | 47138738 | 47138891 | 47142228 | 47142436 | 82232 | 50,54,90,27,24,38,23,45,14 | 0,0,2,10,0,1,0,0,0,0,0,0 | 52,46,28,32,22,15,16,43,14 | 3,4,3,3,3,5,0,0,7,0,3,5,6,4,2 | 298 | 149 | 0.0000000654374961 | 0.0000700705360423611 | 1.0,1.0,0.957,0.57 | 0.897,0.852,0.824, | 0,117 |
| 171729 | ENSG00000173226 | IQCB1 | chr3 | - | 121772556 | 121772736 | 121769760 | 121770574 | 121781742 | 121781874 | 171729 | 55,16,62,42,32,12,23,25,40 | 4,1,4,2,5,1,4,7,1,0,0,0 | 52,36,40,11,29,19,16,27,9, | 0,1,0,0,2,0,0,0,0,2,2,2,0,1,0 | 298 | 149 | 0.0000000659863462 | 0.000070308451957078 | 0.873,0.889,0.886, | 1.0,0.947,1.0,1.0,0 | -0,075 |
| 212633 | ENSG00000117133 | RPF1 | chr1 | + | 84488570 | 84488706 | 84482914 | 84482995 | 84489632 | 84489728 | 212633 | 0,0,5,0,0,0,39,35,1,0,3,0 | 22,19,100,58,68,65,19,40,1 | 2,0,0,3,0,0,0,1,0,0,2,2,1,1,0 | 71,68,39,19,30,26,29,58,25 | 285 | 149 | 0.0000000695655459 | 0.0000737569558081763 | 0.0,0.0,0.025,0.0,0 | 0.015,0.0,0.0,0.0,0.07 | 0,074 |
| 153507 | ENSG00000162086 | ZNF75A | chr16 | + | 3316953 | 3317022 | 3313048 | 3313175 | 3317189 | 3317252 | 153507 | 22,16,33,22,17,20,15,24,12 | 2,2,0,2,0,1,0,0,0,3,0,1 | 19,23,16,15,12,6,12,8,3,6,1 | 1,0,0,3,0,0,4,4,0,3,8,1,0,3,7 | 218 | 149 | 0.0000000706285403 | 0.0000745169184705135 | 0.883,0.845,1.0,0.8 | 0.929,1.0,1.0,0.77 | 0,106 |
| 153505 | ENSG00000162086 | ZNF75A | chr16 | + | 3316938 | 3317022 | 3313048 | 3313175 | 3317189 | 3317252 | 153505 | 24,18,34,23,20,18,13,25,13 | 2,2,0,2,0,1,0,0,0,3,0,1 | 20,28,16,15,14,5,14,7,3,7,1 | 1,0,0,3,0,0,4,4,0,3,8,1,0,3,7 | 233 | 149 | 0.0000000785476192 | 0.000082467720239333 | 0.885,0.852,1.0,0.8 | 0.927,1.0,1.0,0.76 | 0,104 |
| 244404 | ENSG00000099251 | HSD17B7P2 | chr10 | + | 38365909 | 38366011 | 38365433 | 38365629 | 38369952 | 38370009 | 244404 | 5,4,0,2,0,0,0,0,2,0,0,0 | 0,0,12,10,12,2,3,9,2,2,6,6 | 0,0,2,3,17,5,6,10,5,8,5,7,11 | 2,4,3,4,2,4,8,7,0,2,4,2,7,18 | 251 | 149 | 0.0000000803587560 | 0.00008395968656711 | 1.0,1.0,0.0,0.106,0 | 0.0,0.0,0.284,0.30 | -0,241 |
| 13038 | ENSG00000074370 | ATP2A3 | chr17 | - | 3928239 | 3928327 | 3925290 | 3925441 | 3928662 | 3928780 | 13038 | 2,12,28,44,2,9,5,2,14,13,3, | 80,60,111,70,80,62,49,34,6 | 5,4,7,7,11,8,1,2,5,4,1,1,8,3 | 101,125,32,24,62,56,16,71 | 237 | 149 | 0.0000000816950167 | 0.0000848663184802551 | 0.015,0.112,0.137, | 0.03,0.02,0.121,0. | 0,039 |
| 215119 | ENSG00000115355 | CCDC88A | chr2 | - | 55297927 | 55298034 | 55296257 | 55296523 | 55299838 | 55301277 | 215119 | 6,4,55,52,0,1,0,5,13,8,1,0 | 116,69,157,125,39,26,60,3 | 2,4,4,0,20,8,4,0,0,0,0,0,2,0 | 160,126,40,6,15,21,34,45,2 | 256 | 149 | 0.0000000820151104 | 0.0000848663184802551 | 0.029,0.033,0.169, | 0.007,0.018,0.055, | 0,036 |
| 57802 | ENSG00000111674 | ENO2 | chr12 | + | 6922055 | 6922164 | 6921496 | 6921782 | 6922343 | 6922402 | 57802 | 11,11,29,29,30,17,12,2,3,1 | 1,0,1,1,0,3,1,0,0,0,0,3 | 25,23,10,31,5,4,8,21,18,20 | 6,1,0,5,0,0,2,9,1,1,2,2,4,6,4 | 258 | 149 | 0.0000000842345588 | 0.0000867458772395986 | 0.864,1.0,0.944,0. | 0.706,0.93,1.0,0.7 | 0,08 |
| 24995 | ENSG00000167895 | TMC8 | chr17 | + | 78137716 | 78137814 | 78134869 | 78135009 | 78138004 | 78138188 | 24995 | 95,33,36,29,44,21,37,25,55 | 0,3,1,0,0,1,0,0,0,3,0,0 | 31,40,35,101,14,23,28,35,2 | 3,0,1,8,0,0,2,10,2,1,2,5,3,2 | 247 | 149 | 0.0000000870261817 | 0.0000891939624506426 | 1.0,0.869,0.956,1.0 | 0.862,1.0,0.955,0. | 0,049 |
| 130893 | ENSG00000138777 | PPA2 | chr4 | - | 105437949 | 105438036 | 105424195 | 105424322 | 105446382 | 105446502 | 130893 | 48,24,121,100,45,22,28,42 | 4,3,6,4,0,5,1,3,0,0,2,2 | 104,78,48,38,40,33,28,45,3 | 0,0,1,1,0,1,0,2,0,0,2,0,0,3,0 | 236 | 149 | 0.0000000887577482 | 0.000090537530408633 | 0.883,0.835,0.927, | 1.0,1.0,0.968,0.96, | -0,07 |
| 76498 | ENSG00000168256 | NKIRAS2 | chr17 | + | 42022569 | 42022640 | 42022398 | 42022472 | 42023653 | 42024265 | 76498 | 3,7,17,8,4,2,5,3,4,1,1,6 | 2,3,6,6,0,2,2,0,0,0,3,4 | 7,5,2,1,1,3,2,8,1,9,9,9,4,4,4 | 2,2,0,0,1,0,0,2,0,0,2,1,0,2,0 | 220 | 149 | 0.0000000919223835 | 0.0000931348635831161 | 0.504,0.612,0.657, | 0.703,0.629,1.0,1.0 | -0,166 |
| 254149 | ENSG00000123349 | PFDN5 | chr12 | + | 53295838 | 53295891 | 53295290 | 53295639 | 53297849 | 53299100 | 254149 | 3,2,48,5,0,0,0,0,0,0,0,0 | 62,26,120,60,24,30,21,42,2 | 7,0,2,0,2,0,0,1,1,4,0,0,2,8,0 | 50,56,40,22,61,26,27,59,28 | 202 | 149 | 0.0000000921694641 | 0.0000931348635831161 | 0.034,0.054,0.228, | 0.094,0.0,0.036,0.0 | 0,01 |
| 211315 | ENSG00000160695 | VPS11 | chr11 | + | 119073197 | 119073540 | 119070233 | 119070397 | 119073799 | 119073951 | 211315 | 6,4,20,12,2,14,6,4,1,1,8,4 | 2,0,0,0,1,0,2,0,2,0,0,0 | 18,26,8,5,11,3,2,10,9,6,18, | 9,4,2,1,0,0,1,1,0,0,4,0,3,2,1 | 298 | 149 | 0.0000000985471926 | 0.0000991140692365639 | 0.6,1.0,1.0,1.0,0.5, | 0.5,0.765,0.667,0. | 0,066 |
| 198346 | ENSG00000243943 | ZNF512 | chr2 | + | 27599952 | 27600053 | 27599582 | 27599678 | 27600690 | 27600728 | 198346 | 14,19,53,34,15,16,17,17,14 | 1,0,2,0,0,0,1,6,0,2,0,0 | 37,23,14,0,12,21,5,11,6,22 | 10,7,3,2,11,0,0,5,2,0,1,0,1, | 250 | 149 | 0.0000001012820820 | 0.00010082699840125 | 0.893,1.0,0.94,1.0, | 0.688,0.662,0.736, | 0,217 |
| 141570 | ENSG00000214176 | PLEKHM1P1 | chr17 | - | 64814862 | 64815227 | 64804829 | 64805100 | 64821766 | 64822393 | 141570 | 12,8,2,3,9,4,22,21,3,4,7,13 | 27,18,57,42,45,52,12,38,18 | 0,5,1,2,5,1,2,3,1,4,15,15,1, | 31,39,44,59,60,34,24,26,27 | 298 | 149 | 0.0000001013592089 | 0.00010082699840125 | 0.182,0.182,0.017, | 0.0,0.06,0.011,0.0 | 0,102 |
| 130791 | ENSG00000123349 | PFDN5 | chr12 | + | 53295838 | 53295941 | 53295290 | 53295639 | 53297849 | 53297924 | 130791 | 1102,718,800,663,168,216 | 62,26,120,60,24,30,21,42,2 | 475,370,459,272,307,184,1 | 50,56,40,22,61,26,27,59,28 | 252 | 149 | 0.0000001016557031 | 0.00010082699840125 | 0.913,0.942,0.798, | 0.849,0.796,0.872, | 0,065 |
| 93214 | ENSG00000112576 | CCND3 | chr6 | - | 42030008 | 42030190 | 41940409 | 41940585 | 42048500 | 42048686 | 93214 | 4,0,11,1,1,0,1,2,0,0,0,1 | 64,46,53,35,23,45,45,86,47 | 7,6,1,2,0,1,2,2,0,0,29,4,1,0 | 23,18,34,30,11,8,33,66,27, | 298 | 149 | 0.0000001062205173 | 0.00010487132191121 | 0.03,0.0,0.094,0.0 | 0.132,0.143,0.014, | -0,033 |
| 60461 | ENSG00000019582 | CD74 | chr5 | - | 150403120 | 150403312 | 150402562 | 150402625 | 150404679 | 150404767 | 60461 | 677,361,1432,617,962,412 | 4500,3645,6508,6450,4482 | 817,692,832,865,398,219,5 | 3977,4214,2808,3533,2538 | 298 | 149 | 0.0000001082498404 | 0.00010638685572837 | 0.07,0.047,0.099,0 | 0.093,0.076,0.129, | -0,015 |
| 45952 | ENSG00000180329 | CCDC43 | chr17 | - | 44680584 | 44680652 | 44677482 | 44679043 | 44682002 | 44682138 | 45952 | 13,26,15,14,4,4,3,1,9,23,4, | 4,2,5,4,4,2,0,0,1,0,2,0 | 5,6,13,13,13,3,10,7,3,6,4,1 | 0,1,3,1,2,0,0,0,0,0,3,5,0,0,0 | 217 | 149 | 0.0000001099964468 | 0.00010761202386281 | 0.691,0.899,0.673, | 1.0,0.805,0.748,0. | -0,119 |
| 36985 | ENSG00000101161 | PRPF6 | chr20 | + | 64025938 | 64026058 | 64024554 | 64024693 | 64026981 | 64027158 | 36985 | 88,80,160,52,55,56,65,35,8 | 4,0,0,0,0,0,0,1,0,0,0,0 | 53,62,31,56,47,25,40,67,19 | 3,4,2,1,2,0,2,12,0,0,2,3,2,0 | 269 | 149 | 0.0000001105087340 | 0.00010762400605767 | 0.924,1.0,1.0,1.0,1 | 0.907,0.896,0.896, | 0,055 |
| 163012 | ENSG00000112992 | NNT | chr5 | + | 43647923 | 43648178 | 43645356 | 43645510 | 43649146 | 43649308 | 163012 | 3,2,5,7,2,4,15,16,4,4,2,5 | 32,16,75,43,31,38,10,21,34 | 3,1,6,2,1,0,3,1,1,0,3,2,1,0,3 | 17,28,20,10,21,29,13,14,18 | 298 | 149 | 0.0000001137547563 | 0.00011028626113966 | 0.045,0.059,0.032, | 0.081,0.018,0.13,0 | 0,065 |
| 104475 | ENSG00000111412 | SPRING1 | chr12 | - | 116719762 | 116719876 | 116710170 | 116717893 | 116737789 | 116738069 | 104475 | 8,35,60,14,17,24,28,30,14, | 3,1,6,1,0,2,4,3,0,1,2,4 | 26,45,15,8,45,15,26,16,8,1 | 0,8,0,0,0,1,1,0,0,0,0,0,0,4,3 | 263 | 149 | 0.0000001154849770 | 0.00011113409138964 | 0.602,0.952,0.85,0 | 1.0,0.761,1.0,1.0,1 | -0,111 |
| 208362 | ENSG00000005483 | KMT2E | chr7 | + | 105062168 | 105062278 | 105040838 | 105041023 | 105063350 | 105063386 | 208362 | 168,115,249,143,66,102,90 | 4,2,2,1,0,1,1,1,0,0,3,0 | 189,232,125,145,147,78,68 | 7,4,3,24,4,0,2,2,5,2,0,1,6,0 | 259 | 149 | 0.0000001156619467 | 0.00011113409138964 | 0.96,0.971,0.986,0 | 0.94,0.971,0.96,0. | 0,026 |
| 69583 | ENSG00000069248 | NUP133 | chr1 | - | 229477596 | 229477760 | 229475637 | 229475732 | 229484053 | 229484145 | 69583 | 48,41,160,71,70,31,42,50,3 | 2,2,1,0,0,0,0,4,1,0,0,0 | 78,111,30,33,59,30,58,50,7 | 4,1,3,6,2,3,1,3,2,2,4,5,1,1,4 | 298 | 149 | 0.0000001163064494 | 0.00011125668189374 | 0.923,0.911,0.988, | 0.907,0.982,0.833, | 0,072 |
| 41584 | ENSG00000129071 | MBD4 | chr3 | - | 129436442 | 129437308 | 129434061 | 129434136 | 129437719 | 129437950 | 41584 | 192,123,268,198,100,63,69 | 7,0,7,14,2,2,0,0,7,0,0,4 | 182,188,84,64,72,71,28,57 | 0,1,2,1,0,0,2,0,0,2,0,1,0,1,3 | 298 | 149 | 0.0000001227406294 | 0.00011689198413526 | 0.932,1.0,0.95,0.8 | 1.0,0.989,0.955,0. | -0,016 |
| 229389 | ENSG00000162600 | OMA1 | chr1 | - | 58522826 | 58523000 | 58480877 | 58481174 | 58527260 | 58527335 | 229389 | 13,3,7,7,2,10,1,11,9,6,3,3 | 4,4,8,8,0,4,0,6,2,0,2,0 | 17,9,13,11,2,3,2,8,5,1,20,2 | 0,0,0,2,0,0,5,1,0,0,2,3,1,1,0 | 298 | 149 | 0.0000001240777218 | 0.00011764481126378 | 0.619,0.273,0.304, | 1.0,1.0,1.0,0.733,1 | -0,174 |
| 99600 | ENSG00000129103 | SUMF2 | chr7 | + | 56078921 | 56079039 | 56078363 | 56078508 | 56079527 | 56079736 | 99600 | 2,3,8,12,4,13,9,3,5,2,0,6 | 58,40,142,124,79,76,53,59 | 5,3,2,0,2,0,0,2,0,4,6,2,1,0,3 | 74,90,61,28,37,45,44,81,35 | 267 | 149 | 0.0000001265381635 | 0.00011945147135488 | 0.019,0.04,0.03,0.0 | 0.036,0.018,0.018, | 0,022 |
| 235437 | ENSG00000156171 | DRAM2 | chr1 | - | 111119614 | 111119793 | 111118864 | 111118897 | 111119876 | 111119959 | 235437 | 16,1,14,5,8,4,3,13,6,2,1,1 | 70,41,198,95,101,55,57,10 | 9,1,5,6,0,0,1,7,0,0,2,1,4,0,0 | 97,77,75,37,50,48,39,61,54 | 298 | 149 | 0.0000001281506331 | 0.0001204453664359 | 0.103,0.012,0.034, | 0.044,0.006,0.032, | 0,017 |
| 125548 | ENSG00000100225 | FBXO7 | chr22 | + | 32491085 | 32491181 | 32487744 | 32487828 | 32493104 | 32493281 | 125548 | 151,80,229,212,150,110,11 | 3,1,8,3,0,1,0,1,0,0,0,0 | 150,175,142,57,50,68,65,1 | 7,1,9,2,8,4,0,0,0,2,4,0,7,7,4 | 245 | 149 | 0.0000001300279747 | 0.00012167848272685 | 0.968,0.98,0.946,0 | 0.929,0.991,0.906, | 0,043 |
| 55784 | ENSG00000134352 | IL6ST | chr5 | - | 55948897 | 55948982 | 55947564 | 55947589 | 55951463 | 55951506 | 55784 | 0,2,2,0,0,2,1,3,0,1,0,1 | 7,7,12,5,6,9,18,59,13,5,2,1 | 3,2,10,4,3,1,2,7,1,5,8,2,1,0 | 9,25,10,13,17,4,5,13,3,2,26 | 234 | 149 | 0.0000001321937200 | 0.00012316963881804 | 0.0,0.154,0.096,0.0 | 0.175,0.048,0.389, | -0,114 |
| 132953 | ENSG00000290571 | GOLGA2P5 | chr12 | - | 100168451 | 100168633 | 100167549 | 100167639 | 100170564 | 100170667 | 132953 | 36,19,24,10,7,11,12,16,14, | 5,2,3,2,4,3,0,0,5,3,6,3 | 11,12,23,37,6,7,15,24,16,1 | 0,0,1,0,3,1,2,0,0,0,1,0,2,5,2 | 298 | 149 | 0.0000001333659842 | 0.00012372626791984 | 0.783,0.826,0.8,0. | 1.0,1.0,0.92,1.0,0. | -0,151 |
| 76733 | ENSG00000163029 | SMC6 | chr2 | - | 17713384 | 17713580 | 17708638 | 17708753 | 17714860 | 17714877 | 76733 | 2,0,7,5,5,0,3,0,1,0,0,2 | 55,30,138,99,23,10,31,21,2 | 9,8,5,2,2,2,8,10,2,6,0,22,5, | 53,63,38,20,35,22,37,36,35 | 298 | 149 | 0.0000001370943532 | 0.00012663929070487 | 0.018,0.0,0.025,0.0 | 0.078,0.06,0.062,0 | -0,032 |
| 149237 | ENSG00000128881 | TTBK2 | chr15 | - | 42919758 | 42919958 | 42878548 | 42878684 | 42920437 | 42920809 | 149237 | 2,2,5,0,0,1,0,2,0,0,1,1 | 15,4,18,6,2,9,5,8,7,4,5,7 | 3,2,4,0,5,0,0,3,3,8,3,6,1,9,4 | 5,4,7,3,2,3,4,10,5,9,9,5,8,4 | 298 | 149 | 0.0000001386132593 | 0.00012749517271635 | 0.062,0.2,0.122,0.0 | 0.231,0.2,0.222,0.0 | -0,149 |
| 105286 | ENSG00000164074 | ABHD18 | chr4 | + | 127989720 | 127989821 | 127984338 | 127984423 | 128001715 | 128001760 | 105286 | 21,7,44,29,18,25,13,19,14, | 1,1,6,0,0,0,0,2,0,1,0,0 | 32,18,15,9,8,8,9,15,14,30,2 | 5,0,0,3,4,3,0,1,2,0,0,2,5,5,1 | 250 | 149 | 0.0000001408667854 | 0.00012901659191968 | 0.926,0.807,0.814, | 0.792,1.0,1.0,0.64 | 0,102 |

| | | | | | | | | | | | | | | | | | | | | | | |
|---|---|---|---|---|---|---|---|---|---|---|---|---|---|---|---|---|---|---|---|---|---|---|
| 29616 | ENSG00000116459 | ATP5PB | chr1 | + | 111454210 | 111454356 | 111449681 | 111449873 | 111456629 | 111456748 | 29616 | 7,7,30,36,9,5,5,2,7,3,2,9 | 1,2,0,0,2,2,2,0,0,0,0,0 | 16,21,9,1,14,9,3,8,8,14,3,1 | 2,1,2,1,5,1,2,1,2,0,0,6,1,0,2 | 295 | 149 | 0.0000001434614083 | 0.00013083619648957 | 0.78,0.639,1.0,1.0 | 0.802,0.914,0.694, | 0,128 |
| 208880 | ENSG00000124151 | NCOA3 | chr20 | + | 47639007 | 47639196 | 47637647 | 47637783 | 47639576 | 47639822 | 208880 | 193,131,555,357,211,152,1 | 2,1,7,5,2,2,4,8,2,0,1,3 | 320,437,205,241,279,177,2 | 9,12,8,18,12,1,3,10,6,1,10, | 298 | 149 | 0.0000001488374150 | 0.00013516635308254 | 0.98,0.985,0.975,0 | 0.947,0.948,0.928, | 0,038 |
| 60421 | ENSG00000144744 | UBA3 | chr3 | - | 69057255 | 69057309 | 69056778 | 69056815 | 69061813 | 69061927 | 60421 | 26,13,63,47,47,48,17,43,11 | 0,0,6,0,1,0,0,3,0,0,1,0 | 33,17,6,10,21,19,11,33,10, | 1,3,3,2,2,4,3,2,0,3,1,3,3,4,0 | 203 | 149 | 0.0000001613738596 | 0.00014593553435835 | 1.0,1.0,0.885,1.0,0 | 0.96,0.806,0.595,0 | 0,102 |
| 93010 | ENSG00000223501 | VPS52 | chr6 | - | 33269998 | 33270051 | 33269743 | 33269819 | 33270198 | 33270283 | 93010 | 9,5,25,17,11,20,12,21,7,5,9 | 0,0,0,0,2,2,0,5,2,0,0,0 | 19,17,10,12,8,7,10,18,20,1 | 3,3,3,4,2,0,5,3,3,2,6,4,0,0,0 | 202 | 149 | 0.0000001657277498 | 0.00014924581309402 | 1.0,1.0,1.0,1.0,0.8 | 0.824,0.807,0.711, | 0,107 |
| 169636 | ENSG00000136731 | UGGT1 | chr2 | + | 128135100 | 128135240 | 128134875 | 128134961 | 128138716 | 128138852 | 169636 | 11,5,16,3,5,3,0,6,4,0,0,5 | 36,22,49,25,17,16,11,21,18 | 3,3,0,2,0,0,4,2,0,5,1,0,3,6,1 | 44,43,21,24,17,14,11,20,14 | 298 | 149 | 0.0000001740362601 | 0.00015607499297923 | 0.136,0.105,0.144,0 | 0.034,0.035,0.0,0.0 | 0,043 |
| 173763 | ENSG00000156976 | EIF4A2 | chr3 | + | 186788291 | 186788420 | 186787802 | 186787882 | 186789124 | 186789291 | 173763 | 169,65,144,75,45,36,51,73 | 427,137,428,354,181,165,1 | 102,69,66,67,51,39,12,53,3 | 138,51,178,74,110,133,75, | 278 | 149 | 0.0000001840156291 | 0.00016433970076276 | 0.175,0.203,0.153,0 | 0.284,0.42,0.166,0 | -0,074 |
| 9544 | ENSG00000006282 | SPATA20 | chr17 | + | 50547664 | 50547767 | 50547067 | 50547285 | 50548282 | 50548453 | 9544 | 50,21,67,28,7,15,31,27,23, | 7,0,3,4,2,4,4,4,4,4,2,4 | 21,18,28,16,15,4,11,19,14, | 0,1,1,2,0,0,0,2,0,0,5,2,1,1,3 | 252 | 149 | 0.0000001881430420 | 0.00016693043240983 | 0.809,1.0,0.93,0.80 | 1.0,0.914,0.943,0.8 | -0,137 |
| 234887 | ENSG00000101901 | ALG13 | chrX | + | 111728160 | 111728305 | 111727642 | 111727770 | 111730394 | 111730427 | 234887 | 5,4,18,15,23,9,8,13,6,2,4,8 | 0,1,1,1,2,0,2,2,0,0,2,0 | 8,10,12,4,10,7,3,8,16,12,1 | 2,4,0,4,1,2,3,2,2,3,1,1,0,0,2 | 294 | 149 | 0.0000001884677164 | 0.00016693043240983 | 1.0,0.67,0.901,0.8 | 0.67,0.559,1.0,0.3 | 0,111 |
| 206849 | ENSG00000075884 | ARHGAP15 | chr2 | + | 143734644 | 143734826 | 143703418 | 143703524 | 143767988 | 143768352 | 206849 | 2,3,8,3,6,7,4,4,1,1,0,6 | 75,50,155,97,69,55,50,45,4 | 8,12,8,10,8,9,4,8,4,3,10,11 | 94,64,66,18,62,43,14,32,37 | 298 | 149 | 0.0000001949253436 | 0.00017194252720426 | 0.013,0.029,0.025, | 0.041,0.086,0.057, | -0,052 |
| 173767 | ENSG00000156976 | EIF4A2 | chr3 | + | 186788300 | 186788420 | 186787802 | 186787882 | 186789124 | 186789291 | 173767 | 165,65,138,74,45,36,50,73 | 427,137,428,354,181,165,1 | 100,66,66,67,48,38,12,49,3 | 138,51,178,74,110,133,75, | 269 | 149 | 0.0000001957396759 | 0.00017195610688251 | 0.176,0.208,0.152, | 0.286,0.418,0.17,0 | -0,075 |
| 234655 | ENSG00000101199 | ARFGAP1 | chr20 | + | 63282104 | 63282315 | 63281290 | 63281347 | 63282818 | 63283528 | 234655 | 0,2,14,2,4,2,1,1,2,0,2,3 | 16,15,49,25,20,19,15,12,20 | 0,2,0,1,2,1,2,0,0,0,0,0,2,0,0 | 16,21,14,11,17,8,17,20,10, | 298 | 149 | 0.0000001983318107 | 0.00017352501609741 | 0.0,0.062,0.125,0.0 | 0.0,0.0,0.045,0.0,0.04 | 0,038 |
| 164048 | ENSG00000102445 | RUBCNL | chr13 | - | 46369767 | 46370026 | 46368762 | 46368815 | 46371940 | 46372597 | 164048 | 3,1,23,8,2,3,6,8,0,0,4,6 | 118,124,945,696,243,376,4 | 24,27,12,15,12,10,11,18,3, | 540,780,271,253,398,203,2 | 298 | 149 | 0.0000002008711225 | 0.00017503519258945 | 0.013,0.004,0.012, | 0.022,0.017,0.022, | -0,012 |
| 24588 | ENSG00000068650 | ATP11A | chr13 | + | 112858144 | 112858290 | 112857817 | 112857920 | 112859392 | 112859452 | 24588 | 6,2,6,11,13,1,8,15,27,7,4,2 | 0,0,0,1,0,0,0,5,3,1,0,0 | 12,4,22,40,5,9,2,7,6,9,8,8,3 | 0,1,1,8,6,4,0,1,0,0,2,1,4,4,1 | 295 | 149 | 0.0000002046240413 | 0.00017758643971704 | 1.0,1.0,1.0,0.847,1 | 1.0,0.669,0.917,0.1 | 0,172 |
| 47253 | ENSG00000179912 | R3HDM2 | chr12 | - | 57305639 | 57305663 | 57303175 | 57303217 | 57310263 | 57310463 | 47253 | 1,1,4,0,0,1,0,0,2,1,0,0 | 65,36,61,27,13,29,22,42,39 | 2,1,3,2,3,0,3,0,2,1,2,3,11,9 | 35,45,25,37,15,9,37,23,24, | 173 | 149 | 0.0000002069475545 | 0.00017888164301426 | 0.013,0.023,0.053, | 0.047,0.019,0.094, | -0,048 |
| 211313 | ENSG00000160695 | VPS11 | chr11 | + | 119073197 | 119073527 | 119070233 | 119070397 | 119073799 | 119073951 | 211313 | 4,3,16,8,2,14,4,4,1,1,4,3 | 2,0,0,0,1,0,2,0,2,0,0,0 | 16,17,4,4,6,3,2,8,8,4,13,6,1 | 9,4,2,1,0,0,1,1,0,0,4,0,3,2,1 | 298 | 149 | 0.0000002133777933 | 0.00018370206333881 | 0.5,1.0,1.0,1.0,0.5, | 0.471,0.68,0.5,0.66 | 0,102 |
| 203564 | ENSG00000275052 | PPP4R3B | chr2 | - | 55573620 | 55573777 | 55568193 | 55568363 | 55577314 | 55577356 | 203564 | 19,10,34,34,57,15,11,17,15 | 7,2,6,2,5,2,0,0,0,2,1,2 | 40,24,14,6,10,18,18,36,5,2 | 2,0,0,4,0,1,0,0,1,3,0,3,0,1,0 | 298 | 149 | 0.0000002442055581 | 0.00020940484066827 | 0.576,0.714,0.739, | 0.909,1.0,1.0,0.42 | -0,068 |
| 137858 | ENSG00000269352 | PTOV1-AS2 | chr19 | - | 49858581 | 49858626 | 49858346 | 49858427 | 49858842 | 49859026 | 137858 | 1,0,3,0,6,4,4,2,0,0,4,3 | 13,1,7,1,5,8,10,3,15,4,7,4 | 1,1,1,1,0,0,0,2,0,1,2,2,1,2,0 | 2,4,5,7,3,5,6,8,2,0,8,11,5,9 | 194 | 149 | 0.0000002562661589 | 0.00021887468912349 | 0.056,0.0,0.248,0.0 | 0.277,0.161,0.133, | 0,042 |
| 215672 | ENSG00000168887 | C2orf68 | chr2 | - | 85611317 | 85611573 | 85609434 | 85609586 | 85611667 | 85611741 | 215672 | 0,1,2,3,2,0,9,3,3,0,5,3 | 8,4,16,19,16,6,10,8,5,8,8,7 | 1,1,0,0,0,0,2,0,0,1,6,1,3,1,0 | 19,18,1,8,1,6,4,11,3,6,5,24 | 298 | 149 | 0.0000002722501994 | 0.00023160744145345 | 0.0,0.111,0.059,0.0 | 0.026,0.027,0.0,0.0 | 0,043 |
| 255393 | ENSG00000170456 | DENND5B | chr12 | - | 31447598 | 31447769 | 31442774 | 31442925 | 31451939 | 31452476 | 255393 | 20,39,85,61,38,24,23,22,22 | 0,1,2,0,0,0,0,2,0,0,4,0 | 41,55,46,82,56,40,17,55,50 | 3,1,2,5,1,0,0,1,2,5,5,8,1,2,1 | 298 | 149 | 0.0000002755939959 | 0.00023352902103951 | 1.0,0.951,0.955,1.0 | 0.872,0.965,0.92,0 | 0,036 |
| 105827 | ENSG00000104870 | FCGRT | chr19 | + | 49525456 | 49525573 | 49524506 | 49524776 | 49526009 | 49526329 | 105827 | 295,119,292,179,156,60,15 | 5,0,1,1,2,0,2,0,0,0,1,7 | 189,97,65,55,54,52,93,257 | 10,11,1,0,1,0,2,9,1,0,2,1,0, | 266 | 149 | 0.0000002876564448 | 0.00024207416253022 | 0.971,1.0,0.994,0.9 | 0.914,0.832,0.973, | 0,024 |
| 9547 | ENSG00000006282 | SPATA20 | chr17 | + | 50547719 | 50547767 | 50547152 | 50547285 | 50548282 | 50548453 | 9547 | 29,16,38,16,6,14,22,24,17, | 7,0,3,4,2,4,4,4,4,4,2,4 | 13,9,19,8,10,2,8,15,9,25,15 | 0,1,1,2,0,0,0,2,0,0,5,2,1,1,3 | 197 | 149 | 0.0000002879277873 | 0.00024207416253022 | 0.758,1.0,0.905,0.7 | 1.0,0.872,0.935,0.1 | -0,136 |
| 201664 | ENSG00000158019 | BABAM2 | chr2 | + | 28244519 | 28244862 | 28241322 | 28241393 | 28298337 | 28298491 | 201664 | 30,29,21,15,16,20,15,22,33 | 2,6,1,4,0,5,3,2,4,0,2,0 | 20,17,27,27,17,18,8,22,3,1 | 0,3,2,3,0,4,0,2,0,0,0,0,0,0,0 | 298 | 149 | 0.0000002905602657 | 0.00024333687375424 | 0.882,0.707,0.913, | 1.0,0.739,0.871,0.8 | -0,119 |
| 173310 | ENSG00000198586 | TLK1 | chr2 | - | 171091964 | 171092267 | 171082780 | 171082852 | 171117738 | 171117857 | 173310 | 7,11,7,3,1,1,0,0,1,0,0,2 | 121,95,95,87,40,60,31,53,5 | 8,14,8,22,4,1,0,3,7,12,1,7,0 | 42,98,69,126,48,71,21,39,3 | 298 | 149 | 0.0000003035588909 | 0.00025323375335275 | 0.028,0.055,0.036, | 0.087,0.067,0.055, | -0,039 |
| 198049 | ENSG00000112232 | KHDRBS2 | chr6 | - | 61977147 | 61977350 | 61901243 | 61901371 | 61978065 | 61978212 | 198049 | 1,0,20,22,0,0,3,5,9,3,0,0 | 57,30,75,37,29,39,20,30,78 | 0,0,5,0,10,6,0,1,0,0,0,0,0,0 | 55,32,33,33,11,8,14,28,13, | 298 | 149 | 0.0000003212371226 | 0.00026679624723339 | 0.009,0.0,0.118,0.1 | 0.0,0.0,0.07,0.0,0.1 | 0,007 |
| 84790 | ENSG00000162885 | B3GALNT2 | chr1 | - | 235489167 | 235489268 | 235484321 | 235484515 | 235494680 | 235494828 | 84790 | 7,2,11,3,5,8,0,1,1,2,2,3 | 0,0,1,1,2,0,1,0,0,0,1,0 | 3,7,4,3,1,6,1,7,1,0,2,11,15, | 6,3,1,1,4,0,0,1,2,2,0,0,3,3,0 | 250 | 149 | 0.0000003222910467 | 0.00026679624723339 | 1.0,1.0,0.868,0.64 | 0.23,0.582,0.704,0 | 0,161 |
| 130439 | ENSG00000136521 | NDUFB5 | chr3 | + | 179615982 | 179616049 | 179604808 | 179604939 | 179616982 | 179617044 | 130439 | 12,6,43,50,15,4,15,13,13,9 | 5,0,2,2,0,0,1,0,0,0,1,0 | 27,15,11,5,9,7,4,16,16,21,4 | 1,0,1,1,0,3,0,4,2,0,8,7,4,2,4 | 216 | 149 | 0.0000003237290310 | 0.00026695985853246 | 0.623,1.0,0.937,0.9 | 0.949,1.0,0.884,0.1 | 0,125 |
| 107132 | ENSG00000215301 | DDX3X | chrX | + | 41346858 | 41347012 | 41346228 | 41346622 | 41347311 | 41347451 | 107132 | 482,437,848,511,204,253,2 | 16,2,7,7,1,0,2,4,0,0,2,2 | 546,647,363,399,328,237,1 | 9,21,14,30,3,0,5,9,1,3,11,5 | 298 | 149 | 0.0000003419539210 | 0.00028091253582208 | 0.938,0.991,0.984, | 0.968,0.939,0.928, | 0,022 |
| 108629 | ENSG00000116791 | CRYZ | chr1 | - | 74707102 | 74707204 | 74706898 | 74706994 | 74707097 | 74710247 | 108629 | 29,16,43,47,26,17,6,4,16,1 | 0,2,4,8,3,0,0,3,2,3,0,1 | 17,16,5,2,8,4,8,25,12,25,4, | 0,0,0,0,0,0,0,0,0,8,0,1,2,1,1 | 251 | 149 | 0.0000003586031850 | 0.00029346966584974 | 1.0,0.826,0.865,0.7 | 1.0,1.0,1.0,1.0,1.0,1.0, | -0,085 |
| 53573 | ENSG00000291072 | CROCCP2 | chr1 | - | 16619844 | 16620024 | 16618252 | 16619103 | 16625774 | 16626005 | 53573 | 20,18,23,4,19,16,24,15,22, | 3,1,1,2,0,0,0,0,4,0,0,0 | 10,13,12,15,3,2,9,15,7,13,9 | 3,7,2,0,3,0,5,2,0,1,0,1,2,4,5 | 298 | 149 | 0.0000003716390862 | 0.00030298580371127 | 0.769,0.9,0.92,0.5, | 0.625,0.481,0.75,1 | 0,124 |
| 49556 | ENSG00000117543 | DPH5 | chr1 | - | 101025308 | 101025466 | 101021640 | 101021765 | 101025639 | 101025761 | 49556 | 16,13,18,7,8,7,11,7,5,8,7,1 | 0,3,4,10,0,2,0,1,2,3,0,0 | 12,21,11,3,13,3,6,8,9,16,4, | 2,2,3,2,0,0,0,0,0,1,0,2,1,2,0 | 298 | 149 | 0.0000003738518045 | 0.00030363961410321 | 1.0,0.684,0.692,0.1 | 0.75,0.84,0.647,0.4 | -0,119 |
| 13895 | ENSG00000131148 | EMC8 | chr16 | - | 85780370 | 85780473 | 85778623 | 85779867 | 85781210 | 85781258 | 13895 | 22,13,42,44,19,9,14,42,22, | 0,4,5,5,3,3,2,4,0,0,4,5 | 14,34,10,8,13,9,8,26,12,7,1 | 0,1,0,0,0,0,0,0,2,0,3,1,4,1,0 | 244 | 149 | 0.0000003788963303 | 0.00030563269738074 | 1.0,0.665,0.837,0.8 | 1.0,0.954,1.0,1.0,1 | -0,088 |
| 17371 | ENSG00000169375 | SIN3A | chr15 | - | 75381612 | 75381720 | 75380628 | 75380723 | 75384263 | 75384437 | 17371 | 51,52,52,25,19,29,15,22,32 | 2,2,1,2,0,3,0,2,0,1,1,1 | 32,63,31,20,23,9,16,6,10,3 | 4,7,4,4,2,1,4,2,8,0,1,2,2,3,2 | 257 | 149 | 0.0000003791458024 | 0.00030563269738074 | 0.937,0.938,0.968, | 0.823,0.839,0.818, | 0,134 |
| 133808 | ENSG00000228315 | GUSBP11 | chr22 | - | 23640307 | 23640426 | 23638486 | 23638942 | 23683724 | 23683867 | 133808 | 6,5,16,10,5,12,18,14,5,6,6, | 1,0,0,0,0,2,2,0,3,0,2,0 | 20,2,13,11,4,4,4,8,14,7,13, | 2,0,1,5,2,0,2,2,0,1,0,0,2,6,3 | 268 | 149 | 0.0000003813753934 | 0.00030628286311871 | 0.769,1.0,1.0,1.0,1 | 0.848,1.0,0.878,0.9 | 0,088 |
| 126376 | ENSG00000196233 | LCOR | chr10 | + | 96954952 | 96955454 | 96952102 | 96952196 | 96958411 | 96958926 | 126376 | 15,14,20,18,16,21,9,11,12, | 0,0,6,2,0,2,0,0,0,0,0,1 | 33,26,8,15,13,10,15,21,18, | 5,5,2,2,3,2,0,1,0,0,1,0,2,0,2 | 298 | 149 | 0.0000003888701290 | 0.00031098091033231 | 1.0,1.0,0.625,0.81 | 0.767,0.722,0.667, | 0,132 |
| 36784 | ENSG00000101109 | STK4 | chr20 | + | 45053085 | 45053180 | 45024972 | 45025130 | 45075017 | 45075608 | 36784 | 38,37,111,50,20,45,40,65,2 | 216,151,317,174,128,101,1 | 56,161,43,73,58,40,58,33,3 | 213,191,125,127,107,72,10 | 244 | 149 | 0.0000003901167782 | 0.00031098091033231 | 0.097,0.13,0.176,0 | 0.138,0.34,0.174,0 | -0,067 |
| 3541 | ENSG00000204514 | ZNF814 | chr19 | - | 57876915 | 57877042 | 57850314 | 57850561 | 57888766 | 57888780 | 3541 | 20,6,19,9,11,3,2,7,13,4,1,7 | 3,0,5,0,5,0,0,1,3,3,0,1 | 13,19,10,3,4,4,4,4,1,6,5,12 | 2,0,0,3,0,0,0,0,0,0,0,0,1,0,0 | 276 | 149 | 0.0000003923427139 | 0.00031098091033231 | 0.783,1.0,0.672,1.0 | 0.778,1.0,1.0,0.35 | -0,127 |
| 124397 | ENSG00000073921 | PICALM | chr11 | - | 85976622 | 85976682 | 85974707 | 85974812 | 85981128 | 85981228 | 124397 | 138,106,86,52,21,22,23,31 | 0,0,0,1,0,0,0,3,2,0,1,0 | 101,96,42,39,59,19,15,42,3 | 1,6,2,3,0,0,2,0,2,3,0,5,9,5,1 | 209 | 149 | 0.0000003935399400 | 0.00031098091033231 | 1.0,1.0,1.0,0.974,1 | 0.986,0.919,0.937, | 0,041 |
| 198348 | ENSG00000243943 | ZNF512 | chr2 | + | 27599973 | 27600053 | 27599582 | 27599678 | 27600690 | 27600728 | 198348 | 14,19,35,18,15,15,17,17,13 | 1,0,2,0,0,0,1,6,0,2,0,0 | 35,23,14,0,12,22,4,11,6,21 | 10,7,3,2,11,0,0,5,2,0,1,0,1, | 229 | 149 | 0.0000003944496309 | 0.00031098091033231 | 0.901,1.0,0.919,1.0 | 0.695,0.681,0.752, | 0,204 |
| 150554 | ENSG00000166783 | MARF1 | chr16 | - | 15634756 | 15634927 | 15633616 | 15633843 | 15635655 | 15636342 | 150554 | 99,61,95,78,57,74,57,70,59 | 4,1,4,5,1,7,4,11,0,0,3,0 | 84,64,66,68,40,46,51,66,50 | 1,0,0,0,0,2,5,3,2,0,1,1,0,0,3 | 298 | 149 | 0.0000004020129328 | 0.00031518118237209 | 0.925,0.968,0.922, | 0.977,1.0,1.0,1.0,1 | -0,04 |
| 173664 | ENSG00000135870 | RC3H1 | chr1 | - | 173985089 | 173985155 | 173984498 | 173984619 | 173992754 | 173993135 | 173664 | 0,0,23,14,3,6,2,7,1,1,4,9 | 13,12,75,41,45,22,32,60,22 | 3,4,3,3,4,0,3,1,2,1,3,4,2,4 | 66,52,26,35,55,15,20,40,31 | 215 | 149 | 0.0000004050025944 | 0.00031518118237209 | 0.0,0.0,0.175,0.19 | 0.031,0.051,0.074, | 0,054 |
| 54033 | ENSG00000171824 | EXOSC10 | chr1 | - | 11082687 | 11082878 | 11081081 | 11081238 | 11087447 | 11087591 | 54033 | 97,66,154,96,80,44,71,83,4 | 6,8,2,6,4,1,4,2,0,0,0,0 | 96,114,55,50,59,46,53,79,4 | 5,6,11,12,7,4,7,6,5,3,12,9,4 | 298 | 149 | 0.0000004055316356 | 0.00031518118237209 | 0.89,0.805,0.975,0 | 0.906,0.905,0.714, | 0,109 |
| 126380 | ENSG00000196233 | LCOR | chr10 | + | 96954952 | 96955834 | 96952102 | 96952196 | 96958411 | 96958926 | 126380 | 15,14,20,18,16,21,9,11,12, | 0,0,6,2,0,2,0,0,0,0,0,1 | 34,26,8,15,13,10,15,21,18, | 5,5,2,2,3,2,0,1,0,0,1,0,2,0,2 | 298 | 149 | 0.0000004055536622 | 0.00031518118237209 | 1.0,1.0,0.625,0.81 | 0.773,0.722,0.667, | 0,128 |
| 126382 | ENSG00000196233 | LCOR | chr10 | + | 96954952 | 96955957 | 96952102 | 96952196 | 96958411 | 96958877 | 126382 | 15,14,20,18,16,21,7,12,12, | 0,0,6,2,0,2,0,0,0,0,0,1 | 33,26,8,15,13,10,15,21,18, | 5,5,2,2,3,2,0,1,0,0,1,0,2,0,2 | 298 | 149 | 0.0000004074931942 | 0.00031548621475 | 1.0,1.0,0.625,0.81 | 0.767,0.722,0.667, | 0,132 |
| 78226 | ENSG00000075884 | ARHGAP15 | chr2 | + | 143519265 | 143519364 | 143487372 | 143487495 | 143556407 | 143556485 | 78226 | 264,194,429,228,146,153,1 | 6,1,5,5,0,2,5,2,0,0,0,1 | 290,259,193,176,188,136,6 | 3,15,3,15,14,2,0,2,15,3,7,5 | 248 | 149 | 0.0000004105435893 | 0.00031553397679315 | 0.964,0.991,0.981, | 0.983,0.912,0.975, | 0,031 |
| 126378 | ENSG00000196233 | LCOR | chr10 | + | 96954952 | 96955671 | 96952102 | 96952196 | 96958411 | 96958884 | 126378 | 15,14,20,19,16,21,7,11,12, | 0,0,6,2,0,2,0,0,0,0,0,1 | 33,27,8,15,13,10,15,21,18, | 5,5,2,2,3,2,0,1,0,0,1,0,2,0,2 | 298 | 149 | 0.0000004117436155 | 0.00031553397679315 | 1.0,1.0,0.625,0.82 | 0.767,0.73,0.667,0 | 0,132 |
| 116296 | ENSG00000188283 | ZNF383 | chr19 | + | 37224817 | 37224939 | 37217925 | 37218274 | 37230408 | 37230462 | 116296 | 9,10,18,7,7,0,2,4,2,6,7,6 | 0,6,0,3,4,4,4,0,0,5,0,2 | 8,9,5,4,24,7,5,5,7,18,13,10 | 1,1,0,1,6,0,0,1,0,1,1,2,1,0,0 | 271 | 149 | 0.0000004119529597 | 0.00031553397679315 | 1.0,0.478,1.0,0.56 | 0.815,0.832,1.0,0.0 | -0,22 |
| 143735 | ENSG00000177885 | GRB2 | chr17 | - | 75320331 | 75320553 | 75318075 | 75318297 | 75321658 | 75321827 | 143735 | 333,252,566,385,222,137,2 | 6,0,22,11,4,4,14,11,10,0,2, | 299,366,177,64,206,127,11 | 27,14,6,5,4,9,0,11,2,12,9,1 | 298 | 149 | 0.0000004148545529 | 0.00031662964639121 | 0.965,1.0,0.928,0.9 | 0.847,0.929,0.937, | 0,046 |
| 244491 | ENSG00000171634 | BPTF | chr17 | + | 67875555 | 67875744 | 67874783 | 67875020 | 67891696 | 67891962 | 244491 | 38,6,40,8,11,11,7,10,11,8,3 | 2,3,2,0,2,0,0,0,1,1,0,0 | 39,8,10,12,4,7,16,12,5,10,7 | 3,1,3,1,3,0,2,2,2,4,4,0,2,2,0 | 298 | 149 | 0.0000004187021820 | 0.00031843706481274 | 0.905,0.5,0.909,1.0 | 0.867,0.8,0.625,0.8 | 0,119 |
| 189275 | ENSG00000118308 | IRAG2 | chr12 | + | 25081214 | 25081477 | 25079701 | 25079763 | 25083422 | 25083493 | 189275 | 24,14,30,36,9,6,3,0,2,1,2,9 | 118,60,586,422,118,97,172 | 7,23,10,5,11,2,0,2,3,0,6,0,8 | 289,288,122,36,86,87,63,1 | 298 | 149 | 0.0000004286479374 | 0.00032427974651623 | 0.092,0.104,0.025, | 0.012,0.038,0.039, | 0,017 |
| 149080 | ENSG00000169375 | SIN3A | chr15 | - | 75381612 | 75381690 | 75380628 | 75380723 | 75384263 | 75384437 | 149080 | 42,48,45,19,14,20,15,13,22 | 2,2,1,2,0,3,0,2,0,1,1,1 | 29,46,27,19,20,3,13,5,7,32 | 4,7,4,4,2,1,4,2,8,0,1,2,2,3,2 | 227 | 149 | 0.0000004293978458 | 0.00032427974651623 | 0.932,0.94,0.967,0 | 0.826,0.812,0.816, | 0,153 |
| 126814 | ENSG00000150347 | ARID5B | chr10 | + | 62000090 | 62000321 | 61940182 | 61940408 | 62050887 | 62051000 | 126814 | 153,105,222,135,157,188,1 | 11,6,26,12,11,11,13,23,4,6 | 173,78,132,142,103,93,15 | 7,7,5,5,3,1,10,9,4,8,10,3,8, | 298 | 149 | 0.0000004463602242 | 0.00033591103993217 | 0.874,0.897,0.81,0 | 0.925,0.848,0.93,0 | -0,063 |
| 21344 | ENSG00000100902 | PSMA6 | chr14 | + | 35312880 | 35312921 | 35310832 | 35310895 | 35314360 | 35314455 | 21344 | 10,14,31,17,9,8,8,14,7,8,1, | 1,0,3,1,1,2,0,1,0,0,0,3 | 14,15,8,5,8,2,4,8,6,12,5,30 | 0,0,0,2,6,2,1,3,3,2,4,5,0,1, | 190 | 149 | 0.0000004483628079 | 0.00033624242343876 | 0.887,1.0,0.89,0.9 | 1.0,1.0,1.0,0.662,0 | 0,175 |
| 187812 | ENSG00000033030 | ZCCHC8 | chr12 | - | 122478663 | 122478796 | 122478205 | 122478292 | 122480189 | 122481422 | 187812 | 3,0,1,0,0,4,0,1,0,0,0,0 | 18,14,35,36,18,22,9,13,24, | 1,5,5,13,4,3,4,2,0,1,0,0,1,2 | 14,22,25,9,21,9,9,16,5,18,1 | 282 | 149 | 0.0000004572008961 | 0.00034053261819078 | 0.081,0.0,0.015,0.0 | 0.036,0.107,0.096, | -0,059 |
| 51549 | ENSG00000162894 | FCMR | chr1 | - | 206911729 | 206911952 | 206903316 | 206905147 | 206921817 | 206921867 | 51549 | 9,4,4,3,4,10,5,5,8,14,3,3 | 30,8,8,4,7,18,11,37,23,4 | 4,0,6,4,2,0,0,3,0,0,1,2,1,5 | 18,14,16,12,2,10,8,23,5,6,2 | 298 | 149 | 0.0000004572479180 | 0.00034053261819078 | 0.13,0.2,0.2,0.273, | 0.1,0.0,0.158,0.14 | 0,109 |
| 219581 | ENSG00000132424 | PNISR | chr6 | - | 99411928 | 99412318 | 99410940 | 99410964 | 99412550 | 99412739 | 219581 | 59,26,69,23,43,35,29,34,27 | 390,178,238,211,188,206,2 | 79,80,39,16,21,6,20,44,22, | 171,152,190,271,103,126,1 | 298 | 149 | 0.0000004648835694 | 0.00034502536338945 | 0.07,0.068,0.127,0 | 0.188,0.208,0.093, | -0,032 |
| 24994 | ENSG00000167895 | TMC8 | chr17 | + | 78137234 | 78137814 | 78134867 | 78135009 | 78138004 | 78138071 | 24994 | 113,41,73,87,53,40,52,34,6 | 0,3,1,0,0,1,0,0,0,3,0,0 | 49,68,49,133,22,42,27,54,6 | 3,0,1,8,0,0,2,10,2,1,2,5,3,2 | 298 | 149 | 0.0000004673735690 | 0.00034568137678455 | 1.0,0.872,0.973,1.0 | 0.891,1.0,0.961,0.8 | 0,038 |
| 107136 | ENSG00000215301 | DDX3X | chrX | + | 41346858 | 41347451 | 41346228 | 41346622 | 41347639 | 41349851 | 107136 | 533,479,951,641,238,318,2 | 3,1,7,4,3,0,2,0,0,1,0,2 | 614,733,418,472,425,299,2 | 8,5,1,31,0,6,5,8,1,6,10,0,3, | 298 | 149 | 0.0000004744936158 | 0.00034974565928597 | 0.989,0.996,0.985, | 0.975,0.987,0.995, | 0,018 |
| 231062 | ENSG00000111581 | NUP107 | chr12 | + | 68735207 | 68735344 | 68734707 | 68734833 | 68741812 | 68741980 | 231062 | 13,6,35,27,27,18,5,15,5,4,2 | 2,0,1,0,0,2,3,0,0,0,0,2 | 33,21,13,10,16,16,9,33,6,1 | 3,5,0,5,4,0,0,3,1,1,3,2,0,1 | 286 | 149 | 0.0000004967950107 | 0.00036493408521718 | 0.772,1.0,0.948,1.0 | 0.851,0.686,1.0,0.9 | 0,084 |
| 85975 | ENSG00000198721 | ECI2 | chr6 | - | 4125249 | 4125370 | 4119185 | 4119275 | 4127761 | 4127831 | 85975 | 10,7,28,28,7,7,5,4,6,4,2,4 | 0,0,6,2,0,0,0,1,0,0,0,0 | 14,8,12,2,8,4,2,12,3,12,3,1 | 5,7,3,0,1,1,0,1,0,2,2,0,1,2,0 | 270 | 149 | 0.0000005006716643 | 0.00036653082649562 | 1.0,1.0,0.72,0.885, | 0.607,0.387,0.688, | 0,157 |
| 17270 | ENSG00000206149 | HERC2P9 | chr15 | + | 28655494 | 28655691 | 28655272 | 28655420 | 28655795 | 28656026 | 17270 | 4,5,9,7,3,15,2,0,4,1,2,3 | 0,0,5,1,2,4,2,1,4,0,1,2 | 4,6,4,3,4,2,3,3,1,6,6,11,7,8 | 0,0,0,0,2,1,2,4,0,0,0,3,0,0,0 | 298 | 149 | 0.0000005064918520 | 0.00036953473830861 | 1.0,1.0,0.474,0.77 | 1.0,1.0,1.0,1.0,0.5, | -0,257 |
| 242852 | ENSG00000216937 | CCDC7 | chr10 | + | 32481670 | 32481782 | 32473966 | 32474023 | 32491921 | 32491997 | 242852 | 3,0,16,5,2,3,4,5,1,2,4,4 | 5,4,2,1,1,2,0,0,3,2,1,0 | 9,3,4,6,7,6,2,2,7,3,6,6,2,4,2 | 3,2,3,0,0,2,0,0,0,0,1,3,0,1,0 | 261 | 149 | 0.0000005233570740 | 0.00038054954863742 | 0.255,0.0,0.82,0.74 | 0.631,0.461,0.432, | -0,237 |
| 188133 | ENSG00000068097 | HEATR6 | chr17 | - | 60049575 | 60049702 | 60048263 | 60048388 | 60050842 | 60050860 | 188133 | 7,6,57,36,15,20,13,11,3,3,2 | 4,0,3,1,4,7,1,3,0,2,1,2 | 39,25,28,24,17,15,5,20,17, | 0,4,1,3,2,0,0,0,0,3,0,2,0,1,1 | 278 | 149 | 0.0000005266753797 | 0.00038167295844989 | 0.484,1.0,0.911,0.9 | 1.0,0.77,0.938,0.8 | -0,14 |
| 236416 | ENSG00000120899 | PTK2B | chr8 | + | 27404222 | 27404329 | 27397547 | 27397788 | 27419894 | 27420073 | 236416 | 25,8,21,6,0,1,5,14,0,0,1,5 | 127,28,185,160,102,95,84, | 4,3,2,5,0,3,1,3,0,19,9,0,0 | 112,114,88,71,76,65,64,85 | 256 | 149 | 0.0000005424857048 | 0.00039181121050415 | 0.103,0.143,0.062, | 0.02,0.015,0.013,0 | 0,022 |
| 26373 | ENSG00000115760 | BIRC6 | chr2 | + | 32465064 | 32465164 | 32464508 | 32464823 | 32467524 | 32467739 | 26373 | 235,147,210,109,106,128,1 | 4,1,8,3,3,4,7,15,2,3,0,0 | 204,205,132,122,105,73,94 | 8,27,10,26,17,4,2,2,10,4,11 | 249 | 149 | 0.0000005448199196 | 0.00039209896567778 | 0.972,0.989,0.94,0 | 0.938,0.82,0.888,0 | 0,061 |
| 132610 | ENSG00000105953 | OGDH | chr7 | + | 44656317 | 44656362 | 44647443 | 44647534 | 44666735 | 44666851 | 132610 | 1,0,5,7,1,0,5,1,4,0,1,4 | 5,15,34,23,8,14,12,12,5,11 | 0,0,3,2,0,0,0,0,0,2,2,1,1,3 | 9,10,10,5,21,6,9,14,9,11,14 | 194 | 149 | 0.0000005465276363 | 0.00039209896567778 | 0.133,0.0,0.101,0.1 | 0.0,0.0,0.187,0.23 | 0,086 |
| 99010 | ENSG00000136280 | CCM2 | chr7 | + | 45038252 | 45038426 | 45000187 | 45000363 | 45063917 | 45064001 | 99010 | 16,4,29,16,25,17,10,18,36, | 6,5,6,1,6,1,4,0,0,0,0,2 | 22,5,11,4,2,8,5,3,9,44,16,2 | 4,0,0,0,0,0,0,0,3,1,3,2,2,1 | 298 | 149 | 0.0000005677462509 | 0.00040596874859692 | 0.571,0.286,0.707, | 0.733,1.0,1.0,1.0,1 | -0,153 |
| 88891 | ENSG00000073849 | ST6GAL1 | chr3 | + | 186996547 | 186996673 | 186963784 | 186963926 | 187038741 | 187038873 | 88891 | 82,39,110,83,32,18,32,25,4 | 101,51,133,81,66,36,39,47 | 94,46,110,35,56,55,28,52,3 | 98,27,89,42,23,58,34,69,20 | 275 | 149 | 0.0000005698444759 | 0.00040611985563187 | 0.306,0.293,0.309, | 0.342,0.48,0.401,0 | -0,157 |
| 172650 | ENSG00000168216 | LMBRD1 | chr6 | - | 69738401 | 69738492 | 69737941 | 69738015 | 69741788 | 69741877 | 172650 | 11,4,10,9,1,3,3,6,0,0,0,5 | 31,37,138,108,72,60,44,65 | 1,6,1,0,2,0,1,0,0,0,0,2,5,4,2 | 94,100,38,43,69,47,56,58,4 | 240 | 149 | 0.0000005926145222 | 0.00042091615157448 | 0.181,0.063,0.043, | 0.007,0.036,0.016, | 0,026 |
| 169954 | ENSG00000075826 | SEC31B | chr10 | - | 100490007 | 100490136 | 100489702 | 100489761 | 100490705 | 100490883 | 169954 | 30,8,12,17,12,11,13,16,18, | 1,2,4,2,2,1,0,3,2,0,2,3 | 12,4,5,4,8,5,8,10,2,7,4,11,6 | 1,1,2,0,0,1,1,1,0,2,2,1,0,1,0 | 278 | 149 | 0.0000006359577379 | 0.00045025598647828 | 0.941,0.682,0.617, | 0.865,0.682,0.573, | 0,014 |
| 128134 | ENSG00000119541 | VPS4B | chr18 | - | 63397033 | 63397253 | 63393408 | 63393549 | 63400047 | 63400196 | 128134 | 12,7,45,13,10,16,15,13,11, | 0,0,3,0,0,1,0,0,0,0,0,0 | 13,26,7,7,12,11,16,10,9,21 | 0,4,0,3,3,0,3,1,3,1,4,0,3,4,0, | 298 | 149 | 0.0000006382470637 | 0.00045039529529268 | 1.0,1.0,0.882,1.0,1 | 1.0,0.765,1.0,0.53 | 0,143 |
| 73004 | ENSG00000134759 | ELP2 | chr18 | + | 36146249 | 36146381 | 36142825 | 36142966 | 36154849 | 36154999 | 73004 | 42,18,104,43,33,45,22,34,1 | 4,1,3,2,0,2,0,1,0,0,2,0 | 68,70,25,38,28,16,24,63,38 | 4,3,1,17,14,3,2,3,3,5,1,2, | 281 | 149 | 0.0000006589372275 | 0.00046335108790119 | 0.848,0.905,0.948, | 0.9,0.925,0.93,0.5 | 0,097 |
| 32780 | ENSG00000102445 | RUBCNL | chr13 | - | 46345446 | 46345600 | 46344740 | 46344831 | 46349285 | 46349347 | 32780 | 349,355,2123,1456,659,91 | 0,0,0,0,18,28,21,27,0,0,11, | 1314,2021,822,580,901,42 | 0,0,0,0,0,0,21,33,4,24,0,0,0 | 298 | 149 | 0.0000006609121547 | 0.00046335108790119 | 1.0,1.0,1.0,1.0,0.9 | 1.0,1.0,1.0,1.0,1.0, | -0,01 |
| 118053 | ENSG00000291072 | CROCCP2 | chr1 | - | 16619844 | 16620245 | 16618255 | 16619103 | 16625774 | 16626005 | 118053 | 25,14,20,3,15,15,16,16,13, | 3,1,1,2,0,0,0,0,4,0,0,0 | 6,8,7,13,0,2,2,5,9,4,10,8,11, | 3,7,2,0,3,0,5,2,0,1,0,1,2,4,5 | 298 | 149 | 0.0000006666217835 | 0.00046583660096 | 0.806,0.875,0.909, | 0.5,0.364,0.636,1. | 0,165 |
| 108934 | ENSG00000147905 | ZCCHC7 | chr9 | + | 37147361 | 37147445 | 37126311 | 37126942 | 37302187 | 37302231 | 108934 | 20,10,48,13,15,21,16,38,5 | 84,43,152,67,88,72,38,60,8 | 66,46,59,15,14,15,22,20,1 | 151,54,120,29,22,61,33,59 | 233 | 149 | 0.0000006747689487 | 0.00047000386692592 | 0.132,0.129,0.168, | 0.218,0.353,0.239, | -0,092 |
| 181031 | ENSG00000290993 | | chr3 | - | 129399007 | 129399108 | 129397039 | 129397140 | 129399380 | 129399495 | 181031 | 11,3,12,12,6,6,7,5,5,9,2,7 | 2,4,3,2,5,0,2,0,1,1,0,0 | 9,10,2,10,6,1,3,7,6,6,4,7,1 | 0,0,0,1,0,0,0,0,0,0,1,5,1,0,0 | 250 | 149 | 0.0000006783563156 | 0.00047097841348079 | 0.766,0.309,0.704, | 1.0,1.0,1.0,1.0,0.7 | -0,164 |
| 168717 | ENSG00000153064 | BANK1 | chr4 | + | 101847055 | 101847130 | 101829807 | 101830206 | 101855034 | 101855189 | 168717 | 21,10,11,7,6,0,3,8,2,0,3,0 | 251,234,332,207,32,29,53, | 9,14,14,36,2,2,0,2,3,9,6,10 | 99,94,199,294,37,44,15,45 | 224 | 149 | 0.0000006839992580 | 0.00047336927427963 | 0.053,0.028,0.022, | 0.057,0.09,0.045,0 | -0,012 |
| 76526 | ENSG00000129968 | ABHD17A | chr19 | - | 1878663 | 1878904 | 1877507 | 1877687 | 1879920 | 1880017 | 76526 | 3,3,6,0,0,2,1,4,6,0,3,2 | 10,11,22,10,7,3,12,12,16,1 | 0,1,2,1,3,0,0,0,2,1,0,4,3,0,0 | 13,14,5,6,2,1,6,11,2,9,9,59 | 298 | 149 | 0.0000007004059277 | 0.00048317009046409 | 0.13,0.12,0.12,0.0, | 0.0,0.034,0.167,0.0 | 0,035 |
| 22983 | ENSG00000129566 | TEP1 | chr14 | - | 20384599 | 20384713 | 20384037 | 20384232 | 20384984 | 20385109 | 22983 | 16,4,30,22,15,16,5,11,9,8,1 | 2,0,0,0,0,0,0,1,1,1,0,0 | 17,23,14,9,14,11,12,20,26, | 10,6,0,8,0,0,0,1,0,2,1,2,0,0 | 298 | 149 | 0.0000007061358694 | 0.00048504474646968 | 0.819,1.0,1.0,1.0,1 | 0.491,0.685,1.0,0.1 | 0,092 |
| 103321 | ENSG00000231721 | LINC-PINT | chr7 | - | 131002196 | 131002320 | 130984052 | 130984109 | 131052424 | 131052555 | 103321 | 0,2,6,1,16,30,1,1,8,8,0,2 | 132,85,282,117,79,106,96, | 2,3,29,49,35,18,1,7,17,3,2,3, | 176,141,91,124,97,47,76,9 | 273 | 149 | 0.0000007076306405 | 0.00048504474646968 | 0.0,0.013,0.011,0.0 | 0.006,0.011,0.148, | -0,033 |
| 189183 | ENSG00000118308 | IRAG2 | chr12 | + | 25061411 | 25061653 | 25052944 | 25052956 | 25062819 | 25062900 | 189183 | 130,49,231,179,91,65,108, | 241,99,472,328,182,141,16 | 75,105,88,30,39,29,33,70,3 | 221,280,165,85,110,96,48, | 298 | 149 | 0.0000007192104803 | 0.0004914171139856 | 0.212,0.198,0.197, | 0.145,0.158,0.211, | 0,05 |

| | | | | | | | | | | | | | | | | | | | | | |
|---|---|---|---|---|---|---|---|---|---|---|---|---|---|---|---|---|---|---|---|---|---|
| 77485 | ENSG00000162961 | DPY30 | chr2 | - | 31869847 | 31869952 | 31867808 | 31868492 | 31917925 | 31918037 | 77485 | 3,4,11,6,6,3,2,3,4,1,3,7 | 0,0,2,1,0,0,1,1,0,0,0,0 | 5,15,1,4,10,2,2,12,3,7,1,4,2 | 0,7,3,2,2,2,2,3,1,0,0,0,1,0,1 | 254 | 149 | 0.0000007263363452 | 0.00049471549978901 | 1.0,1.0,0.763,0.77 | 1.0,0.557,0.164,0. | 0,209 |
| 151527 | ENSG00000077238 | IL4R | chr16 | + | 27344593 | 27344661 | 27342120 | 27342259 | 27344868 | 27345090 | 151527 | 11,1,63,36,14,8,5,16,21,2,2 | 295,108,1197,554,391,308 | 11,8,7,2,3,3,3,3,2,6,10,1,6, | 532,290,410,100,136,154,1 | 217 | 149 | 0.0000007298613573 | 0.00049554823282023 | 0.025,0.006,0.035, | 0.014,0.019,0.012, | 0,006 |
| 125894 | ENSG00000107938 | EDRF1 | chr10 | + | 125730287 | 125730399 | 125729357 | 125729479 | 125733403 | 125733551 | 125894 | 22,9,52,25,22,24,12,24,7,4 | 0,0,4,1,0,1,0,4,0,1,0,0 | 32,41,15,11,18,16,11,12,14 | 0,0,0,1,9,8,4,2,1,2,0,2,1,0,2 | 261 | 149 | 0.0000007447885583 | 0.00050305801942989 | 1.0,1.0,0.881,0.93 | 1.0,1.0,1.0,0.863,0 | 0,086 |
| 37358 | ENSG00000223547 | ZNF844 | chr19 | + | 12074360 | 12074421 | 12064730 | 12064876 | 12075311 | 12081565 | 37358 | 18,2,62,29,12,19,14,19,4,5 | 2,0,3,3,2,5,5,4,0,0,2,0 | 15,15,28,9,10,9,13,16,12,1 | 2,0,0,0,6,1,0,0,1,0,0,0,0,1,0 | 210 | 149 | 0.0000007455966296 | 0.00050305801942989 | 0.865,1.0,0.936,0. | 0.842,1.0,1.0,1.0,0 | -0,079 |
| 148702 | ENSG00000141646 | SMAD4 | chr18 | + | 51051376 | 51051415 | 51049294 | 51049324 | 51054780 | 51054916 | 148702 | 5,0,4,3,1,8,5,11,2,0,7,2 | 58,56,113,63,55,45,34,47,8 | 7,18,12,18,0,2,5,15,2,10,16 | 47,68,34,23,28,19,37,52,16 | 188 | 149 | 0.0000007563494376 | 0.0005078684881328 | 0.064,0.0,0.027,0. | 0.106,0.173,0.219, | -0,059 |
| 102467 | ENSG00000124486 | USP9X | chrX | + | 41150920 | 41151057 | 41148368 | 41148575 | 41152947 | 41153081 | 102467 | 71,69,247,146,87,129,66,1 | 3,0,7,2,2,3,4,4,2,2,0,0 | 179,176,101,139,99,75,80, | 12,13,7,28,2,0,5,7,4,4,10,1 | 286 | 149 | 0.0000007574456499 | 0.0005078684881328 | 0.925,1.0,0.948,0. | 0.886,0.876,0.883, | 0,045 |
| 163964 | ENSG00000102445 | RUBCNL | chr13 | - | 46345446 | 46345615 | 46344740 | 46344831 | 46349285 | 46349347 | 163964 | 174,164,1098,742,314,464 | 0,0,0,0,18,28,21,27,0,0,11, | 672,961,450,299,474,228,4 | 0,0,0,0,0,0,21,33,4,24,0,0,0 | 298 | 149 | 0.0000007619489394 | 0.000509301342227 | 1.0,1.0,1.0,1.0,0.8 | 1.0,1.0,1.0,1.0,1.0, | -0,02 |
| 27995 | ENSG00000106123 | EPHB6 | chr7 | + | 142863630 | 142863695 | 142862755 | 142863327 | 142863965 | 142864262 | 27995 | 9,3,7,10,5,7,7,8,16,3,2,7 | 4,2,3,2,0,2,2,2,0,4,0,0 | 9,12,2,7,5,3,5,6,4,6,2,30,26 | 2,0,0,1,2,0,0,2,0,0,2,2,1,2,2 | 214 | 149 | 0.0000007854703387 | 0.00052339803865022 | 0.61,0.511,0.619,0 | 0.758,1.0,1.0,0.83, | -0,109 |
| 26353 | ENSG00000115760 | BIRC6 | chr2 | + | 32435495 | 32435585 | 32433643 | 32433804 | 32439507 | 32439686 | 26353 | 24,21,68,42,22,23,11,51,8, | 3,1,2,4,2,0,2,0,0,0,0,0 | 40,48,27,38,25,16,23,41,33 | 5,2,0,15,3,3,0,11,3,2,3,1,0, | 239 | 149 | 0.0000007900981551 | 0.00052485683958419 | 0.833,0.929,0.955, | 0.833,0.937,1.0,0. | 0,079 |
| 158350 | ENSG00000085511 | MAP3K4 | chr6 | + | 161022009 | 161022227 | 160991843 | 160992083 | 161034258 | 161034449 | 158350 | 1,0,2,2,0,0,2,1,1,0,0,3 | 12,6,5,6,4,8,6,5,7,2,6,2 | 4,2,1,3,3,1,4,4,2,1,5,1,2,3,1 | 5,3,10,5,2,5,8,2,1,6,2,7,5,0 | 298 | 149 | 0.0000007929979207 | 0.00052516226303351 | 0.04,0.0,0.167,0.1 | 0.286,0.25,0.048,0 | -0,203 |
| 160560 | ENSG00000113732 | ATP6V0E1 | chr5 | + | 172986599 | 172986760 | 172983830 | 172983964 | 172994774 | 172994822 | 160560 | 3,3,8,6,5,1,3,3,0,0,0,3 | 70,64,115,194,196,99,90,9 | 7,12,3,8,5,3,1,4,4,5,10,8,2, | 121,86,70,21,41,43,48,104 | 298 | 149 | 0.0000008098972802 | 0.00053283286934936 | 0.021,0.023,0.034, | 0.028,0.065,0.021, | -0,027 |
| 174196 | ENSG00000114867 | EIF4G1 | chr3 | + | 184317716 | 184317786 | 184314550 | 184314674 | 184319688 | 184320297 | 174196 | 1,3,2,7,0,1,1,0,4,1,2,2 | 1,0,0,2,2,0,0,4,0,2,0,2 | 1,3,0,2,2,0,2,0,0,2,3,1,1,0,3 | 6,1,1,10,0,1,3,2,1,1,5,4,2,2 | 219 | 149 | 0.0000008109394330 | 0.00053283286934936 | 0.405,1.0,1.0,0.70 | 0.102,0.671,0.0,0. | 0,415 |
| 232081 | ENSG00000104870 | FCGRT | chr19 | + | 49525440 | 49525573 | 49524506 | 49524776 | 49526009 | 49526329 | 232081 | 163,74,126,115,111,41,108 | 5,0,1,1,2,0,2,0,0,0,1,7 | 100,60,37,37,31,29,58,178 | 10,11,1,0,1,0,2,9,1,0,2,1,0, | 282 | 149 | 0.0000008120074763 | 0.00053283286934936 | 0.945,1.0,0.985,0. | 0.841,0.742,0.951, | 0,042 |
| 72999 | ENSG00000134759 | ELP2 | chr18 | + | 36144938 | 36145034 | 36142825 | 36142966 | 36154849 | 36154999 | 72999 | 33,24,50,19,28,26,9,20,18, | 4,1,3,2,0,2,0,1,0,0,2,0 | 44,27,24,21,19,3,11,21,15, | 4,3,1,17,14,3,2,3,3,2,5,1,2, | 245 | 149 | 0.0000008325262917 | 0.00054463667571159 | 0.834,0.936,0.91,0 | 0.87,0.846,0.936,0 | 0,153 |
| 81148 | ENSG00000166986 | MARS1 | chr12 | + | 57489895 | 57489971 | 57489266 | 57489558 | 57490206 | 57490216 | 81148 | 113,23,179,148,77,80,53,6 | 5,3,6,2,1,8,7,3,1,2,1,4 | 83,56,56,22,54,53,37,83,74 | 5,19,9,6,9,1,8,3,4,5,11,6,16 | 225 | 149 | 0.0000008417091754 | 0.00054897547741085 | 0.937,0.835,0.952, | 0.917,0.661,0.805, | 0,077 |
| 132659 | ENSG00000173442 | EHBP1L1 | chr11 | + | 65581538 | 65582016 | 65581210 | 65581373 | 65584240 | 65584398 | 132659 | 40,24,27,13,27,36,42,45,51 | 9,1,1,0,1,1,12,0,2,0,0,0 | 18,14,15,17,16,19,21,42,12 | 0,0,0,0,2,0,0,0,0,2,0,0,0,1,4 | 298 | 149 | 0.0000008541545825 | 0.0005554095013528 | 0.69,0.923,0.931,1 | 1.0,1.0,1.0,1.0,0.8, | -0,02 |
| 189423 | ENSG00000177200 | CHD9 | chr16 | + | 53239747 | 53239881 | 53238342 | 53238586 | 53242839 | 53243016 | 189423 | 4,1,5,0,2,4,1,5,0,1,0,0 | 27,24,88,36,33,43,16,14,10 | 7,13,4,8,7,2,3,2,6,3,1,6,11, | 33,51,22,27,29,23,17,24,42 | 283 | 149 | 0.0000008583756930 | 0.00055646425160451 | 0.072,0.021,0.029, | 0.1,0.118,0.087,0. | -0,049 |
| 141658 | ENSG00000108312 | UBTF | chr17 | - | 44208833 | 44209001 | 44207863 | 44207911 | 44209351 | 44209422 | 141658 | 0,0,1,2,3,0,41,25,1,1,2,1 | 79,39,99,76,54,53,29,38,68 | 0,0,0,0,0,1,2,0,1,2,6,2,2,2,0 | 67,67,45,11,55,41,29,49,19 | 298 | 149 | 0.0000008609475204 | 0.00055646425160451 | 0.0,0.0,0.005,0.01 | 0.0,0.0,0.0,0.0,0.0, | 0,049 |
| 188130 | ENSG00000068097 | HEATR6 | chr17 | - | 60049491 | 60049702 | 60048263 | 60048388 | 60050842 | 60050860 | 188130 | 7,7,68,38,18,22,15,14,7,3,1 | 4,0,3,1,4,7,1,3,0,2,1,2 | 40,28,29,24,19,17,7,28,18, | 0,4,1,3,2,0,0,0,0,3,1,2,0,0,1 | 298 | 149 | 0.0000009231157709 | 0.00059485967213741 | 0.467,1.0,0.919,0. | 1.0,0.778,0.935,0. | -0,132 |
| 28397 | ENSG00000204120 | GIGYF2 | chr2 | + | 232847347 | 232847571 | 232845731 | 232845886 | 232850261 | 232850409 | 28397 | 54,31,65,25,29,28,16,36,44 | 12,6,4,0,0,3,2,5,2,1,0,2 | 28,52,48,44,21,12,23,18,13 | 1,0,1,5,0,0,2,0,5,0,1,1,0,1,2 | 298 | 149 | 0.0000009294213486 | 0.00059650444911857 | 0.692,0.721,0.89,1 | 0.933,1.0,0.96,0.8 | -0,063 |
| 213714 | ENSG00000116793 | PHTF1 | chr1 | - | 113706593 | 113706701 | 113705764 | 113706162 | 113710253 | 113710475 | 213714 | 26,7,37,64,14,18,28,22,21, | 0,0,4,0,1,2,0,3,0,0,0,2 | 19,43,11,6,11,18,11,11,5,2 | 2,5,3,5,1,0,0,8,1,1,4,0,0,0,4 | 257 | 149 | 0.0000009312110936 | 0.00059650444911857 | 1.0,1.0,0.843,1.0,0 | 0.846,0.833,0.68,0 | 0,114 |
| 9006 | ENSG00000291117 | | chr17 | - | 64814862 | 64815245 | 64804841 | 64805100 | 64821766 | 64822052 | 9006 | 12,8,2,2,4,2,22,20,2,1,5,13 | 27,18,57,42,45,52,12,38,18 | 0,2,1,0,3,1,2,2,1,3,15,15,1, | 31,39,44,59,60,34,24,26,27 | 298 | 149 | 0.0000009385635726 | 0.00059813125990882 | 0.182,0.182,0.017, | 0.0,0.025,0.011,0. | 0,086 |
| 459 | ENSG00000183527 | PSMG1 | chr21 | - | 39179923 | 39179986 | 39178448 | 39178647 | 39180284 | 39180436 | 459 | 17,5,27,38,9,12,11,6,11,4,5 | 1,0,0,1,4,0,1,0,3,0,2,0 | 11,5,5,4,7,6,11,4,4,5,6,13,9 | 0,3,1,0,1,0,0,1,3,2,1,4,0,4,3 | 212 | 149 | 0.0000009393087698 | 0.00059813125990882 | 0.923,1.0,1.0,0.96 | 1.0,0.539,0.778,1. | 0,141 |
| 253572 | ENSG00000135968 | GCC2 | chr2 | + | 108474788 | 108475083 | 108472826 | 108472899 | 108475534 | 108475635 | 253572 | 9,8,12,4,1,2,0,3,0,0,5,5 | 65,38,58,37,13,19,6,14,24, | 12,8,14,10,2,5,9,9,4,13,8,3 | 17,16,33,22,13,8,8,20,13,1 | 298 | 149 | 0.0000009778801418 | 0.00061974414752602 | 0.065,0.095,0.094, | 0.261,0.2,0.175,0. | -0,106 |
| 132063 | ENSG00000069493 | CLEC2D | chr12 | + | 9680922 | 9681033 | 9669712 | 9669795 | 9687901 | 9688086 | 132063 | 102,31,410,146,61,39,66,9 | 5,0,17,3,0,0,0,0,0,0,4,8 | 151,106,53,47,55,43,36,78 | 2,0,0,0,0,0,0,0,0,0,1,4,0,1,2 | 260 | 149 | 0.0000009790086472 | 0.00061974414752602 | 0.921,1.0,0.933,0. | 0.977,1.0,1.0,1.0,1 | -0,027 |
| 99367 | ENSG00000176142 | TMEM39A | chr3 | - | 119458017 | 119458240 | 119452446 | 119452530 | 119461961 | 119462008 | 99367 | 10,16,40,10,19,17,21,9,6,3 | 0,1,3,0,0,2,2,0,0,1,0,0 | 32,45,17,6,30,10,13,15,10, | 0,8,1,10,1,0,1,2,0,0,4,9,0,0 | 298 | 149 | 0.0000009888779464 | 0.00062415598030873 | 1.0,0.889,0.87,1.0, | 1.0,0.738,0.895,0. | 0,067 |
| 218269 | ENSG00000134440 | NARS1 | chr18 | - | 57603724 | 57603850 | 57602811 | 57602943 | 57605856 | 57605970 | 218269 | 26,5,5,3,4,14,0,0,7,0,2,3 | 19,7,93,72,39,34,40,51,32, | 6,19,7,8,21,3,7,21,15,1,12, | 46,70,41,9,29,8,28,51,16,3 | 275 | 149 | 0.0000009961218500 | 0.00062688977163282 | 0.426,0.279,0.028, | 0.066,0.128,0.085, | -0,08 |
| 233314 | ENSG00000215301 | DDX3X | chrX | + | 41346836 | 41347012 | 41346504 | 41346622 | 41347311 | 41347451 | 233314 | 207,179,285,149,71,83,90, | 16,2,7,7,1,0,2,4,0,0,2,2 | 174,199,142,115,106,72,40 | 9,21,14,30,3,0,5,9,1,3,11,5 | 298 | 149 | 0.0000010057218731 | 0.00063108607718218 | 0.866,0.978,0.953, | 0.906,0.826,0.835, | 0,064 |
| 233309 | ENSG00000215301 | DDX3X | chrX | + | 41346822 | 41347012 | 41346504 | 41346622 | 41347311 | 41347451 | 233309 | 207,179,285,149,71,83,90, | 16,2,7,7,1,0,2,4,0,0,2,2 | 172,199,142,115,106,72,40 | 9,21,14,30,3,0,5,9,1,3,11,5 | 298 | 149 | 0.0000010088784874 | 0.00063122652827294 | 0.866,0.978,0.953, | 0.905,0.826,0.835, | 0,064 |
| 201918 | ENSG00000134759 | ELP2 | chr18 | + | 36145933 | 36146381 | 36142825 | 36142966 | 36154849 | 36154999 | 201918 | 40,19,110,37,36,42,23,30,1 | 4,1,3,2,0,2,0,1,0,0,2,0 | 67,71,25,30,29,16,25,62,39 | 4,3,1,17,14,3,2,3,3,2,5,1,2, | 298 | 149 | 0.0000010226895982 | 0.00063801305772939 | 0.833,0.905,0.948, | 0.893,0.922,0.926, | 0,103 |
| 183760 | ENSG00000171163 | ZNF692 | chr1 | - | 248856291 | 248856422 | 248855727 | 248855946 | 248856513 | 248856562 | 183760 | 31,14,31,16,21,14,5,9,27,1 | 1,1,5,1,3,2,1,5,0,3,0,2 | 11,14,4,10,10,5,12,12,12,1 | 2,0,0,0,0,0,4,0,0,2,3,0,0,0 | 280 | 149 | 0.0000010350953197 | 0.00064252791789338 | 0.943,0.882,0.767, | 0.745,1.0,1.0,1.0,1 | -0,104 |
| 205873 | ENSG00000128815 | WDFY4 | chr10 | + | 48808288 | 48808365 | 48807858 | 48807958 | 48810529 | 48810735 | 205873 | 8,1,6,4,0,4,2,4,2,2,5,3 | 70,39,99,83,83,71,44,66,56 | 7,19,2,33,3,0,7,5,1,2,10,11 | 121,89,26,41,33,20,69,85,1 | 226 | 149 | 0.0000010358971872 | 0.00064252791789338 | 0.07,0.017,0.038,0 | 0.037,0.123,0.048, | -0,041 |
| 82829 | ENSG00000160695 | VPS11 | chr11 | + | 119073197 | 119073399 | 119070233 | 119070397 | 119073799 | 119073951 | 82829 | 21,16,39,28,12,27,19,17,32 | 2,0,0,0,1,0,2,0,2,0,0,0 | 31,28,20,19,13,6,12,13,13, | 9,4,2,1,0,0,1,1,0,0,4,0,3,2,1 | 298 | 149 | 0.0000010435824047 | 0.00064543472571304 | 0.84,1.0,1.0,1.0,0. | 0.633,0.778,0.833, | 0,081 |
| 79533 | ENSG00000088179 | PTPN4 | chr2 | + | 119926597 | 119926666 | 119920068 | 119920241 | 119932423 | 119932549 | 79533 | 38,22,34,32,15,13,15,14,22 | 3,0,1,1,0,0,3,3,0,0,1,0 | 28,18,8,4,15,12,17,20,2,8,1 | 2,6,1,4,0,1,1,2,4,1,0,9,1,0,3 | 218 | 149 | 0.0000010493690085 | 0.00064715398591885 | 0.896,1.0,0.959,0. | 0.905,0.672,0.845, | 0,13 |
| 132936 | ENSG00000176915 | ANKLE2 | chr12 | - | 132733995 | 132734071 | 132730232 | 132730270 | 132734384 | 132734575 | 132936 | 10,5,17,6,5,9,24,11,8,2,3,5 | 67,61,105,86,48,39,15,43,2 | 0,2,3,4,4,4,0,8,2,8,2,2,9,6,3 | 49,60,26,48,51,26,20,52,36 | 225 | 149 | 0.0000010893149252 | 0.00066939225176332 | 0.09,0.051,0.097,0 | 0.0,0.022,0.071,0. | 0,082 |
| 118052 | ENSG00000291072 | CROCCP2 | chr1 | - | 16619844 | 16620038 | 16618255 | 16619103 | 16625774 | 16626005 | 118052 | 19,10,20,3,13,14,16,13,13, | 3,1,1,2,0,0,0,0,4,0,0,0 | 6,8,6,13,2,2,5,7,4,11,7,11,1 | 3,7,2,0,3,0,5,2,0,1,0,1,2,4,5 | 298 | 149 | 0.0000010916488812 | 0.00066939225176332 | 0.76,0.833,0.909,0 | 0.5,0.364,0.6,1.0,0 | 0,163 |
| 10263 | ENSG00000006740 | ARHGAP44 | chr17 | + | 12928928 | 12929046 | 12919754 | 12919831 | 12941055 | 12941124 | 10263 | 27,2,91,76,44,77,38,54,31, | 0,0,4,10,2,4,6,4,0,2,0,0 | 39,80,9,10,32,24,21,49,23, | 2,0,0,0,0,0,1,0,0,4,7,0,0,0,2 | 267 | 149 | 0.0000011023501858 | 0.0006740339001134 | 1.0,1.0,0.927,0.80 | 0.916,1.0,1.0,1.0,1 | -0,019 |
| 116819 | ENSG00000114999 | TTL | chr2 | + | 112485916 | 112485995 | 112482155 | 112482501 | 112494142 | 112494375 | 116819 | 38,31,22,32,21,10,19,29,24 | 5,2,5,9,2,0,1,0,4,0,1,0 | 26,21,18,8,28,8,16,12,9,27 | 0,0,0,0,1,2,0,1,0,7,2,0,2,2,0 | 228 | 149 | 0.0000011195820127 | 0.00068263103736685 | 0.832,0.91,0.742,0 | 1.0,1.0,1.0,1.0,0.9 | -0,035 |
| 27706 | ENSG00000127603 | MACF1 | chr1 | + | 39315512 | 39315691 | 39310830 | 39311000 | 39316390 | 39316529 | 27706 | 55,27,112,103,58,54,40,20 | 3,0,4,4,1,0,1,1,0,0,0,0 | 103,79,24,35,34,29,17,49,6 | 5,7,0,3,3,0,0,4,2,6,1,8,8,2,1 | 298 | 149 | 0.0000011315365666 | 0.00068797103610438 | 0.902,1.0,0.933,0. | 0.912,0.849,1.0,0. | 0,057 |
| 163185 | ENSG00000247774 | PCED1B-AS1 | chr12 | - | 47209307 | 47209458 | 47206557 | 47206588 | 47210639 | 47210757 | 163185 | 7,11,19,15,21,20,12,9,15,1 | 0,0,0,1,2,0,2,0,1,1,0,0 | 15,15,18,5,5,9,3,8,11,18,4, | 1,4,0,1,0,3,2,2,1,1,0,13,2,1 | 298 | 149 | 0.0000011411495639 | 0.0006918613008409 | 1.0,1.0,1.0,0.882,0 | 0.882,0.652,1.0,0. | 0,129 |
| 217705 | ENSG00000113761 | ZNF346 | chr5 | + | 177023164 | 177023234 | 177022903 | 177022913 | 177041125 | 177041229 | 217705 | 8,1,8,5,1,0,2,1,1,2,6,0 | 10,3,15,3,4,5,11,2,8,14,1,4 | 0,5,0,4,3,0,0,2,2,0,2,0,0,0,0 | 11,9,13,16,5,6,4,4,3,13,5,1 | 219 | 149 | 0.0000011617752272 | 0.00070238776385981 | 0.352,0.185,0.266, | 0.0,0.274,0.0,0.14 | 0,141 |
| 180029 | ENSG00000122224 | LY9 | chr1 | + | 160823506 | 160823649 | 160819320 | 160819374 | 160827747 | 160828261 | 180029 | 18,8,6,9,19,4,16,12,12,5,4, | 0,0,1,0,2,0,0,0,0,0,2,0 | 13,14,16,13,3,7,24,10,6,14 | 2,1,1,1,3,0,0,2,1,0,2,5,4,2,2 | 292 | 149 | 0.0000011663894442 | 0.00070320214698983 | 1.0,1.0,0.754,1.0,0 | 0.768,0.877,0.891, | 0,157 |
| 138030 | ENSG00000198521 | ZNF43 | chr19 | - | 21819094 | 21819407 | 21817887 | 21817986 | 21845391 | 21845506 | 138030 | 24,30,39,36,7,17,11,21,16, | 3,0,1,0,0,0,0,0,1,0,0,0 | 30,24,14,10,9,5,11,25,8,16 | 1,1,0,0,0,1,4,0,1,0,6,3,4,0,2 | 298 | 149 | 0.0000011807192520 | 0.0007098530316643 | 0.8,1.0,0.951,1.0,1 | 0.938,0.923,1.0,1. | 0,147 |
| 130704 | ENSG00000151148 | UBE3B | chr12 | + | 109529889 | 109530072 | 109526357 | 109526416 | 109533465 | 109533558 | 130704 | 21,18,28,6,14,10,8,6,13,2,7 | 1,4,2,2,3,1,6,7,1,1,3,1 | 15,13,9,8,5,11,5,4,4,40,17, | 0,4,0,0,1,1,2,0,1,0,2,2,1,1,0 | 298 | 149 | 0.0000012107021452 | 0.00072585134655582 | 0.913,0.692,0.875, | 1.0,0.619,1.0,1.0,0 | -0,173 |
| 223901 | ENSG00000173200 | PARP15 | chr3 | + | 122620423 | 122620594 | 122619780 | 122619843 | 122621443 | 122621611 | 223901 | 0,4,9,2,6,8,0,2,3,0,0,1 | 335,269,265,198,186,171,1 | 15,13,0,5,15,9,2,0,12,18,8, | 187,244,191,208,238,155,8 | 298 | 149 | 0.0000012178578865 | 0.00072627201218412 | 0.0,0.007,0.017,0. | 0.039,0.026,0.0,0. | -0,014 |
| 152102 | ENSG00000196405 | EVL | chr14 | + | 100072777 | 100073101 | 100065444 | 100065511 | 100084686 | 100085098 | 152102 | 2,0,41,63,0,0,0,7,16,9,0,0 | 443,221,476,369,262,244,2 | 4,0,0,3,12,3,0,0,0,3,0,0,0,0 | 406,236,261,138,276,211,1 | 298 | 149 | 0.0000012181525728 | 0.00072627201218412 | 0.002,0.0,0.041,0. | 0.005,0.0,0.0,0.0,0.01 | 0,011 |
| 167659 | ENSG00000101109 | STK4 | chr20 | + | 44973166 | 44973433 | 44972137 | 44972158 | 44978442 | 44978571 | 167659 | 18,7,16,15,9,15,15,16,1,1,3 | 166,90,299,182,96,74,81,1 | 23,53,16,16,22,10,18,23,28 | 209,231,139,109,89,76,66, | 298 | 149 | 0.0000012337149262 | 0.000733518500788 | 0.051,0.037,0.026, | 0.052,0.103,0.054, | -0,03 |
| 176937 | ENSG00000196204 | RNF216P1 | chr7 | + | 4983699 | 4983773 | 4976364 | 4976436 | 4984909 | 4985075 | 176937 | 22,10,12,10,3,2,8,9,9,3,2,5 | 0,0,2,0,0,0,1,4,0,0,0,2 | 12,7,6,11,9,3,8,10,3,8,10,1 | 0,0,0,8,5,0,1,4,0,0,6,2,1,0,0 | 223 | 149 | 0.0000012712059166 | 0.00075372705411664 | 1.0,1.0,0.8,1.0,1.0, | 1.0,1.0,1.0,0.479,0 | 0,082 |
| 150553 | ENSG00000166783 | MARF1 | chr16 | - | 15634756 | 15634819 | 15633616 | 15633843 | 15635655 | 15636342 | 150553 | 52,28,44,46,21,32,27,35,21 | 4,1,4,5,1,7,4,11,0,0,3,0 | 42,33,22,36,17,23,20,21,26 | 1,0,0,0,0,2,5,3,2,0,1,1,0,0,3 | 212 | 149 | 0.0000012799695605 | 0.00075571122622089 | 0.901,0.952,0.885, | 0.967,1.0,1.0,1.0,1 | -0,047 |
| 187254 | ENSG00000111674 | ENO2 | chr12 | + | 6922037 | 6922164 | 6921580 | 6921782 | 6922343 | 6922402 | 187254 | 4,4,6,16,12,7,3,1,1,2,3,3 | 1,0,1,1,0,3,1,0,0,0,0,3 | 11,16,6,10,3,2,1,10,4,10,7, | 6,1,0,5,0,0,2,9,1,1,2,2,4,6,4 | 276 | 149 | 0.0000012815746689 | 0.00075571122622089 | 0.683,1.0,0.764,0. | 0.497,0.896,1.0,0. | 0,163 |
| 115153 | ENSG00000165525 | NEMF | chr14 | - | 49833422 | 49833496 | 49832206 | 49832277 | 49834362 | 49834449 | 115153 | 67,41,127,66,23,46,31,49,3 | 2,1,3,1,1,1,0,0,5,2,0,1 | 64,76,31,19,49,20,19,44,39 | 5,5,0,4,3,1,2,2,3,9,0,2,7,3,4 | 223 | 149 | 0.0000013509574746 | 0.00079443508753519 | 0.957,0.965,0.966, | 0.895,0.91,1.0,0.7 | 0,064 |
| 150604 | ENSG00000171953 | ATPAF2 | chr17 | - | 18018315 | 18018686 | 18016206 | 18016260 | 18021122 | 18021238 | 150604 | 8,8,12,13,5,13,17,8,7,6,8,8 | 1,0,1,3,0,0,0,0,0,0,0,0 | 9,17,1,8,5,3,8,6,2,5,1,9,14, | 3,4,5,0,2,0,0,1,5,1,0,1,0,0,1 | 298 | 149 | 0.0000013546267829 | 0.00079443508753519 | 0.8,1.0,0.857,0.68 | 0.6,0.68,0.091,1.0, | 0,208 |
| 92639 | ENSG00000124508 | BTN2A2 | chr6 | + | 26385014 | 26385362 | 26383107 | 26383181 | 26388012 | 26388289 | 92639 | 10,9,24,14,11,6,10,11,6,1,5 | 0,3,3,0,0,0,0,0,0,0,0,0 | 16,11,0,2,7,7,5,8,5,10,12,2 | 9,5,2,0,3,0,1,0,2,0,0,5,1,0,2 | 298 | 149 | 0.0000013626376281 | 0.00079634195152373 | 1.0,0.6,0.8,1.0,1.0, | 0.471,0.524,0.0,1. | 0,209 |
| 48244 | ENSG00000067066 | SP100 | chr2 | + | 230464066 | 230464150 | 230462434 | 230462518 | 230466300 | 230466354 | 48244 | 294,235,386,255,159,166,1 | 57,30,72,69,17,16,39,29,3, | 248,205,218,120,174,86,66 | 50,76,27,57,116,31,16,23,5 | 233 | 149 | 0.0000013652781435 | 0.00079634195152373 | 0.767,0.834,0.774, | 0.76,0.633,0.838,0 | 0,129 |
| 255788 | ENSG00000173442 | EHBP1L1 | chr11 | + | 65581538 | 65581641 | 65581210 | 65581373 | 65584240 | 65584398 | 255788 | 40,24,20,12,24,34,40,42,49 | 9,1,1,0,1,1,12,0,2,0,0,0 | 17,13,13,17,15,17,20,36,12 | 0,0,0,0,2,0,0,0,0,2,0,0,0,1,4 | 252 | 149 | 0.0000013897499940 | 0.00080676429706349 | 0.724,0.934,0.922, | 1.0,1.0,1.0,1.0,0.8 | -0,016 |
| 99963 | ENSG00000058091 | CDK14 | chr7 | + | 90709815 | 90709957 | 90604217 | 90604249 | 90726566 | 90726644 | 99963 | 317,149,329,174,171,242,2 | 0,1,18,16,2,0,1,5,0,0,2,4 | 353,254,130,231,193,141,1 | 0,2,0,0,0,0,0,0,0,9,0,2,5,3 | 291 | 149 | 0.0000013906433283 | 0.00080676429706349 | 1.0,0.987,0.903,0. | 1.0,0.985,1.0,1.0,1 | -0,022 |
| 150606 | ENSG00000171953 | ATPAF2 | chr17 | - | 18018379 | 18018686 | 18016206 | 18016260 | 18021122 | 18021238 | 150606 | 8,8,12,13,4,13,17,8,7,6,8,8 | 1,0,1,3,0,0,0,0,0,0,0,0 | 9,17,1,8,5,3,10,6,2,5,1,9,14 | 3,4,5,0,2,0,0,1,5,1,0,1,0,0,1 | 298 | 149 | 0.0000014135547422 | 0.00081785161486925 | 0.8,1.0,0.857,0.68 | 0.6,0.68,0.091,1.0, | 0,208 |
| 12237 | ENSG00000185504 | FAAP100 | chr17 | - | 81546400 | 81546540 | 81545745 | 81545882 | 81546908 | 81547203 | 12237 | 11,3,1,1,3,3,1,2,2,0,0,0 | 13,2,7,8,12,11,5,6,5,6,7,7 | 1,2,0,0,0,0,3,2,0,0,0,0,3,2,2 | 11,4,5,2,7,2,2,11,7,15,12,1 | 289 | 149 | 0.0000014223031188 | 0.00081813271745556 | 0.304,0.436,0.069, | 0.045,0.205,0.0,0. | 0,053 |
| 165949 | ENSG00000172869 | DMXL1 | chr5 | + | 119225675 | 119225756 | 119224708 | 119224769 | 119233339 | 119233467 | 165949 | 5,1,10,0,0,0,1,0,1,1,0,2 | 85,72,92,76,35,39,33,46,49 | 4,10,11,5,4,1,1,8,0,1,4,8,8, | 79,57,70,66,38,33,23,28,41 | 230 | 149 | 0.0000014227294280 | 0.00081813271745556 | 0.037,0.009,0.066, | 0.032,0.102,0.092, | -0,033 |
| 245948 | ENSG00000172977 | KAT5 | chr11 | + | 65717598 | 65717810 | 65716888 | 65716982 | 65718589 | 65718749 | 245948 | 6,5,5,9,0,4,1,4,0,0,1,4 | 25,32,67,63,35,36,21,22,20 | 1,2,1,1,4,0,1,0,0,0,0,1,3,2,0 | 31,36,26,17,50,21,22,28,22 | 298 | 149 | 0.0000014254441462 | 0.00081813271745556 | 0.107,0.072,0.036, | 0.016,0.027,0.019, | 0,033 |
| 233931 | ENSG00000130985 | UBA1 | chrX | + | 47200417 | 47200544 | 47199479 | 47199614 | 47200893 | 47201000 | 233931 | 0,0,13,22,0,0,0,1,3,5,0,0 | 20,24,74,41,27,20,25,21,10 | 0,0,0,0,4,9,0,0,0,3,0,0,0,0,0 | 52,46,15,17,29,28,22,30,29 | 276 | 149 | 0.0000014533888153 | 0.00083195300028147 | 0.0,0.0,0.087,0.22 | 0.0,0.0,0.0,0.0,0.0 | 0,052 |
| 76415 | ENSG00000105983 | LMBR1 | chr7 | - | 156834477 | 156834575 | 156833752 | 156833792 | 156836812 | 156836885 | 76415 | 0,2,6,4,1,12,4,9,4,2,4,3 | 16,10,22,7,19,26,11,35,5,8 | 4,2,0,3,2,0,0,1,1,1,0,2,1,3,1 | 16,20,6,13,7,7,7,5,14,7,18, | 247 | 149 | 0.0000014670114791 | 0.00083718250651575 | 0.0,0.108,0.141,0. | 0.131,0.057,0.0,0. | 0,096 |
| 198120 | ENSG00000061273 | HDAC7 | chr12 | - | 47786608 | 47786703 | 47785751 | 47785885 | 47787711 | 47787809 | 198120 | 9,16,41,40,32,37,22,26,5,1 | 0,0,0,2,1,6,1,3,0,0,0,0 | 31,24,9,9,13,7,10,29,13,21 | 5,4,1,2,2,2,0,5,3,1,1,0,0,2,0 | 244 | 149 | 0.0000014703039407 | 0.00083718250651575 | 1.0,1.0,1.0,0.924,0 | 0.791,0.786,0.846, | 0,087 |
| 187252 | ENSG00000111674 | ENO2 | chr12 | + | 6921954 | 6922164 | 6921580 | 6921782 | 6922343 | 6922402 | 187252 | 4,4,6,16,11,7,3,1,2,2,3,3 | 1,0,1,1,0,3,1,0,0,0,0,3 | 11,16,6,10,3,2,1,10,4,10,7, | 6,1,0,5,0,0,2,9,1,1,2,2,4,6,4 | 298 | 149 | 0.0000014780393871 | 0.00083936647842986 | 0.667,1.0,0.75,0.8 | 0.478,0.889,1.0,0. | 0,169 |
| 13140 | ENSG00000006194 | ZNF263 | chr16 | + | 3288453 | 3288570 | 3286022 | 3286149 | 3289392 | 3290553 | 13140 | 27,6,17,2,14,9,8,5,5,3,2,3 | 1,0,0,2,0,2,0,0,2,0,0,0 | 3,11,3,0,4,4,11,14,3,16,7,1 | 2,6,0,5,1,0,0,1,0,0,3,6,2,0,0 | 266 | 149 | 0.0000014856473340 | 0.000841466740402 | 0.938,1.0,1.0,0.35 | 0.457,0.507,1.0,0. | 0,149 |
| 76310 | ENSG00000076242 | MLH1 | chr3 | + | 37020309 | 37020463 | 37012010 | 37012099 | 37028783 | 37028932 | 76310 | 20,7,25,11,3,11,7,7,7,5,3,1 | 2,4,3,2,0,1,3,5,0,0,4,0 | 14,30,13,9,13,5,20,10,10,1 | 0,1,1,0,0,1,1,1,3,0,5,0,0,0,1 | 298 | 149 | 0.0000014944284636 | 0.00084296580283619 | 0.833,0.467,0.806, | 1.0,0.938,0.867,1. | -0,137 |
| 226626 | ENSG00000058091 | CDK14 | chr7 | + | 90803123 | 90803244 | 90790572 | 90790652 | 90863174 | 90863269 | 226626 | 80,29,34,22,12,19,2,4,27,9 | 618,304,796,604,325,507,3 | 11,9,17,6,5,4,0,6,19,18,36, | 842,571,280,233,370,314,2 | 270 | 149 | 0.0000014961271224 | 0.00084296580283619 | 0.067,0.05,0.023,0 | 0.007,0.009,0.032, | 0,009 |
| 50381 | ENSG00000266094 | RASSF5 | chr1 | + | 206584386 | 206584684 | 206583119 | 206583379 | 206586825 | 206586842 | 50381 | 136,171,100,114,88,75,44, | 0,0,19,20,0,0,0,0,0,0,0,0 | 127,222,45,15,45,33,29,67 | 0,3,0,0,0,0,0,0,1,0,1,2,0,4,0 | 298 | 149 | 0.0000015328444720 | 0.00086139856020192 | 1.0,1.0,0.725,0.74, | 1.0,0.974,1.0,1.0,1 | -0,033 |
| 82359 | ENSG00000145241 | CENPC | chr4 | - | 67474887 | 67474978 | 67468761 | 67472675 | 67489966 | 67490121 | 82359 | 13,19,61,32,19,28,23,33,23 | 0,2,2,1,0,1,0,4,1,0,2,4 | 29,17,18,3,17,15,12,13,6,9 | 7,1,0,12,3,4,0,4,4,3,7,2,3,1 | 240 | 149 | 0.0000015458701589 | 0.0008649342535144 | 1.0,0.855,0.95,0.9 | 0.72,0.913,1.0,0.1 | 0,171 |
| 63003 | ENSG00000225470 | JPX | chrX | + | 73999354 | 74000496 | 73944583 | 73944662 | 74004269 | 74004543 | 63003 | 8,3,15,12,8,5,5,5,1,5,1,3 | 2,0,3,0,0,0,0,1,0,0,1,0 | 6,5,8,6,3,4,3,4,5,2,2,4,3,3,2 | 1,1,5,1,4,3,0,2,2,0,1,2,0,0,0 | 298 | 149 | 0.0000015471734443 | 0.0008649342535144 | 0.667,1.0,0.714,1. | 0.75,0.714,0.444,0 | 0,189 |
| 104949 | ENSG00000137522 | RNF121 | chr11 | + | 71987003 | 71987111 | 71982760 | 71982915 | 71990596 | 71990663 | 104949 | 19,11,36,24,15,10,16,14,10 | 2,0,3,0,0,2,2,3,0,0,0,0 | 18,12,7,8,14,12,5,14,18,20 | 1,0,4,0,4,4,1,2,0,5,0,2,1,1,5 | 257 | 149 | 0.0000015642632289 | 0.00087222263997444 | 0.846,1.0,0.874,1. | 0.913,1.0,0.504,1. | 0,093 |
| 134792 | ENSG00000242259 | C22orf39 | chr22 | - | 19377868 | 19378183 | 19375630 | 19375792 | 19383619 | 19383705 | 134792 | 14,18,22,23,19,8,17,24,14, | 3,0,2,7,0,3,0,1,1,2,1,3 | 14,33,13,21,16,12,16,30,1, | 4,3,0,0,0,0,0,1,2,0,1,2,0,1,0 | 298 | 149 | 0.0000016220370483 | 0.00090209988617898 | 0.7,1.0,0.846,0.62 | 0.636,0.846,1.0,1. | -0,071 |
| 82828 | ENSG00000160695 | VPS11 | chr11 | + | 119071595 | 119071843 | 119070233 | 119070397 | 119073799 | 119073951 | 82828 | 17,14,36,34,22,9,12,14,3,6 | 2,0,0,0,1,0,2,0,2,0,0,0 | 14,25,9,6,15,11,0,3,4,19,11 | 9,4,2,1,0,0,1,1,0,0,4,0,3,2,1 | 298 | 149 | 0.0000016291727742 | 0.0009037332097228 | 0.81,1.0,1.0,1.0,0. | 0.437,0.758,0.692, | 0,159 |
| 48376 | ENSG00000154781 | CCDC174 | chr3 | + | 14668048 | 14668181 | 14666804 | 14666947 | 14669933 | 14670086 | 48376 | 24,35,68,39,23,30,3,36,12, | 1,1,3,2,1,2,4,0,2,0,0,0 | 10,44,16,8,34,10,17,18,18, | 9,8,4,2,3,2,0,2,3,6,6,3,1,2,3 | 282 | 149 | 0.0000016516628941 | 0.0009122423426195 | 0.927,0.949,0.923, | 0.37,0.744,0.679,0 | 0,135 |
| 90708 | ENSG00000082258 | CCNT2 | chr2 | + | 134942611 | 134942674 | 134939001 | 134939062 | 134946100 | 134946146 | 90708 | 50,36,47,57,25,26,19,37,27 | 1,2,1,2,2,2,3,1,0,0,0,1 | 41,72,46,27,35,16,18,34,40 | 2,8,13,13,2,0,1,2,3,3,7,0,2, | 212 | 149 | 0.0000016529891773 | 0.0009122423426195 | 0.972,0.927,0.971, | 0.935,0.863,0.713, | 0,057 |
| 129749 | ENSG00000136021 | SCYL2 | chr12 | + | 100327184 | 100327234 | 100326621 | 100326754 | 100329200 | 100329319 | 129749 | 4,0,3,1,0,2,0,2,0,0,1,0 | 14,10,25,32,9,9,7,8,11,4,1, | 4,2,4,7,4,0,2,9,3,4,1,2,1,1,2 | 19,27,14,6,9,1,7,16,5,16,3, | 199 | 149 | 0.0000017007664984 | 0.00093620888547131 | 0.176,0.0,0.082,0. | 0.136,0.053,0.176, | -0,088 |
| 248031 | ENSG00000149262 | INTS4 | chr11 | - | 77878818 | 77879127 | 77874493 | 77877030 | 77883831 | 77883952 | 248031 | 11,9,32,28,16,16,12,5,9,2,5 | 0,5,1,1,0,0,2,2,0,0,0,0 | 17,11,14,13,13,5,9,15,12,1 | 0,1,1,0,1,3,2,3,2,0,1,2,1,4,6 | 298 | 149 | 0.0000017565428406 | 0.00096295195790704 | 1.0,0.474,0.941,0. | 1.0,0.846,0.875,1. | 0,078 |
| 113482 | ENSG00000156735 | BAG4 | chr8 | + | 38192687 | 38192795 | 38176854 | 38177139 | 38207511 | 38207741 | 113482 | 19,14,30,13,18,16,20,12,19 | 1,1,3,0,0,0,1,0,0,0,3,3 | 23,29,19,26,21,8,2,7,12,16 | 0,6,1,2,1,0,0,1,2,3,2,13,3,2 | 257 | 149 | 0.0000017582974546 | 0.00096295195790704 | 0.917,0.89,0.853,1 | 1.0,0.737,0.917,0. | 0,069 |
| 249673 | ENSG00000100225 | FBXO7 | chr22 | + | 32491085 | 32491452 | 32487287 | 32487828 | 32493104 | 32493281 | 249673 | 40,26,69,66,42,31,19,41,41 | 3,1,8,3,0,1,0,1,0,0,0,0 | 52,47,49,14,11,19,10,31,19 | 7,1,9,2,8,4,0,0,0,2,4,0,7,7,4 | 298 | 149 | 0.0000017853972536 | 0.00097319711317057 | 0.87,0.929,0.812,0 | 0.788,0.959,0.731, | 0,133 |
| 31605 | ENSG00000112992 | NNT | chr5 | + | 43647923 | 43648440 | 43645356 | 43645510 | 43649146 | 43649308 | 31605 | 3,2,6,8,2,7,26,26,4,3,4,5 | 32,16,75,43,31,38,10,21,34 | 4,1,6,2,1,0,3,1,2,2,3,2,3,1,3 | 17,28,20,10,21,29,13,14,18 | 298 | 149 | 0.0000017860478263 | 0.00097319711317057 | 0.045,0.059,0.038, | 0.105,0.018,0.13,0 | 0,082 |

| | | | | | | | | | | | | | | | | | | | | | | |
|---|---|---|---|---|---|---|---|---|---|---|---|---|---|---|---|---|---|---|---|---|---|---|
| 202702 | ENSG00000231424 | | chr1 | - | 171247580 | 171247678 | 171168103 | 171168188 | 171251522 | 171251857 | 202702 | 45,30,103,14,17,54,22,113 | 16,8,108,25,19,47,2,0,19,7 | 95,41,29,37,60,40,0,0,13,2 | 0,0,0,0,0,0,29,50,8,17,0,0, | 247 | 149 | 0.0000018210137827 | 0.00098974398353376 | 0.629,0.693,0.365, | 1.0,1.0,1.0,1.0,1.0, | -0,305 |
| 169593 | ENSG00000047849 | MAP4 | chr3 | - | 47940498 | 47940534 | 47928272 | 47928350 | 47977864 | 47977933 | 169593 | 2,0,9,7,3,9,6,9,0,2,2,4 | 39,27,62,73,48,77,37,45,29 | 2,6,2,1,3,1,0,2,1,0,1,1,1,3, | 29,58,38,50,30,36,32,55,29 | 185 | 149 | 0.0000018265402220 | 0.00099024704920994 | 0.04,0.0,0.105,0.07 | 0.053,0.077,0.041, | 0,037 |
| 142126 | ENSG00000215067 | ALOX12-AS1 | chr17 | - | 7011902 | 7012009 | 7009795 | 7009889 | 7012195 | 7012474 | 142126 | 0,0,4,3,0,1,3,8,4,1,0,2 | 12,2,14,15,9,11,6,8,5,4,2,1 | 2,1,1,0,0,0,0,5,2,0,0,0,0,0, | 7,7,12,8,9,9,8,7,12,15,6,3,1 | 256 | 149 | 0.0000018311845392 | 0.00099027055166045 | 0.0,0.0,0.143,0.104 | 0.143,0.077,0.046, | 0,078 |
| 132955 | ENSG00000290571 | GOLGA2P5 | chr12 | - | 100168923 | 100169199 | 100167549 | 100167639 | 100170564 | 100170667 | 132955 | 42,30,14,15,14,24,17,18,18 | 5,2,3,2,4,3,0,0,5,3,6,3 | 3,14,8,19,16,20,7,20,10,25 | 0,0,1,0,3,1,2,0,0,0,1,0,2,5,2 | 298 | 149 | 0.0000018453765267 | 0.00099544419854177 | 0.808,0.882,0.7,0.7 | 1.0,1.0,0.8,1.0,0.7 | -0,119 |
| 34656 | ENSG00000047188 | YTHDC2 | chr5 | + | 113524980 | 113525177 | 113515271 | 113515362 | 113526585 | 113526703 | 34656 | 48,47,115,38,32,50,43,60,2 | 0,7,5,8,0,6,0,0,8,2,1,4 | 53,44,51,80,46,14,17,45,20 | 1,0,3,0,4,2,0,2,0,2,2,2,0,2,0 | 298 | 149 | 0.0000018749491006 | 0.00100861592499913 | 1.0,0.77,0.92,0.704 | 0.964,1.0,0.895,1.0 | -0,108 |
| 251342 | ENSG00000012779 | ALOX5 | chr10 | + | 45428368 | 45428764 | 45424959 | 45425132 | 45440429 | 45440633 | 251342 | 570,380,366,315,434,279,3 | 1,1,6,0,0,1,0,3,3,1,2,3 | 371,347,250,452,239,179,8 | 12,8,2,7,6,0,5,6,0,1,1,7,0,2 | 298 | 149 | 0.0000018835922633 | 0.00100861592499913 | 0.997,0.995,0.968, | 0.939,0.956,0.984, | 0,02 |
| 163834 | ENSG00000214655 | ZSWIM8 | chr10 | + | 73789366 | 73789615 | 73789095 | 73789190 | 73789716 | 73789824 | 163834 | 15,7,25,19,10,22,12,9,11,7 | 1,0,1,0,0,0,1,1,0,1,1,3 | 16,18,16,9,15,8,6,11,12,18 | 0,4,2,1,2,3,1,2,0,0,2,4,2,1,2 | 298 | 149 | 0.0000018838531710 | 0.00100861592499913 | 0.882,1.0,0.926,1.0 | 1.0,0.692,0.8,0.818 | 0,071 |
| 251340 | ENSG00000012779 | ALOX5 | chr10 | + | 45428365 | 45428764 | 45424959 | 45425132 | 45440429 | 45440633 | 251340 | 570,380,366,314,434,279,3 | 1,1,6,0,0,1,0,3,3,1,2,3 | 371,348,250,451,239,179,8 | 12,8,2,7,6,0,5,6,0,1,1,7,0,2 | 298 | 149 | 0.0000018933431502 | 0.00101048285329268 | 0.997,0.995,0.968, | 0.939,0.956,0.984, | 0,02 |
| 218270 | ENSG00000134440 | NARS1 | chr18 | - | 57603733 | 57603850 | 57602811 | 57602943 | 57605856 | 57605970 | 218270 | 11,5,7,1,5,17,0,0,1,2,4,2 | 19,7,93,72,39,34,40,51,32, | 6,12,3,4,19,5,7,13,13,3,4,2 | 46,70,41,9,29,8,28,51,16,3 | 266 | 149 | 0.0000018967298982 | 0.00101048285329268 | 0.245,0.286,0.04,0 | 0.068,0.088,0.039, | -0,067 |
| 109142 | ENSG00000139218 | SCAF11 | chr12 | - | 45922935 | 45923154 | 45922355 | 45922582 | 45924727 | 45924788 | 109142 | 130,160,174,124,83,41,69, | 0,0,3,6,1,2,3,4,1,0,0,3 | 126,108,95,77,47,50,58,66 | 9,7,5,6,11,4,0,2,2,5,0,5,5,4 | 298 | 149 | 0.0000019214342033 | 0.00102111655558138 | 1.0,1.0,0.967,0.91 | 0.875,0.885,0.905, | 0,055 |
| 198117 | ENSG00000061273 | HDAC7 | chr12 | - | 47786354 | 47786703 | 47785751 | 47785885 | 47787711 | 47787809 | 198117 | 12,20,44,51,39,48,29,29,7, | 0,0,0,2,1,6,1,3,0,0,0,0 | 36,43,13,11,17,9,11,41,18, | 5,4,1,2,2,2,0,5,3,1,1,0,0,2, | 298 | 149 | 0.0000019749908798 | 0.00104699325630256 | 1.0,1.0,1.0,0.927,0 | 0.783,0.843,0.867, | 0,078 |
| 72060 | ENSG00000151353 | TMEM18 | chr2 | - | 675757 | 676238 | 675509 | 675630 | 677288 | 677406 | 72060 | 5,9,1,1,0,1,7,2,5,8,2,4 | 15,15,25,19,2,6,22,12,33,1 | 6,2,0,0,2,1,1,0,0,0,0,3,2,2, | 14,14,6,9,13,0,3,27,2,9,16, | 298 | 149 | 0.0000019809554878 | 0.00104700022318628 | 0.143,0.231,0.02,0 | 0.176,0.067,0.0,0.0 | 0,001 |
| 247600 | ENSG00000149483 | TMEM138 | chr11 | + | 61367427 | 61367656 | 61366044 | 61366216 | 61367922 | 61369161 | 247600 | 0,1,10,1,1,0,3,1,2,1,0,2 | 9,11,38,23,5,4,13,9,13,6,4, | 9,3,1,2,3,1,3,3,0,2,11,11,6, | 18,23,10,9,7,4,6,10,1,2,6,8 | 298 | 149 | 0.0000019847331056 | 0.00104700022318628 | 0.0,0.043,0.116,0.0 | 0.2,0.061,0.048,0.1 | -0,126 |
| 27917 | ENSG00000280987 | MATR3 | chr5 | + | 139279934 | 139280058 | 139279050 | 139279129 | 139307238 | 139307556 | 27917 | 12,10,6,8,6,6,5,14,6,3,7,13 | 0,0,0,2,0,0,1,0,0,0,0,2 | 10,15,5,2,5,5,2,4,4,2,7,23,9 | 3,1,1,0,3,4,0,0,0,2,1,15,0,0 | 273 | 149 | 0.0000019901590633 | 0.00104729566102586 | 1.0,1.0,1.0,0.686,1 | 0.645,0.891,0.732, | 0,177 |
| 99806 | ENSG00000161036 | LRWD1 | chr7 | + | 102465531 | 102465627 | 102465000 | 102465160 | 102465816 | 102466051 | 99806 | 0,0,1,1,0,0,7,24,1,2,0,0 | 6,4,17,10,13,8,11,51,9,13,5 | 1,0,0,0,0,0,0,1,0,0,0,0,0,0,2 | 5,15,1,6,4,5,5,21,3,6,8,8,4, | 245 | 149 | 0.0000020118744041 | 0.00105568461417607 | 0.0,0.0,0.035,0.057 | 0.108,0.0,0.0,0.0,0 | -0,019 |
| 99364 | ENSG00000176142 | TMEM39A | chr3 | - | 119458017 | 119458184 | 119452446 | 119452530 | 119461961 | 119462148 | 99364 | 8,16,36,7,11,14,22,10,5,2,6 | 0,1,3,0,0,2,2,0,0,1,0,0 | 22,33,20,6,19,6,11,11,8,14 | 0,8,1,10,1,0,1,2,0,0,4,9,0,0 | 298 | 149 | 0.0000020159102379 | 0.00105568461417607 | 1.0,0.889,0.857,1.0 | 1.0,0.673,0.909,0.2 | 0,079 |
| 162697 | ENSG00000163297 | ANTXR2 | chr4 | - | 79984118 | 79984271 | 79983877 | 79983970 | 79984818 | 79984863 | 162697 | 12,0,1,2,0,2,1,2,0,2,3,4 | 9,15,12,20,15,19,19,15,18, | 1,10,5,18,12,7,4,2,3,8,16,9 | 21,19,4,14,4,7,17,7,3,6,16, | 298 | 149 | 0.0000021092414285 | 0.00110187898519851 | 0.4,0.0,0.04,0.048, | 0.023,0.208,0.385, | -0,132 |
| 14825 | ENSG00000166747 | AP1G1 | chr16 | - | 71748250 | 71748378 | 71746587 | 71746692 | 71749893 | 71749983 | 14825 | 88,53,124,74,46,88,48,75,4 | 1,0,1,0,0,0,2,0,1,2,1,1 | 90,149,56,82,92,62,55,56,5 | 2,6,0,6,4,1,6,0,1,3,5,0,4,0,3 | 277 | 149 | 0.0000021616085682 | 0.00112650163136323 | 0.979,1.0,0.985,1.0 | 0.96,0.93,1.0,0.88, | 0,029 |
| 163962 | ENSG00000102445 | RUBCNL | chr13 | - | 46345446 | 46345533 | 46344740 | 46344831 | 46349285 | 46349347 | 163962 | 116,107,574,361,155,201,3 | 0,0,0,0,18,28,21,27,0,0,11, | 340,492,242,171,265,119,1 | 0,0,0,0,0,0,21,33,4,24,0,0,0 | 236 | 149 | 0.0000021905561264 | 0.00113606299016563 | 1.0,1.0,1.0,1.0,0.84 | 1.0,1.0,1.0,1.0,1.0, | -0,034 |
| 109533 | ENSG00000147416 | ATP6V1B2 | chr8 | + | 20214817 | 20214968 | 20212781 | 20212905 | 20216412 | 20216495 | 109533 | 18,3,41,42,30,28,17,16,25, | 1,1,1,4,1,1,0,0,0,0,1,0 | 22,24,16,0,17,19,13,14,4,1 | 2,6,3,4,1,0,0,3,1,0,3,0,5,2,0 | 298 | 149 | 0.0000021957095822 | 0.00113606299016563 | 0.9,0.6,0.953,0.84, | 0.846,0.667,0.727, | 0,081 |
| 50322 | ENSG00000132781 | MUTYH | chr1 | - | 45333096 | 45333170 | 45332917 | 45332959 | 45333284 | 45333324 | 50322 | 16,6,9,9,12,25,9,14,19,11,5 | 0,3,1,0,0,1,0,0,2,0,0,0 | 16,5,15,11,3,5,10,16,6,6,3, | 3,2,6,3,2,1,3,0,1,1,1,3,0,1,1 | 223 | 149 | 0.0000022029677505 | 0.00113606299016563 | 1.0,0.572,0.857,1.0 | 0.781,0.626,0.626, | 0,148 |
| 31787 | ENSG00000224032 | EPB41L4A-AS1 | chr5 | + | 112161702 | 112161775 | 112160525 | 112161295 | 112162150 | 112162312 | 31787 | 133,110,210,169,34,55,60, | 3,3,1,3,0,0,2,2,0,0,2,7 | 113,116,98,68,91,64,40,81 | 8,16,6,4,0,2,2,6,0,8,6,9,0,0 | 222 | 149 | 0.0000022031470225 | 0.00113606299016563 | 0.967,0.961,0.993, | 0.905,0.83,0.916,0 | 0,03 |
| 104338 | ENSG00000032389 | EIPR1 | chr2 | - | 3257298 | 3257455 | 3214148 | 3214248 | 3338016 | 3338149 | 104338 | 33,13,68,58,35,35,22,36,35 | 5,0,2,0,3,4,1,0,4,4,2,4 | 66,15,14,4,19,12,21,37,17, | 2,0,1,1,0,0,0,5,0,0,0,5,1,0,0 | 298 | 149 | 0.0000022063472728 | 0.00113606299016563 | 0.767,1.0,0.944,1.0 | 0.943,1.0,0.875,0.6 | -0,085 |
| 202700 | ENSG00000231424 | | chr1 | - | 171247580 | 171247675 | 171168103 | 171168188 | 171251522 | 171251857 | 202700 | 45,29,99,14,16,54,22,112,0 | 16,8,108,25,19,47,2,0,19,7 | 95,41,29,36,59,39,0,0,13,2 | 0,0,0,0,0,0,29,50,8,17,0,0,0 | 244 | 149 | 0.0000022323024995 | 0.00114668424647777 | 0.632,0.689,0.359, | 1.0,1.0,1.0,1.0,1.0, | -0,303 |
| 30583 | ENSG00000181929 | PRKAG1 | chr12 | - | 49005742 | 49005852 | 49005461 | 49005543 | 49013061 | 49013110 | 30583 | 15,10,60,49,22,23,14,11,16 | 1,0,4,1,0,2,2,2,0,0,0,0 | 32,26,22,2,20,13,6,17,9,24 | 2,2,1,1,3,1,5,0,0,0,2,4,3,1,2 | 259 | 149 | 0.0000022439817534 | 0.00114993913516235 | 0.896,1.0,0.896,0.9 | 0.902,0.882,0.927, | 0,112 |
| 202713 | ENSG00000231424 | | chr1 | - | 171247580 | 171247780 | 171168103 | 171168188 | 171251522 | 171251857 | 202713 | 50,32,147,28,24,80,38,147 | 16,8,108,25,19,47,2,0,19,7 | 142,75,34,46,80,60,0,0,20, | 0,0,0,0,0,0,29,50,8,17,0,0,0 | 298 | 149 | 0.0000022858716280 | 0.00116862336433595 | 0.61,0.667,0.405,0 | 1.0,1.0,1.0,1.0,1.0, | -0,283 |
| 204376 | ENSG00000173209 | AHSA2P | chr2 | + | 61185991 | 61186081 | 61185525 | 61185618 | 61186445 | 61186497 | 204376 | 6,2,8,1,0,7,10,15,2,1,1,5 | 65,32,254,161,103,78,48,6 | 11,22,17,0,5,2,3,23,9,10,17 | 61,38,49,35,79,61,76,111,5 | 239 | 149 | 0.0000022913573082 | 0.0011686519545205 | 0.054,0.038,0.019, | 0.101,0.265,0.178, | -0,039 |
| 141571 | ENSG00000214176 | PLEKHM1P1 | chr17 | - | 64814862 | 64815305 | 64804829 | 64805100 | 64821766 | 64822393 | 141571 | 12,8,2,2,4,2,22,20,2,1,5,13 | 27,18,57,42,45,52,12,38,18 | 0,2,2,1,3,1,2,2,1,3,15,15,1, | 31,39,44,59,60,34,24,26,27 | 298 | 149 | 0.0000023044118063 | 0.00117253157801175 | 0.182,0.182,0.017, | 0.0,0.025,0.022,0.0 | 0,085 |
| 64816 | ENSG00000092841 | MYL6 | chr12 | + | 56160625 | 56160670 | 56160242 | 56160320 | 56161386 | 56161550 | 64816 | 11,9,152,112,25,43,17,30,1 | 263,189,911,837,411,375,3 | 38,24,16,5,26,16,19,26,1,1 | 586,439,311,167,324,245,2 | 194 | 149 | 0.0000023190077425 | 0.00117566805365389 | 0.031,0.035,0.114, | 0.047,0.04,0.038,0 | 0,011 |
| 142161 | ENSG00000091592 | NLRP1 | chr17 | - | 5533667 | 5533988 | 5533303 | 5533384 | 5536850 | 5536940 | 142161 | 133,44,82,69,55,56,57,59,6 | 4,2,7,1,0,3,2,1,2,0,2,7 | 35,58,35,54,42,12,24,51,50 | 2,2,4,15,1,0,10,7,1,3,9,7,3, | 298 | 149 | 0.0000023215007262 | 0.00117566805365389 | 0.943,0.917,0.854, | 0.897,0.935,0.814, | 0,138 |
| 159390 | ENSG00000127603 | MACF1 | chr1 | + | 39315521 | 39315691 | 39310830 | 39311000 | 39316390 | 39316529 | 159390 | 30,16,59,44,29,24,26,11,9, | 3,0,4,4,1,0,1,1,0,0,0,0 | 48,47,11,16,13,20,11,29,29 | 5,7,0,3,3,0,0,4,2,6,1,8,8,2,1 | 298 | 149 | 0.0000023319721308 | 0.0011781988114855 | 0.833,1.0,0.881,0.8 | 0.828,0.77,1.0,0.72 | 0,102 |
| 232091 | ENSG00000104870 | FCGRT | chr19 | + | 49525456 | 49525720 | 49524506 | 49524776 | 49526009 | 49526428 | 232091 | 126,42,162,64,34,16,38,38 | 5,0,1,1,2,0,2,0,0,0,1,7 | 88,34,24,13,22,23,27,66,22 | 10,11,1,0,1,0,2,9,1,0,2,1,0, | 298 | 149 | 0.0000023877652645 | 0.00120356230830386 | 0.926,1.0,0.988,0.9 | 0.815,0.607,0.923, | 0,056 |
| 36763 | ENSG00000101109 | STK4 | chr20 | + | 44973326 | 44973433 | 44972077 | 44972158 | 44978442 | 44978571 | 36763 | 16,4,12,14,6,12,12,14,1,1,2 | 166,90,299,182,96,74,81,1 | 16,46,16,14,21,10,16,13,20 | 209,231,139,109,89,76,66, | 256 | 149 | 0.0000024000759216 | 0.00120499404031017 | 0.053,0.025,0.023, | 0.043,0.104,0.063, | -0,032 |
| 202711 | ENSG00000231424 | | chr1 | - | 171247580 | 171247738 | 171168103 | 171168188 | 171251522 | 171251857 | 202711 | 52,32,147,28,24,80,38,147 | 16,8,108,25,19,47,2,0,19,7 | 142,75,34,46,80,60,0,0,20, | 0,0,0,0,0,0,29,50,8,17,0,0,0 | 298 | 149 | 0.0000024018029154 | 0.00120499404031017 | 0.619,0.667,0.405, | 1.0,1.0,1.0,1.0,1.0, | -0,283 |
| 134791 | ENSG00000242259 | C22orf39 | chr22 | - | 19377868 | 19377988 | 19375630 | 19375792 | 19383619 | 19383705 | 134791 | 14,16,17,18,16,6,13,21,13, | 3,0,2,7,0,3,0,1,1,2,1,3 | 13,25,9,20,10,11,11,26,1,1 | 4,3,0,0,0,0,0,1,2,0,1,2,0,1,0 | 269 | 149 | 0.0000024267226853 | 0.00121405700116629 | 0.721,1.0,0.825,0.5 | 0.643,0.822,1.0,1.0 | -0,062 |
| 168597 | ENSG00000005059 | MCUB | chr4 | + | 109684442 | 109684549 | 109682581 | 109682742 | 109685250 | 109685367 | 168597 | 146,67,148,99,53,61,40,50 | 6,1,2,1,0,4,5,0,5,0,0,1 | 71,74,55,45,41,31,27,66,29 | 10,3,8,3,1,3,2,8,3,5,3,10,4, | 256 | 149 | 0.0000024311487076 | 0.00121405700116629 | 0.934,0.975,0.977, | 0.805,0.935,0.8,0.8 | 0,068 |
| 198051 | ENSG00000112232 | KHDRBS2 | chr6 | - | 61977151 | 61977350 | 61901243 | 61901371 | 61978065 | 61978212 | 198051 | 1,0,27,24,0,0,3,5,11,7,0,0 | 57,30,75,37,29,39,20,30,78 | 0,0,5,0,12,9,0,1,0,0,0,1,0,0 | 55,32,33,33,11,8,14,28,13, | 298 | 149 | 0.0000024703679718 | 0.00122852976729097 | 0.009,0.0,0.0.153,0. | 0.0,0.0,0.07,0.0,0.3 | 0,006 |
| 157106 | ENSG00000133103 | COG6 | chr13 | + | 39683798 | 39683883 | 39682170 | 39682264 | 39687502 | 39687631 | 157106 | 5,0,5,3,1,1,0,5,3,0,0,1 | 3,5,17,15,11,5,6,6,5,1,6,8 | 0,0,0,0,1,0,3,0,0,0,0,0,0,1,2 | 9,5,6,5,8,6,5,13,4,4,10,5,8, | 234 | 149 | 0.0000024715463350 | 0.00122852976729097 | 0.515,0.0,0.158,0.1 | 0.0,0.0,0.0,0.0,0.07 | 0,084 |
| 74824 | ENSG00000120451 | SNX19 | chr11 | - | 130880621 | 130880806 | 130879623 | 130879711 | 130903254 | 130903384 | 74824 | 29,27,80,62,34,52,50,75,32 | 0,1,3,0,1,2,0,0,0,0,0,0 | 78,68,24,31,36,26,57,37,25 | 1,0,0,0,8,4,0,0,2,0,0,1,12,4 | 298 | 149 | 0.0000024915997819 | 0.00123564403840259 | 1.0,0.931,0.93,1.0, | 0.975,1.0,1.0,1.0,0 | 0,091 |
| 126549 | ENSG00000136754 | ABI1 | chr10 | - | 26768851 | 26768992 | 26765217 | 26765318 | 26770244 | 26770345 | 126549 | 380,265,719,604,363,326,3 | 0,4,6,5,3,4,5,7,0,4,0,0 | 480,468,299,175,227,174,1 | 2,3,3,19,27,7,7,3,6,0,2,3,1, | 290 | 149 | 0.0000025161475332 | 0.00124494930970683 | 1.0,0.971,0.984,0.9 | 0.992,0.988,0.981, | 0,029 |
| 251344 | ENSG00000012779 | ALOX5 | chr10 | + | 45428558 | 45428764 | 45424959 | 45425132 | 45440429 | 45440633 | 251344 | 582,380,369,318,441,279,3 | 1,1,6,0,0,1,0,3,3,1,2,3 | 379,362,251,455,245,180,8 | 12,8,2,7,6,0,5,6,0,1,1,7,0,2 | 298 | 149 | 0.0000025257387207 | 0.00124595365499746 | 0.997,0.995,0.969, | 0.94,0.958,0.984,0 | 0,019 |
| 200142 | ENSG00000131051 | RBM39 | chr20 | - | 35738968 | 35739017 | 35732059 | 35732135 | 35740823 | 35740837 | 200142 | 57,66,44,24,8,10,9,11,24,1 | 3,4,1,0,0,0,0,0,0,0,0,4 | 9,31,16,33,25,17,5,13,7,22 | 3,1,4,5,1,0,0,0,1,0,9,4,0,1,2 | 198 | 149 | 0.0000025333151059 | 0.00124595365499746 | 0.935,0.925,0.971, | 0.693,0.959,0.751, | 0,05 |
| 185079 | ENSG00000078304 | PPP2R5C | chr14 | + | 101827473 | 101827819 | 101809904 | 101810036 | 101836168 | 101836701 | 185079 | 3,3,2,7,2,0,2,0,0,0,0,2 | 17,16,35,14,10,13,14,10,6, | 7,7,0,8,2,0,3,6,3,1,3,10,4,3 | 12,17,7,6,7,8,8,5,7,18,8,20 | 298 | 149 | 0.0000025355441404 | 0.00124595365499746 | 0.081,0.086,0.028, | 0.226,0.171,0.0,0.4 | -0,1 |
| 209920 | ENSG00000100441 | KHNYN | chr14 | + | 24434099 | 24434200 | 24432935 | 24433032 | 24436071 | 24436179 | 209920 | 2,0,2,1,1,0,2,1,0,0,2,5 | 14,14,27,15,19,16,9,15,7,5 | 1,1,0,2,8,8,1,1,3,10,8,7,8,3 | 24,36,12,22,8,7,10,25,4,16 | 250 | 149 | 0.0000025531748864 | 0.00125175941679899 | 0.078,0.0,0.042,0.0 | 0.024,0.016,0.0,0.0 | -0,096 |
| 12222 | ENSG00000141030 | COPS3 | chr17 | - | 17247479 | 17247560 | 17246615 | 17247151 | 17248925 | 17249039 | 12222 | 128,50,90,67,44,50,40,32,5 | 10,1,12,6,2,0,1,0,2,4,0,3 | 77,99,50,43,83,29,28,35,24 | 5,2,2,4,1,0,0,0,0,0,0,6,0,0,2 | 230 | 149 | 0.0000025662640524 | 0.00125531722336209 | 0.892,0.97,0.829,0 | 0.909,0.97,0.942,0 | -0,033 |
| 251338 | ENSG00000012779 | ALOX5 | chr10 | + | 45428320 | 45428764 | 45424959 | 45425132 | 45440429 | 45440633 | 251338 | 583,389,378,315,445,287,3 | 1,1,6,0,0,1,0,3,3,1,2,3 | 377,353,253,464,245,179,9 | 12,8,2,7,6,0,5,6,0,1,1,7,0,2 | 298 | 149 | 0.0000025794917082 | 0.00125892648492982 | 0.997,0.995,0.969, | 0.94,0.957,0.984,0 | 0,019 |
| 12429 | ENSG00000141568 | FOXK2 | chr17 | + | 82585251 | 82585271 | 82584012 | 82584188 | 82585903 | 82586043 | 12429 | 3,0,5,3,2,0,3,0,9,3,2,3 | 12,11,8,3,16,9,5,10,55,16,6 | 0,1,0,0,0,0,1,0,3,1,2,0,1,3,4 | 6,12,12,6,2,8,4,17,4,11,28, | 169 | 149 | 0.000002634271007 | 0.0012827529029584 | 0.181,0.0,0.355,0.4 | 0.0,0.068,0.0,0.0,0 | 0,115 |
| 118806 | ENSG00000174106 | LEMD3 | chr12 | + | 65245860 | 65245939 | 65245299 | 65245774 | 65246161 | 65246539 | 118806 | 52,42,162,107,65,61,60,15 | 0,0,0,0,8,5,9,15,0,0,0,0 | 89,113,53,35,33,55,45,69,2 | 14,12,5,7,6,16,4,8,9,10,0,1 | 228 | 149 | 0.0000026429895815 | 0.00128409320681745 | 1.0,1.0,1.0,1.0,0.84 | 0.806,0.86,0.874,0 | 0,105 |
| 214848 | ENSG00000055044 | NOP58 | chr2 | + | 202295673 | 202295837 | 202292776 | 202292903 | 202297357 | 202297513 | 214848 | 67,120,145,57,45,53,31,58 | 0,0,16,18,0,0,0,0,0,2,0,0 | 52,137,90,34,82,28,54,53,1 | 0,0,0,1,2,6,0,0,0,0,0,0,0,0,0 | 298 | 149 | 0.0000026750591851 | 0.00129674698982245 | 1.0,1.0,0.819,0.61 | 1.0,1.0,1.0,0.944,0 | -0,043 |
| 108933 | ENSG00000147905 | ZCCHC7 | chr9 | + | 37147361 | 37147445 | 37126311 | 37126942 | 37186711 | 37187054 | 108933 | 20,10,48,12,17,21,16,39,5, | 21,14,20,6,7,6,14,8,6,4,3,1 | 65,47,58,15,14,15,24,20,12 | 6,4,9,17,4,3,6,0,5,4,7,5,1,7 | 233 | 149 | 0.0000027094011230 | 0.00131044295080127 | 0.379,0.314,0.605, | 0.874,0.883,0.805, | -0,246 |
| 189422 | ENSG00000177200 | CHD9 | chr16 | + | 53239741 | 53239881 | 53238342 | 53238586 | 53242839 | 53243016 | 189422 | 5,1,5,0,6,5,1,6,0,1,2,0 | 27,24,88,36,33,43,16,14,10 | 7,18,5,8,11,3,6,2,8,4,1,6,14 | 33,51,22,27,29,23,17,24,42 | 289 | 149 | 0.0000027713444510 | 0.00133739739359145 | 0.087,0.021,0.028, | 0.099,0.154,0.105, | -0,05 |
| 98926 | ENSG00000168090 | COPS6 | chr7 | + | 100090398 | 100090487 | 100089614 | 100089746 | 100090591 | 100090654 | 98926 | 19,11,29,24,18,8,11,15,5,7 | 0,1,0,3,2,0,2,0,0,0,2,0 | 19,18,10,4,5,6,10,12,6,8,9, | 0,1,3,3,0,3,5,0,0,2,5,9,1,1,0 | 238 | 149 | 0.0000027954391961 | 0.00134402265089363 | 1.0,0.873,1.0,0.83 | 1.0,0.918,0.676,0.4 | 0,071 |
| 247599 | ENSG00000149483 | TMEM138 | chr11 | + | 61367427 | 61367620 | 61366044 | 61366216 | 61367922 | 61369161 | 247599 | 0,1,10,1,1,0,3,1,2,1,0,2 | 9,11,38,23,5,4,13,9,13,6,4, | 9,3,1,2,3,1,3,3,0,4,12,9,6,8 | 18,23,10,9,7,4,6,10,1,2,6,8 | 298 | 149 | 0.0000027975623753 | 0.00134402265089363 | 0.0,0.043,0.116,0.0 | 0.2,0.061,0.048,0.1 | -0,133 |
| 141569 | ENSG00000214176 | PLEKHM1P1 | chr17 | - | 64814862 | 64815048 | 64804829 | 64805100 | 64821766 | 64822393 | 141569 | 12,8,2,3,1,3,22,20,5,1,5,13 | 27,18,57,42,45,52,12,38,18 | 0,2,1,0,3,1,2,2,1,3,15,19,1, | 31,39,44,59,60,34,24,26,27 | 298 | 149 | 0.0000028265148315 | 0.00135490782561888 | 0.182,0.182,0.017, | 0.0,0.025,0.011,0.0 | 0,084 |
| 4881 | ENSG00000198521 | ZNF43 | chr19 | - | 21819094 | 21819221 | 21817887 | 21817986 | 21845391 | 21845506 | 4881 | 24,30,37,42,7,18,12,18,16, | 3,0,1,0,0,0,0,0,1,0,0,0 | 32,23,14,10,9,3,11,22,9,16 | 1,1,0,0,0,1,4,0,1,0,6,3,4,0,2 | 276 | 149 | 0.0000029200585286 | 0.00139652633733418 | 0.812,1.0,0.952,1.0 | 0.945,0.925,1.0,1.0 | 0,142 |
| 104207 | ENSG00000145191 | EIF2B5 | chr3 | + | 184135663 | 184135916 | 184135493 | 184135580 | 184136611 | 184136736 | 104207 | 0,0,8,3,0,0,2,3,0,0,2,0 | 32,22,62,46,31,35,26,22,26 | 6,9,3,3,8,0,5,2,1,2,3,2,3,4,2 | 41,42,24,16,12,15,14,30,13 | 298 | 149 | 0.0000029263134870 | 0.00139652633733418 | 0.0,0.0,0.061,0.03 | 0.068,0.097,0.059, | -0,045 |
| 134876 | ENSG00000100138 | SNU13 | chr22 | - | 41681539 | 41681559 | 41680243 | 41680364 | 41682355 | 41682456 | 134876 | 1,3,5,0,1,2,1,0,0,0,0,2 | 14,14,46,41,19,18,14,15,18 | 1,2,4,7,6,0,1,2,3,3,2,9,0,4,1 | 16,22,17,16,15,10,18,28,10 | 169 | 149 | 0.0000029806747197 | 0.00141669235064002 | 0.059,0.159,0.087, | 0.052,0.074,0.172, | -0,072 |
| 90095 | ENSG00000119912 | IDE | chr10 | - | 92559033 | 92559215 | 92537365 | 92537550 | 92573921 | 92574056 | 90095 | 0,2,5,3,2,5,0,2,3,4,0,0 | 9,9,22,7,8,5,10,3,14,5,2,5 | 1,0,0,0,2,4,1,1,1,2,0,0,1,0,1 | 14,11,2,7,8,2,6,4,6,4,20,13 | 298 | 149 | 0.0000029817342057 | 0.00141669235064002 | 0.0,0.1,0.102,0.17 | 0.034,0.0,0.0,0.0,0 | 0,039 |
| 104801 | ENSG00000164062 | APEH | chr3 | + | 49676377 | 49676515 | 49675890 | 49675966 | 49676608 | 49676700 | 104801 | 19,10,47,58,37,16,25,20,33 | 3,0,2,2,3,4,1,2,3,6,2,4 | 34,17,21,6,13,21,13,22,16, | 3,1,0,0,3,0,1,0,0,0,2,2,3,1,2 | 287 | 149 | 0.0000030370326785 | 0.0014397876221174 | 0.767,1.0,0.924,0.9 | 0.855,0.898,1.0,1.0 | -0,091 |
| 24161 | ENSG00000253710 | ALG11 | chr13 | + | 52024005 | 52024937 | 52014564 | 52019143 | 52028318 | 52028749 | 24161 | 20,22,50,15,12,11,18,16,12 | 3,3,1,0,0,2,1,2,0,0,0,0 | 15,28,14,16,21,12,16,9,4,1 | 3,0,0,0,0,3,1,2,0,3,1,10,0,6 | 298 | 149 | 0.0000030557633276 | 0.00144548350936851 | 0.769,0.786,0.962, | 0.714,1.0,1.0,1.0,1 | 0,071 |
| 16296 | ENSG00000197818 | SLC9A8 | chr20 | + | 49849578 | 49849680 | 49845035 | 49845119 | 49855437 | 49855581 | 16296 | 29,28,70,26,13,19,15,10,3, | 0,1,3,0,2,4,1,0,0,0,0,2 | 23,20,18,10,29,10,10,20,13 | 2,2,1,7,2,1,2,1,0,1,8,0,0,3,1 | 251 | 149 | 0.0000030703187813 | 0.00144918373163253 | 1.0,0.943,0.933,1.0 | 0.872,0.856,0.914, | 0,094 |
| 101052 | ENSG00000075142 | SRI | chr7 | - | 88214794 | 88214936 | 88210881 | 88210925 | 88217121 | 88217191 | 101052 | 0,1,5,6,0,3,1,4,0,0,0,2 | 35,15,109,84,32,35,37,47,3 | 4,0,10,6,13,0,3,4,0,0,2,14,2 | 35,34,33,10,29,33,13,42,14 | 291 | 149 | 0.0000031150045861 | 0.00146705810083983 | 0.0,0.033,0.023,0.0 | 0.055,0.0,0.134,0.2 | -0,042 |
| 166822 | ENSG00000084676 | NCOA1 | chr2 | + | 24697758 | 24697798 | 24693251 | 24693347 | 24705085 | 24705233 | 166822 | 9,6,25,23,14,14,8,9,4,2,3,8 | 2,2,0,0,0,2,0,0,1,0,0,0 | 17,15,9,9,7,11,7,16,6,6,6,7 | 1,0,4,4,2,0,1,1,0,0,5,0,1,2,5 | 189 | 149 | 0.0000031472369147 | 0.00147596682390875 | 0.78,0.703,1.0,1.0, | 0.931,1.0,0.639,0.6 | 0,104 |
| 16427 | ENSG00000118900 | UBN1 | chr16 | + | 4877384 | 4877474 | 4876870 | 4877111 | 4880082 | 4880445 | 16427 | 5,6,16,14,5,7,3,11,4,5,7,7 | 0,0,3,0,4,0,2,0,0,2,1,0 | 6,2,2,2,3,5,9,7,1,5,5,9,6,2,3 | 4,6,2,1,1,0,0,1,1,2,2,4,10,0 | 239 | 149 | 0.0000031528853383 | 0.00147596682390875 | 1.0,1.0,0.769,1.0,0 | 0.483,0.172,0.384, | 0,224 |
| 160170 | ENSG00000104320 | NBN | chr8 | - | 89981623 | 89982002 | 89981374 | 89981523 | 89982721 | 89982855 | 160170 | 17,0,1,2,1,1,0,0,0,2,0,0 | 20,8,16,16,12,5,4,5,4,0,1,7 | 6,5,9,5,2,0,2,2,0,3,4,10,2,2 | 16,12,7,3,6,8,6,5,8,18,9,11 | 298 | 149 | 0.0000031544932607 | 0.00147596682390875 | 0.298,0.0,0.03,0.05 | 0.158,0.172,0.391, | -0,055 |
| 128423 | ENSG00000047056 | WDR37 | chr10 | + | 1105125 | 1105267 | 1096169 | 1096246 | 1124217 | 1124352 | 128423 | 18,8,13,8,7,9,8,6,4,4,4,5 | 5,3,0,1,1,3,2,2,2,0,0,0 | 14,4,9,1,2,4,13,15,3,6,3,6,1 | 0,1,2,1,0,0,0,1,0,2,0,0,0,0,1 | 291 | 149 | 0.0000031620594780 | 0.00147620442212419 | 0.648,0.577,1.0,0.8 | 1.0,0.672,0.697,0.3 | -0,112 |
| 149232 | ENSG00000128881 | TTBK2 | chr15 | - | 42871444 | 42871635 | 42840359 | 42840433 | 42872610 | 42872758 | 149232 | 0,0,0,2,2,0,1,0,0,0,0,0 | 12,8,19,6,16,2,4,13,8,6,8,9 | 4,0,1,0,3,1,2,2,0,4,8,2,1,3,0 | 18,8,7,14,5,2,4,18,5,17,15, | 298 | 149 | 0.0000031687184607 | 0.00147620442212419 | 0.0,0.0,0.0,0.143,0 | 0.1,0.0,0.067,0.0,0 | -0,067 |
| 253570 | ENSG00000135968 | GCC2 | chr2 | + | 108474788 | 108474860 | 108472826 | 108472899 | 108475534 | 108475635 | 253570 | 8,2,7,1,0,1,0,2,0,0,2,2 | 65,38,58,37,13,19,6,14,24, | 5,4,12,8,1,2,2,5,2,6,4,3,2,1 | 17,16,33,22,13,8,8,20,13,1 | 221 | 149 | 0.0000032032240585 | 0.0014890564089588 | 0.077,0.034,0.075, | 0.165,0.144,0.197, | -0,077 |
| 235208 | ENSG00000139218 | SCAF11 | chr12 | - | 45922935 | 45923108 | 45922355 | 45922582 | 45924727 | 45925074 | 235208 | 109,124,114,89,67,32,56,6 | 0,0,3,6,1,2,3,4,1,0,0,3 | 93,78,81,55,33,38,37,51,15 | 9,7,5,6,11,4,0,2,2,5,0,5,5,4 | 298 | 149 | 0.0000032594614211 | 0.00151000671093845 | 1.0,1.0,0.95,0.881, | 0.838,0.848,0.89,0 | 0,073 |
| 61722 | ENSG00000144559 | TAMM41 | chr3 | - | 11844890 | 11845096 | 11844028 | 11844211 | 11846501 | 11846885 | 61722 | 8,7,12,3,4,9,11,9,2,0,3,2 | 5,5,12,13,4,5,2,0,4,2,3,2 | 1,9,0,0,0,0,0,6,1,1,0,2,0,0,6 | 4,9,5,4,11,10,1,3,7,3,7,8,5, | 298 | 149 | 0.0000032636773116 | 0.00151000671093845 | 0.444,0.412,0.333, | 0.111,0.333,0.0,0.0 | 0,265 |
| 235207 | ENSG00000139218 | SCAF11 | chr12 | - | 45922935 | 45923090 | 45922355 | 45922582 | 45924727 | 45925074 | 235207 | 109,124,113,89,67,32,56,6 | 0,0,3,6,1,2,3,4,1,0,0,3 | 93,79,81,55,33,38,37,51,15 | 9,7,5,6,11,4,0,2,2,5,0,5,5,4 | 298 | 149 | 0.0000032693391161 | 0.00151000671093845 | 1.0,1.0,0.95,0.881, | 0.838,0.849,0.89,0 | 0,074 |
| 82856 | ENSG00000104218 | CSPP1 | chr8 | + | 67093542 | 67093641 | 67086006 | 67086110 | 67095292 | 67095732 | 82856 | 27,11,35,9,10,8,3,9,9,16,2, | 2,0,4,1,0,0,1,0,2,2,0,0 | 12,10,11,8,10,10,10,7,6,14 | 3,7,3,5,3,1,0,0,3,1,1,2,0,0,3 | 248 | 149 | 0.0000033119788585 | 0.00152642509998675 | 0.89,1.0,0.84,0.84 | 0.706,0.462,0.688, | 0,116 |
| 23845 | ENSG00000173575 | CHD2 | chr15 | + | 92929029 | 92929091 | 92927243 | 92927330 | 92939577 | 92939718 | 23845 | 108,77,123,60,51,65,51,84 | 4,0,2,4,0,0,0,4,0,0,1,2 | 60,96,51,72,69,35,30,46,44 | 0,1,1,10,8,5,1,0,10,2,3,2,0, | 211 | 149 | 0.0000034064842220 | 0.00156662608032422 | 0.95,1.0,0.977,0.91 | 1.0,0.985,0.973,0.8 | 0,039 |
| 107051 | ENSG00000046651 | OFD1 | chrX | + | 13758336 | 13758448 | 13757659 | 13757790 | 13760114 | 13760720 | 107051 | 160,67,396,176,69,108,71, | 3,0,2,0,1,1,0,4,1,1,0,0 | 150,133,96,84,87,90,30,84 | 4,2,1,14,2,4,0,1,6,2,3,4,3,1 | 261 | 149 | 0.0000034249855107 | 0.00157061584621654 | 0.968,1.0,0.991,1.0 | 0.955,0.974,0.982, | 0,034 |
| 99646 | ENSG00000106443 | PHF14 | chr7 | + | 11061790 | 11061841 | 11051611 | 11051780 | 11061963 | 11062085 | 99646 | 34,15,50,28,33,39,21,23,24 | 0,0,2,0,0,0,1,4,0,0,0,0 | 37,24,21,12,33,18,16,18,15 | 3,3,2,4,1,2,0,0,0,0,4,5,2,3,0 | 200 | 149 | 0.0000034297542999 | 0.00157061584621654 | 1.0,1.0,0.949,1.0,1 | 0.902,0.856,0.887, | 0,072 |
| 45184 | ENSG00000138246 | DNAJC13 | chr3 | + | 132446474 | 132446550 | 132434537 | 132434618 | 132447320 | 132447470 | 45184 | 90,22,57,49,32,30,29,33,62 | 7,4,3,2,1,11,1,3,2,2,0,4 | 42,36,42,13,16,28,9,32,28, | 0,4,0,0,1,1,0,6,3,0,1,2,0,0,0 | 225 | 149 | 0.0000034868213169 | 0.00159335889354077 | 0.895,0.785,0.926, | 1.0,0.856,1.0,1.0,0 | -0,052 |
| 225919 | ENSG00000176142 | TMEM39A | chr3 | - | 119458017 | 119458253 | 119452446 | 119452530 | 119461961 | 119462148 | 225919 | 7,13,30,7,11,13,21,8,4,2,8, | 0,1,3,0,0,2,2,0,0,1,0,0 | 21,29,19,6,20,6,9,10,8,14,1 | 0,8,1,10,1,0,1,2,0,0,4,9,0,0 | 298 | 149 | 0.0000035706303687 | 0.00162819988323691 | 1.0,0.867,0.833,1.0 | 1.0,0.644,0.905,0.2 | 0,082 |
| 159392 | ENSG00000127603 | MACF1 | chr1 | + | 39315590 | 39315691 | 39310830 | 39311000 | 39316390 | 39316529 | 159392 | 20,14,46,32,15,13,18,6,8,8 | 3,0,4,4,1,0,1,1,0,0,0,0 | 34,37,7,8,10,17,3,21,18,30 | 5,7,0,3,3,0,0,4,2,6,1,8,8,2,1 | 250 | 149 | 0.0000036520223472 | 0.00166179370361565 | 0.799,1.0,0.873,0.8 | 0.802,0.759,1.0,0.6 | 0,124 |
| 46487 | ENSG00000114446 | IFT57 | chr3 | - | 108165430 | 108165493 | 108163662 | 108163729 | 108167792 | 108167864 | 46487 | 83,42,126,131,64,63,57,73 | 0,0,2,1,4,0,0,1,0,0,0,0 | 99,83,46,35,38,20,55,63,25 | 2,2,0,0,2,0,0,3,1,3,6,14,5,2 | 212 | 149 | 0.0000037757753504 | 0.00171448081109981 | 1.0,1.0,0.978,0.98 | 0.972,0.967,1.0,1.0 | 0,035 |
| 125802 | ENSG00000120438 | TCP1 | chr6 | - | 159780934 | 159781110 | 159780442 | 159780566 | 159783940 | 159784067 | 125802 | 281,283,422,270,149,147,1 | 39,7,3,13,7,12,8,6,4,2,2,2 | 211,358,167,98,227,107,11 | 22,20,21,34,13,2,4,15,10,1 | 298 | 149 | 0.0000038236349172 | 0.00172975074699041 | 0.783,0.953,0.986, | 0.827,0.899,0.799, | 0,066 |

| | | | | | | | | | | | | | | | | | | | | | | |
|---|---|---|---|---|---|---|---|---|---|---|---|---|---|---|---|---|---|---|---|---|---|---|
| 100838 | ENSG00000127914 | AKAP9 | chr7 | + | 92043281 | 92043377 | 92042667 | 92042771 | 92045007 | 92045213 | 100838 | 11,4,6,2,0,6,3,6,0,0,2 | 17,5,71,41,16,40,21,43,19, | 1,1,1,7,3,0,0,0,2,3,4,0,0,1 | 33,53,24,21,18,24,14,21,21 | 245 | 149 | 0.0000038315566051 | 0.00172975074699041 | 0.282,0.327,0.049, | 0.018,0.011,0.025, | 0,05 |
| 4885 | ENSG00000198521 | ZNF43 | chr19 | - | 21843144 | 21843326 | 21817887 | 21817986 | 21845391 | 21845506 | 4885 | 0,2,1,3,6,1,3,12,1,3,1,3 | 3,0,1,0,0,0,0,0,1,0,0,0 | 4,5,10,4,3,2,2,5,0,2,7,9,0,3 | 1,1,0,0,0,1,4,0,1,0,6,3,4,0,2 | 298 | 149 | 0.0000038335142535 | 0.00172975074699041 | 0.0,1.0,0.333,1.0,1 | 0.667,0.714,1.0,1. | 0,223 |
| 155541 | ENSG00000173575 | CHD2 | chr15 | + | 92991475 | 92991833 | 92985497 | 92985673 | 92992858 | 92992998 | 155541 | 24,15,26,14,11,9,10,12,11, | 0,2,0,1,0,0,0,0,0,0,0,3 | 13,42,8,13,25,6,9,12,20,14 | 4,2,2,12,3,0,0,0,1,2,1,0,1,1 | 298 | 149 | 0.0000038555188771 | 0.00173604013274063 | 1.0,0.789,1.0,0.87 | 0.619,0.913,0.667, | 0,103 |
| 41766 | ENSG00000211897 | IGHG3 | chr14 | - | 105770292 | 105770337 | 105770104 | 105770149 | 105770480 | 105770525 | 41766 | 32,11,49,51,47,28,81,84,1 | 2,0,12,12,0,0,18,8,8,5,0,1 | 33,31,32,17,3,2,29,95,3,6, | 8,3,5,1,0,0,0,4,0,0,0,0,0,0,1 | 194 | 149 | 0.0000038685407094 | 0.0017382669842035 | 0.925,1.0,0.758,0. | 0.76,0.888,0.831,0 | -0,045 |
| 224983 | ENSG00000134905 | CARS2 | chr13 | - | 110685519 | 110685598 | 110683050 | 110683134 | 110687720 | 110687826 | 224983 | 16,9,3,1,0,0,2,1,4,3,0,0 | 51,15,95,79,63,46,19,36,2 | 0,2,0,1,3,0,0,3,0,2,1,0,0,1, | 65,34,13,16,41,21,19,29,2 | 228 | 149 | 0.0000039482612268 | 0.00177039210857531 | 0.17,0.282,0.02,0. | 0.0,0.037,0.0,0.03 | 0,055 |
| 240760 | ENSG00000021776 | AQR | chr15 | - | 34963659 | 34963982 | 34960773 | 34960814 | 34964233 | 34964453 | 240760 | 0,2,0,0,0,0,0,2,0,0,0,2 | 24,11,62,31,10,9,13,16,5,3 | 0,8,5,0,4,2,0,3,0,6,5,2,2,0, | 28,28,17,13,14,7,8,16,17,2 | 298 | 149 | 0.0000040046338724 | 0.00179193628480358 | 0.0,0.083,0.0,0.0,0 | 0.0,0.125,0.128,0. | -0,044 |
| 231263 | ENSG00000164062 | APEH | chr3 | + | 49676112 | 49676515 | 49675890 | 49675966 | 49676608 | 49676700 | 231263 | 14,10,45,52,38,17,20,19,2 | 3,0,2,2,3,4,1,2,3,6,2,4 | 36,15,20,7,9,20,14,21,13,1 | 3,1,0,0,3,0,1,0,0,0,2,2,3,1,2 | 298 | 149 | 0.0000041030888317 | 0.0018321823907378 | 0.7,1.0,0.918,0.92 | 0.857,0.882,1.0,1. | -0,097 |
| 244607 | ENSG00000168071 | CCDC88B | chr11 | + | 64351159 | 64351301 | 64349550 | 64349668 | 64351475 | 64351616 | 244607 | 17,13,10,1,15,5,20,6,9,3,5, | 2,0,0,0,3,2,2,1,0,0,0,1 | 1,4,9,3,1,1,3,2,1,3,8,16,2,8 | 0,1,1,6,0,0,0,0,3,1,1,3,2,1,2 | 291 | 149 | 0.0000042183592444 | 0.00187830472767495 | 0.813,1.0,1.0,1.0,0 | 1.0,0.672,0.822,0. | 0,18 |
| 201063 | ENSG00000160908 | ZNF394 | chr7 | - | 99498564 | 99498842 | 99494578 | 99494631 | 99499637 | 99499704 | 201063 | 42,23,53,18,20,29,39,14,2 | 5,3,1,0,5,2,1,4,0,8,2,0 | 35,39,14,51,18,20,23,29,3 | 0,1,2,0,0,1,3,2,0,0,6,2,2,1, | 298 | 149 | 0.0000042318423514 | 0.00187830472767495 | 0.808,0.793,0.964, | 1.0,0.951,0.778,1. | -0,11 |
| 168215 | ENSG00000138639 | ARHGAP24 | chr4 | + | 85604140 | 85604303 | 85570592 | 85570721 | 85721884 | 85721972 | 168215 | 2,3,70,41,8,37,37,108,47,2 | 198,149,408,178,125,188,1 | 83,23,38,49,49,70,38,53,5 | 335,282,171,211,196,103,1 | 298 | 149 | 0.0000042428120511 | 0.00187830472767495 | 0.005,0.01,0.079,0 | 0.11,0.039,0.1,0.1 | -0,049 |
| 68684 | ENSG00000160714 | UBE2Q1 | chr1 | - | 154551396 | 154551492 | 154550937 | 154551004 | 154551770 | 154551819 | 68684 | 166,103,110,130,92,97,60, | 2,2,0,0,8,5,2,0,4,2,0,4 | 87,108,80,39,65,38,50,100 | 6,4,2,8,4,3,3,3,6,1,8,10,3,2 | 245 | 149 | 0.0000042469384551 | 0.00187830472767495 | 0.981,0.969,1.0,1. | 0.898,0.943,0.961, | 0,063 |
| 182362 | ENSG00000182400 | TRAPPC6B | chr14 | - | 39163245 | 39163361 | 39159482 | 39159550 | 39170014 | 39170532 | 182362 | 2,3,11,4,0,2,3,3,1,0,2,0 | 20,9,40,18,25,21,10,22,10, | 4,0,1,0,0,0,1,2,0,2,1,1,0,2, | 10,16,22,8,13,8,15,13,18,2 | 265 | 149 | 0.0000042500123234 | 0.00187830472767495 | 0.053,0.158,0.134, | 0.184,0.0,0.025,0. | 0,048 |
| 155540 | ENSG00000173575 | CHD2 | chr15 | + | 92991475 | 92991780 | 92985497 | 92985673 | 92992858 | 92992998 | 155540 | 11,10,20,7,9,2,6,2,8,2,3,5 | 0,2,0,1,0,0,0,0,0,0,0,3 | 12,24,1,4,13,1,2,11,7,10,4, | 4,2,2,12,3,0,0,0,1,2,1,0,1,1 | 298 | 149 | 0.0000043397441079 | 0.00190547143610708 | 1.0,0.714,1.0,0.77 | 0.6,0.857,0.2,0.14 | 0,157 |
| 155539 | ENSG00000173575 | CHD2 | chr15 | + | 92991475 | 92991763 | 92985497 | 92985673 | 92992858 | 92992998 | 155539 | 12,10,20,7,9,2,5,2,8,2,3,5 | 0,2,0,1,0,0,0,0,0,0,0,3 | 12,24,1,4,13,1,2,11,7,10,4, | 4,2,2,12,3,0,0,0,1,2,1,0,1,1 | 298 | 149 | 0.0000043397529708 | 0.00190547143610708 | 1.0,0.714,1.0,0.77 | 0.6,0.857,0.2,0.14 | 0,157 |
| 241037 | ENSG00000056586 | RC3H2 | chr9 | - | 122892914 | 122893026 | 122890311 | 122890545 | 122897278 | 122897534 | 241037 | 18,20,95,75,13,22,18,40,5, | 1,6,11,1,1,0,3,0,0,0,1,2 | 40,70,34,44,29,29,12,24,3 | 1,0,0,4,2,0,0,1,0,2,0,0,0,0,1 | 261 | 149 | 0.0000043466451741 | 0.00190547143610708 | 0.911,0.656,0.831, | 0.958,1.0,1.0,0.86 | -0,07 |
| 198162 | ENSG00000172500 | FIBP | chr11 | - | 65885949 | 65886422 | 65885529 | 65885663 | 65887599 | 65887726 | 198162 | 33,32,69,62,33,30,17,28,3 | 1,0,4,2,6,0,1,1,4,1,3,9 | 36,23,21,4,25,27,8,33,16,3 | 1,0,0,0,0,3,2,4,0,1,3,0,2,0,1 | 298 | 149 | 0.0000043468946161 | 0.00190547143610708 | 0.943,1.0,0.896,0. | 0.947,1.0,1.0,1.0,1 | -0,064 |
| 193034 | ENSG00000058668 | ATP2B4 | chr1 | + | 203707084 | 203707223 | 203703651 | 203703813 | 203707861 | 203708104 | 193034 | 104,43,139,63,41,26,37,32 | 6,1,3,2,3,1,2,6,0,0,5,2 | 111,64,48,16,12,30,43,38, | 5,0,2,0,3,0,0,0,1,0,2,0,0,1, | 288 | 149 | 0.0000044185279555 | 0.00193293534633033 | 0.9,0.957,0.96,0.9 | 0.92,1.0,0.925,1.0, | -0,054 |
| 3634 | ENSG00000174886 | NDUFA11 | chr19 | - | 5901334 | 5901454 | 5896904 | 5896997 | 5903611 | 5903754 | 3634 | 6,4,7,7,3,3,6,7,1,1,1,7 | 141,90,152,131,57,52,78,7 | 11,5,5,1,8,1,4,14,1,6,19,16 | 77,71,76,39,54,33,61,87,2 | 269 | 149 | 0.0000044279892784 | 0.00193314515292977 | 0.023,0.024,0.025, | 0.073,0.038,0.035, | -0,022 |
| 189186 | ENSG00000118308 | IRAG2 | chr12 | + | 25061588 | 25061653 | 25052944 | 25052956 | 25062819 | 25062900 | 189186 | 129,49,231,179,91,65,108, | 241,99,472,328,182,141,1 | 75,106,88,30,39,29,33,70, | 221,280,165,85,110,96,48, | 214 | 149 | 0.0000044691112003 | 0.00194714832542151 | 0.272,0.256,0.254, | 0.191,0.209,0.271, | 0,06 |
| 139829 | ENSG00000124198 | ARFGEF2 | chr20 | + | 48957130 | 48957300 | 48953555 | 48953790 | 48963829 | 48963898 | 139829 | 2,0,4,2,0,0,0,0,0,0,0,0 | 43,30,75,55,33,32,21,40,2 | 8,5,5,3,1,0,3,8,0,0,10,2,0,1 | 52,57,29,44,41,19,24,31,3 | 298 | 149 | 0.0000045280760249 | 0.00196664866651626 | 0.023,0.0,0.026,0. | 0.071,0.042,0.079, | -0,036 |
| 136108 | ENSG00000105486 | LIG1 | chr19 | - | 48151467 | 48151608 | 48151231 | 48151339 | 48153871 | 48154207 | 136108 | 2,6,3,1,0,1,4,0,6,1,1,4 | 26,18,20,5,27,19,12,15,26, | 0,0,0,0,0,0,0,1,1,0,1,0,0,3, | 14,11,19,13,6,14,4,16,10,1 | 290 | 149 | 0.0000045338832336 | 0.00196664866651626 | 0.038,0.146,0.072, | 0.0,0.0,0.0,0.0,0.0, | 0,042 |
| 241388 | ENSG00000257524 | ST6GALNAC6 | chr9 | - | 127904906 | 127905302 | 127896241 | 127896332 | 127905934 | 127905975 | 241388 | 8,5,13,12,11,4,13,17,17,8, | 5,0,1,3,0,0,2,6,0,4,0,0 | 7,17,5,5,4,8,7,8,3,3,10,8,8, | 0,0,3,0,0,0,2,0,0,1,0,0,0,2, | 298 | 149 | 0.0000045412807042 | 0.00196664866651626 | 0.444,1.0,0.867,0. | 1.0,1.0,0.455,1.0,1 | -0,067 |
| 109679 | ENSG00000197217 | ENTPD4 | chr8 | - | 23442006 | 23442066 | 23441568 | 23441723 | 23443849 | 23443953 | 109679 | 125,80,223,85,78,65,69,10 | 5,0,5,2,1,5,9,4,4,0,1,0 | 98,117,74,111,109,87,61,4 | 5,7,6,51,5,5,7,9,10,4,5,0,4, | 209 | 149 | 0.0000045966509958 | 0.00198663010137678 | 0.947,1.0,0.97,0.9 | 0.933,0.923,0.898, | 0,05 |
| 120687 | ENSG00000102781 | KATNAL1 | chr13 | - | 30255446 | 30255615 | 30240958 | 30241086 | 30280062 | 30280223 | 120687 | 5,4,35,8,11,14,4,11,12,5,2, | 0,1,3,3,7,3,2,4,0,0,0,1 | 28,15,2,7,8,8,1,13,4,6,11,3 | 5,1,2,0,0,0,1,2,0,2,0,4,0,0, | 298 | 149 | 0.0000046313930359 | 0.00199763397700372 | 1.0,0.667,0.854,0. | 0.737,0.882,0.333, | -0,082 |
| 128066 | ENSG00000166224 | SGPL1 | chr10 | + | 70854707 | 70855820 | 70851142 | 70851210 | 70857613 | 70857690 | 128066 | 21,11,21,23,9,6,5,10,16,5, | 3,2,7,1,4,5,0,3,2,0,0,0 | 14,16,10,11,17,7,13,17,7,2 | 0,0,2,1,1,0,0,0,1,4,3,3,1,1, | 298 | 149 | 0.0000046473098267 | 0.00200049028263815 | 0.778,0.733,0.6,0. | 1.0,1.0,0.714,0.84 | -0,088 |
| 146258 | ENSG00000140968 | IRF8 | chr16 | + | 85913130 | 85913213 | 85909486 | 85911658 | 85914472 | 85914520 | 146258 | 340,214,403,290,151,112,1 | 0,4,7,4,6,9,4,4,2,1,1,1 | 186,219,233,134,144,100,1 | 7,11,5,7,4,3,6,1,2,3,9,21,2, | 232 | 149 | 0.0000048407458631 | 0.00207959794984358 | 1.0,0.972,0.974,0. | 0.945,0.927,0.968, | 0,023 |
| 159927 | ENSG00000204120 | GIGYF2 | chr2 | + | 232847347 | 232847658 | 232845731 | 232845886 | 232850261 | 232850409 | 159927 | 47,30,36,20,19,20,13,33,3 | 12,6,4,0,0,3,2,5,2,1,0,2 | 18,26,33,41,18,8,19,15,9,3 | 1,0,1,5,0,0,2,0,5,0,1,1,0,1,2 | 298 | 149 | 0.0000048571236016 | 0.0020824772308899 | 0.662,0.714,0.818, | 0.9,1.0,0.943,0.80 | -0,075 |
| 233928 | ENSG00000130985 | UBA1 | chrX | + | 47200354 | 47200544 | 47199479 | 47199614 | 47200893 | 47201000 | 233928 | 0,0,10,25,0,0,0,1,3,5,0,0 | 20,24,74,41,27,20,25,21,1 | 0,0,0,0,4,10,0,0,0,3,0,0,0,0 | 52,46,15,17,29,28,22,30,2 | 298 | 149 | 0.0000049275876918 | 0.00210801341428842 | 0.0,0.0,0.063,0.23 | 0.0,0.0,0.0,0.0,0.0 | 0,049 |
| 43519 | ENSG00000196455 | PIK3R4 | chr3 | - | 130728462 | 130728684 | 130723413 | 130723587 | 130730307 | 130730442 | 43519 | 8,10,27,18,19,12,16,17,13, | 1,0,0,1,0,3,0,0,0,0,0,4 | 14,28,9,1,5,11,15,4,3,11,1 | 2,3,5,0,1,0,0,2,2,0,1,2,0,6, | 298 | 149 | 0.0000049362720091 | 0.00210801341428842 | 0.8,1.0,1.0,0.9,1.0, | 0.778,0.824,0.474, | 0,119 |
| 13962 | ENSG00000167508 | MVD | chr16 | - | 88657145 | 88657247 | 88656104 | 88656304 | 88657435 | 88657582 | 13962 | 3,1,0,0,1,4,4,6,5,7,2,2 | 5,5,18,8,3,4,8,7,19,14,8,6 | 0,1,0,1,1,0,3,3,0,0,5,1,4,0, | 1,11,4,0,3,1,7,7,1,7,8,45,6, | 251 | 149 | 0.0000049896727943 | 0.00212659854496901 | 0.263,0.106,0.0,0. | 0.0,0.051,0.0,1.0,0 | 0,041 |
| 50225 | ENSG00000122224 | LY9 | chr1 | + | 160823506 | 160823796 | 160819320 | 160819374 | 160827747 | 160828255 | 50225 | 25,12,7,11,25,8,17,12,13,5 | 0,0,1,0,2,0,0,0,0,0,2,0 | 25,17,18,17,6,12,33,22,7,1 | 2,1,1,1,3,0,0,2,1,0,2,5,4,2,2 | 298 | 149 | 0.0000050517171060 | 0.00214855029931991 | 1.0,1.0,0.778,1.0,0 | 0.862,0.895,0.9,0. | 0,108 |
| 153166 | ENSG00000100902 | PSMA6 | chr14 | + | 35312943 | 35313059 | 35310832 | 35310895 | 35314360 | 35314395 | 153166 | 7,10,44,26,8,5,5,7,6,3,2,11 | 1,0,3,1,1,2,0,1,0,0,0,3 | 20,26,11,7,8,16,8,5,6,22,5, | 0,0,0,2,6,2,1,3,3,2,4,5,0,1,3 | 265 | 149 | 0.0000050611436166 | 0.00214855029931991 | 0.797,1.0,0.892,0. | 1.0,1.0,1.0,0.663,0 | 0,162 |
| 11133 | ENSG00000291131 | | chr19 | + | 53200623 | 53200703 | 53197090 | 53197262 | 53204015 | 53204142 | 11133 | 46,8,92,84,26,40,6,13,18,7 | 0,0,0,0,0,0,2,6,3,5,0,0 | 0,0,7,2,7,20,11,35,10,23,2 | 14,4,0,1,0,0,0,0,1,3,3,4,4,2 | 229 | 149 | 0.0000051306516076 | 0.00217377022868616 | 1.0,1.0,1.0,1.0,1.0, | 0.0,0.0,1.0,0.565,1 | 0,244 |
| 101536 | ENSG00000156831 | NSMCE2 | chr8 | + | 125102050 | 125102146 | 125091843 | 125091958 | 125102316 | 125102464 | 101536 | 12,9,33,35,7,35,21,34,5,7, | 1,1,5,10,0,0,0,3,2,2,2,4 | 24,29,9,7,20,8,9,15,13,23,2 | 0,4,0,0,1,1,1,2,0,1,1,2,2,0, | 245 | 149 | 0.0000052412106682 | 0.00221624953504541 | 0.879,0.846,0.801, | 1.0,0.815,1.0,1.0,0 | -0,091 |
| 122690 | ENSG00000261008 | LINC01572 | chr16 | - | 72522552 | 72522623 | 72425520 | 72427457 | 72523331 | 72523415 | 122690 | 30,14,12,10,10,5,2,11,39,2 | 9,5,2,0,4,2,1,0,10,4,0,6 | 4,1,6,5,1,1,1,9,7,16,2,8,1,3 | 0,1,0,0,2,0,0,4,2,2,1,3,0,0, | 220 | 149 | 0.0000052824103741 | 0.00222929111222004 | 0.693,0.655,0.803, | 1.0,0.404,1.0,1.0,0 | -0,061 |
| 47861 | ENSG00000177731 | FLII | chr17 | - | 18253598 | 18253719 | 18253300 | 18253458 | 18254078 | 18254182 | 47861 | 26,12,23,30,17,15,32,8,29, | 1,0,2,3,3,3,3,6,2,0,0,0 | 7,11,16,12,11,8,9,12,4,18, | 0,1,2,0,0,0,3,4,0,0,1,0,0,0,2 | 270 | 149 | 0.0000053024505958 | 0.0022312426920007 | 0.935,1.0,0.864,0. | 1.0,0.859,0.815,1. | -0,044 |
| 158148 | ENSG00000115760 | BIRC6 | chr2 | + | 32436052 | 32436465 | 32433643 | 32433804 | 32439507 | 32439686 | 158148 | 27,10,46,24,22,27,23,23,1 | 3,1,2,4,2,0,2,0,0,0,0,0 | 39,39,21,25,29,19,17,32,2 | 5,2,0,15,3,3,0,11,3,2,3,1,0, | 298 | 149 | 0.0000053077682039 | 0.0022312426920007 | 0.818,0.833,0.92,0 | 0.796,0.907,1.0,0. | 0,129 |
| 134094 | ENSG00000100201 | DDX17 | chr22 | - | 38489222 | 38489423 | 38488025 | 38488115 | 38492055 | 38492115 | 134094 | 19,9,3,1,5,5,5,15,25,12,2,9 | 223,109,242,214,130,97,1 | 15,10,8,31,3,11,6,11,3,10, | 123,52,139,40,64,84,60,14 | 298 | 149 | 0.0000053403466454 | 0.00224056169365857 | 0.041,0.04,0.006,0 | 0.057,0.088,0.028, | -0,028 |
| 32313 | ENSG00000213445 | SIPA1 | chr11 | + | 65642249 | 65642377 | 65640800 | 65641600 | 65642462 | 65642639 | 32313 | 43,29,54,23,26,20,17,25,4 | 0,1,4,1,0,2,0,0,2,3,0,0 | 28,7,10,8,16,7,10,14,12,21 | 7,1,0,0,1,0,1,4,0,3,0,16,3,0 | 277 | 149 | 0.0000054889971528 | 0.00229753735370319 | 1.0,0.94,0.879,0.9 | 0.683,0.79,1.0,1.0, | 0,063 |
| 145442 | ENSG00000074370 | ATP2A3 | chr17 | - | 3928212 | 3928327 | 3925345 | 3925438 | 3928662 | 3928780 | 145442 | 0,2,19,25,1,2,4,0,1,3,0,0 | 24,10,22,13,17,11,19,11,1 | 2,1,1,0,5,7,0,0,1,1,0,0,5,0, | 18,34,11,16,18,16,8,8,14,2 | 264 | 149 | 0.0000054974968157 | 0.00229753735370319 | 0.0,0.101,0.328,0. | 0.059,0.016,0.049, | 0,072 |
| 29457 | ENSG00000114904 | NEK4 | chr3 | - | 52768337 | 52768604 | 52766177 | 52766375 | 52770653 | 52770939 | 29457 | 10,8,36,23,21,16,16,3,11,1 | 1,0,3,1,0,1,0,5,0,0,0,0 | 24,11,18,15,5,8,16,24,8,17 | 2,1,5,0,2,2,0,6,0,2,7,0,2,0, | 298 | 149 | 0.0000055185664569 | 0.00229909747243917 | 0.833,1.0,0.857,0. | 0.857,0.846,0.643, | 0,05 |
| 206821 | ENSG00000075884 | ARHGAP15 | chr2 | + | 143518986 | 143519364 | 143487372 | 143487495 | 143556407 | 143556485 | 206821 | 122,103,164,94,55,56,56,5 | 6,1,5,5,0,2,5,2,0,0,0,1 | 114,91,86,75,91,49,23,41, | 3,15,3,15,14,2,0,2,15,3,7,5 | 298 | 149 | 0.0000055225938329 | 0.00229909747243917 | 0.91,0.981,0.943,0 | 0.95,0.752,0.935,0 | 0,074 |
| 149672 | ENSG00000214013 | GANC | chr15 | + | 42283610 | 42283891 | 42280916 | 42281133 | 42287690 | 42287818 | 149672 | 35,24,44,36,14,30,17,19,2 | 2,3,4,1,0,0,0,0,0,0,1,2 | 12,15,22,17,5,3,6,10,13,25 | 4,1,1,0,4,2,0,2,1,1,3,1,4,2,1 | 298 | 149 | 0.0000056272396795 | 0.00233813981365131 | 0.897,0.8,0.846,0. | 0.6,0.882,0.917,1. | 0,097 |
| 149675 | ENSG00000214013 | GANC | chr15 | + | 42283610 | 42284056 | 42280916 | 42281133 | 42287690 | 42287818 | 149675 | 35,24,43,36,16,30,17,19,2 | 2,3,4,1,0,0,0,0,0,0,1,2 | 15,15,22,19,5,3,6,10,13,25 | 4,1,1,0,4,2,0,2,1,1,3,1,4,2,1 | 298 | 149 | 0.0000057394187360 | 0.00238015574947741 | 0.897,0.8,0.843,0. | 0.652,0.882,0.917, | 0,094 |
| 169376 | ENSG00000157426 | AASDH | chr4 | - | 56371450 | 56371948 | 56355181 | 56355423 | 56378147 | 56378464 | 169376 | 1,4,18,7,20,7,3,8,3,3,5,14 | 2,1,1,2,0,2,0,0,0,1,0,0 | 12,7,2,0,2,5,4,1,5,3,12,13, | 0,0,3,2,4,1,3,4,1,0,1,3,0,0,2 | 298 | 149 | 0.0000057779823960 | 0.00239154024825547 | 0.2,0.667,0.9,0.63 | 1.0,1.0,0.25,0.0,0. | 0,153 |
| 78496 | ENSG00000160352 | ZNF714 | chr19 | + | 21098814 | 21098910 | 21098184 | 21098311 | 21116806 | 21117084 | 78496 | 5,10,10,8,7,7,8,5,6,1,0,3 | 0,0,1,2,1,0,0,1,2,0,2,0 | 10,4,6,4,3,1,6,6,6,5,8,5,4,4 | 0,3,0,1,4,4,4,5,2,0,3,0,0,1, | 245 | 149 | 0.0000059099427267 | 0.00244146426683931 | 1.0,1.0,0.859,0.70 | 1.0,0.448,1.0,0.70 | 0,098 |
| 29687 | ENSG00000065357 | DGKA | chr12 | + | 55932560 | 55932909 | 55932107 | 55932123 | 55936422 | 55936862 | 29687 | 5,0,3,3,1,3,1,2,6,2,1,1 | 4,5,25,14,6,5,6,8,2,2,2,10 | 0,0,0,3,0,1,0,2,2,2,2,0,1,0, | 13,14,2,0,2,5,1,10,8,7,8,1,1 | 298 | 149 | 0.0000059293174814 | 0.00244477572959763 | 0.385,0.0,0.057,0. | 0.0,0.0,0.0,1.0,0.0, | 0,084 |
| 28798 | ENSG00000147050 | KDM6A | chrX | + | 45085864 | 45085979 | 45083459 | 45083608 | 45089742 | 45089930 | 28798 | 63,49,172,96,20,33,29,38, | 2,0,12,4,1,0,1,2,0,0,0,3 | 63,111,46,91,40,33,27,34, | 12,6,6,15,3,5,2,5,2,12,3,5, | 264 | 149 | 0.0000059860674043 | 0.00246345558986462 | 0.947,1.0,0.89,0.9 | 0.748,0.913,0.812, | 0,092 |
| 79915 | ENSG00000138468 | SENP7 | chr3 | - | 101417592 | 101417790 | 101372007 | 101372126 | 101458954 | 101459052 | 79915 | 88,42,135,50,61,76,38,70, | 3,1,8,8,9,4,1,12,3,0,1,4 | 61,46,33,56,65,40,33,52,5 | 5,3,5,1,0,4,1,0,0,4,1,0,0,1,1 | 298 | 149 | 0.0000060447434534 | 0.00248285530196468 | 0.936,0.955,0.894, | 0.859,0.885,0.767, | -0,052 |
| 65664 | ENSG00000196628 | TCF4 | chr18 | - | 55587044 | 55587136 | 55585279 | 55585352 | 55588037 | 55588150 | 65664 | 59,34,91,35,27,27,32,52,5 | 2,6,5,5,0,2,2,1,1,0,2,1 | 45,81,43,56,46,29,20,49,3 | 1,4,3,0,1,2,0,0,0,1,0,0,0,0, | 241 | 149 | 0.0000062158499710 | 0.00254827353356831 | 0.948,0.778,0.918, | 0.965,0.926,0.899, | -0,055 |
| 115715 | ENSG00000021776 | AQR | chr15 | - | 34927034 | 34927138 | 34920331 | 34920434 | 34930257 | 34930371 | 115715 | 50,34,86,52,33,41,34,45,2 | 2,0,0,1,0,0,1,1,0,1,0,0 | 68,63,22,18,46,29,20,49,3 | 0,8,0,4,7,1,2,1,1,0,2,1,0,1, | 253 | 149 | 0.0000062751940368 | 0.0025677115736561 | 0.936,1.0,1.0,0.96 | 1.0,0.823,1.0,0.72 | 0,056 |
| 221842 | ENSG00000068305 | MEF2A | chr15 | + | 99626821 | 99626880 | 99598429 | 99598511 | 99632977 | 99633173 | 221842 | 4,2,1,0,0,2,1,2,0,3,2,2 | 51,39,66,24,15,27,16,34,2 | 4,12,7,11,2,0,2,0,4,0,4,7,1, | 25,28,31,36,22,12,12,11,1 | 208 | 149 | 0.0000063804873623 | 0.00260584188896765 | 0.053,0.035,0.011, | 0.103,0.235,0.139, | -0,035 |
| 31248 | ENSG00000072364 | AFF4 | chr5 | - | 132889078 | 132889173 | 132888096 | 132888160 | 132892163 | 132892404 | 31248 | 139,96,158,88,48,85,60,13 | 14,3,3,1,3,2,3,8,1,0,0,0 | 95,123,58,115,62,27,61,57 | 3,11,1,17,7,4,7,6,3,2,8,3,7, | 244 | 149 | 0.0000064302439579 | 0.00261960209894263 | 0.858,0.951,0.97,0 | 0.951,0.872,0.973, | 0,076 |
| 181161 | ENSG00000162894 | FCMR | chr1 | - | 206912930 | 206913042 | 206911729 | 206911952 | 206921817 | 206921998 | 181161 | 88,26,123,100,97,53,77,64 | 9,3,4,3,4,10,4,4,8,14,3,3 | 100,42,87,31,31,41,47,61, | 4,0,6,4,2,3,2,3,0,0,0,0,2,1, | 261 | 149 | 0.0000064428476165 | 0.00261960209894263 | 0.848,0.832,0.946, | 0.935,1.0,0.892,0. | -0,078 |
| 66788 | ENSG00000180182 | MED14 | chrX | - | 40666719 | 40666851 | 40664306 | 40664489 | 40671860 | 40671972 | 66788 | 52,37,56,36,24,27,32,26,4 | 5,3,2,3,1,1,3,2,14,4,4,3 | 36,30,19,68,28,19,11,19,2 | 1,1,0,1,3,4,3,0,0,4,0,0,2,1, | 281 | 149 | 0.0000064506930341 | 0.00261960209894263 | 0.846,0.867,0.937, | 0.95,0.941,1.0,0.9 | -0,077 |
| 11562 | ENSG00000006025 | OSBPL7 | chr17 | - | 47819728 | 47819782 | 47818985 | 47819099 | 47819970 | 47820096 | 11562 | 20,4,21,30,11,10,8,5,14,8, | 1,0,4,0,3,4,0,2,2,0,2,5 | 12,18,16,5,10,7,4,19,4,6,2, | 1,4,0,0,0,0,0,8,0,0,0,2,2,1, | 203 | 149 | 0.00000652116124 | 0.00264323174170704 | 0.936,1.0,0.794,1. | 0.898,0.768,1.0,1. | -0,134 |
| 198576 | ENSG00000223745 | CCDC18-AS1 | chr1 | - | 93273473 | 93273628 | 93264637 | 93264772 | 93305387 | 93305470 | 198576 | 9,2,7,9,2,3,1,0,4,0,3,0 | 74,43,78,56,21,22,44,34,1 | 2,0,0,0,4,0,0,3,0,2,0,1,0,1,2 | 18,24,22,13,39,25,17,45,2 | 298 | 149 | 0.0000065662921258 | 0.00265399812472519 | 0.057,0.023,0.043, | 0.053,0.0,0.0,0.0,0 | 0,037 |
| 149666 | ENSG00000214013 | GANC | chr15 | + | 42283610 | 42283790 | 42280916 | 42281133 | 42287690 | 42287818 | 149666 | 35,24,43,36,13,30,17,19,2 | 2,3,4,1,0,0,0,0,0,0,1,2 | 13,15,24,17,5,3,6,10,13,25 | 4,1,1,0,4,2,0,2,1,1,3,1,4,2,1 | 298 | 149 | 0.0000065723850211 | 0.00265399812472519 | 0.897,0.8,0.843,0. | 0.619,0.882,0.923, | 0,094 |
| 13036 | ENSG00000074370 | ATP2A3 | chr17 | - | 3928226 | 3928327 | 3925345 | 3925438 | 3928662 | 3928780 | 13036 | 0,2,20,23,1,2,4,0,1,3,0,0 | 24,10,22,13,17,11,19,11,1 | 2,1,1,0,5,7,0,0,1,1,0,0,5,0, | 18,34,11,16,18,16,8,8,14,2 | 250 | 149 | 0.0000066265949239 | 0.00266260321615757 | 0.0,0.107,0.351,0. | 0.062,0.017,0.051, | 0,074 |
| 120235 | ENSG00000168067 | MAP4K2 | chr11 | - | 64801578 | 64801621 | 64801110 | 64801183 | 64801709 | 64801757 | 120235 | 48,54,79,65,35,43,31,55,3 | 0,0,4,5,2,0,0,2,0,0,0,2 | 70,40,28,36,47,24,23,35,2 | 8,4,0,3,0,1,0,2,5,4,7,0,1,1,4 | 192 | 149 | 0.0000066297103175 | 0.00266260321615757 | 1.0,1.0,0.939,0.91, | 0.872,0.886,1.0,0. | 0,051 |
| 253765 | ENSG00000100982 | PCIF1 | chr20 | + | 45947083 | 45947166 | 45946199 | 45946384 | 45947262 | 45947438 | 253765 | 42,40,29,14,13,35,19,17,3 | 2,0,0,2,0,3,2,0,0,4,0,2 | 18,39,20,48,18,9,17,20,8,1 | 4,4,0,1,2,2,7,5,1,2,3,2,1,1,1 | 232 | 149 | 0.0000066496498468 | 0.00266260321615757 | 0.931,1.0,1.0,0.81 | 0.743,0.862,1.0,0. | 0,089 |
| 233930 | ENSG00000130985 | UBA1 | chrX | + | 47200409 | 47200544 | 47199479 | 47199614 | 47200893 | 47201000 | 233930 | 0,0,24,51,0,0,0,2,4,12,0,0 | 20,24,74,41,27,20,25,21,1 | 0,0,0,0,10,14,0,0,0,3,0,0,0, | 52,46,15,17,29,28,22,30,2 | 284 | 149 | 0.0000066533657876 | 0.00266260321615757 | 0.0,0.0,0.145,0.39 | 0.0,0.0,0.0,0.0,0.1 | 0,084 |
| 202289 | ENSG00000136059 | VILL | chr3 | + | 37997886 | 37998168 | 37997482 | 37997685 | 37998265 | 37998364 | 202289 | 10,3,10,11,1,9,4,6,2,2,1,4 | 0,2,0,2,1,0,0,0,0,2,1,0 | 2,2,7,0,1,4,9,9,3,5,6,11,2,4 | 0,5,2,1,3,0,2,1,2,0,2,0,4,0, | 298 | 149 | 0.0000066555492949 | 0.00266260321615757 | 1.0,0.429,1.0,0.73 | 1.0,0.167,0.636,0. | 0,134 |
| 104626 | ENSG00000197157 | SND1 | chr7 | + | 127928059 | 127928186 | 127904625 | 127904819 | 127929187 | 127929329 | 104626 | 2,1,6,0,0,7,3,3,0,0,1,2 | 48,47,153,121,48,49,53,57 | 5,13,4,8,7,2,5,7,10,2,3,1,0, | 60,102,60,23,44,26,49,51,3 | 276 | 149 | 0.0000066880837209 | 0.00267065481881964 | 0.022,0.011,0.021, | 0.043,0.064,0.035, | -0,031 |
| 217002 | ENSG00000135945 | REV1 | chr2 | - | 99424775 | 99424899 | 99424151 | 99424280 | 99429839 | 99429945 | 217002 | 1,1,4,0,1,4,2,0,0,2,0,0 | 15,7,15,6,10,11,12,11,11,3 | 7,6,2,3,2,0,1,2,6,6,6,11,2,5 | 15,22,5,14,16,12,9,22,6,21 | 273 | 149 | 0.0000067327363070 | 0.00268350660758559 | 0.035,0.072,0.127, | 0.203,0.13,0.179,0 | -0,074 |
| 20380 | ENSG00000141956 | PRDM15 | chr21 | - | 41823317 | 41823449 | 41821902 | 41822037 | 41825959 | 41826054 | 20380 | 10,10,17,23,12,8,11,14,12, | 2,0,1,6,3,0,1,1,2,4,0,0 | 15,15,25,13,40,34,10,13,1 | 0,1,1,1,1,1,0,0,3,1,1,0,0,2, | 281 | 149 | 0.0000067571006551 | 0.00268823942902427 | 0.726,1.0,0.9,0.67, | 1.0,0.888,0.93,0.8 | -0,019 |
| 172614 | ENSG00000156931 | VPS8 | chr3 | + | 184944491 | 184944589 | 184940196 | 184940243 | 184957373 | 184957521 | 172614 | 0,4,1,0,2,1,3,3,0,2,3,0 | 20,15,30,24,25,26,20,26,1 | 3,10,1,4,3,2,3,2,7,2,2,5,0,5 | 19,12,8,10,6,6,6,20,8,23,1 | 247 | 149 | 0.0000068022762357 | 0.00270121903411882 | 0.0,0.139,0.02,0.0, | 0.087,0.335,0.07,0 | -0,074 |
| 68057 | ENSG00000174123 | TLR10 | chr4 | - | 38775774 | 38775963 | 38772237 | 38775652 | 38776109 | 38776426 | 68057 | 17,3,55,46,48,9,5,7,31,4,5, | 4,0,9,4,2,1,0,2,2,0,0,0 | 30,14,6,2,4,6,1,4,13,36,1,1 | 2,0,0,0,0,2,0,0,0,0,1,4,0,2, | 298 | 149 | 0.0000068580747189 | 0.00271836147297157 | 0.68,1.0,0.753,0.8 | 0.882,1.0,1.0,1.0,1 | -0,038 |
| 58152 | ENSG00000076770 | MBNL3 | chrX | - | 132439434 | 132440314 | 132406227 | 132406392 | 132489813 | 132489930 | 58152 | 7,12,13,14,7,10,4,13,5,2,1, | 0,0,5,1,0,0,2,7,1,1,0,6 | 8,4,3,9,11,3,3,6,4,1,1,13,1 | 1,0,1,0,1,0,0,0,1,0,4,1,0,1, | 298 | 149 | 0.0000068843840330 | 0.0027237736393781 | 1.0,1.0,0.565,0.87 | 0.8,1.0,0.6,1.0,0.8 | -0,079 |
| 181144 | ENSG00000162894 | FCMR | chr1 | - | 206912899 | 206913042 | 206911729 | 206911952 | 206921817 | 206921998 | 181144 | 95,27,129,107,100,53,79,6 | 9,3,4,3,4,10,4,4,8,14,3,3 | 107,48,89,31,32,46,50,63, | 4,0,6,4,2,3,2,3,0,0,0,0,2,1, | 292 | 149 | 0.0000069182680543 | 0.00273215734238412 | 0.843,0.821,0.943, | 0.932,1.0,0.883,0. | -0,084 |
| 136102 | ENSG00000105486 | LIG1 | chr19 | - | 48151467 | 48151592 | 48151231 | 48151339 | 48153871 | 48154207 | 136102 | 4,6,2,1,2,0,4,0,4,1,1,4 | 26,18,20,5,27,19,12,15,26, | 0,0,0,0,0,0,1,1,1,0,3,0,0,3, | 14,11,19,13,6,14,4,16,10,1 | 274 | 149 | 0.0000069961843719 | 0.00275786768966486 | 0.077,0.153,0.052, | 0.0,0.0,0.0,0.0,0.0, | 0,033 |
| 9521 | ENSG00000029725 | RABEP1 | chr17 | + | 5356507 | 5356573 | 5354400 | 5354490 | 5361207 | 5361629 | 9521 | 25,26,34,8,0,5,17,26,23,13 | 147,67,168,134,73,65,49,5 | 29,24,17,16,21,20,14,18,1 | 85,56,53,30,51,41,50,60,3 | 215 | 149 | 0.0000070292915531 | 0.0027658527427389 | 0.105,0.212,0.123, | 0.191,0.229,0.182, | -0,082 |
| 9500 | ENSG00000129197 | RPAIN | chr17 | + | 5425970 | 5426082 | 5422768 | 5422829 | 5426235 | 5426299 | 9500 | 33,4,106,64,25,28,16,41,1 | 0,0,2,9,6,3,3,2,0,2,0,0 | 28,28,27,8,24,21,11,45,11, | 1,0,0,0,0,0,0,1,0,0,0,9,3,3,2 | 261 | 149 | 0.0000070951679054 | 0.00278304724748281 | 1.0,1.0,0.968,0.80 | 0.941,1.0,1.0,1.0,1 | -0,062 |
| 153980 | ENSG00000012983 | MAP4K5 | chr14 | - | 50461058 | 50461361 | 50456515 | 50456594 | 50462664 | 50462781 | 153980 | 0,0,13,10,1,7,12,9,0,0,1,0 | 45,46,131,67,44,28,31,61,2 | 0,21,12,9,9,5,10,20,4,13,7, | 48,43,26,29,56,32,14,32,3 | 298 | 149 | 0.0000071042248271 | 0.00278304724748281 | 0.0,0.0,0.047,0.06 | 0.0,0.196,0.188,0. | -0,059 |
| 173164 | ENSG00000114127 | XRN1 | chr3 | - | 142376696 | 142376795 | 142376478 | 142376594 | 142380081 | 142380180 | 173164 | 8,1,11,6,1,4,1,2,6,0,0,0 | 26,13,51,34,28,14,18,23,8, | 2,1,0,3,5,2,0,3,0,5,0,1,0,1, | 27,27,13,30,17,17,12,29,1 | 248 | 149 | 0.0000071117821601 | 0.00278304724748281 | 0.156,0.044,0.115, | 0.043,0.022,0.0,0. | 0,047 |
| 167945 | ENSG00000130338 | TULP4 | chr6 | + | 158431583 | 158431663 | 158429735 | 158429897 | 158448995 | 158449176 | 167945 | 0,0,0,0,0,2,0,0,0,0,1,0 | 4,15,11,7,14,15,11,9,14,13 | 1,1,2,2,0,1,2,0,0,2,1,9,0,4,1 | 15,16,10,24,5,8,10,10,6,11 | 229 | 149 | 0.0000071630477629 | 0.00279802165710587 | 0.0,0.0,0.0,0.0,0.0, | 0.042,0.039,0.115, | -0,063 |
| 205914 | ENSG00000128815 | WDFY4 | chr10 | + | 48824680 | 48824785 | 48822379 | 48822537 | 48826670 | 48826909 | 205914 | 0,0,2,0,11,17,0,0,0,0,3,14 | 97,60,97,107,74,48,37,52,8 | 1,1,0,1,0,0,0,0,0,0,0,0,1,0,2 | 65,92,63,46,53,31,34,47,3 | 254 | 149 | 0.0000072530779764 | 0.00282675477757329 | 0.0,0.0,0.012,0.0,0 | 0.009,0.006,0.0,0. | 0,035 |
| 54234 | ENSG00000171163 | ZNF692 | chr1 | - | 248856287 | 248856422 | 248855557 | 248855946 | 248856513 | 248856562 | 54234 | 44,23,42,16,33,33,9,18,44, | 1,1,5,1,3,2,1,5,0,3,0,2 | 19,18,11,12,12,10,18,21,1 | 2,0,0,0,0,0,0,4,0,0,2,3,0,0, | 284 | 149 | 0.0000072628728761 | 0.00282675477757329 | 0.958,0.923,0.815, | 0.833,1.0,1.0,1.0,1 | -0,081 |
| 160912 | ENSG00000164329 | TENT2 | chr5 | + | 79642852 | 79642985 | 79641104 | 79641196 | 79645122 | 79645192 | 160912 | 1,0,5,0,4,1,0,2,0,0,0,0 | 17,6,7,3,5,4,2,5,3,2,6,3 | 8,0,2,1,1,0,1,4,0,4,0,7,1,1, | 6,10,9,15,8,4,3,11,3,6,2,5,4 | 282 | 149 | 0.0000072930891349 | 0.00283339145776583 | 0.03,0.0,0.274,0.0, | 0.413,0.0,0.105,0. | -0,047 |
| 41586 | ENSG00000129071 | MBD4 | chr3 | - | 129436460 | 129437308 | 129434061 | 129434136 | 129437719 | 129437950 | 41586 | 299,224,392,280,171,123,1 | 7,0,7,14,2,2,0,0,7,0,0,4 | 298,327,189,99,107,100,4 | 0,1,2,1,0,0,2,0,0,2,0,1,0,1,3 | 298 | 149 | 0.0000074994236802 | 0.00290830352816278 | 0.955,1.0,0.966,0. | 1.0,0.994,0.979,0. | -0,013 |

| | | | | | | | | | | | | | | | | | | | | | | |
|---|---|---|---|---|---|---|---|---|---|---|---|---|---|---|---|---|---|---|---|---|---|---|
| 21343 | ENSG00000100902 | PSMA6 | chr14 | + | 35312862 | 35313059 | 35310739 | 35310895 | 35314360 | 35314455 | 21343 | 11,10,57,32,10,6,8,8,6,3,4, | 1,0,3,1,1,2,0,1,0,0,0,3 | 28,40,13,9,9,18,9,9,8,22,1 | 0,0,0,2,6,2,1,3,3,2,4,5,0,1, | 298 | 149 | 0.0000075368897339 | 0.00291757610491771 | 0.846,1.0,0.905,0. | 1.0,1.0,1.0,0.692,0 | 0,129 |
| 99760 | ENSG00000165055 | METTL2B | chr7 | + | 128477081 | 128477173 | 128476747 | 128476875 | 128479157 | 128479513 | 99760 | 29,19,50,47,17,10,8,19,3,6 | 0,0,2,0,0,0,0,0,5,7,0,1 | 16,11,18,7,12,13,1,21,7,21 | 9,9,0,0,1,0,10,15,0,0,1,0,4, | 241 | 149 | 0.0000075506849406 | 0.00291766870818085 | 1.0,1.0,0.939,1.0,1 | 0.524,0.43,1.0,1.0, | 0,187 |
| 114166 | ENSG00000179950 | PUF60 | chr8 | - | 143824312 | 143824396 | 143821817 | 143821913 | 143829279 | 143829315 | 114166 | 23,35,32,24,9,10,12,28,12, | 0,3,12,5,0,0,4,1,0,0,0,0 | 19,22,7,2,23,2,6,19,14,18,2 | 2,1,1,1,2,0,0,0,1,0,2,4,0,0,1 | 233 | 149 | 0.0000078172651569 | 0.00301526486917528 | 1.0,0.882,0.63,0.7 | 0.859,0.934,0.817, | -0,001 |
| 253571 | ENSG00000135968 | GCC2 | chr2 | + | 108474788 | 108474864 | 108472826 | 108472899 | 108475534 | 108475635 | 253571 | 8,3,9,2,0,1,0,2,0,0,2,2 | 65,38,58,37,13,19,6,14,24, | 5,5,14,8,1,2,3,5,3,7,5,3,3,1 | 17,16,33,22,13,8,8,20,13,1 | 225 | 149 | 0.0000078627951318 | 0.00302740117894009 | 0.075,0.05,0.093,0 | 0.163,0.171,0.219, | -0,087 |
| 215675 | ENSG00000168887 | C2orf68 | chr2 | - | 85611473 | 85611573 | 85609434 | 85609586 | 85611667 | 85611741 | 215675 | 0,1,0,1,1,2,6,5,3,0,3,3 | 8,4,16,19,16,6,10,8,5,8,8,7 | 1,0,0,0,0,0,2,0,1,3,1,2,1, | 19,18,1,8,1,6,4,11,3,6,5,24 | 249 | 149 | 0.0000080311138583 | 0.00308668690509846 | 0.0,0.13,0.0,0.031, | 0.031,0.0,0.0,0.0,0 | 0,07 |
| 63992 | ENSG00000058668 | ATP2B4 | chr1 | + | 203707008 | 203707223 | 203703651 | 203703813 | 203707861 | 203708104 | 63992 | 123,46,206,99,57,40,57,40 | 6,1,3,2,3,1,2,6,0,0,5,2 | 181,104,59,25,17,38,81,76 | 5,0,2,0,3,0,0,0,1,0,2,0,0,1, | 298 | 149 | 0.0000080888255292 | 0.00310332621656027 | 0.911,0.958,0.972, | 0.948,1.0,0.937,1. | -0,04 |
| 183470 | ENSG00000171824 | EXOSC10 | chr1 | - | 11082687 | 11082768 | 11081081 | 11081238 | 11087447 | 11087591 | 183470 | 39,21,47,24,17,8,6,12,8,3, | 6,8,2,6,4,1,4,2,0,0,0,0 | 20,19,17,13,13,11,9,9,7,19 | 5,6,11,12,7,4,7,6,5,3,12,9, | 230 | 149 | 0.0000081177304865 | 0.00310887411095952 | 0.808,0.63,0.938,0 | 0.722,0.672,0.5,0. | 0,239 |
| 233408 | ENSG00000183354 | KIAA2026 | chr9 | - | 5954015 | 5954128 | 5944870 | 5944960 | 5968018 | 5969436 | 233408 | 25,15,28,12,8,9,11,12,18,1 | 2,3,4,2,1,5,4,2,0,0,1,0 | 12,16,11,26,5,4,1,15,2,8,4, | 0,1,2,1,3,1,0,0,1,0,0,0,0,0, | 262 | 149 | 0.0000082273742207 | 0.00314526817214005 | 0.877,0.74,0.799,0 | 1.0,0.901,0.758,0. | -0,077 |
| 255302 | ENSG00000272917 | | chr12 | + | 9653766 | 9653808 | 9642871 | 9643103 | 9655872 | 9655979 | 255302 | 36,20,120,49,20,34,21,29, | 19,15,0,0,16,3,19,7,14,4,0, | 88,45,14,10,26,5,18,48,42, | 0,0,7,16,29,12,0,0,0,0,0,0, | 191 | 149 | 0.0000083316874278 | 0.00317949896593215 | 0.596,0.51,1.0,1.0, | 1.0,1.0,0.609,0.32 | -0,134 |
| 123458 | ENSG00000149483 | TMEM138 | chr11 | + | 61364432 | 61364518 | 61364249 | 61364315 | 61366044 | 61366216 | 123458 | 13,9,48,28,15,11,7,10,6,2, | 0,5,3,3,0,6,2,4,1,0,0,0 | 21,22,14,3,16,8,5,12,8,5,3, | 2,4,0,2,2,0,0,0,1,0,4,0,1,0, | 235 | 149 | 0.0000084457709139 | 0.00321471671916887 | 1.0,0.533,0.91,0.8 | 0.869,0.777,1.0,0. | -0,047 |
| 6964 | ENSG00000124207 | CSE1L | chr20 | + | 49085282 | 49085386 | 49084025 | 49084162 | 49089246 | 49089397 | 6964 | 29,24,28,17,4,13,5,11,13,1 | 0,0,2,0,0,1,2,1,0,0,0,0 | 36,15,28,3,13,17,12,18,12, | 1,2,0,0,0,1,2,3,1,1,4,2,5,0, | 253 | 149 | 0.0000084538456962 | 0.00321471671916887 | 1.0,1.0,0.892,1.0,1 | 0.955,0.815,1.0,1. | 0,094 |
| 113451 | ENSG00000104325 | DECR1 | chr8 | + | 90017123 | 90017326 | 90001476 | 90001561 | 90018908 | 90018966 | 113451 | 75,50,137,129,66,44,54,46 | 2,0,8,8,0,5,1,0,0,0,0,2 | 59,61,58,23,38,36,16,41,2 | 8,4,4,0,4,1,0,1,5,11,14,27, | 298 | 149 | 0.0000087029047670 | 0.00330358888169531 | 0.949,1.0,0.895,0. | 0.787,0.884,0.879, | 0,102 |
| 128135 | ENSG00000119541 | VPS4B | chr18 | - | 63399241 | 63399323 | 63393408 | 63393549 | 63400047 | 63400196 | 128135 | 10,6,29,26,6,14,6,9,2,5,1,1 | 0,0,3,0,0,1,0,0,0,0,0,0 | 22,34,7,11,12,8,10,13,7,16 | 0,4,0,3,3,0,1,3,1,4,0,3,4,0, | 231 | 149 | 0.0000087757180007 | 0.00332536366375901 | 1.0,1.0,0.862,1.0,1 | 1.0,0.846,1.0,0.70 | 0,122 |
| 30584 | ENSG00000181929 | PRKAG1 | chr12 | - | 49005742 | 49005881 | 49005461 | 49005543 | 49013061 | 49013110 | 30584 | 5,6,41,33,16,12,8,8,3,1,1,1 | 1,0,4,1,0,2,2,2,0,0,0,0 | 15,12,15,1,9,8,8,8,7,17,11, | 2,2,1,1,3,1,5,0,0,0,2,4,3,1, | 288 | 149 | 0.0000088818459743 | 0.00335723832711241 | 0.721,1.0,0.841,0. | 0.795,0.756,0.886, | 0,154 |
| 33805 | ENSG00000087266 | SH3BP2 | chr4 | + | 2823394 | 2823569 | 2822934 | 2823037 | 2824612 | 2824730 | 33805 | 7,3,7,6,11,4,4,0,5,0,4,0 | 82,22,50,35,42,29,26,19,9, | 8,0,2,1,0,0,0,5,1,0,4,2,1,1, | 45,37,29,9,32,19,18,41,11, | 298 | 149 | 0.0000088910326414 | 0.00335723832711241 | 0.041,0.064,0.065, | 0.082,0.0,0.033,0. | 0,047 |
| 91892 | ENSG00000204568 | MRPS18B | chr6 | + | 30622831 | 30622898 | 30619492 | 30619989 | 30624882 | 30624942 | 91892 | 29,11,83,87,43,24,20,36,2 | 0,0,12,7,3,0,0,3,0,0,0,5 | 34,60,29,5,16,35,28,30,10, | 1,0,0,0,3,0,3,0,0,1,1,2,0,2, | 216 | 149 | 0.0000089205108603 | 0.00336247018034594 | 1.0,1.0,0.827,0.89 | 0.959,1.0,1.0,1.0,0 | -0,005 |
| 87976 | ENSG00000135930 | EIF4E2 | chr2 | + | 232574262 | 232574341 | 232567077 | 232567214 | 232580903 | 232581332 | 87976 | 2,0,7,7,0,0,0,0,0,0,0,0 | 9,9,29,13,17,9,11,12,11,7, | 4,0,4,0,5,6,5,1,0,4,0,3,2,6, | 16,17,5,2,3,1,8,3,5,6,10,11 | 228 | 149 | 0.0000089479546362 | 0.00336691822432267 | 0.127,0.0,0.136,0. | 0.14,0.0,0.343,0.0, | -0,166 |
| 189262 | ENSG00000118308 | IRAG2 | chr12 | + | 25076522 | 25076640 | 25069349 | 25069431 | 25079243 | 25079290 | 189262 | 70,30,110,35,12,30,33,46, | 465,206,605,585,254,214, | 50,93,58,46,20,3,13,21,27, | 289,259,273,78,186,115,6 | 267 | 149 | 0.0000090609720566 | 0.00340349402570155 | 0.077,0.075,0.092, | 0.088,0.167,0.106, | -0,04 |
| 233405 | ENSG00000183354 | KIAA2026 | chr9 | - | 5954015 | 5954118 | 5944870 | 5944960 | 5968018 | 5969436 | 233405 | 26,15,29,12,6,9,11,12,18,1 | 2,3,4,2,1,5,4,2,0,0,1,0 | 12,16,11,26,5,4,1,15,2,8,4, | 0,1,2,1,3,1,0,0,1,0,0,0,0,0, | 252 | 149 | 0.0000092132162132 | 0.0034546511128928 | 0.885,0.747,0.811, | 1.0,0.904,0.765,0. | -0,078 |
| 37580 | ENSG00000150961 | SEC24D | chr4 | - | 118766528 | 118766823 | 118764801 | 118764917 | 118768172 | 118768311 | 37580 | 4,2,0,0,0,3,3,0,0,0,0,2 | 12,6,10,23,9,6,11,12,15,6, | 3,2,0,8,3,7,4,1,3,1,2,0,3,0, | 14,15,6,4,8,9,6,11,9,9,8,14 | 298 | 149 | 0.0000092368683296 | 0.00345749636079808 | 0.143,0.143,0.0,0. | 0.097,0.062,0.0,0. | -0,058 |
| 58150 | ENSG00000076770 | MBNL3 | chrX | - | 132439434 | 132439572 | 132406227 | 132406392 | 132489813 | 132489968 | 58150 | 7,12,13,13,7,8,4,13,4,2,1,9 | 0,0,5,1,0,0,2,7,1,1,0,6 | 8,4,3,9,10,3,2,5,4,1,1,13,9, | 1,0,1,0,1,0,0,0,1,0,4,1,0,1, | 287 | 149 | 0.0000093084262738 | 0.00347823245721461 | 1.0,1.0,0.574,0.87 | 0.806,1.0,0.609,1. | -0,073 |
| 32552 | ENSG00000138744 | NAAA | chr4 | - | 75918760 | 75918789 | 75914867 | 75914985 | 75919908 | 75919975 | 32552 | 8,11,29,19,18,14,15,13,18, | 1,1,5,0,11,3,1,3,2,10,1,2 | 15,6,7,2,10,2,7,12,4,10,13, | 0,1,0,0,2,4,0,0,0,4,0,4,0,0, | 178 | 149 | 0.0000094120532472 | 0.00351085898172367 | 0.87,0.902,0.829,1 | 1.0,0.834,1.0,1.0,0 | -0,073 |
| 73818 | ENSG00000231424 | | chr1 | - | 171247580 | 171247727 | 171168103 | 171168188 | 171251522 | 171251832 | 73818 | 152,65,264,72,93,163,91,3 | 16,8,108,25,19,47,2,0,19,7 | 255,150,75,117,168,103,4, | 0,0,0,0,0,0,29,50,8,17,0,0, | 296 | 149 | 0.0000095758885864 | 0.00356174696741827 | 0.827,0.804,0.552, | 1.0,1.0,1.0,1.0,1.0, | -0,144 |
| 183472 | ENSG00000171824 | EXOSC10 | chr1 | - | 11082731 | 11082878 | 11081081 | 11081238 | 11087447 | 11087591 | 183472 | 45,28,69,42,29,27,49,36,2 | 6,8,2,6,4,1,4,2,0,0,0,0 | 47,74,21,20,41,20,20,41,2 | 5,6,11,12,7,4,7,6,5,3,12,9, | 296 | 149 | 0.0000095815727945 | 0.00356174696741827 | 0.791,0.638,0.946, | 0.826,0.861,0.49,0 | 0,186 |
| 187804 | ENSG00000118246 | FASTKD2 | chr2 | + | 206786736 | 206786899 | 206774224 | 206774397 | 206787936 | 206788155 | 187804 | 16,11,41,39,15,25,7,20,8,5 | 0,1,4,4,0,2,2,0,0,0,0,0 | 28,30,22,14,15,9,19,15,12, | 3,0,1,1,8,0,2,2,7,2,0,1,0,1, | 298 | 149 | 0.0000096123052034 | 0.00356688642186505 | 1.0,0.846,0.837,0. | 0.824,1.0,0.917,0. | 0,075 |
| 230354 | ENSG00000150787 | PTS | chr11 | + | 112233431 | 112233486 | 112233162 | 112233233 | 112235848 | 112236034 | 230354 | 2,1,16,7,2,2,2,0,0,0,0,2 | 2,0,6,1,0,0,0,2,4,1,1,3 | 10,10,8,4,5,2,3,2,4,5,7,8,5, | 4,1,3,1,1,0,0,0,0,1,0,0,0,3, | 204 | 149 | 0.0000096285433376 | 0.00356688642186505 | 0.422,1.0,0.661,0. | 0.646,0.88,0.661,0 | -0,345 |
| 156168 | ENSG00000068650 | ATP11A | chr13 | + | 112818487 | 112818602 | 112816082 | 112816211 | 112819303 | 112819407 | 156168 | 0,0,0,0,0,2,1,0,0,0,0,0 | 4,13,5,11,6,14,13,15,13,9, | 0,6,2,8,7,0,0,2,6,0,0,2,0,0, | 4,19,10,20,17,14,6,26,3,11 | 264 | 149 | 0.0000096508754037 | 0.00356901643302072 | 0.0,0.0,0.0,0.0,0.0, | 0.0,0.151,0.101,0. | -0,074 |
| 71640 | ENSG00000156515 | HK1 | chr10 | + | 69398594 | 69398828 | 69394949 | 69395105 | 69400990 | 69401443 | 71640 | 66,28,86,58,62,54,47,39,5 | 10,5,8,2,0,1,4,2,0,1,0,2 | 62,61,22,20,25,32,24,48,1 | 4,4,0,0,0,0,0,0,0,3,2,2,0,0, | 298 | 149 | 0.0000097055843087 | 0.00358309196632106 | 0.767,0.737,0.843, | 0.886,0.884,1.0,1. | -0,045 |
| 230459 | ENSG00000134882 | UBAC2 | chr13 | + | 99279324 | 99279383 | 99244514 | 99244624 | 99314096 | 99314220 | 230459 | 2,4,2,0,2,3,0,5,1,2,4,0 | 36,26,52,18,33,31,37,28,1 | 4,8,6,6,7,2,10,0,4,0,3,7,3,3 | 40,29,30,21,18,33,17,31,1 | 208 | 149 | 0.0000097595200794 | 0.00359683435995487 | 0.038,0.099,0.027, | 0.067,0.165,0.125, | -0,068 |
| 13963 | ENSG00000167508 | MVD | chr16 | - | 88660519 | 88660637 | 88658649 | 88658720 | 88663010 | 88663079 | 13963 | 3,4,1,5,4,0,12,6,5,4,2,0 | 23,9,26,53,36,25,31,24,39, | 3,0,0,0,0,1,0,2,2,1,2,4,4,1, | 10,21,2,7,13,13,6,12,11,25 | 267 | 149 | 0.0000097866648561 | 0.00360010205388961 | 0.068,0.199,0.021, | 0.143,0.0,0.0,0.0,0 | 0,05 |
| 159389 | ENSG00000127603 | MACF1 | chr1 | + | 39315512 | 39315695 | 39310830 | 39311000 | 39316390 | 39316529 | 159389 | 25,11,53,59,26,30,14,9,10, | 3,0,4,4,1,0,1,1,0,0,0,0 | 55,32,13,19,21,11,6,20,32, | 5,7,0,3,3,0,0,4,2,6,1,8,8,2, | 298 | 149 | 0.0000098018399002 | 0.00360010205388961 | 0.806,1.0,0.869,0. | 0.846,0.696,1.0,0. | 0,099 |
| 106965 | ENSG00000086758 | HUWE1 | chrX | - | 53544559 | 53544762 | 53543709 | 53543968 | 53546435 | 53546592 | 106965 | 24,16,26,26,16,14,13,11,1 | 2,0,1,3,1,2,0,0,2,0,1,0 | 24,24,15,8,15,14,13,18,7,2 | 8,2,0,5,4,2,1,0,0,2,3,1,0,0, | 298 | 149 | 0.0000098247348400 | 0.00360236372120986 | 0.857,1.0,0.929,0. | 0.6,0.857,1.0,0.44 | 0,09 |
| 205875 | ENSG00000128815 | WDFY4 | chr10 | + | 48808288 | 48808369 | 48807858 | 48807958 | 48810529 | 48810735 | 205875 | 8,1,6,3,0,4,2,4,2,2,7,4 | 70,39,99,83,83,71,44,66,5 | 7,14,2,34,3,0,8,5,1,2,10,11 | 121,89,26,41,33,20,69,85, | 230 | 149 | 0.0000098450757126 | 0.00360368280733618 | 0.069,0.016,0.038, | 0.036,0.092,0.047, | -0,036 |
| 236747 | ENSG00000067167 | TRAM1 | chr8 | - | 70598133 | 70598207 | 70597890 | 70598011 | 70600018 | 70600082 | 236747 | 153,81,182,85,57,42,41,74 | 1,0,1,1,1,1,1,0,0,0,0,0 | 90,88,73,49,44,49,30,46,2 | 2,2,0,0,1,0,2,1,1,6,0,14,2,4 | 223 | 149 | 0.0000098877218289 | 0.00361314814426322 | 0.99,1.0,0.992,0.9 | 0.968,0.967,1.0,1. | 0,035 |
| 189200 | ENSG00000118308 | IRAG2 | chr12 | + | 25062166 | 25062476 | 25061591 | 25061653 | 25062819 | 25062900 | 189200 | 0,0,33,28,7,20,0,4,8,6,6,6 | 208,78,469,405,181,94,18 | 20,40,42,12,12,4,4,14,12,1 | 169,249,150,55,67,53,65,1 | 298 | 149 | 0.0000099132473735 | 0.0036163358397532 | 0.0,0.0,0.034,0.03 | 0.056,0.074,0.123, | -0,019 |
| 155536 | ENSG00000173575 | CHD2 | chr15 | + | 92991346 | 92991833 | 92985497 | 92985673 | 92992858 | 92992998 | 155536 | 18,5,8,8,2,7,9,11,3,1,1,8 | 0,2,0,1,0,0,0,0,0,0,0,3 | 2,18,7,16,12,5,9,2,13,5,7,4 | 4,2,2,12,3,0,0,0,1,2,1,0,1,1 | 298 | 149 | 0.0000099510185686 | 0.00362397238162156 | 1.0,0.556,1.0,0.8,1 | 0.2,0.818,0.636,0. | 0,148 |
| 206847 | ENSG00000075884 | ARHGAP15 | chr2 | + | 143734644 | 143734731 | 143703418 | 143703524 | 143767988 | 143768352 | 206847 | 2,1,5,6,4,4,4,2,1,0,0,6 | 75,50,155,97,69,55,50,45, | 9,10,6,8,4,7,4,7,1,5,6,6,1,8 | 94,64,66,18,62,43,14,32,3 | 236 | 149 | 0.0000100925826051 | 0.00366931865992952 | 0.017,0.012,0.02,0 | 0.057,0.09,0.054,0 | -0,049 |
| 53333 | ENSG00000196187 | TMEM63A | chr1 | - | 225866573 | 225866682 | 225865896 | 225865967 | 225867887 | 225868030 | 53333 | 25,31,13,18,6,7,9,8,22,19, | 0,0,1,2,3,0,0,5,0,0,0,1 | 11,8,3,7,6,12,12,16,6,10,4, | 4,1,4,6,1,0,0,4,1,1,0,0,1,2, | 258 | 149 | 0.000010178147021 | 0.00369418678157985 | 1.0,1.0,0.882,0.83 | 0.614,0.822,0.302, | 0,122 |
| 198029 | ENSG00000112232 | KHDRBS2 | chr6 | - | 61700735 | 61700777 | 61697194 | 61697253 | 61707329 | 61707403 | 198029 | 13,2,13,11,6,11,1,0,6,3,1,2 | 4,1,2,13,1,0,5,3,2,1,0,0 | 9,1,11,5,0,0,0,1,0,0,3,0,1,0 | 3,3,1,0,5,1,3,5,3,1,8,1,2,3, | 191 | 149 | 0.0000102214443005 | 0.0037036560239746 | 0.717,0.609,0.835, | 0.701,0.206,0.896, | 0,39 |
| 256203 | ENSG00000112297 | CRYBG1 | chr6 | + | 106544756 | 106544933 | 106544583 | 106544697 | 106551851 | 106551978 | 256203 | 78,77,96,41,25,13,30,39,5 | 1,1,1,1,5,0,0,2,2,0,1,0 | 51,81,70,83,55,31,17,32,4 | 0,0,5,7,6,2,2,1,2,2,1,4,0,3, | 298 | 149 | 0.0000103378292957 | 0.00373953165906354 | 0.975,0.975,0.98,0 | 1.0,1.0,0.875,0.85 | 0,039 |
| 102620 | ENSG00000122566 | HNRNPA2B1 | chr7 | - | 26192300 | 26192338 | 26190992 | 26191128 | 26192494 | 26192577 | 102620 | 49,19,35,35,25,18,22,20,2 | 11,5,18,17,12,8,10,28,8,1, | 11,29,6,10,3,25,6,28,3,10, | 23,33,13,20,8,14,7,24,21,1 | 187 | 149 | 0.0000104810357749 | 0.00378497283702601 | 0.78,0.752,0.608,0 | 0.276,0.412,0.269, | 0,248 |
| 185348 | ENSG00000188822 | CNR2 | chr1 | - | 23883211 | 23883444 | 23870514 | 23875662 | 23913245 | 23913362 | 185348 | 11,8,16,2,5,0,5,9,3,2,5,1 | 72,36,191,152,96,74,66,90 | 0,4,2,2,0,3,2,4,3,8,3,3,0,3, | 135,99,48,64,73,71,22,84, | 298 | 149 | 0.0000106478715229 | 0.00383402488769954 | 0.071,0.1,0.04,0.0 | 0.0,0.02,0.02,0.01 | 0,021 |
| 24164 | ENSG00000253710 | ALG11 | chr13 | + | 52024585 | 52024937 | 52018912 | 52019143 | 52028318 | 52028689 | 24164 | 15,21,39,10,11,7,10,13,8,7 | 3,3,1,0,0,2,1,2,0,0,0,0 | 7,16,10,15,14,8,9,9,3,10,1 | 3,0,0,0,0,3,1,2,0,3,1,10,0,6 | 298 | 149 | 0.0000106524937525 | 0.00383402488769954 | 0.714,0.778,0.951, | 0.538,1.0,1.0,1.0,1 | 0,086 |
| 244013 | ENSG00000175216 | CKAP5 | chr11 | - | 46797617 | 46797713 | 46796811 | 46796940 | 46797804 | 46797969 | 244013 | 2,0,8,9,0,1,0,2,4,0,0,1 | 3,1,23,15,4,13,2,12,4,2,3,9 | 0,3,0,3,0,2,2,3,0,0,0,3,0,2, | 30,13,2,1,6,6,6,17,6,4,6,19 | 245 | 149 | 0.0000108457252798 | 0.00389705558880907 | 0.288,0.0,0.175,0. | 0.0,0.123,0.0,0.64 | 0,022 |
| 31886 | ENSG00000247774 | PCED1B-AS1 | chr12 | - | 47209280 | 47209458 | 47205897 | 47206588 | 47210639 | 47210757 | 31886 | 28,20,32,22,39,25,19,17,2 | 0,0,0,1,2,0,2,0,1,1,0,0 | 25,22,30,9,12,14,5,14,15,2 | 1,4,0,1,0,3,2,2,1,1,0,13,2,1 | 298 | 149 | 0.0000108661105703 | 0.00389787307359373 | 1.0,1.0,1.0,0.917,0 | 0.926,0.733,1.0,0. | 0,089 |
| 223426 | ENSG00000106392 | C1GALT1 | chr7 | + | 7215119 | 7215239 | 7182629 | 7182820 | 7234302 | 7235297 | 223426 | 1,0,3,0,0,1,0,2,0,0,0,0 | 18,7,39,15,22,18,8,11,16,6 | 2,2,0,0,3,3,0,0,2,3,3,20,2,0 | 27,27,8,13,10,7,2,11,10,19 | 269 | 149 | 0.0000108948953994 | 0.00390169589302539 | 0.03,0.0,0.041,0.0, | 0.039,0.039,0.0,0. | -0,043 |
| 134095 | ENSG00000100201 | DDX17 | chr22 | - | 38489222 | 38489458 | 38488025 | 38488115 | 38492055 | 38492115 | 134095 | 30,16,16,7,16,10,15,21,40, | 223,109,242,214,130,97,1 | 26,16,20,40,3,13,11,19,9,2 | 123,52,139,40,64,84,60,14 | 298 | 149 | 0.0000109266169103 | 0.00390655595385511 | 0.063,0.068,0.032, | 0.096,0.133,0.067, | -0,035 |
| 218936 | ENSG00000204592 | HLA-E | chr6 | + | 30492403 | 30492429 | 30491536 | 30491653 | 30492540 | 30492583 | 218936 | 179,147,128,135,35,31,60, | 10,5,8,15,4,0,5,9,0,3,2,9 | 73,77,80,55,52,33,36,94,3 | 10,13,4,14,16,5,2,21,5,6,8, | 175 | 149 | 0.0000110486124941 | 0.00394362175077551 | 0.938,0.962,0.932, | 0.861,0.835,0.945, | 0,064 |
| 134161 | ENSG00000100083 | GGA1 | chr22 | + | 37619799 | 37619982 | 37618402 | 37618546 | 37620237 | 37620361 | 134161 | 5,1,10,1,2,8,0,4,13,2,0,1 | 75,49,104,62,43,40,50,56, | 1,2,0,0,2,0,0,0,0,3,3,0,5,4, | 36,49,39,36,37,43,30,34,3 | 298 | 149 | 0.0000110844593678 | 0.00394577837521101 | 0.032,0.01,0.046,0 | 0.014,0.02,0.0,0.0, | 0,021 |
| 164444 | ENSG00000005700 | IBTK | chr6 | - | 82215911 | 82216189 | 82214226 | 82214829 | 82217959 | 82218137 | 164444 | 6,3,13,3,1,8,4,4,1,1,0,1 | 0,0,5,0,0,0,0,1,1,0,2,1 | 8,10,3,1,5,5,0,2,2,0,4,3,3,1 | 2,5,1,5,1,1,2,0,2,2,0,1,0,0, | 298 | 149 | 0.000011104351144 | 0.00394577837521101 | 1.0,1.0,0.565,1.0,1 | 0.667,0.5,0.6,0.09 | 0,144 |
| 201479 | ENSG00000131127 | ZNF141 | chr4 | + | 342271 | 342426 | 337833 | 337986 | 343781 | 343908 | 201479 | 1,0,17,9,0,0,1,4,0,0,0,1 | 8,9,98,59,25,22,11,32,20,8 | 0,3,0,0,0,0,0,0,0,0,1,3,0, | 31,34,9,20,26,17,18,19,13, | 298 | 149 | 0.0000111096528631 | 0.00394577837521101 | 0.059,0.0,0.08,0.0 | 0.0,0.042,0.0,0.0,0 | 0,022 |
| 132133 | ENSG00000170456 | DENND5B | chr12 | - | 31447537 | 31447769 | 31442774 | 31442925 | 31451939 | 31452476 | 132133 | 33,57,145,87,71,30,35,37, | 0,1,2,0,0,0,0,2,0,0,4,0 | 64,80,65,118,81,59,24,85, | 3,1,2,5,1,0,0,1,2,5,5,8,1,2, | 298 | 149 | 0.0000112104520245 | 0.00397501943934952 | 1.0,0.966,0.973,1. | 0.914,0.976,0.942, | 0,021 |
| 149230 | ENSG00000128881 | TTBK2 | chr15 | - | 42871444 | 42871617 | 42840359 | 42840433 | 42872610 | 42872758 | 149230 | 0,0,0,1,2,3,1,0,0,0,0,0 | 12,8,19,6,16,2,4,13,8,6,8,9 | 4,0,1,0,3,1,2,2,0,5,7,2,1,4, | 18,8,7,14,5,2,4,18,5,17,15, | 298 | 149 | 0.0000112779771032 | 0.00399238534522835 | 0.0,0.0,0.0,0.077,0 | 0.1,0.0,0.067,0.0,0 | -0,039 |
| 12166 | ENSG00000063244 | U2AF2 | chr19 | + | 55658861 | 55658981 | 55655108 | 55655153 | 55659209 | 55659345 | 12166 | 8,0,9,12,2,0,5,0,4,1,3,0 | 19,15,31,5,14,10,23,19,15, | 6,0,0,0,1,2,0,5,3,1,3,0,0,0, | 14,15,7,13,12,6,13,7,22,22 | 269 | 149 | 0.0000113881016066 | 0.00402474958438444 | 0.189,0.0,0.139,0. | 0.192,0.0,0.0,0.0,0 | 0,066 |
| 160925 | ENSG00000164329 | TENT2 | chr5 | + | 79679578 | 79679634 | 79668720 | 79669028 | 79681981 | 79682061 | 160925 | 16,17,40,22,10,19,7,17,10, | 2,0,1,0,0,1,0,3,1,0,0,0 | 25,23,13,24,16,9,15,12,6,2 | 2,1,3,10,3,0,0,3,0,2,4,1,1,0 | 205 | 149 | 0.0000117159185372 | 0.00413381780767228 | 0.853,1.0,0.967,1. | 0.901,0.944,0.759, | 0,069 |
| 138218 | ENSG00000130818 | ZNF426 | chr19 | - | 9535425 | 9535520 | 9535187 | 9535279 | 9536207 | 9536569 | 138218 | 4,2,2,0,0,6,3,1,3,0,0,2 | 11,15,54,30,13,21,20,26,1 | 0,9,1,6,3,0,2,9,1,6,3,2,9,1, | 18,48,17,13,15,3,23,19,11, | 244 | 149 | 0.0000117426021619 | 0.0041364517281635 | 0.182,0.075,0.022, | 0.0,0.103,0.035,0. | -0,08 |
| 97961 | ENSG00000166135 | HIF1AN | chr10 | + | 100540633 | 100540782 | 100536410 | 100536661 | 100544950 | 100545096 | 97961 | 54,28,51,50,26,30,30,22,2 | 2,0,6,0,0,0,1,1,2,1,0,2 | 23,47,28,10,22,33,21,27,1 | 2,8,2,1,7,2,6,0,2,2,0,3,0,0, | 298 | 149 | 0.000011838658451 | 0.00416347434161304 | 0.931,1.0,0.81,1.0, | 0.852,0.746,0.875, | 0,087 |
| 234009 | ENSG00000008294 | SPAG9 | chr17 | - | 50996639 | 50996694 | 50995443 | 50995533 | 50998443 | 50998617 | 234009 | 13,6,41,13,6,6,4,16,11,5,9, | 0,0,0,5,0,3,0,0,0,0,1,3 | 24,14,8,6,10,6,4,7,8,11,7,1 | 0,4,0,5,5,1,0,3,0,1,2,0,0,1, | 204 | 149 | 0.0000119539134682 | 0.00419714967320416 | 1.0,1.0,1.0,0.655,1 | 1.0,0.719,1.0,0.46 | 0,071 |
| 239346 | ENSG00000153339 | TRAPPC8 | chr18 | - | 31913403 | 31913522 | 31909666 | 31909760 | 31916271 | 31916446 | 239346 | 24,24,56,47,29,28,18,46,1 | 2,0,10,4,2,6,1,2,0,0,2,5 | 41,40,20,18,28,26,13,30,1 | 2,6,1,1,1,0,1,0,0,2,0,1,2, | 268 | 149 | 0.0000120084486305 | 0.00420943063060447 | 0.87,1.0,0.757,0.8 | 0.919,0.788,0.917, | -0,068 |
| 1748 | ENSG00000284554 | | chr22 | + | 39025069 | 39025349 | 39022821 | 39023014 | 39025556 | 39025671 | 1748 | 12,4,17,27,31,12,15,20,14, | 2,3,2,3,5,3,2,3,0,0,0,0 | 17,13,13,13,15,20,5,8,6,12 | 2,0,2,5,0,0,0,0,0,2,3,2,1, | 298 | 149 | 0.000012060005689 | 0.00422062940560838 | 0.75,0.4,0.81,0.81 | 0.81,1.0,0.765,0.5 | -0,061 |
| 183748 | ENSG00000112531 | QKI | chr6 | + | 163565945 | 163566067 | 163563419 | 163563719 | 163566720 | 163566795 | 183748 | 0,0,3,4,0,2,0,2,1,1,0,0 | 47,22,35,26,37,25,18,37,4 | 2,3,2,4,0,0,3,4,0,0,4,17,3,1 | 52,62,25,39,12,9,38,58,8,3 | 271 | 149 | 0.0000120924234117 | 0.00422510451838247 | 0.0,0.0,0.045,0.07 | 0.021,0.026,0.042, | -0,018 |
| 146260 | ENSG00000140968 | IRF8 | chr16 | + | 85913130 | 85913242 | 85909486 | 85911658 | 85914472 | 85914520 | 146260 | 447,257,493,363,203,158, | 0,4,7,4,6,9,4,4,2,1,1,1 | 237,288,304,171,174,128, | 7,11,5,7,4,3,6,1,2,3,9,21,2, | 261 | 149 | 0.0000121448622605 | 0.00423654918831389 | 1.0,0.973,0.976,0. | 0.951,0.937,0.972, | 0,019 |
| 100822 | ENSG00000127914 | AKAP9 | chr7 | + | 91980872 | 91980958 | 91980288 | 91980333 | 91992157 | 91992211 | 100822 | 4,10,10,0,2,4,1,8,1,0,2,1 | 85,65,54,30,30,25,14,25,4 | 7,14,8,18,3,1,2,7,3,5,10,5, | 40,30,23,43,17,12,12,20,2 | 235 | 149 | 0.0000122733483346 | 0.00427444180486131 | 0.029,0.089,0.105, | 0.1,0.228,0.181,0. | -0,067 |
| 137149 | ENSG00000170889 | RPS9 | chr19 | + | 54206552 | 54206724 | 54206275 | 54206462 | 54207370 | 54207522 | 137149 | 56,32,72,52,19,23,17,41,3 | 0,2,3,5,4,3,0,0,2,2,0,2 | 17,34,27,14,30,22,15,18,2 | 0,0,0,2,0,0,0,0,5,0,0,0,1, | 298 | 149 | 0.0000124166448887 | 0.00431736170604166 | 1.0,0.889,0.923,0. | 1.0,1.0,1.0,0.778,1 | -0,071 |
| 163186 | ENSG00000247774 | PCED1B-AS1 | chr12 | - | 47209307 | 47209458 | 47206557 | 47206588 | 47216164 | 47216251 | 163186 | 2,6,15,7,4,4,1,2,1,7,0,4 | 0,1,1,1,1,2,0,0,0,0,2,3 | 13,9,7,6,12,3,0,6,3,5,1,40, | 2,6,2,2,0,3,3,1,0,4,2,33,4,3 | 298 | 149 | 0.0000125391332811 | 0.00435291966971682 | 1.0,0.75,0.882,0.7 | 0.765,0.429,0.636, | 0,239 |
| 147075 | ENSG00000166747 | AP1G1 | chr16 | - | 71772964 | 71773053 | 71771159 | 71771252 | 71773220 | 71773362 | 147075 | 1,3,4,0,1,4,1,3,1,2,1,0 | 74,85,93,55,39,47,34,43,4 | 5,9,5,4,16,4,0,2,2,1,5,1,1,1 | 34,67,57,64,60,35,34,35,3 | 238 | 149 | 0.0000126178190284 | 0.00437318165426982 | 0.008,0.022,0.026, | 0.084,0.078,0.052, | -0,028 |
| 14586 | ENSG00000168488 | ATXN2L | chr16 | + | 28826861 | 28826986 | 28826239 | 28826390 | 28829400 | 28829492 | 14586 | 87,84,73,78,69,67,46,51,4 | 2,2,8,5,0,1,6,3,0,4,0,3 | 51,45,26,76,22,42,48,35,4 | 10,7,1,15,6,3,5,6,6,2,2,9,1, | 274 | 149 | 0.0000126899504442 | 0.00439111048883538 | 0.959,0.958,0.832, | 0.735,0.778,0.934, | 0,094 |
| 146262 | ENSG00000140968 | IRF8 | chr16 | + | 85913130 | 85913371 | 85909486 | 85911658 | 85914472 | 85914520 | 146262 | 447,258,495,364,205,159, | 0,4,7,4,6,9,4,4,2,1,1,1 | 239,288,307,171,174,128, | 7,11,5,7,4,3,6,1,2,3,9,21,2, | 298 | 149 | 0.0000127175442275 | 0.00439359512300008 | 1.0,0.97,0.972,0.9 | 0.945,0.929,0.968, | 0,021 |
| 111798 | ENSG00000164989 | CCDC171 | chr9 | + | 15594040 | 15594172 | 15591365 | 15591556 | 15623266 | 15623413 | 111798 | 21,18,19,10,6,10,2,7,8,6,2, | 6,0,9,2,3,0,1,4,0,0,0,0 | 4,6,7,3,1,5,6,5,1,17,6,5,8,5 | 0,2,0,0,1,0,0,0,5,0,0,0,3,0, | 281 | 149 | 0.0000127472935026 | 0.00439681526900336 | 0.65,1.0,0.528,0.7 | 1.0,0.614,1.0,1.0,0 | -0,068 |
| 131741 | ENSG00000137955 | RABGGTB | chr1 | + | 75789158 | 75789356 | 75786700 | 75787604 | 75791284 | 75791337 | 131741 | 21,37,64,42,12,13,10,30,1 | 3,2,1,6,0,0,1,0,2,2,0,8 | 21,40,14,9,29,19,7,7,8,20, | 5,1,0,4,1,0,1,0,0,0,0,0,4, | 298 | 149 | 0.0000128734108374 | 0.00442883402206064 | 0.778,0.902,0.97,0 | 0.677,0.952,1.0,0. | -0,023 |
| 146264 | ENSG00000140968 | IRF8 | chr16 | + | 85913130 | 85913478 | 85909486 | 85911658 | 85914472 | 85914520 | 146264 | 447,258,495,364,205,159, | 0,4,7,4,6,9,4,4,2,1,1,1 | 239,288,307,171,174,128, | 7,11,5,7,4,3,6,1,2,3,9,21,2, | 298 | 149 | 0.0000128812768505 | 0.00442883402206064 | 1.0,0.97,0.972,0.9 | 0.945,0.929,0.968, | 0,021 |
| 195104 | ENSG00000185436 | IFNLR1 | chr1 | - | 24159136 | 24159182 | 24157098 | 24157891 | 24159473 | 24159633 | 195104 | 2,0,3,4,5,1,4,7,1,1,1,5 | 29,7,34,49,26,30,31,44,37, | 5,10,7,1,5,2,7,2,0,9,12,2,9, | 22,27,30,16,11,6,12,17,4,1 | 195 | 149 | 0.000012911892166 | 0.00443227984494473 | 0.05,0.0,0.063,0.0 | 0.148,0.221,0.151, | -0,101 |
| 220591 | ENSG00000179344 | HLA-DQB1 | chr6 | - | 32666103 | 32666333 | 32664797 | 32665067 | 32666498 | 32666684 | 220591 | 99,34,0,0,1,9,24,30,6,0,16, | 832,609,117,78,351,145,8 | 21,34,0,0,0,0,0,0,0,0,32,31 | 1034,1216,14,7,7,4,38,52, | 298 | 149 | 0.0000129814769601 | 0.00444907048980777 | 0.056,0.027,0.0,0. | 0.01,0.014,0.0,0.0, | 0,012 |
| 172685 | ENSG00000068745 | IP6K2 | chr3 | - | 48699516 | 48699788 | 48695127 | 48695421 | 48717156 | 48717193 | 172685 | 1,0,1,1,0,0,1,1,0,0,1,2 | 17,8,17,10,5,10,8,4,11,7,2, | 7,4,0,0,4,3,2,3,0,0,0,3,1,1, | 6,8,8,2,4,8,4,2,4,3,16,25,11 | 298 | 149 | 0.0000130652535442 | 0.00447066388803133 | 0.029,0.0,0.029,0. | 0.368,0.2,0.0,0.0,0 | -0,083 |
| 153164 | ENSG00000100902 | PSMA6 | chr14 | + | 35312895 | 35313059 | 35310832 | 35310895 | 35314360 | 35314395 | 153164 | 11,10,50,31,10,6,8,8,6,3,4, | 1,0,3,1,1,2,0,1,0,0,0,3 | 28,40,13,10,9,18,9,9,8,22,1 | 0,0,0,2,6,2,1,3,3,2,4,5,0,1, | 298 | 149 | 0.0000131738791164 | 0.00450067805730458 | 0.846,1.0,0.893,0. | 1.0,1.0,1.0,0.714,0 | 0,124 |
| 101533 | ENSG00000156831 | NSMCE2 | chr8 | + | 125102013 | 125102146 | 125091843 | 125091958 | 125102316 | 125102487 | 101533 | 18,8,37,33,9,29,13,32,3,4, | 1,1,5,10,0,0,0,3,2,2,2,4 | 28,30,10,8,20,9,6,13,21,26 | 0,4,0,0,1,1,1,2,0,1,1,2,2,0, | 282 | 149 | 0.0000132815477123 | 0.0045302706745896 | 0.905,0.809,0.796, | 1.0,0.798,1.0,1.0,0 | -0,127 |
| 189202 | ENSG00000118308 | IRAG2 | chr12 | + | 25062166 | 25062630 | 25061591 | 25061653 | 25062819 | 25062900 | 189202 | 0,0,33,28,7,20,0,4,8,6,6,6 | 208,78,469,405,181,94,18 | 20,40,44,12,12,4,4,11,12,1 | 169,249,150,55,67,53,65,1 | 298 | 149 | 0.0000133175138094 | 0.00453535097264394 | 0.0,0.0,0.034,0.03 | 0.056,0.074,0.128, | -0,019 |
| 241269 | ENSG00000107077 | KDM4C | chr9 | + | 6986464 | 6986666 | 6984280 | 6984404 | 6990415 | 6990524 | 241269 | 52,24,65,38,32,35,24,39,1 | 1,0,0,5,0,1,0,0,0,0,0,1 | 56,35,34,43,40,18,22,33,2 | 1,5,0,3,7,0,2,0,1,0,2,2,2,4, | 298 | 149 | 0.0000134347534396 | 0.00456804963279391 | 0.963,1.0,1.0,0.79 | 0.966,0.778,1.0,0. | 0,062 |
| 247598 | ENSG00000149483 | TMEM138 | chr11 | + | 61367427 | 61367525 | 61366044 | 61366216 | 61367922 | 61369161 | 247598 | 0,1,9,1,0,0,2,1,1,0,0,2 | 9,11,38,23,5,4,13,9,13,6,4, | 6,2,1,1,3,1,2,3,0,1,6,7,6,7, | 18,23,10,9,7,4,6,10,1,2,6,8 | 247 | 149 | 0.0000135549187688 | 0.0046016383620309 | 0.0,0.052,0.125,0. | 0.167,0.05,0.057,0 | -0,109 |
| 31110 | ENSG00000158987 | RAPGEF6 | chr5 | - | 131461704 | 131462088 | 131455800 | 131456012 | 131464040 | 131464281 | 31110 | 24,31,88,73,16,29,21,51,2 | 6,0,6,7,1,2,1,0,0,0,4,0 | 47,51,61,30,39,32,35,34,3 | 3,0,0,3,0,1,0,0,0,4,0,1,1,0, | 298 | 149 | 0.0000136763733852 | 0.00462695507336876 | 0.667,1.0,0.88,0.8 | 0.887,1.0,1.0,0.83 | -0,06 |

| | | | | | | | | | | | | | | | | | | | | | |
|---|---|---|---|---|---|---|---|---|---|---|---|---|---|---|---|---|---|---|---|---|---|
| 110606 | ENSG00000120899 | PTK2B | chr8 | + | 27451042 | 27451078 | 27450748 | 27450895 | 27451484 | 27451509 | 110606 | 63,21,85,69,64,65,51,76,7 | 2,3,11,11,0,1,11,15,5,1,4,5 | 62,34,38,33,44,21,35,70,2 | 0,2,3,3,0,0,2,4,0,0,17,6,0,0 | 185 | 149 | 0.000013682768539 | 0.00462695507336876 | 0.962,0.849,0.862, | 1.0,0.932,0.911,0. | -0,063 |
| 8249 | ENSG00000134444 | RELCH | chr18 | + | 62284238 | 62284361 | 62282315 | 62282444 | 62287350 | 62287467 | 8249 | 9,8,8,3,0,0,0,7,2,2,2,0 | 30,37,43,20,28,27,21,16,1 | 2,4,0,5,0,1,2,0,0,1,2,2,2,4, | 28,43,21,39,22,20,16,12,1 | 272 | 149 | 0.0000136939863762 | 0.00462695507336876 | 0.141,0.106,0.092, | 0.038,0.048,0.0,0. | 0,047 |
| 236855 | ENSG00000033800 | PIAS1 | chr15 | + | 68147118 | 68147208 | 68146661 | 68146700 | 68153589 | 68153961 | 236855 | 9,2,0,1,0,2,0,0,0,0,0,0 | 78,61,133,84,33,41,21,61, | 8,16,2,2,3,4,0,0,4,3,1,0,3,5 | 66,68,40,53,73,21,37,47,3 | 239 | 149 | 0.0000137973852899 | 0.00465458469174054 | 0.067,0.02,0.0,0.0 | 0.07,0.128,0.03,0. | -0,028 |
| 45322 | ENSG00000143612 | C1orf43 | chr1 | - | 154218715 | 154218785 | 154214456 | 154214574 | 154220341 | 154220578 | 45322 | 0,4,2,1,1,1,1,0,2,0,0,0 | 52,54,81,49,25,18,22,35,1 | 3,9,3,0,0,0,0,7,4,0,8,2,0,4, | 29,67,44,17,39,11,20,36,1 | 219 | 149 | 0.0000141966262451 | 0.00478177474704087 | 0.0,0.048,0.017,0. | 0.066,0.084,0.044, | -0,028 |
| 242432 | ENSG00000137145 | DENND4C | chr9 | + | 19357939 | 19358160 | 19356971 | 19357154 | 19360243 | 19360489 | 242432 | 15,11,45,39,16,15,2,9,7,6, | 4,3,2,0,0,2,0,7,1,3,0,0 | 23,11,7,4,3,7,23,16,5,26,1 | 1,1,2,0,0,0,1,1,3,0,3,2,2,0, | 298 | 149 | 0.0000142723598379 | 0.00479977231292352 | 0.652,0.647,0.918, | 0.92,0.846,0.636,1 | -0,061 |
| 172649 | ENSG00000168216 | LMBRD1 | chr6 | - | 69738401 | 69738473 | 69737941 | 69738015 | 69741788 | 69741877 | 172649 | 10,4,8,8,0,2,2,6,0,0,0,5 | 31,37,138,108,72,60,44,65 | 2,9,1,0,2,0,1,0,0,0,0,2,4,2, | 94,100,38,43,69,47,56,58, | 221 | 149 | 0.0000143295169104 | 0.00481147621551061 | 0.179,0.068,0.038, | 0.014,0.057,0.017, | 0,025 |
| 247601 | ENSG00000149483 | TMEM138 | chr11 | + | 61367427 | 61367818 | 61366044 | 61366216 | 61367922 | 61369161 | 247601 | 0,1,10,2,2,0,3,1,2,1,0,2 | 9,11,38,23,5,4,13,9,13,6,4, | 9,3,1,2,3,1,3,3,0,2,11,9,6,8 | 18,23,10,9,7,4,6,10,1,2,6,8 | 298 | 149 | 0.0000145397726928 | 0.0048744701190717 | 0.0,0.043,0.116,0. | 0.2,0.061,0.048,0. | -0,115 |
| 58700 | ENSG00000113532 | ST8SIA4 | chr5 | - | 100856102 | 100856396 | 100806932 | 100812129 | 100886342 | 100886600 | 58700 | 33,32,83,48,32,27,23,30,1 | 6,2,8,5,1,3,2,3,3,0,0,1 | 39,60,29,14,22,6,5,27,9,29 | 6,3,0,0,2,0,0,0,0,2,0,5,2,2, | 298 | 149 | 0.0000147004244827 | 0.00492066417081804 | 0.733,0.889,0.838, | 0.765,0.909,1.0,1. | -0,079 |
| 177509 | ENSG00000137818 | RPLP1 | chr15 | + | 69453333 | 69453721 | 69452883 | 69453020 | 69455169 | 69455287 | 177509 | 1169,845,889,1069,387,40 | 26,16,43,55,17,17,14,10,2 | 650,804,309,421,283,174, | 8,13,2,8,7,5,12,22,6,8,18,2 | 298 | 149 | 0.0000147462515953 | 0.00492626844801265 | 0.957,0.964,0.912, | 0.976,0.969,0.987, | -0,017 |
| 249867 | ENSG00000120438 | TCP1 | chr6 | - | 159780934 | 159781102 | 159780442 | 159780566 | 159783940 | 159784067 | 249867 | 186,167,207,169,88,74,79, | 39,7,3,13,7,12,8,6,4,2,2,2 | 112,200,108,60,127,54,55, | 22,20,21,34,13,2,4,15,10,1 | 298 | 149 | 0.0000147629437626 | 0.00492626844801265 | 0.705,0.923,0.972, | 0.718,0.833,0.72,0 | 0,098 |
| 249869 | ENSG00000120438 | TCP1 | chr6 | - | 159780934 | 159781130 | 159780442 | 159780566 | 159783940 | 159784067 | 249869 | 186,167,207,170,89,73,79, | 39,7,3,13,7,12,8,6,4,2,2,2 | 111,200,108,60,127,54,58, | 22,20,21,34,13,2,4,15,10,1 | 298 | 149 | 0.000014936264776 | 0.00497638886068615 | 0.705,0.923,0.972, | 0.716,0.833,0.72,0 | 0,098 |
| 127209 | ENSG00000135655 | USP15 | chr12 | + | 62325871 | 62325933 | 62321460 | 62321609 | 62349220 | 62349307 | 127209 | 107,94,202,75,82,99,85,11 | 2,0,3,1,0,2,0,2,0,0,1,3 | 109,171,101,103,101,56,8 | 4,6,3,7,0,1,3,2,3,0,8,5,2,3, | 211 | 149 | 0.0000149835271404 | 0.00498441967535615 | 0.974,1.0,0.979,0. | 0.951,0.953,0.96,0 | 0,026 |
| 241263 | ENSG00000107077 | KDM4C | chr9 | + | 6986324 | 6986666 | 6984280 | 6984404 | 6990415 | 6990524 | 241263 | 53,25,65,36,32,35,24,39,2 | 1,0,0,5,0,1,0,0,0,0,0,1 | 56,35,34,43,40,18,22,34,2 | 1,5,0,3,7,0,2,0,1,0,2,2,2,4, | 298 | 149 | 0.0000150456657018 | 0.00499736678188906 | 0.964,1.0,1.0,0.78 | 0.966,0.778,1.0,0. | 0,06 |
| 34817 | ENSG00000172869 | DMXL1 | chr5 | + | 119165182 | 119165280 | 119164506 | 119164676 | 119166615 | 119166781 | 34817 | 33,4,69,47,25,11,19,13,5,4 | 0,0,0,2,1,0,0,3,0,0,0,0 | 47,27,29,25,18,14,6,12,19, | 5,1,0,2,3,1,2,2,2,0,1,0,2,2, | 247 | 149 | 0.0000152423443096 | 0.00505489215423654 | 1.0,1.0,1.0,0.934,0 | 0.85,0.942,1.0,0.8 | 0,103 |
| 237534 | ENSG00000089234 | BRAP | chr12 | - | 111681636 | 111681817 | 111679150 | 111679340 | 111683145 | 111683307 | 237534 | 24,18,29,24,9,18,12,16,27, | 9,1,0,2,1,1,2,3,0,0,2,0 | 21,28,17,12,17,7,7,8,11,11 | 1,8,2,0,2,0,0,0,0,0,0,1,2,1, | 298 | 149 | 0.0000153751829293 | 0.00509110153393256 | 0.571,0.9,1.0,0.85 | 0.913,0.636,0.81,1 | -0,076 |
| 23187 | ENSG00000213983 | AP1G2 | chr14 | - | 23565815 | 23565892 | 23565605 | 23565701 | 23566063 | 23566160 | 23187 | 86,48,129,127,56,51,50,35 | 7,4,8,4,4,4,0,11,6,0,0,3 | 86,58,58,33,34,26,40,55,5 | 3,0,5,2,1,0,3,3,0,4,5,1,0,0, | 226 | 149 | 0.0000156777742023 | 0.00517834529425302 | 0.89,0.888,0.914,0 | 0.95,1.0,0.884,0.9 | -0,042 |
| 249868 | ENSG00000120438 | TCP1 | chr6 | - | 159780934 | 159781108 | 159780442 | 159780566 | 159783940 | 159784067 | 249868 | 186,167,207,169,89,73,79, | 39,7,3,13,7,12,8,6,4,2,2,2 | 111,200,108,60,127,54,58, | 22,20,21,34,13,2,4,15,10,1 | 298 | 149 | 0.0000156867790042 | 0.00517834529425302 | 0.705,0.923,0.972, | 0.716,0.833,0.72,0 | 0,098 |
| 224017 | ENSG00000154978 | VOPP1 | chr7 | - | 55472872 | 55473045 | 55436055 | 55436174 | 55492281 | 55492418 | 224017 | 296,360,380,299,232,287, | 26,21,25,15,12,24,41,79,6, | 187,288,166,117,160,93,1 | 13,24,65,118,51,8,7,21,20, | 298 | 149 | 0.0000159051001636 | 0.00523928976965038 | 0.851,0.896,0.884, | 0.878,0.857,0.561, | 0,118 |
| 77500 | ENSG00000163785 | RYK | chr3 | - | 134177940 | 134178073 | 134175929 | 134176039 | 134183001 | 134183033 | 77500 | 86,86,94,60,45,39,61,89,8 | 6,0,6,5,5,4,2,6,0,0,2,3 | 69,111,53,65,56,25,51,68, | 6,9,5,10,15,11,0,9,8,1,8,5, | 282 | 149 | 0.0000159200835816 | 0.00523928976965038 | 0.883,1.0,0.892,0. | 0.859,0.867,0.849, | 0,091 |
| 93800 | ENSG00000204536 | CCHCR1 | chr6 | - | 31154495 | 31154799 | 31150958 | 31151122 | 31156730 | 31156836 | 93800 | 28,32,32,20,15,10,14,14,1 | 1,0,2,0,0,0,3,2,0,2,2,0 | 30,20,8,11,8,7,14,36,13,20 | 4,2,1,2,6,0,0,0,3,0,0,2,4,3, | 298 | 149 | 0.0000159878750268 | 0.00525356691586746 | 0.933,1.0,0.889,1. | 0.789,0.833,0.8,0. | 0,063 |
| 180028 | ENSG00000122224 | LY9 | chr1 | + | 160823464 | 160823649 | 160819320 | 160819374 | 160827747 | 160828261 | 180028 | 48,16,16,29,40,15,25,10,3 | 0,0,1,0,2,0,0,0,0,0,2,0 | 41,25,25,32,7,21,14,32,10, | 2,1,1,1,3,0,0,2,1,0,2,5,4,2, | 298 | 149 | 0.0000162247235822 | 0.00532326750201293 | 1.0,1.0,0.889,1.0,0 | 0.911,0.926,0.926, | 0,105 |
| 80793 | ENSG00000065600 | PACC1 | chr1 | - | 212377561 | 212377706 | 212375192 | 212375300 | 212379894 | 212380037 | 80793 | 15,7,40,30,21,5,5,15,11,5, | 3,0,1,5,0,0,1,0,0,0,0,3 | 23,14,10,2,9,3,2,11,10,3,6, | 1,1,0,0,1,3,2,0,2,1,4,1,2,2, | 294 | 149 | 0.0000163158156464 | 0.00534500657137034 | 0.717,1.0,0.953,0. | 0.921,0.876,1.0,1. | 0,172 |
| 226599 | ENSG00000058091 | CDK14 | chr7 | + | 90747680 | 90747774 | 90726503 | 90726812 | 90790572 | 90790652 | 226599 | 566,312,470,309,217,296, | 7,1,2,3,2,4,1,5,4,2,3,2 | 429,305,159,191,272,209, | 8,10,7,2,7,1,1,0,7,16,11,5, | 243 | 149 | 0.000016352227845 | 0.00534879384697142 | 0.98,0.995,0.993,0 | 0.97,0.949,0.933,0 | 0,016 |
| 142166 | ENSG00000091592 | NLRP1 | chr17 | - | 5533932 | 5533988 | 5533303 | 5533384 | 5536850 | 5536940 | 142166 | 65,23,44,36,26,35,29,41,4 | 4,2,7,1,0,3,2,1,2,0,2,7 | 20,34,17,36,27,9,18,33,32, | 2,2,4,15,1,0,10,7,1,3,9,7,3, | 205 | 149 | 0.0000164009732297 | 0.00535659767708886 | 0.922,0.893,0.82,0 | 0.879,0.925,0.755, | 0,137 |
| 19822 | ENSG00000185787 | MORF4L1 | chr15 | + | 78884969 | 78885083 | 78880511 | 78880579 | 78886140 | 78886227 | 19822 | 23,18,26,11,2,7,5,7,6,5,1,1 | 189,174,103,76,27,45,22,4 | 15,11,11,23,15,3,3,6,7,13,1 | 42,58,84,63,46,33,24,43,3 | 263 | 149 | 0.0000164775027975 | 0.00537343849183139 | 0.064,0.055,0.125, | 0.168,0.097,0.069, | -0,051 |
| 203559 | ENSG00000275052 | PPP4R3B | chr2 | - | 55573618 | 55573738 | 55568028 | 55568363 | 55577314 | 55577356 | 203559 | 61,33,104,50,26,28,20,19, | 7,2,6,2,5,2,0,0,0,2,1,2 | 94,94,24,17,16,28,9,36,15, | 2,0,0,4,0,1,0,0,1,3,0,3,0,1, | 269 | 149 | 0.0000165393724839 | 0.00538545488514717 | 0.828,0.901,0.906, | 0.963,1.0,1.0,0.70 | -0,069 |
| 226606 | ENSG00000058091 | CDK14 | chr7 | + | 90747680 | 90748068 | 90726503 | 90726812 | 90790572 | 90790652 | 226606 | 582,318,494,326,225,308, | 7,1,2,3,2,4,1,5,4,2,3,2 | 447,320,165,198,285,220, | 8,10,7,2,7,1,1,0,7,16,11,5, | 298 | 149 | 0.0000166021212213 | 0.00538703674854559 | 0.977,0.994,0.992, | 0.965,0.941,0.922, | 0,018 |
| 3056 | ENSG00000129351 | ILF3 | chr19 | + | 10688549 | 10688678 | 10683916 | 10683970 | 10689178 | 10689492 | 3056 | 70,56,74,34,51,83,81,64,4 | 9,6,9,8,0,7,7,1,4,4,2,4 | 34,46,38,27,21,24,29,67,3 | 9,12,17,26,5,6,4,7,6,12,5,9 | 278 | 149 | 0.0000166065889657 | 0.00538703674854559 | 0.807,0.833,0.815, | 0.669,0.673,0.545, | 0,141 |
| 59085 | ENSG00000171853 | TRAPPC12 | chr2 | + | 3421699 | 3421786 | 3401776 | 3401893 | 3421880 | 3421970 | 59085 | 2,0,3,0,2,0,4,2,0,0,2,0 | 21,10,27,23,23,18,14,14,2 | 2,3,2,5,0,0,4,9,2,0,6,1,1,3, | 27,10,12,10,8,9,14,38,9,14 | 236 | 149 | 0.000016619317854 | 0.00538703674854559 | 0.057,0.0,0.066,0. | 0.045,0.159,0.095, | -0,06 |
| 224939 | ENSG00000167081 | PBX3 | chr9 | + | 125876435 | 125876497 | 125748549 | 125748623 | 125929654 | 125929845 | 224939 | 1,0,2,1,1,1,0,0,0,0,1,1 | 4,11,34,13,11,13,19,15,6,2 | 2,5,1,5,3,0,4,4,0,0,0,1,2,1, | 22,19,10,9,2,6,13,17,8,4,1 | 211 | 149 | 0.0000166475618492 | 0.00538807726972205 | 0.15,0.0,0.04,0.05 | 0.06,0.157,0.066,0 | -0,066 |
| 87766 | ENSG00000123106 | CCDC91 | chr12 | + | 28304625 | 28304711 | 28259363 | 28259442 | 28305648 | 28305779 | 87766 | 3,3,5,1,2,2,2,6,3,2,1,2 | 47,31,69,40,53,54,38,43,2 | 12,11,17,15,3,1,1,5,0,0,1,5 | 80,42,29,31,17,19,28,62,1 | 235 | 149 | 0.0000166738513867 | 0.005388483044761 | 0.039,0.058,0.044, | 0.087,0.142,0.271, | -0,044 |
| 233929 | ENSG00000130985 | UBA1 | chrX | + | 47200401 | 47200544 | 47199479 | 47199614 | 47200893 | 47201000 | 233929 | 0,0,9,26,0,0,0,1,3,5,0,0 | 20,24,74,41,27,20,25,21,1 | 0,0,0,0,6,10,0,0,0,3,0,0,0,0 | 52,46,15,17,29,28,22,30,2 | 292 | 149 | 0.0000169870715165 | 0.00547727492380986 | 0.0,0.0,0.058,0.24 | 0.0,0.0,0.0,0.0,0.0 | 0,048 |
| 226596 | ENSG00000058091 | CDK14 | chr7 | + | 90747680 | 90747771 | 90726503 | 90726812 | 90790572 | 90790652 | 226596 | 556,310,467,303,215,291, | 7,1,2,3,2,4,1,5,4,2,3,2 | 421,304,156,188,269,208, | 8,10,7,2,7,1,1,0,7,16,11,5, | 240 | 149 | 0.0000169995012294 | 0.00547727492380986 | 0.98,0.995,0.993,0 | 0.97,0.95,0.933,0. | 0,015 |
| 50983 | ENSG00000130775 | THEMIS2 | chr1 | + | 27879643 | 27880054 | 27876587 | 27876728 | 27885294 | 27885451 | 50983 | 62,88,43,42,25,73,63,85,9 | 8,0,7,4,4,0,2,2,7,3,2,3 | 46,144,30,13,76,35,109,24 | 2,3,0,0,5,7,0,4,2,1,2,0,2,1, | 298 | 149 | 0.0000176862149646 | 0.00568502251677171 | 0.795,1.0,0.754,0. | 0.92,0.96,1.0,1.0,0 | -0,045 |
| 227398 | ENSG00000127914 | AKAP9 | chr7 | + | 92043281 | 92043450 | 92042667 | 92042771 | 92045007 | 92045213 | 227398 | 13,5,9,4,2,7,1,7,0,0,2,3 | 17,5,71,41,16,40,21,43,19, | 5,0,1,12,2,0,0,2,0,1,6,2,0,1 | 33,53,24,21,18,24,14,21,2 | 298 | 149 | 0.0000177093237397 | 0.00568502251677171 | 0.277,0.333,0.06,0 | 0.07,0.0,0.02,0.22 | 0,056 |
| 234656 | ENSG00000101199 | ARFGAP1 | chr20 | + | 63282104 | 63282381 | 63281290 | 63281347 | 63282818 | 63283528 | 234656 | 0,2,13,2,2,0,0,1,2,0,2,4 | 16,15,49,25,20,19,15,12,2 | 0,2,0,1,0,0,2,0,0,0,2,0,1,0, | 16,21,14,11,17,8,17,20,10, | 298 | 149 | 0.0000177248476716 | 0.00568502251677171 | 0.0,0.062,0.117,0. | 0.0,0.045,0.0,0.04 | 0,032 |
| 81744 | ENSG00000169499 | PLEKHA2 | chr8 | + | 38943788 | 38943837 | 38935993 | 38936050 | 38946123 | 38946221 | 81744 | 201,71,569,332,263,272,1 | 4,2,33,22,16,4,10,15,11,6, | 336,179,192,99,79,159,15 | 30,9,10,19,12,5,15,22,7,17 | 198 | 149 | 0.0000177499297558 | 0.00568502251677171 | 0.974,0.964,0.928, | 0.894,0.937,0.935, | 0,037 |
| 12894 | ENSG00000161395 | PGAP3 | chr17 | - | 39676613 | 39676755 | 39674616 | 39674679 | 39684596 | 39684698 | 12894 | 2,0,6,14,4,2,8,5,0,2,1,6 | 22,7,18,23,26,23,21,10,12, | 12,9,6,4,4,3,5,7,5,7,3,5,12, | 11,16,10,8,9,2,8,14,11,13, | 291 | 149 | 0.0000177795929636 | 0.00568606177288052 | 0.044,0.0,0.146,0. | 0.358,0.224,0.235, | -0,153 |
| 42657 | ENSG00000163349 | HIPK1 | chr1 | + | 113954650 | 113954770 | 113940381 | 113941459 | 113955562 | 113955649 | 42657 | 25,17,40,31,14,13,18,17,1 | 0,2,4,5,0,0,0,1,0,0,2,0 | 24,27,15,24,11,19,12,21,1 | 3,8,2,10,2,3,2,3,4,0,0,5,0,2 | 269 | 149 | 0.0000178206878123 | 0.00569074845479101 | 1.0,0.825,0.847,0. | 0.816,0.651,0.806, | 0,099 |
| 136105 | ENSG00000105486 | LIG1 | chr19 | - | 48151467 | 48151600 | 48151231 | 48151339 | 48153871 | 48154207 | 136105 | 2,6,2,1,0,0,6,0,5,1,1,4 | 26,18,20,5,27,19,12,15,26, | 0,0,0,0,0,0,0,1,1,0,1,0,0,3, | 14,11,19,13,6,14,4,16,10,1 | 282 | 149 | 0.0000179841946484 | 0.00572536443611803 | 0.039,0.15,0.05,0. | 0.0,0.0,0.0,0.0,0.0, | 0,043 |
| 13886 | ENSG00000140968 | IRF8 | chr16 | + | 85913130 | 85913236 | 85909486 | 85911658 | 85914472 | 85914520 | 13886 | 643,397,737,542,403,359, | 0,4,7,4,6,9,4,4,2,1,1,1 | 358,490,444,239,266,182, | 7,11,5,7,4,3,6,1,2,3,9,21,2, | 255 | 149 | 0.0000179975235859 | 0.00572536443611803 | 1.0,0.983,0.984,0. | 0.968,0.963,0.981, | 0,013 |
| 168218 | ENSG00000138639 | ARHGAP24 | chr4 | + | 85604218 | 85604303 | 85570592 | 85570721 | 85721884 | 85721972 | 168218 | 2,1,46,30,6,29,26,77,32,12 | 198,149,408,178,125,188, | 64,15,26,29,24,38,29,42,3 | 335,282,171,211,196,103, | 234 | 149 | 0.0000180103320339 | 0.00572536443611803 | 0.006,0.004,0.067, | 0.108,0.033,0.088, | -0,043 |
| 6555 | ENSG00000101084 | RAB5IF | chr20 | + | 36609600 | 36609730 | 36607804 | 36608000 | 36612009 | 36612557 | 6555 | 61,61,60,81,40,34,16,29,5 | 0,0,5,0,0,0,0,0,2,0,0,0 | 66,91,28,22,35,37,13,42,3 | 2,2,0,2,4,1,0,1,0,5,0,9,0,5, | 279 | 149 | 0.0000180354925066 | 0.00572536443611803 | 1.0,1.0,0.865,1.0,1 | 0.946,0.96,1.0,0.8 | 0,049 |
| 153419 | ENSG00000100852 | ARHGAP5 | chr14 | + | 32116747 | 32116823 | 32094037 | 32094386 | 32117139 | 32117287 | 153419 | 3,2,13,6,1,2,6,4,0,0,0,0 | 18,16,93,64,12,31,20,40,1 | 1,1,7,0,0,0,0,1,1,0,4,1,0,0, | 42,62,39,30,20,6,28,53,17, | 225 | 149 | 0.0000181453417002 | 0.00575175263545765 | 0.099,0.076,0.085, | 0.016,0.011,0.106, | 0,029 |
| 3067 | ENSG00000130475 | FCHO1 | chr19 | + | 17754591 | 17754758 | 17754184 | 17754361 | 17755117 | 17755191 | 3067 | 28,11,39,25,20,14,11,7,26, | 6,0,0,1,3,0,0,2,0,0,0,0 | 13,6,13,13,14,9,4,4,7,16,2, | 2,0,4,2,1,0,0,2,1,0,2,0,5,1, | 298 | 149 | 0.0000181759204696 | 0.00575297285087129 | 0.7,1.0,1.0,0.926,0 | 0.765,1.0,0.619,0. | 0,138 |
| 69235 | ENSG00000165029 | ABCA1 | chr9 | - | 104903613 | 104903771 | 104889101 | 104889195 | 104927934 | 104928139 | 69235 | 40,48,56,5,40,47,2,43,27,7 | 2,4,3,4,2,3,0,3,1,0,1,0 | 41,36,6,1,34,13,14,32,17,1 | 0,1,0,0,0,0,1,2,0,0,0,0,0,1, | 298 | 149 | 0.0000182591407915 | 0.00577069727550965 | 0.909,0.857,0.903, | 1.0,0.947,1.0,1.0,1 | -0,067 |
| 141153 | ENSG00000166562 | SEC11C | chr18 | + | 59148064 | 59148231 | 59139898 | 59140035 | 59149512 | 59149622 | 141153 | 0,4,18,7,1,6,5,4,2,5,3,5 | 30,31,99,78,27,24,22,23,2 | 1,0,0,4,0,0,1,2,2,0,0,2,3,4, | 25,25,14,4,11,13,11,18,4,1 | 298 | 149 | 0.0000183075031499 | 0.00577069727550965 | 0.0,0.061,0.083,0. | 0.02,0.0,0.0,0.333, | 0,033 |
| 172368 | ENSG00000143669 | LYST | chr1 | - | 235828944 | 235829036 | 235812970 | 235813061 | 235830225 | 235830424 | 172368 | 5,4,9,4,1,3,2,11,1,0,0,10 | 121,62,64,19,64,49,46,52, | 13,3,4,1,5,3,0,6,4,4,30,22, | 72,28,34,16,5,16,28,30,19, | 241 | 149 | 0.0000183123538857 | 0.00577069727550965 | 0.025,0.038,0.08,0 | 0.1,0.062,0.068,0. | -0,058 |
| 194985 | ENSG00000143384 | MCL1 | chr1 | - | 150578285 | 150578491 | 150577367 | 150577491 | 150578842 | 150578963 | 194985 | 244,280,366,215,277,221, | 2,2,3,4,14,7,14,13,4,1,3,5 | 234,362,173,206,158,124, | 18,10,13,3,1,7,2,4,2,12,8,4 | 298 | 149 | 0.0000183493417603 | 0.00577389938071802 | 0.984,0.986,0.984, | 0.867,0.948,0.869, | 0,029 |
| 53334 | ENSG00000196187 | TMEM63A | chr1 | - | 225867111 | 225867163 | 225865896 | 225865967 | 225867887 | 225868030 | 53334 | 10,9,17,18,9,11,10,10,12,8 | 0,0,1,2,3,0,0,5,0,0,0,1 | 18,12,3,5,7,10,8,10,17,28, | 4,1,4,6,1,0,0,4,1,1,0,0,1,2, | 201 | 149 | 0.000018505305844 | 0.0058064075848767 | 1.0,1.0,0.926,0.87, | 0.769,0.899,0.357, | 0,123 |
| 147386 | ENSG00000103275 | UBE2I | chr16 | + | 1319701 | 1319990 | 1315653 | 1315726 | 1320173 | 1320283 | 147386 | 5,1,5,1,3,5,24,31,2,0,6,4 | 228,233,210,191,126,123, | 4,4,0,3,4,1,8,9,4,0,3,0,2,2, | 144,199,137,104,93,87,73, | 298 | 149 | 0.0000185319474086 | 0.0058064075848767 | 0.011,0.002,0.012, | 0.014,0.01,0.0,0.0 | 0,016 |
| 192687 | ENSG00000117859 | OSBPL9 | chr1 | + | 51784263 | 51784586 | 51783914 | 51784025 | 51785807 | 51785886 | 192687 | 7,3,12,18,13,5,2,8,3,1,3,9 | 4,0,4,0,0,1,1,0,4,1,2,1 | 5,11,2,1,4,3,6,8,4,6,7,20,5, | 1,2,0,0,1,0,1,0,0,2,0,1,0,2, | 298 | 149 | 0.0000185419892705 | 0.0058064075848767 | 0.467,1.0,0.6,1.0,1 | 0.714,0.733,1.0,1. | -0,195 |
| 120569 | ENSG00000107036 | RIC1 | chr9 | + | 5738449 | 5738538 | 5732387 | 5732479 | 5743688 | 5743737 | 120569 | 12,3,18,13,5,5,3,17,4,4,5,4 | 0,0,2,0,0,1,0,0,0,1,0,1 | 18,12,2,11,27,8,1,12,9,12, | 2,1,2,4,3,0,0,1,4,0,0,5,0,2, | 238 | 149 | 0.0000185809627539 | 0.0058064075848767 | 1.0,1.0,0.849,1.0,1 | 0.849,0.883,0.385, | 0,093 |
| 91240 | ENSG00000167842 | MIS12 | chr17 | + | 5488195 | 5488589 | 5486284 | 5486420 | 5488822 | 5488935 | 91240 | 1,2,18,23,8,4,6,9,9,2,4,10 | 0,0,4,2,1,3,0,2,3,0,2,2 | 20,19,5,1,6,3,8,10,3,4,9,5, | 0,0,2,0,0,1,0,4,1,1,0,1,0,0, | 298 | 149 | 0.0000185875400197 | 0.0058064075848767 | 1.0,1.0,0.692,0.85 | 1.0,1.0,0.556,1.0,1 | -0,064 |
| 2461 | ENSG00000198453 | ZNF568 | chr19 | + | 36931552 | 36931645 | 36925199 | 36925258 | 36936745 | 36936872 | 2461 | 1,0,3,4,5,6,0,1,0,0,3,2 | 12,6,23,16,8,5,6,3,15,9,6,4 | 2,1,2,0,1,0,1,2,0,0,0,0,2,1, | 6,9,4,8,8,6,9,6,10,8,9,4,15, | 242 | 149 | 0.0000188791109679 | 0.00588894193149577 | 0.049,0.0,0.074,0. | 0.17,0.064,0.235,0 | 0,075 |
| 93213 | ENSG00000112576 | CCND3 | chr6 | - | 42030008 | 42030190 | 41937174 | 41937394 | 42048500 | 42048686 | 93213 | 4,0,10,3,1,0,1,3,0,0,0,1 | 134,100,83,66,37,83,113,1 | 6,9,1,2,0,1,0,2,0,0,31,4,0,0 | 70,53,59,19,24,48,75,152, | 298 | 149 | 0.0000190803070467 | 0.00594308764973703 | 0.015,0.0,0.057,0. | 0.041,0.078,0.008, | -0,012 |
| 187802 | ENSG00000118246 | FASTKD2 | chr2 | + | 206786682 | 206786899 | 206774224 | 206774397 | 206787936 | 206788155 | 187802 | 11,11,37,37,15,23,7,16,6,5 | 0,1,4,4,0,2,2,0,0,0,0,0 | 27,27,21,14,14,9,20,14,10, | 3,0,1,1,8,0,2,2,7,2,0,1,0,1, | 298 | 149 | 0.0000192595037062 | 0.00599023445403053 | 1.0,0.846,0.822,0. | 0.818,1.0,0.913,0. | 0,083 |
| 116172 | ENSG00000107077 | KDM4C | chr9 | + | 7105401 | 7105500 | 7103684 | 7103870 | 7128065 | 7128236 | 116172 | 13,0,12,8,3,6,0,5,1,0,0,2 | 24,20,31,17,19,12,24,24,1 | 6,3,0,1,2,2,2,4,0,1,3,2,1,2, | 25,27,20,23,10,13,18,21,9, | 248 | 149 | 0.0000194113942762 | 0.00602876450427244 | 0.246,0.0,0.189,0. | 0.126,0.063,0.0,0. | 0,049 |
| 38240 | ENSG00000003056 | M6PR | chr12 | - | 8943800 | 8943910 | 8943295 | 8943535 | 8945417 | 8945479 | 38240 | 232,152,231,239,96,102,9 | 0,1,2,0,1,0,0,0,0,0,0,2 | 175,82,127,39,88,51,38,11 | 5,5,2,2,2,0,1,5,0,2,9,1,4,4, | 259 | 149 | 0.0000195214295207 | 0.00605213884599009 | 1.0,0.989,0.985,1. | 0.953,0.904,0.973, | 0,035 |
| 207089 | ENSG00000160352 | ZNF714 | chr19 | + | 21098459 | 21098910 | 21098184 | 21098311 | 21116806 | 21117084 | 207089 | 1,5,9,8,2,7,8,3,1,1,0,2 | 0,0,1,2,1,0,0,1,2,0,2,0 | 8,1,6,3,2,1,3,3,4,2,4,3,5,4, | 0,3,0,1,4,4,4,5,2,0,3,0,0,1, | 298 | 149 | 0.0000195428934399 | 0.00605213884599009 | 1.0,1.0,0.818,0.66 | 1.0,0.143,1.0,0.6,0 | 0,099 |
| 16960 | ENSG00000130731 | METTL26 | chr16 | - | 635611 | 635774 | 635280 | 635340 | 636093 | 636235 | 16960 | 10,12,22,17,11,5,8,4,20,14 | 3,1,2,0,2,2,0,2,3,2,0,2 | 24,20,5,6,12,1,8,14,4,11,5, | 0,0,0,0,0,0,2,0,0,2,2,1,0,1,1 | 298 | 149 | 0.0000197135854725 | 0.00609566743727893 | 0.625,0.857,0.846, | 1.0,1.0,1.0,1.0,1.0, | -0,121 |
| 193785 | ENSG00000058673 | ZC3H11A | chr1 | + | 203796333 | 203796601 | 203795705 | 203795794 | 203801574 | 203801628 | 193785 | 0,1,3,0,1,0,0,2,1,0,0,0 | 5,7,46,18,26,34,9,14,8,2,1 | 0,3,3,1,0,1,0,3,0,4,12,3,7,2 | 45,29,15,1,9,10,9,19,7,16, | 298 | 149 | 0.000019740094149 | 0.00609566743727893 | 0.0,0.067,0.032,0. | 0.0,0.049,0.091,0. | -0,049 |
| 37618 | ENSG00000005059 | MCUB | chr4 | + | 109684442 | 109684646 | 109682581 | 109682742 | 109685250 | 109685367 | 37618 | 276,150,331,250,142,139, | 6,1,2,1,0,4,5,0,5,0,0,1 | 202,195,144,77,88,83,71,1 | 10,3,8,3,1,3,2,8,3,5,3,10,4, | 298 | 149 | 0.0000199341705518 | 0.00614677859890324 | 0.958,0.987,0.988, | 0.91,0.97,0.9,0.92 | 0,032 |
| 77667 | ENSG00000132953 | XPO4 | chr13 | - | 20787480 | 20787598 | 20786964 | 20787057 | 20790461 | 20790580 | 77667 | 14,9,27,19,15,17,15,19,5,1 | 1,0,0,0,1,2,0,0,0,2,0,0 | 16,17,9,12,17,9,5,15,3,8,2 | 5,1,2,2,0,2,1,0,0,0,2,2,5,0,1 | 267 | 149 | 0.0000199989239115 | 0.00615792330814886 | 0.887,1.0,1.0,1.0,0 | 0.641,0.905,0.715, | 0,066 |
| 184482 | ENSG00000077044 | DGKD | chr2 | + | 233389943 | 233390041 | 233388256 | 233388367 | 233390402 | 233390483 | 184482 | 5,1,22,6,3,6,2,4,0,0,1,3 | 63,43,251,97,46,59,79,124 | 4,1,2,5,2,2,2,0,1,0,0,4,2,1, | 103,107,94,62,89,45,38,57 | 247 | 149 | 0.0000200870050884 | 0.00617620884597345 | 0.046,0.014,0.05,0 | 0.023,0.006,0.013, | 0,011 |
| 61435 | ENSG00000176095 | IP6K1 | chr3 | - | 49747817 | 49748168 | 49738211 | 49738422 | 49786353 | 49786540 | 61435 | 7,27,20,30,16,5,9,6,11,11, | 1,2,6,0,0,0,3,1,0,0,0,2 | 16,26,11,16,16,7,21,15,3,1 | 0,3,2,6,2,1,5,0,0,0,2,1,0,0,1 | 298 | 149 | 0.000020125194389 | 0.00617912369977012 | 0.778,0.871,0.625, | 1.0,0.812,0.733,0. | 0,016 |
| 154178 | ENSG00000213463 | SYNJ2BP | chr14 | - | 70381147 | 70381195 | 70375675 | 70375771 | 70388469 | 70388606 | 154178 | 3,3,2,4,0,1,4,1,2,1,0,3 | 4,8,16,9,10,4,15,6,3,4,6,9 | 2,3,0,1,0,0,0,1,1,0,1,0,0,4, | 12,11,10,4,5,10,1,1,6,8,11, | 197 | 149 | 0.0000204388817701 | 0.00626649709723702 | 0.362,0.221,0.086, | 0.112,0.171,0.0,0. | 0,087 |
| 26169 | ENSG00000175061 | SNHG29 | chr17 | + | 16439580 | 16439692 | 16439326 | 16439414 | 16440184 | 16440253 | 26169 | 15,7,8,20,6,5,12,10,6,0,7,8 | 107,48,427,663,111,100,5 | 70,78,13,3,9,1,4,8,3,9,5,28 | 106,110,84,38,147,106,37, | 261 | 149 | 0.0000206383733057 | 0.00631865963721069 | 0.074,0.077,0.011, | 0.274,0.288,0.081, | -0,019 |
| 189276 | ENSG00000118308 | IRAG2 | chr12 | + | 25081214 | 25081511 | 25079701 | 25079763 | 25083422 | 25083493 | 189276 | 19,14,25,29,7,5,3,0,2,1,2,7 | 118,60,586,422,151,96,17 | 7,24,8,5,9,1,0,2,3,0,4,0,5,0 | 289,288,122,36,86,87,63,1 | 298 | 149 | 0.0000207820719201 | 0.00635361664976568 | 0.075,0.104,0.021, | 0.012,0.04,0.032,0 | 0,014 |
| 103331 | ENSG00000231721 | LINC-PINT | chr7 | - | 131076524 | 131076587 | 131052424 | 131052555 | 131107719 | 131108803 | 103331 | 8,1,5,0,2,1,1,5,0,2,0,1 | 107,73,166,111,52,108,10 | 3,3,6,29,5,2,0,4,9,3,4,3,0,2 | 124,99,93,127,111,37,67,7 | 212 | 149 | 0.0000208482084254 | 0.00636479538667697 | 0.05,0.01,0.021,0. | 0.017,0.021,0.043, | -0,02 |
| 14449 | ENSG00000132600 | PRMT7 | chr16 | + | 68347210 | 68347294 | 68346120 | 68346280 | 68347630 | 68347678 | 14449 | 25,10,41,25,16,11,23,24,4, | 2,0,3,0,0,0,0,0,0,0,0,0 | 19,10,11,6,17,7,7,18,8,23,1 | 1,4,1,1,2,1,7,1,1,0,2,0,0,1, | 233 | 149 | 0.0000209664044307 | 0.00639181330315013 | 0.889,1.0,0.897,1. | 0.924,0.615,0.876, | 0,142 |
| 150808 | ENSG00000138592 | USP8 | chr15 | + | 50481637 | 50482065 | 50477275 | 50477499 | 50484274 | 50484361 | 150808 | 76,64,113,68,36,35,39,77, | 0,0,2,1,0,0,0,4,0,0,0,0 | 58,59,37,41,51,16,36,48,2 | 4,4,0,0,6,1,0,0,2,3,0,2,0,4, | 298 | 149 | 0.0000210224689225 | 0.00639984018197821 | 1.0,1.0,0.966,0.97 | 0.879,0.881,1.0,1. | 0,072 |
| 204377 | ENSG00000173209 | AHSA2P | chr2 | + | 61186016 | 61186081 | 61185525 | 61185618 | 61186445 | 61186497 | 204377 | 6,2,5,0,0,6,9,14,2,1,1,4 | 65,32,254,161,103,78,48,6 | 6,19,12,0,6,2,4,17,9,9,10,3 | 61,38,49,35,79,61,76,111, | 214 | 149 | 0.0000211322104625 | 0.00641853764850812 | 0.06,0.042,0.014,0 | 0.064,0.258,0.146, | -0,033 |
| 35286 | ENSG00000104517 | UBR5 | chr8 | - | 102278825 | 102278939 | 102276971 | 102277192 | 102279035 | 102279208 | 35286 | 103,103,267,130,116,119, | 10,0,8,9,0,1,0,2,0,0,0,2 | 153,161,105,142,107,75,6 | 1,5,3,36,5,2,2,1,9,7,1,10,2, | 263 | 149 | 0.0000211435304059 | 0.00641853764850812 | 0.854,1.0,0.95,0.8 | 0.989,0.948,0.952, | 0,044 |
| 203610 | ENSG00000120451 | SNX19 | chr11 | - | 130907968 | 130908445 | 130906624 | 130906721 | 130910017 | 130910137 | 203610 | 12,10,10,7,3,5,11,15,3,4,2, | 0,2,1,0,0,0,0,1,1,0,0,0 | 7,7,6,8,15,1,6,1,2,4,3,5,8,5 | 0,1,1,0,5,1,2,1,0,0,4,1,2,1,1 | 298 | 149 | 0.0000212577708855 | 0.00644412857106626 | 1.0,0.714,0.833,1. | 1.0,0.778,0.75,1.0, | 0,257 |
| 174949 | ENSG00000096384 | HSP90AB1 | chr6 | + | 44248629 | 44248780 | 44247100 | 44247195 | 44249376 | 44249583 | 174949 | 314,151,812,591,271,306, | 47,12,26,19,11,17,16,10,8, | 528,493,260,71,196,153,1 | 11,11,5,5,4,4,2,11,4,0,32,9 | 298 | 149 | 0.0000214458105374 | 0.00649198769026039 | 0.77,0.863,0.94,0. | 0.96,0.957,0.963,0 | -0,031 |
| 237014 | ENSG00000086102 | NFX1 | chr9 | + | 33291420 | 33291578 | 33290544 | 33290597 | 33294419 | 33295427 | 237014 | 5,6,4,0,0,4,4,2,1,3,1,7 | 11,5,46,18,19,14,10,18,9,4 | 0,0,2,3,2,0,3,1,1,0,2,4,0,0, | 14,19,11,6,12,5,10,11,5,16 | 298 | 149 | 0.0000215198139927 | 0.00650524028857136 | 0.185,0.375,0.042, | 0.0,0.0,0.083,0.2,0 | 0,124 |
| 131986 | ENSG00000089154 | GCN1 | chr12 | - | 120131184 | 120131333 | 120130645 | 120130753 | 120131925 | 120132022 | 131986 | 91,65,116,74,67,77,53,53, | 2,0,1,0,3,5,0,6,0,1,0,0 | 97,87,49,55,61,34,35,43,3 | 6,3,0,4,1,2,4,7,0,2,4,3,7,1, | 298 | 149 | 0.0000215747966085 | 0.00651271395349798 | 0.958,1.0,0.983,1. | 0.89,0.935,1.0,0.8 | 0,049 |
| 233320 | ENSG00000215301 | DDX3X | chrX | + | 41346930 | 41347012 | 41346504 | 41346622 | 41347311 | 41347451 | 233320 | 99,73,146,70,44,33,40,49, | 16,2,7,7,1,0,2,4,0,0,2,2 | 67,84,63,47,51,36,17,39,2 | 9,21,14,30,3,0,5,9,1,3,11,5 | 231 | 149 | 0.0000216775496962 | 0.00653456680484989 | 0.8,0.959,0.931,0. | 0.828,0.721,0.744, | 0,095 |
| 127635 | ENSG00000138134 | STAMBPL1 | chr10 | + | 88908701 | 88908777 | 88905442 | 88905660 | 88910915 | 88911011 | 127635 | 82,30,107,60,25,30,31,43, | 2,0,13,10,4,2,2,4,1,0,2,1 | 27,49,37,27,36,32,12,26,2 | 0,3,2,0,4,0,2,0,2,0,1,0,0,0, | 225 | 149 | 0.0000219363628928 | 0.00660333611437795 | 0.964,1.0,0.845,0. | 1.0,0.915,0.925,1. | -0,035 |

| | | | | | | | | | | | | | | | | | | | | | | |
|---|---|---|---|---|---|---|---|---|---|---|---|---|---|---|---|---|---|---|---|---|---|---|
| 216295 | ENSG00000163219 | ARHGAP25 | chr2 | + | 68806510 | 68806939 | 68787839 | 68787956 | 68807275 | 68807480 | 216295 | 2,0,3,0,0,1,0,3,0,0,6 | 9,27,23,25,24,22,7,17,13,9 | 2,11,0,10,7,0,0,3,4,1,3,2,2, | 29,78,10,4,14,1,6,20,5,12,9 | 298 | 149 | 0.0000220166965414 | 0.00661826202975148 | 0.1,0.0,0.061,0.0,0 | 0.033,0.066,0.0,0.5 | -0,061 |
| 37357 | ENSG00000223547 | ZNF844 | chr19 | + | 12074030 | 12074157 | 12064730 | 12064876 | 12075311 | 12077153 | 37357 | 57,12,124,75,11,58,27,26,2 | 2,0,3,3,2,5,5,4,0,0,2,0 | 59,29,35,11,24,27,10,29,21 | 2,0,0,0,6,1,0,0,1,0,0,0,0,1,0 | 276 | 149 | 0.0000221364704582 | 0.00664498559719504 | 0.939,1.0,0.957,0.9 | 0.941,1.0,1.0,1.0,0 | -0,058 |
| 8884 | ENSG00000282851 | BISPR | chr19 | + | 17413096 | 17413305 | 17406076 | 17406296 | 17414390 | 17415005 | 8884 | 29,11,21,23,10,5,3,27,19,5 | 1,0,6,4,2,0,0,2,0,2,0,3 | 20,12,7,22,10,11,3,22,4,12 | 2,0,0,0,0,0,1,0,0,0,4,0,0,0 | 298 | 149 | 0.0000221854497534 | 0.00664771152422123 | 0.935,1.0,0.636,0.7 | 0.833,1.0,1.0,1.0,1 | -0,114 |
| 80516 | ENSG00000152359 | POC5 | chr5 | - | 75677773 | 75677950 | 75674123 | 75674578 | 75685206 | 75685484 | 80516 | 31,18,28,21,21,22,19,18,30 | 2,0,5,3,0,6,0,0,3,3,0,0 | 24,19,13,7,12,7,13,26,8,27 | 0,1,0,3,4,2,0,2,0,1,0,0,0,2,0 | 298 | 149 | 0.0000222073241583 | 0.00664771152422123 | 0.886,1.0,0.737,0.7 | 1.0,0.905,1.0,0.538 | -0,011 |
| 74805 | ENSG00000275052 | PPP4R3B | chr2 | - | 55573618 | 55573777 | 55568193 | 55568363 | 55577314 | 55577356 | 74805 | 83,49,146,93,90,47,35,40,5 | 7,2,6,2,5,2,0,0,0,2,1,2 | 145,125,39,25,29,46,27,84 | 2,0,0,4,0,1,0,0,1,3,0,3,0,1,0 | 298 | 149 | 0.0000223495930055 | 0.00668100729467607 | 0.856,0.925,0.924, | 0.973,1.0,1.0,0.758 | -0,029 |
| 64547 | ENSG00000114861 | FOXP1 | chr3 | - | 71130497 | 71130584 | 71112535 | 71112637 | 71132090 | 71132099 | 64547 | 174,102,273,219,107,113,1 | 17,10,21,26,10,26,5,19,5,1 | 132,110,94,67,140,129,72, | 5,6,10,7,4,6,7,4,8,3,6,1,3,2 | 236 | 149 | 0.000022387761082 | 0.00668313482030505 | 0.866,0.866,0.891, | 0.943,0.92,0.856,0 | -0,042 |
| 22994 | ENSG00000129566 | TEP1 | chr14 | - | 20400983 | 20401141 | 20396620 | 20396730 | 20401456 | 20401581 | 22994 | 49,64,54,57,50,53,21,33,64 | 0,0,6,4,0,3,9,8,0,1,0,2 | 57,78,46,42,53,38,25,39,55 | 0,1,0,1,2,3,4,0,0,0,2,1,0,2,3 | 298 | 149 | 0.0000226352166658 | 0.00674764587007867 | 1.0,1.0,0.818,0.87 | 1.0,0.975,1.0,0.95 | -0,058 |
| 79388 | ENSG00000123240 | OPTN | chr10 | + | 13123994 | 13124110 | 13122384 | 13122487 | 13125417 | 13125567 | 79388 | 18,15,97,67,38,20,29,41,12 | 5,2,1,0,2,0,0,4,0,0,0,1 | 23,46,27,20,13,17,24,39,18 | 4,3,1,8,3,0,0,1,2,1,4,6,2,0,1 | 265 | 149 | 0.0000227581856473 | 0.00677491985484651 | 0.669,0.808,0.982, | 0.764,0.896,0.938, | 0,069 |
| 130466 | ENSG00000010626 | LRRC23 | chr12 | + | 6912729 | 6913027 | 6909889 | 6910026 | 6913890 | 6914194 | 130466 | 21,14,5,6,10,11,6,1,5,2,4,1 | 3,1,1,0,0,0,0,0,0,0,0,0 | 13,17,10,11,4,5,0,8,3,9,3,1 | 2,0,1,0,2,1,4,2,0,0,1,5,0,3,1 | 298 | 149 | 0.0000230628423117 | 0.00685613068175345 | 0.778,0.875,0.714, | 0.765,1.0,0.833,1.0 | 0,229 |
| 84742 | ENSG00000145425 | RPS3A | chr4 | + | 151104184 | 151104286 | 151102870 | 151103079 | 151104471 | 151104637 | 84742 | 146,107,502,328,72,84,33, | 1,7,6,7,3,2,2,7,6,3,0,5 | 258,320,121,86,172,105,28 | 5,8,2,0,1,0,2,3,0,3,1,3,2,1,1 | 251 | 149 | 0.0000233397003266 | 0.00692886488413026 | 0.989,0.901,0.98,0 | 0.968,0.96,0.973,1 | -0,057 |
| 185248 | ENSG00000072274 | TFRC | chr3 | - | 196061319 | 196061434 | 196060179 | 196060247 | 196062581 | 196064428 | 185248 | 15,2,6,0,5,1,3,4,0,0,0,0 | 39,21,93,42,73,54,48,69,26 | 2,1,14,21,1,5,2,5,3,21,6,9,3 | 52,56,53,27,35,15,21,21,17 | 264 | 149 | 0.0000233901987957 | 0.00693429184159408 | 0.178,0.051,0.035, | 0.021,0.01,0.13,0.3 | -0,06 |
| 219633 | ENSG00000083097 | DOP1A | chr6 | + | 83135669 | 83135878 | 83134186 | 83134287 | 83137172 | 83139162 | 219633 | 17,10,33,6,11,13,9,16,4,3,6 | 0,0,2,0,4,1,0,0,0,0,2,0 | 15,30,7,5,12,17,7,12,17,21 | 0,7,0,2,1,0,1,4,0,2,2,0,2,0,3 | 298 | 149 | 0.0000234725245426 | 0.00694912645093306 | 1.0,1.0,0.892,1.0,0 | 1.0,0.682,1.0,0.556 | 0,091 |
| 213345 | ENSG00000135426 | TESPA1 | chr12 | - | 54983993 | 54984078 | 54974419 | 54974607 | 54984584 | 54984762 | 213345 | 7,4,24,12,5,6,1,6,5,1,4,1 | 41,14,38,23,21,11,9,10,10, | 3,7,1,5,2,2,0,0,4,3,2,2,0,2,2 | 18,23,14,14,13,2,13,14,8,2 | 234 | 149 | 0.0000235881023304 | 0.00697375117125593 | 0.098,0.154,0.287, | 0.096,0.162,0.044, | 0,072 |
| 240154 | ENSG00000165525 | NEMF | chr14 | - | 49833422 | 49833728 | 49832206 | 49832277 | 49834362 | 49834449 | 240154 | 30,14,41,25,9,13,19,27,25, | 2,1,3,1,1,1,0,0,5,2,0,1 | 31,31,10,16,21,13,7,24,21, | 5,5,0,4,3,1,2,2,3,9,0,2,7,3,4 | 298 | 149 | 0.0000237142271616 | 0.007001422258187 | 0.882,0.875,0.872, | 0.756,0.756,1.0,0.0 | 0,144 |
| 228630 | ENSG00000126353 | CCR7 | chr17 | - | 40558892 | 40559164 | 40553768 | 40555818 | 40565399 | 40565472 | 228630 | 88,130,100,64,73,92,66,97 | 3,1,2,3,6,3,1,4,1,0,0,4 | 63,74,42,13,23,22,44,100,2 | 0,0,1,1,1,1,6,1,0,0,0,0,0,0,0 | 298 | 149 | 0.0000237501922591 | 0.00700243510975116 | 0.936,0.985,0.962, | 1.0,1.0,0.955,0.867 | -0,017 |
| 41754 | ENSG00000145012 | LPP | chr3 | + | 188759262 | 188759500 | 188708266 | 188708393 | 188760112 | 188760282 | 41754 | 6,2,6,0,2,0,0,2,1,1,0,0 | 67,24,63,31,29,50,22,47,11 | 7,0,2,10,2,3,3,8,3,3,13,1,0, | 27,28,29,60,47,33,38,38,26 | 298 | 149 | 0.0000238808271122 | 0.00702240412855619 | 0.043,0.04,0.045,0 | 0.115,0.0,0.033,0.0 | -0,026 |
| 22240 | ENSG00000006432 | MAP3K9 | chr14 | - | 70761001 | 70761182 | 70749932 | 70750081 | 70800666 | 70801080 | 22240 | 29,31,53,26,45,28,26,20,30 | 0,0,0,0,3,2,0,0,0,0,0,0 | 27,45,27,24,33,15,23,16,26 | 3,4,4,4,1,2,0,4,1,0,0,0,1,1,1 | 298 | 149 | 0.0000238831758534 | 0.00702240412855619 | 1.0,1.0,1.0,1.0,0.88 | 0.818,0.849,0.771, | 0,094 |
| 107528 | ENSG00000198932 | GPRASP1 | chrX | + | 102652830 | 102652898 | 102652174 | 102652265 | 102653220 | 102653298 | 107528 | 20,7,47,27,12,36,19,25,7,1 | 1,0,3,1,0,1,1,4,0,0,2,0 | 36,27,21,7,10,7,28,44,20,1 | 0,4,0,11,4,0,5,5,0,0,1,0,0,3 | 217 | 149 | 0.0000239636962708 | 0.00703646700144686 | 0.932,1.0,0.915,0.9 | 1.0,0.823,1.0,0.304 | 0,103 |
| 241267 | ENSG00000107077 | KDM4C | chr9 | + | 6986343 | 6986706 | 6984165 | 6984404 | 6990415 | 6990524 | 241267 | 55,23,64,50,23,38,20,46,18 | 1,0,0,5,0,1,0,0,0,0,0,1 | 66,60,35,44,33,39,32,36,31 | 1,5,0,3,7,0,2,0,1,0,2,2,2,4,0 | 298 | 149 | 0.0000240635845744 | 0.00705617080590284 | 0.965,1.0,1.0,0.833 | 0.971,0.857,1.0,0.8 | 0,066 |
| 39558 | ENSG00000266338 | NBPF15 | chr1 | - | 144429699 | 144429863 | 144428605 | 144428657 | 144433772 | 144433824 | 39558 | 20,17,19,12,28,24,10,14,8, | 4,0,5,2,2,1,2,0,0,1,1,1 | 11,16,11,9,9,5,7,14,7,15,5, | 1,0,0,1,1,3,0,1,0,0,0,0,0,0,0 | 298 | 149 | 0.0000241213803921 | 0.00706349499751302 | 0.714,1.0,0.655,0.7 | 0.846,1.0,1.0,0.818 | -0,087 |
| 6476 | ENSG00000170471 | RALGAPB | chr20 | + | 38497479 | 38497516 | 38492929 | 38493132 | 38499446 | 38499633 | 6476 | 9,5,17,12,7,14,3,9,5,1,1,5 | 0,0,2,0,0,2,1,1,0,0,0,3 | 12,15,8,10,13,5,2,5,6,12,12 | 3,3,3,6,4,1,0,4,1,0,1,2,0,0,0 | 186 | 149 | 0.0000242028187857 | 0.00707771316584918 | 1.0,1.0,0.872,1.0,1 | 0.762,0.8,0.681,0.5 | 0,096 |
| 8399 | ENSG00000130813 | SHFL | chr19 | + | 10089110 | 10089378 | 10087250 | 10087300 | 10089656 | 10089695 | 8399 | 15,3,20,12,11,7,9,13,12,7,9 | 22,16,40,31,21,24,16,22,14 | 12,8,3,9,2,5,0,2,4,6,6,4,1,6 | 17,18,22,22,24,8,11,15,7,2 | 298 | 149 | 0.0000243283151146 | 0.00710475928145247 | 0.254,0.086,0.2,0.1 | 0.261,0.182,0.064, | 0,108 |
| 216287 | ENSG00000163219 | ARHGAP25 | chr2 | + | 68788488 | 68788653 | 68787839 | 68787956 | 68807275 | 68807480 | 216287 | 0,0,6,8,0,0,0,3,10,9,0,0 | 9,27,23,25,24,22,7,17,13,9 | 2,0,1,0,16,5,0,0,0,0,0,0,0,0 | 29,78,10,4,14,1,6,20,5,12,9 | 298 | 149 | 0.0000243859139784 | 0.0071119304220664 | 0.0,0.0,0.115,0.138 | 0.033,0.0,0.048,0.0 | 0,006 |
| 255213 | ENSG00000089154 | GCN1 | chr12 | - | 120131184 | 120131351 | 120130645 | 120130753 | 120131925 | 120132022 | 255213 | 64,48,67,41,33,52,28,32,29 | 2,0,1,0,3,5,0,6,0,1,0,0 | 46,48,26,27,34,17,8,23,18, | 6,3,0,4,1,2,4,7,0,2,4,3,7,1,4 | 298 | 149 | 0.0000245116518439 | 0.00713892738567583 | 0.941,1.0,0.971,1.0 | 0.793,0.889,1.0,0.1 | 0,102 |
| 9906 | ENSG00000168961 | LGALS9 | chr17 | + | 27643524 | 27643620 | 27642237 | 27642348 | 27645313 | 27645349 | 9906 | 8,4,30,26,12,2,24,15,28,26 | 82,27,25,42,65,21,18,8,176 | 13,16,38,33,15,11,6,10,17, | 21,15,28,17,3,9,11,39,3,9,1 | 245 | 149 | 0.0000248712526666 | 0.00723387105768241 | 0.056,0.083,0.422, | 0.274,0.393,0.452, | -0,201 |
| 61186 | ENSG00000010270 | STARD3NL | chr7 | + | 38202069 | 38202197 | 38178375 | 38178420 | 38207446 | 38207729 | 61186 | 2,0,3,2,0,4,2,4,2,2,1,0 | 8,7,8,9,2,8,5,1,10,1,1,4 | 0,1,0,1,0,0,0,1,0,0,0,0,0,0,1 | 8,2,7,14,1,1,9,2,1,3,9,17,4, | 277 | 149 | 0.000025058011928 | 0.00727835487891412 | 0.119,0.0,0.168,0.1 | 0.0,0.212,0.0,0.037 | 0,146 |
| 117801 | ENSG00000291015 | | chr22 | - | 38369985 | 38370045 | 38361398 | 38361560 | 38380369 | 38380423 | 117801 | 3,1,7,4,4,5,2,4,0,2,4,0 | 32,22,31,21,22,16,8,18,21, | 2,2,0,1,0,3,0,0,0,2,3,2,2,0,0 | 34,24,11,4,16,15,4,15,13,3 | 209 | 149 | 0.0000252579635618 | 0.00732654549241209 | 0.063,0.031,0.139, | 0.04,0.056,0.0,0.15 | 0,057 |
| 41273 | ENSG00000198843 | SELENOT | chr3 | + | 150618708 | 150618928 | 150603290 | 150603499 | 150622384 | 150622495 | 41273 | 48,29,19,11,12,9,16,23,10, | 52,40,92,80,42,47,34,41,73 | 25,37,36,29,35,9,17,25,14, | 54,68,48,20,45,23,22,50,15 | 298 | 149 | 0.0000255475294449 | 0.00740056569307573 | 0.316,0.266,0.094, | 0.188,0.214,0.273, | -0,083 |
| 224545 | ENSG00000116679 | IVNS1ABP | chr1 | - | 185306447 | 185306625 | 185305535 | 185305643 | 185307013 | 185307139 | 224545 | 2,8,10,2,5,3,5,8,0,0,2,4 | 9,10,10,20,4,12,0,6,4,3,5,4 | 12,15,5,5,10,5,9,4,8,5,12,2 | 13,1,1,0,2,4,9,5,3,14,4,13,3 | 298 | 149 | 0.0000259598492082 | 0.00750989825931464 | 0.1,0.286,0.333,0.0 | 0.316,0.882,0.714, | -0,325 |
| 177510 | ENSG00000137818 | RPLP1 | chr15 | + | 69453649 | 69453721 | 69452883 | 69453020 | 69455169 | 69455287 | 177510 | 1147,837,851,1036,370,38 | 26,16,43,55,17,17,14,10,23 | 617,758,294,415,275,167,2 | 8,13,2,8,7,5,12,22,6,8,18,2 | 221 | 149 | 0.0000260271656021 | 0.00751926561034306 | 0.967,0.972,0.93,0 | 0.981,0.975,0.99,0 | -0,014 |
| 111778 | ENSG00000116604 | MEF2D | chr1 | - | 156476493 | 156476514 | 156475107 | 156475237 | 156477011 | 156477202 | 111778 | 4,8,1,6,1,2,11,2,2,0,2,0 | 17,44,20,15,14,11,4,9,19,1 | 4,4,3,1,1,1,1,0,1,0,0,5,0,0,0 | 13,27,9,10,21,5,12,23,12,1 | 170 | 149 | 0.0000260753616785 | 0.00752309137992658 | 0.171,0.137,0.042, | 0.212,0.115,0.226, | 0,089 |
| 102619 | ENSG00000122566 | HNRNPA2B1 | chr7 | - | 26191421 | 26192338 | 26190992 | 26191128 | 26192494 | 26192577 | 102619 | 152,68,165,126,90,87,92,1 | 11,5,18,17,12,8,10,28,8,1,5 | 80,145,68,79,51,75,43,102 | 23,33,13,20,8,14,7,24,21,1 | 298 | 149 | 0.0000262695063679 | 0.00756895866809837 | 0.874,0.872,0.821, | 0.635,0.687,0.723, | 0,121 |
| 180292 | ENSG00000146409 | SLC18B1 | chr6 | - | 132770889 | 132770932 | 132769369 | 132770336 | 132771035 | 132771129 | 180292 | 38,18,12,10,5,8,1,11,7,4,2, | 3,0,4,1,1,1,0,2,0,0,0,1 | 14,3,12,7,8,8,4,5,7,11,4,2,1 | 2,1,1,10,4,0,0,3,2,0,3,0,3,2 | 192 | 149 | 0.0000263049953556 | 0.00756905141093735 | 0.908,1.0,0.7,0.886 | 0.845,0.7,0.903,0.3 | 0,091 |
| 161378 | ENSG00000165819 | METTL3 | chr14 | - | 21510566 | 21510980 | 21503663 | 21503881 | 21511123 | 21511342 | 161378 | 4,1,4,1,3,3,5,2,4,0,2,4 | 38,12,94,57,70,75,45,57,55 | 4,15,3,3,4,1,7,9,3,5,6,6,3,2 | 63,49,29,7,29,19,42,75,19, | 298 | 149 | 0.0000267319027826 | 0.00768162105180478 | 0.05,0.04,0.021,0.0 | 0.031,0.133,0.049, | -0,036 |
| 243183 | ENSG00000290921 | | chr10 | - | 45965788 | 45965855 | 45962810 | 45962994 | 45972216 | 45972428 | 243183 | 2,0,2,0,2,3,0,4,0,0,1,0 | 11,3,12,5,8,13,7,13,5,3,6,1 | 3,4,2,0,3,0,1,0,2,0,2,4,3,5,1 | 7,12,4,7,6,2,9,3,6,6,4,1,4,5 | 216 | 149 | 0.0000267847152933 | 0.00768653474305633 | 0.111,0.0,0.103,0.0 | 0.228,0.187,0.256, | -0,133 |
| 92222 | ENSG00000124523 | SIRT5 | chr6 | + | 13599031 | 13599155 | 13596962 | 13597016 | 13600833 | 13600949 | 92222 | 15,0,33,34,7,15,4,15,6,2,7, | 2,3,9,8,2,0,2,7,0,0,0,1 | 23,8,6,4,8,7,5,12,8,15,20,1 | 0,0,2,1,0,0,2,2,0,1,1,4,3,0,1 | 273 | 149 | 0.0000270828505947 | 0.00775486398479112 | 0.804,0.0,0.667,0.6 | 1.0,1.0,0.621,0.686 | -0,132 |
| 242435 | ENSG00000137145 | DENND4C | chr9 | + | 19358027 | 19358160 | 19356971 | 19357154 | 19360243 | 19360489 | 242435 | 15,10,46,39,12,14,2,7,7,5,5 | 4,3,2,0,0,2,0,7,1,3,0,0 | 20,11,7,2,3,7,18,15,5,25,14 | 1,1,2,0,0,0,1,1,3,0,3,2,2,0,1 | 282 | 149 | 0.0000270948781382 | 0.00775486398479112 | 0.665,0.638,0.924, | 0.914,0.853,0.649, | -0,064 |
| 168221 | ENSG00000138639 | ARHGAP24 | chr4 | + | 85604225 | 85604303 | 85570592 | 85570721 | 85721884 | 85721972 | 168221 | 2,1,41,27,7,26,18,66,26,11 | 198,149,408,178,125,188,1 | 51,17,21,21,22,33,25,34,27 | 335,282,171,211,196,103,1 | 227 | 149 | 0.0000274400542184 | 0.00784322750263008 | 0.007,0.004,0.062, | 0.091,0.038,0.075, | -0,037 |
| 50662 | ENSG00000146409 | SLC18B1 | chr6 | - | 132770889 | 132770959 | 132769384 | 132770336 | 132771035 | 132771129 | 50662 | 41,18,23,10,7,9,1,15,8,4,4, | 3,0,4,1,1,1,0,2,0,0,0,1 | 18,4,12,9,9,8,6,6,8,22,5,2,2 | 2,1,1,10,4,0,0,3,2,0,3,0,3,2 | 219 | 149 | 0.0000276211623543 | 0.00788452306986517 | 0.903,1.0,0.796,0.8 | 0.86,0.731,0.891,0 | 0,094 |
| 202805 | ENSG00000112033 | PPARD | chr6 | + | 35348913 | 35349003 | 35347066 | 35347150 | 35410986 | 35411217 | 202805 | 7,4,3,2,2,0,1,8,4,1,3,4 | 31,33,19,12,14,10,8,6,14,1 | 1,1,0,1,0,2,1,0,2,3,1,1,3,3,0 | 13,10,21,20,12,3,11,13,4,1 | 239 | 149 | 0.0000277176111942 | 0.00790157506614418 | 0.123,0.07,0.09,0.0 | 0.046,0.059,0.0,0.0 | 0,068 |
| 52244 | ENSG00000102710 | SUPT20H | chr13 | - | 37047345 | 37047408 | 37045246 | 37045373 | 37047534 | 37047601 | 52244 | 5,0,7,0,7,0,3,1,1,0,0,2 | 20,19,45,19,16,11,5,6,22,8 | 1,1,1,0,0,1,0,1,0,1,1,0,0,0,2 | 20,16,10,4,16,13,4,10,4,27 | 212 | 149 | 0.0000281350318252 | 0.0080099616862032 | 0.149,0.0,0.099,0.0 | 0.034,0.042,0.066, | 0,048 |
| 103999 | ENSG00000291151 | NSUN5P1 | chr7 | + | 75415591 | 75415770 | 75415119 | 75415234 | 75415980 | 75416191 | 103999 | 28,17,56,29,55,51,42,48,11 | 1,0,1,1,4,5,1,0,0,0,0,0 | 52,25,14,5,20,29,18,57,18, | 1,0,2,0,1,0,9,2,3,0,0,5,1,0,5 | 298 | 149 | 0.0000283690828313 | 0.00806592611210506 | 0.933,1.0,0.966,0.9 | 0.963,1.0,0.778,1.0 | 0,06 |
| 153311 | ENSG00000092199 | HNRNPC | chr14 | - | 21263619 | 21263829 | 21263310 | 21263336 | 21269297 | 21269389 | 153311 | 553,360,379,322,140,138,2 | 29,16,68,31,7,9,2,10,7,2,2, | 299,324,333,293,156,112,9 | 42,26,21,6,13,19,8,11,8,33 | 298 | 149 | 0.0000285403340988 | 0.00810391114567127 | 0.905,0.918,0.736, | 0.781,0.862,0.888, | 0,065 |
| 45639 | ENSG00000206562 | METTL6 | chr3 | - | 15414729 | 15414914 | 15414020 | 15414162 | 15415497 | 15415583 | 45639 | 2,1,2,6,1,1,0,8,0,3,2,2 | 7,0,7,2,3,0,1,5,5,1,2,2 | 0,2,0,1,1,0,0,7,0,3,0,1,0,0,1 | 2,2,3,2,2,1,4,3,5,1,5,4,2,4,3 | 298 | 149 | 0.0000286458430443 | 0.00811390847045994 | 0.125,1.0,0.125,0.6 | 0.0,0.333,0.0,0.2,0 | 0,259 |
| 20382 | ENSG00000141956 | PRDM15 | chr21 | - | 41825959 | 41826054 | 41821902 | 41822037 | 41828165 | 41828281 | 20382 | 6,3,13,11,6,12,6,7,9,5,5,1 | 0,1,0,2,1,1,1,3,0,3,0,0 | 9,3,18,4,17,19,0,7,10,12,7, | 3,1,0,3,5,1,1,0,2,1,0,3,1,1,2 | 244 | 149 | 0.0000286659183462 | 0.00811390847045994 | 1.0,0.647,1.0,0.77 | 0.647,0.647,1.0,0.4 | 0,121 |
| 225185 | ENSG00000118058 | KMT2A | chr11 | + | 118492994 | 118493230 | 118491743 | 118491928 | 118494287 | 118494398 | 225185 | 93,44,71,49,41,64,38,61,44 | 3,2,8,3,2,7,1,1,2,0,0,2 | 52,66,47,71,42,40,29,23,46 | 0,1,1,4,4,0,2,0,1,0,0,5,0,0,0 | 298 | 149 | 0.0000286886384676 | 0.00811390847045994 | 0.939,0.917,0.816, | 1.0,0.971,0.959,0.8 | -0,038 |
| 26274 | ENSG00000196655 | TRAPPC4 | chr11 | + | 119020149 | 119020253 | 119018765 | 119019317 | 119021759 | 119021886 | 26274 | 22,16,39,71,39,19,30,20,18 | 0,1,3,3,5,1,0,0,0,0,2,0 | 35,31,30,9,19,14,4,5,12,22 | 4,4,2,1,1,0,0,1,0,3,0,4,0,3,3 | 253 | 149 | 0.0000288836802836 | 0.00815835090698098 | 1.0,0.904,0.884,0.9 | 0.837,0.82,0.898,0 | 0,045 |
| 33833 | ENSG00000127419 | TMEM175 | chr4 | + | 947708 | 948154 | 932474 | 932540 | 950420 | 950491 | 33833 | 35,23,10,6,6,12,16,5,19,12 | 0,0,3,0,1,2,2,0,0,1,1,0 | 7,16,10,14,3,1,9,5,7,11,4,5 | 4,0,0,1,3,0,2,1,0,2,0,11,0,0 | 298 | 149 | 0.0000290984320032 | 0.00820823672146886 | 1.0,1.0,0.625,1.0,0 | 0.467,1.0,1.0,0.875 | 0,07 |
| 13117 | ENSG00000177663 | IL17RA | chr22 | + | 17100354 | 17100481 | 17098774 | 17098887 | 17101995 | 17102043 | 13117 | 27,28,36,20,29,21,14,19,49 | 3,2,1,0,4,3,1,0,1,0,0,0 | 23,19,7,15,12,17,16,28,3,3 | 3,5,2,2,2,0,0,2,1,1,6,0,0,1,5 | 276 | 149 | 0.0000291746226905 | 0.0082189570893966 | 0.829,0.883,0.951, | 0.805,0.672,0.654, | 0,094 |
| 76303 | ENSG00000076242 | MLH1 | chr3 | + | 37014431 | 37014544 | 37012010 | 37012099 | 37025636 | 37026007 | 76303 | 12,5,22,23,9,12,7,11,5,1,2, | 0,2,3,0,1,1,0,3,0,1,1,0 | 12,5,11,1,2,9,4,2,2,10,9,6,1 | 0,3,3,0,4,2,4,0,2,1,2,2,1,0,2 | 262 | 149 | 0.0000292952671802 | 0.00824215640583219 | 1.0,0.587,0.807,1.0 | 1.0,0.487,0.676,1.0 | 0,098 |
| 21470 | ENSG00000092199 | HNRNPC | chr14 | - | 21263666 | 21263829 | 21263310 | 21263336 | 21269297 | 21269406 | 21470 | 555,358,377,322,138,135,2 | 29,16,68,31,7,9,2,10,7,2,2, | 300,324,334,291,159,112,9 | 42,26,21,6,13,19,8,11,8,33 | 298 | 149 | 0.0000297005940628 | 0.00834409573035234 | 0.905,0.918,0.735, | 0.781,0.862,0.888, | 0,065 |
| 112875 | ENSG00000147439 | BIN3 | chr8 | - | 22634412 | 22634525 | 22630441 | 22630578 | 22636524 | 22636586 | 112875 | 0,0,7,12,2,9,2,0,0,0,0,0 | 11,2,19,28,13,9,6,13,17,12 | 1,0,0,0,1,0,2,5,0,0,0,3,0,4,3 | 17,8,10,6,11,3,3,6,3,9,12,1 | 262 | 149 | 0.000029735128421 | 0.00834409573035234 | 0.0,0.0,0.173,0.196 | 0.032,0.0,0.0,0.0,0 | 0,011 |
| 243382 | ENSG00000165121 | | chr9 | + | 85829462 | 85829591 | 85822879 | 85823080 | 85837758 | 85837965 | 243382 | 32,22,49,18,6,7,8,5,20,9,8, | 0,0,5,3,0,4,3,0,0,0,0,1 | 13,12,15,28,12,14,2,12,7,2 | 2,4,2,2,1,3,9,1,3,2,5,2,5,1,0 | 278 | 149 | 0.000029802658741 | 0.00835215630662001 | 1.0,1.0,0.84,0.763, | 0.777,0.617,0.801, | 0,173 |
| 74157 | ENSG00000064687 | ABCA7 | chr19 | + | 1054569 | 1054694 | 1054192 | 1054341 | 1054779 | 1054878 | 74157 | 19,15,23,8,14,15,31,15,22, | 0,0,2,0,1,0,2,3,4,8,0,1 | 10,20,3,8,15,3,9,6,6,9,9,54 | 0,0,0,0,0,1,0,1,0,0,2,4,1,2,8 | 274 | 149 | 0.0000298563434022 | 0.00835632073705969 | 1.0,1.0,0.862,1.0,0 | 1.0,1.0,1.0,1.0,1.0, | -0,026 |
| 237882 | ENSG00000211772 | TRBC2 | chr7 | + | 142801904 | 142801961 | 142801040 | 142801427 | 142802104 | 142802211 | 237882 | 11,6,26,9,7,20,10,9,30,14,6 | 1,0,3,1,0,0,0,0,2,0,1,1 | 11,11,6,12,4,2,10,18,5,12,1 | 4,1,1,1,2,2,0,6,0,4,1,1,0,0,4 | 206 | 149 | 0.0000300548993796 | 0.0084009688939879 | 0.888,1.0,0.862,0.8 | 0.665,0.888,0.813, | 0,119 |
| 176711 | ENSG00000165115 | KIF27 | chr9 | - | 83920105 | 83920212 | 83915293 | 83915678 | 83921370 | 83921465 | 176711 | 0,0,5,4,9,3,2,3,2,2,0,3 | 1,3,0,0,2,1,0,1,0,0,1,0 | 4,1,1,1,1,2,1,3,3,10,2,6,5,0 | 1,2,0,2,0,2,7,2,5,7,0,0,0,2,2 | 256 | 149 | 0.0000302952892012 | 0.00845717949424575 | 0.0,0.0,1.0,1.0,0.72 | 0.7,0.225,1.0,0.225 | 0,089 |
| 25431 | ENSG00000125257 | ABCC4 | chr13 | - | 95188452 | 95188542 | 95186700 | 95186892 | 95194835 | 95194937 | 25431 | 12,10,20,12,10,3,6,10,5,14 | 6,2,2,1,0,1,9,2,1,0,0,0 | 2,6,2,5,2,1,4,6,1,9,5,18,7,6 | 0,0,0,0,0,1,0,2,2,0,0,4,0,1,5 | 239 | 149 | 0.0000303938355933 | 0.0084736990007533 | 0.555,0.757,0.862, | 1.0,1.0,1.0,1.0,1.0, | -0,017 |
| 225804 | ENSG00000235621 | LINC00494 | chr20 | + | 48365872 | 48366338 | 48363813 | 48364088 | 48366473 | 48367226 | 225804 | 0,1,5,1,1,2,1,7,0,0,1,4 | 19,6,33,26,23,15,2,25,6,8,1 | 4,5,0,2,6,1,0,8,1,0,15,1,1,2 | 13,6,3,12,10,7,14,11,5,4,11 | 298 | 149 | 0.0000304385899574 | 0.0084751981308165 | 0.0,0.077,0.07,0.01 | 0.133,0.294,0.0,0.0 | -0,077 |
| 240155 | ENSG00000165525 | NEMF | chr14 | - | 49833422 | 49833848 | 49832206 | 49832277 | 49834362 | 49834449 | 240155 | 30,14,39,25,9,13,18,26,23, | 2,1,3,1,1,1,0,0,5,2,0,1 | 34,30,10,14,19,13,7,24,21, | 5,5,0,4,3,1,2,2,3,9,0,2,7,3,4 | 298 | 149 | 0.0000305003841127 | 0.00848143174801102 | 0.882,0.875,0.867, | 0.773,0.75,1.0,0.63 | 0,145 |
| 105669 | ENSG00000133703 | KRAS | chr12 | - | 25213113 | 25213206 | 25205245 | 25209911 | 25225613 | 25225773 | 105669 | 2,0,1,2,0,2,1,2,0,0,0,3 | 3,3,69,50,39,32,21,39,6,4,6 | 3,1,1,14,6,1,0,1,5,5,4,2,0,0 | 22,96,32,55,50,10,22,17,28 | 242 | 149 | 0.000030566821551 | 0.00848893879902358 | 0.291,0.0,0.009,0.0 | 0.077,0.006,0.019, | -0,028 |
| 162420 | ENSG00000146067 | FAM193B | chr5 | - | 177547203 | 177547228 | 177538910 | 177539147 | 177553167 | 177553266 | 162420 | 12,2,2,5,0,2,0,3,9,3,2,3 | 23,9,56,12,18,29,12,23,10, | 2,1,0,4,1,0,3,7,0,1,2,0,1,2,3 | 29,23,19,10,16,9,29,30,21, | 174 | 149 | 0.0000306972393934 | 0.00851341263023713 | 0.309,0.16,0.03,0.2 | 0.056,0.036,0.0,0.2 | 0,097 |
| 142031 | ENSG00000129197 | RPAIN | chr17 | + | 5425970 | 5426071 | 5422768 | 5422829 | 5426235 | 5426299 | 142031 | 18,2,60,34,13,17,10,16,9,9 | 0,0,2,9,6,3,3,2,0,2,0,0 | 18,16,14,6,10,14,6,25,6,19 | 1,0,0,0,0,0,1,0,0,0,9,3,3,2 | 250 | 149 | 0.0000307340560314 | 0.00851341263023713 | 1.0,1.0,0.947,0.69 | 0.915,1.0,1.0,1.0,1 | -0,078 |
| 132574 | ENSG00000112200 | ZNF451 | chr6 | + | 57099468 | 57099568 | 57099060 | 57099141 | 57124733 | 57124859 | 132574 | 0,0,8,0,0,2,0,3,0,0,0,3 | 24,31,37,20,13,30,9,24,12, | 8,9,2,4,3,1,2,1,1,0,7,1,1,0,2 | 16,33,18,30,20,10,14,10,14 | 249 | 149 | 0.0000309400202565 | 0.00854944675432912 | 0.0,0.0,0.115,0.0,0 | 0.23,0.14,0.062,0.0 | -0,044 |
| 238251 | ENSG00000132294 | EFR3A | chr8 | + | 132000768 | 132000901 | 131996389 | 131996497 | 132001758 | 132001807 | 238251 | 5,1,10,3,2,1,1,2,0,0,0,0 | 35,22,44,42,12,17,16,15,64 | 5,9,1,6,2,3,0,5,4,2,0,8,7,6,1 | 32,12,39,8,7,12,9,15,4,37,7 | 282 | 149 | 0.0000309435862939 | 0.00854944675432912 | 0.07,0.023,0.107,0 | 0.076,0.284,0.013, | -0,083 |
| 237075 | ENSG00000173064 | HECTD4 | chr12 | - | 112192306 | 112192362 | 112190785 | 112190965 | 112192559 | 112192765 | 237075 | 0,0,2,0,0,0,0,1,0,0,0,0 | 9,13,30,15,17,25,16,20,9,4 | 4,2,3,1,1,0,5,0,3,0,2,5,3,0,1 | 11,13,4,13,7,3,19,17,18,23 | 205 | 149 | 0.0000311293065295 | 0.00858973304314207 | 0.0,0.0,0.046,0.0,0 | 0.209,0.101,0.353, | -0,087 |
| 211826 | ENSG00000137497 | NUMA1 | chr11 | - | 72011958 | 72012442 | 72010785 | 72010854 | 72012894 | 72012926 | 211826 | 57,33,18,25,11,13,13,13,37 | 67,38,141,80,56,63,46,57,3 | 9,19,8,1,10,11,9,16,11,19,6 | 68,62,48,34,39,27,77,99,36 | 298 | 149 | 0.0000314593814975 | 0.00866969800139362 | 0.298,0.303,0.06,0 | 0.062,0.133,0.077, | 0,07 |
| 69657 | ENSG00000187605 | TET3 | chr2 | + | 74046277 | 74048411 | 74003109 | 74003166 | 74073548 | 74073639 | 69657 | 8,8,19,21,13,9,5,6,6,1,3,8 | 3,2,0,0,1,1,1,0,0,1,0,2 | 15,22,4,8,16,2,6,6,7,11,7,2 | 4,4,0,2,1,0,0,6,2,3,2,0,3,2,2 | 298 | 149 | 0.0000317016405073 | 0.00872528873149192 | 0.571,0.667,1.0,1.0 | 0.652,0.733,1.0,0.6 | 0,057 |
| 93079 | ENSG00000145996 | CDKAL1 | chr6 | + | 21108400 | 21108463 | 21065047 | 21065228 | 21198020 | 21198104 | 93079 | 31,22,60,27,26,10,9,13,30, | 8,2,1,0,0,2,1,4,4,1,0,2 | 26,20,19,11,12,9,10,13,2,1 | 0,3,0,0,1,0,0,1,0,1,1,1,3,0,2 | 212 | 149 | 0.0000318788410031 | 0.00876285418638341 | 0.731,0.885,0.977, | 1.0,0.824,1.0,1.0,0 | -0,073 |
| 170974 | ENSG00000115524 | SF3B1 | chr2 | - | 197398968 | 197399038 | 197398460 | 197398581 | 197400054 | 197400166 | 170974 | 63,46,109,31,41,55,41,65,4 | 702,516,829,512,391,507,3 | 46,90,41,23,43,24,42,77,29 | 497,579,369,255,384,226,3 | 219 | 149 | 0.00003220652582 | 0.00884163617189339 | 0.058,0.057,0.082, | 0.059,0.096,0.07,0 | -0,019 |
| 184325 | ENSG00000004487 | KDM1A | chr1 | + | 23040258 | 23040434 | 23030468 | 23030634 | 23044426 | 23044486 | 184325 | 9,8,4,3,2,6,1,6,2,2,5,7 | 9,6,19,8,14,13,13,14,24,12 | 10,27,7,16,4,4,21,9,7,8,17, | 15,7,6,8,2,2,8,27,1,8,10,17 | 298 | 149 | 0.0000322858013468 | 0.00885210867474281 | 0.333,0.4,0.095,0.1 | 0.25,0.659,0.368,0 | -0,21 |
| 43853 | ENSG00000116095 | PLEKHA3 | chr2 | + | 178488967 | 178489014 | 178485640 | 178485757 | 178490658 | 178490814 | 43853 | 22,8,26,31,2,9,5,8,8,10,7,1 | 15,29,36,44,6,14,12,19,14, | 3,12,8,3,8,3,2,1,4,1,4,14,6, | 21,29,5,9,17,17,12,20,18,2 | 196 | 149 | 0.0000323867967748 | 0.00886850210768063 | 0.527,0.173,0.354, | 0.098,0.239,0.549, | 0,145 |
| 78494 | ENSG00000160352 | ZNF714 | chr19 | + | 21098811 | 21098910 | 21098184 | 21098311 | 21116806 | 21118885 | 78494 | 3,7,11,10,4,5,9,3,1,3,0,2 | 0,0,1,2,1,0,0,1,2,0,2,0 | 8,2,10,2,2,5,3,8,9,2,4,5,6,7 | 0,3,0,1,4,4,4,5,2,0,3,0,0,1,0 | 248 | 149 | 0.0000327260855688 | 0.00895002302802845 | 1.0,1.0,0.869,0.75, | 1.0,0.286,1.0,0.546 | 0,062 |
| 98419 | ENSG00000118058 | KMT2A | chr11 | + | 118493112 | 118493230 | 118491743 | 118491928 | 118494287 | 118494398 | 98419 | 84,41,60,44,34,52,26,52,39 | 3,2,8,3,2,7,1,1,2,0,0,2 | 45,57,46,58,36,33,22,21,35 | 0,1,1,4,4,0,2,0,1,0,0,5,0,0,0 | 267 | 149 | 0.0000329327173899 | 0.00899218417670219 | 0.94,0.92,0.807,0.8 | 1.0,0.97,0.963,0.89 | -0,04 |
| 127503 | ENSG00000107862 | GBF1 | chr10 | + | 102359266 | 102359435 | 102358505 | 102358729 | 102360183 | 102360395 | 127503 | 82,48,50,21,31,34,26,35,73 | 7,2,1,3,0,0,5,2,2,2,2,0 | 25,26,21,57,17,18,10,45,9, | 6,0,0,2,0,0,4,2,0,0,0,0,1,1,0 | 298 | 149 | 0.0000329638077945 | 0.00899218417670219 | 0.854,0.923,0.962, | 0.676,1.0,1.0,0.934 | -0,022 |
| 71414 | ENSG00000112249 | ASCC3 | chr6 | - | 100864063 | 100864214 | 100848003 | 100848707 | 100867907 | 100868032 | 71414 | 215,158,190,74,56,89,64,6 | 5,4,4,4,3,8,3,1,6,1,0,1 | 113,107,73,89,59,65,54,57 | 3,4,8,8,21,2,3,4,2,1,5,9,6,3 | 298 | 149 | 0.0000330424414354 | 0.00900222495263617 | 0.956,0.952,0.96,0 | 0.95,0.93,0.82,0.84 | 0,083 |
| 16496 | ENSG00000153774 | CFDP1 | chr16 | - | 75414577 | 75414695 | 75412534 | 75412754 | 75433288 | 75433461 | 16496 | 97,32,74,73,29,25,29,16,62 | 7,3,5,0,5,4,0,7,6,2,0,1 | 48,13,37,29,16,19,13,25,8, | 0,4,2,2,0,1,1,2,1,1,4,0,0,0,2 | 267 | 149 | 0.0000335860651831 | 0.00913876409029557 | 0.885,0.856,0.892, | 1.0,0.645,0.912,0.8 | -0,042 |
| 103497 | ENSG00000179454 | KLHL28 | chr14 | - | 44931332 | 44931541 | 44924323 | 44929191 | 44934114 | 44934558 | 103497 | 24,14,62,22,19,12,14,22,11 | 1,0,4,2,3,0,0,2,0,0,2,0 | 38,36,27,17,28,10,15,33,16 | 3,1,0,8,1,0,0,2,1,5,1,3,3,0,0 | 298 | 149 | 0.0000339452318475 | 0.00922483105526423 | 0.923,1.0,0.886,0.8 | 0.864,0.947,1.0,0.5 | 0,034 |
| 71260 | ENSG00000131051 | RBM39 | chr20 | - | 35713701 | 35713857 | 35713018 | 35713096 | 35714184 | 35714389 | 71260 | 59,30,79,49,69,128,77,117 | 111,57,424,254,141,147,13 | 98,49,65,31,50,40,39,103,1 | 215,260,132,136,188,104,1 | 298 | 149 | 0.0000342011735994 | 0.00928266430639654 | 0.21,0.208,0.085,0 | 0.186,0.086,0.198, | 0,084 |
| 154179 | ENSG00000213463 | SYNJ2BP | chr14 | - | 70381147 | 70381198 | 70375675 | 70375771 | 70388469 | 70388606 | 154179 | 1,0,3,3,0,2,5,2,2,2,0,3 | 4,8,16,9,10,4,15,6,3,4,6,9 | 1,2,0,1,0,1,0,1,3,0,0,0,0,4,0 | 12,11,10,4,5,10,1,1,6,8,11, | 200 | 149 | 0.0000342506514748 | 0.00928438534958775 | 0.157,0.0,0.123,0.1 | 0.058,0.119,0.0,0.1 | 0,082 |
| 129648 | ENSG00000102225 | CDK16 | chrX | + | 47225771 | 47225866 | 47224987 | 47225102 | 47225964 | 47226027 | 129648 | 30,38,97,34,35,37,32,35,17 | 3,0,0,3,0,0,0,0,0,0,0,0 | 40,40,24,58,65,11,26,50,38 | 2,6,2,4,2,1,2,0,1,3,0,7,0,0,0 | 244 | 149 | 0.0000349536829406 | 0.00946303916099154 | 0.859,1.0,1.0,0.874 | 0.924,0.803,0.88,0 | 0,05 |

| | | | | | | | | | | | | | | | | | | | | | | |
|---|---|---|---|---|---|---|---|---|---|---|---|---|---|---|---|---|---|---|---|---|---|---|
| 163294 | ENSG00000164180 | TMEM161B | chr5 | - | 88236033 | 88236082 | 88228458 | 88228528 | 88240812 | 88240916 | 163294 | 1,3,1,0,1,0,0,0,0,0,3 | 46,36,38,30,17,24,10,12,1 | 3,4,2,0,6,0,0,2,2,0,5,4,1,2, | 17,21,13,9,21,10,10,12,11, | 198 | 149 | 0.0000353615099254 | 0.00956142354617307 | 0.016,0.059,0.019, | 0.117,0.125,0.104, | -0,051 |
| 75824 | ENSG00000149532 | CPSF7 | chr11 | - | 61421191 | 61421257 | 61420469 | 61420573 | 61421389 | 61421608 | 75824 | 2,4,12,18,2,1,5,5,0,0,0,0 | 39,27,82,42,38,29,31,32,1 | 3,2,1,0,2,7,1,3,3,1,2,0,2,3, | 43,50,33,29,38,27,14,55,14 | 215 | 149 | 0.000035705110115 | 0.00964221650584889 | 0.034,0.093,0.092, | 0.046,0.027,0.021, | 0,02 |
| 186812 | ENSG00000116560 | SFPQ | chr1 | - | 35178664 | 35178754 | 35177988 | 35178036 | 35181134 | 35181606 | 186812 | 3,0,2,1,0,3,0,0,0,0,2,1 | 103,13,50,46,34,60,36,60, | 1,5,0,7,4,2,1,3,6,9,1,3,0,2, | 69,33,66,99,17,13,13,74,2 | 239 | 149 | 0.0000358840055953 | 0.00967838397027821 | 0.018,0.0,0.024,0. | 0.009,0.086,0.0,0. | -0,04 |
| 201007 | ENSG00000120733 | KDM3B | chr5 | + | 138428689 | 138428814 | 138427966 | 138428086 | 138429825 | 138429965 | 201007 | 4,6,7,8,1,2,1,4,1,0,0,0 | 73,57,98,65,45,34,49,44,1 | 9,3,7,12,1,2,4,4,3,6,14,2,4, | 45,42,42,51,46,20,28,46,2 | 274 | 149 | 0.0000362152159099 | 0.00975549078285818 | 0.029,0.054,0.037, | 0.098,0.037,0.083, | -0,032 |
| 141469 | ENSG00000290698 | | chr17 | - | 16926423 | 16926489 | 16925845 | 16926140 | 16926868 | 16926937 | 141469 | 13,18,39,27,20,5,16,9,18,1 | 0,0,5,0,0,2,1,2,0,1,0,2 | 46,31,32,68,38,13,8,39,33, | 3,6,1,13,1,1,1,6,1,1,2,4,0,4 | 215 | 149 | 0.0000363922632305 | 0.00978739589737151 | 1.0,1.0,0.844,1.0,1 | 0.914,0.782,0.957, | 0,045 |
| 131987 | ENSG00000089154 | GCN1 | chr12 | - | 120138322 | 120138415 | 120137900 | 120138044 | 120138694 | 120138856 | 131987 | 28,31,53,79,45,64,38,40,3 | 6,0,3,0,2,1,0,4,1,0,0,1 | 40,46,37,30,26,13,41,43,1 | 4,4,1,16,9,0,0,2,1,2,5,4,6,0 | 242 | 149 | 0.0000364684752794 | 0.00978739589737151 | 0.742,1.0,0.916,1. | 0.86,0.876,0.958,0 | 0,061 |
| 251800 | ENSG00000120690 | ELF1 | chr13 | - | 40961442 | 40961561 | 40958835 | 40959016 | 40981982 | 40982282 | 251800 | 37,3,21,15,1,7,4,6,4,6,3,6 | 540,370,520,267,192,189, | 20,19,18,39,15,4,1,17,16,8 | 350,483,303,408,341,138, | 268 | 149 | 0.0000364966607789 | 0.00978739589737151 | 0.037,0.004,0.022, | 0.031,0.021,0.032, | -0,013 |
| 136925 | ENSG00000079805 | DNM2 | chr19 | + | 10797414 | 10797518 | 10795371 | 10795439 | 10798485 | 10798572 | 136925 | 28,26,13,10,12,20,16,25,3 | 1,3,6,1,0,3,2,1,2,0,2,1 | 24,13,16,25,15,3,22,24,3,1 | 0,2,0,0,1,0,2,2,0,0,1,2,4,0, | 253 | 149 | 0.0000365155526183 | 0.00978739589737151 | 0.943,0.836,0.561, | 1.0,0.793,1.0,1.0,0 | -0,101 |
| 83478 | ENSG00000137497 | NUMA1 | chr11 | - | 72012400 | 72012442 | 72010785 | 72010854 | 72012894 | 72016260 | 83478 | 80,65,81,81,51,33,38,28,6 | 67,38,141,80,56,63,46,57, | 50,52,14,5,18,18,25,54,28, | 68,62,48,34,39,27,77,99,3 | 191 | 149 | 0.0000368919998645 | 0.00987599754084104 | 0.482,0.572,0.309, | 0.365,0.396,0.185, | 0,133 |
| 41759 | ENSG00000211897 | IGHG3 | chr14 | - | 105770104 | 105770149 | 105769656 | 105769986 | 105770292 | 105770337 | 41759 | 0,1,189,119,21,15,98,121, | 40,16,88,67,21,34,3,1,1,1, | 78,109,38,22,0,0,13,36,0,2 | 53,68,2,0,11,18,37,140,4,1 | 194 | 149 | 0.0000371821372678 | 0.00994130259165945 | 0.0,0.046,0.623,0. | 0.531,0.552,0.936, | 0,261 |
| 86265 | ENSG00000151687 | ANKAR | chr2 | + | 189676455 | 189677091 | 189674669 | 189674830 | 189689526 | 189689964 | 86265 | 0,5,5,3,0,0,1,5,1,0,0,0 | 11,17,20,13,13,19,11,15,1 | 4,10,2,4,5,2,2,3,5,6,4,1,2,0 | 14,19,14,4,18,3,3,8,10,9,1 | 298 | 149 | 0.0000372595730493 | 0.00994964660914874 | 0.0,0.128,0.111,0. | 0.125,0.208,0.067, | -0,102 |
| 193649 | ENSG00000114861 | FOXP1 | chr3 | - | 71130497 | 71130741 | 71112535 | 71112637 | 71132090 | 71132211 | 193649 | 192,117,305,253,116,121, | 17,10,21,26,10,26,5,19,5,1 | 156,126,101,72,157,138,7 | 5,6,10,7,4,6,7,4,8,3,6,1,3,2 | 298 | 149 | 0.0000375709624338 | 0.0100203665620684 | 0.85,0.854,0.879,0 | 0.94,0.913,0.835,0 | -0,048 |
| 189473 | ENSG00000234127 | TRIM26 | chr6 | - | 30201034 | 30201123 | 30198665 | 30199153 | 30213304 | 30213427 | 189473 | 32,23,27,29,17,14,7,9,10,7 | 0,0,0,3,0,0,1,1,2,4,2,0 | 11,29,4,7,13,4,10,9,2,10,1 | 6,0,0,0,1,4,2,0,0,0,3,8,1,2, | 238 | 149 | 0.0000377099612026 | 0.0100344328497682 | 1.0,1.0,1.0,0.858,1 | 0.534,1.0,1.0,1.0,0 | 0,044 |
| 244915 | ENSG00000136527 | TRA2B | chr3 | - | 185930112 | 185930226 | 185926726 | 185926734 | 185931576 | 185931852 | 244915 | 11,0,10,12,4,0,2,6,3,0,1,1 | 98,34,105,82,65,44,48,52, | 4,4,1,1,0,1,0,4,0,2,2,4,2,4, | 88,78,87,101,35,49,35,66, | 263 | 149 | 0.0000377169467942 | 0.0100344328497682 | 0.06,0.0,0.051,0.0 | 0.025,0.028,0.006, | 0,015 |
| 22630 | ENSG00000032219 | ARID4A | chr14 | + | 58306021 | 58306112 | 58304943 | 58305009 | 58318541 | 58318621 | 22630 | 84,64,112,57,40,41,27,42, | 4,0,7,1,0,0,1,1,2,0,0,3 | 39,122,33,33,61,46,26,33, | 6,8,2,15,0,2,3,0,4,1,2,0,4,3 | 240 | 149 | 0.0000378909240636 | 0.0100564180968835 | 0.929,1.0,0.909,0. | 0.801,0.904,0.911, | 0,065 |
| 129621 | ENSG00000127314 | RAP1B | chr12 | + | 68624674 | 68624873 | 68610924 | 68611043 | 68648698 | 68648781 | 129621 | 0,0,2,7,0,0,0,2,2,2,1,0 | 16,14,21,11,10,12,9,7,17,1 | 2,2,3,1,0,0,1,3,3,0,3,7,4,10 | 10,21,10,9,14,8,9,2,3,14,1 | 298 | 149 | 0.0000379052822235 | 0.0100564180968835 | 0.0,0.0,0.045,0.24 | 0.091,0.045,0.13,0 | -0,074 |
| 33809 | ENSG00000087266 | SH3BP2 | chr4 | + | 2831922 | 2831978 | 2831570 | 2831679 | 2832330 | 2832412 | 33809 | 59,25,69,33,22,9,34,28,6,3 | 5,0,4,3,2,0,2,0,0,0,1,0 | 33,22,12,10,12,6,19,20,15, | 2,1,8,2,5,0,5,1,3,0,2,4,0,1, | 205 | 149 | 0.0000379635934169 | 0.0100564180968835 | 0.896,1.0,0.926,0. | 0.923,0.941,0.522, | 0,102 |
| 136345 | ENSG00000181896 | ZNF101 | chr19 | + | 19678725 | 19678837 | 19677863 | 19677990 | 19679180 | 19683506 | 136345 | 36,51,143,89,27,45,35,58, | 4,1,11,3,1,11,5,10,3,6,4,5 | 89,113,45,68,48,44,25,42, | 13,13,5,33,13,4,6,8,1,6,19, | 261 | 149 | 0.0000380151748899 | 0.0100564180968835 | 0.837,0.967,0.881, | 0.796,0.832,0.837, | 0,105 |
| 68573 | ENSG00000055609 | KMT2C | chr7 | - | 152224434 | 152224616 | 152222572 | 152222682 | 152229922 | 152230027 | 68573 | 19,20,30,19,10,18,10,12,6, | 0,0,2,1,0,4,2,2,0,0,0,0 | 19,23,6,26,8,16,14,17,13,1 | 1,5,1,1,3,0,1,1,0,1,3,0,4,0, | 298 | 149 | 0.0000380332030742 | 0.0100564180968835 | 1.0,1.0,0.882,0.90 | 0.905,0.697,0.75,0 | 0,066 |
| 173217 | ENSG00000174243 | DDX23 | chr12 | - | 48835807 | 48835990 | 48834457 | 48834497 | 48836120 | 48836266 | 173217 | 0,1,14,7,3,3,2,4,3,0,1,1 | 52,26,95,68,17,26,32,25,3 | 5,1,0,0,1,1,1,0,2,1,0,1,2,5, | 48,61,13,13,22,25,22,40,1 | 298 | 149 | 0.0000381949071806 | 0.0100867829047702 | 0.0,0.019,0.069,0. | 0.05,0.008,0.0,0.0, | 0,025 |
| 130437 | ENSG00000136521 | NDUFB5 | chr3 | + | 179614970 | 179615059 | 179604689 | 179604939 | 179616982 | 179617044 | 130437 | 21,12,83,92,18,19,19,26,6, | 5,0,2,2,0,0,1,0,0,1,0,0 | 47,25,22,8,27,21,10,24,11, | 1,0,1,1,0,3,0,4,2,0,8,7,4,2, | 238 | 149 | 0.0000386137027217 | 0.010179996717839 | 0.724,1.0,0.963,0. | 0.967,1.0,0.932,0. | 0,051 |
| 125746 | ENSG00000197857 | ZNF44 | chr19 | - | 12275955 | 12276082 | 12274972 | 12275033 | 12294691 | 12294834 | 125746 | 22,22,71,48,20,22,22,31,2 | 0,0,2,3,2,2,5,1,0,0,0,0 | 21,20,33,9,24,12,5,18,13,1 | 6,0,1,0,4,0,0,0,1,0,6,1,3,2, | 276 | 149 | 0.0000386424693398 | 0.010179996717839 | 1.0,1.0,0.95,0.896, | 0.654,1.0,0.947,1. | 0,07 |
| 236309 | ENSG00000095485 | CWF19L1 | chr10 | - | 100246012 | 100246295 | 100245799 | 100245913 | 100246794 | 100246935 | 236309 | 7,1,7,5,2,4,2,5,6,0,4,3 | 55,24,59,20,17,20,9,24,46, | 5,9,8,19,4,4,2,10,3,15,1,5, | 23,32,14,14,20,13,8,15,18, | 298 | 149 | 0.0000388825831079 | 0.0102307301282352 | 0.06,0.02,0.056,0. | 0.098,0.123,0.222, | -0,064 |
| 51934 | ENSG00000170791 | CHCHD7 | chr8 | + | 56216679 | 56216754 | 56216388 | 56216531 | 56217330 | 56217412 | 51934 | 11,2,14,9,6,6,0,2,1,1,1,0 | 15,5,92,61,45,15,15,21,15, | 17,19,3,6,12,3,7,11,5,3,12, | 43,30,18,3,16,25,11,23,14, | 224 | 149 | 0.000039113809394 | 0.0102717988120912 | 0.328,0.21,0.092,0 | 0.208,0.296,0.1,0. | -0,114 |
| 241266 | ENSG00000107077 | KDM4C | chr9 | + | 6986343 | 6986553 | 6984165 | 6984404 | 6990415 | 6990524 | 241266 | 61,26,70,52,23,38,23,46,1 | 1,0,0,5,0,1,0,0,0,0,0,1 | 72,71,39,50,37,39,30,35,3 | 1,5,0,3,7,0,2,0,1,0,2,2,2,4, | 298 | 149 | 0.0000391341164884 | 0.0102717988120912 | 0.968,1.0,1.0,0.83 | 0.973,0.877,1.0,0. | 0,063 |
| 253139 | ENSG00000111144 | LTA4H | chr12 | - | 96011622 | 96011865 | 96009093 | 96009148 | 96013187 | 96013258 | 253139 | 23,15,47,6,5,3,4,6,1,2,5,7 | 224,107,416,326,157,108, | 4,12,5,6,3,3,6,5,4,11,7,5,3, | 275,164,226,55,136,112,7 | 298 | 149 | 0.0000395782745976 | 0.0103757266990451 | 0.049,0.066,0.053, | 0.007,0.035,0.011, | 0,007 |
| 212991 | ENSG00000145425 | RPS3A | chr4 | + | 151104169 | 151104286 | 151102870 | 151103079 | 151104471 | 151104490 | 212991 | 147,114,515,339,85,101,4 | 1,7,6,7,3,2,2,7,6,3,0,5 | 262,330,124,102,175,111, | 5,8,2,0,1,0,2,3,0,3,1,3,2,1, | 266 | 149 | 0.0000399072598756 | 0.0104352038672132 | 0.988,0.901,0.98,0 | 0.967,0.959,0.972, | -0,042 |
| 145457 | ENSG00000074370 | ATP2A3 | chr17 | - | 3928239 | 3928312 | 3925046 | 3925441 | 3928662 | 3928780 | 145457 | 3,10,5,13,1,5,1,0,7,6,3,4 | 80,60,111,70,80,62,49,34, | 4,4,4,6,2,1,1,2,2,2,1,0,1,2, | 101,125,32,24,62,56,16,71 | 222 | 149 | 0.0000399292427711 | 0.0104352038672132 | 0.025,0.101,0.029, | 0.026,0.021,0.077, | 0,015 |
| 78729 | ENSG00000206530 | CFAP44 | chr3 | - | 113344515 | 113344712 | 113341743 | 113341918 | 113358744 | 113358875 | 78729 | 11,8,13,10,6,10,4,14,8,13, | 1,1,2,2,0,1,1,0,2,2,0,0 | 7,14,7,27,1,9,9,8,7,44,14,5 | 2,7,0,6,0,1,2,1,2,1,0,0,0,1, | 298 | 149 | 0.0000399506018491 | 0.0104352038672132 | 0.846,0.8,0.765,0. | 0.636,0.5,1.0,0.69 | 0,021 |
| 101801 | ENSG00000198874 | TYW1 | chr7 | + | 67055834 | 67055887 | 67049948 | 67050066 | 67067284 | 67067403 | 101801 | 17,11,34,11,23,15,20,19,1 | 6,0,0,6,2,2,0,0,4,2,2,0 | 26,28,8,12,10,7,11,10,8,16 | 0,1,0,0,0,0,0,3,7,0,0,0,0,3, | 202 | 149 | 0.0000401350753652 | 0.0104706817041544 | 0.676,1.0,1.0,0.57 | 1.0,0.954,1.0,1.0,1 | -0,096 |
| 224982 | ENSG00000134905 | CARS2 | chr13 | - | 110685515 | 110685598 | 110683050 | 110683134 | 110687720 | 110687826 | 224982 | 19,7,3,0,0,0,1,0,8,2,0,0 | 51,15,95,79,63,46,19,36,2 | 0,1,0,5,0,0,0,2,0,6,0,0,0,2, | 65,34,13,16,41,21,19,29,2 | 232 | 149 | 0.0000402071511604 | 0.0104767861397143 | 0.193,0.231,0.02,0 | 0.0,0.019,0.0,0.16 | 0,04 |
| 238948 | ENSG00000197265 | GTF2E2 | chr8 | - | 30612920 | 30613011 | 30612469 | 30612481 | 30614607 | 30614715 | 238948 | 5,0,3,0,1,2,0,0,0,0,2,1 | 89,34,78,57,29,30,30,34,4 | 1,8,5,7,8,1,1,0,3,1,0,9,0,4, | 36,44,39,40,44,24,8,28,24, | 240 | 149 | 0.0000403228001955 | 0.0104942159720407 | 0.034,0.0,0.023,0. | 0.017,0.101,0.074, | -0,016 |
| 204117 | ENSG00000241685 | ARPC1A | chr7 | + | 99335452 | 99335506 | 99333324 | 99333417 | 99338180 | 99338285 | 204117 | 2,0,2,1,2,1,0,0,0,0,0,2 | 35,40,90,54,28,27,18,23,6 | 7,6,0,2,3,0,0,4,0,3,5,2,3,1, | 50,33,27,20,26,17,21,24,2 | 203 | 149 | 0.0000403770322183 | 0.0104956389147064 | 0.04,0.0,0.016,0.0 | 0.093,0.118,0.0,0. | -0,028 |
| 11331 | ENSG00000141699 | RETREG3 | chr17 | - | 42587833 | 42587864 | 42586764 | 42586891 | 42592055 | 42592162 | 11331 | 29,23,79,57,36,26,17,25,2 | 9,4,7,9,2,0,0,4,7,1,0,1 | 45,43,13,11,18,35,11,19,1 | 0,2,0,5,2,0,0,5,0,4,2,0,0,3, | 180 | 149 | 0.0000404521040543 | 0.0105024690080953 | 0.727,0.826,0.903, | 1.0,0.947,1.0,0.64 | -0,036 |
| 236308 | ENSG00000095485 | CWF19L1 | chr10 | - | 100246012 | 100246130 | 100245799 | 100245913 | 100246794 | 100246935 | 236308 | 13,6,16,7,6,7,5,9,8,0,4,4 | 55,24,59,20,17,20,9,24,46, | 12,24,16,36,11,8,2,17,4,16 | 23,32,14,14,20,13,8,15,18, | 267 | 149 | 0.0000406040590789 | 0.0105292195657961 | 0.117,0.122,0.131, | 0.226,0.295,0.389, | -0,102 |
| 45892 | ENSG00000169180 | XPO6 | chr16 | - | 28111816 | 28111942 | 28107528 | 28107677 | 28112903 | 28112958 | 45892 | 72,36,79,85,43,39,37,33,2 | 7,4,12,1,2,0,4,4,11,3,2,6 | 64,61,54,39,45,24,22,37,4 | 1,0,0,6,4,1,5,1,2,0,0,6,1,2, | 275 | 149 | 0.0000408017594372 | 0.0105677538934152 | 0.848,0.83,0.781,0 | 0.972,1.0,1.0,0.77 | -0,078 |
| 133549 | ENSG00000183527 | PSMG1 | chr21 | - | 39179923 | 39179966 | 39178448 | 39178647 | 39180284 | 39180436 | 133549 | 6,5,4,11,1,4,3,1,0,2,1,2 | 1,0,0,1,4,0,1,0,3,0,2,0 | 2,2,2,2,2,1,1,0,2,0,1,5,2,2, | 0,3,1,0,1,0,0,1,3,2,1,4,0,4, | 192 | 149 | 0.0000418077382571 | 0.0108152900394398 | 0.823,1.0,1.0,0.89 | 1.0,0.341,0.608,1. | 0,212 |
| 53415 | ENSG00000148843 | PDCD11 | chr10 | + | 103432128 | 103432234 | 103427328 | 103427391 | 103433947 | 103434037 | 53415 | 31,8,46,29,20,15,23,29,26, | 0,0,8,2,0,1,1,2,0,0,0,1 | 23,12,13,3,19,10,12,17,5,1 | 2,0,0,2,3,0,0,8,5,2,7,7,0,0, | 255 | 149 | 0.0000419549812302 | 0.0108403512186761 | 1.0,1.0,0.771,0.89 | 0.87,1.0,1.0,0.467, | 0,144 |
| 228851 | ENSG00000084070 | SMAP2 | chr1 | + | 40375746 | 40375816 | 40373726 | 40374223 | 40406735 | 40406869 | 228851 | 136,46,79,10,17,56,57,93, | 976,732,487,218,186,336, | 65,84,34,36,22,6,42,58,36, | 247,213,376,225,212,139, | 219 | 149 | 0.0000420192045907 | 0.0108439273660203 | 0.087,0.041,0.099, | 0.152,0.212,0.058, | -0,026 |
| 16954 | ENSG00000130731 | METTL26 | chr16 | - | 635280 | 635340 | 634663 | 634797 | 635611 | 635774 | 16954 | 9,9,13,6,5,7,5,8,16,11,1,3 | 2,0,2,2,0,3,2,0,2,1,2,3 | 14,8,4,4,11,2,6,12,1,10,6,2 | 0,1,1,0,0,0,1,0,0,0,1,3,0,0, | 209 | 149 | 0.0000421282945161 | 0.0108590598287362 | 0.762,1.0,0.823,0. | 1.0,0.851,0.74,1.0, | -0,139 |
| 212990 | ENSG00000145425 | RPS3A | chr4 | + | 151104168 | 151104286 | 151102870 | 151103079 | 151104471 | 151104490 | 212990 | 147,114,515,340,85,101,4 | 1,7,6,7,3,2,2,7,6,3,0,5 | 262,330,124,102,175,113, | 5,8,2,0,1,0,2,3,0,3,1,3,2,1, | 267 | 149 | 0.000042236873121 | 0.0108639394189512 | 0.988,0.901,0.98,0 | 0.967,0.958,0.972, | -0,042 |
| 184326 | ENSG00000004487 | KDM1A | chr1 | + | 23040258 | 23040434 | 23030468 | 23030634 | 23050386 | 23050520 | 184326 | 1,2,5,2,2,2,0,0,0,0,1,4 | 8,5,13,14,6,5,2,6,2,8,6,12 | 2,8,0,7,6,0,12,4,2,4,2,3,3,0 | 13,1,4,5,2,1,11,10,2,2,3,6, | 298 | 149 | 0.0000422481765808 | 0.0108639394189512 | 0.059,0.167,0.161, | 0.071,0.8,0.0,0.41 | -0,201 |
| 204752 | ENSG00000196510 | ANAPC7 | chr12 | - | 110388551 | 110388623 | 110387738 | 110387892 | 110395100 | 110395220 | 204752 | 9,2,27,19,12,8,4,7,8,2,1,7 | 0,1,3,0,0,2,2,3,0,0,1,1 | 11,15,4,4,9,2,3,4,9,8,10,10 | 3,2,0,0,2,1,0,0,1,2,1,0,8,5, | 221 | 149 | 0.0000424573789427 | 0.0109047065957235 | 1.0,0.574,0.859,1. | 0.712,0.835,1.0,1. | -0,032 |
| 145552 | ENSG00000177663 | IL17RA | chr22 | + | 17099149 | 17099310 | 17098774 | 17098887 | 17100354 | 17100481 | 145552 | 3,2,3,5,0,3,0,4,2,3,0,1 | 7,23,21,11,13,6,8,4,24,21, | 1,0,0,2,0,0,2,1,0,0,1,0,0,1, | 16,7,4,8,6,7,9,13,2,13,12,3 | 298 | 149 | 0.0000427993575586 | 0.0109794380532718 | 0.176,0.042,0.067, | 0.03,0.0,0.0,0.111, | 0,071 |
| 200083 | ENSG00000131051 | RBM39 | chr20 | - | 35713704 | 35713857 | 35713018 | 35713096 | 35714184 | 35714389 | 200083 | 59,30,76,49,67,127,77,118 | 111,57,424,254,141,147,1 | 98,49,65,30,50,40,41,103, | 215,260,132,136,188,104, | 298 | 149 | 0.0000430730603442 | 0.0110364974415982 | 0.21,0.208,0.082,0 | 0.186,0.086,0.198, | 0,085 |
| 31785 | ENSG00000224032 | EPB41L4A-AS1 | chr5 | + | 112161681 | 112161775 | 112160863 | 112161295 | 112162150 | 112162310 | 31785 | 82,66,97,79,18,24,29,41,3 | 3,3,1,3,0,0,2,2,0,0,2,7 | 68,58,56,41,59,41,13,49,2 | 8,16,6,4,0,2,2,6,0,8,6,9,0,0 | 243 | 149 | 0.0000434194768277 | 0.0111120302224765 | 0.944,0.931,0.983, | 0.839,0.69,0.851,0 | 0,054 |
| 10108 | ENSG00000288894 | | chr17 | + | 63578469 | 63578628 | 63550602 | 63550815 | 63579824 | 63579943 | 10108 | 34,33,53,54,29,17,30,18,1 | 7,4,6,0,0,2,2,1,4,2,2,6 | 23,42,21,16,21,21,7,25,9,2 | 2,0,0,4,5,0,0,2,0,3,4,0,0,0, | 298 | 149 | 0.0000438498032258 | 0.0112088325393018 | 0.708,0.805,0.815, | 0.852,1.0,1.0,0.66 | -0,108 |
| 31786 | ENSG00000224032 | EPB41L4A-AS1 | chr5 | + | 112161685 | 112161775 | 112160871 | 112161295 | 112162150 | 112164257 | 31786 | 82,66,97,79,18,24,29,41,3 | 3,3,1,3,0,0,2,2,0,0,2,7 | 68,58,56,41,59,41,13,49,2 | 8,16,6,4,0,2,2,6,0,8,6,9,0,0 | 239 | 149 | 0.0000439319214616 | 0.011216502239741 | 0.945,0.932,0.984, | 0.841,0.693,0.853, | 0,054 |
| 107126 | ENSG00000215301 | DDX3X | chrX | + | 41343288 | 41343351 | 41342736 | 41342836 | 41343736 | 41343822 | 107126 | 49,30,140,142,57,38,47,60 | 5,0,7,5,3,3,2,5,2,1,0,0 | 109,112,65,47,60,36,31,43 | 9,9,3,17,0,6,2,8,5,8,5,8,8,6 | 212 | 149 | 0.0000440289035083 | 0.0112279442310366 | 0.873,1.0,0.934,0. | 0.895,0.897,0.938, | 0,067 |
| 48689 | ENSG00000072786 | STK10 | chr5 | - | 172096425 | 172096560 | 172090231 | 172090362 | 172105655 | 172105737 | 48689 | 41,29,81,80,28,61,41,34,4 | 2,2,7,0,0,1,3,1,0,0,0,0 | 47,36,18,18,24,24,26,57,2 | 8,3,0,2,2,1,2,2,2,0,2,8,0,2, | 284 | 149 | 0.0000448702990548 | 0.0114245981898465 | 0.915,0.884,0.859, | 0.755,0.863,1.0,0. | 0,052 |
| 155578 | ENSG00000173575 | CHD2 | chr15 | + | 92995752 | 92995926 | 92992858 | 92992998 | 92996956 | 92997095 | 155578 | 18,13,25,17,3,12,4,13,3,15 | 39,40,46,40,36,28,25,24,1 | 5,15,9,9,15,6,4,6,5,9,6,2,1 | 52,75,31,66,45,22,12,36,3 | 298 | 149 | 0.00004490621736 | 0.0114245981898465 | 0.188,0.14,0.214,0 | 0.046,0.091,0.127, | 0,055 |
| 77909 | ENSG00000115421 | PAPOLG | chr2 | + | 60771518 | 60771630 | 60770457 | 60770511 | 60775033 | 60775123 | 77909 | 8,12,32,10,13,4,12,6,15,7, | 2,0,0,3,2,3,0,0,3,0,0,0 | 24,17,7,6,13,2,16,14,4,25, | 1,5,5,0,0,0,4,3,1,0,2,2,0,1, | 261 | 149 | 0.0000451052349527 | 0.0114616822008318 | 0.695,1.0,1.0,0.65 | 0.932,0.66,0.444,1 | 0,029 |
| 95627 | ENSG00000136986 | DERL1 | chr8 | - | 123030604 | 123030716 | 123024985 | 123025050 | 123041969 | 123042122 | 95627 | 51,56,175,93,42,54,66,57, | 2,0,7,10,2,2,5,3,0,1,0,2 | 98,123,51,39,55,30,23,69, | 10,2,2,2,6,6,4,6,7,3,10,7,3, | 261 | 149 | 0.0000453628030227 | 0.0115131210418457 | 0.936,1.0,0.935,0. | 0.848,0.972,0.936, | 0,105 |
| 163173 | ENSG00000157107 | FCHO2 | chr5 | + | 73076465 | 73076609 | 73074741 | 73074853 | 73077337 | 73077493 | 163173 | 1,0,0,0,0,0,4,2,0,0,2,0 | 13,6,43,20,10,21,11,23,11, | 0,0,4,4,0,0,0,0,5,1,8,1,3,4, | 28,15,9,28,16,5,20,8,11,16 | 293 | 149 | 0.0000454146464242 | 0.0115131210418457 | 0.038,0.0,0.0,0.0,0 | 0.0,0.0,0.184,0.06 | -0,049 |
| 200847 | ENSG00000161929 | SCIMP | chr17 | - | 5231141 | 5231553 | 5223353 | 5223456 | 5234734 | 5234860 | 200847 | 6,0,1,0,2,0,0,0,2,0,0,4 | 139,33,107,116,39,36,61,4 | 6,0,2,0,2,2,1,0,0,9,4,0,10,8 | 77,47,33,13,26,48,19,47,3 | 298 | 149 | 0.0000455480721661 | 0.0115333613180963 | 0.021,0.0,0.005,0. | 0.037,0.0,0.029,0. | -0,017 |
| 53947 | ENSG00000132716 | DCAF8 | chr1 | - | 160243959 | 160244034 | 160239221 | 160240370 | 160261284 | 160261325 | 53947 | 206,134,160,109,66,99,83, | 4,3,17,2,0,10,5,3,4,7,0,6 | 98,135,115,87,82,42,28,78 | 12,3,5,3,7,8,4,7,10,12,7,4, | 224 | 149 | 0.0000457158033459 | 0.011550742361639 | 0.972,0.967,0.862, | 0.845,0.968,0.939, | 0,069 |
| 134162 | ENSG00000100083 | GGA1 | chr22 | + | 37619799 | 37620019 | 37618402 | 37618546 | 37620237 | 37620361 | 134162 | 5,5,18,5,3,11,0,5,12,0,2,3 | 75,49,104,62,43,40,50,56, | 1,2,5,0,3,0,1,2,2,6,3,0,7,4, | 36,49,39,36,37,43,30,34,3 | 298 | 149 | 0.0000457240476145 | 0.011550742361639 | 0.032,0.049,0.08,0 | 0.014,0.02,0.06,0. | 0,022 |
| 145045 | ENSG00000011260 | UTP18 | chr17 | + | 51273338 | 51273450 | 51268836 | 51268904 | 51275865 | 51275991 | 145045 | 11,4,20,17,13,10,8,16,3,1, | 0,0,2,2,0,1,1,1,0,0,1,0 | 26,21,20,24,23,4,5,8,6,19, | 2,2,4,7,2,1,0,5,3,1,0,0,1,1, | 261 | 149 | 0.0000460945131868 | 0.0116186060590508 | 1.0,1.0,0.851,0.82 | 0.881,0.857,0.741, | 0,072 |
| 194059 | ENSG00000113318 | MSH3 | chr5 | + | 80719483 | 80719535 | 80678926 | 80679093 | 80725452 | 80725565 | 194059 | 2,0,2,4,0,4,2,4,0,1,3,3 | 19,8,33,27,10,10,18,12,5,6 | 0,3,0,2,0,1,1,1,3,1,0,0,0, | 15,19,14,5,12,6,7,9,9,16,8, | 201 | 149 | 0.0000461006526682 | 0.0116186060590508 | 0.072,0.0,0.043,0. | 0.0,0.105,0.0,0.22 | 0,053 |
| 79914 | ENSG00000138468 | SENP7 | chr3 | - | 101398860 | 101399055 | 101372007 | 101372126 | 101458954 | 101459052 | 79914 | 79,54,101,27,24,38,33,50, | 3,1,8,8,9,4,1,12,3,0,1,4 | 53,45,32,49,28,33,25,26,5 | 5,3,5,1,0,4,1,0,0,4,1,0,0,1, | 298 | 149 | 0.0000466521118668 | 0.0117438370633956 | 0.929,0.964,0.863, | 0.841,0.882,0.762, | -0,084 |
| 120563 | ENSG00000107036 | RIC1 | chr9 | + | 5689958 | 5690038 | 5656582 | 5656690 | 5713895 | 5714003 | 120563 | 110,49,89,55,54,57,28,70, | 2,0,0,0,0,0,4,1,0,0,0,0 | 80,83,49,68,105,38,28,36, | 1,7,0,5,7,0,0,3,0,0,3,7,1,0, | 229 | 149 | 0.0000469629115901 | 0.011808264514543 | 0.973,1.0,1.0,1.0,1 | 0.981,0.885,1.0,0. | 0,033 |
| 91274 | ENSG00000163939 | PBRM1 | chr3 | - | 52563363 | 52563493 | 52561766 | 52561968 | 52564049 | 52564233 | 91274 | 45,12,41,25,15,5,14,13,17, | 5,0,0,3,1,4,3,2,4,2,0,1 | 25,25,15,26,17,17,2,19,27, | 0,0,1,0,4,0,0,0,0,3,0,5,1,0, | 279 | 149 | 0.0000472436434152 | 0.0118447166514133 | 0.828,1.0,1.0,0.81 | 1.0,1.0,0.889,1.0,0 | -0,108 |
| 208184 | ENSG00000231887 | PRH1 | chr12 | - | 11095697 | 11095801 | 11047019 | 11047188 | 11171421 | 11171608 | 208184 | 0,0,9,5,0,2,1,6,2,0,0,4 | 33,17,36,19,18,36,8,24,20, | 3,0,2,1,0,0,0,0,0,0,0,2,2, | 26,15,2,8,16,12,18,21,19,1 | 253 | 149 | 0.0000472897921513 | 0.0118447166514133 | 0.0,0.0,0.128,0.13 | 0.064,0.0,0.371,0. | 0,014 |
| 223881 | ENSG00000173200 | PARP15 | chr3 | + | 122615285 | 122615502 | 122610493 | 122610730 | 122617085 | 122617164 | 223881 | 10,5,2,11,6,12,5,5,5,0,2,8 | 4,0,2,8,3,0,1,3,0,2,0,0 | 10,4,15,8,10,1,1,7,7,3,3,15 | 0,0,3,1,1,0,0,0,4,0,0,2,2,2, | 298 | 149 | 0.0000472960211193 | 0.0118447166514133 | 0.556,1.0,0.333,0. | 1.0,1.0,0.714,0.8,0 | -0,196 |
| 170403 | ENSG00000144645 | OSBPL10 | chr3 | - | 31668641 | 31668824 | 31657911 | 31664232 | 31683779 | 31684114 | 170403 | 137,111,66,72,60,49,46,66 | 0,2,6,0,0,4,1,2,0,0,0,0 | 63,50,106,161,96,99,44,43 | 2,5,3,12,5,4,0,2,2,5,1,2,1,3 | 298 | 149 | 0.0000473280165042 | 0.0118447166514133 | 1.0,0.965,0.846,1. | 0.94,0.833,0.946,0 | 0,036 |
| 176525 | ENSG00000114446 | IFT57 | chr3 | - | 108164856 | 108165245 | 108163662 | 108163729 | 108165430 | 108165493 | 176525 | 22,16,25,39,4,8,5,22,9,8,9, | 163,117,290,258,115,119, | 17,10,8,1,3,3,8,6,3,7,20,10 | 194,159,91,53,79,48,104,1 | 298 | 149 | 0.0000477394330999 | 0.0119222423479601 | 0.063,0.064,0.041, | 0.042,0.03,0.042,0 | 0,016 |
| 94813 | ENSG00000079691 | CARMIL1 | chr6 | + | 25604811 | 25604893 | 25600313 | 25600746 | 25606060 | 25606273 | 94813 | 9,16,32,22,4,12,4,15,26,10 | 1,0,0,0,3,1,0,1,0,0,2,0 | 9,14,3,5,21,10,6,5,6,10,29, | 1,2,0,1,0,0,0,3,2,0,10,0,3,2 | 231 | 149 | 0.0000477485720177 | 0.0119222423479601 | 0.853,1.0,1.0,1.0,0 | 0.853,0.819,1.0,0. | 0,047 |
| 176153 | ENSG00000169180 | XPO6 | chr16 | - | 28111816 | 28112169 | 28107521 | 28107677 | 28112903 | 28113050 | 176153 | 84,44,81,98,56,47,38,39,3 | 7,4,12,1,2,0,4,4,11,3,2,6 | 69,63,57,42,51,28,25,44,4 | 1,0,0,6,4,1,5,1,2,0,0,6,1,2, | 298 | 149 | 0.0000479525220165 | 0.0119592923130166 | 0.857,0.846,0.771, | 0.972,1.0,1.0,0.77 | -0,071 |
| 23684 | ENSG00000100519 | PSMC6 | chr14 | + | 52711409 | 52711524 | 52710545 | 52711168 | 52713880 | 52713902 | 23684 | 71,98,101,67,32,22,28,39, | 0,0,0,3,0,0,1,0,0,0,0,0 | 52,66,43,30,47,16,9,26,24, | 2,3,1,4,0,2,2,2,0,4,2,1,1,3, | 264 | 149 | 0.0000481918615155 | 0.0120050723910215 | 1.0,1.0,1.0,0.926,1 | 0.936,0.925,0.96,0 | 0,08 |
| 36623 | ENSG00000164597 | COG5 | chr7 | - | 107236449 | 107236687 | 107230614 | 107230691 | 107248395 | 107248499 | 36623 | 27,28,73,35,29,34,13,35,1 | 1,4,3,0,1,2,2,3,0,0,0,1 | 33,35,18,15,20,10,20,30,2 | 1,6,1,2,3,2,2,2,5,2,5,3,0,2, | 298 | 149 | 0.0000482880449325 | 0.0120151262414646 | 0.931,0.778,0.924, | 0.943,0.745,0.9,0. | 0,104 |
| 92597 | ENSG00000211450 | SELENOH | chr11 | + | 57742116 | 57742247 | 57741698 | 57741954 | 57742862 | 57743514 | 92597 | 131,92,242,308,120,87,10 | 0,3,5,3,4,0,0,3,0,2,0,0 | 210,149,127,21,84,114,32, | 3,7,8,6,3,2,0,7,4,3,4,3,0,0, | 280 | 149 | 0.0000489336471179 | 0.0121617064697837 | 1.0,0.942,0.963,0. | 0.974,0.919,0.894, | 0,046 |
| 255756 | ENSG00000118454 | ANKRD13C | chr1 | - | 70305924 | 70306290 | 70300763 | 70300908 | 70313744 | 70313790 | 255756 | 26,16,43,21,6,8,4,16,21,3, | 0,0,1,2,0,0,1,0,2,1,0,2 | 18,25,14,15,15,8,4,5,7,16, | 0,4,1,0,2,0,2,3,3,0,4,0,3,0, | 298 | 149 | 0.0000494241131811 | 0.0122694363368873 | 1.0,1.0,0.956,0.84, | 1.0,0.758,0.875,1. | 0,082 |
| 165151 | ENSG00000163738 | MTHFD2L | chr4 | + | 74195738 | 74195838 | 74175280 | 74175403 | 74199793 | 74199946 | 165151 | 4,3,8,14,2,1,6,5,0,5,3,3 | 6,1,13,5,8,12,5,3,3,2,1,5 | 2,4,0,0,2,3,1,0,0,3,0,2,3,1, | 5,6,5,4,4,7,0,2,8,3,7,9,4,9, | 249 | 149 | 0.0000495232206137 | 0.0122798759169438 | 0.285,0.642,0.269, | 0.193,0.285,0.0,0. | 0,181 |
| 25156 | ENSG00000134884 | ARGLU1 | chr13 | - | 106557558 | 106557616 | 106557047 | 106557131 | 106559431 | 106559657 | 25156 | 94,19,84,65,24,22,16,36,3 | 130,49,133,112,66,62,60,6 | 38,25,38,11,11,16,6,46,12, | 104,54,77,34,46,41,18,79, | 207 | 149 | 0.0000500716405788 | 0.0124005292765921 | 0.342,0.218,0.313, | 0.208,0.25,0.262,0 | 0,082 |
| 211828 | ENSG00000137497 | NUMA1 | chr11 | - | 72012198 | 72012442 | 72010785 | 72010854 | 72012894 | 72012926 | 211828 | 57,35,18,25,11,13,14,13,3 | 67,38,141,80,56,63,46,57, | 9,20,8,1,12,13,9,16,12,19, | 68,62,48,34,39,27,77,99,3 | 298 | 149 | 0.0000501417590973 | 0.0124005292765921 | 0.298,0.315,0.06,0 | 0.062,0.139,0.077, | 0,07 |
| 177843 | ENSG00000162601 | MYSM1 | chr1 | - | 58663033 | 58663522 | 58661405 | 58661511 | 58665498 | 58665631 | 177843 | 3,1,4,0,1,4,1,2,0,0,0,1 | 26,27,35,28,18,32,17,25,8, | 3,6,3,8,6,0,4,7,0,8,1,3,5,2, | 34,48,29,26,39,18,12,31,1 | 298 | 149 | 0.0000501826456222 | 0.0124005292765921 | 0.055,0.018,0.054, | 0.042,0.059,0.049, | -0,033 |
| 19134 | ENSG00000137842 | TMEM62 | chr15 | + | 43149028 | 43149151 | 43148754 | 43148879 | 43151789 | 43151890 | 19134 | 35,41,83,36,25,15,27,42,2 | 0,4,5,1,0,1,0,6,0,0,0,0 | 55,73,24,42,36,14,17,25,2 | 7,13,0,8,5,2,1,3,3,0,1,1,2,2 | 272 | 149 | 0.0000502744344841 | 0.0124089642298708 | 1.0,0.849,0.901,0. | 0.811,0.755,1.0,0. | 0,094 |
| 11546 | ENSG00000261371 | PECAM1 | chr17 | - | 64354932 | 64355040 | 64353490 | 64353518 | 64356110 | 64356398 | 11546 | 146,104,179,212,101,29,6 | 17,9,6,4,2,3,5,5,3,5,2,10 | 115,84,66,26,37,66,46,59, | 13,23,7,10,12,2,5,11,8,4,9, | 257 | 149 | 0.0000504267239699 | 0.0124322957924004 | 0.833,0.87,0.945,0 | 0.837,0.679,0.845, | 0,086 |
| 229916 | ENSG00000231721 | LINC-PINT | chr7 | - | 131002196 | 131002318 | 130984052 | 130984109 | 131052424 | 131052555 | 229916 | 0,0,1,1,9,7,1,0,0,5,0,0 | 132,85,282,117,79,106,96, | 2,1,10,23,11,8,6,3,0,0,1,1, | 176,141,91,124,97,47,76,9 | 271 | 149 | 0.0000508462647067 | 0.0125213871843109 | 0.0,0.0,0.002,0.00 | 0.006,0.004,0.057, | -0,013 |
| 10703 | ENSG00000167543 | TP53I13 | chr17 | + | 29571590 | 29571719 | 29569317 | 29569359 | 29571856 | 29572057 | 10703 | 183,68,237,171,162,195,1 | 7,0,13,2,2,8,0,4,5,0,2,6 | 127,95,86,37,120,65,77,13 | 10,5,14,3,9,1,1,5,3,0,8,7,6, | 278 | 149 | 0.0000512910179924 | 0.0126164766783111 | 0.933,1.0,0.907,0. | 0.872,0.911,0.767, | 0,049 |

| | | | | | | | | | | | | | | | | | | | | | | |
|---|---|---|---|---|---|---|---|---|---|---|---|---|---|---|---|---|---|---|---|---|---|---|
| 65997 | ENSG00000143384 | MCL1 | chr1 | - | 150578243 | 150578491 | 150574175 | 150577491 | 150578842 | 150578963 | 65997 | 380,407,693,526,560,412,3 | 2,2,3,4,14,7,14,13,4,1,3,5 | 517,652,305,306,279,243,1 | 18,10,13,3,1,7,2,4,2,12,8,4 | 298 | 149 | 0.0000513772509979 | 0.0126232615405582 | 0.99,0.99,0.991,0.9 | 0.935,0.97,0.921,0 | 0,013 |
| 242147 | ENSG00000172340 | SUCLG2 | chr3 | - | 67408864 | 67409031 | 67400730 | 67400851 | 67495797 | 67495940 | 242147 | 6,5,19,6,8,9,9,5,5,9,0,4 | 35,29,48,34,20,16,23,19,29 | 6,1,4,4,10,4,5,2,5,1,4,1,5,4 | 23,31,31,10,26,24,20,15,17 | 298 | 149 | 0.0000515843706385 | 0.0126596986053535 | 0.079,0.079,0.165, | 0.115,0.016,0.061, | 0,05 |
| 26410 | ENSG00000115760 | BIRC6 | chr2 | + | 32595033 | 32595144 | 32593914 | 32594060 | 32597750 | 32597968 | 26410 | 64,45,103,77,36,28,40,62,3 | 4,5,12,2,4,5,6,10,4,2,0,5 | 101,54,52,25,15,32,29,61,2 | 37,18,5,13,9,3,5,7,5,11,17, | 260 | 149 | 0.0000521732306691 | 0.0127896316744203 | 0.902,0.838,0.831, | 0.61,0.632,0.856,0 | 0,128 |
| 155152 | ENSG00000125746 | EML2 | chr19 | - | 45619623 | 45619697 | 45619059 | 45619191 | 45621206 | 45621332 | 155152 | 2,0,5,4,1,3,1,5,0,0,0,1 | 12,8,19,10,11,18,16,19,12, | 0,3,1,0,0,0,0,0,0,1,1,1,1,1 | 20,21,6,11,7,15,9,20,11,15 | 223 | 149 | 0.0000524677314807 | 0.0128399736063816 | 0.1,0.0,0.15,0.211, | 0.0,0.087,0.1,0.0,0 | 0,04 |
| 169313 | ENSG00000113649 | TCERG1 | chr5 | + | 146507632 | 146507778 | 146507027 | 146507207 | 146507872 | 146507956 | 169313 | 1,3,9,0,2,4,1,2,4,1,0,2 | 92,78,141,35,26,31,32,46,6 | 1,2,3,2,7,8,2,6,3,2,10,5,7,9 | 37,72,42,63,38,28,32,40,26 | 295 | 149 | 0.0000525302072132 | 0.0128399736063816 | 0.005,0.019,0.031, | 0.013,0.014,0.035, | -0,03 |
| 57724 | ENSG00000288701 | PRRC2B | chr9 | + | 131474453 | 131476535 | 131473507 | 131473724 | 131477743 | 131477949 | 57724 | 45,50,137,93,61,71,63,54,2 | 1,4,6,0,1,7,2,9,0,3,0,4 | 57,100,55,79,55,27,51,62,5 | 7,7,4,25,17,5,10,7,7,4,3,5,2 | 298 | 149 | 0.0000525575625594 | 0.0128399736063816 | 0.957,0.862,0.919, | 0.803,0.877,0.873, | 0,093 |
| 97452 | ENSG00000168014 | C2CD3 | chr11 | - | 74161398 | 74161556 | 74139604 | 74139828 | 74168343 | 74168613 | 97452 | 74,39,77,33,30,43,33,40,23 | 2,6,11,2,2,1,2,3,0,0,0,4 | 56,66,45,77,38,24,34,37,24 | 1,5,0,0,0,0,0,1,2,2,3,1,4,3,0 | 298 | 149 | 0.0000529779266871 | 0.0128892005480686 | 0.949,0.765,0.778, | 0.966,0.868,1.0,1.0 | -0,039 |
| 11258 | ENSG00000275700 | AATF | chr17 | + | 37031613 | 37031685 | 37020933 | 37021014 | 37056600 | 37056842 | 11258 | 49,25,96,73,43,42,37,44,61 | 0,1,3,0,5,3,1,4,0,0,0,2 | 50,59,47,31,22,23,24,22,18 | 5,6,2,4,4,1,0,0,3,1,6,1,2,9,1 | 221 | 149 | 0.000052999214626 | 0.0128892005480686 | 1.0,0.944,0.956,1.0 | 0.871,0.869,0.941, | 0,059 |
| 160924 | ENSG00000164329 | TENT2 | chr5 | + | 79679281 | 79679670 | 79668891 | 79669028 | 79681981 | 79682061 | 160924 | 33,21,72,46,21,22,23,36,7, | 2,0,1,0,0,1,0,3,1,0,0,0 | 46,63,31,23,34,19,13,24,39 | 2,1,3,10,3,0,0,3,0,2,4,1,1,0 | 298 | 149 | 0.0000530353397706 | 0.0128892005480686 | 0.892,1.0,0.973,1.0 | 0.92,0.969,0.838,0 | 0,052 |
| 173981 | ENSG00000196455 | PIK3R4 | chr3 | - | 130735868 | 130736024 | 130733547 | 130734130 | 130744485 | 130745264 | 173981 | 3,13,18,28,8,6,6,8,7,8,2,3 | 1,2,3,0,2,0,4,0,0,0,1,0 | 5,18,8,4,11,9,13,8,4,7,5,6,1 | 0,0,2,3,4,2,4,1,2,0,2,0,2,0,1 | 298 | 149 | 0.0000530391581501 | 0.0128892005480686 | 0.6,0.765,0.75,1.0, | 1.0,1.0,0.667,0.4,0 | 0,038 |
| 141208 | ENSG00000067900 | ROCK1 | chr18 | - | 21022444 | 21022564 | 21020150 | 21020239 | 21023619 | 21023680 | 141208 | 6,0,8,0,2,3,2,1,0,0,0,1 | 68,43,84,74,54,53,32,43,40 | 3,4,10,18,1,6,5,0,3,1,3,5,1, | 55,93,33,48,42,34,41,35,48 | 269 | 149 | 0.0000530860422884 | 0.0128892005480686 | 0.047,0.0,0.05,0.0, | 0.029,0.023,0.144, | -0,028 |
| 192519 | ENSG00000196419 | XRCC6 | chr22 | + | 41628161 | 41628230 | 41621989 | 41622086 | 41636112 | 41636251 | 192519 | 116,94,165,101,40,26,38,5 | 4,2,6,4,0,1,0,0,0,0,0,0 | 85,92,82,32,54,33,25,42,29 | 2,7,6,5,11,2,0,0,0,4,3,1,1,1 | 218 | 149 | 0.0000531183746121 | 0.0128892005480686 | 0.952,0.97,0.949,0 | 0.967,0.9,0.903,0.8 | 0,039 |
| 243557 | ENSG00000132109 | TRIM21 | chr11 | - | 4386151 | 4386257 | 4385677 | 4385853 | 4386967 | 4386990 | 243557 | 4,1,19,23,5,9,2,2,15,2,1,5 | 1,0,0,2,1,2,2,0,1,1,0,0 | 9,20,11,8,3,1,3,12,2,5,4,13 | 7,1,0,0,0,2,0,2,0,0,7,5,3,3,0 | 255 | 149 | 0.0000533692143174 | 0.0129244029143555 | 0.7,1.0,1.0,0.87,0.1 | 0.429,0.921,1.0,1.0 | 0,039 |
| 166052 | ENSG00000151247 | EIF4E | chr4 | - | 98895181 | 98895268 | 98891236 | 98891332 | 98901875 | 98901982 | 166052 | 1,0,6,0,0,4,1,0,0,0,0,0 | 46,35,79,113,45,39,33,40,3 | 8,8,6,3,1,2,0,0,2,0,2,3,2,0,2 | 51,41,31,21,43,18,14,30,23 | 236 | 149 | 0.0000533835469373 | 0.0129244029143555 | 0.014,0.0,0.046,0.0 | 0.09,0.11,0.109,0.0 | -0,032 |
| 97596 | ENSG00000004059 | ARF5 | chr7 | + | 127589484 | 127589594 | 127589082 | 127589163 | 127590065 | 127590137 | 97596 | 127,88,171,162,67,93,53,5 | 2,0,1,2,2,1,0,0,2,0,0,2 | 153,110,48,28,98,44,26,74 | 0,3,0,3,2,1,1,3,2,2,6,2,2,2,8 | 259 | 149 | 0.0000537361492465 | 0.0129951518409814 | 0.973,1.0,0.99,0.97 | 1.0,0.955,1.0,0.843 | 0,044 |
| 6965 | ENSG00000124207 | CSE1L | chr20 | + | 49088008 | 49088106 | 49084025 | 49084162 | 49089246 | 49089397 | 6965 | 21,6,48,36,17,22,11,25,20, | 0,0,2,0,0,1,2,1,0,0,0,0 | 22,32,20,9,18,10,7,19,14,2 | 1,2,0,0,0,1,2,3,1,1,4,2,5,0,5 | 247 | 149 | 0.000053985940951 | 0.0130290743470224 | 1.0,1.0,0.935,1.0,1 | 0.93,0.906,1.0,1.0, | 0,09 |
| 30889 | ENSG00000145715 | RASA1 | chr5 | + | 87380508 | 87380595 | 87378395 | 87378538 | 87383712 | 87383780 | 30889 | 11,1,42,28,18,15,4,28,10,6 | 6,0,2,0,0,1,1,3,0,0,0,0 | 14,15,3,1,8,8,11,10,7,12,31 | 0,3,0,6,0,1,0,3,1,2,10,1,2,0 | 236 | 149 | 0.0000539974925431 | 0.0130290743470224 | 0.536,1.0,0.93,1.0, | 1.0,0.759,1.0,0.095 | 0,094 |
| 141151 | ENSG00000166562 | SEC11C | chr18 | + | 59148064 | 59148219 | 59139898 | 59140035 | 59149512 | 59149622 | 141151 | 0,2,11,3,1,3,3,8,0,3,3,3 | 30,31,99,78,27,24,22,23,22 | 0,0,0,1,0,3,1,2,2,0,0,0,1,1,5 | 25,25,14,4,11,13,11,18,4,1 | 298 | 149 | 0.0000542771852149 | 0.0130818956897963 | 0.0,0.031,0.053,0.0 | 0.0,0.0,0.0,0.111,0 | 0,03 |
| 224032 | ENSG00000154978 | VOPP1 | chr7 | - | 55492748 | 55492973 | 55492385 | 55492418 | 55497612 | 55497690 | 224032 | 5,3,10,1,2,5,4,14,14,0,0,3 | 279,248,213,208,203,291,2 | 2,5,10,42,3,3,5,5,6,1,8,2,0, | 156,208,299,279,146,68,79 | 298 | 149 | 0.0000545845926841 | 0.0131293749682543 | 0.009,0.006,0.023, | 0.006,0.012,0.016, | -0,009 |
| 101111 | ENSG00000265681 | RPL17 | chr18 | - | 49491351 | 49491445 | 49490792 | 49490927 | 49491531 | 49491586 | 101111 | 23,13,91,56,27,14,12,23,12 | 7,7,5,1,4,0,3,7,3,1,0,0 | 40,31,24,17,34,19,14,21,21 | 2,3,0,2,1,1,0,0,1,1,4,1,0,6,1 | 243 | 149 | 0.0000545961808317 | 0.0131293749682543 | 0.668,0.532,0.918, | 0.925,0.864,1.0,0.8 | -0,101 |
| 114455 | ENSG00000130749 | ZC3H4 | chr19 | - | 47069091 | 47069343 | 47064186 | 47067869 | 47071777 | 47072121 | 114455 | 34,17,47,27,16,28,15,19,11 | 0,0,3,4,1,2,1,1,1,2,1,1 | 17,24,6,16,4,21,16,17,7,12 | 5,9,0,0,3,2,4,5,4,0,0,4,3,8,4 | 298 | 149 | 0.0000546985182251 | 0.0131336745344217 | 1.0,1.0,0.887,0.77 | 0.63,0.571,1.0,1.0, | 0,128 |
| 212989 | ENSG00000145425 | RPS3A | chr4 | + | 151104043 | 151104286 | 151102870 | 151103079 | 151104471 | 151104490 | 212989 | 154,121,524,341,87,104,46 | 1,7,6,7,3,2,2,7,6,3,0,5 | 263,331,124,103,176,113,3 | 5,8,2,0,1,0,2,3,0,3,1,3,2,1,1 | 298 | 149 | 0.000054736102408 | 0.0131336745344217 | 0.987,0.896,0.978, | 0.963,0.954,0.969, | -0,043 |
| 241718 | ENSG00000148296 | SURF6 | chr9 | - | 133333717 | 133334113 | 133332547 | 133332760 | 133334391 | 133334601 | 241718 | 18,29,21,8,14,5,3,1,19,5,2, | 2,0,3,0,0,0,2,1,0,2,1,0 | 8,12,16,1,8,5,5,5,9,10,2,6,5 | 4,1,1,5,1,0,0,0,0,0,3,4,0,2,2 | 298 | 149 | 0.0000552916880092 | 0.0132431898921253 | 0.818,1.0,0.778,1.0 | 0.5,0.857,0.889,0.0 | 0,063 |
| 239914 | ENSG00000196781 | TLE1 | chr9 | - | 81650234 | 81650372 | 81634096 | 81634301 | 81652183 | 81652288 | 239914 | 2,4,5,4,0,5,2,20,0,0,0,0 | 20,8,92,23,34,49,39,147,31 | 0,2,2,0,1,0,0,6,0,0,2,1,2,0,0 | 7,15,13,2,26,11,28,45,5,13 | 287 | 149 | 0.0000553189673231 | 0.0132431898921253 | 0.049,0.206,0.027, | 0.0,0.065,0.074,0.0 | 0,023 |
| 197386 | ENSG00000163534 | FCRL1 | chr1 | - | 157803844 | 157803996 | 157801914 | 157802193 | 157807101 | 157807122 | 197386 | 17,7,32,2,10,7,7,14,7,2,9,1 | 2,0,6,3,0,0,1,0,0,0,1,0 | 17,11,23,12,3,7,13,4,7,19,1 | 2,1,3,1,3,2,2,0,2,0,0,3,8,6,6 | 298 | 149 | 0.0000553771106528 | 0.0132431898921253 | 0.81,1.0,0.727,0.25 | 0.81,0.846,0.793,0 | 0,109 |
| 129616 | ENSG00000127314 | RAP1B | chr12 | + | 68624507 | 68624873 | 68610907 | 68611043 | 68648698 | 68648757 | 129616 | 0,0,2,7,0,0,0,2,2,2,1,0 | 16,14,21,11,10,12,9,7,17,1 | 2,2,3,0,0,0,1,3,3,0,3,7,4,10 | 10,21,10,9,14,8,9,2,3,14,17 | 298 | 149 | 0.0000560532713572 | 0.013900129272825 | 0.0,0.0,0.045,0.24 | 0.091,0.045,0.13,0 | -0,07 |
| 76921 | ENSG00000259494 | MRPL46 | chr15 | - | 88465586 | 88465773 | 88464702 | 88464876 | 88467149 | 88467388 | 76921 | 17,31,55,24,11,23,5,17,14, | 0,2,0,1,0,1,0,1,2,0,0,0 | 23,19,19,5,17,13,25,13,8,1 | 1,4,3,0,1,0,1,1,0,0,2,4,2,1,6 | 298 | 149 | 0.0000563864790717 | 0.0134546765821298 | 1.0,0.886,1.0,0.92 | 0.92,0.704,0.76,1.0 | 0,097 |
| 109295 | ENSG00000159256 | MORC3 | chr21 | + | 36368987 | 36369876 | 36364092 | 36364259 | 36372373 | 36372531 | 109295 | 336,250,764,537,255,247,1 | 6,0,9,14,5,5,3,5,11,8,1,2 | 318,244,223,240,179,120,1 | 6,14,14,9,14,9,0,3,5,6,6,0,8 | 298 | 149 | 0.0000564606733453 | 0.0134574608912317 | 0.966,1.0,0.977,0.9 | 0.964,0.897,0.888, | 0,031 |
| 112522 | ENSG00000119397 | CNTRL | chr9 | + | 121157749 | 121157880 | 121157469 | 121157600 | 121158854 | 121159019 | 112522 | 9,9,13,7,1,7,3,5,5,4,0,1 | 36,11,20,22,1,16,15,17,9,7 | 0,4,14,1,2,2,0,2,3,0,6,0,7,3 | 22,24,19,38,26,27,8,12,17, | 280 | 149 | 0.0000567513415171 | 0.0135117787456493 | 0.117,0.303,0.257, | 0.0,0.081,0.282,0.0 | 0,114 |
| 3641 | ENSG00000196365 | LONP1 | chr19 | - | 5699026 | 5699205 | 5696248 | 5696757 | 5700788 | 5700927 | 3641 | 28,28,88,36,55,54,37,37,40 | 0,0,0,1,0,3,0,4,0,0,1,1 | 41,50,18,28,37,13,41,48,11 | 1,2,0,10,2,1,4,1,2,0,8,0,0,0 | 298 | 149 | 0.0000569008691961 | 0.0135121306146009 | 1.0,1.0,1.0,0.947,1 | 0.953,0.926,1.0,0.5 | 0,06 |
| 165985 | ENSG00000026297 | RNASET2 | chr6 | - | 166948569 | 166948624 | 166946392 | 166946739 | 166952487 | 166955182 | 165985 | 52,42,58,39,34,24,32,27,49 | 4,0,3,4,0,0,1,3,1,1,0,0 | 76,54,43,11,23,19,13,26,27 | 0,7,4,1,7,4,0,0,3,5,1,8,2,2,1 | 204 | 149 | 0.0000569245138924 | 0.0135121306146009 | 0.905,1.0,0.934,0.8 | 1.0,0.849,0.887,0.8 | 0,057 |
| 51495 | ENSG00000178927 | CYBC1 | chr17 | - | 82447579 | 82447621 | 82446622 | 82446696 | 82449169 | 82449292 | 51495 | 35,19,68,50,20,29,12,20,25 | 0,0,1,4,1,0,5,0,3,0,0,0 | 37,19,15,2,11,9,6,17,8,26,1 | 3,0,0,0,0,6,1,4,0,4,2,5,1,2,1 | 191 | 149 | 0.0000569411584179 | 0.0135121306146009 | 1.0,1.0,0.981,0.90 | 0.906,1.0,1.0,1.0,1 | 0,046 |
| 206844 | ENSG00000075884 | ARHGAP15 | chr2 | + | 143705551 | 143705588 | 143703418 | 143703524 | 143767988 | 143768352 | 206844 | 2,4,8,3,2,5,3,7,1,4,3,2 | 75,50,155,97,69,55,50,45,4 | 5,15,2,8,4,2,7,7,6,4,8,9,8,3 | 94,64,66,18,62,43,14,32,37 | 186 | 149 | 0.0000579312780027 | 0.0137176963297793 | 0.021,0.06,0.04,0.0 | 0.041,0.158,0.024, | -0,056 |
| 71563 | ENSG00000118961 | LDAH | chr2 | - | 20801291 | 20801465 | 20790254 | 20790398 | 20823036 | 20823089 | 71563 | 5,4,30,16,17,5,11,11,5,3,4, | 3,0,4,6,4,2,4,2,3,2,0,2 | 19,10,8,2,6,3,3,14,6,10,5,9 | 2,0,1,0,1,2,0,3,1,1,1,2,0,3,1 | 298 | 149 | 0.0000579348976856 | 0.0137176963297793 | 0.455,1.0,0.789,0.5 | 0.826,1.0,0.8,1.0,0 | -0,17 |
| 136555 | ENSG00000028277 | POU2F2 | chr19 | - | 42124178 | 42124296 | 42122510 | 42122576 | 42132383 | 42132478 | 136555 | 12,2,0,3,1,2,1,0,3,3,0,1 | 41,32,30,20,31,24,14,14,20 | 0,0,0,1,0,1,0,0,0,1,1,0,0,1,2 | 10,3,20,9,3,3,7,7,11,11,3,1 | 267 | 149 | 0.0000582176649581 | 0.0137573608302394 | 0.14,0.034,0.0,0.07 | 0.0,0.0,0.0,0.058,0 | 0,012 |
| 72348 | ENSG00000189077 | TMEM120A | chr7 | - | 75987537 | 75987602 | 75986982 | 75987428 | 75987717 | 75987810 | 72348 | 10,3,16,9,9,5,6,3,7,3,3,4 | 4,6,4,0,2,0,0,1,9,0,0,1 | 5,2,8,6,3,3,2,6,0,2,10,8,2,6 | 0,3,0,1,1,1,0,0,2,0,1,2,3,0,0 | 214 | 149 | 0.0000582302536175 | 0.0137573608302394 | 0.635,0.258,0.736, | 1.0,0.317,1.0,0.807 | -0,013 |
| 68539 | ENSG00000055609 | KMT2C | chr7 | - | 152157760 | 152157854 | 152156204 | 152156346 | 152158862 | 152159072 | 68539 | 14,10,16,20,7,5,9,12,6,3,2, | 4,0,6,0,4,3,2,3,1,0,1,2 | 8,14,10,10,9,7,9,8,3,18,7,1 | 0,0,1,0,1,0,1,2,0,1,5,3,1,2,0 | 243 | 149 | 0.000058850727593 | 0.0138887071826414 | 0.682,1.0,0.621,1.0 | 1.0,1.0,0.86,1.0,0.8 | -0,144 |
| 142436 | ENSG00000178852 | EFCAB13 | chr17 | + | 47344806 | 47344939 | 47344161 | 47344292 | 47345015 | 47345098 | 142436 | 3,5,15,9,2,0,1,0,0,3,2,0 | 37,21,130,92,31,44,34,53,1 | 3,2,0,0,0,2,2,1,2,4,0,1,1,2,1 | 45,44,31,48,36,35,46,57,43 | 282 | 149 | 0.0000592053753148 | 0.0139570998185977 | 0.041,0.112,0.057, | 0.034,0.023,0.0,0.0 | 0,028 |
| 48976 | ENSG00000197548 | ATG7 | chr3 | + | 11331339 | 11331428 | 11315343 | 11315493 | 11340644 | 11340735 | 48976 | 13,3,31,8,18,15,6,15,5,3,5, | 0,2,4,3,0,2,4,1,2,1,0,0 | 14,10,8,8,9,11,2,2,16,9,16, | 0,1,2,0,1,2,0,0,2,0,0,2,0,0,0 | 238 | 149 | 0.0000596561434754 | 0.0140479774795994 | 1.0,0.484,0.829,0.6 | 1.0,0.862,0.715,1.0 | -0,146 |
| 168596 | ENSG00000005059 | MCUB | chr4 | + | 109684424 | 109684646 | 109682581 | 109682742 | 109685250 | 109685367 | 168596 | 95,71,143,121,64,45,59,75 | 6,1,2,1,0,4,5,0,5,0,0,1 | 102,93,72,22,27,43,28,70,3 | 10,3,8,3,1,3,2,8,3,5,3,10,4, | 298 | 149 | 0.0000609882823885 | 0.0143459770565675 | 0.888,0.973,0.973, | 0.836,0.939,0.818, | 0,062 |
| 118773 | ENSG00000165121 | | chr9 | + | 85829462 | 85829606 | 85822879 | 85823080 | 85837758 | 85837965 | 118773 | 44,31,62,25,11,13,11,9,26, | 0,0,5,3,0,4,3,0,0,0,0,1 | 26,18,21,44,15,22,11,20,11 | 2,4,2,2,1,3,9,1,3,2,5,2,5,1,0 | 293 | 149 | 0.0000611048977097 | 0.0143526769602642 | 1.0,1.0,0.863,0.809 | 0.869,0.696,0.842, | 0,119 |
| 136493 | ENSG00000125734 | GPR108 | chr19 | - | 6732986 | 6733056 | 6732471 | 6732549 | 6733167 | 6733301 | 136493 | 16,9,26,30,23,16,27,12,25, | 3,8,2,0,2,2,0,0,0,0,2,2 | 15,16,10,11,6,11,7,17,10,1 | 0,0,0,0,0,0,0,1,0,0,0,1,0,0,0 | 219 | 149 | 0.0000611501353084 | 0.0143526769602642 | 0.784,0.434,0.898, | 1.0,1.0,1.0,1.0,1.0, | -0,13 |
| 151553 | ENSG00000077238 | IL4R | chr16 | + | 27346586 | 27346618 | 27344907 | 27345020 | 27352539 | 27352696 | 151553 | 56,14,181,72,49,25,42,56,6 | 40,17,48,36,11,21,13,26,35 | 70,32,75,11,10,14,25,24,17 | 10,7,15,3,14,6,1,4,1,7,8,7,1 | 181 | 149 | 0.0000612300037696 | 0.0143557679099507 | 0.535,0.404,0.756, | 0.852,0.79,0.805,0 | -0,139 |
| 77767 | ENSG00000106105 | GARS1 | chr7 | + | 30631447 | 30631559 | 30628559 | 30628669 | 30632246 | 30632437 | 77767 | 20,9,29,26,7,14,4,10,2,2,7, | 8,0,2,0,0,0,1,1,0,0,1,0 | 24,29,15,15,21,19,15,8,6,1 | 4,1,1,2,1,0,0,6,1,0,4,2,0,1,6 | 261 | 149 | 0.0000618128305909 | 0.0144766456375517 | 0.588,1.0,0.892,1.0 | 0.774,0.943,0.895, | 0,051 |
| 195649 | ENSG00000182183 | SHISAL2A | chr1 | + | 52648450 | 52648625 | 52642862 | 52643002 | 52667398 | 52667688 | 195649 | 20,11,2,3,2,3,2,0,2,3,3,2 | 19,7,13,19,13,2,15,14,48,3 | 0,1,0,7,0,0,6,1,0,0,2,0,2,0,0 | 13,13,16,28,2,4,0,5,5,13,3, | 298 | 149 | 0.0000619230372318 | 0.0144866926374321 | 0.345,0.44,0.071,0 | 0.0,0.037,0.0,0.11 | 0,041 |
| 228331 | ENSG00000136273 | HUS1 | chr7 | - | 47976786 | 47976837 | 47975654 | 47975687 | 47978416 | 47978534 | 228331 | 9,8,26,13,4,7,15,9,2,7,3,7 | 2,0,2,4,0,2,0,8,0,0,0,2 | 10,15,11,6,13,7,6,7,14,5,14 | 1,0,2,0,2,0,0,0,0,0,1,1,0,0,0 | 200 | 149 | 0.0000620655751785 | 0.0145042734106881 | 0.77,1.0,0.906,0.70 | 0.882,1.0,0.804,1.0 | -0,102 |
| 196491 | ENSG00000166479 | TMX3 | chr18 | - | 68701744 | 68701816 | 68700404 | 68700485 | 68710020 | 68710144 | 196491 | 6,1,22,6,8,5,6,4,5,4,3,2 | 3,0,6,10,0,4,2,2,2,0,1,2 | 20,4,5,7,9,2,1,9,5,11,5,11,1 | 0,3,0,0,3,0,4,1,0,0,0,4,1,0,3 | 221 | 149 | 0.0000622950592427 | 0.0145421126853206 | 0.574,1.0,0.712,0.1 | 1.0,0.473,1.0,1.0,0 | -0,154 |
| 173746 | ENSG00000156976 | EIF4A2 | chr3 | + | 186785035 | 186785101 | 186784563 | 186784696 | 186785882 | 186785913 | 173746 | 352,231,267,229,84,111,86 | 475,304,579,328,154,202,2 | 118,80,126,114,85,102,47, | 280,298,307,267,316,161,8 | 215 | 149 | 0.0000624633739921 | 0.0145656061188447 | 0.339,0.345,0.242, | 0.226,0.157,0.221, | 0,076 |
| 173730 | ENSG00000136560 | TANK | chr2 | + | 161202862 | 161202931 | 161179613 | 161179761 | 161203486 | 161203595 | 173730 | 7,6,9,5,0,5,2,9,3,5,0,0 | 79,95,243,175,102,110,76, | 5,3,0,1,4,0,0,7,2,1,1,0,1,1,0 | 120,137,113,48,117,53,46, | 218 | 149 | 0.0000626781483505 | 0.0145998707225395 | 0.057,0.041,0.025, | 0.028,0.015,0.0,0.0 | 0,023 |
| 73362 | ENSG00000136059 | VILL | chr3 | + | 37998089 | 37998168 | 37997482 | 37997685 | 37998265 | 37998364 | 73362 | 20,2,19,18,4,10,10,10,9,5,0 | 0,2,0,2,1,0,0,0,0,2,1,0 | 5,7,9,3,3,6,11,13,6,7,15,14 | 0,5,2,1,3,0,2,1,2,0,2,0,4,0,0 | 228 | 149 | 0.0000630178775279 | 0.0146548376819253 | 1.0,0.395,1.0,0.85 | 1.0,0.478,0.746,0.0 | 0,02 |
| 222643 | ENSG00000167483 | NIBAN3 | chr19 | + | 17536985 | 17537259 | 17532262 | 17532388 | 17537375 | 17537543 | 222643 | 1,0,0,4,1,0,0,3,1,0,1,3 | 11,9,35,4,17,7,18,19,10,11 | 4,2,5,2,1,1,0,2,0,2,2,4,10,7 | 26,18,7,9,8,5,13,13,7,17,16 | 298 | 149 | 0.0000630905451674 | 0.0146548376819253 | 0.043,0.0,0.0,0.33 | 0.071,0.053,0.263, | -0,03 |
| 213429 | ENSG00000035720 | STAP1 | chr4 | + | 67576837 | 67576943 | 67575384 | 67575498 | 67577202 | 67577259 | 213429 | 0,2,33,8,0,7,0,1,2,0,3,6 | 95,43,180,138,78,65,66,55 | 26,13,17,9,0,0,7,10,0,2,1,0 | 83,59,97,34,67,53,15,55,47 | 255 | 149 | 0.0000631554266852 | 0.0146548376819253 | 0.0,0.026,0.097,0.0 | 0.155,0.114,0.093, | -0,035 |
| 150568 | ENSG00000168970 | JMJD7-PLA2G4 | chr15 | + | 41836780 | 41837219 | 41836474 | 41836551 | 41840157 | 41840230 | 150568 | 17,3,16,7,10,12,7,11,3,6,2, | 0,0,1,0,1,4,0,3,4,3,1,1 | 7,8,6,3,4,1,20,24,3,9,14,6,2 | 1,1,0,0,1,1,0,0,0,0,1,0,2,0,1 | 298 | 149 | 0.0000631864804272 | 0.0146548376819253 | 1.0,1.0,0.889,1.0,0 | 0.778,0.8,1.0,1.0,0 | -0,079 |
| 182980 | ENSG00000175348 | TMEM9B | chr11 | - | 8952989 | 8953337 | 8947500 | 8948475 | 8956189 | 8956298 | 182980 | 27,28,80,45,23,25,15,29,15 | 1,0,1,2,0,1,2,2,0,0,0,0 | 35,44,24,21,17,26,12,35,13 | 5,4,8,0,2,5,0,0,0,1,2,2,1,0,0 | 298 | 149 | 0.0000634833700727 | 0.014701105965543 | 0.931,1.0,0.976,0.9 | 0.778,0.846,0.6,1.0 | 0,056 |
| 154724 | ENSG00000129566 | TEP1 | chr14 | - | 20389637 | 20389740 | 20389237 | 20389297 | 20390680 | 20390758 | 154724 | 31,29,26,25,32,34,18,13,17 | 1,0,1,2,1,3,3,0,0,2,0,1 | 39,34,16,20,34,20,21,19,20 | 0,3,2,5,4,2,4,0,0,0,4,4,6,0,2 | 252 | 149 | 0.0000635689858257 | 0.014701105965543 | 0.948,1.0,0.939,0.8 | 1.0,0.87,0.825,0.70 | 0,056 |
| 32743 | ENSG00000136960 | ENPP2 | chr8 | - | 119572175 | 119572250 | 119570704 | 119570841 | 119580115 | 119580167 | 32743 | 6,2,21,8,0,0,0,5,0,0,0,0 | 24,28,67,62,10,10,7,15,3,2 | 3,0,0,0,2,4,1,0,0,0,0,2,0,0,8 | 6,10,5,1,26,23,1,3,4,2,3,18 | 224 | 149 | 0.0000635908844633 | 0.014701105965543 | 0.143,0.045,0.173, | 0.25,0.0,0.0,0.0,0.0 | -0,015 |
| 1650 | ENSG00000100138 | SNU13 | chr22 | - | 41681225 | 41681559 | 41680243 | 41680364 | 41688793 | 41688981 | 1650 | 5,1,1,0,0,2,3,1,0,0,0,0 | 35,27,61,40,24,23,26,15,20 | 2,5,7,5,2,1,0,0,0,3,6,8,1,0,2 | 29,34,23,10,26,21,11,25,11 | 298 | 149 | 0.0000641310233954 | 0.0148100689875701 | 0.067,0.018,0.008, | 0.033,0.068,0.132, | -0,032 |
| 168594 | ENSG00000005059 | MCUB | chr4 | + | 109684248 | 109684646 | 109682581 | 109682742 | 109685250 | 109685367 | 168594 | 94,71,143,121,64,44,59,75 | 6,1,2,1,0,4,5,0,5,0,0,1 | 101,92,72,22,29,43,28,70,3 | 10,3,8,3,1,3,2,8,3,5,3,10,4, | 298 | 149 | 0.0000644515152942 | 0.0148681287119892 | 0.887,0.973,0.973, | 0.835,0.939,0.818, | 0,061 |
| 160561 | ENSG00000113732 | ATP6V0E1 | chr5 | + | 172986642 | 172986760 | 172983830 | 172983964 | 172994774 | 172994822 | 160561 | 1,5,8,5,1,1,1,3,0,0,0,1 | 70,64,115,194,196,99,90,9 | 6,11,3,4,1,3,1,4,4,3,9,6,1,7 | 121,86,70,21,41,43,48,104 | 267 | 149 | 0.0000648381292551 | 0.0149413012823388 | 0.008,0.042,0.037, | 0.027,0.067,0.023, | -0,02 |
| 209152 | ENSG00000184209 | SNRNP35 | chr12 | + | 123458963 | 123459062 | 123458087 | 123458216 | 123465537 | 123466936 | 209152 | 0,2,6,2,0,0,2,0,3,1,0,1 | 15,9,25,9,11,11,13,17,14,7 | 3,4,2,3,0,0,0,3,0,3,0,8,5,1,3 | 4,9,8,6,5,6,4,13,2,3,8,23,8, | 248 | 149 | 0.0000649483372412 | 0.0149506904521505 | 0.0,0.118,0.126,0.1 | 0.311,0.211,0.131, | -0,078 |
| 238947 | ENSG00000197265 | GTF2E2 | chr8 | - | 30612920 | 30612995 | 30612469 | 30612481 | 30614607 | 30614715 | 238947 | 8,0,3,0,0,6,0,0,0,1,2,1 | 89,34,78,57,29,30,30,34,43 | 3,10,6,8,11,2,1,0,2,1,0,11,0 | 36,44,39,40,44,24,8,28,24, | 224 | 149 | 0.0000657893868582 | 0.0151281148748689 | 0.056,0.0,0.025,0.0 | 0.053,0.131,0.093, | -0,02 |
| 160959 | ENSG00000114904 | NEK4 | chr3 | - | 52768432 | 52768604 | 52766315 | 52766375 | 52770653 | 52770939 | 160959 | 3,5,15,7,3,4,7,2,3,1,2,5 | 1,0,3,1,0,1,0,5,0,0,0,0 | 9,1,7,7,2,4,5,10,5,4,6,3,11, | 2,1,5,0,2,2,0,6,0,2,7,0,2,0,0 | 298 | 149 | 0.0000661381289512 | 0.0151918928737044 | 0.6,1.0,0.714,0.778 | 0.692,0.333,0.412, | 0,124 |
| 191785 | ENSG00000215305 | VPS16 | chr20 | + | 2865387 | 2865495 | 2865147 | 2865317 | 2866211 | 2866315 | 191785 | 37,16,42,26,21,24,12,25,45 | 4,0,4,1,2,3,0,0,8,1,0,4 | 19,15,22,12,12,4,10,10,9,1 | 0,5,1,1,2,0,2,1,0,0,1,2,1,0,3 | 257 | 149 | 0.000066207913895 | 0.0151918928737044 | 0.843,1.0,0.859,0.9 | 1.0,0.635,0.927,0.8 | 0,005 |
| 8398 | ENSG00000130813 | SHFL | chr19 | + | 10089110 | 10089208 | 10087250 | 10087300 | 10089656 | 10089695 | 8398 | 10,1,10,6,3,7,2,7,6,1,4,3 | 22,16,40,31,21,24,16,22,14 | 6,3,2,4,1,3,0,0,1,1,3,3,1,4,2 | 17,18,22,22,24,8,11,15,7,2 | 247 | 149 | 0.000066824183622 | 0.0153169711023926 | 0.215,0.036,0.131, | 0.176,0.091,0.052, | 0,062 |
| 187408 | ENSG00000205323 | SARNP | chr12 | - | 55816933 | 55817169 | 55803628 | 55803728 | 55817665 | 55817724 | 187408 | 0,0,5,2,2,4,5,17,0,1,0,0 | 36,43,77,43,32,35,31,27,45 | 1,0,0,3,0,0,2,0,1,3,2,0,2,0,0 | 25,35,29,22,26,18,18,38,23 | 298 | 149 | 0.0000674149053619 | 0.0154215310686059 | 0.0,0.0,0.0,0.031,0.02 | 0.02,0.0,0.0,0.0,0.064, | 0,023 |
| 26884 | ENSG00000129534 | MIS18BP1 | chr14 | - | 45223917 | 45224746 | 45218281 | 45218454 | 45226742 | 45226836 | 26884 | 72,69,167,108,98,58,42,56 | 5,1,1,2,3,3,5,2,2,0,0,3 | 62,78,43,28,79,85,42,58,60 | 9,8,4,4,16,2,1,1,4,4,2,4,5,4 | 298 | 149 | 0.0000674236552149 | 0.0154215310686059 | 0.878,0.972,0.988, | 0.775,0.83,0.843,0 | 0,054 |
| 196506 | ENSG00000083845 | RPS5 | chr19 | + | 58392988 | 58393487 | 58388136 | 58388245 | 58394496 | 58394595 | 196506 | 2290,1321,2182,1591,1020 | 21,18,27,34,4,5,10,12,2,4,1 | 1046,1367,912,544,709,45 | 21,29,13,5,13,7,14,11,9,18 | 298 | 149 | 0.0000676911356692 | 0.015452898346092 | 0.982,0.973,0.976, | 0.961,0.959,0.972, | 0,012 |
| 86087 | ENSG00000114120 | SLC25A36 | chr3 | + | 140968682 | 140968730 | 140966933 | 140967036 | 140970926 | 140970993 | 86087 | 3,0,12,2,0,0,1,0,0,0,0,1 | 14,10,16,14,2,10,4,12,8,10 | 1,6,2,2,4,0,1,3,0,0,1,2,0,0,1 | 4,13,6,4,4,1,5,6,3,12,3,8,7, | 197 | 149 | 0.0000677043880313 | 0.015452898346092 | 0.139,0.0,0.362,0.0 | 0.159,0.259,0.201, | -0,058 |
| 71120 | ENSG00000151883 | PARP8 | chr5 | + | 50763147 | 50763242 | 50761820 | 50761898 | 50778068 | 50778126 | 71120 | 47,29,108,61,25,34,28,29,3 | 1,0,7,3,0,0,0,2,0,0,1,1 | 51,50,39,38,37,24,23,47,22 | 5,2,1,4,5,1,6,2,1,1,3,2,0,1,1 | 244 | 149 | 0.0000679770947274 | 0.015498705588213 | 0.966,1.0,0.904,0.9 | 0.862,0.939,0.96,0 | 0,058 |
| 240151 | ENSG00000165525 | NEMF | chr14 | - | 49833422 | 49833467 | 49832206 | 49832277 | 49834362 | 49834449 | 240151 | 26,9,30,27,9,10,15,13,13,1 | 2,1,3,1,1,1,0,0,5,2,0,1 | 25,29,12,8,19,7,6,15,14,19 | 5,5,0,4,3,1,2,2,3,9,0,2,7,3,4 | 194 | 149 | 0.0000681433627816 | 0.0155030978987462 | 0.909,0.874,0.885, | 0.793,0.817,1.0,0.0 | 0,119 |
| 196511 | ENSG00000083845 | RPS5 | chr19 | + | 58392993 | 58393487 | 58388136 | 58388245 | 58394496 | 58394595 | 196511 | 2290,1321,2181,1589,1020 | 21,18,27,34,4,5,10,12,2,4,1 | 1046,1367,912,544,709,45 | 21,29,13,5,13,7,14,11,9,18 | 298 | 149 | 0.0000682124494618 | 0.0155030978987462 | 0.982,0.973,0.976, | 0.961,0.959,0.972, | 0,012 |
| 196516 | ENSG00000083845 | RPS5 | chr19 | + | 58393051 | 58393487 | 58388136 | 58388245 | 58394496 | 58394595 | 196516 | 2290,1321,2181,1589,1020 | 21,18,27,34,4,5,10,12,2,4,1 | 1046,1367,912,544,709,45 | 21,29,13,5,13,7,14,11,9,18 | 298 | 149 | 0.0000682124494618 | 0.0155030978987462 | 0.982,0.973,0.976, | 0.961,0.959,0.972, | 0,012 |
| 67658 | ENSG00000083845 | RPS5 | chr19 | + | 58393358 | 58393487 | 58388136 | 58388245 | 58394496 | 58394595 | 67658 | 2281,1317,2167,1578,1009 | 21,18,27,34,4,5,10,12,2,4,1 | 1038,1353,903,538,709,45 | 21,29,13,5,13,7,14,11,9,18 | 278 | 149 | 0.0000683346559294 | 0.0155144897999374 | 0.983,0.975,0.977, | 0.964,0.962,0.974, | 0,011 |
| 209615 | ENSG00000166986 | MARS1 | chr12 | + | 57489895 | 57490058 | 57489266 | 57489558 | 57490206 | 57490379 | 209615 | 43,12,61,56,14,26,21,25,15 | 5,3,6,2,1,8,7,3,1,2,1,4 | 28,20,23,10,15,18,7,24,19, | 5,19,9,6,9,1,8,3,4,5,11,6,16 | 298 | 149 | 0.000068425268748 | 0.0155186923265551 | 0.811,0.667,0.836, | 0.737,0.345,0.561, | 0,2 |
| 173868 | ENSG00000082996 | RNF13 | chr3 | + | 149901770 | 149901854 | 149895472 | 149895570 | 149902071 | 149902162 | 173868 | 4,1,13,2,2,4,1,3,4,0,0,0 | 32,17,57,45,15,20,16,25,19 | 4,2,1,0,3,0,0,2,0,2,3,0,0,1,2 | 41,40,24,13,18,9,25,29,10, | 233 | 149 | 0.0000686990642194 | 0.015551445610465 | 0.074,0.036,0.127, | 0.059,0.031,0.026, | 0,019 |
| 218171 | ENSG00000090975 | PITPNM2 | chr12 | - | 122994800 | 122994928 | 122992498 | 122992669 | 122995388 | 122995660 | 218171 | 76,34,76,18,25,23,48,21,46 | 2,0,2,2,2,2,4,0,4,3,1,0 | 32,57,32,54,38,19,34,24,28 | 0,0,1,0,2,0,0,2,0,0,2,0,1,0,0 | 277 | 149 | 0.0000687765336286 | 0.015551445610465 | 0.953,1.0,0.953,0.8 | 1.0,1.0,0.945,1.0,0 | -0,066 |
| 32432 | ENSG00000113593 | PPWD1 | chr5 | + | 65568917 | 65568994 | 65567512 | 65567615 | 65569631 | 65569732 | 32432 | 6,0,23,19,8,8,3,2,12,6,1,0 | 36,23,58,36,22,17,11,3,46, | 1,0,1,2,3,1,2,3,0,4,1,1,2,3,2 | 29,22,19,22,24,15,15,11,19 | 226 | 149 | 0.0000687864490764 | 0.015551445610465 | 0.099,0.0,0.207,0.2 | 0.022,0.0,0.034,0.0 | 0,074 |
| 32172 | ENSG00000134480 | CCNH | chr5 | - | 87409087 | 87409129 | 87408037 | 87408186 | 87409289 | 87409363 | 32172 | 4,5,4,5,2,1,1,4,1,1,3,0 | 37,26,71,38,21,25,23,54,26 | 5,11,5,2,6,1,7,6,0,3,2,10,5, | 23,59,42,50,31,19,35,33,19 | 191 | 149 | 0.0000689160897563 | 0.0155644059961576 | 0.078,0.13,0.042,0 | 0.145,0.127,0.085, | -0,051 |
| 32913 | ENSG00000083896 | YTHDC1 | chr4 | - | 68330001 | 68330119 | 68324138 | 68324223 | 68332102 | 68332197 | 32913 | 67,76,140,130,51,73,45,14 | 2,0,1,0,0,0,2,0,0,1,0,0 | 84,167,54,127,69,34,56,85 | 1,4,1,13,5,2,2,0,1,0,0,1,1,4 | 267 | 149 | 0.0000692492159372 | 0.0156051992361502 | 0.949,1.0,0.987,1.0 | 0.979,0.959,0.968, | 0,03 |
| 185745 | ENSG00000198160 | MIER1 | chr1 | + | 66975132 | 66975237 | 66972896 | 66972991 | 66976594 | 66976722 | 185745 | 3,0,1,0,0,0,3,2,0,0,1,3 | 25,15,112,80,33,46,27,44,9 | 6,0,4,4,2,7,0,2,6,4,0,2,2,4,2 | 51,91,25,30,43,19,24,51,26 | 254 | 149 | 0.0000692995204595 | 0.0156051992361502 | 0.066,0.0,0.0,0.005,0.0 | 0.065,0.0,0.0,0.086,0.0 | -0,023 |

| | | | | | | | | | | | | | | | | | | | | | | |
|---|---|---|---|---|---|---|---|---|---|---|---|---|---|---|---|---|---|---|---|---|---|---|
| 244681 | ENSG00000168067 | MAP4K2 | chr11 | - | 64801578 | 64801650 | 64801110 | 64801183 | 64801709 | 64801813 | 244681 | 25,20,31,22,10,14,6,18,15, | 0,0,4,5,2,0,0,2,0,0,0,2 | 29,16,14,17,23,8,3,9,8,25,1 | 8,4,0,3,0,1,0,2,5,4,7,0,1,1,4 | 221 | 149 | 0.0000693142273639 | 0.0156051992361502 | 1.0,1.0,0.839,0.748 | 0.71,0.729,1.0,0.79 | 0,129 |
| 162676 | ENSG00000072364 | AFF4 | chr5 | - | 132898262 | 132898365 | 132897203 | 132897240 | 132899103 | 132899141 | 162676 | 1,0,2,0,0,2,1,6,0,0,0,1 | 13,1,7,1,1,3,1,9,3,1,6,3 | 2,5,2,5,5,0,0,3,2,6,4,3,0,1,1 | 5,7,5,31,9,9,5,4,8,5,4,2,2,5 | 252 | 149 | 0.0000697397001249 | 0.0156845824426148 | 0.044,0.0,0.145,0.0 | 0.191,0.297,0.191, | -0,097 |
| 157179 | ENSG00000133112 | TPT1 | chr13 | - | 45339496 | 45339546 | 45338184 | 45338776 | 45339993 | 45340258 | 157179 | 978,588,1752,1299,395,36 | 10,0,7,4,8,6,6,7,0,0,6,6 | 897,812,625,452,462,376,2 | 11,11,8,21,2,9,3,14,5,4,31, | 199 | 149 | 0.0000699066023777 | 0.0157057076579903 | 0.987,1.0,0.995,0.9 | 0.984,0.982,0.983, | 0,007 |
| 160957 | ENSG00000114904 | NEK4 | chr3 | - | 52768376 | 52768604 | 52766315 | 52766375 | 52770653 | 52770939 | 160957 | 2,4,15,7,4,3,7,2,3,1,2,5 | 1,0,3,1,0,1,0,5,0,0,0,0 | 9,1,7,7,2,4,5,8,5,4,6,3,11,4 | 2,1,5,0,2,2,0,6,0,2,7,0,2,0,0 | 298 | 149 | 0.0000704706098568 | 0.0158090430580792 | 0.5,1.0,0.714,0.778 | 0.692,0.333,0.412, | 0,113 |
| 107125 | ENSG00000215301 | DDX3X | chrX | + | 41343220 | 41343351 | 41342736 | 41342836 | 41343736 | 41343822 | 107125 | 78,66,290,285,87,77,83,10 | 5,0,7,5,3,3,2,5,2,1,0,0 | 226,246,130,97,111,68,59, | 9,9,3,17,0,6,2,8,5,8,5,8,8,6 | 280 | 149 | 0.0000705134545477 | 0.0158090430580792 | 0.892,1.0,0.957,0.9 | 0.93,0.936,0.958,0 | 0,041 |
| 80919 | ENSG00000091490 | SEL1L3 | chr4 | - | 25832994 | 25833110 | 25830097 | 25830156 | 25847293 | 25847864 | 80919 | 390,312,453,259,146,149,1 | 6,9,6,3,5,12,4,7,1,3,0,1 | 135,108,208,120,141,105,4 | 7,7,13,2,17,2,2,3,9,2,6,9,6, | 265 | 149 | 0.0000706363828159 | 0.0158201241517679 | 0.973,0.951,0.977, | 0.916,0.897,0.9,0.9 | 0,035 |
| 151808 | ENSG00000166128 | RAB8B | chr15 | + | 63250367 | 63250477 | 63249644 | 63249705 | 63255506 | 63255584 | 151808 | 7,1,1,2,0,1,2,3,4,2,0,1 | 87,85,73,67,39,34,25,29,84 | 5,6,3,15,2,1,0,6,0,2,0,11,0, | 52,69,61,45,34,22,15,30,11 | 259 | 149 | 0.0000707330514736 | 0.0158253070703892 | 0.044,0.007,0.008, | 0.052,0.048,0.028, | -0,023 |
| 61486 | ENSG00000183530 | PRR14L | chr22 | - | 31738386 | 31738911 | 31725537 | 31725610 | 31749992 | 31750132 | 61486 | 34,22,55,23,22,20,9,13,26, | 5,2,0,4,2,2,3,3,2,2,3,0 | 20,15,19,20,11,14,8,16,8,2 | 0,4,0,0,0,0,1,1,1,1,0,0,4,3,3 | 298 | 149 | 0.0000708592423824 | 0.0158370774633503 | 0.773,0.846,1.0,0.7 | 1.0,0.652,1.0,1.0,1 | -0,071 |
| 107218 | ENSG00000125351 | UPF3B | chrX | - | 119840645 | 119840684 | 119838366 | 119838527 | 119841075 | 119841258 | 107218 | 7,4,4,0,6,1,3,0,0,0,2,0 | 23,21,11,7,3,10,6,9,8,6,4,8 | 0,4,0,0,3,1,0,1,0,0,0,3,0,0,1 | 5,19,7,8,11,11,5,7,2,14,5,1 | 188 | 149 | 0.0000709678051199 | 0.0158448876180095 | 0.194,0.131,0.224, | 0.0,0.143,0.0,0.0,0 | 0,087 |
| 51058 | ENSG00000100991 | TRPC4AP | chr20 | - | 35044504 | 35044712 | 35035122 | 35035308 | 35049865 | 35049994 | 51058 | 171,131,266,199,90,166,96 | 3,3,7,1,2,4,2,3,3,2,2,3 | 176,228,107,150,168,108,1 | 4,1,2,1,0,1,0,2,0,0,4,2,1,0,0 | 298 | 149 | 0.0000710733307038 | 0.0158520041872638 | 0.966,0.956,0.95,0 | 0.957,0.991,0.964, | -0,03 |
| 130605 | ENSG00000186815 | TPCN1 | chr12 | + | 113294105 | 113294228 | 113293268 | 113293349 | 113295959 | 113296012 | 130605 | 2,0,4,1,6,2,0,1,0,0,0,2 | 20,17,16,5,10,7,10,9,20,14 | 5,0,3,6,2,2,0,3,1,1,4,5,1,1,4 | 6,2,5,10,5,7,8,9,3,1,5,15,6, | 272 | 149 | 0.0000712951749638 | 0.0158850225700141 | 0.052,0.0,0.12,0.09 | 0.313,0.0,0.247,0.2 | -0,118 |
| 56974 | ENSG00000163466 | ARPC2 | chr2 | + | 218234351 | 218234397 | 218228737 | 218228850 | 218238663 | 218238850 | 56974 | 255,209,571,541,200,108,1 | 5,1,2,3,1,0,0,4,1,0,0,6 | 286,382,179,29,199,130,31 | 7,11,2,3,12,8,1,3,9,3,1,3,1, | 195 | 149 | 0.0000721273532757 | 0.0160538183794025 | 0.975,0.994,0.995, | 0.969,0.964,0.986, | 0,024 |
| 151551 | ENSG00000077238 | IL4R | chr16 | + | 27346476 | 27346618 | 27344907 | 27345020 | 27352539 | 27352696 | 151551 | 1042,307,1116,664,448,34 | 40,17,48,36,11,21,13,26,35 | 555,276,688,133,119,163,1 | 10,7,15,3,14,6,1,4,1,7,8,7,1 | 291 | 149 | 0.0000722742144426 | 0.0160698878602223 | 0.93,0.902,0.923,0 | 0.966,0.953,0.959, | -0,012 |
| 138787 | ENSG00000133243 | BTBD2 | chr19 | - | 2013546 | 2013718 | 1997343 | 1997463 | 2015296 | 2015714 | 138787 | 2,2,15,8,0,2,5,3,6,2,1,4 | 32,28,22,15,19,12,17,5,48, | 2,0,0,8,2,0,2,1,0,4,3,2,2,0,0 | 6,6,9,22,7,6,11,18,4,17,12, | 298 | 149 | 0.0000727115829724 | 0.0161395506421314 | 0.03,0.034,0.254,0 | 0.143,0.0,0.0,0.154 | 0,056 |
| 59805 | ENSG00000162434 | JAK1 | chr1 | - | 64882891 | 64883063 | 64879024 | 64879148 | 64883276 | 64883475 | 59805 | 33,17,26,9,6,9,3,16,7,6,2,6 | 278,204,311,216,177,166,1 | 15,36,12,62,24,12,10,18,6, | 242,263,227,158,177,117,1 | 298 | 149 | 0.0000727603002587 | 0.0161395506421314 | 0.056,0.04,0.04,0.0 | 0.03,0.064,0.026,0 | -0,019 |
| 167545 | ENSG00000112079 | STK38 | chr6 | - | 36499872 | 36500046 | 36498362 | 36498486 | 36506582 | 36506644 | 167545 | 4,4,9,9,9,4,4,4,2,1,1,3 | 1,2,7,6,0,1,0,0,3,2,0,0 | 7,3,3,0,7,5,2,3,6,8,2,7,19,1 | 2,0,1,1,2,0,0,2,2,1,0,0,1,0,0 | 298 | 149 | 0.0000728124836734 | 0.0161395506421314 | 0.667,0.5,0.391,0.4 | 0.636,1.0,0.6,0.0,0 | -0,113 |
| 202292 | ENSG00000136059 | VILL | chr3 | + | 37998073 | 37998168 | 37997482 | 37997685 | 37998265 | 37998364 | 202292 | 7,3,7,10,1,9,3,4,2,2,0,4 | 0,2,0,2,1,0,0,0,0,2,1,0 | 1,1,6,0,1,4,7,10,3,4,6,11,2, | 0,5,2,1,3,0,2,1,2,0,2,0,4,0,0 | 244 | 149 | 0.0000730211217149 | 0.0161691451109255 | 1.0,0.478,1.0,0.753 | 1.0,0.109,0.647,0.0 | 0,108 |
| 97992 | ENSG00000145725 | PPIP5K2 | chr5 | + | 103186319 | 103186439 | 103184671 | 103184744 | 103190841 | 103190982 | 97992 | 11,9,28,15,6,3,4,13,21,7,3, | 23,7,36,17,21,16,12,15,10, | 7,11,3,7,3,4,6,4,0,4,2,6,8,1 | 20,33,17,8,5,8,12,14,8,12,2 | 269 | 149 | 0.0000734536696209 | 0.016248208392781 | 0.209,0.416,0.301, | 0.162,0.156,0.089, | 0,123 |
| 253141 | ENSG00000111144 | LTA4H | chr12 | - | 96011633 | 96011865 | 96009093 | 96009148 | 96013187 | 96013258 | 253141 | 24,15,54,6,5,3,4,8,1,2,5,9 | 224,107,416,326,157,108,1 | 5,13,6,8,4,3,6,8,5,11,8,7,4, | 275,164,226,55,136,112,72 | 298 | 149 | 0.0000737005219489 | 0.0162860749893056 | 0.051,0.066,0.061, | 0.009,0.038,0.013, | 0,006 |
| 213479 | ENSG00000172765 | TMCC1 | chr3 | - | 129857253 | 129857405 | 129848345 | 129848412 | 129880308 | 129880559 | 213479 | 0,3,6,8,5,1,0,4,0,0,2,1 | 5,7,38,15,11,5,10,9,2,8,3,7 | 0,1,1,0,0,3,1,1,0,0,2,0,1,3,2 | 3,11,14,8,13,2,8,14,7,12,12 | 298 | 149 | 0.0000743852186429 | 0.0164089392080688 | 0.0,0.176,0.073,0.1 | 0.0,0.043,0.034,0.0 | 0,03 |
| 82443 | ENSG00000108883 | EFTUD2 | chr17 | - | 44853295 | 44853390 | 44852408 | 44852562 | 44853516 | 44853635 | 82443 | 25,39,53,40,36,22,15,24,42 | 2,3,3,7,0,2,1,2,0,1,0,1 | 37,34,15,24,27,12,9,37,16, | 1,1,0,0,2,0,0,1,0,0,4,0,0,0,0 | 244 | 149 | 0.00007440900552 | 0.0164089392080688 | 0.884,0.888,0.915, | 0.958,0.954,1.0,1.0 | -0,049 |
| 235894 | ENSG00000167302 | TEPSIN | chr17 | - | 81238001 | 81238159 | 81237386 | 81237459 | 81238513 | 81239042 | 235894 | 5,0,1,1,3,0,1,4,0,0,0,3 | 13,6,15,3,9,2,6,3,10,3,1,5 | 2,5,2,1,2,5,2,0,2,2,2,2,4,3,5 | 7,2,0,3,4,1,1,4,3,3,6,0,1,4,2 | 298 | 149 | 0.0000745830677013 | 0.0164304895029872 | 0.161,0.0,0.032,0.1 | 0.125,0.556,1.0,0.1 | -0,305 |
| 115954 | ENSG00000106829 | TLE4 | chr9 | + | 79627373 | 79627448 | 79612655 | 79612718 | 79652589 | 79652629 | 115954 | 74,44,91,79,59,88,54,105,5 | 4,0,5,2,0,4,2,5,0,4,4,0 | 47,84,100,85,51,44,27,34,4 | 0,0,0,2,0,1,0,2,2,0,2,2,0,2,0 | 224 | 149 | 0.0000751439692788 | 0.0165371284783695 | 0.925,1.0,0.924,0.9 | 1.0,1.0,1.0,0.966,1 | -0,016 |
| 72152 | ENSG00000160908 | ZNF394 | chr7 | - | 99498715 | 99498842 | 99493239 | 99494631 | 99499637 | 99499871 | 72152 | 76,48,91,20,25,45,46,26,38 | 5,3,1,0,5,2,1,4,0,8,2,0 | 62,72,34,74,25,22,27,35,36 | 0,1,2,0,0,1,3,2,0,0,6,2,2,1,0 | 276 | 149 | 0.0000755665498391 | 0.0166131400290289 | 0.891,0.896,0.98,1 | 1.0,0.975,0.902,1.0 | -0,068 |
| 184599 | ENSG00000196865 | NHLRC2 | chr10 | + | 113898109 | 113898208 | 113884250 | 113884380 | 113901665 | 113901897 | 184599 | 11,5,18,12,12,8,3,7,8,1,2,8 | 4,5,0,0,0,0,1,2,0,0,0,0 | 20,18,11,12,12,10,11,2,11, | 3,1,2,6,2,0,0,0,5,0,0,1,3,2,1 | 248 | 149 | 0.000075660006027 | 0.0166167130175482 | 0.623,0.375,1.0,1.0 | 0.8,0.915,0.768,0.5 | 0,035 |
| 218320 | ENSG00000163939 | PBRM1 | chr3 | - | 52563276 | 52563493 | 52561766 | 52561968 | 52564049 | 52564233 | 218320 | 48,13,41,26,15,7,18,20,22, | 5,0,0,3,1,4,3,2,4,2,0,1 | 27,26,16,27,18,18,3,21,27, | 0,0,1,0,4,0,0,0,0,3,0,5,1,0,2 | 298 | 149 | 0.0000759121639737 | 0.0166550978228577 | 0.828,1.0,1.0,0.81 | 1.0,1.0,0.889,1.0,0 | -0,087 |
| 6520 | ENSG00000125844 | RRBP1 | chr20 | - | 17616731 | 17616839 | 17615925 | 17616009 | 17618595 | 17618679 | 6520 | 99,52,141,102,56,54,59,82 | 6,0,5,3,0,2,2,0,0,3,5,0 | 108,110,44,70,29,22,120,7 | 6,5,0,10,1,0,12,17,1,0,3,0,1 | 257 | 149 | 0.0000766535402453 | 0.0168006294506478 | 0.905,1.0,0.942,0.9 | 0.913,0.927,1.0,0.8 | 0,036 |
| 236789 | ENSG00000104324 | CPQ | chr8 | + | 97055528 | 97055645 | 97029468 | 97029494 | 97066008 | 97066210 | 236789 | 6,3,2,0,5,10,2,1,7,2,2,3 | 4,4,6,7,2,13,4,4,8,3,5,6 | 1,2,0,5,1,0,0,4,2,0,3,0,0,2,1 | 14,7,3,4,2,2,2,9,3,7,7,23,2, | 266 | 149 | 0.0000784256671734 | 0.0171715511407915 | 0.457,0.296,0.157, | 0.038,0.138,0.0,0.4 | 0,127 |
| 240408 | ENSG00000184465 | WDR27 | chr6 | - | 169694807 | 169695065 | 169689126 | 169689308 | 169701550 | 169701854 | 240408 | 6,0,13,2,0,6,2,8,0,2,4,4 | 8,9,26,15,6,22,8,1,6,7,11,8 | 2,2,1,2,0,1,3,1,0,1,1,1,0,3,0 | 4,3,7,3,6,11,4,12,6,14,14,3 | 298 | 149 | 0.0000788385137974 | 0.0172429291971724 | 0.273,0.0,0.2,0.062 | 0.2,0.25,0.067,0.25 | 0,071 |
| 61207 | ENSG00000178665 | ZNF713 | chr7 | + | 55918041 | 55918297 | 55912634 | 55912723 | 55923161 | 55923288 | 61207 | 1,3,9,2,2,1,2,4,1,6,0,2 | 3,4,19,5,3,9,9,5,3,2,1,4 | 2,0,1,0,5,1,0,1,1,2,2,0,0,1,3 | 3,2,4,3,4,3,4,8,5,9,1,3,2,0,1 | 298 | 149 | 0.0000789118912202 | 0.0172429291971724 | 0.143,0.273,0.191, | 0.25,0.0,0.111,0.0, | 0,029 |
| 167493 | ENSG00000145495 | MARCHF6 | chr5 | + | 10411363 | 10411537 | 10410152 | 10410276 | 10414432 | 10414502 | 167493 | 188,150,192,97,83,105,93, | 0,1,0,2,0,0,4,2,0,0,0,0 | 153,186,123,288,105,53,12 | 0,6,0,14,3,2,1,1,1,3,0,4,1,1 | 298 | 149 | 0.0000792377997142 | 0.0172892833267236 | 1.0,0.987,1.0,0.96, | 1.0,0.939,1.0,0.91 | 0,029 |
| 44617 | ENSG00000159433 | STARD9 | chr15 | + | 42665252 | 42665330 | 42662793 | 42663917 | 42665785 | 42665848 | 44617 | 15,3,14,7,22,21,7,24,14,14 | 5,0,6,0,0,0,1,9,1,0,0,3 | 13,15,6,20,2,5,21,5,5,7,11, | 2,0,0,6,1,1,0,0,0,0,5,0,0,0,0 | 227 | 149 | 0.0000792846878167 | 0.0172892833267236 | 0.663,1.0,0.605,1.0 | 0.81,1.0,1.0,0.686, | -0,004 |
| 186697 | ENSG00000134242 | PTPN22 | chr1 | - | 113825141 | 113825170 | 113819298 | 113819654 | 113829591 | 113829707 | 186697 | 51,30,46,64,7,3,7,2,2,1,2,4 | 4,2,5,5,2,1,4,3,7,3,0,2 | 28,42,19,5,11,12,1,1,12,42 | 0,3,0,1,0,0,1,2,0,2,1,7,1,2,3 | 178 | 149 | 0.0000797084879109 | 0.0173633643080856 | 0.914,0.926,0.885, | 1.0,0.921,1.0,0.807 | -0,157 |
| 59426 | ENSG00000011485 | PPP5C | chr19 | + | 46383410 | 46383476 | 46376452 | 46376574 | 46383779 | 46383878 | 59426 | 21,23,40,24,18,8,3,12,17,3 | 2,0,2,0,2,1,1,1,2,0,0,3 | 15,12,7,2,10,4,4,11,1,11,7, | 2,1,2,0,0,4,0,4,1,1,3,9,2,3,0 | 215 | 149 | 0.0000797857525203 | 0.0173633643080856 | 0.879,1.0,0.933,1.0 | 0.839,0.893,0.708, | 0,094 |
| 63543 | ENSG00000120370 | GORAB | chr1 | + | 170536309 | 170536421 | 170532146 | 170532284 | 170539209 | 170539567 | 63543 | 18,34,25,15,17,9,13,17,2,7 | 0,2,6,3,2,1,0,1,3,1,2,0 | 17,21,3,7,14,5,9,17,10,27,2 | 1,3,0,0,1,0,0,0,0,0,0,0,2,2,2 | 261 | 149 | 0.000080235241063 | 0.0174435466355864 | 1.0,0.907,0.704,0.7 | 0.907,0.8,1.0,1.0,0 | -0,103 |
| 107122 | ENSG00000215301 | DDX3X | chrX | + | 41343215 | 41343351 | 41340005 | 41342836 | 41343736 | 41343822 | 107122 | 367,370,1141,841,248,278 | 5,0,7,5,3,3,2,5,2,1,0,0 | 712,935,459,468,533,285,1 | 9,9,3,17,0,6,2,8,5,8,5,8,8,6 | 285 | 149 | 0.0000810140112729 | 0.0175950824018946 | 0.975,1.0,0.988,0.9 | 0.976,0.982,0.988, | 0,014 |
| 234044 | ENSG00000215021 | PHB2 | chr12 | - | 6968470 | 6968595 | 6967891 | 6968021 | 6969497 | 6969577 | 234044 | 82,78,273,246,89,71,72,89 | 1,2,11,12,3,5,3,2,0,2,0,2 | 150,132,120,42,107,65,37, | 1,5,0,1,0,0,1,5,2,1,5,5,0,4,0 | 274 | 149 | 0.0000816586334729 | 0.0177172069969816 | 0.978,0.955,0.931, | 0.988,0.935,1.0,0.9 | -0,024 |
| 56205 | ENSG00000010322 | NISCH | chr3 | + | 52480878 | 52481037 | 52480183 | 52480295 | 52484512 | 52484637 | 56205 | 0,4,7,0,6,4,0,4,2,0,0,9 | 5,4,8,9,3,8,7,8,6,4,6,9 | 0,4,0,1,1,0,3,5,0,1,0,2,0,2,1 | 7,5,7,7,4,2,8,17,1,2,7,11,5, | 298 | 149 | 0.0000823181585335 | 0.0178243657739675 | 0.0,0.333,0.304,0.0 | 0.0,0.286,0.0,0.067 | 0,083 |
| 56206 | ENSG00000010322 | NISCH | chr3 | + | 52480878 | 52481109 | 52480138 | 52480295 | 52484512 | 52484637 | 56206 | 0,4,7,0,6,4,0,4,2,0,0,9 | 5,4,8,9,3,8,7,8,6,4,6,9 | 0,4,0,1,1,0,3,5,0,1,0,2,0,2,1 | 7,5,7,7,4,2,8,17,1,2,7,11,5, | 298 | 149 | 0.0000823181585335 | 0.0178243657739675 | 0.0,0.333,0.304,0.0 | 0.0,0.286,0.0,0.067 | 0,083 |
| 204021 | ENSG00000144535 | DIS3L2 | chr2 | + | 232015515 | 232015671 | 232014927 | 232014979 | 232024276 | 232024330 | 204021 | 78,49,61,30,27,21,40,29,33 | 3,1,4,3,4,3,3,0,4,3,1,1 | 31,42,32,53,24,36,14,23,23 | 1,0,0,2,0,0,0,0,1,1,0,0,3,2,2 | 298 | 149 | 0.000082788137207 | 0.0178941826355751 | 0.929,0.961,0.884, | 0.939,1.0,1.0,0.93, | -0,07 |
| 63539 | ENSG00000120370 | GORAB | chr1 | + | 170533463 | 170533651 | 170532128 | 170532284 | 170539209 | 170539567 | 63539 | 37,19,19,15,10,12,10,17,3, | 0,2,6,3,2,1,0,1,3,1,2,0 | 19,13,21,31,10,2,1,10,7,16 | 1,3,0,0,1,0,0,0,0,0,0,0,2,2,2 | 298 | 149 | 0.0000828068721747 | 0.0178941826355751 | 1.0,0.826,0.613,0.7 | 0.905,0.684,1.0,1.0 | -0,11 |
| 15354 | ENSG00000103275 | UBE2I | chr16 | + | 1319701 | 1319862 | 1315653 | 1315726 | 1320173 | 1320199 | 15354 | 3,0,3,1,3,5,22,28,2,0,6,2 | 228,233,210,191,126,123,8 | 5,2,1,0,1,1,5,5,3,0,2,0,2,2,2 | 144,199,137,104,93,87,73, | 298 | 149 | 0.0000834747762027 | 0.0180204208193413 | 0.007,0.0,0.007,0.0 | 0.017,0.005,0.004, | 0,018 |
| 130697 | ENSG00000151148 | UBE3B | chr12 | + | 109517916 | 109518013 | 109516764 | 109516884 | 109521147 | 109521324 | 130697 | 0,1,0,5,0,0,0,0,0,0,0,0 | 27,19,34,30,7,16,8,9,8,13,5 | 0,4,8,2,0,3,0,1,3,6,2,0,0,1,1 | 24,13,3,4,18,7,14,6,14,24,1 | 246 | 149 | 0.0000836874162549 | 0.0180482227334252 | 0.0,0.031,0.0,0.092 | 0.0,0.157,0.618,0.1 | -0,106 |
| 74198 | ENSG00000174796 | THAP6 | chr4 | + | 75515432 | 75515532 | 75514437 | 75514520 | 75516771 | 75516789 | 74198 | 33,22,67,36,13,23,28,16,22 | 2,4,6,5,0,4,2,4,1,0,0,0 | 25,33,18,35,37,21,13,20,22 | 0,1,2,2,2,0,5,0,2,0,1,1,0,0,0 | 249 | 149 | 0.0000840317524274 | 0.0181043424491509 | 0.908,0.767,0.87,0 | 1.0,0.952,0.843,0.9 | -0,04 |
| 13907 | ENSG00000047578 | KATNIP | chr16 | + | 27699533 | 27699599 | 27698327 | 27698500 | 27701588 | 27701695 | 13907 | 12,8,21,16,4,15,8,16,5,3,4, | 5,0,1,0,0,0,3,2,4,3,1,1 | 8,11,2,7,6,10,7,14,6,11,8,1 | 0,2,0,0,2,6,1,1,2,0,0,1,0,0,0 | 215 | 149 | 0.0000842260403745 | 0.018128054895844 | 0.625,1.0,0.936,1.0 | 1.0,0.792,1.0,1.0,0 | -0,079 |
| 219672 | ENSG00000211450 | SELENOH | chr11 | + | 57742107 | 57742247 | 57741808 | 57741954 | 57742862 | 57743514 | 219672 | 103,73,172,239,89,66,70,8 | 0,3,5,3,4,0,0,3,0,2,0,0 | 166,107,102,12,59,93,25,8 | 3,7,8,6,3,2,0,7,4,3,4,3,0,0,1 | 289 | 149 | 0.0000846416339257 | 0.0181993042072551 | 1.0,0.926,0.947,0.9 | 0.966,0.887,0.868, | 0,06 |
| 129082 | ENSG00000174718 | RESF1 | chr12 | + | 31987238 | 31987322 | 31980877 | 31985957 | 31992377 | 31992941 | 129082 | 274,171,609,511,275,192,2 | 12,0,40,17,8,13,16,9,7,4,2, | 380,408,415,211,204,180,1 | 5,8,12,12,24,7,1,9,2,1,2,8,6 | 233 | 149 | 0.0000848484265544 | 0.0182255605745809 | 0.936,1.0,0.907,0.9 | 0.98,0.97,0.957,0.9 | -0,016 |
| 85977 | ENSG00000198721 | ECI2 | chr6 | - | 4125249 | 4126237 | 4117307 | 4119275 | 4127761 | 4127831 | 85977 | 28,18,66,73,20,17,7,33,23, | 0,0,6,2,0,0,0,1,0,0,0,0 | 45,11,25,4,26,19,7,21,6,33 | 5,7,3,0,1,1,0,1,0,2,2,0,1,2,0 | 298 | 149 | 0.0000849602361256 | 0.0182313824342463 | 1.0,1.0,0.846,0.948 | 0.818,0.44,0.806,1 | 0,098 |
| 169704 | ENSG00000204463 | BAG6 | chr6 | - | 31640384 | 31640509 | 31640198 | 31640306 | 31640644 | 31640938 | 169704 | 53,33,89,64,42,34,33,33,26 | 2,0,0,0,0,0,1,0,1,2,0,1 | 54,31,24,16,26,16,19,38,20 | 3,1,3,2,1,2,2,0,1,6,2,3,0,0,0 | 274 | 149 | 0.0000852701555571 | 0.0182796622019026 | 0.935,1.0,1.0,1.0,1 | 0.907,0.944,0.813, | 0,072 |
| 44802 | ENSG00000101596 | SMCHD1 | chr18 | + | 2750042 | 2750122 | 2747521 | 2747647 | 2750349 | 2750507 | 44802 | 349,277,650,420,288,338,3 | 38,22,36,23,16,22,34,58,15 | 182,216,318,214,400,227,2 | 15,67,46,67,87,15,16,14,51 | 229 | 149 | 0.0000862793744421 | 0.0184738988788681 | 0.857,0.891,0.922, | 0.888,0.677,0.818, | 0,054 |
| 248474 | ENSG00000110713 | NUP98 | chr11 | - | 3744508 | 3744690 | 3735190 | 3735324 | 3753315 | 3753743 | 248474 | 62,77,84,90,34,57,59,51,41 | 1,2,3,5,0,1,5,0,3,2,3,4 | 69,48,24,56,51,27,38,44,15 | 3,10,6,6,4,1,2,5,3,7,4,3,2,2 | 298 | 149 | 0.0000863478879536 | 0.0184738988788681 | 0.969,0.951,0.933, | 0.92,0.706,0.667,0 | 0,079 |
| 150633 | ENSG00000159459 | UBR1 | chr15 | - | 42976162 | 42976229 | 42970519 | 42970607 | 42976716 | 42976867 | 150633 | 2,2,1,1,0,0,1,2,0,0,0,0 | 5,5,25,9,22,6,12,13,7,2,4,7 | 2,6,1,3,3,3,0,4,1,2,0,0,4,1,0 | 11,15,3,3,10,6,12,19,14,11 | 216 | 149 | 0.0000865608086932 | 0.0185010619819723 | 0.216,0.216,0.027, | 0.111,0.216,0.187, | -0,074 |
| 217003 | ENSG00000135945 | REV1 | chr2 | - | 99424791 | 99424899 | 99424151 | 99424280 | 99429839 | 99429945 | 217003 | 1,1,5,2,1,3,2,2,1,2,0,0 | 15,7,15,6,10,11,12,11,11,3 | 9,5,3,4,2,0,1,3,10,8,10,13,4 | 15,22,5,14,16,12,9,22,6,21 | 257 | 149 | 0.0000866932692276 | 0.018510991100323 | 0.037,0.076,0.162, | 0.258,0.116,0.258, | -0,091 |
| 81982 | ENSG00000145782 | ATG12 | chr5 | - | 115840817 | 115840934 | 115837627 | 115837764 | 115841389 | 115841544 | 81982 | 5,4,2,6,0,1,0,0,0,0,0,0 | 61,77,56,37,14,14,10,18,27 | 4,1,0,0,1,2,1,2,6,8,1,8,7,6,1 | 32,39,37,34,30,20,13,27,15 | 266 | 149 | 0.0000871970129701 | 0.0185817133832962 | 0.044,0.028,0.02,0 | 0.065,0.014,0.0,0.0 | -0,036 |
| 5450 | ENSG00000064607 | SUGP2 | chr19 | - | 18998923 | 18998985 | 18995143 | 18995280 | 19001612 | 19001674 | 5450 | 9,5,11,4,13,9,10,8,4,2,6,5 | 30,32,73,54,39,35,21,35,37 | 11,6,1,4,4,1,4,2,5,3,0,11,6, | 36,40,16,15,28,15,32,41,41 | 211 | 149 | 0.0000871971533707 | 0.0185817133832962 | 0.175,0.099,0.096, | 0.177,0.096,0.042, | 0,06 |
| 68707 | ENSG00000237298 | TTN-AS1 | chr2 | + | 178537360 | 178537447 | 178535731 | 178535828 | 178537603 | 178537691 | 68707 | 1,2,1,1,2,1,3,1,0,0,4,3 | 10,9,13,15,7,16,12,34,8,9,7 | 10,7,2,5,4,0,1,5,0,3,1,3,5,5 | 24,13,9,14,5,9,10,8,6,12,10 | 236 | 149 | 0.0000873610548543 | 0.0185936271291454 | 0.059,0.123,0.046, | 0.208,0.254,0.123, | -0,065 |
| 236417 | ENSG00000120899 | PTK2B | chr8 | + | 27404222 | 27404399 | 27397547 | 27397788 | 27419894 | 27420073 | 236417 | 28,5,30,3,0,0,6,20,0,0,2,4 | 127,28,185,160,102,95,84, | 5,4,3,5,0,1,2,3,3,0,27,16,0, | 112,114,88,71,76,65,64,85 | 298 | 149 | 0.0000874258385395 | 0.0185936271291454 | 0.099,0.082,0.075, | 0.022,0.017,0.017, | 0,014 |
| 178871 | ENSG00000285708 | | chr3 | - | 71288314 | 71288422 | 71198201 | 71198392 | 71299819 | 71299880 | 178871 | 0,0,11,8,0,0,0,0,0,0,0,0 | 194,55,235,123,79,55,92,6 | 0,0,7,8,1,6,2,2,0,0,0,0,22,1 | 188,130,141,121,79,34,64, | 257 | 149 | 0.0000878690315856 | 0.0186667971566883 | 0.0,0.0,0.0,0.026,0.03 | 0.0,0.0,0.0,0.028,0.03 | -0,02 |
| 229298 | ENSG00000122566 | HNRNPA2B1 | chr7 | - | 26191421 | 26191667 | 26190992 | 26191128 | 26192494 | 26192577 | 229298 | 11,0,10,0,0,4,3,7,0,0,1,2 | 11,5,18,17,12,8,10,28,8,1,5 | 0,0,4,4,2,2,0,7,0,1,3,0,6,0,0 | 23,33,13,20,8,14,7,24,21,1 | 298 | 149 | 0.0000879433367725 | 0.0186667971566883 | 0.333,0.0,0.217,0.0 | 0.0,0.0,0.0,0.133,0.09 | 0,039 |
| 13721 | ENSG00000074755 | ZZEF1 | chr17 | - | 4109652 | 4109863 | 4105692 | 4105809 | 4112608 | 4112808 | 13721 | 41,15,99,70,47,58,43,51,14 | 3,1,5,4,0,4,0,3,2,0,0,0 | 70,61,34,55,29,31,68,46,38 | 1,9,1,9,6,2,10,3,8,1,3,2,3,2 | 298 | 149 | 0.0000890730653391 | 0.0188879654443348 | 0.872,0.882,0.908, | 0.972,0.772,0.944, | 0,069 |
| 179005 | ENSG00000091140 | DLD | chr7 | + | 107902328 | 107902393 | 107901737 | 107901817 | 107903477 | 107903547 | 179005 | 9,7,27,25,8,15,5,9,8,4,1,6 | 0,1,4,1,0,2,0,4,1,0,0,0 | 11,23,10,2,14,6,8,11,8,13,8 | 5,3,2,3,2,1,0,4,0,0,5,6,1,1,0 | 214 | 149 | 0.0000892102807245 | 0.0188984428450934 | 1.0,0.83,0.825,0.94 | 0.605,0.842,0.777, | 0,128 |
| 255022 | ENSG00000137955 | RABGGTB | chr1 | + | 75789997 | 75790057 | 75787522 | 75787604 | 75791284 | 75791337 | 255022 | 9,10,24,13,2,3,6,10,2,2,1,6 | 3,2,1,6,0,0,1,0,2,2,0,8 | 9,9,11,13,4,6,1,10,7,6,4,19 | 5,1,0,4,1,0,1,0,0,0,0,0,0,4,2 | 209 | 149 | 0.0000898151487618 | 0.0190078704849076 | 0.681,0.781,0.945, | 0.562,0.865,1.0,0.6 | -0,103 |
| 21358 | ENSG00000197776 | KLHDC1 | chr14 | + | 49709708 | 49709826 | 49709158 | 49709229 | 49710262 | 49710381 | 21358 | 15,15,40,30,6,7,11,13,8,2,1 | 6,2,1,1,0,1,0,5,0,0,2,2 | 36,29,10,4,12,10,7,17,9,16 | 1,1,3,1,2,0,0,0,1,0,0,0,0,2,1 | 267 | 149 | 0.0000899700438042 | 0.0190219474440292 | 0.582,0.807,0.957, | 0.953,0.942,0.65,0 | -0,03 |
| 217292 | ENSG00000152117 | SMPD4BP | chr2 | + | 131502601 | 131502692 | 131501357 | 131501508 | 131506311 | 131506919 | 217292 | 0,1,4,0,1,1,0,0,1,0,0,1 | 11,13,12,6,3,9,2,17,12,5,9, | 1,3,0,2,2,0,1,2,2,4,3,2,0,2,0 | 9,9,7,7,2,8,8,10,3,3,16,20,8 | 240 | 149 | 0.0000904125773356 | 0.0190799515742613 | 0.0,0.046,0.171,0.0 | 0.065,0.171,0.0,0.1 | -0,112 |
| 151841 | ENSG00000125107 | CNOT1 | chr16 | - | 58553781 | 58553853 | 58551588 | 58551819 | 58555250 | 58555537 | 151841 | 16,6,37,24,31,13,23,25,9,5 | 2,3,1,0,0,3,2,0,1,0,2,0 | 43,28,13,12,7,20,22,27,16, | 1,2,2,5,0,1,3,2,4,1,1,5,0,0,0 | 221 | 149 | 0.0000904216892815 | 0.0190799515742613 | 0.844,0.574,0.961, | 0.967,0.904,0.814, | 0,001 |
| 155970 | ENSG00000005810 | MYCBP2 | chr13 | - | 77185870 | 77186075 | 77185102 | 77185377 | 77188950 | 77189047 | 155970 | 38,36,119,78,50,70,45,81,2 | 1,0,0,0,0,0,2,1,0,1,0,0 | 79,82,25,63,39,34,44,47,41 | 3,6,2,8,1,0,0,0,2,1,4,1,1,1,0 | 298 | 149 | 0.0000909029443401 | 0.019162714606527 | 0.95,1.0,1.0,1.0,1.0 | 0.929,0.872,0.862, | 0,033 |
| 97789 | ENSG00000291151 | NSUN5P1 | chr7 | + | 75415591 | 75415778 | 75415119 | 75415234 | 75415980 | 75416030 | 97789 | 16,7,49,27,43,51,29,18,8,6 | 1,0,1,1,4,5,1,0,0,0,0,0 | 47,16,4,5,13,23,14,42,16,2 | 1,0,2,0,1,0,9,2,3,0,0,5,1,0,5 | 298 | 149 | 0.0000912060733345 | 0.019183278484404 | 0.889,1.0,0.961,0.9 | 0.959,1.0,0.5,1.0,0 | 0,086 |
| 218172 | ENSG00000090975 | PITPNM2 | chr12 | - | 122994800 | 122994931 | 122992498 | 122992669 | 122995388 | 122995660 | 218172 | 76,40,84,21,27,22,48,27,48 | 2,0,2,2,2,2,4,0,4,3,1,0 | 29,59,31,48,35,21,37,27,37 | 0,0,1,0,2,0,0,2,0,0,2,0,1,0,0 | 280 | 149 | 0.0000912486670317 | 0.019183278484404 | 0.953,1.0,0.957,0.8 | 1.0,1.0,0.943,1.0,0 | -0,063 |
| 15532 | ENSG00000172775 | PSME3IP1 | chr16 | - | 57178709 | 57178852 | 57173772 | 57173869 | 57185820 | 57186039 | 15532 | 0,0,4,1,0,5,6,0,0,0,0,10 | 12,3,19,12,3,4,9,4,6,7,5,7 | 1,1,7,0,3,2,1,4,5,4,5,4,7,8,6 | 10,3,7,2,0,3,0,2,2,2,3,11,5, | 292 | 149 | 0.0000912678804077 | 0.019183278484404 | 0.0,0.0,0.0,0.097,0.04 | 0.049,0.145,0.338, | -0,304 |
| 68543 | ENSG00000055609 | KMT2C | chr7 | - | 152157760 | 152157922 | 152156204 | 152156346 | 152158862 | 152159072 | 68543 | 22,9,35,31,21,13,10,21,13, | 4,0,6,0,4,3,2,3,1,0,1,2 | 14,16,13,9,17,14,14,12,8,3 | 0,0,1,0,1,0,1,2,0,1,5,3,1,2,0 | 298 | 149 | 0.0000921618120695 | 0.0193510743243491 | 0.733,1.0,0.745,1.0 | 1.0,1.0,0.867,1.0,0 | -0,089 |
| 184818 | ENSG00000142765 | SYTL1 | chr1 | + | 27349971 | 27350296 | 27349397 | 27349765 | 27350388 | 27350485 | 184818 | 11,8,11,3,7,7,7,5,5,4,8,4 | 0,0,0,4,4,1,0,2,0,0,0,0 | 4,9,10,2,2,3,5,4,7,7,6,12,7, | 3,1,0,0,1,0,0,2,5,2,2,4,3,0,0 | 298 | 149 | 0.00009224601594 | 0.0193510743243491 | 1.0,1.0,1.0,0.273,0 | 0.4,0.818,1.0,1.0,0 | 0,096 |
| 231477 | ENSG00000078589 | P2RY10 | chrX | + | 78952192 | 78952567 | 78947814 | 78947863 | 78960507 | 78963727 | 231477 | 85,56,108,85,42,35,54,63,4 | 74,53,104,77,56,70,70,52,6 | 124,136,78,28,37,29,26,10 | 110,66,67,19,44,19,28,60,2 | 298 | 149 | 0.0000928119259553 | 0.0194508311930722 | 0.365,0.346,0.342, | 0.36,0.507,0.368,0 | -0,092 |
| 163393 | ENSG00000182489 | XKRX | chrX | - | 100922888 | 100923061 | 100914298 | 100915083 | 100927969 | 100928917 | 163393 | 19,24,31,9,13,6,16,19,21,1 | 1,0,3,1,0,1,2,0,0,0,0,2 | 12,18,11,3,10,6,5,9,17,20,1 | 3,3,0,0,0,1,0,0,2,4,4,0,2,3,0 | 298 | 149 | 0.0000933935109547 | 0.0195536758329679 | 0.905,1.0,0.838,0.8 | 0.667,0.75,1.0,1.0, | 0,101 |
| 163394 | ENSG00000182489 | XKRX | chrX | - | 100922937 | 100923061 | 100914298 | 100915083 | 100927969 | 100928917 | 163394 | 17,24,28,9,12,6,12,14,18,1 | 1,0,3,1,0,1,2,0,0,0,0,2 | 12,16,7,1,9,6,4,8,15,20,12, | 3,3,0,0,0,1,0,0,2,4,4,0,2,3,0 | 273 | 149 | 0.0000938271588306 | 0.0196253772811165 | 0.903,1.0,0.836,0.8 | 0.686,0.744,1.0,1.0 | 0,095 |
| 99078 | ENSG00000136280 | CCM2 | chr7 | + | 45072725 | 45072783 | 45069825 | 45069961 | 45073459 | 45073519 | 99078 | 36,41,43,77,26,22,32,32,36 | 3,1,4,3,4,0,0,4,2,0,0,3 | 56,50,19,12,32,37,13,27,1 | 4,1,0,0,4,1,0,0,1,0,0,0,0,2,0 | 207 | 149 | 0.0000942982829489 | 0.0197047706188104 | 0.896,0.967,0.886,0.9 | 0.976,0.973,1.0,1.0 | -0,039 |
| 184003 | ENSG00000132549 | VPS13B | chr8 | + | 99274183 | 99274332 | 99192875 | 99193057 | 99275080 | 99275254 | 184003 | 92,47,111,66,39,109,58,84 | 0,0,1,0,0,5,2,0,0,0,0,0 | 86,116,53,99,78,62,72,71, | 2,5,2,4,5,0,0,2,0,0,8,1,3,1 | 298 | 149 | 0.0000947439555513 | 0.019778696699575 | 1.0,1.0,0.982,1.0,1 | 0.956,0.921,0.93,0 | 0,04 |
| 186048 | ENSG00000114423 | CBLB | chr3 | - | 105672336 | 105672429 | 105670232 | 105670352 | 105678430 | 105678571 | 186048 | 1,0,3,0,1,0,2,1,0,0,0,0 | 10,19,23,6,2,1,14,5,2,4,4,4 | 3,4,3,5,2,0,1,1,0,2,1,2,8,0 | 5,18,23,19,23,9,7,9,6,9,12, | 242 | 149 | 0.0000950576449615 | 0.0198249534716169 | 0.058,0.0,0.074,0.0 | 0.27,0.12,0.074,0. | -0,046 |
| 113453 | ENSG00000104325 | DECR1 | chr8 | + | 90017202 | 90017326 | 90001445 | 90001561 | 90018908 | 90018966 | 113453 | 60,42,86,75,35,28,34,31,39 | 2,0,8,8,0,5,1,0,0,0,0,2 | 39,42,41,22,23,25,9,26,16, | 8,4,4,0,4,1,0,1,5,11,14,27,9 | 173 | 149 | 0.0000956514475874 | 0.0199294837518719 | 0.942,1.0,0.854,0.8 | 0.727,0.851,0.848, | 0,117 |
| 215779 | ENSG00000049089 | COL9A2 | chr1 | - | 40312058 | 40312161 | 40311661 | 40311715 | 40312455 | 40312479 | 215779 | 48,26,135,101,91,127,93,8 | 3,0,4,7,3,4,1,1,2,0,0,3 | 28,32,28,18,18,10,43,84,1 | 3,0,0,0,0,2,1,2,0,0,0,0,0,0,0 | 252 | 149 | 0.0000958541651598 | 0.0199524060169332 | 0.904,1.0,0.952,0.8 | 0.847,1.0,1.0,1.0,1 | -0,015 |
| 95402 | ENSG00000111832 | RWDD1 | chr6 | + | 116572857 | 116572985 | 116571408 | 116571655 | 116580294 | 116580360 | 95402 | 4,0,7,6,2,4,9,1,0,0,0,1 | 34,17,20,29,15,14,18,13,55 | 2,9,9,18,9,2,1,4,3,0,3,12,2 | 17,18,16,15,15,8,5,27,4,18 | 277 | 149 | 0.0000961874869774 | 0.0200024434875679 | 0.06,0.0,0.158,0.1 | 0.06,0.212,0.232,0 | -0,082 |

| | | | | | | | | | | | | | | | | | | | | | | |
|---|---|---|---|---|---|---|---|---|---|---|---|---|---|---|---|---|---|---|---|---|---|---|
| 75904 | ENSG00000125637 | PSD4 | chr2 | + | 113192379 | 113192589 | 113185364 | 113186255 | 113193047 | 113193128 | 75904 | 164,148,350,273,222,176,1 | 9,2,11,18,3,8,15,7,1,4,8,9 | 209,123,91,85,63,91,66,14 | 23,18,10,7,16,7,10,13,8,8,9 | 298 | 149 | 0.000096977275398 | 0.0201472161787519 | 0.901,0.974,0.941, | 0.82,0.774,0.82,0.8 | 0,061 |
| 206629 | ENSG00000152061 | RABGAP1L | chr1 | + | 174231146 | 174231355 | 174220971 | 174221164 | 174241482 | 174241657 | 206629 | 58,50,476,318,88,135,112, | 3,1,7,4,0,2,3,5,1,3,0,0 | 211,264,169,118,156,79,34 | 4,7,15,10,4,1,5,3,5,4,9,0,1, | 298 | 149 | 0.0000980761389586 | 0.0203558586925732 | 0.906,0.962,0.971, | 0.963,0.95,0.849,0 | 0,027 |
| 147316 | ENSG00000103222 | ABCC1 | chr16 | + | 16033288 | 16033481 | 16033108 | 16033170 | 16036471 | 16036603 | 147316 | 5,2,4,2,0,0,1,0,1,1,2,7 | 32,26,31,24,23,15,14,11,4, | 13,11,2,4,1,4,2,4,0,0,23,9,2 | 37,38,12,21,19,23,15,13,13 | 298 | 149 | 0.0000982306619551 | 0.0203595454243928 | 0.072,0.037,0.061, | 0.149,0.126,0.077, | -0,054 |
| 46864 | ENSG00000131876 | SNRPA1 | chr15 | - | 101293024 | 101293172 | 101285725 | 101287010 | 101295096 | 101295204 | 46864 | 22,25,28,7,16,19,13,14,8,1 | 0,0,1,0,0,1,2,0,0,2,0,0 | 13,42,17,11,12,3,11,9,5,3,1 | 6,5,2,3,0,0,2,0,0,2,1,0,0,0,4 | 297 | 149 | 0.0000982830897777 | 0.0203595454243928 | 1.0,1.0,0.934,1.0,1 | 0.521,0.808,0.81,0 | 0,102 |
| 158699 | ENSG00000126777 | KTN1 | chr14 | + | 55637197 | 55637358 | 55636448 | 55636536 | 55637778 | 55637847 | 158699 | 147,81,125,66,63,49,36,62 | 5,2,7,3,0,1,0,4,1,0,0,1 | 75,76,97,36,38,42,28,80,16 | 8,7,0,5,4,3,1,7,1,5,3,4,1,5,1 | 298 | 149 | 0.0000987897885973 | 0.0204448317207553 | 0.936,0.953,0.899, | 0.824,0.844,1.0,0.7 | 0,073 |
| 97112 | ENSG00000268750 | | chr19 | + | 57839022 | 57839149 | 57820493 | 57820549 | 57856103 | 57856233 | 97112 | 10,14,15,19,12,3,10,17,8,3 | 1,0,2,1,1,3,0,0,0,0,1,0 | 11,17,4,11,8,7,8,8,6,8,14,2 | 1,2,1,2,1,0,0,2,0,4,3,0,2,0,5 | 276 | 149 | 0.0000989401724063 | 0.020456284579424 | 0.844,1.0,0.802,0.9 | 0.856,0.821,0.683, | 0,082 |
| 76389 | ENSG00000164144 | ARFIP1 | chr4 | + | 152870752 | 152870848 | 152863605 | 152863714 | 152872451 | 152872564 | 76389 | 1,0,3,3,3,1,0,4,1,2,1,6 | 7,6,22,26,13,11,7,7,15,1,2, | 3,0,1,0,0,0,0,0,1,0,2,0,0,3,0 | 12,6,10,2,3,4,6,12,3,10,10, | 245 | 149 | 0.000100575647039 | 0.0207744693741372 | 0.08,0.0,0.077,0.0 | 0.132,0.0,0.057,0.0 | 0,086 |
| 146453 | ENSG00000237440 | ZNF737 | chr19 | - | 20554289 | 20554376 | 20553708 | 20553835 | 20565637 | 20565809 | 146453 | 13,5,3,5,3,4,4,0,34,10,2,0 | 51,32,168,147,46,44,41,64 | 12,10,17,6,0,3,3,14,8,19,0, | 98,44,42,36,11,32,21,41,9, | 236 | 149 | 0.000100831896394 | 0.0207963319886598 | 0.139,0.09,0.011,0 | 0.072,0.125,0.204, | -0,063 |
| 95640 | ENSG00000173692 | PSMD1 | chr2 | + | 231079535 | 231079614 | 231078658 | 231078747 | 231080140 | 231080314 | 95640 | 35,53,74,76,25,58,23,59,42 | 3,1,2,6,3,2,3,0,0,0,1,1 | 65,52,44,35,45,29,32,37,28 | 2,12,3,15,9,0,4,2,8,1,3,2,0, | 228 | 149 | 0.000100891144965 | 0.0207963319886598 | 0.884,0.972,0.96,0 | 0.955,0.739,0.906, | 0,054 |
| 19261 | ENSG00000114062 | UBE3A | chr15 | - | 25409087 | 25409207 | 25405460 | 25405502 | 25438488 | 25438999 | 19261 | 30,30,93,26,21,27,20,58,13 | 0,1,4,1,1,0,1,5,0,0,0,1 | 36,71,23,22,39,14,28,25,20 | 2,1,1,0,1,2,2,3,0,1,3,4,7,5,2 | 269 | 149 | 0.000100971360669 | 0.0207963319886598 | 1.0,0.943,0.928,0.9 | 0.909,0.975,0.927, | 0,077 |
| 31604 | ENSG00000112992 | NNT | chr5 | + | 43647923 | 43648182 | 43645356 | 43645510 | 43649146 | 43649308 | 31604 | 5,3,5,9,5,6,20,23,4,3,4,5 | 32,16,75,43,31,38,10,21,34 | 4,3,6,3,7,0,3,6,0,1,3,3,2,1,4 | 17,28,20,10,21,29,13,14,18 | 298 | 149 | 0.000101373336293 | 0.0208591630436699 | 0.072,0.086,0.032, | 0.105,0.051,0.13,0 | 0,067 |
| 145019 | ENSG00000108984 | MAP2K6 | chr17 | + | 69534383 | 69534516 | 69526569 | 69526709 | 69536114 | 69536160 | 145019 | 4,8,11,10,1,7,0,3,4,0,0,0 | 32,21,137,91,11,21,9,36,22 | 3,1,0,0,0,0,2,5,0,2,2,0,3,3,1 | 14,8,16,2,8,6,8,19,9,26,16, | 282 | 149 | 0.000101511520591 | 0.0208676466937168 | 0.062,0.168,0.041, | 0.102,0.062,0.0,0.0 | 0,015 |
| 33806 | ENSG00000087266 | SH3BP2 | chr4 | + | 2823394 | 2823573 | 2822934 | 2823037 | 2824612 | 2824730 | 33806 | 7,1,6,4,6,0,5,1,0,0,3,0 | 82,22,50,35,42,29,26,19,9, | 0,0,2,0,0,0,3,3,0,2,0,1,1,0,3 | 45,37,29,9,32,19,18,41,11, | 298 | 149 | 0.000102298110664 | 0.021009279252217 | 0.041,0.022,0.057, | 0.0,0.0,0.033,0.0,0 | 0,022 |
| 185923 | ENSG00000118260 | CREB1 | chr2 | + | 207561118 | 207561208 | 207560225 | 207560372 | 207567462 | 207568150 | 185923 | 27,13,22,11,8,11,3,7,25,12 | 57,36,97,46,37,38,15,32,17 | 5,9,10,6,6,3,1,7,3,11,11,7,8 | 57,60,25,20,34,23,13,22,34 | 239 | 149 | 0.000103123481278 | 0.0211585986643901 | 0.228,0.184,0.124, | 0.052,0.086,0.2,0.1 | 0,088 |
| 111470 | ENSG00000035928 | RFC1 | chr4 | - | 39306591 | 39306701 | 39304813 | 39304928 | 39308635 | 39309032 | 111470 | 42,29,85,61,28,19,19,32,36 | 0,2,8,1,2,0,2,1,0,0,0,1 | 48,47,40,12,15,32,9,24,21, | 2,1,3,4,3,6,1,2,3,2,6,0,2,3,2 | 259 | 149 | 0.000103235551363 | 0.0211614199575332 | 1.0,0.893,0.859,0.9 | 0.932,0.964,0.885, | 0,095 |
| 221540 | ENSG00000231925 | TAPBP | chr6 | - | 33303953 | 33304217 | 33301684 | 33301771 | 33304296 | 33304530 | 221540 | 28,37,114,64,43,60,74,35,3 | 0,0,8,3,0,0,0,2,0,0,0,2 | 65,97,29,47,32,31,41,80,39 | 0,2,0,1,4,4,2,5,0,1,0,14,3,0 | 298 | 149 | 0.000103503904754 | 0.0211962406509117 | 1.0,1.0,0.877,0.914 | 1.0,0.96,1.0,0.959, | 0,039 |
| 129618 | ENSG00000127314 | RAP1B | chr12 | + | 68624670 | 68624873 | 68610907 | 68611043 | 68648698 | 68648781 | 129618 | 0,0,2,7,0,0,0,2,2,2,1,1 | 16,14,21,11,10,12,9,7,17,1 | 2,2,3,0,1,0,1,3,3,0,3,8,4,10 | 10,21,10,9,14,8,9,2,3,14,17 | 298 | 149 | 0.000103610985947 | 0.0211980001106072 | 0.0,0.0,0.045,0.24 | 0.091,0.045,0.13,0 | -0,071 |
| 71565 | ENSG00000118961 | LDAH | chr2 | - | 20801309 | 20801465 | 20790254 | 20790398 | 20823036 | 20823101 | 71565 | 44,23,84,26,33,19,35,27,27 | 3,0,4,6,4,2,4,2,3,2,0,2 | 80,29,16,5,14,17,11,39,15, | 2,0,1,0,1,2,0,3,1,1,1,2,0,3,1 | 298 | 149 | 0.000103963594126 | 0.0212499414314654 | 0.88,1.0,0.913,0.68 | 0.952,1.0,0.889,1.0 | -0,066 |
| 142379 | ENSG00000168961 | LGALS9 | chr17 | + | 27647357 | 27647432 | 27647029 | 27647118 | 27648835 | 27649541 | 142379 | 51,29,77,83,29,18,29,15,31 | 11,9,15,22,13,15,11,14,17, | 32,49,24,26,23,29,15,39,34 | 8,10,5,1,3,9,2,3,4,8,0,23,13 | 224 | 149 | 0.000104373461361 | 0.0213018907884223 | 0.755,0.682,0.773, | 0.727,0.765,0.761, | -0,163 |
| 35931 | ENSG00000164190 | NIPBL | chr5 | + | 36962122 | 36962274 | 36961483 | 36961583 | 36970875 | 36971036 | 35931 | 91,83,101,74,70,85,48,136 | 1,0,2,0,0,4,0,0,0,0,1,0 | 127,147,83,173,68,82,83,1 | 0,5,2,9,1,2,1,0,2,1,6,0,3,2,2 | 298 | 149 | 0.000104415696539 | 0.0213018907884223 | 0.978,1.0,0.962,1.0 | 1.0,0.936,0.954,0.9 | 0,031 |
| 155971 | ENSG00000005810 | MYCBP2 | chr13 | - | 77185927 | 77186063 | 77185102 | 77185377 | 77188950 | 77189047 | 155971 | 45,40,106,72,29,43,26,52,4 | 1,0,0,0,0,0,2,1,0,1,0,0 | 73,78,47,55,21,26,31,29,43 | 3,6,2,8,1,0,0,0,2,1,4,1,1,1,0 | 285 | 149 | 0.000104956621726 | 0.0213919684192317 | 0.959,1.0,1.0,1.0,1 | 0.927,0.872,0.925, | 0,035 |
| 88452 | ENSG00000164307 | ERAP1 | chr5 | - | 96788530 | 96788685 | 96786469 | 96786549 | 96790295 | 96790367 | 88452 | 74,78,117,97,63,62,60,88,7 | 2,0,3,4,4,1,1,3,0,0,1,0 | 99,78,40,50,56,40,38,85,47 | 1,2,4,3,2,2,0,2,5,0,11,4,1,3 | 298 | 149 | 0.000105056085398 | 0.021391983269912 | 0.949,1.0,0.951,0.9 | 0.98,0.951,0.833,0 | 0,047 |
| 232789 | ENSG00000104921 | FCER2 | chr19 | - | 7698789 | 7698876 | 7698355 | 7698409 | 7699738 | 7700084 | 232789 | 7,0,11,9,0,0,5,0,1,1,1,4 | 22,13,17,16,8,8,5,5,12,8,0, | 1,3,0,0,2,1,1,1,0,3,0,0,1,1,0 | 10,20,4,8,8,3,1,1,9,1,1,7,1, | 236 | 149 | 0.000105375731341 | 0.0214367902006189 | 0.167,0.0,0.29,0.2 | 0.059,0.087,0.0,0.0 | 0,066 |
| 99564 | ENSG00000106404 | CLDN15 | chr7 | - | 101236746 | 101236886 | 101233768 | 101234442 | 101237364 | 101237750 | 99564 | 2,1,4,2,0,3,1,2,4,3,1,0 | 4,0,10,9,8,5,3,6,1,1,5,1 | 0,3,0,0,1,0,0,2,0,1,0,3,2,1,0 | 1,5,2,2,3,5,4,6,6,9,6,8,3,3,2 | 289 | 149 | 0.000105814919923 | 0.0215058083380049 | 0.205,1.0,0.171,0.1 | 0.0,0.236,0.0,0.0,0 | 0,21 |
| 165547 | ENSG00000109684 | CLNK | chr4 | - | 10590352 | 10590544 | 10584926 | 10584955 | 10597977 | 10598049 | 165547 | 11,4,27,19,6,6,1,12,1,7,2,7 | 132,50,199,85,47,71,95,15 | 2,1,1,11,5,5,1,2,2,2,12,0,3, | 53,62,79,60,73,54,60,68,71 | 298 | 149 | 0.000106293211825 | 0.0215826361078364 | 0.04,0.038,0.064,0 | 0.019,0.008,0.006, | 0,022 |
| 46865 | ENSG00000131876 | SNRPA1 | chr15 | - | 101293024 | 101293172 | 101287655 | 101287702 | 101295096 | 101295204 | 46865 | 22,25,28,6,16,19,13,14,8,1 | 0,2,0,1,0,2,0,3,0,0,0,5 | 13,43,17,8,12,3,11,9,5,3,14 | 3,6,0,0,13,4,1,0,0,2,1,2,0,3 | 297 | 149 | 0.000106859772142 | 0.0216772248990526 | 1.0,0.862,1.0,0.75 | 0.685,0.782,1.0,1.0 | 0,124 |
| 5637 | ENSG00000133243 | BTBD2 | chr19 | - | 2013546 | 2013880 | 1997343 | 1997463 | 2015296 | 2015542 | 5637 | 1,2,13,8,0,2,4,3,2,4,1,2 | 32,28,22,15,19,12,17,5,48, | 2,0,0,3,2,0,2,1,0,4,3,2,2,0,0 | 6,6,9,22,7,6,11,18,4,17,12, | 298 | 149 | 0.000107603743992 | 0.0218075907939192 | 0.015,0.034,0.228, | 0.143,0.0,0.0,0.064 | 0,048 |
| 135915 | ENSG00000081665 | ZNF506 | chr19 | - | 19810662 | 19810693 | 19806941 | 19807068 | 19821600 | 19821751 | 135915 | 22,14,18,5,5,4,2,9,7,2,2,2 | 66,63,226,193,43,50,31,66 | 9,28,14,22,12,1,6,7,9,8,16, | 63,88,46,29,64,30,27,29,28 | 180 | 149 | 0.000107769937296 | 0.0218207256577191 | 0.216,0.155,0.062, | 0.106,0.208,0.201, | -0,061 |
| 213425 | ENSG00000035720 | STAP1 | chr4 | + | 67576822 | 67576943 | 67575384 | 67575498 | 67577202 | 67577259 | 213425 | 0,2,25,4,0,5,0,0,1,0,3,5 | 95,43,180,138,78,65,66,55 | 13,11,9,4,0,0,5,6,0,0,0,0,25 | 83,59,97,34,67,53,15,55,47 | 270 | 149 | 0.00010792530171 | 0.0218216763388431 | 0.0,0.025,0.071,0.0 | 0.08,0.093,0.049,0 | -0,021 |
| 82806 | ENSG00000176978 | DPP7 | chr9 | - | 137113360 | 137113496 | 137112952 | 137113119 | 137113864 | 137114028 | 82806 | 9,8,31,10,6,24,18,8,2,8,5,2 | 2,5,0,0,0,0,2,0,0,0,2,0 | 19,27,16,14,14,5,9,20,6,9,2 | 0,0,5,10,0,0,2,4,0,0,4,4,0,0 | 285 | 149 | 0.000107977407069 | 0.0218216763388431 | 0.702,0.455,1.0,1.0 | 1.0,1.0,0.626,0.42 | -0,002 |
| 239430 | ENSG00000114209 | PDCD10 | chr3 | - | 167698013 | 167698177 | 167697039 | 167697126 | 167704841 | 167704895 | 239430 | 0,0,3,2,2,2,53,47,0,1,2,0 | 29,24,61,69,41,20,5,11,50, | 2,1,1,2,0,0,1,2,4,0,2,0,0,0,1 | 38,32,16,4,21,14,20,13,19, | 298 | 149 | 0.000108387808632 | 0.021884067954666 | 0.0,0.0,0.024,0.01 | 0.026,0.015,0.03,0 | 0,11 |
| 104799 | ENSG00000164062 | APEH | chr3 | + | 49676055 | 49676219 | 49675890 | 49675966 | 49676608 | 49676700 | 104799 | 64,27,74,54,34,35,33,28,40 | 3,0,2,2,3,4,1,2,3,6,2,4 | 36,28,33,13,21,22,16,20,15 | 3,1,0,0,3,0,1,0,0,0,2,2,3,1,2 | 298 | 149 | 0.000108553658684 | 0.0218970126637451 | 0.914,1.0,0.949,0.9 | 0.857,0.933,1.0,1.0 | -0,073 |
| 124741 | ENSG00000119777 | TMEM214 | chr2 | + | 27037814 | 27037892 | 27037560 | 27037702 | 27038145 | 27038237 | 124741 | 3,2,2,8,1,1,0,3,1,0,0,1 | 2,4,15,9,17,7,15,10,6,3,4,7 | 0,0,0,0,0,3,0,0,1,1,0,0,1,2,0 | 11,24,8,4,12,8,23,16,4,8,7, | 227 | 149 | 0.000108732261322 | 0.021912503124153 | 0.496,0.247,0.08,0 | 0.0,0.0,0.0,0.0,0.0, | 0,094 |
| 42655 | ENSG00000163349 | HIPK1 | chr1 | + | 113952765 | 113952889 | 113940381 | 113941459 | 113955562 | 113955649 | 42655 | 16,9,51,42,22,16,8,20,20,9 | 0,2,4,5,0,0,0,1,0,0,2,0 | 35,49,25,43,27,25,20,37,14 | 3,8,2,10,2,3,2,3,4,0,0,5,0,2 | 273 | 149 | 0.00010908963223 | 0.021951629885997 | 1.0,0.711,0.874,0.8 | 0.864,0.77,0.872,0 | 0,017 |
| 240994 | ENSG00000106829 | TLE4 | chr9 | + | 79719647 | 79719734 | 79718721 | 79718971 | 79720045 | 79720293 | 240994 | 6,1,13,3,0,2,1,8,0,4,3,3 | 44,31,99,82,67,68,57,90,61 | 2,10,9,17,3,2,3,4,16,4,10,2 | 41,68,70,34,51,25,38,49,32 | 236 | 149 | 0.000109130394683 | 0.021951629885997 | 0.079,0.02,0.077,0 | 0.03,0.085,0.075,0 | -0,037 |
| 105491 | ENSG00000147586 | MRPS28 | chr8 | - | 80028694 | 80028796 | 80002998 | 80003180 | 80030035 | 80030289 | 105491 | 3,3,13,7,3,5,3,2,2,0,1,5 | 13,5,23,6,4,17,12,27,11,6,2 | 5,2,2,2,0,0,0,2,0,1,5,2,0,1,1 | 15,15,3,11,15,11,8,6,3,2,13 | 251 | 149 | 0.000109565087222 | 0.0220184904648723 | 0.12,0.263,0.251,0 | 0.165,0.073,0.284, | 0,082 |
| 84696 | ENSG00000059588 | TARBP1 | chr1 | - | 234398381 | 234398553 | 234393645 | 234393837 | 234401180 | 234401216 | 84696 | 34,18,73,40,44,43,20,29,4, | 0,0,2,1,0,0,1,0,0,0,1,4 | 29,39,33,17,19,8,10,38,31, | 1,4,2,0,3,0,0,3,4,3,2,3,0,1,1 | 298 | 149 | 0.000110134349164 | 0.0221107462830773 | 1.0,1.0,0.948,0.95 | 0.935,0.83,0.892,1 | 0,039 |
| 98206 | ENSG00000164808 | SPIDR | chr8 | + | 47293866 | 47294030 | 47291032 | 47291137 | 47396375 | 47396626 | 98206 | 142,44,113,93,65,61,38,75 | 12,4,10,6,0,4,1,2,3,4,2,4 | 90,82,51,39,53,26,17,48,52 | 0,1,0,0,8,2,0,1,6,0,2,3,3,0,2 | 298 | 149 | 0.000110229617303 | 0.0221107462830773 | 0.855,0.846,0.85,0 | 1.0,0.976,1.0,1.0,0 | -0,048 |
| 19940 | ENSG00000077238 | IL4R | chr16 | + | 27346466 | 27346618 | 27344868 | 27345020 | 27352539 | 27352696 | 19940 | 1485,449,1739,1057,959,6 | 40,17,48,36,11,21,13,26,35 | 853,442,1128,229,194,288 | 10,7,15,3,14,6,1,4,1,7,8,7,1 | 298 | 149 | 0.000110373823566 | 0.0221190581191189 | 0.949,0.93,0.948,0 | 0.977,0.969,0.974, | -0,006 |
| 26843 | ENSG00000143252 | SDHC | chr1 | + | 161356676 | 161356840 | 161340593 | 161340655 | 161362328 | 161362433 | 26843 | 25,16,70,70,45,44,26,36,15 | 3,2,7,6,1,4,5,1,0,1,0,7 | 47,42,24,13,20,28,28,29,15 | 0,7,0,0,1,1,0,0,2,3,0,4,0,2,0 | 298 | 149 | 0.000110633590212 | 0.0221504913999246 | 0.806,0.8,0.833,0.8 | 1.0,0.75,1.0,1.0,0.9 | -0,062 |
| 148239 | ENSG00000140740 | UQCRC2 | chr16 | + | 21967121 | 21967230 | 21965407 | 21965505 | 21968627 | 21968685 | 148239 | 4,2,9,1,0,5,4,5,0,0,0,0 | 90,41,183,150,66,51,32,55 | 4,12,2,17,4,0,2,10,1,0,5,3,4 | 86,80,70,23,58,60,35,52,49 | 258 | 149 | 0.000110910462863 | 0.0221852879483888 | 0.025,0.027,0.028, | 0.026,0.08,0.016,0 | -0,03 |
| 49217 | ENSG00000126012 | KDM5C | chrX | - | 53218275 | 53218398 | 53217795 | 53217966 | 53220838 | 53220916 | 49217 | 116,67,265,140,71,91,73,7 | 3,0,10,11,0,0,1,2,0,0,1,0 | 115,141,73,137,130,90,57, | 5,2,9,21,8,2,0,1,12,12,1,2,3 | 272 | 149 | 0.00011167426389 | 0.0223173291469903 | 0.955,1.0,0.936,0.8 | 0.926,0.975,0.816, | 0,045 |
| 133550 | ENSG00000183527 | PSMG1 | chr21 | - | 39179923 | 39179971 | 39178448 | 39178647 | 39180284 | 39180436 | 133550 | 13,5,2,14,2,6,4,1,0,2,1,2 | 1,0,0,1,4,0,1,0,3,0,2,0 | 3,1,2,2,3,1,1,0,2,2,2,7,2,2,3 | 0,3,1,0,1,0,0,1,3,2,1,4,0,4,3 | 197 | 149 | 0.000112097724017 | 0.0223811736900769 | 0.908,1.0,1.0,0.91 | 1.0,0.201,0.602,1.0 | 0,176 |
| 200341 | ENSG00000132849 | PATJ | chr1 | + | 62100318 | 62100360 | 62084514 | 62084648 | 62108436 | 62108520 | 200341 | 4,0,7,1,2,2,3,11,3,2,1,2 | 9,5,13,11,8,17,2,10,6,4,2,8 | 3,1,0,0,1,1,1,2,1,2,0,0,1,2,2 | 11,9,7,7,9,7,0,9,1,7,1,9,5,5 | 191 | 149 | 0.000112496980297 | 0.0224400718872137 | 0.257,0.0,0.296,0.0 | 0.175,0.08,0.0,0.0, | 0,055 |
| 200293 | ENSG00000176444 | CLK2 | chr1 | - | 155266728 | 155266907 | 155264644 | 155264774 | 155268009 | 155268126 | 200293 | 4,1,9,3,2,1,5,4,0,0,2,2 | 0,1,2,0,0,1,0,4,1,1,1,0 | 8,8,0,8,6,5,6,4,4,1,7,0,4,1,3 | 3,6,2,0,0,1,5,1,2,3,4,5,2,0,2 | 298 | 149 | 0.000112883717675 | 0.022496366147137 | 1.0,0.333,0.692,1.0 | 0.571,0.4,0.0,1.0,1 | 0,083 |
| 224129 | ENSG00000127980 | PEX1 | chr7 | - | 92504705 | 92504902 | 92503040 | 92503195 | 92506247 | 92506344 | 224129 | 39,15,37,39,25,15,20,31,20 | 0,0,6,0,0,4,2,0,2,0,1,0 | 33,35,24,13,24,8,16,13,17, | 3,4,4,0,1,0,0,3,1,4,4,1,1,0,0 | 298 | 149 | 0.000113233407738 | 0.0225451800007932 | 1.0,1.0,0.755,1.0,1 | 0.846,0.814,0.75,1 | 0,046 |
| 90007 | ENSG00000154473 | BUB3 | chr10 | + | 123160406 | 123160565 | 123157728 | 123157880 | 123162235 | 123162413 | 90007 | 279,205,366,292,174,166,1 | 3,1,1,5,1,6,12,3,2,0,2,3 | 259,257,195,119,280,148,1 | 9,11,5,3,8,6,2,4,7,10,8,2,9, | 298 | 149 | 0.000113641730922 | 0.0226055669021007 | 0.979,0.99,0.995,0 | 0.935,0.921,0.951, | 0,024 |
| 7474 | ENSG00000089057 | SLC23A2 | chr20 | - | 4970792 | 4970915 | 4932454 | 4932716 | 5001405 | 5001510 | 7474 | 111,84,120,86,86,114,93,7 | 0,0,0,3,0,2,0,0,0,0,2,0 | 61,52,47,74,38,27,47,50,46 | 13,13,0,0,0,0,0,2,0,0,2,3,0, | 272 | 149 | 0.000114189165374 | 0.0226934886912386 | 1.0,1.0,1.0,0.94,1.0 | 0.72,0.687,1.0,1.0, | 0,031 |
| 152020 | ENSG00000141956 | PRDM15 | chr21 | - | 41823289 | 41823449 | 41821902 | 41822037 | 41825959 | 41826054 | 152020 | 3,4,4,13,8,3,10,9,2,5,1,6 | 2,0,1,6,3,0,1,1,2,4,0,0 | 5,7,9,8,19,11,2,6,3,16,9,9,5 | 0,1,1,1,1,1,0,0,3,1,1,0,0,2,5 | 298 | 149 | 0.000114436625863 | 0.0227216876578592 | 0.429,1.0,0.667,0.5 | 1.0,0.778,0.818,0.8 | -0,091 |
| 199113 | ENSG00000072501 | SMC1A | chrX | - | 53407426 | 53407474 | 53405770 | 53405956 | 53409061 | 53409269 | 199113 | 0,1,2,1,0,2,1,0,1,0,2,0 | 19,16,35,42,24,28,23,19,27 | 5,3,0,11,1,1,5,1,1,0,2,2,0,1 | 27,22,26,28,16,19,8,17,21, | 197 | 149 | 0.000114736840536 | 0.0227569811385234 | 0.0,0.045,0.041,0.0 | 0.123,0.093,0.0,0.1 | -0,032 |
| 83087 | ENSG00000022840 | RNF10 | chr12 | + | 120539367 | 120539468 | 120534587 | 120534968 | 120546404 | 120546601 | 83087 | 9,2,4,3,1,4,0,1,1,2,0,0 | 31,29,47,33,24,24,20,21,7, | 3,8,2,1,8,4,4,4,0,3,10,5,13, | 18,32,10,21,45,24,12,28,24 | 250 | 149 | 0.000114825845331 | 0.0227569811385234 | 0.148,0.039,0.048, | 0.09,0.13,0.107,0.0 | -0,041 |
| 187092 | ENSG00000163946 | TASOR | chr3 | - | 56623643 | 56623843 | 56622861 | 56623566 | 56624478 | 56624643 | 187092 | 97,66,155,100,93,66,71,92 | 2,0,13,17,5,1,0,0,1,0,0,1 | 105,112,53,58,36,88,42,88 | 5,2,3,0,0,5,2,0,0,3,0,3,0,3,1 | 298 | 149 | 0.00011586280592 | 0.0229413685197493 | 0.96,1.0,0.856,0.7 | 0.913,0.966,0.898, | -0,016 |
| 224611 | ENSG00000223705 | NSUN5P1 | chr7 | + | 75415591 | 75415903 | 75415119 | 75415234 | 75415980 | 75416191 | 224611 | 16,7,50,25,41,48,27,17,5,5 | 1,0,1,1,4,5,1,0,0,0,0,0 | 44,17,3,5,11,25,13,40,16,2 | 1,0,2,0,1,0,9,2,3,0,0,5,1,0,5 | 298 | 149 | 0.0001161922542 | 0.0229854550218016 | 0.889,1.0,0.962,0.9 | 0.957,1.0,0.429,1.0 | 0,092 |
| 9633 | ENSG00000091592 | NLRP1 | chr17 | - | 5533896 | 5533988 | 5532821 | 5532984 | 5536850 | 5536940 | 9633 | 134,41,82,69,54,54,54,59,6 | 2,0,14,7,7,2,9,3,4,3,2,11 | 35,59,35,54,41,12,24,50,51 | 3,4,6,16,3,9,4,15,10,8,9,7,8 | 241 | 149 | 0.000117211366146 | 0.0231657663424883 | 0.976,1.0,0.784,0.8 | 0.878,0.901,0.783, | 0,117 |
| 111472 | ENSG00000035928 | RFC1 | chr4 | - | 39306591 | 39306704 | 39304813 | 39304928 | 39308635 | 39309032 | 111472 | 20,18,60,43,16,18,9,20,24, | 0,2,8,1,2,0,2,1,0,0,0,1 | 27,23,28,5,7,16,7,13,14,27 | 2,1,3,4,3,6,1,2,3,2,6,0,2,3,2 | 262 | 149 | 0.000118052383458 | 0.0233083120133048 | 1.0,0.837,0.81,0.9 | 0.885,0.929,0.841, | 0,134 |
| 163037 | ENSG00000182903 | ZNF721 | chr4 | - | 467417 | 467681 | 444342 | 444432 | 472574 | 472701 | 163037 | 1,0,3,7,0,4,1,3,0,0,0,0 | 82,45,140,64,22,32,24,41,4 | 0,12,4,0,5,1,2,6,0,1,4,7,3,1 | 50,94,57,67,46,33,28,31,29 | 298 | 149 | 0.000118149190435 | 0.0233083120133048 | 0.006,0.0,0.011,0.0 | 0.0,0.06,0.034,0.0, | -0,027 |
| 227625 | ENSG00000186088 | GSAP | chr7 | - | 77321328 | 77321403 | 77320724 | 77320819 | 77323646 | 77323742 | 227625 | 69,34,70,47,24,38,28,38,18 | 12,2,2,0,10,6,0,0,0,0,0,0 | 50,48,47,31,36,20,37,41,27 | 10,4,0,0,0,0,0,0,8,0,0,0,0,0 | 224 | 149 | 0.000118415024814 | 0.0233225284188127 | 0.793,0.919,0.959, | 0.769,0.889,1.0,1.0 | -0,035 |
| 44936 | ENSG00000164073 | MFSD8 | chr4 | - | 127921859 | 127921982 | 127921523 | 127921771 | 127930682 | 127930817 | 44936 | 2,0,7,4,1,0,0,1,0,1,0,1 | 1,7,7,10,9,5,7,10,2,0,4,5 | 9,2,7,1,1,4,5,3,0,4,0,4,6,4,0 | 15,3,1,2,3,1,5,10,4,4,10,8,7 | 272 | 149 | 0.000118437973906 | 0.0233225284188127 | 0.523,0.0,0.354,0.1 | 0.247,0.268,0.793, | -0,065 |
| 115241 | ENSG00000136828 | RALGPS1 | chr9 | + | 127199014 | 127199066 | 127196473 | 127196631 | 127210694 | 127210778 | 115241 | 13,6,4,11,5,14,16,8,12,5,3, | 2,1,3,0,0,0,0,0,0,1,0,2 | 7,3,7,8,5,1,4,5,1,3,6,3,12,9 | 1,2,0,7,1,0,3,0,0,2,0,0,2,3,2 | 201 | 149 | 0.000118848592403 | 0.0233721364616871 | 0.828,0.816,0.497, | 0.838,0.527,1.0,0.4 | 0,087 |
| 59679 | ENSG00000109323 | MANBA | chr4 | - | 102714437 | 102714561 | 102690879 | 102690957 | 102722870 | 102722925 | 59679 | 35,32,48,23,20,31,26,40,17 | 1,3,2,1,0,5,0,2,0,0,2,0 | 48,41,14,14,8,16,12,38,13, | 4,9,1,3,4,0,3,0,4,0,0,8,6,1,3 | 273 | 149 | 0.00011890707856 | 0.0233721364616871 | 0.95,0.853,0.929,0 | 0.868,0.713,0.884, | 0,093 |
| 878 | ENSG00000196588 | MRTFA | chr22 | - | 40423535 | 40423685 | 40420846 | 40421100 | 40424205 | 40424381 | 878 | 31,41,22,18,44,37,36,34,83 | 3,0,0,2,1,0,0,0,3,3,0,0 | 16,19,26,46,14,8,19,46,6,1 | 0,0,0,3,0,3,8,2,1,3,1,0,4,2,1 | 298 | 149 | 0.000120895890256 | 0.0237413716748988 | 0.838,1.0,1.0,0.81 | 1.0,1.0,1.0,0.885,1 | 0,105 |
| 113056 | ENSG00000104529 | EEF1D | chr8 | - | 143592646 | 143592942 | 143589829 | 143590081 | 143597347 | 143597415 | 113056 | 7,3,23,13,4,2,1,9,2,4,3,1 | 2,0,3,0,3,0,0,2,0,0,0,0 | 4,1,3,8,7,3,9,11,1,4,9,14,3, | 2,0,0,0,4,0,0,0,2,0,3,7,8,0,4 | 298 | 149 | 0.000121495614566 | 0.0238373952768047 | 0.636,1.0,0.793,1.0 | 0.5,1.0,1.0,1.0,0.4 | 0,133 |
| 201486 | ENSG00000115446 | UNC50 | chr2 | + | 98610160 | 98610634 | 98609755 | 98610039 | 98610774 | 98610895 | 201486 | 5,1,8,6,2,4,1,7,0,2,0,9 | 13,11,41,18,5,9,10,13,12,7 | 1,2,7,1,2,2,1,0,0,4,4,0,4,3,0 | 23,21,20,4,4,7,8,16,3,11,19 | 298 | 149 | 0.000121759806907 | 0.0238674726779604 | 0.161,0.043,0.089, | 0.021,0.045,0.149, | 0,069 |
| 37771 | ENSG00000227345 | PARG | chr10 | - | 49935075 | 49935142 | 49933176 | 49934163 | 49941508 | 49941824 | 37771 | 13,8,25,9,15,9,11,13,8,7,7, | 4,3,2,0,0,2,2,0,0,1,0,5 | 15,21,10,5,13,4,11,16,4,11 | 0,0,0,0,0,0,0,0,0,0,3,0,0,0 | 216 | 149 | 0.000122126663911 | 0.0239176014560768 | 0.692,0.648,0.896, | 1.0,1.0,1.0,1.0,1.0, | -0,129 |
| 51101 | ENSG00000114439 | BBX | chr3 | + | 107772627 | 107773636 | 107755597 | 107755678 | 107774718 | 107774857 | 51101 | 137,132,143,64,37,61,30,5 | 19,1,13,8,2,9,15,4,0,0,4,2 | 119,81,99,109,42,47,50,56 | 12,19,13,8,12,5,7,14,16,7,1 | 298 | 149 | 0.000122339818255 | 0.0239375649298563 | 0.783,0.985,0.846, | 0.832,0.681,0.792, | 0,118 |
| 170970 | ENSG00000115524 | SF3B1 | chr2 | - | 197392971 | 197393211 | 197388514 | 197392461 | 197396055 | 197396328 | 170970 | 367,231,1002,624,516,667 | 13,1,7,20,15,4,2,1,4,2,2,1 | 668,811,344,228,391,298,3 | 12,7,10,15,9,8,4,10,13,11,1 | 298 | 149 | 0.000123069070751 | 0.0240443286402461 | 0.934,0.991,0.986, | 0.965,0.983,0.945, | 0,018 |
| 70661 | ENSG00000111011 | RSRC2 | chr12 | - | 122526169 | 122526293 | 122522798 | 122522906 | 122526847 | 122526940 | 70661 | 3,0,7,1,0,1,2,0,0,1,2,0 | 54,6,35,27,18,21,26,25,10, | 4,3,3,0,1,0,0,4,1,2,5,5,3,9,0 | 20,14,37,11,6,12,4,15,10,1 | 273 | 149 | 0.000123108893057 | 0.0240443286402461 | 0.029,0.0,0.098,0.0 | 0.098,0.105,0.042, | -0,044 |
| 132930 | ENSG00000145901 | TNIP1 | chr5 | - | 151064959 | 151065131 | 151063612 | 151063747 | 151087084 | 151087158 | 132930 | 79,66,126,94,58,82,71,50,6 | 5,7,2,5,1,0,1,3,4,5,4,1 | 92,88,50,70,42,46,52,88,35 | 6,5,3,2,0,0,0,1,0,1,1,1,0,0,5 | 298 | 149 | 0.000123657673647 | 0.0241296144666523 | 0.888,0.825,0.969, | 0.885,0.898,0.893, | -0,048 |
| 67997 | ENSG00000070814 | TCOF1 | chr5 | + | 150383061 | 150383172 | 150379531 | 150379732 | 150387901 | 150388088 | 67997 | 7,4,4,5,3,11,5,4,5,2,1,0 | 5,2,8,6,4,6,5,6,10,9,2,1 | 1,3,2,0,2,3,2,1,1,9,2,5,1,0,0 | 5,11,1,2,3,6,3,4,4,15,3,25,1 | 260 | 149 | 0.000124325160682 | 0.024237888277851 | 0.445,0.534,0.223, | 0.103,0.135,0.534, | 0,14 |
| 39968 | ENSG00000151576 | QTRT2 | chr3 | + | 114067986 | 114068063 | 114066227 | 114066283 | 114070625 | 114070751 | 39968 | 8,15,31,26,19,19,14,20,21, | 3,0,6,9,0,0,1,1,4,0,1,4 | 18,33,10,9,20,3,7,14,8,21,1 | 1,5,2,1,7,0,2,0,1,0,0,0,0,2,0 | 226 | 149 | 0.000124902194182 | 0.024328347652476 | 0.637,1.0,0.773,0.6 | 0.922,0.813,0.767, | -0,057 |
| 132661 | ENSG00000173442 | EHBP1L1 | chr11 | + | 65583591 | 65583765 | 65581210 | 65581251 | 65584240 | 65584398 | 132661 | 55,39,43,13,46,34,41,62,32 | 5,3,6,1,3,5,6,7,3,0,4,7 | 20,33,23,27,10,9,37,63,19, | 1,4,3,5,0,0,5,1,2,0,1,2,0,0,1 | 298 | 149 | 0.000125755113339 | 0.0244723316447254 | 0.846,0.867,0.782, | 0.909,0.805,0.793, | -0,068 |
| 72561 | ENSG00000115966 | ATF2 | chr2 | - | 175147945 | 175147980 | 175136411 | 175136486 | 175151059 | 175151158 | 72561 | 4,1,13,3,1,1,0,2,4,0,1,3 | 44,18,88,52,33,50,34,51,17 | 6,13,5,5,6,4,3,5,8,4,7,2,2,0 | 65,72,20,32,27,23,21,37,24 | 184 | 149 | 0.000126320249705 | 0.024560102677388 | 0.069,0.043,0.107, | 0.07,0.128,0.168,0 | -0,06 |
| 40608 | ENSG00000115524 | SF3B1 | chr2 | - | 197392971 | 197393186 | 197389791 | 197392461 | 197396055 | 197396328 | 40608 | 369,232,1004,622,522,665 | 13,1,7,20,15,4,2,1,4,2,2,1 | 669,815,348,228,391,296,3 | 12,7,10,15,9,8,4,10,13,11,1 | 298 | 149 | 0.000126788583207 | 0.0246289111482182 | 0.934,0.991,0.986, | 0.965,0.983,0.946, | 0,018 |
| 67594 | ENSG00000166479 | TMX3 | chr18 | - | 68701744 | 68701790 | 68700404 | 68700485 | 68710020 | 68710144 | 67594 | 30,22,64,43,19,20,17,18,24 | 3,0,6,10,0,4,2,2,2,0,1,2 | 40,22,24,13,24,9,11,32,23, | 0,3,0,0,3,0,4,1,0,0,0,4,1,0,3 | 195 | 149 | 0.000127135951098 | 0.0246741189276588 | 0.884,1.0,0.891,0.7 | 1.0,0.849,1.0,1.0,0 | -0,037 |
| 46501 | ENSG00000163611 | SPICE1 | chr3 | - | 113450335 | 113450516 | 113448037 | 113448140 | 113453465 | 113453950 | 46501 | 45,17,63,40,37,23,23,44,21 | 2,5,6,4,10,5,0,3,2,1,1,0 | 32,31,35,38,21,15,33,34,36 | 1,2,0,2,2,1,4,1,3,2,1,2,0,3,0 | 298 | 149 | 0.000129411401208 | 0.0250931038679271 | 0.918,0.63,0.84,0.8 | 0.941,0.886,1.0,0.9 | -0,068 |
| 166338 | ENSG00000104517 | UBR5 | chr8 | - | 102285512 | 102285781 | 102285126 | 102285244 | 102286359 | 102286617 | 166338 | 34,25,86,44,35,35,42,26,37 | 2,2,4,2,3,0,1,0,0,0,0,1 | 48,38,40,63,37,36,29,28,26 | 4,5,2,5,1,0,2,4,0,2,1,5,0,0,5 | 298 | 149 | 0.000130014527409 | 0.0251663958691416 | 0.895,0.862,0.915, | 0.857,0.792,0.909, | 0,042 |
| 255296 | ENSG00000272917 | | chr12 | + | 9650332 | 9650361 | 9642871 | 9643103 | 9653682 | 9653808 | 255296 | 4,2,4,0,5,3,5,10,3,1,0,0 | 37,20,140,62,58,9,58,33,15 | 3,1,0,1,0,0,0,3,0,1,5,1,1,1,2 | 189,133,12,6,48,37,29,90,7 | 178 | 149 | 0.000130023241106 | 0.0251663958691416 | 0.083,0.077,0.023, | 0.013,0.006,0.0,0.1 | 0,004 |
| 201481 | ENSG00000131127 | ZNF141 | chr4 | + | 342307 | 342426 | 337833 | 337986 | 343781 | 343908 | 201481 | 1,0,15,8,0,0,1,3,0,0,0,1 | 8,9,98,59,25,22,11,32,20,8 | 0,3,0,0,0,0,0,0,0,0,1,3,0,0 | 31,34,9,20,26,17,18,19,13, | 268 | 149 | 0.000130711468651 | 0.0252768734134981 | 0.065,0.0,0.078,0.0 | 0.0,0.047,0.0,0.0,0 | 0,021 |
| 34952 | ENSG00000062194 | GPBP1 | chr5 | + | 57237112 | 57237172 | 57235965 | 57236032 | 57246299 | 57246484 | 34952 | 50,42,79,67,37,65,21,43,67 | 39,22,140,90,70,52,20,39,4 | 32,49,27,16,24,25,29,24,13 | 97,66,64,49,42,30,17,78,28 | 209 | 149 | 0.000131556161151 | 0.0254173825140852 | 0.478,0.576,0.287, | 0.19,0.346,0.231,0 | 0,129 |
| 92549 | ENSG00000083097 | DOP1A | chr6 | + | 83113322 | 83113421 | 83110124 | 83110314 | 83118887 | 83118987 | 92549 | 20,18,37,27,16,18,6,11,7,4 | 1,4,4,8,0,1,0,5,1,0,0,1 | 8,21,9,11,10,13,5,12,7,30,7 | 10,8,5,6,11,3,0,7,1,1,3,7,4, | 248 | 149 | 0.000132204182573 | 0.0255196757124389 | 0.923,0.73,0.848,0 | 0.325,0.612,0.52,0 | 0,218 |

| | | | | | | | | | | | | | | | | | | | | | | |
|---|---|---|---|---|---|---|---|---|---|---|---|---|---|---|---|---|---|---|---|---|---|---|
| 60558 | ENSG00000157796 | WDR19 | chr4 | + | 39194543 | 39194659 | 39189655 | 39189781 | 39199477 | 39199593 | 60558 | 42,11,123,48,26,28,22,34, | 7,0,14,8,5,4,1,7,2,4,1,3 | 39,27,37,25,11,9,16,43,30, | 1,2,1,2,0,1,2,5,3,3,3,5,2,2, | 265 | 149 | 0.000132520894756 | 0.0255578895154378 | 0.771,1.0,0.832,0. | 0.956,0.884,0.954, | -0,078 |
| 5136 | ENSG00000090372 | STRN4 | chr19 | - | 46733038 | 46733236 | 46730731 | 46730873 | 46738163 | 46738172 | 5136 | 36,33,25,11,16,15,14,33,2 | 1,3,7,1,1,2,0,2,0,3,4,1 | 21,17,11,17,22,14,18,14,1 | 3,3,4,10,3,1,4,2,5,2,8,5,5,5 | 298 | 149 | 0.000132649731206 | 0.0255598337486111 | 0.947,0.846,0.641, | 0.778,0.739,0.579, | 0,159 |
| 123199 | ENSG00000129255 | MPDU1 | chr17 | + | 7586898 | 7587017 | 7586655 | 7586777 | 7587160 | 7587271 | 123199 | 18,10,60,42,19,18,14,27,3 | 3,0,5,3,1,2,0,1,1,6,0,0 | 24,13,15,7,21,17,10,30,9,1 | 0,0,0,1,4,4,1,0,0,3,2,2,2,1, | 268 | 149 | 0.000134046802631 | 0.0258059278864694 | 0.769,1.0,0.87,0.8 | 1.0,1.0,1.0,0.796,0 | 0,006 |
| 58391 | ENSG00000163257 | DCAF16 | chr4 | - | 17805106 | 17805225 | 17800654 | 17804771 | 17810446 | 17810757 | 58391 | 48,37,106,51,47,19,33,72, | 1,5,5,5,3,3,4,3,13,7,1,11 | 44,51,54,65,84,35,32,72,9, | 3,2,2,6,3,2,1,6,0,2,6,1,1,2, | 268 | 149 | 0.000134343672778 | 0.0258399669666503 | 0.964,0.804,0.922, | 0.891,0.934,0.938, | -0,094 |
| 248917 | ENSG00000154258 | ABCA9 | chr17 | - | 68984879 | 68984979 | 68984055 | 68984175 | 68985052 | 68985128 | 248917 | 2,3,18,5,1,2,8,9,2,5,1,4 | 20,12,47,40,10,7,26,66,25, | 1,9,1,0,4,5,0,2,2,2,2,0,1,1, | 28,63,32,10,30,43,4,12,20, | 249 | 149 | 0.000134580400621 | 0.0258623876839808 | 0.056,0.13,0.186,0 | 0.021,0.079,0.018, | 0,045 |
| 33292 | ENSG00000128739 | SNRPN | chr15 | + | 24967028 | 24967152 | 24962113 | 24962209 | 24967931 | 24968082 | 33292 | 10,4,181,205,10,13,4,25,3 | 126,81,198,89,72,86,83,10 | 9,9,8,0,42,33,6,11,2,12,7,6 | 161,170,110,16,59,53,58,1 | 273 | 149 | 0.000135129774139 | 0.0259447960907325 | 0.042,0.026,0.333, | 0.03,0.028,0.038,0 | 0,085 |
| 227621 | ENSG00000186088 | GSAP | chr7 | - | 77321324 | 77321403 | 77320724 | 77320819 | 77323646 | 77323742 | 227621 | 67,32,70,47,24,38,28,37,1 | 12,2,2,0,10,6,0,0,0,0,0,0 | 50,46,42,31,34,20,35,41,2 | 10,4,0,0,0,0,0,0,8,0,0,0,0,0 | 228 | 149 | 0.000135288569124 | 0.0259521337086699 | 0.785,0.913,0.958, | 0.766,0.883,1.0,1. | -0,036 |
| 111893 | ENSG00000120868 | APAF1 | chr12 | + | 98665552 | 98665791 | 98662674 | 98662806 | 98666189 | 98666357 | 111893 | 5,2,14,10,11,8,2,2,2,3,0,4 | 2,0,7,0,0,0,1,0,0,0,3,0 | 16,10,4,2,3,14,4,2,4,16,3,2 | 0,0,2,3,1,1,0,3,4,2,2,7,1,1, | 298 | 149 | 0.000135767077721 | 0.0260207336642641 | 0.556,1.0,0.5,1.0,1 | 1.0,1.0,0.5,0.25,0. | 0,123 |
| 5322 | ENSG00000269858 | EGLN2 | chr19 | + | 40806554 | 40806674 | 40799544 | 40801415 | 40807137 | 40807274 | 5322 | 21,12,39,33,28,14,19,7,31, | 2,1,0,1,0,0,4,1,1,2,1,7 | 8,18,13,5,7,14,13,8,5,14,6, | 3,0,0,0,1,0,0,1,1,0,3,2,0,1, | 269 | 149 | 0.000136272833115 | 0.0260777623548028 | 0.853,0.869,1.0,0. | 0.596,1.0,1.0,1.0,0 | -0,006 |
| 222228 | ENSG00000152601 | MBNL1 | chr3 | + | 152338467 | 152338673 | 152300197 | 152300367 | 152414940 | 152415111 | 222228 | 50,23,24,10,32,16,12,3,29, | 452,250,444,197,235,233, | 37,20,10,1,7,4,9,9,2,17,0,8 | 365,359,176,132,170,131,1 | 298 | 149 | 0.00013630695694 | 0.0260777623548028 | 0.052,0.044,0.026, | 0.048,0.027,0.028, | 0,012 |
| 102626 | ENSG00000122566 | HNRNPA2B1 | chr7 | - | 26193574 | 26193598 | 26193250 | 26193373 | 26195846 | 26195909 | 102626 | 9,10,23,10,12,12,7,10,6,7, | 122,51,104,76,40,60,59,87 | 10,8,6,2,2,2,2,12,1,7,6,14, | 83,99,103,104,94,25,21,54 | 173 | 149 | 0.000136766682489 | 0.0261347318823768 | 0.06,0.144,0.16,0. | 0.094,0.065,0.048, | 0,062 |
| 122409 | ENSG00000166387 | PPFIBP2 | chr11 | + | 7625776 | 7625891 | 7620934 | 7621027 | 7628284 | 7628346 | 122409 | 74,68,14,4,14,6,5,6,25,11, | 4,7,1,0,0,0,0,0,2,0,0,0 | 15,15,27,27,39,18,12,16,2 | 3,1,2,3,4,1,0,0,2,1,0,5,0,2, | 264 | 149 | 0.000136847586228 | 0.0261347318823768 | 0.913,0.846,0.888, | 0.738,0.894,0.884, | 0,063 |
| 180728 | ENSG00000100991 | TRPC4AP | chr20 | - | 35044504 | 35044606 | 35035122 | 35035308 | 35049865 | 35049994 | 180728 | 70,45,96,50,27,43,24,41,2 | 3,3,7,1,2,4,2,3,3,2,2,3 | 46,55,23,39,45,19,21,43,2 | 4,1,2,1,0,1,0,2,0,0,4,2,1,0, | 251 | 149 | 0.000137013081867 | 0.0261355877061672 | 0.933,0.899,0.891, | 0.872,0.97,0.872,0 | -0,068 |
| 21160 | ENSG00000092148 | HECTD1 | chr14 | - | 31113876 | 31114059 | 31113276 | 31113484 | 31114285 | 31114340 | 21160 | 54,48,140,95,70,59,78,107 | 17,3,11,7,0,12,2,12,2,4,1,0 | 87,114,56,60,84,46,50,76, | 4,8,0,1,3,2,3,4,1,0,7,5,0,3, | 298 | 149 | 0.000137209345426 | 0.0261355877061672 | 0.614,0.889,0.864, | 0.916,0.877,1.0,0. | -0,072 |
| 168156 | ENSG00000287542 | HERC3 | chr4 | + | 88605794 | 88605901 | 88595556 | 88595614 | 88649839 | 88649999 | 168156 | 19,12,42,28,9,26,18,12,13, | 4,5,1,0,2,4,0,6,0,0,0,1 | 33,30,15,33,18,19,15,14,1 | 0,1,3,5,3,1,2,0,2,0,0,1,0,0, | 256 | 149 | 0.00013721635874 | 0.0261355877061672 | 0.734,0.583,0.961, | 1.0,0.946,0.744,0. | -0,038 |
| 122426 | ENSG00000166387 | PPFIBP2 | chr11 | + | 7650839 | 7650965 | 7649531 | 7649654 | 7651655 | 7651844 | 122426 | 78,98,15,13,15,7,6,10,31,1 | 8,4,1,1,0,0,2,0,0,1,0,0 | 15,12,19,36,30,23,12,14,2 | 5,3,1,18,0,4,2,1,1,2,0,1,1,1 | 275 | 149 | 0.000137492841397 | 0.0261650943825974 | 0.841,0.93,0.89,0. | 0.619,0.684,0.911, | 0,08 |
| 227297 | ENSG00000013374 | NUB1 | chr7 | + | 151355767 | 151356221 | 151352811 | 151352882 | 151360140 | 151360914 | 227297 | 97,83,136,95,31,27,31,40, | 0,2,3,4,1,1,2,2,0,5,0,0 | 53,58,40,54,26,17,12,35,2 | 2,0,6,7,3,2,0,0,2,1,7,3,2,0, | 298 | 149 | 0.000137648497873 | 0.0261715758354273 | 1.0,0.954,0.958,0. | 0.93,1.0,0.769,0.7 | 0,071 |
| 47293 | ENSG00000164828 | SUN1 | chr7 | + | 851383 | 851482 | 849519 | 849603 | 851949 | 852043 | 47293 | 21,6,22,11,20,15,6,20,16,9 | 0,0,8,6,0,0,0,4,0,2,0,2 | 19,23,11,4,4,4,4,24,11,13,1 | 0,1,1,0,1,0,2,0,0,0,0,5,0,0, | 248 | 149 | 0.000138386424781 | 0.026288657186266 | 1.0,1.0,0.623,0.52 | 1.0,0.933,0.869,1. | -0,051 |
| 167257 | ENSG00000241258 | CRCP | chr7 | + | 66130743 | 66131069 | 66127703 | 66127740 | 66134279 | 66134489 | 167257 | 15,13,28,26,8,10,16,15,12, | 0,0,2,1,1,0,0,0,0,0,0,1 | 17,25,6,8,10,0,8,9,13,12,1 | 2,1,2,2,0,2,2,1,3,1,0,2,2,2,1 | 298 | 149 | 0.000139048265155 | 0.0263678389053531 | 1.0,1.0,0.875,0.92 | 0.81,0.926,0.6,0.6 | 0,193 |
| 167258 | ENSG00000241258 | CRCP | chr7 | + | 66130743 | 66131132 | 66127703 | 66127740 | 66134279 | 66134489 | 167258 | 15,13,28,26,8,10,16,15,12, | 0,0,2,1,1,0,0,0,0,0,0,1 | 17,25,6,8,10,0,8,9,13,12,1 | 2,1,2,2,0,2,2,1,3,1,0,2,2,2,1 | 298 | 149 | 0.000139048265155 | 0.0263678389053531 | 1.0,1.0,0.875,0.92 | 0.81,0.926,0.6,0.6 | 0,193 |
| 176775 | ENSG00000186469 | GNG2 | chr14 | + | 51877620 | 51877657 | 51860733 | 51860790 | 51950649 | 51950765 | 176775 | 10,10,16,9,17,9,10,14,28,1 | 3,0,3,4,2,0,1,2,0,2,1,1 | 5,7,3,5,7,5,4,6,7,3,5,30,1,6 | 0,2,0,0,0,0,0,1,0,1,0,0,0,0, | 186 | 149 | 0.000139499120795 | 0.0264300486512576 | 0.728,1.0,0.81,0.6 | 1.0,0.737,1.0,1.0,1 | -0,099 |
| 102341 | ENSG00000006576 | PHTF2 | chr7 | + | 77940527 | 77940659 | 77940029 | 77940302 | 77942699 | 77942786 | 102341 | 14,14,49,30,28,7,16,14,19, | 2,0,0,0,4,2,1,0,2,3,0,0 | 30,22,24,14,9,20,5,11,13,2 | 1,3,2,0,1,7,1,1,3,0,0,6,2,3, | 281 | 149 | 0.000141603333455 | 0.0267857475459011 | 0.788,1.0,1.0,1.0,0 | 0.941,0.795,0.864, | 0,061 |
| 170969 | ENSG00000115524 | SF3B1 | chr2 | - | 197392971 | 197393043 | 197388514 | 197392461 | 197396055 | 197396328 | 170969 | 161,118,469,302,280,377, | 13,1,7,20,15,4,2,1,4,2,2,1 | 356,408,184,113,184,136, | 12,7,10,15,9,8,4,10,13,11,1 | 221 | 149 | 0.000141625419699 | 0.0267857475459011 | 0.893,0.988,0.978, | 0.952,0.975,0.925, | 0,022 |
| 238713 | ENSG00000064313 | TAF2 | chr8 | - | 119794830 | 119794944 | 119793365 | 119793451 | 119795531 | 119795631 | 238713 | 0,0,8,9,0,3,0,5,1,0,0,1 | 23,8,36,9,9,22,8,5,12,4,2,1 | 0,1,0,0,2,2,0,0,2,0,0,4,1,1, | 15,16,9,12,12,7,4,14,7,19,1 | 263 | 149 | 0.000142565378733 | 0.0269282686111143 | 0.0,0.0,0.112,0.36 | 0.0,0.034,0.0,0.0,0 | 0,045 |
| 11679 | ENSG00000007202 | KIAA0100 | chr17 | - | 28643163 | 28643311 | 28642868 | 28642987 | 28643609 | 28643659 | 11679 | 90,35,131,123,63,96,58,11 | 2,1,4,3,1,2,2,5,0,2,0,0 | 87,66,51,48,36,56,38,96,2 | 6,4,1,3,6,1,0,3,2,0,2,4,6,2, | 297 | 149 | 0.000142629204049 | 0.0269282686111143 | 0.958,0.946,0.943, | 0.879,0.892,0.962, | 0,05 |
| 245683 | ENSG00000109919 | MTCH2 | chr11 | - | 47628948 | 47629046 | 47627079 | 47627127 | 47630554 | 47630614 | 245683 | 7,9,28,38,13,9,9,24,8,10,4, | 2,0,6,0,0,3,1,0,1,0,0,0 | 21,25,8,1,14,10,12,14,8,10 | 7,5,0,2,3,0,1,2,4,0,0,3,0,4, | 247 | 149 | 0.000142974829931 | 0.0269482265204168 | 0.679,1.0,0.738,1. | 0.644,0.751,1.0,0. | 0,07 |
| 131655 | ENSG00000135457 | TFCP2 | chr12 | - | 51104154 | 51104203 | 51103669 | 51103763 | 51106524 | 51106613 | 131655 | 20,8,21,20,14,18,24,10,31, | 6,0,4,2,2,2,1,1,4,0,1,0 | 16,18,8,23,8,15,11,23,6,13 | 0,4,0,1,0,2,0,1,1,0,1,0,0,1,1 | 198 | 149 | 0.000142985325935 | 0.0269482265204168 | 0.715,1.0,0.798,0. | 1.0,0.772,1.0,0.94 | -0,046 |
| 7476 | ENSG00000089057 | SLC23A2 | chr20 | - | 4970792 | 4970919 | 4932454 | 4932716 | 5001405 | 5001510 | 7476 | 116,85,137,91,85,112,96,7 | 0,0,0,3,0,2,0,0,0,0,2,0 | 65,58,51,79,38,34,51,54,4 | 13,13,0,0,0,0,0,2,0,0,2,3,0, | 276 | 149 | 0.000145028743475 | 0.0273094326219315 | 1.0,1.0,1.0,0.942,1 | 0.73,0.707,1.0,1.0, | 0,03 |
| 173762 | ENSG00000156976 | EIF4A2 | chr3 | + | 186788291 | 186788416 | 186787802 | 186787882 | 186789124 | 186789881 | 173762 | 35,7,48,31,16,21,24,40,10, | 427,137,428,354,181,165,1 | 17,25,36,11,15,7,9,15,11,1 | 138,51,178,74,110,133,75, | 274 | 149 | 0.000145166368697 | 0.027311453409986 | 0.043,0.027,0.057, | 0.063,0.21,0.099,0 | -0,026 |
| 37615 | ENSG00000005059 | MCUB | chr4 | + | 109664289 | 109664394 | 109660194 | 109660365 | 109682581 | 109682742 | 37615 | 233,122,350,237,116,101,1 | 12,8,2,2,0,3,3,0,0,3,4,0 | 237,153,173,97,101,109,6 | 0,1,0,4,1,1,0,0,1,2,2,3,1,0, | 254 | 149 | 0.000146184371131 | 0.0274789592863723 | 0.919,0.899,0.99,0 | 1.0,0.989,1.0,0.93 | -0,033 |
| 2961 | ENSG00000105486 | LIG1 | chr19 | - | 48151467 | 48151734 | 48151231 | 48151339 | 48153871 | 48153967 | 2961 | 3,6,3,1,0,0,4,0,6,1,2,4 | 26,18,20,5,27,19,12,15,26, | 1,0,0,0,0,1,1,1,1,1,3,0,1,3, | 14,11,19,13,6,14,4,16,10,1 | 298 | 149 | 0.000146428267112 | 0.0275007873985889 | 0.055,0.143,0.07,0 | 0.034,0.0,0.0,0.0,0 | 0,017 |
| 92473 | ENSG00000135486 | HNRNPA1 | chr12 | + | 54283078 | 54283234 | 54282572 | 54282665 | 54283811 | 54283967 | 92473 | 7,13,14,24,10,10,2,12,12,4 | 3,0,1,2,0,0,2,0,0,0,3,0 | 7,9,11,3,2,2,0,11,2,8,9,32, | 1,0,1,10,1,1,4,3,0,2,3,2,2,1 | 298 | 149 | 0.000148457445011 | 0.0278575800759918 | 0.538,1.0,0.875,0. | 0.778,1.0,0.846,0. | 0,127 |
| 105530 | ENSG00000136504 | KAT7 | chr17 | + | 49809118 | 49809208 | 49798318 | 49798558 | 49811475 | 49811574 | 105530 | 44,49,40,33,17,16,22,20,2 | 11,0,4,1,0,3,1,2,5,2,0,2 | 28,36,25,75,36,10,14,39,1 | 4,1,0,6,6,2,0,0,2,0,1,0,0,2, | 239 | 149 | 0.000149554427479 | 0.0280389799483734 | 0.714,1.0,0.862,0. | 0.814,0.957,1.0,0. | -0,065 |
| 143066 | ENSG00000167543 | TP53I13 | chr17 | + | 29571590 | 29571669 | 29569317 | 29569359 | 29571856 | 29572057 | 143066 | 62,24,68,42,46,49,48,42,1 | 7,0,13,2,2,8,0,4,5,0,2,6 | 36,30,30,14,26,12,14,42,1 | 10,5,14,3,9,1,1,5,3,0,8,7,6, | 228 | 149 | 0.000151022293888 | 0.0282672274872108 | 0.853,1.0,0.774,0. | 0.702,0.797,0.583, | 0,109 |
| 144072 | ENSG00000007202 | KIAA0100 | chr17 | - | 28643101 | 28643311 | 28642868 | 28642987 | 28643609 | 28643659 | 144072 | 64,29,65,56,33,61,43,76,8 | 2,1,4,3,1,2,2,5,0,2,0,0 | 47,38,31,31,28,40,26,70,1 | 6,4,1,3,6,1,0,3,2,0,2,4,6,2, | 298 | 149 | 0.000151034523885 | 0.0282672274872108 | 0.941,0.935,0.89,0 | 0.797,0.826,0.939, | 0,079 |
| 97026 | ENSG00000173200 | PARP15 | chr3 | + | 122615285 | 122615320 | 122613040 | 122613268 | 122617014 | 122617164 | 97026 | 17,8,33,37,13,12,9,21,8,3,1 | 0,0,2,1,0,0,0,4,1,2,0,3 | 26,19,9,4,14,4,13,15,22,32 | 6,2,0,6,1,1,0,3,0,5,0,0,1,2, | 184 | 149 | 0.000151741424934 | 0.0283748554560988 | 1.0,1.0,0.93,0.968, | 0.778,0.885,1.0,0. | 0,027 |
| 187093 | ENSG00000163946 | TASOR | chr3 | - | 56623728 | 56623843 | 56622861 | 56623566 | 56624478 | 56624643 | 187093 | 89,59,143,84,70,56,57,71, | 2,0,13,17,5,1,0,0,1,0,0,1 | 96,88,48,52,31,78,26,70,4 | 5,2,3,0,0,5,2,0,0,3,0,3,0,3,1 | 264 | 149 | 0.000153908811362 | 0.0287551626547349 | 0.962,1.0,0.861,0. | 0.916,0.961,0.9,1. | -0,015 |
| 122371 | ENSG00000132254 | ARFIP2 | chr11 | - | 6478578 | 6478652 | 6478083 | 6478198 | 6478737 | 6478959 | 122371 | 6,3,3,2,2,3,0,0,0,0,7,4 | 41,37,28,33,9,24,11,16,17, | 6,0,2,0,0,0,1,1,0,1,0,3,0,4, | 25,41,16,28,25,17,6,38,28, | 223 | 149 | 0.000154058999545 | 0.0287582589167996 | 0.089,0.051,0.067, | 0.138,0.0,0.077,0. | 0,052 |
| 144076 | ENSG00000007202 | KIAA0100 | chr17 | - | 28643125 | 28643311 | 28642868 | 28642987 | 28643609 | 28643659 | 144076 | 64,28,64,56,33,61,43,76,8 | 2,1,4,3,1,2,2,5,0,2,0,0 | 47,38,31,31,26,40,26,70,1 | 6,4,1,3,6,1,0,3,2,0,2,4,6,2, | 298 | 149 | 0.000155683319017 | 0.0290203323520412 | 0.941,0.933,0.889, | 0.797,0.826,0.939, | 0,079 |
| 225314 | ENSG00000170265 | ZNF282 | chr7 | + | 149206049 | 149206154 | 149198332 | 149198752 | 149206695 | 149206822 | 225314 | 0,0,1,4,2,4,0,3,0,0,0,1 | 9,4,13,13,7,1,5,7,10,5,6,6 | 3,2,3,4,2,1,3,0,0,3,3,2,0,3, | 9,10,2,3,7,2,5,8,2,2,11,3,8, | 254 | 149 | 0.000155732602955 | 0.0290203323520412 | 0.0,0.0,0.043,0.15 | 0.164,0.105,0.468, | -0,108 |
| 99879 | ENSG00000181090 | EHMT1 | chr9 | + | 137775108 | 137775252 | 137762674 | 137762820 | 137776617 | 137776840 | 99879 | 79,102,182,134,158,116,1 | 3,2,1,0,1,2,0,3,0,1,0,0 | 163,265,89,88,118,74,95,1 | 4,6,0,7,2,0,2,2,2,0,9,1,7,2,1 | 293 | 149 | 0.000156418137364 | 0.0290984821001311 | 0.931,0.963,0.989, | 0.954,0.957,1.0,0. | 0,022 |
| 143059 | ENSG00000167543 | TP53I13 | chr17 | + | 29567863 | 29568060 | 29567179 | 29567411 | 29568277 | 29568508 | 143059 | 0,1,9,4,2,5,1,10,6,5,0,6 | 19,8,31,26,14,25,6,28,37,2 | 1,0,3,0,4,1,0,1,0,1,5,0,4,1, | 19,19,11,1,18,0,15,14,5,13 | 298 | 149 | 0.000156422373124 | 0.0290984821001311 | 0.0,0.059,0.127,0. | 0.026,0.0,0.12,0.0, | -0,049 |
| 55783 | ENSG00000134352 | IL6ST | chr5 | - | 55948101 | 55948982 | 55947492 | 55947589 | 55951463 | 55951604 | 55783 | 3,1,0,1,1,1,1,6,0,6,0,3 | 7,7,12,5,6,9,18,59,13,5,2,1 | 6,6,11,10,4,1,5,12,3,9,13,0 | 9,25,10,13,17,4,5,13,3,2,2 | 298 | 149 | 0.000156625594493 | 0.0291111254994153 | 0.176,0.067,0.0,0. | 0.25,0.107,0.355,0 | -0,08 |
| 85359 | ENSG00000080345 | RIF1 | chr2 | + | 151443528 | 151443709 | 151443258 | 151443329 | 151445337 | 151445445 | 85359 | 47,19,99,53,42,42,28,38,1 | 1,0,13,6,0,2,0,4,0,1,0,0 | 56,67,28,23,30,22,33,43,1 | 3,4,3,1,1,0,0,1,0,0,3,2,0,0, | 298 | 149 | 0.000157046073525 | 0.0291505282093217 | 0.959,1.0,0.792,0. | 0.903,0.893,0.824, | -0,013 |
| 98425 | ENSG00000118058 | KMT2A | chr11 | + | 118507507 | 118507609 | 118502397 | 118506646 | 118509135 | 118509200 | 98425 | 8,7,37,15,10,16,5,7,4,2,3,9 | 2,0,3,0,0,4,1,0,2,0,0,1 | 14,23,8,14,8,8,6,8,12,13,1 | 2,4,0,12,6,1,1,0,1,0,2,2,0,0 | 251 | 149 | 0.000157108468217 | 0.0291505282093217 | 0.704,1.0,0.88,1.0, | 0.806,0.773,1.0,0. | 0,041 |
| 68036 | ENSG00000143119 | CD53 | chr1 | + | 110894326 | 110894401 | 110873147 | 110873248 | 110899123 | 110899922 | 68036 | 15,7,18,12,12,10,11,29,5,9 | 0,0,5,4,0,2,2,0,0,1,0,0 | 10,16,21,11,10,2,4,24,8,5, | 4,5,1,0,3,1,1,0,2,1,3,7,1,0, | 224 | 149 | 0.000157371518708 | 0.0291741854806559 | 1.0,1.0,0.705,0.66 | 0.624,0.68,0.933,1 | 0,079 |
| 159557 | ENSG00000280987 | MATR3 | chr5 | + | 139325295 | 139325340 | 139322597 | 139322967 | 139325439 | 139325662 | 159557 | 9,6,42,15,12,6,6,10,19,9,2, | 696,437,823,592,278,389, | 8,12,9,2,3,3,3,6,5,18,9,6,2, | 536,564,628,543,404,214, | 194 | 149 | 0.000157833297985 | 0.0292346114962216 | 0.01,0.01,0.038,0. | 0.011,0.016,0.011, | 0,007 |
| 52437 | ENSG00000096080 | MRPS18A | chr6 | - | 43675201 | 43675271 | 43671201 | 43671906 | 43675493 | 43675617 | 52437 | 40,31,63,38,23,23,15,31,4 | 0,0,3,1,0,0,2,0,5,0,0,6 | 41,34,28,16,14,14,27,20,4, | 1,3,2,7,2,3,4,2,0,1,2,1,0,3,1 | 219 | 149 | 0.000158019108964 | 0.0292438614285732 | 1.0,1.0,0.935,0.96 | 0.965,0.885,0.905, | 0,056 |
| 155969 | ENSG00000005810 | MYCBP2 | chr13 | - | 77185870 | 77186068 | 77185102 | 77185377 | 77188950 | 77189047 | 155969 | 38,36,119,78,53,70,45,81, | 1,0,0,0,0,0,2,1,0,1,0,0 | 79,82,25,60,39,34,44,47,4 | 3,6,2,8,1,0,0,0,2,1,4,1,1,1, | 298 | 149 | 0.00015824223286 | 0.0292599948631363 | 0.95,1.0,1.0,1.0,1. | 0.929,0.872,0.862, | 0,033 |
| 36638 | ENSG00000112079 | STK38 | chr6 | - | 36499872 | 36499990 | 36498362 | 36498486 | 36506582 | 36506644 | 36638 | 24,14,17,25,15,6,7,5,10,1, | 1,2,7,6,0,1,0,0,3,2,0,0 | 18,7,17,2,10,13,7,6,11,17, | 2,0,1,1,2,0,0,2,2,1,0,0,1,0, | 267 | 149 | 0.000158643969008 | 0.0293090987927561 | 0.931,0.796,0.575, | 0.834,1.0,0.905,0. | -0,086 |
| 27962 | ENSG00000198482 | ZNF808 | chr19 | + | 52532907 | 52533009 | 52527667 | 52527711 | 52543265 | 52543347 | 27962 | 12,12,53,30,14,21,6,12,20, | 2,2,4,3,1,0,2,0,4,0,0,3 | 17,21,18,20,13,5,0,15,8,10 | 1,1,1,0,2,0,2,2,0,0,2,1,0,0, | 251 | 149 | 0.000159503928875 | 0.0294400895980049 | 0.781,0.781,0.887, | 0.91,0.926,0.914,1 | -0,036 |
| 227945 | ENSG00000173715 | C11orf80 | chr11 | + | 66801030 | 66801197 | 66800640 | 66800696 | 66803994 | 66804194 | 227945 | 46,14,91,65,40,48,24,42,1 | 1,0,5,0,0,3,2,5,0,0,0,1 | 64,40,24,19,23,24,19,41,1 | 1,6,2,1,0,0,2,2,1,5,3,0,1,0, | 298 | 149 | 0.000159626561977 | 0.0294400895980049 | 0.958,1.0,0.901,1. | 0.97,0.769,0.857,0 | 0,061 |
| 101341 | ENSG00000173715 | C11orf80 | chr11 | + | 66801030 | 66801161 | 66800640 | 66800696 | 66803994 | 66804194 | 101341 | 44,13,96,65,45,52,28,43,1 | 1,0,5,0,0,3,2,5,0,0,0,1 | 67,39,31,19,21,24,19,39,1 | 1,6,2,1,0,0,2,2,1,5,3,0,1,0, | 280 | 149 | 0.000160634670877 | 0.0296006514105544 | 0.959,1.0,0.911,1. | 0.973,0.776,0.892, | 0,059 |
| 129596 | ENSG00000111144 | LTA4H | chr12 | - | 96011752 | 96011865 | 96009093 | 96009148 | 96013187 | 96013258 | 129596 | 21,12,44,3,7,3,2,4,1,1,4,4 | 224,107,416,326,157,108,1 | 3,11,5,10,3,2,4,1,4,9,4,3,3, | 275,164,226,55,136,112,7 | 262 | 149 | 0.000161092855324 | 0.0296596889172283 | 0.051,0.06,0.057,0 | 0.006,0.037,0.012, | 0,004 |
| 120825 | ENSG00000102043 | MTMR8 | chrX | - | 64345044 | 64345177 | 64343610 | 64343720 | 64348659 | 64348794 | 120825 | 4,7,13,10,10,11,0,3,4,7,1,8 | 2,0,7,2,0,2,2,1,2,2,0,1 | 16,15,3,3,8,6,7,12,4,9,8,6, | 4,2,1,3,1,1,0,0,3,1,1,1,0,0, | 282 | 149 | 0.000162136257375 | 0.0298262810351099 | 0.514,1.0,0.495,0. | 0.679,0.798,0.613, | -0,129 |
| 65168 | ENSG00000146282 | RARS2 | chr6 | - | 87534734 | 87534858 | 87530783 | 87530942 | 87541917 | 87541994 | 65168 | 5,2,3,1,1,0,1,5,3,2,1,3 | 30,22,44,27,17,17,13,26,3 | 8,1,3,13,0,1,7,7,2,0,4,4,2,0 | 17,22,30,13,23,13,6,14,10, | 273 | 149 | 0.000162876131432 | 0.029930264288654 | 0.083,0.047,0.036, | 0.204,0.024,0.052, | -0,076 |
| 115205 | ENSG00000148153 | INIP | chr9 | - | 112689526 | 112689617 | 112683925 | 112687633 | 112694130 | 112694233 | 115205 | 10,11,34,21,13,15,13,10,1 | 0,0,3,2,2,0,2,3,0,2,0,0 | 23,19,23,4,7,7,8,14,5,9,15, | 3,5,2,0,0,0,2,4,2,3,4,4,2,3, | 240 | 149 | 0.000163139827633 | 0.029930264288654 | 1.0,1.0,0.876,0.86 | 0.826,0.702,0.877, | 0,073 |
| 127302 | ENSG00000184047 | DIABLO | chr12 | - | 122209696 | 122209858 | 122208315 | 122208577 | 122216487 | 122216584 | 127302 | 15,5,25,17,8,13,9,6,6,5,2,9 | 25,15,40,32,19,12,18,15,3 | 20,27,16,16,25,8,15,26,12, | 9,21,21,10,22,7,16,14,7,14 | 298 | 149 | 0.000163179795295 | 0.029930264288654 | 0.231,0.143,0.238, | 0.526,0.391,0.276, | -0,123 |
| 8239 | ENSG00000134444 | RELCH | chr18 | + | 62279773 | 62279856 | 62275373 | 62275473 | 62280359 | 62280442 | 8239 | 2,2,0,0,2,3,1,1,0,0,0,2 | 3,5,19,15,11,10,5,3,2,4,1,3 | 1,0,2,5,2,4,0,1,5,0,1,2,6,6, | 17,11,3,2,1,1,7,14,7,6,9,20 | 232 | 149 | 0.000163498597005 | 0.029930264288654 | 0.3,0.204,0.0,0.0,0 | 0.036,0.0,0.3,0.61 | -0,106 |
| 188814 | ENSG00000082512 | TRAF5 | chr1 | + | 211366410 | 211366490 | 211365375 | 211365468 | 211369451 | 211369592 | 188814 | 11,1,26,7,10,6,10,11,7,2,4, | 13,21,56,36,39,25,12,21,2 | 23,12,18,12,5,4,4,8,11,12, | 36,20,15,9,30,8,7,15,21,34 | 229 | 149 | 0.000163557913252 | 0.029930264288654 | 0.355,0.03,0.232,0 | 0.294,0.281,0.438, | -0,118 |
| 116013 | ENSG00000056586 | RC3H2 | chr9 | - | 122892908 | 122893026 | 122890170 | 122890545 | 122897278 | 122897534 | 116013 | 85,65,206,153,31,50,42,59 | 1,6,11,1,1,0,3,0,0,0,1,2 | 81,167,95,127,74,49,34,49 | 1,0,0,4,2,0,0,1,0,2,0,0,0,0,1 | 267 | 149 | 0.000163585757596 | 0.029930264288654 | 0.979,0.858,0.913, | 0.978,1.0,1.0,0.94 | -0,029 |
| 178687 | ENSG00000163564 | PYHIN1 | chr1 | + | 158973989 | 158974065 | 158973646 | 158973771 | 158976700 | 158977059 | 178687 | 6,11,4,0,0,2,3,9,1,4,0,5 | 32,15,16,17,3,6,15,10,2,8, | 1,2,1,1,3,2,0,0,3,5,0,5,3,1, | 2,5,23,12,9,15,7,5,6,16,9,1 | 225 | 149 | 0.000163674940263 | 0.029930264288654 | 0.11,0.327,0.142,0 | 0.249,0.209,0.028, | 0,065 |
| 203614 | ENSG00000120451 | SNX19 | chr11 | - | 130908360 | 130908445 | 130906624 | 130906721 | 130910017 | 130910137 | 203614 | 9,10,10,7,3,5,8,13,4,4,2,7 | 0,2,1,0,0,0,0,1,1,0,0,0 | 7,9,6,8,12,2,6,1,2,4,3,5,8,5 | 0,1,1,0,5,1,2,1,0,0,4,1,2,1,1 | 234 | 149 | 0.000164426167086 | 0.0300278066541061 | 1.0,0.761,0.864,1. | 1.0,0.851,0.793,1. | 0,226 |
| 23892 | ENSG00000126226 | PCID2 | chr13 | - | 113190871 | 113190975 | 113185484 | 113185560 | 113195070 | 113195125 | 23892 | 39,17,53,54,28,30,34,22,4 | 6,1,7,6,2,0,2,6,0,0,0,0 | 31,12,9,7,13,10,9,21,11,18 | 1,1,0,2,4,0,0,3,0,2,0,0,2,0, | 253 | 149 | 0.000164487383533 | 0.0300278066541061 | 0.793,0.909,0.817, | 0.948,0.876,1.0,0. | 0,007 |
| 45894 | ENSG00000169180 | XPO6 | chr16 | - | 28111816 | 28112006 | 28107521 | 28107677 | 28112903 | 28113050 | 45894 | 200,100,152,157,104,92,7 | 7,4,12,1,2,0,4,4,11,3,2,6 | 118,115,115,82,98,67,72,9 | 1,0,0,6,4,1,5,1,2,0,0,6,1,2,1 | 298 | 149 | 0.000164838574952 | 0.0300664163775372 | 0.935,0.926,0.864, | 0.983,1.0,1.0,0.87 | -0,031 |
| 19135 | ENSG00000137842 | TMEM62 | chr15 | + | 43149107 | 43149151 | 43148754 | 43148879 | 43151789 | 43151834 | 19135 | 4,2,20,1,1,10,8,10,1,6,5,5 | 0,4,5,1,0,1,0,6,0,0,0,0 | 8,27,3,4,9,0,12,8,6,4,6,8,9, | 7,13,0,8,5,2,1,3,3,0,1,1,2,2 | 193 | 149 | 0.000165197369648 | 0.0300926058922585 | 1.0,0.279,0.755,0. | 0.469,0.616,1.0,0. | 0,198 |
| 177773 | ENSG00000170854 | RIOX2 | chr3 | - | 97949592 | 97950015 | 97947415 | 97947449 | 97950785 | 97950888 | 177773 | 11,12,8,12,18,4,6,15,6,5,4, | 2,0,2,5,1,4,0,1,3,0,0,0 | 8,1,11,2,4,9,3,5,5,9,4,13,2 | 1,0,0,0,0,0,2,0,2,2,0,1,0,2, | 298 | 149 | 0.000165261789262 | 0.0300926058922585 | 0.733,1.0,0.667,0. | 0.8,1.0,1.0,1.0,1.0, | -0,084 |
| 250550 | ENSG00000148634 | HERC4 | chr10 | - | 67990210 | 67990309 | 67988662 | 67988835 | 67990903 | 67991015 | 250550 | 14,5,29,29,17,11,9,17,8,2,1 | 3,3,5,1,4,2,1,3,0,1,1,0 | 17,16,4,7,18,3,8,11,8,12,1 | 3,2,0,1,1,3,1,1,1,2,3,0,1,0, | 248 | 149 | 0.000165609874236 | 0.0301304977529066 | 0.737,0.5,0.777,0. | 0.773,0.828,1.0,0. | -0,104 |
| 138075 | ENSG00000012061 | ERCC1 | chr19 | - | 45419106 | 45419197 | 45416820 | 45416897 | 45420323 | 45420427 | 138075 | 29,26,95,42,41,31,31,33,2 | 1,2,5,9,0,5,0,2,0,0,0,0 | 42,22,26,16,17,19,16,41,1 | 0,1,0,1,0,0,0,0,0,0,4,3,0,2 | 240 | 149 | 0.000166207353504 | 0.0302136612348137 | 0.947,0.89,0.922,0 | 1.0,0.932,1.0,0.90 | -0,033 |
| 59694 | ENSG00000162511 | LAPTM5 | chr1 | - | 30742455 | 30742549 | 30741639 | 30741716 | 30757658 | 30757774 | 59694 | 2118,1299,3710,2392,107 | 134,31,43,17,6,21,61,58,9, | 2012,2286,1642,394,1012, | 58,25,47,39,23,8,18,96,2,1 | 243 | 149 | 0.000166904092316 | 0.0303147128213207 | 0.906,0.963,0.981, | 0.955,0.982,0.955, | 0,011 |
| 204020 | ENSG00000144535 | DIS3L2 | chr2 | + | 232015513 | 232015821 | 232014834 | 232014979 | 232024276 | 232024330 | 204020 | 99,63,62,40,24,51,20,30,5 | 3,1,4,3,4,3,3,0,4,3,1,1 | 46,38,38,55,42,18,19,24,2 | 1,0,0,2,0,0,0,0,1,1,0,0,3,2, | 298 | 149 | 0.000167692303014 | 0.0304321939882009 | 0.943,0.969,0.886, | 0.958,1.0,1.0,0.93 | -0,065 |
| 228024 | ENSG00000169871 | TRIM56 | chr7 | + | 101087071 | 101087168 | 101085521 | 101085571 | 101087311 | 101087508 | 228024 | 9,9,17,22,15,12,8,12,9,8,3, | 0,2,5,0,0,0,0,0,0,0,2,0 | 7,22,9,5,9,6,2,20,15,8,11,4 | 0,0,1,2,3,0,0,5,0,1,4,9,0,1, | 246 | 149 | 0.00016832635795 | 0.030521525145692 | 1.0,0.732,0.673,1. | 1.0,1.0,0.845,0.60 | 0,047 |
| 74932 | ENSG00000117280 | RAB29 | chr1 | - | 205774832 | 205775086 | 205772495 | 205772567 | 205775373 | 205775449 | 74932 | 15,15,18,23,17,7,5,10,15,1 | 0,1,3,0,2,1,1,0,0,0,0,0 | 11,2,11,5,12,5,4,21,0,24,6, | 2,0,1,0,4,0,0,4,2,2,3,3,0,0,2 | 298 | 149 | 0.00017016770143 | 0.0308294314364312 | 1.0,0.882,0.75,1.0, | 0.733,1.0,0.846,1. | 0,129 |
| 81752 | ENSG00000169499 | PLEKHA2 | chr8 | + | 38952165 | 38952312 | 38950849 | 38950990 | 38952635 | 38952704 | 81752 | 319,108,1065,681,405,391 | 24,2,26,16,9,13,3,0,7,4,6,5 | 617,343,217,211,155,216, | 13,14,14,19,5,4,7,7,2,2,26, | 296 | 149 | 0.000170728661468 | 0.0309050467085106 | 0.87,0.965,0.954,0 | 0.96,0.925,0.886,0 | 0,025 |
| 249058 | ENSG00000080546 | SESN1 | chr6 | - | 108998512 | 108998642 | 108994461 | 108994609 | 109000490 | 109000673 | 249058 | 39,10,100,42,11,31,59,307 | 0,0,2,0,0,2,1,9,2,0,1,1 | 49,62,64,31,11,11,31,23,1 | 4,0,3,1,0,0,0,2,1,0,24,1,0,0 | 279 | 149 | 0.000171307220274 | 0.030983717921675 | 1.0,1.0,0.964,1.0,1 | 0.867,1.0,0.919,0. | 0,027 |
| 249057 | ENSG00000080546 | SESN1 | chr6 | - | 108998512 | 108998637 | 108994461 | 108994609 | 109000490 | 109000673 | 249057 | 33,10,90,38,10,25,47,235, | 0,0,2,0,0,2,1,9,2,0,1,1 | 42,49,48,27,9,10,31,16,16, | 4,0,3,1,0,0,0,2,1,0,24,1,0,0 | 274 | 149 | 0.000172246948952 | 0.0311275256674122 | 1.0,1.0,0.961,1.0,1 | 0.851,1.0,0.897,0. | 0,022 |
| 99837 | ENSG00000130856 | ZNF236 | chr18 | + | 76947520 | 76947652 | 76937155 | 76937343 | 76959686 | 76959816 | 99837 | 11,9,6,4,10,6,5,3,3,9,1,7 | 0,0,1,2,2,2,0,1,0,0,0,4 | 7,7,3,8,8,2,13,13,3,8,18,10 | 0,2,0,0,1,2,5,5,1,0,0,3,0,2, | 281 | 149 | 0.000173239985967 | 0.03121466054667 | 1.0,1.0,0.761,0.51 | 1.0,0.65,1.0,1.0,0. | 0,033 |
| 49747 | ENSG00000159346 | ADIPOR1 | chr1 | - | 202954240 | 202954307 | 202950929 | 202951164 | 202958184 | 202958285 | 49747 | 3,3,10,3,1,2,6,2,0,1,2,1 | 15,4,13,8,3,14,9,5,13,3,3,1 | 8,6,6,9,11,1,2,5,4,8,8,9,0,4 | 6,9,12,6,2,3,6,3,4,2,4,23,6, | 216 | 149 | 0.000173268119106 | 0.03121466054667 | 0.121,0.341,0.347, | 0.479,0.315,0.256, | -0,191 |
| 81945 | ENSG00000140386 | SCAPER | chr15 | - | 76795279 | 76795440 | 76774854 | 76775117 | 76800247 | 76800364 | 81945 | 177,60,133,86,69,99,74,10 | 2,2,6,3,0,5,0,1,0,1,0,3 | 91,71,56,112,38,30,36,64, | 3,2,0,4,4,2,0,7,2,2,6,7,0,4,1 | 298 | 149 | 0.000173298807215 | 0.03121466054667 | 0.978,0.938,0.917, | 0.938,0.947,1.0,0. | 0,048 |
| 245433 | ENSG00000070047 | PHRF1 | chr11 | + | 581961 | 582387 | 581491 | 581606 | 587258 | 587464 | 245433 | 29,38,15,6,5,8,9,7,13,10,7, | 1,3,1,0,0,4,0,2,0,1,0,0 | 10,8,16,18,8,2,8,4,1,6,5,5, | 0,6,1,3,0,0,4,0,1,0,4,7,0,0, | 298 | 149 | 0.000173309232189 | 0.03121466054667 | 0.935,0.864,0.882, | 1.0,0.4,0.889,0.75, | 0,13 |

| | | | | | | | | | | | | | | | | | | | | | | |
|---|---|---|---|---|---|---|---|---|---|---|---|---|---|---|---|---|---|---|---|---|---|---|
| 61567 | ENSG00000048740 | CELF2 | chr10 | + | 11321188 | 11321386 | 11314138 | 11314270 | 11325835 | 11325979 | 61567 | 92,74,62,50,31,46,32,40,7 | 3,2,0,6,0,5,1,1,0,0,0,0 | 55,47,38,22,23,14,15,23,1 | 0,3,1,2,2,0,7,5,0,0,4,1,1,1, | 298 | 149 | 0.000174666522458 | 0.03143281797254 | 0.939,0.949,1.0,0. | 1.0,0.887,0.95,0.8 | 0,073 |
| 227543 | ENSG00000155849 | ELMO1 | chr7 | - | 37430700 | 37430802 | 37429256 | 37429756 | 37448674 | 37449186 | 227543 | 0,0,1,1,0,1,0,2,0,2,0,0 | 9,3,14,3,4,4,3,6,2,3,1,10 | 3,9,0,3,0,3,0,2,0,0,8,0,0,0, | 11,13,4,8,4,0,4,3,1,5,11,10 | 251 | 149 | 0.000175441513343 | 0.0315459084029467 | 0.0,0.0,0.041,0.16 | 0.139,0.291,0.0,0. | -0,099 |
| 199428 | ENSG00000144591 | GMPPA | chr2 | + | 219505231 | 219505362 | 219504082 | 219504213 | 219505457 | 219505555 | 199428 | 41,27,50,25,22,25,10,17,3 | 4,1,3,0,0,3,4,3,2,0,0,1 | 18,24,16,7,10,3,16,18,6,7, | 0,0,0,0,0,0,0,0,0,0,2,0,0, | 280 | 149 | 0.00017658232486 | 0.0317245328563795 | 0.845,0.935,0.899, | 1.0,1.0,1.0,1.0,1.0, | -0,098 |
| 227803 | ENSG00000184863 | RBM33 | chr7 | + | 155706861 | 155707068 | 155700890 | 155700944 | 155711202 | 155711455 | 227803 | 5,0,8,6,7,5,4,7,4,3,5,2 | 2,2,0,0,5,2,2,0,1,0,2,1 | 3,6,3,11,8,2,10,7,4,0,5,8,3, | 0,0,1,2,1,0,0,1,3,1,0,0,1,0, | 298 | 149 | 0.000176817991694 | 0.0317403779568735 | 0.556,0.0,1.0,1.0,0 | 1.0,1.0,0.6,0.733,0 | -0,136 |
| 227805 | ENSG00000184863 | RBM33 | chr7 | + | 155706868 | 155707068 | 155700890 | 155700944 | 155711202 | 155711455 | 227805 | 5,0,8,6,7,5,4,7,4,3,5,2 | 2,2,0,0,5,2,2,0,1,0,2,1 | 3,7,3,11,7,2,10,7,4,0,5,8,3, | 0,0,1,2,1,0,0,1,3,1,0,0,1,0, | 298 | 149 | 0.000177532387431 | 0.0318190692009159 | 0.556,0.0,1.0,1.0,0 | 1.0,1.0,0.6,0.733,0 | -0,135 |
| 133310 | ENSG00000182670 | TTC3 | chr21 | + | 37150820 | 37150884 | 37150077 | 37150170 | 37151892 | 37152029 | 133310 | 28,13,31,20,12,4,4,12,21,5 | 2,0,8,4,0,2,0,3,2,0,0,0 | 9,17,6,24,10,10,3,2,5,16,1 | 0,1,1,0,0,0,0,0,0,1,6,0,0,0, | 213 | 149 | 0.000177552035303 | 0.0318190692009159 | 0.907,1.0,0.731,0. | 1.0,0.922,0.808,1. | -0,064 |
| 163171 | ENSG00000157107 | FCHO2 | chr5 | + | 73076449 | 73076609 | 73074741 | 73074853 | 73077337 | 73077493 | 163171 | 1,0,1,0,0,0,2,1,0,0,2,0 | 13,6,43,20,10,21,11,23,11, | 0,0,3,5,0,0,0,0,4,3,8,0,3,1, | 28,15,9,28,16,5,20,8,11,16 | 298 | 149 | 0.000179223185261 | 0.0320918347644844 | 0.037,0.0,0.011,0. | 0.0,0.0,0.143,0.08 | -0,044 |
| 151547 | ENSG00000077238 | IL4R | chr16 | + | 27346448 | 27346618 | 27344907 | 27345020 | 27352539 | 27352696 | 151547 | 1042,307,1116,664,477,37 | 40,17,48,36,11,21,13,26,3 | 555,276,697,143,119,163,1 | 10,7,15,3,14,6,1,4,1,7,8,7,1 | 298 | 149 | 0.000179599347879 | 0.0321324582238945 | 0.929,0.9,0.921,0. | 0.965,0.952,0.959, | -0,011 |
| 97740 | ENSG00000196511 | TPK1 | chr7 | - | 144591422 | 144591569 | 144548490 | 144548602 | 144623165 | 144623261 | 97740 | 25,20,43,51,23,35,40,36,2 | 2,0,1,4,4,0,2,2,1,1,0,0 | 30,28,13,20,24,20,13,11,2 | 1,1,1,4,2,5,1,2,0,4,0,1,6,2, | 296 | 149 | 0.000179965136934 | 0.0321499220997128 | 0.863,1.0,0.956,0. | 0.938,0.934,0.867, | 0,077 |
| 149435 | ENSG00000273173 | SNURF | chr15 | + | 24967015 | 24967152 | 24962113 | 24962209 | 24967931 | 24968821 | 149435 | 4,0,88,110,1,4,2,12,5,12,1, | 126,81,198,89,72,86,83,10 | 4,3,2,0,24,17,1,6,0,10,2,3,1 | 161,170,110,16,59,53,58,1 | 286 | 149 | 0.00018003510366 | 0.0321499220997128 | 0.016,0.0,0.188,0. | 0.013,0.009,0.009, | 0,037 |
| 144297 | ENSG00000161526 | SAP30BP | chr17 | + | 75703534 | 75703705 | 75703310 | 75703371 | 75703807 | 75703859 | 144297 | 17,1,10,5,8,0,3,17,11,4,2,7 | 50,52,99,76,37,44,42,40,7 | 10,14,4,26,11,4,14,18,3,13 | 74,45,30,22,50,25,33,42,2 | 298 | 149 | 0.000180145081574 | 0.0321499220997128 | 0.145,0.01,0.048,0 | 0.063,0.135,0.062, | -0,05 |
| 88170 | ENSG00000144504 | ANKMY1 | chr2 | - | 240511860 | 240512001 | 240507559 | 240507691 | 240512801 | 240512942 | 88170 | 16,7,17,9,11,11,4,9,5,6,2,1 | 2,4,0,0,0,6,0,11,2,1,0,4 | 4,3,10,4,1,1,4,0,7,4,4,12,5, | 5,1,2,2,1,4,0,8,3,6,4,11,6,4 | 290 | 149 | 0.000181309953195 | 0.032331004586672 | 0.804,0.473,1.0,1. | 0.291,0.607,0.72,0 | 0,287 |
| 116653 | ENSG00000165699 | TSC1 | chr9 | - | 132903650 | 132903802 | 132902604 | 132902787 | 132904410 | 132904454 | 116653 | 21,15,29,8,12,22,19,16,8,1 | 0,1,0,0,0,4,2,0,3,1,0,0 | 25,19,33,28,8,13,10,12,12, | 6,5,1,0,0,1,0,5,0,2,2,0,3,0,1 | 298 | 149 | 0.000182111406457 | 0.0324470365257836 | 1.0,0.882,1.0,1.0,1 | 0.676,0.655,0.943, | 0,029 |
| 107253 | ENSG00000183354 | KIAA2026 | chr9 | - | 5954015 | 5954095 | 5944870 | 5944960 | 5968018 | 5968081 | 107253 | 44,41,113,54,24,24,44,32, | 2,3,4,2,1,5,4,2,0,0,1,0 | 66,48,34,45,31,38,20,40,1 | 0,1,2,1,3,1,0,0,1,0,0,0,0,0, | 229 | 149 | 0.000182594712694 | 0.032506238716164 | 0.935,0.899,0.948, | 1.0,0.969,0.917,0. | -0,038 |
| 127534 | ENSG00000107862 | GBF1 | chr10 | + | 102368738 | 102368832 | 102367477 | 102367560 | 102369210 | 102369387 | 127534 | 10,14,31,15,14,12,8,13,4,6 | 6,1,2,1,0,1,1,0,0,0,1,0 | 15,17,7,8,12,8,6,6,12,11,1 | 3,0,0,0,2,0,0,0,0,0,9,3,0,2, | 243 | 149 | 0.000183195906224 | 0.0325863124731386 | 0.505,0.896,0.905, | 0.754,1.0,1.0,1.0,0 | 0,066 |
| 77416 | ENSG00000115525 | ST3GAL5 | chr2 | - | 85851263 | 85851703 | 85847860 | 85848204 | 85852859 | 85853000 | 77416 | 16,5,31,19,23,14,10,12,7,1 | 0,1,2,0,2,2,0,0,5,0,0,0 | 11,12,6,13,3,5,7,27,9,8,19, | 0,1,0,4,2,1,0,0,8,7,1,0,2,0, | 298 | 149 | 0.000184250080398 | 0.032746762224725 | 1.0,0.714,0.886,1. | 1.0,0.857,1.0,0.61 | 0,111 |
| 212074 | ENSG00000079263 | SP140 | chr2 | + | 230255451 | 230255523 | 230248884 | 230248968 | 230269531 | 230269618 | 212074 | 44,39,42,29,11,14,16,15,3 | 0,0,5,2,1,0,0,0,0,2,0,3 | 19,17,22,19,20,21,6,15,14, | 2,1,0,1,2,0,0,1,2,4,4,4,3,6,1 | 221 | 149 | 0.000184518730139 | 0.0327674511605174 | 1.0,1.0,0.85,0.907, | 0.865,0.92,1.0,0.9 | 0,077 |
| 241717 | ENSG00000148296 | SURF6 | chr9 | - | 133333717 | 133333826 | 133332547 | 133332760 | 133334391 | 133334601 | 241717 | 18,27,21,7,11,5,3,1,19,3,1, | 2,0,3,0,0,0,2,1,0,2,1,0 | 8,11,18,1,9,5,5,5,9,10,2,6, | 4,1,1,5,1,0,0,0,0,0,3,4,0,2, | 258 | 149 | 0.000185744824767 | 0.0329529059610964 | 0.839,1.0,0.802,1. | 0.536,0.864,0.912, | 0,041 |
| 118816 | ENSG00000065923 | SLC9A7 | chrX | - | 46682335 | 46682535 | 46679677 | 46679755 | 46758704 | 46759055 | 118816 | 207,167,457,290,206,263, | 12,4,19,8,6,21,17,29,1,1,5, | 396,336,236,300,215,144,1 | 11,17,4,11,15,4,3,4,4,6,10, | 298 | 149 | 0.000185869265286 | 0.0329529059610964 | 0.896,0.954,0.923, | 0.947,0.908,0.967, | -0,026 |
| 33101 | ENSG00000281649 | EBLN3P | chr9 | + | 37084198 | 37084292 | 37079858 | 37080536 | 37086667 | 37087837 | 33101 | 31,15,55,18,12,10,13,24,1 | 125,93,210,126,63,54,94,1 | 4,11,7,3,7,3,9,10,5,6,18,34 | 156,143,74,65,73,69,35,83 | 243 | 149 | 0.000186074825839 | 0.0329621982223488 | 0.132,0.09,0.138,0 | 0.015,0.045,0.055, | 0,023 |
| 39119 | ENSG00000242086 | MUC20-OT1 | chr3 | + | 195663623 | 195663767 | 195662570 | 195662733 | 195664085 | 195664250 | 39119 | 11,6,0,0,0,0,9,11,6,1,0,0 | 33,8,34,19,15,11,6,7,18,10 | 0,0,1,3,7,7,0,0,1,0,2,1,0,0, | 29,43,9,44,17,7,34,30,12,3 | 293 | 149 | 0.000186265241971 | 0.0329687946502141 | 0.145,0.276,0.0,0. | 0.0,0.0,0.053,0.03 | 0,007 |
| 159857 | ENSG00000055163 | CYFIP2 | chr5 | + | 157360554 | 157360622 | 157360281 | 157360372 | 157361467 | 157361598 | 159857 | 12,5,11,10,2,4,15,20,1,1,4, | 122,110,191,163,145,106,1 | 6,10,8,13,9,8,26,14,12,14, | 121,97,77,85,105,94,64,15 | 217 | 149 | 0.00018646737278 | 0.0329774520220314 | 0.063,0.03,0.038,0 | 0.033,0.066,0.067, | -0,032 |
| 133380 | ENSG00000160305 | DIP2A | chr21 | + | 46562754 | 46562866 | 46561747 | 46561805 | 46563857 | 46563932 | 133380 | 2,0,1,0,1,2,0,1,3,0,0,0 | 3,1,17,10,2,5,2,6,7,2,2,2 | 1,2,3,3,1,1,2,2,2,3,0,0,2,2, | 2,2,1,0,0,4,3,9,1,10,1,8,2,3 | 261 | 149 | 0.00018736475172 | 0.0331089514593164 | 0.276,0.0,0.032,0. | 0.222,0.363,0.631, | -0,243 |
| 27117 | ENSG00000112234 | FBXL4 | chr6 | - | 98880552 | 98880624 | 98875414 | 98875727 | 98899267 | 98899481 | 27117 | 29,10,46,43,21,25,6,17,25, | 1,0,8,2,0,0,0,3,6,2,5,5 | 35,27,14,6,33,8,3,7,12,23,1 | 3,0,1,3,1,1,3,0,0,5,1,1,1,0, | 221 | 149 | 0.000188296293815 | 0.0332462671157475 | 0.951,1.0,0.795,0. | 0.887,1.0,0.904,0. | -0,016 |
| 63614 | ENSG00000117859 | OSBPL9 | chr1 | + | 51784263 | 51784327 | 51783914 | 51784025 | 51785807 | 51785886 | 63614 | 8,3,10,14,14,4,2,8,1,1,3,9 | 4,0,4,0,0,1,1,0,4,1,2,1 | 3,9,2,1,4,3,6,7,4,6,5,18,5,8 | 1,2,0,0,1,0,1,0,0,2,0,1,0,2, | 213 | 149 | 0.000190701134282 | 0.0336432752726632 | 0.583,1.0,0.636,1. | 0.677,0.759,1.0,1. | -0,189 |
| 226491 | ENSG00000171681 | ATF7IP | chr12 | + | 14436366 | 14436510 | 14436194 | 14436251 | 14438129 | 14438267 | 226491 | 13,4,8,10,3,1,0,8,9,2,3,3 | 379,252,496,271,86,125,1 | 7,4,26,21,5,6,2,3,7,10,5,8, | 197,173,324,254,158,171,1 | 293 | 149 | 0.000191063934534 | 0.0336796737868037 | 0.017,0.008,0.008, | 0.018,0.012,0.039, | -0,005 |
| 79836 | ENSG00000005483 | KMT2E | chr7 | + | 105062163 | 105062278 | 105040838 | 105041023 | 105063350 | 105063580 | 79836 | 464,336,500,297,206,281, | 4,2,2,1,0,1,1,1,0,0,3,0 | 381,486,271,443,339,159, | 7,4,3,24,4,0,2,2,5,2,0,1,6,0 | 264 | 149 | 0.000191386512591 | 0.0337089283890945 | 0.985,0.99,0.993,0 | 0.968,0.986,0.981, | 0,012 |
| 245685 | ENSG00000109919 | MTCH2 | chr11 | - | 47628952 | 47629023 | 47627079 | 47627127 | 47630554 | 47630614 | 245685 | 6,7,23,23,9,6,4,17,14,4,5,1 | 2,0,6,0,0,3,1,0,1,0,0,0 | 16,32,8,1,6,5,7,11,5,9,2,26 | 7,5,0,2,3,0,1,2,4,0,0,3,0,4, | 220 | 149 | 0.000191569479856 | 0.0337106885094409 | 0.67,1.0,0.722,1.0, | 0.608,0.813,1.0,0. | 0,077 |
| 80155 | ENSG00000170906 | NDUFA3 | chr19 | + | 54105596 | 54105703 | 54103113 | 54103188 | 54105933 | 54106011 | 80155 | 7,11,5,1,0,2,5,2,4,6,1,1 | 96,67,90,95,62,54,37,61,1 | 0,2,0,5,0,0,2,0,0,3,4,0,3,0, | 61,39,61,32,30,24,21,95,21 | 256 | 149 | 0.000191709757124 | 0.0337106885094409 | 0.041,0.087,0.031, | 0.0,0.029,0.0,0.08 | 0,014 |
| 190514 | ENSG00000009413 | REV3L | chr6 | - | 111364943 | 111365344 | 111363852 | 111363978 | 111367114 | 111367358 | 190514 | 18,17,36,24,9,21,10,12,7,1 | 0,1,1,1,0,2,1,0,0,2,3,1 | 16,26,8,12,19,7,7,13,11,17 | 2,4,0,6,0,1,5,1,1,0,2,5,0,1, | 298 | 149 | 0.000192404731747 | 0.0338048504004543 | 1.0,0.895,0.947,0. | 0.8,0.765,1.0,0.5,1 | 0,01 |
| 247659 | ENSG00000030066 | NUP160 | chr11 | - | 47812866 | 47813047 | 47812301 | 47812429 | 47813315 | 47813415 | 247659 | 20,19,29,15,17,11,13,22,6, | 0,0,6,4,1,2,0,0,0,0,0,0 | 18,26,10,29,16,9,13,7,9,17 | 0,0,0,8,1,0,0,3,0,1,0,7,4,2, | 298 | 149 | 0.00019255937384 | 0.0338048504004543 | 1.0,1.0,0.707,0.65 | 1.0,1.0,1.0,0.644,0 | 0,047 |
| 97450 | ENSG00000168014 | C2CD3 | chr11 | - | 74122987 | 74123135 | 74118227 | 74118382 | 74132843 | 74132972 | 97450 | 24,28,54,31,29,30,32,37,2 | 3,0,1,0,0,4,2,4,0,0,1,1 | 40,26,19,35,21,13,35,45,1 | 1,2,2,11,4,0,1,1,0,1,4,0,0,6 | 297 | 149 | 0.000192806874013 | 0.0338207141814931 | 0.801,1.0,0.964,1. | 0.953,0.867,0.827, | 0,051 |
| 85388 | ENSG00000272752 | STAG3L5P-PVR | chr7 | + | 100351251 | 100351410 | 100349730 | 100349887 | 100352162 | 100352211 | 85388 | 121,52,74,72,81,118,42,71 | 3,0,0,1,0,0,0,1,0,0,0,0 | 42,33,27,41,38,36,21,45,2 | 3,5,1,0,0,0,1,4,0,1,0,3,0,1, | 298 | 149 | 0.00019313830486 | 0.0338512626167123 | 0.953,1.0,1.0,0.97 | 0.875,0.767,0.931, | 0,063 |
| 12236 | ENSG00000264187 | | chr17 | - | 17231793 | 17231882 | 17227989 | 17228161 | 17232787 | 17232901 | 12236 | 24,6,40,18,16,20,9,8,32,8, | 4,0,3,0,2,0,2,0,7,2,0,2 | 22,16,18,10,7,7,17,20,2,14 | 1,0,0,0,2,1,0,0,0,1,2,1,0,3, | 238 | 149 | 0.000194963218687 | 0.034143310432239 | 0.79,1.0,0.893,1.0, | 0.932,1.0,1.0,1.0,0 | -0,042 |
| 195093 | ENSG00000170606 | HSPA4 | chr5 | + | 133091197 | 133091374 | 133089561 | 133089695 | 133092699 | 133092789 | 195093 | 195,184,249,155,107,238, | 4,3,3,0,0,2,14,23,0,0,0,0 | 127,179,157,118,118,74,1 | 0,1,2,2,0,0,0,5,1,0,2,0,0,7, | 298 | 149 | 0.000195122156029 | 0.03414336322299 | 0.961,0.968,0.976, | 1.0,0.989,0.975,0. | -0,011 |
| 90710 | ENSG00000082258 | CCNT2 | chr2 | + | 134945786 | 134945982 | 134942611 | 134942674 | 134946100 | 134946214 | 90710 | 4,2,14,15,2,7,5,7,2,0,3,6 | 26,20,33,28,13,15,8,26,13, | 0,7,0,2,0,5,4,7,0,2,11,4,0,3 | 34,45,37,16,30,6,13,17,30, | 298 | 149 | 0.000195489467058 | 0.0341554742052496 | 0.071,0.048,0.175, | 0.0,0.072,0.0,0.05 | 0,081 |
| 163888 | ENSG00000117091 | CD48 | chr1 | - | 160684767 | 160685189 | 160681201 | 160681468 | 160711681 | 160711803 | 163888 | 278,199,477,632,133,94,1 | 18,12,9,10,0,0,2,2,0,2,2,9 | 270,259,149,90,285,205,6 | 4,4,4,0,4,3,2,2,0,0,4,17,2,5 | 298 | 149 | 0.000195508752089 | 0.0341554742052496 | 0.885,0.892,0.964, | 0.971,0.97,0.949,1 | -0,017 |
| 177994 | ENSG00000142687 | KIAA0319L | chr1 | - | 35435304 | 35435385 | 35434397 | 35435081 | 35441046 | 35443028 | 177994 | 1,0,5,2,1,3,1,2,0,0,0,3 | 9,8,43,30,14,16,19,29,13,5 | 6,0,5,4,4,2,5,1,1,2,5,2,0,5, | 29,18,18,5,6,6,1,31,9,20,1 | 230 | 149 | 0.00019632585869 | 0.0342704062382055 | 0.067,0.0,0.07,0.0 | 0.118,0.0,0.153,0. | -0,118 |
| 255342 | ENSG00000069493 | CLEC2D | chr12 | + | 9693824 | 9694091 | 9692827 | 9692931 | 9694759 | 9699553 | 255342 | 6,0,16,2,0,2,3,8,12,0,3,2 | 105,71,181,91,47,38,51,85 | 3,2,3,1,1,3,2,6,1,2,2,0,6,3, | 159,99,49,47,53,41,30,51, | 298 | 149 | 0.0001965503044 | 0.0342817816582791 | 0.028,0.0,0.042,0. | 0.009,0.01,0.03,0. | 0,008 |
| 58679 | ENSG00000114098 | ARMC8 | chr3 | + | 138237308 | 138237384 | 138235033 | 138235114 | 138237481 | 138237572 | 58679 | 54,35,70,37,26,27,20,27,1 | 1,2,3,3,4,6,2,4,2,4,1,4 | 36,34,24,14,29,13,23,24,1 | 6,13,1,9,6,2,4,3,3,7,8,4,7,3 | 225 | 149 | 0.000196747894273 | 0.0342884583257264 | 0.973,0.921,0.939, | 0.799,0.634,0.941, | 0,101 |
| 145016 | ENSG00000108984 | MAP2K6 | chr17 | + | 69516985 | 69517297 | 69516854 | 69516903 | 69517499 | 69517613 | 145016 | 9,4,7,6,1,2,1,0,4,0,0,0 | 47,53,130,54,17,24,21,40, | 1,0,0,0,0,0,0,0,4,2,0,0,3, | 13,15,34,9,5,8,8,37,11,31,1 | 298 | 149 | 0.000197271958274 | 0.0343519747987632 | 0.087,0.036,0.026, | 0.037,0.0,0.0,0.0,0 | 0,017 |
| 226656 | ENSG00000058091 | CDK14 | chr7 | + | 90908400 | 90908490 | 90899290 | 90899353 | 90917600 | 90917724 | 226656 | 74,25,45,40,15,16,23,53,2 | 730,461,657,466,203,337, | 45,67,18,22,53,15,13,15,5 | 567,453,220,231,330,261,1 | 239 | 149 | 0.000197818715484 | 0.0344058048812449 | 0.059,0.033,0.041, | 0.047,0.084,0.049, | -0,014 |
| 223320 | ENSG00000064419 | TNPO3 | chr7 | - | 129005015 | 129005176 | 129001058 | 129001234 | 129014978 | 129015135 | 223320 | 34,27,113,50,39,48,16,34, | 1,1,4,1,1,4,0,0,1,0,0,4 | 58,55,23,29,54,41,17,27,3 | 1,3,1,6,4,0,4,1,0,1,13,2,0,0 | 298 | 149 | 0.000197900797018 | 0.0344058048812449 | 0.944,0.931,0.934, | 0.967,0.902,0.92,0 | 0,081 |
| 138032 | ENSG00000198521 | ZNF43 | chr19 | - | 21819094 | 21819407 | 21817887 | 21817986 | 21851904 | 21852125 | 138032 | 24,30,40,36,7,17,11,21,16, | 3,0,1,1,1,0,0,0,1,0,0,1 | 30,24,14,10,8,5,11,25,8,16 | 2,3,0,4,4,0,2,1,1,0,3,1,0,3,1 | 298 | 149 | 0.00019827436443 | 0.0344429295646758 | 0.8,1.0,0.952,0.94 | 0.882,0.8,1.0,0.55 | 0,092 |
| 97732 | ENSG00000116679 | IVNS1ABP | chr1 | - | 185306484 | 185306625 | 185305535 | 185305643 | 185307013 | 185307139 | 97732 | 9,12,22,7,10,7,5,11,6,0,2,4 | 9,10,10,20,4,12,0,6,4,3,5,4 | 15,17,12,14,16,8,12,6,12,1 | 13,1,1,0,2,4,9,5,3,14,4,13, | 290 | 149 | 0.000199054163904 | 0.0345505054445337 | 0.339,0.381,0.531, | 0.372,0.897,0.86,1 | -0,287 |
| 46557 | ENSG00000174738 | NR1D2 | chr3 | + | 23967812 | 23968023 | 23964976 | 23965162 | 23977222 | 23977543 | 46557 | 68,94,135,36,24,32,37,71, | 2,1,9,4,5,7,6,9,2,3,9,7 | 62,108,42,86,42,33,33,35, | 14,31,18,11,18,14,4,6,9,11 | 298 | 149 | 0.000200121487573 | 0.0347077742883355 | 0.944,0.979,0.882, | 0.689,0.635,0.538, | 0,14 |
| 195707 | ENSG00000177556 | ATOX1 | chr5 | - | 151751703 | 151751776 | 151742315 | 151746449 | 151758545 | 151758779 | 195707 | 24,13,98,73,25,32,23,22,2 | 2,1,4,0,4,6,0,1,2,1,0,2 | 42,58,31,11,15,26,15,54,8, | 3,0,2,0,1,0,0,2,1,2,0,0,0,2, | 222 | 149 | 0.000201032554166 | 0.0348377114860727 | 0.89,0.897,0.943,1 | 0.904,1.0,0.912,1. | -0,047 |
| 251839 | ENSG00000163399 | ATP1A1 | chr1 | + | 116382508 | 116382593 | 116373268 | 116373523 | 116384013 | 116384124 | 251839 | 4,1,4,2,1,4,0,1,0,0,0,1 | 79,33,42,37,15,4,12,21,12, | 5,6,3,3,6,1,3,3,2,3,4,2,1,1, | 17,33,15,18,5,11,8,30,17,3 | 234 | 149 | 0.000201264649185 | 0.0348498726538509 | 0.031,0.019,0.057, | 0.158,0.104,0.113, | -0,05 |
| 227943 | ENSG00000173715 | C11orf80 | chr11 | + | 66801030 | 66801153 | 66800640 | 66800696 | 66803994 | 66804194 | 227943 | 42,12,80,57,36,41,22,37,1 | 1,0,5,0,0,3,2,5,0,0,0,1 | 62,36,22,16,20,23,15,35,1 | 1,6,2,1,0,0,2,2,1,5,3,0,1,0, | 272 | 149 | 0.000201850115229 | 0.0349231528543834 | 0.958,1.0,0.898,1. | 0.971,0.767,0.858, | 0,063 |
| 185345 | ENSG00000188822 | CNR2 | chr1 | - | 23883206 | 23883444 | 23870514 | 23875662 | 23913245 | 23913362 | 185345 | 11,6,14,2,1,0,6,7,3,2,5,1 | 72,36,191,152,96,74,66,90 | 0,3,2,2,0,3,2,4,2,8,3,3,0,3, | 135,99,48,64,73,71,22,84, | 298 | 149 | 0.000202306272297 | 0.0349739608777153 | 0.071,0.077,0.035, | 0.0,0.015,0.02,0.0 | 0,017 |
| 244028 | ENSG00000151465 | CDC123 | chr10 | + | 12246148 | 12246273 | 12238456 | 12238485 | 12249580 | 12250319 | 244028 | 37,37,103,65,29,39,26,33, | 2,2,3,3,0,2,3,0,1,0,0,0 | 69,72,21,14,51,24,14,11,1 | 1,8,1,4,4,0,0,3,2,0,2,4,3,0, | 274 | 149 | 0.00020313697077 | 0.0350893847157286 | 0.91,0.91,0.949,0. | 0.974,0.83,0.919,0 | 0,065 |
| 195091 | ENSG00000170606 | HSPA4 | chr5 | + | 133091184 | 133091374 | 133089561 | 133089695 | 133092699 | 133092789 | 195091 | 195,184,250,155,107,238, | 4,3,3,0,0,2,14,23,0,0,0,0 | 127,179,156,118,118,74,1 | 0,1,2,2,0,0,0,5,1,0,2,0,0,7, | 298 | 149 | 0.000204920671799 | 0.0353458469395509 | 0.961,0.968,0.977, | 1.0,0.989,0.975,0. | -0,012 |
| 149412 | ENSG00000184056 | VPS33B | chr15 | - | 91016835 | 91017024 | 91014383 | 91014433 | 91017804 | 91017885 | 149412 | 4,7,9,10,9,6,6,7,3,4,5,10 | 0,0,2,2,0,0,0,0,0,0,0,1 | 8,12,8,0,2,10,4,10,3,4,7,4,1 | 0,2,2,3,5,0,2,0,0,0,2,2,1,0,1 | 298 | 149 | 0.000204950109327 | 0.0353458469395509 | 1.0,1.0,0.692,0.71 | 1.0,0.75,0.667,0.0, | 0,203 |
| 195090 | ENSG00000170606 | HSPA4 | chr5 | + | 133091158 | 133091374 | 133089561 | 133089695 | 133092699 | 133092789 | 195090 | 195,184,249,155,107,238, | 4,3,3,0,0,2,14,23,0,0,0,0 | 127,179,156,118,119,74,1 | 0,1,2,2,0,0,0,5,1,0,2,0,0,7, | 298 | 149 | 0.000205116828327 | 0.0353462770837858 | 0.961,0.968,0.976, | 1.0,0.989,0.975,0. | -0,012 |
| 69455 | ENSG00000078140 | UBE2K | chr4 | + | 39777681 | 39777810 | 39774833 | 39774933 | 39778359 | 39778384 | 69455 | 41,29,94,66,37,33,19,45,2 | 4,0,4,2,0,2,0,1,2,0,0,0 | 69,105,38,16,49,35,20,42, | 5,5,1,8,0,1,4,3,0,0,0,2,2,1, | 278 | 149 | 0.00020528200195 | 0.0353464404493604 | 0.846,1.0,0.926,0. | 0.881,0.918,0.953, | 0,051 |
| 88089 | ENSG00000168214 | RBPJ | chr4 | + | 26362445 | 26362628 | 26320786 | 26320815 | 26386352 | 26386391 | 88089 | 43,34,37,16,18,15,20,38,3 | 0,2,0,4,2,4,2,10,0,2,0,2 | 23,23,21,21,12,4,8,5,7,13, | 0,1,0,1,5,0,1,1,0,0,4,2,0,0, | 298 | 149 | 0.000205472295457 | 0.0353509253585176 | 1.0,0.895,1.0,0.66 | 1.0,0.92,1.0,0.913, | -0,065 |
| 116297 | ENSG00000188283 | ZNF383 | chr19 | + | 37224817 | 37224939 | 37219598 | 37219749 | 37230408 | 37230462 | 116297 | 8,16,27,11,7,5,1,12,3,6,6,8 | 0,1,2,3,0,2,0,2,0,2,1,0 | 24,26,11,11,30,6,7,5,8,16,1 | 0,11,1,3,6,0,1,2,0,0,1,0,0,1 | 271 | 149 | 0.000205787399138 | 0.0353768591883953 | 1.0,0.898,0.881,0. | 1.0,0.565,0.858,0. | 0,013 |
| 174121 | ENSG00000160796 | NBEAL2 | chr3 | + | 46991218 | 46991675 | 46989510 | 46989593 | 46991839 | 46991946 | 174121 | 4,6,8,6,9,9,6,4,4,3,1,6 | 2,0,3,0,2,1,4,5,0,1,0,0 | 4,10,4,8,1,4,3,12,3,6,2,11, | 0,0,0,0,2,0,2,1,0,0,0,3,1,0, | 298 | 149 | 0.000207421612964 | 0.035584599524884 | 0.5,1.0,0.571,1.0,0 | 1.0,1.0,1.0,1.0,0.2, | -0,076 |
| 136054 | ENSG00000104856 | RELB | chr19 | + | 45032509 | 45032749 | 45028887 | 45028992 | 45034243 | 45034312 | 136054 | 34,38,74,36,55,39,18,29,5 | 0,9,3,0,0,6,0,1,2,1,0,5 | 24,47,24,34,20,6,36,51,7,7 | 0,0,4,1,1,0,0,0,0,0,2,6,1,1, | 298 | 149 | 0.00020747022543 | 0.035584599524884 | 1.0,0.679,0.925,1. | 1.0,1.0,0.75,0.944, | -0,011 |
| 137661 | ENSG00000104852 | SNRNP70 | chr19 | + | 49103507 | 49103587 | 49102113 | 49102185 | 49104633 | 49104735 | 137661 | 18,6,29,22,5,11,7,4,3,2,9,9 | 14,5,3,3,3,6,4,6,7,4,3,0 | 15,8,4,10,15,10,14,9,9,5,1 | 1,1,3,1,8,3,0,4,0,0,0,3,3,0,1 | 229 | 149 | 0.000207491822292 | 0.035584599524884 | 0.456,0.438,0.863, | 0.907,0.839,0.465, | -0,238 |
| 136057 | ENSG00000104856 | RELB | chr19 | + | 45032555 | 45032749 | 45028887 | 45028992 | 45034243 | 45034312 | 136057 | 34,38,74,36,53,40,19,29,5 | 0,9,3,0,0,6,0,1,2,1,0,5 | 24,47,24,34,20,6,36,50,7,7 | 0,0,4,1,1,0,0,0,0,0,2,6,1,1, | 298 | 149 | 0.000208796962084 | 0.0357799195432336 | 1.0,0.679,0.925,1. | 1.0,1.0,0.75,0.944, | -0,01 |
| 62475 | ENSG00000130939 | UBE4B | chr1 | + | 10158355 | 10158482 | 10151325 | 10151561 | 10161141 | 10161286 | 62475 | 30,24,58,40,26,19,18,25,2 | 5,0,7,7,1,5,3,3,3,2,0,1 | 30,16,19,15,19,14,16,24,7, | 1,0,2,3,2,0,6,3,2,1,0,0,2,0,0 | 276 | 149 | 0.000209111985478 | 0.0358053951841014 | 0.764,1.0,0.817,0. | 0.942,1.0,0.837,0. | -0,054 |
| 136059 | ENSG00000104856 | RELB | chr19 | + | 45032558 | 45032749 | 45028887 | 45028992 | 45034243 | 45034312 | 136059 | 34,39,75,36,53,39,18,29,5 | 0,9,3,0,0,6,0,1,2,1,0,5 | 24,47,24,34,20,6,36,48,7,7 | 0,0,4,1,1,0,0,0,0,0,2,6,1,1, | 298 | 149 | 0.000209687390766 | 0.0358459830260356 | 1.0,0.684,0.926,1. | 1.0,1.0,0.75,0.944, | -0,009 |
| 40609 | ENSG00000115524 | SF3B1 | chr2 | - | 197392971 | 197393188 | 197388514 | 197392461 | 197396055 | 197396328 | 40609 | 998,694,2354,1375,1033,1 | 13,1,7,20,15,4,2,1,4,2,2,1 | 1464,1801,837,551,938,66 | 12,7,10,15,9,8,4,10,13,11,1 | 298 | 149 | 0.000209813342176 | 0.0358459830260356 | 0.975,0.997,0.994, | 0.984,0.992,0.977, | 0,009 |
| 32159 | ENSG00000182489 | XKRX | chrX | - | 100922792 | 100923061 | 100913444 | 100915083 | 100927969 | 100928917 | 32159 | 19,26,41,16,17,8,33,22,28, | 1,0,3,1,0,1,2,0,0,0,0,2 | 13,21,13,3,9,13,4,8,19,26, | 3,3,0,0,0,1,0,0,2,4,4,0,2,3, | 298 | 149 | 0.00020984866777 | 0.0358459830260356 | 0.905,1.0,0.872,0. | 0.684,0.778,1.0,1. | 0,084 |
| 97810 | ENSG00000102390 | PBDC1 | chrX | + | 76176880 | 76176992 | 76175472 | 76175613 | 76177615 | 76178314 | 97810 | 37,41,60,43,16,36,26,22,3 | 3,1,1,0,1,0,0,6,3,0,1,1 | 19,35,21,35,25,21,10,25,1 | 5,2,6,1,0,0,0,0,3,0,0,1,6,3 | 261 | 149 | 0.000211126040493 | 0.0360355819360419 | 0.876,0.959,0.972, | 0.684,0.909,0.666, | 0,019 |
| 204679 | ENSG00000147274 | RBMX | chrX | - | 136873245 | 136873326 | 136872267 | 136872345 | 136875085 | 136875168 | 204679 | 47,32,93,44,31,26,33,43,4 | 6,2,25,18,11,6,9,10,8,6,2,1 | 35,39,32,23,20,9,15,27,9,4 | 17,42,12,13,33,12,5,4,17,2 | 230 | 149 | 0.000211472066748 | 0.0360647034929328 | 0.835,0.912,0.707, | 0.572,0.376,0.633, | 0,176 |
| 204814 | ENSG00000138073 | PREB | chr2 | - | 27132229 | 27132339 | 27131671 | 27132082 | 27132602 | 27132727 | 204814 | 7,5,17,17,9,9,9,4,6,3,5,7 | 0,2,1,0,0,2,0,1,0,1,0,0 | 24,14,8,1,4,2,12,15,1,3,10, | 2,1,2,0,2,0,2,3,0,2,0,2,0,4, | 259 | 149 | 0.000211631784044 | 0.0360647034929328 | 1.0,0.59,0.907,1.0, | 0.873,0.89,0.697,1 | 0,102 |
| 160017 | ENSG00000112624 | BICRAL | chr6 | + | 42822433 | 42822696 | 42822017 | 42822063 | 42822795 | 42822844 | 160017 | 1,1,3,4,0,1,5,4,1,2,0,4 | 19,13,48,21,17,19,38,29,1 | 3,2,2,4,0,4,6,0,4,5,7,8,1,3, | 22,22,10,4,11,16,6,10,11,2 | 298 | 149 | 0.000211853229318 | 0.0360738784804924 | 0.026,0.037,0.03,0 | 0.064,0.043,0.091, | -0,065 |
| 136061 | ENSG00000104856 | RELB | chr19 | + | 45032572 | 45032749 | 45028887 | 45028992 | 45034243 | 45034312 | 136061 | 34,38,74,36,52,39,18,29,5 | 0,9,3,0,0,6,0,1,2,1,0,5 | 24,47,24,34,20,6,36,46,7,7 | 0,0,4,1,1,0,0,0,0,0,2,6,1,1, | 298 | 149 | 0.000212331780679 | 0.0361148670347504 | 1.0,0.679,0.925,1. | 1.0,1.0,0.75,0.944, | -0,011 |
| 94547 | ENSG00000231925 | TAPBP | chr6 | - | 33304127 | 33304217 | 33299860 | 33301771 | 33304296 | 33304530 | 94547 | 28,36,116,68,48,62,74,33, | 0,0,8,3,0,0,0,2,0,0,0,2 | 65,98,29,51,32,31,41,80,4 | 0,2,0,1,4,4,2,5,0,1,0,14,3,0 | 239 | 149 | 0.000212429536944 | 0.0361148670347504 | 1.0,1.0,0.9,0.934,1 | 1.0,0.968,1.0,0.97, | 0,032 |
| 89018 | ENSG00000115392 | FANCL | chr2 | - | 58226727 | 58226784 | 58220593 | 58222042 | 58229813 | 58229874 | 89018 | 17,16,130,92,20,45,16,57, | 0,0,0,0,0,0,0,0,0,0,0,0 | 107,96,50,42,41,37,22,77, | 0,1,0,0,1,0,7,11,3,7,0,0,0,0 | 206 | 149 | 0.000212928714081 | 0.0361711602686407 | 1.0,1.0,1.0,1.0,1.0, | 1.0,0.986,1.0,1.0,0 | 0,075 |
| 187733 | ENSG00000163257 | DCAF16 | chr4 | - | 17805106 | 17805233 | 17800654 | 17804771 | 17810446 | 17810758 | 187733 | 33,20,77,31,23,7,22,56,41, | 1,5,5,5,3,3,4,3,13,7,1,11 | 32,37,38,46,57,31,22,46,3, | 3,2,2,6,3,2,1,6,0,2,6,1,1,2,1 | 276 | 149 | 0.000213189383946 | 0.036179741691857 | 0.947,0.683,0.893, | 0.852,0.909,0.911, | -0,153 |
| 36658 | ENSG00000016864 | GLT8D1 | chr3 | - | 52704723 | 52704952 | 52700444 | 52700496 | 52705446 | 52705764 | 36658 | 3,1,19,27,16,1,2,7,3,6,3,7 | 0,3,7,4,2,9,4,5,0,0,2,1 | 3,10,4,1,5,2,1,2,3,3,8,8,12, | 2,4,0,0,1,0,0,5,0,3,0,2,0,0, | 298 | 149 | 0.000213315424855 | 0.036179741691857 | 1.0,0.143,0.576,0. | 0.429,0.556,1.0,1. | -0,186 |
| 176151 | ENSG00000169180 | XPO6 | chr16 | - | 28111816 | 28111940 | 28107521 | 28107677 | 28112903 | 28113050 | 176151 | 66,32,75,81,40,37,32,30,2 | 7,4,12,1,2,0,4,4,11,3,2,6 | 57,59,48,30,41,20,19,36,4 | 1,0,0,6,4,1,5,1,2,0,0,6,1,2,1 | 273 | 149 | 0.000213891358131 | 0.0362270185988868 | 0.837,0.814,0.773, | 0.969,1.0,1.0,0.73 | -0,083 |
| 198007 | ENSG00000154305 | MIA3 | chr1 | + | 222657921 | 222657972 | 222654654 | 222654793 | 222658721 | 222658823 | 198007 | 25,11,27,11,9,12,2,5,6,9,3, | 86,79,101,54,36,34,25,39, | 29,29,15,22,8,6,4,11,7,12, | 38,34,51,30,36,30,15,27,21 | 200 | 149 | 0.000213930802901 | 0.0362270185988868 | 0.178,0.094,0.166, | 0.362,0.389,0.18,0 | -0,087 |
| 252948 | ENSG00000111647 | UHRF1BP1L | chr12 | - | 100099641 | 100099900 | 100098351 | 100098541 | 100102765 | 100102857 | 252948 | 0,0,1,5,0,3,0,0,1,2,2,0 | 7,12,52,18,9,11,9,14,10,7, | 2,1,0,2,5,7,3,4,3,2,3,1,0,5, | 23,28,12,17,17,7,7,5,12,7,1 | 298 | 149 | 0.000215939622636 | 0.036509972690025 | 0.0,0.0,0.01,0.122, | 0.042,0.018,0.0,0. | -0,05 |
| 32765 | ENSG00000223865 | HLA-DPB1 | chr6 | + | 33080688 | 33080935 | 33076009 | 33076141 | 33084949 | 33085231 | 32765 | 525,357,201,235,141,55,3 | 6,3,15,6,0,0,1,3,1,2,3,0 | 363,398,276,255,269,174,1 | 6,23,5,25,1,1,1,8,5,2,3,19, | 298 | 149 | 0.000215940990073 | 0.036509972690025 | 0.978,0.983,0.87,0 | 0.968,0.896,0.965, | 0,013 |
| 106221 | ENSG00000172534 | HCFC1 | chrX | - | 153956624 | 153956763 | 153956190 | 153956411 | 153956917 | 153957060 | 106221 | 38,18,42,23,33,24,33,16,5 | 6,3,5,1,0,6,0,1,2,0,2,1 | 16,16,17,23,8,14,35,26,7,2 | 5,4,3,5,4,3,4,7,6,2,0,7,3,3, | 288 | 149 | 0.000216316615891 | 0.0365447735909229 | 0.766,0.756,0.813, | 0.623,0.674,0.746, | 0,154 |
| 213424 | ENSG00000035720 | STAP1 | chr4 | + | 67576679 | 67576943 | 67575384 | 67575498 | 67577202 | 67577259 | 213424 | 0,2,26,5,0,7,0,0,1,0,4,7 | 95,43,180,138,78,65,66,55 | 15,14,9,6,0,0,5,7,0,0,0,0,2 | 83,59,97,34,67,53,15,55,4 | 298 | 149 | 0.000216897678214 | 0.0366141993566097 | 0.0,0.023,0.067,0. | 0.083,0.106,0.044, | -0,018 |

| | | | | | | | | | | | | | | | | | | | | | |
|---|---|---|---|---|---|---|---|---|---|---|---|---|---|---|---|---|---|---|---|---|---|
| 18572 | ENSG00000137871 | ZNF280D | chr15 | - | 56707081 | 56707143 | 56704120 | 56704267 | 56707262 | 56707306 | 18572 | 42,17,63,45,10,10,18,25,1 | 3,3,9,1,0,4,1,2,0,0,0,1 | 30,18,15,16,17,14,8,12,24, | 1,0,1,0,0,0,0,0,2,9,1,4,0,1, | 211 | 149 | 0.000217977958166 | 0.0367677225031554 | 0.908,0.8,0.832,0. | 0.955,1.0,0.914,1. | -0,016 |
| 137574 | ENSG00000104980 | TIMM44 | chr19 | - | 7931678 | 7931746 | 7931137 | 7931188 | 7932626 | 7932751 | 137574 | 4,0,1,0,1,1,1,0,0,0,0,2 | 8,2,18,22,13,3,3,13,7,4,3,1 | 6,0,7,2,3,2,3,5,1,0,0,0,0,1,2 | 12,16,9,6,8,3,5,8,2,21,7,21 | 217 | 149 | 0.000218199059493 | 0.0367761955941565 | 0.256,0.0,0.037,0. | 0.256,0.0,0.348,0. | -0,078 |
| 115394 | ENSG00000184465 | WDR27 | chr6 | - | 169694462 | 169695065 | 169689126 | 169689308 | 169701550 | 169701854 | 115394 | 1,0,13,2,0,6,2,9,0,2,4,4 | 8,9,26,15,6,22,8,1,6,7,11,8 | 2,2,1,2,0,1,3,1,2,1,1,1,0,0, | 4,3,7,3,6,11,4,12,6,14,14,3 | 298 | 149 | 0.000219571540544 | 0.0369785620053409 | 0.059,0.0,0.2,0.06 | 0.2,0.25,0.067,0.2 | 0,051 |
| 22506 | ENSG00000072042 | RDH11 | chr14 | - | 67685014 | 67685204 | 67676799 | 67678423 | 67690211 | 67690421 | 22506 | 32,11,39,60,39,45,36,39,3 | 1,3,5,2,0,3,2,4,2,0,0,2 | 39,16,20,3,10,16,10,67,9,5 | 2,0,2,0,3,0,1,6,0,2,2,0,1,0, | 298 | 149 | 0.000220137846175 | 0.036995179750826 | 0.941,0.647,0.796, | 0.907,1.0,0.833,1. | -0,021 |
| 137217 | ENSG00000130303 | BST2 | chr19 | - | 17404370 | 17404409 | 17404128 | 17404189 | 17405290 | 17405630 | 137217 | 2,2,41,27,5,6,5,4,0,9,4,5 | 0,0,5,1,0,0,0,2,2,1,0,2 | 13,12,1,4,3,9,2,8,1,5,7,16, | 4,3,1,3,0,0,0,3,0,0,3,6,2,1,1 | 188 | 149 | 0.000220226908945 | 0.036995179750826 | 1.0,1.0,0.867,0.9 | 0.72,0.76,0.442,0. | 0,028 |
| 91222 | ENSG00000134440 | NARS1 | chr18 | - | 57620568 | 57620651 | 57615640 | 57615975 | 57621707 | 57621776 | 91222 | 133,107,143,86,44,50,43,4 | 4,5,5,3,0,1,0,0,0,1,0,0 | 76,83,72,53,99,52,26,48,3 | 3,3,2,12,8,3,0,2,0,0,1,4,1,3 | 232 | 149 | 0.000220306695702 | 0.036995179750826 | 0.955,0.932,0.948, | 0.942,0.947,0.959, | 0,036 |
| 42661 | ENSG00000163349 | HIPK1 | chr1 | + | 113957123 | 113957286 | 113956626 | 113956811 | 113958065 | 113958291 | 42661 | 38,34,57,47,33,23,31,36,4 | 0,2,1,3,0,0,4,6,0,0,0,1 | 43,40,20,56,42,25,50,83,2 | 3,7,2,0,7,1,4,8,3,2,0,1,2,6, | 298 | 149 | 0.000220357757203 | 0.036995179750826 | 1.0,0.895,0.966,0. | 0.878,0.741,0.833, | 0,064 |
| 15806 | ENSG00000188559 | RALGAPA2 | chr20 | - | 20637362 | 20637501 | 20635350 | 20635617 | 20639784 | 20639900 | 15806 | 15,13,48,28,44,27,16,42,1 | 4,0,4,1,2,1,1,9,1,0,3,4 | 28,15,15,7,9,9,11,10,17,26 | 4,8,4,3,3,1,6,3,9,0,8,10,4,1 | 288 | 149 | 0.000220902517702 | 0.0370577317127975 | 0.66,1.0,0.861,0.9 | 0.784,0.492,0.66,0 | 0,193 |
| 76306 | ENSG00000076242 | MLH1 | chr3 | + | 37017505 | 37017599 | 37012010 | 37012099 | 37025636 | 37026007 | 76306 | 7,2,5,3,8,5,2,2,2,2,2,1 | 0,2,3,0,1,1,0,3,0,1,1,0 | 1,1,5,1,4,0,4,2,2,0,3,1,4,2,2 | 0,3,3,0,4,2,4,0,2,1,2,2,1,0,2 | 243 | 149 | 0.000221387898002 | 0.0371102326128259 | 1.0,0.38,0.505,1.0, | 1.0,0.17,0.505,1.0, | 0,2 |
| 228193 | ENSG00000291136 | | chr7 | - | 66551519 | 66551849 | 66545600 | 66545686 | 66554584 | 66554837 | 228193 | 0,1,2,1,2,2,1,1,0,1,0,0 | 6,0,16,13,6,7,12,10,3,3,7,8 | 2,0,1,0,0,5,2,5,2,0,2,2,4,3,4 | 9,2,0,5,12,6,2,5,3,10,8,9,2, | 298 | 149 | 0.000222216057583 | 0.0372200655950558 | 0.0,1.0,0.059,0.03 | 0.1,0.0,1.0,0.0,0.0, | -0,081 |
| 125162 | ENSG00000165322 | ARHGAP12 | chr10 | - | 31854065 | 31854206 | 31843460 | 31843586 | 31861394 | 31861658 | 125162 | 54,44,57,46,24,58,27,38,3 | 2,3,4,2,0,2,2,7,4,1,1,2 | 45,23,33,41,23,26,14,33,2 | 10,10,0,8,4,3,1,2,7,2,2,4,3, | 290 | 149 | 0.000223727853206 | 0.0374441443027843 | 0.933,0.883,0.88,0 | 0.698,0.542,1.0,0. | 0,113 |
| 136063 | ENSG00000104856 | RELB | chr19 | + | 45032592 | 45032749 | 45028887 | 45028992 | 45034243 | 45034312 | 136063 | 35,38,75,35,50,39,18,29,5 | 0,9,3,0,0,6,0,1,2,1,0,5 | 24,47,24,34,20,6,36,42,7,7 | 0,0,4,1,1,0,0,0,0,0,2,6,1,1,2 | 298 | 149 | 0.000224287177288 | 0.0375085885430253 | 1.0,0.679,0.926,1. | 1.0,1.0,0.75,0.944, | -0,011 |
| 45951 | ENSG00000180329 | CCDC43 | chr17 | - | 44680584 | 44680643 | 44677439 | 44679043 | 44682002 | 44682138 | 45951 | 19,36,46,23,11,7,12,11,13, | 4,2,5,4,4,2,0,0,1,0,2,0 | 11,20,18,22,20,6,16,21,8,1 | 0,1,3,1,2,0,0,0,0,0,3,5,0,0, | 208 | 149 | 0.000224485031527 | 0.0375125293638104 | 0.773,0.928,0.868, | 1.0,0.935,0.811,0. | -0,101 |
| 176634 | ENSG00000174738 | NR1D2 | chr3 | + | 23967778 | 23968023 | 23964976 | 23965162 | 23977222 | 23977543 | 176634 | 37,44,58,26,15,19,27,36,2 | 2,1,9,4,5,7,6,9,2,3,9,7 | 29,53,22,41,21,15,23,25,9, | 14,31,18,11,18,14,4,6,9,11 | 298 | 149 | 0.000224700833932 | 0.0375194609682066 | 0.902,0.957,0.763, | 0.509,0.461,0.379, | 0,201 |
| 9664 | ENSG00000178971 | CTC1 | chr17 | - | 8229890 | 8229922 | 8229141 | 8229446 | 8230293 | 8230348 | 9664 | 3,2,19,10,10,13,10,9,10,6,3 | 2,0,5,4,2,0,0,1,0,0,0,0 | 12,9,7,1,4,4,7,19,1,5,4,4,5, | 3,3,2,12,2,4,0,4,0,0,3,1,0,1 | 181 | 149 | 0.000225418705824 | 0.037610149979229 | 0.553,1.0,0.758,0. | 0.767,0.712,0.742, | 0,138 |
| 139069 | ENSG00000125772 | GPCPD1 | chr20 | - | 5599289 | 5599485 | 5598724 | 5598821 | 5604363 | 5604440 | 139069 | 9,3,15,4,11,10,8,11,2,1,0,5 | 94,47,157,104,69,94,80,10 | 8,16,10,12,29,2,6,6,24,6,9, | 92,87,48,54,93,43,54,70,1 | 298 | 149 | 0.000226084729101 | 0.0376920544764813 | 0.046,0.031,0.046, | 0.042,0.084,0.094, | -0,028 |
| 34688 | ENSG00000125089 | SH3TC1 | chr4 | + | 8209664 | 8209822 | 8205166 | 8205366 | 8212700 | 8212828 | 34688 | 9,21,13,11,9,11,18,8,21,17 | 2,0,1,2,1,2,3,0,1,0,2,5 | 12,19,3,8,12,7,9,14,12,18,7 | 2,1,0,0,0,0,0,1,0,0,4,2,0,0, | 298 | 149 | 0.000226612910632 | 0.0377508694800588 | 0.692,1.0,0.867,0. | 0.75,0.905,1.0,1.0, | -0,115 |
| 48977 | ENSG00000197548 | ATG7 | chr3 | + | 11332971 | 11333093 | 11315343 | 11315493 | 11340644 | 11340735 | 48977 | 11,0,18,27,8,9,3,11,6,6,2,9 | 0,2,4,3,0,2,4,1,2,1,0,0 | 7,9,3,13,1,10,10,2,5,7,7,12 | 0,1,2,0,1,2,0,0,2,0,0,2,0,0, | 271 | 149 | 0.000227162102337 | 0.0377877168340987 | 1.0,0.0,0.712,0.83 | 1.0,0.832,0.452,1. | -0,124 |
| 234342 | ENSG00000116754 | SRSF11 | chr1 | + | 70231094 | 70231566 | 70228421 | 70228555 | 70232267 | 70232377 | 234342 | 18,5,9,8,3,6,2,7,3,5,2,3 | 46,25,40,65,37,29,29,35,5 | 5,13,8,54,8,10,15,14,3,5,5, | 30,46,10,27,36,20,10,40,2 | 298 | 149 | 0.00022718523625 | 0.0377877168340987 | 0.164,0.091,0.101, | 0.077,0.124,0.286, | -0,083 |
| 235105 | ENSG00000063601 | MTMR1 | chrX | + | 150726409 | 150726649 | 150718676 | 150718700 | 150727214 | 150727309 | 235105 | 0,0,11,1,0,3,0,2,1,0,1,1 | 32,58,67,45,26,25,16,32,1 | 3,5,1,8,0,2,4,8,1,2,7,4,5,4,3 | 36,58,17,18,34,29,15,28,1 | 298 | 149 | 0.000227766088268 | 0.0378525826313236 | 0.0,0.0,0.076,0.01 | 0.04,0.041,0.029,0 | -0,037 |
| 246639 | ENSG00000054654 | SYNE2 | chr14 | + | 64098393 | 64098646 | 64097948 | 64098146 | 64098746 | 64098821 | 246639 | 0,3,4,1,1,2,3,7,2,0,0,0 | 110,108,108,92,57,130,84, | 3,4,5,14,3,0,2,6,3,4,1,1,5,4 | 41,85,46,61,87,68,26,82,1 | 298 | 149 | 0.000227926957967 | 0.0378525826313236 | 0.0,0.014,0.018,0. | 0.035,0.023,0.052, | -0,019 |
| 134133 | ENSG00000100101 | | chr22 | + | 37691924 | 37692026 | 37691173 | 37691424 | 37693331 | 37693410 | 134133 | 1,0,5,0,2,0,2,8,3,0,0,1 | 29,24,25,16,26,9,11,10,27, | 2,6,1,0,2,0,2,4,5,6,2,4,4,6,2 | 17,12,21,2,13,10,12,20,8,8 | 251 | 149 | 0.000228462789826 | 0.0379123166669544 | 0.02,0.0,0.106,0.0, | 0.065,0.229,0.027, | -0,056 |
| 218467 | ENSG00000124942 | AHNAK | chr11 | - | 62536217 | 62536567 | 62535944 | 62536098 | 62546659 | 62546796 | 218467 | 83,61,120,39,34,43,53,38, | 2,1,7,1,0,3,1,2,3,2,0,0 | 32,42,30,81,21,12,19,50,2 | 2,0,0,9,1,0,3,3,1,0,3,9,3,2, | 298 | 149 | 0.000228826376846 | 0.0379176549962577 | 0.954,0.968,0.896, | 0.889,1.0,1.0,0.81 | 0,068 |
| 63194 | ENSG00000174501 | ANKRD36C | chr2 | - | 95867439 | 95867581 | 95859860 | 95860074 | 95876438 | 95876509 | 63194 | 22,11,68,39,32,41,16,41,6, | 3,0,12,3,3,0,0,3,4,0,4,1 | 32,58,25,29,28,13,25,39,3 | 2,2,0,1,4,0,2,0,0,2,2,4,1,0, | 291 | 149 | 0.000228847302852 | 0.0379176549962577 | 0.79,1.0,0.744,0.8 | 0.891,0.937,1.0,0. | -0,062 |
| 206076 | ENSG00000101882 | NKAP | chrX | - | 119941564 | 119941756 | 119938729 | 119938810 | 119943219 | 119943751 | 206076 | 4,3,1,1,0,1,0,5,0,2,1,0 | 13,9,42,23,11,16,12,11,9,6 | 1,3,2,9,8,0,1,5,2,1,2,4,2,1, | 14,9,9,11,10,4,7,10,2,9,7,1 | 298 | 149 | 0.00022943204125 | 0.0379852982079067 | 0.133,0.143,0.012, | 0.034,0.143,0.1,0. | -0,061 |
| 232183 | ENSG00000256861 | | chr12 | - | 122209787 | 122209858 | 122207778 | 122208577 | 122216487 | 122216584 | 232183 | 11,2,20,12,5,10,6,5,3,1,2,8 | 25,15,40,32,19,12,18,15,3 | 14,22,16,12,19,5,13,23,9,9 | 9,21,21,10,22,7,16,14,7,14 | 220 | 149 | 0.000229658188868 | 0.0379935139494608 | 0.23,0.083,0.253,0 | 0.513,0.415,0.34,0 | -0,135 |
| 43804 | ENSG00000114867 | EIF4G1 | chr3 | + | 184317320 | 184317497 | 184316071 | 184316218 | 184317716 | 184317773 | 43804 | 21,15,23,45,12,11,3,8,14,1 | 3,4,5,2,0,0,1,0,2,0,1,2 | 13,5,8,8,5,10,1,4,4,18,12,1 | 0,2,4,0,2,0,1,1,0,2,0,3,1,2, | 298 | 149 | 0.000230256344248 | 0.0380632129253773 | 0.778,0.652,0.697, | 1.0,0.556,0.5,1.0,0 | 0,007 |
| 109523 | ENSG00000163428 | LRRC58 | chr3 | - | 120335824 | 120335953 | 120334861 | 120335139 | 120348743 | 120349354 | 109523 | 38,41,73,57,27,37,28,36,1 | 0,1,0,3,2,4,1,1,0,0,2,2 | 35,83,17,18,39,17,15,24,2 | 7,1,3,4,1,0,0,2,2,2,0,4,0,0, | 278 | 149 | 0.000230476817303 | 0.0380704189293492 | 1.0,0.956,1.0,0.91 | 0.728,0.978,0.752, | -0,008 |
| 8211 | ENSG00000074657 | ZNF532 | chr18 | + | 58931645 | 58931755 | 58920329 | 58920633 | 58934432 | 58934578 | 8211 | 6,5,2,0,0,1,0,2,0,0,0,0 | 59,80,22,5,12,21,11,5,7,8,2 | 0,4,2,11,2,0,0,0,2,1,0,6,5,3 | 19,5,24,51,9,4,3,5,1,5,2,35 | 259 | 149 | 0.00023145272589 | 0.0382023018757903 | 0.055,0.035,0.05,0 | 0.0,0.315,0.046,0. | -0,069 |
| 195094 | ENSG00000170606 | HSPA4 | chr5 | + | 133091238 | 133091374 | 133089561 | 133089695 | 133092699 | 133092789 | 195094 | 190,176,238,152,94,218,9 | 4,3,3,0,0,2,14,23,0,0,0,0 | 121,168,148,112,116,72,1 | 0,1,2,2,0,0,0,5,1,0,2,0,0,7, | 285 | 149 | 0.000231804464656 | 0.0382310396416671 | 0.961,0.968,0.976, | 1.0,0.989,0.975,0. | -0,013 |
| 221 | ENSG00000182670 | TTC3 | chr21 | + | 37090541 | 37090589 | 37090232 | 37090286 | 37091292 | 37091413 | 221 | 4,7,2,4,2,6,0,2,8,4,2,3 | 48,17,39,36,25,17,19,14,1 | 2,8,3,2,2,2,5,0,3,0,2,0,1,1,2 | 44,27,29,34,16,23,22,17,3 | 197 | 149 | 0.000233564031259 | 0.0384816077679382 | 0.059,0.237,0.037, | 0.033,0.183,0.073, | 0,07 |
| 211219 | ENSG00000197622 | CDC42SE1 | chr1 | - | 151056312 | 151056786 | 151055676 | 151055993 | 151059478 | 151059615 | 211219 | 66,28,55,56,22,23,17,36,5 | 197,165,295,254,162,112,1 | 53,25,73,107,52,38,20,44,3 | 274,239,156,130,165,98,11 | 298 | 149 | 0.000233681306841 | 0.0384816077679382 | 0.143,0.078,0.085, | 0.088,0.05,0.19,0. | -0,038 |
| 167952 | ENSG00000130338 | TULP4 | chr6 | + | 158501677 | 158501875 | 158498668 | 158498812 | 158506577 | 158511828 | 167952 | 10,11,13,12,7,5,7,17,12,3, | 0,1,2,1,0,0,0,0,1,2,0,0 | 10,9,6,28,9,8,8,8,5,12,14,2 | 1,1,0,2,4,0,0,0,2,0,3,4,0,3, | 298 | 149 | 0.000233863457922 | 0.0384821604831881 | 1.0,0.846,0.765,0. | 0.833,0.818,1.0,0. | 0,108 |
| 107564 | ENSG00000184634 | MED12 | chrX | + | 71120013 | 71120170 | 71119685 | 71119877 | 71120970 | 71121152 | 107564 | 83,49,73,49,48,46,17,33,4 | 0,1,1,0,0,2,3,2,0,3,2,0 | 55,60,38,37,20,23,25,53,4 | 1,1,4,12,1,2,2,4,2,0,2,2,0,0 | 298 | 149 | 0.000234299999228 | 0.0385047171073014 | 1.0,0.961,0.973,1. | 0.965,0.968,0.826, | 0,044 |
| 174167 | ENSG00000060971 | ACAA1 | chr3 | - | 38126161 | 38126352 | 38125825 | 38125881 | 38126509 | 38126700 | 174167 | 24,9,20,12,9,16,11,11,9,5,1 | 3,0,4,1,2,4,2,2,2,0,1,0 | 7,34,8,8,6,9,6,10,6,7,4,18,1 | 2,17,3,3,6,2,1,3,0,1,3,10,2, | 298 | 149 | 0.000234358337835 | 0.0385047171073014 | 0.8,1.0,0.714,0.85 | 0.636,0.5,0.571,0. | 0,21 |
| 225932 | ENSG00000089289 | IGBP1 | chrX | + | 70146744 | 70146828 | 70134522 | 70134816 | 70148760 | 70148840 | 225932 | 28,25,83,74,31,20,26,40,3 | 4,1,3,1,1,3,2,3,1,0,0,2 | 26,29,21,18,34,21,19,12,2 | 7,3,5,0,7,0,2,6,1,0,3,11,0,5 | 233 | 149 | 0.000235234752691 | 0.0386192304015535 | 0.817,0.941,0.947, | 0.704,0.861,0.729, | 0,064 |
| 108669 | ENSG00000120860 | WASHC3 | chr12 | - | 102017719 | 102017805 | 102012941 | 102013192 | 102025973 | 102026038 | 108669 | 11,7,6,2,5,4,2,5,6,1,2,3 | 25,16,64,62,32,22,16,17,4 | 7,9,15,7,4,3,1,8,0,9,2,8,7,1 | 31,21,21,12,12,13,2,13,5,2 | 235 | 149 | 0.00023629732258 | 0.0387520225735599 | 0.218,0.217,0.056, | 0.125,0.214,0.312, | -0,072 |
| 173766 | ENSG00000156976 | EIF4A2 | chr3 | + | 186788300 | 186788416 | 186787802 | 186787882 | 186789124 | 186789881 | 173766 | 31,7,43,30,16,22,24,40,9,1 | 427,137,428,354,181,165,1 | 16,23,36,11,12,7,9,13,11,1 | 138,51,178,74,110,133,75, | 265 | 149 | 0.000236403704109 | 0.0387520225735599 | 0.039,0.028,0.053, | 0.061,0.202,0.102, | -0,024 |
| 138463 | ENSG00000168813 | ZNF507 | chr19 | + | 32359405 | 32359655 | 32356615 | 32356733 | 32360503 | 32360618 | 138463 | 3,0,1,0,3,0,0,1,0,0,0,1 | 4,7,27,9,7,6,2,9,7,4,1,5 | 4,0,0,3,1,4,2,2,4,2,3,1,0,0,2 | 15,15,3,9,2,6,5,7,6,13,5,18 | 298 | 149 | 0.000237690513148 | 0.0389333080938791 | 0.273,0.0,0.018,0. | 0.118,0.0,0.0,0.14 | -0,058 |
| 31252 | ENSG00000072364 | AFF4 | chr5 | - | 132898229 | 132898392 | 132896322 | 132897240 | 132899103 | 132899141 | 31252 | 118,84,214,95,75,130,93,2 | 13,1,7,1,1,3,1,9,3,1,6,3 | 128,146,85,180,99,45,124, | 5,7,5,31,9,9,5,4,8,5,4,2,2,5 | 298 | 149 | 0.000238529756539 | 0.0390410631404148 | 0.819,0.977,0.939, | 0.928,0.912,0.895, | 0,056 |
| 99416 | ENSG00000115677 | HDLBP | chr2 | - | 241255373 | 241255511 | 241255050 | 241255158 | 241256183 | 241256399 | 99416 | 71,36,172,108,47,45,37,60 | 0,0,5,4,4,5,5,5,6,1,0,0 | 78,55,70,27,35,35,31,59,4 | 1,0,0,0,4,1,0,1,2,1,4,1,0,3, | 287 | 149 | 0.000239370807887 | 0.0391489501157499 | 1.0,1.0,0.947,0.93 | 0.976,1.0,1.0,1.0,0 | -0,032 |
| 54376 | ENSG00000138095 | LRPPRC | chr2 | - | 43945331 | 43945417 | 43943686 | 43943894 | 43946112 | 43946243 | 54376 | 63,33,101,100,62,52,44,60 | 1,0,0,1,1,2,0,1,0,0,0,0 | 145,130,60,35,55,43,59,89 | 5,3,0,4,1,0,2,3,4,0,5,2,0,3,1 | 235 | 149 | 0.000239807876609 | 0.0391664926906353 | 0.976,1.0,1.0,0.98 | 0.948,0.965,1.0,0. | 0,04 |
| 187576 | ENSG00000137409 | MTCH1 | chr6 | - | 36977633 | 36977813 | 36975657 | 36977250 | 36978077 | 36978155 | 187576 | 61,53,130,51,24,43,44,44, | 1,0,2,0,1,0,0,0,0,0,0,0 | 57,63,29,43,44,16,28,30,3 | 2,5,1,2,0,0,0,3,3,0,3,2,2,1, | 298 | 149 | 0.000239842017954 | 0.0391664926906353 | 0.968,1.0,0.97,1.0, | 0.934,0.863,0.935, | 0,055 |
| 61574 | ENSG00000048740 | CELF2 | chr10 | + | 11321206 | 11321386 | 11314138 | 11314270 | 11325835 | 11325979 | 61574 | 102,77,77,54,41,50,36,54, | 3,2,0,6,0,5,1,1,0,0,0,0 | 64,55,42,27,28,16,15,28,2 | 0,3,1,2,2,0,7,5,0,0,4,1,1,1, | 298 | 149 | 0.000240084357323 | 0.0391763429196259 | 0.944,0.951,1.0,0. | 1.0,0.902,0.955,0. | 0,061 |
| 218998 | ENSG00000290770 | MFSD14C | chr9 | - | 96962764 | 96962963 | 96958834 | 96958897 | 96972852 | 96972949 | 218998 | 0,0,0,0,1,3,2,3,0,0,0,0 | 22,7,20,14,10,19,14,15,3,1 | 3,2,1,5,3,0,1,4,2,3,0,6,1,0, | 12,14,9,10,17,19,14,14,11, | 298 | 149 | 0.000240273292356 | 0.0391774704447532 | 0.0,0.0,0.0,0.0,0.0 | 0.111,0.067,0.053, | -0,047 |
| 184498 | ENSG00000077044 | DGKD | chr2 | + | 233447725 | 233447905 | 233446711 | 233446796 | 233448086 | 233448181 | 184498 | 20,3,31,19,7,5,10,16,3,4,2, | 20,9,65,49,34,38,20,37,48, | 19,22,32,24,9,3,9,12,10,11 | 20,25,35,20,13,17,33,37,9, | 298 | 149 | 0.000240552977253 | 0.0391933821704318 | 0.333,0.143,0.193, | 0.322,0.306,0.314, | -0,092 |
| 37084 | ENSG00000204525 | HLA-C | chr6 | - | 31269965 | 31270085 | 31269492 | 31269525 | 31270209 | 31270485 | 37084 | 247,272,406,325,630,676, | 50,52,18,10,0,0,7,7,7,10,0, | 491,492,635,1061,1330,93 | 10,17,3,1,0,0,10,35,7,11,0, | 269 | 149 | 0.000240745607262 | 0.0391950966691434 | 0.732,0.743,0.926, | 0.965,0.941,0.992, | -0,013 |
| 210938 | ENSG00000115159 | GPD2 | chr2 | + | 156542916 | 156542944 | 156513332 | 156513496 | 156549607 | 156549772 | 210938 | 4,3,4,4,1,1,3,3,1,2,0,3 | 37,25,102,32,20,9,8,26,27, | 4,5,13,4,8,2,1,2,6,3,2,9,3,2 | 32,11,42,23,25,34,12,24,26 | 177 | 149 | 0.000240985197164 | 0.0392044482016666 | 0.083,0.092,0.032, | 0.095,0.277,0.207, | -0,069 |
| 40545 | ENSG00000119231 | SENP5 | chr3 | + | 196927795 | 196927879 | 196903532 | 196903610 | 196929632 | 196929683 | 40545 | 13,6,29,13,6,7,4,13,14,5,4, | 2,0,3,5,0,0,1,2,0,0,0,2 | 19,14,7,19,10,8,4,9,6,13,9, | 5,2,1,7,4,0,4,3,1,0,3,2,2,2, | 233 | 149 | 0.000242095656999 | 0.0393553552504167 | 0.806,1.0,0.861,0. | 0.708,0.817,0.817, | 0,141 |
| 215729 | ENSG00000135930 | EIF4E2 | chr2 | + | 232574253 | 232574341 | 232567077 | 232567214 | 232580903 | 232581332 | 215729 | 0,0,1,2,0,0,0,0,0,0,0,0 | 9,9,29,13,17,9,11,12,11,7, | 2,0,2,0,2,3,4,2,0,4,0,0,0,6,1 | 16,17,5,2,3,1,8,3,5,6,10,11 | 237 | 149 | 0.000243870671969 | 0.0396139838479697 | 0.0,0.0,0.021,0.08 | 0.073,0.0,0.201,0. | -0,142 |
| 139817 | ENSG00000022277 | RTF2 | chr20 | + | 56516935 | 56516989 | 56513390 | 56513428 | 56517105 | 56517201 | 139817 | 13,7,51,37,16,11,21,12,25, | 2,2,4,13,0,2,0,0,2,2,0,0 | 23,16,16,5,8,10,7,21,5,23,1 | 2,2,0,1,0,2,2,0,0,1,0,2,0,7,2 | 203 | 149 | 0.000244530738451 | 0.0396912483918154 | 0.827,0.72,0.903,0 | 0.894,0.854,1.0,0. | -0,019 |
| 164659 | ENSG00000170242 | USP47 | chr11 | + | 11909066 | 11909118 | 11905398 | 11905548 | 11920155 | 11920251 | 164659 | 0,0,9,1,0,7,1,3,1,0,0,2 | 15,15,55,53,24,22,10,15,1 | 1,0,1,0,1,0,1,4,1,2,0,0,0,0,1 | 45,37,51,35,33,23,19,34,8, | 201 | 149 | 0.000245091137336 | 0.0397423695326544 | 0.0,0.0,0.108,0.01 | 0.016,0.0,0.014,0. | 0,029 |
| 80994 | ENSG00000143933 | CALM2 | chr2 | - | 47172413 | 47172459 | 47170733 | 47170764 | 47176440 | 47176516 | 80994 | 5,2,15,14,6,13,0,0,7,0,2,1 | 98,95,363,296,96,122,96,1 | 5,0,0,2,1,6,4,6,0,1,6,7,4,6,3 | 259,239,130,42,130,55,81, | 195 | 149 | 0.000245214986407 | 0.0397423695326544 | 0.038,0.016,0.031, | 0.015,0.0,0.0,0.03 | 0,011 |
| 146707 | ENSG00000157106 | SMG1 | chr16 | - | 18838013 | 18838262 | 18837252 | 18837443 | 18838356 | 18838466 | 146707 | 72,80,48,23,31,30,42,34,5 | 0,0,8,4,2,0,1,3,4,5,1,0 | 36,54,27,96,39,26,15,25,2 | 1,0,0,0,4,0,2,1,1,0,4,2,0,0,1 | 298 | 149 | 0.000246027429459 | 0.0398440403836644 | 1.0,1.0,0.75,0.742, | 0.947,1.0,1.0,1.0,0 | -0,043 |
| 177382 | ENSG00000115935 | WIPF1 | chr2 | - | 174623398 | 174623508 | 174612606 | 174612771 | 174625820 | 174625960 | 177382 | 9,6,14,4,4,3,2,2,5,6,2,5 | 4,0,1,4,1,0,0,0,0,0,2,0 | 16,20,4,6,5,3,6,2,3,4,4,8,2, | 4,5,0,1,2,0,0,0,2,2,1,0,3,1,1 | 259 | 149 | 0.000246246634009 | 0.0398495558529256 | 0.564,1.0,0.89,0.3 | 0.697,0.697,1.0,0. | 0,1 |
| 175032 | ENSG00000101596 | SMCHD1 | chr18 | + | 2673154 | 2673194 | 2666869 | 2667031 | 2673280 | 2673363 | 175032 | 42,19,53,17,8,13,20,41,6,3 | 514,412,843,542,254,319,3 | 6,31,35,59,70,18,18,14,20, | 231,322,330,168,299,197,1 | 189 | 149 | 0.000247097562915 | 0.0399572168022227 | 0.061,0.035,0.047, | 0.02,0.071,0.077,0 | -0,026 |
| 41127 | ENSG00000144741 | SLC25A26 | chr3 | + | 66346363 | 66346408 | 66263331 | 66263379 | 66362859 | 66362929 | 41127 | 23,12,45,22,22,21,10,28,1 | 1,0,0,1,0,0,0,0,2,0,0,0 | 24,39,11,16,28,6,14,19,1,1 | 4,3,0,3,3,0,0,3,0,2,5,0,0,0,1 | 194 | 149 | 0.000247289414552 | 0.0399582192067879 | 0.946,1.0,1.0,0.94 | 0.822,0.909,1.0,0. | 0,065 |
| 184497 | ENSG00000077044 | DGKD | chr2 | + | 233447671 | 233447905 | 233446711 | 233446796 | 233448086 | 233448181 | 184497 | 20,3,31,19,7,5,10,16,2,6,2, | 20,9,65,49,34,38,20,37,48, | 19,22,32,24,9,3,9,12,10,11 | 20,25,35,20,13,17,33,37,9, | 298 | 149 | 0.000248306453659 | 0.0400924578600752 | 0.333,0.143,0.193, | 0.322,0.306,0.314, | -0,091 |
| 178170 | ENSG00000067066 | SP100 | chr2 | + | 230521306 | 230521400 | 230511124 | 230511166 | 230539266 | 230539382 | 178170 | 10,4,3,1,4,4,6,7,0,2,0,2 | 73,34,112,67,53,53,52,56, | 0,4,9,19,13,1,1,1,1,3,12,4,4 | 51,39,35,45,50,31,22,43,2 | 243 | 149 | 0.000248558777658 | 0.0401031141485075 | 0.077,0.067,0.016, | 0.0,0.059,0.136,0. | -0,031 |
| 228022 | ENSG00000169871 | TRIM56 | chr7 | + | 101086886 | 101087168 | 101085521 | 101085571 | 101087311 | 101087508 | 228022 | 11,9,26,30,22,20,15,26,9,1 | 0,2,5,0,0,0,0,0,0,0,2,0 | 21,33,9,7,11,6,6,27,14,16,1 | 0,0,1,2,3,0,0,5,0,1,4,9,0,1, | 298 | 149 | 0.000249163592008 | 0.0401405835880788 | 1.0,0.692,0.722,1. | 1.0,1.0,0.818,0.63 | 0,045 |
| 161383 | ENSG00000165819 | METTL3 | chr14 | - | 21510831 | 21510980 | 21503663 | 21503881 | 21511123 | 21511342 | 161383 | 4,1,8,2,3,3,5,2,4,0,2,4 | 38,12,94,57,70,75,45,57,5 | 4,16,3,3,4,1,6,9,3,5,6,8,3,2 | 63,49,29,7,29,19,42,75,19, | 298 | 149 | 0.000249233536288 | 0.0401405835880788 | 0.05,0.04,0.041,0. | 0.031,0.14,0.049,0 | -0,034 |
| 151007 | ENSG00000170776 | AKAP13 | chr15 | + | 85522033 | 85522232 | 85521427 | 85521575 | 85533583 | 85536997 | 151007 | 6,2,8,3,1,2,2,1,0,0,1,1 | 105,57,182,121,127,105,7 | 10,8,7,13,10,1,1,3,8,0,3,7,3 | 150,155,83,84,106,56,56,1 | 298 | 149 | 0.000249350512971 | 0.0401405835880788 | 0.028,0.017,0.022, | 0.032,0.025,0.04,0 | -0,013 |
| 120572 | ENSG00000107036 | RIC1 | chr9 | + | 5745930 | 5746083 | 5743688 | 5743737 | 5747301 | 5747505 | 120572 | 102,101,166,111,93,102,6 | 16,0,1,1,0,1,3,4,1,1,3,3 | 128,163,72,81,136,51,58,7 | 9,5,0,28,5,4,1,4,3,5,5,12,4, | 298 | 149 | 0.000251136902408 | 0.0403979421839882 | 0.761,1.0,0.988,0. | 0.877,0.942,1.0,0. | 0,048 |
| 10109 | ENSG00000136485 | DCAF7 | chr17 | + | 63579336 | 63579448 | 63550538 | 63550815 | 63579824 | 63579943 | 10109 | 13,11,37,53,20,11,9,9,6,8, | 7,4,6,0,0,2,2,1,4,2,2,6 | 15,20,13,6,8,5,9,17,4,18,1 | 2,0,0,4,5,0,0,2,0,3,4,0,0,0,1 | 261 | 149 | 0.000252581917424 | 0.0406000438155974 | 0.515,0.611,0.779, | 0.811,1.0,1.0,0.46 | -0,171 |
| 251982 | ENSG00000221817 | PPP3CB-AS1 | chr10 | + | 73498732 | 73498867 | 73496141 | 73496262 | 73499594 | 73499687 | 251982 | 16,8,26,20,4,13,10,17,3,7, | 0,0,2,5,1,1,3,2,0,0,0,0 | 8,2,6,8,7,12,7,16,9,6,8,5,1 | 3,0,4,2,0,2,0,0,0,2,1,4,4,0, | 284 | 149 | 0.000252851696702 | 0.0406130772633344 | 1.0,1.0,0.872,0.67 | 0.583,1.0,0.44,0.6 | 0,127 |
| 85151 | ENSG00000068724 | TTC7A | chr2 | + | 46958494 | 46958552 | 46956838 | 46957007 | 46974972 | 46975103 | 85151 | 6,6,28,11,6,15,10,13,2,0,4, | 48,27,110,72,55,90,37,62, | 8,5,0,1,0,5,4,13,4,9,4,4,15, | 67,108,51,51,45,43,35,166 | 207 | 149 | 0.000253986969721 | 0.0407650033408058 | 0.083,0.138,0.155, | 0.079,0.032,0.0,0. | 0,034 |
| 14834 | ENSG00000166747 | AP1G1 | chr16 | - | 71772959 | 71773053 | 71771155 | 71771252 | 71773220 | 71773362 | 14834 | 7,10,12,3,1,14,6,7,2,4,1,4 | 74,85,93,55,39,47,34,43,4 | 15,25,8,28,16,5,1,6,6,2,10, | 34,67,57,64,60,35,34,35,31 | 243 | 149 | 0.000255154078271 | 0.0409218088080072 | 0.055,0.067,0.073, | 0.213,0.186,0.079, | -0,045 |
| 83304 | ENSG00000131781 | FMO5 | chr1 | - | 147208851 | 147209051 | 147201151 | 147201504 | 147212392 | 147212535 | 83304 | 7,7,25,18,9,6,10,19,8,2,2,6 | 3,5,2,2,0,4,0,2,0,1,0,1 | 21,16,7,11,16,14,2,8,10,7,2 | 4,3,1,0,1,0,0,0,0,3,3,2,2,1, | 298 | 149 | 0.000255392136189 | 0.0409294898466825 | 0.538,0.412,0.862, | 0.724,0.727,0.778, | -0,096 |
| 14726 | ENSG00000125124 | BBS2 | chr16 | - | 56501944 | 56502079 | 56501352 | 56501497 | 56502316 | 56502456 | 14726 | 1,2,3,2,2,1,0,3,0,0,0,1 | 15,8,4,16,11,8,12,8,3,11,9, | 2,2,4,6,3,0,2,2,0,3,7,0,5,1,7 | 12,6,13,7,7,1,12,23,4,9,9,4 | 284 | 149 | 0.000255659694151 | 0.0409418836546234 | 0.034,0.116,0.282, | 0.08,0.149,0.139,0 | -0,044 |
| 225951 | ENSG00000115677 | HDLBP | chr2 | - | 241255373 | 241255496 | 241255050 | 241255158 | 241256183 | 241256399 | 225951 | 69,32,165,105,43,41,34,56 | 0,0,5,4,4,5,5,5,6,1,0,0 | 76,53,65,24,33,30,29,52,4 | 1,0,0,0,4,1,0,1,2,1,4,1,0,3,7 | 272 | 149 | 0.00025593098475 | 0.0409548563410611 | 1.0,1.0,0.948,0.93 | 0.977,1.0,1.0,1.0,0 | -0,033 |
| 164809 | ENSG00000065882 | TBC1D1 | chr4 | + | 38118038 | 38118192 | 38115899 | 38115954 | 38124961 | 38125131 | 164809 | 154,113,315,236,79,84,80, | 6,3,3,4,1,2,6,2,6,2,7,2 | 111,128,132,78,129,100,4 | 0,2,3,0,1,0,3,2,0,4,5,1,0,0,2 | 298 | 149 | 0.000256467339735 | 0.0410089361671609 | 0.928,0.95,0.981,0 | 1.0,0.97,0.957,1.0, | -0,044 |
| 112360 | ENSG00000104365 | IKBKB | chr8 | + | 42290155 | 42290273 | 42272082 | 42272205 | 42305186 | 42305236 | 112360 | 15,9,18,11,3,8,10,3,6,2,3,6 | 2,1,4,1,0,2,2,4,0,4,0,2 | 7,4,4,2,11,1,0,11,5,3,3,1,6, | 2,0,0,0,0,0,5,1,1,1,0,0,3,0,3 | 267 | 149 | 0.000256650004029 | 0.0410089361671609 | 0.807,0.834,0.715, | 0.661,1.0,1.0,1.0,1 | -0,074 |
| 235703 | ENSG00000145041 | DCAF1 | chr3 | - | 51492388 | 51492488 | 51483718 | 51483836 | 51496733 | 51496780 | 235703 | 14,6,21,9,6,11,3,12,8,1,2,1 | 2,3,4,1,1,2,0,3,2,0,0,0 | 11,10,6,9,18,4,6,6,3,5,11,1 | 1,3,2,3,0,1,0,0,0,2,0,2,1,1,4 | 249 | 149 | 0.000256919512111 | 0.0410215456314263 | 0.807,0.545,0.759, | 0.868,0.666,0.642, | 0,033 |
| 163573 | ENSG00000141084 | RANBP10 | chr16 | - | 67770589 | 67770733 | 67744287 | 67744455 | 67772033 | 67772086 | 163573 | 0,0,2,1,0,0,0,0,0,0,0,0 | 3,7,13,14,6,4,4,7,5,6,3,3 | 0,3,0,2,4,2,0,2,2,3,0,3,1,2,2 | 11,3,4,1,3,7,2,11,3,5,1,8,1 | 293 | 149 | 0.000257525180884 | 0.0410259291675761 | 0.0,0.0,0.073,0.03 | 0.0,0.337,0.0,0.50 | -0,153 |
| 44705 | ENSG00000096384 | HSP90AB1 | chr6 | + | 44248629 | 44248776 | 44247100 | 44247195 | 44249376 | 44249583 | 44705 | 1457,818,2417,1908,1202, | 47,12,26,19,11,17,16,10,8, | 1461,1435,979,312,557,48 | 11,11,5,5,4,4,2,11,4,0,32,9 | 296 | 149 | 0.00025763569943 | 0.0410259291675761 | 0.94,0.972,0.979,0 | 0.985,0.985,0.99,0 | -0,006 |
| 121095 | ENSG00000070047 | PHRF1 | chr11 | + | 581964 | 582081 | 581491 | 581606 | 587258 | 587464 | 121095 | 39,34,13,10,5,6,9,22,26,19 | 1,3,1,0,0,4,0,2,0,1,0,0 | 11,17,12,41,15,9,7,16,4,17 | 0,6,1,3,0,0,4,0,1,0,4,7,0,0, | 266 | 149 | 0.000257645699086 | 0.0410259291675761 | 0.956,0.864,0.879, | 1.0,0.613,0.87,0.8 | 0,08 |
| 138736 | ENSG00000127527 | EPS15L1 | chr19 | - | 16395343 | 16395432 | 16393950 | 16394001 | 16402320 | 16402485 | 138736 | 36,32,32,12,22,17,13,8,11, | 3,0,0,0,2,6,1,4,0,2,1,2 | 18,13,16,17,9,14,11,17,12, | 0,0,2,0,1,1,0,1,0,3,0,0,0,2,2 | 238 | 149 | 0.000257709420272 | 0.0410259291675761 | 0.883,1.0,1.0,1.0,0 | 1.0,1.0,0.834,1.0,0 | -0,068 |
| 67338 | ENSG00000117153 | KLHL12 | chr1 | - | 202895521 | 202895717 | 202894590 | 202894749 | 202909009 | 202909124 | 67338 | 27,34,52,44,22,25,14,16,2 | 1,1,1,0,0,0,0,8,0,0,0,2 | 21,34,22,9,23,13,16,18,9,2 | 5,1,0,0,5,0,1,1,1,2,2,8,6,1,1 | 298 | 149 | 0.000259191717646 | 0.0412163882822921 | 0.931,0.944,0.963, | 0.677,0.944,1.0,1. | 0,077 |
| 233549 | ENSG00000143569 | UBAP2L | chr1 | + | 154234590 | 154234714 | 154228614 | 154228725 | 154235195 | 154235294 | 233549 | 121,68,84,45,34,24,17,43,1 | 62,60,23,17,11,26,26,18,2 | 43,30,44,61,20,19,25,39,1 | 31,45,35,67,54,10,20,33,2 | 273 | 149 | 0.000259340364108 | 0.0412163882822921 | 0.516,0.382,0.666, | 0.431,0.267,0.407, | 0,147 |
| 129617 | ENSG00000127314 | RAP1B | chr12 | + | 68624670 | 68624873 | 68610898 | 68611052 | 68648698 | 68648781 | 129617 | 0,0,2,8,0,0,0,2,2,2,1,0 | 7,4,13,13,6,2,1,3,2,2,0,2 | 2,2,3,0,0,0,1,4,4,0,3,7,4,11 | 4,7,6,2,8,2,0,4,0,1,4,9,2,6, | 298 | 149 | 0.000259480307774 | 0.0412163882822921 | 0.0,0.0,0.071,0.23 | 0.2,0.125,0.2,0.0,0 | -0,189 |

| | | | | | | | | | | | | | | | | | | | | | |
|---|---|---|---|---|---|---|---|---|---|---|---|---|---|---|---|---|---|---|---|---|---|
| 116561 | ENSG00000213930 | GALT | chr9 | + | 34647491 | 34647705 | 34647088 | 34647258 | 34648114 | 34648171 | 116561 | 15,12,26,14,24,16,21,13,1 | 0,0,5,4,3,5,1,1,0,1,1,0 | 19,10,9,19,5,3,6,24,12,9,8, | 0,0,0,0,3,0,0,3,0,0,3,1,0,0,2 | 298 | 149 | 0.000261470507645 | 0.0415018870434668 | 1.0,1.0,0.722,0.63 | 1.0,1.0,1.0,1.0,0.4 | -0,044 |
| 155239 | ENSG00000198718 | TOGARAM1 | chr14 | + | 45046900 | 45047024 | 45046544 | 45046703 | 45052435 | 45052562 | 155239 | 5,3,8,0,3,0,0,0,4,0,0,0 | 4,3,21,17,2,6,1,8,3,1,4,3 | 0,0,0,1,0,1,0,0,0,2,0,0,4,2,0 | 6,14,1,4,7,1,6,2,1,4,4,1,8,4 | 273 | 149 | 0.000261987729469 | 0.0415368582050194 | 0.406,0.353,0.172, | 0.0,0.0,0.0,0.12,0.0 | 0,08 |
| 33123 | ENSG00000138182 | KIF20B | chr10 | + | 89709926 | 89710065 | 89709344 | 89709461 | 89710960 | 89711090 | 33123 | 31,17,39,14,28,17,12,24,1 | 0,2,2,1,2,1,0,0,0,0,0,1 | 17,27,23,31,14,8,19,24,13, | 7,2,3,6,1,0,0,0,2,2,0,1,0,1,0 | 288 | 149 | 0.000262076807906 | 0.0415368582050194 | 1.0,0.815,0.91,0.8 | 0.557,0.875,0.799, | 0,073 |
| 63091 | ENSG00000116698 | SMG7 | chr1 | + | 183542075 | 183542364 | 183540983 | 183541103 | 183544352 | 183544497 | 63091 | 18,18,35,10,19,33,15,12,1 | 2,0,1,2,0,0,0,3,2,2,2,0 | 14,47,17,41,8,20,23,16,22, | 1,1,0,6,0,6,0,0,2,5,2,0,0,2,4 | 298 | 149 | 0.000262570852292 | 0.0415845379762027 | 0.818,1.0,0.946,0. | 0.875,0.959,1.0,0. | 0,026 |
| 119472 | ENSG00000183597 | TANGO2 | chr22 | + | 20053436 | 20053551 | 20052464 | 20052584 | 20061529 | 20061681 | 119472 | 6,2,15,23,5,6,5,4,2,6,2,4 | 2,0,6,1,0,0,2,0,0,1,0,2 | 5,3,7,3,4,8,4,6,4,4,3,8,3,3,2 | 2,0,0,1,1,0,1,0,0,1,1,4,7,2,3 | 264 | 149 | 0.000264841837774 | 0.0419133629308351 | 0.629,1.0,0.585,0. | 0.585,1.0,1.0,0.62 | 0,125 |
| 90497 | ENSG00000177879 | AP3S1 | chr5 | + | 115902884 | 115902992 | 115895086 | 115895158 | 115913361 | 115914011 | 90497 | 152,124,127,111,69,54,34, | 15,11,11,9,3,5,7,14,6,0,3,1 | 66,74,64,31,55,71,19,38,1 | 10,21,9,12,25,12,16,13,11, | 257 | 149 | 0.000265219047712 | 0.0419422195871429 | 0.855,0.867,0.87,0 | 0.793,0.671,0.805, | 0,119 |
| 141127 | ENSG00000175354 | PTPN2 | chr18 | - | 12840713 | 12840950 | 12836790 | 12836891 | 12859163 | 12859254 | 141127 | 10,12,8,8,0,8,2,10,14,4,1,2 | 99,84,183,185,73,78,57,93 | 4,19,13,31,4,1,6,14,11,5,8, | 77,52,71,35,45,67,27,103,4 | 298 | 149 | 0.000266939925653 | 0.0421678213165313 | 0.048,0.067,0.021, | 0.025,0.154,0.084, | -0,037 |
| 88898 | ENSG00000073849 | ST6GAL1 | chr3 | + | 187038394 | 187038491 | 186963784 | 186963926 | 187038741 | 187038873 | 88898 | 3,6,23,18,5,7,9,6,6,1,6,8 | 101,51,133,81,66,36,39,47 | 14,19,12,20,5,8,0,12,7,6,5, | 98,27,89,42,23,58,34,69,2 | 246 | 149 | 0.000267037464187 | 0.0421678213165313 | 0.018,0.067,0.095, | 0.08,0.299,0.076,0 | -0,052 |
| 244494 | ENSG00000171634 | BPTF | chr17 | + | 67886127 | 67886316 | 67874783 | 67875020 | 67891846 | 67891962 | 244494 | 21,15,40,24,15,18,15,11,2 | 2,3,2,0,2,0,0,0,1,1,0,0 | 24,5,12,14,12,11,12,22,20, | 3,1,3,1,3,0,2,2,2,4,4,0,2,2,0 | 298 | 149 | 0.000267439183562 | 0.0422002953938657 | 0.84,0.714,0.909,1 | 0.8,0.714,0.667,0. | 0,086 |
| 252711 | ENSG00000174718 | RESF1 | chr12 | + | 31987191 | 31987322 | 31980877 | 31985957 | 31992377 | 31992941 | 252711 | 122,77,327,241,129,119,1 | 12,0,40,17,8,13,16,9,7,4,2, | 197,214,213,119,106,86,8 | 5,8,12,12,24,7,1,9,2,1,2,8,6 | 280 | 149 | 0.000269243635907 | 0.042453902564029 | 0.844,1.0,0.813,0. | 0.954,0.934,0.904, | -0,037 |
| 117075 | ENSG00000125944 | HNRNPR | chr1 | - | 23323555 | 23323732 | 23321527 | 23321663 | 23333517 | 23333631 | 117075 | 148,101,345,265,127,121, | 0,1,6,3,1,1,1,6,6,0,0,1 | 180,210,118,68,85,97,84,1 | 1,11,3,9,8,2,3,2,0,1,3,4,4,5 | 298 | 149 | 0.000270156222955 | 0.0425483475620275 | 1.0,0.981,0.966,0. | 0.989,0.905,0.952, | 0,033 |
| 185067 | ENSG00000078304 | PPP2R5C | chr14 | + | 101824013 | 101824190 | 101809836 | 101810036 | 101836168 | 101836701 | 185067 | 2,0,2,2,0,0,0,0,7,4,0,0 | 17,16,35,14,10,13,14,10,6, | 2,5,5,2,0,0,0,0,0,3,8,0,1,0,4 | 12,17,7,6,7,8,8,5,7,18,8,20 | 298 | 149 | 0.000270349802263 | 0.0425483475620275 | 0.056,0.0,0.028,0. | 0.077,0.128,0.263, | -0,023 |
| 187572 | ENSG00000137409 | MTCH1 | chr6 | - | 36977633 | 36977768 | 36975657 | 36977250 | 36978077 | 36978155 | 187572 | 61,53,129,51,23,41,43,43,2 | 1,0,2,0,1,0,0,0,0,0,0,0 | 54,62,29,43,45,16,23,30,3 | 2,5,1,2,0,0,0,3,3,0,3,2,2,1,0 | 284 | 149 | 0.000270435668955 | 0.0425483475620275 | 0.97,1.0,0.971,1.0, | 0.934,0.867,0.938, | 0,054 |
| 161426 | ENSG00000273294 | C1QTNF3-AMA | chr5 | - | 34023908 | 34023986 | 34020586 | 34020742 | 34028753 | 34028883 | 161426 | 6,8,9,5,3,7,2,14,7,2,4,1 | 2,2,5,1,0,1,1,0,1,0,0,0 | 0,4,3,4,7,2,3,1,4,2,10,4,1,7 | 2,1,0,6,1,0,0,0,5,2,3,3,0,1,2 | 227 | 149 | 0.000271088061372 | 0.0426198353083688 | 0.663,0.724,0.542, | 0.0,0.724,1.0,0.30 | 0,14 |
| 139200 | ENSG00000286235 | | chr20 | + | 6023291 | 6023422 | 6015360 | 6015490 | 6031284 | 6031370 | 139200 | 1,1,8,6,3,5,1,1,2,3,0,4 | 31,12,20,30,11,10,14,13,2 | 2,2,1,1,0,0,0,1,1,2,2,1,1,0,2 | 27,10,7,0,8,17,3,23,4,20,8, | 280 | 149 | 0.000272252558556 | 0.0427716718471288 | 0.017,0.042,0.176, | 0.038,0.096,0.071, | -0,023 |
| 155619 | ENSG00000180776 | ZDHHC20 | chr13 | - | 21400372 | 21400401 | 21391721 | 21391854 | 21401652 | 21401685 | 155619 | 13,9,19,18,3,2,4,5,17,6,2,1 | 1,0,0,4,4,0,0,1,0,0,2,1 | 9,13,6,1,2,5,2,8,1,6,6,8,2,2 | 1,0,1,4,2,2,0,3,2,1,0,1,4,1,2 | 178 | 149 | 0.000272854879972 | 0.0428350318535766 | 0.916,1.0,1.0,0.79, | 0.883,1.0,0.834,0. | 0,111 |
| 30394 | ENSG00000198055 | GRK6 | chr5 | + | 177433346 | 177433410 | 177433146 | 177433239 | 177433535 | 177433676 | 30394 | 128,92,192,274,57,50,68,5 | 9,4,5,1,0,2,0,4,1,0,0,2 | 102,105,59,84,104,60,54,4 | 0,5,12,16,2,0,3,4,7,0,4,3,1, | 213 | 149 | 0.000273855236123 | 0.042960740762383 | 0.909,0.941,0.964, | 1.0,0.936,0.775,0. | 0,036 |
| 240969 | ENSG00000106829 | TLE4 | chr9 | + | 79650902 | 79650986 | 79649819 | 79649858 | 79652592 | 79652799 | 240969 | 4,0,3,4,1,4,0,4,1,2,0,1 | 3,3,4,3,4,10,11,20,2,3,4,3 | 0,3,8,1,4,0,0,0,1,3,0,0,1,0,0 | 6,5,8,11,3,2,4,6,7,3,14,5,0, | 233 | 149 | 0.000275011031529 | 0.0431106331587897 | 0.46,0.0,0.324,0.4 | 0.0,0.277,0.39,0.0 | 0,035 |
| 64812 | ENSG00000092841 | MYL6 | chr12 | + | 56159586 | 56159730 | 56158361 | 56158711 | 56159974 | 56160148 | 64812 | 1212,1021,3172,2605,117 | 38,26,57,25,6,14,8,13,16,1 | 1412,1374,991,529,1049,7 | 11,37,15,54,46,7,0,10,20,8 | 293 | 149 | 0.000275759788652 | 0.0431910711743269 | 0.942,0.952,0.966, | 0.985,0.95,0.971,0 | 0,017 |
| 58183 | ENSG00000137409 | MTCH1 | chr6 | - | 36977633 | 36977796 | 36977198 | 36977250 | 36978077 | 36978155 | 58183 | 61,53,140,58,24,43,44,47,3 | 1,0,2,0,1,0,0,0,0,0,0,0 | 58,64,29,43,45,18,29,30,3 | 2,5,1,2,0,0,0,3,3,0,3,2,2,1,0 | 298 | 149 | 0.00027592550731 4 | 0.0431910711743269 | 0.968,1.0,0.972,1. | 0.935,0.865,0.935, | 0,056 |
| 30834 | ENSG00000058799 | YIPF1 | chr1 | - | 53882942 | 53883031 | 53878641 | 53878722 | 53883112 | 53883276 | 30834 | 4,7,0,1,6,2,6,5,4,0,1,0 | 32,18,30,36,17,20,10,10,2 | 3,0,3,0,4,2,0,1,0,7,0,0,1,1,3 | 17,19,13,21,11,13,18,22,1 | 238 | 149 | 0.000276662784362 | 0.0432750056257395 | 0.073,0.196,0.0,0. | 0.099,0.0,0.126,0. | 0,051 |
| 7580 | ENSG00000141644 | MBD1 | chr18 | - | 50271468 | 50271540 | 50269556 | 50269811 | 50272676 | 50272738 | 7580 | 9,5,10,12,3,11,7,12,9,6,1,1 | 6,1,3,1,2,2,5,0,0,0,0,3 | 16,14,6,8,7,7,9,9,5,8,10,7, | 0,0,0,2,1,0,1,1,0,1,0,4,1,2,3 | 221 | 149 | 0.000277998499021 | 0.0434353236108154 | 0.503,0.771,0.692, | 1.0,1.0,1.0,0.729,0 | -0,059 |
| 154721 | ENSG00000129566 | TEP1 | chr14 | - | 20389609 | 20389765 | 20389237 | 20389297 | 20390680 | 20390758 | 154721 | 34,41,39,32,36,31,19,26,2 | 1,0,1,2,1,3,3,0,0,2,0,1 | 47,64,27,28,44,35,6,32,49, | 0,3,2,5,4,2,4,0,0,0,4,4,6,0,2 | 298 | 149 | 0.000278091334128 | 0.0434353236108154 | 0.944,1.0,0.951,0. | 1.0,0.914,0.871,0. | 0,046 |
| 34964 | ENSG00000151247 | EIF4E | chr4 | - | 98909677 | 98909959 | 98901875 | 98901982 | 98929094 | 98929125 | 34964 | 1,2,6,5,1,0,0,0,0,2,1,2 | 18,13,25,40,17,17,12,14,4, | 3,1,1,36,1,2,3,5,0,1,3,12,3, | 19,13,13,14,13,4,8,8,5,15,2 | 298 | 149 | 0.000279375470183 | 0.0436042507780691 | 0.027,0.071,0.107, | 0.073,0.037,0.037, | -0,052 |
| 233410 | ENSG00000183354 | KIAA2026 | chr9 | - | 5969889 | 5970012 | 5968176 | 5969436 | 5988344 | 5988545 | 233410 | 2,2,7,2,0,2,4,7,2,0,0,1 | 50,42,87,51,21,32,31,34,2 | 15,13,0,3,4,1,2,3,4,4,3,7,4, | 59,30,22,43,36,18,28,27,3 | 272 | 149 | 0.000281307588154 | 0.0438739952941838 | 0.021,0.025,0.042, | 0.122,0.192,0.0,0. | -0,028 |
| 211280 | ENSG00000176978 | DPP7 | chr9 | - | 137113356 | 137113496 | 137112962 | 137113119 | 137113864 | 137114028 | 211280 | 10,7,31,10,6,20,19,8,1,6,4, | 2,5,0,0,0,0,2,0,0,0,2,0 | 19,25,13,12,14,3,6,21,7,10 | 0,0,5,10,0,0,2,4,0,0,4,4,0,0 | 289 | 149 | 0.000282754398799 | 0.0440677132569932 | 0.721,0.419,1.0,1. | 1.0,1.0,0.573,0.38 | 0,003 |
| 28092 | ENSG00000073584 | SMARCE1 | chr17 | - | 40630207 | 40630286 | 40627721 | 40628993 | 40630713 | 40630924 | 28092 | 33,29,24,22,3,6,17,16,14,7 | 166,232,199,151,90,79,83, | 15,44,24,8,16,4,1,18,7,11, | 132,129,119,52,60,36,49,9 | 228 | 149 | 0.000283336032156 | 0.04412640921633 | 0.115,0.076,0.073, | 0.069,0.182,0.116, | -0,023 |
| 161472 | ENSG00000113282 | CLINT1 | chr5 | - | 157789026 | 157789073 | 157787601 | 157787992 | 157789362 | 157790561 | 161472 | 5,0,2,4,3,2,2,4,0,0,2,1 | 104,123,154,107,71,86,10 | 4,6,5,10,9,3,4,10,4,0,13,0, | 111,153,127,414,155,102, | 196 | 149 | 0.000287018925429 | 0.0446676575119372 | 0.035,0.0,0.01,0.0 | 0.027,0.029,0.029, | -0,011 |
| 57419 | ENSG00000170581 | STAT2 | chr12 | - | 56349162 | 56349261 | 56348751 | 56348851 | 56349425 | 56349473 | 57419 | 19,28,36,53,38,29,28,24,1 | 0,1,5,4,1,0,2,2,3,1,0,2 | 43,27,20,4,21,19,23,44,16, | 0,1,2,1,1,0,1,0,2,0,1,0,0,1,0 | 248 | 149 | 0.000288674510657 | 0.0448761387964044 | 1.0,0.944,0.812,0. | 1.0,0.942,0.857,0. | -0,028 |
| 45177 | ENSG00000054967 | RELT | chr11 | + | 73391143 | 73391223 | 73390754 | 73390921 | 73392210 | 73392468 | 45177 | 25,6,16,12,23,5,9,10,4,8,8, | 0,0,2,0,2,0,0,4,0,0,1,1 | 9,14,2,7,7,5,8,11,3,8,3,43,8 | 0,3,0,2,1,4,0,3,1,1,0,11,0,1 | 229 | 149 | 0.000288775558507 | 0.0448761387964044 | 1.0,1.0,0.839,1.0,0 | 1.0,0.752,1.0,0.69 | 0,107 |
| 252710 | ENSG00000174718 | RESF1 | chr12 | + | 31987183 | 31987322 | 31980877 | 31985957 | 31992377 | 31992941 | 252710 | 125,77,339,249,133,123,1 | 12,0,40,17,8,13,16,9,7,4,2, | 200,216,216,119,112,90,8 | 5,8,12,12,24,7,1,9,2,1,2,8,6 | 288 | 149 | 0.000289186847232 | 0.0449076293770495 | 0.843,1.0,0.814,0. | 0.954,0.933,0.903, | -0,038 |
| 26605 | ENSG00000198399 | ITSN2 | chr2 | - | 24284762 | 24284843 | 24275712 | 24275849 | 24286211 | 24286351 | 26605 | 8,0,3,0,3,11,1,8,2,2,0,0 | 255,228,194,117,97,65,64, | 7,12,9,18,6,0,5,6,4,2,7,2,3, | 182,259,127,144,136,77,7 | 230 | 149 | 0.000289646129425 | 0.0449465220492229 | 0.02,0.0,0.01,0.0,0 | 0.024,0.029,0.044, | -0,013 |
| 187528 | ENSG00000076770 | MBNL3 | chrX | - | 132437898 | 132437943 | 132406227 | 132406392 | 132439434 | 132440314 | 187528 | 0,4,6,3,1,1,0,1,4,0,2,0 | 7,12,13,14,7,9,4,13,5,2,1,1 | 2,4,1,2,1,2,2,3,1,3,1,6,5,7, | 8,4,3,9,10,3,3,6,4,1,1,13,9, | 194 | 149 | 0.00029266295613 | 0.0453819457570721 | 0.0,0.204,0.262,0. | 0.161,0.434,0.204, | -0,213 |
| 227940 | ENSG00000173715 | C11orf80 | chr11 | + | 66801030 | 66801097 | 66800640 | 66800696 | 66803994 | 66804194 | 227940 | 41,8,62,43,26,30,19,34,9,2 | 1,0,5,0,0,3,2,5,0,0,0,1 | 52,28,21,14,15,18,12,33,1 | 1,6,2,1,0,0,2,2,1,5,3,0,1,0,8 | 216 | 149 | 0.000292966446566 | 0.0453963004037774 | 0.966,1.0,0.895,1. | 0.973,0.763,0.879, | 0,063 |
| 78063 | ENSG00000152061 | RABGAP1L | chr1 | + | 174231144 | 174231355 | 174220971 | 174221164 | 174241482 | 174241657 | 78063 | 291,184,1029,660,258,336 | 3,1,7,4,0,2,3,5,1,3,0,0 | 493,614,613,372,456,227, | 4,7,15,10,4,1,5,3,5,4,9,0,1, | 298 | 149 | 0.000293203272377 | 0.0454003118827152 | 0.98,0.989,0.987,0 | 0.984,0.978,0.953, | 0,014 |
| 253787 | ENSG00000166908 | PIP4K2C | chr12 | + | 57599399 | 57599869 | 57599064 | 57599211 | 57600323 | 57600437 | 253787 | 1,0,5,3,2,2,0,0,0,0,1,0 | 2,4,10,4,5,7,11,7,4,3,2,5 | 2,1,0,1,3,3,0,0,4,7,1,0,3,0, | 14,12,6,6,16,2,3,6,5,7,13,6 | 298 | 149 | 0.000293532445854 | 0.0454186066524819 | 0.2,0.0,0.2,0.273,0 | 0.067,0.04,0.0,0.0 | -0,019 |
| 73783 | ENSG00000291124 | | chr7 | + | 65754526 | 65754661 | 65751050 | 65751278 | 65755779 | 65755952 | 73783 | 6,11,29,6,5,5,4,10,8,1,11,4 | 3,0,3,1,3,3,0,0,0,2,0,4 | 6,18,6,11,7,3,2,7,6,11,10,4 | 2,4,0,0,0,1,2,1,0,3,0,2,0,0, | 284 | 149 | 0.000294607730809 | 0.0455522388719482 | 0.512,1.0,0.835,0. | 0.611,0.702,1.0,1. | -0,103 |
| 22283 | ENSG00000100485 | SOS2 | chr14 | - | 50215373 | 50215432 | 50204283 | 50204409 | 50231196 | 50231362 | 22283 | 1,3,6,2,0,2,0,3,0,0,0,0 | 6,3,45,21,11,15,19,17,7,5,3 | 0,2,0,2,5,3,2,1,4,5,0,2,0,2,1 | 27,15,20,4,16,6,7,18,12,26 | 208 | 149 | 0.000294931798924 | 0.0455696094861532 | 0.107,0.417,0.087, | 0.0,0.087,0.0,0.26 | -0,046 |
| 142376 | ENSG00000168961 | LGALS9 | chr17 | + | 27647251 | 27647432 | 27647029 | 27647118 | 27648835 | 27649541 | 142376 | 77,41,126,149,60,38,72,42 | 11,9,15,22,13,15,11,14,17, | 57,78,45,51,41,48,21,78,5 | 8,10,5,1,3,9,2,3,4,8,0,23,1 | 298 | 149 | 0.000296408673818 | 0.0457649463949225 | 0.778,0.695,0.808, | 0.781,0.796,0.818, | -0,11 |
| 187570 | ENSG00000137409 | MTCH1 | chr6 | - | 36977633 | 36977759 | 36975657 | 36977250 | 36978077 | 36978155 | 187570 | 61,53,125,45,28,42,44,41,3 | 1,0,2,0,1,0,0,0,0,0,0,0 | 49,67,26,41,46,15,21,33,3 | 2,5,1,2,0,0,0,3,3,0,3,2,2,1,0 | 275 | 149 | 0.000297065489067 | 0.045833478335039 | 0.971,1.0,0.971,1. | 0.93,0.879,0.934,0 | 0,054 |
| 225752 | ENSG00000288044 | | chr14 | + | 24292197 | 24292600 | 24271200 | 24271310 | 24298613 | 24298861 | 225752 | 11,4,7,3,5,1,3,2,3,7,2,1 | 5,2,4,1,0,2,1,0,2,2,2,3 | 8,6,6,2,2,1,4,3,2,7,8,23,8,7 | 2,1,1,1,0,0,0,0,1,1,2,0,4,1,3 | 298 | 149 | 0.000298747996547 | 0.0460351013620358 | 0.524,0.5,0.467,0. | 0.667,0.75,0.75,0. | -0,213 |
| 222189 | ENSG00000152601 | MBNL1 | chr3 | + | 152299404 | 152299631 | 152269102 | 152269407 | 152414940 | 152415111 | 222189 | 34,7,8,4,2,16,5,13,12,20,3, | 2,2,2,1,4,0,6,2,0,0,0,2 | 7,5,4,9,1,0,2,3,4,1,4,13,7,1 | 1,2,1,0,0,3,0,0,1,1,0,0,1,3,0 | 298 | 149 | 0.00029880006413 | 0.0460351013620358 | 0.895,0.636,0.667, | 0.778,0.556,0.667, | -0,007 |
| 163378 | ENSG00000119522 | DENND1A | chr9 | - | 123667175 | 123667288 | 123667025 | 123667079 | 123671290 | 123671371 | 163378 | 2,1,3,0,0,6,1,5,1,0,0,0 | 22,23,40,19,9,19,9,32,13,1 | 2,7,0,22,9,5,3,0,1,1,0,2,1,1 | 22,21,26,55,22,11,21,16,1 | 262 | 149 | 0.00029968651971 | 0.0461386475849092 | 0.049,0.024,0.041, | 0.049,0.159,0.0,0. | -0,029 |
| 66084 | ENSG00000170606 | HSPA4 | chr5 | + | 133091192 | 133091374 | 133089561 | 133089695 | 133092699 | 133092789 | 66084 | 280,251,527,315,206,363, | 4,3,3,0,0,2,14,23,0,0,0,0 | 241,360,264,194,232,138, | 0,1,2,2,0,0,0,5,1,0,2,0,0,7,6 | 298 | 149 | 0.000300700315069 | 0.0462432121736926 | 0.972,0.977,0.989, | 1.0,0.994,0.985,0. | -0,01 |
| 122158 | ENSG00000089597 | GANAB | chr11 | - | 62639929 | 62640069 | 62639626 | 62639731 | 62646561 | 62646606 | 122158 | 0,0,2,5,2,5,2,4,0,2,0,4 | 6,3,20,15,17,7,11,11,3,2,3, | 2,1,2,0,0,0,0,0,1,0,0,4,3,2 | 5,12,4,5,4,4,7,11,8,4,13,6,1 | 289 | 149 | 0.000300795410713 | 0.0462432121736926 | 0.0,0.0,0.049,0.14 | 0.171,0.041,0.205, | 0,036 |
| 187574 | ENSG00000137409 | MTCH1 | chr6 | - | 36977633 | 36977784 | 36975657 | 36977250 | 36978077 | 36978155 | 187574 | 61,54,137,53,24,44,47,44,3 | 1,0,2,0,1,0,0,0,0,0,0,0 | 57,68,31,46,48,16,26,36,3 | 2,5,1,2,0,0,0,3,3,0,3,2,2,1,0 | 298 | 149 | 0.000302389423803 | 0.0464217180268324 | 0.968,1.0,0.972,1. | 0.934,0.872,0.939, | 0,053 |
| 31374 | ENSG00000171206 | TRIM8 | chr10 | + | 102656746 | 102656920 | 102656269 | 102656385 | 102657922 | 102658315 | 31374 | 24,21,73,30,10,31,23,23,9, | 0,2,1,0,2,2,0,1,3,0,0,0 | 22,33,26,8,15,20,15,25,8,2 | 3,1,0,5,0,2,0,2,2,0,2,0,4,2,6 | 298 | 149 | 0.000302473070005 | 0.0464217180268324 | 1.0,0.84,0.973,1.0 | 0.786,0.943,1.0,0. | 0,07 |
| 234238 | ENSG00000135048 | CEMIP2 | chr9 | - | 71690091 | 71690212 | 71685742 | 71685846 | 71694508 | 71694607 | 234238 | 15,22,26,27,12,17,14,37,3, | 5,0,1,2,6,5,4,2,0,0,1,0 | 22,30,9,5,11,1,26,18,1,8,3 | 1,1,3,1,1,0,2,1,0,0,2,5,2,0,0 | 270 | 149 | 0.000302768508038 | 0.0464217180268324 | 0.623,1.0,0.935,0. | 0.924,0.943,0.623, | -0,037 |
| 169708 | ENSG00000204463 | BAG6 | chr6 | - | 31640473 | 31640528 | 31640198 | 31640306 | 31640644 | 31640704 | 169708 | 26,20,58,37,21,18,22,27,2 | 2,0,0,0,0,0,1,0,1,2,0,1 | 41,41,25,13,19,17,20,22,1 | 3,1,3,2,1,2,2,0,1,6,2,3,0,0, | 204 | 149 | 0.000302819259817 | 0.0464217180268324 | 0.905,1.0,1.0,1.0,1 | 0.909,0.968,0.859, | 0,052 |
| 227631 | ENSG00000186088 | GSAP | chr7 | - | 77321332 | 77321806 | 77320724 | 77320819 | 77323646 | 77323742 | 227631 | 105,80,103,59,40,55,35,67 | 12,2,2,0,10,6,0,0,0,0,0,0 | 83,44,56,47,57,33,23,42,3 | 10,4,0,0,0,0,0,0,8,0,0,0,0,0 | 298 | 149 | 0.000303740994727 | 0.0465298776057558 | 0.814,0.952,0.963, | 0.806,0.846,1.0,1. | -0,023 |
| 76685 | ENSG00000159023 | EPB41 | chr1 | + | 29058588 | 29058645 | 29053103 | 29053312 | 29064981 | 29065158 | 76685 | 14,6,20,16,16,16,10,18,21, | 178,160,197,190,182,157, | 9,14,21,9,9,9,2,11,0,13,12, | 184,154,169,147,161,142, | 206 | 149 | 0.000304558478876 | 0.0466219245853203 | 0.054,0.026,0.068, | 0.034,0.062,0.082, | 0,019 |
| 232181 | ENSG00000256861 | | chr12 | - | 122209775 | 122209858 | 122207778 | 122208577 | 122216487 | 122216584 | 232181 | 11,3,21,16,6,11,5,6,3,1,2,8 | 25,15,40,32,19,12,18,15,3 | 16,23,16,14,22,5,14,23,10, | 9,21,21,10,22,7,16,14,7,14 | 232 | 149 | 0.000305694777258 | 0.0467626102373963 | 0.22,0.114,0.252,0 | 0.533,0.413,0.329, | -0,132 |
| 156507 | ENSG00000102763 | VWA8 | chr13 | - | 41837118 | 41837250 | 41833370 | 41833531 | 41865735 | 41865813 | 156507 | 6,1,3,2,1,11,4,3,2,0,0,0 | 8,6,32,12,15,20,21,30,22,2 | 9,3,0,0,0,0,0,2,0,1,1,0,0,1,0 | 11,14,4,22,5,8,18,19,15,12 | 281 | 149 | 0.0003059654941 | 0.0467707807248843 | 0.285,0.081,0.047, | 0.303,0.102,0.0,0. | 0,033 |
| 235872 | ENSG00000110697 | PITPNM1 | chr11 | - | 67503736 | 67503876 | 67502532 | 67502718 | 67504102 | 67504221 | 235872 | 3,2,3,1,5,2,1,0,1,1,4,1 | 12,5,26,5,12,18,15,12,13,1 | 0,0,0,0,1,3,0,0,2,1,1,0,1,1,2 | 12,3,10,3,7,3,8,4,4,13,6,12 | 289 | 149 | 0.000307106982 28 | 0.0469101977927444 | 0.114,0.171,0.056, | 0.0,0.0,0.0,0.0,0.0 | 0,034 |
| 252709 | ENSG00000174718 | RESF1 | chr12 | + | 31987158 | 31987322 | 31980877 | 31985957 | 31992377 | 31992941 | 252709 | 123,77,332,243,133,122,1 | 12,0,40,17,8,13,16,9,7,4,2, | 198,215,213,119,109,88,8 | 5,8,12,12,24,7,1,9,2,1,2,8,6 | 298 | 149 | 0.000307313439457 | 0.0469101977927444 | 0.837,1.0,0.806,0. | 0.952,0.931,0.899, | -0,04 |
| 101457 | ENSG00000087274 | ADD1 | chr4 | + | 2926641 | 2926675 | 2926013 | 2926112 | 2928170 | 2928523 | 101457 | 9,2,4,6,0,0,1,6,8,6,2,3 | 74,48,63,36,29,30,44,15,4 | 0,2,6,7,4,2,0,0,0,0,3,2,2,4,2 | 62,50,37,38,31,9,18,46,10, | 183 | 149 | 0.000308298720807 | 0.0470272444918578 | 0.09,0.033,0.049,0 | 0.0,0.032,0.117,0. | 0,034 |
| 76784 | ENSG00000116984 | MTR | chr1 | + | 236825337 | 236825399 | 236816448 | 236816543 | 236826828 | 236826896 | 76784 | 6,5,24,14,13,28,18,16,6,2,7 | 0,2,2,0,1,0,2,0,3,0,0,0 | 14,27,24,19,17,5,6,10,8,20 | 0,1,3,0,2,0,0,1,5,1,4,4,5,4,0 | 211 | 149 | 0.000309333812191 | 0.0471462519839506 | 1.0,0.638,0.894,1. | 1.0,0.95,0.85,1.0,0 | 0,049 |
| 198976 | ENSG00000054267 | ARID4B | chr1 | - | 235181072 | 235181184 | 235177855 | 235177913 | 235181584 | 235181865 | 198976 | 7,3,26,22,7,13,5,18,0,2,4,1 | 115,97,133,82,44,71,47,48 | 20,21,9,7,34,5,3,10,9,9,11, | 86,76,74,72,55,32,27,51,3 | 261 | 149 | 0.00030951700291 | 0.0471462519839506 | 0.034,0.017,0.1,0. | 0.117,0.136,0.065, | -0,03 |
| 209422 | ENSG00000091490 | SEL1L3 | chr4 | - | 25830097 | 25830273 | 25821995 | 25822128 | 25832994 | 25833110 | 209422 | 27,16,137,49,45,55,42,50,2 | 9,5,5,3,3,7,0,1,3,1,0,2 | 35,16,30,25,25,14,15,17,2 | 4,0,1,2,1,0,4,1,3,1,0,4,1,0,2 | 298 | 149 | 0.000310019944521 | 0.0471894644124465 | 0.6,0.615,0.932,0. | 0.814,1.0,0.938,0. | -0,061 |
| 38770 | ENSG00000163681 | SLMAP | chr3 | + | 57865241 | 57865292 | 57864547 | 57864716 | 57890040 | 57890100 | 38770 | 5,5,2,3,0,1,2,2,2,2,3,3 | 1,2,5,5,3,2,0,0,3,0,4,1 | 0,4,1,0,1,6,3,3,1,0,3,2,1,2 | 1,0,0,1,1,0,2,1,0,6,1,3,0,2,0 | 200 | 149 | 0.000310713941403 | 0.0472616765527273 | 0.788,0.651,0.23,0 | 0.0,1.0,1.0,0.0,0.4 | -0,049 |
| 227630 | ENSG00000186088 | GSAP | chr7 | - | 77321332 | 77321734 | 77320724 | 77320819 | 77323646 | 77323742 | 227630 | 105,80,103,60,40,55,35,67 | 12,2,2,0,10,6,0,0,0,0,0,0 | 83,44,56,47,56,33,23,42,3 | 10,4,0,0,0,0,0,0,8,0,0,0,0,0 | 298 | 149 | 0.000311849106931 | 0.0474008440210918 | 0.814,0.952,0.963, | 0.806,0.846,1.0,1. | -0,023 |
| 142374 | ENSG00000168961 | LGALS9 | chr17 | + | 27647245 | 27647432 | 27647029 | 27647118 | 27648835 | 27649541 | 142374 | 77,39,126,148,61,38,72,42 | 11,9,15,22,13,15,11,14,17, | 57,78,45,51,41,48,21,77,5 | 8,10,5,1,3,9,2,3,4,8,0,23,1 | 298 | 149 | 0.000312274952908 | 0.0474320750806928 | 0.778,0.684,0.808, | 0.781,0.796,0.818, | -0,11 |
| 124222 | ENSG00000149182 | ARFGAP2 | chr11 | - | 47173758 | 47173840 | 47172025 | 47172333 | 47175014 | 47175098 | 124222 | 52,46,72,35,24,36,24,32,3 | 1,3,5,5,2,0,0,4,2,0,1,0 | 40,47,30,28,36,12,12,32,1 | 1,0,1,4,0,0,0,0,0,0,3,0,2,1,0 | 231 | 149 | 0.000314507416866 | 0.0477374794354627 | 0.971,0.908,0.903, | 0.963,1.0,0.951,0. | -0,034 |
| 1730 | ENSG00000243156 | MICAL3 | chr22 | - | 17893807 | 17893904 | 17891484 | 17891632 | 17895283 | 17895288 | 1730 | 58,33,62,17,67,64,23,35,4 | 0,0,0,1,2,2,7,6,1,0,2,3 | 60,29,14,22,20,20,20,33,8, | 0,2,0,0,1,0,5,0,0,0,1,2,0,0,0 | 246 | 149 | 0.000314807061197 | 0.0477492872364281 | 1.0,1.0,1.0,0.911,0 | 1.0,0.898,1.0,1.0,0 | -0,067 |
| 41044 | ENSG00000009790 | TRAF3IP3 | chr1 | + | 209781344 | 209781458 | 209779314 | 209779374 | 209782055 | 209782307 | 41044 | 162,51,133,143,108,73,53, | 11,2,7,4,8,8,0,2,2,0,2,0 | 121,111,64,71,46,68,36,67 | 2,1,0,2,1,1,0,3,2,2,5,2,2,0,2 | 263 | 149 | 0.000315744111657 | 0.047846356889879 | 0.893,0.935,0.915, | 0.972,0.984,1.0,0. | -0,012 |
| 133122 | ENSG00000119801 | YPEL5 | chr2 | + | 30155814 | 30155910 | 30148244 | 30148541 | 30156627 | 30156787 | 133122 | 18,13,25,16,3,7,7,6,11,9,5, | 10,13,14,7,8,3,3,7,8,3,5,3 | 32,22,14,13,5,7,4,36,8,31, | 15,9,5,0,0,1,9,6,1,1,0,5,9,4 | 245 | 149 | 0.000316018298644 | 0.047846356889879 | 0.523,0.378,0.521, | 0.565,0.598,0.63,1 | -0,25 |
| 150998 | ENSG00000170776 | AKAP13 | chr15 | + | 85487197 | 85487369 | 85485709 | 85485753 | 85521427 | 85521575 | 150998 | 1,0,32,22,3,0,0,0,0,0,0,2 | 51,40,109,51,53,84,53,71, | 0,4,2,2,7,4,0,2,2,4,4,3,2,2,5 | 95,65,31,48,62,51,27,43,6 | 298 | 149 | 0.000316113940359 | 0.047846356889879 | 0.01,0.0,0.128,0.1 | 0.0,0.03,0.031,0.0 | 0,006 |
| 166852 | ENSG00000125386 | FAM193A | chr4 | + | 2693201 | 2693382 | 2690697 | 2690970 | 2693585 | 2693874 | 166852 | 2,0,6,0,1,4,2,9,4,0,1,2 | 18,24,5,4,7,10,4,11,8,13,2, | 0,3,1,5,0,0,0,0,3,4,1,1,0,1 | 6,13,1,13,8,3,7,11,2,9,9,5,8 | 298 | 149 | 0.000316418032545 | 0.04785872773204 | 0.053,0.0,0.375,0. | 0.0,0.103,0.333,0. | 0,06 |
| 881 | ENSG00000196588 | MRTFA | chr22 | - | 40424205 | 40424381 | 40423535 | 40423685 | 40429605 | 40429767 | 881 | 40,39,27,21,22,47,40,27,8 | 0,3,2,2,6,0,2,0,5,2,0,2 | 28,20,31,48,19,8,22,43,12, | 0,2,0,2,3,0,0,2,0,1,0,0,2,0,1 | 298 | 149 | 0.000316717091443 | 0.0478703204412699 | 1.0,0.867,0.871,0. | 1.0,0.833,1.0,0.92 | -0,035 |
| 221253 | ENSG00000118873 | RAB3GAP2 | chr1 | - | 220206532 | 220206677 | 220205906 | 220206006 | 220210387 | 220211002 | 221253 | 2,0,4,2,0,0,14,14,0,1,0,0 | 26,23,128,101,34,43,38,34 | 0,0,0,0,0,0,0,0,1,0,0,0,4,0,1 | 45,27,73,63,34,25,14,53,3 | 294 | 149 | 0.000317133739664 | 0.0478996574888578 | 0.038,0.0,0.016,0. | 0.0,0.0,0.0,0.0,0.0, | 0,029 |
| 94926 | ENSG00000172845 | SP3 | chr2 | - | 173910164 | 173910257 | 173907885 | 173907965 | 173913069 | 173913266 | 94926 | 235,191,224,122,71,90,85, | 6,1,17,8,9,3,6,0,1,0,1,6 | 124,212,144,151,121,66,7 | 9,14,6,8,4,3,6,6,6,7,8,17,3, | 242 | 149 | 0.000317789128107 | 0.0479649872591849 | 0.96,0.992,0.89,0. | 0.895,0.903,0.937, | 0,038 |
| 48660 | ENSG00000120008 | WDR11 | chr10 | + | 120867065 | 120867169 | 120866568 | 120866764 | 120871169 | 120871346 | 48660 | 439,289,514,280,249,150,2 | 2,0,5,6,1,3,4,7,3,1,2,4 | 294,379,433,314,287,197, | 6,14,6,11,7,2,6,1,3,9,11,3,4 | 253 | 149 | 0.000321003227985 | 0.0483840200657815 | 0.992,1.0,0.984,0. | 0.967,0.941,0.977, | 0,017 |
| 255826 | ENSG00000182796 | TMEM198B | chr12 | + | 55834517 | 55834726 | 55833446 | 55833541 | 55834850 | 55834988 | 255826 | 2,0,1,1,1,4,1,7,4,1,0,3 | 11,6,6,3,7,10,3,9,8,7,6,3 | 0,1,0,3,1,0,0,3,0,2,1,0,1,2,2 | 9,13,3,24,7,1,8,4,10,7,5,10 | 298 | 149 | 0.000321015005524 | 0.0483840200657815 | 0.083,0.0,0.077,0. | 0.0,0.037,0.0,0.05 | 0,066 |
| 26654 | ENSG00000126214 | KLC1 | chr14 | + | 103692358 | 103692425 | 103686862 | 103687211 | 103700654 | 103700727 | 26654 | 10,6,12,6,7,8,4,1,18,10,1,2 | 2,1,1,6,1,3,3,2,5,2,1,0 | 6,3,5,2,11,7,5,8,12,4,6,12,8 | 0,0,0,1,0,1,1,8,4,1,0,0,0,2,1 | 216 | 149 | 0.000321471504449 | 0.048418917686538 | 0.775,0.805,0.892, | 1.0,1.0,1.0,0.58,1. | -0,163 |
| 116597 | ENSG00000148296 | SURF6 | chr9 | - | 133333717 | 133333806 | 133332547 | 133332760 | 133334391 | 133334601 | 116597 | 18,26,39,21,11,17,9,4,32,8 | 2,0,3,0,0,0,2,1,0,2,1,0 | 14,20,20,5,13,11,9,9,17,17 | 4,1,1,5,1,0,0,0,0,0,3,4,0,2,2 | 238 | 149 | 0.000322142326857 | 0.0484860245816496 | 0.849,1.0,0.891,1. | 0.687,0.926,0.926, | 0,037 |
| 178830 | ENSG00000285708 | | chr3 | - | 71000995 | 71001059 | 70987993 | 70988077 | 71015548 | 71015653 | 178830 | 156,62,317,249,72,83,60,1 | 3,10,3,3,1,2,3,6,15,4,3,3 | 188,134,92,88,115,77,52,1 | 1,0,5,4,2,0,0,4,0,4,6,2,2,0,5 | 213 | 149 | 0.000329227121517 | 0.0495177376598361 | 0.973,0.813,0.987, | 0.992,1.0,0.928,0. | -0,038 |
| 40837 | ENSG00000146232 | NFKBIE | chr6 | - | 44262559 | 44262662 | 44261625 | 44261848 | 44264981 | 44265551 | 40837 | 29,12,61,55,25,34,57,20,2 | 6,3,3,2,1,0,2,8,2,0,3,2 | 45,33,20,10,15,2,2,29,7,27 | 0,0,0,0,2,2,0,4,0,1,2,9,0,1,1 | 252 | 149 | 0.000329717810504 | 0.0495569092685659 | 0.741,0.703,0.923, | 1.0,1.0,1.0,1.0,0.8 | -0,051 |
| 163928 | ENSG00000223865 | HLA-DPB1 | chr6 | + | 33080697 | 33080935 | 33076009 | 33076141 | 33084949 | 33084957 | 163928 | 464,328,201,235,139,55,3 | 6,3,15,6,0,0,1,3,1,2,3,0 | 352,390,268,242,260,173, | 6,23,5,25,1,1,1,8,5,2,3,19,3 | 298 | 149 | 0.000330920272217 | 0.0497029316884418 | 0.975,0.982,0.87,0 | 0.967,0.894,0.964, | 0,014 |
| 14848 | ENSG00000166747 | AP1G1 | chr16 | - | 71808014 | 71808150 | 71789278 | 71789482 | 71808504 | 71808677 | 14848 | 1,2,4,3,5,1,8,4,1,1,2,2 | 0,1,1,0,2,0,1,1,1,0,0,3 | 4,2,2,5,5,4,4,1,3,2,2,4,4,0, | 2,2,0,2,1,0,0,2,1,6,1,2,3,1, | 285 | 149 | 0.000331233100179 | 0.0497152241175916 | 1.0,0.511,0.677,1. | 0.511,0.343,1.0,0. | 0,152 |
| 216180 | ENSG00000073614 | KDM5A | chr12 | - | 321004 | 321109 | 318105 | 318461 | 322416 | 322567 | 216180 | 64,50,120,72,51,39,26,54,4 | 4,0,3,3,4,1,0,2,0,0,1,0 | 83,90,59,55,53,32,32,42,2 | 3,6,5,6,0,1,3,0,0,0,3,1,4,9,0 | 254 | 149 | 0.000331678109894 | 0.0497292988226902 | 0.904,1.0,0.959,0. | 0.942,0.898,0.874, | 0,032 |

| | | | | | | | | | | | | | | | | | | | | | | |
|---|---|---|---|---|---|---|---|---|---|---|---|---|---|---|---|---|---|---|---|---|---|---|
| 215052 | ENSG00000135624 | CCT7 | chr2 | + | 73243996 | 73244423 | 73242972 | 73243129 | 73244543 | 73244715 | 215052 | 35,19,107,71,25,23,17,30, | 4,1,7,5,3,1,3,5,2,0,0,4 | 67,43,32,8,22,29,17,30,18, | 3,5,1,0,3,0,2,2,0,0,1,7,0,0, | 298 | 149 | 0.000331788976074 | 0.0497292988226902 | 0.814,0.905,0.884, | 0.918,0.811,0.941, | -0,065 |
| 26453 | ENSG00000084093 | REST | chr4 | + | 56919786 | 56919870 | 56910629 | 56911536 | 56929840 | 56935822 | 26453 | 51,54,167,105,59,35,40,54 | 3,5,3,1,2,1,3,4,1,0,0,2 | 67,80,49,20,46,36,28,73,2 | 4,4,2,2,4,8,5,2,3,5,2,7,0,6, | 233 | 149 | 0.000334071414802 | 0.0500084205300324 | 0.916,0.874,0.973, | 0.915,0.927,0.94,0 | 0,052 |
| 33103 | ENSG00000281649 | EBLN3P | chr9 | + | 37084198 | 37084292 | 37079873 | 37080034 | 37086667 | 37087837 | 33103 | 22,10,41,13,9,4,8,11,13,2, | 6,1,12,6,2,2,4,1,2,2,0,0 | 2,7,6,1,5,1,5,4,5,6,11,8,12, | 9,3,5,2,2,1,4,4,1,9,6,2,10,8 | 243 | 149 | 0.000334115943903 | 0.0500084205300324 | 0.692,0.86,0.677,0 | 0.12,0.589,0.424,0 | 0,255 |
| 99027 | ENSG00000136280 | CCM2 | chr7 | + | 45063917 | 45064001 | 45027671 | 45027797 | 45069825 | 45069961 | 99027 | 26,10,14,15,5,4,6,9,11,3,1, | 3,2,1,0,0,0,6,0,0,4,1,5 | 13,27,14,2,16,10,1,4,6,24, | 2,4,0,2,5,3,0,0,0,1,0,0,0,1, | 233 | 149 | 0.00033470124822 | 0.050061212199888 | 0.847,0.762,0.9,1. | 0.806,0.812,1.0,0. | -0,126 |
| 104002 | ENSG00000291151 | NSUN5P1 | chr7 | + | 75415595 | 75415770 | 75415119 | 75415234 | 75415980 | 75416191 | 104002 | 14,12,21,6,18,8,20,35,7,15 | 1,0,1,1,4,5,1,0,0,0,0,0 | 14,20,11,4,11,7,7,21,4,14,1 | 1,0,2,0,1,0,9,2,3,0,0,5,1,0, | 298 | 149 | 0.000335815158937 | 0.0501751093055671 | 0.875,1.0,0.913,0. | 0.875,1.0,0.733,1. | 0,08 |
| 206123 | ENSG00000142892 | PIGK | chr1 | - | 77212294 | 77212347 | 77210435 | 77210489 | 77219309 | 77219430 | 206123 | 2,0,1,0,0,2,0,0,0,0,0,0 | 5,9,40,27,14,22,12,11,10,1 | 2,2,1,0,0,0,3,0,3,5,6,0,2,0, | 18,19,10,6,8,26,11,18,12,1 | 202 | 149 | 0.000335928990291 | 0.0501751093055671 | 0.228,0.0,0.018,0. | 0.076,0.072,0.069, | -0,051 |
| 187429 | ENSG00000072858 | SIDT1 | chr3 | + | 113598571 | 113598685 | 113593004 | 113593048 | 113601587 | 113602009 | 187429 | 1,0,12,5,1,0,0,2,3,0,0,0 | 2,3,22,14,17,11,13,9,3,1,5, | 2,1,0,2,0,1,0,0,7,0,2,0,0,1, | 5,6,7,5,2,4,5,4,4,14,3,3,11, | 263 | 149 | 0.000337215288467 | 0.050321133657334 | 0.221,0.0,0.236,0. | 0.185,0.086,0.0,0. | -0,008 |
| 127214 | ENSG00000135655 | USP15 | chr12 | + | 62349220 | 62349307 | 62325871 | 62325933 | 62355330 | 62355475 | 127214 | 128,121,200,78,78,101,87, | 7,9,21,12,8,5,2,9,7,2,1,14 | 114,170,130,114,93,59,70, | 5,15,6,9,17,15,12,24,8,24,1 | 236 | 149 | 0.000337374243801 | 0.050321133657334 | 0.92,0.895,0.857,0 | 0.935,0.877,0.932, | 0,071 |
| 42066 | ENSG00000113971 | NPHP3 | chr3 | - | 132690527 | 132690650 | 132689073 | 132689263 | 132691191 | 132691286 | 42066 | 31,27,64,43,24,16,26,23,1 | 4,3,8,8,2,1,0,5,3,0,2,4 | 21,26,20,8,24,12,13,34,19, | 9,15,4,0,10,1,5,10,4,4,1,3, | 272 | 149 | 0.000339499300482 | 0.050603029045735 | 0.809,0.831,0.814, | 0.561,0.487,0.733, | 0,158 |
| 25458 | ENSG00000172175 | MALT1 | chr18 | + | 58709377 | 58709556 | 58696365 | 58696487 | 58709975 | 58710072 | 25458 | 85,49,100,66,42,30,37,28, | 1,0,3,2,6,0,0,4,0,1,0,1 | 57,49,27,15,23,24,24,48,2 | 2,7,4,2,1,0,0,8,2,1,8,2,2,2, | 298 | 149 | 0.000340493512432 | 0.0507160963143611 | 0.977,1.0,0.943,0. | 0.934,0.778,0.771, | 0,044 |
| 180710 | ENSG00000100991 | TRPC4AP | chr20 | - | 35009031 | 35009178 | 35008663 | 35008747 | 35010186 | 35010288 | 180710 | 1,2,3,3,4,3,4,2,0,2,1,7 | 91,60,118,75,39,68,43,40, | 3,12,7,0,14,0,3,2,9,4,2,11, | 79,65,32,55,63,22,42,46,5 | 296 | 149 | 0.000341798060885 | 0.05087519947603 | 0.006,0.017,0.013, | 0.019,0.085,0.099, | -0,015 |
| 245686 | ENSG00000109919 | MTCH2 | chr11 | - | 47628952 | 47629311 | 47627079 | 47627127 | 47630554 | 47630614 | 245686 | 6,7,33,29,14,9,5,18,14,4,4, | 2,0,6,0,0,3,1,0,1,0,0,0 | 21,40,8,1,11,6,8,13,5,11,4, | 7,5,0,2,3,0,1,2,4,0,0,3,0,4, | 298 | 149 | 0.000344484170896 | 0.0512395802253746 | 0.6,1.0,0.733,1.0,1 | 0.6,0.8,1.0,0.2,0.6 | 0,077 |
| 163925 | ENSG00000223865 | HLA-DPB1 | chr6 | + | 33080673 | 33080935 | 33076009 | 33076141 | 33084949 | 33084957 | 163925 | 462,322,203,235,140,53,3 | 6,3,15,6,0,0,1,3,1,2,3,0 | 352,386,268,241,260,172,1 | 6,23,5,25,1,1,1,8,5,2,3,19, | 298 | 149 | 0.000345259409383 | 0.0512859638096843 | 0.975,0.982,0.871, | 0.967,0.894,0.964, | 0,014 |
| 245569 | ENSG00000109458 | GAB1 | chr4 | + | 143436713 | 143436772 | 143433490 | 143433716 | 143437998 | 143438600 | 245569 | 7,0,0,0,3,4,2,21,0,0,0,0 | 38,35,207,116,89,95,164,3 | 3,4,9,30,1,1,5,7,0,0,6,0,3,2 | 119,102,132,109,83,39,12 | 208 | 149 | 0.000345485122879 | 0.0512859638096843 | 0.117,0.0,0.0,0.0,0 | 0.018,0.027,0.047, | -0,013 |
| 252712 | ENSG00000174718 | RESF1 | chr12 | + | 31987244 | 31987322 | 31980877 | 31985957 | 31992377 | 31992941 | 252712 | 111,70,292,205,114,111,9 | 12,0,40,17,8,13,16,9,7,4,2, | 166,183,185,108,92,71,74, | 5,8,12,12,24,7,1,9,2,1,2,8, | 227 | 149 | 0.000345510858213 | 0.0512859638096843 | 0.859,1.0,0.827,0. | 0.956,0.938,0.91,0 | -0,033 |
| 75159 | ENSG00000124181 | PLCG1 | chr20 | + | 41137520 | 41137858 | 41136959 | 41137035 | 41159605 | 41159758 | 75159 | 34,11,35,17,15,16,13,23,9, | 1,3,3,0,0,0,1,4,0,0,1,1 | 25,36,12,15,12,12,6,13,22, | 2,3,1,3,4,0,0,1,0,0,0,1,0,0, | 298 | 149 | 0.000346261607956 | 0.051361979422452 | 0.944,0.647,0.854, | 0.862,0.857,0.857, | 0,017 |
| 82788 | ENSG00000197622 | CDC42SE1 | chr1 | - | 151056650 | 151056786 | 151055676 | 151055993 | 151059478 | 151059615 | 82788 | 110,32,90,81,43,52,38,63, | 197,165,295,254,162,112,1 | 88,49,108,143,73,66,44,68 | 274,239,156,130,165,98,1 | 285 | 149 | 0.000346960649586 | 0.0514013921226271 | 0.226,0.092,0.138, | 0.144,0.097,0.266, | -0,049 |
| 142123 | ENSG00000215067 | ALOX12-AS1 | chr17 | - | 7011902 | 7011957 | 7009795 | 7009889 | 7012195 | 7012474 | 142123 | 0,0,3,3,0,1,0,8,4,0,0,2 | 12,2,14,15,9,11,6,8,5,4,2,1 | 2,0,0,0,0,0,0,2,0,0,0,0,2,0, | 7,7,12,8,9,9,8,7,12,15,6,3, | 204 | 149 | 0.000347004951676 | 0.0514013921226271 | 0.0,0.0,0.135,0.12 | 0.173,0.0,0.0,0.0,0 | 0,072 |
| 80185 | ENSG00000204681 | GABBR1 | chr6 | - | 29630457 | 29630643 | 29629086 | 29629107 | 29631395 | 29631599 | 80185 | 14,8,18,6,6,12,17,9,5,7,7, | 3,5,4,0,2,1,7,1,0,4,0,0 | 17,6,13,14,4,3,13,10,9,10, | 5,3,1,0,2,0,0,2,0,0,0,3,0,0, | 298 | 149 | 0.000347609713896 | 0.0514340033545577 | 0.7,0.444,0.692,1. | 0.63,0.5,0.867,1.0, | -0,088 |
| 164808 | ENSG00000065882 | TBC1D1 | chr4 | + | 38118032 | 38118504 | 38115709 | 38115954 | 38124961 | 38125131 | 164808 | 152,100,255,198,76,106,9 | 6,3,3,4,1,2,6,2,6,2,7,2 | 97,100,113,68,116,90,34,9 | 0,2,3,0,1,0,3,2,0,4,5,1,0,0, | 298 | 149 | 0.000347904350545 | 0.0514340033545577 | 0.927,0.943,0.977, | 1.0,0.962,0.95,1.0, | -0,038 |
| 171944 | ENSG00000171608 | PIK3CD | chr1 | + | 9710104 | 9710280 | 9630137 | 9630224 | 9710423 | 9710596 | 171944 | 15,6,15,3,10,3,3,3,6,5,7,9 | 0,4,4,0,0,1,2,1,2,0,2,10 | 10,13,11,27,12,6,4,8,15,16 | 1,4,1,2,1,3,0,0,3,1,0,2,0,2, | 298 | 149 | 0.000347942019896 | 0.0514340033545577 | 1.0,0.429,0.652,1. | 0.833,0.619,0.846, | -0,172 |
| 163923 | ENSG00000223865 | HLA-DPB1 | chr6 | + | 33080492 | 33080935 | 33076009 | 33076141 | 33084949 | 33084957 | 163923 | 462,322,201,235,140,53,3 | 6,3,15,6,0,0,1,3,1,2,3,0 | 352,386,268,239,260,172,1 | 6,23,5,25,1,1,1,8,5,2,3,19, | 298 | 149 | 0.000348382325868 | 0.0514567396092526 | 0.975,0.982,0.87,0 | 0.967,0.894,0.964, | 0,014 |
| 121240 | ENSG00000109458 | GAB1 | chr4 | + | 143380611 | 143380739 | 143337177 | 143337260 | 143415476 | 143415771 | 121240 | 3,2,14,3,2,2,12,9,0,0,1,4 | 8,9,82,31,9,22,34,85,12,3, | 5,5,8,5,2,0,4,2,0,2,2,0,0,0, | 26,46,22,30,22,18,25,34,1 | 277 | 149 | 0.000348573980283 | 0.0514567396092526 | 0.168,0.107,0.084, | 0.094,0.055,0.164, | 0,033 |
| 79992 | ENSG00000137075 | RNF38 | chr9 | - | 36390466 | 36390616 | 36375933 | 36376127 | 36400553 | 36400923 | 79992 | 91,56,42,24,15,28,31,43,6 | 2,0,1,1,0,0,4,1,0,2,0,0 | 28,48,52,87,32,9,32,21,32, | 2,4,0,5,0,0,0,3,0,2,1,5,1,3, | 298 | 149 | 0.000350071059028 | 0.0516423194692635 | 0.958,1.0,0.955,0. | 0.875,0.857,1.0,0. | 0,065 |
| 115926 | ENSG00000198771 | RCSD1 | chr1 | + | 167685420 | 167685510 | 167683899 | 167684001 | 167690048 | 167690120 | 115926 | 82,24,148,110,90,72,155,2 | 23,2,28,18,11,14,18,22,20, | 79,26,58,39,31,32,108,162 | 16,10,16,13,7,7,5,33,13,12 | 239 | 149 | 0.000350926133909 | 0.0517330018680602 | 0.69,0.882,0.767,0 | 0.755,0.618,0.693, | 0,094 |
| 129620 | ENSG00000127314 | RAP1B | chr12 | + | 68624674 | 68624873 | 68610907 | 68611052 | 68648698 | 68648781 | 129620 | 0,0,2,7,0,0,0,2,2,2,1,0 | 7,4,13,13,6,2,1,3,2,2,0,2 | 2,2,3,0,0,0,1,3,3,0,3,7,4,10 | 4,7,6,2,8,2,0,4,0,1,4,9,2,6, | 298 | 149 | 0.00035166308868 | 0.0517727482090264 | 0.0,0.0,0.071,0.21 | 0.2,0.125,0.2,0.0,0 | -0,186 |
| 184816 | ENSG00000142765 | SYTL1 | chr1 | + | 27349971 | 27350272 | 27349397 | 27349765 | 27350388 | 27350485 | 184816 | 5,7,10,2,7,7,7,2,5,4,8,4 | 0,0,0,4,4,1,0,2,0,0,0,0 | 3,9,14,2,2,3,5,3,6,8,6,10,9, | 3,1,0,0,1,0,0,2,5,2,2,4,3,0, | 298 | 149 | 0.000351676839682 | 0.0517727482090264 | 1.0,1.0,1.0,0.2,0.4 | 0.333,0.818,1.0,1. | 0,079 |
| 107747 | ENSG00000172725 | CORO1B | chr11 | - | 67440701 | 67440836 | 67440334 | 67440439 | 67441124 | 67441244 | 107747 | 7,3,3,8,8,4,2,13,0,2,0,4 | 20,13,24,17,27,22,11,19,1 | 0,1,1,3,2,3,0,4,0,4,2,6,6,7, | 2,7,18,11,12,15,2,16,13,20 | 284 | 149 | 0.000353021555052 | 0.0519351895525612 | 0.155,0.108,0.062, | 0.0,0.07,0.028,0.1 | 0,051 |
| 52924 | ENSG00000226571 | | chr6 | + | 139283103 | 139283166 | 139271658 | 139271858 | 139286844 | 139286902 | 52924 | 57,25,313,233,193,195,12 | 12,4,0,0,0,0,0,0,0,0,0,0 | 151,120,82,71,106,109,0,0 | 0,0,0,0,0,9,24,15,5,2,0,0, | 212 | 149 | 0.000353384171084 | 0.05195302494985 | 0.77,0.815,1.0,1.0, | 1.0,1.0,1.0,1.0,1.0, | 0,123 |
| 131276 | ENSG00000204852 | TCTN1 | chr12 | + | 110634669 | 110634779 | 110632471 | 110632559 | 110636480 | 110636501 | 131276 | 26,6,38,25,13,8,17,19,10,6 | 14,3,8,6,4,4,2,2,3,1,2,4 | 20,13,44,23,24,13,8,22,4,1 | 12,2,1,2,2,2,2,1,0,0,0,2,3,9 | 259 | 149 | 0.000353929593901 | 0.0519614044580369 | 0.517,0.535,0.732, | 0.489,0.789,0.962, | -0,173 |
| 252197 | ENSG00000198825 | INPP5F | chr10 | + | 119810085 | 119810217 | 119807931 | 119808060 | 119811809 | 119811955 | 252197 | 7,5,32,3,17,38,3,20,9,9,3,1 | 3,2,9,1,4,6,1,4,7,3,4,3 | 7,15,12,7,4,4,16,18,8,7,15, | 0,0,0,0,1,0,6,0,0,1,3,0,0,3, | 281 | 149 | 0.000353975830455 | 0.0519614044580369 | 0.553,0.57,0.653,0 | 1.0,1.0,1.0,1.0,0.6 | -0,222 |
| 181068 | ENSG00000076685 | NT5C2 | chr10 | - | 103177517 | 103177652 | 103174857 | 103174982 | 103181184 | 103181328 | 181068 | 2,0,4,1,0,3,2,0,2,0,0,4 | 14,2,51,29,10,12,8,11,3,5, | 10,0,0,0,2,6,4,2,1,6,3,0,5,7 | 16,9,10,10,15,17,6,20,7,21 | 284 | 149 | 0.000354165433139 | 0.0519614044580369 | 0.07,0.0,0.04,0.01 | 0.247,0.0,0.0,0.0,0 | -0,02 |
| 184996 | ENSG00000108654 | DDX5 | chr17 | - | 64505893 | 64506262 | 64504676 | 64504842 | 64506754 | 64507314 | 184996 | 17,12,11,4,13,9,16,17,29,1 | 19,20,23,8,6,6,13,11,6,6,0, | 4,1,8,0,0,5,0,5,0,4,6,7,9,11 | 13,12,13,9,11,6,9,6,5,6,9,3 | 298 | 149 | 0.000357120002489 | 0.0523591929534809 | 0.309,0.231,0.193, | 0.133,0.04,0.235,0 | 0,238 |
| 102291 | ENSG00000274272 | | chr7 | - | 100577272 | 100577389 | 100577085 | 100577154 | 100577496 | 100577558 | 102291 | 36,25,21,10,6,16,23,16,11, | 0,2,0,0,1,1,0,1,0,2,1,2 | 8,1,5,11,7,2,3,17,7,17,13,2 | 0,2,7,6,0,0,2,0,4,0,0,2,1,0, | 266 | 149 | 0.000357624956436 | 0.0523975336954913 | 1.0,0.875,1.0,1.0,0 | 1.0,0.219,0.286,0. | 0,144 |
| 118791 | ENSG00000291075 | | chr9 | + | 41008880 | 41009006 | 41007439 | 41007609 | 41011616 | 41011730 | 118791 | 74,25,127,96,35,47,30,74, | 1,0,16,11,16,13,5,7,1,0,6,1 | 105,75,85,38,13,17,23,54, | 1,0,0,0,3,4,2,1,0,0,1,20,2,0 | 275 | 149 | 0.000358580923227 | 0.052501857610252 | 0.976,1.0,0.811,0. | 0.983,1.0,1.0,1.0,0 | -0,069 |
| 216299 | ENSG00000163219 | ARHGAP25 | chr2 | + | 68806730 | 68806939 | 68787839 | 68787956 | 68807275 | 68807480 | 216299 | 2,0,2,0,0,2,0,6,1,0,0,12 | 9,27,23,25,24,22,7,17,13,9 | 10,20,1,11,11,0,2,3,4,1,5,5 | 29,78,10,4,14,1,6,20,5,12, | 298 | 149 | 0.000358921805745 | 0.0525160429451408 | 0.1,0.0,0.042,0.0,0 | 0.147,0.114,0.048, | -0,083 |
| 45379 | ENSG00000174953 | DHX36 | chr3 | - | 154306215 | 154306295 | 154305093 | 154305168 | 154309652 | 154309823 | 45379 | 22,19,72,41,15,29,30,53,5, | 0,0,2,0,0,0,0,1,0,0,0,0 | 33,90,20,32,47,29,28,26,2 | 2,3,0,12,0,3,0,2,0,1,3,3,0,0 | 229 | 149 | 0.000359787060292 | 0.0526068809603991 | 1.0,1.0,0.959,1.0,1 | 0.915,0.951,1.0,0. | 0,062 |
| 242626 | ENSG00000132386 | SERPINF1 | chr17 | + | 1771421 | 1771515 | 1771028 | 1771184 | 1771871 | 1772075 | 242626 | 1,1,6,4,7,4,5,11,2,0,2,0 | 12,6,26,22,22,31,21,35,20, | 0,1,1,0,4,3,0,5,1,1,5,0,2,1, | 7,28,5,8,50,21,12,16,35,51 | 243 | 149 | 0.000360382723692 | 0.0526582036679924 | 0.049,0.093,0.124, | 0.0,0.021,0.109,0. | 0,038 |
| 29213 | ENSG00000011243 | AKAP8L | chr19 | - | 15398629 | 15398766 | 15397713 | 15397855 | 15399301 | 15399410 | 29213 | 19,10,23,10,8,18,13,11,23, | 62,60,67,82,41,25,35,53,9 | 12,8,2,0,5,7,6,3,4,8,5,3,18, | 54,75,26,21,35,14,23,43,1 | 286 | 149 | 0.000360908186527 | 0.052699206169873 | 0.138,0.08,0.152,0 | 0.104,0.053,0.039, | 0,048 |
| 196123 | ENSG00000229474 | PATL2 | chr15 | - | 44675212 | 44675685 | 44674149 | 44674230 | 44676474 | 44676824 | 196123 | 0,1,1,2,4,3,0,0,2,1,0,4 | 9,8,4,9,12,3,7,8,2,1,1,8 | 0,0,0,0,4,1,1,0,1,6,2,0,0,1, | 7,4,5,5,6,4,7,3,5,6,2,1,5,2, | 298 | 149 | 0.000362058234579 | 0.0528306356469774 | 0.0,0.059,0.111,0. | 0.0,0.0,0.0,0.0,0.2 | 0,025 |
| 133514 | ENSG00000186866 | POFUT2 | chr21 | - | 45266153 | 45266227 | 45265399 | 45265635 | 45267589 | 45267713 | 133514 | 8,6,5,0,1,1,0,2,1,0,0,3 | 18,4,17,6,5,9,1,2,9,4,3,6 | 0,0,0,1,0,0,2,1,2,1,1,0,3,0, | 12,3,8,7,5,2,4,9,3,6,18,9,7, | 223 | 149 | 0.000362421810432 | 0.0528306356469774 | 0.229,0.501,0.164, | 0.0,0.0,0.0,0.087,0 | 0,054 |
| 57629 | ENSG00000111726 | CMAS | chr12 | + | 22061280 | 22061452 | 22060641 | 22060926 | 22062280 | 22062413 | 57629 | 31,15,54,40,20,11,17,17,1 | 8,1,0,0,0,0,1,2,0,0,0,2 | 22,17,20,5,13,18,7,14,10,1 | 0,3,0,3,0,0,8,0,0,0,4,5,2,2, | 298 | 149 | 0.000362544655977 | 0.0528306356469774 | 0.66,0.882,1.0,1.0, | 1.0,0.739,1.0,0.45 | 0,087 |
| 229549 | ENSG00000179406 | LINC00174 | chr7 | - | 66427109 | 66427157 | 66411614 | 66411702 | 66427948 | 66428040 | 229549 | 0,0,0,0,2,2,2,7,0,0,0,0 | 12,1,11,7,4,15,6,8,2,1,2,9 | 3,1,5,1,1,0,0,0,3,2,4,4,2,1, | 5,7,7,2,2,1,4,7,11,6,7,5,4,6 | 197 | 149 | 0.000365025312049 | 0.0531293940280372 | 0.0,0.0,0.0,0.0,0.2 | 0.312,0.098,0.351, | -0,115 |
| 225187 | ENSG00000118058 | KMT2A | chr11 | + | 118507268 | 118507609 | 118505613 | 118506646 | 118509135 | 118509200 | 225187 | 8,4,37,15,10,16,5,7,4,2,3,9 | 2,0,3,0,0,4,1,0,2,0,0,1 | 14,23,8,14,8,8,6,8,12,13,1 | 2,4,0,12,6,1,1,0,1,0,2,2,0,0 | 298 | 149 | 0.00036508855029 | 0.0531293940280372 | 0.667,1.0,0.86,1.0, | 0.778,0.742,1.0,0. | 0,044 |
| 204813 | ENSG00000138073 | PREB | chr2 | - | 27132227 | 27132403 | 27132009 | 27132082 | 27132602 | 27132620 | 204813 | 53,20,17,25,38,21,9,22,36, | 0,2,1,0,0,2,0,1,0,1,0,0 | 22,28,25,16,9,13,32,27,4,1 | 2,1,2,0,2,0,2,3,0,2,0,2,0,4, | 298 | 149 | 0.000366109555921 | 0.0531819240158175 | 1.0,0.833,0.895,1. | 0.846,0.933,0.862, | 0,069 |
| 213601 | ENSG00000272752 | STAG3L5P-PVR | chr7 | + | 100351251 | 100351423 | 100349730 | 100349887 | 100352162 | 100352489 | 213601 | 104,44,53,49,65,111,36,59 | 3,0,0,1,0,0,0,1,0,0,0,0 | 36,20,26,33,26,17,16,33,2 | 3,5,1,0,0,0,1,4,0,1,0,3,0,1, | 298 | 149 | 0.000366118676375 | 0.0531819240158175 | 0.945,1.0,1.0,0.96 | 0.857,0.667,0.929, | 0,077 |
| 123032 | ENSG00000110330 | BIRC2 | chr11 | + | 102350843 | 102350943 | 102348597 | 102350749 | 102362895 | 102362974 | 123032 | 39,35,107,55,21,19,14,22, | 5,0,2,0,0,0,2,0,0,0,0,0 | 62,72,27,24,49,21,38,20,1 | 2,0,2,0,3,3,4,1,0,0,2,1,2,3, | 249 | 149 | 0.000366190796825 | 0.0531819240158175 | 0.824,1.0,0.97,1.0, | 0.949,1.0,0.89,1.0, | 0,049 |
| 9911 | ENSG00000168961 | LGALS9 | chr17 | + | 27647269 | 27647432 | 27647029 | 27647118 | 27648835 | 27649541 | 9911 | 155,63,226,240,110,62,11 | 11,9,15,22,13,15,11,14,17, | 138,144,86,77,74,69,36,15 | 8,10,5,1,3,9,2,3,4,8,0,23,1 | 298 | 149 | 0.000366536119706 | 0.0531961804318557 | 0.876,0.778,0.883, | 0.896,0.878,0.896, | -0,069 |
| 198273 | ENSG00000125304 | TM9SF2 | chr13 | + | 99546962 | 99547104 | 99543862 | 99543995 | 99549164 | 99549222 | 198273 | 31,24,54,61,41,34,37,31,2 | 2,0,5,8,2,2,0,3,2,0,2,0 | 58,58,27,29,38,24,18,28,2 | 4,1,0,0,4,0,0,0,0,0,1,4,4,3, | 291 | 149 | 0.0003672557135 | 0.0532646997791904 | 0.888,1.0,0.847,0. | 0.881,0.967,1.0,1. | -0,052 |
| 105288 | ENSG00000164074 | ABHD18 | chr4 | + | 128001715 | 128001760 | 127984338 | 127984423 | 128008919 | 128008998 | 105288 | 2,3,11,2,0,0,1,2,0,0,0,0 | 22,8,11,10,3,10,5,12,14,10 | 8,1,0,4,4,4,5,1,3,4,0,2,9,3, | 13,12,2,9,5,7,2,6,6,10,3,5, | 194 | 149 | 0.000367825897832 | 0.0533114719301543 | 0.065,0.224,0.434, | 0.321,0.06,0.0,0.2 | -0,162 |
| 96420 | ENSG00000064419 | TNPO3 | chr7 | - | 129005015 | 129005159 | 129001058 | 129001234 | 129014978 | 129015135 | 96420 | 51,49,131,64,72,82,36,45, | 1,1,4,1,1,4,0,0,1,0,0,4 | 107,125,39,55,58,48,37,56 | 1,3,1,6,4,0,4,1,0,1,13,2,0,0 | 293 | 149 | 0.000368886538345 | 0.0534292183947057 | 0.963,0.961,0.943, | 0.982,0.955,0.952, | 0,051 |
| 91589 | ENSG00000024048 | UBR2 | chr6 | + | 42592150 | 42592229 | 42573733 | 42573993 | 42603587 | 42603718 | 91589 | 14,17,75,37,13,14,12,22,1 | 9,0,2,0,0,2,2,2,0,3,1,0 | 40,33,18,16,21,17,21,27,2 | 0,1,0,1,0,1,0,0,0,0,2,3,0,0, | 228 | 149 | 0.000370000135172 | 0.0535544714816441 | 0.504,1.0,0.961,1. | 1.0,0.956,1.0,0.91 | -0,095 |
| 233198 | ENSG00000046651 | OFD1 | chrX | + | 13736107 | 13736296 | 13734742 | 13735220 | 13736477 | 13736678 | 233198 | 3,2,13,11,21,14,1,11,14,1, | 2,0,4,3,1,2,2,0,0,2,0,3 | 14,11,1,2,2,5,3,13,6,20,1,0 | 1,0,0,1,0,0,0,2,2,3,0,1,1,0, | 298 | 149 | 0.000371106843259 | 0.0536785597993803 | 0.429,1.0,0.619,0. | 0.875,1.0,1.0,0.5,1 | -0,121 |
| 97549 | ENSG00000173875 | ZNF791 | chr19 | + | 12612225 | 12612316 | 12610919 | 12611082 | 12623699 | 12623826 | 97549 | 0,0,16,14,0,2,0,0,0,0,1,0 | 75,43,77,51,27,36,32,30,3 | 1,0,1,0,0,0,0,0,0,0,0,3,0,0,1 | 40,43,34,25,55,30,28,41,2 | 240 | 149 | 0.000371911448458 | 0.0537588132727091 | 0.0,0.0,0.114,0.14 | 0.015,0.0,0.018,0. | 0,022 |
| 109936 | ENSG00000167302 | TEPSIN | chr17 | - | 81232314 | 81232518 | 81231846 | 81232021 | 81233431 | 81233503 | 109936 | 48,36,58,24,35,35,56,21,4 | 2,3,4,0,1,0,2,2,4,1,1,4 | 28,35,30,5,19,11,20,33,19, | 1,3,1,3,5,2,2,6,4,3,9,10,3,0 | 298 | 149 | 0.000372314245411 | 0.0537721811585039 | 0.923,0.857,0.879, | 0.933,0.854,0.938, | 0,116 |
| 66493 | ENSG00000122034 | GTF3A | chr13 | + | 27432730 | 27432804 | 27430532 | 27430621 | 27434138 | 27434150 | 66493 | 196,173,349,267,217,176, | 39,28,52,16,20,10,24,43,1 | 219,237,170,76,123,152,8 | 46,49,31,25,22,31,18,32,1 | 223 | 149 | 0.000372503598958 | 0.0537721811585039 | 0.771,0.805,0.818, | 0.761,0.764,0.786, | 0,062 |
| 193584 | ENSG00000164048 | ZNF589 | chr3 | + | 48256799 | 48257022 | 48247624 | 48247677 | 48260812 | 48260939 | 193584 | 1,0,0,1,1,2,2,3,3,0,0,0 | 2,2,13,8,4,5,3,6,13,7,3,1 | 0,0,0,0,0,0,0,1,0,0,24,2,3 | 3,8,3,1,5,11,4,2,4,9,3,62,1 | 298 | 149 | 0.000373377746851 | 0.0538622425150721 | 0.2,0.0,0.0,0.059,0 | 0.0,0.0,0.0,0.0,0.0, | 0,058 |
| 42828 | ENSG00000174243 | DDX23 | chr12 | - | 48835119 | 48835990 | 48834319 | 48834497 | 48836120 | 48836266 | 42828 | 0,2,13,4,3,3,5,1,3,0,0,1 | 52,26,95,68,17,26,32,25,3 | 3,2,0,0,3,0,1,0,0,0,2,1,2,4, | 48,61,13,13,22,25,22,40,1 | 298 | 149 | 0.000374097792293 | 0.0539299678051003 | 0.0,0.037,0.064,0. | 0.03,0.016,0.0,0.0, | 0,021 |
| 230653 | ENSG00000225648 | SBDSP1 | chr7 | + | 72830690 | 72830821 | 72829655 | 72829783 | 72831601 | 72831798 | 230653 | 10,4,8,2,2,3,9,2,4,5,3,2 | 7,4,6,6,3,2,2,2,5,5,0,1 | 3,6,3,10,7,2,2,7,7,7,15,4,0, | 0,4,0,11,0,0,0,0,3,1,10,1, | 280 | 149 | 0.000374676312397 | 0.0539772137841491 | 0.432,0.347,0.415, | 1.0,0.444,1.0,0.32 | -0,232 |
| 219626 | ENSG00000083097 | DOP1A | chr6 | + | 83113326 | 83113421 | 83110124 | 83110314 | 83118887 | 83118987 | 219626 | 13,9,21,12,9,10,5,3,5,2,1,5 | 1,4,4,8,0,1,0,5,1,0,0,1 | 3,14,8,8,7,6,4,7,7,13,4,7,3, | 10,8,5,6,11,3,0,7,1,1,3,7,4, | 244 | 149 | 0.000375423205397 | 0.0540376210930439 | 0.888,0.579,0.762, | 0.155,0.517,0.494, | 0,274 |
| 39533 | ENSG00000084463 | WBP11 | chr12 | - | 14794970 | 14795104 | 14794536 | 14794736 | 14796806 | 14797003 | 39533 | 183,150,206,166,92,96,11 | 5,2,0,0,0,0,3,1,1,0,0,4 | 117,163,116,178,120,47,1 | 3,1,5,12,0,1,6,9,0,0,3,0,0,0 | 283 | 149 | 0.000375597758479 | 0.0540376210930439 | 0.951,0.975,1.0,1. | 0.954,0.988,0.924, | 0,019 |
| 82790 | ENSG00000197622 | CDC42SE1 | chr1 | - | 151056655 | 151056786 | 151055582 | 151055993 | 151059478 | 151059773 | 82790 | 86,35,85,68,34,48,25,47,6 | 197,165,295,254,162,112,1 | 77,36,95,124,69,48,28,63, | 274,239,156,130,165,98,1 | 280 | 149 | 0.000377119116929 | 0.0542202569510659 | 0.189,0.101,0.133, | 0.13,0.074,0.245,0 | -0,042 |
| 242630 | ENSG00000132386 | SERPINF1 | chr17 | + | 1775951 | 1776054 | 1775062 | 1775200 | 1776531 | 1776742 | 242630 | 0,0,0,1,1,6,21,11,0,0,0,0 | 92,44,40,42,75,40,52,80,1 | 0,0,0,0,0,0,0,0,1,1,4,0,0,0, | 21,27,16,19,118,83,18,40,1 | 252 | 149 | 0.000378470975654 | 0.0543782954345701 | 0.0,0.0,0.0,0.014,0 | 0.0,0.0,0.0,0.0,0.0, | 0,027 |
| 209236 | ENSG00000109466 | KLHL2 | chr4 | + | 165305584 | 165305725 | 165299506 | 165299656 | 165310552 | 165310750 | 209236 | 20,13,31,19,9,15,12,11,8,4 | 1,5,4,0,1,6,2,0,0,1,0,0 | 13,11,6,9,21,4,6,20,2,6,11, | 0,1,0,0,0,0,2,2,1,0,0,2,0,1, | 290 | 149 | 0.000384315421403 | 0.0551811824309467 | 0.911,0.572,0.799, | 1.0,0.85,1.0,1.0,1. | -0,021 |
| 15156 | ENSG00000188603 | CLN3 | chr16 | - | 28479643 | 28479765 | 28477736 | 28477877 | 28482104 | 28482198 | 15156 | 2,3,4,2,2,3,6,1,1,0,2,2 | 12,5,10,10,9,7,11,10,3,4,2, | 1,3,0,1,0,0,3,2,0,0,0,4,0,1,1 | 14,4,10,2,9,1,5,21,3,3,2,26 | 271 | 149 | 0.000385098484999 | 0.0552395066777808 | 0.084,0.248,0.18,0 | 0.038,0.292,0.0,0. | 0,062 |
| 36416 | ENSG00000010256 | UQCRC1 | chr3 | - | 48600481 | 48600567 | 48600062 | 48600151 | 48600679 | 48600840 | 36416 | 32,13,116,72,50,42,24,25, | 8,6,4,11,4,11,3,10,0,4,3,4 | 45,36,11,8,32,19,23,27,8,1 | 7,16,11,19,6,1,8,16,5,6,15, | 235 | 149 | 0.000385368478568 | 0.0552395066777808 | 0.717,0.579,0.948, | 0.803,0.588,0.388, | 0,162 |
| 212057 | ENSG00000079263 | SP140 | chr2 | + | 230250980 | 230251052 | 230248884 | 230248968 | 230253315 | 230253417 | 212057 | 19,13,44,35,38,13,13,20,1 | 1,2,4,5,7,0,2,0,3,5,0,0 | 37,20,12,3,23,17,3,10,13,3 | 4,2,0,1,7,1,0,0,0,1,0,1,1,0,1 | 221 | 149 | 0.000385491583601 | 0.0552395066777808 | 0.928,0.814,0.881, | 0.862,0.871,1.0,0. | -0,06 |
| 238595 | ENSG00000136573 | BLK | chr8 | + | 11555588 | 11555922 | 11555390 | 11555484 | 11556657 | 11556837 | 238595 | 47,21,24,16,24,25,9,5,32,1 | 392,174,307,263,310,194, | 41,48,43,8,11,10,2,11,32,3 | 280,293,166,71,172,94,75, | 298 | 149 | 0.000386630385156 | 0.0553658312890958 | 0.057,0.057,0.038, | 0.068,0.076,0.115, | -0,015 |
| 216696 | ENSG00000073849 | ST6GAL1 | chr3 | + | 187006141 | 187006602 | 186996621 | 186996673 | 187038741 | 187038873 | 216696 | 1,0,2,1,0,0,1,2,1,0,0,1 | 85,39,111,85,36,18,40,26, | 5,8,5,9,3,2,4,2,2,2,0,0,8,4, | 102,48,114,36,57,57,31,56 | 298 | 149 | 0.000387271760798 | 0.0553665881655369 | 0.006,0.0,0.009,0. | 0.024,0.077,0.021, | -0,022 |
| 185467 | ENSG00000109320 | NFKB1 | chr4 | + | 102578880 | 102579128 | 102576875 | 102577039 | 102580534 | 102580639 | 185467 | 15,29,36,30,25,14,16,24,1 | 7,3,0,0,1,0,6,2,6,2,0,0 | 35,16,6,17,10,9,7,29,1,16,1 | 1,0,0,0,4,2,1,8,0,1,2,0,0,1, | 298 | 149 | 0.000387326102305 | 0.0553665881655369 | 0.517,0.829,1.0,1. | 0.946,1.0,1.0,1.0,0 | -0,044 |
| 218969 | ENSG00000204560 | DHX16 | chr6 | - | 30665894 | 30666216 | 30665478 | 30665733 | 30670409 | 30670466 | 218969 | 0,1,6,4,1,4,1,7,0,0,1,2 | 58,26,69,40,22,17,15,18,2 | 1,2,2,2,1,6,2,3,0,3,15,1,0,8 | 15,32,16,10,14,7,15,13,11, | 298 | 149 | 0.000387407398457 | 0.0553665881655369 | 0.0,0.019,0.042,0. | 0.032,0.03,0.059,0 | -0,052 |
| 185163 | ENSG00000160691 | SHC1 | chr1 | - | 154965609 | 154966080 | 154963328 | 154963931 | 154966161 | 154966231 | 185163 | 9,7,11,23,8,9,5,8,8,4,6,5 | 0,0,0,0,2,0,1,1,0,0,0,0 | 7,11,6,6,4,5,11,9,4,9,12,4, | 1,2,0,3,5,0,0,2,1,0,5,2,0,0, | 298 | 149 | 0.000388768107595 | 0.0555241861750361 | 1.0,1.0,1.0,1.0,0.6 | 0.778,0.733,1.0,0. | 0,2 |
| 85939 | ENSG00000197226 | TBC1D9B | chr5 | - | 179875919 | 179876037 | 179874901 | 179875187 | 179878308 | 179878523 | 85939 | 94,63,175,132,69,70,68,75 | 2,0,3,1,0,4,0,0,2,1,3,3 | 82,121,64,35,66,58,38,57, | 6,8,0,2,2,1,1,2,2,0,2,10,2,2 | 267 | 149 | 0.000389565796146 | 0.0556012174206232 | 0.963,1.0,0.97,0.9 | 0.884,0.894,1.0,0. | 0,029 |
| 49691 | ENSG00000171103 | TRMT61B | chr2 | - | 28865016 | 28865119 | 28861117 | 28861308 | 28869578 | 28870295 | 49691 | 7,4,5,6,10,2,1,3,7,8,1,1 | 9,3,10,1,4,1,4,3,1,4,0,1 | 11,15,9,1,3,6,2,6,1,10,6,2, | 3,4,1,7,0,1,0,1,0,0,2,4,0,2, | 252 | 149 | 0.000391060586427 | 0.0557775752665803 | 0.315,0.441,0.228, | 0.684,0.689,0.842, | -0,211 |
| 226553 | ENSG00000058091 | CDK14 | chr7 | + | 90712741 | 90713124 | 90709861 | 90709957 | 90726566 | 90726812 | 226553 | 70,39,53,42,17,28,21,35,2 | 317,149,329,173,171,243, | 32,28,15,13,9,8,7,34,14,46 | 353,254,131,231,193,141,1 | 298 | 149 | 0.000391585903035 | 0.0557819785353677 | 0.099,0.116,0.075, | 0.043,0.052,0.054, | 0,014 |
| 61076 | ENSG00000138614 | INTS14 | chr15 | - | 65607158 | 65607425 | 65605128 | 65605236 | 65610795 | 65611148 | 61076 | 7,4,13,8,4,5,4,9,0,4,5,5 | 0,3,2,1,2,0,2,5,1,1,0,0 | 1,5,9,4,7,4,1,12,4,8,6,16,2, | 0,0,1,1,2,0,0,0,0,4,1,0,0,0 | 298 | 149 | 0.000391763332571 | 0.0557819785353677 | 1.0,0.4,0.765,0.8,0 | 1.0,1.0,0.818,0.66 | -0,202 |
| 240901 | ENSG00000198771 | RCSD1 | chr1 | + | 167640333 | 167640474 | 167638389 | 167638442 | 167640570 | 167640712 | 240901 | 22,15,38,25,0,2,16,33,9,4, | 0,0,2,7,1,1,2,4,3,0,2,2 | 7,17,31,14,2,5,11,21,6,2,3 | 3,2,2,0,0,0,0,0,0,2,0,0,2, | 290 | 149 | 0.00039186897587 | 0.0557819785353677 | 1.0,1.0,0.907,0.64 | 0.545,0.814,0.888, | -0,158 |
| 69536 | ENSG00000107798 | LIPA | chr10 | - | 89222510 | 89222582 | 89215937 | 89216009 | 89223683 | 89223830 | 69536 | 113,88,181,136,56,32,23,4 | 0,0,0,0,0,0,17,27,0,0,0,0 | 122,115,90,22,36,45,41,61 | 0,0,0,0,0,0,0,0,0,0,0,0,0,0,0 | 221 | 149 | 0.000393607450279 | 0.0559916421678599 | 1.0,1.0,1.0,1.0,1.0, | 1.0,1.0,1.0,1.0,1.0, | -0,084 |
| 209090 | ENSG00000204160 | ZDHHC18 | chr1 | + | 26853461 | 26853566 | 26852752 | 26852865 | 26853725 | 26857604 | 209090 | 6,0,3,9,0,0,0,0,3,1,0,2 | 34,21,37,17,19,14,19,11,1 | 2,0,0,0,1,0,0,0,0,0,1,0,0,0, | 17,22,13,7,12,7,11,24,6,23 | 254 | 149 | 0.000393862158528 | 0.0559916421678599 | 0.094,0.0,0.045,0. | 0.065,0.0,0.0,0.0,0 | 0,033 |
| 208435 | ENSG00000138468 | SENP7 | chr3 | - | 101371566 | 101371621 | 101367829 | 101368011 | 101372007 | 101372126 | 208435 | 4,3,0,4,0,0,0,0,0,0,0,0 | 17,5,53,31,23,10,14,17,11, | 2,4,2,3,6,1,2,3,2,1,0,4,1,0, | 29,21,14,9,36,20,3,14,19,3 | 204 | 149 | 0.00039537733963 | 0.0561699407167687 | 0.147,0.305,0.0,0. | 0.048,0.122,0.094, | -0,051 |

| | | | | | | | | | | | | | | | | | | | | | | |
|---|---|---|---|---|---|---|---|---|---|---|---|---|---|---|---|---|---|---|---|---|---|---|
| 90194 | ENSG00000224078 | SNHG14 | chr15 | + | 25106163 | 25106294 | 25101822 | 25103970 | 25107471 | 25107588 | 90194 | 44,16,76,59,14,30,19,36,2 | 4,0,4,4,2,6,6,6,3,0,3,0 | 38,50,76,18,32,21,22,19,2 | 3,6,2,0,3,4,1,0,1,1,0,0,2,2, | 280 | 149 | 0.000396209110985 | 0.0562509783419608 | 0.854,1.0,0.91,0.8 | 0.871,0.816,0.953, | -0,07 |
| 107407 | ENSG00000143569 | UBAP2L | chr1 | + | 154234590 | 154234738 | 154228614 | 154228725 | 154235195 | 154235291 | 107407 | 141,80,90,54,36,32,17,47,1 | 62,60,23,17,11,26,26,18,2 | 51,36,50,79,23,21,35,47,1 | 31,45,35,67,54,10,20,33,2 | 297 | 149 | 0.00039723307464 | 0.0563591772497309 | 0.533,0.401,0.663, | 0.452,0.286,0.417, | 0,141 |
| 52883 | ENSG00000143418 | CERS2 | chr1 | - | 150968082 | 150968201 | 150967819 | 150967877 | 150968394 | 150968478 | 52883 | 129,90,147,138,65,63,60,5 | 0,0,6,3,3,7,3,5,3,1,3,1 | 78,54,67,60,43,37,38,71,2 | 11,8,5,0,1,0,2,7,4,0,9,5,4,5 | 268 | 149 | 0.000397902312284 | 0.0564125082979134 | 1.0,1.0,0.932,0.96 | 0.798,0.79,0.882,1 | 0,042 |
| 46489 | ENSG00000114446 | IFT57 | chr3 | - | 108166853 | 108166985 | 108163662 | 108163729 | 108167792 | 108167864 | 46489 | 102,124,339,321,96,104,9 | 0,0,2,1,4,0,0,1,0,0,0,0 | 212,178,77,74,68,47,92,10 | 2,2,0,0,2,0,0,3,1,3,6,14,5,2 | 281 | 149 | 0.000398325575907 | 0.0564125082979134 | 1.0,1.0,0.989,0.99 | 0.983,0.979,1.0,1. | 0,027 |
| 216178 | ENSG00000073614 | KDM5A | chr12 | - | 321000 | 321109 | 318105 | 318461 | 322416 | 322567 | 216178 | 71,53,120,76,51,43,28,58, | 4,0,3,3,4,1,0,2,0,0,1,0 | 86,96,64,56,60,34,32,44,2 | 3,6,5,6,0,1,3,0,0,0,3,1,4,9, | 258 | 149 | 0.000398466288862 | 0.0564125082979134 | 0.911,1.0,0.959,0. | 0.943,0.902,0.881, | 0,031 |
| 66760 | ENSG00000117419 | ERI3 | chr1 | - | 44339044 | 44339322 | 44319627 | 44319744 | 44352849 | 44352925 | 66760 | 20,16,51,43,25,27,22,37,8, | 0,0,4,3,4,4,0,0,1,0,0,0 | 45,26,28,10,25,17,17,36,1 | 5,1,3,0,0,0,4,1,4,1,2,2,4,0, | 298 | 149 | 0.000398657373339 | 0.0564125082979134 | 1.0,1.0,0.864,0.87 | 0.818,0.929,0.824, | 0,058 |
| 223319 | ENSG00000064419 | TNPO3 | chr7 | - | 129005015 | 129005175 | 129001058 | 129001234 | 129014978 | 129015135 | 223319 | 32,27,92,36,38,48,16,30,1 | 1,1,4,1,1,4,0,0,1,0,0,4 | 58,55,23,29,40,33,17,27,3 | 1,3,1,6,4,0,4,1,0,1,13,2,0,0 | 298 | 149 | 0.000399185732263 | 0.0564501605392232 | 0.941,0.931,0.92,0 | 0.967,0.902,0.92,0 | 0,081 |
| 130326 | ENSG00000111271 | ACAD10 | chr12 | + | 111702161 | 111702310 | 111692696 | 111692896 | 111709525 | 111709684 | 130326 | 47,12,30,15,11,16,20,13,2 | 1,0,1,2,1,0,1,0,0,0,1,0 | 14,13,10,13,14,6,13,14,7,2 | 4,1,1,1,3,0,1,4,1,0,4,3,0,0, | 298 | 149 | 0.000400215635372 | 0.0565586417706835 | 0.959,1.0,0.938,0. | 0.636,0.867,0.833, | 0,11 |
| 181085 | ENSG00000178927 | CYBC1 | chr17 | - | 82446622 | 82446721 | 82445250 | 82445960 | 82450699 | 82450829 | 181085 | 20,17,51,37,22,23,15,41,1 | 1,1,5,3,3,0,0,2,2,0,0,0 | 25,10,10,1,7,13,8,22,6,16, | 1,1,5,0,1,1,1,1,0,6,3,3,1,1, | 248 | 149 | 0.000401056061098 | 0.0566402211851599 | 0.923,0.911,0.86,0 | 0.938,0.857,0.546, | 0,08 |
| 246324 | ENSG00000162144 | CYB561A3 | chr11 | - | 61353373 | 61353562 | 61352985 | 61353139 | 61353783 | 61353992 | 246324 | 42,11,57,46,23,18,21,11,1 | 834,661,652,595,439,196, | 9,10,10,5,18,9,9,6,1,3,1,17 | 525,371,299,119,209,243, | 298 | 149 | 0.000401373244393 | 0.0566478457468523 | 0.025,0.008,0.042, | 0.008,0.013,0.016, | 0,006 |
| 142432 | ENSG00000178852 | EFCAB13 | chr17 | + | 47344750 | 47344939 | 47344161 | 47344292 | 47345015 | 47345098 | 142432 | 3,6,13,9,2,0,1,0,0,3,0,0 | 37,21,130,92,31,44,34,53, | 3,2,0,0,0,2,2,1,2,4,0,1,1,0, | 45,44,31,48,36,35,46,57,4 | 298 | 149 | 0.000402263787736 | 0.0567363285047228 | 0.039,0.125,0.048, | 0.032,0.022,0.0,0. | 0,021 |
| 119098 | ENSG00000110514 | MADD | chr11 | + | 47273826 | 47273976 | 47269673 | 47269750 | 47274562 | 47274726 | 119098 | 44,32,46,38,10,17,21,19,4 | 2,2,1,0,4,3,0,0,3,2,1,2 | 18,19,21,37,8,20,20,10,9,2 | 1,4,0,1,3,2,2,0,0,0,0,2,0,1, | 298 | 149 | 0.000403757893961 | 0.0568882775930028 | 0.917,0.889,0.958, | 0.9,0.704,1.0,0.94 | -0,042 |
| 9746 | ENSG00000125970 | RALY | chr20 | + | 34073818 | 34073866 | 34073562 | 34073635 | 34075873 | 34076039 | 9746 | 50,45,121,89,66,43,49,57, | 7,6,13,4,16,8,6,15,3,1,9,5 | 61,57,38,38,17,27,24,54,1 | 10,8,6,10,6,11,6,18,6,10,4, | 197 | 149 | 0.000403869740707 | 0.0568882775930028 | 0.844,0.85,0.876,0 | 0.822,0.843,0.827, | 0,088 |
| 181837 | ENSG00000123104 | ITPR2 | chr12 | - | 26486816 | 26486895 | 26486103 | 26486360 | 26487067 | 26487251 | 181837 | 10,3,11,4,2,0,0,8,0,0,0,2 | 66,57,98,44,21,60,24,58,4 | 2,3,0,5,6,8,1,6,10,7,6,3,8,9 | 45,33,29,39,32,32,10,34,2 | 228 | 149 | 0.000406763639434 | 0.0572584335376189 | 0.09,0.033,0.068,0 | 0.028,0.056,0.0,0. | -0,049 |
| 82972 | ENSG00000144306 | SCRN3 | chr2 | + | 174398274 | 174398442 | 174397335 | 174397404 | 174399921 | 174400103 | 82972 | 6,10,12,13,3,5,2,1,7,2,3,4 | 4,3,1,7,0,0,0,5,0,0,0,0 | 7,2,3,0,3,6,3,9,2,13,2,4,2,9 | 0,0,2,1,0,0,0,0,2,0,0,0,1, | 298 | 149 | 0.000409048674759 | 0.0575424544555911 | 0.429,0.625,0.857, | 1.0,1.0,0.429,0.0,1 | -0,086 |
| 87133 | ENSG00000135976 | ANKRD36 | chr2 | + | 97198585 | 97198658 | 97198462 | 97198491 | 97200333 | 97200362 | 87133 | 5,5,10,4,1,10,3,7,8,5,4,15 | 7,3,20,34,3,9,4,11,3,5,1,4 | 5,8,7,9,22,19,25,35,5,1,6,1 | 3,4,2,5,5,5,0,0,4,5,2,8,2,6, | 222 | 149 | 0.000411964358662 | 0.0579147621679823 | 0.324,0.528,0.251, | 0.528,0.573,0.701, | -0,179 |
| 219639 | ENSG00000083097 | DOP1A | chr6 | + | 83159795 | 83159842 | 83158566 | 83158622 | 83162789 | 83162919 | 219639 | 9,7,19,2,2,7,2,7,4,1,2,2 | 5,1,1,3,2,3,3,0,0,1,0,0 | 8,9,4,7,15,4,4,7,4,18,2,2,3, | 2,0,0,1,4,0,0,1,2,0,0,1,1,0, | 196 | 149 | 0.000412667996104 | 0.0579508727206447 | 0.578,0.842,0.935, | 0.753,1.0,1.0,0.84 | -0,1 |
| 83723 | ENSG00000079263 | SP140 | chr2 | + | 230250980 | 230251061 | 230248884 | 230248968 | 230269531 | 230269618 | 83723 | 23,16,54,43,44,15,15,27,1 | 0,0,5,2,1,0,0,0,0,2,0,3 | 46,25,16,6,25,22,4,12,14,3 | 2,1,0,1,2,0,0,1,2,4,4,4,3,6,1 | 230 | 149 | 0.00041275972272 | 0.0579508727206447 | 1.0,1.0,0.875,0.93 | 0.937,0.942,1.0,0. | 0,077 |
| 190500 | ENSG00000009413 | REV3L | chr6 | - | 111313351 | 111313456 | 111311068 | 111311259 | 111315266 | 111315637 | 190500 | 15,6,23,16,10,21,6,12,19,4 | 8,0,2,3,2,0,1,5,3,0,0,0 | 20,25,8,7,10,6,10,11,3,11, | 0,1,5,3,0,2,2,0,0,1,0,1,0,3, | 254 | 149 | 0.000413927171289 | 0.0580768963518271 | 0.524,1.0,0.871,0. | 1.0,0.936,0.484,0. | -0,014 |
| 47016 | ENSG00000196204 | RNF216P1 | chr7 | + | 4983670 | 4983773 | 4976300 | 4976436 | 4984909 | 4985075 | 47016 | 29,13,39,21,12,17,15,22,1 | 0,0,2,0,0,0,1,4,0,0,0,2 | 27,25,12,18,23,17,16,25,1 | 0,0,0,8,5,0,1,4,0,0,6,2,1,0, | 252 | 149 | 0.000414665182235 | 0.0581425419137598 | 1.0,1.0,0.92,1.0,1. | 1.0,1.0,1.0,0.571,0 | 0,042 |
| 31018 | ENSG00000081791 | DELE1 | chr5 | + | 141938520 | 141938760 | 141934493 | 141934586 | 141939505 | 141940146 | 31018 | 11,6,6,6,13,3,3,8,10,8,5,3 | 2,2,0,0,2,1,0,1,1,0,0,0 | 10,7,3,4,5,0,2,16,3,2,12,14 | 0,4,0,2,0,1,3,2,0,0,4,0,0,4,1 | 298 | 149 | 0.000415937563519 | 0.0582806797368255 | 0.733,0.6,1.0,1.0,0 | 1.0,0.467,1.0,0.5,1 | 0,167 |
| 15748 | ENSG00000197912 | SPG7 | chr16 | + | 89555920 | 89556085 | 89554485 | 89554563 | 89556886 | 89557343 | 15748 | 14,16,6,3,1,3,0,0,2,6,7,26 | 50,36,67,45,40,34,21,52,3 | 2,0,0,1,2,1,2,5,7,10,6,5,3,5 | 36,38,30,29,34,24,24,41,3 | 298 | 149 | 0.000416191927536 | 0.0582806797368255 | 0.123,0.182,0.043, | 0.027,0.0,0.0,0.0,0.01 | 0,049 |
| 136917 | ENSG00000079805 | DNM2 | chr19 | + | 10796060 | 10796178 | 10795371 | 10795439 | 10798485 | 10798572 | 136917 | 13,5,40,37,13,9,6,6,8,2,3,5 | 1,3,6,1,0,3,2,1,2,0,2,1 | 11,14,5,6,7,18,3,8,11,19,4, | 0,2,0,0,1,0,2,2,0,0,1,2,4,0,1 | 267 | 149 | 0.000416727055492 | 0.0583176728742514 | 0.879,0.482,0.788, | 1.0,0.796,1.0,1.0,0 | -0,112 |
| 47610 | ENSG00000117054 | ACADM | chr1 | + | 75728400 | 75728488 | 75724430 | 75724817 | 75732643 | 75732741 | 47610 | 54,47,94,71,40,35,36,43,5 | 18,8,26,26,6,11,7,20,4,0,0, | 69,60,51,16,46,63,12,57,2 | 13,9,6,2,17,4,0,0,2,6,2,5,2, | 237 | 149 | 0.000417107125215 | 0.0583329328571473 | 0.654,0.787,0.694, | 0.769,0.807,0.842, | -0,097 |
| 200848 | ENSG00000161929 | SCIMP | chr17 | - | 5231440 | 5231553 | 5223353 | 5223456 | 5234734 | 5234860 | 200848 | 5,0,1,0,1,0,0,0,2,0,0,2 | 139,33,107,116,39,36,61,4 | 6,0,2,0,2,2,0,0,0,6,0,1,9,8, | 77,47,33,13,26,48,19,47,3 | 262 | 149 | 0.000418567580733 | 0.0584991681615223 | 0.02,0.0,0.005,0.0, | 0.042,0.0,0.033,0. | -0,018 |
| 35169 | ENSG00000136813 | ECPAS | chr9 | - | 111413894 | 111413986 | 111412013 | 111412148 | 111414428 | 111414651 | 35169 | 89,62,73,59,52,43,34,47,1 | 2,0,1,0,0,1,0,1,0,0,0,0 | 58,48,78,98,51,28,32,62,2 | 2,4,0,14,3,0,0,0,0,2,1,2,0,0 | 241 | 149 | 0.000419018434783 | 0.0585241769868786 | 0.965,1.0,0.978,1. | 0.947,0.881,1.0,0. | 0,032 |
| 116011 | ENSG00000056586 | RC3H2 | chr9 | - | 122865623 | 122865657 | 122865348 | 122865541 | 122877470 | 122877583 | 116011 | 13,8,18,9,5,6,6,7,5,5,9,7 | 1,1,4,0,1,1,2,4,0,0,0,1 | 13,7,8,9,7,5,3,15,4,11,9,1, | 3,4,3,3,0,0,3,2,1,3,2,2,2,1, | 183 | 149 | 0.000421231751869 | 0.0587951564114894 | 0.914,0.867,0.786, | 0.779,0.588,0.685, | 0,158 |
| 76010 | ENSG00000135541 | AHI1 | chr6 | - | 135301407 | 135302841 | 135300499 | 135300558 | 135318518 | 135318616 | 76010 | 33,16,50,21,14,21,11,18,7, | 2,0,3,4,0,3,0,2,0,1,1,1 | 2,9,18,47,9,8,10,22,5,5,11, | 0,1,1,0,2,4,0,5,2,1,2,0,0,1, | 298 | 149 | 0.000423614222728 | 0.0590893796318666 | 0.892,1.0,0.893,0. | 1.0,0.818,0.9,1.0,0 | 0,052 |
| 172543 | ENSG00000113971 | NPHP3 | chr3 | - | 132690527 | 132690646 | 132689073 | 132689263 | 132691191 | 132691286 | 172543 | 23,12,30,20,12,14,16,17,1 | 4,3,8,8,2,1,0,5,3,0,2,4 | 12,16,13,4,9,5,10,23,11,15 | 9,15,4,0,10,1,5,10,4,4,1,3, | 268 | 149 | 0.000424790926335 | 0.0592151398095909 | 0.762,0.69,0.676,0 | 0.426,0.372,0.644, | 0,221 |
| 204476 | ENSG00000125637 | PSD4 | chr2 | + | 113192356 | 113192589 | 113185960 | 113186255 | 113193047 | 113193128 | 204476 | 76,80,159,142,99,89,68,78 | 9,2,11,18,3,8,15,7,1,4,8,9 | 90,56,42,45,30,38,39,70,2 | 23,18,10,7,16,7,10,13,8,8, | 298 | 149 | 0.00042567702157 | 0.0592968416463559 | 0.809,0.952,0.878, | 0.662,0.609,0.677, | 0,104 |
| 178571 | ENSG00000120008 | WDR11 | chr10 | + | 120867068 | 120867169 | 120866568 | 120866764 | 120871169 | 120871346 | 178571 | 207,125,220,123,98,79,10 | 2,0,5,6,1,3,4,7,3,1,2,4 | 137,187,195,159,133,93,3 | 6,14,6,11,7,2,6,1,3,9,11,3, | 250 | 149 | 0.000426007262724 | 0.0592968416463559 | 0.984,1.0,0.963,0. | 0.932,0.888,0.951, | 0,036 |
| 183609 | ENSG00000175309 | PHYKPL | chr5 | - | 178224654 | 178224729 | 178224511 | 178224564 | 178225354 | 178225429 | 183609 | 20,16,34,20,26,21,28,25,2 | 0,3,0,2,3,3,0,0,10,0,0,3 | 29,16,17,8,13,5,16,29,12,1 | 1,3,0,0,1,0,2,0,1,3,0,1,2,0,1 | 224 | 149 | 0.000426203539578 | 0.0592968416463559 | 1.0,0.78,1.0,0.869, | 0.951,0.78,1.0,1.0, | -0,029 |
| 144001 | ENSG00000007202 | KIAA0100 | chr17 | - | 28623204 | 28623292 | 28621398 | 28621498 | 28623759 | 28623966 | 144001 | 3,0,8,2,0,4,1,2,0,1,0,3 | 24,11,89,54,50,40,33,39,2 | 2,13,4,4,9,2,1,2,2,2,1,9,5,0 | 47,35,13,25,34,26,20,43,1 | 237 | 149 | 0.000426680656997 | 0.0593248736990448 | 0.073,0.0,0.053,0. | 0.026,0.189,0.162, | -0,035 |
| 190253 | ENSG00000165671 | NSD1 | chr5 | + | 177136872 | 177136970 | 177135940 | 177136030 | 177191883 | 177192019 | 190253 | 4,1,12,0,0,2,0,9,0,0,1,0 | 61,47,133,81,60,68,45,79, | 7,8,8,8,4,3,3,1,0,0,5,9,5,1, | 89,57,40,41,42,33,28,53,4 | 247 | 149 | 0.000427417818824 | 0.0593890023003927 | 0.038,0.013,0.052, | 0.045,0.078,0.108, | -0,031 |
| 197918 | ENSG00000135905 | DOCK10 | chr2 | - | 224893595 | 224893652 | 224886458 | 224886526 | 224896294 | 224896377 | 197918 | 1,0,3,2,3,2,0,0,1,0,1,0 | 6,6,9,5,3,9,3,6,1,4,6,1 | 10,7,1,1,1,2,0,0,4,2,5,1,1,4 | 15,6,0,6,7,7,6,4,16,16,5,3, | 206 | 149 | 0.000428073424142 | 0.0594417233235528 | 0.108,0.0,0.194,0. | 0.325,0.458,1.0,0. | -0,129 |
| 9594 | ENSG00000091592 | NLRP1 | chr17 | - | 5515472 | 5515517 | 5501398 | 5501872 | 5517745 | 5517887 | 9594 | 54,12,22,11,5,20,15,22,16, | 102,68,92,81,37,69,49,38, | 15,20,21,23,17,10,32,35,1 | 72,50,37,44,24,19,27,36,4 | 194 | 149 | 0.000428420745488 | 0.059451596049083 | 0.289,0.119,0.155, | 0.138,0.235,0.304, | -0,098 |
| 220787 | ENSG00000146414 | SHPRH | chr6 | - | 145940923 | 145941023 | 145940722 | 145940801 | 145941622 | 145941874 | 220787 | 1,2,4,1,4,4,6,6,1,0,3,1 | 22,14,39,20,18,15,11,4,17, | 6,9,4,1,11,1,4,7,6,7,8,5,4,8 | 16,15,10,15,13,5,17,17,11, | 249 | 149 | 0.00043157970359 | 0.0598513731851671 | 0.026,0.079,0.058, | 0.183,0.264,0.193, | -0,07 |
| 244021 | ENSG00000151465 | CDC123 | chr10 | + | 12215739 | 12215849 | 12210289 | 12210322 | 12217360 | 12217467 | 244021 | 45,24,78,87,31,30,38,62,3 | 0,2,6,1,0,2,2,1,2,0,0,1 | 45,79,30,9,42,24,8,26,25,3 | 0,0,6,3,2,4,0,0,1,5,0,13,0,1 | 259 | 149 | 0.000432449557369 | 0.0599153855712368 | 1.0,0.873,0.882,0. | 1.0,1.0,0.742,0.63 | 0,051 |
| 43530 | ENSG00000128513 | POT1 | chr7 | - | 124829253 | 124829342 | 124827200 | 124827305 | 124835278 | 124835414 | 43530 | 3,16,35,39,30,11,17,14,10, | 0,0,3,2,1,1,1,0,0,0,0,0 | 23,20,1,10,3,9,7,8,7,30,15, | 1,4,2,0,2,2,0,2,1,2,2,0,6,1, | 238 | 149 | 0.000432598042 | 0.0599153855712368 | 1.0,1.0,0.88,0.924, | 0.935,0.758,0.238, | 0,156 |
| 133126 | ENSG00000119801 | YPEL5 | chr2 | + | 30155849 | 30155910 | 30148244 | 30148541 | 30156627 | 30156792 | 133126 | 18,12,28,18,1,10,8,7,13,8, | 10,13,14,7,8,3,3,7,8,3,5,3 | 32,24,14,14,5,6,3,38,8,31, | 15,9,5,0,0,1,9,6,1,1,0,5,9,4 | 210 | 149 | 0.00043628952902 | 0.0603876329105479 | 0.561,0.396,0.587, | 0.602,0.654,0.665, | -0,239 |
| 55462 | ENSG00000132185 | FCRLA | chr1 | + | 161711207 | 161711474 | 161710759 | 161710912 | 161711933 | 161712218 | 55462 | 454,372,1110,934,771,320 | 12,5,31,20,13,0,13,16,14,9 | 1022,725,207,161,248,295 | 36,37,3,5,12,5,15,17,0,11, | 298 | 149 | 0.000436623109083 | 0.0603876329105479 | 0.95,0.974,0.947,0 | 0.934,0.907,0.972, | 0,014 |
| 173129 | ENSG00000184470 | TXNRD2 | chr22 | - | 19895047 | 19895200 | 19883324 | 19883461 | 19895406 | 19895581 | 173129 | 1,1,7,0,0,1,4,5,0,0,0,0 | 0,0,11,10,7,9,5,3,3,4,4,4 | 6,6,1,1,6,2,1,6,0,0,5,3,4,2, | 4,0,0,0,2,0,0,5,3,3,1,4,5,5, | 298 | 149 | 0.000436849452178 | 0.0603876329105479 | 1.0,1.0,0.241,0.0,0 | 0.429,1.0,1.0,1.0,0 | -0,237 |
| 113438 | ENSG00000104325 | DECR1 | chr8 | + | 90005447 | 90005731 | 90001476 | 90001561 | 90017123 | 90017326 | 113438 | 1,1,3,0,0,1,0,1,3,0,0,2 | 10,4,40,49,21,13,12,8,7,4, | 3,0,2,2,0,2,0,6,1,5,1,1,2,4, | 17,12,13,0,14,9,4,11,9,13, | 298 | 149 | 0.000437244800433 | 0.0604034888587901 | 0.048,0.111,0.036, | 0.081,0.0,0.071,1. | -0,089 |
| 44761 | ENSG00000107758 | PPP3CB | chr10 | - | 73439871 | 73439901 | 73436432 | 73438420 | 73444724 | 73444822 | 44761 | 25,6,23,21,16,13,11,12,14, | 3,1,1,0,0,1,1,2,1,0,0,0 | 30,16,14,18,12,9,26,19,8,2 | 4,6,4,0,0,1,5,3,2,2,0,2,0,0, | 179 | 149 | 0.000438273059245 | 0.0605067022542403 | 0.874,0.833,0.95,1 | 0.862,0.689,0.744, | 0,044 |
| 41048 | ENSG00000009790 | TRAF3IP3 | chr1 | + | 209781349 | 209781458 | 209779314 | 209779374 | 209782055 | 209782307 | 41048 | 133,50,104,121,82,55,42,3 | 11,2,7,4,8,8,0,2,2,0,2,0 | 93,87,54,56,33,53,31,51,5 | 2,1,0,2,1,1,0,3,2,2,5,2,2,0, | 258 | 149 | 0.000438989444073 | 0.0605667545110743 | 0.875,0.935,0.896, | 0.964,0.98,1.0,0.9 | -0,015 |
| 146386 | ENSG00000167264 | DUS2 | chr16 | + | 68068941 | 68068968 | 68066565 | 68066636 | 68070133 | 68070220 | 146386 | 3,5,3,4,1,5,2,5,1,2,0,0 | 65,63,70,82,72,38,52,32,4 | 7,0,6,2,7,1,3,2,5,2,9,3,2,1, | 36,52,38,8,47,27,14,13,19, | 176 | 149 | 0.00044031209796 | 0.0607103223292945 | 0.038,0.063,0.035, | 0.141,0.0,0.118,0. | -0,056 |
| 250442 | ENSG00000136754 | ABI1 | chr10 | - | 26759085 | 26759235 | 26751597 | 26751783 | 26765217 | 26765318 | 250442 | 31,16,49,59,27,25,23,65,2 | 34,14,49,62,65,45,22,39,1 | 60,60,53,25,21,11,22,34,7, | 42,15,33,3,15,5,20,32,11,3 | 298 | 149 | 0.000440603038039 | 0.0607115444815442 | 0.313,0.364,0.333, | 0.417,0.667,0.445, | -0,133 |
| 242522 | ENSG00000102699 | PARP4 | chr13 | - | 24482146 | 24482280 | 24478092 | 24478276 | 24484652 | 24484748 | 242522 | 6,6,14,3,0,3,3,6,0,1,0,6 | 55,22,124,118,75,53,54,72 | 12,12,9,7,10,4,3,4,5,4,4,7, | 80,66,33,9,31,45,29,67,39, | 283 | 149 | 0.000445814204305 | 0.0613902987887201 | 0.054,0.126,0.056, | 0.073,0.087,0.126, | -0,039 |
| 107409 | ENSG00000143569 | UBAP2L | chr1 | + | 154234590 | 154234792 | 154228614 | 154228725 | 154235195 | 154235291 | 107409 | 154,86,104,57,39,33,18,49 | 62,60,23,17,11,26,26,18,2 | 57,36,53,81,23,21,38,52,1 | 31,45,35,67,54,10,20,33,2 | 298 | 149 | 0.00044713775563 | 0.0615331881598469 | 0.554,0.417,0.693, | 0.479,0.286,0.431, | 0,138 |
| 51488 | ENSG00000178927 | CYBC1 | chr17 | - | 82446622 | 82446692 | 82445863 | 82445960 | 82450699 | 82450728 | 51488 | 18,17,44,31,20,16,13,32,1 | 1,1,5,3,3,0,0,2,2,0,0,0 | 23,6,8,1,7,12,5,19,6,15,14, | 1,1,5,0,1,1,1,1,0,6,3,3,1,1, | 219 | 149 | 0.000447964042362 | 0.0616075072214796 | 0.925,0.92,0.857,0 | 0.94,0.803,0.521,1 | 0,084 |
| 216286 | ENSG00000163219 | ARHGAP25 | chr2 | + | 68788488 | 68788653 | 68787839 | 68787956 | 68807272 | 68807480 | 216286 | 1,0,12,12,0,0,1,7,14,17,0,0 | 145,224,142,169,138,87,7 | 2,1,1,0,30,7,0,0,0,1,0,2,0,0 | 210,364,91,60,101,42,90,8 | 298 | 149 | 0.00044914594467 | 0.0617306071630069 | 0.003,0.0,0.041,0. | 0.005,0.001,0.005, | 0,013 |
| 146253 | ENSG00000140968 | IRF8 | chr16 | + | 85913079 | 85913236 | 85911569 | 85911658 | 85914472 | 85914520 | 146253 | 87,64,175,101,88,90,81,10 | 0,4,7,4,6,9,4,4,2,1,1,1 | 81,121,76,41,54,40,62,72, | 7,11,5,7,4,3,6,1,2,3,9,21,2, | 298 | 149 | 0.00044943738292 | 0.0617312427334107 | 1.0,0.889,0.926,0. | 0.853,0.846,0.884, | 0,051 |
| 210820 | ENSG00000145241 | CENPC | chr4 | - | 67518155 | 67518439 | 67514073 | 67514687 | 67519216 | 67519502 | 210820 | 56,41,94,43,32,39,31,44,6 | 4,0,5,5,1,0,0,2,1,0,0,2 | 39,58,30,41,46,10,22,38,1 | 2,3,3,5,6,1,0,3,0,1,3,2,1,0, | 298 | 149 | 0.000450625521776 | 0.0618549628044453 | 0.875,1.0,0.904,0. | 0.907,0.906,0.833, | 0,068 |
| 192309 | ENSG00000124406 | ATP8A1 | chr4 | - | 42583807 | 42583881 | 42581691 | 42581732 | 42586348 | 42586476 | 192309 | 0,2,9,0,1,1,0,4,0,6,0,1 | 39,15,56,32,30,45,49,77,2 | 3,13,1,3,9,0,1,1,5,0,0,21,3, | 34,41,13,27,20,18,8,53,28, | 223 | 149 | 0.000451203280305 | 0.0618734398383621 | 0.0,0.082,0.097,0. | 0.056,0.175,0.049, | -0,023 |
| 41041 | ENSG00000009790 | TRAF3IP3 | chr1 | + | 209780469 | 209780606 | 209779314 | 209779374 | 209782055 | 209782307 | 41041 | 47,17,75,97,57,27,22,10,3 | 11,2,7,4,8,8,0,2,2,0,2,0 | 42,41,43,20,37,34,6,20,35, | 2,1,0,2,1,1,0,3,2,2,5,2,2,0, | 286 | 149 | 0.000451335079734 | 0.0618734398383621 | 0.69,0.816,0.848,0 | 0.916,0.955,1.0,0. | -0,038 |
| 139490 | ENSG00000170471 | RALGAPB | chr20 | + | 38497392 | 38497516 | 38492929 | 38493132 | 38499446 | 38499633 | 139490 | 16,17,71,41,29,51,33,30,1 | 0,0,2,0,0,2,1,1,0,0,0,3 | 40,56,22,27,37,18,25,22,2 | 3,3,3,6,4,1,0,4,1,0,1,2,0,0, | 273 | 149 | 0.000453740147152 | 0.0621635554498231 | 1.0,1.0,0.951,1.0,1 | 0.879,0.911,0.8,0. | 0,065 |
| 218321 | ENSG00000163939 | PBRM1 | chr3 | - | 52563282 | 52563471 | 52561766 | 52561968 | 52564049 | 52564233 | 218321 | 15,6,25,15,38,30,27,29,49, | 5,0,0,3,1,4,3,2,4,2,0,1 | 23,16,14,12,6,10,14,27,9,1 | 0,0,1,0,4,0,0,0,0,3,0,5,1,0, | 298 | 149 | 0.000455078273663 | 0.0623072213223671 | 0.6,1.0,1.0,0.714,0 | 1.0,1.0,0.875,1.0,0 | -0,066 |
| 146304 | ENSG00000077235 | GTF3C1 | chr16 | - | 27488027 | 27488415 | 27485996 | 27486154 | 27488553 | 27488635 | 146304 | 30,20,37,13,20,25,11,13,2 | 2,1,0,6,0,3,1,2,0,0,0,1 | 24,18,16,14,8,12,5,12,16,2 | 0,6,1,2,2,3,0,4,1,0,2,0,2,2, | 298 | 149 | 0.000456686083923 | 0.0624876049134337 | 0.882,0.909,1.0,0. | 1.0,0.6,0.889,0.77 | 0,051 |
| 67043 | ENSG00000048707 | VPS13D | chr1 | + | 12386184 | 12386334 | 12385259 | 12385373 | 12400180 | 12400330 | 67043 | 48,35,95,52,55,77,43,55,3 | 0,4,2,1,3,1,1,0,0,0,0,0 | 89,71,50,73,35,44,35,38,3 | 5,5,8,4,3,1,0,5,1,0,0,4,1,0, | 298 | 149 | 0.000459062440074 | 0.0627728513593184 | 1.0,0.814,0.96,0.9 | 0.899,0.877,0.758, | 0,037 |
| 52952 | ENSG00000226571 | | chr6 | + | 139600485 | 139600629 | 139598002 | 139598230 | 139600842 | 139600904 | 52952 | 12,5,33,28,12,7,9,6,12,4,5, | 0,2,1,3,2,3,5,1,1,0,0,0 | 16,10,4,3,24,8,8,37,5,13,2 | 1,1,0,0,2,1,0,2,1,0,0,1,0,2, | 293 | 149 | 0.000460923497804 | 0.0629873176862557 | 1.0,0.56,0.944,0.8 | 0.891,0.836,1.0,1. | -0,106 |
| 126785 | ENSG00000066084 | DIP2B | chr12 | + | 50655010 | 50655040 | 50640804 | 50640852 | 50660193 | 50660319 | 126785 | 0,3,5,0,0,1,1,6,3,0,3,0 | 33,30,82,63,30,45,42,44,5 | 1,3,4,2,9,0,3,6,6,0,8,2,2,0, | 50,44,37,45,21,29,24,37,5 | 179 | 149 | 0.00046208230325 | 0.0630798529595353 | 0.0,0.077,0.048,0. | 0.016,0.054,0.083, | -0,033 |
| 227306 | ENSG00000013374 | NUB1 | chr7 | + | 151374058 | 151374201 | 151368829 | 151368887 | 151375847 | 151375943 | 227306 | 4,6,21,18,9,5,4,6,7,3,3,3 | 4,0,1,3,2,2,2,8,0,0,0,0 | 5,4,7,2,2,7,1,14,3,9,11,17, | 1,4,0,0,0,0,1,5,1,0,0,1,0,2, | 292 | 149 | 0.00046218680449 | 0.0630798529595353 | 0.338,1.0,0.915,0. | 0.718,0.338,1.0,1. | -0,055 |
| 34991 | ENSG00000197020 | ZNF100 | chr19 | - | 21744016 | 21744115 | 21722770 | 21727989 | 21744940 | 21745067 | 34991 | 8,12,8,10,7,8,19,21,7,6,4,4 | 3,2,4,3,1,0,2,3,0,4,3,1 | 8,15,3,8,17,12,4,6,4,9,15,6 | 3,2,0,0,2,1,0,0,1,3,2,2,0,0, | 248 | 149 | 0.000462881128357 | 0.063134580568698 | 0.616,0.783,0.546, | 0.616,0.818,1.0,1. | -0,106 |
| 164047 | ENSG00000102445 | RUBCNL | chr13 | - | 46369767 | 46370009 | 46368762 | 46368815 | 46371940 | 46372597 | 164047 | 2,0,21,3,2,3,6,8,0,0,3,5 | 118,124,945,696,243,376, | 22,20,10,15,9,10,7,17,3,6, | 540,780,271,253,398,203, | 298 | 149 | 0.000464283430505 | 0.0632857422615717 | 0.008,0.0,0.011,0. | 0.02,0.013,0.018,0 | -0,011 |
| 68915 | ENSG00000198039 | ZNF273 | chr7 | + | 64917580 | 64917707 | 64903272 | 64903419 | 64927653 | 64929124 | 68915 | 14,9,18,8,2,4,3,4,5,7,0,6 | 5,1,11,0,0,0,4,0,0,0,2,0 | 4,2,1,2,5,2,0,3,8,8,6,21,4,1 | 0,2,4,7,1,1,2,2,0,5,2,0,2,2, | 276 | 149 | 0.000465609678625 | 0.0634263523671756 | 0.602,0.829,0.469, | 1.0,0.351,0.119,0. | 0,245 |
| 116157 | ENSG00000107077 | KDM4C | chr9 | + | 6986343 | 6986666 | 6984165 | 6984404 | 6990415 | 6990524 | 116157 | 107,47,129,86,55,72,44,84 | 1,0,0,5,0,1,0,0,0,0,0,1 | 122,95,69,87,73,57,54,70, | 1,5,0,3,7,0,2,0,1,0,2,2,2,4, | 298 | 149 | 0.000466110668453 | 0.0634544372433951 | 0.982,1.0,1.0,0.89 | 0.984,0.905,1.0,0. | 0,035 |
| 204856 | ENSG00000076242 | MLH1 | chr3 | + | 37020309 | 37020522 | 37012010 | 37012099 | 37025636 | 37026093 | 204856 | 10,5,17,9,1,6,3,2,7,3,2,11 | 0,2,3,0,1,1,0,3,0,1,1,0 | 9,23,9,6,7,5,13,6,6,11,12,1 | 0,3,3,0,4,2,4,0,2,1,2,2,1,0, | 298 | 149 | 0.000468164893791 | 0.0636938042070359 | 1.0,0.556,0.739,1. | 1.0,0.793,0.6,1.0,0 | -0,041 |
| 204736 | ENSG00000196510 | ANAPC7 | chr12 | - | 110377393 | 110377617 | 110376131 | 110376216 | 110381751 | 110381948 | 204736 | 17,10,40,37,35,24,24,15,2 | 0,0,7,2,2,1,0,9,1,0,0,0 | 24,11,13,0,3,8,10,21,7,18, | 3,0,0,1,0,0,0,0,0,2,2,1,0,1 | 298 | 149 | 0.000469414411024 | 0.0638234574220509 | 1.0,1.0,0.741,0.90 | 0.8,1.0,1.0,0.0,1.0, | 0,02 |
| 229959 | ENSG00000026950 | BTN3A1 | chr6 | + | 26408691 | 26408861 | 26407826 | 26407952 | 26409532 | 26409733 | 229959 | 10,7,12,1,3,1,4,3,1,0,1,2 | 61,55,42,42,21,19,8,17,20, | 4,2,6,6,9,1,5,4,4,6,9,9,5,12 | 11,20,9,19,27,16,13,41,12, | 298 | 149 | 0.00046973034742 | 0.063826093690375 | 0.076,0.06,0.125,0 | 0.154,0.048,0.25,0 | -0,053 |
| 176673 | ENSG00000116489 | CAPZA1 | chr1 | + | 112661205 | 112661252 | 112659700 | 112659779 | 112667073 | 112667145 | 176673 | 16,3,15,5,0,9,9,7,1,0,1,1 | 126,99,210,285,115,111,8 | 8,24,2,32,4,1,5,11,9,4,12,1 | 145,104,103,69,79,86,21,4 | 196 | 149 | 0.000470876879153 | 0.0639415151905232 | 0.088,0.023,0.052, | 0.04,0.149,0.015,0 | -0,041 |
| 254172 | ENSG00000123349 | PFDN5 | chr12 | + | 53295842 | 53296275 | 53295585 | 53295639 | 53297849 | 53297924 | 254172 | 204,114,395,315,147,156, | 62,26,120,60,24,30,21,42, | 233,227,123,96,134,60,93, | 50,56,40,22,61,26,27,59,2 | 298 | 149 | 0.000471444461123 | 0.0639782237149839 | 0.622,0.687,0.622, | 0.7,0.67,0.606,0.6 | 0,086 |
| 218322 | ENSG00000163939 | PBRM1 | chr3 | - | 52563282 | 52563474 | 52561766 | 52561968 | 52564049 | 52564233 | 218322 | 15,6,25,15,38,30,27,29,51, | 5,0,0,3,1,4,3,2,4,2,0,1 | 23,16,14,10,6,10,14,27,9,1 | 0,0,1,0,4,0,0,0,0,3,0,5,1,0, | 298 | 149 | 0.000473765874654 | 0.0642527428907719 | 0.6,1.0,1.0,0.714,0 | 1.0,1.0,0.875,1.0,0 | -0,066 |
| 94552 | ENSG00000231925 | TAPBP | chr6 | - | 33304296 | 33304638 | 33302870 | 33303989 | 33304988 | 33305387 | 94552 | 146,143,286,251,230,209, | 1,0,0,3,0,0,2,0,0,0,0,0 | 232,267,104,141,161,107, | 2,6,0,3,4,0,7,8,1,1,2,10,0,0 | 298 | 149 | 0.000477473669449 | 0.0647148207488525 | 0.986,1.0,1.0,0.97 | 0.983,0.957,1.0,0. | 0,022 |
| 103790 | ENSG00000134882 | UBAC2 | chr13 | + | 99201428 | 99201567 | 99200886 | 99200939 | 99238426 | 99238554 | 103790 | 0,1,2,5,2,3,0,0,1,0,0,0 | 4,8,31,12,18,7,15,13,10,12 | 2,1,0,1,0,1,4,0,5,1,6,8,4,4, | 23,22,9,11,8,5,7,13,10,9,8, | 288 | 149 | 0.000480965979607 | 0.0651471294882279 | 0.0,0.061,0.032,0. | 0.043,0.023,0.0,0. | -0,041 |
| 223659 | ENSG00000106100 | NOD1 | chr7 | - | 30450917 | 30451076 | 30448297 | 30448381 | 30451215 | 30453040 | 223659 | 4,2,6,0,0,0,1,5,6,0,0,0 | 14,9,15,3,6,7,3,5,5,10,7,3 | 1,0,0,0,2,0,0,2,2,0,0,1,7,0, | 12,6,1,10,9,10,2,5,4,8,7,17 | 298 | 149 | 0.000482162645054 | 0.06526814355825 | 0.125,0.1,0.167,0. | 0.04,0.0,0.0,0.0,0. | 0,059 |
| 248595 | ENSG00000148925 | BTBD10 | chr11 | - | 13444733 | 13445181 | 13421746 | 13421838 | 13463091 | 13463222 | 248595 | 2,2,14,4,0,0,0,4,0,0,1,0 | 33,22,18,11,5,20,4,14,8,6, | 6,7,0,0,4,1,2,0,0,0,0,1,0,0, | 17,29,6,11,20,5,21,9,9,12, | 298 | 149 | 0.000482972685769 | 0.0653367027848822 | 0.029,0.043,0.28,0 | 0.15,0.108,0.0,0.0, | 0,026 |
| 177511 | ENSG00000137818 | RPLP1 | chr15 | + | 69453674 | 69453721 | 69452883 | 69453020 | 69455169 | 69455287 | 177511 | 926,667,561,759,232,226, | 26,16,43,55,17,17,14,10,2 | 458,493,206,307,201,121, | 8,13,2,8,7,5,12,22,6,8,18,2 | 196 | 149 | 0.000484616257559 | 0.0655178654087193 | 0.964,0.969,0.908, | 0.978,0.966,0.987, | -0,017 |
| 34892 | ENSG00000026297 | RNASET2 | chr6 | - | 166948565 | 166948625 | 166946681 | 166946739 | 166952487 | 166952548 | 34892 | 55,38,32,16,11,10,9,11,7,9 | 4,0,3,4,0,0,1,3,1,1,0,0 | 21,33,27,13,22,17,5,10,8,3 | 0,7,4,1,7,4,0,0,3,5,1,8,2,2, | 209 | 149 | 0.00048514282898 | 0.0655478821244158 | 0.907,1.0,0.884,0. | 1.0,0.771,0.828,0. | 0,062 |
| 19132 | ENSG00000137842 | TMEM62 | chr15 | + | 43146492 | 43146634 | 43135511 | 43135649 | 43148754 | 43148788 | 19132 | 38,24,40,14,9,14,8,21,23,1 | 3,13,40,12,5,10,20,17,2,2, | 27,15,8,11,8,7,8,6,14,15,1 | 18,42,8,26,35,11,13,7,13,8 | 291 | 149 | 0.000486085239365 | 0.0656340101341081 | 0.866,0.486,0.339, | 0.434,0.155,0.339, | 0,208 |
| 48760 | ENSG00000163564 | PYHIN1 | chr1 | + | 158973646 | 158973771 | 158944874 | 158945042 | 158976700 | 158976728 | 48760 | 156,88,31,44,23,13,73,65, | 4,14,20,8,1,5,12,12,7,2,5,2 | 7,10,75,27,35,35,17,36,15, | 2,4,21,1,24,15,3,8,11,24,5, | 274 | 149 | 0.000486812258599 | 0.0656782266522127 | 0.955,0.774,0.457, | 0.656,0.576,0.66,0 | 0,129 |

| | | | | | | | | | | | | | | | | | | | | | | |
|---|---|---|---|---|---|---|---|---|---|---|---|---|---|---|---|---|---|---|---|---|---|---|
| 201842 | ENSG00000175634 | RPS6KB2 | chr11 | + | 67432530 | 67432657 | 67429526 | 67429595 | 67432736 | 67432815 | 201842 | 55,33,37,27,23,23,32,26,4 | 5,4,5,3,0,0,2,5,0,0,0,1 | 23,13,31,26,19,11,16,26,1 | 0,4,4,3,1,0,1,3,0,0,0,4,0,0,1 | 276 | 149 | 0.000487023011262 | 0.0656782266522127 | 0.856,0.817,0.8,0. | 1.0,0.637,0.807,0. | -0,003 |
| 204479 | ENSG00000125637 | PSD4 | chr2 | + | 113192367 | 113192589 | 113185960 | 113186255 | 113193047 | 113193128 | 204479 | 76,80,159,142,96,87,66,80 | 9,2,11,18,3,8,15,7,1,4,8,9 | 90,55,42,45,30,38,39,70,2 | 23,18,10,7,16,7,10,13,8,8, | 298 | 149 | 0.000490342959142 | 0.0660845369061314 | 0.809,0.952,0.878, | 0.662,0.604,0.677, | 0,103 |
| 196828 | ENSG00000143119 | CD53 | chr1 | + | 110877152 | 110877203 | 110873182 | 110873248 | 110891391 | 110891471 | 196828 | 8,4,8,4,1,2,5,4,0,4,0,3 | 250,166,427,384,223,168,1 | 13,11,12,7,3,3,3,6,5,5,8,9, | 299,236,249,120,119,148, | 200 | 149 | 0.000490811603117 | 0.0661063029727628 | 0.023,0.018,0.014, | 0.031,0.034,0.035, | -0,011 |
| 9632 | ENSG00000091592 | NLRP1 | chr17 | - | 5533896 | 5533979 | 5533303 | 5533384 | 5536850 | 5536940 | 9632 | 71,22,48,36,33,37,33,33,3 | 4,2,7,1,0,3,2,1,2,0,2,7 | 48,48,35,23,42,17,20,25,3 | 2,2,4,15,1,0,10,7,1,3,9,7,3, | 232 | 149 | 0.000491206646269 | 0.0661181348862559 | 0.919,0.876,0.815, | 0.939,0.939,0.849, | 0,14 |
| 238706 | ENSG00000064313 | TAF2 | chr8 | - | 119794552 | 119794944 | 119793365 | 119793451 | 119795531 | 119795631 | 238706 | 2,0,8,6,0,2,0,7,3,0,0,1 | 23,8,36,9,9,22,8,5,12,4,2,1 | 0,1,0,0,2,1,0,1,4,0,0,5,1,1, | 15,16,9,12,12,7,4,14,7,19,1 | 298 | 149 | 0.000492302522832 | 0.0662242276822839 | 0.042,0.0,0.1,0.25, | 0.0,0.03,0.0,0.0,0. | 0,039 |
| 189407 | ENSG00000177200 | CHD9 | chr16 | + | 53208264 | 53208347 | 53157163 | 53157541 | 53209481 | 53209541 | 189407 | 8,5,10,7,0,7,5,2,4,1,0,2 | 63,57,85,61,38,40,26,38,3 | 0,8,0,2,0,4,2,2,3,4,0,0,2,6, | 69,78,42,38,79,54,24,27,4 | 232 | 149 | 0.000494448303469 | 0.0664347491632483 | 0.075,0.053,0.07,0 | 0.0,0.062,0.0,0.03 | 0,02 |
| 139068 | ENSG00000125772 | GPCPD1 | chr20 | - | 5599289 | 5599465 | 5598724 | 5598821 | 5604363 | 5604440 | 139068 | 8,3,15,4,11,10,8,11,2,1,0,5 | 94,47,157,104,69,94,80,10 | 8,16,9,12,29,2,6,6,24,5,8,1 | 92,87,48,54,93,43,54,70,1 | 298 | 149 | 0.000494635314838 | 0.0664347491632483 | 0.041,0.031,0.046, | 0.042,0.084,0.086, | -0,026 |
| 172735 | ENSG00000232112 | TMA7 | chr3 | + | 48440540 | 48440912 | 48440283 | 48440458 | 48443847 | 48444208 | 172735 | 102,93,175,103,36,34,48,4 | 0,0,6,2,1,1,0,0,0,0,0,0 | 74,84,83,65,31,39,15,44,1 | 1,5,4,3,2,0,0,2,1,6,1,3,0,0, | 298 | 149 | 0.000494937746729 | 0.0664347491632483 | 1.0,1.0,0.936,0.96 | 0.974,0.894,0.912, | 0,046 |
| 76981 | ENSG00000116127 | ALMS1 | chr2 | + | 73550266 | 73550437 | 73534823 | 73534949 | 73557219 | 73557354 | 76981 | 20,25,40,19,19,23,13,25,1 | 1,1,3,4,1,3,4,2,1,0,0,1 | 41,41,6,32,14,11,25,20,13, | 1,2,0,0,0,0,0,1,0,1,2,0,2,3, | 298 | 149 | 0.000495102181646 | 0.0664347491632483 | 0.909,0.926,0.87,0 | 0.953,0.911,1.0,1. | -0,06 |
| 204473 | ENSG00000125637 | PSD4 | chr2 | + | 113192264 | 113192589 | 113185960 | 113186255 | 113193047 | 113193128 | 204473 | 77,80,159,142,96,87,66,78 | 9,2,11,18,3,8,15,7,1,4,8,9 | 90,55,42,45,30,38,39,70,2 | 23,18,10,7,16,7,10,13,8,8, | 298 | 149 | 0.00049856105025 | 0.0668571921534939 | 0.811,0.952,0.878, | 0.662,0.604,0.677, | 0,103 |
| 139481 | ENSG00000170471 | RALGAPB | chr20 | + | 38497320 | 38497516 | 38492929 | 38493132 | 38499446 | 38499633 | 139481 | 20,18,81,44,42,61,38,51,1 | 0,0,2,0,0,2,1,1,0,0,0,3 | 48,64,23,28,41,22,28,28,2 | 3,3,3,6,4,1,0,4,1,0,1,2,0,0, | 298 | 149 | 0.000500431089198 | 0.0670383099008441 | 1.0,1.0,0.953,1.0,1 | 0.889,0.914,0.793, | 0,064 |
| 91196 | ENSG00000114388 | NPRL2 | chr3 | - | 50349385 | 50349494 | 50348873 | 50349010 | 50349664 | 50349828 | 91196 | 34,9,23,18,20,4,8,15,20,6, | 3,1,2,1,1,0,8,2,0,2,0,1 | 24,10,16,9,14,10,11,6,3,14 | 1,1,2,4,1,0,0,0,0,0,0,2,0,0, | 258 | 149 | 0.000500758865805 | 0.0670383099008441 | 0.867,0.839,0.869, | 0.933,0.852,0.822, | -0,085 |
| 51491 | ENSG00000178927 | CYBC1 | chr17 | - | 82446622 | 82446696 | 82445250 | 82445960 | 82450699 | 82450733 | 51491 | 20,19,56,36,25,20,14,40,1 | 1,1,5,3,3,0,0,2,2,0,0,0 | 26,13,14,1,7,17,6,21,8,21, | 1,1,5,0,1,1,1,1,0,6,3,3,1,1, | 223 | 149 | 0.000500846078495 | 0.0670383099008441 | 0.93,0.927,0.882,0 | 0.946,0.897,0.652, | 0,066 |
| 4884 | ENSG00000198521 | ZNF43 | chr19 | - | 21819094 | 21819221 | 21817887 | 21817986 | 21851904 | 21852068 | 4884 | 24,32,40,42,9,17,13,21,16, | 3,0,1,1,1,0,0,0,1,0,0,1 | 30,23,15,10,9,4,12,22,9,17 | 2,3,0,4,4,0,2,1,1,0,3,1,0,3, | 276 | 149 | 0.000501260835057 | 0.0670521260348994 | 0.812,1.0,0.956,0. | 0.89,0.805,1.0,0.5 | 0,085 |
| 94433 | ENSG00000204351 | SKIV2L | chr6 | + | 31960431 | 31960546 | 31960200 | 31960337 | 31960665 | 31960740 | 94433 | 70,34,100,75,43,45,29,40, | 4,0,3,2,4,1,0,0,0,3,3,2 | 55,66,38,39,40,30,22,61,4 | 0,3,0,0,0,0,1,0,1,0,1,1,0,0, | 264 | 149 | 0.000504036683957 | 0.0673815649222044 | 0.908,1.0,0.95,0.9 | 1.0,0.925,1.0,1.0,1 | -0,044 |
| 155712 | ENSG00000169062 | UPF3A | chr13 | + | 114286258 | 114286400 | 114284093 | 114284358 | 114286518 | 114286629 | 155712 | 55,18,58,33,10,17,18,26,2 | 0,1,2,0,0,1,0,1,4,0,0,0 | 32,24,38,21,25,24,17,19,1 | 2,2,3,0,0,4,1,2,3,4,0,2,0,3,1 | 291 | 149 | 0.000506451997642 | 0.0676624270046464 | 1.0,0.902,0.937,1. | 0.891,0.86,0.866,1 | 0,067 |
| 107139 | ENSG00000215301 | DDX3X | chrX | + | 41347311 | 41347451 | 41346858 | 41347012 | 41347639 | 41347915 | 107139 | 465,400,673,428,181,225, | 23,7,26,30,13,10,10,14,6,1 | 412,484,338,296,309,206,1 | 31,31,15,52,16,17,2,20,13, | 289 | 149 | 0.000512166938975 | 0.0683835002745212 | 0.912,0.967,0.93,0 | 0.873,0.889,0.921, | 0,028 |
| 51446 | ENSG00000076685 | NT5C2 | chr10 | - | 103090610 | 103090787 | 103087184 | 103089908 | 103090935 | 103090996 | 51446 | 126,112,238,166,75,66,81, | 21,12,19,12,2,3,7,5,3,8,0,0 | 87,151,113,63,107,101,73, | 5,3,1,11,1,1,1,9,1,4,7,0,5,3 | 298 | 149 | 0.00051413320407 | 0.0685661120188099 | 0.75,0.824,0.862,0 | 0.897,0.962,0.983, | -0,037 |
| 15860 | ENSG00000140688 | RUSF1 | chr16 | - | 31493466 | 31493524 | 31492977 | 31493048 | 31493602 | 31493781 | 15860 | 24,9,34,35,24,13,11,16,18, | 0,0,2,1,4,3,0,2,4,2,1,0 | 7,8,12,4,8,8,12,8,8,14,11,2 | 2,4,0,1,0,0,4,4,0,0,0,2,2,3, | 207 | 149 | 0.000514400246648 | 0.0685661120188099 | 1.0,1.0,0.924,0.96 | 0.716,0.59,1.0,0.7 | 0,057 |
| 204470 | ENSG00000125637 | PSD4 | chr2 | + | 113192260 | 113192589 | 113185960 | 113186255 | 113193047 | 113193128 | 204470 | 76,80,159,142,96,87,66,78 | 9,2,11,18,3,8,15,7,1,4,8,9 | 90,55,42,45,31,38,39,70,2 | 23,18,10,7,16,7,10,13,8,8, | 298 | 149 | 0.000514583308704 | 0.0685661120188099 | 0.809,0.952,0.878, | 0.662,0.604,0.677, | 0,103 |
| 253573 | ENSG00000135968 | GCC2 | chr2 | + | 108483061 | 108483502 | 108482286 | 108482451 | 108484148 | 108484739 | 253573 | 24,19,45,22,14,14,13,21,2 | 2,5,0,0,0,7,2,2,1,0,0,0 | 19,20,24,29,19,4,13,16,17, | 1,0,0,0,1,0,1,1,0,2,1,0,1,2, | 298 | 149 | 0.000514808912389 | 0.0685661120188099 | 0.857,0.655,1.0,1. | 0.905,1.0,1.0,1.0,0 | -0,068 |
| 165893 | ENSG00000172869 | DMXL1 | chr5 | + | 119081559 | 119081697 | 119071554 | 119071656 | 119097978 | 119098104 | 165893 | 0,2,8,15,2,2,3,6,0,4,0,0 | 101,68,137,64,36,70,52,48 | 0,1,0,0,5,7,0,0,2,2,1,5,1,2, | 65,72,60,73,57,44,22,39,3 | 287 | 149 | 0.000517142009994 | 0.0688342505344167 | 0.0,0.015,0.029,0. | 0.0,0.007,0.0,0.0,0 | 0,019 |
| 156865 | ENSG00000134884 | ARGLU1 | chr13 | - | 106557558 | 106557651 | 106557047 | 106557131 | 106559431 | 106559657 | 156865 | 45,3,49,32,14,9,14,12,12,6 | 130,49,133,112,66,62,60,6 | 13,11,19,9,5,10,0,16,11,10 | 104,54,77,34,46,41,18,79, | 242 | 149 | 0.000517461784616 | 0.0688342505344167 | 0.176,0.036,0.185, | 0.071,0.111,0.132, | 0,04 |
| 191407 | ENSG00000204642 | HLA-F | chr6 | + | 29726402 | 29726529 | 29726010 | 29726043 | 29726882 | 29727296 | 191407 | 32,21,113,174,38,40,28,49 | 4,5,13,8,6,3,3,4,2,2,9,8 | 61,58,40,32,23,22,31,95,4 | 6,0,4,2,1,2,1,8,4,3,1,2,3,3, | 276 | 149 | 0.000520950157619 | 0.0692554807748579 | 0.812,0.694,0.824, | 0.846,1.0,0.844,0. | -0,16 |
| 246239 | ENSG00000172183 | ISG20 | chr15 | + | 88637469 | 88637529 | 88636169 | 88636292 | 88639342 | 88639594 | 246239 | 0,3,0,1,1,3,0,1,4,1,0,6 | 4,7,4,5,2,14,3,14,27,6,3,5 | 0,0,0,0,0,0,0,1,0,0,1,3,1,0, | 4,5,5,2,6,10,1,4,1,11,11,4, | 209 | 149 | 0.000521842567741 | 0.0693312948749773 | 0.0,0.234,0.0,0.12 | 0.0,0.0,0.0,0.0,0.0, | 0,017 |
| 167256 | ENSG00000241258 | CRCP | chr7 | + | 66130743 | 66130847 | 66127703 | 66127740 | 66134279 | 66134489 | 167256 | 15,12,28,26,10,10,16,14,1 | 0,0,2,1,1,0,0,0,0,0,0,1 | 17,24,6,8,10,0,8,9,13,12,1 | 2,1,2,2,0,2,2,1,3,1,0,2,2,2,1 | 253 | 149 | 0.00052351439212 | 0.069510503473399 | 1.0,1.0,0.892,0.93 | 0.833,0.934,0.639, | 0,182 |
| 139487 | ENSG00000170471 | RALGAPB | chr20 | + | 38497362 | 38497516 | 38492929 | 38493132 | 38499446 | 38499633 | 139487 | 18,18,81,43,40,61,38,51,1 | 0,0,2,0,0,2,1,1,0,0,0,3 | 47,64,23,28,45,22,28,28,2 | 3,3,3,6,4,1,0,4,1,0,1,2,0,0, | 298 | 149 | 0.000524010315969 | 0.0695334551888556 | 1.0,1.0,0.953,1.0,1 | 0.887,0.914,0.793, | 0,065 |
| 124283 | ENSG00000110713 | NUP98 | chr11 | - | 3744508 | 3744649 | 3735190 | 3735324 | 3753315 | 3753408 | 124283 | 80,94,154,132,58,91,85,89 | 1,2,3,5,0,1,5,0,3,2,3,4 | 113,96,44,99,81,43,78,77, | 3,10,6,6,4,1,2,5,3,7,4,3,2,2 | 290 | 149 | 0.000526662400593 | 0.0698423137042711 | 0.976,0.96,0.963,0 | 0.951,0.831,0.79,0 | 0,042 |
| 104589 | ENSG00000111581 | NUP107 | chr12 | + | 68735230 | 68735344 | 68734707 | 68734833 | 68741812 | 68741980 | 104589 | 61,46,104,67,43,34,37,61, | 2,0,1,0,0,2,3,0,0,0,0,2 | 81,62,51,35,54,29,25,60,3 | 3,5,0,5,4,0,0,0,3,1,1,3,2,0,1 | 263 | 149 | 0.00052745956686 | 0.0698631542202244 | 0.945,1.0,0.983,1. | 0.939,0.875,1.0,0. | 0,036 |
| 182152 | ENSG00000107099 | DOCK8 | chr9 | + | 314291 | 314397 | 311953 | 312166 | 317042 | 317128 | 182152 | 5,2,14,5,2,9,2,3,5,5,2,2 | 79,24,140,108,59,101,62,7 | 4,8,2,2,1,1,1,8,3,3,1,4,1,2, | 126,112,44,44,75,58,60,70 | 255 | 149 | 0.000527599807284 | 0.0698631542202244 | 0.036,0.046,0.055, | 0.018,0.04,0.026,0 | 0,025 |
| 69025 | ENSG00000153922 | CHD1 | chr5 | - | 98876397 | 98876558 | 98875071 | 98875113 | 98879551 | 98879728 | 69025 | 140,162,122,70,59,45,44,7 | 20,3,9,12,6,9,16,13,12,2,2, | 88,99,67,92,82,33,40,55,4 | 13,13,13,34,18,3,9,10,18,4 | 298 | 149 | 0.000527793341861 | 0.0698631542202244 | 0.778,0.964,0.871, | 0.772,0.792,0.72,0 | 0,095 |
| 255873 | ENSG00000197111 | PCBP2 | chr12 | + | 53467804 | 53468281 | 53467220 | 53467293 | 53468779 | 53468832 | 255873 | 33,14,51,34,18,36,24,35,8, | 0,3,2,2,1,0,1,0,2,0,3,1 | 19,55,15,36,24,16,53,28,2 | 0,4,1,3,4,1,2,6,2,3,3,4,4,4, | 298 | 149 | 0.000530659076615 | 0.0701993139022268 | 1.0,0.7,0.927,0.89 | 1.0,0.873,0.882,0. | 0,045 |
| 186685 | ENSG00000134242 | PTPN22 | chr1 | - | 113819576 | 113819687 | 113813810 | 113814969 | 113825141 | 113825172 | 186685 | 65,43,146,157,43,44,55,21 | 9,2,12,6,0,0,0,0,1,2,1,4 | 71,63,42,7,36,31,20,34,23, | 0,6,0,1,0,0,1,2,4,0,0,2,2,0, | 260 | 149 | 0.000531609385141 | 0.070281830204719 | 0.805,0.925,0.875, | 1.0,0.857,1.0,0.8,1 | -0,029 |
| 205902 | ENSG00000128815 | WDFY4 | chr10 | + | 48817337 | 48817409 | 48813956 | 48814082 | 48820233 | 48820437 | 205902 | 40,34,64,34,45,34,23,22,3 | 0,1,2,2,1,6,0,1,0,0,1,4 | 38,35,11,19,5,8,23,38,23,1 | 2,9,0,5,0,0,1,2,1,0,6,3,0,1, | 221 | 149 | 0.000534090523267 | 0.0705665054716266 | 1.0,0.958,0.956,0. | 0.928,0.724,1.0,0. | 0,021 |
| 121993 | ENSG00000172183 | ISG20 | chr15 | + | 88639342 | 88639594 | 88635669 | 88635700 | 88652109 | 88652310 | 121993 | 374,297,219,269,148,169,1 | 2,2,11,9,1,3,2,1,4,3,0,2 | 184,86,218,95,124,135,81, | 7,0,0,0,0,4,0,0,0,3,3,4,1,1, | 298 | 149 | 0.000535108329715 | 0.0706576079220179 | 0.989,0.987,0.909, | 0.929,1.0,1.0,1.0,1 | -0,017 |
| 15665 | ENSG00000185324 | CDK10 | chr16 | + | 89689251 | 89689324 | 89686688 | 89686797 | 89690552 | 89690557 | 15665 | 28,23,36,23,17,20,10,17,2 | 0,3,4,6,0,1,2,0,10,0,0,0 | 26,13,15,9,11,9,14,24,10,1 | 8,8,4,4,3,3,2,3,2,5,1,5,3,1, | 222 | 149 | 0.000537529943105 | 0.0709338486722454 | 1.0,0.837,0.858,0. | 0.686,0.522,0.716, | 0,171 |
| 228852 | ENSG00000084070 | SMAP2 | chr1 | + | 40375746 | 40375816 | 40373726 | 40374223 | 40408652 | 40408738 | 228852 | 30,10,22,3,5,27,15,28,8,6, | 6,2,1,0,2,4,2,3,2,0,0,0 | 15,26,9,6,6,3,7,24,15,6,23, | 2,0,0,0,4,1,1,1,0,1,2,0,0,0, | 219 | 149 | 0.000538201291964 | 0.07097892295999 | 0.773,0.773,0.937, | 0.836,1.0,1.0,1.0,0 | -0,025 |
| 135660 | ENSG00000270011 | ZNF559-ZNF17 | chr19 | + | 9339138 | 9339319 | 9338493 | 9338582 | 9341101 | 9341184 | 135660 | 15,6,88,40,14,13,3,20,16,1 | 0,2,0,0,0,0,2,4,9,1,2,0 | 39,16,27,3,8,16,12,18,16,3 | 0,0,4,0,0,0,0,1,0,0,0,0,0,0, | 298 | 149 | 0.000538948832932 | 0.0710339842380816 | 1.0,0.6,1.0,1.0,1.0, | 1.0,1.0,0.771,1.0,1 | -0,16 |
| 135659 | ENSG00000270011 | ZNF559-ZNF17 | chr19 | + | 9339049 | 9339319 | 9338493 | 9338582 | 9341101 | 9341184 | 135659 | 15,6,88,40,14,13,3,20,16,1 | 0,2,0,0,0,0,2,4,9,1,2,0 | 39,16,27,3,8,16,12,18,16,3 | 0,0,4,0,0,0,0,1,0,0,0,0,0,0, | 298 | 149 | 0.000539281557048 | 0.0710343383139523 | 1.0,0.6,1.0,1.0,1.0, | 1.0,1.0,0.771,1.0,1 | -0,16 |
| 139484 | ENSG00000170471 | RALGAPB | chr20 | + | 38497356 | 38497516 | 38492929 | 38493132 | 38499446 | 38499633 | 139484 | 18,18,81,43,41,61,38,51,1 | 0,0,2,0,0,2,1,1,0,0,0,3 | 48,64,23,28,41,22,29,28,2 | 3,3,3,6,4,1,0,4,1,0,1,2,0,0, | 298 | 149 | 0.000540094954981 | 0.0710979677403765 | 1.0,1.0,0.953,1.0,1 | 0.889,0.914,0.793, | 0,064 |
| 62615 | ENSG00000121892 | PDS5A | chr4 | - | 39848850 | 39848970 | 39845817 | 39845880 | 39849519 | 39849652 | 62615 | 64,43,179,111,48,54,34,89 | 7,6,12,7,2,0,2,5,0,3,3,6 | 92,75,52,32,40,30,34,56,4 | 6,4,1,0,0,2,3,1,1,1,8,4,3,2,2 | 269 | 149 | 0.000540678048528 | 0.0711312206984902 | 0.835,0.799,0.892, | 0.895,0.912,0.966, | -0,055 |
| 125384 | ENSG00000156650 | KAT6B | chr10 | + | 74985079 | 74985241 | 74981786 | 74981928 | 74989018 | 74989096 | 125384 | 67,42,74,34,26,27,25,29,1 | 3,7,0,3,0,2,0,2,1,4,0,2 | 44,41,25,62,28,16,53,33,2 | 0,0,0,1,0,4,2,1,0,0,0,4,0,0, | 298 | 149 | 0.000541891244887 | 0.0712472782701734 | 0.918,0.75,1.0,0.8 | 1.0,1.0,1.0,0.969,1 | -0,066 |
| 38530 | ENSG00000157426 | AASDH | chr4 | - | 56371450 | 56371643 | 56355181 | 56355423 | 56378147 | 56378464 | 38530 | 15,14,37,15,31,25,9,16,9,8 | 2,1,1,2,0,2,0,0,0,1,0,0 | 30,19,7,6,9,13,8,5,11,13,2 | 0,0,3,2,4,1,3,4,1,0,1,3,0,0,2 | 298 | 149 | 0.000542670742923 | 0.0713062067582785 | 0.789,0.875,0.949, | 1.0,1.0,0.538,0.6,0 | 0,118 |
| 704 | ENSG00000182944 | EWSR1 | chr22 | + | 29288605 | 29288755 | 29286922 | 29287134 | 29292136 | 29292169 | 704 | 92,90,169,129,75,69,64,84 | 13,0,5,13,4,2,5,5,4,5,2,7 | 110,105,68,23,40,54,48,92 | 4,15,10,12,13,13,6,5,2,0,7, | 298 | 149 | 0.000543728555614 | 0.0714016111978992 | 0.78,1.0,0.944,0.8 | 0.932,0.778,0.773, | 0,088 |
| 255870 | ENSG00000197111 | PCBP2 | chr12 | + | 53467804 | 53467827 | 53467220 | 53467293 | 53468779 | 53468832 | 255870 | 19,12,28,17,5,22,12,19,9,1 | 0,3,2,2,1,0,1,0,2,0,3,1 | 12,32,12,23,18,10,23,19,1 | 0,4,1,3,4,1,2,6,2,3,3,4,4,4, | 172 | 149 | 0.00054465736406 | 0.0714799689780475 | 1.0,0.776,0.924,0. | 1.0,0.874,0.912,0. | 0,061 |
| 34805 | ENSG00000172869 | DMXL1 | chr5 | + | 119110150 | 119110283 | 119101934 | 119102006 | 119114474 | 119114541 | 34805 | 79,64,153,65,21,20,34,49, | 2,1,2,0,0,0,0,0,0,0,1,0 | 69,73,84,66,48,26,20,38,3 | 5,8,1,2,3,1,0,0,2,0,2,0,0,1, | 282 | 149 | 0.00054578551615 | 0.0715843768595251 | 0.954,0.971,0.976, | 0.879,0.828,0.978, | 0,044 |
| 199851 | ENSG00000118922 | KLF12 | chr13 | - | 73939089 | 73939263 | 73845826 | 73846373 | 73943980 | 73944070 | 199851 | 7,2,4,1,0,0,1,2,0,0,0,0 | 93,35,67,21,35,47,47,56,3 | 0,4,1,10,1,3,0,2,0,9,0,5,2,3 | 42,25,24,62,43,26,48,43,2 | 298 | 149 | 0.0005469812756 | 0.0716282636335885 | 0.036,0.028,0.029, | 0.0,0.074,0.02,0.0 | -0,022 |
| 222404 | ENSG00000136986 | DERL1 | chr8 | - | 123030604 | 123030740 | 123024985 | 123025050 | 123041969 | 123042302 | 222404 | 10,7,72,39,11,24,19,18,13, | 2,0,7,10,2,2,5,3,0,1,0,2 | 41,43,14,6,29,10,4,18,4,15 | 10,2,2,2,6,6,4,6,7,3,10,7,3, | 285 | 149 | 0.000547100647846 | 0.0716282636335885 | 0.723,1.0,0.843,0. | 0.682,0.918,0.785, | 0,215 |
| 150080 | ENSG00000159322 | ADPGK | chr15 | - | 72780196 | 72780322 | 72774871 | 72775097 | 72783458 | 72783785 | 150080 | 2,4,9,4,4,7,3,1,2,5,1,7 | 16,7,22,17,9,10,12,2,9,10, | 7,19,2,4,8,2,1,6,9,6,3,5,7,6 | 10,14,3,1,11,3,6,4,2,7,4,11 | 275 | 149 | 0.000547118516448 | 0.0716282636335885 | 0.063,0.236,0.181, | 0.275,0.424,0.265, | -0,153 |
| 110150 | ENSG00000102096 | PIM2 | chrX | - | 48914394 | 48914571 | 48913181 | 48914294 | 48915019 | 48915392 | 110150 | 514,1111,1061,801,502,33 | 92,29,75,69,60,44,61,34,6 | 534,994,556,459,566,259, | 89,51,115,59,45,29,19,92, | 298 | 149 | 0.000547962310828 | 0.0716706431510877 | 0.736,0.95,0.876,0 | 0.75,0.907,0.707,0 | 0,035 |
| 132770 | ENSG00000197111 | PCBP2 | chr12 | + | 53467804 | 53467843 | 53467220 | 53467293 | 53468779 | 53468832 | 132770 | 45,28,79,55,31,49,33,47,1 | 0,3,2,2,1,0,1,0,2,0,3,1 | 32,83,24,65,38,27,83,46,4 | 0,4,1,3,4,1,2,6,2,3,3,4,4,4, | 188 | 149 | 0.000548108212231 | 0.0716706431510877 | 1.0,0.881,0.969,0. | 1.0,0.943,0.95,0.9 | 0,022 |
| 225166 | ENSG00000186480 | INSIG1 | chr7 | + | 155302805 | 155302846 | 155298560 | 155298697 | 155308240 | 155308757 | 225166 | 13,18,26,16,5,13,9,8,15,23 | 8,3,7,7,4,3,11,3,5,0,7,4 | 6,8,8,6,4,2,2,7,1,5,6,6,8,5, | 8,12,5,4,8,5,2,2,3,5,10,5,2, | 190 | 149 | 0.000549035732774 | 0.0717083748652497 | 0.56,0.825,0.744,0 | 0.37,0.343,0.556,0 | 0,213 |
| 144636 | ENSG00000277957 | SENP3-EIF4A1 | chr17 | + | 7565716 | 7565782 | 7565439 | 7565587 | 7566926 | 7567004 | 144636 | 13,6,16,12,4,12,5,8,4,5,1,5 | 0,0,2,2,1,0,0,1,0,0,0,2 | 14,12,7,5,7,3,7,9,3,12,7,2, | 2,1,1,1,0,0,5,2,1,2,6,1,1,1, | 215 | 149 | 0.000549063107907 | 0.0717083748652497 | 1.0,1.0,0.847,0.80 | 0.829,0.893,0.829, | 0,102 |
| 131037 | ENSG00000050426 | LETMD1 | chr12 | + | 51052091 | 51052207 | 51048342 | 51049185 | 51055834 | 51056021 | 131037 | 4,4,15,10,2,12,0,1,1,3,1,6 | 3,2,2,5,4,2,5,4,3,2,3,2 | 11,11,4,3,1,5,1,7,1,3,8,13, | 4,1,7,2,0,0,0,3,2,0,2,8,0,0,1 | 265 | 149 | 0.000550219342577 | 0.0718158031062403 | 0.428,0.529,0.808, | 0.607,0.861,0.243, | -0,317 |
| 709 | ENSG00000182944 | EWSR1 | chr22 | + | 29288605 | 29291599 | 29286922 | 29287134 | 29292136 | 29292169 | 709 | 93,92,179,133,78,82,66,87 | 13,0,5,13,4,2,5,5,4,5,2,7 | 120,111,70,27,49,56,49,94 | 4,15,10,12,13,13,6,5,2,0,7, | 298 | 149 | 0.000553691599826 | 0.0722164358577528 | 0.782,1.0,0.947,0. | 0.938,0.787,0.778, | 0,082 |
| 708 | ENSG00000182944 | EWSR1 | chr22 | + | 29288605 | 29288786 | 29286922 | 29287134 | 29292136 | 29292169 | 708 | 92,90,169,129,75,69,64,84 | 13,0,5,13,4,2,5,5,4,5,2,7 | 110,106,68,23,40,54,48,93 | 4,15,10,12,13,13,6,5,2,0,7, | 298 | 149 | 0.000553959864523 | 0.0722164358577528 | 0.78,1.0,0.944,0.8 | 0.932,0.779,0.773, | 0,088 |
| 220594 | ENSG00000179344 | HLA-DQB1 | chr6 | - | 32666221 | 32666333 | 32664797 | 32665067 | 32666498 | 32666684 | 220594 | 92,41,0,0,2,7,24,34,5,0,15, | 832,609,117,78,351,145,8 | 26,49,0,0,0,0,0,0,0,1,29,34 | 1034,1216,14,7,7,4,38,52, | 261 | 149 | 0.000556410987934 | 0.0724920661888758 | 0.059,0.037,0.0,0. | 0.014,0.022,0.0,0. | 0,012 |
| 172124 | ENSG00000145012 | LPP | chr3 | + | 188341655 | 188341719 | 188225404 | 188225527 | 188406111 | 188406313 | 172124 | 89,52,8,0,4,2,2,8,0,8,3,3 | 1,0,38,13,20,38,22,37,33,1 | 30,22,29,29,20,5,12,17,4,3 | 12,13,7,16,16,3,0,4,15,18,1 | 213 | 149 | 0.000559066769342 | 0.0727940107878088 | 0.984,1.0,0.128,0. | 0.636,0.542,0.743, | -0,241 |
| 165608 | ENSG00000047365 | ARAP2 | chr4 | - | 36212395 | 36212484 | 36210389 | 36210743 | 36213242 | 36213319 | 165608 | 21,18,54,37,13,26,13,26,1 | 2,3,0,1,2,3,1,0,0,0,0,1 | 38,15,17,15,14,7,12,28,24, | 1,5,1,11,0,1,2,1,4,2,2,2,3,0 | 238 | 149 | 0.000559987296405 | 0.0728403082787183 | 0.868,0.79,1.0,0.9 | 0.96,0.653,0.914,0 | 0,064 |
| 211086 | ENSG00000118412 | CASP8AP2 | chr6 | + | 89831236 | 89831339 | 89829940 | 89830067 | 89846561 | 89846644 | 211086 | 0,0,6,1,2,1,2,3,3,0,0,0 | 10,4,15,14,3,2,6,4,8,6,2,3 | 2,5,1,1,2,1,3,0,3,6,2,4,2,1, | 11,4,0,8,1,2,1,6,3,2,7,7,1,0 | 252 | 149 | 0.000560099196683 | 0.0728403082787183 | 0.0,0.0,0.191,0.04 | 0.097,0.425,1.0,0. | -0,297 |
| 163589 | ENSG00000113593 | PPWD1 | chr5 | + | 65568541 | 65568994 | 65567512 | 65567615 | 65569631 | 65569732 | 163589 | 7,0,6,5,3,4,0,2,2,0,0,0 | 36,23,58,36,22,17,11,3,46, | 0,0,0,1,2,0,0,0,0,1,1,0,1,0, | 29,22,19,22,24,15,15,11,1 | 298 | 149 | 0.000560454939206 | 0.0728425585871054 | 0.089,0.0,0.049,0. | 0.0,0.0,0.0,0.022,0 | 0,032 |
| 241887 | ENSG00000108424 | KPNB1 | chr17 | + | 47669815 | 47669869 | 47668364 | 47668410 | 47670701 | 47670832 | 241887 | 15,19,54,40,23,23,28,31,4 | 10,5,25,25,10,6,5,2,1,7,6,4 | 38,37,14,14,19,17,14,54,7, | 16,26,14,18,15,4,7,23,13,5 | 203 | 149 | 0.000562318309228 | 0.0729978924850588 | 0.524,0.736,0.613, | 0.635,0.511,0.423, | 0,158 |
| 67792 | ENSG00000144381 | HSPD1 | chr2 | - | 197490937 | 197491042 | 197490196 | 197490296 | 197493323 | 197493492 | 67792 | 19,5,10,8,11,7,4,1,9,0,2,3 | 31,18,84,104,40,39,29,25,1 | 5,9,1,1,0,4,4,2,5,12,4,10,3, | 51,42,36,7,37,25,19,28,23, | 254 | 149 | 0.000562328408734 | 0.0729978924850588 | 0.264,0.14,0.065,0 | 0.054,0.112,0.016, | 0,034 |
| 133785 | ENSG00000182944 | EWSR1 | chr22 | + | 29297826 | 29297925 | 29295525 | 29296368 | 29298732 | 29299851 | 133785 | 97,77,225,119,89,78,56,10 | 1,2,2,3,1,5,5,6,6,2,0,4 | 171,169,73,120,78,47,80,9 | 1,3,2,0,0,2,5,2,0,1,1,2,3,1, | 248 | 149 | 0.000564343246742 | 0.0732152871244891 | 0.983,0.959,0.985, | 0.99,0.971,0.956,1 | -0,033 |
| 168249 | ENSG00000138639 | ARHGAP24 | chr4 | + | 85664004 | 85664157 | 85570592 | 85570721 | 85721884 | 85722386 | 168249 | 0,0,27,10,0,0,18,21,6,2,0,0 | 198,149,408,178,125,188,1 | 0,0,0,0,0,0,0,0,0,0,0,0,0,0, | 335,282,171,211,196,103,1 | 298 | 149 | 0.000565290681927 | 0.0732940233505001 | 0.0,0.0,0.032,0.02 | 0.0,0.0,0.0,0.0,0.0, | 0,015 |
| 184991 | ENSG00000108654 | DDX5 | chr17 | - | 64505891 | 64506262 | 64504676 | 64504842 | 64506754 | 64507314 | 184991 | 17,12,11,3,13,10,15,17,29, | 19,20,23,8,6,6,13,11,6,6,0, | 5,1,8,0,0,5,0,5,0,4,6,7,9,11 | 13,12,13,9,11,6,9,6,5,6,9,3 | 298 | 149 | 0.000568319326492 | 0.0736423461530401 | 0.309,0.231,0.193, | 0.161,0.04,0.235,0 | 0,234 |
| 54960 | ENSG00000159140 | SON | chr21 | + | 33572550 | 33572630 | 33568970 | 33569087 | 33573307 | 33573455 | 54960 | 14,0,22,25,1,6,3,5,7,8,7,4 | 137,106,96,77,54,60,50,40 | 4,2,8,3,0,8,9,6,1,3,2,2,4,0,1 | 41,73,59,53,18,39,54,41,2 | 229 | 149 | 0.000568977448394 | 0.0736832642570933 | 0.062,0.0,0.13,0.1 | 0.06,0.018,0.081,0 | 0,032 |
| 238594 | ENSG00000136573 | BLK | chr8 | + | 11555588 | 11555830 | 11555390 | 11555484 | 11556657 | 11556837 | 238594 | 47,22,24,16,24,25,9,5,32,1 | 392,174,307,263,310,194, | 41,48,43,8,12,9,2,11,31,31 | 280,293,166,71,172,94,75, | 298 | 149 | 0.000570643040393 | 0.0738545232873276 | 0.057,0.059,0.038, | 0.068,0.076,0.115, | -0,014 |
| 204484 | ENSG00000125637 | PSD4 | chr2 | + | 113192424 | 113192589 | 113185960 | 113186255 | 113193047 | 113193128 | 204484 | 76,80,159,142,94,87,66,78 | 9,2,11,18,3,8,15,7,1,4,8,9 | 90,56,42,45,30,38,37,67,2 | 23,18,10,7,16,7,10,13,8,8, | 298 | 149 | 0.000571329648355 | 0.0738989492458504 | 0.809,0.952,0.878, | 0.662,0.609,0.677, | 0,103 |
| 162692 | ENSG00000163297 | ANTXR2 | chr4 | - | 79907563 | 79907664 | 79907404 | 79907467 | 79977620 | 79977701 | 162692 | 1,0,0,0,4,0,0,1,0,0,0,0 | 19,17,21,40,13,14,8,21,12, | 7,3,0,5,0,0,1,4,0,2,2,4,0,3, | 23,13,16,19,16,13,30,13,6, | 250 | 149 | 0.000573218997675 | 0.0740987976508036 | 0.03,0.0,0.0,0.0,0. | 0.154,0.121,0.0,0. | -0,038 |
| 214112 | ENSG00000256206 | | chr11 | - | 14519872 | 14520211 | 14518996 | 14519041 | 14520296 | 14520344 | 214112 | 1,3,6,2,1,0,1,3,5,2,0,2 | 14,13,72,46,26,21,18,22,5, | 3,5,0,13,10,2,0,5,1,2,1,9,2, | 26,39,20,11,27,20,7,22,11, | 298 | 149 | 0.000576320728892 | 0.0744324467576909 | 0.034,0.103,0.04,0 | 0.055,0.06,0.0,0.3 | -0,015 |
| 177745 | ENSG00000177731 | FLII | chr17 | - | 18253544 | 18253719 | 18253426 | 18253458 | 18254078 | 18254178 | 177745 | 30,14,25,39,26,23,44,14,3 | 1,0,2,3,3,3,3,6,2,0,0,0 | 10,16,15,14,14,10,15,21,5, | 0,1,2,0,0,0,3,4,0,0,1,0,0,0, | 298 | 149 | 0.000576491717016 | 0.0744324467576909 | 0.938,1.0,0.862,0. | 1.0,0.889,0.789,1. | -0,04 |
| 230660 | ENSG00000225648 | SBDSP1 | chr7 | + | 72830699 | 72830821 | 72829655 | 72829783 | 72831601 | 72831798 | 230660 | 10,4,7,2,2,3,9,1,4,5,3,2 | 7,4,6,6,3,2,2,2,5,5,0,1 | 2,5,3,10,7,1,2,6,6,6,14,4,0, | 0,4,0,11,0,0,0,0,0,3,1,10,1, | 271 | 149 | 0.000577007297067 | 0.0744543510521747 | 0.44,0.355,0.391,0 | 1.0,0.407,1.0,0.33 | -0,237 |
| 114912 | ENSG00000136937 | NCBP1 | chr9 | + | 97645610 | 97645732 | 97645116 | 97645224 | 97647491 | 97647561 | 114912 | 40,22,111,68,42,62,33,69, | 2,3,7,2,0,4,2,3,3,0,0,6 | 97,104,49,67,66,34,44,32, | 1,2,1,5,4,0,2,0,1,1,0,3,1,1, | 271 | 149 | 0.000577424948857 | 0.0744636004598208 | 0.917,0.801,0.897, | 0.982,0.966,0.964, | -0,053 |
| 40942 | ENSG00000163788 | SNRK | chr3 | + | 43299753 | 43299815 | 43286539 | 43286675 | 43303097 | 43303792 | 40942 | 44,29,27,23,22,13,18,24,3 | 4,2,0,12,3,4,1,1,3,4,1,2 | 26,50,29,27,12,11,15,13,1 | 1,4,0,2,4,0,2,0,1,2,2,0,0,5, | 211 | 149 | 0.000579673300608 | 0.0746999053648772 | 0.886,0.911,1.0,0. | 0.948,0.898,1.0,0. | -0,056 |
| 166333 | ENSG00000104517 | UBR5 | chr8 | - | 102278825 | 102278961 | 102276971 | 102277192 | 102279035 | 102279208 | 166333 | 49,54,95,45,35,44,39,63,5 | 10,0,8,9,0,1,0,2,0,0,0,2 | 49,66,56,57,41,31,20,22,3 | 1,5,3,36,5,2,2,1,9,7,1,10,2, | 285 | 149 | 0.00057995150264 | 0.0746999053648772 | 0.719,1.0,0.861,0. | 0.962,0.873,0.907, | 0,1 |
| 37218 | ENSG00000155189 | AGPAT5 | chr8 | + | 6747669 | 6747828 | 6741660 | 6741751 | 6755050 | 6755174 | 37218 | 80,51,119,115,166,101,58, | 0,0,2,4,5,2,1,4,2,0,4,0 | 136,124,54,12,19,51,49,69 | 3,1,2,0,8,2,5,9,2,4,2,1,2,0,1 | 298 | 149 | 0.000582583969096 | 0.0749940970409696 | 1.0,1.0,0.967,0.93 | 0.958,0.984,0.931, | 0,053 |
| 215919 | ENSG00000115233 | PSMD14 | chr2 | + | 161371995 | 161372153 | 161371171 | 161371322 | 161385463 | 161385571 | 215919 | 4,0,3,0,0,2,4,2,6,0,0,0 | 30,22,47,52,31,22,36,36,3 | 5,4,2,12,3,2,3,2,1,1,2,0,0,5 | 26,28,19,10,26,27,10,24,1 | 298 | 149 | 0.000585440657757 | 0.0753167831498487 | 0.062,0.0,0.031,0. | 0.088,0.067,0.05,0 | -0,044 |
| 79678 | ENSG00000240849 | PEDS1 | chr20 | - | 50130864 | 50130947 | 50129545 | 50129690 | 50143501 | 50143621 | 79678 | 3,3,11,12,2,1,1,1,5,0,1,4 | 8,3,12,1,2,2,6,5,2,5,3,1 | 8,3,1,3,1,3,1,4,1,5,2,4,9,5, | 1,2,0,0,0,4,0,4,0,0,2,0,2,2, | 232 | 149 | 0.000586054418256 | 0.0753507039997952 | 0.194,0.391,0.371, | 0.837,0.491,1.0,1. | -0,347 |
| 114453 | ENSG00000108091 | CCDC6 | chr10 | - | 59794472 | 59794597 | 59788746 | 59793111 | 59804419 | 59804520 | 114453 | 204,85,105,68,65,99,129,1 | 3,0,4,0,4,4,2,1,7,1,1,3 | 71,66,107,67,13,26,37,69, | 0,0,3,1,0,0,0,1,1,0,3,1,0,4, | 274 | 149 | 0.000586684360732 | 0.075386663668483 | 0.974,1.0,0.935,1. | 1.0,1.0,0.951,0.97 | -0,027 |

| | | | | | | | | | | | | | | | | | | | | | | |
|---|---|---|---|---|---|---|---|---|---|---|---|---|---|---|---|---|---|---|---|---|---|---|
| 212916 | ENSG00000152689 | RASGRP3 | chr2 | + | 33558676 | 33559030 | 33558210 | 33558336 | 33562728 | 33564605 | 212916 | 245,116,406,127,115,141,1 | 1,0,13,8,2,12,4,9,4,1,0,0 | 174,182,118,44,72,27,42,1 | 7,5,10,2,3,6,3,14,3,2,5,4,2, | 298 | 149 | 0.000587858254725 | 0.0754924343810957 | 0.992,1.0,0.94,0.8 | 0.926,0.948,0.855, | 0,063 |
| 118905 | ENSG00000132109 | TRIM21 | chr11 | - | 4386156 | 4386257 | 4384896 | 4385853 | 4386967 | 4386990 | 118905 | 25,14,57,65,13,19,8,16,26, | 1,0,0,2,1,2,2,0,1,1,0,0 | 41,50,17,21,20,12,13,28,1 | 7,1,0,0,0,2,0,2,0,0,7,5,3,3,0 | 250 | 149 | 0.000589403845886 | 0.0756457836338042 | 0.937,1.0,1.0,0.95 | 0.777,0.968,1.0,1. | 0,031 |
| 107127 | ENSG00000215301 | DDX3X | chrX | + | 41343736 | 41343822 | 41342639 | 41343351 | 41344029 | 41344109 | 107127 | 195,171,680,530,169,162,1 | 2,0,15,6,5,3,10,4,4,0,5,7 | 430,505,276,267,269,170,1 | 17,8,5,43,1,11,2,12,8,6,13, | 235 | 149 | 0.000591750002151 | 0.0759016357049832 | 0.984,1.0,0.966,0. | 0.941,0.976,0.972, | 0,016 |
| 74792 | ENSG00000163617 | CCDC191 | chr3 | - | 114046590 | 114046732 | 114042702 | 114042846 | 114056376 | 114056549 | 74792 | 18,6,29,16,16,19,4,15,6,4,4 | 2,0,0,3,5,2,0,0,0,2,2,2 | 9,11,6,7,10,3,8,14,8,10,5,1 | 1,0,0,0,0,2,1,0,0,0,0,1,2,0,0 | 291 | 149 | 0.000592730964527 | 0.0759685960722923 | 0.822,1.0,1.0,0.73 | 0.822,1.0,1.0,1.0,1 | -0,095 |
| 139262 | ENSG00000101337 | TM9SF4 | chr20 | + | 32155051 | 32155186 | 32150799 | 32150875 | 32157793 | 32157969 | 139262 | 14,11,13,17,9,7,9,9,16,14,1 | 3,0,1,2,2,0,0,1,0,2,4,2 | 16,7,6,4,3,10,6,7,4,6,11,8,4 | 1,0,0,0,0,0,0,0,0,0,0,0,0,0,2 | 284 | 149 | 0.000592977969723 | 0.0759685960722923 | 0.71,1.0,0.872,0.8 | 0.894,1.0,1.0,1.0,1 | -0,171 |
| 94453 | ENSG00000135535 | CD164 | chr6 | - | 109370410 | 109370467 | 109366513 | 109369017 | 109376073 | 109376112 | 94453 | 91,69,337,251,114,122,90, | 8,7,54,29,15,17,14,24,4,7,7 | 174,162,117,32,65,77,85,1 | 34,46,23,8,24,19,15,19,20, | 206 | 149 | 0.000594423294603 | 0.0761084593222476 | 0.892,0.877,0.819, | 0.787,0.718,0.786, | 0,071 |
| 124594 | ENSG00000110422 | HIPK3 | chr11 | + | 33348521 | 33348818 | 33348165 | 33348228 | 33349146 | 33349287 | 124594 | 122,78,127,71,50,65,57,71 | 0,0,2,6,4,1,6,3,1,0,0,1 | 94,113,71,54,45,56,68,69,3 | 6,1,5,10,1,3,1,3,1,5,12,3,1, | 298 | 149 | 0.000594837084526 | 0.0761161596549438 | 1.0,1.0,0.969,0.85 | 0.887,0.983,0.877, | 0,042 |
| 26321 | ENSG00000196505 | GDAP2 | chr1 | - | 117877775 | 117877841 | 117870183 | 117870616 | 117878008 | 117878152 | 26321 | 1,1,4,2,1,1,4,8,0,0,4,3 | 1,12,17,3,16,13,13,11,9,10 | 5,12,6,0,4,2,5,7,2,4,0,8,4,3 | 6,4,7,6,3,5,2,0,5,18,5,11,3, | 215 | 149 | 0.000596945645689 | 0.0763405872057571 | 0.409,0.055,0.14,0 | 0.366,0.675,0.373, | -0,194 |
| 189008 | ENSG00000162434 | JAK1 | chr1 | - | 64882690 | 64883063 | 64879024 | 64879148 | 64883276 | 64883475 | 189008 | 22,11,17,5,5,4,1,10,6,6,1,4 | 278,204,311,216,177,166,1 | 12,20,10,42,12,8,8,13,5,9,1 | 242,263,227,158,177,117,1 | 298 | 149 | 0.000598200183396 | 0.0764555960050502 | 0.038,0.026,0.027, | 0.024,0.037,0.022, | -0,013 |
| 160565 | ENSG00000290548 | | chr5 | - | 34175388 | 34175537 | 34164697 | 34169153 | 34175821 | 34175957 | 160565 | 15,12,55,16,6,15,13,19,7,4 | 4,2,1,2,0,0,0,2,6,0,0,2 | 10,15,5,10,17,13,3,12,13,4 | 0,0,0,3,2,0,0,0,0,2,0,0,5,0,0 | 298 | 149 | 0.000598689846582 | 0.0764727681719231 | 0.652,0.75,0.965,0 | 1.0,1.0,1.0,0.625,0 | -0,065 |
| 94163 | ENSG00000177728 | TMEM94 | chr17 | + | 75496729 | 75496807 | 75495541 | 75496471 | 75497112 | 75497198 | 94163 | 44,19,26,11,21,26,20,15,4 | 1,3,0,2,1,2,0,2,2,0,2,0 | 18,17,6,9,20,7,15,19,16,24 | 1,0,3,8,0,0,7,3,0,1,1,1,1,0,1 | 227 | 149 | 0.000600021764567 | 0.0765974403377936 | 0.967,0.806,1.0,0. | 0.922,1.0,0.568,0. | 0,044 |
| 204734 | ENSG00000196510 | ANAPC7 | chr12 | - | 110377392 | 110377843 | 110375466 | 110376216 | 110381751 | 110381948 | 204734 | 43,30,70,61,33,52,20,22,2 | 0,0,7,2,2,1,0,9,1,0,0,0 | 49,47,44,31,24,14,13,20,3 | 3,0,0,1,0,0,0,0,0,0,2,2,1,0,1 | 298 | 149 | 0.000600993360083 | 0.0766759940035709 | 1.0,1.0,0.833,0.93 | 0.891,1.0,1.0,0.93 | -0,045 |
| 4750 | ENSG00000213020 | ZNF611 | chr19 | - | 52729905 | 52730005 | 52728729 | 52728831 | 52735000 | 52735011 | 4750 | 12,1,12,7,10,3,8,10,5,3,2,4 | 7,1,2,3,2,1,0,0,4,3,0,0 | 9,10,9,3,6,1,0,4,0,9,3,5,12, | 3,0,0,0,0,0,3,0,2,1,4,0,3,1,0 | 249 | 149 | 0.000602681352496 | 0.0767728410268759 | 0.506,0.374,0.782, | 0.642,1.0,1.0,1.0,1 | -0,067 |
| 255125 | ENSG00000135124 | P2RX4 | chr12 | + | 121221015 | 121221113 | 121217133 | 121217281 | 121221912 | 121221984 | 255125 | 0,0,2,0,0,0,0,1,0,0,2,2 | 22,13,22,16,8,21,18,16,15, | 1,3,2,2,4,0,0,1,0,0,2,3,3,0,3 | 9,13,5,4,8,15,7,14,1,6,9,13 | 247 | 149 | 0.00060273786963 | 0.0767728410268759 | 0.0,0.0,0.052,0.0,0 | 0.063,0.122,0.194, | -0,119 |
| 68511 | ENSG00000163534 | FCRL1 | chr1 | - | 157803844 | 157804111 | 157801914 | 157802193 | 157807101 | 157807122 | 68511 | 163,50,167,10,48,32,61,68 | 2,0,6,3,0,0,1,0,0,0,1,0 | 127,49,101,41,32,18,48,4,2 | 2,1,3,1,3,2,2,0,2,0,0,3,8,6,6 | 298 | 149 | 0.000602822555001 | 0.0767728410268759 | 0.976,1.0,0.933,0. | 0.969,0.961,0.944, | 0,026 |
| 123417 | ENSG00000149187 | CELF1 | chr11 | - | 47475335 | 47475521 | 47473087 | 47473231 | 47478876 | 47478940 | 123417 | 42,29,83,90,49,47,40,34,2 | 0,2,0,2,0,0,0,1,0,1,0,3 | 42,63,16,49,25,36,37,31,2 | 5,8,2,1,0,2,0,2,1,0,2,0,1,0,4 | 298 | 149 | 0.000604193536827 | 0.0768217179105547 | 1.0,0.879,1.0,0.95 | 0.808,0.797,0.8,0. | 0,038 |
| 54107 | ENSG00000175309 | PHYKPL | chr5 | - | 178223319 | 178223451 | 178222851 | 178222934 | 178224447 | 178225010 | 54107 | 3,6,4,3,6,6,0,3,0,0,0,8 | 19,9,30,27,12,29,18,24,17, | 2,1,1,1,0,2,4,3,1,0,1,0,3,5,2 | 27,19,7,5,8,23,13,26,16,24 | 281 | 149 | 0.000604276311582 | 0.0768217179105547 | 0.077,0.261,0.066, | 0.038,0.027,0.07,0 | 0,039 |
| 162479 | ENSG00000081791 | DELE1 | chr5 | + | 141931195 | 141931280 | 141930246 | 141930274 | 141933258 | 141933882 | 162479 | 15,11,7,5,5,2,4,8,8,7,0,0 | 29,29,48,33,18,26,12,23,3 | 10,4,1,1,5,2,1,4,2,3,1,5,5,2 | 38,29,21,10,19,20,25,24,1 | 234 | 149 | 0.000604277118178 | 0.0768217179105547 | 0.248,0.195,0.085, | 0.144,0.081,0.029, | 0,061 |
| 179257 | ENSG00000285565 | | chr6 | + | 31836426 | 31836517 | 31835410 | 31835451 | 31837235 | 31837435 | 179257 | 9,10,38,14,12,10,7,12,11,9 | 141,82,213,111,84,77,76,9 | 10,10,6,4,6,6,2,15,2,6,16,2 | 129,92,62,61,68,35,48,117 | 240 | 149 | 0.000605057412997 | 0.0768755088882865 | 0.038,0.07,0.1,0.0 | 0.046,0.063,0.057, | 0,022 |
| 230668 | ENSG00000225648 | SBDSP1 | chr7 | + | 72830755 | 72830821 | 72829655 | 72829783 | 72831601 | 72831798 | 230668 | 6,1,7,1,1,1,6,3,3,4,1,2 | 7,4,6,6,3,2,2,2,5,5,0,1 | 2,3,3,9,7,1,2,3,2,4,8,2,0,1,1 | 0,4,0,11,0,0,0,0,0,3,1,10,1, | 215 | 149 | 0.000605416501025 | 0.0768757515823668 | 0.373,0.148,0.447, | 1.0,0.342,1.0,0.36 | -0,233 |
| 178683 | ENSG00000163564 | PYHIN1 | chr1 | + | 158973646 | 158974069 | 158944874 | 158945042 | 158976700 | 158977059 | 178683 | 125,74,15,28,20,7,58,56,5 | 4,14,20,8,1,5,12,12,7,2,5,2 | 5,5,53,16,26,20,11,32,9,22 | 2,4,21,1,24,15,3,8,11,24,5, | 298 | 149 | 0.000605990451544 | 0.0769032611298742 | 0.94,0.725,0.273,0 | 0.556,0.385,0.558, | 0,178 |
| 239542 | ENSG00000083223 | TUT7 | chr9 | - | 86298766 | 86298830 | 86287734 | 86288744 | 86301275 | 86301601 | 239542 | 11,4,13,7,7,5,12,2,10,4,1,3 | 46,21,47,30,11,18,22,12,3 | 6,2,2,2,1,1,1,1,1,5,4,11,9,3 | 36,28,35,21,18,17,2,23,18, | 213 | 149 | 0.000607229461742 | 0.077015087967114 | 0.143,0.118,0.162, | 0.104,0.048,0.038, | 0,066 |
| 132726 | ENSG00000182796 | TMEM198B | chr12 | + | 55834568 | 55834726 | 55832958 | 55833541 | 55834850 | 55834988 | 132726 | 2,0,1,1,1,4,1,6,4,1,0,3 | 11,6,6,3,7,10,3,9,8,7,6,3 | 0,1,0,3,1,0,0,3,0,2,1,0,1,2,2 | 9,13,3,24,7,1,8,4,10,7,5,10 | 298 | 149 | 0.000610051969538 | 0.0773275002683353 | 0.083,0.0,0.077,0. | 0.0,0.037,0.0,0.05 | 0,064 |
| 43644 | ENSG00000160796 | NBEAL2 | chr3 | + | 46991218 | 46991304 | 46989510 | 46989593 | 46991839 | 46991946 | 43644 | 4,6,8,6,9,11,7,4,4,3,2,5 | 2,0,3,0,2,1,4,5,0,1,0,0 | 4,10,4,8,1,6,4,15,3,6,2,10,6 | 0,0,0,0,2,0,2,1,0,0,0,3,1,0,2 | 235 | 149 | 0.000611252095631 | 0.0774340198909687 | 0.559,1.0,0.628,1. | 1.0,1.0,1.0,1.0,0.2 | -0,062 |
| 81840 | ENSG00000148248 | SURF4 | chr9 | - | 133364839 | 133365026 | 133361449 | 133363759 | 133365984 | 133366028 | 81840 | 41,39,74,72,36,41,30,57,6 | 1,1,9,13,0,2,4,1,2,4,1,1 | 53,79,56,37,17,28,28,48,2 | 3,5,2,0,1,2,1,4,0,0,0,3,3,0,2 | 298 | 149 | 0.00061350532269 | 0.0776737435928773 | 0.953,0.951,0.804, | 0.898,0.888,0.933, | -0,039 |
| 215387 | ENSG00000116521 | SCAMP3 | chr1 | - | 155261278 | 155261734 | 155260536 | 155260659 | 155262085 | 155262329 | 215387 | 14,12,12,8,9,15,6,3,15,9,3, | 1,3,2,1,2,3,0,8,4,0,0,4 | 7,12,2,4,3,4,3,6,4,4,9,16,9, | 7,3,3,2,4,3,2,4,1,1,3,7,4,4,7 | 298 | 149 | 0.000614119317469 | 0.0776972288403616 | 0.875,0.667,0.75,0 | 0.333,0.667,0.25,0 | 0,274 |
| 153628 | ENSG00000197555 | SIPA1L1 | chr14 | + | 71650334 | 71650508 | 71624047 | 71624236 | 71658332 | 71658436 | 153628 | 81,71,142,62,33,54,50,50,3 | 6,6,7,2,1,2,5,7,7,4,0,2 | 64,60,75,81,93,60,28,36,3 | 0,2,6,5,3,1,2,0,2,3,4,0,2,0,2 | 298 | 149 | 0.000614412809894 | 0.0776972288403616 | 0.871,0.855,0.91,0 | 1.0,0.938,0.862,0. | -0,059 |
| 92557 | ENSG00000083097 | DOP1A | chr6 | + | 83135618 | 83135878 | 83134186 | 83134287 | 83137172 | 83139162 | 92557 | 36,25,84,15,24,36,14,27,1 | 0,0,2,0,4,1,0,0,0,0,2,0 | 34,46,29,26,32,27,18,20,2 | 0,7,0,2,1,0,1,4,0,2,2,0,2,0,3 | 298 | 149 | 0.000615116520979 | 0.0777073503382498 | 1.0,1.0,0.955,1.0,0 | 1.0,0.767,1.0,0.86 | 0,048 |
| 178711 | ENSG00000031698 | SARS1 | chr1 | + | 109236374 | 109236548 | 109235976 | 109236106 | 109237243 | 109237373 | 178711 | 33,31,61,48,51,13,32,29,8 | 2,0,5,3,2,1,0,0,1,0,2,0 | 37,39,38,36,17,17,9,15,10, | 2,0,2,1,4,0,0,7,0,0,3,1,5,0,1 | 298 | 149 | 0.000616018401638 | 0.0777073503382498 | 0.892,1.0,0.859,0. | 0.902,1.0,0.905,0. | 0,06 |
| 111018 | ENSG00000033800 | PIAS1 | chr15 | + | 68141945 | 68142030 | 68086305 | 68086750 | 68146565 | 68146700 | 111018 | 31,21,64,60,21,27,17,27,1 | 4,2,2,1,0,0,1,0,0,0,0,0 | 57,51,10,23,25,12,12,19,2 | 0,0,2,15,0,1,2,0,0,1,10,0,0, | 234 | 149 | 0.000616050949142 | 0.0777073503382498 | 0.832,0.87,0.953,0 | 1.0,1.0,0.761,0.49 | 0,068 |
| 218840 | ENSG00000135049 | AGTPBP1 | chr9 | - | 85621700 | 85621805 | 85621201 | 85621285 | 85632661 | 85633374 | 218840 | 6,0,11,5,0,3,0,2,3,0,0,0 | 20,39,107,60,32,24,24,13,1 | 1,1,1,0,0,3,1,3,0,0,2,2,2,2,0 | 44,45,27,17,43,26,9,32,31, | 254 | 149 | 0.000616091090557 | 0.0777073503382498 | 0.15,0.0,0.057,0.0 | 0.013,0.013,0.021, | 0,025 |
| 37532 | ENSG00000138814 | PPP3CA | chr4 | - | 101063231 | 101063357 | 101061086 | 101061161 | 101080531 | 101080626 | 37532 | 61,50,140,115,43,42,43,41 | 5,0,6,4,4,0,1,6,2,2,0,0 | 71,76,64,48,37,28,56,36,4 | 7,7,3,8,7,2,5,0,2,0,10,5,10, | 275 | 149 | 0.000616298056634 | 0.0777073503382498 | 0.869,1.0,0.927,0. | 0.846,0.855,0.92,0 | 0,069 |
| 53364 | ENSG00000184014 | DENND5A | chr11 | - | 9176771 | 9176902 | 9170626 | 9170777 | 9178131 | 9178366 | 53364 | 4,1,5,1,0,2,0,2,2,0,0,2 | 22,16,18,14,9,0,4,10,9,2,1 | 3,6,7,9,1,2,2,0,1,10,3,1,7,6 | 14,15,2,7,4,7,6,9,4,26,6,41 | 280 | 149 | 0.000617082261023 | 0.0777540755276716 | 0.088,0.032,0.129, | 0.102,0.176,0.651, | -0,049 |
| 136065 | ENSG00000104856 | RELB | chr19 | + | 45032628 | 45032749 | 45028887 | 45028992 | 45034243 | 45034312 | 136065 | 32,33,72,29,38,30,14,22,3 | 0,9,3,0,0,6,0,1,2,1,0,5 | 19,42,19,23,16,5,21,31,6,7 | 0,0,4,1,1,0,0,0,0,0,2,6,1,1,2 | 270 | 149 | 0.000617391152189 | 0.0777540755276716 | 1.0,0.669,0.93,1.0, | 1.0,1.0,0.724,0.92 | -0,003 |
| 114446 | ENSG00000100815 | TRIP11 | chr14 | - | 92011754 | 92011795 | 92007639 | 92007852 | 92014214 | 92014577 | 114446 | 9,8,21,13,8,2,2,5,10,4,2,2 | 9,3,5,3,0,2,3,1,0,2,1,1 | 14,3,6,5,6,1,5,2,3,15,3,5,8, | 0,2,3,0,4,1,1,0,3,0,0,1,0,4,0 | 190 | 149 | 0.000619160202459 | 0.0779312687341831 | 0.44,0.677,0.767,0 | 1.0,0.541,0.611,1. | -0,128 |
| 130705 | ENSG00000151148 | UBE3B | chr12 | + | 109530546 | 109530658 | 109526357 | 109526416 | 109533465 | 109533558 | 130705 | 9,5,11,11,6,13,11,14,6,5,7, | 1,4,2,2,3,1,6,7,1,1,3,1 | 7,1,5,2,2,3,5,8,7,8,5,7,9,10 | 0,4,0,0,1,1,2,0,1,0,2,2,1,1,0 | 261 | 149 | 0.000621454261294 | 0.0781426264350347 | 0.837,0.416,0.758, | 1.0,0.125,1.0,1.0,0 | -0,079 |
| 86165 | ENSG00000115998 | C2orf42 | chr2 | - | 70165091 | 70165192 | 70160624 | 70160787 | 70165527 | 70165635 | 86165 | 27,2,16,16,4,14,6,4,5,5,2,1 | 4,1,3,1,0,0,0,5,0,0,0,0 | 13,16,5,5,8,7,6,7,3,7,4,2,1 | 4,2,1,3,0,3,4,3,2,0,0,1,0,2,0 | 250 | 149 | 0.00062156555727 | 0.0781426264350347 | 0.801,0.544,0.761, | 0.66,0.827,0.749,0 | 0,134 |
| 4245 | ENSG00000197961 | ZNF121 | chr19 | - | 9584185 | 9584348 | 9569001 | 9569082 | 9584460 | 9584497 | 4245 | 2,4,10,4,7,1,6,5,5,4,2,2 | 37,32,41,36,22,22,32,38,2 | 0,3,4,0,0,0,3,2,5,1,3,3,2,0,2 | 25,34,38,30,28,8,22,12,17, | 298 | 149 | 0.000622310299733 | 0.0781905826747421 | 0.026,0.059,0.109, | 0.0,0.042,0.05,0.0, | 0,031 |
| 135270 | ENSG00000054611 | TBC1D22A | chr22 | + | 46978546 | 46978934 | 46974289 | 46974399 | 46997633 | 46997709 | 135270 | 2,6,8,0,2,1,0,2,2,0,2,0 | 87,60,92,100,30,41,43,36,3 | 3,9,5,4,2,2,2,2,3,3,5,9,3,4,3 | 45,62,44,65,48,48,31,36,51 | 298 | 149 | 0.000625756099215 | 0.0785776610210873 | 0.011,0.048,0.042, | 0.032,0.068,0.054, | -0,017 |
| 170341 | ENSG00000100813 | ACIN1 | chr14 | - | 23067303 | 23067404 | 23065965 | 23066008 | 23069475 | 23069617 | 170341 | 9,7,0,7,2,4,3,4,2,3,3,2 | 71,42,74,41,31,41,16,22,9 | 0,2,1,5,1,4,0,0,3,0,3,1,0,2,2 | 56,34,45,25,10,20,36,40,5, | 250 | 149 | 0.000627382666371 | 0.0787359758983654 | 0.07,0.09,0.0,0.09 | 0.0,0.034,0.013,0. | 0,008 |
| 106540 | ENSG00000196924 | FLNA | chrX | - | 154357250 | 154357274 | 154354824 | 154355072 | 154357433 | 154357623 | 106540 | 386,271,188,203,214,182,1 | 53,76,29,32,59,43,57,34,3 | 171,241,90,97,150,126,85, | 43,59,14,33,36,22,40,63,2 | 173 | 149 | 0.000628076130824 | 0.0787770709285433 | 0.862,0.754,0.848, | 0.774,0.779,0.847, | 0,053 |
| 91273 | ENSG00000163939 | PBRM1 | chr3 | - | 52563282 | 52563493 | 52561766 | 52561968 | 52564049 | 52564233 | 91273 | 62,19,66,40,53,37,45,49,7 | 5,0,0,3,1,4,3,2,4,2,0,1 | 50,42,30,37,24,28,17,48,3 | 0,0,1,0,4,0,0,0,0,3,0,5,1,0,2 | 298 | 149 | 0.000628847880564 | 0.0788279313812873 | 0.861,1.0,1.0,0.87, | 1.0,1.0,0.938,1.0,0 | -0,037 |
| 226136 | ENSG00000129103 | SUMF2 | chr7 | + | 56078941 | 56079039 | 56078363 | 56078508 | 56079527 | 56079736 | 226136 | 2,1,2,11,2,10,5,2,2,0,2,4 | 58,40,142,124,79,76,53,59 | 4,0,2,1,4,0,0,0,0,4,2,2,0,2,1 | 74,90,61,28,37,45,44,81,3 | 247 | 149 | 0.000630353384516 | 0.0789706573357178 | 0.02,0.015,0.008,0 | 0.032,0.0,0.019,0. | 0,013 |
| 71753 | ENSG00000053254 | FOXN3 | chr14 | - | 89486568 | 89486671 | 89411933 | 89412490 | 89619027 | 89619137 | 71753 | 20,9,36,21,6,17,11,21,17,1 | 0,0,4,2,5,0,1,2,2,0,0,6 | 39,32,9,6,2,9,13,17,5,11,2 | 3,3,0,0,0,0,2,1,1,2,1,2,3,2,2 | 252 | 149 | 0.000631930078032 | 0.0791221306718472 | 1.0,1.0,0.842,0.86 | 0.885,0.863,1.0,1. | -0,017 |
| 178572 | ENSG00000120008 | WDR11 | chr10 | + | 120867097 | 120867169 | 120866568 | 120866764 | 120871169 | 120871346 | 178572 | 178,97,154,88,76,60,88,65 | 2,0,5,6,1,3,4,7,3,1,2,4 | 102,132,155,133,100,68,3 | 6,14,6,11,7,2,6,1,3,9,11,3,4 | 221 | 149 | 0.000632512994977 | 0.0791490723383109 | 0.984,1.0,0.954,0. | 0.92,0.864,0.946,0 | 0,04 |
| 32726 | ENSG00000117091 | CD48 | chr1 | - | 160684886 | 160685189 | 160680421 | 160681468 | 160711681 | 160711822 | 32726 | 601,416,1024,1257,315,18 | 18,12,9,10,0,0,2,2,0,2,2,9 | 591,562,346,181,599,467,2 | 4,4,4,0,4,3,2,2,0,0,4,17,2,5 | 298 | 149 | 0.000636682860609 | 0.079614228159074 | 0.943,0.945,0.983, | 0.987,0.986,0.977, | -0,009 |
| 230049 | ENSG00000042980 | ADAM28 | chr8 | + | 24339465 | 24339568 | 24335421 | 24335512 | 24341597 | 24341629 | 230049 | 48,5,55,55,35,36,21,19,36, | 5,0,7,3,2,0,0,1,0,1,0,3 | 58,29,32,8,13,6,12,43,7,55 | 3,0,1,3,1,1,2,2,0,0,0,0,0,0,0 | 252 | 149 | 0.00063706935955 | 0.079614228159074 | 0.85,1.0,0.823,0.9 | 0.92,1.0,0.95,0.61 | 0,003 |
| 111263 | ENSG00000214706 | IFRD2 | chr3 | - | 50290387 | 50290472 | 50289928 | 50290294 | 50290559 | 50290679 | 111263 | 16,8,11,32,25,22,17,23,9,7 | 0,0,0,0,1,1,3,3,0,1,0,0 | 24,11,11,9,6,1,14,20,6,9,2 | 5,0,0,1,0,2,2,1,0,0,2,5,5,1,1 | 234 | 149 | 0.000637339951578 | 0.079614228159074 | 1.0,1.0,1.0,1.0,0.9 | 0.753,1.0,1.0,0.85 | 0,094 |
| 78637 | ENSG00000093144 | ECHDC1 | chr6 | - | 127329821 | 127329882 | 127327001 | 127327144 | 127330808 | 127331030 | 78637 | 8,0,1,8,1,1,2,6,0,0,0,0 | 21,7,38,26,25,12,20,13,19, | 3,4,0,0,2,1,0,0,0,2,0,3,0,2,0 | 30,28,16,8,4,20,8,6,5,35,1 | 210 | 149 | 0.000641495839742 | 0.0800868857792984 | 0.213,0.0,0.018,0. | 0.066,0.092,0.0,0. | 0,029 |
| 228959 | ENSG00000144815 | NXPE3 | chr3 | + | 101816794 | 101817091 | 101807052 | 101807126 | 101821403 | 101828231 | 228959 | 9,6,21,26,4,7,3,6,6,5,2,13 | 0,0,2,2,2,0,0,6,2,0,2,3 | 16,18,7,10,7,13,0,4,9,12,8, | 0,2,0,2,0,0,1,1,0,0,1,0,0,2,3 | 298 | 149 | 0.00064304544118 | 0.0802048268276967 | 1.0,1.0,0.84,0.867, | 1.0,0.818,1.0,0.71 | -0,068 |
| 71098 | ENSG00000134215 | VAV3 | chr1 | - | 107777669 | 107777770 | 107777230 | 107777296 | 107779433 | 107779492 | 71098 | 3,0,4,0,1,2,0,1,0,2,0,1 | 34,31,141,48,37,68,11,39,3 | 2,5,6,11,2,0,0,2,1,1,4,7,0,1 | 63,63,51,35,25,13,27,42,1 | 250 | 149 | 0.000643185838028 | 0.0802048268276967 | 0.05,0.0,0.017,0.0, | 0.019,0.045,0.066, | -0,017 |
| 218668 | ENSG00000024048 | UBR2 | chr6 | + | 42634216 | 42634294 | 42632804 | 42632904 | 42635417 | 42635546 | 218668 | 4,2,17,7,2,5,1,5,1,0,1,2 | 31,17,71,47,25,36,23,25,2 | 8,3,5,0,5,2,0,2,1,0,4,1,2,0,1 | 46,45,22,34,46,22,16,28,1 | 227 | 149 | 0.000643733399662 | 0.0802149625577918 | 0.078,0.072,0.136, | 0.102,0.042,0.13,0 | 0,024 |
| 141152 | ENSG00000166562 | SEC11C | chr18 | + | 59148064 | 59148223 | 59139898 | 59140035 | 59149512 | 59149622 | 141152 | 0,2,11,1,1,3,4,5,0,3,3,5 | 30,31,99,78,27,24,22,23,2 | 0,0,0,3,0,0,1,2,2,0,0,0,1,1,8 | 25,25,14,4,11,13,11,18,4,1 | 298 | 149 | 0.000644012504239 | 0.0802149625577918 | 0.0,0.031,0.053,0. | 0.0,0.0,0.0,0.273,0 | 0,023 |
| 39010 | ENSG00000204463 | BAG6 | chr6 | - | 31648910 | 31648964 | 31648676 | 31648751 | 31649198 | 31649395 | 39010 | 97,51,148,93,82,72,52,50,8 | 2,5,5,4,0,3,2,5,7,2,0,2 | 74,72,47,52,60,34,50,78,2 | 9,5,5,0,0,4,5,4,1,1,5,4,8,9,1 | 203 | 149 | 0.000645507242494 | 0.0803166178688411 | 0.973,0.882,0.956, | 0.858,0.914,0.873, | 0,039 |
| 149669 | ENSG00000214013 | GANC | chr15 | + | 42283610 | 42283869 | 42280916 | 42281133 | 42287690 | 42287818 | 149669 | 39,24,48,37,16,30,17,21,2 | 2,3,4,1,0,0,0,0,0,0,1,2 | 12,16,24,19,5,4,8,14,16,27 | 4,1,1,0,4,2,0,2,1,1,3,1,4,2,1 | 298 | 149 | 0.00064582885369 | 0.0803166178688411 | 0.907,0.8,0.857,0. | 0.6,0.889,0.923,1. | 0,103 |
| 91037 | ENSG00000140025 | EFCAB11 | chr14 | - | 89924512 | 89924677 | 89797126 | 89797324 | 89931540 | 89931631 | 91037 | 2,0,5,8,0,4,2,1,0,0,0,1 | 3,2,11,2,6,2,9,2,2,1,1,8 | 1,2,0,0,1,0,1,0,0,0,0,0,0,1,0 | 2,1,6,1,3,2,2,4,7,3,2,1,1,0,4 | 298 | 149 | 0.000645948146554 | 0.0803166178688411 | 0.25,0.0,0.185,0.6 | 0.2,0.5,0.0,0.0,0.1 | 0,021 |
| 193079 | ENSG00000116688 | MFN2 | chr1 | + | 11981969 | 11982113 | 11980208 | 11980484 | 11989164 | 11989343 | 193079 | 9,12,12,7,7,7,7,4,6,4,5,6 | 2,2,4,1,1,2,0,1,0,5,0,0 | 12,5,5,5,5,1,13,8,5,4,10,4,8 | 0,1,2,0,1,1,0,1,0,2,2,0,2,0,0 | 293 | 149 | 0.000646484115137 | 0.0803368490675818 | 0.696,0.753,0.604, | 1.0,0.718,0.56,1.0, | -0,029 |
| 87677 | ENSG00000010244 | ZNF207 | chr17 | + | 32361467 | 32361515 | 32360891 | 32360967 | 32362913 | 32362984 | 87677 | 173,108,313,222,189,166,1 | 2,2,9,15,8,11,11,6,25,8,5,3 | 211,243,155,183,139,98,8 | 3,0,4,3,3,7,4,10,0,7,9,13,8, | 197 | 149 | 0.000650402738637 | 0.0807771678243394 | 0.985,0.976,0.963, | 0.982,1.0,0.967,0. | -0,02 |
| 111712 | ENSG00000116852 | KIF21B | chr1 | - | 200983055 | 200983094 | 200980959 | 200981096 | 200984858 | 200984972 | 111712 | 2,2,0,2,1,1,0,0,2,1,0,1 | 2,5,4,2,8,7,3,3,3,1,3,3 | 5,4,0,0,2,1,2,1,0,1,0,5,0,3,3 | 2,5,1,6,6,4,5,0,2,4,1,8,3,1,8 | 188 | 149 | 0.000651311472448 | 0.0808433791963411 | 0.442,0.241,0.0,0. | 0.665,0.388,0.0,0. | -0,094 |
| 141413 | ENSG00000173821 | RNF213 | chr17 | + | 80278763 | 80278902 | 80273240 | 80273404 | 80287814 | 80288363 | 141413 | 8,6,13,8,12,8,7,15,3,5,2,7 | 64,48,62,34,53,59,48,54,5 | 8,11,4,9,10,6,3,8,9,21,6,32 | 39,38,29,63,23,31,34,41,3 | 288 | 149 | 0.000653852407571 | 0.0811000176393377 | 0.061,0.061,0.098, | 0.096,0.13,0.067,0 | -0,049 |
| 55075 | ENSG00000139631 | CSAD | chr12 | - | 53171325 | 53171441 | 53170422 | 53170502 | 53171881 | 53171988 | 55075 | 7,3,27,17,14,16,5,6,10,4,6, | 0,0,9,0,3,1,1,1,1,0,0,2 | 7,10,10,4,10,5,11,13,8,7,7, | 1,9,0,6,2,2,1,4,4,2,0,8,6,6,3 | 265 | 149 | 0.000654482624491 | 0.0811000176393377 | 1.0,1.0,0.628,1.0,0 | 0.797,0.385,1.0,0. | 0,242 |
| 43408 | ENSG00000176261 | ZBTB8OS | chr1 | - | 32631826 | 32631879 | 32627507 | 32627544 | 32634767 | 32634792 | 43408 | 8,15,31,15,14,13,11,6,2,2,1 | 0,1,2,2,4,0,2,1,0,0,2,0 | 15,9,14,6,5,3,2,10,3,9,6,20 | 0,1,2,0,3,3,0,3,0,1,5,7,3,0,2 | 202 | 149 | 0.000654509483483 | 0.0811000176393377 | 1.0,0.917,0.92,0.8 | 1.0,0.869,0.838,1. | 0,077 |
| 121021 | ENSG00000074266 | EED | chr11 | + | 86266082 | 86266216 | 86263944 | 86264263 | 86276979 | 86277138 | 121021 | 20,30,79,41,16,14,8,12,6,4 | 2,2,8,6,1,1,0,0,0,0,0,5 | 40,85,24,24,43,14,9,18,40, | 2,6,1,0,4,0,0,2,1,3,1,0,1,2,0 | 283 | 149 | 0.000658020039058 | 0.0814730357682571 | 0.84,0.888,0.839,0 | 0.913,0.882,0.927, | -0,02 |
| 146102 | ENSG00000074755 | ZZEF1 | chr17 | - | 4109652 | 4109783 | 4105692 | 4105809 | 4112608 | 4112808 | 146102 | 26,12,49,29,25,34,24,34,7, | 3,1,5,4,0,4,0,3,2,0,0,0 | 37,37,20,30,15,12,30,24,2 | 1,9,1,9,6,2,10,3,8,1,3,2,3,2 | 280 | 149 | 0.000658401038179 | 0.0814730357682571 | 0.822,0.865,0.839, | 0.952,0.686,0.914, | 0,111 |
| 146147 | ENSG00000140983 | RHOT2 | chr16 | + | 668508 | 668569 | 668357 | 668411 | 668655 | 668699 | 146147 | 17,12,25,9,8,8,6,12,5,10,8, | 2,0,9,0,0,2,0,4,0,2,0,2 | 15,10,12,9,12,8,3,6,10,18,1 | 0,2,2,5,1,1,0,3,0,0,0,4,0,0,0 | 210 | 149 | 0.000659353539944 | 0.0814730357682571 | 0.858,1.0,0.663,1. | 1.0,0.78,0.81,0.56 | -0,002 |
| 209419 | ENSG00000091490 | SEL1L3 | chr4 | - | 25830097 | 25830152 | 25821995 | 25822128 | 25832994 | 25833110 | 209419 | 9,9,102,43,31,41,30,42,14, | 9,5,5,3,3,7,0,1,3,1,0,2 | 26,11,16,12,14,8,13,14,10, | 4,0,1,2,1,0,4,1,3,1,0,4,1,0,2 | 204 | 149 | 0.000659606483576 | 0.0814730357682571 | 0.422,0.568,0.937, | 0.826,1.0,0.921,0. | -0,066 |
| 178682 | ENSG00000163564 | PYHIN1 | chr1 | + | 158973646 | 158974065 | 158944874 | 158945042 | 158976700 | 158977059 | 178682 | 127,80,15,28,20,7,61,62,5 | 4,14,20,8,1,5,12,12,7,2,5,2 | 5,6,54,17,27,21,11,32,11,2 | 2,4,21,1,24,15,3,8,11,24,5, | 298 | 149 | 0.000659672608453 | 0.0814730357682571 | 0.941,0.741,0.273, | 0.556,0.429,0.562, | 0,172 |
| 209806 | ENSG00000198589 | LRBA | chr4 | - | 150868243 | 150868305 | 150851884 | 150852943 | 150870524 | 150870606 | 209806 | 145,94,67,47,30,58,27,53,1 | 5,0,1,2,3,3,0,8,0,0,0,1 | 44,34,64,94,26,30,20,33,2 | 3,3,0,2,11,3,0,0,7,0,3,0,4,0 | 211 | 149 | 0.000659791114403 | 0.0814730357682571 | 0.953,1.0,0.979,0. | 0.912,0.889,1.0,0. | 0,047 |
| 169923 | ENSG00000112282 | MED23 | chr6 | - | 131602217 | 131602396 | 131600037 | 131600162 | 131603029 | 131603204 | 169923 | 48,60,42,37,33,33,19,38,4 | 2,3,0,0,0,2,2,7,0,2,0,0 | 31,36,21,36,43,20,28,30,2 | 0,2,2,7,7,0,1,0,1,1,1,2,2,3,3 | 298 | 149 | 0.000661929403617 | 0.0816505758605023 | 0.923,0.909,1.0,1. | 1.0,0.9,0.84,0.72,0 | 0,053 |
| 47906 | ENSG00000170854 | RIOX2 | chr3 | - | 97949843 | 97950012 | 97947360 | 97947449 | 97950785 | 97950888 | 47906 | 6,7,14,14,17,16,8,15,16,6,1 | 2,0,2,5,1,4,0,1,3,0,0,0 | 19,10,7,6,10,16,3,16,5,21,6 | 1,0,0,0,0,0,2,0,2,2,0,1,0,2,0 | 298 | 149 | 0.000662356975814 | 0.0816505758605023 | 0.6,1.0,0.778,0.58 | 0.905,1.0,1.0,1.0,1 | -0,062 |
| 102690 | ENSG00000162600 | OMA1 | chr1 | - | 58530600 | 58530729 | 58527260 | 58527335 | 58533952 | 58534060 | 102690 | 148,61,150,102,140,164,6 | 3,0,6,1,19,6,7,12,1,1,5,9 | 274,162,148,80,83,85,86,1 | 35,20,8,4,3,8,3,17,3,5,25,5 | 278 | 149 | 0.000662366970615 | 0.0816505758605023 | 0.964,1.0,0.931,0. | 0.808,0.813,0.908, | 0,041 |
| 214148 | ENSG00000138380 | CARF | chr2 | + | 202941841 | 202941980 | 202917876 | 202918043 | 202942739 | 202942824 | 214148 | 32,13,34,22,30,6,8,26,26,1 | 2,3,1,3,1,0,2,4,4,0,0,0 | 18,9,10,5,7,7,9,13,14,34,1 | 1,1,0,2,1,0,0,0,0,4,0,0,0,0,2 | 288 | 149 | 0.00066293563433 | 0.0816738978319864 | 0.892,0.692,0.946, | 0.903,0.823,1.0,0. | -0,037 |
| 175030 | ENSG00000101596 | SMCHD1 | chr18 | + | 2673016 | 2673194 | 2666869 | 2667031 | 2673280 | 2673363 | 175030 | 199,132,388,126,66,156,1 | 514,412,843,542,254,319,3 | 132,267,193,365,574,172,1 | 231,322,330,168,299,197,1 | 298 | 149 | 0.000663925279341 | 0.0817490284884684 | 0.162,0.138,0.187, | 0.222,0.293,0.226, | -0,074 |
| 130327 | ENSG00000111271 | ACAD10 | chr12 | + | 111705737 | 111705932 | 111692696 | 111692896 | 111709525 | 111709684 | 130327 | 19,4,27,19,15,13,8,18,14,2 | 1,0,1,2,1,0,1,0,0,0,1,0 | 14,14,12,17,8,7,12,16,13,1 | 4,1,1,1,3,0,1,4,1,0,4,3,0,0,0 | 298 | 149 | 0.000664448182586 | 0.0817666362413764 | 0.905,1.0,0.931,0. | 0.636,0.875,0.857, | 0,101 |
| 46709 | ENSG00000181450 | ZNF678 | chr1 | + | 227646543 | 227646670 | 227563555 | 227563724 | 227654335 | 227654768 | 46709 | 24,18,40,25,14,19,13,20,2 | 0,4,9,3,7,0,2,4,2,0,4,6 | 23,23,12,6,16,20,7,26,10,2 | 1,4,2,0,4,0,0,0,1,1,0,5,1,3,0 | 276 | 149 | 0.000664946998513 | 0.0817812613925437 | 1.0,0.708,0.706,0. | 0.925,0.756,0.764, | -0,137 |
| 209398 | ENSG00000091490 | SEL1L3 | chr4 | - | 25784498 | 25784559 | 25784227 | 25784290 | 25788223 | 25788364 | 209398 | 4,3,5,1,4,1,4,7,0,0,1,2 | 121,103,247,187,127,95,6 | 1,7,9,12,2,1,2,1,7,7,2,8,1,2 | 104,65,92,65,64,64,30,45,6 | 210 | 149 | 0.000668433040652 | 0.0821376671815729 | 0.023,0.02,0.014,0 | 0.007,0.071,0.065, | -0,018 |
| 33674 | ENSG00000065882 | TBC1D1 | chr4 | + | 38118032 | 38118192 | 38115709 | 38115954 | 38124961 | 38125131 | 33674 | 306,213,570,434,169,190,1 | 6,3,3,4,1,2,6,2,6,2,7,2 | 208,228,245,146,245,190,7 | 0,2,3,0,1,0,3,2,0,4,5,1,0,0,2 | 298 | 149 | 0.000668608113618 | 0.0821376671815729 | 0.962,0.973,0.99,0 | 1.0,0.983,0.976,1. | -0,022 |
| 123853 | ENSG00000151702 | FLI1 | chr11 | + | 128805365 | 128805431 | 128781957 | 128782023 | 128809156 | 128809204 | 123853 | 53,13,54,36,28,11,21,30,2 | 0,0,3,0,3,2,1,0,1,0,0,0 | 46,12,17,3,7,16,9,22,13,18 | 3,0,1,0,2,4,1,2,1,1,2,2,4,1,1 | 215 | 149 | 0.000672911465491 | 0.0826191714940636 | 1.0,1.0,0.926,1.0,0 | 0.914,1.0,0.922,1. | 0,059 |
| 5139 | ENSG00000090372 | STRN4 | chr19 | - | 46736822 | 46736901 | 46730731 | 46730873 | 46738163 | 46738172 | 5139 | 11,8,19,18,11,16,11,11,14, | 1,3,7,1,1,2,0,2,0,3,4,1 | 12,12,9,3,16,7,5,10,4,7,14, | 3,3,4,10,3,1,4,2,5,2,8,5,5,5 | 228 | 149 | 0.000675360551783 | 0.0828725923151692 | 0.878,0.635,0.639, | 0.723,0.723,0.595, | 0,185 |
| 247249 | ENSG00000066294 | CD84 | chr1 | - | 160550610 | 160551035 | 160549916 | 160549979 | 160553377 | 160553497 | 247249 | 16,11,30,35,19,21,9,22,14, | 0,2,3,0,1,0,1,0,0,0,0,0 | 35,54,5,6,15,12,10,34,11,1 | 1,4,0,0,2,2,0,3,2,3,1,0,0,2,2 | 298 | 149 | 0.000675814609152 | 0.0828810564913926 | 1.0,0.733,0.833,1. | 0.946,0.871,1.0,1. | 0,072 |

| | | | | | | | | | | | | | | | | | | | | | |
|---|---|---|---|---|---|---|---|---|---|---|---|---|---|---|---|---|---|---|---|---|---|
| 100238 | ENSG00000005471 | ABCB4 | chr7 | - | 87413616 | 87413717 | 87409235 | 87409392 | 87418536 | 87418620 | 100238 | 6,3,14,7,3,5,9,6,6,2,5,4 | 2,0,7,3,1,11,4,0,1,2,1,0 | 9,6,3,1,1,2,3,5,4,8,4,2,6,7, | 6,8,2,8,3,1,3,1,3,3,2,6,6,7, | 250 | 149 | 0.000677149600926 | 0.0829974861941366 | 0.641,1.0,0.544,0. | 0.472,0.309,0.472, | 0,297 |
| 244221 | ENSG00000149761 | NUDT22 | chr11 | + | 64227423 | 64227666 | 64226634 | 64227132 | 64229246 | 64229344 | 244221 | 25,16,21,30,16,9,13,18,36, | 2,0,2,3,0,2,0,0,4,0,2,5 | 17,18,10,9,2,8,2,11,3,12,6, | 0,0,1,0,0,0,1,8,0,0,0,0,3,0, | 298 | 149 | 0.000679883661803 | 0.0832851681351858 | 0.862,1.0,0.84,0.8 | 1.0,1.0,0.833,1.0,1 | -0,069 |
| 111116 | ENSG00000291122 | CASTOR3 | chr7 | - | 100211393 | 100211994 | 100200652 | 100202603 | 100213046 | 100213133 | 111116 | 10,2,14,8,2,7,3,17,7,7,5,9 | 0,3,0,2,2,4,1,11,5,6,0,1 | 4,8,1,6,2,3,3,7,4,4,16,7,7,2 | 3,0,0,2,0,0,0,6,0,0,2,4,1,1,1 | 298 | 149 | 0.000680735481232 | 0.0833420809789787 | 1.0,0.25,1.0,0.667, | 0.4,1.0,1.0,0.6,1.0, | -0,132 |
| 129935 | ENSG00000198270 | TMEM116 | chr12 | - | 111936691 | 111936830 | 111933885 | 111934030 | 111937159 | 111937243 | 129935 | 33,11,41,24,2,7,13,9,9,6,6, | 0,4,3,2,0,0,1,3,4,0,0,2 | 13,12,9,10,17,5,1,7,6,14,8, | 0,2,0,3,1,0,2,0,0,0,1,0,0,0,0 | 288 | 149 | 0.000681315131311 | 0.0833656265077873 | 1.0,0.587,0.876,0.8 | 1.0,0.756,1.0,0.63 | -0,046 |
| 155715 | ENSG00000169062 | UPF3A | chr13 | + | 114286301 | 114286400 | 114284093 | 114284358 | 114286518 | 114286629 | 155715 | 61,13,51,26,9,17,19,24,17, | 0,1,2,0,0,1,0,1,4,0,0,0 | 34,22,38,21,18,15,17,14,1 | 2,2,3,0,0,4,1,2,3,4,0,2,0,3,1 | 248 | 149 | 0.000683082622902 | 0.083488769197465 | 1.0,0.886,0.939,1. | 0.911,0.869,0.884, | 0,067 |
| 241158 | ENSG00000198382 | UVRAG | chr11 | + | 75899346 | 75899424 | 75888828 | 75888903 | 75911953 | 75912039 | 241158 | 0,0,4,0,0,9,21,18,0,0,0,0 | 73,41,311,120,75,205,80,2 | 2,1,3,2,0,0,0,2,1,0,2,1,1,1, | 131,95,76,52,66,63,81,140 | 227 | 149 | 0.00068309733522 | 0.083488769197465 | 0.0,0.0,0.008,0.0,0 | 0.01,0.007,0.025,0 | 0,01 |
| 22417 | ENSG00000075413 | MARK3 | chr14 | + | 103448918 | 103448967 | 103428386 | 103428440 | 103451917 | 103451983 | 22417 | 43,30,124,89,34,34,29,51,1 | 2,0,2,3,0,0,0,0,0,0,0,0 | 59,72,32,20,39,26,17,46,4 | 2,3,1,4,0,0,0,2,4,3,3,2,1,1,2 | 198 | 149 | 0.000687078497798 | 0.0839276913504888 | 0.942,1.0,0.979,0. | 0.957,0.948,0.96,0 | 0,049 |
| 186018 | ENSG00000153898 | MCOLN2 | chr1 | - | 84939625 | 84939702 | 84937980 | 84938082 | 84947032 | 84947132 | 186018 | 1,0,11,5,0,0,1,3,1,0,0,2 | 17,13,39,22,3,3,5,8,1,3,1,3 | 2,2,0,1,0,1,0,1,2,3,2,0,1,3,1 | 17,20,4,3,7,1,7,4,6,3,17,5,1 | 226 | 149 | 0.000688347777287 | 0.084035042798218 | 0.037,0.0,0.157,0. | 0.072,0.062,0.0,0. | -0,043 |
| 100571 | ENSG00000204257 | HLA-DMA | chr6 | - | 32952283 | 32952461 | 32950083 | 32950803 | 32952948 | 32953034 | 100571 | 56,14,42,26,8,5,11,17,10,3 | 1073,624,1591,1596,370,2 | 25,42,27,38,6,9,10,31,3,6,1 | 880,884,520,425,489,430,2 | 298 | 149 | 0.000689058782844 | 0.0840741558335017 | 0.025,0.011,0.013, | 0.014,0.023,0.025, | -0,005 |
| 64943 | ENSG00000164241 | C5orf63 | chr5 | - | 127056396 | 127056506 | 127052612 | 127052669 | 127058881 | 127059002 | 64943 | 3,1,11,15,7,6,9,14,10,3,2,7 | 7,4,22,20,10,10,6,15,4,4,3, | 8,2,3,0,1,2,1,1,0,7,7,7,1,3,1 | 8,11,7,2,7,9,1,5,5,4,10,10, | 259 | 149 | 0.000689568675403 | 0.0840846021833648 | 0.198,0.126,0.223, | 0.365,0.095,0.198, | 0,135 |
| 249915 | ENSG00000135972 | MRPS9 | chr2 | + | 105092199 | 105092326 | 105089919 | 105089995 | 105092400 | 105092569 | 249915 | 3,1,2,1,6,0,0,2,0,0,0,2 | 39,13,14,21,15,15,10,13,2 | 4,4,0,3,0,0,3,5,1,3,4,6,3,1,1 | 17,20,12,8,9,4,6,10,9,17,7, | 276 | 149 | 0.000689925742369 | 0.0840846021833648 | 0.04,0.04,0.072,0. | 0.113,0.097,0.0,0. | -0,056 |
| 21522 | ENSG00000015133 | CCDC88C | chr14 | - | 91285394 | 91285836 | 91283328 | 91283517 | 91289104 | 91289331 | 21522 | 99,87,35,35,34,58,54,50,9 | 4,0,1,3,1,1,2,5,3,2,0,0 | 31,45,53,70,36,22,39,41,1 | 2,1,5,7,0,0,2,3,3,0,13,1,0,0 | 298 | 149 | 0.000691505483281 | 0.0842294378449649 | 0.925,1.0,0.946,0. | 0.886,0.957,0.841, | 0,046 |
| 211221 | ENSG00000197622 | CDC42SE1 | chr1 | - | 151056650 | 151056731 | 151055676 | 151055993 | 151059478 | 151059773 | 211221 | 16,1,22,14,11,13,14,18,5,6 | 197,165,295,254,162,112,1 | 22,12,11,12,14,13,12,14,8, | 274,239,156,130,165,98,11 | 230 | 149 | 0.000693822751284 | 0.0844638939941214 | 0.05,0.004,0.046,0 | 0.049,0.032,0.044, | -0,017 |
| 230649 | ENSG00000225648 | SBDSP1 | chr7 | + | 72830681 | 72830821 | 72829655 | 72829783 | 72831601 | 72831798 | 230649 | 12,4,10,2,2,3,9,2,4,5,3,2 | 7,4,6,6,3,2,2,2,5,5,0,1 | 3,6,3,10,7,2,2,7,7,7,15,4,1, | 0,4,0,11,0,0,0,0,0,3,1,10,1, | 289 | 149 | 0.000694661247081 | 0.0844777596987669 | 0.469,0.34,0.462,0 | 1.0,0.436,1.0,0.31 | -0,249 |
| 4034 | ENSG00000099785 | MARCHF2 | chr19 | + | 8430657 | 8430867 | 8426608 | 8426804 | 8438387 | 8439017 | 4034 | 48,37,60,36,47,59,42,37,3 | 0,5,2,1,6,0,0,0,4,0,0,7 | 30,51,41,13,30,16,16,35,1 | 1,2,0,1,0,4,0,2,0,0,0,0,4,7,2 | 298 | 149 | 0.000695104826486 | 0.0844777596987669 | 1.0,0.787,0.938,0. | 0.938,0.927,1.0,0. | -0,008 |
| 55074 | ENSG00000139631 | CSAD | chr12 | - | 53171272 | 53171441 | 53170422 | 53170502 | 53171881 | 53171988 | 55074 | 4,1,16,11,11,14,6,8,5,5,5,6 | 0,0,9,0,3,1,1,1,1,0,0,2 | 3,5,10,3,9,4,5,13,7,6,4,12,1 | 1,9,0,6,2,2,1,4,4,2,0,8,6,6, | 298 | 149 | 0.000695301081221 | 0.0844777596987669 | 1.0,1.0,0.471,1.0,0 | 0.6,0.217,1.0,0.2,0 | 0,27 |
| 41268 | ENSG00000113811 | SELENOK | chr3 | - | 53885825 | 53885904 | 53885300 | 53885561 | 53886850 | 53886934 | 41268 | 50,36,98,65,11,14,22,29,6, | 1,0,2,1,2,2,2,1,0,0,0,0 | 55,86,55,29,51,15,19,18,2 | 2,1,1,7,1,1,0,2,0,2,8,0,3,1, | 228 | 149 | 0.000696079954604 | 0.0844777596987669 | 0.97,1.0,0.97,0.97 | 0.947,0.983,0.973, | 0,039 |
| 163799 | ENSG00000198130 | HIBCH | chr2 | - | 190294545 | 190294627 | 190290404 | 190290485 | 190296812 | 190296953 | 163799 | 8,5,27,25,9,15,16,12,4,4,3, | 0,0,6,3,0,0,0,0,2,0,0,1 | 12,13,4,1,4,8,5,7,17,9,34,3 | 0,3,0,0,6,0,0,0,1,2,5,3,0,4,1 | 231 | 149 | 0.000696154533986 | 0.0844777596987669 | 1.0,1.0,0.744,0.84 | 1.0,0.737,1.0,1.0,0 | 0,058 |
| 251413 | ENSG00000148429 | USP6NL | chr10 | - | 11489101 | 11489218 | 11485816 | 11485911 | 11490831 | 11490880 | 251413 | 33,21,90,34,12,15,16,36,3, | 4,0,2,0,2,2,0,2,0,0,0,2 | 56,35,21,1,5,10,21,16,20,8 | 5,0,4,2,0,0,1,1,5,0,4,12,3,0 | 266 | 149 | 0.000696291638777 | 0.0844777596987669 | 0.822,1.0,0.962,1. | 0.863,1.0,0.746,0. | 0,07 |
| 242072 | ENSG00000125944 | HNRNPR | chr1 | - | 23323555 | 23323747 | 23321527 | 23321663 | 23333517 | 23333631 | 242072 | 63,46,205,160,55,65,65,64 | 0,1,6,3,1,1,1,6,6,0,0,1 | 108,119,75,38,51,42,52,98 | 1,11,3,9,8,2,3,2,0,1,3,4,4,5 | 298 | 149 | 0.000700853920074 | 0.0849833746881392 | 1.0,0.958,0.945,0. | 0.982,0.844,0.926, | 0,046 |
| 16951 | ENSG00000130731 | METTL26 | chr16 | - | 634888 | 634956 | 634718 | 634797 | 635611 | 635774 | 16951 | 43,20,15,17,13,14,17,24,3 | 2,0,2,2,0,3,2,0,2,1,2,3 | 20,11,15,14,12,8,6,27,4,9,1 | 0,1,1,0,0,0,1,0,0,0,1,3,0,0,2 | 217 | 149 | 0.000710724922244 | 0.0861317768803482 | 0.937,1.0,0.837,0. | 1.0,0.883,0.912,1. | -0,066 |
| 34548 | ENSG00000047365 | ARAP2 | chr4 | - | 36073688 | 36073823 | 36066003 | 36068278 | 36082250 | 36082286 | 34548 | 14,14,32,20,14,23,2,13,14, | 0,0,0,0,1,1,1,2,0,0,0,0 | 31,23,7,11,7,8,18,29,8,21, | 1,4,0,3,2,1,0,1,0,0,5,3,1,1,1 | 284 | 149 | 0.000716157069824 | 0.0867412505882326 | 1.0,1.0,1.0,1.0,0.8 | 0.942,0.751,1.0,0. | 0,049 |
| 121471 | ENSG00000110075 | PPP6R3 | chr11 | + | 68522493 | 68522689 | 68519500 | 68519651 | 68537658 | 68537669 | 121471 | 11,11,16,7,2,5,7,12,8,2,3,1 | 93,68,141,77,85,99,58,111 | 23,18,14,31,9,1,9,16,5,6,1 | 78,107,77,62,57,34,58,78, | 298 | 149 | 0.000717423076117 | 0.0867799938221156 | 0.056,0.075,0.054, | 0.128,0.078,0.083, | -0,023 |
| 23186 | ENSG00000213983 | AP1G2 | chr14 | - | 23565815 | 23565892 | 23565118 | 23565199 | 23566063 | 23566160 | 23186 | 17,12,29,35,5,13,9,5,14,4, | 1,1,2,1,5,6,1,2,0,2,1,0 | 14,16,12,4,14,5,8,11,9,16,1 | 6,1,0,0,0,1,2,0,2,0,0,3,4,0,1 | 226 | 149 | 0.000717433170226 | 0.0867799938221156 | 0.918,0.888,0.905, | 0.606,0.913,1.0,1. | -0,078 |
| 219503 | ENSG00000136044 | APPL2 | chr12 | - | 105229627 | 105229824 | 105229124 | 105229223 | 105235958 | 105236203 | 219503 | 7,3,11,6,3,1,7,8,2,0,1,2 | 12,6,11,12,9,21,15,19,15,6 | 6,7,6,16,8,2,9,11,7,7,13,2, | 7,1,11,15,11,8,7,17,13,17, | 298 | 149 | 0.000717686527514 | 0.0867799938221156 | 0.226,0.2,0.333,0. | 0.3,0.778,0.214,0. | -0,154 |
| 178573 | ENSG00000120008 | WDR11 | chr10 | + | 120867121 | 120867169 | 120866568 | 120866764 | 120871169 | 120871346 | 178573 | 137,71,119,72,67,64,75,70 | 2,0,5,6,1,3,4,7,3,1,2,4 | 85,104,130,125,81,51,28,8 | 6,14,6,11,7,2,6,1,3,9,11,3, | 197 | 149 | 0.000718324838619 | 0.0868084072660337 | 0.981,1.0,0.947,0. | 0.915,0.849,0.942, | 0,042 |
| 55327 | ENSG00000118217 | ATF6 | chr1 | + | 161860206 | 161860277 | 161853223 | 161853323 | 161863197 | 161863312 | 55327 | 87,73,108,75,48,36,30,34, | 3,2,7,3,1,3,0,2,0,3,0,0 | 67,56,32,31,49,32,22,56,3 | 2,6,1,10,10,3,3,0,6,2,0,4,2, | 220 | 149 | 0.000718968979531 | 0.0868374929480565 | 0.952,0.961,0.913, | 0.958,0.863,0.956, | 0,06 |
| 40778 | ENSG00000164327 | RICTOR | chr5 | - | 38962875 | 38963041 | 38962484 | 38962586 | 38964791 | 38964892 | 40778 | 92,63,162,82,60,65,47,113 | 0,0,2,0,0,1,0,2,0,0,0,1 | 104,115,77,92,65,42,41,80 | 4,3,5,5,0,0,0,1,2,0,5,1,3,0, | 298 | 149 | 0.000720009565499 | 0.0869144020145347 | 1.0,1.0,0.976,1.0,1 | 0.929,0.95,0.885,0 | 0,031 |
| 60183 | ENSG00000234127 | TRIM26 | chr6 | - | 30201034 | 30201138 | 30198665 | 30199153 | 30213304 | 30213427 | 60183 | 33,26,33,37,27,18,14,15,4 | 0,0,0,3,0,0,1,1,2,4,2,0 | 20,40,7,13,16,5,9,20,4,12, | 6,0,0,0,1,4,2,0,0,0,3,8,1,2, | 253 | 149 | 0.00072060282345 | 0.0869372568912371 | 1.0,1.0,1.0,0.879,1 | 0.663,1.0,1.0,1.0,0 | 0,074 |
| 146286 | ENSG00000047578 | KATNIP | chr16 | + | 27699479 | 27699599 | 27698327 | 27698500 | 27701588 | 27701695 | 146286 | 2,1,10,5,1,2,5,5,1,0,0,1 | 5,0,1,0,0,0,3,2,4,3,1,1 | 5,2,1,2,3,5,8,7,1,2,6,5,4,6, | 0,2,0,0,2,6,1,1,2,0,0,1,0,0, | 269 | 149 | 0.000722645332512 | 0.0871348333680058 | 0.181,1.0,0.847,1. | 1.0,0.356,1.0,1.0,0 | -0,193 |
| 32719 | ENSG00000204859 | ZBTB48 | chr1 | + | 6588755 | 6588844 | 6588277 | 6588442 | 6588915 | 6589160 | 32719 | 24,9,85,78,34,20,26,77,35, | 0,0,5,0,0,8,1,5,4,1,0,1 | 13,23,6,7,8,29,29,32,23,32 | 0,0,0,1,2,1,0,0,0,0,1,6,0,0, | 238 | 149 | 0.000724519019406 | 0.0873026343902614 | 1.0,1.0,0.914,1.0,1 | 1.0,1.0,1.0,0.814,0 | -0,02 |
| 87971 | ENSG00000135930 | EIF4E2 | chr2 | + | 232557883 | 232558018 | 232556415 | 232556530 | 232564246 | 232564351 | 87971 | 44,59,98,90,41,31,31,43,3 | 18,17,18,6,7,11,5,4,11,9,4, | 63,67,35,10,63,19,10,33,2 | 4,15,9,13,8,1,0,4,0,5,4,2,3, | 284 | 149 | 0.000724927427358 | 0.0873026343902614 | 0.562,0.645,0.741, | 0.892,0.701,0.671, | -0,129 |
| 253822 | ENSG00000111271 | ACAD10 | chr12 | + | 111705784 | 111705932 | 111692696 | 111692896 | 111709525 | 111709684 | 253822 | 16,3,27,16,12,12,8,18,12,2 | 1,0,1,2,1,0,1,0,0,0,1,0 | 14,14,12,17,9,8,11,16,13,1 | 4,1,1,1,3,0,1,4,1,0,4,3,0,0, | 297 | 149 | 0.000725448935815 | 0.0873026343902614 | 0.889,1.0,0.931,0. | 0.637,0.875,0.858, | 0,095 |
| 84024 | ENSG00000145216 | FIP1L1 | chr4 | + | 53379072 | 53379117 | 53377578 | 53377923 | 53379224 | 53379264 | 84024 | 38,28,47,33,21,33,43,18,1 | 10,6,24,15,8,7,9,3,9,3,1,2 | 50,55,30,26,42,24,13,34,1 | 3,7,1,3,6,1,0,7,1,3,10,11,3, | 194 | 149 | 0.000725659467847 | 0.0873026343902614 | 0.745,0.782,0.601, | 0.928,0.858,0.958, | -0,11 |
| 124908 | ENSG00000187147 | RNF220 | chr1 | + | 44635544 | 44635588 | 44632254 | 44632385 | 44636029 | 44636162 | 124908 | 53,52,64,56,29,18,17,15,5 | 2,0,4,1,3,0,4,4,0,0,0,3 | 32,42,18,28,36,26,12,20,3 | 4,2,0,6,2,0,0,2,7,6,3,5,0,0, | 193 | 149 | 0.000727692580392 | 0.087498325011369 | 0.953,1.0,0.925,0. | 0.861,0.942,1.0,0. | 0,027 |
| 173286 | ENSG00000198586 | TLK1 | chr2 | - | 171022871 | 171022956 | 171014850 | 171014948 | 171028338 | 171028405 | 173286 | 6,3,6,1,5,3,2,4,3,2,1,1 | 37,26,82,80,35,42,20,46,1 | 9,12,7,23,2,5,10,0,4,3,0,4, | 47,61,34,70,47,32,55,41,3 | 234 | 149 | 0.000730609918876 | 0.0878000577608042 | 0.094,0.068,0.045, | 0.109,0.111,0.116, | -0,019 |
| 15729 | ENSG00000197912 | SPG7 | chr16 | + | 89545871 | 89546181 | 89542983 | 89544772 | 89546657 | 89546760 | 15729 | 0,1,14,4,3,0,10,3,2,1,2,1 | 36,17,74,42,40,51,18,19,2 | 1,1,0,2,2,3,0,6,0,1,7,3,2,1, | 45,38,15,18,62,22,32,38,4 | 298 | 149 | 0.000731914104178 | 0.0878843709473868 | 0.0,0.029,0.086,0. | 0.011,0.013,0.0,0. | 0,03 |
| 214466 | ENSG00000181904 | C5orf24 | chr5 | + | 134855719 | 134855905 | 134854897 | 134855327 | 134857356 | 134859732 | 214466 | 2,0,9,1,0,0,1,2,4,0,2,0 | 6,1,13,6,1,2,9,4,5,3,4,9 | 1,0,0,4,0,0,0,0,0,0,1,1,0,1, | 6,11,11,4,5,4,1,7,4,3,8,14, | 298 | 149 | 0.000732128165128 | 0.0878843709473868 | 0.143,0.0,0.257,0. | 0.077,0.0,0.0,0.33 | 0,045 |
| 235370 | ENSG00000104361 | NIPAL2 | chr8 | - | 98194367 | 98194546 | 98192757 | 98193090 | 98194727 | 98194822 | 235370 | 2,2,3,1,1,0,0,0,0,0,3,5 | 17,14,63,51,13,14,9,5,15,1 | 6,5,1,1,0,0,2,6,6,4,0,4,1,1, | 37,21,6,5,3,11,14,25,15,20 | 298 | 149 | 0.000733473408004 | 0.0879967754058578 | 0.056,0.067,0.023, | 0.075,0.106,0.077, | -0,033 |
| 117136 | ENSG00000141551 | CSNK1D | chr17 | - | 82255428 | 82255577 | 82253015 | 82253244 | 82265685 | 82265796 | 117136 | 25,37,41,48,26,30,26,54,1 | 0,0,2,4,0,2,2,1,0,0,4,0 | 42,111,36,28,29,28,51,39, | 0,1,0,0,3,0,3,7,0,0,4,0,2,5,1 | 298 | 149 | 0.000737671070932 | 0.0884510764722926 | 1.0,1.0,0.911,0.85 | 1.0,0.982,1.0,1.0,0 | 0,025 |
| 193078 | ENSG00000116688 | MFN2 | chr1 | + | 11981969 | 11982112 | 11980208 | 11980484 | 11989164 | 11989343 | 193078 | 9,12,12,8,7,7,7,4,6,4,5,5 | 2,2,4,1,1,2,0,1,0,5,0,0 | 12,3,5,5,5,1,13,8,5,4,10,4, | 0,1,2,0,1,1,0,1,0,2,2,0,2,0, | 292 | 149 | 0.000742229704374 | 0.0889481300123165 | 0.697,0.754,0.605, | 1.0,0.605,0.561,1. | -0,02 |
| 248597 | ENSG00000148925 | BTBD10 | chr11 | - | 13445023 | 13445148 | 13421641 | 13421838 | 13463091 | 13463297 | 248597 | 9,2,27,11,7,5,6,10,30,14,0, | 33,22,18,11,5,20,4,14,8,6, | 9,7,7,3,5,0,1,7,0,0,3,4,1,0,1 | 17,29,6,11,20,5,21,9,9,12,1 | 274 | 149 | 0.000747221119048 | 0.0894964655947803 | 0.129,0.047,0.449, | 0.224,0.116,0.388, | 0,162 |
| 239884 | ENSG00000148908 | RGS10 | chr10 | - | 119515514 | 119515652 | 119500079 | 119500259 | 119526031 | 119526118 | 239884 | 50,29,102,62,34,16,26,40, | 0,2,6,4,2,0,0,0,4,0,0,3 | 43,32,56,13,36,33,14,25,1 | 0,3,0,0,0,2,0,0,0,0,2,0,0,0,2 | 287 | 149 | 0.00074931767858 | 0.0896976603328432 | 1.0,0.883,0.898,0. | 1.0,0.847,1.0,1.0,1 | -0,034 |
| 239543 | ENSG00000083223 | TUT7 | chr9 | - | 86298766 | 86299198 | 86287734 | 86288744 | 86301275 | 86301601 | 239543 | 4,1,6,4,0,0,5,0,8,3,0,2 | 46,21,47,30,11,18,22,12,3 | 2,0,1,0,3,0,1,1,1,2,1,3,4,2, | 36,28,35,21,18,17,2,23,18, | 298 | 149 | 0.000754323550018 | 0.090246699273999 | 0.042,0.023,0.06,0 | 0.027,0.0,0.014,0. | 0,009 |
| 147764 | ENSG00000197912 | SPG7 | chr16 | + | 89555934 | 89556085 | 89554485 | 89554563 | 89556886 | 89557173 | 147764 | 15,14,6,3,1,3,0,0,2,6,7,22 | 50,36,67,45,40,34,21,52,3 | 2,0,0,1,2,1,2,4,7,10,6,6,3,5 | 36,38,30,29,34,24,24,41,3 | 298 | 149 | 0.000757856255298 | 0.0906189776022466 | 0.13,0.163,0.043,0 | 0.027,0.0,0.0,0.01 | 0,045 |
| 76888 | ENSG00000131778 | CHD1L | chr1 | + | 147252622 | 147252735 | 147242676 | 147242830 | 147254869 | 147254976 | 76888 | 15,8,44,39,23,15,11,27,13, | 0,1,2,2,1,1,0,0,2,0,0,0 | 21,22,9,4,13,13,15,19,11,2 | 5,3,0,0,1,0,0,4,3,0,3,3,0,0, | 262 | 149 | 0.000758993379437 | 0.090704555274628 | 1.0,0.82,0.926,0.9 | 0.705,0.807,1.0,1. | 0,057 |
| 164420 | ENSG00000005700 | IBTK | chr6 | - | 82175903 | 82176026 | 82173366 | 82173438 | 82181878 | 82182028 | 164420 | 1,6,5,6,3,4,0,2,7,1,2,6 | 19,14,42,41,25,23,13,14,1 | 2,3,0,0,1,1,0,0,2,5,0,2,4,3, | 32,20,26,6,26,11,8,16,21,2 | 272 | 149 | 0.000760705451382 | 0.0908587097704768 | 0.028,0.19,0.061,0 | 0.033,0.076,0.0,0. | 0,044 |
| 150538 | ENSG00000166783 | MARF1 | chr16 | - | 15624768 | 15624923 | 15622933 | 15623123 | 15625015 | 15625173 | 150538 | 28,15,34,33,17,27,24,16,2 | 5,0,2,0,2,4,2,4,0,2,0,0 | 21,15,21,10,7,15,25,37,15, | 1,3,3,0,2,0,1,0,0,0,0,0,0,0, | 298 | 149 | 0.000761504697357 | 0.0909037257442288 | 0.737,1.0,0.895,1. | 0.913,0.714,0.778, | -0,037 |
| 102864 | ENSG00000179406 | LINC00174 | chr7 | - | 66467681 | 66467719 | 66450070 | 66450236 | 66476258 | 66476327 | 102864 | 1,2,3,3,2,2,3,1,2,2,1,1 | 0,2,3,8,0,7,4,5,4,0,1,4 | 0,1,1,0,1,1,0,1,2,1,1,1,2,3, | 1,3,3,2,1,1,3,3,0,2,6,0,3,1, | 187 | 149 | 0.00076256197889 | 0.0909794774270918 | 1.0,0.443,0.443,0. | 0.0,0.21,0.21,0.0,0 | 0,137 |
| 240717 | ENSG00000148090 | AUH | chr9 | - | 91342108 | 91342490 | 91325317 | 91325404 | 91355882 | 91355970 | 240717 | 2,0,8,5,0,0,1,5,0,0,0,2 | 11,7,18,17,12,8,10,6,6,5,4, | 0,1,2,1,1,0,1,0,0,0,0,1,2,0, | 17,24,10,6,6,7,5,5,6,17,20, | 298 | 149 | 0.000764457340266 | 0.0911550791151199 | 0.083,0.0,0.182,0. | 0.0,0.02,0.091,0.0 | 0,033 |
| 160352 | ENSG00000166024 | R3HCC1L | chr10 | + | 98205517 | 98205656 | 98186466 | 98186563 | 98208100 | 98209899 | 160352 | 5,1,14,2,0,4,0,5,1,0,0,0 | 0,0,16,6,2,2,2,2,0,5,2,5 | 4,3,6,8,9,0,3,5,3,6,3,4,2,1, | 3,1,1,1,6,4,1,5,0,1,2,1,1,3, | 288 | 149 | 0.000765855792358 | 0.0912712669130702 | 1.0,1.0,0.312,0.14 | 0.408,0.608,0.756, | -0,153 |
| 27022 | ENSG00000126777 | KTN1 | chr14 | + | 55637197 | 55637364 | 55636448 | 55636536 | 55637778 | 55637847 | 27022 | 237,139,228,129,115,123, | 5,2,7,3,0,1,0,4,1,0,0,1 | 145,157,171,57,80,75,87,1 | 8,7,0,5,4,3,1,7,1,5,3,4,1,5,1 | 298 | 149 | 0.000766731978933 | 0.0913165811075826 | 0.96,0.972,0.942,0 | 0.901,0.918,1.0,0. | 0,038 |
| 176636 | ENSG00000174738 | NR1D2 | chr3 | + | 23967949 | 23968023 | 23964976 | 23965162 | 23977222 | 23977543 | 176636 | 22,30,41,11,7,8,22,24,13,7 | 2,1,9,4,5,7,6,9,2,3,9,7 | 20,22,16,28,13,9,9,11,8,10 | 14,31,18,11,18,14,4,6,9,11 | 223 | 149 | 0.000767084567941 | 0.0913165811075826 | 0.88,0.952,0.753,0 | 0.488,0.322,0.373, | 0,186 |
| 70809 | ENSG00000159658 | EFCAB14 | chr1 | - | 46716290 | 46716443 | 46707905 | 46708051 | 46717902 | 46719139 | 70809 | 57,36,125,90,64,57,44,51, | 2,0,0,0,0,0,1,0,0,0,0,0 | 76,99,33,37,31,41,26,68,2 | 3,5,2,2,1,0,2,1,7,2,0,4,0,0, | 298 | 149 | 0.000770126091133 | 0.0916279760755372 | 0.934,1.0,1.0,1.0,1 | 0.927,0.908,0.892, | 0,062 |
| 124909 | ENSG00000187147 | RNF220 | chr1 | + | 44635544 | 44635666 | 44632234 | 44632385 | 44636029 | 44636152 | 124909 | 89,84,125,75,56,32,29,44, | 2,0,4,1,3,0,4,4,0,0,0,3 | 89,66,72,47,64,31,22,43,2 | 4,2,0,6,2,0,0,2,7,6,3,5,0,0, | 271 | 149 | 0.000771842309002 | 0.091781432049066 | 0.961,1.0,0.945,0. | 0.924,0.948,1.0,0. | 0,04 |
| 194358 | ENSG00000060138 | YBX3 | chr12 | - | 10710529 | 10710788 | 10709907 | 10710114 | 10713210 | 10713333 | 194358 | 47,21,80,41,8,21,20,61,33, | 137,70,191,103,66,119,88, | 34,31,23,27,16,16,29,45,1 | 162,111,72,55,33,52,72,20 | 298 | 149 | 0.000773039562578 | 0.091873041465061 | 0.146,0.13,0.173,0 | 0.095,0.123,0.138, | 0,011 |
| 67296 | ENSG00000166377 | ATP9B | chr18 | + | 79348131 | 79348196 | 79347769 | 79347925 | 79359353 | 79359462 | 67296 | 22,4,28,14,12,10,14,8,6,6, | 3,0,0,0,2,0,2,0,0,1,0,2 | 12,3,18,5,5,4,10,16,9,9,14, | 2,0,0,3,0,4,5,4,0,1,0,0,2,4, | 214 | 149 | 0.000776251532773 | 0.0922018362147163 | 0.836,1.0,1.0,1.0,0 | 0.807,1.0,1.0,0.53 | 0,093 |
| 203518 | ENSG00000163617 | CCDC191 | chr3 | - | 114046590 | 114046674 | 114042702 | 114042846 | 114056376 | 114056594 | 203518 | 10,2,14,8,5,9,3,9,2,2,4,2 | 2,0,0,3,5,2,0,0,0,2,2,2 | 6,8,2,4,7,0,3,4,2,4,1,5,4,13 | 1,0,0,0,0,2,1,0,0,0,0,1,2,0, | 233 | 149 | 0.000776662883401 | 0.0922018362147163 | 0.762,1.0,1.0,0.63, | 0.793,1.0,1.0,1.0,1 | -0,06 |
| 89906 | ENSG00000288558 | DUS4L-BCAP29 | chr7 | + | 107571144 | 107571266 | 107567049 | 107567186 | 107575187 | 107575310 | 89906 | 2,3,10,19,5,3,4,7,1,3,1,9 | 1,0,2,4,2,0,0,0,0,0,0,0 | 6,15,4,3,3,4,2,2,1,6,10,8,4, | 0,2,1,2,1,0,0,4,1,2,3,4,0,1, | 271 | 149 | 0.000779149009167 | 0.0924459869856795 | 0.524,1.0,0.733,0. | 1.0,0.805,0.687,0. | 0,173 |
| 255705 | ENSG00000112200 | ZNF451 | chr6 | + | 57150718 | 57151146 | 57147089 | 57148693 | 57152220 | 57152351 | 255705 | 67,60,48,26,32,35,23,39,4 | 0,1,4,2,0,0,0,0,0,0,0,1 | 66,48,36,50,24,24,25,26,3 | 2,2,1,2,2,1,0,4,2,0,5,0,1,1, | 298 | 149 | 0.000780795543549 | 0.0925430990739278 | 1.0,0.968,0.857,0. | 0.943,0.923,0.947, | 0,067 |
| 163547 | ENSG00000059378 | PARP12 | chr7 | - | 140037714 | 140037961 | 140034234 | 140034331 | 140041643 | 140041839 | 163547 | 31,27,60,31,34,30,29,39,3 | 7,1,2,4,5,1,3,1,0,1,0,2 | 28,33,19,22,30,29,10,58,2 | 0,2,2,2,0,0,0,1,0,0,3,2,5,1,1 | 298 | 149 | 0.000780827426896 | 0.0925430990739278 | 0.689,0.931,0.938, | 1.0,0.892,0.826,0. | -0,052 |
| 162101 | ENSG00000241106 | HLA-DOB | chr6 | - | 32813722 | 32813848 | 32813439 | 32813471 | 32814319 | 32814601 | 162101 | 78,55,242,179,74,71,115,7 | 1,0,0,2,0,0,0,1,0,0,0,0 | 157,95,61,19,59,68,17,30, | 2,5,3,2,2,1,0,2,1,0,4,2,2,4,1 | 275 | 149 | 0.000782486028078 | 0.0926427483843569 | 0.977,1.0,1.0,0.98, | 0.977,0.911,0.917, | 0,05 |
| 132647 | ENSG00000118454 | ANKRD13C | chr1 | - | 70306223 | 70306290 | 70300763 | 70300908 | 70313744 | 70313790 | 132647 | 39,30,78,48,11,28,14,29,3 | 0,0,1,2,0,0,1,0,2,1,0,2 | 50,50,16,19,40,13,13,17,1 | 0,4,1,0,2,0,2,3,3,0,4,0,3,0, | 216 | 149 | 0.000782529080675 | 0.0926427483843569 | 1.0,1.0,0.982,0.94 | 1.0,0.896,0.917,1. | 0,037 |
| 76008 | ENSG00000135541 | AHI1 | chr6 | - | 135301407 | 135301539 | 135300499 | 135300558 | 135318518 | 135318616 | 76008 | 34,16,48,20,12,16,9,17,7,2 | 2,0,3,4,0,3,0,2,0,1,1,1 | 4,7,15,43,8,7,7,20,5,5,10,2 | 0,1,1,0,2,4,0,5,2,1,2,0,0,1,2 | 281 | 149 | 0.000783459456504 | 0.0927019033990173 | 0.9,1.0,0.895,0.72 | 1.0,0.788,0.888,1. | 0,065 |
| 15429 | ENSG00000176715 | ACSF3 | chr16 | + | 89114338 | 89114487 | 89112091 | 89112246 | 89120800 | 89120913 | 15429 | 79,35,53,44,54,36,23,31,6 | 19,19,15,9,2,7,3,2,10,7,1,3 | 49,12,31,21,9,42,14,39,20, | 3,2,1,5,4,5,2,9,3,5,3,4,9,5,2 | 298 | 149 | 0.000784623859086 | 0.0927886691290873 | 0.675,0.479,0.639, | 0.891,0.75,0.939,0 | -0,074 |
| 177002 | ENSG00000227372 | TP73-AS1 | chr1 | - | 3742370 | 3742517 | 3740274 | 3740387 | 3742803 | 3742858 | 177002 | 1,0,10,0,5,3,0,3,1,0,0,3 | 13,9,11,14,12,21,9,8,8,2,1 | 0,2,0,0,3,0,1,0,0,1,0,2,3,0, | 7,13,6,8,6,10,3,10,7,2,8,18 | 296 | 149 | 0.000786573361606 | 0.0929427668374019 | 0.037,0.0,0.314,0. | 0.0,0.072,0.0,0.0,0 | 0,014 |
| 25171 | ENSG00000092020 | PPP2R3C | chr14 | - | 35095047 | 35095184 | 35091069 | 35091207 | 35096557 | 35096633 | 25171 | 52,23,66,53,46,37,23,36,1 | 0,1,8,5,0,3,0,2,0,0,1,2 | 51,88,43,17,35,26,23,17,1 | 0,0,4,0,0,0,0,0,0,3,2,1,1,0, | 286 | 149 | 0.000787181664096 | 0.0929427668374019 | 1.0,0.923,0.811,0. | 1.0,1.0,0.848,1.0,1 | -0,057 |
| 44704 | ENSG00000096384 | HSP90AB1 | chr6 | + | 44248616 | 44248776 | 44247086 | 44247195 | 44249376 | 44249583 | 44704 | 1152,668,1609,1324,874,8 | 47,12,26,19,11,17,16,10,8, | 935,945,722,248,363,335, | 11,11,5,5,4,4,2,11,4,0,32,9 | 298 | 149 | 0.000787222398003 | 0.0929427668374019 | 0.925,0.965,0.969, | 0.977,0.977,0.986, | -0,008 |
| 1775 | ENSG00000284431 | | chr22 | + | 40407788 | 40407843 | 40407412 | 40407568 | 40408070 | 40408120 | 1775 | 33,11,40,26,23,29,21,19,6, | 3,0,5,0,0,0,4,0,1,3,0,1 | 31,31,4,12,5,13,14,21,17,2 | 0,4,1,1,0,0,0,1,1,3,2,17,2,0 | 204 | 149 | 0.000791366758572 | 0.0933808436481415 | 0.889,1.0,0.854,1. | 1.0,0.85,0.745,0.8 | 0,024 |
| 188776 | ENSG00000116649 | SRM | chr1 | - | 11055780 | 11055936 | 11054663 | 11055084 | 11056010 | 11056094 | 188776 | 16,17,20,15,8,12,14,8,9,5, | 0,0,4,1,0,0,0,1,2,0,0,1 | 16,20,14,9,8,5,14,16,5,13,1 | 0,3,2,0,0,0,5,3,0,0,5,3,0,1,1 | 298 | 149 | 0.000792964556558 | 0.0935181120397452 | 1.0,1.0,0.714,0.88 | 1.0,0.769,0.778,1. | 0,068 |
| 191771 | ENSG00000130939 | UBE4B | chr1 | + | 10126957 | 10127067 | 10126793 | 10126877 | 10129391 | 10129448 | 191771 | 0,0,4,5,2,1,0,1,0,2,0,0 | 7,6,40,15,10,6,9,13,1,1,2,1 | 5,1,4,5,2,2,4,5,2,1,0,0,1,0, | 13,7,8,3,12,8,6,19,3,33,10, | 259 | 149 | 0.000794410905585 | 0.093634830780498 | 0.0,0.0,0.054,0.16 | 0.181,0.076,0.223, | -0,048 |
| 240687 | ENSG00000136936 | XPA | chr9 | - | 97689533 | 97689660 | 97687095 | 97687261 | 97693648 | 97693759 | 240687 | 26,18,79,78,34,26,33,21,1 | 4,2,2,4,4,1,1,2,2,3,0,0 | 42,34,22,13,32,26,8,11,14, | 3,1,1,3,1,2,2,1,2,2,0,0,1,0, | 276 | 149 | 0.000795104875285 | 0.093634830780498 | 0.778,0.829,0.955, | 0.883,0.948,0.922, | -0,027 |
| 71040 | ENSG00000163605 | PPP4R2 | chr3 | + | 73063672 | 73063747 | 73061022 | 73061060 | 73064002 | 73064146 | 71040 | 43,54,49,31,21,19,13,12,3 | 6,4,18,10,3,0,1,1,6,0,2,6 | 30,38,40,22,25,13,10,14,1 | 11,9,2,9,5,5,5,4,6,7,13,7,5, | 224 | 149 | 0.000795259375586 | 0.093634830780498 | 0.827,0.9,0.644,0. | 0.645,0.737,0.93,0 | 0,161 |
| 163038 | ENSG00000182903 | ZNF721 | chr4 | - | 467471 | 467681 | 444342 | 444432 | 472574 | 472701 | 163038 | 1,0,3,9,0,4,1,3,0,0,0,0 | 82,45,140,64,22,32,24,41, | 0,12,4,0,3,1,2,6,0,1,4,7,3,1 | 50,94,57,67,46,33,28,31,2 | 298 | 149 | 0.00079894587482 | 0.0940089091824457 | 0.006,0.0,0.011,0. | 0.0,0.06,0.034,0.0, | -0,024 |
| 236474 | ENSG00000122882 | ECD | chr10 | - | 73154255 | 73154395 | 73152292 | 73152421 | 73156274 | 73156453 | 236474 | 24,22,51,29,18,34,25,34,1 | 1,0,5,4,0,2,1,0,0,0,2,1 | 26,48,22,25,28,11,24,19,9, | 1,5,6,2,1,2,0,5,2,0,0,5,1,0, | 289 | 149 | 0.000799310061301 | 0.0940089091824457 | 0.925,1.0,0.84,0.7 | 0.931,0.832,0.654, | 0,054 |
| 52280 | ENSG00000123104 | ITPR2 | chr12 | - | 26682573 | 26682673 | 26681873 | 26682034 | 26686480 | 26686632 | 52280 | 107,103,288,133,56,103,9 | 5,6,11,4,2,9,10,8,0,4,2,2 | 140,146,72,89,149,101,59, | 6,23,0,28,12,8,10,13,9,4,1 | 249 | 149 | 0.000800758643147 | 0.0941278446330814 | 0.928,0.911,0.94,0 | 0.933,0.792,1.0,0. | 0,043 |
| 131682 | ENSG00000076513 | ANKRD13A | chr12 | + | 110016387 | 110016433 | 110013124 | 110013249 | 110018344 | 110018488 | 131682 | 273,174,223,133,111,82,9 | 1,0,0,4,2,2,0,1,4,0,1,2 | 138,120,153,84,128,70,31, | 2,0,2,5,3,3,2,2,3,7,0,6,6,8, | 195 | 149 | 0.000802982292756 | 0.094323170230956 | 0.995,1.0,1.0,0.96 | 0.981,1.0,0.983,0. | 0,014 |
| 160562 | ENSG00000113732 | ATP6V0E1 | chr5 | + | 172986695 | 172986760 | 172983830 | 172983964 | 172994774 | 172994822 | 160562 | 3,6,10,9,1,1,4,10,0,4,0,3 | 70,64,115,194,196,99,90,9 | 8,15,6,8,3,4,1,5,6,5,7,7,1,8 | 121,86,70,21,41,43,48,104 | 214 | 149 | 0.000803296788257 | 0.094323170230956 | 0.029,0.061,0.057, | 0.044,0.108,0.056, | -0,028 |
| 167519 | ENSG00000164597 | COG5 | chr7 | - | 107236338 | 107236687 | 107230614 | 107230691 | 107248395 | 107248499 | 167519 | 10,13,34,10,10,15,6,16,4,5 | 1,4,3,0,1,2,2,3,0,0,0,1 | 15,10,4,9,11,3,8,10,12,13,1 | 1,6,1,2,3,2,2,2,5,2,5,3,0,2, | 298 | 149 | 0.000804139610447 | 0.0943540364766949 | 0.833,0.619,0.85,1 | 0.882,0.455,0.667, | 0,197 |
| 184403 | ENSG00000143373 | ZNF687 | chr1 | + | 151283187 | 151283318 | 151282311 | 151282395 | 151286274 | 151288406 | 184403 | 3,0,1,0,0,0,0,3,1,0,2,0 | 9,28,21,14,5,13,16,14,13,1 | 1,3,0,3,0,0,0,3,1,1,0,6,1,3, | 18,16,13,25,6,3,4,14,8,8,1 | 280 | 149 | 0.000804436428464 | 0.0943540364766949 | 0.151,0.0,0.025,0.0 | 0.029,0.091,0.0,0. | -0,022 |

| | | | | | | | | | | | | | | | | | | | | | | |
|---|---|---|---|---|---|---|---|---|---|---|---|---|---|---|---|---|---|---|---|---|---|---|
| 13218 | ENSG00000090565 | RAB11FIP3 | chr16 | + | 491133 | 491268 | 488850 | 489000 | 496823 | 496859 | 13218 | 1,2,2,1,4,0,3,6,2,4,5,3 | 9,5,7,5,11,1,2,3,9,2,4,2 | 6,8,9,3,5,1,3,8,2,6,6,10,0,5 | 1,4,2,3,1,2,0,4,0,0,2,16,1,3 | 284 | 149 | 0.000806215172072 | 0.0945111643247432 | 0.055,0.173,0.13,0 | 0.759,0.512,0.702, | -0,28 |
| 154489 | ENSG00000100888 | CHD8 | chr14 | - | 21412912 | 21412989 | 21409850 | 21409988 | 21414300 | 21414418 | 154489 | 95,62,57,42,19,16,23,31,1 | 2,1,2,1,0,0,0,0,0,0,0,0 | 35,52,22,21,29,18,15,19,1 | 3,8,0,4,4,0,0,0,3,3,0,2,1,0, | 226 | 149 | 0.000807686287682 | 0.0945960843161757 | 0.969,0.976,0.949, | 0.885,0.811,1.0,0. | 0,063 |
| 123204 | ENSG00000149311 | ATM | chr11 | + | 108224783 | 108224871 | 108223115 | 108223186 | 108227594 | 108227696 | 123204 | 13,13,28,19,3,9,6,7,8,5,2,2 | 69,36,113,76,21,30,22,41,4 | 7,9,4,5,9,7,2,6,5,4,12,18,7, | 81,53,26,16,42,44,11,24,42 | 237 | 149 | 0.000807818590134 | 0.0945960843161757 | 0.106,0.185,0.135, | 0.052,0.096,0.088, | 0,033 |
| 203520 | ENSG00000163617 | CCDC191 | chr3 | - | 114046590 | 114046682 | 114042702 | 114042846 | 114056376 | 114056594 | 203520 | 12,4,18,8,7,9,3,9,3,3,4,4 | 2,0,0,3,5,2,0,0,0,2,2,2 | 6,10,4,4,7,0,4,10,3,7,1,6,4, | 1,0,0,0,0,2,1,0,0,0,0,1,2,0,0 | 241 | 149 | 0.000811472225477 | 0.0949722558790866 | 0.788,1.0,1.0,0.62 | 0.788,1.0,1.0,1.0,1 | -0,036 |
| 139526 | ENSG00000125844 | RRBP1 | chr20 | - | 17616722 | 17616839 | 17615925 | 17616009 | 17618595 | 17618679 | 139526 | 63,27,84,62,38,35,34,41,3 | 6,0,5,3,0,2,2,0,0,3,5,0 | 55,51,23,40,16,8,59,33,11, | 6,5,0,10,1,0,12,17,1,0,3,0,1 | 266 | 149 | 0.000812576989154 | 0.0949774021091179 | 0.855,1.0,0.904,0. | 0.837,0.851,1.0,0. | 0,069 |
| 8006 | ENSG00000101546 | RBFA | chr18 | + | 80042134 | 80042219 | 80038504 | 80038617 | 80044211 | 80044285 | 8006 | 36,13,39,45,34,14,13,13,21 | 3,0,3,1,0,0,1,0,5,2,0,1 | 32,18,19,24,15,6,7,38,11,2 | 1,2,0,3,0,2,1,2,2,0,0,5,0,5,4 | 234 | 149 | 0.000812886042814 | 0.0949774021091179 | 0.884,1.0,0.892,0. | 0.953,0.851,1.0,0. | 0,063 |
| 145460 | ENSG00000074370 | ATP2A3 | chr17 | - | 3928239 | 3928376 | 3925046 | 3925441 | 3928662 | 3928780 | 145460 | 2,11,9,15,1,8,3,1,13,6,4,5 | 80,60,111,70,80,62,49,34,6 | 7,7,5,8,1,2,2,3,5,2,2,3,4,3,4 | 101,125,32,24,62,56,16,71 | 286 | 149 | 0.000813121088228 | 0.0949774021091179 | 0.013,0.087,0.041, | 0.035,0.028,0.075, | 0,012 |
| 217885 | ENSG00000205268 | PDE7A | chr8 | - | 65737392 | 65737463 | 65734793 | 65734894 | 65739501 | 65739597 | 217885 | 2,1,2,0,2,4,1,3,2,0,1,5 | 21,20,28,7,69,16,29,44,29, | 4,5,10,6,2,2,2,1,3,2,15,2,1, | 47,34,35,17,16,16,24,41,21 | 220 | 149 | 0.000813281321404 | 0.0949774021091179 | 0.061,0.033,0.046, | 0.054,0.091,0.162, | -0,04 |
| 142164 | ENSG00000091592 | NLRP1 | chr17 | - | 5533896 | 5534361 | 5533291 | 5533384 | 5536850 | 5537044 | 142164 | 77,27,61,47,39,43,36,40,37 | 4,2,7,1,0,3,2,1,2,0,2,7 | 60,61,39,29,48,21,22,31,42 | 2,2,4,15,1,0,10,7,1,3,9,7,3, | 298 | 149 | 0.000814601111891 | 0.0950799413847136 | 0.906,0.871,0.813, | 0.938,0.938,0.83,0 | 0,149 |
| 234167 | ENSG00000169249 | ZRSR2 | chrX | + | 15800208 | 15800329 | 15799882 | 15799953 | 15803687 | 15803796 | 234167 | 2,3,4,7,2,3,0,1,0,0,0,2 | 43,30,50,32,20,26,10,21,27 | 0,7,2,7,9,3,1,2,2,1,0,3,8,2,1 | 16,30,43,37,18,8,10,3,23,2 | 270 | 149 | 0.000816707495046 | 0.0952741305508106 | 0.025,0.052,0.042, | 0.0,0.114,0.025,0. | -0,051 |
| 186695 | ENSG00000134242 | PTPN22 | chr1 | - | 113825133 | 113825172 | 113819576 | 113819654 | 113829591 | 113829707 | 186695 | 6,6,60,25,16,9,15,8,18,9,4, | 4,2,5,5,2,1,4,3,7,3,0,2 | 11,20,5,3,11,2,12,13,4,15,3 | 0,3,0,1,0,0,1,2,0,2,1,7,1,2,3 | 188 | 149 | 0.000818023250331 | 0.0953759275146216 | 0.543,0.704,0.905, | 1.0,0.841,1.0,0.70 | -0,094 |
| 30249 | ENSG00000135074 | ADAM19 | chr5 | - | 157502802 | 157502980 | 157499572 | 157499662 | 157507055 | 157507140 | 30249 | 12,23,14,10,30,17,15,11,8, | 4,0,8,1,1,1,0,2,0,0,0,0 | 30,42,8,14,37,18,12,15,9,3 | 2,2,1,0,1,0,1,8,0,0,0,2,3,0,3 | 298 | 149 | 0.000819302300618 | 0.0954733370137048 | 0.6,1.0,0.467,0.83 | 0.882,0.913,0.8,1. | -0,024 |
| 174939 | ENSG00000096384 | HSP90AB1 | chr6 | + | 44248277 | 44248776 | 44247100 | 44247195 | 44249376 | 44249583 | 174939 | 1151,668,1608,1323,874,8 | 47,12,26,19,11,17,16,10,8, | 934,944,722,248,361,335,4 | 11,11,5,5,4,4,2,11,4,0,32,9 | 298 | 149 | 0.000822367854023 | 0.0957269094587476 | 0.924,0.965,0.969, | 0.977,0.977,0.986, | -0,008 |
| 174943 | ENSG00000096384 | HSP90AB1 | chr6 | + | 44248342 | 44248776 | 44247100 | 44247195 | 44249376 | 44249583 | 174943 | 1151,668,1608,1323,874,8 | 47,12,26,19,11,17,16,10,8, | 934,944,722,248,361,335,4 | 11,11,5,5,4,4,2,11,4,0,32,9 | 298 | 149 | 0.000822367854023 | 0.0957269094587476 | 0.924,0.965,0.969, | 0.977,0.977,0.986, | -0,008 |
| 158038 | ENSG00000196655 | TRAPPC4 | chr11 | + | 119020149 | 119020329 | 119018765 | 119019317 | 119021759 | 119021886 | 158038 | 16,15,28,40,22,10,21,11,16 | 0,1,3,3,5,1,0,0,0,0,2,0 | 27,9,23,7,10,15,2,4,10,18,7 | 4,4,2,1,1,0,0,1,0,3,0,4,0,3,3 | 298 | 149 | 0.000823519636255 | 0.0958091647734053 | 1.0,0.882,0.824,0. | 0.771,0.529,0.852, | 0,082 |
| 63979 | ENSG00000130713 | EXOSC2 | chr9 | + | 130700866 | 130700935 | 130699328 | 130699394 | 130701558 | 130702310 | 63979 | 15,8,15,7,6,9,2,6,8,13,2,1 | 0,3,1,0,0,0,0,4,0,0,0,0 | 13,10,4,12,5,8,7,9,4,9,13,5 | 1,0,0,3,1,0,0,3,0,4,3,6,0,2,1 | 218 | 149 | 0.000824028611763 | 0.0958165867846366 | 1.0,0.646,0.911,1. | 0.899,1.0,1.0,0.73 | 0,097 |
| 103668 | ENSG00000106460 | TMEM106B | chr7 | + | 12231061 | 12231115 | 12230178 | 12230438 | 12231836 | 12232037 | 103668 | 48,19,93,64,30,24,30,22,17 | 3,0,6,1,0,1,1,0,0,0,2,2 | 61,50,18,6,26,30,12,40,21, | 3,2,2,1,4,1,1,1,2,2,4,3,8,2,4 | 203 | 149 | 0.000824628638673 | 0.0958345823597346 | 0.922,1.0,0.919,0. | 0.937,0.948,0.869, | 0,058 |
| 211646 | ENSG00000127081 | ZNF484 | chr9 | - | 92853715 | 92853958 | 92846068 | 92848551 | 92855810 | 92855903 | 211646 | 0,1,1,3,0,1,0,0,0,0,0,1 | 8,6,18,10,6,6,7,7,11,3,3,6 | 1,6,0,3,1,0,5,0,3,3,3,1,1,1,1 | 11,25,22,16,13,6,8,4,10,9,1 | 298 | 149 | 0.000825889502831 | 0.0959293165589957 | 0.0,0.077,0.027,0. | 0.043,0.107,0.0,0.0 | -0,033 |
| 174945 | ENSG00000096384 | HSP90AB1 | chr6 | + | 44248546 | 44248776 | 44247100 | 44247195 | 44249376 | 44249583 | 174945 | 1151,668,1608,1323,874,8 | 47,12,26,19,11,17,16,10,8, | 934,944,722,248,361,335,4 | 11,11,5,5,4,4,2,11,4,0,32,9 | 298 | 149 | 0.000826352867395 | 0.0959313667757785 | 0.924,0.965,0.969, | 0.977,0.977,0.986, | -0,008 |
| 211798 | ENSG00000155368 | DBI | chr2 | + | 119367563 | 119367644 | 119366988 | 119367060 | 119368187 | 119368305 | 211798 | 8,8,14,16,12,4,5,4,2,1,0,5 | 233,188,350,386,203,86,12 | 0,2,7,0,0,1,4,4,2,3,5,0,6,0,3 | 177,174,307,59,44,58,67,1 | 230 | 149 | 0.000826975848928 | 0.09595193473888 | 0.022,0.027,0.025, | 0.0,0.007,0.015,0. | 0,004 |
| 69056 | ENSG00000249915 | PDCD6 | chr5 | + | 304176 | 304221 | 272710 | 272772 | 314416 | 314806 | 69056 | 6,5,19,9,6,8,3,8,9,6,7,10 | 0,0,1,0,0,2,5,0,0,0,0,3 | 25,16,11,2,2,3,0,10,5,8,9,1 | 1,5,0,3,4,1,2,2,1,0,3,3,0,1,0 | 194 | 149 | 0.000827707327956 | 0.0959850624478976 | 1.0,1.0,0.936,1.0,1 | 0.95,0.711,1.0,0.3 | 0,153 |
| 217977 | ENSG00000067248 | DHX29 | chr5 | - | 55289755 | 55289865 | 55289269 | 55289428 | 55290217 | 55290344 | 217977 | 6,0,6,0,0,1,0,1,3,0,0,2 | 8,10,19,19,10,5,8,8,8,4,2,1 | 9,1,2,3,3,0,2,3,4,1,5,7,5,2,0 | 21,21,3,14,10,2,9,7,6,8,11, | 259 | 149 | 0.000829496695656 | 0.0960780054377203 | 0.301,0.0,0.154,0. | 0.198,0.027,0.277, | -0,066 |
| 91284 | ENSG00000163939 | PBRM1 | chr3 | - | 52624896 | 52624941 | 52617261 | 52617538 | 52627272 | 52627370 | 91284 | 5,1,4,7,3,1,0,1,1,1,0,0 | 13,12,59,18,21,17,18,19,10 | 6,0,0,2,0,1,0,1,0,2,0,2,1,0,0 | 33,22,15,10,11,9,22,13,18, | 194 | 149 | 0.000829814522864 | 0.0960780054377203 | 0.228,0.06,0.049,0 | 0.123,0.0,0.0,0.13 | 0,044 |
| 229872 | ENSG00000110497 | AMBRA1 | chr11 | - | 46512457 | 46512813 | 46494123 | 46494204 | 46541944 | 46542981 | 229872 | 14,11,16,2,8,12,3,9,12,11,4 | 0,0,5,1,2,2,0,0,0,0,3,4 | 2,6,7,8,7,3,4,3,4,14,5,5,7,6 | 1,1,0,0,0,2,0,0,0,1,0,1,1,0,0 | 298 | 149 | 0.000829847987087 | 0.0960780054377203 | 1.0,1.0,0.615,0.5,0 | 0.5,0.75,1.0,1.0,1. | -0,099 |
| 243558 | ENSG00000132109 | TRIM21 | chr11 | - | 4386156 | 4386213 | 4384896 | 4385853 | 4386967 | 4386990 | 243558 | 5,8,20,25,5,4,5,6,3,3,2,3 | 1,0,0,2,1,2,2,0,1,1,0,0 | 16,17,5,10,6,2,6,6,6,9,5,8,2 | 7,1,0,0,0,2,0,2,0,0,7,5,3,3,0 | 206 | 149 | 0.000830540808743 | 0.0961065208637444 | 0.783,1.0,1.0,0.9,0 | 0.623,0.925,1.0,1. | 0,053 |
| 114445 | ENSG00000100815 | TRIP11 | chr14 | - | 92010985 | 92011072 | 92007639 | 92007852 | 92014214 | 92014577 | 114445 | 48,17,29,21,27,7,14,7,32,2 | 9,3,5,3,0,2,3,1,0,2,1,1 | 14,14,21,9,10,12,7,18,7,24 | 0,2,3,0,4,1,1,0,3,0,0,1,0,4,0 | 236 | 149 | 0.00083362043956 | 0.0963931460080232 | 0.771,0.782,0.785, | 1.0,0.815,0.815,1. | -0,103 |
| 193138 | ENSG00000213064 | SFT2D2 | chr1 | + | 168235100 | 168235187 | 168231833 | 168231919 | 168236588 | 168236624 | 193138 | 30,36,68,73,36,21,17,30,11 | 3,3,10,2,3,0,3,4,0,1,0,5 | 54,59,16,21,26,24,11,16,26 | 1,4,1,5,5,0,0,2,2,2,0,2,2,1,0 | 236 | 149 | 0.000834019421553 | 0.0963931460080232 | 0.863,0.883,0.811, | 0.972,0.903,0.91,0 | -0,036 |
| 24133 | ENSG00000139496 | NUP58 | chr13 | + | 25309959 | 25310015 | 25309246 | 25309282 | 25312882 | 25312942 | 24133 | 3,2,6,1,0,0,0,0,1,0,1,1 | 20,13,21,18,8,15,18,11,14, | 3,1,9,5,4,1,2,2,1,0,3,4,0,0,1 | 17,16,15,5,16,3,8,8,1,12,16 | 205 | 149 | 0.000834361365291 | 0.0963931460080232 | 0.098,0.101,0.172, | 0.114,0.043,0.304, | -0,084 |
| 116812 | ENSG00000108424 | KPNB1 | chr17 | + | 47669677 | 47669869 | 47668185 | 47668410 | 47670701 | 47670832 | 116812 | 81,87,258,209,118,93,91,1 | 10,5,25,25,10,6,5,2,1,7,6,4 | 194,226,76,62,103,84,88,1 | 16,26,14,18,15,4,7,23,13,5 | 298 | 149 | 0.000836052857891 | 0.0965041070829405 | 0.802,0.897,0.838, | 0.858,0.813,0.731, | 0,072 |
| 72829 | ENSG00000116001 | TIA1 | chr2 | - | 70229058 | 70229091 | 70227734 | 70227822 | 70229263 | 70229318 | 72829 | 36,21,82,29,58,28,25,34,21 | 2,0,2,6,12,5,1,0,2,2,3,2 | 107,67,24,7,8,16,28,36,16, | 6,4,0,0,0,0,2,1,2,3,5,2,1,5,1 | 182 | 149 | 0.000836218573113 | 0.0965041070829405 | 0.936,1.0,0.971,0. | 0.936,0.932,1.0,1. | -0,054 |
| 249674 | ENSG00000100225 | FBXO7 | chr22 | + | 32495492 | 32495534 | 32487744 | 32493281 | 32498143 | 32498829 | 249674 | 26,16,48,25,15,21,8,16,5,5 | 0,2,4,1,2,1,4,0,1,5,0,1 | 30,18,13,12,14,8,7,11,13,1 | 2,0,4,2,0,2,0,1,0,0,0,0,1,0,1 | 191 | 149 | 0.000837285634718 | 0.0965754686205733 | 1.0,0.862,0.903,0. | 0.921,1.0,0.717,0. | -0,079 |
| 87637 | ENSG00000116521 | SCAMP3 | chr1 | - | 155261656 | 155261734 | 155260536 | 155260659 | 155262085 | 155262360 | 87637 | 28,25,21,34,20,22,21,14,48 | 1,3,2,1,2,3,0,8,4,0,0,4 | 18,21,17,18,9,11,11,14,10, | 7,3,3,2,4,3,2,4,1,1,3,7,4,4,7 | 227 | 149 | 0.000842004736769 | 0.0970513497462819 | 0.948,0.845,0.873, | 0.628,0.821,0.788, | 0,136 |
| 92715 | ENSG00000136861 | CDK5RAP2 | chr9 | - | 120529977 | 120530140 | 120528743 | 120528797 | 120536371 | 120536437 | 92715 | 120,83,97,42,44,33,33,59,5 | 6,3,1,4,0,0,4,2,1,1,0,0 | 46,61,62,98,43,40,59,49,28 | 0,0,3,2,3,0,0,0,1,0,1,0,1,0,0 | 298 | 149 | 0.000842313241708 | 0.0970513497462819 | 0.909,0.933,0.98,0 | 1.0,1.0,0.912,0.96 | -0,033 |
| 178487 | ENSG00000138756 | BMP2K | chr4 | + | 78848110 | 78848154 | 78847187 | 78847269 | 78850923 | 78851056 | 178487 | 2,0,16,20,3,2,3,8,0,0,0,7 | 29,17,65,56,39,28,24,31,11 | 5,3,2,2,0,1,5,1,2,1,0,8,1,1,1 | 36,29,20,17,4,15,12,39,7,2 | 193 | 149 | 0.000843123091775 | 0.0970716420253763 | 0.051,0.0,0.16,0.2 | 0.097,0.074,0.072, | 0,007 |
| 167501 | ENSG00000145495 | MARCHF6 | chr5 | + | 10420158 | 10420341 | 10417269 | 10417404 | 10423734 | 10423835 | 167501 | 9,8,4,2,0,1,5,2,2,0,0,1 | 86,76,160,56,62,94,69,86,3 | 3,6,5,25,1,2,5,1,4,5,9,3,1,6 | 105,110,67,80,72,46,73,10 | 298 | 149 | 0.000843391382224 | 0.0970716420253763 | 0.05,0.05,0.012,0. | 0.014,0.027,0.036, | -0,017 |
| 251414 | ENSG00000148429 | USP6NL | chr10 | - | 11489101 | 11489248 | 11485816 | 11485911 | 11490831 | 11490880 | 251414 | 36,22,90,34,14,15,18,36,3, | 4,0,2,0,2,2,0,2,0,0,0,2 | 59,35,22,1,5,10,22,16,20,8 | 5,0,4,2,0,0,1,1,5,0,4,12,3,0 | 296 | 149 | 0.000845635788519 | 0.0972779456968107 | 0.819,1.0,0.958,1. | 0.856,1.0,0.735,0. | 0,075 |
| 89568 | ENSG00000219626 | FAM228B | chr2 | + | 24167626 | 24167683 | 24164197 | 24164335 | 24169355 | 24169496 | 89568 | 24,10,15,17,23,14,6,12,11, | 3,1,10,8,7,1,3,2,3,0,0,2 | 5,4,16,6,9,11,3,19,8,15,3,1 | 10,10,5,1,1,2,2,7,3,14,10,7 | 206 | 149 | 0.000846918816044 | 0.0973734955640845 | 0.853,0.879,0.52,0 | 0.266,0.224,0.698, | 0,231 |
| 162675 | ENSG00000072364 | AFF4 | chr5 | - | 132898229 | 132898365 | 132896322 | 132897240 | 132899103 | 132899141 | 162675 | 86,61,143,60,43,91,59,154 | 13,1,7,1,1,3,1,9,3,1,6,3 | 77,98,58,135,74,29,90,87,4 | 5,7,5,31,9,9,5,4,8,5,4,2,2,5 | 285 | 149 | 0.000849879207777 | 0.0976351986079802 | 0.776,0.97,0.914,0 | 0.89,0.88,0.858,0. | 0,069 |
| 104356 | ENSG00000166579 | NDEL1 | chr17 | + | 8443926 | 8444176 | 8435921 | 8436045 | 8444259 | 8444357 | 104356 | 1,0,12,5,0,2,0,0,2,0,2,4 | 20,4,11,4,6,6,10,14,8,8,3,1 | 2,3,0,2,0,5,0,0,2,0,0,2,0,1,0 | 12,21,6,22,6,1,7,10,8,4,7,2 | 298 | 149 | 0.000850102272402 | 0.0976351986079802 | 0.024,0.0,0.353,0. | 0.077,0.067,0.0,0. | 0,085 |
| 34551 | ENSG00000047365 | ARAP2 | chr4 | - | 36080215 | 36080279 | 36073688 | 36073823 | 36082250 | 36082286 | 34551 | 26,40,51,38,21,30,12,36,29 | 0,0,1,0,0,0,0,2,1,0,0,0 | 51,43,23,35,11,24,16,47,19 | 2,5,0,5,1,0,6,2,1,0,1,2,0,0,1 | 213 | 149 | 0.000852067621868 | 0.0978050666538655 | 1.0,1.0,0.973,1.0,1 | 0.947,0.857,1.0,0. | 0,06 |
| 147687 | ENSG00000185324 | CDK10 | chr16 | + | 89689200 | 89689324 | 89686712 | 89686797 | 89690552 | 89690557 | 147687 | 18,16,27,20,11,18,12,10,11 | 0,3,4,6,0,1,2,0,10,0,0,0 | 22,9,18,12,16,8,11,22,10,1 | 8,8,4,4,3,3,2,3,2,5,1,5,3,1,5 | 273 | 149 | 0.00085282544167 | 0.0978050666538655 | 1.0,0.744,0.787,0. | 0.6,0.38,0.711,0.6 | 0,192 |
| 248437 | ENSG00000139697 | SBNO1 | chr12 | - | 123348028 | 123348129 | 123345257 | 123345570 | 123350309 | 123350441 | 248437 | 57,31,31,17,18,15,15,31,42 | 0,0,0,2,2,0,0,0,0,2,2,2 | 31,33,23,21,19,19,12,17,6, | 1,2,3,2,1,1,0,2,1,5,0,2,5,1,2 | 250 | 149 | 0.000852944557751 | 0.0978050666538655 | 1.0,1.0,1.0,0.835,0 | 0.949,0.908,0.82,0 | 0,046 |
| 217208 | ENSG00000031003 | FAM13B | chr5 | - | 138018954 | 138019153 | 138018301 | 138018514 | 138019880 | 138019977 | 217208 | 43,24,51,36,17,23,12,18,19 | 16,7,9,5,1,1,3,3,0,4,1,1 | 30,24,35,11,20,15,7,25,32, | 5,3,0,0,2,1,0,3,1,0,0,0,4,4,2 | 298 | 149 | 0.000854677726131 | 0.0979516196341327 | 0.573,0.632,0.739, | 0.75,0.8,1.0,1.0,0. | -0,127 |
| 128812 | ENSG00000011258 | MBTD1 | chr17 | - | 51193427 | 51193510 | 51191583 | 51193016 | 51195213 | 51195361 | 128812 | 12,4,32,27,12,20,8,29,4,5,6 | 3,0,7,4,0,2,2,2,0,0,4,1 | 20,31,11,14,11,4,12,12,16, | 11,7,2,14,4,4,5,2,6,0,8,5,3, | 232 | 149 | 0.000857652334473 | 0.0982402179888016 | 0.72,1.0,0.746,0.8 | 0.539,0.74,0.779,0 | 0,174 |
| 147685 | ENSG00000185324 | CDK10 | chr16 | + | 89688946 | 89689324 | 89686712 | 89686797 | 89690552 | 89690557 | 147685 | 21,19,29,21,14,20,12,11,10 | 0,3,4,6,0,1,2,0,10,0,0,0 | 23,9,20,12,20,9,13,23,10,1 | 8,8,4,4,3,3,2,3,2,5,1,5,3,1,5 | 298 | 149 | 0.000859530843501 | 0.0984030228604062 | 1.0,0.76,0.784,0.6 | 0.59,0.36,0.714,0. | 0,187 |
| 51797 | ENSG00000169682 | SPNS1 | chr16 | + | 28981900 | 28982056 | 28981454 | 28981615 | 28982355 | 28982545 | 51797 | 30,24,23,17,28,21,26,29,41 | 1,4,0,1,3,0,2,3,2,2,2,2 | 11,12,8,20,9,4,31,20,5,18,1 | 6,5,0,8,0,0,2,1,1,1,3,4,3,4,6 | 298 | 149 | 0.000862870556119 | 0.0987328509644064 | 0.938,0.75,1.0,0.8 | 0.478,0.545,1.0,0. | 0,106 |
| 173720 | ENSG00000136560 | TANK | chr2 | + | 161177111 | 161177300 | 161136965 | 161137063 | 161179613 | 161180085 | 173720 | 6,1,10,7,0,6,0,5,5,0,2,3 | 191,301,266,174,128,154,8 | 7,7,14,3,4,3,0,14,6,16,2,4,1 | 116,191,136,84,120,41,34, | 298 | 149 | 0.000863643431151 | 0.0987687775398836 | 0.015,0.002,0.018, | 0.029,0.018,0.049, | -0,007 |
| 215423 | ENSG00000010244 | ZNF207 | chr17 | + | 32366664 | 32366713 | 32365329 | 32365487 | 32367771 | 32368433 | 215423 | 32,7,73,50,43,35,38,36,16, | 89,50,53,64,54,64,59,99,13 | 52,65,21,20,38,28,30,34,23 | 78,59,40,69,53,28,24,58,24 | 198 | 149 | 0.000865317656848 | 0.0989076923000807 | 0.213,0.095,0.509, | 0.334,0.453,0.283, | -0,099 |
| 220473 | ENSG00000068028 | RASSF1 | chr3 | - | 50336759 | 50336951 | 50332049 | 50332154 | 50337172 | 50337464 | 220473 | 4,2,3,1,2,1,1,2,0,1,2,3 | 7,11,14,11,4,11,3,3,7,1,1,3 | 1,2,2,3,0,0,0,1,2,1,0,2,2,0,0 | 15,29,5,4,8,1,11,12,3,8,11, | 298 | 149 | 0.000868356038385 | 0.0991573820668916 | 0.222,0.083,0.097, | 0.032,0.033,0.167, | 0,118 |
| 166962 | ENSG00000138443 | ABI2 | chr2 | + | 203383242 | 203383321 | 203380357 | 203380384 | 203391045 | 203391143 | 166962 | 3,0,2,3,0,2,2,2,2,0,1,2 | 9,6,14,10,11,6,10,12,4,4,8, | 1,1,0,0,0,0,0,3,0,2,1,0,0,0,2 | 11,13,8,6,6,3,13,14,6,15,15 | 228 | 149 | 0.000868423531908 | 0.0991573820668916 | 0.179,0.0,0.085,0. | 0.056,0.048,0.0,0. | 0,07 |
| 8749 | ENSG00000101745 | ANKRD12 | chr18 | + | 9204475 | 9204544 | 9182381 | 9182519 | 9208656 | 9208803 | 8749 | 116,61,94,57,20,37,44,40,5 | 4,10,5,5,3,1,2,1,6,0,0,4 | 19,54,52,64,74,25,29,47,27 | 0,1,6,7,4,0,0,0,1,3,1,1,0,0,4 | 218 | 149 | 0.000869911456339 | 0.0992746090452276 | 0.952,0.807,0.928, | 1.0,0.974,0.856,0. | -0,039 |
| 61080 | ENSG00000138614 | INTS14 | chr15 | - | 65607158 | 65607442 | 65605128 | 65605236 | 65610795 | 65611136 | 61080 | 7,4,16,8,5,4,2,13,0,4,5,5 | 0,3,2,1,2,0,2,5,1,1,0,0 | 1,5,9,4,7,6,1,12,5,9,6,17,2, | 0,0,1,1,2,0,0,0,0,0,4,1,0,0,0 | 298 | 149 | 0.00087081545393 | 0.0992755802438761 | 1.0,0.4,0.8,0.8,0.5 | 1.0,1.0,0.818,0.66 | -0,201 |
| 105512 | ENSG00000285171 | | chrX | - | 71110503 | 71110688 | 71109227 | 71109390 | 71111424 | 71111571 | 105512 | 15,18,39,18,6,17,17,13,7,6 | 1,0,6,0,3,4,0,4,2,0,0,0 | 14,39,8,23,24,10,14,20,5,3 | 4,1,1,0,0,0,0,0,1,0,2,1,3,0,0 | 298 | 149 | 0.000870842469256 | 0.0992755802438761 | 0.882,1.0,0.765,1. | 0.636,0.951,0.8,1. | -0,06 |
| 166554 | ENSG00000134851 | TMEM165 | chr4 | + | 55416041 | 55416132 | 55411613 | 55411839 | 55417071 | 55417247 | 166554 | 0,1,0,4,2,1,0,0,0,2,0,0 | 11,7,15,12,7,5,15,9,12,8,4, | 2,4,0,2,0,1,2,3,0,1,2,1,6,0,4 | 14,6,9,3,2,9,0,7,4,15,3,24,1 | 240 | 149 | 0.000876388048793 | 0.0998548841343389 | 0.0,0.081,0.0,0.17 | 0.081,0.293,0.0,0. | -0,139 |
| 107036 | ENSG00000046651 | OFD1 | chrX | + | 13751248 | 13751368 | 13749426 | 13749533 | 13753367 | 13753441 | 107036 | 240,114,331,141,90,97,52, | 50,10,43,34,23,18,12,14,30 | 131,115,143,117,98,57,30, | 19,5,10,13,10,9,6,19,21,44 | 269 | 149 | 0.000879410758517 | 0.10012488518778 | 0.727,0.863,0.81,0 | 0.792,0.927,0.888, | -0,072 |
| 187954 | ENSG00000113532 | ST8SIA4 | chr5 | - | 100856186 | 100856396 | 100806932 | 100812129 | 100886342 | 100886600 | 187954 | 26,26,66,37,18,14,19,21,10 | 6,2,8,5,1,3,2,3,3,0,0,1 | 27,43,22,10,18,5,4,16,8,20 | 6,3,0,0,2,0,0,0,0,2,0,5,2,2,0 | 298 | 149 | 0.000879688139209 | 0.10012488518778 | 0.684,0.867,0.805, | 0.692,0.878,1.0,1. | -0,11 |
| 42138 | ENSG00000163964 | PIGX | chr3 | + | 196726654 | 196726771 | 196722414 | 196722556 | 196727922 | 196728037 | 42138 | 2,7,9,5,0,2,0,0,2,0,0,1 | 5,3,19,22,5,6,2,9,2,2,4,5 | 0,0,0,0,1,0,2,5,1,0,0,2,2,0,4 | 4,17,7,3,3,2,2,2,0,3,4,15,2, | 266 | 149 | 0.000880729519098 | 0.100190430827157 | 0.183,0.567,0.21,0 | 0.0,0.0,0.0,0.0,0.1 | -0,036 |
| 233 | ENSG00000182670 | TTC3 | chr21 | + | 37150819 | 37150884 | 37150077 | 37150170 | 37151892 | 37152029 | 233 | 45,38,65,47,30,16,11,22,29 | 2,0,8,4,0,2,0,3,2,0,0,0 | 43,44,16,35,23,17,14,19,26 | 0,1,1,0,0,0,0,0,0,1,6,0,0,0,0 | 214 | 149 | 0.000883011198864 | 0.100396927280876 | 0.94,1.0,0.85,0.89 | 1.0,0.968,0.918,1. | -0,03 |
| 181725 | ENSG00000143486 | EIF2D | chr1 | - | 206599283 | 206599612 | 206599002 | 206599092 | 206599732 | 206599836 | 181725 | 23,5,25,22,9,18,19,26,11,8 | 0,0,0,0,0,3,0,3,0,0,0,0 | 14,6,12,4,8,7,12,14,7,11,18 | 1,2,1,0,0,0,1,0,0,3,1,7,2,0,2 | 298 | 149 | 0.000887023375112 | 0.100799856414325 | 1.0,1.0,1.0,1.0,1.0, | 0.875,0.6,0.857,1. | 0,097 |
| 95050 | ENSG00000137601 | NEK1 | chr4 | - | 169555719 | 169555851 | 169537808 | 169537911 | 169555931 | 169556095 | 95050 | 8,1,8,1,2,10,3,7,3,3,0,2 | 8,6,13,16,3,10,2,7,7,1,4,4 | 3,4,12,4,3,5,5,14,7,7,5,7,10 | 7,11,0,3,2,3,7,3,2,2,2,1,0,3 | 281 | 149 | 0.000888660000469 | 0.100897494026323 | 0.347,0.081,0.246, | 0.185,0.162,1.0,0. | -0,241 |
| 159499 | ENSG00000038382 | TRIO | chr5 | + | 14399818 | 14399906 | 14399023 | 14399070 | 14400962 | 14401064 | 159499 | 0,1,0,0,2,5,1,11,0,0,0,0 | 7,8,13,3,24,113,15,50,15,4 | 4,3,11,4,0,0,1,8,1,1,0,5,0,2 | 11,10,45,16,13,11,7,28,4,2 | 237 | 149 | 0.000888820145211 | 0.100897494026323 | 0.0,0.073,0.0,0.0,0 | 0.186,0.159,0.133, | -0,068 |
| 117270 | ENSG00000148396 | SEC16A | chr9 | - | 136451255 | 136451408 | 136448083 | 136448161 | 136455600 | 136455793 | 117270 | 18,16,46,25,12,24,15,8,33, | 3,2,2,1,0,0,1,2,0,4,0,2 | 15,16,28,17,13,16,12,12,6, | 1,1,5,10,0,0,0,2,6,1,2,1,3,2 | 298 | 149 | 0.000889645688528 | 0.10093797110573 | 0.75,0.8,0.92,0.92 | 0.882,0.889,0.737, | 0,103 |
| 221885 | ENSG00000172845 | SP3 | chr2 | - | 173910059 | 173910257 | 173907885 | 173907965 | 173913069 | 173913266 | 221885 | 264,215,274,161,99,123,10 | 6,1,17,8,9,3,6,0,1,0,1,6 | 181,272,185,178,157,88,98 | 9,14,6,8,4,3,6,6,6,7,8,17,3, | 298 | 149 | 0.000890249763373 | 0.10095329126477 | 0.957,0.991,0.89,0 | 0.91,0.907,0.939,0 | 0,035 |
| 72964 | ENSG00000176853 | FAM91A1 | chr8 | + | 123786494 | 123786610 | 123786003 | 123786151 | 123787260 | 123787355 | 72964 | 23,24,55,45,20,32,34,30,17 | 0,2,0,1,1,2,0,0,2,0,0,0 | 36,76,21,44,45,22,35,20,38 | 1,5,3,2,1,0,0,0,3,3,0,10,0,0 | 265 | 149 | 0.000892028848054 | 0.101101769876519 | 1.0,0.871,1.0,0.96 | 0.953,0.895,0.797, | 0,014 |
| 150878 | ENSG00000134152 | KATNBL1 | chr15 | - | 34144987 | 34145252 | 34142090 | 34142371 | 34145397 | 34145491 | 150878 | 4,0,4,3,3,4,4,2,0,0,0,1 | 12,7,23,11,21,14,8,11,7,4,5 | 0,1,4,0,0,1,2,1,0,0,2,0,1,1,0 | 18,2,12,2,5,6,1,5,6,16,1,2,4 | 298 | 149 | 0.000896933282499 | 0.101604130171338 | 0.143,0.0,0.08,0.1 | 0.0,0.2,0.143,0.0,0 | -0,034 |
| 50247 | ENSG00000143164 | DCAF6 | chr1 | + | 168038370 | 168038488 | 168022987 | 168023047 | 168043024 | 168043140 | 50247 | 14,23,50,34,13,16,8,15,16, | 2,0,7,3,3,0,4,4,0,0,0,1 | 17,20,16,3,9,1,2,19,11,16,1 | 0,2,2,2,1,0,1,2,1,0,2,1,2,1,1 | 267 | 149 | 0.000900774007644 | 0.101985528900171 | 0.796,1.0,0.799,0. | 1.0,0.848,0.817,0. | 0,025 |
| 83468 | ENSG00000155368 | DBI | chr2 | + | 119367558 | 119367644 | 119366988 | 119367060 | 119368187 | 119368305 | 83468 | 8,13,11,14,13,3,7,5,2,1,0,4 | 233,188,350,386,203,86,12 | 0,6,9,0,0,1,1,3,1,5,2,0,6,0,6 | 177,174,307,59,44,58,67,1 | 235 | 149 | 0.000901914875678 | 0.102061009782887 | 0.021,0.042,0.02,0 | 0.0,0.021,0.018,0. | 0,008 |
| 132401 | ENSG00000196498 | NCOR2 | chr12 | - | 124327546 | 124327633 | 124326179 | 124326370 | 124330844 | 124330898 | 132401 | 6,5,8,4,14,17,3,1,1,3,0,7 | 1,2,1,2,0,1,0,1,0,2,3,6 | 3,7,4,3,4,2,14,18,3,6,4,2,8, | 0,3,1,0,2,0,2,3,1,0,0,0,3,0,1 | 236 | 149 | 0.00090246207826 | 0.102069267244339 | 0.791,0.612,0.835, | 1.0,0.596,0.716,1. | -0,115 |
| 222235 | ENSG00000152601 | MBNL1 | chr3 | + | 152338536 | 152338673 | 152300197 | 152300367 | 152414940 | 152415111 | 222235 | 50,23,23,10,29,13,12,3,24, | 452,250,444,197,235,233,2 | 38,20,9,1,7,3,9,9,2,17,0,7,1 | 365,359,176,132,170,131,1 | 286 | 149 | 0.0009058745745 | 0.10240141310095 | 0.054,0.046,0.026, | 0.051,0.028,0.026, | 0,01 |
| 255706 | ENSG00000112200 | ZNF451 | chr6 | + | 57150718 | 57151178 | 57147089 | 57148693 | 57152220 | 57152351 | 255706 | 67,60,49,26,32,37,23,39,47 | 0,1,4,2,0,0,0,0,0,0,0,1 | 66,50,37,53,24,24,26,27,30 | 2,2,1,2,2,1,0,4,2,0,5,0,1,1,0 | 298 | 149 | 0.000908954333302 | 0.102682495314777 | 1.0,0.968,0.86,0.8 | 0.943,0.926,0.949, | 0,066 |
| 217206 | ENSG00000031003 | FAM13B | chr5 | - | 138018954 | 138019141 | 138018301 | 138018514 | 138019880 | 138019977 | 217206 | 43,24,52,36,17,23,12,18,19 | 16,7,9,5,1,1,3,3,0,4,1,1 | 30,24,35,11,20,15,7,25,32, | 5,3,0,0,2,1,0,3,1,0,0,0,4,4,2 | 298 | 149 | 0.000909315275541 | 0.102682495314777 | 0.573,0.632,0.743, | 0.75,0.8,1.0,1.0,0. | -0,126 |
| 223826 | ENSG00000095015 | MAP3K1 | chr5 | + | 56879020 | 56879101 | 56875031 | 56875310 | 56880710 | 56880802 | 223826 | 40,23,190,83,89,56,43,64,2 | 0,2,4,0,1,0,2,4,2,0,0,2 | 161,164,41,60,64,34,46,69 | 8,11,1,2,0,2,4,0,1,5,2,2,4,3 | 230 | 149 | 0.000910192174951 | 0.1027276203497 | 1.0,0.882,0.969,1. | 0.929,0.906,0.964, | 0,018 |
| 144083 | ENSG00000007202 | KIAA0100 | chr17 | - | 28643163 | 28643332 | 28642868 | 28642987 | 28643609 | 28643659 | 144083 | 26,7,67,67,30,35,15,39,22, | 2,1,4,3,1,2,2,5,0,2,0,0 | 40,28,20,17,10,16,12,25,17 | 6,4,1,3,6,1,0,3,2,0,2,4,6,2,4 | 298 | 149 | 0.000911594092969 | 0.102831922549167 | 0.867,0.778,0.893, | 0.769,0.778,0.909, | 0,078 |
| 31879 | ENSG00000247774 | PCED1B-AS1 | chr12 | - | 47209280 | 47209458 | 47205897 | 47206145 | 47210639 | 47210757 | 31879 | 23,36,55,53,23,21,29,20,46 | 2,0,4,5,0,0,2,0,7,5,0,1 | 30,26,32,14,19,29,11,16,11 | 1,1,1,2,2,4,1,0,0,0,0,10,2,0 | 298 | 149 | 0.000912505410749 | 0.102880802546316 | 0.852,1.0,0.873,0. | 0.938,0.929,0.941, | -0,009 |
| 132660 | ENSG00000173442 | EHBP1L1 | chr11 | + | 65581538 | 65583765 | 65581210 | 65581373 | 65584240 | 65584398 | 132660 | 94,63,70,25,73,69,83,107,8 | 9,1,1,0,1,1,12,0,2,0,0,0 | 38,46,36,43,25,28,58,106,3 | 0,0,0,0,2,0,0,0,0,2,0,0,0,1,4 | 298 | 149 | 0.000913470693722 | 0.102935712502869 | 0.839,0.969,0.972, | 1.0,1.0,1.0,1.0,0. | -0,014 |
| 126970 | ENSG00000099246 | RAB18 | chr10 | + | 27531528 | 27531615 | 27526827 | 27526889 | 27532506 | 27532579 | 126970 | 3,5,5,3,2,7,5,4,0,5,1,2 | 43,38,81,26,22,37,22,32,34 | 5,14,7,17,4,2,4,10,1,5,8,7,1 | 30,53,35,19,33,14,7,24,14, | 236 | 149 | 0.000914576442726 | 0.10300638531887 | 0.042,0.077,0.038, | 0.095,0.143,0.112, | -0,045 |
| 46545 | ENSG00000178694 | NSUN3 | chr3 | + | 94094139 | 94094294 | 94084106 | 94084450 | 94095032 | 94095094 | 46545 | 27,11,30,23,13,17,14,15,10 | 0,0,10,12,0,0,0,0,0,0,0,0 | 25,34,10,25,30,22,4,15,19, | 0,0,0,0,1,0,0,0,0,0,0,0,0,0,0 | 298 | 149 | 0.000915580311818 | 0.103065515738964 | 1.0,1.0,0.6,0.489,1 | 1.0,1.0,1.0,1.0,0.9 | -0,069 |
| 75247 | ENSG00000134313 | KIDINS220 | chr2 | - | 8749370 | 8749421 | 8746504 | 8748000 | 8750111 | 8750335 | 75247 | 2,1,9,0,3,4,7,2,11,1,1,0 | 19,9,41,19,30,11,19,24,23, | 15,7,8,8,0,9,11,6,2,2,5,2,4, | 32,14,8,5,4,6,12,39,4,16,11 | 200 | 149 | 0.000919884431005 | 0.103495894390819 | 0.073,0.076,0.141, | 0.259,0.271,0.427, | -0,136 |
| 240146 | ENSG00000165525 | NEMF | chr14 | - | 49820342 | 49820456 | 49814753 | 49814857 | 49825866 | 49825955 | 240146 | 5,9,19,7,2,10,8,20,7,6,2,9 | 32,13,67,50,28,22,20,36,31 | 10,32,7,15,11,3,9,8,8,7,13, | 34,32,19,9,19,19,9,20,12,1 | 263 | 149 | 0.000929129735249 | 0.10447649172047 | 0.081,0.282,0.138, | 0.143,0.362,0.173, | -0,092 |
| 9776 | ENSG00000198231 | DDX42 | chr17 | + | 63799588 | 63799625 | 63798037 | 63798099 | 63800467 | 63800617 | 9776 | 34,13,61,25,31,36,22,21,16 | 6,5,6,2,6,3,2,2,2,2,0,1 | 32,30,40,18,14,19,18,24,10 | 3,5,0,6,4,0,0,1,0,1,0,2,1,1,2 | 186 | 149 | 0.000929986657493 | 0.10447649172047 | 0.819,0.676,0.891, | 0.895,0.828,1.0,0. | -0,052 |

| | | | | | | | | | | | | | | | | | | | | | | |
|---|---|---|---|---|---|---|---|---|---|---|---|---|---|---|---|---|---|---|---|---|---|---|
| 116957 | ENSG00000137073 | UBAP2 | chr9 | - | 33971650 | 33971754 | 33963725 | 33963791 | 33986759 | 33986837 | 116957 | 14,16,18,17,2,14,4,11,14,1 | 0,1,4,4,4,0,0,1,2,0,0,3 | 13,16,11,5,4,5,6,15,4,11,3, | 2,0,4,0,0,1,2,0,0,1,3,0,0,0, | 253 | 149 | 0.000930056349394 | 0.10447649172047 | 1.0,0.904,0.726,0. | 0.793,1.0,0.618,1. | -0,056 |
| 253112 | ENSG00000124164 | VAPB | chr20 | + | 58439145 | 58439308 | 58438944 | 58439025 | 58440906 | 58441083 | 253112 | 6,3,18,5,6,0,7,7,5,0,3,2 | 22,21,32,22,9,8,9,14,32,12 | 19,14,4,6,10,3,2,3,3,5,5,5, | 20,23,10,9,11,13,8,8,7,18, | 298 | 149 | 0.000931655094034 | 0.104601490633298 | 0.12,0.067,0.22,0. | 0.322,0.233,0.167, | -0,11 |
| 195415 | ENSG00000162889 | MAPKAPK2 | chr1 | + | 206732037 | 206732084 | 206731838 | 206731919 | 206732574 | 206734281 | 195415 | 13,10,18,16,10,11,6,5,8,3, | 48,60,169,90,46,59,50,45, | 10,7,7,2,2,2,7,8,6,10,2,4,3, | 78,101,29,29,72,24,53,59, | 196 | 149 | 0.000933525716694 | 0.10475686836797 | 0.171,0.112,0.075, | 0.089,0.05,0.155,0 | 0,037 |
| 12313 | ENSG00000161956 | SENP3 | chr17 | + | 7565694 | 7565764 | 7565463 | 7565587 | 7566926 | 7567004 | 12313 | 11,7,25,13,11,15,13,9,8,4, | 0,0,2,2,1,0,0,1,0,0,0,2 | 18,13,6,4,4,1,9,18,2,8,9,5, | 2,1,1,1,0,0,5,2,1,2,6,1,1,1, | 219 | 149 | 0.000934958085253 | 0.104862930509165 | 1.0,1.0,0.895,0.81 | 0.86,0.898,0.803,0 | 0,106 |
| 166334 | ENSG00000104517 | UBR5 | chr8 | - | 102278827 | 102278939 | 102276971 | 102277192 | 102279035 | 102279208 | 166334 | 48,48,163,78,69,63,46,68, | 10,0,8,9,0,1,0,2,0,0,0,2 | 93,89,48,76,63,43,37,39,5 | 1,5,3,36,5,2,2,1,9,7,1,10,2, | 261 | 149 | 0.00093580081729 | 0.104886392928098 | 0.733,1.0,0.921,0. | 0.982,0.91,0.901,0 | 0,066 |
| 187173 | ENSG00000288701 | PRRC2B | chr9 | + | 131474453 | 131474496 | 131473507 | 131473724 | 131477743 | 131477949 | 187173 | 8,12,31,15,11,13,10,4,3,9, | 1,4,6,0,1,7,2,9,0,3,0,4 | 8,11,8,10,10,5,5,7,12,10,1 | 7,7,4,25,17,5,10,7,7,4,3,5, | 192 | 149 | 0.000936408360265 | 0.104886392928098 | 0.861,0.7,0.8,1.0,0 | 0.47,0.549,0.608,0 | 0,221 |
| 184651 | ENSG00000139631 | CSAD | chr12 | - | 53171171 | 53171441 | 53170422 | 53170502 | 53171881 | 53171988 | 184651 | 4,1,16,8,11,12,6,8,5,5,3,4 | 0,0,9,0,3,1,1,1,1,0,0,2 | 3,5,10,3,9,4,5,11,5,6,4,10, | 1,9,0,6,2,2,1,4,4,2,0,8,6,6, | 298 | 149 | 0.000936629236531 | 0.104886392928098 | 1.0,1.0,0.471,1.0,0 | 0.6,0.217,1.0,0.2,0 | 0,273 |
| 78933 | ENSG00000196912 | ANKRD36B | chr2 | - | 97560664 | 97560737 | 97558965 | 97558994 | 97560831 | 97560860 | 78933 | 61,26,41,21,19,18,14,32,5, | 21,15,34,15,13,27,9,9,9,12 | 45,41,13,27,48,19,23,41,1 | 6,2,10,8,5,10,0,0,3,17,16,9 | 222 | 149 | 0.000937776955861 | 0.104960307845512 | 0.661,0.538,0.447, | 0.834,0.932,0.466, | -0,188 |
| 212019 | ENSG00000079263 | SP140 | chr2 | + | 230243732 | 230243811 | 230241403 | 230241487 | 230244987 | 230245071 | 212019 | 78,34,63,77,64,36,24,38,7 | 32,20,24,18,8,13,18,24,6,8 | 60,41,51,35,30,30,14,33,3 | 40,96,30,41,46,10,14,6,17, | 228 | 149 | 0.000939352094844 | 0.10508195983647 | 0.614,0.526,0.632, | 0.495,0.218,0.526, | 0,17 |
| 167670 | ENSG00000101109 | STK4 | chr20 | + | 44975323 | 44975441 | 44973326 | 44973433 | 44978442 | 44978571 | 167670 | 5,3,16,9,0,4,4,14,2,3,2,2 | 18,7,14,15,9,14,14,16,1,1, | 9,10,4,5,9,4,1,2,3,5,0,2,0,1 | 23,54,16,16,22,10,18,23,2 | 267 | 149 | 0.000939948225875 | 0.105094024209508 | 0.134,0.193,0.389, | 0.179,0.094,0.122, | 0,161 |
| 234040 | ENSG00000215021 | PHB2 | chr12 | - | 6968410 | 6968542 | 6966925 | 6968021 | 6969497 | 6969577 | 234040 | 167,111,269,197,86,71,89, | 1,2,11,12,3,5,3,2,0,2,0,2 | 129,109,132,60,88,56,62,9 | 1,5,0,1,0,0,1,5,2,1,5,5,0,4, | 281 | 149 | 0.000941631655742 | 0.105227581981831 | 0.989,0.967,0.928, | 0.986,0.92,1.0,0.9 | -0,017 |
| 55414 | ENSG00000160691 | SHC1 | chr1 | - | 154965945 | 154966080 | 154962297 | 154963931 | 154966161 | 154966231 | 55414 | 12,9,11,26,9,9,6,11,8,5,6,5 | 0,0,0,0,2,0,1,1,0,0,0,0 | 8,12,6,8,5,6,11,12,6,8,15,4 | 1,2,0,3,5,0,0,2,1,0,5,2,0,0, | 284 | 149 | 0.00094549463025 | 0.105604439420518 | 1.0,1.0,1.0,1.0,0.7 | 0.808,0.759,1.0,0. | 0,152 |
| 134098 | ENSG00000100201 | DDX17 | chr22 | - | 38489283 | 38489458 | 38488025 | 38488115 | 38492055 | 38492115 | 134098 | 11,7,13,6,11,5,10,6,17,4,4, | 223,109,242,214,130,97,1 | 15,6,14,13,0,2,7,7,6,12,5,1 | 123,52,139,40,64,84,60,14 | 298 | 149 | 0.000948377323626 | 0.105863002159253 | 0.024,0.031,0.026, | 0.057,0.055,0.048, | -0,017 |
| 107935 | ENSG00000215021 | PHB2 | chr12 | - | 6968410 | 6968587 | 6966925 | 6968021 | 6969497 | 6969577 | 107935 | 172,118,269,200,97,80,94, | 1,2,11,12,3,5,3,2,0,2,0,2 | 134,122,135,62,91,58,71,1 | 1,5,0,1,0,0,1,5,2,1,5,5,0,4, | 298 | 149 | 0.000949064359781 | 0.105863002159253 | 0.989,0.967,0.924, | 0.985,0.924,1.0,0. | -0,017 |
| 185080 | ENSG00000078304 | PPP2R5C | chr14 | + | 101827473 | 101827819 | 101809904 | 101810036 | 101856685 | 101856885 | 185080 | 3,3,2,7,2,0,2,0,0,0,0,2 | 112,64,188,136,91,76,109, | 7,6,0,8,3,0,2,6,3,1,3,10,4,3 | 115,112,82,40,129,71,50,8 | 298 | 149 | 0.000949285159514 | 0.105863002159253 | 0.013,0.023,0.005, | 0.03,0.026,0.0,0.0 | -0,017 |
| 76315 | ENSG00000076242 | MLH1 | chr3 | + | 37025636 | 37026007 | 37012010 | 37012099 | 37028783 | 37028847 | 76315 | 36,19,37,30,36,30,24,27,2 | 2,4,3,2,0,1,3,5,0,0,4,0 | 36,36,19,19,30,12,32,26,1 | 0,1,1,0,0,1,1,1,3,0,5,0,0,0, | 298 | 149 | 0.00095128073408 | 0.106022307300906 | 0.9,0.704,0.86,0.8 | 1.0,0.947,0.905,1. | -0,051 |
| 206313 | ENSG00000095002 | MSH2 | chr2 | + | 47482778 | 47482972 | 47480695 | 47480871 | 47512302 | 47513234 | 206313 | 13,3,50,38,20,26,6,20,9,4, | 2,0,6,2,2,2,0,0,0,6,0,1 | 47,28,8,3,3,15,22,41,5,26, | 0,0,0,2,0,2,0,2,3,1,0,2,0,0, | 298 | 149 | 0.000951698861713 | 0.106022307300906 | 0.765,1.0,0.806,0. | 1.0,1.0,1.0,0.429,1 | -0,017 |
| 78089 | ENSG00000152061 | RABGAP1L | chr1 | + | 174957890 | 174958173 | 174957456 | 174957549 | 174969276 | 174969387 | 78089 | 78,90,126,121,28,43,28,44 | 15,2,41,36,13,11,9,13,16,7 | 35,52,79,27,47,18,24,42,2 | 16,25,27,30,21,11,11,33,8, | 298 | 149 | 0.000955528538853 | 0.10639387632999 | 0.722,0.957,0.606, | 0.522,0.51,0.594,0 | 0,138 |
| 85486 | ENSG00000143436 | MRPL9 | chr1 | - | 151762989 | 151763146 | 151762317 | 151762500 | 151763326 | 151763334 | 85486 | 31,35,63,34,35,26,20,27,2 | 3,4,1,9,2,3,6,5,0,0,0,1 | 19,41,32,11,28,12,18,23,1 | 2,0,1,0,2,3,3,2,0,1,2,3,0,0, | 298 | 149 | 0.000956098109793 | 0.106402250397548 | 0.838,0.814,0.969, | 0.826,1.0,0.941,1. | -0,04 |
| 58680 | ENSG00000114098 | ARMC8 | chr3 | + | 138237308 | 138237384 | 138235033 | 138235114 | 138239467 | 138239528 | 58680 | 19,10,15,10,9,7,8,11,4,2,9, | 0,0,3,0,3,0,0,0,2,0,0,0 | 7,20,7,3,8,5,5,12,2,11,10,7 | 1,2,2,4,0,1,2,0,2,2,2,0,2,0, | 225 | 149 | 0.000958235176584 | 0.106584969142817 | 1.0,1.0,0.768,1.0,0 | 0.823,0.869,0.699, | 0,136 |
| 144081 | ENSG00000007202 | KIAA0100 | chr17 | - | 28643163 | 28643309 | 28642868 | 28642987 | 28643609 | 28643659 | 144081 | 26,7,67,67,30,34,14,37,22, | 2,1,4,3,1,2,2,5,0,2,0,0 | 40,28,20,17,10,16,12,25,1 | 6,4,1,3,6,1,0,3,2,0,2,4,6,2, | 295 | 149 | 0.000959114918815 | 0.106627718538983 | 0.868,0.78,0.894,0 | 0.771,0.78,0.91,0. | 0,077 |
| 214628 | ENSG00000170919 | TPT1-AS1 | chr13 | + | 45390575 | 45390902 | 45389734 | 45389820 | 45391031 | 45391619 | 214628 | 22,17,14,12,7,8,9,8,16,8,2, | 7,0,0,0,0,0,4,3,0,3,0,1 | 10,11,12,7,14,6,9,5,16,20, | 0,0,0,0,0,0,1,0,0,0,1,0,0,0, | 298 | 149 | 0.0009614390641 | 0.10683091956908 | 0.611,1.0,1.0,1.0,1 | 1.0,1.0,1.0,1.0,1.0, | -0,125 |
| 250484 | ENSG00000078403 | MLLT10 | chr10 | + | 21673585 | 21673919 | 21670448 | 21670704 | 21681331 | 21681376 | 250484 | 15,3,19,13,13,20,10,9,7,3, | 0,0,4,2,2,1,0,2,0,1,0,1 | 25,11,6,12,5,8,6,14,9,13,9, | 2,2,0,1,1,0,0,2,2,0,3,4,1,1, | 298 | 149 | 0.000963259531255 | 0.106977973256731 | 1.0,1.0,0.704,0.76 | 0.862,0.733,1.0,0. | 0,029 |
| 167343 | ENSG00000058063 | ATP11B | chr3 | + | 182829922 | 182830013 | 182829671 | 182829752 | 182836034 | 182836142 | 167343 | 2,5,17,5,0,7,2,5,3,4,1,2 | 149,67,137,101,42,46,44,7 | 7,20,6,19,11,0,5,7,6,0,10,7 | 71,90,61,45,87,47,14,41,5 | 240 | 149 | 0.000964046548796 | 0.107010161291342 | 0.008,0.044,0.072, | 0.058,0.121,0.058, | -0,033 |
| 107938 | ENSG00000215021 | PHB2 | chr12 | - | 6968410 | 6968708 | 6967891 | 6968021 | 6969497 | 6969577 | 107938 | 170,115,270,200,98,78,94, | 1,2,11,12,3,5,3,2,0,2,0,2 | 134,117,134,62,91,58,71,1 | 1,5,0,1,0,0,1,5,2,1,5,5,0,4, | 298 | 149 | 0.000964863155044 | 0.107045598826431 | 0.988,0.966,0.925, | 0.985,0.921,1.0,0. | -0,017 |
| 206693 | ENSG00000152061 | RABGAP1L | chr1 | + | 174957890 | 174958068 | 174957456 | 174957549 | 174969276 | 174969387 | 206693 | 77,90,123,119,21,43,26,44 | 15,2,41,36,13,11,9,13,16,7 | 35,51,78,26,47,18,18,39,2 | 16,25,27,30,21,11,11,33,8, | 298 | 149 | 0.000967476210441 | 0.107280202086258 | 0.72,0.957,0.6,0.6 | 0.522,0.505,0.591, | 0,143 |
| 32669 | ENSG00000214655 | ZSWIM8 | chr10 | + | 73789366 | 73789539 | 73789095 | 73789190 | 73789716 | 73789824 | 32669 | 34,13,57,46,19,40,26,31,2 | 1,0,1,0,0,0,1,1,0,1,1,3 | 27,47,40,25,27,18,31,25,2 | 0,4,2,1,2,3,1,2,0,0,2,4,2,1, | 298 | 149 | 0.000968269403084 | 0.107307741966181 | 0.944,1.0,0.966,1. | 1.0,0.855,0.909,0. | 0,045 |
| 33104 | ENSG00000281649 | EBLN3P | chr9 | + | 37084198 | 37084292 | 37079916 | 37080030 | 37086667 | 37087837 | 33104 | 22,10,41,14,9,4,8,12,13,2, | 11,8,16,20,2,4,5,3,6,2,2,7 | 2,7,6,1,5,1,5,4,5,6,11,8,12, | 8,23,7,5,11,12,5,10,6,9,9,4 | 243 | 149 | 0.000968721711279 | 0.107307741966181 | 0.551,0.434,0.611, | 0.133,0.157,0.345, | 0,169 |
| 234039 | ENSG00000215021 | PHB2 | chr12 | - | 6968410 | 6968514 | 6966925 | 6968021 | 6969497 | 6969577 | 234039 | 143,101,238,169,76,61,84, | 1,2,11,12,3,5,3,2,0,2,0,2 | 115,95,120,60,76,55,54,82 | 1,5,0,1,0,0,1,5,2,1,5,5,0,4, | 253 | 149 | 0.000972838787469 | 0.107652038006864 | 0.988,0.967,0.927, | 0.985,0.918,1.0,0. | -0,017 |
| 126384 | ENSG00000148688 | RPP30 | chr10 | + | 90895453 | 90895483 | 90894772 | 90894891 | 90895879 | 90895917 | 126384 | 2,0,11,6,11,6,3,2,2,4,0,5 | 2,1,10,4,2,2,1,5,2,0,2,0 | 6,0,1,1,0,2,4,6,0,3,4,0,4,0, | 5,1,1,3,2,5,2,8,6,7,6,6,2,6, | 179 | 149 | 0.00097313106178 | 0.107652038006864 | 0.454,0.0,0.478,0. | 0.5,0.0,0.454,0.21 | 0,25 |
| 186729 | ENSG00000134242 | PTPN22 | chr1 | - | 113851966 | 113852104 | 113848539 | 113848626 | 113854470 | 113854537 | 186729 | 104,67,260,258,44,48,47,4 | 5,2,18,14,1,3,3,5,4,3,4,2 | 165,127,54,19,72,49,28,42 | 13,5,10,2,15,3,3,6,7,3,5,20 | 287 | 149 | 0.000973330356507 | 0.107652038006864 | 0.915,0.946,0.882, | 0.868,0.93,0.737,0 | 0,068 |
| 51138 | ENSG00000111653 | ING4 | chr12 | - | 6655637 | 6655913 | 6653229 | 6653396 | 6656726 | 6656798 | 51138 | 7,3,5,4,4,1,2,8,14,5,0,0 | 42,32,60,73,29,23,37,32,2 | 1,5,1,0,3,3,0,2,2,2,0,3,9,6, | 43,23,36,14,15,34,10,29,11 | 298 | 149 | 0.000974541826884 | 0.10768780893001 | 0.077,0.045,0.04,0 | 0.011,0.098,0.014, | 0,034 |
| 190921 | ENSG00000048740 | CELF2 | chr10 | + | 11228387 | 11228461 | 11217424 | 11217507 | 11249152 | 11249201 | 190921 | 1,1,3,5,1,1,2,8,1,0,0,6 | 155,67,202,131,116,111,9 | 4,20,6,4,2,1,3,5,5,1,2,10,0, | 165,131,91,80,65,45,49,11 | 223 | 149 | 0.000975057804666 | 0.10768780893001 | 0.004,0.01,0.01,0. | 0.016,0.093,0.042, | -0,014 |
| 16046 | ENSG00000090857 | PDPR | chr16 | + | 70131301 | 70131419 | 70130422 | 70130544 | 70132150 | 70132300 | 16046 | 40,17,55,51,22,23,21,35,2 | 0,1,8,3,3,0,4,2,0,2,0,0 | 32,30,16,17,19,18,11,14,2 | 2,0,3,1,6,3,3,1,2,0,2,8,17,1 | 267 | 149 | 0.000975154785345 | 0.10768780893001 | 1.0,0.905,0.793,0. | 0.899,1.0,0.749,0. | 0,112 |
| 55973 | ENSG00000113810 | SMC4 | chr3 | + | 160402675 | 160402867 | 160401914 | 160402093 | 160404327 | 160404336 | 55973 | 22,26,34,48,12,17,14,22,8, | 1,0,0,3,0,1,1,0,0,0,0,0 | 25,34,7,18,25,18,21,53,15, | 3,0,3,1,0,1,0,5,0,3,0,6,2,0, | 298 | 149 | 0.000976312836147 | 0.107760404121413 | 0.917,1.0,1.0,0.88 | 0.806,1.0,0.538,0. | 0,081 |
| 227059 | ENSG00000146833 | TRIM4 | chr7 | - | 99917767 | 99917885 | 99909564 | 99909660 | 99919008 | 99919531 | 227059 | 0,0,4,1,2,1,3,2,3,0,0,2 | 10,4,33,11,11,10,11,10,18, | 3,7,0,7,2,0,0,4,2,0,7,3,0,2, | 10,15,8,10,12,4,4,3,8,16,1 | 267 | 149 | 0.000977096111555 | 0.107791580310658 | 0.0,0.0,0.063,0.04 | 0.143,0.207,0.0,0. | -0,061 |
| 99019 | ENSG00000136280 | CCM2 | chr7 | + | 45038252 | 45038426 | 45027025 | 45027797 | 45069825 | 45069961 | 99019 | 28,14,35,13,21,7,6,19,18,1 | 3,2,1,0,0,0,6,0,0,4,1,5 | 37,12,16,2,7,20,4,9,12,49, | 2,4,0,2,5,3,0,0,0,1,0,0,0,1, | 298 | 149 | 0.000978158811372 | 0.107853534390577 | 0.824,0.778,0.946, | 0.902,0.6,1.0,0.33 | -0,082 |
| 76056 | ENSG00000135541 | AHI1 | chr6 | - | 135467580 | 135467634 | 135463124 | 135466373 | 135490622 | 135490747 | 76056 | 36,18,87,42,35,42,24,30,1 | 2,0,8,1,1,5,1,2,0,1,0,0 | 15,27,24,41,32,13,30,21,5, | 0,0,1,1,1,0,2,0,0,0,2,0,0,0, | 203 | 149 | 0.000980793395025 | 0.108088654994688 | 0.93,1.0,0.889,0.9 | 1.0,1.0,0.946,0.96 | -0,032 |
| 211714 | ENSG00000163516 | ANKZF1 | chr2 | + | 219235473 | 219235585 | 219234825 | 219235293 | 219235707 | 219235875 | 211714 | 29,5,19,18,16,10,11,5,37,2 | 3,3,5,0,2,1,1,0,0,0,0,0 | 11,3,9,1,13,5,4,11,7,21,2,7 | 2,2,0,2,1,1,4,3,0,1,1,1,1,0, | 261 | 149 | 0.000985719031791 | 0.108575891981274 | 0.847,0.488,0.684, | 0.758,0.461,1.0,0. | 0,152 |
| 214321 | ENSG00000129472 | RAB2B | chr14 | - | 21466651 | 21466772 | 21463655 | 21463767 | 21468356 | 21468449 | 214321 | 4,2,3,6,1,3,0,4,0,0,2,4 | 42,17,45,43,26,11,8,3,31,1 | 2,0,0,6,2,0,0,0,1,2,3,0,0,0, | 25,16,13,14,26,16,18,25,1 | 270 | 149 | 0.000989459363037 | 0.108932137169216 | 0.05,0.061,0.035,0 | 0.042,0.0,0.0,0.19 | 0,056 |
| 189261 | ENSG00000118308 | IRAG2 | chr12 | + | 25076522 | 25076640 | 25066364 | 25066512 | 25079243 | 25079290 | 189261 | 65,27,87,32,12,26,32,43,1 | 3,7,22,7,3,11,12,15,4,4,0,0 | 49,85,54,46,18,3,12,17,26, | 8,8,3,2,7,2,2,5,0,0,2,0,6,7, | 267 | 149 | 0.000991458764817 | 0.109096452663767 | 0.924,0.683,0.688, | 0.774,0.856,0.909, | -0,091 |
| 63027 | ENSG00000173039 | RELA | chr11 | - | 65658717 | 65658822 | 65655854 | 65655935 | 65659665 | 65659788 | 63027 | 31,29,62,51,18,34,27,36,2 | 6,1,5,0,0,1,0,4,3,2,0,5 | 34,33,14,38,26,16,21,44,2 | 0,0,1,8,3,0,2,1,0,0,1,4,0,2, | 254 | 149 | 0.000992462433295 | 0.10915108941263 | 0.752,0.944,0.879, | 1.0,1.0,0.891,0.73 | -0,048 |
| 32620 | ENSG00000158290 | CUL4B | chrX | - | 120545443 | 120545517 | 120544480 | 120544643 | 120546546 | 120546616 | 32620 | 73,30,138,85,55,56,25,61, | 0,0,2,0,0,1,0,2,1,0,0,3 | 88,71,47,20,42,44,32,32,2 | 0,3,4,1,5,1,1,0,0,2,2,5,3,1, | 223 | 149 | 0.000993485458392 | 0.109207798107849 | 1.0,1.0,0.979,1.0,1 | 1.0,0.941,0.887,0. | 0,034 |
| 165059 | ENSG00000123933 | MXD4 | chr4 | - | 2255158 | 2255294 | 2252407 | 2252522 | 2257981 | 2258011 | 165059 | 8,2,3,6,1,6,3,3,10,4,0,0 | 8,12,25,24,15,6,13,11,23,1 | 1,0,3,8,0,0,1,2,1,0,5,6,1,3, | 16,29,16,7,13,3,2,8,15,20, | 285 | 149 | 0.001002509881 | 0.110143544766468 | 0.343,0.08,0.059,0 | 0.032,0.0,0.089,0. | 0,042 |
| 248442 | ENSG00000139697 | SBNO1 | chr12 | - | 123348034 | 123348133 | 123345257 | 123345570 | 123350309 | 123350441 | 248442 | 38,28,37,32,25,23,29,46,1 | 0,0,0,2,2,0,0,0,0,2,2,2 | 49,58,27,27,32,16,14,26,2 | 1,2,3,2,1,1,0,2,1,5,0,2,5,1, | 248 | 149 | 0.00100501468931 | 0.110362406425959 | 1.0,1.0,1.0,0.906,0 | 0.967,0.946,0.844, | 0,018 |
| 108537 | ENSG00000179195 | ZNF664 | chr12 | + | 123988045 | 123988138 | 123973885 | 123974020 | 124011380 | 124011441 | 108537 | 28,9,21,14,19,16,13,13,12, | 1,3,6,6,1,2,0,0,1,0,0,0 | 22,8,14,16,7,6,11,23,7,11, | 0,0,0,0,2,2,0,0,0,0,6,0,0,1, | 242 | 149 | 0.00100614733212 | 0.110406079149659 | 0.945,0.649,0.683, | 1.0,1.0,1.0,1.0,0.6 | -0,047 |
| 13249 | ENSG00000167978 | SRRM2 | chr16 | + | 2756333 | 2756606 | 2752637 | 2752846 | 2757471 | 2757574 | 13249 | 107,64,115,54,48,73,77,43 | 0,3,0,0,0,1,0,1,0,0,0,1 | 59,74,42,175,49,38,66,46, | 2,4,0,1,2,2,1,0,0,3,0,2,4,3, | 298 | 149 | 0.00100734098917 | 0.110406079149659 | 1.0,0.914,1.0,1.0,1 | 0.937,0.902,1.0,0. | 0,042 |
| 174048 | ENSG00000134109 | EDEM1 | chr3 | + | 5211119 | 5211199 | 5208092 | 5208263 | 5213318 | 5213522 | 174048 | 13,11,25,21,6,5,5,10,9,8,1, | 0,2,2,2,0,3,0,0,0,0,0,0 | 10,10,3,21,13,3,3,4,7,6,12, | 0,3,2,6,1,1,2,3,0,0,0,2,0,0, | 229 | 149 | 0.00100736647689 | 0.110406079149659 | 1.0,0.782,0.891,0. | 1.0,0.684,0.494,0. | 0,204 |
| 236051 | ENSG00000102096 | PIM2 | chrX | - | 48914394 | 48914564 | 48913181 | 48914294 | 48915019 | 48915735 | 236051 | 330,719,481,393,272,218, | 92,29,75,69,60,44,61,34,6 | 286,527,357,343,287,144, | 89,51,115,59,45,29,19,92, | 298 | 149 | 0.00100798992533 | 0.110406079149659 | 0.642,0.925,0.762, | 0.616,0.838,0.608, | 0,061 |
| 200145 | ENSG00000131051 | RBM39 | chr20 | - | 35738969 | 35739017 | 35732059 | 35732135 | 35740823 | 35740837 | 200145 | 52,58,42,23,7,10,8,11,25,1 | 3,4,1,0,0,0,0,0,0,0,0,4 | 6,29,15,30,23,17,5,13,6,22 | 3,1,4,5,1,0,0,0,1,0,9,4,0,1, | 197 | 149 | 0.00100831292674 | 0.110406079149659 | 0.929,0.916,0.969, | 0.602,0.956,0.739, | 0,056 |
| 15037 | ENSG00000162066 | AMDHD2 | chr16 | + | 2527560 | 2527615 | 2520768 | 2521123 | 2527772 | 2527985 | 15037 | 12,25,24,15,30,13,17,15,1 | 0,1,0,0,1,3,0,4,0,0,0,0 | 17,23,8,7,7,3,14,24,6,11,2 | 2,4,2,3,0,0,2,2,2,1,0,2,3,2, | 204 | 149 | 0.00100849018779 | 0.110406079149659 | 1.0,0.948,1.0,1.0,0 | 0.861,0.808,0.745, | 0,088 |
| 55194 | ENSG00000142765 | SYTL1 | chr1 | + | 27349971 | 27350132 | 27349475 | 27349765 | 27350388 | 27350485 | 55194 | 13,16,19,13,14,11,8,10,8,4 | 0,0,0,4,4,1,0,2,0,0,0,0 | 13,22,13,3,6,10,7,6,8,16,1 | 3,1,0,0,1,0,0,2,5,2,2,4,3,0, | 298 | 149 | 0.00101035735679 | 0.110554257376344 | 1.0,1.0,1.0,0.619,0 | 0.684,0.917,1.0,1. | 0,066 |
| 211225 | ENSG00000197622 | CDC42SE1 | chr1 | - | 151056650 | 151056823 | 151055676 | 151055993 | 151059478 | 151059773 | 211225 | 44,4,37,27,21,30,21,29,31, | 197,165,295,254,162,112,1 | 37,29,36,36,22,32,26,26,1 | 274,239,156,130,165,98,1 | 298 | 149 | 0.00101375569232 | 0.110845190016368 | 0.1,0.012,0.059,0. | 0.063,0.057,0.103, | -0,021 |
| 22874 | ENSG00000168411 | RFWD3 | chr16 | - | 74636345 | 74636577 | 74632522 | 74632673 | 74644361 | 74644453 | 22874 | 24,31,27,32,15,19,15,6,29, | 2,0,7,1,0,1,3,0,0,0,0,2 | 11,22,3,9,10,11,14,28,7,20 | 2,6,5,2,2,3,0,2,0,1,6,2,1,6, | 298 | 149 | 0.00101404620683 | 0.110845190016368 | 0.857,1.0,0.659,0. | 0.733,0.647,0.231, | 0,151 |
| 200275 | ENSG00000112249 | ASCC3 | chr6 | - | 100847625 | 100847733 | 100805759 | 100805880 | 100848147 | 100848707 | 200275 | 3,0,8,4,2,2,0,4,0,3,0,2 | 28,23,66,48,32,34,20,26,1 | 2,0,0,0,0,1,0,0,0,0,0,0,1,0, | 56,56,18,19,32,15,26,18,2 | 257 | 149 | 0.00101504246633 | 0.110897769071407 | 0.058,0.0,0.066,0. | 0.02,0.0,0.0,0.0,0. | 0,047 |
| 106487 | ENSG00000109111 | SUPT6H | chr17 | + | 28690920 | 28691063 | 28690081 | 28690229 | 28693695 | 28693836 | 106487 | 77,24,120,83,54,66,38,52, | 1,0,0,2,0,0,0,1,0,0,0,1 | 78,145,30,72,74,48,56,57, | 1,6,0,7,1,0,0,0,2,0,5,2,1,0, | 292 | 149 | 0.00101703591317 | 0.111059186518261 | 0.975,1.0,1.0,0.95 | 0.975,0.925,1.0,0. | 0,039 |
| 161591 | ENSG00000154822 | PLCL2 | chr3 | + | 17052726 | 17052791 | 17042873 | 17042949 | 17067955 | 17068065 | 161591 | 9,1,16,2,5,5,3,11,8,0,3,2 | 119,77,255,126,56,71,65,1 | 24,32,6,5,4,3,3,6,3,5,23,6, | 175,124,67,36,73,64,43,47 | 214 | 149 | 0.00101895800319 | 0.111212652122001 | 0.05,0.009,0.042,0 | 0.087,0.152,0.059, | -0,018 |
| 236748 | ENSG00000067167 | TRAM1 | chr8 | - | 70598133 | 70598252 | 70597890 | 70598011 | 70600018 | 70600082 | 236748 | 229,127,290,192,98,76,68, | 1,0,1,1,1,1,1,0,0,0,0,0 | 198,161,129,87,102,79,46, | 2,2,0,0,1,0,2,1,1,6,0,14,2,4 | 268 | 149 | 0.001020669943 | 0.11130428665737 | 0.992,1.0,0.994,0. | 0.982,0.978,1.0,1. | 0,023 |
| 83729 | ENSG00000079263 | SP140 | chr2 | + | 230255451 | 230255532 | 230248884 | 230248968 | 230269531 | 230269618 | 83729 | 132,98,196,126,57,42,47,5 | 0,0,5,2,1,0,0,0,0,2,0,3 | 91,87,58,61,105,56,18,44, | 2,1,0,1,2,0,0,1,2,4,4,4,3,6, | 230 | 149 | 0.00102083185908 | 0.11130428665737 | 1.0,1.0,0.962,0.97 | 0.967,0.983,1.0,0. | 0,026 |
| 47523 | ENSG00000160991 | ORAI2 | chr7 | + | 102436201 | 102436333 | 102433529 | 102433661 | 102438943 | 102439181 | 47523 | 37,50,115,114,118,116,13 | 6,2,8,5,4,4,11,30,2,4,7,13 | 75,48,41,43,52,40,49,113, | 5,7,4,9,6,0,1,16,6,7,10,55, | 281 | 149 | 0.00102228830297 | 0.111372990190212 | 0.766,0.93,0.884,0 | 0.888,0.784,0.845, | 0,05 |
| 71922 | ENSG00000161929 | SCIMP | chr17 | - | 5223332 | 5223456 | 5221286 | 5221350 | 5234734 | 5234860 | 71922 | 212,42,191,202,79,67,100, | 3,0,3,13,1,4,0,1,9,3,6,5 | 123,86,64,39,42,86,37,83, | 8,2,10,2,2,4,1,9,6,3,2,3,8,1 | 273 | 149 | 0.00102249689225 | 0.111372990190212 | 0.975,1.0,0.972,0. | 0.894,0.959,0.777, | 0,041 |
| 125227 | ENSG00000170759 | KIF5B | chr10 | - | 32038777 | 32038826 | 32038162 | 32038218 | 32039326 | 32039431 | 125227 | 57,43,113,77,41,50,39,65, | 4,1,8,5,3,1,2,3,0,1,2,0 | 75,101,58,42,43,25,34,60, | 10,11,10,14,1,4,2,1,1,4,2,3 | 198 | 149 | 0.00102455143162 | 0.111496299014528 | 0.915,0.97,0.914,0 | 0.849,0.874,0.814, | 0,037 |
| 26097 | ENSG00000066933 | MYO9A | chr15 | - | 72019038 | 72019095 | 72010349 | 72010447 | 72020917 | 72021017 | 26097 | 36,25,52,20,22,31,47,42,2 | 5,0,6,3,3,4,3,4,6,2,2,4 | 31,39,22,24,16,18,23,44,1 | 7,0,1,0,0,2,0,0,2,1,3,3,0,2, | 206 | 149 | 0.00102466503176 | 0.111496299014528 | 0.839,1.0,0.862,0. | 0.762,1.0,0.941,1. | -0,072 |
| 104729 | ENSG00000214783 | POLR2J4 | chr7 | - | 44014605 | 44014783 | 44013592 | 44013679 | 44016433 | 44016523 | 104729 | 48,31,44,41,33,30,14,46,5 | 1,2,0,3,6,0,3,6,0,0,0,2 | 30,23,12,21,19,10,17,25,1 | 3,2,1,3,0,3,0,4,0,1,1,0,0,0, | 298 | 149 | 0.00102806562398 | 0.111809799047418 | 0.96,0.886,1.0,0.8 | 0.833,0.852,0.857, | 0,017 |
| 33550 | ENSG00000170242 | USP47 | chr11 | + | 11891967 | 11892106 | 11884466 | 11884580 | 11897596 | 11897693 | 33550 | 89,48,264,168,101,110,84, | 1,0,2,2,0,0,2,4,0,0,0,1 | 141,211,68,66,76,41,55,10 | 4,5,0,1,3,0,0,2,4,0,3,1,5,0, | 288 | 149 | 0.00103035097387 | 0.112001752756068 | 0.979,1.0,0.986,0. | 0.948,0.956,1.0,0. | 0,028 |
| 55911 | ENSG00000150753 | CCT5 | chr5 | + | 10254673 | 10254838 | 10254144 | 10254205 | 10255954 | 10256153 | 55911 | 116,74,170,187,125,73,45, | 3,2,8,9,2,1,2,2,0,2,0,4 | 133,111,81,19,81,54,22,34 | 8,8,1,0,10,1,5,5,3,0,8,10,2, | 298 | 149 | 0.00103112398438 | 0.112029200546235 | 0.951,0.949,0.914, | 0.893,0.874,0.976, | 0,069 |
| 75259 | ENSG00000146574 | CCZ1B | chr7 | - | 6819765 | 6819941 | 6814643 | 6814845 | 6822280 | 6822364 | 75259 | 25,29,69,30,30,36,13,32,4, | 0,0,0,0,0,1,0,0,0,0,0,0 | 39,31,46,31,36,32,9,45,17, | 0,0,3,4,7,0,0,1,0,3,0,4,0,3,0 | 298 | 149 | 0.00103226171098 | 0.112085110123689 | 1.0,1.0,1.0,1.0,1.0, | 1.0,1.0,0.885,0.79 | 0,066 |
| 198504 | ENSG00000286192 | | chr7 | - | 2225409 | 2225742 | 2222574 | 2222754 | 2229983 | 2230143 | 198504 | 27,12,30,4,13,7,12,17,10,1 | 1,1,1,0,2,4,1,0,0,2,1,2 | 14,9,2,3,8,3,5,9,16,12,15,3 | 3,3,0,0,1,0,2,1,0,0,0,6,1,0, | 298 | 149 | 0.00103268011288 | 0.112085110123689 | 0.931,0.857,0.938, | 0.7,0.6,1.0,1.0,0.8, | -0,013 |
| 67025 | ENSG00000048707 | VPS13D | chr1 | + | 12319496 | 12319630 | 12318071 | 12318337 | 12321808 | 12321964 | 67025 | 64,29,104,62,53,50,37,47, | 0,0,1,1,0,4,0,0,0,0,1,0 | 73,68,22,36,21,33,37,41,6 | 0,3,1,9,7,3,2,2,2,0,0,0,0,0, | 283 | 149 | 0.00103375574159 | 0.112145303436571 | 1.0,1.0,0.982,0.97, | 1.0,0.923,0.921,0. | 0,059 |
| 222779 | ENSG00000265808 | SEC22B | chr1 | - | 120168839 | 120168990 | 120163209 | 120163370 | 120176306 | 120176520 | 222779 | 52,49,89,81,20,27,26,42,4 | 0,3,1,5,3,1,1,1,1,3,0,1 | 52,59,33,37,24,23,20,42,1 | 1,2,0,1,0,1,0,0,0,0,1,3,2,0, | 298 | 149 | 0.00104319149399 | 0.11311191357328 | 1.0,0.891,0.978,0. | 0.963,0.937,1.0,0. | -0,043 |
| 34658 | ENSG00000047188 | YTHDC2 | chr5 | + | 113526585 | 113526785 | 113515271 | 113515362 | 113532878 | 113533045 | 34658 | 44,26,58,41,18,23,9,18,19, | 31,21,15,11,13,15,5,8,6,1, | 14,19,22,24,29,7,10,10,10, | 26,32,12,11,8,12,14,13,18, | 298 | 149 | 0.00104536210273 | 0.113226407027844 | 0.415,0.382,0.659, | 0.212,0.229,0.478, | 0,177 |
| 94117 | ENSG00000112182 | BACH2 | chr6 | - | 90271848 | 90271941 | 90206568 | 90206676 | 90296479 | 90296742 | 94117 | 42,22,59,12,5,27,10,20,12, | 0,4,3,2,3,2,1,0,2,0,1,6 | 24,16,29,27,5,11,17,7,1,7, | 1,0,0,3,1,0,0,2,0,0,2,0,10,2 | 242 | 149 | 0.00104581711609 | 0.113226407027844 | 1.0,0.772,0.924,0. | 0.937,1.0,1.0,0.84 | -0,087 |
| 55008 | ENSG00000082701 | GSK3B | chr3 | - | 119912703 | 119912810 | 119905463 | 119905852 | 119916043 | 119916174 | 55008 | 47,44,58,39,29,26,17,28,2 | 1,0,2,1,3,6,3,3,0,0,0,4 | 36,34,12,20,18,24,27,36,1 | 2,7,4,3,7,4,3,2,3,1,7,11,3,1 | 256 | 149 | 0.00104582563465 | 0.113226407027844 | 0.965,1.0,0.944,0. | 0.913,0.739,0.636, | 0,096 |
| 2797 | ENSG00000130816 | DNMT1 | chr19 | - | 10180186 | 10180234 | 10177291 | 10177367 | 10180349 | 10180381 | 2797 | 4,2,9,12,3,6,3,8,4,3,0,9 | 16,14,32,18,26,10,13,29,9, | 12,5,4,6,7,6,1,12,2,5,9,13, | 10,9,4,13,5,7,12,36,2,6,11, | 197 | 149 | 0.0010521222237 | 0.113850838777866 | 0.159,0.098,0.175, | 0.476,0.296,0.431, | -0,141 |
| 131354 | ENSG00000004700 | RECQL | chr12 | - | 21501169 | 21501215 | 21499554 | 21499615 | 21501412 | 21501551 | 131354 | 21,12,22,4,13,8,6,28,14,1, | 0,0,2,0,2,2,2,0,1,0,0,1 | 17,24,10,5,16,3,6,11,8,20,1 | 5,1,3,5,3,0,0,2,0,0,1,3,3,0, | 195 | 149 | 0.00105559107406 | 0.114168805256788 | 1.0,1.0,0.894,1.0,0 | 0.722,0.948,0.718, | 0,049 |
| 142343 | ENSG00000083454 | P2RX5 | chr17 | - | 3685203 | 3685580 | 3681895 | 3681978 | 3688011 | 3688105 | 142343 | 11,3,17,19,1,0,4,7,6,3,3,2 | 94,45,101,61,43,18,23,19, | 7,8,0,3,3,0,0,0,1,1,0,10,20, | 33,18,35,19,26,36,13,7,35, | 298 | 149 | 0.00105718349219 | 0.114283606332268 | 0.055,0.032,0.078, | 0.096,0.182,0.0,0. | 0,023 |
| 228973 | ENSG00000113643 | RARS1 | chr5 | + | 168494088 | 168494465 | 168493893 | 168494002 | 168494549 | 168495151 | 228973 | 1,3,9,3,7,1,3,9,1,0,3,0 | 28,16,101,62,25,21,9,13,1 | 2,9,9,7,5,3,8,11,5,2,9,8,4,6 | 52,34,16,19,19,17,14,34,16 | 298 | 149 | 0.0010637581301 | 0.114936609487727 | 0.018,0.086,0.043, | 0.019,0.117,0.22,0 | -0,039 |
| 105477 | ENSG00000165775 | FUNDC2 | chrX | + | 155046508 | 155046584 | 155033402 | 155033553 | 155051669 | 155051732 | 105477 | 81,52,110,98,57,45,61,43, | 6,1,0,0,0,2,0,1,0,0,0,0 | 69,69,43,40,35,18,24,52,2 | 4,0,0,2,3,2,3,0,1,2,4,4,2,1, | 225 | 149 | 0.00106661283393 | 0.11518722873035 | 0.899,0.972,1.0,1. | 0.92,1.0,1.0,0.93,0 | 0,042 |
| 185002 | ENSG00000169155 | ZBTB43 | chr9 | + | 126808882 | 126808915 | 126805039 | 126805132 | 126832486 | 126838210 | 185002 | 0,3,3,2,0,3,0,3,1,1,1,2 | 9,3,7,17,10,12,5,8,0,3,4,8 | 2,3,3,0,0,0,0,1,0,0,0,2,0,0, | 13,19,17,10,12,7,15,9,18,1 | 182 | 149 | 0.00106823963591 | 0.115261274189451 | 0.0,0.45,0.26,0.08 | 0.112,0.114,0.126, | 0,192 |
| 30913 | ENSG00000146067 | FAM193B | chr5 | - | 177523158 | 177523265 | 177521973 | 177522071 | 177523956 | 177524032 | 30913 | 5,1,1,6,3,1,2,4,9,2,0,0 | 43,16,59,27,23,36,30,33,2 | 3,7,3,5,4,7,1,4,1,3,3,8,8,10 | 39,26,27,26,11,7,42,47,17, | 256 | 149 | 0.00106836952859 | 0.115261274189451 | 0.063,0.035,0.01,0 | 0.043,0.135,0.061, | -0,055 |

| | | | | | | | | | | | | | | | | | | | | | | |
|---|---|---|---|---|---|---|---|---|---|---|---|---|---|---|---|---|---|---|---|---|---|---|
| 72080 | ENSG00000120733 | KDM3B | chr5 | + | 138379583 | 138379708 | 138377719 | 138377825 | 138381515 | 138381590 | 72080 | 104,115,146,104,46,35,43, | 8,5,2,0,0,3,4,4,0,4,2,3 | 102,72,59,75,60,55,32,41, | 4,13,0,17,10,0,7,0,3,3,6,3, | 274 | 149 | 0.00106932600867 | 0.115306666418864 | 0.876,0.926,0.975, | 0.933,0.751,1.0,0. | 0,038 |
| 77336 | ENSG00000128815 | WDFY4 | chr10 | + | 48777418 | 48777495 | 48776749 | 48776984 | 48778610 | 48778832 | 77336 | 98,68,219,205,152,144,91, | 9,2,11,3,1,4,10,11,1,4,2,7 | 213,241,37,119,53,59,116, | 15,27,5,17,6,0,13,16,12,0, | 226 | 149 | 0.00107147726643 | 0.115387139528838 | 0.878,0.957,0.929, | 0.903,0.855,0.83,0 | 0,037 |
| 163933 | ENSG00000223865 | HLA-DPB1 | chr6 | + | 33083513 | 33083994 | 33080671 | 33080935 | 33084949 | 33085231 | 163933 | 28,9,0,0,8,0,11,9,11,12,7,2 | 462,322,202,237,140,53,3 | 6,23,33,41,14,3,13,18,8,7, | 352,386,268,239,261,172,1 | 298 | 149 | 0.00107154758141 | 0.115387139528838 | 0.029,0.014,0.0,0. | 0.008,0.029,0.058, | -0,01 |
| 47940 | ENSG00000182362 | YBEY | chr21 | + | 46291333 | 46291462 | 46286869 | 46287123 | 46296161 | 46296230 | 47940 | 42,20,57,51,26,14,13,18,4 | 11,3,14,10,5,4,4,7,14,2,0,8 | 39,28,25,6,23,9,6,16,23,36 | 0,0,4,0,4,4,2,1,2,3,7,6,4,6, | 278 | 149 | 0.00107185285784 | 0.115387139528838 | 0.672,0.781,0.686, | 1.0,1.0,0.77,1.0,0. | -0,106 |
| 823 | ENSG00000290796 | | chr22 | - | 25357254 | 25357426 | 25354595 | 25354812 | 25375812 | 25376008 | 823 | 14,7,29,16,2,8,9,12,24,2,1 | 1,1,5,0,0,0,7,2,1,0,0,0 | 12,27,6,7,6,11,17,18,19,16 | 0,1,0,0,1,1,3,2,0,6,2,0,0,2, | 298 | 149 | 0.0010722167302 | 0.115387139528838 | 0.875,0.778,0.744, | 1.0,0.931,1.0,1.0,0 | 0,001 |
| 31049 | ENSG00000132952 | USPL1 | chr13 | + | 30653147 | 30653305 | 30646931 | 30647057 | 30657473 | 30660766 | 31049 | 56,66,120,55,46,33,21,32, | 2,3,4,11,4,3,3,8,1,0,3,3 | 69,82,52,48,53,17,41,38,1 | 13,4,9,16,5,5,4,2,3,8,14,6, | 298 | 149 | 0.00107412017702 | 0.115534212803694 | 0.933,0.917,0.938, | 0.726,0.911,0.743, | 0,109 |
| 76304 | ENSG00000076242 | MLH1 | chr3 | + | 37014431 | 37014544 | 37012010 | 37012099 | 37028783 | 37028847 | 76304 | 11,5,20,22,8,11,6,11,4,1,1, | 2,4,3,2,0,1,3,5,0,0,4,0 | 9,6,11,1,2,8,8,2,2,9,5,6,9,1 | 0,1,1,0,0,1,1,1,3,0,5,0,0,0, | 262 | 149 | 0.00107481107999 | 0.11555078099766 | 0.758,0.416,0.791, | 1.0,0.773,0.862,1. | -0,04 |
| 185351 | ENSG00000188822 | CNR2 | chr1 | - | 23883278 | 23883444 | 23870514 | 23875662 | 23913245 | 23913362 | 185351 | 11,6,12,2,1,0,5,7,3,2,5,1 | 72,36,191,152,96,74,66,90 | 0,3,2,2,0,4,2,4,2,8,3,3,0,3, | 135,99,48,64,73,71,22,84, | 298 | 149 | 0.00107682834848 | 0.115697032583993 | 0.071,0.077,0.03,0 | 0.0,0.015,0.02,0.0 | 0,016 |
| 72967 | ENSG00000176853 | FAM91A1 | chr8 | + | 123789612 | 123789745 | 123787663 | 123787750 | 123798089 | 123798238 | 72967 | 41,30,64,41,32,50,71,48,4 | 3,3,2,1,0,3,3,0,4,0,0,0 | 38,81,45,53,52,28,29,46,2 | 4,9,1,5,7,0,2,0,1,2,3,10,2,0 | 282 | 149 | 0.00107762989048 | 0.115697032583993 | 0.878,0.841,0.944, | 0.834,0.826,0.96,0 | 0,037 |
| 55073 | ENSG00000139631 | CSAD | chr12 | - | 53171176 | 53171441 | 53170422 | 53170502 | 53171881 | 53171988 | 55073 | 4,1,16,8,11,12,6,8,5,5,3,4 | 0,0,9,0,3,1,1,1,1,0,0,2 | 3,5,10,4,9,4,5,11,5,6,4,10, | 1,9,0,6,2,2,1,4,4,2,0,8,6,6, | 298 | 149 | 0.00107778410327 | 0.115697032583993 | 1.0,1.0,0.471,1.0,0 | 0.6,0.217,1.0,0.25, | 0,269 |
| 82286 | ENSG00000130347 | RTN4IP1 | chr6 | - | 106621424 | 106621493 | 106619201 | 106619326 | 106622817 | 106622969 | 82286 | 15,6,36,18,21,13,8,29,9,1, | 3,2,1,1,0,0,2,0,0,1,1,0 | 25,13,20,7,6,4,5,15,4,13,1 | 0,3,3,0,1,0,0,2,0,4,3,1,4,3, | 218 | 149 | 0.00107941728349 | 0.115811856983882 | 0.774,0.672,0.961, | 1.0,0.748,0.82,1.0, | 0,033 |
| 158810 | ENSG00000075568 | TMEM131 | chr2 | - | 97984455 | 97984536 | 97927425 | 97927487 | 97995475 | 97995948 | 158810 | 1,2,3,0,6,0,5,0,0,0,0,2 | 22,7,25,6,12,26,22,14,10,6 | 1,5,3,6,7,0,0,0,2,0,3,3,1,2, | 11,14,5,12,9,2,4,4,13,8,19, | 230 | 149 | 0.00107992992165 | 0.115811856983882 | 0.029,0.156,0.072, | 0.056,0.188,0.28,0 | -0,035 |
| 116063 | ENSG00000165209 | STRBP | chr9 | - | 123160366 | 123160462 | 123159095 | 123159207 | 123160976 | 123161068 | 116063 | 135,53,318,259,161,119,1 | 0,0,4,0,5,0,4,2,0,0,0,0 | 260,168,167,80,48,75,75,1 | 6,4,0,1,2,2,1,2,2,0,1,3,2,1, | 245 | 149 | 0.00108183999667 | 0.115958916495658 | 1.0,1.0,0.98,1.0,0. | 0.963,0.962,1.0,0. | 0,028 |
| 131030 | ENSG00000050426 | LETMD1 | chr12 | + | 51049033 | 51049185 | 51048300 | 51048478 | 51055834 | 51055848 | 131030 | 14,3,27,26,6,21,15,18,4,4, | 0,4,4,4,1,0,0,4,2,0,0,0 | 14,7,12,8,7,5,1,12,4,10,21, | 0,4,0,0,0,0,0,3,0,0,0,2,0,0, | 298 | 149 | 0.00108266911684 | 0.115990023238721 | 1.0,0.273,0.771,0. | 1.0,0.467,1.0,1.0,1 | -0,102 |
| 223733 | ENSG00000127947 | PTPN12 | chr7 | + | 77583573 | 77583650 | 77581426 | 77581503 | 77585542 | 77585571 | 223733 | 24,8,19,15,14,22,21,57,26, | 3,1,0,0,0,0,2,0,2,0,0,0 | 23,22,5,12,11,22,11,10,10, | 3,5,1,13,1,0,0,0,3,0,2,0,0,0 | 226 | 149 | 0.0010842474031 | 0.116101319809262 | 0.841,0.841,1.0,1. | 0.835,0.744,0.767, | 0,067 |
| 74201 | ENSG00000174796 | THAP6 | chr4 | + | 75521735 | 75521861 | 75516771 | 75516979 | 75539858 | 75539959 | 74201 | 23,9,55,25,27,16,18,35,12, | 3,2,1,3,0,2,5,0,0,0,2,2 | 33,24,16,13,36,20,12,24,2 | 1,4,0,0,1,0,1,0,1,1,1,2,1,1, | 275 | 149 | 0.00108526084372 | 0.116152052041123 | 0.806,0.709,0.968, | 0.947,0.765,1.0,1. | -0,093 |
| 245570 | ENSG00000109458 | GAB1 | chr4 | + | 143436717 | 143436772 | 143433490 | 143433716 | 143437998 | 143438600 | 245570 | 7,0,0,0,3,4,2,19,0,0,0,0 | 38,35,207,116,89,95,164,3 | 3,4,6,28,1,1,5,7,0,0,6,0,3,1 | 119,102,132,109,83,39,127 | 204 | 149 | 0.0010862688522 | 0.116169473985665 | 0.119,0.0,0.0,0.0,0 | 0.018,0.028,0.032, | -0,01 |
| 23688 | ENSG00000100519 | PSMC6 | chr14 | + | 52718228 | 52718352 | 52718080 | 52718142 | 52718976 | 52719038 | 23688 | 36,31,101,58,34,36,36,63, | 8,0,2,9,2,11,6,1,0,0,3,2 | 33,62,38,20,18,15,30,46,8, | 14,1,4,11,9,5,1,10,6,2,6,16 | 273 | 149 | 0.00108650311123 | 0.116169473985665 | 0.711,1.0,0.965,0. | 0.563,0.971,0.838, | 0,129 |
| 194786 | ENSG00000145246 | ATP10D | chr4 | + | 47545402 | 47545511 | 47536730 | 47536938 | 47546623 | 47547564 | 194786 | 1,2,11,9,0,2,2,2,1,0,0,0 | 29,14,76,35,21,10,5,5,16,1 | 10,2,7,6,2,1,3,2,2,4,3,0,3,7 | 32,20,10,6,4,14,10,6,4,10,1 | 258 | 149 | 0.00108929038991 | 0.116409662319126 | 0.02,0.076,0.077,0 | 0.153,0.055,0.288, | -0,078 |
| 210542 | ENSG00000132912 | DCTN4 | chr5 | - | 150755496 | 150755567 | 150753478 | 150753657 | 150756416 | 150756487 | 210542 | 6,0,3,5,7,0,3,5,2,0,0,2 | 25,38,40,15,13,12,12,11,1 | 0,7,0,1,5,0,0,1,1,0,7,0,2,2, | 21,22,14,43,26,9,12,16,16, | 220 | 149 | 0.00109054681144 | 0.116486094676448 | 0.14,0.0,0.048,0.1 | 0.0,0.177,0.0,0.01 | 0,044 |
| 241886 | ENSG00000108424 | KPNB1 | chr17 | + | 47669715 | 47669869 | 47668364 | 47668410 | 47670701 | 47670832 | 241886 | 35,29,135,115,49,49,51,73 | 10,5,25,25,10,6,5,2,1,7,6,4 | 96,102,28,37,42,34,42,103 | 16,26,14,18,15,4,7,23,13,5 | 298 | 149 | 0.00109128558384 | 0.116507186257672 | 0.636,0.744,0.73,0 | 0.75,0.662,0.5,0.5 | 0,113 |
| 129363 | ENSG00000012822 | CALCOCO1 | chr12 | - | 53712774 | 53712846 | 53711118 | 53712121 | 53713099 | 53713206 | 129363 | 7,3,12,8,2,7,2,4,14,2,0,2 | 93,37,81,65,28,31,28,37,7 | 3,4,5,0,4,3,2,4,4,3,0,3,1,3, | 39,56,44,30,39,34,13,59,21 | 221 | 149 | 0.00109692440052 | 0.117051133192028 | 0.048,0.052,0.091, | 0.049,0.046,0.071, | 0,015 |
| 235205 | ENSG00000139218 | SCAF11 | chr12 | - | 45922924 | 45923154 | 45922462 | 45922582 | 45924727 | 45924788 | 235205 | 21,36,61,35,16,9,13,15,24, | 0,0,3,6,1,2,3,4,1,0,0,3 | 33,30,14,22,14,12,20,13,1 | 9,7,5,6,11,4,0,2,2,5,0,5,5,4 | 298 | 149 | 0.00109769633565 | 0.11707546086139 | 1.0,1.0,0.91,0.745, | 0.647,0.682,0.583, | 0,117 |
| 54356 | ENSG00000138095 | LRPPRC | chr2 | - | 43890222 | 43890362 | 43889733 | 43889876 | 43894544 | 43894629 | 54356 | 17,2,19,15,18,7,8,14,5,0,3, | 35,22,68,48,54,34,23,42,3 | 26,50,9,26,19,9,7,15,13,8, | 74,67,47,25,29,16,24,43,2 | 289 | 149 | 0.00109900376326 | 0.117156849415658 | 0.2,0.045,0.126,0. | 0.153,0.278,0.09,0 | -0,074 |
| 176553 | ENSG00000114446 | IFT57 | chr3 | - | 108219353 | 108219572 | 108218534 | 108218653 | 108222110 | 108222360 | 176553 | 205,124,331,261,163,180, | 5,4,2,0,4,2,0,4,0,0,2,6 | 298,245,127,178,101,75,1 | 4,4,2,7,2,0,11,5,0,5,7,1,13, | 298 | 149 | 0.00110021955696 | 0.117228393794088 | 0.953,0.939,0.988, | 0.974,0.968,0.969, | 0,017 |
| 176550 | ENSG00000114446 | IFT57 | chr3 | - | 108219290 | 108219572 | 108218534 | 108218653 | 108222110 | 108222360 | 176550 | 204,124,331,262,162,181, | 5,4,2,0,4,2,0,4,0,0,2,6 | 298,245,127,178,101,75,1 | 4,4,2,7,2,0,11,5,0,5,7,1,13, | 298 | 149 | 0.00110086016181 | 0.117238611324358 | 0.953,0.939,0.988, | 0.974,0.968,0.969, | 0,017 |
| 98410 | ENSG00000118058 | KMT2A | chr11 | + | 118471661 | 118471985 | 118468774 | 118468844 | 118474178 | 118474315 | 98410 | 29,28,82,42,26,39,27,31,1 | 11,3,13,11,12,13,6,4,13,2,0, | 49,48,30,63,21,29,6,20,35, | 13,6,8,2,7,7,0,2,2,2,7,0,2,3 | 298 | 149 | 0.0011073871774 | 0.117875395439653 | 0.569,0.824,0.759, | 0.653,0.8,0.652,0. | -0,139 |
| 87152 | ENSG00000135976 | ANKRD36 | chr2 | + | 97209799 | 97209872 | 97209680 | 97209709 | 97211545 | 97211574 | 87152 | 46,18,36,12,10,22,19,25,1 | 13,15,27,19,8,14,11,13,16, | 18,19,24,45,20,16,9,16,9,2 | 3,3,6,14,7,8,6,5,7,11,7,5,1, | 222 | 149 | 0.00111058476445 | 0.118157325475698 | 0.704,0.446,0.472, | 0.801,0.81,0.729,0 | -0,224 |
| 14584 | ENSG00000168488 | ATXN2L | chr16 | + | 28826861 | 28826923 | 28826239 | 28826390 | 28829400 | 28829492 | 14584 | 31,27,19,25,19,17,9,14,14, | 2,2,8,5,0,1,6,3,0,4,0,3 | 16,14,13,30,5,14,13,9,15,1 | 10,7,1,15,6,3,5,6,6,2,2,9,1, | 211 | 149 | 0.00111182219674 | 0.118195264349863 | 0.916,0.905,0.626, | 0.53,0.585,0.902,0 | 0,15 |
| 154745 | ENSG00000105122 | RASAL3 | chr19 | - | 15452044 | 15452241 | 15451623 | 15451938 | 15452657 | 15452815 | 154745 | 19,22,23,24,5,20,14,21,24, | 0,2,1,1,0,3,2,2,4,3,2,0 | 12,17,7,16,6,9,2,26,15,30, | 0,0,3,2,0,0,0,0,0,0,1,2,0, | 298 | 149 | 0.00111203967044 | 0.118195264349863 | 1.0,0.846,0.92,0.9 | 1.0,1.0,0.538,0.8,1 | -0,063 |
| 236476 | ENSG00000122882 | ECD | chr10 | - | 73154255 | 73154440 | 73152292 | 73152421 | 73156274 | 73156453 | 236476 | 24,22,49,29,18,31,25,36,1 | 1,0,5,4,0,2,1,0,0,0,2,1 | 23,46,23,25,28,11,27,21,8, | 1,5,6,2,1,2,0,5,2,0,0,5,1,0, | 298 | 149 | 0.00111540802455 | 0.118494760381007 | 0.923,1.0,0.831,0. | 0.92,0.821,0.657,0 | 0,058 |
| 86038 | ENSG00000138380 | CARF | chr2 | + | 202941859 | 202941980 | 202917876 | 202918043 | 202942739 | 202942824 | 86038 | 40,19,67,39,46,18,18,37,2 | 2,3,1,3,1,0,2,4,4,0,0,0 | 29,26,12,7,19,12,13,15,22, | 1,1,0,2,1,0,0,0,0,4,0,0,0,0, | 270 | 149 | 0.00111983815574 | 0.118906702564418 | 0.917,0.778,0.974, | 0.941,0.935,1.0,0. | -0,032 |
| 21703 | ENSG00000197555 | SIPA1L1 | chr14 | + | 71413947 | 71414014 | 71321390 | 71321531 | 71512742 | 71512845 | 21703 | 8,15,17,10,11,5,11,15,5,4, | 5,1,9,1,0,1,3,2,0,0,1,0 | 10,18,14,12,15,8,8,6,3,7,4, | 4,0,1,0,0,0,0,0,2,0,0,1,1,2, | 216 | 149 | 0.00112332374321 | 0.119165899614115 | 0.525,0.912,0.566, | 0.633,1.0,0.906,1. | -0,006 |
| 11742 | ENSG00000108587 | GOSR1 | chr17 | + | 30490125 | 30490217 | 30484662 | 30484770 | 30492678 | 30492753 | 11742 | 93,50,84,60,22,38,23,39,2 | 7,0,1,1,0,0,2,0,2,0,6,0 | 40,56,39,23,48,25,32,30,9, | 1,3,0,1,2,0,0,3,3,4,2,4,1,1, | 241 | 149 | 0.00112346755487 | 0.119165899614115 | 0.891,1.0,0.981,0. | 0.961,0.92,1.0,0.9 | 0,026 |
| 202719 | ENSG00000204152 | TIMM23B | chr10 | + | 49943227 | 49943358 | 49942080 | 49942300 | 49945031 | 49945090 | 202719 | 4,4,3,5,1,4,4,1,2,1,3,1 | 33,21,7,4,5,4,4,5,3,4,3,5 | 5,1,0,0,5,1,2,1,0,2,5,2,2,1, | 7,22,10,7,13,5,9,8,6,4,4,10 | 280 | 149 | 0.00112394021408 | 0.119165899614115 | 0.061,0.092,0.186, | 0.275,0.024,0.0,0. | 0,102 |
| 213130 | ENSG00000169045 | HNRNPH1 | chr5 | - | 179616868 | 179616922 | 179616125 | 179616218 | 179617020 | 179617110 | 213130 | 138,73,190,99,91,100,90,1 | 85,63,38,17,26,27,29,23,6 | 143,151,86,99,98,53,41,97 | 26,13,32,30,5,11,6,31,8,33 | 203 | 149 | 0.00112610088729 | 0.119336198952395 | 0.544,0.46,0.786,0 | 0.801,0.895,0.664, | -0,109 |
| 20926 | ENSG00000213928 | IRF9 | chr14 | + | 24163877 | 24163959 | 24163377 | 24163508 | 24164062 | 24164134 | 20926 | 99,107,119,124,78,70,71,8 | 13,5,50,34,10,28,33,22,11, | 78,96,63,47,54,48,40,94,3 | 40,47,13,36,22,24,10,33,1 | 231 | 149 | 0.00112744808694 | 0.119376602204663 | 0.831,0.932,0.606, | 0.557,0.568,0.758, | 0,102 |
| 162477 | ENSG00000081791 | DELE1 | chr5 | + | 141930962 | 141931280 | 141930246 | 141930274 | 141933258 | 141933882 | 162477 | 22,16,12,13,6,5,7,11,10,10 | 29,29,48,33,18,26,12,23,3 | 17,9,2,1,10,2,1,7,3,5,3,11, | 38,29,21,10,19,20,25,24,1 | 298 | 149 | 0.00112759143563 | 0.119376602204663 | 0.275,0.216,0.111, | 0.183,0.134,0.045, | 0,068 |
| 103303 | ENSG00000231721 | LINC-PINT | chr7 | - | 130959868 | 130959970 | 130944168 | 130945835 | 130967388 | 130967517 | 103303 | 6,1,17,2,3,4,4,8,3,0,2,3 | 51,31,72,31,30,30,35,34,2 | 3,9,0,4,3,2,2,0,1,1,1,1,4,3, | 71,82,42,66,37,10,18,24,2 | 251 | 149 | 0.00113464772237 | 0.120064584038062 | 0.065,0.019,0.123, | 0.024,0.061,0.0,0. | 0,011 |
| 156250 | ENSG00000005812 | FBXL3 | chr13 | - | 77001862 | 77002049 | 76992597 | 76992951 | 77015408 | 77015580 | 156250 | 4,1,17,6,1,0,3,5,3,6,5,1 | 0,0,5,1,0,2,0,2,0,2,0,0 | 12,10,11,1,2,0,2,2,4,5,8,11 | 2,2,2,0,7,1,1,0,0,0,3,7,0,0, | 298 | 149 | 0.00113765729545 | 0.12032389059312 | 1.0,1.0,0.63,0.75,1 | 0.75,0.714,0.733,1 | 0,097 |
| 166636 | ENSG00000125247 | TMTC4 | chr13 | - | 100635283 | 100635779 | 100635064 | 100635195 | 100636531 | 100636632 | 166636 | 0,0,3,1,0,3,1,3,0,0,1,2 | 7,11,2,4,3,1,2,7,4,3,5,4 | 1,4,4,2,3,0,0,2,0,0,1,0,4,2, | 7,4,7,1,3,11,2,5,1,5,5,17,3, | 298 | 149 | 0.00114005135423 | 0.120517874765362 | 0.0,0.0,0.429,0.11 | 0.067,0.333,0.222, | -0,013 |
| 43744 | ENSG00000060971 | ACAA1 | chr3 | - | 38126161 | 38126341 | 38125825 | 38125881 | 38126509 | 38126700 | 43744 | 20,9,24,22,12,22,14,14,10, | 3,0,4,1,2,4,2,2,2,0,1,0 | 17,49,15,9,7,15,8,15,7,13, | 2,17,3,3,6,2,1,3,0,1,3,10,2, | 298 | 149 | 0.00114134660251 | 0.120576701877036 | 0.769,1.0,0.75,0.9 | 0.81,0.59,0.714,0. | 0,164 |
| 20553 | ENSG00000100811 | YY1 | chr14 | + | 100276489 | 100276648 | 100274697 | 100274758 | 100277417 | 100277721 | 20553 | 302,242,379,266,157,124, | 1,0,6,4,0,2,0,2,0,0,0,0 | 241,318,146,87,150,98,93, | 7,8,0,2,0,1,0,5,0,1,3,9,1,0, | 298 | 149 | 0.00114172827532 | 0.120576701877036 | 0.993,1.0,0.969,0. | 0.945,0.952,1.0,0. | 0,019 |
| 45293 | ENSG00000005187 | ACSM3 | chr16 | + | 20779796 | 20779907 | 20777372 | 20777580 | 20780713 | 20780857 | 45293 | 5,5,5,0,5,3,1,1,0,1,0,0 | 11,26,12,13,4,16,11,5,13,4 | 0,4,0,0,1,0,0,4,0,0,3,3,1,0, | 9,15,12,44,30,1,14,23,16,8 | 260 | 149 | 0.00115677596203 | 0.122105957373065 | 0.207,0.099,0.193, | 0.0,0.133,0.0,0.0,0 | 0,058 |
| 184331 | ENSG00000004487 | KDM1A | chr1 | + | 23046548 | 23046665 | 23044426 | 23044486 | 23050386 | 23050520 | 184331 | 4,5,9,1,0,7,2,6,0,4,1,2 | 20,12,17,11,18,21,8,17,26, | 7,23,9,12,4,0,5,3,5,5,5,13, | 40,21,8,12,6,5,9,27,5,14,1 | 266 | 149 | 0.00115821316455 | 0.122197734127089 | 0.101,0.189,0.229, | 0.089,0.38,0.387,0 | -0,1 |
| 225093 | ENSG00000136286 | MYO1G | chr7 | - | 44976563 | 44977007 | 44975475 | 44975649 | 44978866 | 44979088 | 225093 | 114,89,50,72,46,37,31,18, | 6,0,0,3,2,2,2,0,2,2,0,2 | 47,49,52,73,40,39,19,14,3 | 0,2,0,0,0,0,0,0,0,0,2,0,2,0, | 298 | 149 | 0.0011602421311 | 0.122351824653985 | 0.905,1.0,1.0,0.92 | 1.0,0.925,1.0,1.0,1 | -0,044 |
| 72993 | ENSG00000134759 | ELP2 | chr18 | + | 36142280 | 36142347 | 36141136 | 36141201 | 36142825 | 36142857 | 72993 | 58,47,104,64,41,43,29,47, | 4,0,0,0,1,3,0,0,0,0,0,0 | 107,72,59,48,49,25,55,77, | 1,2,0,7,1,3,0,4,1,0,5,3,5,0, | 216 | 149 | 0.00116230709324 | 0.122509558268922 | 0.909,1.0,1.0,1.0,0 | 0.987,0.961,1.0,0. | 0,035 |
| 156866 | ENSG00000134884 | ARGLU1 | chr13 | - | 106557558 | 106557663 | 106557047 | 106557131 | 106559431 | 106559657 | 156866 | 46,3,55,34,14,9,14,14,13,5 | 130,49,133,112,66,62,60,6 | 14,12,18,10,5,11,0,21,11,9 | 104,54,77,34,46,41,18,79, | 254 | 149 | 0.00116298635716 | 0.122521153518308 | 0.172,0.035,0.195, | 0.073,0.115,0.121, | 0,035 |
| 173867 | ENSG00000082996 | RNF13 | chr3 | + | 149901770 | 149901854 | 149895472 | 149895560 | 149902071 | 149902162 | 173867 | 6,1,13,2,2,4,1,2,4,0,0,0 | 68,56,111,85,39,37,25,63, | 4,2,1,0,3,0,0,2,0,2,3,0,0,1, | 76,88,34,28,42,12,33,54,1 | 233 | 149 | 0.00117459065699 | 0.123683131944528 | 0.053,0.011,0.07,0 | 0.033,0.014,0.018, | 0,014 |
| 107143 | ENSG00000215301 | DDX3X | chrX | + | 41347314 | 41347451 | 41346858 | 41347012 | 41347639 | 41347694 | 107143 | 371,322,507,353,148,179, | 23,7,26,30,13,10,10,14,6,1 | 290,367,246,245,272,171, | 31,31,15,52,16,17,2,20,13, | 286 | 149 | 0.00117991576507 | 0.124183105150015 | 0.894,0.96,0.91,0. | 0.83,0.86,0.895,0. | 0,032 |
| 66122 | ENSG00000162604 | TM2D1 | chr1 | - | 61723712 | 61723772 | 61709328 | 61709437 | 61724901 | 61725144 | 66122 | 14,5,40,23,9,8,9,6,2,5,8,4 | 2,2,5,3,0,0,0,6,0,1,1,3 | 19,23,12,12,14,13,1,11,6,2 | 1,2,0,2,3,0,2,0,1,1,0,0,1,0, | 209 | 149 | 0.00118112362821 | 0.124249471956631 | 0.833,0.641,0.851, | 0.931,0.891,1.0,0. | -0,035 |
| 230355 | ENSG00000150787 | PTS | chr11 | + | 112233431 | 112233504 | 112233162 | 112233233 | 112235848 | 112236034 | 230355 | 2,2,19,9,2,2,2,0,1,0,0,2 | 2,0,6,1,0,0,0,2,4,1,1,3 | 9,11,10,3,7,2,4,3,4,5,5,11, | 4,1,3,1,1,0,0,0,0,1,0,0,0,3, | 222 | 149 | 0.0011834514176 | 0.124433528119915 | 0.402,1.0,0.68,0.8 | 0.602,0.881,0.691, | -0,326 |
| 248599 | ENSG00000148925 | BTBD10 | chr11 | - | 13445023 | 13445153 | 13421641 | 13421838 | 13463091 | 13463297 | 248599 | 9,2,30,12,7,7,7,10,31,14,0, | 33,22,18,11,5,20,4,14,8,6, | 8,8,7,4,11,0,1,7,0,0,3,4,1,0 | 17,29,6,11,20,5,21,9,9,12,1 | 279 | 149 | 0.00118429945735 | 0.124461892824657 | 0.127,0.046,0.471, | 0.201,0.128,0.384, | 0,163 |
| 107954 | ENSG00000197892 | KIF13B | chr8 | - | 29134039 | 29134210 | 29132307 | 29132465 | 29140062 | 29140191 | 107954 | 37,12,57,34,23,20,14,31,2 | 12,0,1,0,0,8,0,0,2,1,0,3 | 25,28,18,25,17,10,22,30,1 | 0,0,3,6,0,2,1,1,0,0,4,0,2,1,1 | 298 | 149 | 0.00118787943992 | 0.12473084521065 | 0.607,1.0,0.966,1. | 1.0,1.0,0.75,0.676, | -0,016 |
| 188305 | ENSG00000075151 | EIF4G3 | chr1 | - | 20940963 | 20941395 | 20904882 | 20904971 | 20941490 | 20942330 | 188305 | 3,2,12,2,2,7,1,7,2,2,0,7 | 64,20,35,24,19,25,21,26,1 | 12,11,7,4,6,3,10,2,7,2,2,10 | 40,27,21,31,13,15,17,9,15, | 298 | 149 | 0.00118856636925 | 0.12473084521065 | 0.023,0.048,0.146, | 0.13,0.169,0.143,0 | -0,05 |
| 219673 | ENSG00000211450 | SELENOH | chr11 | + | 57742116 | 57742184 | 57741698 | 57741954 | 57742862 | 57743550 | 219673 | 9,8,25,14,8,9,21,3,19,22,3, | 0,3,5,3,4,0,0,3,0,2,0,0 | 9,14,7,3,8,4,4,12,2,5,11,21 | 3,7,8,6,3,2,0,7,4,3,4,3,0,0,1 | 217 | 149 | 0.00118859719802 | 0.12473084521065 | 1.0,0.647,0.774,0. | 0.673,0.579,0.375, | 0,172 |
| 184499 | ENSG00000077044 | DGKD | chr2 | + | 233447857 | 233447905 | 233446711 | 233446796 | 233448086 | 233448181 | 184499 | 5,0,14,5,2,3,3,4,0,2,0,0 | 20,9,65,49,34,38,20,37,48, | 2,5,8,6,1,2,3,3,3,7,9,2,6,7, | 20,25,35,20,13,17,33,37,9, | 197 | 149 | 0.00118978685583 | 0.124794841504457 | 0.159,0.0,0.14,0.0 | 0.07,0.131,0.147,0 | -0,063 |
| 255827 | ENSG00000182796 | TMEM198B | chr12 | + | 55834609 | 55834726 | 55833446 | 55833541 | 55834850 | 55834988 | 255827 | 2,0,1,1,1,5,1,4,3,0,0,3 | 11,6,6,3,7,10,3,9,8,7,6,3 | 0,1,0,2,1,0,0,3,0,2,1,0,1,1, | 9,13,3,24,7,1,8,4,10,7,5,10 | 266 | 149 | 0.00119455062285 | 0.125233475453789 | 0.092,0.0,0.085,0. | 0.0,0.041,0.0,0.04 | 0,068 |
| 55419 | ENSG00000160691 | SHC1 | chr1 | - | 154968203 | 154968257 | 154967979 | 154968031 | 154968494 | 154968594 | 55419 | 33,17,61,46,26,24,25,19,3 | 5,1,5,8,2,2,1,1,1,0,0,0 | 38,22,11,26,23,21,14,17,1 | 1,11,0,12,0,1,6,2,0,3,4,3,3, | 203 | 149 | 0.00119564870221 | 0.125287568561519 | 0.829,0.926,0.9,0. | 0.965,0.595,1.0,0. | 0,099 |
| 91541 | ENSG00000100503 | NIN | chr14 | - | 50756491 | 50758630 | 50754741 | 50754867 | 50759856 | 50760359 | 91541 | 13,3,33,6,11,5,4,8,9,4,5,0 | 32,32,100,60,38,26,24,34, | 15,11,4,17,11,6,7,10,0,9,1, | 35,40,21,22,26,31,11,19,1 | 298 | 149 | 0.00119809151531 | 0.125482450088412 | 0.169,0.045,0.142, | 0.176,0.121,0.087, | -0,051 |
| 26140 | ENSG00000164080 | RAD54L2 | chr3 | + | 51633901 | 51634035 | 51630704 | 51630931 | 51635592 | 51635789 | 26140 | 24,12,28,22,16,14,14,18,9, | 3,0,1,0,4,4,2,0,0,4,0,0 | 15,25,7,14,23,4,9,14,7,12, | 0,0,2,0,0,0,0,0,0,0,0,2,0, | 283 | 149 | 0.00119955500801 | 0.12557462253356 | 0.808,1.0,0.936,1. | 1.0,1.0,0.648,1.0,1 | -0,103 |
| 178713 | ENSG00000031698 | SARS1 | chr1 | + | 109236390 | 109236879 | 109235976 | 109236106 | 109237243 | 109237373 | 178713 | 31,7,55,67,37,22,14,34,31, | 2,0,5,3,2,1,0,0,1,0,2,0 | 32,11,15,10,11,14,7,11,4,2 | 2,0,2,1,4,0,0,7,0,0,3,1,5,0,1 | 298 | 149 | 0.00120089574548 | 0.125653861057562 | 0.886,1.0,0.846,0. | 0.889,1.0,0.789,0. | 0,081 |
| 42765 | ENSG00000114127 | XRN1 | chr3 | - | 142313135 | 142313174 | 142312597 | 142312758 | 142318591 | 142318694 | 42765 | 0,0,4,1,0,2,0,0,1,0,0,0 | 32,19,70,53,43,37,19,38,1 | 2,1,6,0,0,0,0,2,0,4,4,3,0,4, | 20,60,38,30,29,24,30,45,1 | 188 | 149 | 0.00120236595809 | 0.125692624336803 | 0.0,0.0,0.043,0.01 | 0.073,0.013,0.111, | -0,029 |
| 215889 | ENSG00000144504 | ANKMY1 | chr2 | - | 240529036 | 240529523 | 240525684 | 240525849 | 240552913 | 240555560 | 215889 | 1,2,6,6,6,3,3,12,4,3,1,9 | 13,9,9,13,18,22,21,18,15,9 | 2,0,3,3,1,1,3,2,1,1,0,9,3,0, | 18,20,4,15,3,5,8,7,13,7,13, | 298 | 149 | 0.00120243419168 | 0.125692624336803 | 0.037,0.1,0.25,0.1 | 0.053,0.0,0.273,0. | 0,039 |
| 195416 | ENSG00000162889 | MAPKAPK2 | chr1 | + | 206732037 | 206732087 | 206731838 | 206731919 | 206732574 | 206734281 | 195416 | 6,7,9,6,8,4,4,5,7,3,2,8 | 48,60,169,90,46,59,50,45, | 4,3,3,4,1,0,1,5,6,6,2,1,2,2, | 78,101,29,29,72,24,53,59, | 199 | 149 | 0.00120393961155 | 0.125788896375494 | 0.086,0.08,0.038,0 | 0.037,0.022,0.072, | 0,033 |
| 94544 | ENSG00000231925 | TAPBP | chr6 | - | 33303954 | 33303989 | 33299693 | 33301771 | 33304296 | 33304638 | 94544 | 7,5,44,9,9,5,9,9,1,3,1,9 | 0,0,8,3,0,0,0,2,0,0,0,2 | 17,16,6,8,10,9,3,6,9,5,3,12 | 0,2,0,1,4,4,2,5,0,1,0,14,3,0 | 184 | 149 | 0.00121002045455 | 0.126362888138404 | 1.0,1.0,0.817,0.70 | 1.0,0.866,1.0,0.86 | 0,154 |
| 33888 | ENSG00000014824 | SLC30A9 | chr4 | + | 42018388 | 42018505 | 42018110 | 42018170 | 42020415 | 42020515 | 33888 | 17,9,28,7,3,0,14,13,2,0,2,1 | 27,23,82,59,24,41,26,59,2 | 8,11,5,4,10,1,2,0,1,3,5,7,2, | 48,40,35,14,16,21,25,34,1 | 266 | 149 | 0.00121940361281 | 0.127281017938268 | 0.261,0.18,0.161,0 | 0.085,0.133,0.074, | 0,038 |
| 50978 | ENSG00000130775 | THEMIS2 | chr1 | + | 27876587 | 27876728 | 27872542 | 27872665 | 27879643 | 27880054 | 50978 | 78,123,71,65,28,86,78,102 | 1,2,8,0,0,0,3,0,0,0,0,6 | 54,222,41,21,101,41,131,2 | 2,4,0,3,3,0,7,11,2,0,10,6,5, | 290 | 149 | 0.00122134230932 | 0.127421583411174 | 0.976,0.969,0.82,1 | 0.933,0.966,1.0,0. | 0,033 |
| 232579 | ENSG00000134516 | DOCK2 | chr5 | + | 169983898 | 169984051 | 169983107 | 169983166 | 169985827 | 169985922 | 232579 | 1,5,5,5,3,6,5,7,0,1,3,1 | 94,91,156,118,70,90,73,79 | 7,21,1,8,7,4,5,6,12,5,9,7,3, | 104,110,65,44,63,57,45,69 | 298 | 149 | 0.00122317834343 | 0.127551307187395 | 0.005,0.027,0.016, | 0.033,0.087,0.008, | -0,016 |
| 240133 | ENSG00000165525 | NEMF | chr14 | - | 49796208 | 49796328 | 49795790 | 49795944 | 49799474 | 49799524 | 240133 | 1,0,8,2,1,1,3,5,1,0,0,5 | 82,45,91,39,32,35,24,43,6 | 2,16,2,2,2,0,1,1,7,0,7,4,2,2 | 30,41,40,68,39,23,25,30,2 | 269 | 149 | 0.0012265513525 | 0.127725477102635 | 0.007,0.0,0.046,0. | 0.036,0.178,0.027, | -0,026 |
| 167884 | ENSG00000121210 | TMEM131L | chr4 | + | 153617686 | 153617794 | 153612251 | 153612400 | 153620755 | 153620880 | 167884 | 27,10,19,2,3,11,4,12,10,1, | 282,218,270,142,72,92,69, | 12,5,3,3,5,1,9,2,4,3,6,10,2, | 135,112,101,160,73,44,47, | 257 | 149 | 0.00122739582071 | 0.127725477102635 | 0.053,0.026,0.039, | 0.049,0.025,0.017, | 0,009 |
| 139798 | ENSG00000177410 | ZFAS1 | chr20 | + | 49280863 | 49280964 | 49280484 | 49280570 | 49289044 | 49289260 | 139798 | 22,7,66,60,40,32,19,36,10, | 74,29,247,278,49,60,81,92 | 32,32,54,16,19,10,11,18,1 | 63,54,40,35,42,27,23,41,3 | 250 | 149 | 0.00122800024482 | 0.127725477102635 | 0.151,0.126,0.137, | 0.232,0.261,0.446, | -0,054 |
| 204680 | ENSG00000147274 | RBMX | chrX | - | 136873245 | 136873388 | 136872267 | 136872345 | 136875085 | 136875168 | 204680 | 76,53,148,92,66,63,54,117 | 6,2,25,18,11,6,9,10,8,6,2,1 | 77,71,60,46,34,31,44,57,3 | 17,42,12,13,33,12,5,4,17,2 | 292 | 149 | 0.00122814361811 | 0.127725477102635 | 0.866,0.931,0.751, | 0.698,0.463,0.718, | 0,123 |
| 25454 | ENSG00000172175 | MALT1 | chr18 | + | 58700440 | 58700591 | 58696365 | 58696487 | 58709975 | 58710072 | 25454 | 42,23,78,56,53,31,31,37,2 | 1,0,3,2,6,0,0,4,0,1,0,1 | 79,51,34,12,10,33,17,61,2 | 2,7,4,2,1,0,0,8,2,1,8,2,2,2,2 | 298 | 149 | 0.00122861199987 | 0.127725477102635 | 0.955,1.0,0.929,0. | 0.952,0.785,0.81,0 | 0,054 |
| 12798 | ENSG00000141698 | NT5C3B | chr17 | + | 41835587 | 41835682 | 41835202 | 41835272 | 41835858 | 41835957 | 12798 | 9,4,13,9,3,6,1,1,7,7,2,0 | 10,1,11,7,0,3,2,4,5,2,1,5 | 2,0,1,1,2,1,3,3,3,1,1,0,2,0,1 | 2,4,3,0,2,0,2,5,2,8,5,8,0,1,2 | 244 | 149 | 0.00122875833987 | 0.127725477102635 | 0.355,0.71,0.419,0 | 0.379,0.0,0.169,1. | 0,099 |
| 226877 | ENSG00000005471 | ABCB4 | chr7 | - | 87452943 | 87453135 | 87451622 | 87451717 | 87462757 | 87462887 | 226877 | 19,7,36,21,4,24,13,17,7,6 | 1,0,7,5,2,5,0,0,0,0,0,0 | 16,16,6,11,6,1,12,15,9,21,9 | 2,0,0,0,0,1,0,0,0,1,0,3,2,1,0 | 298 | 149 | 0.00122900262081 | 0.127725477102635 | 0.905,1.0,0.72,0.6 | 0.8,1.0,1.0,1.0,1.0, | -0,001 |
| 170499 | ENSG00000182606 | TRAK1 | chr3 | + | 42033837 | 42034050 | 42013801 | 42014117 | 42087103 | 42087400 | 170499 | 7,2,5,2,3,10,8,7,1,1,4,6 | 43,20,17,10,37,26,18,20,4 | 0,8,6,52,17,1,8,4,2,0,7,3,1, | 17,9,14,35,11,7,14,17,2,14 | 298 | 149 | 0.00123402036889 | 0.128152127142641 | 0.075,0.048,0.128,0 | 0.0,0.308,0.176,0. | -0,079 |
| 92928 | ENSG00000111670 | GNPTAB | chr12 | - | 101756419 | 101756566 | 101753371 | 101753539 | 101757211 | 101757310 | 92928 | 0,2,7,7,3,4,5,10,3,1,1,3 | 38,29,140,116,58,59,52,11 | 2,2,4,3,4,9,9,7,8,7,16,8,9,6 | 87,115,23,30,46,29,59,98,3 | 296 | 149 | 0.00123429877465 | 0.128152127142641 | 0.0,0.034,0.025,0.0 | 0.011,0.009,0.08,0 | -0,029 |
| 87699 | ENSG00000198176 | TFDP1 | chr13 | + | 113637817 | 113637896 | 113636533 | 113636700 | 113640119 | 113640684 | 87699 | 19,15,39,33,15,34,18,14,14 | 3,8,14,11,6,1,3,1,7,7,3,12 | 27,19,16,4,9,10,13,7,12,14 | 1,2,0,0,3,2,3,2,2,4,1,26,3,4 | 228 | 149 | 0.00123550038064 | 0.128163448071567 | 0.805,0.551,0.645,0 | 0.946,0.861,1.0,1. | -0,135 |
| 133787 | ENSG00000182944 | EWSR1 | chr22 | + | 29297832 | 29297949 | 29296238 | 29296368 | 29298732 | 29298895 | 133787 | 68,64,181,101,78,74,52,91 | 1,2,2,3,1,5,5,6,6,2,0,4 | 113,173,71,65,49,31,60,10 | 1,3,2,0,0,2,5,2,0,1,1,2,3,1,4 | 266 | 149 | 0.00123559875087 | 0.128163448071567 | 0.974,0.947,0.981,0 | 0.984,0.97,0.952,1 | -0,034 |

| | | | | | | | | | | | | | | | | | | | | | | |
|---|---|---|---|---|---|---|---|---|---|---|---|---|---|---|---|---|---|---|---|---|---|---|
| 36439 | ENSG00000147548 | NSD3 | chr8 | - | 38299443 | 38299590 | 38295795 | 38295952 | 38304586 | 38304757 | 36439 | 39,4,101,33,59,33,35,43,3 | 1,1,5,4,5,4,3,1,1,1,4,2 | 55,43,39,31,15,19,27,32,2 | 5,9,8,8,1,7,0,0,1,5,12,3,14, | 296 | 149 | 0.00123667557646 | 0.128213353081437 | 0.952,0.668,0.91,0 | 0.847,0.706,0.71,0 | 0,063 |
| 190013 | ENSG00000258728 | | chr9 | + | 34647562 | 34647961 | 34647088 | 34647258 | 34648114 | 34648171 | 190013 | 32,13,35,25,29,35,26,17,4 | 0,0,5,4,3,5,1,1,0,1,1,0 | 10,12,16,18,12,9,20,34,7,1 | 0,0,0,0,3,0,0,3,0,0,3,1,0,0, | 298 | 149 | 0.00123919564519 | 0.128371330626117 | 1.0,1.0,0.778,0.75 | 1.0,1.0,1.0,1.0,0.6 | -0,051 |
| 34990 | ENSG00000197020 | ZNF100 | chr19 | - | 21744016 | 21744112 | 21726192 | 21727989 | 21744940 | 21745067 | 34990 | 3,9,12,8,4,5,10,20,6,5,6,4 | 3,2,4,3,1,0,2,3,0,4,3,1 | 6,11,4,5,18,7,3,9,4,7,6,9,4, | 3,2,0,0,2,1,0,0,1,3,2,2,0,0, | 245 | 149 | 0.00123939221135 | 0.128371330626117 | 0.378,0.732,0.646, | 0.549,0.77,1.0,1.0, | -0,109 |
| 40996 | ENSG00000164081 | TEX264 | chr3 | + | 51674270 | 51674562 | 51671174 | 51671288 | 51684412 | 51684467 | 40996 | 29,21,44,40,18,23,19,30,6 | 2,2,0,0,0,1,4,3,12,7,3,2 | 25,13,21,7,9,9,15,35,7,8,3 | 3,1,3,1,0,0,3,3,0,1,1,0,1,2, | 298 | 149 | 0.00124174631751 | 0.128484300412895 | 0.879,0.84,1.0,1.0, | 0.806,0.867,0.778, | -0,047 |
| 83992 | ENSG00000170006 | TMEM154 | chr4 | - | 152647183 | 152647327 | 152644414 | 152644442 | 152652537 | 152652576 | 83992 | 0,3,2,5,3,7,1,8,6,1,0,0 | 59,37,159,144,62,43,29,36 | 2,2,4,10,1,1,1,4,5,9,2,10,1, | 36,59,61,10,38,49,18,29,4 | 293 | 149 | 0.00124199100335 | 0.128484300412895 | 0.0,0.04,0.006,0.0 | 0.027,0.017,0.032, | -0,028 |
| 175024 | ENSG00000101596 | SMCHD1 | chr18 | + | 2672997 | 2673194 | 2666869 | 2667031 | 2673280 | 2673363 | 175024 | 123,74,173,49,21,60,64,15 | 514,412,843,542,254,319, | 31,109,97,177,244,77,66,4 | 231,322,330,168,299,197, | 298 | 149 | 0.00124227378565 | 0.128484300412895 | 0.107,0.082,0.093, | 0.063,0.145,0.128, | -0,042 |
| 208334 | ENSG00000135974 | C2orf49 | chr2 | + | 105341596 | 105341701 | 105339582 | 105339749 | 105342847 | 105343223 | 208334 | 2,0,6,2,0,8,5,9,2,2,0,1 | 24,28,47,35,28,11,28,18,3 | 0,2,3,2,3,0,0,0,4,0,2,3,3,0, | 30,47,31,22,40,21,26,25,2 | 254 | 149 | 0.00124741716323 | 0.128954295609585 | 0.047,0.0,0.07,0.0 | 0.0,0.024,0.054,0. | 0,044 |
| 179174 | ENSG00000126012 | KDM5C | chrX | - | 53218275 | 53218392 | 53217795 | 53217966 | 53220838 | 53220916 | 179174 | 93,40,141,77,46,48,45,50, | 3,0,10,11,0,0,1,2,0,0,1,0 | 78,71,47,89,88,60,43,48,6 | 5,2,9,21,8,2,0,1,12,12,1,2, | 266 | 149 | 0.00124916182957 | 0.129050879731728 | 0.946,1.0,0.888,0. | 0.897,0.952,0.745, | 0,06 |
| 211223 | ENSG00000197622 | CDC42SE1 | chr1 | - | 151056650 | 151056820 | 151055676 | 151055993 | 151059478 | 151059773 | 211223 | 44,4,36,27,21,30,21,29,31, | 197,165,295,254,162,112, | 35,31,36,36,22,32,26,24,1 | 274,239,156,130,165,98,1 | 298 | 149 | 0.00124955063797 | 0.129050879731728 | 0.1,0.012,0.058,0. | 0.06,0.061,0.103,0 | -0,021 |
| 127011 | ENSG00000196683 | TOMM7 | chr7 | - | 22817999 | 22818048 | 22812970 | 22813185 | 22822676 | 22822791 | 127011 | 576,431,1404,1318,435,39 | 4,3,9,14,3,0,6,13,5,0,0,4 | 809,800,354,234,487,278, | 9,14,2,2,6,0,11,13,4,10,12, | 198 | 149 | 0.00125225611833 | 0.129193094538927 | 0.991,0.991,0.992, | 0.985,0.977,0.993, | 0,009 |
| 81862 | ENSG00000168385 | SEPTIN2 | chr2 | + | 241317498 | 241317599 | 241315288 | 241315610 | 241324215 | 241324241 | 81862 | 9,1,38,21,18,17,13,15,3,1, | 0,5,0,0,0,0,2,7,0,0,1,0 | 10,12,4,6,10,4,1,11,4,5,10, | 0,1,4,4,0,3,0,1,3,2,0,2,2,2, | 250 | 149 | 0.0012524114275 | 0.129193094538927 | 1.0,0.107,1.0,1.0,1 | 1.0,0.877,0.373,0. | 0,078 |
| 185947 | ENSG00000118260 | CREB1 | chr2 | + | 207577152 | 207577224 | 207575285 | 207575454 | 207577504 | 207577655 | 185947 | 15,5,22,8,1,6,5,9,5,1,2,4 | 65,19,78,61,49,23,33,31,4 | 8,12,4,1,6,7,5,7,2,9,2,1,1,6 | 57,59,23,38,46,40,42,50,2 | 221 | 149 | 0.00125280860752 | 0.129193094538927 | 0.135,0.151,0.16,0 | 0.086,0.121,0.105, | 0,039 |
| 216176 | ENSG00000073614 | KDM5A | chr12 | - | 320994 | 321105 | 318105 | 318461 | 322416 | 322567 | 216176 | 62,54,149,77,55,41,42,71, | 4,0,3,3,4,1,0,2,0,0,1,0 | 90,114,53,52,72,47,25,57, | 3,6,5,6,0,1,3,0,0,0,3,1,4,9, | 260 | 149 | 0.00125332866268 | 0.129193094538927 | 0.899,1.0,0.966,0. | 0.945,0.916,0.859, | 0,03 |
| 94752 | ENSG00000008083 | JARID2 | chr6 | + | 15410223 | 15410365 | 15374116 | 15374252 | 15452005 | 15452175 | 94752 | 49,48,149,51,59,66,54,82, | 3,0,2,1,9,3,3,4,3,2,1,0 | 83,134,35,59,39,30,31,55, | 8,8,4,1,2,7,2,2,3,5,9,4,2,7, | 291 | 149 | 0.00125423544436 | 0.129224676364312 | 0.893,1.0,0.974,0. | 0.842,0.896,0.818, | 0,059 |
| 116837 | ENSG00000187764 | SEMA4D | chr9 | - | 89418055 | 89418118 | 89405350 | 89405699 | 89455887 | 89455953 | 116837 | 14,6,36,23,8,15,2,23,16,8, | 3,1,3,2,1,1,1,0,2,0,2,2 | 19,27,16,16,9,14,9,7,11,20 | 6,4,2,0,4,1,6,2,2,3,11,2,1,3 | 212 | 149 | 0.00125555444709 | 0.129298679043841 | 0.766,0.808,0.894, | 0.69,0.826,0.849,1 | 0,067 |
| 187810 | ENSG00000033030 | ZCCHC8 | chr12 | - | 122478663 | 122478794 | 122478205 | 122478292 | 122480189 | 122481422 | 187810 | 3,1,0,0,0,3,0,0,0,0,0,0 | 18,14,35,36,18,22,9,13,24, | 0,5,3,12,4,2,3,2,0,1,0,0,1,0 | 14,22,25,9,21,9,9,16,5,18, | 280 | 149 | 0.00125818352532 | 0.129452101175124 | 0.081,0.037,0.0,0. | 0.0,0.108,0.06,0.4 | -0,049 |
| 223658 | ENSG00000106100 | NOD1 | chr7 | - | 30450917 | 30451073 | 30448297 | 30448381 | 30451215 | 30453040 | 223658 | 5,2,6,0,0,0,1,4,4,0,3,1 | 14,9,15,3,6,7,3,5,5,10,7,3 | 1,0,0,0,0,2,0,3,0,1,2,11,0 | 12,6,1,10,9,10,2,5,4,8,7,17 | 298 | 149 | 0.00125824716541 | 0.129452101175124 | 0.152,0.1,0.167,0. | 0.04,0.0,0.0,0.0,0. | 0,058 |
| 159627 | ENSG00000170476 | MZB1 | chr5 | - | 139389256 | 139389436 | 139388460 | 139388667 | 139389679 | 139390081 | 159627 | 0,2,25,21,3,2,3,6,4,3,1,6 | 9,5,9,2,3,4,3,4,2,0,6,10 | 0,3,1,0,6,4,0,3,1,7,5,0,1,0, | 3,8,6,10,0,1,3,39,0,0,18,14 | 298 | 149 | 0.0012602834723 | 0.129599652186623 | 0.0,0.167,0.581,0. | 0.0,0.158,0.077,0. | 0,08 |
| 51497 | ENSG00000178927 | CYBC1 | chr17 | - | 82447579 | 82447646 | 82446622 | 82446696 | 82449169 | 82449292 | 51497 | 10,7,31,28,12,16,6,9,2,1,4, | 0,0,1,4,1,0,5,0,3,0,0,0 | 11,12,8,4,6,4,2,4,5,11,9,22 | 3,0,0,0,0,6,1,4,0,4,2,5,1,2, | 216 | 149 | 0.00126296853837 | 0.129813744738259 | 1.0,1.0,0.955,0.82 | 0.717,1.0,1.0,1.0,1 | 0,076 |
| 105592 | ENSG00000099204 | ABLIM1 | chr10 | - | 114473878 | 114473956 | 114472976 | 114473132 | 114487957 | 114487993 | 105592 | 115,85,86,92,49,35,26,29, | 0,3,7,3,2,0,2,0,0,0,0,3 | 127,71,29,18,69,73,9,36,6 | 6,2,3,0,5,0,0,5,3,2,0,8,5,7, | 227 | 149 | 0.00126933199007 | 0.130405533916351 | 1.0,0.949,0.89,0.9 | 0.933,0.959,0.864, | 0,027 |
| 93658 | ENSG00000169554 | ZEB2 | chr2 | - | 144517277 | 144517419 | 144426512 | 144430026 | 144519938 | 144520006 | 93658 | 103,109,231,105,71,90,78, | 2,1,7,2,0,0,2,2,0,1,1,0 | 182,173,69,102,97,64,64,1 | 4,9,4,2,11,2,1,3,2,1,0,4,0,1 | 291 | 149 | 0.00127618367881 | 0.131046893785284 | 0.963,0.982,0.944, | 0.959,0.908,0.898, | 0,029 |
| 193913 | ENSG00000092841 | MYL6 | chr12 | + | 56159586 | 56159724 | 56158361 | 56158711 | 56159974 | 56160343 | 193913 | 606,515,1388,1172,402,46 | 38,26,57,25,6,14,8,13,16,1 | 573,589,463,236,443,312, | 11,37,15,54,46,7,0,10,20,8 | 287 | 149 | 0.00127806196488 | 0.131177183959508 | 0.892,0.911,0.927, | 0.964,0.892,0.941, | 0,037 |
| 45768 | ENSG00000291075 | | chr9 | + | 41008880 | 41009006 | 40994632 | 40994699 | 41011616 | 41011730 | 45768 | 45,17,85,56,20,28,20,39,3 | 2,0,12,3,18,15,11,19,0,0,2, | 67,44,58,19,5,8,15,29,36,3 | 0,0,0,1,11,4,0,0,0,0,4,16,0, | 275 | 149 | 0.0012822897207 | 0.131548378873204 | 0.924,1.0,0.793,0. | 1.0,1.0,1.0,0.911,0 | -0,09 |
| 125324 | ENSG00000163683 | SMIM14 | chr4 | - | 39556427 | 39556570 | 39546335 | 39552158 | 39572414 | 39572463 | 125324 | 329,142,507,655,286,219, | 5,2,3,3,2,1,0,4,1,0,0,2 | 292,186,231,47,104,156,7 | 1,3,2,2,1,0,0,0,4,6,1,27,6,4 | 292 | 149 | 0.00128318056138 | 0.131561477754099 | 0.971,0.973,0.989, | 0.993,0.969,0.983, | 0,011 |
| 56565 | ENSG00000163870 | TPRA1 | chr3 | - | 127576820 | 127576893 | 127576615 | 127576696 | 127576989 | 127577076 | 56565 | 8,3,6,18,3,2,5,9,5,3,5,6 | 2,0,4,2,4,1,1,0,0,4,0,0 | 13,6,2,3,1,1,3,8,4,4,3,8,4,7 | 1,0,1,2,0,0,0,2,0,1,2,2,0,1, | 222 | 149 | 0.00128363991843 | 0.131561477754099 | 0.729,1.0,0.502,0. | 0.897,1.0,0.573,0. | -0,077 |
| 969 | ENSG00000100099 | HPS4 | chr22 | - | 26471212 | 26471388 | 26470718 | 26470813 | 26472301 | 26472418 | 969 | 4,1,3,11,4,1,5,6,4,0,0,0 | 10,18,23,17,12,17,17,8,21, | 3,2,0,1,1,1,5,4,6,2,2,2,1,0, | 10,23,12,11,14,12,14,32,1 | 298 | 149 | 0.00128605029044 | 0.13174578298985 | 0.167,0.027,0.061, | 0.13,0.042,0.0,0.0 | 0,04 |
| 103567 | ENSG00000107185 | RGP1 | chr9 | + | 35751626 | 35751754 | 35751265 | 35751412 | 35752650 | 35752802 | 103567 | 35,37,35,31,26,19,9,12,11, | 2,3,1,0,0,1,1,1,3,4,1,0 | 22,31,17,14,35,12,12,19,2 | 0,2,0,7,3,4,2,3,0,1,0,0,0,1, | 277 | 149 | 0.00128822650539 | 0.131877378119076 | 0.904,0.869,0.95,1 | 1.0,0.893,1.0,0.51 | -0,014 |
| 175021 | ENSG00000101596 | SMCHD1 | chr18 | + | 2672976 | 2673194 | 2666869 | 2667031 | 2673280 | 2673363 | 175021 | 123,74,173,49,21,60,64,15 | 514,412,843,542,254,319, | 31,109,97,177,244,77,66,4 | 231,322,330,168,299,197, | 298 | 149 | 0.00128856031977 | 0.131877378119076 | 0.107,0.082,0.093, | 0.063,0.145,0.128, | -0,042 |
| 221628 | ENSG00000204599 | TRIM39 | chr6 | + | 30340287 | 30340361 | 30339907 | 30339930 | 30340504 | 30340620 | 221628 | 0,1,10,3,0,2,0,0,0,0,0,1 | 7,11,11,10,10,9,8,6,2,5,17, | 2,5,5,7,0,0,0,2,2,0,2,0,1,3, | 16,25,7,10,10,4,16,12,5,11 | 223 | 149 | 0.00128935990668 | 0.131896493381484 | 0.0,0.057,0.378,0. | 0.077,0.118,0.323, | -0,062 |
| 217376 | ENSG00000154473 | BUB3 | chr10 | + | 123160422 | 123160565 | 123157728 | 123157880 | 123162235 | 123162413 | 217376 | 87,67,145,105,75,70,72,95 | 3,1,1,5,1,6,12,3,2,0,2,3 | 111,107,67,47,101,63,58,7 | 9,11,5,3,8,6,2,4,7,10,8,2,9, | 292 | 149 | 0.00129142212149 | 0.132044691035826 | 0.937,0.972,0.987, | 0.863,0.832,0.872, | 0,05 |
| 156293 | ENSG00000122033 | MTIF3 | chr13 | - | 27441037 | 27441077 | 27440280 | 27440449 | 27445087 | 27445137 | 156293 | 1,0,7,4,2,0,1,5,0,0,0,2 | 6,1,9,15,8,6,4,2,10,5,2,7 | 0,1,2,0,2,0,0,1,0,1,1,0,2,2, | 11,2,10,3,3,11,2,5,6,8,11,9 | 189 | 149 | 0.00129363100872 | 0.13220773771976 | 0.116,0.0,0.38,0.1 | 0.0,0.283,0.136,0. | 0,023 |
| 76300 | ENSG00000076242 | MLH1 | chr3 | + | 37008813 | 37008905 | 37006990 | 37007063 | 37011819 | 37011836 | 76300 | 42,34,108,76,24,44,21,14, | 3,1,5,0,1,0,2,3,0,1,0,0 | 62,45,37,19,52,34,23,47,3 | 5,3,1,7,9,2,1,2,1,2,1,3,0,0, | 241 | 149 | 0.00129528813854 | 0.13231426736882 | 0.896,0.955,0.93,1 | 0.885,0.903,0.958, | 0,032 |
| 190016 | ENSG00000258728 | | chr9 | + | 34647743 | 34647961 | 34647088 | 34647258 | 34648114 | 34648171 | 190016 | 32,13,36,25,29,35,26,17,4 | 0,0,5,4,3,5,1,1,0,1,1,0 | 10,12,16,18,12,9,20,31,7,2 | 0,0,0,0,3,0,0,3,0,0,3,1,0,0, | 298 | 149 | 0.00129655044118 | 0.13238038330437 | 1.0,1.0,0.783,0.75 | 1.0,1.0,1.0,1.0,0.6 | -0,05 |
| 54023 | ENSG00000171824 | EXOSC10 | chr1 | - | 11077329 | 11077443 | 11076841 | 11076948 | 11077600 | 11077651 | 54023 | 4,2,23,12,12,8,8,9,7,7,5,4 | 1,0,4,2,1,2,2,3,0,0,1,0 | 12,6,2,1,12,8,5,4,2,5,5,13, | 1,3,0,2,0,0,4,13,0,0,0,1,0,0 | 263 | 149 | 0.00129946340813 | 0.132614892743114 | 0.694,1.0,0.765,0. | 0.872,0.531,1.0,0. | 0,03 |
| 22767 | ENSG00000100897 | DCAF11 | chr14 | + | 24116971 | 24117044 | 24115325 | 24115749 | 24117265 | 24117330 | 22767 | 74,36,56,60,23,24,14,22,5 | 8,1,4,1,3,2,4,2,1,0,0,1 | 19,23,29,20,30,23,14,26,1 | 6,5,4,2,5,1,2,3,2,2,1,7,0,5, | 222 | 149 | 0.00130400555443 | 0.133015364685082 | 0.861,0.96,0.904,0 | 0.68,0.755,0.83,0. | 0,078 |
| 68340 | ENSG00000110619 | CARS1 | chr11 | - | 3040447 | 3040581 | 3039834 | 3039931 | 3040895 | 3040984 | 68340 | 0,0,5,0,2,3,0,0,0,0,0,0 | 15,9,29,14,8,14,11,9,12,4, | 0,2,4,4,2,3,0,1,4,0,2,2,3,0, | 13,9,6,9,8,8,4,12,8,2,4,12, | 283 | 149 | 0.00130609550802 | 0.133165439264165 | 0.0,0.0,0.083,0.0,0 | 0.0,0.105,0.26,0.1 | -0,077 |
| 21818 | ENSG00000198604 | BAZ1A | chr14 | - | 34786125 | 34786221 | 34785776 | 34786001 | 34792774 | 34792921 | 21818 | 35,12,47,25,27,13,11,16,2 | 2,2,6,5,0,0,0,0,1,0,0,0 | 47,37,22,11,14,13,10,11,1 | 0,8,2,0,1,3,2,2,3,10,0,0,2,2 | 245 | 149 | 0.00130701845471 | 0.133196443667466 | 0.914,0.785,0.827, | 1.0,0.738,0.87,1.0, | 0,095 |
| 159929 | ENSG00000157212 | PAXIP1 | chr7 | - | 154986055 | 154986179 | 154983218 | 154983332 | 154991005 | 154991069 | 159929 | 5,2,6,8,0,0,0,1,0,1,2,2 | 26,15,43,16,5,4,14,13,10,6 | 4,4,0,0,3,2,2,0,0,1,0,0,1,0, | 14,22,9,2,20,5,4,20,7,11,8, | 273 | 149 | 0.00131108286871 | 0.133547409803749 | 0.095,0.068,0.071, | 0.135,0.09,0.0,0.0, | 0,024 |
| 245568 | ENSG00000109458 | GAB1 | chr4 | + | 143436457 | 143436772 | 143433490 | 143433716 | 143437998 | 143438600 | 245568 | 14,6,0,0,8,6,7,49,0,0,4,0 | 38,35,207,116,89,95,164,3 | 3,7,21,52,1,3,19,15,0,0,19, | 119,102,132,109,83,39,12 | 298 | 149 | 0.00131652249286 | 0.134038057076987 | 0.156,0.079,0.0,0. | 0.012,0.033,0.074, | -0,004 |
| 180966 | ENSG00000163346 | PBXIP1 | chr1 | - | 154951794 | 154951948 | 154951470 | 154951535 | 154953670 | 154953757 | 180966 | 165,175,207,218,94,52,10 | 0,1,3,0,3,1,1,1,0,1,0,0 | 201,185,76,118,136,85,11 | 3,4,4,0,1,1,4,9,0,2,6,0,3,1, | 298 | 149 | 0.00131775289269 | 0.134099892598847 | 1.0,0.989,0.972,1. | 0.971,0.959,0.905, | 0,017 |
| 178685 | ENSG00000163564 | PYHIN1 | chr1 | + | 158973649 | 158974065 | 158944874 | 158945042 | 158976700 | 158977059 | 178685 | 5,6,0,0,0,0,3,6,0,3,0,1 | 4,14,20,8,1,5,12,12,7,2,5,2 | 0,1,1,1,1,1,0,0,2,1,0,5,2,1, | 2,4,21,1,24,15,3,8,11,24,5, | 298 | 149 | 0.00131896981638 | 0.134160298936334 | 0.385,0.176,0.0,0. | 0.0,0.111,0.023,0. | 0,061 |
| 175027 | ENSG00000101596 | SMCHD1 | chr18 | + | 2673005 | 2673194 | 2666869 | 2667031 | 2673280 | 2673363 | 175027 | 123,74,173,49,21,60,64,15 | 514,412,843,542,254,319, | 31,109,97,179,244,77,66,4 | 231,322,330,168,299,197, | 298 | 149 | 0.00132398022824 | 0.134606324281683 | 0.107,0.082,0.093, | 0.063,0.145,0.128, | -0,042 |
| 182653 | ENSG00000053900 | ANAPC4 | chr4 | + | 25383578 | 25383644 | 25383260 | 25383393 | 25388499 | 25388574 | 182653 | 9,4,12,1,4,1,2,10,0,2,0,6 | 33,15,47,21,30,44,14,34,1 | 6,13,12,1,9,5,3,4,11,6,13,2 | 32,30,13,6,16,15,5,17,8,27 | 215 | 149 | 0.00132777228641 | 0.134928119441129 | 0.159,0.156,0.15,0 | 0.115,0.231,0.39,0 | -0,104 |
| 51540 | ENSG00000162894 | FCMR | chr1 | - | 206909461 | 206909520 | 206903316 | 206905147 | 206909724 | 206909868 | 51540 | 206,64,118,68,134,75,141, | 10,3,4,1,2,1,4,8,10,3,3,4 | 45,19,138,42,6,25,77,123, | 1,4,3,6,4,2,11,14,0,1,3,3,2, | 208 | 149 | 0.00132901438872 | 0.134990606842187 | 0.937,0.939,0.955, | 0.97,0.773,0.971,0 | 0,065 |
| 200314 | ENSG00000143149 | ALDH9A1 | chr1 | - | 165680486 | 165680613 | 165679441 | 165679582 | 165682106 | 165682241 | 200314 | 35,28,66,49,15,22,23,32,2 | 0,0,0,2,0,2,0,3,0,0,0,0 | 40,47,15,12,30,31,18,25,2 | 2,2,1,0,3,0,0,2,2,2,2,0,2,2, | 276 | 149 | 0.00133165333295 | 0.13519484835102 | 1.0,1.0,1.0,0.93,1. | 0.915,0.927,0.89,1 | 0,061 |
| 165726 | ENSG00000125089 | SH3TC1 | chr4 | + | 8215171 | 8215345 | 8214474 | 8214580 | 8216110 | 8216257 | 165726 | 5,2,0,0,1,0,0,1,0,4,3,9 | 22,16,14,18,17,9,20,24,26, | 4,10,0,1,4,0,8,9,4,1,1,5,6,5 | 23,15,8,3,17,2,9,8,6,20,8,2 | 298 | 149 | 0.00133643778611 | 0.135562513061515 | 0.102,0.059,0.0,0. | 0.08,0.25,0.0,0.14 | -0,051 |
| 10517 | ENSG00000154262 | ABCA6 | chr17 | - | 69097919 | 69098027 | 69096627 | 69096801 | 69100796 | 69100934 | 10517 | 169,93,535,406,134,78,24 | 4,1,11,17,5,2,4,7,0,0,4,2 | 392,311,201,30,162,154,9 | 12,14,0,2,10,4,3,5,12,2,14, | 257 | 149 | 0.00133653448024 | 0.135562513061515 | 0.961,0.982,0.966, | 0.95,0.928,1.0,0.8 | 0,029 |
| 224094 | ENSG00000072609 | CHFR | chr12 | - | 132875497 | 132875620 | 132872284 | 132872394 | 132877554 | 132877654 | 224094 | 0,0,0,3,5,0,0,3,0,0,1,1 | 57,29,26,39,15,14,18,23,1 | 3,2,0,3,2,1,0,1,0,4,5,7,6,2, | 19,28,12,27,17,13,17,6,23, | 272 | 149 | 0.00134068818284 | 0.13591976367444 | 0.0,0.0,0.0,0.04,0. | 0.08,0.038,0.0,0.0 | -0,016 |
| 198332 | ENSG00000107798 | LIPA | chr10 | - | 89222510 | 89222577 | 89215937 | 89216009 | 89223683 | 89223830 | 198332 | 39,34,69,47,15,10,10,20,3 | 0,0,0,0,0,0,17,27,0,0,0,0 | 43,54,37,9,18,13,19,34,10, | 0,0,0,0,0,0,0,0,0,0,0,0,0,0, | 216 | 149 | 0.00134177872218 | 0.135966278791678 | 1.0,1.0,1.0,1.0,1.0, | 1.0,1.0,1.0,1.0,1.0, | -0,114 |
| 1549 | ENSG00000242259 | C22orf39 | chr22 | - | 19377868 | 19378066 | 19375630 | 19375792 | 19383619 | 19383705 | 1549 | 41,31,36,35,22,18,30,49,3 | 3,0,2,7,0,3,0,1,1,2,1,3 | 28,58,21,34,22,22,28,46,7, | 4,3,0,0,0,0,0,1,2,0,1,2,0,1, | 298 | 149 | 0.00135367378598 | 0.137107088296594 | 0.872,1.0,0.9,0.71 | 0.778,0.906,1.0,1. | -0,049 |
| 248372 | ENSG00000132274 | TRIM22 | chr11 | + | 5706593 | 5706715 | 5696166 | 5698545 | 5708172 | 5710760 | 248372 | 53,35,47,37,20,16,16,29,3 | 3,2,0,2,0,4,0,2,0,6,0,2 | 38,37,31,41,20,22,17,21,2 | 0,0,0,0,0,3,0,4,1,1,10,2,1,3 | 271 | 149 | 0.00135960591247 | 0.137623080210439 | 0.907,0.906,1.0,0. | 1.0,1.0,1.0,1.0,1.0, | -0,029 |
| 60927 | ENSG00000165671 | NSD1 | chr5 | + | 177282464 | 177282581 | 177280564 | 177280834 | 177288818 | 177288925 | 60927 | 15,9,29,17,12,16,11,19,2,4 | 4,1,2,4,3,2,3,1,0,0,0,0 | 24,24,7,6,14,11,14,10,13,1 | 0,0,1,6,0,0,0,4,1,0,2,2,0,0, | 266 | 149 | 0.00136004707318 | 0.137623080210439 | 0.677,0.834,0.89,0 | 1.0,1.0,0.797,0.35 | -0,007 |
| 181120 | ENSG00000162894 | FCMR | chr1 | - | 206909461 | 206909514 | 206903316 | 206905147 | 206909724 | 206909945 | 181120 | 204,64,113,66,114,67,116, | 10,3,4,1,2,1,4,8,10,3,3,4 | 45,17,131,34,6,25,66,121, | 1,4,3,6,4,2,11,14,0,1,3,3,2, | 202 | 149 | 0.00136235358104 | 0.137730120213218 | 0.938,0.94,0.954,0 | 0.971,0.758,0.97,0 | 0,068 |
| 28728 | ENSG00000150593 | PDCD4 | chr10 | + | 110875965 | 110876070 | 110871794 | 110872018 | 110881274 | 110881535 | 28728 | 24,12,124,67,30,16,14,25, | 4,1,2,3,0,0,1,2,1,1,0,2 | 35,46,13,13,14,29,10,10,1 | 4,10,0,4,8,2,0,1,0,0,0,3,2,0 | 254 | 149 | 0.0013627145774 | 0.137730120213218 | 0.779,0.876,0.973, | 0.837,0.73,1.0,0.6 | 0,07 |
| 21248 | ENSG00000071537 | SEL1L | chr14 | - | 81492479 | 81492548 | 81490387 | 81490465 | 81495080 | 81495137 | 21248 | 24,25,62,45,14,33,25,47,2 | 2,4,5,0,0,5,1,3,0,4,0,0 | 30,53,17,11,19,14,26,17,2 | 0,2,1,3,0,3,0,1,0,0,0,1,0,0, | 218 | 149 | 0.00136388828991 | 0.137730120213218 | 0.891,0.81,0.894,1 | 1.0,0.948,0.921,0. | -0,027 |
| 21952 | ENSG00000100577 | GSTZ1 | chr14 | + | 77322546 | 77322732 | 77320995 | 77321183 | 77324869 | 77324921 | 21952 | 1,3,2,6,0,5,2,5,0,0,0,0 | 2,0,15,10,12,8,12,0,4,3,3,5 | 0,3,0,0,0,0,0,2,1,0,0,1,2,2, | 14,4,7,2,5,3,2,18,1,2,7,1,1 | 298 | 149 | 0.00136403820015 | 0.137730120213218 | 0.2,1.0,0.062,0.23 | 0.0,0.273,0.0,0.0,0 | 0,159 |
| 161124 | ENSG00000171617 | ENC1 | chr5 | - | 74631709 | 74631802 | 74629928 | 74629992 | 74634683 | 74636498 | 161124 | 2,4,6,0,3,0,0,1,0,0,0,0 | 44,48,34,8,2,8,9,3,8,4,2,5 | 0,3,2,0,4,0,0,2,3,1,3,17,5,5 | 10,52,22,2,30,4,9,14,17,10 | 242 | 149 | 0.00136481232274 | 0.137730120213218 | 0.027,0.049,0.098, | 0.0,0.034,0.053,0. | -0,014 |
| 132873 | ENSG00000120647 | CCDC77 | chr12 | + | 409367 | 409421 | 405510 | 405564 | 411746 | 411978 | 132873 | 0,5,11,8,1,5,9,13,9,5,1,3 | 4,0,8,2,3,3,0,2,5,2,1,2 | 2,9,4,3,9,2,9,9,6,3,5,1,5,6, | 0,0,1,1,0,3,0,1,0,1,7,0,1,5, | 203 | 149 | 0.00136517547821 | 0.137730120213218 | 0.0,1.0,0.502,0.74 | 1.0,1.0,0.746,0.68 | -0,208 |
| 127396 | ENSG00000172661 | WASHC2C | chr10 | + | 45777272 | 45777425 | 45773255 | 45773358 | 45778952 | 45779135 | 127396 | 11,3,16,7,16,13,9,15,19,15 | 7,2,0,0,2,0,1,0,5,0,1,0 | 8,20,10,16,14,7,14,14,3,15 | 0,1,0,0,0,0,0,1,0,0,1,0,0,0, | 298 | 149 | 0.00136621893531 | 0.137730120213218 | 0.44,0.429,1.0,1.0, | 1.0,0.909,1.0,1.0,1 | -0,131 |
| 158172 | ENSG00000115760 | BIRC6 | chr2 | + | 32465040 | 32465164 | 32464508 | 32464823 | 32467524 | 32467739 | 158172 | 56,39,93,49,35,52,35,44,1 | 4,1,8,3,3,4,7,15,2,3,0,0 | 82,84,34,34,45,34,22,76,2 | 8,27,10,26,17,4,2,2,10,4,1 | 273 | 149 | 0.00136777395665 | 0.137730120213218 | 0.884,0.955,0.864, | 0.848,0.629,0.65,0 | 0,12 |
| 204681 | ENSG00000147274 | RBMX | chrX | - | 136873245 | 136873404 | 136872267 | 136872345 | 136875085 | 136875168 | 204681 | 76,53,148,89,71,63,55,118 | 6,2,25,18,11,6,9,10,8,6,2,1 | 78,71,60,46,34,32,48,57,3 | 17,42,12,13,33,12,5,4,17,2 | 298 | 149 | 0.00136804281177 | 0.137730120213218 | 0.864,0.93,0.747,0 | 0.696,0.458,0.714, | 0,124 |
| 167850 | ENSG00000121210 | TMEM131L | chr4 | + | 153583584 | 153583672 | 153583189 | 153583248 | 153584834 | 153584931 | 167850 | 99,52,144,60,36,42,25,60, | 0,2,4,1,1,5,1,6,0,0,1,2 | 62,79,60,43,41,29,37,47,3 | 0,0,0,0,2,0,0,4,0,1,0,0,2,0, | 237 | 149 | 0.00136836240898 | 0.137730120213218 | 1.0,0.942,0.958,0. | 1.0,1.0,1.0,1.0,0.9 | -0,035 |
| 174580 | ENSG00000009844 | VTA1 | chr6 | + | 142211462 | 142211528 | 142203984 | 142204065 | 142218497 | 142224685 | 174580 | 4,0,3,2,5,0,1,0,4,0,3,0 | 5,12,31,39,28,9,8,18,15,2, | 2,2,6,2,4,0,0,4,3,2,7,4,8,1, | 23,31,11,6,15,8,10,9,4,17, | 215 | 149 | 0.00136845400282 | 0.137730120213218 | 0.357,0.0,0.063,0. | 0.057,0.043,0.274, | -0,067 |
| 33751 | ENSG00000204842 | ATXN2 | chr12 | - | 111510384 | 111510582 | 111509890 | 111509998 | 111513356 | 111513539 | 33751 | 25,8,45,28,21,25,18,25,11, | 1,2,6,0,2,2,2,2,0,0,0,0 | 26,7,24,11,14,11,23,35,13, | 4,3,2,3,3,0,5,1,3,0,1,2,5,1, | 298 | 149 | 0.00136975661955 | 0.137730120213218 | 0.926,0.667,0.789, | 0.765,0.538,0.857, | 0,097 |
| 102357 | ENSG00000136247 | ZDHHC4 | chr7 | + | 6580554 | 6580678 | 6578500 | 6578720 | 6581606 | 6581680 | 102357 | 10,22,64,54,20,22,21,31,1 | 0,4,2,2,1,2,1,3,2,0,0,2 | 60,29,12,8,19,11,8,20,20,4 | 5,5,2,0,4,0,1,5,3,2,2,1,1,0, | 273 | 149 | 0.00137012729251 | 0.137730120213218 | 1.0,0.75,0.946,0.9 | 0.868,0.76,0.766,1 | 0,079 |
| 121122 | ENSG00000101017 | CD40 | chr20 | + | 46126639 | 46126701 | 46122609 | 46122756 | 46128137 | 46128224 | 121122 | 19,16,25,23,23,13,18,10,2 | 2,1,7,0,1,0,2,1,0,4,0,3 | 11,18,3,5,3,1,4,19,4,10,5,3 | 0,6,0,2,0,0,0,0,0,0,3,0,0,0, | 211 | 149 | 0.00137080091285 | 0.137730120213218 | 0.87,0.919,0.716,1 | 1.0,0.679,1.0,0.63 | -0,036 |
| 15728 | ENSG00000197912 | SPG7 | chr16 | + | 89545871 | 89546030 | 89543575 | 89544772 | 89546657 | 89546760 | 15728 | 0,1,10,4,1,0,8,3,2,1,0,1 | 36,17,74,42,40,51,18,19,2 | 1,0,0,2,2,2,0,3,0,0,3,3,0,0, | 45,38,15,18,62,22,32,38,4 | 298 | 149 | 0.00137092843175 | 0.137730120213218 | 0.0,0.029,0.063,0. | 0.011,0.0,0.0,0.05 | 0,026 |
| 106592 | ENSG00000128654 | MTX2 | chr2 | + | 176323391 | 176323464 | 176297848 | 176297895 | 176326824 | 176326901 | 106592 | 7,2,23,19,10,14,7,12,4,6,4, | 2,1,1,4,0,1,2,4,1,0,0,0 | 10,12,4,1,14,5,2,10,9,20,4, | 4,5,1,3,1,2,0,2,7,0,3,1,2,5, | 222 | 149 | 0.00137245384703 | 0.137730120213218 | 0.701,0.573,0.939, | 0.627,0.617,0.729, | 0,127 |
| 132441 | ENSG00000187778 | MCRS1 | chr12 | - | 49559452 | 49559535 | 49559213 | 49559301 | 49559728 | 49559780 | 132441 | 32,41,81,71,58,49,48,36,6 | 2,0,0,0,3,0,5,4,0,0,0,1 | 56,65,47,20,38,19,29,25,2 | 4,3,2,2,5,0,3,0,4,1,1,4,1,1, | 232 | 149 | 0.00137272078362 | 0.137730120213218 | 0.911,1.0,1.0,1.0,0 | 0.9,0.933,0.938,0. | 0,038 |
| 172176 | ENSG00000211897 | IGHG3 | chr14 | - | 105770104 | 105770556 | 105769656 | 105769986 | 105770668 | 105770719 | 172176 | 1,0,266,155,1,2,121,142,5 | 18,4,17,52,12,0,12,5,4,7,1 | 154,186,51,65,0,0,3,4,0,0, | 24,17,0,17,3,8,7,39,2,2,33, | 298 | 149 | 0.00137298485238 | 0.137730120213218 | 0.027,0.0,0.887,0. | 0.762,0.845,1.0,0. | 0,287 |
| 116571 | ENSG00000213930 | GALT | chr9 | + | 34647656 | 34647961 | 34647088 | 34647258 | 34648114 | 34648171 | 116571 | 32,13,35,25,29,37,26,19,4 | 0,0,5,4,3,5,1,1,0,1,1,0 | 10,12,16,18,12,9,20,34,7,2 | 0,0,0,0,3,0,0,3,0,0,3,1,0,0, | 298 | 149 | 0.00137333097495 | 0.137730120213218 | 1.0,1.0,0.778,0.75 | 1.0,1.0,1.0,1.0,0.6 | -0,05 |
| 98582 | ENSG00000168137 | SETD5 | chr3 | + | 9452887 | 9452951 | 9448387 | 9448630 | 9453738 | 9453868 | 98582 | 0,1,10,1,0,1,1,2,0,0,0,0 | 12,10,44,19,10,21,16,20,4, | 0,2,1,2,1,3,3,0,0,0,0,11,3,5 | 25,33,7,23,8,16,28,19,14,1 | 213 | 149 | 0.0013739032393 | 0.137730120213218 | 0.0,0.065,0.137,0. | 0.0,0.041,0.091,0. | -0,041 |
| 153472 | ENSG00000182979 | MTA1 | chr14 | + | 105449144 | 105449409 | 105445417 | 105445511 | 105450057 | 105450184 | 153472 | 9,6,15,5,12,9,12,13,7,3,10, | 4,2,2,0,6,2,0,1,0,2,0,4 | 7,7,8,5,5,1,9,15,3,7,6,10,7, | 6,3,0,4,4,0,1,5,5,1,4,6,4,3, | 298 | 149 | 0.00137604012036 | 0.137824717280228 | 0.529,0.6,0.789,1. | 0.368,0.538,1.0,0. | 0,206 |
| 63765 | ENSG00000145860 | RNF145 | chr5 | - | 159158839 | 159159035 | 159157823 | 159158426 | 159161265 | 159161360 | 63765 | 41,25,52,37,30,16,28,41,1 | 4,0,7,1,2,0,0,2,0,0,1,2 | 37,43,14,15,23,17,19,30,2 | 0,0,0,0,0,0,0,3,0,0,1,0,0,0, | 298 | 149 | 0.00137612759052 | 0.137824717280228 | 0.837,1.0,0.788,0. | 1.0,1.0,1.0,1.0,1.0, | -0,067 |
| 165607 | ENSG00000047365 | ARAP2 | chr4 | - | 36212395 | 36212454 | 36210389 | 36210743 | 36213242 | 36213319 | 165607 | 16,19,40,16,16,18,15,22,1 | 2,3,0,1,2,3,1,0,0,0,0,1 | 26,21,12,20,8,3,9,31,31,30 | 1,5,1,11,0,1,2,1,4,2,2,2,3,0 | 208 | 149 | 0.00138000936679 | 0.138088695740295 | 0.851,0.819,1.0,0. | 0.949,0.751,0.896, | 0,053 |
| 162478 | ENSG00000081791 | DELE1 | chr5 | + | 141931110 | 141931280 | 141930246 | 141930274 | 141933258 | 141933882 | 162478 | 22,16,12,13,6,5,6,11,10,10 | 29,29,48,33,18,26,12,23,3 | 17,9,2,1,11,2,1,7,3,5,3,11, | 38,29,21,10,19,20,25,24,1 | 298 | 149 | 0.00138004648279 | 0.138088695740295 | 0.275,0.216,0.111, | 0.183,0.134,0.045, | 0,066 |
| 82801 | ENSG00000176978 | DPP7 | chr9 | - | 137111940 | 137112029 | 137111689 | 137111754 | 137112111 | 137112230 | 82801 | 190,105,115,53,67,64,92,4 | 4,3,11,3,2,3,0,2,2,0,0,4 | 62,63,140,95,50,31,33,89, | 4,2,10,8,5,2,3,12,0,3,5,3,1, | 238 | 149 | 0.00138384431847 | 0.13840436640735 | 0.967,0.956,0.867, | 0.907,0.952,0.898, | 0,037 |
| 81835 | ENSG00000119396 | RAB14 | chr9 | - | 121190553 | 121190731 | 121186952 | 121187019 | 121192170 | 121192224 | 81835 | 23,34,70,46,25,22,11,30,1 | 0,3,2,1,1,4,2,1,4,1,0,1 | 45,48,31,26,16,18,15,15,1 | 2,2,4,2,0,1,0,0,2,4,2,1,4,0, | 298 | 149 | 0.001398886431 | 0.139824042116176 | 1.0,0.85,0.946,0.9 | 0.918,0.923,0.795, | 0,004 |
| 243774 | ENSG00000108854 | SMURF2 | chr17 | - | 64593990 | 64594104 | 64593439 | 64593573 | 64595078 | 64595182 | 243774 | 21,5,18,11,4,10,5,18,5,2,4, | 1,0,4,2,0,4,2,3,0,1,0,2 | 13,12,16,14,11,2,12,9,3,7, | 0,0,4,4,0,0,0,2,0,0,0,3,0,1, | 263 | 149 | 0.001399565162 | 0.139824042116176 | 0.922,1.0,0.718,0. | 1.0,1.0,0.694,0.66 | -0,103 |
| 99369 | ENSG00000089289 | IGBP1 | chrX | + | 70146632 | 70146828 | 70134522 | 70134816 | 70148760 | 70148840 | 99369 | 118,78,334,275,108,95,98, | 4,1,3,1,1,3,2,3,1,0,0,2 | 122,139,111,76,158,113,5 | 7,3,5,0,7,0,2,6,1,0,3,11,0,5 | 298 | 149 | 0.00140001932838 | 0.139824042116176 | 0.937,0.975,0.982, | 0.897,0.959,0.917, | 0,02 |

| | | | | | | | | | | | | | | | | | | | | | | |
|---|---|---|---|---|---|---|---|---|---|---|---|---|---|---|---|---|---|---|---|---|---|---|
| 117530 | ENSG00000090054 | SPTLC1 | chr9 | - | 92070123 | 92070296 | 92067965 | 92068098 | 92080015 | 92080088 | 117530 | 1,1,1,0,1,0,1,4,0,0,0,2 | 72,27,62,40,20,19,14,27,4 | 0,1,0,18,0,3,4,2,0,2,1,1,0,6 | 38,39,31,10,28,12,16,11,5, | 298 | 149 | 0.00140063761634 | 0.139824042116176 | 0.007,0.018,0.008, | 0.0,0.013,0.0,0.47 | -0,042 |
| 62644 | ENSG00000121892 | PDS5A | chr4 | - | 39926774 | 39926861 | 39925835 | 39925933 | 39927960 | 39928164 | 62644 | 37,11,103,119,75,87,36,67 | 3,2,4,0,1,0,1,8,0,0,0,1 | 63,59,35,20,19,21,24,47,3 | 0,4,0,0,10,4,0,3,1,1,3,1,1,7 | 236 | 149 | 0.00140269625672 | 0.139964635155356 | 0.886,0.776,0.942, | 1.0,0.903,1.0,1.0,0 | 0,037 |
| 226652 | ENSG00000058091 | CDK14 | chr7 | + | 90908295 | 90908490 | 90899290 | 90899353 | 90917600 | 90917724 | 226652 | 103,39,60,38,29,28,33,77, | 730,461,657,466,203,337, | 57,95,27,35,68,24,19,25,7 | 567,453,220,231,330,261,1 | 298 | 149 | 0.0014034555238 | 0.139975503170991 | 0.066,0.041,0.044, | 0.048,0.095,0.058, | -0,01 |
| 31348 | ENSG00000164308 | ERAP2 | chr5 | + | 96886654 | 96886789 | 96883791 | 96883930 | 96889184 | 96889305 | 31348 | 123,114,232,118,83,94,19, | 1,1,2,0,1,5,1,1,1,4,1,0 | 112,124,104,101,100,97,1 | 3,1,3,14,2,1,1,5,0,1,3,3,4,0 | 284 | 149 | 0.00140411508496 | 0.13997642142243 | 0.985,0.984,0.984, | 0.951,0.985,0.948, | 0,019 |
| 153428 | ENSG00000100852 | ARHGAP5 | chr14 | + | 32154195 | 32154528 | 32152422 | 32152528 | 32154620 | 32159728 | 153428 | 13,6,22,7,6,4,3,11,2,4,3,4 | 28,22,126,101,40,47,42,65 | 12,11,7,29,1,6,11,8,10,4,8, | 64,60,56,25,19,21,23,29,3 | 298 | 149 | 0.00140595806306 | 0.14009525919929 | 0.188,0.12,0.08,0. | 0.086,0.084,0.059, | -0,037 |
| 60176 | ENSG00000234127 | TRIM26 | chr6 | - | 30189417 | 30189533 | 30189166 | 30189199 | 30190012 | 30190035 | 60176 | 42,32,76,68,26,28,37,34,2 | 0,0,5,1,1,5,1,6,0,2,2,2 | 32,70,24,23,37,26,20,25,9, | 2,1,7,3,5,2,2,4,0,4,8,7,3,3, | 265 | 149 | 0.00140741368131 | 0.140175406775582 | 1.0,1.0,0.895,0.97 | 0.9,0.975,0.658,0. | 0,063 |
| 252131 | ENSG00000241058 | NSUN6 | chr10 | - | 18588514 | 18588672 | 18585948 | 18586093 | 18596207 | 18596327 | 252131 | 5,2,0,3,3,4,1,5,0,6,0,0 | 43,21,64,46,17,13,18,38,3 | 1,3,2,1,0,2,0,0,1,1,0,0,0,0, | 30,33,30,18,28,16,18,28,2 | 298 | 149 | 0.0014108400186 | 0.140451668845188 | 0.055,0.045,0.0,0. | 0.016,0.043,0.032, | 0,04 |
| 143061 | ENSG00000167543 | TP53I13 | chr17 | + | 29567863 | 29568188 | 29567179 | 29567411 | 29568277 | 29568548 | 143061 | 1,1,6,3,2,4,1,8,5,4,0,3 | 19,8,31,26,14,25,6,28,37,2 | 3,0,3,0,3,1,0,1,0,1,3,0,4,0, | 19,19,11,1,18,0,15,14,5,13 | 298 | 149 | 0.00141168101514 | 0.140470419126027 | 0.026,0.059,0.088, | 0.073,0.0,0.12,0.0, | -0,057 |
| 121124 | ENSG00000101017 | CD40 | chr20 | + | 46126639 | 46126741 | 46123125 | 46123219 | 46128137 | 46128224 | 121124 | 12,8,39,24,20,12,14,7,13,4 | 2,4,4,1,2,1,1,1,0,0,0,0 | 18,26,14,2,3,8,13,22,3,9,9, | 3,3,0,0,3,0,1,5,0,2,0,4,0,0, | 251 | 149 | 0.0014197460695 | 0.141207655399517 | 0.781,0.543,0.853, | 0.781,0.837,1.0,1. | 0,003 |
| 103032 | ENSG00000176102 | CSTF3 | chr11 | - | 33099607 | 33099717 | 33099033 | 33099150 | 33102176 | 33102339 | 103032 | 1,2,10,9,6,5,3,8,6,3,3,2 | 4,0,7,3,1,0,0,3,0,0,2,0 | 11,5,4,1,9,5,8,3,2,5,6,4,4,6 | 0,4,2,0,1,2,0,1,2,1,3,0,0,0, | 259 | 149 | 0.0014207207245 | 0.141239326676609 | 0.126,1.0,0.451,0. | 1.0,0.418,0.535,1. | -0,037 |
| 73686 | ENSG00000136068 | FLNB | chr3 | + | 58150104 | 58150227 | 58149087 | 58150002 | 58153374 | 58153641 | 73686 | 11,8,65,86,64,79,32,26,13, | 0,0,4,4,0,3,3,2,0,0,1,0 | 40,44,5,10,17,29,30,151,4 | 5,3,0,0,4,0,1,3,2,1,1,7,2,0, | 272 | 149 | 0.00142714706479 | 0.141812691552085 | 1.0,1.0,0.899,0.92 | 0.814,0.889,1.0,1. | 0,035 |
| 86709 | ENSG00000170919 | TPT1-AS1 | chr13 | + | 45390757 | 45390902 | 45389734 | 45389820 | 45391031 | 45391619 | 86709 | 22,20,18,16,9,8,10,11,16,8 | 7,0,0,0,0,0,4,3,0,3,0,1 | 11,17,12,8,15,5,9,6,19,20, | 0,0,0,0,0,0,1,0,0,0,1,0,0,0, | 294 | 149 | 0.00143101904399 | 0.142131822730508 | 0.614,1.0,1.0,1.0,1 | 1.0,1.0,1.0,1.0,1.0, | -0,116 |
| 1746 | ENSG00000100335 | MIEF1 | chr22 | + | 39509421 | 39509571 | 39504202 | 39504534 | 39511287 | 39511438 | 1746 | 5,0,5,0,1,0,0,0,2,2,1,0 | 9,9,31,21,14,6,6,15,14,3,9, | 3,1,3,2,0,0,5,1,0,1,4,2,6,3, | 18,13,7,8,9,10,8,11,5,8,16, | 298 | 149 | 0.00143426259434 | 0.142265094308406 | 0.217,0.0,0.075,0. | 0.077,0.037,0.176, | -0,047 |
| 234926 | ENSG00000147905 | ZCCHC7 | chr9 | + | 37127094 | 37127263 | 37126311 | 37126942 | 37147361 | 37147445 | 234926 | 8,6,24,19,2,8,9,7,8,1,4,4 | 22,11,57,18,19,21,18,47,5, | 11,6,6,3,7,4,7,9,3,10,18,9, | 66,52,64,15,14,15,24,20,1 | 298 | 149 | 0.00143434200104 | 0.142265094308406 | 0.154,0.214,0.174, | 0.077,0.055,0.045, | 0,081 |
| 179712 | ENSG00000244682 | FCGR2C | chr1 | + | 161591632 | 161591717 | 161591143 | 161591398 | 161592128 | 161592242 | 179712 | 10,9,6,1,10,1,5,6,14,10,3,7 | 3,2,2,1,1,0,3,2,24,21,8,7 | 14,8,0,1,5,2,0,5,4,1,10,5,2 | 18,10,2,11,6,2,5,11,1,1,18, | 234 | 149 | 0.00143434381966 | 0.142265094308406 | 0.68,0.741,0.656,0 | 0.331,0.337,0.0,0. | 0,282 |
| 211269 | ENSG00000176978 | DPP7 | chr9 | - | 137111875 | 137112029 | 137111689 | 137111754 | 137112111 | 137112169 | 211269 | 233,142,161,57,96,69,107, | 4,3,11,3,2,3,0,2,2,0,0,4 | 77,90,174,118,57,35,42,12 | 4,2,10,8,5,2,3,12,0,3,5,3,1, | 298 | 149 | 0.00143695853296 | 0.142458784895216 | 0.967,0.959,0.88,0 | 0.906,0.957,0.897, | 0,036 |
| 152732 | ENSG00000089775 | ZBTB25 | chr14 | - | 64495242 | 64495484 | 64490360 | 64490540 | 64502900 | 64503127 | 152732 | 2,0,6,5,7,8,1,2,5,2,2,0 | 7,6,3,6,9,5,5,1,6,0,0,3 | 5,4,2,2,4,0,3,5,5,7,4,2,1,3, | 10,1,0,1,0,4,0,4,0,3,0,2,0,4 | 298 | 149 | 0.00144122803897 | 0.142729551926914 | 0.125,0.0,0.5,0.29 | 0.2,0.667,1.0,0.5,1 | -0,279 |
| 232106 | ENSG00000139971 | ARMH4 | chr14 | - | 58293765 | 58293926 | 58291634 | 58291707 | 58297954 | 58298139 | 232106 | 16,17,30,23,16,22,12,25,4, | 67,34,129,99,60,80,54,73, | 25,28,18,18,13,14,14,14,9, | 76,42,35,33,53,24,20,71,4 | 298 | 149 | 0.00144140256269 | 0.142729551926914 | 0.107,0.2,0.104,0. | 0.141,0.25,0.205,0 | -0,049 |
| 28290 | ENSG00000055163 | CYFIP2 | chr5 | + | 157300714 | 157300896 | 157296672 | 157296774 | 157302793 | 157302890 | 28290 | 257,192,483,260,307,306, | 4,1,2,3,1,2,4,1,0,0,1,0 | 384,281,155,220,329,218, | 2,5,0,1,11,0,0,2,2,3,4,9,9,2 | 298 | 149 | 0.00144167915351 | 0.142729551926914 | 0.97,0.99,0.992,0. | 0.99,0.966,1.0,0.9 | 0,012 |
| 70571 | ENSG00000144591 | GMPPA | chr2 | + | 219505227 | 219505362 | 219504082 | 219504213 | 219505457 | 219505555 | 70571 | 60,34,77,46,33,46,29,26,5 | 4,1,3,0,0,3,4,3,2,0,0,1 | 23,35,25,16,16,8,21,35,10, | 0,0,0,0,0,0,0,0,0,0,2,0,0, | 284 | 149 | 0.00144248211223 | 0.142743387355115 | 0.887,0.947,0.931, | 1.0,1.0,1.0,1.0,1.0, | -0,062 |
| 156846 | ENSG00000134884 | ARGLU1 | chr13 | - | 106557504 | 106557616 | 106557105 | 106557131 | 106559431 | 106559657 | 156846 | 38,9,20,18,6,10,4,16,16,15 | 130,49,133,112,66,62,60,6 | 16,12,17,2,2,6,5,18,1,9,7,5 | 104,54,77,34,46,41,18,79, | 261 | 149 | 0.00144727315371 | 0.143151676537756 | 0.143,0.095,0.079, | 0.081,0.113,0.112, | 0,038 |
| 162476 | ENSG00000081791 | DELE1 | chr5 | + | 141930950 | 141931280 | 141930246 | 141930274 | 141933258 | 141933882 | 162476 | 32,19,23,25,7,7,13,18,13,1 | 29,29,48,33,18,26,12,23,3 | 18,16,4,2,12,4,4,10,6,8,6,1 | 38,29,21,10,19,20,25,24,1 | 298 | 149 | 0.00145115131601 | 0.143469338032222 | 0.356,0.247,0.193, | 0.191,0.216,0.087, | 0,078 |
| 190003 | ENSG00000258728 | | chr9 | + | 34647474 | 34647961 | 34647088 | 34647258 | 34648114 | 34648171 | 190003 | 34,13,37,25,29,35,28,18,4 | 0,0,5,4,3,5,1,1,0,1,1,0 | 10,12,17,18,14,9,20,34,7,1 | 0,0,0,0,3,0,0,3,0,0,3,1,0,0, | 298 | 149 | 0.00145306001805 | 0.143592084823195 | 1.0,1.0,0.787,0.75 | 1.0,1.0,1.0,1.0,0.7, | -0,05 |
| 14488 | ENSG00000138834 | MAPK8IP3 | chr16 | + | 1729137 | 1729301 | 1724556 | 1724677 | 1729486 | 1729578 | 14488 | 106,51,37,22,20,15,23,31, | 6,2,3,2,2,2,1,7,2,0,0,0 | 33,53,51,64,54,11,16,25,4 | 0,3,1,2,0,0,2,0,0,2,4,1,0,1, | 298 | 149 | 0.00145427580363 | 0.143646276039967 | 0.898,0.927,0.86,0 | 1.0,0.898,0.962,0. | -0,032 |
| 207589 | ENSG00000146192 | FGD2 | chr6 | + | 37026090 | 37026208 | 37025791 | 37025938 | 37027428 | 37029069 | 207589 | 28,25,32,20,10,5,4,6,41,28 | 40,26,95,72,66,39,40,23,6 | 7,6,1,19,5,8,4,6,13,18,5,9, | 37,44,22,17,39,26,24,17,5 | 267 | 149 | 0.00145516860161 | 0.143668529033542 | 0.281,0.349,0.158, | 0.095,0.071,0.025, | 0,053 |
| 143372 | ENSG00000198862 | LTN1 | chr21 | - | 28972215 | 28972258 | 28971270 | 28971444 | 28981118 | 28981299 | 143372 | 4,2,1,3,3,10,0,0,2,0,2,0 | 14,9,41,21,13,10,9,9,12,5, | 1,2,3,0,2,2,2,0,1,2,2,3,0,0, | 25,36,13,13,21,14,16,17,6, | 192 | 149 | 0.00145777211699 | 0.143816556402207 | 0.181,0.147,0.019, | 0.03,0.041,0.152,0 | 0,064 |
| 52392 | ENSG00000267787 | | chr18 | + | 57735150 | 57735592 | 57641558 | 57642052 | 57737761 | 57738044 | 52392 | 31,15,18,8,6,3,20,2,14,17, | 10,2,7,0,1,4,2,0,2,5,0,2 | 13,4,10,9,2,6,19,5,5,7,6,16 | 9,2,8,2,5,10,3,0,2,6,13,11, | 298 | 149 | 0.00145800431197 | 0.143816556402207 | 0.608,0.789,0.562, | 0.419,0.5,0.385,0. | 0,236 |
| 103689 | ENSG00000113013 | HSPA9 | chr5 | - | 138566625 | 138566718 | 138556773 | 138561789 | 138567000 | 138567010 | 103689 | 156,141,347,271,152,139, | 18,3,18,6,4,9,11,16,0,5,2,1 | 186,271,127,77,206,81,95, | 12,34,6,16,28,9,4,12,6,15, | 242 | 149 | 0.00146043980787 | 0.143990801780883 | 0.842,0.967,0.922, | 0.905,0.831,0.929, | 0,039 |
| 155302 | ENSG00000103811 | CTSH | chr15 | - | 78931862 | 78932041 | 78931450 | 78931506 | 78932371 | 78932458 | 155302 | 8,11,9,1,14,41,0,10,1,2,12, | 327,176,243,162,304,494, | 6,8,2,0,3,5,1,9,7,16,1,4,8,7 | 150,135,194,64,120,94,56, | 298 | 149 | 0.00146124961369 | 0.14400467747441 | 0.012,0.03,0.018,0 | 0.02,0.029,0.005,0 | 0,008 |
| 251838 | ENSG00000163399 | ATP1A1 | chr1 | + | 116382414 | 116382593 | 116373268 | 116373523 | 116384013 | 116384124 | 251838 | 14,3,8,5,1,9,0,2,0,0,0,2 | 79,33,42,37,15,4,12,21,12, | 10,15,4,19,11,2,4,7,3,3,5,2 | 17,33,15,18,5,11,8,30,17,3 | 298 | 149 | 0.00146362844143 | 0.144130195245869 | 0.081,0.043,0.087, | 0.227,0.185,0.118, | -0,064 |
| 90841 | ENSG00000155827 | RNF20 | chr9 | + | 101544766 | 101544885 | 101540792 | 101540975 | 101546819 | 101546966 | 90841 | 53,37,71,45,41,32,26,21,4 | 1,2,6,0,0,1,0,0,1,0,4,3 | 64,42,53,56,29,27,20,26,1 | 1,2,2,4,4,2,1,2,0,6,4,10,1,4 | 268 | 149 | 0.00146386257931 | 0.144130195245869 | 0.967,0.911,0.868, | 0.973,0.921,0.936, | 0,053 |
| 136492 | ENSG00000125734 | GPR108 | chr19 | - | 6732955 | 6733062 | 6732475 | 6732549 | 6733167 | 6733301 | 136492 | 65,20,34,27,24,21,22,20,8 | 3,8,2,0,2,2,0,0,0,0,2,2 | 28,15,27,28,18,14,10,19,1 | 0,0,0,0,0,0,1,0,0,0,1,0,0, | 256 | 149 | 0.00146506770654 | 0.144182893253914 | 0.927,0.593,0.908, | 1.0,1.0,1.0,1.0,1.0, | -0,1 |
| 229958 | ENSG00000026950 | BTN3A1 | chr6 | + | 26408678 | 26408861 | 26407826 | 26407952 | 26409532 | 26409733 | 229958 | 6,6,5,1,1,1,3,2,0,0,1,2 | 61,55,42,42,21,19,8,17,20, | 3,1,2,2,9,1,3,3,3,2,6,8,5,5, | 11,20,9,19,27,16,13,41,12, | 298 | 149 | 0.00146583569006 | 0.144188458542174 | 0.047,0.052,0.056, | 0.12,0.024,0.1,0.0 | -0,035 |
| 96986 | ENSG00000095015 | MAP3K1 | chr5 | + | 56887377 | 56887520 | 56885931 | 56886063 | 56888225 | 56888357 | 96986 | 386,355,818,256,222,162, | 43,10,54,26,35,23,15,17,1 | 547,542,322,369,280,151, | 59,41,24,52,23,23,18,26,2 | 292 | 149 | 0.00146646410484 | 0.144188458542174 | 0.821,0.948,0.885, | 0.826,0.871,0.873, | 0,037 |
| 19245 | ENSG00000114062 | UBE3A | chr15 | - | 25407066 | 25407212 | 25405460 | 25405502 | 25409087 | 25409207 | 19245 | 15,6,42,24,9,15,10,19,14,5 | 31,31,96,30,23,39,24,66,1 | 20,11,6,4,6,3,8,6,8,11,20,1 | 40,78,26,27,41,17,36,31,2 | 295 | 149 | 0.00146925908522 | 0.144397306927391 | 0.196,0.089,0.181, | 0.202,0.066,0.104, | 0,058 |
| 146769 | ENSG00000138834 | MAPK8IP3 | chr16 | + | 1729137 | 1729222 | 1724556 | 1724677 | 1729486 | 1729578 | 146769 | 91,48,34,23,18,14,24,29,1 | 6,2,3,2,2,2,1,7,2,0,0,0 | 31,49,42,53,49,11,16,27,3 | 0,3,1,2,0,0,2,0,0,2,4,1,0,1, | 234 | 149 | 0.00147117289114 | 0.144461542895286 | 0.906,0.939,0.878, | 1.0,0.912,0.964,0. | -0,023 |
| 98395 | ENSG00000186480 | INSIG1 | chr7 | + | 155302746 | 155302846 | 155298258 | 155298697 | 155308240 | 155308757 | 98395 | 26,25,85,42,11,20,18,22,2 | 8,3,7,7,4,3,11,3,5,0,7,4 | 11,18,16,7,17,8,8,19,3,11, | 8,12,5,4,8,5,2,2,3,5,10,5,2, | 249 | 149 | 0.0014715633854 | 0.144461542895286 | 0.66,0.833,0.879,0 | 0.451,0.473,0.657, | 0,169 |
| 44871 | ENSG00000103064 | SLC7A6 | chr16 | + | 68290395 | 68290540 | 68274690 | 68275249 | 68291208 | 68291332 | 44871 | 14,6,28,15,8,5,18,9,7,5,4,9 | 1,0,0,0,0,1,4,4,0,0,0,0 | 8,12,11,3,14,9,9,9,13,15,1 | 0,1,0,3,2,4,0,2,4,3,4,1,0,0, | 294 | 149 | 0.00147192627256 | 0.144461542895286 | 0.876,1.0,1.0,1.0,1 | 1.0,0.859,1.0,0.33 | 0,1 |
| 229384 | ENSG00000162600 | OMA1 | chr1 | - | 58506059 | 58506209 | 58480718 | 58481174 | 58522826 | 58522879 | 229384 | 102,43,69,48,35,34,35,45, | 3,0,2,6,0,2,0,10,0,0,0,0 | 82,32,40,37,32,37,18,37,1 | 0,0,0,1,0,0,0,0,0,0,3,1,0,0, | 298 | 149 | 0.00147549420397 | 0.144745712404133 | 0.944,1.0,0.945,0. | 1.0,1.0,1.0,0.949,1 | -0,038 |
| 249554 | ENSG00000155629 | PIK3AP1 | chr10 | - | 96718612 | 96718693 | 96709566 | 96709983 | 96720381 | 96720514 | 249554 | 9,1,27,11,9,6,5,8,7,3,1,5 | 13,11,39,35,15,9,22,4,21,8 | 3,7,1,3,3,3,6,6,0,2,4,6,6,2, | 17,20,9,8,11,10,12,13,7,28 | 230 | 149 | 0.00147695404113 | 0.144822913542802 | 0.31,0.056,0.31,0. | 0.103,0.185,0.067, | 0,092 |
| 227695 | ENSG00000186088 | GSAP | chr7 | - | 77379947 | 77380000 | 77377285 | 77377390 | 77381304 | 77381354 | 227695 | 8,13,11,5,2,12,12,10,2,1,6, | 51,26,156,99,53,60,40,66, | 7,11,5,6,12,7,5,19,5,1,13,1 | 102,45,42,11,27,18,28,56, | 202 | 149 | 0.001478601419 | 0.144842958821365 | 0.104,0.269,0.049, | 0.048,0.153,0.081, | -0,053 |
| 37922 | ENSG00000163138 | PACRGL | chr4 | + | 20724807 | 20724888 | 20713431 | 20713539 | 20727284 | 20727397 | 37922 | 7,5,22,5,5,9,16,3,6,10,2,10 | 2,2,1,0,0,0,0,3,0,0,4,5 | 8,7,3,2,7,8,4,8,7,7,6,8,0,5, | 4,0,1,1,2,4,0,3,1,8,7,3,2,6,1 | 230 | 149 | 0.00147875110905 | 0.144842958821365 | 0.694,0.618,0.934, | 0.564,1.0,0.66,0.5 | 0,188 |
| 5480 | ENSG00000167232 | ZNF91 | chr19 | - | 23373741 | 23373837 | 23357695 | 23362725 | 23395324 | 23395457 | 5480 | 12,12,17,6,4,5,8,3,7,1,2,8 | 2,3,3,7,0,3,2,0,0,0,2,1 | 15,5,15,11,8,1,1,12,6,30,9, | 5,0,0,5,3,2,0,0,0,2,0,3,0,0, | 245 | 149 | 0.0014791773652 | 0.144842958821365 | 0.785,0.709,0.775, | 0.646,1.0,1.0,0.57 | -0,097 |
| 158170 | ENSG00000115760 | BIRC6 | chr2 | + | 32465036 | 32465164 | 32464508 | 32464823 | 32467524 | 32467739 | 158170 | 56,39,91,50,35,51,35,44,1 | 4,1,8,3,3,4,7,15,2,3,0,0 | 82,84,34,34,45,34,22,76,2 | 8,27,10,26,17,4,2,2,10,4,11 | 277 | 149 | 0.00148225580047 | 0.145032294615822 | 0.883,0.955,0.86,0 | 0.846,0.626,0.647, | 0,121 |
| 149140 | ENSG00000169375 | SIN3A | chr15 | - | 75446232 | 75446395 | 75430186 | 75430408 | 75451422 | 75451746 | 149140 | 0,2,2,1,0,2,2,2,0,1,2,2 | 12,18,15,6,6,7,8,6,12,10,2, | 6,7,3,8,1,2,4,5,4,0,0,4,1,3, | 9,24,7,17,19,7,16,9,11,12,1 | 298 | 149 | 0.00148245860566 | 0.145032294615822 | 0.0,0.053,0.062,0. | 0.25,0.127,0.176,0 | -0,022 |
| 210999 | ENSG00000068878 | PSME4 | chr2 | - | 53913870 | 53913931 | 53910074 | 53910130 | 53919150 | 53919246 | 210999 | 1,1,0,2,1,1,2,2,0,0,3,0 | 4,9,22,12,10,21,11,15,2,7, | 6,2,3,5,3,4,8,8,3,0,2,11,2,0 | 39,71,13,32,28,15,26,20,1 | 210 | 149 | 0.00148554747265 | 0.145268454377979 | 0.151,0.073,0.0,0. | 0.098,0.02,0.141,0 | 0,028 |
| 116030 | ENSG00000011454 | RABGAP1 | chr9 | + | 123071411 | 123071501 | 123070349 | 123070424 | 123073551 | 123073677 | 116030 | 1,1,3,1,7,2,2,3,0,0,0,0 | 17,8,46,28,6,12,3,5,7,11,2, | 3,14,3,4,0,3,3,2,4,8,1,2,3,1 | 13,13,18,8,8,8,20,8,4,16,9, | 239 | 149 | 0.00148711928409 | 0.145356117454121 | 0.035,0.072,0.039, | 0.126,0.402,0.094, | -0,071 |
| 126315 | ENSG00000157625 | TAB3 | chrX | - | 30845932 | 30846071 | 30842987 | 30843049 | 30846550 | 30846644 | 126315 | 2,1,4,7,1,0,1,3,3,0,0,0 | 14,13,31,22,5,10,6,13,10,6 | 4,1,2,2,5,2,2,1,6,5,0,5,1,2, | 7,8,11,13,8,4,5,9,5,15,19,1 | 288 | 149 | 0.00148943669415 | 0.145516545219518 | 0.069,0.038,0.063, | 0.228,0.061,0.086, | -0,077 |
| 15747 | ENSG00000197912 | SPG7 | chr16 | + | 89555877 | 89556085 | 89554485 | 89554563 | 89556886 | 89557701 | 15747 | 14,16,6,5,1,3,0,2,2,6,8,27 | 50,36,67,45,40,34,21,52,3 | 4,0,0,1,4,1,2,5,8,11,6,5,3,5 | 36,38,30,29,34,24,24,41,3 | 298 | 149 | 0.00149086911452 | 0.145585059159707 | 0.123,0.182,0.043, | 0.053,0.0,0.0,0.01 | 0,05 |
| 111024 | ENSG00000033800 | PIAS1 | chr15 | + | 68145815 | 68145906 | 68086305 | 68086750 | 68146565 | 68146700 | 111024 | 39,19,65,62,25,28,19,41,1 | 4,2,2,1,0,0,1,0,0,0,0,0 | 76,60,23,23,41,28,21,31,2 | 0,0,2,15,0,1,2,0,0,1,10,0,0, | 240 | 149 | 0.00149149079569 | 0.145585059159707 | 0.858,0.855,0.953, | 1.0,1.0,0.877,0.48 | 0,044 |
| 192829 | ENSG00000096060 | FKBP5 | chr6 | - | 35643276 | 35643348 | 35642719 | 35642843 | 35688803 | 35688942 | 192829 | 0,0,0,0,0,2,0,9,0,0,0,3 | 11,5,9,12,3,11,30,37,13,1, | 2,6,3,5,2,0,0,1,0,1,10,1,1,0 | 6,14,4,0,8,6,16,13,5,15,43, | 221 | 149 | 0.00149304331417 | 0.145670537421633 | 0.0,0.0,0.0,0.0,0.0, | 0.183,0.224,0.336, | -0,118 |
| 34620 | ENSG00000136709 | WDR33 | chr2 | - | 127709489 | 127709582 | 127708676 | 127708892 | 127709692 | 127709856 | 34620 | 53,31,105,49,41,40,40,43, | 2,0,0,0,0,1,1,4,2,0,0,0 | 64,54,32,35,22,21,43,48,2 | 5,0,0,0,0,0,0,10,2,0,5,1,1,2 | 242 | 149 | 0.00149680176902 | 0.145971065495217 | 0.942,1.0,1.0,1.0,1 | 0.887,1.0,1.0,1.0,1 | 0,031 |
| 187827 | ENSG00000139645 | ANKRD52 | chr12 | - | 56257518 | 56257728 | 56257282 | 56257361 | 56257827 | 56257911 | 187827 | 0,1,2,2,3,0,0,1,0,0,0,2 | 11,8,18,25,5,19,10,11,2,2, | 5,0,0,6,0,2,4,2,1,2,1,1,5,0, | 15,16,7,10,17,10,18,10,14, | 298 | 149 | 0.00149931796392 | 0.146150228574486 | 0.0,0.059,0.053,0. | 0.143,0.0,0.0,0.23 | -0,027 |
| 109098 | ENSG00000075539 | FRYL | chr4 | - | 48539970 | 48540068 | 48535656 | 48535827 | 48540352 | 48540960 | 109098 | 44,59,71,57,23,24,27,37,3 | 2,4,1,2,2,8,2,2,1,1,3,3 | 46,55,39,34,34,34,37,36,2 | 3,12,5,10,5,2,2,4,6,8,5,2,7, | 247 | 149 | 0.00151030915094 | 0.147101415731919 | 0.93,0.899,0.977,0 | 0.902,0.734,0.825, | 0,053 |
| 73621 | ENSG00000165119 | HNRNPK | chr9 | - | 83973901 | 83973973 | 83973285 | 83973399 | 83974516 | 83974589 | 73621 | 504,398,1187,706,408,475 | 49,32,56,50,14,25,17,24,2 | 868,780,614,517,420,314, | 31,25,17,8,15,19,26,19,21, | 221 | 149 | 0.00151044286728 | 0.147101415731919 | 0.874,0.893,0.935, | 0.95,0.955,0.961,0 | -0,018 |
| 237400 | ENSG00000179526 | SHARPIN | chr8 | - | 144100742 | 144100998 | 144099928 | 144100069 | 144103050 | 144103225 | 237400 | 8,0,9,1,0,2,0,6,0,1,0,1 | 35,24,64,38,29,20,33,31,4 | 2,9,0,9,4,0,1,1,2,2,5,7,6,4, | 27,35,19,19,21,13,22,49,1 | 298 | 149 | 0.00151472901769 | 0.147452121757773 | 0.103,0.0,0.066,0. | 0.036,0.114,0.0,0. | -0,032 |
| 11433 | ENSG00000125447 | GGA3 | chr17 | - | 75248839 | 75249008 | 75246711 | 75246796 | 75261547 | 75261600 | 11433 | 0,0,17,3,2,2,0,6,4,0,0,2 | 34,20,17,14,7,13,10,8,13,1 | 3,3,0,0,2,1,1,0,1,6,1,1,0,2, | 13,16,8,4,11,10,3,13,11,13 | 298 | 149 | 0.00151563766079 | 0.147473874036841 | 0.0,0.0,0.333,0.09 | 0.103,0.086,0.0,0. | 0,041 |
| 253319 | ENSG00000136021 | SCYL2 | chr12 | + | 100296761 | 100296982 | 100291502 | 100291660 | 100298030 | 100298175 | 253319 | 10,5,7,0,3,10,6,9,0,0,3,9 | 35,24,62,54,19,15,27,34,1 | 16,23,10,11,10,3,17,6,4,5, | 54,34,28,29,18,16,21,31,1 | 298 | 149 | 0.00151758532044 | 0.147572970928941 | 0.125,0.094,0.053, | 0.129,0.253,0.152, | -0,048 |
| 64471 | ENSG00000204469 | PRRC2A | chr6 | + | 31625791 | 31625871 | 31625459 | 31625611 | 31626019 | 31626162 | 64471 | 5,2,45,30,15,16,9,10,6,3,4, | 2,0,9,4,0,0,0,2,0,0,0,0 | 8,15,2,3,13,4,14,20,5,8,11, | 1,5,2,1,4,3,1,1,3,1,3,6,3,1, | 229 | 149 | 0.00151802741072 | 0.147572970928941 | 0.619,1.0,0.765,0. | 0.839,0.661,0.394, | 0,201 |
| 30392 | ENSG00000198055 | GRK6 | chr5 | + | 177431994 | 177432107 | 177430871 | 177430967 | 177432232 | 177432257 | 30392 | 70,38,84,65,40,35,56,18,1 | 16,9,7,6,4,12,7,2,1,1,2,0 | 32,64,41,35,39,28,27,31,1 | 15,20,7,9,13,4,7,9,7,2,11,1 | 262 | 149 | 0.00151984832635 | 0.147651002683817 | 0.713,0.706,0.872, | 0.548,0.645,0.769, | 0,102 |
| 125471 | ENSG00000085274 | MYNN | chr3 | + | 169774030 | 169774097 | 169773426 | 169773462 | 169774264 | 169774561 | 125471 | 11,4,15,20,4,4,4,6,7,3,2,2 | 18,14,45,30,13,16,8,8,8,4, | 1,7,4,1,3,8,1,5,2,2,7,3,10,5 | 16,37,13,11,28,18,13,12,1 | 216 | 149 | 0.00152020211748 | 0.147651002683817 | 0.297,0.165,0.187, | 0.041,0.115,0.175, | 0,106 |
| 189238 | ENSG00000118308 | IRAG2 | chr12 | + | 25069349 | 25069423 | 25066364 | 25066512 | 25079243 | 25079290 | 189238 | 207,102,387,267,124,108, | 3,7,22,7,3,11,12,15,4,4,0,0 | 146,198,142,46,85,68,37,6 | 8,8,3,2,7,2,2,5,0,0,2,0,6,7, | 223 | 149 | 0.0015214263525 | 0.147703254521844 | 0.979,0.907,0.922, | 0.924,0.943,0.969, | -0,022 |
| 64555 | ENSG00000114861 | FOXP1 | chr3 | - | 71335159 | 71335277 | 71299819 | 71299880 | 71359149 | 71359244 | 64555 | 2,0,6,5,1,0,1,0,2,0,0,0 | 61,10,145,69,52,9,51,65,2 | 6,5,4,6,0,1,0,0,8,5,2,11,2,4 | 155,105,63,69,33,12,43,47 | 267 | 149 | 0.00152329432867 | 0.147817926805218 | 0.018,0.0,0.023,0. | 0.021,0.026,0.034, | -0,015 |
| 181088 | ENSG00000178927 | CYBC1 | chr17 | - | 82447579 | 82447638 | 82446622 | 82446696 | 82449169 | 82449292 | 181088 | 9,5,31,28,12,13,4,8,2,1,4,5 | 0,0,1,4,1,0,5,0,3,0,0,0 | 12,12,10,3,6,4,2,3,5,11,6,2 | 3,0,0,0,0,6,1,4,0,4,2,5,1,2, | 208 | 149 | 0.00152435836858 | 0.147854518264012 | 1.0,1.0,0.957,0.83 | 0.741,1.0,1.0,1.0,1 | 0,078 |
| 183987 | ENSG00000132549 | VPS13B | chr8 | + | 99099967 | 99100022 | 99096311 | 99096432 | 99102952 | 99103120 | 183987 | 3,0,1,1,1,3,1,3,0,0,0,0 | 10,14,82,43,23,55,27,33,8, | 3,7,0,4,1,3,0,0,5,1,0,4,3,2, | 24,45,10,19,36,22,20,31,2 | 204 | 149 | 0.00153364398021 | 0.148688165542603 | 0.18,0.0,0.009,0.0 | 0.084,0.102,0.0,0. | -0,024 |
| 70456 | ENSG00000168421 | RHOH | chr4 | + | 40197100 | 40197300 | 40191010 | 40191338 | 40242713 | 40242834 | 70456 | 294,139,485,273,146,157, | 0,3,13,12,5,6,2,18,2,4,0,2 | 355,405,185,196,173,69,2 | 4,2,3,0,3,1,1,0,1,2,6,3,14,1 | 298 | 149 | 0.00154006374704 | 0.149243340990169 | 1.0,0.959,0.949,0. | 0.978,0.99,0.969,1 | -0,021 |
| 83422 | ENSG00000196247 | ZNF107 | chr7 | + | 64679176 | 64679336 | 64666129 | 64666285 | 64691247 | 64691374 | 83422 | 105,104,64,105,45,62,64,9 | 31,31,69,44,13,26,48,30,2 | 80,66,32,155,34,25,41,68, | 58,39,35,23,30,19,19,28,2 | 298 | 149 | 0.00154264585352 | 0.149426286993233 | 0.629,0.627,0.317, | 0.408,0.458,0.314, | 0,135 |
| 232005 | ENSG00000139083 | ETV6 | chr12 | + | 11826401 | 11826603 | 11752449 | 11752579 | 11839139 | 11839304 | 232005 | 24,7,11,4,1,14,23,14,16,15 | 21,22,47,51,29,20,25,19,2 | 8,10,1,18,13,1,9,4,5,3,10,0 | 36,34,23,16,14,48,20,25,2 | 298 | 149 | 0.00154469092848 | 0.149557072976914 | 0.364,0.137,0.105, | 0.1,0.128,0.021,0. | 0,079 |
| 115426 | ENSG00000149177 | PTPRJ | chr11 | + | 48121002 | 48121266 | 48112746 | 48112983 | 48123612 | 48123870 | 115426 | 103,130,259,221,173,72,9 | 18,10,23,17,2,10,9,9,2,0,2, | 197,238,67,100,122,79,63, | 21,25,4,5,39,17,9,10,15,15 | 298 | 149 | 0.00154671500736 | 0.149685709419559 | 0.741,0.867,0.849, | 0.824,0.826,0.893, | 0,069 |
| 92917 | ENSG00000198315 | ZKSCAN8 | chr6 | + | 28148315 | 28148824 | 28141882 | 28142029 | 28149482 | 28149624 | 92917 | 47,35,64,36,38,48,28,43,3 | 4,3,3,5,0,3,2,5,0,1,0,2 | 55,47,18,72,36,12,33,45,2 | 4,8,12,3,7,2,6,2,3,0,4,2,0,3 | 298 | 149 | 0.0015477010538 | 0.149691382812748 | 0.855,0.854,0.914, | 0.873,0.746,0.429, | 0,102 |
| 64662 | ENSG00000058673 | ZC3H11A | chr1 | + | 203802956 | 203803016 | 203795653 | 203795794 | 203816925 | 203817124 | 64662 | 26,24,75,25,20,30,29,65,4 | 10,8,33,16,9,16,15,16,4,8, | 37,43,31,29,20,13,25,47,1 | 28,38,27,26,29,9,15,18,6,7 | 209 | 149 | 0.00154850999035 | 0.149691382812748 | 0.65,0.681,0.618,0 | 0.485,0.447,0.45,0 | 0,114 |
| 63542 | ENSG00000120370 | GORAB | chr1 | + | 170533463 | 170536421 | 170532165 | 170532284 | 170539209 | 170539567 | 63542 | 64,58,47,29,31,23,24,35,1 | 0,2,6,3,2,1,0,1,3,1,2,0 | 40,38,25,47,26,6,12,28,21, | 1,3,0,0,1,0,0,0,0,0,0,2,2, | 298 | 149 | 0.00154886010623 | 0.149691382812748 | 1.0,0.935,0.797,0. | 0.952,0.864,1.0,1. | -0,057 |
| 130261 | ENSG00000100982 | PCIF1 | chr20 | + | 45947072 | 45947166 | 45946199 | 45946384 | 45947262 | 45947438 | 130261 | 70,60,81,44,33,72,44,37,5 | 2,0,0,2,0,3,2,0,0,4,0,2 | 30,89,30,89,50,25,54,49,2 | 4,4,0,1,2,2,7,5,1,2,3,2,1,1, | 243 | 149 | 0.00154991556001 | 0.149726156147447 | 0.955,1.0,1.0,0.93 | 0.821,0.932,1.0,0. | 0,04 |
| 143595 | ENSG00000185813 | PCYT2 | chr17 | - | 81908875 | 81909035 | 81908567 | 81908634 | 81909513 | 81909602 | 143595 | 35,22,30,43,15,9,15,20,29, | 2,0,6,2,0,2,0,0,0,0,0,0 | 10,6,10,4,7,6,3,26,2,13,15, | 3,0,0,0,0,0,1,6,1,3,5,3,0,0, | 298 | 149 | 0.00155105362056 | 0.149768874745065 | 0.897,1.0,0.714,0. | 0.625,1.0,1.0,1.0,1 | 0,108 |
| 70461 | ENSG00000168421 | RHOH | chr4 | + | 40197120 | 40197300 | 40191010 | 40191338 | 40242713 | 40242834 | 70461 | 294,139,486,273,143,157, | 0,3,13,12,5,6,2,18,2,4,0,2 | 355,405,185,196,173,69,2 | 4,2,3,0,3,1,1,0,1,2,6,3,14,1 | 298 | 149 | 0.00155456426465 | 0.150039557058925 | 1.0,0.959,0.949,0. | 0.978,0.99,0.969,1 | -0,021 |
| 81301 | ENSG00000198589 | LRBA | chr4 | - | 150868181 | 150868305 | 150851884 | 150852943 | 150870524 | 150870606 | 81301 | 165,111,143,96,53,89,39,8 | 5,0,1,2,3,3,0,8,0,0,0,1 | 74,64,77,123,51,48,29,53, | 3,3,0,2,11,3,0,0,7,0,3,0,4,0 | 273 | 149 | 0.00155525111066 | 0.150039557058925 | 0.947,1.0,0.987,0. | 0.931,0.921,1.0,0. | 0,034 |
| 27501 | ENSG00000137177 | KIF13A | chr6 | - | 17771113 | 17771218 | 17763692 | 17764946 | 17771907 | 17772059 | 27501 | 1,2,3,3,1,2,2,3,2,0,1,0 | 14,4,25,11,10,20,3,7,6,1,2, | 0,0,0,0,2,1,0,0,0,5,2,0,1,0, | 7,6,15,10,8,4,5,6,3,8,6,13, | 254 | 149 | 0.00155990306775 | 0.150407522694506 | 0.04,0.227,0.066,0 | 0.0,0.0,0.0,0.0,0.1 | 0,074 |
| 63086 | ENSG00000116698 | SMG7 | chr1 | + | 183527625 | 183527731 | 183526595 | 183526767 | 183527955 | 183528027 | 63086 | 4,2,9,4,0,3,3,3,4,0,0,2 | 29,10,22,16,18,30,14,16,1 | 4,4,1,4,2,0,3,1,0,0,2,0,1,4, | 25,31,15,28,14,14,20,33,2 | 255 | 149 | 0.00156069592189 | 0.150407522694506 | 0.075,0.105,0.193, | 0.085,0.07,0.037,0 | 0,042 |
| 231462 | ENSG00000078589 | P2RY10 | chrX | + | 78947814 | 78947862 | 78945390 | 78945495 | 78960507 | 78963727 | 231462 | 2,8,26,16,2,7,5,4,3,4,1,2 | 152,92,179,96,49,38,68,52 | 11,10,13,2,4,2,4,6,5,4,7,3, | 87,78,106,43,60,24,28,65, | 197 | 149 | 0.00156129010892 | 0.150407522694506 | 0.01,0.062,0.099,0 | 0.087,0.088,0.085, | -0,034 |
| 85271 | ENSG00000172765 | TMCC1 | chr3 | - | 129857249 | 129857405 | 129848342 | 129848412 | 129880308 | 129880559 | 85271 | 0,9,9,12,8,7,1,5,9,2,2,3 | 5,7,38,15,11,5,10,9,2,8,3,7 | 0,2,6,0,0,7,1,3,0,1,4,0,2,4, | 3,11,14,8,13,2,8,14,7,12,1 | 298 | 149 | 0.00156240822012 | 0.150407522694506 | 0.0,0.391,0.106,0. | 0.0,0.083,0.176,0. | 0,119 |

| | | | | | | | | | | | | | | | | | | | | | | |
|---|---|---|---|---|---|---|---|---|---|---|---|---|---|---|---|---|---|---|---|---|---|---|
| 11132 | ENSG00000291131 | | chr19 | + | 53200623 | 53200703 | 53197090 | 53197262 | 53202551 | 53202702 | 11132 | 24,5,54,43,16,20,1,5,17,5,1 | 0,0,0,0,0,0,2,1,4,0,0,0 | 0,0,5,2,3,10,7,25,6,21,14,5 | 8,2,0,0,0,0,0,0,0,3,2,2,1,0,3 | 229 | 149 | 0.00156255939314 | 0.150407522694506 | 1.0,1.0,1.0,1.0,1.0, | 0.0,0.0,1.0,1.0,1.0, | 0,219 |
| 173745 | ENSG00000156976 | EIF4A2 | chr3 | + | 186784911 | 186785101 | 186784563 | 186784696 | 186785882 | 186785913 | 173745 | 498,337,599,452,203,257,1 | 475,304,579,328,154,202,2 | 293,197,238,175,156,196,1 | 280,298,307,267,316,161,8 | 298 | 149 | 0.0015650479644 | 0.150579722139373 | 0.344,0.357,0.341, | 0.343,0.248,0.279, | 0,059 |
| 73780 | ENSG00000291124 | | chr7 | + | 65752163 | 65752310 | 65751063 | 65751278 | 65754526 | 65754661 | 73780 | 7,5,9,1,2,3,4,4,2,8,2,3 | 0,0,0,1,2,1,3,2,4,1,1,2 | 14,4,2,5,9,2,5,11,4,2,12,6,7 | 0,2,0,4,0,0,0,0,1,0,1,1,0,1,2 | 296 | 149 | 0.00156798870261 | 0.150795253106101 | 1.0,1.0,1.0,0.335,0 | 1.0,0.502,1.0,0.38 | -0,19 |
| 225810 | ENSG00000235621 | LINC00494 | chr20 | + | 48366127 | 48366338 | 48363813 | 48364088 | 48366473 | 48367226 | 225810 | 0,0,2,1,1,1,1,7,0,0,1,4 | 19,6,33,26,23,15,2,25,6,8,1 | 4,2,0,1,3,0,1,8,1,0,13,1,1,0 | 13,6,3,12,10,7,14,11,5,4,11 | 298 | 149 | 0.00158199715107 | 0.152074510416234 | 0.0,0.0,0.029,0.01 | 0.133,0.143,0.0,0. | -0,048 |
| 107042 | ENSG00000046651 | OFD1 | chrX | + | 13752703 | 13753441 | 13751248 | 13751368 | 13756577 | 13756767 | 107042 | 12,16,78,17,10,42,4,25,21, | 2,0,1,0,1,0,0,0,0,0,0,0 | 11,12,34,17,18,9,9,20,21,2 | 1,1,4,0,1,1,2,2,0,3,0,2,3,0,2 | 298 | 149 | 0.00158626953236 | 0.152417132910435 | 0.75,1.0,0.975,1.0, | 0.846,0.857,0.81,1 | 0,095 |
| 1228 | ENSG00000100077 | GRK3 | chr22 | + | 25718244 | 25718381 | 25714407 | 25714570 | 25721283 | 25721397 | 1228 | 111,121,95,45,36,26,24,26 | 24,7,8,8,3,4,5,2,1,2,4,4 | 70,71,33,28,31,23,16,19,1 | 14,13,11,15,6,6,0,7,5,3,8,2 | 286 | 149 | 0.00158848741589 | 0.152562130749407 | 0.707,0.9,0.861,0. | 0.723,0.74,0.61,0. | 0,089 |
| 206332 | ENSG00000171132 | PRKCE | chr2 | + | 45697940 | 45698009 | 45652400 | 45652448 | 45842999 | 45843063 | 206332 | 0,1,0,1,2,0,1,0,0,0,5,0 | 86,57,110,49,54,52,62,69,5 | 8,17,0,9,5,0,0,1,1,1,0,2,3,0 | 102,110,38,127,68,32,73,1 | 218 | 149 | 0.00159649232864 | 0.15326255146544 | 0.0,0.012,0.0,0.01 | 0.051,0.096,0.0,0. | -0,007 |
| 71418 | ENSG00000176444 | CLK2 | chr1 | - | 155265859 | 155265954 | 155264644 | 155264774 | 155266728 | 155266895 | 71418 | 57,42,80,56,40,60,20,49,4 | 2,1,9,3,2,1,5,4,0,0,2,2 | 31,47,29,38,39,29,24,36,2 | 8,8,0,8,6,5,6,4,4,1,7,0,4,1,3 | 244 | 149 | 0.00159735781089 | 0.153277271063605 | 0.946,0.962,0.844, | 0.703,0.782,1.0,0. | 0,077 |
| 182716 | ENSG00000263001 | GTF2I | chr7 | + | 74714859 | 74714916 | 74711031 | 74711109 | 74716893 | 74716950 | 182716 | 1,8,2,4,2,1,1,3,2,3,0,2 | 108,83,110,57,63,61,45,57 | 1,5,1,4,6,1,4,4,5,4,3,22,2,6 | 92,76,81,50,63,84,21,45,4 | 206 | 149 | 0.00160194576579 | 0.153649014757909 | 0.007,0.065,0.013, | 0.008,0.045,0.009, | -0,021 |
| 247653 | ENSG00000030066 | NUP160 | chr11 | - | 47787922 | 47788305 | 47786452 | 47786554 | 47788500 | 47788611 | 247653 | 23,14,20,26,11,14,2,16,5,5 | 1,0,4,2,0,2,1,2,0,0,1,0 | 12,22,10,17,36,11,8,15,17, | 0,2,0,2,6,0,0,0,1,0,3,5,0,2,4 | 298 | 149 | 0.00160367662179 | 0.153746513578835 | 0.92,1.0,0.714,0.8 | 1.0,0.846,1.0,0.81, | 0,056 |
| 19243 | ENSG00000114062 | UBE3A | chr15 | - | 25407066 | 25407137 | 25405460 | 25405502 | 25409087 | 25409207 | 19243 | 7,1,23,7,3,5,3,5,1,1,2,2 | 31,31,96,30,23,39,24,66,14 | 6,4,1,0,2,1,0,0,5,8,5,4,4,1,1 | 40,78,26,27,41,17,36,31,2 | 220 | 149 | 0.00161168872842 | 0.154445848934357 | 0.133,0.021,0.14,0 | 0.092,0.034,0.025, | 0,022 |
| 206834 | ENSG00000075884 | ARHGAP15 | chr2 | + | 143614122 | 143614281 | 143556407 | 143556485 | 143624132 | 143624267 | 206834 | 7,2,0,3,6,4,1,10,2,0,3,7 | 129,105,173,109,58,67,71, | 16,10,2,11,14,4,1,3,2,0,9,1 | 115,116,96,53,91,51,37,51 | 298 | 149 | 0.00161567127081 | 0.15475858624286 | 0.026,0.009,0.0,0. | 0.065,0.041,0.01,0 | -0,008 |
| 6925 | ENSG00000101191 | DIDO1 | chr20 | - | 62891986 | 62892076 | 62887213 | 62891155 | 62892808 | 62892962 | 6925 | 33,15,70,47,35,27,23,56,2 | 3,2,2,2,0,0,0,1,0,0,0,0 | 39,47,8,12,21,29,25,31,8,4 | 1,2,0,0,1,0,1,5,0,5,0,3,3,2,1 | 239 | 149 | 0.00161880786673 | 0.154990051585482 | 0.873,0.824,0.956, | 0.96,0.936,1.0,1.0, | 0,033 |
| 42581 | ENSG00000150995 | ITPR1 | chr3 | + | 4795064 | 4795187 | 4787946 | 4788139 | 4800424 | 4800600 | 42581 | 92,140,116,74,53,94,97,11 | 20,12,4,3,10,6,6,17,12,6,4, | 153,103,50,63,59,56,74,12 | 7,8,4,3,7,2,2,8,1,1,7,12,0,4 | 272 | 149 | 0.00162419650147 | 0.155415320281264 | 0.716,0.865,0.941, | 0.923,0.876,0.873, | -0,049 |
| 165055 | ENSG00000123933 | MXD4 | chr4 | - | 2255006 | 2255294 | 2252407 | 2252522 | 2257981 | 2258011 | 165055 | 7,2,6,5,1,5,3,1,10,5,0,1 | 8,12,25,24,15,6,13,11,23,1 | 1,2,2,7,0,0,1,3,1,0,6,5,1,3,0 | 16,29,16,7,13,3,2,8,15,20,5 | 298 | 149 | 0.00162545269178 | 0.155415320281264 | 0.304,0.077,0.107, | 0.03,0.033,0.059,0 | 0,031 |
| 158565 | ENSG00000155111 | CDK19 | chr6 | - | 110623289 | 110623350 | 110622814 | 110622912 | 110626775 | 110626845 | 158565 | 23,10,12,4,8,7,11,5,8,17,2, | 9,0,0,0,1,0,0,3,6,0,0,1 | 4,10,3,3,7,6,5,5,5,7,8,6,3,9 | 4,2,1,11,0,3,0,3,0,0,2,1,2,0 | 210 | 149 | 0.00162612048267 | 0.155415320281264 | 0.645,1.0,1.0,1.0,0 | 0.415,0.78,0.68,0. | 0,058 |
| 724 | ENSG00000291299 | | chr22 | - | 23683724 | 23683867 | 23658093 | 23660000 | 23690231 | 23693185 | 724 | 53,19,0,0,25,26,34,58,13,5 | 0,0,19,10,4,0,0,0,1,0,0,0 | 38,38,25,31,11,11,40,42,2 | 0,0,0,0,4,1,0,0,2,4,0,5,0,0,0 | 292 | 149 | 0.00162689982206 | 0.155415320281264 | 1.0,1.0,0.0,0.0,0.7 | 1.0,1.0,1.0,1.0,0.5 | -0,113 |
| 254891 | ENSG00000089248 | ERP29 | chr12 | + | 112016052 | 112016190 | 112013479 | 112013609 | 112022149 | 112023449 | 254891 | 0,2,14,5,2,0,0,2,0,0,1,2 | 28,13,121,67,43,24,29,67,3 | 11,8,1,7,5,0,0,8,0,5,4,4,6,3 | 44,81,42,14,42,13,13,19,34 | 287 | 149 | 0.00162725047929 | 0.155415320281264 | 0.0,0.074,0.057,0. | 0.115,0.049,0.012, | -0,032 |
| 247258 | ENSG00000066294 | CD84 | chr1 | - | 160550937 | 160551039 | 160549916 | 160549979 | 160553377 | 160553497 | 247258 | 29,12,49,39,18,16,22,5,8,4 | 0,2,3,0,1,0,1,0,0,0,0,0 | 39,48,4,6,22,14,18,24,13,1 | 1,4,0,0,2,2,0,3,2,3,1,0,0,2,2 | 251 | 149 | 0.00162773160461 | 0.155415320281264 | 1.0,0.781,0.907,1. | 0.959,0.877,1.0,1. | 0,06 |
| 118793 | ENSG00000291075 | | chr9 | + | 41008919 | 41009006 | 41007439 | 41007609 | 41011616 | 41011730 | 118793 | 34,13,57,41,17,19,14,24,2 | 1,0,16,11,16,13,5,7,1,0,6,1 | 49,29,45,13,5,3,16,19,20,2 | 1,0,0,0,3,4,2,1,0,0,1,20,2,0 | 236 | 149 | 0.00162830422771 | 0.155415320281264 | 0.955,1.0,0.692,0. | 0.969,1.0,1.0,1.0,0 | -0,085 |
| 180963 | ENSG00000163346 | PBXIP1 | chr1 | - | 154951794 | 154951915 | 154951470 | 154951535 | 154953670 | 154953757 | 180963 | 155,164,193,193,86,44,98, | 0,1,3,0,3,1,1,1,0,1,0,0 | 190,177,74,108,123,77,10 | 3,4,4,0,1,1,4,9,0,2,6,0,3,1, | 270 | 149 | 0.00162919765945 | 0.155418442484834 | 1.0,0.989,0.973,1. | 0.972,0.961,0.911, | 0,016 |
| 238445 | ENSG00000254995 | STX16-NPEPL1 | chr20 | + | 58691404 | 58691596 | 58673630 | 58673711 | 58691723 | 58692206 | 238445 | 0,4,5,1,1,2,3,0,3,0,0,3 | 67,28,22,9,16,24,12,10,13, | 5,2,5,11,2,0,0,2,9,0,1,7,3,3 | 29,42,18,20,24,15,15,9,7,2 | 298 | 149 | 0.00162979721971 | 0.155418442484834 | 0.0,0.067,0.102,0. | 0.079,0.023,0.122, | -0,035 |
| 79480 | ENSG00000000938 | FGR | chr1 | - | 27617192 | 27617296 | 27616856 | 27617006 | 27621558 | 27621657 | 79480 | 484,234,302,153,209,73,1 | 21,5,6,2,3,2,0,3,6,1,3,4 | 213,91,88,65,99,142,46,12 | 9,5,11,7,7,7,1,7,4,5,11,10, | 253 | 149 | 0.00163067811841 | 0.155418442484834 | 0.931,0.965,0.967, | 0.933,0.915,0.825, | 0,041 |
| 246635 | ENSG00000054654 | SYNE2 | chr14 | + | 64082431 | 64082719 | 64081442 | 64081580 | 64087670 | 64087924 | 246635 | 2,1,7,11,5,0,1,6,4,0,0,4 | 34,67,92,80,51,100,74,121 | 0,0,1,5,2,0,1,0,9,5,1,0,1,1,1 | 66,86,38,60,97,89,27,62,9 | 298 | 149 | 0.00163122534195 | 0.155418442484834 | 0.029,0.007,0.037, | 0.0,0.0,0.013,0.04, | 0,016 |
| 199604 | ENSG00000159658 | EFCAB14 | chr1 | - | 46716294 | 46716446 | 46707235 | 46708051 | 46717902 | 46719146 | 199604 | 48,37,177,124,70,65,56,66 | 2,0,0,0,0,0,1,0,0,0,0,0 | 104,72,44,39,53,28,53,77,3 | 3,5,2,2,1,0,2,1,7,2,0,4,0,0,0 | 298 | 149 | 0.00163272906545 | 0.155492880303482 | 0.923,1.0,1.0,1.0,1 | 0.945,0.878,0.917, | 0,05 |
| 197919 | ENSG00000135905 | DOCK10 | chr2 | - | 224893595 | 224893652 | 224886458 | 224886531 | 224896294 | 224896377 | 197919 | 1,0,3,2,3,8,0,0,1,0,1,1 | 117,90,244,152,107,106,81 | 13,9,2,1,4,2,0,0,7,2,5,3,2,5 | 220,167,71,98,115,65,42,4 | 206 | 149 | 0.00163741433378 | 0.155812667381136 | 0.006,0.0,0.009,0. | 0.041,0.038,0.02,0 | -0,009 |
| 64369 | ENSG00000159592 | GPBP1L1 | chr1 | - | 45634095 | 45634236 | 45630481 | 45630606 | 45640209 | 45640403 | 64369 | 59,43,138,85,45,45,30,46,6 | 4,2,6,2,0,1,0,2,0,0,0,3 | 63,66,32,24,39,24,33,36,2 | 0,7,4,0,4,1,2,0,0,2,0,2,3,0,1 | 290 | 149 | 0.00163762960698 | 0.155812667381136 | 0.883,0.917,0.922, | 1.0,0.829,0.804,1. | 0,038 |
| 78957 | ENSG00000143870 | PDIA6 | chr2 | - | 10784933 | 10785030 | 10783390 | 10784326 | 10787280 | 10787439 | 78957 | 49,58,256,234,47,55,53,90 | 19,3,24,17,13,19,11,4,11,5 | 226,136,56,21,32,25,46,82 | 31,48,11,11,6,9,13,19,2,5,2 | 246 | 149 | 0.00163825873728 | 0.155812667381136 | 0.61,0.921,0.866,0 | 0.815,0.632,0.755, | 0,086 |
| 247260 | ENSG00000066294 | CD84 | chr1 | - | 160550937 | 160551043 | 160549916 | 160549979 | 160553377 | 160553497 | 247260 | 31,12,49,39,18,16,21,5,8,4 | 0,2,3,0,1,0,1,0,0,0,0,0 | 40,49,4,6,22,14,18,24,13,1 | 1,4,0,0,2,2,0,3,2,3,1,0,0,2,2 | 255 | 149 | 0.00163946567702 | 0.155858585305076 | 1.0,0.778,0.905,1. | 0.959,0.877,1.0,1. | 0,058 |
| 83703 | ENSG00000079263 | SP140 | chr2 | + | 230243730 | 230243811 | 230241403 | 230241487 | 230244987 | 230245071 | 83703 | 131,76,233,206,133,72,45, | 32,20,24,18,8,13,18,24,6,8 | 158,110,94,60,85,93,31,55 | 40,96,30,41,46,10,14,6,17, | 230 | 149 | 0.00164538248351 | 0.156350703254586 | 0.726,0.711,0.863, | 0.719,0.426,0.67,0 | 0,111 |
| 154746 | ENSG00000105122 | RASAL3 | chr19 | - | 15452044 | 15452250 | 15451623 | 15451938 | 15452657 | 15452815 | 154746 | 23,22,24,24,5,20,18,23,25, | 0,2,1,1,0,3,2,2,4,3,2,0 | 13,17,6,17,7,9,2,29,15,31,1 | 0,0,3,2,0,0,0,0,0,0,1,2,0,5 | 298 | 149 | 0.00164609509585 | 0.156350703254586 | 1.0,0.846,0.923,0. | 1.0,1.0,0.5,0.81,1. | -0,054 |
| 188794 | ENSG00000082512 | TRAF5 | chr1 | + | 211359878 | 211360146 | 211356366 | 211356468 | 211360701 | 211360779 | 188794 | 11,1,16,2,5,6,14,3,6,6,1,7 | 2,0,0,2,0,1,0,4,0,0,0,0 | 9,14,2,8,10,5,2,5,5,4,16,5,7 | 1,0,0,2,2,4,1,2,5,4,0,0,2,0,0 | 298 | 149 | 0.00164808891956 | 0.15643657121342 | 0.733,1.0,1.0,0.33 | 0.818,1.0,1.0,0.66 | 0,094 |
| 111256 | ENSG00000214706 | IFRD2 | chr3 | - | 50289254 | 50289360 | 50288798 | 50288937 | 50289446 | 50289601 | 111256 | 28,3,28,36,21,15,7,11,41,1 | 2,0,3,1,1,0,1,3,1,0,0,3 | 20,21,22,4,8,5,13,22,6,24,8 | 4,0,0,5,0,2,4,1,2,1,0,4,0,1,3 | 255 | 149 | 0.00164845279496 | 0.15643657121342 | 0.891,1.0,0.845,0. | 0.745,1.0,1.0,0.31 | 0,066 |
| 194859 | ENSG00000158711 | ELK4 | chr1 | - | 205618697 | 205618822 | 205616477 | 205616644 | 205618956 | 205619073 | 194859 | 4,9,3,3,0,1,6,5,5,1,0,1 | 119,109,176,86,78,85,96,1 | 5,6,9,9,5,0,5,6,0,9,9,14,2,5 | 161,128,91,99,84,76,70,12 | 274 | 149 | 0.00165462336319 | 0.156952948912625 | 0.018,0.043,0.009, | 0.017,0.025,0.051, | -0,013 |
| 137185 | ENSG00000104957 | YJU2B | chr19 | + | 13747515 | 13747954 | 13732174 | 13732285 | 13751607 | 13751811 | 137185 | 10,7,19,18,13,10,10,13,14, | 1,3,2,1,0,4,1,1,4,5,1,0 | 12,10,6,14,6,2,8,9,2,8,18,7 | 1,0,0,0,1,0,1,2,0,2,2,8,0,1,0 | 298 | 149 | 0.00166101098647 | 0.157489451818909 | 0.833,0.538,0.826, | 0.857,1.0,1.0,1.0,0 | -0,066 |
| 60587 | ENSG00000213930 | GALT | chr9 | + | 34647831 | 34647961 | 34647088 | 34647258 | 34648114 | 34648171 | 60587 | 31,12,39,26,28,35,26,17,41 | 0,0,5,4,3,5,1,1,0,1,1,0 | 10,12,16,18,14,8,20,30,7,1 | 0,0,0,0,3,0,0,3,0,0,3,1,0,0,2 | 279 | 149 | 0.0016628592783 | 0.157595273151822 | 1.0,1.0,0.806,0.77 | 1.0,1.0,1.0,1.0,0.7 | -0,052 |
| 142441 | ENSG00000178852 | EFCAB13 | chr17 | + | 47344830 | 47344939 | 47344161 | 47344292 | 47345015 | 47345098 | 142441 | 6,5,29,19,6,6,4,3,3,4,0,2 | 37,21,130,92,31,44,34,53,1 | 4,4,2,4,2,2,9,5,7,11,3,4,2,2 | 45,44,31,48,36,35,46,57,4 | 258 | 149 | 0.00166384968018 | 0.157619731740679 | 0.086,0.121,0.114, | 0.049,0.05,0.036,0 | 0,032 |
| 204815 | ENSG00000138073 | PREB | chr2 | - | 27132229 | 27132356 | 27131671 | 27132082 | 27132602 | 27132727 | 204815 | 9,5,18,20,11,12,13,5,7,4,5, | 0,2,1,0,0,2,0,1,0,1,0,0 | 29,16,9,1,4,2,19,16,1,4,11, | 2,1,2,0,2,0,2,3,0,2,0,2,0,4,2 | 276 | 149 | 0.00166469890806 | 0.157630801003371 | 1.0,0.574,0.907,1. | 0.887,0.896,0.708, | 0,093 |
| 51358 | ENSG00000163346 | PBXIP1 | chr1 | - | 154951794 | 154951919 | 154951470 | 154951535 | 154953670 | 154953757 | 51358 | 157,165,199,202,89,48,99, | 0,1,3,0,3,1,1,1,0,1,0,0 | 194,178,74,110,126,78,11 | 3,4,4,0,1,1,4,9,0,2,6,0,3,1, | 274 | 149 | 0.00167496244905 | 0.158499420282859 | 1.0,0.989,0.973,1. | 0.972,0.96,0.91,1. | 0,017 |
| 135141 | ENSG00000100239 | PPP6R2 | chr22 | + | 50418866 | 50418989 | 50416091 | 50416157 | 50419348 | 50419462 | 135141 | 22,11,46,33,20,16,9,25,18, | 0,1,2,0,0,1,0,0,3,1,0,0 | 29,22,6,24,15,10,17,29,14, | 1,3,1,6,0,0,1,1,2,0,0,2,1,1,2 | 272 | 149 | 0.00167534500673 | 0.158499420282859 | 1.0,0.858,0.926,1. | 0.941,0.801,0.767, | 0,061 |
| 131687 | ENSG00000076513 | ANKRD13A | chr12 | + | 110021107 | 110021301 | 110019138 | 110019328 | 110024045 | 110024112 | 131687 | 6,0,17,3,7,2,0,5,0,2,4,4 | 353,341,216,100,90,67,67, | 5,7,3,5,3,3,2,3,4,12,7,11,1 | 136,119,176,110,120,58,2 | 298 | 149 | 0.00168686062639 | 0.159518760755073 | 0.008,0.0,0.038,0. | 0.018,0.029,0.008, | -0,004 |
| 23226 | ENSG00000100461 | RBM23 | chr14 | - | 22913723 | 22913845 | 22911327 | 22911403 | 22918998 | 22919149 | 23226 | 0,2,6,4,1,1,1,0,2,1,1,3 | 31,46,66,56,14,17,8,14,27, | 5,6,2,2,1,3,5,4,1,3,2,2,0,5, | 34,26,17,21,28,8,10,24,8,2 | 271 | 149 | 0.00168766905444 | 0.159525120007104 | 0.0,0.023,0.048,0. | 0.075,0.113,0.061, | -0,039 |
| 78660 | ENSG00000158435 | CNOT11 | chr2 | + | 101257790 | 101257955 | 101252885 | 101253478 | 101262538 | 101262691 | 78660 | 40,28,32,35,23,22,9,17,60, | 2,2,6,0,2,1,1,5,1,2,0,0 | 31,19,16,16,8,19,4,10,3,35 | 1,1,1,1,2,1,2,2,1,1,2,1,0,1,2 | 298 | 149 | 0.00168932846526 | 0.159611876605081 | 0.909,0.875,0.727, | 0.939,0.905,0.889, | 0,045 |
| 184255 | ENSG00000169905 | TOR1AIP2 | chr1 | - | 179848224 | 179848388 | 179847534 | 179847636 | 179850844 | 179851363 | 184255 | 2,0,0,0,1,6,0,6,0,4,1,1 | 19,15,21,9,15,16,7,11,13,2 | 1,2,0,1,1,0,0,0,8,3,1,1,0,4,1 | 7,11,12,3,4,9,6,14,6,9,7,6,1 | 298 | 149 | 0.00169114278018 | 0.159713186362844 | 0.05,0.0,0.0,0.0,0. | 0.067,0.083,0.0,0. | 0,011 |
| 185502 | ENSG00000134352 | IL6ST | chr5 | - | 55946196 | 55946283 | 55942669 | 55942751 | 55947492 | 55947594 | 185502 | 0,1,2,2,0,0,0,1,0,0,0,4 | 9,9,17,9,9,6,30,74,19,9,4,6 | 1,4,2,12,0,0,0,2,1,1,5,0,1,0 | 26,40,27,18,28,7,21,24,10, | 236 | 149 | 0.00170403781641 | 0.160860422484097 | 0.0,0.066,0.069,0. | 0.024,0.059,0.045, | -0,006 |
| 190008 | ENSG00000258728 | | chr9 | + | 34647540 | 34647961 | 34647088 | 34647258 | 34648114 | 34648171 | 190008 | 32,14,43,26,29,36,26,17,4 | 0,0,5,4,3,5,1,1,0,1,1,0 | 14,12,16,18,12,9,20,38,7,1 | 0,0,0,0,3,0,0,3,0,0,3,1,0,0,2 | 298 | 149 | 0.00170938075534 | 0.161294050571058 | 1.0,1.0,0.811,0.76 | 1.0,1.0,1.0,1.0,0.6 | -0,049 |
| 192518 | ENSG00000196419 | XRCC6 | chr22 | + | 41628120 | 41628230 | 41621989 | 41622086 | 41636112 | 41636251 | 192518 | 194,158,260,187,92,65,55, | 4,2,6,4,0,1,0,0,0,0,0,0 | 172,156,154,56,113,67,47, | 2,7,6,5,11,2,0,0,0,4,3,1,1,1 | 259 | 149 | 0.00171081249337 | 0.16135840655588 | 0.965,0.978,0.961, | 0.98,0.928,0.937,0 | 0,024 |
| 34690 | ENSG00000125089 | SH3TC1 | chr4 | + | 8209747 | 8209822 | 8205166 | 8205366 | 8212700 | 8212828 | 34690 | 12,14,11,4,14,6,11,8,13,19 | 2,0,1,2,1,2,3,0,1,0,2,5 | 8,5,3,1,6,10,10,6,10,16,3,2 | 2,1,0,0,0,0,0,1,0,0,4,2,0,0,0 | 224 | 149 | 0.00171322779582 | 0.161515432204176 | 0.8,1.0,0.88,0.571, | 0.727,0.769,1.0,1. | -0,074 |
| 150340 | ENSG00000247556 | OIP5-AS1 | chr15 | + | 41297836 | 41297929 | 41286010 | 41286100 | 41298040 | 41306534 | 150340 | 1,0,6,5,7,4,1,5,0,0,0,0 | 29,11,80,66,68,66,35,46,1 | 1,1,1,0,5,1,1,10,2,8,1,10,2, | 31,45,19,3,16,29,11,31,6,4 | 242 | 149 | 0.00171597456945 | 0.16170355628603 | 0.021,0.0,0.044,0. | 0.019,0.013,0.031, | -0,043 |
| 208581 | ENSG00000196504 | PRPF40A | chr2 | - | 152693414 | 152693476 | 152693037 | 152693068 | 152694541 | 152694622 | 208581 | 7,0,16,4,2,8,1,5,0,0,3,3 | 36,36,77,68,40,37,36,40,2 | 5,4,3,1,1,0,1,7,2,1,1,2,1,2,3 | 53,79,32,27,45,24,31,52,1 | 211 | 149 | 0.00172760157522 | 0.16269060689435 | 0.121,0.0,0.128,0. | 0.062,0.035,0.062, | 0,021 |
| 37186 | ENSG00000168310 | IRF2 | chr4 | - | 184447676 | 184449050 | 184428977 | 184429070 | 184474378 | 184474558 | 37186 | 0,1,1,4,2,0,0,2,2,0,0,0 | 61,23,88,35,58,44,27,25,5 | 1,2,0,1,0,2,1,2,3,4,15,1,0,0 | 61,25,17,14,11,27,13,16,2 | 298 | 149 | 0.00172870802166 | 0.16269060689435 | 0.0,0.021,0.006,0. | 0.008,0.038,0.0,0. | -0,02 |
| 58584 | ENSG00000079277 | MKNK1 | chr1 | - | 46585873 | 46585974 | 46583227 | 46583329 | 46594112 | 46594206 | 58584 | 0,3,10,3,1,0,2,0,0,0,0,3 | 30,12,14,20,11,12,7,4,18,1 | 0,1,1,5,0,0,0,0,0,0,1,0,0,0 | 8,10,15,3,8,13,1,9,7,14,2,8 | 250 | 149 | 0.00172897796448 | 0.16269060689435 | 0.0,0.13,0.299,0.0 | 0.0,0.056,0.038,0. | 0,032 |
| 182092 | ENSG00000121310 | ECHDC2 | chr1 | - | 52898579 | 52898754 | 52897436 | 52897484 | 52899173 | 52899224 | 182092 | 6,4,9,0,4,3,5,5,2,0,0,3 | 12,9,13,2,16,7,6,9,1,2,9,10 | 3,3,1,1,0,0,2,5,1,1,0,0,4,1,3 | 10,6,6,7,3,2,9,8,7,6,5,10,7, | 298 | 149 | 0.00172947256006 | 0.16269060689435 | 0.2,0.182,0.257,0. | 0.13,0.2,0.077,0.0 | 0,081 |
| 136478 | ENSG00000213793 | ZNF888 | chr19 | - | 52917607 | 52917704 | 52915195 | 52915322 | 52917858 | 52917997 | 136478 | 3,1,3,0,0,1,1,0,0,0,2,0 | 7,6,13,10,5,2,4,4,2,3,1,5 | 3,6,4,0,0,2,0,4,3,0,1,0,0,0, | 5,2,2,5,4,1,4,8,3,1,7,12,2,4 | 246 | 149 | 0.00173572645137 | 0.163124108136447 | 0.206,0.092,0.123, | 0.267,0.645,0.548, | -0,074 |
| 113204 | ENSG00000165923 | AGBL2 | chr11 | - | 47705863 | 47705917 | 47704542 | 47704728 | 47710376 | 47710511 | 113204 | 2,4,15,11,5,2,6,5,2,1,2,7 | 1,0,0,5,1,0,2,2,0,0,1,2 | 6,4,2,4,9,5,5,2,7,6,6,4,5,7,1 | 0,2,4,2,0,2,0,0,2,5,0,3,2,3, | 203 | 149 | 0.00173583884374 | 0.163124108136447 | 0.595,1.0,1.0,0.61 | 1.0,0.595,0.268,0. | 0,08 |
| 182371 | ENSG00000143418 | CERS2 | chr1 | - | 150967984 | 150968201 | 150967819 | 150967877 | 150968394 | 150968478 | 182371 | 114,80,77,68,29,46,30,22, | 0,0,6,3,3,7,3,5,3,1,3,1 | 40,31,60,51,28,25,25,40,1 | 11,8,5,0,1,0,2,7,4,0,9,5,4,5 | 298 | 149 | 0.00173635457597 | 0.163124108136447 | 1.0,1.0,0.865,0.91 | 0.645,0.66,0.857,1 | 0,077 |
| 221112 | ENSG00000120278 | PLEKHG1 | chr6 | + | 150638084 | 150638125 | 150632942 | 150633304 | 150650727 | 150650786 | 221112 | 1,3,8,6,3,3,3,4,5,1,1,5 | 13,10,60,56,17,24,10,10,2 | 6,10,5,5,4,5,1,5,6,3,4,2,7,4 | 19,19,11,20,15,18,8,23,8,1 | 190 | 149 | 0.00174448214702 | 0.163816159243133 | 0.057,0.19,0.095,0 | 0.198,0.292,0.263, | -0,087 |
| 166167 | ENSG00000077232 | DNAJC10 | chr2 | + | 182737377 | 182737463 | 182736248 | 182736386 | 182739544 | 182740388 | 166167 | 47,24,82,19,25,23,10,28,2 | 160,163,209,50,85,47,45,3 | 50,87,38,24,27,13,13,13,1 | 110,103,63,36,40,12,37,47 | 235 | 149 | 0.00174873382947 | 0.164143798888207 | 0.157,0.085,0.199, | 0.224,0.349,0.277, | -0,069 |
| 46498 | ENSG00000114446 | IFT57 | chr3 | - | 108219409 | 108219572 | 108218534 | 108218653 | 108222110 | 108222360 | 46498 | 295,165,795,608,286,305, | 5,4,2,0,4,2,0,4,0,0,2,6 | 619,516,196,292,216,163, | 4,4,2,7,2,0,11,5,0,5,7,1,13, | 298 | 149 | 0.00175006476413 | 0.164190932360541 | 0.967,0.954,0.995, | 0.987,0.985,0.98,0 | 0,009 |
| 47290 | ENSG00000164828 | SUN1 | chr7 | + | 849922 | 850033 | 849519 | 849603 | 851949 | 852043 | 47290 | 17,15,18,3,23,23,11,13,9,0 | 0,0,8,6,0,0,0,4,0,2,0,2 | 17,3,16,5,2,1,10,20,9,14,1 | 0,1,1,0,1,0,2,0,0,0,0,5,0,0, | 260 | 149 | 0.00175076169217 | 0.164190932360541 | 1.0,1.0,0.563,0.22 | 1.0,0.632,0.902,1. | -0,134 |
| 179150 | ENSG00000144357 | UBR3 | chr2 | + | 170067038 | 170067090 | 170061317 | 170061443 | 170073427 | 170073607 | 179150 | 0,1,0,0,0,0,0,4,0,0,0,3 | 19,4,19,16,21,13,10,20,11, | 2,3,2,6,1,0,6,0,0,1,3,0,1,2, | 13,21,16,25,10,13,21,10,1 | 201 | 149 | 0.00175296632801 | 0.164270453633465 | 0.0,0.156,0.0,0.0,0 | 0.102,0.096,0.085, | -0,032 |
| 68031 | ENSG00000143119 | CD53 | chr1 | + | 110892344 | 110892533 | 110873147 | 110873248 | 110899123 | 110899514 | 68031 | 8,6,28,26,21,5,17,16,8,6,7, | 0,0,5,4,0,2,2,0,0,1,0,0 | 13,12,15,15,5,10,4,5,1,13,2 | 4,5,1,0,3,1,1,0,2,1,3,7,1,0, | 298 | 149 | 0.00175313608168 | 0.164270453633465 | 1.0,1.0,0.737,0.76 | 0.619,0.545,0.882, | 0,117 |
| 38768 | ENSG00000163681 | SLMAP | chr3 | + | 57864806 | 57864857 | 57864547 | 57864716 | 57890040 | 57890100 | 38768 | 14,10,11,14,11,4,5,3,3,3,2, | 1,2,5,5,3,2,0,0,3,0,4,1 | 8,5,2,2,4,2,4,3,4,4,14,4,9,6 | 1,0,0,1,1,0,2,1,0,6,1,3,0,2, | 200 | 149 | 0.00175901615771 | 0.164749698276798 | 0.913,0.788,0.621, | 0.856,1.0,1.0,0.59 | -0,103 |
| 147329 | ENSG00000103222 | ABCC1 | chr16 | + | 16083463 | 16083542 | 16079351 | 16079478 | 16086823 | 16086991 | 147329 | 3,2,16,18,4,12,10,10,2,4,2, | 5,0,3,0,0,1,0,0,0,1,1,7 | 22,23,10,9,10,7,7,7,14,18, | 1,0,0,0,1,1,2,0,4,4,0,0,3,0,3 | 228 | 149 | 0.00176480584129 | 0.165220063517481 | 0.282,1.0,0.777,1. | 0.935,1.0,1.0,1.0,0 | -0,064 |
| 142077 | ENSG00000029725 | RABEP1 | chr17 | + | 5356507 | 5356555 | 5354400 | 5354490 | 5361207 | 5361675 | 142077 | 1,2,14,3,0,1,4,5,12,2,0,8 | 147,67,168,134,73,65,49,5 | 7,10,3,6,3,4,7,6,5,10,5,7,2, | 85,56,53,30,51,41,50,60,3 | 197 | 149 | 0.00177094453205 | 0.165722679381588 | 0.005,0.022,0.059, | 0.059,0.119,0.041, | -0,034 |
| 211067 | ENSG00000115825 | PRKD3 | chr2 | - | 37292921 | 37292976 | 37290867 | 37290999 | 37293132 | 37293271 | 211067 | 5,4,17,17,5,6,0,1,0,0,2,0 | 18,24,89,39,25,39,19,31,1 | 17,7,3,2,1,0,1,0,1,3,0,5,2,0 | 73,79,12,11,34,23,20,21,1 | 204 | 149 | 0.00177266100629 | 0.165811212970362 | 0.169,0.109,0.122, | 0.145,0.061,0.154, | 0,031 |
| 205874 | ENSG00000128815 | WDFY4 | chr10 | + | 48808288 | 48808365 | 48807858 | 48807958 | 48811538 | 48811708 | 205874 | 8,1,6,2,0,4,2,4,2,2,5,3 | 9,4,10,3,8,21,13,11,11,4,7, | 7,15,2,34,3,0,7,5,1,2,10,13 | 15,17,2,19,4,1,8,7,4,3,17,7 | 226 | 149 | 0.00177551765435 | 0.16600627292068 | 0.369,0.142,0.283, | 0.235,0.368,0.397, | -0,102 |
| 71275 | ENSG00000131051 | RBM39 | chr20 | - | 35734185 | 35735081 | 35731940 | 35732135 | 35738967 | 35739017 | 71275 | 37,19,45,22,18,13,7,21,12, | 1004,870,810,476,304,322 | 11,21,23,22,19,19,4,14,10, | 410,524,509,502,430,198,1 | 298 | 149 | 0.00178159683065 | 0.166459202625875 | 0.018,0.011,0.027, | 0.013,0.02,0.022,0 | -0,007 |
| 207637 | ENSG00000245937 | SLC12A2-DT | chr5 | - | 128023928 | 128024298 | 127966849 | 127966928 | 128061143 | 128061244 | 207637 | 1,0,14,3,5,1,7,4,9,2,1,2 | 5,4,14,9,1,0,1,0,6,0,2,4 | 7,5,2,5,3,2,3,6,4,3,4,7,8,9, | 4,3,1,0,0,2,0,1,2,2,0,5,0,3, | 298 | 149 | 0.00178205126316 | 0.166459202625875 | 0.091,0.0,0.333,0. | 0.467,0.455,0.5,1. | -0,225 |
| 41387 | ENSG00000173226 | IQCB1 | chr3 | - | 121795456 | 121795566 | 121790072 | 121790215 | 121799195 | 121799374 | 41387 | 42,28,40,18,11,14,17,15,2 | 2,0,1,2,0,0,0,0,0,0,0,0 | 35,21,25,14,13,17,11,14,1 | 3,1,0,1,6,1,1,0,0,2,0,3,2,0, | 259 | 149 | 0.00178268215105 | 0.166459202625875 | 0.924,1.0,0.958,0. | 0.87,0.924,1.0,0.8 | 0,103 |
| 113725 | ENSG00000075856 | SART3 | chr12 | - | 108535358 | 108535468 | 108531203 | 108531280 | 108536513 | 108536572 | 113725 | 17,22,38,28,20,24,10,18,1 | 1,1,4,1,0,3,0,5,1,2,0,0 | 23,57,9,6,36,6,21,16,13,15 | 0,0,0,0,0,1,0,0,0,0,2,2,0,1,1 | 259 | 149 | 0.00178464108078 | 0.166569854491483 | 0.907,0.927,0.845, | 1.0,1.0,1.0,1.0,1.0, | -0,071 |
| 240946 | ENSG00000106829 | TLE4 | chr9 | + | 79574479 | 79574528 | 79573688 | 79573786 | 79574872 | 79575029 | 240946 | 3,1,3,2,0,3,0,6,1,0,0,1 | 73,34,165,92,67,109,59,18 | 1,6,4,10,7,0,1,0,3,0,4,1,5,0 | 72,78,125,98,78,51,36,70,8 | 198 | 149 | 0.00179428944859 | 0.167397794672507 | 0.03,0.022,0.013,0 | 0.01,0.055,0.024,0 | -0,029 |
| 8147 | ENSG00000124155 | PIGT | chr20 | + | 45418771 | 45418979 | 45416312 | 45416343 | 45419294 | 45419395 | 8147 | 29,9,29,18,10,19,21,14,35, | 2,0,0,0,0,0,3,2,0,0,2,1 | 13,17,11,12,14,7,16,15,8,1 | 0,1,0,3,0,1,2,5,0,0,3,7,0,0,2 | 298 | 149 | 0.00180946884013 | 0.168740809328432 | 0.879,1.0,1.0,1.0,1 | 1.0,0.895,1.0,0.66 | 0,049 |
| 19504 | ENSG00000170776 | AKAP13 | chr15 | + | 85662387 | 85662453 | 85658536 | 85658590 | 85664562 | 85664755 | 19504 | 11,6,21,18,16,13,9,13,6,5,5 | 62,29,67,25,36,34,15,26,24 | 13,10,9,0,8,9,5,4,10,11,1,1 | 64,82,18,35,48,16,26,26,4 | 215 | 149 | 0.00181249450541 | 0.168949763921135 | 0.109,0.125,0.178, | 0.123,0.078,0.257, | 0,069 |
| 95422 | ENSG00000147251 | DOCK11 | chrX | + | 118590209 | 118590302 | 118588412 | 118588478 | 118593213 | 118593337 | 95422 | 140,78,225,154,120,134,1 | 3,5,13,4,5,4,3,6,0,1,2,4 | 166,94,100,75,50,74,73,10 | 13,8,1,5,15,5,1,5,6,3,5,9,7, | 242 | 149 | 0.00181755403354 | 0.169337701796541 | 0.966,0.906,0.914, | 0.887,0.879,0.984, | 0,042 |
| 38919 | ENSG00000066654 | THUMPD1 | chr16 | - | 20738087 | 20738269 | 20737707 | 20737956 | 20738896 | 20739071 | 38919 | 12,2,14,8,11,4,4,9,9,2,0,8 | 15,11,27,24,13,26,24,16,15 | 6,4,6,6,9,3,5,3,2,1,8,9,4,6,4 | 35,42,25,18,9,6,20,21,12,2 | 298 | 149 | 0.00181822985003 | 0.169337701796541 | 0.286,0.083,0.206, | 0.079,0.045,0.107, | 0,05 |
| 91634 | ENSG00000196262 | PPIA | chr7 | + | 44799391 | 44799874 | 44797972 | 44799277 | 44801286 | 44801647 | 91634 | 1143,912,1559,1101,787,6 | 17,4,19,20,3,9,11,7,7,2,4,1 | 835,818,753,314,671,450,3 | 14,17,4,24,16,7,1,19,9,0,8, | 298 | 149 | 0.00181919651152 | 0.169354448257336 | 0.971,0.991,0.976, | 0.968,0.96,0.989,0 | 0,015 |
| 110605 | ENSG00000120899 | PTK2B | chr8 | + | 27445793 | 27445919 | 27444205 | 27444271 | 27450748 | 27450895 | 110605 | 20,3,12,9,3,4,8,19,6,6,2,4 | 68,19,66,54,58,53,46,47,9 | 6,6,19,17,12,12,3,8,4,1,13, | 29,27,40,8,27,24,26,38,28, | 275 | 149 | 0.00182145665311 | 0.169470910671848 | 0.137,0.079,0.09,0 | 0.101,0.107,0.205, | -0,069 |
| 128487 | ENSG00000198825 | INPP5F | chr10 | + | 119781060 | 119781242 | 119757894 | 119757977 | 119781634 | 119781771 | 128487 | 0,0,10,6,3,3,0,3,2,0,0,0 | 11,1,39,16,20,26,8,8,1,1,1 | 3,0,2,0,0,0,0,1,0,4,0,0,0,2, | 13,14,17,4,5,4,10,16,2,7,1 | 298 | 149 | 0.00182202232622 | 0.169470910671848 | 0.0,0.0,0.0,0.114,0.15 | 0.103,0.0,0.0,0.056,0. | 0,053 |
| 225289 | ENSG00000198198 | SZT2 | chr1 | + | 43432706 | 43432799 | 43432516 | 43432604 | 43432988 | 43433095 | 225289 | 6,6,9,3,9,14,5,7,2,3,5,5 | 2,0,3,1,0,0,3,1,2,0,0,1 | 8,4,6,5,4,7,6,5,5,9,4,3,9,5,3 | 1,7,2,0,1,0,9,3,2,0,2,1,0,1, | 242 | 149 | 0.0018240239968 | 0.169583805121063 | 0.649,1.0,0.649,0. | 0.831,0.26,0.649,1 | 0,045 |

| | | | | | | | | | | | | | | | | | | | | | | |
|---|---|---|---|---|---|---|---|---|---|---|---|---|---|---|---|---|---|---|---|---|---|---|
| 97757 | ENSG00000196511 | TPK1 | chr7 | - | 144835559 | 144835618 | 144765879 | 144765951 | 144835966 | 144836053 | 97757 | 26,14,45,26,12,23,21,35,8, | 5,0,2,0,1,0,0,0,0,0,0,0 | 19,25,18,6,37,29,7,14,18,1 | 5,3,2,0,6,0,0,0,0,3,2,0,0,1, | 208 | 149 | 0.00182997899643 | 0.170003157133088 | 0.788,1.0,0.942,1. | 0.731,0.857,0.866, | 0,048 |
| 46860 | ENSG00000131876 | SNRPA1 | chr15 | - | 101287655 | 101287702 | 101285725 | 101287010 | 101295096 | 101295203 | 46860 | 7,13,20,12,5,15,3,15,2,2,3, | 0,0,1,0,0,1,2,0,0,2,0,0 | 8,26,10,7,26,12,11,6,12,9, | 6,5,2,3,0,0,2,0,0,2,1,0,0,0, | 196 | 149 | 0.00183011422647 | 0.170003157133088 | 1.0,1.0,0.938,1.0,1 | 0.503,0.798,0.792, | 0,048 |
| 236749 | ENSG00000067167 | TRAM1 | chr8 | - | 70598133 | 70598274 | 70597890 | 70598011 | 70600018 | 70600082 | 236749 | 244,130,306,206,117,84,8 | 1,0,1,1,1,1,1,0,0,0,0,0 | 220,186,145,92,111,85,55, | 2,2,0,0,1,0,2,1,1,6,0,14,2,4 | 290 | 149 | 0.00183110498487 | 0.170019766308051 | 0.992,1.0,0.994,0. | 0.983,0.98,1.0,1.0, | 0,023 |
| 239076 | ENSG00000066117 | SMARCD1 | chr12 | + | 50096849 | 50097337 | 50094436 | 50094572 | 50098713 | 50098815 | 239076 | 43,40,102,58,21,33,29,35, | 2,0,0,4,1,2,2,0,6,1,0,2 | 76,104,50,33,53,29,19,32, | 2,0,2,0,0,2,0,3,0,0,0,2,3,0, | 298 | 149 | 0.00183187290896 | 0.170019766308051 | 0.915,1.0,1.0,0.87 | 0.95,1.0,0.926,1.0, | -0,049 |
| 96314 | ENSG00000164654 | MIOS | chr7 | + | 7568025 | 7568123 | 7567482 | 7567688 | 7572435 | 7572667 | 96314 | 13,14,46,25,11,18,7,14,7,1 | 2,1,0,2,3,2,1,1,0,0,1,2 | 32,23,28,7,10,5,5,8,17,23,1 | 2,3,5,1,3,2,4,7,0,0,8,6,0,0,1 | 247 | 149 | 0.00183294603352 | 0.1700460378192 | 0.797,0.894,1.0,0. | 0.906,0.822,0.772, | 0,081 |
| 19249 | ENSG00000114062 | UBE3A | chr15 | - | 25407066 | 25407262 | 25405460 | 25405502 | 25409087 | 25409207 | 19249 | 17,7,49,32,12,15,11,21,22, | 31,31,96,30,23,39,24,66,1 | 20,11,10,4,6,3,11,8,8,16,2 | 40,78,26,27,41,17,36,31,2 | 298 | 149 | 0.00183969868727 | 0.170536246217857 | 0.215,0.101,0.203, | 0.2,0.066,0.161,0. | 0,068 |
| 104977 | ENSG00000148985 | PGAP2 | chr11 | + | 3823882 | 3824135 | 3808293 | 3808315 | 3824269 | 3824343 | 104977 | 4,0,12,3,4,5,4,11,9,2,3,2 | 7,1,2,0,4,2,0,0,0,2,1,2 | 6,6,5,1,1,2,3,5,4,0,5,20,3,3 | 2,4,5,0,2,0,1,2,0,1,0,0,0,0,1 | 298 | 149 | 0.00183981472798 | 0.170536246217857 | 0.222,0.0,0.75,1.0, | 0.6,0.429,0.333,1. | -0,123 |
| 116562 | ENSG00000213930 | GALT | chr9 | + | 34647491 | 34647961 | 34647088 | 34647258 | 34648114 | 34648171 | 116562 | 47,25,61,39,53,50,47,30,5 | 0,0,5,4,3,5,1,1,0,1,1,0 | 29,22,25,34,17,12,26,58,1 | 0,0,0,0,3,0,0,3,0,0,3,1,0,0, | 298 | 149 | 0.00184204540764 | 0.170669511464384 | 1.0,1.0,0.859,0.83, | 1.0,1.0,1.0,1.0,0.7 | -0,028 |
| 217742 | ENSG00000115145 | STAM2 | chr2 | - | 152146538 | 152146669 | 152144887 | 152144957 | 152147161 | 152147308 | 217742 | 1,0,0,2,0,0,0,0,0,0,0,0 | 24,18,45,40,26,26,19,16,2 | 0,4,3,9,5,1,0,0,1,1,4,0,5,0, | 31,37,16,5,17,12,11,23,10, | 280 | 149 | 0.0018445295018 | 0.170826131326126 | 0.022,0.0,0.0,0.02 | 0.0,0.054,0.091,0. | -0,063 |
| 158566 | ENSG00000155111 | CDK19 | chr6 | - | 110623289 | 110623478 | 110622814 | 110622912 | 110626775 | 110626845 | 158566 | 31,11,18,8,10,8,13,6,10,15 | 9,0,0,0,1,0,0,3,6,0,0,1 | 8,13,6,3,8,7,5,9,5,9,8,8,6,1 | 4,2,1,11,0,3,0,3,0,0,2,1,2,0 | 298 | 149 | 0.0018553462571 | 0.17175399151049 | 0.633,1.0,1.0,1.0,0 | 0.5,0.765,0.75,0.1 | 0,041 |
| 164238 | ENSG00000143624 | INTS3 | chr1 | + | 153730941 | 153731176 | 153728552 | 153728784 | 153740650 | 153740734 | 164238 | 4,3,2,0,0,1,0,5,1,0,0,1 | 16,7,23,14,14,24,13,19,19, | 3,0,0,9,2,2,3,0,4,2,3,3,2,0, | 16,5,11,14,3,11,9,29,6,11,1 | 298 | 149 | 0.00185650692396 | 0.171787550193824 | 0.111,0.176,0.042, | 0.086,0.0,0.0,0.24 | -0,054 |
| 42731 | ENSG00000157601 | MX1 | chr21 | + | 41449136 | 41449295 | 41445999 | 41446141 | 41451166 | 41451243 | 42731 | 85,40,156,208,62,42,49,63 | 1,2,9,8,4,1,1,0,4,0,0,1 | 117,107,41,55,42,141,24,8 | 2,4,2,5,8,10,0,6,2,3,0,11,1 | 298 | 149 | 0.001857755771 | 0.171829236075677 | 0.977,0.909,0.897, | 0.967,0.93,0.911,0 | 0,045 |
| 4405 | ENSG00000196268 | ZNF493 | chr19 | + | 21405128 | 21405255 | 21397118 | 21397267 | 21422912 | 21423040 | 4405 | 0,3,7,23,8,7,2,6,10,7,4,2 | 7,3,5,0,0,0,1,3,4,2,1,0 | 13,6,6,0,6,7,2,8,10,3,8,8,1 | 12,10,8,12,8,2,6,10,6,2,14, | 276 | 149 | 0.00185957535043 | 0.171843624620178 | 0.0,0.351,0.43,1.0, | 0.369,0.245,0.288, | 0,254 |
| 112151 | ENSG00000070501 | POLB | chr8 | + | 42369848 | 42369988 | 42368935 | 42369335 | 42371562 | 42371765 | 112151 | 31,16,27,38,24,14,3,12,27, | 0,0,3,3,2,2,1,0,2,0,0,4 | 28,21,10,7,14,15,13,34,4,2 | 1,1,3,4,0,3,0,0,0,0,5,2,0,1, | 289 | 149 | 0.00186019284185 | 0.171843624620178 | 1.0,1.0,0.823,0.86 | 0.935,0.915,0.632, | 0,03 |
| 149084 | ENSG00000169375 | SIN3A | chr15 | - | 75393351 | 75393507 | 75392241 | 75392815 | 75394679 | 75394863 | 149084 | 9,3,8,0,0,0,8,2,0,0,2,2 | 77,92,69,45,33,33,23,32,5 | 4,7,3,23,3,2,3,4,5,5,4,2,5,1 | 52,85,40,63,32,28,34,33,1 | 298 | 149 | 0.00186030657928 | 0.171843624620178 | 0.055,0.016,0.055, | 0.037,0.04,0.036,0 | -0,018 |
| 160189 | ENSG00000151292 | CSNK1G3 | chr5 | + | 123550227 | 123550302 | 123545633 | 123545841 | 123553106 | 123553147 | 160189 | 9,5,17,44,7,16,5,12,10,0,2, | 38,11,150,76,64,97,33,73,2 | 7,9,8,4,2,6,3,19,2,7,0,21,6, | 144,86,30,30,41,19,29,82,2 | 224 | 149 | 0.00187148703726 | 0.172774036246474 | 0.136,0.232,0.07,0 | 0.031,0.065,0.151, | 0,04 |
| 119883 | ENSG00000109971 | HSPA8 | chr11 | - | 123060592 | 123060798 | 123059813 | 123060268 | 123061119 | 123061258 | 119883 | 612,415,1331,1480,537,43 | 1,4,35,20,6,4,6,3,6,2,2,1 | 689,606,704,203,536,392,2 | 20,12,3,19,5,4,8,1,4,5,26,1 | 298 | 149 | 0.00187198429839 | 0.172774036246474 | 0.997,0.981,0.95,0 | 0.945,0.962,0.992, | 0,022 |
| 31162 | ENSG00000288643 | | chr5 | + | 62526141 | 62526257 | 62515387 | 62515501 | 62536701 | 62536781 | 31162 | 3,2,8,8,10,7,4,7,2,2,1,4 | 0,0,4,0,2,6,0,1,0,1,2,3 | 17,7,2,7,7,5,3,9,8,9,10,5,3, | 1,3,0,2,0,0,0,0,0,0,0,4,0,2 | 265 | 149 | 0.00187364562214 | 0.172853245134511 | 1.0,1.0,0.529,1.0,0 | 0.905,0.567,1.0,0. | -0,12 |
| 101064 | ENSG00000186088 | GSAP | chr7 | - | 77311840 | 77311940 | 77310750 | 77311449 | 77312100 | 77312202 | 101064 | 193,109,331,131,154,123,2 | 28,7,31,27,22,18,20,21,14, | 209,131,136,52,82,78,83,1 | 35,35,28,34,38,15,15,23,21 | 249 | 149 | 0.00187640554609 | 0.173033694126177 | 0.805,0.903,0.865, | 0.781,0.691,0.744, | 0,07 |
| 9828 | ENSG00000108433 | GOSR2 | chr17 | + | 46948942 | 46949347 | 46938598 | 46938704 | 46966533 | 46966626 | 9828 | 0,0,0,0,4,2,7,9,0,0,0,1 | 2,2,6,6,2,0,4,5,3,9,2,1 | 11,3,2,5,0,0,0,0,0,2,5,5,2,3 | 0,0,0,0,2,1,6,5,2,2,2,8,1,3, | 298 | 149 | 0.00187754890153 | 0.173064979711008 | 0.0,0.0,0.0,0.0,0.5, | 1.0,1.0,1.0,1.0,0.0, | -0,225 |
| 180606 | ENSG00000130775 | THEMIS2 | chr1 | + | 27876572 | 27876728 | 27872557 | 27872665 | 27879643 | 27880054 | 180606 | 59,88,42,42,25,73,63,85,9 | 1,2,8,0,0,0,3,0,0,0,0,6 | 44,140,30,13,72,35,104,23 | 2,4,0,3,3,0,7,11,2,0,10,6,5, | 298 | 149 | 0.00187919564754 | 0.173142619184795 | 0.967,0.957,0.724, | 0.917,0.946,1.0,0. | 0,046 |
| 137080 | ENSG00000257103 | LSM14A | chr19 | + | 34215092 | 34215300 | 34208928 | 34209051 | 34215595 | 34215661 | 137080 | 75,43,114,122,39,43,25,39 | 16,16,31,9,17,7,11,20,2,5,3 | 62,50,52,32,38,41,17,29,3 | 21,51,21,19,37,4,11,15,17, | 298 | 149 | 0.00188254979635 | 0.173377439117761 | 0.701,0.573,0.648, | 0.596,0.329,0.553, | 0,137 |
| 155591 | ENSG00000126226 | PCID2 | chr13 | - | 113190567 | 113190975 | 113185484 | 113185560 | 113195070 | 113195125 | 155591 | 26,13,32,30,18,15,18,21,2 | 6,1,7,6,2,0,2,6,0,0,0,0 | 11,12,5,10,12,8,6,15,5,16,2 | 1,1,0,2,4,0,0,3,0,2,0,0,2,0, | 298 | 149 | 0.00188710132517 | 0.173722286277872 | 0.684,0.867,0.696, | 0.846,0.857,1.0,0. | 0,006 |
| 193869 | ENSG00000139173 | TMEM117 | chr12 | + | 43924838 | 43925000 | 43844850 | 43844928 | 43944209 | 43944342 | 193869 | 2,0,13,1,0,2,0,4,2,2,0,0 | 11,5,23,14,6,10,4,9,7,6,4,5 | 2,6,0,2,1,0,0,2,0,2,3,2,3,3, | 20,14,6,3,7,6,9,13,6,14,5,3 | 298 | 149 | 0.00189649588378 | 0.174449800194443 | 0.083,0.0,0.22,0.0 | 0.048,0.176,0.0,0. | -0,013 |
| 73098 | ENSG00000099194 | SCD | chr10 | + | 100356531 | 100356764 | 100354426 | 100354632 | 100360733 | 100364826 | 73098 | 10,12,15,18,7,12,9,20,7,11 | 4,0,1,0,2,1,0,3,0,0,0,1 | 8,6,2,10,11,3,7,23,6,7,7,11 | 2,2,1,0,4,0,0,3,1,0,4,2,0,0, | 298 | 149 | 0.00189662517228 | 0.174449800194443 | 0.556,1.0,0.882,1. | 0.667,0.6,0.5,1.0,0 | 0,124 |
| 121180 | ENSG00000181924 | COA4 | chr11 | - | 73876478 | 73876575 | 73872996 | 73873394 | 73876756 | 73876843 | 121180 | 1,1,2,5,0,0,0,4,0,0,0,1 | 9,4,35,24,21,11,9,7,7,9,4,1 | 2,2,0,4,0,0,0,4,2,3,10,6,3,0 | 14,7,13,7,16,7,13,14,6,8,1 | 246 | 149 | 0.00190736234017 | 0.175362453582712 | 0.063,0.132,0.033, | 0.08,0.148,0.0,0.2 | -0,042 |
| 70936 | ENSG00000279457 | WASH9P | chr1 | - | 188129 | 188266 | 187375 | 187577 | 188438 | 188584 | 70936 | 36,17,22,15,12,5,28,18,20, | 3,0,0,1,2,4,4,4,2,4,1,2 | 11,20,15,18,14,3,16,15,3,7 | 2,0,4,2,0,0,0,1,0,3,0,0,0,1, | 286 | 149 | 0.00190917824601 | 0.175454459037992 | 0.862,1.0,1.0,0.88 | 0.741,1.0,0.661,0. | -0,081 |
| 198391 | ENSG00000069248 | NUP133 | chr1 | - | 229488741 | 229488845 | 229487465 | 229487613 | 229489954 | 229490102 | 198391 | 5,4,14,6,1,4,4,4,2,0,2,0 | 35,21,53,42,27,18,16,31,2 | 0,4,8,5,1,0,3,2,1,1,5,3,0,0,1 | 66,55,13,21,17,13,19,31,4 | 253 | 149 | 0.00191566186162 | 0.175975167792716 | 0.078,0.101,0.135, | 0.0,0.041,0.266,0. | 0,019 |
| 161882 | ENSG00000198055 | GRK6 | chr5 | + | 177432010 | 177432107 | 177430871 | 177430967 | 177432232 | 177432257 | 161882 | 40,21,41,37,17,21,28,11,3 | 16,9,7,6,4,12,7,2,1,1,2,0 | 16,28,16,19,24,13,14,19,7, | 15,20,7,9,13,4,7,9,7,2,11,1 | 246 | 149 | 0.00192099036824 | 0.176389367724686 | 0.602,0.586,0.78,0 | 0.392,0.459,0.581, | 0,162 |
| 139177 | ENSG00000088812 | ATRN | chr20 | + | 3559188 | 3559483 | 3549169 | 3549338 | 3560661 | 3560905 | 139177 | 12,14,17,15,5,10,9,13,25,6 | 2,0,0,0,0,0,0,0,0,0,0,0,0 | 31,12,6,9,7,8,9,6,14,13,13, | 2,0,2,1,2,1,4,2,0,0,3,0,0,0, | 298 | 149 | 0.00192724048615 | 0.176887802590427 | 0.75,1.0,1.0,1.0,1. | 0.886,1.0,0.6,0.81 | 0,17 |
| 20257 | ENSG00000125107 | CNOT1 | chr16 | - | 58623281 | 58623409 | 58599262 | 58599511 | 58629727 | 58629885 | 20257 | 67,38,92,73,29,45,21,44,3, | 43,38,72,39,33,28,21,24,2 | 57,81,41,56,69,33,13,27,2 | 38,25,26,9,8,17,12,25,13,4 | 277 | 149 | 0.00192833438447 | 0.176912761254843 | 0.456,0.35,0.407,0 | 0.447,0.635,0.459, | -0,145 |
| 114281 | ENSG00000143748 | NVL | chr1 | - | 224301690 | 224301771 | 224300561 | 224300663 | 224303722 | 224303857 | 114281 | 4,0,7,5,0,2,0,0,1,1,0,0 | 8,8,16,9,10,20,7,23,11,5,9, | 1,2,2,0,2,3,2,2,3,3,5,3,4,6, | 7,11,5,11,10,3,6,13,12,19,1 | 230 | 149 | 0.00194233456417 | 0.178121265692745 | 0.245,0.0,0.221,0. | 0.085,0.105,0.206, | -0,074 |
| 89675 | ENSG00000225830 | ERCC6 | chr10 | - | 49524032 | 49524777 | 49504991 | 49506012 | 49528416 | 49528525 | 89675 | 29,23,44,25,14,19,28,11,2 | 2,3,2,2,5,4,1,0,0,2,0,2 | 11,23,22,9,20,14,10,12,15, | 4,3,0,1,2,3,0,3,0,0,2,0,2,2, | 298 | 149 | 0.00194662792228 | 0.17843895840726 | 0.879,0.793,0.917, | 0.579,0.793,1.0,0. | 0,001 |
| 17103 | ENSG00000186812 | ZNF397 | chr18 | + | 35242390 | 35242884 | 35241056 | 35241690 | 35243151 | 35243293 | 17103 | 13,10,43,31,26,19,14,12,1 | 2,1,2,2,0,0,0,3,1,6,1,0 | 13,30,9,12,13,6,12,12,4,13 | 0,1,0,0,2,0,0,0,0,0,0,0,0,0, | 298 | 149 | 0.00194970792895 | 0.178645205302613 | 0.765,0.833,0.915, | 1.0,0.938,1.0,1.0,0 | -0,136 |
| 197130 | ENSG00000198836 | OPA1 | chr3 | + | 193631754 | 193631848 | 193631611 | 193631665 | 193635417 | 193635522 | 197130 | 3,0,5,5,0,3,4,4,0,4,0,2 | 27,24,82,51,26,20,18,31,1 | 7,17,5,6,4,0,2,2,4,1,4,6,3,1 | 51,60,23,17,34,19,14,22,2 | 243 | 149 | 0.00195150520511 | 0.178733794383417 | 0.064,0.0,0.036,0. | 0.078,0.148,0.118, | -0,032 |
| 66197 | ENSG00000100038 | TOP3B | chr22 | - | 21968618 | 21968775 | 21967602 | 21967716 | 21971876 | 21971951 | 66197 | 24,12,36,21,7,19,11,23,19, | 0,7,1,0,1,5,1,2,0,1,0,3 | 15,35,15,12,12,11,7,11,22, | 2,2,2,1,0,1,0,0,0,0,2,0,0,1, | 298 | 149 | 0.00196378655182 | 0.179782111158984 | 1.0,0.462,0.947,1. | 0.789,0.897,0.789, | -0,04 |
| 33920 | ENSG00000120137 | PANK3 | chr5 | - | 168563888 | 168564065 | 168561392 | 168561516 | 168566012 | 168566266 | 33920 | 103,114,106,90,58,59,49,9 | 11,5,13,13,4,2,1,9,1,4,0,1 | 91,68,56,66,40,24,50,48,3 | 6,15,12,15,9,1,3,12,5,3,6,2 | 298 | 149 | 0.00196530160515 | 0.17984431538182 | 0.824,0.919,0.803, | 0.883,0.694,0.7,0. | 0,094 |
| 149767 | ENSG00000128928 | IVD | chr15 | + | 40418129 | 40418215 | 40416289 | 40416362 | 40419179 | 40419373 | 149767 | 3,2,10,17,12,6,6,1,1,1,4,4 | 1,0,1,1,2,1,1,0,0,0,0,0 | 4,9,4,2,4,6,3,6,6,5,3,1,4,8, | 0,8,1,0,0,0,0,4,0,0,0,0,5,0, | 235 | 149 | 0.0019677238567 | 0.17998944895937 | 0.655,1.0,0.864,0. | 1.0,0.416,0.717,1. | 0,059 |
| 164169 | ENSG00000083896 | YTHDC1 | chr4 | - | 68330201 | 68330699 | 68330001 | 68330119 | 68332102 | 68332197 | 164169 | 52,58,123,73,41,62,54,75, | 5,0,0,6,1,2,2,1,1,0,0,1 | 84,115,73,57,48,44,39,53, | 1,8,0,13,4,3,0,4,0,2,0,1,0,0 | 298 | 149 | 0.00196905221845 | 0.180034442663217 | 0.839,1.0,1.0,0.85 | 0.977,0.878,1.0,0. | 0,018 |
| 16492 | ENSG00000153774 | CFDP1 | chr16 | - | 75391379 | 75391508 | 75305023 | 75305182 | 75395089 | 75395136 | 16492 | 4,0,4,1,3,3,0,4,1,2,0,1 | 27,19,43,57,13,21,14,11,2 | 5,4,3,1,3,7,1,1,1,5,2,1,7,2,2 | 16,13,8,2,13,7,13,20,11,13 | 278 | 149 | 0.00197153894156 | 0.180119905588274 | 0.074,0.0,0.047,0. | 0.143,0.142,0.167, | -0,07 |
| 212015 | ENSG00000079263 | SP140 | chr2 | + | 230243700 | 230243811 | 230241403 | 230241487 | 230244987 | 230245071 | 212015 | 87,36,68,82,66,39,27,38,7 | 32,20,24,18,8,13,18,24,6,8 | 64,45,54,36,46,36,14,35,3 | 40,96,30,41,46,10,14,6,17, | 260 | 149 | 0.00197217500349 | 0.180119905588274 | 0.609,0.508,0.619, | 0.478,0.212,0.508, | 0,163 |
| 32134 | ENSG00000139278 | GLIPR1 | chr12 | + | 75490405 | 75490518 | 75481833 | 75482079 | 75495576 | 75495662 | 32134 | 73,44,43,66,51,17,20,21,2 | 4,0,1,0,0,0,0,0,2,0,0,2 | 90,48,2,10,47,29,24,2,17,4 | 1,3,0,0,4,0,0,0,6,4,0,4,6,4, | 262 | 149 | 0.00197291551685 | 0.180119905588274 | 0.912,1.0,0.961,1. | 0.981,0.901,1.0,1. | 0,038 |
| 109339 | ENSG00000104361 | NIPAL2 | chr8 | - | 98193384 | 98193412 | 98189832 | 98193090 | 98194727 | 98194822 | 109339 | 1,0,4,0,1,1,1,9,0,0,0,0 | 17,14,63,51,13,14,9,5,15,1 | 2,2,1,2,3,0,0,0,4,1,4,9,2,2, | 37,21,6,5,3,11,14,25,15,20 | 177 | 149 | 0.00197365984337 | 0.180119905588274 | 0.047,0.0,0.051,0. | 0.044,0.074,0.123, | -0,043 |
| 203568 | ENSG00000275052 | PPP4R3B | chr2 | - | 55574368 | 55574413 | 55573618 | 55573777 | 55577314 | 55577356 | 203568 | 1,0,16,24,1,2,0,6,6,9,0,2 | 17,10,34,34,58,15,11,17,1 | 2,4,2,6,12,2,0,3,0,3,1,0,0,0 | 40,24,14,6,10,18,18,37,5,2 | 194 | 149 | 0.00197417127311 | 0.180119905588274 | 0.043,0.0,0.265,0. | 0.037,0.113,0.099, | 0,05 |
| 210323 | ENSG00000148248 | SURF4 | chr9 | - | 133364839 | 133365025 | 133361449 | 133363759 | 133365984 | 133366077 | 210323 | 37,24,45,40,23,31,18,45,5 | 1,1,9,13,0,2,4,1,2,4,1,1 | 32,57,34,28,13,19,20,35,1 | 3,5,2,0,1,2,1,4,0,0,0,3,3,0, | 298 | 149 | 0.00197797216291 | 0.180324665672502 | 0.949,0.923,0.714, | 0.842,0.851,0.895, | -0,055 |
| 33544 | ENSG00000038219 | BOD1L1 | chr4 | - | 13623389 | 13626312 | 13619942 | 13620067 | 13627344 | 13627701 | 33544 | 6,5,4,2,8,1,8,5,0,0,5,5 | 26,24,51,29,13,12,12,30,1 | 0,1,2,5,4,7,2,4,3,6,2,5,0,2, | 33,44,24,18,35,18,15,22,1 | 298 | 149 | 0.00197846543606 | 0.180324665672502 | 0.103,0.094,0.038, | 0.0,0.011,0.04,0.1 | 0,05 |
| 156858 | ENSG00000134884 | ARGLU1 | chr13 | - | 106557519 | 106557616 | 106557105 | 106557131 | 106559431 | 106559657 | 156858 | 38,10,20,17,6,10,3,14,15,1 | 130,49,133,112,66,62,60,6 | 14,12,16,2,2,5,5,18,1,9,7,5 | 104,54,77,34,46,41,18,79, | 246 | 149 | 0.00197965159533 | 0.180324665672502 | 0.15,0.11,0.083,0. | 0.075,0.119,0.112, | 0,039 |
| 41410 | ENSG00000112149 | CD83 | chr6 | + | 14131519 | 14131748 | 14117949 | 14118065 | 14133648 | 14133755 | 41410 | 614,556,277,126,79,113,9 | 25,13,19,12,2,4,9,6,1,2,12, | 126,233,111,47,102,21,11 | 9,16,16,5,4,0,4,18,8,0,5,72 | 298 | 149 | 0.00197976678538 | 0.180324665672502 | 0.925,0.955,0.879, | 0.875,0.879,0.776, | 0,04 |
| 131255 | ENSG00000111229 | ARPC3 | chr12 | - | 110436664 | 110436683 | 110436177 | 110436204 | 110437083 | 110437152 | 131255 | 82,53,99,93,43,30,35,55,6 | 82,40,42,42,27,8,19,7,10,8 | 64,87,78,42,55,47,18,30,2 | 30,34,18,10,3,14,3,7,6,8,7, | 168 | 149 | 0.00198077811505 | 0.180340462978141 | 0.47,0.54,0.676,0. | 0.654,0.694,0.794, | -0,113 |
| 238646 | ENSG00000164933 | SLC25A32 | chr8 | - | 103413422 | 103413531 | 103407633 | 103407784 | 103414783 | 103415189 | 238646 | 14,9,11,7,8,2,4,7,14,8,3,4 | 69,46,50,36,16,22,20,24,3 | 12,16,12,7,11,3,3,5,10,15, | 33,21,21,25,23,13,6,20,11, | 258 | 149 | 0.00198630153235 | 0.180718310705203 | 0.105,0.102,0.113, | 0.174,0.306,0.248, | -0,062 |
| 180601 | ENSG00000130775 | THEMIS2 | chr1 | + | 27876272 | 27876728 | 27872557 | 27872665 | 27879643 | 27880054 | 180601 | 59,88,41,42,25,73,63,85,9 | 1,2,8,0,0,0,3,0,0,0,0,6 | 44,140,30,13,72,35,106,24 | 2,4,0,3,3,0,7,11,2,0,10,6,5, | 298 | 149 | 0.00198660751996 | 0.180718310705203 | 0.967,0.957,0.719, | 0.917,0.946,1.0,0. | 0,045 |
| 150034 | ENSG00000159322 | ADPGK | chr15 | - | 72756250 | 72756344 | 72755555 | 72755654 | 72771782 | 72771845 | 150034 | 13,12,33,13,19,24,19,13,4, | 0,0,3,2,0,2,1,3,2,2,0,2 | 13,26,20,15,5,13,4,14,15,1 | 3,2,2,2,3,1,1,2,1,3,3,0,1,1,1 | 243 | 149 | 0.00198803498576 | 0.180771760887246 | 1.0,1.0,0.871,0.79 | 0.727,0.889,0.86,0 | 0,042 |
| 26519 | ENSG00000107201 | DDX58 | chr9 | - | 32467761 | 32467932 | 32466289 | 32466441 | 32472974 | 32473065 | 26519 | 40,9,62,32,30,19,21,27,26, | 2,0,5,1,2,2,2,0,0,2,0,3 | 55,17,22,29,2,19,7,23,16,3 | 2,0,0,4,1,5,0,5,1,1,0,5,2,0, | 298 | 149 | 0.00198957449085 | 0.180805921481351 | 0.909,1.0,0.861,0. | 0.932,1.0,1.0,0.78 | 0,005 |
| 55235 | ENSG00000010292 | NCAPD2 | chr12 | + | 6522827 | 6523002 | 6521797 | 6522037 | 6525582 | 6525716 | 55235 | 24,13,31,36,21,21,17,20,1 | 3,0,2,0,0,1,0,0,0,1,0,0 | 26,13,8,3,8,9,11,16,9,19,2 | 0,0,0,1,1,2,2,2,0,4,3,5,5,0, | 298 | 149 | 0.00199037000178 | 0.180805921481351 | 0.8,1.0,0.886,1.0,1 | 1.0,1.0,1.0,0.6,0.8, | 0,094 |
| 117535 | ENSG00000090054 | SPTLC1 | chr9 | - | 92079433 | 92079566 | 92067965 | 92068098 | 92080015 | 92080088 | 117535 | 6,4,10,2,1,1,4,3,2,0,2,6 | 72,27,62,40,20,19,14,27,4 | 6,9,9,19,8,2,6,5,0,1,6,2,0,9 | 38,39,31,10,28,12,16,11,5, | 282 | 149 | 0.00199093083204 | 0.180805921481351 | 0.042,0.073,0.079, | 0.077,0.109,0.133, | -0,045 |
| 227693 | ENSG00000186088 | GSAP | chr7 | - | 77379847 | 77380000 | 77377285 | 77377390 | 77381304 | 77381354 | 227693 | 21,15,51,27,16,50,32,44,1 | 51,26,156,99,53,60,40,66, | 43,57,33,21,20,17,17,44,2 | 102,45,42,11,27,18,28,56,2 | 298 | 149 | 0.00199612907148 | 0.181201542042898 | 0.171,0.224,0.14,0 | 0.174,0.388,0.282, | -0,078 |
| 69986 | ENSG00000115207 | GTF3C2 | chr2 | - | 27335597 | 27335706 | 27333973 | 27333999 | 27335916 | 27336028 | 69986 | 25,22,56,41,32,29,22,36,1 | 0,0,5,2,0,5,1,2,0,0,3,0 | 54,62,13,14,30,5,42,53,15, | 8,5,1,3,3,0,2,5,0,0,2,1,2,1,1 | 258 | 149 | 0.00200183206493 | 0.181642629497027 | 1.0,1.0,0.866,0.92 | 0.796,0.877,0.882, | 0,026 |
| 124899 | ENSG00000080546 | SESN1 | chr6 | - | 108998512 | 108998755 | 108994461 | 108994609 | 109000490 | 109000673 | 124899 | 44,14,144,52,21,41,85,495 | 0,0,2,0,0,2,1,9,2,0,1,1 | 76,96,82,37,17,12,50,31,2 | 4,0,3,1,0,0,0,2,1,0,24,1,0,0 | 298 | 149 | 0.00200820250756 | 0.182143882808532 | 1.0,1.0,0.973,1.0,1 | 0.905,1.0,0.932,0. | 0,018 |
| 89778 | ENSG00000152056 | AP1S3 | chr2 | - | 223775900 | 223776009 | 223765176 | 223765350 | 223777690 | 223777810 | 89778 | 78,119,65,64,32,36,11,20, | 5,4,7,9,1,2,3,2,4,0,2,1 | 53,88,24,22,52,23,17,24,3 | 1,2,5,5,2,1,0,1,0,1,1,0,2,0, | 258 | 149 | 0.00201192140962 | 0.182404320519765 | 0.9,0.945,0.843,0. | 0.968,0.962,0.735, | -0,053 |
| 18515 | ENSG00000159322 | ADPGK | chr15 | - | 72774871 | 72775097 | 72771782 | 72771845 | 72783245 | 72783731 | 18515 | 108,90,127,114,78,93,50,8 | 8,7,6,1,5,3,6,1,5,0,3,6 | 102,113,79,50,75,35,58,71 | 2,11,8,10,23,2,3,6,10,3,3,1 | 298 | 149 | 0.00201495758536 | 0.182602667812471 | 0.871,0.865,0.914, | 0.962,0.837,0.832, | 0,059 |
| 35627 | ENSG00000125247 | TMTC4 | chr13 | - | 100670359 | 100670569 | 100668578 | 100668794 | 100674743 | 100674795 | 35627 | 3,4,10,5,4,4,5,12,2,3,3,7 | 3,3,2,1,1,4,0,3,5,2,0,0 | 11,18,4,1,1,3,3,4,4,9,7,25,2 | 1,0,0,1,4,0,0,1,0,0,2,2,5,1, | 298 | 149 | 0.0020188434687 | 0.182808242422707 | 0.333,0.4,0.714,0. | 0.846,1.0,1.0,0.33 | -0,093 |
| 145325 | ENSG00000161395 | PGAP3 | chr17 | - | 39676664 | 39676755 | 39674616 | 39674679 | 39684596 | 39684698 | 145325 | 0,0,3,6,2,1,2,1,0,2,0,1 | 22,7,18,23,26,23,21,10,12, | 4,4,4,0,2,0,2,3,2,4,0,2,1,1, | 11,16,10,8,9,2,8,14,11,13, | 240 | 149 | 0.00201892474708 | 0.182808242422707 | 0.0,0.0,0.094,0.13 | 0.184,0.134,0.199, | -0,055 |
| 114913 | ENSG00000136937 | NCBP1 | chr9 | + | 97651309 | 97651373 | 97650502 | 97650600 | 97653797 | 97653908 | 114913 | 49,32,71,46,39,35,48,62,3 | 3,2,4,2,1,7,2,1,3,1,0,4 | 49,62,42,73,26,11,27,54,4 | 4,14,5,8,8,2,2,2,3,4,6,4,0,3 | 213 | 149 | 0.00202044502893 | 0.182868967208424 | 0.92,0.918,0.925,0 | 0.895,0.756,0.855, | 0,052 |
| 118792 | ENSG00000291075 | | chr9 | + | 41008919 | 41009006 | 40994632 | 40994699 | 41011616 | 41011730 | 118792 | 31,12,54,41,16,17,14,25,2 | 2,0,12,3,18,15,11,19,0,0,2, | 48,31,43,10,5,3,15,21,20,1 | 0,0,0,1,11,4,0,0,0,0,4,16,0, | 236 | 149 | 0.00202449062451 | 0.183158109123124 | 0.907,1.0,0.74,0.8 | 1.0,1.0,1.0,0.863,0 | -0,099 |
| 189944 | ENSG00000138688 | KIAA1109 | chr4 | + | 122337210 | 122337260 | 122336880 | 122337020 | 122339206 | 122339408 | 189944 | 2,3,8,2,0,5,4,2,5,2,3,2 | 36,29,53,31,20,20,16,25,2 | 5,24,5,6,4,0,3,2,4,3,4,4,5,2 | 38,18,14,10,14,11,15,16,8, | 199 | 149 | 0.00202821215373 | 0.183417701705656 | 0.04,0.072,0.102,0 | 0.09,0.5,0.211,0.3 | -0,063 |
| 66006 | ENSG00000137776 | SLTM | chr15 | - | 58890280 | 58890461 | 58889429 | 58889554 | 58892896 | 58893060 | 66006 | 238,128,182,170,123,86,7 | 17,13,53,32,22,36,18,29,1 | 81,132,135,89,52,80,47,12 | 28,56,11,20,25,9,17,12,31, | 298 | 149 | 0.00202917378892 | 0.183427595028576 | 0.875,0.831,0.632, | 0.591,0.541,0.86,0 | 0,1 |
| 84267 | ENSG00000064666 | CNN2 | chr19 | + | 1036878 | 1037000 | 1036415 | 1036562 | 1037624 | 1038309 | 84267 | 7,1,6,8,1,3,1,3,3,1,1,5 | 153,112,138,139,166,58,1 | 7,10,2,8,8,2,5,5,6,6,6,10,2, | 119,111,69,71,60,62,53,97 | 271 | 149 | 0.00204156067249 | 0.184469834215237 | 0.025,0.005,0.023, | 0.031,0.047,0.016, | -0,014 |
| 164085 | ENSG00000102445 | RUBCNL | chr13 | - | 46372268 | 46372597 | 46368762 | 46368815 | 46387133 | 46387486 | 164085 | 41,46,172,95,65,59,102,10 | 3,2,10,2,2,6,5,6,0,2,0,1 | 76,142,69,28,55,51,51,69,2 | 11,12,2,2,4,2,1,3,2,1,8,4,9, | 298 | 149 | 0.00205176564484 | 0.185294459359393 | 0.872,0.92,0.896,0 | 0.776,0.855,0.945, | 0,053 |
| 134390 | ENSG00000100077 | GRK3 | chr22 | + | 25666632 | 25666720 | 25663629 | 25663704 | 25667738 | 25667800 | 134390 | 7,7,2,1,1,3,0,5,0,0,0,7 | 68,53,100,60,21,18,9,13,2 | 11,15,1,4,4,3,0,3,5,2,2,8,0, | 50,42,21,24,21,14,9,13,24, | 237 | 149 | 0.00205240876599 | 0.185294459359393 | 0.061,0.077,0.012, | 0.122,0.183,0.029, | -0,027 |
| 77951 | ENSG00000146535 | GNA12 | chr7 | - | 2733450 | 2733501 | 2728104 | 2731750 | 2794927 | 2795143 | 77951 | 71,50,81,37,41,71,28,109, | 1,1,6,1,0,1,2,0,6,2,0,0 | 54,87,47,50,39,29,37,42,2 | 0,4,0,16,3,0,2,0,0,2,5,6,0,0 | 200 | 149 | 0.00205344810297 | 0.185310561278967 | 0.981,0.974,0.91,0 | 1.0,0.942,1.0,0.7,0 | 0,012 |
| 31873 | ENSG00000247774 | PCED1B-AS1 | chr12 | - | 47206510 | 47206588 | 47205897 | 47206169 | 47209280 | 47209458 | 31873 | 14,4,7,6,16,3,3,5,4,1,3,1 | 2,0,4,0,1,0,0,0,3,1,2,0 | 7,5,9,3,7,2,2,1,4,9,2,43,10, | 3,2,1,0,1,0,0,2,0,5,2,6,3,0, | 227 | 149 | 0.00205431232305 | 0.185310853144331 | 0.821,1.0,0.535,1. | 0.605,0.621,0.855, | 0,057 |
| 255215 | ENSG00000089154 | GCN1 | chr12 | - | 120144634 | 120144835 | 120144305 | 120144448 | 120144922 | 120145061 | 255215 | 53,33,51,50,27,42,24,26,3 | 5,0,7,6,0,8,2,1,0,2,0,0 | 23,26,21,35,22,13,17,24,1 | 0,10,4,12,7,2,4,5,0,2,4,2,4, | 298 | 149 | 0.00205604626936 | 0.185389566234027 | 0.841,1.0,0.785,0. | 1.0,0.565,0.724,0. | 0,125 |
| 6473 | ENSG00000170471 | RALGAPB | chr20 | + | 38497352 | 38497516 | 38492929 | 38493132 | 38499446 | 38499633 | 6473 | 80,75,155,97,81,108,77,83 | 0,0,2,0,0,2,1,1,0,0,0,3 | 106,127,72,73,77,36,59,61 | 3,3,3,6,4,1,0,4,1,0,1,2,0,0, | 298 | 149 | 0.00206545170935 | 0.186154633546808 | 1.0,1.0,0.975,1.0,1 | 0.946,0.955,0.923, | 0,03 |
| 218895 | ENSG00000291118 | ZNF767P | chr7 | - | 149558607 | 149558833 | 149552800 | 149553016 | 149619923 | 149620037 | 218895 | 10,3,8,15,16,6,7,12,39,11, | 0,1,2,0,2,2,2,1,2,2,0,4 | 3,5,7,15,3,3,6,6,2,10,3,2,1 | 0,2,2,2,2,0,1,0,1,0,1,0,0,0, | 298 | 149 | 0.00206626099188 | 0.186154633546808 | 1.0,0.6,0.667,1.0,0 | 1.0,0.556,0.636,0. | -0,062 |
| 104004 | ENSG00000291151 | NSUN5P1 | chr7 | + | 75415595 | 75415778 | 75415119 | 75415234 | 75415980 | 75416191 | 104004 | 2,2,14,4,6,8,7,5,4,3,4,5 | 1,0,1,1,4,5,1,0,0,0,0,0 | 9,11,1,4,4,1,3,7,2,9,3,9,9,6 | 1,0,2,0,1,0,9,2,3,0,0,5,1,0, | 298 | 149 | 0.00207066578518 | 0.186461757849206 | 0.5,1.0,0.875,0.66 | 0.818,1.0,0.2,1.0,0 | 0,137 |
| 175312 | ENSG00000186001 | LRCH3 | chr3 | + | 197831959 | 197832056 | 197830769 | 197830863 | 197832196 | 197832317 | 175312 | 0,0,8,6,1,4,2,5,0,4,4,4 | 47,38,77,55,28,33,35,55,2 | 5,3,0,3,3,1,3,1,0,1,3,0,0,0, | 37,25,21,12,18,15,27,40,1 | 246 | 149 | 0.00207140264654 | 0.186461757849206 | 0.0,0.0,0.059,0.06 | 0.076,0.068,0.0,0. | 0,035 |
| 154722 | ENSG00000129566 | TEP1 | chr14 | - | 20389620 | 20389740 | 20389237 | 20389297 | 20390680 | 20390758 | 154722 | 34,31,33,31,31,37,19,15,1 | 1,0,1,2,1,3,3,0,0,2,0,1 | 51,41,19,24,40,24,22,20,2 | 0,3,2,5,4,2,4,0,0,0,4,4,6,0, | 269 | 149 | 0.00207263742799 | 0.186494910645366 | 0.95,1.0,0.948,0.8 | 1.0,0.883,0.84,0.7 | 0,049 |
| 128100 | ENSG00000065613 | SLK | chr10 | + | 104010815 | 104010908 | 104008176 | 104008356 | 104018159 | 104018289 | 128100 | 5,4,11,5,0,16,2,16,0,0,6,2 | 30,33,56,37,24,26,25,33,4, | 2,3,8,2,3,3,2,4,0,3,3,3,0,4, | 27,31,28,34,29,9,16,28,11, | 242 | 149 | 0.00207696516573 | 0.186806222141761 | 0.093,0.069,0.108, | 0.044,0.056,0.15,0 | 0,025 |
| 152734 | ENSG00000089775 | ZBTB25 | chr14 | - | 64495242 | 64495559 | 64490360 | 64490540 | 64502900 | 64503127 | 152734 | 2,0,8,6,9,10,1,2,5,2,2,0 | 7,6,3,6,9,5,5,1,6,0,0,3 | 4,6,2,2,4,0,3,5,5,9,4,2,1,3, | 10,1,0,1,0,4,0,4,0,3,0,2,0,4 | 298 | 149 | 0.00207793644202 | 0.186815513096243 | 0.125,0.0,0.571,0. | 0.167,0.75,1.0,0.5, | -0,267 |
| 82481 | ENSG00000115159 | GPD2 | chr2 | + | 156495561 | 156495691 | 156476097 | 156476207 | 156496043 | 156496215 | 82481 | 4,0,5,7,0,3,3,4,0,0,0,3 | 30,14,74,23,7,15,9,9,8,8,2, | 5,6,4,8,12,4,0,2,4,0,8,3,2,5 | 25,10,24,9,16,20,9,9,16,23 | 279 | 149 | 0.00208033194395 | 0.186940295786393 | 0.066,0.0,0.035,0. | 0.097,0.243,0.082, | -0,087 |

| | | | | | | | | | | | | | | | | | | | | | | |
|---|---|---|---|---|---|---|---|---|---|---|---|---|---|---|---|---|---|---|---|---|---|---|
| 40140 | ENSG00000291102 | GUSBP1 | chr5 | + | 21466618 | 21466748 | 21461782 | 21461866 | 21491320 | 21491508 | 40140 | 2,2,4,2,3,0,2,6,0,0,0,1 | 8,1,18,8,7,6,2,7,3,1,3,3 | 0,1,2,3,0,0,0,0,0,3,1,2,1,1, | 9,7,15,8,7,3,9,7,5,11,7,3,5, | 279 | 149 | 0.00208189736496 | 0.186940295786393 | 0.118,0.516,0.106, | 0.0,0.071,0.066,0. | 0,078 |
| 228964 | ENSG00000111667 | USP5 | chr12 | + | 6856304 | 6856455 | 6856016 | 6856150 | 6856706 | 6856891 | 228964 | 32,39,47,54,31,35,39,22,2 | 0,0,2,0,0,2,0,2,0,0,0,0 | 29,45,21,15,20,10,8,36,12, | 0,1,5,1,1,1,0,6,0,0,3,3,2,3, | 298 | 149 | 0.00208193006119 | 0.186940295786393 | 1.0,1.0,0.922,1.0,1 | 1.0,0.957,0.677,0. | 0,07 |
| 203556 | ENSG00000275052 | PPP4R3B | chr2 | - | 55569655 | 55569840 | 55568193 | 55568363 | 55573618 | 55573777 | 203556 | 8,13,5,7,1,2,4,14,6,9,7,8 | 66,39,112,59,33,32,24,23, | 17,38,11,44,13,11,14,12,1 | 105,101,25,19,19,28,11,49 | 298 | 149 | 0.00208600728985 | 0.187192464341327 | 0.057,0.143,0.022, | 0.075,0.158,0.18,0 | -0,056 |
| 743 | ENSG00000228315 | GUSBP11 | chr22 | - | 23659864 | 23660000 | 23638486 | 23638942 | 23683724 | 23683867 | 743 | 60,19,12,9,28,24,30,51,13, | 1,0,0,0,0,2,2,0,3,0,2,0 | 35,38,28,45,14,17,40,34,3 | 2,0,1,5,2,0,2,2,0,1,0,0,2,6, | 285 | 149 | 0.00208647788634 | 0.187192464341327 | 0.969,1.0,1.0,1.0,1 | 0.901,1.0,0.936,0. | 0,02 |
| 7407 | ENSG00000062598 | ELMO2 | chr20 | - | 46374535 | 46374640 | 46374331 | 46374440 | 46375235 | 46375370 | 7407 | 22,17,41,51,17,22,16,31,1 | 6,2,4,1,1,0,1,2,0,0,1,0 | 24,31,10,21,13,13,18,35,2 | 0,9,1,3,1,1,0,6,0,4,2,1,0,1, | 254 | 149 | 0.00209056445735 | 0.187411380958816 | 0.683,0.833,0.857, | 1.0,0.669,0.854,0. | 0,044 |
| 47460 | ENSG00000115935 | WIPF1 | chr2 | - | 174630381 | 174630696 | 174629665 | 174629868 | 174634517 | 174634536 | 47460 | 12,2,28,16,4,3,3,15,5,4,4,8 | 0,0,16,8,0,1,0,0,1,0,0,0 | 16,42,5,10,18,4,7,5,14,6,1 | 2,0,2,0,0,1,4,1,3,0,0,2,1,2, | 298 | 149 | 0.00209065945743 | 0.187411380958816 | 1.0,1.0,0.467,0.5,1 | 0.8,1.0,0.556,1.0,1 | 0,044 |
| 227610 | ENSG00000186088 | GSAP | chr7 | - | 77311840 | 77311958 | 77310750 | 77311449 | 77312100 | 77312202 | 227610 | 130,69,181,69,84,51,132,1 | 28,7,31,27,22,18,20,21,14, | 112,65,74,29,39,41,44,65, | 35,35,28,34,38,15,15,23,2 | 267 | 149 | 0.00210512751815 | 0.188629767218544 | 0.722,0.846,0.765, | 0.641,0.509,0.596, | 0,11 |
| 156852 | ENSG00000134884 | ARGLU1 | chr13 | - | 106557508 | 106557616 | 106557105 | 106557131 | 106559431 | 106559657 | 156852 | 38,9,20,17,7,10,4,14,15,15 | 130,49,133,112,66,62,60,6 | 14,12,16,3,2,5,5,18,1,9,7,5 | 104,54,77,34,46,41,18,79, | 257 | 149 | 0.00211074627935 | 0.189054528693625 | 0.145,0.096,0.08,0 | 0.072,0.114,0.108, | 0,037 |
| 87956 | ENSG00000162645 | GBP2 | chr1 | - | 89108022 | 89108291 | 89106131 | 89107052 | 89109676 | 89109870 | 87956 | 174,147,200,259,69,100,1 | 9,7,6,4,7,7,3,7,4,8,0,6 | 80,135,76,31,54,46,56,169 | 11,9,9,6,5,1,5,15,1,6,14,2, | 298 | 149 | 0.00211430581758 | 0.189294573803478 | 0.906,0.913,0.943, | 0.784,0.882,0.809, | 0,05 |
| 159630 | ENSG00000170476 | MZB1 | chr5 | - | 139389326 | 139389436 | 139388460 | 139388667 | 139389679 | 139390081 | 159630 | 0,2,26,21,3,4,3,6,4,3,1,2 | 9,5,9,2,3,4,3,4,2,0,6,10 | 0,4,1,0,6,4,0,3,1,7,4,0,1,0, | 3,8,6,10,0,1,3,39,0,0,18,14 | 259 | 149 | 0.00211584979241 | 0.189354040195508 | 0.0,0.187,0.624,0. | 0.0,0.223,0.087,0. | 0,09 |
| 175963 | ENSG00000276291 | FRG1HP | chr9 | + | 41008855 | 41009006 | 40994632 | 40994699 | 41011616 | 41011730 | 175963 | 41,18,74,53,18,27,20,42,3 | 2,0,12,3,18,15,11,19,0,0,2, | 65,44,58,19,6,7,15,24,33,2 | 0,0,0,1,11,4,0,0,0,0,4,16,0, | 298 | 149 | 0.00211841938891 | 0.189505205109929 | 0.911,1.0,0.755,0. | 1.0,1.0,1.0,0.905,0 | -0,098 |
| 226895 | ENSG00000105968 | H2AZ2 | chr7 | - | 44835528 | 44835696 | 44831977 | 44834562 | 44840898 | 44841012 | 226895 | 39,44,115,124,52,38,37,21 | 1,2,1,9,0,1,0,3,0,2,0,0 | 70,68,43,25,29,43,16,34,4 | 3,2,0,0,3,2,6,0,3,1,0,2,1,7, | 298 | 149 | 0.0021193476692 | 0.1895094799292 | 0.951,0.917,0.983, | 0.921,0.944,1.0,1. | 0,013 |
| 5781 | ENSG00000291100 | | chr20 | - | 28601656 | 28601839 | 28600391 | 28600461 | 28602608 | 28602732 | 5781 | 13,6,10,4,8,4,5,6,8,0,0,3 | 36,20,19,21,22,16,22,30,2 | 0,2,1,0,1,3,2,1,2,3,6,7,1,1, | 12,16,4,1,22,15,2,3,1,12,31 | 298 | 149 | 0.00212337976793 | 0.189791175594411 | 0.153,0.13,0.208,0 | 0.0,0.059,0.111,0. | -0,004 |
| 48690 | ENSG00000072786 | STK10 | chr5 | - | 172096425 | 172096560 | 172093411 | 172093960 | 172105655 | 172105737 | 48690 | 104,61,156,141,87,133,99, | 0,2,2,0,2,0,0,0,0,1,1,0 | 118,88,67,57,70,48,75,167 | 5,1,0,1,2,0,1,9,0,0,0,7,4,0, | 284 | 149 | 0.00212709140445 | 0.189863884735104 | 1.0,0.941,0.976,1. | 0.925,0.979,1.0,0. | 0,017 |
| 114168 | ENSG00000179950 | PUF60 | chr8 | - | 143824312 | 143824399 | 143821817 | 143821913 | 143829279 | 143829315 | 114168 | 27,35,43,29,13,10,14,32,1 | 0,3,12,5,0,0,4,1,0,0,0,0 | 22,29,8,2,23,2,11,21,15,18 | 2,1,1,1,2,0,0,0,1,0,2,4,0,0, | 236 | 149 | 0.00212729729382 | 0.189863884735104 | 1.0,0.88,0.693,0.7 | 0.874,0.948,0.835, | 0,003 |
| 221884 | ENSG00000172845 | SP3 | chr2 | - | 173909944 | 173910257 | 173907885 | 173907965 | 173913069 | 173913266 | 221884 | 301,217,299,171,113,127, | 6,1,17,8,9,3,6,0,1,0,1,6 | 217,294,211,219,173,102, | 9,14,6,8,4,3,6,6,6,7,8,17,3, | 298 | 149 | 0.00212745938059 | 0.189863884735104 | 0.962,0.991,0.898, | 0.923,0.913,0.946, | 0,027 |
| 151375 | ENSG00000185787 | MORF4L1 | chr15 | + | 78894057 | 78894196 | 78893538 | 78893627 | 78894819 | 78894908 | 151375 | 97,68,133,147,49,63,68,89 | 8,4,3,0,4,1,0,1,2,0,0,3 | 73,142,67,102,58,39,30,61 | 9,4,0,6,0,1,0,10,0,4,7,7,5,2 | 288 | 149 | 0.00212793036657 | 0.189863884735104 | 0.863,0.898,0.958, | 0.808,0.948,1.0,0. | 0,038 |
| 163000 | ENSG00000112992 | NNT | chr5 | + | 43614499 | 43614619 | 43612907 | 43613137 | 43615847 | 43616065 | 163000 | 2,5,8,9,3,2,4,6,5,0,2,2 | 6,5,37,35,6,12,5,17,13,12, | 3,7,3,0,0,0,0,3,0,2,0,2,5,3, | 10,22,6,3,8,12,2,19,7,20,8, | 269 | 149 | 0.00212892427514 | 0.189863884735104 | 0.156,0.356,0.107, | 0.142,0.15,0.217,0 | 0,123 |
| 215680 | ENSG00000162645 | GBP2 | chr1 | - | 89108021 | 89108291 | 89106131 | 89107052 | 89109676 | 89109870 | 215680 | 167,145,193,245,66,93,10 | 9,7,6,4,7,7,3,7,4,8,0,6 | 79,126,68,29,48,42,50,156 | 11,9,9,6,5,1,5,15,1,6,14,2, | 298 | 149 | 0.00212948607659 | 0.189863884735104 | 0.903,0.912,0.941, | 0.782,0.875,0.791, | 0,053 |
| 116168 | ENSG00000107077 | KDM4C | chr9 | + | 7049091 | 7049200 | 7046861 | 7046917 | 7076433 | 7077263 | 116168 | 63,27,46,19,24,24,23,28,2 | 0,2,5,4,1,1,0,0,5,4,0,2 | 37,38,23,41,24,24,19,15,1 | 1,1,1,3,0,1,0,0,0,2,0,3,1,0, | 258 | 149 | 0.00213407176401 | 0.190193954384943 | 1.0,0.886,0.842,0. | 0.955,0.956,0.93,0 | -0,05 |
| 31791 | ENSG00000001084 | GCLC | chr6 | - | 53516108 | 53516217 | 53514438 | 53514497 | 53520777 | 53520960 | 31791 | 36,21,69,78,22,18,22,16,5 | 1,0,3,8,0,1,5,2,3,1,0,2 | 34,29,29,3,24,13,4,12,16,3 | 0,3,2,2,2,0,0,1,2,1,2,1,1,1, | 258 | 149 | 0.00213789520893 | 0.190367834302209 | 0.954,1.0,0.93,0.8 | 1.0,0.848,0.893,0. | 0,038 |
| 150735 | ENSG00000114062 | UBE3A | chr15 | - | 25407066 | 25407384 | 25405460 | 25405502 | 25409087 | 25409412 | 150735 | 15,6,40,24,11,15,10,20,14, | 31,31,96,30,23,39,24,66,1 | 20,11,6,4,6,4,10,6,8,11,20, | 40,78,26,27,41,17,36,31,2 | 298 | 149 | 0.00213817466732 | 0.190367834302209 | 0.195,0.088,0.172, | 0.2,0.066,0.103,0. | 0,057 |
| 170479 | ENSG00000111880 | RNGTT | chr6 | - | 88803697 | 88803809 | 88801563 | 88801632 | 88844356 | 88844521 | 170479 | 6,2,13,3,0,5,1,9,0,0,2,6 | 28,6,55,37,16,30,16,25,9,5 | 6,7,0,1,0,2,0,2,3,0,5,0,0,0, | 27,21,20,9,36,16,5,16,24,2 | 261 | 149 | 0.00213867622853 | 0.190367834302209 | 0.109,0.16,0.119,0 | 0.113,0.16,0.0,0.0 | 0,012 |
| 215683 | ENSG00000162645 | GBP2 | chr1 | - | 89108028 | 89108291 | 89106131 | 89107052 | 89109676 | 89109870 | 215683 | 169,145,193,245,66,93,10 | 9,7,6,4,7,7,3,7,4,8,0,6 | 79,126,68,29,48,42,50,156 | 11,9,9,6,5,1,5,15,1,6,14,2, | 298 | 149 | 0.0021421576208 | 0.19059889494932 | 0.904,0.912,0.941, | 0.782,0.875,0.791, | 0,053 |
| 1919 | ENSG00000093000 | NUP50 | chr22 | + | 45175893 | 45176080 | 45171599 | 45171683 | 45181285 | 45181367 | 1919 | 27,11,35,23,33,15,8,25,22, | 2,5,5,4,0,1,1,1,0,0,0,0 | 25,10,9,5,7,21,8,18,3,11,1 | 3,5,6,2,5,0,4,0,3,2,0,0,3,3, | 298 | 149 | 0.00214342264836 | 0.190632644640153 | 0.871,0.524,0.778, | 0.806,0.5,0.429,0. | 0,186 |
| 154719 | ENSG00000129566 | TEP1 | chr14 | - | 20389472 | 20389740 | 20389237 | 20389297 | 20390680 | 20390758 | 154719 | 36,37,32,31,34,42,22,18,2 | 1,0,1,2,1,3,3,0,0,2,0,1 | 49,46,24,28,41,26,24,25,2 | 0,3,2,5,4,2,4,0,0,0,4,4,6,0, | 298 | 149 | 0.00214530128891 | 0.190720917684175 | 0.947,1.0,0.941,0. | 1.0,0.885,0.857,0. | 0,045 |
| 19821 | ENSG00000185787 | MORF4L1 | chr15 | + | 78884966 | 78885083 | 78880511 | 78880579 | 78886140 | 78886227 | 19821 | 43,26,51,17,3,11,9,20,12,1 | 189,174,103,76,27,45,22,4 | 24,14,17,39,25,4,9,12,12,1 | 42,58,84,63,46,33,24,43,3 | 266 | 149 | 0.00214747316291 | 0.190835176022412 | 0.113,0.077,0.217, | 0.242,0.119,0.102, | -0,045 |
| 136927 | ENSG00000079805 | DNM2 | chr19 | + | 10797431 | 10797518 | 10795371 | 10795439 | 10798485 | 10798572 | 136927 | 28,23,10,9,12,19,15,22,30, | 1,3,6,1,0,3,2,1,2,0,2,1 | 19,13,14,22,14,2,23,23,3,1 | 0,2,0,0,1,0,2,2,0,0,1,2,4,0, | 236 | 149 | 0.00214929825216 | 0.190904836469016 | 0.946,0.829,0.513, | 1.0,0.804,1.0,1.0,0 | -0,128 |
| 92596 | ENSG00000211450 | SELENOH | chr11 | + | 57742116 | 57742227 | 57741808 | 57741954 | 57742862 | 57743038 | 92596 | 26,20,66,65,24,19,25,14,3 | 0,3,5,3,4,0,0,3,0,2,0,0 | 44,38,26,9,21,18,9,25,9,27 | 3,7,8,6,3,2,0,7,4,3,4,3,0,0, | 260 | 149 | 0.00215003100669 | 0.190904836469016 | 1.0,0.793,0.883,0. | 0.894,0.757,0.651, | 0,11 |
| 169508 | ENSG00000174227 | PIGG | chr4 | + | 525170 | 525575 | 523831 | 523913 | 527038 | 527540 | 169508 | 7,1,4,2,2,4,0,2,2,2,1,0 | 5,3,9,5,9,2,6,0,11,1,7,10 | 0,1,0,4,0,3,4,0,0,0,2,2,0,0, | 3,8,1,0,1,5,0,3,2,9,10,10,1, | 298 | 149 | 0.00215650328416 | 0.191365358412457 | 0.412,0.143,0.182, | 0.0,0.059,0.0,1.0,0 | 0,108 |
| 206489 | ENSG00000125962 | ARMCX5 | chrX | + | 102600919 | 102601044 | 102599514 | 102599847 | 102601463 | 102601509 | 206489 | 3,0,3,1,2,4,0,4,4,2,0,0 | 17,6,23,14,14,8,11,13,6,3, | 2,3,0,0,1,2,0,0,1,5,4,5,0,0, | 21,12,11,1,14,4,9,10,2,4,1 | 274 | 149 | 0.00215699578364 | 0.191365358412457 | 0.088,0.0,0.066,0. | 0.049,0.12,0.0,0.0, | 0,002 |
| 134073 | ENSG00000100055 | CYTH4 | chr22 | + | 37300069 | 37300188 | 37299225 | 37299306 | 37300906 | 37301019 | 134073 | 9,0,6,5,6,1,4,0,0,1,0,0 | 57,19,54,47,65,49,45,32,6 | 2,0,2,0,2,1,3,0,0,0,3,1,2,3, | 55,33,29,14,12,25,24,56,1 | 268 | 149 | 0.00216263663711 | 0.19178675156235 | 0.081,0.0,0.058,0. | 0.02,0.0,0.037,0.0, | 0,005 |
| 239667 | ENSG00000108669 | CYTH1 | chr17 | - | 78702537 | 78702603 | 78702121 | 78702240 | 78708196 | 78708261 | 239667 | 276,178,159,159,66,58,83, | 106,52,56,20,16,44,37,56, | 134,102,114,72,76,51,16,5 | 73,103,31,161,65,21,17,35 | 215 | 149 | 0.00216688914286 | 0.192084726979778 | 0.643,0.703,0.663, | 0.56,0.407,0.718,0 | 0,121 |
| 27738 | ENSG00000127603 | MACF1 | chr1 | + | 39480919 | 39481030 | 39479797 | 39480009 | 39484600 | 39484730 | 27738 | 14,8,6,21,4,11,8,5,12,2,4,1 | 17,8,21,27,25,10,10,8,18,1 | 9,5,9,1,1,5,5,3,0,10,0,26,8, | 28,30,19,10,9,9,8,7,16,10, | 260 | 149 | 0.00216886236528 | 0.192180492277307 | 0.321,0.364,0.141, | 0.156,0.087,0.214, | 0,109 |
| 18191 | ENSG00000128928 | IVD | chr15 | + | 40414888 | 40414982 | 40412990 | 40413087 | 40415400 | 40415482 | 18191 | 8,5,38,28,20,29,9,15,9,15, | 2,2,4,2,0,6,1,6,3,0,0,2 | 25,14,9,4,6,14,3,7,7,13,7,4 | 3,4,3,4,4,1,2,10,1,3,1,1,6,4 | 243 | 149 | 0.00217040411392 | 0.192237962075356 | 0.71,0.605,0.853,0 | 0.836,0.682,0.648, | 0,167 |
| 173980 | ENSG00000196455 | PIK3R4 | chr3 | - | 130735868 | 130735932 | 130733547 | 130734130 | 130744485 | 130745264 | 173980 | 0,2,13,10,3,4,3,2,3,2,1,2 | 1,2,3,0,2,0,4,0,0,0,1,0 | 9,11,3,2,7,3,3,4,2,4,3,4,5,1 | 0,0,2,3,4,2,4,1,2,0,2,0,2,0, | 213 | 149 | 0.00218358616742 | 0.193325970547089 | 0.0,0.412,0.752,1. | 1.0,1.0,0.512,0.31 | 0,008 |
| 215678 | ENSG00000162645 | GBP2 | chr1 | - | 89108010 | 89108291 | 89106131 | 89107052 | 89109676 | 89109870 | 215678 | 172,145,196,245,66,94,10 | 9,7,6,4,7,7,3,7,4,8,0,6 | 79,127,68,31,48,43,50,161 | 11,9,9,6,5,1,5,15,1,6,14,2, | 298 | 149 | 0.00219034025689 | 0.193844212101436 | 0.905,0.912,0.942, | 0.782,0.876,0.791, | 0,052 |
| 129649 | ENSG00000102225 | CDK16 | chrX | + | 47226278 | 47226402 | 47225964 | 47226027 | 47226582 | 47226703 | 129649 | 14,18,20,13,14,9,15,12,18, | 0,1,10,0,0,4,2,0,0,0,0,3 | 18,18,14,13,21,6,7,7,10,26 | 4,4,4,9,4,0,6,8,3,2,0,5,1,3, | 273 | 149 | 0.00219682959066 | 0.194338606505278 | 1.0,0.908,0.522,1. | 0.711,0.711,0.656, | 0,172 |
| 205250 | ENSG00000116984 | MTR | chr1 | + | 236827033 | 236827274 | 236826828 | 236826896 | 236829188 | 236829268 | 205250 | 6,0,2,2,2,6,0,6,0,0,0,2 | 22,8,28,31,25,35,30,20,11, | 2,2,2,0,0,0,0,2,3,1,0,0,0,0, | 35,57,27,15,29,11,13,31,7, | 298 | 149 | 0.00219779648004 | 0.19434426220283 | 0.12,0.0,0.034,0.0 | 0.028,0.017,0.036, | 0,012 |
| 22986 | ENSG00000129566 | TEP1 | chr14 | - | 20389609 | 20389740 | 20389237 | 20389297 | 20390680 | 20390758 | 22986 | 70,75,70,62,70,75,39,40,4 | 1,0,1,2,1,3,3,0,0,2,0,1 | 93,109,46,53,85,60,30,54, | 0,3,2,5,4,2,4,0,0,0,4,4,6,0, | 280 | 149 | 0.00220276757742 | 0.194703847415065 | 0.974,1.0,0.974,0. | 1.0,0.951,0.924,0. | 0,023 |
| 221542 | ENSG00000231925 | TAPBP | chr6 | - | 33303954 | 33304208 | 33299693 | 33301771 | 33304296 | 33305387 | 221542 | 12,8,48,15,19,8,9,14,3,5,3, | 0,0,8,3,0,0,0,2,0,0,0,2 | 21,20,6,10,9,12,5,7,14,7,5, | 0,2,0,1,4,4,2,5,0,1,0,14,3,0 | 298 | 149 | 0.0022087946339 | 0.195156435898576 | 1.0,1.0,0.75,0.714, | 1.0,0.833,1.0,0.83 | 0,169 |
| 59987 | ENSG00000118308 | IRAG2 | chr12 | + | 25069349 | 25069431 | 25066364 | 25066512 | 25079243 | 25079290 | 59987 | 665,311,991,845,369,326, | 3,7,22,7,3,11,12,15,4,4,0,0 | 443,465,412,125,276,185, | 8,8,3,2,7,2,2,5,0,0,2,0,6,7, | 231 | 149 | 0.00221303931868 | 0.195451237422575 | 0.993,0.966,0.967, | 0.973,0.974,0.989, | -0,009 |
| 131990 | ENSG00000089154 | GCN1 | chr12 | - | 120144638 | 120144835 | 120144305 | 120144448 | 120144922 | 120145061 | 131990 | 82,53,96,87,55,66,54,73,6 | 5,0,7,6,0,8,2,1,0,2,0,0 | 57,50,50,50,31,33,33,52,2 | 0,10,4,12,7,2,4,5,0,2,4,2,4, | 298 | 149 | 0.00221739596934 | 0.195755681655873 | 0.891,1.0,0.873,0. | 1.0,0.714,0.862,0. | 0,073 |
| 215685 | ENSG00000162645 | GBP2 | chr1 | - | 89108188 | 89108291 | 89106131 | 89107052 | 89109676 | 89109870 | 215685 | 125,108,170,206,45,60,72, | 9,7,6,4,7,7,3,7,4,8,0,6 | 56,89,46,14,33,31,18,97,4 | 11,9,9,6,5,1,5,15,1,6,14,2, | 252 | 149 | 0.00221851604594 | 0.19577426284695 | 0.891,0.901,0.944, | 0.751,0.854,0.751, | 0,069 |
| 255214 | ENSG00000089154 | GCN1 | chr12 | - | 120144589 | 120144835 | 120144305 | 120144448 | 120144922 | 120145061 | 255214 | 53,33,51,50,27,42,24,24,3 | 5,0,7,6,0,8,2,1,0,2,0,0 | 23,26,21,35,22,13,17,24,1 | 0,10,4,12,7,2,4,5,0,2,4,2,4, | 298 | 149 | 0.00222059271976 | 0.195807561919213 | 0.841,1.0,0.785,0. | 1.0,0.565,0.724,0. | 0,124 |
| 226654 | ENSG00000058091 | CDK14 | chr7 | + | 90908317 | 90908490 | 90899290 | 90899353 | 90917600 | 90917724 | 226654 | 202,78,122,85,64,49,62,13 | 730,461,657,466,203,337, | 125,180,46,81,128,46,38,5 | 567,453,220,231,330,261,1 | 298 | 149 | 0.00222071290216 | 0.195807561919213 | 0.122,0.078,0.085, | 0.099,0.166,0.095, | -0,014 |
| 22325 | ENSG00000092108 | SCFD1 | chr14 | + | 30628208 | 30628279 | 30622290 | 30622399 | 30630476 | 30630565 | 22325 | 21,11,72,39,22,46,37,39,1 | 0,0,2,2,4,0,3,6,2,0,0,1 | 27,32,15,22,33,10,13,21,4 | 6,2,1,5,5,1,0,2,0,2,0,0,0,5, | 220 | 149 | 0.00222285660804 | 0.19591631883909 | 1.0,1.0,0.961,0.93, | 0.753,0.916,0.91,0 | 0,044 |
| 168241 | ENSG00000138639 | ARHGAP24 | chr4 | + | 85653063 | 85653122 | 85570592 | 85570721 | 85721884 | 85722386 | 168241 | 16,9,20,3,8,22,13,20,5,3,0, | 198,149,408,178,125,188, | 12,54,6,26,26,3,8,11,21,23 | 335,282,171,211,196,103,1 | 208 | 149 | 0.00222379971766 | 0.195919212866017 | 0.055,0.041,0.034, | 0.025,0.121,0.025, | -0,017 |
| 84677 | ENSG00000152689 | RASGRP3 | chr2 | + | 33558671 | 33559030 | 33558210 | 33558336 | 33562728 | 33564605 | 84677 | 339,178,621,221,201,242, | 1,0,13,8,2,12,4,9,4,1,0,0 | 287,321,180,64,112,49,86, | 7,5,10,2,3,6,3,14,3,2,5,4,2, | 298 | 149 | 0.00222508343384 | 0.195952100060891 | 0.994,1.0,0.96,0.9 | 0.953,0.97,0.9,0.9 | 0,039 |
| 126156 | ENSG00000099290 | WASHC2A | chr10 | + | 50080757 | 50080931 | 50078674 | 50078737 | 50084071 | 50084165 | 126156 | 28,13,38,41,26,30,32,21,3 | 0,0,1,0,0,0,0,0,0,0,0,0 | 28,16,12,14,18,13,9,20,20, | 0,1,4,4,2,0,0,1,0,1,3,0,0,2, | 298 | 149 | 0.00222963957432 | 0.196273028720028 | 1.0,1.0,0.95,1.0,1. | 1.0,0.889,0.6,0.63 | 0,123 |
| 45577 | ENSG00000100578 | KIAA0586 | chr14 | + | 58540070 | 58540136 | 58512521 | 58512627 | 58547780 | 58548036 | 45577 | 6,5,29,16,6,9,5,8,6,4,3,14 | 4,1,3,1,2,3,0,1,1,0,2,0 | 10,11,8,6,11,9,16,12,9,15, | 3,1,0,0,1,5,0,1,0,4,2,0,3,1, | 215 | 149 | 0.0022316324014 | 0.196368141204302 | 0.51,0.776,0.87,0. | 0.698,0.884,1.0,1. | -0,048 |
| 58899 | ENSG00000068097 | HEATR6 | chr17 | - | 60073745 | 60074159 | 60073163 | 60073279 | 60076129 | 60076185 | 58899 | 33,12,76,55,33,31,28,25,2 | 0,1,0,7,2,0,4,2,0,0,1,2 | 34,18,39,56,14,11,15,19,1 | 2,2,1,1,0,1,1,0,0,1,8,6,1,4, | 298 | 149 | 0.00223435635206 | 0.196468487105803 | 1.0,0.857,1.0,0.79 | 0.895,0.818,0.951, | 0,046 |
| 137642 | ENSG00000104852 | SNRNP70 | chr19 | + | 49098814 | 49098963 | 49098641 | 49098704 | 49101389 | 49101668 | 137642 | 28,7,26,13,12,9,11,18,10,5 | 480,291,503,287,327,340, | 10,13,7,1,5,9,7,10,2,20,16, | 361,387,238,132,296,173,1 | 298 | 149 | 0.00223459843812 | 0.196468487105803 | 0.028,0.012,0.025, | 0.014,0.017,0.014, | 0,008 |
| 194356 | ENSG00000060138 | YBX3 | chr12 | - | 10709774 | 10710114 | 10704059 | 10704148 | 10713210 | 10713333 | 194356 | 137,70,192,103,64,117,88, | 219,157,296,162,92,240,2 | 162,111,72,55,33,52,72,19 | 186,182,139,92,119,51,14 | 298 | 149 | 0.00223725695814 | 0.196621907863385 | 0.238,0.182,0.245, | 0.303,0.234,0.206, | -0,037 |
| 93377 | ENSG00000197323 | TRIM33 | chr1 | - | 114430797 | 114430912 | 114427747 | 114427894 | 114433616 | 114433733 | 93377 | 178,127,174,98,81,80,75,7 | 7,5,8,1,1,6,2,2,0,1,0,0 | 107,150,106,132,99,76,29, | 4,5,6,5,15,4,3,3,2,4,2,5,4,4 | 264 | 149 | 0.00223958236874 | 0.19674593992093 | 0.935,0.935,0.925, | 0.938,0.944,0.909, | 0,043 |
| 156840 | ENSG00000134884 | ARGLU1 | chr13 | - | 106557491 | 106557616 | 106557105 | 106557131 | 106559431 | 106559657 | 156840 | 37,9,20,17,7,10,4,15,16,15 | 130,49,133,112,66,62,60,6 | 14,12,16,2,2,6,5,18,1,10,8, | 104,54,77,34,46,41,18,79, | 274 | 149 | 0.00224573850327 | 0.197206260219219 | 0.134,0.091,0.076, | 0.068,0.108,0.102, | 0,036 |
| 141504 | ENSG00000182446 | NPLOC4 | chr17 | - | 81563916 | 81563979 | 81559395 | 81559416 | 81565504 | 81565607 | 141504 | 0,0,9,6,4,3,5,1,1,7,3,0 | 13,12,62,57,16,6,53,34,46, | 2,5,0,0,0,0,2,1,1,1,0,0,3,1, | 14,24,14,6,7,12,10,18,8,17 | 212 | 149 | 0.00225004331134 | 0.19750369981363 | 0.0,0.0,0.093,0.06 | 0.091,0.128,0.0,0. | 0,031 |
| 15800 | ENSG00000188559 | RALGAPA2 | chr20 | - | 20611314 | 20611426 | 20605174 | 20605412 | 20616042 | 20616191 | 15800 | 41,26,57,39,24,16,21,29,1 | 0,2,3,4,0,2,4,0,1,3,0,0 | 36,40,26,37,14,19,2,22,21, | 1,0,2,1,3,0,2,0,3,0,1,1,0,2, | 261 | 149 | 0.00225417508277 | 0.197785714325181 | 1.0,0.881,0.916,0. | 0.954,1.0,0.881,0. | 0,018 |
| 156868 | ENSG00000134884 | ARGLU1 | chr13 | - | 106557558 | 106557759 | 106557047 | 106557131 | 106559431 | 106559657 | 156868 | 58,5,64,40,18,12,21,18,23, | 130,49,133,112,66,62,60,6 | 16,18,28,11,8,14,1,29,12,1 | 104,54,77,34,46,41,18,79, | 298 | 149 | 0.00226067906762 | 0.198275556806406 | 0.182,0.049,0.194, | 0.071,0.143,0.154, | 0,035 |
| 253317 | ENSG00000136021 | SCYL2 | chr12 | + | 100296761 | 100296871 | 100291502 | 100291660 | 100298030 | 100298175 | 253317 | 8,5,4,0,3,7,5,2,0,0,3,5 | 35,24,62,54,19,15,27,34,1 | 13,15,6,8,5,1,10,3,3,4,7,1, | 54,34,28,29,18,16,21,31,1 | 259 | 149 | 0.00226258614305 | 0.19833004193219 | 0.116,0.107,0.036, | 0.122,0.202,0.11,0 | -0,031 |
| 86059 | ENSG00000138380 | CARF | chr2 | + | 202974333 | 202974496 | 202971504 | 202971738 | 202977268 | 202977332 | 86059 | 43,37,70,52,36,25,33,49,3 | 1,0,4,6,0,4,4,5,3,1,4,0 | 40,26,43,13,22,17,15,34,2 | 5,0,0,2,3,0,0,3,1,5,1,1,0,1, | 298 | 149 | 0.00226314324138 | 0.19833004193219 | 0.956,1.0,0.897,0. | 0.8,1.0,1.0,0.765,0 | -0,025 |
| 49812 | ENSG00000138709 | LARP1B | chr4 | + | 128091000 | 128091144 | 128082164 | 128082305 | 128098185 | 128098330 | 49812 | 37,21,35,21,10,15,13,8,13, | 2,0,3,1,3,3,1,0,2,0,0,2 | 9,17,11,10,37,12,8,4,7,21,1 | 2,0,0,1,2,1,2,0,0,0,0,0,0,0, | 293 | 149 | 0.00226470446655 | 0.198386083451373 | 0.904,1.0,0.856,0. | 0.696,1.0,1.0,0.83 | -0,075 |
| 233028 | ENSG00000005889 | ZFX | chrX | + | 24208899 | 24209099 | 24208217 | 24208370 | 24210192 | 24216255 | 233028 | 31,22,91,46,21,14,24,20,6, | 4,3,8,3,0,2,4,0,5,2,0,0 | 55,46,22,10,39,24,13,12,2 | 9,4,2,0,0,2,1,0,2,1,0,2,1,0, | 298 | 149 | 0.00226575489432 | 0.198397352180386 | 0.795,0.786,0.85,0 | 0.753,0.852,0.846, | -0,085 |
| 217225 | ENSG00000152056 | AP1S3 | chr2 | - | 223775900 | 223775977 | 223765176 | 223765350 | 223777690 | 223777869 | 217225 | 4,10,28,21,3,8,1,7,6,4,5,7 | 5,4,7,9,1,2,3,2,4,0,2,1 | 21,30,2,5,11,4,8,9,7,10,6,1 | 1,2,5,5,2,1,0,1,0,1,1,0,2,0, | 226 | 149 | 0.00226961281546 | 0.198654345621908 | 0.345,0.622,0.725, | 0.933,0.908,0.209, | -0,164 |
| 59335 | ENSG00000082068 | WDR70 | chr5 | + | 37396374 | 37396570 | 37391999 | 37392120 | 37437921 | 37437981 | 59335 | 59,25,122,82,67,87,47,70, | 3,2,1,2,1,1,5,0,0,0,0,1 | 60,44,49,48,47,34,35,71,4 | 6,6,0,1,1,0,0,3,5,3,3,6,2,0, | 298 | 149 | 0.00227328331709 | 0.198894732366097 | 0.908,0.862,0.984, | 0.833,0.786,1.0,0. | 0,031 |
| 105010 | ENSG00000069509 | FUNDC1 | chrX | - | 44527236 | 44527365 | 44523638 | 44524275 | 44538466 | 44538542 | 105010 | 34,23,72,71,21,26,9,7,24,5 | 0,0,17,2,1,0,1,1,0,0,0,0 | 36,37,29,8,18,10,10,15,21, | 1,0,0,0,0,1,0,0,0,0,0,0,0,0, | 278 | 149 | 0.00228171315262 | 0.199551159508962 | 1.0,1.0,0.694,0.95, | 0.951,1.0,1.0,1.0,1 | -0,035 |
| 249548 | ENSG00000155629 | PIK3AP1 | chr10 | - | 96669530 | 96669752 | 96656797 | 96656934 | 96709566 | 96709983 | 249548 | 10,1,9,14,4,0,1,1,2,8,0,3 | 161,53,180,152,58,73,41,4 | 2,2,0,0,5,1,2,4,3,4,0,5,0,2, | 101,67,64,12,42,45,33,69, | 298 | 149 | 0.00228784811025 | 0.199930417560849 | 0.03,0.009,0.024,0 | 0.01,0.015,0.0,0.0, | 0,007 |
| 143415 | ENSG00000126858 | RHOT1 | chr17 | + | 32191271 | 32191330 | 32183170 | 32183272 | 32192200 | 32192299 | 143415 | 2,0,3,1,0,0,0,1,0,0,0,0 | 13,4,22,26,12,12,8,10,2,2, | 1,3,6,1,2,0,1,1,0,0,3,3,0,1, | 10,11,14,10,7,9,3,17,7,21,1 | 208 | 149 | 0.00228790749684 | 0.199930417560849 | 0.099,0.0,0.089,0. | 0.067,0.163,0.235, | -0,079 |
| 151195 | ENSG00000174485 | DENND4A | chr15 | - | 65732751 | 65732772 | 65731641 | 65731700 | 65737706 | 65737945 | 151195 | 6,12,27,18,1,8,6,8,3,4,1,6 | 0,0,3,0,0,3,4,2,0,0,0,0 | 16,11,7,9,8,2,4,8,12,4,11,1 | 2,2,0,2,0,1,1,8,1,0,4,0,0,0, | 170 | 149 | 0.0022947173463 | 0.20044411897788 | 1.0,1.0,0.887,1.0,1 | 0.875,0.828,1.0,0. | 0,05 |
| 245949 | ENSG00000172977 | KAT5 | chr11 | + | 65717715 | 65717810 | 65716888 | 65716982 | 65718589 | 65718749 | 245949 | 5,4,3,6,0,3,4,0,0,1,0 | 25,32,67,63,35,36,21,22,2 | 0,1,1,1,2,0,1,0,0,0,0,1,1,1, | 31,36,26,17,50,21,22,28,2 | 244 | 149 | 0.00229879604198 | 0.200718933432616 | 0.109,0.071,0.027, | 0.0,0.017,0.023,0. | 0,025 |
| 78617 | ENSG00000136715 | SAP130 | chr2 | - | 127977984 | 127978089 | 127954985 | 127955344 | 127986784 | 127986959 | 78617 | 17,17,27,13,6,13,8,9,20,24 | 10,5,1,0,0,1,0,0,6,2,4,2 | 5,7,17,32,9,4,9,8,2,10,7,10 | 1,0,0,5,1,1,1,1,0,2,1,2,0,0, | 254 | 149 | 0.00231071443971 | 0.201677769494413 | 0.499,0.666,0.941, | 0.746,1.0,1.0,0.79, | -0,055 |
| 193122 | ENSG00000196591 | HDAC2 | chr6 | - | 113953276 | 113953386 | 113946007 | 113949260 | 113956012 | 113956602 | 193122 | 12,16,34,22,12,7,3,8,7,7,2, | 6,0,3,1,1,2,3,1,0,0,0,2 | 28,16,10,10,11,22,5,9,3,17 | 2,1,0,3,0,0,1,1,1,1,0,2,0,0, | 259 | 149 | 0.00231661003522 | 0.202110374742779 | 0.535,1.0,0.867,0. | 0.89,0.902,1.0,0.6 | -0,053 |
| 21506 | ENSG00000151332 | MBIP | chr14 | - | 36308091 | 36308189 | 36300784 | 36300823 | 36311572 | 36311590 | 21506 | 23,30,50,37,18,49,12,21,3 | 0,0,0,3,1,4,2,4,4,0,0,1 | 23,55,29,19,21,8,20,26,12, | 0,2,2,0,1,4,0,4,0,2,3,7,4,1, | 247 | 149 | 0.00232621056736 | 0.20286573201923 | 1.0,1.0,1.0,0.882,0 | 1.0,0.943,0.897,1. | 0,063 |
| 107408 | ENSG00000143569 | UBAP2L | chr1 | + | 154234590 | 154234759 | 154228614 | 154228725 | 154235195 | 154235291 | 107408 | 225,130,150,94,71,66,35,6 | 62,60,23,17,11,26,26,18,2 | 79,54,75,122,48,55,43,87, | 31,45,35,67,54,10,20,33,2 | 298 | 149 | 0.00233214616019 | 0.203300992387142 | 0.645,0.52,0.765,0 | 0.56,0.375,0.517,0 | 0,113 |
| 147542 | ENSG00000172775 | PSME3IP1 | chr16 | - | 57178755 | 57178852 | 57173780 | 57173869 | 57185820 | 57186031 | 147542 | 0,0,3,0,0,2,3,0,0,0,0,4 | 12,3,19,12,3,4,9,4,6,7,5,7 | 3,0,0,0,1,1,1,4,0,0,3,2,4,4, | 10,3,7,2,0,3,0,2,2,2,3,11,5, | 246 | 149 | 0.00233966583279 | 0.203873933950293 | 0.0,0.0,0.087,0.0,0 | 0.154,0.0,0.0,0.0,0.1 | -0,224 |
| 167065 | ENSG00000174652 | ZNF266 | chr19 | - | 9418761 | 9419167 | 9418504 | 9418631 | 9420064 | 9420218 | 167065 | 18,9,32,47,17,7,16,15,25,1 | 4,0,3,4,0,0,3,2,0,0,2,5 | 38,21,10,21,15,7,11,24,19, | 1,3,0,2,3,0,0,0,0,0,6,0,0,0, | 298 | 149 | 0.00234729361685 | 0.204455828590952 | 0.692,1.0,0.842,0. | 0.95,0.778,1.0,0.8 | -0,12 |
| 133301 | ENSG00000182670 | TTC3 | chr21 | + | 37099623 | 37099748 | 37096580 | 37096643 | 37108391 | 37108446 | 133301 | 8,0,4,4,0,5,0,2,3,0,0,1 | 37,9,46,45,32,26,34,29,23, | 2,1,0,0,0,0,0,0,0,0,1,0,1,0, | 58,32,37,30,40,21,32,61,1 | 274 | 149 | 0.00234952993067 | 0.204567830302603 | 0.105,0.0,0.045,0. | 0.018,0.017,0.0,0. | 0,032 |
| 218682 | ENSG00000024048 | UBR2 | chr6 | + | 42662512 | 42662578 | 42662183 | 42662277 | 42663257 | 42663419 | 218682 | 6,0,4,1,1,1,2,1,0,0,2,0 | 10,15,26,23,22,9,15,7,4,2, | 3,0,0,1,0,2,5,2,3,1,3,0,3,0, | 11,41,11,8,9,9,6,15,6,13,4, | 215 | 149 | 0.00235252943175 | 0.204746163414874 | 0.294,0.0,0.096,0. | 0.159,0.0,0.0,0.08, | -0,041 |
| 107194 | ENSG00000134982 | APC | chr5 | + | 112834950 | 112835165 | 112828855 | 112828972 | 112837552 | 112837722 | 107194 | 45,25,61,30,25,26,18,26,2 | 2,0,5,2,3,3,1,3,0,0,0,0 | 44,35,13,47,10,18,23,34,6, | 1,0,2,0,0,1,1,1,0,1,3,0,1,3, | 298 | 149 | 0.00235395884509 | 0.204787759170398 | 0.918,1.0,0.859,0. | 0.957,1.0,0.765,1. | -0,022 |
| 10041 | ENSG00000141349 | G6PC3 | chr17 | + | 44074968 | 44075087 | 44074679 | 44074770 | 44075309 | 44075451 | 10041 | 14,9,27,30,26,20,10,17,18, | 2,0,0,3,0,0,3,1,1,0,0,4 | 25,9,15,4,12,4,19,36,4,7,9, | 5,4,1,2,0,0,0,5,3,1,6,0,0,0, | 268 | 149 | 0.002370822443 | 0.206171509183569 | 0.796,1.0,1.0,0.84 | 0.735,0.556,0.893, | 0,066 |

| | | | | | | | | | | | | | | | | | | | | | | |
|---|---|---|---|---|---|---|---|---|---|---|---|---|---|---|---|---|---|---|---|---|---|---|
| 24292 | ENSG00000196199 | MPHOSPH8 | chr13 | + | 19670839 | 19670964 | 19670235 | 19670363 | 19671205 | 19671289 | 24292 | 18,6,22,14,3,5,6,7,30,7,3,8 | 15,11,57,30,25,25,26,21,18 | 9,4,5,7,3,1,5,4,3,13,4,9,0,1 | 30,29,10,28,11,17,17,23,3, | 274 | 149 | 0.0023759124013 | 0.206530695494427 | 0.395,0.229,0.173, | 0.14,0.07,0.214,0.1 | 0,085 |
| 8800 | ENSG00000154889 | MPPE1 | chr18 | - | 11886712 | 11886781 | 11886114 | 11886621 | 11886916 | 11887025 | 8800 | 7,6,7,14,5,7,3,5,5,2,1,6 | 2,1,1,7,1,1,0,4,2,0,0,0 | 8,8,4,2,6,6,5,8,2,11,2,7,16, | 5,4,0,0,1,0,0,1,1,0,0,0,3,0,0 | 218 | 149 | 0.00238090653892 | 0.206881265756274 | 0.705,0.804,0.827, | 0.522,0.578,1.0,1.0 | -0,044 |
| 233578 | ENSG00000143569 | UBAP2L | chr1 | + | 154263102 | 154263436 | 154261591 | 154261697 | 154266500 | 154266568 | 233578 | 31,21,29,22,8,13,12,27,14, | 17,15,15,19,19,24,13,22,40 | 22,29,20,30,24,10,24,20,10 | 18,0,30,15,9,9,11,15,5,13,1 | 298 | 149 | 0.00238577740172 | 0.207196358931863 | 0.477,0.412,0.492, | 0.379,1.0,0.25,0.5, | -0,171 |
| 226489 | ENSG00000171681 | ATF7IP | chr12 | + | 14436362 | 14436510 | 14436194 | 14436251 | 14438129 | 14438267 | 226489 | 15,4,21,20,3,6,1,16,9,6,5,5 | 379,252,496,271,86,125,12 | 8,5,31,26,10,6,10,2,11,19,4 | 197,173,324,254,158,171,1 | 297 | 149 | 0.00238645814865 | 0.207196358931863 | 0.019,0.008,0.021, | 0.02,0.014,0.046,0 | -0,003 |
| 247325 | ENSG00000115084 | SLC35F5 | chr2 | - | 113739961 | 113740032 | 113735776 | 113735858 | 113742691 | 113742879 | 247325 | 5,0,5,4,0,2,0,9,2,1,3,2 | 28,13,38,23,28,23,6,11,20, | 7,6,5,2,1,1,4,8,4,2,5,2,1,2,2 | 10,11,10,6,7,15,9,11,5,16,1 | 220 | 149 | 0.00238812201204 | 0.207257213215073 | 0.108,0.0,0.082,0.1 | 0.322,0.27,0.253,0 | -0,089 |
| 58469 | ENSG00000015532 | XYLT2 | chr17 | + | 50354853 | 50355056 | 50354407 | 50354583 | 50355500 | 50355581 | 58469 | 44,14,22,17,16,21,14,11,22 | 2,3,0,3,4,0,1,1,0,1,1,2 | 15,2,11,8,1,7,3,16,2,18,4,2 | 2,2,0,1,1,0,0,0,0,1,4,1,5,1,1 | 298 | 149 | 0.00239444142361 | 0.207694110205672 | 0.917,0.7,1.0,0.739 | 0.789,0.333,1.0,0.8 | 0,057 |
| 3886 | ENSG00000079805 | DNM2 | chr19 | + | 10797379 | 10797518 | 10795371 | 10795439 | 10798485 | 10798572 | 3886 | 34,31,40,29,28,36,24,42,38 | 1,3,6,1,0,3,2,1,2,0,2,1 | 31,35,24,42,33,8,31,36,5,1 | 0,2,0,0,1,0,2,2,0,0,1,2,4,0,1 | 288 | 149 | 0.00239562122211 | 0.207694110205672 | 0.946,0.842,0.775, | 1.0,0.901,1.0,1.0,0 | -0,053 |
| 173771 | ENSG00000156976 | EIF4A2 | chr3 | + | 186788309 | 186788412 | 186787802 | 186787882 | 186789124 | 186789897 | 173771 | 244,140,187,105,70,112,10 | 427,137,428,354,181,165,1 | 179,122,168,159,61,52,37, | 138,51,178,74,110,133,75, | 252 | 149 | 0.00239605110621 | 0.207694110205672 | 0.253,0.377,0.205, | 0.434,0.586,0.358, | -0,073 |
| 252929 | ENSG00000152256 | PDK1 | chr2 | + | 172584602 | 172584637 | 172570725 | 172570824 | 172586277 | 172586388 | 252929 | 1,1,5,2,0,5,1,5,0,0,0,0 | 2,5,11,8,17,11,7,9,5,14,3,7 | 0,3,0,0,2,2,0,0,3,1,0,1,0,0,0 | 15,20,5,5,6,6,6,3,3,5,8,30,2 | 184 | 149 | 0.00240128882663 | 0.207986789807779 | 0.288,0.139,0.269, | 0.0,0.108,0.0,0.0,0 | 0,057 |
| 225753 | ENSG00000288044 | | chr14 | + | 24292237 | 24292600 | 24271200 | 24271310 | 24298613 | 24298861 | 225753 | 6,2,7,3,5,1,3,1,3,6,2,1 | 5,2,4,1,0,2,1,0,2,2,2,3 | 4,4,2,2,1,1,4,2,2,3,6,14,8,7 | 2,1,1,1,0,0,0,0,1,1,2,0,4,1,3 | 298 | 149 | 0.00240230041399 | 0.207986789807779 | 0.375,0.333,0.467, | 0.5,0.667,0.5,0.5,1 | -0,19 |
| 206845 | ENSG00000075884 | ARHGAP15 | chr2 | + | 143705551 | 143705651 | 143703418 | 143703524 | 143767988 | 143768352 | 206845 | 7,5,23,9,5,5,12,14,3,3,3,8 | 75,50,155,97,69,55,50,45,4 | 24,19,5,13,9,6,6,11,15,11,6 | 94,64,66,18,62,43,14,32,37 | 249 | 149 | 0.00240232661402 | 0.207986789807779 | 0.053,0.056,0.082, | 0.133,0.151,0.043, | -0,047 |
| 80786 | ENSG00000004455 | AK2 | chr1 | - | 33021366 | 33021461 | 33008896 | 33014594 | 33021592 | 33021703 | 80786 | 102,64,84,63,51,45,36,38,1 | 5,2,6,1,2,5,8,4,9,3,0,1 | 55,39,43,24,32,26,41,43,17 | 5,6,3,2,7,2,3,5,5,4,4,11,1,5 | 244 | 149 | 0.00240412923108 | 0.207998883644293 | 0.926,0.951,0.895, | 0.87,0.799,0.897,0 | 0,077 |
| 195661 | ENSG00000177051 | FBXO46 | chr19 | - | 45723472 | 45723885 | 45712943 | 45713573 | 45730848 | 45730896 | 195661 | 2,6,3,10,7,3,5,4,8,2,5,3 | 3,2,2,3,7,5,7,2,12,4,4,3 | 3,7,3,5,3,2,0,6,2,6,4,11,7,3 | 0,2,1,2,1,0,2,3,0,0,9,4,0,0,5 | 298 | 149 | 0.00240439909914 | 0.207998883644293 | 0.25,0.6,0.429,0.62 | 1.0,0.636,0.6,0.556 | -0,285 |
| 162095 | ENSG00000124201 | ZNFX1 | chr20 | - | 49263333 | 49263564 | 49260462 | 49260577 | 49264715 | 49264864 | 162095 | 26,27,57,41,10,8,14,19,27, | 0,2,4,1,2,1,0,1,0,0,0,5 | 29,40,16,37,15,12,7,22,9,1 | 4,4,1,4,5,0,1,1,1,1,5,1,2,1,1 | 298 | 149 | 0.00240770603132 | 0.208193339469406 | 1.0,0.871,0.877,0.9 | 0.784,0.833,0.889, | 0,092 |
| 198067 | ENSG00000112232 | KHDRBS2 | chr6 | - | 62187049 | 62187216 | 62177184 | 62177312 | 62285857 | 62286225 | 198067 | 0,0,12,4,1,2,6,0,6,0,0,2 | 30,9,123,57,46,61,30,43,21 | 1,4,0,0,0,1,0,0,0,0,0,1,0,0,2 | 69,55,31,36,28,20,18,28,31 | 298 | 149 | 0.00240864193375 | 0.208193339469406 | 0.0,0.0,0.047,0.034 | 0.007,0.035,0.0,0.0 | 0,022 |
| 34290 | ENSG00000109762 | SNX25 | chr4 | + | 185320732 | 185320864 | 185310634 | 185310816 | 185323527 | 185323800 | 34290 | 85,49,125,59,60,43,30,34,5 | 0,3,0,1,1,8,2,2,1,0,3,0 | 98,91,19,36,61,36,14,20,51 | 1,0,0,0,3,1,0,0,1,0,0,2,0,2,1 | 281 | 149 | 0.00240954885039 | 0.208193339469406 | 1.0,0.896,1.0,0.969 | 0.981,1.0,1.0,1.0,0 | -0,045 |
| 225013 | ENSG00000164808 | SPIDR | chr8 | + | 47293877 | 47294030 | 47291052 | 47291137 | 47396375 | 47396626 | 225013 | 61,17,52,37,23,19,20,25,39 | 12,4,10,6,0,4,1,2,3,4,2,4 | 32,21,25,16,24,13,8,21,23, | 0,1,0,0,8,2,0,1,6,0,2,3,3,0,2 | 298 | 149 | 0.00241370688208 | 0.208468918915313 | 0.718,0.68,0.722,0 | 1.0,0.913,1.0,1.0,0 | -0,098 |
| 173769 | ENSG00000156976 | EIF4A2 | chr3 | + | 186788309 | 186788410 | 186787802 | 186787882 | 186789124 | 186789897 | 173769 | 233,132,180,94,66,107,104 | 427,137,428,354,181,165,1 | 175,120,165,150,57,49,36, | 138,51,178,74,110,133,75, | 250 | 149 | 0.00242541311644 | 0.209395944831327 | 0.245,0.365,0.2,0.1 | 0.43,0.584,0.356,0 | -0,072 |
| 87678 | ENSG00000010244 | ZNF207 | chr17 | + | 32366664 | 32366757 | 32365329 | 32365487 | 32367771 | 32368014 | 87678 | 376,202,320,304,208,200,2 | 89,50,53,64,54,64,59,99,13 | 302,358,264,201,175,143,1 | 78,59,40,69,53,28,24,58,24 | 242 | 149 | 0.00242719451696 | 0.209465718957425 | 0.722,0.713,0.788, | 0.704,0.789,0.803, | -0,053 |
| 219442 | ENSG00000213676 | ATF6B | chr6 | - | 32125163 | 32125273 | 32121262 | 32121348 | 32126116 | 32126252 | 219442 | 2,1,2,0,1,5,1,7,0,0,1,1 | 42,36,80,57,60,13,24,44,63 | 6,7,4,13,1,2,2,1,1,1,2,1,0,2 | 49,26,22,30,18,24,24,44,27 | 259 | 149 | 0.00243299013449 | 0.209824956175236 | 0.027,0.016,0.014, | 0.066,0.134,0.095, | -0,015 |
| 124255 | ENSG00000139697 | SBNO1 | chr12 | - | 123348028 | 123348133 | 123345257 | 123345570 | 123350309 | 123350441 | 124255 | 104,65,79,51,45,41,55,84,5 | 0,0,0,2,2,0,0,0,0,2,2,2 | 85,98,53,56,65,38,29,47,33 | 1,2,3,2,1,1,0,2,1,5,0,2,5,1,2 | 254 | 149 | 0.00243330696142 | 0.209824956175236 | 1.0,1.0,1.0,0.937,0 | 0.98,0.966,0.912,0 | 0,016 |
| 3991 | ENSG00000257103 | LSM14A | chr19 | + | 34215123 | 34215300 | 34208195 | 34209051 | 34215595 | 34215661 | 3991 | 162,113,238,246,111,99,80 | 16,16,31,9,17,7,11,20,2,5,3 | 132,152,137,90,81,100,51, | 21,51,21,19,37,4,11,15,17, | 298 | 149 | 0.00243795879126 | 0.210131931762433 | 0.835,0.779,0.793, | 0.759,0.598,0.765, | 0,079 |
| 107936 | ENSG00000215021 | PHB2 | chr12 | - | 6968410 | 6968595 | 6967170 | 6968021 | 6969497 | 6969577 | 107936 | 264,208,565,460,227,169,1 | 1,2,11,12,3,5,3,2,0,2,0,2 | 298,257,269,113,204,128,1 | 1,5,0,1,0,0,1,5,2,1,5,5,0,4,0 | 298 | 149 | 0.00243881952666 | 0.210131931762433 | 0.992,0.981,0.963, | 0.993,0.963,1.0,0.9 | -0,009 |
| 191305 | ENSG00000054118 | THRAP3 | chr1 | + | 36296582 | 36296670 | 36293850 | 36293935 | 36300885 | 36301084 | 191305 | 139,75,213,192,46,84,71,6 | 4,1,8,2,8,3,4,6,1,5,3,0 | 107,156,55,50,61,43,25,50 | 1,3,1,0,1,3,0,6,0,1,4,0,1,1,4 | 237 | 149 | 0.00244189692632 | 0.210312892096351 | 0.956,0.979,0.944, | 0.985,0.97,0.972,1 | -0,031 |
| 224621 | ENSG00000128607 | KLHDC10 | chr7 | + | 130117518 | 130117834 | 130116444 | 130116666 | 130120748 | 130120903 | 224621 | 4,0,3,0,2,0,1,0,0,0,0,0 | 27,7,40,20,10,8,18,15,20,1 | 3,2,0,1,3,0,3,0,1,4,3,3,3,2,2 | 7,17,12,19,4,14,16,19,6,22 | 298 | 149 | 0.00244423239651 | 0.210429833173298 | 0.069,0.0,0.036,0.0 | 0.176,0.056,0.0,0.0 | -0,049 |
| 233321 | ENSG00000215301 | DDX3X | chrX | + | 41346930 | 41347012 | 41346504 | 41346622 | 41347314 | 41347451 | 233321 | 51,40,71,39,21,29,12,23,19 | 3,0,1,0,0,0,0,0,0,0,2,0 | 34,49,22,38,46,28,10,9,19, | 3,4,5,8,0,2,0,2,0,3,0,0,0,0,0 | 231 | 149 | 0.00244669695747 | 0.21055779002528 | 0.916,1.0,0.979,1.0 | 0.88,0.888,0.739,0 | 0,049 |
| 15749 | ENSG00000197912 | SPG7 | chr16 | + | 89555958 | 89556085 | 89554485 | 89554563 | 89556886 | 89557488 | 15749 | 12,13,6,4,1,3,0,0,2,6,6,15 | 50,36,67,45,40,34,21,52,37 | 2,0,0,1,2,1,2,2,7,11,5,4,3,5 | 36,38,30,29,34,24,24,41,35 | 276 | 149 | 0.00245612419068 | 0.211284598594823 | 0.115,0.163,0.046, | 0.029,0.0,0.0,0.018 | 0,041 |
| 39009 | ENSG00000204463 | BAG6 | chr6 | - | 31647590 | 31647826 | 31646393 | 31646523 | 31648676 | 31648751 | 39009 | 186,117,152,129,116,76,10 | 7,5,8,1,4,5,7,7,3,3,1,1 | 97,80,38,85,58,46,104,102 | 3,5,6,12,5,4,5,2,3,2,4,13,5, | 298 | 149 | 0.00246069257856 | 0.211593018128665 | 0.93,0.921,0.905,0 | 0.942,0.889,0.76,0 | 0,05 |
| 41469 | ENSG00000144468 | RHBDD1 | chr2 | + | 226838072 | 226838154 | 226837547 | 226837770 | 226839408 | 226839627 | 41469 | 5,2,12,6,0,5,4,3,2,4,1,3 | 1,2,1,5,1,2,2,1,2,0,0,0 | 7,3,0,2,2,4,1,1,4,5,3,1,1,3,3 | 0,3,1,0,0,0,0,0,1,1,1,1,1,0,3 | 231 | 149 | 0.00246619179553 | 0.211981200616501 | 0.763,0.392,0.886, | 1.0,0.392,0.0,1.0,1 | -0,09 |
| 5563 | ENSG00000127527 | EPS15L1 | chr19 | - | 16395343 | 16395467 | 16393950 | 16394001 | 16402320 | 16402485 | 5563 | 60,45,53,24,45,19,18,31,21 | 3,0,0,0,2,6,1,4,0,2,1,2 | 36,27,32,30,17,23,17,35,26 | 0,0,2,0,1,1,0,1,0,3,0,0,0,2,2 | 273 | 149 | 0.00248166980758 | 0.213226456828443 | 0.916,1.0,1.0,1.0,0 | 1.0,1.0,0.897,1.0,0 | -0,055 |
| 88900 | ENSG00000073849 | ST6GAL1 | chr3 | + | 187038394 | 187038491 | 187021855 | 187022004 | 187038741 | 187038873 | 88900 | 3,10,23,17,4,7,9,6,6,1,5,7 | 79,106,124,65,70,51,23,25 | 12,22,10,18,5,8,0,14,6,6,7, | 81,112,9,35,120,33,14,19,1 | 246 | 149 | 0.00248500650266 | 0.213427946757388 | 0.022,0.054,0.101, | 0.082,0.106,0.402, | -0,091 |
| 184474 | ENSG00000077044 | DGKD | chr2 | + | 233389848 | 233390041 | 233388256 | 233388367 | 233390402 | 233390483 | 184474 | 7,4,26,13,9,7,3,5,2,2,3,3 | 63,43,251,97,46,59,79,124 | 5,1,6,10,3,4,3,5,1,2,4,5,4,2 | 103,107,94,62,89,45,38,57 | 298 | 149 | 0.00248840479626 | 0.21363456430149 | 0.053,0.044,0.049, | 0.024,0.005,0.031, | 0,014 |
| 61335 | ENSG00000132155 | RAF1 | chr3 | - | 12606200 | 12606299 | 12604135 | 12604289 | 12608765 | 12608923 | 61335 | 126,86,185,102,112,131,91 | 45,20,36,26,10,18,20,11,20 | 120,159,109,86,70,54,100, | 5,20,20,23,18,1,6,13,3,11,2 | 248 | 149 | 0.00248948510935 | 0.213642093130187 | 0.627,0.721,0.755, | 0.935,0.827,0.766, | -0,07 |
| 81944 | ENSG00000140386 | SCAPER | chr15 | - | 76795240 | 76795440 | 76774854 | 76775117 | 76800247 | 76800267 | 81944 | 111,41,69,48,45,57,36,61,6 | 2,2,6,3,0,5,0,1,0,1,0,3 | 47,36,35,80,18,14,26,40,21 | 3,2,0,4,4,2,0,7,2,2,6,7,0,4,1 | 298 | 149 | 0.00249269465886 | 0.213832269478317 | 0.965,0.911,0.852, | 0.887,0.9,1.0,0.909 | 0,078 |
| 151006 | ENSG00000170776 | AKAP13 | chr15 | + | 85522033 | 85522110 | 85521427 | 85521575 | 85533583 | 85536997 | 151006 | 5,1,13,4,3,5,3,7,0,2,1,2 | 105,57,182,121,127,105,77 | 7,20,3,14,5,3,6,6,9,4,4,7,4, | 150,155,83,84,106,56,56,1 | 226 | 149 | 0.00249834730279 | 0.214231788178006 | 0.03,0.011,0.045,0 | 0.03,0.078,0.023,0 | -0,019 |
| 44868 | ENSG00000103064 | SLC7A6 | chr16 | + | 68287745 | 68287871 | 68274690 | 68275249 | 68291208 | 68291332 | 44868 | 3,6,30,23,17,13,21,12,9,6,5 | 1,0,0,0,0,1,4,4,0,0,0,0 | 14,34,6,2,9,10,6,16,7,20,12 | 0,1,0,3,2,4,0,2,4,3,4,1,0,0,0 | 275 | 149 | 0.00250324366432 | 0.214481829318545 | 0.619,1.0,1.0,1.0,1 | 1.0,0.949,1.0,0.265 | 0,093 |
| 212040 | ENSG00000079263 | SP140 | chr2 | + | 230247248 | 230247727 | 230245862 | 230245940 | 230247915 | 230248065 | 212040 | 26,9,7,7,0,5,7,11,8,3,4,6 | 259,138,178,134,120,71,56 | 9,19,7,18,13,5,3,4,10,4,5,2 | 139,166,121,130,140,122,3 | 298 | 149 | 0.00250325629323 | 0.214481829318545 | 0.048,0.032,0.019, | 0.031,0.054,0.028, | -0,012 |
| 3189 | ENSG00000104904 | OAZ1 | chr19 | + | 2271384 | 2271530 | 2269514 | 2269744 | 2271781 | 2271954 | 3189 | 838,560,1746,1265,680,61 | 12,17,31,8,10,4,11,9,11,4,5 | 994,605,370,113,401,396,2 | 14,23,9,8,13,1,12,9,0,4,32, | 295 | 149 | 0.00250562340493 | 0.214507417022602 | 0.972,0.943,0.966, | 0.973,0.93,0.954,0 | 0,016 |
| 233315 | ENSG00000215301 | DDX3X | chrX | + | 41346836 | 41347012 | 41346504 | 41346622 | 41347314 | 41347451 | 233315 | 123,109,137,85,42,42,29,6 | 3,0,1,0,0,0,0,0,0,0,2,0 | 66,95,64,77,83,46,17,26,42 | 3,4,5,8,0,2,0,2,0,3,0,0,0,0,0 | 298 | 149 | 0.00250623098177 | 0.214507417022602 | 0.953,1.0,0.986,1.0 | 0.917,0.922,0.865, | 0,029 |
| 128692 | ENSG00000151490 | PTPRO | chr12 | + | 15560192 | 15560276 | 15557454 | 15557523 | 15565592 | 15565628 | 128692 | 11,7,24,20,9,9,7,8,3,2,2,7 | 5,1,8,1,3,0,1,0,0,2,1,0 | 5,5,1,2,5,5,4,7,3,16,3,5,17, | 5,3,4,2,3,0,5,1,1,1,0,2,8,8,3 | 233 | 149 | 0.00250654484629 | 0.214507417022602 | 0.585,0.817,0.657, | 0.39,0.516,0.138,0 | 0,187 |
| 233310 | ENSG00000215301 | DDX3X | chrX | + | 41346822 | 41347012 | 41346504 | 41346622 | 41347314 | 41347451 | 233310 | 123,109,137,85,42,42,29,6 | 3,0,1,0,0,0,0,0,0,0,2,0 | 64,95,64,77,83,46,17,26,42 | 3,4,5,8,0,2,0,2,0,3,0,0,0,0,0 | 298 | 149 | 0.00250900461969 | 0.21463258080306 | 0.953,1.0,0.986,1.0 | 0.914,0.922,0.865, | 0,03 |
| 73262 | ENSG00000153207 | AHCTF1 | chr1 | - | 246916141 | 246916395 | 246913231 | 246913412 | 246918249 | 246918377 | 73262 | 16,1,32,16,11,24,10,14,9,6 | 2,4,0,4,0,6,3,9,1,0,2,6 | 2,20,5,16,12,6,15,16,9,3,12 | 6,8,8,9,12,3,5,11,17,3,4,3,2 | 298 | 149 | 0.00251579206898 | 0.215127708700292 | 0.8,0.111,1.0,0.667 | 0.143,0.556,0.238, | 0,207 |
| 141965 | ENSG00000108559 | NUP88 | chr17 | - | 5394890 | 5394975 | 5390353 | 5391662 | 5399551 | 5399650 | 141965 | 98,39,148,60,71,81,71,105 | 2,0,3,1,0,6,4,3,2,0,1,2 | 97,95,121,62,45,47,46,80,2 | 0,0,0,1,1,0,0,0,1,1,0,0,0,3,4 | 234 | 149 | 0.00251752885916 | 0.215157925317715 | 0.969,1.0,0.969,0.9 | 1.0,1.0,1.0,0.975,0 | -0,02 |
| 19253 | ENSG00000114062 | UBE3A | chr15 | - | 25408571 | 25408684 | 25407066 | 25407262 | 25409087 | 25409207 | 19253 | 32,15,82,49,16,10,14,15,23 | 2,1,9,8,1,0,1,1,8,0,1,1 | 31,27,16,9,5,3,12,11,5,19,1 | 0,0,4,0,0,0,1,2,0,5,2,3,1,0,3 | 262 | 149 | 0.00251814475552 | 0.215157925317715 | 0.901,0.895,0.838, | 1.0,1.0,0.695,1.0,1 | -0,028 |
| 1832 | ENSG00000100014 | SPECC1L | chr22 | + | 24411587 | 24411704 | 24369217 | 24369320 | 24412647 | 24412707 | 1832 | 40,32,63,36,40,21,23,39,30 | 0,0,1,0,1,2,0,3,0,3,1,0 | 26,27,37,13,20,17,14,30,15 | 8,2,1,0,0,0,7,1,1,0,3,1,1,0,2 | 266 | 149 | 0.0025204763043 | 0.215271680734442 | 1.0,1.0,0.972,1.0,0 | 0.645,0.883,0.954, | 0,046 |
| 179958 | ENSG00000204252 | HLA-DOA | chr6 | - | 33007311 | 33007592 | 33007079 | 33007215 | 33008012 | 33008054 | 179958 | 265,125,284,208,133,77,27 | 0,5,4,3,0,1,3,4,1,1,0,1 | 129,132,169,97,82,47,96,1 | 5,4,0,1,0,1,0,3,0,2,2,4,4,3,5 | 298 | 149 | 0.00252740624288 | 0.2157779345741 | 1.0,0.926,0.973,0.9 | 0.928,0.943,1.0,0.9 | 0,017 |
| 11614 | ENSG00000196267 | ZNF836 | chr19 | - | 52169647 | 52169781 | 52168057 | 52168152 | 52171335 | 52171621 | 11614 | 34,26,52,39,20,26,9,18,19, | 0,1,5,4,4,4,2,2,0,3,0,0 | 36,40,16,11,35,21,13,15,9, | 2,2,0,2,2,0,0,1,2,2,4,0,0,3,0 | 283 | 149 | 0.00253057161776 | 0.215962513823197 | 1.0,0.932,0.846,0.8 | 0.905,0.913,1.0,0.1 | -0,022 |
| 56014 | ENSG00000143106 | PSMA5 | chr1 | - | 109413067 | 109413135 | 109412076 | 109412184 | 109415236 | 109415363 | 56014 | 63,47,128,138,83,46,57,58 | 4,1,13,5,4,3,7,5,3,0,0,2 | 77,97,52,28,45,55,39,45,43 | 3,13,7,9,12,3,8,7,6,1,6,11,5 | 217 | 149 | 0.00253508301472 | 0.2162617726283 | 0.915,0.97,0.871,0 | 0.946,0.837,0.836, | 0,067 |
| 163049 | ENSG00000182903 | ZNF721 | chr4 | - | 472574 | 473025 | 439987 | 444432 | 473980 | 474167 | 163049 | 81,45,139,64,21,26,21,40,4 | 2,1,1,2,6,0,2,4,0,0,2,2 | 50,94,57,68,45,33,21,31,29 | 0,0,0,7,1,0,1,0,1,0,1,0,0,0,1 | 298 | 149 | 0.00253685785954 | 0.216327438180132 | 0.953,0.957,0.986, | 1.0,1.0,1.0,0.829,0 | -0,088 |
| 211229 | ENSG00000197622 | CDC42SE1 | chr1 | - | 151056655 | 151056786 | 151055582 | 151055993 | 151059482 | 151059574 | 211229 | 22,7,32,14,12,25,8,13,6,5,5 | 2,0,3,3,3,0,1,2,4,3,0,1 | 24,13,23,20,20,13,5,21,6,1 | 1,1,3,0,3,0,0,0,0,0,2,1,2,3,1 | 280 | 149 | 0.00254197412375 | 0.216628647538589 | 0.854,1.0,0.85,0.71 | 0.927,0.874,0.803, | -0,122 |
| 163047 | ENSG00000182903 | ZNF721 | chr4 | - | 472574 | 472743 | 439987 | 444432 | 473980 | 474167 | 163047 | 81,45,139,64,21,26,21,40,4 | 2,1,1,2,6,0,2,4,0,0,2,2 | 50,94,57,67,45,33,21,32,29 | 0,0,0,7,1,0,1,0,1,0,1,0,0,0,1 | 298 | 149 | 0.00254253826575 | 0.216628647538589 | 0.953,0.957,0.986, | 1.0,1.0,1.0,0.827,0 | -0,088 |
| 204482 | ENSG00000125637 | PSD4 | chr2 | + | 113192379 | 113192509 | 113185364 | 113186255 | 113193047 | 113193128 | 204482 | 80,66,184,124,96,69,63,84 | 9,2,11,18,3,8,15,7,1,4,8,9 | 116,60,48,38,32,49,23,54,4 | 23,18,10,7,16,7,10,13,8,8,9 | 279 | 149 | 0.00254340960331 | 0.216628647538589 | 0.826,0.946,0.899, | 0.729,0.64,0.719,0 | 0,095 |
| 100873 | ENSG00000127914 | AKAP9 | chr7 | + | 92107292 | 92107422 | 92105677 | 92105763 | 92108493 | 92108633 | 100873 | 63,51,115,105,53,46,54,81 | 2,1,2,2,4,4,3,7,2,0,0,1 | 65,66,48,42,42,38,40,56,29 | 0,8,2,1,4,5,1,7,3,3,2,8,0,4,2 | 279 | 149 | 0.00254818595174 | 0.216874190979147 | 0.944,0.965,0.968, | 1.0,0.815,0.928,0.9 | 0,054 |
| 49212 | ENSG00000126012 | KDM5C | chrX | - | 53210675 | 53210857 | 53208890 | 53210576 | 53211496 | 53211655 | 49212 | 96,81,266,168,95,110,88,1 | 1,7,13,9,2,7,1,3,0,1,0,2 | 105,144,82,201,186,76,119 | 2,6,8,19,19,5,9,6,7,5,6,5,7, | 298 | 149 | 0.00254830776694 | 0.216874190979147 | 0.98,0.853,0.911,0 | 0.963,0.923,0.837, | 0,041 |
| 199127 | ENSG00000242372 | EIF6 | chr20 | - | 35284183 | 35284261 | 35280073 | 35280118 | 35284380 | 35284492 | 199127 | 18,20,56,56,17,15,21,24,26 | 2,0,11,3,1,0,0,4,1,0,0,3 | 32,13,31,8,25,11,8,22,10,2 | 0,1,5,0,2,0,0,2,0,1,1,4,0,0,0 | 227 | 149 | 0.00254932221356 | 0.21687477049278 | 0.855,1.0,0.77,0.92 | 1.0,0.895,0.803,1.0 | -0,022 |
| 108262 | ENSG00000116754 | SRSF11 | chr1 | + | 70231094 | 70231203 | 70228421 | 70228555 | 70232267 | 70232377 | 108262 | 18,3,6,7,2,6,5,6,2,5,2,2 | 46,25,40,65,37,29,29,35,58 | 3,11,9,50,7,10,11,13,0,2,3, | 30,46,10,27,36,20,10,40,20 | 258 | 149 | 0.00255276507975 | 0.217081857321087 | 0.184,0.065,0.08,0 | 0.055,0.121,0.342, | -0,074 |
| 39332 | ENSG00000075826 | SEC31B | chr10 | - | 100497134 | 100497280 | 100496257 | 100496431 | 100497666 | 100497793 | 39332 | 62,28,65,54,28,17,44,28,20 | 0,0,1,3,1,7,0,3,1,0,1,5 | 18,22,14,27,21,29,18,25,43 | 10,16,3,7,8,1,0,1,1,0,4,4,0, | 295 | 149 | 0.00255831618076 | 0.2174679896924 | 1.0,1.0,0.97,0.901, | 0.476,0.41,0.702,0 | 0,109 |
| 145059 | ENSG00000108960 | MMD | chr17 | - | 55402213 | 55402303 | 55401468 | 55401538 | 55403766 | 55403868 | 145059 | 1,2,11,2,0,1,4,0,0,0,0,2 | 24,20,34,3,14,3,4,17,2,3,1, | 8,2,0,0,0,3,0,0,0,4,0,6,1,4,0 | 46,19,8,2,7,9,5,9,5,12,4,54 | 239 | 149 | 0.00256300092032 | 0.217780201769204 | 0.025,0.059,0.168, | 0.098,0.062,0.0,0.0 | 0,062 |
| 138791 | ENSG00000141968 | VAV1 | chr19 | + | 6825279 | 6825406 | 6825052 | 6825121 | 6826611 | 6826711 | 138791 | 68,41,72,53,32,19,19,23,43 | 7,2,3,0,1,2,0,2,1,1,2,0 | 37,28,28,11,39,34,8,10,31, | 4,1,0,2,0,1,3,0,0,0,0,0,1,0,2 | 276 | 149 | 0.00256745765756 | 0.217988710343153 | 0.84,0.917,0.928,1 | 0.833,0.938,1.0,0.1 | -0,018 |
| 253216 | ENSG00000122965 | RBM19 | chr12 | - | 113959211 | 113959398 | 113957781 | 113958050 | 113959864 | 113959903 | 253216 | 58,33,60,31,45,54,51,51,73 | 0,2,1,0,0,0,2,0,0,2,0,2 | 50,33,50,50,20,14,22,39,22 | 1,0,0,6,4,0,0,1,2,0,1,0,4,2,2 | 298 | 149 | 0.00256748043135 | 0.217988710343153 | 1.0,0.892,0.968,1.0 | 0.962,1.0,1.0,0.806 | 0,042 |
| 239093 | ENSG00000219481 | NBPF1 | chr1 | - | 16597571 | 16597680 | 16594930 | 16595013 | 16608507 | 16608779 | 239093 | 0,3,3,2,2,0,0,1,0,1,0,0 | 18,5,16,22,6,12,7,8,8,6,8,3 | 1,4,0,0,2,2,2,2,3,3,1,0,2,3,5 | 14,8,10,8,6,6,11,16,13,20,4 | 258 | 149 | 0.00257084726666 | 0.218188496865338 | 0.0,0.257,0.098,0.0 | 0.04,0.224,0.0,0.0, | -0,037 |
| 226501 | ENSG00000171681 | ATF7IP | chr12 | + | 14457206 | 14457249 | 14456560 | 14456634 | 14460494 | 14461133 | 226501 | 62,32,112,80,23,43,33,57,3 | 35,7,36,20,7,12,14,18,22,1 | 51,53,63,57,47,45,22,34,35 | 11,16,13,17,11,5,3,13,6,13 | 192 | 149 | 0.00257237286349 | 0.218231921080732 | 0.579,0.78,0.707,0 | 0.783,0.72,0.79,0.1 | -0,112 |
| 154603 | ENSG00000100897 | DCAF11 | chr14 | + | 24116918 | 24117044 | 24115542 | 24115749 | 24117265 | 24117306 | 154603 | 96,54,81,105,37,42,20,33,8 | 8,1,4,1,3,2,4,2,1,0,0,1 | 30,43,37,21,44,30,27,40,22 | 6,5,4,2,5,1,2,3,2,2,1,7,0,5,3 | 275 | 149 | 0.00257590459724 | 0.218445438285486 | 0.867,0.967,0.916, | 0.73,0.823,0.834,0 | 0,057 |
| 176554 | ENSG00000114446 | IFT57 | chr3 | - | 108219409 | 108219570 | 108218392 | 108218653 | 108222110 | 108222435 | 176554 | 91,41,464,347,119,125,117 | 5,4,2,0,4,2,0,4,0,0,2,6 | 321,271,69,114,115,88,76, | 4,4,2,7,2,0,11,5,0,5,7,1,13, | 298 | 149 | 0.00258192779744 | 0.218869988881768 | 0.901,0.837,0.991, | 0.976,0.971,0.945, | 0,013 |
| 64136 | ENSG00000213064 | SFT2D2 | chr1 | + | 168235100 | 168235182 | 168231833 | 168231919 | 168236588 | 168236624 | 64136 | 50,69,125,131,74,58,39,65 | 3,3,10,2,3,0,3,4,0,1,0,5 | 116,135,29,38,54,43,21,46 | 1,4,1,5,5,0,0,2,2,2,0,2,2,1,0 | 231 | 149 | 0.00258806269674 | 0.219303670189782 | 0.915,0.937,0.89,0 | 0.987,0.956,0.949, | -0,029 |
| 34803 | ENSG00000172869 | DMXL1 | chr5 | + | 119105179 | 119105258 | 119101934 | 119102006 | 119114474 | 119114541 | 34803 | 116,96,84,54,31,42,51,43,4 | 2,1,2,0,0,0,0,0,0,0,1,0 | 56,73,87,57,44,39,21,35,21 | 5,8,1,2,3,1,0,0,2,0,2,0,0,1,2 | 228 | 149 | 0.00258950247192 | 0.21933931780158 | 0.974,0.984,0.965, | 0.88,0.856,0.983,0 | 0,039 |
| 230905 | ENSG00000166579 | NDEL1 | chr17 | + | 8454778 | 8454887 | 8450780 | 8450953 | 8460008 | 8460160 | 230905 | 39,14,42,34,23,25,21,21,20 | 1,2,0,0,0,0,0,0,2,0,0,0 | 22,38,20,35,28,15,8,13,24, | 0,1,6,2,4,0,1,2,1,0,1,3,2,0,0 | 258 | 149 | 0.00259391112925 | 0.219626312454605 | 0.957,0.802,1.0,1.0 | 1.0,0.956,0.658,0.9 | 0,069 |
| 253217 | ENSG00000122965 | RBM19 | chr12 | - | 113959211 | 113959401 | 113957781 | 113958050 | 113959864 | 113959903 | 253217 | 58,33,60,31,45,54,51,51,73 | 0,2,1,0,0,0,2,0,0,2,0,2 | 50,33,50,50,20,14,22,39,23 | 1,0,0,6,4,0,0,1,2,0,1,0,4,2,2 | 298 | 149 | 0.00259754810629 | 0.219847768959852 | 1.0,0.892,0.968,1.0 | 0.962,1.0,1.0,0.806 | 0,042 |
| 39828 | ENSG00000205981 | DNAJC19 | chr3 | - | 180988177 | 180988229 | 180984564 | 180988096 | 180989599 | 180989673 | 39828 | 15,4,16,11,6,4,4,1,5,1,6,6 | 7,0,2,0,2,2,0,2,2,0,0,2 | 14,8,14,10,3,4,2,5,3,18,12, | 3,0,1,0,0,0,2,0,1,0,2,1,0,0,2 | 201 | 149 | 0.00261081802439 | 0.220884030741936 | 0.614,1.0,0.856,1.0 | 0.776,1.0,0.912,1.0 | -0,091 |
| 244676 | ENSG00000168067 | MAP4K2 | chr11 | - | 64801552 | 64801621 | 64801110 | 64801183 | 64801709 | 64801757 | 244676 | 1,4,10,4,3,5,5,9,5,1,3,8 | 0,0,4,5,2,0,0,2,0,0,0,2 | 5,3,3,4,1,2,7,7,0,4,8,3,5,3,1 | 8,4,0,3,0,1,0,2,5,4,7,0,1,1,4 | 218 | 149 | 0.00261298612743 | 0.220980596146517 | 1.0,1.0,0.631,0.353 | 0.299,0.339,1.0,0.4 | 0,219 |
| 241232 | ENSG00000107077 | KDM4C | chr9 | + | 6825306 | 6825737 | 6814630 | 6814745 | 6849506 | 6849700 | 241232 | 5,0,8,2,0,1,1,3,0,1,0,6 | 67,26,90,51,46,66,40,56,25 | 7,7,4,8,3,3,0,3,6,7,4,7,5,0,2 | 73,77,37,59,51,41,19,39,36 | 298 | 149 | 0.00261875865464 | 0.22138179261462 | 0.036,0.0,0.043,0.0 | 0.046,0.043,0.051, | -0,018 |
| 30188 | ENSG00000115561 | CHMP3 | chr2 | - | 86542251 | 86542312 | 86529217 | 86529397 | 86563303 | 86563374 | 30188 | 91,62,171,105,57,38,31,50 | 7,4,9,2,2,6,2,4,1,8,0,0 | 103,108,61,29,79,44,38,35 | 9,13,10,8,6,1,8,17,2,1,10,8 | 210 | 149 | 0.00262573003242 | 0.221858600381826 | 0.902,0.917,0.931, | 0.89,0.855,0.812,0 | 0,055 |
| 161498 | ENSG00000113282 | CLINT1 | chr5 | - | 157790559 | 157790651 | 157789362 | 157789513 | 157791702 | 157791841 | 161498 | 1,2,5,2,2,1,1,1,1,0,1,0 | 77,63,57,52,40,25,29,142,1 | 0,5,8,6,3,0,0,2,1,4,6,3,2,7,1 | 60,85,75,104,93,44,30,52,4 | 241 | 149 | 0.00262720688585 | 0.221858600381826 | 0.008,0.019,0.051, | 0.0,0.035,0.062,0.0 | -0,01 |
| 178095 | ENSG00000067066 | SP100 | chr2 | + | 230464046 | 230464150 | 230462466 | 230462518 | 230466300 | 230466354 | 178095 | 83,61,157,95,52,60,50,81,3 | 57,30,72,69,17,16,39,29,3, | 99,75,63,40,70,30,15,57,46 | 50,76,27,57,116,31,16,23,5 | 253 | 149 | 0.00262749126461 | 0.221858600381826 | 0.462,0.545,0.562, | 0.538,0.368,0.579, | 0,16 |
| 191628 | ENSG00000186470 | BTN3A2 | chr6 | + | 26373953 | 26374199 | 26373386 | 26373413 | 26374326 | 26374373 | 191628 | 8,0,13,1,2,2,0,0,0,0,0,0 | 24,11,37,39,12,12,18,19,15 | 0,0,1,0,0,3,0,0,0,2,0,0,3,0,0 | 13,10,15,4,3,3,7,36,13,16,1 | 298 | 149 | 0.00262919136565 | 0.22190338806744 | 0.143,0.0,0.149,0.0 | 0.0,0.0,0.032,0.0,0 | 0,007 |
| 151192 | ENSG00000174485 | DENND4A | chr15 | - | 65725887 | 65726098 | 65722847 | 65722948 | 65729071 | 65729247 | 151192 | 2,0,7,4,2,15,4,5,0,1,0,3 | 71,47,234,139,69,78,59,92 | 6,18,5,13,12,4,1,7,10,8,1,6 | 114,141,55,105,104,67,52, | 298 | 149 | 0.00263145269028 | 0.22190338806744 | 0.014,0.0,0.015,0.0 | 0.026,0.06,0.043,0 | -0,013 |
| 127650 | ENSG00000176155 | CCDC57 | chr17 | - | 82179026 | 82179189 | 82178473 | 82178605 | 82183773 | 82183932 | 127650 | 27,18,24,5,14,8,6,15,11,6,1 | 5,1,3,0,3,0,1,4,0,1,0,0 | 6,4,10,13,4,4,14,32,3,8,10, | 1,1,0,0,0,1,0,0,0,2,0,1,0,0,1 | 298 | 149 | 0.00263194017756 | 0.22190338806744 | 0.73,0.9,0.8,1.0,0.1 | 0.75,0.667,1.0,1.0, | -0,028 |
| 169724 | ENSG00000204463 | BAG6 | chr6 | - | 31647590 | 31647871 | 31646393 | 31646523 | 31648676 | 31648751 | 169724 | 117,73,93,84,52,38,57,32,5 | 7,5,8,1,4,5,7,7,3,3,1,1 | 39,42,15,50,28,27,64,44,21 | 3,5,6,12,5,4,5,2,3,2,4,13,5, | 298 | 149 | 0.0026321456934 | 0.22190338806744 | 0.893,0.88,0.853,0 | 0.867,0.808,0.556, | 0,09 |
| 48896 | ENSG00000285708 | | chr3 | - | 71335159 | 71335281 | 71299819 | 71299880 | 71359149 | 71359244 | 48896 | 6,0,8,3,3,0,5,1,2,0,2,0 | 61,10,145,69,52,9,51,65,26 | 14,7,5,9,1,1,1,6,8,5,5,10,5, | 155,105,63,69,33,12,43,47 | 271 | 149 | 0.00263546337462 | 0.22209609145765 | 0.051,0.0,0.0,0.029,0.0 | 0.047,0.035,0.042, | -0,018 |
| 15079 | ENSG00000100266 | PACSIN2 | chr22 | - | 42876887 | 42877010 | 42876136 | 42876333 | 42879047 | 42879169 | 15079 | 16,13,34,33,29,32,25,21,46 | 0,0,19,21,4,7,3,10,4,1,2,0 | 16,13,36,8,8,15,20,34,6,19 | 9,11,8,11,10,2,14,8,2,0,9,8 | 272 | 149 | 0.00263977862717 | 0.222372678553591 | 1.0,1.0,0.495,0.463 | 0.493,0.393,0.711, | 0,21 |

| | | | | | | | | | | | | | | | | | | | | | |
|---|---|---|---|---|---|---|---|---|---|---|---|---|---|---|---|---|---|---|---|---|---|
| 40831 | ENSG00000136819 | C9orf78 | chr9 | - | 129834706 | 129834766 | 129833657 | 129833709 | 129835138 | 129835275 | 40831 | 200,124,125,76,58,39,25,5 | 0,1,2,1,1,0,8,17,0,0,0,0 | 83,92,114,72,87,21,31,69,2 | 0,1,0,0,1,0,1,1,0,0,7,1,0,0,3 | 209 | 149 | 0.00264152645255 | 0.222432855989354 | 1.0,0.989,0.978,0.9 | 1.0,0.985,1.0,1.0,0 | -0,037 |
| 121519 | ENSG00000110075 | PPP6R3 | chr11 | + | 68567013 | 68567166 | 68564302 | 68564432 | 68569747 | 68569897 | 121519 | 181,112,305,126,147,156,1 | 26,7,20,4,0,8,24,18,1,5,2,8 | 220,180,148,93,107,96,107 | 23,15,33,28,12,3,11,22,8,4 | 298 | 149 | 0.00265092605778 | 0.223094281541047 | 0.777,0.889,0.884, | 0.827,0.857,0.692, | 0,06 |
| 218948 | ENSG00000204569 | PPP1R10 | chr6 | - | 30604865 | 30605094 | 30604587 | 30604735 | 30605922 | 30605998 | 218948 | 29,15,38,23,10,9,8,14,10,6 | 2,0,4,5,0,3,0,0,2,0,0,1 | 14,18,15,8,11,4,19,17,0,12 | 3,1,0,1,2,0,2,1,2,2,8,0,0,0,0 | 298 | 149 | 0.00265145435454 | 0.223094281541047 | 0.879,1.0,0.826,0.6 | 0.7,0.9,1.0,0.8,0.73 | 0,054 |
| 122189 | ENSG00000165916 | PSMC3 | chr11 | - | 47424108 | 47424183 | 47422829 | 47422973 | 47424428 | 47424491 | 122189 | 27,38,59,22,12,22,9,24,24, | 1,0,4,5,2,2,2,1,0,0,0,2 | 32,26,31,15,22,14,4,10,7,2 | 1,1,0,2,0,0,0,0,0,0,0,2,0,0,0 | 224 | 149 | 0.00265338779931 | 0.223169718418636 | 0.947,1.0,0.908,0.1 | 0.955,0.945,1.0,0.8 | -0,081 |
| 190578 | ENSG00000177463 | NR2C2 | chr3 | + | 15040205 | 15040345 | 15039121 | 15039227 | 15042833 | 15049273 | 190578 | 2,2,1,0,0,0,0,1,2,0,0,0 | 16,26,44,23,20,21,21,29,23 | 2,2,0,5,1,2,1,0,0,3,4,6,1,0,1 | 24,30,22,27,21,13,21,24,16 | 289 | 149 | 0.00265481315324 | 0.223202378822265 | 0.061,0.038,0.012, | 0.041,0.033,0.0,0.0 | -0,023 |
| 70826 | ENSG00000047410 | TPR | chr1 | - | 186353687 | 186353880 | 186352014 | 186352110 | 186355409 | 186355558 | 70826 | 331,295,218,94,101,118,61 | 19,3,6,3,0,3,4,0,3,6,0,6 | 126,121,167,161,105,58,52 | 1,6,2,8,0,2,3,3,0,2,0,5,1,1,1 | 298 | 149 | 0.00265743592512 | 0.223335646856502 | 0.897,0.98,0.948,0 | 0.984,0.91,0.977,0 | -0,019 |
| 181152 | ENSG00000162894 | FCMR | chr1 | - | 206912928 | 206912968 | 206911729 | 206911952 | 206921817 | 206921998 | 181152 | 230,82,160,113,136,121,20 | 9,3,4,3,4,10,4,4,8,14,3,3 | 116,47,107,64,25,33,84,19 | 4,0,6,4,2,3,2,3,0,0,0,0,2,1,5 | 189 | 149 | 0.00265988266659 | 0.223454022721636 | 0.953,0.956,0.969, | 0.958,1.0,0.934,0.9 | -0,014 |
| 234084 | ENSG00000068308 | OTUD5 | chrX | - | 48936190 | 48936355 | 48934953 | 48935018 | 48944189 | 48944283 | 234084 | 30,12,17,13,4,2,10,9,7,3,1, | 169,118,125,76,49,64,33,5 | 17,24,12,78,4,5,7,8,4,3,7,2 | 57,65,65,31,53,46,35,54,32 | 298 | 149 | 0.00266137985814 | 0.223492566620105 | 0.082,0.048,0.064, | 0.13,0.156,0.085,0 | -0,049 |
| 156867 | ENSG00000134884 | ARGLU1 | chr13 | - | 106557558 | 106557704 | 106557047 | 106557131 | 106559431 | 106559657 | 156867 | 58,4,65,40,17,12,23,15,23, | 130,49,133,112,66,62,60,6 | 16,14,28,11,8,14,1,26,13,1 | 104,54,77,34,46,41,18,79,3 | 295 | 149 | 0.00267120866369 | 0.224230464857513 | 0.184,0.04,0.198,0 | 0.072,0.116,0.155, | 0,035 |
| 141012 | ENSG00000130813 | SHFL | chr19 | + | 10089110 | 10089523 | 10087250 | 10087300 | 10089656 | 10089695 | 141012 | 12,1,11,7,3,6,3,8,11,3,5,3 | 22,16,40,31,21,24,16,22,14 | 7,4,3,6,2,5,0,1,3,3,4,4,1,4,2 | 17,18,22,22,24,8,11,15,7,2 | 298 | 149 | 0.00267535131512 | 0.224438286514172 | 0.214,0.03,0.121,0 | 0.171,0.1,0.064,0.1 | 0,051 |
| 30643 | ENSG00000112983 | BRD8 | chr5 | - | 138167933 | 138168078 | 138165827 | 138166108 | 138169221 | 138169358 | 30643 | 12,3,24,35,21,8,7,6,9,8,7,6 | 4,1,1,3,0,1,0,2,1,0,0,1 | 16,14,10,1,4,3,17,17,0,11,5 | 3,1,1,0,4,2,0,3,4,0,0,1,2,2,7 | 294 | 149 | 0.00267576995505 | 0.224438286514172 | 0.603,0.603,0.924, | 0.73,0.876,0.835,1 | 0,094 |
| 3154 | ENSG00000268027 | | chr19 | + | 41553705 | 41553746 | 41549519 | 41549552 | 41554950 | 41555099 | 3154 | 22,3,7,7,10,11,11,13,40,20 | 6,0,1,2,3,3,4,2,5,6,1,0 | 16,3,12,6,3,7,1,4,2,12,7,5,7 | 4,1,1,0,0,1,1,3,0,0,3,2,0,2,1 | 190 | 149 | 0.00268347329965 | 0.224996743964538 | 0.742,1.0,0.846,0.1 | 0.758,0.702,0.904, | -0,025 |
| 21642 | ENSG00000182979 | MTA1 | chr14 | + | 105449358 | 105449409 | 105445417 | 105445511 | 105450057 | 105450184 | 21642 | 17,14,38,10,22,17,24,19,11 | 4,2,2,0,6,2,0,1,0,2,0,4 | 18,9,8,6,10,3,21,22,13,21,1 | 6,3,0,4,4,0,1,5,5,1,4,6,4,3,4 | 200 | 149 | 0.00268490101675 | 0.225028789227461 | 0.76,0.839,0.934,1 | 0.691,0.691,1.0,0.5 | 0,082 |
| 209616 | ENSG00000166986 | MARS1 | chr12 | + | 57489898 | 57489971 | 57489529 | 57489558 | 57490206 | 57490216 | 209616 | 43,10,65,52,29,26,16,18,27 | 5,3,6,2,1,8,7,3,1,2,1,4 | 28,22,20,6,26,24,16,31,33, | 5,19,9,6,9,1,8,3,4,5,11,6,16 | 222 | 149 | 0.00268874742352 | 0.225263447532749 | 0.852,0.691,0.879, | 0.79,0.437,0.599,0 | 0,105 |
| 41567 | ENSG00000171608 | PIK3CD | chr1 | + | 9691466 | 9691571 | 9629972 | 9630224 | 9710423 | 9710596 | 41567 | 51,28,57,40,36,32,22,14,16 | 0,4,4,0,0,1,2,1,2,0,2,10 | 40,41,40,41,20,15,30,45,17 | 1,4,1,2,1,3,0,0,3,1,0,2,0,2,0 | 254 | 149 | 0.00268982902923 | 0.225266378128483 | 1.0,0.804,0.893,1.0 | 0.959,0.857,0.959, | -0,053 |
| 102957 | ENSG00000152457 | DCLRE1C | chr10 | - | 14928015 | 14928152 | 14922980 | 14923069 | 14932853 | 14932955 | 102957 | 75,32,54,31,30,17,25,37,25 | 2,2,5,3,0,0,1,4,1,0,0,1 | 43,63,20,17,29,21,9,17,19, | 0,1,1,0,1,0,0,1,0,0,1,1,1,0,2 | 286 | 149 | 0.00269674136454 | 0.22575742537196 | 0.951,0.893,0.849, | 1.0,0.97,0.912,1.0, | -0,017 |
| 240924 | ENSG00000198771 | RCSD1 | chr1 | + | 167685420 | 167685502 | 167683899 | 167684001 | 167690048 | 167690120 | 240924 | 41,10,65,34,46,40,99,150,7 | 23,2,28,18,11,14,18,22,20, | 27,7,28,21,14,20,64,105,11 | 16,10,16,13,7,7,5,33,13,12 | 231 | 149 | 0.00269803930289 | 0.225778264852379 | 0.535,0.763,0.6,0.5 | 0.521,0.311,0.53,0 | 0,132 |
| 102461 | ENSG00000124486 | USP9X | chrX | + | 41140655 | 41140771 | 41136803 | 41137022 | 41140965 | 41141217 | 102461 | 67,44,248,150,81,103,51,1 | 5,0,7,2,1,4,4,8,0,0,1,0 | 156,151,81,131,97,61,44,6 | 14,10,1,34,7,1,2,4,4,0,0,6,2 | 265 | 149 | 0.0027006653637 | 0.225910185345711 | 0.883,1.0,0.952,0.9 | 0.862,0.895,0.979, | 0,024 |
| 128757 | ENSG00000100354 | TNRC6B | chr22 | + | 40300909 | 40301005 | 40300454 | 40300586 | 40301149 | 40301333 | 128757 | 69,44,103,72,45,72,54,63,3 | 3,0,8,2,3,2,2,9,0,3,4,0 | 53,60,36,73,28,38,51,70,46 | 1,2,2,1,1,0,3,0,0,2,2,1,3,0,1 | 245 | 149 | 0.0027018049801 | 0.225917710828245 | 0.933,1.0,0.887,0.9 | 0.97,0.948,0.916,0 | -0,026 |
| 96801 | ENSG00000127947 | PTPN12 | chr7 | + | 77563935 | 77564195 | 77537553 | 77537645 | 77571077 | 77571186 | 96801 | 3,0,1,0,2,0,4,0,0,0,0,0 | 38,17,21,20,11,15,11,24,13 | 6,3,0,2,0,6,0,2,4,2,0,2,0,0,2 | 14,30,16,6,7,21,6,7,10,12,2 | 298 | 149 | 0.00270316980844 | 0.225944054772951 | 0.038,0.0,0.023,0.0 | 0.176,0.048,0.0,0.1 | -0,035 |
| 240569 | ENSG00000152795 | HNRNPDL | chr4 | - | 82425565 | 82425667 | 82424630 | 82424883 | 82426036 | 82426129 | 240569 | 9,7,14,8,2,3,4,8,15,2,1,9 | 68,36,370,202,166,186,95, | 10,8,4,2,3,3,4,0,5,5,5,8,8,1 | 272,222,114,64,104,80,61, | 251 | 149 | 0.00270447004709 | 0.225956562485642 | 0.073,0.103,0.022, | 0.021,0.021,0.02,0 | 0,037 |
| 12996 | ENSG00000109046 | WSB1 | chr17 | + | 27301787 | 27301956 | 27294092 | 27294435 | 27303366 | 27303411 | 12996 | 77,48,260,203,64,87,68,10 | 4,3,37,11,32,20,10,15,5,2,0 | 82,89,19,17,27,48,31,139,3 | 15,20,23,21,20,11,3,12,11, | 298 | 149 | 0.00270541911493 | 0.225956562485642 | 0.906,0.889,0.778, | 0.732,0.69,0.292,0 | 0,146 |
| 199626 | ENSG00000047410 | TPR | chr1 | - | 186353687 | 186353870 | 186351951 | 186352110 | 186355409 | 186355558 | 199626 | 329,291,210,97,101,116,60 | 19,3,6,3,0,3,4,0,3,6,0,6 | 121,121,165,160,107,56,52 | 1,6,2,8,0,2,3,3,0,2,0,5,1,1,1 | 298 | 149 | 0.00270851573074 | 0.226091325752523 | 0.896,0.98,0.946,0 | 0.984,0.91,0.976,0 | -0,019 |
| 230370 | ENSG00000113013 | HSPA9 | chr5 | - | 138559863 | 138560059 | 138558552 | 138558657 | 138561579 | 138561789 | 230370 | 53,37,208,136,56,58,36,52 | 3,0,8,11,2,3,1,3,3,0,0,3 | 97,150,46,33,77,40,35,55,1 | 3,7,1,0,1,1,0,2,0,1,2,6,0,1,2 | 298 | 149 | 0.00270913357795 | 0.226091325752523 | 0.898,1.0,0.929,0.8 | 0.942,0.915,0.958, | -0,032 |
| 45783 | ENSG00000116191 | RALGPS2 | chr1 | + | 178811314 | 178811404 | 178808044 | 178808128 | 178821611 | 178821704 | 45783 | 612,390,830,454,424,242,3 | 8,2,5,1,1,3,2,1,5,1,2,2 | 846,712,350,274,316,324,1 | 16,12,4,15,7,3,3,1,10,3,2,6 | 239 | 149 | 0.00271308202288 | 0.226333084056777 | 0.979,0.992,0.99,0 | 0.971,0.974,0.982, | 0,007 |
| 221491 | ENSG00000112695 | COX7A2 | chr6 | - | 75240300 | 75240384 | 75237674 | 75237988 | 75241175 | 75241265 | 221491 | 210,106,236,202,102,92,96 | 19,14,26,17,4,8,16,15,5,6,6 | 136,111,126,57,85,83,63,9 | 8,34,24,8,13,4,9,30,6,14,17 | 233 | 149 | 0.0027149627619 | 0.226381573689727 | 0.876,0.829,0.853, | 0.916,0.676,0.77,0 | 0,047 |
| 26869 | ENSG00000150867 | PIP4K2A | chr10 | - | 22541803 | 22542047 | 22539970 | 22540074 | 22550658 | 22550772 | 26869 | 127,94,88,74,44,23,43,39,6 | 5,1,1,7,2,0,0,0,2,2,0,2 | 71,37,73,52,31,45,14,26,27 | 0,1,2,0,0,1,0,0,3,2,0,3,0,2,0 | 298 | 149 | 0.00271660641953 | 0.226381573689727 | 0.927,0.979,0.978, | 1.0,0.949,0.948,1.0 | -0,018 |
| 95453 | ENSG00000152601 | MBNL1 | chr3 | + | 152335110 | 152335230 | 152300197 | 152300367 | 152414940 | 152415109 | 95453 | 10,7,14,7,8,6,9,7,9,15,5,7 | 452,250,444,197,235,233,2 | 9,7,1,1,0,3,1,8,2,7,7,4,4,10 | 365,359,176,132,170,131,1 | 269 | 149 | 0.00271681869638 | 0.226381573689727 | 0.012,0.015,0.017, | 0.013,0.011,0.003, | 0,009 |
| 178047 | ENSG00000067066 | SP100 | chr2 | + | 230436380 | 230436607 | 230417590 | 230417665 | 230442936 | 230443099 | 178047 | 26,8,39,27,0,0,0,0,0,0,0,0 | 261,213,313,243,99,136,13 | 0,4,0,0,5,0,11,15,17,17,0,0 | 224,216,191,114,279,122,4 | 298 | 149 | 0.00272086400203 | 0.226630913320789 | 0.047,0.018,0.059, | 0.0,0.009,0.0,0.0,0 | -0,009 |
| 188792 | ENSG00000082512 | TRAF5 | chr1 | + | 211359878 | 211360005 | 211356366 | 211356468 | 211360701 | 211360779 | 188792 | 11,1,16,1,3,5,10,3,6,4,1,8 | 2,0,0,2,0,1,0,4,0,0,0,0 | 9,14,2,8,10,5,2,5,5,4,15,3,7 | 1,0,0,2,2,4,1,2,5,4,0,0,2,0,0 | 276 | 149 | 0.00272733764056 | 0.227082246698402 | 0.748,1.0,1.0,0.21 | 0.829,1.0,1.0,0.68 | 0,076 |
| 162090 | ENSG00000124201 | ZNFX1 | chr20 | - | 49263333 | 49263471 | 49260462 | 49260577 | 49264715 | 49264864 | 162090 | 23,27,57,42,9,8,14,19,27,2 | 0,2,4,1,2,1,0,1,0,0,0,5 | 29,39,16,41,15,12,7,21,9,1 | 4,4,1,4,5,0,1,1,1,1,5,1,2,1,1 | 287 | 149 | 0.00273150220532 | 0.227278207295394 | 1.0,0.875,0.881,0.9 | 0.79,0.835,0.893,0 | 0,089 |
| 44775 | ENSG00000101596 | SMCHD1 | chr18 | + | 2690665 | 2690761 | 2688692 | 2688747 | 2694526 | 2694693 | 44775 | 31,18,11,5,7,15,13,21,7,3,2 | 673,562,474,275,227,306,2 | 1,2,11,7,13,1,11,0,2,17,12, | 165,277,370,259,410,172,2 | 245 | 149 | 0.00273180314301 | 0.227278207295394 | 0.027,0.019,0.014, | 0.004,0.004,0.018, | 0,006 |
| 99071 | ENSG00000136280 | CCM2 | chr7 | + | 45071749 | 45071872 | 45064602 | 45064646 | 45072725 | 45072786 | 99071 | 7,0,4,2,0,0,0,5,3,0,0,0 | 9,2,10,6,2,4,5,11,9,6,5,2 | 6,2,2,4,4,3,3,3,2,4,1,6,2,1,1 | 4,5,4,2,2,0,1,1,2,4,9,5,3,5,4 | 272 | 149 | 0.00273798757337 | 0.227691489393459 | 0.299,0.0,0.18,0.15 | 0.451,0.18,0.215,0 | -0,279 |
| 167073 | ENSG00000174652 | ZNF266 | chr19 | - | 9418765 | 9419167 | 9418504 | 9418631 | 9420064 | 9420218 | 167073 | 29,24,86,56,32,6,42,66,41, | 4,0,3,4,0,0,3,2,0,0,2,5 | 81,38,41,56,38,27,28,50,40 | 1,3,0,2,3,0,0,0,0,0,6,0,0,0,1 | 298 | 149 | 0.00273888643383 | 0.227691489393459 | 0.784,1.0,0.935,0.8 | 0.976,0.864,1.0,0.9 | -0,09 |
| 57197 | ENSG00000134242 | PTPN22 | chr1 | - | 113852026 | 113852104 | 113848539 | 113848626 | 113854470 | 113854537 | 57197 | 264,139,672,628,180,120,1 | 5,2,18,14,1,3,3,5,4,3,4,2 | 449,404,141,54,176,157,74 | 13,5,10,2,15,3,3,6,7,3,5,20 | 227 | 149 | 0.00274445890854 | 0.228001613196516 | 0.972,0.979,0.961, | 0.958,0.981,0.902, | 0,023 |
| 147756 | ENSG00000197912 | SPG7 | chr16 | + | 89545871 | 89546011 | 89542983 | 89544772 | 89546657 | 89547635 | 147756 | 0,1,10,4,1,0,7,3,1,1,0,1 | 36,17,74,42,40,51,18,19,23 | 1,0,0,2,2,2,0,3,0,0,2,4,0,0,0 | 45,38,15,18,62,22,32,38,46 | 289 | 149 | 0.00274473556222 | 0.228001613196516 | 0.0,0.029,0.065,0.0 | 0.011,0.0,0.0,0.0,0.05 | 0,024 |
| 26460 | ENSG00000055211 | GINM1 | chr6 | + | 149572506 | 149572603 | 149572284 | 149572344 | 149578821 | 149578973 | 26460 | 55,47,150,95,47,38,32,35,5 | 0,3,10,2,2,3,0,0,0,0,0,2 | 61,77,59,44,63,42,23,42,29 | 0,0,0,0,3,1,0,0,1,1,0,0,1,0 | 246 | 149 | 0.00275154387724 | 0.228478989290989 | 1.0,0.905,0.901,0.9 | 1.0,1.0,1.0,1.0,1.0, | -0,016 |
| 101378 | ENSG00000008282 | SYPL1 | chr7 | - | 106092563 | 106092669 | 106091456 | 106091939 | 106092948 | 106093137 | 101378 | 11,4,7,6,0,2,3,0,0,2,1,0 | 93,78,154,90,77,69,53,48,3 | 0,2,5,1,10,2,16,2,3,0,9,25,4 | 77,98,69,71,63,53,24,51,48 | 255 | 149 | 0.00275617923537 | 0.228775631703787 | 0.065,0.029,0.026, | 0.0,0.012,0.041,0.0 | -0,026 |
| 138027 | ENSG00000198521 | ZNF43 | chr19 | - | 21810205 | 21810271 | 21809680 | 21809807 | 21817887 | 21817986 | 138027 | 3,3,9,3,0,4,2,4,2,0,0,3 | 18,19,29,22,10,9,7,6,18,11 | 11,7,4,9,3,1,1,6,0,4,6,5,3,0 | 23,18,9,23,14,4,9,10,8,8,24 | 215 | 149 | 0.00275966222124 | 0.228894966237612 | 0.104,0.099,0.177, | 0.249,0.212,0.235, | -0,082 |
| 107948 | ENSG00000197892 | KIF13B | chr8 | - | 29092744 | 29092878 | 29075280 | 29075343 | 29099132 | 29099241 | 107948 | 29,10,19,17,15,15,18,31,21 | 2,0,3,0,1,0,1,3,2,0,0,4 | 13,7,17,14,5,2,12,25,7,6,14 | 3,3,1,2,3,0,0,3,1,2,0,0,0,2,0 | 283 | 149 | 0.00275974389092 | 0.228894966237612 | 0.884,1.0,0.769,1.0 | 0.695,0.551,0.9,0.1 | 0,062 |
| 205305 | ENSG00000176407 | KCMF1 | chr2 | + | 85035636 | 85035811 | 85035015 | 85035155 | 85043563 | 85043665 | 205305 | 4,0,10,13,2,0,3,3,1,1,2,2 | 60,55,158,144,72,46,35,51 | 18,18,4,11,1,0,2,5,2,2,11,5 | 116,170,55,61,76,33,20,49 | 298 | 149 | 0.0027621466483 | 0.229006003567125 | 0.032,0.0,0.031,0.0 | 0.072,0.05,0.035,0 | -0,012 |
| 223214 | ENSG00000125779 | PANK2 | chr20 | + | 3918483 | 3918796 | 3916926 | 3917050 | 3923243 | 3923631 | 223214 | 19,6,41,34,16,12,6,9,31,1,3 | 6,0,2,2,6,2,1,1,0,0,0,0 | 19,20,20,8,10,10,4,11,6,15 | 0,0,2,2,0,0,1,1,0,3,1,1,0,1,0 | 298 | 149 | 0.00276528280916 | 0.2291391181031 | 0.613,1.0,0.911,0.8 | 1.0,1.0,0.833,0.667 | -0,013 |
| 169722 | ENSG00000204463 | BAG6 | chr6 | - | 31647590 | 31647844 | 31646393 | 31646523 | 31648676 | 31648751 | 169722 | 117,73,93,84,52,38,57,32,5 | 7,5,8,1,4,5,7,7,3,3,1,1 | 39,42,15,50,28,27,63,44,21 | 3,5,6,12,5,4,5,2,3,2,4,13,5, | 298 | 149 | 0.002766189698 | 0.2291391181031 | 0.893,0.88,0.853,0 | 0.867,0.808,0.556, | 0,089 |
| 137152 | ENSG00000170889 | RPS9 | chr19 | + | 54206586 | 54206724 | 54206275 | 54206462 | 54207397 | 54207522 | 137152 | 41,30,80,64,23,18,28,31,33 | 970,517,1906,1616,681,79 | 14,35,58,23,50,27,13,24,15 | 749,634,603,319,436,414,5 | 287 | 149 | 0.00276694606237 | 0.2291391181031 | 0.021,0.029,0.021, | 0.01,0.028,0.048,0 | -0,007 |
| 150068 | ENSG00000159322 | ADPGK | chr15 | - | 72774871 | 72775095 | 72771782 | 72771845 | 72783245 | 72783731 | 150068 | 87,81,121,88,66,81,36,78,8 | 8,7,6,1,5,3,6,1,5,0,3,6 | 83,97,70,43,62,32,55,62,42 | 2,11,8,10,23,2,3,6,10,3,3,1 | 298 | 149 | 0.00277997608842 | 0.230129628264125 | 0.845,0.853,0.91,0 | 0.954,0.815,0.814, | 0,065 |
| 148394 | ENSG00000153774 | CFDP1 | chr16 | - | 75391397 | 75391508 | 75305023 | 75305182 | 75395089 | 75395136 | 148394 | 1,0,3,1,1,2,0,4,2,1,0,1 | 27,19,43,57,13,21,14,11,20 | 5,4,1,1,1,6,0,1,1,2,1,3,4,1,2 | 16,13,8,2,13,7,13,20,11,13 | 260 | 149 | 0.00279932840591 | 0.231602901663135 | 0.021,0.0,0.038,0.0 | 0.152,0.15,0.067,0 | -0,056 |
| 167100 | ENSG00000174652 | ZNF266 | chr19 | - | 9419216 | 9419289 | 9418504 | 9418631 | 9420064 | 9420218 | 167100 | 4,2,74,60,15,11,38,34,9,4,7 | 4,0,3,4,0,0,3,2,0,0,2,5 | 43,28,14,22,12,9,14,28,12, | 1,3,0,2,3,0,0,0,0,0,6,0,0,0,1 | 222 | 149 | 0.00279992542955 | 0.231602901663135 | 0.402,1.0,0.943,0.9 | 0.967,0.862,1.0,0.8 | -0,087 |
| 199125 | ENSG00000242372 | EIF6 | chr20 | - | 35284170 | 35284261 | 35280073 | 35280118 | 35284380 | 35284492 | 199125 | 21,23,69,71,24,18,24,31,31 | 2,0,11,3,1,0,0,4,1,0,0,3 | 39,21,38,11,36,15,13,27,11 | 0,1,5,0,2,0,0,2,0,1,1,4,0,0,0 | 240 | 149 | 0.00280654407637 | 0.23206119404579 | 0.867,1.0,0.796,0.9 | 1.0,0.929,0.825,1.0 | -0,018 |
| 176158 | ENSG00000169180 | XPO6 | chr16 | - | 28155711 | 28155975 | 28152929 | 28152964 | 28156073 | 28156332 | 176158 | 17,10,10,2,5,14,5,13,9,9,5, | 3,1,4,2,0,1,1,5,0,0,0,2 | 18,20,7,5,19,7,7,8,9,19,9,8 | 1,2,0,0,2,0,2,2,0,1,1,3,0,1,0 | 298 | 149 | 0.00280929028031 | 0.232199061567359 | 0.739,0.833,0.556, | 0.9,0.833,1.0,1.0,0 | -0,095 |
| 151885 | ENSG00000125107 | CNOT1 | chr16 | - | 58623246 | 58623409 | 58599373 | 58599511 | 58629727 | 58629847 | 151885 | 65,39,68,66,33,50,24,52,2, | 43,38,72,39,33,28,21,24,21 | 61,74,37,56,63,32,11,31,19 | 38,25,26,9,8,17,12,25,13,4 | 298 | 149 | 0.00281132354561 | 0.232203647871025 | 0.43,0.339,0.321,0 | 0.445,0.597,0.416, | -0,137 |
| 184478 | ENSG00000077044 | DGKD | chr2 | + | 233389928 | 233390041 | 233388256 | 233388367 | 233390402 | 233390483 | 184478 | 6,3,35,17,8,9,7,10,18,8,1,5 | 63,43,251,97,46,59,79,124 | 10,6,4,12,4,2,11,8,1,3,2,7,5 | 103,107,94,62,89,45,38,57 | 262 | 149 | 0.00281150348394 | 0.232203647871025 | 0.051,0.038,0.073, | 0.052,0.031,0.024, | 0,019 |
| 61572 | ENSG00000048740 | CELF2 | chr10 | + | 11321206 | 11321386 | 11314138 | 11314258 | 11325835 | 11325979 | 61572 | 119,102,116,90,60,69,66,7 | 8,3,3,3,0,2,3,2,0,4,0,1 | 118,104,60,43,37,27,48,47 | 2,3,2,1,0,0,1,6,0,2,0,0,0,0,0 | 298 | 149 | 0.00281454668553 | 0.232365821892331 | 0.881,0.944,0.951, | 0.967,0.945,0.938, | -0,014 |
| 202073 | ENSG00000181852 | RNF41 | chr12 | - | 56212995 | 56213093 | 56210247 | 56210568 | 56213957 | 56214070 | 202073 | 21,17,63,29,6,15,13,12,8,1 | 38,81,146,71,50,52,34,59,2 | 52,69,10,9,22,12,23,27,18, | 77,161,20,7,68,27,71,65,21 | 247 | 149 | 0.00281832506403 | 0.232588543656534 | 0.25,0.112,0.207,0 | 0.289,0.205,0.232, | -0,073 |
| 179166 | ENSG00000126012 | KDM5C | chrX | - | 53210675 | 53210781 | 53208890 | 53210576 | 53211496 | 53211906 | 179166 | 44,40,112,82,37,61,42,77,4 | 1,7,13,9,2,7,1,3,0,1,0,2 | 46,58,49,106,76,45,72,57,3 | 2,6,8,19,19,5,9,6,7,5,6,5,7, | 255 | 149 | 0.00282036444233 | 0.232667634836002 | 0.963,0.77,0.834,0 | 0.931,0.85,0.782,0 | 0,069 |
| 202173 | ENSG00000153207 | AHCTF1 | chr1 | - | 246843794 | 246843869 | 246842693 | 246842776 | 246849614 | 246851442 | 202173 | 7,11,7,3,3,4,1,0,11,3,1,1 | 23,19,12,14,6,10,10,14,13, | 2,0,2,7,4,0,1,4,1,2,4,3,1,3,2 | 6,25,14,17,19,11,2,15,12,1 | 224 | 149 | 0.00282391578622 | 0.232804905276227 | 0.168,0.278,0.28,0 | 0.181,0.0,0.087,0.1 | 0,056 |
| 121531 | ENSG00000110075 | PPP6R3 | chr11 | + | 68583042 | 68583129 | 68575957 | 68576043 | 68587944 | 68588024 | 121531 | 34,21,99,49,32,38,27,49,18 | 0,1,4,1,2,2,2,0,2,0,0,0 | 69,61,30,30,28,11,33,37,27 | 5,0,2,2,4,2,0,0,3,2,2,8,3,2,2 | 236 | 149 | 0.00282419171809 | 0.232804905276227 | 1.0,0.93,0.94,0.96 | 0.897,1.0,0.904,0.9 | 0,041 |
| 36231 | ENSG00000155229 | MMS19 | chr10 | - | 97468965 | 97469104 | 97468251 | 97468406 | 97469645 | 97469723 | 36231 | 48,44,59,50,45,42,49,42,74 | 11,7,12,14,6,9,6,11,5,5,6,2 | 40,36,27,15,25,12,24,27,20 | 6,16,6,7,12,0,9,7,5,6,15,15 | 288 | 149 | 0.00282530900473 | 0.232807841652773 | 0.693,0.765,0.718, | 0.775,0.538,0.7,0.5 | 0,11 |
| 15788 | ENSG00000188559 | RALGAPA2 | chr20 | - | 20524405 | 20524543 | 20520916 | 20521100 | 20524829 | 20524898 | 15788 | 23,6,37,26,23,20,18,31,18, | 9,5,5,1,2,5,3,2,0,0,3,4 | 20,19,10,13,19,5,7,18,12,3 | 3,4,2,3,2,0,2,3,0,2,0,0,0,1,0 | 287 | 149 | 0.00283215611036 | 0.233282737003021 | 0.57,0.384,0.793,0 | 0.776,0.711,0.722, | -0,089 |
| 169723 | ENSG00000204463 | BAG6 | chr6 | - | 31647590 | 31647868 | 31646393 | 31646523 | 31648676 | 31648751 | 169723 | 117,73,93,84,52,38,57,32,5 | 7,5,8,1,4,5,7,7,3,3,1,1 | 39,42,15,50,30,27,63,44,21 | 3,5,6,12,5,4,5,2,3,2,4,13,5, | 298 | 149 | 0.00284348646346 | 0.234092687584215 | 0.893,0.88,0.853,0 | 0.867,0.808,0.556, | 0,089 |
| 64457 | ENSG00000115947 | ORC4 | chr2 | - | 147981927 | 147982081 | 147975901 | 147975975 | 148020607 | 148020760 | 64457 | 2,2,28,5,2,2,6,2,4,0,1,1 | 0,0,8,3,4,0,0,2,1,1,0,2 | 11,9,2,5,2,1,3,5,1,5,11,7,3, | 0,0,2,0,0,2,0,0,2,1,1,2,2,0,1 | 298 | 149 | 0.00284432502044 | 0.234092687584215 | 1.0,1.0,0.636,0.45 | 1.0,1.0,0.333,1.0,1 | -0,127 |
| 193594 | ENSG00000083720 | OXCT1 | chr5 | - | 41740464 | 41740558 | 41739389 | 41739491 | 41749526 | 41749607 | 193594 | 8,3,3,3,1,3,2,6,4,0,0,0 | 54,14,39,33,39,30,22,13,15 | 1,0,2,0,2,0,1,2,2,0,2,0,6,2,2 | 30,23,28,8,6,16,6,10,19,29 | 243 | 149 | 0.00284525217427 | 0.234092687584215 | 0.083,0.116,0.045, | 0.02,0.0,0.042,0.0, | 0,017 |
| 151889 | ENSG00000125107 | CNOT1 | chr16 | - | 58623257 | 58623409 | 58599373 | 58599511 | 58629727 | 58629847 | 151889 | 63,39,71,66,33,50,24,52,2, | 43,38,72,39,33,28,21,24,21 | 61,74,37,56,63,32,11,31,19 | 38,25,26,9,8,17,12,25,13,4 | 298 | 149 | 0.00285097612418 | 0.234473993956204 | 0.423,0.339,0.33,0 | 0.445,0.597,0.416, | -0,137 |
| 229330 | ENSG00000242588 | | chr7 | + | 128532123 | 128532338 | 128531706 | 128531800 | 128533644 | 128533730 | 229330 | 1,0,4,3,0,3,2,2,0,4,3,4 | 1,6,11,3,2,11,2,5,8,0,3,15 | 0,4,2,2,0,0,0,2,0,2,2,1,0,1,1 | 4,7,0,11,1,2,4,0,5,3,7,3,2,3 | 298 | 149 | 0.00285283037358 | 0.234536873237585 | 0.333,0.0,0.154,0.3 | 0.0,0.222,1.0,0.08 | 0,043 |
| 196518 | ENSG00000083845 | RPS5 | chr19 | + | 58393358 | 58393486 | 58388136 | 58388245 | 58394496 | 58394595 | 196518 | 5,7,4,2,0,4,2,2,2,4,0,3 | 21,18,27,34,4,5,10,12,2,4,1 | 2,1,5,0,2,3,2,1,0,0,0,9,0,3,6 | 21,29,13,5,13,7,14,11,9,18 | 277 | 149 | 0.00285715179201 | 0.234802457940475 | 0.114,0.173,0.074, | 0.049,0.018,0.171, | 0,075 |
| 106659 | ENSG00000005889 | ZFX | chrX | + | 24173606 | 24173723 | 24172714 | 24172800 | 24179182 | 24179770 | 106659 | 9,12,40,17,2,11,6,7,3,1,2,4 | 16,9,87,41,17,18,8,27,5,4,9 | 21,11,14,9,4,12,10,5,2,5,6, | 58,71,25,17,32,13,13,20,29 | 266 | 149 | 0.00285892268044 | 0.23485831581442 | 0.24,0.428,0.205,0 | 0.169,0.08,0.239,0 | 0,067 |
| 105826 | ENSG00000104870 | FCGRT | chr19 | + | 49525456 | 49525573 | 49514210 | 49514486 | 49526009 | 49526329 | 105826 | 167,74,126,117,111,45,110 | 6,0,1,2,6,2,0,3,3,2,2,5 | 101,60,37,37,31,29,56,178 | 13,2,9,2,2,1,2,4,1,4,4,4,1,2 | 266 | 149 | 0.00286050326506 | 0.234898503716951 | 0.94,1.0,0.986,0.97 | 0.813,0.944,0.697, | 0,04 |
| 192748 | ENSG00000110046 | ATG2A | chr11 | - | 64908990 | 64909145 | 64907747 | 64907890 | 64909270 | 64909367 | 192748 | 8,14,14,11,14,8,9,13,11,6,0 | 1,1,0,0,0,0,3,0,0,0,1,0 | 8,4,5,7,6,5,1,10,9,9,8,1,9,4 | 1,2,0,2,3,0,0,8,1,0,1,1,1,0,1 | 298 | 149 | 0.002867905852 | 0.235416569195962 | 0.8,0.875,1.0,1.0,1 | 0.8,0.5,1.0,0.636,0 | 0,131 |
| 91637 | ENSG00000196262 | PPIA | chr7 | + | 44799701 | 44799874 | 44799246 | 44799277 | 44801286 | 44801630 | 91637 | 1011,747,1474,1034,745,6 | 17,4,19,20,3,9,11,7,7,2,4,1 | 786,755,691,280,589,406,3 | 14,17,4,24,16,7,1,19,9,0,8, | 298 | 149 | 0.00287171900265 | 0.235503335024648 | 0.967,0.989,0.975, | 0.966,0.957,0.989, | 0,016 |
| 11142 | ENSG00000073536 | NLE1 | chr17 | - | 35137542 | 35137640 | 35137000 | 35137193 | 35137813 | 35137890 | 11142 | 7,2,20,12,4,6,6,3,11,4,2,10 | 0,0,4,5,0,0,0,0,0,0,2,2 | 6,12,7,0,9,3,2,11,1,2,4,20,9 | 2,0,0,1,1,0,2,2,1,0,2,4,3,2,2 | 247 | 149 | 0.00287197295025 | 0.235503335024648 | 1.0,1.0,0.751,0.59 | 0.644,1.0,1.0,0.0,0 | 0,155 |
| 20280 | ENSG00000138613 | APH1B | chr15 | + | 63302344 | 63302472 | 63279160 | 63279331 | 63305613 | 63305735 | 20280 | 13,16,82,40,32,16,20,46,21 | 0,1,5,1,0,0,2,7,1,1,1,0 | 43,54,30,20,19,19,32,17,21 | 3,2,3,6,3,0,5,4,2,1,9,0,0,2,1 | 277 | 149 | 0.00287224542208 | 0.235503335024648 | 1.0,0.896,0.898,0.9 | 0.885,0.936,0.843, | 0,053 |
| 201463 | ENSG00000115966 | ATF2 | chr2 | - | 175074999 | 175075363 | 175074504 | 175074835 | 175080659 | 175080765 | 201463 | 13,0,17,5,1,4,2,6,2,2,0,0 | 21,13,58,46,37,20,26,20,21 | 3,2,3,6,2,2,2,6,0,6,1,6,1,1,1 | 33,32,30,15,25,18,15,29,18 | 298 | 149 | 0.00288016675093 | 0.236062897931993 | 0.236,0.0,0.128,0.0 | 0.043,0.03,0.048,0 | 0,015 |
| 65039 | ENSG00000103657 | HERC1 | chr15 | - | 63635960 | 63636142 | 63634732 | 63634888 | 63637504 | 63637643 | 65039 | 34,40,65,49,35,32,26,30,35 | 10,4,8,1,1,1,5,8,0,0,2,4 | 54,66,17,23,38,20,39,43,25 | 8,13,6,10,10,1,13,9,8,2,3,1 | 298 | 149 | 0.0028817320266 | 0.236101281239872 | 0.63,0.833,0.802,0 | 0.771,0.717,0.586, | 0,115 |
| 47039 | ENSG00000116731 | PRDM2 | chr1 | + | 13715540 | 13715614 | 13700187 | 13700300 | 13730999 | 13731117 | 47039 | 13,9,33,18,15,18,4,15,2,4,2 | 2,4,0,0,0,2,10,0,0,0,3,1 | 19,24,10,15,17,13,8,17,8,8 | 0,1,1,1,2,0,0,0,1,1,2,3,0,0,2 | 223 | 149 | 0.00289329689523 | 0.23695859362909 | 0.813,0.601,1.0,1.0 | 1.0,0.941,0.87,0.9 | -0,099 |
| 218731 | ENSG00000196262 | PPIA | chr7 | + | 44799392 | 44799874 | 44799246 | 44799277 | 44801286 | 44801630 | 218731 | 1010,747,1474,1034,749,6 | 17,4,19,20,3,9,11,7,7,2,4,1 | 786,755,690,280,587,406,3 | 14,17,4,24,16,7,1,19,9,0,8, | 298 | 149 | 0.00289943177108 | 0.237370711114994 | 0.967,0.989,0.975, | 0.966,0.957,0.989, | 0,016 |
| 164043 | ENSG00000102445 | RUBCNL | chr13 | - | 46368443 | 46368627 | 46368041 | 46368249 | 46368732 | 46368815 | 164043 | 0,0,27,27,3,3,7,17,6,7,2,4 | 51,81,407,326,73,59,37,71 | 11,9,1,0,10,2,1,8,9,3,9,0,6, | 161,205,114,41,153,102,83 | 298 | 149 | 0.00290591180938 | 0.237793883024142 | 0.0,0.0,0.032,0.04, | 0.033,0.021,0.004, | 0,017 |
| 210579 | ENSG00000144566 | RAB5A | chr3 | + | 19983222 | 19983312 | 19978309 | 19978403 | 19983707 | 19985175 | 210579 | 8,10,5,5,0,0,2,1,2,0,0,3 | 36,58,50,39,27,17,17,21,31 | 7,10,1,8,1,4,7,5,8,1,3,6,7,8 | 33,65,37,35,33,9,15,22,17, | 239 | 149 | 0.00290681038622 | 0.237793883024142 | 0.122,0.097,0.059, | 0.117,0.088,0.017, | -0,068 |
| 124637 | ENSG00000135387 | CAPRIN1 | chr11 | + | 34091905 | 34092056 | 34090528 | 34090678 | 34096478 | 34096673 | 124637 | 251,188,223,189,140,125,8 | 17,8,25,11,4,13,16,14,7,7,6 | 192,245,169,70,138,93,57, | 23,24,18,15,19,7,2,19,9,9,1 | 298 | 149 | 0.0029099472267 | 0.237960049980953 | 0.881,0.922,0.817, | 0.807,0.836,0.824, | 0,049 |
| 184480 | ENSG00000077044 | DGKD | chr2 | + | 233389931 | 233390041 | 233388256 | 233388367 | 233390402 | 233390483 | 184480 | 5,1,22,8,5,5,3,5,2,2,1,3 | 63,43,251,97,46,59,79,124 | 4,1,3,6,2,2,5,6,2,0,1,5,2,1,4 | 103,107,94,62,89,45,38,57 | 259 | 149 | 0.00291183461003 | 0.238023955545525 | 0.044,0.013,0.048, | 0.022,0.005,0.018, | 0,008 |
| 121126 | ENSG00000168002 | POLR2G | chr11 | + | 62762541 | 62762742 | 62761794 | 62761904 | 62762866 | 62763026 | 121126 | 11,1,29,13,0,10,9,5,7,6,2,3 | 39,42,90,111,36,20,31,15,3 | 18,16,9,1,6,7,7,11,9,9,14,7 | 37,51,31,10,33,26,9,16,16, | 298 | 149 | 0.00291350348859 | 0.238034356818142 | 0.124,0.012,0.139, | 0.196,0.136,0.127, | -0,059 |
| 9287 | ENSG00000174231 | PRPF8 | chr17 | - | 1660992 | 1661162 | 1660697 | 1660827 | 1661270 | 1661329 | 9287 | 223,110,340,251,136,163,1 | 0,0,6,6,1,1,2,3,0,0,4,0 | 168,312,126,229,126,145,2 | 6,2,4,3,5,9,4,2,5,0,4,4,4,0,1 | 298 | 149 | 0.00291498162304 | 0.238034356818142 | 1.0,1.0,0.966,0.95 | 0.933,0.987,0.94,0 | 0,016 |

| | | | | | | | | | | | | | | | | | | | | | |
|---|---|---|---|---|---|---|---|---|---|---|---|---|---|---|---|---|---|---|---|---|---|
| 167072 | ENSG00000174652 | ZNF266 | chr19 | - | 9418765 | 9419163 | 9418504 | 9418631 | 9420064 | 9420218 | 167072 | 27,23,80,51,30,6,42,66,33, | 4,0,3,4,0,0,3,2,0,0,2,5 | 77,35,40,50,28,27,24,50,3 | 1,3,0,2,3,0,0,0,0,0,6,0,0,0, | 298 | 149 | 0.0029152796975 | 0.238034356818142 | 0.771,1.0,0.93,0.8 | 0.975,0.854,1.0,0. | -0,093 |
| 167071 | ENSG00000174652 | ZNF266 | chr19 | - | 9418765 | 9419143 | 9418504 | 9418631 | 9420064 | 9420218 | 167071 | 27,23,80,51,30,6,42,66,33, | 4,0,3,4,0,0,3,2,0,0,2,5 | 77,35,40,50,28,27,24,50,3 | 1,3,0,2,3,0,0,0,0,0,6,0,0,0, | 298 | 149 | 0.00291646974545 | 0.238041221002256 | 0.771,1.0,0.93,0.8 | 0.975,0.854,1.0,0. | -0,093 |
| 253445 | ENSG00000092531 | SNAP23 | chr15 | + | 42513173 | 42513247 | 42512954 | 42512996 | 42513398 | 42513447 | 253445 | 24,6,13,8,2,5,8,2,3,0,0,3 | 87,73,139,168,74,38,35,52 | 17,14,11,7,6,1,0,11,4,8,5,7 | 79,70,49,17,30,43,9,34,24, | 223 | 149 | 0.00291881989695 | 0.238142731326932 | 0.156,0.052,0.059, | 0.126,0.118,0.13,0 | -0,04 |
| 80674 | ENSG00000109466 | KLHL2 | chr4 | + | 165305607 | 165305725 | 165299506 | 165299656 | 165310552 | 165310750 | 80674 | 35,19,47,28,12,23,20,25,1 | 1,5,4,0,1,6,2,0,0,1,0,0 | 25,21,10,10,32,6,14,31,3,1 | 0,1,0,0,0,0,2,2,1,0,0,2,0,1, | 267 | 149 | 0.00292100206889 | 0.238230464679524 | 0.951,0.68,0.868,1 | 1.0,0.921,1.0,1.0,1 | -0,021 |
| 6120 | ENSG00000286235 | | chr20 | + | 6015360 | 6015490 | 6009774 | 6009912 | 6031284 | 6031370 | 6120 | 33,16,33,48,18,12,23,15,3 | 3,0,6,3,0,2,2,2,0,0,0,2 | 36,17,9,0,10,23,5,24,5,24, | 2,0,0,2,0,0,0,0,2,0,0,1,1,0, | 279 | 149 | 0.00292681839459 | 0.238614412835606 | 0.855,1.0,0.746,0. | 0.906,1.0,1.0,0.0,1 | 0,003 |
| 25691 | ENSG00000136758 | YME1L1 | chr10 | - | 27142386 | 27142485 | 27136275 | 27136385 | 27145427 | 27145590 | 25691 | 99,75,232,161,99,82,93,10 | 12,2,10,3,3,6,1,3,0,0,0,4 | 145,186,90,59,93,55,65,13 | 4,4,2,27,20,3,2,7,6,9,8,2,4, | 248 | 149 | 0.00292939759309 | 0.238734257235272 | 0.832,0.958,0.933, | 0.956,0.965,0.964, | 0,057 |
| 213749 | ENSG00000116793 | PHTF1 | chr1 | - | 113757991 | 113758083 | 113757698 | 113757755 | 113758658 | 113759489 | 213749 | 2,0,4,10,0,4,0,3,0,0,2,1 | 18,13,71,41,35,31,33,22,2 | 1,1,2,0,0,0,0,0,0,0,1,0,0, | 38,46,15,38,23,15,14,25,1 | 241 | 149 | 0.00293110353158 | 0.238782870630392 | 0.064,0.0,0.034,0. | 0.016,0.013,0.076, | 0,038 |
| 234979 | ENSG00000147905 | ZCCHC7 | chr9 | + | 37186711 | 37186774 | 37126311 | 37126942 | 37302187 | 37302231 | 234979 | 7,4,10,3,4,2,6,2,1,2,0,3 | 84,43,152,67,88,72,38,60, | 1,2,4,4,5,0,2,0,1,1,2,1,1,3, | 151,54,120,29,22,61,33,59 | 212 | 149 | 0.00293608749773 | 0.23900405594556 | 0.055,0.061,0.044, | 0.005,0.025,0.023, | 0,005 |
| 199598 | ENSG00000159658 | EFCAB14 | chr1 | - | 46692090 | 46692191 | 46691826 | 46691937 | 46696550 | 46696649 | 199598 | 6,1,3,0,1,2,2,0,0,0,2,0 | 78,74,159,107,61,40,56,66 | 6,4,3,13,2,1,1,3,0,2,3,3,1,1 | 81,59,68,30,32,28,37,69,2 | 250 | 149 | 0.00293614951416 | 0.23900405594556 | 0.044,0.008,0.011, | 0.042,0.039,0.026, | -0,018 |
| 28926 | ENSG00000132842 | AP3B1 | chr5 | - | 78121711 | 78121873 | 78116125 | 78116234 | 78128029 | 78128160 | 28926 | 7,1,3,0,1,1,2,0,2,5,2,2 | 50,29,48,37,22,20,24,15,3 | 0,0,0,3,0,0,0,2,0,0,2,2,0,3, | 24,34,17,32,15,12,11,26,1 | 298 | 149 | 0.00293714998293 | 0.23900405594556 | 0.065,0.017,0.03,0 | 0.0,0.0,0.0,0.045,0 | 0,017 |
| 234085 | ENSG00000068308 | OTUD5 | chrX | - | 48936190 | 48936361 | 48934953 | 48935018 | 48944189 | 48944283 | 234085 | 31,11,17,13,4,2,10,9,6,3,1, | 169,118,125,76,49,64,33,5 | 17,24,11,76,4,5,7,8,4,3,8,2 | 57,65,65,31,53,46,35,54,3 | 298 | 149 | 0.00293980536336 | 0.239129723416983 | 0.084,0.045,0.064, | 0.13,0.156,0.078,0 | -0,048 |
| 249514 | ENSG00000163683 | SMIM14 | chr4 | - | 39556430 | 39556570 | 39551838 | 39552158 | 39572414 | 39572463 | 249514 | 87,53,179,228,88,85,44,55 | 5,2,3,3,2,1,0,4,1,0,0,2 | 119,74,81,10,47,54,24,68, | 1,3,2,2,1,0,0,0,4,6,1,27,6,4 | 289 | 149 | 0.00294791398561 | 0.23969870609627 | 0.9,0.932,0.969,0. | 0.984,0.927,0.954, | 0,026 |
| 234086 | ENSG00000068308 | OTUD5 | chrX | - | 48936190 | 48936379 | 48934953 | 48935018 | 48944189 | 48944283 | 234086 | 30,11,18,13,4,2,10,9,6,3,1, | 169,118,125,76,49,64,33,5 | 17,24,11,76,4,5,7,8,4,3,7,2 | 57,65,65,31,53,46,35,54,3 | 298 | 149 | 0.00296218317689 | 0.240749190088876 | 0.082,0.045,0.067, | 0.13,0.156,0.078,0 | -0,048 |
| 42700 | ENSG00000168297 | PXK | chr3 | + | 58391772 | 58391847 | 58382513 | 58382700 | 58394997 | 58395102 | 42700 | 18,13,52,34,6,8,15,12,10,2 | 8,0,2,2,0,2,4,0,1,1,1,0 | 18,12,14,2,17,15,5,10,16,1 | 1,0,1,8,2,0,0,0,0,1,0,8,0,0, | 224 | 149 | 0.00296380202391 | 0.240749190088876 | 0.599,1.0,0.945,0. | 0.923,1.0,0.903,0. | -0,057 |
| 227017 | ENSG00000223482 | NUTM2A-AS1 | chr10 | - | 87241181 | 87245855 | 87240175 | 87240531 | 87246114 | 87246212 | 227017 | 8,3,1,1,12,7,3,8,7,4,1,4 | 1,0,2,0,1,0,0,3,0,0,0,0 | 1,2,0,3,3,4,2,5,5,4,0,1,1,7, | 3,2,1,0,0,0,0,0,0,1,3,3,2,2, | 298 | 149 | 0.00296418895854 | 0.240749190088876 | 0.8,1.0,0.2,1.0,0.8 | 0.143,0.333,0.0,1. | 0,236 |
| 12688 | ENSG00000108848 | LUC7L3 | chr17 | + | 50736959 | 50737026 | 50719564 | 50719831 | 50740305 | 50740345 | 12688 | 72,47,191,109,131,55,96,1 | 7,5,8,7,0,3,3,3,1,3,0,0 | 105,102,51,44,75,68,44,64 | 6,1,6,8,9,1,2,6,1,5,7,7,4,12 | 216 | 149 | 0.00297130125433 | 0.241200463834004 | 0.876,0.866,0.943, | 0.923,0.986,0.854, | 0,032 |
| 68292 | ENSG00000198836 | OPA1 | chr3 | + | 193628508 | 193628616 | 193618868 | 193618936 | 193631611 | 193631665 | 68292 | 1,0,5,3,2,1,0,2,4,1,0,2 | 15,5,38,32,15,9,13,17,4,5, | 5,5,4,2,3,0,2,3,2,6,3,8,2,4, | 36,29,13,3,18,8,11,7,11,22 | 257 | 149 | 0.00297198651722 | 0.241200463834004 | 0.037,0.0,0.071,0. | 0.075,0.091,0.151, | -0,053 |
| 912 | ENSG00000100201 | DDX17 | chr22 | - | 38489227 | 38489458 | 38488025 | 38488115 | 38492055 | 38492115 | 912 | 11,14,14,8,11,6,12,7,17,3, | 223,109,242,214,130,97,1 | 13,14,13,10,0,2,6,9,7,12,5, | 123,52,139,40,64,84,60,14 | 298 | 149 | 0.00297457448996 | 0.241319503222232 | 0.024,0.06,0.028,0 | 0.05,0.119,0.045,0 | -0,016 |
| 67135 | ENSG00000229474 | PATL2 | chr15 | - | 44668341 | 44668482 | 44667105 | 44667203 | 44668979 | 44669139 | 67135 | 16,12,33,58,29,13,28,17,1 | 1,0,1,1,2,5,2,0,3,0,0,0 | 11,23,12,18,31,19,4,18,24, | 2,1,0,1,1,1,0,2,1,3,0,0,3,5, | 290 | 149 | 0.00297947123829 | 0.241625687297813 | 0.892,1.0,0.944,0. | 0.739,0.922,1.0,0. | 0,012 |
| 225403 | ENSG00000006634 | DBF4 | chr7 | + | 87886831 | 87886894 | 87884978 | 87885158 | 87887328 | 87887398 | 225403 | 10,8,5,9,1,5,5,1,2,1,2,5 | 3,0,3,3,0,0,6,1,2,0,1,1 | 6,5,5,4,6,2,0,1,0,4,6,2,1,1, | 5,5,1,7,6,2,1,5,3,2,5,5,1,0, | 212 | 149 | 0.00298170577799 | 0.241715825349366 | 0.701,1.0,0.539,0. | 0.458,0.413,0.778, | 0,249 |
| 150074 | ENSG00000159322 | ADPGK | chr15 | - | 72780196 | 72780272 | 72774871 | 72775097 | 72783458 | 72783785 | 150074 | 2,0,6,2,1,3,3,1,1,1,0,5 | 16,7,22,17,9,10,12,2,9,10, | 4,13,1,2,2,0,1,4,2,2,3,3,3,1 | 10,14,3,1,11,3,6,4,2,7,4,11 | 225 | 149 | 0.00299940114025 | 0.243058775157059 | 0.076,0.0,0.153,0. | 0.209,0.381,0.181, | -0,118 |
| 103673 | ENSG00000150787 | PTS | chr11 | + | 112230207 | 112230230 | 112228593 | 112228673 | 112230625 | 112230682 | 103673 | 2,1,10,0,2,0,1,1,1,1,0,0 | 2,9,38,24,2,4,8,5,3,2,2,5 | 3,4,2,3,5,2,0,1,2,1,2,6,3,0, | 14,23,6,14,19,5,3,8,10,11, | 172 | 149 | 0.00300334141943 | 0.243286479881573 | 0.464,0.088,0.186, | 0.157,0.131,0.224, | 0,024 |
| 47598 | ENSG00000135148 | TRAFD1 | chr12 | + | 112142088 | 112142295 | 112140818 | 112141224 | 112145585 | 112145662 | 47598 | 23,23,87,84,40,30,15,22,4 | 2,0,2,4,3,4,2,1,1,2,1,1 | 58,62,26,8,30,25,16,28,15, | 1,3,0,0,1,1,1,4,2,0,3,1,0,0, | 298 | 149 | 0.00300596073263 | 0.24340704832381 | 0.852,1.0,0.956,0. | 0.967,0.912,1.0,1. | -0,043 |
| 50631 | ENSG00000152223 | EPG5 | chr18 | - | 45916437 | 45916582 | 45916008 | 45916206 | 45923267 | 45923387 | 50631 | 5,11,17,10,6,13,7,6,3,5,5,5 | 1,0,1,0,0,0,0,0,0,0,0,1 | 23,19,13,16,9,8,6,9,4,10,1 | 3,1,0,1,1,2,0,0,2,2,0,2,2,3, | 294 | 149 | 0.0030099321579 | 0.243541983190567 | 0.717,1.0,0.896,1. | 0.795,0.906,1.0,0. | 0,134 |
| 244950 | ENSG00000107036 | RIC1 | chr9 | + | 5710415 | 5710474 | 5689958 | 5690038 | 5713895 | 5714003 | 244950 | 2,0,3,6,1,1,1,1,1,1,1,3 | 12,1,45,33,35,29,5,34,7,4, | 8,11,3,6,1,3,1,4,0,0,4,7,3,2 | 27,26,6,9,40,15,17,21,20,2 | 208 | 149 | 0.00301058868478 | 0.243541983190567 | 0.107,0.0,0.046,0. | 0.175,0.233,0.264, | -0,043 |
| 198041 | ENSG00000112232 | KHDRBS2 | chr6 | - | 61939411 | 61939528 | 61901243 | 61901371 | 61978065 | 61978212 | 198041 | 4,2,3,2,0,3,10,11,5,1,5,11 | 57,30,75,37,29,39,20,30,7 | 10,9,2,2,12,4,6,7,3,1,13,7, | 55,32,33,33,11,8,14,28,13, | 266 | 149 | 0.0030114746254 | 0.243541983190567 | 0.038,0.036,0.022, | 0.092,0.136,0.033, | -0,039 |
| 196517 | ENSG00000083845 | RPS5 | chr19 | + | 58393358 | 58393485 | 58388136 | 58388245 | 58394496 | 58394595 | 196517 | 5,7,4,2,0,4,2,2,2,4,0,3 | 21,18,27,34,4,5,10,12,2,4, | 2,1,3,0,2,3,2,1,0,0,2,9,0,3, | 21,29,13,5,13,7,14,11,9,18 | 276 | 149 | 0.00301322802546 | 0.243541983190567 | 0.114,0.174,0.074, | 0.049,0.018,0.111, | 0,075 |
| 135202 | ENSG00000100150 | DEPDC5 | chr22 | + | 31815415 | 31815806 | 31814991 | 31815212 | 31819021 | 31819225 | 135202 | 24,3,9,7,8,6,0,0,16,15,2,8 | 32,48,74,42,29,32,43,21,3 | 0,0,2,5,10,6,1,6,6,10,1,4,2, | 52,41,12,13,27,24,11,23,4 | 298 | 149 | 0.00301328480208 | 0.243541983190567 | 0.273,0.03,0.057,0 | 0.0,0.0,0.077,0.16 | 0,023 |
| 92919 | ENSG00000198315 | ZKSCAN8 | chr6 | + | 28148315 | 28148838 | 28141909 | 28142029 | 28149482 | 28149624 | 92919 | 18,14,28,23,19,22,12,18,1 | 4,3,3,5,0,3,2,5,0,1,0,2 | 26,18,9,26,12,7,20,13,7,16 | 4,8,12,3,7,2,6,2,3,0,4,2,0,3 | 298 | 149 | 0.00301794046295 | 0.24382670562357 | 0.692,0.7,0.824,0. | 0.765,0.529,0.273, | 0,162 |
| 227309 | ENSG00000013374 | NUB1 | chr7 | + | 151374096 | 151374158 | 151368734 | 151368887 | 151375847 | 151375943 | 227309 | 16,21,20,19,8,15,4,9,26,15 | 4,0,1,3,2,2,2,8,0,0,0,0 | 14,7,12,6,2,2,0,6,6,9,12,19 | 1,4,0,0,0,0,1,5,1,0,0,1,0,2, | 211 | 149 | 0.00302739973701 | 0.244499164276322 | 0.739,1.0,0.934,0. | 0.908,0.553,1.0,1. | 0,009 |
| 56692 | ENSG00000085465 | OVGP1 | chr1 | - | 111416322 | 111416458 | 111414318 | 111415344 | 111419609 | 111419726 | 56692 | 30,23,16,15,30,13,25,18,7, | 5,0,2,1,0,2,0,1,0,3,1,0 | 24,32,10,5,33,15,12,13,8,1 | 5,4,2,3,2,0,0,1,0,2,0,7,1,2, | 285 | 149 | 0.00302942875295 | 0.24457126028739 | 0.758,1.0,0.807,0. | 0.715,0.807,0.723, | 0,074 |
| 185236 | ENSG00000132185 | FCRLA | chr1 | + | 161711318 | 161711474 | 161710759 | 161710912 | 161711933 | 161712218 | 185236 | 165,144,454,409,375,191, | 12,5,31,20,13,0,13,16,14,9 | 449,310,97,48,101,131,14 | 36,37,3,5,12,5,15,17,0,11, | 298 | 149 | 0.0030307762781 | 0.244588304878793 | 0.873,0.935,0.88,0 | 0.862,0.807,0.942, | 0,029 |
| 137219 | ENSG00000130303 | BST2 | chr19 | - | 17404370 | 17404516 | 17404128 | 17404189 | 17405290 | 17405630 | 137219 | 12,6,69,56,16,26,15,12,6,1 | 0,0,5,1,0,0,0,2,2,1,0,2 | 22,22,7,16,6,16,6,25,3,15, | 4,3,1,3,0,0,0,3,0,0,3,6,2,1, | 295 | 149 | 0.00303267431225 | 0.244649746963973 | 1.0,1.0,0.875,0.96 | 0.735,0.787,0.78,0 | 0,035 |
| 179167 | ENSG00000126012 | KDM5C | chrX | - | 53210675 | 53210806 | 53208890 | 53210576 | 53211496 | 53211906 | 179167 | 60,43,130,96,47,70,53,87, | 1,7,13,9,2,7,1,3,0,1,0,2 | 54,70,59,131,100,56,79,68 | 2,6,8,19,19,5,9,6,7,5,6,5,7, | 280 | 149 | 0.00303757788086 | 0.244863360798877 | 0.97,0.766,0.842,0 | 0.935,0.861,0.797, | 0,06 |
| 16240 | ENSG00000103472 | RRN3P2 | chr16 | + | 29084442 | 29084532 | 29077900 | 29077957 | 29084638 | 29084768 | 16240 | 28,13,74,33,19,48,40,51,1 | 2,0,4,2,0,3,0,6,0,0,0,2 | 28,41,21,59,34,23,23,26,3 | 2,2,0,1,0,0,0,3,2,0,1,1,0,0, | 239 | 149 | 0.00303759761992 | 0.244863360798877 | 0.897,1.0,0.92,0.9 | 0.897,0.927,1.0,0. | -0,025 |
| 76670 | ENSG00000159023 | EPB41 | chr1 | + | 28997214 | 28997319 | 28993329 | 28993542 | 29011864 | 29011907 | 76670 | 208,131,431,239,217,192, | 2,2,4,11,3,0,3,15,2,0,0,4 | 224,209,207,236,211,191, | 11,8,5,5,13,5,5,2,9,10,3,0, | 254 | 149 | 0.00304190337177 | 0.24511864642809 | 0.984,0.975,0.984, | 0.923,0.939,0.96,0 | 0,021 |
| 20201 | ENSG00000166128 | RAB8B | chr15 | + | 63223041 | 63223073 | 63189559 | 63189748 | 63244755 | 63244816 | 20201 | 9,8,10,11,9,8,11,7,8,3,1,7 | 97,123,73,68,59,32,51,42, | 19,15,14,7,7,1,8,6,6,1,6,20 | 75,72,38,40,44,21,15,26,1 | 181 | 149 | 0.00304664640111 | 0.245328501849899 | 0.071,0.051,0.101, | 0.173,0.146,0.233, | -0,051 |
| 109020 | ENSG00000198947 | DMD | chrX | - | 31247693 | 31247768 | 31223046 | 31223121 | 31260954 | 31261016 | 109020 | 16,10,13,9,0,13,16,8,0,3,2, | 127,52,147,110,50,50,51,8 | 0,4,6,8,5,2,5,8,4,1,8,4,0,2, | 7,4,14,26,8,2,40,36,4,2,50, | 224 | 149 | 0.00304713102663 | 0.245328501849899 | 0.077,0.113,0.056, | 0.0,0.399,0.222,0. | -0,126 |
| 11378 | ENSG00000213339 | QTRT1 | chr19 | + | 10706709 | 10706825 | 10702115 | 10702254 | 10707301 | 10707380 | 11378 | 5,1,7,2,10,1,4,5,3,0,2,7 | 53,11,54,35,23,28,22,34,1 | 0,2,1,1,5,0,1,4,2,6,4,0,0,4, | 23,37,33,26,14,7,24,23,8,2 | 265 | 149 | 0.00304792717567 | 0.245328501849899 | 0.05,0.049,0.068,0 | 0.0,0.029,0.017,0. | 0,024 |
| 88895 | ENSG00000073849 | ST6GAL1 | chr3 | + | 187038391 | 187038491 | 187021854 | 187022004 | 187038741 | 187038873 | 88895 | 3,7,25,19,4,11,9,10,6,1,5,1 | 79,106,124,65,70,51,23,25 | 13,20,10,18,10,8,0,12,6,8, | 81,112,9,35,120,33,14,19, | 249 | 149 | 0.00305826698926 | 0.246068733594549 | 0.022,0.038,0.108, | 0.088,0.097,0.399, | -0,09 |
| 108002 | ENSG00000128731 | HERC2 | chr15 | - | 28163093 | 28163285 | 28152676 | 28152830 | 28167686 | 28167827 | 108002 | 10,8,22,19,36,30,5,6,1,7,5, | 0,0,2,2,2,10,0,1,0,0,0,2 | 5,12,2,3,10,3,3,6,8,21,12,3 | 0,1,2,0,2,0,0,2,0,0,1,0,0,0, | 298 | 149 | 0.00306842807477 | 0.246794036980875 | 1.0,1.0,0.846,0.82 | 1.0,0.857,0.333,1. | 0,018 |
| 70825 | ENSG00000047410 | TPR | chr1 | - | 186353687 | 186353850 | 186351975 | 186352110 | 186355409 | 186355558 | 70825 | 422,375,368,167,187,198, | 19,3,6,3,0,3,4,0,3,6,0,6 | 233,245,236,223,157,82,8 | 1,6,2,8,0,2,3,3,0,2,0,5,1,1, | 298 | 149 | 0.00307064317172 | 0.246820221254958 | 0.917,0.984,0.968, | 0.991,0.953,0.983, | -0,016 |
| 155899 | ENSG00000196199 | MPHOSPH8 | chr13 | + | 19661962 | 19662081 | 19661697 | 19661838 | 19663039 | 19663126 | 155899 | 8,0,13,4,2,4,2,1,0,2,2,5 | 26,18,91,52,45,23,25,30,4 | 2,2,3,4,5,2,0,0,0,0,3,1,4,2, | 67,48,18,24,13,7,28,62,22, | 268 | 149 | 0.0030710471657 | 0.246820221254958 | 0.146,0.0,0.074,0. | 0.016,0.023,0.085, | 0,009 |
| 202875 | ENSG00000089127 | OAS1 | chr12 | + | 112917479 | 112917700 | 112916508 | 112916738 | 112919486 | 112919615 | 202875 | 9,8,11,12,14,4,5,0,2,4,1,5 | 3,1,1,5,3,1,3,1,0,0,1,0 | 2,4,5,5,2,2,4,0,2,4,2,4,12,1 | 2,0,0,0,1,0,0,1,2,2,0,3,2,2, | 298 | 149 | 0.00307400594786 | 0.246965798493414 | 0.6,0.8,0.846,0.54 | 0.333,1.0,1.0,1.0,0 | -0,062 |
| 157824 | ENSG00000255073 | ZFP91-CNTF | chr11 | + | 58616018 | 58616130 | 58614228 | 58614343 | 58616715 | 58616815 | 157824 | 4,0,7,3,0,0,2,6,6,0,0,1 | 39,47,78,69,19,30,36,34,2 | 2,0,0,1,0,1,0,2,0,0,8,2,0,0, | 51,60,36,41,36,22,16,29,2 | 261 | 149 | 0.00307675958653 | 0.247094792003149 | 0.055,0.0,0.049,0. | 0.022,0.0,0.0,0.01 | 0,019 |
| 167064 | ENSG00000174652 | ZNF266 | chr19 | - | 9418761 | 9419163 | 9418504 | 9418631 | 9420064 | 9420218 | 167064 | 16,8,26,42,15,7,16,15,17,1 | 4,0,3,4,0,0,3,2,0,0,2,5 | 34,18,9,15,5,7,7,24,17,47, | 1,3,0,2,3,0,0,0,0,0,6,0,0,0, | 298 | 149 | 0.00307832445907 | 0.247123892824736 | 0.667,1.0,0.812,0. | 0.944,0.75,1.0,0.7 | -0,104 |
| 36107 | ENSG00000174652 | ZNF266 | chr19 | - | 9418761 | 9418895 | 9418504 | 9418631 | 9420064 | 9420218 | 36107 | 16,8,25,41,15,6,15,15,17,1 | 4,0,3,4,0,0,3,2,0,0,2,5 | 34,18,9,16,5,7,7,24,17,44, | 1,3,0,2,3,0,0,0,0,0,6,0,0,0, | 283 | 149 | 0.00308050293737 | 0.247123892824736 | 0.678,1.0,0.814,0. | 0.947,0.76,1.0,0.8 | -0,105 |
| 10778 | ENSG00000108389 | MTMR4 | chr17 | - | 58514991 | 58515083 | 58512851 | 58512941 | 58516579 | 58516618 | 10778 | 2,0,5,3,0,4,0,0,0,2,3,0 | 3,6,12,3,7,4,2,5,1,2,3,11 | 2,1,2,0,0,3,0,0,0,1,0,0,1,0, | 2,10,5,3,1,2,4,10,4,5,10,11 | 241 | 149 | 0.00308156214555 | 0.247123892824736 | 0.292,0.0,0.205,0. | 0.382,0.058,0.198, | 0,08 |
| 167063 | ENSG00000174652 | ZNF266 | chr19 | - | 9418761 | 9419143 | 9418504 | 9418631 | 9420064 | 9420218 | 167063 | 16,8,26,42,15,7,16,15,17,1 | 4,0,3,4,0,0,3,2,0,0,2,5 | 34,18,9,15,5,7,7,24,17,46, | 1,3,0,2,3,0,0,0,0,0,6,0,0,0, | 298 | 149 | 0.00308171466165 | 0.247123892824736 | 0.667,1.0,0.812,0. | 0.944,0.75,1.0,0.7 | -0,104 |
| 219969 | ENSG00000111670 | GNPTAB | chr12 | - | 101789164 | 101789383 | 101788547 | 101788589 | 101789937 | 101790057 | 219969 | 1,3,1,3,1,2,3,34,4,0,1,0 | 35,14,97,135,112,100,74,1 | 1,3,1,4,0,0,1,6,4,1,5,4,1,1, | 79,82,16,15,45,33,46,145, | 298 | 149 | 0.00308427621289 | 0.247237189413977 | 0.014,0.097,0.005, | 0.006,0.018,0.03,0 | 0,004 |
| 155593 | ENSG00000126226 | PCID2 | chr13 | - | 113190867 | 113190975 | 113185484 | 113185560 | 113195070 | 113195125 | 155593 | 22,11,31,27,17,13,15,18,1 | 6,1,7,6,2,0,2,6,0,0,0,0 | 9,10,5,7,9,6,7,13,3,13,27,1 | 1,1,0,2,4,0,0,3,0,2,0,0,2,0, | 257 | 149 | 0.00308855965576 | 0.247488377985436 | 0.68,0.864,0.72,0. | 0.839,0.853,1.0,0. | 0,027 |
| 232080 | ENSG00000104870 | FCGRT | chr19 | + | 49525440 | 49525573 | 49514210 | 49514486 | 49526009 | 49526329 | 232080 | 163,74,127,115,111,41,10 | 6,0,1,2,6,2,0,3,3,2,2,5 | 100,60,37,37,31,29,58,178 | 13,2,9,2,2,1,2,4,1,4,4,4,1,2 | 282 | 149 | 0.00309751928459 | 0.248113946089166 | 0.935,1.0,0.985,0. | 0.803,0.941,0.685, | 0,042 |
| 109313 | ENSG00000167862 | MRPL58 | chr17 | + | 75018585 | 75018694 | 75017077 | 75017114 | 75019699 | 75019759 | 109313 | 5,1,9,3,0,1,1,3,2,1,1,2 | 8,5,16,5,5,5,2,3,17,2,1,0 | 3,2,0,0,1,0,1,1,2,2,2,3,1,1, | 11,9,2,4,9,4,2,10,2,8,7,17, | 258 | 149 | 0.00309975863495 | 0.248200948943052 | 0.265,0.104,0.245, | 0.136,0.114,0.0,0. | 0,168 |
| 41272 | ENSG00000198843 | SELENOT | chr3 | + | 150618708 | 150618803 | 150603327 | 150603499 | 150622384 | 150622495 | 41272 | 6,9,5,5,5,4,5,5,3,4,2,0 | 52,40,92,80,42,47,34,41,7 | 10,9,13,6,8,2,3,3,3,4,8,12, | 54,68,48,20,45,23,22,50,1 | 244 | 149 | 0.00311007934514 | 0.248934729465908 | 0.066,0.121,0.032, | 0.102,0.075,0.142, | -0,028 |
| 207894 | ENSG00000123240 | OPTN | chr10 | + | 13124045 | 13124110 | 13122384 | 13122487 | 13125417 | 13125567 | 207894 | 7,4,36,21,16,4,12,12,4,3,3, | 5,2,1,0,2,0,0,4,0,0,0,1 | 7,23,11,10,4,5,8,14,4,15,2, | 4,3,1,8,3,0,0,1,2,1,4,6,2,0, | 214 | 149 | 0.00311343288576 | 0.249110510570636 | 0.494,0.582,0.962, | 0.549,0.842,0.885, | 0,137 |
| 245728 | ENSG00000134809 | TIMM10 | chr11 | - | 57530344 | 57530538 | 57530118 | 57530234 | 57530657 | 57530754 | 245728 | 0,0,2,0,0,0,0,2,3,0,0,1 | 9,19,31,22,10,14,8,13,7,4, | 0,0,1,1,2,2,5,3,7,8,0,0,0,1, | 10,33,14,5,9,5,5,8,7,9,12,4 | 298 | 149 | 0.00311501953014 | 0.249144841505597 | 0.0,0.0,0.031,0.0,0 | 0.0,0.0,0.034,0.09 | -0,072 |
| 37214 | ENSG00000155189 | AGPAT5 | chr8 | + | 6741660 | 6741751 | 6732560 | 6732650 | 6747669 | 6747828 | 37214 | 68,43,104,103,135,63,53,5 | 7,2,7,5,7,4,2,4,0,0,1,8 | 115,113,32,7,18,41,41,71, | 7,15,7,0,7,8,4,2,7,6,4,2,6,3 | 240 | 149 | 0.00311919778773 | 0.249386351801975 | 0.858,0.93,0.902,0 | 0.911,0.824,0.739, | 0,076 |
| 66798 | ENSG00000159377 | PSMB4 | chr1 | + | 151400763 | 151400845 | 151400441 | 151400588 | 151401238 | 151401355 | 66798 | 127,103,269,322,173,151, | 3,2,4,9,7,5,1,0,0,0,2,2 | 216,175,158,44,140,105,8 | 3,9,2,15,1,1,4,9,1,0,8,8,5,2 | 231 | 149 | 0.00314629842274 | 0.251459693956462 | 0.965,0.971,0.977, | 0.979,0.926,0.981, | 0,035 |
| 22670 | ENSG00000100888 | CHD8 | chr14 | - | 21431810 | 21431858 | 21428963 | 21429335 | 21437193 | 21437245 | 22670 | 5,3,8,8,3,3,2,2,4,3,0,2 | 0,0,2,0,1,1,2,0,0,3,1,0 | 4,13,4,3,5,2,4,6,3,1,1,2,8,4 | 0,6,3,1,0,0,0,2,0,0,1,2,0,2, | 197 | 149 | 0.00315050463613 | 0.25152126426307 | 1.0,1.0,0.752,1.0,0 | 1.0,0.621,0.502,0. | 0,012 |
| 141045 | ENSG00000186814 | ZSCAN30 | chr18 | - | 35263475 | 35263657 | 35251067 | 35254381 | 35263944 | 35264046 | 141045 | 10,3,20,15,12,4,12,8,7,1,5, | 0,0,8,1,1,0,2,3,3,0,0,0 | 8,7,1,5,2,2,2,14,5,8,1,6,17, | 3,1,1,0,1,1,0,0,0,1,0,0,1,0, | 298 | 149 | 0.00315167069459 | 0.25152126426307 | 1.0,1.0,0.556,0.88 | 0.571,0.778,0.333, | 0,028 |
| 27198 | ENSG00000130449 | ZSWIM6 | chr5 | + | 61539595 | 61539759 | 61538813 | 61538971 | 61541883 | 61541965 | 27198 | 4,6,5,15,5,2,5,0,10,1,1,4 | 3,0,1,4,2,2,0,1,0,3,2,0 | 4,3,7,5,1,6,4,4,6,2,9,6,10,5 | 1,2,0,2,0,0,0,2,2,2,1,0,2,1, | 298 | 149 | 0.0031528375548 | 0.25152126426307 | 0.4,1.0,0.714,0.65 | 0.667,0.429,1.0,0. | -0,188 |
| 117286 | ENSG00000148396 | SEC16A | chr9 | - | 136478708 | 136478830 | 136474048 | 136477684 | 136482937 | 136483040 | 117286 | 1,0,0,2,2,0,0,0,0,0,0,2 | 26,12,28,15,17,17,7,19,26, | 3,3,4,6,0,3,5,0,0,1,0,3,0,0, | 8,14,8,54,15,5,14,12,12,9, | 271 | 149 | 0.00315341348772 | 0.25152126426307 | 0.021,0.0,0.0,0.06 | 0.171,0.105,0.216, | -0,056 |
| 252274 | ENSG00000066651 | TRMT11 | chr6 | + | 126008625 | 126008745 | 126008391 | 126008472 | 126011252 | 126011274 | 252274 | 0,3,2,2,1,1,0,0,1,0,0,2 | 1,0,3,7,9,5,2,6,1,5,3,0 | 3,6,2,1,3,3,0,1,0,3,3,1,3,2, | 2,4,5,5,4,3,3,0,4,1,3,5,4,6, | 269 | 149 | 0.0031536676433 | 0.25152126426307 | 0.0,1.0,0.27,0.137, | 0.454,0.454,0.181, | -0,092 |
| 34693 | ENSG00000125089 | SH3TC1 | chr4 | + | 8215171 | 8215242 | 8214474 | 8214580 | 8216110 | 8216257 | 34693 | 4,0,2,1,2,1,0,3,0,0,2,5 | 22,16,14,18,17,9,20,24,26, | 1,7,2,4,6,0,4,1,2,3,0,2,1,1, | 23,15,8,3,17,2,9,8,6,20,8,2 | 220 | 149 | 0.00315408046353 | 0.25152126426307 | 0.11,0.0,0.088,0.0 | 0.029,0.24,0.145,0 | -0,04 |
| 177184 | ENSG00000164828 | SUN1 | chr7 | + | 872469 | 872656 | 868473 | 869516 | 873214 | 874934 | 177184 | 69,32,68,22,24,30,25,27,4 | 2,0,1,0,0,4,4,2,2,0,0,4 | 20,23,23,22,6,5,11,23,10,3 | 0,0,1,0,1,0,0,0,0,0,0,2,0,0, | 298 | 149 | 0.0031559017661 | 0.251556249317336 | 0.945,1.0,0.971,1. | 1.0,1.0,0.92,1.0,0. | -0,053 |
| 138654 | ENSG00000197372 | ZNF675 | chr19 | - | 23658602 | 23658672 | 23654545 | 23654706 | 23662113 | 23662209 | 138654 | 0,3,0,1,0,0,0,3,0,0,0,2 | 26,58,65,37,7,11,11,15,11, | 2,6,3,3,1,2,0,1,1,1,2,0,1,2, | 16,25,19,6,12,10,6,6,4,18, | 219 | 149 | 0.00315685672327 | 0.251556249317336 | 0.0,0.034,0.0,0.01 | 0.078,0.14,0.097,0 | -0,053 |
| 78237 | ENSG00000119787 | ATL2 | chr2 | - | 38310308 | 38310447 | 38309378 | 38309506 | 38313150 | 38313243 | 78237 | 27,13,77,46,18,24,22,25,1 | 0,0,2,0,0,0,0,3,0,0,0,0 | 55,45,27,9,22,30,25,34,29, | 0,5,0,1,2,2,3,0,0,1,0,1,2,0, | 288 | 149 | 0.00315851883065 | 0.251595546424734 | 1.0,1.0,0.952,1.0,1 | 1.0,0.823,1.0,0.82 | 0,089 |
| 123638 | ENSG00000133943 | DGLUCY | chr14 | + | 91204705 | 91204825 | 91199756 | 91199905 | 91215404 | 91215556 | 123638 | 68,46,74,79,44,21,37,28,2 | 2,1,7,3,0,1,0,4,1,0,0,2 | 45,33,43,17,28,37,21,49,3 | 0,1,0,2,5,2,2,5,6,0,2,4,2,4, | 269 | 149 | 0.00316198935663 | 0.251778812880811 | 0.95,0.962,0.854,0 | 1.0,0.948,1.0,0.82 | 0,059 |
| 109598 | ENSG00000104219 | ZDHHC2 | chr8 | + | 17205654 | 17205775 | 17198380 | 17198413 | 17207959 | 17208092 | 109598 | 22,14,108,71,60,36,36,57, | 3,1,16,12,5,8,2,5,6,2,5,1 | 52,37,58,28,26,24,24,54,1 | 1,1,7,3,3,3,1,1,3,5,1,3,1,2, | 270 | 149 | 0.00316465427126 | 0.251897819326021 | 0.802,0.885,0.788, | 0.966,0.953,0.821, | -0,057 |
| 240166 | ENSG00000165525 | NEMF | chr14 | - | 49840717 | 49840816 | 49838138 | 49838206 | 49846139 | 49846265 | 240166 | 39,41,67,50,25,39,20,41,2 | 1,2,4,1,0,0,3,4,0,0,0,0 | 42,46,29,58,48,24,19,22,3 | 2,4,7,4,1,2,2,1,0,2,1,2,2,2, | 248 | 149 | 0.00317037807728 | 0.252260127153808 | 0.959,0.925,0.91,0 | 0.927,0.874,0.713, | 0,057 |
| 77332 | ENSG00000128815 | WDFY4 | chr10 | + | 48775711 | 48775806 | 48774457 | 48774672 | 48776749 | 48776984 | 77332 | 349,270,303,278,315,308, | 3,1,4,1,3,4,3,1,1,2,2,2 | 331,422,131,273,120,97,2 | 10,10,2,10,2,0,6,5,3,0,3,3, | 244 | 149 | 0.0031728598799 | 0.252364303329918 | 0.986,0.994,0.979, | 0.953,0.963,0.976, | 0,01 |
| 80688 | ENSG00000162735 | PEX19 | chr1 | - | 160282416 | 160282502 | 160282038 | 160282200 | 160282943 | 160283109 | 80688 | 40,46,157,89,54,54,28,57, | 2,0,5,5,3,2,0,0,0,2,3,0 | 83,71,46,7,43,35,23,65,36, | 4,4,0,2,5,3,0,9,4,0,5,3,0,1, | 235 | 149 | 0.00318192947785 | 0.252992191890334 | 0.927,1.0,0.952,0. | 0.929,0.918,1.0,0. | 0,038 |
| 88169 | ENSG00000144504 | ANKMY1 | chr2 | - | 240511860 | 240512001 | 240500451 | 240500565 | 240512801 | 240512942 | 88169 | 9,8,10,4,5,7,4,9,5,6,2,1 | 2,4,2,2,2,2,2,2,0,0,2,3 | 3,1,2,2,1,2,0,0,1,2,3,6,3,7, | 5,1,0,3,0,1,2,2,6,0,1,2,4,5, | 290 | 149 | 0.00319544710088 | 0.25397314437574 | 0.698,0.507,0.72,0 | 0.236,0.339,1.0,0. | 0,147 |
| 109302 | ENSG00000147421 | HMBOX1 | chr8 | + | 28924639 | 28924755 | 28890394 | 28890678 | 28963810 | 28963890 | 109302 | 0,1,1,3,2,4,6,8,0,2,0,2 | 10,5,12,13,9,9,6,1,6,1,2,5 | 4,3,0,1,2,0,1,0,0,3,2,0,1,3, | 7,6,3,6,0,5,5,3,11,19,6,7,3, | 265 | 149 | 0.00319947208032 | 0.254184487635602 | 0.0,0.101,0.045,0. | 0.243,0.219,0.0,0. | 0,066 |
| 41389 | ENSG00000173226 | IQCB1 | chr3 | - | 121797117 | 121797227 | 121790072 | 121790215 | 121799195 | 121799374 | 41389 | 18,11,86,37,19,12,17,14,1 | 2,0,1,2,0,0,0,0,0,0,0,0 | 37,23,10,22,14,15,4,9,15,3 | 3,1,0,1,6,1,1,0,0,2,0,3,2,0, | 259 | 149 | 0.00320046815511 | 0.254184487635602 | 0.838,1.0,0.98,0.9 | 0.876,0.93,1.0,0.9 | 0,1 |
| 209452 | ENSG00000091490 | SEL1L3 | chr4 | - | 25860600 | 25860688 | 25847342 | 25847864 | 25862674 | 25862988 | 209452 | 1,0,2,2,2,2,0,0,0,0,0,0 | 15,5,87,26,19,6,9,7,11,4,4, | 1,3,3,0,4,2,0,0,4,2,3,1,2,0, | 11,16,12,8,12,3,4,5,13,16, | 237 | 149 | 0.00320236736625 | 0.254241508891683 | 0.04,0.0,0.014,0.0 | 0.054,0.105,0.136, | -0,051 |
| 251929 | ENSG00000171862 | PTEN | chr10 | + | 87927547 | 87927599 | 87925441 | 87925557 | 87930216 | 87930255 | 251929 | 2,5,12,2,0,4,2,4,2,1,4,3 | 3,1,5,2,5,1,0,0,0,3,0,0 | 6,4,1,1,2,1,2,2,9,3,3,13,1,5 | 0,1,0,6,0,0,0,2,1,0,1,6,0,1, | 201 | 149 | 0.00321333406149 | 0.255018105969231 | 0.331,0.788,0.64,0 | 1.0,0.748,1.0,0.11, | -0,088 |
| 55163 | ENSG00000122674 | CCZ1 | chr7 | + | 5905093 | 5905269 | 5902595 | 5902744 | 5910034 | 5910116 | 55163 | 38,16,55,28,28,25,28,47,1 | 4,0,0,0,2,2,3,6,0,1,0,3 | 33,29,2,1,13,7,14,23,7,16, | 3,3,0,0,0,0,2,0,0,0,1,1,1,0, | 298 | 149 | 0.00321804176706 | 0.25529758480136 | 0.826,1.0,1.0,1.0,0 | 0.846,0.829,1.0,1. | 0,022 |
| 8152 | ENSG00000124155 | PIGT | chr20 | + | 45418851 | 45418979 | 45416122 | 45416343 | 45419294 | 45419395 | 8152 | 29,12,29,18,10,19,21,16,3 | 2,0,0,0,0,0,3,2,0,0,2,1 | 13,17,13,15,14,7,16,15,8,1 | 0,1,0,3,0,1,2,5,0,0,3,7,0,0, | 277 | 149 | 0.00322501079242 | 0.255526350189303 | 0.886,1.0,1.0,1.0,1 | 1.0,0.901,1.0,0.72 | 0,044 |
| 55609 | ENSG00000148660 | CAMK2G | chr10 | - | 73819531 | 73819645 | 73817478 | 73817554 | 73821681 | 73821730 | 55609 | 3,0,4,6,1,1,0,6,0,3,1,1 | 19,12,18,9,10,8,3,12,9,7,1, | 4,1,2,0,2,0,2,2,0,2,2,2,0,2, | 12,13,9,3,8,4,1,6,7,18,7,26 | 263 | 149 | 0.00322534312249 | 0.255526350189303 | 0.082,0.0,0.112,0. | 0.159,0.042,0.112, | 0,014 |

| | | | | | | | | | | | | | | | | | | | | | | |
|---|---|---|---|---|---|---|---|---|---|---|---|---|---|---|---|---|---|---|---|---|---|---|
| 62640 | ENSG00000121892 | PDS5A | chr4 | - | 39910243 | 39910338 | 39908394 | 39908540 | 39913610 | 39913726 | 62640 | 121,91,293,158,123,139,97 | 7,2,11,12,0,0,3,4,6,0,2,0 | 199,149,106,69,73,60,91,9 | 6,12,6,13,20,2,1,3,11,2,1,3 | 244 | 149 | 0.00322543119615 | 0.255526350189303 | 0.913,0.965,0.942, | 0.953,0.883,0.915, | 0,043 |
| 185234 | ENSG00000132185 | FCRLA | chr1 | + | 161711225 | 161711474 | 161710759 | 161710912 | 161711933 | 161712218 | 185234 | 165,148,456,409,402,191,2 | 12,5,31,20,13,0,13,16,14,9 | 449,310,97,48,101,133,18 | 36,37,3,5,12,5,15,17,0,11,1 | 298 | 149 | 0.00322567424518 | 0.255526350189303 | 0.873,0.937,0.88,0 | 0.862,0.807,0.942, | 0,027 |
| 116166 | ENSG00000107077 | KDM4C | chr9 | + | 7015852 | 7015929 | 7013787 | 7014001 | 7046861 | 7046917 | 116166 | 76,22,60,44,33,22,24,28,2 | 5,0,10,1,0,0,4,0,0,2,0,3 | 47,35,41,24,26,31,24,16,1 | 4,9,2,9,5,2,0,2,0,0,10,2,3,0 | 226 | 149 | 0.00323180042202 | 0.255917452771077 | 0.909,1.0,0.798,0. | 0.886,0.719,0.931, | 0,091 |
| 214884 | ENSG00000184007 | PTP4A2 | chr1 | - | 31915890 | 31915987 | 31911695 | 31911826 | 31918969 | 31919109 | 214884 | 89,66,234,173,75,84,72,81 | 10,5,16,12,5,1,1,3,13,9,4,7 | 144,151,93,67,85,74,47,89 | 6,5,5,8,5,6,3,8,2,7,2,13,4,4 | 246 | 149 | 0.00323408668599 | 0.255930695637584 | 0.844,0.889,0.899, | 0.936,0.948,0.918, | -0,065 |
| 40592 | ENSG00000074935 | TUBE1 | chr6 | - | 112079632 | 112079754 | 112076321 | 112076509 | 112081091 | 112081203 | 40592 | 45,30,40,25,15,21,11,13,1 | 0,0,4,4,4,2,2,0,0,0,0,0 | 16,6,19,4,11,5,6,14,10,35, | 6,1,2,2,8,1,2,0,2,2,2,0,0,1, | 271 | 149 | 0.00323434585229 | 0.255930695637584 | 1.0,1.0,0.846,0.77 | 0.595,0.767,0.839, | 0,123 |
| 139012 | ENSG00000198276 | UCKL1 | chr20 | - | 63945082 | 63945503 | 63944544 | 63944734 | 63945650 | 63945722 | 139012 | 6,2,10,7,4,1,1,1,1,1,2,3 | 38,32,93,19,43,53,30,34,2 | 2,2,1,0,5,2,2,2,0,0,0,2,5,3, | 59,34,20,24,30,23,41,63,3 | 298 | 149 | 0.00324054519202 | 0.256305064094097 | 0.073,0.03,0.051,0 | 0.017,0.029,0.024, | 0,02 |
| 96672 | ENSG00000106100 | NOD1 | chr7 | - | 30428503 | 30428609 | 30424529 | 30425710 | 30429373 | 30429457 | 96672 | 4,4,2,0,2,0,0,0,4,2,1,0 | 13,9,27,11,5,3,11,3,6,4,3,8 | 0,1,0,2,0,0,0,0,1,0,1,0,0,1, | 22,8,2,7,7,7,9,18,5,8,11,19 | 255 | 149 | 0.00324215572551 | 0.256305064094097 | 0.152,0.206,0.041, | 0.0,0.068,0.0,0.14 | 0,08 |
| 10126 | ENSG00000276293 | PIP4K2B | chr17 | - | 38784242 | 38784339 | 38780451 | 38780604 | 38786822 | 38786920 | 10126 | 42,30,66,36,33,32,14,46,4 | 3,0,5,2,0,2,2,8,0,3,2,2 | 43,17,25,12,14,13,15,36,1 | 1,1,0,0,0,0,2,3,0,1,1,6,0,1, | 246 | 149 | 0.00324264947674 | 0.256305064094097 | 0.895,1.0,0.889,0. | 0.963,0.911,1.0,1. | -0,057 |
| 152749 | ENSG00000089775 | ZBTB25 | chr14 | - | 64502652 | 64502815 | 64490360 | 64490540 | 64502900 | 64503127 | 152749 | 7,7,28,4,10,8,15,13,2,10,4, | 7,6,3,6,9,5,5,1,6,0,0,3 | 12,8,8,16,3,4,9,16,13,17,5, | 10,1,0,1,0,4,0,4,0,3,0,2,0,4 | 298 | 149 | 0.00326202623494 | 0.257653893722878 | 0.333,0.368,0.824, | 0.375,0.8,1.0,0.88 | -0,22 |
| 164385 | ENSG00000124203 | ZNF831 | chr20 | + | 59173333 | 59173455 | 59146230 | 59146374 | 59190983 | 59194757 | 164385 | 5,2,5,0,0,0,1,11,4,0,0,0 | 10,3,7,4,7,22,8,18,8,2,11,6 | 4,6,0,0,0,0,6,1,1,0,2,0,0,2, | 7,4,2,1,2,4,13,0,6,11,36,1,4 | 271 | 149 | 0.00326210843417 | 0.257653893722878 | 0.216,0.268,0.282, | 0.239,0.452,0.0,0. | -0,037 |
| 27653 | ENSG00000165097 | KDM1B | chr6 | + | 18201485 | 18201657 | 18200438 | 18200576 | 18205536 | 18205664 | 27653 | 7,10,34,31,21,11,10,9,19,9 | 1,0,8,1,2,2,1,2,2,0,0,0 | 18,18,11,11,12,7,8,5,9,28, | 1,2,0,0,8,1,2,0,1,3,4,0,0,0, | 298 | 149 | 0.00326363556081 | 0.257679950619478 | 0.778,1.0,0.68,0.9 | 0.9,0.818,1.0,1.0,0 | 0,009 |
| 160760 | ENSG00000006459 | KDM7A | chr7 | - | 140126635 | 140126823 | 140120441 | 140120529 | 140127441 | 140127583 | 160760 | 15,2,23,23,4,8,9,6,5,6,3,11 | 40,12,23,19,10,16,12,12,9, | 11,14,6,4,0,18,5,3,6,22,4,3 | 30,51,30,34,31,15,4,23,21, | 298 | 149 | 0.00326756173119 | 0.257895335154292 | 0.158,0.077,0.333, | 0.155,0.121,0.091, | 0,069 |
| 248436 | ENSG00000139697 | SBNO1 | chr12 | - | 123348028 | 123348129 | 123345257 | 123345567 | 123350309 | 123350441 | 248436 | 47,45,47,29,17,14,28,32,4 | 4,0,4,2,0,0,1,1,3,2,0,2 | 26,48,40,27,33,14,15,27,1 | 0,1,3,0,3,0,0,0,0,1,1,2,0,0, | 250 | 149 | 0.0032697401605 | 0.257972670265607 | 0.875,1.0,0.875,0. | 1.0,0.966,0.888,1. | -0,027 |
| 12695 | ENSG00000108848 | LUC7L3 | chr17 | + | 50738140 | 50738199 | 50736959 | 50737026 | 50740305 | 50740345 | 12695 | 8,5,17,9,3,2,4,7,0,1,5,1 | 59,33,146,87,96,38,64,87, | 8,18,6,10,6,6,4,4,5,8,4,9,8, | 78,65,31,24,51,50,31,43,3 | 208 | 149 | 0.00327833993804 | 0.25855638812909 | 0.089,0.098,0.077, | 0.068,0.166,0.122, | -0,037 |
| 188810 | ENSG00000082512 | TRAF5 | chr1 | + | 211366360 | 211366490 | 211365375 | 211365468 | 211369451 | 211369592 | 188810 | 16,2,30,12,23,10,16,15,9,3 | 13,21,56,36,39,25,12,21,2 | 36,12,19,20,7,5,5,19,15,21 | 36,20,15,9,30,8,7,15,21,34 | 279 | 149 | 0.00327991707651 | 0.25858601915543 | 0.397,0.048,0.222, | 0.348,0.243,0.404, | -0,097 |
| 190894 | ENSG00000133805 | AMPD3 | chr11 | + | 10478529 | 10478730 | 10461514 | 10461740 | 10482062 | 10482225 | 190894 | 10,3,9,3,16,8,1,2,1,0,7,14 | 2,2,4,0,2,0,0,1,1,2,0,4 | 18,16,2,3,5,6,2,2,13,19,6,3 | 0,2,0,0,0,2,0,2,3,0,0,6,1,0, | 298 | 149 | 0.00328434929021 | 0.258840637891318 | 0.714,0.429,0.529, | 1.0,0.8,1.0,1.0,1.0, | -0,217 |
| 50761 | ENSG00000151092 | NGLY1 | chr3 | - | 25732318 | 25732483 | 25729132 | 25729318 | 25733871 | 25733982 | 50761 | 89,39,121,93,44,31,25,42, | 32,9,46,40,20,23,22,29,0,9 | 51,31,33,36,30,46,15,33,2 | 39,67,21,27,33,26,10,16,3 | 298 | 149 | 0.00328558148004 | 0.258842967617309 | 0.582,0.684,0.568, | 0.395,0.188,0.44,0 | 0,137 |
| 13378 | ENSG00000038532 | CLEC16A | chr16 | + | 11166387 | 11166552 | 11125978 | 11126146 | 11178334 | 11182182 | 13378 | 10,15,10,2,10,1,16,9,14,7, | 2,0,1,2,2,0,0,1,0,0,0,2 | 3,2,7,6,1,0,5,6,1,2,13,21,7, | 4,3,2,0,0,3,0,4,0,0,1,2,1,0, | 298 | 149 | 0.0032903805813 | 0.259126199375697 | 0.714,1.0,0.833,0. | 0.273,0.25,0.636,1 | 0,121 |
| 192943 | ENSG00000158006 | PAFAH2 | chr1 | - | 25985753 | 25985945 | 25984459 | 25984528 | 25988230 | 25988327 | 192943 | 6,0,2,1,2,2,5,2,0,0,3,4 | 4,3,3,4,15,2,6,2,6,4,4,3 | 0,0,0,4,2,1,4,2,0,0,1,1,0,1, | 4,2,4,3,2,5,0,1,2,4,3,1,9,2, | 298 | 149 | 0.00329176670761 | 0.259140541421217 | 0.429,0.0,0.25,0.1 | 0.0,0.0,0.0,0.4,0.3 | -0,002 |
| 217759 | ENSG00000082258 | CCNT2 | chr2 | + | 134942596 | 134942674 | 134939001 | 134939062 | 134946100 | 134946146 | 217759 | 16,7,13,19,5,5,4,9,5,2,2,1 | 1,2,1,2,2,2,3,1,0,0,0,1 | 12,24,19,7,20,1,6,6,13,12, | 2,8,13,13,2,0,1,2,3,3,7,0,2, | 227 | 149 | 0.00329680797245 | 0.259442514339446 | 0.913,0.697,0.895, | 0.798,0.663,0.49,0 | 0,069 |
| 160755 | ENSG00000006459 | KDM7A | chr7 | - | 140124620 | 140125024 | 140120441 | 140120529 | 140127441 | 140127583 | 160755 | 28,10,38,12,12,7,15,6,15,1 | 40,12,23,19,10,16,12,12,9, | 26,8,4,7,8,8,4,15,11,22,5,1 | 30,51,30,34,31,15,4,23,21, | 298 | 149 | 0.00330394901438 | 0.259817964149925 | 0.259,0.294,0.452, | 0.302,0.073,0.062, | 0,094 |
| 175956 | ENSG00000276291 | FRG1HP | chr9 | + | 41007439 | 41007592 | 40994632 | 40994699 | 41011616 | 41011730 | 175956 | 22,9,30,10,9,22,9,8,10,6,6, | 2,0,12,3,18,15,11,19,0,0,2, | 19,11,13,7,6,7,10,15,11,26 | 0,0,0,1,11,4,0,0,0,0,4,16,0, | 298 | 149 | 0.00330399323461 | 0.259817964149925 | 0.846,1.0,0.556,0. | 1.0,1.0,1.0,0.778,0 | -0,127 |
| 150339 | ENSG00000247556 | OIP5-AS1 | chr15 | + | 41297833 | 41297929 | 41286010 | 41286100 | 41298040 | 41306534 | 150339 | 1,0,7,5,7,4,1,5,0,0,0,0 | 29,11,80,66,68,66,35,46,1 | 1,2,1,0,4,1,1,8,4,8,1,10,2,8 | 31,45,19,3,16,29,11,31,6,4 | 245 | 149 | 0.00330594094487 | 0.25986005555218 | 0.021,0.0,0.051,0. | 0.019,0.026,0.031, | -0,047 |
| 129777 | ENSG00000089022 | MAPKAPK5 | chr12 | + | 111883580 | 111883768 | 111880446 | 111880527 | 111885915 | 111886036 | 129777 | 82,71,122,87,74,71,57,73, | 5,7,6,6,1,1,5,1,6,0,2,4 | 80,55,56,26,50,44,39,64,1 | 8,3,2,5,9,6,6,5,2,0,5,11,7,6 | 298 | 149 | 0.0033069787437 | 0.25986005555218 | 0.891,0.835,0.91,0 | 0.833,0.902,0.933, | 0,073 |
| 214413 | ENSG00000144224 | UBXN4 | chr2 | + | 135775463 | 135775539 | 135772446 | 135772547 | 135776248 | 135776351 | 214413 | 3,2,3,2,2,2,0,3,0,0,0,1 | 25,13,41,31,20,26,25,15,2 | 2,6,5,2,4,1,1,3,5,4,3,3,5,3, | 21,81,29,40,38,22,30,36,1 | 225 | 149 | 0.00330815055551 | 0.25986005555218 | 0.074,0.092,0.046, | 0.059,0.047,0.102, | -0,027 |
| 167498 | ENSG00000145495 | MARCHF6 | chr5 | + | 10417269 | 10417400 | 10415472 | 10415669 | 10423734 | 10423835 | 167498 | 135,89,132,93,75,114,74,7 | 0,0,1,2,0,1,0,0,0,0,0,1 | 113,135,68,141,64,40,99,9 | 2,5,2,4,3,0,5,2,1,0,2,1,0,1, | 280 | 149 | 0.00331110837156 | 0.259997506719894 | 1.0,1.0,0.986,0.96 | 0.968,0.935,0.948, | 0,028 |
| 192237 | ENSG00000131408 | NR1H2 | chr19 | + | 50378157 | 50378439 | 50377732 | 50377870 | 50378521 | 50378678 | 192237 | 12,8,19,4,12,10,3,2,14,5,1, | 3,4,2,0,2,2,0,2,1,1,0,1 | 8,11,6,12,5,2,4,9,4,4,7,5,3, | 1,3,0,1,0,1,0,0,2,2,0,6,0,0, | 298 | 149 | 0.00331768979339 | 0.260419289540891 | 0.667,0.5,0.826,1. | 0.8,0.647,1.0,0.85 | -0,004 |
| 44623 | ENSG00000159433 | STARD9 | chr15 | + | 42665785 | 42665848 | 42665252 | 42665330 | 42669157 | 42669337 | 44623 | 16,4,15,6,26,18,5,25,10,11 | 0,0,1,0,0,1,1,2,0,2,0,0 | 12,10,4,23,1,4,17,8,7,6,14, | 3,4,1,1,2,0,2,2,0,0,2,1,0,2, | 212 | 149 | 0.0033197852297 | 0.260478104051986 | 1.0,1.0,0.913,1.0,1 | 0.738,0.637,0.738, | 0,123 |
| 60334 | ENSG00000163479 | SSR2 | chr1 | - | 156019084 | 156019266 | 156018131 | 156018368 | 156020012 | 156020167 | 60334 | 3,2,9,13,2,1,5,2,2,4,1,0 | 212,136,392,288,128,95,1 | 7,9,6,3,6,3,0,11,4,9,5,11,5, | 161,171,173,76,128,94,71, | 298 | 149 | 0.00332085953008 | 0.260478104051986 | 0.007,0.007,0.011, | 0.021,0.026,0.017, | -0,009 |
| 138015 | ENSG00000142453 | CARM1 | chr19 | + | 10908549 | 10908794 | 10908038 | 10908145 | 10909102 | 10909207 | 138015 | 7,4,6,9,0,13,4,0,6,3,2,6 | 3,7,12,24,6,11,3,2,4,12,7,9 | 2,3,3,4,1,4,9,5,0,1,0,5,4,2, | 9,7,7,8,7,10,6,22,6,13,10,2 | 298 | 149 | 0.00332814906141 | 0.260850386600117 | 0.538,0.222,0.2,0. | 0.1,0.176,0.176,0. | 0,11 |
| 114710 | ENSG00000108669 | CYTH1 | chr17 | - | 78702537 | 78702604 | 78702121 | 78702240 | 78708196 | 78708261 | 114710 | 638,469,473,492,258,195, | 106,52,56,20,16,44,37,56, | 390,355,353,239,248,183, | 73,103,31,161,65,21,17,35 | 216 | 149 | 0.00332922270187 | 0.260850386600117 | 0.806,0.862,0.854, | 0.787,0.704,0.887, | 0,053 |
| 244243 | ENSG00000166266 | CUL5 | chr11 | + | 108061039 | 108061172 | 108054874 | 108054955 | 108070095 | 108070189 | 244243 | 7,4,18,9,6,0,1,0,0,4,1,4 | 13,14,55,37,16,15,10,9,3,4 | 5,1,0,2,4,2,1,0,0,2,2,7,2,4, | 25,33,8,12,25,8,7,5,13,10, | 282 | 149 | 0.00333101956977 | 0.260850386600117 | 0.221,0.131,0.147, | 0.096,0.016,0.0,0. | 0,046 |
| 143057 | ENSG00000167543 | TP53I13 | chr17 | + | 29567863 | 29567985 | 29567179 | 29567411 | 29568277 | 29568548 | 143057 | 0,1,6,3,2,4,1,8,5,4,0,3 | 19,8,31,26,14,25,6,28,37,2 | 1,0,3,0,3,1,0,1,0,1,8,0,4,0, | 19,19,11,1,18,0,15,14,5,13 | 271 | 149 | 0.0033314543562 | 0.260850386600117 | 0.0,0.064,0.096,0. | 0.028,0.0,0.13,0.0, | -0,069 |
| 213427 | ENSG00000035720 | STAP1 | chr4 | + | 67576828 | 67576943 | 67575384 | 67575498 | 67577202 | 67577259 | 213427 | 0,2,27,9,0,5,0,0,1,0,3,5 | 95,43,180,138,78,65,66,55 | 14,11,11,4,0,0,6,6,0,0,0,0, | 83,59,97,34,67,53,15,55,4 | 264 | 149 | 0.00333166557217 | 0.260850386600117 | 0.0,0.026,0.078,0. | 0.087,0.095,0.06,0 | -0,02 |
| 127390 | ENSG00000172661 | WASHC2C | chr10 | + | 45751481 | 45751553 | 45750750 | 45750838 | 45752587 | 45752706 | 127390 | 42,20,100,65,37,43,51,50, | 6,3,5,0,4,0,3,6,6,4,0,0 | 46,50,26,24,36,23,25,47,2 | 1,0,0,1,0,0,2,6,1,1,5,4,1,6, | 221 | 149 | 0.00334530921853 | 0.261757996038747 | 0.825,0.818,0.931, | 0.969,1.0,1.0,0.94 | -0,05 |
| 114291 | ENSG00000143748 | NVL | chr1 | - | 224313033 | 224313151 | 224311799 | 224311857 | 224317693 | 224317793 | 114291 | 0,7,0,0,2,0,0,6,0,1,0,3 | 7,6,22,16,9,4,9,9,2,3,3,14 | 1,1,3,0,0,0,0,1,5,5,1,3,1,0, | 7,12,6,5,6,1,5,10,7,2,3,21, | 267 | 149 | 0.00334569019845 | 0.261757996038747 | 0.0,0.394,0.0,0.0,0 | 0.074,0.044,0.218, | -0,02 |
| 375 | ENSG00000159267 | HLCS | chr21 | - | 36948165 | 36948287 | 36938831 | 36938994 | 36962035 | 36962170 | 375 | 3,0,15,1,4,6,4,5,2,3,0,2 | 15,6,22,18,12,13,5,14,8,0, | 3,3,0,3,0,1,2,5,0,0,0,4,2,4, | 11,17,15,13,16,13,5,7,2,15 | 271 | 149 | 0.00335215803775 | 0.262155643601676 | 0.099,0.0,0.273,0. | 0.13,0.088,0.0,0.1 | 0,121 |
| 19510 | ENSG00000170776 | AKAP13 | chr15 | + | 85715787 | 85715923 | 85710442 | 85710645 | 85717289 | 85717348 | 19510 | 422,289,357,306,189,190, | 2,2,2,5,2,1,3,1,3,1,0,0 | 271,402,310,228,235,188, | 5,3,8,7,2,5,0,3,2,4,5,3,5,0, | 285 | 149 | 0.0033544068645 | 0.262155643601676 | 0.991,0.987,0.989, | 0.966,0.986,0.953, | 0,012 |
| 180759 | ENSG00000117533 | VAMP4 | chr1 | - | 171728034 | 171728082 | 171719170 | 171719221 | 171728523 | 171728570 | 180759 | 2,8,0,7,1,0,1,4,0,2,2,3 | 35,27,20,12,13,16,7,17,29, | 7,9,4,3,2,4,2,0,1,4,5,2,2,2, | 14,17,11,23,8,13,4,11,5,16 | 197 | 149 | 0.00335442683665 | 0.262155643601676 | 0.041,0.183,0.0,0. | 0.274,0.286,0.216, | -0,063 |
| 130796 | ENSG00000123349 | PFDN5 | chr12 | + | 53296218 | 53296275 | 53295524 | 53295639 | 53297849 | 53297924 | 130796 | 157,91,320,253,90,119,75, | 62,26,120,60,24,30,21,42, | 172,171,105,75,105,51,68, | 50,56,40,22,61,26,27,59,2 | 206 | 149 | 0.00336105202082 | 0.262578071685339 | 0.647,0.717,0.659, | 0.713,0.688,0.655, | 0,065 |
| 49563 | ENSG00000133789 | SWAP70 | chr11 | + | 9724657 | 9724885 | 9694145 | 9694286 | 9728052 | 9728071 | 49563 | 145,103,314,209,128,47,5 | 5,8,2,4,3,5,2,0,0,0,1,3 | 175,153,48,56,78,98,28,42 | 9,10,7,1,6,2,2,3,0,0,1,10,2, | 298 | 149 | 0.00337116090065 | 0.263272253921553 | 0.935,0.866,0.987, | 0.907,0.884,0.774, | 0,032 |
| 204618 | ENSG00000158985 | CDC42SE2 | chr5 | + | 131256076 | 131256192 | 131255094 | 131255229 | 131315975 | 131316144 | 204618 | 29,62,50,61,23,21,5,26,10, | 0,4,6,2,2,0,1,0,0,0,0,0 | 19,37,14,14,35,11,3,11,12, | 2,5,1,2,1,2,0,0,2,5,2,3,2,2, | 265 | 149 | 0.0033739098264 | 0.263375591342709 | 1.0,0.897,0.824,0. | 0.842,0.806,0.887, | 0,09 |
| 230328 | ENSG00000158636 | EMSY | chr11 | + | 76538283 | 76538402 | 76537794 | 76537950 | 76539598 | 76539640 | 230328 | 8,0,0,1,0,0,1,4,2,0,4,0 | 21,10,12,2,11,6,7,8,16,3,1 | 0,0,0,0,0,0,0,2,0,0,1,0,0, | 14,7,7,6,4,4,3,22,8,7,7,10, | 268 | 149 | 0.0033749314965 | 0.263375591342709 | 0.175,0.0,0.0,0.21 | 0.0,0.0,0.0,0.0,0.0, | 0,059 |
| 163045 | ENSG00000182903 | ZNF721 | chr4 | - | 472574 | 472713 | 439987 | 444432 | 473980 | 474167 | 163045 | 81,45,139,64,21,26,21,40, | 2,1,1,2,6,0,2,4,0,0,2,2 | 50,96,57,67,45,33,21,31,2 | 0,0,0,7,1,0,1,0,1,0,1,0,0,0, | 288 | 149 | 0.00337931842018 | 0.263622356974904 | 0.954,0.959,0.986, | 1.0,1.0,1.0,0.832,0 | -0,086 |
| 157264 | ENSG00000120686 | UFM1 | chr13 | + | 38350149 | 38350227 | 38349998 | 38350055 | 38354238 | 38354296 | 157264 | 1,0,3,1,2,5,2,0,0,2,0,0 | 16,14,83,48,20,19,12,29,2, | 2,4,2,0,7,0,0,3,4,3,3,7,2,1, | 41,36,17,11,36,11,15,29,7, | 227 | 149 | 0.00338177436538 | 0.263718361751849 | 0.039,0.0,0.023,0. | 0.031,0.068,0.072, | -0,006 |
| 83027 | ENSG00000170266 | GLB1 | chr3 | - | 33058088 | 33058269 | 33053490 | 33053549 | 33065462 | 33065557 | 83027 | 70,55,79,56,57,67,44,36,4 | 1,0,2,4,3,6,3,0,1,2,1,0 | 53,65,27,20,30,25,14,37,1 | 1,2,2,0,4,2,0,3,0,0,0,0,0,0, | 298 | 149 | 0.00338910458832 | 0.264194266442847 | 0.972,1.0,0.952,0. | 0.964,0.942,0.871, | -0,024 |
| 165456 | ENSG00000132466 | ANKRD17 | chr4 | - | 73101110 | 73101173 | 73098072 | 73098520 | 73102375 | 73102547 | 165456 | 0,0,5,3,0,2,3,6,2,0,0,0 | 25,22,45,25,18,24,8,20,19, | 2,1,1,0,0,0,3,1,1,1,0,0,0,2, | 26,18,10,24,15,20,13,17,1 | 212 | 149 | 0.00339554229035 | 0.264600276138422 | 0.0,0.0,0.072,0.07 | 0.051,0.038,0.066, | 0,026 |
| 154492 | ENSG00000100888 | CHD8 | chr14 | - | 21430800 | 21430976 | 21428963 | 21429335 | 21431810 | 21431858 | 154492 | 28,16,38,43,42,17,17,14,4 | 8,7,13,14,7,7,6,10,5,3,0,5 | 22,27,17,8,15,19,15,27,10, | 12,21,10,7,10,5,9,24,7,5,6, | 298 | 149 | 0.00339781666194 | 0.264681678597904 | 0.636,0.533,0.594, | 0.478,0.391,0.459, | 0,165 |
| 34007 | ENSG00000073331 | ALPK1 | chr4 | + | 112302338 | 112302485 | 112297368 | 112297469 | 112315800 | 112315852 | 34007 | 0,0,3,3,6,4,0,0,2,1,0,2 | 6,8,10,11,4,4,9,5,8,0,2,1 | 2,2,1,0,0,0,0,0,0,0,1,1,2, | 10,8,8,11,1,6,2,1,9,5,7,11, | 296 | 149 | 0.00340444502069 | 0.265102064489193 | 0.0,0.0,0.131,0.12 | 0.091,0.112,0.059, | 0,167 |
| 200811 | ENSG00000123728 | RAP2C | chrX | - | 132213902 | 132213955 | 132205448 | 132205587 | 132214133 | 132214446 | 200811 | 1,5,8,1,1,0,0,1,1,0,0,0 | 83,65,199,104,61,50,48,63 | 7,5,0,3,3,4,3,3,3,3,5,3,3,3, | 103,130,53,60,103,50,47,5 | 202 | 149 | 0.00340641989903 | 0.265159913666592 | 0.009,0.054,0.029, | 0.048,0.028,0.0,0. | -0,019 |
| 107419 | ENSG00000143569 | UBAP2L | chr1 | + | 154260891 | 154261109 | 154259735 | 154260029 | 154261591 | 154261697 | 107419 | 93,78,97,59,57,59,41,70,4 | 7,0,1,0,0,2,0,6,2,0,3,0 | 65,71,60,92,64,30,77,69,4 | 2,1,3,3,3,2,0,3,0,0,6,6,2,1, | 298 | 149 | 0.00340993567747 | 0.265337623209525 | 0.869,1.0,0.98,1.0, | 0.942,0.973,0.909, | 0,026 |
| 91355 | ENSG00000094880 | CDC23 | chr5 | - | 138212990 | 138213063 | 138205870 | 138206684 | 138213151 | 138213215 | 91355 | 13,22,49,36,21,22,17,15,1 | 9,0,1,0,0,2,8,0,0,0,0,1 | 20,46,17,8,32,13,8,19,5,12 | 9,4,1,1,8,2,1,0,3,0,2,6,4,1, | 222 | 149 | 0.00341197174862 | 0.265400105322454 | 0.492,1.0,0.97,1.0, | 0.599,0.885,0.919, | 0,089 |
| 110247 | ENSG00000166169 | POLL | chr10 | - | 101585861 | 101586156 | 101584610 | 101584919 | 101587245 | 101587382 | 110247 | 4,1,14,23,1,5,7,8,0,1,8,10 | 1,2,2,0,1,2,0,2,2,2,0,0 | 5,3,5,1,3,3,5,5,3,12,6,3,21, | 0,0,0,0,0,0,0,0,0,0,1,0,0, | 298 | 149 | 0.00341497207571 | 0.265537519807492 | 0.667,0.2,0.778,1. | 1.0,1.0,1.0,1.0,1.0, | -0,343 |
| 148715 | ENSG00000090905 | TNRC6A | chr16 | + | 24730252 | 24730300 | 24640883 | 24641279 | 24750725 | 24750813 | 148715 | 4,1,4,3,3,5,3,1,5,4,2,6 | 4,0,5,1,2,0,0,1,0,0,1,4 | 3,3,3,8,3,2,4,4,3,4,3,3,1,6, | 0,1,2,7,0,0,3,5,0,0,0,0,0,0, | 197 | 149 | 0.00342301176412 | 0.266066538462734 | 0.431,1.0,0.377,0. | 1.0,0.694,0.532,0. | -0,102 |
| 36111 | ENSG00000174652 | ZNF266 | chr19 | - | 9418765 | 9418895 | 9418504 | 9418631 | 9420064 | 9420218 | 36111 | 27,22,78,50,28,5,41,65,33, | 4,0,3,4,0,0,3,2,0,0,2,5 | 73,35,40,50,28,27,20,42,3 | 1,3,0,2,3,0,0,0,0,6,0,0,0,1 | 279 | 149 | 0.0034261104997 | 0.26618039971122 | 0.783,1.0,0.933,0. | 0.975,0.862,1.0,0. | -0,09 |
| 33818 | ENSG00000161013 | MGAT4B | chr5 | - | 179800483 | 179800597 | 179800183 | 179800259 | 179800906 | 179800939 | 33818 | 57,40,191,135,104,96,103, | 1,0,2,1,0,0,0,0,1,0,0 | 30,47,40,18,40,27,34,130, | 2,3,1,0,3,0,3,4,1,0,0,4,1,3, | 263 | 149 | 0.00342695005645 | 0.26618039971122 | 0.97,1.0,0.982,0.9 | 0.895,0.899,0.958, | 0,051 |
| 100139 | ENSG00000177189 | RPS6KA3 | chrX | - | 20167588 | 20167747 | 20164898 | 20165060 | 20169401 | 20169491 | 100139 | 75,67,181,105,86,60,45,51 | 1,3,1,6,1,1,2,2,1,0,0,0 | 101,130,39,15,36,55,34,87 | 0,7,4,2,4,0,0,2,0,5,4,4,2,1, | 298 | 149 | 0.0034309452959 | 0.266389203241215 | 0.974,0.918,0.989, | 1.0,0.903,0.83,0.7 | 0,05 |
| 129797 | ENSG00000120805 | ARL1 | chr12 | - | 101405843 | 101405981 | 101402864 | 101402946 | 101407641 | 101407746 | 129797 | 45,18,77,95,45,30,25,30,4 | 0,5,9,3,0,7,1,1,2,0,0,2 | 29,17,22,11,22,22,22,37,1 | 1,12,2,0,9,2,1,2,4,1,7,2,5,4 | 287 | 149 | 0.00343211368524 | 0.266389203241215 | 1.0,0.651,0.816,0. | 0.938,0.424,0.851, | 0,106 |
| 167588 | ENSG00000182150 | ERCC6L2 | chr9 | + | 95907669 | 95907761 | 95907122 | 95907271 | 95915667 | 95915829 | 167588 | 0,1,2,0,4,2,0,0,2,0,0,0 | 16,11,44,21,7,9,19,26,10,6 | 4,0,4,8,1,1,0,4,4,0,3,0,0,0, | 25,17,19,14,12,10,13,22,12 | 241 | 149 | 0.0034451378298 | 0.267303698718704 | 0.0,0.053,0.027,0. | 0.09,0.0,0.115,0.2 | -0,022 |
| 77222 | ENSG00000165630 | PRPF18 | chr10 | + | 13613740 | 13613881 | 13611614 | 13611683 | 13614014 | 13614060 | 77222 | 133,79,161,116,65,62,62,7 | 3,1,8,4,5,2,0,6,7,0,0,1 | 98,108,68,40,74,55,32,48, | 3,1,10,1,1,0,0,2,1,0,2,3,0,4 | 290 | 149 | 0.00344669363121 | 0.267328042140166 | 0.958,0.976,0.912, | 0.944,0.982,0.777, | -0,017 |
| 243405 | ENSG00000291075 | | chr9 | + | 41008880 | 41009261 | 41007439 | 41007609 | 41011616 | 41011907 | 243405 | 31,9,52,42,14,19,8,30,15,1 | 1,0,16,11,16,13,5,7,1,0,6,1 | 42,35,30,20,8,10,8,22,21,2 | 1,0,0,0,3,4,2,1,0,0,1,20,2,0 | 298 | 149 | 0.00345173409979 | 0.267622543959619 | 0.939,1.0,0.619,0. | 0.955,1.0,1.0,1.0,0 | -0,131 |
| 250507 | ENSG00000120616 | EPC1 | chr10 | - | 32273013 | 32273114 | 32272025 | 32272167 | 32284697 | 32285050 | 250507 | 8,3,9,12,4,5,4,8,9,1,2,4 | 0,0,1,0,0,1,0,0,0,0,0,0 | 8,10,4,13,4,2,9,7,2,5,5,6,5, | 2,0,1,0,6,1,2,2,2,1,2,0,0,0, | 250 | 149 | 0.0034576094445 | 0.267896229046444 | 1.0,1.0,0.843,1.0,1 | 0.704,1.0,0.704,1. | 0,193 |
| 124456 | ENSG00000148925 | BTBD10 | chr11 | - | 13445023 | 13445181 | 13421641 | 13421838 | 13463091 | 13463222 | 124456 | 11,4,38,16,7,6,6,14,30,16, | 33,22,18,11,5,20,4,14,8,6, | 14,15,7,5,9,1,5,7,0,0,3,5,1, | 17,29,6,11,20,5,21,9,9,12,1 | 298 | 149 | 0.00345775341048 | 0.267896229046444 | 0.143,0.083,0.514, | 0.292,0.205,0.368, | 0,155 |
| 210495 | ENSG00000140386 | SCAPER | chr15 | - | 76795376 | 76795440 | 76774931 | 76775117 | 76800247 | 76800267 | 210495 | 22,6,18,15,11,9,6,12,6,6,5, | 2,2,6,3,0,5,0,1,0,1,0,3 | 9,9,9,4,2,2,1,7,2,9,4,6,2,4, | 3,2,0,4,4,2,0,7,2,2,6,7,0,4,1 | 213 | 149 | 0.00346047964468 | 0.268010973157294 | 0.885,0.677,0.677, | 0.677,0.759,1.0,0. | 0,196 |
| 235871 | ENSG00000110697 | PITPNM1 | chr11 | - | 67501102 | 67501264 | 67500094 | 67500421 | 67501861 | 67502086 | 235871 | 4,2,6,3,2,2,0,3,3,2,0,1 | 49,25,65,33,41,26,36,28,3 | 0,0,0,0,0,4,0,2,1,1,3,3,0, | 19,23,24,6,12,11,17,16,33, | 298 | 149 | 0.00346933880332 | 0.268590766219973 | 0.039,0.038,0.044, | 0.0,0.0,0.0,0.0,0.0, | 0,014 |
| 56229 | ENSG00000118260 | CREB1 | chr2 | + | 207577109 | 207577224 | 207575271 | 207575454 | 207577504 | 207577655 | 56229 | 24,9,34,11,5,11,10,12,7,2, | 65,19,78,61,49,23,33,31,4 | 16,20,6,3,9,8,11,8,5,17,7,1 | 57,59,23,38,46,40,42,50,2 | 264 | 149 | 0.00347046160106 | 0.268590766219973 | 0.172,0.211,0.197, | 0.137,0.161,0.128, | 0,042 |
| 25000 | ENSG00000171295 | ZNF440 | chr19 | + | 11830282 | 11830409 | 11814272 | 11814450 | 11830616 | 11830677 | 25000 | 1,5,13,3,4,12,2,5,1,4,4,8 | 3,0,3,0,0,1,1,0,0,0,0,2 | 7,12,5,7,8,6,2,2,6,9,4,10,1 | 0,1,0,0,2,2,0,2,0,1,0,3,2,0, | 276 | 149 | 0.0034731876002 | 0.268705118755804 | 0.153,1.0,0.701,1. | 1.0,0.866,1.0,1.0,0 | 0,05 |
| 185238 | ENSG00000132185 | FCRLA | chr1 | + | 161711327 | 161711474 | 161710759 | 161710912 | 161711933 | 161712218 | 185238 | 165,144,454,411,364,185, | 12,5,31,20,13,0,13,16,14,9 | 449,310,97,47,101,131,14 | 36,37,3,5,12,5,15,17,0,11,1 | 296 | 149 | 0.00348747781706 | 0.269713739864765 | 0.874,0.935,0.881, | 0.863,0.808,0.942, | 0,029 |
| 207591 | ENSG00000146192 | FGD2 | chr6 | + | 37026090 | 37026437 | 37025791 | 37025938 | 37027428 | 37029069 | 207591 | 40,34,40,23,20,5,6,7,53,29 | 40,26,95,72,66,39,40,23,6 | 10,9,1,22,6,12,4,12,20,24, | 37,44,22,17,39,26,24,17,51 | 298 | 149 | 0.00348975361939 | 0.269792802174903 | 0.333,0.395,0.174, | 0.119,0.093,0.022, | 0,056 |
| 256293 | ENSG00000111450 | STX2 | chr12 | - | 130804162 | 130804540 | 130801414 | 130801488 | 130806981 | 130807090 | 256293 | 0,0,0,0,1,5,2,3,0,0,0,0 | 1,1,10,13,11,13,6,8,5,1,4,7 | 0,2,0,0,1,5,1,1,2,0,4,9,4,2, | 3,0,1,14,1,2,2,10,2,5,5,16,1 | 298 | 149 | 0.00350200386211 | 0.270642654666355 | 0.0,0.0,0.0,0.0,0.0 | 0.0,1.0,0.0,0.0,0.3 | -0,174 |
| 95841 | ENSG00000167483 | NIBAN3 | chr19 | + | 17546685 | 17546797 | 17543523 | 17543631 | 17549443 | 17549527 | 95841 | 474,330,515,342,254,179, | 27,3,9,1,7,3,3,2,9,6,10,2 | 346,385,349,158,327,166, | 1,7,6,3,5,0,0,7,3,3,2,8,17,9 | 261 | 149 | 0.00350586061981 | 0.270764095137826 | 0.909,0.984,0.97,0 | 0.995,0.969,0.971, | -0,016 |
| 74931 | ENSG00000117280 | RAB29 | chr1 | - | 205774832 | 205775086 | 205772495 | 205772567 | 205775272 | 205775460 | 74931 | 14,16,20,23,17,7,9,8,15,11 | 1,0,2,1,0,0,0,0,0,2,0,0 | 13,0,11,5,12,5,4,19,0,25,6, | 2,3,3,1,0,0,1,0,3,1,0,0,2,3, | 298 | 149 | 0.00350719853971 | 0.270764095137826 | 0.875,1.0,0.833,0. | 0.765,0.0,0.647,0. | 0,183 |
| 40843 | ENSG00000004534 | RBM6 | chr3 | + | 49975322 | 49975392 | 49962575 | 49962685 | 50048244 | 50048319 | 40843 | 4,2,14,5,3,0,3,7,3,7,1,0 | 2,2,11,13,10,3,3,5,5,9,0,4 | 3,4,5,9,7,2,1,4,5,9,4,3,7,4, | 4,1,5,1,2,2,3,0,1,8,2,3,7,3, | 219 | 149 | 0.00350734930026 | 0.270764095137826 | 0.576,0.405,0.464, | 0.338,0.731,0.405, | -0,213 |
| 205229 | ENSG00000163029 | SMC6 | chr2 | - | 17728319 | 17728586 | 17726388 | 17726469 | 17731077 | 17731139 | 205229 | 2,0,5,0,0,2,0,5,0,0,0,1 | 87,53,125,59,21,15,23,27, | 2,5,2,6,1,0,0,2,1,0,5,13,2,2 | 52,71,28,21,37,26,16,34,2 | 298 | 149 | 0.00351631786263 | 0.271359128681146 | 0.011,0.0,0.02,0.0, | 0.019,0.034,0.034, | -0,023 |
| 243966 | ENSG00000136270 | TBRG4 | chr7 | - | 45104175 | 45104256 | 45103332 | 45103443 | 45104537 | 45104709 | 243966 | 11,8,9,16,5,18,4,18,15,16, | 22,4,13,9,5,12,6,21,3,5,1,7 | 11,6,12,6,7,1,6,6,1,8,10,50 | 11,17,8,8,19,1,13,16,2,8,2 | 230 | 149 | 0.00352006679406 | 0.271551073889723 | 0.245,0.564,0.31,0 | 0.393,0.186,0.493, | 0,162 |
| 53844 | ENSG00000121897 | LIAS | chr4 | + | 39462195 | 39462289 | 39460789 | 39460962 | 39463524 | 39463605 | 53844 | 10,9,26,20,6,11,3,11,6,4,2, | 1,1,3,5,0,1,0,1,0,1,2,3 | 13,18,9,5,8,2,3,19,6,10,11, | 0,0,0,0,1,0,2,4,0,0,0,1,0,4,1 | 243 | 149 | 0.00352251627936 | 0.271642673351104 | 0.86,0.847,0.842,0 | 1.0,1.0,1.0,1.0,0.8 | -0,072 |
| 221794 | ENSG00000112159 | MDN1 | chr6 | - | 89668247 | 89668443 | 89668013 | 89668151 | 89670918 | 89671080 | 221794 | 1,1,0,1,0,0,0,1,0,0,0,2 | 48,24,28,22,11,17,10,11,2 | 3,4,0,4,9,1,0,4,2,0,1,2,0,1, | 23,23,17,7,12,6,10,6,10,19 | 298 | 149 | 0.00353335782449 | 0.272381138224501 | 0.01,0.02,0.0,0.02 | 0.061,0.08,0.0,0.2 | -0,057 |
| 185751 | ENSG00000067225 | PKM | chr15 | - | 72231115 | 72231184 | 72218943 | 72219110 | 72231323 | 72231624 | 185751 | 232,185,108,126,75,61,84, | 8,5,2,3,5,5,3,5,19,5,0,11 | 81,120,54,46,55,48,27,52, | 2,0,5,0,0,1,0,3,0,1,3,10,1,8 | 218 | 149 | 0.00354003783954 | 0.272798383187266 | 0.952,0.962,0.974, | 0.965,1.0,0.881,1. | -0,023 |
| 153640 | ENSG00000197555 | SIPA1L1 | chr14 | + | 71684420 | 71684758 | 71672347 | 71672622 | 71685361 | 71685631 | 153640 | 8,0,11,4,0,2,3,3,2,0,0,1 | 85,77,113,42,27,34,48,47, | 0,9,14,9,8,3,1,0,2,0,2,1,0,0 | 44,43,44,66,41,41,28,23,1 | 298 | 149 | 0.00354262287502 | 0.272899879747111 | 0.045,0.0,0.046,0. | 0.0,0.095,0.137,0. | -0,018 |
| 82132 | ENSG00000184922 | FMNL1 | chr17 | + | 45246504 | 45246604 | 45246209 | 45246330 | 45246866 | 45247318 | 82132 | 0,4,10,13,1,2,6,9,5,2,2,5 | 44,44,43,22,64,59,37,41,3 | 7,4,0,5,5,0,0,2,3,2,0,0,3,3, | 32,28,15,33,16,12,41,29,34 | 249 | 149 | 0.00354438745557 | 0.272938123953412 | 0.0,0.052,0.122,0. | 0.116,0.079,0.0,0. | 0,029 |

| | | | | | | | | | | | | | | | | | | | | | | |
|---|---|---|---|---|---|---|---|---|---|---|---|---|---|---|---|---|---|---|---|---|---|---|
| 107247 | ENSG00000183354 | KIAA2026 | chr9 | - | 5944870 | 5944960 | 5929043 | 5929159 | 5968018 | 5968081 | 107247 | 11,9,24,13,9,7,8,6,4,1,6,6 | 4,1,1,2,1,0,2,1,2,0,2,0 | 9,8,7,7,13,8,8,1,7,1,4,4,8,4 | 2,2,0,0,0,1,1,1,0,3,0,0,0,0, | 239 | 149 | 0.00354681408571 | 0.272939156206116 | 0.632,0.849,0.937, | 0.737,0.714,1.0,1. | -0,04 |
| 139978 | ENSG00000101265 | RASSF2 | chr20 | - | 4803400 | 4803586 | 4800971 | 4801062 | 4822328 | 4822403 | 139978 | 3,11,5,3,3,0,2,2,2,2,2,2 | 24,14,38,41,74,26,33,34,1 | 35,10,2,0,1,0,2,0,1,6,2,20, | 87,37,13,16,16,10,5,7,16,2 | 298 | 149 | 0.00354693710436 | 0.272939156206116 | 0.059,0.282,0.062, | 0.167,0.119,0.071, | -0,007 |
| 99072 | ENSG00000136280 | CCM2 | chr7 | + | 45071749 | 45071872 | 45069825 | 45069961 | 45072725 | 45072786 | 99072 | 8,0,4,4,3,7,0,6,2,0,0,2 | 21,31,29,51,12,9,19,20,11, | 14,7,4,6,6,5,4,7,2,4,3,6,7,2 | 37,26,7,6,19,25,6,17,5,30, | 272 | 149 | 0.00356214518243 | 0.273929639780166 | 0.173,0.0,0.07,0.0 | 0.172,0.129,0.238, | -0,074 |
| 237757 | ENSG00000184445 | KNTC1 | chr12 | + | 122601850 | 122601991 | 122601535 | 122601625 | 122602568 | 122602740 | 237757 | 1,3,11,5,0,0,4,3,0,0,0,1 | 30,14,83,54,29,26,33,36,2 | 0,1,0,3,0,0,0,0,3,5,0,1,1,2, | 36,43,54,21,25,25,19,38,2 | 290 | 149 | 0.00356235422288 | 0.273929639780166 | 0.017,0.099,0.064, | 0.0,0.012,0.0,0.06 | 0,013 |
| 172849 | ENSG00000211456 | SACM1L | chr3 | + | 45713726 | 45713807 | 45713140 | 45713196 | 45714045 | 45714082 | 172849 | 6,0,7,3,0,1,5,1,0,0,1,1 | 16,1,23,17,12,1,9,8,10,9,4, | 0,4,3,1,1,0,1,0,0,3,1,2,0,1, | 5,10,4,10,6,6,0,3,7,9,3,14,1 | 230 | 149 | 0.00356944548783 | 0.274283368187355 | 0.195,0.0,0.165,0. | 0.0,0.206,0.327,0. | -0,029 |
| 179168 | ENSG00000126012 | KDM5C | chrX | - | 53210675 | 53210866 | 53208890 | 53210576 | 53211496 | 53211906 | 179168 | 70,48,139,104,58,80,57,96 | 1,7,13,9,2,7,1,3,0,1,0,2 | 57,79,61,146,113,58,96,77 | 2,6,8,19,19,5,9,6,7,5,6,5,7, | 298 | 149 | 0.00356950306551 | 0.274283368187355 | 0.972,0.774,0.842, | 0.934,0.868,0.792, | 0,056 |
| 133075 | ENSG00000112297 | CRYBG1 | chr6 | + | 106544787 | 106544933 | 106544570 | 106544697 | 106551851 | 106551978 | 133075 | 136,125,222,119,46,36,61, | 1,1,1,1,5,0,0,2,2,0,1,0 | 119,195,149,172,140,86,3 | 0,0,5,7,6,2,2,1,2,2,1,4,0,3,1 | 295 | 149 | 0.00357660773248 | 0.27473121301549 | 0.986,0.984,0.991, | 1.0,1.0,0.938,0.92 | 0,02 |
| 232609 | ENSG00000176871 | WSB2 | chr12 | - | 118034985 | 118035353 | 118032686 | 118034358 | 118036337 | 118036510 | 232609 | 85,52,102,121,49,79,81,73 | 1,2,1,0,0,1,0,5,0,0,0,0 | 62,157,66,32,56,23,18,49, | 1,2,1,2,0,2,0,2,0,1,5,0,2,5, | 298 | 149 | 0.00357858449623 | 0.274784987409233 | 0.977,0.929,0.981, | 0.969,0.975,0.971, | 0,035 |
| 47527 | ENSG00000160991 | ORAI2 | chr7 | + | 102436229 | 102436333 | 102433569 | 102433661 | 102438943 | 102439181 | 47527 | 31,44,84,84,116,114,124,2 | 6,2,8,5,4,4,11,30,2,4,7,13 | 70,46,37,40,36,29,49,106, | 5,7,4,9,6,0,1,16,6,7,10,55, | 253 | 149 | 0.00358566717639 | 0.275230646234521 | 0.753,0.928,0.861, | 0.892,0.795,0.845, | 0,045 |
| 218666 | ENSG00000024048 | UBR2 | chr6 | + | 42633912 | 42634294 | 42632804 | 42632904 | 42635417 | 42635546 | 218666 | 4,2,17,5,2,4,1,5,1,0,1,2 | 31,17,71,47,25,36,23,25,2 | 8,3,5,0,5,2,0,2,1,0,4,2,2,0,1 | 46,45,22,34,46,22,16,28,1 | 298 | 149 | 0.0035994664546 | 0.276174672610669 | 0.061,0.056,0.107, | 0.08,0.032,0.102,0 | 0,016 |
| 49617 | ENSG00000144218 | AFF3 | chr2 | - | 100011496 | 100011571 | 100008811 | 100008932 | 100142483 | 100142747 | 49617 | 16,5,30,22,8,14,10,8,10,12 | 0,4,1,7,0,0,6,0,0,1,0,0 | 6,15,10,40,27,5,9,4,13,10, | 2,0,0,0,9,0,0,0,1,1,1,4,0,0, | 224 | 149 | 0.0036005321322 | 0.276174672610669 | 1.0,0.454,0.952,0. | 0.666,1.0,1.0,1.0,0 | -0,022 |
| 191554 | ENSG00000159363 | ATP13A2 | chr1 | - | 16989886 | 16990044 | 16989690 | 16989770 | 16990126 | 16990287 | 191554 | 12,0,10,10,4,15,10,1,1,0,2, | 0,1,2,5,2,2,3,0,1,2,0,1 | 4,1,2,1,1,7,6,10,5,5,4,11,1, | 1,0,1,0,2,0,2,8,0,0,0,2,0,0, | 298 | 149 | 0.00360204170788 | 0.276192033782943 | 1.0,0.0,0.714,0.5,0 | 0.667,1.0,0.5,1.0,0 | -0,176 |
| 200491 | ENSG00000156515 | HK1 | chr10 | + | 69398594 | 69398826 | 69394949 | 69395105 | 69400990 | 69401884 | 200491 | 17,7,47,31,25,9,20,13,15,1 | 10,5,8,2,0,1,4,2,0,1,0,2 | 27,33,5,5,11,13,6,12,6,22,1 | 4,4,0,0,0,0,0,0,0,3,2,2,0,0, | 298 | 149 | 0.00360452189856 | 0.276283779469006 | 0.459,0.412,0.746, | 0.771,0.805,1.0,1. | -0,119 |
| 103622 | ENSG00000158636 | EMSY | chr11 | + | 76539598 | 76539640 | 76537794 | 76537950 | 76542215 | 76542367 | 103622 | 16,8,6,2,6,5,4,5,15,3,7,5 | 2,2,4,0,1,0,0,1,1,6,0,2 | 13,7,5,5,4,5,0,24,8,7,7,4,7, | 0,1,0,1,0,0,1,4,0,0,0,5,1,1, | 191 | 149 | 0.00361007058111 | 0.276572779739241 | 0.862,0.757,0.539, | 1.0,0.845,1.0,0.79 | -0,004 |
| 68256 | ENSG00000143110 | C1orf162 | chr1 | + | 111477333 | 111477428 | 111476797 | 111476867 | 111477725 | 111477775 | 68256 | 79,23,71,85,17,1,32,15,11 | 9,0,4,3,0,0,0,0,2,3,0,1 | 29,14,65,29,29,42,3,5,19,8 | 1,3,1,4,5,1,0,0,1,3,2,2,2,3,1 | 244 | 149 | 0.00361086233427 | 0.276572779739241 | 0.843,1.0,0.916,0. | 0.947,0.74,0.975,0 | 0,08 |
| 94136 | ENSG00000112303 | VNN2 | chr6 | - | 132753417 | 132753491 | 132752713 | 132752749 | 132755842 | 132756035 | 94136 | 1,0,4,1,5,2,4,9,2,1,0,0 | 11,8,20,11,21,30,21,7,9,3, | 2,0,0,0,4,1,4,0,1,3,0,6,3,0, | 10,9,4,10,5,11,6,8,7,13,4,2 | 223 | 149 | 0.00361366484393 | 0.276688971193133 | 0.057,0.0,0.118,0. | 0.118,0.0,0.0,0.0,0 | 0,021 |
| 59970 | ENSG00000118308 | IRAG2 | chr12 | + | 25062819 | 25062900 | 25052270 | 25052956 | 25063719 | 25063816 | 59970 | 419,164,1059,701,400,304 | 12,7,61,38,8,15,10,22,8,3, | 458,609,372,145,213,195, | 5,7,14,9,8,6,7,10,3,11,11,3 | 230 | 149 | 0.00362152295203 | 0.277192036446788 | 0.958,0.938,0.918, | 0.983,0.983,0.945, | -0,007 |
| 171196 | ENSG00000136819 | C9orf78 | chr9 | - | 129834308 | 129834766 | 129833657 | 129833709 | 129835138 | 129835254 | 171196 | 93,43,51,23,9,5,5,11,28,26 | 0,1,2,1,1,0,8,17,0,0,0,0 | 18,35,42,26,26,4,10,16,6,1 | 0,1,0,0,1,0,1,1,0,0,7,1,0,0, | 298 | 149 | 0.00365168893234 | 0.279401585707953 | 1.0,0.956,0.927,0. | 1.0,0.946,1.0,1.0,0 | -0,075 |
| 132589 | ENSG00000112200 | ZNF451 | chr6 | + | 57150718 | 57150862 | 57147089 | 57148693 | 57152220 | 57152351 | 132589 | 86,64,95,45,59,58,37,66,6 | 0,1,4,2,0,0,0,0,0,0,0,1 | 109,91,62,66,48,33,41,57, | 2,2,1,2,2,1,0,4,2,0,5,0,1,1, | 293 | 149 | 0.00365406708298 | 0.279422078543486 | 1.0,0.97,0.924,0.9 | 0.965,0.959,0.969, | 0,041 |
| 103687 | ENSG00000113013 | HSPA9 | chr5 | - | 138559863 | 138560091 | 138558552 | 138558657 | 138561579 | 138561789 | 103687 | 236,197,412,319,152,155, | 3,0,8,11,2,3,1,3,3,0,0,3 | 251,348,195,120,182,110, | 3,7,1,0,1,1,0,2,0,1,2,6,0,1, | 298 | 149 | 0.00365579625265 | 0.279422078543486 | 0.975,1.0,0.963,0. | 0.977,0.961,0.99,1 | -0,012 |
| 238928 | ENSG00000104325 | DECR1 | chr8 | + | 90017296 | 90017326 | 90001516 | 90001561 | 90018908 | 90018966 | 238928 | 7,8,21,22,10,7,5,12,7,1,1,9 | 2,0,8,8,0,5,1,0,0,0,0,2 | 8,15,10,5,5,7,1,2,2,18,6,21 | 8,4,4,0,4,1,0,1,5,11,14,27, | 179 | 149 | 0.00365585149527 | 0.279422078543486 | 0.744,1.0,0.686,0. | 0.454,0.757,0.675, | 0,189 |
| 135034 | ENSG00000284431 | | chr22 | + | 40364569 | 40365056 | 40364275 | 40364365 | 40366435 | 40366498 | 135034 | 47,14,74,74,37,29,20,27,3 | 5,1,9,9,4,1,2,7,6,1,0,7 | 35,42,32,8,22,27,7,11,15,3 | 2,10,4,3,11,7,2,0,4,13,2,9,1 | 298 | 149 | 0.00366836145765 | 0.280132885451736 | 0.825,0.875,0.804, | 0.897,0.677,0.8,0. | 0,131 |
| 155892 | ENSG00000196199 | MPHOSPH8 | chr13 | + | 19660151 | 19660269 | 19659200 | 19659289 | 19661697 | 19661838 | 155892 | 11,1,7,2,2,4,7,3,1,0,0,2 | 37,25,36,28,32,28,20,31,31 | 7,8,0,3,0,0,2,0,1,0,1,0,5,1, | 28,22,17,14,7,11,9,31,13,2 | 267 | 149 | 0.00366932843174 | 0.280132885451736 | 0.142,0.022,0.098, | 0.122,0.169,0.0,0. | 0,008 |
| 58897 | ENSG00000068097 | HEATR6 | chr17 | - | 60073745 | 60073886 | 60073163 | 60073279 | 60076129 | 60076237 | 58897 | 25,11,79,45,36,32,23,18,2 | 0,1,0,7,2,0,4,2,0,0,1,2 | 42,14,38,57,11,11,13,13,1 | 2,2,1,1,0,1,1,0,0,1,8,6,1,4, | 290 | 149 | 0.00366952269078 | 0.280132885451736 | 1.0,0.85,1.0,0.768, | 0.915,0.782,0.951, | 0,039 |
| 91859 | ENSG00000204592 | HLA-E | chr6 | + | 30492403 | 30492436 | 30491536 | 30491653 | 30492540 | 30492583 | 91859 | 667,612,1017,1044,393,31 | 10,5,8,15,4,0,5,9,0,3,2,9 | 670,789,508,340,436,291, | 10,13,4,14,16,5,2,21,5,6,8, | 182 | 149 | 0.0036709749318 | 0.280132885451736 | 0.982,0.99,0.99,0. | 0.982,0.98,0.99,0. | 0,008 |
| 63158 | ENSG00000124406 | ATP8A1 | chr4 | - | 42594288 | 42594362 | 42588259 | 42588329 | 42600477 | 42600518 | 63158 | 29,8,62,35,18,32,39,55,21, | 1,0,3,2,2,4,3,0,1,2,0,1 | 35,64,24,24,9,3,13,33,17,3 | 1,4,0,0,3,4,0,4,3,3,0,31,4,6 | 223 | 149 | 0.00367165914696 | 0.280132885451736 | 0.951,1.0,0.932,0. | 0.959,0.914,1.0,1. | 0,071 |
| 69727 | ENSG00000177225 | GATD1 | chr11 | - | 773427 | 773629 | 772426 | 772521 | 774007 | 774113 | 69727 | 24,9,17,15,19,13,14,16,46, | 1,1,2,1,2,4,1,5,0,1,1,0 | 13,3,15,19,7,6,17,22,3,13, | 0,1,1,1,0,0,2,0,0,1,4,10,0,0 | 298 | 149 | 0.00367587725372 | 0.280286529818995 | 0.923,0.818,0.81,0 | 1.0,0.6,0.882,0.90 | -0,004 |
| 98268 | ENSG00000136286 | MYO1G | chr7 | - | 44976563 | 44976657 | 44975475 | 44975649 | 44978866 | 44979069 | 98268 | 114,90,51,73,53,40,34,18, | 6,0,0,3,2,2,2,0,2,2,0,2 | 49,50,53,74,40,37,18,15,3 | 0,2,0,0,0,0,0,0,0,0,2,0,2, | 243 | 149 | 0.00367627745854 | 0.280286529818995 | 0.921,1.0,1.0,0.93 | 1.0,0.939,1.0,1.0,1 | -0,035 |
| 42465 | ENSG00000078618 | NRDC | chr1 | - | 51792045 | 51792098 | 51791577 | 51791661 | 51792376 | 51792424 | 42465 | 79,48,90,58,33,34,27,52,1 | 0,0,3,0,6,0,0,1,0,1,2,0 | 51,49,30,24,28,40,37,44,3 | 4,2,2,6,5,3,0,2,2,2,2,3,0,0,1 | 202 | 149 | 0.00367768970147 | 0.280294912229848 | 1.0,1.0,0.957,1.0,0 | 0.904,0.948,0.917, | 0,028 |
| 232608 | ENSG00000176871 | WSB2 | chr12 | - | 118034985 | 118035314 | 118032686 | 118034358 | 118036337 | 118036510 | 232608 | 85,52,102,121,49,79,81,71 | 1,2,1,0,0,1,0,5,0,0,0,0 | 62,157,66,32,56,23,18,49, | 1,2,1,2,0,2,0,2,0,1,5,0,2,5, | 298 | 149 | 0.00368285590811 | 0.28040583476697 | 0.977,0.929,0.981, | 0.969,0.975,0.971, | 0,035 |
| 203606 | ENSG00000120451 | SNX19 | chr11 | - | 130907952 | 130908423 | 130906624 | 130906721 | 130910017 | 130910137 | 203606 | 24,12,53,30,2,22,20,23,14, | 0,2,1,0,0,0,0,1,1,0,0,0 | 28,17,17,7,25,18,19,34,39, | 0,1,1,0,5,1,2,1,0,0,4,1,2,1,1 | 298 | 149 | 0.00368286954948 | 0.28040583476697 | 1.0,0.75,0.964,1.0, | 1.0,0.895,0.895,1. | 0,079 |
| 208655 | ENSG00000152520 | PAN3 | chr13 | + | 28239578 | 28240037 | 28220230 | 28220378 | 28256291 | 28256539 | 208655 | 5,1,5,9,3,11,7,10,2,0,0,1 | 58,50,74,37,49,47,74,98,4 | 10,11,9,9,6,5,11,8,5,6,2,1, | 46,66,49,77,37,39,40,50,3 | 298 | 149 | 0.00368321672302 | 0.28040583476697 | 0.041,0.01,0.033,0 | 0.098,0.077,0.084, | -0,027 |
| 87436 | ENSG00000036257 | CUL3 | chr2 | - | 224556294 | 224556420 | 224535527 | 224535641 | 224557658 | 224557820 | 87436 | 2,2,6,1,0,0,0,6,0,0,0,4 | 28,34,106,66,39,59,33,59, | 7,7,4,2,0,1,2,4,2,8,4,4,1,1, | 69,99,35,70,49,20,49,35,61 | 275 | 149 | 0.00368435634607 | 0.28040583476697 | 0.037,0.031,0.03,0 | 0.052,0.037,0.058, | -0,021 |
| 232610 | ENSG00000176871 | WSB2 | chr12 | - | 118034985 | 118035475 | 118032686 | 118034358 | 118036337 | 118036510 | 232610 | 85,52,102,121,49,79,81,71 | 1,2,1,0,0,1,0,5,0,0,0,0 | 63,157,66,32,56,23,18,49, | 1,2,1,2,0,2,0,2,0,1,5,0,2,5, | 298 | 149 | 0.0036861464906 | 0.280442911035111 | 0.977,0.929,0.981, | 0.969,0.975,0.971, | 0,035 |
| 192102 | ENSG00000266028 | SRGAP2 | chr1 | + | 206449746 | 206449805 | 206446074 | 206446299 | 206450385 | 206450465 | 192102 | 1,0,1,6,0,0,1,9,0,3,0,3 | 13,13,9,8,3,15,3,7,7,7,2,6 | 2,4,0,6,0,0,2,1,1,0,0,3,2,0, | 8,15,3,3,9,4,15,8,4,11,7,18 | 208 | 149 | 0.00369448973013 | 0.280938659365788 | 0.052,0.0,0.074,0. | 0.152,0.16,0.0,0.5 | 0,042 |
| 203710 | ENSG00000117280 | RAB29 | chr1 | - | 205774832 | 205775091 | 205772495 | 205772567 | 205775272 | 205775482 | 203710 | 12,15,18,23,17,7,5,8,15,11 | 1,0,2,1,0,0,0,0,0,2,0,0 | 11,0,11,5,12,5,4,19,0,26,4, | 2,3,3,1,0,0,1,0,3,1,0,0,2,3,1 | 298 | 149 | 0.00369532235264 | 0.280938659365788 | 0.857,1.0,0.818,0. | 0.733,0.0,0.647,0. | 0,18 |
| 203704 | ENSG00000117280 | RAB29 | chr1 | - | 205774832 | 205774958 | 205772495 | 205772567 | 205775272 | 205775482 | 203704 | 12,13,17,21,13,5,5,8,15,11 | 1,0,2,1,0,0,0,0,0,2,0,0 | 10,0,11,4,11,4,3,16,0,23,3, | 2,3,3,1,0,0,1,0,3,1,0,0,2,3,1 | 275 | 149 | 0.0036965784823 | 0.280938659365788 | 0.867,1.0,0.822,0. | 0.73,0.0,0.665,0.6 | 0,189 |
| 84265 | ENSG00000064666 | CNN2 | chr19 | + | 1036129 | 1036246 | 1032558 | 1032696 | 1036415 | 1036472 | 84265 | 377,231,331,360,366,174, | 4,6,9,3,11,0,2,0,2,5,2,0 | 310,311,193,155,163,136, | 6,0,6,7,1,4,9,6,3,3,0,14,15, | 266 | 149 | 0.00369879526824 | 0.281007908358123 | 0.981,0.956,0.954, | 0.967,1.0,0.947,0. | 0,019 |
| 89533 | ENSG00000163754 | GYG1 | chr3 | + | 149024052 | 149024272 | 149009275 | 149009402 | 149026759 | 149027632 | 89533 | 12,11,54,61,17,26,16,31,7, | 0,1,5,2,0,1,0,0,0,2,0,0 | 36,31,27,18,37,17,14,10,1 | 3,4,0,0,0,2,0,1,0,8,0,4,0,4,1 | 298 | 149 | 0.0037012051596 | 0.281091774066996 | 1.0,0.846,0.844,0. | 0.857,0.795,1.0,1. | 0,052 |
| 198064 | ENSG00000112232 | KHDRBS2 | chr6 | - | 62152193 | 62152312 | 62047877 | 62047994 | 62177184 | 62177312 | 198064 | 8,0,16,4,6,8,9,2,6,2,4,1 | 57,24,105,56,44,53,34,57, | 2,0,1,7,4,0,3,1,1,2,4,4,1,3, | 71,60,43,32,41,27,11,27,3 | 268 | 149 | 0.00370482504338 | 0.281267442296903 | 0.072,0.0,0.078,0. | 0.015,0.0,0.013,0. | 0,029 |
| 206937 | ENSG00000048991 | R3HDM1 | chr2 | + | 135633695 | 135633828 | 135631860 | 135632001 | 135635889 | 135635998 | 206937 | 13,4,4,5,8,1,4,5,0,4,0,0 | 34,19,66,38,7,21,13,22,9,9 | 2,1,2,1,2,1,4,2,0,1,3,2,2,5, | 23,24,12,16,12,11,18,13,2 | 282 | 149 | 0.00372194499873 | 0.282381725499843 | 0.168,0.1,0.031,0. | 0.044,0.022,0.081, | 0,045 |
| 83257 | ENSG00000130706 | ADRM1 | chr20 | + | 62308367 | 62308470 | 62308020 | 62308178 | 62308654 | 62308860 | 83257 | 59,40,48,38,43,15,23,23,1 | 31,37,38,11,8,13,24,15,24, | 23,26,27,8,31,23,10,10,7,3 | 20,26,15,23,25,6,18,18,8,1 | 252 | 149 | 0.00372212625153 | 0.282381725499843 | 0.529,0.39,0.428,0 | 0.405,0.372,0.516, | 0,159 |
| 254221 | ENSG00000134283 | PPHLN1 | chr12 | + | 42351884 | 42352241 | 42335882 | 42335974 | 42355160 | 42355678 | 254221 | 51,22,54,59,18,28,22,36,4 | 48,30,63,46,31,35,18,39,41 | 33,60,24,34,33,25,6,30,23, | 45,19,21,24,23,10,4,26,13, | 298 | 149 | 0.00372714354505 | 0.282662731622501 | 0.347,0.268,0.3,0. | 0.268,0.612,0.364, | -0,105 |
| 30877 | ENSG00000145715 | RASA1 | chr5 | + | 87363347 | 87363504 | 87362550 | 87362671 | 87369812 | 87369900 | 30877 | 57,27,113,112,57,50,45,75 | 6,0,11,8,4,3,3,8,0,2,2,1 | 85,50,32,10,37,41,36,34,4 | 8,8,11,10,4,0,2,2,6,6,21,6,1 | 298 | 149 | 0.00373427866058 | 0.283104096652094 | 0.826,1.0,0.837,0. | 0.842,0.758,0.593, | 0,118 |
| 245837 | ENSG00000110075 | PPP6R3 | chr11 | + | 68583034 | 68583129 | 68575957 | 68576043 | 68587944 | 68588024 | 245837 | 9,6,22,13,3,3,1,4,4,1,1,3 | 0,1,4,1,2,2,2,0,2,0,0,0 | 14,10,2,7,7,2,5,2,8,8,18,14 | 5,0,2,2,4,2,0,0,3,2,2,8,3,2, | 244 | 149 | 0.00373724278016 | 0.283177826979873 | 1.0,0.786,0.771,0. | 0.631,1.0,0.379,0. | 0,09 |
| 241284 | ENSG00000107077 | KDM4C | chr9 | + | 7015784 | 7015929 | 7013787 | 7014001 | 7046861 | 7046917 | 241284 | 55,19,27,18,23,14,19,11,2 | 5,0,10,1,0,0,4,0,0,2,0,3 | 30,9,25,17,17,13,11,6,8,26 | 4,9,2,9,5,2,0,2,0,0,10,2,3,0 | 294 | 149 | 0.00373788258406 | 0.283177826979873 | 0.848,1.0,0.578,0. | 0.792,0.336,0.864, | 0,159 |
| 13155 | ENSG00000180747 | SMG1P3 | chr16 | - | 21471653 | 21471751 | 21469746 | 21469919 | 21472672 | 21472745 | 13155 | 17,5,22,8,17,25,17,12,6,9, | 0,0,0,0,12,9,0,0,0,0,0,0 | 18,25,17,30,16,15,7,19,27, | 0,0,0,1,0,0,0,0,0,0,0,0,0,0, | 247 | 149 | 0.00374700910187 | 0.283769358200064 | 1.0,1.0,1.0,1.0,0.4 | 1.0,1.0,1.0,0.948,1 | -0,073 |
| 135143 | ENSG00000100239 | PPP6R2 | chr22 | + | 50418868 | 50418979 | 50416091 | 50416157 | 50419348 | 50419462 | 135143 | 52,29,50,46,24,33,18,27,3 | 0,1,2,0,0,1,0,0,3,1,0,0 | 33,31,28,24,24,27,26,36,2 | 1,3,1,6,0,0,1,1,2,0,0,2,1,1, | 260 | 149 | 0.00374998603381 | 0.283894915245501 | 1.0,0.943,0.935,1. | 0.95,0.856,0.941,0 | 0,05 |
| 63268 | ENSG00000119004 | CYP20A1 | chr2 | + | 203285611 | 203285732 | 203280058 | 203280113 | 203289764 | 203289876 | 63268 | 49,31,85,77,68,46,42,60,3 | 0,0,8,3,0,3,1,2,2,2,0,0 | 63,57,25,30,48,31,27,80,2 | 5,5,0,0,6,0,0,0,3,2,2,4,1,0, | 270 | 149 | 0.00375208756916 | 0.283954134879703 | 1.0,1.0,0.854,0.93 | 0.874,0.863,1.0,1. | 0,024 |
| 172676 | ENSG00000068745 | IP6K2 | chr3 | - | 48693269 | 48693318 | 48693058 | 48693179 | 48695089 | 48695588 | 172676 | 4,4,1,3,0,0,1,4,0,0,2,4 | 49,51,86,47,16,17,20,33,1 | 0,0,1,2,1,1,1,2,0,0,0,0,3,0,1 | 44,67,27,15,39,23,15,21,1 | 198 | 149 | 0.00375722561874 | 0.284154922522639 | 0.058,0.056,0.009, | 0.0,0.0,0.027,0.09 | 0,015 |
| 245345 | ENSG00000130764 | LRRC47 | chr1 | - | 3783769 | 3783841 | 3782660 | 3782763 | 3783995 | 3784175 | 245345 | 1,2,7,0,0,4,6,0,0,3,1,0 | 5,4,24,15,18,20,26,25,13,6 | 0,0,0,0,2,1,0,2,0,1,5,0,3,0, | 19,22,14,6,11,7,19,21,13,2 | 221 | 149 | 0.00375738118347 | 0.284154922522639 | 0.119,0.252,0.164, | 0.0,0.0,0.0,0.0,0.1 | 0,048 |
| 419 | ENSG00000186866 | POFUT2 | chr21 | - | 45283382 | 45283527 | 45282348 | 45282459 | 45285677 | 45285691 | 419 | 13,16,31,21,11,26,19,15,1 | 1,2,4,3,1,2,1,0,5,5,0,0 | 15,6,13,6,7,10,22,14,7,16,1 | 1,5,0,4,0,0,1,0,0,0,0,3,0,2,1 | 294 | 149 | 0.00376463524772 | 0.284603515631199 | 0.868,0.802,0.797, | 0.884,0.378,1.0,0. | -0,027 |
| 102712 | ENSG00000126746 | ZNF384 | chr12 | - | 6672349 | 6672532 | 6670759 | 6670838 | 6673215 | 6673440 | 102712 | 19,5,27,19,15,16,17,7,23,1 | 42,21,33,20,32,18,30,39,1 | 11,10,22,4,6,4,4,19,6,5,7,1 | 36,29,24,20,23,12,15,29,1 | 298 | 149 | 0.00377408368431 | 0.285205547512607 | 0.184,0.106,0.29,0 | 0.133,0.147,0.314, | 0,09 |
| 45029 | ENSG00000186001 | LRCH3 | chr3 | + | 197858833 | 197858905 | 197854391 | 197854445 | 197865422 | 197865471 | 45029 | 33,19,54,43,22,23,26,33,3 | 2,2,4,8,5,6,3,2,2,0,1,2 | 29,36,20,22,20,17,31,51,2 | 1,2,1,0,2,0,0,2,0,9,1,2,3,1, | 221 | 149 | 0.0037752489412 | 0.285205547512607 | 0.918,0.865,0.901, | 0.951,0.924,0.931, | -0,054 |
| 184586 | ENSG00000198265 | HELZ | chr17 | - | 67190159 | 67190355 | 67189588 | 67189696 | 67193966 | 67194042 | 184586 | 26,24,43,34,13,26,13,24,7, | 0,3,1,4,0,2,0,4,0,2,0,3 | 46,30,35,29,19,21,15,16,1 | 3,0,1,5,1,0,4,0,1,1,1,2,2,3, | 298 | 149 | 0.00378011655105 | 0.285473075578611 | 1.0,0.8,0.956,0.81, | 0.885,1.0,0.946,0. | 0,021 |
| 161415 | ENSG00000164187 | LMBRD2 | chr5 | - | 36118062 | 36118156 | 36117734 | 36117916 | 36122279 | 36122463 | 161415 | 7,2,2,0,0,1,1,3,0,5,0,0 | 15,5,19,14,8,14,9,6,1,4,6,6 | 0,0,0,1,1,0,1,1,2,0,1,1,1,0, | 11,12,4,2,11,2,0,11,7,3,3,1 | 243 | 149 | 0.00378234562629 | 0.285541224655217 | 0.222,0.197,0.061, | 0.0,0.0,0.0,0.235,0 | -0,017 |
| 167214 | ENSG00000155229 | MMS19 | chr10 | - | 97468965 | 97469126 | 97468251 | 97468406 | 97469645 | 97469723 | 167214 | 33,33,39,25,30,34,36,22,5 | 11,7,12,14,6,9,6,11,5,5,6,2 | 17,26,24,6,12,9,22,14,9,10 | 6,16,6,7,12,0,9,7,5,6,15,15 | 298 | 149 | 0.00379167596936 | 0.286145235119678 | 0.6,0.702,0.619,0. | 0.586,0.448,0.667, | 0,135 |
| 136550 | ENSG00000028277 | POU2F2 | chr19 | - | 42122686 | 42122752 | 42122510 | 42122576 | 42132383 | 42132478 | 136550 | 5,2,0,6,1,5,5,4,2,2,0,0 | 41,32,30,20,31,24,14,14,2 | 0,0,0,1,0,0,0,2,0,0,3,2,2,0, | 10,3,20,9,3,3,7,7,11,11,3,1 | 215 | 149 | 0.00379346511378 | 0.286179912339286 | 0.078,0.042,0.0,0. | 0.0,0.0,0.0,0.071,0 | 0,01 |
| 102231 | ENSG00000144815 | NXPE3 | chr3 | + | 101816794 | 101817001 | 101807052 | 101807126 | 101821403 | 101822072 | 102231 | 23,8,44,47,18,27,11,12,22, | 0,0,2,2,2,0,0,6,2,0,2,3 | 34,53,13,18,9,18,7,22,24,3 | 0,2,0,2,0,0,1,1,0,0,1,0,0,2, | 298 | 149 | 0.00379783439192 | 0.286409142959823 | 1.0,1.0,0.917,0.92 | 1.0,0.93,1.0,0.818, | -0,062 |
| 122942 | ENSG00000135365 | PHF21A | chr11 | - | 46117921 | 46118114 | 46092182 | 46092223 | 46118741 | 46118800 | 122942 | 9,11,21,12,6,6,4,9,2,3,2,2 | 0,5,7,0,6,6,2,2,4,4,5,0 | 11,5,10,17,1,5,3,7,14,6,6,5 | 2,1,0,1,0,3,0,0,1,0,7,0,3,4, | 298 | 149 | 0.00380552507456 | 0.286888605016681 | 1.0,0.524,0.6,1.0,0 | 0.733,0.714,1.0,0. | -0,219 |
| 169145 | ENSG00000003056 | M6PR | chr12 | - | 8941821 | 8941940 | 8940362 | 8940869 | 8942415 | 8942542 | 169145 | 62,32,51,54,45,39,40,50,6 | 11,7,25,13,6,8,4,14,1,6,4,1 | 51,15,25,8,7,23,14,52,25,4 | 20,19,10,11,12,8,10,12,7,4 | 268 | 149 | 0.00380977862203 | 0.287035062885176 | 0.758,0.718,0.531, | 0.586,0.305,0.582, | 0,139 |
| 119121 | ENSG00000110514 | MADD | chr11 | + | 47293882 | 47293983 | 47290609 | 47290816 | 47295495 | 47295567 | 119121 | 36,30,73,59,39,33,26,31,2 | 7,1,5,4,1,0,0,0,2,2,0,0 | 33,25,27,27,20,20,21,29,1 | 0,4,0,0,0,0,0,2,0,0,0,4,0,1,1 | 250 | 149 | 0.00381013503939 | 0.287035062885176 | 0.754,0.947,0.897, | 1.0,0.788,1.0,1.0,1 | -0,026 |
| 167500 | ENSG00000145495 | MARCHF6 | chr5 | + | 10420134 | 10420341 | 10417269 | 10417404 | 10423734 | 10423835 | 167500 | 9,8,4,2,0,1,5,4,2,0,0,1 | 86,76,160,56,62,94,69,86, | 3,6,4,24,1,2,5,1,4,5,9,3,1,6 | 105,110,67,80,72,46,73,10 | 298 | 149 | 0.00381284103548 | 0.287138414593211 | 0.05,0.05,0.012,0. | 0.014,0.027,0.029, | -0,015 |
| 21878 | ENSG00000258289 | CHURC1 | chr14 | + | 64926009 | 64926080 | 64923990 | 64924126 | 64943298 | 64944056 | 21878 | 11,7,42,30,6,13,12,7,4,4,5, | 1,1,1,2,0,0,1,2,6,2,1,1 | 21,35,16,6,15,14,10,9,15,2 | 0,1,1,1,2,1,3,0,0,0,1,0,1,1, | 220 | 149 | 0.00381653504671 | 0.287276386128067 | 0.882,0.826,0.966, | 1.0,0.96,0.916,0.8 | -0,115 |
| 93007 | ENSG00000223501 | VPS52 | chr6 | - | 33267681 | 33267739 | 33266556 | 33266712 | 33267864 | 33267997 | 93007 | 14,10,33,22,11,11,8,11,12, | 0,0,2,0,1,0,0,0,1,3,0,0 | 8,12,22,13,18,16,12,7,13,1 | 4,5,2,1,2,0,1,2,0,0,2,1,0,0, | 207 | 149 | 0.0038174420745 | 0.287276386128067 | 1.0,1.0,0.922,1.0,0 | 0.59,0.633,0.888,0 | 0,082 |
| 66108 | ENSG00000171791 | BCL2 | chr18 | - | 63301661 | 63301865 | 63280886 | 63281214 | 63318081 | 63318666 | 66108 | 9,2,4,0,5,2,1,12,8,2,0,4 | 2,0,4,5,7,4,5,6,11,2,3,3 | 6,6,3,6,1,3,4,5,4,1,7,4,2,7, | 7,0,1,0,0,4,0,11,3,1,8,3,0,3 | 298 | 149 | 0.00381867733139 | 0.287276386128067 | 0.692,1.0,0.333,0. | 0.3,1.0,0.6,1.0,1.0, | -0,282 |
| 224599 | ENSG00000239713 | APOBEC3G | chr22 | + | 39078985 | 39079085 | 39077274 | 39077378 | 39080932 | 39081194 | 224599 | 93,51,261,207,117,40,35,3 | 0,1,9,7,10,2,2,2,3,0,1,0 | 96,43,80,16,33,42,3,48,15, | 4,6,6,0,2,3,1,5,3,4,1,5,3,6, | 249 | 149 | 0.00382296563323 | 0.287498503216536 | 1.0,0.968,0.946,0. | 0.935,0.811,0.889, | 0,063 |
| 146540 | ENSG00000179583 | CIITA | chr16 | + | 10935117 | 10935200 | 10923227 | 10923325 | 10941185 | 10943021 | 146540 | 6,1,4,1,5,1,5,6,0,0,1,2 | 37,12,59,13,28,22,21,13,7, | 1,2,1,0,1,0,2,1,4,2,0,6,2,0, | 32,37,11,35,23,9,22,35,34, | 232 | 149 | 0.00382882003943 | 0.287838199757792 | 0.094,0.051,0.042, | 0.02,0.034,0.055,0 | 0,023 |
| 123679 | ENSG00000179532 | DNHD1 | chr11 | + | 6520237 | 6520289 | 6519964 | 6520102 | 6528521 | 6528787 | 123679 | 5,4,16,21,4,17,20,14,11,4, | 3,1,6,3,1,5,4,6,2,2,0,6 | 11,10,18,25,12,3,24,26,18, | 0,0,6,3,0,0,2,5,4,1,3,1,1,2, | 201 | 149 | 0.00383064207675 | 0.287874624588331 | 0.553,0.748,0.664, | 1.0,1.0,0.69,0.861, | -0,15 |
| 40832 | ENSG00000136819 | C9orf78 | chr9 | - | 129834711 | 129834766 | 129833657 | 129833709 | 129835138 | 129835254 | 40832 | 90,42,51,24,6,4,6,10,27,24 | 0,1,2,1,1,0,8,17,0,0,0,0 | 16,32,43,23,26,3,7,19,6,17 | 0,1,0,0,1,0,1,1,0,0,7,1,0,0, | 204 | 149 | 0.0038338504938 | 0.288015174740338 | 1.0,0.968,0.949,0. | 1.0,0.959,1.0,1.0,0 | -0,068 |
| 89539 | ENSG00000148690 | FRA10AC1 | chr10 | - | 93692008 | 93692093 | 93687403 | 93687449 | 93692645 | 93692729 | 89539 | 21,7,13,13,9,11,6,18,29,8, | 8,4,2,4,6,1,4,1,2,2,0,5 | 33,15,22,4,4,7,11,17,6,14, | 2,0,1,2,3,3,0,0,0,2,2,4,0,0, | 234 | 149 | 0.00384138558354 | 0.288480551476238 | 0.626,0.527,0.805, | 0.913,1.0,0.933,0. | -0,1 |
| 159495 | ENSG00000038382 | TRIO | chr5 | + | 14311294 | 14311495 | 14304460 | 14304592 | 14316512 | 14316743 | 159495 | 0,0,1,0,6,5,0,3,2,0,0,2 | 8,16,20,1,52,246,46,147,1 | 0,2,4,0,0,0,0,3,0,4,0,18,2,8 | 11,8,77,69,29,21,9,84,14,2 | 298 | 149 | 0.00384581279437 | 0.288712289342536 | 0.0,0.0,0.024,0.0,0 | 0.0,0.111,0.025,0. | -0,014 |
| 121956 | ENSG00000053372 | MRTO4 | chr1 | + | 19257453 | 19257521 | 19257063 | 19257145 | 19257832 | 19257984 | 121956 | 12,9,42,30,19,17,6,17,21,1 | 0,1,1,2,4,1,3,0,4,0,2,1 | 10,14,10,2,15,7,7,8,3,7,17, | 7,6,2,1,2,2,3,3,1,1,1,9,4,2, | 217 | 149 | 0.00386545452034 | 0.290085649186645 | 1.0,0.861,0.966,0. | 0.495,0.616,0.774, | 0,118 |
| 143794 | ENSG00000125450 | NUP85 | chr17 | + | 75218127 | 75218306 | 75215800 | 75215823 | 75225102 | 75225237 | 143794 | 22,35,45,17,28,21,29,23,2 | 4,0,6,2,5,1,2,2,2,0,2,4 | 33,23,12,10,16,9,20,21,22, | 1,4,1,1,0,1,1,1,1,2,2,2,1,1, | 298 | 149 | 0.0038725364221 | 0.290515819681075 | 0.733,1.0,0.789,0. | 0.943,0.742,0.857, | -0,068 |
| 76305 | ENSG00000076242 | MLH1 | chr3 | + | 37017505 | 37017599 | 37012010 | 37012099 | 37020309 | 37020463 | 76305 | 19,19,37,23,12,12,8,7,13,1 | 10,5,17,9,1,6,3,2,7,3,2,11 | 16,11,17,5,12,7,11,13,9,6, | 8,23,9,6,7,5,13,7,6,11,14,1 | 243 | 149 | 0.00387933526987 | 0.290861843473332 | 0.538,0.7,0.572,0. | 0.551,0.227,0.537, | 0,167 |
| 119921 | ENSG00000095370 | SH2D3C | chr9 | - | 127766900 | 127767271 | 127761610 | 127761650 | 127773989 | 127774467 | 119921 | 1,0,9,4,0,1,2,1,4,3,0,0 | 7,6,46,41,41,27,22,19,49,2 | 0,3,8,6,2,0,0,7,2,2,1,2,0,4, | 34,20,23,6,3,18,6,29,7,23,1 | 298 | 149 | 0.0038798516599 | 0.290861843473332 | 0.067,0.0,0.089,0. | 0.0,0.07,0.148,0.3 | -0,057 |
| 43352 | ENSG00000082996 | RNF13 | chr3 | + | 149813924 | 149814026 | 149812769 | 149813353 | 149846010 | 149846140 | 43352 | 4,2,5,8,4,10,5,7,6,1,0,6 | 30,29,68,71,31,35,42,41,2 | 4,12,6,9,8,1,3,5,8,7,5,2,8,8 | 48,60,14,13,18,14,18,40,1 | 251 | 149 | 0.0038833410605 | 0.291022068172868 | 0.073,0.039,0.042, | 0.047,0.106,0.203, | -0,057 |
| 124395 | ENSG00000073921 | PICALM | chr11 | - | 85960674 | 85960777 | 85958754 | 85959060 | 85974707 | 85974812 | 124395 | 2,2,1,0,0,0,1,0,1,0,1,1 | 40,31,75,37,20,11,6,14,18, | 1,3,2,2,4,0,2,3,6,1,0,5,2,0, | 66,65,39,45,46,16,29,31,1 | 252 | 149 | 0.00388584942139 | 0.291108686674275 | 0.029,0.037,0.008, | 0.009,0.027,0.029, | -0,016 |
| 55818 | ENSG00000165240 | ATP7A | chrX | + | 77922998 | 77923042 | 77910708 | 77910835 | 77971620 | 77971761 | 55818 | 4,3,9,0,1,0,0,0,2,0,0,2 | 11,7,14,8,3,2,4,3,2,1,4,0 | 7,8,2,2,8,0,0,5,4,1,3,1,4,1, | 11,3,6,6,4,2,11,2,2,9,6,2,8, | 193 | 149 | 0.00389563824846 | 0.291710574410825 | 0.219,0.249,0.332, | 0.329,0.673,0.205, | -0,128 |
| 196835 | ENSG00000143119 | CD53 | chr1 | + | 110891397 | 110891471 | 110873182 | 110873248 | 110899123 | 110899514 | 196835 | 9,5,16,15,8,9,16,8,8,5,7,13 | 0,0,5,4,0,2,2,0,0,1,0,0 | 14,34,13,5,8,0,5,5,5,6,19,1 | 4,5,1,0,3,1,1,0,2,1,3,7,1,0, | 223 | 149 | 0.00389732307577 | 0.291710574410825 | 1.0,1.0,0.681,0.71 | 0.7,0.82,0.897,1.0, | 0,118 |

| | | | | | | | | | | | | | | | | | | | | | | |
|---|---|---|---|---|---|---|---|---|---|---|---|---|---|---|---|---|---|---|---|---|---|---|
| 196481 | ENSG00000166479 | TMX3 | chr18 | - | 68689947 | 68689981 | 68688416 | 68689054 | 68691294 | 68691361 | 196481 | 4,1,3,4,2,3,4,22,13,12,1,7 | 1,0,2,2,0,2,0,8,2,2,0,0 | 2,2,9,1,9,9,1,4,9,2,4,2,6,5,6 | 0,0,0,0,0,1,0,2,0,3,0,1,0,3,0 | 183 | 149 | 0.00389794970058 | 0.291710574410825 | 0.765,1.0,0.55,0.62 | 1.0,1.0,1.0,1.0,1.0, | -0,039 |
| 24835 | ENSG00000102763 | VWA8 | chr13 | - | 41690165 | 41690275 | 41689353 | 41689508 | 41691319 | 41691445 | 24835 | 4,5,26,15,16,19,20,19,8,3,6 | 0,2,0,2,0,2,0,4,0,2,0,2 | 10,15,7,13,16,6,19,33,6,11 | 2,6,0,3,6,0,0,0,0,0,3,5,0,1,3 | 259 | 149 | 0.00390698136132 | 0.292284847194392 | 1.0,0.59,1.0,0.812, | 0.742,0.59,1.0,0.71 | 0,021 |
| 55749 | ENSG00000109320 | NFKB1 | chr4 | + | 102578880 | 102579039 | 102576875 | 102577039 | 102580534 | 102580588 | 55749 | 47,43,67,57,46,32,25,48,27 | 7,3,0,0,1,0,6,2,6,2,0,0 | 64,38,18,27,22,18,19,49,15 | 1,0,0,0,4,2,1,8,0,1,2,0,0,1,0 | 298 | 149 | 0.00391298717294 | 0.292632433015653 | 0.77,0.878,1.0,1.0, | 0.97,1.0,1.0,1.0,0.7 | -0,036 |
| 178091 | ENSG00000067066 | SP100 | chr2 | + | 230463756 | 230464150 | 230462466 | 230462518 | 230466300 | 230466354 | 178091 | 87,64,165,99,53,65,53,82,3 | 57,30,72,69,17,16,39,29,3, | 102,78,63,43,74,33,19,66,4 | 50,76,27,57,116,31,16,23,5 | 298 | 149 | 0.00391688200554 | 0.292821962811525 | 0.433,0.516,0.534, | 0.505,0.339,0.538, | 0,158 |
| 45129 | ENSG00000269343 | ZNF587B | chr19 | + | 57820493 | 57820549 | 57819740 | 57819945 | 57839022 | 57839149 | 45129 | 15,16,24,24,26,23,15,23,14 | 0,1,4,0,0,0,0,2,1,1,1,2 | 20,31,12,20,14,8,13,31,9,6 | 0,2,4,0,5,0,4,2,1,3,0,5,1,1,2 | 205 | 149 | 0.00392409164754 | 0.293259086594334 | 1.0,0.921,0.813,1.0 | 1.0,0.918,0.686,1.0 | 0,05 |
| 47525 | ENSG00000160991 | ORAI2 | chr7 | + | 102436224 | 102436333 | 102433548 | 102433661 | 102438943 | 102439047 | 47525 | 40,56,94,89,145,133,151,3 | 6,2,8,5,4,4,11,30,2,4,7,13 | 79,56,47,48,43,35,62,128,3 | 5,7,4,9,6,0,1,16,6,7,10,55,8 | 258 | 149 | 0.00393854023522 | 0.294147734376659 | 0.794,0.942,0.872, | 0.901,0.822,0.872, | 0,039 |
| 124237 | ENSG00000149182 | ARFGAP2 | chr11 | - | 47175850 | 47175923 | 47175153 | 47175313 | 47176515 | 47176634 | 124237 | 87,99,89,94,60,98,46,55,69 | 7,4,6,3,5,3,6,2,2,3,0,0 | 52,71,47,26,72,27,40,62,33 | 1,7,4,8,3,5,3,3,2,4,8,4,6,1,4 | 222 | 149 | 0.00393871593996 | 0.294147734376659 | 0.893,0.943,0.909, | 0.972,0.872,0.887, | 0,055 |
| 132990 | ENSG00000166037 | CEP57 | chr11 | + | 95821870 | 95821978 | 95818826 | 95818904 | 95827785 | 95828027 | 132990 | 19,5,27,26,12,10,3,5,17,2,1 | 0,2,3,4,0,1,3,0,0,0,0,1 | 23,10,15,2,3,19,9,23,7,9,7, | 0,2,0,1,3,4,0,0,4,0,0,2,1,8,4 | 257 | 149 | 0.00394183358314 | 0.294278454364483 | 1.0,0.592,0.839,0.7 | 1.0,0.744,1.0,0.53 | 0,066 |
| 209618 | ENSG00000166986 | MARS1 | chr12 | + | 57489902 | 57489971 | 57489529 | 57489558 | 57490206 | 57490216 | 209618 | 40,9,55,44,27,20,13,17,27, | 5,3,6,2,1,8,7,3,1,2,1,4 | 25,19,18,6,23,23,13,27,27, | 5,19,9,6,9,1,8,3,4,5,11,6,16 | 218 | 149 | 0.00394337258013 | 0.294291270386255 | 0.845,0.672,0.862, | 0.774,0.406,0.578, | 0,107 |
| 29092 | ENSG00000290548 | | chr5 | - | 34175444 | 34175537 | 34164697 | 34169153 | 34175821 | 34175957 | 29092 | 21,15,54,19,10,16,12,27,8, | 4,2,1,2,0,0,0,2,6,0,0,2 | 13,12,8,9,17,8,3,12,11,17,1 | 0,0,0,3,2,0,0,0,0,2,0,0,5,0,0 | 242 | 149 | 0.00394568328263 | 0.294361649429372 | 0.764,0.822,0.971, | 1.0,1.0,1.0,0.649,0 | -0,055 |
| 151483 | ENSG00000077238 | IL4R | chr16 | + | 27315360 | 27315566 | 27313991 | 27314020 | 27330065 | 27330198 | 151483 | 4,0,23,7,6,0,4,1,3,2,4,0 | 91,44,257,89,74,72,81,76,7 | 2,2,4,0,0,0,1,2,2,1,1,0,3,3,9 | 89,44,59,14,13,31,27,28,21 | 298 | 149 | 0.00394711172147 | 0.294366182579248 | 0.022,0.0,0.043,0.0 | 0.011,0.022,0.033, | 0,006 |
| 216298 | ENSG00000163219 | ARHGAP25 | chr2 | + | 68806730 | 68806939 | 68787839 | 68787956 | 68807272 | 68807480 | 216298 | 9,2,3,2,0,2,3,9,7,0,0,7 | 145,224,142,169,138,87,73 | 16,19,2,17,11,1,2,0,1,0,16, | 210,364,91,60,101,42,90,8 | 298 | 149 | 0.00395351612829 | 0.294741679878069 | 0.03,0.004,0.01,0.0 | 0.037,0.025,0.011, | -0,009 |
| 33102 | ENSG00000281649 | EBLN3P | chr9 | + | 37084198 | 37084292 | 37079870 | 37080038 | 37086667 | 37087837 | 33102 | 22,10,42,12,9,4,8,16,13,2,2 | 1,6,18,3,4,2,9,6,0,0,2,5 | 2,7,6,1,5,1,5,4,5,7,11,8,13, | 8,9,6,1,2,1,1,4,3,12,8,6,5,1 | 243 | 149 | 0.00395663575968 | 0.294872116063603 | 0.931,0.505,0.589, | 0.133,0.323,0.38,0 | 0,201 |
| 231901 | ENSG00000099204 | ABLIM1 | chr10 | - | 114447883 | 114448020 | 114445311 | 114445403 | 114451623 | 114451671 | 231901 | 58,46,23,39,43,18,19,7,39, | 7,14,9,7,0,2,4,2,2,4,0,5 | 26,25,15,9,12,17,2,11,13,3 | 8,10,6,1,11,6,7,3,17,6,0,25 | 286 | 149 | 0.00396507822385 | 0.29539901391397 | 0.812,0.631,0.571, | 0.629,0.566,0.566, | 0,154 |
| 30645 | ENSG00000112983 | BRD8 | chr5 | - | 138168469 | 138168688 | 138165827 | 138166108 | 138169221 | 138169349 | 30645 | 20,12,11,30,29,19,22,27,40 | 4,1,1,3,0,1,0,2,1,0,0,1 | 15,18,17,12,4,11,12,33,5,9 | 3,1,1,0,4,2,0,3,4,0,0,1,2,2,7 | 298 | 149 | 0.00397477804007 | 0.296019187661698 | 0.714,0.857,0.846, | 0.714,0.9,0.895,1.0 | 0,097 |
| 6829 | ENSG00000124198 | ARFGEF2 | chr20 | + | 48941863 | 48941987 | 48921710 | 48922010 | 48951322 | 48951469 | 6829 | 19,14,60,39,18,28,20,22,11 | 1,1,0,2,0,0,0,0,2,4,0,0 | 62,37,27,28,19,16,9,22,14, | 5,0,0,4,2,0,2,4,0,1,2,2,3,2,2 | 273 | 149 | 0.00397654994134 | 0.296048709935853 | 0.912,0.884,1.0,0.9 | 0.871,1.0,1.0,0.79 | 0,03 |
| 115860 | ENSG00000109689 | STIM2 | chr4 | + | 27021493 | 27021612 | 27017710 | 27017984 | 27022518 | 27022840 | 115860 | 8,12,74,44,16,22,22,22,5,0 | 28,11,149,73,17,35,41,44,8 | 24,21,13,15,13,6,4,13,10,7 | 95,72,49,19,23,21,25,54,22 | 268 | 149 | 0.00398131878937 | 0.296301253234749 | 0.137,0.378,0.216, | 0.123,0.14,0.129,0 | 0,054 |
| 83145 | ENSG00000095059 | DHPS | chr19 | - | 12677107 | 12677211 | 12676016 | 12676142 | 12677290 | 12677396 | 83145 | 78,52,85,66,44,35,55,52,60 | 1,2,8,8,4,2,8,0,3,4,1,5 | 35,29,36,30,15,25,36,53,31 | 3,2,3,3,5,4,5,8,2,2,3,14,8,3 | 253 | 149 | 0.00399143717242 | 0.296951612187048 | 0.979,0.939,0.862, | 0.873,0.895,0.876, | 0,072 |
| 173979 | ENSG00000196455 | PIK3R4 | chr3 | - | 130731607 | 130731734 | 130730307 | 130730442 | 130733547 | 130734130 | 173979 | 2,0,4,0,0,2,1,1,0,0,0,0 | 5,9,26,18,12,25,10,14,2,7,1 | 4,1,0,7,0,0,2,2,0,0,2,2,2,2,2 | 10,12,5,15,17,9,6,11,8,8,10 | 276 | 149 | 0.00400048903175 | 0.297522202761778 | 0.178,0.0,0.077,0.0 | 0.178,0.043,0.0,0.1 | -0,039 |
| 125160 | ENSG00000165322 | ARHGAP12 | chr10 | - | 31852516 | 31852597 | 31843460 | 31843586 | 31861394 | 31861658 | 125160 | 57,54,78,61,26,29,24,40,27 | 2,3,4,2,0,2,2,7,4,1,1,2 | 70,83,51,20,46,17,13,49,48 | 10,10,0,8,4,3,1,2,7,2,2,4,3, | 230 | 149 | 0.00400365612174 | 0.297654891446709 | 0.949,0.921,0.927, | 0.819,0.843,1.0,0.6 | 0,058 |
| 121094 | ENSG00000070047 | PHRF1 | chr11 | + | 581961 | 582081 | 581491 | 581606 | 587258 | 587464 | 121094 | 59,64,27,15,10,15,17,19,24 | 1,3,1,0,0,4,0,2,0,1,0,0 | 21,20,27,47,21,8,15,19,4,2 | 0,6,1,3,0,0,4,0,1,0,4,7,0,0,0 | 269 | 149 | 0.00400764549759 | 0.297848600860426 | 0.97,0.922,0.937,1 | 1.0,0.649,0.937,0.8 | 0,076 |
| 238066 | ENSG00000119397 | CNTRL | chr9 | + | 121158043 | 121158109 | 121157749 | 121157880 | 121158854 | 121159019 | 238066 | 29,14,44,24,14,19,4,8,11,1 | 9,9,13,7,1,7,3,5,5,4,0,2 | 19,21,18,22,15,18,3,7,14,2 | 0,4,14,1,2,0,0,5,3,0,6,0,7,3 | 215 | 149 | 0.00401267164999 | 0.298119203278908 | 0.691,0.519,0.701, | 1.0,0.784,0.471,0.9 | -0,129 |
| 83950 | ENSG00000130038 | CRACR2A | chr12 | - | 3678914 | 3679098 | 3673445 | 3673592 | 3680237 | 3680349 | 83950 | 14,28,45,18,18,20,10,22,24 | 0,0,3,1,1,7,3,2,2,0,1,0 | 21,23,19,23,21,18,18,39,17 | 1,1,1,1,1,4,2,1,0,0,2,0,1,0,0 | 298 | 149 | 0.00401826158976 | 0.298431490761088 | 1.0,1.0,0.882,0.9,0 | 0.913,0.92,0.905,0 | -0,044 |
| 97895 | ENSG00000122482 | ZNF644 | chr1 | - | 90937484 | 90938090 | 90918051 | 90918154 | 90982309 | 90982370 | 97895 | 23,24,122,47,42,39,26,65,1 | 5,2,16,2,7,3,6,2,0,0,0,2 | 92,97,23,44,47,32,22,37,26 | 3,11,0,0,0,2,0,1,4,1,7,2,0,1 | 298 | 149 | 0.00402268746464 | 0.298657138910635 | 0.697,0.857,0.792, | 0.939,0.815,1.0,1.0 | -0,058 |
| 155222 | ENSG00000198718 | TOGARAM1 | chr14 | + | 44975172 | 44975255 | 44962218 | 44964467 | 44995745 | 44997404 | 155222 | 2,1,5,2,1,3,0,4,0,0,2,0 | 10,9,47,25,11,22,6,13,8,2,6 | 0,0,1,2,1,0,0,0,0,0,3,0,0,0,1 | 16,15,22,11,7,10,7,5,3,13,1 | 232 | 149 | 0.00402521684321 | 0.29874187806239 | 0.114,0.067,0.064, | 0.0,0.0,0.028,0.105 | 0,03 |
| 15247 | ENSG00000103222 | ABCC1 | chr16 | + | 16068155 | 16068302 | 16056091 | 16056295 | 16071641 | 16071729 | 15247 | 78,50,98,57,72,56,57,60,49 | 1,7,4,2,2,0,2,7,4,0,1,0 | 76,110,41,87,54,45,69,91,8 | 3,2,1,2,3,4,2,0,0,1,0,0,1,4,2 | 296 | 149 | 0.0040270874359 | 0.29877768214967 | 0.975,0.782,0.925, | 0.927,0.965,0.954, | -0,028 |
| 89125 | ENSG00000168958 | MFF | chr2 | + | 227342744 | 227342819 | 227340291 | 227340380 | 227355676 | 227355761 | 89125 | 9,4,13,4,1,2,4,3,1,1,1,1 | 58,40,90,62,32,25,29,27,25 | 7,7,4,1,6,6,4,6,1,5,3,10,7,4 | 42,41,35,9,46,14,7,37,7,21 | 224 | 149 | 0.00402958435099 | 0.298859913662277 | 0.094,0.062,0.088, | 0.1,0.102,0.071,0.0 | -0,056 |
| 47076 | ENSG00000116731 | PRDM2 | chr1 | + | 13778417 | 13782831 | 13749360 | 13749487 | 13816426 | 13816570 | 47076 | 21,3,4,3,0,6,1,2,2,3,0,2 | 1,3,3,1,2,0,1,1,6,0,1,1 | 4,2,5,6,4,5,5,5,2,4,2,0,3,2,0 | 3,1,0,0,0,0,0,6,0,2,0,2,2,0,1 | 298 | 149 | 0.00403418514226 | 0.299098071771878 | 0.913,0.333,0.4,0.6 | 0.4,0.5,1.0,1.0,1.0, | -0,168 |
| 55006 | ENSG00000082701 | GSK3B | chr3 | - | 119912703 | 119912800 | 119905754 | 119905852 | 119916043 | 119916174 | 55006 | 23,15,27,14,15,9,6,11,9,2,5 | 1,0,2,1,3,6,3,3,0,0,0,4 | 15,6,7,9,9,7,9,13,6,13,6,27 | 2,7,4,3,7,4,3,2,3,1,7,11,3,1 | 246 | 149 | 0.00404434984806 | 0.299666499888736 | 0.933,1.0,0.891,0.8 | 0.82,0.342,0.515,0 | 0,175 |
| 76259 | ENSG00000138073 | PREB | chr2 | - | 27132229 | 27132403 | 27131671 | 27132082 | 27132602 | 27132620 | 76259 | 64,25,35,48,51,32,25,30,47 | 0,2,1,0,0,2,0,1,0,1,0,0 | 54,45,34,17,13,17,50,47,5, | 2,1,2,0,2,0,2,3,0,2,0,2,0,4,2 | 298 | 149 | 0.00404463660986 | 0.299666499888736 | 1.0,0.862,0.946,1.0 | 0.931,0.957,0.895, | 0,043 |
| 154134 | ENSG00000100934 | SEC23A | chr14 | - | 39037469 | 39037608 | 39033203 | 39033328 | 39039030 | 39041433 | 154134 | 2,0,8,7,4,1,2,6,0,0,1,0 | 4,3,55,23,15,22,9,24,9,5,7, | 2,10,3,6,6,3,6,1,1,5,10,3,2, | 22,26,14,4,13,11,6,14,9,14 | 288 | 149 | 0.00405332245196 | 0.300206691210531 | 0.206,0.0,0.07,0.13 | 0.045,0.166,0.1,0.4 | -0,079 |
| 11863 | ENSG00000108395 | TRIM37 | chr17 | - | 59001597 | 59001714 | 58998201 | 58999459 | 59012327 | 59012446 | 11863 | 19,19,30,24,11,13,4,4,1,3,3 | 5,2,3,4,2,3,0,0,0,1,0,4 | 20,12,11,16,4,9,12,11,5,14 | 7,5,8,4,3,1,4,4,0,0,5,7,2,1,5 | 266 | 149 | 0.00406558080279 | 0.300943222207592 | 0.68,0.842,0.849,0 | 0.615,0.573,0.435, | 0,188 |
| 223539 | ENSG00000124422 | USP22 | chr17 | - | 21015751 | 21015794 | 21012829 | 21012935 | 21017941 | 21018111 | 223539 | 28,14,28,19,20,23,37,27,42 | 0,2,4,0,2,1,0,2,5,3,0,0 | 13,16,9,22,11,6,21,26,13,1 | 0,1,1,1,0,0,0,3,0,0,0,0,0,0,2 | 192 | 149 | 0.00406606339319 | 0.300943222207592 | 1.0,0.845,0.845,1.0 | 1.0,0.925,0.875,0.9 | -0,038 |
| 183814 | ENSG00000162591 | MEGF6 | chr1 | - | 3508552 | 3508689 | 3507794 | 3507923 | 3509074 | 3509245 | 183814 | 46,16,93,38,16,13,23,14,6, | 5,5,3,0,2,3,0,2,0,0,0,2 | 49,27,19,31,21,5,23,21,36, | 2,2,1,3,0,0,0,1,1,0,2,0,2,1,2 | 286 | 149 | 0.00407542369559 | 0.301512880672926 | 0.827,0.625,0.942, | 0.927,0.876,0.908, | -0,049 |
| 115160 | ENSG00000165525 | NEMF | chr14 | - | 49840717 | 49840866 | 49838138 | 49838206 | 49846139 | 49846265 | 115160 | 65,42,156,89,56,55,32,69,3 | 1,2,4,1,0,0,3,4,0,0,0,0 | 82,91,39,73,71,55,30,43,37 | 2,4,7,4,1,2,2,1,0,2,1,2,2,2,2 | 298 | 149 | 0.00407656184638 | 0.301512880672926 | 0.97,0.913,0.951,0 | 0.953,0.919,0.736, | 0,038 |
| 12693 | ENSG00000108848 | LUC7L3 | chr17 | + | 50738136 | 50738199 | 50736959 | 50737026 | 50740305 | 50740345 | 12693 | 8,5,18,10,3,2,4,7,1,1,5,2 | 59,33,146,87,96,38,64,87,4 | 8,18,6,10,6,6,4,6,5,9,4,10,9 | 78,65,31,24,51,50,31,43,36 | 212 | 149 | 0.004084092027 | 0.301966063573767 | 0.087,0.096,0.08,0 | 0.067,0.163,0.12,0 | -0,036 |
| 123825 | ENSG00000149262 | INTS4 | chr11 | - | 77918030 | 77918264 | 77907716 | 77907810 | 77918820 | 77918978 | 123825 | 0,0,2,4,1,1,4,1,0,2,0,2 | 31,16,56,45,19,22,24,25,16 | 2,5,3,2,0,2,2,0,2,2,6,3,3,4,0 | 25,23,21,8,24,15,12,19,9,2 | 298 | 149 | 0.00409979921988 | 0.303023312463596 | 0.0,0.0,0.018,0.04 | 0.038,0.098,0.067, | -0,016 |
| 76059 | ENSG00000135541 | AHI1 | chr6 | - | 135495312 | 135495898 | 135492227 | 135492291 | 135497187 | 135497249 | 76059 | 7,8,4,5,3,1,0,0,0,1,4,6 | 3,2,24,7,10,5,10,6,1,0,3,11 | 0,1,0,3,4,3,2,3,0,3,1,0,1,0,0 | 10,8,11,9,7,1,2,7,2,1,8,2,8, | 298 | 149 | 0.004102933753 | 0.303150887261223 | 0.538,0.667,0.077, | 0.0,0.059,0.0,0.14 | 0,141 |
| 177449 | ENSG00000160991 | ORAI2 | chr7 | + | 102436183 | 102436333 | 102433575 | 102433661 | 102438943 | 102439047 | 177449 | 40,50,100,98,120,119,136, | 6,2,8,5,4,4,11,30,2,4,7,13 | 77,48,43,42,45,36,49,118,3 | 5,7,4,9,6,0,1,16,6,7,10,55,8 | 298 | 149 | 0.00410477867692 | 0.303183122653455 | 0.769,0.926,0.862, | 0.885,0.774,0.843, | 0,049 |
| 135743 | ENSG00000291230 | | chr19 | - | 50336505 | 50336690 | 50335058 | 50335181 | 50337016 | 50337135 | 135743 | 265,244,276,686,332,289,2 | 14,3,5,13,2,4,8,2,10,5,1,0 | 243,155,194,76,168,147,10 | 6,4,2,1,4,2,2,3,7,5,7,15,8,1 | 298 | 149 | 0.00410637359617 | 0.303196876664585 | 0.904,0.976,0.965, | 0.953,0.951,0.98,0 | 0,021 |
| 87041 | ENSG00000184007 | PTP4A2 | chr1 | - | 31911695 | 31911826 | 31910037 | 31910112 | 31915894 | 31915987 | 87041 | 599,414,547,460,252,189,1 | 14,4,9,1,0,3,4,1,2,4,3,4 | 376,384,296,342,216,214,1 | 10,8,2,8,8,0,0,8,2,6,3,11,7, | 280 | 149 | 0.00411185606146 | 0.303408687532184 | 0.958,0.982,0.97,0 | 0.952,0.962,0.987, | 0,011 |
| 30709 | ENSG00000124357 | NAGK | chr2 | + | 71071685 | 71071827 | 71068315 | 71068712 | 71072640 | 71072690 | 30709 | 58,28,133,89,77,70,44,61,3 | 1,1,2,4,9,5,2,3,0,1,1,2 | 144,97,32,20,22,30,21,64,4 | 1,4,1,0,0,0,0,1,0,0,1,4,4,2,5 | 291 | 149 | 0.00411206165251 | 0.303408687532184 | 0.967,0.935,0.971, | 0.987,0.925,0.942, | -0,039 |
| 233295 | ENSG00000215301 | DDX3X | chrX | + | 41343701 | 41343822 | 41343288 | 41343351 | 41344029 | 41344109 | 233295 | 119,109,383,249,82,92,100 | 2,0,15,6,5,3,10,4,4,0,5,7 | 203,267,156,179,168,104,7 | 17,8,5,43,1,11,2,12,8,6,13, | 270 | 149 | 0.00411488296041 | 0.303512808242634 | 0.97,1.0,0.934,0.9 | 0.868,0.949,0.945, | 0,027 |
| 178088 | ENSG00000067066 | SP100 | chr2 | + | 230463747 | 230464150 | 230462466 | 230462518 | 230466300 | 230466354 | 178088 | 87,64,160,97,53,65,54,82,3 | 57,30,72,69,17,16,39,29,3, | 102,78,63,43,74,33,19,66,4 | 50,76,27,57,116,31,16,23,5 | 298 | 149 | 0.00411909457168 | 0.303719370934312 | 0.433,0.516,0.526, | 0.505,0.339,0.538, | 0,157 |
| 103235 | ENSG00000110497 | AMBRA1 | chr11 | - | 46512726 | 46512813 | 46494123 | 46494204 | 46541944 | 46542981 | 103235 | 14,11,18,4,9,13,2,9,12,9,4, | 0,0,5,1,2,2,0,0,0,0,3,4 | 4,6,7,9,9,3,4,4,7,14,5,5,9,6 | 1,1,0,0,0,2,0,0,0,1,0,1,1,0,0 | 236 | 149 | 0.00412489332232 | 0.304042778991868 | 1.0,1.0,0.694,0.71 | 0.716,0.791,1.0,1.0 | -0,075 |
| 210639 | ENSG00000134262 | AP4B1 | chr1 | - | 113899083 | 113899352 | 113898717 | 113898801 | 113899903 | 113900008 | 210639 | 10,3,14,4,1,1,4,6,3,0,3,1 | 19,10,45,23,17,14,24,13,10 | 5,1,3,3,3,0,3,1,1,5,3,2,1,4,0 | 23,22,7,11,25,15,4,10,11,1 | 298 | 149 | 0.0041287826785 | 0.304225273767968 | 0.208,0.13,0.135,0 | 0.098,0.022,0.176, | 0,022 |
| 3542 | ENSG00000204514 | ZNF814 | chr19 | - | 57876915 | 57877042 | 57858814 | 57858983 | 57888766 | 57888780 | 3542 | 8,3,16,6,10,3,2,6,4,4,2,2 | 3,2,8,1,0,0,2,0,0,1,0,2 | 9,16,3,3,3,2,4,2,1,3,5,9,7,6 | 0,0,0,0,0,0,2,0,0,0,1,0,0,2,5 | 276 | 149 | 0.00414133137291 | 0.305045480055713 | 0.59,0.447,0.519,0 | 1.0,1.0,1.0,1.0,1.0, | -0,112 |
| 135745 | ENSG00000291230 | | chr19 | - | 50336505 | 50336734 | 50335058 | 50335181 | 50337016 | 50337135 | 135745 | 265,244,276,686,340,289,2 | 14,3,5,13,2,4,8,2,10,5,1,0 | 243,155,194,76,168,147,10 | 6,4,2,1,4,2,2,3,7,5,7,15,8,1 | 298 | 149 | 0.00414574812028 | 0.305266340634959 | 0.904,0.976,0.965, | 0.953,0.951,0.98,0 | 0,021 |
| 48771 | ENSG00000031698 | SARS1 | chr1 | + | 109236390 | 109236548 | 109235976 | 109236106 | 109237243 | 109237373 | 48771 | 61,38,116,115,95,33,46,63 | 2,0,5,3,2,1,0,0,1,0,2,0 | 67,46,53,45,28,31,16,30,14 | 2,0,2,1,4,0,0,7,0,0,3,1,5,0,1 | 298 | 149 | 0.00415535861471 | 0.305821251904686 | 0.938,1.0,0.921,0.9 | 0.944,1.0,0.93,0.95 | 0,047 |
| 163591 | ENSG00000113593 | PPWD1 | chr5 | + | 65568917 | 65569078 | 65567512 | 65567615 | 65569631 | 65569732 | 163591 | 2,0,7,7,2,2,3,0,0,2,0,0 | 36,23,58,36,22,17,11,3,46, | 0,0,0,0,0,1,2,0,0,2,0,1,0,0,5 | 29,22,19,22,24,15,15,11,19 | 298 | 149 | 0.0041561260312 | 0.305821251904686 | 0.027,0.0,0.057,0.0 | 0.0,0.0,0.0,0.0,0.0, | 0,021 |
| 99627 | ENSG00000106443 | PHF14 | chr7 | + | 10982371 | 10983159 | 10974834 | 10974945 | 10990702 | 10990847 | 99627 | 64,73,82,30,41,44,31,46,30 | 25,8,8,6,11,3,7,1,4,7,2,9 | 42,42,41,38,30,25,14,25,29 | 10,12,8,7,9,6,9,2,15,4,7,31 | 298 | 149 | 0.00417234944647 | 0.306910096962811 | 0.561,0.82,0.837,0 | 0.677,0.636,0.719, | 0,118 |
| 130100 | ENSG00000141376 | BCAS3 | chr17 | + | 60688019 | 60688195 | 60683981 | 60684036 | 60689685 | 60689761 | 130100 | 4,2,1,0,0,1,2,1,0,0,0,0 | 30,23,57,20,24,35,8,37,19, | 3,4,1,16,0,0,0,2,0,0,4,0,0,3 | 34,29,25,50,27,28,7,17,20, | 298 | 149 | 0.00418107281489 | 0.307446697308367 | 0.062,0.042,0.009, | 0.042,0.065,0.02,0 | -0,009 |
| 30549 | ENSG00000124201 | ZNFX1 | chr20 | - | 49263333 | 49263483 | 49260462 | 49260577 | 49264715 | 49264864 | 30549 | 38,38,96,89,16,19,19,33,42 | 0,2,4,1,2,1,0,1,0,0,0,5 | 66,74,29,49,31,22,13,39,20 | 4,4,1,4,5,0,1,1,1,1,5,1,2,1,1 | 298 | 149 | 0.00419684283996 | 0.308500915740243 | 1.0,0.905,0.923,0.9 | 0.892,0.902,0.935, | 0,044 |
| 92529 | ENSG00000132424 | PNISR | chr6 | - | 99416348 | 99416428 | 99414571 | 99414686 | 99425214 | 99425270 | 92529 | 89,44,115,53,18,31,28,30,1 | 7,3,4,0,4,2,2,0,3,2,3,0 | 87,59,54,46,36,23,11,20,49 | 1,2,0,0,0,0,3,4,1,3,3,0,1,4,3 | 229 | 149 | 0.00420408525524 | 0.308920783039155 | 0.892,0.905,0.949, | 0.983,0.95,1.0,1.0, | -0,055 |
| 132399 | ENSG00000196498 | NCOR2 | chr12 | - | 124327408 | 124327633 | 124326190 | 124326370 | 124330844 | 124330898 | 132399 | 5,6,6,5,7,6,3,3,2,2,0,5 | 1,2,1,2,0,1,0,1,0,2,3,6 | 3,5,3,7,5,2,13,5,5,3,2,3,4,7 | 0,3,1,0,2,0,2,3,1,0,0,0,3,0,1 | 298 | 149 | 0.00420542530725 | 0.308920783039155 | 0.714,0.6,0.75,0.5 | 1.0,0.455,0.6,1.0,0 | -0,1 |
| 143599 | ENSG00000183684 | ALYREF | chr17 | - | 81888875 | 81888963 | 81888519 | 81888583 | 81889181 | 81889329 | 143599 | 13,3,7,4,3,3,2,4,3,0,4,7 | 189,149,184,97,64,77,56,4 | 3,1,0,2,1,3,2,0,0,2,7,11,1,4 | 114,138,106,32,62,39,62,9 | 237 | 149 | 0.00421601809338 | 0.309519477027926 | 0.041,0.012,0.023, | 0.016,0.005,0.0,0.0 | 0,014 |
| 15444 | ENSG00000213380 | COG8 | chr16 | - | 69332713 | 69332882 | 69330812 | 69331095 | 69334520 | 69335348 | 15444 | 37,14,83,72,43,42,32,41,31 | 5,0,1,0,2,2,0,5,6,0,0,0 | 62,51,18,14,15,25,28,59,15 | 0,2,0,0,0,0,0,0,0,0,0,3,0,0 | 298 | 149 | 0.00421645165727 | 0.309519477027926 | 0.787,1.0,0.976,1.0 | 1.0,0.927,1.0,1.0,1 | -0,063 |
| 63604 | ENSG00000117859 | OSBPL9 | chr1 | + | 51756319 | 51756358 | 51750144 | 51750195 | 51760689 | 51760764 | 63604 | 6,7,18,12,4,8,3,7,7,1,0,2 | 15,15,45,25,18,16,14,16,14 | 23,8,5,2,3,3,3,13,9,12,10,2 | 16,14,17,2,23,5,14,19,11,2 | 188 | 149 | 0.00422925617802 | 0.310353575332909 | 0.241,0.27,0.241,0 | 0.533,0.312,0.189, | -0,13 |
| 79221 | ENSG00000143924 | EML4 | chr2 | + | 42304483 | 42304551 | 42303314 | 42303446 | 42315961 | 42316050 | 79221 | 99,80,106,61,83,72,28,58,3 | 2,0,2,0,1,1,0,0,0,0,0,2 | 137,135,49,112,85,57,63,1 | 1,6,1,7,1,2,0,1,3,3,0,8,1,5,0 | 217 | 149 | 0.00423164355837 | 0.310422929349534 | 0.971,1.0,0.973,1.0 | 0.989,0.939,0.971, | 0,016 |
| 15769 | ENSG00000072864 | NDE1 | chr16 | + | 15696708 | 15696860 | 15694164 | 15694256 | 15724190 | 15725119 | 15769 | 21,12,66,57,33,40,34,31,12 | 3,1,3,10,1,6,1,1,5,2,0,0 | 28,48,15,30,37,18,11,25,28 | 3,4,0,2,5,0,2,2,0,2,0,3,0,2,0 | 298 | 149 | 0.00423710340104 | 0.310717547567032 | 0.778,0.857,0.917, | 0.824,0.857,1.0,0.8 | -0,06 |
| 244361 | ENSG00000109971 | HSPA8 | chr11 | - | 123060592 | 123060773 | 123059813 | 123060268 | 123061119 | 123061332 | 244361 | 244,146,585,703,275,238,2 | 1,4,35,20,6,4,6,3,6,2,2,1 | 348,289,267,88,235,173,92 | 20,12,3,19,5,4,8,1,4,5,26,1 | 298 | 149 | 0.00424249644011 | 0.311007067881919 | 0.992,0.948,0.893, | 0.897,0.923,0.978, | 0,041 |
| 234135 | ENSG00000112305 | SMAP1 | chr6 | + | 70699274 | 70699512 | 70667882 | 70668141 | 70732377 | 70732511 | 234135 | 4,1,3,5,2,3,1,0,2,0,0,1 | 30,10,26,27,15,19,10,24,4, | 3,2,0,7,4,1,2,7,3,2,3,1,0,7,6 | 22,18,9,9,10,21,8,11,10,13 | 298 | 149 | 0.00424509249554 | 0.311091420806118 | 0.062,0.048,0.055, | 0.064,0.053,0.0,0.1 | -0,051 |
| 28843 | ENSG00000166888 | STAT6 | chr12 | - | 57109940 | 57110110 | 57108162 | 57108299 | 57111128 | 57111413 | 28843 | 44,33,63,63,23,19,15,28,26 | 55,45,164,114,97,88,46,56 | 26,13,28,15,8,8,9,22,11,16 | 102,83,36,30,38,49,20,65,3 | 298 | 149 | 0.00425636304937 | 0.311811189747772 | 0.286,0.268,0.161, | 0.113,0.073,0.28,0 | 0,033 |
| 148322 | ENSG00000167984 | NLRC3 | chr16 | - | 3575063 | 3575415 | 3567242 | 3567324 | 3577148 | 3577403 | 148322 | 0,0,3,2,3,0,2,0,0,1,0,0 | 14,5,21,17,8,8,9,5,6,1,13,6 | 0,8,0,0,2,0,1,3,0,0,2,6,0,2,6 | 17,5,8,2,8,2,10,16,2,8,7,15 | 298 | 149 | 0.00426358403911 | 0.312233908241471 | 0.0,0.0,0.067,0.056 | 0.0,0.444,0.0,0.0,0 | -0,021 |
| 56130 | ENSG00000055917 | PUM2 | chr2 | - | 20331716 | 20331937 | 20327309 | 20327378 | 20350596 | 20350702 | 56130 | 6,3,4,1,0,2,3,1,5,4,0,1 | 17,19,7,16,4,8,6,13,4,5,3,2 | 5,2,2,2,1,0,0,3,1,0,2,2,0,0,2 | 14,29,7,15,2,6,7,9,10,12,11 | 298 | 149 | 0.00426698896396 | 0.312376973368053 | 0.15,0.073,0.222,0 | 0.152,0.033,0.125, | 0,073 |
| 176850 | ENSG00000036054 | TBC1D23 | chr3 | + | 100302075 | 100302237 | 100299238 | 100299331 | 100304845 | 100304888 | 176850 | 8,10,23,11,8,12,9,12,8,2,4, | 2,0,4,3,2,1,1,1,0,0,0,0 | 16,20,3,14,27,8,5,17,13,17 | 1,3,1,5,2,1,0,8,0,0,2,0,2,0,0 | 298 | 149 | 0.0042722180272 | 0.312576758951014 | 0.667,1.0,0.742,0.6 | 0.889,0.769,0.6,0.5 | 0,05 |
| 51773 | ENSG00000153914 | SREK1 | chr5 | + | 66156064 | 66158932 | 66153462 | 66153596 | 66159218 | 66159334 | 51773 | 104,40,72,43,15,39,36,78,5 | 18,10,11,12,4,7,17,7,25,2,6 | 51,68,59,158,53,36,37,32,3 | 7,7,9,8,3,9,3,3,3,5,4,7,10,7 | 298 | 149 | 0.0042726225536 | 0.312576758951014 | 0.743,0.667,0.766, | 0.785,0.829,0.766, | -0,081 |
| 112340 | ENSG00000156738 | MS4A1 | chr11 | + | 60461071 | 60461160 | 60455846 | 60455945 | 60462358 | 60462533 | 112340 | 157,94,338,76,31,2,21,10,1 | 145,107,343,247,78,2,46,1 | 113,70,133,48,156,99,8,0,7 | 94,71,135,42,85,99,10,1,64 | 238 | 149 | 0.00427690690835 | 0.312783877265062 | 0.404,0.355,0.382, | 0.429,0.382,0.381, | -0,075 |
| 246872 | ENSG00000173933 | RBM4 | chr11 | + | 66646026 | 66646364 | 66639699 | 66640123 | 66649735 | 66649930 | 246872 | 12,11,32,34,23,10,17,9,15, | 2,4,1,6,1,2,0,1,2,0,0,0 | 19,30,15,9,9,9,6,16,4,13,7, | 0,0,0,2,2,0,0,0,0,1,0,1,1,2,0 | 298 | 149 | 0.00428098039647 | 0.312975438761085 | 0.75,0.579,0.941,0 | 1.0,1.0,1.0,0.692,0 | -0,061 |
| 181823 | ENSG00000123104 | ITPR2 | chr12 | - | 26445062 | 26445130 | 26443542 | 26443650 | 26475295 | 26475418 | 181823 | 5,6,16,0,2,6,5,16,0,1,1,1 | 20,16,91,38,19,24,21,54,11 | 11,23,5,13,22,6,0,13,3,12,9 | 48,44,23,34,23,41,14,23,29 | 217 | 149 | 0.00429006786479 | 0.313533309543843 | 0.147,0.205,0.108, | 0.136,0.264,0.13,0 | -0,067 |
| 11914 | ENSG00000108469 | RECQL5 | chr17 | - | 75630618 | 75630692 | 75630183 | 75630277 | 75630778 | 75630837 | 11914 | 10,9,21,15,13,11,8,13,11,7 | 3,1,8,2,0,0,2,0,0,0,2,1 | 15,4,8,8,7,11,2,4,3,19,11,6 | 1,0,0,0,0,0,0,3,1,0,2,1,0,0,1 | 223 | 149 | 0.00429309439598 | 0.313647997264485 | 0.69,0.857,0.637,0 | 0.909,1.0,1.0,1.0,1 | -0,078 |
| 68546 | ENSG00000055609 | KMT2C | chr7 | - | 152157760 | 152157925 | 152156204 | 152156346 | 152158862 | 152159072 | 68546 | 33,19,55,45,30,18,16,22,21 | 4,0,6,0,4,3,2,3,1,0,1,2 | 34,15,12,12,23,20,21,18,9, | 0,0,1,0,1,0,1,2,0,1,5,3,1,2,0 | 298 | 149 | 0.00429983787154 | 0.314034070216975 | 0.805,1.0,0.821,1.0 | 1.0,1.0,0.857,1.0,0 | -0,066 |
| 12412 | ENSG00000154265 | ABCA5 | chr17 | - | 69285387 | 69285420 | 69283952 | 69284072 | 69285897 | 69286037 | 12412 | 17,4,25,5,16,13,10,13,2,2,4 | 9,12,51,18,13,25,9,11,6,1,3 | 14,5,4,2,2,3,8,9,6,7,6,2,4,2 | 13,15,17,5,6,6,17,11,28,24 | 182 | 149 | 0.00430504072744 | 0.314307401902184 | 0.607,0.214,0.286, | 0.469,0.214,0.162, | 0,16 |
| 60952 | ENSG00000037474 | NSUN2 | chr5 | - | 6605272 | 6605408 | 6604604 | 6604685 | 6606819 | 6606912 | 60952 | 25,29,59,47,18,19,10,21,30 | 6,2,7,5,2,2,3,0,0,0,0,0 | 33,32,45,20,20,9,19,24,9,2 | 2,1,0,0,1,1,0,3,2,4,0,0,2,0,3 | 285 | 149 | 0.00430719544869 | 0.31435808193184 | 0.685,0.883,0.815, | 0.896,0.944,1.0,1.0 | -0,029 |
| 178082 | ENSG00000067066 | SP100 | chr2 | + | 230463679 | 230464150 | 230462466 | 230462518 | 230466300 | 230466354 | 178082 | 87,64,160,97,53,65,53,82,3 | 57,30,72,69,17,16,39,29,3, | 102,78,63,43,74,33,19,66,4 | 50,76,27,57,116,31,16,23,5 | 298 | 149 | 0.00431637288648 | 0.314921102620331 | 0.433,0.516,0.526, | 0.505,0.339,0.538, | 0,157 |
| 192235 | ENSG00000131408 | NR1H2 | chr19 | + | 50378032 | 50378439 | 50377732 | 50377870 | 50378521 | 50378678 | 192235 | 14,10,21,4,12,10,3,2,14,9,1 | 3,4,2,0,2,2,0,2,1,1,0,1 | 8,10,10,16,5,3,4,9,4,4,11,5 | 1,3,0,1,0,1,0,0,2,2,0,6,0,0,0 | 298 | 149 | 0.00432173502846 | 0.31520547336851 | 0.7,0.556,0.84,1.0, | 0.8,0.625,1.0,0.889 | 0,006 |
| 51045 | ENSG00000143772 | ITPKB | chr1 | - | 226647166 | 226647380 | 226641920 | 226642125 | 226648671 | 226648771 | 51045 | 165,58,75,55,60,38,36,23,4 | 1,0,1,5,1,0,0,0,1,0,0,0 | 57,25,31,10,16,50,24,58,20 | 2,2,0,0,0,0,0,8,2,4,0,2,6,0,0 | 298 | 149 | 0.00432471512119 | 0.315315975694053 | 0.988,1.0,0.974,0.8 | 0.934,0.862,1.0,1.0 | 0,038 |
| 213428 | ENSG00000035720 | STAP1 | chr4 | + | 67576828 | 67577259 | 67575384 | 67575498 | 67581304 | 67581471 | 213428 | 95,41,113,85,38,50,40,46,9 | 3,0,4,1,1,0,3,4,1,1,2,2 | 58,46,71,34,51,30,19,38,20 | 2,3,3,11,3,0,2,2,2,0,0,0,3,4 | 298 | 149 | 0.00435213490562 | 0.317207703309007 | 0.941,1.0,0.934,0.9 | 0.935,0.885,0.922, | 0,016 |
| 1794 | ENSG00000100330 | MTMR3 | chr22 | + | 30023456 | 30023483 | 30022608 | 30022697 | 30025629 | 30026037 | 1794 | 4,2,8,5,6,8,6,17,3,3,0,1 | 32,14,47,18,25,16,27,23,22 | 7,19,6,6,3,1,22,10,4,2,9,14 | 17,27,6,27,27,11,19,26,6,2 | 176 | 149 | 0.00435714536438 | 0.317465387244709 | 0.096,0.108,0.126, | 0.258,0.373,0.458, | -0,108 |
| 79552 | ENSG00000137817 | PARP6 | chr15 | - | 72250844 | 72250954 | 72250019 | 72250092 | 72251206 | 72251255 | 79552 | 60,42,94,56,41,42,48,57,52 | 8,5,5,3,9,6,6,7,0,3,0,5 | 51,73,45,35,29,33,42,52,32 | 13,2,4,5,4,1,6,23,12,8,13,6 | 259 | 149 | 0.00437753863088 | 0.318836301104145 | 0.812,0.829,0.915, | 0.693,0.955,0.866, | 0,076 |

| | | | | | | | | | | | | | | | | | | | | | | |
|---|---|---|---|---|---|---|---|---|---|---|---|---|---|---|---|---|---|---|---|---|---|---|
| 240638 | ENSG00000130653 | PNPLA7 | chr9 | - | 137541416 | 137541580 | 137540641 | 137540722 | 137542641 | 137542801 | 240638 | 0,2,1,2,2,2,0,1,1,2,0,1 | 8,6,15,7,7,12,10,8,2,2,1,3 | 0,0,2,6,0,0,2,0,10,7,1,1,3,4 | 9,13,8,9,4,4,5,8,4,9,1,4,4,4 | 298 | 149 | 0.00437977765657 | 0.318836301104145 | 0.0,0.143,0.032,0. | 0.0,0.0,0.111,0.25, | -0,048 |
| 169509 | ENSG00000174227 | PIGG | chr4 | + | 525170 | 525634 | 523831 | 523913 | 527038 | 527230 | 169509 | 6,1,5,2,2,4,0,2,2,2,1,0 | 5,3,9,5,9,2,6,0,11,1,7,10 | 0,0,2,4,0,3,4,0,0,0,1,4,0,0, | 3,8,1,0,1,5,0,3,2,9,10,10,1, | 298 | 149 | 0.00438040497124 | 0.318836301104145 | 0.375,0.143,0.217, | 0.0,0.0,0.5,1.0,0.0, | 0,079 |
| 169087 | ENSG00000100412 | ACO2 | chr22 | + | 41515766 | 41515857 | 41515376 | 41515535 | 41517526 | 41517631 | 169087 | 28,20,35,34,19,31,17,23,1 | 4,0,0,1,1,4,0,10,0,0,0,0 | 19,13,15,10,5,7,5,14,15,15 | 0,2,1,2,1,1,1,0,0,0,0,2,0,0, | 240 | 149 | 0.00438552321936 | 0.319100929014899 | 0.813,1.0,1.0,0.95 | 1.0,0.801,0.903,0. | 0,006 |
| 71281 | ENSG00000131051 | RBM39 | chr20 | - | 35740524 | 35740597 | 35731940 | 35732135 | 35740823 | 35740887 | 71281 | 46,18,99,56,41,41,30,54,2 | 3,4,1,0,0,0,0,0,0,0,0,4 | 58,67,35,9,25,18,18,18,13, | 3,1,4,5,1,0,0,0,1,0,9,4,0,1, | 222 | 149 | 0.0043882091262 | 0.319124773980792 | 0.911,0.751,0.985, | 0.928,0.978,0.854, | 0,046 |
| 17751 | ENSG00000184056 | VPS33B | chr15 | - | 91006651 | 91006729 | 91006371 | 91006445 | 91006949 | 91007046 | 17751 | 11,9,29,28,14,21,13,6,13,6 | 3,1,10,3,3,2,1,6,6,0,3,4 | 6,8,11,3,10,12,1,6,1,5,13,3 | 14,8,8,3,2,1,0,5,0,3,14,6,4, | 227 | 149 | 0.00438881634608 | 0.319124773980792 | 0.706,0.855,0.656, | 0.22,0.396,0.474,0 | 0,157 |
| 163039 | ENSG00000182903 | ZNF721 | chr4 | - | 467545 | 467681 | 444342 | 444432 | 472574 | 472701 | 163039 | 1,1,3,8,0,4,1,4,0,0,0,2 | 82,45,140,64,22,32,24,41, | 0,12,3,0,4,1,2,6,0,1,3,7,4,3 | 50,94,57,67,46,33,28,31,2 | 285 | 149 | 0.00439045208702 | 0.319135897717461 | 0.006,0.011,0.011, | 0.0,0.063,0.027,0. | -0,019 |
| 129775 | ENSG00000089022 | MAPKAPK5 | chr12 | + | 111882789 | 111882853 | 111880446 | 111880527 | 111883580 | 111883768 | 129775 | 15,3,21,13,3,5,7,8,6,0,3,7 | 46,37,60,42,29,35,37,32,5 | 6,14,4,16,11,6,10,14,1,8,1 | 43,23,36,13,22,18,18,32,8, | 213 | 149 | 0.00439939677767 | 0.319678112037371 | 0.186,0.054,0.197, | 0.089,0.299,0.072, | -0,083 |
| 8425 | ENSG00000186814 | ZSCAN30 | chr18 | - | 35263512 | 35263657 | 35251061 | 35254381 | 35263944 | 35264455 | 8425 | 10,5,22,16,12,5,12,10,6,1, | 0,0,8,1,1,0,2,3,3,0,0,0 | 9,10,1,5,2,3,2,14,5,9,1,7,1 | 3,1,1,0,1,1,0,0,0,1,0,0,1,0, | 294 | 149 | 0.00440415924807 | 0.319916165751385 | 1.0,1.0,0.582,0.89, | 0.603,0.835,0.336, | 0,026 |
| 22661 | ENSG00000100888 | CHD8 | chr14 | - | 21430800 | 21431643 | 21428963 | 21429335 | 21431810 | 21431858 | 22661 | 28,16,38,43,42,17,17,14,4 | 8,7,13,14,7,7,6,10,5,3,0,5 | 22,27,17,8,15,19,15,29,10, | 12,21,10,7,10,5,9,24,7,5,6, | 298 | 149 | 0.00441297347549 | 0.32034020037207 | 0.636,0.533,0.594, | 0.478,0.391,0.459, | 0,159 |
| 22662 | ENSG00000100888 | CHD8 | chr14 | - | 21430800 | 21431724 | 21428963 | 21429335 | 21431810 | 21431858 | 22662 | 28,16,38,43,42,17,17,14,4 | 8,7,13,14,7,7,6,10,5,3,0,5 | 22,27,17,8,15,19,15,29,10, | 12,21,10,7,10,5,9,24,7,5,6, | 298 | 149 | 0.00441297347549 | 0.32034020037207 | 0.636,0.533,0.594, | 0.478,0.391,0.459, | 0,159 |
| 135004 | ENSG00000100335 | MIEF1 | chr22 | + | 39509443 | 39509571 | 39504202 | 39504534 | 39511287 | 39511422 | 135004 | 4,0,3,0,0,0,0,0,2,1,1,0 | 9,9,31,21,14,6,6,15,14,3,9, | 1,0,2,1,0,0,4,0,0,0,6,5,4,3, | 18,13,7,8,9,10,8,11,5,8,16, | 277 | 149 | 0.00441631025635 | 0.32047433337305 | 0.193,0.0,0.049,0. | 0.029,0.0,0.133,0. | -0,041 |
| 178085 | ENSG00000067066 | SP100 | chr2 | + | 230463744 | 230464150 | 230462466 | 230462518 | 230466300 | 230466354 | 178085 | 87,64,160,97,53,65,53,82, | 57,30,72,69,17,16,39,29,3, | 102,78,63,43,74,33,19,67, | 50,76,27,57,116,31,16,23, | 298 | 149 | 0.00441876083842 | 0.320544089657558 | 0.433,0.516,0.526, | 0.505,0.339,0.538, | 0,156 |
| 178094 | ENSG00000067066 | SP100 | chr2 | + | 230463777 | 230464150 | 230462466 | 230462518 | 230466300 | 230466354 | 178094 | 87,64,160,97,53,65,53,82, | 57,30,72,69,17,16,39,29,3, | 102,78,63,43,74,33,19,66, | 50,76,27,57,116,31,16,23, | 298 | 149 | 0.00442872589857 | 0.321158727720728 | 0.433,0.516,0.526, | 0.505,0.339,0.538, | 0,156 |
| 242748 | ENSG00000175938 | ORAI3 | chr16 | + | 30953184 | 30953460 | 30949067 | 30949517 | 30956199 | 30956461 | 242748 | 12,10,21,19,17,13,16,18,2 | 0,3,4,1,1,0,1,3,2,2,0,3 | 13,22,10,18,8,7,25,35,3,11 | 3,1,3,4,0,2,0,0,1,1,0,1,0,0, | 298 | 149 | 0.00443074021513 | 0.321196580411804 | 1.0,0.625,0.724,0. | 0.684,0.917,0.625, | 0,011 |
| 141632 | ENSG00000160551 | TAOK1 | chr17 | + | 29403725 | 29403847 | 29390955 | 29391024 | 29451454 | 29451707 | 141632 | 7,2,3,1,5,0,0,3,0,0,1,2 | 22,6,22,14,10,13,16,6,17,6 | 2,0,0,2,1,0,0,1,0,0,1,0,0,0, | 16,5,3,5,4,11,7,18,3,21,13, | 271 | 149 | 0.0044372131037 | 0.321480471894996 | 0.149,0.155,0.07,0 | 0.064,0.0,0.0,0.18, | 0,115 |
| 9607 | ENSG00000091592 | NLRP1 | chr17 | - | 5517745 | 5517887 | 5514295 | 5515517 | 5521523 | 5521786 | 9607 | 136,56,138,91,43,58,54,77 | 1,2,0,1,0,1,0,2,0,0,0,2 | 85,86,72,76,67,28,76,84,5 | 1,4,2,7,3,0,2,4,1,1,1,0,1,0, | 291 | 149 | 0.00443764365728 | 0.321480471894996 | 0.986,0.935,1.0,0. | 0.978,0.917,0.949, | 0,021 |
| 9433 | ENSG00000175106 | TVP23C | chr17 | - | 15540432 | 15540561 | 15502263 | 15503232 | 15545784 | 15545916 | 9433 | 0,0,20,10,19,18,8,5,15,4,3, | 12,4,2,1,0,0,2,0,0,0,0,1 | 7,18,14,4,8,6,3,7,10,19,36, | 3,0,0,1,0,0,0,0,0,0,0,0,0,0, | 278 | 149 | 0.00444867128088 | 0.322170918053527 | 0.0,0.0,0.843,0.84 | 0.556,1.0,1.0,0.68 | -0,14 |
| 23504 | ENSG00000198718 | TOGARAM1 | chr14 | + | 45008912 | 45009145 | 45006007 | 45006267 | 45011974 | 45012075 | 23504 | 7,9,55,17,5,7,1,12,11,3,5,9 | 1,0,10,2,3,4,0,10,2,3,5,2 | 15,24,18,14,10,11,3,11,2,8 | 2,3,0,0,2,1,6,4,1,0,1,1,2,0, | 298 | 149 | 0.00446058204142 | 0.322924834630632 | 0.778,1.0,0.733,0. | 0.789,0.8,1.0,1.0,0 | -0,134 |
| 88893 | ENSG00000073849 | ST6GAL1 | chr3 | + | 187038391 | 187038491 | 186963784 | 186963926 | 187038741 | 187038873 | 88893 | 1,7,25,19,7,12,9,10,8,4,7,7 | 101,51,133,81,66,36,39,47 | 12,17,13,22,5,8,0,13,6,6,1 | 98,27,89,42,23,58,34,69,2 | 249 | 149 | 0.00446448543429 | 0.323098743950125 | 0.006,0.076,0.101, | 0.068,0.274,0.08,0 | -0,038 |
| 98397 | ENSG00000186480 | INSIG1 | chr7 | + | 155303824 | 155303895 | 155302746 | 155302846 | 155308240 | 155308960 | 98397 | 16,18,104,69,2,18,12,21,1 | 26,26,97,53,18,26,28,27,2 | 19,24,22,9,21,4,8,30,5,13, | 16,21,16,8,19,5,11,30,4,11 | 220 | 149 | 0.00447000528592 | 0.323326087685268 | 0.294,0.319,0.421, | 0.446,0.436,0.482, | -0,127 |
| 31852 | ENSG00000188295 | ZNF669 | chr1 | - | 247101986 | 247102113 | 247101050 | 247101319 | 247104196 | 247104289 | 31852 | 13,16,36,28,6,18,10,21,14, | 2,0,1,1,0,0,1,2,0,0,0,0 | 34,37,13,13,29,5,19,12,17, | 3,0,0,1,0,1,2,0,1,0,10,2,2,2 | 276 | 149 | 0.00447063126107 | 0.323326087685268 | 0.778,1.0,0.951,0. | 0.86,1.0,1.0,0.875, | 0,068 |
| 198727 | ENSG00000223745 | CCDC18-AS1 | chr1 | - | 93335423 | 93335607 | 93325818 | 93325895 | 93337376 | 93339274 | 198727 | 3,3,2,4,0,1,9,9,2,0,2,7 | 26,13,45,34,17,15,16,24,1 | 1,1,2,1,2,0,0,4,3,2,3,0,0,3, | 6,15,13,6,38,8,14,21,16,42 | 298 | 149 | 0.0044746238643 | 0.323506136693703 | 0.055,0.103,0.022, | 0.077,0.032,0.071, | 0,021 |
| 240062 | ENSG00000160563 | MED27 | chr9 | - | 131996452 | 131996600 | 131939380 | 131939474 | 132014336 | 132014467 | 240062 | 0,0,0,4,0,4,1,0,0,0,0,0 | 2,4,17,8,8,7,4,3,5,3,2,3 | 4,4,2,6,1,0,2,1,0,0,3,3,0,0, | 8,6,2,2,6,1,3,7,2,3,11,9,2,5 | 297 | 149 | 0.00448542361115 | 0.324178042058907 | 0.0,0.0,0.0,0.201,0 | 0.201,0.251,0.334, | -0,083 |
| 238592 | ENSG00000136573 | BLK | chr8 | + | 11555585 | 11555922 | 11555390 | 11555484 | 11556657 | 11556837 | 238592 | 73,34,31,15,30,19,9,13,35, | 392,174,307,263,310,194, | 39,53,34,21,25,3,4,10,40,5 | 280,293,166,71,172,94,75, | 298 | 149 | 0.00449283042948 | 0.32460435923713 | 0.085,0.089,0.048, | 0.065,0.083,0.093, | -0,016 |
| 60667 | ENSG00000156983 | BRPF1 | chr3 | + | 9744223 | 9744508 | 9743577 | 9743901 | 9745007 | 9745155 | 60667 | 12,19,17,5,15,10,12,9,7,7, | 1,0,0,3,2,2,2,3,0,2,0,0 | 23,25,19,16,13,4,8,14,2,10 | 4,3,1,1,0,0,6,2,2,4,7,3,3,2, | 298 | 149 | 0.00451129190529 | 0.325781083861416 | 0.857,1.0,1.0,0.45 | 0.742,0.806,0.905, | 0,102 |
| 185339 | ENSG00000188822 | CNR2 | chr1 | - | 23879503 | 23879604 | 23870514 | 23875662 | 23913245 | 23913362 | 185339 | 5,4,14,18,11,9,12,10,12,1, | 72,36,191,152,96,74,66,90 | 11,2,5,11,1,8,0,1,9,6,7,7,8, | 135,99,48,64,73,71,22,84, | 250 | 149 | 0.00451279969171 | 0.325781083861416 | 0.04,0.062,0.042,0 | 0.046,0.012,0.058, | 0,017 |
| 195662 | ENSG00000177051 | FBXO46 | chr19 | - | 45723472 | 45723890 | 45712943 | 45713573 | 45730848 | 45730896 | 195662 | 2,6,4,9,7,2,3,3,8,2,5,3 | 3,2,2,3,7,5,7,2,12,4,4,3 | 3,3,3,5,3,2,0,6,2,6,3,9,7,3, | 0,2,1,2,1,0,2,3,0,0,9,4,0,0, | 298 | 149 | 0.00451365831165 | 0.325781083861416 | 0.25,0.6,0.5,0.6,0. | 1.0,0.429,0.6,0.55 | -0,281 |
| 52389 | ENSG00000267787 | | chr18 | + | 57734025 | 57734096 | 57641558 | 57642052 | 57735150 | 57735592 | 52389 | 6,1,1,2,0,1,1,1,1,0,2,1 | 16,3,7,6,5,1,11,2,2,1,0,6 | 2,3,4,0,2,1,6,2,0,1,5,6,2,5, | 3,3,2,6,1,2,8,2,4,5,5,9,3,7, | 220 | 149 | 0.00451912535475 | 0.326066332292389 | 0.203,0.184,0.088, | 0.311,0.404,0.575, | -0,052 |
| 195301 | ENSG00000134744 | TUT4 | chr1 | - | 52471951 | 52472089 | 52468180 | 52468267 | 52474831 | 52475535 | 195301 | 70,58,109,46,17,29,30,29, | 0,0,9,2,2,2,1,3,0,1,0,0 | 61,48,19,25,26,20,21,37,2 | 3,0,0,4,0,0,0,0,0,0,0,0,0,0, | 287 | 149 | 0.00453609622088 | 0.327181141325812 | 1.0,1.0,0.863,0.92 | 0.913,1.0,1.0,0.76 | -0,06 |
| 135035 | ENSG00000284431 | | chr22 | + | 40364912 | 40365056 | 40364275 | 40364365 | 40366435 | 40366498 | 135035 | 47,14,75,74,34,29,20,27,3 | 5,1,9,9,4,1,2,7,6,1,0,7 | 35,42,32,8,22,27,7,11,15,3 | 2,10,4,3,11,7,2,0,4,13,2,9, | 293 | 149 | 0.00454341555574 | 0.327599287597144 | 0.827,0.877,0.809, | 0.899,0.681,0.803, | 0,129 |
| 215413 | ENSG00000010244 | ZNF207 | chr17 | + | 32361467 | 32361547 | 32360891 | 32360967 | 32362913 | 32362984 | 215413 | 28,23,94,53,32,42,46,55,2 | 2,2,9,15,8,11,11,6,25,8,5,3 | 66,69,35,35,39,31,24,47,3 | 3,0,4,3,3,7,4,10,0,7,9,13,8, | 229 | 149 | 0.00455177225277 | 0.3280919269042 | 0.901,0.882,0.872, | 0.935,1.0,0.851,0. | -0,09 |
| 12995 | ENSG00000109046 | WSB1 | chr17 | + | 27301766 | 27301956 | 27294075 | 27294435 | 27303366 | 27303635 | 12995 | 57,31,168,128,42,42,33,64 | 4,3,37,11,32,20,10,15,5,2, | 44,66,17,9,20,30,19,86,17, | 15,20,23,21,20,11,3,12,11, | 298 | 149 | 0.00455621411834 | 0.328302149616484 | 0.877,0.838,0.694, | 0.595,0.623,0.27,0 | 0,166 |
| 197983 | ENSG00000153922 | CHD1 | chr5 | - | 98904896 | 98905094 | 98903791 | 98903908 | 98926333 | 98926534 | 197983 | 133,109,83,31,37,60,51,61 | 3,2,0,3,0,3,3,1,0,0,2,2 | 80,113,77,182,85,41,33,59 | 6,3,4,0,4,3,2,4,2,1,0,13,1,0 | 298 | 149 | 0.00456951851285 | 0.329105752877441 | 0.957,0.965,1.0,0. | 0.87,0.95,0.906,1. | 0,027 |
| 101582 | ENSG00000291136 | | chr7 | - | 66560578 | 66560675 | 66554584 | 66554837 | 66573384 | 66573550 | 101582 | 6,11,14,8,16,16,15,7,28,14 | 20,7,17,22,17,20,5,9,12,6, | 12,10,6,3,5,11,3,7,1,12,8,5 | 18,19,11,1,2,7,13,19,6,9,1 | 246 | 149 | 0.0045704247778 | 0.329105752877441 | 0.154,0.488,0.333, | 0.288,0.242,0.248, | 0,123 |
| 213426 | ENSG00000035720 | STAP1 | chr4 | + | 67576822 | 67577259 | 67575384 | 67575498 | 67581304 | 67581471 | 213426 | 95,41,111,80,38,50,40,46, | 3,0,4,1,1,0,3,4,1,1,2,2 | 57,45,70,34,51,30,19,38,2 | 2,3,3,11,3,0,2,2,2,0,0,0,3,4 | 298 | 149 | 0.00457894222743 | 0.329608800853507 | 0.941,1.0,0.933,0. | 0.934,0.882,0.921, | 0,016 |
| 105708 | ENSG00000167548 | KMT2D | chr12 | - | 49054306 | 49054416 | 49053977 | 49054140 | 49054527 | 49054751 | 105708 | 21,20,19,19,12,43,8,17,5,2 | 2,1,1,1,0,1,0,0,0,2,0,0 | 14,38,8,85,18,17,39,21,14, | 2,1,0,12,0,0,4,1,0,0,0,0,1,1 | 259 | 149 | 0.00458788793271 | 0.33014232953698 | 0.858,0.92,0.916,0 | 0.801,0.956,1.0,0. | 0,03 |
| 88865 | ENSG00000143207 | COP1 | chr1 | - | 176136487 | 176136547 | 176135009 | 176135086 | 176149005 | 176149074 | 88865 | 78,67,104,81,43,68,30,69, | 11,6,4,1,8,12,8,13,2,1,0,1 | 59,67,48,82,45,27,24,44,4 | 14,14,7,3,2,6,3,15,11,8,6,4 | 209 | 149 | 0.00459612207682 | 0.330621894647174 | 0.835,0.888,0.949, | 0.75,0.773,0.83,0. | 0,075 |
| 95720 | ENSG00000196295 | GARS1-DT | chr7 | - | 30561465 | 30562128 | 30550635 | 30550781 | 30563730 | 30563998 | 95720 | 91,31,68,60,38,44,45,57,1 | 0,0,2,4,0,0,0,1,0,0,3,0 | 27,41,15,34,27,31,38,63,2 | 2,0,3,1,1,0,1,5,2,0,0,8,0,2, | 298 | 149 | 0.00459762455538 | 0.330621894647174 | 1.0,1.0,0.944,0.88 | 0.871,1.0,0.714,0. | 0,036 |
| 105779 | ENSG00000104870 | FCGRT | chr19 | + | 49513386 | 49513473 | 49506815 | 49506924 | 49513881 | 49514133 | 105779 | 32,6,18,15,16,6,6,3,2,2,6,1 | 10,3,2,0,0,0,1,2,0,0,2,1 | 11,15,3,5,5,3,11,13,1,9,10, | 2,1,0,0,2,0,0,0,0,0,2,2,2,1, | 236 | 149 | 0.00460340451008 | 0.330926972648306 | 0.669,0.558,0.85,1 | 0.776,0.904,1.0,1. | -0,059 |
| 103323 | ENSG00000231721 | LINC-PINT | chr7 | - | 131009708 | 131009778 | 130984052 | 130984109 | 131052424 | 131052555 | 103323 | 1,1,11,1,3,7,3,3,6,1,0,4 | 132,85,282,117,79,106,96, | 8,15,5,9,4,1,2,4,7,8,7,0,1,2 | 176,141,91,124,97,47,76,9 | 219 | 149 | 0.00460696287602 | 0.33107219591608 | 0.005,0.008,0.026, | 0.03,0.067,0.036,0 | -0,011 |
| 175927 | ENSG00000173889 | PHC3 | chr3 | - | 170149085 | 170149211 | 170136365 | 170136665 | 170171372 | 170171450 | 175927 | 5,4,1,6,1,0,0,3,1,0,0,1 | 31,10,12,2,12,5,6,13,16,6, | 0,5,3,0,0,1,1,0,4,2,2,3,0,3, | 18,12,19,17,12,19,8,5,7,10 | 275 | 149 | 0.00461512606884 | 0.331548130481476 | 0.08,0.178,0.043,0 | 0.0,0.184,0.079,0. | 0,012 |
| 167404 | ENSG00000147548 | NSD3 | chr8 | - | 38299452 | 38299590 | 38295795 | 38295952 | 38304586 | 38304757 | 167404 | 22,2,41,14,32,13,14,19,29, | 1,1,5,4,5,4,3,1,1,1,4,2 | 30,18,16,9,7,9,10,11,9,25, | 5,9,8,8,1,7,0,0,1,5,12,3,14, | 287 | 149 | 0.00461963745689 | 0.331761491319283 | 0.919,0.509,0.81,0 | 0.757,0.509,0.509, | 0,113 |
| 140957 | ENSG00000141627 | DYM | chr18 | - | 49390705 | 49390812 | 49379745 | 49379758 | 49391592 | 49391645 | 140957 | 1,0,1,0,0,0,0,1,3,0,0,0 | 41,13,67,41,25,30,23,19,1 | 1,3,0,7,4,0,1,0,2,3,2,0,3,3, | 34,32,24,25,10,17,25,26,2 | 256 | 149 | 0.00463327837016 | 0.332523627136891 | 0.014,0.0,0.009,0. | 0.017,0.052,0.0,0. | -0,034 |
| 177447 | ENSG00000160991 | ORAI2 | chr7 | + | 102436164 | 102436333 | 102433575 | 102433661 | 102438943 | 102439047 | 177447 | 37,50,100,98,118,117,133, | 6,2,8,5,4,4,11,30,2,4,7,13 | 75,48,41,42,45,36,49,115, | 5,7,4,9,6,0,1,16,6,7,10,55, | 298 | 149 | 0.0046333397967 | 0.332523627136891 | 0.755,0.926,0.862, | 0.882,0.774,0.837, | 0,047 |
| 31902 | ENSG00000247774 | PCED1B-AS1 | chr12 | - | 47210639 | 47210761 | 47205897 | 47206145 | 47216164 | 47216225 | 31902 | 12,9,35,46,27,19,11,5,15,1 | 2,2,4,22,1,2,3,4,5,4,0,1 | 27,15,10,6,18,22,11,8,17,3 | 2,1,2,0,3,1,1,0,0,0,2,16,5,6 | 271 | 149 | 0.00463902069942 | 0.332820354718955 | 0.767,0.712,0.828, | 0.881,0.892,0.733, | -0,127 |
| 201301 | ENSG00000055070 | SZRD1 | chr1 | + | 16391381 | 16391424 | 16367264 | 16367308 | 16393227 | 16393385 | 201301 | 12,7,10,5,3,5,5,7,19,9,3,5 | 27,19,30,12,10,12,29,7,24, | 5,7,3,1,1,1,2,12,1,2,6,10,5, | 22,26,15,9,7,3,16,32,1,13,1 | 192 | 149 | 0.00464369691844 | 0.333044828874628 | 0.256,0.222,0.206, | 0.15,0.173,0.134,0 | 0,081 |
| 58827 | ENSG00000065243 | PKN2 | chr1 | + | 88804390 | 88804534 | 88786103 | 88786213 | 88804845 | 88804894 | 58827 | 86,55,131,70,37,66,48,84, | 3,2,3,5,0,5,2,1,0,0,0,3 | 63,85,51,81,96,44,50,55,5 | 5,20,6,2,4,0,0,3,1,3,1,5,3,0 | 293 | 149 | 0.00465440249149 | 0.333592404761136 | 0.936,0.933,0.957, | 0.865,0.684,0.812, | 0,046 |
| 152794 | ENSG00000092148 | HECTD1 | chr14 | - | 31113923 | 31114059 | 31113276 | 31113484 | 31114285 | 31114340 | 152794 | 21,25,68,46,41,35,51,59,1 | 17,3,11,7,0,12,2,12,2,4,1,0 | 46,59,34,28,45,27,24,39,1 | 4,8,0,1,3,2,3,4,1,0,7,5,0,3, | 285 | 149 | 0.00465443171057 | 0.333592404761136 | 0.392,0.813,0.764, | 0.857,0.794,1.0,0. | -0,102 |
| 106319 | ENSG00000176871 | WSB2 | chr12 | - | 118035213 | 118035324 | 118032686 | 118034358 | 118036337 | 118036510 | 106319 | 15,7,71,79,17,45,46,59,26, | 1,2,1,0,0,1,0,5,0,0,0,0 | 48,66,25,5,17,16,6,28,10,2 | 1,2,1,2,0,2,0,2,0,1,5,0,2,5, | 260 | 149 | 0.00465936695717 | 0.333779089436595 | 0.896,0.667,0.976, | 0.965,0.95,0.935,0 | 0,048 |
| 204044 | ENSG00000144535 | DIS3L2 | chr2 | + | 232259639 | 232259761 | 232249339 | 232249346 | 232263206 | 232263440 | 204044 | 4,4,2,4,0,1,2,3,2,2,2,0 | 32,21,16,18,20,28,20,27,2 | 2,3,2,2,0,2,2,7,5,6,2,7,3,2, | 15,23,6,38,17,5,7,27,11,10 | 271 | 149 | 0.00466013800873 | 0.333779089436595 | 0.064,0.095,0.064, | 0.068,0.067,0.155, | -0,066 |
| 69661 | ENSG00000232956 | SNHG15 | chr7 | - | 44984356 | 44984426 | 44983022 | 44984046 | 44986020 | 44986097 | 69661 | 24,8,39,36,21,10,13,19,14, | 0,2,3,3,2,1,0,2,0,2,1,3 | 21,21,11,10,10,4,20,3,6,17 | 0,6,0,1,3,0,1,1,0,0,0,15,0,1 | 219 | 149 | 0.00466868739526 | 0.334280191872656 | 1.0,0.731,0.898,0. | 1.0,0.704,1.0,0.87 | -0,06 |
| 30087 | ENSG00000113282 | CLINT1 | chr5 | - | 157790599 | 157790651 | 157789362 | 157789513 | 157791702 | 157791841 | 30087 | 2,9,3,2,6,1,1,11,3,1,3,4 | 77,63,57,52,40,25,29,142, | 9,6,11,17,6,5,0,3,2,10,6,11 | 60,85,75,104,93,44,30,52, | 201 | 149 | 0.00467104070673 | 0.334337466694448 | 0.019,0.096,0.038, | 0.1,0.05,0.098,0.1 | -0,018 |
| 236124 | ENSG00000166169 | POLL | chr10 | - | 101585865 | 101586156 | 101584877 | 101584919 | 101587245 | 101587382 | 236124 | 4,1,14,17,3,5,7,8,0,1,8,10 | 1,2,2,0,1,2,0,2,2,2,0,0 | 5,3,5,1,3,3,5,5,3,12,4,3,23, | 0,0,0,0,0,0,0,0,0,0,1,0,0, | 298 | 149 | 0.00467325984629 | 0.334385103050812 | 0.667,0.2,0.778,1. | 1.0,1.0,1.0,1.0,1.0, | -0,318 |
| 172365 | ENSG00000143669 | LYST | chr1 | - | 235808078 | 235808180 | 235805742 | 235806772 | 235808454 | 235810534 | 172365 | 10,1,4,1,0,0,2,9,6,0,0,1 | 34,21,52,12,14,16,23,20,1 | 5,6,2,2,0,0,2,0,2,0,3,1,0,0, | 44,22,15,9,6,8,12,9,20,22, | 251 | 149 | 0.00468175451419 | 0.33488159050968 | 0.149,0.027,0.044, | 0.063,0.139,0.073, | 0,023 |
| 108150 | ENSG00000135048 | CEMIP2 | chr9 | - | 71690091 | 71690246 | 71685742 | 71685846 | 71694508 | 71694607 | 108150 | 31,37,71,62,29,50,43,84,1 | 5,0,1,2,6,5,4,2,0,0,1,0 | 49,64,23,12,22,3,81,37,5,1 | 1,1,3,1,1,0,2,1,0,0,2,5,2,0, | 298 | 149 | 0.00468540377393 | 0.335031275636787 | 0.756,1.0,0.973,0. | 0.961,0.97,0.793,0 | -0,026 |
| 194983 | ENSG00000143384 | MCL1 | chr1 | - | 150578243 | 150578342 | 150574175 | 150577491 | 150578842 | 150579610 | 194983 | 106,95,234,200,171,129,9 | 2,2,3,4,14,7,14,13,4,1,3,5 | 200,190,95,71,82,80,32,74 | 18,10,13,3,1,7,2,4,2,12,8,4 | 248 | 149 | 0.00468769835903 | 0.335084027071533 | 0.97,0.966,0.979,0 | 0.87,0.919,0.814,0 | 0,02 |
| 197389 | ENSG00000163534 | FCRL1 | chr1 | - | 157803844 | 157803996 | 157801914 | 157802664 | 157820006 | 157820120 | 197389 | 299,121,354,37,107,60,17 | 0,1,14,0,2,2,4,5,1,0,0,2 | 203,142,222,103,43,33,10 | 7,4,4,3,2,3,0,0,0,4,6,6,11,2 | 298 | 149 | 0.00469291020048 | 0.335345204302626 | 1.0,0.984,0.927,1. | 0.935,0.947,0.965, | 0,029 |
| 63456 | ENSG00000111481 | COPZ1 | chr12 | + | 54343224 | 54343316 | 54342205 | 54342287 | 54345459 | 54345475 | 63456 | 89,55,269,222,74,66,74,75 | 6,3,1,1,1,0,0,0,0,0,0,1 | 120,86,99,30,70,65,35,69, | 5,4,2,1,2,2,0,1,0,4,0,5,1,1, | 241 | 149 | 0.00469473948319 | 0.335358732237936 | 0.902,0.919,0.994, | 0.937,0.93,0.968,0 | 0,017 |
| 76688 | ENSG00000159023 | EPB41 | chr1 | + | 29060421 | 29060484 | 29053103 | 29053312 | 29064981 | 29065158 | 76688 | 10,5,15,17,11,9,6,13,20,9, | 178,160,197,190,182,157, | 13,8,8,11,7,11,2,5,0,12,7,1 | 184,154,169,147,161,142,1 | 212 | 149 | 0.00469716917082 | 0.335358732237936 | 0.038,0.021,0.051, | 0.047,0.035,0.032, | 0,011 |
| 219922 | ENSG00000198315 | ZKSCAN8 | chr6 | + | 28148315 | 28148770 | 28141882 | 28142029 | 28149482 | 28149624 | 219922 | 16,13,21,19,17,15,11,12,1 | 4,3,3,5,0,3,2,5,0,1,0,2 | 23,12,6,25,7,5,15,13,5,8,1 | 4,8,12,3,7,2,6,2,3,0,4,2,0,3 | 298 | 149 | 0.00469777391592 | 0.335358732237936 | 0.667,0.684,0.778, | 0.742,0.429,0.2,0. | 0,172 |
| 232434 | ENSG00000197713 | RPE | chr2 | + | 210016178 | 210016257 | 210015972 | 210016112 | 210016506 | 210016597 | 232434 | 0,0,9,1,1,4,0,5,0,1,0,1 | 14,13,36,19,25,23,14,27,1 | 0,0,1,0,1,0,0,0,4,0,0,1,1,1, | 27,22,21,10,7,9,12,22,10,1 | 228 | 149 | 0.00472180839561 | 0.336962713128493 | 0.0,0.0,0.14,0.033, | 0.0,0.0,0.03,0.0,0. | 0,012 |
| 48645 | ENSG00000120008 | WDR11 | chr10 | + | 120856081 | 120856130 | 120852523 | 120852635 | 120858642 | 120858796 | 48645 | 19,21,76,26,29,6,9,5,9,6,2, | 169,180,400,232,157,101, | 29,42,20,14,39,40,6,22,47, | 216,290,278,248,208,166, | 198 | 149 | 0.00473795510518 | 0.338002921857142 | 0.078,0.081,0.125, | 0.092,0.098,0.051, | -0,034 |
| 113441 | ENSG00000104325 | DECR1 | chr8 | + | 90005447 | 90005845 | 90001476 | 90001561 | 90017123 | 90017326 | 113441 | 1,1,5,1,0,1,0,1,3,0,0,2 | 10,4,40,49,21,13,12,8,7,4, | 3,0,2,2,0,2,0,6,1,5,1,1,2,4, | 17,12,13,0,14,9,4,11,9,13, | 298 | 149 | 0.00474923907681 | 0.338695651338931 | 0.048,0.111,0.059, | 0.081,0.0,0.071,1. | -0,086 |
| 156268 | ENSG00000102805 | CLN5 | chr13 | + | 76998042 | 76998291 | 76995901 | 76996127 | 77000457 | 77005117 | 156268 | 1,0,1,8,3,3,0,3,0,0,0,2 | 9,13,30,14,11,10,7,13,8,5, | 2,1,0,0,1,0,1,0,0,0,0,0,2,0, | 15,16,8,4,9,8,9,12,1,6,14,6 | 298 | 149 | 0.00476085744367 | 0.339411761662318 | 0.053,0.0,0.016,0. | 0.062,0.03,0.0,0.0, | 0,031 |
| 190397 | ENSG00000164300 | SERINC5 | chr5 | - | 80205206 | 80205360 | 80202885 | 80203053 | 80255895 | 80256048 | 190397 | 0,0,3,0,0,8,0,0,0,0,2,7 | 6,2,14,5,3,7,15,8,11,2,11,2 | 1,0,5,9,0,0,0,2,3,2,6,1,2,0, | 2,4,7,23,13,4,13,27,21,11, | 298 | 149 | 0.00476268188554 | 0.339429398975715 | 0.0,0.0,0.097,0.0,0 | 0.2,0.0,0.263,0.16 | -0,021 |
| 131178 | ENSG00000188906 | LRRK2 | chr12 | + | 40316277 | 40316403 | 40315211 | 40315300 | 40319987 | 40320175 | 131178 | 0,2,2,2,4,0,2,4,0,0,0,2 | 18,16,99,45,14,13,30,21,1 | 4,2,4,0,4,1,2,1,3,6,2,1,5,5, | 38,37,19,13,19,23,8,14,16, | 275 | 149 | 0.00476866206373 | 0.339743099847293 | 0.0,0.063,0.011,0. | 0.054,0.028,0.102, | -0,022 |
| 201175 | ENSG00000181722 | ZBTB20 | chr3 | - | 114741555 | 114741708 | 114713975 | 114714015 | 114801100 | 114801177 | 201175 | 2,0,3,1,1,4,1,0,2,0,0,1 | 6,7,7,4,14,18,7,9,7,2,3,4 | 0,0,6,5,2,2,0,1,2,9,3,1,0,2, | 3,10,7,3,11,2,8,4,6,2,5,9,7, | 298 | 149 | 0.00477396250952 | 0.340008181630211 | 0.143,0.0,0.176,0. | 0.0,0.0,0.3,0.455,0 | -0,117 |
| 174063 | ENSG00000134109 | EDEM1 | chr3 | + | 5213886 | 5214191 | 5213318 | 5213522 | 5215828 | 5219958 | 174063 | 3,17,14,6,2,1,2,10,5,5,5,4 | 135,91,256,170,104,122,1 | 2,16,5,28,6,8,4,11,2,7,8,1, | 116,163,82,144,82,67,79,1 | 298 | 149 | 0.00478289237792 | 0.340531495002349 | 0.011,0.085,0.027, | 0.009,0.047,0.03,0 | -0,004 |
| 130716 | ENSG00000137343 | ATAT1 | chr6 | + | 30640534 | 30640603 | 30640376 | 30640422 | 30642175 | 30642247 | 130716 | 14,8,4,10,7,9,15,6,7,6,1,12 | 2,0,4,0,0,1,3,1,2,4,0,0 | 8,1,1,8,4,2,8,12,4,5,9,3,9,8 | 0,0,0,0,0,0,0,2,0,0,2,2,0,0, | 218 | 149 | 0.00479004244936 | 0.340927786513956 | 0.827,1.0,0.406,1. | 1.0,1.0,1.0,1.0,1.0, | -0,095 |
| 54355 | ENSG00000138095 | LRPPRC | chr2 | - | 43890222 | 43890274 | 43889733 | 43889876 | 43894544 | 43894629 | 54355 | 7,0,5,5,3,1,3,3,3,0,1,0 | 35,22,68,48,54,34,23,42,3 | 5,14,3,9,3,2,6,5,3,3,6,8,1,1 | 74,67,47,25,29,16,24,43,2 | 201 | 149 | 0.00479211094846 | 0.340962258362973 | 0.129,0.0,0.052,0. | 0.048,0.134,0.045, | -0,034 |
| 9658 | ENSG00000178971 | CTC1 | chr17 | - | 8228726 | 8228892 | 8228502 | 8228629 | 8229141 | 8229206 | 9658 | 35,15,56,35,38,33,31,45,3 | 7,0,3,1,0,0,3,2,1,5,2,10 | 40,30,13,20,15,19,28,37,1 | 6,12,6,15,3,1,3,14,2,0,9,0, | 298 | 149 | 0.00480508332705 | 0.341772270179874 | 0.714,1.0,0.903,0. | 0.769,0.556,0.52,0 | 0,113 |
| 47799 | ENSG00000163714 | U2SURP | chr3 | + | 143023974 | 143024018 | 143022852 | 143023064 | 143027148 | 143027253 | 47799 | 103,52,123,81,57,54,52,81 | 26,21,18,4,7,2,9,4,5,4,6,6 | 103,58,89,44,25,34,17,68, | 5,21,3,65,12,3,13,25,5,9,6, | 193 | 149 | 0.00480712184963 | 0.341804308826467 | 0.754,0.657,0.841, | 0.941,0.681,0.958, | 0,097 |
| 241673 | ENSG00000175893 | ZDHHC21 | chr9 | - | 14652737 | 14652835 | 14639895 | 14640012 | 14658748 | 14658887 | 241673 | 2,7,8,7,2,1,0,0,4,1,0,0 | 95,93,131,120,46,55,57,41 | 7,10,4,0,0,0,0,2,0,0,2,21,0, | 54,68,48,16,56,58,55,101, | 247 | 149 | 0.0048188299803 | 0.342473202017307 | 0.013,0.043,0.036, | 0.073,0.081,0.048, | -0,015 |
| 98694 | ENSG00000197694 | SPTAN1 | chr9 | + | 128592982 | 128593042 | 128591476 | 128591625 | 128594174 | 128594373 | 98694 | 134,78,161,173,68,64,53,5 | 0,0,4,3,0,0,1,8,5,2,0,3 | 92,82,64,35,62,63,34,51,4 | 4,2,1,14,5,0,1,5,2,2,0,2,5,0 | 209 | 149 | 0.00482103789908 | 0.342473202017307 | 1.0,1.0,0.966,0.97 | 0.943,0.967,0.979, | 0,026 |
| 138095 | ENSG00000187912 | CLEC17A | chr19 | + | 14585611 | 14585830 | 14583356 | 14583434 | 14587613 | 14587691 | 138095 | 3,4,3,5,5,2,7,0,0,3,1,2 | 21,2,31,24,73,24,44,31,25, | 6,5,4,3,2,2,1,1,4,3,0,8,2,0, | 16,14,4,4,11,5,8,13,6,7,4,4 | 298 | 149 | 0.0048213027032 | 0.342473202017307 | 0.067,0.5,0.046,0. | 0.158,0.152,0.333, | -0,042 |
| 123083 | ENSG00000066294 | CD84 | chr1 | - | 160550937 | 160551035 | 160549916 | 160549979 | 160553377 | 160553497 | 123083 | 39,22,65,64,28,28,27,8,15, | 0,2,3,0,1,0,1,0,0,0,0,0 | 59,67,7,7,23,15,26,42,15,2 | 1,4,0,0,2,2,0,3,2,3,1,0,0,2, | 247 | 149 | 0.00482713128031 | 0.342774098512835 | 1.0,0.869,0.929,1. | 0.973,0.91,1.0,1.0, | 0,053 |
| 99417 | ENSG00000115677 | HDLBP | chr2 | - | 241255373 | 241255580 | 241255050 | 241255158 | 241256183 | 241256399 | 99417 | 109,66,271,189,79,70,68,9 | 0,0,5,4,4,5,5,5,6,1,0,0 | 121,90,99,35,59,63,59,104 | 1,0,0,0,4,1,0,1,2,1,4,1,0,3, | 298 | 149 | 0.00483767685634 | 0.343409639665869 | 1.0,1.0,0.964,0.95 | 0.984,1.0,1.0,1.0,0 | -0,023 |
| 248801 | ENSG00000110422 | HIPK3 | chr11 | + | 33348576 | 33348818 | 33348165 | 33348228 | 33349146 | 33349287 | 248801 | 80,45,104,52,35,44,51,46, | 0,0,2,6,4,1,6,3,1,0,0,1 | 68,65,45,38,34,45,52,53,3 | 6,1,5,10,1,3,1,3,1,5,12,3,1, | 298 | 149 | 0.00484287533163 | 0.343665315035297 | 1.0,1.0,0.963,0.81 | 0.85,0.97,0.818,0. | 0,058 |
| 114756 | ENSG00000147162 | OGT | chrX | + | 71537828 | 71538072 | 71533086 | 71533336 | 71544566 | 71544624 | 114756 | 208,146,654,489,176,218, | 2,2,3,0,0,4,0,4,1,0,0,0 | 446,490,262,151,390,303, | 8,8,7,1,6,0,0,3,4,2,9,0,0,1, | 298 | 149 | 0.00484714373465 | 0.343854842832055 | 0.981,0.973,0.991, | 0.965,0.968,0.949, | 0,013 |
| 69803 | ENSG00000223745 | CCDC18-AS1 | chr1 | - | 93315148 | 93315269 | 93305387 | 93305470 | 93324634 | 93324691 | 69803 | 2,2,1,2,1,7,3,3,0,0,3,2 | 18,9,27,25,15,8,16,20,13,7 | 0,1,1,4,2,5,7,2,3,10,4,4,3,4 | 10,9,3,2,16,10,4,15,3,10,1 | 270 | 149 | 0.00485491741207 | 0.344292826859057 | 0.058,0.109,0.02,0 | 0.0,0.058,0.155,0. | -0,097 |

| | | | | | | | | | | | | | | | | | | | | | | |
|---|---|---|---|---|---|---|---|---|---|---|---|---|---|---|---|---|---|---|---|---|---|---|
| 72307 | ENSG00000055070 | SZRD1 | chr1 | + | 16393230 | 16393482 | 16367264 | 16367308 | 16395037 | 16395139 | 72307 | 103,101,150,126,100,89,8 | 0,0,2,0,1,3,2,2,0,0,0,1 | 106,93,72,95,77,31,41,134 | 2,0,2,2,0,1,3,2,0,3,2,8,1,0,5 | 298 | 149 | 0.00485655094529 | 0.344295229415584 | 1.0,1.0,0.974,1.0,0 | 0.964,1.0,0.947,0. | 0,022 |
| 28432 | ENSG00000185404 | SP140L | chr2 | + | 230400954 | 230401063 | 230400126 | 230400242 | 230401365 | 230401443 | 28432 | 14,11,43,20,12,8,5,12,3,2, | 1,0,2,4,0,1,1,3,0,2,0,0 | 29,28,14,8,12,6,2,9,10,14,2 | 1,4,0,7,2,1,0,3,0,0,0,1,0,4,3 | 258 | 149 | 0.00485928492899 | 0.344375618884243 | 0.89,1.0,0.925,0.74 | 0.944,0.802,1.0,0.3 | 0,003 |
| 151298 | ENSG00000167491 | GATAD2A | chr19 | + | 19457006 | 19457229 | 19436130 | 19436219 | 19465339 | 19465614 | 151298 | 1,6,2,5,0,0,1,1,0,0,1,1 | 11,7,10,7,6,1,3,4,10,4,1,2 | 0,0,0,2,2,4,0,2,0,1,0,2,2,9,3 | 4,9,10,8,5,1,2,2,2,4,6,3,6,3 | 298 | 149 | 0.00486895143983 | 0.344812114934584 | 0.043,0.3,0.091,0.2 | 0.0,0.0,0.0,0.111,0 | -0,063 |
| 175931 | ENSG00000173889 | PHC3 | chr3 | - | 170149103 | 170149244 | 170145422 | 170145521 | 170171372 | 170171450 | 175931 | 80,84,105,53,53,51,54,49,1 | 31,12,35,23,4,31,22,16,17, | 51,66,52,48,41,46,34,56,20 | 34,46,25,29,26,14,18,29,18 | 290 | 149 | 0.00487011717451 | 0.344812114934584 | 0.57,0.782,0.607,0 | 0.435,0.424,0.517, | 0,116 |
| 250719 | ENSG00000119977 | TCTN3 | chr10 | - | 95680471 | 95680575 | 95659822 | 95664300 | 95682650 | 95683629 | 250719 | 10,8,21,23,8,14,3,4,6,5,2,0 | 4,3,4,0,0,3,0,0,0,0,0,2 | 11,4,6,2,4,4,7,9,6,5,7,9,9,5 | 0,1,0,3,2,2,8,2,1,3,1,0,0,1,1 | 253 | 149 | 0.00487025023998 | 0.344812114934584 | 0.596,0.611,0.756, | 1.0,0.702,1.0,0.28 | 0,113 |
| 142026 | ENSG00000129197 | RPAIN | chr17 | + | 5425945 | 5426082 | 5422768 | 5422829 | 5426235 | 5426299 | 142026 | 12,2,37,24,5,4,2,18,6,0,2,6 | 0,0,2,9,6,3,3,2,0,2,0,0 | 7,7,9,1,10,5,4,10,3,7,9,15,1 | 1,0,0,0,0,0,0,1,0,0,0,9,3,3,2 | 286 | 149 | 0.00487589873997 | 0.344939707803966 | 1.0,1.0,0.906,0.58 | 0.785,1.0,1.0,1.0,1 | -0,2 |
| 43784 | ENSG00000159147 | DONSON | chr21 | - | 33583487 | 33583666 | 33582164 | 33582246 | 33584589 | 33584726 | 43784 | 14,3,17,8,4,3,8,12,8,9,2,6 | 2,1,1,1,1,0,0,5,3,2,0,1 | 5,8,14,6,6,0,3,2,4,8,3,8,2,1 | 0,3,0,3,0,1,1,1,2,1,0,3,2,2,0 | 298 | 149 | 0.00487682331635 | 0.344939707803966 | 0.778,0.6,0.895,0.8 | 1.0,0.571,1.0,0.5,1 | 0,114 |
| 212037 | ENSG00000079263 | SP140 | chr2 | + | 230247248 | 230247496 | 230245862 | 230245940 | 230247915 | 230248065 | 212037 | 26,9,7,7,0,5,7,13,8,3,4,6 | 259,138,178,134,120,71,56 | 9,19,7,18,13,5,3,4,10,4,5,2 | 139,166,121,130,140,122,3 | 298 | 149 | 0.00487686035398 | 0.344939707803966 | 0.048,0.032,0.019, | 0.031,0.054,0.028, | -0,011 |
| 12399 | ENSG00000141298 | SSH2 | chr17 | - | 29848848 | 29848929 | 29793615 | 29793937 | 29929937 | 29930167 | 12399 | 9,4,0,1,6,2,5,0,0,2,0,1 | 4,1,5,0,0,2,1,3,3,5,2,2 | 1,6,10,4,5,1,2,5,3,5,3,5,5,2 | 0,1,2,2,3,2,0,2,0,0,3,4,4,2,2 | 230 | 149 | 0.00488551737118 | 0.345438498461381 | 0.593,0.722,0.0,1.0 | 1.0,0.795,0.764,0.5 | -0,228 |
| 253299 | ENSG00000130177 | CDC16 | chr13 | + | 114265144 | 114265240 | 114262878 | 114263014 | 114272183 | 114272481 | 253299 | 89,39,72,46,30,17,10,28,33 | 6,5,19,11,9,8,9,11,0,1,8,8 | 31,25,39,19,8,28,11,15,33, | 6,23,13,22,19,13,5,7,15,23 | 245 | 149 | 0.00489416458804 | 0.345885095355063 | 0.9,0.826,0.697,0.7 | 0.759,0.398,0.646, | 0,133 |
| 90930 | ENSG00000111679 | PTPN6 | chr12 | + | 6955150 | 6955267 | 6954804 | 6954994 | 6955371 | 6955394 | 90930 | 947,502,1043,753,610,268 | 31,6,36,11,8,13,5,13,13,7,0 | 600,420,259,55,147,248,11 | 19,10,4,3,6,2,2,7,1,4,1,26,1 | 266 | 149 | 0.00489504764858 | 0.345885095355063 | 0.945,0.979,0.942, | 0.946,0.959,0.973, | -0,008 |
| 173215 | ENSG00000174243 | DDX23 | chr12 | - | 48833606 | 48833721 | 48833276 | 48833519 | 48834319 | 48834897 | 173215 | 7,3,5,3,0,2,0,1,0,0,0,2 | 76,50,85,53,35,49,40,60,56 | 4,4,6,5,1,0,3,8,0,0,7,0,10,1 | 68,67,30,20,27,21,31,42,26 | 264 | 149 | 0.00489883295543 | 0.345985808855127 | 0.049,0.033,0.032, | 0.032,0.033,0.101, | -0,024 |
| 156863 | ENSG00000134884 | ARGLU1 | chr13 | - | 106557558 | 106557614 | 106557047 | 106557131 | 106559431 | 106559657 | 156863 | 31,3,31,27,8,7,7,9,8,4,5,5 | 130,49,133,112,66,62,60,6 | 9,5,15,6,5,6,0,13,9,8,10,15 | 104,54,77,34,46,41,18,79,3 | 205 | 149 | 0.00489968798821 | 0.345985808855127 | 0.148,0.043,0.145, | 0.059,0.063,0.124, | 0,024 |
| 149692 | ENSG00000214013 | GANC | chr15 | + | 42327855 | 42328119 | 42327362 | 42327442 | 42329305 | 42329504 | 149692 | 2,5,25,18,17,7,13,19,9,3,5, | 24,29,27,39,35,31,20,18,15 | 29,36,13,22,23,16,17,12,24 | 33,21,16,9,20,19,20,14,15, | 298 | 149 | 0.00491091795904 | 0.34666506501874 | 0.04,0.079,0.316,0 | 0.305,0.462,0.289, | -0,075 |
| 225140 | ENSG00000068885 | IFT80 | chr3 | - | 160366042 | 160366181 | 160357484 | 160357578 | 160375811 | 160375880 | 225140 | 26,8,52,43,20,27,13,18,8,9 | 4,2,7,0,0,2,1,1,1,0,1,0 | 24,14,15,15,15,20,16,29,17 | 5,3,5,2,3,1,0,0,1,2,1,4,0,4,4 | 288 | 149 | 0.00492316584786 | 0.347415707737953 | 0.771,0.674,0.794, | 0.713,0.707,0.608, | 0,046 |
| 90855 | ENSG00000205268 | PDE7A | chr8 | - | 65780843 | 65780947 | 65779719 | 65779803 | 65782782 | 65782843 | 90855 | 8,6,7,1,0,3,7,6,1,3,1,13 | 57,36,33,15,81,58,45,67,30 | 13,18,9,13,6,1,7,6,9,5,3,5,3 | 72,27,23,27,32,23,35,43,32 | 253 | 149 | 0.00492690029562 | 0.347565282702913 | 0.076,0.089,0.111, | 0.096,0.282,0.187, | -0,041 |
| 181568 | ENSG00000125703 | ATG4C | chr1 | + | 62861984 | 62862064 | 62841427 | 62841547 | 62863991 | 62865516 | 181568 | 1,0,6,2,3,3,0,2,3,0,0,1 | 12,9,60,21,12,18,7,12,5,4,5 | 0,0,0,2,0,0,1,0,0,1,4,1,0,0,0 | 15,13,10,6,8,9,8,17,14,11,4 | 229 | 149 | 0.00493014917664 | 0.347680516853671 | 0.051,0.0,0.061,0.0 | 0.0,0.0,0.0,0.178,0 | 0,016 |
| 69060 | ENSG00000249915 | PDCD6 | chr5 | + | 306601 | 306754 | 304176 | 304221 | 314416 | 314969 | 69060 | 62,32,80,77,36,42,25,23,75 | 3,1,4,5,0,3,0,7,8,2,5,6 | 49,26,30,5,15,17,15,35,22, | 16,13,7,1,2,1,0,2,4,1,8,7,8, | 298 | 149 | 0.0049421604151 | 0.348274855640819 | 0.912,0.941,0.909, | 0.605,0.5,0.682,0.7 | 0,084 |
| 10142 | ENSG00000270647 | TAF15 | chr17 | + | 35824077 | 35824198 | 35822639 | 35822833 | 35834565 | 35834598 | 10142 | 20,12,27,24,13,24,14,11,11 | 2,0,7,4,2,4,0,2,0,0,0,2 | 23,39,16,7,7,23,13,14,13,1 | 8,11,1,10,6,4,2,1,4,0,3,4,2, | 270 | 149 | 0.00494298605516 | 0.348274855640819 | 0.847,1.0,0.68,0.76 | 0.613,0.662,0.898, | 0,174 |
| 23524 | ENSG00000203546 | | chr14 | - | 31345086 | 31345276 | 31343895 | 31344065 | 31347747 | 31347907 | 23524 | 17,1,29,20,7,7,15,14,4,1,1, | 3,2,1,1,2,4,0,2,0,0,0,0 | 17,24,5,5,9,9,9,16,3,16,4,1 | 4,0,1,2,0,2,0,5,0,1,0,1,1,0,1 | 298 | 149 | 0.00494343140153 | 0.348274855640819 | 0.739,0.2,0.935,0.9 | 0.68,1.0,0.714,0.5 | -0,015 |
| 168225 | ENSG00000138639 | ARHGAP24 | chr4 | + | 85647372 | 85647510 | 85604152 | 85604303 | 85721884 | 85722386 | 168225 | 0,0,2,0,0,0,11,8,0,0,0,0 | 3,3,70,41,8,36,37,108,47,2 | 0,1,0,2,7,2,0,1,35,17,2,0,0, | 85,24,38,48,50,70,38,54,59 | 287 | 149 | 0.00494935228031 | 0.348509947860393 | 0.0,0.0,0.015,0.0,0 | 0.0,0.021,0.0,0.02 | -0,014 |
| 170473 | ENSG00000111880 | RNGTT | chr6 | - | 88635705 | 88635796 | 88614271 | 88614395 | 88678352 | 88678419 | 170473 | 3,0,3,1,0,2,1,2,3,0,0,6 | 10,5,27,16,7,7,15,11,6,5,4, | 1,1,3,2,5,0,0,2,0,5,0,0,5,1,3 | 15,9,9,5,10,4,15,23,3,5,11, | 240 | 149 | 0.00495000678624 | 0.348509947860393 | 0.157,0.0,0.065,0.0 | 0.04,0.065,0.171,0 | -0,015 |
| 113724 | ENSG00000075856 | SART3 | chr12 | - | 108532221 | 108532334 | 108531203 | 108531280 | 108536513 | 108536572 | 113724 | 23,24,59,50,14,12,20,24,16 | 1,1,4,1,0,3,0,5,1,2,0,0 | 23,48,30,20,35,21,8,21,17, | 0,0,0,0,0,1,0,0,0,0,2,2,0,1,1 | 262 | 149 | 0.00495248477515 | 0.348570389352619 | 0.929,0.932,0.893, | 1.0,1.0,1.0,1.0,1.0, | -0,066 |
| 146566 | ENSG00000149932 | TMEM219 | chr16 | + | 29972045 | 29972140 | 29971407 | 29971578 | 29972949 | 29973048 | 146566 | 10,3,4,4,1,1,2,4,8,2,0,2 | 48,21,74,59,36,34,16,40,23 | 8,2,2,6,1,0,0,1,1,1,1,1,4,1,4 | 64,45,21,29,21,15,19,62,20 | 244 | 149 | 0.00496035080217 | 0.349009893266378 | 0.113,0.08,0.032,0 | 0.071,0.026,0.055, | 0,024 |
| 197397 | ENSG00000163534 | FCRL1 | chr1 | - | 157803844 | 157804010 | 157801914 | 157802664 | 157820006 | 157820120 | 197397 | 296,121,339,36,109,59,172 | 0,1,14,0,2,2,4,5,1,0,0,2 | 199,140,216,102,44,33,106 | 7,4,4,3,2,3,0,0,0,4,6,6,11,2 | 298 | 149 | 0.00496643892083 | 0.349324057310184 | 1.0,0.984,0.924,1.0 | 0.934,0.946,0.964, | 0,029 |
| 119881 | ENSG00000109971 | HSPA8 | chr11 | - | 123060592 | 123060675 | 123060115 | 123060268 | 123061119 | 123061329 | 119881 | 172,109,397,440,130,123,1 | 1,4,35,20,6,4,6,3,6,2,2,1 | 223,186,179,63,150,105,47 | 20,12,3,19,5,4,8,1,4,5,26,1 | 232 | 149 | 0.00498676805089 | 0.350507200680206 | 0.991,0.946,0.879, | 0.877,0.909,0.975, | 0,048 |
| 80299 | ENSG00000135632 | SMYD5 | chr2 | + | 73224865 | 73224960 | 73223946 | 73224003 | 73225630 | 73225701 | 80299 | 24,20,68,40,29,36,16,27,29 | 3,0,3,2,1,4,1,4,2,0,0,3 | 39,47,36,21,8,13,15,25,5,1 | 5,7,4,6,0,1,2,4,4,1,3,6,0,2,5 | 244 | 149 | 0.00498747735064 | 0.350507200680206 | 0.83,1.0,0.933,0.92 | 0.826,0.804,0.846, | 0,102 |
| 142039 | ENSG00000129197 | RPAIN | chr17 | + | 5425975 | 5426082 | 5422768 | 5422829 | 5426235 | 5426299 | 142039 | 12,2,36,23,5,3,2,16,6,0,2,5 | 0,0,2,9,6,3,3,2,0,2,0,0 | 6,8,9,1,10,6,3,10,2,7,9,14,1 | 1,0,0,0,0,0,0,1,0,0,0,9,3,3,2 | 256 | 149 | 0.00498814555377 | 0.350507200680206 | 1.0,1.0,0.913,0.598 | 0.777,1.0,1.0,1.0,1 | -0,205 |
| 46355 | ENSG00000127481 | UBR4 | chr1 | - | 19146022 | 19146106 | 19145792 | 19145933 | 19146825 | 19147000 | 46355 | 0,2,5,6,5,1,0,1,1,2,2,3 | 18,26,58,50,25,32,18,24,13 | 3,8,4,6,10,1,5,3,3,2,9,5,3,1 | 29,35,21,25,24,18,21,34,36 | 233 | 149 | 0.00499301085872 | 0.350676819802193 | 0.0,0.047,0.052,0.0 | 0.062,0.128,0.109, | -0,038 |
| 213430 | ENSG00000035720 | STAP1 | chr4 | + | 67576837 | 67577259 | 67575384 | 67575498 | 67581304 | 67581471 | 213430 | 95,41,117,81,38,50,40,47,9 | 3,0,4,1,1,0,3,4,1,1,2,2 | 65,46,75,37,51,30,20,41,20 | 2,3,3,11,3,0,2,2,2,0,0,0,3,4 | 298 | 149 | 0.00499381804988 | 0.350676819802193 | 0.941,1.0,0.936,0.9 | 0.942,0.885,0.926, | 0,013 |
| 175862 | ENSG00000069849 | ATP1B3 | chr3 | + | 141913651 | 141913735 | 141907166 | 141907274 | 141915969 | 141916020 | 175862 | 11,6,41,58,17,12,2,14,14,3 | 0,2,2,3,0,1,4,4,0,1,0,1 | 45,27,12,6,6,15,4,21,9,8,17 | 1,2,0,0,0,0,0,0,0,1,0,0,0,0 | 233 | 149 | 0.00499715051506 | 0.350796380465388 | 1.0,0.657,0.929,0.9 | 0.966,0.896,1.0,1.0 | -0,168 |
| 226213 | ENSG00000136193 | SCRN1 | chr7 | - | 29940786 | 29940876 | 29936555 | 29936721 | 29943976 | 29944179 | 226213 | 22,1,61,59,13,8,17,32,4,7,3 | 0,0,6,5,0,4,4,1,2,0,0,0 | 27,32,16,11,4,10,10,15,11, | 0,1,2,1,2,0,4,3,0,0,2,2,0,1,2 | 239 | 149 | 0.00500647597857 | 0.351249430873235 | 1.0,1.0,0.864,0.88, | 1.0,0.952,0.833,0.8 | -0,005 |
| 8895 | ENSG00000173821 | RNF213 | chr17 | + | 80278763 | 80278910 | 80273240 | 80273404 | 80287814 | 80288363 | 8895 | 21,17,20,18,25,12,30,23,15 | 64,48,62,34,53,59,48,54,58 | 14,15,13,26,18,13,6,25,20, | 39,38,29,63,23,31,34,41,33 | 296 | 149 | 0.00500686822028 | 0.351249430873235 | 0.142,0.151,0.14,0 | 0.153,0.166,0.184, | -0,053 |
| 26398 | ENSG00000115760 | BIRC6 | chr2 | + | 32499546 | 32500109 | 32493539 | 32493667 | 32501712 | 32501888 | 26398 | 45,38,135,90,110,118,52,1 | 4,6,4,5,0,7,2,8,0,0,2,3 | 163,140,65,59,66,41,97,10 | 8,26,7,8,10,4,5,8,4,3,6,4,1, | 298 | 149 | 0.00500998342127 | 0.351353451203442 | 0.849,0.76,0.944,0 | 0.911,0.729,0.823, | 0,047 |
| 203580 | ENSG00000275052 | PPP4R3B | chr2 | - | 55591516 | 55591695 | 55588878 | 55588956 | 55598415 | 55599039 | 203580 | 4,5,10,9,2,8,2,8,8,1,3,4 | 106,86,85,71,52,39,36,64,8 | 3,17,6,12,6,5,3,7,6,6,4,7,3, | 91,103,35,82,25,43,39,65,2 | 298 | 149 | 0.00501818446091 | 0.351813960816326 | 0.019,0.028,0.056, | 0.016,0.076,0.079, | -0,03 |
| 136551 | ENSG00000028277 | POU2F2 | chr19 | - | 42122686 | 42122810 | 42122510 | 42122576 | 42132383 | 42132478 | 136551 | 12,5,5,13,2,6,6,5,6,3,1,1 | 41,32,30,20,31,24,14,14,20 | 0,0,0,1,0,0,0,7,0,0,5,2,2,0,8 | 10,3,20,9,3,3,7,7,11,11,3,1 | 273 | 149 | 0.0050371194014 | 0.353026455839376 | 0.138,0.079,0.083, | 0.0,0.0,0.0,0.057,0 | 0,034 |
| 101225 | ENSG00000126653 | NSRP1 | chr17 | + | 30163066 | 30163162 | 30118079 | 30118173 | 30172541 | 30172598 | 101225 | 5,0,1,1,4,5,2,0,0,4,1,2 | 8,6,17,10,13,8,7,3,6,3,2,5 | 4,1,1,3,3,0,5,2,0,2,0,0,0,3,0 | 6,17,1,6,7,5,7,9,4,4,11,4,2, | 245 | 149 | 0.00504104963594 | 0.353186899151368 | 0.275,0.0,0.035,0.0 | 0.288,0.035,0.378, | 0,025 |
| 205304 | ENSG00000176407 | KCMF1 | chr2 | + | 85035636 | 85035799 | 85035015 | 85035155 | 85043563 | 85043665 | 205304 | 4,0,10,13,2,0,4,3,1,1,2,2 | 60,55,158,144,72,46,35,51 | 18,17,4,11,1,0,2,5,2,2,11,5 | 116,170,55,61,76,33,20,49 | 298 | 149 | 0.00504532363751 | 0.353371315270067 | 0.032,0.0,0.031,0.0 | 0.072,0.048,0.035, | -0,011 |
| 6803 | ENSG00000101247 | NDUFAF5 | chr20 | + | 13801817 | 13801873 | 13801542 | 13801683 | 13808841 | 13808902 | 6803 | 1,0,8,5,1,1,0,5,0,0,2,2 | 10,7,40,33,10,17,14,15,12, | 2,13,2,2,7,5,2,11,6,0,5,6,3, | 10,50,18,18,30,6,8,11,16,1 | 205 | 149 | 0.00505338540976 | 0.353820818193902 | 0.068,0.0,0.127,0.0 | 0.127,0.159,0.075, | -0,09 |
| 36530 | ENSG00000145495 | MARCHF6 | chr5 | + | 10417269 | 10417410 | 10415487 | 10415669 | 10423734 | 10423835 | 36530 | 148,98,144,96,80,119,82,7 | 0,0,1,2,0,1,0,0,0,0,0,1 | 129,141,74,152,68,43,103, | 2,5,2,4,3,0,5,2,1,0,2,1,0,1,0 | 290 | 149 | 0.00505597626976 | 0.353887098704623 | 1.0,1.0,0.987,0.96 | 0.971,0.935,0.95,0 | 0,028 |
| 171074 | ENSG00000164463 | CREBRF | chr5 | + | 173080584 | 173080798 | 173056351 | 173056479 | 173086500 | 173086626 | 171074 | 78,68,161,95,41,39,28,49,1 | 3,0,5,3,0,2,1,0,0,0,1,1 | 87,92,79,77,74,36,38,64,35 | 4,0,0,5,7,3,0,2,2,3,7,0,3,3,0 | 298 | 149 | 0.00506244242027 | 0.354224494329367 | 0.929,1.0,0.942,0.9 | 0.916,1.0,1.0,0.885 | 0,026 |
| 177995 | ENSG00000142687 | KIAA0319L | chr1 | - | 35435304 | 35435417 | 35434397 | 35435081 | 35441046 | 35443028 | 177995 | 2,0,6,2,1,3,3,5,0,0,1,3 | 9,8,43,30,14,16,19,29,13,5 | 6,1,5,4,4,3,7,1,1,5,5,3,0,5,4 | 29,18,18,5,6,6,1,31,9,20,17 | 262 | 149 | 0.00506670052066 | 0.354407221242175 | 0.112,0.0,0.074,0.0 | 0.105,0.031,0.136, | -0,106 |
| 59967 | ENSG00000118308 | IRAG2 | chr12 | + | 25061591 | 25061653 | 25052297 | 25052956 | 25062819 | 25062900 | 59967 | 299,102,752,589,256,158,2 | 241,99,472,328,182,141,16 | 299,374,234,82,98,87,82,1 | 221,280,165,85,110,96,48, | 211 | 149 | 0.00507486563492 | 0.354754047938802 | 0.467,0.421,0.529, | 0.489,0.485,0.5,0.4 | 0,045 |
| 99147 | ENSG00000067596 | DHX8 | chr17 | + | 43520750 | 43520879 | 43517166 | 43520267 | 43521368 | 43521565 | 99147 | 34,50,75,58,25,38,17,31,25 | 3,0,2,0,3,1,1,2,0,0,0,2 | 41,79,25,31,54,17,21,49,14 | 0,2,4,1,2,1,2,1,2,2,3,9,2,0,3 | 278 | 149 | 0.00507495534381 | 0.354754047938802 | 0.859,1.0,0.953,1.0 | 1.0,0.955,0.77,0.94 | 0,042 |
| 36724 | ENSG00000144043 | TEX261 | chr2 | - | 70991829 | 70991983 | 70989576 | 70989816 | 70993695 | 70993775 | 36724 | 20,13,60,38,39,22,47,42,22 | 1,0,6,0,2,0,1,6,1,0,1,1 | 73,53,35,26,17,23,26,45,23 | 5,0,0,2,3,0,1,3,2,3,8,1,2,0,2 | 298 | 149 | 0.00507840659014 | 0.354880041818968 | 0.909,1.0,0.833,1.0 | 0.88,1.0,1.0,0.867, | 0,036 |
| 137155 | ENSG00000170889 | RPS9 | chr19 | + | 54206611 | 54206724 | 54206275 | 54206462 | 54207397 | 54207522 | 137155 | 22,20,54,39,12,10,17,19,17 | 970,517,1906,1616,681,79 | 13,26,34,19,31,9,8,11,7,29 | 749,634,603,319,436,414,5 | 262 | 149 | 0.00508431140273 | 0.355177353950334 | 0.013,0.022,0.016, | 0.01,0.023,0.031,0 | -0,005 |
| 38420 | ENSG00000113649 | TCERG1 | chr5 | + | 146481149 | 146481200 | 146480027 | 146480094 | 146482591 | 146482727 | 38420 | 0,4,7,0,1,2,2,1,1,2,0,2 | 24,13,27,18,18,11,12,12,10 | 2,3,2,1,0,3,0,1,3,2,2,3,4,2,1 | 16,24,17,12,16,3,4,16,8,7,2 | 200 | 149 | 0.00508868508326 | 0.355367546773242 | 0.0,0.186,0.162,0.0 | 0.085,0.085,0.081, | -0,005 |
| 150730 | ENSG00000114062 | UBE3A | chr15 | - | 25356690 | 25356742 | 25355891 | 25356056 | 25360382 | 25360527 | 150730 | 5,5,59,24,8,6,9,14,7,2,5,5 | 0,2,2,3,0,1,0,3,0,0,0,0 | 12,18,8,4,6,12,5,6,2,8,16,9 | 5,1,1,1,3,0,0,4,1,1,0,0,0,2,0 | 201 | 149 | 0.00509262868413 | 0.355483349816839 | 1.0,0.65,0.956,0.85 | 0.64,0.93,0.856,0.7 | 0,105 |
| 220477 | ENSG00000068028 | RASSF1 | chr3 | - | 50337172 | 50337448 | 50332049 | 50332154 | 50337904 | 50338023 | 220477 | 7,11,14,11,4,11,3,3,7,1,1,4 | 14,3,10,5,13,3,7,8,8,5,5,10 | 15,29,5,4,8,1,11,12,3,8,11, | 8,6,5,2,4,2,3,6,1,5,2,4,17,8 | 298 | 149 | 0.00509364659754 | 0.355483349816839 | 0.2,0.647,0.412,0.5 | 0.484,0.707,0.333, | -0,171 |
| 37535 | ENSG00000138814 | PPP3CA | chr4 | - | 101171083 | 101171226 | 101108953 | 101109078 | 101195915 | 101195928 | 37535 | 2,1,1,0,3,0,4,3,0,0,0,2 | 114,64,68,62,42,29,31,50,6 | 1,5,3,4,4,0,2,2,1,3,3,4,3,3,0 | 47,31,48,47,23,18,18,40,19 | 292 | 149 | 0.00510927174814 | 0.356344496948403 | 0.009,0.008,0.007, | 0.011,0.076,0.031, | -0,017 |
| 4797 | ENSG00000071626 | DAZAP1 | chr19 | + | 1433738 | 1433827 | 1432513 | 1432690 | 1434739 | 1435687 | 4797 | 2,2,3,0,2,5,2,1,2,4,0,0 | 14,5,5,3,2,3,8,4,12,8,3,6 | 0,2,1,0,2,0,1,3,1,0,2,3,2,1,0 | 8,2,6,14,5,3,5,4,4,3,3,19,6, | 238 | 149 | 0.00510929706958 | 0.356344496948403 | 0.082,0.2,0.273,0.0 | 0.0,0.385,0.094,0.0 | 0,048 |
| 74121 | ENSG00000186432 | KPNA4 | chr3 | - | 160509799 | 160509871 | 160508106 | 160508269 | 160514076 | 160514181 | 74121 | 150,155,146,134,80,77,56, | 9,2,7,4,0,9,4,3,2,2,4,0 | 101,185,79,54,78,56,69,90 | 7,5,5,14,10,1,0,6,3,4,4,2,7, | 221 | 149 | 0.00512038156894 | 0.357001893574514 | 0.918,0.981,0.934, | 0.907,0.961,0.914, | 0,037 |
| 80528 | ENSG00000152359 | POC5 | chr5 | - | 75712853 | 75712951 | 75707736 | 75707875 | 75717305 | 75717344 | 80528 | 4,7,19,17,12,1,4,6,4,7,5,6 | 0,0,3,1,0,0,0,0,0,0,0,0 | 15,14,6,0,6,4,3,9,7,8,11,10 | 0,4,0,1,3,0,0,0,3,0,2,0,0,0,0 | 247 | 149 | 0.0051223521728 | 0.357023633582875 | 1.0,1.0,0.793,0.91 | 1.0,0.679,1.0,0.0,0 | 0,182 |
| 220038 | ENSG00000223501 | VPS52 | chr6 | - | 33267187 | 33267307 | 33266556 | 33266712 | 33267864 | 33267997 | 220038 | 22,15,35,33,15,18,19,18,14 | 0,0,2,0,1,0,0,0,1,3,0,0 | 21,16,11,9,22,15,8,24,22,3 | 4,5,2,1,2,0,1,2,0,0,2,1,0,0,0 | 269 | 149 | 0.00513527252732 | 0.357808300850635 | 1.0,1.0,0.906,1.0,0 | 0.744,0.639,0.753, | 0,046 |
| 148665 | ENSG00000069345 | DNAJA2 | chr16 | - | 46960217 | 46960380 | 46959274 | 46959419 | 46964610 | 46964807 | 148665 | 18,2,19,7,1,16,9,18,2,3,1,3 | 52,45,126,91,47,32,33,44,2 | 23,6,20,31,11,6,5,10,8,14,2 | 72,87,33,24,75,27,38,52,23 | 298 | 149 | 0.00514714881368 | 0.35851973667222 | 0.148,0.022,0.07,0 | 0.138,0.033,0.233, | -0,045 |
| 248723 | ENSG00000197860 | SGTB | chr5 | - | 65705328 | 65705445 | 65704278 | 65704378 | 65708488 | 65708558 | 248723 | 1,0,4,3,0,6,0,4,0,0,3,0 | 24,19,61,19,28,30,27,30,16 | 4,5,2,5,3,6,4,0,1,3,2,4,2,1,0 | 18,44,9,17,46,7,18,11,12,1 | 266 | 149 | 0.00515567703603 | 0.35899758141112 | 0.023,0.0,0.035,0.0 | 0.111,0.06,0.111,0 | -0,052 |
| 40845 | ENSG00000004534 | RBM6 | chr3 | + | 49999439 | 49999513 | 49962575 | 49962685 | 50048244 | 50048319 | 40845 | 20,22,27,15,9,11,15,14,23, | 2,2,11,13,10,3,3,5,5,9,0,4 | 18,10,18,14,4,3,16,15,13,1 | 4,1,5,1,2,2,3,0,1,8,2,3,7,3,2 | 223 | 149 | 0.00515809238722 | 0.359049606272234 | 0.87,0.88,0.621,0.4 | 0.75,0.87,0.706,0.9 | -0,14 |
| 155576 | ENSG00000173575 | CHD2 | chr15 | + | 92995752 | 92995926 | 92991475 | 92991517 | 92996956 | 92997095 | 155576 | 12,10,14,6,3,9,2,8,3,12,0,6 | 0,2,2,1,1,1,6,1,1,0,1,0 | 5,13,6,3,12,4,3,4,2,7,6,2,7, | 0,0,0,0,1,0,1,1,0,2,0,0,0,1,1 | 298 | 149 | 0.00516559867042 | 0.359340853851074 | 1.0,0.714,0.778,0.7 | 1.0,1.0,1.0,1.0,0.85 | -0,156 |
| 161881 | ENSG00000198055 | GRK6 | chr5 | + | 177431917 | 177432107 | 177430871 | 177430967 | 177432232 | 177432257 | 161881 | 49,26,44,38,24,24,34,15,51 | 16,9,7,6,4,12,7,2,1,1,2,0 | 20,33,21,21,30,20,19,26,11 | 15,20,7,9,13,4,7,9,7,2,11,1 | 298 | 149 | 0.00516561555638 | 0.359340853851074 | 0.605,0.591,0.759, | 0.4,0.452,0.6,0.538 | 0,144 |
| 92545 | ENSG00000083097 | DOP1A | chr6 | + | 83096730 | 83096823 | 83067665 | 83067779 | 83096924 | 83097115 | 92545 | 9,15,24,6,1,7,10,12,0,0,9,7 | 13,6,11,9,3,4,0,7,4,4,0,3 | 18,31,6,9,17,11,4,9,10,13,2 | 6,8,2,2,2,2,2,3,4,6,2,1,1,3,4 | 242 | 149 | 0.00517346122754 | 0.359681768149792 | 0.299,0.606,0.573, | 0.649,0.705,0.649, | -0,214 |
| 7283 | ENSG00000154930 | ACSS1 | chr20 | - | 25012811 | 25012939 | 25012600 | 25012664 | 25013535 | 25013662 | 7283 | 39,19,58,49,35,20,33,23,24 | 2,3,12,4,5,3,0,8,5,3,0,4 | 50,54,14,15,16,24,6,49,22, | 0,3,0,4,1,1,7,2,3,0,1,3,2,2,3 | 277 | 149 | 0.00517385857145 | 0.359681768149792 | 0.913,0.773,0.722, | 1.0,0.906,1.0,0.66 | -0,055 |
| 29288 | ENSG00000006459 | KDM7A | chr7 | - | 140124620 | 140124783 | 140120441 | 140120529 | 140127441 | 140127583 | 29288 | 33,10,40,12,12,7,16,6,15,1 | 40,12,23,19,10,16,12,12,9, | 28,11,4,7,9,8,4,15,13,22,5, | 30,51,30,34,31,15,4,23,21, | 298 | 149 | 0.00517800684752 | 0.359853920503254 | 0.292,0.294,0.465, | 0.318,0.097,0.062, | 0,09 |
| 28742 | ENSG00000150593 | PDCD4 | chr10 | + | 110894411 | 110894522 | 110894090 | 110894198 | 110895947 | 110896087 | 28742 | 323,294,1076,689,315,266 | 8,4,23,16,3,6,5,3,0,0,0,1 | 473,606,235,247,295,230,8 | 8,17,6,2,21,3,2,1,16,11,5,0 | 260 | 149 | 0.00518209568759 | 0.359910360309099 | 0.959,0.977,0.964, | 0.971,0.953,0.957, | 0,022 |
| 86095 | ENSG00000140612 | SEC11A | chr15 | - | 84680712 | 84680832 | 84670724 | 84670782 | 84687624 | 84687774 | 86095 | 194,183,462,411,141,125,1 | 11,2,12,11,2,8,1,7,7,1,8,3 | 179,296,120,65,182,159,67 | 9,12,2,12,12,4,5,13,4,7,10, | 269 | 149 | 0.00518216338073 | 0.359910360309099 | 0.907,0.981,0.955, | 0.917,0.932,0.971, | 0,025 |
| 253109 | ENSG00000124164 | VAPB | chr20 | + | 58439110 | 58439308 | 58438944 | 58439025 | 58440906 | 58441083 | 253109 | 4,1,11,3,2,0,3,3,4,0,1,1 | 22,21,32,22,9,8,9,14,32,12 | 10,6,2,2,1,1,0,2,2,3,2,4,6,1 | 20,23,10,9,11,13,8,8,7,18,1 | 298 | 149 | 0.00518556373248 | 0.360030344421098 | 0.083,0.023,0.147, | 0.2,0.115,0.091,0.1 | -0,061 |
| 108302 | ENSG00000134294 | SLC38A2 | chr12 | - | 46363020 | 46363145 | 46362493 | 46362638 | 46363725 | 46363826 | 108302 | 29,32,261,186,81,93,49,15 | 10,0,19,15,7,9,7,20,1,0,1,4 | 105,197,60,60,40,36,41,70 | 25,13,9,15,0,4,0,5,7,14,35, | 274 | 149 | 0.00520468648472 | 0.360921934054326 | 0.612,1.0,0.882,0.8 | 0.695,0.892,0.784, | 0,064 |
| 64814 | ENSG00000092841 | MYL6 | chr12 | + | 56159622 | 56159730 | 56158683 | 56158711 | 56159974 | 56160148 | 64814 | 485,416,1269,1037,520,54 | 38,26,57,25,6,14,8,13,16,1 | 582,542,405,229,422,313,1 | 11,37,15,54,46,7,0,10,20,8 | 257 | 149 | 0.00520472167585 | 0.360921934054326 | 0.881,0.903,0.928, | 0.968,0.895,0.94,0 | 0,029 |
| 61772 | ENSG00000132704 | FCRL2 | chr1 | - | 157769865 | 157770150 | 157768413 | 157768701 | 157770408 | 157770666 | 61772 | 169,104,131,18,51,26,67,3 | 20,6,11,0,11,1,4,0,2,0,4,7 | 61,51,118,18,19,25,36,2,35 | 6,2,6,0,0,1,0,0,1,1,2,7,2,2,9 | 298 | 149 | 0.00520475036797 | 0.360921934054326 | 0.809,0.897,0.856, | 0.836,0.927,0.908, | -0,05 |
| 37873 | ENSG00000086061 | DNAJA1 | chr9 | + | 33026812 | 33026990 | 33026474 | 33026616 | 33029884 | 33029989 | 37873 | 129,192,192,139,62,58,29, | 13,4,11,24,3,4,5,8,4,2,4,2 | 77,99,90,47,68,57,44,90,22 | 22,8,10,9,17,2,0,2,8,20,2,2 | 298 | 149 | 0.00520511303346 | 0.360921934054326 | 0.832,0.96,0.897,0 | 0.636,0.861,0.818, | 0,057 |
| 57163 | ENSG00000071564 | TCF3 | chr19 | - | 1615284 | 1615515 | 1612206 | 1612433 | 1615685 | 1615821 | 57163 | 42,10,52,18,23,13,19,22,35 | 45,34,66,71,24,53,32,30,60 | 19,15,8,7,14,6,20,16,2,8,5, | 35,38,20,21,10,9,25,85,11, | 298 | 149 | 0.00521135389003 | 0.36116671619688 | 0.318,0.128,0.283, | 0.213,0.165,0.167, | 0,046 |
| 153644 | ENSG00000197555 | SIPA1L1 | chr14 | + | 71730179 | 71730301 | 71724669 | 71724835 | 71733665 | 71733812 | 153644 | 51,32,78,33,19,17,46,36,63 | 5,0,2,0,1,0,5,5,0,5,0,0 | 46,33,22,29,48,23,20,18,8, | 0,0,0,0,2,1,0,6,0,0,2,0,0,0,3 | 271 | 149 | 0.00521199929614 | 0.36116671619688 | 0.849,1.0,0.955,1.0 | 1.0,1.0,1.0,1.0,0.93 | -0,017 |
| 193800 | ENSG00000058673 | ZC3H11A | chr1 | + | 203802568 | 203803016 | 203795705 | 203795794 | 203816925 | 203817124 | 193800 | 84,48,182,88,87,84,80,157 | 10,8,33,16,9,16,15,16,4,8,8 | 110,139,79,66,66,49,88,12 | 28,38,27,26,29,9,15,18,6,7 | 298 | 149 | 0.00522143553672 | 0.361685871088488 | 0.808,0.75,0.734,0 | 0.663,0.647,0.594, | 0,076 |
| 58535 | ENSG00000118246 | FASTKD2 | chr2 | + | 206786732 | 206786899 | 206774224 | 206774397 | 206787936 | 206788155 | 58535 | 26,28,83,63,33,53,29,53,30 | 0,1,4,4,0,2,2,0,0,0,0,0 | 51,83,45,28,30,15,28,43,23 | 3,0,1,1,8,0,2,2,7,2,0,1,0,1,0 | 298 | 149 | 0.00522285213256 | 0.361685871088488 | 1.0,0.933,0.912,0.8 | 0.895,1.0,0.957,0.9 | 0,053 |
| 234368 | ENSG00000010671 | BTK | chrX | - | 101354629 | 101354690 | 101353869 | 101353988 | 101356051 | 101356349 | 234368 | 18,13,107,88,32,24,28,20,4 | 0,0,5,3,0,0,3,0,0,0,0,0 | 52,34,10,2,25,35,10,19,9,4 | 1,4,0,0,1,2,1,6,0,0,1,4,6,1,1 | 210 | 149 | 0.00523022103163 | 0.362079672839742 | 1.0,1.0,0.938,0.95 | 0.974,0.858,1.0,1.0 | 0,067 |
| 237326 | ENSG00000120910 | PPP3CC | chr8 | + | 22539731 | 22539936 | 22539468 | 22539498 | 22540614 | 22541142 | 237326 | 35,17,53,54,22,14,11,8,24, | 91,45,100,64,53,29,69,74,9 | 22,26,9,11,10,31,6,24,7,25 | 64,54,74,56,43,28,28,67,30 | 298 | 149 | 0.00523828233014 | 0.362521139613621 | 0.161,0.159,0.209, | 0.147,0.194,0.057, | 0,043 |
| 241687 | ENSG00000175893 | ZDHHC21 | chr9 | - | 14677297 | 14677461 | 14674186 | 14674385 | 14680032 | 14680162 | 241687 | 19,31,68,57,4,13,17,28,28, | 12,10,23,19,5,5,8,9,2,2,1,7 | 14,8,10,3,13,14,6,38,5,18,3 | 11,21,6,0,9,5,2,13,16,19,15 | 298 | 149 | 0.00524120416572 | 0.362606754674407 | 0.442,0.608,0.596, | 0.389,0.16,0.455,1 | 0,118 |
| 35969 | ENSG00000205937 | RNPS1 | chr16 | - | 2267174 | 2267218 | 2264572 | 2264760 | 2268054 | 2268094 | 35969 | 6,4,1,1,1,3,2,0,1,3,0,0 | 50,62,33,25,14,11,7,19,21, | 0,1,2,0,3,0,0,0,2,0,1,1,2,0,0 | 20,22,15,21,23,9,12,10,9,2 | 193 | 149 | 0.00524438121899 | 0.362647959217961 | 0.085,0.047,0.023, | 0.0,0.034,0.093,0.0 | 0,02 |
| 61665 | ENSG00000142252 | GEMIN7 | chr19 | + | 45079906 | 45080029 | 45079259 | 45079377 | 45090106 | 45090537 | 61665 | 8,5,25,18,26,12,14,16,2,3,8 | 1,3,1,4,2,1,0,2,1,0,0,3 | 3,5,9,6,21,14,8,8,15,20,8,2 | 3,2,0,1,1,0,2,0,1,0,0,0,0,1,4 | 272 | 149 | 0.00524673274657 | 0.362647959217961 | 0.814,0.477,0.932, | 0.354,0.578,1.0,0.7 | -0,052 |
| 254466 | ENSG00000166598 | HSP90B1 | chr12 | + | 103932276 | 103932731 | 103931520 | 103931623 | 103932825 | 103932942 | 254466 | 29,45,344,162,55,55,56,95 | 2,2,4,3,0,2,0,2,0,0,0,2 | 115,148,58,32,77,41,13,72 | 7,4,1,3,2,3,0,2,0,2,2,5,1,4,0 | 298 | 149 | 0.00524796687291 | 0.362647959217961 | 0.879,0.918,0.977, | 0.891,0.949,0.967, | 0,023 |
| 91138 | ENSG00000090975 | PITPNM2 | chr12 | - | 122994800 | 122994979 | 122992498 | 122992669 | 122995388 | 122995660 | 91138 | 155,114,157,57,64,61,97,6 | 2,0,2,2,2,2,4,0,4,3,1,0 | 69,103,78,143,73,46,68,62 | 0,0,1,0,2,0,0,2,0,0,2,0,1,0,0 | 298 | 149 | 0.0052485394435 | 0.362647959217961 | 0.975,1.0,0.975,0.9 | 1.0,1.0,0.975,1.0,0 | -0,029 |

| | | | | | | | | | | | | | | | | | | | | | | |
|---|---|---|---|---|---|---|---|---|---|---|---|---|---|---|---|---|---|---|---|---|---|---|
| 178057 | ENSG00000067066 | SP100 | chr2 | + | 230450149 | 230450255 | 230449568 | 230449710 | 230461261 | 230461328 | 178057 | 104,69,270,192,72,105,93, | 9,4,5,8,1,1,5,8,4,0,0,2 | 135,138,141,93,184,68,54, | 2,6,10,1,4,1,2,2,0,2,6,2,0,0 | 255 | 149 | 0.00525388840941 | 0.362901045008255 | 0.871,0.91,0.969,0 | 0.975,0.931,0.892, | -0,025 |
| 163260 | ENSG00000113391 | FAM172A | chr5 | - | 94057078 | 94057207 | 94053125 | 94053227 | 94074645 | 94074771 | 163260 | 7,4,6,1,4,3,8,11,3,3,1,4 | 42,26,40,10,28,32,19,15,22 | 4,2,3,1,2,2,2,3,1,3,1,2,3,3, | 27,25,18,17,33,7,13,14,18, | 278 | 149 | 0.00525560797017 | 0.362903355478877 | 0.082,0.076,0.074, | 0.074,0.041,0.082, | 0,033 |
| 101880 | ENSG00000122515 | ZMIZ2 | chr7 | + | 44760150 | 44760228 | 44759280 | 44759460 | 44760424 | 44760593 | 101880 | 22,10,8,3,9,9,6,10,4,8,3,4 | 8,0,9,1,3,0,0,2,0,0,0,3 | 4,7,2,12,4,9,3,7,4,5,0,9,1,5 | 2,2,3,14,1,2,0,3,5,1,1,8,0,2 | 227 | 149 | 0.00526524832257 | 0.363452425181226 | 0.644,1.0,0.368,0. | 0.568,0.697,0.304, | 0,219 |
| 145463 | ENSG00000074370 | ATP2A3 | chr17 | - | 3928239 | 3928479 | 3925046 | 3925441 | 3928662 | 3928780 | 145463 | 2,11,9,15,1,7,3,2,13,7,4,5 | 80,60,111,70,80,62,49,34, | 7,7,5,8,1,2,4,5,5,2,3,3,4,3, | 101,125,32,24,62,56,16,71 | 298 | 149 | 0.00526998334292 | 0.363662643437004 | 0.012,0.084,0.039, | 0.033,0.027,0.072, | 0,007 |
| 161055 | ENSG00000140853 | NLRC5 | chr16 | + | 57062350 | 57062484 | 57061617 | 57061701 | 57065211 | 57065298 | 161055 | 3,0,3,1,0,1,2,2,3,0,0,2 | 17,28,11,12,7,6,13,14,2,2,1 | 4,3,2,9,1,1,5,2,2,9,3,9,3,1, | 25,27,8,6,15,10,5,42,24,28 | 283 | 149 | 0.00527310758375 | 0.363761608448108 | 0.085,0.0,0.126,0. | 0.078,0.055,0.116, | -0,021 |
| 151191 | ENSG00000174485 | DENND4A | chr15 | - | 65725887 | 65725956 | 65722847 | 65722948 | 65729071 | 65729247 | 151191 | 1,0,5,4,0,7,1,2,0,1,0,2 | 71,47,234,139,69,78,59,92 | 2,9,3,3,12,1,1,3,7,4,1,3,4,0 | 114,141,55,105,104,67,52, | 218 | 149 | 0.00527674673684 | 0.363807223566658 | 0.01,0.0,0.014,0.0 | 0.012,0.042,0.036, | -0,01 |
| 195304 | ENSG00000134744 | TUT4 | chr1 | - | 52471951 | 52472124 | 52468180 | 52468267 | 52474831 | 52475535 | 195304 | 74,60,114,48,21,30,37,36, | 0,0,9,2,2,2,1,3,0,1,0,0 | 64,51,20,31,28,21,25,46,3 | 3,0,0,4,0,0,0,0,0,0,0,0,0,0, | 298 | 149 | 0.00527714944397 | 0.363807223566658 | 1.0,1.0,0.864,0.92 | 0.914,1.0,1.0,0.79 | -0,056 |
| 255359 | ENSG00000069493 | CLEC2D | chr12 | + | 9693850 | 9694091 | 9693033 | 9693115 | 9694759 | 9699553 | 255359 | 3,0,10,1,0,2,3,6,4,0,2,2 | 48,24,69,43,13,16,12,41,2 | 2,2,2,0,0,3,2,4,1,1,1,0,6,2, | 44,26,27,24,21,8,5,24,27,1 | 298 | 149 | 0.00527983895839 | 0.363876086728542 | 0.03,0.0,0.068,0.0 | 0.022,0.037,0.036, | 0,005 |
| 202427 | ENSG00000154222 | CC2D1B | chr1 | - | 52356869 | 52357126 | 52356383 | 52356442 | 52357525 | 52357698 | 202427 | 14,9,42,19,1,3,14,14,13,7, | 7,1,7,1,3,2,2,2,2,5,0,7 | 10,23,15,7,11,15,8,15,6,17 | 0,1,0,6,3,0,2,5,3,0,2,0,0,0, | 298 | 149 | 0.00528215322826 | 0.363909663428236 | 0.5,0.818,0.75,0.9 | 1.0,0.92,1.0,0.368, | -0,162 |
| 148807 | ENSG00000005339 | CREBBP | chr16 | - | 3745439 | 3745591 | 3745276 | 3745354 | 3749626 | 3749683 | 148807 | 4,0,5,3,0,0,4,4,0,2,1,4 | 26,20,40,21,10,19,17,9,9,2 | 4,3,6,22,2,3,7,6,6,3,0,7,8,6 | 20,54,12,22,15,13,32,28,2 | 298 | 149 | 0.00528458467969 | 0.363909663428236 | 0.071,0.0,0.059,0. | 0.091,0.027,0.2,0. | -0,021 |
| 215446 | ENSG00000198176 | TFDP1 | chr13 | + | 113637817 | 113637884 | 113636533 | 113636700 | 113640119 | 113641473 | 215446 | 10,3,18,21,1,9,4,5,4,1,6,18 | 3,8,14,11,6,1,3,1,7,7,3,12 | 4,6,3,1,9,2,10,2,3,5,3,62,1 | 1,2,0,0,3,2,3,2,2,4,1,26,3,4 | 216 | 149 | 0.00528539851544 | 0.363909663428236 | 0.697,0.206,0.47,0 | 0.734,0.674,1.0,1. | -0,201 |
| 161486 | ENSG00000113282 | CLINT1 | chr5 | - | 157789358 | 157789513 | 157787601 | 157787992 | 157791702 | 157791724 | 161486 | 77,63,57,52,35,25,29,143, | 0,2,0,0,0,0,0,0,0,2,0,1 | 60,85,75,104,93,44,28,52, | 1,3,0,13,0,0,0,0,2,0,6,0,0,0 | 298 | 149 | 0.00529206230185 | 0.364251954361841 | 1.0,0.94,1.0,1.0,1. | 0.968,0.934,1.0,0. | 0,017 |
| 98466 | ENSG00000198198 | SZT2 | chr1 | + | 43419733 | 43419944 | 43416534 | 43416641 | 43420152 | 43420323 | 98466 | 70,29,86,43,29,38,52,45,2 | 2,1,5,1,1,5,1,14,0,2,2,2 | 49,58,37,56,21,20,50,34,4 | 1,3,1,5,8,0,0,1,2,0,1,3,0,1, | 298 | 149 | 0.00529500969232 | 0.364338309171268 | 0.946,0.935,0.896, | 0.961,0.906,0.949, | -0,031 |
| 64657 | ENSG00000058673 | ZC3H11A | chr1 | + | 203801574 | 203803016 | 203795622 | 203795794 | 203816925 | 203817124 | 64657 | 88,55,223,99,110,115,84,1 | 10,8,33,16,9,16,15,16,4,8, | 149,162,92,67,73,58,93,14 | 28,38,27,26,29,9,15,18,6,7 | 298 | 149 | 0.00530644509331 | 0.365008464007096 | 0.815,0.775,0.772, | 0.727,0.681,0.63,0 | 0,069 |
| 98596 | ENSG00000006634 | DBF4 | chr7 | + | 87886843 | 87886894 | 87884978 | 87885158 | 87887328 | 87887398 | 98596 | 36,28,20,25,3,6,7,8,12,9,4, | 3,0,3,3,0,0,6,1,2,0,1,1 | 15,20,21,13,12,11,6,7,9,13 | 5,5,1,7,6,2,1,5,3,2,5,5,1,0, | 200 | 149 | 0.00531643776641 | 0.365578983035844 | 0.899,1.0,0.832,0. | 0.691,0.749,0.94,0 | 0,104 |
| 173712 | ENSG00000136560 | TANK | chr2 | + | 161177111 | 161177187 | 161136965 | 161137063 | 161179613 | 161180085 | 173712 | 5,1,4,2,2,3,0,4,1,0,0,1 | 191,301,266,174,128,154, | 6,3,12,3,0,2,0,9,2,10,2,1,2, | 116,191,136,84,120,41,34, | 225 | 149 | 0.00531903410286 | 0.365640699135312 | 0.017,0.002,0.01,0 | 0.033,0.01,0.055,0 | -0,014 |
| 161531 | ENSG00000131446 | MGAT1 | chr5 | - | 180795671 | 180795875 | 180793061 | 180793097 | 180803601 | 180803904 | 161531 | 4,0,8,9,3,2,0,1,5,0,1,5 | 2,1,12,17,20,3,6,9,22,12,1, | 6,7,0,3,2,3,7,1,7,4,6,7,4,7, | 5,25,2,4,1,3,7,11,6,6,9,14,1 | 298 | 149 | 0.00532474199243 | 0.365901895839935 | 0.5,0.0,0.25,0.209, | 0.375,0.123,0.0,0. | -0,093 |
| 232580 | ENSG00000134516 | DOCK2 | chr5 | + | 169983898 | 169984382 | 169983107 | 169983166 | 169985827 | 169985922 | 232580 | 3,2,2,2,2,3,4,4,0,0,1,0 | 94,91,156,118,70,90,73,79 | 6,16,1,5,5,1,0,5,4,5,7,3,4,0 | 104,110,65,44,63,57,45,69 | 298 | 149 | 0.00532751704036 | 0.365901895839935 | 0.016,0.011,0.006, | 0.028,0.068,0.008, | -0,011 |
| 167632 | ENSG00000205339 | IPO7 | chr11 | + | 9416912 | 9416978 | 9414254 | 9414411 | 9417058 | 9417148 | 167632 | 0,3,5,5,2,5,2,15,2,4,0,0 | 25,12,77,75,45,43,35,34,2 | 1,5,10,4,2,0,0,4,5,10,10,7,1 | 63,53,18,23,15,18,22,34,2 | 215 | 149 | 0.00532793390154 | 0.365901895839935 | 0.0,0.148,0.043,0. | 0.011,0.061,0.278, | -0,039 |
| 110062 | ENSG00000177565 | TBL1XR1 | chr3 | - | 177131362 | 177131449 | 177098465 | 177098541 | 177197120 | 177197163 | 110062 | 0,6,5,0,2,1,1,2,4,0,0,0 | 10,7,28,10,8,10,11,23,2,3, | 0,0,2,4,1,0,0,1,0,0,0,1,1,0, | 26,14,8,13,6,3,7,12,1,13,1 | 236 | 149 | 0.0053354095854 | 0.366298418014427 | 0.0,0.351,0.101,0. | 0.0,0.0,0.136,0.16 | 0,076 |
| 210493 | ENSG00000140386 | SCAPER | chr15 | - | 76795279 | 76795438 | 76774854 | 76775117 | 76800247 | 76800364 | 210493 | 67,19,66,38,22,43,38,43,3 | 2,2,6,3,0,5,0,1,0,1,0,3 | 44,35,23,32,20,16,11,24,1 | 3,2,0,4,4,2,0,7,2,2,6,7,0,4,1 | 298 | 149 | 0.00534750833035 | 0.367011978778559 | 0.944,0.826,0.846, | 0.88,0.897,1.0,0.8, | 0,086 |
| 48203 | ENSG00000142687 | KIAA0319L | chr1 | - | 35506611 | 35507135 | 35478965 | 35479212 | 35554349 | 35554510 | 48203 | 64,39,100,87,62,68,54,65, | 4,0,6,2,4,1,0,1,3,0,0,1 | 51,73,43,61,30,44,44,46,3 | 3,0,1,0,0,0,0,0,1,2,0,4,4,0, | 298 | 149 | 0.00535203992602 | 0.367205899049159 | 0.889,1.0,0.893,0. | 0.895,1.0,0.956,1. | -0,018 |
| 149694 | ENSG00000214013 | GANC | chr15 | + | 42327922 | 42328119 | 42327362 | 42327442 | 42329305 | 42329504 | 149694 | 2,5,25,19,17,7,13,19,9,3,5, | 24,29,27,39,35,31,20,18,1 | 25,37,13,22,23,17,16,12,2 | 33,21,16,9,20,19,20,14,15, | 298 | 149 | 0.00535854909319 | 0.367535334568635 | 0.04,0.079,0.316,0 | 0.275,0.468,0.289, | -0,074 |
| 161883 | ENSG00000198055 | GRK6 | chr5 | + | 177433315 | 177433410 | 177433146 | 177433239 | 177433535 | 177433676 | 161883 | 59,32,72,121,22,30,26,26, | 9,4,5,1,0,2,0,4,1,0,0,2 | 43,42,16,18,30,24,15,18,3 | 0,5,12,16,2,0,3,4,7,0,4,3,1, | 244 | 149 | 0.00536400889954 | 0.367792608938163 | 0.8,0.83,0.898,0.9 | 1.0,0.837,0.449,0. | 0,096 |
| 142627 | ENSG00000100425 | BRD1 | chr22 | - | 49820833 | 49820986 | 49804203 | 49804360 | 49822950 | 49823205 | 142627 | 2,0,16,10,0,3,4,8,1,1,1,2 | 21,20,34,12,15,7,17,12,11, | 2,3,4,4,4,7,1,4,2,4,2,0,0,1, | 14,25,7,18,23,11,12,21,8,2 | 298 | 149 | 0.00537547373728 | 0.368353766681131 | 0.045,0.0,0.19,0.2 | 0.067,0.057,0.222, | 0,037 |
| 201772 | ENSG00000162664 | ZNF326 | chr1 | + | 90007633 | 90007750 | 90005132 | 90005244 | 90010087 | 90010286 | 201772 | 55,24,20,20,11,20,18,27,1 | 8,2,1,2,5,1,1,0,0,1,0,0 | 30,30,45,64,12,15,24,25,2 | 0,3,2,3,0,1,4,0,0,0,2,0,1,0, | 266 | 149 | 0.0053756158785 | 0.368353766681131 | 0.794,0.87,0.918,0 | 1.0,0.849,0.926,0. | -0,059 |
| 132503 | ENSG00000171792 | RHNO1 | chr12 | + | 2885282 | 2885534 | 2877222 | 2877282 | 2887910 | 2887934 | 132503 | 108,82,63,47,45,42,29,38, | 3,2,2,7,0,3,3,2,1,0,0,2 | 14,34,48,35,32,15,30,49,1 | 7,1,1,0,3,2,1,4,2,0,1,3,4,4, | 298 | 149 | 0.00537952034985 | 0.368503992494769 | 0.947,0.953,0.94,0 | 0.5,0.944,0.96,1.0, | 0,062 |
| 106574 | ENSG00000147140 | NONO | chrX | + | 71284983 | 71285097 | 71284396 | 71284453 | 71290628 | 71290791 | 106574 | 13,3,13,8,0,3,0,4,0,4,1,4 | 114,74,124,77,99,115,93,1 | 11,12,11,4,1,1,3,9,5,4,7,11 | 95,83,81,61,52,28,64,108, | 263 | 149 | 0.00538192786236 | 0.368551611754249 | 0.061,0.022,0.056, | 0.062,0.076,0.071, | -0,016 |
| 72908 | ENSG00000117625 | RCOR3 | chr1 | + | 211312719 | 211312835 | 211304082 | 211304140 | 211313423 | 211313672 | 72908 | 4,2,2,8,4,2,2,1,4,0,2,1 | 44,12,17,19,22,31,24,34,7 | 8,4,2,1,0,2,0,3,2,14,0,5,4,1 | 24,19,16,13,18,7,11,17,2,1 | 265 | 149 | 0.00538382387 | 0.368564184276072 | 0.049,0.086,0.062, | 0.158,0.106,0.066, | -0,051 |
| 95905 | ENSG00000290554 | ADCY10P1 | chr6 | + | 41124484 | 41124820 | 41122976 | 41123249 | 41132293 | 41132513 | 95905 | 6,6,7,2,6,5,5,9,1,2,1,2 | 1,0,1,0,4,7,0,0,0,0,2,0 | 12,8,3,24,2,2,10,6,1,6,12,2 | 0,2,1,7,2,1,2,3,1,0,1,0,2,0, | 298 | 149 | 0.00538664230095 | 0.368616238328973 | 0.75,1.0,0.778,1.0, | 1.0,0.667,0.6,0.63 | 0,099 |
| 97021 | ENSG00000173200 | PARP15 | chr3 | + | 122613040 | 122613268 | 122610493 | 122610730 | 122617085 | 122617164 | 97021 | 36,26,99,88,60,31,36,47,2 | 4,0,2,8,3,0,1,3,0,2,0,0 | 57,101,19,34,40,31,16,49, | 0,0,3,1,1,0,0,0,4,0,0,2,2,2,1 | 298 | 149 | 0.0053889294686 | 0.368616238328973 | 0.818,1.0,0.961,0. | 1.0,1.0,0.76,0.944, | -0,027 |
| 196237 | ENSG00000166377 | ATP9B | chr18 | + | 79077981 | 79078153 | 79069284 | 79069529 | 79096475 | 79096649 | 196237 | 0,4,2,1,0,0,4,4,0,1,0,0 | 17,1,9,8,10,11,9,9,4,4,2,13 | 3,5,1,4,4,0,0,1,1,0,4,4,3,4, | 5,5,12,22,6,9,4,7,11,3,7,15 | 298 | 149 | 0.00539019254359 | 0.368616238328973 | 0.0,0.667,0.1,0.05 | 0.231,0.333,0.04,0 | -0,008 |
| 68210 | ENSG00000147099 | HDAC8 | chrX | - | 72494195 | 72494381 | 72490928 | 72491006 | 72495155 | 72495268 | 68210 | 5,0,5,4,0,2,0,3,1,2,1,4 | 9,5,12,13,7,16,8,6,2,6,2,14 | 3,0,1,4,1,0,2,4,1,2,0,1,0,0, | 5,6,5,3,4,6,3,9,4,10,6,7,6,9 | 298 | 149 | 0.00539143486886 | 0.368616238328973 | 0.217,0.0,0.172,0. | 0.231,0.0,0.091,0. | 0,025 |
| 141981 | ENSG00000108559 | NUP88 | chr17 | - | 5404098 | 5404113 | 5399551 | 5399650 | 5405056 | 5405243 | 141981 | 2,2,7,4,0,2,0,1,0,0,1,0 | 7,3,49,35,9,21,9,25,5,10,2, | 8,2,3,0,2,2,0,1,0,2,1,2,3,3, | 40,23,16,1,14,9,15,23,8,10 | 164 | 149 | 0.00539378623634 | 0.368659893754746 | 0.206,0.377,0.115, | 0.154,0.073,0.146, | 0,01 |
| 27030 | ENSG00000126777 | KTN1 | chr14 | + | 55663954 | 55664041 | 55661273 | 55661612 | 55667240 | 55667330 | 27030 | 36,21,57,24,21,16,14,30,1 | 50,36,58,36,25,30,28,47,6 | 37,29,37,13,19,15,12,26,2 | 36,37,21,12,22,16,24,41,2 | 236 | 149 | 0.00540452046016 | 0.369218882341199 | 0.313,0.269,0.383, | 0.394,0.331,0.527, | -0,088 |
| 32860 | ENSG00000102445 | RUBCNL | chr13 | - | 46377889 | 46378005 | 46368732 | 46368815 | 46387133 | 46387230 | 32860 | 5,9,42,23,20,31,37,25,28,1 | 3,2,10,2,2,6,5,6,0,2,0,1 | 23,30,11,12,14,3,13,18,14, | 11,12,2,2,4,2,1,3,2,1,8,4,9, | 265 | 149 | 0.00540539559012 | 0.369218882341199 | 0.484,0.717,0.703, | 0.54,0.584,0.756,0 | 0,125 |
| 69035 | ENSG00000153922 | CHD1 | chr5 | - | 98904896 | 98905098 | 98903791 | 98903908 | 98926333 | 98926534 | 69035 | 151,144,138,56,60,81,66,8 | 3,2,0,3,0,3,3,1,0,0,2,2 | 111,166,95,211,120,54,61, | 6,3,4,0,4,3,2,4,2,1,0,13,1,0 | 298 | 149 | 0.00541302527043 | 0.369622729054543 | 0.962,0.973,1.0,0. | 0.902,0.965,0.922, | 0,023 |
| 151858 | ENSG00000125107 | CNOT1 | chr16 | - | 58564810 | 58564908 | 58560211 | 58560362 | 58574608 | 58574760 | 151858 | 0,0,7,0,0,2,2,8,0,1,1,0 | 53,22,166,105,75,69,72,66 | 4,3,2,4,1,0,6,2,3,2,10,2,7,1 | 116,75,47,64,33,56,61,60, | 247 | 149 | 0.00542503344161 | 0.370288000603561 | 0.0,0.0,0.025,0.0,0 | 0.02,0.024,0.025,0 | -0,014 |
| 13091 | ENSG00000182628 | SKA2 | chr17 | - | 59119318 | 59119495 | 59109856 | 59112345 | 59131280 | 59131367 | 13091 | 18,11,42,27,5,9,10,2,9,2,2, | 6,3,14,9,1,7,1,1,3,2,1,0 | 7,16,5,22,13,6,8,2,4,21,8,2 | 1,2,4,1,1,1,1,0,0,2,0,6,3,1, | 298 | 149 | 0.00542620883564 | 0.370288000603561 | 0.6,0.647,0.6,0.6,0 | 0.778,0.8,0.385,0. | -0,177 |
| 63092 | ENSG00000116698 | SMG7 | chr1 | + | 183542075 | 183542502 | 183540983 | 183541103 | 183544352 | 183544497 | 63092 | 17,10,27,16,10,18,19,24,5, | 2,0,1,2,0,0,0,3,2,2,2,0 | 18,43,9,36,6,22,14,24,13,1 | 1,1,0,6,0,6,0,0,2,5,2,0,0,2, | 298 | 149 | 0.00544031196283 | 0.371013981096909 | 0.81,1.0,0.931,0.8, | 0.9,0.956,1.0,0.75, | 0,004 |
| 195270 | ENSG00000134744 | TUT4 | chr1 | - | 52463676 | 52463777 | 52461711 | 52461769 | 52465069 | 52465173 | 195270 | 7,2,20,1,3,10,2,2,2,0,0,5 | 44,16,80,45,18,22,18,8,3,4 | 3,6,5,9,3,0,10,17,5,3,2,3,9, | 29,30,8,20,26,8,11,14,19,1 | 250 | 149 | 0.00544185141543 | 0.371013981096909 | 0.087,0.069,0.13,0 | 0.058,0.107,0.271, | -0,066 |
| 160162 | ENSG00000104320 | NBN | chr8 | - | 89957964 | 89958022 | 89955282 | 89955555 | 89958724 | 89958854 | 160162 | 12,7,15,20,3,11,2,8,3,2,4,9 | 12,19,40,36,15,13,22,10,3, | 14,36,18,7,11,14,6,22,7,10 | 20,15,12,11,13,12,7,13,5,2 | 207 | 149 | 0.00544375820697 | 0.371013981096909 | 0.419,0.21,0.213,0 | 0.335,0.633,0.519, | -0,103 |
| 172474 | ENSG00000081307 | UBA5 | chr3 | + | 132675247 | 132675514 | 132671776 | 132672177 | 132675604 | 132675680 | 172474 | 27,11,41,22,12,19,5,16,13, | 7,2,14,5,5,0,0,3,1,1,0,2 | 21,10,6,15,9,12,5,5,2,27,3, | 4,10,4,5,6,1,5,3,3,2,3,4,3,2 | 298 | 149 | 0.00544479615223 | 0.371013981096909 | 0.659,0.733,0.594, | 0.724,0.333,0.429, | 0,233 |
| 23003 | ENSG00000129566 | TEP1 | chr14 | - | 20406232 | 20406400 | 20405450 | 20405585 | 20407872 | 20408463 | 23003 | 61,47,67,70,39,51,34,47,6 | 3,0,4,4,5,1,3,1,0,0,0,1 | 89,75,39,30,48,32,20,26,4 | 1,7,1,5,8,0,1,2,1,2,2,5,4,0, | 298 | 149 | 0.00544546634214 | 0.371013981096909 | 0.91,1.0,0.893,0.8 | 0.978,0.843,0.951, | 0,033 |
| 227919 | ENSG00000173715 | C11orf80 | chr11 | + | 66796477 | 66796598 | 66796280 | 66796364 | 66800640 | 66800696 | 227919 | 1,0,6,4,3,1,3,5,1,1,6,5 | 48,19,131,75,57,67,30,62,1 | 12,0,5,5,1,2,3,5,4,10,5,0,1, | 77,61,23,25,28,28,32,65,1 | 270 | 149 | 0.00544948669187 | 0.371170401955023 | 0.011,0.0,0.025,0. | 0.079,0.0,0.107,0. | -0,018 |
| 253111 | ENSG00000124164 | VAPB | chr20 | + | 58439138 | 58439308 | 58438944 | 58439025 | 58440906 | 58441083 | 253111 | 4,1,11,3,2,0,3,3,4,0,1,1 | 22,21,32,22,9,8,9,14,32,12 | 10,6,1,2,1,1,0,2,2,3,2,4,6,1 | 20,23,10,9,11,13,8,8,7,18,1 | 298 | 149 | 0.00545287792502 | 0.371283887592528 | 0.083,0.023,0.147, | 0.2,0.115,0.048,0. | -0,061 |
| 220670 | ENSG00000137185 | ZSCAN9 | chr6 | + | 28230317 | 28230488 | 28227689 | 28227837 | 28232561 | 28233015 | 220670 | 3,0,5,0,5,0,0,0,6,2,0,1 | 11,27,14,11,5,5,9,10,2,6,2, | 1,2,1,1,0,0,1,0,0,1,1,0,0,4, | 4,8,12,3,10,1,2,8,2,20,9,2,1 | 298 | 149 | 0.00546824482847 | 0.372212461314493 | 0.12,0.0,0.152,0.0, | 0.111,0.111,0.04,0 | 0,046 |
| 41770 | ENSG00000211897 | IGHG3 | chr14 | - | 105770480 | 105770525 | 105770104 | 105770149 | 105770668 | 105770719 | 41770 | 28,18,66,74,11,26,45,38,2 | 0,0,12,7,0,0,15,10,4,9,0,0 | 64,75,25,4,10,7,20,54,3,2,1 | 15,12,4,22,0,0,0,0,0,0,0,0, | 194 | 149 | 0.00547289557946 | 0.372411251489964 | 1.0,1.0,0.809,0.89, | 0.766,0.828,0.828, | -0,015 |
| 141289 | ENSG00000101745 | ANKRD12 | chr18 | + | 9254210 | 9254297 | 9221851 | 9221999 | 9263789 | 9263888 | 141289 | 11,10,36,23,8,15,10,13,3,1 | 0,5,3,0,0,4,0,1,1,0,1,4 | 9,37,9,16,25,14,4,17,13,11 | 0,0,6,1,1,0,0,2,4,0,2,0,1,4, | 236 | 149 | 0.00547523105201 | 0.372452419265223 | 1.0,0.558,0.883,1. | 1.0,1.0,0.486,0.91, | -0,069 |
| 31016 | ENSG00000081791 | DELE1 | chr5 | + | 141937197 | 141937357 | 141934493 | 141934586 | 141939505 | 141940146 | 31016 | 17,4,30,25,11,9,11,14,9,3, | 2,2,0,0,2,1,0,1,1,0,0,0 | 27,5,12,1,10,6,5,7,7,12,17, | 0,4,0,2,0,1,3,2,0,0,4,0,0,4, | 298 | 149 | 0.0054801857637 | 0.37267167839081 | 0.81,0.5,1.0,1.0,0. | 1.0,0.385,1.0,0.2,1 | 0,094 |
| 179377 | ENSG00000116406 | EDEM3 | chr1 | - | 184723760 | 184723856 | 184721288 | 184721386 | 184726254 | 184726389 | 179377 | 43,22,31,23,6,10,11,6,29,1 | 2,1,4,1,1,7,0,0,0,0,2,1 | 21,17,13,6,5,10,4,7,4,23,8, | 2,2,3,5,3,0,0,4,5,4,2,3,0,0, | 245 | 149 | 0.00548551787248 | 0.372916455215333 | 0.929,0.93,0.825,0 | 0.865,0.838,0.725, | 0,052 |
| 46963 | ENSG00000130023 | ERMARD | chr6 | + | 169766637 | 169766667 | 169762428 | 169762531 | 169768102 | 169768171 | 46963 | 63,21,78,42,20,21,16,36,1 | 4,1,13,7,5,17,6,8,2,3,6,6 | 41,38,19,31,31,15,10,24,1 | 2,5,3,8,6,3,0,3,3,2,4,4,4,1,1 | 179 | 149 | 0.00548955704619 | 0.372993312443528 | 0.929,0.946,0.833, | 0.945,0.864,0.841, | -0,147 |
| 178757 | ENSG00000070190 | DAPP1 | chr4 | + | 99865502 | 99865836 | 99863769 | 99863855 | 99866033 | 99866121 | 178757 | 31,25,30,4,8,2,5,11,4,4,4,0 | 33,29,44,28,20,13,14,19,2 | 18,15,8,7,15,8,3,6,5,22,6,2 | 18,26,10,9,16,12,2,2,5,15, | 298 | 149 | 0.00549011440648 | 0.372993312443528 | 0.32,0.301,0.254,0 | 0.333,0.224,0.286, | -0,129 |
| 8236 | ENSG00000134444 | RELCH | chr18 | + | 62264728 | 62264852 | 62263988 | 62264145 | 62266700 | 62266749 | 8236 | 43,22,111,57,45,38,33,51, | 4,3,9,4,3,5,6,0,5,0,2,2 | 79,82,31,24,43,31,34,46,2 | 2,4,1,1,0,4,1,0,0,1,4,1,5,7,1 | 273 | 149 | 0.00550177855165 | 0.373667813016782 | 0.854,0.8,0.871,0. | 0.956,0.918,0.944, | -0,056 |
| 23614 | ENSG00000103811 | CTSH | chr15 | - | 78931868 | 78932041 | 78931450 | 78931506 | 78932371 | 78932458 | 23614 | 13,11,12,1,23,55,0,11,0,2,1 | 327,176,243,162,304,494,1 | 10,8,5,0,3,8,1,11,10,24,2,5 | 150,135,194,64,120,94,56, | 298 | 149 | 0.00551056565774 | 0.374068175837684 | 0.019,0.03,0.024,0 | 0.032,0.029,0.013, | 0,006 |
| 209576 | ENSG00000162191 | UBXN1 | chr11 | - | 62676902 | 62677005 | 62676612 | 62676639 | 62677517 | 62677563 | 209576 | 53,29,103,75,41,27,35,49, | 6,1,0,0,1,1,1,7,3,0,0,2 | 62,63,17,23,38,23,28,57,3 | 0,8,2,4,3,0,2,2,0,3,10,4,0,0 | 252 | 149 | 0.00551435142812 | 0.374068175837684 | 0.839,0.945,1.0,1. | 1.0,0.823,0.834,0. | 0,039 |
| 129002 | ENSG00000133316 | WDR74 | chr11 | - | 62833617 | 62833674 | 62832910 | 62833131 | 62833791 | 62833937 | 129002 | 38,45,61,53,30,60,45,43,4 | 8,3,12,4,1,2,1,5,2,2,4,7 | 51,58,19,15,40,13,32,57,8, | 4,4,5,13,0,0,16,9,6,7,15,5,1 | 206 | 149 | 0.00551471130321 | 0.374068175837684 | 0.775,0.916,0.786, | 0.902,0.913,0.733, | 0,096 |
| 186308 | ENSG00000085465 | OVGP1 | chr1 | - | 111421275 | 111421486 | 111419609 | 111419726 | 111421564 | 111421673 | 186308 | 25,14,5,15,14,11,16,5,8,7, | 6,0,5,0,1,2,0,2,1,0,3,0 | 17,14,9,9,15,8,2,3,4,10,1,6 | 5,6,1,3,5,4,0,0,3,1,2,2,1,0, | 298 | 149 | 0.00551725765701 | 0.374068175837684 | 0.676,1.0,0.333,1. | 0.63,0.538,0.818,0 | 0,208 |
| 250718 | ENSG00000119977 | TCTN3 | chr10 | - | 95680471 | 95680564 | 95659822 | 95664300 | 95682650 | 95683629 | 250718 | 10,6,19,21,8,11,3,4,4,5,2,0 | 4,3,4,0,0,3,0,0,0,0,0,2 | 10,3,8,2,4,4,5,8,6,4,7,5,9,5 | 0,1,0,3,2,2,8,2,1,3,1,0,0,1,1 | 242 | 149 | 0.00551940968794 | 0.374068175837684 | 0.606,0.552,0.745, | 1.0,0.649,1.0,0.29 | 0,12 |
| 61864 | ENSG00000001461 | NIPAL3 | chr1 | + | 24457733 | 24457833 | 24456165 | 24456273 | 24458887 | 24458976 | 61864 | 6,0,4,4,3,1,1,0,0,2,0,2 | 11,4,42,23,13,6,12,5,17,7, | 4,8,0,0,2,2,0,1,4,2,2,3,6,14 | 13,13,5,3,6,6,5,5,3,4,7,11,1 | 249 | 149 | 0.00551997854743 | 0.374068175837684 | 0.246,0.0,0.054,0. | 0.155,0.269,0.0,0. | -0,083 |
| 237242 | ENSG00000120910 | PPP3CC | chr8 | + | 22477646 | 22477800 | 22475591 | 22475624 | 22498000 | 22498112 | 237242 | 15,1,13,17,7,4,11,10,7,4,0, | 128,43,231,130,93,73,75,1 | 2,3,4,5,0,3,2,4,3,11,12,9,1 | 163,146,77,70,91,99,42,10 | 298 | 149 | 0.00552154109142 | 0.374068175837684 | 0.055,0.011,0.027, | 0.006,0.01,0.025,0 | 0,013 |
| 170982 | ENSG00000115524 | SF3B1 | chr2 | - | 197409862 | 197410007 | 197408368 | 197408581 | 197416740 | 197416911 | 170982 | 867,597,646,443,363,501, | 24,26,9,5,15,8,9,15,11,11,1 | 459,494,512,377,298,126, | 22,26,17,7,10,3,11,15,9,11 | 294 | 149 | 0.00552256109202 | 0.374068175837684 | 0.948,0.921,0.973, | 0.914,0.906,0.939, | 0,013 |
| 27787 | ENSG00000170144 | HNRNPA3 | chr2 | + | 177217704 | 177217845 | 177215977 | 177216940 | 177219036 | 177219156 | 27787 | 1,2,11,10,11,9,10,7,3,4,2,2 | 3,1,6,4,1,5,3,3,0,0,2,2 | 11,4,2,2,4,3,1,12,4,4,2,9,2, | 9,15,4,0,4,6,0,8,1,4,5,8,1,3 | 290 | 149 | 0.00552331524182 | 0.374068175837684 | 0.146,0.507,0.485, | 0.386,0.121,0.204, | 0,168 |
| 178061 | ENSG00000067066 | SP100 | chr2 | + | 230450171 | 230450251 | 230449374 | 230449710 | 230461261 | 230461414 | 178061 | 234,135,359,267,117,133, | 9,4,5,8,1,1,5,8,4,0,0,2 | 176,206,204,124,229,115, | 2,6,10,1,4,1,2,2,0,2,6,2,0,0 | 229 | 149 | 0.00552613509419 | 0.37414142260384 | 0.944,0.956,0.979, | 0.983,0.957,0.93,0 | -0,013 |
| 141451 | ENSG00000173821 | RNF213 | chr17 | + | 80382259 | 80382454 | 80381546 | 80381727 | 80382978 | 80383070 | 141451 | 6,3,6,2,0,0,0,3,2,2,2,1 | 55,33,29,22,25,14,15,26,2 | 2,7,7,4,4,5,2,4,0,3,4,2,1,2, | 10,32,25,17,17,17,5,13,28, | 298 | 149 | 0.005534856884 | 0.374614082390001 | 0.052,0.043,0.094, | 0.091,0.099,0.123, | -0,051 |
| 224493 | ENSG00000088451 | TGDS | chr13 | - | 94590852 | 94590971 | 94583093 | 94583236 | 94592240 | 94592309 | 224493 | 5,11,8,8,7,3,5,4,5,4,2,7 | 5,0,6,3,0,0,2,2,0,0,1,0 | 6,10,2,2,0,0,11,2,1,3,1,7,1, | 1,0,4,0,3,4,2,0,1,6,2,1,2,0, | 268 | 149 | 0.00554865021049 | 0.375330797969959 | 0.357,1.0,0.426,0. | 0.769,1.0,0.218,1. | 0,24 |
| 252958 | ENSG00000188342 | GTF2F2 | chr13 | + | 45195341 | 45195515 | 45151686 | 45151831 | 45207423 | 45207505 | 252958 | 4,1,6,4,1,3,2,3,0,0,0,1 | 37,19,67,68,17,29,11,22,1 | 0,1,0,1,0,2,0,2,0,0,0,3,2,1, | 42,28,16,10,22,6,13,38,15, | 298 | 149 | 0.00554964250332 | 0.375330797969959 | 0.051,0.026,0.043, | 0.0,0.018,0.0,0.04 | 0,012 |
| 28802 | ENSG00000147050 | KDM6A | chrX | + | 45106541 | 45106649 | 45090816 | 45090864 | 45107409 | 45107536 | 28802 | 1,3,9,1,0,1,0,4,0,0,3,2 | 28,17,66,28,6,18,14,23,9,1 | 3,5,3,11,2,2,0,1,0,7,5,7,4,1 | 37,35,38,27,31,15,10,11,2 | 257 | 149 | 0.00555311767918 | 0.375330797969959 | 0.02,0.093,0.073,0 | 0.045,0.076,0.044, | -0,035 |
| 64112 | ENSG00000196591 | HDAC2 | chr6 | - | 113953276 | 113953418 | 113946007 | 113949260 | 113956012 | 113956151 | 64112 | 44,50,80,58,26,28,13,16,5 | 6,0,3,1,1,2,3,1,0,0,0,2 | 67,37,28,19,25,50,7,30,14, | 2,1,0,3,0,0,1,1,1,1,0,2,0,0, | 291 | 149 | 0.00555353617107 | 0.375330797969959 | 0.79,1.0,0.932,0.9 | 0.945,0.95,1.0,0.7 | -0,034 |
| 29148 | ENSG00000134644 | PUM1 | chr1 | - | 30967098 | 30967310 | 30966146 | 30966278 | 30968353 | 30968492 | 29148 | 34,22,120,57,55,43,29,39, | 4,0,7,0,2,0,1,0,13,6,2,2 | 48,63,24,48,35,20,25,57,1 | 2,1,1,1,1,3,1,2,0,0,2,1,0,1, | 298 | 149 | 0.00555416548515 | 0.375330797969959 | 0.81,1.0,0.896,1.0, | 0.923,0.969,0.923, | -0,027 |
| 59615 | ENSG00000170248 | PDCD6IP | chr3 | + | 33827621 | 33827739 | 33826479 | 33826580 | 33828852 | 33828969 | 59615 | 2,0,3,0,1,1,0,1,0,0,0,2 | 41,32,88,49,44,53,32,34,2 | 2,4,7,1,1,0,1,4,0,6,2,0,4,2, | 55,44,34,21,19,27,22,32,1 | 267 | 149 | 0.00558172509078 | 0.377074787512138 | 0.027,0.0,0.019,0. | 0.02,0.048,0.103,0 | -0,022 |
| 242870 | ENSG00000216937 | CCDC7 | chr10 | + | 32571858 | 32571893 | 32567669 | 32567891 | 32583167 | 32583307 | 242870 | 26,13,44,17,8,10,20,22,5,8 | 0,1,8,2,3,6,3,7,2,0,3,2 | 18,8,15,7,11,4,5,6,6,15,18, | 4,4,12,2,4,2,4,1,1,2,13,6,3, | 184 | 149 | 0.00558410161672 | 0.377116967388849 | 1.0,0.913,0.817,0. | 0.785,0.618,0.503, | 0,09 |
| 249388 | ENSG00000170759 | KIF5B | chr10 | - | 32050309 | 32050523 | 32048463 | 32048551 | 32055847 | 32056425 | 249388 | 6,1,8,2,0,2,2,7,2,0,1,0 | 53,34,134,69,55,51,39,42, | 6,1,5,12,3,2,8,3,3,5,3,11,4, | 70,81,40,50,50,39,36,51,2 | 298 | 149 | 0.00558954995253 | 0.377311379103245 | 0.054,0.014,0.029, | 0.041,0.006,0.059, | -0,017 |
| 79188 | ENSG00000112078 | KCTD20 | chr6 | + | 36465600 | 36465698 | 36443006 | 36443111 | 36470051 | 36470257 | 79188 | 0,0,0,0,2,0,3,1,0,0,0,2 | 5,3,18,13,8,2,5,5,6,5,2,7 | 2,3,2,0,0,2,0,0,0,2,4,1,2,3, | 11,11,4,6,9,2,5,2,7,6,16,25 | 247 | 149 | 0.00559150190765 | 0.377311379103245 | 0.0,0.0,0.0,0.0,0.1 | 0.099,0.141,0.232, | -0,051 |
| 181044 | ENSG00000076685 | NT5C2 | chr10 | - | 103100662 | 103100964 | 103099925 | 103100019 | 103101044 | 103101102 | 181044 | 2,0,5,4,0,1,2,4,0,2,1,2 | 49,53,127,96,26,25,44,34, | 0,1,6,5,8,4,1,1,0,6,7,0,0,4, | 49,55,35,25,48,49,8,42,43, | 298 | 149 | 0.00559395348721 | 0.377311379103245 | 0.02,0.0,0.019,0.0 | 0.0,0.009,0.079,0. | -0,018 |
| 137146 | ENSG00000170889 | RPS9 | chr19 | + | 54206548 | 54206724 | 54206275 | 54206462 | 54207397 | 54207522 | 137146 | 41,30,80,64,27,22,32,32,4 | 970,517,1906,1616,681,79 | 14,35,59,23,51,26,16,28,1 | 749,634,603,319,436,414, | 298 | 149 | 0.00559399255088 | 0.377311379103245 | 0.021,0.028,0.021, | 0.009,0.027,0.047, | -0,006 |
| 12877 | ENSG00000108599 | AKAP10 | chr17 | - | 19931804 | 19931978 | 19924407 | 19924517 | 19936285 | 19936430 | 12877 | 53,19,69,30,25,27,13,31,1 | 4,4,1,6,0,2,1,4,0,0,0,0 | 37,36,22,23,18,9,19,27,16, | 0,1,0,0,1,1,0,2,2,0,2,0,3,1,1 | 298 | 149 | 0.00560327709961 | 0.377813494524708 | 0.869,0.704,0.972, | 1.0,0.947,1.0,1.0,0 | -0,004 |
| 234629 | ENSG00000104613 | INTS10 | chr8 | + | 19824805 | 19824972 | 19823894 | 19824044 | 19826425 | 19826559 | 234629 | 36,9,89,62,37,21,21,27,12, | 2,1,9,3,2,0,1,0,2,1,3,2 | 56,31,18,7,12,15,16,26,9,3 | 4,2,1,0,0,0,0,6,1,1,3,1,6,0, | 298 | 149 | 0.0056066916228 | 0.377813494524708 | 0.9,0.818,0.832,0. | 0.875,0.886,0.9,1. | -0,044 |
| 208157 | ENSG00000123552 | USP45 | chr6 | - | 99469776 | 99469891 | 99468536 | 99468618 | 99476142 | 99476230 | 208157 | 1,2,1,1,0,0,0,4,0,0,0,2 | 30,22,58,34,35,44,13,26,2 | 5,0,3,2,1,1,2,0,2,1,2,2,2,3, | 40,37,24,17,34,14,17,19,1 | 264 | 149 | 0.00560835036796 | 0.377813494524708 | 0.018,0.049,0.01,0 | 0.066,0.0,0.066,0. | -0,026 |
| 188561 | ENSG00000082068 | WDR70 | chr5 | + | 37396460 | 37396570 | 37391999 | 37392120 | 37437921 | 37437981 | 188561 | 22,8,53,30,23,21,13,22,12, | 3,2,1,2,1,1,5,0,0,0,0,1 | 20,15,15,21,19,16,7,18,16, | 6,6,0,1,1,0,0,3,5,3,3,6,2,0, | 259 | 149 | 0.00560845842377 | 0.377813494524708 | 0.808,0.697,0.968, | 0.657,0.59,1.0,0.9 | 0,062 |

| | | | | | | | | | | | | | | | | | | | | | | |
|---|---|---|---|---|---|---|---|---|---|---|---|---|---|---|---|---|---|---|---|---|---|---|
| 103394 | ENSG00000042980 | ADAM28 | chr8 | + | 24313387 | 24313580 | 24310162 | 24310241 | 24320235 | 24320307 | 103394 | 175,74,270,120,117,123,1 | 20,8,6,4,0,2,7,1,3,1,0,1 | 184,157,75,64,54,22,42,10 | 4,1,3,2,2,0,0,3,5,7,0,4,2,2, | 298 | 149 | 0.00561210631313 | 0.377940943016672 | 0.814,0.822,0.957, | 0.958,0.987,0.926, | -0,013 |
| 12422 | ENSG00000141568 | FOXK2 | chr17 | + | 82582740 | 82582934 | 82571723 | 82571870 | 82585903 | 82586200 | 12422 | 6,4,10,8,1,10,1,10,4,1,4,3 | 7,11,1,4,6,2,4,3,2,2,1,3 | 13,2,4,4,1,3,4,7,5,11,8,19, | 1,0,2,1,2,0,0,10,0,1,6,5,2,1 | 298 | 149 | 0.00563410510229 | 0.379181931064173 | 0.3,0.154,0.833,0. | 0.867,1.0,0.5,0.66 | -0,249 |
| 126720 | ENSG00000225484 | NUTM2B-AS1 | chr10 | - | 79808758 | 79808832 | 79806038 | 79806122 | 79814612 | 79814716 | 126720 | 26,35,28,20,26,27,11,16,1 | 2,1,3,6,0,3,1,4,5,0,3,3 | 28,32,26,24,13,11,15,15,1 | 1,2,0,0,4,1,0,1,1,3,2,4,1,2, | 223 | 149 | 0.00563470793628 | 0.379181931064173 | 0.897,0.959,0.862, | 0.949,0.914,1.0,1. | -0,064 |
| 155706 | ENSG00000169062 | UPF3A | chr13 | + | 114284115 | 114284358 | 114282836 | 114282943 | 114286301 | 114286400 | 155706 | 9,0,2,0,3,2,3,3,1,0,0,0 | 6,4,40,17,4,3,7,11,11,2,2,7 | 6,3,4,3,1,0,0,0,0,4,0,1,6,1, | 13,9,11,8,17,5,10,4,10,16, | 298 | 149 | 0.00563581917788 | 0.379181931064173 | 0.429,0.0,0.024,0. | 0.188,0.143,0.154, | 0,044 |
| 84263 | ENSG00000064666 | CNN2 | chr19 | + | 1036066 | 1036246 | 1032558 | 1032696 | 1036415 | 1036562 | 84263 | 204,112,130,135,183,98,1 | 4,6,9,3,11,0,2,0,2,5,2,0 | 157,164,126,69,56,47,56,1 | 6,0,6,7,1,4,9,6,3,3,0,14,15, | 298 | 149 | 0.00565526085714 | 0.380371077982219 | 0.962,0.903,0.878, | 0.929,1.0,0.913,0. | 0,041 |
| 5923 | ENSG00000174365 | SNHG11 | chr20 | + | 38447929 | 38448093 | 38446886 | 38447623 | 38448661 | 38448730 | 5923 | 0,0,9,8,0,0,2,0,4,0,0,0 | 10,2,27,10,2,7,4,8,8,5,6,9 | 0,1,4,0,0,0,0,0,4,0,0,0,1,1, | 4,11,3,7,11,3,7,6,2,1,2,16, | 298 | 149 | 0.00568794381762 | 0.382449808125639 | 0.0,0.0,0.143,0.28 | 0.0,0.043,0.4,0.0,0 | -0,009 |
| 201773 | ENSG00000162664 | ZNF326 | chr1 | + | 90007646 | 90007750 | 90005132 | 90005244 | 90010087 | 90010286 | 201773 | 52,22,18,18,10,15,18,21,8, | 8,2,1,2,5,1,1,0,0,1,0,0 | 28,28,43,52,8,14,20,20,22, | 0,3,2,3,0,1,4,0,0,0,2,0,1,0, | 253 | 149 | 0.0057064713634 | 0.383575745788865 | 0.793,0.866,0.914, | 1.0,0.846,0.927,0. | -0,064 |
| 86708 | ENSG00000170919 | TPT1-AS1 | chr13 | + | 45390713 | 45390902 | 45387584 | 45389820 | 45391031 | 45391676 | 86708 | 27,19,28,19,11,16,12,21,1 | 7,0,0,0,0,0,4,3,0,3,0,1 | 18,21,15,14,17,11,9,12,22, | 0,0,0,0,0,0,1,0,0,0,1,0,0,0, | 298 | 149 | 0.00570938753459 | 0.383637235366082 | 0.659,1.0,1.0,1.0,1 | 1.0,1.0,1.0,1.0,1.0, | -0,092 |
| 196941 | ENSG00000128699 | ORMDL1 | chr2 | - | 189773886 | 189774114 | 189771310 | 189771902 | 189775564 | 189775716 | 196941 | 4,2,9,4,9,7,12,18,6,0,1,5 | 92,61,206,194,101,102,75, | 5,4,0,3,5,4,0,7,3,7,7,3,4,6, | 143,176,75,20,81,63,51,92 | 298 | 149 | 0.00571095103453 | 0.383637235366082 | 0.021,0.016,0.021, | 0.017,0.011,0.0,0. | 0,008 |
| 118111 | ENSG00000216937 | CCDC7 | chr10 | + | 32623718 | 32623817 | 32584231 | 32584304 | 32685969 | 32686080 | 118111 | 5,5,22,15,10,6,13,18,13,1, | 3,1,6,2,1,5,6,1,0,0,2,2 | 7,5,9,6,15,8,1,4,5,8,4,8,6,4 | 3,3,0,1,2,2,0,0,0,2,1,1,0,3, | 248 | 149 | 0.00571374475009 | 0.383689742636389 | 0.5,0.75,0.688,0.8 | 0.584,0.5,1.0,0.78 | -0,092 |
| 218659 | ENSG00000024048 | UBR2 | chr6 | + | 42612172 | 42612291 | 42606588 | 42606651 | 42615070 | 42615178 | 218659 | 54,22,73,36,32,57,39,57,1 | 1,0,1,3,0,1,0,2,2,0,0,0 | 67,67,47,38,43,14,24,55,3 | 1,1,5,2,1,0,0,0,4,0,3,6,0,2, | 268 | 149 | 0.00571529805136 | 0.383689742636389 | 0.968,1.0,0.976,0. | 0.974,0.974,0.839, | 0,024 |
| 126460 | ENSG00000148840 | PPRC1 | chr10 | + | 102146671 | 102147392 | 102144249 | 102144307 | 102148371 | 102148521 | 126460 | 5,20,6,11,10,10,10,3,7,14, | 0,0,0,0,2,0,0,0,0,0,0,1 | 6,3,1,14,8,2,2,11,3,0,13,25 | 0,4,0,0,0,0,0,0,2,0,2,2,2, | 298 | 149 | 0.00572586238516 | 0.384206238791062 | 1.0,1.0,1.0,1.0,0.7 | 1.0,0.273,1.0,1.0,1 | 0,188 |
| 70148 | ENSG00000185344 | ATP6V0A2 | chr12 | + | 123735892 | 123736540 | 123735530 | 123735624 | 123737058 | 123737271 | 70148 | 0,0,13,4,2,2,10,14,4,4,0,3 | 51,38,51,29,40,31,20,19,21 | 2,5,0,1,2,0,2,0,6,8,3,2,0,4, | 19,42,17,20,31,21,10,28,1 | 298 | 149 | 0.00572656175942 | 0.384206238791062 | 0.0,0.0,0.113,0.06 | 0.05,0.056,0.0,0.0 | 0,033 |
| 28256 | ENSG00000112851 | ERBIN | chr5 | + | 66075023 | 66075230 | 66053405 | 66054951 | 66076315 | 66076408 | 28256 | 139,136,296,188,151,122, | 20,11,43,22,14,5,24,17,7,5 | 164,146,146,96,111,86,83, | 41,50,18,17,27,9,8,10,14,5 | 298 | 149 | 0.00573918523878 | 0.384933181093131 | 0.777,0.861,0.775, | 0.667,0.593,0.802, | 0,059 |
| 60507 | ENSG00000138688 | KIAA1109 | chr4 | + | 122192228 | 122192379 | 122189999 | 122190079 | 122196634 | 122196791 | 60507 | 154,119,205,77,81,113,69, | 8,1,4,5,0,0,4,3,1,1,0,3 | 110,144,87,127,74,43,49,6 | 0,0,0,0,8,1,0,2,0,0,9,3,0,0, | 298 | 149 | 0.00575329397059 | 0.385759256879768 | 0.906,0.983,0.962, | 1.0,1.0,1.0,1.0,0.8 | -0,021 |
| 173123 | ENSG00000157601 | MX1 | chr21 | + | 41449218 | 41449295 | 41445999 | 41446141 | 41451166 | 41451243 | 173123 | 25,15,41,57,20,24,15,18,2 | 1,2,9,8,4,1,1,0,4,0,0,1 | 21,26,12,16,12,38,8,29,21, | 2,4,2,5,8,10,0,6,2,3,0,11,1 | 226 | 149 | 0.00575662460364 | 0.385862369998767 | 0.943,0.832,0.75,0 | 0.874,0.811,0.798, | 0,1 |
| 146795 | ENSG00000167377 | ZNF23 | chr16 | - | 71450593 | 71450715 | 71449656 | 71449885 | 71453242 | 71453350 | 146795 | 6,2,2,2,2,2,0,1,2,1,0,0 | 28,18,19,15,20,6,10,8,11,9 | 2,2,5,1,0,0,0,0,1,1,0,2,0,0, | 15,26,11,9,14,8,9,12,4,8,2 | 271 | 149 | 0.00575890420735 | 0.385894991112126 | 0.105,0.058,0.055, | 0.068,0.041,0.2,0. | 0,016 |
| 44976 | ENSG00000140262 | TCF12 | chr15 | + | 57232290 | 57232430 | 57231151 | 57231257 | 57232711 | 57232856 | 44976 | 117,92,149,97,82,77,60,91 | 7,7,8,6,10,4,10,13,2,1,2,3 | 102,84,77,75,60,30,63,94, | 7,2,3,2,5,2,6,3,1,1,6,2,9,2, | 289 | 149 | 0.00576512727673 | 0.386191755025794 | 0.896,0.871,0.906, | 0.883,0.956,0.93,0 | -0,033 |
| 255349 | ENSG00000069493 | CLEC2D | chr12 | + | 9693826 | 9694091 | 9693033 | 9693115 | 9694759 | 9699553 | 255349 | 3,0,10,2,0,5,4,6,4,0,2,2 | 48,24,69,43,13,16,12,41,2 | 2,2,3,0,0,3,2,4,2,2,1,0,6,2, | 44,26,27,24,21,8,5,24,27,1 | 298 | 149 | 0.00577449552885 | 0.386698956804578 | 0.03,0.0,0.068,0.0 | 0.022,0.037,0.053, | 0,01 |
| 70157 | ENSG00000185344 | ATP6V0A2 | chr12 | + | 123756814 | 123756986 | 123752282 | 123752402 | 123757926 | 123758015 | 70157 | 6,9,39,31,26,6,13,12,3,1,1 | 10,1,5,3,0,0,3,0,0,0,0,0 | 16,20,5,7,17,7,6,11,14,23, | 1,0,0,2,1,0,1,1,1,1,0,3,0,2, | 298 | 149 | 0.00578027905565 | 0.386965860474838 | 0.231,0.818,0.796, | 0.889,1.0,1.0,0.63 | -0,018 |
| 3305 | ENSG00000102575 | ACP5 | chr19 | - | 11578584 | 11578764 | 11577592 | 11577679 | 11578843 | 11578961 | 3305 | 4,2,13,8,19,6,13,7,12,6,2,1 | 6,4,6,3,15,0,23,7,16,13,5,5 | 12,8,4,2,17,9,6,12,9,8,11,1 | 1,12,0,0,2,4,3,4,0,3,2,167, | 298 | 149 | 0.00578264323855 | 0.387003758357076 | 0.25,0.2,0.52,0.57 | 0.857,0.25,1.0,1.0, | -0,236 |
| 142171 | ENSG00000091592 | NLRP1 | chr17 | - | 5536850 | 5536939 | 5532821 | 5532984 | 5539414 | 5542027 | 142171 | 1,0,9,4,3,2,5,2,3,2,1,6 | 4,2,0,0,0,0,2,1,0,2,2,0 | 3,1,5,9,2,5,2,8,5,6,8,5,7,7, | 0,2,0,0,1,1,0,0,1,1,2,0,0,0, | 238 | 149 | 0.00578466233736 | 0.387018545083099 | 0.135,0.0,1.0,1.0,1 | 1.0,0.238,1.0,1.0,0 | -0,203 |
| 100885 | ENSG00000140396 | NCOA2 | chr8 | - | 70144641 | 70144848 | 70141183 | 70141399 | 70148272 | 70148483 | 100885 | 99,55,114,62,42,49,61,80, | 14,7,31,25,9,14,10,12,11,5 | 46,45,59,72,32,37,35,74,2 | 8,16,13,29,23,13,13,16,9,2 | 298 | 149 | 0.00578932508806 | 0.387210139225681 | 0.78,0.797,0.648,0 | 0.742,0.584,0.694, | 0,093 |
| 13251 | ENSG00000167978 | SRRM2 | chr16 | + | 2759746 | 2760008 | 2759568 | 2759661 | 2760300 | 2760499 | 13251 | 54,52,32,29,35,61,31,45,1 | 33,49,64,29,29,30,24,26,1 | 22,50,19,47,36,30,31,20,5 | 40,60,20,96,37,25,42,62,3 | 298 | 149 | 0.00579840210696 | 0.387696764176175 | 0.45,0.347,0.2,0.3 | 0.216,0.294,0.322, | 0,09 |
| 151059 | ENSG00000170776 | AKAP13 | chr15 | + | 85715771 | 85715923 | 85710581 | 85710645 | 85717289 | 85717348 | 151059 | 245,174,212,175,120,110, | 2,2,2,5,2,1,3,1,3,1,0,0 | 162,247,199,147,162,124, | 5,3,8,7,2,5,0,3,2,4,5,3,5,0, | 298 | 149 | 0.00580171134028 | 0.387708444827019 | 0.984,0.978,0.981, | 0.942,0.976,0.926, | 0,017 |
| 27688 | ENSG00000167881 | SRP68 | chr17 | - | 76050427 | 76050526 | 76047905 | 76047970 | 76057402 | 76057543 | 27688 | 42,29,44,41,16,12,10,20,2 | 3,2,0,0,0,5,5,0,0,0,2,4 | 32,83,23,42,23,26,22,36,1 | 2,4,0,4,0,2,0,2,2,1,0,1,2,0, | 248 | 149 | 0.00580217952241 | 0.387708444827019 | 0.894,0.897,1.0,1. | 0.906,0.926,1.0,0. | -0,039 |
| 198118 | ENSG00000061273 | HDAC7 | chr12 | - | 47786584 | 47786698 | 47785450 | 47785885 | 47787711 | 47788169 | 198118 | 12,15,57,42,29,35,18,24,3, | 0,0,0,2,1,6,1,3,0,0,0,0 | 31,50,15,7,18,9,12,29,18,3 | 5,4,1,2,2,2,0,5,3,1,1,0,0,2, | 263 | 149 | 0.00580992226442 | 0.388105331748411 | 1.0,1.0,1.0,0.922,0 | 0.778,0.876,0.895, | 0,067 |
| 245913 | ENSG00000166405 | RIC3 | chr11 | - | 8128218 | 8128291 | 8126658 | 8126804 | 8137377 | 8137471 | 245913 | 0,0,2,0,0,1,1,2,0,0,1,1 | 45,19,67,33,46,36,45,29,2 | 1,0,1,3,0,1,1,2,2,3,0,2,1,10 | 54,46,20,33,17,5,20,19,5,2 | 222 | 149 | 0.00582028628412 | 0.388648470262203 | 0.0,0.0,0.02,0.0,0. | 0.012,0.0,0.032,0. | -0,044 |
| 10668 | ENSG00000132388 | UBE2G1 | chr17 | - | 4296716 | 4296814 | 4289229 | 4289408 | 4307020 | 4307123 | 10668 | 92,47,271,186,87,41,65,53 | 4,4,8,3,0,2,0,3,0,0,1,7 | 78,149,98,74,137,92,38,34 | 10,10,2,4,14,6,5,2,2,3,0,3, | 247 | 149 | 0.005821664482 | 0.388648470262203 | 0.933,0.876,0.953, | 0.825,0.9,0.967,0. | 0,026 |
| 86224 | ENSG00000158417 | EIF5B | chr2 | + | 99361147 | 99361820 | 99360464 | 99360549 | 99363644 | 99363862 | 86224 | 60,70,36,43,10,15,14,20,5 | 4,1,0,2,1,2,2,3,0,1,0,0 | 19,46,30,44,24,25,5,15,14, | 3,8,2,1,6,3,0,2,1,0,0,2,0,3, | 298 | 149 | 0.00582479152719 | 0.388736652771669 | 0.882,0.972,1.0,0. | 0.76,0.742,0.882,0 | 0,04 |
| 178554 | ENSG00000120008 | WDR11 | chr10 | + | 120860116 | 120860282 | 120858703 | 120858796 | 120862734 | 120862910 | 178554 | 660,633,365,231,241,162, | 6,7,12,4,2,8,5,13,8,12,2,9 | 231,271,582,547,198,157, | 7,24,11,25,12,0,10,10,4,2, | 298 | 149 | 0.00583165497265 | 0.389074064295856 | 0.982,0.978,0.938, | 0.943,0.85,0.964,0 | 0,015 |
| 37290 | ENSG00000251022 | THAP9-AS1 | chr4 | - | 82897988 | 82898062 | 82893008 | 82894847 | 82900076 | 82900223 | 37290 | 7,15,10,11,5,5,7,7,13,2,2,5 | 1,2,10,0,0,0,0,0,2,2,2,2 | 16,3,10,7,4,1,2,7,3,12,8,2, | 3,2,4,1,3,3,0,1,6,0,0,6,6,8, | 223 | 149 | 0.00583963180164 | 0.389485525968013 | 0.824,0.834,0.401, | 0.781,0.501,0.626, | 0,175 |
| 62953 | ENSG00000225470 | JPX | chrX | + | 73946998 | 73947374 | 73944326 | 73944662 | 74004269 | 74004542 | 62953 | 9,3,29,28,4,13,8,4,4,4,0,9 | 2,0,3,0,0,0,0,1,0,0,1,0 | 9,15,13,7,27,5,7,13,12,8,1 | 1,1,5,1,4,3,0,2,2,0,1,2,0,0, | 298 | 149 | 0.00584260257329 | 0.389562947475768 | 0.692,1.0,0.829,1. | 0.818,0.882,0.565, | 0,026 |
| 226454 | ENSG00000181090 | EHMT1 | chr9 | + | 137782759 | 137782805 | 137782337 | 137782397 | 137790847 | 137790970 | 226454 | 3,6,4,5,2,0,5,1,4,0,0,0 | 48,33,91,67,71,56,66,57,5 | 1,2,0,2,0,0,0,1,0,3,1,2,0,1, | 57,65,63,47,40,16,31,54,5 | 195 | 149 | 0.00584891329727 | 0.389862947626113 | 0.046,0.122,0.032, | 0.013,0.023,0.0,0. | 0,021 |
| 50329 | ENSG00000132781 | MUTYH | chr1 | - | 45333412 | 45333617 | 45333284 | 45333324 | 45334390 | 45334477 | 50329 | 21,11,18,9,14,23,10,16,19, | 1,2,3,0,0,2,2,0,1,4,0,3 | 19,14,16,15,8,8,7,10,7,7,6, | 1,1,0,3,2,0,0,0,0,0,0,3,0,1, | 298 | 149 | 0.00585178557453 | 0.389933641173891 | 0.913,0.733,0.75,1 | 0.905,0.875,1.0,0. | -0,069 |
| 151058 | ENSG00000170776 | AKAP13 | chr15 | + | 85715726 | 85715923 | 85710581 | 85710645 | 85717289 | 85717348 | 151058 | 245,174,212,175,120,110, | 2,2,2,5,2,1,3,1,3,1,0,0 | 162,247,199,149,161,124, | 5,3,8,7,2,5,0,3,2,4,5,3,5,0, | 298 | 149 | 0.00585613716156 | 0.38998496669443 | 0.984,0.978,0.981, | 0.942,0.976,0.926, | 0,017 |
| 231476 | ENSG00000078589 | P2RY10 | chrX | + | 78952192 | 78952567 | 78945420 | 78945495 | 78960507 | 78963727 | 231476 | 59,30,106,70,19,26,32,33, | 152,92,179,96,49,38,68,52 | 54,38,51,21,29,24,17,23,1 | 87,78,106,43,60,24,28,65, | 298 | 149 | 0.00585617969696 | 0.38998496669443 | 0.163,0.14,0.228,0 | 0.237,0.196,0.194, | -0,044 |
| 167089 | ENSG00000174652 | ZNF266 | chr19 | - | 9419050 | 9419143 | 9418504 | 9418631 | 9419216 | 9419299 | 167089 | 3,0,16,14,3,3,7,10,8,2,0,0 | 3,8,98,86,25,11,62,61,26,5 | 12,15,7,8,6,6,2,3,3,20,0,0, | 69,29,16,38,18,12,17,50,1 | 242 | 149 | 0.00586450377992 | 0.390337063513649 | 0.381,0.0,0.091,0. | 0.097,0.242,0.212, | -0,033 |
| 144884 | ENSG00000108510 | MED13 | chr17 | - | 61953368 | 61953463 | 61952964 | 61953113 | 61955381 | 61955567 | 144884 | 9,5,12,3,6,5,4,11,3,2,0,6 | 59,38,83,41,30,39,29,49,2 | 9,11,7,0,4,3,1,7,4,0,4,5,6,3 | 81,77,60,51,46,31,25,48,3 | 244 | 149 | 0.00586616143829 | 0.390337063513649 | 0.085,0.074,0.081, | 0.064,0.08,0.067,0 | 0,027 |
| 69062 | ENSG00000249915 | PDCD6 | chr5 | + | 306601 | 306760 | 304176 | 304221 | 314416 | 314974 | 69062 | 65,34,81,77,36,42,25,23,7 | 3,1,4,5,0,3,0,7,8,2,5,6 | 55,31,30,6,15,17,17,36,22, | 16,13,7,1,2,1,0,2,4,1,8,7,8, | 298 | 149 | 0.00586690765023 | 0.390337063513649 | 0.915,0.944,0.91,0 | 0.632,0.544,0.682, | 0,077 |
| 100316 | ENSG00000128891 | CCDC32 | chr15 | - | 40553594 | 40554127 | 40531330 | 40532302 | 40557215 | 40557372 | 100316 | 78,41,67,72,37,48,46,31,9 | 1,3,4,9,0,0,1,4,0,1,2,2 | 45,44,42,45,30,38,13,41,3 | 1,0,5,1,0,2,2,3,4,24,0,6,4,4 | 298 | 149 | 0.00587301847319 | 0.39059785142091 | 0.975,0.872,0.893, | 0.957,1.0,0.808,0. | 0,043 |
| 133525 | ENSG00000186866 | POFUT2 | chr21 | - | 45283382 | 45283484 | 45282348 | 45282459 | 45285677 | 45285928 | 133525 | 6,12,13,6,3,11,9,7,9,3,2,3 | 1,2,4,3,1,2,1,0,5,5,0,0 | 4,4,3,2,2,2,4,7,2,8,3,1,4,2, | 1,5,0,4,0,0,1,0,0,0,0,3,0,2, | 251 | 149 | 0.00587445695578 | 0.39059785142091 | 0.781,0.781,0.659, | 0.704,0.322,1.0,0. | -0,004 |
| 176275 | ENSG00000173193 | PARP14 | chr3 | + | 122721174 | 122721653 | 122720254 | 122720388 | 122727811 | 122727986 | 176275 | 10,9,13,11,2,5,6,6,1,1,6,3 | 29,34,27,20,19,7,12,25,17, | 19,4,1,5,1,0,2,2,6,3,3,2,20, | 27,22,29,9,25,19,11,27,16, | 298 | 149 | 0.00588107346288 | 0.390917023622336 | 0.147,0.117,0.194, | 0.26,0.083,0.017,0 | 0,069 |
| 248802 | ENSG00000110422 | HIPK3 | chr11 | + | 33348681 | 33348818 | 33348165 | 33348228 | 33349146 | 33349287 | 248802 | 76,44,102,52,30,40,46,44, | 0,0,2,6,4,1,6,3,1,0,0,1 | 66,65,42,38,33,44,43,50,3 | 6,1,5,10,1,3,1,3,1,5,12,3,1, | 286 | 149 | 0.00589388654328 | 0.391544740617147 | 1.0,1.0,0.964,0.81 | 0.851,0.971,0.814, | 0,058 |
| 109659 | ENSG00000221914 | PPP2R2A | chr8 | + | 26339902 | 26339973 | 26338889 | 26338987 | 26354467 | 26354633 | 109659 | 0,3,8,6,3,1,0,1,1,1,0,1 | 13,4,32,29,13,11,3,5,5,9,7, | 4,2,0,0,0,0,0,0,4,0,2,2,3, | 16,24,17,7,12,6,9,11,8,18, | 220 | 149 | 0.00589415539397 | 0.391544740617147 | 0.0,0.337,0.145,0. | 0.145,0.053,0.0,0. | 0,027 |
| 152793 | ENSG00000092148 | HECTD1 | chr14 | - | 31113876 | 31114086 | 31113276 | 31113484 | 31114285 | 31115472 | 152793 | 33,23,73,48,27,23,27,44,1 | 17,3,11,7,0,12,2,12,2,4,1,0 | 40,55,23,32,39,19,23,34,2 | 4,8,0,1,3,2,3,4,1,0,7,5,0,3, | 298 | 149 | 0.00589726419167 | 0.391630382362643 | 0.493,0.793,0.768, | 0.833,0.775,1.0,0. | -0,105 |
| 45857 | ENSG00000124596 | OARD1 | chr6 | - | 41068840 | 41068953 | 41064771 | 41067437 | 41070075 | 41070134 | 45857 | 24,17,54,39,23,23,18,10,2 | 1,1,10,2,0,2,1,3,3,1,0,2 | 24,30,13,3,8,18,11,12,15,2 | 2,8,1,7,3,1,0,3,0,1,5,8,2,3, | 262 | 149 | 0.00590442485109 | 0.391984967651126 | 0.932,0.906,0.754, | 0.872,0.681,0.881, | 0,137 |
| 175119 | ENSG00000101596 | SMCHD1 | chr18 | + | 2751360 | 2751393 | 2750429 | 2750507 | 2752487 | 2752552 | 175119 | 26,16,52,35,24,18,23,55,1 | 1,0,2,0,0,1,3,2,0,0,0,0 | 13,31,30,22,32,12,17,26,1 | 2,6,0,4,3,0,0,2,2,2,2,1,0,0, | 182 | 149 | 0.00590836811638 | 0.392125802669314 | 0.955,1.0,0.955,1. | 0.842,0.809,1.0,0. | 0,04 |
| 24629 | ENSG00000122026 | RPL21 | chr13 | + | 27255241 | 27255354 | 27253764 | 27253843 | 27256183 | 27256307 | 24629 | 18,16,46,24,9,5,8,10,4,3,1, | 2,0,6,0,1,0,0,0,0,0,0,0 | 14,23,13,27,6,21,8,7,9,24,1 | 0,4,3,2,1,2,3,0,0,0,0,1,3,6, | 262 | 149 | 0.00591048565758 | 0.39214541879365 | 0.837,1.0,0.813,1. | 1.0,0.766,0.711,0. | 0,082 |
| 100820 | ENSG00000127914 | AKAP9 | chr7 | + | 91979435 | 91979574 | 91973710 | 91973968 | 91980288 | 91980333 | 100820 | 9,0,3,5,3,4,1,4,3,3,0,2 | 141,109,120,72,46,53,38,8 | 1,9,3,12,4,0,2,6,0,8,3,9,1,2 | 71,70,47,81,58,20,36,36,5 | 288 | 149 | 0.00591939331241 | 0.392615390454027 | 0.032,0.0,0.013,0. | 0.007,0.062,0.032, | -0,014 |
| 43288 | ENSG00000156976 | EIF4A2 | chr3 | + | 186788309 | 186788416 | 186787802 | 186787882 | 186789124 | 186789881 | 43288 | 320,168,245,143,92,148,1 | 427,137,428,354,181,165, | 213,159,234,192,80,61,54, | 138,51,178,74,110,133,75, | 256 | 149 | 0.00593973724183 | 0.393760282240011 | 0.304,0.416,0.25,0 | 0.473,0.645,0.433, | -0,074 |
| 50996 | ENSG00000114796 | KLHL24 | chr3 | + | 183643479 | 183643542 | 183635618 | 183635793 | 183650295 | 183650590 | 50996 | 43,33,90,36,21,37,27,39,1 | 2,3,3,1,3,1,5,0,4,2,0,2 | 36,72,47,29,24,23,32,26,2 | 1,3,0,5,2,0,0,1,3,2,0,2,2,0, | 212 | 149 | 0.0059414531834 | 0.393760282240011 | 0.938,0.885,0.955, | 0.962,0.944,1.0,0. | -0,058 |
| 133123 | ENSG00000119801 | YPEL5 | chr2 | + | 30155849 | 30155910 | 30146940 | 30147062 | 30156627 | 30156792 | 133123 | 29,21,38,23,7,15,11,9,18,1 | 115,111,144,121,51,54,75, | 39,42,18,16,9,12,12,45,10, | 148,206,88,88,80,54,71,89 | 210 | 149 | 0.00594214307751 | 0.393760282240011 | 0.152,0.118,0.158, | 0.158,0.126,0.127, | -0,032 |
| 60505 | ENSG00000138688 | KIAA1109 | chr4 | + | 122189999 | 122190079 | 122187894 | 122188088 | 122196634 | 122196791 | 60505 | 68,45,54,21,20,23,26,20,2 | 4,3,3,0,0,2,3,0,0,0,1,4 | 30,27,28,39,38,15,15,14,2 | 0,1,0,0,3,1,0,1,1,0,1,2,0,0, | 229 | 149 | 0.00594493408271 | 0.393823978933751 | 0.917,0.907,0.921, | 1.0,0.946,1.0,1.0,0 | -0,018 |
| 226927 | ENSG00000128891 | CCDC32 | chr15 | - | 40553661 | 40554127 | 40531340 | 40532302 | 40557215 | 40557372 | 226927 | 78,41,65,72,37,48,46,31,9 | 1,3,4,9,0,0,1,4,0,1,2,2 | 45,44,42,46,30,38,13,41,3 | 1,0,5,1,0,2,2,3,4,24,0,6,4,4 | 298 | 149 | 0.00595035473962 | 0.393979626294789 | 0.975,0.872,0.89,0 | 0.957,1.0,0.808,0. | 0,043 |
| 27379 | ENSG00000012660 | ELOVL5 | chr6 | - | 53290400 | 53290535 | 53276178 | 53276256 | 53291775 | 53291963 | 27379 | 23,4,21,16,15,9,10,13,6,4, | 274,131,314,306,154,82,1 | 17,34,16,9,10,2,7,15,13,10 | 166,184,189,64,148,88,54, | 284 | 149 | 0.00595094463662 | 0.393979626294789 | 0.042,0.016,0.034, | 0.051,0.088,0.043, | -0,016 |
| 333 | ENSG00000155313 | USP25 | chr21 | + | 15808808 | 15808885 | 15805120 | 15805258 | 15811136 | 15811210 | 333 | 44,29,62,40,35,37,20,18,3 | 7,3,8,9,6,4,1,7,3,0,3,2 | 57,35,18,14,7,15,17,27,18, | 12,13,7,7,9,3,2,6,9,5,10,8, | 226 | 149 | 0.00595489467797 | 0.394119906652571 | 0.806,0.864,0.836, | 0.758,0.64,0.629,0 | 0,12 |
| 153024 | ENSG00000140563 | MCTP2 | chr15 | + | 94314792 | 94314864 | 94314281 | 94314344 | 94315528 | 94315637 | 153024 | 1,1,6,2,2,1,1,4,3,0,0,2 | 49,39,141,68,73,99,91,76, | 3,1,5,10,2,4,2,2,3,7,2,6,1,1 | 86,89,52,98,127,61,60,88, | 221 | 149 | 0.00596306544088 | 0.394539359946524 | 0.014,0.017,0.028, | 0.023,0.008,0.061, | -0,014 |
| 92325 | ENSG00000204619 | PPP1R11 | chr6 | + | 30068278 | 30068353 | 30067204 | 30067479 | 30068589 | 30068698 | 92325 | 20,2,13,4,4,4,4,4,7,5,1,3 | 70,50,88,58,35,37,24,18,4 | 9,15,7,7,9,5,4,7,5,6,8,2,9,2 | 30,22,30,20,18,34,30,32,1 | 224 | 149 | 0.00596709673965 | 0.394684756721453 | 0.16,0.026,0.089,0 | 0.166,0.312,0.134, | -0,055 |
| 1914 | ENSG00000093000 | NUP50 | chr22 | + | 45170987 | 45171152 | 45168167 | 45168246 | 45171599 | 45171639 | 1914 | 4,10,9,2,2,9,3,3,3,1,3,0 | 11,10,13,10,10,8,16,10,25, | 13,8,5,1,3,6,3,4,5,10,6,7,0, | 7,10,5,5,6,6,5,5,4,4,7,7,11, | 298 | 149 | 0.0059959481251 | 0.396471247592442 | 0.154,0.333,0.257, | 0.481,0.286,0.333, | -0,105 |
| 180059 | ENSG00000132781 | MUTYH | chr1 | - | 45333412 | 45333633 | 45332676 | 45333324 | 45334390 | 45334511 | 180059 | 21,11,18,9,14,23,10,16,21, | 1,2,3,0,0,2,2,0,1,4,0,3 | 19,18,16,15,8,8,7,10,7,7,6, | 1,1,0,3,2,0,0,0,0,0,0,3,0,1, | 298 | 149 | 0.00600204670477 | 0.39675261496141 | 0.913,0.733,0.75,1 | 0.905,0.9,1.0,0.71 | -0,071 |
| 93534 | ENSG00000288721 | | chr6 | - | 41909268 | 41909522 | 41907015 | 41907287 | 41916784 | 41916939 | 93534 | 7,4,12,17,17,6,6,11,7,4,5,5 | 0,0,4,2,1,3,1,0,0,0,2,0 | 9,9,4,2,6,5,7,5,2,4,2,22,17, | 0,1,1,3,2,0,0,0,3,0,0,6,0,0, | 298 | 149 | 0.00601044397579 | 0.397185713034467 | 1.0,1.0,0.6,0.81,0. | 1.0,0.818,0.667,0. | 0,102 |
| 143160 | ENSG00000109118 | PHF12 | chr17 | - | 28919142 | 28919301 | 28917284 | 28917449 | 28921687 | 28921808 | 143160 | 24,10,42,43,28,22,20,26,1 | 0,2,8,1,0,5,0,0,4,0,0,0 | 24,37,8,18,17,23,12,19,12, | 2,6,0,0,0,0,0,8,0,1,0,0,1,2, | 298 | 149 | 0.00601893143932 | 0.397590979244276 | 1.0,0.714,0.724,0. | 0.857,0.755,1.0,1. | -0,032 |
| 7544 | ENSG00000089063 | TMEM230 | chr20 | - | 5111763 | 5111808 | 5109331 | 5109445 | 5112960 | 5113103 | 7544 | 0,3,1,5,0,2,1,4,3,3,0,3 | 20,33,33,23,21,26,33,34,2 | 4,1,0,0,3,0,1,0,0,0,1,0,3,0, | 17,29,8,4,4,8,17,21,8,20,3 | 194 | 149 | 0.00602027124976 | 0.397590979244276 | 0.0,0.065,0.023,0. | 0.153,0.026,0.0,0. | 0,004 |
| 137261 | ENSG00000104814 | MAP4K1 | chr19 | - | 38605174 | 38605249 | 38601440 | 38601525 | 38605408 | 38605491 | 137261 | 0,2,0,2,0,0,1,0,0,2,0,0 | 3,5,14,10,14,13,5,9,11,17, | 0,4,0,4,5,1,1,2,0,1,2,2,2,0, | 11,11,5,8,6,4,6,10,4,14,9,2 | 224 | 149 | 0.00603162622252 | 0.398218694324909 | 0.0,0.21,0.0,0.117, | 0.0,0.195,0.0,0.25, | -0,049 |
| 21883 | ENSG00000079134 | THOC1 | chr18 | - | 254272 | 254355 | 252538 | 252612 | 259179 | 259275 | 21883 | 47,34,58,24,24,29,26,51,1 | 1,1,3,3,3,7,0,9,4,0,1,0 | 17,13,22,10,36,8,20,46,14, | 1,4,1,11,7,1,2,2,5,3,12,3,4, | 232 | 149 | 0.00605200775811 | 0.399441791409314 | 0.968,0.956,0.925, | 0.916,0.676,0.934, | 0,107 |
| 124086 | ENSG00000084234 | APLP2 | chr11 | + | 130123611 | 130123779 | 130122304 | 130122513 | 130126699 | 130126830 | 124086 | 78,63,453,245,153,168,11 | 34,27,141,57,57,80,34,100 | 148,127,149,122,81,67,10 | 47,41,20,19,24,28,43,29,4, | 298 | 149 | 0.00605496726843 | 0.399468722966091 | 0.534,0.538,0.616, | 0.612,0.608,0.788, | -0,073 |
| 50226 | ENSG00000122224 | LY9 | chr1 | + | 160824180 | 160824249 | 160823464 | 160823796 | 160827747 | 160828255 | 50226 | 45,16,52,39,60,21,36,37,3 | 7,4,1,2,6,4,0,0,1,0,5,12 | 58,29,38,25,13,26,36,47,2 | 12,3,2,4,3,5,3,11,1,3,11,29 | 218 | 149 | 0.0060561278024 | 0.399468722966091 | 0.815,0.732,0.973, | 0.768,0.869,0.929, | 0,048 |
| 82773 | ENSG00000100243 | CYB5R3 | chr22 | - | 42636478 | 42636598 | 42631377 | 42631450 | 42636714 | 42636846 | 82773 | 19,12,17,4,2,9,0,4,4,9,7,10 | 97,127,49,24,41,27,55,64, | 17,13,20,58,6,3,17,6,0,4,3 | 33,20,46,128,64,40,26,74, | 269 | 149 | 0.00605978110411 | 0.39958723860867 | 0.098,0.05,0.161,0 | 0.222,0.265,0.194, | -0,037 |
| 719 | ENSG00000182944 | EWSR1 | chr22 | + | 29297826 | 29297949 | 29295525 | 29296368 | 29298732 | 29298895 | 719 | 184,161,489,256,212,189, | 1,2,2,3,1,5,5,6,6,2,0,4 | 337,432,173,219,155,89,1 | 1,3,2,0,0,2,5,2,0,1,1,2,3,1, | 272 | 149 | 0.00606775624627 | 0.399990580288189 | 0.99,0.978,0.993,0 | 0.995,0.987,0.979, | -0,015 |
| 125412 | ENSG00000267699 | | chr18 | + | 51051353 | 51051415 | 51049294 | 51049324 | 51054780 | 51054993 | 125412 | 9,0,16,15,5,12,9,21,5,1,6,2 | 58,56,113,63,55,45,34,47, | 5,23,15,21,4,2,5,21,1,21,1 | 47,68,34,23,28,19,37,52,1 | 211 | 149 | 0.00607338929735 | 0.400239330023863 | 0.099,0.0,0.091,0. | 0.07,0.193,0.238,0 | -0,04 |
| 193797 | ENSG00000058673 | ZC3H11A | chr1 | + | 203802546 | 203803016 | 203795705 | 203795794 | 203816925 | 203817124 | 193797 | 84,48,182,83,87,84,80,157 | 10,8,33,16,9,16,15,16,4,8, | 110,139,79,66,66,49,88,12 | 28,38,27,26,29,9,15,18,6,7 | 298 | 149 | 0.0060780444417 | 0.400423502672645 | 0.808,0.75,0.734,0 | 0.663,0.647,0.594, | 0,074 |
| 121120 | ENSG00000101017 | CD40 | chr20 | + | 46123125 | 46123219 | 46122609 | 46122756 | 46128137 | 46128224 | 121120 | 29,33,58,71,33,38,19,16,1 | 2,1,7,0,1,0,2,1,0,4,0,3 | 50,58,26,6,19,17,9,23,11,2 | 0,6,0,2,0,0,0,0,0,3,0,0,0, | 243 | 149 | 0.0060815127175 | 0.400529395257112 | 0.899,0.953,0.836, | 1.0,0.856,1.0,0.64 | -0,057 |
| 167342 | ENSG00000058063 | ATP11B | chr3 | + | 182829922 | 182830002 | 182829671 | 182829752 | 182836034 | 182836142 | 167342 | 5,13,32,10,1,13,4,18,3,10, | 149,67,137,101,42,46,44,7 | 23,39,17,47,21,1,9,16,8,0, | 71,90,61,45,87,47,14,41,5 | 229 | 149 | 0.00608416849338 | 0.400581728050985 | 0.021,0.112,0.132, | 0.174,0.22,0.153,0 | -0,054 |
| 85835 | ENSG00000114956 | DGUOK | chr2 | + | 73932597 | 73932683 | 73926896 | 73927052 | 73958145 | 73958245 | 85835 | 1,3,2,3,0,7,0,3,1,0,1,2 | 4,5,11,2,7,3,1,0,5,2,4,3 | 2,6,2,3,2,4,2,6,1,4,3,7,6,0, | 5,10,4,0,5,2,4,1,0,2,2,3,2,3 | 235 | 149 | 0.00609118841398 | 0.400921276308663 | 0.137,0.276,0.103, | 0.202,0.276,0.241, | -0,217 |
| 66531 | ENSG00000145287 | PLAC8 | chr4 | - | 83104895 | 83105020 | 83092860 | 83094791 | 83114665 | 83114736 | 66531 | 517,328,103,94,36,1,31,14 | 68,41,4,1,1,0,4,0,4,0,5,10 | 155,42,173,209,93,162,52, | 2,0,9,20,8,4,2,2,10,14,11,7 | 274 | 149 | 0.00610400396103 | 0.401641967758009 | 0.805,0.813,0.933, | 0.977,1.0,0.913,0. | -0,011 |
| 189206 | ENSG00000118308 | IRAG2 | chr12 | + | 25062821 | 25062900 | 25052944 | 25052956 | 25063719 | 25063816 | 189206 | 198,63,598,381,207,153,1 | 12,7,61,38,8,15,10,22,8,3, | 246,337,200,62,107,100,4 | 5,7,14,9,8,6,7,10,3,11,11,3 | 228 | 149 | 0.00611208293338 | 0.401953589008364 | 0.915,0.855,0.865, | 0.97,0.969,0.903,0 | -0,016 |
| 238591 | ENSG00000136573 | BLK | chr8 | + | 11555585 | 11555830 | 11555390 | 11555484 | 11556657 | 11556837 | 238591 | 73,35,31,15,30,19,9,13,35, | 392,174,307,263,310,194, | 39,53,34,21,26,2,4,10,39,5 | 280,293,166,71,172,94,75, | 298 | 149 | 0.006112474954 | 0.401953589008364 | 0.085,0.091,0.048, | 0.065,0.083,0.093, | -0,015 |
| 63876 | ENSG00000158006 | PAFAH2 | chr1 | - | 25972557 | 25972712 | 25959766 | 25962083 | 25974479 | 25974650 | 63876 | 22,7,21,8,13,11,5,4,13,0,1, | 1,0,4,0,4,0,0,0,0,8,0,0 | 9,17,2,4,6,8,6,13,6,12,6,4, | 2,1,0,0,0,0,1,0,0,0,0,1,2,0, | 298 | 149 | 0.00611747978849 | 0.402159832729533 | 0.917,1.0,0.724,1. | 0.692,0.895,1.0,1. | -0,045 |
| 216822 | ENSG00000153201 | RANBP2 | chr2 | + | 108768349 | 108768388 | 108758412 | 108758548 | 108771700 | 108771871 | 216822 | 17,15,39,23,9,19,13,19,14, | 0,0,3,6,2,5,5,0,0,0,0,0 | 21,29,13,21,21,4,7,11,11,2 | 3,2,0,1,0,0,0,0,1,1,2,0,0,0, | 188 | 149 | 0.00613787605563 | 0.403377466054748 | 1.0,1.0,0.912,0.75 | 0.847,0.92,1.0,0.9 | -0,027 |

| | | | | | | | | | | | | | | | | | | | | | | |
|---|---|---|---|---|---|---|---|---|---|---|---|---|---|---|---|---|---|---|---|---|---|---|
| 238710 | ENSG00000064313 | TAF2 | chr8 | - | 119794779 | 119794944 | 119793365 | 119793451 | 119795531 | 119795631 | 238710 | 1,2,21,10,2,4,0,7,6,4,1,2 | 23,8,36,9,9,22,8,5,12,4,2,1 | 4,1,2,1,3,2,0,3,4,3,3,9,2,2, | 15,16,9,12,12,7,4,14,7,19,1 | 298 | 149 | 0.00614986744925 | 0.40404216146811 | 0.021,0.111,0.226, | 0.118,0.03,0.1,0.0 | 0,079 |
| 189210 | ENSG00000118308 | IRAG2 | chr12 | + | 25062829 | 25062900 | 25052944 | 25052956 | 25063719 | 25063816 | 189210 | 169,55,536,325,181,129,14 | 12,7,61,38,8,15,10,22,8,3, | 221,300,169,51,90,87,38,1 | 5,7,14,9,8,6,7,10,3,11,11,3 | 220 | 149 | 0.00615463202628 | 0.404212046258156 | 0.905,0.842,0.856, | 0.968,0.967,0.891, | -0,018 |
| 200841 | ENSG00000161929 | SCIMP | chr17 | - | 5223353 | 5223430 | 5221286 | 5221350 | 5234734 | 5234860 | 200841 | 21,6,16,20,6,15,10,9,4,11, | 3,0,3,13,1,4,0,1,9,3,6,5 | 6,0,7,6,3,5,4,4,3,12,6,9,13, | 8,2,10,2,2,4,1,9,6,3,2,3,8,1 | 226 | 149 | 0.00615779941896 | 0.404212046258156 | 0.822,1.0,0.779,0. | 0.331,0.0,0.316,0. | 0,216 |
| 8739 | ENSG00000152242 | C18orf25 | chr18 | + | 46240004 | 46240187 | 46215810 | 46216595 | 46253696 | 46253823 | 8739 | 11,4,23,8,4,5,4,10,5,1,0,9 | 17,9,63,28,18,17,23,24,7,3 | 9,9,0,3,6,4,2,6,9,7,13,3,1,2 | 30,78,28,24,36,21,9,10,11, | 298 | 149 | 0.00615808735582 | 0.404212046258156 | 0.244,0.182,0.154, | 0.13,0.055,0.0,0.0 | 0,044 |
| 58338 | ENSG00000065413 | ANKRD44 | chr2 | - | 197086679 | 197086748 | 197083368 | 197083509 | 197099815 | 197099930 | 58338 | 242,134,122,82,63,109,71, | 10,0,6,0,0,0,1,4,5,0,0,0 | 124,81,90,78,79,49,64,56, | 4,0,0,30,4,1,4,3,0,0,3,0,3,0 | 218 | 149 | 0.006162018274 | 0.404345506471796 | 0.943,1.0,0.933,1. | 0.955,1.0,1.0,0.64, | 0,025 |
| 162518 | ENSG00000132952 | USPL1 | chr13 | + | 30619681 | 30619740 | 30617936 | 30618056 | 30621763 | 30621892 | 162518 | 2,0,0,2,2,3,1,0,0,1,0,0 | 21,9,42,25,16,15,13,28,10, | 3,3,0,0,2,2,2,2,0,0,10,1,0,1 | 17,27,16,14,24,11,15,12,2 | 208 | 149 | 0.00616662056899 | 0.404345506471796 | 0.064,0.0,0.0,0.05 | 0.112,0.074,0.0,0. | -0,011 |
| 68514 | ENSG00000163534 | FCRL1 | chr1 | - | 157803844 | 157804111 | 157802376 | 157802664 | 157820006 | 157820060 | 68514 | 306,127,343,37,119,62,17 | 0,1,14,0,2,2,4,5,1,0,0,2 | 211,147,228,107,44,33,12 | 7,4,4,3,2,3,0,0,0,4,6,6,11,2 | 298 | 149 | 0.00616708903717 | 0.404345506471796 | 1.0,0.984,0.925,1. | 0.938,0.948,0.966, | 0,027 |
| 160169 | ENSG00000104320 | NBN | chr8 | - | 89981374 | 89981550 | 89980733 | 89980893 | 89982721 | 89982855 | 160169 | 24,12,39,34,23,13,9,15,12, | 0,0,0,0,0,0,1,0,0,0,0,2 | 25,15,20,7,23,18,5,23,16,3 | 0,1,3,6,2,1,0,1,3,0,0,3,1,0, | 298 | 149 | 0.00616763522795 | 0.404345506471796 | 1.0,1.0,1.0,1.0,1.0, | 1.0,0.882,0.769,0. | 0,084 |
| 250266 | ENSG00000057608 | GDI2 | chr10 | - | 5794987 | 5795019 | 5786005 | 5786050 | 5796762 | 5796862 | 250266 | 54,26,72,75,23,17,23,26,2 | 1,0,9,13,3,2,1,2,1,1,9,3 | 52,46,12,17,18,13,6,13,12, | 15,11,1,5,3,1,1,3,3,3,8,19, | 181 | 149 | 0.00617281369414 | 0.404561773508967 | 0.978,1.0,0.868,0. | 0.741,0.775,0.908, | 0,068 |
| 71944 | ENSG00000116539 | ASH1L | chr1 | - | 155415743 | 155415923 | 155395458 | 155395553 | 155438326 | 155439068 | 71944 | 104,59,165,107,80,93,95,1 | 1,1,4,3,1,8,0,3,0,0,0,2 | 94,78,76,113,69,51,63,93, | 6,3,5,1,5,0,5,3,2,0,4,4,0,4, | 298 | 149 | 0.00617546187624 | 0.40461212635769 | 0.981,0.967,0.954, | 0.887,0.929,0.884, | 0,034 |
| 105484 | ENSG00000147586 | MRPS28 | chr8 | - | 80002998 | 80003180 | 79918716 | 79919148 | 80030035 | 80030259 | 105484 | 27,23,61,50,19,22,18,48,3 | 0,0,3,2,0,0,0,1,0,0,0,0 | 32,37,13,15,26,22,19,16,6, | 1,0,1,2,4,1,0,1,0,0,0,4,0,0, | 298 | 149 | 0.00618893132046 | 0.405371234642096 | 1.0,1.0,0.91,0.926, | 0.941,1.0,0.867,0. | 0,072 |
| 158227 | ENSG00000115760 | BIRC6 | chr2 | + | 32595036 | 32595144 | 32593914 | 32594060 | 32597750 | 32597968 | 158227 | 28,15,46,37,18,10,24,24,1 | 4,5,12,2,4,5,6,10,4,2,0,5 | 53,18,33,7,7,16,13,32,10,1 | 37,18,5,13,9,3,5,7,5,11,17, | 257 | 149 | 0.00619404080649 | 0.405582475455324 | 0.802,0.635,0.69,0 | 0.454,0.367,0.793, | 0,145 |
| 50328 | ENSG00000132781 | MUTYH | chr1 | - | 45333412 | 45333603 | 45333284 | 45333324 | 45334390 | 45334511 | 50328 | 21,11,19,10,14,23,10,16,1 | 1,2,3,0,0,2,2,0,1,4,0,3 | 20,14,16,15,8,8,7,10,7,7,6, | 1,1,0,3,2,0,0,0,0,0,3,0,1, | 298 | 149 | 0.00622169843701 | 0.40726957916548 | 0.913,0.733,0.76,1 | 0.909,0.875,1.0,0. | -0,069 |
| 58769 | ENSG00000154710 | RABGEF1 | chr7 | + | 66770322 | 66770467 | 66740732 | 66740792 | 66771882 | 66772035 | 58769 | 3,2,4,1,2,0,3,2,0,2,1,2 | 7,8,23,13,1,6,1,1,3,0,2,5 | 0,0,0,0,0,4,0,0,0,1,0,8,0,1, | 5,15,4,1,1,4,4,5,2,4,4,14,3, | 294 | 149 | 0.0062293562649 | 0.407646876938489 | 0.178,0.112,0.081, | 0.0,0.0,0.0,0.0,0.0, | 0,216 |
| 12998 | ENSG00000109046 | WSB1 | chr17 | + | 27301817 | 27301956 | 27294086 | 27294435 | 27303366 | 27303422 | 12998 | 59,31,165,125,38,39,28,59 | 4,3,37,11,32,20,10,15,5,2, | 44,65,17,9,20,30,16,73,17, | 15,20,23,21,20,11,3,12,11, | 288 | 149 | 0.00623679572838 | 0.407846843814717 | 0.884,0.842,0.698, | 0.603,0.627,0.277, | 0,16 |
| 203207 | ENSG00000109381 | ELF2 | chr4 | - | 139064579 | 139064891 | 139061936 | 139062057 | 139067683 | 139067770 | 203207 | 3,5,11,2,1,1,3,4,0,5,0,2 | 13,16,98,42,15,15,19,33,9, | 8,2,7,16,3,3,2,0,4,5,2,4,3,3 | 31,63,18,10,16,12,3,12,11, | 298 | 149 | 0.00623845823798 | 0.407846843814717 | 0.103,0.135,0.053, | 0.114,0.016,0.163, | -0,054 |
| 48422 | ENSG00000135241 | PNPLA8 | chr7 | - | 108521475 | 108521521 | 108514435 | 108515574 | 108526028 | 108526135 | 48422 | 15,9,66,21,21,17,22,29,4,1 | 1,0,15,1,0,0,0,0,0,0,2,3 | 33,28,16,22,28,12,15,10,1 | 1,1,0,0,1,2,0,2,0,0,1,2,2,0,1 | 195 | 149 | 0.0062414513907 | 0.407846843814717 | 0.92,1.0,0.771,0.9 | 0.962,0.955,1.0,1. | -0,002 |
| 194998 | ENSG00000137776 | SLTM | chr15 | - | 58894433 | 58894582 | 58894188 | 58894193 | 58897114 | 58897120 | 194998 | 180,110,111,81,45,47,57,5 | 3,1,5,13,5,10,0,8,0,1,0,2 | 62,64,89,65,56,44,32,43,5 | 2,17,2,3,4,1,2,4,2,3,4,16,16 | 298 | 149 | 0.00624205858066 | 0.407846843814717 | 0.968,0.982,0.917, | 0.939,0.653,0.957, | 0,058 |
| 198389 | ENSG00000069248 | NUP133 | chr1 | - | 229488725 | 229488845 | 229487465 | 229487613 | 229489954 | 229490102 | 198389 | 4,7,18,6,1,6,4,5,3,0,5,0 | 35,21,53,42,27,18,16,31,2 | 1,12,8,5,3,0,5,3,1,1,5,3,1,0 | 66,55,13,21,17,13,19,31,4 | 269 | 149 | 0.0062431447939 | 0.407846843814717 | 0.06,0.156,0.158,0 | 0.008,0.108,0.254, | 0,027 |
| 110280 | ENSG00000165392 | WRN | chr8 | + | 31090464 | 31090532 | 31088889 | 31088965 | 31090833 | 31090942 | 110280 | 13,4,30,13,15,24,9,18,8,3, | 0,0,0,2,0,0,3,0,0,0,2,0 | 14,29,4,15,10,1,15,7,11,19 | 1,4,0,3,0,0,2,0,1,1,1,7,4,1, | 217 | 149 | 0.00624378156664 | 0.407846843814717 | 1.0,1.0,1.0,0.817,1 | 0.906,0.833,1.0,0. | 0,038 |
| 189302 | ENSG00000118308 | IRAG2 | chr12 | + | 25102419 | 25102693 | 25102197 | 25102241 | 25103836 | 25103899 | 189302 | 69,4,57,26,24,6,12,34,17,1 | 109,85,524,396,147,91,147 | 29,38,18,50,16,19,23,31,4, | 162,227,139,35,87,35,49,1 | 298 | 149 | 0.00624906977242 | 0.408068427241423 | 0.24,0.023,0.052,0 | 0.082,0.077,0.061, | -0,041 |
| 175195 | ENSG00000145016 | RUBCN | chr3 | - | 197699316 | 197699673 | 197696953 | 197697049 | 197700612 | 197701146 | 175195 | 7,7,8,0,0,4,0,3,0,0,0,0 | 45,52,48,41,32,28,28,22,2 | 2,9,1,21,4,3,2,5,2,4,5,1,4,3 | 34,35,13,54,44,23,40,31,2 | 298 | 149 | 0.00625721838996 | 0.408458867343068 | 0.072,0.063,0.077, | 0.029,0.114,0.037, | -0,022 |
| 193647 | ENSG00000114861 | FOXP1 | chr3 | - | 71130400 | 71130584 | 71112590 | 71112637 | 71132090 | 71132099 | 193647 | 1,2,8,1,2,4,3,1,0,1,0,2 | 17,10,21,26,10,26,5,19,5,1 | 6,6,8,0,5,5,2,4,1,4,2,0,0,1, | 5,6,10,7,4,6,7,4,8,3,6,1,3,2 | 298 | 149 | 0.00625884442528 | 0.408458867343068 | 0.029,0.091,0.16,0 | 0.375,0.333,0.286, | -0,152 |
| 245443 | ENSG00000101017 | CD40 | chr20 | + | 46121819 | 46122050 | 46118270 | 46118394 | 46122232 | 46122358 | 245443 | 26,21,54,11,19,7,12,17,18, | 2,0,1,2,0,0,3,0,0,0,0,2 | 18,17,4,7,6,9,5,12,3,7,13,4 | 3,0,1,12,2,0,0,1,0,0,0,7,0,0 | 298 | 149 | 0.00626381575433 | 0.408645667750938 | 0.867,1.0,0.964,0. | 0.75,1.0,0.667,0.2 | 0,07 |
| 126621 | ENSG00000148634 | HERC4 | chr10 | - | 67990210 | 67990400 | 67988662 | 67988835 | 67990903 | 67991015 | 126621 | 41,17,100,95,44,34,31,57, | 3,3,5,1,4,2,1,3,0,1,1,0 | 67,63,16,16,48,19,19,40,2 | 3,2,0,1,1,3,1,1,1,2,3,0,1,0, | 298 | 149 | 0.00626550405647 | 0.408645667750938 | 0.872,0.739,0.909, | 0.918,0.94,1.0,0.8 | -0,045 |
| 30059 | ENSG00000153113 | CAST | chr5 | + | 96730779 | 96730860 | 96729589 | 96729725 | 96736171 | 96736187 | 30059 | 143,105,219,223,133,193, | 11,15,20,6,1,9,8,17,5,6,2,1 | 87,126,138,113,105,64,92, | 6,19,13,20,16,6,4,6,8,6,17, | 230 | 149 | 0.00626892993796 | 0.408729142287839 | 0.894,0.819,0.876, | 0.904,0.811,0.873, | 0,033 |
| 64587 | ENSG00000144401 | METTL21A | chr2 | - | 207620640 | 207620694 | 207612722 | 207613443 | 207621805 | 207621917 | 64587 | 0,4,3,5,3,1,2,1,4,0,0,0 | 4,10,16,14,10,5,3,4,13,14, | 2,2,0,0,0,1,0,3,2,0,2,1,1,3, | 12,15,9,2,4,4,6,10,3,10,6,2 | 203 | 149 | 0.00627127677884 | 0.408729142287839 | 0.0,0.227,0.121,0. | 0.109,0.089,0.0,0. | 0,033 |
| 105583 | ENSG00000099204 | ABLIM1 | chr10 | - | 114447879 | 114448020 | 114445311 | 114445403 | 114451623 | 114451671 | 105583 | 100,76,78,79,61,28,41,21, | 7,14,9,7,0,2,4,2,2,4,0,5 | 62,49,29,24,36,42,12,16,3 | 8,10,6,1,11,6,7,3,17,6,0,25 | 290 | 149 | 0.00627248099473 | 0.408729142287839 | 0.88,0.736,0.817,0 | 0.799,0.716,0.713, | 0,102 |
| 28259 | ENSG00000112851 | ERBIN | chr5 | + | 66075101 | 66075230 | 66053405 | 66054951 | 66076315 | 66076408 | 28259 | 60,86,188,82,70,56,65,82, | 20,11,43,22,14,5,24,17,7,5 | 77,72,68,46,67,40,33,57,3 | 41,50,18,17,27,9,8,10,14,5 | 278 | 149 | 0.00628063613495 | 0.409136681586387 | 0.617,0.807,0.701, | 0.502,0.436,0.669, | 0,087 |
| 128372 | ENSG00000088298 | EDEM2 | chr20 | - | 35134737 | 35134949 | 35131641 | 35131783 | 35137879 | 35138005 | 128372 | 34,15,62,47,37,36,31,18,3 | 2,5,4,0,6,0,2,0,1,0,0,1 | 46,58,24,3,16,28,28,35,14, | 4,1,2,1,2,0,3,2,0,4,1,0,5,1, | 298 | 149 | 0.00629132182597 | 0.409708770930514 | 0.895,0.6,0.886,1. | 0.852,0.967,0.857, | 0,025 |
| 8730 | ENSG00000141425 | RPRD1A | chr18 | - | 36062977 | 36063075 | 36033751 | 36033837 | 36067253 | 36067576 | 8730 | 10,5,4,2,0,3,0,2,2,0,0,3 | 10,19,48,36,13,13,12,25,1 | 0,0,0,3,3,5,0,0,2,2,3,6,1,4, | 40,27,17,10,5,12,17,26,5,1 | 247 | 149 | 0.00631048779372 | 0.410832606875423 | 0.376,0.137,0.048, | 0.0,0.0,0.0,0.153,0 | 0,007 |
| 71441 | ENSG00000143149 | ALDH9A1 | chr1 | - | 165680486 | 165680683 | 165679441 | 165679582 | 165682106 | 165682241 | 71441 | 53,47,113,106,47,48,48,51 | 0,0,0,2,0,2,0,3,0,0,0,0 | 75,91,32,21,57,54,31,46,3 | 2,2,1,0,3,0,0,2,2,2,2,0,2,2, | 298 | 149 | 0.00631883348807 | 0.411251542325611 | 1.0,1.0,1.0,0.964,1 | 0.949,0.958,0.941, | 0,035 |
| 19825 | ENSG00000185787 | MORF4L1 | chr15 | + | 78894057 | 78894230 | 78893538 | 78893627 | 78894819 | 78894904 | 19825 | 183,142,419,389,129,185, | 8,4,3,0,4,1,0,1,2,0,0,3 | 245,424,159,249,176,139, | 9,4,0,6,0,1,0,10,0,4,7,7,5,2 | 298 | 149 | 0.00632802424755 | 0.411701537515644 | 0.92,0.947,0.986,1 | 0.932,0.981,1.0,0. | 0,011 |
| 39327 | ENSG00000075826 | SEC31B | chr10 | - | 100490007 | 100490322 | 100489702 | 100489761 | 100490705 | 100490883 | 39327 | 35,16,29,38,16,17,24,27,2 | 1,2,4,2,2,1,0,3,2,0,2,3 | 19,9,10,8,16,9,12,24,8,13, | 1,1,2,0,0,1,1,1,0,2,2,1,0,1, | 298 | 149 | 0.0063295732847 | 0.411701537515644 | 0.946,0.8,0.784,0. | 0.905,0.818,0.714, | -0,003 |
| 187941 | ENSG00000114098 | ARMC8 | chr3 | + | 138237308 | 138237377 | 138235033 | 138235114 | 138237481 | 138237572 | 187941 | 17,7,12,10,10,7,8,11,4,2,8, | 1,2,3,3,4,6,2,4,2,4,1,4 | 9,19,5,3,8,5,5,12,2,7,12,7, | 6,13,1,9,6,2,4,3,3,7,8,4,7,3 | 218 | 149 | 0.00633352800293 | 0.411821155350463 | 0.921,0.705,0.732, | 0.506,0.5,0.774,0. | 0,156 |
| 82800 | ENSG00000176978 | DPP7 | chr9 | - | 137111872 | 137112029 | 137111689 | 137111754 | 137112111 | 137112169 | 82800 | 297,202,264,99,169,120,2 | 4,3,11,3,2,3,0,2,2,0,0,4 | 149,159,229,166,92,49,76, | 4,2,10,8,5,2,3,12,0,3,5,3,1, | 298 | 149 | 0.00633523909365 | 0.411821155350463 | 0.974,0.971,0.923, | 0.949,0.975,0.92,0 | 0,023 |
| 209395 | ENSG00000091490 | SEL1L3 | chr4 | - | 25784227 | 25784559 | 25782241 | 25782418 | 25788223 | 25788364 | 209395 | 102,76,252,106,87,99,65,7 | 6,2,2,8,1,0,0,1,2,0,1,0 | 83,48,97,68,64,37,16,30,7 | 2,2,0,1,0,0,0,0,0,0,1,2,0,0, | 298 | 149 | 0.00633993155521 | 0.41200175379209 | 0.895,0.95,0.984,0 | 0.954,0.923,1.0,0. | -0,023 |
| 181051 | ENSG00000076685 | NT5C2 | chr10 | - | 103100778 | 103100964 | 103099925 | 103100019 | 103101044 | 103101102 | 181051 | 4,0,6,8,0,1,2,5,0,2,1,2 | 49,53,127,96,26,25,44,34, | 0,1,6,7,11,4,1,3,2,7,11,0,1, | 49,55,35,25,48,49,8,42,43, | 298 | 149 | 0.00635140257528 | 0.41262261626323 | 0.039,0.0,0.023,0. | 0.0,0.009,0.079,0. | -0,023 |
| 110642 | ENSG00000122882 | ECD | chr10 | - | 73154255 | 73154448 | 73152292 | 73152421 | 73156274 | 73156453 | 110642 | 98,76,123,58,35,61,54,69, | 1,0,5,4,0,2,1,0,0,0,2,1 | 58,114,45,67,91,23,65,61, | 1,5,6,2,1,2,0,5,2,0,0,5,1,0, | 298 | 149 | 0.00636385633335 | 0.413306928932786 | 0.98,1.0,0.925,0.8 | 0.967,0.919,0.789, | 0,019 |
| 50219 | ENSG00000122224 | LY9 | chr1 | + | 160816593 | 160816863 | 160799752 | 160800082 | 160818217 | 160818319 | 50219 | 31,7,7,6,7,5,13,5,14,6,2,7 | 0,0,0,0,1,0,0,0,0,0,0,0 | 13,4,11,15,8,1,7,11,12,16, | 3,2,0,0,1,4,4,0,0,1,1,1,0,2, | 298 | 149 | 0.00637472026885 | 0.413887607295582 | 1.0,1.0,1.0,1.0,0.7 | 0.684,0.5,1.0,1.0,0 | 0,195 |
| 16058 | ENSG00000090857 | PDPR | chr16 | + | 70153390 | 70153573 | 70148463 | 70148553 | 70156474 | 70156614 | 16058 | 31,12,53,48,35,41,17,16,1 | 7,4,22,11,9,6,6,12,2,0,0,0 | 33,45,12,9,22,26,33,32,29, | 11,10,0,3,2,3,0,3,0,0,5,8,1, | 298 | 149 | 0.00638160279736 | 0.414209514981481 | 0.689,0.6,0.546,0. | 0.6,0.692,1.0,0.6,0 | -0,104 |
| 121424 | ENSG00000109919 | MTCH2 | chr11 | - | 47628952 | 47629046 | 47627079 | 47627127 | 47630554 | 47630614 | 121424 | 15,16,73,78,33,26,16,54,2 | 2,0,6,0,0,3,1,0,1,0,0,0 | 52,66,20,3,28,16,22,32,17, | 7,5,0,2,3,0,1,2,4,0,0,3,0,4, | 243 | 149 | 0.00638572393308 | 0.414352049394254 | 0.821,1.0,0.882,1. | 0.82,0.89,1.0,0.47 | 0,048 |
| 13562 | ENSG00000102893 | PHKB | chr16 | + | 47463881 | 47463993 | 47461294 | 47461426 | 47497398 | 47497488 | 13562 | 25,0,51,29,31,27,21,34,17, | 0,2,1,1,0,4,2,4,2,0,0,3 | 20,33,11,18,22,35,12,20,3 | 5,5,0,0,2,6,2,0,0,1,0,2,1,0, | 261 | 149 | 0.00640429535244 | 0.41530960399795 | 1.0,0.0,0.967,0.94 | 0.695,0.79,1.0,1.0, | -0,066 |
| 161479 | ENSG00000113282 | CLINT1 | chr5 | - | 157789026 | 157789513 | 157787601 | 157787992 | 157791702 | 157791724 | 161479 | 78,63,57,55,37,25,31,147, | 0,2,0,0,0,0,0,0,0,2,0,1 | 61,86,76,106,95,44,30,57, | 1,3,0,13,0,0,0,0,2,0,6,0,0,0 | 298 | 149 | 0.00640457446691 | 0.41530960399795 | 1.0,0.94,1.0,1.0,1. | 0.968,0.935,1.0,0. | 0,016 |
| 105101 | ENSG00000185753 | CXorf38 | chrX | - | 40636532 | 40636712 | 40626920 | 40630773 | 40637006 | 40637156 | 105101 | 48,35,216,133,32,35,36,36 | 3,1,25,13,2,3,8,3,3,0,2,2 | 70,85,34,44,61,56,16,39,3 | 5,14,8,11,11,2,5,3,9,11,12, | 298 | 149 | 0.00640626993915 | 0.41530960399795 | 0.889,0.946,0.812, | 0.875,0.752,0.68,0 | 0,104 |
| 170609 | ENSG00000076108 | BAZ2A | chr12 | - | 56630775 | 56630841 | 56617394 | 56617532 | 56636181 | 56636379 | 170609 | 6,5,6,7,2,4,8,10,4,6,2,5 | 11,13,20,7,8,19,7,13,4,9,3, | 6,6,1,5,4,4,0,11,2,5,1,2,5,1 | 10,35,11,32,23,11,5,10,12, | 215 | 149 | 0.0064087566504 | 0.415345709913352 | 0.274,0.21,0.172,0 | 0.294,0.106,0.059, | 0,124 |
| 224326 | ENSG00000131023 | LATS1 | chr6 | - | 149676254 | 149676366 | 149661696 | 149662238 | 149676554 | 149676737 | 224326 | 8,11,26,12,7,15,10,20,13,3 | 1,1,0,0,4,0,1,1,1,2,0,0 | 18,26,6,17,19,9,9,14,4,21, | 3,1,0,2,2,1,1,1,2,1,4,1,1,3, | 261 | 149 | 0.00641076957054 | 0.4153510973621 | 0.82,0.863,1.0,1.0, | 0.774,0.937,1.0,0. | 0,032 |
| 41950 | ENSG00000137502 | RAB30 | chr11 | - | 82994038 | 82994122 | 82987400 | 82987770 | 82997223 | 82997324 | 41950 | 83,126,140,108,26,17,25,4 | 5,3,6,8,0,1,3,3,2,3,0,4 | 116,182,110,94,118,39,78, | 3,9,5,13,19,9,14,1,7,1,3,8, | 233 | 149 | 0.00641415247296 | 0.415445215440161 | 0.914,0.964,0.937, | 0.961,0.928,0.934, | 0,037 |
| 254249 | ENSG00000138777 | PPA2 | chr4 | - | 105383817 | 105383885 | 105370836 | 105370873 | 105386566 | 105386636 | 254249 | 7,0,3,6,0,0,6,0,2,0,0,0 | 9,6,31,35,20,21,8,15,6,9,2, | 1,0,2,3,0,4,0,3,0,0,0,0,4, | 21,8,10,8,10,12,9,15,10,27 | 217 | 149 | 0.00641668206953 | 0.415484024821604 | 0.348,0.0,0.062,0. | 0.032,0.0,0.121,0. | 0,039 |
| 249501 | ENSG00000181192 | DHTKD1 | chr10 | + | 12094205 | 12094271 | 12091512 | 12091684 | 12097683 | 12097958 | 249501 | 18,3,34,22,19,13,9,9,8,2,7, | 4,0,3,1,0,0,2,3,2,4,0,0 | 13,7,6,10,9,10,7,3,11,25,1 | 0,0,1,2,0,1,0,5,0,1,1,0,2,2, | 215 | 149 | 0.00643068336909 | 0.416144366923181 | 0.757,1.0,0.887,0. | 1.0,1.0,0.806,0.77 | -0,048 |
| 52508 | ENSG00000181135 | ZNF707 | chr8 | + | 143692244 | 143692330 | 143691599 | 143691713 | 143693670 | 143693703 | 52508 | 2,2,2,1,4,4,1,0,4,1,1,0 | 13,6,14,6,1,7,7,5,3,6,5,2 | 5,1,1,0,2,0,2,0,1,0,0,4,0,0, | 1,14,0,5,0,6,2,6,7,6,5,14,2, | 235 | 149 | 0.00643096747282 | 0.416144366923181 | 0.089,0.174,0.083, | 0.76,0.043,1.0,0.0, | -0,033 |
| 239469 | ENSG00000143437 | ARNT | chr1 | - | 150817683 | 150818030 | 150817393 | 150817433 | 150823193 | 150823345 | 239469 | 56,56,77,37,25,33,31,42,5 | 0,0,3,4,9,3,0,3,6,2,0,2 | 44,54,26,19,37,38,20,30,1 | 0,0,2,2,3,0,2,2,3,1,0,0,2,1, | 298 | 149 | 0.00643268074187 | 0.416144366923181 | 1.0,1.0,0.928,0.82 | 1.0,1.0,0.867,0.82 | -0,026 |
| 232985 | ENSG00000005889 | ZFX | chrX | + | 24161714 | 24161844 | 24152719 | 24152830 | 24172714 | 24172914 | 232985 | 2,1,11,5,1,2,0,0,2,0,0,3 | 12,7,34,15,14,7,1,8,6,2,1,2 | 8,0,0,0,3,2,0,0,1,4,0,3,3,1, | 16,5,9,9,9,5,1,4,2,11,11,13 | 279 | 149 | 0.00643636741013 | 0.416224444162348 | 0.082,0.071,0.147, | 0.211,0.0,0.0,0.0,0 | 0,013 |
| 87915 | ENSG00000166164 | BRD7 | chr16 | - | 50323586 | 50323698 | 50321981 | 50322038 | 50325747 | 50325883 | 87915 | 92,85,160,74,64,61,29,55, | 22,13,26,11,11,20,7,7,0,0, | 86,36,59,31,57,51,33,63,1 | 18,26,9,21,8,8,5,13,5,8,10, | 261 | 149 | 0.00643778626042 | 0.416224444162348 | 0.705,0.789,0.778, | 0.732,0.441,0.789, | 0,115 |
| 64646 | ENSG00000058673 | ZC3H11A | chr1 | + | 203796292 | 203796496 | 203795653 | 203795794 | 203816925 | 203817124 | 64646 | 39,26,29,22,27,24,34,47,2 | 10,8,33,16,9,16,15,16,4,8, | 17,26,25,17,15,8,21,25,12, | 28,38,27,26,29,9,15,18,6,7 | 298 | 149 | 0.00644114946162 | 0.416266484907537 | 0.661,0.619,0.305, | 0.233,0.255,0.316, | 0,134 |
| 190317 | ENSG00000037474 | NSUN2 | chr5 | - | 6625581 | 6625669 | 6623213 | 6623285 | 6631872 | 6631977 | 190317 | 7,19,12,15,10,8,5,15,10,3, | 0,2,3,2,5,0,0,1,0,2,2,2 | 6,16,12,11,16,3,8,7,12,11, | 0,3,2,0,3,3,0,2,1,0,0,2,0,2, | 237 | 149 | 0.00644230459937 | 0.416266484907537 | 1.0,0.857,0.715,0. | 1.0,0.77,0.79,1.0,0 | -0,025 |
| 80657 | ENSG00000109466 | KLHL2 | chr4 | + | 165223937 | 165224057 | 165219933 | 165220059 | 165228806 | 165228913 | 80657 | 2,2,1,0,3,0,0,0,0,1,0,0 | 45,19,55,32,15,16,10,32,2 | 3,4,0,0,2,0,4,0,2,0,0,2,3,4, | 15,30,15,17,14,10,12,28,1 | 269 | 149 | 0.00645666233369 | 0.417068994820658 | 0.024,0.055,0.01,0 | 0.1,0.069,0.0,0.0,0 | -0,04 |
| 175302 | ENSG00000213413 | PVRIG | chr7 | + | 100220113 | 100220395 | 100219712 | 100220028 | 100220546 | 100220673 | 175302 | 13,14,26,14,1,19,25,28,16, | 1,5,0,0,0,1,3,1,0,4,2,3 | 16,15,1,6,5,5,12,10,12,13, | 0,0,1,1,0,0,4,0,0,2,1,4,1,0, | 298 | 149 | 0.00646711014545 | 0.417618536968302 | 0.867,0.583,1.0,1. | 1.0,1.0,0.333,0.75, | 0,011 |
| 107146 | ENSG00000215301 | DDX3X | chrX | + | 41347314 | 41347454 | 41346858 | 41347012 | 41347639 | 41349884 | 107146 | 123,109,137,85,40,42,29,5 | 23,7,26,30,13,10,10,14,6,1 | 64,97,64,76,83,46,14,25,4 | 31,31,15,52,16,17,2,20,13, | 289 | 149 | 0.0064699903222 | 0.417679210269175 | 0.734,0.889,0.731, | 0.516,0.617,0.687, | 0,084 |
| 164595 | ENSG00000185246 | PRPF39 | chr14 | + | 45108424 | 45108522 | 45107450 | 45107616 | 45109615 | 45109780 | 164595 | 47,31,91,35,14,23,14,21,1 | 0,2,1,1,1,2,0,0,0,1,0,0 | 34,50,37,20,15,21,11,16,2 | 0,4,2,2,3,0,0,0,4,4,0,0,2,2, | 247 | 149 | 0.00647292387936 | 0.417743292197461 | 1.0,0.903,0.982,0. | 1.0,0.883,0.918,0. | 0,032 |
| 27146 | ENSG00000129515 | SNX6 | chr14 | - | 34581560 | 34581600 | 34575755 | 34575842 | 34586229 | 34586305 | 27146 | 28,17,84,79,35,29,19,26,3 | 1,2,1,0,0,4,2,3,2,1,0,1 | 42,48,27,16,32,25,17,29,2 | 3,1,2,4,1,2,0,1,4,10,3,2,3,0 | 189 | 149 | 0.00647610552214 | 0.41782334161742 | 0.957,0.87,0.985,1 | 0.917,0.974,0.914, | 0,023 |
| 97779 | ENSG00000239713 | APOBEC3G | chr22 | + | 39078931 | 39079085 | 39077274 | 39077378 | 39080932 | 39081194 | 97779 | 265,151,614,487,319,162, | 0,1,9,7,10,2,2,2,3,0,1,0 | 261,143,225,58,70,103,23, | 4,6,6,0,2,3,1,5,3,4,1,5,3,6, | 298 | 149 | 0.00648947466251 | 0.418560419864156 | 1.0,0.987,0.972,0. | 0.97,0.923,0.949,1 | 0,025 |
| 173122 | ENSG00000157601 | MX1 | chr21 | + | 41449136 | 41449435 | 41445999 | 41446141 | 41451166 | 41451827 | 173122 | 41,15,92,117,29,14,21,34, | 1,2,9,8,4,1,1,0,4,0,0,1 | 73,62,22,20,26,71,16,36,1 | 2,4,2,5,8,10,0,6,2,3,0,11,1 | 298 | 149 | 0.00649345357458 | 0.418691583675982 | 0.953,0.789,0.836, | 0.948,0.886,0.846, | 0,073 |
| 218469 | ENSG00000124942 | AHNAK | chr11 | - | 62536468 | 62536844 | 62535944 | 62536098 | 62546659 | 62546833 | 218469 | 285,191,394,260,327,359, | 2,1,7,1,0,3,1,2,3,2,0,0 | 232,186,130,293,181,91,8 | 2,0,0,9,1,0,3,3,1,0,3,9,3,2, | 298 | 149 | 0.00649922552422 | 0.418938247620064 | 0.986,0.99,0.966,0 | 0.983,1.0,1.0,0.94 | 0,014 |
| 82366 | ENSG00000145241 | CENPC | chr4 | - | 67518155 | 67518368 | 67514073 | 67514687 | 67519216 | 67519502 | 82366 | 123,99,185,81,59,75,56,88 | 4,0,5,5,1,0,0,2,1,0,0,2 | 88,109,65,92,67,35,45,71, | 2,3,3,5,6,1,0,3,0,1,3,2,1,0, | 298 | 149 | 0.00650464264867 | 0.419161898777213 | 0.939,1.0,0.949,0. | 0.957,0.948,0.915, | 0,039 |
| 234937 | ENSG00000147905 | ZCCHC7 | chr9 | + | 37144634 | 37144708 | 37126311 | 37126942 | 37147361 | 37147445 | 234937 | 13,2,10,5,10,4,1,14,0,0,1,5 | 22,11,57,18,19,21,18,47,5, | 14,4,2,5,2,3,2,2,2,7,6,5,5,8 | 66,52,64,15,14,15,24,20,1 | 223 | 149 | 0.00651731621237 | 0.419852884078003 | 0.283,0.108,0.105, | 0.124,0.049,0.02,0 | 0,025 |
| 253361 | ENSG00000089022 | MAPKAPK5 | chr12 | + | 111890471 | 111890639 | 111890039 | 111890144 | 111892966 | 111902222 | 253361 | 5,0,6,2,2,1,3,7,2,0,2,4 | 31,13,57,37,33,23,11,19,1 | 0,0,0,1,3,2,1,1,2,1,0,2,5,1, | 35,23,12,11,8,13,12,17,14, | 298 | 149 | 0.00654222322205 | 0.421331312479067 | 0.075,0.0,0.05,0.0 | 0.0,0.0,0.0,0.043,0 | 0,022 |
| 8139 | ENSG00000124155 | PIGT | chr20 | + | 45416516 | 45416654 | 45416106 | 45416343 | 45419294 | 45419395 | 8139 | 26,29,50,19,11,19,22,26,1 | 2,0,0,0,0,0,3,2,0,0,2,1 | 27,25,17,7,24,15,13,22,10, | 0,1,0,3,0,1,2,5,0,0,3,7,0,0, | 287 | 149 | 0.00655851447269 | 0.422254151502106 | 0.871,1.0,1.0,1.0,1 | 1.0,0.928,1.0,0.54 | 0,06 |
| 175194 | ENSG00000145016 | RUBCN | chr3 | - | 197699316 | 197699463 | 197696953 | 197697049 | 197700612 | 197701146 | 175194 | 7,4,3,0,0,0,0,3,0,0,0,0 | 45,52,48,41,32,28,28,22,2 | 2,1,1,15,4,0,2,5,0,1,3,0,4,3 | 34,35,13,54,44,23,40,31,2 | 296 | 149 | 0.00656474288303 | 0.422528760603298 | 0.073,0.037,0.031, | 0.029,0.014,0.037, | -0,014 |
| 173428 | ENSG00000145545 | SRD5A1 | chr5 | + | 6653446 | 6653636 | 6651841 | 6652008 | 6656077 | 6656179 | 173428 | 4,0,1,3,3,3,2,1,4,1,0,4 | 4,3,35,25,16,10,1,9,11,4,2, | 0,3,0,0,0,0,2,0,1,2,2,4,1,3, | 12,5,7,6,7,4,0,7,2,2,11,10,1 | 298 | 149 | 0.00656816533501 | 0.422622658660549 | 0.333,0.0,0.014,0. | 0.0,0.231,0.0,0.0,0 | -0,012 |
| 222650 | ENSG00000167483 | NIBAN3 | chr19 | + | 17537053 | 17537259 | 17533586 | 17533701 | 17537375 | 17537543 | 222650 | 1,0,1,4,4,0,1,3,1,0,1,4 | 173,90,161,87,108,74,74,8 | 6,3,5,3,1,1,1,3,2,2,4,6,14,1 | 128,111,120,73,88,55,73,8 | 298 | 149 | 0.00658738162184 | 0.423732436894873 | 0.003,0.0,0.003,0. | 0.023,0.013,0.02,0 | -0,009 |
| 13067 | ENSG00000185722 | ANKFY1 | chr17 | - | 4209823 | 4209947 | 4207932 | 4208082 | 4216982 | 4217118 | 13067 | 44,11,59,51,18,24,6,26,28, | 3,0,1,5,1,4,0,6,0,0,0,3 | 37,40,38,29,21,20,20,21,2 | 1,2,2,3,1,4,2,2,0,0,0,1,0,0, | 273 | 149 | 0.00659172324576 | 0.423867521731466 | 0.889,1.0,0.97,0.8 | 0.953,0.916,0.912, | -0,024 |
| 185898 | ENSG00000010322 | NISCH | chr3 | + | 52476542 | 52476599 | 52473733 | 52473829 | 52477573 | 52477642 | 185898 | 43,28,38,19,13,17,15,12,1 | 0,0,0,1,0,2,0,0,2,0,0,0 | 25,29,16,17,30,12,10,28,2 | 2,0,5,2,1,0,0,1,1,4,4,0,0,2,1 | 206 | 149 | 0.00659342038441 | 0.423867521731466 | 1.0,1.0,1.0,0.932,1 | 0.9,1.0,0.698,0.86, | 0,06 |
| 19082 | ENSG00000166783 | MARF1 | chr16 | - | 15624768 | 15624927 | 15622933 | 15623123 | 15625015 | 15625173 | 19082 | 71,36,114,106,60,71,44,45 | 5,0,2,0,2,4,2,4,0,2,0,0 | 63,69,63,40,46,36,42,73,5 | 1,3,3,0,2,0,1,0,0,0,0,0,0,0, | 298 | 149 | 0.0066153660232 | 0.425060067053847 | 0.877,1.0,0.966,1. | 0.969,0.92,0.913,1 | -0,025 |
| 165319 | ENSG00000258315 | C17orf49 | chr17 | + | 7016632 | 7016685 | 7016494 | 7016525 | 7016909 | 7017014 | 165319 | 7,1,14,16,25,13,18,22,7,6, | 2,2,1,2,3,1,2,2,2,0,1,3 | 9,1,3,2,10,3,6,28,7,5,5,13, | 0,4,2,1,0,0,4,6,2,3,4,12,0,4 | 202 | 149 | 0.0066159206835 | 0.425060067053847 | 0.721,0.269,0.912, | 1.0,0.156,0.525,0. | 0,128 |
| 109817 | ENSG00000205581 | HMGN1 | chr21 | - | 39347378 | 39347483 | 39345145 | 39345274 | 39348291 | 39348339 | 109817 | 6,0,6,1,3,0,0,1,1,2,0,0 | 23,15,23,16,6,8,6,12,4,10, | 0,4,1,0,0,0,0,0,2,0,1,2,1,0, | 13,9,2,5,16,6,3,6,5,20,9,39 | 254 | 149 | 0.00663313625746 | 0.426038958468927 | 0.133,0.0,0.133,0. | 0.0,0.207,0.227,0. | 0,021 |
| 149171 | ENSG00000140400 | MAN2C1 | chr15 | - | 75363209 | 75363263 | 75362641 | 75362748 | 75363998 | 75364188 | 149171 | 2,0,4,5,1,1,4,1,2,0,0,0 | 7,7,44,26,13,16,12,29,3,1, | 1,0,1,0,0,0,0,0,0,0,6,1,0,0, | 12,15,12,7,7,1,15,17,13,4, | 203 | 149 | 0.00665916513964 | 0.427496586227064 | 0.173,0.0,0.063,0. | 0.058,0.0,0.058,0. | 0,059 |
| 90933 | ENSG00000111679 | PTPN6 | chr12 | + | 6957918 | 6958073 | 6957653 | 6957785 | 6959926 | 6959994 | 90933 | 668,305,670,539,709,337, | 14,2,17,7,12,5,0,4,4,0,7,1 | 549,365,244,55,109,209,1 | 18,7,1,3,4,2,2,8,5,2,2,18,2 | 298 | 149 | 0.00665980297271 | 0.427496586227064 | 0.96,0.987,0.952,0 | 0.938,0.963,0.992, | 0,016 |
| 192942 | ENSG00000158006 | PAFAH2 | chr1 | - | 25985753 | 25985926 | 25984459 | 25984528 | 25988230 | 25988327 | 192942 | 6,0,2,1,2,2,5,2,0,0,3,4 | 4,3,3,4,15,2,6,2,6,4,4,3 | 0,0,0,5,2,1,4,2,0,0,1,1,0,1, | 4,2,4,3,2,5,0,1,2,4,3,1,9,2, | 298 | 149 | 0.00666443978598 | 0.427666678466387 | 0.429,0.0,0.25,0.1 | 0.0,0.0,0.0,0.455,0 | -0,005 |
| 163306 | ENSG00000118965 | WDR35 | chr2 | - | 19960553 | 19960604 | 19953833 | 19953978 | 19966723 | 19966909 | 163306 | 6,9,14,6,1,6,4,0,4,0,4,2 | 8,0,7,3,0,0,2,1,1,3,0,5 | 2,4,1,1,0,2,1,5,1,3,1,2,4,11 | 5,3,4,9,3,0,1,6,2,2,2,3,6,4, | 200 | 149 | 0.00667241761319 | 0.428051003071385 | 0.358,1.0,0.598,0. | 0.23,0.498,0.157,0 | 0,182 |

| | | | | | | | | | | | | | | | | | | | | | | |
|---|---|---|---|---|---|---|---|---|---|---|---|---|---|---|---|---|---|---|---|---|---|---|
| 251384 | ENSG00000108055 | SMC3 | chr10 | + | 110602473 | 110602591 | 110601965 | 110602178 | 110602824 | 110603002 | 251384 | 101,96,83,50,21,24,16,26,6 | 0,0,0,3,1,2,4,0,0,0,0,0 | 32,66,44,49,40,33,20,24,7, | 1,1,2,6,0,2,2,0,0,0,1,9,0,3,0 | 267 | 149 | 0.00667457124871 | 0.428061574621902 | 1.0,1.0,1.0,0.903,0 | 0.947,0.974,0.925, | 0,029 |
| 8574 | ENSG00000175354 | PTPN2 | chr18 | - | 12821335 | 12821425 | 12817155 | 12817365 | 12825809 | 12825944 | 8574 | 2,3,11,7,0,1,2,7,0,2,0,0 | 17,12,75,77,30,24,21,26,43 | 2,6,4,0,11,7,3,7,2,8,2,4,8,5 | 24,17,22,11,27,31,8,36,15, | 239 | 149 | 0.00667730177362 | 0.428109126612453 | 0.068,0.135,0.084, | 0.049,0.18,0.102,0 | -0,038 |
| 93535 | ENSG00000124641 | MED20 | chr6 | - | 41909344 | 41909522 | 41906803 | 41907287 | 41916784 | 41916939 | 93535 | 4,4,8,11,11,6,6,8,3,4,4,2 | 0,0,4,2,1,3,1,0,0,0,2,0 | 5,7,4,4,3,3,6,4,2,3,1,9,10,7 | 0,1,1,3,2,0,0,0,3,0,0,6,0,0,4 | 298 | 149 | 0.00670599611555 | 0.429820801055968 | 1.0,1.0,0.5,0.733,0 | 1.0,0.778,0.667,0. | 0,122 |
| 185240 | ENSG00000132185 | FCRLA | chr1 | + | 161711352 | 161711474 | 161710759 | 161710912 | 161711933 | 161712218 | 185240 | 135,113,378,327,302,153,1 | 12,5,31,20,13,0,13,16,14,9 | 369,249,73,39,86,97,120,1 | 36,37,3,5,12,5,15,17,0,11,1 | 271 | 149 | 0.00671289887895 | 0.430135140998895 | 0.861,0.926,0.87,0 | 0.849,0.787,0.93,0 | 0,029 |
| 99584 | ENSG00000129103 | SUMF2 | chr7 | + | 56074173 | 56074218 | 56072996 | 56073111 | 56074585 | 56074639 | 99584 | 61,28,191,131,69,62,57,89 | 0,1,18,7,2,4,0,0,0,2,4,1 | 80,123,62,32,48,28,49,79,3 | 0,1,1,2,2,4,1,0,0,1,2,3,0,7,4 | 194 | 149 | 0.00673997308782 | 0.431741412995413 | 1.0,0.956,0.891,0. | 1.0,0.99,0.979,0.9 | -0,009 |
| 172897 | ENSG00000189079 | ARID2 | chr12 | + | 45755930 | 45756048 | 45731216 | 45731314 | 45811417 | 45811551 | 172897 | 0,0,8,2,2,0,2,4,2,0,0,2 | 35,45,55,32,31,39,24,32,7, | 6,6,5,9,5,0,0,2,2,5,2,2,8,2,0 | 31,60,16,39,47,23,21,30,37 | 267 | 149 | 0.00674884095221 | 0.432180835163675 | 0.0,0.0,0.075,0.03 | 0.097,0.053,0.148, | -0,019 |
| 218898 | ENSG00000291118 | ZNF767P | chr7 | - | 149558621 | 149558839 | 149552800 | 149553016 | 149619923 | 149620037 | 218898 | 14,4,38,13,24,19,17,11,4,1 | 0,1,2,0,2,2,2,1,2,2,0,4 | 13,9,6,17,8,6,10,27,8,10,2, | 0,2,2,2,2,0,1,0,1,0,1,0,0,0,2 | 298 | 149 | 0.00675843843399 | 0.432666705111868 | 1.0,0.667,0.905,1. | 1.0,0.692,0.6,0.81, | -0,073 |
| 108827 | ENSG00000101901 | ALG13 | chrX | + | 111716952 | 111717004 | 111713224 | 111713297 | 111717845 | 111717927 | 108827 | 1,3,5,1,2,3,1,2,0,0,3,2 | 10,5,40,11,11,12,6,7,2,2,3, | 0,0,0,2,0,0,2,0,4,1,0,2,5,0,0 | 10,15,4,3,10,6,7,2,15,17,3, | 201 | 149 | 0.00678563229184 | 0.434278448945886 | 0.069,0.308,0.085, | 0.0,0.0,0.0,0.331,0 | 0,067 |
| 201488 | ENSG00000115446 | UNC50 | chr2 | + | 98610207 | 98610634 | 98609755 | 98610039 | 98610774 | 98610895 | 201488 | 4,1,8,6,2,2,1,5,0,0,0,6 | 13,11,41,18,5,9,10,13,12,7 | 1,2,3,1,2,2,1,1,0,5,4,0,4,2,0 | 23,21,20,4,4,7,8,16,3,11,19 | 298 | 149 | 0.00678930741828 | 0.43438449017355 | 0.133,0.043,0.089, | 0.021,0.045,0.07,0 | 0,039 |
| 200062 | ENSG00000197343 | ZNF655 | chr7 | + | 99563862 | 99563933 | 99561887 | 99562494 | 99571259 | 99571571 | 200062 | 49,18,27,12,15,12,23,30,14 | 0,0,2,0,4,3,0,3,0,0,1,0 | 15,22,17,45,17,6,13,15,10, | 3,7,1,0,0,0,0,2,2,2,3,2,3,0,3 | 220 | 149 | 0.00681644638431 | 0.435965583098071 | 1.0,1.0,0.901,1.0,0 | 0.772,0.68,0.92,1. | 0,048 |
| 172825 | ENSG00000186448 | ZNF197 | chr3 | + | 44641899 | 44642009 | 44632472 | 44632599 | 44646490 | 44648471 | 172825 | 19,6,16,14,5,4,6,5,7,4,3,0 | 3,0,5,3,0,3,0,1,0,0,0,1 | 6,12,3,2,5,5,6,6,2,5,10,6,4, | 0,0,0,1,2,0,4,7,0,1,2,3,2,2,4 | 259 | 149 | 0.00681807059721 | 0.435965583098071 | 0.785,1.0,0.648,0. | 1.0,1.0,1.0,0.535,0 | 0,047 |
| 9494 | ENSG00000072849 | DERL2 | chr17 | - | 5480386 | 5480459 | 5480053 | 5480144 | 5481100 | 5481142 | 9494 | 12,9,44,30,12,7,17,15,8,2,6 | 1,0,3,0,0,0,3,0,0,2,2,0 | 16,34,15,18,24,5,15,6,7,8,1 | 4,2,4,0,1,2,0,0,0,2,0,7,1,0,0 | 222 | 149 | 0.00682196573937 | 0.436085092976045 | 0.89,1.0,0.908,1.0, | 0.729,0.919,0.716, | 0,013 |
| 202423 | ENSG00000154222 | CC2D1B | chr1 | - | 52356760 | 52357126 | 52356383 | 52356442 | 52357525 | 52357698 | 202423 | 16,9,45,22,1,3,14,17,13,7,7 | 7,1,7,1,3,2,2,2,2,5,0,7 | 10,24,16,8,11,15,8,17,6,20 | 0,1,0,6,3,0,2,5,3,0,2,0,0,0,3 | 298 | 149 | 0.00682685979271 | 0.436268366996664 | 0.533,0.818,0.763, | 1.0,0.923,1.0,0.4,0 | -0,158 |
| 170468 | ENSG00000052802 | MSMO1 | chr4 | + | 165340721 | 165340813 | 165340220 | 165340375 | 165341750 | 165343164 | 170468 | 2,0,2,4,0,0,3,1,1,0,0,2 | 3,11,39,44,11,18,20,22,4,6 | 4,0,0,3,2,1,5,11,0,0,1,3,0,1 | 17,29,3,10,10,5,23,15,4,16 | 241 | 149 | 0.00684885231585 | 0.437543880318406 | 0.292,0.0,0.031,0. | 0.127,0.0,0.0,0.15 | -0,015 |
| 129729 | ENSG00000130177 | CDC16 | chr13 | + | 114265149 | 114265240 | 114262878 | 114263014 | 114272183 | 114272481 | 129729 | 129,65,204,103,52,31,29,6 | 6,5,19,11,9,8,9,11,0,1,8,8 | 82,65,63,45,24,52,51,45,74 | 6,23,13,22,19,13,5,7,15,23 | 240 | 149 | 0.00686017547014 | 0.438137218579734 | 0.93,0.89,0.87,0.8 | 0.895,0.637,0.751, | 0,07 |
| 136204 | ENSG00000105397 | TYK2 | chr19 | - | 10369756 | 10369874 | 10368294 | 10368418 | 10378213 | 10378426 | 136204 | 3,5,15,5,4,6,4,3,3,0,2,4 | 47,25,66,60,61,53,28,45,33 | 0,7,0,0,6,0,2,0,4,7,5,5,3,3,3 | 43,55,24,25,52,28,30,56,26 | 267 | 149 | 0.00686222891074 | 0.438138353808211 | 0.034,0.1,0.113,0. | 0.0,0.066,0.0,0.0,0 | 0,021 |
| 46225 | ENSG00000188612 | SUMO2 | chr17 | - | 75181056 | 75181188 | 75174751 | 75174823 | 75182813 | 75182956 | 46225 | 424,485,785,643,261,229,2 | 13,3,13,20,4,13,3,15,7,1,2, | 447,349,184,97,282,192,13 | 10,16,12,9,14,10,1,3,8,11,1 | 281 | 149 | 0.00686599959006 | 0.438249097796027 | 0.945,0.988,0.97,0 | 0.96,0.92,0.89,0.8 | 0,023 |
| 209396 | ENSG00000091490 | SEL1L3 | chr4 | - | 25784227 | 25784562 | 25782241 | 25782418 | 25788223 | 25788364 | 209396 | 103,76,258,108,87,99,66,7 | 6,2,2,8,1,0,0,1,2,0,1,0 | 84,49,97,68,66,37,16,30,70 | 2,2,0,1,0,0,0,0,0,0,1,2,0,0,2 | 298 | 149 | 0.00687077410159 | 0.438423830613495 | 0.896,0.95,0.985,0 | 0.955,0.925,1.0,0. | -0,023 |
| 160311 | ENSG00000166888 | STAT6 | chr12 | - | 57109940 | 57110108 | 57108162 | 57108299 | 57111128 | 57111433 | 160311 | 15,16,34,23,19,19,7,20,16, | 55,45,164,114,97,88,46,56 | 15,4,24,7,3,3,2,14,5,9,4,16 | 102,83,36,30,38,49,20,65,3 | 298 | 149 | 0.00687807717231 | 0.438759759298593 | 0.12,0.151,0.094,0 | 0.068,0.024,0.25,0 | 0,022 |
| 179577 | ENSG00000144218 | AFF3 | chr2 | - | 100137861 | 100137937 | 100129223 | 100129306 | 100142483 | 100142595 | 179577 | 10,13,117,37,12,31,19,12,4 | 109,73,157,70,52,96,29,17 | 22,55,21,18,10,5,5,13,12,9 | 107,123,100,137,63,34,27, | 225 | 149 | 0.00688108330002 | 0.438821463628624 | 0.057,0.105,0.33,0 | 0.12,0.228,0.122,0 | 0,06 |
| 9515 | ENSG00000029725 | RABEP1 | chr17 | + | 5335183 | 5335344 | 5331948 | 5332152 | 5338018 | 5338138 | 9515 | 513,181,373,247,114,98,77 | 8,1,21,11,7,5,11,16,0,1,1,5 | 177,139,194,195,130,103,7 | 22,16,9,9,9,8,0,3,9,4,15,2,6 | 298 | 149 | 0.00688441802604 | 0.438904080616888 | 0.97,0.989,0.899,0 | 0.801,0.813,0.915, | 0,019 |
| 39911 | ENSG00000168827 | GFM1 | chr3 | + | 158649040 | 158649157 | 158646742 | 158646947 | 158652095 | 158652246 | 39911 | 14,2,36,31,20,10,11,14,7,8 | 3,4,20,7,3,2,4,8,1,1,1,3 | 24,6,7,6,2,13,10,17,7,11,15 | 5,17,11,1,11,5,4,7,3,5,11,1 | 266 | 149 | 0.00689130356603 | 0.438932525637813 | 0.723,0.219,0.502, | 0.729,0.165,0.263, | 0,161 |
| 118790 | ENSG00000291075 | | chr9 | + | 41007439 | 41007609 | 40994632 | 40994699 | 41011616 | 41011730 | 118790 | 23,9,46,21,25,35,14,15,12, | 2,0,12,3,18,15,11,19,0,0,2, | 20,11,13,7,9,11,14,16,11,2 | 0,0,0,1,11,4,0,0,0,0,4,16,0, | 298 | 149 | 0.00689190766286 | 0.438932525637813 | 0.852,1.0,0.657,0. | 1.0,1.0,1.0,0.778,0 | -0,09 |
| 1771 | ENSG00000239900 | ADSL | chr22 | + | 40364879 | 40365056 | 40364275 | 40364365 | 40366435 | 40366489 | 1771 | 65,42,118,114,67,52,40,57 | 5,1,9,9,4,1,2,7,6,1,0,7 | 59,74,61,19,38,49,12,39,27 | 2,10,4,3,11,7,2,0,4,13,2,9,1 | 298 | 149 | 0.00689205181457 | 0.438932525637813 | 0.867,0.955,0.868, | 0.937,0.787,0.884, | 0,085 |
| 178071 | ENSG00000067066 | SP100 | chr2 | + | 230457702 | 230457743 | 230450171 | 230450255 | 230461261 | 230461414 | 178071 | 22,4,20,10,8,15,19,26,3,6,4 | 151,106,402,277,155,187,1 | 15,24,25,37,11,4,15,16,13, | 223,221,204,143,281,119,1 | 190 | 149 | 0.00689302162168 | 0.438932525637813 | 0.103,0.029,0.038, | 0.05,0.078,0.088,0 | -0,017 |
| 237267 | ENSG00000120910 | PPP3CC | chr8 | + | 22511171 | 22511231 | 22498000 | 22498112 | 22513292 | 22513432 | 237267 | 53,20,98,64,32,19,27,34,22 | 1,0,3,0,2,4,0,0,1,0,0,2 | 52,47,23,31,36,40,11,30,23 | 4,1,5,2,0,1,2,0,5,1,2,1,2,4,1 | 209 | 149 | 0.00689759311829 | 0.439093719148972 | 0.974,1.0,0.959,1. | 0.903,0.971,0.766, | 0,044 |
| 69069 | ENSG00000249915 | PDCD6 | chr5 | + | 313700 | 313850 | 311292 | 311402 | 314416 | 314954 | 69069 | 1,4,2,6,0,3,3,5,0,1,2,3 | 30,13,82,81,45,45,34,27,34 | 0,2,4,0,2,0,2,2,1,0,5,0,0,0,1 | 49,24,15,11,15,10,17,52,8, | 298 | 149 | 0.00689986834485 | 0.439108682356715 | 0.016,0.133,0.012, | 0.0,0.04,0.118,0.0, | 0,013 |
| 149688 | ENSG00000214013 | GANC | chr15 | + | 42327817 | 42328119 | 42327362 | 42327442 | 42329305 | 42329504 | 149688 | 2,5,25,18,19,7,13,19,9,3,5, | 24,29,27,39,35,31,20,18,15 | 25,36,13,22,23,16,17,12,24 | 33,21,16,9,20,19,20,14,15, | 298 | 149 | 0.00690311357903 | 0.439185349904879 | 0.04,0.079,0.316,0 | 0.275,0.462,0.289, | -0,072 |
| 197349 | ENSG00000163534 | FCRL1 | chr1 | - | 157802376 | 157802739 | 157801914 | 157802193 | 157807101 | 157807122 | 197349 | 125,48,313,15,108,48,145, | 2,0,6,3,0,0,1,0,0,0,1,0 | 207,172,195,88,37,31,131, | 2,1,3,1,3,2,2,0,2,0,0,3,8,6,6 | 298 | 149 | 0.00690921294962 | 0.439443502470349 | 0.969,1.0,0.963,0. | 0.981,0.989,0.97,0 | 0,009 |
| 233748 | ENSG00000130741 | EIF2S3 | chrX | + | 24074337 | 24074397 | 24073090 | 24073263 | 24076721 | 24078810 | 233748 | 1,0,21,30,0,0,0,3,3,4,0,0 | 260,155,437,371,113,109,9 | 1,1,0,0,4,15,0,0,2,4,0,0,0,0 | 239,202,233,136,133,122,6 | 209 | 149 | 0.00691229055682 | 0.439449227466838 | 0.003,0.0,0.033,0. | 0.003,0.004,0.0,0. | 0,002 |
| 241391 | ENSG00000257524 | ST6GALNAC6 | chr9 | - | 127905206 | 127905326 | 127896241 | 127896332 | 127905934 | 127906022 | 241391 | 60,47,18,36,31,21,33,14,63 | 5,0,1,3,0,0,2,6,0,4,0,0 | 30,44,20,25,34,13,5,14,20, | 0,0,3,0,0,0,2,0,0,1,0,0,0,2,0 | 269 | 149 | 0.00691338647408 | 0.439449227466838 | 0.869,1.0,0.909,0.8 | 1.0,1.0,0.787,1.0,1 | -0,023 |
| 163613 | ENSG00000105875 | WDR91 | chr7 | - | 135197043 | 135197119 | 135196143 | 135196337 | 135197992 | 135201343 | 163613 | 8,4,11,5,15,5,4,3,2,3,5,8 | 14,14,44,37,36,35,14,15,33 | 9,17,8,4,5,1,5,11,4,7,1,0,6, | 21,28,8,6,7,3,21,25,2,11,20 | 225 | 149 | 0.00691898230939 | 0.439675075710753 | 0.275,0.159,0.142, | 0.221,0.287,0.398, | -0,095 |
| 47363 | ENSG00000182179 | UBA7 | chr3 | - | 49809389 | 49809464 | 49808975 | 49809159 | 49809541 | 49809725 | 47363 | 51,40,40,22,27,35,37,43,27 | 26,13,26,9,7,8,8,11,3,5,14, | 20,42,35,23,10,6,29,64,21, | 10,15,29,24,5,6,11,35,10,1 | 224 | 149 | 0.00693137102064 | 0.440332324717641 | 0.566,0.672,0.506, | 0.571,0.651,0.445, | 0,111 |
| 98471 | ENSG00000198198 | SZT2 | chr1 | + | 43422479 | 43422632 | 43422082 | 43422225 | 43422768 | 43422883 | 98471 | 61,38,49,25,31,46,29,28,39 | 0,0,3,0,0,0,3,2,0,0,1,0 | 17,25,31,48,40,22,35,51,38 | 0,1,4,7,3,1,2,1,0,0,0,3,0,1,0 | 298 | 149 | 0.00693425932539 | 0.440340939024856 | 1.0,1.0,0.891,1.0,1 | 1.0,0.926,0.795,0. | 0,038 |
| 201770 | ENSG00000162664 | ZNF326 | chr1 | + | 90007608 | 90007750 | 90005132 | 90005244 | 90010087 | 90010286 | 201770 | 60,28,24,20,10,19,20,24,12 | 8,2,1,2,5,1,1,0,0,1,0,0 | 30,29,46,60,11,18,24,24,26 | 0,3,2,3,0,1,4,0,0,0,2,0,1,0,0 | 291 | 149 | 0.00693594741748 | 0.440340939024856 | 0.793,0.878,0.925, | 1.0,0.832,0.922,0. | -0,069 |
| 158775 | ENSG00000075568 | TMEM131 | chr2 | - | 97775590 | 97775726 | 97772296 | 97772424 | 97775842 | 97776018 | 158775 | 6,8,6,3,1,3,1,0,1,0,0,3 | 38,50,36,20,31,22,8,9,18,5 | 6,4,5,3,9,3,1,2,0,3,2,13,3,5 | 33,25,19,8,11,11,12,10,17, | 285 | 149 | 0.00693764431812 | 0.440340939024856 | 0.076,0.077,0.08,0 | 0.087,0.077,0.121, | -0,05 |
| 40774 | ENSG00000164327 | RICTOR | chr5 | - | 38946467 | 38946552 | 38945490 | 38945724 | 38947263 | 38947441 | 40774 | 40,33,73,40,24,37,38,41,22 | 5,0,2,1,2,0,4,1,0,0,3,0 | 46,63,30,46,40,22,13,39,28 | 1,5,4,6,6,1,0,3,1,2,2,4,3,2,1 | 234 | 149 | 0.00695105443876 | 0.440861045817849 | 0.836,1.0,0.959,0.9 | 0.967,0.889,0.827, | 0,03 |
| 66008 | ENSG00000137776 | SLTM | chr15 | - | 58894432 | 58894582 | 58894089 | 58894193 | 58897114 | 58897120 | 66008 | 247,161,220,163,87,79,90, | 3,1,5,13,5,10,0,8,0,1,0,2 | 130,139,148,115,100,67,58 | 2,17,2,3,4,1,2,4,2,3,4,16,16 | 298 | 149 | 0.00695123644647 | 0.440861045817849 | 0.976,0.988,0.957, | 0.97,0.803,0.974,0 | 0,033 |
| 231714 | ENSG00000100883 | SRP54 | chr14 | + | 35022915 | 35023080 | 35018965 | 35019074 | 35028087 | 35028183 | 231714 | 26,32,60,31,38,17,20,28,48 | 1,2,3,5,0,1,2,1,0,0,4,0 | 42,40,38,1,5,9,16,20,8,23,8 | 1,0,1,0,1,1,0,0,0,0,0,2,0,0,1 | 298 | 149 | 0.00695198363389 | 0.440861045817849 | 0.929,0.889,0.909, | 0.955,1.0,0.95,1.0, | -0,074 |
| 4792 | ENSG00000071626 | DAZAP1 | chr19 | + | 1425877 | 1425960 | 1422347 | 1422396 | 1428841 | 1428995 | 4792 | 199,110,129,104,70,60,65, | 15,2,6,2,7,5,6,3,12,3,0,2 | 78,62,89,51,29,18,41,64,20 | 11,5,3,4,4,1,1,6,4,9,2,12,9, | 232 | 149 | 0.00696056852395 | 0.441259192609783 | 0.895,0.972,0.932, | 0.82,0.888,0.95,0.8 | 0,061 |
| 248805 | ENSG00000110422 | HIPK3 | chr11 | + | 33348697 | 33348818 | 33348165 | 33348228 | 33349146 | 33349287 | 248805 | 72,44,88,47,22,31,43,38,38 | 0,0,2,6,4,1,6,3,1,0,0,1 | 61,58,40,35,28,41,34,42,26 | 6,1,5,10,1,3,1,3,1,5,12,3,1, | 270 | 149 | 0.00696236238322 | 0.441259192609783 | 1.0,1.0,0.96,0.812, | 0.849,0.97,0.815,0 | 0,064 |
| 235004 | ENSG00000147905 | ZCCHC7 | chr9 | + | 37260454 | 37260565 | 37126311 | 37126942 | 37302187 | 37302231 | 235004 | 2,2,8,3,0,1,2,3,3,2,1,0 | 84,43,152,67,88,72,38,60,8 | 8,3,5,2,0,1,1,3,2,5,6,8,4,3,4 | 151,54,120,29,22,61,33,59 | 260 | 149 | 0.00696670866611 | 0.441404672627472 | 0.013,0.026,0.029, | 0.029,0.031,0.023, | -0,013 |
| 197341 | ENSG00000163534 | FCRL1 | chr1 | - | 157802376 | 157802695 | 157801914 | 157802193 | 157807101 | 157807122 | 197341 | 125,48,313,15,108,48,145, | 2,0,6,3,0,0,1,0,0,0,1,0 | 207,172,194,88,37,31,131, | 2,1,3,1,3,2,2,0,2,0,0,3,8,6,6 | 298 | 149 | 0.0069748308685 | 0.441789235626287 | 0.969,1.0,0.963,0. | 0.981,0.989,0.97,0 | 0,009 |
| 119509 | ENSG00000148950 | IMMP1L | chr11 | - | 31456259 | 31456386 | 31433459 | 31433570 | 31509518 | 31509563 | 119509 | 4,5,15,9,3,4,0,6,12,2,3,12 | 0,2,14,8,2,0,1,0,4,0,0,4 | 2,4,3,3,1,1,0,0,6,3,6,4,12,7 | 13,4,4,0,3,2,1,2,0,2,2,5,2,3 | 276 | 149 | 0.00699001567277 | 0.442528087822559 | 1.0,0.574,0.366,0. | 0.077,0.351,0.288, | 0,244 |
| 127231 | ENSG00000122884 | P4HA1 | chr10 | - | 73043886 | 73043957 | 73030266 | 73030370 | 73046924 | 73047101 | 127231 | 23,14,27,18,11,8,11,10,18, | 2,1,4,1,0,2,0,1,0,0,0,1 | 28,10,3,5,2,5,11,20,1,19,8, | 2,1,0,5,0,2,1,0,0,2,3,0,0,2,4 | 220 | 149 | 0.00699060775909 | 0.442528087822559 | 0.886,0.905,0.821, | 0.905,0.871,1.0,0. | 0,075 |
| 48397 | ENSG00000143297 | FCRL5 | chr1 | - | 157546942 | 157547197 | 157544830 | 157545082 | 157549559 | 157549580 | 48397 | 325,254,619,210,310,152,2 | 6,4,28,21,12,7,9,12,4,2,0,7 | 584,259,409,111,66,88,126 | 19,2,3,2,2,5,6,1,9,4,0,6,6,2 | 298 | 149 | 0.00699514667461 | 0.442685214326076 | 0.964,0.969,0.917, | 0.939,0.985,0.986, | 0,003 |
| 246335 | ENSG00000162144 | CYB561A3 | chr11 | - | 61357202 | 61357382 | 61356529 | 61356728 | 61357732 | 61361264 | 246335 | 13,7,19,13,12,8,9,1,12,13,1 | 62,31,122,114,89,45,44,39 | 3,7,5,1,6,6,2,9,4,10,1,11,12 | 70,43,39,7,33,45,17,17,44, | 298 | 149 | 0.00699782927923 | 0.442724807054072 | 0.095,0.101,0.072, | 0.021,0.075,0.06,0 | 0,051 |
| 217591 | ENSG00000133872 | SARAF | chr8 | - | 30073944 | 30074051 | 30069873 | 30070059 | 30082846 | 30083124 | 217591 | 5,4,8,3,2,4,5,2,0,2,0,4 | 22,10,58,11,17,12,31,33,7, | 2,0,0,5,0,3,0,1,0,4,2,5,2,1,2 | 21,23,14,13,23,10,10,22,7, | 256 | 149 | 0.00700400055064 | 0.442985025716955 | 0.117,0.189,0.074, | 0.053,0.0,0.0,0.18 | 0,037 |
| 43277 | ENSG00000156976 | EIF4A2 | chr3 | + | 186784961 | 186785101 | 186784563 | 186784696 | 186785882 | 186785913 | 43277 | 705,488,1012,726,359,414 | 475,304,579,328,154,202,2 | 436,313,408,253,266,299,1 | 280,298,307,267,316,161,8 | 289 | 149 | 0.00700816373946 | 0.443074485861472 | 0.433,0.453,0.474, | 0.445,0.351,0.407, | 0,061 |
| 139622 | ENSG00000132670 | PTPRA | chr20 | + | 3034900 | 3035086 | 3027706 | 3027841 | 3035584 | 3035710 | 139622 | 7,3,5,1,0,2,3,2,8,0,0,4 | 62,33,70,64,20,27,15,26,37 | 5,0,10,9,3,2,4,2,3,5,4,10,4, | 31,38,22,22,33,22,18,34,16 | 298 | 149 | 0.00701093337758 | 0.443074485861472 | 0.053,0.043,0.034, | 0.075,0.0,0.185,0. | -0,029 |
| 124417 | ENSG00000073921 | PICALM | chr11 | - | 86023415 | 86023541 | 86022377 | 86022469 | 86026291 | 86026367 | 124417 | 19,7,24,8,5,1,3,4,5,3,0,1 | 172,139,150,72,26,30,38,4 | 12,17,10,25,6,5,4,11,1,2,14 | 87,83,74,96,54,40,50,82,30 | 275 | 149 | 0.00701159079707 | 0.443074485861472 | 0.056,0.027,0.08,0 | 0.07,0.1,0.068,0.1 | -0,02 |
| 183271 | ENSG00000198408 | OGA | chr10 | - | 101795364 | 101795480 | 101792838 | 101792943 | 101797979 | 101798154 | 183271 | 27,16,41,13,5,13,10,17,8,2 | 46,11,34,12,30,15,18,12,15 | 29,27,23,50,19,5,17,23,8,9 | 10,11,22,19,18,9,5,16,11,9 | 265 | 149 | 0.00701505281315 | 0.443163144123008 | 0.248,0.45,0.404,0 | 0.62,0.58,0.37,0.5 | -0,129 |
| 32289 | ENSG00000167766 | ZNF83 | chr19 | - | 52616717 | 52616815 | 52612407 | 52614797 | 52618935 | 52619062 | 32289 | 76,41,82,54,19,36,39,33,24 | 12,10,31,21,7,13,2,5,7,3,1, | 46,50,47,16,38,27,10,23,31 | 15,15,2,1,5,2,0,2,0,2,3,4,8, | 247 | 149 | 0.00702513395271 | 0.443559755825551 | 0.793,0.712,0.615, | 0.649,0.668,0.934, | -0,082 |
| 95769 | ENSG00000166295 | ANAPC16 | chr10 | + | 72223887 | 72224056 | 72216011 | 72216138 | 72230365 | 72230440 | 95769 | 288,137,389,297,148,138,1 | 48,26,42,56,20,26,19,18,53 | 248,159,186,106,90,86,90, | 33,11,25,9,15,19,5,14,12,3 | 298 | 149 | 0.00702545268855 | 0.443559755825551 | 0.75,0.725,0.822,0 | 0.79,0.878,0.788,0 | -0,042 |
| 3464 | ENSG00000028277 | POU2F2 | chr19 | - | 42110700 | 42110756 | 42099715 | 42099821 | 42117249 | 42117366 | 3464 | 8,2,3,3,0,4,1,2,0,1,0,4 | 125,62,41,36,49,26,36,33,6 | 0,2,0,0,0,0,0,3,0,0,3,0,0,0,0 | 26,31,36,33,10,10,22,26,22 | 205 | 149 | 0.00704309157422 | 0.444476677693085 | 0.044,0.023,0.05,0 | 0.0,0.045,0.0,0.0,0 | 0,027 |
| 147692 | ENSG00000185324 | CDK10 | chr16 | + | 89689251 | 89689516 | 89686688 | 89686797 | 89690552 | 89690624 | 147692 | 5,6,3,7,2,0,0,2,2,1,1,1 | 0,3,4,6,0,1,2,0,10,0,0,0 | 5,1,0,0,2,1,0,2,0,2,0,1,4,7,0 | 8,8,4,4,3,3,2,3,2,5,1,5,3,1,5 | 298 | 149 | 0.00704410585655 | 0.444476677693085 | 1.0,0.5,0.273,0.36 | 0.238,0.059,0.0,0. | 0,392 |
| 88017 | ENSG00000049089 | COL9A2 | chr1 | - | 40315237 | 40315465 | 40314351 | 40314387 | 40315589 | 40315664 | 88017 | 9,0,14,10,11,9,3,14,2,0,4,3 | 8,3,26,7,40,64,31,19,10,6,6 | 9,14,1,3,3,1,8,18,5,6,8,4,6, | 0,4,18,6,1,2,21,25,0,8,19,1 | 298 | 149 | 0.00705630378901 | 0.445115861902817 | 0.36,0.0,0.212,0.4 | 1.0,0.636,0.027,0. | -0,224 |
| 167219 | ENSG00000155229 | MMS19 | chr10 | - | 97469448 | 97469548 | 97468251 | 97468406 | 97469645 | 97469723 | 167219 | 9,8,3,2,4,4,7,3,1,1,0,2 | 11,7,12,14,6,9,6,11,5,5,6,2 | 1,4,3,4,3,1,3,1,0,1,2,0,1,4,1 | 6,16,6,7,12,0,9,7,5,6,15,15 | 249 | 149 | 0.007067413157 | 0.445686024375701 | 0.329,0.406,0.13,0 | 0.091,0.13,0.23,0. | 0,025 |
| 4923 | ENSG00000012061 | ERCC1 | chr19 | - | 45419410 | 45419489 | 45419097 | 45419197 | 45420323 | 45420382 | 4923 | 1,0,3,3,2,3,2,5,6,1,0,2 | 37,30,114,56,63,43,46,46,4 | 0,4,0,1,3,0,1,2,0,1,1,0,1,1,0 | 57,26,37,23,23,24,30,65,18 | 228 | 149 | 0.00707708632258 | 0.446108010020353 | 0.017,0.0,0.017,0. | 0.0,0.091,0.0,0.0,0.02 | 0,011 |
| 102974 | ENSG00000152457 | DCLRE1C | chr10 | - | 14945434 | 14945565 | 14945104 | 14945189 | 14949035 | 14949087 | 102974 | 5,1,7,6,0,8,0,2,3,4,1,4 | 21,15,51,28,17,21,12,18,15 | 15,4,3,3,3,0,0,4,7,7,6,5,9,8 | 32,37,6,11,12,7,2,17,28,18 | 280 | 149 | 0.0070790412493 | 0.446108010020353 | 0.112,0.034,0.068, | 0.2,0.054,0.21,0.1 | -0,039 |
| 217310 | ENSG00000168016 | TRANK1 | chr3 | - | 36850153 | 36850363 | 36847199 | 36847346 | 36851718 | 36851856 | 217310 | 2,6,6,8,0,3,0,1,2,0,0,1 | 16,11,39,40,12,26,10,12,8, | 1,3,0,6,0,0,0,4,1,0,0,0,1,2,0 | 12,11,12,4,7,24,4,21,4,19,6 | 298 | 149 | 0.00708032282631 | 0.446108010020353 | 0.059,0.214,0.071, | 0.04,0.12,0.0,0.42 | -0,004 |
| 116280 | ENSG00000158122 | PRXL2C | chr9 | - | 96645892 | 96646024 | 96639576 | 96641886 | 96651389 | 96651495 | 116280 | 13,5,27,40,37,11,20,10,13, | 19,5,60,36,20,22,17,16,7,5 | 26,12,15,7,1,5,7,17,5,15,20 | 17,44,27,36,20,7,14,20,19, | 281 | 149 | 0.0070837020829 | 0.446190308166418 | 0.266,0.347,0.193, | 0.448,0.126,0.228, | 0,148 |
| 73750 | ENSG00000138078 | PREPL | chr2 | - | 44359496 | 44360038 | 44346267 | 44346390 | 44361379 | 44361494 | 73750 | 33,16,46,22,11,13,13,17,16 | 1,4,5,4,4,2,2,5,10,0,6,4 | 38,16,19,16,17,13,20,25,5, | 3,1,4,3,0,2,0,2,0,2,4,0,1,8,4 | 298 | 149 | 0.00709463736119 | 0.446748359826298 | 0.943,0.667,0.821, | 0.864,0.889,0.704, | -0,133 |
| 82699 | ENSG00000239779 | WBP1 | chr2 | + | 74459477 | 74459562 | 74458831 | 74459117 | 74459642 | 74459745 | 82699 | 15,4,22,23,5,11,10,14,8,3,2 | 0,3,6,1,0,1,0,0,1,0,0,3 | 10,18,7,8,9,11,8,12,5,4,5,9 | 2,2,6,3,1,0,0,2,0,0,1,2,3,0,1 | 234 | 149 | 0.00710092603041 | 0.447013574276448 | 1.0,0.459,0.7,0.93 | 0.761,0.851,0.426, | 0,038 |
| 139667 | ENSG00000124243 | BCAS4 | chr20 | + | 50851941 | 50852120 | 50841765 | 50841900 | 50875996 | 50876093 | 139667 | 3,4,0,2,2,5,5,11,0,0,0,2 | 10,8,18,6,10,7,13,11,5,5,3, | 3,1,1,3,2,0,0,1,1,6,0,2,2,1,1 | 2,9,2,4,4,5,4,12,3,5,13,8,5, | 298 | 149 | 0.0071044938348 | 0.447107401332701 | 0.13,0.2,0.0,0.143, | 0.429,0.053,0.2,0. | -0,018 |
| 167667 | ENSG00000101109 | STK4 | chr20 | + | 44975247 | 44975441 | 44972137 | 44972158 | 44978442 | 44978571 | 167667 | 5,3,16,8,0,2,4,9,2,3,3,4 | 166,90,299,182,96,74,81,1 | 5,11,4,5,9,6,1,2,1,3,0,0,0,0 | 209,231,139,109,89,76,66, | 298 | 149 | 0.00712505285475 | 0.44827016982774 | 0.015,0.016,0.026, | 0.012,0.023,0.014, | 0,008 |
| 198009 | ENSG00000154305 | MIA3 | chr1 | + | 222657921 | 222658189 | 222654654 | 222654793 | 222658721 | 222658823 | 198009 | 10,3,16,5,4,4,5,1,4,6,3,4 | 86,79,101,54,36,34,25,39,9 | 16,11,10,12,6,7,0,7,3,2,1,1 | 38,34,51,30,36,30,15,27,21 | 298 | 149 | 0.00713459470966 | 0.448739320267338 | 0.055,0.019,0.073, | 0.174,0.139,0.089, | -0,027 |
| 130850 | ENSG00000134283 | PPHLN1 | chr12 | + | 42351884 | 42352049 | 42335882 | 42335974 | 42355160 | 42355222 | 130850 | 112,67,131,131,67,61,53,7 | 48,30,63,46,31,35,18,39,41 | 78,122,52,74,70,56,19,62,5 | 45,19,21,24,23,10,4,26,13, | 298 | 149 | 0.0071404320826 | 0.448975266599498 | 0.538,0.528,0.51,0 | 0.464,0.763,0.553, | -0,101 |
| 54445 | ENSG00000132549 | VPS13B | chr8 | + | 99274197 | 99274332 | 99192875 | 99193057 | 99275080 | 99275254 | 54445 | 139,89,238,129,75,182,99, | 0,0,1,0,0,5,2,0,0,0,0,0 | 149,205,102,163,140,104,1 | 2,5,2,4,5,0,0,2,0,0,8,1,3,1,0 | 284 | 149 | 0.00715064888926 | 0.44930976578818 | 1.0,1.0,0.992,1.0,1 | 0.975,0.956,0.964, | 0,023 |
| 47727 | ENSG00000072803 | FBXW11 | chr5 | - | 171914342 | 171914405 | 171910571 | 171910797 | 172006457 | 172006873 | 47727 | 26,25,74,49,14,11,22,15,24 | 2,1,14,6,2,7,7,3,3,2,1,5 | 47,52,24,36,22,13,4,13,19, | 5,1,3,4,6,4,0,3,1,5,0,2,1,1,2 | 212 | 149 | 0.00715131242691 | 0.44930976578818 | 0.901,0.946,0.788, | 0.869,0.973,0.849, | -0,089 |
| 17008 | ENSG00000005339 | CREBBP | chr16 | - | 3791980 | 3792094 | 3782683 | 3782926 | 3793385 | 3793626 | 17008 | 113,98,106,80,98,104,82,1 | 14,10,7,8,4,7,4,7,1,6,0,12 | 93,102,62,158,60,53,125,9 | 10,27,3,30,15,7,13,8,9,10,1 | 263 | 149 | 0.00715201461495 | 0.44930976578818 | 0.821,0.847,0.896, | 0.84,0.682,0.921,0 | 0,049 |
| 163565 | ENSG00000141084 | RANBP10 | chr16 | - | 67735635 | 67735758 | 67734968 | 67735042 | 67738012 | 67738035 | 163565 | 4,1,3,0,0,1,2,0,0,0,0,3 | 16,16,19,19,19,7,8,5,13,17 | 2,2,1,8,1,0,1,3,4,0,1,3,1,1,0 | 17,9,12,7,8,11,2,10,10,21,4 | 272 | 149 | 0.0071678464149 | 0.450172965195606 | 0.12,0.033,0.08,0. | 0.061,0.109,0.044, | -0,051 |
| 242148 | ENSG00000172340 | SUCLG2 | chr3 | - | 67408864 | 67409034 | 67400730 | 67400851 | 67495797 | 67495940 | 242148 | 8,6,17,13,6,9,14,12,4,12,2, | 35,29,48,34,20,16,23,19,29 | 15,6,7,10,9,1,9,2,7,2,4,4,6, | 23,31,31,10,26,24,20,15,17 | 298 | 149 | 0.00717351409689 | 0.450273761945024 | 0.103,0.094,0.15,0 | 0.246,0.088,0.101, | 0,047 |
| 2075 | ENSG00000054611 | TBC1D22A | chr22 | + | 46788979 | 46789037 | 46762686 | 46762848 | 46792519 | 46792576 | 2075 | 0,0,3,2,1,2,0,1,0,1,0,2 | 94,59,55,56,50,61,34,38,48 | 0,0,1,3,3,4,0,0,2,6,2,3,6,2,0 | 42,39,31,55,44,26,32,46,42 | 207 | 149 | 0.0071736354415 | 0.450273761945024 | 0.0,0.0,0.038,0.02 | 0.0,0.0,0.023,0.03 | -0,021 |
| 172962 | ENSG00000136100 | VPS36 | chr13 | - | 52449930 | 52450253 | 52442376 | 52442445 | 52450498 | 52450634 | 172962 | 2,0,3,2,0,0,0,2,4,3,3,1 | 31,18,58,49,27,15,25,17,18 | 5,3,5,0,0,6,0,0,2,6,4,4,6,2,0 | 35,44,17,10,24,27,13,16,15 | 298 | 149 | 0.00718361296237 | 0.450768571867014 | 0.031,0.0,0.025,0. | 0.067,0.033,0.128, | -0,011 |
| 198697 | ENSG00000223745 | CCDC18-AS1 | chr1 | - | 93326601 | 93326983 | 93325768 | 93325895 | 93337376 | 93339274 | 198697 | 7,8,25,11,4,6,7,11,4,0,2,8 | 26,13,45,34,17,15,16,24,11 | 5,6,7,3,7,4,2,9,7,3,6,5,2,4,1 | 6,15,13,6,38,8,14,21,16,42 | 298 | 149 | 0.00718588824027 | 0.450779921842481 | 0.119,0.235,0.217, | 0.294,0.167,0.212, | 0,02 |
| 154740 | ENSG00000129566 | TEP1 | chr14 | - | 20406232 | 20406378 | 20405450 | 20405585 | 20407872 | 20408463 | 154740 | 52,34,42,44,34,38,27,31,49 | 3,0,4,4,5,1,3,1,0,0,0,1 | 65,57,31,22,38,21,14,16,30 | 1,7,1,5,8,0,1,2,1,2,2,5,4,0,2 | 295 | 149 | 0.00719820143512 | 0.45131597279041 | 0.897,1.0,0.841,0.8 | 0.97,0.804,0.94,0. | 0,046 |
| 12792 | ENSG00000141698 | NT5C3B | chr17 | - | 41828789 | 41828952 | 41827425 | 41827626 | 41830800 | 41830890 | 12792 | 102,41,101,125,69,69,19,4 | 7,2,2,6,0,1,2,1,2,1,3,1 | 68,53,47,44,20,11,32,50,31 | 1,4,0,4,0,0,1,3,0,0,0,0,0,1,4 | 298 | 149 | 0.00719862721722 | 0.45131597279041 | 0.879,0.911,0.962, | 0.971,0.869,1.0,0.8 | -0,038 |
| 245832 | ENSG00000110075 | PPP6R3 | chr11 | + | 68583018 | 68583129 | 68575957 | 68576043 | 68587944 | 68588024 | 245832 | 9,6,24,13,3,3,1,4,4,2,1,3 | 0,1,4,1,2,2,2,0,2,0,0,0 | 14,10,2,7,7,2,5,2,8,8,18,14 | 5,0,2,2,4,2,0,0,3,2,2,8,3,2,2 | 260 | 149 | 0.00720716937457 | 0.451719939329667 | 1.0,0.775,0.775,0.8 | 0.616,1.0,0.364,0. | 0,095 |
| 200290 | ENSG00000176444 | CLK2 | chr1 | - | 155265859 | 155266033 | 155264644 | 155264774 | 155266728 | 155266895 | 200290 | 16,29,38,29,17,25,6,14,20, | 2,1,9,3,2,1,5,4,0,0,2,2 | 11,21,16,24,16,14,9,8,8,21 | 8,8,0,8,6,5,6,4,4,1,7,0,4,1,3 | 298 | 149 | 0.00721023802746 | 0.451727108007384 | 0.8,0.935,0.679,0.8 | 0.407,0.568,1.0,0. | 0,138 |

| | | | | | | | | | | | | | | | | | | | | | | |
|---|---|---|---|---|---|---|---|---|---|---|---|---|---|---|---|---|---|---|---|---|---|---|
| 111866 | ENSG00000111530 | CAND1 | chr12 | + | 67304604 | 67304746 | 67302322 | 67302615 | 67307396 | 67307492 | 111866 | 106,83,73,57,66,52,53,56,8 | 0,0,2,2,0,2,3,5,0,0,0,1 | 62,69,58,106,24,30,39,39,2 | 0,0,3,7,2,0,0,2,1,3,3,1,6,2,0 | 291 | 149 | 0.00721148135312 | 0.451727108007384 | 1.0,1.0,0.949,0.93 | 1.0,1.0,0.908,0.88 | 0,026 |
| 12999 | ENSG00000109046 | WSB1 | chr17 | + | 27301880 | 27301956 | 27294113 | 27294435 | 27303366 | 27303635 | 12999 | 38,18,97,81,18,24,13,39,11 | 4,3,37,11,32,20,10,15,5,2,0 | 26,38,14,6,14,22,4,41,9,20 | 15,20,23,21,20,11,3,12,11, | 225 | 149 | 0.00721920088655 | 0.452079088162072 | 0.863,0.799,0.635, | 0.534,0.557,0.287, | 0,185 |
| 173119 | ENSG00000157601 | MX1 | chr21 | + | 41449114 | 41449295 | 41445999 | 41446141 | 41451166 | 41451243 | 173119 | 44,25,64,91,34,28,28,29,45 | 1,2,9,8,4,1,1,0,4,0,0,1 | 44,47,19,35,16,70,7,52,32, | 2,4,2,5,8,10,0,6,2,3,0,11,16 | 298 | 149 | 0.00722141796597 | 0.452086390411195 | 0.957,0.862,0.78,0 | 0.917,0.855,0.826, | 0,077 |
| 207689 | ENSG00000112078 | KCTD20 | chr6 | + | 36475027 | 36475062 | 36470051 | 36470257 | 36479120 | 36479223 | 207689 | 1,1,6,3,1,1,1,1,0,0,0,2 | 5,8,17,7,8,7,2,9,2,7,3,2 | 4,2,5,0,1,2,0,0,1,2,0,3,4,4,0 | 21,17,5,2,3,0,2,3,5,0,5,9,16 | 184 | 149 | 0.0072245176045 | 0.452148923679599 | 0.139,0.092,0.222, | 0.134,0.087,0.447, | -0,085 |
| 91633 | ENSG00000196262 | PPIA | chr7 | + | 44799391 | 44799480 | 44796978 | 44799277 | 44801286 | 44801624 | 91633 | 135,163,89,69,42,27,31,53 | 17,4,19,20,3,9,11,7,7,2,4,1 | 48,70,62,38,87,43,18,21,40 | 14,17,4,24,16,7,1,19,9,0,8, | 238 | 149 | 0.00723398574568 | 0.45248201664764 | 0.833,0.962,0.746, | 0.682,0.721,0.907, | 0,085 |
| 130706 | ENSG00000151148 | UBE3B | chr12 | + | 109530546 | 109530658 | 109529889 | 109530072 | 109533465 | 109533558 | 130706 | 21,8,30,26,25,20,17,25,29, | 4,1,1,1,0,1,4,0,4,0,0,1 | 9,4,16,10,4,7,8,21,12,17,17 | 0,4,0,0,0,0,2,0,0,0,1,2,1,2,4 | 261 | 149 | 0.00723404444222 | 0.45248201664764 | 0.75,0.82,0.945,0.9 | 1.0,0.363,1.0,1.0,1 | 0,029 |
| 130798 | ENSG00000123349 | PFDN5 | chr12 | + | 53296243 | 53296275 | 53295508 | 53295639 | 53297849 | 53297924 | 130798 | 134,77,276,221,79,106,68, | 62,26,120,60,24,30,21,42,2 | 157,146,92,71,84,43,58,13 | 50,56,40,22,61,26,27,59,28 | 181 | 149 | 0.00723647252369 | 0.452502387491668 | 0.64,0.709,0.654,0 | 0.721,0.682,0.654, | 0,062 |
| 121338 | ENSG00000163131 | CTSS | chr1 | - | 150748164 | 150748285 | 150747776 | 150747879 | 150751780 | 150752008 | 121338 | 0,1,4,5,6,2,0,1,1,1,0,1 | 19,22,10,8,10,0,5,3,5,5,1,0 | 3,6,10,2,8,0,1,3,8,6,2,6,6,1 | 8,22,11,5,8,6,6,3,6,6,9,15,2 | 270 | 149 | 0.00724768953406 | 0.453072165583871 | 0.0,0.024,0.181,0.2 | 0.171,0.131,0.334, | 0,075 |
| 155138 | ENSG00000171858 | RPS21 | chr20 | + | 62387875 | 62387914 | 62387610 | 62387674 | 62388308 | 62388364 | 155138 | 41,16,575,385,9,4,6,7,1,3,0 | 29,9,22,14,15,6,11,12,6,2,5 | 277,336,52,40,99,107,1,1,8 | 11,19,15,19,8,4,1,17,6,13,6 | 188 | 149 | 0.00725465312807 | 0.453375797737408 | 0.528,0.585,0.954, | 0.952,0.933,0.733, | -0,21 |
| 86873 | ENSG00000119820 | YIPF4 | chr2 | + | 32298233 | 32298311 | 32292176 | 32292348 | 32301381 | 32301495 | 86873 | 14,15,55,40,15,20,11,19,8, | 7,2,17,7,3,2,7,3,0,6,1,3 | 50,36,12,7,9,6,23,24,9,15,1 | 5,4,6,1,1,2,3,1,0,1,2,7,0,0,2 | 227 | 149 | 0.00726411617707 | 0.453835410422918 | 0.568,0.831,0.68,0 | 0.868,0.855,0.568, | -0,076 |
| 95768 | ENSG00000166295 | ANAPC16 | chr10 | + | 72223887 | 72224056 | 72215999 | 72216072 | 72230365 | 72230440 | 95768 | 271,122,359,270,129,117,1 | 3,8,20,16,7,1,9,4,0,1,1,2 | 211,130,156,96,83,63,82,2 | 14,4,3,2,9,4,2,9,4,16,15,10 | 298 | 149 | 0.00726925400349 | 0.454024610686348 | 0.978,0.884,0.9,0.8 | 0.883,0.942,0.963, | 0,033 |
| 97679 | ENSG00000104299 | INTS9 | chr8 | - | 28812326 | 28812461 | 28796543 | 28796655 | 28813491 | 28813612 | 97679 | 18,6,36,22,22,19,18,21,13, | 2,0,5,1,0,1,2,7,0,0,0,1 | 16,6,8,6,8,5,12,21,8,15,13, | 0,4,0,0,0,2,0,1,0,0,1,0,3,0,0 | 284 | 149 | 0.00727221414041 | 0.454077726038464 | 0.825,1.0,0.791,0.9 | 1.0,0.44,1.0,1.0,1.0 | -0,015 |
| 190254 | ENSG00000165671 | NSD1 | chr5 | + | 177136872 | 177136974 | 177135940 | 177136030 | 177191883 | 177192019 | 190254 | 2,1,7,0,0,2,1,10,0,0,0,0 | 61,47,133,81,60,68,45,79,1 | 7,7,5,8,4,1,0,1,0,0,3,5,5,0,1 | 89,57,40,41,42,33,28,53,47 | 251 | 149 | 0.00727558029133 | 0.454156154780524 | 0.019,0.012,0.03,0 | 0.045,0.068,0.069, | -0,021 |
| 1191 | ENSG00000138942 | RNF185 | chr22 | + | 31187046 | 31187270 | 31160181 | 31160303 | 31192683 | 31192702 | 1191 | 15,30,42,25,14,14,5,3,20,1 | 4,0,9,4,4,7,1,3,2,2,0,3 | 21,23,18,9,13,11,11,11,1,1 | 2,0,2,1,1,1,3,7,1,0,0,2,0,1,0 | 298 | 149 | 0.00728378472818 | 0.454536465883128 | 0.652,1.0,0.7,0.75 | 0.84,1.0,0.818,0.8 | -0,11 |
| 203644 | ENSG00000231365 | WARS2-AS1 | chr1 | + | 119146358 | 119146836 | 119141746 | 119141813 | 119146952 | 119147069 | 203644 | 9,1,10,12,3,5,3,3,8,2,4,4 | 1,0,6,3,1,3,0,2,0,0,0,1 | 9,5,6,9,10,4,1,8,5,6,1,2,4,3 | 2,2,3,3,1,1,0,1,3,2,0,0,0,0,0 | 298 | 149 | 0.00728895128976 | 0.45458817289468 | 0.818,1.0,0.455,0.6 | 0.692,0.556,0.5,0.6 | -0,036 |
| 211938 | ENSG00000164985 | PSIP1 | chr9 | - | 15470975 | 15471231 | 15469937 | 15469993 | 15472631 | 15472750 | 211938 | 8,3,8,2,2,1,1,6,9,2,0,0 | 51,20,59,33,59,32,29,36,30 | 5,6,0,3,1,3,7,6,3,4,6,5,5,9,4 | 35,20,18,1,13,25,13,27,6,2 | 298 | 149 | 0.0072898470956 | 0.45458817289468 | 0.073,0.07,0.063,0 | 0.067,0.13,0.0,0.6, | -0,08 |
| 110238 | ENSG00000166169 | POLL | chr10 | - | 101585315 | 101585478 | 101584601 | 101584919 | 101587245 | 101587382 | 110238 | 9,1,12,5,9,13,3,10,4,1,5,4 | 1,2,2,0,1,2,0,2,2,2,0,0 | 4,3,4,4,5,1,2,6,1,2,8,10,2,7 | 0,0,0,0,0,0,0,0,0,0,0,1,0,0,0 | 298 | 149 | 0.00729094959756 | 0.45458817289468 | 0.818,0.2,0.75,1.0, | 1.0,1.0,1.0,1.0,1.0, | -0,241 |
| 71754 | ENSG00000167528 | ZNF641 | chr12 | - | 48345344 | 48345474 | 48344598 | 48344712 | 48347251 | 48347343 | 71754 | 17,8,15,8,8,4,8,1,13,12,4,6 | 13,10,6,4,2,3,4,1,5,1,2,2 | 10,15,6,11,2,5,4,5,7,20,8,1 | 4,4,7,0,0,2,5,2,0,0,0,3,8,7,2 | 279 | 149 | 0.0072979496374 | 0.454892846338615 | 0.411,0.299,0.572, | 0.572,0.667,0.314, | -0,195 |
| 61818 | ENSG00000054118 | THRAP3 | chr1 | + | 36296582 | 36296770 | 36293850 | 36293935 | 36300885 | 36301084 | 61818 | 312,214,689,557,235,252,2 | 4,1,8,2,8,3,4,6,1,5,3,0 | 366,462,170,110,160,147,1 | 1,3,1,0,1,3,0,6,0,1,4,0,1,1,4 | 298 | 149 | 0.00730686110155 | 0.455316450998178 | 0.975,0.991,0.977, | 0.995,0.987,0.988, | -0,012 |
| 45733 | ENSG00000173889 | PHC3 | chr3 | - | 170149085 | 170149244 | 170136365 | 170136665 | 170171372 | 170171450 | 45733 | 89,89,106,59,58,58,59,55,1 | 31,10,12,2,12,5,6,13,16,6,6 | 51,71,55,52,41,48,36,58,24 | 18,12,19,17,12,19,8,5,7,10 | 298 | 149 | 0.00731076542905 | 0.455427888295184 | 0.589,0.817,0.815, | 0.586,0.747,0.591, | 0,086 |
| 180112 | ENSG00000266094 | RASSF5 | chr1 | + | 206584549 | 206584684 | 206583271 | 206583379 | 206586825 | 206586842 | 180112 | 31,36,22,17,21,19,7,13,11, | 0,0,19,20,0,0,0,0,0,0,0,0 | 41,61,2,2,11,6,2,15,12,12,1 | 0,3,0,0,0,0,0,0,1,0,1,2,0,4,0 | 284 | 149 | 0.00731401927553 | 0.455498750778819 | 1.0,1.0,0.378,0.30 | 1.0,0.914,1.0,1.0,1 | -0,068 |
| 8586 | ENSG00000175354 | PTPN2 | chr18 | - | 12840713 | 12840962 | 12836790 | 12836891 | 12859163 | 12859254 | 8586 | 15,14,9,14,3,8,1,14,17,5,1, | 99,84,183,185,73,78,57,93 | 7,16,16,31,5,3,8,16,9,5,9,1 | 77,52,71,35,45,67,27,103,4 | 298 | 149 | 0.00732121908694 | 0.455815245965051 | 0.07,0.077,0.024,0 | 0.043,0.133,0.101, | -0,034 |
| 221793 | ENSG00000112159 | MDN1 | chr6 | - | 89668247 | 89668411 | 89668013 | 89668151 | 89670918 | 89671080 | 221793 | 1,1,0,0,0,0,0,1,0,0,0,2 | 48,24,28,22,11,17,10,11,21 | 3,5,0,4,7,0,1,3,2,0,1,2,0,0,0 | 23,23,17,7,12,6,10,6,10,19 | 298 | 149 | 0.0073296766287 | 0.456209841084942 | 0.01,0.02,0.0,0.0,0 | 0.061,0.098,0.0,0. | -0,05 |
| 166272 | ENSG00000138698 | RAP1GDS1 | chr4 | + | 98404455 | 98404602 | 98391951 | 98392080 | 98416744 | 98416888 | 166272 | 15,17,78,40,7,18,19,23,8,6 | 2,1,0,0,0,0,0,0,0,0,0,4 | 45,45,28,15,30,21,6,8,22,1 | 0,3,0,0,3,2,0,1,0,5,3,0,1,2,0 | 296 | 149 | 0.00734547535903 | 0.45706100231263 | 0.791,0.895,1.0,1.0 | 1.0,0.883,1.0,1.0,0 | 0,06 |
| 248377 | ENSG00000132274 | TRIM22 | chr11 | + | 5709052 | 5709126 | 5708576 | 5708603 | 5734181 | 5734235 | 248377 | 90,63,163,110,55,70,51,13 | 1,1,11,0,3,6,2,3,4,2,0,2 | 102,108,77,107,68,61,67,9 | 19,10,2,5,4,10,6,10,3,3,8,7 | 223 | 149 | 0.0073526726231 | 0.45737661310475 | 0.984,0.977,0.908, | 0.782,0.878,0.963, | 0,032 |
| 162670 | ENSG00000072364 | AFF4 | chr5 | - | 132889078 | 132889167 | 132888096 | 132888160 | 132892163 | 132892404 | 162670 | 20,17,71,27,16,32,16,40,6, | 14,3,3,1,3,2,3,8,1,0,0,0 | 39,52,6,38,27,7,22,15,30,2 | 3,11,1,17,7,4,7,6,3,2,8,3,7, | 238 | 149 | 0.00736162080102 | 0.457733762842256 | 0.472,0.78,0.937,0 | 0.891,0.747,0.79,0 | 0,122 |
| 46668 | ENSG00000136631 | VPS45 | chr1 | + | 150098849 | 150099080 | 150093526 | 150093648 | 150110495 | 150110627 | 46668 | 1,5,7,8,4,1,0,3,2,3,1,8 | 20,13,49,37,13,18,26,11,14 | 6,4,9,3,5,0,4,8,2,4,4,4,8,4,4 | 23,9,12,8,5,19,3,17,10,20,9 | 298 | 149 | 0.00736266749195 | 0.457733762842256 | 0.024,0.161,0.067, | 0.115,0.182,0.273, | -0,076 |
| 102628 | ENSG00000122566 | HNRNPA2B1 | chr7 | - | 26193574 | 26193694 | 26193250 | 26193373 | 26195846 | 26195909 | 102628 | 253,219,360,208,125,124,1 | 122,51,104,76,40,60,59,87 | 155,218,111,74,55,160,21, | 83,99,103,104,94,25,21,54 | 269 | 149 | 0.00736613244585 | 0.457816936890771 | 0.535,0.704,0.657, | 0.508,0.549,0.374, | 0,106 |
| 32180 | ENSG00000131503 | ANKHD1 | chr5 | + | 140472238 | 140472295 | 140464666 | 140464776 | 140482579 | 140482667 | 32180 | 36,17,42,37,21,32,14,30,25 | 11,8,23,14,19,12,15,7,7,3,2 | 37,33,23,32,30,7,26,23,33, | 9,38,22,43,23,12,24,6,14,1 | 206 | 149 | 0.00737018021436 | 0.457833750473045 | 0.703,0.606,0.569, | 0.748,0.386,0.431, | 0,121 |
| 252144 | ENSG00000241058 | NSUN6 | chr10 | - | 18643500 | 18643640 | 18642475 | 18642555 | 18648489 | 18648645 | 252144 | 5,4,11,16,3,2,3,8,2,3,4,6 | 39,24,44,33,21,11,11,10,18 | 1,5,2,4,7,0,1,2,5,6,11,5,1,4 | 22,18,29,11,15,22,19,17,27 | 289 | 149 | 0.00737065731883 | 0.457833750473045 | 0.062,0.079,0.114, | 0.023,0.125,0.034, | 0,05 |
| 237805 | ENSG00000182986 | ZNF320 | chr19 | - | 52883639 | 52883704 | 52881953 | 52881983 | 52888126 | 52888253 | 237805 | 1,1,7,6,3,2,2,0,1,1,1,2 | 4,0,7,8,0,3,3,8,15,10,0,9 | 4,0,1,0,1,1,0,1,0,1,5,0,0,4,1 | 6,14,3,2,0,3,3,2,2,6,2,1,1,0 | 214 | 149 | 0.00738068616602 | 0.458324427062507 | 0.148,1.0,0.41,0.3 | 0.317,0.0,0.188,0.0 | 0,151 |
| 162943 | ENSG00000126456 | IRF3 | chr19 | - | 49661449 | 49661567 | 49660712 | 49660828 | 49661947 | 49662329 | 162943 | 7,1,14,14,7,5,2,2,6,0,0,1 | 30,8,47,34,41,31,17,19,14, | 2,6,1,3,6,2,4,0,2,9,9,0,5,3,1 | 19,36,23,16,13,17,35,66,11 | 267 | 149 | 0.0073950563683 | 0.459084331469737 | 0.115,0.065,0.143, | 0.055,0.085,0.024, | 0,013 |
| 36732 | ENSG00000196850 | PPTC7 | chr12 | - | 110538143 | 110538273 | 110533244 | 110537095 | 110539821 | 110539945 | 36732 | 36,45,41,46,19,35,23,19,17 | 21,2,9,11,10,7,4,1,4,0,1,3 | 32,75,22,30,26,27,26,19,29 | 5,2,2,15,0,3,1,8,0,1,1,2,10, | 279 | 149 | 0.0074036578582 | 0.459485779838017 | 0.478,0.923,0.709, | 0.774,0.952,0.855, | -0,089 |
| 45829 | ENSG00000179262 | RAD23A | chr19 | + | 12948496 | 12948552 | 12948176 | 12948358 | 12948685 | 12948813 | 45829 | 75,55,75,57,35,37,50,26,34 | 10,6,5,1,3,6,6,5,1,0,2,3 | 33,24,28,34,19,15,34,46,24 | 3,0,3,1,4,0,1,0,7,1,2,7,2,1,0 | 205 | 149 | 0.00740907899761 | 0.459689674757739 | 0.845,0.869,0.916, | 0.889,1.0,0.872,0.9 | -0,041 |
| 83721 | ENSG00000079263 | SP140 | chr2 | + | 230250980 | 230251061 | 230248884 | 230248968 | 230253315 | 230253417 | 83721 | 38,23,84,84,76,25,20,30,31 | 1,2,4,5,7,0,2,0,3,5,0,0 | 76,34,25,7,42,43,4,16,23,4 | 4,2,0,1,7,1,0,0,0,1,0,1,1,0,1 | 230 | 149 | 0.00741122968409 | 0.459690598310194 | 0.961,0.882,0.932, | 0.925,0.917,1.0,0.8 | -0,024 |
| 239545 | ENSG00000083223 | TUT7 | chr9 | - | 86301739 | 86301843 | 86301275 | 86301601 | 86303085 | 86303201 | 239545 | 40,16,12,3,6,14,7,7,8,7,5,5 | 40,47,52,35,19,23,20,14,55 | 10,18,22,21,8,7,5,6,5,17,12 | 21,16,25,15,15,11,8,32,3,1 | 253 | 149 | 0.00742084972683 | 0.460035183916485 | 0.371,0.167,0.12,0 | 0.219,0.399,0.341, | -0,098 |
| 100580 | ENSG00000257923 | CUX1 | chr7 | + | 102198801 | 102198867 | 102196633 | 102197305 | 102200070 | 102200172 | 100580 | 1,5,10,5,4,6,5,13,14,1,4,10 | 1,0,3,2,0,4,0,2,3,2,0,1 | 6,19,2,5,4,10,10,4,3,7,9,12 | 1,1,0,4,3,1,0,3,0,1,0,0,0,1,3 | 215 | 149 | 0.00742105997072 | 0.460035183916485 | 0.409,1.0,0.698,0.6 | 0.806,0.929,1.0,0.4 | -0,061 |
| 176079 | ENSG00000179262 | RAD23A | chr19 | + | 12949122 | 12949159 | 12948688 | 12948813 | 12949274 | 12949408 | 176079 | 9,3,12,5,0,3,4,2,2,5,1,3 | 1,0,5,1,2,3,2,1,0,0,0,2 | 6,7,3,6,1,2,1,6,2,3,3,6,11,4 | 2,3,0,0,0,0,0,4,0,0,4,0,0,0,0 | 186 | 149 | 0.00742556055566 | 0.460181636612513 | 0.878,1.0,0.658,0.8 | 0.706,0.651,1.0,1.0 | -0,179 |
| 189301 | ENSG00000118308 | IRAG2 | chr12 | + | 25102419 | 25102678 | 25102197 | 25102241 | 25103836 | 25103899 | 189301 | 69,4,56,26,24,6,14,37,17,1 | 109,85,524,396,147,91,147 | 33,40,18,51,16,19,20,31,4, | 162,227,139,35,87,35,49,1 | 298 | 149 | 0.00743382847743 | 0.460561409621686 | 0.24,0.023,0.051,0 | 0.092,0.081,0.061, | -0,039 |
| 89654 | ENSG00000144233 | AMMECR1L | chr2 | - | 127869453 | 127869544 | 127866899 | 127866996 | 127873827 | 127874272 | 89654 | 9,7,7,9,4,2,0,4,2,5,1,2 | 2,1,1,2,4,0,1,0,1,1,1,2 | 4,3,0,0,2,2,5,3,5,12,1,6,3,2 | 1,0,1,2,1,0,3,0,0,1,0,1,0,0,2 | 240 | 149 | 0.00744055908799 | 0.46084574764523 | 0.736,0.813,0.813, | 0.713,1.0,0.0,0.0,0 | -0,124 |
| 8729 | ENSG00000141425 | RPRD1A | chr18 | - | 36040806 | 36040865 | 36033707 | 36033837 | 36067253 | 36067530 | 8729 | 20,9,7,9,16,7,4,12,11,2,1,6 | 10,19,48,36,13,13,12,25,11 | 7,9,5,4,7,0,2,2,1,2,2,18,8,1 | 40,27,17,10,5,12,17,26,5,1 | 208 | 149 | 0.00744534746512 | 0.461009660605651 | 0.589,0.253,0.095, | 0.111,0.193,0.174, | 0,097 |
| 27830 | ENSG00000187688 | TRPV2 | chr17 | + | 16431783 | 16431850 | 16428816 | 16428982 | 16431965 | 16432300 | 27830 | 8,7,36,46,18,13,10,16,9,17 | 3,0,1,4,0,1,0,0,2,0,1,2 | 21,16,8,3,3,15,8,14,12,16,5 | 3,0,0,0,1,0,1,3,2,0,6,3,0,0,2 | 216 | 149 | 0.00744874935565 | 0.461087653887232 | 0.648,1.0,0.961,0.8 | 0.828,1.0,1.0,1.0,0 | 0,017 |
| 73543 | ENSG00000143222 | UFC1 | chr1 | + | 161158120 | 161158211 | 161157616 | 161157693 | 161158411 | 161158856 | 73543 | 188,139,262,229,77,83,63, | 19,17,48,41,9,14,13,5,13,1 | 91,148,78,24,92,81,36,109 | 17,35,21,29,27,14,5,21,13, | 240 | 149 | 0.00746755373321 | 0.462118762953571 | 0.86,0.835,0.772,0 | 0.769,0.724,0.698, | 0,081 |
| 100241 | ENSG00000005471 | ABCB4 | chr7 | - | 87417311 | 87417515 | 87409235 | 87409392 | 87418536 | 87418620 | 100241 | 6,4,47,27,9,7,7,17,6,8,7,10 | 2,0,7,3,1,11,4,0,1,2,1,0 | 16,16,7,10,5,1,7,11,2,22,13 | 6,8,2,8,3,1,3,1,3,3,2,6,6,7,5 | 298 | 149 | 0.00747089955935 | 0.462192924132929 | 0.6,1.0,0.77,0.818, | 0.571,0.5,0.636,0.3 | 0,202 |
| 14319 | ENSG00000125814 | NAPB | chr20 | - | 23395138 | 23395185 | 23394921 | 23394999 | 23402992 | 23403072 | 14319 | 4,2,10,3,2,1,1,0,0,2,0,2 | 4,1,10,1,7,6,2,2,2,0,1,4 | 2,1,2,2,0,2,1,0,2,1,1,2,1,1,1 | 13,10,0,3,5,4,0,5,4,12,0,7,1 | 196 | 149 | 0.00747332501084 | 0.462210119370145 | 0.432,0.603,0.432, | 0.105,0.071,1.0,0.3 | 0,04 |
| 101315 | ENSG00000173715 | C11orf80 | chr11 | + | 66748378 | 66748517 | 66744735 | 66744919 | 66756352 | 66756505 | 101315 | 28,1,37,24,20,24,20,8,18,6 | 2,2,8,4,13,6,4,4,3,2,1,4 | 24,25,11,8,14,7,13,33,12,1 | 2,3,3,4,0,0,1,3,0,3,3,1,1,1,5 | 288 | 149 | 0.00747736760914 | 0.462327293272856 | 0.879,0.206,0.705, | 0.861,0.812,0.655, | -0,162 |
| 112523 | ENSG00000119397 | CNTRL | chr9 | + | 121157963 | 121158109 | 121157739 | 121157880 | 121158854 | 121159019 | 112523 | 113,87,104,59,27,32,16,22 | 9,9,13,7,1,7,3,5,5,4,0,2 | 47,52,70,61,40,36,14,29,34 | 0,4,14,1,2,0,0,5,3,0,6,0,7,3 | 295 | 149 | 0.0074814091642 | 0.462362887292961 | 0.864,0.83,0.802,0 | 1.0,0.868,0.716,0.9 | -0,072 |
| 69808 | ENSG00000223745 | CCDC18-AS1 | chr1 | - | 93315148 | 93315269 | 93309752 | 93309790 | 93324634 | 93324691 | 69808 | 3,1,1,0,1,4,2,1,0,0,3,1 | 1,4,10,5,3,4,10,11,3,1,7,3 | 0,0,0,3,2,4,5,5,2,10,7,2,0,3 | 1,4,2,1,8,1,2,6,3,6,9,0,3,3,0 | 270 | 149 | 0.00748223971659 | 0.462362887292961 | 0.623,0.121,0.052, | 0.0,0.0,0.0,0.623,0 | -0,244 |
| 188009 | ENSG00000135040 | NAA35 | chr9 | + | 85966595 | 85966674 | 85962012 | 85962180 | 85974966 | 85975033 | 188009 | 10,7,16,3,6,12,6,14,2,0,1,9 | 15,14,27,16,13,15,3,6,7,4,6 | 4,4,11,6,2,2,4,3,4,3,8,4,1,0 | 17,12,15,19,5,6,10,9,13,12 | 228 | 149 | 0.00749077104376 | 0.462757216567023 | 0.303,0.246,0.279, | 0.133,0.179,0.324, | 0,099 |
| 158713 | ENSG00000126777 | KTN1 | chr14 | + | 55668646 | 55668803 | 55667240 | 55667330 | 55670728 | 55670809 | 158713 | 5,3,5,5,6,6,3,14,0,4,3,3 | 118,74,108,72,54,55,56,71 | 4,6,5,6,1,1,1,4,3,4,0,2,4,2,2 | 80,79,63,28,50,36,33,67,42 | 298 | 149 | 0.00749472391271 | 0.462868557376323 | 0.021,0.02,0.023,0 | 0.024,0.037,0.038, | 0,015 |
| 39122 | ENSG00000242086 | MUC20-OT1 | chr3 | + | 195685817 | 195685939 | 195683749 | 195683865 | 195688532 | 195688797 | 39122 | 4,0,16,9,0,0,1,2,0,0,0,0 | 30,18,6,0,4,13,7,6,11,15,8, | 1,1,4,4,0,0,4,4,8,14,0,0,3,2 | 9,21,9,14,8,8,8,12,2,7,2,4,8 | 271 | 149 | 0.00752120960514 | 0.464371045474437 | 0.068,0.0,0.595,1.0 | 0.058,0.026,0.196, | -0,009 |
| 52893 | ENSG00000117523 | PRRC2C | chr1 | + | 171512031 | 171512200 | 171485529 | 171485735 | 171512994 | 171513172 | 52893 | 248,191,258,143,105,141,8 | 3,2,7,4,2,13,2,1,2,0,0,5 | 149,175,85,272,136,113,11 | 6,12,2,17,8,2,0,4,15,0,1,3,1 | 298 | 149 | 0.00752935143705 | 0.464616729458779 | 0.976,0.979,0.949, | 0.925,0.879,0.955, | 0,023 |
| 197767 | ENSG00000066027 | PPP2R5A | chr1 | + | 212347957 | 212348021 | 212347346 | 212347406 | 212348388 | 212348497 | 197767 | 10,5,4,8,4,0,3,2,2,1,1,3 | 11,14,47,10,31,10,18,4,9,2 | 3,6,1,1,3,2,0,2,2,1,2,1,1,1,2 | 28,17,8,10,4,5,13,5,7,16,16 | 213 | 149 | 0.00752950621589 | 0.464616729458779 | 0.389,0.2,0.056,0.3 | 0.07,0.198,0.08,0.0 | 0,056 |
| 215415 | ENSG00000010244 | ZNF207 | chr17 | + | 32361467 | 32361777 | 32360891 | 32360967 | 32362913 | 32362984 | 215415 | 31,23,95,53,31,43,52,61,27 | 2,2,9,15,8,11,11,6,25,8,5,3 | 67,71,35,36,43,33,24,49,35 | 3,0,4,3,3,7,4,10,0,7,9,13,8, | 298 | 149 | 0.00753229927612 | 0.46465586285428 | 0.886,0.852,0.841, | 0.918,1.0,0.814,0.8 | -0,101 |
| 85405 | ENSG00000168763 | CNNM3 | chr2 | + | 96825057 | 96825201 | 96816267 | 96817502 | 96826832 | 96826982 | 85405 | 52,45,44,27,36,44,33,29,12 | 1,1,3,0,1,3,0,0,0,0,0,0 | 36,35,29,28,63,35,19,47,35 | 1,0,2,2,7,2,0,1,2,0,1,0,1,3,2 | 293 | 149 | 0.00753567825591 | 0.464731107936322 | 0.964,0.958,0.882, | 0.948,1.0,0.881,0.8 | 0,041 |
| 34621 | ENSG00000136709 | WDR33 | chr2 | - | 127713582 | 127714021 | 127709692 | 127709856 | 127717154 | 127717263 | 34621 | 27,25,83,30,39,38,35,49,30 | 3,4,13,3,0,6,3,12,0,0,1,1 | 31,19,18,21,10,7,37,71,16, | 5,2,3,0,1,1,0,5,0,1,5,4,1,1,2 | 298 | 149 | 0.0075387418125 | 0.464786863089713 | 0.818,0.758,0.761, | 0.756,0.826,0.75,1 | -0,026 |
| 194076 | ENSG00000121486 | TRMT1L | chr1 | - | 185137441 | 185137506 | 185128668 | 185128747 | 185137605 | 185137796 | 194076 | 0,0,2,2,1,1,0,0,0,0,2,1 | 2,1,18,28,7,6,9,3,7,1,3,6 | 5,3,0,1,2,0,0,1,0,3,5,1,2,0,0 | 8,11,7,3,14,7,14,4,3,6,10,7 | 214 | 149 | 0.00754239171034 | 0.464878725718267 | 0.0,0.0,0.072,0.04 | 0.303,0.16,0.0,0.1 | -0,043 |
| 38772 | ENSG00000163681 | SLMAP | chr3 | + | 57865241 | 57865292 | 57864806 | 57864857 | 57871635 | 57871698 | 38772 | 11,16,29,5,9,5,6,8,17,4,2,2 | 7,3,5,5,3,4,4,1,10,0,0,1 | 15,12,9,11,10,12,4,7,6,6,9, | 2,2,3,0,0,0,2,0,2,5,1,1,3,3,1 | 200 | 149 | 0.00754789667523 | 0.465084840912232 | 0.539,0.799,0.812, | 0.848,0.817,0.691, | -0,123 |
| 89079 | ENSG00000153201 | RANBP2 | chr2 | + | 108762100 | 108762195 | 108758412 | 108758548 | 108771700 | 108771871 | 89079 | 48,26,20,27,31,22,19,27,31 | 0,0,3,6,2,5,5,0,0,0,0,0 | 36,48,35,18,20,4,8,24,9,18 | 3,2,0,1,0,0,0,0,1,1,2,0,0,0,4 | 244 | 149 | 0.00755084424834 | 0.465133302351021 | 1.0,1.0,0.803,0.73 | 0.88,0.936,1.0,0.9 | -0,032 |
| 72748 | ENSG00000168710 | AHCYL1 | chr1 | + | 110019050 | 110019119 | 110018551 | 110018650 | 110019547 | 110019626 | 72748 | 42,15,41,28,39,17,27,31,64 | 3,3,7,0,0,3,0,1,0,3,1,2 | 38,16,18,15,10,27,33,31,5, | 2,6,0,3,2,0,1,5,0,0,3,0,1,2,1 | 218 | 149 | 0.00755337246299 | 0.46515591089608 | 0.905,0.774,0.8,1.0 | 0.929,0.646,1.0,0.7 | 0,004 |
| 105752 | ENSG00000290683 | | chr1 | - | 247202015 | 247202095 | 247200066 | 247200193 | 247210690 | 247210856 | 105752 | 28,11,80,49,32,42,13,32,44 | 5,2,4,5,0,4,0,0,2,2,3,2 | 45,30,25,30,34,20,14,33,43 | 1,4,1,1,6,1,4,1,2,0,0,2,0,3,0 | 229 | 149 | 0.00756000174725 | 0.465430988576191 | 0.785,0.782,0.929, | 0.967,0.83,0.942,0 | -0,023 |
| 215871 | ENSG00000148832 | PAOX | chr10 | + | 133383959 | 133384302 | 133381459 | 133381659 | 133388955 | 133389068 | 215871 | 9,6,23,11,10,2,5,11,11,2,1, | 1,1,1,0,3,8,1,2,0,0,0,4 | 6,2,7,0,4,9,9,10,2,4,3,14,8, | 2,4,1,3,8,2,2,0,3,2,4,4,1,4,2 | 298 | 149 | 0.00757304058531 | 0.465970877662761 | 0.818,0.75,0.92,1.0 | 0.6,0.2,0.778,0.0,0 | 0,226 |
| 82662 | ENSG00000115825 | PRKD3 | chr2 | - | 37268300 | 37268372 | 37267429 | 37267536 | 37269614 | 37269687 | 82662 | 1,4,4,5,0,3,0,2,0,0,0,0 | 15,17,50,42,25,18,6,19,35, | 4,5,3,2,4,0,4,2,0,3,5,2,6,2,4 | 65,68,18,9,10,10,10,25,4,2 | 221 | 149 | 0.0075731011335 | 0.465970877662761 | 0.043,0.137,0.051, | 0.04,0.047,0.101,0 | -0,045 |
| 150545 | ENSG00000166783 | MARF1 | chr16 | - | 15624851 | 15624927 | 15622933 | 15623123 | 15625015 | 15625173 | 150545 | 32,12,38,49,25,33,9,18,25, | 5,0,2,0,2,4,2,4,0,2,0,0 | 31,36,32,19,29,14,1,9,15,3 | 1,3,3,0,2,0,1,0,0,0,0,0,0,0,0 | 225 | 149 | 0.00758183275759 | 0.466374806015677 | 0.809,1.0,0.926,1.0 | 0.954,0.888,0.876, | -0,018 |
| 124156 | ENSG00000167323 | STIM1 | chr11 | + | 4084672 | 4084765 | 4083262 | 4083498 | 4086476 | 4086543 | 124156 | 0,0,2,0,0,0,0,2,0,0,1,0 | 21,10,39,30,21,18,17,22,11 | 0,4,0,0,0,1,0,7,0,0,3,6,0,2,5 | 16,24,13,30,14,22,14,31,14 | 242 | 149 | 0.00759078992435 | 0.46672896243203 | 0.0,0.0,0.031,0.0,0 | 0.0,0.093,0.0,0.0,0 | -0,021 |
| 184736 | ENSG00000184677 | ZBTB40 | chr1 | + | 22501491 | 22501549 | 22491399 | 22491533 | 22502298 | 22502441 | 184736 | 28,35,19,17,3,3,3,14,3,1,1, | 0,0,2,0,0,1,0,4,0,0,0,0 | 13,22,15,4,16,8,7,4,15,10,1 | 2,1,2,2,4,0,1,2,0,0,2,1,0,1,2 | 207 | 149 | 0.0075919272664 | 0.46672896243203 | 1.0,1.0,0.872,1.0,1 | 0.824,0.941,0.844, | 0,11 |
| 109057 | ENSG00000177485 | ZBTB33 | chrX | + | 120252668 | 120252770 | 120250811 | 120250977 | 120253413 | 120258398 | 109057 | 2,2,6,4,0,3,0,0,1,0,0,3 | 8,5,22,13,4,6,12,8,2,4,2,8 | 1,3,5,4,0,0,0,4,2,1,2,3,0,1,2 | 29,23,3,8,7,5,4,5,10,10,13, | 251 | 149 | 0.00759612199305 | 0.466763298526583 | 0.129,0.192,0.139, | 0.02,0.072,0.497,0 | 0,005 |
| 190706 | ENSG00000132155 | RAF1 | chr3 | - | 12606043 | 12606299 | 12604135 | 12604289 | 12608765 | 12608923 | 190706 | 67,42,88,48,37,46,29,39,20 | 45,20,36,26,10,18,20,11,20 | 57,88,43,34,32,19,32,40,21 | 5,20,20,23,18,1,6,13,3,11,2 | 298 | 149 | 0.00759707192662 | 0.466763298526583 | 0.427,0.512,0.55,0 | 0.851,0.688,0.518, | -0,13 |
| 240085 | ENSG00000186350 | RXRA | chr9 | + | 134429107 | 134429220 | 134421675 | 134421805 | 134431904 | 134431996 | 240085 | 22,15,51,6,21,34,23,45,17, | 4,0,4,1,2,0,4,3,3,0,0,0 | 14,12,17,31,4,5,26,58,3,17 | 2,1,2,12,0,0,2,2,0,0,0,0,0,0 | 262 | 149 | 0.00760269421576 | 0.466763298526583 | 0.758,1.0,0.879,0.1 | 0.799,0.872,0.829, | -0,036 |
| 127029 | ENSG00000130640 | TUBGCP2 | chr10 | - | 133282222 | 133282342 | 133281272 | 133281436 | 133283077 | 133283184 | 127029 | 47,29,65,27,23,32,42,23,42 | 4,1,3,2,2,0,0,0,0,2,2,0 | 42,35,19,17,17,11,19,19,27 | 5,1,4,6,4,0,2,3,1,4,1,1,0,1,1 | 269 | 149 | 0.00760399678124 | 0.466763298526583 | 0.867,0.941,0.923, | 0.823,0.951,0.725, | 0,055 |
| 193963 | ENSG00000109184 | DCUN1D4 | chr4 | + | 51874720 | 51874824 | 51874270 | 51874385 | 51877762 | 51877972 | 193963 | 0,3,4,2,3,3,1,1,4,1,0,0 | 16,10,47,39,11,12,11,14,2, | 1,0,1,0,0,0,0,0,0,3,1,0,2,2 | 45,17,14,9,17,16,6,24,5,30 | 253 | 149 | 0.00760469900743 | 0.466763298526583 | 0.0,0.15,0.048,0.0 | 0.013,0.0,0.04,0.0, | 0,083 |
| 18558 | ENSG00000137871 | ZNF280D | chr15 | - | 56666394 | 56666535 | 56658423 | 56658486 | 56666799 | 56666986 | 18558 | 59,24,46,34,14,13,15,21,9, | 3,5,11,8,0,5,5,7,0,0,0,5 | 32,14,20,4,13,27,7,22,12,2 | 2,2,1,3,2,1,0,3,0,2,1,0,0,1,2 | 290 | 149 | 0.00760549775791 | 0.466763298526583 | 0.91,0.712,0.682,0 | 0.892,0.782,0.911, | -0,063 |
| 98340 | ENSG00000068885 | IFT80 | chr3 | - | 160366092 | 160366152 | 160357488 | 160357578 | 160375811 | 160375880 | 98340 | 46,30,43,15,17,6,3,7,32,13 | 4,2,7,0,0,2,1,1,1,0,1,0 | 11,6,13,12,12,12,11,25,14, | 5,3,5,2,3,1,0,0,1,2,1,4,0,4,4 | 209 | 149 | 0.0076282941603 | 0.468028899776377 | 0.891,0.914,0.814, | 0.611,0.588,0.65,0 | 0,071 |
| 121279 | ENSG00000149743 | TRPT1 | chr11 | - | 64224800 | 64224970 | 64224542 | 64224717 | 64225498 | 64225580 | 121279 | 43,15,55,42,27,19,34,31,52 | 2,0,4,2,2,6,0,4,0,3,0,0 | 43,25,34,24,19,27,13,41,11 | 0,0,0,2,0,1,2,2,1,0,0,5,1,1,3 | 298 | 149 | 0.00764124962769 | 0.46869016774504 | 0.915,1.0,0.873,0.9 | 1.0,1.0,1.0,0.857,1 | -0,034 |
| 46337 | ENSG00000127481 | UBR4 | chr1 | - | 19094905 | 19095025 | 19093948 | 19094139 | 19095544 | 19095652 | 46337 | 66,51,77,64,47,66,42,47,53 | 2,1,6,6,0,2,1,7,1,0,0,6 | 50,61,30,35,53,30,51,38,85 | 2,14,2,9,7,5,2,3,5,2,5,3,3,2 | 269 | 149 | 0.00764725206229 | 0.468924703310182 | 0.948,0.966,0.877, | 0.933,0.707,0.893, | 0,067 |
| 142378 | ENSG00000168961 | LGALS9 | chr17 | + | 27647269 | 27647437 | 27647029 | 27647118 | 27648835 | 27649560 | 142378 | 78,24,100,93,50,24,47,47,8 | 11,9,15,22,13,15,11,14,17, | 81,66,41,26,33,21,15,79,24 | 8,10,5,1,3,9,2,3,4,8,0,23,13 | 298 | 149 | 0.00767368027674 | 0.470411244557968 | 0.78,0.571,0.769,0 | 0.835,0.767,0.804, | -0,098 |
| 191881 | ENSG00000121892 | PDS5A | chr4 | - | 39848850 | 39848903 | 39845817 | 39845880 | 39849519 | 39849652 | 191881 | 24,12,41,28,7,14,13,18,21, | 7,6,12,7,2,0,2,5,0,3,3,6 | 27,22,16,8,16,12,10,12,7,2 | 6,4,1,0,0,2,3,1,1,1,8,4,3,2,2 | 202 | 149 | 0.00767739610954 | 0.470425959542306 | 0.717,0.596,0.716, | 0.768,0.802,0.922, | -0,107 |
| 188812 | ENSG00000082512 | TRAF5 | chr1 | + | 211366390 | 211366490 | 211365375 | 211365468 | 211369451 | 211369592 | 188812 | 21,3,42,20,25,15,18,20,17, | 13,21,56,36,39,25,12,21,23 | 46,13,25,30,7,7,6,23,14,32 | 36,20,15,9,30,8,7,15,21,34 | 249 | 149 | 0.00767829167672 | 0.470425959542306 | 0.492,0.079,0.31,0 | 0.433,0.28,0.499,0 | -0,109 |
| 206653 | ENSG00000152061 | RABGAP1L | chr1 | + | 174393994 | 174394073 | 174370978 | 174371072 | 174637374 | 174637488 | 206653 | 45,45,157,101,42,47,39,37 | 1,2,6,0,0,3,0,5,0,0,0,0 | 66,92,91,96,67,29,26,40,58 | 2,5,6,5,2,0,0,3,1,1,3,1,1,4,3 | 228 | 149 | 0.0076899879487 | 0.471008479279069 | 0.967,0.936,0.945, | 0.956,0.923,0.908, | 0,028 |
| 91847 | ENSG00000291118 | ZNF767P | chr7 | - | 149558607 | 149558839 | 149552800 | 149553016 | 149619923 | 149620037 | 91847 | 24,7,46,28,41,25,23,22,43, | 0,1,2,0,2,2,2,1,2,2,0,4 | 16,14,13,32,11,9,17,35,10, | 0,2,2,2,2,0,1,0,1,0,1,0,0,0,2 | 298 | 149 | 0.00770082316544 | 0.471537943306064 | 1.0,0.778,0.92,1.0, | 1.0,0.778,0.765,0.8 | -0,021 |

| | | | | | | | | | | | | | | | | | | | | | | |
|---|---|---|---|---|---|---|---|---|---|---|---|---|---|---|---|---|---|---|---|---|---|---|
| 175998 | ENSG00000116191 | RALGPS2 | chr1 | + | 178808889 | 178809123 | 178808044 | 178808128 | 178811314 | 178811404 | 175998 | 8,6,21,4,20,10,12,9,13,3,3, | 242,127,468,225,230,129,1 | 21,28,13,15,5,18,2,10,20,2 | 470,387,162,140,164,181,9 | 298 | 149 | 0.00770502120987 | 0.471660813430469 | 0.016,0.023,0.022, | 0.022,0.035,0.039, | -0,006 |
| 171751 | ENSG00000173226 | IQCB1 | chr3 | - | 121828469 | 121828532 | 121826050 | 121826180 | 121828860 | 121828972 | 171751 | 54,27,43,30,13,15,5,16,4,1 | 2,0,2,0,2,1,1,7,2,2,1,2 | 23,17,20,14,16,18,13,12,9, | 0,2,1,6,1,0,2,0,1,3,0,0,2,1,2 | 212 | 149 | 0.00771235669128 | 0.4719756164404 | 0.95,1.0,0.938,1.0, | 1.0,0.857,0.934,0. | -0,127 |
| 180110 | ENSG00000266094 | RASSF5 | chr1 | + | 206584252 | 206584684 | 206583271 | 206583379 | 206586825 | 206586842 | 180110 | 32,38,23,17,22,19,7,13,11, | 0,0,19,20,0,0,0,0,0,0,0,0 | 40,65,2,2,13,6,3,19,11,12,1 | 0,3,0,0,0,0,0,0,1,0,1,2,0,4,0 | 298 | 149 | 0.00772268076938 | 0.472473082624908 | 1.0,1.0,0.377,0.29 | 1.0,0.915,1.0,1.0,1 | -0,067 |
| 200867 | ENSG00000106028 | SSBP1 | chr7 | + | 141741748 | 141742095 | 141739123 | 141739190 | 141742168 | 141742229 | 200867 | 0,0,20,4,1,3,3,6,0,0,1,0 | 29,29,163,132,62,38,24,21 | 7,8,5,1,2,5,5,4,3,6,2,9,3,2,5 | 72,46,49,6,37,23,18,30,31, | 298 | 149 | 0.00772723790547 | 0.472617545220862 | 0.0,0.0,0.058,0.01 | 0.046,0.08,0.049,0 | -0,026 |
| 122082 | ENSG00000162144 | CYB561A3 | chr11 | - | 61357202 | 61357371 | 61356529 | 61356728 | 61357732 | 61357828 | 122082 | 17,7,22,16,19,10,9,1,14,9,1 | 62,31,122,114,89,45,44,39 | 3,8,6,1,6,9,2,9,5,11,3,11,1 | 70,43,39,7,33,45,17,17,44, | 298 | 149 | 0.00773528549329 | 0.47297535000179 | 0.121,0.101,0.083, | 0.021,0.085,0.071, | 0,046 |
| 14392 | ENSG00000157106 | SMG1 | chr16 | - | 18899986 | 18900044 | 18896792 | 18896956 | 18925949 | 18926158 | 14392 | 8,8,18,5,3,11,8,2,1,3,3,14 | 10,5,13,7,11,4,10,8,1,2,4,6 | 14,24,9,13,21,7,1,6,2,1,9,2 | 6,5,2,2,5,2,1,4,0,14,7,19,3, | 207 | 149 | 0.00774086137859 | 0.473181861793611 | 0.365,0.535,0.499, | 0.627,0.776,0.764, | -0,195 |
| 57603 | ENSG00000173230 | GOLGB1 | chr3 | - | 121702480 | 121702591 | 121699811 | 121699885 | 121714860 | 121714971 | 57603 | 26,36,29,22,14,16,6,18,15, | 0,0,3,2,2,4,1,2,0,0,1,0 | 7,29,20,54,21,19,22,23,20, | 2,2,1,3,1,1,0,6,6,1,0,1,0,0, | 260 | 149 | 0.00774323134905 | 0.473192341421743 | 1.0,1.0,0.847,0.86 | 0.667,0.893,0.92,0 | -0,027 |
| 215077 | ENSG00000170802 | FOXN2 | chr2 | + | 48334496 | 48334678 | 48328560 | 48328702 | 48346200 | 48346751 | 215077 | 4,4,4,0,2,0,0,6,0,0,0,2 | 13,12,45,33,26,20,10,23,8, | 0,0,0,2,2,1,0,2,0,2,0,2,0,1, | 63,68,15,13,22,18,10,26,9, | 298 | 149 | 0.00774689959922 | 0.473251879420802 | 0.133,0.143,0.043, | 0.0,0.0,0.0,0.071,0 | 0,026 |
| 132665 | ENSG00000234608 | MAPKAPK5-AS1 | chr12 | - | 111840332 | 111840477 | 111839762 | 111840003 | 111841326 | 111841470 | 132665 | 10,6,37,20,10,5,8,16,8,6,5, | 1,3,11,6,3,3,5,1,0,0,1,5 | 18,15,13,2,13,7,11,8,3,13,1 | 8,9,3,2,7,4,6,8,6,5,3,13,6,6 | 294 | 149 | 0.00774860323596 | 0.473251879420802 | 0.835,0.503,0.63,0 | 0.533,0.458,0.687, | 0,197 |
| 149409 | ENSG00000184056 | VPS33B | chr15 | - | 91006651 | 91006744 | 91006371 | 91006445 | 91006949 | 91007046 | 149409 | 7,6,15,12,7,12,9,4,10,3,2,4 | 3,1,10,3,3,2,1,6,6,0,3,4 | 5,5,6,2,6,10,1,2,1,3,6,2,5,7 | 14,8,8,3,2,1,0,5,0,3,14,6,4, | 242 | 149 | 0.00775091873271 | 0.473259004187207 | 0.59,0.787,0.48,0. | 0.18,0.278,0.316,0 | 0,152 |
| 192704 | ENSG00000158863 | FHIP2B | chr8 | + | 22094609 | 22094750 | 22094439 | 22094518 | 22096336 | 22097034 | 192704 | 1,1,2,0,4,4,2,3,1,0,2,2 | 3,8,9,3,10,4,3,13,3,7,3,11 | 3,2,2,1,1,0,1,0,2,1,0,0,0,1, | 4,4,3,5,2,2,9,5,4,6,9,4,3,9,1 | 290 | 149 | 0.00775762501278 | 0.473534143257417 | 0.146,0.06,0.102,0 | 0.278,0.204,0.255, | 0,045 |
| 18727 | ENSG00000103994 | ZNF106 | chr15 | - | 42449770 | 42451954 | 42446588 | 42446658 | 42466052 | 42466114 | 18727 | 11,7,12,10,3,1,8,6,4,1,4,2 | 0,0,1,0,0,0,1,2,0,1,0,0 | 7,10,3,4,8,5,7,2,2,2,2,5,6,7 | 1,0,0,0,3,3,0,4,2,0,0,2,0,1, | 298 | 149 | 0.00776434029147 | 0.473809675439008 | 1.0,1.0,0.857,1.0,1 | 0.778,1.0,1.0,1.0,0 | 0,113 |
| 3053 | ENSG00000129351 | ILF3 | chr19 | + | 10684415 | 10684476 | 10683916 | 10683970 | 10687345 | 10687708 | 3053 | 8,20,20,6,4,17,5,12,2,2,2,9 | 9,1,8,5,14,14,3,4,0,2,0,3 | 20,20,9,13,11,4,8,17,6,18,1 | 6,2,1,1,4,3,2,2,1,3,5,5,4,6,1 | 210 | 149 | 0.00777444921674 | 0.474292085988709 | 0.387,0.934,0.639, | 0.703,0.876,0.865, | -0,14 |
| 87698 | ENSG00000198176 | TFDP1 | chr13 | + | 113637619 | 113637896 | 113636533 | 113636700 | 113640119 | 113640310 | 87698 | 3,9,21,13,10,19,8,6,7,6,3,1 | 3,8,14,11,6,1,3,1,7,7,3,12 | 16,9,7,3,7,7,2,3,9,9,5,52,7, | 1,2,0,0,3,2,3,2,2,4,1,26,3,4 | 298 | 149 | 0.0077801273742 | 0.474372675712695 | 0.333,0.36,0.429,0 | 0.889,0.692,1.0,1. | -0,182 |
| 42697 | ENSG00000168297 | PXK | chr3 | + | 58390581 | 58390659 | 58382513 | 58382700 | 58394997 | 58395102 | 42697 | 21,12,60,51,10,2,12,13,7,5 | 8,0,2,2,0,2,4,0,1,1,1,0 | 29,16,9,4,18,19,4,4,11,14,7 | 1,0,1,8,2,0,0,0,0,1,0,8,0,0, | 227 | 149 | 0.00778017825158 | 0.474372675712695 | 0.633,1.0,0.952,0. | 0.95,1.0,0.855,0.2 | -0,084 |
| 193251 | ENSG00000070610 | GBA2 | chr9 | - | 35740571 | 35740628 | 35740208 | 35740362 | 35740824 | 35741064 | 193251 | 20,11,27,20,18,17,13,16,1 | 10,0,12,2,1,0,1,3,2,5,0,7 | 9,8,7,2,4,4,3,14,13,17,6,18 | 5,6,2,2,4,1,3,3,4,1,6,7,5,4, | 206 | 149 | 0.00778614547086 | 0.474602060560371 | 0.591,1.0,0.619,0. | 0.566,0.491,0.717, | 0,191 |
| 202996 | ENSG00000239305 | RNF103 | chr2 | - | 86616719 | 86616791 | 86612158 | 86612274 | 86620329 | 86620469 | 202996 | 5,0,4,2,1,0,0,2,0,0,0,1 | 12,12,42,17,3,8,4,10,14,4,4 | 5,5,2,6,1,3,2,0,0,0,4,2,1,0, | 19,38,7,8,18,6,20,12,7,6,1 | 221 | 149 | 0.00779139598807 | 0.47478764153689 | 0.219,0.0,0.06,0.0 | 0.151,0.081,0.162, | -0,052 |
| 29290 | ENSG00000006459 | KDM7A | chr7 | - | 140126636 | 140126823 | 140120441 | 140120529 | 140127441 | 140127583 | 29290 | 16,2,23,27,4,8,11,5,7,6,3,1 | 40,12,23,19,10,16,12,12,9, | 12,16,6,4,0,18,5,5,6,24,4,3 | 30,51,30,34,31,15,4,23,21, | 298 | 149 | 0.00781088407492 | 0.475776280206112 | 0.167,0.077,0.333, | 0.167,0.136,0.091, | 0,066 |
| 35815 | ENSG00000084676 | NCOA1 | chr2 | + | 24697657 | 24697798 | 24693251 | 24693347 | 24705085 | 24705233 | 35815 | 98,68,109,106,63,42,36,69 | 2,2,0,0,0,2,0,0,1,0,0,0 | 122,98,85,108,52,43,62,13 | 1,0,4,4,2,0,1,1,0,0,5,0,1,2, | 290 | 149 | 0.00781204089675 | 0.475776280206112 | 0.962,0.946,1.0,1. | 0.984,1.0,0.916,0. | 0,029 |
| 164988 | ENSG00000178950 | GAK | chr4 | - | 851689 | 851974 | 850935 | 851084 | 859605 | 859722 | 164988 | 34,22,33,16,18,24,23,13,2 | 2,3,6,1,0,1,0,1,0,6,0,1 | 15,11,12,12,18,11,23,24,6, | 0,2,1,4,0,0,0,2,0,1,1,5,0,1, | 298 | 149 | 0.00782841996983 | 0.476638941591649 | 0.895,0.786,0.733, | 1.0,0.733,0.857,0. | -0,018 |
| 29408 | ENSG00000164329 | TENT2 | chr5 | + | 79679578 | 79679670 | 79668720 | 79669028 | 79681981 | 79682061 | 29408 | 88,72,178,101,53,66,53,86 | 2,0,1,0,0,1,0,3,1,0,0,0 | 109,147,69,72,77,42,47,58 | 2,1,3,10,3,0,0,3,0,2,4,1,1,0 | 241 | 149 | 0.00783575684393 | 0.47695072999884 | 0.965,1.0,0.991,1. | 0.971,0.989,0.934, | 0,025 |
| 245600 | ENSG00000149743 | TRPT1 | chr11 | - | 64224800 | 64224938 | 64223876 | 64224717 | 64225498 | 64225580 | 245600 | 30,14,38,34,11,11,27,26,3 | 2,0,4,2,2,6,0,4,0,3,0,0 | 31,23,26,21,18,23,8,31,10, | 0,0,0,2,0,1,2,2,1,0,0,5,1,1, | 287 | 149 | 0.00783997478774 | 0.477072551184639 | 0.886,1.0,0.831,0. | 1.0,1.0,1.0,0.845,1 | -0,052 |
| 102466 | ENSG00000124486 | USP9X | chrX | + | 41150920 | 41151057 | 41148368 | 41148575 | 41152932 | 41153081 | 102466 | 47,27,174,112,75,120,66,8 | 0,0,1,3,1,1,0,2,0,0,0,0 | 139,143,60,92,85,72,68,81 | 5,4,1,4,6,0,0,1,3,0,1,0,2,0, | 286 | 149 | 0.00784481778847 | 0.477232328273088 | 1.0,1.0,0.989,0.95 | 0.935,0.949,0.969, | 0,025 |
| 32851 | ENSG00000102445 | RUBCNL | chr13 | - | 46371940 | 46372597 | 46368732 | 46368815 | 46387133 | 46387486 | 32851 | 159,170,1117,791,308,435 | 3,2,10,2,2,6,5,6,0,2,0,1 | 616,922,340,281,453,254, | 11,12,2,2,4,2,1,3,2,1,8,4,9, | 298 | 149 | 0.00784906394964 | 0.477347629278641 | 0.964,0.977,0.982, | 0.966,0.975,0.988, | 0,007 |
| 202344 | ENSG00000008869 | HEATR5B | chr2 | - | 37020686 | 37020836 | 37019808 | 37019877 | 37027922 | 37028174 | 202344 | 61,35,82,66,40,19,37,36,2 | 2,0,0,3,0,0,5,0,0,2,0,0 | 99,82,31,17,30,31,30,49,3 | 0,6,0,0,0,2,2,1,2,0,2,1,8,2,1 | 298 | 149 | 0.00785350276429 | 0.477347629278641 | 0.938,1.0,1.0,0.9 | 1.0,0.872,1.0,1.0,1 | 0,019 |
| 255789 | ENSG00000173442 | EHBP1L1 | chr11 | + | 65581538 | 65583765 | 65580927 | 65581326 | 65584240 | 65584398 | 255789 | 54,39,43,12,46,33,41,66,3 | 0,4,5,5,6,4,3,8,9,0,3,0 | 20,32,23,26,9,9,37,64,19,2 | 1,2,4,1,3,0,3,3,3,1,4,1,2,1, | 298 | 149 | 0.00785441322438 | 0.477347629278641 | 1.0,0.83,0.811,0.5 | 0.909,0.889,0.742, | -0,049 |
| 153622 | ENSG00000197555 | SIPA1L1 | chr14 | + | 71619204 | 71619388 | 71618756 | 71618887 | 71624047 | 71624236 | 153622 | 4,0,6,2,0,2,0,3,1,0,1,0 | 49,53,146,81,43,43,32,70, | 2,11,5,4,2,3,3,2,3,0,3,2,0,0 | 59,70,77,59,83,50,28,25,4 | 298 | 149 | 0.00785558447856 | 0.477347629278641 | 0.039,0.0,0.02,0.0 | 0.017,0.073,0.031, | -0,018 |
| 205812 | ENSG00000128815 | WDFY4 | chr10 | + | 48775715 | 48775806 | 48774457 | 48774672 | 48776749 | 48776984 | 205812 | 103,64,92,66,102,100,53,5 | 3,1,4,1,3,4,3,1,1,2,2,2 | 95,161,33,70,33,24,63,73, | 10,10,2,10,2,0,6,5,3,0,3,3,1 | 240 | 149 | 0.00787393031087 | 0.47832737644337 | 0.955,0.975,0.935, | 0.855,0.909,0.911, | 0,027 |
| 1921 | ENSG00000093000 | NUP50 | chr22 | + | 45178237 | 45178900 | 45171599 | 45171683 | 45181285 | 45181367 | 1921 | 13,14,39,30,29,11,14,29,3 | 2,5,5,4,0,1,1,1,0,0,0,0 | 19,25,9,11,11,14,13,10,8,2 | 3,5,6,2,5,0,4,0,3,2,0,0,3,3, | 298 | 149 | 0.00787693914076 | 0.478375137755337 | 0.765,0.583,0.796, | 0.76,0.714,0.429,0 | 0,138 |
| 81269 | ENSG00000101104 | PABPC1L | chr20 | + | 44938673 | 44938748 | 44938044 | 44938191 | 44939125 | 44939184 | 81269 | 11,20,49,28,11,22,23,15,4, | 3,0,11,2,0,1,4,3,1,0,0,2 | 23,21,18,20,12,4,19,15,14, | 2,2,1,2,2,0,2,0,2,3,1,0,0,0, | 224 | 149 | 0.00787933875809 | 0.478385884412544 | 0.709,1.0,0.748,0. | 0.884,0.875,0.923, | -0,037 |
| 63699 | ENSG00000110046 | ATG2A | chr11 | - | 64908990 | 64909150 | 64907747 | 64907890 | 64909270 | 64909367 | 63699 | 21,19,23,15,23,15,14,18,1 | 1,1,0,0,0,0,3,0,0,0,1,0 | 14,4,6,12,10,10,13,14,15,2 | 1,2,0,2,3,0,0,8,1,0,1,1,1,0,1 | 298 | 149 | 0.00788535780694 | 0.478506285634543 | 0.913,0.905,1.0,1. | 0.875,0.5,1.0,0.75, | 0,101 |
| 184737 | ENSG00000184677 | ZBTB40 | chr1 | + | 22501491 | 22501554 | 22491399 | 22491533 | 22502298 | 22502441 | 184737 | 27,36,19,16,4,3,5,14,3,2,1, | 0,0,2,0,0,1,0,4,0,0,0,0 | 15,20,17,5,17,8,8,4,14,9,1 | 2,1,2,2,4,0,1,2,0,0,2,1,0,1, | 212 | 149 | 0.00788761952457 | 0.478506285634543 | 1.0,1.0,0.87,1.0,1. | 0.841,0.934,0.857, | 0,101 |
| 74332 | ENSG00000109381 | ELF2 | chr4 | - | 139067683 | 139067770 | 139061864 | 139062057 | 139071865 | 139072003 | 74332 | 47,44,210,92,34,42,36,68, | 0,1,19,1,4,2,5,9,3,0,0,1 | 87,123,51,48,40,28,14,33, | 9,12,5,11,6,3,1,2,0,12,5,7,4 | 236 | 149 | 0.00788799151345 | 0.478506285634543 | 1.0,0.965,0.875,0. | 0.859,0.866,0.866, | 0,058 |
| 238078 | ENSG00000119397 | CNTRL | chr9 | + | 121171407 | 121171548 | 121169610 | 121169816 | 121173356 | 121173509 | 238078 | 68,43,112,58,34,36,35,42, | 4,0,6,6,0,0,0,0,3,3,0,0 | 59,40,65,56,43,21,8,48,14, | 0,0,0,2,3,3,0,0,0,1,0,0,0,0, | 290 | 149 | 0.00789154720513 | 0.478587093972199 | 0.897,1.0,0.906,0. | 1.0,1.0,1.0,0.935,0 | -0,029 |
| 126524 | ENSG00000150093 | ITGB1 | chr10 | - | 32932505 | 32932600 | 32929821 | 32930044 | 32935491 | 32935558 | 126524 | 215,222,80,64,34,30,74,62 | 4,5,12,2,2,0,12,17,2,0,0,3 | 34,47,1,28,47,36,15,37,43, | 9,3,0,3,7,4,2,4,4,6,7,12,5,1 | 244 | 149 | 0.00790542473074 | 0.479293653583634 | 0.97,0.964,0.803,0 | 0.698,0.905,1.0,0. | 0,06 |
| 145581 | ENSG00000180747 | SMG1P3 | chr16 | - | 21471653 | 21471683 | 21469746 | 21469919 | 21472672 | 21472745 | 145581 | 7,1,8,2,6,5,8,4,3,4,7,6 | 0,0,0,0,12,9,0,0,0,0,0,0 | 8,12,3,12,3,5,1,6,9,15,14,5 | 0,0,0,1,0,0,0,0,0,0,0,0,0,0, | 179 | 149 | 0.00792278871068 | 0.480211134043471 | 1.0,1.0,1.0,1.0,0.2 | 1.0,1.0,1.0,0.909,1 | -0,11 |
| 181631 | ENSG00000072210 | ALDH3A2 | chr17 | + | 19655278 | 19655387 | 19652546 | 19652632 | 19656365 | 19656574 | 181631 | 7,4,11,2,2,11,3,6,1,1,2,0 | 33,15,28,15,12,5,6,3,16,4, | 8,5,21,5,1,7,5,0,3,9,3,2,18, | 7,6,6,1,2,5,6,8,13,12,2,7,1 | 258 | 149 | 0.00792719920283 | 0.480343190209545 | 0.109,0.133,0.185, | 0.398,0.325,0.669, | -0,135 |
| 27329 | ENSG00000189091 | SF3B3 | chr16 | + | 70561629 | 70561784 | 70560349 | 70560591 | 70563875 | 70564033 | 27329 | 149,148,206,256,128,101, | 20,18,5,7,4,5,7,10,12,9,3,5 | 123,114,136,69,74,73,69,8 | 8,17,17,23,16,3,5,8,10,8,3, | 298 | 149 | 0.00794942752308 | 0.481554527222075 | 0.788,0.804,0.954, | 0.885,0.77,0.8,0.6, | 0,056 |
| 126525 | ENSG00000150093 | ITGB1 | chr10 | - | 32932510 | 32932600 | 32929821 | 32930044 | 32935491 | 32935558 | 126525 | 216,222,80,64,34,30,74,62 | 4,5,12,2,2,0,12,17,2,0,0,3 | 34,47,1,28,47,36,15,37,43, | 9,3,0,3,7,4,2,4,4,6,7,12,5,1 | 239 | 149 | 0.00796767042931 | 0.482523825033996 | 0.971,0.965,0.806, | 0.702,0.907,1.0,0. | 0,059 |
| 26554 | ENSG00000085511 | MAP3K4 | chr6 | + | 161108950 | 161109041 | 161108742 | 161108859 | 161109754 | 161109914 | 26554 | 5,1,15,11,4,3,3,3,5,2,5,4 | 21,3,16,11,7,9,3,3,4,8,1,8 | 11,16,4,11,5,8,2,14,2,16,8, | 1,4,7,1,7,5,4,2,5,13,7,8,9,6 | 240 | 149 | 0.00797105799758 | 0.482593188151094 | 0.129,0.171,0.368, | 0.872,0.713,0.262, | -0,151 |
| 102986 | ENSG00000134250 | NOTCH2 | chr1 | - | 119959391 | 119959502 | 119955039 | 119955232 | 119963573 | 119963807 | 102986 | 64,55,18,17,60,55,46,58,4 | 7,2,1,0,2,4,3,3,0,0,1,2 | 43,48,11,14,52,24,10,15,3 | 11,4,0,3,4,1,2,1,3,7,7,21,0, | 260 | 149 | 0.0079796550275 | 0.482977820929092 | 0.84,0.94,0.912,1. | 0.691,0.873,1.0,0. | 0,061 |
| 53330 | ENSG00000196187 | TMEM63A | chr1 | - | 225860859 | 225860997 | 225859195 | 225859349 | 225862217 | 225862351 | 53330 | 207,155,67,69,85,61,56,84 | 3,1,5,5,3,1,5,0,5,1,0,2 | 92,75,37,130,67,68,90,154 | 4,7,2,4,3,0,11,8,0,0,2,1,3,0 | 287 | 149 | 0.00800058703759 | 0.484108616442939 | 0.973,0.988,0.874, | 0.923,0.848,0.906, | 0,02 |
| 21040 | ENSG00000100697 | DICER1 | chr14 | - | 95141153 | 95141187 | 95140658 | 95140759 | 95141682 | 95141748 | 21040 | 5,12,10,2,4,5,2,8,5,2,2,1 | 15,21,38,12,11,5,9,16,11,4 | 4,9,4,4,4,0,1,5,2,0,2,5,4,2, | 23,41,17,13,27,7,3,13,17,1 | 183 | 149 | 0.00800550957335 | 0.484270329112337 | 0.213,0.318,0.176, | 0.124,0.152,0.161, | 0,085 |
| 91590 | ENSG00000024048 | UBR2 | chr6 | + | 42594190 | 42594304 | 42573733 | 42573993 | 42603587 | 42603718 | 91590 | 101,144,112,75,23,45,46,3 | 9,0,2,0,0,2,2,2,0,3,1,0 | 68,86,57,61,71,35,31,41,2 | 0,1,0,1,0,1,0,0,0,0,2,3,0,0,1 | 263 | 149 | 0.0080206445936 | 0.48504955226893 | 0.864,1.0,0.969,1. | 1.0,0.98,1.0,0.972, | -0,034 |
| 64661 | ENSG00000058673 | ZC3H11A | chr1 | + | 203802956 | 203803016 | 203795653 | 203795775 | 203816925 | 203817124 | 64661 | 26,24,75,25,20,30,29,65,4 | 3,0,2,1,1,0,4,4,0,5,0,2 | 37,43,31,29,19,13,25,46,1 | 7,0,3,11,1,2,0,2,4,0,4,1,2,1 | 209 | 149 | 0.00802469330385 | 0.485158079910376 | 0.861,1.0,0.964,0. | 0.79,1.0,0.88,0.65 | 0,029 |
| 21735 | ENSG00000197555 | SIPA1L1 | chr14 | + | 71650334 | 71650509 | 71624047 | 71624236 | 71658332 | 71658436 | 21735 | 211,167,236,120,58,84,10 | 6,6,7,2,1,2,5,7,7,4,0,2 | 120,116,167,195,171,83,6 | 0,2,6,5,3,1,2,0,2,3,4,0,2,0, | 298 | 149 | 0.00804132944549 | 0.486027345656349 | 0.946,0.933,0.944, | 1.0,0.967,0.933,0. | -0,018 |
| 88159 | ENSG00000148832 | PAOX | chr10 | + | 133383959 | 133384212 | 133381459 | 133381659 | 133388955 | 133389068 | 88159 | 9,6,29,15,17,6,11,13,13,2, | 1,1,1,0,3,8,1,2,0,0,0,4 | 10,2,7,0,4,9,12,10,3,7,9,20 | 2,4,1,3,8,2,2,0,3,2,4,4,1,4, | 298 | 149 | 0.00804805219264 | 0.48629711439475 | 0.818,0.75,0.935,1 | 0.714,0.2,0.778,0. | 0,201 |
| 172061 | ENSG00000163539 | CLASP2 | chr3 | - | 33700301 | 33700417 | 33696854 | 33696933 | 33717807 | 33718356 | 172061 | 0,0,3,0,0,0,0,6,0,0,1,0 | 13,16,24,8,20,16,19,18,14, | 3,4,3,2,1,1,0,0,2,1,3,3,1,1, | 21,21,10,14,14,8,10,14,13, | 265 | 149 | 0.00805128113042 | 0.486346341328001 | 0.0,0.0,0.066,0.0,0 | 0.074,0.097,0.144, | -0,028 |
| 38185 | ENSG00000100412 | ACO2 | chr22 | + | 41515766 | 41515992 | 41515376 | 41515535 | 41517526 | 41517631 | 38185 | 35,33,46,46,29,44,30,34,2 | 4,0,0,1,1,4,0,10,0,0,0,0 | 32,23,23,18,14,13,14,21,2 | 0,2,1,2,1,1,1,0,0,0,0,2,0,0, | 298 | 149 | 0.00805580730155 | 0.486346341328001 | 0.814,1.0,1.0,0.95 | 1.0,0.852,0.92,0.8 | -0,016 |
| 149436 | ENSG00000273173 | SNURF | chr15 | + | 24967015 | 24967240 | 24962113 | 24962209 | 24967931 | 24968821 | 149436 | 6,0,12,21,0,0,0,2,2,7,0,0 | 126,81,198,89,72,86,83,10 | 1,2,5,0,4,5,0,0,1,0,0,0,0,5, | 161,170,110,16,59,53,58,1 | 298 | 149 | 0.00805739433956 | 0.486346341328001 | 0.023,0.0,0.029,0. | 0.003,0.006,0.022, | 0,012 |
| 62485 | ENSG00000215305 | VPS16 | chr20 | + | 2865398 | 2865495 | 2865147 | 2865317 | 2866211 | 2866315 | 62485 | 75,27,90,53,52,50,30,44,6 | 4,0,4,1,2,3,0,0,8,1,0,4 | 41,35,31,18,21,18,24,28,2 | 0,5,1,1,2,0,2,1,0,0,1,2,1,0, | 246 | 149 | 0.00805790547447 | 0.486346341328001 | 0.919,1.0,0.932,0. | 1.0,0.809,0.949,0. | -0,003 |
| 90712 | ENSG00000082258 | CCNT2 | chr2 | + | 134945888 | 134945975 | 134942611 | 134942674 | 134946100 | 134946146 | 90712 | 26,9,38,25,15,11,10,19,12, | 26,20,33,28,13,15,8,26,13, | 24,6,9,12,6,6,27,16,6,3,27, | 34,45,37,16,30,6,13,17,30, | 236 | 149 | 0.00806689521623 | 0.486570233578997 | 0.387,0.221,0.421, | 0.308,0.078,0.133, | 0,094 |
| 24212 | ENSG00000132963 | POMP | chr13 | + | 28659818 | 28659919 | 28659177 | 28659187 | 28662409 | 28662507 | 24212 | 5,0,3,0,0,3,0,0,2,0,1,0 | 12,15,39,29,16,9,13,18,7,5 | 3,4,0,4,6,0,0,0,4,3,3,4,2,1,1 | 39,30,17,13,23,11,7,23,9,1 | 250 | 149 | 0.00806992927233 | 0.486570233578997 | 0.199,0.0,0.044,0. | 0.044,0.074,0.0,0. | -0,016 |
| 27838 | ENSG00000187688 | TRPV2 | chr17 | + | 16434889 | 16434969 | 16433231 | 16433698 | 16436788 | 16437003 | 27838 | 27,12,50,62,43,39,29,16,4 | 1,0,3,3,5,1,2,0,0,1,4,3 | 38,31,5,10,17,14,5,19,19,2 | 3,7,0,6,0,2,0,2,0,6,3,1,5,4, | 229 | 149 | 0.00807034604665 | 0.486570233578997 | 0.946,1.0,0.916,0. | 0.892,0.742,1.0,0. | 0,066 |
| 9270 | ENSG00000177427 | MIEF2 | chr17 | + | 18262713 | 18262867 | 18260596 | 18260737 | 18263085 | 18263248 | 9270 | 8,3,4,0,6,9,9,2,6,2,0,12 | 2,1,0,1,0,2,1,0,4,0,2,2 | 3,3,4,2,2,3,3,13,2,8,6,7,1,3 | 0,0,0,1,0,0,0,2,0,0,0,0,0,2,1 | 298 | 149 | 0.00807065772996 | 0.486570233578997 | 0.667,0.6,1.0,0.0,1 | 1.0,1.0,1.0,0.5,1.0, | -0,203 |
| 104266 | ENSG00000214021 | TTLL3 | chr3 | + | 9818821 | 9818920 | 9817644 | 9817759 | 9820545 | 9820741 | 104266 | 10,2,12,5,13,13,9,9,2,1,1,6 | 1,1,0,0,1,1,1,0,0,0,0,0 | 15,6,5,8,7,5,3,10,12,13,9,6 | 2,1,1,0,0,1,0,8,0,0,3,0,1,3, | 248 | 149 | 0.00807494687098 | 0.486608172603365 | 0.857,0.546,1.0,1. | 0.818,0.783,0.75,1 | 0,076 |
| 207615 | ENSG00000149499 | EML3 | chr11 | - | 62610635 | 62610768 | 62609628 | 62609696 | 62610878 | 62611007 | 207615 | 9,0,10,4,11,4,7,3,0,2,2,2 | 29,43,31,16,21,25,33,22,2 | 5,1,0,3,1,0,0,3,3,3,3,0,8,3, | 23,21,19,12,9,9,12,22,19,4 | 282 | 149 | 0.00807709018477 | 0.486608172603365 | 0.141,0.0,0.146,0. | 0.103,0.025,0.0,0. | 0,032 |
| 232045 | ENSG00000185101 | ANO9 | chr11 | - | 423941 | 424187 | 421140 | 421198 | 428087 | 428199 | 232045 | 6,1,14,20,9,6,1,8,6,2,5,5 | 15,6,36,22,11,10,17,11,4,4 | 1,8,2,0,1,1,2,0,12,7,2,17,2, | 11,16,6,20,18,7,3,8,21,22, | 298 | 149 | 0.00807871533932 | 0.486608172603365 | 0.167,0.077,0.163, | 0.043,0.2,0.143,0. | 0,078 |
| 191882 | ENSG00000121892 | PDS5A | chr4 | - | 39848850 | 39849131 | 39845817 | 39845880 | 39849519 | 39849652 | 191882 | 37,24,74,45,15,22,19,29,3 | 7,6,12,7,2,0,2,5,0,3,3,6 | 39,30,26,14,16,17,23,22,1 | 6,4,1,0,0,2,3,1,1,1,8,4,3,2, | 298 | 149 | 0.00808033047695 | 0.486608172603365 | 0.725,0.667,0.755, | 0.765,0.789,0.929, | -0,093 |
| 73370 | ENSG00000136059 | VILL | chr3 | + | 38002395 | 38002575 | 38001701 | 38001860 | 38004254 | 38004399 | 73370 | 27,11,33,11,28,27,21,22,2 | 3,5,6,5,2,9,5,2,5,0,1,5 | 22,20,15,10,9,11,12,22,10, | 11,5,5,2,5,2,9,11,1,1,11,8, | 298 | 149 | 0.00808442295516 | 0.486683189322522 | 0.818,0.524,0.733, | 0.5,0.667,0.6,0.71 | 0,125 |
| 38481 | ENSG00000025293 | PHF20 | chr20 | + | 35800093 | 35800173 | 35772025 | 35772079 | 35801490 | 35801605 | 38481 | 19,18,14,10,5,4,12,12,3,0, | 4,13,12,9,9,6,5,10,17,4,4,9 | 9,18,24,36,16,2,0,11,21,8, | 6,1,5,0,2,1,5,13,7,6,6,11,3, | 229 | 149 | 0.00808609858718 | 0.486683189322522 | 0.756,0.474,0.432, | 0.494,0.921,0.757, | -0,19 |
| 12702 | ENSG00000108848 | LUC7L3 | chr17 | + | 50749218 | 50749344 | 50746541 | 50746702 | 50750500 | 50750550 | 12702 | 61,39,44,31,36,26,35,49,4 | 160,115,279,183,198,161,1 | 38,25,20,29,14,27,17,31,1 | 194,220,120,89,123,143,3 | 275 | 149 | 0.00810738328818 | 0.487827848056547 | 0.171,0.155,0.079, | 0.096,0.058,0.083, | 0,024 |
| 179256 | ENSG00000285565 | | chr6 | + | 31836412 | 31836517 | 31835410 | 31835451 | 31837235 | 31837435 | 179256 | 10,11,40,17,12,8,9,12,11,9 | 141,82,213,111,84,77,76,9 | 11,11,9,4,8,7,2,16,2,6,18,2 | 129,92,62,61,68,35,48,117 | 254 | 149 | 0.00811228143725 | 0.487986150369132 | 0.04,0.073,0.099,0 | 0.048,0.066,0.078, | 0,016 |
| 143997 | ENSG00000007202 | KIAA0100 | chr17 | - | 28620508 | 28620573 | 28619859 | 28620015 | 28620987 | 28621170 | 143997 | 25,19,36,29,19,16,16,26,2 | 3,2,2,1,0,3,2,2,1,0,0,2 | 24,39,17,22,35,19,11,34,1 | 0,2,0,0,0,1,3,0,0,0,1,0,0,1, | 214 | 149 | 0.00811960436737 | 0.488290183736634 | 0.853,0.869,0.926, | 1.0,0.931,1.0,1.0,1 | -0,059 |
| 153096 | ENSG00000140563 | MCTP2 | chr15 | + | 94476052 | 94476135 | 94470332 | 94470442 | 94476695 | 94476793 | 153096 | 2,0,3,1,3,3,1,1,3,0,1,2 | 11,18,34,31,8,17,7,20,5,5, | 6,4,1,6,1,3,2,3,0,1,0,3,0,1,1 | 17,23,12,4,18,12,10,22,8,2 | 232 | 149 | 0.00812200764075 | 0.488298275565995 | 0.105,0.0,0.054,0. | 0.185,0.1,0.051,0. | 0 |
| 201788 | ENSG00000134186 | PRPF38B | chr1 | + | 108693589 | 108693636 | 108692326 | 108692867 | 108695701 | 108695770 | 201788 | 4,5,11,6,0,0,3,2,0,1,1,5 | 61,46,56,22,37,21,26,42,1 | 5,7,4,4,3,4,3,3,2,5,6,6,2,4, | 32,39,19,22,13,10,17,25,1 | 196 | 149 | 0.00813741226014 | 0.489016185971526 | 0.047,0.076,0.13,0 | 0.106,0.12,0.138,0 | -0,06 |
| 226586 | ENSG00000058091 | CDK14 | chr7 | + | 90735077 | 90735229 | 90726566 | 90726812 | 90790572 | 90790652 | 226586 | 20,8,8,13,12,21,6,19,2,11, | 7,1,2,3,2,4,1,5,4,2,3,2 | 14,4,4,6,13,12,3,14,5,11,6, | 8,10,7,2,7,1,1,0,7,16,11,5, | 298 | 149 | 0.00813931926915 | 0.489016185971526 | 0.588,0.8,0.667,0. | 0.467,0.167,0.222, | 0,15 |
| 167664 | ENSG00000101109 | STK4 | chr20 | + | 44975157 | 44975441 | 44972137 | 44972158 | 44978442 | 44978571 | 167664 | 5,3,16,8,0,2,4,9,2,3,3,4 | 166,90,299,182,96,74,81,1 | 5,11,4,5,10,6,1,2,1,3,0,0,0, | 209,231,139,109,89,76,66, | 298 | 149 | 0.00814299841556 | 0.489016185971526 | 0.015,0.016,0.026, | 0.012,0.023,0.014, | 0,008 |
| 220347 | ENSG00000149657 | LSM14B | chr20 | + | 62126105 | 62126439 | 62124616 | 62124780 | 62129784 | 62129952 | 220347 | 19,3,31,16,14,15,11,8,16,1 | 0,0,3,3,1,2,0,0,0,0,0,1 | 15,7,9,7,10,11,6,11,12,11,1 | 3,1,2,3,0,0,0,2,0,5,2,2,3,0, | 298 | 149 | 0.00814303706493 | 0.489016185971526 | 1.0,1.0,0.838,0.72 | 0.714,0.778,0.692, | 0,109 |
| 254250 | ENSG00000138777 | PPA2 | chr4 | - | 105383821 | 105383885 | 105370836 | 105370873 | 105386566 | 105386636 | 254250 | 4,0,7,6,0,2,7,3,2,2,0,1 | 9,6,31,35,20,21,8,15,6,9,2, | 2,1,1,6,0,4,0,2,2,1,3,1,0,3, | 21,8,10,8,10,12,9,15,10,27 | 213 | 149 | 0.00816339246355 | 0.490101847509716 | 0.237,0.0,0.136,0. | 0.062,0.08,0.065,0 | 0,041 |
| 131684 | ENSG00000076513 | ANKRD13A | chr12 | + | 110018344 | 110018488 | 110013124 | 110013249 | 110019138 | 110019328 | 131684 | 172,115,158,143,66,58,40, | 7,4,7,2,3,9,3,2,2,1,2,2 | 88,77,73,42,61,46,37,44,7 | 4,3,5,10,9,4,4,4,6,3,7,10,6, | 293 | 149 | 0.00819243514555 | 0.491708312552112 | 0.926,0.936,0.92,0 | 0.918,0.929,0.881, | 0,052 |
| 185160 | ENSG00000160691 | SHC1 | chr1 | - | 154965609 | 154965781 | 154963328 | 154963931 | 154965945 | 154966077 | 185160 | 9,2,9,13,0,16,2,3,13,3,3,11 | 3,2,0,3,1,0,1,3,0,1,0,0 | 6,11,1,4,5,2,3,3,3,10,5,15, | 1,1,0,2,1,1,0,3,3,1,3,0,0,2, | 298 | 149 | 0.00819524730351 | 0.491739969997703 | 0.6,0.333,1.0,0.68 | 0.75,0.846,1.0,0.5, | -0,032 |
| 34483 | ENSG00000132466 | ANKRD17 | chr4 | - | 73139530 | 73140283 | 73135116 | 73135265 | 73141740 | 73141792 | 34483 | 49,27,61,61,45,52,27,36,3 | 34,30,26,19,20,16,19,20,3 | 51,25,31,52,25,31,41,44,2 | 11,21,13,8,17,7,7,11,8,15,1 | 298 | 149 | 0.00819863451246 | 0.491806105003143 | 0.419,0.31,0.54,0. | 0.699,0.373,0.544, | -0,113 |
| 73876 | ENSG00000120798 | NR2C1 | chr12 | - | 95059905 | 95059984 | 95057730 | 95058489 | 95062507 | 95062738 | 73876 | 47,48,64,32,36,34,38,42,2 | 4,4,3,1,0,3,2,6,0,0,0,4 | 41,38,36,21,39,25,22,32,2 | 8,5,3,4,4,4,0,4,3,7,5,6,2,2, | 228 | 149 | 0.00820709673305 | 0.492176549721673 | 0.885,0.887,0.933, | 0.77,0.832,0.887,0 | 0,074 |
| 85269 | ENSG00000172765 | TMCC1 | chr3 | - | 129857249 | 129857405 | 129848342 | 129848412 | 129860939 | 129861012 | 85269 | 0,6,6,9,3,6,1,3,9,2,0,2 | 19,8,18,13,2,11,10,16,9,9, | 0,4,7,0,2,7,0,2,0,2,2,0,1,1,1 | 15,17,19,28,12,10,13,13,6, | 298 | 149 | 0.00821934385737 | 0.492773704112981 | 0.0,0.273,0.143,0. | 0.0,0.105,0.156,0. | 0,056 |
| 88597 | ENSG00000009954 | BAZ1B | chr7 | - | 73476867 | 73478569 | 73470344 | 73470483 | 73489193 | 73489391 | 88597 | 79,65,117,109,103,83,56,6 | 6,1,7,5,4,6,5,6,2,0,2,2 | 99,63,60,45,42,39,28,96,2 | 10,15,11,6,6,0,2,19,11,2,3, | 298 | 149 | 0.0082220495987 | 0.492798651400111 | 0.868,0.97,0.893,0 | 0.832,0.677,0.732, | 0,069 |
| 8731 | ENSG00000141425 | RPRD1A | chr18 | - | 36062977 | 36063075 | 36040806 | 36040865 | 36067253 | 36067576 | 8731 | 4,3,3,2,0,0,0,2,0,0,0,3 | 7,3,4,7,14,4,4,7,3,4,1,5 | 0,1,0,2,0,2,2,0,1,3,1,4,0,2, | 4,5,4,3,3,0,2,2,1,1,1,13,5,4 | 247 | 149 | 0.00822870532666 | 0.493036625035841 | 0.256,0.376,0.311, | 0.0,0.108,0.0,0.28 | -0,116 |
| 73747 | ENSG00000138078 | PREPL | chr2 | - | 44359496 | 44359750 | 44346267 | 44346390 | 44361379 | 44361806 | 73747 | 34,17,49,24,14,14,13,18,1 | 1,4,5,4,4,2,2,5,10,0,6,4 | 41,12,21,16,15,12,23,25,6, | 3,1,4,3,0,2,0,2,0,2,4,0,1,8, | 298 | 149 | 0.00823060151072 | 0.493036625035841 | 0.944,0.68,0.831,0 | 0.872,0.857,0.724, | -0,134 |
| 230655 | ENSG00000225648 | SBDSP1 | chr7 | + | 72830691 | 72830819 | 72829655 | 72829783 | 72831601 | 72831798 | 230655 | 29,12,42,27,8,23,9,21,8,4, | 7,4,6,6,3,2,2,2,5,5,0,1 | 27,23,15,32,17,8,9,14,7,7, | 0,4,0,11,0,0,0,0,0,3,1,10,1, | 277 | 149 | 0.00823386105216 | 0.493094643327059 | 0.69,0.617,0.79,0. | 1.0,0.756,1.0,0.61, | -0,134 |
| 143141 | ENSG00000108506 | INTS2 | chr17 | - | 61925451 | 61925573 | 61924960 | 61925099 | 61926351 | 61926662 | 143141 | 1,3,3,0,0,0,0,1,0,2,0,0 | 9,0,20,9,4,10,5,9,4,0,2,10 | 4,3,0,3,1,2,2,0,1,5,2,2,2,0, | 9,14,5,4,7,7,2,5,4,2,9,15,4, | 271 | 149 | 0.00824499885589 | 0.493624297288473 | 0.058,1.0,0.076,0. | 0.196,0.105,0.0,0. | 0,032 |

| | | | | | | | | | | | | | | | | | | | | | | |
|---|---|---|---|---|---|---|---|---|---|---|---|---|---|---|---|---|---|---|---|---|---|---|
| 230656 | ENSG00000225648 | SBDSP1 | chr7 | + | 72830691 | 72830821 | 72829655 | 72829783 | 72831601 | 72831798 | 230656 | 39,16,50,29,10,26,18,23,12 | 7,4,6,6,3,2,2,2,5,5,0,1 | 30,29,18,42,24,9,11,20,13, | 0,4,0,11,0,0,0,0,0,3,1,10,1, | 279 | 149 | 0.00826548174656 | 0.494712987150683 | 0.748,0.681,0.817, | 1.0,0.795,1.0,0.67 | -0,106 |
| 209153 | ENSG00000184209 | SNRNP35 | chr12 | + | 123458963 | 123459206 | 123458087 | 123458216 | 123465537 | 123466936 | 209153 | 2,6,14,8,1,0,6,3,8,6,0,3 | 15,9,25,9,11,11,13,17,14,7 | 9,6,6,7,0,2,1,6,1,10,7,19,13 | 4,9,8,6,5,6,4,13,2,3,8,23,8, | 298 | 149 | 0.00828099689662 | 0.495503820699588 | 0.062,0.25,0.219,0 | 0.529,0.25,0.273,0 | -0,118 |
| 64771 | ENSG00000271601 | LIX1L | chr1 | - | 145937603 | 145937699 | 145936907 | 145936985 | 145942712 | 145942853 | 64771 | 22,15,40,40,16,18,8,16,13, | 2,4,6,0,0,1,2,1,0,1,0,0 | 30,15,7,8,4,9,1,17,2,15,9,7 | 0,1,0,0,0,1,0,1,0,1,0,2,1,1,1 | 245 | 149 | 0.0083101274683 | 0.497052392548263 | 0.87,0.695,0.802,1 | 1.0,0.901,1.0,1.0,1 | -0,041 |
| 174411 | ENSG00000173145 | NOC3L | chr10 | - | 94361323 | 94361441 | 94358082 | 94358215 | 94361664 | 94361872 | 174411 | 5,3,3,2,0,3,0,0,0,2,0,0 | 17,11,27,29,7,7,5,2,18,1,3, | 4,1,6,2,0,0,0,3,1,8,4,1,4,0,1 | 12,11,4,2,5,5,3,3,4,14,5,12 | 267 | 149 | 0.008311972674 | 0.497052392548263 | 0.141,0.132,0.058, | 0.157,0.048,0.456, | -0,054 |
| 228642 | ENSG00000066422 | ZBTB11 | chr3 | - | 101654072 | 101654403 | 101652779 | 101652938 | 101654703 | 101655332 | 228642 | 4,0,1,2,0,0,0,0,0,0,0,1 | 21,23,37,19,15,8,5,13,13,4 | 2,7,0,5,2,0,0,0,0,3,3,2,4,0,0 | 36,37,11,20,13,10,19,16,11 | 298 | 149 | 0.00831560599672 | 0.497052392548263 | 0.087,0.0,0.013,0.0 | 0.027,0.086,0.0,0.1 | -0,026 |
| 165727 | ENSG00000125089 | SH3TC1 | chr4 | + | 8215171 | 8215390 | 8214474 | 8214580 | 8216110 | 8216257 | 165727 | 5,2,0,0,1,0,0,3,0,4,3,10 | 22,16,14,18,17,9,20,24,26, | 4,10,0,1,4,0,8,9,4,1,1,5,4,5 | 23,15,8,3,17,2,9,8,6,20,8,2 | 298 | 149 | 0.00831611461902 | 0.497052392548263 | 0.102,0.059,0.0,0.0 | 0.08,0.25,0.0,0.14 | -0,043 |
| 12120 | ENSG00000184381 | PLA2G6 | chr22 | - | 38127406 | 38127458 | 38126370 | 38126449 | 38128268 | 38128430 | 12120 | 4,1,1,3,0,1,1,3,2,0,1,3 | 5,1,7,11,5,6,5,0,4,1,5,5 | 3,2,1,0,1,0,0,2,2,1,0,4,0,0,3 | 1,1,1,14,1,3,6,6,1,7,9,11,3, | 201 | 149 | 0.00831905099396 | 0.49708985687979 | 0.372,0.426,0.096, | 0.69,0.597,0.426,0 | 0,015 |
| 164593 | ENSG00000185246 | PRPF39 | chr14 | + | 45108414 | 45108520 | 45107450 | 45107616 | 45109615 | 45109780 | 164593 | 95,63,79,49,36,27,20,26,41 | 0,2,1,1,1,2,0,0,0,1,0,0 | 59,76,45,44,54,25,24,26,40 | 0,4,2,2,3,0,0,0,4,4,0,0,2,2,1 | 255 | 149 | 0.008323084707 | 0.497115216214091 | 1.0,0.948,0.979,0.9 | 1.0,0.917,0.929,0.9 | 0,019 |
| 201172 | ENSG00000181722 | ZBTB20 | chr3 | - | 114741555 | 114741708 | 114693527 | 114693575 | 114801100 | 114801177 | 201172 | 2,3,2,1,3,3,1,0,4,0,0,3 | 35,40,17,14,18,22,26,27,26 | 0,1,4,5,2,2,4,1,4,7,3,3,0,2,1 | 17,27,13,13,13,8,11,12,14, | 298 | 149 | 0.00832485613129 | 0.497115216214091 | 0.028,0.036,0.056, | 0.0,0.018,0.133,0.1 | -0,054 |
| 109939 | ENSG00000167302 | TEPSIN | chr17 | - | 81237550 | 81237660 | 81237386 | 81237459 | 81238513 | 81238873 | 109939 | 11,5,14,14,6,8,4,4,1,4,8,8 | 13,6,15,3,9,2,6,3,10,3,1,5 | 3,15,2,15,9,7,0,4,5,6,8,6,14 | 7,2,0,3,4,1,1,4,3,3,6,0,1,4,2 | 259 | 149 | 0.00832640444198 | 0.497115216214091 | 0.327,0.324,0.349, | 0.198,0.812,1.0,0.1 | -0,174 |
| 95845 | ENSG00000167483 | NIBAN3 | chr19 | + | 17549443 | 17549527 | 17543523 | 17543631 | 17551785 | 17552024 | 95845 | 113,63,121,81,109,47,66,9 | 11,4,13,7,2,0,2,4,0,2,8,4 | 130,103,54,38,87,45,29,95 | 3,2,3,7,3,3,0,1,6,2,4,6,7,6,1 | 233 | 149 | 0.00833408729365 | 0.497435924098609 | 0.868,0.91,0.856,0 | 0.965,0.971,0.92,0 | -0,015 |
| 28411 | ENSG00000185404 | SP140L | chr2 | + | 230328756 | 230328831 | 230327183 | 230327301 | 230357804 | 230357967 | 28411 | 67,41,67,51,10,37,30,43,27 | 2,1,10,3,1,0,2,1,0,0,0,4 | 65,65,44,18,37,25,16,31,37 | 6,3,1,1,4,2,0,6,9,2,2,2,1,4,3 | 224 | 149 | 0.00833895221264 | 0.497518154130113 | 0.957,0.965,0.817, | 0.878,0.935,0.967, | 0,038 |
| 2520 | ENSG00000291230 | | chr19 | - | 50336505 | 50336705 | 50335058 | 50335181 | 50337016 | 50337135 | 2520 | 500,460,544,1308,608,452 | 14,3,5,13,2,4,8,2,10,5,1,0 | 473,317,316,124,318,305,1 | 6,4,2,1,4,2,2,3,7,5,7,15,8,1 | 298 | 149 | 0.0083419185157 | 0.497518154130113 | 0.947,0.987,0.982, | 0.975,0.975,0.988, | 0,011 |
| 158776 | ENSG00000075568 | TMEM131 | chr2 | - | 97775590 | 97776018 | 97766477 | 97766602 | 97792385 | 97792984 | 158776 | 25,27,40,15,30,26,20,23,22 | 4,2,3,2,0,2,0,0,2,2,0,0 | 34,24,33,27,32,19,12,5,12, | 0,0,1,0,3,2,0,0,0,0,0,0,0,0,2 | 298 | 149 | 0.00834239964622 | 0.497518154130113 | 0.758,0.871,0.87,0 | 1.0,1.0,0.943,1.0,0 | -0,064 |
| 86081 | ENSG00000114120 | SLC25A36 | chr3 | + | 140966933 | 140967036 | 140963126 | 140963227 | 140970926 | 140970993 | 86081 | 13,9,15,14,0,5,2,10,5,10,3, | 16,12,27,28,13,15,19,7,15, | 2,6,5,6,4,2,4,4,3,11,2,6,4,2 | 16,32,15,45,16,7,17,19,22, | 252 | 149 | 0.00835698075308 | 0.498249674367358 | 0.325,0.307,0.247, | 0.069,0.1,0.165,0.0 | 0,059 |
| 43190 | ENSG00000168591 | TMUB2 | chr17 | + | 44187675 | 44187743 | 44186969 | 44187109 | 44187906 | 44188009 | 43190 | 19,7,39,19,3,10,7,18,6,3,1, | 0,6,13,3,0,1,0,3,0,0,0,3 | 12,15,17,8,17,3,2,4,8,18,6, | 6,10,7,0,1,1,2,1,2,6,9,0,5,1 | 217 | 149 | 0.00836290331825 | 0.498464703431256 | 1.0,0.445,0.673,0.8 | 0.579,0.507,0.625, | 0,205 |
| 55445 | ENSG00000132185 | FCRLA | chr1 | + | 161710474 | 161710492 | 161706971 | 161707343 | 161710759 | 161710912 | 55445 | 82,48,149,145,171,123,8,1 | 345,256,627,562,206,101,3 | 27,25,35,14,33,44,11,8,44, | 666,397,175,131,141,184,1 | 167 | 149 | 0.00837455448237 | 0.499020967828067 | 0.175,0.143,0.175, | 0.035,0.053,0.151, | 0,079 |
| 121569 | ENSG00000131626 | PPFIA1 | chr11 | + | 70350993 | 70351023 | 70349922 | 70350039 | 70354300 | 70354452 | 121569 | 17,10,7,4,3,11,8,15,2,0,3,3 | 3,0,1,0,0,1,2,1,2,4,2,0 | 8,5,8,3,2,3,2,11,3,7,13,11,1 | 0,0,0,0,1,0,0,0,0,0,0,3,0,0 | 179 | 149 | 0.00838723252126 | 0.499638096535652 | 0.825,1.0,0.854,1.0 | 1.0,1.0,1.0,1.0,0.62 | -0,154 |
| 105823 | ENSG00000104870 | FCGRT | chr19 | + | 49524506 | 49525573 | 49513194 | 49514486 | 49526009 | 49526331 | 105823 | 215,100,192,133,148,72,16 | 6,0,1,2,6,2,0,3,3,2,2,5 | 135,105,61,49,36,40,108,2 | 13,2,9,2,2,1,2,4,1,4,4,4,1,2 | 298 | 149 | 0.00839262879408 | 0.49982121969525 | 0.947,1.0,0.99,0.97 | 0.839,0.963,0.772, | 0,03 |
| 153309 | ENSG00000092199 | HNRNPC | chr14 | - | 21263619 | 21263824 | 21263310 | 21263336 | 21269297 | 21269494 | 153309 | 5,2,10,5,0,4,2,4,1,0,1,4 | 29,16,68,31,7,9,2,10,7,2,2, | 1,2,1,2,0,1,1,1,3,4,5,2,0,7,1 | 42,26,21,6,13,19,8,11,8,33 | 298 | 149 | 0.0083989981427 | 0.499928162006692 | 0.079,0.059,0.068, | 0.012,0.037,0.023, | 0,059 |
| 240992 | ENSG00000106829 | TLE4 | chr9 | + | 79719572 | 79719734 | 79718721 | 79718971 | 79720045 | 79720293 | 240992 | 10,3,20,12,1,5,2,25,4,4,2,9 | 44,31,99,82,67,68,57,90,61 | 5,15,15,28,9,2,6,6,18,9,20, | 41,68,70,34,51,25,38,49,32 | 298 | 149 | 0.00839906999371 | 0.499928162006692 | 0.102,0.046,0.092, | 0.057,0.099,0.097, | -0,035 |
| 40530 | ENSG00000075711 | DLG1 | chr3 | - | 197161639 | 197161738 | 197149742 | 197149796 | 197194424 | 197194589 | 40530 | 55,35,29,12,21,20,19,41,31 | 36,14,38,25,28,18,23,22,20 | 31,18,15,38,13,9,12,20,7,1 | 27,42,28,25,21,11,16,25,19 | 248 | 149 | 0.00844005293988 | 0.502228652005339 | 0.479,0.6,0.314,0.1 | 0.408,0.205,0.243, | 0,092 |
| 240083 | ENSG00000186350 | RXRA | chr9 | + | 134429107 | 134429203 | 134421675 | 134421805 | 134431904 | 134431996 | 240083 | 20,15,44,4,20,32,21,39,16, | 4,0,4,1,2,0,4,3,3,0,0,0 | 14,9,16,23,4,5,22,52,1,15,1 | 2,1,2,12,0,0,2,2,0,0,0,0,0,0 | 245 | 149 | 0.00845270595702 | 0.50276869961333 | 0.753,1.0,0.87,0.70 | 0.81,0.846,0.83,0.5 | -0,038 |
| 63147 | ENSG00000124406 | ATP8A1 | chr4 | - | 42485495 | 42485668 | 42464892 | 42465076 | 42503449 | 42503514 | 63147 | 53,24,88,54,52,53,31,61,23 | 7,2,9,3,1,2,7,8,0,2,0,2 | 50,87,24,19,9,22,19,52,28, | 3,14,6,10,3,1,4,9,6,5,7,27,6 | 298 | 149 | 0.0084538004465 | 0.50276869961333 | 0.791,0.857,0.83,0 | 0.893,0.757,0.667, | 0,104 |
| 211647 | ENSG00000127081 | ZNF484 | chr9 | - | 92853913 | 92853958 | 92846068 | 92848551 | 92855810 | 92855903 | 211647 | 3,5,12,5,0,2,2,3,0,0,1,3 | 8,6,18,10,6,6,7,7,11,3,3,6 | 6,18,5,10,7,2,11,9,3,3,14,1 | 11,25,22,16,13,6,8,4,10,9,1 | 194 | 149 | 0.00846290224658 | 0.503170970561785 | 0.224,0.39,0.339,0 | 0.295,0.356,0.149, | -0,133 |
| 133605 | ENSG00000160214 | RRP1 | chr21 | + | 43798223 | 43798391 | 43797906 | 43798100 | 43799569 | 43799649 | 133605 | 2,1,3,1,0,4,5,1,0,3,0,0 | 10,6,22,11,14,8,8,16,8,10,6 | 1,1,0,1,1,0,1,1,0,1,0,1,0,0,2 | 4,1,6,4,13,6,9,17,4,12,10,3 | 298 | 149 | 0.0084756210542 | 0.503788013012019 | 0.091,0.077,0.064, | 0.111,0.333,0.0,0.1 | 0,012 |
| 255790 | ENSG00000173442 | EHBP1L1 | chr11 | + | 65583591 | 65583765 | 65580927 | 65581326 | 65584240 | 65584398 | 255790 | 56,39,44,12,46,33,41,62,34 | 0,4,5,5,6,4,3,8,9,0,3,0 | 20,32,21,26,9,9,37,63,19,2 | 1,2,4,1,3,0,3,3,3,1,4,1,2,1,3 | 298 | 149 | 0.00848349130402 | 0.50411659769617 | 1.0,0.83,0.815,0.54 | 0.909,0.889,0.724, | -0,048 |
| 224430 | ENSG00000184903 | IMMP2L | chr7 | - | 111329393 | 111329451 | 110963499 | 110963565 | 111487237 | 111487341 | 224430 | 8,1,5,4,3,12,5,5,2,3,1,1 | 7,15,70,30,39,45,31,40,13, | 4,6,0,1,6,6,2,3,3,2,1,3,0,2,4 | 43,51,26,16,31,17,15,31,37 | 207 | 149 | 0.00849479014731 | 0.504648682913519 | 0.451,0.046,0.049, | 0.063,0.078,0.0,0.0 | 0,053 |
| 181844 | ENSG00000123104 | ITPR2 | chr12 | - | 26510476 | 26510631 | 26495151 | 26495260 | 26550246 | 26550355 | 181844 | 1,0,5,0,0,6,4,6,0,2,0,0 | 60,38,72,43,15,31,21,46,37 | 1,2,4,3,12,2,1,0,3,3,3,4,5,5 | 34,39,18,45,37,23,19,17,22 | 298 | 149 | 0.00850060802707 | 0.504854957581209 | 0.008,0.0,0.034,0.0 | 0.014,0.025,0.1,0.0 | -0,02 |
| 186997 | ENSG00000181666 | ZNF875 | chr19 | + | 37322437 | 37322479 | 37322184 | 37322232 | 37323532 | 37323570 | 186997 | 4,6,3,2,1,2,1,1,0,2,0,0 | 13,2,21,18,6,4,6,11,7,4,2,3 | 0,2,2,0,0,0,2,0,1,0,1,3,2,2,0 | 3,11,4,3,4,3,2,1,4,7,5,5,2,6 | 191 | 149 | 0.00851456098212 | 0.505544130963495 | 0.194,0.701,0.1,0.0 | 0.0,0.124,0.281,0.0 | 0,03 |
| 179680 | ENSG00000136450 | SRSF1 | chr17 | - | 58005397 | 58005617 | 58004922 | 58005041 | 58005800 | 58005973 | 179680 | 3,0,10,7,1,2,0,0,0,3,2,0 | 51,23,46,28,22,14,16,32,32 | 0,0,6,0,0,4,0,3,0,0,0,0,2,0,0 | 36,40,21,51,25,9,6,26,15,4 | 298 | 149 | 0.00853471155454 | 0.506555557373108 | 0.029,0.0,0.098,0.1 | 0.0,0.0,0.125,0.0,0 | 0,03 |
| 34699 | ENSG00000125089 | SH3TC1 | chr4 | + | 8232407 | 8233065 | 8231975 | 8232156 | 8233362 | 8233513 | 34699 | 10,13,5,3,9,9,13,6,12,5,5,8 | 32,58,15,11,12,25,18,22,20 | 2,2,7,3,9,7,2,8,2,0,2,15,2,6 | 12,28,14,4,18,13,3,11,27,4 | 298 | 149 | 0.00853630288663 | 0.506555557373108 | 0.135,0.101,0.143, | 0.077,0.034,0.2,0.2 | 0,038 |
| 54328 | ENSG00000143458 | GABPB2 | chr1 | + | 151073038 | 151073157 | 151070577 | 151070934 | 151088189 | 151088297 | 54328 | 3,2,1,3,2,2,2,2,0,0,0,2 | 30,8,21,12,11,6,14,9,11,1,2 | 10,2,0,0,0,1,0,2,2,2,6,8,3,4 | 16,8,11,14,4,7,11,10,4,6,13 | 268 | 149 | 0.00854673812463 | 0.507035003666549 | 0.053,0.122,0.026, | 0.258,0.122,0.0,0.0 | -0,032 |
| 116248 | ENSG00000257524 | ST6GALNAC6 | chr9 | - | 127905206 | 127905302 | 127896241 | 127896332 | 127905934 | 127905975 | 116248 | 65,46,32,48,48,24,44,30,77 | 5,0,1,3,0,0,2,6,0,4,0,0 | 38,55,24,32,37,21,16,24,18 | 0,0,3,0,0,0,2,0,0,1,0,0,0,2,0 | 245 | 149 | 0.00855345046743 | 0.507293385934259 | 0.888,1.0,0.951,0.9 | 1.0,1.0,0.83,1.0,1.0 | -0,019 |
| 149690 | ENSG00000214013 | GANC | chr15 | + | 42327847 | 42328119 | 42327362 | 42327442 | 42329305 | 42329504 | 149690 | 2,5,30,21,23,10,16,24,11,3 | 24,29,27,39,35,31,20,18,15 | 28,45,15,25,26,18,18,17,27 | 33,21,16,9,20,19,20,14,15, | 298 | 149 | 0.00855605399905 | 0.507308005032928 | 0.04,0.079,0.357,0 | 0.298,0.517,0.319, | -0,079 |
| 193909 | ENSG00000092841 | MYL6 | chr12 | + | 56159470 | 56159730 | 56158683 | 56158711 | 56159974 | 56160148 | 193909 | 605,502,1779,1425,745,77 | 38,26,57,25,6,14,8,13,16,1 | 836,784,523,287,605,431,2 | 11,37,15,54,46,7,0,10,20,8 | 298 | 149 | 0.00856743579151 | 0.507842955891625 | 0.888,0.906,0.94,0 | 0.974,0.914,0.946, | 0,023 |
| 206205 | ENSG00000132953 | XPO4 | chr13 | - | 20855626 | 20856002 | 20843769 | 20843886 | 20862716 | 20862858 | 206205 | 10,8,43,32,17,24,26,29,10, | 2,0,8,5,1,2,0,2,0,0,0,1 | 34,57,10,7,34,12,12,20,16, | 17,6,3,13,8,3,0,2,1,0,0,1,9, | 298 | 149 | 0.00857540028156 | 0.508175104075011 | 0.714,1.0,0.729,0.7 | 0.5,0.826,0.625,0.2 | 0,135 |
| 220438 | ENSG00000196367 | TRRAP | chr7 | + | 98950875 | 98950987 | 98950063 | 98950262 | 98953166 | 98953433 | 220438 | 25,26,35,20,36,59,28,42,23 | 0,3,0,1,1,7,2,8,2,0,0,0 | 25,31,15,45,21,12,20,27,23 | 0,4,0,3,0,0,4,2,0,0,0,2,1,2,0 | 261 | 149 | 0.00858531054993 | 0.508622343785297 | 1.0,0.832,1.0,0.919 | 1.0,0.816,1.0,0.89 | -0,013 |
| 7177 | ENSG00000126001 | CEP250 | chr20 | + | 35507737 | 35507851 | 35502389 | 35505005 | 35508034 | 35508190 | 7177 | 37,19,38,24,20,14,13,12,16 | 1,1,9,2,2,0,1,0,0,4,2,3 | 28,25,12,15,6,15,12,14,10, | 5,2,4,4,1,1,0,5,2,8,2,12,3,6 | 263 | 149 | 0.00859430464277 | 0.509015075004961 | 0.954,0.915,0.705, | 0.76,0.876,0.63,0.6 | 0,097 |
| 98396 | ENSG00000186480 | INSIG1 | chr7 | + | 155302746 | 155302846 | 155302250 | 155302417 | 155308240 | 155308757 | 98396 | 79,78,235,175,43,50,68,91 | 0,0,5,9,1,3,9,5,0,0,0,5 | 39,51,58,29,51,23,35,53,10 | 5,6,4,4,6,5,2,5,5,0,0,1,0,3,3 | 249 | 149 | 0.0085976839805 | 0.509068097726181 | 1.0,1.0,0.966,0.92 | 0.824,0.836,0.897, | 0,071 |
| 42801 | ENSG00000114127 | XRN1 | chr3 | - | 142426743 | 142426841 | 142425428 | 142425538 | 142432660 | 142432893 | 42801 | 116,96,140,72,50,92,46,10 | 2,8,11,3,2,5,6,7,0,2,1,3 | 72,93,62,96,50,43,44,74,68 | 6,10,5,17,9,3,1,3,4,0,18,14 | 247 | 149 | 0.00859993032292 | 0.509068097726181 | 0.972,0.879,0.885, | 0.879,0.849,0.882, | 0,042 |
| 232849 | ENSG00000083750 | RRAGB | chrX | + | 55755067 | 55755265 | 55753391 | 55753514 | 55755840 | 55755932 | 232849 | 18,18,21,16,3,3,3,13,9,7,2, | 3,1,14,14,2,3,4,2,4,0,2,5 | 16,8,9,6,3,1,6,5,0,4,0,9,9,8 | 5,2,9,0,3,4,5,8,1,11,2,8,15, | 298 | 149 | 0.00860556151687 | 0.509261372240156 | 0.75,0.9,0.429,0.36 | 0.615,0.667,0.333, | 0,203 |
| 210934 | ENSG00000115159 | GPD2 | chr2 | + | 156495140 | 156495367 | 156476097 | 156476207 | 156496043 | 156496215 | 210934 | 3,0,2,1,0,0,1,0,0,0,0,0 | 30,14,74,23,7,15,9,9,8,8,2, | 0,2,1,3,4,4,2,0,1,4,0,3,1,0,1 | 25,10,24,9,16,20,9,9,16,23 | 298 | 149 | 0.00861518639533 | 0.509611578999869 | 0.048,0.0,0.013,0.0 | 0.0,0.091,0.02,0.14 | -0,039 |
| 208373 | ENSG00000005483 | KMT2E | chr7 | + | 105068478 | 105068545 | 105066726 | 105066807 | 105073618 | 105073677 | 208373 | 6,0,2,3,0,2,1,0,2,0,3,1 | 173,135,196,130,48,71,68, | 9,7,12,3,4,3,1,1,3,2,5,4,0,1 | 150,174,110,186,119,54,76 | 216 | 149 | 0.00861621483885 | 0.509611578999869 | 0.023,0.0,0.007,0.0 | 0.04,0.027,0.07,0.0 | -0,008 |
| 193073 | ENSG00000120942 | UBIAD1 | chr1 | + | 11278637 | 11278693 | 11273933 | 11274060 | 11285643 | 11288434 | 193073 | 0,0,1,2,0,0,3,0,0,0,0,1 | 28,23,30,26,21,31,25,22,29 | 1,5,2,1,0,1,1,2,0,0,0,5,0,3,2 | 27,28,23,5,13,5,19,20,11,2 | 205 | 149 | 0.00862336913897 | 0.509741070993441 | 0.0,0.0,0.024,0.05 | 0.026,0.115,0.059, | -0,04 |
| 127687 | ENSG00000138286 | FAM149B1 | chr10 | + | 73235192 | 73235318 | 73234816 | 73234940 | 73239311 | 73239384 | 127687 | 29,18,42,20,12,16,22,22,27 | 1,1,1,7,2,2,0,1,0,0,0,2 | 9,14,10,12,4,4,7,23,3,21,11 | 1,0,0,0,1,0,0,0,0,1,0,0,0,0,1 | 275 | 149 | 0.00862474818454 | 0.509741070993441 | 0.94,0.907,0.958,0 | 0.83,1.0,1.0,1.0,0.6 | -0,062 |
| 49430 | ENSG00000115685 | PPP1R7 | chr2 | + | 241159212 | 241159343 | 241158465 | 241158549 | 241160331 | 241160421 | 49430 | 60,45,142,104,38,39,28,25 | 0,0,2,2,1,0,0,0,0,0,0,0 | 98,56,34,13,29,46,12,22,18 | 2,1,0,2,3,1,7,0,0,0,6,2,0,2,1 | 280 | 149 | 0.00862550924615 | 0.509741070993441 | 1.0,1.0,0.974,0.96 | 0.963,0.968,1.0,0.1 | 0,082 |
| 178340 | ENSG00000135241 | PNPLA8 | chr7 | - | 108485693 | 108485762 | 108479183 | 108479379 | 108487758 | 108487953 | 178340 | 34,24,43,23,9,21,14,21,5,5 | 13,14,31,29,15,8,4,19,8,5,2 | 27,29,27,26,15,5,16,10,11, | 25,46,29,23,48,10,21,16,10 | 218 | 149 | 0.00863743298609 | 0.510263461293489 | 0.641,0.54,0.487,0 | 0.425,0.301,0.389, | 0,133 |
| 154261 | ENSG00000092108 | SCFD1 | chr14 | + | 30670272 | 30670395 | 30653488 | 30653588 | 30673256 | 30673347 | 154261 | 97,103,85,72,34,37,25,33,4 | 3,1,2,0,0,2,0,0,0,0,1,0 | 53,52,70,48,41,28,21,37,28 | 3,4,1,5,4,0,0,0,1,2,2,2,0,0,0 | 272 | 149 | 0.00863909033993 | 0.510263461293489 | 0.947,0.983,0.959, | 0.906,0.877,0.975, | 0,033 |
| 121890 | ENSG00000166333 | ILK | chr11 | + | 6609070 | 6609156 | 6608693 | 6608967 | 6609298 | 6609393 | 121890 | 17,12,85,93,42,22,21,20,35 | 3,0,5,0,2,3,0,4,0,0,5,3 | 35,17,11,1,6,19,10,24,7,31 | 2,0,0,3,1,0,0,5,3,1,2,1,0,0,1 | 235 | 149 | 0.0086441062485 | 0.510419652118218 | 0.782,1.0,0.915,1.0 | 0.917,1.0,1.0,0.17 | 0,011 |
| 105543 | ENSG00000241769 | EOLA1-DT | chrX | - | 149538723 | 149538906 | 149538114 | 149538172 | 149540694 | 149540813 | 105543 | 6,4,10,2,5,6,2,9,1,0,7,5 | 2,5,5,5,4,7,2,4,5,2,0,3 | 6,6,5,7,6,0,4,20,1,4,12,6,8, | 3,0,0,3,7,2,0,2,4,1,3,1,0,2,2 | 298 | 149 | 0.00865010097784 | 0.51063353910079 | 0.6,0.286,0.5,0.16 | 0.5,1.0,1.0,0.538,0 | -0,223 |
| 48687 | ENSG00000072786 | STK10 | chr5 | - | 172093411 | 172093960 | 172090231 | 172090362 | 172105655 | 172105737 | 48687 | 76,42,112,99,44,52,33,71,8 | 2,2,7,0,0,1,3,1,0,0,0,0 | 62,89,54,40,52,47,37,63,32 | 8,3,0,2,2,1,2,2,2,0,2,8,0,2,0 | 298 | 149 | 0.00865657187151 | 0.51087541005675 | 0.95,0.913,0.889,1 | 0.795,0.937,1.0,0.9 | 0,028 |
| 198045 | ENSG00000112232 | KHDRBS2 | chr6 | - | 61968727 | 61968919 | 61901243 | 61901371 | 61978065 | 61978212 | 198045 | 3,0,12,6,0,3,0,0,7,2,1,2 | 57,30,75,37,29,39,20,30,78 | 3,12,2,2,6,0,2,0,3,6,8,3,0,3 | 55,32,33,33,11,8,14,28,13, | 298 | 149 | 0.00865971323979 | 0.510920707322709 | 0.026,0.0,0.074,0.0 | 0.027,0.158,0.029, | -0,036 |
| 163487 | ENSG00000167766 | ZNF83 | chr19 | - | 52616717 | 52616835 | 52612377 | 52614797 | 52618935 | 52619062 | 163487 | 90,45,90,65,28,35,50,37,28 | 12,10,31,21,7,13,2,5,7,3,1, | 59,61,50,17,49,30,11,27,39 | 15,15,2,1,5,2,0,2,0,2,3,4,8, | 267 | 149 | 0.008665863082 | 0.511109015848654 | 0.807,0.715,0.618, | 0.687,0.694,0.933, | -0,075 |
| 244297 | ENSG00000130309 | COLGALT1 | chr19 | + | 17571865 | 17571991 | 17568508 | 17568713 | 17572482 | 17572602 | 244297 | 1,0,3,3,6,1,2,1,0,0,0,0 | 29,14,35,25,16,22,22,21,21 | 2,4,6,1,0,3,1,0,5,5,1,3,6,2,0 | 40,21,14,11,25,13,6,26,14, | 275 | 149 | 0.00866765432418 | 0.511109015848654 | 0.018,0.0,0.044,0.0 | 0.026,0.094,0.188, | -0,033 |
| 118292 | ENSG00000116701 | NCF2 | chr1 | - | 183574486 | 183574621 | 183573184 | 183573292 | 183577598 | 183577707 | 118292 | 37,19,62,100,44,12,28,16,3 | 2,4,4,1,0,2,2,0,0,0,2,0 | 53,52,42,19,31,24,14,26,52 | 0,12,2,2,3,2,0,3,7,0,0,4,1,0 | 284 | 149 | 0.00868059182165 | 0.511731705934142 | 0.907,0.714,0.89,0 | 1.0,0.695,0.917,0.8 | -0,009 |
| 36934 | ENSG00000121210 | TMEM131L | chr4 | + | 153591052 | 153591194 | 153588889 | 153589007 | 153593798 | 153593871 | 36934 | 116,77,189,88,58,50,52,73 | 3,3,9,7,4,5,5,5,1,0,2,3 | 94,84,69,54,44,30,41,63,45 | 14,11,2,2,10,1,0,11,5,1,7,1 | 291 | 149 | 0.00868361574846 | 0.51176979768806 | 0.952,0.929,0.915, | 0.775,0.796,0.946, | 0,046 |
| 171748 | ENSG00000173226 | IQCB1 | chr3 | - | 121819172 | 121819230 | 121808915 | 121809009 | 121826050 | 121826180 | 171748 | 0,0,2,10,0,7,1,7,0,2,1,2 | 28,19,49,48,37,20,25,15,13 | 6,2,4,1,0,3,0,2,2,1,1,0,2,0,0 | 47,46,16,8,20,22,24,25,12, | 207 | 149 | 0.00869151668262 | 0.512095216840127 | 0.0,0.0,0.029,0.13, | 0.084,0.03,0.153,0 | 0,036 |
| 146450 | ENSG00000237440 | ZNF737 | chr19 | - | 20552474 | 20552553 | 20537991 | 20545976 | 20553708 | 20553835 | 146450 | 35,22,156,129,52,37,28,54 | 1,0,5,11,0,1,0,1,4,0,0,5 | 95,65,53,30,23,29,15,33,27 | 0,0,2,0,2,11,0,2,0,0,0,0,0,0 | 228 | 149 | 0.00869869411375 | 0.512377841487829 | 0.958,1.0,0.953,0.8 | 1.0,1.0,0.945,1.0,0 | -0,009 |
| 189303 | ENSG00000118308 | IRAG2 | chr12 | + | 25102419 | 25102719 | 25102197 | 25102241 | 25103836 | 25103899 | 189303 | 67,4,56,28,24,6,12,36,17,1 | 109,85,524,396,147,91,147 | 29,38,18,50,16,19,20,31,4, | 162,227,139,35,87,35,49,1 | 298 | 149 | 0.00870617455589 | 0.512678154812246 | 0.235,0.023,0.051, | 0.082,0.077,0.061, | -0,039 |
| 43290 | ENSG00000156976 | EIF4A2 | chr3 | + | 186788309 | 186788420 | 186787802 | 186787882 | 186789124 | 186789291 | 43290 | 561,275,353,194,146,186,1 | 427,137,428,354,181,165,1 | 298,199,314,288,120,91,64 | 138,51,178,74,110,133,75, | 260 | 149 | 0.00871339459356 | 0.512824375737296 | 0.43,0.535,0.321,0 | 0.553,0.691,0.503, | -0,078 |
| 126527 | ENSG00000150093 | ITGB1 | chr10 | - | 32932514 | 32932600 | 32929821 | 32930044 | 32935491 | 32935558 | 126527 | 554,457,261,245,133,130,2 | 4,5,12,2,2,0,12,17,2,0,0,3 | 154,148,5,74,134,121,64,1 | 9,3,0,3,7,4,2,4,4,6,7,12,5,1 | 235 | 149 | 0.00871342298308 | 0.512824375737296 | 0.989,0.983,0.932, | 0.916,0.969,1.0,0.9 | 0,019 |
| 52569 | ENSG00000121310 | ECHDC2 | chr1 | - | 52897436 | 52897484 | 52895909 | 52896597 | 52899173 | 52899224 | 52569 | 13,10,23,4,18,18,14,14,2,3 | 4,0,1,0,1,0,2,1,0,0,0,3 | 18,10,10,13,3,3,9,13,14,10 | 2,2,2,0,0,0,2,0,8,3,0,0,0,3,3 | 197 | 149 | 0.00872153905032 | 0.513135668671336 | 0.711,1.0,0.946,1.0 | 0.872,0.791,0.791, | 0,049 |
| 88761 | ENSG00000168918 | INPP5D | chr2 | + | 233139841 | 233139929 | 233130507 | 233130648 | 233146161 | 233146242 | 88761 | 198,191,263,220,156,151,1 | 8,1,3,2,3,6,8,20,0,2,0,1 | 185,215,118,88,99,101,81, | 2,1,2,5,7,0,2,2,2,3,10,3,0,0 | 237 | 149 | 0.00872348040788 | 0.513135668671336 | 0.94,0.992,0.982,0 | 0.983,0.993,0.974, | -0,004 |
| 144079 | ENSG00000007202 | KIAA0100 | chr17 | - | 28643163 | 28643226 | 28642868 | 28642987 | 28643609 | 28643659 | 144079 | 16,6,32,31,16,14,7,21,21,4 | 2,1,4,3,1,2,2,5,0,2,0,0 | 22,19,11,9,5,8,6,16,10,11,1 | 6,4,1,3,6,1,0,3,2,0,2,4,6,2,4 | 212 | 149 | 0.00872878917941 | 0.513307656796064 | 0.849,0.808,0.849, | 0.72,0.77,0.885,0.6 | 0,098 |
| 178414 | ENSG00000160271 | RALGDS | chr9 | - | 133105239 | 133105370 | 133104262 | 133104331 | 133105931 | 133106016 | 178414 | 2,0,32,28,1,2,4,13,1,3,1,2 | 22,20,21,17,19,33,10,17,12 | 1,2,3,4,11,6,1,6,0,2,1,4,0,0 | 16,23,20,15,14,8,17,28,8,1 | 280 | 149 | 0.00873963049236 | 0.513804810298043 | 0.046,0.0,0.448,0.4 | 0.032,0.044,0.074, | 0,079 |
| 254727 | ENSG00000150990 | DHX37 | chr12 | - | 124975418 | 124975554 | 124972502 | 124972599 | 124977341 | 124977490 | 254727 | 13,6,17,17,6,19,14,7,2,8,5, | 0,2,1,2,1,2,0,6,0,0,0,4 | 24,19,2,11,2,2,4,13,4,6,13, | 0,4,1,4,1,2,0,0,0,0,1,0,1,2,0 | 285 | 149 | 0.00874274736816 | 0.513847694919838 | 1.0,0.611,0.899,0.8 | 1.0,0.713,0.511,0.5 | -0,007 |
| 88531 | ENSG00000163219 | ARHGAP25 | chr2 | + | 68787839 | 68787956 | 68782232 | 68782320 | 68807275 | 68807480 | 88531 | 156,265,230,214,134,103,7 | 3,1,8,9,1,1,4,3,8,3,0,3 | 226,534,88,57,155,46,70,8 | 3,23,4,15,12,1,6,2,3,3,2,11 | 266 | 149 | 0.00875778526614 | 0.514484546883483 | 0.967,0.993,0.942, | 0.977,0.929,0.925, | 0,022 |
| 205862 | ENSG00000128815 | WDFY4 | chr10 | + | 48782278 | 48782374 | 48779940 | 48780119 | 48786638 | 48786870 | 205862 | 5,4,17,8,9,9,7,6,0,2,5,0 | 72,54,195,189,152,174,85, | 8,14,0,7,0,0,1,5,4,1,0,8,5,5 | 210,191,38,88,69,50,79,13 | 245 | 149 | 0.00875836371053 | 0.514484546883483 | 0.041,0.043,0.05,0 | 0.023,0.043,0.0,0.0 | 0,013 |
| 157858 | ENSG00000066933 | MYO9A | chr15 | - | 71902676 | 71903063 | 71901190 | 71901340 | 71903928 | 71904039 | 157858 | 14,4,24,12,12,14,8,21,13,9 | 0,0,2,2,0,1,1,0,0,0,2,0 | 18,13,14,13,11,7,16,14,3,1 | 2,1,0,1,0,0,4,1,1,0,2,2,2,4,0 | 298 | 149 | 0.00877001128161 | 0.515028185034707 | 1.0,1.0,0.857,0.75, | 0.818,0.867,1.0,0.8 | 0,073 |
| 101377 | ENSG00000008282 | SYPL1 | chr7 | - | 106092547 | 106092669 | 106091406 | 106091939 | 106092948 | 106093137 | 101377 | 6,2,7,3,1,4,3,5,0,3,0,3 | 93,78,154,90,77,69,53,48,3 | 4,2,2,4,11,3,11,3,2,1,3,32,3 | 77,98,69,71,63,53,24,51,48 | 271 | 149 | 0.00878984010222 | 0.516051848074444 | 0.034,0.014,0.024, | 0.028,0.011,0.016, | -0,012 |
| 255190 | ENSG00000135297 | MTO1 | chr6 | + | 73471459 | 73471561 | 73466488 | 73466606 | 73473364 | 73473654 | 255190 | 13,3,4,5,3,3,2,7,3,6,1,0 | 39,48,62,40,28,32,24,27,39 | 4,2,8,2,3,2,0,1,0,2,2,2,3,0,3 | 47,62,45,35,32,14,22,19,24 | 251 | 149 | 0.0088029571774 | 0.516681013430319 | 0.165,0.036,0.037, | 0.048,0.019,0.095, | 0,02 |
| 115375 | ENSG00000184465 | WDR27 | chr6 | - | 169663562 | 169663878 | 169662303 | 169662424 | 169664165 | 169664286 | 115375 | 11,6,20,7,5,20,12,22,13,10 | 1,0,14,5,2,4,0,2,1,2,0,2 | 8,2,9,1,6,2,10,12,5,8,16,9,9 | 3,0,0,1,0,1,3,3,0,0,7,2,0,0,0 | 298 | 149 | 0.00880818888867 | 0.516726142436647 | 0.846,1.0,0.417,0.4 | 0.571,1.0,1.0,0.33 | -0,041 |
| 98581 | ENSG00000168137 | SETD5 | chr3 | + | 9449479 | 9449592 | 9448558 | 9448630 | 9453738 | 9453868 | 98581 | 4,0,0,7,3,8,7,2,2,0,2,5 | 12,10,44,19,10,21,16,20,4, | 3,1,1,9,2,2,0,5,0,3,4,6,1,0,0 | 25,33,7,23,8,16,28,19,14,1 | 262 | 149 | 0.00880852765912 | 0.516726142436647 | 0.159,0.0,0.0,0.17 | 0.064,0.017,0.075, | 0,036 |
| 199659 | ENSG00000114933 | INO80D | chr2 | - | 206063776 | 206063980 | 206063150 | 206063244 | 206085900 | 206086303 | 199659 | 0,1,3,1,0,2,0,0,1,0,1,0 | 8,5,19,10,9,27,13,15,12,7,1 | 2,6,1,12,4,1,0,1,1,2,2,0,1,0 | 8,5,4,27,12,13,8,3,7,2,12,1 | 298 | 149 | 0.00882622886986 | 0.517342016033097 | 0.0,0.091,0.073,0.0 | 0.111,0.375,0.111, | -0,109 |
| 125957 | ENSG00000148700 | ADD3 | chr10 | + | 110079239 | 110079297 | 110008208 | 110008299 | 110100624 | 110100704 | 125957 | 42,22,20,9,4,1,14,4,0,0,6,3 | 165,97,58,73,20,10,21,17,1 | 20,2,10,6,20,14,0,3,23,27,1 | 48,34,24,5,62,34,2,18,37,7 | 207 | 149 | 0.00882654842603 | 0.517342016033097 | 0.155,0.14,0.199,0 | 0.231,0.041,0.231, | -0,077 |
| 76205 | ENSG00000196510 | ANAPC7 | chr12 | - | 110377392 | 110377617 | 110375476 | 110376216 | 110381751 | 110381948 | 76205 | 56,40,109,95,69,71,44,37,5 | 0,0,7,2,2,1,0,9,1,0,0,0 | 73,56,57,31,27,22,22,41,46 | 3,0,0,1,0,0,0,0,0,2,2,1,0,1 | 298 | 149 | 0.00882833291214 | 0.517342016033097 | 1.0,1.0,0.886,0.96, | 0.924,1.0,1.0,0.93 | -0,028 |
| 43267 | ENSG00000136560 | TANK | chr2 | + | 161230970 | 161231551 | 161224630 | 161224746 | 161235341 | 161236187 | 43267 | 193,211,459,300,169,179,1 | 4,0,6,0,0,0,1,3,2,0,1,2 | 243,248,155,100,167,133,8 | 1,5,0,2,3,2,0,8,2,0,2,6,3,2,1 | 298 | 149 | 0.00882864097128 | 0.517342016033097 | 0.96,1.0,0.975,1.0, | 0.992,0.961,1.0,0.9 | 0,014 |
| 21028 | ENSG00000100697 | DICER1 | chr14 | - | 95113091 | 95113224 | 95112171 | 95112247 | 95115666 | 95115821 | 21028 | 127,94,238,133,105,124,89 | 6,10,8,3,2,12,5,7,1,0,1,2 | 188,160,93,127,99,67,97,9 | 5,12,8,18,8,2,4,4,2,4,7,8,5, | 282 | 149 | 0.00883835167493 | 0.517653747557553 | 0.918,0.832,0.94,0 | 0.952,0.876,0.86,0 | 0,038 |
| 104924 | ENSG00000189376 | C8orf76 | chr8 | - | 123226499 | 123226632 | 123219966 | 123220297 | 123231299 | 123231592 | 104924 | 4,14,11,12,8,5,2,4,9,2,2,4 | 0,0,1,5,1,1,0,4,1,0,0,0 | 16,10,5,1,10,8,3,3,3,7,7,40 | 0,5,3,0,0,2,2,0,1,0,0,1,4,0,0 | 282 | 149 | 0.00883877100545 | 0.517653747557553 | 1.0,1.0,0.853,0.55 | 1.0,0.514,0.468,1.0 | 0,07 |

| | | | | | | | | | | | | | | | | | | | | | |
|---|---|---|---|---|---|---|---|---|---|---|---|---|---|---|---|---|---|---|---|---|---|
| 83696 | ENSG00000079263 | SP140 | chr2 | + | 230238212 | 230238381 | 230237082 | 230237260 | 230241403 | 230241487 | 83696 | 383,269,615,465,271,151,1 | 3,2,10,4,3,3,5,3,1,0,3,2 | 430,496,241,234,341,244,1 | 6,0,0,0,1,1,1,1,2,0,3,4,1,1,1 | 298 | 149 | 0.00884959668425 | 0.518146774740972 | 0.985,0.985,0.969, | 0.973,1.0,1.0,1.0,0 | -0,016 |
| 7860 | ENSG00000166401 | SERPINB8 | chr18 | + | 63983578 | 63983721 | 63981720 | 63981838 | 63985092 | 63985245 | 7860 | 37,46,99,99,33,31,21,41,7 | 2,2,3,5,1,2,2,0,0,1,1,4 | 53,99,31,14,16,11,13,17,1 | 0,2,1,0,2,0,1,0,1,0,1,2,0,0,4 | 292 | 149 | 0.00885601110337 | 0.518379976306455 | 0.904,0.921,0.944, | 1.0,0.962,0.941,1. | -0,04 |
| 254965 | ENSG00000076513 | ANKRD13A | chr12 | + | 110021107 | 110021296 | 110019138 | 110019328 | 110024045 | 110024112 | 254965 | 5,8,19,3,8,1,0,10,6,4,5,2 | 353,341,216,100,90,67,67, | 6,13,5,7,1,4,2,2,6,13,8,8,2 | 136,119,176,110,120,58,2 | 298 | 149 | 0.00885950357441 | 0.518379976306455 | 0.007,0.012,0.042, | 0.022,0.052,0.014, | -0,004 |
| 135181 | ENSG00000093000 | NUP50 | chr22 | + | 45178590 | 45178900 | 45171599 | 45171683 | 45181285 | 45181367 | 135181 | 6,10,33,28,18,10,10,21,23, | 2,5,5,4,0,1,1,1,0,0,0,0 | 18,17,5,5,8,14,13,7,7,20,1 | 3,5,6,2,5,0,4,0,3,2,0,0,3,3, | 298 | 149 | 0.00886080505518 | 0.518379976306455 | 0.6,0.5,0.767,0.77 | 0.75,0.63,0.294,0. | 0,153 |
| 101236 | ENSG00000146909 | NOM1 | chr7 | + | 156958605 | 156958856 | 156954111 | 156954298 | 156959850 | 156960118 | 101236 | 5,5,6,1,2,3,8,4,1,1,2,3 | 15,12,27,21,8,10,6,7,12,8, | 1,8,0,3,0,1,0,0,1,1,3,5,3,4,1 | 15,21,7,12,9,12,5,5,9,14,6, | 298 | 149 | 0.00886515847842 | 0.518476935384088 | 0.143,0.172,0.1,0. | 0.032,0.16,0.0,0.1 | 0,062 |
| 126768 | ENSG00000119977 | TCTN3 | chr10 | - | 95680471 | 95680609 | 95659822 | 95664300 | 95682650 | 95682804 | 126768 | 26,12,38,49,18,27,9,16,16, | 4,3,4,0,0,3,0,0,0,0,0,2 | 27,9,10,3,10,8,11,21,8,14,1 | 0,1,0,3,2,2,8,2,1,3,1,0,0,1,1 | 287 | 149 | 0.00886728026701 | 0.518476935384088 | 0.771,0.675,0.831, | 1.0,0.824,1.0,0.34 | 0,118 |
| 162911 | ENSG00000173926 | MARCHF3 | chr5 | - | 126917636 | 126917676 | 126914930 | 126915134 | 126917983 | 126918228 | 162911 | 6,0,3,1,2,0,0,0,5,1,0,0 | 87,49,133,63,45,35,53,53, | 3,0,8,2,3,0,4,4,5,1,0,2,2,2, | 27,45,80,70,50,34,60,93,31 | 189 | 149 | 0.00888055242093 | 0.519111944081799 | 0.052,0.0,0.017,0. | 0.081,0.0,0.073,0. | -0,027 |
| 133133 | ENSG00000111450 | STX2 | chr12 | - | 130804304 | 130804543 | 130801414 | 130801488 | 130806981 | 130807090 | 133133 | 1,0,2,0,4,7,11,22,0,0,0,0 | 1,1,10,13,11,13,6,8,5,1,4,7 | 5,7,0,0,9,2,8,10,1,3,3,17,6, | 3,0,1,14,1,2,2,10,2,5,5,16,1 | 298 | 149 | 0.00889082862871 | 0.519571527718133 | 0.333,0.0,0.091,0. | 0.455,1.0,0.0,0.0,0 | -0,207 |
| 240563 | ENSG00000198000 | NOL8 | chr9 | - | 92310905 | 92311259 | 92310552 | 92310675 | 92314266 | 92316138 | 240563 | 46,30,31,27,10,18,13,10,2 | 5,0,7,3,2,1,2,1,2,0,0,1 | 17,28,25,34,13,17,13,13,2 | 0,0,0,1,2,1,1,2,1,3,1,4,2,0, | 298 | 149 | 0.00891485616692 | 0.520634152178538 | 0.821,1.0,0.689,0. | 1.0,1.0,1.0,0.944,0 | 0,008 |
| 184476 | ENSG00000077044 | DGKD | chr2 | + | 233389894 | 233390041 | 233388256 | 233388367 | 233390402 | 233390483 | 184476 | 8,4,29,13,7,11,3,5,2,2,2,3 | 63,43,251,97,46,59,79,124 | 5,1,12,11,2,3,4,5,1,1,6,5,3, | 103,107,94,62,89,45,38,57 | 296 | 149 | 0.00891538868792 | 0.520634152178538 | 0.06,0.045,0.055,0 | 0.024,0.005,0.06,0 | 0,012 |
| 208574 | ENSG00000196504 | PRPF40A | chr2 | - | 152679055 | 152679141 | 152677450 | 152677475 | 152679351 | 152679472 | 208574 | 13,12,42,19,4,6,4,5,3,4,4,4 | 7,4,5,8,2,4,7,8,8,3,0,4 | 25,32,14,11,12,12,4,5,9,13 | 1,1,1,4,9,8,0,0,4,12,3,5,1,5 | 235 | 149 | 0.00891626896186 | 0.520634152178538 | 0.541,0.655,0.842, | 0.941,0.953,0.899, | -0,184 |
| 163617 | ENSG00000105875 | WDR91 | chr7 | - | 135197043 | 135197299 | 135196143 | 135196337 | 135197992 | 135201343 | 163617 | 10,7,20,9,16,6,5,13,6,6,5,1 | 14,14,44,37,36,35,14,15,3 | 11,23,7,5,5,1,8,13,10,12,6, | 21,28,8,6,7,3,21,25,2,11,2 | 298 | 149 | 0.00892182744713 | 0.520817424267219 | 0.263,0.2,0.185,0. | 0.208,0.291,0.304, | -0,088 |
| 116831 | ENSG00000187764 | SEMA4D | chr9 | - | 89388635 | 89388792 | 89387385 | 89387608 | 89388871 | 89389047 | 116831 | 61,48,90,88,47,62,39,65,6 | 6,0,4,1,2,2,3,6,2,2,0,1 | 48,58,34,75,28,22,64,52,4 | 0,1,0,1,1,1,10,0,2,0,1,0,0,2 | 298 | 149 | 0.00892850374504 | 0.520886865134991 | 0.836,1.0,0.918,0. | 1.0,0.967,1.0,0.97 | -0,027 |
| 140244 | ENSG00000154930 | ACSS1 | chr20 | - | 25012801 | 25012939 | 25012600 | 25012664 | 25013535 | 25013662 | 140244 | 31,12,29,27,16,15,22,5,14, | 2,3,12,4,5,3,0,8,5,3,0,4 | 27,40,11,7,13,10,3,24,18,2 | 0,3,0,4,1,1,7,2,3,0,1,3,2,2, | 287 | 149 | 0.00892941154435 | 0.520886865134991 | 0.889,0.675,0.556, | 1.0,0.874,1.0,0.47 | -0,099 |
| 112525 | ENSG00000119397 | CNTRL | chr9 | + | 121157982 | 121158109 | 121157739 | 121157880 | 121158854 | 121159019 | 112525 | 199,149,164,84,69,59,37,3 | 9,9,13,7,1,7,3,5,5,4,0,2 | 102,79,135,91,61,54,30,43 | 0,4,14,1,2,0,0,5,3,0,6,0,7,3 | 276 | 149 | 0.00893027738731 | 0.520886865134991 | 0.923,0.899,0.872, | 1.0,0.914,0.839,0. | -0,039 |
| 230657 | ENSG00000225648 | SBDSP1 | chr7 | + | 72830691 | 72830825 | 72829655 | 72829783 | 72831601 | 72831798 | 230657 | 29,12,44,27,8,23,9,21,10,4 | 7,4,6,6,3,2,2,2,5,5,0,1 | 27,23,15,32,17,8,9,13,7,7,2 | 0,4,0,11,0,0,0,0,3,1,10,1, | 283 | 149 | 0.00894267483777 | 0.521468666488235 | 0.686,0.612,0.794, | 1.0,0.752,1.0,0.60 | -0,131 |
| 218462 | ENSG00000124942 | AHNAK | chr11 | - | 62533660 | 62534074 | 62491810 | 62491831 | 62535002 | 62535190 | 218462 | 473,306,900,642,661,750,6 | 11,6,11,8,7,8,17,25,3,2,1,3 | 481,486,267,568,266,232,4 | 13,17,9,0,11,3,7,8,6,4,15,2 | 298 | 149 | 0.00894612224277 | 0.521528395567072 | 0.956,0.962,0.976, | 0.949,0.935,0.937, | 0,014 |
| 148183 | ENSG00000177455 | CD19 | chr16 | + | 28936150 | 28936201 | 28933233 | 28933509 | 28936511 | 28936600 | 148183 | 33,16,28,11,36,30,53,58,3 | 6,2,1,0,0,0,4,0,0,0,1,4 | 16,18,9,6,12,11,30,39,10,1 | 0,2,0,1,1,0,2,1,1,0,3,14,2,2 | 200 | 149 | 0.00895393145055 | 0.521842301660798 | 0.804,0.856,0.954, | 1.0,0.87,1.0,0.817, | 0,046 |
| 159628 | ENSG00000170476 | MZB1 | chr5 | - | 139389326 | 139389436 | 139388460 | 139388585 | 139389679 | 139390081 | 159628 | 2,3,31,22,4,5,3,6,6,3,4,4 | 13,20,30,0,13,20,15,12,11, | 1,8,3,2,6,4,0,5,7,8,4,2,3,1, | 35,58,10,16,0,0,6,83,11,17 | 259 | 149 | 0.00896928258548 | 0.522470487551497 | 0.081,0.079,0.373, | 0.016,0.074,0.147, | 0,052 |
| 220378 | ENSG00000014123 | UFL1 | chr6 | + | 96522862 | 96522977 | 96521594 | 96521950 | 96523145 | 96523291 | 220378 | 6,0,3,1,0,0,4,1,0,0,0,0 | 9,9,34,22,19,15,15,6,11,6, | 2,5,0,3,2,4,0,0,3,3,8,0,1,3, | 18,36,12,9,23,16,8,8,19,34 | 264 | 149 | 0.00897163803466 | 0.522470487551497 | 0.273,0.0,0.047,0. | 0.059,0.073,0.0,0. | -0,025 |
| 57578 | ENSG00000173230 | GOLGB1 | chr3 | - | 121664925 | 121665031 | 121663198 | 121664614 | 121667475 | 121667610 | 57578 | 60,42,86,59,34,44,22,52,3 | 2,0,0,1,0,2,0,0,0,0,2,0 | 50,57,36,62,34,23,38,49,1 | 5,1,0,7,0,0,0,4,4,0,1,0,2,1,1 | 255 | 149 | 0.00897199251962 | 0.522470487551497 | 0.946,1.0,1.0,0.97 | 0.854,0.971,1.0,0. | 0,034 |
| 237293 | ENSG00000120910 | PPP3CC | chr8 | + | 22535251 | 22535369 | 22532920 | 22533018 | 22540614 | 22541142 | 237293 | 9,6,2,0,1,1,0,3,4,0,3,1 | 32,26,40,20,11,4,11,4,12,6 | 3,2,4,1,3,0,0,0,1,0,3,1,0,0, | 19,19,10,5,16,7,12,11,9,14 | 267 | 149 | 0.00898215963543 | 0.522841692001999 | 0.136,0.114,0.027, | 0.081,0.055,0.182, | 0,037 |
| 12359 | ENSG00000266967 | AARSD1 | chr17 | - | 42953723 | 42953778 | 42951799 | 42951894 | 42954875 | 42954967 | 12359 | 9,6,10,3,5,11,4,11,10,13,3, | 22,8,8,0,3,4,2,4,14,7,3,5 | 2,0,10,1,2,1,10,10,0,5,2,12 | 9,2,6,4,4,3,0,12,1,7,8,7,8,1 | 204 | 149 | 0.00898322535798 | 0.522841692001999 | 0.23,0.354,0.477,1 | 0.14,0.0,0.549,0.1 | 0,237 |
| 222517 | ENSG00000166295 | ANAPC16 | chr10 | + | 72223892 | 72224056 | 72216028 | 72216072 | 72230365 | 72230440 | 222517 | 252,119,331,247,115,106,1 | 3,8,20,16,7,1,9,4,0,1,1,2 | 193,126,149,91,78,60,70,1 | 14,4,3,2,9,4,2,9,4,16,15,10 | 298 | 149 | 0.00899026484707 | 0.523109946823391 | 0.977,0.881,0.892, | 0.873,0.94,0.961,0 | 0,034 |
| 241691 | ENSG00000175893 | ZDHHC21 | chr9 | - | 14683566 | 14683729 | 14680032 | 14680162 | 14690336 | 14690385 | 241691 | 10,6,5,1,5,5,4,4,6,4,0,1 | 11,25,27,19,4,14,7,7,1,3,1, | 2,8,2,2,2,1,0,0,2,4,0,15,4,5 | 22,32,20,2,13,12,7,9,12,30 | 298 | 149 | 0.00899482538064 | 0.523233854459602 | 0.312,0.107,0.085, | 0.043,0.111,0.048, | 0,134 |
| 203607 | ENSG00000120451 | SNX19 | chr11 | - | 130907952 | 130908445 | 130906624 | 130906721 | 130910017 | 130910137 | 203607 | 33,22,63,37,5,27,29,34,17, | 0,2,1,0,0,0,0,1,1,0,0,0 | 35,24,23,15,37,19,25,35,4 | 0,1,1,0,5,1,2,1,0,0,4,1,2,1,1 | 298 | 149 | 0.00900330373011 | 0.523585535027102 | 1.0,0.846,0.969,1. | 1.0,0.923,0.92,1.0, | 0,07 |
| 211829 | ENSG00000137497 | NUMA1 | chr11 | - | 72012400 | 72012433 | 72010785 | 72010854 | 72012894 | 72016260 | 211829 | 4,5,11,4,7,4,4,0,3,2,8,3 | 67,38,141,80,56,63,46,57,3 | 5,6,0,1,1,1,2,7,0,3,2,2,2,5,5 | 68,62,48,34,39,27,77,99,3 | 182 | 149 | 0.00901285222611 | 0.523999242970795 | 0.047,0.097,0.06,0 | 0.057,0.073,0.0,0. | 0,03 |
| 121116 | ENSG00000101017 | CD40 | chr20 | + | 46121723 | 46121898 | 46118343 | 46118394 | 46122232 | 46122358 | 121116 | 81,96,83,58,42,27,39,44,4 | 2,0,1,2,0,0,3,0,0,0,0,2 | 36,51,64,19,18,14,9,19,22, | 3,0,1,12,2,0,0,1,0,0,0,7,0,0 | 298 | 149 | 0.00902487854221 | 0.524556746831866 | 0.953,1.0,0.976,0. | 0.857,1.0,0.97,0.4 | 0,047 |
| 176947 | ENSG00000214106 | PAXIP1-AS2 | chr7 | + | 154945007 | 154945125 | 154944026 | 154944620 | 154946359 | 154946493 | 176947 | 0,1,2,6,4,0,2,0,1,0,5,2 | 6,7,17,19,16,6,1,2,6,5,2,7 | 1,0,0,2,0,0,0,6,0,3,5,0,0,0, | 8,8,9,11,5,5,5,4,2,3,1,5,13, | 267 | 149 | 0.009039822494 | 0.525157956385979 | 0.0,0.074,0.062,0. | 0.065,0.0,0.0,0.09 | 0,028 |
| 35435 | ENSG00000122008 | POLK | chr5 | + | 75534827 | 75534964 | 75511890 | 75511914 | 75547009 | 75547157 | 35435 | 7,17,29,10,6,9,10,11,7,6,1, | 7,8,7,5,8,5,1,3,2,3,0,1 | 2,13,29,13,10,6,4,5,12,7,2, | 2,1,4,0,4,1,1,2,3,2,2,2,2,3, | 286 | 149 | 0.00904010216191 | 0.525157956385979 | 0.343,0.525,0.683, | 0.343,0.871,0.791, | -0,095 |
| 25488 | ENSG00000136152 | COG3 | chr13 | + | 45480124 | 45480290 | 45479004 | 45479066 | 45481229 | 45481304 | 25488 | 36,33,86,41,39,24,24,39,4 | 2,1,2,0,5,2,5,2,0,0,3,0 | 43,42,33,29,32,17,20,48,1 | 2,2,4,0,2,1,0,2,1,1,2,0,3,2,4 | 298 | 149 | 0.00906242174371 | 0.526312491721653 | 0.9,0.943,0.956,1. | 0.915,0.913,0.805, | 0,003 |
| 220344 | ENSG00000149657 | LSM14B | chr20 | + | 62126032 | 62126439 | 62124616 | 62124780 | 62129784 | 62129952 | 220344 | 19,3,30,16,14,15,11,8,16,1 | 0,0,3,3,1,2,0,0,0,0,0,1 | 15,7,9,7,10,11,6,11,12,11,1 | 3,1,2,3,0,0,0,2,0,5,2,2,3,0, | 298 | 149 | 0.00906519431282 | 0.5263314910015 | 1.0,1.0,0.833,0.72 | 0.714,0.778,0.692, | 0,107 |
| 147248 | ENSG00000100266 | PACSIN2 | chr22 | - | 42876883 | 42877010 | 42876136 | 42876333 | 42879047 | 42879169 | 147248 | 8,7,15,14,9,16,12,9,22,7,7, | 0,0,19,21,4,7,3,10,4,1,2,0 | 7,4,16,3,4,9,8,16,4,7,6,50,1 | 9,11,8,11,10,2,14,8,2,0,9,8 | 276 | 149 | 0.00907712189675 | 0.526881883214509 | 1.0,1.0,0.299,0.26 | 0.296,0.164,0.519, | 0,226 |
| 37288 | ENSG00000251022 | THAP9-AS1 | chr4 | - | 82895691 | 82895774 | 82893008 | 82894847 | 82900076 | 82900223 | 37288 | 9,4,9,11,11,4,1,4,4,0,2,2 | 1,2,10,0,0,0,0,0,2,2,2,2 | 4,7,3,0,3,0,4,5,1,6,4,8,5,7,4 | 3,2,4,1,3,3,0,1,6,0,0,6,6,8,2 | 232 | 149 | 0.0090829954764 | 0.527080668476961 | 0.853,0.562,0.366, | 0.461,0.692,0.325, | 0,168 |
| 215201 | ENSG00000115355 | CCDC88A | chr2 | - | 55355578 | 55355772 | 55348440 | 55349599 | 55362307 | 55362448 | 215201 | 366,240,716,427,86,109,1 | 12,4,10,9,0,0,11,1,0,0,0,3 | 421,480,127,59,124,92,11 | 10,6,0,0,3,2,2,2,0,3,0,0,6,6 | 298 | 149 | 0.009093691251 | 0.52755910017358 | 0.938,0.968,0.973, | 0.955,0.976,1.0,1. | -0,01 |
| 179957 | ENSG00000204252 | HLA-DOA | chr6 | - | 33007310 | 33007608 | 33007079 | 33007215 | 33008012 | 33008261 | 179957 | 99,46,379,361,99,74,161,1 | 0,5,4,3,0,1,3,4,1,1,0,1 | 136,177,73,27,53,56,39,12 | 5,4,0,1,0,1,0,3,0,2,2,4,4,3, | 298 | 149 | 0.0091211264317 | 0.529008128003563 | 1.0,0.821,0.979,0. | 0.932,0.957,1.0,0. | 0,009 |
| 158067 | ENSG00000048405 | ZNF800 | chr7 | - | 127377185 | 127377329 | 127373083 | 127373479 | 127386059 | 127386154 | 158067 | 103,92,171,106,52,43,66,7 | 4,1,11,1,0,2,0,6,0,0,0,0 | 89,142,118,81,81,41,15,57 | 1,8,4,1,5,0,1,3,2,2,3,4,4,0, | 293 | 149 | 0.00912732302169 | 0.529119199283555 | 0.929,0.979,0.888, | 0.978,0.9,0.938,0. | 0,05 |
| 256294 | ENSG00000111450 | STX2 | chr12 | - | 130804162 | 130804543 | 130801414 | 130801488 | 130806981 | 130807090 | 256294 | 1,0,2,0,5,7,9,22,0,0,0,0 | 1,1,10,13,11,13,6,8,5,1,4,7 | 5,7,0,0,9,2,8,10,1,3,3,17,6, | 3,0,1,14,1,2,2,10,2,5,5,16,1 | 298 | 149 | 0.00912795827246 | 0.529119199283555 | 0.333,0.0,0.091,0. | 0.455,1.0,0.0,0.0,0 | -0,209 |
| 124220 | ENSG00000149182 | ARFGAP2 | chr11 | - | 47172709 | 47172751 | 47172280 | 47172333 | 47173425 | 47173482 | 124220 | 3,1,11,5,5,2,6,5,1,2,1,3 | 26,34,61,55,21,19,27,30,2 | 1,3,3,0,6,0,2,6,1,1,1,2,3,0,1 | 31,51,19,20,39,10,26,30,1 | 191 | 149 | 0.00914562052818 | 0.530000283225824 | 0.083,0.022,0.123, | 0.025,0.044,0.11,0 | 0,031 |
| 72396 | ENSG00000020922 | MRE11 | chr11 | - | 94476288 | 94476403 | 94471573 | 94471759 | 94478734 | 94478876 | 72396 | 22,13,95,60,20,13,14,26,2 | 3,3,2,0,2,2,1,0,0,0,2,0 | 52,43,14,22,33,26,25,43,4 | 2,10,1,0,2,2,5,0,2,4,3,5,1,1 | 264 | 149 | 0.00915038271337 | 0.530133518649082 | 0.805,0.71,0.964,1 | 0.936,0.708,0.888, | -0,002 |
| 224866 | ENSG00000177600 | RPLP2 | chr11 | + | 810233 | 810356 | 809966 | 810039 | 811596 | 811839 | 224866 | 295,268,501,137,150,99,1 | 4,0,4,3,2,1,3,6,2,1,0,0 | 174,253,107,136,196,140,1 | 2,3,1,1,0,1,0,0,1,4,0,32,4,3 | 272 | 149 | 0.00916162029513 | 0.530515698624621 | 0.976,1.0,0.986,0. | 0.979,0.979,0.983, | 0,001 |
| 131380 | ENSG00000130921 | MTRFR | chr12 | + | 123233739 | 123233924 | 123233461 | 123233531 | 123253646 | 123253956 | 131380 | 8,3,2,0,0,0,4,5,0,0,0,2 | 10,10,25,15,16,17,7,25,4,9 | 2,1,1,1,0,1,0,5,0,2,2,4,1,0, | 6,17,11,5,11,1,7,7,3,3,6,10 | 298 | 149 | 0.00916190907345 | 0.530515698624621 | 0.286,0.13,0.038,0 | 0.143,0.029,0.043, | -0,036 |
| 152712 | ENSG00000100697 | DICER1 | chr14 | - | 95113150 | 95113224 | 95112171 | 95112247 | 95115666 | 95115821 | 152712 | 51,33,71,39,36,37,28,43,5 | 6,10,8,3,2,12,5,7,1,0,1,2 | 62,41,27,47,26,30,26,21,1 | 5,12,8,18,8,2,4,4,2,4,7,8,5, | 223 | 149 | 0.0091667467779 | 0.530573962415757 | 0.85,0.688,0.856,0 | 0.892,0.695,0.693, | 0,083 |
| 225442 | ENSG00000163492 | CCDC141 | chr2 | - | 178915596 | 178916033 | 178905328 | 178905501 | 178918712 | 178918907 | 225442 | 20,10,31,20,7,4,11,13,19,1 | 225,164,311,178,80,112,1 | 9,21,11,7,16,11,1,15,10,6, | 126,172,129,96,162,91,49, | 298 | 149 | 0.00916784555303 | 0.530573962415757 | 0.043,0.03,0.047,0 | 0.034,0.058,0.041, | 0,015 |
| 97260 | ENSG00000127980 | PEX1 | chr7 | - | 92504731 | 92504902 | 92503040 | 92503195 | 92506247 | 92506344 | 97260 | 48,26,72,57,36,33,32,55,2 | 0,0,6,0,0,4,2,0,2,0,1,0 | 58,54,35,24,40,14,30,27,2 | 3,4,4,0,1,0,0,3,1,4,4,1,1,0, | 298 | 149 | 0.00917625355377 | 0.530917803395557 | 1.0,1.0,0.857,1.0,1 | 0.906,0.871,0.814, | 0,036 |
| 22740 | ENSG00000196497 | IPO4 | chr14 | - | 24187666 | 24187796 | 24187399 | 24187579 | 24188215 | 24188257 | 22740 | 17,10,16,15,11,8,15,11,15, | 0,1,2,0,3,1,2,1,0,0,0,4 | 12,8,8,4,7,2,3,7,4,8,3,38,7, | 0,0,1,2,2,0,4,5,0,0,2,1,1,0, | 279 | 149 | 0.00918783801449 | 0.53138098691692 | 1.0,0.842,0.81,1.0, | 1.0,1.0,0.81,0.516, | 0,068 |
| 217958 | ENSG00000111679 | PTPN6 | chr12 | + | 6957922 | 6958073 | 6957653 | 6957785 | 6959926 | 6959994 | 217958 | 294,136,294,208,320,155,1 | 14,2,17,7,12,5,0,4,4,0,7,1 | 233,164,94,24,50,102,68,1 | 18,7,1,3,4,2,2,8,5,2,2,18,24 | 298 | 149 | 0.00918919687826 | 0.53138098691692 | 0.913,0.971,0.896, | 0.866,0.921,0.979, | 0,034 |
| 97279 | ENSG00000161405 | IKZF3 | chr17 | - | 39777884 | 39778010 | 39777650 | 39777767 | 39788257 | 39788374 | 97279 | 53,60,113,78,68,50,27,24,3 | 167,127,340,225,133,121, | 153,151,41,26,63,53,5,45,5 | 255,239,100,81,80,103,11 | 275 | 149 | 0.00919302221996 | 0.531459405163634 | 0.147,0.204,0.153, | 0.245,0.255,0.182, | -0,05 |
| 34260 | ENSG00000258315 | C17orf49 | chr17 | + | 7016949 | 7017014 | 7016494 | 7016685 | 7017256 | 7017304 | 34260 | 16,3,10,10,11,15,5,23,16,5 | 27,8,15,8,19,10,7,6,14,6,2, | 14,12,15,5,8,3,6,11,2,5,8,1 | 8,3,4,21,1,3,10,7,0,1,7,11,2 | 214 | 149 | 0.00924693700128 | 0.534403868979703 | 0.292,0.207,0.317, | 0.549,0.736,0.723, | -0,204 |
| 218118 | ENSG00000064933 | PMS1 | chr2 | + | 189818737 | 189818829 | 189818126 | 189818180 | 189843963 | 189844080 | 218118 | 1,0,6,2,1,2,3,8,5,0,1,1 | 43,25,62,64,30,30,23,25,7 | 2,2,1,1,2,0,1,2,3,0,3,1,0,0,1 | 42,36,15,11,15,20,7,24,5,3 | 241 | 149 | 0.00924892051772 | 0.534403868979703 | 0.014,0.0,0.056,0. | 0.029,0.033,0.04,0 | 0,003 |
| 28016 | ENSG00000131732 | ZCCHC9 | chr5 | + | 81304740 | 81305141 | 81301623 | 81301670 | 81308560 | 81308711 | 28016 | 12,9,10,16,13,6,10,8,6,4,4, | 0,0,4,3,1,1,2,2,0,0,0,2 | 4,10,8,1,1,7,7,11,1,14,2,21 | 1,1,0,2,1,0,1,2,0,0,1,0,0,0,1 | 298 | 149 | 0.00926920799596 | 0.535432341969529 | 1.0,1.0,0.556,0.72 | 0.667,0.833,1.0,0. | 0,037 |
| 164740 | ENSG00000065882 | TBC1D1 | chr4 | + | 37983144 | 37983257 | 37902002 | 37902512 | 38014508 | 38014973 | 164740 | 1,4,0,0,2,0,2,1,1,2,0,0 | 33,11,46,27,19,12,8,13,19, | 4,0,2,3,1,1,0,0,3,5,0,3,5,0,1 | 29,14,9,14,5,20,16,9,15,23 | 262 | 149 | 0.00927538682653 | 0.535645500955428 | 0.017,0.171,0.0,0. | 0.073,0.0,0.112,0. | -0,032 |
| 71272 | ENSG00000131051 | RBM39 | chr20 | - | 35729461 | 35729551 | 35729311 | 35729365 | 35731940 | 35732135 | 71272 | 160,93,338,153,87,125,79, | 4,0,8,4,1,5,0,1,4,5,0,0 | 170,209,133,102,139,75,6 | 6,0,0,0,0,2,2,1,1,12,13,5,7, | 239 | 149 | 0.00929103852182 | 0.536405448521953 | 0.961,1.0,0.963,0. | 0.946,1.0,1.0,1.0,1 | 0,012 |
| 101318 | ENSG00000173715 | C11orf80 | chr11 | + | 66756352 | 66756505 | 66744735 | 66744919 | 66759042 | 66759084 | 101318 | 36,14,88,56,46,36,33,40,1 | 18,7,32,3,11,12,11,9,2,3,5, | 43,43,30,28,19,13,23,41,2 | 6,5,3,3,3,0,4,4,3,13,3,10,1, | 298 | 149 | 0.00929648864948 | 0.536576172838892 | 0.5,0.5,0.579,0.90 | 0.782,0.811,0.833, | -0,132 |
| 194750 | ENSG00000276087 | | chr2 | + | 24146747 | 24146826 | 24139369 | 24139450 | 24146929 | 24147086 | 194750 | 32,20,43,28,24,27,6,21,5,8 | 4,3,5,5,0,0,0,0,4,0,1,0 | 61,31,20,7,12,12,13,48,3,1 | 0,0,0,0,2,0,1,3,1,4,2,0,0,0,2 | 228 | 149 | 0.00929939429364 | 0.536599982095022 | 0.839,0.813,0.849, | 1.0,1.0,1.0,1.0,0.7 | -0,03 |
| 50327 | ENSG00000132781 | MUTYH | chr1 | - | 45333412 | 45333594 | 45333284 | 45333324 | 45334390 | 45334511 | 50327 | 21,11,19,10,16,23,10,16,1 | 1,2,3,0,0,2,2,0,1,4,0,3 | 21,14,16,16,8,8,7,10,7,7,6, | 1,1,0,3,2,0,0,0,0,0,3,0,1,2 | 298 | 149 | 0.00930641267716 | 0.536861031068835 | 0.913,0.733,0.76,1 | 0.913,0.875,1.0,0. | -0,066 |
| 250856 | ENSG00000165983 | PTER | chr10 | + | 16484336 | 16484816 | 16467131 | 16467210 | 16486351 | 16486617 | 250856 | 16,9,45,27,13,8,14,13,8,6, | 0,4,7,2,0,1,1,0,0,1,0,2 | 30,28,11,5,9,14,9,15,8,13,2 | 1,2,2,0,0,0,0,0,1,0,0,0,0,0 | 298 | 149 | 0.00931907282478 | 0.537447310597595 | 1.0,0.529,0.763,0. | 0.938,0.875,0.733, | -0,107 |
| 44576 | ENSG00000075790 | BCAP29 | chr7 | + | 107613653 | 107613732 | 107613331 | 107613432 | 107618327 | 107618664 | 44576 | 3,0,3,5,2,0,2,0,6,2,0,0 | 14,4,32,21,23,16,15,26,18, | 6,4,4,0,6,1,0,2,0,2,0,4,3,6,1 | 36,14,25,10,14,7,18,18,8,1 | 228 | 149 | 0.00934789924043 | 0.538965363358422 | 0.123,0.0,0.058,0. | 0.098,0.157,0.095, | -0,019 |
| 88107 | ENSG00000168214 | RBPJ | chr4 | + | 26414048 | 26414155 | 26406174 | 26406270 | 26415474 | 26415640 | 88107 | 0,1,3,0,0,4,0,4,0,0,1,2 | 45,26,48,33,43,53,49,75,4 | 3,4,3,3,3,0,3,1,0,0,13,0,0,0 | 42,60,17,9,51,21,44,48,18, | 256 | 149 | 0.00936204011855 | 0.539636115895992 | 0.0,0.022,0.035,0. | 0.04,0.037,0.093,0 | -0,016 |
| 3874 | ENSG00000079432 | CIC | chr19 | + | 42292299 | 42292466 | 42292085 | 42292207 | 42292565 | 42292859 | 3874 | 20,2,20,7,5,21,26,9,32,22,2 | 1,2,3,0,0,2,2,0,1,2,2,2 | 5,13,18,18,8,16,19,10,11,9 | 0,2,0,2,0,1,0,0,0,1,0,0,0,0,5 | 298 | 149 | 0.00936869469619 | 0.539875107939938 | 0.909,0.333,0.769, | 1.0,0.765,1.0,0.81 | -0,126 |
| 102717 | ENSG00000126746 | ZNF384 | chr12 | - | 6678662 | 6678710 | 6678126 | 6678460 | 6678945 | 6679183 | 102717 | 17,11,27,28,20,15,20,21,1 | 2,1,2,4,6,5,0,2,0,1,0,2 | 26,12,20,12,11,12,10,23,7, | 0,0,0,3,0,0,2,1,1,0,0,4,4,2, | 197 | 149 | 0.00938379011338 | 0.540600248900667 | 0.865,0.893,0.911, | 1.0,1.0,1.0,0.752,1 | -0,04 |
| 199266 | ENSG00000166526 | ZNF3 | chr7 | - | 100076277 | 100076550 | 100075541 | 100075630 | 100077302 | 100077839 | 199266 | 3,3,0,1,0,0,2,2,0,1,1,2 | 8,17,16,15,7,5,2,6,6,4,2,8 | 2,2,2,3,4,2,0,5,0,3,1,1,0,1, | 7,15,4,7,14,3,5,12,2,1,11,1 | 298 | 149 | 0.00939581981914 | 0.541148433367225 | 0.158,0.081,0.0,0. | 0.125,0.062,0.2,0. | -0,086 |
| 79926 | ENSG00000129292 | PHF20L1 | chr8 | + | 132798771 | 132798860 | 132794409 | 132794581 | 132803818 | 132804032 | 79926 | 16,16,30,13,7,9,7,11,3,9,7, | 2,2,1,7,0,0,2,1,2,0,0,0 | 7,5,13,4,10,10,11,7,9,13,6, | 3,1,0,0,2,0,0,0,2,1,3,4,1,0, | 238 | 149 | 0.00941649610896 | 0.542194187808339 | 0.834,0.834,0.949, | 0.594,0.758,1.0,1. | -0,008 |
| 169970 | ENSG00000075826 | SEC31B | chr10 | - | 100496939 | 100497280 | 100496257 | 100496431 | 100497666 | 100497793 | 169970 | 34,16,36,33,14,7,21,12,12, | 0,0,1,3,1,7,0,3,1,0,1,5 | 13,13,3,13,16,15,7,13,21,1 | 10,16,3,7,8,1,0,1,1,0,4,4,0, | 298 | 149 | 0.00942650644326 | 0.542625410080046 | 1.0,1.0,0.947,0.84 | 0.394,0.289,0.333, | 0,133 |
| 85529 | ENSG00000123607 | TTC21B | chr2 | - | 165936920 | 165937067 | 165932972 | 165933057 | 165941026 | 165941184 | 85529 | 12,0,9,5,7,0,4,1,4,0,0,5 | 43,32,36,18,11,19,14,19,4 | 12,15,3,10,9,3,3,1,6,3,1,2,1 | 23,14,13,31,7,17,13,18,8,2 | 296 | 149 | 0.00943725553811 | 0.542966373144936 | 0.123,0.0,0.112,0. | 0.208,0.35,0.104,0 | -0,074 |
| 127834 | ENSG00000107679 | PLEKHA1 | chr10 | + | 122428275 | 122428420 | 122426941 | 122427031 | 122429623 | 122432354 | 127834 | 0,0,6,5,0,0,1,5,3,0,0,2 | 26,29,32,4,11,3,16,2,11,7,1 | 2,1,0,0,1,0,0,0,0,0,2,0,0,0 | 10,8,12,4,6,4,10,3,1,10,6,2 | 294 | 149 | 0.00943843604497 | 0.542966373144936 | 0.0,0.0,0.087,0.38 | 0.092,0.06,0.0,0.0, | 0,09 |
| 162672 | ENSG00000072364 | AFF4 | chr5 | - | 132889078 | 132889193 | 132888096 | 132888160 | 132892163 | 132892404 | 162672 | 22,17,90,31,17,32,16,42,6, | 14,3,3,1,3,2,3,8,1,0,0,0 | 45,60,7,42,32,10,22,15,34, | 3,11,1,17,7,4,7,6,3,2,8,3,7, | 264 | 149 | 0.00943999780844 | 0.542966373144936 | 0.47,0.762,0.944,0 | 0.894,0.755,0.798, | 0,112 |
| 88777 | ENSG00000123684 | LPGAT1 | chr1 | - | 211787631 | 211787727 | 211783228 | 211783502 | 211793071 | 211793190 | 88777 | 211,141,138,90,60,69,72,7 | 0,3,0,1,0,0,0,0,0,0,0,5 | 107,114,100,139,76,64,43, | 2,2,1,7,0,1,0,0,2,3,2,2,5,1,4 | 245 | 149 | 0.00945331920539 | 0.543587321906304 | 1.0,0.966,1.0,0.98 | 0.97,0.972,0.984,0 | 0,017 |
| 9685 | ENSG00000178971 | CTC1 | chr17 | - | 8238030 | 8238242 | 8237374 | 8237519 | 8238391 | 8238578 | 9685 | 96,32,154,99,90,101,83,14 | 3,0,5,4,1,9,2,12,3,6,1,0 | 113,153,60,86,88,48,64,10 | 23,13,7,6,3,6,9,4,4,0,4,2,5, | 298 | 149 | 0.00946016003029 | 0.543835390886578 | 0.941,1.0,0.939,0. | 0.711,0.855,0.811, | 0,042 |
| 45045 | ENSG00000111642 | CHD4 | chr12 | - | 6582136 | 6582281 | 6581648 | 6581814 | 6582614 | 6582748 | 45045 | 168,138,268,180,90,122,7 | 14,4,1,1,1,1,1,2,0,4,1,3 | 151,181,81,49,127,76,31,8 | 3,5,6,4,1,1,6,3,1,2,8,1,1,4,5 | 294 | 149 | 0.00947886265955 | 0.544765043812445 | 0.859,0.946,0.993, | 0.962,0.948,0.872, | 0,038 |
| 164118 | ENSG00000102445 | RUBCNL | chr13 | - | 46377646 | 46378005 | 46371940 | 46372597 | 46387676 | 46387831 | 164118 | 2,0,9,1,3,3,1,10,0,0,0,0 | 26,13,33,18,7,16,10,27,12, | 2,7,0,0,1,1,3,0,2,1,1,0,4,0,1 | 24,44,12,27,24,7,19,24,6,4 | 298 | 149 | 0.009493854379 | 0.54548098554366 | 0.037,0.0,0.12,0.0 | 0.04,0.074,0.0,0.0, | 0,006 |
| 254701 | ENSG00000079332 | SAR1A | chr10 | - | 70162689 | 70162817 | 70161857 | 70161931 | 70170412 | 70170523 | 254701 | 22,20,44,26,14,16,16,23,7, | 2,6,9,9,11,2,1,5,3,5,5,2 | 25,52,35,47,44,16,20,34,1 | 5,8,2,0,7,3,4,0,2,3,1,8,5,2,1 | 277 | 149 | 0.00949838445769 | 0.545595619218862 | 0.855,0.642,0.724, | 0.729,0.778,0.904, | -0,138 |
| 32445 | ENSG00000105875 | WDR91 | chr7 | - | 135197043 | 135197423 | 135196143 | 135196337 | 135197992 | 135198151 | 32445 | 10,7,20,9,16,6,5,13,6,6,5,1 | 14,14,44,37,36,35,14,15,3 | 11,22,7,5,5,1,8,13,10,12,6, | 21,28,8,6,7,3,21,25,2,11,2 | 298 | 149 | 0.00950863387208 | 0.545892978640077 | 0.263,0.2,0.185,0. | 0.208,0.282,0.304, | -0,088 |
| 163618 | ENSG00000105875 | WDR91 | chr7 | - | 135197043 | 135197404 | 135196143 | 135196337 | 135197992 | 135201343 | 163618 | 10,7,20,9,16,6,5,13,6,6,5,1 | 14,14,44,37,36,35,14,15,3 | 11,22,7,5,5,1,8,13,10,12,6, | 21,28,8,6,7,3,21,25,2,11,2 | 298 | 149 | 0.00950863387208 | 0.545892978640077 | 0.263,0.2,0.185,0. | 0.208,0.282,0.304, | -0,088 |
| 210935 | ENSG00000115159 | GPD2 | chr2 | + | 156495140 | 156495373 | 156476097 | 156476207 | 156496043 | 156496215 | 210935 | 3,0,2,1,0,0,1,0,0,0,0,0 | 30,14,74,23,7,15,9,9,8,8,2, | 0,3,1,3,3,4,2,0,1,4,0,3,1,0,1 | 25,10,24,9,16,20,9,9,16,23 | 298 | 149 | 0.00951933973516 | 0.546361869476859 | 0.048,0.0,0.013,0. | 0.0,0.13,0.02,0.14 | -0,04 |
| 197895 | ENSG00000135905 | DOCK10 | chr2 | - | 224861181 | 224861219 | 224856859 | 224856982 | 224862663 | 224862746 | 197895 | 2,0,11,3,1,6,1,5,4,4,1,4 | 272,202,313,188,141,103,1 | 7,5,9,15,5,1,4,4,7,11,3,1,3, | 211,223,149,177,176,85,4 | 187 | 149 | 0.0095295441639 | 0.546759765162672 | 0.006,0.0,0.027,0. | 0.026,0.018,0.046, | -0,007 |
| 64257 | ENSG00000119778 | ATAD2B | chr2 | - | 23757417 | 23758101 | 23754646 | 23754774 | 23762208 | 23762346 | 64257 | 66,60,73,14,22,19,24,22,2 | 1,1,3,2,0,1,0,0,0,0,0,0 | 46,51,28,52,21,13,22,28,2 | 1,8,0,0,1,0,0,2,0,0,0,0,4,4,1 | 298 | 149 | 0.00953135300626 | 0.546759765162672 | 0.971,0.968,0.924, | 0.958,0.761,1.0,1. | 0,039 |
| 90866 | ENSG00000085721 | RRN3 | chr16 | - | 15083512 | 15083582 | 15079997 | 15080096 | 15084641 | 15084705 | 90866 | 21,20,72,45,16,19,23,16,7, | 9,0,8,4,1,5,7,6,5,1,0,0 | 32,63,22,23,24,13,20,16,1 | 10,17,2,12,10,2,4,5,5,16,1 | 219 | 149 | 0.00953671760922 | 0.546784215606483 | 0.614,1.0,0.86,0.8 | 0.685,0.716,0.882, | 0,108 |
| 208573 | ENSG00000196504 | PRPF40A | chr2 | - | 152679055 | 152679125 | 152677450 | 152677475 | 152679351 | 152679472 | 208573 | 14,12,42,18,4,6,4,5,3,4,4,4 | 7,4,5,8,2,4,7,8,8,3,0,4 | 25,32,14,10,12,12,4,5,9,13 | 1,1,1,4,9,8,0,0,4,12,3,5,1,5 | 219 | 149 | 0.00953757686985 | 0.546784215606483 | 0.576,0.671,0.851, | 0.944,0.956,0.905, | -0,18 |
| 127740 | ENSG00000107669 | ATE1 | chr10 | - | 121898840 | 121898969 | 121870005 | 121870038 | 121899865 | 121899994 | 127740 | 6,4,23,27,11,9,17,21,9,5,1 | 4,0,17,10,9,9,5,2,6,0,3,11 | 13,21,6,5,8,5,4,14,5,3,13,2 | 14,8,2,4,6,5,7,11,6,12,12,1 | 278 | 149 | 0.00953940059565 | 0.546784215606483 | 0.446,1.0,0.42,0.5 | 0.332,0.585,0.617, | 0,219 |

| | | | | | | | | | | | | | | | | | | | | | |
|---|---|---|---|---|---|---|---|---|---|---|---|---|---|---|---|---|---|---|---|---|---|
| 3162 | ENSG00000105699 | LSR | chr19 | + | 35262545 | 35262692 | 35258944 | 35259064 | 35266358 | 35266532 | 3162 | 137,50,70,35,56,41,71,56,1 | 17,13,3,3,4,7,22,7,22,6,3,1 | 34,27,55,56,40,23,68,94,12 | 2,1,7,5,7,0,5,15,0,6,1,0,4,1 | 296 | 149 | 0.00955701794501 | 0.547590408917274 | 0.802,0.659,0.922, | 0.895,0.931,0.798, | -0,085 |
| 160310 | ENSG00000166888 | STAT6 | chr12 | - | 57109940 | 57110104 | 57108162 | 57108299 | 57111128 | 57111433 | 160310 | 14,16,32,23,17,17,5,18,16, | 55,45,164,114,97,88,46,56 | 15,4,20,7,4,3,2,14,5,9,4,16 | 102,83,36,30,38,49,20,65,3 | 298 | 149 | 0.00955855414091 | 0.547590408917274 | 0.113,0.151,0.089, | 0.068,0.024,0.217, | 0,018 |
| 138376 | ENSG00000157540 | DYRK1A | chr21 | + | 37488198 | 37488351 | 37486466 | 37486614 | 37490174 | 37490461 | 138376 | 6,6,11,10,2,5,1,11,4,5,1,5 | 66,50,199,91,61,84,55,85,5 | 7,14,3,3,2,2,2,6,1,9,4,3,0,3 | 137,148,83,79,52,59,63,85 | 298 | 149 | 0.00959086857219 | 0.549295432054557 | 0.043,0.057,0.027, | 0.025,0.045,0.018, | 0,018 |
| 61151 | ENSG00000009413 | REV3L | chr6 | - | 111313351 | 111313489 | 111311068 | 111311259 | 111315266 | 111315381 | 61151 | 34,25,62,43,16,36,18,33,3 | 8,0,2,3,2,0,1,5,3,0,0,0 | 36,59,31,34,32,16,21,23,2 | 0,1,5,3,0,2,2,0,0,1,0,1,0,3,0 | 287 | 149 | 0.00959461921807 | 0.549364056643901 | 0.688,1.0,0.942,0. | 1.0,0.968,0.763,0. | -0,028 |
| 43951 | ENSG00000154767 | XPC | chr3 | - | 14168256 | 14168380 | 14167168 | 14167253 | 14170437 | 14170460 | 43951 | 51,57,99,94,67,31,34,63,6 | 8,4,7,5,2,4,0,3,0,0,2,0 | 63,65,33,39,58,49,36,51,3 | 0,1,2,3,3,3,3,2,0,0,2,1,0,2,0 | 273 | 149 | 0.00960415342936 | 0.54976370924324 | 0.777,0.886,0.885, | 1.0,0.973,0.9,0.87 | -0,029 |
| 129553 | ENSG00000090006 | LTBP4 | chr19 | + | 40616888 | 40617020 | 40614191 | 40614446 | 40617099 | 40617225 | 129553 | 6,10,27,9,25,9,15,8,9,9,9,1 | 2,0,3,2,4,3,2,0,1,0,1,0 | 12,16,9,4,15,6,4,16,7,6,10, | 0,0,3,2,0,2,0,2,0,2,0,4,1,1,0 | 281 | 149 | 0.00961017934017 | 0.549962379570361 | 0.614,1.0,0.827,0. | 1.0,1.0,0.614,0.51 | 0,003 |
| 96317 | ENSG00000164654 | MIOS | chr7 | + | 7583117 | 7583372 | 7574097 | 7574196 | 7588497 | 7588563 | 96317 | 51,23,42,18,18,17,14,10,3 | 2,8,5,3,0,9,0,3,0,3,2,1 | 17,12,25,10,9,1,17,5,12,22 | 4,8,5,3,3,0,7,6,3,1,7,5,3,1,7 | 298 | 149 | 0.0096217539982 | 0.550478398401538 | 0.927,0.59,0.808,0 | 0.68,0.429,0.714,0 | 0,114 |
| 143803 | ENSG00000125450 | NUP85 | chr17 | + | 75226744 | 75227015 | 75226050 | 75226157 | 75231339 | 75231423 | 143803 | 9,2,1,5,2,0,2,7,10,0,0,0 | 30,18,32,12,24,28,11,12,1 | 0,0,2,4,0,0,0,0,13,7,1,0,1,2 | 30,36,7,3,12,12,19,10,6,19 | 298 | 149 | 0.00962914072299 | 0.550754607214951 | 0.13,0.053,0.015,0 | 0.0,0.0,0.125,0.4,0 | 0 |
| 212889 | ENSG00000152689 | RASGRP3 | chr2 | + | 33520552 | 33520788 | 33519951 | 33520014 | 33521954 | 33522102 | 212889 | 140,38,354,144,105,138,1 | 8,3,18,11,9,8,4,4,6,0,5,3 | 187,216,120,27,70,48,65,1 | 10,10,1,3,0,1,0,3,0,0,6,9,1, | 298 | 149 | 0.00963177150369 | 0.550758717457679 | 0.897,0.864,0.908, | 0.903,0.915,0.984, | -0,036 |
| 35178 | ENSG00000170876 | TMEM43 | chr3 | + | 14128854 | 14129085 | 14125014 | 14125205 | 14129411 | 14129561 | 35178 | 15,4,35,13,11,18,7,20,4,2,8 | 29,18,34,44,16,21,9,30,24, | 11,15,12,15,10,3,12,16,6,9 | 29,18,3,1,4,18,4,28,5,27,21 | 298 | 149 | 0.00965111580658 | 0.55162256637633 | 0.205,0.1,0.34,0.1 | 0.159,0.294,0.667, | -0,164 |
| 83847 | ENSG00000014641 | MDH1 | chr2 | + | 63595422 | 63595519 | 63594487 | 63594586 | 63597398 | 63597538 | 83847 | 91,68,240,216,156,123,83, | 0,1,3,4,1,1,1,3,0,0,0,1 | 150,182,113,18,65,53,88,1 | 3,9,0,0,4,0,1,3,0,3,1,2,3,3,2 | 246 | 149 | 0.0096520045206 | 0.55162256637633 | 1.0,0.976,0.98,0.9 | 0.968,0.925,1.0,1. | 0,016 |
| 224410 | ENSG00000184903 | IMMP2L | chr7 | - | 110835354 | 110835735 | 110663425 | 110663721 | 110886592 | 110886695 | 224410 | 0,0,3,0,2,0,5,9,3,0,4,0 | 5,6,27,37,20,13,15,43,11,6 | 4,0,2,0,0,0,0,0,0,0,2,1,0,0,4 | 30,28,17,11,18,35,16,22,12 | 298 | 149 | 0.00966624582125 | 0.552258272610215 | 0.0,0.0,0.053,0.0,0 | 0.062,0.0,0.056,0. | 0,031 |
| 19913 | ENSG00000196547 | MAN2A2 | chr15 | + | 90916585 | 90916660 | 90916122 | 90916256 | 90918193 | 90918388 | 19913 | 8,1,10,0,1,1,4,2,2,0,0,0 | 42,22,20,4,17,12,22,13,18, | 8,2,1,3,2,1,4,4,1,5,6,10,6,0 | 21,18,8,6,12,3,20,44,10,16 | 224 | 149 | 0.00966825954995 | 0.552258272610215 | 0.112,0.029,0.25,0 | 0.202,0.069,0.077, | -0,071 |
| 58096 | ENSG00000122483 | CCDC18 | chr1 | + | 93217737 | 93217869 | 93216635 | 93216746 | 93221608 | 93221743 | 58096 | 27,17,12,11,8,7,12,4,9,7,0, | 2,1,3,4,2,1,0,0,1,2,2,0 | 9,20,7,2,8,5,2,13,8,13,2,26 | 0,0,0,2,2,0,2,1,0,1,0,0,2,0,2 | 281 | 149 | 0.00968204458005 | 0.552821756467582 | 0.877,0.9,0.68,0.5 | 1.0,1.0,1.0,0.347,0 | -0,078 |
| 60359 | ENSG00000074603 | DPP8 | chr15 | - | 65454262 | 65454415 | 65451959 | 65452102 | 65456224 | 65456371 | 60359 | 23,23,55,34,26,25,23,36,1 | 4,3,9,9,5,1,5,2,3,0,2,4 | 42,67,27,16,22,17,11,20,1 | 3,6,6,0,2,0,0,0,1,5,4,1,0,6,2 | 298 | 149 | 0.00968326134192 | 0.552821756467582 | 0.742,0.793,0.753, | 0.875,0.848,0.692, | -0,106 |
| 225427 | ENSG00000163492 | CCDC141 | chr2 | - | 178855156 | 178855207 | 178853440 | 178853624 | 178855346 | 178855541 | 225427 | 12,8,0,0,0,1,0,0,10,2,0,0 | 203,143,207,121,49,78,11 | 10,12,12,8,0,0,0,0,7,11,7,0 | 70,137,125,65,156,83,49,1 | 200 | 149 | 0.00970046354528 | 0.553656979399141 | 0.042,0.04,0.0,0.0, | 0.096,0.061,0.067, | -0,015 |
| 254118 | ENSG00000100599 | RIN3 | chr14 | + | 92676474 | 92676632 | 92659160 | 92659469 | 92684986 | 92688049 | 254118 | 54,27,54,37,28,18,21,17,2 | 1,5,4,2,0,0,4,1,2,0,0,2 | 26,58,19,13,14,19,20,21,3 | 2,4,0,0,8,0,3,0,1,2,2,1,1,2,1 | 298 | 149 | 0.00970374894583 | 0.553697664199347 | 0.964,0.73,0.871,0 | 0.867,0.879,1.0,1. | 0,012 |
| 16251 | ENSG00000177455 | CD19 | chr16 | + | 28936152 | 28936201 | 28933233 | 28933509 | 28936511 | 28936600 | 16251 | 33,16,27,12,36,30,53,57,3 | 6,2,1,0,0,0,4,0,0,0,1,4 | 16,18,9,6,12,10,30,39,10,1 | 0,2,0,1,1,0,2,1,1,0,3,14,2,2 | 198 | 149 | 0.00970741469428 | 0.553760024400896 | 0.805,0.858,0.953, | 1.0,0.871,1.0,0.81 | 0,045 |
| 53016 | ENSG00000100258 | LMF2 | chr22 | - | 50505056 | 50505153 | 50504801 | 50504984 | 50505228 | 50505257 | 53016 | 56,48,61,33,39,63,63,35,2 | 5,3,2,0,3,2,2,3,4,3,1,5 | 24,50,31,26,11,15,39,63,1 | 1,3,2,0,1,0,0,3,0,1,4,8,0,0,0 | 246 | 149 | 0.00972668769996 | 0.554712432525196 | 0.872,0.906,0.949, | 0.936,0.91,0.904,1 | -0,051 |
| 13619 | ENSG00000102977 | ACD | chr16 | - | 67658554 | 67658641 | 67657746 | 67658362 | 67658719 | 67658816 | 13619 | 48,25,38,6,39,20,28,21,77, | 5,2,0,0,0,2,0,2,6,2,0,1 | 24,9,22,20,4,7,7,27,17,24,3 | 0,0,2,2,0,0,1,1,0,0,3,0,1,0,5 | 236 | 149 | 0.0097299733768 | 0.554752821155508 | 0.858,0.888,1.0,1. | 1.0,1.0,0.874,0.86 | -0,016 |
| 88016 | ENSG00000049089 | COL9A2 | chr1 | - | 40315237 | 40315409 | 40314351 | 40314387 | 40315589 | 40315664 | 88016 | 9,0,14,10,9,9,3,12,2,0,4,3 | 8,3,26,7,40,64,31,19,10,6,6 | 9,14,1,3,3,1,8,17,3,4,8,4,6, | 0,4,18,6,1,2,21,25,0,8,19,1 | 298 | 149 | 0.00973346996633 | 0.55480521036101 | 0.36,0.0,0.212,0.4 | 1.0,0.636,0.027,0. | -0,223 |
| 184123 | ENSG00000180098 | TRNAU1AP | chr1 | + | 28560849 | 28561260 | 28560632 | 28560732 | 28561345 | 28561398 | 184123 | 20,6,32,17,16,11,10,18,8,3 | 13,4,22,15,14,9,9,4,10,5,0, | 12,7,1,3,10,1,2,9,3,14,10,9 | 11,10,6,5,10,4,7,9,2,7,5,30 | 298 | 149 | 0.00974635452427 | 0.55532020001906 | 0.435,0.429,0.421, | 0.353,0.259,0.077, | 0,19 |
| 232783 | ENSG00000104921 | FCER2 | chr19 | - | 7698789 | 7698852 | 7698355 | 7698409 | 7699738 | 7700084 | 232783 | 6,0,3,2,0,1,2,0,1,1,1,5 | 22,13,17,16,8,8,5,5,12,8,0, | 1,2,0,0,1,0,1,1,1,3,0,0,1,1,0 | 10,20,4,8,8,3,1,1,9,1,1,7,1, | 212 | 149 | 0.00974766513965 | 0.55532020001906 | 0.161,0.0,0.11,0.0 | 0.066,0.066,0.0,0. | 0,039 |
| 92398 | ENSG00000137692 | DCUN1D5 | chr11 | - | 103082747 | 103082839 | 103066458 | 103066567 | 103083255 | 103083326 | 92398 | 29,26,44,27,12,4,5,11,7,9,3 | 2,0,1,0,0,0,0,1,0,1,0,0 | 27,47,17,4,25,23,4,13,20,1 | 1,2,1,0,0,0,0,2,5,0,0,2,2,4,3 | 241 | 149 | 0.00975329057089 | 0.555493644578784 | 0.9,1.0,0.965,1.0,1 | 0.943,0.936,0.913, | 0,052 |
| 152951 | ENSG00000257621 | PSMA3-AS1 | chr14 | - | 58293765 | 58294062 | 58291634 | 58291707 | 58297954 | 58298134 | 152951 | 12,15,21,17,11,15,8,13,2,5 | 67,34,129,99,60,80,54,73,5 | 14,23,10,18,12,8,10,12,4,1 | 76,42,35,33,53,24,20,71,41 | 298 | 149 | 0.0097701816441 | 0.556308456465949 | 0.082,0.181,0.075, | 0.084,0.215,0.125, | -0,029 |
| 221652 | ENSG00000188994 | ZNF292 | chr6 | + | 87233615 | 87233650 | 87233426 | 87233527 | 87243474 | 87243611 | 221652 | 3,1,14,2,0,3,2,0,0,0,0,0 | 0,16,47,37,11,15,2,5,9,3,4, | 3,14,4,4,6,3,5,5,3,2,0,0,1,0 | 42,46,10,15,24,15,5,7,15,2 | 184 | 149 | 0.00980051229422 | 0.557887876645666 | 1.0,0.048,0.194,0. | 0.055,0.198,0.245, | 0,026 |
| 205226 | ENSG00000163029 | SMC6 | chr2 | - | 17728175 | 17728586 | 17726388 | 17726469 | 17731077 | 17731139 | 205226 | 2,0,5,0,0,2,0,5,0,0,0,1 | 87,53,125,59,21,15,23,27,5 | 2,4,2,6,1,0,0,2,1,0,5,14,2,0 | 52,71,28,21,37,26,16,34,27 | 298 | 149 | 0.00982564247855 | 0.559032987206698 | 0.011,0.0,0.02,0.0, | 0.019,0.027,0.034, | -0,023 |
| 45831 | ENSG00000179262 | RAD23A | chr19 | + | 12948513 | 12948552 | 12948176 | 12948358 | 12948685 | 12948813 | 45831 | 8,6,21,10,7,16,2,7,3,6,5,9 | 10,6,5,1,3,6,6,5,1,0,2,3 | 6,8,9,5,8,2,9,9,6,5,19,24,9, | 3,0,3,1,4,0,1,0,7,1,2,7,2,1,0 | 188 | 149 | 0.00982582337397 | 0.559032987206698 | 0.388,0.442,0.769, | 0.613,1.0,0.704,0. | -0,162 |
| 223420 | ENSG00000106392 | C1GALT1 | chr7 | + | 7215041 | 7215234 | 7182629 | 7182820 | 7234302 | 7235297 | 223420 | 2,0,4,0,7,4,4,2,4,4,0,5 | 18,7,39,15,22,18,8,11,16,6 | 1,1,0,2,0,0,1,1,1,5,0,11,2,5 | 27,27,8,13,10,7,2,11,10,19 | 298 | 149 | 0.00983438236772 | 0.559245438656847 | 0.053,0.0,0.049,0. | 0.018,0.018,0.0,0. | 0,044 |
| 209172 | ENSG00000115317 | HTRA2 | chr2 | + | 74532598 | 74532714 | 74531855 | 74531925 | 74532819 | 74532934 | 209172 | 25,16,48,43,14,23,21,20,1 | 4,1,3,0,1,3,2,2,2,0,0,4 | 26,30,18,10,17,14,3,24,10, | 2,0,0,4,0,0,0,0,0,0,0,10,1,0 | 265 | 149 | 0.00983676164131 | 0.559245438656847 | 0.778,0.9,0.9,1.0,0 | 0.88,1.0,1.0,0.584, | -0,053 |
| 184994 | ENSG00000108654 | DDX5 | chr17 | - | 64505893 | 64506259 | 64504676 | 64504842 | 64506754 | 64507562 | 184994 | 4,0,2,1,2,0,3,2,2,2,1,2 | 19,20,23,8,6,6,13,11,6,6,0, | 0,0,0,0,0,2,0,0,0,2,4,0,0,3 | 13,12,13,9,11,6,9,6,5,6,9,3 | 298 | 149 | 0.0098396241761 | 0.559245438656847 | 0.095,0.0,0.042,0. | 0.0,0.0,0.0,0.0,0.0, | 0,127 |
| 30710 | ENSG00000124357 | NAGK | chr2 | + | 71071685 | 71071827 | 71070740 | 71070839 | 71072640 | 71072690 | 30710 | 83,56,246,215,172,137,83, | 0,0,5,3,2,3,3,1,4,0,0,0 | 229,170,55,37,40,51,37,11 | 1,3,0,0,0,0,0,13,0,2,2,4,0,7 | 291 | 149 | 0.00983995091875 | 0.559245438656847 | 1.0,1.0,0.962,0.97 | 0.992,0.967,1.0,1. | 0,021 |
| 29151 | ENSG00000134644 | PUM1 | chr1 | - | 30967166 | 30967310 | 30965981 | 30966278 | 30968353 | 30968492 | 29151 | 78,52,148,91,93,55,43,76,1 | 4,0,7,0,2,0,1,0,13,6,2,2 | 78,75,66,61,41,37,38,83,2 | 2,1,1,1,1,3,1,2,0,0,2,1,0,1,5 | 293 | 149 | 0.00984607749881 | 0.559445909753531 | 0.908,1.0,0.915,1. | 0.952,0.974,0.971, | 0,002 |
| 3885 | ENSG00000079805 | DNM2 | chr19 | + | 10796060 | 10796199 | 10795371 | 10795439 | 10798485 | 10798572 | 3885 | 142,51,154,135,75,54,66,5 | 1,3,6,1,0,3,2,1,2,0,2,1 | 73,65,30,40,50,69,27,54,9 | 0,2,0,0,1,0,2,2,0,0,1,2,4,0,1 | 288 | 149 | 0.0098505078426 | 0.559549921739414 | 0.987,0.898,0.93,0 | 1.0,0.944,1.0,1.0,0 | -0,016 |
| 965 | ENSG00000100099 | HPS4 | chr22 | - | 26466050 | 26466104 | 26465454 | 26465551 | 26466225 | 26466262 | 965 | 7,3,14,13,2,0,4,4,0,0,1,5 | 5,6,10,10,9,11,2,4,5,6,0,6 | 9,10,1,3,0,0,1,14,3,4,5,8,8, | 5,2,3,8,1,2,2,2,5,6,4,2,4,5,1 | 203 | 149 | 0.00985496739122 | 0.559655537356114 | 0.507,0.268,0.507, | 0.569,0.786,0.197, | -0,126 |
| 94481 | ENSG00000112695 | COX7A2 | chr6 | - | 75240300 | 75240385 | 75237674 | 75237988 | 75241175 | 75241265 | 94481 | 289,156,454,348,199,169,1 | 19,14,26,17,4,8,16,15,5,6,6 | 274,257,201,101,165,139,1 | 8,34,24,8,13,4,9,30,6,14,17 | 234 | 149 | 0.00987721036495 | 0.56077073702415 | 0.906,0.876,0.917, | 0.956,0.828,0.842, | 0,021 |
| 255343 | ENSG00000069493 | CLEC2D | chr12 | + | 9693824 | 9694091 | 9693033 | 9693115 | 9694759 | 9699553 | 255343 | 4,0,14,4,0,3,5,7,5,0,2,2 | 48,24,69,43,13,16,12,41,2 | 2,4,2,0,0,3,2,5,1,2,4,0,8,2,0 | 44,26,27,24,21,8,5,24,27,1 | 298 | 149 | 0.00989735387069 | 0.561766184320274 | 0.04,0.0,0.092,0.0 | 0.022,0.071,0.036, | 0,01 |
| 255511 | ENSG00000117151 | CTBS | chr1 | - | 84564207 | 84564304 | 84563734 | 84563832 | 84565840 | 84566012 | 255511 | 7,6,40,15,0,3,6,6,2,0,0,0 | 20,12,9,27,13,9,12,8,4,2,3, | 6,5,4,5,7,2,4,3,0,2,0,1,3,0,1 | 13,19,7,14,16,15,14,11,9,1 | 246 | 149 | 0.00990677019237 | 0.562081384736757 | 0.175,0.232,0.729, | 0.218,0.137,0.257, | 0,085 |
| 30063 | ENSG00000153113 | CAST | chr5 | + | 96740744 | 96740783 | 96740037 | 96740118 | 96741265 | 96741317 | 30063 | 45,44,96,97,37,40,20,25,2 | 0,0,93,103,33,50,19,41,15, | 69,98,29,21,38,16,30,27,6 | 0,0,24,30,48,23,22,34,1,0,0 | 188 | 149 | 0.00990813021215 | 0.562081384736757 | 1.0,1.0,0.45,0.427, | 1.0,1.0,0.489,0.35 | -0,093 |
| 95820 | ENSG00000167483 | NIBAN3 | chr19 | + | 17530754 | 17530885 | 17527237 | 17527395 | 17532262 | 17532388 | 95820 | 185,186,376,162,247,149,1 | 45,36,41,14,40,27,53,39,5 | 180,268,167,59,182,101,1 | 26,38,25,17,12,6,18,37,12, | 280 | 149 | 0.00991735403936 | 0.562456397166138 | 0.686,0.733,0.83,0 | 0.787,0.79,0.78,0. | -0,059 |
| 41462 | ENSG00000144468 | RHBDD1 | chr2 | + | 226837547 | 226837770 | 226836055 | 226836087 | 226864603 | 226864856 | 41462 | 7,3,10,13,5,9,3,7,2,2,3,3 | 6,1,7,1,1,2,2,7,2,2,3,4 | 2,7,0,1,10,2,0,6,3,4,3,3,3,4 | 9,13,7,2,3,4,2,4,5,3,5,1,6,1 | 298 | 149 | 0.0099217002343 | 0.562554653089732 | 0.368,0.6,0.417,0. | 0.1,0.212,0.0,0.2,0 | 0,163 |
| 241899 | ENSG00000114999 | TTL | chr2 | + | 112521221 | 112521341 | 112520281 | 112520425 | 112528680 | 112541739 | 241899 | 4,0,4,3,1,0,0,0,5,2,0,1 | 29,23,9,16,18,10,7,8,15,6,5 | 0,1,0,1,0,1,0,0,0,3,0,3,0,0,4 | 14,7,12,5,7,6,4,14,4,15,4,2 | 269 | 149 | 0.00992692700021 | 0.562702771973189 | 0.071,0.0,0.198,0. | 0.0,0.073,0.0,0.1,0 | 0,026 |
| 102408 | ENSG00000130429 | ARPC1B | chr7 | + | 99388654 | 99388843 | 99388038 | 99388261 | 99389904 | 99390012 | 102408 | 9,2,9,3,5,3,15,10,2,3,0,4 | 77,80,208,215,139,96,104, | 8,4,3,2,5,1,3,3,0,0,0,5,5,5,6 | 110,124,68,22,85,59,41,73 | 298 | 149 | 0.00995507401619 | 0.56402946427822 | 0.055,0.012,0.021, | 0.035,0.016,0.022, | 0,006 |
| 27781 | ENSG00000132780 | NASP | chr1 | + | 45617462 | 45617591 | 45616625 | 45616703 | 45618060 | 45618213 | 27781 | 291,160,186,95,91,60,44,4 | 8,2,2,2,0,2,4,4,2,0,0,5 | 147,92,103,55,41,53,42,50 | 5,8,2,7,1,0,3,4,0,4,2,5,2,1,6 | 278 | 149 | 0.00995557301129 | 0.56402946427822 | 0.951,0.977,0.98,0 | 0.94,0.86,0.965,0. | 0,027 |
| 174446 | ENSG00000157020 | SEC13 | chr3 | - | 10301252 | 10301374 | 10300949 | 10301147 | 10304025 | 10304172 | 174446 | 74,63,69,81,20,26,29,23,6 | 7,4,6,11,4,3,1,5,0,3,3,3 | 43,45,45,40,22,18,26,34,1 | 3,12,5,2,7,4,4,0,3,1,5,11,6, | 271 | 149 | 0.00996372490696 | 0.564234056421765 | 0.853,0.896,0.863, | 0.887,0.673,0.832, | 0,066 |
| 113261 | ENSG00000225889 | | chr2 | + | 64229844 | 64229958 | 64229496 | 64229640 | 64251118 | 64251983 | 113261 | 6,2,8,3,3,2,13,8,13,6,2,0 | 92,53,119,64,106,59,138,6 | 3,1,9,1,33,11,3,0,2,2,0,0,3, | 130,232,101,86,148,58,66, | 263 | 149 | 0.00996442728259 | 0.564234056421765 | 0.036,0.021,0.037, | 0.013,0.002,0.048, | 0,012 |
| 164331 | ENSG00000137076 | TLN1 | chr9 | - | 35718355 | 35718406 | 35717618 | 35717786 | 35718811 | 35718910 | 164331 | 35,16,23,21,16,19,14,9,15, | 77,57,91,91,67,71,34,66,8 | 13,14,4,6,17,10,9,13,1,12,8 | 84,87,37,29,42,34,76,59,3 | 200 | 149 | 0.00997635088336 | 0.564760646232629 | 0.253,0.173,0.158, | 0.103,0.107,0.075, | 0,032 |
| 44330 | ENSG00000150779 | TIMM8B | chr11 | - | 112085977 | 112086310 | 112084799 | 112085462 | 112086639 | 112086798 | 44330 | 2,2,13,4,4,2,3,1,4,2,0,2 | 11,7,14,18,5,3,2,2,8,5,3,5 | 6,14,3,4,3,3,2,6,4,8,0,19,7, | 9,11,4,0,1,3,1,4,1,9,13,46,9 | 298 | 149 | 0.0099815627352 | 0.564907107299456 | 0.083,0.125,0.317, | 0.25,0.389,0.273,1 | -0,193 |
| 142971 | ENSG00000108840 | HDAC5 | chr17 | - | 44085021 | 44085168 | 44084554 | 44084675 | 44086571 | 44086755 | 142971 | 38,13,55,40,43,27,31,23,3 | 6,0,1,2,2,1,3,1,0,0,2,2 | 44,24,24,7,10,16,30,31,12, | 2,0,0,3,0,1,1,0,1,1,2,0,0,0,0 | 296 | 149 | 0.0100134412186 | 0.566498469831783 | 0.761,1.0,0.965,0. | 0.917,1.0,1.0,0.54, | -0,042 |
| 217059 | ENSG00000148690 | FRA10AC1 | chr10 | - | 93692008 | 93692325 | 93687403 | 93687449 | 93692645 | 93692729 | 217059 | 15,3,8,6,8,4,4,20,14,4,2,4 | 8,4,2,4,6,1,4,1,2,2,0,5 | 16,7,13,3,1,2,9,9,4,9,3,4,2, | 2,0,1,2,3,3,0,0,0,2,2,4,0,0,0 | 298 | 149 | 0.0100171677979 | 0.566498469831783 | 0.484,0.273,0.667, | 0.8,1.0,0.867,0.42 | -0,155 |
| 149764 | ENSG00000128928 | IVD | chr15 | + | 40414873 | 40414982 | 40412990 | 40413087 | 40415400 | 40415461 | 149764 | 3,2,15,11,5,16,5,10,1,2,2,4 | 2,2,4,2,0,6,1,6,3,0,0,2 | 12,7,2,0,1,3,3,2,4,3,6,1,8,3 | 3,4,3,4,4,1,2,10,1,3,1,1,6,4 | 258 | 149 | 0.0100201335921 | 0.566498469831783 | 0.464,0.366,0.684, | 0.698,0.503,0.278, | 0,229 |
| 110173 | ENSG00000164024 | METAP1 | chr4 | + | 99037766 | 99037941 | 99035399 | 99035460 | 99039373 | 99039465 | 110173 | 7,0,8,4,3,9,1,3,4,0,2,3 | 1,2,16,28,10,8,4,4,9,5,2,1 | 3,1,3,4,4,1,0,2,2,0,2,10,1,1 | 16,10,2,3,10,6,2,8,3,15,13, | 298 | 149 | 0.0100202093316 | 0.566498469831783 | 0.778,0.0,0.2,0.06 | 0.086,0.048,0.429, | 0,129 |
| 68519 | ENSG00000163534 | FCRL1 | chr1 | - | 157807101 | 157807122 | 157803844 | 157804111 | 157820006 | 157820060 | 68519 | 149,46,162,10,42,28,51,60 | 10,6,4,1,3,3,6,5,0,0,2,0 | 119,40,94,35,29,16,41,4,1 | 12,7,12,5,1,0,2,0,4,3,9,3,9, | 170 | 149 | 0.0100420432702 | 0.567583774970697 | 0.929,0.87,0.973,0 | 0.897,0.834,0.873, | 0,034 |
| 19814 | ENSG00000137845 | ADAM10 | chr15 | - | 58698279 | 58698404 | 58682251 | 58682314 | 58717576 | 58717727 | 19814 | 1,2,1,0,1,0,0,6,0,1,1,0 | 45,26,30,24,21,28,22,18,7, | 4,2,0,2,1,0,4,0,5,0,2,4,3,2,3 | 30,38,10,25,15,27,2,20,18, | 274 | 149 | 0.0100513039476 | 0.567958046717221 | 0.012,0.04,0.018,0 | 0.068,0.028,0.0,0. | -0,022 |
| 217058 | ENSG00000148690 | FRA10AC1 | chr10 | - | 93692008 | 93692280 | 93687403 | 93687449 | 93692645 | 93692729 | 217058 | 15,3,7,5,8,4,4,18,14,4,4,4 | 8,4,2,4,6,1,4,1,2,2,0,5 | 16,7,13,3,0,2,9,9,4,9,3,4,2, | 2,0,1,2,3,3,0,0,0,2,2,4,0,0,0 | 298 | 149 | 0.0100568209771 | 0.568083658654738 | 0.484,0.273,0.636, | 0.8,1.0,0.867,0.42 | -0,153 |
| 11864 | ENSG00000108395 | TRIM37 | chr17 | - | 59001597 | 59001741 | 58999380 | 58999459 | 59012327 | 59012446 | 11864 | 12,12,18,19,6,10,1,6,1,13,1 | 5,2,3,4,2,3,0,0,0,1,0,4 | 14,9,8,12,8,12,8,6,6,9,2,8,6 | 7,5,8,4,3,1,4,4,0,0,5,7,2,1,5 | 293 | 149 | 0.0100588057628 | 0.568083658654738 | 0.55,0.753,0.753,0 | 0.504,0.478,0.337, | 0,233 |
| 6995 | ENSG00000101265 | RASSF2 | chr20 | - | 4822328 | 4822403 | 4800971 | 4801062 | 4823556 | 4823608 | 6995 | 24,13,40,39,69,22,29,33,1 | 0,4,0,0,0,1,0,2,0,2,0,1 | 92,35,13,15,16,9,5,5,17,24 | 7,6,0,0,1,0,0,0,0,1,0,6,0,1,3 | 224 | 149 | 0.0100673694826 | 0.568418153491469 | 1.0,0.684,1.0,1.0,1 | 0.897,0.795,1.0,1. | -0,009 |
| 91073 | ENSG00000215375 | MYL5 | chr4 | + | 677921 | 678029 | 676072 | 676209 | 678657 | 678765 | 91073 | 4,7,5,0,1,3,4,2,3,5,2,4 | 6,3,10,6,1,2,1,0,1,2,0,2 | 2,4,4,2,0,1,1,0,3,1,0,8,2,1,1 | 5,5,5,2,2,2,0,10,1,2,1,24,1, | 257 | 149 | 0.0100722755127 | 0.568536185549982 | 0.279,0.575,0.225, | 0.188,0.317,0.317, | 0,135 |
| 44537 | ENSG00000187514 | PTMA | chr2 | + | 231712442 | 231712516 | 231711347 | 231711983 | 231712803 | 231712856 | 44537 | 1027,717,1087,758,865,77 | 19,6,10,8,9,3,12,18,7,7,2,8 | 806,637,579,416,454,264,4 | 9,12,12,15,13,2,7,11,2,7,29 | 223 | 149 | 0.0100747430049 | 0.568536185549982 | 0.973,0.988,0.986, | 0.984,0.973,0.97,0 | 0,007 |
| 7574 | ENSG00000141644 | MBD1 | chr18 | - | 50271468 | 50271540 | 50267516 | 50267663 | 50272676 | 50272738 | 7574 | 8,5,7,10,1,11,7,12,8,6,1,10 | 1,0,1,0,1,0,0,0,1,3,4,2 | 15,12,5,8,5,5,7,4,5,8,9,7,1 | 6,0,1,1,0,0,1,3,0,2,0,6,3,6,0 | 221 | 149 | 0.010079233464 | 0.56864049742862 | 0.844,1.0,0.825,1. | 0.628,1.0,0.771,0. | -0,019 |
| 53601 | ENSG00000105856 | HBP1 | chr7 | + | 107174447 | 107174745 | 107168989 | 107169027 | 107179878 | 107180062 | 53601 | 0,3,11,0,1,1,2,0,0,0,0,0 | 14,9,35,13,8,18,10,13,11,2 | 1,1,4,16,0,4,2,4,0,3,6,3,0,0 | 25,29,23,12,19,4,9,6,12,19 | 298 | 149 | 0.0100827138143 | 0.568687782223691 | 0.0,0.143,0.136,0. | 0.02,0.017,0.08,0. | -0,052 |
| 237264 | ENSG00000120910 | PPP3CC | chr8 | + | 22511168 | 22511231 | 22498000 | 22498112 | 22513292 | 22513432 | 237264 | 58,22,105,67,36,22,30,38,2 | 1,0,3,0,2,4,0,0,1,0,0,2 | 59,52,28,34,41,47,12,35,2 | 4,1,5,2,0,1,2,0,5,1,2,1,2,4,1 | 212 | 149 | 0.0100858020551 | 0.568712932177424 | 0.976,1.0,0.961,1. | 0.912,0.973,0.797, | 0,04 |
| 79839 | ENSG00000005483 | KMT2E | chr7 | + | 105064027 | 105064116 | 105063350 | 105063580 | 105066726 | 105066807 | 79839 | 5,2,10,7,5,1,3,4,10,3,1,1 | 188,152,202,123,62,90,56, | 4,4,3,2,2,0,3,7,0,4,2,6,0,3,0 | 154,227,110,179,121,54,62 | 238 | 149 | 0.0100932047173 | 0.568981284575484 | 0.016,0.008,0.03,0 | 0.016,0.011,0.017, | 0,011 |
| 103752 | ENSG00000122545 | SEPTIN7 | chr7 | + | 35838563 | 35838657 | 35832797 | 35832900 | 35863551 | 35863658 | 103752 | 15,4,38,15,8,10,2,3,4,1,6,5 | 309,194,603,457,280,276,2 | 35,27,15,9,11,3,8,11,6,12,1 | 472,477,287,161,299,260,1 | 243 | 149 | 0.0101037048381 | 0.56942406284606 | 0.029,0.012,0.037, | 0.043,0.034,0.031, | -0,007 |
| 102824 | ENSG00000123066 | MED13L | chr12 | - | 116236820 | 116236890 | 116111427 | 116111512 | 116237467 | 116237569 | 102824 | 5,4,2,3,2,1,2,1,1,0,3,4 | 80,46,86,44,21,33,54,82,2 | 9,3,2,1,5,2,4,14,2,0,10,15,8 | 83,98,22,53,54,31,65,104,6 | 219 | 149 | 0.0101097880709 | 0.56961774771934 | 0.041,0.056,0.016, | 0.069,0.02,0.058,0 | -0,008 |
| 236606 | ENSG00000104783 | KCNN4 | chr19 | - | 43773478 | 43773891 | 43771999 | 43772135 | 43774191 | 43774619 | 236606 | 21,5,18,6,7,8,4,9,22,5,3,12 | 259,98,204,103,120,137,1 | 9,5,3,11,6,6,2,8,3,5,13,0,3, | 141,123,107,155,97,66,79, | 298 | 149 | 0.0101273025104 | 0.570455233346219 | 0.039,0.025,0.042, | 0.031,0.02,0.014,0 | 0,007 |
| 188528 | ENSG00000138802 | SEC24B | chr4 | + | 109481676 | 109481938 | 109473003 | 109473186 | 109491326 | 109491407 | 188528 | 14,3,6,8,0,4,1,9,9,1,1,2 | 51,48,41,22,38,33,27,34,4 | 14,5,11,13,3,0,0,8,6,7,8,1,5 | 30,35,36,46,22,22,21,40,1 | 298 | 149 | 0.0101424266295 | 0.571157672918345 | 0.121,0.03,0.068,0 | 0.189,0.067,0.133, | -0,042 |
| 155385 | ENSG00000100519 | PSMC6 | chr14 | + | 52718239 | 52718352 | 52718080 | 52718142 | 52718976 | 52719038 | 155385 | 23,20,49,34,17,18,21,26,8, | 8,0,2,9,2,11,6,1,0,0,3,2 | 20,29,28,13,11,9,20,30,3,1 | 14,1,4,11,9,5,1,10,6,2,6,16 | 262 | 149 | 0.0101489042294 | 0.57137295479937 | 0.62,1.0,0.933,0.6 | 0.448,0.943,0.799, | 0,152 |
| 36423 | ENSG00000182093 | GET1 | chr21 | + | 39393165 | 39393280 | 39391768 | 39391836 | 39396865 | 39397116 | 36423 | 1,6,20,16,7,7,8,23,1,5,0,8 | 0,1,5,1,2,0,3,2,2,0,1,3 | 13,6,9,4,1,4,1,22,2,17,8,26 | 0,2,2,0,0,0,0,0,1,0,2,4,0,1,2 | 264 | 149 | 0.0101611310517 | 0.571834232637709 | 1.0,0.772,0.693,0. | 1.0,0.629,0.717,1. | -0,149 |
| 28310 | ENSG00000055163 | CYFIP2 | chr5 | + | 157327972 | 157328049 | 157325599 | 157326267 | 157330741 | 157330850 | 28310 | 150,110,249,158,191,165,2 | 6,1,2,1,5,5,9,4,1,1,1,1 | 179,160,103,134,173,110,1 | 2,4,1,18,8,4,2,6,0,0,7,8,2,4 | 226 | 149 | 0.0101640154184 | 0.571834232637709 | 0.943,0.986,0.988, | 0.983,0.963,0.985, | 0,014 |
| 193923 | ENSG00000092841 | MYL6 | chr12 | + | 56159594 | 56159730 | 56158683 | 56158711 | 56159974 | 56160148 | 193923 | 572,475,1650,1311,622,62 | 38,26,57,25,6,14,8,13,16,1 | 771,713,492,270,560,399,2 | 11,37,15,54,46,7,0,10,20,8 | 285 | 149 | 0.0101671306183 | 0.571834232637709 | 0.887,0.905,0.938, | 0.973,0.91,0.945,0 | 0,024 |
| 68837 | ENSG00000126698 | DNAJC8 | chr1 | - | 28230130 | 28230252 | 28228921 | 28229023 | 28232920 | 28233020 | 68837 | 4,0,19,18,0,10,5,10,4,0,2,5 | 18,21,69,43,21,19,27,31,1 | 8,6,6,5,6,3,0,5,2,1,2,8,3,4,2 | 29,40,19,6,17,9,8,31,9,25,4 | 271 | 149 | 0.0101677249481 | 0.571834232637709 | 0.109,0.0,0.131,0. | 0.132,0.076,0.148, | 0,015 |
| 235236 | ENSG00000139218 | SCAF11 | chr12 | - | 45974189 | 45974338 | 45964106 | 45964188 | 45990352 | 45990584 | 235236 | 2,3,10,12,2,2,2,4,2,4,2,0 | 110,76,118,98,66,72,63,68 | 1,0,1,6,2,1,1,0,1,3,1,9,3,8,2 | 119,103,45,70,52,53,25,74 | 298 | 149 | 0.0101736053469 | 0.572015478688253 | 0.009,0.019,0.041, | 0.004,0.0,0.011,0. | 0,011 |
| 185714 | ENSG00000143106 | PSMA5 | chr1 | - | 109413067 | 109413134 | 109412076 | 109412184 | 109415236 | 109415365 | 185714 | 32,22,38,42,28,13,13,15,2 | 4,1,13,5,4,3,7,5,3,0,0,2 | 20,29,23,8,16,17,15,14,15, | 3,13,7,9,12,3,8,7,6,1,6,11,5 | 216 | 149 | 0.0101836237792 | 0.57242923207652 | 0.847,0.938,0.668, | 0.821,0.606,0.694, | 0,123 |
| 240061 | ENSG00000160563 | MED27 | chr9 | - | 131996332 | 131996600 | 131939380 | 131939474 | 132014336 | 132014467 | 240061 | 0,0,0,4,0,4,1,0,1,1,0,0 | 2,4,17,8,8,7,4,3,5,3,2,3 | 4,4,2,5,1,0,2,1,0,0,3,5,0,0,0 | 8,6,2,2,6,1,3,7,2,3,11,9,2,5 | 298 | 149 | 0.0101879438791 | 0.572522545441925 | 0.0,0.0,0.0,0.0,0.2,0.0, | 0.2,0.25,0.333,0.5 | -0,065 |
| 52064 | ENSG00000112972 | HMGCS1 | chr5 | - | 43298008 | 43298134 | 43297001 | 43297166 | 43298517 | 43298975 | 52064 | 7,10,11,26,14,13,11,23,18, | 8,3,9,5,0,2,2,3,2,1,4,0 | 5,33,8,3,16,4,10,8,0,8,4,13 | 6,2,2,8,5,1,6,4,4,3,4,14,6,3 | 275 | 149 | 0.0101993439922 | 0.573013575250639 | 0.322,0.644,0.398, | 0.311,0.899,0.684, | 0,189 |
| 51617 | ENSG00000163565 | IFI16 | chr1 | + | 159053532 | 159053724 | 159051678 | 159052098 | 159054820 | 159055124 | 51617 | 344,149,456,335,186,133,9 | 17,7,31,19,8,21,14,6,4,5,2, | 271,283,127,72,125,178,4 | 32,28,6,2,6,9,8,21,4,11,17, | 298 | 149 | 0.0102093783913 | 0.573427641058948 | 0.91,0.914,0.88,0. | 0.809,0.835,0.914, | 0,044 |
| 160239 | ENSG00000150593 | PDCD4 | chr10 | + | 110894302 | 110894522 | 110894090 | 110894198 | 110895947 | 110896087 | 160239 | 241,194,603,393,151,143,7 | 8,4,23,16,3,6,5,3,0,0,0,1 | 292,344,153,172,177,127,4 | 8,17,6,2,21,3,2,1,16,11,5,0 | 298 | 149 | 0.0102286367615 | 0.574359436163425 | 0.938,0.96,0.929,0 | 0.948,0.91,0.927,0 | 0,038 |
| 71924 | ENSG00000161929 | SCIMP | chr17 | - | 5223353 | 5223456 | 5221286 | 5221350 | 5234734 | 5234823 | 71924 | 146,36,102,107,34,46,55,4 | 3,0,3,13,1,4,0,1,9,3,6,5 | 65,42,35,19,23,50,25,45,3 | 8,2,10,2,2,4,1,9,6,3,2,3,8,1 | 252 | 149 | 0.0102342806934 | 0.574474697726517 | 0.966,1.0,0.953,0. | 0.828,0.925,0.674, | 0,033 |
| 176546 | ENSG00000114446 | IFT57 | chr3 | - | 108215277 | 108215392 | 108213930 | 108214021 | 108218534 | 108218653 | 176546 | 7,3,22,8,3,5,9,28,4,5,5,4 | 187,146,512,396,163,167,1 | 19,24,7,20,16,6,7,6,8,7,17, | 338,271,127,137,117,95,11 | 264 | 149 | 0.0102360276437 | 0.574474697726517 | 0.021,0.011,0.024, | 0.031,0.048,0.03,0 | -0,011 |

| | | | | | | | | | | | | | | | | | | | | | | |
|---|---|---|---|---|---|---|---|---|---|---|---|---|---|---|---|---|---|---|---|---|---|---|
| 135723 | ENSG00000291230 | | chr19 | - | 50334306 | 50334386 | 50333795 | 50334101 | 50334828 | 50334973 | 135723 | 166,121,187,379,156,104,9 | 0,0,0,1,2,0,2,2,0,0,0,0 | 120,101,103,50,96,112,46, | 6,6,0,8,0,0,0,0,4,2,1,8,0,2,0 | 229 | 149 | 0.0102450760025 | 0.574705595619376 | 1.0,1.0,1.0,0.996,0 | 0.929,0.916,1.0,0.8 | 0,026 |
| 58006 | ENSG00000072858 | SIDT1 | chr3 | + | 113576921 | 113576967 | 113567539 | 113567710 | 113583408 | 113583496 | 58006 | 8,10,19,11,7,9,5,7,12,3,2,8 | 0,0,0,0,1,0,0,1,0,0,0,0 | 14,11,10,2,5,5,8,4,8,11,15, | 2,1,0,0,5,0,0,0,4,2,3,2,0,0,1 | 195 | 149 | 0.0102454821582 | 0.574705595619376 | 1.0,1.0,1.0,1.0,0.84 | 0.842,0.894,1.0,1.0 | 0,097 |
| 3936 | ENSG00000161091 | MFSD12 | chr19 | - | 3542775 | 3542977 | 3538274 | 3538756 | 3544808 | 3544939 | 3936 | 15,8,9,8,2,2,5,10,6,10,6,11 | 2,2,0,0,0,0,1,0,0,0,0,0 | 6,4,6,7,11,5,1,12,1,6,8,6,6, | 0,1,1,0,0,0,1,2,1,0,2,4,3,0,1 | 298 | 149 | 0.0102534045232 | 0.574962714351332 | 0.789,0.667,1.0,1.0 | 1.0,0.667,0.75,1.0, | 0,205 |
| 70767 | ENSG00000204475 | NCR3 | chr6 | - | 31589974 | 31590126 | 31589781 | 31589899 | 31592678 | 31592749 | 70767 | 34,5,22,12,54,15,26,16,22, | 3,0,1,0,2,6,2,2,3,0,0,0 | 47,54,9,3,13,17,8,29,25,32 | 2,1,0,0,1,0,0,1,7,4,2,2,0,0,1 | 298 | 149 | 0.0102559413758 | 0.574962714351332 | 0.85,1.0,0.917,1.0, | 0.922,0.964,1.0,1.0 | -0,024 |
| 180330 | ENSG00000291073 | | chr16 | - | 21495170 | 21495427 | 21487577 | 21487616 | 21519827 | 21520444 | 180330 | 9,4,12,10,11,13,22,17,13,1 | 44,31,120,36,32,95,69,60,4 | 4,7,9,11,9,0,15,17,10,11,9, | 63,101,38,130,55,17,27,61 | 298 | 149 | 0.0102580800308 | 0.574962714351332 | 0.093,0.061,0.048, | 0.031,0.033,0.106, | 0,033 |
| 192058 | ENSG00000087206 | UIMC1 | chr5 | - | 176969365 | 176969706 | 176958092 | 176958154 | 176970741 | 176970793 | 192058 | 19,11,56,27,35,30,27,25,7, | 4,1,14,9,0,6,6,11,4,1,0,7 | 38,35,10,32,25,22,19,27,13 | 13,13,2,12,19,13,2,2,4,11,5 | 298 | 149 | 0.0102707569957 | 0.575523379052722 | 0.704,0.846,0.667, | 0.594,0.574,0.714, | 0,124 |
| 158058 | ENSG00000196505 | GDAP2 | chr1 | - | 117884136 | 117884249 | 117883487 | 117883627 | 117886576 | 117886653 | 158058 | 1,0,0,0,1,1,4,5,0,2,0,2 | 14,10,20,18,11,8,9,4,4,7,2, | 4,3,1,1,5,0,1,4,3,6,2,1,3,1,1 | 15,15,7,5,12,8,6,9,8,5,13,8 | 262 | 149 | 0.0102851516764 | 0.576179979298086 | 0.039,0.0,0.0,0.0,0 | 0.132,0.102,0.075, | -0,039 |
| 101776 | ENSG00000164896 | FASTK | chr7 | - | 151078841 | 151079021 | 151077620 | 151078092 | 151079499 | 151079922 | 101776 | 58,39,59,43,40,62,24,32,54 | 2,7,8,8,1,6,0,5,6,0,0,7 | 21,14,16,11,16,21,12,35,18 | 7,7,2,1,2,1,5,1,2,2,10,7,6,6 | 298 | 149 | 0.0102933806702 | 0.576490922463652 | 0.935,0.736,0.787, | 0.6,0.5,0.8,0.846,0 | 0,118 |
| 127504 | ENSG00000107862 | GBF1 | chr10 | + | 102359269 | 102359435 | 102358505 | 102358729 | 102360183 | 102360395 | 127504 | 94,64,73,40,53,63,39,60,90 | 7,2,1,3,0,0,5,2,2,2,2,0 | 39,37,36,67,45,28,13,64,19 | 6,0,0,2,0,0,4,2,0,0,0,0,1,1,0 | 298 | 149 | 0.0103106613661 | 0.577308521458655 | 0.87,0.941,0.973,0 | 0.765,1.0,1.0,0.94 | -0,014 |
| 28522 | ENSG00000170234 | PWWP2A | chr5 | - | 160108544 | 160108614 | 160093757 | 160094065 | 160118804 | 160118833 | 28522 | 2,2,6,6,4,4,3,4,7,3,1,2 | 2,2,12,2,7,3,5,4,2,0,7,9 | 5,4,1,0,2,3,1,4,0,3,1,2,2,7,1 | 9,12,8,3,3,1,5,7,1,7,13,10,5 | 219 | 149 | 0.0103201605376 | 0.577690109926706 | 0.405,0.405,0.254, | 0.274,0.185,0.078, | 0,217 |
| 174878 | ENSG00000127483 | HP1BP3 | chr1 | - | 20780344 | 20780558 | 20779811 | 20779911 | 20787194 | 20787323 | 174878 | 73,52,141,112,60,48,62,55 | 15,10,15,15,5,3,3,1,11,7,2, | 68,85,49,23,44,41,25,52,25 | 10,8,8,6,4,3,3,4,1,2,17,8,1, | 298 | 149 | 0.0103308634707 | 0.578138865226789 | 0.709,0.722,0.825, | 0.773,0.842,0.754, | -0,059 |
| 161884 | ENSG00000198055 | GRK6 | chr5 | + | 177433350 | 177433410 | 177433146 | 177433239 | 177433535 | 177433676 | 161884 | 34,19,69,108,20,30,23,22,1 | 9,4,5,1,0,2,0,4,1,0,0,2 | 35,41,17,20,28,24,15,17,28 | 0,5,12,16,2,0,3,4,7,0,4,3,1, | 209 | 149 | 0.0103376674168 | 0.578369247669686 | 0.729,0.772,0.908, | 1.0,0.854,0.502,0.4 | 0,076 |
| 93126 | ENSG00000135317 | SNX14 | chr6 | - | 85538837 | 85538864 | 85536791 | 85536924 | 85541984 | 85542043 | 93126 | 69,35,57,54,34,23,17,22,55 | 73,59,67,54,44,24,34,30,74 | 39,14,20,25,14,13,12,27,4, | 37,41,35,35,27,15,16,23,23 | 176 | 149 | 0.0103417897985 | 0.578449521860955 | 0.445,0.334,0.419, | 0.472,0.224,0.326, | 0,066 |
| 68496 | ENSG00000163534 | FCRL1 | chr1 | - | 157802376 | 157802664 | 157801914 | 157802193 | 157807101 | 157807122 | 68496 | 143,49,359,21,114,55,154, | 2,0,6,3,0,0,1,0,0,0,1,0 | 234,186,226,89,49,35,136, | 2,1,3,1,3,2,2,0,2,0,0,3,8,6,6 | 298 | 149 | 0.0103471420716 | 0.578501795792214 | 0.973,1.0,0.968,0.7 | 0.983,0.989,0.974, | 0,011 |
| 30714 | ENSG00000124357 | NAGK | chr2 | + | 71075554 | 71075642 | 71072640 | 71073594 | 71076603 | 71076641 | 30714 | 25,22,188,157,87,73,50,58 | 3,5,8,1,3,4,5,8,0,1,1,1 | 99,68,21,18,22,39,20,74,17 | 3,12,5,2,3,5,2,6,5,2,1,8,6,5 | 237 | 149 | 0.0103511378729 | 0.578501795792214 | 0.84,0.734,0.937,0 | 0.954,0.781,0.725, | 0,048 |
| 45735 | ENSG00000173889 | PHC3 | chr3 | - | 170149085 | 170149244 | 170145422 | 170145521 | 170171372 | 170171450 | 45735 | 220,178,326,184,145,142,1 | 31,12,35,23,4,31,22,16,17, | 256,177,156,134,112,122,9 | 34,46,25,29,26,14,18,29,18 | 298 | 149 | 0.0103511899037 | 0.578501795792214 | 0.78,0.881,0.823,0 | 0.79,0.658,0.757,0 | 0,06 |
| 32843 | ENSG00000102445 | RUBCNL | chr13 | - | 46371940 | 46372210 | 46368732 | 46368815 | 46387133 | 46387252 | 32843 | 122,124,948,699,249,376,4 | 3,2,10,2,2,6,5,6,0,2,0,1 | 542,784,275,254,399,203,2 | 11,12,2,2,4,2,1,3,2,1,8,4,9, | 298 | 149 | 0.0103543896924 | 0.578501795792214 | 0.953,0.969,0.979, | 0.961,0.97,0.986,0 | 0,008 |
| 119596 | ENSG00000254536 | | chr10 | + | 133401528 | 133401590 | 133399528 | 133399619 | 133402148 | 133402245 | 119596 | 16,10,24,19,10,9,20,10,12, | 5,0,9,10,5,13,6,6,2,3,3,6 | 10,10,6,3,6,8,8,12,3,3,2,31 | 0,14,1,2,2,4,0,2,0,2,1,4,3,4 | 211 | 149 | 0.0103561634671 | 0.578501795792214 | 0.693,1.0,0.653,0.5 | 1.0,0.335,0.809,0.5 | -0,136 |
| 23004 | ENSG00000105122 | RASAL3 | chr19 | - | 15452044 | 15452108 | 15451623 | 15451938 | 15452657 | 15452815 | 23004 | 51,35,45,47,27,35,19,27,40 | 0,2,1,1,0,3,2,2,4,3,2,0 | 18,26,15,29,12,13,11,49,20 | 0,0,3,2,0,0,0,0,0,0,1,2,0,5 | 213 | 149 | 0.0103798602565 | 0.579675065611508 | 1.0,0.924,0.969,0.9 | 1.0,1.0,0.778,0.91, | -0,04 |
| 102338 | ENSG00000006576 | PHTF2 | chr7 | + | 77937709 | 77937838 | 77929108 | 77929327 | 77940029 | 77940302 | 102338 | 62,35,118,103,50,40,55,52 | 3,0,4,2,2,0,3,1,4,0,0,2 | 129,143,74,69,84,45,22,23 | 9,2,0,16,5,1,0,0,0,5,2,0,3,4 | 278 | 149 | 0.0103951295217 | 0.580377204172507 | 0.917,1.0,0.941,0.9 | 0.885,0.975,1.0,0.6 | 0,003 |
| 6172 | ENSG00000171456 | ASXL1 | chr20 | + | 32429900 | 32430053 | 32429337 | 32429431 | 32431320 | 32431484 | 6172 | 42,60,50,19,22,29,32,35,34 | 3,1,10,5,1,10,6,11,9,5,0,0 | 18,49,47,66,22,10,7,34,28, | 6,2,3,0,1,1,3,7,1,2,1,3,3,1,5 | 298 | 149 | 0.0104072921586 | 0.580905575359864 | 0.875,0.968,0.714, | 0.6,0.925,0.887,1.0 | -0,057 |
| 91542 | ENSG00000100503 | NIN | chr14 | - | 50759856 | 50760359 | 50754741 | 50754867 | 50761789 | 50761911 | 91542 | 180,118,216,141,108,80,77 | 14,3,22,6,5,10,3,10,0,1,2,1 | 91,102,102,105,65,62,44,9 | 8,10,2,7,9,6,8,4,3,2,12,21,1 | 298 | 149 | 0.01041151043 | 0.580990355550773 | 0.865,0.952,0.831, | 0.85,0.836,0.962,0 | 0,052 |
| 44939 | ENSG00000164073 | MFSD8 | chr4 | - | 127938782 | 127938836 | 127932421 | 127933093 | 127939852 | 127939997 | 44939 | 10,8,9,4,6,9,4,10,15,1,5,10 | 6,0,3,0,3,0,3,0,0,0,2,0 | 13,13,7,3,5,1,3,10,5,8,6,10 | 0,1,3,4,0,1,2,0,0,0,2,0,1,1,0 | 203 | 149 | 0.0104194352587 | 0.581281874070829 | 0.55,1.0,0.688,1.0, | 1.0,0.905,0.631,0.3 | 0,019 |
| 144334 | ENSG00000215041 | NEURL4 | chr17 | - | 7318275 | 7318356 | 7318064 | 7318172 | 7318494 | 7318613 | 144334 | 9,2,7,3,2,9,3,3,10,8,5,7 | 0,0,4,1,1,0,0,3,0,2,2,4 | 6,1,2,4,1,2,2,0,1,8,13,1,6,9 | 0,2,4,0,0,0,0,2,0,0,0,0,0,1,0 | 230 | 149 | 0.0104390944246 | 0.582227709795564 | 1.0,1.0,0.531,0.66, | 1.0,0.245,0.245,1.0 | -0,02 |
| 207590 | ENSG00000146192 | FGD2 | chr6 | + | 37026090 | 37026325 | 37025791 | 37025938 | 37027428 | 37029069 | 207590 | 55,39,59,45,39,9,8,15,80,3 | 40,26,95,72,66,39,40,23,66 | 16,14,6,30,11,17,7,19,28,3 | 37,44,22,17,39,26,24,17,51 | 298 | 149 | 0.0104446027803 | 0.582384015804857 | 0.407,0.429,0.237, | 0.178,0.137,0.12,0 | 0,061 |
| 45833 | ENSG00000179262 | RAD23A | chr19 | + | 12949080 | 12949159 | 12948685 | 12948813 | 12949277 | 12949408 | 45833 | 23,16,43,40,22,18,30,15,22 | 2,0,1,4,2,4,2,2,0,2,0,2 | 41,20,16,9,10,13,16,39,21, | 2,10,0,0,1,2,2,2,0,0,4,10,0, | 228 | 149 | 0.010449262213 | 0.582492917732764 | 0.883,1.0,0.966,0.8 | 0.931,0.567,1.0,1.0 | -0,016 |
| 184491 | ENSG00000077044 | DGKD | chr2 | + | 233434768 | 233434812 | 233434379 | 233434484 | 233435817 | 233435924 | 184491 | 48,32,54,30,23,42,29,38,40 | 3,8,2,2,2,4,1,1,4,2,0,0 | 23,12,33,54,23,17,21,20,15 | 4,7,3,0,0,1,3,5,2,3,5,1,2,8,2 | 193 | 149 | 0.0104556750065 | 0.582699478851373 | 0.925,0.755,0.954, | 0.816,0.57,0.895,1 | 0,066 |
| 174771 | ENSG00000187514 | PTMA | chr2 | + | 231711886 | 231711947 | 231711347 | 231711419 | 231712442 | 231712675 | 174771 | 95,74,50,47,27,28,26,18,65 | 72,29,55,23,35,27,46,37,24 | 32,23,37,22,25,20,12,22,13 | 59,49,18,53,37,8,14,42,18, | 210 | 149 | 0.0104629606468 | 0.582954564579708 | 0.484,0.644,0.392, | 0.278,0.25,0.593,0 | 0,103 |
| 136922 | ENSG00000079805 | DNM2 | chr19 | + | 10796163 | 10796199 | 10795371 | 10795439 | 10798485 | 10798562 | 136922 | 48,17,21,24,14,15,28,16,35 | 1,3,6,1,0,3,2,1,2,0,2,1 | 13,10,12,13,11,12,11,17,23 | 0,2,0,0,1,0,2,2,0,0,1,2,4,0,1 | 185 | 149 | 0.010465924428 | 0.582968783789562 | 0.975,0.82,0.738,0 | 1.0,0.801,1.0,1.0,0 | -0,022 |
| 101386 | ENSG00000169871 | TRIM56 | chr7 | + | 101087005 | 101087168 | 101085480 | 101085571 | 101087311 | 101088128 | 101386 | 14,13,53,48,30,28,19,32,11 | 0,2,5,0,0,0,0,0,0,0,2,0 | 33,45,13,10,19,16,8,37,27, | 0,0,1,2,3,0,0,5,0,1,4,9,0,1,0 | 298 | 149 | 0.0104691034011 | 0.582994979074296 | 1.0,0.765,0.841,1.0 | 1.0,1.0,0.867,0.714 | 0,04 |
| 22766 | ENSG00000100897 | DCAF11 | chr14 | + | 24116916 | 24117044 | 24115373 | 24115749 | 24117265 | 24117306 | 22766 | 114,70,145,158,72,61,35,5 | 8,1,4,1,3,2,4,2,1,0,0,1 | 55,77,65,27,69,47,37,62,37 | 6,5,4,2,5,1,2,3,2,2,1,7,0,5,3 | 277 | 149 | 0.0104794936224 | 0.583282465955978 | 0.885,0.974,0.951, | 0.831,0.892,0.897, | 0,034 |
| 63784 | ENSG00000145860 | RNF145 | chr5 | - | 159203433 | 159203656 | 159194715 | 159194824 | 159209230 | 159209510 | 63784 | 18,25,48,32,20,16,26,39,15 | 0,1,1,0,1,0,4,5,0,0,0,4 | 29,33,14,7,23,5,10,24,16,1 | 8,0,1,3,0,2,2,3,0,1,7,2,4,2,2 | 298 | 149 | 0.0104796859925 | 0.583282465955978 | 1.0,0.926,0.96,1.0, | 0.644,1.0,0.875,0.5 | 0,099 |
| 234209 | ENSG00000170430 | MGMT | chr10 | + | 129646761 | 129646834 | 129536240 | 129536377 | 129707894 | 129708043 | 234209 | 4,2,9,5,0,0,0,0,0,0,2,0 | 16,7,15,23,13,22,21,36,15, | 0,0,3,0,0,0,0,0,0,0,0,0,2,0 | 32,23,26,5,15,8,12,28,8,20 | 222 | 149 | 0.0104927904837 | 0.583860855376741 | 0.144,0.161,0.287, | 0.0,0.0,0.072,0.0,0 | 0,061 |
| 59848 | ENSG00000129084 | PSMA1 | chr11 | - | 14519904 | 14520211 | 14518996 | 14519041 | 14520296 | 14520445 | 59848 | 4,2,7,6,0,3,4,3,3,2,2,2 | 14,13,72,46,26,21,18,22,5, | 3,3,3,11,1,9,2,7,0,1,9,10,7, | 26,39,20,11,27,20,7,22,11, | 298 | 149 | 0.0105119923571 | 0.584598671983346 | 0.125,0.071,0.046, | 0.055,0.037,0.07,0 | -0,025 |
| 122157 | ENSG00000089597 | GANAB | chr11 | - | 62639929 | 62640066 | 62639626 | 62639731 | 62646561 | 62646617 | 122157 | 0,0,2,4,0,5,1,4,0,2,0,3 | 6,3,20,15,17,7,11,11,3,2,3, | 2,1,2,0,0,0,0,0,0,0,0,4,1,2 | 5,12,4,5,4,4,7,11,8,4,13,6,1 | 286 | 149 | 0.0105135687005 | 0.584598671983346 | 0.0,0.0,0.05,0.122, | 0.172,0.042,0.207, | 0,039 |
| 58681 | ENSG00000114098 | ARMC8 | chr3 | + | 138237481 | 138237572 | 138235033 | 138235114 | 138239467 | 138239528 | 58681 | 47,42,41,27,12,13,9,29,17, | 0,0,3,0,3,0,0,0,2,0,0,0 | 43,37,20,16,37,13,10,16,10 | 1,2,2,4,0,1,2,0,2,2,2,0,2,0,3 | 240 | 149 | 0.0105141985088 | 0.584598671983346 | 1.0,1.0,0.895,1.0,0 | 0.964,0.92,0.861,0 | 0,057 |
| 225148 | ENSG00000068885 | IFT80 | chr3 | - | 160373614 | 160373672 | 160366092 | 160366152 | 160375811 | 160375880 | 225148 | 5,4,6,4,6,1,2,1,2,2,0,1 | 72,41,61,44,41,15,10,21,43 | 2,3,1,4,7,3,2,9,7,5,3,3,5,3,3 | 15,17,21,19,27,28,16,42,21 | 207 | 149 | 0.0105460454659 | 0.586217952394402 | 0.048,0.066,0.066, | 0.088,0.113,0.033, | -0,041 |
| 154370 | ENSG00000072042 | RDH11 | chr14 | - | 67692437 | 67692846 | 67691139 | 67691244 | 67692933 | 67693052 | 154370 | 3,10,19,23,17,2,15,25,12,4 | 1,0,3,3,2,1,1,1,1,0,0,2 | 19,9,8,4,2,17,5,28,5,16,13, | 3,1,3,1,2,0,0,2,0,1,2,0,2,2,4 | 298 | 149 | 0.010553379635 | 0.586461081896963 | 0.6,1.0,0.76,0.793, | 0.76,0.818,0.571,0 | 0,02 |
| 193926 | ENSG00000092841 | MYL6 | chr12 | + | 56160625 | 56160846 | 56160242 | 56160320 | 56161386 | 56161587 | 193926 | 2,4,56,41,4,11,7,6,6,3,5,2 | 263,189,911,837,411,375,3 | 9,5,5,0,9,7,7,3,1,11,4,2,14, | 586,439,311,167,324,245,2 | 298 | 149 | 0.010555868956 | 0.586461081896963 | 0.004,0.01,0.03,0.0 | 0.008,0.006,0.008, | 0,003 |
| 235887 | ENSG00000167302 | TEPSIN | chr17 | - | 81232306 | 81232518 | 81231861 | 81232021 | 81233431 | 81233503 | 235887 | 2,7,7,1,2,1,23,9,15,6,8,13 | 2,3,4,0,1,0,2,2,4,1,1,4 | 2,0,4,1,1,2,4,6,0,13,22,25,9 | 1,3,1,3,5,2,2,6,4,3,9,10,3,0 | 298 | 149 | 0.0105691357437 | 0.586983250029177 | 0.333,0.538,0.467, | 0.5,0.0,0.667,0.14 | 0,231 |
| 230205 | ENSG00000179454 | KLHL28 | chr14 | - | 44931332 | 44931400 | 44924323 | 44929191 | 44934114 | 44934558 | 230205 | 4,5,19,7,5,1,5,4,2,1,1,4 | 1,0,4,2,3,0,0,2,0,0,2,0 | 5,6,7,0,5,3,7,7,4,1,4,7,0,2,6 | 3,1,0,8,1,0,0,2,1,5,1,3,3,0,0 | 217 | 149 | 0.010570722048 | 0.586983250029177 | 0.733,1.0,0.765,0.7 | 0.534,0.805,1.0,0.0 | 0,109 |
| 13137 | ENSG00000006194 | ZNF263 | chr16 | + | 3286022 | 3286149 | 3283441 | 3284205 | 3288453 | 3288570 | 13137 | 15,11,11,5,4,8,6,2,9,5,1,8 | 3,3,5,5,3,2,0,2,4,4,2,1 | 9,10,7,7,5,2,5,3,4,10,12,7,5 | 1,1,0,0,1,0,3,3,0,7,0,3,3,2,0 | 276 | 149 | 0.010574797298 | 0.5870580854387 | 0.73,0.664,0.543,0 | 0.829,0.844,1.0,1.0 | -0,19 |
| 134824 | ENSG00000128159 | TUBGCP6 | chr22 | - | 50221779 | 50221874 | 50219956 | 50220015 | 50222027 | 50222102 | 134824 | 48,29,49,20,22,32,27,29,24 | 6,4,5,0,9,3,3,10,7,0,3,2 | 31,20,9,15,13,9,15,20,17,2 | 3,10,1,4,9,1,1,8,2,4,5,9,4,6 | 244 | 149 | 0.0105794391376 | 0.587164328268382 | 0.83,0.816,0.857,1 | 0.863,0.55,0.846,0 | 0,111 |
| 17341 | ENSG00000101266 | CSNK2A1 | chr20 | - | 527932 | 528049 | 508450 | 508660 | 543671 | 543712 | 17341 | 2,2,11,11,8,13,4,2,7,2,2,4 | 15,15,57,35,24,31,11,34,18 | 9,6,7,3,0,12,3,9,8,6,4,5,20, | 21,29,13,6,28,5,15,25,7,19 | 266 | 149 | 0.0105973363618 | 0.588006007343794 | 0.069,0.069,0.098, | 0.194,0.104,0.232, | -0,084 |
| 101762 | ENSG00000164896 | FASTK | chr7 | - | 151076912 | 151077027 | 151076623 | 151076832 | 151077100 | 151077236 | 101762 | 20,14,55,40,25,20,36,27,19 | 0,0,7,0,0,2,0,3,0,0,3,3 | 18,42,7,2,12,13,14,24,16,2 | 0,1,0,0,0,2,0,0,0,0,1,2,1,0,0 | 264 | 149 | 0.0106075077899 | 0.588418687919579 | 1.0,1.0,0.816,1.0,1 | 1.0,0.96,1.0,1.0,1.0 | -0,044 |
| 39137 | ENSG00000242086 | MUC20-OT1 | chr3 | + | 195689338 | 195689438 | 195688366 | 195688797 | 195699029 | 195699170 | 39137 | 60,35,29,30,25,24,12,33,23 | 10,2,4,3,4,3,0,1,5,5,2,0 | 60,60,38,51,16,12,39,60,8, | 5,0,1,5,0,0,1,8,5,0,1,0,1,0,4 | 249 | 149 | 0.0106174753465 | 0.588819849601273 | 0.782,0.913,0.813, | 0.878,1.0,0.958,0.8 | -0,05 |
| 160920 | ENSG00000164329 | TENT2 | chr5 | + | 79656957 | 79657005 | 79648861 | 79649190 | 79668891 | 79669028 | 160920 | 41,26,22,18,13,18,12,22,25 | 0,0,6,5,3,2,0,10,0,0,2,5 | 22,17,20,7,11,5,14,18,5,17 | 3,7,7,1,5,3,0,3,3,3,4,2,7,2,3 | 197 | 149 | 0.0106314654672 | 0.58927980240006 | 1.0,1.0,0.735,0.73 | 0.847,0.647,0.684, | 0,07 |
| 163262 | ENSG00000113391 | FAM172A | chr5 | - | 94057080 | 94057203 | 94053125 | 94053227 | 94111510 | 94111699 | 163262 | 8,3,4,1,4,9,2,3,0,4,0,1 | 45,16,55,33,15,35,21,9,19, | 2,2,3,2,0,2,0,4,1,0,0,1,3,3,1 | 18,9,7,16,18,3,9,7,12,15,31 | 272 | 149 | 0.0106333719216 | 0.58927980240006 | 0.089,0.093,0.038, | 0.057,0.109,0.19,0 | -0,002 |
| 69753 | ENSG00000172531 | PPP1CA | chr11 | - | 67401067 | 67401199 | 67398939 | 67400919 | 67401727 | 67401788 | 69753 | 238,176,405,337,193,175,1 | 2,4,10,4,3,3,1,1,4,4,0,0 | 229,226,137,57,119,109,65 | 3,0,0,3,0,0,0,1,2,3,2,1,2,4,2 | 281 | 149 | 0.0106339828023 | 0.58927980240006 | 0.984,0.959,0.956, | 0.976,1.0,1.0,0.91, | -0,005 |
| 241689 | ENSG00000175893 | ZDHHC21 | chr9 | - | 14677297 | 14677500 | 14674186 | 14674385 | 14680032 | 14680162 | 241689 | 18,31,60,47,4,13,15,24,25, | 12,10,23,19,5,5,8,9,2,2,1,7 | 14,8,10,3,10,16,6,38,5,18,3 | 11,21,6,0,9,5,2,13,16,19,15 | 298 | 149 | 0.0106490918515 | 0.589965170731067 | 0.429,0.608,0.566, | 0.389,0.16,0.455,1 | 0,11 |
| 208572 | ENSG00000196504 | PRPF40A | chr2 | - | 152679055 | 152679116 | 152677450 | 152677475 | 152679351 | 152679472 | 208572 | 15,12,42,18,4,5,4,5,5,4,3,4 | 7,4,5,8,2,4,7,8,8,3,0,4 | 26,32,14,11,12,12,4,5,9,11 | 1,1,1,4,9,8,0,0,4,12,3,5,1,5 | 210 | 149 | 0.0106698854341 | 0.590965031360468 | 0.603,0.68,0.856,0 | 0.949,0.958,0.909, | -0,171 |
| 54108 | ENSG00000175309 | PHYKPL | chr5 | - | 178223319 | 178223451 | 178222851 | 178222934 | 178224615 | 178224729 | 54108 | 5,4,1,3,9,4,0,4,0,0,0,5 | 8,5,20,13,18,7,28,12,10,3,8 | 4,1,2,1,0,2,4,3,0,0,3,0,3,3,2 | 13,13,7,1,12,9,11,14,2,1,7, | 281 | 149 | 0.0106727020678 | 0.590968957744961 | 0.249,0.298,0.026, | 0.14,0.039,0.132,0 | 0,023 |
| 90167 | ENSG00000224078 | SNHG14 | chr15 | + | 25085189 | 25085322 | 25082767 | 25083527 | 25091589 | 25091719 | 90167 | 10,2,35,22,1,10,9,14,8,3,8, | 5,1,0,0,2,1,0,1,0,0,2,4 | 13,15,21,1,5,2,10,11,3,12,1 | 0,4,0,0,0,0,0,4,0,0,0,0,2,0,1 | 282 | 149 | 0.010686037861 | 0.591531667029993 | 0.514,0.514,1.0,1.0 | 1.0,0.665,1.0,1.0,1 | -0,179 |
| 57184 | ENSG00000134242 | PTPN22 | chr1 | - | 113825141 | 113825172 | 113819298 | 113819654 | 113829591 | 113829707 | 57184 | 208,120,332,310,128,91,11 | 4,2,5,5,2,1,4,3,7,3,0,2 | 197,244,107,32,74,66,46,6 | 0,3,0,1,0,0,1,2,0,2,1,7,1,2,3 | 180 | 149 | 0.010688361124 | 0.591531667029993 | 0.977,0.98,0.982,0 | 1.0,0.985,1.0,0.964 | -0,009 |
| 104476 | ENSG00000111412 | SPRING1 | chr12 | - | 116720295 | 116720447 | 116719762 | 116719876 | 116723066 | 116723223 | 104476 | 30,70,69,58,39,36,44,61,89 | 3,6,2,4,0,2,1,3,1,4,0,0 | 57,71,54,23,78,26,40,73,13 | 1,2,0,0,1,0,1,1,0,0,2,7,0,2,3 | 298 | 149 | 0.010701189203 | 0.592060089962853 | 0.833,0.854,0.945, | 0.966,0.947,1.0,1.0 | -0,028 |
| 205876 | ENSG00000128815 | WDFY4 | chr10 | + | 48808288 | 48808369 | 48807858 | 48807958 | 48811538 | 48811708 | 205876 | 8,1,6,3,0,4,2,5,2,2,7,4 | 9,4,10,3,8,21,13,11,11,4,7, | 7,14,2,34,3,0,7,5,1,2,10,11 | 15,17,2,19,4,1,8,7,4,3,17,7 | 230 | 149 | 0.0107034108007 | 0.592060089962853 | 0.365,0.139,0.28,0 | 0.232,0.348,0.393, | -0,079 |
| 10578 | ENSG00000073605 | GSDMB | chr17 | - | 39906960 | 39906987 | 39906110 | 39906271 | 39909755 | 39909924 | 10578 | 4,6,1,2,2,2,3,7,2,1,4,0 | 1,0,6,4,0,1,1,1,0,0,0,2 | 4,0,5,2,0,3,2,0,1,1,1,1,2,4,1 | 0,1,1,2,4,2,0,1,0,1,0,2,4,10 | 176 | 149 | 0.010707711676 | 0.592145809798858 | 0.772,1.0,0.124,0.2 | 1.0,0.0,0.809,0.45 | 0,163 |
| 43540 | ENSG00000128513 | POT1 | chr7 | - | 124870910 | 124871041 | 124863349 | 124863640 | 124892265 | 124892380 | 43540 | 29,25,66,58,26,17,14,35,21 | 2,7,5,2,2,1,0,0,0,0,1,0 | 29,31,31,12,17,21,11,19,20 | 0,4,2,0,0,3,2,2,0,0,0,0,3,2,0 | 280 | 149 | 0.0107329443497 | 0.59338873499365 | 0.885,0.655,0.875, | 1.0,0.805,0.892,1.0 | -0,02 |
| 163863 | ENSG00000140471 | LINS1 | chr15 | - | 100576544 | 100576702 | 100575043 | 100575128 | 100580262 | 100580352 | 163863 | 1,1,3,3,1,2,0,3,2,0,0,2 | 5,2,15,14,12,12,3,7,3,4,6,5 | 3,3,0,1,1,1,5,3,0,0,2,0,2,2,1 | 13,10,7,1,12,3,8,11,9,4,11, | 298 | 149 | 0.0107465708527 | 0.593989520081529 | 0.091,0.2,0.091,0.0 | 0.103,0.13,0.0,0.3 | -0,074 |
| 210053 | ENSG00000151779 | NBAS | chr2 | - | 15536417 | 15536496 | 15534542 | 15534641 | 15539222 | 15539356 | 210053 | 11,7,21,5,4,10,14,8,8,3,6,5 | 2,0,0,1,2,0,0,3,0,0,7,0 | 23,13,6,6,8,10,13,17,4,7,10 | 7,5,0,6,3,2,4,2,1,0,0,2,1,0,1 | 228 | 149 | 0.0107527024478 | 0.594175840960832 | 0.782,1.0,1.0,0.76 | 0.682,0.63,1.0,0.39 | 0,052 |
| 178311 | ENSG00000143297 | FCRL5 | chr1 | - | 157547639 | 157547695 | 157544830 | 157545082 | 157549559 | 157549580 | 178311 | 2,3,11,0,2,3,0,3,1,0,2,0 | 6,4,28,21,12,7,9,12,4,2,0,7 | 2,2,0,1,1,0,1,0,0,0,1,0,4,2,1 | 19,2,3,2,2,5,6,1,9,4,0,6,6,2 | 205 | 149 | 0.0107815599296 | 0.59561753727098 | 0.195,0.353,0.222, | 0.071,0.421,0.0,0. | 0,018 |
| 162671 | ENSG00000072364 | AFF4 | chr5 | - | 132889078 | 132889177 | 132888096 | 132888160 | 132892163 | 132892404 | 162671 | 22,17,86,28,17,31,16,40,6, | 14,3,3,1,3,2,3,8,1,0,0,0 | 43,59,7,42,28,9,22,15,34,2 | 3,11,1,17,7,4,7,6,3,2,8,3,7, | 248 | 149 | 0.0107856080595 | 0.595688275148639 | 0.486,0.773,0.945, | 0.896,0.763,0.808, | 0,109 |
| 135959 | ENSG00000196214 | ZNF766 | chr19 | + | 52283284 | 52283667 | 52282110 | 52282237 | 52290065 | 52296046 | 135959 | 55,41,66,38,36,27,28,35,16 | 0,1,0,0,0,1,0,2,0,0,0,0 | 80,53,43,25,22,9,12,38,10, | 6,1,0,0,3,1,2,0,0,0,0,4,1,3,0 | 298 | 149 | 0.0107889096267 | 0.595717754711203 | 1.0,0.953,1.0,1.0,1 | 0.87,0.964,1.0,1.0, | 0,071 |
| 201388 | ENSG00000148187 | MRRF | chr9 | + | 122265461 | 122265700 | 122264886 | 122264938 | 122270863 | 122271075 | 201388 | 2,1,8,3,1,1,0,1,0,0,0,1 | 6,1,6,2,1,5,2,2,2,4,1,0 | 3,3,2,7,4,1,0,2,1,2,1,3,1,2,1 | 2,6,4,5,5,2,2,2,5,3,3,5,3,0,4 | 298 | 149 | 0.0107962074008 | 0.595967816127619 | 0.143,0.333,0.4,0.4 | 0.429,0.2,0.2,0.41 | -0,049 |
| 7529 | ENSG00000088876 | ZNF343 | chr20 | - | 2486640 | 2486763 | 2481822 | 2484656 | 2492698 | 2492825 | 7529 | 0,0,3,0,0,0,1,0,5,0,0,1 | 5,5,13,17,6,7,11,2,2,2,2,6 | 3,0,1,0,1,2,0,2,1,0,3,1,4,5,0 | 12,9,4,2,2,3,1,9,0,4,0,4,9,1 | 272 | 149 | 0.0108098662999 | 0.596568803485584 | 0.0,0.0,0.112,0.0,0 | 0.12,0.0,0.12,0.0,0 | -0,173 |
| 142861 | ENSG00000154262 | ABCA6 | chr17 | - | 69097919 | 69097962 | 69096627 | 69096801 | 69100796 | 69100934 | 142861 | 11,12,59,44,11,7,54,57,17, | 4,1,11,17,5,2,4,7,0,0,4,2 | 38,31,22,6,19,9,13,28,9,20 | 12,14,0,2,10,4,3,5,12,2,14, | 192 | 149 | 0.0108203606851 | 0.596994886084275 | 0.681,0.903,0.806, | 0.711,0.632,1.0,0.7 | 0,101 |
| 196413 | ENSG00000143575 | HAX1 | chr1 | + | 154273733 | 154273961 | 154273479 | 154273598 | 154274949 | 154275001 | 196413 | 77,75,88,85,57,67,46,68,11 | 2,2,7,3,0,3,1,3,2,1,1,0 | 65,40,46,23,25,25,38,57,17 | 2,0,0,2,1,4,0,0,1,0,4,0,0,0,4 | 298 | 149 | 0.0108566238148 | 0.59884213231246 | 0.951,0.949,0.863, | 0.942,1.0,1.0,0.85 | -0,003 |
| 216880 | ENSG00000115866 | DARS1 | chr2 | - | 135932782 | 135932838 | 135924386 | 135924498 | 135933909 | 135933990 | 216880 | 16,7,38,35,14,19,18,22,17, | 1,1,1,1,0,2,1,0,0,0,0,4 | 26,30,12,2,12,8,6,13,12,13 | 0,1,2,2,9,0,0,0,0,1,6,1,0,0,2 | 205 | 149 | 0.0108728552036 | 0.599385092685039 | 0.921,0.836,0.965, | 1.0,0.956,0.813,0.4 | 0,078 |
| 127927 | ENSG00000095585 | BLNK | chr10 | - | 96215320 | 96215389 | 96209837 | 96209907 | 96216652 | 96216734 | 127927 | 157,101,186,131,122,91,98 | 12,3,12,8,7,1,5,1,25,16,5,5 | 148,62,130,50,42,76,30,12 | 5,4,6,5,4,1,3,4,2,4,0,12,11, | 218 | 149 | 0.0108731929683 | 0.599385092685039 | 0.899,0.958,0.914, | 0.953,0.914,0.937, | -0,015 |
| 115960 | ENSG00000106829 | TLE4 | chr9 | + | 79649819 | 79649858 | 79627373 | 79627448 | 79652589 | 79652798 | 115960 | 37,32,45,28,10,46,41,80,12 | 10,7,11,33,32,34,23,41,13, | 23,69,54,77,60,14,21,30,27 | 13,7,11,7,9,11,11,12,3,9,26 | 188 | 149 | 0.0108748218748 | 0.599385092685039 | 0.746,0.784,0.764, | 0.584,0.887,0.796, | -0,114 |
| 85059 | ENSG00000023516 | AKAP11 | chr13 | + | 42317527 | 42317688 | 42313893 | 42313940 | 42319087 | 42323261 | 85059 | 74,58,156,88,52,37,35,48,5 | 1,1,1,6,4,3,4,2,1,3,0,4 | 63,68,54,32,45,52,34,43,57 | 2,6,7,0,6,1,2,0,2,4,1,5,6,8,1 | 298 | 149 | 0.0108808072492 | 0.599561450346279 | 0.974,0.967,0.987, | 0.94,0.85,0.794,1.0 | 0,027 |
| 246189 | ENSG00000171204 | TMEM126B | chr11 | + | 85631913 | 85631989 | 85631686 | 85631808 | 85634085 | 85635087 | 246189 | 9,3,3,3,0,1,0,0,2,0,0,1 | 14,2,28,24,3,7,5,3,8,2,5,8 | 1,3,0,1,3,0,0,0,2,2,1,2,0,1,2 | 16,24,5,5,4,5,2,16,9,17,14, | 225 | 149 | 0.0109096878725 | 0.600998203315395 | 0.299,0.498,0.066, | 0.04,0.076,0.0,0.1 | 0,037 |
| 96807 | ENSG00000127947 | PTPN12 | chr7 | + | 77583554 | 77583650 | 77581426 | 77581503 | 77585542 | 77585571 | 96807 | 90,41,101,64,50,45,49,127 | 3,1,0,0,0,0,2,0,2,0,0,0 | 67,111,38,50,49,74,34,22,3 | 3,5,1,13,1,0,0,0,3,0,2,0,0,0 | 245 | 149 | 0.0109124660414 | 0.600998203315395 | 0.948,0.961,1.0,1.0 | 0.931,0.931,0.959, | 0,024 |
| 32291 | ENSG00000167766 | ZNF83 | chr19 | - | 52616717 | 52616841 | 52612397 | 52614797 | 52618935 | 52619062 | 32291 | 99,51,113,72,34,38,63,43,3 | 12,10,31,21,7,13,2,5,7,3,1, | 67,68,58,17,53,32,17,33,44 | 15,15,2,1,5,2,0,2,0,2,3,4,8, | 273 | 149 | 0.0109210882976 | 0.601319200660206 | 0.818,0.736,0.665, | 0.709,0.712,0.941, | -0,065 |
| 159131 | ENSG00000065491 | TBC1D22B | chr6 | + | 37282184 | 37282504 | 37279303 | 37279611 | 37282881 | 37282952 | 159131 | 15,28,21,14,15,13,4,8,17,7 | 0,1,5,4,0,2,0,1,0,1,0,2 | 8,17,2,9,3,2,8,21,4,11,15,2 | 1,1,0,0,4,0,2,0,0,0,1,0,0,0,1 | 298 | 149 | 0.0109266087766 | 0.601428273375155 | 1.0,0.933,0.677,0.6 | 0.8,0.895,1.0,1.0,0 | -0,027 |
| 141987 | ENSG00000108559 | NUP88 | chr17 | - | 5405341 | 5405573 | 5405056 | 5405243 | 5408732 | 5408909 | 141987 | 6,0,12,3,3,5,4,16,2,5,2,2 | 60,23,150,79,109,88,79,99 | 1,4,4,5,3,0,8,6,1,1,8,4,1,0,0 | 97,101,115,46,55,29,56,85 | 298 | 149 | 0.0109286579404 | 0.601428273375155 | 0.048,0.0,0.038,0.0 | 0.005,0.019,0.017, | 0,008 |
| 3037 | ENSG00000105397 | TYK2 | chr19 | - | 10369756 | 10369877 | 10368294 | 10368418 | 10378213 | 10378426 | 3037 | 3,7,15,5,5,10,6,4,3,1,3,5 | 47,25,66,60,61,53,28,45,33 | 4,7,2,6,5,0,3,0,4,8,4,5,4,5,3 | 43,55,24,25,52,28,30,56,26 | 270 | 149 | 0.0109484269907 | 0.602257579086088 | 0.034,0.134,0.111, | 0.049,0.066,0.044, | 0,022 |
| 73622 | ENSG00000165119 | HNRNPK | chr9 | - | 83975461 | 83975505 | 83974516 | 83974589 | 83976994 | 83977051 | 73622 | 427,331,956,679,385,384,3 | 8,3,16,24,5,18,13,8,4,4,4,5 | 680,763,591,453,371,334,2 | 21,20,31,5,11,4,8,10,4,14,1 | 193 | 149 | 0.0109503002973 | 0.602257579086088 | 0.976,0.988,0.979, | 0.962,0.967,0.936, | 0,008 |
| 246433 | ENSG00000081237 | PTPRC | chr1 | + | 198739063 | 198739210 | 198735126 | 198735252 | 198741868 | 198742026 | 246433 | 36,15,25,5,1,14,8,13,20,19 | 356,273,494,275,229,137,1 | 17,10,12,13,8,2,5,9,4,3,15, | 306,268,183,165,257,140,9 | 296 | 149 | 0.0109521219738 | 0.602257579086088 | 0.048,0.027,0.025, | 0.027,0.018,0.032, | 0,006 |
| 210113 | ENSG00000136536 | MARCHF7 | chr2 | + | 159745769 | 159745934 | 159743060 | 159743253 | 159747804 | 159748903 | 210113 | 99,65,257,170,82,83,67,89 | 7,3,3,4,5,5,2,6,17,0,2,3 | 104,173,78,59,93,75,44,85 | 7,5,4,2,0,2,3,6,0,7,0,2,1,3,3 | 298 | 149 | 0.0109837674025 | 0.60384348449744 | 0.876,0.915,0.977, | 0.881,0.945,0.907, | -0,044 |

| | | | | | | | | | | | | | | | | | | | | | | |
|---|---|---|---|---|---|---|---|---|---|---|---|---|---|---|---|---|---|---|---|---|---|---|
| 213437 | ENSG00000035720 | STAP1 | chr4 | + | 67592139 | 67592271 | 67590883 | 67590953 | 67606295 | 67607337 | 213437 | 4,2,4,4,1,1,6,4,1,0,0,1 | 3,1,18,8,1,5,5,3,0,1,2,0 | 2,7,4,1,1,0,1,0,0,1,2,2,0,0,0 | 1,3,5,0,5,2,0,1,5,9,3,3,1,3,2 | 281 | 149 | 0.0109877426403 | 0.6039077722713 | 0.414,0.515,0.105, | 0.515,0.553,0.298, | 0,1 |
| 77911 | ENSG00000115421 | PAPOLG | chr2 | + | 60787510 | 60787620 | 60786946 | 60787066 | 60791760 | 60791882 | 77911 | 49,46,39,34,6,17,8,29,26,2 | 1,2,2,7,1,0,0,2,1,2,0,4 | 47,64,20,14,27,22,13,31,12 | 4,11,1,6,3,6,0,6,0,6,4,5,4,0 | 259 | 149 | 0.0109927878767 | 0.604030821416905 | 0.966,0.93,0.918,0 | 0.871,0.77,0.92,0. | 0,072 |
| 28602 | ENSG00000290835 | | chr7 | + | 6730633 | 6730727 | 6730181 | 6730301 | 6731896 | 6731955 | 28602 | 3,5,9,2,2,4,0,2,1,0,2,0 | 3,5,9,7,5,1,1,3,2,2,2,5 | 5,5,2,0,4,0,4,1,2,3,5,4,5,4,4 | 5,1,0,1,2,1,0,1,0,4,3,5,1,2,2 | 243 | 149 | 0.0109970183149 | 0.604109047711649 | 0.38,0.38,0.38,0.14 | 0.38,0.754,1.0,0.0, | -0,253 |
| 148821 | ENSG00000005339 | CREBBP | chr16 | - | 3782683 | 3782827 | 3781203 | 3781306 | 3791980 | 3792094 | 148821 | 77,56,78,28,45,51,36,70,51 | 0,1,3,0,0,3,1,0,1,11,0,3 | 50,79,46,116,37,28,64,54,4 | 1,1,0,1,0,1,0,1,0,0,5,2,0,2,0 | 293 | 149 | 0.0110163326182 | 0.605015638108651 | 1.0,0.966,0.93,1.0, | 0.962,0.976,1.0,0.9 | -0,012 |
| 186654 | ENSG00000071564 | TCF3 | chr19 | - | 1615284 | 1615480 | 1612206 | 1612433 | 1615685 | 1615821 | 186654 | 42,10,53,18,23,12,19,22,35 | 45,34,66,71,24,53,32,30,60 | 19,16,8,7,14,6,20,13,5,9,7, | 35,38,20,21,10,9,25,85,11, | 298 | 149 | 0.0110315437669 | 0.605696478697361 | 0.318,0.128,0.286, | 0.213,0.174,0.167, | 0,035 |
| 75985 | ENSG00000077157 | PPP1R12B | chr1 | + | 202445045 | 202445228 | 202442446 | 202442572 | 202448988 | 202449097 | 75985 | 17,10,8,13,6,11,7,11,16,9,6 | 2,0,2,7,0,5,2,4,4,3,0,7 | 5,5,8,12,5,2,9,9,6,17,7,10,2 | 11,5,0,7,1,2,7,7,0,2,9,3,2,8 | 298 | 149 | 0.0110494251473 | 0.606523546003195 | 0.81,1.0,0.667,0.48 | 0.185,0.333,1.0,0.4 | 0,201 |
| 152522 | ENSG00000258643 | BCL2L2-PABPN | chr14 | + | 23323978 | 23324052 | 23323376 | 23323483 | 23324137 | 23324289 | 152522 | 226,129,227,130,69,98,101 | 9,3,3,7,5,2,3,6,16,6,1,7 | 147,127,95,46,96,72,36,12 | 8,1,4,0,4,1,0,6,4,1,4,6,2,3,4 | 223 | 149 | 0.0110538976708 | 0.606614342066281 | 0.944,0.966,0.981, | 0.925,0.988,0.941, | -0,028 |
| 203333 | ENSG00000115806 | GORASP2 | chr2 | + | 170956478 | 170956559 | 170951327 | 170951458 | 170961662 | 170961749 | 203333 | 62,61,46,53,29,27,28,21,13 | 4,1,0,4,0,3,8,1,0,0,0,3 | 19,47,37,47,29,22,9,40,11, | 1,4,3,3,4,2,0,2,4,4,6,1,1,0,3 | 230 | 149 | 0.0110681922945 | 0.607156255803439 | 0.909,0.975,1.0,0.8 | 0.925,0.884,0.889, | 0,05 |
| 235015 | ENSG00000147905 | ZCCHC7 | chr9 | + | 37304187 | 37304231 | 37302187 | 37302231 | 37305543 | 37305714 | 235015 | 29,16,57,37,23,21,21,34,35 | 5,0,0,1,0,1,0,2,1,0,0,0 | 55,29,55,15,15,20,10,21,18 | 5,2,2,3,1,0,0,1,0,3,2,2,2,0,0 | 193 | 149 | 0.0110694144792 | 0.607156255803439 | 0.817,1.0,1.0,0.96 | 0.895,0.918,0.955, | 0,031 |
| 227690 | ENSG00000186088 | GSAP | chr7 | - | 77379846 | 77380000 | 77377285 | 77377390 | 77381304 | 77381354 | 227690 | 73,61,178,60,53,97,80,118 | 51,26,156,99,53,60,40,66,5 | 109,136,106,71,58,44,36,7 | 102,45,42,11,27,18,28,56,2 | 298 | 149 | 0.0110825141808 | 0.607719900547201 | 0.417,0.54,0.363,0 | 0.348,0.602,0.558, | -0,098 |
| 71789 | ENSG00000124256 | ZBP1 | chr20 | - | 57616243 | 57616468 | 57615511 | 57615580 | 57620218 | 57620478 | 71789 | 5,7,11,19,11,10,3,10,6,7,5, | 1,5,2,6,5,5,2,4,9,2,4,0 | 9,21,3,5,21,11,4,2,12,10,2, | 10,3,2,2,0,7,1,0,1,1,1,7,0,0 | 298 | 149 | 0.0111145776526 | 0.609322889135698 | 0.714,0.412,0.733, | 0.31,0.778,0.429,0 | -0,161 |
| 126268 | ENSG00000057608 | GDI2 | chr10 | - | 5794884 | 5795019 | 5785851 | 5786050 | 5796762 | 5796862 | 126268 | 524,226,1345,1249,492,30 | 1,0,9,13,3,2,1,2,1,1,9,3 | 829,546,343,189,320,334,1 | 15,11,1,5,3,1,1,3,3,3,8,19,8 | 284 | 149 | 0.0111207822863 | 0.60944083721401 | 0.996,1.0,0.987,0.9 | 0.967,0.963,0.994, | 0,004 |
| 161186 | ENSG00000065357 | DGKA | chr12 | + | 55938646 | 55938698 | 55938552 | 55938560 | 55938914 | 55938942 | 161186 | 1,0,3,14,1,1,2,0,5,2,1,1 | 54,27,134,90,64,54,75,65,2 | 2,0,0,0,1,4,0,0,0,2,2,0,1,3,0 | 72,86,60,26,70,35,32,58,78 | 201 | 149 | 0.0111223922603 | 0.60944083721401 | 0.014,0.0,0.016,0.1 | 0.02,0.0,0.0,0.0,0.0 | 0,021 |
| 169971 | ENSG00000075826 | SEC31B | chr10 | - | 100497134 | 100497489 | 100496257 | 100496431 | 100497666 | 100497793 | 169971 | 29,12,29,21,14,11,23,16,8, | 0,0,1,3,1,7,0,3,1,0,1,5 | 5,9,11,14,5,14,8,12,22,20,3 | 10,16,3,7,8,1,0,1,1,0,4,4,0, | 298 | 149 | 0.0111325937689 | 0.609738188768769 | 1.0,1.0,0.935,0.778 | 0.2,0.22,0.647,0.5, | 0,164 |
| 168680 | ENSG00000227345 | PARG | chr10 | - | 49859213 | 49859329 | 49857305 | 49857453 | 49861587 | 49861663 | 168680 | 2,0,1,0,1,0,1,3,0,0,1,0 | 8,7,17,14,8,16,9,10,6,2,5,8 | 4,2,2,3,1,0,1,3,3,1,2,0,0,1,0 | 13,9,7,12,15,5,6,9,10,13,9, | 265 | 149 | 0.0111334848691 | 0.609738188768769 | 0.123,0.0,0.032,0.0 | 0.147,0.111,0.138, | -0,028 |
| 67481 | ENSG00000143575 | HAX1 | chr1 | + | 154273782 | 154273961 | 154273335 | 154273598 | 154274949 | 154275001 | 67481 | 77,75,88,85,58,67,46,68,11 | 2,2,7,3,0,3,1,3,2,1,1,0 | 65,39,46,23,25,25,38,57,17 | 2,0,0,2,1,4,0,0,1,0,4,0,0,0,4 | 298 | 149 | 0.0111394390964 | 0.609909085768829 | 0.951,0.949,0.863, | 0.942,1.0,1.0,0.85 | -0,003 |
| 211598 | ENSG00000183283 | DAZAP2 | chr12 | + | 51242329 | 51242518 | 51238907 | 51241116 | 51246060 | 51246717 | 211598 | 311,223,499,326,215,211,2 | 8,7,6,8,2,0,2,0,8,0,1,1 | 359,386,227,184,290,150,1 | 1,7,3,0,3,2,2,3,0,3,0,4,2,8,0 | 298 | 149 | 0.0111636782632 | 0.611080782366938 | 0.951,0.941,0.977, | 0.994,0.965,0.974, | -0,012 |
| 167883 | ENSG00000121210 | TMEM131L | chr4 | + | 153617578 | 153617794 | 153612251 | 153612400 | 153620755 | 153620880 | 167883 | 16,7,9,2,0,7,1,4,10,0,0,3 | 282,218,270,142,72,92,69, | 6,1,2,2,2,1,3,0,3,2,2,7,1,1,0 | 135,112,101,160,73,44,47, | 298 | 149 | 0.0111799358036 | 0.611503655790379 | 0.028,0.016,0.016, | 0.022,0.004,0.01,0 | 0,005 |
| 115915 | ENSG00000198771 | RCSD1 | chr1 | + | 167640410 | 167640474 | 167638389 | 167638442 | 167640570 | 167640712 | 115915 | 16,15,47,23,0,5,19,36,7,3,1 | 0,0,2,7,1,1,2,4,3,0,2,2 | 11,22,35,10,2,4,7,25,6,3,32 | 3,2,2,0,0,0,0,0,0,0,2,0,0,2,3 | 213 | 149 | 0.0111805933772 | 0.611503655790379 | 1.0,1.0,0.943,0.69 | 0.719,0.885,0.924, | -0,161 |
| 216995 | ENSG00000135945 | REV1 | chr2 | - | 99406324 | 99406366 | 99405909 | 99406106 | 99408028 | 99408403 | 216995 | 5,2,6,5,4,3,3,9,6,2,3,8 | 1,1,3,9,0,0,3,2,2,0,0,1 | 4,7,3,1,0,1,1,6,1,4,3,4,3,3,4 | 4,0,0,1,3,0,1,1,0,0,0,0,0,3,2 | 191 | 149 | 0.0111823097727 | 0.611503655790379 | 0.796,0.609,0.609, | 0.438,1.0,1.0,0.43 | 0,037 |
| 120372 | ENSG00000189057 | FAM111B | chr11 | + | 59108667 | 59108712 | 59107235 | 59107296 | 59124178 | 59126436 | 120372 | 7,15,106,19,12,4,9,1,2,3,6, | 0,0,4,0,0,1,0,0,0,0,0,0 | 115,148,61,26,62,11,24,11 | 4,8,1,3,5,2,0,0,0,0,0,0,0,2,0 | 194 | 149 | 0.0111827682313 | 0.611503655790379 | 1.0,1.0,0.953,1.0,1 | 0.957,0.934,0.979, | 0,022 |
| 115912 | ENSG00000198771 | RCSD1 | chr1 | + | 167640410 | 167640474 | 167630265 | 167630429 | 167640570 | 167640712 | 115912 | 14,15,25,19,0,2,13,28,7,3,2 | 3,2,5,3,4,2,4,15,6,2,3,12 | 6,16,28,12,2,4,6,19,6,2,25, | 1,0,8,6,0,0,3,3,1,0,2,0,3,0,1 | 213 | 149 | 0.0111857099363 | 0.611509152984451 | 0.766,0.84,0.778,0 | 0.808,1.0,0.71,0.58 | -0,231 |
| 237869 | ENSG00000104365 | IKBKB | chr8 | + | 42319559 | 42319646 | 42319269 | 42319421 | 42320734 | 42320844 | 237869 | 41,15,18,14,6,13,15,7,32,1 | 4,13,2,2,2,1,2,1,4,1,0,3 | 15,7,7,11,8,2,10,11,9,9,2,1 | 0,4,3,12,6,2,0,3,2,11,2,7,0, | 236 | 149 | 0.0112105415903 | 0.612711040381376 | 0.866,0.421,0.85,0 | 1.0,0.525,0.596,0.3 | 0,174 |
| 86626 | ENSG00000119707 | RBM25 | chr14 | + | 73088369 | 73088474 | 73088000 | 73088161 | 73096914 | 73097100 | 86626 | 13,13,7,7,4,11,6,16,7,2,0,4 | 312,256,212,117,75,115,71 | 2,15,9,10,7,7,12,15,0,27,8, | 133,175,150,250,108,60,86 | 254 | 149 | 0.0112226836537 | 0.613123569502604 | 0.024,0.029,0.019, | 0.009,0.048,0.034, | -0,013 |
| 191020 | ENSG00000270629 | NBPF14 | chr1 | - | 148607068 | 148607159 | 148598612 | 148598695 | 148607409 | 148607745 | 191020 | 21,13,14,9,14,17,9,7,8,7,9, | 1,0,1,2,4,0,1,1,3,2,0,0 | 2,8,6,15,12,7,14,6,11,17,8, | 2,0,1,0,0,1,0,2,2,3,1,7,0,0,0 | 240 | 149 | 0.0112241844904 | 0.613123569502604 | 0.929,1.0,0.897,0.7 | 0.383,1.0,0.788,1.0 | 0,009 |
| 49613 | ENSG00000144218 | AFF3 | chr2 | - | 100011496 | 100011571 | 100008811 | 100008932 | 100063125 | 100063249 | 49613 | 16,5,32,22,10,14,10,8,10,1 | 4,0,9,8,5,5,0,5,6,0,1,3 | 8,15,13,41,26,6,9,4,13,10,8 | 6,2,0,10,2,0,4,1,0,0,2,3,0,0 | 224 | 149 | 0.0112266355098 | 0.613123569502604 | 0.727,1.0,0.703,0.6 | 0.47,0.833,1.0,0.7 | -0,09 |
| 27092 | ENSG00000075568 | TMEM131 | chr2 | - | 97775842 | 97776018 | 97766477 | 97766602 | 97792385 | 97792984 | 27092 | 31,33,41,19,31,26,20,23,22 | 4,2,3,2,0,2,0,0,2,2,0,0 | 33,26,39,26,35,20,12,3,11, | 0,0,1,0,3,2,0,0,0,0,0,0,0,0,2 | 298 | 149 | 0.0112303008383 | 0.613123877074837 | 0.795,0.892,0.872, | 1.0,1.0,0.951,1.0,0 | -0,057 |
| 127454 | ENSG00000108055 | SMC3 | chr10 | + | 110602473 | 110602665 | 110601965 | 110602178 | 110602824 | 110603002 | 127454 | 128,120,168,112,58,53,30, | 0,0,0,3,1,2,4,0,0,0,0,0 | 70,129,68,63,87,54,54,66,1 | 1,1,2,6,0,2,2,0,0,0,1,9,0,3,0 | 298 | 149 | 0.0112323384982 | 0.613123877074837 | 1.0,1.0,1.0,0.949,0 | 0.972,0.985,0.944, | 0,017 |
| 36765 | ENSG00000101109 | STK4 | chr20 | + | 44975323 | 44975410 | 44973326 | 44973433 | 44978442 | 44978571 | 36765 | 1,0,17,7,4,4,7,7,6,1,1,0 | 18,7,14,15,9,14,14,16,1,1,3 | 10,5,5,2,4,1,2,3,2,6,1,1,4,1 | 23,54,16,16,22,10,18,23,28 | 236 | 149 | 0.0112468984934 | 0.61376298418686 | 0.034,0.0,0.434,0.1 | 0.215,0.055,0.165, | 0,137 |
| 122117 | ENSG00000081237 | PTPRC | chr1 | + | 198702386 | 198702530 | 198699563 | 198699704 | 198703297 | 198703346 | 122117 | 934,1263,579,994,431,498 | 317,161,311,27,227,156,21 | 452,683,491,680,393,293,1 | 283,104,173,75,161,86,95, | 293 | 149 | 0.0112544307294 | 0.613820345332885 | 0.6,0.8,0.486,0.949 | 0.448,0.77,0.591,0 | 0,056 |
| 206640 | ENSG00000152061 | RABGAP1L | chr1 | + | 174273576 | 174273661 | 174272413 | 174272480 | 174275832 | 174275935 | 206640 | 1,2,15,7,0,9,6,2,7,5,2,1 | 173,129,384,235,96,104,12 | 0,4,10,5,2,1,0,3,3,5,2,1,2 | 173,260,263,141,160,105,5 | 234 | 149 | 0.0112563092395 | 0.613820345332885 | 0.004,0.01,0.024,0 | 0.0,0.01,0.024,0.0 | 0,009 |
| 173740 | ENSG00000136560 | TANK | chr2 | + | 161228622 | 161228752 | 161224630 | 161224746 | 161230970 | 161231551 | 173740 | 20,6,15,10,14,8,5,19,6,4,1, | 86,97,204,129,74,76,41,59 | 18,18,12,7,6,8,1,8,10,19,14 | 89,110,87,35,72,51,35,54,3 | 279 | 149 | 0.0112565053502 | 0.613820345332885 | 0.11,0.032,0.038,0 | 0.097,0.08,0.069,0 | -0,017 |
| 131888 | ENSG00000028203 | VEZT | chr12 | + | 95257149 | 95257239 | 95256549 | 95256622 | 95266356 | 95266384 | 131888 | 5,1,18,14,16,11,3,6,7,4,2,7 | 0,0,3,2,0,2,0,2,0,0,0,0 | 8,0,4,5,3,0,8,3,2,16,4,7,13, | 0,4,0,0,3,2,4,1,2,0,1,0,0,0,2 | 239 | 149 | 0.011267342419 | 0.61425566772639 | 1.0,1.0,0.789,0.814 | 1.0,0.0,1.0,1.0,0.38 | 0,209 |
| 67556 | ENSG00000204217 | BMPR2 | chr2 | + | 202530793 | 202530954 | 202520086 | 202520201 | 202542310 | 202542447 | 67556 | 88,62,142,98,41,56,37,120 | 2,5,3,0,0,3,4,1,0,0,0,1 | 132,116,44,32,66,46,36,50 | 6,6,2,4,6,1,4,0,2,4,6,0,0,0,2 | 298 | 149 | 0.0112782961614 | 0.614697128669102 | 0.957,0.861,0.959, | 0.917,0.906,0.917, | 0,026 |
| 99753 | ENSG00000132256 | TRIM5 | chr11 | - | 5665980 | 5666081 | 5663616 | 5665682 | 5678203 | 5678434 | 99753 | 8,14,7,15,5,5,4,2,29,13,2,4 | 1,0,0,0,0,0,0,0,1,0,0,0 | 13,3,8,6,3,5,2,3,2,16,4,3,8, | 1,0,0,4,0,1,0,0,1,0,3,0,1,3,4 | 250 | 149 | 0.011283494924 | 0.61482478379429 | 0.827,1.0,1.0,1.0,1 | 0.886,1.0,1.0,0.47 | 0,175 |
| 23960 | ENSG00000088387 | DOCK9 | chr13 | - | 98825871 | 98825975 | 98824397 | 98824504 | 98826829 | 98826887 | 23960 | 2,2,4,0,2,2,3,2,0,1,3,0 | 7,9,14,3,3,15,1,4,18,2,3,7 | 0,5,0,1,1,2,0,2,4,0,0,0,1,0,1 | 3,9,3,8,17,5,2,8,6,8,11,7,10 | 253 | 149 | 0.0112997074509 | 0.61555234987716 | 0.144,0.116,0.144, | 0.0,0.247,0.0,0.069 | 0,087 |
| 6528 | ENSG00000125844 | RRBP1 | chr20 | - | 17642978 | 17643127 | 17636576 | 17636729 | 17658595 | 17660507 | 6528 | 30,15,53,51,46,47,52,41,23 | 0,0,0,0,2,0,0,1,2,1,0,2 | 46,37,7,45,9,13,71,77,14,2 | 0,0,1,2,0,1,5,1,0,2,2,0,0,2,2 | 298 | 149 | 0.0113127047931 | 0.61610444466693 | 1.0,1.0,1.0,1.0,0.9 | 1.0,1.0,0.778,0.918 | 0,044 |
| 159793 | ENSG00000112851 | ERBIN | chr5 | + | 66075003 | 66075230 | 66054456 | 66054951 | 66076315 | 66076408 | 159793 | 61,92,197,89,81,61,70,90,7 | 20,11,43,22,14,5,24,17,7,5 | 88,78,73,51,69,44,43,66,38 | 41,50,18,17,27,9,8,10,14,5 | 298 | 149 | 0.0113314275861 | 0.616967996656688 | 0.604,0.807,0.696, | 0.518,0.438,0.67,0 | 0,082 |
| 75514 | ENSG00000121940 | CLCC1 | chr1 | - | 108938165 | 108938216 | 108937076 | 108937418 | 108939635 | 108939782 | 75514 | 6,3,4,5,1,0,1,1,1,0,0,0 | 62,26,28,18,9,25,14,30,31, | 0,4,0,0,1,0,0,0,2,2,1,2,0,1,0 | 36,22,49,38,14,15,7,13,6,3 | 200 | 149 | 0.0113529757256 | 0.617984906018365 | 0.067,0.079,0.096, | 0.0,0.119,0.0,0.0,0 | 0,005 |
| 105459 | ENSG00000164162 | ANAPC10 | chr4 | - | 144999256 | 144999294 | 144995160 | 144995603 | 145064571 | 145064692 | 105459 | 0,2,9,2,2,1,1,4,2,0,0,0 | 12,10,38,20,12,15,7,17,24, | 1,4,2,0,5,0,2,3,1,3,1,2,7,1,4 | 14,17,16,8,19,10,9,15,8,12 | 187 | 149 | 0.0113981558671 | 0.620287354091479 | 0.0,0.137,0.159,0.0 | 0.054,0.158,0.091, | -0,033 |
| 3107 | ENSG00000076928 | ARHGEF1 | chr19 | + | 41897945 | 41898113 | 41896376 | 41896482 | 41898441 | 41898587 | 3107 | 9,6,22,3,18,6,13,15,4,2,3,2 | 5,2,8,2,9,9,5,7,19,5,4,9 | 6,13,6,0,8,0,9,12,5,8,9,61,3 | 4,9,9,13,7,3,8,11,11,5,7,52 | 298 | 149 | 0.0114067882341 | 0.620600211934676 | 0.474,0.6,0.579,0.4 | 0.429,0.419,0.25,0 | 0,15 |
| 121779 | ENSG00000122359 | ANXA11 | chr10 | - | 80180940 | 80181455 | 80176106 | 80176155 | 80205342 | 80205522 | 121779 | 4,0,17,6,1,0,1,14,3,3,2,4 | 11,10,35,12,16,13,21,22,18 | 2,6,0,2,4,6,21,11,7,4,4,5,11 | 28,18,9,8,8,7,12,17,7,10,11 | 298 | 149 | 0.0114134919043 | 0.62080800506808 | 0.154,0.0,0.195,0.1 | 0.034,0.143,0.0,0.1 | -0,086 |
| 163615 | ENSG00000105875 | WDR91 | chr7 | - | 135197043 | 135197147 | 135196143 | 135196337 | 135197992 | 135201343 | 163615 | 8,7,16,6,15,6,2,5,2,5,5,9 | 14,14,44,37,36,35,14,15,33 | 8,16,6,4,4,1,8,8,8,11,2,0,11 | 21,28,8,6,7,3,21,25,2,11,20 | 253 | 149 | 0.011423836401 | 0.621213676711377 | 0.252,0.227,0.176, | 0.183,0.252,0.306, | -0,092 |
| 107788 | ENSG00000204116 | CHIC1 | chrX | + | 73578703 | 73578785 | 73577406 | 73577461 | 73584416 | 73584572 | 107788 | 6,2,3,4,0,2,1,0,0,0,0,0 | 16,8,13,16,13,16,11,11,7,3 | 2,1,0,0,1,0,0,1,1,1,0,2,1,2,2 | 24,17,13,2,16,9,9,21,3,9,17 | 231 | 149 | 0.0114331045037 | 0.621445784329753 | 0.195,0.139,0.13,0 | 0.051,0.037,0.0,0.0 | 0,014 |
| 43259 | ENSG00000136560 | TANK | chr2 | + | 161202862 | 161202979 | 161179613 | 161179761 | 161203486 | 161203595 | 43259 | 7,6,15,8,0,7,1,10,7,8,1,0 | 79,95,243,175,102,110,76, | 16,4,4,1,4,2,0,5,1,2,7,0,2,3 | 120,137,113,48,117,53,46, | 266 | 149 | 0.0114340595079 | 0.621445784329753 | 0.047,0.034,0.033, | 0.069,0.016,0.019, | 0,019 |
| 158701 | ENSG00000126777 | KTN1 | chr14 | + | 55651253 | 55651375 | 55650568 | 55650637 | 55651889 | 55651927 | 158701 | 5,7,14,13,6,3,3,9,2,1,2,5 | 120,56,110,64,59,63,46,49 | 7,3,2,3,2,5,1,5,3,5,3,7,3,7,1 | 89,59,67,42,44,41,34,47,47 | 271 | 149 | 0.0114367667842 | 0.621445784329753 | 0.022,0.064,0.065, | 0.041,0.027,0.016, | 0,019 |
| 96229 | ENSG00000144580 | CNOT9 | chr2 | + | 218584611 | 218584721 | 218582970 | 218583086 | 218587585 | 218587695 | 96229 | 55,27,43,52,53,27,29,33,35 | 2,6,24,7,10,14,7,14,0,1,7,4 | 42,20,22,11,18,21,15,37,10 | 17,20,16,11,14,3,4,11,5,5,1 | 259 | 149 | 0.011450718308 | 0.62203343550934 | 0.941,0.721,0.508, | 0.587,0.365,0.442, | 0,125 |
| 166556 | ENSG00000134851 | TMEM165 | chr4 | + | 55416041 | 55416157 | 55411613 | 55411839 | 55417071 | 55417247 | 166556 | 3,1,4,10,0,1,3,1,0,3,1,1 | 11,7,15,12,7,5,15,9,12,8,4, | 10,14,0,4,2,1,4,5,2,4,4,4,8, | 14,6,9,3,2,9,0,7,4,15,3,24,1 | 265 | 149 | 0.0114533617598 | 0.62203343550934 | 0.133,0.074,0.13,0 | 0.287,0.567,0.0,0.4 | -0,188 |
| 110367 | ENSG00000166946 | CCNDBP1 | chr15 | + | 43190054 | 43190151 | 43189198 | 43189280 | 43190324 | 43190398 | 110367 | 82,81,271,243,105,95,92,7 | 26,9,61,39,17,22,28,38,7,8 | 78,87,91,15,62,92,31,68,52 | 58,43,22,31,20,14,25,30,22 | 246 | 149 | 0.0114661027917 | 0.622568307255394 | 0.656,0.845,0.729, | 0.449,0.551,0.715, | 0,1 |
| 152002 | ENSG00000171606 | ZNF274 | chr19 | + | 58210208 | 58210430 | 58209960 | 58210073 | 58211559 | 58211686 | 152002 | 16,10,25,5,5,0,7,12,3,5,5,3 | 50,26,59,28,33,32,22,34,83 | 11,10,17,45,5,7,10,9,2,6,20 | 36,56,52,25,25,9,15,32,5,6 | 298 | 149 | 0.0114770938212 | 0.623007914307868 | 0.138,0.161,0.175, | 0.133,0.082,0.14,0 | -0,066 |
| 202147 | ENSG00000167258 | CDK12 | chr17 | + | 39530572 | 39530953 | 39525866 | 39526316 | 39544248 | 39544360 | 202147 | 4,1,2,1,0,0,6,1,0,2,1,0 | 7,1,10,14,8,13,9,4,8,0,5,1 | 0,1,6,3,2,0,4,2,3,1,2,4,3,0,5 | 5,16,0,3,12,10,3,8,7,11,5,1 | 298 | 149 | 0.0114830212285 | 0.623040117046827 | 0.222,0.333,0.091, | 0.0,0.03,1.0,0.333, | -0,044 |
| 58684 | ENSG00000114098 | ARMC8 | chr3 | + | 138241782 | 138241983 | 138238019 | 138239528 | 138245087 | 138245183 | 58684 | 85,89,149,90,67,54,51,68,3 | 2,2,6,7,2,6,0,5,6,2,1,3 | 100,90,58,63,92,39,49,50,3 | 6,11,5,14,17,4,0,6,1,5,2,7,0 | 298 | 149 | 0.0114834765639 | 0.623040117046827 | 0.955,0.957,0.925, | 0.893,0.804,0.853, | 0,035 |
| 101070 | ENSG00000186088 | GSAP | chr7 | - | 77321332 | 77321403 | 77320724 | 77320819 | 77323646 | 77323742 | 101070 | 207,133,220,133,96,132,10 | 12,2,2,0,10,6,0,0,0,0,0,0 | 172,140,136,85,125,73,75, | 10,4,0,0,0,0,0,0,8,0,0,0,0,0 | 220 | 149 | 0.011503073705 | 0.623946082812715 | 0.921,0.978,0.987, | 0.921,0.96,1.0,1.0, | -0,01 |
| 89832 | ENSG00000272030 | | chr1 | - | 153633035 | 153633157 | 153632033 | 153632240 | 153633957 | 153634110 | 89832 | 8,1,1,3,1,0,2,2,8,2,1,1 | 6,3,10,13,4,6,6,9,4,0,2,3 | 0,7,0,0,2,1,2,0,0,5,2,2,0,0,2 | 2,5,2,3,6,5,0,1,4,3,0,5,3,11 | 271 | 149 | 0.0115146612341 | 0.624417246680921 | 0.423,0.155,0.052, | 0.0,0.435,0.0,0.0,0 | 0,029 |
| 80737 | ENSG00000181381 | DDX60L | chr4 | - | 168377727 | 168377843 | 168375376 | 168375524 | 168378353 | 168378475 | 80737 | 7,0,10,2,2,0,4,11,0,3,2,3 | 11,3,8,5,6,7,0,1,8,2,6,8 | 7,1,5,4,4,4,6,5,1,3,6,0,13,1 | 6,7,0,0,3,2,2,3,0,3,11,11,1, | 265 | 149 | 0.0115177469178 | 0.624427251099499 | 0.264,0.0,0.413,0.1 | 0.396,0.074,1.0,1.0 | -0,229 |
| 193918 | ENSG00000092841 | MYL6 | chr12 | + | 56159588 | 56159730 | 56158683 | 56158711 | 56159974 | 56160148 | 193918 | 607,502,1779,1425,670,68 | 38,26,57,25,6,14,8,13,16,1 | 836,784,519,283,605,431,2 | 11,37,15,54,46,7,0,10,20,8 | 291 | 149 | 0.0115267422777 | 0.624757559096361 | 0.891,0.908,0.941, | 0.975,0.916,0.947, | 0,023 |
| 24468 | ENSG00000120697 | ALG5 | chr13 | - | 36994988 | 36995035 | 36993603 | 36993672 | 36995424 | 36995596 | 24468 | 47,43,91,42,15,28,27,25,32 | 4,0,3,1,1,0,0,3,3,1,1,2 | 42,55,26,11,32,17,13,19,17 | 1,6,7,3,1,3,5,0,0,1,3,1,4,2,2 | 196 | 149 | 0.0115423023941 | 0.625440424151606 | 0.899,1.0,0.958,0.9 | 0.97,0.875,0.738,0 | 0,04 |
| 97742 | ENSG00000196511 | TPK1 | chr7 | - | 144606928 | 144607050 | 144591422 | 144591569 | 144623165 | 144623261 | 97742 | 3,0,4,4,3,2,0,4,0,3,0,2 | 23,13,21,27,14,11,22,30,16 | 1,5,2,3,5,1,1,4,2,9,7,0,2,4,0 | 16,15,9,16,14,13,6,2,13,20 | 271 | 149 | 0.0115451529062 | 0.625440424151606 | 0.067,0.0,0.095,0.0 | 0.033,0.155,0.109, | -0,048 |
| 213757 | ENSG00000123607 | TTC21B | chr2 | - | 165894570 | 165894824 | 165890837 | 165890988 | 165898685 | 165898767 | 213757 | 2,5,7,1,4,6,5,9,4,3,5,2 | 17,10,21,22,2,8,4,9,3,6,4,1 | 15,18,4,21,11,6,10,7,1,1,6, | 15,11,3,7,2,6,7,6,9,6,11,4,4 | 298 | 149 | 0.0115792377147 | 0.627129067079163 | 0.056,0.2,0.143,0.0 | 0.333,0.45,0.4,0.6, | -0,071 |
| 223505 | ENSG00000171316 | CHD7 | chr8 | + | 60846499 | 60846632 | 60845249 | 60845409 | 60848514 | 60848604 | 223505 | 0,0,8,3,5,2,1,3,0,0,1,0 | 4,12,46,15,22,17,10,16,6,1 | 8,9,2,6,3,0,2,2,0,2,3,4,0,4,4 | 33,40,26,16,10,3,20,32,1,2 | 282 | 149 | 0.0116026310332 | 0.628185897475501 | 0.0,0.0,0.084,0.09 | 0.114,0.106,0.039, | -0,024 |
| 231035 | ENSG00000244055 | | chr7 | + | 92489788 | 92490142 | 92462957 | 92463026 | 92491242 | 92491579 | 231035 | 1,0,3,0,0,2,2,3,6,0,1,2 | 14,4,15,11,6,8,7,11,2,1,3,3 | 2,3,0,1,0,0,0,0,1,3,0,2,0,1,0 | 10,6,5,9,3,2,3,15,3,9,11,7,1 | 298 | 149 | 0.0116045882255 | 0.628185897475501 | 0.034,0.0,0.091,0.0 | 0.091,0.2,0.0,0.05 | 0,071 |
| 220833 | ENSG00000172890 | NADSYN1 | chr11 | + | 71481189 | 71481404 | 71480754 | 71480879 | 71481922 | 71482025 | 220833 | 21,4,34,16,13,6,12,2,21,17 | 4,0,4,1,11,4,3,3,0,2,4,1 | 7,9,9,12,5,8,7,12,8,8,8,11,7 | 0,2,1,0,1,0,1,0,2,5,0,2,1,3,1 | 298 | 149 | 0.0116283170378 | 0.629312120785198 | 0.724,1.0,0.81,0.88 | 1.0,0.692,0.818,1.0 | -0,124 |
| 240898 | ENSG00000198771 | RCSD1 | chr1 | + | 167640333 | 167640474 | 167630265 | 167630429 | 167640570 | 167640712 | 240898 | 22,15,36,25,0,2,16,32,9,3,2 | 3,2,5,3,4,2,4,15,6,2,3,12 | 7,17,31,14,2,5,11,21,6,2,32 | 1,0,8,6,0,0,3,3,1,0,2,0,3,0,1 | 290 | 149 | 0.0116475699663 | 0.63019560870204 | 0.79,0.794,0.787,0 | 0.782,1.0,0.666,0.5 | -0,246 |
| 76302 | ENSG00000076242 | MLH1 | chr3 | + | 37014431 | 37014544 | 37012010 | 37012099 | 37020309 | 37020463 | 76302 | 9,5,24,24,10,15,9,14,4,1,1, | 10,5,17,9,1,6,3,2,7,3,2,11 | 12,6,11,3,2,9,5,3,2,13,5,6,1 | 8,23,9,6,7,5,13,7,6,11,14,1 | 262 | 149 | 0.0116558053081 | 0.630482692200973 | 0.339,0.363,0.445, | 0.46,0.129,0.41,0.3 | 0,148 |
| 73050 | ENSG00000125633 | CCDC93 | chr2 | - | 117985968 | 117986069 | 117977993 | 117978030 | 117995445 | 117995502 | 73050 | 69,56,73,30,33,43,25,26,38 | 1,1,2,0,2,0,0,2,0,1,0,0 | 45,48,27,41,35,22,42,37,14 | 0,4,0,1,8,0,0,0,2,0,0,3,1,0,1 | 250 | 149 | 0.0116626723497 | 0.630610406636622 | 0.976,0.971,0.956, | 1.0,0.877,1.0,0.96 | 0,031 |
| 149482 | ENSG00000140367 | UBE2Q2 | chr15 | + | 75874824 | 75875223 | 75873427 | 75873568 | 75876186 | 75876271 | 149482 | 6,1,10,7,1,7,3,6,6,2,2,3 | 37,65,79,48,18,20,11,11,57 | 3,1,0,0,4,6,1,1,0,1,4,5,5,6,1 | 43,36,41,16,23,15,18,44,26 | 298 | 149 | 0.0116640262268 | 0.630610406636622 | 0.075,0.008,0.06,0 | 0.034,0.014,0.0,0.0 | 0,032 |
| 104464 | ENSG00000149970 | CNKSR2 | chrX | + | 21561470 | 21561560 | 21531855 | 21532067 | 21563237 | 21563452 | 104464 | 17,0,6,6,5,2,2,2,8,3,1,4 | 7,2,14,9,4,0,1,8,14,4,3,5 | 2,1,2,5,2,4,0,0,2,3,0,6,0,3,1 | 7,4,5,13,5,2,1,4,5,8,7,10,2, | 239 | 149 | 0.0116670531535 | 0.630615649744088 | 0.602,0.0,0.211,0.1 | 0.151,0.135,0.2,0.1 | 0,204 |
| 180031 | ENSG00000122224 | LY9 | chr1 | + | 160824180 | 160824289 | 160823464 | 160823796 | 160827747 | 160828261 | 180031 | 32,13,24,18,27,10,16,16,15 | 7,4,1,2,6,4,0,0,1,0,5,12 | 29,9,17,8,7,12,18,28,12,31 | 12,3,2,4,3,5,3,11,1,3,11,29 | 258 | 149 | 0.0116765161264 | 0.630803396741129 | 0.725,0.652,0.933, | 0.583,0.634,0.831, | 0,125 |
| 218924 | ENSG00000204592 | HLA-E | chr6 | + | 30491508 | 30491653 | 30491136 | 30491412 | 30492403 | 30492436 | 218924 | 368,373,662,703,250,215,4 | 11,6,11,3,1,5,6,1,0,2,3,8 | 453,510,335,207,303,196,2 | 2,4,8,13,4,0,4,17,1,0,12,18 | 294 | 149 | 0.0116793059791 | 0.630803396741129 | 0.944,0.969,0.968, | 0.991,0.985,0.955, | 0,01 |
| 45390 | ENSG00000109861 | CTSC | chr11 | - | 88328111 | 88328196 | 88312387 | 88312554 | 88334936 | 88335082 | 45390 | 13,6,26,19,1,2,2,8,7,7,4,2 | 25,21,52,60,11,1,7,14,5,4,2 | 4,11,8,4,2,6,0,2,9,11,3,4,3, | 41,30,16,11,18,17,10,18,15 | 234 | 149 | 0.0116793191316 | 0.630803396741129 | 0.249,0.154,0.241, | 0.058,0.189,0.241, | 0,151 |
| 218725 | ENSG00000196262 | PPIA | chr7 | + | 44799391 | 44799880 | 44796978 | 44799277 | 44801286 | 44803117 | 218725 | 133,165,86,67,38,28,30,53 | 17,4,19,20,3,9,11,7,7,2,4,1 | 51,63,63,34,84,44,18,19,36 | 14,17,4,24,16,7,1,19,9,0,8, | 298 | 149 | 0.0117058390584 | 0.632077131555065 | 0.796,0.954,0.694, | 0.646,0.649,0.887, | 0,097 |
| 16764 | ENSG00000122299 | ZC3H7A | chr16 | - | 11761467 | 11761511 | 11758430 | 11758539 | 11761909 | 11762043 | 16764 | 11,8,15,6,2,9,5,13,4,0,5,7 | 7,0,7,6,4,6,4,6,2,2,2,2 | 9,21,5,8,12,9,5,5,11,14,6,1 | 1,2,0,0,6,1,0,3,6,1,3,0,2,0,6 | 193 | 149 | 0.0117087973694 | 0.632078296115709 | 0.548,1.0,0.623,0.4 | 0.874,0.89,1.0,1.0, | -0,207 |
| 24999 | ENSG00000171295 | ZNF440 | chr19 | + | 11830273 | 11830409 | 11814343 | 11814450 | 11830616 | 11830677 | 24999 | 1,4,10,2,2,9,2,5,2,4,4,8 | 3,0,3,0,0,1,1,0,0,0,0,2 | 7,11,6,7,6,5,2,3,7,5,4,6,10, | 0,1,0,0,2,2,0,2,0,1,0,3,2,0,3 | 285 | 149 | 0.011715884792 | 0.63230230683725 | 0.148,1.0,0.635,1.0 | 1.0,0.852,1.0,1.0,0 | 0,059 |
| 159328 | ENSG00000259040 | BLOC1S5-TXND | chr6 | - | 7889539 | 7889581 | 7888704 | 7888848 | 7891620 | 7891736 | 159328 | 7,18,62,32,14,27,57,100,46 | 0,1,4,0,0,3,4,6,2,0,2,4 | 21,30,101,72,26,27,10,29,3 | 1,0,0,2,1,2,6,2,2,1,2,0,0,0,1 | 191 | 149 | 0.0117194146964 | 0.632334255332381 | 1.0,0.934,0.924,1.0 | 0.942,1.0,1.0,0.96 | 0,005 |
| 162124 | ENSG00000181929 | PRKAG1 | chr12 | - | 49003841 | 49003922 | 49003143 | 49003290 | 49004506 | 49004633 | 162124 | 5,6,28,12,16,4,1,9,17,7,2,7 | 1,1,1,2,0,1,0,0,0,1,0,1 | 11,22,8,1,13,6,1,4,1,8,8,12 | 2,10,2,0,0,2,1,0,1,0,0,2,1,1 | 230 | 149 | 0.011734028581 | 0.632964086595792 | 0.764,0.795,0.948, | 0.781,0.588,0.722, | 0,092 |
| 218724 | ENSG00000196262 | PPIA | chr7 | + | 44799391 | 44799871 | 44796978 | 44799277 | 44801286 | 44803117 | 218724 | 133,165,85,68,38,28,30,53 | 17,4,19,20,3,9,11,7,7,2,4,1 | 49,63,63,34,85,44,18,19,36 | 14,17,4,24,16,7,1,19,9,0,8, | 298 | 149 | 0.0117683313483 | 0.634655405769471 | 0.796,0.954,0.691, | 0.636,0.649,0.887, | 0,096 |
| 140284 | ENSG00000197586 | ENTPD6 | chr20 | + | 25207075 | 25207522 | 25195692 | 25195867 | 25209848 | 25209925 | 140284 | 1,2,3,0,0,0,7,1,2,2,0,1 | 11,9,11,5,2,5,4,5,2,1,7,5 | 0,0,1,0,0,0,1,2,0,0,1,0,0,0,0 | 6,8,9,2,3,1,3,3,1,17,1,9,10, | 298 | 149 | 0.0117776395334 | 0.634998280163631 | 0.043,0.1,0.12,0.0, | 0.0,0.0,0.053,0.0,0 | 0,09 |
| 99276 | ENSG00000102531 | FNDC3A | chr13 | + | 49191014 | 49191120 | 49188514 | 49188633 | 49191208 | 49191384 | 99276 | 27,19,62,59,36,39,38,41,19 | 2,1,2,13,4,4,2,4,2,0,0,3 | 36,54,29,29,19,10,33,23,9, | 10,16,3,7,0,1,1,0,3,9,7,4,2, | 255 | 149 | 0.0118158482144 | 0.636898779622721 | 0.887,0.917,0.948, | 0.678,0.664,0.85,0 | 0,079 |
| 122197 | ENSG00000103429 | BFAR | chr16 | + | 14649803 | 14649973 | 14644287 | 14644609 | 14655065 | 14655210 | 122197 | 17,9,33,31,19,22,9,15,23,1 | 2,2,3,0,2,1,4,1,0,0,1,0 | 23,26,15,5,14,7,6,14,21,21 | 2,1,0,0,0,0,0,0,0,0,0,0,0,2,0 | 298 | 149 | 0.0118289637512 | 0.637446093423767 | 0.81,0.692,0.846,1 | 0.852,0.929,1.0,1.0 | -0,109 |
| 90913 | ENSG00000093167 | LRRFIP2 | chr3 | - | 37154613 | 37154656 | 37148893 | 37149038 | 37174538 | 37174605 | 90913 | 1,3,1,0,0,0,0,0,0,1,0,1 | 15,9,28,9,5,10,3,14,5,6,8,4 | 1,8,1,2,0,0,0,2,1,0,2,0,2,2,2 | 14,14,11,13,25,7,8,16,8,8,1 | 192 | 149 | 0.0118392788051 | 0.637789611615714 | 0.049,0.206,0.027, | 0.053,0.307,0.066, | -0,037 |

| | | | | | | | | | | | | | | | | | | | | | | |
|---|---|---|---|---|---|---|---|---|---|---|---|---|---|---|---|---|---|---|---|---|---|---|
| 252084 | ENSG00000107854 | TNKS2 | chr10 | + | 91830033 | 91830094 | 91828284 | 91828406 | 91830922 | 91831014 | 252084 | 2,2,18,6,1,10,7,6,4,0,4,3 | 26,14,65,46,15,17,14,14,1 | 4,5,10,11,4,1,4,2,2,6,4,4,1, | 42,55,21,30,28,17,12,33,2 | 210 | 149 | 0.0118412649108 | 0.637789611615714 | 0.052,0.092,0.164, | 0.063,0.061,0.253, | 0,052 |
| 190348 | ENSG00000163781 | TOPBP1 | chr3 | - | 133618212 | 133618429 | 133617159 | 133617326 | 133620154 | 133620347 | 190348 | 52,38,22,22,28,23,19,23,5 | 15,1,7,3,11,14,5,16,20,6,5, | 24,21,24,20,14,8,6,21,3,12 | 13,2,15,14,11,9,2,10,5,7,8, | 298 | 149 | 0.0118468244424 | 0.637801339448766 | 0.634,0.95,0.611,0 | 0.48,0.84,0.444,0. | 0,149 |
| 29552 | ENSG00000140853 | NLRC5 | chr16 | + | 57055435 | 57055519 | 57055031 | 57055094 | 57058064 | 57058148 | 29552 | 14,13,33,42,5,11,6,22,6,6, | 4,3,1,2,0,2,6,0,1,2,0,5 | 11,35,10,13,15,23,6,35,11, | 5,5,9,3,3,1,1,10,4,3,7,12,1 | 233 | 149 | 0.0118474093189 | 0.637801339448766 | 0.691,0.735,0.955, | 0.585,0.817,0.415, | 0,097 |
| 116114 | ENSG00000198382 | UVRAG | chr11 | + | 76012926 | 76013102 | 76008806 | 76008867 | 76016814 | 76016980 | 116114 | 3,4,24,5,5,9,4,19,0,0,2,1 | 82,44,290,142,77,152,56,1 | 2,1,2,0,2,2,1,0,3,7,4,5,11,1 | 119,119,68,71,53,63,76,11 | 298 | 149 | 0.0118986428839 | 0.640399301461536 | 0.018,0.043,0.04,0 | 0.008,0.004,0.014, | 0,003 |
| 88901 | ENSG00000073849 | ST6GAL1 | chr3 | + | 187038394 | 187038491 | 187025482 | 187025517 | 187038741 | 187038873 | 88901 | 1,6,26,18,4,7,9,7,6,1,6,9 | 1,2,32,24,6,6,8,3,6,3,2,4 | 11,18,10,18,5,8,0,12,6,7,5, | 15,2,8,1,0,6,3,3,7,4,1,0,7,1 | 246 | 149 | 0.0119183843496 | 0.641301445487189 | 0.377,0.645,0.33,0 | 0.308,0.845,0.431, | -0,179 |
| 183681 | ENSG00000105483 | CARD8 | chr19 | - | 48229502 | 48229616 | 48223718 | 48223868 | 48230407 | 48230700 | 183681 | 0,0,1,1,0,2,3,3,2,0,0,0 | 9,8,15,14,8,2,12,5,7,5,2,7 | 0,0,0,1,0,2,0,4,4,0,5,0,6,6, | 13,12,6,5,8,8,2,6,5,13,0,13 | 263 | 149 | 0.0119228611175 | 0.641358477089147 | 0.0,0.0,0.036,0.03 | 0.0,0.0,0.0,0.102,0 | -0,091 |
| 214879 | ENSG00000184007 | PTP4A2 | chr1 | - | 31911695 | 31911841 | 31910037 | 31910112 | 31915894 | 31915987 | 214879 | 311,190,276,254,126,91,8 | 14,4,9,1,0,3,4,1,2,4,3,4 | 208,218,153,154,116,129,6 | 10,8,2,8,8,0,0,8,2,6,3,11,7, | 295 | 149 | 0.011925403986 | 0.641358477089147 | 0.918,0.96,0.939,0 | 0.913,0.932,0.975, | 0,017 |
| 198722 | ENSG00000223745 | CCDC18-AS1 | chr1 | - | 93335353 | 93335607 | 93326601 | 93326662 | 93337376 | 93339274 | 198722 | 1,2,1,2,0,0,7,6,2,1,2,6 | 13,9,32,12,1,10,6,16,5,1,2, | 0,0,1,1,1,0,0,1,3,4,3,0,0,2,2 | 14,4,14,0,8,6,4,8,5,11,10,5 | 298 | 149 | 0.0119352402668 | 0.641584311869452 | 0.037,0.1,0.015,0. | 0.0,0.0,0.034,1.0,0 | 0,005 |
| 115877 | ENSG00000070061 | ELP1 | chr9 | - | 108919252 | 108919349 | 108917546 | 108917670 | 108922841 | 108922927 | 115877 | 27,22,15,19,16,17,7,7,29,1 | 1,0,2,1,0,4,2,4,1,0,0,5 | 18,8,19,17,3,2,12,12,10,20 | 5,1,2,0,8,1,4,3,2,3,1,5,0,0,4 | 246 | 149 | 0.0119355649731 | 0.641584311869452 | 0.942,1.0,0.82,0.9 | 0.686,0.829,0.852, | 0,141 |
| 217065 | ENSG00000148690 | FRA10AC1 | chr10 | - | 93701506 | 93701795 | 93700029 | 93700106 | 93702374 | 93702592 | 217065 | 0,4,2,1,6,0,2,3,0,0,0,0 | 3,6,6,5,7,7,3,0,3,6,2,10 | 4,0,0,0,0,1,0,1,1,0,1,0,0,0,0 | 3,2,4,2,2,5,4,3,1,3,3,5,3,7,8 | 298 | 149 | 0.0119687877762 | 0.643209528554133 | 0.0,0.25,0.143,0.0 | 0.4,0.0,0.0,0.0,0.0, | 0,1 |
| 89434 | ENSG00000135945 | REV1 | chr2 | - | 99406324 | 99406490 | 99405909 | 99406106 | 99408028 | 99408131 | 89434 | 48,18,32,29,22,19,28,33,28 | 1,1,3,9,0,0,3,2,2,0,0,1 | 32,29,30,21,12,19,16,30,1 | 4,0,0,1,3,0,1,1,0,0,0,0,0,3,2 | 298 | 149 | 0.0119948999717 | 0.64445190110059 | 0.96,0.9,0.842,0.6 | 0.8,1.0,1.0,0.913,0 | -0,009 |
| 152998 | ENSG00000071537 | SEL1L | chr14 | - | 81484224 | 81484290 | 81479277 | 81479740 | 81485671 | 81485746 | 152998 | 19,15,19,12,9,11,5,18,8,6, | 4,2,2,1,2,0,1,2,0,2,0,0 | 16,17,6,9,8,3,14,11,2,15,1 | 1,1,0,4,0,2,0,1,0,0,0,0,0,0,0 | 215 | 149 | 0.0120052307109 | 0.644845972332847 | 0.767,0.839,0.868, | 0.917,0.922,1.0,0. | -0,058 |
| 215831 | ENSG00000188647 | PTAR1 | chr9 | - | 69740685 | 69740776 | 69734149 | 69734254 | 69741791 | 69741858 | 215831 | 9,7,14,4,0,3,1,4,0,0,4,1 | 39,5,62,44,58,36,26,36,9,1 | 12,8,10,5,8,0,5,0,5,6,4,13,1 | 32,41,19,25,34,19,11,33,2 | 240 | 149 | 0.0120166312435 | 0.645297295202033 | 0.125,0.465,0.123, | 0.189,0.108,0.246, | -0,017 |
| 141878 | ENSG00000118046 | STK11 | chr19 | + | 1216203 | 1216398 | 1207090 | 1207203 | 1218416 | 1218500 | 141878 | 10,13,7,1,0,1,1,10,3,0,1,5 | 34,29,77,74,33,41,25,27,2 | 3,1,5,3,0,1,0,3,0,8,4,2,5,4,0 | 23,25,12,16,11,21,18,50,1 | 298 | 149 | 0.0120289712103 | 0.645798828277396 | 0.128,0.183,0.043, | 0.061,0.02,0.172,0 | 0,018 |
| 211520 | ENSG00000164548 | TRA2A | chr7 | - | 23522126 | 23522432 | 23521706 | 23521840 | 23531788 | 23531980 | 211520 | 175,79,69,22,21,13,52,57,3 | 14,7,72,52,14,24,4,4,4,10,4, | 44,40,59,42,22,21,4,15,9,4 | 36,49,23,11,10,15,5,17,19, | 298 | 149 | 0.0120470837035 | 0.646498156322494 | 0.862,0.849,0.324, | 0.379,0.29,0.562,0 | 0,155 |
| 189679 | ENSG00000074603 | DPP8 | chr15 | - | 65454278 | 65454415 | 65452043 | 65452102 | 65456224 | 65456371 | 189679 | 7,6,29,10,16,13,14,20,7,4,3 | 4,3,9,9,5,1,5,2,3,0,2,4 | 14,31,14,9,8,10,6,9,4,12,8, | 3,6,6,0,2,0,0,0,1,5,4,1,0,6,2 | 286 | 149 | 0.0120480047252 | 0.646498156322494 | 0.477,0.51,0.627,0 | 0.709,0.729,0.549, | -0,163 |
| 188468 | ENSG00000143294 | PRCC | chr1 | + | 156791066 | 156791368 | 156786890 | 156787174 | 156791696 | 156791792 | 188468 | 1,0,1,4,2,2,0,6,2,1,0,1 | 9,14,1,9,8,3,3,2,3,1,1,4 | 3,0,0,3,4,1,1,0,0,0,0,1,0,4,3 | 1,9,3,3,9,2,2,9,2,4,3,3,2,3,2 | 298 | 149 | 0.0120838356228 | 0.648186644692726 | 0.053,0.0,0.333,0. | 0.6,0.0,0.0,0.333,0 | 0,03 |
| 38594 | ENSG00000155508 | CNOT8 | chr5 | + | 154870660 | 154870822 | 154865191 | 154865385 | 154871729 | 154871831 | 38594 | 88,115,256,151,56,100,66, | 1,0,1,4,2,1,1,7,6,1,0,0 | 172,236,72,97,144,79,60,6 | 1,1,0,0,1,1,0,0,2,0,1,2,0,1,0 | 298 | 149 | 0.0120854902139 | 0.648186644692726 | 0.978,1.0,0.992,0.9 | 0.989,0.992,1.0,1.0 | -0,027 |
| 79143 | ENSG00000155097 | ATP6V1C1 | chr8 | + | 103064713 | 103064811 | 103063134 | 103063228 | 103066320 | 103066447 | 79143 | 19,19,45,28,15,14,11,21,1 | 2,1,0,3,1,1,3,1,2,2,0,0,0 | 43,22,16,14,11,8,22,32, | 4,3,4,3,2,0,0,3,1,2,0,3,7,1,1 | 247 | 149 | 0.0121148101689 | 0.649597087060916 | 0.92,1.0,0.9,0.944, | 0.866,0.816,0.707, | 0,038 |
| 171869 | ENSG00000152102 | FAM168B | chr2 | - | 131072021 | 131072254 | 131071854 | 131071938 | 131082576 | 131082657 | 171869 | 0,0,0,0,0,2,0,0,0,0,0,0 | 25,16,17,23,16,15,19,19,4 | 1,5,3,6,1,1,1,2,0,0,3,0,0,0,4 | 16,26,13,17,10,5,19,18,3,2 | 298 | 149 | 0.0121315000681 | 0.650329985344267 | 0.0,0.0,0.0,0.0,0.0, | 0.03,0.088,0.103,0 | -0,052 |
| 10895 | ENSG00000108819 | PPP1R9B | chr17 | - | 50140092 | 50140228 | 50139428 | 50139581 | 50141268 | 50141373 | 10895 | 398,305,304,253,186,121,9 | 3,0,10,4,2,1,0,0,1,2,0,1 | 178,231,142,130,85,101,3 | 3,6,0,3,0,2,1,0,0,7,0,12,3,2 | 285 | 149 | 0.0121518197295 | 0.651257049850601 | 0.986,1.0,0.941,0.9 | 0.969,0.953,1.0,0.9 | 0,016 |
| 40146 | ENSG00000291102 | GUSBP1 | chr5 | + | 21491320 | 21491508 | 21461782 | 21461866 | 21495097 | 21495182 | 40146 | 101,41,164,83,94,155,128, | 1,2,3,4,1,1,1,4,0,3,0,2 | 39,44,49,83,43,20,126,137 | 1,4,2,6,10,1,0,2,1,0,1,0,5,3 | 298 | 149 | 0.0121604166076 | 0.651465423362075 | 0.981,0.911,0.965, | 0.951,0.846,0.925, | 0,066 |
| 158225 | ENSG00000115760 | BIRC6 | chr2 | + | 32594886 | 32595144 | 32593914 | 32594060 | 32597750 | 32597968 | 158225 | 29,15,47,37,18,10,26,22,1 | 4,5,12,2,4,5,6,10,4,2,0,5 | 58,18,34,8,7,16,14,33,11,2 | 37,18,5,13,9,3,5,7,5,11,17, | 298 | 149 | 0.0121617614148 | 0.651465423362075 | 0.784,0.6,0.662,0.9 | 0.439,0.333,0.773, | 0,14 |
| 206949 | ENSG00000048991 | R3HDM1 | chr2 | + | 135715554 | 135715694 | 135710058 | 135710231 | 135721923 | 135722006 | 206949 | 2,6,12,9,3,4,8,5,6,4,4,8 | 0,0,1,2,1,1,0,2,0,0,0,1 | 8,7,13,1,1,6,4,8,6,4,9,8,6,5 | 1,2,2,3,4,1,0,0,3,0,0,0,0,1,2 | 289 | 149 | 0.0121663824052 | 0.651550796579647 | 1.0,1.0,0.861,0.69 | 0.805,0.643,0.77,0 | 0,09 |
| 255509 | ENSG00000117151 | CTBS | chr1 | - | 84564206 | 84564304 | 84563734 | 84563832 | 84565840 | 84566012 | 255509 | 7,6,40,15,0,3,6,6,2,0,0,0 | 20,12,9,27,13,9,12,8,4,2,3, | 6,5,4,5,8,2,4,3,0,2,0,1,3,0,1 | 13,19,7,14,16,15,14,11,9,1 | 247 | 149 | 0.0121782226419 | 0.652022646129049 | 0.174,0.232,0.728, | 0.218,0.137,0.256, | 0,083 |
| 158464 | ENSG00000135506 | OS9 | chr12 | + | 57720100 | 57720263 | 57719092 | 57719182 | 57720405 | 57720518 | 158464 | 37,29,96,101,31,31,33,30,5 | 49,24,134,85,53,41,41,55,3 | 52,44,38,4,14,20,18,26,18, | 67,85,51,24,37,28,28,36,3 | 298 | 149 | 0.0122008088609 | 0.652909640621549 | 0.274,0.377,0.264, | 0.28,0.206,0.271,0 | 0,065 |
| 228593 | ENSG00000088930 | XRN2 | chr20 | + | 21387431 | 21387516 | 21386867 | 21387006 | 21389272 | 21389825 | 228593 | 0,0,5,5,1,3,6,1,5,0,1,2 | 40,30,76,42,47,41,32,53,6 | 9,7,2,5,8,3,4,6,4,6,0,1,4,0,0 | 56,45,42,54,26,22,20,58,2 | 234 | 149 | 0.0122008566358 | 0.652909640621549 | 0.0,0.0,0.04,0.07,0 | 0.093,0.09,0.029,0 | -0,026 |
| 36516 | ENSG00000145495 | MARCHF6 | chr5 | + | 10411332 | 10411537 | 10410138 | 10410276 | 10414432 | 10414502 | 36516 | 307,222,312,160,146,184,1 | 0,1,0,2,0,0,4,2,0,0,0,0 | 251,298,190,439,168,107,1 | 0,6,0,14,3,2,1,1,1,3,0,4,1,1 | 298 | 149 | 0.0122145440607 | 0.653397254464873 | 1.0,0.991,1.0,0.97 | 1.0,0.961,1.0,0.94, | 0,019 |
| 124058 | ENSG00000110721 | CHKA | chr11 | - | 68061492 | 68061830 | 68053514 | 68054047 | 68061952 | 68062034 | 124058 | 6,4,5,3,0,1,4,0,0,0,2,5 | 7,15,54,32,17,16,5,10,10,3 | 2,9,0,4,3,3,1,4,3,6,1,4,3,9,2 | 10,13,4,2,3,4,6,15,3,22,7,1 | 298 | 149 | 0.0122160402171 | 0.653397254464873 | 0.3,0.118,0.044,0.0 | 0.091,0.257,0.0,0. | -0,072 |
| 200344 | ENSG00000132849 | PATJ | chr1 | + | 62100318 | 62100407 | 62084514 | 62084648 | 62108436 | 62108520 | 200344 | 3,0,3,0,1,1,2,4,0,2,1,1 | 9,5,13,11,8,17,2,10,6,4,2,8 | 4,4,1,0,0,1,2,1,4,2,0,0,0,0,1 | 11,9,7,7,9,7,0,9,1,7,1,9,5,5 | 238 | 149 | 0.0122230943375 | 0.65361212853527 | 0.173,0.0,0.126,0.0 | 0.185,0.218,0.082, | -0,042 |
| 83127 | ENSG00000164548 | TRA2A | chr7 | - | 23522131 | 23522432 | 23521706 | 23521840 | 23531788 | 23532041 | 83127 | 179,80,77,24,26,14,57,58,3 | 14,7,72,52,14,24,4,4,4,10,4, | 44,43,59,43,25,23,6,17,9,4 | 36,49,23,11,10,15,5,17,19, | 298 | 149 | 0.0122264075149 | 0.653626904083319 | 0.865,0.851,0.348, | 0.379,0.305,0.562, | 0,148 |
| 5812 | ENSG00000118705 | RPN2 | chr20 | + | 37232295 | 37232391 | 37229972 | 37230059 | 37234019 | 37234095 | 5812 | 155,83,151,134,117,139,14 | 4,0,7,8,6,5,3,11,2,3,1,2 | 117,123,105,106,50,41,85, | 13,14,6,6,7,1,3,4,0,2,5,8,5, | 245 | 149 | 0.0122367749203 | 0.654018699496173 | 0.959,1.0,0.929,0.9 | 0.846,0.842,0.914, | 0,027 |
| 137657 | ENSG00000104852 | SNRNP70 | chr19 | + | 49102113 | 49102198 | 49101389 | 49101471 | 49104633 | 49104735 | 137657 | 93,53,171,113,106,120,89, | 60,48,65,32,32,23,36,20,6 | 185,159,66,33,81,60,72,13 | 25,28,37,33,32,15,11,30,1 | 234 | 149 | 0.0122465975951 | 0.654381193393984 | 0.497,0.413,0.626, | 0.825,0.783,0.532, | -0,111 |
| 215141 | ENSG00000115355 | CCDC88A | chr2 | - | 55302960 | 55303152 | 55301871 | 55302072 | 55308808 | 55309023 | 215141 | 144,78,243,129,35,21,52,4 | 88,80,222,151,22,17,41,42 | 177,154,74,7,23,36,53,65,2 | 106,57,30,5,26,26,33,45,3 | 298 | 149 | 0.012273210858 | 0.655565376263991 | 0.45,0.328,0.354,0 | 0.455,0.575,0.552, | -0,086 |
| 90163 | ENSG00000224078 | SNHG14 | chr15 | + | 25079825 | 25080079 | 25078861 | 25078995 | 25081090 | 25081183 | 90163 | 4,2,13,9,0,4,1,5,4,1,2,4 | 6,3,16,12,2,16,3,4,6,8,2,9 | 8,10,13,2,13,4,2,4,5,2,11,2 | 2,3,12,3,2,5,3,5,3,6,7,4,4,3 | 298 | 149 | 0.0122791548111 | 0.655565376263991 | 0.25,0.25,0.289,0. | 0.667,0.625,0.351, | -0,152 |
| 18348 | ENSG00000130703 | OSBPL2 | chr20 | + | 62259980 | 62260125 | 62256056 | 62256221 | 62263615 | 62263691 | 18348 | 105,58,94,58,24,37,34,42,9 | 9,0,3,2,0,4,3,5,0,0,0,3 | 68,64,43,67,46,30,21,46,3 | 6,13,2,4,12,3,2,0,6,6,5,4,1, | 298 | 149 | 0.0122803357409 | 0.655565376263991 | 0.855,1.0,0.941,0.9 | 0.852,0.714,0.916, | 0,056 |
| 9963 | ENSG00000178852 | EFCAB13 | chr17 | + | 47391436 | 47391580 | 47379181 | 47379253 | 47394024 | 47394099 | 9963 | 73,32,145,124,52,39,46,49 | 1,3,8,4,3,4,3,12,1,2,1,1 | 65,58,40,41,40,34,40,110,3 | 0,5,1,1,5,5,3,4,0,0,10,1,1,2 | 293 | 149 | 0.0122809427875 | 0.655565376263991 | 0.974,0.844,0.902, | 1.0,0.855,0.953,0. | -0,049 |
| 236889 | ENSG00000110344 | UBE4A | chr11 | + | 118364147 | 118364278 | 118359643 | 118359674 | 118365039 | 118365201 | 236889 | 0,0,3,4,1,3,2,2,0,0,1,2 | 11,9,52,23,11,8,10,12,10,4 | 6,8,2,6,4,0,0,2,3,6,2,2,1,0 | 16,17,8,2,7,4,3,10,4,14,12, | 280 | 149 | 0.0122910565978 | 0.655942574411379 | 0.0,0.0,0.03,0.085, | 0.166,0.2,0.117,0. | -0,063 |
| 252877 | ENSG00000179889 | PDXDC1 | chr16 | + | 15009498 | 15009561 | 15008778 | 15008847 | 15009680 | 15009759 | 252877 | 3,3,6,5,1,3,0,0,0,2,0,1 | 73,48,120,99,50,58,69,56,3 | 2,4,8,8,0,0,2,6,0,0,5,6,1,4,1 | 74,57,29,50,47,27,37,94,3 | 212 | 149 | 0.0123054229285 | 0.656546475538915 | 0.028,0.042,0.034, | 0.071,0.036,0.046, | -0,015 |
| 179981 | ENSG00000122224 | LY9 | chr1 | + | 160796889 | 160797275 | 160796210 | 160796311 | 160799752 | 160800227 | 179981 | 6,3,6,3,11,1,1,5,1,0,0,2 | 93,93,62,41,60,24,46,18,3 | 0,2,1,10,0,2,2,3,2,1,6,11,0, | 49,70,55,55,59,27,72,81,2 | 298 | 149 | 0.0123184973341 | 0.657081164737466 | 0.031,0.016,0.046, | 0.0,0.014,0.009,0. | 0,011 |
| 94358 | ENSG00000137265 | IRF4 | chr6 | + | 398827 | 398935 | 397107 | 397252 | 401423 | 401777 | 94358 | 62,66,162,123,72,53,67,1 | 22,7,14,4,3,2,1,11,4,5,11,7 | 130,111,85,9,42,44,27,13 | 12,10,14,8,6,2,5,14,1,3,44, | 257 | 149 | 0.0123313125136 | 0.657446813788816 | 0.62,0.845,0.87,0.9 | 0.863,0.866,0.779, | 0,064 |
| 211936 | ENSG00000164985 | PSIP1 | chr9 | - | 15470975 | 15471176 | 15469937 | 15469993 | 15472631 | 15472750 | 211936 | 17,4,9,4,4,3,6,12,4,1,2,8 | 51,20,59,33,59,32,29,36,3 | 3,7,4,6,6,1,9,10,2,13,13,15 | 35,20,18,1,13,25,13,27,6,2 | 298 | 149 | 0.0123314614868 | 0.657446813788816 | 0.143,0.091,0.071, | 0.041,0.149,0.1,0. | -0,086 |
| 247400 | ENSG00000149311 | ATM | chr11 | + | 108244848 | 108245026 | 108243961 | 108244118 | 108246963 | 108247127 | 247400 | 52,25,268,148,28,41,19,3 | 4,0,2,0,0,0,0,0,2,0,1,0 | 119,100,20,16,66,55,18,45 | 9,10,0,0,4,0,0,0,0,1,0,0,0,0 | 298 | 149 | 0.0123423933622 | 0.657866682946921 | 0.867,1.0,0.985,1. | 0.869,0.833,1.0,1. | 0,008 |
| 227858 | ENSG00000173715 | C11orf80 | chr11 | + | 66752637 | 66752757 | 66744891 | 66744919 | 66756352 | 66756505 | 227858 | 13,12,7,11,9,17,11,12,13,6 | 2,2,8,4,13,6,4,4,3,2,1,4 | 12,13,12,2,7,2,6,14,9,14,3, | 2,3,3,4,0,0,1,3,0,3,3,1,1,5 | 269 | 149 | 0.0123498829147 | 0.658032206516785 | 0.783,0.769,0.326, | 0.769,0.706,0.689, | -0,135 |
| 239915 | ENSG00000196781 | TLE1 | chr9 | - | 81650234 | 81650372 | 81634096 | 81634301 | 81652213 | 81652288 | 239915 | 2,3,2,4,0,2,2,17,2,0,0,0 | 7,4,29,7,11,25,59,99,69,2 | 1,0,2,0,1,0,0,4,0,0,2,1,2,0, | 0,10,8,5,18,7,8,21,2,8,48,1 | 287 | 149 | 0.0123516134494 | 0.658032206516785 | 0.129,0.28,0.035,0 | 1.0,0.0,0.115,0.0,0 | -0,028 |
| 107840 | ENSG00000029363 | BCLAF1 | chr6 | - | 136279362 | 136274189 | 136273081 | 136273187 | 136275531 | 136275701 | 107840 | 8,3,41,6,14,4,3,14,5,2,10,3 | 140,81,176,151,70,118,75, | 10,24,4,6,4,4,9,7,0,7,7,8,6, | 129,150,105,105,94,49,79, | 298 | 149 | 0.0123640085981 | 0.658529555698506 | 0.028,0.018,0.104, | 0.037,0.074,0.019, | 0,014 |
| 82127 | ENSG00000184922 | FMNL1 | chr17 | + | 45234831 | 45234849 | 45234071 | 45234200 | 45236135 | 45236244 | 82127 | 6,4,5,10,5,3,3,0,1,0,2,1 | 49,49,71,49,69,71,57,54,2 | 3,4,0,0,3,3,4,2,0,2,1,4,1,1,2 | 36,82,36,32,54,24,57,92,3 | 167 | 149 | 0.0123677409356 | 0.658565375880783 | 0.098,0.068,0.059, | 0.069,0.042,0.0,0. | 0,019 |
| 139779 | ENSG00000101343 | CRNKL1 | chr20 | - | 20039969 | 20040079 | 20039608 | 20039848 | 20040685 | 20040766 | 139779 | 1,0,5,1,1,3,2,2,1,0,0,2 | 27,25,46,21,16,16,18,19,1 | 5,4,11,3,0,1,1,0,3,3,0,0,2,4 | 41,50,17,7,21,7,16,22,8,19 | 259 | 149 | 0.012373087 2927 | 0.65868710143337 | 0.021,0.0,0.059,0. | 0.066,0.044,0.271, | -0,043 |
| 34950 | ENSG00000062194 | GPBP1 | chr5 | + | 57221375 | 57221396 | 57214073 | 57214193 | 57230845 | 57230969 | 34950 | 35,13,41,28,20,22,22,41,2 | 55,50,160,95,29,68,35,61,4 | 41,41,33,33,35,10,7,55,19, | 74,53,42,51,40,19,27,52,2 | 170 | 149 | 0.0123956685458 | 0.659698057220563 | 0.358,0.186,0.183, | 0.327,0.404,0.408, | -0,076 |
| 44655 | ENSG00000127483 | HP1BP3 | chr1 | - | 20780344 | 20780540 | 20779811 | 20779911 | 20787194 | 20787307 | 44655 | 96,61,158,131,72,69,81,7 | 15,10,15,15,5,3,3,1,11,7,2, | 87,95,63,24,46,44,36,66,3 | 10,8,8,6,4,3,3,4,1,2,17,8,1, | 298 | 149 | 0.0123982076999 | 0.659698057220563 | 0.762,0.753,0.84,0 | 0.813,0.856,0.797, | -0,043 |
| 216994 | ENSG00000135945 | REV1 | chr2 | - | 99406324 | 99406361 | 99405909 | 99406106 | 99408028 | 99408403 | 216994 | 5,2,5,4,3,3,1,6,6,2,3,8 | 1,1,3,9,0,0,3,2,2,0,0,1 | 4,9,3,1,0,1,1,4,2,3,1,5,1,3,4 | 4,0,0,1,3,0,1,1,0,0,0,0,0,3,2 | 186 | 149 | 0.0124128061641 | 0.660311587618736 | 0.8,0.616,0.572,0. | 0.445,1.0,1.0,0.44 | 0,009 |
| 131182 | ENSG00000188906 | LRRK2 | chr12 | + | 40340293 | 40340454 | 40334966 | 40335157 | 40346752 | 40346923 | 131182 | 90,90,144,108,44,28,48,3 | 0,1,1,3,0,4,0,2,0,2,0,0 | 103,126,72,47,72,51,43,50 | 0,5,3,3,8,1,0,0,0,1,2,5,0,3,1 | 298 | 149 | 0.0124204918233 | 0.660557172132612 | 1.0,0.978,0.986,0.9 | 1.0,0.926,0.923,0. | 0,008 |
| 202586 | ENSG00000138078 | PREPL | chr2 | - | 44359496 | 44359709 | 44346267 | 44346390 | 44361379 | 44361806 | 202586 | 32,15,44,20,9,13,13,16,15, | 1,4,5,4,4,2,2,5,10,0,6,4 | 31,11,19,15,15,11,18,25,5, | 3,1,4,3,0,2,0,2,0,2,4,0,1,8,4 | 298 | 149 | 0.0124474556341 | 0.661606057022127 | 0.941,0.652,0.815, | 0.838,0.846,0.704, | -0,141 |
| 100457 | ENSG00000223482 | NUTM2A-AS1 | chr10 | + | 87330895 | 87331096 | 87326200 | 87326672 | 87336785 | 87336889 | 100457 | 7,6,5,3,1,1,4,4,4,1,2,0 | 31,26,38,19,28,34,17,21,1 | 2,9,1,1,4,2,0,3,10,4,6,3,6 | 15,20,17,11,13,9,10,18,7,2 | 298 | 149 | 0.0124487140954 | 0.661606057022127 | 0.101,0.103,0.062, | 0.062,0.184,0.029, | -0,042 |
| 30713 | ENSG00000124357 | NAGK | chr2 | + | 71075554 | 71075610 | 71073481 | 71073594 | 71076603 | 71076701 | 30713 | 12,10,69,51,32,24,22,21,9 | 3,5,8,1,3,4,5,8,0,1,1,1 | 38,25,9,8,11,9,12,21,4,11, | 3,12,5,2,3,5,2,6,5,2,1,8,6,5 | 205 | 149 | 0.0124494358663 | 0.661606057022127 | 0.744,0.592,0.862, | 0.902,0.602,0.567, | 0,101 |
| 152711 | ENSG00000100697 | DICER1 | chr14 | - | 95113092 | 95113224 | 95112171 | 95112247 | 95115666 | 95115821 | 152711 | 66,50,120,62,56,78,52,78, | 6,10,8,3,2,12,5,7,1,0,1,2 | 119,80,58,77,51,41,47,44, | 5,12,8,18,8,2,4,4,2,4,7,8,5, | 281 | 149 | 0.012459660581 | 0.661722305635328 | 0.854,0.726,0.888, | 0.927,0.779,0.794, | 0,06 |
| 215607 | ENSG00000166164 | BRD7 | chr16 | - | 50326283 | 50326385 | 50325747 | 50325883 | 50328668 | 50328744 | 215607 | 23,15,45,33,12,12,4,12,17, | 3,6,5,4,2,3,2,2,6,2,0,0 | 28,26,15,4,12,13,7,17,9,20 | 4,4,3,1,1,4,1,2,0,2,1,1,0,1,1 | 251 | 149 | 0.0124601527868 | 0.661722305635328 | 0.82,0.597,0.842,0 | 0.806,0.794,0.748, | -0,059 |
| 134830 | ENSG00000128159 | TUBGCP6 | chr22 | - | 50226991 | 50227077 | 50226734 | 50226842 | 50227906 | 50228028 | 134830 | 15,8,18,11,19,32,15,9,11,1 | 4,2,1,1,1,4,1,2,0,0,0 | 21,8,9,2,5,8,7,5,7,5,14,8,1 | 4,2,0,4,0,0,0,0,1,1,0,3,2,0 | 235 | 149 | 0.0124608467402 | 0.661722305635328 | 0.704,0.717,0.919, | 0.769,0.717,1.0,0. | 0,029 |
| 199477 | ENSG00000111011 | RSRC2 | chr12 | - | 122513768 | 122513874 | 122511110 | 122511188 | 122515104 | 122515227 | 199477 | 19,17,20,8,6,12,4,15,7,11,6 | 42,90,167,99,39,46,37,59,1 | 9,18,11,16,6,2,8,11,8,9,6,4 | 51,130,59,58,98,34,22,46,3 | 255 | 149 | 0.0124913127591 | 0.663176550679293 | 0.209,0.099,0.065, | 0.093,0.075,0.098, | 0,041 |
| 120570 | ENSG00000107036 | RIC1 | chr9 | + | 5742868 | 5743013 | 5732387 | 5732479 | 5743688 | 5743737 | 120570 | 46,38,72,47,47,45,34,75,7 | 0,0,2,0,0,1,0,0,0,1,0,1 | 90,100,39,78,115,39,19,45 | 2,1,2,4,3,0,0,1,4,0,0,5,0,2,0 | 298 | 149 | 0.0124983365129 | 0.663385811592597 | 1.0,1.0,0.948,1.0,1 | 0.958,0.981,0.908, | 0,012 |
| 44532 | ENSG00000187514 | PTMA | chr2 | + | 231711886 | 231711983 | 231711347 | 231711419 | 231712442 | 231712516 | 44532 | 700,516,701,492,477,449,4 | 72,29,55,23,35,27,46,37,2 | 545,371,348,241,290,168,2 | 59,49,18,53,37,8,14,42,18, | 246 | 149 | 0.0125064771112 | 0.663654234497211 | 0.855,0.915,0.885, | 0.848,0.821,0.921, | 0,025 |
| 19325 | ENSG00000138592 | USP8 | chr15 | + | 50481480 | 50482065 | 50477275 | 50477499 | 50484274 | 50484361 | 19325 | 131,122,268,156,89,87,10 | 0,0,2,1,0,0,0,4,0,0,0,0 | 138,188,111,109,139,59,8 | 4,4,0,0,6,1,0,0,2,3,0,2,0,4,5 | 298 | 149 | 0.0125171001389 | 0.664054222330684 | 1.0,1.0,0.985,0.98 | 0.945,0.959,1.0,1.0 | 0,03 |
| 223317 | ENSG00000064419 | TNPO3 | chr7 | - | 128999222 | 128999290 | 128997388 | 128997535 | 129000428 | 129000567 | 223317 | 2,5,11,2,0,5,0,0,0,0,0,0 | 43,53,60,44,21,22,25,18,6, | 2,1,3,2,1,2,0,0,0,0,0,4,2,0,0 | 42,43,25,22,30,16,15,11,9, | 217 | 149 | 0.0125301694494 | 0.664400845026664 | 0.031,0.061,0.112, | 0.032,0.016,0.076, | -0,007 |
| 145190 | ENSG00000141968 | NT5C3B | chr17 | - | 41828751 | 41828952 | 41827425 | 41827626 | 41830800 | 41830880 | 145190 | 52,18,43,66,42,38,8,27,16, | 7,2,2,6,0,1,2,1,2,1,3,1 | 35,23,19,23,8,6,21,29,8,26 | 1,4,0,4,0,0,1,3,0,0,0,0,0,1,4 | 298 | 149 | 0.0125320571676 | 0.664400845026664 | 0.788,0.818,0.915, | 0.946,0.742,1.0,0. | -0,067 |
| 216731 | ENSG00000073849 | ST6GAL1 | chr3 | + | 187064519 | 187064644 | 187051248 | 187051346 | 187072848 | 187072947 | 216731 | 17,3,25,5,22,8,4,10,4,2,0,4 | 245,187,357,279,185,130,1 | 19,13,11,15,6,5,8,10,3,10,5 | 222,155,178,106,113,131,7 | 274 | 149 | 0.0125328945682 | 0.664400845026664 | 0.036,0.009,0.037, | 0.044,0.044,0.033, | -0,013 |
| 29551 | ENSG00000140853 | NLRC5 | chr16 | + | 57055432 | 57055519 | 57055031 | 57055094 | 57058064 | 57058148 | 29551 | 11,9,35,37,7,14,8,21,6,4,7, | 4,3,1,2,0,2,6,0,1,2,0,5 | 7,42,13,12,16,17,6,26,10,3 | 5,5,9,3,3,1,1,10,4,3,7,12,1 | 236 | 149 | 0.01253839 4536 | 0.664432776421829 | 0.635,0.654,0.957, | 0.469,0.841,0.477, | 0,093 |
| 154131 | ENSG00000100934 | SEC23A | chr14 | - | 39037465 | 39037608 | 39033203 | 39033328 | 39039030 | 39041433 | 154131 | 2,0,4,12,5,2,1,3,0,0,1,0 | 4,3,55,23,15,22,9,24,9,5,7, | 0,8,3,7,7,0,2,1,1,5,5,4,6,3, | 22,26,14,4,13,11,6,14,9,14 | 292 | 149 | 0.012539671041 | 0.664432776421829 | 0.203,0.0,0.036,0. | 0.0,0.136,0.099,0. | -0,07 |
| 29617 | ENSG00000116459 | ATP5PB | chr1 | + | 111456085 | 111456249 | 111449839 | 111449873 | 111456629 | 111456748 | 29617 | 88,69,219,181,92,107,103, | 1,2,0,0,2,2,2,0,0,0,0,0 | 125,128,107,38,106,77,39, | 2,1,2,1,5,1,2,1,2,0,0,6,1,0,2 | 298 | 149 | 0.0125651123136 | 0.665616955050072 | 0.978,0.945,1.0,1. | 0.969,0.985,0.964, | 0,02 |
| 52302 | ENSG00000163166 | IWS1 | chr2 | - | 127498139 | 127498237 | 127495997 | 127496148 | 127502814 | 127502872 | 52302 | 26,19,43,30,17,17,9,20,36, | 5,3,6,3,10,5,3,14,8,5,4,1 | 19,8,9,11,4,12,8,12,2,7,26, | 6,5,12,2,5,6,10,11,4,5,1,6,1 | 247 | 149 | 0.0125867095546 | 0.666596969524142 | 0.758,0.793,0.812, | 0.656,0.491,0.311, | 0,153 |
| 56053 | ENSG00000145388 | METTL14 | chr4 | + | 118706510 | 118706821 | 118703934 | 118704051 | 118709997 | 118715430 | 56053 | 63,45,103,50,32,40,29,44,2 | 3,2,15,9,2,9,13,5,6,0,1,6 | 41,83,45,43,67,20,29,58,2 | 15,26,8,7,9,5,5,8,6,7,8,22,0 | 298 | 149 | 0.0126342223739 | 0.668826501922837 | 0.913,0.918,0.774, | 0.577,0.615,0.738, | 0,112 |
| 161360 | ENSG00000165819 | METTL3 | chr14 | - | 21500498 | 21500682 | 21499763 | 21499802 | 21500912 | 21501129 | 161360 | 62,17,91,66,58,54,35,54,3 | 8,4,15,3,10,5,2,12,2,7,1,4 | 80,33,31,13,22,28,25,42,3 | 6,10,2,1,0,2,2,1,0,0,4,7,2,2 | 298 | 149 | 0.0126350226353 | 0.668826501922837 | 0.795,0.68,0.752,0 | 0.87,0.623,0.886,0 | -0,074 |
| 60506 | ENSG00000138688 | KIAA1109 | chr4 | + | 122192229 | 122192379 | 122187894 | 122188088 | 122196634 | 122196791 | 60506 | 33,19,97,34,39,43,27,49,3 | 4,3,3,0,0,2,3,0,0,0,1,4 | 64,84,40,40,33,15,15,28,3 | 0,1,0,0,3,1,0,1,1,0,1,2,0,0,1 | 298 | 149 | 0.0126567400524 | 0.669811364204107 | 0.805,0.76,0.942,1 | 1.0,0.977,1.0,1.0,0 | -0,071 |
| 225164 | ENSG00000186480 | INSIG1 | chr7 | + | 155302729 | 155302846 | 155302250 | 155302417 | 155308240 | 155308757 | 225164 | 24,25,85,40,11,20,18,23,2 | 0,0,5,9,1,3,9,5,0,0,0,5 | 11,18,14,7,17,5,7,19,3,11,7 | 5,6,4,4,6,5,2,5,5,0,0,1,0,3, | 298 | 149 | 0.0126601138071 | 0.669825210131745 | 1.0,1.0,0.905,0.71 | 0.552,0.627,0.662, | 0,155 |
| 73105 | ENSG00000179335 | CLK3 | chr15 | + | 74622119 | 74622216 | 74619966 | 74620225 | 74624901 | 74625018 | 73105 | 12,11,14,9,2,3,7,7,3,6,3 | 4,5,3,1,3,7,6,2,2,6,2,3 | 11,23,5,8,9,3,6,19,3,16,9,1 | 6,4,1,2,4,0,0,3,4,8,0,4,1,2,5 | 246 | 149 | 0.0126750421994 | 0.670450235345063 | 0.645,0.571,0.739, | 0.526,0.777,0.752, | -0,193 |
| 121278 | ENSG00000149743 | TRPT1 | chr11 | - | 64224794 | 64224970 | 64224542 | 64224717 | 64225498 | 64225580 | 121278 | 13,1,19,8,12,8,4,3,17,8,1,1 | 2,0,4,2,2,6,0,4,0,3,0,0 | 12,9,3,2,3,3,7,1,9,9,11,22 | 0,0,0,2,0,1,2,2,1,0,0,5,1,1,3 | 298 | 149 | 0.012682725045 | 0.670691792177001 | 0.765,1.0,0.704,0. | 1.0,1.0,1.0,0.429,1 | -0,013 |
| 247398 | ENSG00000149311 | ATM | chr11 | + | 108244768 | 108245026 | 108243961 | 108244118 | 108246963 | 108247127 | 247398 | 52,25,269,148,30,41,19,30 | 4,0,2,0,0,0,0,0,2,0,1,0 | 119,100,20,16,64,55,20,46 | 9,10,0,0,4,0,0,0,0,1,0,0,0,0 | 298 | 149 | 0.0127429541717 | 0.673640437155276 | 0.867,1.0,0.985,1. | 0.869,0.833,1.0,1. | 0,009 |
| 227699 | ENSG00000186088 | GSAP | chr7 | - | 77379948 | 77380000 | 77377285 | 77377390 | 77381304 | 77381354 | 227699 | 7,4,10,4,2,11,12,10,2,1,9,4 | 51,26,156,99,53,60,40,66, | 6,9,5,6,7,6,4,16,4,1,10,15, | 102,45,42,11,27,18,28,56,2 | 201 | 149 | 0.0127447433692 | 0.673640437155276 | 0.092,0.102,0.045, | 0.042,0.129,0.081, | -0,038 |
| 227965 | ENSG00000173715 | C11orf80 | chr11 | + | 66828715 | 66828943 | 66828265 | 66828364 | 66838366 | 66838445 | 227965 | 3,0,9,3,1,8,0,1,3,3,0,5 | 101,38,109,54,42,52,59,69 | 0,4,0,1,0,2,1,1,0,9,2,0,0,0,3 | 66,36,45,52,36,13,43,56,3 | 298 | 149 | 0.012753829295 | 0.67395177628091 | 0.015,0.0,0.04,0.0 | 0.0,0.053,0.0,0.01, | 0,011 |
| 16304 | ENSG00000167397 | VKORC1 | chr16 | - | 31093311 | 31093421 | 31090853 | 31091342 | 31094556 | 31094797 | 16304 | 14,6,34,34,20,15,19,17,15, | 0,1,3,2,0,0,0,2,2,0,0,6 | 16,11,16,6,0,20,6,18,2,11, | 0,0,0,0,2,0,2,4,1,3,2,5,3,0,1 | 259 | 149 | 0.0127642879911 | 0.67434228488278 | 0.775,0.867,0.9 | 1.0,1.0,1.0,1.0,0.0, | 0,099 |
| 155662 | ENSG00000088387 | DOCK9 | chr13 | - | 98825871 | 98825963 | 98824397 | 98824504 | 98826829 | 98826887 | 155662 | 2,2,4,1,1,2,3,2,0,1,3,0 | 7,9,14,3,3,15,1,4,18,2,3,7 | 0,5,0,1,1,2,0,2,4,0,0,0,1,0,1 | 3,9,3,8,17,5,2,8,6,8,11,7,1 | 241 | 149 | 0.0127772013341 | 0.674858851617099 | 0.15,0.121,0.15,0. | 0.0,0.256,0.0,0.07 | 0,094 |

| | | | | | | | | | | | | | | | | | | | | | | |
|---|---|---|---|---|---|---|---|---|---|---|---|---|---|---|---|---|---|---|---|---|---|---|
| 173413 | ENSG00000180370 | PAK2 | chr3 | + | 196827177 | 196827333 | 196820370 | 196820567 | 196828318 | 196832647 | 173413 | 171,131,211,196,97,123,1 | 3,1,2,0,0,0,0,1,0,1,0,1 | 145,167,104,112,89,80,67, | 2,5,0,2,1,0,2,4,2,0,3,0,1,1, | 298 | 149 | 0.0127998819229 | 0.675789817905163 | 0.966,0.985,0.981, | 0.973,0.944,1.0,0. | 0,023 |
| 225163 | ENSG00000186480 | INSIG1 | chr7 | + | 155302728 | 155302846 | 155302250 | 155302417 | 155308240 | 155308757 | 225163 | 25,25,87,44,11,20,18,23,2 | 0,0,5,9,1,3,9,5,0,0,0,5 | 11,18,14,7,17,7,7,20,3,11, | 5,6,4,4,6,5,2,5,5,0,0,1,0,3, | 267 | 149 | 0.0128011071249 | 0.675789817905163 | 1.0,1.0,0.907,0.73 | 0.551,0.626,0.661, | 0,151 |
| 120211 | ENSG00000068831 | RASGRP2 | chr11 | - | 64740367 | 64740449 | 64740012 | 64740163 | 64740947 | 64741079 | 120211 | 31,24,73,42,35,12,18,21,9, | 101,57,221,144,107,73,87, | 38,31,12,13,49,36,14,22,3 | 160,99,46,30,104,58,42,76 | 231 | 149 | 0.0128120471185 | 0.676201499107865 | 0.165,0.214,0.176, | 0.133,0.168,0.144, | -0,043 |
| 161347 | ENSG00000165819 | METTL3 | chr14 | - | 21500238 | 21500682 | 21499763 | 21499802 | 21500912 | 21501129 | 161347 | 58,14,86,65,56,53,32,52,2 | 8,4,15,3,10,5,2,12,2,7,1,4 | 76,32,30,13,22,26,24,37,2 | 6,10,2,1,0,2,2,1,0,0,4,7,2,2 | 298 | 149 | 0.0128256828995 | 0.676755223373936 | 0.784,0.636,0.741, | 0.864,0.615,0.882, | -0,084 |
| 202809 | ENSG00000112033 | PPARD | chr6 | + | 35383576 | 35383891 | 35347066 | 35347150 | 35410986 | 35411217 | 202809 | 1,1,2,0,3,0,0,6,8,0,0,3 | 31,33,19,12,14,10,8,6,14,1 | 1,4,0,0,1,3,2,0,0,3,0,1,0,0, | 13,10,21,20,12,3,11,13,4,1 | 298 | 149 | 0.0128366562932 | 0.677168228098463 | 0.016,0.015,0.05,0 | 0.037,0.167,0.0,0. | 0,02 |
| 32837 | ENSG00000102445 | RUBCNL | chr13 | - | 46371940 | 46372078 | 46368732 | 46368815 | 46387133 | 46387457 | 32837 | 117,124,939,690,219,347, | 3,2,10,2,2,6,5,6,0,2,0,1 | 538,768,271,251,395,202, | 11,12,2,2,4,2,1,3,2,1,8,4,9, | 287 | 149 | 0.0128459212404 | 0.677490927344409 | 0.953,0.97,0.98,0. | 0.962,0.971,0.986, | 0,008 |
| 28224 | ENSG00000056277 | ZNF280C | chrX | - | 130230509 | 130230727 | 130228976 | 130229134 | 130236213 | 130236320 | 28224 | 18,12,23,16,10,11,6,10,13, | 7,2,4,7,3,5,7,4,3,0,4,2 | 6,11,6,22,4,15,6,24,8,17,1 | 2,0,11,7,0,1,2,9,0,0,2,5,1,0 | 298 | 149 | 0.0128503475074 | 0.677558340118866 | 0.562,0.75,0.742,0 | 0.6,1.0,0.214,0.61 | -0,147 |
| 37032 | ENSG00000173020 | GRK2 | chr11 | + | 67283706 | 67283773 | 67283127 | 67283228 | 67283853 | 67283949 | 37032 | 141,127,324,278,170,131, | 3,6,7,4,1,4,2,6,6,0,2,4 | 184,216,83,77,150,73,87,2 | 2,7,5,18,7,0,4,6,1,0,4,4,5,1 | 216 | 149 | 0.0128559064999 | 0.677685430291447 | 0.97,0.936,0.97,0. | 0.984,0.955,0.92,0 | 0,019 |
| 202430 | ENSG00000154222 | CC2D1B | chr1 | - | 52357000 | 52357146 | 52356383 | 52356442 | 52357525 | 52357698 | 202430 | 19,14,21,12,0,12,8,9,14,12 | 7,1,7,1,3,2,2,2,2,5,0,7 | 16,21,13,13,7,1,10,9,7,8,6, | 0,1,0,6,3,0,2,5,3,0,2,0,0,0, | 295 | 149 | 0.012860426605 | 0.677757707791566 | 0.578,0.876,0.602, | 1.0,0.914,1.0,0.52 | -0,155 |
| 102846 | ENSG00000179406 | LINC00174 | chr7 | - | 66400321 | 66400437 | 66399691 | 66399795 | 66400922 | 66401027 | 102846 | 9,5,8,10,2,1,0,1,6,2,4,2 | 2,0,10,2,0,0,1,4,4,0,2,4 | 3,7,1,0,0,1,3,0,6,2,1,3,3,4, | 6,7,1,2,3,1,4,6,3,0,2,2,0,2, | 265 | 149 | 0.0128733556322 | 0.678273000262922 | 0.717,1.0,0.31,0.7 | 0.219,0.36,0.36,0. | 0,218 |
| 152077 | ENSG00000154719 | MRPL39 | chr21 | - | 25589227 | 25589347 | 25588834 | 25588882 | 25592811 | 25592965 | 152077 | 3,0,3,10,0,2,5,4,0,0,0,0 | 31,26,25,27,30,22,11,22,1 | 2,0,0,0,7,7,1,1,0,0,2,2,1,0, | 29,23,17,26,30,24,8,25,10, | 269 | 149 | 0.0128838397039 | 0.67865925191143 | 0.051,0.0,0.062,0. | 0.037,0.0,0.0,0.0,0 | 0,022 |
| 189670 | ENSG00000074603 | DPP8 | chr15 | - | 65454262 | 65454405 | 65451959 | 65452102 | 65456224 | 65456371 | 189670 | 16,17,26,24,8,12,9,12,5,10 | 4,3,9,9,5,1,5,2,3,0,2,4 | 29,37,13,7,14,7,5,7,10,15, | 3,6,6,0,2,0,0,0,1,5,4,1,0,6, | 292 | 149 | 0.0129136422404 | 0.680062670184373 | 0.671,0.743,0.596, | 0.831,0.759,0.525, | -0,201 |
| 54062 | ENSG00000163568 | AIM2 | chr1 | - | 159073237 | 159073519 | 159068567 | 159068701 | 159076632 | 159076766 | 54062 | 81,48,201,300,187,125,19 | 4,0,5,4,0,1,0,3,1,0,4,2 | 126,69,90,23,109,231,25,7 | 3,0,5,0,7,11,0,1,2,4,2,9,0,2 | 298 | 149 | 0.0129217966764 | 0.680325641012292 | 0.91,1.0,0.953,0.9 | 0.955,1.0,0.9,1.0,0 | -0,005 |
| 24517 | ENSG00000136147 | PHF11 | chr13 | + | 49525772 | 49525937 | 49524084 | 49524216 | 49526386 | 49526458 | 24517 | 19,2,13,1,9,18,20,45,9,3,0, | 102,43,103,47,55,50,26,57 | 27,21,0,13,13,7,10,4,21,18 | 87,47,67,54,20,19,13,27,4 | 298 | 149 | 0.0129428800741 | 0.681269019620596 | 0.085,0.023,0.059, | 0.134,0.183,0.0,0. | -0,051 |
| 121777 | ENSG00000122359 | ANXA11 | chr10 | - | 80180940 | 80181136 | 80176106 | 80176155 | 80205342 | 80205572 | 121777 | 3,0,13,5,1,0,1,12,3,2,2,4 | 11,10,35,12,16,13,21,22,1 | 2,5,0,2,4,5,18,9,6,4,4,3,11, | 28,18,9,8,8,7,12,17,7,10,1 | 298 | 149 | 0.0129532513022 | 0.681648222744207 | 0.12,0.0,0.157,0.1 | 0.034,0.122,0.0,0. | -0,079 |
| 158085 | ENSG00000165025 | SYK | chr9 | + | 90846003 | 90846057 | 90845433 | 90845594 | 90862205 | 90862344 | 158085 | 6,2,12,6,5,11,3,13,2,6,10,9 | 403,397,525,457,224,204, | 7,12,3,4,34,18,2,3,7,1,6,4, | 395,368,378,301,175,149, | 203 | 149 | 0.0129590768386 | 0.681712405458602 | 0.011,0.004,0.017, | 0.013,0.023,0.006, | -0,01 |
| 37016 | ENSG00000130338 | TULP4 | chr6 | + | 158501677 | 158504178 | 158498668 | 158498812 | 158506577 | 158511828 | 37016 | 18,24,20,21,8,15,13,21,18, | 0,1,2,1,0,0,0,0,1,2,0,0 | 14,16,5,47,11,12,15,12,6,1 | 1,1,0,2,4,0,0,0,2,0,3,4,0,3, | 298 | 149 | 0.0129608056606 | 0.681712405458602 | 1.0,0.923,0.833,0. | 0.875,0.889,1.0,0. | 0,096 |
| 39647 | ENSG00000184203 | PPP1R2 | chr3 | - | 195527757 | 195527902 | 195524818 | 195524896 | 195529793 | 195529901 | 39647 | 10,4,14,12,6,17,21,8,12,8, | 157,100,244,175,84,84,80, | 23,11,8,0,4,5,0,11,3,14,11, | 189,155,125,82,104,78,57, | 294 | 149 | 0.0129743681792 | 0.682259036789004 | 0.031,0.02,0.028,0 | 0.058,0.035,0.031, | 0,018 |
| 74505 | ENSG00000115806 | GORASP2 | chr2 | + | 170956435 | 170956559 | 170951327 | 170951458 | 170961662 | 170961749 | 74505 | 93,104,80,70,51,45,43,37, | 4,1,0,4,0,3,8,1,0,0,0,3 | 28,80,78,63,35,37,13,61,1 | 1,4,3,3,4,2,0,2,4,4,6,1,1,0, | 273 | 149 | 0.0129872687308 | 0.682770599950859 | 0.927,0.983,1.0,0. | 0.939,0.916,0.934, | 0,039 |
| 189567 | ENSG00000138138 | ATAD1 | chr10 | - | 87759700 | 87759800 | 87756788 | 87756922 | 87767672 | 87767723 | 189567 | 2,1,27,31,2,3,3,11,9,5,0,2 | 20,11,26,11,16,15,8,10,8,3 | 2,12,2,0,24,8,0,3,6,6,3,2,0, | 31,40,6,24,14,8,4,12,18,10 | 249 | 149 | 0.0129922103716 | 0.682863597189216 | 0.056,0.052,0.383, | 0.037,0.152,0.166, | 0,105 |
| 215778 | ENSG00000049089 | COL9A2 | chr1 | - | 40310506 | 40310767 | 40310289 | 40310317 | 40311092 | 40311146 | 215778 | 38,18,157,83,58,102,41,75 | 0,5,3,2,1,1,0,8,0,0,0,5 | 22,41,15,10,15,16,32,55,1 | 2,0,0,0,0,1,1,8,0,0,0,2,0,0, | 298 | 149 | 0.0130021215474 | 0.683217681339954 | 1.0,0.643,0.963,0. | 0.846,1.0,1.0,1.0,1 | -0,022 |
| 35679 | ENSG00000169762 | TAPT1 | chr4 | - | 16170652 | 16170729 | 16166632 | 16166793 | 16174203 | 16174272 | 35679 | 47,43,181,33,25,94,38,181 | 4,13,15,9,8,19,12,31,3,4,4, | 33,47,106,39,26,16,53,89, | 10,26,7,19,2,5,11,23,5,7,1 | 226 | 149 | 0.0130269443092 | 0.684354954994734 | 0.886,0.686,0.888, | 0.685,0.544,0.909, | 0,084 |
| 201302 | ENSG00000055070 | SZRD1 | chr1 | + | 16391381 | 16391424 | 16367264 | 16367308 | 16393230 | 16393482 | 201302 | 8,4,3,4,1,4,2,4,10,9,0,2 | 34,38,22,18,30,20,24,23,2 | 1,2,6,0,1,1,3,1,1,2,1,7,2,2, | 18,28,14,22,14,7,13,30,5,1 | 192 | 149 | 0.0130484590945 | 0.685303503793392 | 0.154,0.076,0.096, | 0.041,0.053,0.25,0 | 0,029 |
| 6589 | ENSG00000240849 | PEDS1 | chr20 | - | 50130855 | 50130947 | 50129545 | 50129690 | 50143501 | 50143621 | 6589 | 39,21,25,39,18,9,8,14,22,1 | 8,3,12,1,2,2,6,5,2,5,3,1 | 27,16,4,6,7,12,6,9,1,22,8,2 | 1,2,0,0,0,4,0,4,0,0,2,0,2,2, | 241 | 149 | 0.0130513683533 | 0.685303503793392 | 0.751,0.812,0.563, | 0.943,0.832,1.0,1. | -0,123 |
| 66963 | ENSG00000074201 | CLNS1A | chr11 | - | 77622499 | 77622673 | 77619605 | 77619695 | 77624962 | 77625070 | 66963 | 213,78,345,231,120,177,8 | 0,0,0,2,0,2,1,5,2,0,0,1 | 257,153,114,85,77,53,84,1 | 0,6,0,2,1,0,0,2,2,1,7,4,0,3, | 298 | 149 | 0.0130690698883 | 0.686065605153341 | 1.0,1.0,1.0,0.983,1 | 1.0,0.927,1.0,0.95 | 0,016 |
| 83660 | ENSG00000039650 | PNKP | chr19 | - | 49862370 | 49862463 | 49861816 | 49862281 | 49862537 | 49862608 | 83660 | 40,30,58,27,31,46,37,37,1 | 0,0,0,0,0,0,0,0,0,0,0,0 | 31,42,24,18,10,7,17,41,19, | 0,0,0,0,0,0,0,0,0,0,12,11,0, | 242 | 149 | 0.0130915056923 | 0.687075801428779 | 1.0,1.0,1.0,1.0,1.0, | 1.0,1.0,1.0,1.0,1.0, | 0,095 |
| 110069 | ENSG00000177565 | TBL1XR1 | chr3 | - | 177160870 | 177160988 | 177098465 | 177098541 | 177197120 | 177197225 | 110069 | 17,11,17,16,3,5,8,5,9,7,1,8 | 10,7,28,10,8,10,11,23,2,3, | 4,3,6,2,0,1,3,2,1,4,4,17,11, | 26,14,8,13,6,3,7,12,1,13,1 | 267 | 149 | 0.0131091819212 | 0.687835771350999 | 0.487,0.467,0.253, | 0.079,0.107,0.295, | 0,132 |
| 251500 | ENSG00000138134 | STAMBPL1 | chr10 | + | 88913420 | 88913458 | 88910915 | 88911011 | 88914533 | 88914658 | 251500 | 7,3,9,10,3,4,5,1,0,3,1,2 | 48,2,40,31,14,30,13,23,4,1 | 2,2,1,2,2,2,4,4,1,1,4,1,1,2, | 24,39,21,8,24,7,9,13,35,18 | 187 | 149 | 0.0131125085609 | 0.687842634674889 | 0.104,0.544,0.152, | 0.062,0.039,0.037, | 0,054 |
| 94129 | ENSG00000112303 | VNN2 | chr6 | - | 132752460 | 132752577 | 132751144 | 132751518 | 132755842 | 132756035 | 94129 | 18,25,24,26,24,14,17,17,4 | 5,6,0,2,1,2,1,0,4,1,0,0 | 10,12,24,12,9,19,7,9,20,21 | 0,3,2,2,2,0,0,0,0,0,0,1,2,1, | 266 | 149 | 0.0131229972056 | 0.688180842307904 | 0.668,0.7,1.0,0.87 | 1.0,0.691,0.87,0.7 | -0,01 |
| 181876 | ENSG00000123104 | ITPR2 | chr12 | - | 26682573 | 26682685 | 26681873 | 26682034 | 26686480 | 26686632 | 181876 | 63,44,141,67,27,33,39,125 | 5,6,11,4,2,9,10,8,0,4,2,2 | 79,54,28,34,62,38,29,34,4 | 6,23,0,28,12,8,10,13,9,4,1 | 261 | 149 | 0.0131253507054 | 0.688180842307904 | 0.878,0.807,0.88,0 | 0.883,0.573,1.0,0. | 0,078 |
| 184567 | ENSG00000198265 | HELZ | chr17 | - | 67135969 | 67136055 | 67128494 | 67128855 | 67137930 | 67138114 | 184567 | 27,4,21,6,13,12,8,6,11,7,2, | 2,1,9,2,5,14,1,8,9,0,2,1 | 7,12,9,12,3,3,3,3,3,10,5,9, | 5,6,4,9,10,3,3,7,3,2,10,7,5, | 235 | 149 | 0.0131418911619 | 0.688819767597886 | 0.895,0.717,0.597, | 0.47,0.559,0.588,0 | 0,188 |
| 202488 | ENSG00000138496 | PARP9 | chr3 | - | 122546175 | 122546368 | 122545431 | 122545489 | 122550583 | 122550802 | 202488 | 0,0,9,6,0,4,1,1,1,0,1,3 | 34,35,76,76,17,16,15,16,4 | 4,4,2,3,0,0,0,1,7,3,5,11,4,2 | 28,34,38,15,7,30,4,15,27,5 | 298 | 149 | 0.0131439373767 | 0.688819767597886 | 0.0,0.0,0.056,0.03 | 0.067,0.056,0.026, | 0,004 |
| 188861 | ENSG00000109323 | MANBA | chr4 | - | 102635864 | 102635938 | 102634787 | 102635045 | 102639712 | 102639857 | 188861 | 32,20,12,10,10,5,8,9,7,13, | 2,0,6,4,9,3,2,5,0,2,0,0 | 9,11,9,2,10,5,5,5,18,6,3,9, | 4,4,0,8,6,2,2,1,5,1,6,10,6,4 | 223 | 149 | 0.0131616982222 | 0.689582636334549 | 0.914,1.0,0.572,0. | 0.601,0.648,1.0,0. | 0,158 |
| 199249 | ENSG00000076321 | KLHL20 | chr1 | + | 173725427 | 173725858 | 173716002 | 173716066 | 173733712 | 173734513 | 199249 | 11,3,5,3,0,6,1,4,2,1,5,1 | 11,6,28,14,13,16,5,8,1,7,9, | 12,0,3,5,5,0,0,4,2,2,0,0,0,0 | 11,26,2,5,22,5,11,20,1,11, | 298 | 149 | 0.0131840138829 | 0.690583716726807 | 0.333,0.2,0.082,0. | 0.353,0.0,0.429,0. | 0,044 |
| 207073 | ENSG00000119866 | BCL11A | chr2 | - | 60538380 | 60538549 | 60468739 | 60468833 | 60545970 | 60546283 | 207073 | 10,0,4,2,0,0,0,0,0,0,0,2 | 52,21,52,46,43,21,20,15,4 | 0,9,0,2,0,2,7,0,0,4,1,4,11,3 | 57,32,28,26,18,28,59,42,1 | 298 | 149 | 0.0131992357476 | 0.691125887501602 | 0.088,0.0,0.037,0. | 0.0,0.123,0.0,0.03 | -0,015 |
| 239546 | ENSG00000083223 | TUT7 | chr9 | - | 86301739 | 86301885 | 86301275 | 86301601 | 86303085 | 86303201 | 239546 | 40,16,12,3,6,14,6,7,8,7,6,5 | 40,47,52,35,19,23,20,14,5 | 10,16,24,21,8,6,5,6,5,15,1 | 21,16,25,15,15,11,8,32,3,1 | 295 | 149 | 0.013200786706 | 0.691125887501602 | 0.336,0.147,0.104, | 0.194,0.336,0.327, | -0,082 |
| 137641 | ENSG00000104852 | SNRNP70 | chr19 | + | 49098814 | 49098959 | 49098641 | 49098704 | 49101389 | 49101668 | 137641 | 19,5,18,12,6,10,12,14,7,2, | 480,291,503,287,327,340, | 8,12,6,1,4,9,5,8,1,13,14,5, | 361,387,238,132,296,173, | 294 | 149 | 0.0132051505307 | 0.691186224190927 | 0.02,0.009,0.018,0 | 0.011,0.015,0.013, | 0,005 |
| 159682 | ENSG00000112081 | SRSF3 | chr6 | + | 36600111 | 36600276 | 36598848 | 36598983 | 36601151 | 36601190 | 159682 | 77,21,129,30,35,24,31,43, | 117,111,314,232,133,144, | 48,49,21,31,42,41,36,38,1 | 208,166,92,21,79,51,74,17 | 298 | 149 | 0.0132178615815 | 0.691628583663338 | 0.248,0.086,0.17,0 | 0.103,0.129,0.102, | -0,057 |
| 150637 | ENSG00000159459 | UBR1 | chr15 | - | 42976173 | 42976229 | 42970519 | 42970607 | 42976716 | 42976867 | 150637 | 2,1,2,0,0,0,0,1,0,0,0,0 | 5,5,25,9,22,6,12,13,7,2,4,7 | 2,5,1,3,2,3,0,2,1,0,0,0,2,1, | 11,15,3,3,10,6,12,19,14,11 | 205 | 149 | 0.0132200286817 | 0.691628583663338 | 0.225,0.127,0.055, | 0.117,0.195,0.195, | -0,07 |
| 62797 | ENSG00000087206 | UIMC1 | chr5 | - | 176969600 | 176969706 | 176958092 | 176958154 | 176970741 | 176970793 | 62797 | 18,12,68,32,29,29,26,22,7, | 4,1,14,9,0,6,6,11,4,1,0,7 | 42,42,13,26,24,28,23,24,1 | 13,13,2,12,19,13,2,2,4,11, | 255 | 149 | 0.0132255020101 | 0.691746785695221 | 0.724,0.875,0.739, | 0.654,0.654,0.792, | 0,108 |
| 255032 | ENSG00000137955 | RABGGTB | chr1 | + | 75790151 | 75790512 | 75789951 | 75790057 | 75791284 | 75791337 | 255032 | 67,60,32,18,12,18,18,36,2 | 29,19,44,34,8,11,12,21,11, | 15,35,18,19,26,9,5,10,14,1 | 28,24,14,24,7,11,3,20,12,1 | 298 | 149 | 0.01324490636 | 0.692593401547415 | 0.536,0.612,0.267, | 0.211,0.422,0.391, | 0,132 |
| 45797 | ENSG00000116191 | RALGPS2 | chr1 | + | 178833423 | 178833550 | 178821611 | 178821704 | 178877497 | 178877635 | 45797 | 514,356,928,546,486,233, | 28,20,46,30,38,14,35,17,3, | 990,825,428,265,333,275, | 27,30,24,9,30,16,5,5,30,19 | 276 | 149 | 0.0132824163365 | 0.694386142949048 | 0.908,0.906,0.916, | 0.952,0.937,0.906, | -0,008 |
| 79515 | ENSG00000088179 | PTPN4 | chr2 | + | 119885794 | 119885882 | 119882502 | 119882623 | 119900717 | 119900806 | 79515 | 16,25,53,28,20,13,11,16,5, | 7,0,6,1,2,2,1,1,0,0,0,0 | 30,31,17,11,14,14,11,26,1 | 3,5,3,6,3,1,0,1,2,0,2,1,0,2, | 237 | 149 | 0.0132873293667 | 0.694474304741187 | 0.59,1.0,0.847,0.9 | 0.863,0.796,0.781, | 0,042 |
| 115373 | ENSG00000184465 | WDR27 | chr6 | - | 169663562 | 169663649 | 169662303 | 169662424 | 169664165 | 169664286 | 115373 | 11,2,13,6,10,25,7,10,5,6,2, | 1,0,14,5,2,4,0,2,1,2,0,2 | 5,7,3,1,6,3,5,6,4,8,5,11,6,5 | 3,0,0,1,0,1,3,3,0,0,7,2,0,0, | 236 | 149 | 0.0133113354132 | 0.695560095246043 | 0.874,1.0,0.37,0.4 | 0.513,1.0,1.0,0.38 | -0,028 |
| 28844 | ENSG00000166888 | STAT6 | chr12 | - | 57110042 | 57110110 | 57108162 | 57108299 | 57111128 | 57111374 | 28844 | 17,8,26,39,4,2,11,10,14,11 | 55,45,164,114,97,88,46,56 | 9,7,12,4,6,5,8,9,5,5,3,13,1 | 102,83,36,30,38,49,20,65, | 217 | 149 | 0.0133161016191 | 0.69562361321758 | 0.175,0.109,0.098, | 0.057,0.055,0.186, | 0,015 |
| 251726 | ENSG00000095585 | BLNK | chr10 | - | 96238924 | 96239164 | 96230793 | 96230834 | 96242734 | 96242784 | 251726 | 0,1,3,2,1,4,3,10,2,0,1,3 | 81,58,161,113,99,71,57,84 | 6,7,1,4,3,6,1,3,1,3,2,6,3,5, | 148,64,76,42,30,54,54,74, | 298 | 149 | 0.0133210193535 | 0.69562361321758 | 0.0,0.009,0.009,0. | 0.02,0.052,0.007,0 | -0,012 |
| 13856 | ENSG00000169189 | NSMCE1 | chr16 | - | 27268162 | 27268374 | 27257434 | 27257581 | 27268705 | 27268723 | 13856 | 4,0,14,3,2,5,4,0,2,3,0,0 | 26,16,52,54,17,20,22,11,4 | 1,0,2,0,0,1,0,1,2,2,3,4,2,0, | 42,10,23,6,8,9,8,18,8,29,1 | 298 | 149 | 0.0133222469518 | 0.69562361321758 | 0.071,0.0,0.119,0. | 0.012,0.0,0.042,0. | 0,006 |
| 73189 | ENSG00000155545 | MIER3 | chr5 | - | 56938295 | 56938338 | 56937577 | 56937698 | 56938882 | 56938971 | 73189 | 3,0,3,3,1,1,3,0,0,0,3,1 | 10,11,27,22,15,13,7,6,7,3, | 6,13,3,4,2,1,3,1,2,2,2,2,3,1 | 16,30,19,14,13,6,9,16,9,11 | 192 | 149 | 0.0133280298251 | 0.695698118120176 | 0.189,0.0,0.079,0. | 0.225,0.252,0.109, | -0,036 |
| 132451 | ENSG00000011523 | CEP68 | chr2 | + | 65069398 | 65069801 | 65056353 | 65056524 | 65071453 | 65072569 | 132451 | 62,39,68,34,43,24,21,30,2 | 0,3,4,2,1,2,1,2,0,0,0,0 | 46,107,29,20,19,9,56,80,1 | 0,11,0,0,3,2,3,5,2,0,1,2,1,0 | 298 | 149 | 0.0133301384983 | 0.695698118120176 | 1.0,0.867,0.895,0. | 1.0,0.829,1.0,1.0,0 | 0,06 |
| 209007 | ENSG00000163728 | TTC14 | chr3 | + | 180604455 | 180604607 | 180604224 | 180604309 | 180604851 | 180604912 | 209007 | 33,14,84,68,65,71,24,39,5 | 1,0,3,4,3,1,0,1,5,0,0,0 | 83,70,25,20,18,58,8,42,24, | 2,1,0,0,0,2,0,0,3,1,0,0,1,0, | 298 | 149 | 0.0133409023182 | 0.696091090145092 | 0.943,1.0,0.933,0. | 0.954,0.972,1.0,1. | -0,008 |
| 79396 | ENSG00000116809 | ZBTB17 | chr1 | - | 15943398 | 15943519 | 15943063 | 15943194 | 15943598 | 15943715 | 79396 | 34,46,68,28,34,39,46,33,6 | 1,9,3,0,0,0,7,4,2,0,0,4 | 17,28,32,21,23,5,23,28,19, | 1,5,0,10,2,0,0,9,0,3,5,6,0,1 | 270 | 149 | 0.0133467722452 | 0.696228583884305 | 0.949,0.738,0.926, | 0.904,0.756,1.0,0. | 0,074 |
| 122126 | ENSG00000081237 | PTPRC | chr1 | + | 198729633 | 198729765 | 198729136 | 198729171 | 198731616 | 198731726 | 122126 | 70,32,75,37,13,48,37,44,7, | 244,157,721,381,221,142, | 65,105,46,110,62,28,17,24 | 415,397,147,101,287,180, | 281 | 149 | 0.0133819516205 | 0.697894555180963 | 0.132,0.098,0.052, | 0.077,0.123,0.142, | -0,026 |
| 200838 | ENSG00000161929 | SCIMP | chr17 | - | 5223332 | 5223430 | 5221286 | 5221350 | 5234734 | 5234860 | 200838 | 60,6,64,69,28,19,18,43,31, | 3,0,3,13,1,4,0,1,9,3,6,5 | 37,30,24,16,11,30,14,33,1 | 8,2,10,2,2,4,1,9,6,3,2,3,8,1 | 247 | 149 | 0.0134098344441 | 0.699179282276668 | 0.923,1.0,0.928,0. | 0.736,0.9,0.591,0. | 0,102 |
| 254703 | ENSG00000079332 | SAR1A | chr10 | - | 70162689 | 70162876 | 70161857 | 70161931 | 70170412 | 70170523 | 254703 | 17,20,41,24,11,16,15,23,7, | 2,6,9,9,11,2,1,5,3,5,5,2 | 22,45,32,45,39,15,17,33,1 | 5,8,2,0,7,3,4,0,2,3,1,8,5,2, | 298 | 149 | 0.0134297013374 | 0.700045543363996 | 0.81,0.625,0.695,0 | 0.688,0.738,0.889, | -0,136 |
| 7172 | ENSG00000126001 | CEP250 | chr20 | + | 35496576 | 35496715 | 35494523 | 35494657 | 35497718 | 35498067 | 7172 | 50,48,57,31,30,23,16,26,4 | 5,4,2,1,0,0,1,0,0,0,0,0 | 36,31,21,19,20,15,10,21,1 | 1,2,2,1,3,4,0,2,0,2,0,4,0,0, | 288 | 149 | 0.0134351289175 | 0.700051361566639 | 0.838,0.861,0.936, | 0.949,0.889,0.845, | 0,063 |
| 134448 | ENSG00000100299 | ARSA | chr22 | - | 50625604 | 50625681 | 50625362 | 50625464 | 50625935 | 50626063 | 134448 | 21,15,22,10,5,12,12,9,14,1 | 0,0,0,0,1,0,0,1,0,0,0,0 | 18,8,10,3,5,6,13,8,10,20,1 | 0,2,6,2,6,2,0,0,0,0,0,1,0,0, | 226 | 149 | 0.0134363180705 | 0.700051361566639 | 1.0,1.0,1.0,1.0,0.7 | 1.0,0.725,0.524,0. | 0,112 |
| 152944 | ENSG00000257621 | PSMA3-AS1 | chr14 | - | 58293065 | 58293547 | 58291634 | 58291707 | 58297954 | 58298134 | 152944 | 12,4,3,2,1,0,3,7,0,0,4,1 | 67,34,129,99,60,80,54,73, | 6,5,5,3,3,3,2,5,2,11,6,2,2,8 | 76,42,35,33,53,24,20,71,4 | 298 | 149 | 0.0134440678343 | 0.700285615693181 | 0.082,0.056,0.011, | 0.038,0.056,0.067, | -0,009 |
| 141258 | ENSG00000101745 | ANKRD12 | chr18 | + | 9195552 | 9195698 | 9182385 | 9182519 | 9208656 | 9208797 | 141258 | 282,154,259,213,57,100,8 | 4,10,5,5,3,1,2,1,6,0,0,4 | 111,121,125,119,86,73,67, | 0,1,6,7,4,0,0,0,1,3,1,1,0,0, | 295 | 149 | 0.0134542581454 | 0.700618470281602 | 0.973,0.886,0.963, | 1.0,0.984,0.913,0. | -0,017 |
| 182652 | ENSG00000053900 | ANAPC4 | chr4 | + | 25383546 | 25383644 | 25383260 | 25383393 | 25388499 | 25388574 | 182652 | 6,1,5,1,0,1,1,7,0,1,0,3 | 33,15,47,21,30,44,14,34,1 | 1,9,11,0,5,3,1,2,6,2,1,0,3,0 | 32,30,13,6,16,15,5,17,8,27 | 247 | 149 | 0.0134569683556 | 0.700618470281602 | 0.099,0.039,0.06,0 | 0.019,0.153,0.338, | -0,049 |
| 99696 | ENSG00000092820 | EZR | chr6 | - | 158818081 | 158818166 | 158789287 | 158789371 | 158819316 | 158819349 | 99696 | 647,462,937,642,231,408, | 18,0,3,1,1,10,17,14,0,3,1,3 | 437,887,433,428,705,221, | 11,17,11,15,12,7,0,9,8,3,9, | 234 | 149 | 0.0134622867819 | 0.700725863689267 | 0.958,1.0,0.995,0. | 0.962,0.971,0.962, | 0,01 |
| 82216 | ENSG00000143390 | RFX5 | chr1 | - | 151345927 | 151345961 | 151344988 | 151345188 | 151346204 | 151346333 | 82216 | 141,108,159,125,49,59,60, | 0,9,5,1,1,0,3,3,0,1,2,0 | 82,101,39,57,54,45,40,88, | 5,5,2,10,1,3,4,1,0,0,2,11,1, | 183 | 149 | 0.0134857655161 | 0.701750562238463 | 1.0,0.907,0.963,0. | 0.93,0.943,0.941,0 | 0,018 |
| 81700 | ENSG00000136536 | MARCHF7 | chr2 | + | 159745769 | 159745937 | 159743060 | 159743253 | 159747804 | 159748903 | 81700 | 290,209,557,375,208,250, | 7,3,3,4,5,5,2,6,17,0,2,3 | 268,485,228,120,197,157, | 7,5,4,2,0,2,3,6,0,7,0,2,1,3, | 298 | 149 | 0.0134910201229 | 0.701750562238463 | 0.954,0.972,0.989, | 0.95,0.98,0.966,0. | -0,017 |
| 205345 | ENSG00000018699 | TTC27 | chr2 | + | 32812503 | 32812839 | 32811023 | 32811221 | 32817456 | 32817557 | 205345 | 3,3,10,11,12,3,3,17,5,5,2,4 | 4,0,0,1,0,0,0,0,0,0,0,0 | 20,6,6,2,6,4,3,10,4,5,4,6,9, | 0,0,0,4,0,2,0,7,0,0,0,2,4,0, | 298 | 149 | 0.0134917545639 | 0.701750562238463 | 0.273,1.0,1.0,0.84 | 1.0,1.0,1.0,0.2,1.0, | 0,14 |
| 172963 | ENSG00000136100 | VPS36 | chr13 | - | 52449930 | 52450256 | 52442376 | 52442445 | 52450498 | 52450634 | 172963 | 2,0,3,2,0,0,0,2,2,3,5,1 | 31,18,58,49,27,15,25,17,1 | 5,4,5,0,0,5,0,0,2,6,1,5,6,2, | 35,44,17,10,24,27,13,16,1 | 298 | 149 | 0.0135235540899 | 0.70323461471932 | 0.031,0.0,0.025,0. | 0.067,0.043,0.128, | -0,004 |
| 142076 | ENSG00000029725 | RABEP1 | chr17 | + | 5356502 | 5356573 | 5354400 | 5354490 | 5361207 | 5361629 | 142076 | 17,14,16,3,0,1,8,14,3,5,0,7 | 147,67,168,134,73,65,49,5 | 12,8,6,6,10,4,4,5,8,10,12,2 | 85,56,53,30,51,41,50,60,3 | 220 | 149 | 0.0135315641662 | 0.703481180448162 | 0.073,0.124,0.061, | 0.087,0.088,0.071, | -0,03 |
| 215244 | ENSG00000036257 | CUL3 | chr2 | - | 224556294 | 224556444 | 224535527 | 224535641 | 224557658 | 224557856 | 215244 | 2,2,6,1,0,0,0,6,0,0,0,4 | 28,34,106,66,39,59,33,59, | 10,7,0,2,0,1,1,4,2,7,6,3,1,1 | 69,99,35,70,49,20,49,35,61 | 298 | 149 | 0.0135394774942 | 0.703524601259182 | 0.034,0.029,0.028, | 0.068,0.034,0.0,0. | -0,014 |
| 153499 | ENSG00000162086 | ZNF75A | chr16 | + | 3316550 | 3317022 | 3312676 | 3312768 | 3317189 | 3317252 | 153499 | 33,23,61,40,32,34,23,35,2 | 3,3,7,9,5,3,3,1,1,2,0,2 | 27,39,26,30,22,8,21,11,10, | 4,4,8,4,5,3,4,5,3,5,6,2,3,1, | 298 | 149 | 0.0135443500818 | 0.703524601259182 | 0.846,0.793,0.813, | 0.771,0.83,0.619,0 | 0,118 |
| 211985 | ENSG00000039650 | PNKP | chr19 | - | 49862370 | 49862446 | 49861816 | 49862281 | 49862537 | 49862608 | 211985 | 17,6,18,10,16,15,11,12,3,3 | 0,0,0,0,0,0,0,0,0,0,0,0 | 6,9,11,10,2,2,9,13,2,7,3,4, | 0,0,0,0,0,0,0,0,0,12,11,0, | 225 | 149 | 0.0135451017227 | 0.703524601259182 | 1.0,1.0,1.0,1.0,1.0, | 1.0,1.0,1.0,1.0,1.0, | 0,104 |
| 91543 | ENSG00000100503 | NIN | chr14 | - | 50759993 | 50760359 | 50754741 | 50754867 | 50761789 | 50761867 | 91543 | 148,86,118,81,70,54,53,73 | 14,3,22,6,5,10,3,10,0,1,2,1 | 56,62,81,85,42,31,33,78,2 | 8,10,2,7,9,6,8,4,3,2,12,21,1 | 298 | 149 | 0.0135454741539 | 0.703524601259182 | 0.841,0.935,0.728, | 0.778,0.756,0.953, | 0,068 |
| 143037 | ENSG00000132388 | UBE2G1 | chr17 | - | 4322174 | 4322231 | 4307020 | 4307123 | 4366270 | 4366675 | 143037 | 0,2,12,3,1,4,3,3,0,0,2,4 | 38,26,104,42,20,19,16,20,1 | 2,9,9,2,7,3,7,4,1,0,1,5,5,2, | 25,40,40,30,45,34,9,17,32, | 206 | 149 | 0.013561244398 | 0.704173749825317 | 0.0,0.053,0.077,0. | 0.055,0.14,0.14,0. | -0,038 |
| 236547 | ENSG00000034677 | RNF19A | chr8 | - | 100287939 | 100288267 | 100274952 | 100275161 | 100309866 | 100309953 | 236547 | 31,12,81,37,17,43,50,137, | 1,0,0,0,0,1,2,6,1,0,0,0 | 37,26,40,50,22,14,38,59,3 | 0,0,1,2,1,0,2,2,4,0,8,4,0,0, | 298 | 149 | 0.0135752198874 | 0.704503809346729 | 0.939,1.0,1.0,1.0,1 | 1.0,1.0,0.952,0.92 | 0,018 |
| 51567 | ENSG00000162894 | FCMR | chr1 | - | 206913861 | 206914094 | 206911729 | 206911952 | 206921817 | 206921941 | 51567 | 1167,447,1219,948,777,48 | 9,3,4,3,4,10,4,4,8,14,3,3 | 857,366,531,338,204,325, | 4,0,6,4,2,3,2,3,0,0,0,0,2,1, | 298 | 149 | 0.0135773632966 | 0.704503809346729 | 0.985,0.987,0.993, | 0.991,1.0,0.978,0. | -0,012 |
| 119487 | ENSG00000136270 | TBRG4 | chr7 | - | 45104098 | 45104256 | 45103030 | 45103443 | 45104537 | 45104709 | 119487 | 25,16,25,38,20,25,20,29,3 | 22,4,13,9,5,12,6,21,3,5,1,7 | 19,21,25,14,32,8,19,14,6,2 | 11,17,8,8,19,1,13,16,2,8,2 | 298 | 149 | 0.0135774205443 | 0.704503809346729 | 0.362,0.667,0.49,0 | 0.463,0.382,0.61,0 | 0,138 |
| 174772 | ENSG00000187514 | PTMA | chr2 | + | 231711886 | 231711977 | 231711347 | 231711419 | 231712442 | 231712675 | 174772 | 107,83,64,52,42,34,33,22, | 72,29,55,23,35,27,46,37,2 | 41,28,43,27,31,25,16,27,1 | 59,49,18,53,37,8,14,42,18, | 240 | 149 | 0.0135958857877 | 0.705291900210282 | 0.48,0.64,0.419,0. | 0.301,0.262,0.597, | 0,096 |
| 36829 | ENSG00000163110 | PDLIM5 | chr4 | + | 94584985 | 94585003 | 94575942 | 94576034 | 94585564 | 94585737 | 36829 | 2,1,5,2,1,0,2,1,0,0,0,1 | 17,8,15,18,6,16,7,5,10,5,4, | 5,0,2,1,6,0,2,2,0,0,4,0,0,0, | 15,9,1,14,11,7,10,4,5,17,9, | 167 | 149 | 0.0136049364683 | 0.70551174799836 | 0.095,0.1,0.229,0. | 0.229,0.0,0.641,0. | -0,043 |
| 232846 | ENSG00000083750 | RRAGB | chrX | + | 55755067 | 55755141 | 55753391 | 55753514 | 55755840 | 55755932 | 232846 | 17,18,18,14,2,2,3,12,7,7,2, | 3,1,14,14,2,3,4,2,4,0,2,5 | 15,7,8,6,3,1,4,4,0,4,0,8,8,8 | 5,2,9,0,3,4,5,8,1,11,2,8,15, | 223 | 149 | 0.0136066796416 | 0.70551174799836 | 0.791,0.923,0.462, | 0.667,0.7,0.373,1. | 0,191 |
| 243848 | ENSG00000166889 | PATL1 | chr11 | - | 59655955 | 59656010 | 59655522 | 59655740 | 59656498 | 59656600 | 243848 | 3,0,0,0,6,0,4,2,3,5,1,3 | 19,14,10,15,7,12,6,10,10,1 | 0,0,3,4,1,0,1,1,4,2,1,1,2,2, | 5,6,6,15,6,3,13,8,9,8,8,8,9, | 204 | 149 | 0.0136250067694 | 0.706185956190816 | 0.103,0.0,0.0,0.0,0 | 0.0,0.0,0.268,0.16 | 0,029 |
| 227413 | ENSG00000127914 | AKAP9 | chr7 | + | 92107271 | 92107422 | 92105677 | 92105763 | 92108493 | 92108633 | 227413 | 29,33,63,63,32,26,34,51,1 | 2,1,2,2,4,4,3,7,2,0,0,1 | 41,28,22,21,29,21,23,27,1 | 0,8,2,1,4,5,1,7,3,3,2,8,0,4, | 298 | 149 | 0.0136262447141 | 0.706185956190816 | 0.879,0.943,0.94,0 | 1.0,0.636,0.846,0. | 0,09 |
| 134200 | ENSG00000100395 | L3MBTL2 | chr22 | + | 41217515 | 41217965 | 41217122 | 41217202 | 41219418 | 41219536 | 134200 | 3,1,5,2,0,0,1,2,1,0,0,5 | 18,6,14,22,11,14,8,8,10,12 | 1,3,0,0,1,0,1,1,1,2,2,0,2,0, | 22,3,13,3,4,3,5,13,1,6,6,13 | 298 | 149 | 0.0136527746721 | 0.707319505423787 | 0.077,0.077,0.152, | 0.022,0.333,0.0,0. | -0,004 |
| 115696 | ENSG00000173611 | SCAI | chr9 | - | 125065989 | 125066058 | 125055875 | 125056007 | 125142632 | 125142677 | 115696 | 18,17,20,3,4,4,2,5,1,9,2,1 | 2,0,6,4,1,4,5,2,6,0,5,1 | 14,16,5,2,5,6,2,2,20,1,4,12 | 3,0,2,0,0,1,0,2,1,7,0,3,3,3, | 218 | 149 | 0.0136546898218 | 0.707319505423787 | 0.86,1.0,0.695,0.3 | 0.761,1.0,0.631,1. | -0,204 |

| | | | | | | | | | | | | | | | | | | | | | | |
|---|---|---|---|---|---|---|---|---|---|---|---|---|---|---|---|---|---|---|---|---|---|---|
| 178415 | ENSG00000160271 | RALGDS | chr9 | - | 133105239 | 133105514 | 133104262 | 133104331 | 133105931 | 133106016 | 178415 | 1,5,29,16,1,5,5,12,1,1,2,2 | 22,20,21,17,19,33,10,17,1 | 2,4,8,4,7,3,3,10,0,2,1,4,0,0 | 16,23,20,15,14,8,17,28,8,1 | 298 | 149 | 0.0136784003106 | 0.708377232254752 | 0.022,0.111,0.408, | 0.059,0.08,0.167,0 | 0,059 |
| 32292 | ENSG00000167766 | ZNF83 | chr19 | - | 52616717 | 52616875 | 52614664 | 52614797 | 52618935 | 52619062 | 32292 | 100,52,117,78,40,41,68,47 | 12,10,31,21,7,13,2,5,7,3,1, | 72,69,59,20,54,35,17,35,4 | 15,15,2,1,5,2,0,2,0,2,3,4,8, | 298 | 149 | 0.0137130444616 | 0.71000054667179 | 0.806,0.722,0.654, | 0.706,0.697,0.937, | -0,066 |
| 204741 | ENSG00000196510 | ANAPC7 | chr12 | - | 110377704 | 110377843 | 110377580 | 110377617 | 110381751 | 110381948 | 204741 | 7,0,9,3,2,8,3,6,2,0,1,10 | 17,10,40,36,36,24,24,15,2 | 2,13,3,28,7,0,9,1,4,3,4,7,7, | 24,11,13,0,3,8,10,22,7,18, | 288 | 149 | 0.0137198498228 | 0.71018205801132 | 0.176,0.0,0.104,0. | 0.041,0.379,0.107, | -0,127 |
| 112720 | ENSG00000132294 | EFR3A | chr8 | + | 131996405 | 131996497 | 131987574 | 131987702 | 132001758 | 132001807 | 112720 | 39,25,60,53,15,22,20,21,6 | 4,1,2,3,0,0,1,3,2,5,0,1 | 40,13,43,10,10,14,11,17,4, | 6,8,1,2,7,1,0,2,0,0,1,8,3,1, | 241 | 149 | 0.0137403514051 | 0.711072270562894 | 0.858,0.939,0.949, | 0.805,0.501,0.964, | 0,091 |
| 221647 | ENSG00000188994 | ZNF292 | chr6 | + | 87222769 | 87222894 | 87218595 | 87218731 | 87233324 | 87233777 | 221647 | 2,1,25,12,3,6,4,12,1,0,1,3 | 23,29,74,34,30,12,11,18,8, | 16,40,5,5,6,4,2,14,7,4,14,1 | 44,33,27,21,32,18,8,22,11, | 274 | 149 | 0.0137570092277 | 0.711763185838245 | 0.045,0.018,0.155, | 0.165,0.397,0.091, | -0,056 |
| 233324 | ENSG00000215301 | DDX3X | chrX | + | 41347317 | 41347451 | 41346858 | 41347012 | 41347639 | 41347694 | 233324 | 242,211,360,260,106,137, | 23,7,26,30,13,10,10,14,6,1 | 222,272,175,164,187,126, | 31,31,15,52,16,17,2,20,13, | 283 | 149 | 0.0137615598641 | 0.711827515287216 | 0.847,0.941,0.879, | 0.79,0.822,0.86,0. | 0,032 |
| 231528 | ENSG00000262919 | CCNQ | chrX | - | 153592505 | 153592703 | 153587931 | 153588394 | 153594546 | 153594679 | 231528 | 9,24,21,6,10,14,4,10,25,10 | 1,0,4,1,1,3,1,2,0,1,2,2 | 4,8,2,1,4,5,1,8,4,2,7,34,4,8 | 1,0,0,0,1,0,2,1,1,1,0,3,4,1, | 298 | 149 | 0.0137724348457 | 0.711933322868889 | 0.818,1.0,0.724,0. | 0.667,1.0,1.0,1.0,0 | 0,012 |
| 4987 | ENSG00000127452 | FBXL12 | chr19 | - | 9814137 | 9814339 | 9811155 | 9811717 | 9818544 | 9818813 | 4987 | 1,0,0,1,4,3,2,2,6,0,1,0 | 1,2,9,1,1,4,4,5,1,6,2,5 | 2,0,0,2,0,0,2,2,1,2,2,0,0,1, | 5,5,3,3,4,1,7,2,0,2,3,2,1,5, | 298 | 149 | 0.0137727295939 | 0.711933322868889 | 0.333,0.0,0.0,0.33 | 0.167,0.0,0.0,0.25, | 0,072 |
| 168138 | ENSG00000155189 | AGPAT5 | chr8 | + | 6741682 | 6741751 | 6732560 | 6732650 | 6747669 | 6747828 | 168138 | 23,15,37,23,46,19,18,22,2 | 7,2,7,5,7,4,2,4,0,0,1,8 | 32,43,15,2,3,13,11,19,12,5 | 7,15,7,0,7,8,4,2,7,6,4,2,6,3 | 218 | 149 | 0.0137735287037 | 0.711933322868889 | 0.692,0.837,0.783, | 0.758,0.662,0.594, | 0,132 |
| 67444 | ENSG00000186283 | TOR3A | chr1 | + | 179085627 | 179085893 | 179082069 | 179082387 | 179087910 | 179088083 | 67444 | 9,31,45,27,19,20,30,30,13, | 0,0,0,0,0,2,4,1,0,1,0,0 | 52,52,29,26,24,30,10,30,1 | 0,1,2,0,1,2,0,1,0,0,1,7,1,2, | 298 | 149 | 0.0137964997575 | 0.71294944521164 | 1.0,1.0,1.0,1.0,1.0, | 1.0,0.963,0.879,1. | 0,023 |
| 81129 | ENSG00000136710 | CCDC115 | chr2 | - | 130339559 | 130339762 | 130338242 | 130339232 | 130340907 | 130341039 | 81129 | 6,4,18,9,8,6,10,0,8,0,3,10 | 36,13,97,85,50,35,40,46,3 | 3,7,5,3,2,6,0,2,3,3,13,3,5,7 | 66,71,31,26,40,19,22,43,1 | 298 | 149 | 0.013814278252 | 0.713570164439018 | 0.077,0.133,0.085, | 0.022,0.047,0.075, | 0,019 |
| 212071 | ENSG00000079263 | SP140 | chr2 | + | 230253333 | 230253417 | 230250980 | 230251061 | 230255451 | 230255532 | 212071 | 46,26,61,35,32,18,13,13,2 | 17,6,10,15,3,2,7,9,0,1,7,5 | 59,39,32,20,38,29,5,9,20,5 | 7,2,1,1,0,1,0,2,4,7,6,13,2,7 | 233 | 149 | 0.0138151422203 | 0.713570164439018 | 0.634,0.735,0.796, | 0.844,0.926,0.953, | -0,116 |
| 16763 | ENSG00000122299 | ZC3H7A | chr16 | - | 11761405 | 11761511 | 11758430 | 11758539 | 11761909 | 11762043 | 16763 | 83,73,106,67,32,45,35,86, | 7,0,7,6,4,6,4,6,2,2,2,2 | 54,146,56,64,63,32,48,56, | 1,2,0,0,6,1,0,3,6,1,3,0,2,0, | 255 | 149 | 0.0138387991101 | 0.714620578518698 | 0.874,1.0,0.898,0. | 0.969,0.977,1.0,1. | -0,039 |
| 208650 | ENSG00000152520 | PAN3 | chr13 | + | 28239578 | 28239696 | 28220230 | 28220378 | 28256291 | 28256539 | 208650 | 10,8,16,17,7,17,16,26,2,3, | 58,50,74,37,49,47,74,98,4 | 13,17,17,23,12,11,17,13,6, | 46,66,49,77,37,39,40,50,3 | 267 | 149 | 0.0138473067991 | 0.714862141448925 | 0.088,0.082,0.108, | 0.136,0.126,0.162, | -0,037 |
| 69868 | ENSG00000223745 | CCDC18-AS1 | chr1 | - | 93339648 | 93339679 | 93338130 | 93339536 | 93345723 | 93345799 | 69868 | 33,21,70,43,17,26,21,20,1 | 0,1,8,0,0,1,0,8,4,2,1,0 | 15,15,12,15,36,24,11,19,2 | 4,1,1,0,4,3,1,2,5,0,6,1,2,1, | 180 | 149 | 0.0138522261254 | 0.714862141448925 | 1.0,0.946,0.879,1. | 0.756,0.925,0.909, | 0,01 |
| 135415 | ENSG00000099917 | MED15 | chr22 | + | 20518877 | 20518997 | 20507581 | 20507746 | 20537116 | 20537215 | 135415 | 6,1,3,2,0,1,4,2,0,3,0,0 | 33,24,28,16,13,8,11,12,10, | 1,9,2,2,2,3,4,3,4,2,1,3,6,6, | 23,24,13,39,17,14,8,8,23,1 | 269 | 149 | 0.0138534411492 | 0.714862141448925 | 0.091,0.023,0.056, | 0.024,0.172,0.079, | -0,043 |
| 92768 | ENSG00000081377 | CDC14B | chr9 | - | 96515648 | 96515792 | 96503564 | 96503789 | 96522505 | 96522603 | 92768 | 17,2,13,11,1,7,7,11,6,2,1,6 | 0,0,5,1,4,5,0,0,0,0,2,1 | 10,4,7,8,2,4,2,4,6,1,6,0,11, | 4,0,0,2,2,0,4,0,2,6,4,1,3,2, | 293 | 149 | 0.0138615524382 | 0.715109250437734 | 1.0,1.0,0.569,0.84 | 0.56,1.0,1.0,0.67,0 | 0,167 |
| 237874 | ENSG00000104365 | IKBKB | chr8 | + | 42319590 | 42319646 | 42319269 | 42319421 | 42320734 | 42320844 | 237874 | 24,11,11,9,6,10,11,7,26,13 | 4,13,2,2,2,1,2,1,4,1,0,3 | 11,3,5,10,5,4,7,11,4,7,2,7, | 0,4,3,12,6,2,0,3,2,11,2,7,0, | 205 | 149 | 0.0138972033091 | 0.716776651191206 | 0.813,0.381,0.8,0. | 1.0,0.353,0.548,0. | 0,177 |
| 141262 | ENSG00000101745 | ANKRD12 | chr18 | + | 9195557 | 9195698 | 9182385 | 9182519 | 9208656 | 9208797 | 141262 | 281,154,257,213,55,95,75, | 4,10,5,5,3,1,2,1,6,0,0,4 | 111,121,122,119,86,71,65, | 0,1,6,7,4,0,0,0,1,3,1,1,0,0, | 290 | 149 | 0.0139103302885 | 0.717189641471838 | 0.973,0.888,0.964, | 1.0,0.984,0.913,0. | -0,017 |
| 76063 | ENSG00000135541 | AHI1 | chr6 | - | 135495813 | 135495898 | 135492227 | 135492291 | 135497187 | 135497249 | 76063 | 8,6,21,15,8,9,2,2,3,2,4,9 | 3,2,24,7,10,5,10,6,1,0,3,11 | 1,1,1,5,10,1,4,6,2,5,3,2,3,6 | 10,8,11,9,7,1,2,7,2,1,8,2,8, | 234 | 149 | 0.0139118749304 | 0.717189641471838 | 0.629,0.656,0.358, | 0.06,0.074,0.055,0 | 0,187 |
| 414 | ENSG00000186866 | POFUT2 | chr21 | - | 45277232 | 45277773 | 45277016 | 45277142 | 45278102 | 45278169 | 414 | 6,5,3,9,6,2,7,9,1,3,4,3 | 7,7,14,5,10,4,7,11,5,7,6,5 | 8,5,3,0,0,1,1,4,1,1,1,0,1,5, | 6,6,0,8,8,2,2,24,5,8,8,3,7,5 | 298 | 149 | 0.0139167851844 | 0.717270975101436 | 0.3,0.263,0.097,0. | 0.4,0.294,1.0,0.0,0 | 0,023 |
| 203043 | ENSG00000064687 | ABCA7 | chr19 | + | 1057943 | 1058269 | 1057313 | 1057429 | 1058617 | 1058732 | 203043 | 52,24,37,40,37,31,55,44,5 | 0,0,2,1,0,3,1,3,0,0,2,0 | 33,36,18,33,21,25,17,32,1 | 1,0,0,2,1,2,0,2,3,0,4,1,2,2, | 298 | 149 | 0.01392034346 | 0.717282605515743 | 1.0,1.0,0.902,0.95 | 0.943,1.0,1.0,0.89 | 0,041 |
| 70255 | ENSG00000095787 | WAC | chr10 | + | 28535909 | 28536059 | 28535561 | 28535757 | 28583398 | 28583505 | 70255 | 30,18,26,18,6,13,6,15,2,9, | 261,169,200,188,132,122, | 19,23,33,50,21,3,7,10,16,1 | 193,195,133,125,118,88,64 | 298 | 149 | 0.0139321636177 | 0.717719843849016 | 0.054,0.051,0.061, | 0.047,0.056,0.11,0 | -0,014 |
| 230445 | ENSG00000062725 | APPBP2 | chr17 | - | 60455305 | 60455420 | 60454301 | 60454492 | 60456295 | 60456381 | 230445 | 6,0,13,7,2,1,2,4,0,0,4,4 | 21,8,40,30,28,10,15,31,15, | 1,2,3,2,1,1,2,6,0,3,6,2,2,0, | 25,37,23,27,13,16,12,29,8, | 264 | 149 | 0.0139590427905 | 0.718932457248649 | 0.139,0.0,0.155,0. | 0.022,0.03,0.069,0 | 0,038 |
| 51564 | ENSG00000162894 | FCMR | chr1 | - | 206913758 | 206914094 | 206911729 | 206911952 | 206921817 | 206921867 | 51564 | 1167,447,1219,947,784,48 | 9,3,4,3,4,10,4,4,8,14,3,3 | 859,366,530,338,204,325, | 4,0,6,4,2,3,2,3,0,0,0,0,2,1, | 298 | 149 | 0.0140243259716 | 0.722121938563024 | 0.985,0.987,0.993, | 0.991,1.0,0.978,0. | -0,012 |
| 142767 | ENSG00000173812 | EIF1 | chr17 | + | 41689777 | 41690008 | 41688884 | 41689069 | 41690087 | 41690620 | 142767 | 68,45,178,164,47,79,64,85 | 3,0,7,7,2,2,1,2,0,1,1,2 | 138,272,48,112,103,89,43, | 5,13,3,9,2,6,1,5,6,7,2,3,4,0 | 298 | 149 | 0.0140361650958 | 0.722558682069871 | 0.919,1.0,0.927,0. | 0.932,0.913,0.889, | 0,026 |
| 47627 | ENSG00000117054 | ACADM | chr1 | + | 75742167 | 75745914 | 75739979 | 75740110 | 75749418 | 75749559 | 47627 | 41,20,113,67,32,36,20,24, | 2,0,3,1,1,5,3,4,0,1,1,3 | 56,29,31,17,29,23,16,37,1 | 0,2,0,0,4,2,0,1,2,0,0,0,3,3, | 298 | 149 | 0.0140477107773 | 0.72298011437328 | 0.911,1.0,0.95,0.9 | 1.0,0.879,1.0,1.0,0 | -0,038 |
| 203047 | ENSG00000064687 | ABCA7 | chr19 | + | 1057973 | 1058269 | 1057313 | 1057429 | 1058617 | 1058732 | 203047 | 52,24,38,41,37,31,55,44,5 | 0,0,2,1,0,3,1,3,0,0,2,0 | 33,36,18,35,21,25,17,32,1 | 1,0,0,2,1,2,0,2,3,0,4,1,2,2, | 298 | 149 | 0.0140602708544 | 0.723453539627867 | 1.0,1.0,0.905,0.95 | 0.943,1.0,1.0,0.89 | 0,041 |
| 53796 | ENSG00000137074 | APTX | chr9 | - | 32988082 | 32988124 | 32987543 | 32987731 | 32989758 | 32989895 | 53796 | 17,11,7,2,4,9,4,2,2,1,4,8 | 2,3,0,1,0,0,0,1,3,0,2,2 | 9,8,4,4,5,2,0,5,6,9,2,6,1,2, | 0,6,5,1,0,0,3,2,0,1,1,0,2,1, | 191 | 149 | 0.0140658959108 | 0.723568279659362 | 0.869,0.741,1.0,0. | 1.0,0.51,0.384,0.7 | 0,11 |
| 51406 | ENSG00000158864 | NDUFS2 | chr1 | + | 161203436 | 161203543 | 161201665 | 161202480 | 161206406 | 161206593 | 51406 | 35,31,94,74,37,37,26,31,2 | 4,3,1,2,1,0,0,0,0,0,0,0 | 43,53,27,20,29,13,22,44,2 | 0,2,2,0,3,0,0,0,3,1,3,2,3,2, | 256 | 149 | 0.0140692244629 | 0.723568279659362 | 0.836,0.857,0.982, | 1.0,0.939,0.887,1. | 0,035 |
| 30887 | ENSG00000145715 | RASA1 | chr5 | + | 87379734 | 87379850 | 87378395 | 87378538 | 87383712 | 87383780 | 30887 | 39,15,102,100,30,52,42,81 | 6,0,2,0,0,1,1,3,0,0,0,0 | 67,68,35,15,27,36,18,25,4 | 0,3,0,6,0,1,0,3,1,2,10,1,2,0 | 265 | 149 | 0.0140835436344 | 0.724131672234961 | 0.785,1.0,0.966,1. | 1.0,0.927,1.0,0.58 | 0,051 |
| 238065 | ENSG00000119397 | CNTRL | chr9 | + | 121157982 | 121158281 | 121157739 | 121157880 | 121158854 | 121159019 | 238065 | 90,64,61,28,29,25,20,16,4 | 9,9,13,7,1,7,3,5,5,4,0,2 | 55,30,64,32,24,21,14,9,19, | 0,4,14,1,2,0,0,5,3,0,6,0,7,3 | 298 | 149 | 0.0140962973459 | 0.724614323872797 | 0.833,0.78,0.701,0 | 1.0,0.789,0.696,0. | -0,071 |
| 226007 | ENSG00000155096 | AZIN1 | chr8 | - | 102849998 | 102850127 | 102843550 | 102843747 | 102858012 | 102858150 | 226007 | 28,20,102,65,29,28,28,38, | 69,36,283,241,104,98,75,8 | 56,86,21,13,35,23,11,35,2 | 161,132,81,38,133,74,34,7 | 278 | 149 | 0.0141035764507 | 0.724704804190414 | 0.179,0.229,0.162, | 0.157,0.259,0.122, | -0,024 |
| 18429 | ENSG00000198690 | FAN1 | chr15 | + | 30905724 | 30905897 | 30904511 | 30904590 | 30908117 | 30908258 | 18429 | 7,4,36,21,26,11,5,8,5,1,5,1 | 0,0,1,4,0,0,0,0,0,0,0,0 | 25,20,11,16,7,3,7,11,11,20 | 1,3,0,3,1,0,0,2,2,2,0,2,0,1, | 298 | 149 | 0.0141050645263 | 0.724704804190414 | 1.0,1.0,0.947,0.72 | 0.926,0.769,1.0,0. | 0,112 |
| 91848 | ENSG00000291118 | ZNF767P | chr7 | - | 149563519 | 149563603 | 149558785 | 149558839 | 149619923 | 149620037 | 91848 | 5,1,8,3,0,1,1,2,4,0,2,0 | 14,4,38,13,25,19,17,10,4,1 | 5,0,4,0,2,0,0,1,0,4,0,0,4,3, | 13,9,6,17,8,6,11,29,8,10,2, | 233 | 149 | 0.0141093766811 | 0.724704804190414 | 0.186,0.138,0.119, | 0.197,0.0,0.299,0. | 0,034 |
| 181981 | ENSG00000291153 | STAG3L4 | chr7 | + | 67320102 | 67320409 | 67309563 | 67309715 | 67330262 | 67330891 | 181981 | 19,40,42,22,9,10,19,23,20, | 0,0,1,1,4,0,3,6,0,0,0,3 | 21,20,15,14,13,12,6,16,14, | 0,0,2,0,0,0,0,2,0,0,0,0,1,4, | 298 | 149 | 0.0141115259157 | 0.724704804190414 | 1.0,1.0,0.955,0.91 | 1.0,1.0,0.789,1.0,1 | -0,052 |
| 73830 | ENSG00000204152 | TIMM23B | chr10 | + | 49943234 | 49943358 | 49942048 | 49942300 | 49945031 | 49945090 | 73830 | 9,7,5,7,3,9,11,2,2,2,4,2 | 33,21,7,4,5,4,4,5,3,4,3,5 | 7,1,4,0,6,4,2,4,5,5,5,2,5,3, | 7,22,10,7,13,5,9,8,6,4,4,10 | 273 | 149 | 0.0141307818178 | 0.72552058717245 | 0.13,0.154,0.28,0. | 0.353,0.024,0.179, | 0,117 |
| 26391 | ENSG00000115760 | BIRC6 | chr2 | + | 32487646 | 32487801 | 32485642 | 32485759 | 32488587 | 32488714 | 26391 | 138,81,190,121,123,127,9 | 0,0,0,2,0,0,0,0,0,0,2,1 | 164,191,89,89,74,32,111,1 | 4,6,2,4,0,0,0,5,0,1,1,0,5,0, | 298 | 149 | 0.0141357260059 | 0.725601346047189 | 1.0,1.0,1.0,0.968,1 | 0.953,0.941,0.957, | 0,021 |
| 236629 | ENSG00000104783 | KCNN4 | chr19 | - | 43776370 | 43776636 | 43774218 | 43774619 | 43780702 | 43780973 | 236629 | 97,44,93,75,62,105,89,78, | 0,0,5,3,5,1,0,4,3,3,0,0 | 70,74,105,80,43,40,32,88, | 0,3,0,1,0,0,4,0,4,1,2,1,0,0, | 298 | 149 | 0.0141461091318 | 0.725766696008325 | 1.0,1.0,0.903,0.92 | 1.0,0.925,1.0,0.97 | -0,006 |
| 9886 | ENSG00000083454 | P2RX5 | chr17 | - | 3689491 | 3689630 | 3688625 | 3688676 | 3690069 | 3690150 | 9886 | 93,34,134,94,43,10,28,26, | 11,3,12,9,5,0,2,3,0,0,0,2 | 58,25,47,10,31,45,6,13,21, | 12,5,8,2,8,3,2,0,4,1,1,21,1 | 288 | 149 | 0.0141479733389 | 0.725766696008325 | 0.814,0.854,0.852, | 0.714,0.721,0.752, | 0,134 |
| 28909 | ENSG00000132842 | AP3B1 | chr5 | - | 78003458 | 78003494 | 78002335 | 78003055 | 78015409 | 78015548 | 28909 | 6,6,31,11,0,8,1,4,4,0,2,3 | 79,70,206,139,24,37,15,42 | 7,10,11,1,5,9,2,11,4,6,8,28 | 29,42,70,26,81,102,33,59, | 185 | 149 | 0.0141502798402 | 0.725766696008325 | 0.058,0.065,0.108, | 0.163,0.161,0.112, | -0,04 |
| 38035 | ENSG00000101844 | ATG4A | chrX | + | 108126792 | 108127094 | 108126076 | 108126187 | 108128780 | 108128852 | 38035 | 0,1,7,4,1,0,1,4,1,1,0,1 | 9,7,10,10,9,5,15,15,2,2,4,8 | 0,0,2,8,3,0,4,6,1,1,2,0,0,1, | 19,7,5,1,6,5,6,10,6,10,3,19 | 298 | 149 | 0.014152435398 | 0.725766696008325 | 0.0,0.067,0.259,0. | 0.0,0.0,0.167,0.8,0 | -0,036 |
| 109202 | ENSG00000068400 | GRIPAP1 | chrX | - | 49000860 | 49000959 | 48999437 | 48999504 | 49002187 | 49002211 | 109202 | 1,0,0,0,0,0,0,0,0,0,0,1 | 9,5,18,14,9,6,3,8,2,5,2,3 | 1,5,1,0,1,0,4,5,0,1,3,2,0,2, | 14,19,4,4,5,4,1,5,4,15,7,12 | 248 | 149 | 0.0141973705232 | 0.727897631033554 | 0.063,0.0,0.0,0.0,0 | 0.041,0.137,0.131, | -0,101 |
| 48146 | ENSG00000142687 | KIAA0319L | chr1 | - | 35442245 | 35442336 | 35438541 | 35441138 | 35442905 | 35443028 | 48146 | 11,14,32,25,14,10,4,18,9,4 | 7,4,9,18,8,12,4,5,4,5,1,3 | 20,17,15,8,2,9,8,16,13,20, | 9,10,9,14,8,7,6,18,5,8,18,1 | 240 | 149 | 0.0142349703544 | 0.729551090202091 | 0.494,0.685,0.688, | 0.58,0.513,0.509,0 | 0,148 |
| 189902 | ENSG00000138688 | KIAA1109 | chr4 | + | 122192228 | 122192333 | 122189999 | 122190079 | 122196634 | 122196791 | 189902 | 87,74,77,31,22,49,27,47,2 | 8,1,4,5,0,0,4,3,1,1,0,3 | 31,37,39,62,31,17,27,31,4 | 0,0,0,0,8,1,0,2,0,0,9,3,0,0, | 254 | 149 | 0.0142363998627 | 0.729551090202091 | 0.864,0.977,0.919, | 1.0,1.0,1.0,1.0,0.6 | -0,033 |
| 22268 | ENSG00000100485 | SOS2 | chr14 | - | 50138611 | 50138784 | 50134122 | 50134239 | 50139941 | 50140059 | 22268 | 135,83,132,102,69,91,49,8 | 4,0,15,0,0,4,0,9,0,2,0,4 | 80,103,87,54,42,44,51,95, | 6,5,3,0,2,0,2,1,1,1,0,0,2,1, | 298 | 149 | 0.0142506759527 | 0.730108839794233 | 0.944,1.0,0.815,1. | 0.87,0.912,0.935,1 | -0,015 |
| 167180 | ENSG00000170088 | TMEM192 | chr4 | - | 165100627 | 165100861 | 165088467 | 165088602 | 165102949 | 165103096 | 167180 | 6,3,14,25,9,11,9,5,2,5,3,9 | 1,0,4,1,0,1,2,0,0,0,0,0 | 11,16,13,2,18,4,14,26,1,7, | 3,0,1,2,3,0,0,0,0,0,2,7,1,0, | 298 | 149 | 0.0142607380946 | 0.730450480923097 | 0.75,1.0,0.636,0.9 | 0.647,1.0,0.867,0. | 0,114 |
| 140823 | ENSG00000134265 | NAPG | chr18 | + | 10538859 | 10539187 | 10534465 | 10534496 | 10539761 | 10539871 | 140823 | 13,6,4,7,5,3,1,8,0,0,2,1 | 8,14,32,30,12,13,1,12,9,2, | 3,7,7,3,8,1,3,5,3,2,16,3,5,7 | 3,9,4,1,6,4,2,6,6,5,12,10,1 | 298 | 149 | 0.0142653239746 | 0.730511526142549 | 0.448,0.176,0.059, | 0.333,0.28,0.467,0 | -0,121 |
| 101200 | ENSG00000184863 | RBM33 | chr7 | + | 155684932 | 155685052 | 155680589 | 155680908 | 155700772 | 155701490 | 101200 | 20,8,21,3,4,9,4,13,5,8,1,3 | 85,72,119,68,39,53,36,52, | 14,34,12,33,8,10,6,9,13,17 | 57,121,38,89,96,49,46,51, | 269 | 149 | 0.0143003859213 | 0.731961867088184 | 0.115,0.058,0.089, | 0.12,0.135,0.149,0 | -0,016 |
| 23188 | ENSG00000213983 | AP1G2 | chr14 | - | 23566561 | 23566686 | 23566277 | 23566419 | 23567110 | 23567315 | 23188 | 195,118,241,256,137,141, | 13,13,20,2,9,13,8,11,8,8,3, | 180,140,89,33,92,77,72,11 | 8,11,13,7,10,3,8,10,4,11,7, | 274 | 149 | 0.0143004476637 | 0.731961867088184 | 0.891,0.832,0.868, | 0.924,0.874,0.788, | 0,042 |
| 54562 | ENSG00000198925 | ATG9A | chr2 | - | 219227921 | 219228079 | 219227769 | 219227813 | 219228433 | 219228485 | 54562 | 6,7,22,12,8,5,8,10,2,2,4,8 | 0,1,2,3,0,0,3,1,0,0,2,0 | 9,23,14,10,6,10,13,8,6,5,9, | 2,2,2,2,3,0,2,1,0,0,0,3,2,4, | 298 | 149 | 0.0143149091836 | 0.732361305276808 | 1.0,0.778,0.846,0. | 0.692,0.852,0.778, | 0,058 |
| 152710 | ENSG00000100697 | DICER1 | chr14 | - | 95113004 | 95113224 | 95112171 | 95112247 | 95115666 | 95115821 | 152710 | 66,50,124,62,57,79,51,81, | 6,10,8,3,2,12,5,7,1,0,1,2 | 121,82,58,77,51,41,47,45, | 5,12,8,18,8,2,4,4,2,4,7,8,5, | 298 | 149 | 0.0143150568984 | 0.732361305276808 | 0.846,0.714,0.886, | 0.924,0.774,0.784, | 0,061 |
| 24341 | ENSG00000005810 | MYCBP2 | chr13 | - | 77129155 | 77129230 | 77126317 | 77126542 | 77139195 | 77139336 | 24341 | 15,19,26,16,10,7,16,24,5,6 | 25,9,86,55,36,30,13,15,13, | 22,4,12,7,13,3,2,14,4,8,18, | 50,53,27,21,20,15,20,21,1 | 224 | 149 | 0.0143324595506 | 0.733077376790682 | 0.285,0.584,0.167, | 0.226,0.048,0.228, | 0,098 |
| 159680 | ENSG00000112081 | SRSF3 | chr6 | + | 36599865 | 36600276 | 36598848 | 36598983 | 36601151 | 36601190 | 159680 | 77,21,130,30,38,24,31,43, | 117,111,314,232,133,144, | 48,50,21,31,42,41,37,42,1 | 208,166,92,21,79,51,74,17 | 298 | 149 | 0.0143378062902 | 0.733137525893303 | 0.248,0.086,0.172, | 0.103,0.131,0.102, | -0,054 |
| 185733 | ENSG00000198160 | MIER1 | chr1 | + | 66948923 | 66949159 | 66946149 | 66946295 | 66958058 | 66958220 | 185733 | 3,2,20,13,1,2,1,5,0,1,2,5 | 116,96,121,111,46,66,49,7 | 4,8,3,1,1,6,2,1,4,3,1,2,1,0, | 105,113,95,120,67,41,38,5 | 298 | 149 | 0.0143420962959 | 0.733137525893303 | 0.013,0.01,0.076,0 | 0.019,0.034,0.016, | 0,006 |
| 203038 | ENSG00000064687 | ABCA7 | chr19 | + | 1057865 | 1058269 | 1057313 | 1057429 | 1058617 | 1058732 | 203038 | 53,24,39,41,39,31,57,47,5 | 0,0,2,1,0,3,1,3,0,0,2,0 | 34,39,18,34,22,25,17,34,1 | 1,0,0,2,1,2,0,2,3,0,4,1,2,2, | 298 | 149 | 0.0143458400755 | 0.733137525893303 | 1.0,1.0,0.907,0.95 | 0.944,1.0,1.0,0.89 | 0,043 |
| 37446 | ENSG00000197142 | ACSL5 | chr10 | + | 112426787 | 112426859 | 112426257 | 112426359 | 112427217 | 112428363 | 37446 | 125,87,99,100,42,31,36,39 | 2,0,3,0,1,0,0,1,2,0,2,8 | 57,63,55,39,21,38,30,37,3 | 0,0,0,0,0,0,0,0,0,0,2,0,7,3, | 221 | 149 | 0.0143535075328 | 0.733137525893303 | 0.977,1.0,0.957,1. | 1.0,1.0,1.0,1.0,1.0, | 0,005 |
| 190166 | ENSG00000084733 | RAB10 | chr2 | + | 26123982 | 26124047 | 26109767 | 26109906 | 26127143 | 26127278 | 190166 | 14,5,7,17,1,6,4,6,2,4,0,3 | 51,23,63,48,36,26,15,30,6 | 11,10,8,5,3,3,4,17,4,11,8,5 | 48,16,36,18,15,28,30,40,1 | 214 | 149 | 0.0143557767427 | 0.733137525893303 | 0.16,0.131,0.072,0 | 0.138,0.303,0.134, | -0,05 |
| 50568 | ENSG00000127463 | EMC1 | chr1 | - | 19231260 | 19231422 | 19230843 | 19230963 | 19232623 | 19232773 | 50568 | 9,17,26,23,21,27,13,28,27, | 2,0,13,4,2,2,2,0,1,0,2,0 | 20,4,20,9,9,5,13,21,6,15,1 | 5,6,1,0,5,0,3,6,0,1,2,7,2,2, | 298 | 149 | 0.0143570925994 | 0.733137525893303 | 0.692,1.0,0.5,0.74 | 0.667,0.25,0.909,1 | 0,116 |
| 32152 | ENSG00000005156 | LIG3 | chr17 | + | 35001908 | 35002104 | 35001256 | 35001403 | 35002667 | 35002789 | 32152 | 30,18,30,12,29,24,4,21,22, | 0,2,5,0,0,0,1,1,0,0,0,0 | 24,23,25,25,17,14,22,29,9, | 3,3,3,3,1,0,1,0,0,1,2,2,2,1, | 298 | 149 | 0.0143584029027 | 0.733137525893303 | 1.0,0.818,0.75,1.0, | 0.8,0.793,0.806,0. | 0,034 |
| 66088 | ENSG00000185436 | IFNLR1 | chr1 | - | 24159051 | 24159182 | 24154167 | 24157891 | 24159473 | 24159633 | 66088 | 50,8,80,41,27,25,23,51,37, | 29,7,34,49,26,30,31,44,37, | 46,45,24,15,23,12,47,21,7, | 22,27,30,16,11,6,12,17,4,1 | 280 | 149 | 0.0143608857886 | 0.733137525893303 | 0.478,0.378,0.556, | 0.527,0.47,0.299,0 | -0,102 |
| 65155 | ENSG00000146282 | RARS2 | chr6 | - | 87519582 | 87519707 | 87518823 | 87518891 | 87520179 | 87520252 | 65155 | 26,18,31,31,22,2,6,13,27,9 | 14,2,11,11,12,6,13,10,3,2, | 23,21,43,10,17,11,16,24,1 | 2,3,4,4,5,1,2,4,4,3,9,2,6,9, | 274 | 149 | 0.0143906513575 | 0.734308117731254 | 0.502,0.83,0.605,0 | 0.862,0.792,0.854, | -0,141 |
| 185959 | ENSG00000186517 | ARHGAP30 | chr1 | - | 161056387 | 161056528 | 161054622 | 161054705 | 161059613 | 161059716 | 185959 | 53,24,97,97,49,39,33,32,4 | 4,2,7,8,2,6,1,0,1,2,2,5 | 88,62,41,21,39,51,18,34,3 | 5,3,0,2,2,2,0,1,3,4,0,2,5,2, | 290 | 149 | 0.0143931316717 | 0.734308117731254 | 0.872,0.86,0.877,0 | 0.9,0.914,1.0,0.84 | -0,048 |
| 49033 | ENSG00000091140 | DLD | chr7 | + | 107902324 | 107902393 | 107901737 | 107901817 | 107903477 | 107903547 | 49033 | 43,40,68,58,39,50,23,20,3 | 0,1,4,1,0,2,0,4,1,0,0,0 | 38,103,42,14,42,27,17,49, | 5,3,2,3,2,1,0,4,0,0,5,6,1,1, | 218 | 149 | 0.014397183536 | 0.734308117731254 | 1.0,0.965,0.921,0. | 0.839,0.959,0.935, | 0,035 |
| 54879 | ENSG00000077044 | DGKD | chr2 | + | 233438216 | 233438379 | 233437376 | 233437479 | 233441886 | 233441995 | 54879 | 109,67,352,143,121,135,1 | 7,2,27,11,5,16,12,14,5,4,1 | 144,145,122,104,125,56,8 | 9,15,2,7,13,1,6,4,4,5,13,4, | 298 | 149 | 0.0143974625255 | 0.734308117731254 | 0.886,0.944,0.867, | 0.889,0.829,0.968, | -0,043 |
| 11641 | ENSG00000007202 | KIAA0100 | chr17 | - | 28628264 | 28628559 | 28624232 | 28624390 | 28631478 | 28631681 | 11641 | 196,99,226,166,109,102,6 | 12,5,22,7,4,6,5,6,0,4,4,5 | 119,87,56,71,88,79,63,133 | 8,12,3,6,18,1,3,17,7,6,9,9, | 298 | 149 | 0.0144155179118 | 0.734898681127719 | 0.891,0.908,0.837, | 0.881,0.784,0.903, | 0,039 |
| 9451 | ENSG00000108406 | DHX40 | chr17 | + | 59573739 | 59573967 | 59573115 | 59573235 | 59574187 | 59574254 | 9451 | 26,51,110,34,36,34,19,34, | 3,1,4,3,0,1,2,1,0,0,2,2 | 51,71,50,67,49,22,28,42,1 | 6,4,3,12,1,2,3,1,1,0,2,6,2,0 | 298 | 149 | 0.0144158705378 | 0.734898681127719 | 0.812,0.962,0.932, | 0.81,0.899,0.893,0 | 0,023 |
| 215134 | ENSG00000115355 | CCDC88A | chr2 | - | 55301620 | 55302072 | 55301205 | 55301277 | 55308808 | 55309023 | 215134 | 88,80,223,151,22,21,41,42 | 7,6,5,6,0,0,2,0,2,0,0,0 | 106,57,30,5,26,26,32,45,3 | 6,3,3,0,0,0,2,6,1,1,0,1,8,6, | 298 | 149 | 0.0144271088457 | 0.73517233852545 | 0.863,0.87,0.957,0 | 0.898,0.905,0.833, | 0,053 |
| 73500 | ENSG00000154222 | CC2D1B | chr1 | - | 52357000 | 52357174 | 52356383 | 52356442 | 52357525 | 52357698 | 73500 | 19,14,21,11,0,12,8,10,12,1 | 7,1,7,1,3,2,2,2,2,5,0,7 | 16,21,13,13,7,1,10,6,7,8,6, | 0,1,0,6,3,0,2,5,3,0,2,0,0,0, | 298 | 149 | 0.0144280701104 | 0.73517233852545 | 0.576,0.875,0.6,0. | 1.0,0.913,1.0,0.52, | -0,151 |
| 149763 | ENSG00000128928 | IVD | chr15 | + | 40414822 | 40414982 | 40412990 | 40413087 | 40415400 | 40415461 | 149763 | 3,4,15,20,3,16,6,10,2,1,2,6 | 2,2,4,2,0,6,1,6,3,0,0,2 | 12,10,3,0,1,3,3,3,9,3,6,3,9, | 3,4,3,4,4,1,2,10,1,3,1,1,6,4 | 298 | 149 | 0.0144402339192 | 0.735617985009073 | 0.429,0.5,0.652,0. | 0.667,0.556,0.333, | 0,233 |
| 15727 | ENSG00000197912 | SPG7 | chr16 | + | 89545871 | 89545995 | 89543689 | 89544772 | 89546657 | 89546760 | 15727 | 0,1,10,3,1,0,6,3,0,1,0,1 | 36,17,74,42,40,51,18,19,2 | 1,0,0,2,2,2,0,3,0,1,2,3,0,0, | 45,38,15,18,62,22,32,38,4 | 273 | 149 | 0.0144472301709 | 0.7358002358999 | 0.0,0.031,0.069,0. | 0.012,0.0,0.0,0.05 | 0,02 |
| 190311 | ENSG00000037474 | NSUN2 | chr5 | - | 6605286 | 6605408 | 6604604 | 6604685 | 6606819 | 6606912 | 190311 | 14,15,27,18,4,9,4,10,16,22 | 6,2,7,5,2,2,3,0,0,0,0,0 | 10,20,19,14,14,2,16,12,5,7 | 2,1,0,0,1,1,0,3,2,4,0,0,2,0, | 271 | 149 | 0.0144639472075 | 0.73633352167839 | 0.562,0.805,0.68,0 | 0.733,0.917,1.0,1. | -0,03 |
| 124414 | ENSG00000073921 | PICALM | chr11 | - | 86022366 | 86022469 | 86014869 | 86014963 | 86026291 | 86026367 | 124414 | 250,220,263,136,33,53,58, | 9,1,5,2,2,1,2,4,4,4,2,2 | 145,166,136,155,105,80,7 | 4,11,2,10,7,1,3,3,4,0,6,7,3, | 252 | 149 | 0.0144645433495 | 0.73633352167839 | 0.943,0.992,0.969, | 0.955,0.899,0.976, | 0,011 |
| 211948 | ENSG00000164985 | PSIP1 | chr9 | - | 15470999 | 15471231 | 15469937 | 15469993 | 15472631 | 15472750 | 211948 | 6,3,8,2,2,1,1,6,9,2,0,0 | 51,20,59,33,59,32,29,36,3 | 5,4,0,2,1,3,5,6,2,2,8,5,5,9, | 35,20,18,1,13,25,13,27,6,2 | 298 | 149 | 0.0144714445881 | 0.73651063848223 | 0.056,0.07,0.063,0 | 0.067,0.091,0.0,0. | -0,065 |
| 80440 | ENSG00000290927 | | chr16 | + | 31701662 | 31701858 | 31700557 | 31700738 | 31702534 | 31702681 | 80440 | 27,16,26,12,3,10,6,16,5,6, | 16,13,29,10,7,11,5,8,2,13, | 13,8,11,11,8,14,7,19,14,13 | 8,8,4,9,8,1,0,1,4,6,3,3,13,5 | 298 | 149 | 0.0144790565313 | 0.73672383363788 | 0.458,0.381,0.31,0 | 0.448,0.333,0.579, | -0,164 |
| 107046 | ENSG00000046651 | OFD1 | chrX | + | 13753367 | 13753441 | 13751248 | 13751368 | 13756577 | 13756767 | 107046 | 33,16,91,37,11,27,12,28,8, | 2,0,1,0,1,0,0,0,0,0,0,0 | 26,23,16,15,18,8,7,15,18,6 | 1,1,4,0,1,1,2,2,0,3,0,2,3,0, | 223 | 149 | 0.0144860957906 | 0.736907795581808 | 0.917,1.0,0.984,1. | 0.946,0.939,0.728, | 0,086 |
| 157877 | ENSG00000066933 | MYO9A | chr15 | - | 72061558 | 72061603 | 72046343 | 72046634 | 72117679 | 72118150 | 157877 | 0,0,1,0,2,0,0,0,2,6,6,3 | 24,13,9,4,9,15,5,9,20,12,4, | 2,1,3,0,1,0,0,0,0,0,1,0,0,2, | 4,3,8,10,5,3,3,6,1,10,15,11 | 194 | 149 | 0.0144933863672 | 0.737104452079117 | 0.0,0.0,0.079,0.0,0 | 0.277,0.204,0.224, | 0,022 |
| 143028 | ENSG00000132388 | UBE2G1 | chr17 | - | 4296716 | 4296809 | 4289229 | 4289408 | 4307020 | 4307123 | 143028 | 39,26,141,90,33,18,31,24, | 4,4,8,3,0,2,0,3,0,0,1,7 | 48,84,48,37,57,55,21,18,2 | 10,10,2,4,14,6,5,2,2,3,0,3, | 242 | 149 | 0.0145118064414 | 0.73786690578525 | 0.857,0.8,0.916,0. | 0.747,0.838,0.937, | 0,044 |
| 165897 | ENSG00000172869 | DMXL1 | chr5 | + | 119099081 | 119099416 | 119097978 | 119098104 | 119101934 | 119102210 | 165897 | 4,1,8,7,5,2,4,3,1,5,0,4 | 160,127,129,92,50,78,67,7 | 2,1,3,1,1,2,2,0,0,1,6,5,5,0, | 91,115,119,74,73,59,41,65 | 298 | 149 | 0.0145327342819 | 0.738611982427275 | 0.012,0.004,0.03,0 | 0.011,0.004,0.012, | 0,011 |
| 38184 | ENSG00000100412 | ACO2 | chr22 | + | 41515766 | 41515917 | 41515376 | 41515535 | 41517526 | 41517631 | 38184 | 76,63,105,96,64,73,61,65, | 4,0,0,1,1,4,0,10,0,0,0,0 | 69,72,48,33,35,27,44,51,5 | 0,2,1,2,1,1,1,0,0,0,0,2,0,0, | 298 | 149 | 0.0145333234784 | 0.738611982427275 | 0.905,1.0,1.0,0.98, | 1.0,0.947,0.96,0.8 | -0,016 |

| | | | | | | | | | | | | | | | | | | | | | | |
|---|---|---|---|---|---|---|---|---|---|---|---|---|---|---|---|---|---|---|---|---|---|---|
| 96461 | ENSG00000168944 | CEP120 | chr5 | - | 123422466 | 123422575 | 123418358 | 123418515 | 123422949 | 123423324 | 96461 | 4,0,5,2,1,0,3,5,0,0,0,4 | 11,6,27,18,15,20,14,21,9,7 | 9,6,2,3,7,2,1,3,2,3,2,4,2,0, | 21,8,4,11,6,4,9,26,8,10,36, | 258 | 149 | 0.0145375423975 | 0.738651980112446 | 0.174,0.0,0.097,0. | 0.198,0.302,0.224, | -0,083 |
| 164890 | ENSG00000204084 | INPP5B | chr1 | - | 37866665 | 37866751 | 37866458 | 37866543 | 37868500 | 37868614 | 164890 | 0,0,1,0,1,3,0,1,0,0,0,0 | 24,23,30,26,22,30,18,25,1 | 0,0,9,4,1,1,2,0,0,4,0,1,1,2, | 29,17,38,7,16,11,8,27,12,1 | 235 | 149 | 0.0145495030479 | 0.73908522315378 | 0.0,0.0,0.021,0.0,0 | 0.0,0.0,0.131,0.26 | -0,047 |
| 253360 | ENSG00000089022 | MAPKAPK5 | chr12 | + | 111890439 | 111890639 | 111890039 | 111890144 | 111892966 | 111902222 | 253360 | 5,0,2,2,0,1,1,6,0,0,2,4 | 31,13,57,37,33,23,11,19,1 | 0,0,0,1,0,2,1,0,2,0,0,1,2,0, | 35,23,12,11,8,13,12,17,14, | 298 | 149 | 0.0145560073528 | 0.739241155863732 | 0.075,0.0,0.017,0. | 0.0,0.0,0.0,0.043,0 | 0,027 |
| 106223 | ENSG00000172534 | HCFC1 | chrX | - | 153958568 | 153958766 | 153958024 | 153958249 | 153959330 | 153959491 | 106223 | 36,24,58,35,21,31,35,30,4 | 4,1,1,1,2,2,4,2,5,6,0,2 | 15,37,21,22,22,29,38,48,1 | 0,8,4,16,3,0,7,7,3,4,6,8,6,0 | 298 | 149 | 0.0145634550665 | 0.739444915644695 | 0.818,0.923,0.967, | 1.0,0.698,0.724,0. | 0,075 |
| 58586 | ENSG00000247828 | TMEM161B-DT | chr5 | + | 88269467 | 88269617 | 88269041 | 88269119 | 88270525 | 88270585 | 58586 | 1,1,12,6,5,3,0,0,0,0,1,1 | 1,3,8,9,0,0,1,3,2,2,0,4 | 6,1,0,2,8,1,0,2,4,3,6,3,0,5, | 3,5,3,1,0,0,4,1,4,1,0,1,5,3, | 298 | 149 | 0.0145780640939 | 0.740012102121271 | 0.333,0.143,0.429, | 0.5,0.091,0.0,0.5,1 | -0,149 |
| 4643 | ENSG00000269352 | PTOV1-AS2 | chr19 | - | 49858553 | 49858626 | 49858346 | 49858427 | 49858842 | 49859097 | 4643 | 10,0,6,3,15,10,12,10,8,2,1 | 13,1,7,1,5,8,10,3,15,4,7,4 | 3,1,1,3,3,4,3,9,2,3,6,3,5,7, | 2,4,5,7,3,5,6,8,2,0,8,11,5,9 | 222 | 149 | 0.0145938349939 | 0.740637986223082 | 0.34,0.0,0.365,0.6 | 0.502,0.144,0.118, | 0,097 |
| 181252 | ENSG00000163565 | IFI16 | chr1 | + | 159053532 | 159053677 | 159051678 | 159052098 | 159054820 | 159055155 | 181252 | 284,121,225,170,79,67,45, | 17,7,31,19,8,21,14,6,4,5,2, | 133,146,69,40,66,106,15,4 | 32,28,6,2,6,9,8,21,4,11,17, | 294 | 149 | 0.0146007627892 | 0.740814892947267 | 0.894,0.898,0.786, | 0.678,0.725,0.854, | 0,074 |
| 162513 | ENSG00000132952 | USPL1 | chr13 | + | 30619678 | 30619740 | 30617936 | 30618056 | 30621763 | 30621892 | 162513 | 5,2,10,9,2,6,5,2,1,1,0,0 | 21,9,42,25,16,15,13,28,10, | 12,8,7,4,4,4,4,8,2,6,18,4,8, | 17,27,16,14,24,11,15,12,2 | 211 | 149 | 0.0146143905271 | 0.741331578491223 | 0.144,0.136,0.144, | 0.333,0.173,0.236, | -0,077 |
| 252290 | ENSG00000066651 | TRMT11 | chr6 | + | 126010924 | 126011054 | 126008391 | 126008472 | 126011252 | 126011417 | 252290 | 5,1,5,0,2,0,4,10,0,0,0,2 | 1,0,3,7,9,5,2,6,1,5,3,0 | 4,5,1,3,3,5,0,10,3,5,9,1,7,2 | 2,4,5,5,4,3,3,0,4,1,3,5,4,6, | 279 | 149 | 0.0146559919948 | 0.743266685445994 | 0.728,1.0,0.471,0. | 0.516,0.4,0.097,0. | -0,054 |
| 243748 | ENSG00000214413 | BBIP1 | chr10 | - | 110907696 | 110907836 | 110901537 | 110901612 | 110918120 | 110918213 | 243748 | 2,5,6,4,6,2,13,6,2,0,0,2 | 23,17,23,18,12,14,1,2,6,4, | 5,2,2,1,0,0,2,5,0,7,3,0,1,2, | 18,20,13,1,3,5,11,24,2,15,1 | 289 | 149 | 0.0146709386214 | 0.743849420594239 | 0.043,0.132,0.119, | 0.125,0.049,0.073, | 0,107 |
| 149971 | ENSG00000198690 | FAN1 | chr15 | + | 30905740 | 30905897 | 30904511 | 30904590 | 30908117 | 30908258 | 149971 | 7,4,36,21,26,11,5,8,5,1,5,1 | 0,0,1,4,0,0,0,0,0,0,0,0 | 25,20,11,16,7,3,7,11,11,22 | 1,3,0,3,1,0,0,2,2,2,0,2,0,1, | 298 | 149 | 0.014687182273 | 0.74449762783798 | 1.0,1.0,0.947,0.72 | 0.926,0.769,1.0,0. | 0,111 |
| 149283 | ENSG00000247982 | LINC00926 | chr15 | + | 57306865 | 57307150 | 57300364 | 57300924 | 57307541 | 57307769 | 149283 | 8,2,7,2,2,5,3,2,4,2,0,2 | 80,66,36,12,48,24,17,12,6 | 0,4,2,0,1,2,2,0,0,3,4,2,3,3, | 42,13,4,6,20,12,3,9,2,32,1 | 298 | 149 | 0.0147164730699 | 0.745806737769637 | 0.048,0.015,0.089, | 0.0,0.133,0.2,0.0,0 | -0,005 |
| 49351 | ENSG00000138069 | RAB1A | chr2 | - | 65090982 | 65091078 | 65088938 | 65089070 | 65097970 | 65098066 | 49351 | 27,33,108,62,30,30,29,51, | 6,2,7,0,1,4,4,4,1,2,1,1 | 92,104,44,58,46,23,27,27, | 9,11,2,4,6,4,0,4,8,5,4,5,4,9 | 245 | 149 | 0.0147487477973 | 0.747266416469086 | 0.732,0.909,0.904, | 0.861,0.852,0.93,0 | 0,037 |
| 43422 | ENSG00000176261 | ZBTB8OS | chr1 | - | 32633644 | 32633727 | 32627507 | 32627544 | 32634767 | 32634792 | 43422 | 7,6,9,22,4,4,7,7,5,1,0,2 | 0,1,2,2,4,0,2,1,0,0,2,0 | 9,11,10,5,3,0,4,7,1,7,14,5, | 0,1,2,0,3,3,0,3,0,1,5,7,3,0, | 232 | 149 | 0.0147649336314 | 0.747910433141881 | 1.0,0.794,0.743,0. | 1.0,0.876,0.763,1. | 0,065 |
| 87642 | ENSG00000268362 | | chr19 | + | 24036252 | 24036382 | 24033432 | 24033621 | 24046183 | 24046279 | 87642 | 4,1,2,2,2,3,2,1,6,1,0,0 | 23,19,45,30,19,37,16,56,1 | 0,0,2,0,0,0,0,2,0,2,4,4,0,0, | 51,53,22,54,20,17,25,22,4 | 279 | 149 | 0.0147815013441 | 0.748542368060022 | 0.085,0.027,0.023, | 0.0,0.0,0.0,0.046,0.0,0 | 0,023 |
| 247660 | ENSG00000030066 | NUP160 | chr11 | - | 47812883 | 47813047 | 47812301 | 47812429 | 47813315 | 47813415 | 247660 | 11,3,21,14,12,7,9,15,3,3,5, | 0,0,6,4,1,2,0,0,0,0,0,0 | 17,20,8,13,12,4,7,2,6,11,1 | 0,0,0,8,1,0,0,3,0,1,0,7,4,2, | 298 | 149 | 0.0147843647366 | 0.748542368060022 | 1.0,1.0,0.636,0.63 | 1.0,1.0,1.0,0.448,0 | 0,075 |
| 189231 | ENSG00000118308 | IRAG2 | chr12 | + | 25068155 | 25068328 | 25066364 | 25066512 | 25069349 | 25069431 | 189231 | 9,4,24,26,4,5,3,20,5,0,1,3 | 295,145,614,469,233,194, | 14,16,27,7,2,1,8,3,7,4,14,4 | 254,318,226,76,135,100,6 | 298 | 149 | 0.0148084326352 | 0.749449541220511 | 0.015,0.014,0.019, | 0.027,0.025,0.056, | -0,011 |
| 12263 | ENSG00000121067 | SPOP | chr17 | - | 49675894 | 49676016 | 49622732 | 49622876 | 49677932 | 49678000 | 12263 | 0,0,7,6,2,1,0,2,2,0,2,1 | 6,1,13,17,10,4,6,4,7,13,2,9 | 6,1,1,1,0,2,3,2,1,0,5,5,4,0, | 12,10,7,7,5,5,4,6,6,11,32,1 | 271 | 149 | 0.0148092463391 | 0.749449541220511 | 0.0,0.0,0.228,0.16 | 0.216,0.052,0.073, | 0,008 |
| 34547 | ENSG00000047365 | ARAP2 | chr4 | - | 36073688 | 36073823 | 36066003 | 36068278 | 36080215 | 36080279 | 34547 | 33,45,66,49,26,47,9,31,31, | 0,1,4,0,1,2,2,2,2,0,0,0 | 68,40,25,31,16,30,27,60,2 | 1,2,1,2,1,0,3,4,1,1,5,5,0,1, | 284 | 149 | 0.0148340870293 | 0.750530179925545 | 1.0,0.959,0.896,1. | 0.973,0.913,0.929, | 0,02 |
| 15843 | ENSG00000103274 | NUBP1 | chr16 | + | 10767948 | 10768032 | 10761756 | 10761859 | 10769046 | 10769313 | 15843 | 40,33,90,69,28,28,35,55,6 | 6,5,5,2,3,0,3,5,1,0,2,9 | 35,55,35,20,33,10,6,16,6,2 | 4,2,1,2,1,0,0,3,0,1,5,1,2,0, | 233 | 149 | 0.014846355561 | 0.750952064580773 | 0.81,0.808,0.92,0. | 0.848,0.946,0.957, | -0,047 |
| 47630 | ENSG00000117054 | ACADM | chr1 | + | 75745834 | 75745914 | 75739979 | 75740110 | 75749418 | 75749559 | 47630 | 34,16,76,59,28,27,17,22,2 | 2,0,3,1,1,5,3,4,0,1,1,3 | 46,25,24,17,24,18,13,36,1 | 0,2,0,0,4,2,0,1,2,0,0,0,3,3, | 229 | 149 | 0.0148494036029 | 0.750952064580773 | 0.917,1.0,0.943,0. | 1.0,0.891,1.0,1.0,0 | -0,038 |
| 81299 | ENSG00000198589 | LRBA | chr4 | - | 150867670 | 150867863 | 150851884 | 150852943 | 150868181 | 150868305 | 81299 | 134,78,173,101,96,122,79, | 1,1,7,4,0,6,1,8,0,1,0,1 | 134,110,51,102,71,66,48,7 | 2,6,1,3,9,1,2,2,10,0,5,7,3,2 | 298 | 149 | 0.0148648539102 | 0.751556817464472 | 0.985,0.975,0.925, | 0.971,0.902,0.962, | 0,037 |
| 5673 | ENSG00000160633 | SAFB | chr19 | + | 5648988 | 5649499 | 5648015 | 5648043 | 5649925 | 5649975 | 5673 | 71,40,93,50,38,22,35,36,4 | 4,1,5,5,0,0,4,0,2,1,2,1 | 43,50,41,53,30,36,33,34,1 | 7,3,4,4,0,3,1,3,4,2,2,3,1,0, | 298 | 149 | 0.0148901971484 | 0.752661348625477 | 0.899,0.952,0.903, | 0.754,0.893,0.837, | 0,041 |
| 135733 | ENSG00000291230 | | chr19 | - | 50336309 | 50336705 | 50335058 | 50335181 | 50337016 | 50337135 | 135733 | 264,223,287,640,286,175, | 14,3,5,13,2,4,8,2,10,5,1,0 | 242,175,152,55,157,163,9 | 6,4,2,1,4,2,2,3,7,5,7,15,8,1 | 298 | 149 | 0.0148962912958 | 0.752792597296626 | 0.904,0.974,0.966, | 0.953,0.956,0.974, | 0,018 |
| 49655 | ENSG00000144218 | AFF3 | chr2 | - | 100129223 | 100129306 | 100112385 | 100112524 | 100142483 | 100142583 | 49655 | 62,28,137,41,41,73,17,11, | 31,19,36,6,16,29,21,8,7,7, | 71,55,57,88,23,14,28,34,7 | 7,12,17,34,12,12,15,11,17, | 232 | 149 | 0.0148998168008 | 0.752794007007743 | 0.562,0.486,0.71,0 | 0.867,0.746,0.683, | -0,1 |
| 108841 | ENSG00000101901 | ALG13 | chrX | + | 111728184 | 111728305 | 111727613 | 111727770 | 111730394 | 111730427 | 108841 | 42,29,39,29,34,28,15,26,1 | 0,1,1,1,2,0,2,2,0,0,2,0 | 23,26,29,21,25,24,8,19,37, | 2,4,0,4,1,2,3,2,2,3,1,1,0,0, | 270 | 149 | 0.0149265375261 | 0.753967049584611 | 1.0,0.941,0.956,0. | 0.864,0.782,1.0,0. | 0,04 |
| 210135 | ENSG00000155252 | PI4K2A | chr10 | + | 97654038 | 97654109 | 97650940 | 97651141 | 97656284 | 97656416 | 210135 | 2,4,14,2,2,10,0,3,0,0,1,0 | 40,75,81,62,25,38,51,21,1 | 2,4,1,10,6,0,0,8,4,2,0,0,0,0 | 34,104,63,65,101,36,26,37 | 220 | 149 | 0.0149653686713 | 0.75575111790065 | 0.033,0.035,0.105, | 0.038,0.025,0.011, | 0,014 |
| 164237 | ENSG00000143624 | INTS3 | chr1 | + | 153730941 | 153731051 | 153728552 | 153728784 | 153740650 | 153740734 | 164237 | 2,2,1,0,0,0,1,0,0,0,1,0 | 16,7,23,14,14,24,13,19,19, | 2,0,0,5,2,2,2,0,2,1,1,2,2,0, | 16,5,11,14,3,11,9,29,6,11,1 | 259 | 149 | 0.014996346123 | 0.757137830729396 | 0.067,0.141,0.024, | 0.067,0.0,0.0,0.17, | -0,048 |
| 63428 | ENSG00000060339 | CCAR1 | chr10 | + | 68737844 | 68737889 | 68736875 | 68737048 | 68740628 | 68740661 | 63428 | 197,114,93,36,83,73,55,11 | 14,1,6,3,2,2,6,6,23,1,1,3 | 71,74,93,117,69,41,51,110 | 4,6,0,3,4,0,2,2,4,6,7,1,8,5, | 194 | 149 | 0.0150043161054 | 0.757362560900879 | 0.915,0.989,0.923, | 0.932,0.905,1.0,0. | -0,015 |
| 10746 | ENSG00000258315 | C17orf49 | chr17 | + | 7016909 | 7017014 | 7016494 | 7016685 | 7017256 | 7017375 | 10746 | 44,13,77,75,59,63,46,71,5 | 27,8,15,8,19,10,7,6,14,6,2, | 65,46,25,10,49,31,29,81,1 | 8,3,4,21,1,3,10,7,0,1,7,11, | 254 | 149 | 0.0150203812188 | 0.75799570225171 | 0.489,0.488,0.751, | 0.827,0.9,0.786,0. | -0,077 |
| 56493 | ENSG00000134248 | LAMTOR5 | chr1 | - | 110407032 | 110407128 | 110406317 | 110406379 | 110407585 | 110407668 | 56493 | 32,21,39,32,8,17,17,22,23, | 31,35,67,85,30,49,29,46,2 | 26,43,21,46,33,11,13,30,2 | 44,49,31,22,33,28,17,26,2 | 245 | 149 | 0.0150339838591 | 0.758504308480533 | 0.386,0.267,0.261, | 0.264,0.348,0.292, | -0,086 |
| 103986 | ENSG00000159352 | PSMD4 | chr1 | + | 151265165 | 151265234 | 151264831 | 151264918 | 151265393 | 151265609 | 103986 | 86,35,256,200,115,82,60,1 | 1,4,14,4,2,1,2,0,2,0,0,3 | 116,99,64,29,58,40,62,92, | 3,15,4,1,4,0,2,8,0,3,3,3,2,0 | 218 | 149 | 0.0150517330048 | 0.759071580652264 | 0.983,0.857,0.926, | 0.964,0.819,0.916, | 0,02 |
| 74927 | ENSG00000117280 | RAB29 | chr1 | - | 205772495 | 205772567 | 205771471 | 205771653 | 205774832 | 205775086 | 74927 | 36,37,43,54,25,16,7,15,19, | 9,18,9,9,1,2,4,2,4,4,3,1 | 24,17,25,15,18,21,7,49,9,5 | 1,1,6,0,6,2,2,3,4,2,4,3,4,10 | 221 | 149 | 0.0150522810665 | 0.759071580652264 | 0.729,0.581,0.763, | 0.942,0.92,0.737,1 | -0,057 |
| 112359 | ENSG00000104365 | IKBKB | chr8 | + | 42288633 | 42288728 | 42271462 | 42272205 | 42290155 | 42290273 | 112359 | 25,16,41,41,36,21,30,22,2 | 15,9,17,11,3,8,10,4,6,2,3,8 | 37,21,27,13,21,21,6,36,23, | 7,4,3,2,11,1,2,9,4,4,4,4,3,3 | 244 | 149 | 0.0150596538088 | 0.759265483467283 | 0.504,0.521,0.596, | 0.763,0.762,0.846, | -0,078 |
| 76668 | ENSG00000159023 | EPB41 | chr1 | + | 28997214 | 28997319 | 28987430 | 28987905 | 29011864 | 29011907 | 76668 | 129,80,142,78,89,89,69,10 | 4,4,11,3,4,6,2,4,1,8,0,5 | 68,61,114,132,58,70,56,83 | 4,7,7,6,21,4,8,3,11,11,0,1, | 254 | 149 | 0.0150753453328 | 0.759877837143759 | 0.95,0.921,0.883,0 | 0.909,0.836,0.905, | 0,041 |
| 238247 | ENSG00000132294 | EFR3A | chr8 | + | 131996387 | 131996497 | 131987574 | 131987702 | 132001758 | 132001807 | 238247 | 30,20,33,28,7,9,13,11,51,1 | 4,1,2,3,0,0,1,3,2,5,0,1 | 24,9,35,7,3,10,4,9,4,27,3,2 | 6,8,1,2,7,1,0,2,0,0,1,8,3,1, | 259 | 149 | 0.0150788605844 | 0.759877837143759 | 0.812,0.92,0.905,0 | 0.697,0.393,0.953, | 0,142 |
| 223851 | ENSG00000120314 | WDR55 | chr5 | + | 140667043 | 140667113 | 140664903 | 140665103 | 140668233 | 140668502 | 223851 | 1,3,2,3,1,5,2,1,0,1,2,2 | 11,17,33,25,21,14,17,8,9,1 | 2,6,2,1,3,2,2,2,0,4,5,5,3,0, | 13,18,7,12,10,8,8,18,1,7,1 | 219 | 149 | 0.0150830763215 | 0.759912359492689 | 0.058,0.107,0.04,0 | 0.095,0.185,0.163, | -0,057 |
| 41399 | ENSG00000033327 | GAB2 | chr11 | - | 78220318 | 78220444 | 78215292 | 78219415 | 78221676 | 78221779 | 41399 | 127,74,285,126,112,173,1 | 3,0,4,4,0,2,2,10,2,2,0,0 | 77,115,159,216,117,85,11 | 1,2,0,3,0,0,2,2,0,2,4,0,0,0, | 275 | 149 | 0.0150953232443 | 0.760351396488166 | 0.958,1.0,0.975,0. | 0.977,0.969,1.0,0. | -0,009 |
| 242944 | ENSG00000106948 | AKNA | chr9 | - | 114359593 | 114359793 | 114357920 | 114358167 | 114359895 | 114360062 | 242944 | 51,35,46,46,54,36,40,33,2 | 10,0,7,5,3,1,4,3,0,0,0,0 | 38,37,31,37,19,24,30,28,2 | 5,2,2,3,7,1,2,3,6,2,4,15,0,0 | 298 | 149 | 0.0150996843797 | 0.760393113880957 | 0.718,1.0,0.767,0. | 0.792,0.902,0.886, | 0,098 |
| 63791 | ENSG00000177272 | KCNA3 | chr1 | - | 110669769 | 110669829 | 110668222 | 110668344 | 110671475 | 110674940 | 63791 | 3,0,3,4,4,2,1,2,15,4,0,0 | 73,41,85,39,52,31,59,59,8 | 4,0,0,0,1,3,1,4,0,8,2,3,0,2, | 92,82,73,94,46,34,59,63,4 | 209 | 149 | 0.0151033871529 | 0.760401665568613 | 0.028,0.0,0.025,0. | 0.03,0.0,0.0,0.0,0. | 0 |
| 83190 | ENSG00000183283 | DAZAP2 | chr12 | + | 51240870 | 51241039 | 51240342 | 51240461 | 51242329 | 51242816 | 83190 | 742,559,531,383,255,241, | 7,6,14,10,3,9,6,7,0,0,3,0 | 399,474,435,384,336,171, | 5,12,5,7,9,1,10,14,1,5,23,1 | 298 | 149 | 0.0151123657556 | 0.760675770332915 | 0.981,0.979,0.95,0 | 0.976,0.952,0.978, | 0,011 |
| 251368 | ENSG00000012779 | ALOX5 | chr10 | + | 45445013 | 45445386 | 45444115 | 45444286 | 45445507 | 45446119 | 251368 | 7,3,8,10,6,12,8,5,0,1,3,3 | 159,87,178,160,174,105,1 | 7,9,6,6,2,1,0,3,7,2,5,7,1,2, | 160,151,119,127,152,65,5 | 298 | 149 | 0.0151295778524 | 0.761364080137925 | 0.022,0.017,0.022, | 0.021,0.029,0.025, | 0,008 |
| 21987 | ENSG00000125952 | MAX | chr14 | - | 65077562 | 65077643 | 65077384 | 65077428 | 65077912 | 65078036 | 21987 | 0,6,5,2,1,4,0,0,2,0,0,0 | 23,5,26,17,6,0,6,8,7,1,12,1 | 2,1,0,0,0,0,0,0,0,0,9,0,0,0, | 6,9,7,1,3,3,1,12,12,9,8,3,1 | 230 | 149 | 0.0151354030137 | 0.761479178597865 | 0.0,0.437,0.111,0. | 0.178,0.067,0.0,0. | 0,095 |
| 81568 | ENSG00000168769 | TET2 | chr4 | + | 105241338 | 105241429 | 105233896 | 105237351 | 105242833 | 105242927 | 81568 | 50,49,93,59,11,28,10,13,2 | 1,2,1,3,1,0,0,3,2,0,0,0 | 48,49,28,35,33,33,14,32,2 | 1,6,0,1,4,3,0,3,3,3,0,0,2,0, | 240 | 149 | 0.0151608641498 | 0.762581900403273 | 0.969,0.938,0.983, | 0.968,0.835,1.0,0. | 0,02 |
| 169395 | ENSG00000124535 | WRNIP1 | chr6 | + | 2772978 | 2773141 | 2770194 | 2770361 | 2779262 | 2779492 | 169395 | 4,1,4,2,0,1,0,1,0,0,0,2 | 11,2,53,24,29,12,17,4,8,9, | 4,1,4,0,1,1,4,6,0,1,3,2,3,0, | 27,19,5,4,8,11,6,14,4,23,2 | 298 | 149 | 0.0151803692882 | 0.763384593987985 | 0.154,0.2,0.036,0. | 0.069,0.026,0.286, | -0,024 |
| 245444 | ENSG00000101017 | CD40 | chr20 | + | 46121829 | 46121898 | 46118347 | 46118394 | 46122232 | 46122358 | 245444 | 74,86,66,51,38,21,35,35,4 | 2,0,1,2,0,0,3,0,0,0,0,2 | 29,46,58,17,16,12,7,17,19, | 3,0,1,12,2,0,0,1,0,0,0,7,0,0 | 218 | 149 | 0.0151864360511 | 0.763511286548541 | 0.962,1.0,0.978,0. | 0.869,1.0,0.975,0. | 0,044 |
| 237243 | ENSG00000120910 | PPP3CC | chr8 | + | 22477646 | 22477810 | 22475591 | 22475624 | 22498000 | 22498112 | 237243 | 15,1,13,15,7,4,11,10,7,4,0, | 128,43,231,130,93,73,75,1 | 2,3,4,7,0,3,2,5,3,11,12,9,1 | 163,146,77,70,91,99,42,10 | 298 | 149 | 0.015192475443 | 0.763636544155145 | 0.055,0.011,0.027, | 0.006,0.01,0.025,0 | 0,012 |
| 185489 | ENSG00000116120 | FARSB | chr2 | - | 222646800 | 222646992 | 222642850 | 222643005 | 222648739 | 222648795 | 185489 | 3,1,3,1,1,4,1,0,2,2,0,2 | 6,2,26,12,8,17,0,9,0,3,6,6 | 2,0,2,0,0,3,2,0,0,1,0,1,0,0, | 17,6,12,1,3,4,1,12,5,9,1,12 | 298 | 149 | 0.0151985745179 | 0.763734498465315 | 0.2,0.2,0.055,0.04, | 0.056,0.0,0.077,0. | 0,184 |
| 233316 | ENSG00000215301 | DDX3X | chrX | + | 41346836 | 41347012 | 41346504 | 41346622 | 41347639 | 41347694 | 233316 | 23,7,26,30,13,10,10,14,6,1 | 3,1,7,4,3,0,2,0,0,1,0,2 | 33,31,15,52,16,17,2,19,13, | 8,5,1,31,0,6,5,8,1,6,10,0,3, | 298 | 149 | 0.0152015211165 | 0.763734498465315 | 0.793,0.778,0.65,0 | 0.673,0.756,0.882, | 0,159 |
| 33097 | ENSG00000281649 | EBLN3P | chr9 | + | 37080341 | 37080536 | 37079873 | 37080034 | 37086667 | 37087995 | 33097 | 125,93,210,126,62,54,94,1 | 6,1,12,6,2,2,4,1,2,2,0,0 | 157,143,74,65,73,70,30,81 | 9,3,5,2,2,1,4,4,1,9,6,2,10,8 | 298 | 149 | 0.0152156436339 | 0.764265623096366 | 0.912,0.979,0.897, | 0.897,0.96,0.881,0 | 0,031 |
| 157907 | ENSG00000164080 | RAD54L2 | chr3 | + | 51572821 | 51572915 | 51541588 | 51541650 | 51590366 | 51590559 | 157907 | 0,0,1,1,3,0,1,0,0,0,0,0 | 13,8,33,17,2,14,12,19,12,3 | 3,4,2,1,1,0,0,4,1,2,3,2,2,0, | 23,11,3,19,5,5,14,22,9,0,1 | 243 | 149 | 0.0152207982627 | 0.764284749125007 | 0.0,0.0,0.018,0.03 | 0.074,0.182,0.29,0 | -0,087 |
| 218934 | ENSG00000204592 | HLA-E | chr6 | + | 30492385 | 30492436 | 30491536 | 30491653 | 30492540 | 30492583 | 218934 | 149,115,292,252,74,61,78, | 10,5,8,15,4,0,5,9,0,3,2,9 | 193,238,116,85,92,81,88,2 | 10,13,4,14,16,5,2,21,5,6,8, | 200 | 149 | 0.015227560651 | 0.764284749125007 | 0.917,0.945,0.965, | 0.935,0.932,0.956, | 0,026 |
| 28887 | ENSG00000108039 | XPNPEP1 | chr10 | - | 109870730 | 109870904 | 109869952 | 109870029 | 109871791 | 109871861 | 28887 | 31,24,58,27,32,19,16,23,1 | 6,8,9,2,2,2,6,2,0,1,1,5 | 29,16,21,10,23,13,15,22,1 | 4,15,5,10,3,3,1,7,4,3,12,8, | 298 | 149 | 0.0152276037808 | 0.764284749125007 | 0.721,0.6,0.763,0. | 0.784,0.348,0.677, | 0,12 |
| 189775 | ENSG00000019582 | CD74 | chr5 | - | 150403120 | 150403251 | 150402562 | 150402625 | 150404679 | 150404767 | 189775 | 287,170,475,219,376,167, | 4500,3645,6508,6450,4482 | 283,207,384,464,131,75,2 | 3977,4214,2808,3533,2538 | 280 | 149 | 0.0152302284011 | 0.764284749125007 | 0.033,0.024,0.037, | 0.036,0.025,0.068, | -0,004 |
| 172792 | ENSG00000131370 | SH3BP5 | chr3 | - | 15330446 | 15330566 | 15304102 | 15304231 | 15332270 | 15332476 | 172792 | 29,16,77,41,31,18,44,35,8, | 6,2,2,2,1,5,5,1,0,0,0,2 | 35,24,18,11,37,23,7,14,23, | 0,3,2,3,0,0,1,1,3,4,2,5,0,2,1 | 269 | 149 | 0.015234507315 | 0.764321268977801 | 0.728,0.816,0.955, | 1.0,0.816,0.833,0. | 0,017 |
| 162012 | ENSG00000174989 | FBXW8 | chr12 | + | 116918530 | 116918726 | 116910964 | 116911157 | 116928022 | 116928127 | 162012 | 0,2,10,5,7,4,6,5,6,2,0,2 | 1,0,6,2,2,0,0,2,0,2,2,0 | 7,2,5,4,1,4,0,5,2,2,4,0,5,3, | 1,1,2,0,0,2,2,1,2,2,1,8,1,6, | 298 | 149 | 0.0152433201549 | 0.764585187662382 | 0.0,1.0,0.455,0.55 | 0.778,0.5,0.556,1. | 0,135 |
| 196222 | ENSG00000168010 | ATG16L2 | chr11 | + | 72827194 | 72827293 | 72826702 | 72826823 | 72828358 | 72828508 | 196222 | 64,41,39,36,10,40,10,27,9, | 0,2,0,0,0,4,0,0,2,0,0,2 | 59,20,13,17,20,20,21,36,2 | 1,2,5,2,1,1,2,5,0,0,1,1,0,0,1 | 248 | 149 | 0.0152513046127 | 0.764807442473447 | 1.0,0.925,1.0,1.0,1 | 0.973,0.857,0.61,0 | 0,043 |
| 199726 | ENSG00000279457 | WASH9P | chr1 | - | 188129 | 188275 | 187375 | 187577 | 188438 | 188584 | 199726 | 17,5,18,9,2,3,18,12,10,4,2, | 3,0,0,1,2,4,4,4,2,4,1,2 | 11,9,10,15,7,2,9,11,3,2,6,3 | 2,0,4,2,0,0,0,1,0,3,0,0,0,1, | 295 | 149 | 0.0152893491617 | 0.766335627663455 | 0.741,1.0,1.0,0.82, | 0.735,1.0,0.558,0. | -0,158 |
| 233203 | ENSG00000046651 | OFD1 | chrX | + | 13736187 | 13736296 | 13735263 | 13735346 | 13736477 | 13736678 | 233203 | 3,2,8,8,19,13,1,7,13,1,2,3 | 62,32,139,79,55,63,36,61, | 8,9,0,2,1,4,2,10,3,14,1,0,6, | 80,73,48,74,37,34,28,78,4 | 258 | 149 | 0.0152906453656 | 0.766335627663455 | 0.027,0.035,0.032, | 0.055,0.066,0.0,0. | 0,02 |
| 150141 | ENSG00000156232 | WHAMM | chr15 | + | 82817537 | 82817702 | 82816691 | 82816842 | 82817919 | 82818089 | 150141 | 10,3,9,2,2,2,4,1,0,2,0,0 | 92,37,79,38,33,43,39,42,1 | 3,2,5,3,6,0,2,4,1,3,0,0,0,0,1 | 62,69,29,85,46,18,25,59,3 | 298 | 149 | 0.015293759134 | 0.766335627663455 | 0.052,0.039,0.054, | 0.024,0.014,0.079, | 0,008 |
| 225162 | ENSG00000186480 | INSIG1 | chr7 | + | 155302521 | 155302846 | 155302250 | 155302417 | 155308240 | 155308757 | 225162 | 32,27,92,45,12,23,26,27,2 | 0,0,5,9,1,3,9,5,0,0,0,5 | 12,19,17,14,17,8,13,22,3,1 | 5,6,4,4,6,5,2,5,5,0,0,1,0,3, | 298 | 149 | 0.0152960208169 | 0.766335627663455 | 1.0,1.0,0.902,0.71 | 0.545,0.613,0.68,0 | 0,139 |
| 31775 | ENSG00000106624 | AEBP1 | chr7 | + | 44107808 | 44107931 | 44107628 | 44107700 | 44108006 | 44108084 | 31775 | 4,13,36,9,11,28,18,47,18,1 | 3,3,1,4,3,6,7,14,2,2,2,4 | 19,16,0,6,8,10,23,27,12,8,1 | 7,2,1,0,5,2,6,0,2,0,4,0,0,0, | 272 | 149 | 0.0153136532657 | 0.767040471498691 | 0.422,0.704,0.952, | 0.598,0.814,0.0,1. | -0,088 |
| 92807 | ENSG00000112739 | PRPF4B | chr6 | + | 4061014 | 4061147 | 4060412 | 4060686 | 4061654 | 4061700 | 92807 | 10,11,30,17,4,8,15,15,8,4, | 0,0,7,0,1,5,0,5,0,0,3,4 | 33,18,14,7,19,4,6,10,23,44 | 4,5,2,0,1,0,1,0,1,5,0,1,1,0, | 282 | 149 | 0.0153233524392 | 0.767347712620162 | 1.0,1.0,0.694,1.0,0 | 0.813,0.655,0.787, | -0,111 |
| 87337 | ENSG00000115355 | CCDC88A | chr2 | - | 55299838 | 55299919 | 55296257 | 55296523 | 55301205 | 55301277 | 87337 | 187,117,326,241,57,39,85, | 5,4,10,12,0,0,3,7,0,0,0,1 | 275,241,71,8,27,36,52,92, | 13,6,7,0,0,2,0,12,2,1,4,0,9, | 230 | 149 | 0.0153581500513 | 0.768911373270842 | 0.96,0.95,0.955,0. | 0.932,0.963,0.868, | 0,029 |
| 76456 | ENSG00000144848 | ATG3 | chr3 | - | 112553279 | 112553329 | 112550191 | 112550262 | 112558375 | 112558417 | 76456 | 59,25,45,33,29,27,18,13,3 | 4,5,6,4,2,4,1,2,0,3,0,1 | 15,29,14,17,16,14,8,13,10, | 3,10,2,2,12,3,0,2,0,1,7,7,2, | 199 | 149 | 0.015364963589 | 0.768927476804968 | 0.917,0.789,0.849, | 0.789,0.685,0.84,0 | 0,072 |
| 140638 | ENSG00000134046 | MBD2 | chr18 | - | 54187713 | 54187963 | 54166075 | 54166166 | 54188873 | 54189011 | 140638 | 6,1,11,8,7,4,6,15,6,2,2,7 | 190,106,237,155,109,92,8 | 12,8,11,24,2,10,7,9,2,14,8, | 141,140,145,49,94,86,45,8 | 298 | 149 | 0.0153672600369 | 0.768927476804968 | 0.016,0.005,0.023, | 0.041,0.028,0.037, | -0,015 |
| 23208 | ENSG00000100461 | RBM23 | chr14 | - | 22908332 | 22908380 | 22906027 | 22906368 | 22909482 | 22909595 | 23208 | 80,57,57,35,27,29,26,27,7 | 21,5,49,28,6,16,10,8,14,6, | 35,31,37,26,21,12,20,18,1 | 26,20,7,5,7,6,13,12,9,13,1 | 197 | 149 | 0.0153691894068 | 0.768927476804968 | 0.742,0.896,0.468, | 0.504,0.54,0.8,0.7 | 0,107 |
| 137337 | ENSG00000196684 | HSH2D | chr19 | + | 16153031 | 16153204 | 16152551 | 16152641 | 16154398 | 16154863 | 137337 | 21,18,68,50,49,49,19,32,2 | 0,0,2,1,0,0,0,3,3,4,0,5 | 29,44,14,16,30,16,5,13,11, | 0,1,0,0,0,0,0,1,1,0,0,0,0,0, | 298 | 149 | 0.0153790021388 | 0.769239602448539 | 1.0,1.0,0.944,0.96 | 1.0,0.957,1.0,1.0,1 | -0,048 |
| 211591 | ENSG00000183283 | DAZAP2 | chr12 | + | 51240870 | 51241284 | 51240342 | 51240461 | 51242329 | 51243933 | 211591 | 743,559,531,383,266,241, | 7,6,14,10,3,9,6,7,0,0,3,0 | 399,474,435,384,336,171, | 5,12,5,7,9,1,10,14,1,5,23,1 | 298 | 149 | 0.0153908948456 | 0.769655596773544 | 0.982,0.979,0.95,0 | 0.976,0.952,0.978, | 0,011 |
| 50239 | ENSG00000143164 | DCAF6 | chr1 | + | 168015780 | 168015951 | 168004532 | 168004793 | 168022987 | 168023047 | 50239 | 10,4,9,6,4,3,7,1,0,2,5,0 | 2,2,26,19,9,5,3,9,1,4,4,3 | 15,13,7,7,9,3,0,7,6,3,13,15 | 7,3,1,1,6,2,1,11,5,7,6,4,3,8 | 298 | 149 | 0.0153991365209 | 0.769888862376267 | 0.714,0.5,0.148,0. | 0.517,0.684,0.778, | -0,15 |
| 201572 | ENSG00000173473 | SMARCC1 | chr3 | - | 47728646 | 47729094 | 47720665 | 47720735 | 47736033 | 47736126 | 201572 | 18,7,93,51,11,22,13,26,7,3 | 0,2,4,0,1,5,1,0,0,1,0,2 | 28,40,9,14,15,21,6,10,9,14 | 0,3,2,4,8,1,2,0,6,0,1,0,0,2, | 298 | 149 | 0.0154404200836 | 0.771773584536301 | 1.0,0.636,0.921,1. | 1.0,0.87,0.692,0.6 | 0,032 |
| 20583 | ENSG00000140105 | WARS1 | chr14 | - | 100374135 | 100374179 | 100369086 | 100369258 | 100375282 | 100375347 | 20583 | 4,1,0,0,10,5,0,2,0,0,0,0 | 47,19,27,20,24,12,14,28,4, | 6,2,0,0,1,3,0,2,0,0,0,0,0,2, | 15,21,14,2,10,13,17,30,4,6 | 193 | 149 | 0.0154627805702 | 0.772617057110444 | 0.062,0.039,0.0,0. | 0.236,0.068,0.0,0. | 0,006 |
| 225168 | ENSG00000186480 | INSIG1 | chr7 | + | 155302808 | 155302846 | 155302250 | 155302417 | 155308240 | 155308757 | 225168 | 13,18,26,14,5,12,8,8,15,22 | 0,0,5,9,1,3,9,5,0,0,0,5 | 6,8,8,6,4,2,2,7,1,5,6,6,8,5, | 5,6,4,4,6,5,2,5,5,0,0,1,0,3, | 187 | 149 | 0.0154644743649 | 0.772617057110444 | 1.0,1.0,0.806,0.55 | 0.489,0.515,0.614, | 0,188 |
| 30215 | ENSG00000131503 | ANKHD1 | chr5 | + | 140535361 | 140535538 | 140528183 | 140529796 | 140537388 | 140537589 | 30215 | 93,59,56,34,58,35,39,51,3 | 0,0,4,0,0,3,3,3,0,0,0,1 | 44,75,39,73,35,31,53,53,2 | 0,8,0,3,3,1,1,0,0,1,0,4,0,2,1 | 298 | 149 | 0.0154703834751 | 0.772676564691145 | 1.0,1.0,0.875,1.0,1 | 1.0,0.824,1.0,0.92 | 0,024 |
| 110144 | ENSG00000147649 | MTDH | chr8 | + | 97706625 | 97706750 | 97699753 | 97699852 | 97713661 | 97713769 | 110144 | 227,157,514,426,134,134, | 10,0,13,8,3,2,4,5,4,4,4,3 | 127,181,182,168,67,83,10 | 0,3,2,8,1,3,0,4,1,0,4,10,0,0 | 274 | 149 | 0.0154728454257 | 0.772676564691145 | 0.925,1.0,0.956,0. | 1.0,0.97,0.98,0.91 | -0,021 |
| 135726 | ENSG00000291230 | | chr19 | - | 50334306 | 50334468 | 50333795 | 50334101 | 50334828 | 50334973 | 135726 | 235,166,240,505,214,155, | 0,0,0,1,2,0,2,2,0,0,0,0 | 153,128,132,69,133,148,5 | 6,6,0,8,0,0,0,0,4,2,1,8,0,2, | 298 | 149 | 0.015483031699 | 0.773005890885518 | 1.0,1.0,1.0,0.996,0 | 0.927,0.914,1.0,0. | 0,026 |
| 193648 | ENSG00000114861 | FOXP1 | chr3 | - | 71130400 | 71130741 | 71112590 | 71112637 | 71132090 | 71132211 | 193648 | 1,2,10,1,2,4,3,1,0,1,0,2 | 17,10,21,26,10,26,5,19,5,1 | 5,6,7,0,5,5,2,4,1,4,2,0,0,1, | 5,6,10,7,4,6,7,4,8,3,6,1,3,2 | 298 | 149 | 0.0155035835914 | 0.77380423192427 | 0.029,0.091,0.192, | 0.333,0.333,0.259, | -0,145 |
| 246980 | ENSG00000261008 | LINC01572 | chr16 | - | 72391070 | 72391215 | 72368652 | 72368722 | 72427389 | 72427457 | 246980 | 46,21,34,9,19,8,13,19,78,2 | 3,3,2,0,3,0,0,1,0,2,2,2 | 9,6,8,5,12,4,2,7,12,24,1,18 | 0,0,0,2,1,0,0,0,0,0,0,0,2, | 294 | 149 | 0.015506212638 | 0.77380423192427 | 0.886,0.78,0.896,1 | 1.0,1.0,1.0,0.559,0 | -0,087 |
| 126526 | ENSG00000150093 | ITGB1 | chr10 | - | 32932514 | 32932590 | 32929821 | 32930044 | 32935491 | 32935558 | 126526 | 232,151,115,108,51,56,93, | 4,5,12,2,2,0,12,17,2,0,0,3 | 68,59,1,40,60,48,28,61,64, | 9,3,0,3,7,4,2,4,4,6,7,12,5,1 | 225 | 149 | 0.0155401378759 | 0.775317435133944 | 0.975,0.952,0.864, | 0.833,0.929,1.0,0. | 0,038 |
| 201232 | ENSG00000181722 | ZBTB20 | chr3 | - | 115071218 | 115071414 | 114974365 | 114974416 | 115110086 | 115110200 | 201232 | 18,2,24,4,5,15,14,15,7,4,1, | 1,1,5,0,3,2,1,6,3,0,0,0 | 17,19,3,7,11,8,4,12,14,10,1 | 3,1,1,0,1,1,0,2,3,0,2,1,2,1, | 298 | 149 | 0.0155452061503 | 0.775390559660537 | 0.9,0.5,0.706,1.0,0 | 0.739,0.905,0.6,1. | -0,066 |

| | | | | | | | | | | | | | | | | | | | | | | |
|---|---|---|---|---|---|---|---|---|---|---|---|---|---|---|---|---|---|---|---|---|---|---|
| 124253 | ENSG00000139697 | SBNO1 | chr12 | - | 123348028 | 123348133 | 123345257 | 123345567 | 123350309 | 123350441 | 124253 | 94,80,98,73,47,45,65,83,5 | 4,0,4,2,0,0,1,1,3,2,0,2 | 85,122,76,68,73,37,32,60,4 | 0,1,3,0,3,0,0,0,0,1,1,2,0,0,0 | 254 | 149 | 0.0155625732406 | 0.775905337148691 | 0.932,1.0,0.935,0.9 | 1.0,0.986,0.937,1.0 | -0,018 |
| 92553 | ENSG00000083097 | DOP1A | chr6 | + | 83130122 | 83130397 | 83128886 | 83129508 | 83132175 | 83132328 | 92553 | 29,22,50,31,28,21,21,31,14 | 6,1,5,1,0,6,0,0,0,0,0,0 | 52,35,14,11,14,27,17,14,12 | 3,1,1,1,0,0,0,0,1,0,0,0,0,0,0 | 298 | 149 | 0.0155627365039 | 0.775905337148691 | 0.707,0.917,0.833, | 0.897,0.946,0.875, | -0,044 |
| 103758 | ENSG00000122545 | SEPTIN7 | chr7 | + | 35879595 | 35879719 | 35873640 | 35873775 | 35879822 | 35879940 | 103758 | 13,4,40,21,5,8,7,17,11,4,4, | 568,311,697,470,348,270,2 | 6,16,7,13,9,10,9,13,9,12,16 | 510,431,303,177,222,173,1 | 273 | 149 | 0.0155718751629 | 0.776181163313138 | 0.012,0.007,0.03,0 | 0.006,0.02,0.012,0 | -0,008 |
| 79554 | ENSG00000137817 | PARP6 | chr15 | - | 72250844 | 72250957 | 72250019 | 72250092 | 72251206 | 72251255 | 79554 | 59,27,75,39,34,28,33,46,41 | 8,5,5,3,9,6,6,7,0,3,0,5 | 46,66,37,42,30,28,37,40,27 | 13,2,4,5,4,1,6,23,12,8,13,6 | 262 | 149 | 0.0155892238543 | 0.776865996616079 | 0.807,0.754,0.895, | 0.668,0.949,0.84,0 | 0,079 |
| 91882 | ENSG00000204569 | PPP1R10 | chr6 | - | 30604993 | 30605094 | 30604587 | 30604735 | 30605922 | 30606035 | 91882 | 44,32,64,38,28,18,17,31,42 | 2,0,4,5,0,3,0,0,2,0,0,1 | 16,26,30,15,21,6,28,32,2,2 | 3,1,0,1,2,0,2,1,2,2,8,0,0,0,0 | 250 | 149 | 0.0155998550314 | 0.777215832931309 | 0.929,1.0,0.905,0.8 | 0.761,0.939,1.0,0.8 | 0,05 |
| 195742 | ENSG00000159377 | PSMB4 | chr1 | + | 151400713 | 151400845 | 151400441 | 151400588 | 151401238 | 151401355 | 195742 | 36,47,84,73,49,48,46,55,93 | 3,2,4,9,7,5,1,0,0,0,2,2 | 66,57,70,21,33,27,27,41,23 | 3,9,2,15,1,1,4,9,1,0,8,8,5,2 | 281 | 149 | 0.0156167026949 | 0.777875153373298 | 0.864,0.926,0.918, | 0.921,0.771,0.949, | 0,067 |
| 81578 | ENSG00000143622 | RIT1 | chr1 | - | 155910655 | 155910804 | 155910449 | 155910506 | 155911242 | 155911349 | 81578 | 28,41,81,68,50,35,32,41,27 | 0,0,4,10,5,5,7,0,2,5,0,1 | 71,75,30,28,35,18,22,45,25 | 2,2,1,5,1,2,0,3,0,1,2,2,2,4,4 | 298 | 149 | 0.0156444567351 | 0.779077294667355 | 1.0,1.0,0.91,0.773, | 0.947,0.949,0.938, | -0,041 |
| 81130 | ENSG00000162191 | UBXN1 | chr11 | - | 62676812 | 62677005 | 62676498 | 62676639 | 62677517 | 62677634 | 81130 | 233,151,327,278,208,113,1 | 6,1,0,0,1,1,1,7,3,0,0,2 | 230,223,144,123,127,140,1 | 0,8,2,4,3,0,2,2,0,3,10,4,0,0 | 298 | 149 | 0.0156629314103 | 0.779816884193911 | 0.951,0.987,1.0,1.0 | 1.0,0.933,0.973,0.9 | 0,014 |
| 100727 | ENSG00000013374 | NUB1 | chr7 | + | 151374096 | 151374201 | 151368734 | 151368887 | 151375847 | 151375943 | 100727 | 27,30,57,48,23,33,15,23,33 | 4,0,1,3,2,2,2,8,0,0,0,0 | 25,14,28,9,3,11,5,32,12,24 | 1,4,0,0,0,0,1,5,1,0,0,1,0,2,3 | 254 | 149 | 0.0157011851345 | 0.781438957896929 | 0.798,1.0,0.971,0.9 | 0.936,0.672,1.0,1.0 | -0,017 |
| 151998 | ENSG00000171606 | ZNF274 | chr19 | + | 58209960 | 58210430 | 58206719 | 58207202 | 58211559 | 58211686 | 151998 | 35,19,68,27,20,27,17,25,18 | 5,3,3,0,0,0,0,2,2,2,0,0 | 38,68,35,42,28,13,21,22,12 | 1,1,0,4,0,0,0,0,0,0,1,0,0,0,0 | 298 | 149 | 0.0157027728018 | 0.781438957896929 | 0.778,0.76,0.919,1 | 0.95,0.971,1.0,0.84 | -0,07 |
| 208447 | ENSG00000138468 | SENP7 | chr3 | - | 101417592 | 101417746 | 101398860 | 101399055 | 101458954 | 101459052 | 208447 | 39,21,38,22,12,13,12,13,23 | 7,2,2,4,5,1,6,8,7,2,3,6 | 26,12,17,24,16,21,6,15,31, | 5,0,0,5,0,3,2,1,0,5,2,0,9,6,2 | 298 | 149 | 0.0157071819906 | 0.781477690018222 | 0.736,0.84,0.905,0 | 0.722,1.0,1.0,0.706 | -0,118 |
| 205287 | ENSG00000165487 | MICU2 | chr13 | - | 21547836 | 21547912 | 21539656 | 21539688 | 21566796 | 21566944 | 205287 | 10,11,2,6,1,5,2,2,2,0,1,0 | 51,45,69,69,43,28,32,39,38 | 0,4,2,4,5,1,0,1,2,2,3,2,0,2,0 | 42,39,32,29,40,28,16,28,10 | 225 | 149 | 0.0157442573073 | 0.783141262885945 | 0.115,0.139,0.019, | 0.0,0.064,0.04,0.08 | 0,01 |
| 95837 | ENSG00000167483 | NIBAN3 | chr19 | + | 17543523 | 17543631 | 17543316 | 17543433 | 17549443 | 17549527 | 95837 | 152,112,197,122,97,62,58, | 2,1,4,2,0,2,0,2,4,1,0,2 | 112,129,86,37,79,45,38,81 | 2,2,1,2,0,3,2,3,4,2,2,3,8,4,1 | 257 | 149 | 0.0157520018542 | 0.783223355677105 | 0.978,0.985,0.966, | 0.97,0.974,0.98,0.9 | 0,028 |
| 181415 | ENSG00000169682 | SPNS1 | chr16 | + | 28981747 | 28982056 | 28981469 | 28981615 | 28982355 | 28982545 | 181415 | 21,14,17,11,14,11,17,18,23 | 1,4,0,1,3,0,2,3,2,2,2,2 | 9,8,8,16,6,3,14,13,5,14,6,6 | 6,5,0,8,0,0,2,1,1,1,3,4,3,4,6 | 298 | 149 | 0.0157531856783 | 0.783223355677105 | 0.913,0.636,1.0,0.8 | 0.429,0.444,1.0,0.5 | 0,097 |
| 171836 | ENSG00000118855 | MFSD1 | chr3 | + | 158818660 | 158818699 | 158813964 | 158814067 | 158819648 | 158819747 | 171836 | 0,0,4,1,0,4,0,7,0,0,1,7 | 12,8,66,43,33,29,11,15,9,7 | 4,1,2,0,3,1,2,0,1,2,0,1,0,0,0 | 25,27,13,6,12,16,5,20,12,1 | 188 | 149 | 0.0157629354332 | 0.783378089792059 | 0.0,0.0,0.046,0.018 | 0.113,0.029,0.109, | 0,015 |
| 226977 | ENSG00000146872 | TLK2 | chr17 | + | 62574232 | 62574311 | 62573214 | 62573367 | 62576708 | 62576775 | 226977 | 5,0,9,0,0,2,0,2,1,0,0,0 | 125,60,69,24,35,35,45,25,6 | 4,0,3,1,0,0,0,6,8,1,3,3,0,5,1 | 39,53,82,70,31,16,15,29,33 | 228 | 149 | 0.0157635773048 | 0.783378089792059 | 0.025,0.0,0.079,0.0 | 0.063,0.0,0.023,0.0 | -0,02 |
| 110698 | ENSG00000034677 | RNF19A | chr8 | - | 100261541 | 100261755 | 100257059 | 100259246 | 100264670 | 100264785 | 110698 | 52,13,49,25,14,11,8,63,13, | 2,0,0,0,4,3,0,1,2,0,2,1 | 28,18,21,97,6,5,16,56,16,1 | 0,4,0,1,0,0,0,6,1,3,11,3,0,0 | 298 | 149 | 0.0157691602678 | 0.783474638411556 | 0.929,1.0,1.0,1.0,0 | 1.0,0.692,1.0,0.98, | -0,007 |
| 59659 | ENSG00000109323 | MANBA | chr4 | - | 102635864 | 102636007 | 102634787 | 102635045 | 102639712 | 102639857 | 59659 | 61,47,60,35,33,25,39,27,9, | 2,0,6,4,9,3,2,5,0,2,0,0 | 45,44,28,6,18,12,14,18,34, | 4,4,0,8,6,2,2,1,5,1,6,10,6,4 | 292 | 149 | 0.0157744847542 | 0.78355830328438 | 0.94,1.0,0.836,0.81 | 0.852,0.849,1.0,0.2 | 0,119 |
| 87926 | ENSG00000131508 | UBE2D2 | chr5 | + | 139562243 | 139562407 | 139561166 | 139561815 | 139600371 | 139600435 | 87926 | 19,10,49,50,13,23,17,31,5, | 42,11,68,35,26,26,41,22,41 | 32,49,12,17,21,8,7,17,12,1 | 38,20,20,12,11,12,5,38,6,3 | 298 | 149 | 0.0157790058812 | 0.783602033875532 | 0.184,0.312,0.265, | 0.296,0.551,0.231, | -0,097 |
| 226243 | ENSG00000092820 | EZR | chr6 | - | 158787107 | 158787194 | 158785308 | 158785583 | 158789287 | 158789371 | 226243 | 588,444,505,339,240,306,3 | 86,8,26,9,8,8,27,30,9,8,27, | 307,609,397,728,486,151,4 | 15,33,24,78,27,10,14,30,13 | 236 | 149 | 0.0157917082368 | 0.784051938988397 | 0.812,0.972,0.925, | 0.928,0.921,0.913, | 0,006 |
| 163489 | ENSG00000167766 | ZNF83 | chr19 | - | 52616717 | 52616921 | 52612377 | 52614797 | 52618935 | 52619062 | 163489 | 100,52,117,80,40,41,68,47 | 12,10,31,21,7,13,2,5,7,3,1, | 72,69,59,20,54,35,18,35,48 | 15,15,2,1,5,2,0,2,0,2,3,4,8, | 298 | 149 | 0.0157980383136 | 0.784185328476578 | 0.806,0.722,0.654, | 0.706,0.697,0.937, | -0,064 |
| 2756 | ENSG00000167555 | ZNF528 | chr19 | + | 52402428 | 52402545 | 52401946 | 52402028 | 52405906 | 52405931 | 2756 | 10,5,11,5,5,2,4,2,9,4,0,1 | 21,14,43,39,22,18,14,23,12 | 3,2,4,7,2,3,2,0,0,1,0,2,1,5,5 | 29,26,16,17,22,12,3,20,6,6 | 266 | 149 | 0.0158333393752 | 0.785694557819304 | 0.211,0.167,0.125, | 0.055,0.041,0.123, | 0,02 |
| 61566 | ENSG00000048740 | CELF2 | chr10 | + | 11321188 | 11321386 | 11314138 | 11314258 | 11325835 | 11325979 | 61566 | 103,79,80,63,34,49,35,43,7 | 8,3,3,3,0,2,3,2,0,4,0,1 | 57,50,42,23,25,23,18,25,18 | 2,3,2,1,0,0,1,6,0,2,0,0,0,0,0 | 298 | 149 | 0.0158371622982 | 0.785694557819304 | 0.866,0.929,0.93,0 | 0.934,0.893,0.913, | -0,006 |
| 57181 | ENSG00000134242 | PTPN22 | chr1 | - | 113819576 | 113819654 | 113813810 | 113814969 | 113825141 | 113825172 | 57181 | 266,157,424,441,157,126,1 | 9,2,12,6,0,0,0,0,1,2,1,4 | 256,286,145,37,104,96,52, | 0,6,0,1,0,0,1,2,4,0,0,2,2,0,3 | 227 | 149 | 0.0158393943548 | 0.785694557819304 | 0.951,0.981,0.959, | 1.0,0.969,1.0,0.96, | -0,005 |
| 251103 | ENSG00000096717 | SIRT1 | chr10 | + | 67906165 | 67906636 | 67891483 | 67891554 | 67906789 | 67906937 | 251103 | 8,2,2,3,0,1,0,0,2,0,3,5 | 9,9,48,30,17,8,9,5,11,2,2,1 | 1,1,5,0,1,0,1,1,3,1,3,1,0,2,0 | 18,35,7,1,10,3,15,13,0,14,1 | 298 | 149 | 0.0158578474284 | 0.786428654346074 | 0.308,0.1,0.02,0.04 | 0.027,0.014,0.263, | -0,011 |
| 119319 | ENSG00000183431 | SF3A3 | chr1 | - | 37978987 | 37979055 | 37978719 | 37978827 | 37979464 | 37979533 | 119319 | 83,64,122,93,46,29,36,35,6 | 2,1,2,1,0,0,0,0,0,0,0,3 | 72,47,39,21,31,26,32,52,24 | 2,8,1,5,0,0,3,3,1,0,0,0,2,1,0 | 217 | 149 | 0.0158652289356 | 0.786458253902869 | 0.966,0.978,0.977, | 0.961,0.801,0.964, | 0,038 |
| 73948 | ENSG00000174446 | SNAPC5 | chr15 | - | 66495329 | 66495419 | 66493467 | 66494552 | 66497641 | 66497750 | 73948 | 41,37,69,36,9,27,15,28,34, | 14,9,8,12,3,8,7,6,13,2,0,4 | 30,26,19,34,20,7,19,37,9,1 | 10,0,6,0,14,2,0,0,0,4,3,0,1, | 239 | 149 | 0.0158665690533 | 0.786458253902869 | 0.646,0.719,0.843, | 0.652,1.0,0.664,1.0 | -0,074 |
| 90594 | ENSG00000113761 | ZNF346 | chr5 | + | 177023159 | 177023234 | 177022734 | 177022913 | 177041125 | 177041229 | 90594 | 8,1,13,16,2,0,2,1,2,3,7,2 | 10,3,15,3,4,5,11,2,8,14,1,4 | 1,9,2,4,5,1,1,3,2,0,4,2,2,0,2 | 11,9,13,16,5,6,4,4,3,13,5,1 | 224 | 149 | 0.0158735067347 | 0.786458253902869 | 0.347,0.181,0.366, | 0.057,0.399,0.093, | 0,129 |
| 133686 | ENSG00000156273 | BACH1 | chr21 | + | 29305203 | 29305322 | 29299415 | 29299598 | 29321220 | 29321355 | 133686 | 97,46,82,33,11,15,28,16,5, | 2,5,5,2,2,0,3,0,0,0,0,3 | 54,91,33,41,41,27,27,55,33 | 3,4,1,2,8,0,2,4,0,4,1,2,2,0,0 | 268 | 149 | 0.0158759777545 | 0.786458253902869 | 0.964,0.836,0.901, | 0.909,0.927,0.948, | 0,007 |
| 233311 | ENSG00000215301 | DDX3X | chrX | + | 41346822 | 41347012 | 41346504 | 41346622 | 41347639 | 41347694 | 233311 | 23,7,26,30,13,10,10,14,6,1 | 3,1,7,4,3,0,2,0,0,1,0,2 | 31,31,15,52,16,17,2,19,13, | 8,5,1,31,0,6,5,8,1,6,10,0,3, | 298 | 149 | 0.0158780882376 | 0.786458253902869 | 0.793,0.778,0.65,0 | 0.66,0.756,0.882,0 | 0,156 |
| 229395 | ENSG00000162600 | OMA1 | chr1 | - | 58530600 | 58530754 | 58527260 | 58527335 | 58533952 | 58534060 | 229395 | 78,38,73,48,52,73,33,33,62 | 3,0,6,1,19,6,7,12,1,1,5,9 | 140,80,62,47,40,38,41,62,3 | 35,20,8,4,3,8,3,17,3,5,25,5 | 298 | 149 | 0.0158831571291 | 0.786458253902869 | 0.929,1.0,0.859,0.9 | 0.667,0.667,0.795, | 0,064 |
| 74683 | ENSG00000166912 | MTMR10 | chr15 | - | 30967919 | 30968010 | 30960880 | 30961073 | 30974313 | 30974456 | 74683 | 24,13,38,8,7,16,7,19,6,8,3, | 1,0,3,7,0,0,2,0,0,0,0,5 | 13,39,10,12,8,6,4,12,11,16 | 0,0,0,0,3,2,0,0,2,0,0,2,0,2,1 | 240 | 149 | 0.0158840224211 | 0.786458253902869 | 0.937,1.0,0.887,0.4 | 1.0,1.0,1.0,1.0,0.62 | -0,044 |
| 148662 | ENSG00000069345 | DNAJA2 | chr16 | - | 46960217 | 46960306 | 46959274 | 46959419 | 46964610 | 46964807 | 148662 | 5,0,14,6,2,6,2,12,3,1,2,3 | 52,45,126,91,47,32,33,44,2 | 14,9,5,13,4,3,1,7,6,10,12,3 | 72,87,33,24,75,27,38,52,23 | 238 | 149 | 0.0158893798024 | 0.78654257227469 | 0.057,0.0,0.065,0.0 | 0.109,0.061,0.087, | -0,025 |
| 218692 | ENSG00000024048 | UBR2 | chr6 | + | 42688229 | 42688386 | 42684793 | 42684871 | 42689568 | 42689670 | 218692 | 58,75,89,44,27,26,29,30,39 | 21,4,9,5,0,8,2,6,8,1,0,2 | 41,50,29,44,23,20,20,32,22 | 7,10,10,8,18,11,0,5,8,4,8,2 | 298 | 149 | 0.0158975076594 | 0.786763962069515 | 0.58,0.904,0.832,0 | 0.745,0.714,0.592, | 0,111 |
| 252840 | ENSG00000133639 | BTG1 | chr12 | - | 92145059 | 92145178 | 92144117 | 92144447 | 92145387 | 92145846 | 252840 | 44,8,13,11,4,1,0,1,5,5,4,6 | 622,314,1047,517,252,330 | 18,12,15,2,8,9,6,39,16,15,3 | 870,696,481,249,392,310,3 | 268 | 149 | 0.0159167544791 | 0.787461513305207 | 0.038,0.014,0.007, | 0.011,0.009,0.017, | -0,004 |
| 47521 | ENSG00000072736 | NFATC3 | chr16 | + | 68221195 | 68221284 | 68190767 | 68191775 | 68226349 | 68229258 | 47521 | 2,1,2,1,3,0,1,1,9,0,0,3 | 30,7,59,46,12,17,19,18,55, | 2,4,0,3,0,0,4,0,1,2,4,0,3,7,8 | 32,25,7,11,9,14,23,27,15,1 | 238 | 149 | 0.0159189198786 | 0.787461513305207 | 0.04,0.082,0.021,0 | 0.038,0.091,0.0,0.1 | -0,029 |
| 213577 | ENSG00000080345 | RIF1 | chr2 | + | 151469053 | 151469107 | 151468640 | 151468756 | 151469710 | 151469864 | 213577 | 0,2,12,2,3,2,0,2,2,0,0,0 | 7,10,67,45,29,41,21,29,10, | 4,6,2,1,10,1,0,2,2,0,8,1,2,2 | 35,37,25,22,31,25,7,44,35, | 203 | 149 | 0.0159429569549 | 0.788469340156269 | 0.0,0.128,0.116,0.0 | 0.077,0.106,0.055, | -0,019 |
| 100813 | ENSG00000097033 | SH3GLB1 | chr1 | + | 86728402 | 86728489 | 86724312 | 86724405 | 86734601 | 86734691 | 100813 | 5,3,8,5,0,4,5,5,0,0,3,0 | 33,34,91,66,53,27,39,25,16 | 3,10,4,10,9,6,3,4,5,2,10,2,1 | 59,68,40,19,44,20,18,34,27 | 236 | 149 | 0.0159644227631 | 0.789349569429078 | 0.087,0.053,0.053, | 0.031,0.085,0.059, | -0,024 |
| 50635 | ENSG00000152223 | EPG5 | chr18 | - | 45922340 | 45922600 | 45916008 | 45916206 | 45923267 | 45923387 | 50635 | 20,10,23,10,9,10,5,7,8,1,1, | 1,0,1,0,0,0,0,0,0,0,0,1 | 14,5,13,20,1,1,8,11,6,11,10 | 3,1,0,1,1,2,0,0,2,2,0,2,2,3,1 | 298 | 149 | 0.0159715705071 | 0.789521610430326 | 0.909,1.0,0.92,1.0, | 0.7,0.714,1.0,0.909 | 0,215 |
| 165439 | ENSG00000166913 | YWHAB | chr20 | + | 44902897 | 44903315 | 44901610 | 44901833 | 44903992 | 44904116 | 165439 | 1,2,12,2,2,4,0,2,0,0,0,3 | 74,56,102,117,130,102,47, | 6,12,4,7,0,0,8,4,0,3,7,8,1,1 | 111,108,39,47,43,36,43,46 | 298 | 149 | 0.015998073528 | 0.790650140873701 | 0.007,0.018,0.056, | 0.026,0.053,0.049, | -0,013 |
| 255551 | ENSG00000128585 | MKLN1 | chr7 | + | 131429071 | 131429145 | 131414644 | 131414710 | 131437784 | 131437997 | 255551 | 8,2,32,7,8,15,2,13,6,0,4,8 | 1,0,4,0,0,0,0,6,2,2,2,0 | 27,14,9,10,8,5,11,10,9,3,16 | 0,4,0,2,1,0,0,2,1,0,1,0,0,1,2 | 223 | 149 | 0.0160098549157 | 0.791050753755975 | 0.842,1.0,0.842,1.0 | 1.0,0.7,1.0,0.77,0.8 | -0,11 |
| 70179 | ENSG00000242372 | EIF6 | chr20 | - | 35280653 | 35280829 | 35279941 | 35280118 | 35284380 | 35284492 | 70179 | 22,21,89,63,40,15,30,25,39 | 2,0,11,3,1,0,0,4,1,0,0,3 | 41,34,10,3,21,22,12,18,15, | 0,1,5,0,2,0,0,2,0,1,1,4,0,0,0 | 298 | 149 | 0.0160250176097 | 0.791613540089914 | 0.846,1.0,0.802,0.9 | 1.0,0.944,0.5,1.0,0 | 0 |
| 102579 | ENSG00000198286 | CARD11 | chr7 | - | 2980519 | 2980580 | 2958499 | 2958631 | 3043666 | 3043845 | 102579 | 14,12,42,16,13,18,16,21,21 | 35,24,26,8,25,26,26,15,48, | 14,4,11,17,4,3,15,16,10,14 | 26,46,33,41,19,15,15,20,21 | 210 | 149 | 0.0160286009344 | 0.791613540089914 | 0.221,0.262,0.534, | 0.276,0.058,0.191, | 0,095 |
| 65752 | ENSG00000112339 | HBS1L | chr6 | - | 135037810 | 135038026 | 135002620 | 135002842 | 135039572 | 135039767 | 65752 | 8,7,22,25,14,13,7,11,5,1,3, | 31,7,80,58,19,38,21,32,12, | 5,22,16,1,12,11,19,13,12,1 | 19,35,19,10,23,8,19,35,14, | 298 | 149 | 0.0160639795012 | 0.793110602851454 | 0.114,0.333,0.121, | 0.116,0.239,0.296, | -0,058 |
| 47069 | ENSG00000116731 | PRDM2 | chr1 | + | 13769038 | 13769173 | 13749435 | 13749487 | 13773077 | 13773188 | 47069 | 5,7,22,17,3,2,2,8,2,1,0,8 | 63,135,103,56,35,44,40,52 | 10,19,19,9,20,5,2,12,14,10 | 64,127,71,116,98,35,47,71 | 284 | 149 | 0.0160662833348 | 0.793110602851454 | 0.04,0.026,0.101,0 | 0.076,0.073,0.123, | -0,027 |
| 180030 | ENSG00000122224 | LY9 | chr1 | + | 160824180 | 160824252 | 160823464 | 160823796 | 160827747 | 160828261 | 180030 | 26,13,24,18,29,9,17,12,17, | 7,4,1,2,6,4,0,0,1,0,5,12 | 29,9,15,9,6,12,19,24,12,30 | 12,3,2,4,3,5,3,11,1,3,11,29 | 221 | 149 | 0.0160774110914 | 0.793341752150651 | 0.715,0.687,0.942, | 0.62,0.669,0.835,0 | 0,102 |
| 248422 | ENSG00000149182 | ARFGAP2 | chr11 | - | 47175014 | 47175052 | 47173758 | 47173840 | 47175181 | 47175413 | 248422 | 11,4,25,18,17,18,9,11,7,5,3 | 2,2,1,3,0,1,5,2,0,0,0,0 | 14,27,9,7,8,4,4,20,7,15,8,1 | 0,0,2,0,0,0,2,0,0,0,0,1,2,0,0 | 187 | 149 | 0.0160783377993 | 0.793341752150651 | 0.814,0.614,0.952, | 1.0,1.0,0.782,1.0,1 | -0,07 |
| 253354 | ENSG00000089022 | MAPKAPK5 | chr12 | + | 111883582 | 111883768 | 111880446 | 111880527 | 111885915 | 111886036 | 253354 | 36,34,63,45,45,36,20,41,16 | 5,7,6,6,1,1,5,1,6,0,2,4 | 37,32,20,13,28,26,21,32,11 | 8,3,2,5,9,6,6,5,2,0,5,11,7,6 | 298 | 149 | 0.0160880287501 | 0.7936379821024 | 0.783,0.708,0.84,0 | 0.698,0.842,0.833, | 0,096 |
| 244373 | ENSG00000134910 | STT3A | chr11 | + | 125609709 | 125609780 | 125609433 | 125609589 | 125611427 | 125611519 | 244373 | 0,1,3,7,6,9,4,4,0,2,3,4 | 2,1,17,21,6,5,7,11,13,3,4,9 | 9,13,5,2,3,1,1,2,8,3,6,2,3,8 | 6,6,3,0,0,1,3,9,3,3,4,15,1,8 | 220 | 149 | 0.0160946465454 | 0.793782509306367 | 0.0,0.404,0.107,0.1 | 0.504,0.595,0.53,1 | -0,217 |
| 94220 | ENSG00000234745 | HLA-B | chr6 | - | 31355106 | 31355223 | 31354632 | 31354665 | 31355316 | 31355592 | 94220 | 5077,4965,8805,7752,3210 | 18,0,57,44,1,0,0,2,1,0,2,0 | 5868,6727,3114,5033,3483 | 22,24,1,12,3,3,34,103,61,4 | 266 | 149 | 0.0161049772167 | 0.793953533721905 | 0.994,1.0,0.989,0.9 | 0.993,0.994,0.999, | 0,009 |
| 131945 | ENSG00000135297 | MTO1 | chr6 | + | 73471459 | 73471557 | 73466488 | 73466606 | 73473364 | 73473654 | 131945 | 23,17,12,13,12,7,4,11,16,1 | 39,48,62,40,28,32,24,27,39 | 11,6,16,5,15,2,7,2,8,10,5,8 | 47,62,45,35,32,14,22,19,24 | 247 | 149 | 0.0161076066016 | 0.793953533721905 | 0.262,0.176,0.105, | 0.124,0.055,0.177, | 0,039 |
| 195739 | ENSG00000159377 | PSMB4 | chr1 | + | 151400691 | 151400845 | 151400441 | 151400588 | 151401238 | 151401355 | 195739 | 36,47,84,72,50,49,47,55,93 | 3,2,4,9,7,5,1,0,0,0,2,2 | 66,56,70,21,33,27,27,42,23 | 3,9,2,15,1,1,4,9,1,0,8,8,5,2 | 298 | 149 | 0.0161163153931 | 0.793953533721905 | 0.857,0.922,0.913, | 0.917,0.757,0.946, | 0,07 |
| 182600 | ENSG00000227232 | WASH7P | chr1 | - | 17525 | 17742 | 16857 | 17055 | 17914 | 18061 | 182600 | 3,5,10,3,3,7,7,7,0,0,2,10 | 0,3,0,2,5,0,5,4,2,1,3,5 | 2,4,0,2,0,1,2,9,0,1,4,6,1,3,1 | 3,2,3,4,1,4,3,0,1,1,2,4,3,2,1 | 298 | 149 | 0.0161176631895 | 0.793953533721905 | 1.0,0.455,1.0,0.429 | 0.25,0.5,0.0,0.2,0.0 | 0,201 |
| 213330 | ENSG00000135426 | TESPA1 | chr12 | - | 54956753 | 54956804 | 54950353 | 54950390 | 54961167 | 54961267 | 213330 | 14,2,3,6,2,3,4,4,1,1,0,2 | 25,14,15,11,5,8,12,14,59,1 | 1,2,2,0,0,0,0,1,2,3,1,3,0,2,3 | 9,12,14,1,4,2,6,7,0,9,2,8,12 | 200 | 149 | 0.0161191939012 | 0.793953533721905 | 0.294,0.096,0.13,0 | 0.076,0.11,0.096,0 | -0,03 |
| 123621 | ENSG00000133943 | DGLUCY | chr14 | + | 91147780 | 91147858 | 91114685 | 91114994 | 91157638 | 91157690 | 123621 | 6,5,3,17,10,4,4,6,1,3,4,4 | 6,3,2,11,1,3,5,18,6,3,3,4 | 3,3,8,1,1,1,3,7,4,2,2,8,1,3,4 | 5,7,2,7,1,2,7,19,3,6,4,9,7,4 | 227 | 149 | 0.0161202472802 | 0.793953533721905 | 0.396,0.522,0.496, | 0.283,0.22,0.724,0 | 0,149 |
| 94292 | ENSG00000104343 | UBE2W | chr8 | - | 73870268 | 73870301 | 73830380 | 73830472 | 73878807 | 73878862 | 94292 | 6,4,6,0,1,4,2,0,0,2,0,2 | 16,21,16,14,13,5,0,5,5,9,3, | 1,2,0,2,3,2,1,0,2,1,0,0,2,0,3 | 12,20,5,5,10,10,0,5,14,5,10 | 182 | 149 | 0.0162057515859 | 0.797982182471938 | 0.235,0.135,0.235, | 0.064,0.076,0.0,0.2 | 0,04 |
| 36490 | ENSG00000151612 | ZNF827 | chr4 | - | 145870244 | 145870478 | 145849321 | 145849561 | 145885677 | 145885769 | 36490 | 22,19,41,25,7,19,17,20,7,6 | 3,5,8,1,1,5,2,2,0,2,3,1 | 23,34,13,52,23,6,22,27,33, | 4,12,2,2,3,0,0,0,1,2,2,0,0,1 | 298 | 149 | 0.0162132962748 | 0.798171082003998 | 0.786,0.655,0.719, | 0.742,0.586,0.765, | -0,093 |
| 93554 | ENSG00000065427 | KARS1 | chr16 | - | 75642752 | 75642835 | 75641563 | 75641723 | 75644282 | 75644462 | 93554 | 3,3,2,5,0,1,0,6,3,1,2,1 | 12,8,28,14,21,8,9,11,8,7,5, | 0,2,0,0,0,0,4,5,0,0,5,1,1,2,0 | 14,11,6,6,9,11,15,15,8,12,1 | 232 | 149 | 0.0162535338531 | 0.799851473322711 | 0.138,0.194,0.044, | 0.0,0.105,0.0,0.0,0 | 0,06 |
| 149703 | ENSG00000214013 | GANC | chr15 | + | 42329361 | 42329449 | 42327362 | 42327442 | 42330575 | 42330672 | 149703 | 70,64,64,42,29,27,33,22,23 | 6,2,4,0,9,2,0,2,0,0,0,0 | 32,31,29,48,28,19,30,32,43 | 2,5,3,5,3,0,0,2,1,2,0,2,1,3,3 | 237 | 149 | 0.016254862656 | 0.799851473322711 | 0.88,0.953,0.91,1.0 | 0.91,0.796,0.859,0 | 0,014 |
| 82130 | ENSG00000184922 | FMNL1 | chr17 | + | 45245631 | 45245733 | 45245252 | 45245416 | 45245877 | 45245973 | 82130 | 60,51,86,76,84,67,54,53,58 | 9,4,10,5,8,9,8,1,1,3,2,9 | 49,56,30,58,41,26,49,70,51 | 4,12,3,15,10,1,17,21,8,3,7, | 251 | 149 | 0.0162620630347 | 0.800022877490632 | 0.798,0.883,0.836, | 0.879,0.735,0.856, | 0,053 |
| 235760 | ENSG00000197217 | ENTPD4 | chr8 | - | 23441392 | 23441487 | 23439748 | 23439915 | 23441592 | 23441723 | 235760 | 13,6,14,4,1,9,1,5,3,2,2,6 | 20,22,34,18,6,15,12,32,7,7 | 13,19,6,15,18,4,2,2,5,5,12, | 16,16,10,14,18,6,6,20,17,2 | 244 | 149 | 0.0162687364915 | 0.800168286974871 | 0.284,0.143,0.201, | 0.332,0.42,0.268,0 | -0,089 |
| 162158 | ENSG00000149418 | ST14 | chr11 | + | 130192409 | 130192531 | 130190453 | 130190694 | 130194148 | 130194288 | 162158 | 0,0,3,3,3,14,3,1,0,0,0,1 | 18,40,33,46,41,37,40,34,47 | 0,0,0,0,0,0,0,0,0,0,9,2,3,0 | 28,25,34,43,43,24,28,34,32 | 271 | 149 | 0.0162827389802 | 0.800674021806586 | 0.0,0.0,0.048,0.035 | 0.0,0.0,0.0,0.0,0.0, | 0,022 |
| 170816 | ENSG00000131374 | TBC1D5 | chr3 | - | 17562018 | 17562140 | 17508473 | 17508605 | 17623848 | 17623916 | 170816 | 4,2,16,4,2,13,4,3,5,2,0,4 | 29,31,71,38,28,29,31,41,21 | 14,5,4,12,2,5,3,7,5,8,10,10 | 41,41,11,27,26,25,15,12,24 | 271 | 149 | 0.0162942761778 | 0.801058327095494 | 0.07,0.034,0.11,0.0 | 0.158,0.063,0.167, | -0,045 |
| 47459 | ENSG00000115935 | WIPF1 | chr2 | - | 174630381 | 174630625 | 174629665 | 174629868 | 174634517 | 174634579 | 47459 | 12,2,29,17,4,3,3,17,5,4,4,9 | 0,0,16,8,0,1,0,0,1,0,0,0 | 16,44,5,10,18,4,4,5,14,6,14 | 2,0,2,0,0,1,4,1,3,0,0,2,1,2,0 | 298 | 149 | 0.0163017057711 | 0.801240565156343 | 1.0,1.0,0.475,0.515 | 0.8,1.0,0.556,1.0,1 | 0,054 |
| 6793 | ENSG00000173418 | NAA20 | chr20 | + | 20032507 | 20032653 | 20026783 | 20026919 | 20033101 | 20033629 | 6793 | 33,21,51,51,26,15,15,13,32 | 7,2,15,8,3,3,4,13,0,0,3,4 | 20,23,30,5,14,20,5,10,8,26 | 8,13,3,2,4,4,7,10,2,4,10,10 | 295 | 149 | 0.016310138705 | 0.801472023656588 | 0.704,0.841,0.632, | 0.558,0.472,0.835, | 0,14 |
| 84224 | ENSG00000183486 | MX2 | chr21 | + | 41363268 | 41363389 | 41362006 | 41362055 | 41376835 | 41377155 | 84224 | 2,0,4,0,2,0,0,6,0,0,0,4 | 14,6,23,20,4,15,3,5,5,3,7,2 | 5,0,2,3,0,2,0,4,2,2,10,1,5,3 | 8,7,11,8,2,4,2,5,7,0,18,8,12 | 270 | 149 | 0.0163164400358 | 0.801595895329597 | 0.073,0.0,0.088,0.0 | 0.256,0.0,0.091,0.1 | -0,083 |
| 189958 | ENSG00000157796 | WDR19 | chr4 | + | 39218267 | 39218443 | 39217982 | 39218105 | 39224883 | 39225033 | 189958 | 5,1,3,2,0,0,1,1,2,2,0,1 | 23,16,37,30,8,25,15,10,15, | 3,10,1,2,0,0,4,3,0,0,4,6,0,2 | 13,15,13,8,8,2,8,23,9,15,13 | 298 | 149 | 0.0163232696353 | 0.801595895329597 | 0.098,0.03,0.039,0 | 0.103,0.25,0.037,0 | 0,04 |
| 144434 | ENSG00000065559 | MAP2K4 | chr17 | + | 12125620 | 12125737 | 12125293 | 12125371 | 12129138 | 12129287 | 144434 | 1,3,3,5,5,0,2,0,2,1,0,3 | 8,1,26,14,15,2,9,10,0,0,1,6 | 5,12,8,4,3,4,1,4,5,4,2,1,1,6 | 10,11,7,3,9,3,3,10,1,11,13, | 266 | 149 | 0.0163238325763 | 0.801595895329597 | 0.065,0.627,0.061, | 0.219,0.379,0.39,0 | 0,023 |
| 160209 | ENSG00000150593 | PDCD4 | chr10 | + | 110875965 | 110876237 | 110871794 | 110872018 | 110881274 | 110881535 | 160209 | 23,9,104,42,22,13,6,17,7,0 | 4,1,2,3,0,0,1,2,1,1,0,2 | 27,39,8,12,13,23,6,12,10,2 | 4,10,0,4,8,2,0,1,0,0,0,3,2,0 | 298 | 149 | 0.0163597961072 | 0.803178666959116 | 0.742,0.818,0.963, | 0.771,0.661,1.0,0.6 | 0,004 |
| 252613 | ENSG00000133316 | WDR74 | chr11 | - | 62833617 | 62833673 | 62832910 | 62833131 | 62833791 | 62833937 | 252613 | 10,16,15,14,3,13,9,12,20,6 | 8,3,12,4,1,2,1,5,2,2,4,7 | 11,13,5,3,11,2,10,9,1,13,5, | 4,4,5,13,0,0,16,9,6,7,15,5,1 | 205 | 149 | 0.0163688848742 | 0.803325146109696 | 0.476,0.795,0.476, | 0.667,0.703,0.421, | 0,143 |
| 238319 | ENSG00000147457 | CHMP7 | chr8 | + | 23247935 | 23248197 | 23246665 | 23246994 | 23249209 | 23249381 | 238319 | 0,1,5,2,0,4,0,3,0,2,1,3 | 14,3,27,15,13,16,11,19,29, | 3,4,4,0,0,0,0,1,0,0,0,3,0,0,0 | 7,22,14,5,16,8,7,10,10,23,1 | 298 | 149 | 0.0163702444854 | 0.803325146109696 | 0.0,0.143,0.085,0.0 | 0.176,0.083,0.125, | 0,027 |
| 243439 | ENSG00000174106 | LEMD3 | chr12 | + | 65240655 | 65240809 | 65240135 | 65240238 | 65240908 | 65241087 | 243439 | 6,5,18,4,3,5,7,11,2,0,5,6 | 10,16,89,68,23,33,35,63,11 | 5,5,2,7,6,8,3,9,3,3,11,0,2,2 | 62,63,20,19,33,20,21,29,11 | 298 | 149 | 0.0163803941036 | 0.803639982519246 | 0.231,0.135,0.092, | 0.039,0.038,0.048, | 0,03 |
| 142279 | ENSG00000125970 | RALY | chr20 | + | 34073818 | 34074147 | 34073562 | 34073635 | 34075873 | 34076229 | 142279 | 31,24,38,26,17,7,12,11,24, | 7,6,13,4,16,8,6,15,3,1,9,5 | 14,15,31,28,3,6,1,5,5,14,11 | 10,8,6,10,6,11,6,18,6,10,4, | 298 | 149 | 0.0164469002031 | 0.806700024473865 | 0.689,0.667,0.594, | 0.412,0.484,0.721, | 0,166 |
| 212289 | ENSG00000170006 | TMEM154 | chr4 | - | 152647173 | 152647327 | 152644414 | 152644442 | 152652537 | 152652576 | 212289 | 0,4,2,5,3,4,1,7,7,1,0,0 | 59,37,159,144,62,43,29,36 | 2,0,4,10,1,0,1,4,6,6,2,9,1,1 | 36,59,61,10,38,49,18,29,41 | 298 | 149 | 0.0164502623108 | 0.806700024473865 | 0.0,0.051,0.006,0.0 | 0.027,0.0,0.032,0.3 | -0,024 |
| 47458 | ENSG00000115935 | WIPF1 | chr2 | - | 174630381 | 174630560 | 174629665 | 174629868 | 174634517 | 174634605 | 47458 | 16,3,31,20,5,3,3,16,5,4,5,8 | 0,0,16,8,0,1,0,0,1,0,0,0 | 16,45,5,12,18,4,2,5,17,6,14 | 2,0,2,0,0,1,4,1,3,0,0,2,1,2,0 | 298 | 149 | 0.0164686691133 | 0.807418706588764 | 1.0,1.0,0.492,0.556 | 0.8,1.0,0.556,1.0,1 | 0,062 |
| 68946 | ENSG00000104643 | MTMR9 | chr8 | + | 11297899 | 11297951 | 11295193 | 11295302 | 11300022 | 11300148 | 68946 | 0,2,4,1,2,4,4,3,0,0,1,1 | 14,4,22,9,7,8,18,6,7,9,2,3 | 1,3,1,0,0,0,2,2,0,2,2,0,1,5,0 | 17,21,7,2,12,7,7,10,10,15,9 | 201 | 149 | 0.0164836730204 | 0.807945127706571 | 0.0,0.27,0.119,0.07 | 0.042,0.096,0.096, | 0,083 |
| 82827 | ENSG00000160695 | VPS11 | chr11 | + | 119071595 | 119071843 | 119070233 | 119070397 | 119073197 | 119073399 | 82827 | 39,24,88,55,48,26,32,31,29 | 4,2,16,8,2,14,4,4,1,1,4,3 | 31,55,18,25,41,26,7,6,6,35 | 16,17,4,4,6,1,2,8,8,4,13,6,1 | 298 | 149 | 0.0164869140639 | 0.807945127706571 | 0.83,0.857,0.733,0 | 0.492,0.618,0.692, | 0,142 |
| 212914 | ENSG00000152689 | RASGRP3 | chr2 | + | 33558671 | 33559025 | 33558210 | 33558336 | 33562728 | 33564750 | 212914 | 101,62,226,97,94,103,105, | 1,0,13,8,2,12,4,9,4,1,0,0 | 120,144,64,22,43,24,48,17 | 7,5,10,2,3,6,3,14,3,2,5,4,2, | 298 | 149 | 0.0164953391076 | 0.808139639898504 | 0.981,1.0,0.897,0.8 | 0.896,0.935,0.762, | 0,077 |
| 138378 | ENSG00000157540 | DYRK1A | chr21 | + | 37488246 | 37488351 | 37486466 | 37486614 | 37490174 | 37490461 | 138378 | 6,6,11,10,2,5,1,11,4,4,1,4 | 66,50,199,91,61,84,55,85,5 | 7,16,3,3,2,2,0,6,1,9,4,3,0,2 | 137,148,83,79,52,59,63,85 | 254 | 149 | 0.0165000153214 | 0.808139639898504 | 0.051,0.066,0.031, | 0.029,0.06,0.021,0 | 0,019 |
| 16799 | ENSG00000166123 | GPT2 | chr16 | + | 46885466 | 46885577 | 46884693 | 46884958 | 46897647 | 46897737 | 16799 | 13,17,11,1,6,1,7,5,2,1,4,5 | 19,25,26,5,12,14,17,22,8,1 | 6,4,3,5,5,4,6,7,4,10,18,2,18 | 11,20,11,5,2,1,5,31,6,12,21 | 260 | 149 | 0.0165021475408 | 0.808139639898504 | 0.282,0.28,0.195,0 | 0.238,0.103,0.135, | -0,164 |

| | | | | | | | | | | | | | | | | | | | | | | |
|---|---|---|---|---|---|---|---|---|---|---|---|---|---|---|---|---|---|---|---|---|---|---|
| 68913 | ENSG00000198039 | ZNF273 | chr7 | + | 64917029 | 64917118 | 64903287 | 64903419 | 64917580 | 64917707 | 68913 | 13,5,26,18,10,8,6,8,9,3,2,6 | 1,5,9,4,2,3,2,0,4,6,0,3 | 15,18,7,4,12,5,6,6,9,10,17, | 2,0,1,0,5,1,0,3,1,4,1,2,2,1, | 238 | 149 | 0.0165215703518 | 0.808906758050106 | 0.891,0.385,0.644, | 0.824,1.0,0.814,1. | -0,13 |
| 139644 | ENSG00000124243 | BCAS4 | chr20 | + | 50818210 | 50818365 | 50794893 | 50795173 | 50830278 | 50830380 | 139644 | 1,1,7,2,5,6,3,7,9,9,3,7 | 2,0,0,0,1,2,3,4,0,0,2,0 | 2,4,4,1,1,2,1,3,0,2,10,8,3,4 | 0,0,0,0,1,0,0,0,1,2,0,2,0,0, | 298 | 149 | 0.0165367918819 | 0.809467876627978 | 0.2,1.0,1.0,1.0,0.7 | 1.0,1.0,1.0,1.0,0.3 | -0,097 |
| 217876 | ENSG00000155827 | RNF20 | chr9 | + | 101544778 | 101544885 | 101540792 | 101540975 | 101546819 | 101546966 | 217876 | 30,20,32,15,8,10,15,10,28, | 1,2,6,0,0,1,0,0,1,0,4,3 | 27,17,23,24,14,16,7,16,7,3 | 1,2,2,4,4,2,1,2,0,6,4,10,1,4 | 256 | 149 | 0.0165586261181 | 0.810352355167071 | 0.946,0.853,0.756, | 0.94,0.832,0.87,0. | 0,114 |
| 222686 | ENSG00000167483 | NIBAN3 | chr19 | + | 17546662 | 17546797 | 17543523 | 17543631 | 17549443 | 17549527 | 222686 | 264,154,228,161,125,77,1 | 27,3,9,1,7,3,3,2,9,6,10,2 | 143,176,180,88,176,98,53, | 1,7,6,3,5,0,0,7,3,3,2,8,17,9 | 284 | 149 | 0.0165839954925 | 0.811409396191468 | 0.837,0.964,0.93,0 | 0.987,0.93,0.94,0. | -0,03 |
| 110980 | ENSG00000047249 | ATP6V1H | chr8 | - | 53771988 | 53772167 | 53769617 | 53769743 | 53801798 | 53801896 | 110980 | 8,6,54,21,15,13,8,15,6,2,6, | 2,0,7,3,0,1,0,0,0,2,0,0 | 21,38,5,8,23,10,5,7,15,12, | 2,3,4,0,0,1,0,2,0,0,0,4,1,2, | 298 | 149 | 0.0165898318567 | 0.811492529965722 | 0.667,1.0,0.794,0. | 0.84,0.864,0.385,1 | 0,009 |
| 78020 | ENSG00000138029 | HADHB | chr2 | + | 26285406 | 26285571 | 26284882 | 26284957 | 26289917 | 26290412 | 78020 | 119,78,200,154,107,47,67, | 4,3,15,10,4,12,7,7,6,0,0,1 | 165,133,104,31,84,77,64,9 | 15,23,14,4,19,5,9,5,6,2,8,9 | 298 | 149 | 0.0165932352885 | 0.811492529965722 | 0.937,0.929,0.87,0 | 0.846,0.743,0.788, | 0,062 |
| 164688 | ENSG00000005436 | GCFC2 | chr2 | - | 75674478 | 75674550 | 75673443 | 75673520 | 75680192 | 75680314 | 164688 | 1,1,8,2,0,1,1,3,0,0,0,0 | 17,7,25,27,10,2,4,12,10,3, | 3,4,1,0,1,3,2,2,3,1,3,0,0,1, | 21,9,5,5,8,8,7,3,1,8,4,33,4, | 221 | 149 | 0.0166079160081 | 0.812025981449198 | 0.038,0.088,0.177, | 0.088,0.231,0.119, | -0,077 |
| 44076 | ENSG00000111731 | C2CD5 | chr12 | - | 22472285 | 22472347 | 22471966 | 22472065 | 22472743 | 22472807 | 44076 | 26,18,33,16,28,21,11,17,8, | 6,0,5,2,8,3,4,8,3,1,1,3 | 28,21,16,13,5,9,10,17,11,1 | 1,13,0,22,11,0,3,10,19,5,11 | 211 | 149 | 0.0166176954734 | 0.812319603551069 | 0.754,1.0,0.823,0. | 0.952,0.533,1.0,0. | 0,135 |
| 86761 | ENSG00000123600 | METTL8 | chr2 | - | 171392042 | 171392197 | 171360421 | 171360513 | 171414602 | 171414726 | 86761 | 14,20,24,28,25,15,14,9,11, | 0,0,1,0,2,5,2,0,2,1,0,3 | 6,7,4,11,7,1,8,10,13,34,5,1 | 2,1,0,0,3,0,0,0,2,3,0,0,4,0, | 298 | 149 | 0.0166466203586 | 0.813330554333839 | 1.0,1.0,0.923,1.0,0 | 0.6,0.778,1.0,1.0,0 | -0,017 |
| 232254 | ENSG00000196776 | CD47 | chr3 | - | 108054252 | 108054715 | 108051938 | 108051970 | 108057476 | 108057569 | 232254 | 6,2,15,4,3,0,7,4,5,0,0,5 | 125,111,377,215,127,145, | 9,5,5,17,3,2,11,3,5,5,5,0,8, | 156,155,104,59,68,83,82,1 | 298 | 149 | 0.0166468815599 | 0.813330554333839 | 0.023,0.009,0.02,0 | 0.028,0.016,0.023, | -0,014 |
| 167087 | ENSG00000174652 | ZNF266 | chr19 | - | 9419050 | 9419131 | 9418504 | 9418631 | 9419216 | 9419299 | 167087 | 2,0,15,8,1,2,5,6,6,2,0,0 | 3,8,98,86,25,11,62,61,26,5 | 9,8,1,5,6,4,0,3,2,14,0,0,13, | 69,29,16,38,18,12,17,50,18 | 230 | 149 | 0.016649713203 | 0.813330554333839 | 0.302,0.0,0.09,0.0 | 0.078,0.152,0.039, | -0,013 |
| 102320 | ENSG00000106245 | BUD31 | chr7 | + | 99410036 | 99410169 | 99408968 | 99409004 | 99411063 | 99411186 | 102320 | 146,90,166,135,53,48,58,5 | 4,4,2,1,0,2,1,2,2,0,0,1 | 73,117,87,56,92,74,24,54, | 2,0,1,0,2,2,0,1,0,0,0,2,0,2,1 | 282 | 149 | 0.0166547674735 | 0.813392842770338 | 0.951,0.922,0.978, | 0.951,1.0,0.979,1. | -0,015 |
| 133904 | ENSG00000228315 | GUSBP11 | chr22 | - | 23700725 | 23700831 | 23694118 | 23695517 | 23705428 | 23705541 | 133904 | 3,5,11,7,3,1,12,11,0,3,4,5 | 12,3,6,1,5,4,7,13,7,0,3,10 | 2,6,2,2,4,3,3,0,1,4,2,1,8,3, | 12,10,9,6,0,1,2,3,12,4,9,8,1 | 255 | 149 | 0.0166733790339 | 0.81411706961101 | 0.127,0.493,0.517, | 0.089,0.26,0.115,0 | 0,151 |
| 58634 | ENSG00000263956 | NBPF11 | chr1 | - | 148137710 | 148137809 | 148135671 | 148135813 | 148143414 | 148143686 | 58634 | 11,5,27,15,12,8,7,9,6,5,3,3 | 2,2,1,4,5,0,0,0,0,1,0,0 | 11,15,2,3,15,3,7,15,4,10,4, | 2,0,0,0,0,1,0,0,0,1,0,0,0,2, | 248 | 149 | 0.0166784447202 | 0.814179708680736 | 0.768,0.6,0.942,0. | 0.768,1.0,1.0,1.0,1 | -0,07 |
| 251483 | ENSG00000138134 | STAMBPL1 | chr10 | + | 88893870 | 88893944 | 88880223 | 88880638 | 88901655 | 88901738 | 251483 | 6,2,16,5,3,6,4,8,1,0,1,2 | 35,15,46,31,22,23,26,16,4 | 6,10,10,1,3,2,0,5,0,6,6,4,2, | 16,30,16,17,19,10,11,19,1 | 223 | 149 | 0.0166862426962 | 0.814375669330119 | 0.103,0.082,0.189, | 0.2,0.182,0.295,0. | -0,06 |
| 232949 | ENSG00000124275 | MTRR | chr5 | + | 7890276 | 7890412 | 7889094 | 7889275 | 7891371 | 7891414 | 232949 | 12,7,11,13,0,2,6,17,0,5,0,3 | 20,27,19,23,17,13,15,16,3 | 10,11,14,12,6,7,6,11,4,14, | 18,16,19,8,9,11,6,19,5,28,1 | 285 | 149 | 0.0167131998057 | 0.815506394781369 | 0.239,0.119,0.232, | 0.225,0.264,0.278, | -0,087 |
| 182823 | ENSG00000196187 | TMEM63A | chr1 | - | 225860859 | 225860986 | 225859195 | 225859349 | 225862217 | 225862351 | 182823 | 130,112,29,31,46,30,32,47 | 3,1,5,5,3,1,5,0,5,1,0,2 | 31,34,23,73,37,22,57,83,1 | 4,7,2,4,3,0,11,8,0,0,2,1,3,0 | 276 | 149 | 0.016739306426 | 0.816595118171896 | 0.959,0.984,0.758, | 0.807,0.724,0.861, | 0,037 |
| 39568 | ENSG00000266338 | NBPF15 | chr1 | - | 144450771 | 144450870 | 144440140 | 144440295 | 144456536 | 144456805 | 39568 | 9,5,21,11,10,10,7,20,3,1,2, | 1,1,5,1,2,0,0,0,2,0,0,3 | 18,6,3,3,4,2,2,11,7,10,3,4, | 0,0,0,1,0,0,0,0,0,0,0,2,0, | 248 | 149 | 0.0167809493275 | 0.818441084230037 | 0.844,0.75,0.716,0 | 1.0,1.0,1.0,0.643,1 | -0,101 |
| 206638 | ENSG00000152061 | RABGAP1L | chr1 | + | 174252739 | 174252773 | 174252479 | 174252590 | 174272413 | 174272480 | 206638 | 17,11,16,2,4,1,3,5,7,0,0,0 | 153,106,352,204,73,89,98, | 15,23,17,4,5,1,7,1,15,10,5, | 175,189,222,132,120,89,3 | 183 | 149 | 0.0167937776955 | 0.818881188758532 | 0.083,0.078,0.036, | 0.065,0.09,0.059,0 | -0,022 |
| 231778 | ENSG00000164162 | ANAPC10 | chr4 | - | 145033041 | 145033371 | 144995455 | 144995603 | 145064571 | 145064692 | 231778 | 2,1,7,3,0,0,0,2,2,1,0,2 | 12,10,38,20,12,15,7,17,24, | 0,1,0,2,0,0,0,0,1,1,3,0,0,1 | 14,17,16,8,19,10,9,15,8,12 | 298 | 149 | 0.0167984135583 | 0.818921709754579 | 0.077,0.048,0.084, | 0.0,0.029,0.0,0.11 | 0,03 |
| 163614 | ENSG00000105875 | WDR91 | chr7 | - | 135197043 | 135197123 | 135196143 | 135196337 | 135197992 | 135201343 | 163614 | 8,4,11,7,15,5,2,3,2,3,5,7 | 14,14,44,37,36,35,14,15,3 | 8,18,7,2,3,1,6,7,5,7,1,0,7,9 | 21,28,8,6,7,3,21,25,2,11,2 | 229 | 149 | 0.0168040821432 | 0.818945417224294 | 0.271,0.157,0.14,0 | 0.199,0.295,0.363, | -0,085 |
| 16684 | ENSG00000159593 | NAE1 | chr16 | - | 66827934 | 66828077 | 66826676 | 66826780 | 66830846 | 66830976 | 16684 | 6,0,16,5,0,0,0,1,1,0,0,0 | 7,5,31,27,8,13,6,17,7,3,12, | 1,1,2,2,4,4,0,3,0,0,1,0,0,2,1 | 13,18,11,12,6,6,5,11,13,16 | 292 | 149 | 0.0168065097866 | 0.818945417224294 | 0.304,0.0,0.208,0. | 0.038,0.028,0.085, | -0,005 |
| 169392 | ENSG00000124535 | WRNIP1 | chr6 | + | 2772937 | 2773141 | 2770194 | 2770361 | 2779262 | 2779492 | 169392 | 2,1,4,2,0,1,0,1,2,0,0,2 | 11,2,53,24,29,12,17,4,8,9, | 4,1,4,0,1,1,4,6,0,1,3,2,3,0, | 27,19,5,4,8,11,6,14,4,23,2 | 298 | 149 | 0.0168244879945 | 0.819635893061165 | 0.083,0.2,0.036,0. | 0.069,0.026,0.286, | -0,02 |
| 225685 | ENSG00000136280 | CCM2 | chr7 | + | 45070388 | 45070572 | 45069825 | 45069961 | 45072725 | 45073288 | 225685 | 4,2,3,0,0,2,4,2,0,0,0,2 | 21,31,29,51,12,9,19,20,11, | 11,10,0,1,0,0,0,2,4,9,0,2,1, | 37,26,7,6,19,25,6,17,5,30, | 298 | 149 | 0.0168306194184 | 0.819749049115558 | 0.087,0.031,0.049, | 0.129,0.161,0.0,0. | -0,022 |
| 215803 | ENSG00000119812 | FAM98A | chr2 | - | 33595219 | 33595330 | 33592079 | 33592214 | 33595488 | 33595637 | 215803 | 6,4,2,2,0,0,2,1,2,0,2,0 | 8,4,13,13,14,7,2,3,9,2,0,2 | 3,4,0,0,1,3,1,6,2,0,1,1,0,0,1 | 9,16,2,5,14,8,14,11,4,6,3,7 | 260 | 149 | 0.0168460886249 | 0.820316855390464 | 0.301,0.364,0.081, | 0.16,0.125,0.0,0.0, | 0,125 |
| 73051 | ENSG00000125633 | CCDC93 | chr2 | - | 118000820 | 118000932 | 117996263 | 117996362 | 118006721 | 118006816 | 73051 | 64,37,95,63,40,52,45,51,3 | 2,4,4,2,0,3,1,7,0,1,0,0 | 71,64,38,56,58,36,31,36,2 | 5,4,2,13,1,0,6,2,4,2,4,5,0,4 | 261 | 149 | 0.0168590608106 | 0.820702354698287 | 0.948,0.841,0.931, | 0.89,0.901,0.916,0 | 0,042 |
| 23872 | ENSG00000173575 | CHD2 | chr15 | + | 92991692 | 92991759 | 92991475 | 92991517 | 92996956 | 92997095 | 23872 | 30,16,20,25,1,8,10,8,7,8,1, | 0,2,2,1,1,1,6,1,1,0,1,0 | 10,16,3,19,7,8,1,4,7,11,13, | 0,0,0,0,1,0,1,1,0,2,0,0,0,1,1 | 216 | 149 | 0.0168616315144 | 0.820702354698287 | 1.0,0.847,0.873,0. | 1.0,1.0,1.0,1.0,0.8 | -0,089 |
| 239883 | ENSG00000148908 | RGS10 | chr10 | - | 119515508 | 119515745 | 119499816 | 119500259 | 119526031 | 119526118 | 239883 | 69,33,190,154,80,48,48,52 | 0,2,6,4,2,0,0,0,4,0,0,3 | 69,77,50,11,38,46,26,43,4 | 0,3,0,0,0,2,0,0,0,0,2,0,0,0, | 298 | 149 | 0.0168754542607 | 0.821189440647688 | 1.0,0.892,0.941,0. | 1.0,0.928,1.0,1.0,1 | -0,02 |
| 122021 | ENSG00000171940 | ZNF217 | chr20 | - | 53571720 | 53571853 | 53567070 | 53569264 | 53575726 | 53577280 | 122021 | 109,85,119,74,46,42,54,86 | 9,8,17,23,6,5,8,7,4,2,8,11 | 127,139,71,62,37,52,35,48 | 17,25,7,12,9,8,7,7,10,23,1 | 282 | 149 | 0.0168910460879 | 0.821762373540869 | 0.865,0.849,0.787, | 0.798,0.746,0.843, | 0,061 |
| 161994 | ENSG00000124177 | CHD6 | chr20 | - | 41549764 | 41550205 | 41533155 | 41533570 | 41551304 | 41551360 | 161994 | 2,6,6,0,3,4,1,12,0,0,0,0 | 33,22,57,22,19,37,28,39,1 | 3,3,0,2,4,0,0,3,2,0,0,2,0,2, | 33,28,22,20,17,18,15,17,1 | 298 | 149 | 0.0169027941806 | 0.822148089103891 | 0.029,0.12,0.05,0. | 0.043,0.051,0.0,0. | 0,007 |
| 2123 | ENSG00000198089 | SFI1 | chr22 | + | 31528689 | 31528863 | 31508254 | 31508376 | 31546860 | 31546971 | 2123 | 30,22,68,53,27,41,18,28,3 | 5,0,7,4,4,2,1,1,2,1,0,1 | 29,30,13,18,35,25,16,26,3 | 4,1,3,3,5,1,1,1,4,8,6,8,2,9, | 298 | 149 | 0.016922525518 | 0.822921845857356 | 0.75,1.0,0.829,0.8 | 0.784,0.938,0.684, | 0,074 |
| 204403 | ENSG00000149532 | CPSF7 | chr11 | - | 61413943 | 61413986 | 61411768 | 61411937 | 61415665 | 61416519 | 204403 | 0,0,3,5,2,1,2,2,1,0,1,2 | 27,14,58,31,31,17,12,29,2 | 3,3,3,2,3,2,2,5,0,0,6,1,3,0, | 17,47,24,11,19,29,20,25,6, | 192 | 149 | 0.0169327672216 | 0.823114390450731 | 0.0,0.0,0.039,0.11 | 0.12,0.047,0.088,0 | -0,034 |
| 56699 | ENSG00000085465 | OVGP1 | chr1 | - | 111421275 | 111421461 | 111419609 | 111419726 | 111421564 | 111421673 | 56699 | 31,19,14,21,21,15,18,13,9, | 6,0,5,0,1,2,0,2,1,0,3,0 | 20,20,10,14,23,12,6,7,9,12 | 5,6,1,3,5,4,0,0,3,1,2,2,1,0, | 298 | 149 | 0.0169363321899 | 0.823114390450731 | 0.721,1.0,0.583,1. | 0.667,0.625,0.833, | 0,205 |
| 151404 | ENSG00000048471 | SNX29 | chr16 | + | 12083344 | 12083394 | 12078832 | 12078915 | 12126632 | 12126696 | 151404 | 1,0,11,1,2,8,3,2,0,0,4,0 | 64,29,236,116,74,84,39,62 | 0,5,2,1,0,1,0,5,4,0,0,0,0,2, | 108,102,56,34,63,29,42,75 | 199 | 149 | 0.0169379579861 | 0.823114390450731 | 0.012,0.0,0.034,0. | 0.0,0.035,0.026,0. | 0,014 |
| 225167 | ENSG00000186480 | INSIG1 | chr7 | + | 155302805 | 155302846 | 155302250 | 155302417 | 155308240 | 155308757 | 225167 | 14,18,26,15,7,13,11,8,15,2 | 0,0,5,9,1,3,9,5,0,0,0,5 | 6,9,9,6,5,2,2,7,1,5,6,6,8,6, | 5,6,4,4,6,5,2,5,5,0,0,1,0,3, | 190 | 149 | 0.0169531883572 | 0.82366855153691 | 1.0,1.0,0.803,0.56 | 0.485,0.541,0.638, | 0,178 |
| 185250 | ENSG00000072274 | TFRC | chr3 | - | 196061370 | 196061434 | 196060179 | 196060247 | 196062581 | 196064428 | 185250 | 2,1,1,0,3,2,2,3,0,0,0,0 | 39,21,93,42,73,54,48,69,2 | 2,0,6,6,0,0,1,0,3,15,0,2,0,0 | 52,56,53,27,35,15,21,21,1 | 213 | 149 | 0.0169743592337 | 0.824511016075036 | 0.035,0.032,0.007, | 0.026,0.0,0.073,0. | -0,028 |
| 154934 | ENSG00000213983 | AP1G2 | chr14 | - | 23566554 | 23566686 | 23566338 | 23566419 | 23567110 | 23567315 | 154934 | 158,86,128,138,92,92,99,8 | 13,13,20,2,9,13,8,11,8,8,3, | 95,92,66,22,56,40,53,78,5 | 8,11,13,7,10,3,8,10,4,11,7, | 281 | 149 | 0.0169995649767 | 0.825549045464828 | 0.866,0.778,0.772, | 0.863,0.816,0.729, | 0,058 |
| 167124 | ENSG00000174652 | ZNF266 | chr19 | - | 9433667 | 9433786 | 9419951 | 9420218 | 9434109 | 9434235 | 167124 | 13,16,90,76,55,18,63,41,6 | 0,0,15,5,1,0,2,4,2,0,0,0 | 103,53,45,16,21,11,16,57, | 1,2,2,1,4,1,4,2,2,1,3,0,0,2, | 268 | 149 | 0.0170231548313 | 0.826192686896691 | 1.0,1.0,0.769,0.89 | 0.983,0.936,0.926, | 0,052 |
| 160492 | ENSG00000186532 | SMYD4 | chr17 | - | 1783615 | 1784085 | 1783359 | 1783476 | 1784325 | 1784461 | 160492 | 0,1,7,1,2,2,1,5,2,6,0,5 | 8,0,32,9,4,5,3,5,10,0,1,9 | 1,0,0,2,0,0,0,1,0,3,2,1,2,2, | 5,2,5,3,3,1,1,5,6,5,8,13,5,1 | 298 | 149 | 0.0170249622743 | 0.826192686896691 | 0.0,1.0,0.099,0.05 | 0.091,0.0,0.0,0.25, | 0,136 |
| 18835 | ENSG00000185418 | TARS3 | chr15 | - | 101684074 | 101684237 | 101675599 | 101675737 | 101685895 | 101686062 | 18835 | 63,40,105,83,16,26,41,34, | 3,3,3,0,0,1,0,2,0,2,4,1 | 38,56,26,19,28,27,18,59,2 | 0,0,1,3,1,0,0,0,4,1,0,0,0,0, | 298 | 149 | 0.017025529584 | 0.826192686896691 | 0.913,0.87,0.946,1 | 1.0,1.0,0.929,0.76, | -0,043 |
| 29901 | ENSG00000165819 | METTL3 | chr14 | - | 21500494 | 21500682 | 21499763 | 21499802 | 21500912 | 21501129 | 29901 | 273,104,179,134,131,109, | 8,4,15,3,10,5,2,12,2,7,1,4 | 146,90,89,39,72,72,50,137 | 6,10,2,1,0,2,2,1,0,0,4,7,2,2 | 298 | 149 | 0.0170281732607 | 0.826192686896691 | 0.945,0.929,0.856, | 0.924,0.818,0.957, | -0,025 |
| 139052 | ENSG00000125772 | GPCPD1 | chr20 | - | 5582763 | 5582885 | 5580007 | 5580131 | 5584280 | 5586037 | 139052 | 5,0,7,6,3,5,5,14,0,1,3,3 | 66,42,188,141,83,96,108,8 | 3,8,21,10,10,3,3,1,4,4,1,18 | 98,123,58,46,83,45,43,66, | 271 | 149 | 0.0170381305391 | 0.826489491562099 | 0.04,0.0,0.02,0.02 | 0.017,0.035,0.166, | -0,015 |
| 54364 | ENSG00000138095 | LRPPRC | chr2 | - | 43896633 | 43896708 | 43894544 | 43894629 | 43899218 | 43899334 | 54364 | 11,9,62,62,39,24,24,26,9,8 | 7,3,0,1,5,0,1,1,0,0,3,2 | 58,54,19,5,11,10,35,55,21, | 7,8,7,12,3,1,3,2,1,2,2,8,6,2 | 224 | 149 | 0.0170432187369 | 0.82655002522796 8 | 0.511,0.666,1.0,0. | 0.846,0.818,0.644, | 0,063 |
| 149256 | ENSG00000247982 | LINC00926 | chr15 | + | 57302710 | 57303013 | 57300364 | 57300924 | 57307541 | 57307769 | 149256 | 0,0,1,0,11,18,8,8,0,2,3,3 | 80,66,36,12,48,24,17,12,6 | 0,0,2,0,0,0,2,8,1,0,0,0,3,4, | 42,13,4,6,20,12,3,9,2,32,1 | 298 | 149 | 0.017088274626 | 0.828548419920839 | 0.0,0.0,0.014,0.0,0 | 0.0,0.0,0.2,0.0,0.0, | 0,008 |
| 32856 | ENSG00000102445 | RUBCNL | chr13 | - | 46376491 | 46376584 | 46371940 | 46372597 | 46387133 | 46387500 | 32856 | 2,1,7,8,2,0,0,8,0,0,0,2 | 41,46,173,95,65,63,102,10 | 2,5,4,3,9,3,0,2,1,2,4,3,7,3, | 76,142,69,28,55,51,49,70, | 242 | 149 | 0.0170930058352 | 0.828591157413498 | 0.029,0.013,0.024, | 0.016,0.021,0.034, | -0,019 |
| 186104 | ENSG00000065308 | TRAM2 | chr6 | - | 52519615 | 52519807 | 52516627 | 52516737 | 52535782 | 52535846 | 186104 | 6,0,12,2,6,0,6,1,8,2,2,0 | 20,16,43,28,22,16,16,14,4, | 2,0,1,6,1,1,0,4,0,1,0,7,4,4, | 17,12,7,12,17,15,4,19,14,1 | 298 | 149 | 0.0171069940258 | 0.829082510958559 | 0.13,0.0,0.122,0.0 | 0.056,0.0,0.067,0. | 0,035 |
| 227775 | ENSG00000196305 | IARS1 | chr9 | - | 92257803 | 92257951 | 92256679 | 92256800 | 92258853 | 92258998 | 227775 | 12,3,12,6,0,6,8,5,2,4,1,4 | 39,28,52,55,28,35,25,28,3 | 3,19,8,15,7,3,8,11,8,5,9,4, | 30,44,50,28,18,15,13,26,1 | 297 | 149 | 0.0171209805294 | 0.829573561531583 | 0.134,0.051,0.104, | 0.048,0.178,0.074, | -0,046 |
| 51152 | ENSG00000197111 | PCBP2 | chr12 | + | 53464759 | 53464852 | 53462480 | 53462567 | 53467220 | 53467285 | 51152 | 66,30,92,71,72,103,81,108 | 13,10,24,9,14,10,3,15,2,5, | 72,86,46,140,65,31,164,82 | 14,17,5,21,8,11,35,20,8,5, | 242 | 149 | 0.0171365929365 | 0.830143154246192 | 0.758,0.649,0.702, | 0.76,0.757,0.85,0. | 0,056 |
| 184328 | ENSG00000004487 | KDM1A | chr1 | + | 23040258 | 23040646 | 23030468 | 23030634 | 23044426 | 23044486 | 184328 | 1,2,4,2,2,2,0,0,0,0,1,2 | 9,6,19,8,14,13,13,14,24,12 | 2,4,0,2,2,0,9,2,2,2,1,3,2,0, | 15,7,6,8,2,2,8,27,1,8,10,17 | 298 | 149 | 0.017148890414 | 0.830551942550773 | 0.053,0.143,0.095, | 0.062,0.222,0.0,0. | -0,073 |
| 229668 | ENSG00000152457 | DCLRE1C | chr10 | - | 14920379 | 14920814 | 14919737 | 14919832 | 14922980 | 14923675 | 229668 | 8,0,10,1,1,1,2,4,4,1,1,1 | 42,40,51,30,33,28,19,30,3 | 6,2,1,0,0,2,1,0,1,6,0,2,2,1, | 35,20,13,21,15,5,13,19,14, | 298 | 149 | 0.0171552987025 | 0.83067538696013 | 0.087,0.0,0.089,0. | 0.079,0.048,0.037, | 0,008 |
| 29601 | ENSG00000198612 | COPS8 | chr2 | + | 237087463 | 237087563 | 237087126 | 237087197 | 237088604 | 237088653 | 29601 | 1,3,3,1,1,3,3,5,1,0,1,1 | 11,6,32,48,26,13,19,7,16,1 | 0,3,7,3,3,0,1,9,3,0,3,0,2,5, | 19,16,21,5,19,19,10,18,4,2 | 249 | 149 | 0.0171592864825 | 0.830681598946234 | 0.052,0.23,0.053,0 | 0.0,0.101,0.166,0. | -0,038 |
| 143546 | ENSG00000291131 | | chr19 | + | 53204951 | 53205127 | 53204015 | 53204142 | 53210980 | 53213415 | 143546 | 14,1,26,21,7,10,6,8,7,2,1,1 | 15,2,17,19,7,1,6,7,9,5,2,0 | 4,2,4,1,3,3,2,6,2,5,13,2,0,4 | 31,0,3,0,7,8,1,5,4,5,11,3,5, | 298 | 149 | 0.0171746506228 | 0.831238414255873 | 0.318,0.2,0.433,0. | 0.061,1.0,0.4,1.0,0 | 0,059 |
| 182059 | ENSG00000172432 | GTPBP2 | chr6 | - | 43628516 | 43628589 | 43626921 | 43626948 | 43628976 | 43629264 | 182059 | 4,0,0,1,2,1,2,2,1,2,1,1 | 6,2,3,6,9,10,5,7,23,9,7,7 | 2,0,1,2,1,0,0,0,1,2,0,1,2,0, | 3,2,4,2,1,3,1,7,2,4,7,15,3,1 | 222 | 149 | 0.0172070734241 | 0.832620418197497 | 0.309,0.0,0.0,0.10 | 0.309,0.0,0.144,0. | -0,027 |
| 222683 | ENSG00000167483 | NIBAN3 | chr19 | + | 17546591 | 17546797 | 17543523 | 17543631 | 17549443 | 17549527 | 222683 | 263,154,228,161,127,78,1 | 27,3,9,1,7,3,3,2,9,6,10,2 | 143,177,180,88,176,98,54, | 1,7,6,3,5,0,0,7,3,3,2,8,17,9 | 298 | 149 | 0.0172230521188 | 0.83320627794593 | 0.83,0.963,0.927,0 | 0.986,0.927,0.938, | -0,031 |
| 53278 | ENSG00000168175 | MAPK1IP1L | chr14 | + | 55061679 | 55061701 | 55051646 | 55051803 | 55062617 | 55063325 | 53278 | 19,11,16,9,5,5,5,10,18,6,2, | 13,9,6,14,7,2,11,9,7,2,2,5 | 3,14,25,4,5,3,9,8,6,4,10,11 | 9,16,8,20,13,6,5,8,5,7,7,20 | 171 | 149 | 0.0172348485784 | 0.83358959424216 | 0.56,0.516,0.699,0 | 0.225,0.433,0.731, | 0,144 |
| 51932 | ENSG00000170791 | CHCHD7 | chr8 | + | 56212761 | 56212909 | 56211755 | 56211837 | 56214597 | 56214667 | 51932 | 10,7,40,34,8,6,5,20,7,8,6,8 | 10,2,26,17,10,7,3,3,9,2,6,1 | 28,22,19,6,13,6,9,15,11,32 | 14,9,4,1,3,3,0,0,6,14,8,21, | 297 | 149 | 0.0172511682601 | 0.834116079026259 | 0.334,0.637,0.436, | 0.501,0.551,0.704, | -0,165 |
| 154931 | ENSG00000213983 | AP1G2 | chr14 | - | 23566553 | 23566686 | 23566338 | 23566419 | 23567110 | 23567315 | 154931 | 158,86,128,138,92,93,99,8 | 13,13,20,2,9,13,8,11,8,8,3, | 97,92,67,22,56,40,53,78,5 | 8,11,13,7,10,3,8,10,4,11,7, | 282 | 149 | 0.0172534847853 | 0.834116079026259 | 0.865,0.778,0.772, | 0.865,0.815,0.731, | 0,058 |
| 49085 | ENSG00000166938 | DIS3L | chr15 | + | 66323492 | 66323585 | 66322686 | 66322934 | 66325830 | 66325988 | 49085 | 16,18,89,38,28,32,16,28,1 | 0,0,11,5,0,3,2,4,0,1,2,5 | 37,16,19,20,6,24,13,30,15, | 6,8,0,1,3,2,2,5,4,5,2,4,3,0, | 242 | 149 | 0.0172700007613 | 0.834615696320074 | 1.0,1.0,0.833,0.82 | 0.792,0.552,1.0,0. | 0,105 |
| 76887 | ENSG00000018699 | TTC27 | chr2 | + | 32812503 | 32812615 | 32811023 | 32811221 | 32817456 | 32817557 | 76887 | 23,12,31,36,27,17,7,29,25, | 4,0,0,1,0,0,0,0,0,0,0,0 | 37,22,15,5,17,8,18,34,10,2 | 0,0,0,4,0,2,0,7,0,0,0,2,4,0, | 261 | 149 | 0.0172715747797 | 0.834615696320074 | 0.766,1.0,1.0,0.95 | 1.0,1.0,1.0,0.416,1 | 0,085 |
| 148920 | ENSG00000140521 | POLG | chr15 | - | 89317375 | 89317552 | 89316304 | 89316827 | 89318540 | 89318749 | 148920 | 46,13,76,38,24,32,29,31,1 | 2,0,1,3,0,1,0,2,2,0,0,3 | 33,48,18,9,38,16,13,32,11, | 1,5,2,6,0,0,2,4,1,2,1,0,2,2, | 298 | 149 | 0.0172894964122 | 0.835294186822496 | 0.92,1.0,0.974,0.8 | 0.943,0.828,0.818, | 0,063 |
| 248330 | ENSG00000167323 | STIM1 | chr11 | + | 3945804 | 3946068 | 3856181 | 3856409 | 3967551 | 3968721 | 248330 | 1,0,2,6,2,6,0,5,3,0,1,0 | 22,21,69,57,17,23,20,45,1 | 0,3,2,5,0,2,1,8,3,5,9,2,3,6, | 53,41,18,30,25,25,27,49,2 | 298 | 149 | 0.0172942587415 | 0.835336760141783 | 0.022,0.0,0.014,0. | 0.0,0.035,0.053,0. | -0,019 |
| 204733 | ENSG00000196510 | ANAPC7 | chr12 | - | 110375628 | 110375686 | 110374245 | 110374333 | 110376065 | 110376216 | 204733 | 3,0,4,0,1,4,0,1,1,0,0,1 | 27,12,25,26,17,24,14,12,2 | 1,4,2,7,5,0,1,3,2,2,0,5,0,1, | 23,16,25,14,10,5,18,17,14, | 207 | 149 | 0.0173026856297 | 0.835556277937169 | 0.074,0.0,0.103,0. | 0.03,0.153,0.054,0 | -0,031 |
| 204411 | ENSG00000149532 | CPSF7 | chr11 | - | 61420464 | 61420573 | 61420057 | 61420094 | 61421389 | 61421608 | 204411 | 34,27,74,34,28,25,31,27,1 | 2,0,6,3,2,0,1,0,2,1,3,4 | 39,45,33,29,37,23,10,44,1 | 6,5,2,3,0,1,1,3,0,0,10,4,3,2 | 258 | 149 | 0.0173103468395 | 0.835738730509741 | 0.908,1.0,0.877,0. | 0.79,0.839,0.905,0 | 0,031 |
| 184330 | ENSG00000004487 | KDM1A | chr1 | + | 23044426 | 23044486 | 23040258 | 23040434 | 23050386 | 23050520 | 184330 | 16,10,7,7,4,10,2,10,5,4,5,6 | 0,0,1,0,0,0,0,0,0,0,0,2 | 30,24,4,11,6,6,12,12,7,5,1 | 0,4,0,6,4,0,3,2,0,2,1,0,1,0, | 209 | 149 | 0.0173210500848 | 0.835998341128995 | 1.0,1.0,0.833,1.0,1 | 1.0,0.811,1.0,0.56 | 0,089 |
| 202585 | ENSG00000138078 | PREPL | chr2 | - | 44359496 | 44359596 | 44346267 | 44346390 | 44361379 | 44361806 | 202585 | 24,10,37,15,5,7,7,8,16,3,4, | 1,4,5,4,4,2,2,5,10,0,6,4 | 21,8,15,9,14,10,8,11,3,16, | 3,1,4,3,0,2,0,2,0,2,4,0,1,8, | 249 | 149 | 0.0173281076078 | 0.835998341128995 | 0.935,0.599,0.816, | 0.807,0.827,0.692, | -0,168 |
| 19098 | ENSG00000168970 | JMJD7-PLA2G4 | chr15 | + | 41843125 | 41843223 | 41842192 | 41842591 | 41843675 | 41843811 | 19098 | 3,4,12,4,4,4,9,2,7,2,4,5 | 3,0,1,4,5,1,0,1,2,1,0,3 | 5,2,9,1,1,1,5,16,2,3,7,1,3,7 | 2,3,2,5,0,1,6,4,1,8,0,2,4,6, | 247 | 149 | 0.0173348373796 | 0.835998341128995 | 0.376,1.0,0.879,0. | 0.601,0.287,0.731, | 0,159 |
| 64500 | ENSG00000164048 | ZNF589 | chr3 | + | 48256710 | 48257022 | 48247624 | 48247677 | 48260812 | 48260939 | 64500 | 0,0,1,0,1,2,2,2,2,0,0,0 | 2,2,13,8,4,5,3,6,13,7,3,1 | 0,1,0,0,0,0,2,0,1,0,0,24,2,3 | 3,8,3,1,5,11,4,2,4,9,3,62,1 | 298 | 149 | 0.0173356068836 | 0.835998341128995 | 0.0,0.0,0.037,0.0,0 | 0.0,0.059,0.0,0.0,0 | 0,016 |
| 88391 | ENSG00000136699 | SMPD4 | chr2 | - | 130161090 | 130161272 | 130157250 | 130157396 | 130164373 | 130164445 | 88391 | 14,7,1,5,4,11,12,13,19,7,1, | 14,2,3,5,4,6,17,8,19,13,4,1 | 2,3,5,13,3,0,2,4,7,6,1,2,1,0 | 7,1,0,9,3,5,0,13,2,16,5,14,1 | 298 | 149 | 0.0173367961382 | 0.835998341128995 | 0.333,0.636,0.143, | 0.125,0.6,1.0,0.41 | 0,02 |
| 102610 | ENSG00000122566 | HNRNPA2B1 | chr7 | - | 26190992 | 26191128 | 26173593 | 26174782 | 26192494 | 26192577 | 102610 | 10,5,12,19,11,6,3,17,7,1,2, | 3,0,4,0,1,9,2,6,1,3,1,1 | 22,24,12,12,4,8,5,14,14,7, | 1,2,3,2,3,2,0,1,0,1,0,2,0,0, | 285 | 149 | 0.0173390292049 | 0.835998341128995 | 0.635,1.0,0.611,1. | 0.92,0.863,0.677,0 | -0,142 |
| 157079 | ENSG00000102580 | DNAJC3 | chr13 | + | 95724360 | 95724461 | 95723241 | 95723366 | 95725177 | 95725252 | 157079 | 15,7,0,0,0,0,0,0,1,0,0,0 | 11,9,31,15,15,17,11,27,5,3 | 1,0,0,0,0,0,0,0,0,0,0,0,0,0, | 36,38,12,19,10,7,17,20,25, | 250 | 149 | 0.0173488403113 | 0.836284042338502 | 0.448,0.317,0.0,0. | 0.016,0.0,0.0,0.0,0 | 0,072 |
| 74175 | ENSG00000064687 | ABCA7 | chr19 | + | 1058145 | 1058269 | 1057313 | 1057429 | 1058617 | 1058732 | 74175 | 56,24,37,41,37,32,56,45,5 | 0,0,2,1,0,3,1,3,0,0,2,0 | 33,38,19,33,21,25,16,32,1 | 1,0,0,2,1,2,0,2,3,0,4,1,2,2, | 273 | 149 | 0.0173568912262 | 0.836484786275471 | 1.0,1.0,0.91,0.957, | 0.947,1.0,1.0,0.9,0 | 0,04 |
| 212192 | ENSG00000014641 | MDH1 | chr2 | + | 63594487 | 63594596 | 63588608 | 63589046 | 63597398 | 63598186 | 212192 | 9,8,54,51,17,25,14,14,4,9, | 0,0,3,0,2,2,0,2,0,0,1,0 | 38,41,12,0,12,5,7,17,7,27,1 | 3,5,0,2,2,1,0,2,0,0,3,5,1,1, | 258 | 149 | 0.0173725774533 | 0.837053328375019 | 1.0,1.0,0.912,1.0,0 | 0.88,0.826,1.0,0.0, | 0,072 |
| 43857 | ENSG00000115129 | TP53I3 | chr2 | - | 24079443 | 24079640 | 24077432 | 24077761 | 24080818 | 24081031 | 43857 | 28,12,53,70,24,10,14,14,1 | 0,1,4,12,0,4,1,2,2,0,0,0 | 27,53,8,2,26,23,11,6,59,66 | 5,14,4,0,3,3,2,1,5,5,2,1,2,5 | 298 | 149 | 0.017388676138 | 0.837641484748854 | 1.0,0.857,0.869,0. | 0.73,0.654,0.5,1.0, | 0,091 |
| 20258 | ENSG00000125107 | CNOT1 | chr16 | - | 58624652 | 58624794 | 58623281 | 58623409 | 58629727 | 58629815 | 20258 | 15,15,27,19,9,17,5,10,17,1 | 63,39,68,66,33,51,24,52,2, | 15,16,4,13,12,7,18,9,4,7,1 | 62,74,37,56,63,33,13,31,1 | 291 | 149 | 0.0173949334761 | 0.837755409933873 | 0.109,0.165,0.169, | 0.11,0.1,0.052,0.1 | 0,095 |
| 46749 | ENSG00000015479 | MATR3 | chr5 | + | 139307240 | 139308327 | 139293717 | 139293772 | 139314674 | 139314736 | 46749 | 141,108,351,296,159,182, | 3,2,6,11,0,9,9,6,2,0,3,2 | 260,228,134,115,64,59,13 | 5,14,2,5,2,3,11,5,4,2,10,11 | 298 | 149 | 0.0174103617445 | 0.838135924374531 | 0.959,0.964,0.967, | 0.963,0.891,0.971, | 0,023 |
| 241688 | ENSG00000175893 | ZDHHC21 | chr9 | - | 14677297 | 14677472 | 14674186 | 14674385 | 14680032 | 14680162 | 241688 | 18,34,80,69,4,14,18,30,32, | 12,10,23,19,5,5,8,9,2,2,1,7 | 16,8,14,6,13,20,8,43,5,20, | 11,21,6,0,9,5,2,13,16,19,1 | 298 | 149 | 0.0174106226235 | 0.838135924374531 | 0.429,0.63,0.635,0 | 0.421,0.16,0.538,1 | 0,098 |
| 151374 | ENSG00000185787 | MORF4L1 | chr15 | + | 78894029 | 78894230 | 78893538 | 78893627 | 78894819 | 78894904 | 151374 | 86,74,285,238,72,114,97,1 | 8,4,3,0,4,1,0,1,2,0,0,3 | 171,281,90,144,118,100,7 | 9,4,0,6,0,1,0,10,0,4,7,7,5,2 | 298 | 149 | 0.0174262395882 | 0.838514122185707 | 0.843,0.902,0.979, | 0.905,0.972,1.0,0. | 0,008 |
| 214830 | ENSG00000055044 | NOP58 | chr2 | + | 202277951 | 202278002 | 202275161 | 202275189 | 202282350 | 202282472 | 214830 | 225,234,87,29,16,21,15,23 | 14,7,7,4,3,3,3,2,0,0,1,5 | 28,53,56,30,58,11,13,25,2 | 7,4,4,12,12,6,0,2,1,2,4,2,1, | 200 | 149 | 0.0174262706977 | 0.838514122185707 | 0.923,0.961,0.903, | 0.749,0.908,0.913, | 0,03 |
| 127532 | ENSG00000107862 | GBF1 | chr10 | + | 102368217 | 102368454 | 102367477 | 102367560 | 102369210 | 102369387 | 127532 | 42,44,52,29,10,22,28,23,2 | 6,1,2,1,0,1,1,0,0,0,1,0 | 33,24,18,54,31,14,23,23,2 | 3,0,0,0,2,0,0,0,0,9,3,0,2, | 298 | 149 | 0.017435460473 | 0.838768795946416 | 0.778,0.957,0.929, | 0.846,1.0,1.0,1.0,0 | 0,073 |
| 3303 | ENSG00000102575 | ACP5 | chr19 | - | 11578584 | 11578657 | 11577592 | 11577679 | 11578843 | 11578954 | 3303 | 3,4,12,5,13,7,13,4,6,5,0,15 | 6,4,6,3,15,0,23,7,16,13,5,5 | 9,3,5,2,10,6,1,7,7,10,10,15 | 1,12,0,0,2,4,3,4,0,3,2,167,1 | 222 | 149 | 0.0174490786044 | 0.839236343486842 | 0.251,0.402,0.573, | 0.858,0.144,1.0,1. | -0,209 |

| | | | | | | | | | | | | | | | | | | | | | | |
|---|---|---|---|---|---|---|---|---|---|---|---|---|---|---|---|---|---|---|---|---|---|---|
| 247857 | ENSG00000179532 | DNHD1 | chr11 | + | 6538352 | 6538510 | 6533923 | 6534229 | 6538611 | 6538810 | 247857 | 8,8,15,4,7,14,11,9,9,7,3,12 | 1,1,2,3,0,2,0,1,0,0,4,0 | 9,7,15,21,4,1,3,12,9,20,10, | 0,0,0,1,0,0,0,0,2,0,0,0,0,0,0 | 298 | 149 | 0.0174680798365 | 0.839808132265694 | 0.8,0.8,0.789,0.4,1 | 1.0,1.0,1.0,0.913,1 | -0,129 |
| 246607 | ENSG00000054654 | SYNE2 | chr14 | + | 64025959 | 64026075 | 64025129 | 64025421 | 64026578 | 64026730 | 246607 | 12,4,15,4,4,17,7,20,3,0,0,1 | 121,154,200,132,98,184,12 | 6,18,11,33,16,4,1,11,13,11 | 74,149,88,154,129,99,30,1 | 265 | 149 | 0.0174713946539 | 0.839808132265694 | 0.053,0.014,0.04,0 | 0.044,0.064,0.066, | -0,016 |
| 242860 | ENSG00000216937 | CCDC7 | chr10 | + | 32558985 | 32559146 | 32544246 | 32544301 | 32565557 | 32565620 | 242860 | 9,5,16,5,1,6,3,3,5,2,4,3 | 50,36,78,48,13,23,27,49,52 | 5,11,11,8,4,6,5,7,0,1,4,17,1 | 21,14,25,18,19,11,11,13,17 | 298 | 149 | 0.0174740936577 | 0.839808132265694 | 0.083,0.065,0.093, | 0.106,0.282,0.18,0 | -0,014 |
| 252219 | ENSG00000198825 | INPP5F | chr10 | + | 119823846 | 119823902 | 119823070 | 119823199 | 119826630 | 119826702 | 252219 | 67,22,76,39,43,67,29,33,41 | 1,0,1,2,5,2,3,0,0,0,0,3 | 56,41,79,26,11,24,48,35,17 | 7,2,2,0,0,0,0,0,4,4,4,0,0,1,5 | 205 | 149 | 0.0174765745846 | 0.839808132265694 | 0.98,1.0,0.982,0.93 | 0.853,0.937,0.966, | 0,033 |
| 241907 | ENSG00000187764 | SEMA4D | chr9 | - | 89388635 | 89388791 | 89387171 | 89387608 | 89388871 | 89389047 | 241907 | 39,32,42,47,25,41,26,48,56 | 6,0,4,1,2,2,3,6,2,2,0,1 | 22,23,31,59,12,15,48,22,19 | 0,1,0,1,1,1,10,0,2,0,1,0,0,2 | 298 | 149 | 0.0174909004809 | 0.840308928884953 | 0.765,1.0,0.84,0.95 | 1.0,0.92,1.0,0.967, | -0,031 |
| 164647 | ENSG00000038219 | BOD1L1 | chr4 | - | 13623389 | 13623447 | 13619942 | 13620067 | 13627344 | 13627725 | 164647 | 1,3,4,1,2,1,6,1,0,0,0,1 | 26,24,51,29,13,12,12,30,17 | 0,1,0,2,1,3,0,1,3,2,0,2,0,0,0 | 33,44,24,18,35,18,15,22,14 | 207 | 149 | 0.0174955739757 | 0.840345878679734 | 0.027,0.083,0.053, | 0.0,0.016,0.0,0.074 | 0,028 |
| 224867 | ENSG00000177600 | RPLP2 | chr11 | + | 810233 | 810360 | 809966 | 810039 | 811596 | 811839 | 224867 | 301,275,524,141,163,115,2 | 4,0,4,3,2,1,3,6,2,1,0,0 | 190,268,117,142,216,144,2 | 2,3,1,1,0,1,0,0,1,4,0,32,4,3 | 276 | 149 | 0.0175135364363 | 0.841020964016351 | 0.976,1.0,0.986,0.9 | 0.981,0.98,0.984,0 | 0,001 |
| 43800 | ENSG00000114867 | EIF4G1 | chr3 | + | 184317320 | 184317497 | 184314541 | 184314674 | 184317716 | 184317773 | 43800 | 21,15,24,41,12,11,3,8,14,1 | 7,9,11,8,5,4,10,5,1,7,8,3 | 13,4,8,8,5,10,1,4,4,15,10,1 | 10,15,8,14,11,3,9,5,3,4,6,7 | 298 | 149 | 0.0175303641805 | 0.841641269893642 | 0.6,0.455,0.522,0.7 | 0.394,0.118,0.333, | 0,148 |
| 126528 | ENSG00000150093 | ITGB1 | chr10 | - | 32932514 | 32933583 | 32929821 | 32930044 | 32935491 | 32935558 | 126528 | 277,179,141,129,64,72,113 | 4,5,12,2,2,0,12,17,2,0,0,3 | 81,74,1,41,69,59,38,77,79, | 9,3,0,3,7,4,2,4,4,6,7,12,5,1 | 298 | 149 | 0.017541812539 | 0.842003089781782 | 0.972,0.947,0.855, | 0.818,0.925,1.0,0.8 | 0,041 |
| 247576 | ENSG00000149187 | CELF1 | chr11 | - | 47536619 | 47536730 | 47488982 | 47489024 | 47552991 | 47553132 | 247576 | 4,4,11,5,3,2,15,3,8,2,5,5 | 6,11,15,11,9,5,22,12,5,13,4 | 1,0,7,1,3,2,0,0,6,5,0,6,4,5,4 | 15,10,3,9,7,3,7,1,12,3,11,2 | 260 | 149 | 0.0176146964881 | 0.845312985469398 | 0.276,0.172,0.296, | 0.037,0.0,0.572,0.0 | 0,064 |
| 106571 | ENSG00000147140 | NONO | chrX | + | 71284983 | 71285097 | 71283589 | 71283721 | 71290628 | 71290791 | 106571 | 8,2,5,3,0,1,0,1,0,4,1,2 | 34,11,12,6,7,18,13,24,14,1 | 7,4,9,3,1,1,3,7,2,3,3,5,3,2,6 | 13,12,23,45,14,6,21,27,2,9 | 263 | 149 | 0.0176223336086 | 0.845445903034644 | 0.118,0.093,0.191, | 0.234,0.159,0.181, | -0,063 |
| 30492 | ENSG00000124177 | CHD6 | chr20 | - | 41564010 | 41564151 | 41551304 | 41551360 | 41618339 | 41618375 | 30492 | 3,1,6,0,1,2,2,0,0,1,0,6 | 10,9,27,6,5,17,14,12,4,9,1, | 3,0,2,3,0,0,0,0,2,1,0,1,0,2,0 | 7,12,8,7,9,11,8,5,4,7,5,21,6 | 290 | 149 | 0.0176253224067 | 0.845445903034644 | 0.134,0.054,0.102, | 0.18,0.0,0.114,0.18 | 0,014 |
| 192034 | ENSG00000070961 | ATP2B1 | chr12 | - | 89677621 | 89677816 | 89655678 | 89656107 | 89709218 | 89709366 | 192034 | 0,0,2,3,0,0,0,0,0,0,0,1 | 3,3,28,10,3,14,6,10,3,1,2,8 | 0,1,0,2,0,3,4,0,0,0,2,1,4,6,1 | 11,13,5,3,10,4,1,13,8,9,12, | 298 | 149 | 0.0176367061334 | 0.845803453163506 | 0.0,0.0,0.034,0.13, | 0.0,0.037,0.0,0.25, | -0,142 |
| 3605 | ENSG00000196646 | ZNF136 | chr19 | + | 12164465 | 12164874 | 12163069 | 12163206 | 12185784 | 12185911 | 3605 | 0,1,0,0,0,0,3,9,2,0,0,4 | 16,12,34,13,5,5,3,15,5,4,1, | 1,0,0,0,2,0,0,1,2,0,2,0,0,0,0 | 17,23,3,17,4,6,4,5,5,10,20, | 298 | 149 | 0.0176491065095 | 0.846209588582356 | 0.0,0.04,0.0,0.0,0.0 | 0.029,0.0,0.0,0.0,0 | 0,052 |
| 124592 | ENSG00000110422 | HIPK3 | chr11 | + | 33348521 | 33348818 | 33347851 | 33348013 | 33349146 | 33349287 | 124592 | 126,73,152,82,54,54,70,63 | 7,0,5,7,3,0,3,0,0,1,0,0 | 100,84,65,59,55,54,59,76,4 | 0,2,8,1,5,5,2,1,0,0,4,1,0,4,2 | 298 | 149 | 0.0176702759356 | 0.846775264470639 | 0.9,1.0,0.938,0.854 | 1.0,0.955,0.802,0.9 | 0,029 |
| 19563 | ENSG00000166411 | IDH3A | chr15 | + | 78160091 | 78160206 | 78156907 | 78157631 | 78161580 | 78161633 | 19563 | 43,22,57,47,38,34,40,34,27 | 2,0,2,3,0,1,0,0,0,2,7,0 | 35,42,19,16,29,31,23,25,23 | 3,4,2,0,5,0,4,2,2,2,1,1,3,2,0 | 264 | 149 | 0.0176714741403 | 0.846775264470639 | 0.924,1.0,0.941,0.8 | 0.868,0.856,0.843, | 0,03 |
| 34901 | ENSG00000026297 | RNASET2 | chr6 | - | 166955494 | 166955627 | 166952487 | 166952548 | 166956096 | 166956611 | 34901 | 3,4,10,1,12,2,5,0,9,14,0,5 | 62,37,49,29,22,11,22,13,26 | 8,1,2,8,4,1,6,3,1,3,9,4,8,2,2 | 22,46,47,27,30,25,13,16,14 | 282 | 149 | 0.0176727074074 | 0.846775264470639 | 0.025,0.054,0.097, | 0.161,0.011,0.022, | 0,011 |
| 217001 | ENSG00000135945 | REV1 | chr2 | - | 99424772 | 99424899 | 99424151 | 99424280 | 99429839 | 99429945 | 217001 | 3,1,11,2,2,4,4,0,0,2,0,0 | 15,7,15,6,10,11,12,11,11,3 | 9,7,4,3,4,1,3,4,9,10,7,16,2, | 15,22,5,14,16,12,9,22,6,21 | 276 | 149 | 0.0176987295327 | 0.847833353227811 | 0.097,0.072,0.284, | 0.245,0.147,0.302, | -0,077 |
| 218447 | ENSG00000112343 | TRIM38 | chr6 | + | 25974789 | 25974930 | 25973172 | 25973285 | 25983163 | 25991231 | 218447 | 0,0,28,28,0,1,0,7,5,0,4,2 | 47,39,99,87,47,42,24,29,19 | 4,7,4,0,13,15,1,2,3,4,2,2,1, | 53,87,31,21,48,39,20,37,47 | 290 | 149 | 0.0177108954519 | 0.848227356254537 | 0.0,0.0,0.127,0.14 | 0.037,0.04,0.062,0 | 0,01 |
| 252217 | ENSG00000198825 | INPP5F | chr10 | + | 119823825 | 119823902 | 119823070 | 119823199 | 119826630 | 119826702 | 252217 | 93,26,125,61,70,89,37,53,5 | 1,0,1,2,5,2,3,0,0,0,0,3 | 74,67,112,33,17,33,67,54,2 | 7,2,2,0,0,0,0,0,4,4,4,0,0,1,5 | 226 | 149 | 0.0177154277924 | 0.848255670564192 | 0.984,1.0,0.988,0.9 | 0.875,0.957,0.974, | 0,029 |
| 135739 | ENSG00000291230 | | chr19 | - | 50336404 | 50336705 | 50335058 | 50335181 | 50337016 | 50337135 | 135739 | 235,216,268,622,258,163,1 | 14,3,5,13,2,4,8,2,10,5,1,0 | 230,162,124,48,150,158,77 | 6,4,2,1,4,2,2,3,7,5,7,15,8,1 | 298 | 149 | 0.0177427507099 | 0.849365910417672 | 0.894,0.973,0.964, | 0.95,0.953,0.969,0 | 0,019 |
| 97493 | ENSG00000131023 | LATS1 | chr6 | - | 149676259 | 149676366 | 149658152 | 149662238 | 149676554 | 149676737 | 97493 | 26,30,66,27,24,29,15,35,20 | 1,1,0,0,4,0,1,1,1,2,0,0 | 36,42,13,27,29,13,25,40,11 | 3,1,0,2,2,1,1,1,2,1,4,1,1,3,0 | 256 | 149 | 0.0177465072371 | 0.849365910417672 | 0.938,0.946,1.0,1.0 | 0.875,0.961,1.0,0.8 | 0,032 |
| 189905 | ENSG00000138688 | KIAA1109 | chr4 | + | 122192228 | 122192387 | 122189999 | 122190079 | 122196634 | 122196791 | 189905 | 121,100,108,45,41,70,44,5 | 8,1,4,5,0,0,4,3,1,1,0,3 | 48,60,47,87,41,28,34,40,50 | 0,0,0,0,8,1,0,2,0,0,9,3,0,0,0 | 298 | 149 | 0.0177944158829 | 0.851469525320687 | 0.883,0.98,0.931,0 | 1.0,1.0,1.0,1.0,0.7 | -0,026 |
| 156770 | ENSG00000139746 | RBM26 | chr13 | - | 79348706 | 79348789 | 79344668 | 79344794 | 79354438 | 79354570 | 156770 | 9,1,9,0,0,1,1,2,1,1,0,1 | 86,72,113,83,47,70,53,106 | 3,12,1,8,7,4,1,2,3,7,0,8,0,1 | 72,125,60,83,91,57,43,75,8 | 232 | 149 | 0.01781554526 | 0.85226002483397 | 0.063,0.009,0.049, | 0.026,0.058,0.011, | -0,012 |
| 153506 | ENSG00000162086 | ZNF75A | chr16 | + | 3316953 | 3317022 | 3312676 | 3312768 | 3317189 | 3317252 | 153506 | 22,18,34,24,17,18,14,24,12 | 3,3,7,9,5,3,3,1,1,2,0,2 | 20,23,16,15,12,7,12,7,3,6,1 | 4,4,8,4,5,3,4,5,3,5,6,2,3,1,6 | 218 | 149 | 0.0178188556098 | 0.85226002483397 | 0.834,0.804,0.769, | 0.774,0.797,0.578, | 0,139 |
| 47026 | ENSG00000214106 | PAXIP1-AS2 | chr7 | + | 154928955 | 154929024 | 154928459 | 154928767 | 154944026 | 154944398 | 47026 | 21,17,26,32,26,9,12,15,22, | 0,2,2,0,2,0,0,2,5,0,0,2 | 22,21,8,10,10,5,12,9,6,22,9 | 0,3,0,0,0,3,4,3,0,3,0,1,2,0,0 | 218 | 149 | 0.0178232907662 | 0.85228275825372 | 1.0,0.853,0.899,1.0 | 1.0,0.827,1.0,1.0,1 | 0,055 |
| 250200 | ENSG00000035403 | VCL | chr10 | + | 74063685 | 74063797 | 74043090 | 74043153 | 74070669 | 74070820 | 250200 | 3,1,2,0,3,4,2,1,0,0,1,2 | 4,6,9,19,8,4,3,3,2,4,2,5 | 4,0,1,9,2,1,0,3,3,0,0,2,4,0,2 | 10,5,0,5,0,9,3,2,5,4,6,11,14 | 261 | 149 | 0.0178333057466 | 0.852411467365405 | 0.3,0.087,0.113,0.0 | 0.186,0.0,1.0,0.507 | -0,108 |
| 58950 | ENSG00000160055 | TMEM234 | chr1 | - | 32217436 | 32217577 | 32217258 | 32217351 | 32221130 | 32221197 | 58950 | 13,4,29,5,1,12,5,6,5,2,4,1 | 16,14,23,21,10,9,3,9,12,5,5 | 9,10,5,19,5,1,7,11,8,8,9,9,6 | 16,3,7,7,5,4,4,6,11,6,6,8,12 | 290 | 149 | 0.0178339032832 | 0.852411467365405 | 0.295,0.128,0.393, | 0.224,0.631,0.268, | -0,126 |
| 115560 | ENSG00000152795 | HNRNPDL | chr4 | - | 82425562 | 82425667 | 82424628 | 82424883 | 82426036 | 82426129 | 115560 | 11,9,22,11,9,6,11,12,15,2,2 | 68,36,370,202,166,186,95, | 19,14,11,6,4,5,5,1,9,10,9,1 | 272,222,114,64,104,80,61, | 254 | 149 | 0.0178503195662 | 0.852940921078119 | 0.087,0.128,0.034, | 0.039,0.036,0.054, | 0,035 |
| 198032 | ENSG00000112232 | KHDRBS2 | chr6 | - | 61749520 | 61749683 | 61732681 | 61732764 | 61894634 | 61894833 | 198032 | 13,9,13,14,4,10,10,3,11,1,1 | 14,11,27,19,11,10,11,16,34 | 17,18,7,23,5,3,0,3,2,4,14,9 | 26,8,5,9,2,0,9,5,4,4,9,6,3,4 | 298 | 149 | 0.0178561968078 | 0.852940921078119 | 0.317,0.29,0.194,0 | 0.246,0.529,0.412, | -0,157 |
| 50215 | ENSG00000122224 | LY9 | chr1 | + | 160814419 | 160814761 | 160799752 | 160800082 | 160818217 | 160818319 | 50215 | 9,7,10,13,14,4,15,7,10,2,2 | 0,0,0,0,1,0,0,0,0,0,0,0 | 20,14,7,3,2,7,6,21,4,22,11, | 3,2,0,0,1,4,4,0,0,1,1,1,0,2,0 | 298 | 149 | 0.0178568690866 | 0.852940921078119 | 1.0,1.0,1.0,1.0,0.87 | 0.769,0.778,1.0,1.0 | 0,156 |
| 9673 | ENSG00000178971 | CTC1 | chr17 | - | 8235830 | 8235959 | 8234454 | 8235285 | 8236082 | 8236342 | 9673 | 42,18,46,21,22,32,22,24,25 | 7,7,22,10,12,21,18,9,4,9,4, | 22,9,11,10,19,22,23,27,8,2 | 14,17,7,20,10,5,12,20,13,8 | 278 | 149 | 0.0178704130658 | 0.853398463404748 | 0.763,0.58,0.528,0 | 0.457,0.221,0.457, | 0,114 |
| 8143 | ENSG00000124155 | PIGT | chr20 | + | 45416516 | 45416694 | 45416096 | 45416343 | 45419294 | 45419395 | 8143 | 35,40,54,20,16,25,30,31,19 | 2,0,0,0,0,0,3,2,0,0,2,1 | 31,32,19,12,31,15,15,25,16 | 0,1,0,3,0,1,2,5,0,0,3,7,0,0,2 | 298 | 149 | 0.0178835084702 | 0.853834385880571 | 0.897,1.0,1.0,1.0,1 | 1.0,0.941,1.0,0.667 | 0,05 |
| 32764 | ENSG00000223865 | HLA-DPB1 | chr6 | + | 33080671 | 33080935 | 33075989 | 33076141 | 33084949 | 33084957 | 32764 | 2271,1665,1755,1929,410, | 6,3,15,6,0,0,1,3,1,2,3,0 | 1104,1284,804,781,918,61 | 6,23,5,25,1,1,1,8,5,2,3,19,3 | 298 | 149 | 0.0179006014781 | 0.854147743981758 | 0.995,0.996,0.983, | 0.989,0.965,0.988, | 0,009 |
| 213364 | ENSG00000068724 | TTC7A | chr2 | + | 46958371 | 46958552 | 46956838 | 46957007 | 46974972 | 46975103 | 213364 | 3,3,6,5,1,19,6,13,2,2,0,4 | 48,27,110,72,55,90,37,62,4 | 5,5,0,0,3,4,3,5,4,6,3,3,8,5,6 | 67,108,51,51,45,43,35,166 | 298 | 149 | 0.0179025989558 | 0.854147743981758 | 0.03,0.053,0.027,0 | 0.036,0.023,0.0,0.0 | 0,009 |
| 252215 | ENSG00000198825 | INPP5F | chr10 | + | 119823500 | 119823902 | 119823070 | 119823199 | 119826630 | 119826702 | 252215 | 225,89,189,93,120,149,60, | 1,0,1,2,5,2,3,0,0,0,0,3 | 129,111,201,83,29,53,99,9 | 7,2,2,0,0,0,0,0,4,4,4,0,0,1,5 | 298 | 149 | 0.0179034956145 | 0.854147743981758 | 0.991,1.0,0.99,0.95 | 0.902,0.965,0.98,1 | 0,017 |
| 237789 | ENSG00000176623 | RMDN1 | chr8 | - | 86474050 | 86474358 | 86472352 | 86472479 | 86474819 | 86474914 | 237789 | 6,4,67,61,17,11,10,19,2,2,3 | 0,0,0,0,1,0,0,0,0,0,0,0 | 21,22,18,5,19,14,4,14,6,19 | 4,2,2,7,0,0,0,5,0,0,0,0,0,0,0 | 298 | 149 | 0.0179059458017 | 0.854147743981758 | 1.0,1.0,1.0,1.0,0.89 | 0.724,0.846,0.818, | 0,102 |
| 5479 | ENSG00000167232 | ZNF91 | chr19 | - | 23373741 | 23373837 | 23357695 | 23362725 | 23374637 | 23374764 | 5479 | 61,54,74,48,17,42,28,63,43 | 4,2,7,1,1,1,1,1,0,4,2,0 | 52,43,53,55,24,11,20,36,29 | 0,7,5,7,8,6,1,1,1,0,1,5,2,2,5 | 245 | 149 | 0.0179615168817 | 0.856527402014374 | 0.903,0.943,0.865, | 1.0,0.789,0.866,0.8 | 0,046 |
| 232787 | ENSG00000104921 | FCER2 | chr19 | - | 7698789 | 7698872 | 7698355 | 7698409 | 7699738 | 7700084 | 232787 | 5,0,3,3,0,0,3,0,1,1,1,4 | 22,13,17,16,8,8,5,5,12,8,0, | 1,3,0,0,0,0,1,1,0,3,0,0,1,1,0 | 10,20,4,8,8,3,1,1,9,1,1,7,1, | 232 | 149 | 0.0179637909627 | 0.856527402014374 | 0.127,0.0,0.102,0.1 | 0.06,0.088,0.0,0.0, | 0,044 |
| 112558 | ENSG00000196150 | ZNF250 | chr8 | - | 144889580 | 144889694 | 144886839 | 144886902 | 144889932 | 144890059 | 112558 | 19,26,31,11,12,29,15,4,14, | 3,0,3,0,2,4,3,2,0,0,0,2 | 18,19,19,15,24,17,4,30,16, | 5,0,0,2,0,0,0,0,4,1,4,0,0,0,1 | 263 | 149 | 0.0179779819909 | 0.856923005099187 | 0.782,1.0,0.854,1.0 | 0.671,1.0,1.0,0.809 | -0,061 |
| 153023 | ENSG00000140563 | MCTP2 | chr15 | + | 94314731 | 94314864 | 94314281 | 94314344 | 94315528 | 94315637 | 153023 | 3,1,4,7,3,2,3,7,4,0,1,4 | 49,39,141,68,73,99,91,76,4 | 4,3,6,11,4,9,1,2,4,11,4,6,1, | 86,89,52,98,127,61,60,88,9 | 282 | 149 | 0.0179800506945 | 0.856923005099187 | 0.031,0.013,0.015, | 0.024,0.017,0.057, | -0,011 |
| 230080 | ENSG00000042980 | ADAM28 | chr8 | + | 24353518 | 24353832 | 24351986 | 24352052 | 24354383 | 24354472 | 230080 | 16,6,111,38,2,6,6,11,8,2,1, | 0,0,5,3,5,5,3,3,1,0,4,3 | 35,26,14,6,14,3,4,9,18,13,2 | 3,0,3,1,1,1,1,5,0,2,0,1,2,1,2 | 298 | 149 | 0.01803710809 | 0.859049013582338 | 1.0,1.0,0.917,0.864 | 0.854,1.0,0.7,0.75, | -0,127 |
| 226267 | ENSG00000071462 | BUD23 | chr7 | + | 73693991 | 73694029 | 73693623 | 73693669 | 73697604 | 73697694 | 226267 | 15,16,25,18,3,7,6,8,10,7,2, | 0,2,0,0,0,1,1,0,0,0,0,0 | 14,27,10,9,8,11,8,7,8,6,6,2 | 3,2,0,10,2,0,3,4,0,0,0,0,0,0,0 | 187 | 149 | 0.0180396080255 | 0.859049013582338 | 1.0,0.864,1.0,1.0,1 | 0.788,0.915,1.0,0.4 | 0,078 |
| 212258 | ENSG00000130038 | CRACR2A | chr12 | - | 3678925 | 3679098 | 3673445 | 3673592 | 3680237 | 3680349 | 212258 | 4,7,25,14,9,12,4,14,11,2,9, | 0,0,3,1,1,7,3,2,2,0,1,0 | 12,7,8,5,11,6,8,15,12,12,24 | 1,1,1,1,1,4,2,1,0,0,2,0,1,0,0 | 298 | 149 | 0.018044408153 | 0.859049013582338 | 1.0,1.0,0.806,0.875 | 0.857,0.778,0.8,0.7 | -0,016 |
| 67557 | ENSG00000204217 | BMPR2 | chr2 | + | 202532584 | 202532732 | 202520086 | 202520201 | 202542310 | 202542447 | 67557 | 77,38,152,76,44,54,46,106 | 2,5,3,0,0,3,4,1,0,0,0,1 | 92,133,52,36,53,33,30,48,7 | 6,6,2,4,6,1,4,0,2,4,6,0,0,0,2 | 297 | 149 | 0.018045641763 | 0.859049013582338 | 0.951,0.792,0.962, | 0.885,0.917,0.929, | 0,023 |
| 32884 | ENSG00000102445 | RUBCNL | chr13 | - | 46386415 | 46386725 | 46368732 | 46368815 | 46387133 | 46387500 | 32884 | 49,40,21,8,6,15,21,41,52,7 | 3,2,10,2,2,6,5,6,0,2,0,1 | 22,26,24,39,3,7,23,20,0,9,4 | 11,12,2,2,4,2,1,3,2,1,8,4,9, | 298 | 149 | 0.0180474732341 | 0.859049013582338 | 0.891,0.909,0.512, | 0.5,0.52,0.857,0.90 | 0,152 |
| 175538 | ENSG00000100151 | PICK1 | chr22 | + | 38070241 | 38070413 | 38069032 | 38069122 | 38070837 | 38070891 | 175538 | 5,5,4,3,5,0,2,1,1,0,0,3 | 21,12,14,7,3,8,4,6,15,6,3,1 | 0,1,5,2,2,0,1,1,1,1,9,0,2,2,2 | 9,7,11,7,3,7,6,2,5,3,11,6,6, | 298 | 149 | 0.0180486065642 | 0.859049013582338 | 0.106,0.172,0.125, | 0.0,0.067,0.185,0.1 | 0,009 |
| 92371 | ENSG00000137312 | FLOT1 | chr6 | - | 30741613 | 30741704 | 30741189 | 30741333 | 30742146 | 30742203 | 92371 | 29,16,42,32,16,17,11,14,12 | 1,0,3,2,0,0,0,2,4,1,0,0 | 21,26,8,12,11,7,9,28,4,20,1 | 4,0,2,0,2,0,1,3,0,2,2,0,2,0,3 | 240 | 149 | 0.0180782496193 | 0.860212475791703 | 0.947,1.0,0.897,0.9 | 0.765,1.0,0.713,1.0 | 0,054 |
| 183865 | ENSG00000143458 | GABPB2 | chr1 | + | 151073040 | 151073157 | 151070803 | 151070934 | 151088189 | 151088297 | 183865 | 3,2,0,1,2,2,0,2,0,0,0,2 | 30,8,21,12,11,6,14,9,11,1,2 | 6,1,0,0,0,1,0,2,2,2,5,0,3,3,4 | 16,8,11,14,4,7,11,10,4,6,13 | 266 | 149 | 0.0180810442756 | 0.860212475791703 | 0.053,0.123,0.0,0.0 | 0.174,0.065,0.0,0.0 | -0,022 |
| 238877 | ENSG00000171988 | JMJD1C | chr10 | - | 63264650 | 63264785 | 63219876 | 63219983 | 63380317 | 63380482 | 238877 | 52,31,108,76,16,28,30,31,1 | 1,0,4,1,0,1,1,5,0,1,0,1 | 66,67,32,82,49,38,29,25,42 | 0,3,4,2,4,0,2,3,1,1,4,3,2,1,3 | 284 | 149 | 0.0180951016636 | 0.8606910113057 | 0.965,1.0,0.934,0.9 | 1.0,0.921,0.808,0.9 | 0,008 |
| 149963 | ENSG00000213614 | HEXA | chr15 | - | 72362322 | 72362466 | 72356524 | 72356617 | 72375719 | 72375972 | 149963 | 2,8,2,1,1,1,0,8,0,1,2,9 | 20,14,90,73,47,44,31,35,18 | 0,5,0,4,3,0,0,0,1,4,4,1,3,8,0 | 72,77,20,16,38,26,19,55,27 | 293 | 149 | 0.0181013966754 | 0.860800200583963 | 0.048,0.225,0.011, | 0.0,0.032,0.0,0.113 | 0,025 |
| 43295 | ENSG00000114648 | KLHL18 | chr3 | + | 47297561 | 47297672 | 47282916 | 47283094 | 47319652 | 47319783 | 43295 | 1,0,3,9,2,3,0,1,0,0,0,2 | 7,12,9,13,5,10,5,6,4,2,4,12 | 1,1,0,0,6,1,0,0,1,2,0,1,0,5,0 | 13,8,5,7,9,7,6,11,6,15,6,20 | 260 | 149 | 0.0181377586719 | 0.862338841774179 | 0.076,0.0,0.16,0.28 | 0.042,0.067,0.0,0.0 | 0,032 |
| 146504 | ENSG00000179583 | CIITA | chr16 | + | 10901797 | 10902184 | 10901513 | 10901571 | 10902657 | 10902722 | 146504 | 381,98,179,163,168,66,164 | 6,2,2,0,5,2,4,0,1,1,0,4 | 172,164,76,153,59,59,53,1 | 1,2,0,2,2,0,0,0,0,1,1,2,0,0,1 | 298 | 149 | 0.018154082638 | 0.862924328679191 | 0.969,0.961,0.978, | 0.989,0.976,1.0,0.9 | -0,009 |
| 10897 | ENSG00000108819 | PPP1R9B | chr17 | - | 50140092 | 50140359 | 50139459 | 50139581 | 50141268 | 50141373 | 10897 | 236,177,201,140,118,84,49 | 3,0,10,4,2,1,0,0,1,2,0,1 | 91,155,72,82,52,56,16,73,4 | 3,6,0,3,0,2,1,0,0,7,0,12,3,2 | 298 | 149 | 0.0181663322771 | 0.863190663360845 | 0.975,1.0,0.91,0.94 | 0.938,0.928,1.0,0.9 | 0,026 |
| 203045 | ENSG00000064687 | ABCA7 | chr19 | + | 1057973 | 1058059 | 1057313 | 1057429 | 1058145 | 1058269 | 203045 | 8,17,27,16,15,13,16,19,15, | 5,0,0,2,1,2,1,1,1,1,1,1 | 14,21,11,10,10,12,2,11,12, | 0,2,1,1,0,0,0,1,0,0,2,1,0,1,0 | 235 | 149 | 0.0181713139661 | 0.863190663360845 | 0.504,1.0,1.0,0.835 | 1.0,0.869,0.875,0.8 | -0,085 |
| 169179 | ENSG00000100422 | CERK | chr22 | - | 46691869 | 46692022 | 46691571 | 46691777 | 46693426 | 46693503 | 169179 | 17,6,4,1,4,2,3,1,2,5,2,5 | 15,18,34,59,17,8,11,8,5,6,2 | 7,0,2,1,1,0,1,1,2,7,4,5,6,2,4 | 28,19,18,2,4,4,21,34,9,9,7, | 298 | 149 | 0.0181717173441 | 0.863190663360845 | 0.362,0.143,0.056, | 0.111,0.0,0.053,0.2 | 0,043 |
| 55033 | ENSG00000148399 | DPH7 | chr9 | - | 137577469 | 137577603 | 137576308 | 137576402 | 137578624 | 137578889 | 55033 | 24,19,24,14,21,27,13,13,30 | 1,0,2,0,1,2,3,2,0,4,0,1 | 10,19,27,37,6,7,6,13,15,39 | 0,0,0,2,0,0,0,0,0,0,0,0,0,0,0 | 283 | 149 | 0.0181935049092 | 0.864034919486325 | 0.927,1.0,0.863,1.0 | 1.0,1.0,1.0,0.907,1 | -0,104 |
| 3055 | ENSG00000129351 | ILF3 | chr19 | + | 10687345 | 10687708 | 10683916 | 10683970 | 10689178 | 10689846 | 3055 | 20,13,19,8,21,16,9,12,5,4,4 | 9,6,9,8,0,7,7,1,4,4,2,4 | 12,8,10,15,7,4,18,7,1,3,19, | 9,12,17,26,5,6,4,7,6,12,5,9 | 298 | 149 | 0.0182048931328 | 0.864385032840432 | 0.526,0.52,0.514,0 | 0.4,0.25,0.227,0.22 | 0,148 |
| 120267 | ENSG00000166532 | RIMKLB | chr12 | + | 8699784 | 8700614 | 8697921 | 8698297 | 8713810 | 8714041 | 120267 | 1,0,7,2,0,4,0,8,0,2,1,0 | 0,4,45,7,2,10,10,19,16,7,2, | 3,3,0,2,2,0,5,10,2,2,2,0,6,6 | 11,6,2,9,6,3,8,14,12,16,8,5 | 298 | 149 | 0.0182347807152 | 0.865613164559597 | 1.0,0.0,0.072,0.125 | 0.12,0.2,0.0,0.1,0.1 | 0,041 |
| 211024 | ENSG00000177646 | ACAD9 | chr3 | + | 128883966 | 128884104 | 128879653 | 128879841 | 128884652 | 128884746 | 211024 | 9,0,1,2,0,1,1,2,0,1,0,1 | 10,6,18,1,8,2,7,10,3,3,1,2 | 0,0,0,2,1,0,0,1,1,0,0,0,0,0,0 | 11,3,4,3,1,4,3,3,3,2,9,7,10, | 287 | 149 | 0.0182446289958 | 0.865889689833524 | 0.318,0.0,0.028,0.5 | 0.0,0.0,0.0,0.257,0 | 0,076 |
| 81981 | ENSG00000145782 | ATG12 | chr5 | - | 115840496 | 115840612 | 115837627 | 115837764 | 115841389 | 115841837 | 81981 | 8,9,8,4,6,0,1,2,0,0,0,2 | 61,77,56,37,14,14,10,18,27 | 1,0,1,0,0,0,0,2,1,1,1,5,6,10 | 32,39,37,34,30,20,13,27,15 | 265 | 149 | 0.0182554565939 | 0.866034742153004 | 0.069,0.062,0.074, | 0.017,0.0,0.015,0.0 | 0,016 |
| 1792 | ENSG00000100330 | MTMR3 | chr22 | + | 30022028 | 30022139 | 30019479 | 30020884 | 30022608 | 30022697 | 1792 | 28,20,27,8,20,15,17,25,20, | 3,13,26,9,9,14,11,7,2,0,1,7 | 17,12,11,11,10,10,19,21,4, | 8,14,3,24,13,3,18,21,4,6,6, | 260 | 149 | 0.0182587595473 | 0.866034742153004 | 0.842,0.469,0.373, | 0.549,0.329,0.678, | 0,158 |
| 68585 | ENSG00000055609 | KMT2C | chr7 | - | 152238706 | 152238826 | 152235816 | 152235933 | 152247901 | 152248620 | 68585 | 91,71,122,62,46,59,30,51,7 | 6,2,2,3,0,2,4,1,2,0,0,3 | 68,72,42,64,43,43,34,49,48 | 0,5,0,18,2,0,2,0,0,1,1,1,3,4 | 269 | 149 | 0.0182643627034 | 0.866034742153004 | 0.894,0.952,0.971, | 1.0,0.889,1.0,0.663 | 0,013 |
| 195467 | ENSG00000172667 | ZMAT3 | chr3 | - | 179030692 | 179030999 | 179027756 | 179027812 | 179067482 | 179067809 | 195467 | 4,11,14,11,3,0,0,3,4,1,1,5 | 4,0,1,1,0,2,2,2,0,0,0,0 | 7,11,7,7,9,7,2,9,11,15,8,25 | 0,0,0,3,1,2,0,0,1,0,0,1,2,0,0 | 298 | 149 | 0.0182674171568 | 0.866034742153004 | 0.333,1.0,0.875,0.8 | 1.0,1.0,1.0,0.538,0 | -0,204 |
| 142345 | ENSG00000083454 | P2RX5 | chr17 | - | 3689491 | 3689624 | 3688625 | 3688676 | 3690069 | 3690150 | 142345 | 30,10,34,20,9,2,11,4,7,7,1, | 11,3,12,9,5,0,2,3,0,0,0,2 | 18,7,14,2,13,13,5,3,4,9,0,5 | 12,5,8,2,8,3,2,0,4,1,1,21,11 | 282 | 149 | 0.0182678040309 | 0.866034742153004 | 0.59,0.638,0.6,0.54 | 0.442,0.425,0.48,0 | 0,254 |
| 37215 | ENSG00000155189 | AGPAT5 | chr8 | + | 6741660 | 6741751 | 6732560 | 6732650 | 6755050 | 6755174 | 37215 | 25,12,42,35,43,24,18,24,22 | 11,6,15,20,21,23,14,18,4,1 | 34,36,8,3,18,14,11,40,13,4 | 36,24,19,7,44,7,17,26,10,8 | 240 | 149 | 0.0182764854798 | 0.866156277205558 | 0.585,0.554,0.635, | 0.37,0.482,0.207,0 | 0,148 |
| 84084 | ENSG00000198799 | LRIG2 | chr1 | + | 113110241 | 113110562 | 113107593 | 113107757 | 113112478 | 113112760 | 84084 | 17,7,43,18,20,15,7,15,3,6,5 | 0,2,9,2,1,2,3,6,0,0,0,2 | 10,9,8,7,2,9,7,13,11,16,7,6 | 0,4,3,10,0,7,7,4,0,3,14,3,2, | 298 | 149 | 0.0182784162647 | 0.866156277205558 | 1.0,0.636,0.705,0.8 | 1.0,0.529,0.571,0.2 | 0,157 |
| 120076 | ENSG00000171634 | BPTF | chr17 | + | 67891843 | 67892034 | 67874816 | 67875020 | 67893369 | 67893725 | 120076 | 67,82,94,86,49,70,38,55,32 | 8,5,13,2,2,4,7,6,4,0,0,6 | 60,64,69,57,44,29,27,52,44 | 11,15,16,11,10,1,2,3,4,4,2, | 298 | 149 | 0.0183088605502 | 0.867407960616354 | 0.807,0.891,0.783, | 0.732,0.681,0.683, | 0,06 |
| 181558 | ENSG00000075914 | EXOSC7 | chr3 | + | 44990014 | 44990295 | 44989549 | 44989644 | 44997086 | 44997252 | 181558 | 0,0,1,2,0,1,2,1,0,0,0,0 | 29,27,33,46,21,26,18,12,13 | 0,1,2,2,0,0,0,0,0,8,8,3,0,3 | 28,32,13,7,26,8,12,16,11,1 | 298 | 149 | 0.0183144294345 | 0.867480845426248 | 0.0,0.0,0.015,0.02 | 0.0,0.015,0.071,0.1 | -0,042 |
| 223833 | ENSG00000095015 | MAP3K1 | chr5 | + | 56888225 | 56888268 | 56885931 | 56886063 | 56893530 | 56896152 | 223833 | 23,8,20,15,9,2,7,1,2,1,3,3 | 2,1,1,0,0,0,2,0,0,1,0,4 | 20,15,11,21,5,13,5,12,9,9,4 | 5,2,0,4,1,4,0,0,2,2,0,1,1,4,2 | 192 | 149 | 0.0183240979577 | 0.867747838841304 | 0.899,0.861,0.939, | 0.756,0.853,1.0,0.8 | 0,007 |
| 205664 | ENSG00000122912 | SLC25A16 | chr10 | - | 68496433 | 68496478 | 68493448 | 68493570 | 68503631 | 68503695 | 205664 | 5,0,3,0,3,2,1,0,1,0,0,1 | 14,20,39,22,13,6,22,11,15, | 2,0,0,2,1,2,0,2,4,6,0,5,3,1,2 | 6,17,9,12,10,9,5,10,4,12,22 | 194 | 149 | 0.0183561057487 | 0.86904067730064 | 0.215,0.0,0.056,0.0 | 0.204,0.0,0.0,0.113 | -0,053 |
| 184341 | ENSG00000137996 | RTCA | chr1 | + | 100267165 | 100267245 | 100266523 | 100266624 | 100268136 | 100268295 | 184341 | 1,4,1,0,3,2,0,2,0,0,2,1 | 7,8,4,7,0,8,1,4,5,5,0,4 | 4,1,0,0,0,0,0,1,0,0,0,0,0,4,1 | 3,11,7,3,6,2,1,7,1,2,8,2,5,8 | 229 | 149 | 0.018359474052 | 0.86904067730064 | 0.085,0.245,0.14,0 | 0.465,0.056,0.0,0.0 | 0,172 |
| 211943 | ENSG00000164985 | PSIP1 | chr9 | - | 15470994 | 15471231 | 15469937 | 15469993 | 15472631 | 15472750 | 211943 | 6,3,8,2,2,1,1,6,9,2,2,1 | 51,20,59,33,59,32,29,36,30 | 5,5,0,4,1,3,5,6,2,2,6,6,5,9,4 | 35,20,18,1,13,25,13,27,6,2 | 298 | 149 | 0.0183998736493 | 0.87076147876264 | 0.056,0.07,0.063,0 | 0.067,0.111,0.0,0.0 | -0,067 |
| 67095 | ENSG00000111605 | CPSF6 | chr12 | + | 69258034 | 69258145 | 69257731 | 69257905 | 69258589 | 69259094 | 67095 | 29,23,28,13,17,17,5,17,8,1 | 53,30,72,83,82,63,47,79,69 | 44,28,24,19,6,7,11,23,13,1 | 92,71,25,23,17,23,26,122,2 | 260 | 149 | 0.0184170858156 | 0.871384435519324 | 0.239,0.305,0.182, | 0.215,0.184,0.355, | -0,051 |
| 197536 | ENSG00000132300 | PTCD3 | chr2 | + | 86132928 | 86133063 | 86132317 | 86132424 | 86133177 | 86133256 | 197536 | 3,4,2,1,4,8,5,8,6,0,0,1 | 46,19,75,59,51,53,31,49,11 | 10,14,2,0,1,0,0,5,1,3,8,0,4, | 85,83,37,21,40,31,15,55,83 | 284 | 149 | 0.018424590749 | 0.871439842498444 | 0.033,0.099,0.014, | 0.058,0.081,0.028, | 0,032 |
| 159858 | ENSG00000055163 | CYFIP2 | chr5 | + | 157360559 | 157360622 | 157360281 | 157360372 | 157361467 | 157361598 | 159858 | 5,3,10,11,8,8,9,13,3,0,2,5 | 122,110,191,163,145,106,1 | 6,8,3,8,6,3,23,14,9,13,14,0 | 121,97,77,85,105,94,64,15 | 212 | 149 | 0.0184283274582 | 0.871439842498444 | 0.028,0.019,0.035, | 0.034,0.055,0.027, | -0,021 |
| 8240 | ENSG00000134444 | RELCH | chr18 | + | 62279773 | 62279856 | 62275373 | 62275473 | 62280645 | 62280709 | 8240 | 2,1,0,0,2,4,1,1,0,0,0,3 | 11,5,12,9,8,19,2,9,3,3,2,3 | 1,0,4,3,0,4,0,1,3,0,1,2,7,2,4 | 6,13,14,2,4,3,6,8,6,5,3,20,1 | 232 | 149 | 0.0184304034412 | 0.871439842498444 | 0.105,0.114,0.0,0.0 | 0.097,0.0,0.155,0.4 | -0,052 |
| 197034 | ENSG00000147099 | HDAC8 | chrX | - | 72568805 | 72568884 | 72567888 | 72568030 | 72572056 | 72572104 | 197034 | 5,0,12,3,1,2,8,3,4,2,4,5 | 0,2,3,0,0,2,0,6,1,2,2,0 | 7,4,1,4,3,2,0,0,1,1,5,6,6,5,3 | 4,1,2,1,4,2,1,1,3,2,2,0,1,1,4 | 228 | 149 | 0.0184374991844 | 0.871583875896683 | 1.0,0.0,0.723,1.0,1 | 0.534,0.723,0.246, | 0,216 |
| 149085 | ENSG00000169375 | SIN3A | chr15 | - | 75393351 | 75393704 | 75392241 | 75392815 | 75394679 | 75394863 | 149085 | 10,3,11,0,0,2,10,3,2,0,2,3 | 77,92,69,45,33,33,23,32,57 | 4,9,3,26,3,4,3,5,6,5,4,2,6,2 | 52,85,40,63,32,28,34,33,19 | 298 | 149 | 0.018464058018 | 0.872647709985103 | 0.061,0.016,0.074, | 0.037,0.05,0.036,0 | -0,014 |
| 46746 | ENSG00000015479 | MATR3 | chr5 | + | 139307238 | 139308327 | 139293741 | 139293772 | 139314674 | 139314736 | 46746 | 141,108,351,296,163,191,1 | 3,2,6,11,0,9,9,6,2,0,3,2 | 260,228,134,115,65,60,132 | 5,14,2,5,2,3,11,5,4,2,10,11 | 298 | 149 | 0.0185195732738 | 0.874988917381142 | 0.959,0.964,0.967, | 0.963,0.891,0.971, | 0,023 |

| | | | | | | | | | | | | | | | | | | | | | | |
|---|---|---|---|---|---|---|---|---|---|---|---|---|---|---|---|---|---|---|---|---|---|---|
| 168073 | ENSG00000146350 | TBC1D32 | chr6 | - | 121256083 | 121256299 | 121255327 | 121255410 | 121279120 | 121279245 | 168073 | 3,3,8,1,7,2,5,8,5,0,3,4 | 11,4,5,2,2,4,2,1,2,4,2,0 | 9,3,9,12,3,1,6,10,3,3,5,11,2 | 4,5,4,3,2,1,0,0,3,0,6,5,0,0,0 | 298 | 149 | 0.0185239232626 | 0.874988917381142 | 0.12,0.273,0.444,0 | 0.529,0.231,0.529, | -0,182 |
| 147579 | ENSG00000172775 | PSME3IP1 | chr16 | - | 57181553 | 57181659 | 57178525 | 57178852 | 57185820 | 57186055 | 147579 | 11,12,33,19,9,9,9,12,6,3,3, | 1,0,3,1,0,5,6,1,0,0,0,10 | 5,5,5,9,13,4,15,13,15,11,14 | 1,0,0,0,3,2,1,4,5,4,3,4,5,8,7 | 255 | 149 | 0.0185257908782 | 0.874988917381142 | 0.865,1.0,0.865,0. | 0.745,1.0,1.0,1.0,0 | 0,137 |
| 218927 | ENSG00000204592 | HLA-E | chr6 | + | 30491538 | 30491653 | 30491136 | 30491412 | 30492403 | 30492436 | 218927 | 364,373,655,698,242,208,3 | 11,6,11,3,1,5,6,1,0,2,3,8 | 452,506,331,204,303,196,2 | 2,4,8,13,4,0,4,17,1,0,12,18 | 264 | 149 | 0.0185759686199 | 0.87716636727286 | 0.949,0.972,0.971, | 0.992,0.986,0.959, | 0,009 |
| 60274 | ENSG00000138138 | ATAD1 | chr10 | - | 87778520 | 87778731 | 87776320 | 87776427 | 87784469 | 87784670 | 60274 | 3,12,21,12,4,4,2,5,0,2,4,1 | 17,36,46,33,35,33,28,32,12 | 2,13,3,4,12,3,0,0,11,2,1,5,5 | 54,58,29,39,30,23,24,31,28 | 298 | 149 | 0.0185855330818 | 0.877425503559749 | 0.081,0.143,0.186, | 0.018,0.101,0.049, | 0,037 |
| 8178 | ENSG00000124155 | PIGT | chr20 | + | 45424460 | 45424579 | 45424269 | 45424381 | 45425573 | 45426189 | 8178 | 55,48,42,18,13,25,15,19,29 | 1,1,0,1,0,1,1,0,4,1,2,1 | 33,20,42,19,19,9,20,17,4,1 | 2,3,2,2,5,0,3,0,1,0,0,1,1,0,3 | 268 | 149 | 0.0185967551346 | 0.877762764117345 | 0.968,0.964,1.0,0. | 0.902,0.788,0.921, | -0,014 |
| 4185 | ENSG00000104814 | MAP4K1 | chr19 | - | 38589177 | 38589275 | 38587640 | 38587817 | 38593281 | 38593337 | 4185 | 0,0,8,4,2,1,3,4,1,0,1,2 | 44,28,95,64,56,41,62,54,48 | 8,2,2,1,2,7,1,8,2,0,4,7,7,5,2 | 77,53,35,38,63,34,45,58,37 | 247 | 149 | 0.0186080229227 | 0.8780866742109 | 0.0,0.0,0.048,0.03 | 0.059,0.022,0.033, | -0,017 |
| 150983 | ENSG00000129003 | VPS13C | chr15 | - | 62032770 | 62032874 | 62023410 | 62023520 | 62033440 | 62033542 | 150983 | 7,2,16,7,4,10,1,18,2,0,3,7 | 107,54,181,76,71,102,69,1 | 13,22,9,31,6,1,6,19,5,5,13, | 144,121,85,131,74,47,53,7 | 253 | 149 | 0.0186117771499 | 0.8780866742109 | 0.037,0.021,0.049, | 0.05,0.097,0.059,0 | -0,017 |
| 251631 | ENSG00000107957 | SH3PXD2A | chr10 | - | 103660771 | 103660864 | 103627088 | 103627202 | 103660982 | 103661114 | 251631 | 0,0,13,3,0,0,2,2,0,0,0,2 | 5,10,82,39,14,25,15,22,1,2 | 0,0,0,0,0,0,0,0,0,0,0,0,0,0 | 17,23,13,20,4,2,6,6,11,8,31 | 242 | 149 | 0.0186174965154 | 0.878164013260148 | 0.0,0.0,0.089,0.04 | 0.0,0.0,0.0,0.0,0.0, | 0,028 |
| 120550 | ENSG00000149179 | CSTPP1 | chr11 | + | 46987207 | 46987310 | 46936696 | 46936851 | 47052387 | 47052518 | 120550 | 19,9,26,28,10,26,8,10,7,8,5 | 0,2,0,5,0,2,0,1,0,0,0,2 | 14,6,13,11,6,15,2,12,8,12,1 | 0,1,1,3,1,2,3,1,0,0,0,8,3,1,0 | 252 | 149 | 0.0186836731204 | 0.8809673409983 | 1.0,0.727,1.0,0.76 | 1.0,0.78,0.885,0.6 | 0,086 |
| 99918 | ENSG00000171681 | ATF7IP | chr12 | + | 14457206 | 14457295 | 14456560 | 14456634 | 14460494 | 14461133 | 99918 | 245,142,530,386,158,230,2 | 35,7,36,20,7,12,14,18,22,1 | 257,242,304,239,176,181,1 | 11,16,13,17,11,5,3,13,6,13 | 238 | 149 | 0.0186908485495 | 0.8809673409983 | 0.814,0.927,0.902, | 0.936,0.904,0.936, | -0,018 |
| 28013 | ENSG00000113558 | SKP1 | chr5 | - | 134174461 | 134174514 | 134173925 | 134174022 | 134176854 | 134176950 | 28013 | 4,5,23,8,2,6,4,14,3,0,2,2 | 107,95,370,249,86,104,70, | 7,20,11,2,10,3,8,1,2,0,6,27 | 160,118,75,57,89,53,43,10 | 202 | 149 | 0.0186944148262 | 0.8809673409983 | 0.027,0.037,0.044, | 0.031,0.111,0.098, | -0,02 |
| 194573 | ENSG00000158161 | EYA3 | chr1 | - | 28080331 | 28080455 | 28057993 | 28058094 | 28088523 | 28088637 | 194573 | 7,0,5,6,0,2,0,2,2,0,0,0 | 9,2,24,16,10,7,7,17,8,1,1,2 | 2,0,0,0,1,1,0,1,1,3,0,0,1,6,3 | 10,19,2,3,8,16,10,11,5,20,6 | 273 | 149 | 0.0187011920462 | 0.8809673409983 | 0.298,0.0,0.102,0. | 0.098,0.0,0.0,0.0,0.0 | 0,015 |
| 202299 | ENSG00000136059 | VILL | chr3 | + | 38002395 | 38002491 | 38001701 | 38001860 | 38004254 | 38004399 | 202299 | 16,7,14,5,22,14,16,9,11,18 | 3,5,6,5,2,9,5,2,5,0,1,5 | 10,12,8,5,2,7,8,14,6,8,6,5,1 | 11,5,5,2,5,2,9,11,1,1,11,8,5 | 245 | 149 | 0.0187019479625 | 0.8809673409983 | 0.764,0.46,0.587,0 | 0.356,0.593,0.493, | 0,134 |
| 163104 | ENSG00000157916 | RER1 | chr1 | + | 2393269 | 2393355 | 2391889 | 2391958 | 2395783 | 2396367 | 163104 | 2,1,22,10,5,1,0,10,0,0,1,14 | 1,2,23,8,6,5,1,4,8,4,5,6 | 5,2,7,7,4,3,4,6,4,6,14,4,7,7 | 4,3,4,0,3,1,4,2,1,5,7,17,9,2 | 235 | 149 | 0.0187041861167 | 0.8809673409983 | 0.559,0.241,0.378, | 0.442,0.297,0.526, | -0,22 |
| 161044 | ENSG00000140853 | NLRC5 | chr16 | + | 57055432 | 57055514 | 57055031 | 57055094 | 57058064 | 57058148 | 161044 | 4,2,18,11,5,8,6,7,0,4,3,5 | 4,3,1,2,0,2,6,0,1,2,0,5 | 8,11,6,6,10,6,3,6,5,15,10,8 | 5,5,9,3,3,1,1,10,4,3,7,12,10 | 231 | 149 | 0.0187055802759 | 0.8809673409983 | 0.392,0.301,0.921, | 0.508,0.587,0.301, | 0,139 |
| 113559 | ENSG00000134014 | ELP3 | chr8 | + | 28107912 | 28107976 | 28106712 | 28106783 | 28110369 | 28110422 | 113559 | 46,18,57,57,42,18,26,39,45 | 3,3,2,3,0,5,2,2,0,2,0,3 | 53,38,35,8,10,19,5,41,30,4 | 0,1,5,4,1,1,0,0,5,3,6,0,4,1,0 | 213 | 149 | 0.0187316805556 | 0.882003574198719 | 0.915,0.808,0.952, | 1.0,0.964,0.83,0.5 | -0,001 |
| 242398 | ENSG00000090054 | SPTLC1 | chr9 | - | 92070123 | 92070426 | 92067965 | 92068098 | 92080015 | 92080088 | 242398 | 1,1,3,0,1,0,1,4,0,0,0,2 | 72,27,62,40,20,19,14,27,49 | 0,0,0,18,0,3,4,2,0,0,1,1,0,6 | 38,39,31,10,28,12,16,11,5, | 298 | 149 | 0.0187481211668 | 0.882463255451597 | 0.007,0.018,0.024, | 0.0,0.0,0.0,0.474,0 | -0,039 |
| 140487 | ENSG00000141644 | MBD1 | chr18 | - | 50273123 | 50273426 | 50272847 | 50272955 | 50273563 | 50273863 | 140487 | 2,1,9,6,1,7,1,3,4,0,3,4 | 30,22,28,15,14,20,10,11,21 | 1,5,0,1,2,2,0,5,3,5,4,1,0,3,1 | 17,11,14,43,14,6,12,24,16, | 298 | 149 | 0.0187496432539 | 0.882463255451597 | 0.032,0.022,0.138, | 0.029,0.185,0.0,0.0 | 0,027 |
| 1945 | ENSG00000100150 | DEPDC5 | chr22 | + | 31822692 | 31822790 | 31819021 | 31819225 | 31833914 | 31833980 | 1945 | 29,37,18,20,15,11,11,11,15 | 15,22,6,5,8,9,4,2,33,29,4,1 | 26,17,16,9,21,12,10,9,34,3 | 9,5,2,2,6,2,4,7,10,10,0,9,8, | 247 | 149 | 0.0187549801794 | 0.882521455835689 | 0.538,0.504,0.644, | 0.635,0.672,0.828, | -0,121 |
| 175934 | ENSG00000173889 | PHC3 | chr3 | - | 170149212 | 170149244 | 170145422 | 170145521 | 170171372 | 170171450 | 175934 | 40,34,19,10,19,6,14,12,53, | 31,12,35,23,4,31,22,16,17, | 15,15,21,21,6,13,11,13,5,2 | 34,46,25,29,26,14,18,29,18 | 181 | 149 | 0.0187640836607 | 0.882756828497513 | 0.515,0.7,0.309,0. | 0.266,0.212,0.409, | 0,13 |
| 47608 | ENSG00000117054 | ACADM | chr1 | + | 75728388 | 75728488 | 75724779 | 75724817 | 75732643 | 75732741 | 47608 | 24,19,59,35,20,12,19,22,27 | 18,8,26,26,6,11,7,20,4,0,0, | 30,39,29,8,28,26,6,31,9,29 | 13,9,6,2,17,4,0,0,2,6,2,5,2, | 249 | 149 | 0.0187702602499 | 0.882854432658703 | 0.444,0.587,0.576, | 0.58,0.722,0.743,0 | -0,119 |
| 102245 | ENSG00000111667 | USP5 | chr12 | + | 6856304 | 6856450 | 6856016 | 6856150 | 6856706 | 6856891 | 102245 | 83,69,119,125,73,71,65,55 | 0,0,2,0,0,2,0,2,0,0,0,0 | 80,91,61,25,35,27,22,67,23 | 0,1,5,1,1,1,0,6,0,0,3,3,2,3,0 | 295 | 149 | 0.0187789046815 | 0.883068043151393 | 1.0,1.0,0.968,1.0,1 | 1.0,0.979,0.86,0.9 | 0,036 |
| 113737 | ENSG00000075856 | SART3 | chr12 | - | 108538064 | 108538203 | 108537487 | 108537595 | 108538933 | 108539089 | 113737 | 39,48,113,88,42,57,37,57,5 | 2,1,2,2,2,2,1,2,2,0,0,1 | 42,66,30,18,71,44,36,50,27 | 2,0,1,4,1,1,0,2,1,0,1,1,0,1,0 | 288 | 149 | 0.0187865255026 | 0.883233436096571 | 0.91,0.961,0.967,0 | 0.916,1.0,0.939,0. | 0 |
| 20241 | ENSG00000125107 | CNOT1 | chr16 | - | 58558472 | 58558674 | 58556846 | 58556993 | 58560211 | 58560362 | 20241 | 183,155,190,132,150,115,1 | 14,5,7,3,3,9,12,3,3,1,4,2 | 145,112,130,100,36,81,104 | 16,6,2,15,3,1,5,7,4,5,13,6,5 | 298 | 149 | 0.0187928388886 | 0.883337302212768 | 0.867,0.939,0.931, | 0.819,0.903,0.97,0 | 0,027 |
| 75943 | ENSG00000143442 | POGZ | chr1 | - | 151429602 | 151429711 | 151428122 | 151428413 | 151430665 | 151430841 | 75943 | 77,52,79,38,40,36,36,65,85 | 5,3,0,2,1,0,6,2,3,2,1,1 | 62,70,55,82,33,38,31,63,30 | 3,8,1,17,2,4,3,3,0,3,3,1,0,0 | 258 | 149 | 0.0187974916701 | 0.883363085075828 | 0.899,0.909,1.0,0. | 0.923,0.835,0.969, | 0,013 |
| 205072 | ENSG00000118482 | PHF3 | chr6 | + | 63706706 | 63706876 | 63706028 | 63706224 | 63709150 | 63709214 | 205072 | 78,47,264,133,50,80,59,11 | 3,0,7,1,4,6,6,4,0,0,2,0 | 132,149,59,65,80,45,60,80 | 4,6,2,4,0,2,4,8,3,4,8,1,4,9,3 | 298 | 149 | 0.0188078713375 | 0.883450926188043 | 0.929,1.0,0.95,0.9 | 0.943,0.925,0.937, | 0,031 |
| 154928 | ENSG00000213983 | AP1G2 | chr14 | - | 23566517 | 23566686 | 23566338 | 23566419 | 23567110 | 23567315 | 154928 | 158,86,128,138,92,92,99,8 | 13,13,20,2,9,13,8,11,8,8,3, | 95,92,66,22,56,40,53,78,57 | 8,11,13,7,10,3,8,10,4,11,7, | 298 | 149 | 0.0188090495868 | 0.883450926188043 | 0.859,0.768,0.762, | 0.856,0.807,0.717, | 0,059 |
| 240719 | ENSG00000148090 | AUH | chr9 | - | 91342108 | 91342543 | 91325317 | 91325404 | 91355882 | 91355970 | 240719 | 2,0,6,5,0,0,1,5,0,0,0,2 | 11,7,18,17,12,8,10,6,6,5,4, | 0,1,0,3,1,0,1,0,0,0,0,0,1,2,0 | 17,24,10,6,6,7,5,5,6,17,20, | 298 | 149 | 0.0188116748736 | 0.883450926188043 | 0.083,0.0,0.143,0. | 0.0,0.02,0.0,0.2,0.0 | 0,028 |
| 226269 | ENSG00000071462 | BUD23 | chr7 | + | 73693991 | 73694168 | 73693623 | 73693669 | 73697604 | 73697694 | 226269 | 24,34,42,28,6,17,12,20,15, | 0,2,0,0,0,1,1,0,0,0,0,0 | 30,41,19,18,17,15,15,15,11 | 3,2,0,10,2,0,3,4,0,0,0,0,0,0 | 298 | 149 | 0.0188420423871 | 0.884684036871274 | 1.0,0.895,1.0,1.0,1 | 0.833,0.911,1.0,0. | 0,071 |
| 54560 | ENSG00000198925 | ATG9A | chr2 | - | 219227921 | 219228053 | 219227769 | 219227813 | 219228433 | 219228482 | 54560 | 14,11,29,14,14,5,12,17,3,2 | 0,1,2,3,0,0,3,1,0,0,2,0 | 15,35,16,16,6,13,12,14,7,6 | 2,2,2,2,3,0,2,1,0,0,0,3,2,4,0 | 281 | 149 | 0.0188680280716 | 0.885547502888146 | 1.0,0.854,0.885,0. | 0.799,0.903,0.809, | 0,046 |
| 222518 | ENSG00000166295 | ANAPC16 | chr10 | + | 72223892 | 72224056 | 72220053 | 72220379 | 72230365 | 72230440 | 222518 | 252,119,330,247,115,106,1 | 2,6,11,8,3,2,5,1,12,1,1,3 | 193,126,149,91,78,60,70,1 | 5,1,1,2,0,1,1,3,1,6,6,3,7,2,3 | 298 | 149 | 0.0188723883003 | 0.885547502888146 | 0.984,0.908,0.938, | 0.951,0.984,0.987, | -0,018 |
| 127095 | ENSG00000141759 | TXNL4A | chr18 | - | 79976717 | 79976787 | 79973524 | 79973856 | 79977597 | 79977701 | 127095 | 2,2,4,3,2,0,0,3,4,0,0,0 | 86,59,75,71,30,22,28,30,51 | 0,10,0,0,2,2,1,0,5,0,0,16,2, | 70,31,57,11,47,36,18,24,26 | 219 | 149 | 0.0188727757421 | 0.885547502888146 | 0.016,0.023,0.035, | 0.0,0.18,0.0,0.0,0.0 | -0,017 |
| 55411 | ENSG00000160691 | SHC1 | chr1 | - | 154965542 | 154965781 | 154962297 | 154963931 | 154966161 | 154966231 | 55411 | 13,7,16,26,19,6,4,7,11,10,3 | 0,0,0,0,2,0,1,1,0,0,0,0 | 28,20,5,4,14,12,8,15,4,11,1 | 1,2,0,3,5,0,0,2,1,0,5,2,0,0,0 | 298 | 149 | 0.0189041897868 | 0.886828176112195 | 1.0,1.0,1.0,1.0,0.8 | 0.933,0.833,1.0,0. | 0,103 |
| 89886 | ENSG00000140750 | ARHGAP17 | chr16 | - | 24927683 | 24927732 | 24919394 | 24920260 | 24930783 | 24931160 | 89886 | 4,4,3,7,6,5,4,5,5,0,0,4 | 16,7,41,14,12,19,17,10,22, | 1,1,2,2,2,0,2,2,0,7,5,6,4,1,3 | 20,23,20,10,9,9,7,7,17,25,1 | 198 | 149 | 0.0189232546408 | 0.887529095575076 | 0.158,0.301,0.052, | 0.036,0.032,0.07,0 | 0,037 |
| 183273 | ENSG00000198408 | OGA | chr10 | - | 101795373 | 101795480 | 101793912 | 101793998 | 101797979 | 101798154 | 183273 | 27,16,39,13,5,13,8,14,8,2,1 | 81,75,183,75,45,28,41,43,7 | 30,24,20,46,21,4,17,23,8,9 | 113,106,69,36,83,32,48,45 | 256 | 149 | 0.0189320984696 | 0.887750432616662 | 0.162,0.11,0.11,0.0 | 0.134,0.116,0.144, | -0,035 |
| 121895 | ENSG00000159063 | ALG8 | chr11 | - | 78102873 | 78102948 | 78100931 | 78101195 | 78103979 | 78104052 | 121895 | 6,2,11,3,4,2,2,4,0,0,0,7 | 6,3,14,8,8,11,9,9,3,2,7,9 | 5,4,1,0,0,1,2,1,0,2,6,2,0,3,4 | 14,5,14,1,2,8,10,11,3,8,7,2 | 224 | 149 | 0.0189479575081 | 0.888300553784768 | 0.399,0.307,0.343, | 0.192,0.347,0.045, | 0,064 |
| 175143 | ENSG00000101596 | SMCHD1 | chr18 | + | 2778186 | 2778239 | 2777805 | 2777915 | 2784449 | 2784621 | 175143 | 201,169,168,86,58,63,67,6 | 115,73,107,95,26,44,54,72 | 39,67,105,81,78,55,36,53,5 | 23,76,62,109,168,43,26,42 | 202 | 149 | 0.0189629966442 | 0.888706773661058 | 0.563,0.631,0.537, | 0.556,0.394,0.555, | 0,075 |
| 174773 | ENSG00000187514 | PTMA | chr2 | + | 231711886 | 231712015 | 231711347 | 231711419 | 231712442 | 231712675 | 174773 | 117,88,67,60,48,36,35,23,7 | 72,29,55,23,35,27,46,37,24 | 43,31,47,28,32,26,18,30,15 | 59,49,18,53,37,8,14,42,18, | 278 | 149 | 0.0189648805768 | 0.888706773661058 | 0.466,0.619,0.395, | 0.281,0.253,0.583, | 0,089 |
| 92540 | ENSG00000112237 | CCNC | chr6 | - | 99550217 | 99550309 | 99549507 | 99549575 | 99550992 | 99551028 | 92540 | 21,16,46,34,23,24,12,13,23 | 5,3,5,4,2,1,4,2,0,1,0,1 | 35,50,37,18,16,9,17,35,8,3 | 2,0,0,0,3,0,1,3,2,0,0,5,3,2,1 | 241 | 149 | 0.0190099636659 | 0.890625487543605 | 0.722,0.767,0.85,0 | 0.915,1.0,1.0,1.0,0 | -0,048 |
| 151190 | ENSG00000174485 | DENND4A | chr15 | - | 65725887 | 65725934 | 65722847 | 65722948 | 65729071 | 65729247 | 151190 | 5,4,13,13,0,8,1,18,0,3,0,5 | 71,47,234,139,69,78,59,92 | 1,24,5,11,46,3,2,7,21,4,6,9 | 114,141,55,105,104,67,52, | 196 | 149 | 0.0190207685702 | 0.890892771155432 | 0.051,0.061,0.041, | 0.007,0.115,0.065, | -0,018 |
| 94247 | ENSG00000118873 | RAB3GAP2 | chr1 | - | 220196249 | 220196398 | 220195297 | 220195377 | 220202275 | 220202374 | 94247 | 50,37,217,199,91,74,116,1 | 5,6,20,6,6,7,11,13,1,3,0,4 | 85,50,70,43,51,46,22,55,67 | 7,11,17,17,10,6,4,2,10,6,4, | 298 | 149 | 0.0190239471834 | 0.890892771155432 | 0.833,0.755,0.844, | 0.859,0.694,0.673, | 0,063 |
| 93162 | ENSG00000135317 | SNX14 | chr6 | - | 85567533 | 85567577 | 85565331 | 85565419 | 85572136 | 85572215 | 93162 | 43,34,107,58,48,44,27,44,2 | 4,0,13,2,4,3,4,8,2,1,0,6 | 74,45,28,23,23,35,20,37,23 | 7,21,0,10,11,2,5,4,0,2,10,0 | 193 | 149 | 0.0190357537043 | 0.89125175234505 | 0.892,1.0,0.864,0. | 0.891,0.623,1.0,0.0 | 0,061 |
| 85959 | ENSG00000240344 | PPIL3 | chr2 | - | 200882341 | 200882435 | 200881420 | 200881488 | 200885261 | 200885367 | 85959 | 12,7,34,23,6,11,4,11,8,2,9, | 1,0,6,2,2,0,1,0,0,1,2,0 | 8,13,10,4,5,10,4,12,6,9,19, | 0,0,0,0,0,3,0,3,0,1,0,3,5,1,4 | 243 | 149 | 0.0190894705051 | 0.893572385011566 | 0.88,1.0,0.777,0.8 | 1.0,1.0,1.0,1.0,1.0, | -0,022 |
| 226386 | ENSG00000130856 | ZNF236 | chr18 | + | 76928973 | 76929146 | 76927926 | 76928106 | 76937155 | 76937343 | 226386 | 4,0,2,2,2,2,1,6,1,0,0,2 | 14,20,20,25,20,22,13,31,6, | 0,1,0,2,0,0,0,0,0,0,2,5,0,6,1 | 28,21,31,23,8,8,11,16,7,15 | 298 | 149 | 0.0190998770779 | 0.893865110535659 | 0.125,0.0,0.048,0.0 | 0.0,0.023,0.0,0.04 | 0,02 |
| 17031 | ENSG00000125877 | ITPA | chr20 | + | 3213168 | 3213226 | 3209497 | 3209617 | 3213984 | 3214058 | 17031 | 18,8,23,18,9,7,4,4,14,1,3,7 | 3,0,10,7,2,3,5,5,0,4,0,1 | 16,11,8,2,13,2,11,4,4,10,6, | 0,0,3,0,1,3,0,0,2,1,0,10,3,1 | 207 | 149 | 0.019120763578 | 0.894391677286736 | 0.812,1.0,0.623,0. | 1.0,1.0,0.657,1.0,0 | -0,129 |
| 43294 | ENSG00000165782 | PIP4P1 | chr14 | - | 20459627 | 20459733 | 20459387 | 20459440 | 20460191 | 20460298 | 43294 | 48,57,148,101,37,46,44,52 | 4,3,9,5,0,1,0,3,2,1,2,0 | 75,171,49,45,103,49,47,46 | 6,12,4,18,3,2,1,6,2,0,5,2,2, | 255 | 149 | 0.0191222169397 | 0.894391677286736 | 0.875,0.917,0.906, | 0.88,0.893,0.877,0 | 0,036 |
| 212033 | ENSG00000079263 | SP140 | chr2 | + | 230247248 | 230247440 | 230245862 | 230245940 | 230247915 | 230248065 | 212033 | 37,27,11,8,6,5,10,12,8,5,5, | 259,138,178,134,120,71,56 | 10,30,8,23,17,14,4,5,13,6,6 | 139,166,121,130,140,122,3 | 298 | 149 | 0.0191243510361 | 0.894391677286736 | 0.067,0.089,0.03,0 | 0.035,0.083,0.032, | -0,011 |
| 221143 | ENSG00000120278 | PLEKHG1 | chr6 | + | 150821113 | 150821233 | 150819678 | 150819774 | 150823653 | 150823676 | 221143 | 11,12,46,37,13,14,13,7,1,2 | 2,0,0,4,0,4,1,1,0,0,2,1 | 23,21,19,14,15,13,3,9,12,1 | 4,0,0,8,3,0,1,0,1,2,4,7,1,3,2 | 269 | 149 | 0.0191277506054 | 0.894391677286736 | 0.753,1.0,1.0,0.83 | 0.761,1.0,1.0,0.49 | 0,065 |
| 207785 | ENSG00000144909 | OSBPL11 | chr3 | - | 125552550 | 125552679 | 125538450 | 125538633 | 125560378 | 125560519 | 207785 | 24,20,37,59,17,9,12,12,17, | 4,0,1,2,1,0,0,0,2,0,3,0 | 34,49,6,17,14,23,14,11,11, | 1,3,2,2,0,1,0,3,3,2,2,0,2,1,0 | 278 | 149 | 0.0191426642974 | 0.894894608903931 | 0.763,1.0,0.952,0. | 0.948,0.897,0.617, | 0,035 |
| 159428 | ENSG00000132780 | NASP | chr1 | + | 45617441 | 45617591 | 45616700 | 45616703 | 45618060 | 45618213 | 159428 | 153,88,87,32,48,39,23,21,8 | 8,2,2,2,0,2,4,4,2,0,0,5 | 87,45,63,32,26,29,13,20,15 | 5,8,2,7,1,0,3,4,0,4,2,5,2,1,6 | 298 | 149 | 0.0191582836783 | 0.894994782522425 | 0.905,0.957,0.956, | 0.897,0.738,0.94,0 | 0,052 |
| 98600 | ENSG00000006634 | DBF4 | chr7 | + | 87904291 | 87904416 | 87900763 | 87900878 | 87907187 | 87907219 | 98600 | 125,79,68,49,11,27,35,39,3 | 5,4,3,8,3,2,3,2,2,1,0,1 | 38,112,48,83,47,16,27,23,3 | 7,6,5,8,3,2,2,7,2,4,3,7,0,3,6 | 274 | 149 | 0.0191683997092 | 0.894994782522425 | 0.931,0.915,0.925, | 0.747,0.91,0.839,0 | 0,046 |
| 44534 | ENSG00000187514 | PTMA | chr2 | + | 231711889 | 231711983 | 231711347 | 231711419 | 231712442 | 231712516 | 44534 | 896,636,917,639,630,563,5 | 72,29,55,23,35,27,46,37,24 | 685,560,470,343,411,227,3 | 59,49,18,53,37,8,14,42,18, | 243 | 149 | 0.0191692333004 | 0.894994782522425 | 0.884,0.931,0.911, | 0.877,0.875,0.941, | 0,016 |
| 181619 | ENSG00000134255 | CEPT1 | chr1 | + | 111161189 | 111161296 | 111159379 | 111159527 | 111174878 | 111174963 | 181619 | 7,10,35,27,25,8,3,4,14,4,7, | 0,0,2,4,0,3,0,2,2,0,0,2 | 20,2,9,2,3,11,2,7,2,14,4,21 | 3,0,0,0,0,1,2,0,2,0,0,4,1,3,2 | 256 | 149 | 0.0191748727033 | 0.894994782522425 | 1.0,1.0,0.911,0.79 | 0.795,1.0,1.0,1.0,1 | 0,049 |
| 18100 | ENSG00000137815 | RTF1 | chr15 | + | 41452900 | 41453048 | 41438320 | 41438431 | 41457671 | 41457876 | 18100 | 83,89,107,35,44,45,32,51,7 | 6,2,2,1,2,2,0,1,1,0,0,2 | 66,82,52,34,33,28,36,67,37 | 1,2,1,0,1,0,1,2,5,2,0,1,0,0,0 | 297 | 149 | 0.019177665622 | 0.894994782522425 | 0.874,0.957,0.964, | 0.971,0.954,0.963, | -0,007 |
| 95771 | ENSG00000166295 | ANAPC16 | chr10 | + | 72223887 | 72224056 | 72220053 | 72220379 | 72230365 | 72230440 | 95771 | 258,125,349,268,133,130,1 | 2,6,11,8,3,2,5,1,12,1,1,3 | 205,134,159,93,88,68,91,2 | 5,1,1,2,0,1,1,3,1,6,6,3,7,2,3 | 298 | 149 | 0.0191809882307 | 0.894994782522425 | 0.985,0.912,0.941, | 0.953,0.985,0.988, | -0,016 |
| 74136 | ENSG00000239305 | RNF103 | chr2 | - | 86612158 | 86612274 | 86603397 | 86605418 | 86620329 | 86620469 | 74136 | 56,43,118,44,13,40,24,79,2 | 4,2,8,3,0,1,1,2,1,0,1,0 | 83,134,40,44,49,19,44,26,3 | 5,1,4,8,5,0,5,0,1,2,5,1,0,2,1 | 265 | 149 | 0.0191841513506 | 0.894994782522425 | 0.887,0.924,0.892, | 0.903,0.987,0.849, | 0,025 |
| 178037 | ENSG00000067066 | SP100 | chr2 | + | 230417590 | 230417856 | 230415941 | 230416328 | 230442936 | 230443099 | 178037 | 53,49,86,33,15,31,46,50,15 | 0,0,5,2,2,0,2,6,3,0,2,0 | 45,59,45,37,46,23,13,24,34 | 0,1,0,4,6,0,0,2,1,3,0,0,0,0,2 | 298 | 149 | 0.0191856600259 | 0.894994782522425 | 1.0,1.0,0.896,0.89 | 1.0,0.967,1.0,0.82 | -0,057 |
| 123507 | ENSG00000030066 | NUP160 | chr11 | - | 47788181 | 47788305 | 47786452 | 47786554 | 47788500 | 47788611 | 123507 | 35,22,50,43,18,28,6,28,9,7 | 1,0,4,2,0,2,1,2,0,0,1,0 | 24,33,10,19,38,18,23,19,28 | 0,2,0,2,6,0,0,0,1,0,3,5,0,2,4 | 273 | 149 | 0.0191859374398 | 0.894994782522425 | 0.95,1.0,0.872,0.9 | 1.0,0.9,1.0,0.838,0 | 0,048 |
| 8679 | ENSG00000175387 | SMAD2 | chr18 | - | 47920071 | 47920169 | 47896520 | 47896809 | 47930799 | 47931146 | 8679 | 21,13,5,2,4,6,5,3,1,4,2,5 | 12,9,10,2,10,2,2,7,10,6,2,7 | 7,9,2,4,11,1,12,3,8,8,3,8,5, | 4,10,3,8,4,0,0,4,1,2,3,4,10, | 247 | 149 | 0.0191863900951 | 0.894994782522425 | 0.514,0.466,0.232, | 0.514,0.352,0.287, | -0,184 |
| 207989 | ENSG00000162924 | REL | chr2 | + | 60900503 | 60900884 | 60894396 | 60894545 | 60900991 | 60901083 | 207989 | 6,3,4,0,1,2,1,5,0,0,0,0 | 108,105,175,97,86,58,70,1 | 8,10,2,9,0,3,2,2,2,1,3,6,0,1 | 145,153,56,79,91,26,89,59 | 298 | 149 | 0.0191949947915 | 0.895202150372554 | 0.027,0.014,0.011, | 0.027,0.032,0.018, | -0,013 |
| 720 | ENSG00000182944 | EWSR1 | chr22 | + | 29299194 | 29299280 | 29298732 | 29298895 | 29299598 | 29299851 | 720 | 16,11,11,5,8,5,5,2,10,6,7,5 | 0,0,2,1,1,0,1,0,0,0,2,0 | 8,11,9,8,8,2,3,3,3,7,10,5,2, | 4,0,0,9,0,0,2,3,0,1,0,0,0,0,0 | 235 | 149 | 0.0192057782338 | 0.895511017122835 | 1.0,1.0,0.777,0.76, | 0.559,1.0,1.0,0.36, | 0,069 |
| 88225 | ENSG00000091009 | RBM27 | chr5 | + | 146261509 | 146261806 | 146260744 | 146260898 | 146263490 | 146263631 | 88225 | 179,124,181,99,56,75,74,8 | 2,11,16,8,1,6,2,7,5,2,3,2 | 130,148,90,103,122,52,73, | 16,22,3,8,2,2,4,5,8,9,5,8,15 | 298 | 149 | 0.0192222949885 | 0.896087020288032 | 0.978,0.849,0.85,0 | 0.802,0.771,0.938, | 0,039 |
| 132209 | ENSG00000110888 | CAPRIN2 | chr12 | - | 30719078 | 30720915 | 30714958 | 30716676 | 30723258 | 30723314 | 132209 | 92,67,61,45,27,63,51,48,46 | 15,1,1,2,3,5,3,4,0,4,1,7 | 42,34,37,38,29,22,28,61,30 | 3,2,1,2,0,1,1,5,1,3,4,0,2,5,4 | 298 | 149 | 0.0192449718869 | 0.896949879642567 | 0.754,0.971,0.968, | 0.875,0.895,0.949, | -0,036 |
| 80924 | ENSG00000091490 | SEL1L3 | chr4 | - | 25835196 | 25835323 | 25830097 | 25830156 | 25847293 | 25847864 | 80924 | 134,82,432,200,119,70,72, | 6,9,6,3,5,12,4,7,1,3,0,1 | 112,64,115,56,79,63,43,37 | 7,7,13,2,17,2,2,3,9,2,6,9,6, | 276 | 149 | 0.0192494583984 | 0.89696475006409 | 0.923,0.831,0.975, | 0.896,0.832,0.827, | 0,03 |
| 31859 | ENSG00000247774 | PCED1B-AS1 | chr12 | - | 47206510 | 47206585 | 47205897 | 47206069 | 47209280 | 47209458 | 31859 | 10,13,18,10,17,8,3,3,14,3,1 | 25,9,9,11,17,17,3,6,18,21,0 | 20,14,9,5,14,4,8,6,9,17,16, | 16,7,9,0,7,11,6,13,6,16,15, | 224 | 149 | 0.0192698275216 | 0.897707796853489 | 0.21,0.49,0.571,0. | 0.454,0.571,0.399, | -0,11 |
| 219586 | ENSG00000132424 | PNISR | chr6 | - | 99416328 | 99416428 | 99414571 | 99414686 | 99425214 | 99425258 | 219586 | 46,29,81,36,2,6,6,5,2,1,1,3 | 7,3,4,0,4,2,2,0,3,2,3,0 | 46,34,22,32,23,12,1,2,31,4 | 1,2,0,0,0,0,3,4,1,3,3,0,1,4,3 | 249 | 149 | 0.0192754425716 | 0.897707796853489 | 0.797,0.853,0.924, | 0.965,0.91,1.0,1.0, | -0,116 |
| 216996 | ENSG00000135945 | REV1 | chr2 | - | 99406324 | 99406599 | 99405909 | 99406106 | 99408028 | 99408403 | 216996 | 41,15,23,20,18,14,20,26,24 | 1,1,3,9,0,0,3,2,2,0,0,1 | 24,25,24,16,9,14,13,21,14, | 4,0,0,1,3,0,1,1,0,0,0,0,0,3,2 | 298 | 149 | 0.0192779173867 | 0.897707796853489 | 0.953,0.882,0.793, | 0.75,1.0,1.0,0.889, | -0,006 |
| 171076 | ENSG00000164463 | CREBRF | chr5 | + | 173080588 | 173080798 | 173056351 | 173056479 | 173086500 | 173086626 | 171076 | 16,22,56,30,6,16,10,23,4,5 | 3,0,5,3,0,2,1,0,0,0,1,1 | 29,39,29,21,22,11,11,19,14 | 4,0,0,5,7,3,0,2,2,3,7,0,3,3,0 | 298 | 149 | 0.0193278576025 | 0.899460666268002 | 0.727,1.0,0.848,0. | 0.784,1.0,1.0,0.67 | 0,06 |
| 162253 | ENSG00000124357 | NAGK | chr2 | + | 71075554 | 71075606 | 71072640 | 71073594 | 71076603 | 71076701 | 162253 | 7,7,50,42,26,20,13,14,6,8,4 | 3,5,8,1,3,4,5,8,0,1,1,1 | 24,22,8,5,9,6,10,12,3,8,7,1 | 3,12,5,2,3,5,2,6,5,2,1,8,6,5 | 201 | 149 | 0.0193346092468 | 0.899460666268002 | 0.634,0.509,0.822, | 0.856,0.576,0.543, | 0,111 |
| 33790 | ENSG00000172795 | DCP2 | chr5 | + | 112981041 | 112981171 | 112976812 | 112976986 | 112985834 | 112985986 | 33790 | 4,0,3,4,1,1,3,0,0,0,2,1 | 20,19,54,39,24,20,23,19,18 | 7,3,2,1,3,1,0,0,4,10,0,6,1,2 | 35,33,17,22,16,15,7,12,21, | 279 | 149 | 0.0193350772464 | 0.899460666268002 | 0.097,0.0,0.029,0.0 | 0.097,0.046,0.059, | -0,009 |
| 235650 | ENSG00000188938 | FAM120AOS | chr9 | - | 93448129 | 93448489 | 93447634 | 93447697 | 93450478 | 93450587 | 235650 | 8,5,16,20,5,9,6,7,13,4,12,4 | 20,10,53,20,14,8,13,17,15, | 11,8,11,6,9,1,4,8,2,9,8,12,0 | 17,16,37,13,14,11,7,13,11, | 298 | 149 | 0.0193354834556 | 0.899460666268002 | 0.167,0.2,0.131,0. | 0.244,0.2,0.129,0. | 0,088 |
| 82243 | ENSG00000004897 | CDC27 | chr17 | - | 47169916 | 47170042 | 47158205 | 47158303 | 47181561 | 47181605 | 82243 | 40,26,58,28,34,22,26,38,19 | 2,1,3,0,3,0,2,1,0,0,0,3 | 49,46,34,25,21,14,15,37,10 | 0,3,3,0,2,1,1,1,0,2,5,7,4,0,4 | 275 | 149 | 0.0193364547989 | 0.899460666268002 | 0.916,0.934,0.913, | 1.0,0.893,0.86,1.0, | 0,042 |
| 130883 | ENSG00000120837 | NFYB | chr12 | - | 104131753 | 104131838 | 104128423 | 104128517 | 104135447 | 104135532 | 130883 | 6,6,24,13,5,6,7,7,2,3,5,10 | 21,3,64,45,27,35,11,22,16, | 16,18,8,2,11,5,9,7,7,12,5,1 | 38,12,22,4,21,13,10,18,13, | 234 | 149 | 0.0193721976835 | 0.900928582458381 | 0.154,0.56,0.193,0 | 0.211,0.489,0.188, | -0,067 |
| 99047 | ENSG00000136280 | CCM2 | chr7 | + | 45068442 | 45068579 | 45027025 | 45027797 | 45069825 | 45069961 | 99047 | 32,19,55,70,45,32,35,28,22 | 3,2,1,0,0,0,6,0,0,4,1,5 | 41,49,26,15,27,31,9,25,16, | 2,4,0,2,5,3,0,0,0,1,0,0,0,1,0 | 286 | 149 | 0.0194026847078 | 0.902151486788616 | 0.847,0.832,0.966, | 0.914,0.865,1.0,0. | -0,041 |
| 188306 | ENSG00000075151 | EIF4G3 | chr1 | - | 20981047 | 20981218 | 20980333 | 20980448 | 21002712 | 21002808 | 188306 | 56,56,43,34,27,25,40,58,37 | 1,2,3,0,4,3,2,2,0,5,0,0 | 48,28,26,47,13,19,11,25,5, | 3,2,0,2,1,0,0,0,4,0,6,2,4,1,2 | 298 | 149 | 0.0194086928696 | 0.902235934128915 | 0.966,0.933,0.878, | 0.889,0.875,1.0,0. | 0,033 |
| 202029 | ENSG00000115267 | IFIH1 | chr2 | - | 162276479 | 162276601 | 162273794 | 162273944 | 162276686 | 162276946 | 202029 | 2,2,7,6,0,1,0,6,3,0,0,0 | 9,5,8,13,12,4,12,0,3,4,2,6 | 1,2,0,3,0,0,0,0,0,1,4,3,4,1,0 | 19,9,5,10,1,13,8,7,2,6,4,26 | 271 | 149 | 0.0194189216124 | 0.902516501308241 | 0.109,0.18,0.325,0 | 0.028,0.109,0.0,0.0. | 0,123 |
| 53293 | ENSG00000134684 | YARS1 | chr1 | - | 32810590 | 32810766 | 32806481 | 32806611 | 32817187 | 32818031 | 53293 | 54,44,71,37,12,20,42,21,95 | 1,2,4,5,0,3,3,0,0,2,0,2 | 39,50,33,27,35,31,9,17,38, | 2,5,0,0,4,0,1,2,2,0,0,0,2,0,0 | 298 | 149 | 0.0194246956941 | 0.902589956592581 | 0.964,0.917,0.899, | 0.907,0.833,1.0,1.0 | -0,033 |
| 190309 | ENSG00000037474 | NSUN2 | chr5 | - | 6605272 | 6605375 | 6604604 | 6604685 | 6606819 | 6606926 | 190309 | 6,11,15,23,5,3,2,7,8,2,1,4 | 6,2,7,5,2,2,3,0,0,0,0,0 | 11,12,11,3,4,3,1,9,1,3,2,13 | 2,1,0,0,1,1,0,3,2,4,0,0,2,0,3 | 252 | 149 | 0.0194533644702 | 0.903726977829833 | 0.372,0.765,0.559, | 0.765,0.876,1.0,1.0 | -0,06 |
| 208580 | ENSG00000196504 | PRPF40A | chr2 | - | 152693391 | 152693476 | 152693037 | 152693068 | 152694541 | 152694622 | 208580 | 9,0,18,7,2,10,1,6,0,2,4,3 | 36,36,77,68,40,37,36,40,22 | 7,9,4,3,2,0,1,9,2,1,2,3,3,2,4 | 53,79,32,27,45,24,31,52,16 | 234 | 149 | 0.0194593260105 | 0.90380884690676 | 0.137,0.0,0.13,0.0 | 0.078,0.068,0.074, | 0,015 |
| 56007 | ENSG00000163931 | TKT | chr3 | - | 53237796 | 53237921 | 53234982 | 53235174 | 53240250 | 53240348 | 56007 | 2,3,6,0,3,3,2,1,6,0,3,3 | 19,15,23,45,21,20,29,11,31 | 0,1,3,0,1,0,1,10,1,2,0,2,2,3 | 23,22,25,9,18,18,16,21,15, | 274 | 149 | 0.0194760862781 | 0.904392130684302 | 0.054,0.098,0.124, | 0.0,0.024,0.061,0.0 | 0,025 |

| | | | | | | | | | | | | | | | | | | | | | | |
|---|---|---|---|---|---|---|---|---|---|---|---|---|---|---|---|---|---|---|---|---|---|---|
| 169562 | ENSG00000174839 | DENND6A | chr3 | - | 57630409 | 57630523 | 57628810 | 57628885 | 57630714 | 57630824 | 169562 | 41,31,51,21,11,10,15,28,33 | 2,5,11,0,1,1,1,9,1,2,0,4 | 31,40,33,24,19,9,10,15,7,2 | 0,6,1,0,7,4,7,13,7,2,11,0,5, | 263 | 149 | 0.0194809983044 | 0.90442509621534 | 0.921,0.778,0.724, | 1.0,0.791,0.949,1.0 | 0,147 |
| 227062 | ENSG00000198742 | SMURF1 | chr7 | - | 99032771 | 99032912 | 99027440 | 99030683 | 99033036 | 99033121 | 227062 | 3,0,0,1,0,1,1,0,0,0,0,0 | 9,12,26,10,4,10,2,9,3,3,7,9 | 0,5,2,12,5,0,0,0,0,1,0,0,1,0 | 9,33,10,10,22,4,25,12,3,5,7 | 290 | 149 | 0.0195098406845 | 0.905433379054521 | 0.146,0.0,0.0,0.0,0.049 | 0.0,0.072,0.093,0.3 | -0,013 |
| 96814 | ENSG00000127947 | PTPN12 | chr7 | + | 77604878 | 77604935 | 77600663 | 77600806 | 77607234 | 77607301 | 96814 | 8,0,2,2,1,1,0,4,0,2,1,0 | 34,24,46,47,23,17,21,43,29 | 0,2,0,2,4,1,2,0,0,0,4,2,0,0,0 | 34,56,36,26,18,38,29,12,36 | 206 | 149 | 0.0195111299583 | 0.905433379054521 | 0.145,0.0,0.03,0.03 | 0.0,0.025,0.0,0.053 | 0,022 |
| 187940 | ENSG00000114098 | ARMC8 | chr3 | + | 138237308 | 138237362 | 138235033 | 138235114 | 138237481 | 138237572 | 187940 | 14,7,10,9,7,6,5,7,4,2,5,5 | 1,2,3,3,4,6,2,4,2,4,1,4 | 7,19,5,3,8,5,5,13,3,7,7,4,11 | 6,13,1,9,6,2,4,3,3,7,8,4,7,3 | 203 | 149 | 0.0195453568156 | 0.906826189432723 | 0.911,0.72,0.71,0.6 | 0.461,0.518,0.786, | 0,137 |
| 194712 | ENSG00000184588 | PDE4B | chr1 | + | 66355527 | 66355620 | 66332510 | 66332620 | 66361614 | 66361793 | 194712 | 264,207,276,98,66,67,100, | 13,7,2,4,0,0,7,3,0,0,5,7 | 216,334,250,140,279,96,29 | 0,1,9,6,4,3,5,14,1,1,5,1,6,3 | 242 | 149 | 0.0195636781271 | 0.907480604951263 | 0.926,0.948,0.988, | 1.0,0.995,0.945,0.9 | -0,012 |
| 184989 | ENSG00000108654 | DDX5 | chr17 | - | 64505891 | 64506259 | 64504676 | 64504842 | 64506754 | 64507562 | 184989 | 4,0,2,0,2,1,2,2,2,2,1,2 | 19,20,23,8,6,6,13,11,6,6,0, | 1,0,0,0,0,0,2,0,0,0,2,4,0,0,3 | 13,12,13,9,11,6,9,6,5,6,9,3 | 298 | 149 | 0.0195782017484 | 0.907958616787305 | 0.095,0.0,0.042,0.0 | 0.037,0.0,0.0,0.0,0.0 | 0,123 |
| 88897 | ENSG00000073849 | ST6GAL1 | chr3 | + | 187038391 | 187038491 | 187025482 | 187025517 | 187038741 | 187038873 | 88897 | 1,7,24,18,4,9,9,8,10,2,7,8 | 1,2,32,24,6,6,8,3,6,3,2,4 | 13,17,10,18,5,10,0,12,6,6,5 | 15,2,8,1,0,6,3,3,7,4,1,0,7,1 | 249 | 149 | 0.0195930057255 | 0.908282625269006 | 0.374,0.677,0.31,0 | 0.342,0.836,0.428, | -0,157 |
| 168423 | ENSG00000197142 | ACSL5 | chr10 | + | 112410257 | 112410339 | 112409532 | 112409685 | 112410462 | 112410495 | 168423 | 0,0,2,3,1,6,2,2,1,0,1,0 | 18,12,29,38,22,16,26,46,21 | 7,3,1,2,3,1,4,0,0,0,3,1,2,2,3 | 20,13,9,2,6,13,10,7,8,17,12 | 231 | 149 | 0.0195936283766 | 0.908282625269006 | 0.0,0.0,0.043,0.048 | 0.184,0.13,0.067,0 | -0,068 |
| 86330 | ENSG00000151687 | ANKAR | chr2 | + | 189738564 | 189738682 | 189728659 | 189728821 | 189741341 | 189741451 | 86330 | 9,9,36,18,9,7,11,18,4,10,10 | 2,1,5,2,2,0,0,0,2,1,2,1 | 11,24,12,7,12,4,2,4,10,21,7 | 3,0,0,0,4,0,0,0,0,4,0,0,1,5,1 | 267 | 149 | 0.0196083551785 | 0.908752827083933 | 0.715,0.834,0.801, | 0.672,1.0,1.0,1.0,0 | -0,063 |
| 220350 | ENSG00000149657 | LSM14B | chr20 | + | 62126339 | 62126439 | 62124616 | 62124780 | 62129784 | 62129952 | 220350 | 19,3,31,17,16,17,11,8,14,1 | 0,0,3,3,1,2,0,0,0,0,0,1 | 11,13,8,7,10,11,7,8,12,11,1 | 3,1,2,3,0,0,0,2,0,5,2,2,3,0,0 | 249 | 149 | 0.0196122160925 | 0.908752827083933 | 1.0,1.0,0.861,0.772 | 0.687,0.886,0.705, | 0,109 |
| 128972 | ENSG00000129315 | CCNT1 | chr12 | - | 48695998 | 48696162 | 48695758 | 48695829 | 48698137 | 48698183 | 128972 | 23,32,37,26,13,7,16,15,21, | 59,55,36,34,10,10,21,17,12 | 31,20,23,35,39,9,9,28,14,2 | 26,30,18,24,9,10,11,17,14, | 298 | 149 | 0.0196269519404 | 0.909239882282444 | 0.163,0.225,0.339, | 0.373,0.25,0.39,0.4 | -0,076 |
| 152610 | ENSG00000213928 | IRF9 | chr14 | + | 24163862 | 24163959 | 24163382 | 24163508 | 24164062 | 24164134 | 152610 | 39,30,40,39,17,21,17,31,19 | 13,5,50,34,10,28,33,22,11, | 27,39,16,15,14,20,11,32,10 | 40,47,13,36,22,24,10,33,17 | 246 | 149 | 0.0196339300672 | 0.909367420334307 | 0.645,0.784,0.326, | 0.29,0.334,0.427,0 | 0,121 |
| 246247 | ENSG00000172183 | ISG20 | chr15 | + | 88638959 | 88639076 | 88637361 | 88637529 | 88639342 | 88639531 | 246247 | 84,63,41,18,14,29,34,33,10 | 6,3,5,3,4,8,0,1,8,2,2,7 | 15,28,58,47,21,10,23,30,13 | 0,0,3,0,0,0,2,5,2,2,7,10,5,1 | 266 | 149 | 0.019639376388 | 0.909423971464206 | 0.887,0.922,0.821, | 1.0,1.0,0.915,1.0,1 | -0,07 |
| 178301 | ENSG00000143297 | FCRL5 | chr1 | - | 157546942 | 157547165 | 157544830 | 157545082 | 157549559 | 157549580 | 178301 | 203,175,401,129,162,81,14 | 6,4,28,21,12,7,9,12,4,2,0,7 | 328,151,232,67,39,59,61,2 | 19,2,3,2,2,5,6,1,9,4,0,6,6,2 | 298 | 149 | 0.0196650377404 | 0.910416377264795 | 0.944,0.956,0.877, | 0.896,0.974,0.975, | 0,007 |
| 181617 | ENSG00000134255 | CEPT1 | chr1 | + | 111161184 | 111161296 | 111159379 | 111159527 | 111174878 | 111174963 | 181617 | 7,10,35,27,27,8,4,4,17,4,9, | 0,0,2,4,0,3,0,2,2,0,0,2 | 20,4,9,3,3,11,2,7,2,14,4,21 | 3,0,0,0,0,1,2,0,2,0,0,4,1,3,2 | 261 | 149 | 0.0196848657697 | 0.911138353651248 | 1.0,1.0,0.909,0.794 | 0.792,1.0,1.0,1.0,1 | 0,045 |
| 146507 | ENSG00000179583 | CIITA | chr16 | + | 10902037 | 10902184 | 10901513 | 10901571 | 10902657 | 10902801 | 146507 | 390,99,186,166,166,64,163 | 6,2,2,0,5,2,4,0,1,1,0,4 | 174,166,79,154,59,60,56,1 | 1,2,0,2,2,0,0,0,0,1,1,2,0,0,1 | 296 | 149 | 0.0196937045065 | 0.911351475948936 | 0.97,0.961,0.979,1 | 0.989,0.977,1.0,0.9 | -0,009 |
| 101154 | ENSG00000144840 | RABL3 | chr3 | - | 120738526 | 120738667 | 120730695 | 120730787 | 120742461 | 120742533 | 101154 | 12,5,12,8,2,10,2,4,1,2,1,5 | 7,7,14,6,5,4,5,8,1,1,1,3 | 7,9,9,9,14,0,2,2,3,2,5,7,8,6 | 3,6,0,0,5,4,0,1,2,7,3,8,1,6,2 | 290 | 149 | 0.0197017688979 | 0.911511442365915 | 0.468,0.268,0.306, | 0.545,0.435,1.0,1.0 | -0,206 |
| 77487 | ENSG00000162959 | MEMO1 | chr2 | - | 31883385 | 31883462 | 31869847 | 31869952 | 31891991 | 31892134 | 77487 | 37,23,59,40,18,11,7,15,34, | 1,4,4,4,0,4,1,3,7,1,1,0 | 36,22,17,9,17,17,7,27,6,33 | 8,2,2,6,1,2,5,5,2,8,5,1,5,2,3 | 226 | 149 | 0.0197056313511 | 0.911511442365915 | 0.961,0.791,0.907, | 0.748,0.879,0.849, | 0,091 |
| 227967 | ENSG00000173715 | C11orf80 | chr11 | + | 66831254 | 66831364 | 66828265 | 66828364 | 66838366 | 66838445 | 227967 | 8,1,4,1,3,7,6,0,0,0,0,1 | 101,38,109,54,42,52,59,69 | 0,5,0,1,0,1,5,0,0,1,0,2,0,0,2 | 66,36,45,52,36,13,43,56,33 | 259 | 149 | 0.0197258130324 | 0.912248918086911 | 0.044,0.015,0.021, | 0.0,0.074,0.0,0.01 | 0,005 |
| 83348 | ENSG00000115649 | CNPPD1 | chr2 | - | 219174145 | 219174207 | 219173349 | 219173467 | 219174777 | 219174906 | 83348 | 42,58,171,109,62,31,63,32 | 8,3,13,5,0,3,3,0,0,0,1,2 | 90,183,44,25,66,16,26,44,2 | 2,11,7,1,7,1,11,10,1,3,6,6,6 | 211 | 149 | 0.0197496329545 | 0.913154296547796 | 0.788,0.932,0.903, | 0.969,0.922,0.816, | 0,05 |
| 218448 | ENSG00000112343 | TRIM38 | chr6 | + | 25974804 | 25974930 | 25973172 | 25973285 | 25983163 | 25991231 | 218448 | 0,0,24,17,0,1,0,2,3,0,4,2 | 47,39,99,87,47,42,24,29,19 | 2,5,6,0,6,15,1,1,2,3,2,3,0,4 | 53,87,31,21,48,39,20,37,47 | 275 | 149 | 0.0197579739918 | 0.913343750049206 | 0.0,0.0,0.0,0.116,0.096 | 0.02,0.03,0.095,0.0 | 0,002 |
| 98920 | ENSG00000175324 | LSM1 | chr8 | - | 38169801 | 38169917 | 38163338 | 38163840 | 38171964 | 38172033 | 98920 | 40,47,114,81,28,31,26,36,3 | 2,0,4,12,3,1,2,4,0,8,4,0 | 48,82,33,22,59,28,20,28,16 | 9,16,2,1,14,3,1,2,5,5,1,19,2 | 265 | 149 | 0.0197782938351 | 0.914086742628818 | 0.918,1.0,0.941,0.7 | 0.75,0.742,0.903,0 | 0,031 |
| 238915 | ENSG00000164823 | OSGIN2 | chr8 | + | 89915212 | 89915263 | 89914554 | 89914746 | 89921079 | 89921319 | 238915 | 0,0,2,3,2,1,0,3,0,1,0,2 | 4,11,27,24,14,18,12,12,2,4 | 0,6,2,0,0,2,0,1,4,2,1,6,1,1,1 | 8,28,14,4,24,4,6,12,10,9,15 | 200 | 149 | 0.0198041035126 | 0.915083083538087 | 0.0,0.0,0.0,0.052,0.085 | 0.0,0.138,0.096,0.0 | -0,046 |
| 25422 | ENSG00000083520 | DIS3 | chr13 | - | 72781288 | 72781356 | 72780845 | 72781003 | 72781604 | 72781925 | 25422 | 8,2,6,4,2,6,3,0,6,2,3,3 | 13,7,26,14,10,6,15,15,14,1 | 0,3,9,0,2,0,3,0,3,4,5,4,0,1,1 | 25,27,19,11,13,10,7,10,1,1 | 217 | 149 | 0.0198908777809 | 0.918620026411106 | 0.297,0.164,0.137, | 0.0,0.071,0.245,0.0 | 0,046 |
| 27305 | ENSG00000189091 | SF3B3 | chr16 | + | 70539103 | 70539207 | 70538322 | 70538460 | 70544437 | 70544533 | 27305 | 46,41,74,76,32,13,12,30,17 | 7,1,8,0,1,0,4,1,0,0,0,0 | 57,67,39,9,31,23,15,23,15, | 5,4,1,2,1,2,5,1,2,0,0,2,1,0,4 | 253 | 149 | 0.0198925315356 | 0.918620026411106 | 0.795,0.96,0.845,1 | 0.87,0.908,0.958,0 | 0,044 |
| 194163 | ENSG00000103657 | HERC1 | chr15 | - | 63636987 | 63637111 | 63635960 | 63636142 | 63637504 | 63637643 | 194163 | 1,0,8,2,1,1,0,1,0,1,2,3 | 23,21,30,26,21,20,14,16,25 | 1,12,3,3,2,1,3,1,4,0,2,4,2,2 | 30,31,12,12,22,12,15,22,15 | 273 | 149 | 0.0198985199847 | 0.918620026411106 | 0.023,0.0,0.127,0.0 | 0.018,0.174,0.12,0 | -0,029 |
| 203551 | ENSG00000275052 | PPP4R3B | chr2 | - | 55565155 | 55565211 | 55564901 | 55565041 | 55568193 | 55568363 | 203551 | 36,36,66,14,12,28,13,34,6, | 18,8,50,60,28,20,18,19,19, | 51,104,21,53,26,18,24,35,2 | 37,50,8,9,7,19,9,42,7,22,21 | 205 | 149 | 0.0199050722758 | 0.918620026411106 | 0.592,0.766,0.49,0 | 0.5,0.602,0.656,0.8 | -0,145 |
| 212975 | ENSG00000145425 | RPS3A | chr4 | + | 151100484 | 151100587 | 151099627 | 151099714 | 151102870 | 151104205 | 212975 | 935,662,1191,1166,289,27 | 42,14,67,40,25,11,10,28,5, | 854,760,671,409,618,357,1 | 24,18,11,27,22,11,14,21,6, | 252 | 149 | 0.0199103750446 | 0.918620026411106 | 0.929,0.965,0.913, | 0.955,0.961,0.973, | -0,006 |
| 84732 | ENSG00000145425 | RPS3A | chr4 | + | 151100484 | 151100588 | 151099627 | 151099714 | 151102870 | 151104205 | 84732 | 940,662,1203,1167,290,27 | 42,14,67,40,25,11,10,28,5, | 862,766,676,409,620,357,1 | 24,18,11,27,22,11,14,21,6, | 253 | 149 | 0.0199137735048 | 0.918620026411106 | 0.929,0.965,0.914, | 0.955,0.962,0.973, | -0,005 |
| 252218 | ENSG00000198825 | INPP5F | chr10 | + | 119823835 | 119823902 | 119823070 | 119823199 | 119826630 | 119826702 | 252218 | 91,26,113,55,53,75,31,36,5 | 1,0,1,2,5,2,3,0,0,0,0,3 | 72,64,105,31,15,33,49,45,2 | 7,2,2,0,0,0,0,0,4,4,4,0,0,1,5 | 216 | 149 | 0.019914564342 | 0.918620026411106 | 0.984,1.0,0.987,0.9 | 0.876,0.957,0.973, | 0,028 |
| 84188 | ENSG00000085365 | SCAMP1 | chr5 | + | 78449932 | 78450034 | 78421800 | 78421960 | 78459244 | 78459362 | 84188 | 34,13,27,24,20,11,9,12,24, | 0,1,4,3,3,0,2,2,2,0,0,3 | 29,11,17,4,7,6,9,15,2,16,11 | 8,7,0,1,2,0,4,6,3,1,3,2,3,0,2 | 251 | 149 | 0.0199187199149 | 0.918620026411106 | 1.0,0.885,0.8,0.826 | 0.683,0.483,1.0,0.1 | 0,137 |
| 56236 | ENSG00000186517 | ARHGAP30 | chr1 | - | 161056387 | 161056532 | 161054622 | 161054705 | 161059613 | 161059716 | 56236 | 98,37,171,183,123,83,71,8 | 4,2,7,8,2,6,1,0,1,2,2,5 | 146,131,88,37,60,108,37,6 | 5,3,0,2,2,2,0,1,3,4,0,2,5,2,1 | 294 | 149 | 0.0199190621391 | 0.918620026411106 | 0.925,0.904,0.925, | 0.937,0.957,1.0,0.9 | -0,023 |
| 230372 | ENSG00000113013 | HSPA9 | chr5 | - | 138566625 | 138566714 | 138556773 | 138561789 | 138567000 | 138567163 | 230372 | 39,53,162,117,53,48,50,71 | 18,3,18,6,4,9,11,16,0,5,2,1 | 89,110,67,39,71,30,37,64,3 | 12,34,6,16,28,9,4,12,6,15,1 | 238 | 149 | 0.0199300746215 | 0.918930996328206 | 0.576,0.917,0.849, | 0.823,0.669,0.875, | 0,068 |
| 24136 | ENSG00000139496 | NUP58 | chr13 | + | 25319325 | 25319350 | 25315356 | 25315467 | 25320529 | 25320595 | 24136 | 7,3,19,10,2,0,2,5,6,4,2,6 | 7,5,18,5,11,10,6,5,2,0,5,6 | 1,5,0,2,4,7,0,1,0,4,5,5,5,1,2 | 9,7,4,5,14,2,3,9,1,4,2,6,12, | 174 | 149 | 0.0199518098679 | 0.919535508258595 | 0.461,0.339,0.475, | 0.087,0.38,0.0,0.25 | 0,182 |
| 102609 | ENSG00000122566 | HNRNPA2B1 | chr7 | - | 26185198 | 26185279 | 26173570 | 26174782 | 26192494 | 26192577 | 102609 | 1,1,3,5,1,5,0,0,4,0,0,4 | 3,0,4,0,1,9,2,6,1,3,1,1 | 5,4,2,2,2,1,1,0,1,5,2,0,2,2,2 | 1,2,3,2,3,2,0,1,0,1,0,2,0,0,4 | 230 | 149 | 0.019953528158 | 0.919535508258595 | 0.178,1.0,0.327,1.0 | 0.764,0.564,0.302, | -0,188 |
| 68794 | ENSG00000141524 | TMC6 | chr17 | - | 78121555 | 78121711 | 78120176 | 78121164 | 78122604 | 78122749 | 68794 | 69,53,30,6,46,17,45,22,44, | 0,1,0,0,0,1,2,0,2,4,0,0 | 14,19,27,25,7,11,20,51,14, | 5,6,1,0,0,0,0,0,1,2,0,5,1,2,0 | 298 | 149 | 0.0199560024303 | 0.919535508258595 | 1.0,0.964,1.0,1.0,1 | 0.583,0.613,0.931, | 0,056 |
| 178036 | ENSG00000067066 | SP100 | chr2 | + | 230417590 | 230417679 | 230415941 | 230416328 | 230442936 | 230443099 | 178036 | 53,46,83,30,17,31,43,46,15 | 0,0,5,2,2,0,2,6,3,0,2,0 | 39,58,44,33,44,23,12,24,32 | 0,1,0,4,6,0,0,2,1,3,0,0,0,0,2 | 238 | 149 | 0.019975793775 | 0.920095143998726 | 1.0,1.0,0.912,0.904 | 1.0,0.973,1.0,0.838 | -0,05 |
| 77766 | ENSG00000106105 | GARS1 | chr7 | + | 30631447 | 30631541 | 30627379 | 30628669 | 30632246 | 30632437 | 77766 | 61,39,85,82,45,47,26,55,76 | 8,0,2,0,0,0,1,1,0,0,1,0 | 59,73,44,76,51,55,38,47,13 | 4,1,1,2,1,0,0,6,1,0,4,2,0,1,6 | 243 | 149 | 0.0199786170228 | 0.920095143998726 | 0.824,1.0,0.963,1.0 | 0.9,0.978,0.964,0.9 | 0,021 |
| 206197 | ENSG00000132953 | XPO4 | chr13 | - | 20807483 | 20807634 | 20800830 | 20800990 | 20808435 | 20808581 | 206197 | 17,11,47,29,17,14,13,27,8, | 1,2,1,3,2,2,1,3,1,0,0,0 | 28,47,20,31,25,22,3,20,26, | 0,2,4,0,1,2,2,0,0,0,0,2,0,1,0 | 298 | 149 | 0.019981510339 | 0.920095143998726 | 0.895,0.733,0.959, | 1.0,0.922,0.714,1.0 | -0,037 |
| 112372 | ENSG00000104365 | IKBKB | chr8 | + | 42319584 | 42319646 | 42319269 | 42319421 | 42320734 | 42320844 | 112372 | 104,39,70,53,24,34,34,20,8 | 4,13,2,2,2,1,2,1,4,1,0,3 | 45,29,33,28,32,34,29,44,28 | 0,4,3,12,6,2,0,3,2,11,2,7,0, | 211 | 149 | 0.0199852474699 | 0.920095143998726 | 0.948,0.679,0.961, | 1.0,0.837,0.886,0.0 | 0,054 |
| 237642 | ENSG00000147601 | TERF1 | chr8 | + | 73012855 | 73012920 | 73008863 | 73009205 | 73013894 | 73013990 | 237642 | 4,3,0,1,2,1,0,0,0,0,0,5 | 2,5,22,17,8,8,3,8,6,2,6,6 | 1,6,0,0,0,0,0,0,2,1,0,11,0,0 | 19,14,4,2,5,2,1,12,3,10,13, | 214 | 149 | 0.0199955083359 | 0.920295865134834 | 0.582,0.295,0.0,0.0 | 0.035,0.23,0.0,0.0, | 0,057 |
| 183316 | ENSG00000198408 | OGA | chr10 | - | 101811941 | 101812380 | 101810183 | 101810314 | 101813029 | 101813127 | 183316 | 2,1,10,1,8,3,1,12,0,0,5,1 | 97,107,295,175,100,114,10 | 8,10,7,13,5,0,3,10,4,8,7,13 | 189,261,104,119,180,75,54 | 298 | 149 | 0.019998159007 | 0.920295865134834 | 0.01,0.005,0.017,0 | 0.021,0.019,0.033, | -0,006 |
| 78220 | ENSG00000075884 | ARHGAP15 | chr2 | + | 143436912 | 143437042 | 143435313 | 143435699 | 143487372 | 143487479 | 78220 | 676,445,667,363,233,293,2 | 9,8,43,16,6,26,4,8,5,2,3,14 | 540,564,426,492,451,295,1 | 9,13,3,24,24,14,12,14,37,1 | 279 | 149 | 0.020021929288 | 0.921085363701276 | 0.976,0.967,0.892, | 0.97,0.959,0.987,0 | 0,025 |
| 74632 | ENSG00000115020 | PIKFYVE | chr2 | + | 208285725 | 208285933 | 208276711 | 208276830 | 208288728 | 208288818 | 74632 | 21,15,54,33,26,28,20,20,13 | 1,0,2,1,0,3,0,6,0,2,0,1 | 40,43,14,27,30,14,14,37,18 | 4,8,0,2,5,0,0,0,1,0,1,2,0,2,0 | 298 | 149 | 0.0200238739622 | 0.921085363701276 | 0.913,1.0,0.931,0.9 | 0.833,0.729,1.0,0.8 | -0,009 |
| 77917 | ENSG00000115421 | PAPOLG | chr2 | + | 60795289 | 60795336 | 60794963 | 60795020 | 60797061 | 60798466 | 77917 | 5,4,5,3,5,8,4,8,2,0,0,0 | 23,10,29,21,20,3,4,20,13,4 | 1,1,3,2,0,5,5,13,3,0,2,2,2,5 | 49,53,21,10,29,11,21,22,11 | 196 | 149 | 0.0200289278004 | 0.921120974232456 | 0.142,0.233,0.116, | 0.015,0.014,0.098, | 0,079 |
| 141083 | ENSG00000175354 | PTPN2 | chr18 | - | 12793481 | 12793958 | 12785480 | 12785844 | 12794383 | 12794485 | 141083 | 5,3,9,6,1,6,2,2,0,3,0,2 | 51,52,137,124,74,55,43,68 | 0,2,3,0,0,5,1,2,0,2,3,0,1,4,0 | 61,49,62,24,29,55,23,79,42 | 298 | 149 | 0.0200464463752 | 0.921664957641911 | 0.047,0.028,0.032, | 0.0,0.02,0.024,0.0, | 0,012 |
| 198651 | ENSG00000223745 | CCDC18-AS1 | chr1 | - | 93315148 | 93315266 | 93311279 | 93311608 | 93324634 | 93324729 | 198651 | 1,0,1,0,0,2,1,4,0,0,0,5 | 9,2,17,8,7,9,15,16,5,4,7,5 | 0,2,1,0,0,0,0,2,4,7,4,2,1,5,0 | 5,5,6,2,8,8,6,18,7,18,15,14 | 267 | 149 | 0.0200493206447 | 0.921664957641911 | 0.058,0.0,0.032,0.0 | 0.0,0.182,0.085,0.0 | -0,025 |
| 183317 | ENSG00000198408 | OGA | chr10 | - | 101812247 | 101812380 | 101810183 | 101810314 | 101813029 | 101813127 | 183317 | 2,3,18,2,11,10,8,16,2,1,8,2 | 97,107,295,175,100,114,10 | 11,11,15,23,9,1,3,14,14,11 | 189,261,104,119,180,75,54 | 282 | 149 | 0.0200580149413 | 0.921867737311753 | 0.011,0.015,0.031, | 0.03,0.022,0.071,0 | -0,009 |
| 46413 | ENSG00000198625 | MDM4 | chr1 | + | 204544534 | 204544684 | 204542783 | 204542944 | 204546796 | 204546832 | 46413 | 322,278,446,227,148,249,2 | 12,9,10,13,6,13,14,11,5,1,5 | 254,323,204,400,245,151,1 | 8,16,10,39,30,10,5,4,13,15 | 298 | 149 | 0.0200888434497 | 0.923087503100423 | 0.931,0.939,0.957, | 0.941,0.91,0.911,0 | 0,013 |
| 236526 | ENSG00000034677 | RNF19A | chr8 | - | 100261541 | 100261762 | 100259853 | 100259997 | 100264033 | 100264580 | 236526 | 82,96,194,60,47,64,30,233 | 5,8,19,8,0,20,6,41,2,1,3,8 | 73,115,85,126,61,34,73,96 | 0,14,8,0,7,1,2,12,4,3,35,6,6 | 298 | 149 | 0.0201024987937 | 0.923367297758144 | 0.891,0.857,0.836, | 1.0,0.804,0.842,1.0 | -0,052 |
| 222976 | ENSG00000104133 | SPG11 | chr15 | - | 44598630 | 44598747 | 44597384 | 44598373 | 44600466 | 44600632 | 222976 | 43,15,53,75,39,24,16,23,27 | 1,0,7,9,2,4,0,3,6,1,0,1 | 31,15,14,20,17,31,9,18,23, | 12,8,3,0,2,2,0,4,3,9,4,4,3,0 | 266 | 149 | 0.0201070830405 | 0.923367297758144 | 0.96,1.0,0.809,0.82 | 0.591,0.512,0.723, | 0,099 |
| 82661 | ENSG00000115825 | PRKD3 | chr2 | - | 37267429 | 37267536 | 37260222 | 37260384 | 37269614 | 37269687 | 82661 | 34,27,115,92,46,38,17,41,4 | 1,0,6,3,0,3,5,6,0,0,2,2 | 142,128,47,23,27,23,18,47 | 3,12,12,4,1,5,0,3,4,0,1,11,4 | 256 | 149 | 0.0201078028936 | 0.923367297758144 | 0.952,1.0,0.918,0.9 | 0.965,0.861,0.695, | 0,04 |
| 67684 | ENSG00000133226 | SRRM1 | chr1 | + | 24663182 | 24663224 | 24662659 | 24662804 | 24666814 | 24666845 | 67684 | 69,40,18,22,22,19,33,32,38 | 29,7,14,10,3,16,16,8,28,16 | 23,40,31,90,22,12,35,35,8, | 11,8,12,19,2,6,4,11,5,2,5,3 | 191 | 149 | 0.0201225895789 | 0.923626327127798 | 0.65,0.817,0.501,0 | 0.62,0.796,0.668,0 | -0,074 |
| 80593 | ENSG00000115317 | HTRA2 | chr2 | + | 74532618 | 74532714 | 74530910 | 74531925 | 74532819 | 74532934 | 80593 | 30,21,87,77,36,44,43,44,21 | 4,1,3,0,1,3,2,2,2,0,0,4 | 60,54,34,19,28,16,21,50,12 | 2,0,0,4,0,0,0,0,0,0,10,1,0 | 245 | 149 | 0.0201247380933 | 0.923626327127798 | 0.82,0.927,0.946,1 | 0.948,1.0,1.0,0.743 | -0,035 |
| 155547 | ENSG00000173575 | CHD2 | chr15 | + | 92991692 | 92991763 | 92991475 | 92991517 | 92992858 | 92992998 | 155547 | 31,17,21,26,1,9,11,10,7,8,1 | 33,33,44,27,36,24,17,22,23 | 10,15,3,18,7,8,2,6,7,13,15, | 35,83,16,22,33,18,10,23,26 | 220 | 149 | 0.0201269623027 | 0.923626327127798 | 0.389,0.259,0.244, | 0.162,0.109,0.113, | 0,065 |
| 129067 | ENSG00000174718 | RESF1 | chr12 | + | 31960776 | 31960871 | 31959369 | 31959491 | 31970188 | 31970246 | 129067 | 135,73,458,278,170,142,16 | 17,6,26,12,3,5,8,25,7,6,2,5 | 284,275,210,134,161,122,7 | 7,8,10,14,6,3,2,15,0,2,15,1 | 244 | 149 | 0.0201306089762 | 0.923626327127798 | 0.829,0.881,0.915, | 0.961,0.955,0.928, | -0,024 |
| 252006 | ENSG00000234420 | ZNF37BP | chr10 | - | 42525539 | 42525639 | 42523611 | 42523738 | 42550556 | 42552808 | 252006 | 6,1,11,3,5,9,0,4,9,1,1,1 | 27,20,35,15,25,28,25,37,23 | 4,3,0,1,1,1,6,13,0,5,1,0,3,0 | 37,30,22,16,18,8,8,14,20,3 | 249 | 149 | 0.02014229244 | 0.92396541861757 | 0.117,0.029,0.158, | 0.061,0.056,0.0,0.0 | 0,013 |
| 10942 | ENSG00000159842 | ABR | chr17 | - | 1079329 | 1079390 | 1073624 | 1073677 | 1091664 | 1091850 | 10942 | 47,43,80,34,61,73,31,46,46 | 3,1,7,1,0,2,1,0,0,0,1,0 | 48,40,30,50,38,25,37,61,34 | 3,1,1,4,0,0,3,5,4,1,4,4,0,0,1 | 210 | 149 | 0.0201496338644 | 0.924105230400315 | 0.917,0.968,0.89,0 | 0.919,0.966,0.955, | 0,028 |
| 252216 | ENSG00000198825 | INPP5F | chr10 | + | 119823770 | 119823902 | 119823070 | 119823199 | 119826630 | 119826702 | 252216 | 225,89,187,91,109,134,57, | 1,0,1,2,5,2,3,0,0,0,0,3 | 128,110,199,81,29,53,94,9 | 7,2,2,0,0,0,0,0,4,4,4,0,0,1,5 | 281 | 149 | 0.0201744651936 | 0.92498127653574 | 0.992,1.0,0.99,0.96 | 0.907,0.967,0.981, | 0,017 |
| 52133 | ENSG00000092529 | CAPN3 | chr15 | + | 42411746 | 42411979 | 42411286 | 42411345 | 42412085 | 42412316 | 52133 | 74,58,34,35,47,21,63,37,49 | 3,0,5,6,10,1,1,6,2,1,2,4 | 28,27,27,35,17,24,26,38,21 | 2,6,0,3,11,2,0,4,6,6,6,9,7,3 | 298 | 149 | 0.0201773308368 | 0.92498127653574 | 0.925,1.0,0.773,0.7 | 0.875,0.692,1.0,0.8 | 0,088 |
| 196788 | ENSG00000265491 | RNF115 | chr1 | - | 145793834 | 145794196 | 145788907 | 145788966 | 145823771 | 145824095 | 196788 | 6,3,10,1,2,3,3,4,3,6,0,0 | 26,18,34,12,12,18,14,12,19 | 7,3,0,3,3,0,2,2,2,4,1,2,1,0,2 | 15,27,15,15,13,16,10,16,7, | 298 | 149 | 0.0201961801432 | 0.925335530170238 | 0.103,0.077,0.128, | 0.189,0.053,0.0,0.0 | 0,027 |
| 101785 | ENSG00000196792 | STRN3 | chr14 | - | 30913523 | 30913657 | 30912006 | 30912182 | 30918965 | 30919106 | 101785 | 12,14,22,8,7,19,6,15,8,10,2 | 2,3,7,3,0,0,0,2,2,0,0,0 | 11,17,6,1,10,6,7,16,1,3,7,9 | 1,0,1,10,2,6,2,4,2,2,2,0,1,2 | 283 | 149 | 0.0201961894992 | 0.925335530170238 | 0.76,0.711,0.623,0 | 0.853,1.0,0.76,0.05 | 0,224 |
| 197964 | ENSG00000153922 | CHD1 | chr5 | - | 98876397 | 98876553 | 98875071 | 98875113 | 98879551 | 98879728 | 197964 | 41,31,43,17,26,20,18,39,17 | 20,3,9,12,6,9,16,13,12,2,2, | 34,41,20,29,29,12,13,21,11 | 13,13,13,34,18,3,9,10,18,4 | 298 | 149 | 0.0201979562458 | 0.925335530170238 | 0.506,0.838,0.705, | 0.567,0.612,0.435, | 0,127 |
| 189559 | ENSG00000143398 | PIP5K1A | chr1 | + | 151242119 | 151242567 | 151239954 | 151240039 | 151246919 | 151246965 | 189559 | 14,15,34,33,16,22,17,25,9, | 1,3,1,7,0,0,2,0,1,2,0,2 | 35,46,18,12,17,10,6,19,9,2 | 0,0,1,0,0,0,0,1,0,2,4,0,2,2,0 | 298 | 149 | 0.0202055074933 | 0.925484482504884 | 0.875,0.714,0.944, | 1.0,1.0,0.9,1.0,1.0, | -0,061 |
| 79696 | ENSG00000161813 | LARP4 | chr12 | + | 50453459 | 50453672 | 50441589 | 50441643 | 50454313 | 50454417 | 79696 | 19,4,35,32,7,5,10,7,12,2,5, | 2,1,6,1,1,2,2,1,0,0,0,2 | 21,15,3,8,8,15,4,11,10,12,6 | 3,0,1,0,0,2,1,2,0,0,1,0,4,2,0 | 298 | 149 | 0.0202161233888 | 0.925773713424428 | 0.826,0.667,0.745, | 0.778,1.0,0.6,1.0,1 | -0,065 |
| 171473 | ENSG00000147854 | UHRF2 | chr9 | + | 6465743 | 6465846 | 6460572 | 6460791 | 6481642 | 6481766 | 171473 | 36,10,86,31,12,9,11,23,11, | 7,0,8,1,0,1,1,3,0,0,0,2 | 35,44,11,17,18,11,11,11,11 | 1,2,3,0,4,0,2,0,2,1,1,0,0,0,2 | 252 | 149 | 0.0202384620039 | 0.926599535324697 | 0.753,1.0,0.864,0.9 | 0.954,0.929,0.684, | 0,008 |
| 121516 | ENSG00000110075 | PPP6R3 | chr11 | + | 68566998 | 68567166 | 68564302 | 68564432 | 68569747 | 68569897 | 121516 | 114,52,162,97,59,87,63,11 | 26,7,20,4,0,8,24,18,1,5,2,8 | 128,121,100,56,78,65,63,7 | 23,15,33,28,12,3,11,22,8,4 | 298 | 149 | 0.0202510343216 | 0.926977960032388 | 0.687,0.788,0.802, | 0.736,0.801,0.602, | 0,074 |
| 113390 | ENSG00000171988 | JMJD1C | chr10 | - | 63264650 | 63264764 | 63219876 | 63219983 | 63380317 | 63380482 | 113390 | 70,47,132,83,26,38,45,46,2 | 1,0,4,1,0,1,1,5,0,1,0,1 | 82,77,40,101,73,53,35,37,5 | 0,3,4,2,4,0,2,3,1,1,4,3,2,1,3 | 263 | 149 | 0.0202626258885 | 0.927311340124972 | 0.975,1.0,0.949,0.9 | 1.0,0.936,0.85,0.96 | 0,012 |
| 82099 | ENSG00000134262 | AP4B1 | chr1 | - | 113896257 | 113896465 | 113895756 | 113896038 | 113898717 | 113898801 | 82099 | 16,4,25,18,11,9,7,7,5,4,10, | 0,0,6,4,0,2,2,1,0,2,0,0 | 16,13,10,6,3,5,12,21,5,18,9 | 2,5,1,2,4,2,2,4,0,1,3,13,2,0 | 298 | 149 | 0.0202828561939 | 0.928039842999424 | 1.0,1.0,0.676,0.692 | 0.8,0.565,0.833,0.0 | 0,176 |
| 148819 | ENSG00000005339 | CREBBP | chr16 | - | 3782679 | 3782926 | 3781203 | 3781306 | 3791980 | 3792094 | 148819 | 58,51,57,44,57,58,41,65,39 | 0,1,3,0,0,3,1,0,1,11,0,3 | 47,58,41,73,36,39,57,54,37 | 1,1,0,1,0,1,0,1,0,0,5,2,0,2,0 | 298 | 149 | 0.0202872162006 | 0.928042046773929 | 1.0,0.962,0.905,1.0 | 0.959,0.967,1.0,0.9 | -0,017 |
| 185557 | ENSG00000141446 | ESCO1 | chr18 | - | 21540636 | 21540700 | 21539919 | 21540009 | 21560858 | 21560981 | 185557 | 0,2,7,4,1,0,0,0,3,1,0,0 | 17,6,40,22,18,12,11,17,11, | 2,2,1,0,0,0,0,0,2,0,0,2,0,0 | 20,22,12,6,9,6,5,12,8,18,15 | 213 | 149 | 0.020323463002 | 0.929502606328828 | 0.0,0.189,0.109,0.1 | 0.065,0.06,0.055,0 | 0,036 |
| 163616 | ENSG00000105875 | WDR91 | chr7 | - | 135197043 | 135197150 | 135196143 | 135196337 | 135197992 | 135201343 | 163616 | 8,7,16,7,15,6,2,6,2,5,5,9 | 14,14,44,37,36,35,14,15,33 | 9,16,6,3,3,1,8,8,8,11,2,0,11 | 21,28,8,6,7,3,21,25,2,11,20 | 256 | 149 | 0.0203315684173 | 0.929675759937093 | 0.25,0.225,0.175,0 | 0.2,0.25,0.304,0.2 | -0,083 |
| 240273 | ENSG00000197329 | PELI1 | chr2 | - | 64106205 | 64106299 | 64104700 | 64104830 | 64108239 | 64108379 | 240273 | 27,8,51,23,6,8,23,16,14,3,3 | 198,141,190,167,28,24,77, | 16,63,38,70,22,3,9,30,29,1 | 110,268,192,123,192,35,10 | 243 | 149 | 0.0203506160881 | 0.930349076307955 | 0.077,0.034,0.141, | 0.082,0.126,0.108, | -0,02 |
| 162677 | ENSG00000072364 | AFF4 | chr5 | - | 132898262 | 132898392 | 132897203 | 132897240 | 132899103 | 132899141 | 162677 | 30,15,67,31,17,28,16,47,13 | 13,1,7,1,1,3,1,9,3,1,6,3 | 46,40,18,28,25,12,15,17,28 | 5,7,5,31,9,9,5,4,8,5,4,2,2,5 | 279 | 149 | 0.0203581458786 | 0.930495666935009 | 0.552,0.889,0.836, | 0.831,0.753,0.658, | 0,115 |
| 240710 | ENSG00000148090 | AUH | chr9 | - | 91327499 | 91327703 | 91325317 | 91325404 | 91355882 | 91355970 | 240710 | 0,5,2,3,0,3,0,2,0,0,0,0 | 11,7,18,17,12,8,10,6,6,5,4, | 1,0,5,4,2,3,3,1,0,2,9,0,1,1,1 | 17,24,10,6,6,7,5,5,6,17,20, | 298 | 149 | 0.0203708530355 | 0.930878783372336 | 0.0,0.263,0.053,0.0 | 0.029,0.0,0.2,0.25, | -0,047 |
| 155607 | ENSG00000180776 | ZDHHC20 | chr13 | - | 21390194 | 21390413 | 21387507 | 21387634 | 21391721 | 21391854 | 155607 | 2,3,10,4,2,2,0,14,4,0,3,2 | 96,49,170,155,52,41,49,72 | 5,4,0,2,0,1,3,2,0,1,5,4,1,6,2 | 67,110,58,67,79,41,46,81,4 | 298 | 149 | 0.0203784416068 | 0.931027884838287 | 0.01,0.03,0.029,0.0 | 0.036,0.018,0.0,0.0 | 0,014 |
| 236524 | ENSG00000034677 | RNF19A | chr8 | - | 100261541 | 100261753 | 100259853 | 100259997 | 100264033 | 100264580 | 236524 | 82,96,194,60,47,64,30,233 | 5,8,19,8,0,20,6,41,2,1,3,8 | 73,115,85,126,61,34,72,96 | 0,14,8,0,7,1,2,12,4,3,35,6,6 | 298 | 149 | 0.020395651027 | 0.931616376526366 | 0.891,0.857,0.836, | 1.0,0.804,0.842,1.0 | -0,052 |
| 182010 | ENSG00000163125 | RPRD2 | chr1 | + | 150441880 | 150441957 | 150440922 | 150441023 | 150443230 | 150443283 | 182010 | 24,13,38,20,15,18,10,20,15 | 9,5,15,24,9,12,7,9,11,7,0,4 | 20,21,14,22,10,11,10,6,3,1 | 24,17,15,18,18,6,5,10,5,8,1 | 226 | 149 | 0.0204133699461 | 0.932163530759092 | 0.637,0.632,0.625, | 0.355,0.449,0.381, | 0,144 |
| 235652 | ENSG00000188938 | FAM120AOS | chr9 | - | 93448368 | 93448485 | 93446904 | 93447697 | 93450478 | 93450599 | 235652 | 36,20,46,43,13,18,20,21,35 | 20,10,53,20,14,8,13,17,15, | 15,22,37,41,17,3,15,29,6,2 | 17,16,37,13,14,11,7,13,11, | 266 | 149 | 0.0204191634848 | 0.932163530759092 | 0.502,0.528,0.327, | 0.331,0.435,0.359, | 0,098 |
| 243446 | ENSG00000065923 | SLC9A7 | chrX | - | 46631381 | 46631450 | 46620976 | 46621059 | 46631585 | 46631649 | 243446 | 24,10,24,9,5,4,8,4,22,2,1,7 | 176,139,189,124,125,139,1 | 4,14,16,6,9,1,9,10,0,2,28,4 | 158,152,130,142,108,51,96 | 218 | 149 | 0.0204206227148 | 0.932163530759092 | 0.085,0.047,0.08,0 | 0.017,0.059,0.078, | 0,011 |

| | | | | | | | | | | | | | | | | | | | | | | |
|---|---|---|---|---|---|---|---|---|---|---|---|---|---|---|---|---|---|---|---|---|---|---|
| 31867 | ENSG00000247774 | PCED1B-AS1 | chr12 | - | 47206510 | 47206588 | 47205892 | 47206145 | 47216164 | 47216264 | 31867 | 10,5,12,11,5,6,0,1,14,4,2,3 | 2,2,4,22,1,2,3,4,5,4,0,1 | 5,9,2,2,2,11,1,1,3,8,2,57,6, | 2,1,2,0,3,1,1,0,0,0,2,16,5,6 | 227 | 149 | 0.0204269895085 | 0.932256441667507 | 0.766,0.621,0.663, | 0.621,0.855,0.396, | -0,166 |
| 61412 | ENSG00000095564 | BTAF1 | chr10 | + | 91964076 | 91964201 | 91962537 | 91962678 | 91980453 | 91980558 | 61412 | 204,87,255,148,108,101,85 | 2,2,3,1,1,0,1,2,0,0,0,1 | 192,215,168,105,80,62,65, | 4,11,1,0,7,3,0,0,4,0,1,0,0,1 | 274 | 149 | 0.0204406494794 | 0.932682092028989 | 0.982,0.959,0.979, | 0.963,0.914,0.989, | 0,017 |
| 225444 | ENSG00000163492 | CCDC141 | chr2 | - | 178915620 | 178915807 | 178905328 | 178905501 | 178918712 | 178918907 | 225444 | 19,7,35,17,7,5,7,7,9,4,0,12 | 225,164,311,178,80,112,11 | 11,19,20,15,14,5,10,19,15, | 126,172,129,96,162,91,49, | 298 | 149 | 0.0204549985708 | 0.933138999023284 | 0.041,0.021,0.053, | 0.042,0.052,0.072, | -0,014 |
| 15232 | ENSG00000121281 | ADCY7 | chr16 | + | 50315013 | 50315138 | 50314291 | 50314406 | 50315358 | 50315738 | 15232 | 81,46,126,111,70,65,44,70 | 4,0,14,6,2,10,5,3,0,0,0,0 | 79,70,34,49,31,30,42,58,65 | 2,6,2,0,2,2,1,4,2,4,1,1,5,6,2 | 274 | 149 | 0.0204679424102 | 0.93353161959944 | 0.917,1.0,0.83,0.91 | 0.956,0.864,0.902, | -0,001 |
| 193680 | ENSG00000163867 | ZMYM6 | chr1 | - | 35012667 | 35012747 | 35012430 | 35012581 | 35014696 | 35014888 | 193680 | 4,3,17,2,0,1,1,2,0,0,0,0 | 4,2,22,22,15,3,9,6,2,4,4,11 | 1,3,1,1,2,2,1,3,2,1,2,0,2,7,2 | 17,21,5,5,7,4,7,10,10,10,3, | 229 | 149 | 0.0204815348051 | 0.933953647804339 | 0.394,0.494,0.335, | 0.037,0.085,0.115, | 0,023 |
| 251249 | ENSG00000130803 | ZNF317 | chr19 | + | 9157974 | 9158075 | 9157267 | 9157394 | 9158825 | 9158908 | 251249 | 8,9,14,10,8,12,4,13,6,3,5,1 | 19,16,28,12,13,7,9,9,4,9,4, | 11,20,4,2,4,1,3,3,0,4,10,6,4 | 11,25,8,12,9,8,8,16,7,8,21, | 250 | 149 | 0.0204969604885 | 0.9344590770812 | 0.201,0.251,0.23,0 | 0.373,0.323,0.23,0 | 0,112 |
| 178307 | ENSG00000143297 | FCRL5 | chr1 | - | 157546942 | 157547236 | 157544830 | 157545082 | 157549559 | 157549580 | 178307 | 198,171,381,122,156,74,14 | 6,4,28,21,12,7,9,12,4,2,0,7 | 314,140,223,67,39,59,61,2 | 19,2,3,2,2,5,6,1,9,4,0,6,6,2 | 298 | 149 | 0.0205132228608 | 0.935002428960789 | 0.943,0.955,0.872, | 0.892,0.972,0.974, | 0,005 |
| 183274 | ENSG00000198408 | OGA | chr10 | - | 101795387 | 101795480 | 101793912 | 101793998 | 101797979 | 101798154 | 183274 | 25,15,33,10,3,11,7,12,8,2,1 | 81,75,183,75,45,28,41,43,7 | 25,22,15,46,21,4,14,18,7,8 | 113,106,69,36,83,32,48,45 | 242 | 149 | 0.0205190368871 | 0.935069411019991 | 0.16,0.11,0.1,0.076 | 0.12,0.113,0.118,0 | -0,033 |
| 234325 | ENSG00000116754 | SRSF11 | chr1 | + | 70222557 | 70222798 | 70221723 | 70221839 | 70228421 | 70228555 | 234325 | 9,3,13,8,7,1,5,14,1,2,8,7 | 41,43,50,39,50,37,25,35,19 | 3,8,2,8,15,11,5,11,2,3,3,7,5 | 44,84,46,43,31,31,31,48,20 | 298 | 149 | 0.0205527564758 | 0.936407774987915 | 0.099,0.034,0.115, | 0.033,0.045,0.021, | 0,023 |
| 63953 | ENSG00000138674 | SEC31A | chr4 | - | 82862533 | 82862572 | 82857688 | 82857764 | 82863317 | 82863392 | 63953 | 14,4,28,28,2,4,7,9,5,1,2,9 | 29,30,57,31,11,17,24,21,13 | 18,9,11,5,5,3,8,10,1,4,4,5,4 | 38,40,29,29,20,3,17,28,14, | 188 | 149 | 0.0205753999524 | 0.937241038551324 | 0.277,0.096,0.28,0 | 0.273,0.151,0.231, | 0,03 |
| 29862 | ENSG00000127022 | CANX | chr5 | + | 179708977 | 179709059 | 179708238 | 179708380 | 179709872 | 179710065 | 29862 | 344,228,506,364,178,211,2 | 22,20,29,7,5,8,15,37,4,6,3, | 302,372,288,143,144,153,1 | 5,22,11,11,6,4,13,14,11,5,8 | 231 | 149 | 0.0205939391671 | 0.937887033617034 | 0.91,0.88,0.918,0.9 | 0.975,0.916,0.944, | -0,007 |
| 187485 | ENSG00000122483 | CCDC18 | chr1 | + | 93220297 | 93220470 | 93216635 | 93216746 | 93221608 | 93221743 | 187485 | 20,13,12,6,3,0,6,6,2,2,2,5 | 2,1,3,4,2,1,0,0,1,2,2,0 | 2,10,2,2,1,2,0,6,4,6,3,1,6,4 | 0,0,0,2,2,0,2,1,0,1,0,0,2,0,2 | 298 | 149 | 0.0206198188701 | 0.938651123191572 | 0.833,0.867,0.667, | 1.0,1.0,1.0,0.333,0 | -0,075 |
| 232105 | ENSG00000139971 | ARMH4 | chr14 | - | 58293065 | 58293544 | 58291634 | 58291707 | 58297954 | 58298139 | 232105 | 13,4,3,3,1,0,3,7,0,1,4,1 | 67,34,129,99,60,80,54,73,5 | 7,5,7,3,5,3,2,5,2,11,6,2,4,8 | 76,42,35,33,53,24,20,71,41 | 298 | 149 | 0.0206228572458 | 0.938651123191572 | 0.088,0.056,0.011, | 0.044,0.056,0.091, | -0,007 |
| 241118 | ENSG00000165209 | STRBP | chr9 | - | 123213588 | 123213918 | 123184131 | 123184298 | 123268435 | 123268586 | 241118 | 2,0,2,5,1,0,0,0,0,0,1,0 | 19,13,21,14,12,6,13,10,9,5 | 1,0,7,3,0,1,0,0,0,0,2,0,4,3,3 | 14,12,11,12,5,2,2,2,3,11,8, | 298 | 149 | 0.0206238002963 | 0.938651123191572 | 0.05,0.0,0.045,0.15 | 0.034,0.0,0.241,0.1 | -0,028 |
| 215620 | ENSG00000131508 | UBE2D2 | chr5 | + | 139562243 | 139562631 | 139561166 | 139561815 | 139600371 | 139600435 | 215620 | 5,9,20,7,4,1,2,10,3,0,10,5 | 42,11,68,35,26,26,41,22,41 | 11,19,17,2,6,2,9,3,3,3,12,7 | 38,20,20,12,11,12,5,38,6,3 | 298 | 149 | 0.0206373587306 | 0.938992504520001 | 0.056,0.29,0.128,0 | 0.126,0.322,0.298, | -0,053 |
| 75837 | ENSG00000257093 | DENND11 | chr7 | - | 141685477 | 141685636 | 141674066 | 141674220 | 141686558 | 141686658 | 75837 | 116,92,131,86,137,147,145 | 6,7,5,3,6,10,10,4,4,0,8,4 | 122,133,100,69,71,48,99,1 | 2,10,2,2,4,1,2,3,4,7,5,1,1,0 | 298 | 149 | 0.0206413160517 | 0.938992504520001 | 0.906,0.868,0.929, | 0.968,0.869,0.962, | -0,021 |
| 164696 | ENSG00000005436 | GCFC2 | chr2 | - | 75680210 | 75680314 | 75673443 | 75673520 | 75687826 | 75687977 | 164696 | 24,6,32,33,10,7,6,24,9,6,6, | 0,0,6,0,7,0,0,3,1,2,0,0 | 30,23,8,1,7,8,6,6,1,4,13,19 | 3,3,0,0,1,0,2,0,2,0,0,2,0,0,1 | 253 | 149 | 0.0206443891976 | 0.938992504520001 | 1.0,1.0,0.759,1.0,0 | 0.855,0.819,1.0,1.0 | 0,008 |
| 32494 | ENSG00000048028 | USP28 | chr11 | - | 113835206 | 113835327 | 113834248 | 113834335 | 113840597 | 113840757 | 32494 | 13,8,14,7,2,1,4,1,9,2,3,2 | 33,18,39,29,21,11,11,16,42 | 29,7,14,28,5,10,7,12,8,2,20 | 50,13,24,7,33,25,4,18,23,4 | 270 | 149 | 0.020660818553 | 0.939383772608399 | 0.179,0.197,0.165, | 0.242,0.229,0.244, | -0,099 |
| 14731 | ENSG00000125124 | BBS2 | chr16 | - | 56509356 | 56509494 | 56506119 | 56506224 | 56509956 | 56510034 | 14731 | 3,2,4,11,2,4,1,3,4,0,1,4 | 33,14,37,18,17,15,16,16,14 | 4,2,6,4,1,0,4,0,4,3,5,0,1,3,3 | 29,31,17,12,24,9,24,30,15, | 287 | 149 | 0.0206617205678 | 0.939383772608399 | 0.045,0.069,0.053, | 0.067,0.032,0.155, | 0,041 |
| 20928 | ENSG00000213928 | IRF9 | chr14 | + | 24163883 | 24163959 | 24163377 | 24163508 | 24164062 | 24164134 | 20928 | 38,29,38,38,15,18,17,30,18 | 13,5,50,34,10,28,33,22,11, | 26,36,16,12,12,15,10,30,9, | 40,47,13,36,22,24,10,33,17 | 225 | 149 | 0.0206705827794 | 0.939389318956008 | 0.659,0.793,0.335, | 0.301,0.337,0.449, | 0,122 |
| 24357 | ENSG00000005810 | MYCBP2 | chr13 | - | 77185870 | 77186063 | 77185102 | 77185377 | 77188950 | 77189047 | 24357 | 87,77,230,156,91,121,71,1 | 1,0,0,0,0,0,2,1,0,1,0,0 | 152,161,79,120,60,61,87,8 | 3,6,2,8,1,0,0,0,2,1,4,1,1,1,0 | 298 | 149 | 0.0206712302573 | 0.939389318956008 | 0.978,1.0,1.0,1.0,1 | 0.962,0.931,0.952, | 0,019 |
| 194858 | ENSG00000158711 | ELK4 | chr1 | - | 205618697 | 205618820 | 205616477 | 205616644 | 205618956 | 205619073 | 194858 | 8,15,11,3,4,4,15,9,7,3,3,6 | 119,109,176,86,78,85,96,1 | 15,12,12,14,5,0,12,16,2,13 | 161,128,91,99,84,76,70,12 | 272 | 149 | 0.0206749362494 | 0.939389318956008 | 0.036,0.07,0.033,0 | 0.049,0.049,0.067, | -0,012 |
| 47201 | ENSG00000291147 | | chr2 | + | 242116182 | 242116232 | 242114638 | 242114741 | 242118964 | 242119057 | 47201 | 15,10,13,6,4,6,4,2,4,11,2,1 | 4,4,2,0,1,0,0,0,0,0,0,0 | 5,9,2,3,2,9,3,5,2,17,12,14,3 | 5,2,1,1,1,5,1,0,0,1,2,4,0,1,0 | 199 | 149 | 0.0206838104129 | 0.939439906631491 | 0.737,0.652,0.83,1 | 0.428,0.771,0.6,0.6 | 0,137 |
| 224591 | ENSG00000133606 | MKRN1 | chr7 | - | 140469970 | 140470216 | 140459928 | 140459936 | 140471882 | 140472011 | 224591 | 1,0,12,2,0,5,3,12,5,0,0,1 | 37,24,170,128,86,59,67,93 | 2,3,6,2,2,0,0,0,3,1,1,2,2,5,0 | 106,140,78,42,52,46,32,59 | 298 | 149 | 0.0206847792257 | 0.939439906631491 | 0.013,0.0,0.034,0.0 | 0.009,0.011,0.037, | 0,005 |
| 151906 | ENSG00000125107 | CNOT1 | chr16 | - | 58625558 | 58625656 | 58623281 | 58623409 | 58629727 | 58629885 | 151906 | 2,5,8,2,2,4,0,2,2,0,4,3 | 63,39,68,66,33,51,24,52,2, | 2,2,1,3,4,1,0,0,1,2,4,4,1,3,2 | 62,74,37,56,63,33,13,31,19 | 247 | 149 | 0.0206918560638 | 0.939563053263236 | 0.019,0.072,0.066, | 0.019,0.016,0.016, | 0,045 |
| 69818 | ENSG00000223745 | CCDC18-AS1 | chr1 | - | 93318167 | 93318228 | 93310127 | 93311608 | 93324634 | 93324691 | 69818 | 2,0,2,0,0,2,0,1,0,0,2,0 | 9,2,17,8,7,9,15,16,5,4,7,5 | 2,3,4,1,2,0,0,0,3,0,2,1,1,0,1 | 5,5,6,2,8,8,6,18,7,18,15,14 | 210 | 149 | 0.020711584129 | 0.940260485903564 | 0.136,0.0,0.077,0.0 | 0.221,0.299,0.321, | -0,087 |
| 98339 | ENSG00000068885 | IFT80 | chr3 | - | 160366042 | 160366152 | 160357484 | 160357578 | 160375811 | 160375880 | 98339 | 96,49,109,86,58,41,21,39,5 | 4,2,7,0,0,2,1,1,1,0,1,0 | 39,31,33,30,41,48,31,70,38 | 5,3,5,2,3,1,0,0,1,2,1,4,0,4,4 | 259 | 149 | 0.0207280558391 | 0.940809824189231 | 0.932,0.934,0.9,1.0 | 0.818,0.856,0.792, | 0,039 |
| 102582 | ENSG00000131931 | THAP1 | chr8 | - | 42839185 | 42839381 | 42836673 | 42838336 | 42843023 | 42843201 | 102582 | 22,23,51,47,41,33,27,30,48 | 1,1,3,2,5,0,2,7,1,1,0,0 | 35,46,24,18,17,14,13,36,9, | 1,1,0,0,1,0,1,1,0,0,1,0,6,4,3 | 298 | 149 | 0.0207616865636 | 0.942137583970102 | 0.917,0.92,0.895,0 | 0.946,0.958,1.0,1.0 | -0,022 |
| 95587 | ENSG00000146143 | PRIM2 | chr6 | + | 57398004 | 57398122 | 57382030 | 57382168 | 57507386 | 57507454 | 95587 | 5,0,3,2,2,0,4,2,0,0,1,0 | 15,10,13,17,12,12,14,8,12, | 0,0,2,0,2,3,0,0,0,0,0,0,1,0,0 | 18,4,7,13,6,3,2,11,8,11,7,5 | 267 | 149 | 0.0207791498471 | 0.942701553130071 | 0.157,0.0,0.114,0.0 | 0.0,0.0,0.138,0.0,0 | 0,003 |
| 185713 | ENSG00000143106 | PSMA5 | chr1 | - | 109413067 | 109413122 | 109412076 | 109412184 | 109415236 | 109415365 | 185713 | 28,18,22,27,15,8,8,6,15,11 | 4,1,13,5,4,3,7,5,3,0,0,2 | 10,16,16,5,12,8,11,11,10,1 | 3,13,7,9,12,3,8,7,6,1,6,11,5 | 204 | 149 | 0.0207828745376 | 0.942701553130071 | 0.836,0.929,0.553, | 0.709,0.473,0.625, | 0,134 |
| 92706 | ENSG00000136861 | CDK5RAP2 | chr9 | - | 120460571 | 120460667 | 120458449 | 120458622 | 120467859 | 120467997 | 92706 | 82,48,80,43,23,32,34,29,32 | 6,1,4,1,3,3,0,3,3,0,0,0 | 50,46,60,83,52,27,47,28,33 | 1,1,5,2,3,4,1,1,5,2,0,4,4,9,1 | 245 | 149 | 0.0208114605088 | 0.943642792603602 | 0.893,0.967,0.924, | 0.968,0.965,0.879, | 0,019 |
| 22908 | ENSG00000100664 | EIF5 | chr14 | + | 103336035 | 103336117 | 103334948 | 103335932 | 103336676 | 103336728 | 22908 | 159,91,419,295,117,203,17 | 6,2,7,15,7,6,16,2,2,0,1,1 | 294,312,188,73,225,69,120 | 15,11,6,19,0,8,2,25,5,0,0,1 | 231 | 149 | 0.0208123938303 | 0.943642792603602 | 0.945,0.967,0.975, | 0.927,0.948,0.953, | 0,033 |
| 4562 | ENSG00000267120 | | chr19 | - | 35740379 | 35740564 | 35739251 | 35739626 | 35741023 | 35741228 | 4562 | 16,5,22,14,18,10,14,13,4,2 | 29,15,26,20,14,16,8,18,25, | 3,13,11,0,10,7,4,4,12,16,2, | 12,32,16,14,12,2,20,70,13, | 298 | 149 | 0.0208343144988 | 0.944437730389895 | 0.216,0.143,0.297, | 0.111,0.169,0.256, | 0,075 |
| 106462 | ENSG00000085224 | ATRX | chrX | - | 77693823 | 77693937 | 77688817 | 77688924 | 77696576 | 77696704 | 106462 | 99,64,105,35,34,26,25,47,5 | 5,2,4,2,10,3,6,9,3,2,2,7 | 60,46,39,76,36,25,10,34,37 | 5,6,7,6,9,4,3,10,8,5,6,5,9,5 | 263 | 149 | 0.0208505668142 | 0.944975436089088 | 0.918,0.948,0.937, | 0.872,0.813,0.759, | 0,073 |
| 209138 | ENSG00000162923 | WDR26 | chr1 | - | 224418334 | 224418416 | 224411426 | 224411565 | 224419517 | 224419615 | 209138 | 30,25,53,30,25,22,21,30,12 | 2,4,0,3,0,1,1,0,0,0,0,0 | 45,53,16,58,28,21,28,11,18 | 1,2,1,3,2,0,1,2,3,2,0,0,4,2,4 | 231 | 149 | 0.0208579039162 | 0.945108951113188 | 0.906,0.801,1.0,0.8 | 0.967,0.945,0.912, | 0,062 |
| 130260 | ENSG00000100982 | PCIF1 | chr20 | + | 45945710 | 45945883 | 45944867 | 45945030 | 45946028 | 45946115 | 130260 | 11,27,56,29,21,28,21,19,20 | 7,5,6,3,4,5,7,2,0,0,0,2 | 17,67,13,21,23,14,27,22,5, | 1,10,2,14,5,0,2,14,2,1,17,9 | 298 | 149 | 0.0208816339057 | 0.945985044655381 | 0.44,0.73,0.824,0.8 | 0.895,0.77,0.765,0 | 0,11 |
| 57030 | ENSG00000077097 | TOP2B | chr3 | - | 25604759 | 25604870 | 25601099 | 25601225 | 25606042 | 25606122 | 57030 | 173,111,274,151,104,89,61 | 11,4,19,8,7,7,5,12,9,4,5,6 | 153,120,95,77,62,58,53,97 | 10,19,1,19,19,6,8,15,4,3,7, | 260 | 149 | 0.0209307647203 | 0.947751667018299 | 0.9,0.941,0.892,0.9 | 0.898,0.784,0.982, | 0,035 |
| 241686 | ENSG00000175893 | ZDHHC21 | chr9 | - | 14677297 | 14677422 | 14674186 | 14674385 | 14680032 | 14680162 | 241686 | 22,31,57,44,2,7,13,21,18,1 | 12,10,23,19,5,5,8,9,2,2,1,7 | 14,8,9,3,9,14,6,31,3,17,27, | 11,21,6,0,9,5,2,13,16,19,15 | 274 | 149 | 0.0209352507194 | 0.947751667018299 | 0.499,0.628,0.574, | 0.409,0.172,0.449, | 0,095 |
| 94130 | ENSG00000112303 | VNN2 | chr6 | - | 132752460 | 132752749 | 132751144 | 132751518 | 132755842 | 132756035 | 94130 | 32,36,44,42,53,46,40,30,57 | 5,6,0,2,1,2,1,0,4,1,0,0 | 20,24,29,22,17,36,15,17,28 | 0,3,2,2,2,0,0,0,0,0,0,1,2,1,0 | 298 | 149 | 0.0209359369261 | 0.947751667018299 | 0.762,0.75,1.0,0.91 | 1.0,0.8,0.879,0.84 | -0,01 |
| 138094 | ENSG00000187912 | CLEC17A | chr19 | + | 14585475 | 14585830 | 14583356 | 14583434 | 14587613 | 14587691 | 138094 | 3,4,6,5,5,1,9,0,0,3,1,2 | 21,2,31,24,73,24,44,31,25, | 6,6,4,3,2,2,0,1,3,1,4,8,1,0,7 | 16,14,4,4,11,5,8,13,6,7,4,4 | 298 | 149 | 0.0209382439178 | 0.947751667018299 | 0.067,0.5,0.088,0.0 | 0.158,0.176,0.333, | -0,043 |
| 253036 | ENSG00000091039 | OSBPL8 | chr12 | - | 76459858 | 76459940 | 76450850 | 76450988 | 76487509 | 76487618 | 253036 | 17,10,10,6,2,3,1,8,16,2,1,3 | 23,31,35,36,20,12,20,20,22 | 5,4,3,2,4,2,3,2,4,3,4,4,5,9,5 | 25,34,12,15,18,13,9,17,8,1 | 231 | 149 | 0.0209629075267 | 0.948668534457352 | 0.323,0.172,0.156, | 0.114,0.071,0.139, | 0,035 |
| 140822 | ENSG00000134265 | NAPG | chr18 | + | 10538859 | 10538974 | 10534465 | 10534496 | 10539761 | 10539871 | 140822 | 10,5,5,7,2,2,1,7,2,0,2,0 | 8,14,32,30,12,13,1,12,9,2,6 | 2,7,6,4,6,2,2,4,2,3,14,2,6,7 | 3,9,4,1,6,4,2,6,6,5,12,10,17 | 264 | 149 | 0.0209693725852 | 0.948761620955472 | 0.414,0.168,0.081, | 0.273,0.305,0.458, | -0,119 |
| 120625 | ENSG00000107290 | SETX | chr9 | - | 132266126 | 132266249 | 132261355 | 132264985 | 132269614 | 132269702 | 120625 | 7,7,9,1,0,12,2,1,3,5,1,4 | 103,41,126,113,114,57,76, | 4,6,2,0,1,1,3,0,3,2,8,0,4,1,6 | 144,98,76,36,19,57,36,77,3 | 272 | 149 | 0.0209968886464 | 0.949806923130164 | 0.036,0.086,0.038, | 0.015,0.032,0.014, | 0,015 |
| 168784 | ENSG00000055955 | ITIH4 | chr3 | - | 52818650 | 52818727 | 52818525 | 52818536 | 52819392 | 52819598 | 168784 | 2,2,5,2,8,5,7,3,5,1,1,2 | 1,0,2,5,5,9,4,13,6,1,5,9 | 1,4,9,2,2,4,3,8,3,4,2,5,2,1,5 | 3,5,1,3,1,1,0,4,0,1,7,0,3,1,6 | 226 | 149 | 0.0210126914008 | 0.950010368236221 | 0.569,1.0,0.622,0.2 | 0.18,0.345,0.856,0 | -0,165 |
| 82805 | ENSG00000176978 | DPP7 | chr9 | - | 137113205 | 137113287 | 137112816 | 137113119 | 137113864 | 137114028 | 82805 | 104,55,75,58,48,47,74,61,7 | 2,5,0,0,0,0,2,0,0,0,2,0 | 57,60,94,70,43,9,36,79,47, | 0,0,5,10,0,0,2,4,0,0,4,4,0,0 | 231 | 149 | 0.0210151794227 | 0.950010368236221 | 0.971,0.876,1.0,1.0 | 1.0,1.0,0.924,0.81 | 0,013 |
| 26685 | ENSG00000135506 | OS9 | chr12 | + | 57720098 | 57720263 | 57718992 | 57719182 | 57720405 | 57720518 | 26685 | 89,67,205,209,79,78,72,81 | 49,24,134,85,53,41,41,55,3 | 111,113,61,21,44,48,41,69 | 67,85,51,24,37,28,28,36,35 | 298 | 149 | 0.0210169281001 | 0.950010368236221 | 0.476,0.583,0.433, | 0.453,0.399,0.374, | 0,067 |
| 57162 | ENSG00000071564 | TCF3 | chr19 | - | 1615284 | 1615485 | 1612206 | 1612433 | 1615685 | 1615798 | 57162 | 49,10,56,19,24,14,22,24,35 | 45,34,66,71,24,53,32,30,60 | 19,15,8,8,14,6,28,15,2,10,1 | 35,38,20,21,10,9,25,85,11, | 298 | 149 | 0.0210229485636 | 0.950010368236221 | 0.353,0.128,0.298, | 0.213,0.165,0.167, | 0,04 |
| 5649 | ENSG00000141968 | VAV1 | chr19 | + | 6825302 | 6825406 | 6825052 | 6825121 | 6826611 | 6826711 | 5649 | 118,58,103,74,53,29,33,44 | 7,2,3,0,1,2,0,2,1,1,2,0 | 50,53,52,10,57,46,15,19,36 | 4,1,0,2,0,1,3,0,0,0,0,0,1,0,2 | 253 | 149 | 0.0210234556542 | 0.950010368236221 | 0.908,0.945,0.953, | 0.88,0.969,1.0,0.74 | -0,003 |
| 4060 | ENSG00000170889 | RPS9 | chr19 | + | 54206552 | 54206724 | 54206275 | 54206462 | 54207397 | 54207522 | 4060 | 97,61,152,116,47,45,49,73 | 970,517,1906,1616,681,79 | 31,69,86,37,81,48,33,50,36 | 749,634,603,319,436,414,5 | 298 | 149 | 0.0210333086355 | 0.950256098011608 | 0.048,0.056,0.038, | 0.02,0.052,0.067,0 | -0,007 |
| 195182 | ENSG00000162604 | TM2D1 | chr1 | - | 61723712 | 61724031 | 61709328 | 61709437 | 61724901 | 61725144 | 195182 | 17,11,43,22,13,9,8,6,1,5,3, | 2,2,5,3,0,0,0,6,0,1,1,3 | 23,17,11,14,8,7,1,6,5,22,22 | 1,2,0,2,3,0,2,0,1,1,0,0,1,0,2 | 298 | 149 | 0.0210480376219 | 0.950721969653548 | 0.81,0.733,0.811,0 | 0.92,0.81,1.0,0.778 | -0,025 |
| 37555 | ENSG00000113522 | RAD50 | chr5 | + | 132592763 | 132592965 | 132591876 | 132592034 | 132594868 | 132595044 | 37555 | 0,1,4,1,4,1,2,1,1,0,0,0 | 9,1,16,15,8,6,5,13,10,2,2,8 | 2,4,4,0,3,3,3,0,1,0,2,6,0,4,2 | 12,12,13,2,4,5,4,12,10,13,5 | 298 | 149 | 0.0211028612555 | 0.95299830694136 | 0.0,0.333,0.111,0.0 | 0.077,0.143,0.133, | -0,025 |
| 123181 | ENSG00000151498 | ACAD8 | chr11 | + | 134257087 | 134257257 | 134256547 | 134256648 | 134258514 | 134258624 | 123181 | 27,16,44,17,28,18,15,18,9, | 3,0,3,1,2,3,2,0,1,1,2,2 | 20,21,21,6,7,10,11,29,9,23 | 4,1,0,1,2,1,0,9,3,1,4,2,0,3,1 | 298 | 149 | 0.0211124508981 | 0.953231365481217 | 0.818,1.0,0.88,0.89 | 0.714,0.913,1.0,0.7 | 0,019 |
| 14265 | ENSG00000168802 | CHTF8 | chr16 | - | 69121052 | 69121170 | 69118009 | 69120649 | 69121435 | 69121492 | 14265 | 22,15,45,37,21,16,16,15,20 | 10,6,9,2,7,2,3,5,1,0,2,4 | 38,10,15,4,10,25,2,29,12,3 | 0,3,0,5,2,2,3,3,0,2,2,6,5,4,0 | 267 | 149 | 0.021119677022 | 0.95335763530245 | 0.551,0.582,0.736, | 1.0,0.65,1.0,0.309, | -0,049 |
| 227719 | ENSG00000186088 | GSAP | chr7 | - | 77399467 | 77399537 | 77397345 | 77397415 | 77404558 | 77404615 | 227719 | 23,20,79,32,19,23,36,32,5, | 70,53,212,82,65,61,71,141 | 46,55,12,11,19,13,18,19,27 | 137,72,61,29,57,20,49,58,3 | 219 | 149 | 0.0211584421872 | 0.954731568255641 | 0.183,0.204,0.202, | 0.186,0.342,0.118, | -0,043 |
| 221645 | ENSG00000188994 | ZNF292 | chr6 | + | 87222739 | 87222894 | 87218595 | 87218731 | 87233324 | 87233777 | 221645 | 0,0,8,4,0,1,2,5,0,0,0,2 | 23,29,74,34,30,12,11,18,8, | 0,2,0,1,0,0,0,0,2,0,5,2,0,1,0 | 44,33,27,21,32,18,8,22,11, | 298 | 149 | 0.0211589853719 | 0.954731568255641 | 0.0,0.0,0.051,0.05 | 0.0,0.029,0.0,0.02 | 0,014 |
| 32253 | ENSG00000231074 | HCG18 | chr6 | - | 30314268 | 30314482 | 30296131 | 30296237 | 30326353 | 30326389 | 32253 | 1,1,0,1,0,2,2,1,4,0,0,2 | 17,1,16,10,7,9,2,9,5,2,3,12 | 8,1,3,0,2,1,0,5,4,1,1,4,0,0,2 | 6,2,5,2,2,5,2,10,6,9,4,16,1, | 298 | 149 | 0.021168513025 | 0.954961271616805 | 0.029,0.333,0.0,0.0 | 0.4,0.2,0.231,0.0,0 | -0,022 |
| 236618 | ENSG00000104783 | KCNN4 | chr19 | - | 43773785 | 43773891 | 43771999 | 43772135 | 43774191 | 43774619 | 236618 | 6,3,17,10,4,2,3,4,17,5,1,2 | 259,98,204,103,120,137,16 | 1,0,5,3,6,11,1,8,1,3,6,0,0,5 | 141,123,107,155,97,66,79, | 255 | 149 | 0.0211913092139 | 0.955769472334242 | 0.013,0.018,0.046, | 0.004,0.0,0.027,0.0 | 0,003 |
| 43205 | ENSG00000254087 | LYN | chr8 | + | 55941854 | 55941991 | 55879834 | 55880103 | 55946447 | 55946493 | 43205 | 289,248,547,277,150,133,1 | 2,7,19,10,0,7,0,0,1,0,1,0 | 289,276,158,117,208,135,8 | 9,19,3,2,7,4,2,3,3,3,0,11,3, | 286 | 149 | 0.0211953096508 | 0.955769472334242 | 0.987,0.949,0.937, | 0.944,0.883,0.965, | 0,019 |
| 19229 | ENSG00000114062 | UBE3A | chr15 | - | 25354527 | 25354683 | 25352281 | 25354426 | 25355891 | 25356056 | 19229 | 142,106,353,123,61,63,81, | 11,7,27,0,1,4,5,8,6,2,7,3 | 81,111,70,48,104,62,54,70 | 5,14,14,6,5,4,1,9,0,3,17,5,5 | 298 | 149 | 0.0212183433437 | 0.956607720194364 | 0.866,0.883,0.867, | 0.89,0.799,0.714,0 | 0,043 |
| 73412 | ENSG00000008869 | HEATR5B | chr2 | - | 37020654 | 37020836 | 37019808 | 37019877 | 37027922 | 37028174 | 73412 | 109,65,145,126,72,51,74,7 | 2,0,0,3,0,0,5,0,0,2,0,0 | 151,143,48,38,43,60,61,98 | 0,6,0,0,0,2,2,1,2,0,2,1,8,2,1 | 298 | 149 | 0.0212763542694 | 0.958844986497136 | 0.965,1.0,1.0,0.95 | 1.0,0.923,1.0,1.0,1 | 0,014 |
| 189563 | ENSG00000143398 | PIP5K1A | chr1 | + | 151242179 | 151242567 | 151239954 | 151240039 | 151246919 | 151246965 | 189563 | 14,15,34,33,18,22,17,25,9, | 1,3,1,7,0,0,2,0,1,2,0,2 | 35,46,18,10,17,10,6,19,9,2 | 0,0,1,0,0,0,0,1,0,2,4,0,2,2,0 | 298 | 149 | 0.0212768776594 | 0.958844986497136 | 0.875,0.714,0.944, | 1.0,1.0,0.9,1.0,1.0, | -0,061 |
| 48073 | ENSG00000118007 | STAG1 | chr3 | - | 136568764 | 136568861 | 136542118 | 136542195 | 136604308 | 136604473 | 48073 | 68,29,116,45,30,57,23,38,1 | 0,3,2,2,3,1,5,0,0,0,0,6 | 74,60,42,52,41,24,19,42,39 | 2,0,1,1,1,0,0,1,0,0,2,1,0,0,4 | 246 | 149 | 0.021295578631 | 0.959486850393293 | 1.0,0.854,0.972,0.9 | 0.957,1.0,0.962,0.9 | -0,04 |
| 88463 | ENSG00000073614 | KDM5A | chr12 | - | 286125 | 286207 | 285455 | 285662 | 292758 | 292925 | 88463 | 11,12,15,18,12,6,6,16,16,1 | 72,47,117,87,63,50,53,88,3 | 16,18,8,5,10,1,9,3,1,13,8,1 | 75,99,51,47,74,35,45,68,42 | 231 | 149 | 0.0213016963222 | 0.959561615973928 | 0.09,0.141,0.076,0 | 0.121,0.105,0.092, | 0,034 |
| 117289 | ENSG00000080298 | RFX3 | chr9 | - | 3270370 | 3270525 | 3266207 | 3266305 | 3271002 | 3271118 | 117289 | 16,12,74,28,14,38,24,41,9, | 4,6,14,5,3,6,5,13,2,2,0,2 | 19,45,26,43,18,22,52,17,17 | 7,13,8,17,15,8,9,7,6,7,11,7 | 298 | 149 | 0.0213061876829 | 0.959563105498692 | 0.667,0.5,0.725,0.7 | 0.576,0.634,0.619, | 0,104 |
| 207252 | ENSG00000158435 | CNOT11 | chr2 | + | 101257837 | 101257955 | 101252885 | 101253478 | 101262538 | 101262691 | 207252 | 36,27,26,27,19,19,7,15,56, | 2,2,6,0,2,1,1,5,1,2,0,0 | 28,17,15,15,6,12,4,9,3,31,2 | 1,1,1,1,2,1,2,2,1,1,2,1,0,1,2 | 267 | 149 | 0.0213444127173 | 0.961083534216987 | 0.909,0.883,0.707, | 0.94,0.905,0.893,0 | 0,05 |
| 118194 | ENSG00000106948 | AKNA | chr9 | - | 114359593 | 114359794 | 114357920 | 114358167 | 114359895 | 114360062 | 118194 | 68,74,75,79,74,55,52,55,45 | 10,0,7,5,3,1,4,3,0,0,0,0 | 52,61,43,84,35,40,37,46,39 | 5,2,2,3,7,1,2,3,6,2,4,15,0,0 | 298 | 149 | 0.0213529780017 | 0.961268104640011 | 0.773,1.0,0.843,0.8 | 0.839,0.938,0.915, | 0,06 |
| 148507 | ENSG00000159593 | NAE1 | chr16 | - | 66827884 | 66828077 | 66826676 | 66826780 | 66830846 | 66830976 | 148507 | 6,0,15,5,0,0,0,1,1,0,0,0 | 7,5,31,27,8,13,6,17,7,3,12, | 1,1,2,2,4,4,0,3,0,0,1,0,0,2,1 | 13,18,11,12,6,6,5,11,13,16 | 298 | 149 | 0.0213578022281 | 0.961284218184063 | 0.3,0.0,0.195,0.085 | 0.037,0.027,0.083, | -0,005 |
| 118193 | ENSG00000106948 | AKNA | chr9 | - | 114356862 | 114356969 | 114355924 | 114356136 | 114357920 | 114358167 | 118193 | 183,148,117,104,134,112,8 | 15,0,6,2,0,2,8,6,4,3,0,3 | 120,163,121,126,90,75,65, | 3,3,5,22,10,0,3,5,6,5,1,4,2, | 256 | 149 | 0.0213689430477 | 0.961584566401739 | 0.877,1.0,0.919,0.9 | 0.959,0.969,0.934, | 0,024 |
| 113495 | ENSG00000197265 | GTF2E2 | chr8 | - | 30636929 | 30636960 | 30635031 | 30635123 | 30653432 | 30653602 | 113495 | 1,1,1,4,0,0,2,3,0,0,0,0 | 43,25,45,44,29,21,26,14,27 | 0,4,2,3,3,2,0,0,2,0,1,4,0,2,1 | 26,19,23,28,21,17,14,23,17 | 180 | 149 | 0.0213739628937 | 0.961609407937697 | 0.019,0.032,0.018, | 0.0,0.148,0.067,0.0 | -0,025 |
| 30573 | ENSG00000181929 | PRKAG1 | chr12 | - | 49003557 | 49003595 | 49003143 | 49003290 | 49004506 | 49004633 | 30573 | 10,7,14,14,13,5,5,10,8,7,4, | 1,1,1,2,0,1,0,0,0,1,0,1 | 6,6,7,4,3,2,9,11,15,4,9,23,1 | 2,10,2,0,0,2,1,0,1,0,0,2,1,1 | 187 | 149 | 0.0214325668363 | 0.964044470792829 | 0.888,0.848,0.918, | 0.705,0.323,0.736, | 0,08 |
| 54544 | ENSG00000180098 | TRNAU1AP | chr1 | + | 28561112 | 28561260 | 28560632 | 28560732 | 28561345 | 28561398 | 54544 | 24,7,37,20,23,12,13,18,14, | 13,4,22,15,14,9,9,4,10,5,0, | 16,10,4,3,11,2,3,15,4,17,14 | 11,10,6,5,10,4,7,9,2,7,5,30 | 297 | 149 | 0.021438504654 | 0.964110070034491 | 0.481,0.468,0.458, | 0.422,0.334,0.251, | 0,165 |
| 248572 | ENSG00000073921 | PICALM | chr11 | - | 86023419 | 86023541 | 86022377 | 86022469 | 86026291 | 86026367 | 248572 | 11,5,12,6,1,0,1,1,2,0,0,0 | 172,139,150,72,26,30,38,4 | 4,12,2,14,3,0,1,7,1,1,7,3,4, | 87,83,74,96,54,40,50,82,30 | 271 | 149 | 0.0214768679299 | 0.965484214865777 | 0.034,0.019,0.042, | 0.025,0.074,0.015, | -0,015 |
| 210199 | ENSG00000169499 | PLEKHA2 | chr8 | + | 38943788 | 38943934 | 38935993 | 38936050 | 38946123 | 38946221 | 210199 | 43,11,133,75,43,60,33,70,4 | 4,2,33,22,16,4,10,15,11,6,7 | 68,52,38,12,20,43,36,50,29 | 30,9,10,19,12,5,15,22,7,17 | 295 | 149 | 0.0214780325361 | 0.965484214865777 | 0.844,0.735,0.671, | 0.534,0.745,0.657, | 0,089 |
| 725 | ENSG00000291299 | | chr22 | - | 23683724 | 23683867 | 23658093 | 23660000 | 23690235 | 23690297 | 725 | 50,13,0,0,23,19,31,57,9,2,1 | 0,0,9,5,1,0,0,0,0,3,0,0 | 27,35,20,30,12,12,22,42,28 | 0,0,0,0,1,1,0,2,4,2,0,6,0,0,0 | 292 | 149 | 0.0214988160663 | 0.966216680051329 | 1.0,1.0,0.0,0.0,0.92 | 1.0,1.0,1.0,1.0,0.8 | -0,162 |
| 236548 | ENSG00000034677 | RNF19A | chr8 | - | 100303159 | 100303355 | 100287906 | 100288267 | 100309866 | 100310148 | 236548 | 2,0,11,10,5,14,6,27,0,0,4,0 | 31,13,81,37,17,43,52,137,9 | 0,13,6,13,2,7,7,25,4,2,71,1 | 39,26,40,50,22,14,38,59,30 | 298 | 149 | 0.021512311246 | 0.966435933394214 | 0.031,0.0,0.064,0.1 | 0.0,0.2,0.07,0.115, | -0,039 |
| 79261 | ENSG00000144909 | OSBPL11 | chr3 | - | 125552180 | 125552679 | 125538450 | 125538633 | 125560378 | 125560519 | 79261 | 27,21,40,60,18,12,12,14,19 | 4,0,1,2,1,0,0,0,2,0,3,0 | 39,50,6,17,14,22,15,11,11, | 1,3,2,2,0,1,0,3,3,2,2,0,2,1,0 | 298 | 149 | 0.021515284237 | 0.966435933394214 | 0.771,1.0,0.952,0.9 | 0.951,0.893,0.6,0.8 | 0,041 |
| 172850 | ENSG00000211456 | SACM1L | chr3 | + | 45713729 | 45713807 | 45713140 | 45713196 | 45714045 | 45714082 | 172850 | 8,0,11,6,0,4,3,2,0,0,1,0 | 16,1,23,17,12,1,9,8,10,9,4, | 3,4,4,8,6,3,4,0,4,6,2,1,2,1,5 | 5,10,4,10,6,6,0,3,7,9,3,14,1 | 227 | 149 | 0.0215171652449 | 0.966435933394214 | 0.247,0.0,0.239,0.1 | 0.283,0.208,0.396, | -0,143 |
| 53828 | ENSG00000198408 | OGA | chr10 | - | 101793912 | 101793998 | 101792838 | 101792943 | 101797979 | 101798154 | 53828 | 127,131,325,167,84,55,78, | 46,11,34,12,30,15,18,12,15 | 204,196,121,109,125,58,78 | 10,11,22,19,18,9,5,16,11,9 | 235 | 149 | 0.0215395373653 | 0.967203083481522 | 0.636,0.883,0.858, | 0.928,0.919,0.777, | -0,046 |
| 35402 | ENSG00000001629 | ANKIB1 | chr7 | + | 92389970 | 92390116 | 92387974 | 92388041 | 92391165 | 92391344 | 35402 | 75,32,94,91,36,57,45,56,68 | 0,2,0,0,4,7,5,4,0,2,1,0 | 39,55,79,54,26,42,13,95,36 | 0,2,7,0,3,4,0,6,8,3,1,1,3,2,0 | 295 | 149 | 0.0215432330018 | 0.967203083481522 | 1.0,0.89,1.0,1.0,0.8 | 1.0,0.933,0.851,1.0 | 0,002 |
| 252431 | ENSG00000011258 | MBTD1 | chr17 | - | 51208972 | 51209430 | 51206887 | 51207005 | 51217333 | 51217416 | 252431 | 2,2,12,1,1,0,2,2,3,1,2,1 | 18,8,24,22,16,19,11,10,7,2 | 0,7,0,1,1,2,2,0,0,3,2,0,3,1,3 | 14,28,7,10,21,16,11,14,4,2 | 298 | 149 | 0.0215703361863 | 0.968217941128996 | 0.053,0.111,0.2,0.0 | 0.0,0.111,0.0,0.048 | 0,053 |

| | | | | | | | | | | | | | | | | | | | | | | |
|---|---|---|---|---|---|---|---|---|---|---|---|---|---|---|---|---|---|---|---|---|---|---|
| 233025 | ENSG00000005889 | ZFX | chrX | + | 24208545 | 24209040 | 24208217 | 24208370 | 24210192 | 24211986 | 233025 | 42,48,156,67,39,36,26,49,2 | 4,3,8,3,0,2,4,0,5,2,0,0 | 81,96,41,36,63,45,15,50,32 | 9,4,2,0,0,2,1,0,2,1,0,2,1,0,1 | 298 | 149 | 0.021579871438 | 0.968443976120137 | 0.84,0.889,0.907,0 | 0.818,0.923,0.911, | -0,044 |
| 42987 | ENSG00000183726 | TMEM50A | chr1 | + | 25342960 | 25343073 | 25340473 | 25340579 | 25351625 | 25351693 | 42987 | 38,44,173,123,56,69,53,75 | 1,1,7,7,2,0,2,6,1,0,0,4 | 107,155,53,48,43,48,29,57 | 1,6,2,2,2,5,7,3,1,4,7,6,3,0,6 | 262 | 149 | 0.0215903736868 | 0.968713303936554 | 0.956,0.962,0.934, | 0.984,0.936,0.938, | 0,046 |
| 227991 | ENSG00000224137 | LINC01857 | chr2 | + | 207674949 | 207675077 | 207667498 | 207667564 | 207678541 | 207679116 | 227991 | 31,23,57,43,20,7,19,16,26, | 51,22,55,29,22,7,24,4,8,9,1 | 28,20,2,6,7,3,1,5,4,12,0,92 | 19,16,9,0,13,2,4,3,11,29,9, | 277 | 149 | 0.0216100399273 | 0.969393602249418 | 0.246,0.36,0.358,0 | 0.442,0.402,0.107, | 0,098 |
| 82645 | ENSG00000031081 | ARHGAP31 | chr3 | + | 119368371 | 119368516 | 119365315 | 119365418 | 119380903 | 119380986 | 82645 | 10,23,12,25,13,12,7,5,12,3 | 5,9,2,1,0,0,0,0,0,0,1,2 | 16,6,8,1,11,15,3,8,9,21,3,4 | 0,2,1,1,1,2,0,1,1,1,0,2,1,2,2 | 294 | 149 | 0.0216442285798 | 0.970724934665333 | 0.503,0.564,0.753, | 1.0,0.603,0.802,0.3 | 0,097 |
| 165396 | ENSG00000138767 | CNOT6L | chr4 | - | 77730249 | 77730332 | 77728876 | 77729081 | 77731386 | 77731881 | 165396 | 2,2,1,2,0,1,0,4,0,0,0,0 | 23,25,77,53,29,34,16,34,24 | 2,1,6,5,0,1,0,1,0,2,4,0,6,3,0 | 38,58,15,31,45,12,50,32,14 | 232 | 149 | 0.0216627786951 | 0.971354483609389 | 0.053,0.049,0.008, | 0.033,0.011,0.204, | -0,029 |
| 38223 | ENSG00000134852 | CLOCK | chr4 | - | 55442431 | 55442634 | 55438281 | 55438537 | 55443686 | 55443896 | 38223 | 46,21,39,24,15,21,10,18,12 | 3,0,4,4,0,0,2,0,0,1,2,0 | 26,19,17,24,19,17,25,35,15 | 3,6,1,1,1,0,0,0,2,2,0,0,3,0,4 | 298 | 149 | 0.0216714784378 | 0.971542173640102 | 0.885,1.0,0.83,0.75 | 0.812,0.613,0.895, | -0,039 |
| 243450 | ENSG00000065923 | SLC9A7 | chrX | - | 46635588 | 46635646 | 46631585 | 46631649 | 46643235 | 46643389 | 243450 | 65,55,55,27,20,23,20,46,23 | 181,139,132,102,86,90,127 | 52,50,32,32,33,19,9,30,25, | 118,79,161,172,55,42,66,1 | 207 | 149 | 0.0216951459084 | 0.972400655770687 | 0.205,0.222,0.231, | 0.241,0.313,0.125, | -0,064 |
| 37855 | ENSG00000164038 | SLC9B2 | chr4 | - | 103057800 | 103057971 | 103050239 | 103050382 | 103066326 | 103066507 | 37855 | 16,15,33,35,11,29,23,17,16 | 2,0,18,4,5,5,0,6,0,1,0,5 | 22,25,20,7,16,12,7,22,13,1 | 7,10,1,2,5,2,1,12,2,2,5,12,4 | 298 | 149 | 0.0217295239065 | 0.973738738271893 | 0.8,1.0,0.478,0.814 | 0.611,0.556,0.909, | 0,107 |
| 94716 | ENSG00000146247 | PHIP | chr6 | - | 78946537 | 78946607 | 78946000 | 78946260 | 78946710 | 78946874 | 94716 | 23,11,23,8,7,20,6,24,4,5,5, | 89,52,67,26,37,41,27,38,32 | 20,36,17,22,17,6,10,36,16, | 41,40,70,55,58,30,21,59,29 | 219 | 149 | 0.0217344958674 | 0.973758801006738 | 0.15,0.126,0.189,0 | 0.249,0.38,0.142,0 | -0,054 |
| 180306 | ENSG00000146409 | SLC18B1 | chr6 | - | 132787433 | 132787528 | 132783932 | 132784089 | 132789763 | 132789837 | 180306 | 23,14,27,21,28,15,6,22,9,4 | 4,2,10,7,6,2,1,5,2,4,0,6 | 45,7,8,6,5,5,3,10,6,4,14,11 | 10,6,8,2,2,0,3,3,3,7,10,2,10 | 244 | 149 | 0.0217497679973 | 0.974240232222034 | 0.778,0.81,0.622,0 | 0.733,0.416,0.379, | 0,14 |
| 204157 | ENSG00000142655 | PEX14 | chr1 | + | 10576777 | 10576899 | 10536212 | 10536297 | 10599237 | 10599366 | 204157 | 1,0,1,1,0,0,0,0,2,0,0,0 | 23,16,11,13,3,11,4,13,17,6 | 4,5,1,1,0,2,0,2,0,4,0,1,1,2,2 | 8,12,8,4,3,1,2,14,11,9,4,12 | 271 | 149 | 0.0217620909212 | 0.974472368179741 | 0.023,0.0,0.048,0.0 | 0.216,0.186,0.064, | -0,101 |
| 22664 | ENSG00000100888 | CHD8 | chr14 | - | 21430800 | 21431858 | 21428963 | 21429335 | 21437193 | 21437273 | 22664 | 28,16,38,45,42,17,17,14,40 | 0,0,2,0,1,1,2,0,0,3,1,0 | 22,27,17,8,17,19,15,29,12, | 0,6,3,1,0,0,0,2,0,0,1,2,0,2,2 | 298 | 149 | 0.0217640055282 | 0.974472368179741 | 1.0,1.0,0.905,1.0,0 | 1.0,0.692,0.739,0.8 | 0,04 |
| 165191 | ENSG00000115073 | ACTR1B | chr2 | - | 97658447 | 97658643 | 97658223 | 97658316 | 97658878 | 97659003 | 165191 | 12,11,26,22,32,19,4,18,14, | 5,0,4,2,0,4,0,0,0,1,0,2 | 12,17,5,2,4,12,6,23,8,14,31 | 0,0,0,1,1,0,0,2,0,2,0,2,0,6,0 | 298 | 149 | 0.0217799543763 | 0.974983644002792 | 0.545,1.0,0.765,0.8 | 1.0,1.0,1.0,0.5,0.66 | -0,019 |
| 221084 | ENSG00000196664 | TLR7 | chrX | + | 12877443 | 12877566 | 12867478 | 12867581 | 12885511 | 12890361 | 221084 | 0,0,11,5,4,4,0,2,0,0,0,1 | 5,4,28,59,49,15,17,22,18,1 | 4,2,2,2,5,6,1,1,0,2,0,2,5,7,4 | 28,55,21,13,13,15,5,18,4,2 | 272 | 149 | 0.0217948931606 | 0.975449500904367 | 0.0,0.0,0.177,0.04 | 0.073,0.02,0.05,0.0 | -0,037 |
| 85125 | ENSG00000135426 | TESPA1 | chr12 | - | 54961167 | 54961267 | 54948014 | 54950390 | 54962430 | 54963238 | 85125 | 43,28,54,41,25,16,24,23,98 | 4,1,0,0,1,0,0,2,6,1,0,4 | 25,32,22,5,20,16,7,13,9,23 | 0,0,0,2,2,0,0,0,1,0,3,0,0,0,0 | 249 | 149 | 0.0218219279036 | 0.976456416760859 | 0.865,0.944,1.0,1.0 | 1.0,1.0,1.0,0.599,0 | 0,004 |
| 151661 | ENSG00000156958 | GALK2 | chr15 | + | 49281986 | 49282101 | 49239219 | 49239367 | 49283565 | 49283718 | 151661 | 6,3,27,11,7,9,2,10,2,0,0,11 | 3,2,3,1,2,4,3,0,2,3,1,3 | 12,4,4,4,7,8,3,6,2,4,11,15,1 | 2,0,1,5,5,0,1,0,1,0,0,2,4,0,1 | 264 | 149 | 0.0218329174998 | 0.976745097983674 | 0.53,0.458,0.836,0 | 0.772,1.0,0.693,0.3 | -0,216 |
| 216486 | ENSG00000254999 | BRK1 | chr3 | + | 10118905 | 10118970 | 10115674 | 10115819 | 10125625 | 10125708 | 216486 | 23,24,72,36,14,14,22,20,13 | 113,97,196,234,96,69,80,5 | 43,75,34,10,34,8,13,15,17, | 127,114,112,31,130,92,33, | 214 | 149 | 0.0218787915004 | 0.97855885972618 | 0.124,0.147,0.204, | 0.191,0.314,0.174, | -0,031 |
| 25044 | ENSG00000139746 | RBM26 | chr13 | - | 79353152 | 79353224 | 79344668 | 79344794 | 79354438 | 79354561 | 25044 | 1,1,3,1,3,4,2,5,1,2,0,0 | 86,72,113,83,47,70,53,106 | 7,7,5,5,1,0,1,5,3,12,0,1,3,0 | 72,125,60,83,91,57,43,75,8 | 221 | 149 | 0.021882553126 | 0.97855885972618 | 0.008,0.009,0.018, | 0.062,0.036,0.053, | -0,008 |
| 228756 | ENSG00000160218 | TRAPPC10 | chr21 | + | 44063569 | 44063785 | 44059102 | 44059214 | 44074323 | 44074470 | 228756 | 50,62,90,80,42,43,53,53,57 | 7,0,5,3,3,5,2,3,0,0,5,2 | 66,70,40,66,68,27,43,52,34 | 2,0,2,2,5,1,2,4,2,0,5,0,1,2,0 | 298 | 149 | 0.021891421371 | 0.978630257453631 | 0.781,1.0,0.9,0.93, | 0.943,1.0,0.909,0.9 | -0,036 |
| 251472 | ENSG00000137752 | CASP1 | chr11 | - | 105029116 | 105029267 | 105026841 | 105026951 | 105029664 | 105029899 | 251472 | 20,4,53,58,8,6,7,10,2,6,4,5 | 1,1,4,2,0,0,0,0,0,0,0,0 | 9,9,4,7,12,21,3,12,14,18,9, | 0,1,1,0,0,0,0,1,0,0,2,7,7,0,0 | 298 | 149 | 0.0218932434902 | 0.978630257453631 | 0.909,0.667,0.869, | 1.0,0.818,0.667,1.0 | 0,054 |
| 41515 | ENSG00000118855 | MFSD1 | chr3 | + | 158805361 | 158805474 | 158804230 | 158804371 | 158807039 | 158807082 | 41515 | 38,16,76,43,37,24,20,27,31 | 7,3,1,9,1,5,2,4,1,2,0,0 | 45,37,19,7,35,15,5,17,13,3 | 5,7,3,1,5,2,2,3,0,5,4,25,2,3 | 262 | 149 | 0.0219026119663 | 0.978845738396743 | 0.755,0.752,0.977, | 0.837,0.75,0.783,0 | 0,075 |
| 67479 | ENSG00000143575 | HAX1 | chr1 | + | 154273773 | 154273961 | 154272606 | 154273598 | 154274949 | 154275001 | 67479 | 122,103,188,160,110,126,1 | 2,2,7,3,0,3,1,3,2,1,1,0 | 107,68,85,40,53,45,73,106 | 2,0,0,2,1,4,0,0,1,0,4,0,0,0,4 | 298 | 149 | 0.0219249302168 | 0.979639745794495 | 0.968,0.963,0.931, | 0.964,1.0,1.0,0.909 | 0,001 |
| 231960 | ENSG00000133703 | KRAS | chr12 | - | 25213113 | 25213153 | 25205245 | 25209911 | 25225613 | 25225773 | 231960 | 2,0,2,1,0,3,2,2,0,0,0,2 | 3,3,69,50,39,32,21,39,6,4,6 | 0,8,1,3,2,0,0,2,5,1,2,2,0,1,4 | 22,96,32,55,50,10,22,17,28 | 189 | 149 | 0.0219361181488 | 0.979881829567065 | 0.345,0.0,0.022,0.0 | 0.0,0.062,0.024,0.0 | 0 |
| 37115 | ENSG00000146350 | TBC1D32 | chr6 | - | 121256083 | 121256285 | 121255327 | 121255410 | 121279120 | 121279245 | 37115 | 5,7,16,4,7,5,7,12,14,0,5,6 | 11,4,5,2,2,4,2,1,2,4,2,0 | 18,7,13,16,7,3,6,14,10,3,8, | 4,5,4,3,2,1,0,0,3,0,6,5,0,0,0 | 298 | 149 | 0.0219395666422 | 0.979881829567065 | 0.185,0.467,0.615, | 0.692,0.412,0.619, | -0,153 |
| 83125 | ENSG00000164548 | TRA2A | chr7 | - | 23522120 | 23522432 | 23521706 | 23521840 | 23531788 | 23531980 | 83125 | 192,82,87,34,32,17,56,68,3 | 14,7,72,52,14,24,4,4,10,4,6 | 65,53,64,45,28,29,17,21,12 | 36,49,23,11,10,15,5,17,19, | 298 | 149 | 0.0219440062933 | 0.979881829567065 | 0.873,0.854,0.377, | 0.474,0.351,0.582, | 0,12 |
| 185146 | ENSG00000151466 | SCLT1 | chr4 | - | 128943001 | 128943188 | 128936654 | 128936851 | 128946006 | 128946152 | 185146 | 8,7,17,12,6,8,7,6,6,3,1,8 | 9,10,6,14,5,5,1,2,9,2,2,5 | 19,11,10,9,10,4,3,8,6,15,1, | 4,10,5,2,8,1,1,0,2,4,0,13,8, | 298 | 149 | 0.0219725727229 | 0.98095390991962 | 0.308,0.259,0.586, | 0.704,0.355,0.5,0.6 | -0,195 |
| 43594 | ENSG00000134109 | EDEM1 | chr3 | + | 5211119 | 5211216 | 5208092 | 5208263 | 5213318 | 5213522 | 43594 | 68,54,142,106,41,49,28,47 | 0,2,2,2,0,3,0,0,0,0,0,0 | 51,52,49,40,38,44,21,42,34 | 0,3,2,6,1,1,2,3,0,0,0,2,0,0,2 | 246 | 149 | 0.0220137451528 | 0.98258821712615 | 1.0,0.942,0.977,0.9 | 1.0,0.913,0.937,0.8 | 0,054 |
| 253184 | ENSG00000088992 | TESC | chr12 | - | 117048680 | 117048813 | 117046776 | 117046838 | 117049018 | 117049158 | 253184 | 7,7,12,9,0,3,1,4,4,0,5,2 | 19,35,129,59,96,99,39,69,4 | 2,2,2,5,4,5,7,1,0,2,0,2,3,9,2 | 79,67,53,23,30,20,37,66,18 | 282 | 149 | 0.0220318192977 | 0.983191063500574 | 0.163,0.096,0.047, | 0.013,0.016,0.02,0 | 0,013 |
| 69406 | ENSG00000117335 | CD46 | chr1 | + | 207767606 | 207767651 | 207767012 | 207767195 | 207767778 | 207767823 | 69406 | 8,6,5,1,5,3,7,4,0,0,4,5 | 43,34,61,62,42,62,37,40,26 | 1,12,0,3,0,0,3,3,2,1,0,1,0,2 | 25,41,41,22,15,14,11,26,15 | 194 | 149 | 0.0220681460882 | 0.984608032899953 | 0.125,0.119,0.059, | 0.03,0.184,0.0,0.09 | 0,018 |
| 199467 | ENSG00000111011 | RSRC2 | chr12 | - | 122513728 | 122513874 | 122511110 | 122511188 | 122515104 | 122515227 | 199467 | 18,15,21,9,4,12,4,15,7,11,6 | 42,90,167,99,39,46,37,59,1 | 8,15,13,16,6,3,8,11,7,6,4,4 | 51,130,59,58,98,34,22,46,3 | 295 | 149 | 0.0220981564216 | 0.985742653839874 | 0.178,0.078,0.06,0 | 0.073,0.055,0.1,0.1 | 0,034 |
| 222724 | ENSG00000167483 | NIBAN3 | chr19 | + | 17549664 | 17549755 | 17549454 | 17549527 | 17551785 | 17553645 | 222724 | 15,4,27,25,15,12,10,10,4,5 | 167,98,173,131,155,69,103 | 14,29,15,9,30,3,10,16,13,9 | 162,132,91,59,133,72,49,1 | 240 | 149 | 0.0221094031655 | 0.98603998191333 | 0.053,0.025,0.088, | 0.051,0.12,0.093,0 | -0,015 |
| 106085 | ENSG00000111203 | ITFG2 | chr12 | + | 2830216 | 2830885 | 2824089 | 2824252 | 2836744 | 2836956 | 106085 | 22,7,14,7,15,9,30,39,9,12,3 | 2,4,1,1,0,0,2,1,0,0,0,1 | 8,7,5,10,6,7,8,11,4,4,8,11,1 | 0,2,0,3,2,1,0,0,0,4,0,0,4,7,1 | 298 | 149 | 0.0221359137186 | 0.986880260550217 | 0.846,0.467,0.875, | 1.0,0.636,1.0,0.625 | 0,096 |
| 42340 | ENSG00000131375 | CAPN7 | chr3 | + | 15240753 | 15240853 | 15240472 | 15240617 | 15242177 | 15242253 | 42340 | 32,21,55,25,18,21,15,35,30 | 0,2,4,2,0,0,1,6,2,3,0,6 | 28,44,14,21,9,12,25,38,8,1 | 1,6,5,0,1,0,1,6,4,3,5,5,5,1,1 | 249 | 149 | 0.0221374146751 | 0.986880260550217 | 1.0,0.863,0.892,0.8 | 0.944,0.814,0.626, | 0,09 |
| 157229 | ENSG00000152484 | USP12 | chr13 | - | 27072776 | 27072907 | 27071070 | 27071149 | 27075190 | 27075388 | 157229 | 1,0,3,0,0,0,1,3,0,0,0,0 | 24,18,57,33,26,12,18,18,7, | 0,4,1,2,0,1,0,0,0,2,6,6,2,1 | 15,23,12,7,15,6,16,13,5,6,1 | 280 | 149 | 0.0221519342286 | 0.987323039129386 | 0.022,0.0,0.027,0.0 | 0.0,0.085,0.042,0.1 | -0,032 |
| 232690 | ENSG00000196092 | PAX5 | chr9 | - | 36923354 | 36923477 | 36882003 | 36882105 | 36966548 | 36966724 | 232690 | 79,47,105,27,44,43,53,46,5 | 7,0,1,0,2,0,4,1,0,2,0,3 | 50,56,59,26,17,18,30,44,19 | 0,1,1,0,2,0,0,0,0,0,0,1,3,0,0 | 272 | 149 | 0.0221604196154 | 0.987496744149515 | 0.861,1.0,0.983,1.0 | 1.0,0.968,0.97,1.0, | -0,025 |
| 51945 | ENSG00000125703 | ATG4C | chr1 | + | 62829039 | 62829176 | 62821138 | 62821209 | 62834037 | 62834116 | 51945 | 56,23,119,70,39,22,30,23,2 | 1,0,10,2,0,0,3,3,0,0,0,0 | 60,36,15,12,8,27,15,17,12, | 0,2,1,1,1,1,0,0,0,0,0,1,0,1,2 | 286 | 149 | 0.0221690077975 | 0.987674956999322 | 0.967,1.0,0.861,0.9 | 1.0,0.904,0.887,0.8 | 0 |
| 242184 | ENSG00000182158 | CREB3L2 | chr7 | - | 137882367 | 137882628 | 137874978 | 137880551 | 137884994 | 137885121 | 242184 | 18,21,110,67,14,38,35,66,1 | 3,4,3,0,2,1,2,2,1,0,2,1 | 65,120,25,62,44,37,49,56,2 | 2,4,2,15,2,1,5,0,3,0,8,0,0,1 | 298 | 149 | 0.022177371382 | 0.987688556489837 | 0.75,0.724,0.948,1 | 0.942,0.938,0.862, | -0,032 |
| 203204 | ENSG00000109381 | ELF2 | chr4 | - | 139064579 | 139064715 | 139061936 | 139062057 | 139067683 | 139067770 | 203204 | 3,5,11,2,1,1,3,4,0,5,0,2 | 13,16,98,42,15,15,19,33,9, | 8,2,8,15,3,3,2,0,4,5,2,3,3,3 | 31,63,18,10,16,12,3,12,11, | 285 | 149 | 0.0221784909865 | 0.987688556489837 | 0.108,0.14,0.055,0 | 0.119,0.016,0.189, | -0,053 |
| 40650 | ENSG00000111224 | PARP11 | chr12 | - | 3863835 | 3863957 | 3829889 | 3830018 | 3873211 | 3873355 | 40650 | 8,5,29,16,16,16,5,8,2,5,6,6 | 12,11,13,3,9,11,5,6,5,2,2,6 | 12,7,10,15,9,5,5,9,11,19,11 | 4,6,5,8,6,5,5,5,4,1,5,1,8,7,2 | 271 | 149 | 0.0221843760492 | 0.987746264262591 | 0.268,0.2,0.551,0.7 | 0.623,0.391,0.524, | -0,139 |
| 187723 | ENSG00000018189 | RUFY3 | chr4 | + | 70796662 | 70796746 | 70794794 | 70794894 | 70800140 | 70800205 | 187723 | 9,3,2,0,3,4,3,4,4,0,2,4 | 11,3,7,4,5,9,7,5,6,2,1,9 | 0,3,1,1,3,3,0,2,5,0,0,2,1,0,6 | 7,2,3,3,1,4,11,7,2,4,2,29,6, | 233 | 149 | 0.0221958767017 | 0.988008216019349 | 0.343,0.39,0.154,0 | 0.0,0.49,0.176,0.17 | 0,05 |
| 138936 | ENSG00000118705 | RPN2 | chr20 | + | 37232295 | 37232395 | 37229972 | 37230059 | 37234019 | 37234095 | 138936 | 102,51,75,58,45,66,60,71,1 | 4,0,7,8,6,5,3,11,2,3,1,2 | 62,59,63,52,20,22,35,51,15 | 13,14,6,6,7,1,3,4,0,2,5,8,5, | 249 | 149 | 0.0221994402882 | 0.988008216019349 | 0.938,1.0,0.865,0.8 | 0.741,0.716,0.863, | 0,047 |
| 107919 | ENSG00000102103 | PQBP1 | chrX | + | 48902232 | 48902517 | 48901929 | 48902042 | 48902731 | 48902795 | 107919 | 50,49,89,91,39,37,30,36,96 | 5,5,5,5,3,1,3,3,4,5,0,8 | 33,34,49,27,18,25,12,74,15 | 7,9,2,2,8,2,2,5,1,1,10,27,4, | 298 | 149 | 0.0222040424237 | 0.988008735765014 | 0.833,0.831,0.899, | 0.702,0.654,0.925, | 0,07 |
| 36626 | ENSG00000164597 | COG5 | chr7 | - | 107253047 | 107253076 | 107248395 | 107248499 | 107256731 | 107256794 | 36626 | 6,0,2,0,4,1,0,2,4,2,0,2 | 29,20,36,21,19,25,12,23,18 | 7,7,4,4,2,2,7,0,4,2,0,0,2,0,3 | 24,17,8,24,12,17,9,22,12,2 | 178 | 149 | 0.0222166168385 | 0.988363922853905 | 0.148,0.0,0.044,0.0 | 0.196,0.256,0.295, | -0,068 |
| 115392 | ENSG00000184465 | WDR27 | chr6 | - | 169670878 | 169670957 | 169670568 | 169670693 | 169672254 | 169672396 | 115392 | 0,0,7,2,2,4,0,0,0,0,2,0 | 5,3,15,6,7,16,7,12,4,9,3,6 | 0,1,0,0,1,0,1,2,0,0,0,0,0,2,0 | 4,1,6,7,6,4,7,8,3,5,7,7,6,10 | 228 | 149 | 0.022229987093 | 0.988754360821137 | 0.0,0.0,0.234,0.179 | 0.0,0.395,0.0,0.0,0 | 0,032 |
| 17174 | ENSG00000140521 | POLG | chr15 | - | 89317375 | 89317476 | 89316329 | 89316827 | 89318540 | 89318749 | 17174 | 33,10,60,33,19,26,19,16,11 | 2,0,1,3,0,1,0,2,2,0,0,3 | 25,36,7,8,29,12,9,25,9,27,1 | 1,5,2,6,0,0,2,4,1,2,1,0,2,2,0 | 250 | 149 | 0.0222372162641 | 0.98887154829308 | 0.908,1.0,0.973,0.8 | 0.937,0.811,0.676, | 0,063 |
| 135182 | ENSG00000093000 | NUP50 | chr22 | + | 45178799 | 45178900 | 45171599 | 45171683 | 45181285 | 45181367 | 135182 | 3,6,25,22,9,5,6,12,20,1,2,4 | 2,5,5,4,0,1,1,1,0,0,0,0 | 16,17,7,5,4,10,7,6,5,13,5,9 | 3,5,6,2,5,0,4,0,3,2,0,0,3,3,5 | 250 | 149 | 0.0222549710203 | 0.989327650184225 | 0.472,0.417,0.749, | 0.761,0.67,0.41,0.5 | 0,131 |
| 20560 | ENSG00000171469 | ZNF561 | chr19 | - | 9617620 | 9617758 | 9617044 | 9617171 | 9618090 | 9618179 | 20560 | 10,2,0,0,0,1,0,0,2,0,0,0 | 14,11,35,16,7,9,5,13,20,8,1 | 2,0,1,0,0,5,1,3,2,5,0,1,1,2,2 | 10,9,6,5,8,3,7,12,2,5,7,5,3, | 287 | 149 | 0.0222566660109 | 0.989327650184225 | 0.271,0.086,0.0,0.0 | 0.094,0.0,0.08,0.0, | -0,09 |
| 90622 | ENSG00000113761 | ZNF346 | chr5 | + | 177050702 | 177050936 | 177041777 | 177041870 | 177062057 | 177062151 | 90622 | 5,7,13,22,10,7,15,10,9,8,3, | 0,1,4,1,0,3,0,4,1,0,0,2 | 14,11,6,13,3,7,13,8,6,9,1,1 | 3,3,0,0,0,1,0,1,0,0,2,0,2,1,4 | 298 | 149 | 0.0222651856512 | 0.989501997293708 | 1.0,0.778,0.619,0.9 | 0.7,0.647,1.0,1.0,1 | 0,002 |
| 99970 | ENSG00000058091 | CDK14 | chr7 | + | 90716339 | 90716394 | 90709397 | 90709610 | 90726566 | 90726812 | 99970 | 31,10,13,6,9,11,11,12,4,2,4 | 3,3,5,1,3,6,2,10,0,0,0,3 | 14,9,4,1,4,6,5,9,9,15,20,4,1 | 7,4,0,5,5,4,4,8,1,4,12,5,6,4 | 204 | 149 | 0.0222839145395 | 0.989943771466884 | 0.883,0.709,0.655, | 0.594,0.622,1.0,0.1 | 0,184 |
| 231732 | ENSG00000139436 | GIT2 | chr12 | - | 109959846 | 109959949 | 109952679 | 109953234 | 109961277 | 109961375 | 231732 | 132,113,127,88,39,53,56,4 | 1,9,5,2,2,0,0,0,3,0,0,0 | 71,89,55,34,64,33,32,48,48 | 3,0,0,2,2,1,0,0,0,2,0,4,0,1,0 | 252 | 149 | 0.0222843250868 | 0.989943771466884 | 0.987,0.881,0.938, | 0.933,1.0,1.0,0.91, | -0,001 |
| 132939 | ENSG00000176915 | ANKLE2 | chr12 | - | 132741808 | 132741891 | 132741418 | 132741485 | 132743153 | 132743276 | 132939 | 12,14,9,17,4,13,5,5,2,9,5,8 | 101,70,84,58,32,25,19,18,2 | 6,3,2,5,5,2,3,9,3,4,8,8,12,3 | 53,38,28,30,32,16,14,17,21 | 232 | 149 | 0.022289727767 | 0.98997944635146 | 0.071,0.114,0.064, | 0.068,0.048,0.044, | 0,054 |
| 62407 | ENSG00000158773 | USF1 | chr1 | - | 161041650 | 161041846 | 161041323 | 161041411 | 161042115 | 161042217 | 62407 | 390,364,341,223,180,147,1 | 21,9,23,5,11,13,5,6,17,18,1 | 229,300,186,252,214,95,10 | 26,36,5,18,13,5,7,7,7,22,11 | 298 | 149 | 0.0223041850716 | 0.990417177046738 | 0.903,0.953,0.881, | 0.815,0.806,0.949, | 0,023 |
| 183066 | ENSG00000215908 | CROCCP2 | chr1 | - | 16632606 | 16632769 | 16632274 | 16632371 | 16633094 | 16633346 | 183066 | 4,0,9,13,0,0,14,4,0,0,4,1 | 12,27,35,22,23,16,14,21,13 | 8,3,1,2,1,3,5,8,14,19,3,2,5, | 21,19,11,18,10,4,12,17,8,2 | 298 | 149 | 0.0223375832362 | 0.991695622423796 | 0.143,0.0,0.114,0.1 | 0.16,0.073,0.043,0 | -0,065 |
| 34470 | ENSG00000132466 | ANKRD17 | chr4 | - | 73085248 | 73085446 | 73078641 | 73078890 | 73090666 | 73092300 | 34470 | 75,69,150,116,63,89,33,93 | 2,1,8,3,1,6,0,3,0,0,0,0 | 109,89,72,42,38,63,48,86,3 | 1,0,0,2,0,2,3,8,0,2,4,0,2,5,0 | 298 | 149 | 0.0223505325776 | 0.992048107678395 | 0.949,0.972,0.904, | 0.982,1.0,1.0,0.913 | 0,019 |
| 200836 | ENSG00000161929 | SCIMP | chr17 | - | 5223058 | 5223456 | 5221286 | 5221350 | 5234734 | 5234823 | 200836 | 127,33,100,104,29,29,51,4 | 3,0,3,13,1,4,0,1,9,3,6,5 | 70,45,29,11,24,44,17,35,32 | 8,2,10,2,2,4,1,9,6,3,2,3,8,1 | 298 | 149 | 0.0223563240881 | 0.992048107678395 | 0.955,1.0,0.943,0.8 | 0.814,0.918,0.592, | 0,043 |
| 120947 | ENSG00000110274 | CEP164 | chr11 | + | 117363428 | 117363506 | 117362403 | 117362538 | 117371079 | 117371466 | 120947 | 19,2,17,8,11,7,5,11,8,8,3,4 | 2,3,3,3,5,4,0,5,5,2,3,6 | 17,2,9,12,5,2,1,3,3,9,3,9,13 | 5,3,5,6,8,6,6,5,2,3,1,22,4,4 | 227 | 149 | 0.0223623689378 | 0.992048107678395 | 0.862,0.304,0.788, | 0.691,0.304,0.542, | 0,153 |
| 97988 | ENSG00000145725 | PPIP5K2 | chr5 | + | 103176865 | 103176910 | 103173857 | 103173972 | 103177666 | 103177774 | 97988 | 1,6,25,3,4,9,9,16,2,1,5,6 | 12,12,35,24,26,17,20,10,16 | 16,21,12,9,12,3,10,3,2,9,10 | 18,22,7,2,10,10,9,12,2,20,1 | 194 | 149 | 0.0223639597384 | 0.992048107678395 | 0.06,0.277,0.354,0 | 0.406,0.423,0.568, | -0,127 |
| 205073 | ENSG00000118482 | PHF3 | chr6 | + | 63706733 | 63706876 | 63706028 | 63706224 | 63709150 | 63709214 | 205073 | 78,47,260,132,47,74,55,11 | 3,0,7,1,4,6,6,4,0,0,2,0 | 132,149,59,66,80,45,55,77 | 4,6,2,4,0,2,4,8,3,4,8,1,4,9,3 | 292 | 149 | 0.0223769307348 | 0.992418953014988 | 0.93,1.0,0.95,0.985 | 0.944,0.927,0.938, | 0,029 |
| 249272 | ENSG00000136770 | DNAJC1 | chr10 | - | 21879743 | 21879883 | 21805979 | 21806099 | 21882281 | 21882439 | 249272 | 0,0,1,0,0,3,3,0,0,0,2,1 | 2,3,26,15,20,12,15,9,11,4,1 | 0,2,4,4,3,0,2,5,0,2,0,0,0,0,1 | 22,16,15,24,5,6,9,20,3,13,8 | 289 | 149 | 0.0223881476421 | 0.992711867564241 | 0.0,0.0,0.019,0.0,0 | 0.0,0.061,0.121,0.0 | -0,028 |
| 28431 | ENSG00000185404 | SP140L | chr2 | + | 230400333 | 230400413 | 230400126 | 230400242 | 230401365 | 230401443 | 28431 | 6,12,7,5,9,1,0,9,1,1,2,1 | 1,0,2,4,0,1,1,3,0,2,0,0 | 6,10,9,1,5,2,6,2,5,5,2,3,8,6 | 1,4,0,7,2,1,0,3,0,0,0,1,0,4,3 | 229 | 149 | 0.0224144737661 | 0.993674480566729 | 0.796,1.0,0.695,0.4 | 0.796,0.619,1.0,0.0 | 0,034 |
| 241439 | ENSG00000158122 | PRXL2C | chr9 | - | 96645892 | 96645934 | 96639576 | 96641886 | 96651389 | 96651495 | 241439 | 1,0,5,5,7,0,4,2,5,4,2,0 | 19,5,60,36,20,22,17,16,7,5 | 2,2,3,0,1,0,1,3,2,3,4,2,2,3,2 | 17,44,27,36,20,7,14,20,19, | 191 | 149 | 0.0224371442073 | 0.99447466739732 | 0.039,0.0,0.061,0.0 | 0.084,0.034,0.08,0 | 0,108 |
| 87560 | ENSG00000146463 | ZMYM4 | chr1 | + | 35370371 | 35370627 | 35361618 | 35361789 | 35381258 | 35381433 | 87560 | 14,8,11,10,2,3,3,17,9,0,4,5 | 6,1,6,5,4,1,5,7,2,6,0,3 | 6,5,6,12,4,3,8,9,1,8,4,13,6, | 2,4,3,2,2,1,1,5,1,2,0,0,2,1,1 | 298 | 149 | 0.0224465723337 | 0.994687710511547 | 0.538,0.8,0.478,0.5 | 0.6,0.385,0.5,0.75, | -0,136 |
| 251494 | ENSG00000138134 | STAMBPL1 | chr10 | + | 88908712 | 88908777 | 88905442 | 88905660 | 88910915 | 88911011 | 251494 | 25,6,36,12,4,9,7,13,9,8,4,2 | 2,0,13,10,4,2,2,4,1,0,2,1 | 4,11,11,7,12,5,2,6,5,15,12, | 0,3,2,0,4,0,2,0,2,0,1,0,0,0,2 | 214 | 149 | 0.0224865364402 | 0.996253545607387 | 0.897,1.0,0.658,0.4 | 1.0,0.719,0.793,1.0 | -0,086 |
| 138190 | ENSG00000152454 | ZNF256 | chr19 | - | 57943933 | 57944015 | 57940832 | 57942647 | 57947441 | 57947706 | 138190 | 6,8,8,11,9,11,10,12,0,6,1,9 | 2,0,8,5,5,2,9,4,3,4,3,2 | 11,20,3,3,12,11,4,6,5,2,16, | 5,2,4,2,1,2,2,2,0,0,4,1,0,0,3 | 231 | 149 | 0.0225280965596 | 0.997688517425194 | 0.659,1.0,0.392,0.5 | 0.587,0.866,0.326, | -0,186 |
| 220599 | ENSG00000124782 | RREB1 | chr6 | + | 7107979 | 7108060 | 7107596 | 7107768 | 7176654 | 7176773 | 220599 | 19,10,25,13,15,12,11,12,5, | 1,1,4,3,0,0,0,0,1,0,2,0 | 23,22,7,28,11,4,7,10,14,7,8 | 1,4,2,2,1,0,0,1,1,3,2,1,1,1,0 | 230 | 149 | 0.0225281961924 | 0.997688517425194 | 0.925,0.866,0.802, | 0.937,0.781,0.694, | 0,062 |
| 202807 | ENSG00000112033 | PPARD | chr6 | + | 35352928 | 35353050 | 35347066 | 35347150 | 35410986 | 35411217 | 202807 | 5,3,3,2,1,0,4,5,3,0,1,0 | 31,33,19,12,14,10,8,6,14,1 | 9,5,3,3,3,1,0,0,1,5,0,2,1,3,4 | 13,10,21,20,12,3,11,13,4,1 | 271 | 149 | 0.0225542763095 | 0.998638025996707 | 0.081,0.048,0.08,0 | 0.276,0.216,0.073, | -0,012 |
| 65477 | ENSG00000198624 | CCDC69 | chr5 | - | 151185446 | 151185541 | 151184343 | 151184441 | 151186022 | 151186124 | 65477 | 42,48,122,96,57,33,53,51,3 | 1,0,3,0,0,0,0,2,0,0,0,2 | 87,97,32,39,44,25,30,55,34 | 0,8,0,1,2,1,1,0,1,0,3,3,2,2,1 | 244 | 149 | 0.0225709164542 | 0.999169255317548 | 0.962,1.0,0.961,1.0 | 1.0,0.881,1.0,0.96, | 0,037 |
| 165406 | ENSG00000138767 | CNOT6L | chr4 | - | 77818251 | 77818334 | 77776270 | 77776392 | 77819232 | 77819368 | 165406 | 2,0,0,2,2,1,0,2,0,0,0,0 | 21,19,19,19,21,5,6,12,13,1 | 6,2,1,4,0,1,1,0,0,0,2,1,0,2,0 | 10,19,13,10,12,9,7,10,3,18 | 232 | 149 | 0.0226067095561 | 1 | 0.058,0.0,0.0,0.063 | 0.278,0.063,0.047, | -0,046 |
| 235660 | ENSG00000168300 | PCMTD1 | chr8 | - | 51839446 | 51839664 | 51833517 | 51833689 | 51845660 | 51845763 | 235660 | 2,0,19,11,1,0,0,8,3,0,0,0 | 25,14,162,93,68,53,51,69,3 | 2,1,2,2,2,4,4,4,0,3,3,5,3,1,1 | 120,113,64,37,27,47,41,81 | 298 | 149 | 0.0226096318252 | 1 | 0.038,0.0,0.055,0.0 | 0.008,0.004,0.015, | -0,001 |
| 5031 | ENSG00000205744 | DENND1C | chr19 | - | 6479986 | 6480051 | 6479858 | 6479902 | 6481678 | 6481787 | 5031 | 52,25,17,19,24,29,22,22,29 | 2,0,2,0,0,4,0,1,7,2,0,0 | 15,22,10,12,10,6,5,20,7,11 | 0,0,0,1,0,0,2,4,0,0,0,2,0,0,0 | 214 | 149 | 0.0226231375036 | 1 | 0.948,1.0,0.855,1.0 | 1.0,1.0,1.0,0.893,1 | -0,016 |
| 137655 | ENSG00000104852 | SNRNP70 | chr19 | + | 49102113 | 49102168 | 49101389 | 49101471 | 49104633 | 49104735 | 137655 | 64,28,123,87,81,91,67,110 | 60,48,65,32,32,23,36,20,65 | 136,117,50,22,61,47,55,10 | 25,28,37,33,32,15,11,30,14 | 204 | 149 | 0.0226281916311 | 1 | 0.438,0.299,0.58,0 | 0.799,0.753,0.497, | -0,114 |
| 170693 | ENSG00000278259 | MYO19 | chr17 | - | 36515112 | 36515195 | 36514445 | 36514548 | 36515857 | 36515990 | 170693 | 12,8,21,6,5,4,4,3,5,5,4,4 | 0,2,2,2,0,0,0,0,1,0,0,0 | 10,14,2,10,12,5,10,11,2,4,6 | 0,3,5,0,2,0,0,0,2,0,0,4,0,0,2 | 232 | 149 | 0.0226498648345 | 1 | 1.0,0.72,0.871,0.65 | 1.0,0.75,0.204,1.0, | 0,079 |
| 228753 | ENSG00000160218 | TRAPPC10 | chr21 | + | 44063510 | 44063785 | 44059102 | 44059214 | 44074323 | 44074470 | 228753 | 50,62,90,79,42,43,53,53,57 | 7,0,5,3,3,5,2,3,0,0,5,2 | 63,70,40,66,68,27,43,56,34 | 2,0,2,2,5,1,2,4,2,0,5,0,1,2,0 | 298 | 149 | 0.0226624696336 | 1 | 0.781,1.0,0.9,0.929 | 0.94,1.0,0.909,0.94 | -0,036 |
| 191292 | ENSG00000054118 | THRAP3 | chr1 | + | 36259384 | 36259484 | 36224442 | 36224505 | 36282532 | 36282700 | 191292 | 46,35,225,146,77,80,77,87 | 2,2,2,2,0,4,0,0,4,6,0,1 | 112,181,36,33,62,31,58,73 | 0,0,2,1,1,0,0,3,0,0,0,2,2,1,1 | 249 | 149 | 0.0226717001296 | 1 | 0.932,0.913,0.985, | 1.0,1.0,0.915,0.952 | -0,028 |
| 208624 | ENSG00000152520 | PAN3 | chr13 | + | 28191250 | 28191307 | 28177864 | 28177935 | 28197184 | 28197346 | 208624 | 7,7,2,0,3,6,6,1,1,2,1,0 | 21,22,53,22,25,32,32,57,18 | 1,7,7,9,11,3,3,1,3,2,2,5,1,4 | 29,32,35,37,27,22,30,31,23 | 206 | 149 | 0.0226864197255 | 1 | 0.194,0.187,0.027, | 0.024,0.137,0.126, | -0,04 |
| 210212 | ENSG00000169499 | PLEKHA2 | chr8 | + | 38952221 | 38952312 | 38950849 | 38950990 | 38952635 | 38952704 | 210212 | 77,23,324,195,71,60,59,12 | 24,2,26,16,9,13,3,0,7,4,6,5 | 195,89,58,35,29,50,50,65,4 | 13,14,14,19,5,4,7,7,2,2,26, | 240 | 149 | 0.0226865944388 | 1 | 0.666,0.877,0.886, | 0.903,0.798,0.72,0 | 0,061 |
| 193964 | ENSG00000109184 | DCUN1D4 | chr4 | + | 51874720 | 51874966 | 51874270 | 51874385 | 51877762 | 51877972 | 193964 | 0,3,4,5,3,3,1,0,2,1,0,0 | 16,10,47,39,11,12,11,14,2, | 0,1,0,1,0,0,0,0,1,0,2,3,0,2,1 | 45,17,14,9,17,16,6,24,5,30 | 298 | 149 | 0.0227042853809 | 1 | 0.0,0.13,0.041,0.06 | 0.0,0.029,0.0,0.053 | 0,055 |
| 124901 | ENSG00000080546 | SESN1 | chr6 | - | 109000490 | 109000673 | 108994461 | 108994609 | 109001287 | 109001488 | 124901 | 16,7,79,40,22,46,48,440,34 | 1,1,0,3,1,3,0,18,2,0,2,0 | 43,40,42,30,12,5,16,29,13, | 3,0,5,4,2,0,0,8,0,1,30,0,1,0 | 298 | 149 | 0.0227162491797 | 1 | 0.889,0.778,1.0,0.8 | 0.878,1.0,0.808,0.7 | 0,024 |
| 234682 | ENSG00000121390 | PSPC1 | chr13 | - | 19759528 | 19759693 | 19759346 | 19759418 | 19772241 | 19772543 | 234682 | 14,2,10,7,5,10,4,26,2,2,2,1 | 64,35,130,85,66,67,64,91,4 | 5,8,10,24,12,2,5,12,9,11,20 | 50,77,45,47,62,34,42,50,39 | 298 | 149 | 0.0227393845041 | 1 | 0.099,0.028,0.037, | 0.048,0.049,0.1,0.2 | -0,023 |
| 137459 | ENSG00000167604 | NFKBID | chr19 | - | 35889619 | 35889704 | 35888240 | 35888612 | 35889889 | 35890054 | 137459 | 10,8,7,9,1,5,4,1,4,1,7,0 | 41,117,34,9,29,32,57,28,34 | 2,1,5,11,2,2,0,0,0,0,8,4,0,0 | 28,29,3,47,17,2,15,32,7,5,3 | 234 | 149 | 0.0227445325041 | 1 | 0.134,0.042,0.116, | 0.044,0.021,0.515, | 0,008 |

| | | | | | | | | | | | | | | | | | | | | | | |
|---|---|---|---|---|---|---|---|---|---|---|---|---|---|---|---|---|---|---|---|---|---|---|
| 189561 | ENSG00000143398 | PIP5K1A | chr1 | + | 151242127 | 151242567 | 151239954 | 151240039 | 151246919 | 151246965 | 189561 | 16,15,35,33,16,23,18,29,9, | 1,3,1,7,0,0,2,0,1,2,0,2 | 36,49,18,12,17,10,8,19,11, | 0,0,1,0,0,0,0,1,0,2,4,0,2,2,0 | 298 | 149 | 0.0227665060098 | | 1 | 0.889,0.714,0.946, | 1.0,1.0,0.9,1.0,1.0, | -0,06 |
| 211946 | ENSG00000164985 | PSIP1 | chr9 | - | 15470999 | 15471176 | 15469937 | 15469993 | 15472631 | 15472750 | 211946 | 15,4,9,4,4,3,6,12,4,1,2,8 | 51,20,59,33,59,32,29,36,30 | 3,5,4,5,6,1,7,10,1,11,15,15 | 35,20,18,1,13,25,13,27,6,2 | 298 | 149 | 0.0227800215901 | | 1 | 0.128,0.091,0.071, | 0.041,0.111,0.1,0. | -0,076 |
| 27091 | ENSG00000075568 | TMEM131 | chr2 | - | 97772296 | 97772424 | 97766477 | 97766602 | 97792385 | 97792984 | 27091 | 59,52,47,17,22,20,13,23,19 | 4,2,3,2,0,2,0,0,2,2,0,0 | 30,32,34,41,14,26,15,7,15, | 0,0,1,0,3,2,0,0,0,0,0,0,0,2 | 277 | 149 | 0.022804942923 | | 1 | 0.888,0.933,0.894, | 1.0,1.0,0.948,1.0,0 | -0,075 |
| 64310 | ENSG00000135596 | MICAL1 | chr6 | - | 109447680 | 109447722 | 109447356 | 109447440 | 109448202 | 109448393 | 64310 | 40,9,21,19,17,21,29,23,10, | 3,0,3,1,1,7,2,2,2,2,0,0 | 23,15,3,8,13,11,17,16,19,1 | 6,0,3,3,1,0,1,1,0,2,0,4,7,0,1 | 191 | 149 | 0.0228107392224 | | 1 | 0.912,1.0,0.845,0. | 0.749,1.0,0.438,0. | 0,034 |
| 135310 | ENSG00000198089 | SFI1 | chr22 | + | 31580250 | 31580364 | 31578381 | 31578452 | 31583874 | 31583877 | 135310 | 23,10,26,7,4,12,12,14,10,5 | 2,3,7,4,4,5,3,7,5,5,5,8 | 10,11,6,3,10,10,4,15,16,30 | 2,1,4,3,0,0,5,4,4,6,1,2,5,1, | 263 | 149 | 0.022820951576 | | 1 | 0.867,0.654,0.678, | 0.739,0.862,0.459, | -0,156 |
| 197965 | ENSG00000153922 | CHD1 | chr5 | - | 98876397 | 98876833 | 98875071 | 98875113 | 98879551 | 98879728 | 197965 | 38,30,41,17,26,20,18,39,17 | 20,3,9,12,6,9,16,13,12,2,2, | 34,41,20,29,30,12,14,21,11 | 13,13,13,34,18,3,9,10,18,4 | 298 | 149 | 0.0228212840794 | | 1 | 0.487,0.833,0.695, | 0.567,0.612,0.435, | 0,125 |
| 253110 | ENSG00000124164 | VAPB | chr20 | + | 58439135 | 58439308 | 58438944 | 58439025 | 58440906 | 58441083 | 253110 | 4,4,11,3,2,0,3,3,6,0,1,1 | 22,21,32,22,9,8,9,14,32,12 | 10,7,2,2,1,1,0,2,2,3,2,4,6,1 | 20,23,10,9,11,13,8,8,7,18,1 | 298 | 149 | 0.02285949069 | | 1 | 0.083,0.087,0.147, | 0.2,0.132,0.091,0. | -0,055 |
| 235197 | ENSG00000075539 | FRYL | chr4 | - | 48687005 | 48687160 | 48684672 | 48684795 | 48710518 | 48710698 | 235197 | 4,5,4,5,0,0,0,0,2,2,1,1 | 16,16,62,37,24,33,21,27,4, | 0,3,4,2,1,0,0,0,0,0,0,0,1,2 | 41,36,6,15,19,12,13,21,30, | 298 | 149 | 0.0228654720586 | | 1 | 0.111,0.135,0.031, | 0.0,0.04,0.25,0.06 | 0,039 |
| 52260 | ENSG00000123104 | ITPR2 | chr12 | - | 26486816 | 26486933 | 26486103 | 26486360 | 26487067 | 26487251 | 52260 | 30,21,41,15,10,0,9,30,0,4,5 | 66,57,98,44,21,60,24,58,47 | 11,15,7,30,14,23,3,11,22,1 | 45,33,29,39,32,32,10,34,23 | 266 | 149 | 0.0228661873811 | | 1 | 0.203,0.171,0.19,0 | 0.12,0.203,0.119,0 | -0,057 |
| 39476 | ENSG00000162736 | NCSTN | chr1 | + | 160355862 | 160355958 | 160354117 | 160354290 | 160356259 | 160356347 | 39476 | 59,42,48,25,24,22,30,36,42 | 2,0,2,3,6,1,0,2,3,0,0,5 | 40,36,31,7,21,15,24,36,13, | 1,3,1,10,4,1,1,1,3,1,2,8,2,1 | 245 | 149 | 0.0228762715201 | | 1 | 0.947,1.0,0.936,0.8 | 0.961,0.879,0.95,0 | 0,089 |
| 178448 | ENSG00000143514 | TP53BP2 | chr1 | - | 223809106 | 223809164 | 223806910 | 223806947 | 223810430 | 223810513 | 178448 | 3,0,7,3,1,3,1,6,0,0,3,0 | 8,22,27,16,13,7,13,26,1,6,3 | 4,3,3,17,1,1,1,9,2,4,6,2,5,4 | 20,23,11,8,21,11,7,18,7,12 | 207 | 149 | 0.0229006199721 | | 1 | 0.213,0.0,0.157,0. | 0.126,0.086,0.164, | -0,05 |
| 207467 | ENSG00000182973 | CNOT10 | chr3 | + | 32715991 | 32716134 | 32713226 | 32713369 | 32716224 | 32716311 | 207467 | 4,3,28,7,5,2,6,4,2,0,3,6 | 35,21,68,30,36,13,15,23,18 | 12,7,1,3,6,2,2,6,7,10,9,5,3, | 23,15,14,2,9,10,5,18,11,32 | 292 | 149 | 0.0229132199029 | | 1 | 0.055,0.068,0.174, | 0.21,0.192,0.035,0 | -0,082 |
| 73271 | ENSG00000165689 | ENTR1 | chr9 | - | 136408998 | 136409067 | 136407825 | 136407938 | 136410327 | 136410593 | 73271 | 4,5,11,1,10,3,5,7,7,2,9,8 | 9,10,14,8,13,1,8,0,19,12,2, | 7,1,2,0,0,4,0,0,2,4,4,2,6,2,6 | 5,4,6,4,5,4,2,10,4,7,2,10,4, | 218 | 149 | 0.0229228885774 | | 1 | 0.233,0.255,0.349, | 0.489,0.146,0.186, | 0,157 |
| 217586 | ENSG00000133872 | SARAF | chr8 | - | 30073889 | 30074051 | 30069873 | 30070059 | 30082846 | 30083124 | 217586 | 6,6,9,3,4,6,6,5,0,2,0,9 | 22,10,58,11,17,12,31,33,7, | 4,1,1,6,2,4,0,2,0,8,3,5,3,5,4 | 21,23,14,13,23,10,10,22,7, | 298 | 149 | 0.0229263487189 | | 1 | 0.12,0.231,0.072,0 | 0.087,0.021,0.034, | 0,032 |
| 217875 | ENSG00000155827 | RNF20 | chr9 | + | 101544752 | 101544885 | 101540792 | 101540975 | 101546819 | 101546966 | 217875 | 33,22,37,25,9,10,15,11,29, | 1,2,6,0,0,1,0,0,1,0,4,3 | 29,19,30,29,18,19,11,16,7, | 1,2,2,4,4,2,1,2,0,6,4,10,1,4 | 282 | 149 | 0.0229371731416 | | 1 | 0.946,0.853,0.765, | 0.939,0.834,0.888, | 0,101 |
| 215127 | ENSG00000115355 | CCDC88A | chr2 | - | 55299831 | 55299919 | 55296257 | 55296523 | 55301205 | 55301277 | 215127 | 73,48,178,119,23,14,28,27 | 5,4,10,12,0,0,3,7,0,0,0,1 | 120,122,35,2,12,17,19,52,2 | 13,6,7,0,0,2,0,12,2,1,4,0,9, | 237 | 149 | 0.0229414406802 | | 1 | 0.902,0.883,0.918, | 0.853,0.927,0.759, | 0,06 |
| 195264 | ENSG00000134744 | TUT4 | chr1 | - | 52458335 | 52458371 | 52446589 | 52446667 | 52461133 | 52461223 | 195264 | 10,6,15,19,3,10,3,5,1,2,2,8 | 24,36,36,21,12,16,18,29,10 | 13,10,4,7,11,8,10,13,10,15 | 10,48,20,32,21,18,20,12,11 | 185 | 149 | 0.0229415720505 | | 1 | 0.251,0.118,0.251, | 0.511,0.144,0.139, | -0,111 |
| 222521 | ENSG00000166295 | ANAPC16 | chr10 | + | 72223942 | 72224056 | 72220053 | 72220379 | 72230365 | 72230440 | 222521 | 235,114,296,221,93,93,98, | 2,6,11,8,3,2,5,1,12,1,1,3 | 182,119,131,85,72,54,55,1 | 5,1,1,2,0,1,1,3,1,6,6,3,7,2,3 | 263 | 149 | 0.0229544686264 | | 1 | 0.985,0.915,0.938, | 0.954,0.985,0.987, | -0,016 |
| 123288 | ENSG00000130935 | NOL11 | chr17 | + | 67736665 | 67736754 | 67735899 | 67736023 | 67737070 | 67737145 | 123288 | 52,72,136,92,35,48,31,62,4 | 13,2,11,10,11,6,7,9,1,2,5,3 | 49,107,70,39,62,30,48,40,1 | 15,11,5,12,10,5,8,13,5,7,15 | 238 | 149 | 0.0229812848321 | | 1 | 0.715,0.958,0.886, | 0.672,0.859,0.898, | 0,056 |
| 86084 | ENSG00000114120 | SLC25A36 | chr3 | + | 140966999 | 140967036 | 140963126 | 140963227 | 140970926 | 140970993 | 86084 | 5,8,9,9,0,2,1,8,3,4,2,1 | 16,12,27,28,13,15,19,7,15, | 1,3,3,2,1,1,1,0,3,6,1,5,4,2,5 | 16,32,15,45,16,7,17,19,22, | 186 | 149 | 0.0229981828672 | | 1 | 0.2,0.348,0.211,0. | 0.048,0.07,0.138,0 | 0,039 |
| 214711 | ENSG00000101452 | DHX35 | chr20 | + | 38970638 | 38970765 | 38969080 | 38969214 | 38972558 | 38972651 | 214711 | 0,3,2,0,2,2,1,13,1,0,0,1 | 12,18,12,4,4,6,1,6,3,1,4,8 | 4,4,4,15,3,2,4,0,5,1,3,2,3,2 | 9,4,6,5,8,6,2,5,9,7,5,9,2,16 | 276 | 149 | 0.0230268859105 | | 1 | 0.0,0.083,0.083,0. | 0.194,0.351,0.265, | -0,095 |
| 161109 | ENSG00000198612 | COPS8 | chr2 | + | 237087517 | 237087563 | 237087126 | 237087197 | 237088604 | 237088653 | 161109 | 0,0,2,0,1,2,2,4,0,0,0,1 | 11,6,32,48,26,13,19,7,16,1 | 0,1,5,0,0,0,1,7,1,0,1,0,1,3,1 | 19,16,21,5,19,19,10,18,4,2 | 195 | 149 | 0.0230330912237 | | 1 | 0.0,0.0,0.046,0.0,0 | 0.0,0.046,0.154,0. | -0,027 |
| 242854 | ENSG00000216937 | CCDC7 | chr10 | + | 32481670 | 32481782 | 32473966 | 32474023 | 32543299 | 32543386 | 242854 | 3,0,18,6,2,4,3,6,1,2,4,4 | 1,7,10,5,3,8,9,1,2,4,2,4 | 9,3,4,6,7,7,2,2,8,3,7,4,2,5,2 | 2,2,6,1,6,1,1,0,3,2,8,3,0,6,6 | 261 | 149 | 0.0230785766178 | | 1 | 0.631,0.0,0.507,0.4 | 0.72,0.461,0.276,0 | -0,22 |
| 217327 | ENSG00000140750 | ARHGAP17 | chr16 | - | 24959670 | 24959771 | 24954602 | 24954730 | 24959910 | 24959981 | 217327 | 11,2,69,37,8,16,6,7,6,5,4,6 | 0,0,0,0,0,2,0,0,0,0,1,0 | 27,24,8,3,7,6,6,12,19,28,14 | 1,3,0,7,2,0,0,0,1,3,2,0,1,2,0 | 250 | 149 | 0.0230859418402 | | 1 | 1.0,1.0,1.0,1.0,1.0, | 0.941,0.827,1.0,0. | 0,086 |
| 134620 | ENSG00000099991 | CABIN1 | chr22 | + | 24054978 | 24055159 | 24050824 | 24050974 | 24056191 | 24056304 | 134620 | 31,27,47,33,21,37,15,24,12 | 3,9,1,3,0,3,0,6,0,1,0,0 | 17,27,19,36,23,6,17,17,11, | 2,2,0,3,4,0,2,0,2,0,0,0,0,4,0 | 298 | 149 | 0.0230918603453 | | 1 | 0.838,0.6,0.959,0.8 | 0.81,0.871,1.0,0.8 | -0,013 |
| 250064 | ENSG00000270099 | | chr10 | + | 95861250 | 95861361 | 95860547 | 95860582 | 95864723 | 95864861 | 250064 | 0,0,3,6,1,0,0,2,0,2,0,0 | 69,35,53,69,42,18,16,21,25 | 0,10,3,2,0,0,4,1,5,2,4,2,4,2 | 46,81,58,30,36,38,35,35,51 | 260 | 149 | 0.0230925029291 | | 1 | 0.0,0.0,0.031,0.04 | 0.0,0.066,0.029,0. | -0,01 |
| 189668 | ENSG00000074603 | DPP8 | chr15 | - | 65454262 | 65454342 | 65451959 | 65452102 | 65456224 | 65456371 | 189668 | 9,8,16,16,5,8,8,9,3,5,0,9 | 4,3,9,9,5,1,5,2,3,0,2,4 | 16,19,6,4,9,3,4,4,6,9,4,10,1 | 3,6,6,0,2,0,0,0,1,5,4,1,0,6,2 | 229 | 149 | 0.0230991246128 | | 1 | 0.594,0.634,0.536, | 0.776,0.673,0.394, | -0,191 |
| 208259 | ENSG00000081320 | STK17B | chr2 | - | 196147237 | 196147415 | 196145910 | 196146055 | 196156438 | 196156651 | 208259 | 17,3,1,0,0,2,1,7,0,0,6,1 | 145,71,124,98,57,31,53,42 | 0,0,0,2,0,1,3,0,1,0,0,0,2,0,0 | 110,142,68,56,110,57,49,4 | 298 | 149 | 0.0231200240534 | | 1 | 0.055,0.021,0.004, | 0.0,0.0,0.0,0.018,0 | 0,019 |
| 56503 | ENSG00000132581 | SDF2 | chr17 | - | 28651485 | 28651577 | 28648924 | 28649276 | 28655286 | 28655483 | 56503 | 6,2,18,19,8,6,5,3,5,2,3,4 | 13,15,38,44,14,19,24,8,17, | 11,12,11,3,8,4,9,14,3,6,2,1 | 13,31,6,0,12,10,8,15,7,14,1 | 241 | 149 | 0.0231421189092 | | 1 | 0.222,0.076,0.227, | 0.343,0.193,0.531, | -0,143 |
| 252761 | ENSG00000082898 | XPO1 | chr2 | - | 61525041 | 61525333 | 61522610 | 61522683 | 61526419 | 61526521 | 252761 | 10,5,13,12,1,14,8,9,2,2,0,9 | 99,29,98,58,70,49,36,41,42 | 24,15,8,4,16,8,5,12,5,5,4,9 | 106,44,55,35,31,20,31,54,1 | 298 | 149 | 0.0231446965797 | | 1 | 0.048,0.079,0.062, | 0.102,0.146,0.068, | -0,029 |
| 139623 | ENSG00000132670 | PTPRA | chr20 | + | 3034900 | 3035107 | 3027706 | 3027841 | 3035584 | 3035710 | 139623 | 7,3,4,1,0,2,3,2,7,0,0,4 | 62,33,70,64,20,27,15,26,37 | 5,0,10,11,3,2,4,2,3,3,3,9,3, | 31,38,22,22,33,22,18,34,16 | 298 | 149 | 0.0231456634776 | | 1 | 0.053,0.043,0.028, | 0.075,0.0,0.185,0. | -0,028 |
| 148040 | ENSG00000090857 | PDPR | chr16 | + | 70153390 | 70153699 | 70148463 | 70148553 | 70156474 | 70162537 | 148040 | 17,6,30,34,23,26,9,10,2,9,7 | 7,4,22,11,9,6,6,12,2,0,0,0 | 23,29,8,5,14,15,15,23,17,2 | 11,10,0,3,2,3,0,3,0,0,5,8,1, | 298 | 149 | 0.0231677837085 | | 1 | 0.548,0.429,0.405, | 0.511,0.592,1.0,0. | -0,147 |
| 16037 | ENSG00000090857 | PDPR | chr16 | + | 70114820 | 70114930 | 70114331 | 70114440 | 70120460 | 70120719 | 16037 | 2,2,10,2,0,0,3,1,0,0,1,2 | 2,0,5,13,2,4,0,0,1,1,1,5 | 6,2,2,1,6,2,0,1,2,6,2,2,5,6,2 | 4,5,1,0,3,5,5,2,2,5,0,4,8,1,0 | 259 | 149 | 0.0231692264488 | | 1 | 0.365,1.0,0.535,0.0 | 0.463,0.187,0.535, | -0,093 |
| 233026 | ENSG00000005889 | ZFX | chrX | + | 24208893 | 24209040 | 24208217 | 24208370 | 24210192 | 24211986 | 233026 | 42,48,160,66,39,36,26,49,2 | 4,3,8,3,0,2,4,0,5,2,0,0 | 81,96,41,36,63,45,14,50,32 | 9,4,2,0,0,2,1,0,2,1,0,2,1,0,1 | 296 | 149 | 0.0231755072239 | | 1 | 0.841,0.89,0.91,0.9 | 0.819,0.924,0.912, | -0,042 |
| 2144 | ENSG00000198089 | SFI1 | chr22 | + | 31580271 | 31580364 | 31578381 | 31578452 | 31583874 | 31583877 | 2144 | 34,22,54,33,27,27,24,37,21 | 2,3,7,4,4,5,3,7,5,5,5,8 | 21,31,12,16,19,26,14,34,26 | 2,1,4,3,0,0,5,4,4,6,1,2,5,1,9 | 242 | 149 | 0.0231917126646 | | 1 | 0.913,0.819,0.826, | 0.866,0.95,0.649,0 | -0,073 |
| 219328 | ENSG00000132436 | FIGNL1 | chr7 | - | 50447776 | 50448055 | 50447182 | 50447297 | 50448180 | 50448289 | 219328 | 1,0,3,7,7,2,3,5,0,0,2,0 | 13,4,17,9,5,3,4,4,7,1,1,7 | 2,1,0,0,0,0,1,2,1,2,2,0,1,4,5 | 6,13,6,10,4,4,0,4,0,7,0,15,5 | 298 | 149 | 0.0232096113721 | | 1 | 0.037,0.0,0.081,0. | 0.143,0.037,0.0,0. | -0,12 |
| 154938 | ENSG00000213983 | AP1G2 | chr14 | - | 23566600 | 23566686 | 23566338 | 23566419 | 23567110 | 23567315 | 154938 | 127,57,91,96,72,60,72,46,8 | 13,13,20,2,9,13,8,11,8,8,3, | 69,63,55,19,42,29,30,47,41 | 8,11,13,7,10,3,8,10,4,11,7, | 235 | 149 | 0.0232198038062 | | 1 | 0.861,0.735,0.743, | 0.845,0.784,0.728, | 0,059 |
| 108052 | ENSG00000124222 | STX16 | chr20 | + | 58669301 | 58669453 | 58667986 | 58668127 | 58670511 | 58670603 | 108052 | 34,15,28,14,4,9,15,15,9,11 | 22,5,9,7,12,13,10,8,3,5,6,3 | 9,27,10,36,12,11,16,22,13, | 18,25,11,25,12,13,12,12,23 | 298 | 149 | 0.0232207177445 | | 1 | 0.436,0.6,0.609,0.5 | 0.2,0.351,0.312,0. | 0,128 |
| 208087 | ENSG00000088179 | PTPN4 | chr2 | + | 119915324 | 119915369 | 119915178 | 119915242 | 119920068 | 119920241 | 208087 | 3,6,1,0,0,2,0,1,0,0,0,0 | 8,6,24,14,3,6,5,13,6,4,3,3 | 4,0,1,2,3,2,1,0,0,0,2,3,3,0,1 | 11,5,0,2,1,6,8,10,2,4,5,19,2 | 194 | 149 | 0.0232227860791 | | 1 | 0.224,0.434,0.031, | 0.218,0.0,1.0,0.43 | -0,146 |
| 171777 | ENSG00000112149 | CD83 | chr6 | + | 14131377 | 14131748 | 14117949 | 14118065 | 14133648 | 14133755 | 171777 | 366,354,191,99,48,74,58,4 | 25,13,19,12,2,4,9,6,1,2,12, | 90,160,85,26,58,15,71,72,1 | 9,16,16,5,4,0,4,18,8,0,5,72 | 298 | 149 | 0.023231348133 | | 1 | 0.88,0.932,0.834,0 | 0.833,0.833,0.726, | 0,054 |
| 237290 | ENSG00000120910 | PPP3CC | chr8 | + | 22535226 | 22535369 | 22532920 | 22533018 | 22539468 | 22539498 | 237290 | 12,0,4,1,0,3,3,3,0,2,0,2 | 91,54,107,61,44,26,57,62,8 | 4,1,1,16,2,3,1,8,9,1,4,0,9,1 | 68,61,67,41,40,31,30,66,28 | 292 | 149 | 0.0232625582927 | | 1 | 0.063,0.0,0.019,0.0 | 0.029,0.008,0.008, | -0,017 |
| 119474 | ENSG00000183597 | TANGO2 | chr22 | + | 20055942 | 20056013 | 20052464 | 20052584 | 20061529 | 20061576 | 119474 | 11,15,17,22,6,7,2,7,8,11,4, | 2,0,6,1,0,0,2,0,0,1,0,2 | 2,6,1,5,6,2,1,1,2,9,9,20,6,6 | 2,0,0,1,1,0,1,0,0,1,1,4,7,2,3 | 220 | 149 | 0.0232892696036 | | 1 | 0.788,1.0,0.657,0.9 | 0.404,1.0,1.0,0.77 | 0,059 |
| 4920 | ENSG00000012061 | ERCC1 | chr19 | - | 45419097 | 45419197 | 45416490 | 45416897 | 45420323 | 45420427 | 4920 | 100,48,201,103,94,68,76,7 | 1,2,5,9,0,5,0,2,0,0,0,0 | 115,64,76,40,48,47,62,136 | 0,1,0,1,0,0,0,0,0,0,0,4,3,0,2 | 249 | 149 | 0.0233125706741 | | 1 | 0.984,0.935,0.96,0 | 1.0,0.975,1.0,0.96, | -0,02 |
| 209802 | ENSG00000198589 | LRBA | chr4 | - | 150867670 | 150867860 | 150851884 | 150852943 | 150868181 | 150868305 | 209802 | 103,53,100,58,57,76,53,92 | 1,1,7,4,0,6,1,8,0,1,0,1 | 86,70,33,67,47,46,28,40,56 | 2,6,1,3,9,1,2,2,10,0,5,7,3,2 | 298 | 149 | 0.023334028905 | | 1 | 0.981,0.964,0.877, | 0.956,0.854,0.943, | 0,052 |
| 255750 | ENSG00000118454 | ANKRD13C | chr1 | - | 70293503 | 70293614 | 70292387 | 70292549 | 70296129 | 70296261 | 255750 | 8,4,12,7,4,3,0,3,2,1,1,3 | 42,24,85,68,22,24,19,35,27 | 6,8,3,2,1,0,5,0,4,1,2,2,2,0,8 | 55,88,27,39,32,19,12,33,19 | 260 | 149 | 0.0233353538385 | | 1 | 0.098,0.087,0.075, | 0.059,0.05,0.06,0. | 0,012 |
| 101703 | ENSG00000141380 | SS18 | chr18 | - | 26035830 | 26035923 | 26035004 | 26035127 | 26038554 | 26038659 | 101703 | 51,17,73,58,31,26,34,30,39 | 15,14,21,24,31,11,28,17,34 | 49,39,33,12,29,21,12,25,19 | 9,20,6,12,12,10,12,9,0,7,17 | 242 | 149 | 0.0233544678798 | | 1 | 0.677,0.428,0.682, | 0.77,0.546,0.772,0 | -0,129 |
| 87332 | ENSG00000115355 | CCDC88A | chr2 | - | 55299827 | 55299919 | 55296257 | 55296523 | 55301205 | 55301277 | 87332 | 76,49,182,125,24,14,28,27 | 5,4,10,12,0,0,3,7,0,0,0,1 | 124,129,34,2,13,18,20,53,2 | 13,6,7,0,0,2,0,12,2,1,4,0,9, | 241 | 149 | 0.0233546892394 | | 1 | 0.904,0.883,0.918, | 0.855,0.93,0.75,1. | 0,057 |
| 211941 | ENSG00000164985 | PSIP1 | chr9 | - | 15470994 | 15471176 | 15469937 | 15469993 | 15472631 | 15472750 | 211941 | 15,4,9,4,4,3,6,12,4,1,4,9 | 51,20,59,33,59,32,29,36,30 | 3,6,4,7,6,1,7,10,1,11,13,16 | 35,20,18,1,13,25,13,27,6,2 | 298 | 149 | 0.0233615038137 | | 1 | 0.128,0.091,0.071, | 0.041,0.13,0.1,0.7 | -0,075 |
| 37152 | ENSG00000203875 | SNHG5 | chr6 | - | 85677794 | 85677899 | 85677423 | 85677492 | 85677953 | 85678032 | 37152 | 19,8,75,42,3,1,1,3,0,0,5,5 | 1,5,25,43,14,8,6,7,2,3,10,1 | 16,74,25,13,13,10,3,8,24,3 | 11,9,2,4,15,10,11,15,4,12,2 | 254 | 149 | 0.0233695327198 | | 1 | 0.918,0.484,0.638, | 0.46,0.828,0.88,0. | -0,173 |
| 84449 | ENSG00000115053 | NCL | chr2 | - | 231456009 | 231456136 | 231454750 | 231455624 | 231456630 | 231456764 | 84449 | 601,419,647,456,256,343,2 | 39,8,29,24,6,13,16,27,8,9,2 | 380,569,314,135,253,227,2 | 15,16,17,24,12,13,8,17,4,1 | 276 | 149 | 0.0233740677157 | | 1 | 0.893,0.966,0.923, | 0.932,0.95,0.909,0 | 0,032 |
| 116066 | ENSG00000165209 | STRBP | chr9 | - | 123168188 | 123168258 | 123160976 | 123161068 | 123169901 | 123170046 | 116066 | 4,1,20,14,14,12,11,9,3,0,2, | 67,24,174,123,123,78,60,9 | 14,8,3,3,1,1,8,10,3,1,5,7,10 | 179,135,73,44,42,36,51,46 | 219 | 149 | 0.0234016767282 | | 1 | 0.039,0.028,0.073, | 0.051,0.039,0.027, | 0,01 |
| 211233 | ENSG00000197622 | CDC42SE1 | chr1 | - | 151056655 | 151056823 | 151055582 | 151055993 | 151059478 | 151059773 | 211233 | 20,7,33,14,13,25,9,16,6,5,5 | 197,165,295,254,162,112,1 | 28,14,25,20,18,13,5,23,6,1 | 274,239,156,130,165,98,11 | 298 | 149 | 0.0234016816146 | | 1 | 0.048,0.021,0.053, | 0.049,0.028,0.074, | -0,009 |
| 29289 | ENSG00000006459 | KDM7A | chr7 | - | 140126636 | 140126823 | 140113490 | 140113582 | 140127441 | 140127583 | 29289 | 15,2,23,23,4,8,9,6,5,6,3,11 | 5,0,0,0,0,0,0,0,0,0,0,0 | 11,14,6,4,1,18,5,3,6,22,4,3 | 1,3,2,3,2,1,0,1,0,0,1,0,2,0,0 | 298 | 149 | 0.0234019295677 | | 1 | 0.6,1.0,1.0,1.0,1.0, | 0.846,0.7,0.6,0.4,0 | 0,197 |
| 151159 | ENSG00000092439 | TRPM7 | chr15 | - | 50614122 | 50614220 | 50613706 | 50613841 | 50619744 | 50619798 | 151159 | 35,25,58,40,32,32,20,34,26 | 3,4,3,2,0,2,0,0,2,1,0,1 | 34,22,27,13,28,25,17,37,18 | 2,0,0,1,3,0,0,0,4,0,1,1,0,0,2 | 247 | 149 | 0.0234147631243 | | 1 | 0.876,0.79,0.921,0 | 0.911,1.0,1.0,0.88 | -0,019 |
| 222180 | ENSG00000147251 | DOCK11 | chrX | + | 118676825 | 118676945 | 118676590 | 118676737 | 118680481 | 118680692 | 222180 | 0,0,14,8,1,0,0,0,0,0,8,3 | 69,54,175,89,68,44,33,75,2 | 0,2,7,24,0,0,0,0,0,2,3,16,0, | 77,71,39,26,58,45,30,28,49 | 269 | 149 | 0.0234684205809 | | 1 | 0.0,0.0,0.042,0.04 | 0.0,0.015,0.09,0.3 | -0,007 |
| 95819 | ENSG00000167483 | NIBAN3 | chr19 | + | 17530754 | 17530816 | 17527290 | 17527395 | 17532262 | 17532388 | 95819 | 19,22,66,20,34,13,22,18,14 | 45,36,41,14,40,27,53,39,53 | 27,48,16,3,28,18,14,16,16, | 26,38,25,17,12,6,18,37,12, | 211 | 149 | 0.0235365018911 | | 1 | 0.23,0.301,0.532,0 | 0.423,0.471,0.311, | -0,097 |
| 239657 | ENSG00000108669 | CYTH1 | chr17 | - | 78696006 | 78696059 | 78692416 | 78692493 | 78698268 | 78698380 | 239657 | 0,0,3,0,0,2,3,0,5,2,0,0 | 4,1,2,3,2,0,1,2,2,1,2,1 | 2,3,2,1,0,0,1,0,0,0,4,0,1,0,0 | 1,6,2,16,1,5,3,2,5,2,7,2,3,1 | 202 | 149 | 0.0235577145565 | | 1 | 0.0,0.0,0.525,0.0,0 | 0.596,0.269,0.425, | 0,162 |
| 97240 | ENSG00000172115 | CYCS | chr7 | - | 25123950 | 25124127 | 25118655 | 25123849 | 25125199 | 25125260 | 97240 | 197,132,240,180,54,97,95, | 29,1,5,7,0,2,6,0,2,0,0,0 | 139,192,147,186,150,76,79 | 2,3,4,14,2,0,0,4,5,0,5,4,7,0 | 298 | 149 | 0.023574768539 | | 1 | 0.773,0.985,0.96,0 | 0.972,0.97,0.948,0 | -0,005 |
| 69766 | ENSG00000223745 | CCDC18-AS1 | chr1 | - | 93269682 | 93269758 | 93264637 | 93264772 | 93324634 | 93324729 | 69766 | 11,4,5,4,2,3,4,1,6,0,1,2 | 7,16,27,4,5,9,3,7,2,2,8,7 | 0,1,2,4,6,0,0,0,2,5,2,6,1,0,4 | 3,9,4,0,6,8,5,12,14,8,15,7,2 | 225 | 149 | 0.023575072133 | | 1 | 0.51,0.142,0.109,0 | 0.0,0.069,0.249,1. | 0,062 |
| 109483 | ENSG00000147475 | ERLIN2 | chr8 | + | 37741771 | 37741818 | 37740364 | 37740446 | 37744354 | 37744416 | 109483 | 10,5,17,35,8,11,9,1,4,8,3,3 | 0,2,7,1,0,1,0,3,2,4,0,0 | 8,9,10,1,2,1,4,15,2,8,6,15,8 | 3,1,0,4,2,0,0,3,0,0,1,0,3,0,0 | 196 | 149 | 0.023597276101 | | 1 | 1.0,0.655,0.649,0.9 | 0.67,0.872,1.0,0.1 | -0,031 |
| 170694 | ENSG00000278259 | MYO19 | chr17 | - | 36515112 | 36515287 | 36514445 | 36514548 | 36515857 | 36515990 | 170694 | 12,8,22,6,8,4,4,3,5,6,4,4 | 0,2,2,2,0,0,0,0,1,0,0,0 | 10,16,3,11,13,5,10,11,2,4,5 | 0,3,5,0,2,0,0,0,2,0,0,4,0,0,2 | 298 | 149 | 0.0236112983758 | | 1 | 1.0,0.667,0.846,0. | 1.0,0.727,0.231,1. | 0,074 |
| 120318 | ENSG00000005801 | ZNF195 | chr11 | - | 3361742 | 3361889 | 3358993 | 3360565 | 3370974 | 3371070 | 120318 | 3,7,8,6,15,4,3,7,9,2,1,6 | 1,3,10,9,1,1,0,0,0,2,1,5 | 16,7,5,2,1,5,5,2,9,8,2,7,10, | 14,10,2,2,7,2,2,1,2,4,6,4,3, | 296 | 149 | 0.0236197734292 | | 1 | 0.602,0.54,0.287,0 | 0.365,0.261,0.557, | 0,184 |
| 227201 | ENSG00000157191 | NECAP2 | chr1 | + | 16441165 | 16441298 | 16440735 | 16440853 | 16443631 | 16443732 | 227201 | 7,15,6,5,1,3,8,5,4,6,2,4 | 66,124,79,40,25,41,36,29,5 | 0,7,1,7,0,0,2,4,0,4,5,3,2,0,9 | 55,63,55,17,30,26,23,58,26 | 282 | 149 | 0.0236245304715 | | 1 | 0.053,0.06,0.039,0 | 0.0,0.055,0.01,0.1 | 0,014 |
| 252904 | ENSG00000135720 | DYNC1LI2 | chr16 | - | 66724669 | 66724748 | 66723605 | 66723822 | 66725827 | 66725944 | 252904 | 1,0,1,2,0,2,0,0,0,2,0,0 | 22,23,51,25,14,18,23,20,5, | 0,1,0,0,2,3,3,1,2,0,8,1,4,1,0 | 21,50,25,20,42,13,15,20,11 | 228 | 149 | 0.0236299705025 | | 1 | 0.029,0.0,0.013,0.0 | 0.0,0.013,0.0,0.0,0 | -0,023 |
| 35013 | ENSG00000131507 | NDFIP1 | chr5 | + | 142135729 | 142135817 | 142132211 | 142132342 | 142140562 | 142140629 | 35013 | 55,40,157,124,56,53,31,40 | 0,0,3,5,0,4,1,1,0,1,2,0 | 55,78,70,31,37,25,23,56,25 | 1,9,9,0,6,0,0,1,4,0,0,0,0,0,0 | 237 | 149 | 0.0236875541129 | | 1 | 1.0,1.0,0.971,0.94, | 0.972,0.845,0.83,1 | 0,013 |
| 219614 | ENSG00000083097 | DOP1A | chr6 | + | 83096711 | 83096823 | 83067670 | 83067779 | 83096924 | 83097115 | 219614 | 8,11,23,9,1,8,9,9,1,0,9,7 | 13,6,11,9,3,4,0,7,4,4,0,3 | 12,25,5,10,14,10,4,6,10,10 | 6,8,2,2,2,2,2,3,4,6,2,1,1,3,4 | 261 | 149 | 0.0237218016949 | | 1 | 0.26,0.511,0.544,0 | 0.533,0.641,0.588, | -0,166 |
| 34899 | ENSG00000026297 | RNASET2 | chr6 | - | 166955466 | 166955627 | 166952487 | 166952548 | 166956096 | 166956182 | 34899 | 5,9,11,3,16,5,8,2,9,14,0,8 | 62,37,49,29,22,11,22,13,26 | 9,4,3,8,3,3,9,3,2,5,13,6,11, | 22,46,47,27,30,25,13,16,14 | 298 | 149 | 0.0237254529392 | | 1 | 0.039,0.108,0.101, | 0.17,0.042,0.031,0 | 0,026 |
| 113727 | ENSG00000075856 | SART3 | chr12 | - | 108537487 | 108537595 | 108536707 | 108536785 | 108538064 | 108538203 | 113727 | 39,53,96,75,44,44,35,39,36 | 10,9,13,8,0,3,6,6,0,0,0,0 | 54,57,23,14,68,26,30,44,11 | 4,12,5,5,7,2,7,3,2,3,11,12,5 | 257 | 149 | 0.0237354348025 | | 1 | 0.693,0.773,0.811, | 0.887,0.734,0.727, | 0,096 |
| 211345 | ENSG00000116750 | UCHL5 | chr1 | - | 193044048 | 193044329 | 193029531 | 193029657 | 193049745 | 193049851 | 211345 | 0,2,4,5,0,0,1,1,0,0,2,2 | 7,2,16,20,7,4,13,5,3,1,0,2 | 6,2,1,2,1,0,1,0,2,4,0,4,2,2,2 | 8,15,10,5,1,4,2,9,9,10,3,7,1 | 298 | 149 | 0.0237396801459 | | 1 | 0.0,0.333,0.111,0. | 0.273,0.062,0.048, | 0,02 |
| 181319 | ENSG00000162739 | SLAMF6 | chr1 | - | 160496060 | 160496246 | 160491124 | 160491388 | 160523143 | 160523262 | 181319 | 79,25,208,224,157,128,83, | 2,0,16,8,6,4,4,7,3,3,6,11 | 258,86,68,52,42,75,40,80,2 | 7,3,0,0,3,2,1,8,2,4,7,2,9,4,3 | 298 | 149 | 0.0237585207205 | | 1 | 0.952,1.0,0.867,0.9 | 0.949,0.935,1.0,1. | -0,019 |
| 196675 | ENSG00000104946 | TBC1D17 | chr19 | + | 49879451 | 49879504 | 49878497 | 49878572 | 49880278 | 49880402 | 196675 | 2,0,8,2,2,4,2,1,0,0,2,0 | 10,5,17,21,32,14,20,15,6,6 | 1,1,0,2,1,0,2,2,0,3,2,1,1,2,0 | 7,16,3,6,7,7,4,18,3,21,7,9,1 | 202 | 149 | 0.0237585850049 | | 1 | 0.129,0.0,0.258,0. | 0.095,0.044,0.0,0. | 0,005 |
| 150106 | ENSG00000137871 | ZNF280D | chr15 | - | 56732680 | 56733024 | 56707262 | 56707306 | 56733457 | 56733577 | 150106 | 0,0,0,2,0,2,0,0,0,0,0,0 | 29,11,56,28,5,16,13,12,9,4 | 0,1,0,2,1,2,0,2,0,1,0,8,4,2,3 | 26,14,5,7,9,6,4,11,12,21,10 | 298 | 149 | 0.0237882012349 | | 1 | 0.0,0.0,0.0,0.034,0 | 0.0,0.034,0.0,0.12 | -0,048 |
| 183213 | ENSG00000198952 | SMG5 | chr1 | - | 156272319 | 156272449 | 156268289 | 156268415 | 156273360 | 156273450 | 183213 | 16,15,40,38,6,11,7,7,5,8,3, | 2,1,0,0,0,0,3,1,0,0,0,1 | 29,22,17,12,17,16,7,10,6,2 | 0,2,0,2,0,1,3,2,0,1,5,1,0,1,2 | 279 | 149 | 0.023801696651 | | 1 | 0.81,0.889,1.0,1.0, | 1.0,0.855,1.0,0.76 | 0,042 |
| 125326 | ENSG00000163683 | SMIM14 | chr4 | - | 39611965 | 39612110 | 39605070 | 39605180 | 39638738 | 39638902 | 125326 | 0,1,11,5,0,1,2,0,0,0,1,2 | 10,1,47,36,6,4,1,10,6,6,6,1 | 2,5,1,2,1,1,0,0,1,0,2,8,3,3,5 | 14,9,9,1,2,4,2,8,6,19,2,11,1 | 294 | 149 | 0.0238069010888 | | 1 | 0.0,0.336,0.106,0. | 0.068,0.22,0.053,0 | -0,055 |
| 205276 | ENSG00000165487 | MICU2 | chr13 | - | 21495475 | 21495842 | 21495160 | 21495318 | 21496051 | 21496652 | 205276 | 2,2,1,0,4,1,0,0,0,1,0,0 | 28,11,110,48,34,32,17,38,3 | 2,3,1,7,3,1,1,3,0,2,3,2,0,0,0 | 27,37,32,16,36,10,9,28,10, | 298 | 149 | 0.0238118123004 | | 1 | 0.034,0.083,0.005, | 0.036,0.039,0.015, | -0,025 |
| 222199 | ENSG00000152601 | MBNL1 | chr3 | + | 152300880 | 152300973 | 152300197 | 152300367 | 152301918 | 152302449 | 222199 | 12,0,33,12,11,12,6,9,5,2,0, | 39,20,96,41,47,36,33,36,12 | 13,20,4,1,5,5,2,5,3,3,6,4,12 | 85,83,23,30,25,19,7,25,43, | 242 | 149 | 0.0238140599716 | | 1 | 0.159,0.0,0.175,0. | 0.086,0.129,0.097, | 0,045 |
| 20927 | ENSG00000213928 | IRF9 | chr14 | + | 24163880 | 24163959 | 24163377 | 24163508 | 24164062 | 24164134 | 20927 | 38,30,39,38,15,19,17,30,18 | 13,5,50,34,10,28,33,22,11, | 27,37,16,13,12,17,10,31,10 | 40,47,13,36,22,24,10,33,17 | 228 | 149 | 0.0238402979374 | | 1 | 0.656,0.797,0.338, | 0.306,0.34,0.446,0 | 0,119 |
| 82103 | ENSG00000134262 | AP4B1 | chr1 | - | 113897839 | 113897943 | 113895756 | 113896038 | 113898717 | 113898763 | 82103 | 22,6,38,28,14,12,13,9,21,8 | 0,0,6,4,0,2,2,1,0,2,0,0 | 43,43,6,17,20,7,1,10,13,9,7 | 2,5,1,2,4,2,2,4,0,1,3,13,2,0 | 253 | 149 | 0.0238709760795 | | 1 | 1.0,1.0,0.789,0.80 | 0.927,0.835,0.779, | 0,17 |
| 70805 | ENSG00000159658 | EFCAB14 | chr1 | - | 46688352 | 46688544 | 46684490 | 46686870 | 46689586 | 46689691 | 70805 | 131,97,231,171,86,68,98,7 | 24,36,62,62,57,34,22,32,34 | 116,60,66,48,35,39,38,105 | 50,59,47,40,44,21,20,24,23 | 298 | 149 | 0.0238907577783 | | 1 | 0.732,0.574,0.651, | 0.537,0.337,0.413, | 0,083 |
| 126116 | ENSG00000197535 | MYO5A | chr15 | - | 52376787 | 52376945 | 52376346 | 52376558 | 52379624 | 52379733 | 126116 | 3,13,12,8,6,7,8,9,5,1,0,7 | 27,19,33,41,14,20,28,18,25 | 8,10,2,8,5,1,3,7,6,3,5,14,3, | 39,35,18,31,32,18,14,23,22 | 298 | 149 | 0.0238945716325 | | 1 | 0.053,0.255,0.154, | 0.093,0.125,0.053, | 0,042 |

| | | | | | | | | | | | | | | | | | | | | | | | |
|---|---|---|---|---|---|---|---|---|---|---|---|---|---|---|---|---|---|---|---|---|---|---|---|
| 76698 | ENSG00000159023 | EPB41 | chr1 | + | 29097806 | 29097935 | 29064956 | 29065158 | 29109335 | 29109346 | 76698 | 433,329,433,412,426,399, | 4,14,8,6,2,5,4,12,24,12,2,3 | 360,326,353,358,342,301, | 5,8,7,10,2,2,1,5,4,8,5,11,2, | 278 | 149 | 0.0239061944427 | | 1 | 0.983,0.926,0.967, | 0.975,0.956,0.964, | -0,008 |
| 192469 | ENSG00000157873 | TNFRSF14 | chr1 | + | 2561493 | 2561674 | 2560553 | 2560714 | 2561776 | 2561815 | 192469 | 23,18,15,6,5,13,8,14,7,10, | 2,1,10,6,3,1,2,5,3,1,2,3 | 10,9,8,14,5,5,10,25,4,7,6,2 | 7,6,0,7,2,1,1,4,2,6,6,12,7,3 | 298 | 149 | 0.023907997143 | | 1 | 0.852,0.9,0.429,0. | 0.417,0.429,1.0,0. | 0,129 |
| 199362 | ENSG00000168421 | RHOH | chr4 | + | 40209462 | 40209837 | 40197060 | 40197300 | 40242713 | 40243187 | 199362 | 26,7,17,4,20,11,13,13,2,0, | 294,139,484,273,154,157, | 17,6,10,45,2,1,6,18,4,3,7,1 | 355,405,185,196,173,69,2 | 298 | 149 | 0.023909954349 | | 1 | 0.042,0.025,0.017, | 0.023,0.007,0.026, | 0,009 |
| 247337 | ENSG00000151498 | ACAD8 | chr11 | + | 134263120 | 134263555 | 134262519 | 134262622 | 134264907 | 134265850 | 247337 | 1,4,5,2,11,3,2,3,1,0,5,2 | 26,7,27,14,14,7,18,4,15,4, | 3,1,0,0,0,1,2,4,4,2,1,1,4,3, | 10,15,4,8,5,10,3,20,6,17,1 | 298 | 149 | 0.0239264781101 | | 1 | 0.019,0.222,0.085, | 0.13,0.032,0.0,0.0, | 0,029 |
| 9338 | ENSG00000163597 | SNHG16 | chr17 | + | 76558944 | 76559043 | 76557734 | 76557857 | 76561287 | 76561398 | 9338 | 11,18,20,22,6,15,8,13,5,17 | 23,11,63,78,15,23,21,25,2, | 14,20,14,3,8,7,2,11,4,10,9, | 50,28,29,7,41,23,10,38,12, | 248 | 149 | 0.0239331839744 | | 1 | 0.223,0.496,0.16,0 | 0.144,0.3,0.225,0. | 0,123 |
| 47021 | ENSG00000196204 | RNF216P1 | chr7 | + | 4989062 | 4989177 | 4984909 | 4985075 | 4996609 | 4998165 | 47021 | 14,5,30,11,5,10,9,17,4,2,5, | 1,0,3,0,0,2,0,2,5,0,1,1 | 21,20,5,13,12,11,5,10,10,1 | 5,2,4,6,1,1,0,2,0,0,2,0,0,2, | 264 | 149 | 0.0239368857932 | | 1 | 0.888,1.0,0.849,1. | 0.703,0.849,0.414, | 0,005 |
| 240868 | ENSG00000109689 | STIM2 | chr4 | + | 27021478 | 27021612 | 27017710 | 27017984 | 27022518 | 27022661 | 240868 | 8,12,74,34,18,25,23,26,6,0 | 28,11,149,73,17,35,41,44, | 30,24,18,15,14,5,6,18,11,1 | 95,72,49,19,23,21,25,54,2 | 283 | 149 | 0.023945711191 | | 1 | 0.131,0.365,0.207, | 0.143,0.149,0.162, | 0,044 |
| 73066 | ENSG00000131165 | CHMP1A | chr16 | - | 89653903 | 89653923 | 89651568 | 89651646 | 89657581 | 89657607 | 73066 | 16,13,43,43,22,17,21,18,2 | 0,1,3,4,2,0,0,1,1,2,2,2 | 19,25,11,12,9,8,14,24,13,2 | 1,0,0,1,1,1,0,4,0,0,0,0,0,1, | 169 | 149 | 0.0239485736644 | | 1 | 1.0,0.92,0.927,0.9 | 0.944,1.0,1.0,0.91 | -0,03 |
| 83938 | ENSG00000130038 | CRACR2A | chr12 | - | 3644594 | 3644640 | 3641731 | 3641838 | 3648541 | 3648613 | 83938 | 32,11,9,9,12,12,1,10,18,9, | 7,1,2,0,0,1,0,2,0,2,1,0 | 6,17,14,18,9,8,8,9,10,17,4, | 0,0,1,0,0,0,0,0,0,0,0,0,0,0, | 195 | 149 | 0.0239548235805 | | 1 | 0.777,0.894,0.775, | 1.0,1.0,0.915,1.0,1 | -0,11 |
| 158399 | ENSG00000198399 | ITSN2 | chr2 | - | 24264861 | 24265092 | 24261560 | 24261742 | 24270670 | 24270768 | 158399 | 33,38,33,14,11,10,12,12,1 | 43,47,99,71,54,25,22,89,1 | 23,29,10,17,7,11,4,27,5,14 | 72,117,50,53,37,38,46,60, | 298 | 149 | 0.0239699670589 | | 1 | 0.277,0.288,0.143, | 0.138,0.11,0.091,0 | 0,059 |
| 134826 | ENSG00000128159 | TUBGCP6 | chr22 | - | 50225128 | 50225182 | 50224510 | 50224592 | 50225793 | 50225943 | 134826 | 0,0,6,5,1,5,1,4,1,1,0,0 | 84,48,67,34,42,56,60,36,3 | 1,1,1,0,1,1,1,3,1,2,0,1,0,0, | 51,29,26,30,26,23,24,57,3 | 203 | 149 | 0.0239746760181 | | 1 | 0.0,0.0,0.062,0.09 | 0.014,0.025,0.027, | 0,012 |
| 119006 | ENSG00000149428 | HYOU1 | chr11 | - | 119055749 | 119055843 | 119055492 | 119055571 | 119056069 | 119056167 | 119006 | 28,44,46,52,38,39,41,36,3 | 2,6,4,4,0,0,5,3,0,1,0,4 | 28,46,20,20,15,9,8,25,5,28 | 0,1,0,2,6,2,0,0,0,0,2,4,4,0, | 243 | 149 | 0.0240165837376 | | 1 | 0.896,0.818,0.876, | 1.0,0.966,1.0,0.86, | -0,011 |
| 235181 | ENSG00000075539 | FRYL | chr4 | - | 48608268 | 48608336 | 48606437 | 48606606 | 48608986 | 48609067 | 235181 | 1,3,7,8,0,4,0,6,1,0,2,3 | 25,47,108,37,28,33,33,46, | 2,1,2,0,2,5,2,0,2,1,0,5,3,2, | 45,43,25,31,41,23,15,22,3 | 217 | 149 | 0.0240237838583 | | 1 | 0.027,0.042,0.043, | 0.03,0.016,0.052,0 | 0,011 |
| 94118 | ENSG00000112182 | BACH2 | chr6 | - | 90271848 | 90271941 | 90206568 | 90206681 | 90296479 | 90296742 | 94118 | 74,33,115,27,22,41,19,61, | 21,13,115,37,16,38,21,31, | 57,27,70,74,21,21,50,27,8, | 33,12,30,53,5,16,32,22,7,1 | 242 | 149 | 0.0240277408785 | | 1 | 0.685,0.61,0.381,0 | 0.515,0.581,0.59,0 | -0,097 |
| 38805 | ENSG00000174839 | DENND6A | chr3 | - | 57630420 | 57630523 | 57628810 | 57628885 | 57630714 | 57630824 | 38805 | 82,44,101,52,23,27,35,56, | 2,5,11,0,1,1,1,9,1,2,0,4 | 66,78,63,50,28,20,31,38,3 | 0,6,1,0,7,4,7,13,7,2,11,0,5, | 252 | 149 | 0.0240401743172 | | 1 | 0.96,0.839,0.844,1 | 1.0,0.885,0.974,1. | 0,067 |
| 220239 | ENSG00000124788 | ATXN1 | chr6 | - | 16432677 | 16433096 | 16326393 | 16328470 | 16485971 | 16486109 | 220239 | 4,1,10,5,4,3,4,17,0,2,2,8 | 66,36,75,34,40,36,40,33,3 | 2,6,1,0,6,0,3,4,0,0,7,2,0,0, | 20,43,17,8,5,10,21,40,2,1, | 298 | 149 | 0.0240548106693 | | 1 | 0.029,0.014,0.062, | 0.048,0.065,0.029, | -0,001 |
| 208446 | ENSG00000138468 | SENP7 | chr3 | - | 101417592 | 101417746 | 101372007 | 101372126 | 101458954 | 101459052 | 208446 | 9,8,24,12,8,14,1,10,4,3,4,4 | 3,1,8,8,9,4,1,12,3,0,1,4 | 7,9,1,5,7,6,4,13,8,9,11,4,7, | 5,3,5,1,0,4,1,0,0,4,1,0,0,1,1 | 298 | 149 | 0.0240580262127 | | 1 | 0.6,0.8,0.6,0.429,0 | 0.412,0.6,0.091,0. | -0,126 |
| 106669 | ENSG00000005889 | ZFX | chrX | + | 24208899 | 24209040 | 24208217 | 24208370 | 24210192 | 24211986 | 106669 | 73,70,247,112,62,50,50,69 | 4,3,8,3,0,2,4,0,5,2,0,0 | 136,142,63,46,102,69,29,6 | 9,4,2,0,0,2,1,0,2,1,0,2,1,0,1 | 290 | 149 | 0.0240633556683 | | 1 | 0.904,0.923,0.941, | 0.886,0.948,0.942, | -0,034 |
| 183272 | ENSG00000198408 | OGA | chr10 | - | 101795364 | 101795480 | 101793912 | 101793998 | 101797979 | 101798154 | 183272 | 27,16,42,12,5,14,10,17,8,2 | 81,75,183,75,45,28,41,43, | 29,27,24,50,19,5,17,23,8,9 | 113,106,69,36,83,32,48,45 | 265 | 149 | 0.0240989174019 | | 1 | 0.158,0.107,0.114, | 0.126,0.125,0.164, | -0,033 |
| 30535 | ENSG00000133835 | HSD17B4 | chr5 | + | 119531265 | 119531404 | 119529893 | 119529980 | 119536422 | 119536550 | 30535 | 131,61,263,183,133,65,71, | 30,7,15,14,26,8,18,19,20,9 | 116,35,108,23,36,70,25,54 | 25,17,10,6,9,11,7,15,24,14 | 288 | 149 | 0.0241319271905 | | 1 | 0.693,0.818,0.901, | 0.706,0.516,0.848, | 0,069 |
| 23867 | ENSG00000173575 | CHD2 | chr15 | + | 92991475 | 92991517 | 92985497 | 92985673 | 92992858 | 92992998 | 23867 | 39,38,56,34,43,25,18,24,2 | 0,2,0,1,0,0,0,0,0,0,0,3 | 45,104,17,26,44,19,12,28, | 4,2,2,12,3,0,0,0,1,2,1,0,1,1 | 191 | 149 | 0.0241447531868 | | 1 | 1.0,0.937,1.0,0.96 | 0.898,0.976,0.869, | 0,037 |
| 242523 | ENSG00000102699 | PARP4 | chr13 | - | 24482146 | 24482288 | 24478092 | 24478276 | 24484652 | 24484748 | 242523 | 14,11,33,16,1,5,12,27,3,9, | 55,22,124,118,75,53,54,72 | 24,36,15,20,26,12,8,13,9,7 | 80,66,33,9,31,45,29,67,39, | 291 | 149 | 0.0241556844667 | | 1 | 0.115,0.204,0.12,0 | 0.133,0.218,0.189, | -0,049 |
| 138019 | ENSG00000127445 | PIN1 | chr19 | + | 9836657 | 9836875 | 9835368 | 9835402 | 9838435 | 9838648 | 138019 | 3,0,7,4,0,0,0,4,0,1,2,1 | 14,10,34,11,17,7,6,9,11,12 | 0,5,2,6,0,0,3,7,0,0,2,5,2,0, | 20,14,12,3,3,1,3,13,4,3,11, | 298 | 149 | 0.0241715005585 | | 1 | 0.097,0.0,0.093,0. | 0.0,0.152,0.077,0. | 0,023 |
| 13144 | ENSG00000103495 | MAZ | chr16 | + | 29809539 | 29809764 | 29808569 | 29808741 | 29810076 | 29811164 | 13144 | 19,9,10,11,12,5,4,13,7,18, | 5,3,11,5,5,5,2,1,1,2,6,4 | 6,6,5,3,5,2,5,10,3,5,4,64,1 | 10,12,3,0,3,1,0,5,2,2,8,41, | 298 | 149 | 0.0241759162798 | | 1 | 0.655,0.6,0.312,0. | 0.231,0.2,0.455,1. | 0,115 |
| 167797 | ENSG00000254870 | ATP6V1G2-DD | chr6 | - | 31534485 | 31534854 | 31532779 | 31532911 | 31535366 | 31535485 | 167797 | 3,4,3,8,5,3,0,0,1,4,2,1 | 38,12,94,77,42,21,32,29,4 | 0,4,3,18,2,4,11,3,2,2,12,1, | 79,46,30,5,20,35,27,41,12, | 298 | 149 | 0.0241826953799 | | 1 | 0.038,0.143,0.016, | 0.0,0.042,0.048,0. | -0,044 |
| 74171 | ENSG00000064687 | ABCA7 | chr19 | + | 1057914 | 1058059 | 1056906 | 1057429 | 1058145 | 1058269 | 74171 | 59,41,54,57,80,41,67,71,5 | 5,0,0,2,1,2,1,1,1,1,1,1 | 72,78,28,34,38,42,21,40,4 | 0,2,1,1,0,0,0,1,0,0,2,1,0,1, | 294 | 149 | 0.0242474223973 | | 1 | 0.857,1.0,1.0,0.93 | 1.0,0.952,0.934,0. | -0,026 |
| 241684 | ENSG00000175893 | ZDHHC21 | chr9 | - | 14677297 | 14677371 | 14674186 | 14674385 | 14680032 | 14680162 | 241684 | 24,27,39,30,3,6,12,17,15,1 | 12,10,23,19,5,5,8,9,2,2,1,7 | 11,8,7,4,5,10,5,26,3,14,27, | 11,21,6,0,9,5,2,13,16,19,1 | 223 | 149 | 0.0242538310059 | | 1 | 0.572,0.643,0.531, | 0.401,0.203,0.438, | 0,093 |
| 78173 | ENSG00000075292 | ZNF638 | chr2 | + | 71348752 | 71350271 | 71331781 | 71331875 | 71355718 | 71355780 | 78173 | 44,8,90,67,58,36,42,44,31, | 15,17,8,10,15,9,5,10,5,4,0, | 66,40,41,37,23,25,22,49,2 | 8,15,7,5,5,1,2,1,5,8,8,6,3,4 | 298 | 149 | 0.0242886014554 | | 1 | 0.595,0.19,0.849,0 | 0.805,0.571,0.745, | -0,107 |
| 98922 | ENSG00000175324 | LSM1 | chr8 | - | 38170942 | 38171012 | 38169801 | 38169917 | 38171964 | 38172033 | 98922 | 0,0,1,3,0,0,0,0,2,1,0,0 | 25,29,58,42,17,11,18,15,2 | 2,0,4,2,2,0,0,0,2,5,0,0,2,2, | 32,50,20,12,35,20,8,13,11, | 219 | 149 | 0.0243091294419 | | 1 | 0.0,0.0,0.012,0.04 | 0.041,0.0,0.12,0.1 | -0,043 |
| 169932 | ENSG00000112282 | MED23 | chr6 | - | 131615906 | 131616122 | 131610045 | 131610246 | 131618406 | 131618519 | 169932 | 25,21,23,17,6,7,3,13,10,10 | 2,1,9,7,6,4,3,7,3,1,0,0 | 12,10,11,3,11,7,2,2,7,11,1 | 7,7,2,26,11,6,0,6,2,3,8,0,5, | 298 | 149 | 0.0243149522903 | | 1 | 0.862,0.913,0.561, | 0.462,0.417,0.733, | 0,126 |
| 42933 | ENSG00000164258 | NDUFS4 | chr5 | + | 53604723 | 53604895 | 53603451 | 53603530 | 53646232 | 53646405 | 42933 | 12,1,3,1,3,4,1,4,2,3,0,2 | 48,34,80,71,20,33,23,31,3 | 6,4,2,9,4,2,2,0,2,6,3,9,0,6, | 30,46,16,22,24,23,1,35,10, | 298 | 149 | 0.0243256087704 | | 1 | 0.111,0.014,0.018, | 0.091,0.042,0.059, | -0,046 |
| 193744 | ENSG00000077254 | USP33 | chr1 | - | 77720278 | 77720586 | 77718595 | 77718641 | 77721171 | 77721738 | 193744 | 10,9,41,22,5,16,9,14,1,4,6, | 8,8,34,24,25,20,14,15,6,12 | 22,35,7,34,15,10,17,9,15,4 | 18,20,11,8,8,7,9,19,4,12,2 | 298 | 149 | 0.0243350871345 | | 1 | 0.385,0.36,0.376,0 | 0.379,0.467,0.241, | -0,114 |
| 143033 | ENSG00000132388 | UBE2G1 | chr17 | - | 4296716 | 4296839 | 4289229 | 4289408 | 4307020 | 4307123 | 143033 | 39,22,134,84,33,16,29,23, | 4,4,8,3,0,2,0,3,0,0,1,7 | 46,81,44,34,59,51,21,16,2 | 10,10,2,4,14,6,5,2,2,3,0,3, | 272 | 149 | 0.0243392044046 | | 1 | 0.842,0.751,0.902, | 0.716,0.816,0.923, | 0,046 |
| 13492 | ENSG00000187555 | USP7 | chr16 | - | 8921156 | 8921295 | 8920358 | 8920447 | 8923214 | 8923413 | 13492 | 367,298,640,279,159,173, | 5,0,1,0,7,1,5,0,0,0,5,0 | 311,407,219,339,305,118, | 6,1,7,6,1,1,0,0,5,2,3,11,4,1 | 288 | 149 | 0.0243413131025 | | 1 | 0.974,1.0,0.997,1. | 0.964,0.995,0.942, | 0,001 |
| 21975 | ENSG00000172590 | MRPL52 | chr14 | + | 22830963 | 22831037 | 22830186 | 22830254 | 22833417 | 22833468 | 21975 | 14,4,20,5,4,7,4,10,12,13,0, | 54,26,85,66,42,32,52,11,4 | 2,4,3,4,2,3,2,6,3,9,5,8,4,6, | 60,50,23,10,22,18,14,28,17 | 223 | 149 | 0.024360033551 | | 1 | 0.148,0.093,0.136, | 0.022,0.051,0.08,0 | 0,028 |
| 80549 | ENSG00000162923 | WDR26 | chr1 | - | 224418259 | 224418416 | 224411426 | 224411565 | 224419517 | 224419615 | 80549 | 139,146,188,120,72,90,61, | 2,4,0,3,0,1,1,0,0,0,0,0 | 132,195,101,223,142,65,7 | 1,2,1,3,2,0,1,2,3,2,0,0,4,2, | 298 | 149 | 0.0243621408451 | | 1 | 0.972,0.948,1.0,0. | 0.985,0.98,0.981,0 | 0,019 |
| 237378 | ENSG00000121022 | COPS5 | chr8 | - | 67060496 | 67060545 | 67059210 | 67059445 | 67061845 | 67062024 | 237378 | 0,2,0,2,1,1,0,1,0,0,2,0 | 5,4,11,3,7,6,5,4,6,3,4,1 | 1,1,4,1,3,2,0,0,2,1,1,1,1,0, | 1,0,3,4,2,0,2,2,0,1,4,5,2,4, | 198 | 149 | 0.0243759873864 | | 1 | 0.0,0.273,0.0,0.33 | 0.429,1.0,0.501,0. | -0,247 |
| 70773 | ENSG00000068120 | COASY | chr17 | + | 42564982 | 42565047 | 42563306 | 42564577 | 42565226 | 42565311 | 70773 | 4,6,24,4,11,7,3,4,9,10,5,11 | 0,0,4,7,2,1,2,1,1,1,1,1 | 16,5,1,5,5,6,5,8,2,4,8,5,13, | 4,10,3,1,1,2,0,7,2,0,1,8,3,0 | 214 | 149 | 0.0243818268231 | | 1 | 1.0,1.0,0.807,0.28 | 0.736,0.258,0.188, | 0,168 |
| 106471 | ENSG00000083750 | RRAGB | chrX | + | 55727306 | 55727390 | 55722185 | 55722285 | 55729293 | 55729360 | 106471 | 0,0,4,3,2,2,4,1,0,0,2,2 | 45,31,32,31,13,14,18,13,2 | 0,5,5,3,2,1,1,5,1,4,5,1,0,0, | 13,15,16,7,10,6,4,17,8,30,1 | 233 | 149 | 0.024409908954 | | 1 | 0.0,0.0,0.074,0.05 | 0.0,0.176,0.167,0. | -0,043 |
| 152999 | ENSG00000071537 | SEL1L | chr14 | - | 81484224 | 81484334 | 81479277 | 81479740 | 81485671 | 81485746 | 152999 | 24,17,36,27,13,15,9,25,9,9 | 4,2,2,1,2,0,1,2,0,2,0,0 | 23,26,15,15,20,6,21,21,3,2 | 1,1,0,4,0,2,0,1,0,0,0,0,0,0, | 259 | 149 | 0.024423698024 | | 1 | 0.775,0.83,0.912,0 | 0.93,0.937,1.0,0.6 | -0,054 |
| 228758 | ENSG00000160218 | TRAPPC10 | chr21 | + | 44063573 | 44063785 | 44059102 | 44059214 | 44074323 | 44074470 | 228758 | 50,62,90,79,46,43,53,53,5 | 7,0,5,3,3,5,2,3,0,0,5,2 | 62,70,40,66,68,27,43,52,3 | 2,0,2,2,5,1,2,4,2,0,5,0,1,2, | 298 | 149 | 0.0244237069922 | | 1 | 0.781,1.0,0.9,0.92 | 0.939,1.0,0.909,0. | -0,035 |
| 76214 | ENSG00000196510 | ANAPC7 | chr12 | - | 110388511 | 110388623 | 110387738 | 110387892 | 110395100 | 110395220 | 76214 | 27,6,79,53,30,28,16,19,13, | 0,1,3,0,0,2,2,3,0,0,1,1 | 53,40,31,11,25,8,17,32,14, | 3,2,0,0,2,1,0,0,1,2,1,0,8,5, | 261 | 149 | 0.0244672656628 | | 1 | 1.0,0.774,0.938,1. | 0.91,0.919,1.0,1.0, | -0,009 |
| 107405 | ENSG00000143569 | UBAP2L | chr1 | + | 154225083 | 154225213 | 154220848 | 154220975 | 154227281 | 154227359 | 107405 | 137,86,133,67,38,59,49,50 | 2,0,1,3,0,0,0,3,1,0,0,0 | 75,71,54,52,63,42,27,46,3 | 1,2,1,6,3,1,2,3,0,0,1,0,0,1, | 279 | 149 | 0.0244746141586 | | 1 | 0.973,1.0,0.986,0. | 0.976,0.95,0.966,0 | 0,03 |
| 135727 | ENSG00000291230 | | chr19 | - | 50334306 | 50334557 | 50333795 | 50334101 | 50334828 | 50334973 | 135727 | 255,173,256,513,218,156, | 0,0,0,1,2,0,2,2,0,0,0,0 | 160,135,134,73,141,148,5 | 6,6,0,8,0,0,0,0,4,2,1,8,0,2, | 298 | 149 | 0.0244975874887 | | 1 | 1.0,1.0,1.0,0.996,0 | 0.93,0.918,1.0,0.8 | 0,025 |
| 196289 | ENSG00000117153 | KLHL12 | chr1 | - | 202893282 | 202893425 | 202891119 | 202892659 | 202894590 | 202894749 | 196289 | 6,1,1,0,0,1,0,3,4,0,0,1 | 6,9,10,6,1,3,2,6,1,4,2,0 | 0,2,0,0,3,0,5,3,0,0,4,6,2,1, | 7,6,4,6,2,1,3,12,1,5,11,6,1, | 292 | 149 | 0.0245099920764 | | 1 | 0.338,0.054,0.049, | 0.0,0.145,0.0,0.0,0 | 0,035 |
| 27152 | ENSG00000129515 | SNX6 | chr14 | - | 34608029 | 34608140 | 34605486 | 34605717 | 34609637 | 34609742 | 27152 | 170,97,300,263,93,121,69, | 5,2,5,7,4,2,2,4,2,2,0,0 | 153,211,82,51,104,101,46, | 8,5,1,6,8,5,2,3,2,8,5,7,0,1, | 260 | 149 | 0.0245111890541 | | 1 | 0.951,0.965,0.972, | 0.916,0.96,0.979,0 | 0,021 |
| 91345 | ENSG00000146063 | TRIM41 | chr5 | + | 181233412 | 181233435 | 181232658 | 181232889 | 181233635 | 181233763 | 91345 | 15,6,10,1,4,1,4,6,5,5,3,11 | 0,0,6,2,0,0,2,0,2,5,0,2 | 13,16,9,8,2,7,10,17,2,9,7,3 | 0,1,1,0,0,3,0,0,0,0,0,2,6,0, | 172 | 149 | 0.0245361718102 | | 1 | 1.0,1.0,0.591,0.30 | 1.0,0.933,0.886,1. | -0,117 |
| 177069 | ENSG00000151422 | FER | chr5 | + | 108897962 | 108898132 | 108897658 | 108897848 | 108946129 | 108946222 | 177069 | 5,0,16,2,4,4,2,3,1,1,2,2 | 6,3,8,10,10,12,5,5,4,0,2,5 | 1,2,0,5,0,1,2,3,0,1,2,3,4,1, | 13,20,5,2,5,6,6,11,6,4,7,17 | 298 | 149 | 0.0245541248496 | | 1 | 0.294,0.0,0.5,0.09 | 0.037,0.048,0.0,0. | 0,162 |
| 251813 | ENSG00000120690 | ELF1 | chr13 | - | 41014158 | 41014221 | 40981982 | 40982282 | 41019227 | 41019316 | 251813 | 5,0,14,4,2,8,6,4,0,2,0,1 | 55,42,200,127,48,44,53,65 | 3,2,0,4,2,4,1,2,0,3,4,2,9,8, | 127,171,49,53,126,59,33,6 | 212 | 149 | 0.0245683100933 | | 1 | 0.06,0.0,0.047,0.0 | 0.016,0.008,0.0,0. | 0,012 |
| 91753 | ENSG00000135049 | AGTPBP1 | chr9 | - | 85588297 | 85588478 | 85586830 | 85586960 | 85589527 | 85589681 | 91753 | 52,37,138,118,28,33,33,45 | 2,1,9,10,1,3,0,7,3,0,2,0 | 53,73,36,33,68,39,22,37,2 | 10,14,5,3,7,0,0,4,5,1,8,9,6, | 298 | 149 | 0.0245723061899 | | 1 | 0.929,0.949,0.885, | 0.726,0.723,0.783, | 0,056 |
| 79929 | ENSG00000129292 | PHF20L1 | chr8 | + | 132798771 | 132798860 | 132794732 | 132794817 | 132803818 | 132804032 | 79929 | 67,43,63,34,15,14,13,31,2 | 8,4,7,8,0,2,1,1,0,0,0,3 | 23,22,21,14,18,22,15,18,1 | 2,1,3,11,4,1,1,4,1,2,3,8,4,7 | 238 | 149 | 0.0245896824408 | | 1 | 0.84,0.871,0.849,0 | 0.878,0.932,0.814, | 0,09 |
| 225365 | ENSG00000168137 | SETD5 | chr3 | + | 9434562 | 9434630 | 9434333 | 9434485 | 9434823 | 9434882 | 225365 | 1,0,0,0,1,2,0,1,0,1,0,0 | 22,10,28,17,11,14,16,17,7, | 2,0,2,1,1,2,2,2,1,0,1,1,2,1, | 10,12,5,10,13,4,5,12,6,20,1 | 217 | 149 | 0.0245963045434 | | 1 | 0.03,0.0,0.0,0.0,0. | 0.121,0.0,0.215,0. | -0,068 |
| 185205 | ENSG00000132185 | FCRLA | chr1 | + | 161710477 | 161710492 | 161707294 | 161707343 | 161710759 | 161710863 | 185205 | 7,1,18,6,21,14,1,0,0,0,0,7 | 345,256,627,562,206,101, | 2,1,4,1,5,9,1,1,3,23,5,6,0,0 | 666,397,175,131,141,184,1 | 164 | 149 | 0.0245996758942 | | 1 | 0.018,0.004,0.025, | 0.003,0.002,0.02,0 | 0,008 |
| 118234 | ENSG00000107164 | FUBP3 | chr9 | + | 130631160 | 130631293 | 130630627 | 130630788 | 130631556 | 130631630 | 118234 | 2,1,0,4,2,10,0,0,1,0,2,7 | 25,14,24,17,21,27,17,19,2 | 2,3,12,0,2,1,4,0,1,3,6,1,0,6 | 10,18,19,16,10,1,18,23,3,1 | 282 | 149 | 0.0246535559452 | | 1 | 0.041,0.036,0.0,0. | 0.096,0.081,0.25,0 | -0,053 |
| 21079 | ENSG00000089775 | ZBTB25 | chr14 | - | 64499440 | 64499563 | 64490360 | 64490540 | 64502900 | 64503127 | 21079 | 7,3,12,9,3,5,7,12,4,2,2,0 | 7,6,3,6,9,5,5,1,6,0,0,3 | 3,8,2,1,8,10,4,0,10,8,4,0,4, | 10,1,0,1,0,4,0,4,0,3,0,2,0,4 | 272 | 149 | 0.0246557452851 | | 1 | 0.354,0.215,0.687, | 0.141,0.814,1.0,0. | -0,14 |
| 28727 | ENSG00000150593 | PDCD4 | chr10 | + | 110875965 | 110876070 | 110871794 | 110872018 | 110881232 | 110881535 | 28727 | 80,59,255,204,130,67,33,7 | 17,15,30,17,3,10,11,28,5,5 | 92,91,49,26,45,75,20,47,3 | 16,48,16,63,37,7,0,8,8,19,1 | 254 | 149 | 0.024660318084 | | 1 | 0.734,0.698,0.833, | 0.771,0.527,0.642, | 0,08 |
| 2985 | ENSG00000268400 | | chr19 | + | 7630572 | 7630659 | 7629811 | 7629860 | 7630822 | 7630846 | 2985 | 17,4,40,18,3,4,4,6,6,6,5,9 | 4,2,6,2,0,6,3,11,2,0,1,3 | 15,7,12,4,12,8,1,5,5,6,8,15 | 6,12,4,7,1,2,3,14,0,8,10,18 | 236 | 149 | 0.0246620319053 | | 1 | 0.729,0.558,0.808, | 0.612,0.269,0.654, | 0,174 |
| 55454 | ENSG00000132185 | FCRLA | chr1 | + | 161710759 | 161710912 | 161706971 | 161707343 | 161713084 | 161713705 | 55454 | 348,258,631,564,186,101, | 10,2,4,2,0,4,2,0,6,12,0,6 | 668,404,176,131,144,189, | 3,4,0,2,0,0,2,2,0,0,2,7,2,2, | 298 | 149 | 0.0246648383942 | | 1 | 0.946,0.985,0.987, | 0.991,0.981,1.0,0. | -0,012 |
| 39097 | ENSG00000164944 | VIRMA | chr8 | - | 94534838 | 94535007 | 94530962 | 94531085 | 94537102 | 94537151 | 39097 | 2,11,18,17,8,10,17,17,11,2 | 1,0,1,0,0,3,0,3,2,0,0,0 | 20,33,8,9,11,10,4,8,11,17, | 0,1,1,1,0,2,3,3,1,0,0,10,1,0 | 298 | 149 | 0.0247129638627 | | 1 | 0.5,1.0,0.9,1.0,1.0, | 1.0,0.943,0.8,0.81 | 0,051 |
| 200831 | ENSG00000161929 | SCIMP | chr17 | - | 5223053 | 5223456 | 5221286 | 5221350 | 5234734 | 5234823 | 200831 | 127,32,99,103,29,29,50,42 | 3,0,3,13,1,4,0,1,9,3,6,5 | 70,44,29,11,24,44,17,35,3 | 8,2,10,2,2,4,1,9,6,3,2,3,8,1 | 298 | 149 | 0.0247270830438 | | 1 | 0.955,1.0,0.943,0. | 0.814,0.917,0.592, | 0,042 |
| 24410 | ENSG00000150403 | TMCO3 | chr13 | + | 113534044 | 113534240 | 113520615 | 113520733 | 113539356 | 113539507 | 24410 | 17,9,33,14,21,4,13,9,5,13, | 0,0,3,0,4,3,0,0,4,0,0,0 | 19,6,9,5,4,1,6,24,4,15,20,2 | 0,0,0,0,0,0,0,0,0,0,0,3,0, | 298 | 149 | 0.024742787656 | | 1 | 1.0,1.0,0.846,1.0,0 | 1.0,1.0,1.0,1.0,1.0, | -0,119 |
| 9661 | ENSG00000178971 | CTC1 | chr17 | - | 8229141 | 8229206 | 8228726 | 8228892 | 8229301 | 8229446 | 9661 | 18,7,19,17,13,8,15,9,15,11 | 1,1,3,2,0,0,0,1,2,0,2,6 | 15,7,8,4,6,8,13,14,8,9,12,5 | 1,1,0,0,1,0,2,4,0,2,1,1,0,2, | 214 | 149 | 0.0247446383116 | | 1 | 0.926,0.83,0.815,0 | 0.913,0.83,1.0,1.0, | -0,013 |
| 254928 | ENSG00000111300 | NAA25 | chr12 | - | 112078386 | 112078541 | 112078187 | 112078266 | 112078633 | 112078741 | 254928 | 1,0,3,0,0,0,0,4,2,1,0,2 | 47,45,35,29,28,30,11,14,2 | 0,6,2,2,4,0,0,0,0,5,3,3,1, | 25,14,17,19,26,3,18,14,14, | 298 | 149 | 0.024752218231 | | 1 | 0.011,0.0,0.041,0. | 0.0,0.176,0.056,0. | -0,017 |
| 48978 | ENSG00000197548 | ATG7 | chr3 | + | 11332971 | 11333093 | 11331339 | 11331428 | 11340644 | 11340735 | 48978 | 17,5,39,31,15,17,10,18,9,9 | 0,0,4,0,0,8,0,1,3,0,0,0 | 19,10,13,11,3,13,15,6,8,13 | 3,2,0,0,0,2,0,0,2,0,0,4,0,2,1 | 271 | 149 | 0.024769100584 | | 1 | 1.0,1.0,0.843,1.0,1 | 0.777,0.733,1.0,1. | 0,04 |
| 146448 | ENSG00000237440 | ZNF737 | chr19 | - | 20552474 | 20552549 | 20537991 | 20545976 | 20553708 | 20553835 | 146448 | 34,23,160,129,50,36,29,58 | 1,0,5,11,0,1,0,1,4,0,0,5 | 93,73,50,33,22,30,15,32,2 | 0,0,2,0,2,11,0,2,0,0,0,0,0,0 | 224 | 149 | 0.0247855361182 | | 1 | 0.958,1.0,0.955,0. | 1.0,1.0,0.943,1.0,0 | -0,009 |
| 48474 | ENSG00000264364 | DYNLL2 | chr17 | + | 58087081 | 58087222 | 58083418 | 58083683 | 58089141 | 58095542 | 48474 | 141,86,140,114,66,52,44,6 | 0,0,5,0,2,2,0,1,5,0,0,0 | 108,127,105,117,45,46,41, | 1,1,2,3,1,0,2,4,0,0,7,3,2,2, | 290 | 149 | 0.0247894602856 | | 1 | 1.0,1.0,0.935,1.0,0 | 0.982,0.985,0.964, | 0,024 |
| 243452 | ENSG00000065923 | SLC9A7 | chrX | - | 46635588 | 46635653 | 46631585 | 46631649 | 46643235 | 46643389 | 243452 | 70,60,55,28,21,27,22,49,2 | 181,139,132,102,86,90,12 | 52,54,32,33,35,19,12,31,2 | 118,79,161,172,55,42,66,1 | 214 | 149 | 0.024829562721 | | 1 | 0.212,0.231,0.225, | 0.235,0.322,0.122, | -0,061 |
| 176103 | ENSG00000124596 | OARD1 | chr6 | - | 41068840 | 41068944 | 41064771 | 41067437 | 41070075 | 41070134 | 176103 | 11,3,27,18,9,10,14,5,9,6,2, | 1,1,10,2,0,2,1,3,3,1,0,2 | 13,18,5,0,3,10,7,8,10,10,9, | 2,8,1,7,3,1,0,3,0,1,5,8,2,3, | 253 | 149 | 0.0248393052172 | | 1 | 0.866,0.639,0.614, | 0.793,0.57,0.746,0 | 0,154 |
| 141421 | ENSG00000173821 | RNF213 | chr17 | + | 80294120 | 80294573 | 80291627 | 80291827 | 80294719 | 80295003 | 141421 | 1,0,10,0,4,0,4,9,5,2,2,0 | 77,40,91,44,44,38,23,61,4 | 0,2,0,1,3,1,0,2,2,0,4,2,2,8, | 49,53,25,48,33,37,30,36,4 | 298 | 149 | 0.0248428257749 | | 1 | 0.006,0.0,0.052,0. | 0.0,0.019,0.0,0.01, | 0,011 |
| 174785 | ENSG00000187514 | PTMA | chr2 | + | 231711910 | 231711983 | 231711347 | 231711419 | 231712803 | 231712856 | 174785 | 4,4,2,4,4,1,4,7,1,0,0,5 | 1,0,3,0,5,2,5,3,1,2,1,0 | 4,4,6,3,6,1,6,6,1,1,10,20,6, | 0,0,1,5,0,0,1,4,0,0,2,4,5,2, | 222 | 149 | 0.0248480048684 | | 1 | 0.729,1.0,0.309,1. | 1.0,1.0,0.801,0.28 | -0,248 |
| 158592 | ENSG00000150867 | PIP4K2A | chr10 | - | 22591628 | 22591767 | 22573310 | 22573457 | 22607926 | 22608023 | 158592 | 42,23,36,61,19,21,21,24,2 | 2,6,5,0,1,0,0,2,0,2,2,5 | 41,31,29,18,16,14,7,31,22, | 2,4,0,1,1,1,0,2,1,2,1,5,0,1,1 | 288 | 149 | 0.0248562034717 | | 1 | 0.916,0.665,0.788, | 0.914,0.8,1.0,0.90 | -0,065 |
| 16979 | ENSG00000102858 | MGRN1 | chr16 | + | 4680031 | 4680097 | 4677462 | 4677572 | 4681549 | 4681580 | 16979 | 12,11,26,8,12,2,12,12,7,8, | 3,1,0,0,3,0,2,0,2,0,0,1 | 5,16,15,8,7,4,7,3,5,10,10,1 | 2,1,0,9,4,1,1,0,0,3,0,1,1,0, | 215 | 149 | 0.0248647743525 | | 1 | 0.735,0.884,1.0,1. | 0.634,0.917,1.0,0. | 0,056 |
| 122350 | ENSG00000149292 | TTC12 | chr11 | + | 113324604 | 113324682 | 113323993 | 113324015 | 113325523 | 113325645 | 122350 | 43,17,74,43,14,16,17,16,2 | 0,0,3,1,3,1,0,0,0,0,0,0 | 45,38,18,36,31,25,16,22,2 | 2,2,0,2,0,0,0,0,3,10,1,0,0 | 227 | 149 | 0.024871319055 | | 1 | 1.0,1.0,0.942,0.96 | 0.937,0.926,1.0,0. | 0,033 |
| 211216 | ENSG00000197622 | CDC42SE1 | chr1 | - | 151055937 | 151055993 | 151055015 | 151055126 | 151059478 | 151059574 | 211216 | 116,100,106,69,49,31,63,5 | 10,5,4,4,5,4,4,3,6,2,0,3 | 54,45,88,68,40,29,35,45,2 | 1,3,1,12,7,0,2,2,2,2,5,12,2, | 205 | 149 | 0.0248821962552 | | 1 | 0.894,0.936,0.951, | 0.975,0.916,0.985, | 0,03 |
| 186029 | ENSG00000153898 | MCOLN2 | chr1 | - | 84956387 | 84956584 | 84952445 | 84952530 | 84958528 | 84958702 | 186029 | 112,87,198,109,55,70,67,1 | 8,3,41,20,1,3,10,5,3,4,3,4 | 101,132,55,86,52,37,36,62 | 10,11,6,8,26,5,4,20,10,14,1 | 298 | 149 | 0.0248930476856 | | 1 | 0.875,0.935,0.707, | 0.835,0.857,0.821, | 0,061 |
| 146610 | ENSG00000168802 | CHTF8 | chr16 | - | 69121052 | 69121184 | 69118009 | 69120649 | 69121435 | 69121493 | 146610 | 17,11,32,29,20,9,7,10,10,8 | 10,6,9,2,7,2,3,5,1,0,2,4 | 34,7,13,4,8,18,0,23,10,25, | 0,3,0,5,2,2,3,3,0,2,2,6,5,4, | 281 | 149 | 0.0249032678965 | | 1 | 0.474,0.493,0.653, | 1.0,0.553,1.0,0.29 | -0,076 |
| 217067 | ENSG00000148690 | FRA10AC1 | chr10 | - | 93701675 | 93701795 | 93700029 | 93700106 | 93702374 | 93702592 | 217067 | 4,1,2,1,4,2,1,2,0,0,0,2 | 3,6,6,5,7,7,3,0,3,6,2,10 | 4,2,1,2,0,0,0,1,0,2,0,0,0,3, | 3,2,4,2,2,5,4,3,1,3,3,5,3,7, | 269 | 149 | 0.0249399659008 | | 1 | 0.425,0.085,0.156, | 0.425,0.356,0.122, | 0,054 |
| 89751 | ENSG00000031003 | FAM13B | chr5 | - | 138019880 | 138019977 | 138018478 | 138018514 | 138021030 | 138021197 | 89751 | 17,11,15,12,2,4,3,7,2,9,1,1 | 22,9,34,11,12,11,8,7,8,2,6, | 10,7,5,3,8,3,0,4,1,1,0,2,4,4 | 26,20,14,7,11,10,10,23,5,1 | 246 | 149 | 0.0250100357084 | | 1 | 0.319,0.425,0.211, | 0.189,0.175,0.178, | 0,112 |
| 239117 | ENSG00000075856 | SART3 | chr12 | - | 108538003 | 108538203 | 108537487 | 108537595 | 108538933 | 108539089 | 239117 | 20,22,64,41,16,34,18,36,2 | 2,1,2,2,2,2,1,2,2,0,0,1 | 23,35,13,13,27,24,20,32,2 | 2,0,1,4,1,1,0,2,1,0,1,1,0,1, | 298 | 149 | 0.0250119754487 | | 1 | 0.833,0.917,0.941, | 0.852,1.0,0.867,0. | -0,013 |
| 43266 | ENSG00000136560 | TANK | chr2 | + | 161223919 | 161223991 | 161204674 | 161204793 | 161224630 | 161224746 | 43266 | 93,82,124,78,33,36,16,30, | 4,1,10,6,2,0,1,0,1,1,2,1 | 59,70,42,31,42,28,11,36,1 | 5,1,0,2,0,2,0,0,0,0,1,3,3,2, | 221 | 149 | 0.0250525417648 | | 1 | 0.94,0.982,0.893,0 | 0.888,0.979,1.0,0. | -0,014 |

| | | | | | | | | | | | | | | | | | | | | | | | |
|---|---|---|---|---|---|---|---|---|---|---|---|---|---|---|---|---|---|---|---|---|---|---|---|
| 62562 | ENSG00000138101 | DTNB | chr2 | - | 25383835 | 25383889 | 25379289 | 25379323 | 25387288 | 25387378 | 62562 | 12,3,9,6,6,3,4,4,6,4,1,9 | 1,5,10,2,6,3,7,3,3,11,3,4 | 4,10,8,4,9,3,8,8,3,6,2,3,0,0 | 2,0,3,2,1,1,4,1,0,3,2,6,3,2,1 | 203 | 149 | 0.0250935532491 | | 1 | 0.898,0.306,0.398, | 0.595,1.0,0.662,0. | -0,155 |
| 155299 | ENSG00000103811 | CTSH | chr15 | - | 78929411 | 78929481 | 78923780 | 78927781 | 78931450 | 78935082 | 155299 | 182,158,151,95,161,274,88 | 10,3,7,0,3,14,2,8,5,0,0,2 | 85,78,122,45,56,77,20,134 | 0,1,1,1,3,0,0,3,6,1,6,7,5,0,1 | 219 | 149 | 0.0250977694653 | | 1 | 0.925,0.973,0.936, | 1.0,0.982,0.988,0. | 0 |
| 231859 | ENSG00000136504 | KAT7 | chr17 | + | 49817825 | 49818011 | 49815802 | 49815913 | 49821336 | 49821426 | 231859 | 47,34,117,43,34,33,23,52,1 | 1,0,1,2,0,1,0,2,0,0,1,0 | 37,44,30,43,48,21,22,26,14 | 2,4,0,1,2,0,0,2,2,0,2,6,1,0,0 | 298 | 149 | 0.0251016121019 | | 1 | 0.959,1.0,0.983,0. | 0.902,0.846,1.0,0. | 0,034 |
| 140120 | ENSG00000126001 | CEP250 | chr20 | + | 35507737 | 35507792 | 35502389 | 35505005 | 35508034 | 35508190 | 140120 | 11,6,11,2,1,5,7,3,3,2,1,0 | 1,1,9,2,2,0,1,0,0,4,2,3 | 5,3,2,5,1,2,2,3,2,6,2,9,3,6,2 | 5,2,4,4,1,1,0,5,2,8,2,12,3,6 | 204 | 149 | 0.025140322924 | | 1 | 0.889,0.814,0.472, | 0.422,0.523,0.268, | 0,133 |
| 83844 | ENSG00000014641 | MDH1 | chr2 | + | 63594487 | 63594586 | 63588608 | 63589046 | 63597398 | 63597574 | 83844 | 9,8,57,54,17,26,15,15,4,5,2 | 0,0,3,0,2,2,0,2,0,0,1,0 | 41,49,12,0,16,5,6,20,7,28,1 | 3,5,0,2,2,1,0,2,0,0,3,5,1,1,0 | 248 | 149 | 0.0251506236851 | | 1 | 1.0,1.0,0.919,1.0,0 | 0.891,0.855,1.0,0. | 0,064 |
| 185897 | ENSG00000010322 | NISCH | chr3 | + | 52476450 | 52476599 | 52473733 | 52473829 | 52477573 | 52477642 | 185897 | 69,31,75,40,34,41,31,38,33 | 0,0,0,1,0,2,0,0,2,0,0,0 | 47,66,32,33,46,19,26,60,35 | 2,0,5,2,1,0,0,1,1,4,4,0,0,2,1 | 298 | 149 | 0.0251600704282 | | 1 | 1.0,1.0,1.0,0.952,1 | 0.922,1.0,0.762,0.8 | 0,045 |
| 73507 | ENSG00000154222 | CC2D1B | chr1 | - | 52359704 | 52359820 | 52359458 | 52359534 | 52360073 | 52360233 | 73507 | 8,9,20,9,10,14,10,11,13,5,5 | 0,2,2,0,2,10,2,6,1,0,0,3 | 15,13,5,10,7,5,10,22,5,7,8, | 4,4,2,5,4,1,2,5,1,0,6,2,3,1,8 | 265 | 149 | 0.0251621654615 | | 1 | 1.0,0.717,0.849,1. | 0.678,0.646,0.584, | 0,148 |
| 33255 | ENSG00000005700 | IBTK | chr6 | - | 82216075 | 82216250 | 82214226 | 82214829 | 82217959 | 82218137 | 33255 | 27,17,156,106,38,27,25,35 | 0,0,5,0,0,0,0,1,1,0,2,1 | 81,59,47,12,44,40,32,38,40 | 2,5,1,5,1,1,2,0,2,2,0,1,0,0,0 | 298 | 149 | 0.0252250060403 | | 1 | 1.0,1.0,0.94,1.0,1. | 0.953,0.855,0.959, | 0,027 |
| 59831 | ENSG00000009780 | FAM76A | chr1 | + | 27749067 | 27749154 | 27744653 | 27744811 | 27755194 | 27755330 | 59831 | 11,8,65,32,16,11,20,16,18, | 4,1,5,7,0,4,5,1,3,3,2,4 | 26,20,10,7,11,9,14,12,6,17 | 11,8,5,2,4,0,1,8,5,3,4,16,6, | 236 | 149 | 0.0252277606208 | | 1 | 0.635,0.835,0.891, | 0.599,0.612,0.558, | 0,078 |
| 238996 | ENSG00000134014 | ELP3 | chr8 | + | 28183171 | 28183264 | 28162020 | 28162078 | 28189648 | 28190087 | 238996 | 3,1,4,3,2,0,0,0,3,3,0,2 | 23,4,21,22,28,20,20,12,25, | 1,0,0,0,2,0,0,1,1,0,0,0,0,0,0 | 30,8,21,8,1,13,13,10,14,28 | 242 | 149 | 0.0252283525067 | | 1 | 0.074,0.133,0.105, | 0.02,0.0,0.0,0.0,0. | 0,008 |
| 107889 | ENSG00000008294 | SPAG9 | chr17 | - | 50996564 | 50996694 | 50995443 | 50995533 | 50998443 | 50998617 | 107889 | 27,24,139,65,21,45,27,56,3 | 0,0,0,5,0,3,0,0,0,0,1,3 | 78,57,35,25,51,20,34,35,21 | 0,4,0,5,5,1,0,3,0,1,2,0,0,1,5 | 279 | 149 | 0.0252318689547 | | 1 | 1.0,1.0,1.0,0.874,1 | 1.0,0.884,1.0,0.72 | 0,038 |
| 247387 | ENSG00000149311 | ATM | chr11 | + | 108235669 | 108236168 | 108229177 | 108229323 | 108243952 | 108244118 | 247387 | 105,86,373,223,49,53,40,4 | 2,6,15,7,2,1,0,3,1,2,0,2 | 180,199,38,39,144,107,25, | 12,3,0,1,4,0,1,0,1,4,2,7,1,6 | 298 | 149 | 0.0252710457073 | | 1 | 0.963,0.878,0.926, | 0.882,0.971,1.0,0. | -0,023 |
| 46799 | ENSG00000138757 | G3BP2 | chr4 | - | 75653982 | 75654081 | 75648638 | 75648741 | 75655065 | 75655246 | 46799 | 8,3,14,15,5,4,8,8,8,5,0,4 | 110,112,251,203,93,129,92 | 14,18,15,9,16,4,5,2,6,8,5,1 | 183,322,118,149,169,59,87 | 248 | 149 | 0.025284812739 | | 1 | 0.042,0.016,0.032, | 0.044,0.032,0.071, | -0,009 |
| 173738 | ENSG00000136560 | TANK | chr2 | + | 161223887 | 161223991 | 161204674 | 161204793 | 161224630 | 161224637 | 173738 | 111,97,134,89,38,50,21,42 | 4,1,10,6,2,0,1,0,1,1,2,1 | 69,83,54,37,53,33,12,44,24 | 5,1,0,2,0,2,0,0,0,0,1,3,3,2,0 | 253 | 149 | 0.0252936367794 | | 1 | 0.942,0.983,0.888, | 0.89,0.98,1.0,0.91 | -0,014 |
| 55785 | ENSG00000134352 | IL6ST | chr5 | - | 55951463 | 55951604 | 55947492 | 55947589 | 55951928 | 55952075 | 55785 | 16,13,24,16,12,15,34,155,3 | 1,0,3,3,0,2,4,2,0,0,0,1 | 31,61,33,48,35,12,23,25,12 | 0,1,2,8,0,1,2,2,0,0,6,2,0,0,1 | 290 | 149 | 0.0253414618537 | | 1 | 0.892,1.0,0.804,0. | 1.0,0.969,0.894,0. | 0,007 |
| 130269 | ENSG00000123329 | ARHGAP9 | chr12 | - | 57473602 | 57473691 | 57472265 | 57472688 | 57474041 | 57474176 | 130269 | 42,27,83,76,34,13,20,28,28 | 38,11,18,14,10,11,13,14,28 | 72,44,20,15,34,29,19,27,39 | 13,13,23,9,4,10,5,9,2,15,4, | 238 | 149 | 0.025349056185 | | 1 | 0.409,0.606,0.743, | 0.776,0.679,0.352, | -0,104 |
| 227692 | ENSG00000186088 | GSAP | chr7 | - | 77379846 | 77380018 | 77377285 | 77377390 | 77381304 | 77381354 | 227692 | 52,46,127,33,35,47,48,74,2 | 51,26,156,99,53,60,40,66,5 | 66,79,73,50,38,27,18,28,49 | 102,45,42,11,27,18,28,56,2 | 298 | 149 | 0.0253586792074 | | 1 | 0.338,0.469,0.289, | 0.244,0.467,0.465, | -0,086 |
| 949 | ENSG00000100083 | GGA1 | chr22 | + | 37625796 | 37625949 | 37623551 | 37623633 | 37629461 | 37629526 | 949 | 9,10,9,4,16,16,7,5,10,3,17, | 3,4,4,0,2,4,1,0,6,0,0,3 | 6,1,11,6,10,3,6,13,2,7,6,24 | 2,3,1,2,2,1,3,9,2,3,3,10,3,1 | 298 | 149 | 0.0253697590788 | | 1 | 0.6,0.556,0.529,1. | 0.6,0.143,0.846,0. | 0,202 |
| 138217 | ENSG00000130818 | ZNF426 | chr19 | - | 9535401 | 9535520 | 9535187 | 9535279 | 9536207 | 9536569 | 138217 | 4,0,2,0,0,4,3,6,1,0,0,2 | 11,15,54,30,13,21,20,26,19 | 0,8,0,5,0,0,0,5,1,1,3,3,8,0,6 | 18,48,17,13,15,3,23,19,11, | 268 | 149 | 0.0253723013211 | | 1 | 0.168,0.0,0.02,0.0, | 0.0,0.085,0.0,0.17 | -0,034 |
| 77408 | ENSG00000115525 | ST3GAL5 | chr2 | - | 85847381 | 85847562 | 85846376 | 85846424 | 85847860 | 85847984 | 77408 | 7,6,16,5,5,1,6,7,2,2,6,10 | 0,1,17,4,3,1,0,1,3,0,2,6 | 15,15,0,4,8,4,0,3,5,22,16,1 | 8,7,9,1,4,3,2,10,3,2,3,4,2,7 | 298 | 149 | 0.0253834886439 | | 1 | 1.0,0.75,0.32,0.38 | 0.484,0.517,0.0,0. | 0,22 |
| 192468 | ENSG00000157873 | TNFRSF14 | chr1 | + | 2561493 | 2561672 | 2560553 | 2560714 | 2561776 | 2561815 | 192468 | 23,18,15,6,5,13,8,14,7,10,6 | 2,1,10,6,3,1,2,5,3,1,2,3 | 10,9,8,14,5,5,12,22,4,7,6,2 | 7,6,0,7,2,1,1,4,2,6,6,12,7,3 | 298 | 149 | 0.0254207382017 | | 1 | 0.852,0.9,0.429,0. | 0.417,0.429,1.0,0. | 0,128 |
| 39271 | ENSG00000168026 | TTC21A | chr3 | + | 39130247 | 39130358 | 39130078 | 39130151 | 39130700 | 39130839 | 39271 | 11,8,15,8,9,3,5,8,7,5,4,3 | 1,0,1,3,4,0,0,0,0,1,0,4 | 9,4,3,6,5,4,8,8,7,16,11,9,5, | 2,2,0,0,0,0,0,0,0,2,0,1,1,0,0 | 260 | 149 | 0.0254322952438 | | 1 | 0.863,1.0,0.896,0. | 0.721,0.534,1.0,1. | -0,085 |
| 179681 | ENSG00000136450 | SRSF1 | chr17 | - | 58005413 | 58005617 | 58004587 | 58005041 | 58005800 | 58005973 | 179681 | 9,6,20,17,7,5,2,4,11,5,4,3 | 51,23,46,28,22,14,16,32,32 | 0,8,13,19,5,6,2,3,7,6,0,3,5, | 36,40,21,51,25,9,6,26,15,4 | 298 | 149 | 0.0254481033373 | | 1 | 0.081,0.115,0.179, | 0.0,0.091,0.236,0. | 0,044 |
| 203288 | ENSG00000092871 | RFFL | chr17 | - | 35025182 | 35025337 | 35021621 | 35021781 | 35026373 | 35026568 | 203288 | 0,1,2,5,1,4,0,4,0,0,0,1 | 8,1,19,20,6,7,5,16,5,6,2,12 | 3,5,1,0,2,0,0,1,0,0,7,0,5,1,0 | 10,6,1,5,6,8,7,13,3,10,2,14 | 298 | 149 | 0.0254494721083 | | 1 | 0.0,0.333,0.05,0.1 | 0.13,0.294,0.333,0 | -0,05 |
| 70315 | ENSG00000166526 | ZNF3 | chr7 | - | 100079535 | 100079656 | 100077017 | 100077433 | 100081634 | 100081703 | 70315 | 10,21,56,21,14,28,7,28,20, | 0,0,4,5,0,1,4,0,0,0,2,1 | 20,32,26,7,26,5,14,14,16,5 | 0,1,0,0,3,0,0,2,0,0,2,4,5,0,0 | 270 | 149 | 0.0254688793502 | | 1 | 1.0,1.0,0.885,0.69 | 1.0,0.946,1.0,1.0,0 | -0,045 |
| 149586 | ENSG00000140598 | EFL1 | chr15 | - | 82215582 | 82215668 | 82214782 | 82214855 | 82219651 | 82219818 | 149586 | 0,2,7,1,0,4,3,4,2,0,2,2 | 14,6,20,9,23,10,11,8,6,3,3, | 4,0,2,0,1,1,1,2,0,2,0,0,0,2,2 | 9,4,3,4,0,3,10,14,2,6,22,9,6 | 235 | 149 | 0.0254986231271 | | 1 | 0.0,0.174,0.182,0. | 0.22,0.0,0.297,0.0, | -0,021 |
| 177998 | ENSG00000142687 | KIAA0319L | chr1 | - | 35435304 | 35435654 | 35434397 | 35435081 | 35441046 | 35443028 | 177998 | 3,0,5,4,3,5,4,8,0,0,2,3 | 9,8,43,30,14,16,19,29,13,5 | 7,2,5,5,4,3,9,5,2,6,7,4,0,5,6 | 29,18,18,5,6,6,1,31,9,20,17 | 298 | 149 | 0.0254993912973 | | 1 | 0.143,0.0,0.055,0. | 0.108,0.053,0.122, | -0,099 |
| 231858 | ENSG00000136504 | KAT7 | chr17 | + | 49817822 | 49818011 | 49815802 | 49815913 | 49821336 | 49821426 | 231858 | 48,34,117,43,34,33,23,52,1 | 1,0,1,2,0,1,0,2,0,0,1,0 | 37,44,29,43,48,21,22,26,14 | 2,4,0,1,2,0,0,2,2,0,2,6,1,0,0 | 298 | 149 | 0.0255085036232 | | 1 | 0.96,1.0,0.983,0.9 | 0.902,0.846,1.0,0. | 0,034 |
| 85904 | ENSG00000115970 | THADA | chr2 | - | 43556344 | 43556555 | 43552067 | 43552339 | 43566697 | 43566821 | 85904 | 33,17,30,10,20,27,11,23,9, | 3,0,5,4,2,0,0,2,0,0,0,0 | 48,41,21,29,28,18,19,38,17 | 3,4,1,0,2,0,0,0,1,4,0,0,4,3,3 | 298 | 149 | 0.0255427030916 | | 1 | 0.846,1.0,0.75,0.5 | 0.889,0.837,0.913, | 0,036 |
| 165188 | ENSG00000115073 | ACTR1B | chr2 | - | 97658426 | 97658632 | 97658223 | 97658316 | 97658878 | 97659003 | 165188 | 41,27,55,50,38,35,28,22,31 | 5,0,4,2,0,4,0,0,0,1,0,2 | 28,37,22,9,20,15,31,36,20, | 0,0,0,1,1,0,0,2,0,2,0,2,0,6,0 | 298 | 149 | 0.0255483737235 | | 1 | 0.804,1.0,0.873,0. | 1.0,1.0,1.0,0.818,0 | -0,01 |
| 243339 | ENSG00000197136 | PCNX3 | chr11 | + | 65635930 | 65636031 | 65635528 | 65635803 | 65636173 | 65636307 | 243339 | 0,0,1,4,0,4,3,4,2,2,0,1 | 8,6,10,4,12,6,13,4,10,8,5,5 | 2,6,0,1,3,2,0,2,0,0,0,3,3,0,0 | 5,6,3,9,3,1,17,9,1,2,7,7,6,6 | 250 | 149 | 0.0255497462146 | | 1 | 0.0,0.0,0.0,0.056,0.37 | 0.193,0.373,0.0,0.0 | -0,002 |
| 215877 | ENSG00000144504 | ANKMY1 | chr2 | - | 240507530 | 240507691 | 240500543 | 240500565 | 240512801 | 240512942 | 215877 | 2,4,0,0,0,6,4,13,2,1,0,2 | 2,4,2,2,2,2,2,2,0,0,2,3 | 5,1,2,2,1,4,0,7,3,7,5,10,7,4 | 5,1,0,3,0,1,2,2,6,0,1,2,4,5,5 | 298 | 149 | 0.0255817271509 | | 1 | 0.333,0.333,0.0,0.0 | 0.333,0.333,1.0,0. | -0,172 |
| 124187 | ENSG00000103479 | RBL2 | chr16 | + | 53447041 | 53447106 | 53442657 | 53442858 | 53451702 | 53451831 | 124187 | 28,16,155,105,77,75,33,47 | 1,1,2,7,2,6,1,6,0,0,2,5 | 88,87,40,17,60,53,34,72,37 | 10,12,0,5,10,3,0,1,6,2,3,5,2 | 214 | 149 | 0.0255835323449 | | 1 | 0.951,0.918,0.982, | 0.86,0.835,1.0,0.7 | 0,026 |
| 191603 | ENSG00000023318 | ERP44 | chr9 | - | 100007563 | 100007689 | 100006505 | 100006647 | 100016321 | 100016438 | 191603 | 23,8,74,47,13,15,16,27,7,3 | 0,0,0,0,0,1,0,1,0,2,0,0 | 33,31,34,26,12,15,8,28,12, | 2,3,4,1,2,0,0,0,0,0,0,1,1,4,2 | 275 | 149 | 0.0256271320073 | | 1 | 1.0,1.0,1.0,1.0,1.0, | 0.899,0.848,0.822, | 0,022 |
| 172826 | ENSG00000186448 | ZNF197 | chr3 | + | 44641899 | 44642217 | 44632472 | 44632599 | 44646490 | 44648471 | 172826 | 29,12,27,17,7,5,8,5,11,9,5, | 3,0,5,3,0,3,0,1,0,0,0,1 | 7,15,5,3,9,6,6,12,6,5,19,12 | 0,0,0,1,2,0,4,7,0,1,2,3,2,2,4 | 298 | 149 | 0.0256437031276 | | 1 | 0.829,1.0,0.73,0.7 | 1.0,1.0,1.0,0.6,0.6 | 0,051 |
| 117705 | ENSG00000167985 | SDHAF2 | chr11 | + | 61438003 | 61438113 | 61430041 | 61430182 | 61445940 | 61446342 | 117705 | 19,16,49,38,13,14,16,15,22 | 3,3,17,7,2,5,6,8,0,0,0,8 | 43,38,13,2,16,12,16,38,4,1 | 1,5,2,0,1,5,2,6,0,3,3,4,4,4,2 | 259 | 149 | 0.0256502081704 | | 1 | 0.785,0.754,0.624, | 0.961,0.814,0.789, | -0,069 |
| 4859 | ENSG00000142453 | CARM1 | chr19 | + | 10908549 | 10908668 | 10908038 | 10908145 | 10909102 | 10909207 | 4859 | 5,1,8,6,1,13,5,0,2,2,0,2 | 3,7,12,24,6,11,3,2,4,12,7,9 | 2,3,3,4,2,3,6,3,0,1,0,2,3,1,3 | 9,7,7,8,7,10,6,22,6,13,10,2 | 268 | 149 | 0.0256618345164 | | 1 | 0.481,0.074,0.27,0 | 0.11,0.192,0.192,0 | 0,078 |
| 109617 | ENSG00000188938 | FAM120AOS | chr9 | - | 93448368 | 93448489 | 93446904 | 93447697 | 93450478 | 93450587 | 109617 | 45,26,60,65,23,29,28,33,51 | 20,10,53,20,14,8,13,17,15, | 27,33,51,51,29,4,21,40,10, | 17,16,37,13,14,11,7,13,11, | 270 | 149 | 0.0256738158943 | | 1 | 0.554,0.589,0.385, | 0.467,0.532,0.432, | 0,106 |
| 33737 | ENSG00000204842 | ATXN2 | chr12 | - | 111464661 | 111464715 | 111457213 | 111457359 | 111470107 | 111470240 | 33737 | 12,12,8,8,6,12,6,7,15,7,6,8 | 1,0,2,0,4,0,0,0,7,2,0,0 | 4,8,5,8,5,5,2,17,2,5,4,9,3,2 | 1,0,2,0,0,1,2,7,1,3,1,0,2,5,3 | 203 | 149 | 0.02569738637 | | 1 | 0.898,1.0,0.746,1. | 0.746,1.0,0.647,1. | 0,152 |
| 59249 | ENSG00000119950 | MXI1 | chr10 | + | 110244827 | 110244857 | 110227642 | 110228321 | 110279179 | 110279294 | 59249 | 93,64,63,29,37,26,44,35,46 | 0,3,4,0,0,0,3,2,3,0,1,2 | 31,24,43,37,26,12,32,57,17 | 2,0,4,2,0,0,2,0,0,0,1,0,0,0,0 | 179 | 149 | 0.0257179970266 | | 1 | 1.0,0.947,0.929,1. | 0.928,1.0,0.899,0. | -0,017 |
| 185825 | ENSG00000055917 | PUM2 | chr2 | - | 20311493 | 20311655 | 20308313 | 20308584 | 20312235 | 20312423 | 185825 | 34,21,111,68,52,62,25,67,6 | 1,2,1,7,0,2,4,2,0,2,4,2 | 88,192,21,90,24,44,81,65,5 | 5,6,1,5,2,0,5,0,0,4,0,4,0,5,0 | 298 | 149 | 0.0257667700669 | | 1 | 0.944,0.84,0.982,0 | 0.898,0.941,0.913, | -0,062 |
| 52360 | ENSG00000154358 | OBSCN | chr1 | + | 228292522 | 228292786 | 228291897 | 228292161 | 228293352 | 228293616 | 52360 | 13,7,5,5,21,17,8,0,6,1,2,9 | 1,2,1,0,2,2,0,2,1,0,3,0 | 22,21,19,23,14,7,8,15,13,1 | 5,7,0,5,1,4,0,4,1,11,0,2,1,2 | 298 | 149 | 0.0257829444312 | | 1 | 0.867,0.636,0.714, | 0.688,0.6,1.0,0.69 | 0,06 |
| 124943 | ENSG00000165629 | ATP5F1C | chr10 | + | 7801512 | 7801569 | 7800051 | 7800091 | 7802269 | 7802425 | 124943 | 4,4,19,9,0,3,5,7,1,0,1,3 | 178,140,228,219,100,94,88 | 9,12,5,0,0,1,5,18,1,4,10,2,5 | 123,114,111,52,72,56,71,1 | 206 | 149 | 0.0258067898271 | | 1 | 0.016,0.02,0.057,0 | 0.05,0.071,0.032,0 | -0,012 |
| 79260 | ENSG00000144909 | OSBPL11 | chr3 | - | 125547405 | 125547592 | 125538450 | 125538633 | 125560378 | 125560519 | 79260 | 73,43,46,32,29,21,13,18,13 | 4,0,1,2,1,0,0,0,2,0,3,0 | 26,49,9,15,26,35,10,12,24, | 1,3,2,2,0,1,0,3,3,2,2,0,2,1,0 | 298 | 149 | 0.0258285482548 | | 1 | 0.901,1.0,0.958,0. | 0.929,0.891,0.692, | 0,033 |
| 79995 | ENSG00000137075 | RNF38 | chr9 | - | 36390466 | 36390616 | 36375933 | 36376127 | 36424612 | 36424683 | 79995 | 92,57,42,39,16,34,33,40,64 | 4,6,6,7,4,1,6,1,12,1,4,0 | 34,60,56,83,34,10,31,25,33 | 2,2,3,1,2,1,0,1,0,10,3,1,7,2 | 298 | 149 | 0.0258594189573 | | 1 | 0.92,0.826,0.778,0 | 0.895,0.938,0.903, | -0,064 |
| 226250 | ENSG00000092820 | EZR | chr6 | - | 158818081 | 158818165 | 158789287 | 158789371 | 158819316 | 158819368 | 226250 | 557,389,731,505,167,318,2 | 18,0,3,1,1,10,17,14,0,3,1,3 | 357,736,359,389,612,198,3 | 11,17,11,15,12,7,0,9,8,3,9, | 233 | 149 | 0.0258629831127 | | 1 | 0.952,1.0,0.994,0. | 0.954,0.965,0.954, | 0,011 |
| 222769 | ENSG00000196821 | ILRUN | chr6 | - | 34688840 | 34688933 | 34654624 | 34654779 | 34696445 | 34696859 | 222769 | 1,5,12,8,14,10,5,3,5,2,0,9 | 132,127,237,159,73,74,58, | 7,6,8,5,6,6,2,0,0,2,2,4,6,1,1 | 91,123,80,47,71,50,20,89,3 | 242 | 149 | 0.0258836826536 | | 1 | 0.005,0.024,0.03,0 | 0.045,0.029,0.058, | 0,009 |
| 4972 | ENSG00000152454 | ZNF256 | chr19 | - | 57943933 | 57944060 | 57940832 | 57942647 | 57947441 | 57947706 | 4972 | 16,19,27,15,18,17,19,24,12 | 2,0,8,5,5,2,9,4,3,4,3,2 | 23,36,10,8,26,14,8,12,7,6,2 | 5,2,4,2,1,2,2,2,0,0,4,1,0,0,3 | 276 | 149 | 0.0258870937954 | | 1 | 0.812,1.0,0.646,0. | 0.713,0.907,0.574, | -0,097 |
| 232847 | ENSG00000083750 | RRAGB | chrX | + | 55755067 | 55755206 | 55753391 | 55753514 | 55755840 | 55755932 | 232847 | 20,18,27,19,3,3,3,12,9,7,2, | 3,1,14,14,2,3,4,2,4,0,2,5 | 17,10,9,6,4,1,5,5,1,4,0,9,9, | 5,2,9,0,3,4,5,8,1,11,2,8,15, | 288 | 149 | 0.0259005089407 | | 1 | 0.775,0.903,0.499, | 0.638,0.721,0.341, | 0,158 |
| 157992 | ENSG00000124767 | GLO1 | chr6 | - | 38688346 | 38688469 | 38686891 | 38686974 | 38702970 | 38703145 | 157992 | 1,0,6,5,0,2,4,9,9,2,0,1 | 32,36,99,41,45,135,128,16 | 2,0,1,2,2,2,3,5,0,1,60,0,0,0 | 102,90,66,31,30,41,72,96,1 | 272 | 149 | 0.0259247398393 | | 1 | 0.017,0.0,0.032,0. | 0.011,0.0,0.008,0. | 0,003 |
| 75036 | ENSG00000143393 | PI4KB | chr1 | - | 151324744 | 151324917 | 151315572 | 151316509 | 151326171 | 151326373 | 75036 | 24,14,44,33,19,15,9,9,20,1 | 64,32,21,22,11,25,12,9,39, | 17,24,6,9,4,14,3,18,8,19,18 | 22,39,27,61,19,8,22,15,17, | 298 | 149 | 0.025925029471 | | 1 | 0.158,0.179,0.512, | 0.279,0.235,0.1,0. | 0,095 |
| 87937 | ENSG00000042493 | CAPG | chr2 | - | 85398689 | 85398782 | 85398019 | 85398152 | 85399135 | 85399285 | 87937 | 113,78,320,492,366,231,12 | 2,0,1,5,6,4,0,1,0,7,1,2 | 389,347,67,22,92,58,77,11 | 2,15,0,2,2,0,4,2,0,2,6,8,1,4 | 242 | 149 | 0.0259263802502 | | 1 | 0.972,1.0,0.995,0. | 0.992,0.934,1.0,0. | 0,015 |
| 248277 | ENSG00000084234 | APLP2 | chr11 | + | 130123611 | 130123690 | 130122304 | 130122513 | 130126699 | 130126830 | 248277 | 21,11,141,56,34,50,31,44,4 | 34,27,141,57,57,80,34,100 | 40,32,45,38,16,15,26,33,6, | 47,41,20,19,24,28,43,29,4, | 228 | 149 | 0.0259291574157 | | 1 | 0.288,0.21,0.395,0 | 0.357,0.338,0.595, | -0,082 |
| 227985 | ENSG00000224137 | LINC01857 | chr2 | + | 207668105 | 207668224 | 207667498 | 207667564 | 207678541 | 207679116 | 227985 | 22,6,7,10,7,4,2,3,2,3,2,4 | 51,22,55,29,22,7,24,4,8,9,1 | 16,2,5,2,2,2,0,1,1,0,0,34,2, | 19,16,9,0,13,2,4,3,11,29,9, | 268 | 149 | 0.0259512986645 | | 1 | 0.193,0.132,0.066, | 0.319,0.065,0.236, | -0,01 |
| 113039 | ENSG00000104529 | EEF1D | chr8 | - | 143590607 | 143590735 | 143589694 | 143590081 | 143592646 | 143592893 | 113039 | 7,8,29,17,4,13,7,6,7,2,5,8 | 7,3,21,13,4,1,1,4,0,0,3,1 | 10,3,6,3,7,6,0,2,4,4,10,16,1 | 4,0,3,8,6,3,9,9,1,4,7,10,1,1 | 277 | 149 | 0.0259563707254 | | 1 | 0.35,0.589,0.426,0 | 0.574,1.0,0.518,0. | 0,184 |
| 4858 | ENSG00000142453 | CARM1 | chr19 | + | 10908549 | 10908633 | 10908038 | 10908145 | 10909102 | 10909207 | 4858 | 4,0,4,4,0,4,4,0,0,2,0,2 | 3,7,12,24,6,11,3,2,4,12,7,9 | 1,1,2,3,1,2,3,0,0,1,0,1,3,1,2 | 9,7,7,8,7,10,6,22,6,13,10,2 | 233 | 149 | 0.0259646818496 | | 1 | 0.46,0.0,0.176,0.0 | 0.066,0.084,0.154, | 0,049 |
| 218186 | ENSG00000090975 | PITPNM2 | chr12 | - | 123000331 | 123000447 | 122997437 | 122997572 | 123000777 | 123000848 | 218186 | 0,0,7,1,4,0,2,1,0,1,1,3 | 20,32,69,34,25,32,19,7,16, | 1,4,2,7,0,1,5,1,2,3,11,0,3,1 | 19,31,12,22,32,15,18,46,17 | 265 | 149 | 0.0259885753445 | | 1 | 0.0,0.0,0.054,0.01 | 0.029,0.068,0.086, | -0,02 |
| 19596 | ENSG00000092439 | TRPM7 | chr15 | - | 50628148 | 50628249 | 50624165 | 50624300 | 50631416 | 50631489 | 19596 | 34,14,110,70,44,53,27,58,1 | 5,0,14,6,3,13,1,3,0,0,0,2 | 58,46,23,25,26,15,33,66,28 | 3,8,8,5,2,2,2,5,6,3,11,9,7,6 | 250 | 149 | 0.0259898146707 | | 1 | 0.802,1.0,0.824,0. | 0.92,0.774,0.631,0 | 0,095 |
| 53800 | ENSG00000137074 | APTX | chr9 | - | 32988082 | 32988129 | 32987543 | 32987731 | 32989758 | 32989895 | 53800 | 21,20,10,6,5,10,5,8,11,5,5, | 2,3,0,1,0,0,0,1,3,0,2,2 | 6,11,3,6,10,4,0,5,4,9,4,5,5, | 0,6,5,1,0,0,3,2,0,1,1,0,2,1,2 | 196 | 149 | 0.0260090540111 | | 1 | 0.889,0.835,1.0,0. | 1.0,0.582,0.313,0. | 0,154 |
| 229532 | ENSG00000179406 | LINC00174 | chr7 | - | 66400321 | 66400441 | 66399691 | 66399795 | 66400922 | 66401027 | 229532 | 8,5,7,7,2,1,0,1,2,1,3,0 | 2,0,10,2,0,0,1,4,4,0,2,4 | 2,5,0,0,0,1,3,0,3,2,0,2,3,1,1 | 6,7,1,2,3,1,4,6,3,0,2,2,0,2,1 | 269 | 149 | 0.0260203563896 | | 1 | 0.689,1.0,0.279,0. | 0.156,0.283,0.0,0. | 0,24 |
| 126760 | ENSG00000138185 | ENTPD1 | chr10 | + | 95842343 | 95842494 | 95839690 | 95839808 | 95844475 | 95844635 | 126760 | 145,98,99,142,74,28,30,33 | 11,4,1,4,2,4,3,4,7,1,5,7 | 109,113,178,100,62,71,45, | 8,10,7,4,13,5,1,4,13,8,8,21 | 298 | 149 | 0.0260319565944 | | 1 | 0.868,0.925,0.98,0 | 0.872,0.85,0.927,0 | 0,017 |
| 13687 | ENSG00000289733 | | chr16 | + | 81931496 | 81931654 | 81928557 | 81928624 | 81936168 | 81936378 | 13687 | 89,85,133,155,72,46,59,48 | 3,1,5,3,3,7,2,1,0,0,1,1 | 87,105,37,62,63,47,46,37,4 | 6,7,0,3,1,0,0,1,2,0,4,16,0,0 | 298 | 149 | 0.0260532081417 | | 1 | 0.937,0.977,0.93,0 | 0.879,0.882,1.0,0. | 0,007 |
| 81649 | ENSG00000151779 | NBAS | chr2 | - | 15536417 | 15536551 | 15534542 | 15534641 | 15539222 | 15539356 | 81649 | 42,13,85,31,14,31,37,47,23 | 2,0,0,1,2,0,0,3,0,0,7,0 | 99,65,20,19,23,22,37,65,25 | 7,5,0,6,3,2,4,2,1,0,0,2,1,0,1 | 283 | 149 | 0.0260714111426 | | 1 | 0.917,1.0,1.0,0.94 | 0.882,0.873,1.0,0. | 0,02 |
| 180559 | ENSG00000138668 | HNRNPD | chr4 | - | 82356536 | 82356686 | 82353846 | 82355401 | 82356795 | 82356895 | 180559 | 16,16,51,27,35,30,30,43,7, | 25,14,68,35,24,50,30,44,15 | 65,89,23,96,53,25,35,35,64 | 50,99,19,62,40,27,45,34,34 | 298 | 149 | 0.0260779154934 | | 1 | 0.242,0.364,0.273, | 0.394,0.31,0.377,0 | -0,059 |
| 158544 | ENSG00000127990 | SGCE | chr7 | - | 94621612 | 94621995 | 94618807 | 94618956 | 94623324 | 94623397 | 158544 | 4,0,7,4,0,2,2,6,4,0,0,0 | 15,19,54,26,9,21,18,31,20, | 2,0,2,2,0,0,2,1,0,1,5,0,2,1,1 | 7,17,13,4,4,7,11,21,5,9,13, | 298 | 149 | 0.0260779741041 | | 1 | 0.118,0.0,0.061,0. | 0.125,0.0,0.071,0. | -0,012 |
| 251670 | ENSG00000134452 | FBH1 | chr10 | + | 5913783 | 5913839 | 5910937 | 5911128 | 5914177 | 5914269 | 251670 | 81,35,116,58,37,51,35,82,4 | 2,1,4,6,1,1,2,6,0,0,0,0 | 75,54,68,64,26,28,27,87,30 | 3,0,1,0,5,0,0,0,1,0,0,1,1,6,0 | 205 | 149 | 0.0260855552348 | | 1 | 0.967,0.962,0.955, | 0.948,1.0,0.98,1.0, | -0,004 |
| 39920 | ENSG00000168827 | GFM1 | chr3 | + | 158690162 | 158690323 | 158681994 | 158682157 | 158691138 | 158691192 | 39920 | 14,13,22,20,30,18,3,4,5,12 | 3,4,9,8,2,4,2,4,5,7,7,2 | 11,7,11,1,5,10,10,5,4,6,11, | 10,11,10,7,3,0,7,2,7,2,5,5,9 | 298 | 149 | 0.0261251896648 | | 1 | 0.7,0.619,0.55,0.5 | 0.355,0.241,0.355, | 0,113 |
| 254091 | ENSG00000139428 | MMAB | chr12 | - | 109568254 | 109568393 | 109565118 | 109565176 | 109568769 | 109568863 | 254091 | 2,2,1,0,1,0,0,3,0,2,0,1 | 5,2,22,14,15,2,6,5,6,0,6,3 | 3,1,2,2,2,0,3,5,0,0,2,3,1,0,3 | 7,2,12,2,0,3,2,7,4,1,5,12,6, | 288 | 149 | 0.0261283871217 | | 1 | 0.171,0.341,0.023, | 0.181,0.206,0.079, | -0,055 |
| 169898 | ENSG00000168026 | TTC21A | chr3 | + | 39136906 | 39137060 | 39136356 | 39136507 | 39137194 | 39137253 | 169898 | 11,11,23,16,2,11,4,13,6,1,3 | 3,0,2,0,0,0,0,1,0,0,0,0 | 8,7,10,7,1,5,10,5,4,19,10,2 | 0,1,0,1,1,0,0,1,2,0,1,2,1,6,0 | 298 | 149 | 0.0261591276431 | | 1 | 0.647,1.0,0.852,1. | 1.0,0.778,1.0,0.77 | 0,207 |
| 247848 | ENSG00000179532 | DNHD1 | chr11 | + | 6533896 | 6534173 | 6533680 | 6533806 | 6538382 | 6538510 | 247848 | 5,7,27,12,12,14,14,25,7,1,7 | 2,1,0,3,2,2,0,0,0,0,2,2 | 12,20,28,22,13,15,7,21,6,1 | 1,0,0,0,0,1,1,2,0,0,0,2,0,1,1 | 298 | 149 | 0.0261910070364 | | 1 | 0.556,0.778,1.0,0. | 0.857,1.0,1.0,1.0,1 | -0,124 |
| 6597 | ENSG00000087460 | GNAS | chr20 | + | 58898940 | 58899162 | 58895611 | 58895684 | 58903530 | 58903585 | 6597 | 38,27,65,43,15,17,22,60,32 | 127,64,176,105,79,81,70,9 | 26,27,21,41,21,13,14,23,19 | 110,114,73,94,79,48,60,66 | 298 | 149 | 0.0261923457753 | | 1 | 0.13,0.174,0.156,0 | 0.106,0.106,0.126, | 0,022 |
| 235852 | ENSG00000174197 | MGA | chr15 | + | 41743055 | 41743172 | 41740056 | 41740203 | 41748636 | 41748927 | 235852 | 11,6,21,14,13,9,14,12,9,0,2 | 13,5,11,16,6,15,8,2,8,11,3, | 12,8,5,4,4,5,6,12,4,4,10,3,5 | 10,29,9,4,15,7,8,6,9,7,4,6,5 | 266 | 149 | 0.0262652898228 | | 1 | 0.322,0.402,0.517, | 0.402,0.134,0.237, | 0,126 |
| 163541 | ENSG00000059378 | PARP12 | chr7 | - | 140033974 | 140034081 | 140028612 | 140028688 | 140034234 | 140034331 | 163541 | 4,2,7,3,2,0,4,7,3,1,1,6 | 40,31,50,42,29,34,22,28,52 | 10,4,4,17,2,0,0,5,7,0,6,0,5, | 29,17,23,19,24,17,9,54,13, | 256 | 149 | 0.0262765045257 | | 1 | 0.055,0.036,0.075, | 0.167,0.12,0.092,0 | -0,03 |
| 254071 | ENSG00000289963 | | chr22 | + | 39068256 | 39068309 | 39066279 | 39066358 | 39068708 | 39069975 | 254071 | 0,0,2,1,3,2,1,3,1,1,0,2 | 7,1,12,9,6,3,7,6,7,3,3,2 | 0,2,0,0,1,1,0,2,0,2,0,2,2,3,2 | 18,5,6,1,0,10,6,12,1,1,8,14 | 202 | 149 | 0.0263461513531 | | 1 | 0.0,0.0,0.109,0.07 | 0.0,0.228,0.0,0.0,1 | -0,014 |
| 99036 | ENSG00000136280 | CCM2 | chr7 | + | 45064462 | 45064646 | 45027025 | 45027797 | 45072725 | 45072783 | 99036 | 9,2,10,6,2,4,5,11,9,6,4,2 | 0,0,3,1,0,1,1,1,1,2,3,1 | 4,7,4,2,2,0,1,1,2,4,9,5,3,5,4 | 1,1,0,0,3,2,1,2,0,0,1,0,0,4,2 | 298 | 149 | 0.0263492947518 | | 1 | 1.0,1.0,0.625,0.75, | 0.667,0.778,1.0,1. | 0,06 |
| 228660 | ENSG00000146757 | ZNF92 | chr7 | + | 65387901 | 65388254 | 65373798 | 65374000 | 65398340 | 65401136 | 228660 | 46,55,308,197,50,46,66,94 | 6,4,28,23,0,2,2,17,5,0,2,2 | 186,146,71,94,131,128,28, | 13,20,12,2,22,18,6,4,4,23,2 | 298 | 149 | 0.0263566494876 | | 1 | 0.793,0.873,0.846, | 0.877,0.785,0.747, | 0,069 |
| 169552 | ENSG00000163681 | SLMAP | chr3 | + | 57896219 | 57896371 | 57890047 | 57890100 | 57896510 | 57896591 | 169552 | 4,2,8,5,3,11,5,4,2,0,1,5 | 13,5,15,16,11,4,3,2,8,1,4,3 | 9,10,3,4,4,3,6,4,6,3,9,4,6,2 | 12,0,0,0,6,4,5,4,2,11,6,8,8, | 298 | 149 | 0.0263599799411 | | 1 | 0.133,0.167,0.211, | 0.273,1.0,1.0,1.0,0 | -0,203 |
| 50782 | ENSG00000151092 | NGLY1 | chr3 | - | 25737333 | 25737455 | 25736308 | 25736400 | 25739576 | 25739799 | 50782 | 13,11,56,39,34,9,17,21,21, | 2,0,2,0,0,0,0,1,0,0,0,1 | 27,13,17,4,3,30,12,12,12,3 | 2,2,0,0,1,0,2,4,0,0,3,1,0,2,2 | 271 | 149 | 0.0263723770804 | | 1 | 0.781,1.0,0.939,1. | 0.881,0.781,1.0,1. | 0,095 |
| 112055 | ENSG00000147601 | TERF1 | chr8 | + | 73038705 | 73038795 | 73032041 | 73032133 | 73039115 | 73039219 | 112055 | 14,10,19,0,0,6,3,10,4,4,5,2 | 18,11,26,20,11,18,8,14,13, | 9,10,14,17,6,2,7,9,0,2,16,2 | 10,14,4,10,8,10,8,4,5,9,8,2 | 239 | 149 | 0.0263945509803 | | 1 | 0.327,0.362,0.313, | 0.359,0.308,0.686, | -0,113 |
| 231733 | ENSG00000139436 | GIT2 | chr12 | - | 109959846 | 109959953 | 109952679 | 109953234 | 109961277 | 109961375 | 231733 | 143,115,130,94,44,55,60,5 | 1,9,5,2,2,0,0,0,3,0,0,0 | 74,94,55,35,67,34,35,49,50 | 3,0,0,2,2,1,0,0,0,2,0,4,0,1,0 | 256 | 149 | 0.0264072445789 | | 1 | 0.988,0.881,0.938, | 0.935,1.0,1.0,0.91 | -0,001 |

| | | | | | | | | | | | | | | | | | | | | | | |
|---|---|---|---|---|---|---|---|---|---|---|---|---|---|---|---|---|---|---|---|---|---|---|
| 245781 | ENSG00000110075 | PPP6R3 | chr11 | + | 68522522 | 68522689 | 68519634 | 68519651 | 68537658 | 68537669 | 245781 | 11,6,12,3,0,5,6,9,5,1,2,7 | 93,68,141,77,85,99,58,111 | 11,10,11,20,9,1,7,9,2,4,9,2 | 78,107,77,62,57,34,58,78, | 298 | 149 | 0.0264142510406 | | 1 | 0.056,0.042,0.041, | 0.066,0.045,0.067, | -0,013 |
| 234768 | ENSG00000102054 | RBBP7 | chrX | - | 16852550 | 16852621 | 16852045 | 16852122 | 16852748 | 16852970 | 234768 | 33,19,83,52,22,20,12,18,2 | 2,4,5,2,0,0,0,0,0,4,0,0 | 59,34,13,9,37,24,18,26,4,2 | 1,5,0,12,5,2,0,4,0,0,0,1,0,0 | 220 | 149 | 0.0264345985893 | | 1 | 0.918,0.763,0.918, | 0.976,0.822,1.0,0. | 0,029 |
| 57035 | ENSG00000077097 | TOP2B | chr3 | - | 25618417 | 25618509 | 25615430 | 25615586 | 25618653 | 25618849 | 57035 | 126,78,191,124,87,79,61,1 | 3,1,4,1,0,3,0,1,1,2,1,3 | 92,86,58,40,60,59,54,54,3 | 1,9,3,2,7,2,1,2,0,2,0,3,1,3, | 241 | 149 | 0.0264401004442 | | 1 | 0.963,0.98,0.967,0 | 0.983,0.855,0.923, | 0,017 |
| 199960 | ENSG00000151883 | PARP8 | chr5 | + | 50821300 | 50821338 | 50815431 | 50815524 | 50822334 | 50822400 | 199960 | 0,2,7,5,4,1,1,5,0,0,0,0 | 5,0,8,6,2,2,0,0,1,2,2,4 | 4,4,4,4,0,0,2,2,1,6,6,5,8,5, | 2,1,5,15,4,2,1,5,2,0,5,0,3,0 | 187 | 149 | 0.026451939996 | | 1 | 0.0,1.0,0.411,0.39 | 0.614,0.761,0.389, | -0,092 |
| 102087 | ENSG00000160218 | TRAPPC10 | chr21 | + | 44063537 | 44063785 | 44059102 | 44059214 | 44074323 | 44074470 | 102087 | 95,94,154,127,83,60,78,87 | 7,0,5,3,3,5,2,3,0,0,5,2 | 109,119,72,98,102,44,60,9 | 2,0,2,2,5,1,2,4,2,0,5,0,1,2, | 298 | 149 | 0.0264569754636 | | 1 | 0.872,1.0,0.939,0. | 0.965,1.0,0.947,0. | -0,024 |
| 42017 | ENSG00000081307 | UBA5 | chr3 | + | 132675247 | 132675383 | 132671776 | 132672177 | 132675604 | 132675680 | 42017 | 51,32,67,27,20,24,17,25,1 | 7,2,14,5,5,0,0,3,1,1,0,2 | 38,15,16,29,15,26,14,14,7, | 4,10,4,5,6,1,5,3,3,2,3,4,3,2 | 285 | 149 | 0.0264928554825 | | 1 | 0.792,0.893,0.714, | 0.832,0.44,0.677,0 | 0,146 |
| 11964 | ENSG00000215041 | NEURL4 | chr17 | - | 7327164 | 7327230 | 7326710 | 7327009 | 7327439 | 7327500 | 11964 | 24,6,11,20,16,17,10,4,18,4 | 2,2,4,2,1,3,2,3,4,3,3,8 | 8,8,6,3,5,2,11,12,5,22,9,11 | 0,6,0,4,2,1,2,1,2,1,1,0,1,1, | 215 | 149 | 0.0265099425298 | | 1 | 0.893,0.675,0.656, | 1.0,0.48,1.0,0.342, | -0,061 |
| 200301 | ENSG00000172071 | EIF2AK3 | chr2 | - | 88593912 | 88593996 | 88593271 | 88593405 | 88595468 | 88595663 | 200301 | 0,2,4,3,1,1,3,0,2,0,1,0 | 18,7,36,30,6,23,7,16,4,5,7, | 3,4,2,7,2,0,4,2,2,2,2,5,1,1, | 12,45,14,17,28,1,16,18,8,4 | 233 | 149 | 0.0265655871996 | | 1 | 0.0,0.154,0.066,0. | 0.138,0.054,0.084, | -0,013 |
| 218932 | ENSG00000204592 | HLA-E | chr6 | + | 30491984 | 30492436 | 30491536 | 30491653 | 30492540 | 30492583 | 218932 | 197,132,346,291,101,78,9 | 10,5,8,15,4,0,5,9,0,3,2,9 | 237,266,142,146,116,108, | 10,13,4,14,16,5,2,21,5,6,8, | 298 | 149 | 0.026565921242 | | 1 | 0.908,0.93,0.956,0 | 0.922,0.911,0.947, | 0,027 |
| 184568 | ENSG00000198265 | HELZ | chr17 | - | 67135969 | 67136165 | 67128494 | 67128855 | 67137930 | 67138114 | 184568 | 32,6,39,17,27,16,11,8,17,1 | 2,1,9,2,5,14,1,8,9,0,2,1 | 18,16,12,13,4,6,9,8,3,23,2 | 5,6,4,9,10,3,3,7,3,2,10,7,5, | 298 | 149 | 0.0266359749958 | | 1 | 0.889,0.75,0.684,0 | 0.643,0.571,0.6,0. | 0,167 |
| 55097 | ENSG00000106462 | EZH2 | chr7 | - | 148818977 | 148819059 | 148816683 | 148818117 | 148819595 | 148819687 | 55097 | 9,2,9,5,1,4,0,2,3,0,7,6 | 11,16,7,13,2,8,10,3,6,6,2,1 | 0,5,0,3,3,0,1,5,0,0,1,9,2,0, | 3,8,1,6,5,2,1,4,4,5,0,13,2,8 | 231 | 149 | 0.0266363530993 | | 1 | 0.345,0.075,0.453, | 0.0,0.287,0.0,0.24 | 0,07 |
| 240557 | ENSG00000198000 | NOL8 | chr9 | - | 92300501 | 92300806 | 92299889 | 92300016 | 92301550 | 92301822 | 240557 | 0,0,2,3,1,4,2,0,0,0,0,0 | 11,9,31,39,14,12,6,20,10,6 | 4,4,4,5,1,0,0,0,0,0,2,8,0,0, | 11,20,30,17,18,8,6,13,21,1 | 298 | 149 | 0.0266377231872 | | 1 | 0.0,0.0,0.031,0.03 | 0.154,0.091,0.062, | -0,012 |
| 104400 | ENSG00000119321 | FKBP15 | chr9 | - | 113211476 | 113211592 | 113206926 | 113207296 | 113221190 | 113221275 | 104400 | 56,18,43,21,20,20,22,24,2 | 7,0,3,0,1,3,2,1,0,0,0,0 | 25,6,32,15,6,9,14,18,14,44 | 0,1,2,0,0,0,0,0,0,0,0,1,0,0, | 265 | 149 | 0.0266609634881 | | 1 | 0.818,1.0,0.89,1.0, | 1.0,0.771,0.9,1.0,1 | -0,026 |
| 126113 | ENSG00000197535 | MYO5A | chr15 | - | 52359967 | 52360069 | 52353870 | 52354014 | 52364553 | 52364702 | 126113 | 55,51,59,39,25,22,19,19,2 | 7,0,0,0,1,0,1,1,0,0,0,0 | 49,62,31,49,43,18,31,17,1 | 1,3,0,3,2,2,1,1,1,0,0,4,2,1, | 251 | 149 | 0.0266628117684 | | 1 | 0.823,1.0,1.0,1.0,0 | 0.967,0.925,1.0,0. | 0,04 |
| 41529 | ENSG00000136141 | LRCH1 | chr13 | + | 46715559 | 46715664 | 46712524 | 46712597 | 46723220 | 46723330 | 41529 | 1,1,7,6,0,3,1,4,0,0,2,1 | 5,7,20,20,8,9,11,18,11,7,4, | 1,1,0,0,1,1,2,4,3,1,3,2,2,0, | 13,19,3,11,12,13,12,15,11, | 254 | 149 | 0.0266767910278 | | 1 | 0.105,0.077,0.17,0 | 0.043,0.03,0.0,0.0, | 0,028 |
| 235919 | ENSG00000176731 | RBIS | chr8 | - | 85219322 | 85219377 | 85217385 | 85217502 | 85220305 | 85220421 | 235919 | 15,8,6,10,8,3,5,3,4,1,4,3 | 56,25,52,55,27,22,24,13,2 | 2,6,9,4,2,0,1,4,1,4,3,0,1,3, | 37,16,41,7,25,22,5,6,13,43 | 204 | 149 | 0.0266852047899 | | 1 | 0.164,0.189,0.078, | 0.038,0.215,0.138, | 0,059 |
| 134447 | ENSG00000100299 | ARSA | chr22 | - | 50625578 | 50625798 | 50622753 | 50625464 | 50625935 | 50626063 | 134447 | 24,17,43,31,16,13,14,12,7, | 0,0,0,0,1,0,0,1,0,0,0,0 | 23,15,12,8,14,8,3,23,14,12 | 0,2,6,2,6,2,0,0,0,0,0,1,0,0, | 298 | 149 | 0.0266858318606 | | 1 | 1.0,1.0,1.0,1.0,0.8 | 1.0,0.789,0.5,0.66 | 0,097 |
| 58083 | ENSG00000122483 | CCDC18 | chr1 | + | 93183977 | 93184146 | 93183359 | 93183495 | 93186344 | 93186398 | 58083 | 25,17,25,26,13,16,5,25,15, | 2,3,3,6,0,4,2,2,2,0,0,0 | 17,26,10,13,12,16,9,21,17, | 2,1,0,2,4,2,1,4,1,0,1,1,5,2,1 | 298 | 149 | 0.026687280639 | | 1 | 0.862,0.739,0.806, | 0.81,0.929,1.0,0.7 | -0,003 |
| 27385 | ENSG00000065491 | TBC1D22B | chr6 | + | 37282184 | 37282364 | 37279303 | 37279611 | 37282881 | 37282952 | 27385 | 23,36,29,25,17,19,7,9,24,1 | 0,1,5,4,0,2,0,1,0,1,0,2 | 18,33,6,10,11,4,8,30,9,14,1 | 1,1,0,0,4,0,2,0,0,0,1,0,0,0,1 | 298 | 149 | 0.0267464900887 | | 1 | 1.0,0.947,0.744,0. | 0.9,0.943,1.0,1.0,0 | -0,028 |
| 132570 | ENSG00000112200 | ZNF451 | chr6 | + | 57090810 | 57090894 | 57090029 | 57090274 | 57099060 | 57099141 | 132570 | 16,16,39,21,12,29,17,26,9, | 6,7,17,8,1,8,8,12,1,5,1,3 | 20,18,15,14,13,6,12,18,10, | 15,22,6,21,8,6,7,10,3,6,14, | 233 | 149 | 0.0267904824268 | | 1 | 0.63,0.594,0.595,0 | 0.46,0.343,0.615,0 | 0,127 |
| 228516 | ENSG00000105778 | AVL9 | chr7 | + | 32562557 | 32562913 | 32558928 | 32559464 | 32570019 | 32570154 | 228516 | 3,4,17,17,1,4,3,9,4,7,0,5 | 26,6,15,13,18,26,23,20,14, | 1,8,2,13,11,2,12,5,1,0,10,1 | 23,29,25,3,15,8,14,18,10,3 | 298 | 149 | 0.0267956803934 | | 1 | 0.055,0.25,0.362,0 | 0.021,0.121,0.038, | 0,038 |
| 115919 | ENSG00000198771 | RCSD1 | chr1 | + | 167640570 | 167640712 | 167630248 | 167630429 | 167683899 | 167684001 | 115919 | 3,3,5,3,5,4,6,15,6,2,3,14 | 91,38,133,49,79,69,170,22 | 1,0,8,6,0,1,3,5,1,0,2,0,3,0,1 | 41,28,78,48,20,20,89,160,1 | 291 | 149 | 0.0268041983609 | | 1 | 0.017,0.039,0.019, | 0.012,0.0,0.05,0.0 | 0,008 |
| 181322 | ENSG00000162739 | SLAMF6 | chr1 | - | 160496060 | 160496303 | 160491124 | 160491388 | 160523143 | 160523262 | 181322 | 72,25,186,203,139,116,71, | 2,0,16,8,6,4,4,7,3,3,6,11 | 235,75,61,48,39,59,40,77,2 | 7,3,0,0,3,2,1,8,2,4,7,2,9,4,3 | 298 | 149 | 0.0268065693289 | | 1 | 0.947,1.0,0.853,0. | 0.944,0.926,1.0,1. | -0,019 |
| 152899 | ENSG00000257621 | PSMA3-AS1 | chr14 | - | 58286639 | 58286827 | 58285585 | 58285756 | 58291634 | 58291707 | 152899 | 22,6,8,2,10,6,18,11,22,7,4, | 32,7,29,26,18,15,16,51,12, | 23,5,7,8,6,5,4,22,8,13,7,11 | 19,19,11,13,10,4,11,15,3,8 | 298 | 149 | 0.026812476057 | | 1 | 0.256,0.3,0.121,0. | 0.377,0.116,0.241, | -0,091 |
| 195745 | ENSG00000159377 | PSMB4 | chr1 | + | 151400763 | 151401120 | 151400441 | 151400588 | 151401238 | 151401630 | 195745 | 55,32,119,165,71,55,47,49 | 3,2,4,9,7,5,1,0,0,0,2,2 | 95,77,55,23,62,43,28,38,2 | 3,9,2,15,1,1,4,9,1,0,8,8,5,2 | 298 | 149 | 0.0268187007685 | | 1 | 0.902,0.889,0.937, | 0.941,0.811,0.932, | 0,069 |
| 205896 | ENSG00000128815 | WDFY4 | chr10 | + | 48817254 | 48817409 | 48813956 | 48814082 | 48820233 | 48820437 | 205896 | 86,71,98,70,98,85,53,66,1 | 0,1,2,2,1,6,0,1,0,0,1,4 | 81,80,21,53,19,31,71,85,4 | 2,9,0,5,0,0,1,2,1,0,6,3,0,1, | 298 | 149 | 0.0268248664002 | | 1 | 1.0,0.973,0.961,0. | 0.953,0.816,1.0,0. | 0,016 |
| 14686 | ENSG00000102984 | ZNF821 | chr16 | - | 71867911 | 71868037 | 71864898 | 71865048 | 71879906 | 71880023 | 14686 | 22,11,54,31,21,8,18,12,16, | 7,1,2,1,0,0,0,2,0,0,0,3 | 50,25,13,7,16,5,7,10,12,24 | 0,0,0,3,1,0,1,0,0,1,1,0,0,1, | 275 | 149 | 0.0268303157352 | | 1 | 0.63,0.856,0.936,0 | 1.0,1.0,1.0,0.558,0 | -0,019 |
| 210638 | ENSG00000134262 | AP4B1 | chr1 | - | 113899083 | 113899209 | 113898717 | 113898801 | 113899903 | 113900400 | 210638 | 1,1,6,0,1,0,4,7,0,0,1,0 | 19,10,45,23,17,14,24,13,1 | 0,1,0,1,2,0,2,0,0,1,0,2,0,0, | 23,22,7,11,25,15,4,10,11,1 | 275 | 149 | 0.0268541856398 | | 1 | 0.028,0.051,0.067, | 0.0,0.024,0.0,0.04 | 0,02 |
| 233225 | ENSG00000046651 | OFD1 | chrX | + | 13752703 | 13753104 | 13751248 | 13751368 | 13753367 | 13755242 | 233225 | 12,16,81,18,10,43,7,25,21, | 74,45,143,58,39,47,23,41, | 15,12,36,21,19,9,9,20,21,2 | 53,39,34,35,26,14,9,33,39, | 298 | 149 | 0.0268991074827 | | 1 | 0.075,0.151,0.221, | 0.124,0.133,0.346, | -0,075 |
| 249497 | ENSG00000181192 | DHTKD1 | chr10 | + | 12094072 | 12094256 | 12091512 | 12091684 | 12097683 | 12097996 | 249497 | 17,7,34,41,22,30,10,39,18, | 4,0,3,1,0,0,2,3,2,4,0,0 | 37,15,18,9,14,17,8,21,4,17 | 0,0,1,2,0,1,0,5,0,1,1,0,2,2, | 298 | 149 | 0.0269007030188 | | 1 | 0.68,1.0,0.85,0.95 | 1.0,1.0,0.9,0.692,1 | -0,017 |
| 225690 | ENSG00000136280 | CCM2 | chr7 | + | 45071753 | 45071872 | 45069825 | 45069961 | 45072725 | 45072783 | 225690 | 7,0,4,4,0,0,0,5,1,0,0,0 | 21,31,29,51,12,9,19,20,11, | 6,2,2,3,4,3,3,3,2,4,1,5,2,1,1 | 37,26,7,6,19,25,6,17,5,30,5 | 268 | 149 | 0.0269275912857 | | 1 | 0.156,0.0,0.071,0. | 0.083,0.041,0.137, | -0,056 |
| 147571 | ENSG00000172775 | PSME3IP1 | chr16 | - | 57181553 | 57181651 | 57178525 | 57178852 | 57185820 | 57186055 | 147571 | 11,9,25,16,9,8,9,11,5,2,3,7 | 1,0,3,1,0,5,6,1,0,0,0,10 | 5,4,5,9,15,4,13,11,14,8,12, | 1,0,0,0,3,2,1,4,5,4,3,4,5,8, | 247 | 149 | 0.0269499159507 | | 1 | 0.869,1.0,0.834,0. | 0.751,1.0,1.0,1.0,0 | 0,135 |
| 105097 | ENSG00000131196 | NFATC1 | chr18 | + | 79486247 | 79486630 | 79467449 | 79467582 | 79527527 | 79529323 | 105097 | 43,24,31,25,28,30,34,16,3 | 1,2,2,0,0,0,1,3,3,0,1,2 | 12,14,25,20,16,15,14,28,3, | 1,0,1,0,4,0,2,0,1,2,2,3,0,2, | 298 | 149 | 0.0269522076408 | | 1 | 0.956,0.857,0.886, | 0.857,1.0,0.926,1. | 0,045 |
| 181614 | ENSG00000134255 | CEPT1 | chr1 | + | 111161099 | 111161296 | 111159379 | 111159527 | 111174878 | 111174963 | 181614 | 10,12,41,31,30,8,4,5,17,5, | 0,0,2,4,0,3,0,2,2,0,0,2 | 26,6,10,3,3,13,2,9,2,19,5,3 | 3,0,0,0,0,1,2,0,2,0,0,4,1,3, | 298 | 149 | 0.0269690345915 | | 1 | 1.0,1.0,0.911,0.79 | 0.812,1.0,1.0,1.0,1 | 0,036 |
| 253782 | ENSG00000123329 | ARHGAP9 | chr12 | - | 57474874 | 57474968 | 57474625 | 57474703 | 57475290 | 57475411 | 253782 | 68,35,68,54,39,21,26,19,3 | 2,1,3,3,0,3,0,0,0,0,2,0 | 52,32,42,14,18,28,15,6,23, | 6,3,0,2,4,0,0,4,1,0,0,0,1,1, | 243 | 149 | 0.0269722146338 | | 1 | 0.954,0.955,0.933, | 0.842,0.867,1.0,0. | 0,052 |
| 102050 | ENSG00000146757 | ZNF92 | chr7 | + | 65387901 | 65388028 | 65373798 | 65374000 | 65398340 | 65401135 | 102050 | 57,58,325,205,54,43,82,10 | 6,4,28,23,0,2,2,17,5,0,2,2 | 201,154,76,109,150,143,2 | 13,20,12,2,22,18,6,4,4,23, | 276 | 149 | 0.0269804478671 | | 1 | 0.837,0.887,0.862, | 0.893,0.806,0.774, | 0,061 |
| 189722 | ENSG00000115756 | HPCAL1 | chr2 | + | 10408558 | 10408734 | 10396834 | 10396920 | 10419733 | 10420135 | 189722 | 2,2,7,13,2,1,7,4,9,4,4,4 | 19,14,34,16,20,26,37,12,1 | 5,4,1,0,0,0,7,5,0,0,1,2,6,0, | 22,20,6,5,10,15,20,19,3,17 | 298 | 149 | 0.0269932246055 | | 1 | 0.05,0.067,0.093,0 | 0.102,0.091,0.077, | 0,054 |
| 185896 | ENSG00000010322 | NISCH | chr3 | + | 52476432 | 52476599 | 52473733 | 52473829 | 52477573 | 52477642 | 185896 | 69,31,78,43,37,43,32,40,3 | 0,0,0,1,0,2,0,0,2,0,0,0 | 47,67,31,33,46,19,27,65,3 | 2,0,5,2,1,0,0,1,1,4,4,0,0,2,1 | 298 | 149 | 0.0269996603316 | | 1 | 1.0,1.0,1.0,0.956,1 | 0.922,1.0,0.756,0. | 0,046 |
| 97914 | ENSG00000054523 | KIF1B | chr1 | + | 10337370 | 10337533 | 10337073 | 10337203 | 10339768 | 10339859 | 97914 | 8,8,12,7,7,4,7,1,5,0,3,3 | 2,1,2,0,0,2,1,2,0,1,0,1 | 21,2,7,3,1,4,1,0,9,10,2,5,1 | 0,6,1,4,1,1,1,6,0,0,1,0,0,2, | 298 | 149 | 0.0270260736645 | | 1 | 0.667,0.8,0.75,1.0, | 1.0,0.143,0.778,0. | 0,036 |
| 73508 | ENSG00000154222 | CC2D1B | chr1 | - | 52359704 | 52359836 | 52359458 | 52359534 | 52360073 | 52360233 | 73508 | 20,11,41,21,13,23,18,19,3 | 0,2,2,0,2,10,2,6,1,0,0,3 | 30,30,6,11,19,8,17,30,8,14 | 4,4,2,5,4,1,2,5,1,0,6,2,3,1, | 281 | 149 | 0.027044924136 | | 1 | 1.0,0.745,0.916,1. | 0.799,0.799,0.614, | 0,096 |
| 79122 | ENSG00000130348 | QRSL1 | chr6 | + | 106654729 | 106654922 | 106652466 | 106652582 | 106655614 | 106655732 | 79122 | 54,42,209,225,99,62,45,71 | 4,8,10,11,8,5,2,2,1,2,2,0 | 103,74,55,31,34,86,21,68, | 5,2,5,2,0,0,0,2,4,11,3,0,0,0 | 298 | 149 | 0.0270739573585 | | 1 | 0.871,0.724,0.913, | 0.912,0.949,0.846, | -0,027 |
| 168525 | ENSG00000138814 | PPP3CA | chr4 | - | 101183217 | 101183271 | 101108953 | 101109078 | 101195915 | 101196116 | 168525 | 9,1,4,3,2,2,3,3,0,1,0,0 | 114,64,68,62,42,29,31,50, | 7,5,2,2,8,1,5,2,2,1,1,9,0,1, | 47,31,48,47,23,18,18,40,1 | 203 | 149 | 0.0270758672498 | | 1 | 0.055,0.011,0.041, | 0.099,0.106,0.03,0 | -0,032 |
| 248279 | ENSG00000084234 | APLP2 | chr11 | + | 130123625 | 130123779 | 130122304 | 130122513 | 130126699 | 130126830 | 248279 | 23,26,190,106,66,68,54,94 | 34,27,141,57,57,80,34,100 | 57,52,54,38,32,29,35,59,8, | 47,41,20,19,24,28,43,29,4, | 298 | 149 | 0.0270771104622 | | 1 | 0.253,0.325,0.403, | 0.377,0.388,0.574, | -0,073 |
| 171035 | ENSG00000111224 | PARP11 | chr12 | - | 3863835 | 3864223 | 3829889 | 3830018 | 3873211 | 3873448 | 171035 | 6,4,18,10,9,12,4,6,1,1,3,3 | 12,11,13,3,9,11,5,6,5,2,2,6 | 6,5,9,10,6,3,5,4,8,14,8,4,1 | 4,6,5,8,6,5,5,5,4,1,5,1,8,7, | 298 | 149 | 0.0270818961319 | | 1 | 0.2,0.154,0.409,0. | 0.429,0.294,0.474, | -0,143 |
| 229393 | ENSG00000162600 | OMA1 | chr1 | - | 58530600 | 58530725 | 58527260 | 58527335 | 58533952 | 58534060 | 229393 | 70,39,72,47,52,62,29,30,4 | 3,0,6,1,19,6,7,12,1,1,5,9 | 135,76,58,41,42,38,39,56, | 35,20,8,4,3,8,3,17,3,5,25,5 | 274 | 149 | 0.0271259296645 | | 1 | 0.927,1.0,0.867,0. | 0.677,0.674,0.798, | 0,057 |
| 89581 | ENSG00000104907 | TRMT1 | chr19 | - | 13109914 | 13110001 | 13109768 | 13109838 | 13110157 | 13110306 | 89581 | 6,4,23,11,10,14,19,24,11,7 | 3,3,6,2,8,20,8,10,5,6,3,10 | 18,21,7,2,2,5,4,17,6,8,29,5 | 2,6,8,3,4,2,0,4,3,10,2,12,3, | 236 | 149 | 0.0271357364133 | | 1 | 0.558,0.457,0.708, | 0.85,0.688,0.356,0 | -0,104 |
| 134000 | ENSG00000100387 | RBX1 | chr22 | + | 40955286 | 40955446 | 40953554 | 40953633 | 40964046 | 40964411 | 134000 | 2,3,12,9,6,14,5,2,2,2,3,0 | 32,33,149,125,62,51,40,61 | 6,4,11,0,9,3,8,3,2,10,15,13 | 82,66,42,18,35,28,34,43,1 | 298 | 149 | 0.0271460605234 | | 1 | 0.03,0.043,0.039,0 | 0.035,0.029,0.116, | -0,018 |
| 41369 | ENSG00000095951 | HIVEP1 | chr6 | + | 12129758 | 12129892 | 12119889 | 12125870 | 12130766 | 12130942 | 41369 | 81,101,261,118,102,158,6 | 4,8,19,6,9,26,3,6,3,1,1,10 | 170,192,83,127,94,57,151, | 12,8,8,12,16,1,15,13,4,15, | 283 | 149 | 0.0271474493175 | | 1 | 0.914,0.869,0.879, | 0.882,0.927,0.845, | 0,055 |
| 111880 | ENSG00000089234 | BRAP | chr12 | - | 111681636 | 111681835 | 111679150 | 111679340 | 111683145 | 111683307 | 111880 | 49,42,71,62,31,41,34,34,5 | 9,1,0,2,1,1,2,3,0,0,2,0 | 41,54,51,35,51,12,33,41,2 | 1,8,2,0,2,0,0,0,0,0,1,2,1, | 298 | 149 | 0.027158184047 | | 1 | 0.731,0.955,1.0,0. | 0.953,0.771,0.927, | -0,038 |
| 219917 | ENSG00000137193 | PIM1 | chr6 | + | 37172995 | 37173050 | 37171124 | 37171491 | 37173933 | 37175428 | 219917 | 23,13,201,53,38,39,36,37, | 5,3,5,3,4,5,3,5,0,0,2,1 | 40,68,40,13,56,21,20,64,1 | 0,3,1,0,4,1,0,10,2,2,5,2,0,3 | 204 | 149 | 0.0271931414159 | | 1 | 0.771,0.76,0.967,0 | 1.0,0.943,0.967,1. | -0,041 |
| 237598 | ENSG00000104408 | EIF3E | chr8 | - | 108233320 | 108233672 | 108229069 | 108229195 | 108234997 | 108235102 | 237598 | 2,2,6,12,16,8,8,17,10,0,1,1 | 296,169,599,433,165,197,1 | 11,8,7,14,16,3,5,7,2,2,13,5 | 264,297,308,156,149,84,1 | 298 | 149 | 0.0272038799045 | | 1 | 0.003,0.006,0.005, | 0.02,0.013,0.011,0 | -0,003 |
| 103369 | ENSG00000026950 | BTN3A1 | chr6 | + | 26409893 | 26409914 | 26409532 | 26409733 | 26410005 | 26410032 | 103369 | 59,58,27,20,18,17,18,9,17, | 1,6,7,1,1,0,0,5,1,0,2,1 | 18,9,20,26,17,9,14,32,14,1 | 1,2,0,1,0,2,0,3,2,0,3,7,0,0, | 170 | 149 | 0.027213920736 | | 1 | 0.981,0.894,0.772, | 0.94,0.798,1.0,0.9 | -0,039 |
| 32689 | ENSG00000112685 | EXOC2 | chr6 | - | 657162 | 657250 | 637700 | 637861 | 693018 | 693111 | 32689 | 4,1,1,0,0,4,1,0,1,0,0,3 | 17,19,20,13,27,31,24,21,1 | 4,2,2,6,0,4,1,0,0,2,3,4,2,1, | 17,15,5,12,16,14,16,17,11, | 237 | 149 | 0.0272146385023 | | 1 | 0.129,0.032,0.03,0 | 0.129,0.077,0.201, | -0,036 |
| 128359 | ENSG00000088298 | EDEM2 | chr20 | - | 35118597 | 35118719 | 35115363 | 35115933 | 35123889 | 35124034 | 128359 | 54,22,78,69,37,19,35,33,4 | 17,10,11,7,10,18,9,9,5,6,5, | 36,38,27,5,10,21,21,24,17, | 21,9,4,12,11,2,16,17,7,11, | 271 | 149 | 0.0272409504152 | | 1 | 0.636,0.547,0.796, | 0.485,0.699,0.788, | 0,1 |
| 181918 | ENSG00000163166 | IWS1 | chr2 | - | 127496260 | 127496361 | 127495997 | 127496148 | 127502814 | 127502872 | 181918 | 1,3,4,4,1,0,0,2,0,1,0,0 | 5,3,6,3,10,5,3,14,8,5,4,1 | 0,1,1,1,0,0,0,0,0,3,0,0,0, | 6,5,12,2,5,6,10,11,4,5,1,6,1 | 250 | 149 | 0.0272587147168 | | 1 | 0.107,0.373,0.284, | 0.0,0.107,0.047,0. | 0,042 |
| 52937 | ENSG00000226571 | | chr6 | + | 139436336 | 139436460 | 139429080 | 139429613 | 139451557 | 139451669 | 52937 | 79,42,119,57,39,49,46,47, | 3,8,11,12,13,6,3,1,2,2,1,4 | 60,64,11,39,79,21,33,59,5 | 3,1,2,2,11,1,0,4,2,2,7,3,0,4 | 273 | 149 | 0.0272723892316 | | 1 | 0.935,0.741,0.855, | 0.916,0.972,0.75,0 | -0,038 |
| 34671 | ENSG00000047188 | YTHDC2 | chr5 | + | 113581416 | 113581709 | 113579585 | 113579695 | 113584301 | 113584479 | 34671 | 40,21,71,60,35,40,26,43,1 | 2,0,2,1,4,1,0,7,0,0,0,0 | 30,58,31,27,32,21,24,46,3 | 0,3,6,11,0,2,0,1,1,0,0,0,2,0 | 298 | 149 | 0.0272791721203 | | 1 | 0.909,1.0,0.947,0. | 1.0,0.906,0.721,0. | 0,039 |
| 11241 | ENSG00000185813 | PCYT2 | chr17 | - | 81907203 | 81907257 | 81906759 | 81906898 | 81907553 | 81907598 | 11241 | 1,1,15,12,1,4,4,3,2,5,3,3 | 7,2,28,20,13,8,13,12,19,1, | 2,7,2,0,3,1,1,2,1,4,1,0,2,1, | 6,5,3,5,4,1,2,20,8,15,5,15, | 203 | 149 | 0.0273247573691 | | 1 | 0.095,0.268,0.282, | 0.197,0.507,0.329, | 0,078 |
| 142287 | ENSG00000108592 | FTSJ3 | chr17 | - | 63825047 | 63825104 | 63824824 | 63824929 | 63825241 | 63825436 | 142287 | 6,7,12,13,4,5,2,3,14,1,4,1 | 0,0,4,1,0,2,0,0,0,0,0,2 | 20,22,8,1,6,2,4,4,3,12,3,2, | 0,3,1,2,5,2,3,0,0,0,4,0,0,0, | 206 | 149 | 0.0273260276152 | | 1 | 1.0,1.0,0.685,0.90 | 1.0,0.841,0.853,0. | 0,128 |
| 13131 | ENSG00000197006 | METTL9 | chr16 | + | 21612644 | 21612835 | 21599557 | 21599898 | 21617864 | 21618009 | 13131 | 86,74,227,122,112,76,80,8 | 1,2,12,10,8,3,8,3,1,0,2,5 | 120,114,76,51,45,74,36,98 | 7,4,3,4,3,1,5,3,0,3,3,6,2,1, | 298 | 149 | 0.0273286169986 | | 1 | 0.977,0.949,0.904, | 0.896,0.934,0.927, | -0,009 |
| 34657 | ENSG00000047188 | YTHDC2 | chr5 | + | 113524980 | 113525177 | 113515271 | 113515362 | 113532878 | 113533045 | 34657 | 16,15,46,9,11,24,17,23,10, | 31,21,15,11,13,15,5,8,6,1, | 22,12,18,19,8,3,3,14,7,8,3 | 26,32,12,11,8,12,14,13,18, | 298 | 149 | 0.027337374349 | | 1 | 0.205,0.263,0.605, | 0.297,0.158,0.429, | 0,139 |
| 159561 | ENSG00000113648 | MACROH2A1 | chr5 | - | 135343259 | 135343362 | 135334380 | 135335141 | 135345967 | 135347606 | 159561 | 41,27,59,76,35,28,17,27,1 | 0,5,1,1,1,0,0,3,0,0,0,0 | 26,47,19,10,27,19,17,26,2 | 0,4,1,0,7,2,0,2,1,0,0,3,0,0, | 252 | 149 | 0.0273399497258 | | 1 | 1.0,0.761,0.972,0. | 1.0,0.874,0.918,1. | 0,029 |
| 69794 | ENSG00000223745 | CCDC18-AS1 | chr1 | - | 93311489 | 93311608 | 93305387 | 93305470 | 93324634 | 93324729 | 69794 | 14,5,24,13,9,11,19,15,7,4, | 18,9,27,25,15,8,16,20,13,7 | 8,8,12,7,9,12,7,22,5,20,22, | 10,9,3,2,16,10,4,15,3,10,17 | 268 | 149 | 0.0273561989095 | | 1 | 0.302,0.236,0.331, | 0.308,0.331,0.69,0 | -0,118 |
| 170129 | ENSG00000290763 | SDHAP1 | chr3 | - | 195963291 | 195963421 | 195959920 | 195960086 | 195965428 | 195965542 | 170129 | 12,2,7,3,2,3,6,14,9,2,2,22 | 5,3,5,3,3,1,6,2,3,0,2,0 | 4,1,6,10,5,1,16,14,10,11,5, | 1,6,2,1,2,0,4,1,1,0,1,1,0,2, | 279 | 149 | 0.0273920582434 | | 1 | 0.562,0.263,0.428, | 0.681,0.082,0.616, | -0,188 |
| 170246 | ENSG00000167515 | TRAPPC2L | chr16 | + | 88859654 | 88859750 | 88858641 | 88858789 | 88859892 | 88859972 | 170246 | 143,147,214,148,57,55,66, | 4,0,21,7,0,3,4,10,8,4,0,4 | 120,157,39,7,60,46,51,103 | 8,9,2,2,5,0,5,14,3,2,7,4,7,3 | 245 | 149 | 0.0274005539964 | | 1 | 0.956,1.0,0.861,0. | 0.901,0.914,0.922, | 0,056 |
| 94728 | ENSG00000146247 | PHIP | chr6 | - | 78970048 | 78970173 | 78969834 | 78969917 | 78978591 | 78978711 | 94728 | 24,17,70,25,15,15,17,29,1 | 3,0,3,6,1,2,3,1,0,0,0,0 | 33,43,16,33,22,14,14,22,1 | 0,2,0,13,1,0,0,1,2,0,3,0,0,0 | 274 | 149 | 0.0274210981599 | | 1 | 0.813,1.0,0.927,0. | 1.0,0.921,1.0,0.58, | -0,031 |
| 253717 | ENSG00000078674 | PCM1 | chr8 | + | 17980569 | 17980755 | 17972672 | 17972687 | 17985446 | 17985619 | 253717 | 49,21,85,29,25,31,21,42,3 | 51,16,92,48,38,27,40,75,1 | 40,26,44,15,13,21,11,29,1 | 70,94,41,60,49,16,27,21,3 | 298 | 149 | 0.0274366644893 | | 1 | 0.325,0.396,0.316, | 0.222,0.121,0.349, | 0,062 |
| 81072 | ENSG00000204387 | SNHG32 | chr6 | + | 31836298 | 31836517 | 31835410 | 31835451 | 31837235 | 31837435 | 81072 | 10,12,40,18,12,8,9,13,11,9 | 141,82,213,111,84,77,76,9 | 11,12,11,6,11,10,2,17,2,8, | 129,92,62,61,68,35,48,117 | 298 | 149 | 0.0274372170538 | | 1 | 0.034,0.068,0.086, | 0.041,0.061,0.081, | 0,01 |
| 140636 | ENSG00000134046 | MBD2 | chr18 | - | 54187713 | 54187854 | 54166075 | 54166166 | 54188873 | 54189011 | 140636 | 9,1,14,10,9,4,6,15,6,2,2,6 | 190,106,237,155,109,92,8 | 12,10,11,24,2,12,8,12,2,16 | 141,140,145,49,94,86,45,8 | 290 | 149 | 0.027441543308 | | 1 | 0.024,0.005,0.029, | 0.042,0.035,0.038, | -0,016 |
| 104007 | ENSG00000291152 | SBDSP1 | chr7 | + | 72830691 | 72830813 | 72829464 | 72829783 | 72831601 | 72831798 | 104007 | 177,78,249,125,44,72,72,1 | 7,4,6,6,3,2,2,2,5,5,0,1 | 119,130,112,134,118,70,3 | 0,4,0,11,0,0,0,0,3,1,10,1, | 271 | 149 | 0.0274940548723 | | 1 | 0.933,0.915,0.958, | 1.0,0.947,1.0,0.87, | -0,028 |
| 89078 | ENSG00000153201 | RANBP2 | chr2 | + | 108762100 | 108762195 | 108758412 | 108758548 | 108763236 | 108768388 | 89078 | 71,52,68,55,63,63,36,59,7 | 8,10,11,3,3,15,2,14,5,2,1,9 | 72,91,74,39,31,20,22,44,1 | 2,16,2,52,14,1,7,8,7,4,16,1 | 244 | 149 | 0.0275326631749 | | 1 | 0.844,0.761,0.791, | 0.956,0.776,0.958, | 0,103 |
| 217945 | ENSG00000111679 | PTPN6 | chr12 | + | 6955150 | 6955274 | 6954804 | 6954994 | 6955371 | 6955485 | 217945 | 402,208,468,300,236,116, | 31,6,36,11,8,13,5,13,13,7, | 255,186,124,17,63,110,37, | 19,10,4,3,6,2,2,7,1,4,1,26,1 | 273 | 149 | 0.0275484278175 | | 1 | 0.876,0.95,0.876,0 | 0.88,0.91,0.944,0. | -0,011 |
| 180611 | ENSG00000130775 | THEMIS2 | chr1 | + | 27876587 | 27876733 | 27872542 | 27872665 | 27879643 | 27880054 | 180611 | 19,35,30,23,4,16,19,21,23, | 1,2,8,0,0,0,3,0,0,0,0,6 | 10,83,12,10,29,6,31,52,14, | 2,4,0,3,3,0,7,11,2,0,10,6,5, | 295 | 149 | 0.0275614661236 | | 1 | 0.906,0.898,0.654, | 0.716,0.913,1.0,0. | 0,095 |
| 137701 | ENSG00000142546 | NOSIP | chr19 | - | 49576758 | 49576856 | 49560621 | 49560692 | 49580514 | 49580558 | 137701 | 5,4,5,5,1,1,1,1,1,3,0,2 | 14,6,40,34,12,13,17,19,4,5 | 5,1,2,0,0,0,0,1,2,0,2,1,8,1, | 23,12,5,7,5,3,6,19,3,14,9,2 | 247 | 149 | 0.0275717794682 | | 1 | 0.177,0.287,0.07,0 | 0.116,0.048,0.194, | 0,023 |
| 215842 | ENSG00000188647 | PTAR1 | chr9 | - | 69758259 | 69758704 | 69741791 | 69741858 | 69759852 | 69759959 | 215842 | 13,9,24,10,5,7,12,25,15,6, | 7,2,7,12,7,5,4,7,0,0,2,6 | 17,23,14,2,6,2,6,7,7,8,18,1 | 13,14,8,8,15,4,1,4,0,9,5,15 | 298 | 149 | 0.0275827251185 | | 1 | 0.481,0.692,0.632, | 0.395,0.451,0.467, | 0,156 |
| 225443 | ENSG00000163492 | CCDC141 | chr2 | - | 178915596 | 178916051 | 178905328 | 178905501 | 178918712 | 178918907 | 225443 | 22,10,31,22,7,4,9,14,19,12 | 225,164,311,178,80,112,1 | 7,21,16,9,16,11,1,15,14,6, | 126,172,129,96,162,91,49, | 298 | 149 | 0.0276092891016 | | 1 | 0.047,0.03,0.047,0 | 0.027,0.058,0.058, | 0,013 |
| 240275 | ENSG00000197329 | PELI1 | chr2 | - | 64106231 | 64106299 | 64104700 | 64104830 | 64108239 | 64108379 | 240275 | 16,4,38,12,4,3,11,6,7,1,1,0 | 198,141,190,167,28,24,77, | 11,38,23,41,6,2,3,15,20,7, | 110,268,192,123,192,35,1 | 217 | 149 | 0.0276391825994 | | 1 | 0.053,0.019,0.121, | 0.064,0.089,0.076, | -0,019 |
| 239125 | ENSG00000075856 | SART3 | chr12 | - | 108538074 | 108538203 | 108537487 | 108537595 | 108538933 | 108539089 | 239125 | 21,24,64,39,16,32,18,37,2 | 2,1,2,2,2,2,1,2,2,0,0,1 | 24,29,15,13,25,22,15,30,2 | 2,0,1,4,1,1,0,2,1,0,1,1,0,1, | 278 | 149 | 0.0276439424585 | | 1 | 0.849,0.928,0.945, | 0.865,1.0,0.889,0. | -0,011 |
| 110703 | ENSG00000034677 | RNF19A | chr8 | - | 100264033 | 100264195 | 100257059 | 100259246 | 100264670 | 100264785 | 110703 | 24,48,88,32,18,33,18,73,1 | 2,0,0,0,4,3,0,1,2,0,2,1 | 32,50,34,16,23,6,25,40,9,1 | 0,4,0,1,0,0,0,6,1,3,11,3,0,0 | 298 | 149 | 0.0276543613792 | | 1 | 0.857,1.0,1.0,1.0,0 | 1.0,0.862,1.0,0.88 | -0,011 |
| 50326 | ENSG00000132781 | MUTYH | chr1 | - | 45333412 | 45333564 | 45333284 | 45333324 | 45334390 | 45334511 | 50326 | 24,14,20,9,15,26,11,17,23, | 1,2,3,0,0,2,2,0,1,4,0,3 | 20,15,20,17,8,8,7,11,8,7,6, | 1,1,0,3,2,0,0,0,0,0,0,3,0,1, | 298 | 149 | 0.0276725471307 | | 1 | 0.923,0.778,0.769, | 0.909,0.882,1.0,0. | -0,053 |

| | | | | | | | | | | | | | | | | | | | | | | | |
|---|---|---|---|---|---|---|---|---|---|---|---|---|---|---|---|---|---|---|---|---|---|---|---|
| 51317 | ENSG00000042429 | MED17 | chr11 | + | 93809716 | 93809876 | 93806100 | 93807635 | 93811852 | 93811888 | 51317 | 50,47,70,53,13,26,8,15,18, | 4,0,3,5,1,0,0,0,0,0,1,3 | 49,66,36,27,41,15,35,27,2 | 2,3,2,4,1,3,0,4,0,6,1,6,5,0, | 298 | 149 | 0.0276830303575 | | 1 | 0.862,1.0,0.921,0. | 0.925,0.917,0.9,0. | 0,054 |
| 127941 | ENSG00000095585 | BLNK | chr10 | - | 96238258 | 96238371 | 96230793 | 96230834 | 96242734 | 96242784 | 127941 | 3,2,3,3,5,8,0,5,2,0,0,1 | 81,58,161,113,99,71,57,84 | 0,5,8,2,6,2,5,4,2,5,4,2,9,3, | 148,64,76,42,30,54,54,74, | 262 | 149 | 0.0276838470321 | | 1 | 0.021,0.019,0.01,0 | 0.0,0.043,0.056,0. | -0,016 |
| 66664 | ENSG00000152683 | SLC30A6 | chr2 | + | 32197343 | 32197392 | 32193888 | 32193983 | 32197706 | 32197784 | 66664 | 10,1,26,16,14,4,1,8,4,1,3,4 | 2,2,4,0,0,2,1,2,0,0,2,0 | 18,18,4,4,6,6,9,10,5,8,12,7 | 6,3,0,0,2,3,3,3,4,6,1,1,0,1, | 198 | 149 | 0.0276921785562 | | 1 | 0.79,0.273,0.83,1. | 0.693,0.819,1.0,1. | -0,03 |
| 1899 | ENSG00000100239 | PPP6R2 | chr22 | + | 50418866 | 50418979 | 50416091 | 50416157 | 50419348 | 50419462 | 1899 | 79,40,97,80,46,52,35,54,6 | 0,1,2,0,0,1,0,0,3,1,0,0 | 65,55,38,48,41,39,48,73,3 | 1,3,1,6,0,0,1,1,2,0,0,2,1,1, | 262 | 149 | 0.027708645225 | | 1 | 1.0,0.958,0.965,1. | 0.974,0.912,0.956, | 0,028 |
| 185826 | ENSG00000055917 | PUM2 | chr2 | - | 20311493 | 20311678 | 20308313 | 20308584 | 20312235 | 20312423 | 185826 | 34,21,114,74,52,62,25,67, | 1,2,1,7,0,2,4,2,0,2,4,2 | 88,192,21,90,24,44,82,65, | 5,6,1,5,2,0,5,0,0,4,0,4,0,5, | 298 | 149 | 0.0277144439411 | | 1 | 0.944,0.84,0.983,0 | 0.898,0.941,0.913, | -0,061 |
| 129681 | ENSG00000122965 | RBM19 | chr12 | - | 113959211 | 113959404 | 113957781 | 113958050 | 113959864 | 113959903 | 129681 | 88,45,129,72,72,100,75,90 | 0,2,1,0,0,0,2,0,0,2,0,2 | 83,73,78,63,41,31,45,73,4 | 1,0,0,6,4,0,0,1,2,0,1,0,4,2, | 298 | 149 | 0.0277171447532 | | 1 | 1.0,0.918,0.985,1. | 0.976,1.0,1.0,0.84, | 0,021 |
| 130870 | ENSG00000139168 | ZCRB1 | chr12 | - | 42324067 | 42324104 | 42322417 | 42322446 | 42325923 | 42326087 | 130870 | 7,1,6,3,6,3,1,1,1,2,1,3 | 0,0,2,2,0,2,0,0,0,0,0,2 | 5,5,3,3,3,0,2,10,0,4,2,11,1 | 0,5,2,5,4,1,1,0,1,0,0,0,0,0, | 186 | 149 | 0.0277338961665 | | 1 | 1.0,1.0,0.706,0.54 | 1.0,0.445,0.546,0. | 0,17 |
| 127371 | ENSG00000130803 | ZNF317 | chr19 | + | 9157833 | 9158075 | 9157267 | 9157394 | 9158825 | 9158908 | 127371 | 13,10,12,11,10,12,4,17,8,3 | 19,16,28,12,13,7,9,9,4,9,4, | 15,21,5,3,5,2,4,4,1,6,12,7, | 11,25,8,12,9,8,8,16,7,8,21, | 298 | 149 | 0.0277373467126 | | 1 | 0.255,0.238,0.176, | 0.405,0.296,0.238, | 0,098 |
| 157353 | ENSG00000120685 | PROSER1 | chr13 | - | 39025421 | 39025522 | 39024502 | 39024556 | 39026276 | 39026472 | 157353 | 0,1,0,0,0,2,0,4,2,0,0,0 | 18,14,14,14,4,16,8,13,11,1 | 1,2,1,10,0,0,0,0,3,0,1,4,0,1 | 4,32,15,37,16,3,21,5,9,12, | 250 | 149 | 0.0277516184064 | | 1 | 0.0,0.041,0.0,0.0,0 | 0.13,0.036,0.038,0 | -0,036 |
| 195566 | ENSG00000122034 | GTF3A | chr13 | + | 27432543 | 27432804 | 27430537 | 27430621 | 27434138 | 27434150 | 195566 | 70,57,135,108,106,70,104, | 39,28,52,16,20,10,24,43,1 | 90,93,81,37,57,57,44,77,2 | 46,49,31,25,22,31,18,32,1 | 298 | 149 | 0.0277542391284 | | 1 | 0.473,0.504,0.565, | 0.495,0.487,0.566, | 0,087 |
| 106569 | ENSG00000147140 | NONO | chrX | + | 71284401 | 71284453 | 71283693 | 71283721 | 71290628 | 71290791 | 106569 | 57,41,33,33,55,55,47,68,5 | 34,11,12,6,7,18,13,24,14,1 | 31,28,37,31,23,14,31,57,8, | 13,12,23,45,14,6,21,27,2,9 | 201 | 149 | 0.0277594341592 | | 1 | 0.554,0.734,0.671, | 0.639,0.634,0.544, | 0,079 |
| 162818 | ENSG00000164308 | ERAP2 | chr5 | + | 96902642 | 96902842 | 96902273 | 96902353 | 96903376 | 96903560 | 162818 | 0,0,0,0,0,0,1,0,0,0,0,0 | 80,56,86,73,97,51,15,30,4 | 0,0,0,0,1,0,12,17,0,0,0,0,0, | 61,66,52,46,29,42,6,14,19, | 298 | 149 | 0.0277837819715 | | 1 | 0.0,0.0,0.0,0.0,0.0, | 0.0,0.0,0.0,0.0,0.0 | -0,053 |
| 77434 | ENSG00000115525 | ST3GAL5 | chr2 | - | 85852859 | 85853099 | 85847860 | 85848204 | 85861180 | 85861292 | 77434 | 0,1,6,1,2,2,1,0,5,0,0,0 | 24,42,47,39,22,12,8,10,33, | 0,3,0,4,2,3,0,5,8,8,1,0,3,0, | 34,41,29,11,19,20,4,34,2,2 | 298 | 149 | 0.0278038163612 | | 1 | 0.0,0.012,0.06,0.0 | 0.0,0.035,0.0,0.15 | -0,062 |
| 190067 | ENSG00000109436 | TBC1D9 | chr4 | - | 140687763 | 140687918 | 140686343 | 140686462 | 140701503 | 140701614 | 190067 | 3,5,0,1,0,2,3,1,8,0,0,0 | 146,194,62,74,64,85,36,49 | 3,7,3,12,9,1,0,1,4,1,5,10,4, | 138,144,34,136,182,122,4 | 298 | 149 | 0.0278125757531 | | 1 | 0.01,0.013,0.0,0.0 | 0.011,0.024,0.042, | -0,009 |
| 142583 | ENSG00000277791 | PSMB3 | chr17 | + | 38760430 | 38760571 | 38755882 | 38755990 | 38762410 | 38762505 | 142583 | 34,47,146,155,75,55,54,54 | 20,11,20,21,7,10,6,11,10,8 | 98,66,66,14,52,58,39,59,1 | 1,9,5,8,7,4,4,11,3,5,20,16,1 | 290 | 149 | 0.0278343724992 | | 1 | 0.466,0.687,0.79,0 | 0.981,0.79,0.871,0 | -0,062 |
| 231957 | ENSG00000285547 | | chrX | - | 72568753 | 72568879 | 72567888 | 72568030 | 72572056 | 72572109 | 231957 | 28,11,34,18,8,11,10,17,19, | 0,2,3,0,0,2,0,6,1,2,2,0 | 10,21,16,17,29,10,9,13,12, | 4,1,2,1,4,2,1,1,3,2,2,0,1,1, | 275 | 149 | 0.0278353699189 | | 1 | 1.0,0.749,0.86,1.0, | 0.575,0.919,0.813, | 0,029 |
| 126267 | ENSG00000057608 | GDI2 | chr10 | - | 5794884 | 5794989 | 5785851 | 5786050 | 5796762 | 5796862 | 126267 | 188,77,626,564,232,163,1 | 1,0,9,13,3,2,1,2,1,1,9,3 | 405,256,173,76,142,171,7 | 15,11,1,5,3,1,1,3,3,3,8,19, | 254 | 149 | 0.027838485546 | | 1 | 0.991,1.0,0.976,0. | 0.941,0.932,0.99,0 | 0,008 |
| 50292 | ENSG00000125347 | IRF1 | chr5 | - | 132486250 | 132486373 | 132485666 | 132485716 | 132486556 | 132486639 | 50292 | 150,226,217,85,40,101,77, | 12,7,9,2,0,5,4,4,1,3,2,10 | 149,266,162,290,78,43,84, | 13,7,12,42,0,4,4,17,5,7,27, | 272 | 149 | 0.0278403780555 | | 1 | 0.873,0.946,0.93,0 | 0.863,0.954,0.881, | 0,023 |
| 135358 | ENSG00000244625 | MIATNB | chr22 | + | 26717256 | 26717459 | 26693692 | 26693836 | 26774155 | 26774217 | 135358 | 10,5,6,4,2,0,1,3,13,5,0,2 | 0,0,1,0,1,6,1,3,0,0,1,1 | 13,3,0,1,8,5,3,0,8,9,2,11,1, | 3,3,2,0,3,0,2,1,2,0,1,10,1,0 | 298 | 149 | 0.0278712984947 | | 1 | 1.0,1.0,0.75,1.0,0. | 0.684,0.333,0.0,1. | 0,001 |
| 248332 | ENSG00000167323 | STIM1 | chr11 | + | 3945812 | 3946068 | 3856181 | 3856409 | 3967551 | 3968721 | 248332 | 1,0,2,6,2,6,0,5,3,0,1,0 | 22,21,69,57,17,23,20,45,1 | 0,3,2,5,0,1,1,8,3,5,9,2,3,6, | 53,41,18,30,25,25,27,49,2 | 298 | 149 | 0.0278774499915 | | 1 | 0.022,0.0,0.014,0. | 0.0,0.035,0.053,0. | -0,017 |
| 126883 | ENSG00000266412 | NCOA4 | chr10 | - | 46016539 | 46016694 | 46015125 | 46015266 | 46030525 | 46030623 | 126883 | 60,32,221,169,102,62,61,7 | 2,0,20,17,9,0,1,7,0,1,0,2 | 138,100,54,26,81,63,56,70 | 3,4,5,2,3,1,1,3,0,2,9,5,3,2, | 298 | 149 | 0.0278802363137 | | 1 | 0.938,1.0,0.847,0. | 0.958,0.926,0.844, | 0,006 |
| 195720 | ENSG00000117419 | ERI3 | chr1 | - | 44339044 | 44339243 | 44319627 | 44319744 | 44352849 | 44352925 | 195720 | 15,13,37,29,20,25,13,22,7, | 0,0,4,3,4,4,0,0,1,0,0,0 | 29,17,16,4,22,11,12,21,7,1 | 5,1,3,0,0,0,4,1,4,1,2,2,4,0, | 298 | 149 | 0.0279331340943 | | 1 | 1.0,1.0,0.822,0.82 | 0.744,0.895,0.727, | 0,095 |
| 7670 | ENSG00000263753 | LINC00667 | chr18 | + | 5240647 | 5240819 | 5238099 | 5239049 | 5241698 | 5241889 | 7670 | 1,0,13,5,5,3,1,2,5,2,1,1 | 7,4,47,31,7,7,10,16,5,2,2,3 | 2,0,0,0,1,1,3,7,0,6,2,1,1,1, | 9,12,6,2,6,10,2,15,1,20,11, | 298 | 149 | 0.0279511774312 | | 1 | 0.067,0.0,0.121,0. | 0.1,0.0,0.0,0.0,0.0 | 0,059 |
| 254780 | ENSG00000121749 | TBC1D15 | chr12 | + | 71884810 | 71885017 | 71880468 | 71880607 | 71893221 | 71893324 | 254780 | 26,19,50,33,12,8,8,21,13,6 | 0,2,2,0,1,1,0,3,2,0,0,0 | 20,50,15,30,24,17,14,19,1 | 0,5,0,0,1,3,1,0,0,0,5,1,3,1, | 298 | 149 | 0.0279557990527 | | 1 | 1.0,0.826,0.926,1. | 1.0,0.833,1.0,1.0,0 | 0,046 |
| 109305 | ENSG00000147421 | HMBOX1 | chr8 | + | 28980070 | 28980156 | 28970042 | 28970519 | 29009071 | 29009182 | 109305 | 109,43,89,49,40,44,40,32, | 16,3,0,5,1,4,3,4,5,2,0,2 | 59,60,43,47,40,48,33,39,5 | 2,4,3,4,1,0,0,3,8,3,2,0,4,1, | 235 | 149 | 0.0279655753933 | | 1 | 0.812,0.901,1.0,0. | 0.949,0.905,0.901, | -0,019 |
| 13831 | ENSG00000103522 | IL21R | chr16 | + | 27403079 | 27403220 | 27402173 | 27402618 | 27430055 | 27430120 | 13831 | 0,1,19,6,4,1,4,5,11,0,4,1 | 8,8,22,12,8,5,13,15,7,11,7, | 4,1,1,0,1,1,3,2,2,5,17,0,0,1 | 8,4,5,1,2,0,15,14,3,7,41,32 | 290 | 149 | 0.0279874584195 | | 1 | 0.0,0.06,0.307,0.2 | 0.204,0.114,0.093, | -0,021 |
| 52410 | ENSG00000291153 | STAG3L4 | chr7 | + | 67320102 | 67320336 | 67309563 | 67309715 | 67330262 | 67330394 | 52410 | 21,40,40,24,13,10,19,24,2 | 0,0,1,1,4,0,3,6,0,0,0,3 | 21,23,15,14,14,12,7,17,15, | 0,0,2,0,0,0,0,2,0,0,0,0,1,4, | 298 | 149 | 0.0280204242173 | | 1 | 1.0,1.0,0.952,0.92 | 1.0,1.0,0.789,1.0,1 | -0,046 |
| 235108 | ENSG00000063601 | MTMR1 | chrX | + | 150734923 | 150735228 | 150732541 | 150732730 | 150735463 | 150735499 | 235108 | 5,2,6,2,0,2,2,5,2,0,1,7 | 14,3,16,8,4,6,2,11,7,5,0,2 | 4,14,3,21,4,2,7,6,2,7,21,0, | 6,4,11,13,6,3,7,8,7,1,12,12 | 298 | 149 | 0.0280402372189 | | 1 | 0.152,0.25,0.158,0 | 0.25,0.636,0.12,0. | -0,045 |
| 17489 | ENSG00000140326 | CDAN1 | chr15 | - | 42726096 | 42726160 | 42725488 | 42725670 | 42726309 | 42726417 | 17489 | 15,1,19,4,4,4,3,5,7,4,6,12 | 1,2,1,0,0,1,0,0,0,0,0,2 | 2,10,2,1,3,8,3,5,6,15,3,2,1, | 2,1,0,0,0,0,0,1,1,2,3,0,4,0, | 213 | 149 | 0.0280638628134 | | 1 | 0.913,0.259,0.93,1 | 0.412,0.875,1.0,1. | 0,093 |
| 86911 | ENSG00000166801 | FAM111A | chr11 | + | 59143496 | 59143695 | 59143082 | 59143304 | 59145826 | 59145856 | 86911 | 25,13,37,36,11,15,11,29,2 | 10,7,11,17,1,7,9,12,6,1,3,1 | 13,8,6,6,7,13,3,18,10,43,1 | 7,9,5,6,1,7,4,16,4,9,14,1,6, | 298 | 149 | 0.0280684279416 | | 1 | 0.556,0.481,0.627, | 0.481,0.308,0.375, | 0,148 |
| 15025 | ENSG00000153066 | TXNDC11 | chr16 | - | 11733981 | 11734079 | 11730644 | 11730774 | 11736016 | 11736233 | 15025 | 59,53,86,54,39,38,35,52,6 | 2,0,2,0,3,2,2,0,0,3,0,5 | 63,58,30,29,31,20,24,87,1 | 3,2,4,5,3,0,2,3,2,1,3,3,0,0,1 | 247 | 149 | 0.0281508293806 | | 1 | 0.947,1.0,0.963,1. | 0.927,0.946,0.819, | 0,031 |
| 106049 | ENSG00000015285 | WAS | chrX | + | 48690149 | 48690267 | 48689319 | 48689434 | 48691106 | 48691395 | 106049 | 2,0,2,2,0,2,1,7,2,0,0,0 | 49,23,38,16,27,24,22,17,3 | 8,3,0,5,1,2,2,1,4,0,2,0,1,3, | 23,10,17,15,10,2,16,18,16, | 267 | 149 | 0.0281545416642 | | 1 | 0.022,0.0,0.029,0. | 0.163,0.143,0.0,0. | -0,048 |
| 181250 | ENSG00000163565 | IFI16 | chr1 | + | 159053532 | 159053626 | 159051678 | 159052098 | 159054820 | 159055155 | 181250 | 235,103,174,136,61,63,34, | 17,7,31,19,8,21,14,6,4,5,2, | 105,114,52,36,56,90,10,38 | 32,28,6,2,6,9,8,21,4,11,17, | 243 | 149 | 0.0281582363191 | | 1 | 0.894,0.9,0.775,0. | 0.668,0.714,0.842, | 0,073 |
| 214958 | ENSG00000135976 | ANKRD36 | chr2 | + | 97143042 | 97143086 | 97142762 | 97142835 | 97144517 | 97144546 | 214958 | 0,0,0,0,0,0,0,0,0,0,0,0 | 27,15,85,57,16,46,21,35,1 | 0,0,0,1,0,0,0,0,0,0,0,0,0,1 | 39,30,19,35,28,14,14,26,2 | 193 | 149 | 0.0281586186277 | | 1 | 0.0,0.0,0.0,0.0,0.0, | 0.0,0.0,0.0,0.022,0 | -0,058 |
| 97795 | ENSG00000128607 | KLHDC10 | chr7 | + | 130120748 | 130120903 | 130116444 | 130116666 | 130122053 | 130122202 | 97795 | 59,33,108,66,39,36,50,52, | 2,0,5,2,6,0,0,4,0,0,1,0 | 40,63,50,41,17,26,40,74,1 | 0,0,1,0,2,2,0,0,2,1,1,0,2,3, | 298 | 149 | 0.0281604899569 | | 1 | 0.937,1.0,0.915,0. | 1.0,1.0,0.962,1.0,0 | 0,002 |
| 54486 | ENSG00000132549 | VPS13B | chr8 | + | 99854409 | 99854519 | 99853450 | 99854256 | 99859303 | 99859480 | 54486 | 3,3,2,1,2,1,0,1,2,1,0,0 | 9,17,24,22,21,10,6,15,8,7, | 3,4,0,6,3,0,1,0,5,0,3,3,0,0, | 8,18,1,8,10,6,12,8,6,15,7,2 | 259 | 149 | 0.0281749249503 | | 1 | 0.161,0.092,0.046, | 0.177,0.113,0.0,0. | -0,04 |
| 241710 | ENSG00000148308 | GTF3C5 | chr9 | + | 133054943 | 133055220 | 133054711 | 133054809 | 133056011 | 133056094 | 241710 | 1,2,23,10,3,4,2,17,2,0,4,1 | 16,10,29,23,18,6,7,6,15,3, | 5,12,6,6,7,10,1,8,6,15,6,3, | 12,12,15,15,6,9,22,10,4,16 | 298 | 149 | 0.028192091543 | | 1 | 0.03,0.091,0.284,0 | 0.172,0.333,0.167, | -0,094 |
| 108536 | ENSG00000179195 | ZNF664 | chr12 | + | 123988042 | 123988138 | 123973122 | 123974020 | 124011380 | 124011441 | 108536 | 36,22,55,26,35,18,21,27,3 | 1,3,6,6,1,2,0,0,1,0,0,0 | 37,19,11,26,10,4,31,35,14, | 0,0,0,0,2,2,0,0,0,0,6,0,0,1, | 245 | 149 | 0.0282193391591 | | 1 | 0.956,0.817,0.848, | 1.0,1.0,1.0,1.0,0.7 | -0,005 |
| 217947 | ENSG00000111679 | PTPN6 | chr12 | + | 6955166 | 6955267 | 6954953 | 6954994 | 6955371 | 6955394 | 217947 | 417,220,420,297,223,79,1 | 31,6,36,11,8,13,5,13,13,7, | 243,177,92,28,55,93,46,70 | 19,10,4,3,6,2,2,7,1,4,1,26,1 | 250 | 149 | 0.0282489434887 | | 1 | 0.889,0.956,0.874, | 0.884,0.913,0.932, | -0,016 |
| 104369 | ENSG00000004961 | HCCS | chrX | + | 11120906 | 11120993 | 11118500 | 11118620 | 11121611 | 11121879 | 104369 | 8,8,25,18,9,3,9,7,12,1,1,12 | 1,0,0,2,0,0,0,1,0,2,0,0 | 17,32,19,12,23,10,6,8,9,15 | 0,0,0,1,0,0,2,0,8,2,2,4,2,0, | 236 | 149 | 0.0282906330218 | | 1 | 0.835,1.0,1.0,0.85, | 1.0,1.0,1.0,0.883,1 | 0,016 |
| 198035 | ENSG00000112232 | KHDRBS2 | chr6 | - | 61816610 | 61817005 | 61732681 | 61732764 | 61894634 | 61894833 | 198035 | 2,0,5,5,7,6,2,9,13,2,1,2 | 14,11,27,19,11,10,11,16,3 | 6,5,4,3,3,1,1,5,3,2,7,2,1,2, | 26,8,5,9,2,0,9,5,4,4,9,6,3,4 | 298 | 149 | 0.0282965015799 | | 1 | 0.067,0.0,0.085,0. | 0.103,0.238,0.286, | -0,164 |
| 35151 | ENSG00000064726 | BTBD1 | chr15 | - | 83050072 | 83050178 | 83041727 | 83041925 | 83056388 | 83056545 | 35151 | 86,87,199,142,61,70,65,73 | 1,0,7,3,0,1,3,2,0,0,0,0 | 159,160,95,47,76,51,29,45 | 1,9,3,1,7,0,0,0,6,0,2,1,2,4, | 255 | 149 | 0.0283062069603 | | 1 | 0.98,1.0,0.943,0.9 | 0.989,0.912,0.949, | 0,02 |
| 3057 | ENSG00000129351 | ILF3 | chr19 | + | 10688549 | 10688678 | 10685927 | 10687708 | 10689178 | 10689492 | 3057 | 117,97,117,53,101,120,13 | 11,12,11,3,7,2,6,8,5,2,4,0 | 64,62,68,43,31,39,50,125, | 6,6,9,14,3,1,16,5,0,0,14,14 | 278 | 149 | 0.0283756828321 | | 1 | 0.851,0.812,0.851, | 0.851,0.847,0.802, | 0,049 |
| 39604 | ENSG00000290763 | SDHAP1 | chr3 | - | 195963299 | 195963421 | 195959920 | 195960086 | 195965428 | 195965542 | 39604 | 20,3,18,12,5,5,9,19,12,2,4, | 5,3,5,3,3,1,6,2,3,0,2,0 | 6,6,13,15,9,2,25,20,21,22, | 1,6,2,1,2,0,4,1,1,0,1,1,0,2, | 271 | 149 | 0.0284025883951 | | 1 | 0.687,0.355,0.664, | 0.767,0.355,0.781, | -0,14 |
| 18490 | ENSG00000159322 | ADPGK | chr15 | - | 72760406 | 72760527 | 72751368 | 72752892 | 72771782 | 72771845 | 18490 | 42,47,76,41,41,37,22,36,4 | 2,5,10,4,1,5,1,4,0,0,0,5 | 42,69,51,24,21,28,26,38,3 | 4,6,1,2,12,3,4,4,4,6,2,8,4,1 | 270 | 149 | 0.0284044090529 | | 1 | 0.921,0.838,0.807, | 0.853,0.864,0.966, | 0,083 |
| 227175 | ENSG00000204257 | HLA-DMA | chr6 | - | 32950518 | 32950666 | 32949610 | 32949889 | 32952948 | 32953108 | 227175 | 407,214,947,925,248,145, | 8,2,14,11,6,4,0,5,0,0,2,0 | 559,567,308,203,266,231, | 2,0,2,4,1,0,2,1,2,1,7,6,0,2,1 | 297 | 149 | 0.0284077798142 | | 1 | 0.962,0.982,0.971, | 0.993,1.0,0.987,0. | -0,005 |
| 188960 | ENSG00000122678 | POLM | chr7 | - | 44075483 | 44075633 | 44074397 | 44074530 | 44076508 | 44076629 | 188960 | 0,0,10,0,5,0,5,5,1,0,1,2 | 7,5,37,18,6,16,8,12,12,10, | 4,1,0,1,1,1,1,3,4,2,0,1,5,2, | 3,8,3,3,1,7,7,6,4,5,14,18,1 | 298 | 149 | 0.0284098043838 | | 1 | 0.0,0.0,0.119,0.0,0 | 0.4,0.059,0.0,0.14 | -0,054 |
| 158710 | ENSG00000126777 | KTN1 | chr14 | + | 55663954 | 55664332 | 55661273 | 55661612 | 55667240 | 55667330 | 158710 | 17,13,23,11,7,9,4,13,7,0,2, | 50,36,58,36,25,30,28,47,6 | 15,12,16,5,6,10,6,11,8,10, | 36,37,21,12,22,16,24,41,2 | 298 | 149 | 0.028431168026 | | 1 | 0.145,0.153,0.165, | 0.172,0.14,0.276,0 | -0,051 |
| 181395 | ENSG00000153914 | SREK1 | chr5 | + | 66156064 | 66156202 | 66153462 | 66153596 | 66158808 | 66158932 | 181395 | 24,4,12,13,3,10,15,14,12,4 | 10,4,19,11,2,3,15,6,2,2,3,4 | 18,29,15,13,21,13,8,3,6,6, | 6,9,2,9,6,3,2,10,1,4,4,7,1,5 | 287 | 149 | 0.028432473753 | | 1 | 0.555,0.342,0.247, | 0.609,0.626,0.796, | -0,145 |
| 170050 | ENSG00000162736 | NCSTN | chr1 | + | 160355631 | 160355958 | 160354222 | 160354290 | 160356259 | 160356347 | 170050 | 59,41,51,26,22,23,30,38,4 | 2,0,2,3,6,1,0,2,3,0,0,5 | 37,40,32,5,21,15,22,37,16, | 1,3,1,10,4,1,1,1,3,1,2,8,2,1 | 298 | 149 | 0.0284386979279 | | 1 | 0.937,1.0,0.927,0. | 0.949,0.87,0.941,0 | 0,095 |
| 169345 | ENSG00000025293 | PHF20 | chr20 | + | 35812622 | 35812719 | 35801490 | 35801605 | 35842572 | 35842744 | 169345 | 6,4,6,0,5,4,1,0,1,0,0,4 | 41,28,55,41,36,35,24,29,4 | 4,6,6,5,0,4,0,2,0,2,0,8,10,6 | 34,32,22,28,21,17,17,29,2 | 246 | 149 | 0.0284578913191 | | 1 | 0.081,0.08,0.062,0 | 0.067,0.102,0.142, | -0,023 |
| 88530 | ENSG00000163219 | ARHGAP25 | chr2 | + | 68787839 | 68787956 | 68782232 | 68782320 | 68807272 | 68807480 | 88530 | 286,456,349,355,243,167, | 35,45,31,31,6,14,10,13,3,1 | 404,818,169,113,242,85,1 | 38,77,14,31,29,9,15,7,9,15 | 266 | 149 | 0.0284798474308 | | 1 | 0.821,0.85,0.863,0 | 0.856,0.856,0.871, | 0,036 |
| 19310 | ENSG00000138592 | USP8 | chr15 | + | 50441348 | 50441493 | 50439008 | 50439177 | 50449399 | 50449440 | 19310 | 106,111,289,164,60,83,12 | 1,3,1,1,1,7,3,3,10,7,2,2 | 138,232,88,120,177,65,85, | 3,7,4,2,2,2,0,1,4,2,3,1,0,2, | 294 | 149 | 0.0285058245995 | | 1 | 0.982,0.949,0.993, | 0.959,0.944,0.918, | -0,041 |
| 64472 | ENSG00000204469 | PRRC2A | chr6 | + | 31626771 | 31626862 | 31625753 | 31626162 | 31626981 | 31627074 | 64472 | 76,36,120,57,79,92,49,73, | 0,2,5,0,2,0,0,0,0,0,0,0 | 80,69,36,57,43,40,85,79,3 | 1,0,5,1,2,0,3,2,0,2,1,0,2,2, | 240 | 149 | 0.0285267198051 | | 1 | 1.0,0.918,0.937,1. | 0.98,1.0,0.817,0.9 | 0,03 |
| 29992 | ENSG00000273294 | C1QTNF3-AMA | chr5 | - | 34023908 | 34024008 | 34020586 | 34020742 | 34028753 | 34028883 | 29992 | 11,12,29,11,2,13,9,21,8,6, | 2,2,5,1,0,1,1,0,1,0,0,0 | 1,13,12,10,11,8,10,5,7,4,1 | 2,1,0,6,1,0,0,0,5,2,3,3,0,1, | 249 | 149 | 0.0285345845145 | | 1 | 0.767,0.782,0.776, | 0.23,0.886,1.0,0.4 | 0,117 |
| 104479 | ENSG00000111412 | SPRING1 | chr12 | - | 116723066 | 116723223 | 116720295 | 116720447 | 116737789 | 116738069 | 104479 | 30,37,66,43,21,19,24,34,2 | 0,2,0,1,0,0,3,1,0,0,0,7 | 37,34,19,12,42,15,22,41,9, | 1,4,0,0,0,2,2,4,0,0,3,2,4,1, | 298 | 149 | 0.0285423654388 | | 1 | 1.0,0.902,1.0,0.95 | 0.949,0.81,1.0,1.0, | 0,035 |
| 172388 | ENSG00000137502 | RAB30 | chr11 | - | 83013686 | 83013770 | 82997223 | 82997324 | 83034171 | 83034633 | 172388 | 0,1,2,0,1,2,0,2,0,0,1,0 | 8,12,15,16,0,5,2,6,3,5,2,7 | 2,9,0,0,4,0,3,0,2,0,4,2,1,0, | 14,36,20,11,22,3,14,8,13,3 | 233 | 149 | 0.0285739965104 | | 1 | 0.0,0.051,0.079,0. | 0.084,0.138,0.0,0. | 0,079 |
| 170616 | ENSG00000076108 | BAZ2A | chr12 | - | 56630775 | 56630925 | 56617394 | 56617532 | 56636181 | 56636379 | 170616 | 7,6,12,7,2,5,8,13,6,4,1,9 | 11,13,20,7,8,19,7,13,4,9,3, | 7,8,4,8,8,7,0,13,5,7,1,2,5,1 | 10,35,11,32,23,11,5,10,12, | 298 | 149 | 0.028619066227 | | 1 | 0.241,0.188,0.231, | 0.259,0.103,0.154, | 0,086 |
| 81071 | ENSG00000204387 | SNHG32 | chr6 | + | 31836294 | 31836517 | 31835410 | 31835451 | 31837235 | 31837435 | 81071 | 10,11,42,17,12,8,9,12,11,9 | 141,82,213,111,84,77,76,9 | 11,12,11,4,12,7,2,17,2,10,1 | 129,92,62,61,68,35,48,117 | 298 | 149 | 0.0286427797784 | | 1 | 0.034,0.063,0.09,0 | 0.041,0.061,0.081, | 0,01 |
| 66758 | ENSG00000117419 | ERI3 | chr1 | - | 44339044 | 44339316 | 44319627 | 44319744 | 44352849 | 44352925 | 66758 | 15,13,38,29,18,25,15,22,7, | 0,0,4,3,4,4,0,0,1,0,0,0 | 30,17,17,5,22,13,13,21,7,1 | 5,1,3,0,0,0,4,1,4,1,2,2,4,0, | 298 | 149 | 0.0286466481665 | | 1 | 1.0,1.0,0.826,0.82 | 0.75,0.895,0.739,1 | 0,091 |
| 190255 | ENSG00000165671 | NSD1 | chr5 | + | 177136872 | 177136980 | 177135940 | 177136030 | 177191883 | 177192019 | 190255 | 2,1,7,0,2,2,1,10,0,0,0,0 | 61,47,133,81,60,68,45,79,1 | 7,7,4,8,4,1,0,1,0,0,3,5,5,0,1 | 89,57,40,41,42,33,28,53,4 | 257 | 149 | 0.0286553970459 | | 1 | 0.019,0.012,0.03,0 | 0.044,0.066,0.055, | -0,018 |
| 180807 | ENSG00000257379 | | chr12 | + | 53464711 | 53464852 | 53462492 | 53462567 | 53465931 | 53465973 | 180807 | 38,32,93,71,38,84,44,87,3 | 59,28,91,64,39,48,61,68,4 | 55,109,47,111,63,32,133,7 | 56,80,24,66,44,31,112,73, | 290 | 149 | 0.0286796475145 | | 1 | 0.249,0.37,0.344,0 | 0.335,0.412,0.502, | -0,055 |
| 21598 | ENSG00000256053 | COA8 | chr14 | + | 103574106 | 103574170 | 103571622 | 103571820 | 103587273 | 103587364 | 21598 | 23,16,28,28,20,10,9,11,13, | 3,0,0,0,0,0,2,0,2,0,0,0 | 30,13,11,8,19,9,12,13,9,28 | 3,4,0,0,2,2,1,0,1,0,0,1,3,2, | 213 | 149 | 0.0286808084651 | | 1 | 0.843,1.0,1.0,1.0,1 | 0.875,0.695,1.0,1. | 0,052 |
| 237700 | ENSG00000184445 | KNTC1 | chr12 | + | 122528861 | 122529121 | 122527248 | 122527351 | 122530021 | 122530192 | 237700 | 7,3,11,0,0,0,2,0,1,5,0,0 | 36,16,100,65,33,50,17,40, | 0,1,3,0,0,2,0,3,3,6,2,0,0,0, | 39,38,48,32,15,22,11,26,3 | 298 | 149 | 0.0287147808127 | | 1 | 0.089,0.086,0.052, | 0.0,0.013,0.03,0.0, | 0,026 |
| 157191 | ENSG00000133112 | TPT1 | chr13 | - | 45340712 | 45340785 | 45339993 | 45340184 | 45341041 | 45341086 | 157191 | 471,331,1777,1399,366,45 | 31,14,25,12,14,8,10,8,8,13 | 842,710,313,227,455,285, | 17,19,6,23,8,6,9,10,6,7,17, | 222 | 149 | 0.0287258215074 | | 1 | 0.911,0.941,0.979, | 0.971,0.962,0.972, | -0,011 |
| 184569 | ENSG00000198265 | HELZ | chr17 | - | 67135969 | 67136170 | 67128494 | 67128855 | 67137930 | 67138114 | 184569 | 32,6,39,17,27,16,11,7,17,1 | 2,1,9,2,5,14,1,8,9,0,2,1 | 18,16,11,14,4,6,9,8,3,23,2 | 5,6,4,9,10,3,3,7,3,2,10,7,5, | 298 | 149 | 0.0287389516305 | | 1 | 0.889,0.75,0.684,0 | 0.643,0.571,0.579, | 0,165 |
| 30401 | ENSG00000115661 | STK16 | chr2 | + | 219247661 | 219247757 | 219247414 | 219247535 | 219248192 | 219248314 | 30401 | 7,5,15,16,16,7,11,9,5,6,6,1 | 4,2,6,1,2,0,1,5,2,1,2,2 | 6,8,2,0,1,6,2,12,2,12,4,9,1 | 5,9,0,2,2,6,2,8,2,3,4,2,1,2, | 245 | 149 | 0.0287410036751 | | 1 | 0.516,0.603,0.603, | 0.422,0.351,1.0,0. | 0,18 |
| 130898 | ENSG00000138777 | PPA2 | chr4 | - | 105456247 | 105456291 | 105453597 | 105453642 | 105456680 | 105456745 | 130898 | 0,0,1,2,0,1,3,3,1,1,1,2 | 12,13,39,32,5,21,10,26,15, | 4,3,3,2,0,0,0,2,1,1,2,4,3,4, | 30,23,19,15,30,19,12,26,1 | 193 | 149 | 0.0287459301093 | | 1 | 0.0,0.0,0.019,0.04 | 0.093,0.091,0.109, | 0,009 |
| 199587 | ENSG00000068120 | COASY | chr17 | + | 42564824 | 42565047 | 42564445 | 42564577 | 42565226 | 42565311 | 199587 | 4,5,16,3,7,9,5,3,7,9,3,11 | 0,0,4,7,2,1,2,1,1,1,1,1 | 10,3,2,5,5,6,1,8,2,4,8,4,14, | 4,10,3,1,1,2,0,7,2,0,1,8,3,0 | 298 | 149 | 0.0287486101445 | | 1 | 1.0,1.0,0.667,0.17 | 0.556,0.13,0.25,0. | 0,16 |
| 252902 | ENSG00000135720 | DYNC1LI2 | chr16 | - | 66724661 | 66724780 | 66723600 | 66723822 | 66725827 | 66725944 | 252902 | 2,4,8,2,0,3,2,3,2,2,0,0 | 22,23,51,25,14,18,23,20,5, | 1,4,0,0,2,1,2,5,0,0,7,0,1,0, | 21,50,25,20,42,13,15,20,1 | 268 | 149 | 0.0287517188837 | | 1 | 0.048,0.088,0.08,0 | 0.026,0.043,0.0,0. | 0,034 |
| 217326 | ENSG00000140750 | ARHGAP17 | chr16 | - | 24959670 | 24959745 | 24954602 | 24954730 | 24959910 | 24959981 | 217326 | 11,2,63,34,8,13,5,6,6,3,4,6 | 0,0,0,0,0,2,0,0,0,0,1,0 | 26,24,6,3,7,5,6,12,15,27,1 | 1,3,0,7,2,0,0,0,1,3,2,0,1,2, | 224 | 149 | 0.0287567481446 | | 1 | 1.0,1.0,1.0,1.0,1.0, | 0.945,0.842,1.0,0. | 0,083 |
| 81276 | ENSG00000198589 | LRBA | chr4 | - | 150277852 | 150278004 | 150264434 | 150265812 | 150282449 | 150282643 | 81276 | 69,57,98,69,49,32,33,53,5 | 7,1,6,3,0,2,1,2,0,1,4,9 | 67,70,47,36,25,27,53,51,2 | 6,4,5,6,1,4,2,2,0,0,1,1,1,2,1 | 298 | 149 | 0.0287625528161 | | 1 | 0.831,0.966,0.891, | 0.848,0.897,0.825, | -0,013 |
| 117051 | ENSG00000178177 | LCORL | chr4 | - | 17873387 | 17878213 | 17841186 | 17845901 | 17886067 | 17886161 | 117051 | 14,2,3,4,2,1,4,0,4,1,2,4 | 1,2,4,0,0,0,4,2,0,0,0,1 | 4,2,2,6,2,5,1,1,4,5,6,3,2,7, | 1,2,3,5,4,0,0,4,0,3,0,9,0,1, | 298 | 149 | 0.0288287474761 | | 1 | 0.875,0.333,0.273, | 0.667,0.333,0.25,0 | 0,062 |
| 21877 | ENSG00000258289 | CHURC1 | chr14 | + | 64926009 | 64926080 | 64923990 | 64924126 | 64932137 | 64935368 | 21877 | 17,26,112,67,55,58,52,76, | 14,8,25,13,6,8,13,17,20,10 | 81,128,44,10,29,39,35,49, | 8,14,19,8,13,0,12,8,4,13,7, | 220 | 149 | 0.0288378503177 | | 1 | 0.451,0.688,0.752, | 0.873,0.861,0.611, | -0,092 |
| 109385 | ENSG00000156171 | DRAM2 | chr1 | - | 111119614 | 111119681 | 111118864 | 111118897 | 111119876 | 111119959 | 109385 | 25,1,28,12,8,7,3,21,13,7,3, | 70,41,198,95,101,55,57,10 | 19,9,14,17,3,3,6,14,0,4,3,1 | 97,77,75,37,50,48,39,61,5 | 216 | 149 | 0.0288516732801 | | 1 | 0.198,0.017,0.089, | 0.119,0.075,0.114, | 0,011 |
| 180815 | ENSG00000257379 | | chr12 | + | 53464764 | 53464852 | 53462492 | 53462567 | 53465931 | 53465973 | 180815 | 36,32,78,59,36,69,36,80,3 | 59,28,91,64,39,48,61,68,4 | 47,95,39,102,56,31,113,70 | 56,80,24,66,44,31,112,73, | 237 | 149 | 0.028866490158 | | 1 | 0.277,0.418,0.35,0 | 0.345,0.427,0.505, | -0,055 |
| 4453 | ENSG00000104852 | SNRNP70 | chr19 | + | 49102113 | 49102185 | 49101389 | 49101471 | 49104633 | 49104735 | 4453 | 84,36,137,95,102,113,82,1 | 60,48,65,32,32,23,36,20,6 | 154,129,55,29,80,57,67,13 | 25,28,37,33,32,15,11,30,1 | 221 | 149 | 0.028883747236 | | 1 | 0.486,0.336,0.587, | 0.806,0.756,0.501, | -0,101 |
| 4130 | ENSG00000130303 | BST2 | chr19 | - | 17404370 | 17404437 | 17404128 | 17404189 | 17405290 | 17405630 | 4130 | 72,56,212,253,65,79,32,47 | 0,0,5,1,0,0,0,2,2,1,0,2 | 80,71,42,36,25,64,35,117, | 4,3,1,3,0,0,0,3,0,0,3,6,2,1,1 | 216 | 149 | 0.0288904003092 | | 1 | 1.0,1.0,0.967,0.99 | 0.932,0.942,0.967, | 0,02 |
| 138377 | ENSG00000157540 | DYRK1A | chr21 | + | 37488243 | 37488351 | 37486466 | 37486614 | 37490174 | 37490461 | 138377 | 8,6,14,16,4,5,2,13,4,4,2,6 | 66,50,199,91,61,84,55,85, | 17,18,4,4,5,3,0,6,1,11,8,4,1 | 137,148,83,79,52,59,63,85 | 257 | 149 | 0.0289210071841 | | 1 | 0.066,0.065,0.039, | 0.067,0.066,0.027, | 0,021 |
| 8823 | ENSG00000154889 | MPPE1 | chr18 | - | 11905737 | 11905953 | 11897525 | 11897620 | 11906202 | 11906309 | 8823 | 11,6,11,6,1,2,8,10,9,10,4,1 | 4,2,13,6,2,3,1,5,2,5,0,3 | 0,0,1,0,9,3,1,14,3,10,4,10, | 8,8,0,1,10,2,1,3,6,4,6,7,1,1 | 298 | 149 | 0.0289262338803 | | 1 | 0.579,0.6,0.297,0. | 0.0,0.0,1.0,0.0,0.3 | 0,139 |

| | | | | | | | | | | | | | | | | | | | | | | | |
|---|---|---|---|---|---|---|---|---|---|---|---|---|---|---|---|---|---|---|---|---|---|---|---|
| 211231 | ENSG00000197622 | CDC42SE1 | chr1 | - | 151056655 | 151056820 | 151055582 | 151055993 | 151059478 | 151059773 | 211231 | 20,7,32,14,13,25,9,16,6,5, | 197,165,295,254,162,112, | 26,16,25,20,18,13,5,21,6,1 | 274,239,156,130,165,98,1 | 298 | 149 | 0.0289526220573 | | 1 | 0.048,0.021,0.051, | 0.045,0.032,0.074, | -0,009 |
| 137844 | ENSG00000007255 | TRAPPC6A | chr19 | - | 45165126 | 45165185 | 45164724 | 45164970 | 45178134 | 45178237 | 137844 | 1,7,13,15,9,5,6,10,4,1,3,5 | 2,0,4,4,7,1,2,2,1,5,4,5 | 9,4,8,4,3,3,1,7,5,4,4,17,7,6 | 0,0,0,2,1,0,0,4,3,8,2,17,2,0 | 208 | 149 | 0.028970302002 | | 1 | 0.264,1.0,0.7,0.72 | 1.0,1.0,1.0,0.589,0 | -0,185 |
| 94399 | ENSG00000072121 | ZFYVE26 | chr14 | - | 67784333 | 67784436 | 67782779 | 67783525 | 67785058 | 67785277 | 94399 | 10,15,40,19,18,13,26,27,6, | 0,1,0,0,0,3,0,0,3,0,0,0 | 13,29,10,11,16,9,5,26,27,3 | 2,0,0,3,0,2,0,5,3,0,0,1,0,4, | 252 | 149 | 0.0289794448956 | | 1 | 1.0,0.899,1.0,1.0,1 | 0.794,1.0,1.0,0.684 | 0,032 |
| 167119 | ENSG00000174652 | ZNF266 | chr19 | - | 9433667 | 9433783 | 9419951 | 9420218 | 9434109 | 9434235 | 167119 | 14,13,79,73,56,18,60,37,5 | 0,0,15,5,1,0,2,4,2,0,0,0 | 97,42,37,14,19,13,18,50,3 | 1,2,2,1,4,1,4,2,2,1,3,0,0,2, | 265 | 149 | 0.0289957358517 | | 1 | 1.0,1.0,0.748,0.89 | 0.982,0.922,0.912, | 0,053 |
| 233682 | ENSG00000101974 | ATP11C | chrX | - | 139802037 | 139802097 | 139800059 | 139800110 | 139802235 | 139802339 | 233682 | 4,2,2,0,1,2,0,2,4,2,0,0 | 73,33,44,34,31,24,22,22,2 | 5,2,2,3,3,0,1,1,2,9,1,7,2,0, | 19,17,45,28,21,19,11,26,1 | 209 | 149 | 0.0290280305693 | | 1 | 0.038,0.041,0.031, | 0.158,0.077,0.031, | -0,023 |
| 172062 | ENSG00000163539 | CLASP2 | chr3 | - | 33700301 | 33700421 | 33696854 | 33696933 | 33717807 | 33718356 | 172062 | 0,0,3,0,0,0,0,6,2,0,1,0 | 13,16,24,8,20,16,19,18,14, | 3,4,3,2,1,1,0,0,2,0,3,3,1,1, | 21,21,10,14,14,8,10,14,13, | 269 | 149 | 0.0290338466036 | | 1 | 0.0,0.0,0.065,0.0,0 | 0.073,0.095,0.142, | -0,02 |
| 101795 | ENSG00000196792 | STRN3 | chr14 | - | 30929200 | 30929311 | 30918965 | 30919106 | 30935162 | 30935304 | 101795 | 17,15,28,9,3,11,6,10,13,5, | 0,0,2,1,2,1,0,0,0,0,0,1 | 14,5,6,22,4,10,6,15,4,2,11, | 4,0,0,2,0,0,6,4,0,1,1,0,1,0, | 260 | 149 | 0.0290515493665 | | 1 | 1.0,1.0,0.889,0.83 | 0.667,1.0,1.0,0.86 | 0,086 |
| 180560 | ENSG00000138668 | HNRNPD | chr4 | - | 82356536 | 82356705 | 82353846 | 82355401 | 82356795 | 82356895 | 180560 | 16,16,51,27,35,30,30,44,7, | 25,14,68,35,24,50,30,44,1 | 65,89,23,96,51,25,35,35,6 | 50,99,19,62,40,27,45,34,3 | 298 | 149 | 0.0290598946984 | | 1 | 0.242,0.364,0.273, | 0.394,0.31,0.377,0 | -0,058 |
| 224645 | ENSG00000001631 | KRIT1 | chr7 | - | 92240748 | 92240847 | 92237666 | 92237759 | 92240992 | 92241152 | 224645 | 0,1,0,0,2,5,0,1,0,0,1,2 | 9,2,23,23,9,14,11,20,7,4,1 | 3,1,1,0,2,5,0,4,2,0,1,1,0,4, | 16,14,8,10,10,10,10,18,6,1 | 248 | 149 | 0.0290630361011 | | 1 | 0.0,0.231,0.0,0.0,0 | 0.101,0.041,0.07,0 | -0,013 |
| 138928 | ENSG00000118705 | RPN2 | chr20 | + | 37213801 | 37213865 | 37210046 | 37210165 | 37223877 | 37223969 | 138928 | 112,111,71,71,46,45,36,66 | 10,1,5,8,2,6,3,4,4,2,0,4 | 58,69,65,50,31,21,21,48,1 | 7,7,0,5,4,5,3,5,2,6,2,9,3,7, | 213 | 149 | 0.0291035148613 | | 1 | 0.887,0.987,0.909, | 0.853,0.873,1.0,0. | 0,05 |
| 215016 | ENSG00000003402 | CFLAR | chr2 | + | 201162995 | 201163106 | 201160431 | 201160942 | 201163834 | 201176687 | 215016 | 10,11,49,32,11,20,9,18,10, | 62,40,139,85,64,66,69,106 | 14,21,14,39,13,18,9,35,20, | 68,102,28,26,46,25,41,105 | 260 | 149 | 0.0291102019846 | | 1 | 0.085,0.136,0.168, | 0.106,0.106,0.223, | -0,05 |
| 83961 | ENSG00000155438 | NIFK | chr2 | - | 121732095 | 121732204 | 121730382 | 121731104 | 121735612 | 121735654 | 83961 | 43,40,121,68,25,41,31,45, | 3,0,2,3,0,0,0,0,0,0,2,0 | 47,100,44,13,67,35,39,15, | 5,3,2,0,0,0,0,1,0,1,2,4,0,1, | 258 | 149 | 0.0291297761646 | | 1 | 0.892,1.0,0.972,0. | 0.844,0.951,0.927, | 0,025 |
| 50324 | ENSG00000132781 | MUTYH | chr1 | - | 45333412 | 45333497 | 45332676 | 45333324 | 45334390 | 45334511 | 50324 | 18,17,16,10,13,23,8,15,19, | 1,2,3,0,0,2,2,0,1,4,0,3 | 20,9,17,18,11,4,5,14,7,12, | 1,1,0,3,2,0,0,0,0,0,0,3,0,1, | 234 | 149 | 0.0291308567818 | | 1 | 0.92,0.844,0.773,1 | 0.927,0.851,1.0,0. | -0,061 |
| 98168 | ENSG00000134905 | CARS2 | chr13 | - | 110687946 | 110688018 | 110683050 | 110683134 | 110701437 | 110701555 | 98168 | 22,13,47,31,18,22,25,27,1 | 2,0,6,0,0,2,5,3,2,0,2,0 | 23,15,7,12,23,17,6,8,12,17 | 3,2,0,4,0,0,0,5,4,4,3,5,3,2, | 221 | 149 | 0.0291387813666 | | 1 | 0.881,1.0,0.841,1. | 0.838,0.835,1.0,0. | 0,071 |
| 220414 | ENSG00000197323 | TRIM33 | chr1 | - | 114430797 | 114430877 | 114427747 | 114427894 | 114433616 | 114433733 | 220414 | 44,42,53,30,16,26,21,27,9, | 7,5,8,1,1,6,2,2,0,1,0,0 | 33,43,31,39,30,20,7,32,16, | 4,5,6,5,15,4,3,3,2,4,2,5,4,4 | 229 | 149 | 0.029179118365 | | 1 | 0.804,0.845,0.812, | 0.843,0.848,0.771, | 0,096 |
| 149663 | ENSG00000214013 | GANC | chr15 | + | 42280921 | 42281133 | 42278481 | 42278590 | 42287690 | 42287818 | 149663 | 2,3,4,1,0,0,0,0,0,0,1,2 | 58,28,55,42,38,39,34,25,1 | 4,1,1,0,4,3,0,2,1,1,3,1,4,2, | 38,32,22,14,24,26,10,21,2 | 298 | 149 | 0.0291839218801 | | 1 | 0.017,0.051,0.035, | 0.05,0.015,0.022,0 | -0,015 |
| 41892 | ENSG00000143669 | LYST | chr1 | - | 235753043 | 235753274 | 235746335 | 235746527 | 235755477 | 235755647 | 41892 | 43,32,108,17,57,34,17,60, | 0,1,2,4,2,0,0,3,0,2,2,2 | 54,29,21,13,8,12,15,16,12, | 0,0,0,0,1,2,0,0,0,1,0,1,2,3, | 298 | 149 | 0.0292009394949 | | 1 | 1.0,0.941,0.964,0. | 1.0,1.0,1.0,1.0,0.8, | -0,031 |
| 37572 | ENSG00000154144 | TBRG1 | chr11 | + | 124626472 | 124626609 | 124625670 | 124625903 | 124626903 | 124627050 | 37572 | 12,6,19,4,13,6,5,4,15,2,3,5 | 6,0,6,12,5,4,10,4,2,0,2,0 | 8,4,2,2,3,4,4,2,3,4,4,3,13,1 | 4,11,3,3,3,3,0,3,2,10,4,4,8, | 286 | 149 | 0.0292230055613 | | 1 | 0.51,1.0,0.623,0.1 | 0.51,0.159,0.258,0 | 0,209 |
| 254305 | ENSG00000111596 | CNOT2 | chr12 | + | 70329408 | 70329570 | 70319297 | 70319364 | 70332766 | 70332846 | 254305 | 13,18,10,7,0,4,3,11,8,6,3,1 | 1,3,0,0,1,3,0,1,0,0,0,0 | 4,5,9,4,5,5,6,1,6,3,14,3,21, | 3,1,2,0,3,0,2,0,0,0,0,6,1, | 298 | 149 | 0.0292261205214 | | 1 | 0.867,0.75,1.0,1.0, | 0.4,0.714,0.692,1. | 0,043 |
| 186554 | ENSG00000077097 | TOP2B | chr3 | - | 25618169 | 25618509 | 25615472 | 25615586 | 25618653 | 25618849 | 186554 | 72,35,81,68,44,46,34,36,2 | 3,1,4,1,0,3,0,1,1,2,1,3 | 41,40,27,23,33,21,30,32,2 | 1,9,3,2,7,2,1,2,0,2,0,3,1,3,1 | 298 | 149 | 0.0292500863822 | | 1 | 0.923,0.946,0.91,0 | 0.953,0.69,0.818,0 | 0,034 |
| 51565 | ENSG00000162894 | FCMR | chr1 | - | 206913758 | 206914094 | 206912928 | 206913042 | 206921817 | 206921867 | 51565 | 2178,737,2172,1648,1412, | 117,37,147,119,123,70,10 | 1552,785,1185,672,403,66 | 120,56,126,41,34,55,64,72 | 298 | 149 | 0.0292632313078 | | 1 | 0.903,0.909,0.881, | 0.866,0.875,0.825, | -0,013 |
| 79195 | ENSG00000112078 | KCTD20 | chr6 | + | 36474788 | 36475062 | 36470051 | 36470257 | 36479590 | 36479711 | 79195 | 19,4,38,37,21,10,9,13,22,1 | 4,4,14,11,0,1,3,4,3,0,1,6 | 14,26,8,9,7,9,10,8,19,25,7, | 14,15,6,4,1,2,5,5,3,4,18,12 | 298 | 149 | 0.0292787842284 | | 1 | 0.704,0.333,0.576, | 0.333,0.464,0.4,0. | 0,107 |
| 201325 | ENSG00000157450 | RNF111 | chr15 | + | 59085221 | 59085330 | 59084128 | 59084254 | 59085658 | 59085785 | 201325 | 9,0,3,1,4,0,0,1,5,0,0,2 | 8,15,13,12,15,12,9,7,17,10 | 0,3,0,4,0,0,0,2,1,1,1,0,4,0, | 7,6,9,4,5,10,11,13,5,6,8,17 | 258 | 149 | 0.029282259549 | | 1 | 0.394,0.0,0.118,0. | 0.0,0.224,0.0,0.36 | 0 |
| 173586 | ENSG00000168591 | TMUB2 | chr17 | + | 44187566 | 44187743 | 44187049 | 44187109 | 44187906 | 44188009 | 173586 | 34,19,50,22,12,17,5,23,11, | 0,6,13,3,0,1,0,3,0,0,0,3 | 21,22,32,5,25,7,0,12,15,22 | 6,10,7,0,1,1,2,1,2,6,9,0,5,1 | 298 | 149 | 0.0293156573135 | | 1 | 1.0,0.613,0.658,0. | 0.636,0.524,0.696, | 0,184 |
| 217367 | ENSG00000128708 | HAT1 | chr2 | + | 171922921 | 171923117 | 171922461 | 171922507 | 171925536 | 171925641 | 217367 | 2,0,4,1,0,1,0,2,4,3,0,0 | 2,1,25,11,10,11,6,7,2,2,4,8 | 4,3,2,0,0,3,3,0,1,2,6,6,1,0, | 6,7,5,2,8,7,1,6,5,1,2,6,11,5 | 298 | 149 | 0.0293210540219 | | 1 | 0.333,0.0,0.074,0. | 0.25,0.176,0.167,0 | -0,055 |
| 31866 | ENSG00000247774 | PCED1B-AS1 | chr12 | - | 47206510 | 47206588 | 47205892 | 47206145 | 47210639 | 47210761 | 31866 | 10,5,12,11,6,4,1,1,15,5,1,3 | 2,0,4,5,0,0,2,0,7,5,0,1 | 3,8,1,2,2,11,0,1,4,8,2,44,7, | 1,1,1,2,2,4,1,0,0,0,0,10,2,0 | 227 | 149 | 0.0293299081558 | | 1 | 0.766,1.0,0.663,0. | 0.663,0.84,0.396,0 | 0,044 |
| 150050 | ENSG00000159322 | ADPGK | chr15 | - | 72760394 | 72760527 | 72751371 | 72752892 | 72771782 | 72771845 | 150050 | 46,45,63,45,42,40,24,37,4 | 2,5,10,4,1,5,1,4,0,0,0,5 | 41,61,54,24,22,29,26,40,3 | 4,6,1,2,12,3,4,4,4,6,2,8,4,1 | 282 | 149 | 0.0293318374947 | | 1 | 0.924,0.826,0.769, | 0.844,0.843,0.966, | 0,087 |
| 79223 | ENSG00000143924 | EML4 | chr2 | + | 42315961 | 42316050 | 42304483 | 42304551 | 42325466 | 42325554 | 79223 | 35,26,34,25,38,22,12,21,1 | 0,4,1,0,0,1,0,0,0,0,2,0 | 55,61,25,43,29,17,27,60,1 | 0,5,0,5,3,0,2,1,0,2,0,4,2,1, | 238 | 149 | 0.0293369794729 | | 1 | 1.0,0.803,0.955,1. | 1.0,0.884,1.0,0.84 | -0,004 |
| 131367 | ENSG00000079332 | SAR1A | chr10 | - | 70162689 | 70162759 | 70161857 | 70161931 | 70170412 | 70170523 | 131367 | 13,18,21,26,6,10,11,19,7,7 | 2,6,9,9,11,2,1,5,3,5,5,2 | 15,34,21,30,24,12,11,16,1 | 5,8,2,0,7,3,4,0,2,3,1,8,5,2,1 | 219 | 149 | 0.0293440524929 | | 1 | 0.816,0.671,0.614, | 0.671,0.743,0.877, | -0,134 |
| 243441 | ENSG00000174106 | LEMD3 | chr12 | + | 65240658 | 65240809 | 65240135 | 65240238 | 65240908 | 65241087 | 243441 | 6,5,19,5,3,5,7,10,2,0,5,6 | 10,16,89,68,23,33,35,63,1 | 5,5,2,7,8,8,3,9,3,3,12,0,2,2 | 62,63,20,19,33,20,21,29,1 | 298 | 149 | 0.0293609787468 | | 1 | 0.231,0.135,0.096, | 0.039,0.038,0.048, | 0,027 |
| 76701 | ENSG00000159023 | EPB41 | chr1 | + | 29109335 | 29109437 | 29096231 | 29097935 | 29112367 | 29112448 | 76701 | 402,318,455,334,309,261, | 3,5,9,10,0,3,4,0,4,2,0,2 | 307,261,323,354,344,270, | 2,3,8,2,7,5,2,3,2,17,0,4,3,3 | 251 | 149 | 0.0293701659119 | | 1 | 0.988,0.974,0.968, | 0.989,0.981,0.96,0 | 0,01 |
| 114634 | ENSG00000135018 | UBQLN1 | chr9 | - | 83666349 | 83666433 | 83665029 | 83665145 | 83669184 | 83669327 | 114634 | 131,112,125,82,78,55,41,4 | 12,13,9,3,1,3,7,1,3,3,0,0 | 96,125,95,151,79,62,90,66 | 7,16,2,28,13,7,8,3,3,4,0,4,1 | 233 | 149 | 0.0293830397358 | | 1 | 0.875,0.846,0.899, | 0.898,0.833,0.968, | 0,037 |
| 224289 | ENSG00000168014 | C2CD3 | chr11 | - | 74108041 | 74108143 | 74106370 | 74106493 | 74109033 | 74109152 | 224289 | 3,1,2,2,1,0,3,1,0,0,1,3 | 12,3,9,14,5,9,3,6,7,1,10,10 | 0,0,0,1,1,0,5,0,0,0,1,3,0,2, | 15,8,8,13,5,7,8,9,7,4,9,2,8, | 251 | 149 | 0.0294091531634 | | 1 | 0.129,0.165,0.117, | 0.0,0.0,0.0,0.044,0 | 0,038 |
| 253194 | ENSG00000122965 | RBM19 | chr12 | - | 113822199 | 113822646 | 113816741 | 113818186 | 113823208 | 113823321 | 253194 | 7,3,9,4,5,2,3,5,2,2,1,1 | 3,0,13,8,11,18,14,5,9,3,2,2 | 13,5,10,4,2,2,2,1,5,9,5,4,3, | 6,2,1,0,2,5,5,14,0,1,5,8,0,8 | 298 | 149 | 0.0294184558364 | | 1 | 0.538,1.0,0.257,0. | 0.52,0.556,0.833,1 | -0,184 |
| 243469 | ENSG00000065923 | SLC9A7 | chrX | - | 46682076 | 46682535 | 46679677 | 46679755 | 46758704 | 46759055 | 243469 | 94,85,189,121,79,115,87,1 | 12,4,19,8,6,21,17,29,1,1,5, | 172,113,69,111,78,51,58,1 | 11,17,4,11,15,4,3,4,4,6,10, | 298 | 149 | 0.0294517799355 | | 1 | 0.797,0.914,0.833, | 0.887,0.769,0.896, | -0,037 |
| 5153 | ENSG00000157540 | DYRK1A | chr21 | + | 37420298 | 37420384 | 37366772 | 37367628 | 37472683 | 37472880 | 5153 | 31,21,32,15,19,28,21,34,4 | 2,1,2,3,2,4,3,0,5,0,1,0 | 39,37,32,42,19,17,23,32,9, | 2,2,0,0,3,0,3,0,0,0,2,0,3,4, | 235 | 149 | 0.0294610641164 | | 1 | 0.908,0.93,0.91,0. | 0.925,0.921,1.0,1. | -0,03 |
| 155549 | ENSG00000173575 | CHD2 | chr15 | + | 92991692 | 92991780 | 92991475 | 92991517 | 92992858 | 92992998 | 155549 | 33,17,27,26,2,11,13,12,8,9 | 33,33,44,27,36,24,17,22,2 | 14,15,3,19,11,9,4,9,11,14, | 35,83,16,22,33,18,10,23,2 | 237 | 149 | 0.0294788576573 | | 1 | 0.386,0.245,0.278, | 0.201,0.102,0.105, | 0,057 |
| 132499 | ENSG00000171792 | RHNO1 | chr12 | + | 2885262 | 2885534 | 2877198 | 2877282 | 2887910 | 2888137 | 132499 | 79,62,58,47,38,30,24,31,5 | 3,2,2,7,0,3,3,2,1,0,0,2 | 6,24,32,27,25,12,22,35,9,3 | 7,1,1,0,3,2,1,4,2,0,1,3,4,4, | 298 | 149 | 0.0294792915292 | | 1 | 0.929,0.939,0.935, | 0.3,0.923,0.941,1. | 0,089 |
| 52025 | ENSG00000072210 | ALDH3A2 | chr17 | + | 19655280 | 19655387 | 19652546 | 19652632 | 19656365 | 19656574 | 52025 | 5,4,9,1,0,8,3,2,1,1,2,0 | 33,15,28,15,12,5,6,3,16,4, | 4,5,18,4,1,4,5,0,0,4,1,0,11, | 7,6,6,1,2,5,6,8,13,12,2,7,1 | 256 | 149 | 0.0294846209902 | | 1 | 0.081,0.134,0.158, | 0.25,0.327,0.636,0 | -0,103 |
| 76584 | ENSG00000118482 | PHF3 | chr6 | + | 63706728 | 63706876 | 63706028 | 63706224 | 63709150 | 63709214 | 76584 | 112,77,390,194,95,122,10 | 3,0,7,1,4,6,6,4,0,0,2,0 | 200,235,110,111,111,87,1 | 4,6,2,4,0,2,4,8,3,4,8,1,4,9, | 297 | 149 | 0.0294978695537 | | 1 | 0.949,1.0,0.965,0. | 0.962,0.952,0.965, | 0,018 |
| 110094 | ENSG00000064115 | TM7SF3 | chr12 | - | 26979783 | 26979936 | 26975495 | 26975658 | 26980565 | 26980646 | 110094 | 6,3,35,14,19,11,3,13,9,5,6, | 0,0,5,1,2,2,2,2,0,0,0,1 | 17,12,4,5,4,3,8,16,5,10,10, | 1,0,2,0,0,0,1,1,1,1,0,5,2,4, | 298 | 149 | 0.0295003759973 | | 1 | 1.0,1.0,0.778,0.87 | 0.895,1.0,0.5,1.0,1 | -0,018 |
| 131036 | ENSG00000050426 | LETMD1 | chr12 | + | 51052091 | 51052207 | 51048342 | 51049185 | 51053811 | 51053860 | 131036 | 3,4,15,10,2,13,0,3,1,5,1,6 | 3,3,2,3,8,0,1,6,7,7,0,1 | 10,11,6,3,1,4,2,7,1,4,8,13, | 1,3,3,1,1,2,0,0,2,1,3,5,5,0, | 265 | 149 | 0.0295210677455 | | 1 | 0.36,0.428,0.808,0 | 0.849,0.673,0.529, | -0,186 |
| 5014 | ENSG00000125731 | SH2D3A | chr19 | - | 6760637 | 6760987 | 6757329 | 6759670 | 6763679 | 6763816 | 5014 | 44,17,68,47,64,41,18,30,2 | 6,0,1,1,2,2,4,4,0,2,1,0 | 44,24,16,17,19,9,27,22,21, | 1,4,0,3,1,0,0,1,0,0,0,2,2,0, | 298 | 149 | 0.0295228825264 | | 1 | 0.786,1.0,0.971,0. | 0.957,0.75,1.0,0.7 | -0,037 |
| 71231 | ENSG00000197343 | ZNF655 | chr7 | + | 99562348 | 99562494 | 99560532 | 99560695 | 99571687 | 99571717 | 71231 | 2,3,34,19,12,14,14,16,7,2, | 4,0,1,5,5,1,0,2,0,0,0,0 | 32,25,10,8,7,1,8,23,11,12, | 1,0,0,2,0,0,0,2,1,4,0,3,4,2, | 295 | 149 | 0.0295423951144 | | 1 | 0.202,1.0,0.945,0. | 0.942,1.0,1.0,0.66 | -0,04 |
| 87435 | ENSG00000036257 | CUL3 | chr2 | - | 224556294 | 224556389 | 224535606 | 224535641 | 224557658 | 224557856 | 87435 | 2,1,8,0,1,0,0,6,0,0,0,2 | 28,34,106,66,39,59,33,59, | 6,8,0,4,0,1,1,7,0,5,4,2,1,1, | 69,99,35,70,49,20,49,35,61 | 244 | 149 | 0.0295696679894 | | 1 | 0.042,0.018,0.044, | 0.05,0.047,0.0,0.0 | -0,016 |
| 235157 | ENSG00000075539 | FRYL | chr4 | - | 48539970 | 48540024 | 48535656 | 48535827 | 48540352 | 48540960 | 235157 | 6,8,18,20,4,7,7,14,3,5,3,8 | 2,4,1,2,2,8,2,2,1,1,3,3 | 12,10,18,7,7,6,2,8,4,11,10, | 3,12,5,10,5,2,2,4,6,8,5,2,7, | 203 | 149 | 0.0295764289678 | | 1 | 0.688,0.595,0.93,0 | 0.746,0.38,0.725,0 | 0,105 |
| 231102 | ENSG00000178761 | FAM219B | chr15 | - | 74901949 | 74902129 | 74899991 | 74900137 | 74902603 | 74902786 | 231102 | 4,1,14,9,12,2,2,4,1,0,1,0 | 0,0,11,6,4,2,2,3,0,2,0,2 | 14,4,4,2,2,9,0,7,2,4,4,5,6,9 | 4,0,0,1,2,0,7,1,0,0,3,3,2,0, | 298 | 149 | 0.0295765285615 | | 1 | 1.0,1.0,0.389,0.42 | 0.636,1.0,1.0,0.5,0 | -0,154 |
| 93282 | ENSG00000149657 | LSM14B | chr20 | + | 62126225 | 62126439 | 62124616 | 62124780 | 62129784 | 62129952 | 93282 | 19,5,30,16,14,17,13,10,16, | 0,0,3,3,1,2,0,0,0,0,0,1 | 17,7,12,9,10,11,6,21,12,11 | 3,1,2,3,0,0,0,2,0,5,2,2,3,0, | 298 | 149 | 0.0296102944426 | | 1 | 1.0,1.0,0.833,0.72 | 0.739,0.778,0.75,0 | 0,092 |
| 171870 | ENSG00000152102 | FAM168B | chr2 | - | 131072021 | 131072295 | 131071854 | 131071938 | 131082576 | 131082657 | 171870 | 0,0,0,0,0,2,0,0,0,0,2,0 | 25,16,17,23,16,15,19,19,4 | 0,5,3,6,1,1,1,2,0,0,3,0,0,0, | 16,26,13,17,10,5,19,18,3,2 | 298 | 149 | 0.0296236190694 | | 1 | 0.0,0.0,0.0,0.0,0.0, | 0.0,0.088,0.103,0. | -0,043 |
| 177016 | ENSG00000256771 | ZNF253 | chr19 | + | 19872127 | 19872383 | 19865917 | 19865999 | 19891473 | 19894674 | 177016 | 2,3,8,9,3,4,8,5,0,0,0,2 | 24,13,35,20,6,3,13,12,8,6, | 6,9,0,2,4,3,0,6,3,1,1,11,3,1 | 8,7,5,6,9,7,14,8,3,12,14,2, | 298 | 149 | 0.0296289178013 | | 1 | 0.04,0.103,0.103,0 | 0.273,0.391,0.0,0. | -0,111 |
| 205532 | ENSG00000134686 | PHC2 | chr1 | - | 33325794 | 33326149 | 33323625 | 33325019 | 33328869 | 33329146 | 205532 | 4,1,1,4,0,2,0,4,2,0,1,0 | 11,2,25,29,6,7,11,14,17,5, | 1,7,0,0,0,0,0,2,0,0,2,0,0,0, | 13,28,6,4,13,6,9,23,8,21,1 | 298 | 149 | 0.0296610048614 | | 1 | 0.154,0.2,0.02,0.0 | 0.037,0.111,0.0,0. | 0,042 |
| 49562 | ENSG00000133789 | SWAP70 | chr11 | + | 9713465 | 9713639 | 9694145 | 9694286 | 9728052 | 9728071 | 49562 | 119,73,239,176,80,53,62,5 | 5,8,2,4,3,5,2,0,0,0,1,3 | 129,154,87,63,81,78,30,28 | 9,10,7,1,6,2,2,3,0,0,1,10,2, | 298 | 149 | 0.029668095091 | | 1 | 0.922,0.82,0.984,0 | 0.878,0.885,0.861, | 0,032 |
| 20984 | ENSG00000100906 | NFKBIA | chr14 | - | 35402770 | 35402859 | 35401551 | 35402663 | 35403149 | 35403360 | 20984 | 526,308,256,194,469,531, | 18,6,0,0,2,5,5,2,7,2,0,0 | 219,242,114,144,43,43,52, | 2,4,1,0,0,0,4,7,0,0,3,0,0,0, | 238 | 149 | 0.02967229606 | | 1 | 0.948,0.97,1.0,1.0, | 0.986,0.974,0.986, | -0,003 |
| 232042 | ENSG00000185101 | ANO9 | chr11 | - | 423941 | 424067 | 421140 | 421198 | 428087 | 428199 | 232042 | 1,0,5,8,2,2,0,3,0,0,1,3 | 15,6,36,22,11,10,17,11,4,4 | 1,2,2,0,0,1,0,0,4,3,2,3,1,1, | 11,16,6,20,18,7,3,8,21,22, | 275 | 149 | 0.0297058365193 | | 1 | 0.035,0.0,0.07,0.1 | 0.047,0.063,0.153, | 0,004 |
| 29118 | ENSG00000273841 | TAF9 | chr5 | - | 69366502 | 69366595 | 69364742 | 69365754 | 69369656 | 69370013 | 29118 | 90,120,88,101,28,27,18,51 | 0,0,1,0,0,0,0,0,0,0,0,0 | 66,79,96,34,63,28,9,24,31, | 0,0,0,19,2,0,4,0,2,0,1,0,0,1 | 242 | 149 | 0.029715096618 | | 1 | 1.0,1.0,0.982,1.0,1 | 1.0,1.0,1.0,0.524,0 | 0,07 |
| 79321 | ENSG00000138795 | LEF1 | chr4 | - | 108081585 | 108081669 | 108079491 | 108079614 | 108083355 | 108083446 | 79321 | 260,69,456,376,187,223,2 | 91,12,122,90,22,33,49,66, | 193,168,188,79,94,97,136, | 50,37,112,57,32,32,50,97, | 233 | 149 | 0.0297191557895 | | 1 | 0.646,0.786,0.705, | 0.712,0.744,0.518, | 0,1 |
| 186214 | ENSG00000132581 | SDF2 | chr17 | - | 28661725 | 28661939 | 28655286 | 28655483 | 28662045 | 28662189 | 186214 | 6,10,14,26,11,5,5,6,26,7,1, | 5,17,30,35,5,15,9,8,2,2,5,6 | 6,16,8,4,2,1,5,3,9,5,4,12,2, | 24,15,14,1,18,6,10,18,7,0,1 | 298 | 149 | 0.0297347260978 | | 1 | 0.375,0.227,0.189, | 0.111,0.348,0.222, | 0,086 |
| 192125 | ENSG00000117682 | DHDDS | chr1 | + | 26447558 | 26447682 | 26446315 | 26446432 | 26457790 | 26457905 | 192125 | 19,16,30,39,20,11,14,9,19, | 0,1,2,2,0,1,0,1,4,1,0,0 | 18,14,14,6,16,7,4,22,9,23, | 1,2,1,0,0,3,1,0,1,1,0,2,3,2, | 273 | 149 | 0.0297396321246 | | 1 | 1.0,0.897,0.891,0. | 0.908,0.793,0.884, | 0,064 |
| 112524 | ENSG00000119397 | CNTRL | chr9 | + | 121157982 | 121158109 | 121157457 | 121157600 | 121158854 | 121159019 | 112524 | 111,87,103,55,27,32,16,22 | 36,11,20,22,1,16,15,17,9,7 | 47,49,70,61,37,34,14,29,3 | 22,24,19,38,26,27,8,12,17, | 276 | 149 | 0.0297440664674 | | 1 | 0.625,0.81,0.735,0 | 0.536,0.524,0.665, | 0,116 |
| 170614 | ENSG00000076108 | BAZ2A | chr12 | - | 56630775 | 56630893 | 56617394 | 56617532 | 56636181 | 56636379 | 170614 | 7,6,12,7,2,5,8,13,6,4,1,7 | 11,13,20,7,8,19,7,13,4,9,3, | 5,8,4,8,8,7,0,13,5,7,1,2,5,1 | 10,35,11,32,23,11,5,10,12, | 267 | 149 | 0.0297551918327 | | 1 | 0.262,0.205,0.251, | 0.218,0.113,0.169, | 0,092 |
| 127639 | ENSG00000138134 | STAMBPL1 | chr10 | + | 88913100 | 88913458 | 88910915 | 88911011 | 88914533 | 88914658 | 127639 | 91,37,119,108,42,37,38,32 | 48,2,40,31,14,30,13,23,4,1 | 35,36,45,30,29,36,18,51,2 | 24,39,21,8,24,7,9,13,35,18 | 298 | 149 | 0.029838517425 | | 1 | 0.487,0.902,0.598, | 0.422,0.316,0.517, | 0,108 |
| 145170 | ENSG00000159210 | SNF8 | chr17 | - | 48936854 | 48936954 | 48936040 | 48936242 | 48937019 | 48937239 | 145170 | 2,1,10,4,3,1,1,4,1,0,0,3 | 32,15,52,47,26,14,34,31,1 | 7,4,5,2,6,1,2,2,1,2,8,4,1,1, | 24,29,19,16,18,16,10,36,1 | 249 | 149 | 0.0298654740454 | | 1 | 0.036,0.038,0.103, | 0.149,0.076,0.136, | -0,043 |
| 2176 | ENSG00000244625 | MIATNB | chr22 | + | 26717252 | 26717459 | 26693680 | 26693836 | 26774155 | 26774217 | 2176 | 11,5,14,5,15,6,6,7,18,13,5, | 0,0,1,0,1,6,1,3,0,0,1,1 | 25,17,0,1,13,14,9,4,13,16, | 3,3,2,0,3,0,2,1,2,0,1,10,1,0 | 298 | 149 | 0.0298706800287 | | 1 | 1.0,1.0,0.875,1.0,0 | 0.806,0.739,0.0,1. | 0,039 |
| 243471 | ENSG00000065923 | SLC9A7 | chrX | - | 46682139 | 46682535 | 46679677 | 46679755 | 46758704 | 46759055 | 243471 | 94,85,189,121,79,115,87,1 | 12,4,19,8,6,21,17,29,1,1,5, | 172,113,69,108,78,50,58,1 | 11,17,4,11,15,4,3,4,4,6,10, | 298 | 149 | 0.0298954977109 | | 1 | 0.797,0.914,0.833, | 0.887,0.769,0.896, | -0,037 |
| 196932 | ENSG00000162971 | TYW5 | chr2 | - | 199939900 | 199939958 | 199938932 | 199939070 | 199940088 | 199940133 | 196932 | 4,0,11,8,3,7,1,5,0,0,3,5 | 7,10,17,9,2,5,4,7,4,1,4,3 | 7,12,3,4,9,3,4,9,6,1,6,0,8,4 | 7,20,9,3,9,5,0,1,10,1,4,2,4, | 207 | 149 | 0.0299138938474 | | 1 | 0.291,0.0,0.318,0. | 0.419,0.302,0.194, | -0,2 |
| 103490 | ENSG00000012124 | CD22 | chr19 | + | 35338167 | 35338431 | 35337754 | 35338021 | 35340880 | 35341138 | 103490 | 399,249,260,200,302,173, | 3,4,5,3,11,3,2,2,0,2,1,1 | 264,129,82,128,52,87,55,7 | 9,6,1,4,2,0,0,2,2,2,2,28,9,2 | 298 | 149 | 0.0299246815106 | | 1 | 0.985,0.969,0.963, | 0.936,0.915,0.976, | 0,012 |
| 126570 | ENSG00000078403 | MLLT10 | chr10 | + | 21617111 | 21617207 | 21614830 | 21614924 | 21651672 | 21651768 | 126570 | 28,17,20,20,13,12,5,16,30, | 4,0,13,4,2,8,7,5,3,1,1,0 | 13,9,9,19,3,15,3,16,7,18,1 | 5,11,4,13,6,0,2,10,7,0,8,6,2 | 245 | 149 | 0.0299511420404 | | 1 | 0.81,1.0,0.483,0.7 | 0.613,0.332,0.578, | 0,155 |
| 107080 | ENSG00000147133 | TAF1 | chrX | + | 71459149 | 71459251 | 71458240 | 71458366 | 71459551 | 71459708 | 107080 | 9,6,23,26,13,8,7,7,10,5,5,8 | 17,23,67,37,20,19,17,23,5, | 7,12,5,7,9,8,4,8,4,9,7,3,11, | 29,50,19,40,28,28,13,7,12, | 251 | 149 | 0.0299759402944 | | 1 | 0.239,0.134,0.169, | 0.125,0.125,0.135, | 0,09 |
| 210485 | ENSG00000140386 | SCAPER | chr15 | - | 76739301 | 76739437 | 76733228 | 76733384 | 76753807 | 76753948 | 210485 | 1,0,14,3,0,0,0,6,4,0,0,0 | 86,35,69,29,25,60,19,43,4 | 4,12,1,3,4,1,0,1,2,0,6,1,2,1 | 39,39,38,43,28,24,13,27,2 | 285 | 149 | 0.029981971883 | | 1 | 0.006,0.0,0.096,0. | 0.051,0.139,0.014, | -0,023 |
| 160587 | ENSG00000236935 | | chr11 | - | 64338970 | 64339402 | 64327483 | 64327536 | 64339899 | 64340154 | 160587 | 4,5,22,5,16,5,6,6,4,6,1,5 | 2,2,2,4,0,0,0,4,6,2,2,6 | 8,3,2,5,1,7,4,3,3,5,6,7,2,2,1 | 1,5,0,0,0,0,0,2,0,2,2,13,2,0 | 298 | 149 | 0.0300094925568 | | 1 | 0.5,0.556,0.846,0. | 0.8,0.231,1.0,1.0,1 | -0,172 |
| 10747 | ENSG00000161939 | RNASEK-C17orf | chr17 | + | 7016909 | 7017014 | 7016494 | 7016685 | 7017260 | 7017502 | 10747 | 62,24,128,116,119,105,84, | 9,6,3,0,2,4,1,2,1,0,5,1 | 80,70,34,18,69,49,57,175, | 1,1,6,5,0,0,4,3,0,1,0,1,0,1, | 254 | 149 | 0.0300117568384 | | 1 | 0.802,0.701,0.962, | 0.979,0.976,0.769, | -0,023 |
| 131112 | ENSG00000061936 | SFSWAP | chr12 | + | 131764455 | 131764686 | 131756472 | 131756644 | 131766117 | 131766308 | 131112 | 99,78,117,62,89,61,63,79, | 2,2,4,2,10,5,8,1,1,3,1,5 | 77,94,75,148,50,24,72,82, | 3,11,1,0,2,0,1,3,3,2,0,3,6,8 | 298 | 149 | 0.0300266337935 | | 1 | 0.961,0.951,0.936, | 0.928,0.81,0.974,1 | -0,035 |
| 75710 | ENSG00000173209 | AHSA2P | chr2 | + | 61185979 | 61186081 | 61185501 | 61185618 | 61186445 | 61186497 | 75710 | 25,19,18,6,2,16,27,39,6,5, | 65,32,254,161,103,78,48,6 | 28,50,39,10,19,4,8,38,31,1 | 61,38,49,35,79,61,76,111, | 251 | 149 | 0.0300285608247 | | 1 | 0.186,0.261,0.04,0 | 0.214,0.439,0.321, | -0,039 |
| 151660 | ENSG00000156958 | GALK2 | chr15 | + | 49281986 | 49282089 | 49239219 | 49239367 | 49283565 | 49283718 | 151660 | 6,3,27,11,7,8,2,11,2,0,2,11 | 3,2,3,1,2,4,3,0,2,3,1,3 | 12,6,4,4,7,7,3,6,2,4,11,15, | 2,0,1,5,5,0,1,0,1,0,0,2,4,0,1 | 252 | 149 | 0.0300315650664 | | 1 | 0.542,0.47,0.842,0 | 0.78,1.0,0.703,0.3 | -0,172 |
| 236481 | ENSG00000122882 | ECD | chr10 | - | 73163732 | 73163952 | 73160433 | 73160551 | 73167865 | 73168095 | 236481 | 56,26,102,46,21,27,33,40, | 8,7,4,4,0,2,0,2,0,2,0,3 | 59,76,37,48,71,27,28,31,3 | 3,4,3,0,2,2,2,4,2,1,0,0,0,1, | 298 | 149 | 0.0300350911729 | | 1 | 0.778,0.65,0.927,0 | 0.908,0.905,0.86,1 | -0,033 |
| 10615 | ENSG00000108840 | HDAC5 | chr17 | - | 44085021 | 44085155 | 44084554 | 44084675 | 44086571 | 44086737 | 10615 | 52,22,122,70,74,53,62,41, | 6,0,1,2,2,1,3,1,0,0,2,2 | 64,38,39,13,17,34,52,70,2 | 2,0,0,3,0,1,1,0,1,1,2,0,0,0, | 283 | 149 | 0.0300454936298 | | 1 | 0.82,1.0,0.985,0.9 | 0.944,1.0,1.0,0.69 | -0,018 |
| 9751 | ENSG00000108604 | SMARCD2 | chr17 | - | 63833295 | 63833420 | 63832132 | 63833170 | 63833586 | 63833722 | 9751 | 128,135,99,58,45,55,42,44 | 5,0,3,0,0,3,5,3,2,0,2,0 | 58,95,142,70,58,46,32,37, | 3,12,3,1,3,5,3,6,0,0,1,0,0,3 | 274 | 149 | 0.0300721663348 | | 1 | 0.933,1.0,0.947,1. | 0.913,0.812,0.963, | 0,02 |
| 184479 | ENSG00000077044 | DGKD | chr2 | + | 233389928 | 233390066 | 233388256 | 233388367 | 233390402 | 233390483 | 184479 | 0,1,12,9,1,4,3,5,15,6,0,2 | 63,43,251,97,46,59,79,124 | 6,5,1,3,2,0,6,2,0,3,1,0,2,7, | 103,107,94,62,89,45,38,57 | 287 | 149 | 0.0300876473269 | | 1 | 0.0,0.012,0.024,0. | 0.029,0.024,0.005, | 0,011 |
| 205887 | ENSG00000128815 | WDFY4 | chr10 | + | 48817145 | 48817409 | 48813956 | 48814082 | 48820233 | 48820437 | 205887 | 81,70,95,67,95,83,50,64,9 | 0,1,2,2,1,6,0,1,0,0,1,4 | 79,75,21,50,19,31,71,85,4 | 2,9,0,5,0,0,1,2,1,0,6,3,0,1, | 298 | 149 | 0.0301036643835 | | 1 | 1.0,0.972,0.96,0.9 | 0.952,0.806,1.0,0. | 0,017 |
| 65558 | ENSG00000158161 | EYA3 | chr1 | - | 27998283 | 27998351 | 27997319 | 27997378 | 27999959 | 28000049 | 65558 | 12,2,21,18,5,8,4,6,4,2,5,11 | 18,4,20,19,10,3,5,6,4,13,6, | 14,4,7,3,3,11,7,9,6,15,12,4 | 6,10,8,0,14,3,0,1,0,5,9,6,1 | 217 | 149 | 0.0301628775051 | | 1 | 0.314,0.256,0.419, | 0.616,0.215,0.375, | -0,192 |

| | | | | | | | | | | | | | | | | | | | | | | | |
|---|---|---|---|---|---|---|---|---|---|---|---|---|---|---|---|---|---|---|---|---|---|---|---|
| 22354 | ENSG00000092108 | SCFD1 | chr14 | + | 30670255 | 30670395 | 30653488 | 30653588 | 30673256 | 30673347 | 22354 | 127,143,185,130,66,72,55, | 3,1,2,0,0,2,0,0,0,0,1,0 | 88,107,87,78,85,52,40,77, | 3,4,1,5,4,0,0,0,1,2,2,2,0,0, | 289 | 149 | 0.0301766447816 | | 1 | 0.956,0.987,0.979, | 0.938,0.932,0.978, | 0,02 |
| 238240 | ENSG00000132294 | EFR3A | chr8 | + | 131954700 | 131954776 | 131953817 | 131953967 | 131955767 | 131955905 | 238240 | 4,4,8,1,0,1,1,4,0,0,0,0 | 30,12,74,72,24,31,22,44,14 | 3,6,8,7,0,1,4,2,8,6,3,1,2,0,2 | 41,52,22,30,41,24,15,25,40 | 225 | 149 | 0.0301934611276 | | 1 | 0.081,0.181,0.067, | 0.046,0.071,0.194, | -0,027 |
| 241157 | ENSG00000198382 | UVRAG | chr11 | + | 75892128 | 75892393 | 75888828 | 75888903 | 75911953 | 75912039 | 241157 | 4,3,14,7,3,8,6,14,3,1,0,9 | 73,41,311,120,75,205,80,2 | 6,2,13,10,2,6,2,11,8,12,9,1 | 131,95,76,52,66,63,81,140 | 298 | 149 | 0.0302134855831 | | 1 | 0.027,0.035,0.022, | 0.022,0.01,0.079,0 | -0,009 |
| 167107 | ENSG00000174652 | ZNF266 | chr19 | - | 9419236 | 9419299 | 9418504 | 9418631 | 9420064 | 9420218 | 167107 | 18,7,140,81,52,18,80,65,4 | 4,0,3,4,0,0,3,2,0,0,2,5 | 74,86,36,16,49,35,26,62,3 | 1,3,0,2,3,0,0,0,0,0,6,0,0,0,1 | 212 | 149 | 0.0302358829847 | | 1 | 0.76,1.0,0.97,0.93 | 0.981,0.953,1.0,0. | -0,053 |
| 183397 | ENSG00000177409 | SAMD9L | chr7 | - | 93147087 | 93147232 | 93145384 | 93146040 | 93148193 | 93148385 | 183397 | 0,1,9,7,4,4,5,2,0,1,1,5 | 4,4,25,26,5,10,20,5,4,1,4,4 | 11,2,7,4,4,2,7,12,5,5,0,12, | 11,4,8,2,1,6,0,14,13,20,14, | 294 | 149 | 0.0302651148886 | | 1 | 0.0,0.112,0.154,0. | 0.336,0.202,0.307, | -0,146 |
| 66897 | ENSG00000198837 | DENND4B | chr1 | - | 153942186 | 153942356 | 153941868 | 153942113 | 153942555 | 153942625 | 66897 | 11,6,17,20,14,7,15,11,5,7, | 0,3,2,2,2,2,0,0,0,0,4,0 | 3,19,2,10,10,10,5,6,14,13, | 1,0,5,0,0,1,0,0,2,0,1,2,0,0, | 298 | 149 | 0.0302697243466 | | 1 | 1.0,0.5,0.81,0.833, | 0.6,1.0,0.167,1.0,1 | 0,012 |
| 65018 | ENSG00000165637 | VDAC2 | chr10 | + | 75217887 | 75217991 | 75214020 | 75214070 | 75219062 | 75219215 | 65018 | 25,6,20,18,10,7,9,10,1,9,2, | 7,12,22,22,13,14,5,12,19,9 | 20,20,10,9,13,8,4,12,4,13, | 6,4,4,4,7,3,5,6,5,8,10,23,1 | 253 | 149 | 0.0303298408931 | | 1 | 0.678,0.227,0.349, | 0.663,0.746,0.596, | -0,122 |
| 76573 | ENSG00000118482 | PHF3 | chr6 | + | 63691736 | 63692043 | 63679999 | 63680161 | 63694580 | 63694690 | 76573 | 112,48,239,82,63,52,44,91 | 4,1,8,8,0,3,3,6,2,0,5,1 | 91,113,65,72,55,44,41,68, | 8,7,5,16,14,2,3,3,2,1,4,2,3, | 298 | 149 | 0.0303325584623 | | 1 | 0.933,0.96,0.937,0 | 0.85,0.89,0.867,0. | 0,03 |
| 100187 | ENSG00000051825 | MPHOSPH9 | chr12 | - | 123202163 | 123202319 | 123198246 | 123198334 | 123202623 | 123203084 | 100187 | 27,15,63,28,19,17,18,12,3 | 7,6,8,11,1,2,6,2,0,0,1,5 | 29,20,21,13,19,20,12,16,3 | 3,4,3,1,7,0,0,1,0,1,3,4,3,11 | 298 | 149 | 0.0303508465488 | | 1 | 0.659,0.556,0.797, | 0.829,0.714,0.778, | -0,07 |
| 205890 | ENSG00000128815 | WDFY4 | chr10 | + | 48817151 | 48817409 | 48813956 | 48814082 | 48820233 | 48820437 | 205890 | 80,70,95,67,95,81,50,64,9 | 0,1,2,2,1,6,0,1,0,0,1,4 | 79,75,21,50,19,31,71,85,4 | 2,9,0,5,0,0,1,2,1,0,6,3,0,1, | 298 | 149 | 0.0303658754404 | | 1 | 1.0,0.972,0.96,0.9 | 0.952,0.806,1.0,0. | 0,017 |
| 57592 | ENSG00000173230 | GOLGB1 | chr3 | - | 121682916 | 121682961 | 121681686 | 121681865 | 121690669 | 121692581 | 57592 | 3,1,5,7,1,1,1,3,2,0,0,0 | 15,10,47,41,16,12,21,17,9, | 2,5,0,1,0,0,4,2,7,0,0,3,6,7, | 15,21,22,35,10,10,8,15,16, | 194 | 149 | 0.0303730360198 | | 1 | 0.133,0.071,0.076, | 0.093,0.155,0.0,0. | -0,022 |
| 156911 | ENSG00000185989 | RASA3 | chr13 | - | 114040999 | 114041089 | 114029810 | 114029887 | 114052051 | 114052155 | 156911 | 12,9,10,5,8,5,12,2,3,2,4,7 | 1,3,3,1,2,2,3,0,1,0,0,0 | 12,17,4,18,10,6,4,19,5,8,7, | 0,2,3,3,3,1,4,3,3,1,1,0,2,0, | 239 | 149 | 0.030374442938 | | 1 | 0.882,0.652,0.675, | 1.0,0.841,0.454,0. | 0,015 |
| 244387 | ENSG00000095370 | SH2D3C | chr9 | - | 127774962 | 127775060 | 127774346 | 127774467 | 127778590 | 127778666 | 244387 | 2,0,8,6,3,1,5,3,9,1,0,1 | 49,8,76,72,62,40,41,39,66, | 0,2,4,0,2,1,2,1,0,6,2,0,3,3, | 49,26,46,26,10,19,17,29,8, | 247 | 149 | 0.0303762045415 | | 1 | 0.024,0.0,0.06,0.0 | 0.0,0.044,0.05,0.0, | 0,002 |
| 203071 | ENSG00000242247 | ARFGAP3 | chr22 | - | 42829478 | 42829531 | 42826939 | 42826999 | 42831548 | 42831652 | 203071 | 2,1,3,3,0,2,0,5,2,1,0,0 | 18,11,36,34,16,14,13,7,18, | 1,1,0,0,1,0,4,0,2,0,0,2,0,0,1 | 20,13,19,11,15,12,15,19,4, | 202 | 149 | 0.0303915118455 | | 1 | 0.076,0.063,0.058, | 0.036,0.054,0.0,0. | 0,029 |
| 89702 | ENSG00000284634 | | chr12 | + | 9656567 | 9656649 | 9655872 | 9655979 | 9656920 | 9657302 | 89702 | 83,31,79,35,15,18,25,15,2 | 95,54,423,153,71,103,91,1 | 122,49,20,15,27,28,50,92, | 369,254,94,51,101,62,109, | 231 | 149 | 0.0304036606065 | | 1 | 0.36,0.27,0.108,0. | 0.176,0.111,0.121, | -0,021 |
| 64004 | ENSG00000143771 | CNIH4 | chr1 | + | 224371282 | 224371423 | 224365878 | 224365991 | 224375794 | 224375998 | 64004 | 62,19,81,73,31,27,20,30,2 | 1,3,4,2,0,3,0,2,0,0,0,3 | 40,58,38,20,38,29,18,32,1 | 0,0,0,0,2,0,2,0,0,0,0,3,0,1, | 290 | 149 | 0.0304136661187 | | 1 | 0.97,0.765,0.912,0 | 1.0,1.0,1.0,1.0,0.9 | -0,018 |
| 3492 | ENSG00000214046 | SMIM7 | chr19 | - | 16654034 | 16654125 | 16646147 | 16647261 | 16659394 | 16659447 | 3492 | 67,68,164,138,53,47,59,92 | 4,0,4,6,1,1,2,5,7,0,1,2 | 85,123,46,27,87,47,45,65, | 5,5,8,0,3,2,0,7,5,6,0,6,2,0, | 240 | 149 | 0.0304210491775 | | 1 | 0.912,1.0,0.962,0. | 0.913,0.939,0.781, | 0,021 |
| 28680 | ENSG00000116906 | GNPAT | chr1 | + | 231258221 | 231258309 | 231250960 | 231251143 | 231260506 | 231260683 | 28680 | 0,0,6,0,0,2,0,1,0,0,0,2 | 45,32,66,42,25,23,17,28,2 | 3,7,2,0,0,0,0,1,2,1,8,0,2,2, | 39,35,32,22,25,9,17,32,15, | 237 | 149 | 0.0304270802233 | | 1 | 0.0,0.0,0.054,0.0,0 | 0.046,0.112,0.038, | -0,017 |
| 175517 | ENSG00000010404 | IDS | chrX | - | 149503311 | 149503445 | 149500948 | 149501037 | 149504156 | 149504293 | 175517 | 52,29,57,50,38,40,24,39,1 | 9,5,7,2,2,16,11,4,3,6,1,7 | 61,46,21,47,27,14,34,59,3 | 3,7,6,8,7,3,1,1,3,0,5,3,3,8,1 | 283 | 149 | 0.0304771023988 | | 1 | 0.753,0.753,0.811, | 0.915,0.776,0.648, | -0,069 |
| 127230 | ENSG00000122884 | P4HA1 | chr10 | - | 73043886 | 73043957 | 73030266 | 73030370 | 73044980 | 73045051 | 127230 | 15,11,11,4,7,6,7,3,8,3,1,5 | 24,25,28,25,8,12,4,8,6,10, | 12,2,0,0,1,0,2,9,0,10,1,10, | 19,9,10,10,3,9,0,17,5,16,2, | 220 | 149 | 0.0304830617495 | | 1 | 0.297,0.23,0.21,0. | 0.3,0.131,0.0,0.0,0 | 0,064 |
| 179960 | ENSG00000204252 | HLA-DOA | chr6 | - | 33007353 | 33007608 | 33007079 | 33007215 | 33008012 | 33008261 | 179960 | 9,3,28,27,9,0,16,24,3,6,1,6 | 0,5,4,3,0,1,3,4,1,1,0,1 | 2,11,11,5,2,7,3,9,3,4,5,13,1 | 5,4,0,1,0,1,0,3,0,2,2,4,4,3, | 298 | 149 | 0.0304897608533 | | 1 | 1.0,0.231,0.778,0. | 0.167,0.579,1.0,0. | 0,068 |
| 25886 | ENSG00000137210 | TMEM14B | chr6 | + | 10751132 | 10751234 | 10749621 | 10749698 | 10755141 | 10755232 | 25886 | 15,15,88,74,32,20,32,23,7, | 1,0,4,8,1,2,2,3,1,0,0,5 | 58,54,11,17,30,9,16,40,15, | 5,9,0,0,0,2,2,3,4,4,6,3,3,7, | 251 | 149 | 0.0305151180649 | | 1 | 0.899,1.0,0.929,0. | 0.873,0.781,1.0,1. | 0,064 |
| 36935 | ENSG00000121210 | TMEM131L | chr4 | + | 153592474 | 153592584 | 153588889 | 153589007 | 153593798 | 153593871 | 36935 | 45,31,131,85,40,80,44,54, | 3,3,9,7,4,5,5,5,1,0,2,3 | 68,95,51,54,39,21,51,60,4 | 14,11,2,2,10,1,0,11,5,1,7,1 | 259 | 149 | 0.0305182586699 | | 1 | 0.896,0.856,0.893, | 0.736,0.832,0.936, | 0,041 |
| 21945 | ENSG00000270580 | PKD1P6-NPIPP | chr16 | - | 15114442 | 15114571 | 15110721 | 15110821 | 15118925 | 15119054 | 21945 | 16,13,57,28,8,21,13,17,12, | 0,0,0,0,0,0,0,0,0,0,6,10 | 43,9,16,15,4,11,16,27,13,1 | 0,0,0,0,0,0,0,0,0,0,0,0,0,0, | 278 | 149 | 0.0305529825342 | | 1 | 1.0,1.0,1.0,1.0,1.0, | 1.0,1.0,1.0,1.0,1.0, | -0,092 |
| 133299 | ENSG00000182670 | TTC3 | chr21 | + | 37097914 | 37098160 | 37096580 | 37096643 | 37108391 | 37108446 | 133299 | 0,0,4,4,2,2,0,5,0,0,0,2 | 37,9,46,45,32,26,34,29,23, | 6,8,2,8,2,3,0,1,4,2,1,1,4,0, | 58,32,37,30,40,21,32,61,1 | 298 | 149 | 0.0305875253653 | | 1 | 0.0,0.0,0.042,0.04 | 0.049,0.111,0.026, | -0,018 |
| 113726 | ENSG00000075856 | SART3 | chr12 | - | 108535358 | 108535468 | 108532221 | 108532334 | 108536513 | 108536572 | 113726 | 44,66,90,63,47,47,33,36,5 | 4,5,5,5,4,5,2,0,2,0,0,0 | 62,97,54,14,99,21,32,34,2 | 1,2,5,1,7,1,1,2,1,0,5,1,1,1, | 259 | 149 | 0.0305934775862 | | 1 | 0.864,0.884,0.912, | 0.973,0.965,0.861, | -0,015 |
| 25748 | ENSG00000145088 | EAF2 | chr3 | + | 121836644 | 121836794 | 121835276 | 121835391 | 121844452 | 121844547 | 25748 | 0,1,2,2,0,0,0,0,4,1,0,2 | 63,7,116,78,35,11,14,13,1 | 3,3,3,0,1,0,2,2,0,8,0,6,1,5, | 79,14,71,4,18,33,6,31,17,6 | 298 | 149 | 0.0305964356158 | | 1 | 0.0,0.067,0.009,0. | 0.019,0.097,0.021, | -0,009 |
| 178758 | ENSG00000070190 | DAPP1 | chr4 | + | 99865725 | 99865836 | 99863769 | 99863855 | 99866033 | 99866121 | 178758 | 34,23,28,4,8,7,4,11,5,4,3,0 | 33,29,44,28,20,13,14,19,2 | 21,14,9,5,9,8,2,2,4,18,5,28 | 18,26,10,9,16,12,2,2,5,15, | 260 | 149 | 0.0306255348915 | | 1 | 0.371,0.312,0.267, | 0.401,0.236,0.34,0 | -0,095 |
| 250950 | ENSG00000196683 | TOMM7 | chr7 | - | 22817999 | 22818047 | 22812970 | 22813185 | 22822676 | 22822849 | 250950 | 61,51,335,243,49,50,59,87 | 4,3,9,14,3,0,6,13,5,0,0,4 | 183,141,42,27,82,55,33,82 | 9,14,2,2,6,0,11,13,4,10,12, | 197 | 149 | 0.0306301462767 | | 1 | 0.92,0.928,0.966,0 | 0.939,0.884,0.941, | 0,025 |
| 189322 | ENSG00000118308 | IRAG2 | chr12 | + | 25104417 | 25104829 | 25104008 | 25104058 | 25106942 | 25107050 | 189322 | 3,0,10,1,0,1,0,5,1,0,1,3 | 5,1,6,14,6,5,6,0,5,1,2,2 | 4,1,3,2,2,2,4,2,2,4,0,1,5,5, | 7,3,2,1,0,3,0,2,4,6,3,0,6,6, | 298 | 149 | 0.0306472989155 | | 1 | 0.231,0.0,0.455,0. | 0.222,0.143,0.429, | -0,192 |
| 153504 | ENSG00000162086 | ZNF75A | chr16 | + | 3316938 | 3317022 | 3312676 | 3312768 | 3317189 | 3317252 | 153504 | 24,18,34,23,18,18,13,25,1 | 3,3,7,9,5,3,3,1,1,2,0,2 | 20,27,16,15,15,6,14,7,3,7,1 | 4,4,8,4,5,3,4,5,3,5,6,2,3,1, | 233 | 149 | 0.0306650989828 | | 1 | 0.836,0.793,0.756, | 0.762,0.812,0.561, | 0,13 |
| 247629 | ENSG00000152404 | CWF19L2 | chr11 | - | 107416208 | 107416297 | 107392778 | 107392895 | 107418193 | 107418287 | 247629 | 46,43,64,34,12,9,4,9,25,15 | 0,0,0,3,0,0,0,0,1,0,0,0 | 24,30,34,31,24,14,7,10,24, | 2,0,2,10,4,1,0,1,2,0,0,0,1,1 | 238 | 149 | 0.030666237557 | | 1 | 1.0,1.0,1.0,0.876,1 | 0.883,1.0,0.914,0. | 0,102 |
| 93044 | ENSG00000112511 | PHF1 | chr6 | + | 33414965 | 33415144 | 33414724 | 33414829 | 33415589 | 33415670 | 93044 | 7,7,16,6,10,10,6,15,9,7,5,7 | 1,2,0,0,0,2,2,1,0,0,1,0 | 4,14,5,17,4,2,7,27,2,2,10,5 | 0,1,0,4,1,0,1,4,0,5,3,0,0,0, | 298 | 149 | 0.030686244326 | | 1 | 0.778,0.636,1.0,1. | 1.0,0.875,1.0,0.68, | 0,041 |
| 253785 | ENSG00000123329 | ARHGAP9 | chr12 | - | 57476845 | 57476963 | 57476589 | 57476651 | 57477155 | 57477269 | 253785 | 113,69,53,54,60,35,65,44, | 8,0,9,10,2,2,4,2,8,3,3,3 | 60,61,52,29,30,26,29,45,3 | 7,4,3,2,6,5,4,7,6,3,3,3,10,4 | 267 | 149 | 0.0306917105708 | | 1 | 0.887,1.0,0.767,0. | 0.827,0.895,0.906, | 0,055 |
| 45216 | ENSG00000010404 | IDS | chrX | - | 149503311 | 149503489 | 149500948 | 149501037 | 149504156 | 149504219 | 45216 | 134,65,111,89,101,100,75, | 9,5,7,2,2,16,11,4,3,6,1,7 | 96,95,82,135,55,31,87,137 | 3,7,6,8,7,3,1,1,3,0,5,3,3,8,1 | 298 | 149 | 0.0306940165917 | | 1 | 0.882,0.867,0.888, | 0.941,0.872,0.872, | -0,022 |
| 58669 | ENSG00000114098 | ARMC8 | chr3 | + | 138188454 | 138188530 | 138187266 | 138187599 | 138209816 | 138209893 | 58669 | 15,11,29,24,12,5,5,18,5,3, | 12,4,10,19,4,11,6,9,2,5,2,1 | 19,18,8,9,22,6,7,14,5,15,1 | 14,13,12,8,16,14,13,16,15, | 225 | 149 | 0.0307237082981 | | 1 | 0.453,0.646,0.658, | 0.473,0.478,0.306, | 0,129 |
| 108523 | ENSG00000121390 | PSPC1 | chr13 | - | 19699379 | 19699398 | 19677723 | 19677823 | 19730238 | 19730316 | 108523 | 3,1,8,1,5,5,0,3,2,0,2,4 | 6,9,24,18,14,9,15,15,9,3,4, | 3,2,3,0,2,2,0,3,0,0,1,0,1,7, | 19,20,5,14,12,4,6,16,7,22, | 168 | 149 | 0.0307257756657 | | 1 | 0.307,0.09,0.228,0 | 0.123,0.081,0.347, | 0,08 |
| 202435 | ENSG00000154222 | CC2D1B | chr1 | - | 52359704 | 52359826 | 52359458 | 52359534 | 52360073 | 52360233 | 202435 | 13,9,24,9,12,15,10,11,13,5 | 0,2,2,0,2,10,2,6,1,0,0,3 | 16,14,5,10,11,8,10,25,5,9, | 4,4,2,5,4,1,2,5,1,0,6,2,3,1, | 271 | 149 | 0.0307925357944 | | 1 | 1.0,0.712,0.868,1. | 0.687,0.658,0.579, | 0,135 |
| 175895 | ENSG00000173889 | PHC3 | chr3 | - | 170106831 | 170106937 | 170102801 | 170102934 | 170113359 | 170113519 | 175895 | 52,24,99,65,30,43,34,49,1 | 2,0,4,0,0,1,0,1,0,2,0,0 | 60,68,37,43,55,44,40,44,2 | 1,3,2,2,4,0,0,2,2,0,2,0,0,4, | 255 | 149 | 0.0308125654826 | | 1 | 0.938,1.0,0.935,1. | 0.972,0.93,0.915,0 | 0,022 |
| 197025 | ENSG00000147099 | HDAC8 | chrX | - | 72494195 | 72494363 | 72490928 | 72491006 | 72495155 | 72495268 | 197025 | 2,0,4,4,0,0,0,2,0,2,1,3 | 9,5,12,13,7,16,8,6,2,6,2,14 | 3,0,0,2,1,0,2,2,1,2,0,1,0,0, | 5,6,5,3,4,6,3,9,4,10,6,7,6,9 | 298 | 149 | 0.0308646926659 | | 1 | 0.1,0.0,0.143,0.13 | 0.231,0.0,0.0,0.25, | 0,004 |
| 235508 | ENSG00000139197 | PEX5 | chr12 | + | 7202261 | 7202351 | 7201750 | 7201841 | 7202611 | 7202704 | 235508 | 6,8,1,3,3,0,1,0,6,2,1,1 | 25,22,31,11,10,11,15,16,2 | 3,0,1,2,0,1,0,3,2,0,1,0,0,10 | 6,24,7,10,12,4,7,15,6,11,1 | 239 | 149 | 0.0308661949152 | | 1 | 0.13,0.185,0.02,0. | 0.238,0.0,0.082,0. | 0,006 |
| 229303 | ENSG00000122566 | HNRNPA2B1 | chr7 | - | 26193574 | 26193710 | 26193250 | 26193373 | 26195846 | 26195909 | 229303 | 163,148,168,112,66,60,65, | 122,51,104,76,40,60,59,87 | 89,104,70,36,36,85,8,94,1 | 83,99,103,104,94,25,21,54 | 285 | 149 | 0.0308755332165 | | 1 | 0.411,0.603,0.458, | 0.359,0.355,0.262, | 0,078 |
| 123088 | ENSG00000066294 | CD84 | chr1 | - | 160552704 | 160552737 | 160550937 | 160551035 | 160553377 | 160553497 | 123088 | 6,1,1,10,3,3,3,1,0,3,0,4 | 10,11,20,30,10,12,6,4,9,4, | 6,14,0,1,3,1,4,10,2,0,4,0,3, | 24,20,3,2,3,1,9,21,2,7,4,3, | 182 | 149 | 0.0308806296837 | | 1 | 0.329,0.069,0.039, | 0.17,0.364,0.0,0.2 | -0,12 |
| 217944 | ENSG00000111679 | PTPN6 | chr12 | + | 6955150 | 6955263 | 6954804 | 6954994 | 6955371 | 6955485 | 217944 | 392,206,460,294,227,114, | 31,6,36,11,8,13,5,13,13,7, | 250,182,117,17,59,107,34, | 19,10,4,3,6,2,2,7,1,4,1,26,1 | 262 | 149 | 0.0308967397944 | | 1 | 0.878,0.951,0.879, | 0.882,0.912,0.943, | -0,01 |
| 204884 | ENSG00000215790 | SLC35E2A | chr1 | - | 1737551 | 1737809 | 1734689 | 1734835 | 1738320 | 1738448 | 204884 | 10,10,16,15,9,9,16,18,25,1 | 1,0,4,3,0,0,1,2,0,0,0,0 | 14,9,3,3,17,14,15,7,4,23,8, | 5,4,0,0,1,0,0,0,0,0,1,0,2,7, | 298 | 149 | 0.0309059275223 | | 1 | 0.833,1.0,0.667,0. | 0.583,0.529,1.0,1. | 0,075 |
| 54936 | ENSG00000198265 | HELZ | chr17 | - | 67122969 | 67123160 | 67120404 | 67120612 | 67123962 | 67124014 | 54936 | 72,55,88,80,73,29,52,52,5 | 1,0,0,4,3,1,0,1,2,4,0,0 | 45,61,49,75,44,49,33,56,2 | 1,3,3,8,2,2,0,0,1,1,1,0,3,2, | 298 | 149 | 0.0309110210264 | | 1 | 0.973,1.0,1.0,0.90 | 0.957,0.91,0.891,0 | 0,023 |
| 123598 | ENSG00000149806 | FAU | chr11 | - | 65120980 | 65121036 | 65120629 | 65120806 | 65121499 | 65121539 | 123598 | 655,474,1199,955,409,463 | 6,9,8,6,3,6,5,8,6,7,0,1 | 810,771,432,295,414,310, | 4,11,2,38,2,1,4,17,2,1,17,1 | 205 | 149 | 0.0309439682594 | | 1 | 0.988,0.975,0.991, | 0.993,0.981,0.994, | 0,006 |
| 124726 | ENSG00000103544 | VPS35L | chr16 | + | 19650381 | 19650459 | 19647783 | 19647882 | 19651975 | 19652090 | 124726 | 31,18,43,23,26,38,27,30,2 | 0,2,7,3,0,6,2,1,1,2,0,4 | 21,16,25,13,12,28,22,22,8, | 2,1,4,0,5,1,0,0,0,1,0,2,5,0,1 | 227 | 149 | 0.030948993604 | | 1 | 1.0,0.855,0.801,0. | 0.873,0.913,0.804, | -0,023 |
| 205893 | ENSG00000128815 | WDFY4 | chr10 | + | 48817234 | 48817409 | 48813956 | 48814082 | 48820233 | 48820437 | 205893 | 80,70,95,67,94,81,50,64,9 | 0,1,2,2,1,6,0,1,0,0,1,4 | 79,75,21,50,19,31,71,85,4 | 2,9,0,5,0,0,1,2,1,0,6,3,0,1, | 298 | 149 | 0.0309707615063 | | 1 | 1.0,0.972,0.96,0.9 | 0.952,0.806,1.0,0. | 0,016 |
| 59746 | ENSG00000122678 | POLM | chr7 | - | 44074130 | 44074233 | 44073782 | 44074025 | 44074397 | 44074530 | 59746 | 26,16,47,25,25,29,26,37,2 | 4,5,6,6,2,4,13,17,11,6,7,9 | 11,18,21,24,16,20,17,32,1 | 2,8,2,5,2,4,2,7,2,6,9,7,1,2, | 252 | 149 | 0.0309986448237 | | 1 | 0.794,0.654,0.822, | 0.765,0.571,0.861, | -0,09 |
| 36320 | ENSG00000136169 | SETDB2 | chr13 | + | 49451552 | 49451909 | 49444273 | 49444857 | 49460106 | 49460232 | 36320 | 29,15,108,38,44,32,30,68, | 4,0,0,0,0,2,0,0,0,0,0,0 | 36,49,31,6,24,16,3,19,20,5 | 2,3,0,0,4,2,0,0,2,2,3,1,0,1, | 298 | 149 | 0.0309991615127 | | 1 | 0.784,1.0,1.0,1.0,1 | 0.9,0.891,1.0,1.0,0 | 0,049 |
| 123618 | ENSG00000133943 | DGLUCY | chr14 | + | 91145220 | 91145313 | 91114685 | 91114994 | 91157638 | 91157690 | 123618 | 13,2,8,12,4,0,2,7,8,3,5,3 | 6,3,2,11,1,3,5,18,6,3,3,4 | 2,6,3,5,6,3,2,5,4,4,3,1,2,4, | 5,7,2,7,1,2,7,19,3,6,4,9,7,4 | 242 | 149 | 0.0310490320161 | | 1 | 0.572,0.291,0.711, | 0.198,0.345,0.48,0 | 0,111 |
| 97009 | ENSG00000221829 | FANCG | chr9 | - | 35078140 | 35078343 | 35077263 | 35077399 | 35078604 | 35078736 | 97009 | 12,6,16,21,6,3,5,3,8,2,0,6 | 2,2,2,0,0,0,0,1,2,0,1,3 | 10,7,4,3,1,5,2,9,1,4,6,22,8, | 1,2,0,0,0,0,0,0,1,1,2,0,2, | 298 | 149 | 0.0310516912934 | | 1 | 0.75,0.6,0.8,1.0,1. | 0.833,0.636,1.0,1. | -0,153 |
| 154768 | ENSG00000100603 | SNW1 | chr14 | - | 77723180 | 77723275 | 77720710 | 77720828 | 77730987 | 77731129 | 154768 | 59,42,49,23,23,23,16,33,2 | 21,10,10,3,2,6,1,7,4,3,1,4 | 47,61,65,23,25,24,28,30,1 | 6,6,2,12,6,2,4,1,4,1,5,5,3,4 | 244 | 149 | 0.0310578565795 | | 1 | 0.632,0.719,0.75,0 | 0.827,0.861,0.952, | -0,043 |
| 250951 | ENSG00000196683 | TOMM7 | chr7 | - | 22817999 | 22818054 | 22812970 | 22813185 | 22822676 | 22822849 | 250951 | 61,52,338,243,52,54,62,93 | 4,3,9,14,3,0,6,13,5,0,0,4 | 184,142,45,28,82,56,35,90 | 9,14,2,2,6,0,11,13,4,10,12, | 204 | 149 | 0.0310619978639 | | 1 | 0.918,0.927,0.965, | 0.937,0.881,0.943, | 0,025 |
| 12813 | ENSG00000186566 | GPATCH8 | chr17 | - | 44464471 | 44464544 | 44436477 | 44436545 | 44474828 | 44474903 | 12813 | 39,32,89,51,24,29,16,28,2 | 10,3,7,3,1,9,7,18,0,0,1,5 | 39,44,35,28,38,24,21,25,1 | 10,6,12,21,5,10,5,11,11,2, | 222 | 149 | 0.0310648904038 | | 1 | 0.724,0.877,0.895, | 0.724,0.831,0.662, | 0,086 |
| 73509 | ENSG00000154222 | CC2D1B | chr1 | - | 52359704 | 52359883 | 52359458 | 52359534 | 52360073 | 52360196 | 73509 | 38,19,56,26,35,27,25,31,3 | 0,2,2,0,2,10,2,6,1,0,0,3 | 38,40,16,21,25,19,27,74,1 | 4,4,2,5,4,1,2,5,1,0,6,2,3,1, | 298 | 149 | 0.0310717983849 | | 1 | 1.0,0.826,0.933,1. | 0.826,0.833,0.8,0. | 0,068 |
| 172369 | ENSG00000143669 | LYST | chr1 | - | 235828944 | 235829039 | 235812970 | 235813061 | 235830225 | 235830424 | 172369 | 10,1,10,7,15,12,7,33,3,1,1 | 121,62,64,19,64,49,46,52, | 11,6,3,2,13,2,13,9,6,3,24,2 | 72,28,34,16,5,16,28,30,19, | 244 | 149 | 0.031080462232 | | 1 | 0.048,0.01,0.087,0 | 0.085,0.116,0.051, | -0,059 |
| 159937 | ENSG00000185404 | SP140L | chr2 | + | 230328733 | 230328831 | 230327269 | 230327301 | 230357804 | 230357967 | 159937 | 49,23,50,30,6,17,19,27,18, | 2,1,10,3,1,0,2,1,0,0,0,4 | 36,37,23,13,26,14,11,23,1 | 6,3,1,1,4,2,0,6,9,2,2,2,1,4, | 247 | 149 | 0.0310838587222 | | 1 | 0.937,0.933,0.751, | 0.784,0.882,0.933, | 0,079 |
| 201489 | ENSG00000115446 | UNC50 | chr2 | + | 98610523 | 98610634 | 98609755 | 98610039 | 98610774 | 98610895 | 201489 | 3,2,7,4,1,1,1,2,0,0,0,6 | 13,11,41,18,5,9,10,13,12,7 | 1,2,3,1,2,2,1,0,0,4,2,0,4,2, | 23,21,20,4,4,7,8,16,3,11,1 | 260 | 149 | 0.0310981403995 | | 1 | 0.117,0.094,0.089, | 0.024,0.052,0.079, | 0,026 |
| 104404 | ENSG00000196998 | WDR45 | chrX | - | 49075135 | 49075281 | 49074432 | 49074912 | 49075363 | 49075378 | 104404 | 55,37,102,91,43,33,30,29, | 2,0,3,5,0,0,0,4,2,0,0,0 | 44,57,30,33,48,34,30,65,2 | 2,1,0,1,2,2,0,3,1,0,8,3,2,0, | 295 | 149 | 0.0311108878939 | | 1 | 0.933,1.0,0.945,0. | 0.917,0.966,1.0,0. | 0,027 |
| 227776 | ENSG00000196305 | IARS1 | chr9 | - | 92257803 | 92257986 | 92256679 | 92256800 | 92258853 | 92258998 | 227776 | 13,3,14,6,0,6,7,7,4,4,3,4 | 39,28,52,55,28,35,25,28,3 | 3,19,10,16,6,3,10,15,8,5,9, | 30,44,50,28,18,15,13,26,1 | 298 | 149 | 0.0311265262261 | | 1 | 0.143,0.051,0.119, | 0.048,0.178,0.091, | -0,041 |
| 123357 | ENSG00000153827 | TRIP12 | chr2 | - | 229787504 | 229787661 | 229785756 | 229785855 | 229788797 | 229788940 | 123357 | 199,113,231,144,102,83,7 | 1,0,6,5,2,2,0,4,0,4,0,0 | 145,159,142,153,125,85,6 | 0,3,0,2,0,0,0,2,0,0,1,0,2,2, | 298 | 149 | 0.0311312518403 | | 1 | 0.99,1.0,0.951,0.9 | 1.0,0.964,1.0,0.97 | -0,021 |
| 109243 | ENSG00000185015 | CA13 | chr8 | + | 85224954 | 85225295 | 85220926 | 85221125 | 85230084 | 85230200 | 109243 | 11,8,24,12,18,20,28,20,4,7 | 2,1,2,1,2,2,2,3,0,0,1,1 | 40,15,9,5,6,11,3,14,10,24,1 | 8,0,0,2,0,0,0,0,2,4,1,0,0,0, | 298 | 149 | 0.0311491799731 | | 1 | 0.733,0.8,0.857,0. | 0.714,1.0,1.0,0.55 | -0,058 |
| 175296 | ENSG00000174891 | RSRC1 | chr3 | + | 158455610 | 158455766 | 158354856 | 158354908 | 158460934 | 158461983 | 175296 | 17,2,1,0,4,5,1,3,0,2,0,3 | 53,25,41,23,17,23,19,34,3 | 0,5,0,9,3,0,2,0,2,0,2,1,0,1,1 | 24,21,22,10,14,19,13,13,8, | 298 | 149 | 0.0311507741312 | | 1 | 0.138,0.038,0.012, | 0.0,0.106,0.0,0.31, | -0,004 |
| 250519 | ENSG00000120616 | EPC1 | chr10 | - | 32345562 | 32345623 | 32305771 | 32305931 | 32346762 | 32347121 | 250519 | 6,0,4,3,1,2,6,6,0,0,0,0 | 53,28,80,27,29,35,27,43,3 | 2,2,1,0,1,1,1,1,2,4,11,14,0, | 24,24,42,38,30,26,23,31,2 | 210 | 149 | 0.031167481517 | | 1 | 0.074,0.0,0.034,0. | 0.056,0.056,0.017, | -0,034 |
| 227215 | ENSG00000105939 | ZC3HAV1 | chr7 | - | 139080580 | 139080780 | 139079469 | 139080243 | 139083779 | 139084032 | 227215 | 10,7,16,14,14,5,8,23,12,4, | 55,58,114,102,65,43,43,49 | 14,13,5,9,8,2,3,8,12,11,7,1 | 87,114,49,20,58,25,72,80, | 298 | 149 | 0.0312045788448 | | 1 | 0.083,0.057,0.066, | 0.074,0.054,0.049, | 0,017 |
| 201976 | ENSG00000125633 | CCDC93 | chr2 | - | 118000820 | 118000930 | 117996263 | 117996362 | 118006721 | 118006816 | 201976 | 34,11,67,24,19,25,17,24,1 | 2,4,4,2,0,3,1,7,0,1,0,0 | 38,27,13,23,33,19,13,17,1 | 5,4,2,13,1,0,6,2,4,2,4,5,0,4 | 259 | 149 | 0.0312321103464 | | 1 | 0.907,0.613,0.906, | 0.814,0.795,0.789, | 0,073 |
| 150654 | ENSG00000159459 | UBR1 | chr15 | - | 43053093 | 43053140 | 43048391 | 43048491 | 43054741 | 43054899 | 150654 | 0,4,8,4,3,6,0,0,0,0,0,1 | 20,20,44,21,22,27,19,26,6, | 7,1,0,0,2,4,0,1,4,1,0,0,0,3, | 26,43,23,10,20,22,19,17,1 | 196 | 149 | 0.0312486311797 | | 1 | 0.0,0.132,0.121,0. | 0.17,0.017,0.0,0.0, | 0,006 |
| 187944 | ENSG00000114098 | ARMC8 | chr3 | + | 138241782 | 138241956 | 138238019 | 138239528 | 138245087 | 138246887 | 187944 | 50,50,73,45,34,26,28,46,1 | 2,2,6,7,2,6,0,5,6,2,1,3 | 53,34,32,27,44,17,14,27,2 | 6,11,5,14,17,4,0,6,1,5,2,7, | 298 | 149 | 0.0312868626653 | | 1 | 0.926,0.926,0.859, | 0.815,0.607,0.762, | 0,058 |
| 200042 | ENSG00000197343 | ZNF655 | chr7 | + | 99562274 | 99562494 | 99560532 | 99560695 | 99571687 | 99571717 | 200042 | 2,3,34,19,11,14,14,19,7,2, | 4,0,1,5,5,1,0,2,0,0,0,0 | 30,25,10,8,7,1,10,24,11,12 | 1,0,0,2,0,0,0,2,1,4,0,3,4,2, | 298 | 149 | 0.0313082584135 | | 1 | 0.2,1.0,0.944,0.65 | 0.938,1.0,1.0,0.66 | -0,04 |
| 165044 | ENSG00000120137 | PANK3 | chr5 | - | 168563888 | 168564032 | 168561392 | 168561516 | 168566012 | 168566266 | 165044 | 29,33,45,38,16,23,22,37,1 | 11,5,13,13,4,2,1,9,1,4,0,1 | 33,22,9,18,13,9,18,15,12,2 | 6,15,12,15,9,1,3,12,5,3,6,2 | 293 | 149 | 0.0313082924621 | | 1 | 0.573,0.77,0.638,0 | 0.737,0.427,0.276, | 0,135 |
| 248434 | ENSG00000139697 | SBNO1 | chr12 | - | 123347783 | 123348133 | 123345257 | 123345567 | 123350309 | 123350441 | 248434 | 37,29,41,32,24,23,31,46,1 | 4,0,4,2,0,0,1,1,3,2,0,2 | 51,58,28,31,37,19,15,21,2 | 0,1,3,0,3,0,0,0,0,1,1,2,0,0, | 298 | 149 | 0.0313129891464 | | 1 | 0.822,1.0,0.837,0. | 1.0,0.967,0.824,1. | -0,063 |
| 113041 | ENSG00000104529 | EEF1D | chr8 | - | 143590607 | 143590738 | 143588990 | 143590081 | 143592646 | 143592797 | 113041 | 7,8,32,18,4,13,7,9,7,2,5,8 | 7,3,21,13,4,1,1,4,0,0,3,1 | 10,3,6,3,9,5,0,5,4,4,14,17,1 | 4,0,3,8,6,3,9,9,1,4,7,10,1,1 | 280 | 149 | 0.0313163298872 | | 1 | 0.347,0.587,0.448, | 0.571,1.0,0.516,0. | 0,181 |
| 99031 | ENSG00000136280 | CCM2 | chr7 | + | 45063917 | 45064001 | 45038252 | 45038426 | 45069825 | 45069961 | 99031 | 9,4,17,12,8,9,6,7,13,2,3,3 | 4,1,0,0,6,0,1,4,0,0,0,0 | 14,5,8,3,1,5,2,1,3,23,10,13 | 0,0,3,0,0,1,0,0,0,1,2,0,0,0,0 | 233 | 149 | 0.0313240398685 | | 1 | 0.59,0.719,1.0,1.0, | 1.0,1.0,0.63,1.0,1. | -0,102 |
| 26719 | ENSG00000196182 | STK40 | chr1 | - | 36358736 | 36358837 | 36358238 | 36358382 | 36361220 | 36361340 | 26719 | 6,3,21,12,14,12,5,1,5,1,7,6 | 4,0,4,0,1,2,1,2,2,0,0,0 | 11,6,4,6,1,4,4,9,5,9,2,10,1 | 2,1,0,0,0,0,0,2,1,2,0,0,1,0,0 | 250 | 149 | 0.0313701173759 | | 1 | 0.472,1.0,0.758,1. | 0.766,0.781,1.0,1. | -0,092 |
| 191915 | ENSG00000121892 | PDS5A | chr4 | - | 39910243 | 39910356 | 39908394 | 39908540 | 39913610 | 39913726 | 191915 | 56,48,134,63,50,66,57,36, | 7,2,11,12,0,0,3,4,6,0,2,0 | 91,63,45,29,34,33,29,41,3 | 6,12,6,13,20,2,1,3,11,2,1,3 | 262 | 149 | 0.0313867808215 | | 1 | 0.82,0.932,0.874,0 | 0.896,0.749,0.81,0 | 0,072 |
| 158396 | ENSG00000198399 | ITSN2 | chr2 | - | 24264750 | 24265092 | 24261560 | 24261742 | 24270670 | 24270768 | 158396 | 33,38,33,14,11,10,12,12,1 | 43,47,99,71,54,25,22,89,1 | 23,30,10,18,7,11,4,27,5,12 | 72,117,50,53,37,38,46,60, | 298 | 149 | 0.0313983451371 | | 1 | 0.277,0.288,0.143, | 0.138,0.114,0.091, | 0,057 |
| 72685 | ENSG00000058272 | PPP1R12A | chr12 | - | 79806165 | 79806297 | 79805591 | 79805768 | 79807225 | 79807330 | 72685 | 21,8,20,13,6,8,11,13,22,9, | 0,0,4,1,4,2,0,0,5,0,0,2 | 11,19,19,17,14,9,5,15,14,1 | 1,0,0,1,0,1,1,0,1,2,2,0,3,3, | 281 | 149 | 0.0314121862771 | | 1 | 1.0,1.0,0.726,0.87 | 0.854,1.0,1.0,0.9,1 | -0,089 |
| 27944 | ENSG00000113648 | MACROH2A1 | chr5 | - | 135350822 | 135350913 | 135345967 | 135346057 | 135360496 | 135360604 | 27944 | 22,16,18,6,5,13,4,6,10,4,3, | 0,1,1,1,0,2,5,1,0,0,0,0 | 16,15,8,6,9,3,6,4,6,9,4,14, | 1,4,1,1,2,2,3,0,0,2,0,2,1,0,1 | 240 | 149 | 0.0314250595843 | | 1 | 1.0,0.909,0.918,0. | 0.909,0.7,0.832,0. | 0,054 |

| | | | | | | | | | | | | | | | | | | | | | | | |
|---|---|---|---|---|---|---|---|---|---|---|---|---|---|---|---|---|---|---|---|---|---|---|---|
| 89433 | ENSG00000135945 | REV1 | chr2 | - | 99406324 | 99406431 | 99405927 | 99406106 | 99408028 | 99408131 | 89433 | 39,16,21,13,12,13,16,21,1 | 1,1,3,9,0,0,3,2,2,0,0,1 | 17,19,14,13,9,11,11,19,13, | 4,0,0,1,3,0,1,1,0,0,0,0,0,3, | 256 | 149 | 0.0314446069124 | | 1 | 0.958,0.903,0.803, | 0.712,1.0,1.0,0.88 | -0,014 |
| 104783 | ENSG00000165813 | CCDC186 | chr10 | - | 114173140 | 114173265 | 114161607 | 114163329 | 114174014 | 114174202 | 104783 | 11,14,27,27,13,14,16,10,1 | 30,24,30,38,25,25,24,30,2 | 29,41,14,15,14,9,15,8,13,1 | 24,38,14,28,25,27,20,17,2 | 274 | 149 | 0.031447284384 | | 1 | 0.166,0.241,0.329, | 0.397,0.37,0.352,0 | -0,075 |
| 124773 | ENSG00000154258 | ABCA9 | chr17 | - | 68984884 | 68984979 | 68984055 | 68984175 | 68985052 | 68985128 | 124773 | 11,16,59,24,7,7,21,38,19,1 | 20,12,47,40,10,7,26,66,25, | 10,29,5,8,19,16,5,11,7,11,1 | 28,63,32,10,30,43,4,12,20, | 244 | 149 | 0.0314475960267 | | 1 | 0.251,0.449,0.434, | 0.179,0.219,0.087, | 0,067 |
| 236604 | ENSG00000104783 | KCNN4 | chr19 | - | 43773478 | 43773641 | 43771999 | 43772135 | 43774191 | 43774619 | 236604 | 22,6,17,5,7,8,6,10,19,5,4,1 | 259,98,204,103,120,137,1 | 9,5,5,11,7,4,4,10,3,8,11,2, | 141,123,107,155,97,66,79, | 298 | 149 | 0.0314509953598 | | 1 | 0.041,0.03,0.04,0. | 0.031,0.02,0.023,0 | 0,006 |
| 74852 | ENSG00000140307 | GTF2A2 | chr15 | - | 59652205 | 59652326 | 59650668 | 59650773 | 59657405 | 59657515 | 74852 | 52,57,130,115,36,52,30,48 | 1,0,0,2,5,0,0,1,0,0,0,0 | 58,95,30,27,44,43,45,52,2 | 3,2,0,0,3,2,0,2,0,0,3,0,5,3, | 270 | 149 | 0.0314564879498 | | 1 | 0.966,1.0,1.0,0.96 | 0.914,0.963,1.0,1. | 0,026 |
| 213647 | ENSG00000168763 | CNNM3 | chr2 | + | 96825057 | 96825169 | 96816267 | 96817502 | 96826832 | 96826982 | 213647 | 18,27,22,12,14,17,13,8,1,4 | 1,1,3,0,1,3,0,0,0,0,0,0 | 19,15,10,8,34,16,7,18,15,1 | 1,0,2,2,7,2,0,1,2,0,1,0,1,3, | 261 | 149 | 0.0314847650406 | | 1 | 0.911,0.939,0.807, | 0.916,1.0,0.741,0. | 0,092 |
| 39296 | ENSG00000112282 | MED23 | chr6 | - | 131615906 | 131616002 | 131610045 | 131610246 | 131618406 | 131618519 | 39296 | 40,31,68,34,31,24,18,29,1 | 2,1,9,7,6,4,3,7,3,1,0,0 | 38,48,18,14,33,15,11,15,1 | 7,7,2,26,11,6,0,6,2,3,8,0,5, | 245 | 149 | 0.0315290149579 | | 1 | 0.924,0.95,0.821,0 | 0.768,0.807,0.846, | 0,092 |
| 114491 | ENSG00000083223 | TUT7 | chr9 | - | 86326370 | 86326491 | 86325333 | 86325514 | 86328339 | 86328492 | 114491 | 23,9,44,22,19,25,14,13,6,1 | 27,14,67,59,35,34,27,27,1 | 11,27,13,7,8,17,9,10,14,7,1 | 35,59,43,31,24,20,22,15,1 | 270 | 149 | 0.0315623837043 | | 1 | 0.32,0.262,0.266,0 | 0.148,0.202,0.143, | 0,068 |
| 175991 | ENSG00000116191 | RALGPS2 | chr1 | + | 178796621 | 178796769 | 178785556 | 178785607 | 178808044 | 178808128 | 175991 | 3,4,5,3,4,4,6,7,0,2,0,6 | 180,65,513,269,214,119,1 | 12,25,10,5,8,7,0,1,8,1,5,9, | 516,398,125,132,166,150, | 297 | 149 | 0.0316276375655 | | 1 | 0.008,0.03,0.005,0 | 0.012,0.031,0.039, | -0,003 |
| 66104 | ENSG00000171791 | BCL2 | chr18 | - | 63280886 | 63281214 | 63127776 | 63128759 | 63318081 | 63318666 | 66104 | 5,2,8,6,8,4,5,6,15,2,4,3 | 103,84,240,145,124,163,9 | 8,0,1,1,0,6,0,12,3,4,11,4,0, | 162,94,84,52,40,77,126,28 | 298 | 149 | 0.0316376633733 | | 1 | 0.024,0.012,0.016, | 0.024,0.0,0.006,0. | 0,008 |
| 167495 | ENSG00000145495 | MARCHF6 | chr5 | + | 10413274 | 10413450 | 10411332 | 10411537 | 10414432 | 10414502 | 167495 | 9,4,11,3,6,11,6,6,4,2,4,6 | 188,150,192,97,84,105,93, | 1,6,5,11,4,3,4,9,7,0,6,2,5,2 | 151,186,123,288,105,53,1 | 298 | 149 | 0.0316453081383 | | 1 | 0.023,0.013,0.028, | 0.003,0.016,0.02,0 | 0,009 |
| 32655 | ENSG00000083312 | TNPO1 | chr5 | + | 72897055 | 72897151 | 72895458 | 72896556 | 72900005 | 72900081 | 32655 | 153,123,182,114,70,83,72, | 0,1,4,0,0,1,2,1,2,0,0,1 | 123,176,110,69,98,52,35,6 | 1,3,0,4,3,2,0,0,1,2,0,0,10,3 | 245 | 149 | 0.0316483804123 | | 1 | 1.0,0.987,0.965,1. | 0.987,0.973,1.0,0. | 0,017 |
| 64736 | ENSG00000135778 | NTPCR | chr1 | + | 232955556 | 232955719 | 232950610 | 232950744 | 232956346 | 232956443 | 64736 | 7,12,63,61,15,16,29,24,16, | 0,2,2,2,0,1,0,0,1,2,0,3 | 29,12,8,8,9,16,4,2,9,21,8,3 | 0,0,2,0,1,0,0,3,2,0,0,8,1,1, | 298 | 149 | 0.0316587971829 | | 1 | 1.0,0.75,0.94,0.93 | 1.0,1.0,0.667,1.0,0 | 0,083 |
| 110700 | ENSG00000034677 | RNF19A | chr8 | - | 100261541 | 100261755 | 100259853 | 100259997 | 100264033 | 100264195 | 110700 | 119,148,307,90,77,99,49,3 | 5,8,19,8,0,20,6,41,2,1,3,8 | 92,165,116,133,81,47,136, | 0,14,8,0,7,1,2,12,4,3,35,6, | 298 | 149 | 0.0316688114807 | | 1 | 0.922,0.902,0.89,0 | 1.0,0.855,0.879,1. | -0,031 |
| 33610 | ENSG00000111885 | MAN1A1 | chr6 | - | 119248259 | 119248354 | 119204758 | 119204882 | 119290682 | 119290763 | 33610 | 76,95,159,131,70,78,61,88 | 5,8,11,8,2,5,8,10,1,6,2,9 | 193,228,48,28,99,95,149,3 | 12,28,2,18,22,5,22,34,11,1 | 244 | 149 | 0.0316899872245 | | 1 | 0.903,0.879,0.898, | 0.908,0.833,0.936, | 0,045 |
| 132991 | ENSG00000166037 | CEP57 | chr11 | + | 95822498 | 95822576 | 95818826 | 95818904 | 95827785 | 95828027 | 132991 | 6,5,18,15,3,7,2,3,7,3,6,13 | 0,2,3,4,0,1,3,0,0,0,0,1 | 9,8,10,0,4,3,5,9,10,8,8,11, | 0,2,0,1,3,4,0,0,4,0,0,2,1,8, | 227 | 149 | 0.0317260423576 | | 1 | 1.0,0.621,0.798,0. | 1.0,0.724,1.0,0.0,0 | 0,1 |
| 161351 | ENSG00000165819 | METTL3 | chr14 | - | 21500494 | 21500627 | 21499763 | 21499802 | 21500912 | 21501129 | 161351 | 204,89,86,63,56,47,52,63, | 8,4,15,3,10,5,2,12,2,7,1,4 | 66,54,55,23,47,46,22,92,3 | 6,10,2,1,0,2,2,1,0,0,4,7,2,2 | 282 | 149 | 0.0317465361531 | | 1 | 0.931,0.922,0.752, | 0.853,0.74,0.936,0 | -0,041 |
| 240476 | ENSG00000080802 | CNOT4 | chr7 | - | 135486525 | 135486560 | 135438157 | 135438423 | 135509888 | 135510127 | 240476 | 1,0,4,2,0,3,2,2,0,2,1,0 | 10,12,41,22,24,13,23,18,8, | 0,1,2,2,4,0,2,2,6,0,1,2,1,0, | 16,11,8,17,16,4,8,23,6,15, | 184 | 149 | 0.0317490294292 | | 1 | 0.075,0.0,0.073,0. | 0.0,0.069,0.168,0. | 0,01 |
| 80746 | ENSG00000181381 | DDX60L | chr4 | - | 168419301 | 168419397 | 168416681 | 168416797 | 168420260 | 168420380 | 80746 | 13,3,29,17,9,7,7,10,5,2,4,1 | 1,1,5,1,0,2,4,1,0,0,0,0 | 32,14,10,4,4,6,1,3,2,14,12, | 2,2,0,0,1,3,1,1,0,0,6,0,0,0, | 245 | 149 | 0.0317904310565 | | 1 | 0.888,0.646,0.779, | 0.907,0.81,1.0,1.0, | 0,041 |
| 161353 | ENSG00000165819 | METTL3 | chr14 | - | 21500494 | 21500634 | 21499763 | 21499802 | 21500912 | 21501129 | 161353 | 216,90,93,78,61,48,53,66, | 8,4,15,3,10,5,2,12,2,7,1,4 | 70,58,59,25,52,46,24,92,4 | 6,10,2,1,0,2,2,1,0,0,4,7,2,2 | 289 | 149 | 0.0317931026779 | | 1 | 0.933,0.921,0.762, | 0.857,0.749,0.938, | -0,038 |
| 68618 | ENSG00000132300 | PTCD3 | chr2 | + | 86134838 | 86134987 | 86134291 | 86134377 | 86136520 | 86136562 | 68618 | 111,80,171,142,126,106,6 | 6,1,6,6,0,4,2,5,0,0,0,1 | 234,184,80,82,98,60,61,12 | 3,2,0,4,2,0,0,5,2,1,2,0,1,0, | 298 | 149 | 0.0318037180676 | | 1 | 0.902,0.976,0.934, | 0.975,0.979,1.0,0. | -0,007 |
| 165996 | ENSG00000026297 | RNASET2 | chr6 | - | 166955497 | 166955627 | 166952487 | 166952548 | 166956096 | 166956182 | 165996 | 2,3,10,1,10,3,4,0,9,14,0,4 | 62,37,49,29,22,11,22,13,2 | 8,1,2,8,4,1,6,3,1,3,9,4,8,2, | 22,46,47,27,30,25,13,16,1 | 279 | 149 | 0.0318130060933 | | 1 | 0.017,0.042,0.098, | 0.163,0.011,0.022, | 0,01 |
| 246499 | ENSG00000165916 | PSMC3 | chr11 | - | 47424045 | 47424255 | 47422829 | 47422973 | 47424428 | 47424491 | 246499 | 61,57,125,86,31,27,33,42, | 1,0,4,5,2,2,2,1,0,0,0,2 | 59,58,41,25,60,23,23,45,2 | 1,1,0,2,0,0,0,0,0,0,2,0,0,0 | 298 | 149 | 0.0318257592546 | | 1 | 0.968,1.0,0.94,0.8 | 0.967,0.967,1.0,0. | -0,04 |
| 167212 | ENSG00000155229 | MMS19 | chr10 | - | 97468965 | 97469095 | 97468251 | 97468406 | 97469645 | 97469723 | 167212 | 28,26,32,23,19,22,23,9,12, | 11,7,12,14,6,9,6,11,5,5,6,2 | 14,20,14,3,8,7,14,10,9,8,1 | 6,16,6,7,12,0,9,7,5,6,15,15 | 279 | 149 | 0.0318262146085 | | 1 | 0.576,0.665,0.587, | 0.555,0.4,0.555,0. | 0,099 |
| 211267 | ENSG00000176978 | DPP7 | chr9 | - | 137111872 | 137111997 | 137111689 | 137111754 | 137112111 | 137112604 | 211267 | 60,54,91,41,54,44,67,66,5 | 4,3,11,3,2,3,0,2,2,0,0,4 | 67,65,50,43,28,13,29,88,2 | 4,2,10,8,5,2,3,12,0,3,5,3,1, | 274 | 149 | 0.0318355129176 | | 1 | 0.891,0.907,0.818, | 0.901,0.946,0.731, | 0,058 |
| 195143 | ENSG00000171791 | BCL2 | chr18 | - | 63242001 | 63242110 | 63128211 | 63128759 | 63318081 | 63320128 | 195143 | 1,0,6,3,2,5,3,9,0,0,2,6 | 103,84,240,145,124,163,9 | 4,11,2,8,2,0,3,7,2,3,12,7,2, | 162,94,84,52,40,77,126,28 | 258 | 149 | 0.031842673642 | | 1 | 0.006,0.0,0.014,0. | 0.014,0.063,0.014, | -0,011 |
| 145995 | ENSG00000289733 | | chr16 | + | 81851940 | 81852126 | 81785942 | 81786182 | 81854443 | 81854587 | 145995 | 4,0,2,0,5,2,3,1,1,0,3,1 | 65,64,127,85,70,38,56,67, | 0,3,7,5,2,3,1,4,3,2,6,5,2,1, | 111,96,19,28,57,34,46,25, | 298 | 149 | 0.0318757997325 | | 1 | 0.03,0.0,0.008,0.0, | 0.0,0.015,0.156,0. | -0,024 |
| 167499 | ENSG00000145495 | MARCHF6 | chr5 | + | 10418675 | 10418831 | 10417269 | 10417404 | 10423734 | 10423835 | 167499 | 3,6,9,9,6,4,6,3,0,0,0,4 | 86,76,160,56,62,94,69,86, | 2,7,0,9,4,1,2,4,0,1,4,3,1,1, | 105,110,67,80,72,46,73,10 | 298 | 149 | 0.0318763114746 | | 1 | 0.017,0.038,0.027, | 0.009,0.031,0.0,0. | 0,008 |
| 41255 | ENSG00000114650 | SCAP | chr3 | - | 47423932 | 47424045 | 47422441 | 47422536 | 47425484 | 47425611 | 41255 | 9,5,47,25,19,19,18,6,19,3, | 4,2,3,3,8,4,3,8,4,0,1,7 | 16,4,15,6,7,7,8,14,3,10,16, | 4,4,4,2,4,3,3,14,0,2,8,8,15, | 262 | 149 | 0.0318835723442 | | 1 | 0.561,0.587,0.899, | 0.695,0.363,0.681, | 0,107 |
| 136298 | ENSG00000104904 | OAZ1 | chr19 | + | 2271384 | 2271474 | 2269514 | 2269744 | 2271781 | 2272262 | 136298 | 183,126,375,211,190,178, | 12,17,31,8,10,4,11,9,11,4, | 219,147,78,18,81,86,76,17 | 14,23,9,8,13,1,12,9,0,4,32, | 239 | 149 | 0.031891489011 | | 1 | 0.905,0.822,0.883, | 0.907,0.799,0.844, | 0,045 |
| 92576 | ENSG00000083097 | DOP1A | chr6 | + | 83159795 | 83159960 | 83158566 | 83158622 | 83162789 | 83162919 | 92576 | 18,11,38,17,16,15,7,20,3,5 | 5,1,1,3,2,3,3,0,0,1,0,0 | 24,20,11,9,23,13,10,13,12, | 2,0,0,1,4,0,0,1,2,0,0,1,1,0, | 298 | 149 | 0.031893794604 | | 1 | 0.643,0.846,0.95,0 | 0.857,1.0,1.0,0.81 | -0,065 |
| 131267 | ENSG00000204852 | TCTN1 | chr12 | + | 110619835 | 110619956 | 110614026 | 110614402 | 110626361 | 110626382 | 131267 | 43,14,68,42,19,27,32,41,3 | 5,3,0,3,4,0,1,2,2,0,1,2 | 37,25,65,55,22,20,28,29,7, | 0,0,5,5,1,4,2,0,0,3,4,5,0,1, | 270 | 149 | 0.0318949418338 | | 1 | 0.826,0.72,1.0,0.8 | 1.0,1.0,0.878,0.85 | -0,002 |
| 234950 | ENSG00000147905 | ZCCHC7 | chr9 | + | 37147361 | 37147445 | 37126311 | 37126942 | 37247069 | 37247103 | 234950 | 20,10,48,12,14,21,17,38,5, | 6,2,10,2,1,10,6,8,3,1,3,2 | 65,46,59,15,14,15,22,20,1 | 7,6,4,7,0,0,1,10,4,5,13,3,1, | 233 | 149 | 0.0319115058106 | | 1 | 0.681,0.762,0.754, | 0.856,0.831,0.904, | -0,109 |
| 173392 | ENSG00000183726 | TMEM50A | chr1 | + | 25344977 | 25345231 | 25342960 | 25343073 | 25351625 | 25351693 | 173392 | 2,3,4,0,3,2,0,7,2,0,0,5 | 23,36,102,80,39,43,35,48, | 1,3,2,0,2,0,0,0,1,2,2,2,1,2, | 54,87,22,27,28,22,27,35,1 | 298 | 149 | 0.0319213019288 | | 1 | 0.042,0.04,0.019,0 | 0.009,0.017,0.043, | 0,011 |
| 167383 | ENSG00000185033 | SEMA4B | chr15 | + | 90221959 | 90222082 | 90221749 | 90221765 | 90223558 | 90223792 | 167383 | 2,4,0,2,3,3,2,0,4,0,0,4 | 54,32,45,24,45,22,19,44,4 | 0,0,1,0,1,1,5,1,2,2,0,0,2,2, | 42,67,15,32,32,25,45,38,2 | 272 | 149 | 0.0319229346829 | | 1 | 0.02,0.064,0.0,0.0 | 0.0,0.0,0.035,0.0,0 | 0,015 |
| 217805 | ENSG00000124789 | NUP153 | chr6 | - | 17633510 | 17633578 | 17632649 | 17632844 | 17637152 | 17637770 | 217805 | 9,6,15,7,4,10,9,10,2,1,2,6 | 14,18,24,12,15,14,7,6,8,9, | 10,20,5,10,17,12,6,5,10,16 | 12,15,2,14,13,8,15,15,9,5, | 217 | 149 | 0.0319266411946 | | 1 | 0.306,0.186,0.3,0. | 0.364,0.478,0.632, | -0,122 |
| 122283 | ENSG00000054654 | SYNE2 | chr14 | + | 64107490 | 64107607 | 64101931 | 64102042 | 64113340 | 64113571 | 122283 | 157,244,258,206,129,225, | 1,6,2,5,4,5,13,14,4,0,2,4 | 100,210,106,184,235,137, | 0,9,3,7,3,1,0,6,8,2,0,2,2,8, | 266 | 149 | 0.0319344085909 | | 1 | 0.989,0.958,0.986, | 1.0,0.929,0.952,0. | -0,019 |
| 210606 | ENSG00000177479 | ARIH2 | chr3 | + | 48933565 | 48933692 | 48927778 | 48927813 | 48961611 | 48961679 | 210606 | 8,6,13,11,7,6,15,8,0,3,2,4 | 27,18,30,20,21,18,18,35,2 | 15,19,1,10,10,5,7,16,6,5,5, | 20,47,27,15,12,16,14,27,7, | 276 | 149 | 0.0319948126842 | | 1 | 0.138,0.153,0.19,0 | 0.288,0.179,0.02,0 | -0,06 |
| 36764 | ENSG00000101109 | STK4 | chr20 | + | 44975323 | 44975410 | 44972077 | 44972158 | 44978442 | 44978571 | 36764 | 2,0,34,10,5,5,9,11,11,8,2,3 | 166,90,299,182,96,74,81,1 | 14,14,12,4,8,1,4,5,1,11,2,5 | 209,231,139,109,89,76,66, | 236 | 149 | 0.0320101012736 | | 1 | 0.008,0.0,0.067,0. | 0.041,0.037,0.052, | 0,01 |
| 144280 | ENSG00000161526 | SAP30BP | chr17 | + | 75668515 | 75668624 | 75667337 | 75667478 | 75671815 | 75672156 | 144280 | 142,110,143,78,27,42,51,4 | 1,6,10,3,0,4,0,2,3,0,4,0 | 67,68,39,61,84,56,26,26,2 | 4,3,3,7,4,0,2,8,2,0,5,8,1,0, | 258 | 149 | 0.0320207768573 | | 1 | 0.988,0.914,0.892, | 0.906,0.929,0.882, | 0,047 |
| 34005 | ENSG00000109189 | USP46 | chr4 | - | 52658173 | 52658281 | 52631063 | 52631144 | 52659114 | 52659275 | 34005 | 0,0,1,7,1,2,2,2,1,1,0,0 | 4,6,4,11,5,7,4,8,2,4,1,3 | 0,3,1,0,2,0,0,0,3,1,0,0,1,2, | 1,10,3,3,10,3,2,10,0,6,3,7,1 | 257 | 149 | 0.0320266769467 | | 1 | 0.0,0.0,0.127,0.27, | 0.0,0.148,0.162,0. | -0,039 |
| 253318 | ENSG00000136021 | SCYL2 | chr12 | + | 100296761 | 100296897 | 100291502 | 100291660 | 100298030 | 100298175 | 253318 | 13,7,14,4,8,17,8,10,0,0,5,9 | 35,24,62,54,19,15,27,34,1 | 25,31,10,16,10,3,19,11,4,5 | 54,34,28,29,18,16,21,31,1 | 285 | 149 | 0.0320295772019 | | 1 | 0.163,0.132,0.106, | 0.195,0.323,0.157, | -0,041 |
| 54135 | ENSG00000162777 | DENND2D | chr1 | - | 111194577 | 111194726 | 111192139 | 111192317 | 111195915 | 111196056 | 54135 | 121,29,86,121,83,70,53,78 | 2,0,2,6,5,0,4,5,4,0,0,2 | 81,41,49,21,37,55,23,66,3 | 1,0,2,8,8,1,4,2,0,0,7,0,2,8, | 298 | 149 | 0.0320561231322 | | 1 | 0.968,1.0,0.956,0. | 0.976,1.0,0.925,0. | 0,061 |
| 16863 | ENSG00000069329 | VPS35 | chr16 | - | 46668929 | 46669052 | 46662982 | 46663162 | 46671704 | 46671860 | 16863 | 37,36,264,151,93,78,55,69 | 0,3,6,1,1,1,3,0,5,0,0,3 | 93,97,50,37,44,45,40,49,4 | 1,6,1,1,7,4,0,1,5,0,3,7,2,3, | 272 | 149 | 0.0320641628169 | | 1 | 1.0,0.868,0.96,0.9 | 0.981,0.899,0.965, | 0,016 |
| 197542 | ENSG00000132300 | PTCD3 | chr2 | + | 86134844 | 86134987 | 86134295 | 86134377 | 86136520 | 86136562 | 197542 | 43,22,72,49,44,34,27,44,1 | 6,1,6,6,0,4,2,5,0,0,0,1 | 92,74,28,24,36,19,21,45,6 | 3,2,0,4,2,0,0,5,2,1,2,0,1,0, | 292 | 149 | 0.0320648350462 | | 1 | 0.785,0.918,0.86,0 | 0.94,0.95,1.0,0.75 | -0,011 |
| 161045 | ENSG00000140853 | NLRC5 | chr16 | + | 57055432 | 57055889 | 57055031 | 57055094 | 57058064 | 57058148 | 161045 | 5,4,16,11,4,9,7,7,0,4,3,5 | 4,3,1,2,0,2,6,0,1,2,0,5 | 6,12,6,8,11,8,3,5,4,21,10,1 | 5,5,9,3,3,1,1,10,4,3,7,12,1 | 298 | 149 | 0.0320656696696 | | 1 | 0.385,0.4,0.889,0. | 0.375,0.545,0.25,0 | 0,158 |
| 19257 | ENSG00000114062 | UBE3A | chr15 | - | 25408619 | 25408684 | 25407066 | 25407262 | 25409087 | 25409207 | 19257 | 31,21,120,63,18,11,14,21, | 2,1,9,8,1,0,1,1,8,0,1,1 | 39,28,20,15,9,9,14,17,10,2 | 0,0,4,0,0,0,1,2,0,5,2,3,1,0, | 214 | 149 | 0.0320722784258 | | 1 | 0.915,0.936,0.903, | 1.0,1.0,0.777,1.0,1 | -0,02 |
| 170105 | ENSG00000266338 | NBPF15 | chr1 | - | 144458846 | 144459032 | 144456536 | 144456805 | 144459365 | 144459488 | 170105 | 7,8,6,4,9,1,5,7,4,0,4,4 | 22,16,21,10,23,16,10,13,1 | 2,0,3,3,1,4,2,5,3,5,0,4,2,3, | 14,31,11,13,8,4,6,12,14,15 | 298 | 149 | 0.0321471052337 | | 1 | 0.137,0.2,0.125,0. | 0.067,0.0,0.12,0.1 | 0,02 |
| 207473 | ENSG00000120675 | DNAJC15 | chr13 | + | 43037764 | 43037849 | 43023585 | 43023734 | 43065685 | 43065737 | 207473 | 2,0,8,0,1,0,0,1,0,0,0,0 | 41,29,89,75,36,45,20,33,2 | 5,2,5,0,6,0,1,1,2,0,2,2,0,0, | 53,55,27,9,39,27,24,24,36, | 234 | 149 | 0.0321517974589 | | 1 | 0.03,0.0,0.054,0.0, | 0.057,0.023,0.105, | -0,023 |
| 253786 | ENSG00000123329 | ARHGAP9 | chr12 | - | 57476851 | 57476963 | 57476589 | 57476651 | 57477155 | 57477269 | 253786 | 115,69,54,53,60,35,65,44, | 8,0,9,10,2,2,4,2,8,3,3,3 | 60,60,52,29,31,26,28,47,3 | 7,4,3,2,6,5,4,7,6,3,3,3,10,4 | 261 | 149 | 0.0321558786201 | | 1 | 0.891,1.0,0.774,0. | 0.83,0.895,0.908,0 | 0,053 |
| 4860 | ENSG00000142453 | CARM1 | chr19 | + | 10908549 | 10908747 | 10908038 | 10908145 | 10909102 | 10909207 | 4860 | 7,1,6,6,0,11,4,0,2,3,2,4 | 3,7,12,24,6,11,3,2,4,12,7,9 | 2,3,3,4,1,4,7,3,0,1,0,4,4,2, | 9,7,7,8,7,10,6,22,6,13,10,2 | 298 | 149 | 0.0321602565697 | | 1 | 0.538,0.067,0.2,0. | 0.1,0.176,0.176,0. | 0,072 |
| 135747 | ENSG00000291230 | | chr19 | - | 50336561 | 50336705 | 50335058 | 50335181 | 50337016 | 50337135 | 135747 | 235,216,268,622,228,152, | 14,3,5,13,2,4,8,2,10,5,1,0 | 230,162,119,45,150,158,6 | 6,4,2,1,4,2,2,3,7,5,7,15,8,1 | 293 | 149 | 0.0321975628838 | | 1 | 0.895,0.973,0.965, | 0.951,0.954,0.968, | 0,017 |
| 123977 | ENSG00000162241 | SLC25A45 | chr11 | - | 65377332 | 65377423 | 65376817 | 65377076 | 65379375 | 65379561 | 123977 | 0,0,4,2,1,3,4,2,1,0,0,1 | 13,6,22,10,7,11,16,7,14,3, | 1,1,0,2,1,0,2,0,0,0,1,2,2,0, | 12,11,8,5,4,6,1,9,2,7,6,11, | 240 | 149 | 0.032220045715 | | 1 | 0.0,0.0,0.101,0.11, | 0.049,0.053,0.0,0. | -0,013 |
| 17043 | ENSG00000059145 | UNKL | chr16 | - | 1401567 | 1401701 | 1399139 | 1399509 | 1403167 | 1403190 | 17043 | 18,15,20,12,33,12,29,26,1 | 0,3,5,0,2,1,0,2,2,0,0,1 | 17,19,14,22,10,2,31,27,19, | 0,1,0,4,1,0,4,3,3,0,0,2,3,1, | 283 | 149 | 0.0322415540258 | | 1 | 1.0,0.725,0.678,1. | 1.0,0.909,1.0,0.74 | 0,046 |
| 95898 | ENSG00000265808 | SEC22B | chr1 | - | 120168839 | 120168949 | 120163209 | 120163370 | 120176306 | 120176520 | 95898 | 114,123,163,153,38,46,46, | 0,3,1,5,3,1,1,1,1,3,0,1 | 105,112,72,76,49,34,33,72 | 1,2,0,1,0,1,0,0,0,0,1,3,2,0, | 259 | 149 | 0.0322491816351 | | 1 | 1.0,0.959,0.989,0. | 0.984,0.97,1.0,0.9 | -0,02 |
| 85134 | ENSG00000135426 | TESPA1 | chr12 | - | 54983993 | 54984166 | 54974419 | 54974607 | 54984584 | 54984731 | 85134 | 1,1,50,38,12,1,3,11,6,4,4,1 | 41,14,38,23,21,11,9,10,10, | 8,13,0,2,1,13,2,9,5,4,0,1,8, | 18,23,14,14,13,2,13,14,8,2 | 298 | 149 | 0.0322529217644 | | 1 | 0.012,0.034,0.397, | 0.182,0.22,0.0,0.0 | 0,053 |
| 76666 | ENSG00000159023 | EPB41 | chr1 | + | 28993329 | 28993542 | 28987430 | 28987905 | 29011864 | 29011907 | 76666 | 186,160,296,241,165,209, | 4,4,11,3,4,6,2,4,1,8,0,5 | 170,146,232,136,199,190, | 4,7,7,6,21,4,8,3,11,11,0,1, | 298 | 149 | 0.0322846484161 | | 1 | 0.959,0.952,0.931, | 0.955,0.912,0.943, | 0,014 |
| 35440 | ENSG00000122008 | POLK | chr5 | + | 75568676 | 75568790 | 75552471 | 75552591 | 75569339 | 75569369 | 35440 | 11,6,10,2,2,14,4,7,1,1,0,2 | 32,49,75,59,28,32,20,24,2 | 10,14,14,17,11,1,1,12,7,4, | 40,66,46,41,55,24,11,30,2 | 263 | 149 | 0.0322925271641 | | 1 | 0.163,0.065,0.07,0 | 0.124,0.107,0.147, | -0,033 |
| 62686 | ENSG00000138768 | USO1 | chr4 | + | 75793689 | 75793901 | 75790642 | 75790797 | 75799621 | 75799732 | 62686 | 28,10,77,95,28,39,11,29,1 | 0,0,0,2,0,0,0,1,0,2,1,0 | 49,19,19,9,14,26,12,21,12, | 0,0,2,0,1,0,2,4,4,0,1,1,3,1, | 298 | 149 | 0.0323018001858 | | 1 | 1.0,1.0,1.0,0.96,1. | 1.0,1.0,0.826,1.0,0 | 0,059 |
| 127440 | ENSG00000012779 | ALOX5 | chr10 | + | 45444115 | 45444286 | 45443727 | 45443828 | 45445507 | 45446117 | 127440 | 259,146,258,192,339,220, | 19,29,21,14,23,17,9,11,10, | 214,223,206,280,190,99,9 | 13,6,11,10,3,9,2,16,10,6,1 | 298 | 149 | 0.0323040954242 | | 1 | 0.872,0.716,0.86,0 | 0.892,0.949,0.904, | -0,026 |
| 115361 | ENSG00000184465 | WDR27 | chr6 | - | 169638538 | 169638660 | 169634427 | 169634525 | 169643696 | 169643786 | 115361 | 6,2,10,12,8,10,10,7,6,4,5,1 | 2,0,6,1,0,0,0,2,2,2,0,2 | 5,1,7,4,0,1,2,8,2,0,9,4,5,6, | 1,2,1,0,1,1,4,0,0,2,2,1,4,2, | 271 | 149 | 0.0323167742934 | | 1 | 0.623,1.0,0.478,0. | 0.733,0.216,0.794, | 0,254 |
| 72481 | ENSG00000165661 | QSOX2 | chr9 | - | 136221795 | 136221941 | 136219029 | 136219164 | 136223762 | 136223853 | 72481 | 102,86,227,84,73,76,67,51 | 6,2,14,5,0,3,2,0,2,0,3,0 | 136,56,77,31,49,39,48,52, | 12,6,7,4,9,0,5,1,6,4,1,2,2,1 | 295 | 149 | 0.0323419711332 | | 1 | 0.896,0.956,0.891, | 0.851,0.825,0.847, | 0,052 |
| 163823 | ENSG00000083312 | TNPO1 | chr5 | + | 72891802 | 72891896 | 72889785 | 72889957 | 72893138 | 72893246 | 163823 | 51,58,75,39,30,30,34,42,3 | 8,0,0,2,0,2,0,1,0,0,2,2 | 39,71,44,24,41,18,17,35,1 | 0,4,1,6,2,1,0,3,2,1,6,1,2,2, | 243 | 149 | 0.0323512174079 | | 1 | 0.796,1.0,1.0,0.92 | 1.0,0.916,0.964,0. | 0,026 |
| 228393 | ENSG00000164896 | FASTK | chr7 | - | 151077157 | 151077236 | 151076715 | 151076832 | 151077309 | 151077401 | 228393 | 33,7,16,23,18,12,12,6,20,1 | 0,2,11,2,0,1,6,3,0,2,0,0 | 16,9,14,7,4,2,8,17,2,10,9,6 | 7,4,9,0,1,2,0,2,4,0,2,2,4,4, | 228 | 149 | 0.0323551823953 | | 1 | 1.0,0.696,0.487,0. | 0.599,0.595,0.504, | 0,123 |
| 35284 | ENSG00000104517 | UBR5 | chr8 | - | 102271133 | 102271245 | 102270038 | 102270183 | 102272528 | 102272772 | 35284 | 120,115,117,85,75,58,56,7 | 19,2,13,13,2,11,12,15,4,2, | 83,78,71,40,69,60,39,66,4 | 13,12,14,26,10,6,2,11,18,1 | 261 | 149 | 0.032368801872 | | 1 | 0.783,0.97,0.837,0 | 0.785,0.788,0.743, | 0,066 |
| 177997 | ENSG00000142687 | KIAA0319L | chr1 | - | 35435304 | 35435640 | 35434397 | 35435081 | 35441046 | 35443028 | 177997 | 3,0,5,2,5,5,4,8,0,0,4,3 | 9,8,43,30,14,16,19,29,13,5 | 8,2,6,5,5,3,9,5,2,6,7,4,0,5, | 29,18,18,5,6,6,1,31,9,20,1 | 298 | 149 | 0.0324017781762 | | 1 | 0.143,0.0,0.055,0. | 0.121,0.053,0.143, | -0,095 |
| 4561 | ENSG00000267120 | | chr19 | - | 35739709 | 35740088 | 35739251 | 35739626 | 35741023 | 35741228 | 4561 | 153,98,163,166,77,56,122, | 29,15,26,20,14,16,8,18,25, | 100,134,59,23,74,38,24,74 | 12,32,16,14,12,2,20,70,13, | 298 | 149 | 0.0324103295468 | | 1 | 0.725,0.766,0.758, | 0.806,0.677,0.648, | 0,09 |
| 141170 | ENSG00000175387 | SMAD2 | chr18 | - | 47868051 | 47868097 | 47865058 | 47865133 | 47868322 | 47868457 | 141170 | 11,4,11,6,3,2,1,6,6,3,0,2 | 72,47,173,105,67,67,79,64 | 3,13,1,4,5,2,1,3,3,3,3,0,3,5 | 105,86,69,42,64,39,46,83, | 195 | 149 | 0.0324454323063 | | 1 | 0.105,0.061,0.046, | 0.021,0.104,0.011, | 0,03 |
| 240867 | ENSG00000109689 | STIM2 | chr4 | + | 27021459 | 27021612 | 27017710 | 27017984 | 27022518 | 27022661 | 240867 | 8,12,78,36,22,27,26,29,6,0 | 28,11,149,73,17,35,41,44, | 31,24,18,15,15,6,6,21,11,1 | 95,72,49,19,23,21,25,54,2 | 298 | 149 | 0.0324727421659 | | 1 | 0.125,0.353,0.207, | 0.14,0.143,0.155,0 | 0,043 |
| 147815 | ENSG00000188559 | RALGAPA2 | chr20 | - | 20619276 | 20619474 | 20616042 | 20616191 | 20620462 | 20620630 | 147815 | 6,4,19,13,4,16,4,5,6,2,7,13 | 0,0,0,0,0,0,2,0,0,0,0,1 | 16,17,6,9,12,2,1,2,9,14,8,6 | 0,1,1,1,6,0,1,0,3,0,0,3,2,0, | 298 | 149 | 0.0324831282914 | | 1 | 1.0,1.0,1.0,1.0,1.0, | 1.0,0.895,0.75,0.8 | 0,129 |
| 170975 | ENSG00000115524 | SF3B1 | chr2 | - | 197398968 | 197399064 | 197398460 | 197398581 | 197400054 | 197400166 | 170975 | 17,16,26,6,9,11,7,13,18,5, | 702,516,829,512,391,507, | 12,22,20,4,10,10,15,14,8,1 | 497,579,369,255,384,226, | 245 | 149 | 0.0324927587727 | | 1 | 0.015,0.019,0.019, | 0.014,0.023,0.032, | -0,004 |
| 104195 | ENSG00000175582 | RAB6A | chr11 | - | 73718612 | 73718718 | 73716250 | 73716362 | 73718784 | 73718890 | 104195 | 108,82,124,73,27,30,27,59 | 17,7,47,34,9,3,1,7,16,5,2,1 | 56,53,52,30,36,13,19,23,1 | 18,6,14,5,7,10,6,20,5,26,1 | 255 | 149 | 0.0325150463642 | | 1 | 0.788,0.873,0.607, | 0.645,0.838,0.685, | 0,091 |
| 151377 | ENSG00000185787 | MORF4L1 | chr15 | + | 78894170 | 78894230 | 78893538 | 78893627 | 78894819 | 78894904 | 151377 | 25,13,93,67,22,34,26,55,1 | 8,4,3,0,4,1,0,1,2,0,0,3 | 44,73,28,36,37,20,21,34,1 | 9,4,0,6,0,1,0,10,0,4,7,7,5,2 | 209 | 149 | 0.0325355799413 | | 1 | 0.69,0.699,0.957,1 | 0.777,0.929,1.0,0. | 0,017 |
| 87691 | ENSG00000198176 | TFDP1 | chr13 | + | 113623179 | 113623286 | 113610995 | 113611062 | 113631622 | 113631744 | 87691 | 32,61,55,83,45,38,37,25,5 | 2,0,0,1,1,1,0,0,2,0,4,0 | 16,54,34,7,26,14,22,10,19, | 2,2,0,0,0,0,1,0,1,2,0,8,0,4, | 256 | 149 | 0.0325488862094 | | 1 | 0.903,1.0,1.0,0.98, | 0.823,0.94,1.0,1.0, | 0,013 |
| 193049 | ENSG00000143771 | CNIH4 | chr1 | + | 224371329 | 224371423 | 224365878 | 224365991 | 224375794 | 224375998 | 193049 | 29,4,20,31,14,13,11,9,10,8 | 1,3,4,2,0,3,0,2,0,0,0,3 | 13,24,20,7,16,17,9,15,6,10 | 0,0,0,0,2,0,2,0,0,0,0,3,0,1, | 243 | 149 | 0.0325914979054 | | 1 | 0.947,0.45,0.754,0 | 1.0,1.0,1.0,1.0,0.8 | -0,066 |
| 153626 | ENSG00000197555 | SIPA1L1 | chr14 | + | 71650148 | 71650509 | 71624047 | 71624236 | 71658332 | 71658436 | 153626 | 131,96,97,58,23,30,56,83, | 6,6,7,2,1,2,5,7,7,4,0,2 | 56,58,96,116,81,25,37,37, | 0,2,6,5,3,1,2,0,2,3,4,0,2,0, | 298 | 149 | 0.0326047561765 | | 1 | 0.916,0.889,0.874, | 1.0,0.935,0.889,0. | -0,026 |
| 105528 | ENSG00000136504 | KAT7 | chr17 | + | 49805362 | 49805445 | 49798318 | 49798558 | 49811475 | 49811574 | 105528 | 73,72,133,79,22,50,34,54, | 11,0,4,1,0,3,1,2,5,2,0,2 | 51,53,59,73,55,21,22,20,3 | 4,1,0,6,6,2,0,0,2,0,1,0,0,2, | 232 | 149 | 0.0326676355799 | | 1 | 0.81,1.0,0.955,0.9 | 0.891,0.971,1.0,0. | -0,03 |
| 192230 | ENSG00000116698 | SMG7 | chr1 | + | 183542122 | 183542364 | 183540983 | 183541103 | 183544352 | 183544497 | 192230 | 13,15,17,0,17,22,4,4,11,18 | 2,0,1,2,0,0,0,3,2,2,2,0 | 4,26,11,24,8,9,18,8,13,8,4, | 1,1,0,6,0,6,0,0,2,5,2,0,0,2, | 298 | 149 | 0.0327343380679 | | 1 | 0.765,1.0,0.895,0. | 0.667,0.929,1.0,0. | -0,002 |
| 209208 | ENSG00000142188 | TMEM50B | chr21 | - | 33436882 | 33437084 | 33432484 | 33432802 | 33439213 | 33439297 | 209208 | 8,1,15,12,14,5,1,10,7,4,4,6 | 18,7,18,7,27,19,9,10,0,2,8, | 19,12,5,21,2,1,6,5,3,11,15, | 11,14,1,11,4,1,0,14,6,5,16, | 298 | 149 | 0.0327505203395 | | 1 | 0.182,0.067,0.294, | 0.463,0.3,0.714,0. | -0,128 |
| 25445 | ENSG00000102543 | CDADC1 | chr13 | + | 49265878 | 49266003 | 49259345 | 49259523 | 49267489 | 49267525 | 25445 | 1,6,6,4,0,3,0,5,0,0,1,4 | 6,4,17,9,3,4,2,2,2,3,1,3 | 6,12,6,7,3,1,0,6,4,2,4,3,1,2 | 8,7,3,1,0,0,1,1,2,5,5,7,3,10 | 274 | 149 | 0.0327922047841 | | 1 | 0.083,0.449,0.161, | 0.29,0.482,0.521,0 | -0,221 |

| | | | | | | | | | | | | | | | | | | | | | | | |
|---|---|---|---|---|---|---|---|---|---|---|---|---|---|---|---|---|---|---|---|---|---|---|---|
| 234697 | ENSG00000179195 | ZNF664 | chr12 | + | 124004553 | 124004935 | 123988045 | 123988138 | 124011380 | 124011441 | 234697 | 4,4,10,4,12,7,3,6,10,4,1,3 | 28,18,28,18,25,11,16,17,24 | 4,9,1,2,2,3,1,9,4,4,1,3,3,4,3 | 26,8,10,18,7,5,19,22,8,16,1 | 298 | 149 | 0.0327976130569 | | 1 | 0.067,0.1,0.152,0.1 | 0.071,0.36,0.048,0 | 0,029 |
| 156476 | ENSG00000121741 | ZMYM2 | chr13 | + | 20082015 | 20082151 | 20067238 | 20067390 | 20082780 | 20083032 | 156476 | 40,11,87,54,18,32,28,46,7, | 0,0,2,2,0,0,2,0,0,0,0,0 | 37,61,16,23,32,23,15,36,17 | 3,5,0,1,2,2,0,2,2,0,1,1,0,2,0 | 285 | 149 | 0.0328142604466 | | 1 | 1.0,1.0,0.958,0.934 | 0.866,0.864,1.0,0.9 | 0,047 |
| 19326 | ENSG00000138592 | USP8 | chr15 | + | 50481480 | 50482065 | 50477275 | 50477499 | 50489800 | 50489881 | 19326 | 81,87,173,114,78,75,72,13 | 4,1,0,1,0,0,3,2,5,2,0,3 | 96,159,83,95,117,53,73,57 | 0,2,0,0,9,3,1,2,8,1,3,2,0,5,1 | 298 | 149 | 0.0328181754519 | | 1 | 0.91,0.978,1.0,0.98 | 1.0,0.975,1.0,1.0,0 | 0,022 |
| 192228 | ENSG00000116698 | SMG7 | chr1 | + | 183542101 | 183542364 | 183540983 | 183541103 | 183544352 | 183544497 | 192228 | 13,15,17,0,16,22,4,4,11,18 | 2,0,1,2,0,0,0,3,2,2,2,0 | 4,25,11,23,9,9,18,8,13,8,4, | 1,1,0,6,0,6,0,0,2,5,2,0,0,2,4 | 298 | 149 | 0.0328232954717 | | 1 | 0.765,1.0,0.895,0.0 | 0.667,0.926,1.0,0.0 | -0,001 |
| 113456 | ENSG00000104325 | DECR1 | chr8 | + | 90036840 | 90036940 | 90020908 | 90021056 | 90042727 | 90042748 | 113456 | 127,70,156,135,80,76,40,4 | 1,0,0,3,1,0,0,1,1,1,0,1 | 79,89,83,31,69,38,10,38,42 | 0,1,0,3,4,0,1,4,1,2,4,5,0,2,0 | 249 | 149 | 0.0328257455581 | | 1 | 0.987,1.0,1.0,0.964 | 1.0,0.982,1.0,0.861 | 0,038 |
| 5908 | ENSG00000125772 | GPCPD1 | chr20 | - | 5586193 | 5586269 | 5584280 | 5584322 | 5593326 | 5593411 | 5908 | 86,61,274,168,85,107,83,1 | 17,9,22,34,16,7,23,10,7,7,6 | 100,159,93,32,107,52,59,7 | 34,15,23,15,17,9,2,5,20,30 | 225 | 149 | 0.0328334366114 | | 1 | 0.77,0.818,0.892,0 | 0.661,0.875,0.728, | 0,04 |
| 124799 | ENSG00000154258 | ABCA9 | chr17 | - | 69035659 | 69035801 | 69035245 | 69035431 | 69043488 | 69043715 | 124799 | 20,13,98,70,14,20,87,135,1 | 0,0,8,3,1,1,2,1,2,0,0,1 | 51,95,25,8,46,31,17,33,33, | 2,0,0,0,0,1,0,0,0,2,0,0,0,0,0 | 291 | 149 | 0.0328339476319 | | 1 | 1.0,1.0,0.862,0.923 | 0.929,1.0,1.0,1.0,1 | -0,05 |
| 188062 | ENSG00000143553 | SNAPIN | chr1 | + | 153659118 | 153659184 | 153658713 | 153658886 | 153659447 | 153659566 | 188062 | 6,2,11,8,8,3,2,5,4,2,2,2 | 0,1,3,0,3,0,0,1,3,2,0,3 | 9,3,6,1,2,1,1,2,3,2,3,1,13,3 | 0,1,0,0,0,0,0,2,0,0,7,0,3,0,2 | 215 | 149 | 0.0328741535064 | | 1 | 1.0,0.581,0.718,1.0 | 1.0,0.675,1.0,1.0,1 | -0,095 |
| 246244 | ENSG00000172183 | ISG20 | chr15 | + | 88638947 | 88639178 | 88636169 | 88636292 | 88639342 | 88639594 | 246244 | 4,1,2,2,1,0,2,3,5,2,0,0 | 4,7,4,5,2,14,3,14,27,6,3,5 | 3,4,1,1,0,1,1,0,0,4,4,4,0,0,2 | 4,5,5,2,6,10,1,4,1,11,11,4,5 | 298 | 149 | 0.0328974576655 | | 1 | 0.333,0.067,0.2,0.1 | 0.273,0.286,0.091, | -0,063 |
| 53254 | ENSG00000263001 | GTF2I | chr7 | + | 74714856 | 74714916 | 74711031 | 74711109 | 74716893 | 74716950 | 53254 | 8,8,13,10,6,7,6,6,2,1,0,8 | 108,83,110,57,63,61,45,57 | 6,18,4,12,9,4,3,6,11,6,9,26 | 92,76,81,50,63,84,21,45,45 | 209 | 149 | 0.0329456794438 | | 1 | 0.05,0.064,0.078,0 | 0.044,0.144,0.034, | -0,025 |
| 11245 | ENSG00000185813 | PCYT2 | chr17 | - | 81909161 | 81909321 | 81908875 | 81909037 | 81909513 | 81909602 | 11245 | 1,0,5,2,1,9,0,5,1,0,0,0 | 21,17,15,18,9,11,4,10,10,1 | 4,2,0,1,1,0,0,2,0,0,4,3,1,1,4 | 8,1,6,4,9,4,10,4,2,4,2,20,11 | 298 | 149 | 0.0329512918867 | | 1 | 0.023,0.0,0.143,0.0 | 0.2,0.5,0.0,0.111,0 | -0,047 |
| 215600 | ENSG00000166164 | BRD7 | chr16 | - | 50323586 | 50323817 | 50321981 | 50322038 | 50325747 | 50325883 | 215600 | 41,29,88,36,22,28,10,29,29 | 22,13,26,11,11,20,7,7,0,0,2 | 51,23,15,7,32,24,14,25,5,4 | 18,26,9,21,8,8,5,13,5,8,10, | 298 | 149 | 0.0329569914461 | | 1 | 0.482,0.527,0.629, | 0.586,0.307,0.455, | 0,155 |
| 236285 | ENSG00000108239 | TBC1D12 | chr10 | + | 94447634 | 94447670 | 94441895 | 94442019 | 94474667 | 94474783 | 236285 | 0,4,1,2,1,2,2,2,2,3,0,4 | 12,3,32,9,11,8,6,11,8,1,6,9 | 0,1,0,0,2,6,0,3,3,0,0,4,0,0,2 | 13,10,9,10,14,3,2,11,11,14 | 185 | 149 | 0.0329656873219 | | 1 | 0.0,0.518,0.025,0.1 | 0.0,0.075,0.0,0.0,0 | 0,089 |
| 218532 | ENSG00000059758 | CDK17 | chr12 | - | 96323788 | 96323844 | 96313409 | 96313454 | 96323947 | 96324112 | 218532 | 11,1,10,4,4,4,4,6,3,0,1,0 | 23,18,82,32,38,29,41,49,22 | 4,9,5,16,13,2,11,6,3,0,7,6,5 | 30,31,23,28,29,22,22,54,29 | 205 | 149 | 0.0329713652349 | | 1 | 0.258,0.039,0.081, | 0.088,0.174,0.136, | -0,047 |
| 155372 | ENSG00000154001 | PPP2R5E | chr14 | - | 63453434 | 63453885 | 63421992 | 63422094 | 63539528 | 63539692 | 155372 | 13,13,40,30,14,13,7,13,5,6 | 1,0,11,4,1,9,3,4,0,2,4,4 | 27,36,13,17,8,14,12,17,15, | 4,9,0,0,1,1,1,6,1,4,3,2,4,1,0 | 298 | 149 | 0.0329872442942 | | 1 | 0.867,1.0,0.645,0.1 | 0.771,0.667,1.0,1.0 | -0,123 |
| 95953 | ENSG00000186376 | ZNF75D | chrX | - | 135295767 | 135296039 | 135292280 | 135292473 | 135341767 | 135344109 | 95953 | 4,1,9,6,11,1,2,2,6,1,2,0 | 0,0,2,3,0,1,0,6,0,0,0,2 | 7,1,2,2,0,6,1,3,1,1,3,6,4,6,3 | 1,0,0,0,1,2,0,0,0,0,0,0,2,2,0 | 298 | 149 | 0.0329931148227 | | 1 | 1.0,1.0,0.692,0.5,1 | 0.778,1.0,1.0,1.0,0 | -0,12 |
| 217180 | ENSG00000031003 | FAM13B | chr5 | - | 137946227 | 137946322 | 137945901 | 137945997 | 137948954 | 137949184 | 217180 | 17,8,19,10,9,7,15,1,0,1,2,4 | 7,7,11,12,6,10,4,4,8,8,2,1 | 7,5,19,10,12,3,10,9,8,6,5,7 | 7,11,3,1,2,2,1,8,8,4,2,5,2,7 | 244 | 149 | 0.0330324052096 | | 1 | 0.597,0.411,0.513, | 0.379,0.217,0.795, | -0,156 |
| 188022 | ENSG00000121481 | RNF2 | chr1 | + | 185094520 | 185094570 | 185093060 | 185093276 | 185098071 | 185098344 | 188022 | 10,0,2,0,1,0,0,0,0,0,0,1 | 42,46,87,54,37,31,18,16,39 | 1,4,1,0,4,1,0,0,1,4,5,2,0,5,0 | 35,29,30,15,46,20,10,16,15 | 199 | 149 | 0.0330565759358 | | 1 | 0.151,0.0,0.017,0.0 | 0.021,0.094,0.024, | -0,024 |
| 175896 | ENSG00000173889 | PHC3 | chr3 | - | 170106831 | 170106944 | 170102801 | 170102934 | 170113359 | 170113519 | 175896 | 59,25,109,75,33,50,34,55,1 | 2,0,4,0,0,1,0,1,0,2,0,0 | 68,79,38,45,58,46,50,46,34 | 1,3,2,2,4,0,0,2,2,0,2,0,0,4,0 | 262 | 149 | 0.0330613195257 | | 1 | 0.944,1.0,0.939,1.0 | 0.975,0.937,0.915, | 0,02 |
| 52882 | ENSG00000143418 | CERS2 | chr1 | - | 150968082 | 150968174 | 150967819 | 150967877 | 150968394 | 150968512 | 52882 | 10,9,55,53,27,14,25,21,3,5 | 0,0,6,3,3,7,3,5,3,1,3,1 | 27,16,5,8,10,7,8,21,10,18,2 | 11,8,5,0,1,0,2,7,4,0,9,5,4,5 | 241 | 149 | 0.0330903659984 | | 1 | 1.0,1.0,0.85,0.916, | 0.603,0.553,0.382, | 0,062 |
| 240384 | ENSG00000184465 | WDR27 | chr6 | - | 169663562 | 169663801 | 169662303 | 169662424 | 169664165 | 169664632 | 240384 | 12,5,12,7,7,18,14,10,7,10,8 | 1,0,14,5,2,4,0,2,1,2,0,2 | 9,1,8,2,5,2,6,12,1,4,11,10,1 | 3,0,0,1,0,1,3,3,0,0,7,2,0,0,0 | 298 | 149 | 0.0331023614793 | | 1 | 0.857,1.0,0.3,0.41 | 0.6,1.0,1.0,0.5,1.0 | -0,073 |
| 68544 | ENSG00000055609 | KMT2C | chr7 | - | 152157760 | 152157925 | 152156204 | 152156274 | 152158862 | 152159072 | 68544 | 21,17,35,25,14,15,8,13,12, | 0,0,0,0,0,1,0,0,2,2,0,1 | 26,17,5,15,19,12,11,11,4,2 | 0,5,2,2,1,0,3,2,0,0,0,2,2,0,0 | 298 | 149 | 0.0331564291716 | | 1 | 1.0,1.0,1.0,1.0,1.0 | 1.0,0.63,0.556,0.78 | 0,083 |
| 509 | ENSG00000160216 | AGPAT3 | chr21 | + | 43978045 | 43978121 | 43971387 | 43971490 | 43980988 | 43981187 | 509 | 16,26,62,51,29,39,14,27,7, | 4,0,2,2,0,1,0,3,0,0,0,2 | 30,21,20,20,19,17,4,10,27, | 0,1,0,2,4,2,0,6,2,4,0,2,0,0,0 | 225 | 149 | 0.033161984224 | | 1 | 0.726,1.0,0.954,0.9 | 1.0,0.933,1.0,0.869 | 0,047 |
| 222498 | ENSG00000166295 | ANAPC16 | chr10 | + | 72220053 | 72220270 | 72216028 | 72216072 | 72223887 | 72224056 | 222498 | 10,3,5,3,1,3,1,0,2,0,1,0 | 19,3,29,23,13,11,24,21,12, | 10,3,4,6,4,1,4,4,1,5,5,0,1,4 | 18,4,7,5,5,3,11,31,3,8,34,2 | 298 | 149 | 0.0331759302 | | 1 | 0.208,0.333,0.079, | 0.217,0.273,0.222, | -0,055 |
| 51153 | ENSG00000197111 | PCBP2 | chr12 | + | 53465931 | 53465973 | 53462480 | 53462567 | 53467220 | 53467293 | 51153 | 89,51,153,107,62,94,69,11 | 13,10,24,9,14,10,3,15,2,5,0 | 85,133,49,136,69,51,185,1 | 14,17,5,21,8,11,35,20,8,5,6 | 191 | 149 | 0.0332128283659 | | 1 | 0.842,0.799,0.833, | 0.826,0.859,0.884, | 0,038 |
| 98826 | ENSG00000179818 | PCBP1-AS1 | chr2 | - | 70055655 | 70055910 | 70048708 | 70051305 | 70059642 | 70059679 | 98826 | 2,1,2,0,7,0,1,2,2,0,1,4 | 4,3,4,2,3,5,0,5,4,2,2,1 | 10,4,5,2,3,7,0,8,1,5,2,5,12, | 7,13,9,2,5,2,2,3,4,11,5,3,7, | 298 | 149 | 0.0332153099712 | | 1 | 0.2,0.143,0.2,0.0,0 | 0.417,0.133,0.217, | -0,068 |
| 99064 | ENSG00000136280 | CCM2 | chr7 | + | 45069825 | 45069961 | 45064462 | 45064646 | 45072725 | 45072783 | 99064 | 26,40,42,55,18,13,18,24,23 | 9,2,10,6,2,4,5,11,9,6,5,2 | 43,36,7,7,25,30,5,17,6,34,1 | 4,5,4,2,2,0,1,1,2,4,9,5,3,5,4 | 285 | 149 | 0.0332187709299 | | 1 | 0.602,0.913,0.687, | 0.849,0.79,0.478,0 | -0,083 |
| 152982 | ENSG00000165355 | FBXO33 | chr14 | - | 39401697 | 39401861 | 39399565 | 39399787 | 39402400 | 39402511 | 152982 | 82,46,78,55,26,21,23,35,52 | 1,0,3,2,3,0,3,3,1,0,0,0 | 50,81,31,21,36,23,32,48,22 | 2,2,0,2,0,0,0,0,3,0,4,12,1,1 | 298 | 149 | 0.0332198437897 | | 1 | 0.976,1.0,0.929,0.9 | 0.926,0.953,1.0,0.8 | 0,03 |
| 3160 | ENSG00000105699 | LSR | chr19 | + | 35261924 | 35261981 | 35258944 | 35259064 | 35262545 | 35262692 | 3160 | 12,2,7,8,9,2,8,12,3,6,0,4 | 51,7,46,14,29,16,24,24,27, | 4,3,1,0,4,2,5,13,1,3,1,0,7,3 | 22,15,12,16,23,13,32,42,5, | 206 | 149 | 0.033224861352 | | 1 | 0.145,0.171,0.099, | 0.116,0.126,0.057, | 0,051 |
| 150666 | ENSG00000196705 | ZNF431 | chr19 | + | 21161545 | 21161637 | 21143550 | 21143643 | 21166334 | 21166461 | 150666 | 0,1,3,6,0,2,5,1,1,0,1,0 | 13,12,31,33,12,24,11,32,5, | 3,1,1,0,1,0,0,0,5,0,0,0,0,3,0 | 27,31,19,15,19,8,9,16,10,5 | 241 | 149 | 0.0332301939297 | | 1 | 0.0,0.049,0.056,0.1 | 0.064,0.02,0.032,0 | 0,024 |
| 249784 | ENSG00000095574 | IKZF5 | chr10 | - | 123006239 | 123006347 | 122998492 | 122998671 | 123007025 | 123007176 | 249784 | 4,3,21,14,2,3,5,3,0,0,2,11 | 9,5,2,10,0,6,2,0,1,1,1,0 | 4,9,10,13,5,1,1,0,4,8,0,0,0, | 9,19,5,0,11,6,6,2,5,2,1,8,5, | 257 | 149 | 0.033238570095 | | 1 | 0.205,0.258,0.859, | 0.205,0.215,0.537, | 0,188 |
| 21513 | ENSG00000100938 | GMPR2 | chr14 | + | 24234106 | 24234241 | 24233478 | 24233598 | 24235736 | 24235820 | 21513 | 0,1,4,5,2,6,2,0,1,0,2,0 | 5,7,29,36,18,20,6,12,8,2,6, | 4,2,0,2,1,3,0,0,0,1,2,2,2,0,0 | 22,12,17,1,8,3,1,16,9,8,43, | 284 | 149 | 0.0332394494311 | | 1 | 0.0,0.07,0.067,0.00 | 0.087,0.08,0.0,0.5 | -0,019 |
| 142709 | ENSG00000108474 | PIGL | chr17 | + | 16284020 | 16284138 | 16233970 | 16234070 | 16299887 | 16300187 | 142709 | 6,0,13,9,7,8,16,10,4,3,4,6 | 17,4,20,9,7,4,15,15,7,4,7,9 | 4,1,7,2,3,3,0,3,5,8,11,0,9,5 | 6,16,8,4,17,8,12,9,5,9,11,4 | 267 | 149 | 0.0332585810252 | | 1 | 0.165,0.0,0.266,0.1 | 0.271,0.034,0.328, | 0,073 |
| 200893 | ENSG00000116539 | ASH1L | chr1 | - | 155415743 | 155415821 | 155395458 | 155395553 | 155438326 | 155439068 | 200893 | 9,6,46,24,11,22,7,25,1,4,4, | 1,1,4,3,1,8,0,3,0,0,0,2 | 22,22,6,11,12,11,8,9,8,9,14 | 6,3,5,1,5,0,5,3,2,0,4,4,0,4,3 | 227 | 149 | 0.0332972120946 | | 1 | 0.855,0.798,0.883, | 0.706,0.828,0.441, | 0,132 |
| 222194 | ENSG00000152601 | MBNL1 | chr3 | + | 152300204 | 152300367 | 152268946 | 152269092 | 152414940 | 152415111 | 222194 | 452,251,444,197,225,233,2 | 74,49,24,11,25,26,41,14,35 | 365,359,176,132,170,131,1 | 26,23,12,1,13,10,13,12,11, | 298 | 149 | 0.0333067738226 | | 1 | 0.753,0.719,0.902, | 0.875,0.886,0.88,0 | -0,037 |
| 3609 | ENSG00000196646 | ZNF136 | chr19 | + | 12185784 | 12185911 | 12163095 | 12163206 | 12186113 | 12186157 | 3609 | 60,34,85,33,19,23,24,45,15 | 13,10,12,5,5,6,2,3,0,0,0,2 | 34,61,16,38,24,18,16,21,19 | 0,5,0,3,5,0,0,2,4,4,3,12,1,1 | 276 | 149 | 0.0333238601381 | | 1 | 0.714,0.647,0.793, | 1.0,0.868,1.0,0.87 | -0,052 |
| 235515 | ENSG00000111364 | DDX55 | chr12 | + | 123603847 | 123604038 | 123602142 | 123602256 | 123605930 | 123606159 | 235515 | 2,0,0,1,2,2,0,2,0,0,4,1 | 1,6,5,3,3,4,2,1,5,2,0,2 | 0,0,0,0,0,0,0,1,0,0,2,0,0,0 | 3,3,1,2,1,2,2,4,1,4,1,5,3,1,1 | 298 | 149 | 0.0333270435476 | | 1 | 0.5,0.0,0.0,0.143,0 | 0.0,0.0,0.0,0.0,0.0 | 0,202 |
| 39699 | ENSG00000167515 | TRAPPC2L | chr16 | + | 88859572 | 88859750 | 88858641 | 88858789 | 88859892 | 88859972 | 39699 | 158,154,225,159,70,69,75, | 4,0,21,7,0,3,4,10,8,4,0,4 | 134,172,40,8,65,54,62,115 | 8,9,2,2,5,0,5,14,3,2,7,4,7,3 | 298 | 149 | 0.0333366639602 | | 1 | 0.952,1.0,0.843,0.9 | 0.893,0.905,0.909, | 0,057 |
| 31790 | ENSG00000001084 | GCLC | chr6 | - | 53508594 | 53508711 | 53507479 | 53507618 | 53509175 | 53509250 | 31790 | 80,18,205,178,71,49,51,43 | 2,0,2,1,0,0,0,0,4,0,2,2 | 85,101,70,18,54,53,22,46,3 | 0,6,2,3,0,0,0,5,0,0,4,1,0,2,8 | 266 | 149 | 0.0333779115394 | | 1 | 0.957,1.0,0.983,0.9 | 1.0,0.904,0.951,0.7 | 0,023 |
| 195383 | ENSG00000134744 | TUT4 | chr1 | - | 52529767 | 52529873 | 52526019 | 52526373 | 52552930 | 52553100 | 195383 | 0,3,22,11,1,2,1,1,0,0,0,3 | 5,17,75,19,20,17,24,33,6,6 | 7,13,0,2,9,2,0,2,2,2,4,9,6,4 | 15,25,15,9,14,16,17,31,15, | 255 | 149 | 0.03337967327 | | 1 | 0.0,0.093,0.146,0.1 | 0.214,0.233,0.0,0.1 | -0,059 |
| 193296 | ENSG00000119778 | ATAD2B | chr2 | - | 23816636 | 23816739 | 23810315 | 23810502 | 23819746 | 23819882 | 193296 | 3,1,5,1,0,4,1,2,0,1,0,1 | 28,16,43,19,17,30,15,28,9, | 5,11,1,3,1,1,1,3,0,2,3,5,3,1 | 39,31,15,23,26,15,14,47,13 | 252 | 149 | 0.03341084218 | | 1 | 0.06,0.036,0.064,0 | 0.07,0.173,0.038,0 | -0,03 |
| 152964 | ENSG00000257621 | PSMA3-AS1 | chr14 | - | 58295745 | 58295813 | 58291634 | 58291707 | 58297954 | 58298134 | 152964 | 6,5,0,0,3,9,0,2,0,0,0,0 | 67,34,129,99,60,80,54,73,5 | 0,0,2,4,0,0,0,0,2,6,16,7,2,7 | 76,42,35,33,53,24,20,71,41 | 217 | 149 | 0.0334121416574 | | 1 | 0.058,0.092,0.0,0.0 | 0.0,0.0,0.038,0.07 | -0,013 |
| 67968 | ENSG00000265491 | RNF115 | chr1 | - | 145751437 | 145751510 | 145750406 | 145750500 | 145752977 | 145753049 | 67968 | 34,18,82,53,36,31,41,56,26 | 1,0,2,1,0,6,2,2,0,2,0,1 | 36,60,30,23,45,14,26,33,29 | 1,2,1,1,13,2,0,2,0,1,4,1,4,0 | 222 | 149 | 0.0334214695338 | | 1 | 0.958,1.0,0.965,0.9 | 0.96,0.953,0.953,0 | 0,029 |
| 162521 | ENSG00000132952 | USPL1 | chr13 | + | 30621072 | 30621238 | 30617919 | 30618056 | 30621763 | 30621892 | 162521 | 2,1,6,1,0,0,0,0,2,0,0,8 | 21,9,42,25,16,15,13,28,10, | 2,3,0,2,2,0,0,1,0,0,3,0,0,0,0 | 17,27,16,14,24,11,15,12,20 | 298 | 149 | 0.033438528186 | | 1 | 0.045,0.053,0.067, | 0.056,0.053,0.0,0.0 | 0,018 |
| 30208 | ENSG00000131503 | ANKHD1 | chr5 | + | 140496519 | 140497278 | 140486957 | 140487060 | 140504820 | 140504966 | 30208 | 50,27,133,76,68,95,46,45,2 | 16,13,27,6,16,11,11,11,4,2 | 103,112,54,155,66,43,136, | 12,31,1,5,30,8,10,5,6,16,3, | 298 | 149 | 0.033440764267 | | 1 | 0.61,0.509,0.711,0 | 0.811,0.644,0.964, | -0,075 |
| 117203 | ENSG00000182158 | CREB3L2 | chr7 | - | 137882411 | 137882628 | 137874978 | 137880551 | 137884994 | 137885121 | 117203 | 93,92,166,129,35,82,60,15 | 3,4,3,0,2,1,2,2,1,0,2,1 | 110,202,72,147,62,57,97,9 | 2,4,2,15,2,1,5,0,3,0,8,0,0,1 | 298 | 149 | 0.0334472618322 | | 1 | 0.939,0.92,0.965,1 | 0.965,0.962,0.947, | 0,003 |
| 131807 | ENSG00000089094 | KDM2B | chr12 | - | 121443679 | 121443793 | 121442991 | 121443030 | 121444011 | 121444272 | 131807 | 84,43,68,45,49,30,25,43,57 | 2,1,5,3,4,2,1,1,6,0,0,3 | 38,27,18,13,23,26,29,48,13 | 3,1,0,0,0,1,2,6,2,1,2,2,0,3,0 | 263 | 149 | 0.0335088287923 | | 1 | 0.96,0.961,0.885,0 | 0.878,0.939,1.0,1.0 | -0,013 |
| 80642 | ENSG00000142188 | TMEM50B | chr21 | - | 33436882 | 33436992 | 33432484 | 33432802 | 33439213 | 33439297 | 80642 | 9,1,23,17,21,13,6,14,5,5,2, | 18,7,18,7,27,19,9,10,0,2,8, | 28,13,5,24,5,1,7,8,8,21,15, | 11,14,1,11,4,1,0,14,6,5,16, | 259 | 149 | 0.0335199454704 | | 1 | 0.223,0.076,0.424, | 0.594,0.348,0.742, | -0,132 |
| 214578 | ENSG00000119707 | RBM25 | chr14 | + | 73111289 | 73111382 | 73110830 | 73111155 | 73111527 | 73111802 | 214578 | 4,1,10,4,2,1,3,4,0,0,0,5 | 70,44,149,103,58,55,60,98 | 1,4,4,0,1,6,3,6,1,0,0,0,0,4,0 | 96,154,65,73,62,44,40,64,4 | 242 | 149 | 0.0335336389127 | | 1 | 0.034,0.014,0.04,0 | 0.006,0.016,0.037, | 0,003 |
| 99966 | ENSG00000058091 | CDK14 | chr7 | + | 90709815 | 90709957 | 90709250 | 90709610 | 90726566 | 90726644 | 99966 | 319,149,329,173,175,242,2 | 3,3,5,1,3,6,2,10,0,0,0,3 | 355,256,132,234,197,144,1 | 7,4,0,5,5,4,4,8,1,4,12,5,6,4 | 291 | 149 | 0.0335366699141 | | 1 | 0.982,0.962,0.971, | 0.963,0.97,1.0,0.9 | 0,017 |
| 71468 | ENSG00000100075 | SLC25A1 | chr22 | - | 19176577 | 19176693 | 19176420 | 19176494 | 19176845 | 19176950 | 71468 | 69,58,108,82,54,48,51,66,5 | 4,1,7,4,1,3,1,0,2,2,0,1 | 37,34,50,34,25,24,27,62,35 | 2,0,1,8,7,0,1,2,0,1,8,5,0,0,0 | 265 | 149 | 0.0335460424026 | | 1 | 0.907,0.97,0.897,0 | 0.912,1.0,0.966,0.7 | 0,037 |
| 201280 | ENSG00000197956 | S100A6 | chr1 | - | 153535788 | 153535933 | 153535201 | 153535360 | 153536116 | 153536215 | 201280 | 0,0,5,3,2,8,0,3,2,1,2,0 | 6,3,9,8,7,6,4,2,4,2,0,13 | 0,2,2,0,0,0,7,3,0,0,7,2,1,0,0 | 7,8,0,1,2,4,1,3,1,3,3,10,5,5 | 294 | 149 | 0.0335521004904 | | 1 | 0.0,0.0,0.22,0.16,0 | 0.0,0.112,1.0,0.0,0 | 0,044 |
| 136676 | ENSG00000196646 | ZNF136 | chr19 | + | 12164625 | 12164874 | 12163115 | 12163206 | 12185784 | 12185911 | 136676 | 0,1,0,0,0,2,3,6,2,0,0,4 | 16,12,34,13,5,5,3,15,5,4,1, | 0,0,0,0,2,0,0,0,2,0,2,0,0,0,0 | 17,23,3,17,4,6,4,5,5,10,20, | 298 | 149 | 0.0335533132304 | | 1 | 0.0,0.04,0.0,0.0,0.0 | 0.0,0.0,0.0,0.0,0.2, | 0,068 |
| 51774 | ENSG00000153914 | SREK1 | chr5 | + | 66158808 | 66158932 | 66153462 | 66153596 | 66159218 | 66159334 | 51774 | 89,39,71,38,11,29,33,63,43 | 18,10,11,12,4,7,17,7,25,2,6 | 34,46,41,143,37,24,26,38,2 | 7,7,9,8,3,9,3,3,3,5,4,7,10,7 | 273 | 149 | 0.0336264457835 | | 1 | 0.73,0.68,0.779,0.6 | 0.726,0.782,0.713, | -0,067 |
| 28554 | ENSG00000136153 | LMO7 | chr13 | + | 75809153 | 75809183 | 75804226 | 75804541 | 75817160 | 75817278 | 28554 | 27,20,84,40,47,30,42,48,33 | 3,7,28,15,18,20,9,23,10,6,4 | 46,46,10,10,16,11,21,56,5, | 20,17,5,17,36,21,4,18,16,6 | 179 | 149 | 0.0336303400148 | | 1 | 0.882,0.704,0.714, | 0.657,0.693,0.625, | 0,123 |
| 210196 | ENSG00000169499 | PLEKHA2 | chr8 | + | 38943788 | 38943834 | 38935993 | 38936050 | 38946123 | 38946221 | 210196 | 38,11,121,66,29,45,29,55,3 | 4,2,33,22,16,4,10,15,11,6,7 | 63,42,35,9,19,41,27,40,25, | 30,9,10,19,12,5,15,22,7,17 | 195 | 149 | 0.0336544636358 | | 1 | 0.879,0.808,0.737, | 0.616,0.781,0.728, | 0,076 |
| 32740 | ENSG00000231113 | LINC02976 | chr6 | + | 42941509 | 42941615 | 42940772 | 42940929 | 42943797 | 42943970 | 32740 | 9,9,9,15,4,4,9,8,7,3,2,5 | 7,5,13,5,3,4,16,8,3,6,0,5 | 6,6,4,4,7,5,1,3,5,2,8,4,2,3,5 | 14,11,2,6,13,10,3,8,2,7,12, | 255 | 149 | 0.0336563977884 | | 1 | 0.429,0.513,0.288, | 0.2,0.242,0.539,0.1 | 0,14 |
| 95842 | ENSG00000167483 | NIBAN3 | chr19 | + | 17546685 | 17546797 | 17543523 | 17543631 | 17551785 | 17551960 | 95842 | 206,171,270,192,124,97,94 | 11,4,13,7,2,0,2,4,0,2,8,4 | 192,188,165,65,160,63,56, | 3,2,3,7,3,3,0,1,6,2,4,6,7,6,1 | 261 | 149 | 0.0336608025982 | | 1 | 0.914,0.961,0.922, | 0.973,0.982,0.969, | -0,011 |
| 102108 | ENSG00000160218 | TRAPPC10 | chr21 | + | 44098356 | 44098534 | 44094062 | 44094233 | 44102777 | 44102972 | 102108 | 58,45,127,78,43,60,56,41,4 | 3,2,1,0,0,6,5,3,1,1,0,1 | 89,74,21,18,76,35,30,31,27 | 2,0,0,1,3,0,0,3,1,0,0,5,0,0,2 | 298 | 149 | 0.033721790499 | | 1 | 0.906,0.918,0.984, | 0.957,1.0,1.0,0.9,0 | -0,016 |
| 28659 | ENSG00000104320 | NBN | chr8 | - | 89981374 | 89981523 | 89980733 | 89980893 | 89982721 | 89982855 | 28659 | 44,20,55,50,35,18,13,20,16 | 0,0,0,0,0,0,1,0,0,0,0,2 | 41,25,27,10,29,26,11,28,24 | 0,1,3,6,2,1,0,1,3,0,0,3,1,0,0 | 298 | 149 | 0.0337292293282 | | 1 | 1.0,1.0,1.0,1.0,1.0, | 1.0,0.926,0.818,0.4 | 0,068 |
| 68399 | ENSG00000144674 | GOLGA4 | chr3 | + | 37337142 | 37337163 | 37335052 | 37335166 | 37337665 | 37337734 | 68399 | 56,54,60,40,24,41,20,32,20 | 1,0,1,3,0,1,0,3,0,0,0,0 | 26,45,26,44,36,17,20,17,18 | 3,1,1,2,0,0,2,1,2,4,0,0,0,4,3 | 170 | 149 | 0.0337426031275 | | 1 | 0.98,1.0,0.981,0.92 | 0.884,0.975,0.958, | 0,038 |
| 222496 | ENSG00000166295 | ANAPC16 | chr10 | + | 72220053 | 72220264 | 72216028 | 72216072 | 72223887 | 72224056 | 222496 | 10,3,5,3,1,3,1,0,2,0,1,0 | 19,3,29,23,13,11,24,21,12, | 10,2,4,6,4,1,4,4,1,5,5,1,1,4 | 18,4,7,5,5,3,11,31,3,8,34,2 | 298 | 149 | 0.0337486269504 | | 1 | 0.208,0.333,0.079, | 0.217,0.2,0.222,0.1 | -0,051 |
| 217181 | ENSG00000031003 | FAM13B | chr5 | - | 137946231 | 137946311 | 137945901 | 137945997 | 137948954 | 137949184 | 217181 | 8,5,22,11,6,8,7,8,6,1,5,2 | 7,7,11,12,6,10,4,4,8,8,2,1 | 15,23,4,12,9,4,5,11,4,5,3,5 | 7,11,3,1,2,2,1,8,8,4,2,5,2,7 | 229 | 149 | 0.0337490484772 | | 1 | 0.426,0.317,0.565, | 0.582,0.576,0.465, | -0,129 |
| 103234 | ENSG00000110497 | AMBRA1 | chr11 | - | 46508190 | 46508370 | 46494123 | 46494204 | 46541944 | 46542981 | 103234 | 30,33,41,29,28,32,24,24,17 | 0,0,5,1,2,2,0,0,0,0,3,4 | 30,41,22,21,24,19,25,25,14 | 1,1,0,0,0,2,0,0,0,1,0,1,1,0,0 | 298 | 149 | 0.0337612759462 | | 1 | 1.0,1.0,0.804,0.93 | 0.938,0.953,1.0,1.0 | -0,074 |
| 207654 | ENSG00000130348 | QRSL1 | chr6 | + | 106654781 | 106654922 | 106652466 | 106652582 | 106655614 | 106655732 | 207654 | 36,28,126,146,52,39,25,42 | 4,8,10,11,8,5,2,2,1,2,2,0 | 64,55,28,16,26,63,15,39,11 | 5,2,5,2,0,0,0,2,4,11,3,0,0,0 | 290 | 149 | 0.0337973244679 | | 1 | 0.822,0.643,0.866, | 0.868,0.934,0.742, | -0,033 |
| 74627 | ENSG00000115020 | PIKFYVE | chr2 | + | 208277536 | 208277708 | 208276711 | 208276830 | 208288728 | 208288818 | 74627 | 36,17,32,23,24,26,17,25,11 | 1,0,2,1,0,3,0,6,0,2,0,1 | 47,31,9,10,15,6,18,20,8,27 | 4,8,0,2,5,0,0,0,1,0,1,2,0,2,0 | 298 | 149 | 0.0338090542092 | | 1 | 0.947,1.0,0.889,0.9 | 0.855,0.66,1.0,0.71 | 0,022 |
| 162910 | ENSG00000173926 | MARCHF3 | chr5 | - | 126917510 | 126917676 | 126914930 | 126915134 | 126917983 | 126918228 | 162910 | 2,0,11,2,4,2,0,2,1,0,0,2 | 87,49,133,63,45,35,53,53,8 | 5,2,3,8,2,0,0,4,1,8,0,3,5,1,1 | 27,45,80,70,50,34,60,93,31 | 298 | 149 | 0.0338399119607 | | 1 | 0.011,0.0,0.04,0.0 | 0.085,0.022,0.018, | -0,02 |
| 171775 | ENSG00000112149 | CD83 | chr6 | + | 14131361 | 14131748 | 14117949 | 14118065 | 14133648 | 14133755 | 171775 | 365,351,186,93,48,74,58,4 | 25,13,19,12,2,4,9,6,1,2,12, | 90,157,85,26,58,15,71,72,1 | 9,16,16,5,4,0,4,18,8,0,5,72 | 298 | 149 | 0.0338432087476 | | 1 | 0.88,0.931,0.83,0.1 | 0.833,0.831,0.726, | 0,049 |
| 222604 | ENSG00000167483 | NIBAN3 | chr19 | + | 17530754 | 17530884 | 17527237 | 17527395 | 17532262 | 17532388 | 222604 | 65,74,168,71,103,64,77,72 | 45,36,41,14,40,27,53,39,53 | 77,124,83,14,69,47,59,64,5 | 26,38,25,17,12,6,18,37,12, | 279 | 149 | 0.0338612218271 | | 1 | 0.435,0.523,0.686, | 0.613,0.635,0.639, | -0,082 |
| 221779 | ENSG00000166788 | SAAL1 | chr11 | - | 18093701 | 18093965 | 18092244 | 18092324 | 18096770 | 18096854 | 221779 | 0,0,2,1,3,2,2,0,1,2,0,2 | 16,13,12,12,4,15,10,11,4,6 | 8,3,1,5,0,0,4,1,0,3,2,2,1,0,0 | 19,15,15,6,14,6,12,5,11,12 | 298 | 149 | 0.0338922437673 | | 1 | 0.0,0.0,0.077,0.04, | 0.174,0.091,0.032, | -0,004 |
| 74222 | ENSG00000237877 | LINC01473 | chr2 | - | 186058756 | 186058969 | 186039499 | 186039555 | 186081329 | 186081416 | 74222 | 1,1,7,2,2,0,2,4,0,0,1,2 | 11,0,19,10,2,6,8,21,2,3,1,1 | 3,5,7,5,2,0,0,1,0,3,2,1,2,0,0 | 8,5,7,4,5,3,2,8,6,6,5,4,2,9,4 | 298 | 149 | 0.0339120716993 | | 1 | 0.043,1.0,0.156,0.0 | 0.158,0.333,0.333, | 0,03 |
| 44866 | ENSG00000103064 | SLC7A6 | chr16 | + | 68287322 | 68287435 | 68274690 | 68275249 | 68287745 | 68287871 | 44866 | 2,0,6,18,4,2,2,5,3,1,5,2 | 2,6,20,20,13,16,20,12,6,7,6 | 3,1,4,2,5,6,10,13,6,5,8,14,6 | 13,25,4,2,7,7,5,13,3,16,8,3 | 262 | 149 | 0.0339350367077 | | 1 | 0.363,0.0,0.146,0.1 | 0.116,0.022,0.363, | -0,109 |
| 23927 | ENSG00000180776 | ZDHHC20 | chr13 | - | 21421060 | 21421164 | 21413651 | 21413772 | 21425651 | 21425678 | 23927 | 121,59,182,129,36,61,47,6 | 3,0,0,4,0,0,0,0,4,0,0,1 | 81,88,49,45,67,58,32,51,24 | 0,1,6,0,1,0,3,7,1,1,3,3,1,0,0 | 253 | 149 | 0.0339364003948 | | 1 | 0.96,1.0,1.0,0.95,1 | 1.0,0.981,0.828,1.0 | 0,026 |
| 102954 | ENSG00000152457 | DCLRE1C | chr10 | - | 14926842 | 14926897 | 14922980 | 14923069 | 14928015 | 14928152 | 102954 | 50,38,77,62,32,26,14,27,31 | 5,2,8,2,3,2,3,4,4,3,2,3 | 62,33,12,25,16,11,11,33,16 | 3,1,1,2,0,0,0,2,3,10,1,1,4,4 | 204 | 149 | 0.0339472133935 | | 1 | 0.88,0.933,0.875,0 | 0.938,0.96,0.898,0 | -0,06 |
| 232373 | ENSG00000111203 | ITFG2 | chr12 | + | 2830698 | 2830885 | 2824089 | 2824252 | 2836744 | 2836956 | 232373 | 22,7,11,6,13,7,23,23,7,11,3 | 2,4,1,1,0,0,2,1,0,0,0,1 | 8,7,4,10,6,4,6,8,4,3,5,9,12, | 0,2,0,3,2,1,0,0,0,4,0,0,4,7,1 | 298 | 149 | 0.0339546255755 | | 1 | 0.846,0.467,0.846, | 1.0,0.636,1.0,0.62 | 0,095 |
| 28610 | ENSG00000143774 | GUK1 | chr1 | + | 228141625 | 228141829 | 228141123 | 228141288 | 228145510 | 228145624 | 28610 | 2,2,6,2,5,5,2,2,0,0,2,0 | 7,13,20,9,13,11,7,5,7,11,4, | 4,4,0,1,0,0,4,0,0,2,0,2,1,0,2 | 17,8,10,6,3,3,2,9,4,8,9,18,4 | 298 | 149 | 0.0339558255961 | | 1 | 0.125,0.071,0.13,0 | 0.105,0.2,0.0,0.07 | 0,013 |
| 105100 | ENSG00000131196 | NFATC1 | chr18 | + | 79486247 | 79486937 | 79467449 | 79467582 | 79527527 | 79527741 | 105100 | 47,27,32,31,32,39,38,17,38 | 1,2,2,0,0,0,1,3,3,0,1,2 | 12,29,24,25,16,16,10,28,8, | 1,0,1,0,4,0,2,0,1,2,2,3,0,2,4 | 298 | 149 | 0.0339570768253 | | 1 | 0.959,0.871,0.889, | 0.857,1.0,0.923,1.0 | 0,03 |
| 178313 | ENSG00000143297 | FCRL5 | chr1 | - | 157547639 | 157547695 | 157546942 | 157547197 | 157549559 | 157549580 | 178313 | 3,3,17,1,2,5,1,9,5,0,2,1 | 127,83,238,88,154,78,115, | 8,6,2,1,1,0,1,0,0,3,1,0,5,5,3 | 270,119,186,44,27,29,65,2 | 205 | 149 | 0.0339629308705 | | 1 | 0.017,0.026,0.049, | 0.021,0.035,0.008, | 0,005 |
| 93517 | ENSG00000124782 | RREB1 | chr6 | + | 7246423 | 7247221 | 7228996 | 7231907 | 7248510 | 7249274 | 93517 | 33,40,45,37,24,22,14,17,21 | 6,0,1,9,5,2,3,8,4,6,1,0 | 14,29,17,44,32,9,21,35,12, | 3,5,8,9,4,2,3,8,2,7,3,8,4,8,6 | 298 | 149 | 0.0339719811612 | | 1 | 0.733,1.0,0.957,0.0 | 0.7,0.744,0.515,0.1 | 0,095 |
| 21751 | ENSG00000197555 | SIPA1L1 | chr14 | + | 71730054 | 71730301 | 71724669 | 71724835 | 71733665 | 71733812 | 21751 | 154,123,186,76,52,57,89,9 | 5,0,2,0,1,0,5,5,0,5,0,0 | 103,77,115,82,107,76,55,5 | 0,0,0,0,2,1,0,6,0,0,2,0,0,0,3 | 298 | 149 | 0.0340641915358 | | 1 | 0.939,1.0,0.979,1.0 | 1.0,1.0,1.0,1.0,0.9 | -0,012 |

| | | | | | | | | | | | | | | | | | | | | | |
|---|---|---|---|---|---|---|---|---|---|---|---|---|---|---|---|---|---|---|---|---|---|
| 32854 | ENSG00000102445 | RUBCNL | chr13 | - | 46376491 | 46376584 | 46371940 | 46372597 | 46377889 | 46378005 | 32854 | 2,1,21,13,2,1,0,10,0,0,0,1 | 64,29,180,136,85,90,113,1 | 2,9,4,10,11,3,0,4,2,2,6,3,1 | 105,116,58,59,57,55,70,95 | 242 | 149 | 0.0340641992559 | | 1 | 0.019,0.021,0.067, | 0.012,0.046,0.041, | -0,026 |
| 253783 | ENSG00000123329 | ARHGAP9 | chr12 | - | 57474874 | 57475087 | 57474625 | 57474703 | 57475290 | 57475411 | 253783 | 76,39,73,58,46,22,26,22,41 | 2,1,3,3,0,3,0,0,0,0,2,0 | 57,42,46,15,24,36,16,6,28, | 6,3,0,2,4,0,0,4,1,0,0,0,1,1, | 298 | 149 | 0.0341063421461 | | 1 | 0.95,0.951,0.924,0 | 0.826,0.875,1.0,0. | 0,052 |
| 158709 | ENSG00000126777 | KTN1 | chr14 | + | 55663954 | 55664044 | 55661273 | 55661612 | 55667240 | 55667330 | 158709 | 17,13,24,11,7,8,4,13,7,0,2, | 50,36,58,36,25,30,28,47,62 | 15,14,18,5,6,8,6,11,8,10,7, | 36,37,21,12,22,16,24,41,27 | 239 | 149 | 0.0341402683658 | | 1 | 0.175,0.184,0.205, | 0.206,0.191,0.348, | -0,058 |
| 97842 | ENSG00000001631 | KRIT1 | chr7 | - | 92244001 | 92244149 | 92242033 | 92242137 | 92245426 | 92245914 | 97842 | 25,11,59,28,17,24,23,29,8, | 3,4,8,2,2,6,0,4,4,1,2,10 | 25,22,25,27,18,24,17,46,1 | 1,3,0,0,1,3,1,1,2,15,3,4,7,4 | 297 | 149 | 0.0341602960468 | | 1 | 0.807,0.58,0.787,0 | 0.926,0.786,1.0,1. | -0,096 |
| 177090 | ENSG00000179912 | R3HDM2 | chr12 | - | 57269235 | 57269449 | 57268921 | 57269082 | 57269751 | 57269994 | 177090 | 79,45,33,31,39,33,25,27,4 | 10,2,10,0,2,8,2,2,0,4,3,0 | 19,14,23,27,12,12,24,35,9, | 11,5,2,3,6,2,8,6,0,1,5,4,4,3 | 298 | 149 | 0.0341623483434 | | 1 | 0.798,0.918,0.623, | 0.463,0.583,0.852, | 0,09 |
| 213878 | ENSG00000175198 | PCCA | chr13 | + | 100333572 | 100333779 | 100330560 | 100330671 | 100340156 | 100340259 | 213878 | 1,3,0,0,0,2,0,0,0,2,2,1 | 13,20,24,20,13,13,9,12,3,3 | 0,4,1,4,1,4,2,0,4,2,0,1,0,0, | 14,21,16,14,16,7,7,19,10,1 | 298 | 149 | 0.0341936197373 | | 1 | 0.037,0.07,0.0,0.0, | 0.0,0.087,0.03,0.1 | -0,035 |
| 59265 | ENSG00000117616 | RSRP1 | chr1 | - | 25244453 | 25244498 | 25243549 | 25243633 | 25245149 | 25245301 | 59265 | 12,3,45,39,34,14,22,24,22, | 3,3,4,2,0,0,1,3,0,0,3,1 | 57,44,23,6,25,32,3,43,34,5 | 2,4,0,3,0,1,0,3,2,0,0,0,1,0, | 194 | 149 | 0.0342142979124 | | 1 | 0.754,0.434,0.896, | 0.956,0.894,1.0,0. | -0,074 |
| 230679 | ENSG00000225648 | SBDSP1 | chr7 | + | 72833763 | 72833889 | 72831601 | 72831798 | 72836576 | 72836701 | 230679 | 90,28,198,118,40,69,90,95 | 8,0,6,1,0,1,0,9,4,0,2,0 | 116,105,114,119,95,74,29, | 6,0,4,1,3,0,0,1,3,2,4,0,1,0, | 275 | 149 | 0.0342196252255 | | 1 | 0.859,1.0,0.947,0. | 0.913,1.0,0.939,0. | -0,023 |
| 131622 | ENSG00000110851 | PRDM4 | chr12 | - | 107754828 | 107754860 | 107754025 | 107754109 | 107756831 | 107756965 | 131622 | 10,5,10,1,1,3,9,5,1,3,3,5 | 38,42,45,32,24,20,48,32,3 | 4,9,5,1,7,4,6,11,2,6,13,17, | 25,38,24,20,54,21,39,47,2 | 181 | 149 | 0.0342853826535 | | 1 | 0.178,0.089,0.155, | 0.116,0.163,0.146, | -0,032 |
| 112469 | ENSG00000119397 | CNTRL | chr9 | + | 121094887 | 121095018 | 121090274 | 121090405 | 121096421 | 121096563 | 112469 | 77,46,192,85,51,69,67,108 | 1,1,6,7,1,2,1,2,0,6,0,0 | 87,76,71,98,62,30,48,51,4 | 7,0,0,0,0,4,0,5,4,5,0,0,2,4, | 280 | 149 | 0.0342985775517 | | 1 | 0.976,0.961,0.945, | 0.869,1.0,1.0,1.0,1 | -0,007 |
| 2698 | ENSG00000161551 | ZNF577 | chr19 | - | 51886820 | 51886912 | 51880678 | 51880877 | 51890832 | 51890950 | 2698 | 2,0,15,12,16,9,4,19,32,17, | 30,21,32,16,17,16,3,11,4,1 | 8,0,41,39,16,10,0,0,2,13,0, | 16,13,7,8,4,5,11,21,0,11,1 | 241 | 149 | 0.034315059043 | | 1 | 0.04,0.0,0.225,0.3 | 0.236,0.0,0.784,0. | 0,004 |
| 116169 | ENSG00000107077 | KDM4C | chr9 | + | 7049091 | 7049200 | 7046861 | 7046917 | 7103684 | 7103870 | 116169 | 103,46,62,25,40,46,30,55, | 1,0,1,2,1,1,4,0,0,0,0,2 | 57,39,33,37,38,49,27,32,2 | 0,3,3,3,2,2,0,2,0,2,0,2,0,0, | 258 | 149 | 0.0343166585698 | | 1 | 0.983,1.0,0.973,0. | 1.0,0.882,0.864,0. | 0,019 |
| 59695 | ENSG00000163249 | CCNYL1 | chr2 | + | 207733946 | 207734047 | 207726841 | 207726876 | 207737410 | 207737446 | 59695 | 17,18,40,20,2,17,7,9,10,3, | 0,2,1,0,0,0,0,1,0,0,0,1 | 10,22,14,7,7,9,1,4,6,6,3,5, | 0,0,0,0,6,0,1,2,1,3,1,0,1,1, | 250 | 149 | 0.0344040491002 | | 1 | 1.0,0.843,0.96,1.0, | 1.0,1.0,1.0,1.0,0.4 | 0,146 |
| 126140 | ENSG00000213047 | DENND1B | chr1 | - | 197658299 | 197658369 | 197652234 | 197652315 | 197672036 | 197672156 | 126140 | 17,28,34,17,10,13,12,22,3, | 4,1,1,2,3,3,0,3,0,1,0,1 | 14,42,20,42,15,7,6,10,13,2 | 6,2,7,11,0,2,1,3,4,1,1,1,8,1 | 219 | 149 | 0.0344781095947 | | 1 | 0.743,0.95,0.959,0 | 0.614,0.935,0.66,0 | 0,077 |
| 232411 | ENSG00000164754 | RAD21 | chr8 | - | 116859967 | 116860080 | 116858351 | 116858458 | 116861840 | 116861940 | 232411 | 10,2,5,2,3,7,3,8,0,6,0,4 | 150,96,331,232,120,111,1 | 11,7,3,4,2,6,1,10,2,8,14,8, | 173,185,122,92,91,72,89,1 | 262 | 149 | 0.0344873436604 | | 1 | 0.037,0.012,0.009, | 0.035,0.021,0.014, | -0,004 |
| 133440 | ENSG00000155313 | USP25 | chr21 | + | 15808808 | 15809238 | 15805120 | 15805258 | 15811136 | 15811210 | 133440 | 31,25,38,20,15,15,7,9,16,6 | 7,3,8,9,6,4,1,7,3,0,3,2 | 30,20,12,11,8,11,11,11,10, | 12,13,7,7,9,3,2,6,9,5,10,8, | 298 | 149 | 0.0345041857317 | | 1 | 0.689,0.806,0.704, | 0.556,0.435,0.462, | 0,134 |
| 37062 | ENSG00000100650 | SRSF5 | chr14 | + | 69768797 | 69768896 | 69768603 | 69768674 | 69769181 | 69769251 | 37062 | 1346,777,1560,1157,878,7 | 39,11,20,18,3,23,12,2,17,3 | 1117,928,1006,631,656,55 | 4,13,10,37,8,9,1,30,15,12, | 248 | 149 | 0.0345569078044 | | 1 | 0.954,0.977,0.979, | 0.994,0.977,0.984, | 0,011 |
| 250263 | ENSG00000057608 | GDI2 | chr10 | - | 5794884 | 5795014 | 5785851 | 5786050 | 5796762 | 5796862 | 250263 | 203,85,700,640,262,175,1 | 1,0,9,13,3,2,1,2,1,1,9,3 | 445,293,190,88,166,190,8 | 15,11,1,5,3,1,1,3,3,3,8,19, | 279 | 149 | 0.0345609996105 | | 1 | 0.991,1.0,0.976,0. | 0.941,0.934,0.99,0 | 0,006 |
| 239732 | ENSG00000147162 | OGT | chrX | + | 71544733 | 71545037 | 71544566 | 71544635 | 71547906 | 71548023 | 239732 | 5,3,3,2,2,1,6,3,5,0,1,5 | 32,35,69,115,17,15,5,15,5 | 2,15,0,1,3,6,2,2,3,5,2,13,6, | 41,34,28,4,36,25,2,13,8,31 | 298 | 149 | 0.0345647353682 | | 1 | 0.072,0.041,0.021, | 0.024,0.181,0.0,0. | -0,005 |
| 112811 | ENSG00000147457 | CHMP7 | chr8 | + | 23247986 | 23248197 | 23246658 | 23246994 | 23249209 | 23249381 | 112811 | 9,5,13,7,2,11,10,21,3,7,3,2 | 14,3,27,15,13,16,11,19,29, | 8,24,13,17,3,0,4,7,1,5,8,0, | 7,22,14,5,16,8,7,10,10,23,1 | 298 | 149 | 0.0345803445092 | | 1 | 0.243,0.455,0.194, | 0.364,0.353,0.317, | 0,057 |
| 155228 | ENSG00000198718 | TOGARAM1 | chr14 | + | 45023502 | 45023581 | 45011974 | 45012075 | 45025782 | 45025872 | 155228 | 0,0,6,3,1,0,0,3,4,3,0,1 | 6,1,14,10,4,2,2,9,5,0,2,5 | 5,0,0,1,0,0,0,0,1,2,0,0,0,0, | 3,2,10,4,5,3,3,6,0,0,3,7,1,1 | 228 | 149 | 0.034590916789 | | 1 | 0.0,0.0,0.219,0.16 | 0.521,0.0,0.0,0.14, | 0,014 |
| 77850 | ENSG00000171132 | PRKCE | chr2 | + | 45840379 | 45840551 | 45701382 | 45701491 | 45842999 | 45843063 | 77850 | 4,5,15,4,1,5,5,4,9,2,4,4 | 3,2,2,1,0,0,4,0,0,5,0,0 | 11,6,1,14,1,7,2,10,2,0,8,7, | 2,6,5,7,3,2,0,2,1,2,0,0,4,0, | 298 | 149 | 0.0345934714616 | | 1 | 0.4,0.556,0.789,0. | 0.733,0.333,0.091, | 0,183 |
| 171780 | ENSG00000112149 | CD83 | chr6 | + | 14131520 | 14131748 | 14117949 | 14118065 | 14133648 | 14133755 | 171780 | 365,351,185,93,48,74,58,4 | 25,13,19,12,2,4,9,6,1,2,12, | 90,157,85,26,59,15,71,72, | 9,16,16,5,4,0,4,18,8,0,5,72 | 298 | 149 | 0.0345952829508 | | 1 | 0.88,0.931,0.83,0. | 0.833,0.831,0.726, | 0,049 |
| 85502 | ENSG00000116793 | PHTF1 | chr1 | - | 113713278 | 113713438 | 113711939 | 113712113 | 113724758 | 113724893 | 85502 | 44,29,141,107,31,19,39,33 | 3,2,7,4,3,2,0,0,0,1,1,1 | 42,66,31,13,14,23,5,20,14, | 2,1,2,7,3,0,2,0,0,4,0,0,0,3, | 298 | 149 | 0.0346013243919 | | 1 | 0.88,0.879,0.91,0. | 0.913,0.971,0.886, | 0,05 |
| 185342 | ENSG00000188822 | CNR2 | chr1 | - | 23883144 | 23883444 | 23870514 | 23875662 | 23913245 | 23913362 | 185342 | 18,9,33,24,6,8,28,28,3,3,1 | 72,36,191,152,96,74,66,90 | 10,23,12,7,7,9,8,17,15,29, | 135,99,48,64,73,71,22,84, | 298 | 149 | 0.0346031972185 | | 1 | 0.111,0.111,0.08,0 | 0.036,0.104,0.111, | 0,011 |
| 154654 | ENSG00000166887 | VPS39 | chr15 | - | 42180040 | 42180089 | 42178449 | 42178570 | 42184516 | 42184700 | 154654 | 6,1,3,1,0,7,2,2,4,0,2,1 | 67,42,56,38,20,22,37,32,4 | 4,2,0,0,0,1,0,1,0,0,4,1,0,6, | 31,59,19,28,49,14,25,27,2 | 198 | 149 | 0.0346271285357 | | 1 | 0.063,0.018,0.039, | 0.089,0.025,0.0,0. | 0,016 |
| 143219 | ENSG00000108819 | PPP1R9B | chr17 | - | 50140092 | 50140155 | 50139428 | 50139581 | 50141268 | 50141373 | 143219 | 88,60,92,60,39,28,15,14,1 | 3,0,10,4,2,1,0,0,1,2,0,1 | 47,53,29,20,18,22,2,23,13, | 3,6,0,3,0,2,1,0,0,7,0,12,3,2 | 212 | 149 | 0.0346716136182 | | 1 | 0.954,1.0,0.866,0. | 0.917,0.861,1.0,0. | 0,059 |
| 136218 | ENSG00000129351 | ILF3 | chr19 | + | 10687345 | 10687441 | 10683916 | 10683970 | 10689178 | 10692400 | 136218 | 9,1,9,3,9,10,3,3,0,2,0,3 | 9,6,9,8,0,7,7,1,4,4,2,4 | 6,2,1,1,3,3,2,2,1,2,5,5,3,6, | 9,12,17,26,5,6,4,7,6,12,5,9 | 245 | 149 | 0.0346725922704 | | 1 | 0.378,0.092,0.378, | 0.288,0.092,0.035, | 0,126 |
| 167978 | ENSG00000173020 | GRK2 | chr11 | + | 67277795 | 67277951 | 67277271 | 67277348 | 67279199 | 67279273 | 167978 | 5,3,27,11,3,7,0,12,7,3,1,1 | 153,107,203,137,99,93,76, | 3,12,8,1,11,8,2,3,2,4,5,2,4, | 144,144,46,55,134,63,50,9 | 298 | 149 | 0.0347344313381 | | 1 | 0.016,0.014,0.062, | 0.01,0.04,0.08,0.0 | 0,004 |
| 141657 | ENSG00000108312 | UBTF | chr17 | - | 44208618 | 44209004 | 44207863 | 44207911 | 44209351 | 44209541 | 141657 | 6,2,1,3,1,0,42,22,6,2,0,1 | 79,39,99,76,54,53,29,38,6 | 4,2,1,4,6,4,1,5,1,5,3,3,1,5, | 67,67,45,11,55,41,29,49,1 | 298 | 149 | 0.034743226416 | | 1 | 0.037,0.025,0.005, | 0.029,0.015,0.011, | 0,028 |
| 156472 | ENSG00000121741 | ZMYM2 | chr13 | + | 20081988 | 20082130 | 20067238 | 20067390 | 20082780 | 20083032 | 156472 | 66,27,143,61,45,32,33,72, | 0,0,2,2,0,0,2,0,0,0,0,0 | 80,92,48,39,67,26,20,62,5 | 3,5,0,1,2,2,0,2,2,0,1,1,0,2, | 291 | 149 | 0.0348003512024 | | 1 | 1.0,1.0,0.973,0.94, | 0.932,0.904,1.0,0. | 0,026 |
| 218680 | ENSG00000024048 | UBR2 | chr6 | + | 42662509 | 42662578 | 42662183 | 42662277 | 42663257 | 42663419 | 218680 | 2,0,4,1,3,1,3,3,4,0,6,0 | 10,15,26,23,22,9,15,7,4,2, | 5,1,4,0,2,1,0,4,1,1,4,0,3,0, | 11,41,11,8,9,9,6,15,6,13,4, | 218 | 149 | 0.0348167196973 | | 1 | 0.12,0.0,0.095,0.0 | 0.237,0.016,0.199, | 0,025 |
| 39415 | ENSG00000120709 | FAM53C | chr5 | + | 138341808 | 138341866 | 138338014 | 138338307 | 138344824 | 138345215 | 39415 | 11,6,12,8,2,3,19,13,19,6,3, | 5,0,3,0,0,0,0,1,3,2,1,0 | 9,16,18,6,2,1,7,4,1,6,8,14, | 0,2,4,9,0,1,0,0,0,0,1,1,0,0, | 207 | 149 | 0.0348235582334 | | 1 | 0.613,1.0,0.742,1. | 1.0,0.852,0.764,0. | 0,018 |
| 234982 | ENSG00000147905 | ZCCHC7 | chr9 | + | 37188338 | 37188458 | 37126311 | 37126942 | 37302187 | 37302231 | 234982 | 14,0,4,3,0,6,3,3,8,5,0,3 | 84,43,152,67,88,72,38,60, | 2,4,7,2,1,3,1,2,0,5,1,3,6,3, | 151,54,120,29,22,61,33,59 | 269 | 149 | 0.0348270019983 | | 1 | 0.085,0.0,0.014,0. | 0.007,0.039,0.031, | 0,007 |
| 4454 | ENSG00000104852 | SNRNP70 | chr19 | + | 49102113 | 49103587 | 49101389 | 49101471 | 49104633 | 49104735 | 4454 | 259,146,273,180,187,215, | 60,48,65,32,32,23,36,20,6 | 298,241,157,94,141,106,1 | 25,28,37,33,32,15,11,30,1 | 298 | 149 | 0.0348588548507 | | 1 | 0.683,0.603,0.677, | 0.856,0.811,0.68,0 | -0,045 |
| 238753 | ENSG00000171988 | JMJD1C | chr10 | - | 63174437 | 63174477 | 63168434 | 63168566 | 63176296 | 63176555 | 238753 | 3,2,14,6,0,6,2,5,2,0,0,0 | 116,122,290,174,75,78,79, | 11,4,3,1,1,0,2,0,0,1,1,2,3,0 | 190,272,111,120,163,60,7 | 189 | 149 | 0.0349456414438 | | 1 | 0.02,0.013,0.037,0 | 0.044,0.011,0.021, | 0,006 |
| 34701 | ENSG00000125089 | SH3TC1 | chr4 | + | 8232706 | 8233065 | 8231975 | 8232156 | 8233362 | 8233513 | 34701 | 3,1,3,0,7,1,6,3,4,0,3,0 | 32,58,15,11,12,25,18,22,2 | 1,0,2,0,3,0,2,2,8,2,0,4,1,1, | 12,28,14,4,18,13,3,11,27,4 | 298 | 149 | 0.0349575168821 | | 1 | 0.045,0.009,0.091, | 0.04,0.0,0.067,0.0, | 0,039 |
| 217447 | ENSG00000224078 | SNHG14 | chr15 | + | 25106185 | 25106294 | 25103858 | 25103970 | 25107471 | 25107588 | 217447 | 31,7,34,33,7,19,9,23,6,10, | 4,0,4,4,2,6,6,6,3,0,3,0 | 17,24,36,10,13,12,6,10,15, | 3,6,2,0,3,4,1,0,1,1,0,0,2,2, | 258 | 149 | 0.0349590419291 | | 1 | 0.817,1.0,0.831,0. | 0.766,0.698,0.912, | -0,095 |
| 225937 | ENSG00000089289 | IGBP1 | chrX | + | 70162345 | 70162464 | 70150209 | 70150322 | 70165832 | 70166324 | 225937 | 12,5,9,6,5,3,2,2,7,2,1,14 | 106,48,153,133,87,88,91,8 | 2,8,6,12,14,2,4,9,8,9,10,16 | 67,91,105,50,103,73,45,78 | 268 | 149 | 0.0349786270783 | | 1 | 0.059,0.055,0.032, | 0.016,0.047,0.031, | -0,014 |
| 98486 | ENSG00000198198 | SZT2 | chr1 | + | 43432061 | 43432180 | 43431715 | 43431901 | 43432271 | 43432439 | 98486 | 7,1,4,3,0,3,2,3,2,0,5,10 | 20,7,20,12,7,22,11,12,15,7 | 3,2,0,3,3,0,5,2,5,6,2,0,3,1, | 13,17,11,16,16,1,28,11,9,2 | 268 | 149 | 0.0349917910206 | | 1 | 0.163,0.074,0.1,0. | 0.114,0.061,0.0,0. | 0,037 |
| 134009 | ENSG00000100387 | RBX1 | chr22 | + | 40967798 | 40967883 | 40964046 | 40964117 | 40972475 | 40973309 | 134009 | 26,19,94,72,46,29,36,48,3 | 0,0,1,2,0,2,0,2,0,0,0,0 | 49,46,29,15,31,16,15,26,1 | 0,1,2,0,2,1,2,7,0,0,0,3,2,0, | 234 | 149 | 0.0349966100366 | | 1 | 1.0,1.0,0.984,0.95 | 1.0,0.967,0.902,1. | 0,048 |
| 178592 | ENSG00000120008 | WDR11 | chr10 | + | 120880362 | 120880733 | 120878352 | 120878459 | 120880825 | 120880901 | 178592 | 5,1,14,6,1,0,3,1,3,0,1,1 | 154,137,314,171,134,71,8 | 4,8,7,8,5,5,1,5,9,7,5,3,0,1, | 173,168,204,140,128,109, | 298 | 149 | 0.0350005897373 | | 1 | 0.016,0.004,0.022, | 0.011,0.023,0.017, | -0,01 |
| 101339 | ENSG00000173715 | C11orf80 | chr11 | + | 66801030 | 66801102 | 66800640 | 66800696 | 66803994 | 66804168 | 101339 | 81,32,128,85,57,61,65,81, | 1,0,5,0,0,3,2,5,0,0,0,1 | 89,74,47,32,39,38,41,80,2 | 1,6,2,1,0,0,2,2,1,5,3,0,1,0, | 221 | 149 | 0.0350169370959 | | 1 | 0.982,1.0,0.945,1. | 0.984,0.893,0.941, | 0,032 |
| 120590 | ENSG00000107036 | RIC1 | chr9 | + | 5765413 | 5765572 | 5763139 | 5763868 | 5768969 | 5769106 | 120590 | 57,45,73,61,36,47,41,88,7 | 0,0,2,2,0,2,4,4,0,0,0,0 | 59,61,46,56,72,28,16,52,3 | 6,4,2,1,6,0,1,2,1,2,0,1,0,0, | 298 | 149 | 0.0350564339165 | | 1 | 1.0,1.0,0.948,0.93 | 0.831,0.884,0.92,0 | 0,023 |
| 193736 | ENSG00000162384 | CZIB | chr1 | - | 53218413 | 53218490 | 53218171 | 53218203 | 53218866 | 53219920 | 193736 | 22,10,25,12,7,11,6,13,23,8 | 0,0,2,4,3,5,0,1,0,0,0,0 | 12,11,9,6,3,5,8,13,3,17,12, | 0,0,0,0,0,0,0,0,0,0,0,0,0,0, | 226 | 149 | 0.0350692661769 | | 1 | 1.0,1.0,0.892,0.66 | 1.0,1.0,1.0,1.0,1.0, | -0,113 |
| 175313 | ENSG00000186001 | LRCH3 | chr3 | + | 197831959 | 197832062 | 197830769 | 197830863 | 197832196 | 197832317 | 175313 | 0,0,6,4,0,1,2,3,0,2,2,5 | 47,38,77,55,28,33,35,55,2 | 5,1,0,3,2,0,2,0,0,0,2,0,0,0, | 37,25,21,12,18,15,27,40,1 | 252 | 149 | 0.0350901155826 | | 1 | 0.0,0.0,0.044,0.04 | 0.074,0.023,0.0,0. | 0,022 |
| 218739 | ENSG00000196262 | PPIA | chr7 | + | 44799701 | 44799880 | 44799391 | 44799480 | 44801286 | 44803117 | 218739 | 187,117,517,348,216,205, | 2,1,5,4,5,2,3,3,0,0,1,0 | 267,254,166,40,193,102,9 | 6,11,0,4,8,2,0,3,5,0,7,6,0,0 | 298 | 149 | 0.0350980295369 | | 1 | 0.979,0.983,0.981, | 0.957,0.92,1.0,0.8 | 0,02 |
| 112501 | ENSG00000119397 | CNTRL | chr9 | + | 121141429 | 121141588 | 121140640 | 121140786 | 121142090 | 121142172 | 112501 | 77,43,83,52,43,33,31,30,6 | 13,9,24,6,11,16,20,9,4,4,7, | 68,51,35,55,24,30,25,40,1 | 10,13,24,18,5,8,7,21,12,23 | 298 | 149 | 0.0351231910387 | | 1 | 0.748,0.705,0.634, | 0.773,0.662,0.422, | 0,091 |
| 88488 | ENSG00000215717 | TMEM167B | chr1 | + | 109092889 | 109093021 | 109090763 | 109090882 | 109094416 | 109094989 | 88488 | 38,40,99,74,37,38,31,22,2 | 1,4,4,1,3,0,4,4,0,0,1,1 | 93,101,54,44,59,45,51,67, | 0,2,0,0,3,2,1,2,0,0,6,4,1,0, | 281 | 149 | 0.0351580965876 | | 1 | 0.953,0.841,0.929, | 1.0,0.964,1.0,1.0,0 | -0,046 |
| 218735 | ENSG00000196262 | PPIA | chr7 | + | 44799701 | 44799871 | 44799391 | 44799480 | 44801286 | 44803117 | 218735 | 187,117,516,349,214,205, | 2,1,5,4,5,2,3,3,0,0,1,0 | 265,254,166,40,194,102,9 | 6,11,0,4,8,2,0,3,5,0,7,6,0,0 | 298 | 149 | 0.0351713511655 | | 1 | 0.979,0.983,0.981, | 0.957,0.92,1.0,0.8 | 0,02 |
| 32048 | ENSG00000140350 | ANP32A | chr15 | - | 68787412 | 68787535 | 68784270 | 68784595 | 68787769 | 68787829 | 32048 | 354,265,506,412,214,197, | 34,41,48,34,15,20,11,52,1 | 275,326,299,105,148,185, | 44,89,31,19,17,14,22,32,7, | 272 | 149 | 0.0351784137994 | | 1 | 0.851,0.78,0.852,0 | 0.774,0.667,0.841, | 0,04 |
| 1400 | ENSG00000099991 | CABIN1 | chr22 | + | 24054872 | 24055159 | 24050824 | 24050974 | 24056191 | 24056304 | 1400 | 57,42,84,49,44,65,42,57,4 | 3,9,1,3,0,3,0,6,0,1,0,0 | 32,58,28,69,34,11,53,36,2 | 2,2,0,3,4,0,2,0,2,0,0,0,0,4, | 298 | 149 | 0.0352184866644 | | 1 | 0.905,0.7,0.977,0. | 0.889,0.935,1.0,0. | -0,012 |
| 152158 | ENSG00000196405 | EVL | chr14 | + | 100138847 | 100138897 | 100137739 | 100137802 | 100141179 | 100141229 | 152158 | 29,6,20,18,3,6,9,13,8,3,2,5 | 90,41,211,149,113,85,82,1 | 7,7,5,9,6,3,2,1,3,7,9,21,8,5 | 123,87,62,32,77,57,71,85, | 199 | 149 | 0.0352540883363 | | 1 | 0.194,0.099,0.066, | 0.041,0.057,0.057, | 0,013 |
| 241111 | ENSG00000165209 | STRBP | chr9 | - | 123173676 | 123173856 | 123169901 | 123170046 | 123179006 | 123179227 | 241111 | 355,152,482,378,204,165, | 11,10,19,9,10,12,10,9,3,8, | 343,292,293,151,142,114, | 16,16,10,26,13,3,2,3,7,1,8, | 298 | 149 | 0.0352616343149 | | 1 | 0.942,0.884,0.927, | 0.915,0.901,0.936, | 0,019 |
| 253784 | ENSG00000123329 | ARHGAP9 | chr12 | - | 57474874 | 57475178 | 57474625 | 57474703 | 57475290 | 57475411 | 253784 | 77,39,73,58,44,22,26,22,41 | 2,1,3,3,0,3,0,0,0,0,2,0 | 57,42,45,15,24,36,16,6,28, | 6,3,0,2,4,0,0,4,1,0,0,0,1,1, | 298 | 149 | 0.0352620549048 | | 1 | 0.951,0.951,0.924, | 0.826,0.875,1.0,0. | 0,054 |
| 126792 | ENSG00000066084 | DIP2B | chr12 | + | 50692948 | 50693013 | 50686572 | 50686682 | 50695266 | 50695360 | 126792 | 49,39,27,18,10,2,11,13,35, | 10,5,6,2,1,2,3,10,2,0,0,1 | 26,21,16,3,13,7,5,8,7,22,2 | 1,8,5,7,7,3,2,12,7,5,4,0,0,0 | 214 | 149 | 0.0352761259826 | | 1 | 0.773,0.844,0.758, | 0.948,0.646,0.69,0 | 0,122 |
| 247778 | ENSG00000110324 | IL10RA | chr11 | + | 117986683 | 117986751 | 117986390 | 117986534 | 117988381 | 117988502 | 247778 | 11,18,0,0,2,2,0,1,4,0,1,7 | 42,117,47,26,30,18,38,19, | 0,4,3,4,5,0,1,2,2,3,0,10,3,4 | 53,75,55,50,73,12,28,27,2 | 217 | 149 | 0.0352862521366 | | 1 | 0.152,0.096,0.0,0. | 0.0,0.035,0.036,0. | 0,01 |
| 33976 | ENSG00000248672 | LY75-CD302 | chr2 | - | 159840728 | 159840955 | 159835479 | 159835645 | 159842244 | 159842374 | 33976 | 67,67,191,128,51,46,61,49 | 1,2,9,6,5,3,2,1,0,2,0,0 | 92,53,40,29,32,46,35,58,2 | 4,0,0,1,3,0,0,0,6,4,5,7,0,0, | 298 | 149 | 0.0352904606875 | | 1 | 0.971,0.944,0.914, | 0.92,1.0,1.0,0.935, | 0,014 |
| 126097 | ENSG00000028116 | VRK2 | chr2 | + | 58154753 | 58154802 | 58146315 | 58146474 | 58159348 | 58159871 | 126097 | 26,18,29,32,18,12,15,23,1 | 25,19,37,26,6,18,16,21,7,5 | 43,26,10,22,16,16,6,21,10, | 31,46,26,13,36,20,4,25,18, | 198 | 149 | 0.0353411480227 | | 1 | 0.439,0.416,0.371, | 0.511,0.298,0.224, | 0,091 |
| 216219 | ENSG00000215717 | TMEM167B | chr1 | + | 109092889 | 109093194 | 109090763 | 109090882 | 109094416 | 109096934 | 216219 | 22,25,57,52,6,12,15,8,8,5, | 1,4,4,1,3,0,4,4,0,0,1,1 | 48,54,28,22,40,25,32,20,1 | 0,2,0,0,3,2,1,2,0,0,6,4,1,0, | 298 | 149 | 0.0353484749375 | | 1 | 0.917,0.758,0.877, | 1.0,0.931,1.0,1.0,0 | -0,124 |
| 228442 | ENSG00000198874 | TYW1 | chr7 | + | 67014431 | 67014561 | 67009582 | 67009684 | 67017852 | 67018143 | 228442 | 50,33,46,20,13,17,15,22,3 | 0,1,1,3,3,0,0,0,0,0,0,1 | 13,14,13,42,15,21,17,13,1 | 3,1,1,7,0,0,1,0,2,1,2,0,0,0, | 279 | 149 | 0.035381934791 | | 1 | 1.0,0.946,0.961,0. | 0.698,0.882,0.874, | 0,05 |
| 21030 | ENSG00000100697 | DICER1 | chr14 | - | 95129471 | 95129632 | 95126579 | 95126748 | 95130057 | 95130192 | 21030 | 91,57,207,100,76,56,67,10 | 3,3,10,9,5,4,1,5,1,0,4,3 | 169,116,62,70,101,34,55,8 | 7,28,5,2,4,2,5,5,3,0,6,7,2,4 | 298 | 149 | 0.0354016612728 | | 1 | 0.938,0.905,0.912, | 0.923,0.674,0.861, | 0,036 |
| 163309 | ENSG00000118965 | WDR35 | chr2 | - | 19977924 | 19978167 | 19975529 | 19975663 | 19978750 | 19978879 | 163309 | 7,2,9,3,3,2,0,0,5,0,0,2 | 32,20,43,18,7,7,10,21,15,1 | 1,3,2,4,3,1,2,0,2,2,2,0,3,2, | 34,24,26,14,12,13,5,20,10, | 298 | 149 | 0.0354276921526 | | 1 | 0.099,0.048,0.095, | 0.014,0.059,0.037, | 0,012 |
| 184888 | ENSG00000144554 | FANCD2 | chr3 | + | 10034694 | 10034811 | 10034468 | 10034536 | 10035172 | 10035233 | 184888 | 30,8,30,16,15,15,16,14,5,0 | 1,0,6,1,0,0,1,1,2,2,0,0 | 28,28,21,18,14,8,10,10,13, | 0,0,0,1,2,3,2,0,1,6,5,2,0,0, | 266 | 149 | 0.0354666168364 | | 1 | 0.944,1.0,0.737,0. | 1.0,1.0,1.0,0.91,0. | -0,035 |
| 23835 | ENSG00000173575 | CHD2 | chr15 | + | 92901166 | 92901299 | 92892003 | 92892072 | 92924320 | 92924552 | 23835 | 147,62,186,88,72,108,66,1 | 5,7,9,1,0,5,6,8,2,0,2,5 | 110,126,75,99,117,59,46,6 | 4,7,2,3,4,1,4,2,0,1,3,0,1,5, | 282 | 149 | 0.0354739802457 | | 1 | 0.94,0.824,0.916,0 | 0.936,0.905,0.952, | -0,024 |
| 222603 | ENSG00000167483 | NIBAN3 | chr19 | + | 17530754 | 17530883 | 17527237 | 17527395 | 17532262 | 17532388 | 222603 | 65,74,167,70,103,63,76,72 | 45,36,41,14,40,27,53,39,5 | 77,123,83,14,69,47,59,64, | 26,38,25,17,12,6,18,37,12, | 278 | 149 | 0.035478837477 | | 1 | 0.436,0.524,0.686, | 0.613,0.634,0.64,0 | -0,081 |
| 215422 | ENSG00000010244 | ZNF207 | chr17 | + | 32366331 | 32366757 | 32365329 | 32365487 | 32367771 | 32368014 | 215422 | 282,161,160,168,101,100, | 89,50,53,64,54,64,59,99,1 | 155,209,182,127,102,76,7 | 78,59,40,69,53,28,24,58,2 | 298 | 149 | 0.0354938476331 | | 1 | 0.613,0.617,0.602, | 0.498,0.639,0.695, | -0,059 |
| 136480 | ENSG00000213793 | ZNF888 | chr19 | - | 52917607 | 52917723 | 52915195 | 52915322 | 52917858 | 52917997 | 136480 | 6,0,2,0,0,1,3,0,0,0,0,0 | 7,6,13,10,5,2,4,4,2,3,1,5 | 2,7,4,0,0,0,0,4,3,0,2,0,0,0, | 5,2,2,5,4,1,4,8,3,1,7,12,2,4 | 265 | 149 | 0.0355047899622 | | 1 | 0.325,0.0,0.08,0.0, | 0.184,0.663,0.529, | -0,077 |
| 230169 | ENSG00000012124 | CD22 | chr19 | + | 35338167 | 35338429 | 35337754 | 35338021 | 35340880 | 35341843 | 230169 | 253,155,132,105,142,91,5 | 3,4,5,3,11,3,2,2,0,2,1,1 | 137,70,54,68,33,49,23,26, | 9,6,1,4,2,0,0,2,2,2,2,28,9,2 | 298 | 149 | 0.0355242605377 | | 1 | 0.977,0.951,0.93,0 | 0.884,0.854,0.964, | 0,023 |
| 251701 | ENSG00000095585 | BLNK | chr10 | - | 96215313 | 96215389 | 96209837 | 96209907 | 96216652 | 96216734 | 251701 | 60,36,82,66,45,39,39,28,7 | 12,3,12,8,7,1,5,1,25,16,5,5 | 61,37,48,22,23,32,5,44,15, | 5,4,6,5,4,1,3,4,2,4,0,12,11, | 225 | 149 | 0.035545113743 | | 1 | 0.768,0.888,0.819, | 0.89,0.86,0.841,0. | -0,039 |
| 55364 | ENSG00000078304 | PPP2R5C | chr14 | + | 101831700 | 101831804 | 101809868 | 101810036 | 101836168 | 101836432 | 55364 | 8,3,2,5,0,3,1,5,5,4,2,4 | 17,16,35,14,10,13,14,10,6, | 7,14,6,9,5,2,3,5,2,1,9,7,3,7 | 12,17,7,6,7,8,8,5,7,18,8,20 | 253 | 149 | 0.0355622129601 | | 1 | 0.217,0.099,0.033, | 0.256,0.327,0.335, | -0,067 |
| 168072 | ENSG00000146350 | TBC1D32 | chr6 | - | 121256083 | 121256277 | 121255327 | 121255410 | 121279120 | 121279245 | 168072 | 3,4,10,2,8,2,6,9,5,0,3,3 | 11,4,5,2,2,4,2,1,2,4,2,0 | 9,3,7,13,3,1,6,10,3,3,5,13, | 4,5,4,3,2,1,0,0,3,0,6,5,0,0, | 298 | 149 | 0.0355718919326 | | 1 | 0.12,0.333,0.5,0.3 | 0.529,0.231,0.467, | -0,154 |
| 151849 | ENSG00000125107 | CNOT1 | chr16 | - | 58559803 | 58559894 | 58558472 | 58558674 | 58560211 | 58560362 | 151849 | 11,5,18,2,5,5,2,4,6,8,4,3 | 99,93,86,59,86,61,68,101, | 9,11,5,3,8,1,1,4,5,4,5,10,7, | 62,59,64,62,21,33,47,77,3 | 240 | 149 | 0.0355789852624 | | 1 | 0.065,0.032,0.115, | 0.083,0.104,0.046, | -0,02 |
| 214951 | ENSG00000135976 | ANKRD36 | chr2 | + | 97124459 | 97124676 | 97122886 | 97122993 | 97127066 | 97127134 | 214951 | 2,0,16,12,3,16,3,1,2,2,3,5 | 3,2,6,12,4,8,3,4,1,0,10,11 | 12,14,5,11,9,3,2,3,7,3,6,8, | 5,7,0,6,5,3,4,4,9,1,2,4,3,2, | 298 | 149 | 0.0355797693341 | | 1 | 0.25,0.0,0.571,0.3 | 0.545,0.5,1.0,0.47 | -0,141 |
| 227323 | ENSG00000145375 | SPATA5 | chr4 | + | 123019161 | 123019249 | 122979231 | 122979386 | 123028185 | 123028395 | 227323 | 11,7,5,1,3,3,2,3,2,0,0,0 | 40,16,30,22,13,30,11,29,1 | 11,2,1,0,4,1,1,2,0,3,3,0,0,4 | 26,18,7,18,15,15,10,26,8,1 | 237 | 149 | 0.0356458408296 | | 1 | 0.147,0.216,0.095, | 0.21,0.065,0.082,0 | 0,002 |
| 161665 | ENSG00000254996 | ANKHD1-EIF4E | chr5 | + | 140512808 | 140512923 | 140510018 | 140510181 | 140513362 | 140513479 | 161665 | 33,21,42,24,7,24,17,10,9,1 | 0,0,4,0,2,0,0,2,1,2,0,5 | 25,24,34,17,11,12,21,7,23, | 0,10,0,6,2,2,2,3,2,0,1,0,0,0 | 264 | 149 | 0.0356752389287 | | 1 | 1.0,1.0,0.856,1.0,0 | 1.0,0.575,1.0,0.61 | 0,01 |
| 129068 | ENSG00000174718 | RESF1 | chr12 | + | 31960776 | 31960871 | 31959369 | 31959491 | 31980877 | 31981060 | 129068 | 79,44,201,114,48,48,62,11 | 54,30,17,33,7,8,15,15,19,8 | 88,107,96,51,57,23,21,51, | 11,20,22,44,20,5,1,7,3,5,1 | 244 | 149 | 0.035677025349 | | 1 | 0.472,0.472,0.878, | 0.83,0.766,0.727,0 | -0,083 |
| 200114 | ENSG00000131051 | RBM39 | chr20 | - | 35729461 | 35729506 | 35729311 | 35729365 | 35731940 | 35732806 | 200114 | 101,51,172,89,59,91,54,88 | 4,0,8,4,1,5,0,1,4,5,0,0 | 96,110,80,64,70,44,47,87, | 6,0,0,0,0,2,2,1,1,12,13,5,7, | 194 | 149 | 0.0356885242941 | | 1 | 0.951,1.0,0.943,0. | 0.925,1.0,1.0,1.0,1 | 0,017 |
| 57164 | ENSG00000071564 | TCF3 | chr19 | - | 1615284 | 1615520 | 1612206 | 1612433 | 1615685 | 1615804 | 57164 | 70,32,104,66,76,57,63,86, | 45,34,66,71,24,53,32,30,6 | 49,54,20,35,27,17,75,54,1 | 35,38,20,21,10,9,25,85,11, | 298 | 149 | 0.0356993348983 | | 1 | 0.438,0.32,0.441,0 | 0.412,0.415,0.333, | 0,071 |
| 224690 | ENSG00000122482 | ZNF644 | chr1 | - | 90920236 | 90920306 | 90918051 | 90918154 | 90937484 | 90938090 | 224690 | 4,0,4,1,1,1,0,2,0,0,0,0 | 21,24,121,47,42,39,25,65, | 9,4,1,0,0,0,0,2,1,1,4,7,1,4, | 90,97,23,44,47,31,22,35,2 | 219 | 149 | 0.0357385914416 | | 1 | 0.115,0.0,0.022,0. | 0.064,0.027,0.029, | -0,008 |

| | | | | | | | | | | | | | | | | | | | | | | |
|---|---|---|---|---|---|---|---|---|---|---|---|---|---|---|---|---|---|---|---|---|---|---|
| 40436 | ENSG00000131374 | TBC1D5 | chr3 | - | 17622647 | 17622765 | 17586351 | 17586466 | 17623848 | 17623913 | 40436 | 20,10,3,10,8,19,3,6,22,9,3, | 0,0,4,7,0,0,4,0,2,0,2,0 | 10,5,9,6,4,8,3,6,7,8,18,22,1 | 0,0,1,0,2,1,1,1,0,0,2,4,1,0,2 | 267 | 149 | 0.0357465177718 | | 1 | 1.0,1.0,0.295,0.44 | 1.0,1.0,0.834,1.0,0 | -0,084 |
| 152980 | ENSG00000165355 | FBXO33 | chr14 | - | 39401605 | 39401861 | 39399565 | 39399787 | 39402400 | 39402511 | 152980 | 80,46,76,55,28,20,23,35,5 | 1,0,3,2,3,0,3,3,1,0,0,0 | 49,81,31,21,35,23,35,48,2 | 2,2,0,2,0,0,0,0,3,0,4,12,1,1 | 298 | 149 | 0.0357472690363 | | 1 | 0.976,1.0,0.927,0. | 0.925,0.953,1.0,0. | 0,029 |
| 46708 | ENSG00000181450 | ZNF678 | chr1 | + | 227582513 | 227582606 | 227563542 | 227563724 | 227654335 | 227654353 | 46708 | 20,10,21,28,18,11,13,22,1 | 0,4,9,3,7,0,2,4,2,0,4,6 | 20,10,8,7,9,15,16,11,3,13,2 | 1,4,2,0,4,0,0,0,1,1,0,5,1,3,0 | 242 | 149 | 0.0357506180523 | | 1 | 1.0,0.606,0.59,0.8 | 0.925,0.606,0.711, | -0,084 |
| 252260 | ENSG00000019995 | ZRANB1 | chr10 | + | 124981224 | 124981328 | 124974199 | 124974398 | 124981708 | 124981829 | 252260 | 5,6,3,0,0,0,0,2,6,2,0,0 | 62,25,69,31,17,20,23,27,4 | 0,1,0,1,3,0,2,0,1,0,2,1,1,0,0 | 43,55,29,51,28,13,27,38,13 | 253 | 149 | 0.035753607453 | | 1 | 0.045,0.124,0.025, | 0.0,0.011,0.0,0.01 | 0,015 |
| 17176 | ENSG00000140521 | POLG | chr15 | - | 89317375 | 89317536 | 89316304 | 89316827 | 89318540 | 89318749 | 17176 | 62,23,124,57,48,46,44,59,3 | 2,0,1,3,0,1,0,2,2,0,0,3 | 57,69,29,19,51,30,23,50,2 | 1,5,2,6,0,0,2,4,1,2,1,0,2,2,0 | 298 | 149 | 0.0357567940908 | | 1 | 0.939,1.0,0.984,0. | 0.966,0.873,0.879, | 0,042 |
| 155827 | ENSG00000134900 | TPP2 | chr13 | + | 102634660 | 102634830 | 102633949 | 102634098 | 102635586 | 102635702 | 155827 | 3,0,5,0,0,3,2,0,0,0,0,0 | 39,34,86,33,27,39,36,47,1 | 0,3,1,15,5,0,2,0,0,0,1,2,4,0 | 56,68,29,36,55,36,20,58,40 | 298 | 149 | 0.0357613777529 | | 1 | 0.037,0.0,0.028,0.0 | 0.0,0.022,0.017,0. | -0,02 |
| 169798 | ENSG00000242086 | MUC20-OT1 | chr3 | + | 195680794 | 195680961 | 195677783 | 195677848 | 195681189 | 195681301 | 169798 | 0,0,0,3,2,2,2,0,0,0,0,0 | 19,14,19,3,6,20,15,15,11,4 | 0,3,0,2,2,0,4,2,3,1,0,0,3,0,3 | 2,18,6,16,12,6,0,12,4,11,1, | 298 | 149 | 0.0357797710097 | | 1 | 0.0,0.0,0.0,0.333,0 | 0.0,0.077,0.0,0.05 | -0,064 |
| 251248 | ENSG00000130803 | ZNF317 | chr19 | + | 9157886 | 9158075 | 9157267 | 9157394 | 9158825 | 9158908 | 251248 | 11,10,12,12,10,12,4,16,8,3 | 19,16,28,12,13,7,9,9,4,9,4, | 15,20,5,3,5,2,3,5,1,5,13,7,5 | 11,25,8,12,9,8,8,16,7,8,21, | 298 | 149 | 0.0358065474854 | | 1 | 0.224,0.238,0.176, | 0.405,0.286,0.238, | 0,096 |
| 117753 | ENSG00000156990 | RPUSD3 | chr3 | - | 9843464 | 9843601 | 9842198 | 9842243 | 9843889 | 9843986 | 117753 | 8,1,12,7,4,5,3,3,8,7,0,7 | 0,0,4,5,0,0,1,0,1,1,1,1 | 10,2,11,4,6,2,1,8,4,9,1,15,9 | 4,1,1,0,0,0,2,2,1,1,2,5,1,0,0 | 286 | 149 | 0.0358242792752 | | 1 | 1.0,1.0,0.61,0.422, | 0.566,0.51,0.851,1 | 0,046 |
| 91738 | ENSG00000169826 | CSGALNACT2 | chr10 | + | 43163865 | 43164044 | 43160493 | 43160595 | 43167003 | 43167098 | 91738 | 10,22,31,16,18,12,12,7,2,2 | 0,1,0,0,7,1,0,2,2,2,0,0 | 15,35,11,10,21,7,29,33,7,1 | 1,2,1,0,2,0,0,0,0,0,0,0,0,0 | 298 | 149 | 0.0358243121877 | | 1 | 1.0,0.917,1.0,1.0,0 | 0.882,0.897,0.846, | -0,163 |
| 37571 | ENSG00000154144 | TBRG1 | chr11 | + | 124626472 | 124626609 | 124625670 | 124625903 | 124626812 | 124627050 | 37571 | 13,4,8,4,1,3,3,4,11,1,2,5 | 4,1,8,2,3,4,2,2,0,3,3,2 | 4,4,2,1,1,3,3,1,1,5,2,2,7,7,0 | 4,6,5,12,7,1,1,3,2,2,2,7,1,1 | 286 | 149 | 0.035846150974 | | 1 | 0.629,0.676,0.343, | 0.343,0.258,0.172, | 0,08 |
| 242364 | ENSG00000137078 | SIT1 | chr9 | - | 35650501 | 35650592 | 35650356 | 35650415 | 35650753 | 35650931 | 242364 | 15,15,48,44,23,13,17,19,2 | 2,0,3,4,2,1,1,3,4,1,2,0 | 16,21,11,6,10,8,4,6,14,15,1 | 0,1,5,1,0,0,0,0,3,2,2,1,3,5,3 | 240 | 149 | 0.0358674259423 | | 1 | 0.823,1.0,0.909,0.8 | 1.0,0.929,0.577,0. | 0,017 |
| 115283 | ENSG00000143409 | MINDY1 | chr1 | - | 151000456 | 151000615 | 150999861 | 150999964 | 151001249 | 151001314 | 115283 | 6,5,15,22,6,1,6,16,2,8,8,5 | 1,0,5,2,0,2,0,1,1,0,0,0 | 10,7,9,3,6,6,5,6,3,16,12,4,1 | 5,2,2,0,0,0,3,2,0,1,0,0,3,1,0 | 298 | 149 | 0.0358741606723 | | 1 | 0.75,1.0,0.6,0.846, | 0.5,0.636,0.692,1.0 | -0,001 |
| 148851 | ENSG00000059145 | UNKL | chr16 | - | 1397949 | 1398139 | 1397177 | 1397295 | 1399373 | 1399509 | 148851 | 0,0,8,0,2,4,0,4,0,1,0,2 | 5,5,12,2,5,4,15,15,11,11,2, | 5,2,0,0,0,0,0,2,2,1,3,0,1,0,1 | 3,1,2,2,9,2,8,15,1,6,9,5,2,6 | 298 | 149 | 0.0358749032371 | | 1 | 0.0,0.0,0.25,0.0,0.1 | 0.455,0.5,0.0,0.0,0.0 | -0,042 |
| 251372 | ENSG00000012779 | ALOX5 | chr10 | + | 45445256 | 45445386 | 45444115 | 45444286 | 45445507 | 45446119 | 251372 | 7,3,8,10,5,6,10,5,0,1,3,3 | 159,87,178,160,174,105,1 | 7,10,4,4,2,1,1,2,8,2,5,7,1,2 | 160,151,119,127,152,65,50 | 279 | 149 | 0.0359273015065 | | 1 | 0.023,0.018,0.023, | 0.023,0.034,0.018, | 0,006 |
| 206620 | ENSG00000152061 | RABGAP1L | chr1 | + | 174220351 | 174220703 | 174219124 | 174219295 | 174220971 | 174221164 | 206620 | 7,6,28,18,3,5,7,3,10,7,0,1 | 120,83,280,158,70,126,10 | 7,19,4,9,10,5,5,3,11,10,0,0 | 120,165,237,196,147,55,5 | 298 | 149 | 0.03593241061 | | 1 | 0.028,0.035,0.048, | 0.028,0.054,0.008, | 0,005 |
| 158941 | ENSG00000178385 | PLEKHM3 | chr2 | - | 208020468 | 208020688 | 208001029 | 208001957 | 208025388 | 208025527 | 158941 | 1,1,1,2,0,3,3,0,5,0,0,2 | 11,9,13,6,15,19,8,19,10,4,3 | 0,0,1,1,0,0,1,0,0,0,0,2,1,0,0 | 7,23,7,9,3,8,9,11,8,13,9,16 | 298 | 149 | 0.0359387260777 | | 1 | 0.043,0.053,0.037, | 0.0,0.0,0.067,0.05 | 0,051 |
| 64619 | ENSG00000077254 | USP33 | chr1 | - | 77714613 | 77714783 | 77713199 | 77713281 | 77717866 | 77718047 | 64619 | 28,18,75,30,28,23,12,34,9, | 1,1,8,6,2,1,0,4,0,0,0,0 | 46,28,14,8,8,18,10,23,19,3 | 3,3,2,3,9,3,0,2,6,0,3,4,4,0,1 | 298 | 149 | 0.0359401510095 | | 1 | 0.933,0.9,0.824,0.7 | 0.885,0.824,0.778, | 0,128 |
| 238033 | ENSG00000119397 | CNTRL | chr9 | + | 121141389 | 121141588 | 121140640 | 121140786 | 121142090 | 121142172 | 238033 | 77,43,83,52,43,33,31,30,6 | 13,9,24,6,11,16,20,9,4,4,7, | 68,51,35,55,24,30,25,37,1 | 10,13,24,18,5,8,7,21,12,23 | 298 | 149 | 0.0359404331109 | | 1 | 0.748,0.705,0.634, | 0.773,0.662,0.422, | 0,09 |
| 185744 | ENSG00000198160 | MIER1 | chr1 | + | 66972898 | 66972991 | 66971654 | 66971736 | 66976594 | 66976722 | 185744 | 15,12,77,47,14,24,14,26,5, | 3,0,0,2,0,2,3,2,0,0,0,1 | 31,58,18,15,27,12,11,25,1 | 2,1,0,1,0,4,3,6,0,0,5,0,1,3,0 | 242 | 149 | 0.035951024481 | | 1 | 0.755,1.0,1.0,0.93 | 0.905,0.973,1.0,0. | 0,026 |
| 72228 | ENSG00000181722 | ZBTB20 | chr3 | - | 114710267 | 114710313 | 114693527 | 114693575 | 114713975 | 114713997 | 72228 | 20,13,21,4,16,21,13,16,12, | 2,3,0,2,2,4,2,2,2,0,0,0 | 11,15,8,15,14,4,7,13,25,10 | 0,0,1,3,1,0,2,0,1,2,0,0,0,2,2 | 195 | 149 | 0.035965178359 | | 1 | 0.884,0.768,1.0,0.6 | 1.0,1.0,0.859,0.79 | -0,04 |
| 120609 | ENSG00000140299 | BNIP2 | chr15 | - | 59668102 | 59668138 | 59662855 | 59664120 | 59668891 | 59668949 | 120609 | 7,4,21,8,8,8,5,4,16,5,0,2 | 141,129,240,198,103,94,1 | 11,22,10,3,5,0,14,14,2,14,6 | 186,282,125,106,155,92,65 | 185 | 149 | 0.0359712069872 | | 1 | 0.038,0.024,0.066, | 0.045,0.059,0.061, | -0,015 |
| 21729 | ENSG00000197555 | SIPA1L1 | chr14 | + | 71576534 | 71576587 | 71529311 | 71529370 | 71587570 | 71587694 | 21729 | 22,31,49,13,17,23,9,38,7,1 | 28,14,138,61,25,16,43,58,2 | 18,35,20,39,28,25,2,4,7,10 | 58,34,36,40,61,54,23,25,26 | 202 | 149 | 0.0359875834561 | | 1 | 0.367,0.62,0.208,0 | 0.186,0.432,0.291, | 0,093 |
| 146240 | ENSG00000140968 | IRF8 | chr16 | + | 85909483 | 85909514 | 85908989 | 85909173 | 85911569 | 85911658 | 146240 | 183,44,88,67,16,27,25,44,8 | 373,252,464,442,261,184,2 | 37,34,72,79,48,9,18,14,30, | 261,254,341,168,145,148,1 | 180 | 149 | 0.0360019033131 | | 1 | 0.289,0.126,0.136, | 0.105,0.1,0.149,0.2 | 0,014 |
| 31600 | ENSG00000112992 | NNT | chr5 | + | 43619031 | 43619119 | 43615847 | 43616065 | 43624031 | 43624120 | 31600 | 47,21,78,72,33,25,17,30,4 | 10,2,11,5,5,0,2,6,1,2,2,6 | 35,29,21,9,20,28,7,18,10,4 | 3,3,1,1,3,1,1,4,0,1,3,6,4,3,0 | 237 | 149 | 0.0360306125689 | | 1 | 0.747,0.868,0.817, | 0.88,0.859,0.93,0.8 | -0,029 |
| 156311 | ENSG00000124193 | SRSF6 | chr20 | + | 43459152 | 43459317 | 43458360 | 43458509 | 43459770 | 43459895 | 156311 | 14,0,17,10,5,6,12,16,6,1,5, | 19,19,33,10,18,20,20,14,6 | 11,6,15,28,16,13,6,14,12,1 | 18,22,24,9,8,11,19,13,6,16 | 298 | 149 | 0.0360387052735 | | 1 | 0.269,0.0,0.205,0.3 | 0.234,0.12,0.238,0 | -0,076 |
| 210701 | ENSG00000244754 | N4BP2L2 | chr13 | - | 32535768 | 32536233 | 32526818 | 32527532 | 32538682 | 32538825 | 210701 | 50,36,157,88,81,58,54,75,6 | 2,0,4,1,2,2,0,2,2,1,0,0 | 99,55,35,23,30,31,36,75,3 | 0,0,0,0,0,1,0,4,0,0,0,2,0,0,2 | 298 | 149 | 0.0360786574008 | | 1 | 0.926,1.0,0.952,0.9 | 1.0,1.0,1.0,1.0,1.0, | -0,022 |
| 32050 | ENSG00000140350 | ANP32A | chr15 | - | 68787769 | 68787919 | 68784384 | 68784595 | 68820697 | 68820868 | 32050 | 86,73,214,133,92,88,68,12 | 8,3,1,1,2,0,4,8,2,2,4,3 | 182,196,74,41,53,56,56,11 | 3,1,1,2,3,2,1,6,0,3,1,3,4,1,2 | 298 | 149 | 0.036089271202 | | 1 | 0.843,0.924,0.991, | 0.968,0.99,0.974,0 | -0,03 |
| 255990 | ENSG00000145901 | TNIP1 | chr5 | - | 151033607 | 151033811 | 151032286 | 151032383 | 151035001 | 151035067 | 255990 | 16,9,27,13,21,12,17,20,12, | 21,7,11,14,12,12,6,11,18,8 | 11,10,5,3,5,2,2,22,2,10,12, | 6,17,14,6,4,0,2,16,4,10,6,1 | 298 | 149 | 0.0361179167591 | | 1 | 0.276,0.391,0.551, | 0.478,0.227,0.152, | 0,065 |
| 60416 | ENSG00000213397 | HAUS7 | chrX | - | 153464987 | 153465055 | 153462609 | 153462671 | 153469145 | 153469261 | 60416 | 11,8,19,16,20,5,8,10,12,7,4 | 4,2,5,0,0,1,1,0,1,1,0,1 | 8,7,4,2,2,6,5,10,4,15,10,29 | 4,5,1,3,6,0,0,4,0,0,1,1,0,0,2 | 217 | 149 | 0.0361422929611 | | 1 | 0.654,0.733,0.723, | 0.579,0.49,0.733,0 | 0,117 |
| 207652 | ENSG00000130348 | QRSL1 | chr6 | + | 106654696 | 106654922 | 106652466 | 106652582 | 106655614 | 106655732 | 207652 | 36,24,125,144,49,39,25,43 | 4,8,10,11,8,5,2,2,1,2,2,0 | 61,48,28,16,24,60,14,39,1 | 5,2,5,2,0,0,0,2,4,11,3,0,0,0 | 298 | 149 | 0.0361607245847 | | 1 | 0.818,0.6,0.862,0.8 | 0.859,0.923,0.737, | -0,037 |
| 67065 | ENSG00000142197 | DOP1B | chr21 | + | 36214080 | 36214190 | 36211973 | 36212097 | 36214441 | 36214556 | 67065 | 9,17,50,26,13,16,13,15,10, | 5,0,9,3,2,10,4,4,0,2,0,2 | 12,24,12,10,14,14,7,5,7,19 | 4,8,1,11,18,9,0,3,8,12,1,0,5 | 259 | 149 | 0.0361666925678 | | 1 | 0.509,1.0,0.762,0.8 | 0.633,0.633,0.873, | 0,107 |
| 251801 | ENSG00000120690 | ELF1 | chr13 | - | 40961442 | 40961690 | 40958835 | 40959016 | 40981982 | 40982282 | 251801 | 22,2,14,12,1,4,3,3,4,2,3,3 | 540,370,520,267,192,189,1 | 6,9,14,22,9,3,0,10,10,5,10, | 350,483,303,408,341,138,3 | 298 | 149 | 0.0362126387755 | | 1 | 0.02,0.003,0.013,0 | 0.008,0.009,0.023, | -0,006 |
| 221057 | ENSG00000112182 | BACH2 | chr6 | - | 90249688 | 90249767 | 90206568 | 90206681 | 90252512 | 90252590 | 221057 | 7,1,44,8,6,13,8,7,9,2,5,10 | 39,10,120,17,24,24,23,36,2 | 12,9,6,14,0,4,2,6,1,3,6,3,2 | 72,34,36,42,8,10,20,21,16, | 228 | 149 | 0.0362499131875 | | 1 | 0.105,0.061,0.193, | 0.098,0.147,0.098, | 0,069 |
| 227397 | ENSG00000127914 | AKAP9 | chr7 | + | 92042058 | 92042186 | 92040703 | 92040898 | 92042667 | 92042771 | 227397 | 7,4,43,21,8,14,6,22,1,2,5,1 | 4,5,18,9,5,12,4,8,4,0,9,0 | 12,33,19,14,14,8,12,11,10, | 4,9,3,0,13,1,4,1,1,0,1,21,7, | 277 | 149 | 0.0362559602446 | | 1 | 0.485,0.301,0.562, | 0.617,0.664,0.773, | -0,155 |
| 123514 | ENSG00000030066 | NUP160 | chr11 | - | 47812881 | 47813047 | 47812063 | 47812429 | 47813315 | 47813415 | 123514 | 35,56,74,35,25,22,16,31,1 | 0,0,6,4,1,2,0,0,0,0,0,0 | 37,45,24,54,27,17,21,14,2 | 0,0,0,8,1,0,0,3,0,1,0,7,4,2,0 | 298 | 149 | 0.0362589617842 | | 1 | 1.0,1.0,0.86,0.814, | 1.0,1.0,1.0,0.771,0 | 0,031 |
| 38986 | ENSG00000204463 | BAG6 | chr6 | - | 31640384 | 31640528 | 31640198 | 31640306 | 31640644 | 31640704 | 38986 | 119,83,225,171,107,89,80, | 2,0,0,0,0,0,1,0,1,2,0,1 | 138,108,75,46,73,52,78,11 | 3,1,3,2,1,2,2,0,1,6,2,3,0,0,0 | 293 | 149 | 0.0362852378553 | | 1 | 0.968,1.0,1.0,1.0,1 | 0.959,0.982,0.927, | 0,028 |
| 132979 | ENSG00000166037 | CEP57 | chr11 | + | 95795515 | 95795559 | 95790512 | 95790743 | 95799231 | 95799388 | 132979 | 0,0,8,6,0,0,0,2,0,1,4,3 | 14,16,55,34,14,13,6,15,9,8 | 4,7,0,0,3,0,8,10,0,0,6,13,4, | 36,22,9,12,16,11,15,22,10, | 193 | 149 | 0.0362855805477 | | 1 | 0.0,0.0,0.101,0.12, | 0.079,0.197,0.0,0.0 | -0,036 |
| 165189 | ENSG00000115073 | ACTR1B | chr2 | - | 97658426 | 97658762 | 97658223 | 97658316 | 97658878 | 97659003 | 165189 | 41,27,55,50,40,35,28,23,3 | 5,0,4,2,0,4,0,0,0,1,0,2 | 27,37,22,9,20,15,31,36,20, | 0,0,0,1,1,0,0,2,0,2,0,2,0,6,0 | 298 | 149 | 0.0363033193345 | | 1 | 0.804,1.0,0.873,0.9 | 1.0,1.0,1.0,0.818,0 | -0,01 |
| 21734 | ENSG00000197555 | SIPA1L1 | chr14 | + | 71650289 | 71650509 | 71624047 | 71624236 | 71658332 | 71658436 | 21734 | 130,96,94,58,23,30,56,77,1 | 6,6,7,2,1,2,5,7,7,4,0,2 | 56,56,92,114,78,24,34,37,2 | 0,2,6,5,3,1,2,0,2,3,4,0,2,0,2 | 298 | 149 | 0.0363046028238 | | 1 | 0.915,0.889,0.87,0 | 1.0,0.933,0.885,0.9 | -0,026 |
| 24808 | ENSG00000121741 | ZMYM2 | chr13 | + | 20027984 | 20028064 | 20027202 | 20027318 | 20031318 | 20031379 | 24808 | 0,0,13,4,2,9,2,1,3,0,3,1 | 30,10,96,30,28,38,20,33,5, | 2,1,1,0,1,7,5,2,3,4,2,4,0,1,2 | 67,90,25,30,36,21,24,31,28 | 229 | 149 | 0.0363100688564 | | 1 | 0.0,0.0,0.081,0.08, | 0.019,0.007,0.025, | 0,025 |
| 144282 | ENSG00000161526 | SAP30BP | chr17 | + | 75668515 | 75668629 | 75667337 | 75667478 | 75671815 | 75672156 | 144282 | 144,112,146,79,29,43,51,4 | 1,6,10,3,0,4,0,2,3,0,4,0 | 70,69,41,61,84,58,26,28,2 | 4,3,3,7,4,0,2,8,2,0,5,8,1,0,2 | 263 | 149 | 0.0363114165818 | | 1 | 0.988,0.914,0.892, | 0.908,0.929,0.886, | 0,046 |
| 137357 | ENSG00000196684 | HSH2D | chr19 | + | 16155666 | 16155763 | 16154401 | 16154491 | 16157209 | 16157248 | 137357 | 27,8,19,11,17,21,18,25,24, | 135,148,162,110,120,130,1 | 11,10,15,16,9,6,3,9,6,8,4,1 | 69,124,78,82,129,45,43,67 | 246 | 149 | 0.0363361821402 | | 1 | 0.108,0.032,0.066, | 0.088,0.047,0.104, | 0,009 |
| 111969 | ENSG00000104408 | EIF3E | chr8 | - | 108242111 | 108242256 | 108241798 | 108241913 | 108248612 | 108248717 | 111969 | 3,0,10,2,1,8,5,9,0,3,0,2 | 111,48,450,246,90,76,75,1 | 6,10,1,8,2,3,1,3,4,1,7,29,0, | 180,169,117,34,78,28,65,1 | 294 | 149 | 0.0363907998474 | | 1 | 0.014,0.0,0.011,0.0 | 0.017,0.029,0.004, | -0,005 |
| 177719 | ENSG00000143643 | TTC13 | chr1 | - | 230928055 | 230928171 | 230925516 | 230925647 | 230928936 | 230929093 | 177719 | 1,0,4,0,2,0,0,2,0,0,0,0 | 19,16,33,35,23,13,19,11,9, | 3,1,2,1,0,0,2,5,0,4,2,0,0,3,2 | 18,18,9,7,15,9,7,8,7,40,3,2 | 265 | 149 | 0.0364160465153 | | 1 | 0.029,0.0,0.064,0.0 | 0.086,0.03,0.111,0 | -0,054 |
| 163657 | ENSG00000048028 | USP28 | chr11 | - | 113835206 | 113835314 | 113834248 | 113834335 | 113840597 | 113840757 | 163657 | 3,0,2,2,0,0,3,1,6,0,1,2 | 33,18,39,29,21,11,11,16,4 | 6,0,3,5,1,5,2,2,4,1,9,5,1,0,0 | 50,13,24,7,33,25,4,18,23,4 | 257 | 149 | 0.0364255721271 | | 1 | 0.05,0.0,0.029,0.0 | 0.065,0.0,0.068,0.2 | -0,046 |
| 130790 | ENSG00000123349 | PFDN5 | chr12 | + | 53295838 | 53295941 | 53295290 | 53295639 | 53296453 | 53296551 | 130790 | 1094,718,804,657,167,216 | 10,2,8,5,3,9,1,11,3,1,1,6 | 469,364,459,271,307,185,1 | 7,10,2,1,9,2,3,7,1,5,12,7,5, | 252 | 149 | 0.0364488737907 | | 1 | 0.985,0.995,0.983, | 0.975,0.956,0.993, | 0,005 |
| 156470 | ENSG00000121741 | ZMYM2 | chr13 | + | 20081754 | 20082130 | 20067238 | 20067390 | 20082780 | 20083032 | 156470 | 66,27,143,61,45,32,33,72,2 | 0,0,2,2,0,0,2,0,0,0,0,0 | 80,92,48,39,67,26,20,62,5 | 3,5,0,1,2,2,0,2,2,0,1,1,0,2,0 | 298 | 149 | 0.0364545806676 | | 1 | 1.0,1.0,0.973,0.93 | 0.93,0.902,1.0,0.9 | 0,027 |
| 194440 | ENSG00000103365 | GGA2 | chr16 | - | 23495138 | 23495170 | 23494302 | 23494378 | 23495693 | 23495778 | 194440 | 35,9,39,28,3,26,17,15,15,2 | 74,57,140,186,92,85,65,54 | 43,41,9,21,36,21,13,22,15, | 124,97,48,43,69,70,58,86,5 | 181 | 149 | 0.036456651592 | | 1 | 0.28,0.115,0.187,0 | 0.222,0.258,0.134, | -0,045 |
| 51800 | ENSG00000114395 | CYB561D2 | chr3 | + | 50352008 | 50352046 | 50350857 | 50351560 | 50353240 | 50354032 | 51800 | 8,12,25,23,5,7,7,13,2,3,6,1 | 1,1,7,6,0,0,1,2,2,2,0,2 | 21,9,4,1,2,8,1,5,4,20,7,15,2 | 11,4,2,0,3,1,0,4,4,5,6,2,4,2 | 187 | 149 | 0.0364874513072 | | 1 | 0.864,0.905,0.74,0 | 0.603,0.642,0.614, | 0,103 |
| 80651 | ENSG00000142188 | TMEM50B | chr21 | - | 33469530 | 33469683 | 33468786 | 33468926 | 33479837 | 33479974 | 80651 | 2,0,3,8,1,0,1,0,0,0,0,0 | 10,26,54,25,25,17,10,13,5, | 0,2,0,10,5,0,0,0,2,1,0,2,0,6 | 20,16,11,27,11,5,3,8,5,12,2 | 298 | 149 | 0.0365259547366 | | 1 | 0.091,0.0,0.027,0.1 | 0.0,0.059,0.0,0.15 | -0,033 |
| 199935 | ENSG00000134215 | VAV3 | chr1 | - | 107782987 | 107783086 | 107779433 | 107779492 | 107874900 | 107875017 | 199935 | 0,2,1,1,0,6,0,2,0,0,0,0 | 24,30,124,51,26,66,14,35,2 | 1,6,8,2,1,1,4,0,0,3,0,0,2,2,1 | 59,61,55,43,17,12,20,43,25 | 248 | 149 | 0.0365429517297 | | 1 | 0.0,0.039,0.005,0.0 | 0.01,0.056,0.08,0.0 | -0,02 |
| 190575 | ENSG00000177463 | NR2C2 | chr3 | + | 15032378 | 15032427 | 15030274 | 15030452 | 15034669 | 15034809 | 190575 | 22,24,13,15,14,10,15,9,16, | 2,0,4,1,0,2,1,4,0,0,0,0 | 17,17,13,17,4,5,8,15,6,9,1 | 3,3,0,2,2,0,0,0,1,0,3,2,0,4,0 | 198 | 149 | 0.0365562267265 | | 1 | 0.892,1.0,0.71,0.9 | 0.81,0.81,1.0,0.86 | 0,03 |
| 22273 | ENSG00000100485 | SOS2 | chr14 | - | 50158564 | 50158646 | 50155891 | 50157121 | 50159430 | 50160086 | 22273 | 92,82,165,111,57,82,58,64 | 0,3,16,6,14,9,9,5,0,0,2,5 | 83,76,95,25,53,35,44,50,4 | 6,10,3,15,3,2,14,9,11,10,1 | 231 | 149 | 0.0365565635701 | | 1 | 1.0,0.946,0.869,0.9 | 0.899,0.831,0.953, | 0,058 |
| 750 | ENSG00000228315 | GUSBP11 | chr22 | - | 23683724 | 23683867 | 23659864 | 23660000 | 23686837 | 23686995 | 750 | 39,14,0,0,20,22,32,57,10,0 | 0,1,3,1,4,0,0,0,2,1,0,0 | 29,33,19,28,9,11,22,42,27, | 0,0,0,0,0,0,0,0,0,7,2,0,0,0,0 | 292 | 149 | 0.0365690721753 | | 1 | 1.0,0.877,0.0,0.0,0 | 1.0,1.0,1.0,1.0,1.0, | -0,27 |
| 23666 | ENSG00000154001 | PPP2R5E | chr14 | - | 63453688 | 63453885 | 63421992 | 63422094 | 63539528 | 63539692 | 23666 | 48,43,94,70,44,32,30,41,2 | 1,0,11,4,1,9,3,4,0,2,4,4 | 69,81,44,30,28,29,24,35,3 | 4,9,0,0,1,1,1,6,1,4,3,2,4,1,0 | 298 | 149 | 0.0365713349412 | | 1 | 0.96,1.0,0.81,0.89 | 0.896,0.818,1.0,1.0 | -0,08 |
| 214828 | ENSG00000055044 | NOP58 | chr2 | + | 202277783 | 202278002 | 202275161 | 202275189 | 202282350 | 202282472 | 214828 | 230,244,93,28,20,27,16,27 | 14,7,7,4,3,3,3,2,0,0,1,5 | 35,57,72,42,59,12,14,29,2 | 7,4,4,12,12,6,0,2,1,2,4,2,1, | 298 | 149 | 0.0365831245501 | | 1 | 0.891,0.946,0.869, | 0.714,0.877,0.9,0.6 | 0,033 |
| 199824 | ENSG00000174606 | ANGEL2 | chr1 | - | 213011207 | 213011458 | 213008209 | 213008466 | 213013092 | 213013418 | 199824 | 0,0,9,3,0,3,3,5,1,0,0,2 | 38,40,73,44,39,34,29,33,2 | 2,1,0,2,0,4,0,3,0,1,0,0,0,1,0 | 39,49,21,34,24,15,15,36,2 | 298 | 149 | 0.0366191816577 | | 1 | 0.0,0.0,0.058,0.03 | 0.025,0.01,0.0,0.0 | 0,009 |
| 242627 | ENSG00000132386 | SERPINF1 | chr17 | + | 1771424 | 1771515 | 1771028 | 1771184 | 1771871 | 1772075 | 242627 | 0,1,2,4,3,3,6,5,2,0,1,0 | 12,6,26,22,22,31,21,35,20, | 1,2,1,2,4,2,0,1,1,4,1,0,1,0,4 | 7,28,5,8,50,21,12,16,35,51 | 240 | 149 | 0.0366209867704 | | 1 | 0.0,0.094,0.046,0.1 | 0.081,0.042,0.11,0 | 0,016 |
| 18350 | ENSG00000130703 | OSBPL2 | chr20 | + | 62264628 | 62264718 | 62263615 | 62263691 | 62272124 | 62272259 | 18350 | 9,0,3,1,1,3,5,4,2,2,0,0 | 46,29,51,38,24,33,27,28,2 | 0,2,3,1,0,0,0,2,1,6,4,1,0,3,0 | 28,35,17,31,22,14,13,35,28 | 239 | 149 | 0.0366258577628 | | 1 | 0.109,0.0,0.035,0.0 | 0.0,0.034,0.099,0.0 | 0,014 |
| 45654 | ENSG00000069849 | ATP1B3 | chr3 | + | 141913651 | 141913836 | 141907166 | 141907274 | 141915969 | 141916010 | 45654 | 18,13,83,142,40,36,25,49,1 | 0,2,2,3,0,1,4,4,0,1,0,1 | 106,88,28,18,11,31,17,51,2 | 1,2,0,0,0,0,0,0,0,0,1,0,0,0,0 | 298 | 149 | 0.0366342338576 | | 1 | 1.0,0.765,0.954,0.9 | 0.981,0.957,1.0,1.0 | -0,089 |
| 122845 | ENSG00000175224 | ATG13 | chr11 | + | 46668793 | 46668910 | 46668498 | 46668576 | 46669403 | 46669441 | 122845 | 65,53,105,65,34,37,31,36,3 | 3,0,4,1,0,2,1,2,2,0,2,0 | 59,85,32,84,53,39,24,21,1 | 5,0,1,13,1,0,5,0,0,3,0,6,1,0 | 266 | 149 | 0.0366381090149 | | 1 | 0.924,1.0,0.936,0.9 | 0.869,1.0,0.947,0.7 | 0,01 |
| 7251 | ENSG00000188785 | ZNF548 | chr19 | + | 57390721 | 57390845 | 57390004 | 57390114 | 57394187 | 57394223 | 7251 | 3,2,9,0,1,1,4,4,2,0,0,1 | 19,12,21,9,21,15,17,19,20, | 5,1,7,7,0,2,1,2,1,0,0,6,3,3,3 | 13,30,12,12,5,10,21,20,8,6 | 273 | 149 | 0.0366382055369 | | 1 | 0.079,0.083,0.19,0 | 0.173,0.018,0.241, | -0,042 |
| 184198 | ENSG00000143033 | MTF2 | chr1 | + | 93115491 | 93115618 | 93114987 | 93115088 | 93118344 | 93118440 | 184198 | 8,9,38,17,12,4,14,25,11,6,2 | 1,0,0,1,3,1,2,0,1,0,1,0 | 21,27,7,1,15,8,4,18,7,6,15, | 3,0,3,2,0,0,0,2,0,0,11,3,1,0 | 276 | 149 | 0.0366397539636 | | 1 | 0.812,1.0,1.0,0.90 | 0.791,1.0,0.557,0.2 | 0,016 |
| 237863 | ENSG00000104365 | IKBKB | chr8 | + | 42288640 | 42288728 | 42272082 | 42272205 | 42290155 | 42290273 | 237863 | 11,4,18,15,18,13,12,4,13,7 | 15,9,17,11,3,8,10,4,6,2,3,8 | 15,11,10,9,4,10,3,12,13,17 | 7,4,3,2,11,1,2,9,4,4,4,4,3,3 | 237 | 149 | 0.0366428723155 | | 1 | 0.316,0.218,0.4,0.4 | 0.574,0.634,0.677, | -0,123 |
| 35858 | ENSG00000198707 | CEP290 | chr12 | - | 88092680 | 88092832 | 88089031 | 88090839 | 88093769 | 88093975 | 35858 | 38,12,43,40,29,17,30,22,8, | 0,0,2,0,0,3,3,3,0,2,0,2 | 19,23,18,19,20,20,13,34,1 | 0,0,0,3,0,0,1,0,0,0,0,0,0,0,2 | 298 | 149 | 0.0366489299363 | | 1 | 1.0,1.0,0.915,1.0,1 | 1.0,1.0,1.0,0.76,1.0 | -0,111 |
| 224577 | ENSG00000133606 | MKRN1 | chr7 | - | 140455789 | 140455855 | 140455094 | 140455233 | 140456651 | 140456919 | 224577 | 45,61,71,50,22,21,19,33,6 | 3,0,0,2,0,2,3,2,1,2,0,1 | 39,42,26,15,20,13,12,20,1 | 3,1,5,9,1,0,0,0,1,2,0,0,1,0,6 | 215 | 149 | 0.0366623024571 | | 1 | 0.912,1.0,1.0,0.94 | 0.9,0.967,0.783,0.5 | 0,027 |
| 75477 | ENSG00000141504 | SAT2 | chr17 | - | 7626942 | 7627044 | 7626706 | 7626793 | 7627142 | 7627226 | 75477 | 24,5,13,13,7,5,15,6,19,6,4, | 10,7,21,12,14,11,1,3,0,2,4, | 11,3,6,2,4,6,1,5,4,34,7,6,1 | 8,7,6,9,7,6,17,29,3,11,8,4,4 | 251 | 149 | 0.0366668815599 | | 1 | 0.588,0.298,0.269, | 0.449,0.203,0.372, | 0,144 |
| 179411 | ENSG00000133789 | SWAP70 | chr11 | + | 9713465 | 9713582 | 9694145 | 9694286 | 9728052 | 9728199 | 179411 | 104,62,202,158,60,38,51,4 | 5,8,2,4,3,5,2,0,0,0,1,3 | 108,129,77,56,61,63,19,23 | 9,10,7,1,6,2,2,3,0,0,1,10,2, | 266 | 149 | 0.0366980198219 | | 1 | 0.921,0.813,0.983, | 0.87,0.878,0.86,0.9 | 0,036 |
| 222732 | ENSG00000167483 | NIBAN3 | chr19 | + | 17549664 | 17549790 | 17549454 | 17549527 | 17551785 | 17553645 | 222732 | 17,8,30,30,15,14,15,12,4,5 | 167,98,173,131,155,69,10 | 18,31,16,13,31,4,13,18,15, | 162,132,91,59,133,72,49,1 | 275 | 149 | 0.0367078003763 | | 1 | 0.052,0.042,0.086, | 0.057,0.113,0.087, | -0,013 |
| 18824 | ENSG00000069943 | PIGB | chr15 | + | 55350698 | 55350912 | 55341737 | 55341802 | 55354797 | 55354978 | 18824 | 45,33,49,26,23,12,10,22,1 | 7,0,4,2,1,0,2,0,3,6,0,1 | 27,55,12,6,24,13,23,5,27,9 | 0,1,0,1,1,0,1,2,3,5,0,1,0,2,0 | 298 | 149 | 0.0367424829557 | | 1 | 0.763,1.0,0.86,0.8 | 1.0,0.965,1.0,0.75, | -0,024 |
| 197540 | ENSG00000132300 | PTCD3 | chr2 | + | 86134805 | 86134987 | 86134295 | 86134377 | 86136520 | 86136562 | 197540 | 43,22,71,49,49,34,25,45,2 | 6,1,6,6,0,4,2,5,0,0,0,1 | 92,73,27,25,35,19,21,47,6 | 3,2,0,4,2,0,0,5,2,1,2,0,1,0,5 | 298 | 149 | 0.0367521487347 | | 1 | 0.782,0.917,0.855, | 0.939,0.948,1.0,0.7 | -0,014 |
| 74172 | ENSG00000064687 | ABCA7 | chr19 | + | 1057968 | 1058059 | 1057313 | 1057429 | 1058145 | 1058269 | 74172 | 11,18,32,19,22,14,24,20,1 | 5,0,0,2,1,2,1,1,1,1,1,1 | 20,22,11,10,13,18,2,12,18, | 0,2,1,1,0,0,0,1,0,0,2,1,0,1,0 | 240 | 149 | 0.0367958664646 | | 1 | 0.577,1.0,1.0,0.85 | 1.0,0.872,0.872,0.8 | -0,076 |
| 242557 | ENSG00000167985 | SDHAF2 | chr11 | + | 61437668 | 61438113 | 61430143 | 61430182 | 61445940 | 61446342 | 242557 | 18,13,48,39,13,16,17,15,2 | 3,3,17,7,2,5,6,8,0,0,0,8 | 39,37,13,2,15,12,15,38,4,1 | 1,5,2,0,1,5,2,6,0,3,3,4,4,4,2 | 298 | 149 | 0.036804509209 | | 1 | 0.75,0.684,0.585,0 | 0.951,0.787,0.765, | -0,072 |
| 36358 | ENSG00000136169 | SETDB2 | chr13 | + | 49490821 | 49490910 | 49488289 | 49488630 | 49491731 | 49495002 | 36358 | 29,11,71,29,24,10,21,43,2 | 2,0,3,0,2,0,2,4,0,0,0,0 | 33,36,20,4,9,12,3,14,24,56 | 0,2,0,2,0,0,0,1,6,0,5,2,1,2,5 | 238 | 149 | 0.0368047186761 | | 1 | 0.901,1.0,0.937,1.0 | 1.0,0.918,1.0,0.55 | 0,059 |
| 111657 | ENSG00000165102 | HGSNAT | chr8 | + | 43178073 | 43178234 | 43172309 | 43172386 | 43182144 | 43182260 | 111657 | 15,4,33,25,15,7,2,11,1,0,9, | 0,1,3,1,0,1,0,1,0,2,3,2 | 23,9,2,2,10,7,3,7,5,5,6,6,1 | 1,2,0,0,1,0,0,1,3,0,1,0,0,0,0 | 298 | 149 | 0.0368696160678 | | 1 | 1.0,0.667,0.846,0.9 | 0.92,0.692,1.0,1.0, | -0,124 |
| 233979 | ENSG00000029363 | BCLAF1 | chr6 | - | 136269140 | 136269192 | 136268161 | 136268339 | 136269436 | 136269959 | 233979 | 5,4,24,8,1,6,8,12,7,6,0,6 | 122,84,225,146,80,89,101, | 5,13,5,12,3,3,7,5,3,5,8,6,2, | 127,137,109,94,71,65,65,8 | 201 | 149 | 0.0369081249627 | | 1 | 0.029,0.034,0.073, | 0.028,0.066,0.033, | 0,008 |
| 143608 | ENSG00000275700 | AATF | chr17 | + | 37031399 | 37031685 | 37020937 | 37021014 | 37056600 | 37056842 | 143608 | 9,5,29,25,14,13,8,12,8,4,4, | 0,1,3,0,5,3,1,4,0,0,0,2 | 23,22,11,5,8,8,6,7,7,20,14, | 5,6,2,4,4,1,0,0,3,1,6,1,2,9,1 | 298 | 149 | 0.0369451329313 | | 1 | 1.0,0.714,0.829,1.0 | 0.697,0.647,0.733, | 0,132 |
| 216218 | ENSG00000215717 | TMEM167B | chr1 | + | 109092889 | 109093094 | 109090763 | 109090882 | 109094416 | 109096934 | 216218 | 22,25,57,52,6,12,15,9,8,5,1 | 1,4,4,1,3,0,4,4,0,0,1,1 | 48,54,28,22,40,25,32,20,1 | 0,2,0,0,3,2,1,2,0,0,6,4,1,0,2 | 298 | 149 | 0.0369803389032 | | 1 | 0.917,0.758,0.877, | 1.0,0.931,1.0,1.0,0 | -0,121 |
| 205160 | ENSG00000159023 | EPB41 | chr1 | + | 28997214 | 28997574 | 28993329 | 28993542 | 29011864 | 29011907 | 205160 | 73,45,264,149,107,97,74,1 | 2,2,4,11,3,0,3,15,2,0,0,4 | 148,135,91,93,138,115,38, | 11,8,5,5,13,5,5,2,9,10,3,0,4 | 298 | 149 | 0.0369884095283 | | 1 | 0.948,0.918,0.971, | 0.871,0.894,0.901, | 0,031 |

| | | | | | | | | | | | | | | | | | | | | | |
|---|---|---|---|---|---|---|---|---|---|---|---|---|---|---|---|---|---|---|---|---|---|
| 59356 | ENSG00000166046 | TCP11L2 | chr12 | + | 106311040 | 106311232 | 106302732 | 106302941 | 106314357 | 106314493 | 59356 | 103,78,189,104,81,71,61,7 | 5,0,18,0,2,1,0,0,0,0,0,2 | 116,207,58,48,150,66,46,8 | 0,4,0,0,7,3,3,4,3,0,0,0,5,3, | 298 | 149 | 0.0370104273436 | | 1 | 0.912,1.0,0.84,1.0, | 1.0,0.963,1.0,1.0,0 | 0,012 |
| 197769 | ENSG00000066027 | PPP2R5A | chr1 | + | 212347976 | 212348021 | 212347346 | 212347406 | 212348388 | 212348497 | 197769 | 11,5,7,11,4,2,6,2,1,0,1,1 | 11,14,47,10,31,10,18,4,9,2 | 1,3,1,2,4,1,0,4,2,3,5,1,1,2, | 28,17,8,10,4,5,13,5,7,16,1 | 194 | 149 | 0.0370474314356 | | 1 | 0.434,0.215,0.103, | 0.027,0.119,0.088, | 0,028 |
| 105062 | ENSG00000078589 | P2RY10 | chrX | + | 78952192 | 78952335 | 78947814 | 78947863 | 78960507 | 78961080 | 105062 | 344,182,343,207,122,109, | 74,53,104,77,56,70,70,52, | 315,306,247,123,104,71,1 | 110,66,67,19,44,19,28,60, | 292 | 149 | 0.0370772644383 | | 1 | 0.703,0.637,0.627, | 0.594,0.703,0.653, | -0,05 |
| 28984 | ENSG00000214253 | FIS1 | chr7 | - | 101244006 | 101244139 | 101240829 | 101240906 | 101251894 | 101252133 | 28984 | 113,89,161,105,33,37,30,6 | 2,3,2,2,4,1,2,1,5,0,0,2 | 90,81,105,45,50,36,22,53, | 0,6,1,0,4,3,2,0,3,3,0,15,1,4 | 282 | 149 | 0.0371091689468 | | 1 | 0.968,0.94,0.977,0 | 1.0,0.877,0.982,1. | 0,025 |
| 144718 | ENSG00000141298 | SSH2 | chr17 | - | 29637485 | 29637612 | 29635967 | 29636802 | 29648143 | 29648344 | 144718 | 17,10,15,9,12,10,11,6,10,9 | 109,79,95,92,86,75,57,78, | 5,8,5,8,5,1,0,5,3,14,10,9,1 | 83,48,33,38,65,50,73,86,4 | 276 | 149 | 0.0371122872687 | | 1 | 0.078,0.064,0.079, | 0.031,0.083,0.076, | 0,013 |
| 37209 | ENSG00000155189 | AGPAT5 | chr8 | + | 6724869 | 6724939 | 6708641 | 6708887 | 6730710 | 6730743 | 37209 | 25,23,53,40,61,39,20,36,1 | 5,0,6,5,8,6,2,0,1,0,2,8 | 43,61,23,2,39,21,20,23,25, | 4,5,2,0,3,2,3,0,3,2,2,1,5,1, | 219 | 149 | 0.0371150606483 | | 1 | 0.773,1.0,0.857,0. | 0.88,0.892,0.887,1 | -0,018 |
| 106536 | ENSG00000196924 | FLNA | chrX | - | 154352552 | 154352675 | 154352180 | 154352447 | 154352771 | 154352783 | 106536 | 433,257,303,309,352,218, | 14,16,7,5,1,4,11,14,3,6,2,8 | 268,421,108,114,256,192, | 7,12,9,14,25,2,1,11,8,7,4,5 | 272 | 149 | 0.0371262893311 | | 1 | 0.944,0.898,0.96,0 | 0.954,0.951,0.868, | 0,016 |
| 253034 | ENSG00000091039 | OSBPL8 | chr12 | - | 76459858 | 76459934 | 76450850 | 76450988 | 76487509 | 76487618 | 253034 | 17,10,10,6,2,3,2,8,16,2,1,3 | 23,31,35,36,20,12,20,20,2 | 5,4,3,3,4,2,3,4,4,3,4,4,6,9, | 25,34,12,15,18,13,9,17,8,1 | 225 | 149 | 0.0371306459203 | | 1 | 0.329,0.176,0.159, | 0.117,0.072,0.142, | 0,031 |
| 245939 | ENSG00000166405 | RIC3 | chr11 | - | 8137384 | 8137471 | 8135786 | 8135844 | 8138271 | 8138347 | 245939 | 60,25,85,39,47,39,37,39,3 | 1,0,2,0,0,0,1,0,0,0,3,0 | 56,61,60,64,26,14,17,29,1 | 6,1,2,6,0,0,0,3,0,0,4,1,1,0, | 236 | 149 | 0.0371878684768 | | 1 | 0.974,1.0,0.964,1. | 0.855,0.975,0.95,0 | 0,029 |
| 152247 | ENSG00000140943 | MBTPS1 | chr16 | - | 84058217 | 84058358 | 84056004 | 84056135 | 84059301 | 84061631 | 152247 | 1,3,0,4,2,0,5,3,2,0,0,0 | 26,15,81,81,25,27,23,28,2 | 2,4,0,2,3,0,6,4,1,1,10,1,0,2 | 40,42,15,9,20,20,15,30,13, | 290 | 149 | 0.0372451944763 | | 1 | 0.019,0.093,0.0,0. | 0.025,0.047,0.0,0. | -0,021 |
| 192988 | ENSG00000138674 | SEC31A | chr4 | - | 82861630 | 82861708 | 82857688 | 82857764 | 82862595 | 82862652 | 192988 | 23,13,34,23,17,18,6,23,3,6 | 0,0,0,3,0,4,5,0,0,0,0,4 | 24,20,14,14,11,8,8,21,13,2 | 0,0,1,4,1,3,2,2,0,0,4,7,1,2, | 227 | 149 | 0.0372485573879 | | 1 | 1.0,1.0,1.0,0.834,1 | 1.0,1.0,0.902,0.69 | 0,034 |
| 181625 | ENSG00000134255 | CEPT1 | chr1 | + | 111173113 | 111173258 | 111161154 | 111161296 | 111174878 | 111174963 | 181625 | 19,10,83,30,48,26,17,29,9, | 10,12,41,31,30,8,4,5,17,5, | 60,54,23,32,24,16,3,17,38, | 26,6,10,3,3,13,2,10,2,19,5, | 294 | 149 | 0.0372614536462 | | 1 | 0.491,0.297,0.506, | 0.539,0.82,0.538,0 | -0,126 |
| 97410 | ENSG00000083168 | KAT6A | chr8 | - | 42049575 | 42049707 | 42048377 | 42049302 | 42051900 | 42051984 | 97410 | 23,18,70,33,25,17,21,32,2 | 16,18,25,18,20,31,24,14,1 | 50,37,27,82,30,17,26,33,1 | 19,21,18,22,25,7,9,12,8,15 | 281 | 149 | 0.0372693517805 | | 1 | 0.433,0.347,0.598, | 0.583,0.483,0.443, | -0,093 |
| 19872 | ENSG00000140319 | SRP14 | chr15 | - | 40037308 | 40037355 | 40036985 | 40037018 | 40038281 | 40038394 | 19872 | 23,0,12,6,4,10,1,8,19,13,2, | 814,539,440,340,197,165, | 3,4,3,2,7,6,1,10,0,1,5,2,4,2 | 266,343,299,169,175,119, | 196 | 149 | 0.0372694475393 | | 1 | 0.021,0.0,0.02,0.0 | 0.009,0.009,0.008, | 0,004 |
| 119742 | ENSG00000149761 | NUDT22 | chr11 | + | 64227567 | 64227666 | 64226634 | 64227132 | 64229246 | 64229344 | 119742 | 30,19,30,34,36,15,28,24,5 | 2,0,2,3,0,2,0,0,4,0,2,5 | 22,31,15,16,2,9,5,10,5,18, | 0,0,1,0,0,0,1,8,0,0,0,0,3,0, | 248 | 149 | 0.0372729685022 | | 1 | 0.9,1.0,0.9,0.872,1 | 1.0,1.0,0.9,1.0,1.0, | -0,04 |
| 178369 | ENSG00000108523 | RNF167 | chr17 | + | 4943158 | 4943519 | 4942850 | 4942941 | 4944557 | 4944638 | 178369 | 83,19,81,65,29,24,26,18,3 | 3,0,2,3,0,4,2,5,1,0,0,1 | 39,25,28,20,23,14,23,36,1 | 4,6,0,1,2,0,1,4,0,3,3,7,0,2, | 298 | 149 | 0.0373056622833 | | 1 | 0.933,1.0,0.953,0. | 0.83,0.676,1.0,0.9 | 0,049 |
| 36511 | ENSG00000145495 | MARCHF6 | chr5 | + | 10390331 | 10390500 | 10386924 | 10387066 | 10391541 | 10391574 | 36511 | 331,250,252,117,71,114,8 | 12,11,15,11,0,8,7,5,7,0,0,7 | 143,242,165,274,97,96,10 | 3,24,9,19,23,5,3,4,23,5,6,7 | 298 | 149 | 0.0373250534198 | | 1 | 0.932,0.919,0.894, | 0.96,0.834,0.902,0 | 0,039 |
| 205823 | ENSG00000128815 | WDFY4 | chr10 | + | 48777447 | 48777495 | 48776749 | 48776984 | 48778610 | 48778832 | 205823 | 26,13,44,36,24,33,14,22,1 | 9,2,11,3,1,4,10,11,1,4,2,7 | 44,59,3,26,6,14,25,30,15,1 | 15,27,5,17,6,0,13,16,12,0, | 197 | 149 | 0.0373398839091 | | 1 | 0.686,0.831,0.752, | 0.689,0.623,0.312, | 0,112 |
| 36779 | ENSG00000101109 | STK4 | chr20 | + | 44997168 | 44997306 | 44995089 | 44995257 | 45000391 | 45000520 | 36779 | 579,411,740,493,243,254, | 6,6,6,0,0,1,5,3,0,3,2,0 | 480,621,354,373,366,246, | 2,12,3,12,4,4,4,4,2,3,2,2,1, | 287 | 149 | 0.0373493279127 | | 1 | 0.98,0.973,0.985,1 | 0.992,0.964,0.984, | 0,007 |
| 191914 | ENSG00000121892 | PDS5A | chr4 | - | 39910243 | 39910306 | 39908394 | 39908540 | 39913610 | 39913726 | 191914 | 26,21,87,35,30,34,40,20,1 | 7,2,11,12,0,0,3,4,6,0,2,0 | 52,39,20,17,17,22,13,24,1 | 6,12,6,13,20,2,1,3,11,2,1,3 | 212 | 149 | 0.0373509506167 | | 1 | 0.723,0.881,0.848, | 0.859,0.696,0.701, | 0,092 |
| 231794 | ENSG00000139146 | SINHCAF | chr12 | - | 31304981 | 31305468 | 31298076 | 31298224 | 31326023 | 31326199 | 231794 | 16,15,58,22,10,18,17,43,1 | 20,16,82,33,4,29,18,44,38, | 35,39,51,19,56,14,36,26,1 | 32,19,22,37,75,29,27,26,2 | 298 | 149 | 0.037352946947 | | 1 | 0.286,0.319,0.261, | 0.354,0.506,0.537, | -0,059 |
| 190601 | ENSG00000115942 | ORC2 | chr2 | - | 200962137 | 200962176 | 200959391 | 200959440 | 200963489 | 200963680 | 190601 | 0,1,6,4,0,2,1,0,0,3,0,1 | 32,12,34,12,18,14,6,12,12, | 0,2,0,0,0,1,0,0,0,0,0,0,0,0 | 8,8,9,4,9,8,6,14,8,14,5,7,7, | 188 | 149 | 0.0373596602602 | | 1 | 0.0,0.062,0.123,0. | 0.0,0.165,0.0,0.0,0 | 0,055 |
| 217581 | ENSG00000133872 | SARAF | chr8 | - | 30073877 | 30074051 | 30069873 | 30070059 | 30082846 | 30083124 | 217581 | 6,6,9,3,4,6,6,5,0,2,0,9 | 22,10,58,11,17,12,31,33,7, | 3,1,1,6,3,4,0,2,0,8,3,5,3,5, | 21,23,14,13,23,10,10,22,7, | 298 | 149 | 0.0373875177025 | | 1 | 0.12,0.231,0.072,0 | 0.067,0.021,0.034, | 0,033 |
| 178651 | ENSG00000163608 | NEPRO | chr3 | - | 113008647 | 113008805 | 113008170 | 113008430 | 113010642 | 113010704 | 178651 | 6,0,0,1,0,4,0,7,3,2,0,0 | 27,33,55,36,23,18,17,29,1 | 1,0,0,3,1,0,0,2,0,1,0,4,1,0, | 37,39,16,8,23,12,23,25,15, | 298 | 149 | 0.0374034174859 | | 1 | 0.1,0.0,0.0,0.014,0 | 0.013,0.0,0.0,0.15 | 0,021 |
| 12730 | ENSG00000073584 | SMARCE1 | chr17 | - | 40632194 | 40632367 | 40630713 | 40631693 | 40635930 | 40636098 | 12730 | 246,212,416,280,138,139, | 1,0,1,2,2,5,6,15,0,0,1,0 | 237,269,139,102,136,96,5 | 1,1,4,0,4,0,6,0,0,0,2,1,0,0, | 298 | 149 | 0.0375223747355 | | 1 | 0.992,1.0,0.995,0. | 0.992,0.993,0.946, | -0,012 |
| 230835 | ENSG00000214021 | TTLL3 | chr3 | + | 9818769 | 9818920 | 9817689 | 9817759 | 9820545 | 9820616 | 230835 | 34,15,21,15,18,22,12,14,1 | 1,1,0,0,1,1,1,0,0,0,0,0 | 21,16,15,35,9,7,11,29,12,2 | 2,1,1,0,0,1,0,8,0,0,3,0,1,3, | 298 | 149 | 0.0375264251235 | | 1 | 0.944,0.882,1.0,1. | 0.84,0.889,0.882,1 | 0,07 |
| 70064 | ENSG00000054267 | ARID4B | chr1 | - | 235181070 | 235181184 | 235177855 | 235177913 | 235181584 | 235181865 | 70064 | 54,41,62,50,23,20,24,34,2 | 115,97,133,82,44,71,47,48 | 40,66,25,47,65,22,9,21,27, | 86,76,74,72,55,32,27,51,31 | 263 | 149 | 0.0375325287337 | | 1 | 0.21,0.193,0.209,0 | 0.209,0.33,0.161,0 | -0,038 |
| 18388 | ENSG00000213614 | HEXA | chr15 | - | 72353067 | 72353178 | 72351131 | 72351234 | 72355558 | 72355624 | 18388 | 53,46,115,53,69,38,46,66, | 1,7,0,6,4,6,1,1,3,3,0,1 | 74,80,52,43,51,27,27,41,4 | 1,6,3,1,1,2,0,7,1,1,2,0,3,9, | 260 | 149 | 0.0375453435495 | | 1 | 0.968,0.79,1.0,0.8 | 0.977,0.884,0.909, | -0,02 |
| 19851 | ENSG00000169612 | RAMAC | chr15 | + | 82989009 | 82989188 | 82987335 | 82987384 | 82989880 | 82991057 | 19851 | 66,41,122,60,41,27,28,36, | 5,0,1,1,0,0,0,2,0,0,0,0 | 67,68,46,28,41,27,26,43,3 | 0,0,0,0,3,0,0,3,1,0,2,1,0,3, | 298 | 149 | 0.0375492378103 | | 1 | 0.868,1.0,0.984,0. | 1.0,1.0,1.0,1.0,0.8 | 0,029 |
| 249545 | ENSG00000155629 | PIK3AP1 | chr10 | - | 96669530 | 96669621 | 96656797 | 96656934 | 96709566 | 96709983 | 249545 | 8,1,9,14,4,0,0,1,1,2,0,3 | 161,53,180,152,58,73,41,4 | 2,2,2,0,5,1,2,4,3,4,0,3,0,2, | 101,67,64,12,42,45,33,69, | 240 | 149 | 0.0375506466971 | | 1 | 0.03,0.012,0.03,0. | 0.012,0.018,0.019, | 0,001 |
| 223728 | ENSG00000127947 | PTPN12 | chr7 | + | 77569657 | 77569794 | 77537553 | 77537645 | 77571077 | 77571186 | 223728 | 5,3,3,2,1,7,15,12,2,4,0,0 | 38,17,21,20,11,15,11,24,1 | 0,12,11,15,12,8,3,2,12,0,1 | 14,30,16,6,7,21,6,7,10,12, | 286 | 149 | 0.0375518384929 | | 1 | 0.064,0.084,0.069, | 0.0,0.172,0.264,0. | -0,081 |
| 230677 | ENSG00000225648 | SBDSP1 | chr7 | + | 72833763 | 72833851 | 72831601 | 72831798 | 72836576 | 72836701 | 230677 | 58,22,160,73,31,46,66,69, | 8,0,6,1,0,1,0,9,4,0,2,0 | 78,71,80,82,80,56,22,73,5 | 6,0,4,1,3,0,0,1,3,2,4,0,1,0, | 237 | 149 | 0.0376037317927 | | 1 | 0.82,1.0,0.944,0.9 | 0.891,1.0,0.926,0. | -0,029 |
| 116618 | ENSG00000160325 | CACFD1 | chr9 | + | 133465321 | 133465447 | 133463482 | 133463555 | 133467920 | 133468028 | 116618 | 20,13,31,28,16,18,13,8,5,6 | 3,4,2,2,2,0,0,2,3,1,3,0 | 14,8,10,7,7,7,14,13,4,8,1,1 | 0,1,0,0,0,0,4,5,0,3,0,2,1,0, | 275 | 149 | 0.0376311033618 | | 1 | 0.783,0.638,0.894, | 1.0,0.813,1.0,1.0,1 | -0,079 |
| 246237 | ENSG00000172183 | ISG20 | chr15 | + | 88637361 | 88637529 | 88636169 | 88636292 | 88639342 | 88639594 | 246237 | 23,11,21,12,10,24,2,9,11,2 | 4,7,4,5,2,14,3,14,27,6,3,5 | 2,0,7,3,0,0,3,7,3,22,15,13, | 4,5,5,2,6,10,1,4,1,11,11,4, | 298 | 149 | 0.0376617834475 | | 1 | 0.742,0.44,0.724,0 | 0.2,0.0,0.412,0.42 | 0,079 |
| 214681 | ENSG00000169967 | MAP3K2 | chr2 | - | 127332320 | 127332370 | 127330391 | 127330505 | 127335869 | 127335969 | 214681 | 1,1,4,4,0,1,1,1,2,0,0,2 | 30,35,53,36,23,23,11,31,2 | 4,1,1,2,1,0,0,1,1,0,0,1,2,0, | 37,73,43,51,39,24,14,27,2 | 199 | 149 | 0.0376656354347 | | 1 | 0.024,0.021,0.053, | 0.075,0.01,0.017,0 | 0,018 |
| 225277 | ENSG00000198198 | SZT2 | chr1 | + | 43429764 | 43429844 | 43428239 | 43428486 | 43430010 | 43430103 | 225277 | 5,2,19,10,4,22,6,7,2,0,9,7 | 4,2,2,0,0,0,1,2,0,1,2,0 | 12,5,2,3,9,3,13,12,8,15,7,6 | 3,1,2,6,2,0,0,0,0,1,2,1,0,1, | 229 | 149 | 0.0376675248275 | | 1 | 0.449,0.394,0.861, | 0.722,0.765,0.394, | -0,044 |
| 223204 | ENSG00000125779 | PANK2 | chr20 | + | 3908671 | 3908905 | 3907925 | 3908278 | 3910576 | 3910830 | 223204 | 5,0,19,1,4,6,1,7,6,3,4,1 | 28,8,69,42,21,21,22,18,23, | 6,4,5,1,6,4,3,0,1,2,1,2,3,2, | 23,39,38,9,23,21,3,30,13,3 | 298 | 149 | 0.0376719192262 | | 1 | 0.082,0.0,0.121,0. | 0.115,0.049,0.062, | 0,015 |
| 191921 | ENSG00000121892 | PDS5A | chr4 | - | 39926774 | 39926838 | 39925835 | 39925933 | 39927960 | 39928164 | 191921 | 17,1,21,38,25,43,19,33,7,6 | 3,2,4,0,1,0,1,8,0,0,0,1 | 21,19,17,10,6,7,10,17,14,2 | 0,4,0,0,10,4,0,3,1,1,3,1,1,7 | 213 | 149 | 0.0377076952922 | | 1 | 0.799,0.259,0.786, | 1.0,0.769,1.0,1.0,0 | 0,054 |
| 252598 | ENSG00000129315 | CCNT1 | chr12 | - | 48695998 | 48696141 | 48695758 | 48695829 | 48698137 | 48698183 | 252598 | 18,28,26,11,7,5,6,10,17,2, | 59,55,36,34,10,10,21,17,1 | 22,10,12,27,26,4,2,17,11,1 | 26,30,18,24,9,10,11,17,14, | 292 | 149 | 0.0377080831624 | | 1 | 0.135,0.206,0.269, | 0.302,0.145,0.254, | -0,056 |
| 134416 | ENSG00000100077 | GRK3 | chr22 | + | 25718244 | 25718385 | 25714407 | 25714570 | 25721283 | 25721397 | 134416 | 55,63,38,23,9,14,11,12,42, | 24,7,8,8,3,4,5,2,1,2,4,4 | 30,40,21,11,16,12,10,14,1 | 14,13,11,15,6,6,0,7,5,3,8,2 | 290 | 149 | 0.0377114244738 | | 1 | 0.541,0.822,0.709, | 0.524,0.613,0.495, | 0,079 |
| 211308 | ENSG00000160695 | VPS11 | chr11 | + | 119071595 | 119071960 | 119070233 | 119070397 | 119073197 | 119073399 | 211308 | 17,14,36,34,22,9,12,13,3,6 | 4,2,16,8,2,14,4,4,1,1,4,3 | 14,21,8,6,15,10,0,4,4,19,1 | 16,17,4,4,6,1,2,8,8,4,13,6,1 | 298 | 149 | 0.0377253361827 | | 1 | 0.68,0.778,0.529,0 | 0.304,0.382,0.5,0. | 0,159 |
| 240257 | ENSG00000197329 | PELI1 | chr2 | - | 64104553 | 64105051 | 64100397 | 64100499 | 64108239 | 64108379 | 240257 | 1,1,4,2,1,0,2,3,0,0,2,2 | 54,12,11,12,7,13,22,6,6,7, | 1,1,6,2,5,0,3,2,0,0,2,4,1,1, | 5,30,45,109,26,7,15,21,8,1 | 298 | 149 | 0.0377516523255 | | 1 | 0.009,0.04,0.154,0 | 0.091,0.016,0.062, | 0,06 |
| 191856 | ENSG00000141456 | PELP1 | chr17 | - | 4676035 | 4676151 | 4675273 | 4675884 | 4676356 | 4676507 | 191856 | 8,1,18,11,12,14,15,8,11,3, | 0,0,3,0,2,0,0,2,2,0,0,2 | 1,4,7,3,3,3,16,6,5,6,10,2,8, | 3,0,0,2,0,0,3,3,0,4,1,2,1,0, | 265 | 149 | 0.0377772742925 | | 1 | 1.0,1.0,0.771,1.0,0 | 0.158,1.0,1.0,0.45 | 0,157 |
| 249997 | ENSG00000023191 | RNH1 | chr11 | - | 504823 | 505022 | 502061 | 502181 | 507112 | 507300 | 249997 | 10,7,13,34,10,14,16,13,8,1 | 18,25,22,12,23,26,21,14,1 | 6,6,8,11,5,4,2,8,3,2,6,17,5, | 12,12,10,20,16,4,12,24,5,2 | 298 | 149 | 0.0377945632737 | | 1 | 0.217,0.123,0.228, | 0.2,0.2,0.286,0.21 | 0,084 |
| 201087 | ENSG00000132680 | KHDC4 | chr1 | - | 155926675 | 155926863 | 155925631 | 155925843 | 155927103 | 155927156 | 201087 | 27,13,41,43,21,17,13,32,2 | 13,2,10,12,2,3,0,2,3,2,1,3 | 25,17,6,8,15,12,2,15,22,39 | 19,4,4,1,3,2,1,3,7,2,8,5,4,1 | 298 | 149 | 0.0378134246662 | | 1 | 0.509,0.765,0.672, | 0.397,0.68,0.429,0 | 0,143 |
| 73154 | ENSG00000181852 | RNF41 | chr12 | - | 56212995 | 56213077 | 56210296 | 56210568 | 56213957 | 56214070 | 73154 | 17,12,53,20,5,11,11,8,9,1, | 38,81,146,71,50,52,34,59, | 40,53,7,7,13,8,19,21,13,8, | 77,161,20,7,68,27,71,65,21 | 231 | 149 | 0.0378203115146 | | 1 | 0.224,0.087,0.19,0 | 0.251,0.175,0.184, | -0,049 |
| 53502 | ENSG00000197386 | HTT | chr4 | + | 3154292 | 3154419 | 3148004 | 3148207 | 3157071 | 3157199 | 53502 | 38,31,72,47,39,60,78,61,1 | 0,3,0,0,0,0,4,0,4,0,0,0 | 42,65,34,62,53,37,22,42,4 | 3,5,0,7,2,0,0,0,1,2,0,1,1,0,1 | 276 | 149 | 0.0378821472409 | | 1 | 1.0,0.848,1.0,1.0,1 | 0.883,0.875,1.0,0. | 0,013 |
| 110737 | ENSG00000153317 | ASAP1 | chr8 | - | 130091972 | 130092143 | 130079901 | 130079971 | 130112093 | 130112322 | 110737 | 8,11,5,6,4,4,3,5,4,8,0,7 | 2,1,2,3,2,1,1,7,0,1,1,1 | 8,6,3,5,2,0,2,5,0,5,1,9,0,2, | 2,3,0,2,0,2,3,5,1,1,0,5,3,1, | 298 | 149 | 0.037893792461 | | 1 | 0.667,0.846,0.556, | 0.667,0.5,1.0,0.55 | 0,114 |
| 101600 | ENSG00000196313 | POM121 | chr7 | + | 72913764 | 72913828 | 72890932 | 72891110 | 72926261 | 72926477 | 101600 | 4,4,7,6,5,5,4,12,1,1,4,6 | 5,5,14,9,9,7,1,4,2,0,8,6 | 5,15,0,3,1,3,4,0,3,1,2,3,8,4 | 5,9,5,4,6,3,12,13,7,4,5,18,1 | 213 | 149 | 0.0379182608926 | | 1 | 0.359,0.359,0.259, | 0.412,0.538,0.0,0. | 0,191 |
| 49618 | ENSG00000144218 | AFF3 | chr2 | - | 100063125 | 100063249 | 100008886 | 100008932 | 100104401 | 100104525 | 49618 | 4,0,11,8,7,6,0,6,7,0,1,4 | 158,101,137,61,85,85,70,6 | 7,1,2,12,2,0,7,3,0,0,8,3,1,0 | 116,94,80,144,87,55,43,64 | 273 | 149 | 0.0379601775175 | | 1 | 0.014,0.0,0.042,0. | 0.032,0.006,0.013, | 0,01 |
| 232375 | ENSG00000111203 | ITFG2 | chr12 | + | 2830714 | 2830885 | 2824089 | 2824252 | 2836744 | 2836956 | 232375 | 22,7,11,5,11,7,23,23,7,11, | 2,4,1,1,0,0,2,1,0,0,0,1 | 8,7,4,11,6,4,6,8,4,3,5,9,12, | 0,2,0,3,2,1,0,0,0,4,0,0,4,7,1 | 298 | 149 | 0.0379672558486 | | 1 | 0.846,0.467,0.846, | 1.0,0.636,1.0,0.64 | 0,091 |
| 207653 | ENSG00000130348 | QRSL1 | chr6 | + | 106654765 | 106654922 | 106652466 | 106652582 | 106655614 | 106655732 | 207653 | 36,24,127,144,49,38,25,43 | 4,8,10,11,8,5,2,2,1,2,2,0 | 61,48,28,16,24,60,14,38,1 | 5,2,5,2,0,0,0,2,4,11,3,0,0,0 | 298 | 149 | 0.0379767007577 | | 1 | 0.818,0.6,0.864,0. | 0.859,0.923,0.737, | -0,036 |
| 11809 | ENSG00000007168 | PAFAH1B1 | chr17 | + | 2665371 | 2665456 | 2638098 | 2638320 | 2666015 | 2666090 | 11809 | 109,65,324,187,87,90,93,1 | 1,1,4,1,1,0,0,0,1,2,2,0 | 123,148,102,80,135,70,52, | 4,2,0,3,2,0,2,3,2,1,2,0,1,3, | 234 | 149 | 0.037986608906 | | 1 | 0.986,0.976,0.981, | 0.951,0.979,1.0,0. | 0,012 |
| 14904 | ENSG00000273611 | ZNHIT3 | chr17 | + | 36486934 | 36486966 | 36486680 | 36486785 | 36492812 | 36492899 | 14904 | 55,30,104,90,20,26,32,32, | 1,2,8,8,1,0,1,2,0,1,0,6 | 49,45,39,14,42,17,13,23,1 | 3,5,1,0,2,2,0,1,4,6,0,7,2,1, | 181 | 149 | 0.0379887227185 | | 1 | 0.978,0.925,0.915, | 0.931,0.881,0.97,1 | 0,028 |
| 147608 | ENSG00000141002 | TCF25 | chr16 | + | 89878460 | 89878631 | 89873643 | 89873859 | 89883350 | 89884063 | 147608 | 9,2,9,3,4,6,9,3,4,0,0,3 | 170,76,79,67,70,42,67,43, | 5,7,2,18,2,0,7,2,7,8,2,3,5,2 | 52,56,39,120,65,33,31,52, | 298 | 149 | 0.0380159070766 | | 1 | 0.026,0.013,0.054, | 0.046,0.059,0.025, | -0,014 |
| 199265 | ENSG00000166526 | ZNF3 | chr7 | - | 100076277 | 100076455 | 100075541 | 100075630 | 100077302 | 100077839 | 199265 | 3,3,0,2,0,0,2,2,0,1,1,2 | 8,17,16,15,7,5,2,6,6,4,2,8 | 2,2,2,2,4,2,0,5,0,3,1,1,0,1, | 7,15,4,7,14,3,5,12,2,1,11,1 | 298 | 149 | 0.0380537979659 | | 1 | 0.158,0.081,0.0,0. | 0.125,0.062,0.2,0. | -0,08 |
| 112721 | ENSG00000132294 | EFR3A | chr8 | + | 132000768 | 132000998 | 131996389 | 131996497 | 132001758 | 132001807 | 112721 | 26,3,32,11,3,3,4,6,0,1,0,1 | 35,22,44,42,12,17,16,15,6 | 8,15,11,14,2,2,6,3,3,11,9,2 | 32,12,39,8,7,12,9,15,4,37, | 298 | 149 | 0.0380710700512 | | 1 | 0.271,0.064,0.267, | 0.111,0.385,0.124, | -0,085 |
| 203040 | ENSG00000064687 | ABCA7 | chr19 | + | 1057914 | 1058157 | 1056906 | 1057429 | 1058617 | 1058747 | 203040 | 33,14,18,27,40,18,32,40,2 | 0,0,2,1,0,3,1,3,0,0,2,0 | 50,40,6,17,20,19,14,21,25, | 1,0,0,2,1,2,0,2,3,0,4,1,2,2, | 298 | 149 | 0.0381659078259 | | 1 | 1.0,1.0,0.818,0.93 | 0.962,1.0,1.0,0.81, | 0,033 |
| 227993 | ENSG00000224137 | LINC01857 | chr2 | + | 207674949 | 207675410 | 207667498 | 207667564 | 207678541 | 207679116 | 227993 | 26,21,51,40,20,7,15,16,26, | 51,22,55,29,22,7,24,4,8,9, | 28,20,2,6,7,3,1,5,4,11,0,89 | 19,16,9,0,13,2,4,3,11,29,9, | 298 | 149 | 0.038175022193 | | 1 | 0.203,0.323,0.317, | 0.424,0.385,0.1,1. | 0,081 |
| 124766 | ENSG00000273784 | | chr13 | + | 52621988 | 52622264 | 52617524 | 52617682 | 52627437 | 52627578 | 124766 | 38,28,89,65,38,28,30,45,3 | 0,3,1,7,0,0,2,9,0,0,4,0 | 47,70,39,50,54,16,34,36,3 | 2,8,2,1,3,5,0,0,2,0,0,0,0,0, | 298 | 149 | 0.0382166763873 | | 1 | 1.0,0.824,0.978,0. | 0.922,0.814,0.907, | -0,037 |
| 242111 | ENSG00000141551 | CSNK1D | chr17 | - | 82255428 | 82255558 | 82253015 | 82253244 | 82265685 | 82265796 | 242111 | 12,30,20,29,12,15,20,31,7, | 0,0,2,4,0,2,2,1,0,0,4,0 | 23,77,16,22,15,20,31,20,1 | 0,1,0,0,3,0,3,7,0,0,4,0,2,5, | 279 | 149 | 0.0382663672826 | | 1 | 1.0,1.0,0.842,0.79 | 1.0,0.976,1.0,1.0,0 | 0,03 |
| 28626 | ENSG00000143774 | GUK1 | chr1 | + | 228148370 | 228148539 | 228147620 | 228147699 | 228148664 | 228148808 | 28626 | 37,35,53,25,16,21,24,9,29, | 4,2,5,2,0,1,2,4,3,1,0,6 | 22,48,24,8,9,13,15,23,12,1 | 3,6,4,6,1,1,4,2,1,2,0,16,7,5 | 298 | 149 | 0.0382674996942 | | 1 | 0.822,0.897,0.841, | 0.786,0.8,0.75,0.4, | 0,086 |
| 227424 | ENSG00000140396 | NCOA2 | chr8 | - | 70126605 | 70127047 | 70124687 | 70124865 | 70128432 | 70128510 | 227424 | 60,60,113,73,44,59,37,59, | 1,0,3,2,1,2,3,0,0,1,0,2 | 47,52,45,38,63,30,50,58,3 | 0,4,5,8,2,0,0,0,0,1,1,1,2,2, | 298 | 149 | 0.03828867674 | | 1 | 0.968,1.0,0.95,0.9 | 1.0,0.867,0.818,0. | 0,028 |
| 123425 | ENSG00000149187 | CELF1 | chr11 | - | 47477296 | 47477425 | 47476845 | 47476956 | 47478876 | 47478940 | 123425 | 55,49,68,57,60,40,35,32,6 | 3,1,3,7,0,2,0,4,2,0,0,0 | 65,63,30,24,26,37,27,42,1 | 1,2,0,0,1,1,0,0,4,0,0,0,1,3, | 278 | 149 | 0.038336868899 | | 1 | 0.908,0.963,0.924, | 0.972,0.944,1.0,1. | -0,007 |
| 51851 | ENSG00000198793 | MTOR | chr1 | - | 11117003 | 11117086 | 11115395 | 11115468 | 11121245 | 11121368 | 51851 | 30,13,20,30,19,17,12,16,1 | 9,3,5,8,4,2,9,4,8,0,2,2 | 18,4,7,1,10,16,8,19,3,11,1 | 7,4,8,10,9,0,10,5,3,5,6,4,5, | 232 | 149 | 0.0383451359329 | | 1 | 0.682,0.736,0.72,0 | 0.623,0.391,0.36,0 | 0,139 |
| 183518 | ENSG00000163568 | AIM2 | chr1 | - | 159073237 | 159073387 | 159068567 | 159068701 | 159076632 | 159076766 | 183518 | 34,24,92,109,74,67,100,10 | 4,0,5,4,0,1,0,3,1,0,4,2 | 49,32,46,9,47,110,5,42,50, | 3,0,5,0,7,11,0,1,2,4,2,9,0,2 | 298 | 149 | 0.038347828279 | | 1 | 0.81,1.0,0.902,0.9 | 0.891,1.0,0.821,1. | -0,006 |
| 251272 | ENSG00000172661 | WASHC2C | chr10 | + | 45741976 | 45742073 | 45740190 | 45740246 | 45743389 | 45743483 | 251272 | 5,1,7,2,2,3,5,2,0,0,0,2 | 19,4,8,10,8,6,4,9,14,9,4,9 | 3,1,4,4,9,2,2,2,5,3,0,3,5,4, | 4,14,6,4,2,6,6,12,4,4,5,9,7, | 246 | 149 | 0.0383549907218 | | 1 | 0.137,0.132,0.346, | 0.312,0.041,0.288, | -0,102 |
| 103612 | ENSG00000158636 | EMSY | chr11 | + | 76454748 | 76454790 | 76453313 | 76453388 | 76458182 | 76458358 | 103612 | 11,1,12,8,7,2,4,11,3,1,8,2 | 2,2,9,16,15,2,8,7,5,2,0,5 | 12,10,2,4,12,1,10,11,9,6,1 | 7,3,1,4,7,2,4,4,1,3,3,3,4,7, | 191 | 149 | 0.0383684575573 | | 1 | 0.811,0.281,0.51,0 | 0.572,0.722,0.609, | -0,137 |
| 185001 | ENSG00000169155 | ZBTB43 | chr9 | + | 126808795 | 126808915 | 126805039 | 126805132 | 126832486 | 126838210 | 185001 | 1,14,10,18,3,4,5,6,8,6,3,8 | 9,3,7,17,10,12,5,8,0,3,4,8 | 6,12,9,3,5,1,4,9,8,2,3,9,6,1 | 13,19,17,10,12,7,15,9,18,1 | 269 | 149 | 0.0383706447229 | | 1 | 0.058,0.721,0.442, | 0.204,0.259,0.227, | 0,148 |
| 178373 | ENSG00000108523 | RNF167 | chr17 | + | 4943243 | 4943519 | 4942850 | 4942941 | 4944557 | 4944638 | 178373 | 83,19,79,65,29,24,26,18,3 | 3,0,2,3,0,4,2,5,1,0,0,1 | 39,25,28,20,23,14,23,36,1 | 4,6,0,1,2,0,1,4,0,3,3,7,0,2, | 298 | 149 | 0.0383738726405 | | 1 | 0.933,1.0,0.952,0. | 0.83,0.676,1.0,0.9 | 0,049 |
| 101787 | ENSG00000196792 | STRN3 | chr14 | - | 30913523 | 30913657 | 30912006 | 30912182 | 30935162 | 30935304 | 101787 | 16,16,26,8,10,19,4,11,6,5, | 7,1,9,1,2,0,3,8,1,3,7,0 | 11,17,7,0,8,5,5,12,2,5,6,7, | 9,11,6,5,3,4,6,1,5,1,3,0,9,8 | 283 | 149 | 0.0383766747456 | | 1 | 0.546,0.894,0.603, | 0.392,0.449,0.381, | 0,154 |
| 213433 | ENSG00000035720 | STAP1 | chr4 | + | 67588331 | 67588552 | 67583573 | 67583702 | 67590883 | 67590953 | 213433 | 2,0,9,14,4,6,6,7,6,2,4,5 | 147,67,196,143,88,62,67,5 | 7,5,4,5,1,0,5,12,0,0,6,26,8, | 96,40,67,21,74,66,26,49,4 | 298 | 149 | 0.0384125647136 | | 1 | 0.007,0.0,0.022,0. | 0.035,0.059,0.029, | -0,01 |
| 39292 | ENSG00000112282 | MED23 | chr6 | - | 131602217 | 131602381 | 131600037 | 131600162 | 131603029 | 131603204 | 39292 | 111,87,125,89,65,96,44,90 | 2,3,0,0,0,2,2,7,0,2,0,0 | 85,86,51,79,90,37,71,75,4 | 0,2,2,7,7,0,1,0,1,1,1,2,2,3, | 298 | 149 | 0.0384190938875 | | 1 | 0.965,0.935,1.0,1. | 1.0,0.956,0.927,0. | 0,023 |
| 252670 | ENSG00000161970 | RPL26 | chr17 | - | 8379914 | 8379936 | 8377678 | 8377692 | 8382142 | 8382309 | 252670 | 261,102,588,453,94,102,1 | 10,2,4,2,0,0,2,1,7,1,0,1 | 332,261,195,87,228,122,5 | 0,3,6,7,3,4,0,5,1,3,6,18,4,2 | 171 | 149 | 0.0384210057404 | | 1 | 0.958,0.978,0.992, | 1.0,0.987,0.966,0. | 0,006 |
| 226443 | ENSG00000181090 | EHMT1 | chr9 | + | 137778239 | 137778734 | 137777969 | 137778055 | 137779634 | 137779717 | 226443 | 9,9,17,3,3,10,9,20,6,2,2,15 | 83,60,101,85,67,48,80,65, | 14,23,7,9,10,0,17,8,11,6,7, | 77,62,60,77,27,32,26,50,4 | 298 | 149 | 0.0384400874224 | | 1 | 0.051,0.07,0.078,0 | 0.083,0.156,0.055, | -0,025 |
| 167686 | ENSG00000101109 | STK4 | chr20 | + | 44997137 | 44997306 | 44995089 | 44995257 | 45000391 | 45000520 | 167686 | 289,211,332,242,104,100, | 6,6,6,0,0,1,5,3,0,3,2,0 | 205,274,162,151,180,125, | 2,12,3,12,4,4,4,4,2,3,2,2,1, | 298 | 149 | 0.038450471323 | | 1 | 0.96,0.946,0.965,1 | 0.981,0.919,0.964, | 0,013 |
| 10186 | ENSG00000100425 | BRD1 | chr22 | - | 49787389 | 49787494 | 49777677 | 49777813 | 49794033 | 49794294 | 10186 | 10,5,17,11,16,5,13,7,16,3, | 0,1,2,0,1,2,1,0,5,1,0,0 | 6,5,3,6,3,1,11,9,6,8,5,6,13, | 1,2,1,0,0,2,3,3,0,0,4,2,0,2, | 254 | 149 | 0.0384562946079 | | 1 | 1.0,0.746,0.833,1. | 0.779,0.595,0.638, | 0,109 |
| 143593 | ENSG00000185813 | PCYT2 | chr17 | - | 81906986 | 81907257 | 81906759 | 81906898 | 81907553 | 81907598 | 143593 | 2,0,6,2,1,4,3,2,2,0,2,1 | 7,2,28,20,13,8,13,12,19,1, | 3,3,0,0,0,3,3,0,0,3,3,0,4,1, | 6,5,3,5,4,1,2,20,8,15,5,15,1 | 298 | 149 | 0.0384752699769 | | 1 | 0.125,0.0,0.097,0. | 0.2,0.231,0.0,0.0,0 | -0,039 |
| 116428 | ENSG00000060237 | WNK1 | chr12 | + | 871264 | 871348 | 862082 | 862270 | 878214 | 878361 | 116428 | 39,17,45,52,15,31,23,18,3 | 7,10,12,10,8,14,4,15,10,5, | 17,11,14,33,6,15,24,29,17, | 19,22,13,13,8,6,9,15,13,7,1 | 233 | 149 | 0.038491788208 | | 1 | 0.781,0.521,0.706, | 0.364,0.242,0.408, | 0,108 |
| 170298 | ENSG00000114416 | FXR1 | chr3 | + | 180970266 | 180970358 | 180968050 | 180968254 | 180976121 | 180976418 | 170298 | 25,28,59,36,16,16,11,17,2 | 0,0,7,1,1,0,0,0,0,0,0,0 | 30,31,21,16,27,20,12,14,1 | 0,2,2,0,3,0,0,0,0,3,5,2,0,1, | 241 | 149 | 0.0385016106354 | | 1 | 1.0,1.0,0.839,0.95 | 1.0,0.906,0.867,1. | 0,057 |

| | | | | | | | | | | | | | | | | | | | | | | | |
|---|---|---|---|---|---|---|---|---|---|---|---|---|---|---|---|---|---|---|---|---|---|---|---|
| 124274 | ENSG00000110713 | NUP98 | chr11 | - | 3700609 | 3700839 | 3698928 | 3699348 | 3702462 | 3702892 | 124274 | 65,56,118,87,48,74,54,64,3 | 8,12,10,4,4,1,3,5,2,3,0,3 | 95,112,59,76,65,33,43,59,3 | 6,2,5,15,0,1,2,4,6,2,3,2,1,2 | 298 | 149 | 0.0385040539996 | | 1 | 0.802,0.7,0.855,0.9 | 0.888,0.966,0.855, | -0,032 |
| 177996 | ENSG00000142687 | KIAA0319L | chr1 | - | 35435304 | 35435637 | 35434397 | 35435081 | 35441046 | 35443028 | 177996 | 3,0,8,4,3,6,4,8,0,3,2,3 | 9,8,43,30,14,16,19,29,13,5 | 7,3,5,5,4,3,9,7,3,7,7,5,1,5,6 | 29,18,18,5,6,6,1,31,9,20,17 | 298 | 149 | 0.0385042210674 | | 1 | 0.143,0.0,0.085,0.0 | 0.108,0.077,0.122, | -0,089 |
| 240053 | ENSG00000106771 | TMEM245 | chr9 | - | 109090910 | 109091155 | 109087172 | 109087342 | 109093474 | 109093591 | 240053 | 26,15,11,9,17,10,7,29,15,1 | 1,2,3,0,0,0,0,2,0,0,0,0 | 6,11,4,12,6,7,16,39,4,11,12 | 2,1,2,0,4,0,2,1,3,0,2,0,0,1,0 | 298 | 149 | 0.0385170336816 | | 1 | 0.929,0.789,0.647, | 0.6,0.846,0.5,1.0,0 | 0,12 |
| 100443 | ENSG00000223482 | NUTM2A-AS1 | chr10 | - | 87307956 | 87308060 | 87287958 | 87288495 | 87326629 | 87326672 | 100443 | 4,9,11,3,8,16,0,3,3,0,3,2 | 34,27,38,18,17,27,16,21,9, | 4,5,9,1,7,7,1,2,5,13,4,11,5, | 17,29,14,14,12,10,12,18,7, | 253 | 149 | 0.038611073531 | | 1 | 0.065,0.164,0.146, | 0.122,0.092,0.275, | -0,077 |
| 227427 | ENSG00000140396 | NCOA2 | chr8 | - | 70126887 | 70127047 | 70124687 | 70124865 | 70128432 | 70128510 | 227427 | 56,59,109,69,39,56,34,55,3 | 1,0,3,2,1,2,3,0,0,1,0,2 | 47,50,45,38,61,30,46,47,32 | 0,4,5,8,2,0,0,0,0,1,1,1,2,2,3 | 298 | 149 | 0.0386290860869 | | 1 | 0.966,1.0,0.948,0.9 | 1.0,0.862,0.818,0.7 | 0,026 |
| 32872 | ENSG00000102445 | RUBCNL | chr13 | - | 46377889 | 46378005 | 46371940 | 46372597 | 46387676 | 46387831 | 32872 | 61,24,178,128,74,84,98,13 | 26,13,33,18,7,16,10,27,12, | 99,120,54,56,53,53,61,80,8 | 24,44,12,27,24,7,19,24,6,4 | 265 | 149 | 0.0386700688297 | | 1 | 0.569,0.509,0.752, | 0.699,0.605,0.717, | 0,081 |
| 32624 | ENSG00000158290 | CUL4B | chrX | - | 120554907 | 120554953 | 120547135 | 120547239 | 120557923 | 120558039 | 32624 | 10,2,4,2,0,3,0,6,0,0,1,2 | 31,35,94,42,24,32,15,41,22 | 2,5,5,0,3,1,0,0,2,0,0,3,3,1,1 | 55,60,28,28,41,26,25,31,21 | 195 | 149 | 0.0386726481561 | | 1 | 0.198,0.042,0.031, | 0.027,0.06,0.12,0.0 | 0,016 |
| 10437 | ENSG00000173812 | EIF1 | chr17 | + | 41689777 | 41689941 | 41688884 | 41689069 | 41690087 | 41690189 | 10437 | 323,204,680,630,161,303,2 | 3,0,7,7,2,2,1,2,0,1,1,2 | 585,1131,239,541,447,343 | 5,13,3,9,2,6,1,5,6,7,2,3,4,0 | 298 | 149 | 0.0386755814878 | | 1 | 0.982,1.0,0.98,0.97 | 0.983,0.978,0.976, | 0,006 |
| 151017 | ENSG00000170776 | AKAP13 | chr15 | + | 85577762 | 85577871 | 85575130 | 85575329 | 85578929 | 85582107 | 151017 | 71,54,88,27,21,54,23,55,26 | 303,208,227,146,147,150,1 | 40,82,39,95,118,38,17,26,4 | 194,205,147,158,186,128,5 | 258 | 149 | 0.0386939052819 | | 1 | 0.119,0.13,0.183,0 | 0.106,0.188,0.133, | -0,019 |
| 201085 | ENSG00000132680 | KHDC4 | chr1 | - | 155926675 | 155926836 | 155925631 | 155925843 | 155927103 | 155927156 | 201085 | 27,13,41,43,21,17,14,30,27 | 13,2,10,12,2,3,0,2,3,2,1,3 | 25,17,6,8,15,12,2,15,22,39 | 19,4,4,1,3,2,1,3,7,2,8,5,4,1 | 298 | 149 | 0.0386966575035 | | 1 | 0.509,0.765,0.672, | 0.397,0.68,0.429,0 | 0,142 |
| 88082 | ENSG00000188647 | PTAR1 | chr9 | - | 69750780 | 69750950 | 69741515 | 69741858 | 69759852 | 69759955 | 88082 | 61,19,116,72,81,72,67,67,3 | 7,2,7,12,7,5,4,7,0,0,2,6 | 80,88,54,55,62,29,25,59,52 | 13,14,8,8,15,4,1,4,0,9,5,15 | 298 | 149 | 0.0387153971178 | | 1 | 0.813,0.826,0.892, | 0.755,0.759,0.771, | 0,054 |
| 163600 | ENSG00000105875 | WDR91 | chr7 | - | 135188432 | 135188604 | 135186971 | 135187169 | 135189343 | 135189452 | 163600 | 31,18,50,50,20,26,12,21,18 | 3,1,15,12,2,1,3,9,2,0,1,8 | 29,52,14,4,14,3,10,34,9,27 | 13,4,0,16,1,0,1,0,0,0,0,1,10 | 298 | 149 | 0.0387265524058 | | 1 | 0.838,0.9,0.625,0.6 | 0.527,0.867,1.0,0.1 | -0,04 |
| 31372 | ENSG00000171206 | TRIM8 | chr10 | + | 102654652 | 102654748 | 102644478 | 102645187 | 102655079 | 102655313 | 31372 | 77,55,138,76,42,37,45,36,4 | 2,0,5,0,1,2,0,7,3,0,0,0 | 44,47,40,56,42,22,24,47,28 | 7,0,3,3,2,1,4,1,2,1,3,2,2,2,3 | 245 | 149 | 0.0387339544707 | | 1 | 0.959,1.0,0.944,1.0 | 0.793,1.0,0.89,0.91 | 0,036 |
| 71790 | ENSG00000124256 | ZBP1 | chr20 | - | 57616243 | 57616468 | 57615511 | 57615580 | 57620261 | 57620394 | 71790 | 12,10,27,32,20,16,10,15,13 | 13,10,15,10,18,21,5,15,15, | 16,33,4,4,37,21,7,3,30,27,4 | 13,11,6,4,5,9,12,9,3,2,3,44 | 298 | 149 | 0.0387485929038 | | 1 | 0.316,0.333,0.474, | 0.381,0.6,0.25,0.33 | -0,113 |
| 58151 | ENSG00000076770 | MBNL3 | chrX | - | 132439434 | 132440314 | 132406227 | 132406392 | 132488850 | 132489038 | 58151 | 22,19,17,18,9,12,10,19,8,1 | 7,6,7,21,21,18,13,13,7,12,2 | 12,14,12,13,12,7,4,8,5,7,1, | 15,20,8,3,6,6,8,18,5,13,19, | 298 | 149 | 0.0387940789601 | | 1 | 0.611,0.613,0.548, | 0.286,0.259,0.429, | 0,084 |
| 220440 | ENSG00000196367 | TRRAP | chr7 | + | 98950913 | 98951004 | 98950063 | 98950262 | 98953166 | 98953433 | 220440 | 29,45,24,6,52,69,52,58,79, | 0,3,0,1,1,7,2,8,2,0,0,0 | 7,19,39,37,19,21,24,37,9,3 | 0,4,0,3,0,0,4,2,0,0,0,2,1,2,0 | 240 | 149 | 0.0388612881549 | | 1 | 1.0,0.903,1.0,0.788 | 1.0,0.747,1.0,0.884 | -0,002 |
| 177108 | ENSG00000179912 | R3HDM2 | chr12 | - | 57297327 | 57297356 | 57296410 | 57296551 | 57298089 | 57298168 | 177108 | 9,1,14,2,2,2,4,0,1,0,0,1 | 11,1,10,3,2,5,0,5,0,1,3,5 | 5,6,3,6,5,3,3,2,1,7,0,4,15,1 | 6,6,2,3,0,7,0,8,3,5,6,0,5,4,0 | 178 | 149 | 0.0388668793361 | | 1 | 0.406,0.456,0.54,0 | 0.411,0.456,0.557, | -0,158 |
| 75030 | ENSG00000143393 | PI4KB | chr1 | - | 151310210 | 151310255 | 151307573 | 151307801 | 151315572 | 151316509 | 75030 | 28,17,32,31,13,12,10,17,10 | 32,39,72,56,26,30,25,23,36 | 13,16,8,3,3,9,12,13,2,12,21 | 54,46,34,31,31,19,15,38,17 | 194 | 149 | 0.0389139411498 | | 1 | 0.402,0.251,0.254, | 0.156,0.211,0.153, | 0,043 |
| 205899 | ENSG00000128815 | WDFY4 | chr10 | + | 48817280 | 48817409 | 48813956 | 48814082 | 48820233 | 48820437 | 205899 | 74,65,93,60,73,68,41,46,59 | 0,1,2,2,1,6,0,1,0,0,1,4 | 74,71,19,29,17,30,54,71,37 | 2,9,0,5,0,0,1,2,1,0,6,3,0,1,4 | 278 | 149 | 0.0389444459536 | | 1 | 1.0,0.972,0.961,0.9 | 0.952,0.809,1.0,0.7 | 0,02 |
| 8353 | ENSG00000141627 | DYM | chr18 | - | 49417934 | 49418057 | 49391619 | 49391645 | 49430254 | 49430447 | 8353 | 2,3,2,0,3,4,0,5,0,0,0,0 | 52,23,84,39,32,32,28,34,18 | 7,2,4,5,2,0,2,5,0,3,1,1,0,5,5 | 38,44,30,57,40,19,26,32,36 | 272 | 149 | 0.0389467443218 | | 1 | 0.021,0.067,0.013, | 0.092,0.024,0.068, | -0,017 |
| 28517 | ENSG00000170234 | PWWP2A | chr5 | - | 160080650 | 160080770 | 160075881 | 160078168 | 160093100 | 160094065 | 28517 | 12,4,25,4,9,6,4,10,13,6,2,2 | 0,0,2,6,1,0,0,2,3,0,0,2 | 9,8,10,2,3,6,3,3,2,2,6,6,10, | 2,2,0,0,4,1,0,1,0,2,2,0,2,4,5 | 269 | 149 | 0.0390197856364 | | 1 | 1.0,1.0,0.874,0.27, | 0.714,0.689,1.0,1.0 | 0,132 |
| 226889 | ENSG00000136451 | VEZF1 | chr17 | - | 57985253 | 57985465 | 57982698 | 57983393 | 57988078 | 57988259 | 226889 | 4,2,3,3,0,2,1,3,4,0,3,1 | 7,7,8,11,5,13,7,9,7,1,4,5 | 2,1,1,2,2,0,0,4,0,1,7,0,2,2,0 | 17,4,9,13,5,2,8,10,2,5,9,8,6 | 298 | 149 | 0.0390207596807 | | 1 | 0.222,0.125,0.158, | 0.056,0.111,0.053, | 0,049 |
| 151294 | ENSG00000167491 | GATAD2A | chr19 | + | 19456921 | 19457229 | 19436130 | 19436219 | 19465339 | 19465614 | 151294 | 1,6,3,2,0,0,1,1,0,0,1,1 | 11,7,10,7,6,1,3,4,10,4,1,2 | 1,0,0,1,2,4,0,2,0,2,0,2,4,7,1 | 4,9,10,8,5,1,2,2,2,4,6,3,6,3 | 298 | 149 | 0.0390558557889 | | 1 | 0.043,0.3,0.13,0.12 | 0.111,0.0,0.0,0.059 | -0,066 |
| 129415 | ENSG00000091039 | OSBPL8 | chr12 | - | 76459858 | 76459895 | 76450900 | 76450979 | 76487509 | 76487618 | 129415 | 3,2,7,6,3,4,1,2,10,1,1,1 | 29,11,14,16,10,16,0,1,16,5 | 3,3,2,2,2,5,5,2,2,14,6,8,15, | 13,15,7,7,3,8,0,8,8,12,9,9,1 | 186 | 149 | 0.0391506617597 | | 1 | 0.077,0.127,0.286, | 0.156,0.138,0.186, | -0,09 |
| 105061 | ENSG00000078589 | P2RY10 | chrX | + | 78952192 | 78952335 | 78945420 | 78945495 | 78960507 | 78961080 | 105061 | 318,156,341,192,102,103,1 | 152,92,179,96,49,38,68,52 | 245,208,222,117,96,66,90, | 87,78,106,43,60,24,28,65,2 | 292 | 149 | 0.0391512275984 | | 1 | 0.516,0.464,0.493, | 0.59,0.576,0.517,0 | -0,052 |
| 215935 | ENSG00000091009 | RBM27 | chr5 | + | 146261509 | 146261795 | 146260744 | 146260898 | 146263490 | 146263631 | 215935 | 144,97,105,50,28,37,51,59 | 2,11,16,8,1,6,2,7,5,2,3,2 | 73,100,67,64,78,40,48,50,3 | 16,22,3,8,2,2,4,5,8,9,5,8,15 | 298 | 149 | 0.0391607906673 | | 1 | 0.973,0.815,0.766, | 0.695,0.694,0.918, | 0,044 |
| 158261 | ENSG00000084093 | REST | chr4 | + | 56919786 | 56919874 | 56910629 | 56911536 | 56929840 | 56935844 | 158261 | 10,11,90,43,24,13,14,23,6, | 3,5,3,1,2,1,3,4,1,0,0,2 | 32,40,24,6,18,14,11,23,10, | 4,4,2,2,4,8,5,2,3,5,2,7,0,6,6 | 237 | 149 | 0.0391649719076 | | 1 | 0.677,0.58,0.95,0.9 | 0.834,0.863,0.883, | 0,059 |
| 159124 | ENSG00000012660 | ELOVL5 | chr6 | - | 53292057 | 53292153 | 53291775 | 53291963 | 53295641 | 53295707 | 159124 | 40,2,30,33,7,3,9,9,12,15,1, | 226,150,155,141,80,49,84, | 9,9,12,12,6,7,4,5,3,6,1,16,4 | 81,104,140,19,87,44,19,48 | 245 | 149 | 0.0391719583036 | | 1 | 0.097,0.008,0.105, | 0.063,0.05,0.05,0.1 | -0,002 |
| 63341 | ENSG00000290842 | CROCCP3 | chr1 | - | 16491009 | 16491316 | 16490864 | 16490926 | 16499202 | 16499257 | 63341 | 3,0,4,2,1,2,3,6,7,0,1,3 | 14,1,2,3,3,1,2,2,3,4,0,0 | 4,13,0,1,1,2,2,8,2,4,6,2,0,3 | 1,1,1,2,1,1,0,5,0,3,2,1,3,0,5 | 298 | 149 | 0.0392082893412 | | 1 | 0.097,0.0,0.5,0.25, | 0.667,0.867,0.0,0.1 | -0,142 |
| 208578 | ENSG00000196504 | PRPF40A | chr2 | - | 152693157 | 152693232 | 152693037 | 152693068 | 152694541 | 152694622 | 208578 | 8,0,4,4,1,1,3,3,0,0,1,0 | 36,36,77,68,40,37,36,40,22 | 7,3,2,6,1,0,3,12,2,1,6,0,1,5 | 53,79,32,27,45,24,31,52,16 | 224 | 149 | 0.0392251256883 | | 1 | 0.129,0.0,0.033,0.0 | 0.081,0.025,0.04,0 | -0,021 |
| 59118 | ENSG00000164615 | CAMLG | chr5 | + | 134743986 | 134744052 | 134738582 | 134738792 | 134750758 | 134751377 | 59118 | 18,7,48,19,10,9,6,20,11,1,3 | 2,2,3,0,1,1,9,11,1,1,2,2 | 14,18,4,7,12,5,6,16,6,11,13 | 4,4,5,5,5,0,0,3,0,2,9,11,3,1 | 215 | 149 | 0.0392313155026 | | 1 | 0.862,0.708,0.917, | 0.708,0.757,0.357, | 0,058 |
| 109668 | ENSG00000221914 | PPP2R2A | chr8 | + | 26363720 | 26363890 | 26362683 | 26362848 | 26366314 | 26366406 | 109668 | 29,8,69,44,24,27,25,34,25, | 3,1,5,3,1,3,5,3,0,0,2,2 | 48,39,20,27,22,12,28,23,20 | 6,12,2,7,2,1,1,3,1,5,0,4,1,3 | 298 | 149 | 0.0392514473806 | | 1 | 0.829,0.8,0.873,0.8 | 0.8,0.619,0.833,0.6 | 0,047 |
| 14868 | ENSG00000040199 | PHLPP2 | chr16 | - | 71702597 | 71702731 | 71690518 | 71690709 | 71714511 | 71714801 | 14868 | 20,12,13,21,9,5,8,9,10,8,3, | 1,1,2,2,0,4,1,5,0,1,0,0 | 15,13,13,12,14,11,7,20,9,8 | 0,3,0,0,2,0,3,0,1,0,4,0,1,3,0 | 283 | 149 | 0.0392537316929 | | 1 | 0.913,0.863,0.774, | 1.0,0.695,1.0,1.0,0 | 0,006 |
| 216174 | ENSG00000073614 | KDM5A | chr12 | - | 309519 | 309571 | 307453 | 308005 | 309802 | 309964 | 216174 | 1,1,7,4,3,4,3,1,1,0,2,7 | 75,76,92,38,31,27,42,57,66 | 6,3,8,3,4,2,2,3,5,5,9,6,6,1,8 | 83,83,51,81,43,33,38,34,19 | 201 | 149 | 0.0392731494523 | | 1 | 0.01,0.01,0.053,0.0 | 0.051,0.026,0.104, | -0,017 |
| 105480 | ENSG00000147586 | MRPS28 | chr8 | - | 80002998 | 80003120 | 79918939 | 79919148 | 80030035 | 80030289 | 105480 | 11,17,36,34,10,4,3,18,16,1 | 0,0,3,2,0,0,0,1,0,0,0,0 | 14,19,10,2,11,9,9,7,3,8,10, | 1,0,1,2,4,1,0,1,0,0,0,4,0,0,2 | 271 | 149 | 0.0392900544916 | | 1 | 1.0,1.0,0.868,0.903 | 0.885,1.0,0.846,0.3 | 0,132 |
| 117079 | ENSG00000125944 | HNRNPR | chr1 | - | 23333517 | 23333631 | 23323555 | 23323732 | 23337753 | 23337861 | 117079 | 215,143,287,199,115,114,7 | 5,5,26,12,3,3,9,12,5,2,1,14 | 146,193,119,85,88,98,70,1 | 4,12,6,8,6,2,2,9,0,4,5,10,2, | 263 | 149 | 0.0392969959992 | | 1 | 0.961,0.942,0.862, | 0.954,0.901,0.918, | -0,023 |
| 232919 | ENSG00000147140 | NONO | chrX | + | 71294017 | 71294089 | 71291778 | 71291972 | 71294226 | 71295154 | 232919 | 7,3,14,8,5,8,6,10,8,0,0,2 | 185,89,272,220,141,97,109 | 10,15,1,5,8,1,5,1,6,3,1,2,1, | 219,165,118,82,112,77,75, | 221 | 149 | 0.0392972069251 | | 1 | 0.025,0.022,0.034, | 0.03,0.058,0.006,0 | 0,004 |
| 190709 | ENSG00000132155 | RAF1 | chr3 | - | 12606200 | 12606320 | 12604135 | 12604289 | 12608765 | 12609335 | 190709 | 49,30,76,46,44,52,44,42,40 | 45,20,36,26,10,18,20,11,20 | 43,45,50,35,25,27,34,67,22 | 5,20,20,23,18,1,6,13,3,11,2 | 269 | 149 | 0.0393272675984 | | 1 | 0.376,0.454,0.539, | 0.826,0.555,0.581, | -0,089 |
| 121816 | ENSG00000101972 | STAG2 | chrX | + | 124021366 | 124021431 | 123961705 | 123961856 | 124022530 | 124022671 | 121816 | 20,16,36,18,18,20,10,21,6, | 17,13,12,22,13,24,10,13,17 | 23,42,23,31,8,6,7,25,5,18,1 | 12,5,5,6,8,6,3,7,3,3,13,13,1 | 214 | 149 | 0.0394084689329 | | 1 | 0.45,0.461,0.676,0 | 0.572,0.854,0.762, | -0,088 |
| 163537 | ENSG00000059378 | PARP12 | chr7 | - | 140027275 | 140027322 | 140026195 | 140026348 | 140028612 | 140029142 | 163537 | 8,4,9,6,6,4,3,4,12,2,4,4 | 1,2,6,2,0,3,0,0,0,0,0,0 | 5,7,9,6,5,3,11,22,4,4,5,8,6, | 0,0,2,0,0,0,0,0,1,0,0,0,0,0,0 | 196 | 149 | 0.039480914077 | | 1 | 0.859,0.603,0.533, | 1.0,1.0,0.774,1.0,1 | -0,121 |
| 60265 | ENSG00000143398 | PIP5K1A | chr1 | + | 151242437 | 151242567 | 151239954 | 151240039 | 151246919 | 151246965 | 60265 | 14,14,36,35,16,24,23,31,9, | 1,3,1,7,0,0,2,0,1,2,0,2 | 40,51,18,12,25,10,6,19,13, | 0,0,1,0,0,0,0,1,0,2,4,0,2,2,0 | 279 | 149 | 0.0395038644677 | | 1 | 0.882,0.714,0.951, | 1.0,1.0,0.906,1.0,1 | -0,053 |
| 1798 | ENSG00000100030 | MAPK1 | chr22 | - | 21788693 | 21788808 | 21788256 | 21788388 | 21799011 | 21799128 | 1798 | 113,81,198,194,101,114,82 | 3,3,2,1,1,3,8,3,0,1,0,0 | 153,110,75,47,60,70,56,10 | 9,2,3,4,7,3,0,1,1,6,6,3,0,2,0 | 264 | 149 | 0.0395377501394 | | 1 | 0.955,0.938,0.982, | 0.906,0.969,0.934, | 0,018 |
| 48442 | ENSG00000108523 | RNF167 | chr17 | + | 4943425 | 4943519 | 4942850 | 4942941 | 4944557 | 4944638 | 48442 | 74,17,72,62,28,19,21,15,35 | 3,0,2,3,0,4,2,5,1,0,0,1 | 36,21,29,19,20,14,21,32,13 | 4,6,0,1,2,0,1,4,0,3,3,7,0,2,2 | 243 | 149 | 0.0396073925839 | | 1 | 0.938,1.0,0.957,0.9 | 0.847,0.682,1.0,0.9 | 0,042 |
| 196352 | ENSG00000153187 | HNRNPU | chr1 | - | 244858201 | 244858274 | 244857597 | 244857717 | 244858728 | 244858835 | 196352 | 105,57,264,164,59,66,51,9 | 11,0,4,4,2,6,3,3,5,0,2,0 | 153,136,65,88,58,71,43,56 | 8,1,4,12,2,0,0,9,0,9,7,5,6,0 | 222 | 149 | 0.0396579841553 | | 1 | 0.865,1.0,0.978,0.9 | 0.928,0.989,0.916, | 0,019 |
| 143106 | ENSG00000161939 | RNASEK-C17orf | chr17 | + | 7016909 | 7017164 | 7016494 | 7016685 | 7017256 | 7017520 | 143106 | 26,8,52,57,37,38,29,31,32, | 27,8,15,8,19,10,7,6,14,6,2, | 35,28,8,5,32,26,18,53,14,2 | 8,3,4,21,1,3,10,7,0,1,7,11,2 | 298 | 149 | 0.0396602298635 | | 1 | 0.325,0.333,0.634, | 0.686,0.824,0.5,0.1 | -0,103 |
| 210211 | ENSG00000169499 | PLEKHA2 | chr8 | + | 38952183 | 38952312 | 38950849 | 38950990 | 38952635 | 38952704 | 210211 | 89,27,427,262,104,94,91,1 | 24,2,26,16,9,13,3,0,7,4,6,5 | 260,143,69,48,42,70,73,11 | 13,14,14,19,5,4,7,7,2,2,26, | 278 | 149 | 0.0396618420953 | | 1 | 0.665,0.879,0.898, | 0.915,0.846,0.725, | 0,047 |
| 221408 | ENSG00000072121 | ZFYVE26 | chr14 | - | 67784333 | 67784494 | 67782779 | 67783525 | 67785058 | 67785277 | 221408 | 6,5,19,10,13,10,14,12,5,7,3 | 0,1,0,0,0,3,0,0,3,0,0,0 | 6,11,8,3,10,3,5,10,13,19,5, | 2,0,0,3,0,2,0,5,3,0,0,1,0,4,0 | 298 | 149 | 0.0396621554408 | | 1 | 1.0,0.714,1.0,1.0,1 | 0.6,1.0,1.0,0.333,1 | 0,102 |
| 255993 | ENSG00000145901 | TNIP1 | chr5 | - | 151033607 | 151033839 | 151032286 | 151032383 | 151035001 | 151035067 | 255993 | 16,9,27,13,21,12,17,20,9,2 | 21,7,11,14,12,12,6,11,18,8 | 10,8,5,3,5,2,2,22,2,10,12,1 | 6,17,14,6,4,0,2,16,4,10,6,1 | 298 | 149 | 0.0397067538664 | | 1 | 0.276,0.391,0.551, | 0.455,0.19,0.152,0 | 0,064 |
| 4734 | ENSG00000213020 | ZNF611 | chr19 | - | 52711206 | 52711304 | 52706655 | 52706864 | 52714014 | 52714141 | 4734 | 3,0,6,2,3,0,2,2,1,0,0,0 | 13,4,26,16,2,7,3,16,4,2,2,5 | 2,2,2,0,1,1,0,0,0,0,1,0,0,0,0 | 17,32,8,7,8,6,2,6,1,8,8,8,9, | 247 | 149 | 0.0397655503854 | | 1 | 0.122,0.0,0.122,0.0 | 0.066,0.036,0.131, | 0,073 |
| 195158 | ENSG00000171791 | BCL2 | chr18 | - | 63280886 | 63281214 | 63242001 | 63242110 | 63318081 | 63318510 | 195158 | 2,0,4,6,7,5,5,6,11,2,3,3 | 1,1,5,8,3,6,3,10,0,0,4,6 | 8,0,1,1,0,4,0,13,3,1,9,7,0,3 | 5,9,2,9,2,0,3,6,2,3,15,9,2,0 | 298 | 149 | 0.0398013398961 | | 1 | 0.5,0.0,0.286,0.273 | 0.444,0.0,0.2,0.053 | 0,124 |
| 161561 | ENSG00000196670 | ZFP62 | chr5 | - | 180860527 | 180860654 | 180850914 | 180851493 | 180861218 | 180861285 | 161561 | 7,10,15,7,0,3,5,14,8,4,2,7 | 0,2,7,6,3,5,3,3,6,6,0,3 | 16,1,6,5,1,2,4,9,7,8,13,7,6, | 3,3,5,3,0,2,0,2,0,3,1,1,1,1,2 | 276 | 149 | 0.0398181649881 | | 1 | 1.0,0.73,0.536,0.38 | 0.742,0.153,0.393, | -0,147 |
| 176555 | ENSG00000163611 | SPICE1 | chr3 | - | 113450446 | 113450516 | 113448037 | 113448140 | 113453465 | 113453950 | 176555 | 13,2,13,10,3,4,3,11,3,1,2,5 | 2,5,6,4,10,5,0,3,2,1,1,0 | 8,7,8,9,2,2,11,9,8,8,2,1,5,7 | 1,2,0,2,2,1,4,1,3,2,1,2,0,3,0 | 219 | 149 | 0.0398255147013 | | 1 | 0.816,0.214,0.596, | 0.845,0.704,1.0,0.7 | -0,082 |
| 173828 | ENSG00000175455 | CCDC14 | chr3 | - | 123949383 | 123949850 | 123948895 | 123949132 | 123955842 | 123958900 | 173828 | 10,8,10,6,7,3,6,10,7,3,3,11 | 98,34,91,37,54,83,63,94,74 | 8,5,9,0,0,0,8,8,4,8,4,12,12, | 48,52,27,13,25,12,25,59,16 | 298 | 149 | 0.0398326786556 | | 1 | 0.049,0.105,0.052, | 0.077,0.046,0.143, | -0,017 |
| 48742 | ENSG00000160679 | CHTOP | chr1 | + | 153642245 | 153642429 | 153638083 | 153638448 | 153645063 | 153646302 | 48742 | 29,17,40,24,24,20,10,13,10 | 0,4,0,1,0,3,4,1,1,0,1,1 | 28,28,32,10,9,10,4,23,10,1 | 0,4,2,2,2,0,0,0,1,1,3,3,0,7,0 | 298 | 149 | 0.0398834551747 | | 1 | 1.0,0.68,1.0,0.923, | 1.0,0.778,0.889,0.7 | -0,015 |
| 153000 | ENSG00000071537 | SEL1L | chr14 | - | 81484224 | 81484421 | 81479277 | 81479740 | 81485671 | 81485746 | 153000 | 29,20,54,44,23,20,10,29,11 | 4,2,2,1,2,0,1,2,0,2,0,0 | 33,40,17,22,30,7,33,30,5,3 | 1,1,0,4,0,2,0,1,0,0,0,0,0,0,0 | 298 | 149 | 0.039890776536 | | 1 | 0.784,0.833,0.931, | 0.943,0.952,1.0,0.7 | -0,053 |
| 97745 | ENSG00000196511 | TPK1 | chr7 | - | 144667725 | 144667846 | 144648806 | 144648879 | 144682908 | 144682978 | 97745 | 6,2,1,2,3,3,5,8,0,0,6,0 | 16,11,27,15,18,21,20,26,14 | 7,3,2,0,7,6,2,2,4,4,3,2,4,3,6 | 10,15,9,14,28,14,5,4,17,16 | 270 | 149 | 0.0399196098876 | | 1 | 0.171,0.091,0.02,0 | 0.279,0.099,0.109, | -0,069 |
| 239075 | ENSG00000066117 | SMARCD1 | chr12 | + | 50096814 | 50096972 | 50094505 | 50094572 | 50098713 | 50098815 | 239075 | 39,55,98,56,44,29,36,22,20 | 2,0,0,4,1,2,2,0,6,1,0,2 | 52,56,49,57,79,29,22,33,36 | 2,0,2,0,0,2,0,3,0,0,0,2,3,0,0 | 298 | 149 | 0.0399594702439 | | 1 | 0.907,1.0,1.0,0.875 | 0.929,1.0,0.925,1.0 | -0,052 |
| 15726 | ENSG00000197912 | SPG7 | chr16 | + | 89545871 | 89545979 | 89544647 | 89544772 | 89546657 | 89546760 | 15726 | 0,1,10,2,1,0,6,3,0,1,0,1 | 36,17,74,42,40,51,18,19,23 | 0,0,0,1,1,2,0,3,0,0,2,3,0,0,0 | 45,38,15,18,62,22,32,38,46 | 257 | 149 | 0.0399602853479 | | 1 | 0.0,0.033,0.073,0.0 | 0.0,0.0,0.0,0.031,0 | 0,025 |
| 232891 | ENSG00000196924 | FLNA | chrX | - | 154352552 | 154352655 | 154352180 | 154352447 | 154352771 | 154352924 | 232891 | 190,133,125,117,158,108,1 | 14,16,7,5,1,4,11,14,3,6,2,8 | 130,170,59,55,114,83,79,1 | 7,12,9,14,25,2,1,11,8,7,4,5 | 252 | 149 | 0.0399637278204 | | 1 | 0.889,0.831,0.913, | 0.917,0.893,0.795, | 0,031 |
| 14799 | ENSG00000186153 | WWOX | chr16 | + | 78114975 | 78115154 | 78109777 | 78109835 | 78164182 | 78164289 | 14799 | 27,16,22,13,11,21,13,17,8, | 0,0,0,0,0,0,0,0,0,0,0,0 | 22,16,11,12,12,9,6,16,6,14 | 0,0,0,0,0,0,0,0,0,0,6,12,0,0 | 298 | 149 | 0.0399725189451 | | 1 | 1.0,1.0,1.0,1.0,1.0, | 1.0,1.0,1.0,1.0,1.0, | 0,087 |
| 180224 | ENSG00000185104 | FAF1 | chr1 | - | 50752437 | 50752523 | 50744683 | 50744775 | 50787999 | 50788205 | 180224 | 3,2,1,3,0,3,1,1,0,5,0,0 | 28,21,52,46,42,43,21,23,19 | 3,1,2,4,3,1,4,1,1,0,4,9,0,0,4 | 52,36,30,32,17,15,30,24,22 | 235 | 149 | 0.039987057854 | | 1 | 0.064,0.057,0.012, | 0.035,0.017,0.041, | -0,008 |
| 230249 | ENSG00000147065 | MSN | chrX | + | 65727861 | 65727909 | 65716817 | 65716901 | 65729437 | 65729589 | 230249 | 64,30,130,120,67,51,72,68 | 10,4,7,9,1,0,0,3,1,7,0,0 | 100,108,54,21,50,28,35,53 | 4,1,2,6,2,1,5,3,1,1,2,1,0,1,3 | 197 | 149 | 0.039987958192 | | 1 | 0.829,0.85,0.934,0 | 0.95,0.988,0.953,0 | -0,004 |
| 187736 | ENSG00000163257 | DCAF16 | chr4 | - | 17809536 | 17809692 | 17800654 | 17804771 | 17810446 | 17810711 | 187736 | 15,6,14,4,5,9,5,20,5,7,4,6 | 1,5,5,5,3,3,4,3,13,7,1,11 | 3,9,9,16,4,2,2,10,2,6,25,22 | 3,2,2,6,3,2,1,6,0,2,6,1,1,2,1 | 298 | 149 | 0.0400101339197 | | 1 | 0.882,0.375,0.583, | 0.333,0.692,0.692, | -0,122 |
| 98413 | ENSG00000118058 | KMT2A | chr11 | + | 118484861 | 118484975 | 118484182 | 118484314 | 118488613 | 118488760 | 98413 | 69,68,123,62,54,53,27,34,6 | 5,3,14,4,4,2,1,0,3,1,5,6 | 46,24,34,67,15,34,29,19,31 | 4,16,13,22,3,0,3,1,2,3,10,4 | 263 | 149 | 0.0400213862471 | | 1 | 0.887,0.928,0.833, | 0.867,0.459,0.597, | 0,063 |
| 27864 | ENSG00000138303 | ASCC1 | chr10 | - | 72215781 | 72216109 | 72213186 | 72213331 | 72216206 | 72216275 | 27864 | 4,3,0,0,0,4,0,4,0,0,0,1 | 6,7,17,19,10,3,15,15,7,10,2 | 1,4,5,2,0,1,0,1,0,1,0,2,4,6,2 | 6,13,10,1,4,4,1,4,2,22,6,7,7 | 298 | 149 | 0.0400352308726 | | 1 | 0.25,0.176,0.0,0.0, | 0.077,0.133,0.2,0.5 | -0,032 |
| 80449 | ENSG00000290927 | | chr16 | + | 31705671 | 31705792 | 31705348 | 31705472 | 31705889 | 31705994 | 80449 | 79,48,52,50,40,35,26,46,40 | 4,2,4,7,2,1,0,1,0,0,0,4 | 35,29,30,41,23,17,28,48,13 | 6,9,0,4,4,2,0,2,6,0,4,2,0,0,4 | 270 | 149 | 0.0400428804379 | | 1 | 0.916,0.93,0.878,0 | 0.763,0.64,1.0,0.85 | 0,072 |
| 86271 | ENSG00000151687 | ANKAR | chr2 | + | 189694980 | 189695161 | 189689526 | 189689964 | 189696149 | 189696369 | 86271 | 1,0,8,4,6,0,5,6,0,0,2,2 | 15,10,26,23,18,19,15,8,10, | 4,7,2,1,1,2,4,5,2,5,6,6,1,3,6 | 16,15,22,6,11,14,9,5,8,12,9 | 298 | 149 | 0.0400652888379 | | 1 | 0.032,0.0,0.133,0.0 | 0.111,0.189,0.043, | 0,005 |
| 122252 | ENSG00000054654 | SYNE2 | chr14 | + | 64009965 | 64010116 | 64007042 | 64007222 | 64016472 | 64016631 | 122252 | 254,285,400,286,173,264,2 | 4,1,7,4,6,10,4,10,4,1,6,3 | 165,289,172,265,285,224,4 | 4,3,4,13,12,3,0,3,7,3,3,2,0, | 298 | 149 | 0.0400992609879 | | 1 | 0.969,0.993,0.966, | 0.954,0.98,0.956,0 | -0,025 |
| 165530 | ENSG00000109684 | CLNK | chr4 | - | 10554420 | 10554459 | 10542254 | 10542280 | 10558406 | 10558452 | 165530 | 2,2,4,2,1,1,3,5,0,3,1,4 | 115,68,109,68,38,54,78,74 | 0,3,5,5,5,3,2,0,5,2,5,0,0,0,7 | 31,40,76,69,43,22,33,60,28 | 188 | 149 | 0.0401223053944 | | 1 | 0.014,0.023,0.028, | 0.0,0.056,0.05,0.05 | -0,014 |
| 18731 | ENSG00000103994 | ZNF106 | chr15 | - | 42456957 | 42457158 | 42448071 | 42448705 | 42466052 | 42466114 | 18731 | 35,23,51,46,27,18,33,17,68 | 12,4,12,7,19,9,1,3,4,2,2,8 | 29,31,26,9,8,20,8,13,4,23,1 | 4,21,17,2,8,3,7,3,7,8,7,8,16 | 298 | 149 | 0.0401229271805 | | 1 | 0.593,0.742,0.68,0 | 0.784,0.425,0.433, | 0,108 |
| 133104 | ENSG00000109685 | NSD2 | chr4 | + | 1939653 | 1939778 | 1935143 | 1935262 | 1951071 | 1951180 | 133104 | 15,10,18,15,12,13,11,12,3, | 0,0,2,3,1,0,0,0,0,1,0,0 | 11,8,2,4,6,7,8,19,8,16,18,3 | 0,5,1,0,0,0,0,3,2,0,0,9,0,0,0 | 274 | 149 | 0.0401231372477 | | 1 | 1.0,1.0,0.83,0.731, | 1.0,0.465,0.521,1.0 | 0,066 |
| 176077 | ENSG00000179262 | RAD23A | chr19 | + | 12949080 | 12949191 | 12948685 | 12948813 | 12949274 | 12949408 | 176077 | 12,3,19,25,10,5,9,10,5,6,4, | 1,0,5,1,2,3,2,1,0,0,0,2 | 11,10,4,5,3,6,3,17,7,9,13,3 | 2,3,0,0,0,0,0,4,0,0,4,0,0,0,0 | 260 | 149 | 0.0401320086087 | | 1 | 0.873,1.0,0.685,0.9 | 0.759,0.656,1.0,1.0 | -0,087 |
| 160985 | ENSG00000043143 | JADE2 | chr5 | + | 134536653 | 134536844 | 134535857 | 134535915 | 134537988 | 134538083 | 160985 | 10,6,11,8,0,9,7,10,2,2,2,1 | 77,76,110,84,64,61,46,60,6 | 4,9,9,15,12,2,5,11,3,7,5,2,2 | 57,62,47,51,37,32,34,86,39 | 298 | 149 | 0.0401349821425 | | 1 | 0.061,0.038,0.048, | 0.034,0.068,0.087, | -0,017 |
| 1943 | ENSG00000100150 | DEPDC5 | chr22 | + | 31821501 | 31821637 | 31819021 | 31819225 | 31833914 | 31833980 | 1943 | 78,60,90,34,23,34,35,17,41 | 15,22,6,5,8,9,4,2,33,29,4,1 | 41,52,19,37,40,19,8,21,40, | 9,5,2,2,6,2,4,7,10,10,0,9,8, | 285 | 149 | 0.0401372033322 | | 1 | 0.731,0.588,0.887, | 0.704,0.845,0.832, | -0,082 |
| 135297 | ENSG00000198089 | SFI1 | chr22 | + | 31550977 | 31551124 | 31550253 | 31550348 | 31556941 | 31557059 | 135297 | 2,0,2,0,1,2,0,4,0,0,1,0 | 54,26,50,31,27,34,31,36,24 | 0,1,1,0,3,3,1,1,3,3,0,0,3,6,0 | 24,25,19,18,26,22,16,24,34 | 296 | 149 | 0.040153233561 | | 1 | 0.018,0.0,0.02,0.0, | 0.0,0.02,0.026,0.0, | -0,016 |
| 49195 | ENSG00000144357 | UBR3 | chr2 | + | 170079813 | 170080023 | 170073403 | 170073607 | 170080544 | 170080684 | 49195 | 24,7,42,16,17,23,4,29,5,3,9 | 0,0,1,1,1,1,1,1,0,0,2,0 | 17,32,9,40,15,12,24,17,13, | 0,1,1,3,0,0,0,4,4,0,3,1,1,1,1 | 298 | 149 | 0.0401806936693 | | 1 | 1.0,1.0,0.955,0.889 | 1.0,0.941,0.818,0.8 | 0,023 |
| 165340 | ENSG00000109762 | SNX25 | chr4 | + | 185318906 | 185318939 | 185310634 | 185310816 | 185320732 | 185320864 | 165340 | 1,2,1,0,0,1,0,1,0,0,0,0 | 52,24,64,32,24,25,13,22,19 | 6,2,0,0,3,1,0,0,5,2,2,1,0,1,2 | 64,42,10,18,32,19,6,7,33,3 | 182 | 149 | 0.0402427314484 | | 1 | 0.015,0.064,0.013, | 0.071,0.038,0.0,0.0 | -0,022 |
| 234137 | ENSG00000112305 | SMAP1 | chr6 | + | 70699274 | 70699694 | 70667882 | 70668141 | 70732377 | 70732511 | 234137 | 3,1,4,5,2,3,1,3,2,0,0,1 | 30,10,26,27,15,19,10,24,4, | 7,2,0,4,2,1,2,7,2,0,3,1,2,8,5 | 22,18,9,9,10,21,8,11,10,13 | 298 | 149 | 0.0402743388941 | | 1 | 0.048,0.048,0.071, | 0.137,0.053,0.0,0.1 | -0,035 |
| 201685 | ENSG00000168710 | AHCYL1 | chr1 | + | 110020650 | 110020851 | 110019547 | 110019626 | 110021673 | 110022045 | 201685 | 18,18,17,31,14,10,16,23,21 | 9,0,4,1,6,2,2,4,5,1,0,3 | 16,5,7,4,1,19,2,23,1,15,19, | 5,6,1,2,0,1,6,6,0,4,5,6,1,1,8 | 298 | 149 | 0.0402960808992 | | 1 | 0.5,1.0,0.68,0.939, | 0.615,0.294,0.778, | 0,143 |
| 85905 | ENSG00000115970 | THADA | chr2 | - | 43560233 | 43560385 | 43552203 | 43552339 | 43566697 | 43566821 | 85905 | 28,13,48,20,11,14,11,16,9, | 3,0,5,4,2,0,0,2,0,0,0,0 | 23,40,5,18,7,8,25,21,13,7,9 | 3,4,1,0,2,0,0,0,1,4,0,0,4,3,3 | 298 | 149 | 0.040301134506 | | 1 | 0.824,1.0,0.828,0.7 | 0.793,0.833,0.714, | 0,141 |

| | | | | | | | | | | | | | | | | | | | | | | |
|---|---|---|---|---|---|---|---|---|---|---|---|---|---|---|---|---|---|---|---|---|---|---|
| 28625 | ENSG00000143774 | GUK1 | chr1 | + | 228148370 | 228148456 | 228147465 | 228147699 | 228148664 | 228148691 | 28625 | 91,87,171,97,67,80,56,48,5 | 4,2,5,2,0,1,2,4,3,1,0,6 | 95,134,79,28,44,37,50,84,4 | 3,6,4,6,1,1,4,2,1,2,0,16,7,5 | 235 | 149 | 0.0403214832184 | 1 | 0.935,0.965,0.956, | 0.953,0.934,0.926, | 0,024 |
| 145328 | ENSG00000161395 | PGAP3 | chr17 | - | 39685190 | 39685308 | 39684601 | 39684749 | 39685921 | 39686019 | 145328 | 2,1,3,0,2,1,6,1,0,2,8,2 | 16,10,23,9,27,14,13,10,29, | 2,3,2,0,2,0,0,2,0,0,4,2,3,0,1 | 15,20,12,11,9,8,18,24,4,20 | 267 | 149 | 0.0403235501775 | 1 | 0.065,0.053,0.068, | 0.069,0.077,0.085, | 0,02 |
| 240256 | ENSG00000197329 | PELI1 | chr2 | - | 64104553 | 64104800 | 64100397 | 64100499 | 64108239 | 64108379 | 240256 | 1,1,4,2,1,0,2,3,0,0,2,1 | 54,12,11,12,7,13,22,6,6,7,0 | 1,1,6,2,5,0,3,2,0,0,2,4,1,1,1 | 5,30,45,109,26,7,15,21,8,1 | 298 | 149 | 0.0403524929176 | 1 | 0.009,0.04,0.154,0 | 0.091,0.016,0.062, | 0,053 |
| 110956 | ENSG00000170633 | RNF34 | chr12 | + | 121402741 | 121402807 | 121400118 | 121400218 | 121416158 | 121416377 | 110956 | 9,5,11,1,9,7,7,10,8,6,2,7 | 9,15,27,14,11,12,17,13,32, | 5,9,5,2,10,1,3,7,1,1,6,5,9,7 | 22,22,8,25,19,6,6,19,6,10,2 | 215 | 149 | 0.0403603840105 | 1 | 0.409,0.188,0.22,0 | 0.136,0.221,0.302, | 0,073 |
| 70601 | ENSG00000136717 | BIN1 | chr2 | - | 127050801 | 127050912 | 127050381 | 127050522 | 127059010 | 127059155 | 70601 | 14,3,28,33,15,13,9,12,6,4,1 | 2,2,3,0,1,0,2,2,2,0,0,1 | 28,19,5,1,14,14,5,13,14,13 | 0,2,0,0,4,0,0,1,3,1,2,6,1,2,5 | 260 | 149 | 0.0403729257167 | 1 | 0.8,0.462,0.842,1.0 | 1.0,0.845,1.0,1.0,0 | -0,014 |
| 247326 | ENSG00000115084 | SLC35F5 | chr2 | - | 113739961 | 113740106 | 113735776 | 113735858 | 113742691 | 113742879 | 247326 | 4,0,1,0,0,2,0,6,2,0,2,0 | 28,13,38,23,28,23,6,11,20, | 5,6,4,1,1,0,1,2,0,2,6,0,0,0,0 | 10,11,10,6,7,15,9,11,5,16,1 | 294 | 149 | 0.0404012937214 | 1 | 0.068,0.0,0.013,0.0 | 0.202,0.217,0.169, | -0,021 |
| 189618 | ENSG00000162869 | PPP1R21 | chr2 | + | 48462195 | 48462272 | 48461137 | 48461232 | 48464936 | 48464989 | 189618 | 2,1,5,0,2,2,1,1,0,0,1,0 | 19,18,15,19,17,16,5,12,2,6 | 5,8,0,3,0,0,2,0,0,0,0,0,2,5,3 | 17,23,11,4,11,11,4,13,1,9,7 | 226 | 149 | 0.0404138840269 | 1 | 0.065,0.035,0.18,0 | 0.162,0.187,0.0,0.3 | -0,04 |
| 144305 | ENSG00000161526 | SAP30BP | chr17 | + | 75705610 | 75705664 | 75704755 | 75704814 | 75706007 | 75706092 | 144305 | 0,0,9,8,2,4,5,7,6,7,2,2 | 31,42,39,28,28,24,26,33,58 | 8,1,1,2,5,1,3,7,0,2,2,1,0,2,1 | 22,41,13,24,32,5,19,33,7,2 | 203 | 149 | 0.0404218778109 | 1 | 0.0,0.0,0.145,0.173 | 0.211,0.018,0.053, | 0,02 |
| 55358 | ENSG00000078304 | PPP2R5C | chr14 | + | 101824013 | 101824169 | 101809836 | 101810036 | 101836168 | 101836701 | 55358 | 5,0,0,2,0,0,1,0,11,4,1,2 | 17,16,35,14,10,13,14,10,6, | 2,7,4,2,0,0,4,6,0,5,9,0,1,2,7 | 12,17,7,6,7,8,8,5,7,18,8,20 | 298 | 149 | 0.0404321692121 | 1 | 0.128,0.0,0.0,0.067 | 0.077,0.171,0.222, | -0,038 |
| 132498 | ENSG00000171792 | RHNO1 | chr12 | + | 2885161 | 2885534 | 2877198 | 2877282 | 2887910 | 2888057 | 132498 | 79,61,45,41,38,30,24,31,50 | 3,2,2,7,0,3,3,2,1,0,0,2 | 6,24,32,27,24,12,22,35,9,3 | 7,1,1,0,3,2,1,4,2,0,1,3,4,4,6 | 298 | 149 | 0.0404428895906 | 1 | 0.929,0.938,0.918, | 0.3,0.923,0.941,1.0 | 0,085 |
| 135730 | ENSG00000291230 | | chr19 | - | 50336309 | 50336505 | 50335058 | 50335181 | 50337016 | 50337135 | 135730 | 37,9,28,27,31,14,24,24,55, | 14,3,5,13,2,4,8,2,10,5,1,0 | 18,17,34,7,9,5,14,25,16,23 | 6,4,2,1,4,2,2,3,7,5,7,15,8,1 | 298 | 149 | 0.0404475969297 | 1 | 0.569,0.6,0.737,0.5 | 0.6,0.68,0.895,0.77 | 0,117 |
| 130279 | ENSG00000123329 | ARHGAP9 | chr12 | - | 57476870 | 57476963 | 57476589 | 57476651 | 57477155 | 57477269 | 130279 | 147,102,101,79,107,60,124 | 8,0,9,10,2,2,4,2,8,3,3,3 | 100,90,76,38,54,47,39,70,5 | 7,4,3,2,6,5,4,7,6,3,3,3,10,4 | 242 | 149 | 0.0404525516609 | 1 | 0.919,1.0,0.874,0.8 | 0.898,0.933,0.94,0 | 0,032 |
| 255985 | ENSG00000145901 | TNIP1 | chr5 | - | 151033607 | 151033792 | 151032286 | 151032383 | 151035001 | 151035067 | 255985 | 16,9,28,13,21,12,17,20,9,2 | 21,7,11,14,12,12,6,11,18,8 | 10,8,5,3,5,2,2,22,2,10,12,1 | 6,17,14,6,4,0,2,16,4,10,6,1 | 298 | 149 | 0.0404751396042 | 1 | 0.276,0.391,0.56,0 | 0.455,0.19,0.152,0 | 0,064 |
| 253770 | ENSG00000123329 | ARHGAP9 | chr12 | - | 57473602 | 57473755 | 57472263 | 57472688 | 57474041 | 57474176 | 253770 | 56,37,115,97,47,23,33,38,3 | 38,11,18,14,10,11,13,14,28 | 85,54,39,19,43,33,29,39,45 | 13,13,23,9,4,10,5,9,2,15,4, | 298 | 149 | 0.0404837450667 | 1 | 0.424,0.627,0.762, | 0.766,0.675,0.459, | -0,091 |
| 144476 | ENSG00000184381 | PLA2G6 | chr22 | - | 38122199 | 38122285 | 38120758 | 38120909 | 38123094 | 38123258 | 144476 | 0,0,2,1,2,5,0,0,0,0,1,3 | 7,18,20,17,9,9,13,9,10,7,4, | 3,2,2,0,4,2,1,0,1,4,2,4,1,3,0 | 17,17,11,10,12,8,2,15,15,1 | 235 | 149 | 0.0404846633137 | 1 | 0.0,0.0,0.06,0.036, | 0.101,0.069,0.103, | -0,02 |
| 207483 | ENSG00000068697 | LAPTM4A | chr2 | - | 20037537 | 20037636 | 20037315 | 20037438 | 20040890 | 20041011 | 207483 | 42,32,52,17,25,30,35,31,12 | 0,0,2,5,2,1,2,1,0,0,0,3 | 39,118,27,21,46,25,41,50,1 | 6,6,2,0,6,4,0,4,4,1,0,1,2,0,2 | 248 | 149 | 0.0405048158 | 1 | 1.0,1.0,0.94,0.671, | 0.796,0.922,0.89,1 | 0,021 |
| 196081 | ENSG00000229474 | PATL2 | chr15 | - | 44668341 | 44668441 | 44667105 | 44667203 | 44668979 | 44669139 | 196081 | 8,9,12,20,13,3,11,8,3,5,8,9 | 1,0,1,1,2,5,2,0,3,0,0,0 | 6,8,4,4,7,4,4,6,6,25,6,3,13, | 2,1,0,1,1,1,0,2,1,3,0,0,3,5,4 | 249 | 149 | 0.0405166746373 | 1 | 0.827,1.0,0.878,0.9 | 0.642,0.827,1.0,0.7 | 0,039 |
| 1631 | ENSG00000183765 | CHEK2 | chr22 | - | 28730455 | 28730548 | 28725242 | 28725367 | 28734402 | 28734727 | 1631 | 1,2,4,2,1,1,0,3,0,0,1,0 | 12,8,15,10,7,4,1,2,4,4,0,10 | 5,0,1,4,2,1,0,0,1,1,0,4,4,5,3 | 11,5,2,0,3,6,3,6,13,18,5,17 | 242 | 149 | 0.0405364363768 | 1 | 0.049,0.133,0.141, | 0.219,0.0,0.235,1.0 | 0,011 |
| 245627 | ENSG00000163131 | CTSS | chr1 | - | 150750005 | 150750165 | 150747776 | 150747879 | 150751780 | 150752008 | 245627 | 295,258,809,550,305,191,2 | 19,22,10,8,10,0,5,3,5,5,1,0 | 435,381,297,143,240,184,1 | 8,22,11,5,8,6,6,3,6,6,9,15,2 | 298 | 149 | 0.0405433264162 | 1 | 0.886,0.854,0.976, | 0.965,0.896,0.931, | 0,015 |
| 179643 | ENSG00000163444 | TMEM183A | chr1 | + | 203017819 | 203017919 | 203015959 | 203016140 | 203018480 | 203018561 | 179643 | 4,3,20,9,4,4,3,8,3,1,4,2 | 1,2,3,3,3,4,1,5,4,4,3,14 | 12,15,3,2,6,6,3,6,3,6,8,8,7, | 7,7,1,0,0,2,2,3,0,2,3,12,2,1 | 249 | 149 | 0.0405506878933 | 1 | 0.705,0.473,0.8,0.6 | 0.506,0.562,0.642, | -0,187 |
| 237300 | ENSG00000120910 | PPP3CC | chr8 | + | 22539468 | 22539940 | 22532920 | 22533018 | 22540614 | 22541142 | 237300 | 19,13,36,16,4,6,9,3,11,2,3, | 32,26,40,20,11,4,11,4,12,6 | 18,13,9,8,12,14,11,10,12,1 | 19,19,10,5,16,7,12,11,9,14 | 298 | 149 | 0.0405541057483 | 1 | 0.229,0.2,0.31,0.28 | 0.321,0.255,0.31,0 | -0,087 |
| 227107 | ENSG00000114503 | NCBP2 | chr3 | - | 196937171 | 196937282 | 196937022 | 196937082 | 196937509 | 196937648 | 227107 | 2,1,2,4,1,3,0,0,0,0,2,2 | 29,14,49,32,21,21,6,19,31, | 2,6,1,1,3,0,0,3,2,1,2,4,3,2,2 | 21,22,15,3,20,18,13,16,4,1 | 260 | 149 | 0.0405903455502 | 1 | 0.038,0.039,0.023, | 0.052,0.135,0.037, | -0,031 |
| 105815 | ENSG00000104870 | FCGRT | chr19 | + | 49521694 | 49521813 | 49513881 | 49514486 | 49524506 | 49524568 | 105815 | 15,3,7,3,7,7,7,6,2,2,3,1 | 51,26,66,18,37,31,56,36,29 | 2,3,3,3,3,0,5,3,4,1,7,2,5,3,1 | 35,45,24,12,5,11,50,112,6, | 268 | 149 | 0.0405989680622 | 1 | 0.141,0.06,0.056,0 | 0.031,0.036,0.065, | 0,009 |
| 73723 | ENSG00000015171 | ZMYND11 | chr10 | + | 247397 | 247466 | 246765 | 246973 | 248335 | 248531 | 73723 | 180,69,163,91,95,96,62,65 | 5,0,8,11,8,0,1,6,4,8,1,2 | 97,65,68,53,20,24,64,122,2 | 4,0,1,1,6,2,0,6,0,0,6,1,3,3,2 | 218 | 149 | 0.0406042653516 | 1 | 0.961,1.0,0.933,0.8 | 0.943,1.0,0.979,0.9 | -0,004 |
| 189229 | ENSG00000118308 | IRAG2 | chr12 | + | 25068155 | 25068267 | 25066364 | 25066512 | 25069349 | 25069431 | 189229 | 8,4,22,23,3,4,3,16,5,0,0,3 | 295,145,614,469,233,194,2 | 13,15,18,7,3,1,7,3,5,4,13,4 | 254,318,226,76,135,100,65 | 261 | 149 | 0.0406284794251 | 1 | 0.015,0.016,0.02,0 | 0.028,0.026,0.043, | -0,012 |
| 160367 | ENSG00000020633 | RUNX3 | chr1 | - | 24907258 | 24907524 | 24899510 | 24902666 | 24919239 | 24919344 | 160367 | 75,77,19,19,34,27,27,16,47 | 8,2,3,1,0,1,2,0,2,0,3,0 | 11,21,16,22,2,24,9,52,6,37 | 0,2,1,1,3,1,10,3,2,0,2,7,3,5 | 298 | 149 | 0.0406492239654 | 1 | 0.824,0.951,0.76,0 | 1.0,0.84,0.889,0.9 | 0,119 |
| 51562 | ENSG00000162894 | FCMR | chr1 | - | 206913758 | 206914094 | 206903316 | 206905147 | 206921817 | 206921867 | 51562 | 1167,447,1219,947,786,48 | 30,8,8,4,7,18,11,11,37,23,4 | 857,366,530,338,204,325,5 | 18,14,16,12,2,10,8,23,5,6,2 | 298 | 149 | 0.0406979467543 | 1 | 0.951,0.965,0.987, | 0.96,0.929,0.943,0 | 0,009 |
| 217260 | ENSG00000015676 | NUDCD3 | chr7 | - | 44481291 | 44481461 | 44427570 | 44427703 | 44484967 | 44485284 | 217260 | 6,2,14,6,1,9,3,7,0,3,5,1 | 29,19,55,34,24,46,14,19,11 | 2,2,0,2,0,1,4,7,4,4,1,13,6,3 | 30,43,17,25,26,12,16,19,13 | 298 | 149 | 0.0407123358186 | 1 | 0.094,0.05,0.113,0 | 0.032,0.023,0.0,0.0 | 0,018 |
| 48338 | ENSG00000213782 | DDX47 | chr12 | + | 12821207 | 12821396 | 12814130 | 12814224 | 12821654 | 12821726 | 48338 | 32,56,95,72,32,49,33,79,28 | 4,2,15,11,9,8,7,12,1,0,0,0 | 44,99,27,12,55,15,60,20,19 | 9,4,7,11,7,5,2,19,4,4,7,10,8 | 298 | 149 | 0.0407636752541 | 1 | 0.8,0.933,0.76,0.76 | 0.71,0.925,0.659,0 | 0,119 |
| 108542 | ENSG00000179195 | ZNF664 | chr12 | + | 124004877 | 124004935 | 123988042 | 123988138 | 124011380 | 124011441 | 108542 | 9,16,19,13,14,10,11,17,18, | 28,18,28,18,25,11,16,17,24 | 7,20,2,0,6,7,8,9,7,9,5,7,11, | 26,8,10,18,7,5,19,22,8,16,1 | 207 | 149 | 0.0408027408581 | 1 | 0.188,0.39,0.328,0 | 0.162,0.643,0.126, | 0,085 |
| 147759 | ENSG00000197912 | SPG7 | chr16 | + | 89549554 | 89549713 | 89549080 | 89549336 | 89550493 | 89550609 | 147759 | 6,0,2,1,5,0,0,0,3,3,1,2 | 19,7,23,8,6,7,9,17,9,8,3,20 | 1,0,4,0,5,2,1,0,3,1,0,0,4,2,0 | 8,15,2,16,11,2,8,6,4,7,12,7 | 298 | 149 | 0.0408390163237 | 1 | 0.136,0.0,0.042,0.0 | 0.059,0.0,0.5,0.0,0 | -0,093 |
| 239544 | ENSG00000083223 | TUT7 | chr9 | - | 86301739 | 86301828 | 86301275 | 86301601 | 86303085 | 86303201 | 239544 | 87,41,54,21,15,25,20,14,29 | 40,47,52,35,19,23,20,14,55 | 33,42,47,52,13,10,12,25,22 | 21,16,25,15,15,11,8,32,3,1 | 238 | 149 | 0.040854466918 | 1 | 0.577,0.353,0.394, | 0.496,0.622,0.541, | -0,091 |
| 125744 | ENSG00000197857 | ZNF44 | chr19 | - | 12272486 | 12274063 | 12250241 | 12250368 | 12274972 | 12275033 | 125744 | 12,10,49,22,12,20,24,39,3, | 4,4,6,2,7,1,1,6,2,0,3,4 | 22,12,25,20,24,12,4,7,16,1 | 0,0,7,4,4,1,0,2,0,4,1,4,0,1,0 | 298 | 149 | 0.0408575580731 | 1 | 0.6,0.556,0.803,0.8 | 1.0,1.0,0.641,0.714 | -0,166 |
| 123022 | ENSG00000149196 | HIKESHI | chr11 | + | 86326512 | 86326676 | 86306244 | 86306482 | 86337378 | 86337530 | 123022 | 6,1,15,18,8,6,0,0,1,0,2,2 | 17,8,47,46,10,11,6,3,16,12 | 7,1,1,1,3,6,2,3,2,0,3,21,0,0 | 38,17,11,3,9,13,11,29,11,3 | 298 | 149 | 0.0408631461938 | 1 | 0.15,0.059,0.138,0 | 0.084,0.029,0.043, | 0,015 |
| 254082 | ENSG00000151148 | UBE3B | chr12 | + | 109508349 | 109508719 | 109507563 | 109507735 | 109509595 | 109509714 | 254082 | 0,2,2,2,2,0,1,0,3,0,2,2 | 9,7,18,21,9,6,9,5,5,3,4,4 | 4,3,0,0,0,4,1,1,6,1,8,0,1,3,0 | 17,6,5,1,6,7,6,8,10,13,8,6,2 | 298 | 149 | 0.0408642828655 | 1 | 0.0,0.125,0.053,0.0 | 0.105,0.2,0.0,0.0,0 | -0,015 |
| 44352 | ENSG00000163755 | HPS3 | chr3 | + | 149157349 | 149157531 | 149155106 | 149155215 | 149158665 | 149158846 | 44352 | 87,37,92,33,62,58,36,55,49 | 5,0,3,0,0,0,3,1,1,4,0,4 | 92,51,56,17,22,53,31,53,45 | 0,0,0,0,0,1,0,2,0,1,2,4,3,0,2 | 298 | 149 | 0.0408951003635 | 1 | 0.897,1.0,0.939,1.0 | 1.0,1.0,1.0,1.0,1.0, | -0,013 |
| 104736 | ENSG00000188419 | CHM | chrX | - | 85981736 | 85981809 | 85978655 | 85978891 | 86027490 | 86027557 | 104736 | 14,16,69,37,10,13,15,28,4, | 6,2,0,1,1,0,4,0,0,0,2,0 | 18,17,11,9,18,16,9,36,14,2 | 0,0,0,2,0,0,0,0,0,0,0,0,0,0,0 | 222 | 149 | 0.0409363447001 | 1 | 0.61,0.843,1.0,0.96 | 1.0,1.0,1.0,0.751,1 | -0,089 |
| 237288 | ENSG00000120910 | PPP3CC | chr8 | + | 22535202 | 22535369 | 22532920 | 22533018 | 22539468 | 22539498 | 237288 | 12,1,4,1,0,4,3,5,0,2,0,4 | 91,54,107,61,44,26,57,62,8 | 5,1,1,16,2,3,2,9,9,1,5,0,9,1 | 68,61,67,41,40,31,30,66,28 | 298 | 149 | 0.0409544006485 | 1 | 0.062,0.009,0.018, | 0.035,0.008,0.007, | -0,015 |
| 239479 | ENSG00000143437 | ARNT | chr1 | - | 150839400 | 150839654 | 150836423 | 150836493 | 150846262 | 150846307 | 239479 | 6,1,16,11,13,6,5,4,1,1,5,7 | 0,0,3,2,0,0,1,0,5,2,0,0 | 7,10,3,3,8,2,4,9,6,12,5,7,14 | 1,0,1,0,5,1,0,0,0,0,0,0,0,0,0 | 298 | 149 | 0.0409589943287 | 1 | 1.0,1.0,0.727,0.73 | 0.778,1.0,0.6,1.0,0 | -0,093 |
| 230830 | ENSG00000214021 | TTLL3 | chr3 | + | 9818551 | 9818920 | 9817689 | 9817759 | 9820545 | 9820616 | 230830 | 34,17,20,9,24,22,12,16,15, | 1,1,0,0,1,1,1,0,0,0,0,0 | 21,16,15,34,9,8,11,32,12,2 | 2,1,1,0,0,1,0,8,0,0,3,0,1,3,0 | 298 | 149 | 0.0409771099288 | 1 | 0.944,0.895,1.0,1.0 | 0.84,0.889,0.882,1 | 0,07 |
| 178089 | ENSG00000067066 | SP100 | chr2 | + | 230463756 | 230463911 | 230462466 | 230462518 | 230464066 | 230464150 | 178089 | 1,0,20,14,0,0,8,14,0,0,0,0 | 215,177,239,165,109,108,1 | 0,2,0,0,0,0,7,3,1,3,6,0,2,6,5 | 155,135,167,83,112,60,51, | 298 | 149 | 0.0409882043053 | 1 | 0.002,0.0,0.04,0.04 | 0.0,0.007,0.0,0.0,0 | 0,001 |
| 41738 | ENSG00000145012 | LPP | chr3 | + | 188341662 | 188341719 | 188225404 | 188225527 | 188406111 | 188406313 | 41738 | 47,28,15,0,7,8,2,16,5,10,5, | 1,0,38,13,20,38,22,37,33,1 | 16,13,18,18,14,7,9,9,8,5,11 | 12,13,7,16,16,3,0,4,15,18,1 | 206 | 149 | 0.0410067634788 | 1 | 0.971,1.0,0.222,0.0 | 0.491,0.42,0.65,0.4 | -0,154 |
| 225227 | ENSG00000198198 | SZT2 | chr1 | + | 43403600 | 43403690 | 43403176 | 43403302 | 43404379 | 43404550 | 225227 | 10,1,23,16,8,9,7,14,0,3,8,1 | 0,4,0,0,0,0,3,4,2,0,0,0 | 31,19,5,10,9,7,11,7,8,21,18 | 1,4,1,2,0,0,0,2,0,0,0,0,0,1,1 | 239 | 149 | 0.0410141589084 | 1 | 1.0,0.135,1.0,1.0,1 | 0.951,0.748,0.757, | -0,112 |
| 120581 | ENSG00000107036 | RIC1 | chr9 | + | 5753535 | 5753646 | 5753199 | 5753238 | 5754840 | 5754930 | 120581 | 46,29,90,50,41,30,27,57,48 | 2,3,3,8,7,8,5,7,1,1,2,2 | 49,86,41,39,53,38,12,57,42 | 11,11,5,11,11,7,6,5,5,3,5,5 | 260 | 149 | 0.0410142557028 | 1 | 0.929,0.847,0.945, | 0.719,0.818,0.825, | 0,055 |
| 1097 | ENSG00000093072 | ADA2 | chr22 | - | 17203853 | 17203889 | 17203562 | 17203773 | 17207070 | 17207290 | 1097 | 2,7,7,4,4,4,2,7,3,0,1,1 | 45,32,67,42,42,51,28,53,17 | 5,2,1,5,3,3,1,5,2,3,10,4,2,2 | 95,74,27,55,51,40,111,68,2 | 185 | 149 | 0.0410156212105 | 1 | 0.035,0.15,0.078,0 | 0.041,0.021,0.029, | 0,027 |
| 93555 | ENSG00000065427 | KARS1 | chr16 | - | 75642752 | 75642835 | 75641563 | 75641723 | 75647577 | 75647650 | 93555 | 4,4,5,5,0,1,0,8,3,1,2,2 | 40,35,62,32,29,25,20,29,49 | 0,3,2,0,0,0,6,2,0,0,7,7,0,2,0 | 37,26,35,19,35,22,8,29,19, | 232 | 149 | 0.0410255954318 | 1 | 0.06,0.068,0.049,0 | 0.0,0.069,0.035,0.0 | 0,008 |
| 226846 | ENSG00000005471 | ABCB4 | chr7 | - | 87413646 | 87413717 | 87409235 | 87409392 | 87417311 | 87417515 | 226846 | 15,14,8,15,3,5,5,6,13,13,3, | 0,3,15,7,2,1,5,3,1,5,1,2 | 6,9,16,2,1,1,0,4,3,7,0,7,6,1 | 4,4,2,5,4,0,2,5,1,2,5,4,4,4,3 | 220 | 149 | 0.0410283953738 | 1 | 1.0,0.76,0.265,0.59 | 0.504,0.604,0.844, | 0,162 |
| 184053 | ENSG00000132549 | VPS13B | chr8 | + | 99507769 | 99507885 | 99507136 | 99507203 | 99511103 | 99511512 | 184053 | 9,0,5,2,2,5,1,4,1,2,0,1 | 51,27,56,48,18,29,23,24,23 | 3,1,0,1,0,0,5,3,2,3,4,1,2,2,1 | 34,50,34,57,56,33,23,26,25 | 265 | 149 | 0.0410707738292 | 1 | 0.09,0.0,0.048,0.02 | 0.047,0.011,0.0,0.0 | 0,005 |
| 139645 | ENSG00000124243 | BCAS4 | chr20 | + | 50818210 | 50818368 | 50794893 | 50795173 | 50830278 | 50830380 | 139645 | 2,3,16,11,9,17,5,11,9,9,4,8 | 2,0,0,0,1,2,3,4,0,0,2,0 | 10,5,8,6,1,4,1,5,0,6,15,9,3, | 0,0,0,0,1,0,0,0,1,2,0,2,0,0,1 | 298 | 149 | 0.0411083760077 | 1 | 0.333,1.0,1.0,1.0,0 | 1.0,1.0,1.0,1.0,0.33 | -0,052 |
| 190117 | ENSG00000026508 | CD44 | chr11 | + | 35221716 | 35221732 | 35219343 | 35219387 | 35229128 | 35229164 | 190117 | 18,15,9,11,6,10,12,5,9,8,4, | 2,0,0,0,0,2,0,1,0,0,0,0 | 8,10,8,6,6,4,4,6,3,8,6,11,2, | 0,2,0,1,6,0,2,1,0,1,0,4,2,0,0 | 165 | 149 | 0.0411322029574 | 1 | 0.89,1.0,1.0,1.0,1.0 | 1.0,0.819,1.0,0.84 | 0,105 |
| 82098 | ENSG00000134262 | AP4B1 | chr1 | - | 113896257 | 113896465 | 113895756 | 113896038 | 113897839 | 113897943 | 82098 | 27,12,50,33,30,22,15,10,12 | 4,1,6,7,3,0,1,1,0,4,5,4 | 33,32,14,14,13,9,18,31,12, | 12,24,2,7,7,5,1,4,6,3,2,3,0, | 298 | 149 | 0.0411672397302 | 1 | 0.771,0.857,0.806, | 0.579,0.4,0.778,0.5 | 0,111 |
| 134418 | ENSG00000100077 | GRK3 | chr22 | + | 25718244 | 25718417 | 25714407 | 25714570 | 25721283 | 25721397 | 134418 | 55,63,38,23,9,14,11,12,42, | 24,7,8,8,3,4,5,2,1,2,4,4 | 30,40,21,11,16,12,10,13,10 | 14,13,11,15,6,6,0,7,5,3,8,2 | 298 | 149 | 0.0411688074304 | 1 | 0.534,0.818,0.704, | 0.517,0.606,0.488, | 0,078 |
| 186786 | ENSG00000122188 | LAX1 | chr1 | + | 203765516 | 203765654 | 203765176 | 203765383 | 203770827 | 203770937 | 186786 | 49,39,122,62,33,29,23,58,2 | 19,10,38,22,7,9,5,9,7,2,1,8 | 78,88,50,29,72,42,68,75,30 | 23,14,7,3,7,5,14,14,8,6,21, | 287 | 149 | 0.041170344328 | 1 | 0.572,0.669,0.625, | 0.638,0.765,0.788, | -0,055 |
| 20584 | ENSG00000140105 | WARS1 | chr14 | - | 100374135 | 100374179 | 100369086 | 100369258 | 100376259 | 100376805 | 20584 | 3,1,0,0,10,5,0,2,0,0,0,0 | 22,14,37,26,17,14,16,15,4, | 6,3,0,0,1,3,0,2,0,0,0,0,0,2,0 | 16,25,21,3,17,15,13,6,3,3,7 | 193 | 149 | 0.0411714232166 | 1 | 0.095,0.052,0.0,0.0 | 0.225,0.085,0.0,0.0 | 0,012 |
| 9418 | ENSG00000170004 | CHD3 | chr17 | + | 7907600 | 7907702 | 7907352 | 7907488 | 7907893 | 7908019 | 9418 | 57,13,48,39,30,28,20,18,32 | 20,1,32,9,8,7,15,5,5,9,7,14 | 31,33,13,0,7,5,20,21,9,26,1 | 11,20,5,16,15,4,15,8,11,11 | 251 | 149 | 0.0412175442619 | 1 | 0.629,0.885,0.471, | 0.626,0.495,0.607, | 0,118 |
| 226725 | ENSG00000089006 | SNX5 | chr20 | - | 17955364 | 17955498 | 17953995 | 17954117 | 17956932 | 17957037 | 226725 | 37,25,137,78,47,33,27,41,3 | 0,0,2,4,0,0,3,0,0,0,0,1 | 65,75,31,23,38,29,19,40,34 | 0,4,0,2,1,0,0,4,1,0,0,5,6,1,0 | 283 | 149 | 0.0412676114377 | 1 | 1.0,1.0,0.973,0.91 | 1.0,0.908,1.0,0.858 | 0,023 |
| 253771 | ENSG00000123329 | ARHGAP9 | chr12 | - | 57473602 | 57473938 | 57472263 | 57472688 | 57474041 | 57474176 | 253771 | 56,37,115,97,47,23,33,38,3 | 38,11,18,14,10,11,13,14,28 | 85,54,39,19,43,33,29,37,45 | 13,13,23,9,4,10,5,9,2,15,4, | 298 | 149 | 0.0413196182576 | 1 | 0.424,0.627,0.762, | 0.766,0.675,0.459, | -0,09 |
| 4597 | ENSG00000213096 | ZNF254 | chr19 | + | 24091900 | 24092015 | 24087175 | 24087337 | 24105939 | 24106066 | 4597 | 1,5,12,8,2,7,0,3,3,3,1,0 | 5,3,24,15,9,4,9,19,16,6,3,6 | 1,6,2,0,4,0,0,1,3,0,2,2,4,1,3 | 12,12,2,6,18,8,1,8,5,17,3,8 | 264 | 149 | 0.04136920487 | 1 | 0.101,0.485,0.22,0 | 0.045,0.22,0.361,0 | 0,065 |
| 180814 | ENSG00000257379 | | chr12 | + | 53464759 | 53465031 | 53462492 | 53462567 | 53467220 | 53467528 | 180814 | 59,35,89,71,65,96,78,99,78 | 13,10,24,9,14,10,3,15,2,5,0 | 71,83,44,154,66,34,156,85 | 14,17,5,21,8,11,35,20,8,5,6 | 298 | 149 | 0.0413871825986 | 1 | 0.694,0.636,0.65,0 | 0.717,0.709,0.815, | 0,059 |
| 49640 | ENSG00000144218 | AFF3 | chr2 | - | 100112385 | 100112524 | 100008856 | 100008932 | 100142483 | 100142583 | 49640 | 31,19,36,6,14,28,21,6,7,8,4 | 0,4,1,7,0,0,6,0,0,1,0,0 | 7,12,17,34,12,12,15,11,17, | 2,0,0,0,9,0,0,0,1,1,1,4,0,0,0 | 288 | 149 | 0.041397091158 | 1 | 1.0,0.711,0.949,0.3 | 0.644,1.0,1.0,1.0,0 | 0,007 |
| 246103 | ENSG00000122359 | ANXA11 | chr10 | - | 80180940 | 80181023 | 80176106 | 80176155 | 80205342 | 80205572 | 246103 | 1,0,5,4,0,1,1,3,1,2,1,3 | 11,10,35,12,16,13,21,22,18 | 0,3,0,0,2,2,12,3,3,4,2,3,5,3 | 28,18,9,8,8,7,12,17,7,10,11 | 232 | 149 | 0.0414286231352 | 1 | 0.055,0.0,0.084,0.1 | 0.0,0.097,0.0,0.0,0 | -0,048 |
| 225228 | ENSG00000198198 | SZT2 | chr1 | + | 43403600 | 43403731 | 43403176 | 43403302 | 43404379 | 43404550 | 225228 | 15,5,37,25,17,14,11,19,0,7 | 0,4,0,0,0,0,3,4,2,0,0,0 | 42,36,10,23,12,10,17,15,16 | 1,4,1,2,0,0,0,2,0,0,0,0,0,1,1 | 280 | 149 | 0.0414633043559 | 1 | 1.0,0.399,1.0,1.0,1 | 0.957,0.827,0.842, | -0,115 |
| 51566 | ENSG00000162894 | FCMR | chr1 | - | 206913861 | 206914094 | 206904893 | 206905147 | 206921817 | 206921941 | 51566 | 1167,447,1219,947,777,48 | 30,8,8,4,7,18,11,11,37,23,4 | 857,366,530,338,204,325,5 | 18,14,16,12,2,10,8,23,5,6,2 | 298 | 149 | 0.0414791795683 | 1 | 0.951,0.965,0.987, | 0.96,0.929,0.943,0 | 0,009 |
| 35114 | ENSG00000077232 | DNAJC10 | chr2 | + | 182732482 | 182732542 | 182731029 | 182731107 | 182736248 | 182736386 | 35114 | 43,32,96,30,43,34,14,17,15 | 4,0,1,3,3,4,3,0,1,1,2,4 | 65,46,23,11,13,22,6,19,13, | 4,0,3,0,0,1,5,2,1,1,4,0,1,3,4 | 209 | 149 | 0.0414996194225 | 1 | 0.885,1.0,0.986,0.8 | 0.921,1.0,0.845,1.0 | 0,007 |
| 173291 | ENSG00000198586 | TLK1 | chr2 | - | 171022871 | 171023282 | 171014850 | 171014948 | 171028338 | 171028405 | 173291 | 16,5,14,6,10,7,3,7,4,3,1,1 | 37,26,82,80,35,42,20,46,16 | 18,14,8,30,11,9,8,0,6,5,7,8 | 47,61,34,70,47,32,55,41,37 | 298 | 149 | 0.0415135258741 | 1 | 0.178,0.088,0.079, | 0.161,0.103,0.105, | -0,02 |
| 70128 | ENSG00000133657 | ATP13A3 | chr3 | - | 194457093 | 194457174 | 194455892 | 194455962 | 194459470 | 194459541 | 70128 | 52,29,66,28,19,27,19,38,26 | 0,0,5,2,0,0,0,0,2,0,0,2 | 33,31,16,13,24,18,16,34,17 | 1,1,1,0,1,1,0,2,0,2,2,3,0,2,2 | 230 | 149 | 0.0415749282156 | 1 | 1.0,1.0,0.895,0.90 | 0.955,0.953,0.912, | 0,036 |
| 116534 | ENSG00000175893 | ZDHHC21 | chr9 | - | 14683570 | 14683729 | 14680032 | 14680162 | 14690336 | 14690385 | 116534 | 8,4,3,1,3,3,5,6,3,1,0,0 | 11,25,27,19,4,14,7,7,1,3,1, | 2,10,2,1,2,1,0,0,2,1,0,5,6,3 | 22,32,20,2,13,12,7,9,12,30 | 298 | 149 | 0.0415774094735 | 1 | 0.267,0.074,0.053, | 0.043,0.135,0.048, | 0,1 |
| 40244 | ENSG00000278259 | MYO19 | chr17 | - | 36512657 | 36512779 | 36511364 | 36511455 | 36513428 | 36513457 | 40244 | 1,0,4,5,1,3,6,2,4,0,0,0 | 5,1,14,9,8,19,9,6,3,2,1,4 | 3,0,0,0,3,3,1,2,2,0,5,0,1,0,0 | 10,11,4,6,6,5,3,3,7,12,4,9,8 | 271 | 149 | 0.0415840838172 | 1 | 0.099,0.0,0.136,0.1 | 0.142,0.0,0.0,0.0,0 | 0,013 |
| 123476 | ENSG00000152404 | CWF19L2 | chr11 | - | 107416208 | 107416298 | 107392778 | 107392895 | 107418193 | 107418287 | 123476 | 95,75,142,70,29,33,13,31,4 | 0,0,0,3,0,0,0,0,1,0,0,0 | 50,70,54,63,56,28,14,26,52 | 2,0,2,10,4,1,0,1,2,0,0,0,1,1 | 239 | 149 | 0.0416039952556 | 1 | 1.0,1.0,1.0,0.936,1 | 0.94,1.0,0.944,0.79 | 0,041 |
| 17481 | ENSG00000128881 | TTBK2 | chr15 | - | 42872610 | 42872758 | 42840359 | 42840433 | 42878548 | 42878684 | 17481 | 22,9,30,15,24,11,9,18,11,9 | 0,0,4,1,0,3,2,1,1,0,0,1 | 22,12,19,23,9,6,7,22,6,22,2 | 0,2,2,1,1,2,0,3,1,3,0,3,3,1,0 | 297 | 149 | 0.0416132805192 | 1 | 1.0,1.0,0.79,0.883, | 1.0,0.751,0.827,0.9 | 0,05 |
| 146424 | ENSG00000167261 | DPEP2 | chr16 | - | 67991837 | 67992005 | 67991114 | 67991184 | 67992063 | 67992193 | 146424 | 24,7,45,37,41,10,33,21,12, | 3,3,16,1,6,9,4,6,6,2,2,3 | 20,32,25,8,22,17,11,30,24, | 9,2,1,1,4,2,4,3,6,2,4,3,0,0,8 | 298 | 149 | 0.041664207444 | 1 | 0.8,0.538,0.584,0.9 | 0.526,0.889,0.926, | -0,107 |
| 170110 | ENSG00000168936 | TMEM129 | chr4 | - | 1717515 | 1717733 | 1715951 | 1717428 | 1718151 | 1718626 | 170110 | 7,8,20,11,13,5,5,5,8,4,3,4 | 0,1,0,2,3,0,0,0,0,0,0,0 | 4,9,5,3,15,7,5,18,3,11,5,15 | 1,1,0,0,1,1,2,1,0,1,3,4,0,0,3 | 298 | 149 | 0.0416895672874 | 1 | 1.0,0.8,1.0,0.733,0 | 0.667,0.818,1.0,1.0 | 0,11 |
| 94070 | ENSG00000048828 | FAM120A | chr9 | + | 93515666 | 93515767 | 93498789 | 93498886 | 93527154 | 93527242 | 94070 | 34,26,57,54,24,35,18,45,22 | 6,2,10,5,7,2,3,4,2,1,6,3 | 51,46,45,27,17,25,16,23,19 | 3,3,3,6,4,3,1,4,1,2,1,8,3,1,4 | 250 | 149 | 0.0417078347971 | 1 | 0.772,0.886,0.773, | 0.91,0.901,0.899,0 | -0,052 |
| 254215 | ENSG00000111639 | MRPL51 | chr12 | - | 6493057 | 6493219 | 6492861 | 6492972 | 6493458 | 6493840 | 254215 | 37,41,91,106,51,31,31,23,6 | 4,4,7,3,3,0,0,2,6,1,0,3 | 24,26,35,7,27,22,13,23,11, | 4,2,3,0,3,1,2,1,5,0,3,16,4,5 | 298 | 149 | 0.0417444701489 | 1 | 0.822,0.837,0.867, | 0.75,0.867,0.854,1 | 0,066 |
| 182002 | ENSG00000096080 | MRPS18A | chr6 | - | 43675201 | 43675252 | 43671201 | 43671906 | 43675493 | 43675617 | 182002 | 5,3,18,6,2,2,5,8,6,11,4,2 | 0,0,3,1,0,0,2,0,5,0,0,6 | 15,11,6,5,2,2,6,4,1,7,2,13,1 | 1,3,2,7,2,3,4,2,0,1,2,1,0,3,1 | 200 | 149 | 0.0417453316466 | 1 | 1.0,1.0,0.817,0.81 | 0.918,0.732,0.691, | 0,159 |
| 175646 | ENSG00000187862 | TTC24 | chr1 | + | 156585948 | 156586366 | 156585712 | 156585826 | 156586468 | 156587719 | 175646 | 11,7,5,5,1,3,1,5,1,2,2,3 | 5,1,2,6,2,0,0,2,2,0,0,0 | 6,3,4,2,1,0,2,8,5,7,4,2,9,13 | 2,4,7,2,2,1,3,6,1,2,3,0,3,1,5 | 298 | 149 | 0.0417686006636 | 1 | 0.524,0.778,0.556, | 0.6,0.273,0.222,0.3 | 0,243 |
| 212895 | ENSG00000152689 | RASGRP3 | chr2 | + | 33531051 | 33531099 | 33527202 | 33527412 | 33534322 | 33534400 | 212895 | 11,1,29,20,4,9,2,7,14,4,2,8 | 283,181,375,172,186,130,2 | 10,11,7,0,3,0,3,10,3,2,3,1,0 | 223,246,153,37,88,50,77,2 | 197 | 149 | 0.0417785933715 | 1 | 0.029,0.004,0.055, | 0.033,0.033,0.033, | 0,008 |
| 144160 | ENSG00000153944 | MSI2 | chr17 | + | 57596867 | 57596949 | 57529675 | 57529724 | 57615969 | 57616631 | 144160 | 34,11,235,152,63,56,36,45 | 3,0,2,4,0,1,0,1,2,0,0,3 | 124,161,32,58,70,73,29,82 | 5,5,1,3,2,0,2,1,8,2,1,0,5,0,1 | 231 | 149 | 0.0417807053762 | 1 | 0.88,1.0,0.987,0.96 | 0.941,0.954,0.954, | -0,007 |
| 247190 | ENSG00000149196 | HIKESHI | chr11 | + | 86303356 | 86303644 | 86302283 | 86302478 | 86306244 | 86306482 | 247190 | 1,3,13,2,1,1,0,3,2,2,0,2 | 33,21,59,40,14,13,6,6,19,1 | 9,3,2,0,5,0,1,3,1,2,4,5,11,3 | 26,16,12,7,11,11,10,7,9,25 | 298 | 149 | 0.0417980235845 | 1 | 0.015,0.067,0.099, | 0.148,0.086,0.077, | -0,027 |
| 176438 | ENSG00000127481 | UBR4 | chr1 | - | 19168026 | 19168150 | 19167021 | 19167231 | 19169434 | 19169532 | 176438 | 43,32,59,71,32,32,23,32,12 | 1,1,0,0,0,0,1,0,0,0,0,0 | 47,64,31,49,46,21,25,30,52 | 0,5,0,5,0,1,1,1,2,0,1,2,2,0,2 | 273 | 149 | 0.0418716563414 | 1 | 0.959,0.946,1.0,1.0 | 1.0,0.875,1.0,0.842 | 0,043 |

| | | | | | | | | | | | | | | | | | | | | | | | |
|---|---|---|---|---|---|---|---|---|---|---|---|---|---|---|---|---|---|---|---|---|---|---|---|
| 84648 | ENSG00000152689 | RASGRP3 | chr2 | + | 33520552 | 33520684 | 33519951 | 33520014 | 33521954 | 33522102 | 84648 | 450,222,813,341,257,308,3 | 8,3,18,11,9,8,4,4,6,0,5,3 | 410,428,278,94,177,129,1 | 10,10,1,3,0,1,0,3,0,0,6,9,1, | 281 | 149 | 0.0419007518716 | | 1 | 0.968,0.975,0.96,0 | 0.956,0.958,0.993, | -0,01 |
| 83726 | ENSG00000079263 | SP140 | chr2 | + | 230253315 | 230253417 | 230250980 | 230251061 | 230255451 | 230255532 | 83726 | 54,31,78,77,57,27,18,19,3 | 17,6,10,15,3,2,7,9,0,1,7,5 | 74,48,29,17,56,53,1,7,27,5 | 7,2,1,1,0,1,0,2,4,7,6,13,2,7 | 251 | 149 | 0.0419143000257 | | 1 | 0.653,0.754,0.822, | 0.863,0.934,0.945, | -0,089 |
| 31956 | ENSG00000164180 | TMEM161B | chr5 | - | 88225768 | 88225866 | 88220562 | 88220719 | 88228444 | 88228528 | 31956 | 13,6,52,32,21,23,10,11,8,1 | 0,0,4,5,3,2,1,1,1,0,2,4 | 34,28,12,6,18,8,10,17,14,2 | 3,3,1,0,0,1,2,4,0,0,0,3,3,0,1 | 247 | 149 | 0.0419756359032 | | 1 | 1.0,1.0,0.887,0.79 | 0.872,0.849,0.879, | -0,03 |
| 209043 | ENSG00000290927 | | chr16 | + | 31702534 | 31702676 | 31700562 | 31700738 | 31703155 | 31703367 | 209043 | 17,13,30,13,7,13,4,7,2,10,0 | 15,15,19,12,5,2,7,5,2,8,7,7 | 8,8,4,9,8,1,0,1,4,6,4,3,13,5 | 12,8,3,13,9,4,5,7,2,8,5,9,20 | 291 | 149 | 0.0420067562666 | | 1 | 0.367,0.307,0.447, | 0.254,0.339,0.406, | 0,099 |
| 247860 | ENSG00000179532 | DNHD1 | chr11 | + | 6538435 | 6538510 | 6533680 | 6533806 | 6538611 | 6538810 | 247860 | 3,2,7,2,4,8,8,5,3,6,1,6 | 3,0,0,3,0,0,0,6,0,0,2,5 | 5,2,7,9,0,0,1,1,3,7,4,0,2,0,5 | 0,0,2,1,2,2,0,0,3,2,3,1,5,1,0 | 224 | 149 | 0.0420179169548 | | 1 | 0.399,1.0,1.0,0.30 | 1.0,1.0,0.7,0.857,0 | 0,2 |
| 155212 | ENSG00000053770 | AP5M1 | chr14 | + | 57274243 | 57274437 | 57268923 | 57269388 | 57280194 | 57280422 | 155212 | 40,25,80,80,40,35,37,47,21 | 0,0,0,0,2,0,2,0,0,0,1,1 | 55,77,33,26,35,30,22,46,41 | 1,2,0,0,0,0,0,2,0,0,0,2,3,5,4 | 298 | 149 | 0.0420473981545 | | 1 | 1.0,1.0,1.0,1.0,0.9 | 0.965,0.951,1.0,1. | 0,022 |
| 78380 | ENSG00000048991 | R3HDM1 | chr2 | + | 135715549 | 135715694 | 135710058 | 135710231 | 135721923 | 135722006 | 78380 | 17,19,21,16,8,10,13,12,20, | 0,0,1,2,1,1,0,2,0,0,0,1 | 18,11,24,22,5,10,8,21,11,1 | 1,2,2,3,4,1,0,0,3,0,0,0,0,1,2 | 294 | 149 | 0.0420630940602 | | 1 | 1.0,1.0,0.914,0.80 | 0.901,0.736,0.859, | 0,045 |
| 241117 | ENSG00000165209 | STRBP | chr9 | - | 123213588 | 123213918 | 123184131 | 123184298 | 123236829 | 123236966 | 241117 | 2,0,2,5,1,0,1,0,0,0,0,0 | 47,21,52,63,21,21,12,20,44 | 1,0,11,3,0,1,0,0,0,0,2,1,4,3 | 38,23,28,19,15,15,7,11,10, | 298 | 149 | 0.0420925794334 | | 1 | 0.021,0.0,0.019,0.0 | 0.013,0.0,0.164,0.0 | -0,033 |
| 159849 | ENSG00000055163 | CYFIP2 | chr5 | + | 157327972 | 157328045 | 157325599 | 157326267 | 157330741 | 157330850 | 159849 | 23,23,55,34,24,24,33,43,13 | 6,1,2,1,5,5,9,4,1,1,1,1 | 31,19,13,24,25,14,14,34,8, | 2,4,1,18,8,4,2,6,0,0,7,8,2,4 | 222 | 149 | 0.0421389858499 | | 1 | 0.72,0.939,0.949,0 | 0.912,0.761,0.897, | 0,058 |
| 3489 | ENSG00000214046 | SMIM7 | chr19 | - | 16652604 | 16652961 | 16645660 | 16647261 | 16654034 | 16654125 | 3489 | 8,5,10,7,3,5,4,8,3,2,4,5 | 51,50,115,93,43,33,36,66,4 | 2,5,6,1,3,5,2,9,5,3,1,3,6,2,4 | 47,89,32,20,55,26,34,57,27 | 298 | 149 | 0.0421489109068 | | 1 | 0.073,0.048,0.042, | 0.021,0.027,0.086, | 0,02 |
| 20802 | ENSG00000258643 | BCL2L2-PABPN | chr14 | + | 23323964 | 23324052 | 23323376 | 23323483 | 23324137 | 23324289 | 20802 | 636,458,606,320,269,325,3 | 9,3,3,7,5,2,3,6,16,6,1,7 | 391,403,358,249,270,189,1 | 8,1,4,0,4,1,0,6,4,1,4,6,2,3,4 | 237 | 149 | 0.0422552404978 | | 1 | 0.978,0.99,0.992,0 | 0.968,0.996,0.983, | -0,005 |
| 188109 | ENSG00000126107 | HECTD3 | chr1 | - | 45008603 | 45008701 | 45008239 | 45008321 | 45009143 | 45009226 | 188109 | 5,3,24,15,8,6,4,9,11,2,2,7 | 1,0,7,0,1,6,0,2,0,0,0,1 | 8,7,6,1,6,10,5,5,6,6,5,12,8, | 2,6,2,8,1,0,1,1,0,4,4,2,4,0,2 | 247 | 149 | 0.0422653701798 | | 1 | 0.751,1.0,0.674,1. | 0.707,0.413,0.644, | 0,186 |
| 4619 | ENSG00000007255 | TRAPPC6A | chr19 | - | 45165126 | 45165194 | 45164724 | 45164970 | 45178092 | 45178211 | 4619 | 24,17,27,33,28,18,12,29,18 | 0,3,2,2,1,0,0,2,3,0,1,0 | 17,18,13,8,20,5,6,14,14,15 | 0,0,0,0,1,0,4,0,0,0,1,13,5,1 | 217 | 149 | 0.0423014422504 | | 1 | 1.0,0.796,0.903,0. | 1.0,1.0,1.0,1.0,0.9 | 0,029 |
| 128775 | ENSG00000149100 | EIF3M | chr11 | + | 32586933 | 32587144 | 32583838 | 32583929 | 32588593 | 32588732 | 128775 | 69,46,215,182,63,48,49,50 | 3,2,10,9,2,1,1,4,0,4,0,1 | 127,136,87,40,96,44,24,42 | 5,4,3,2,8,0,0,3,0,1,2,7,0,0,2 | 298 | 149 | 0.042304757631 | | 1 | 0.92,0.92,0.915,0. | 0.927,0.944,0.935, | -0,014 |
| 188797 | ENSG00000082512 | TRAF5 | chr1 | + | 211359911 | 211360005 | 211356366 | 211356468 | 211360701 | 211360779 | 188797 | 33,30,70,52,22,32,26,28,19 | 2,0,0,2,0,1,0,4,0,0,0,0 | 38,28,25,26,32,26,14,18,35 | 1,0,0,2,2,4,1,2,5,4,0,0,2,0,0 | 243 | 149 | 0.0423050985727 | | 1 | 0.91,1.0,1.0,0.941, | 0.959,1.0,1.0,0.88 | 0,033 |
| 159091 | ENSG00000012660 | ELOVL5 | chr6 | - | 53270299 | 53270727 | 53267403 | 53269270 | 53273219 | 53273344 | 159091 | 207,95,248,206,93,71,95,7 | 5,2,5,0,2,1,3,3,1,3,0,0 | 133,128,111,43,81,52,65,5 | 5,5,3,2,4,0,4,0,0,0,2,6,0,2,7 | 298 | 149 | 0.0423272848646 | | 1 | 0.954,0.96,0.961,1 | 0.93,0.928,0.949,0 | 0,017 |
| 130088 | ENSG00000139350 | NEDD1 | chr12 | + | 96924832 | 96924942 | 96919984 | 96920125 | 96934975 | 96935205 | 130088 | 10,0,18,9,1,2,2,7,3,4,0,3 | 18,17,11,18,15,14,3,13,8,3 | 13,9,8,5,6,4,6,14,0,7,12,7,4 | 21,24,12,4,12,7,8,23,2,15,1 | 259 | 149 | 0.0423304780235 | | 1 | 0.242,0.0,0.485,0. | 0.263,0.177,0.277, | -0,077 |
| 15525 | ENSG00000172775 | PSME3IP1 | chr16 | - | 57178501 | 57178852 | 57173727 | 57173869 | 57185820 | 57186039 | 15525 | 20,22,28,9,6,11,16,11,4,6,4 | 12,3,19,12,3,4,9,4,6,7,5,7 | 8,5,3,14,9,4,5,15,10,13,19, | 10,3,7,2,0,3,0,2,2,2,3,11,5, | 298 | 149 | 0.0423603529902 | | 1 | 0.455,0.786,0.424, | 0.286,0.455,0.176, | -0,158 |
| 92011 | ENSG00000139163 | ETNK1 | chr12 | + | 22656431 | 22656745 | 22643762 | 22644022 | 22659013 | 22659154 | 92011 | 2,3,1,4,0,0,3,2,0,0,0,0 | 14,5,35,23,21,22,7,10,10,8 | 1,2,6,5,6,1,2,1,0,1,1,0,2,3,0 | 32,8,10,1,6,9,3,26,5,10,7,1 | 298 | 149 | 0.0424014533567 | | 1 | 0.067,0.231,0.014, | 0.015,0.111,0.231, | -0,068 |
| 14067 | ENSG00000237440 | ZNF737 | chr19 | - | 20552474 | 20552570 | 20537991 | 20545976 | 20553708 | 20553835 | 14067 | 73,51,312,274,141,81,61,1 | 1,0,5,11,0,1,0,1,4,0,0,5 | 197,146,110,77,62,53,33,9 | 0,0,2,0,2,11,0,2,0,0,0,0,0,0 | 245 | 149 | 0.0424349016333 | | 1 | 0.978,1.0,0.974,0. | 1.0,1.0,0.971,1.0,0 | -0,003 |
| 160351 | ENSG00000166024 | R3HCC1L | chr10 | + | 98205517 | 98205656 | 98163292 | 98163397 | 98208100 | 98209899 | 160351 | 6,3,13,5,2,6,0,5,1,0,0,0 | 13,18,30,6,3,11,8,6,3,5,4,3 | 4,4,11,9,9,0,3,4,6,10,4,4,2, | 4,6,8,4,27,10,4,4,6,6,7,10,3 | 288 | 149 | 0.0424535882265 | | 1 | 0.193,0.079,0.183, | 0.341,0.256,0.416, | -0,112 |
| 19251 | ENSG00000114062 | UBE3A | chr15 | - | 25408571 | 25408684 | 25405460 | 25405502 | 25409087 | 25409207 | 19251 | 32,15,83,49,16,10,14,15,20 | 31,31,96,30,23,39,24,66,14 | 31,27,16,9,5,3,12,11,5,17,1 | 40,78,26,27,41,17,36,31,23 | 262 | 149 | 0.0425119689595 | | 1 | 0.37,0.216,0.33,0. | 0.306,0.164,0.259, | 0,059 |
| 241309 | ENSG00000119314 | PTBP3 | chr9 | - | 112262434 | 112262607 | 112252677 | 112252788 | 112268048 | 112268195 | 241309 | 67,63,160,116,69,68,62,83 | 2,0,4,1,2,2,2,4,1,0,0,0 | 113,155,85,64,61,45,35,81 | 3,6,3,5,1,1,5,4,1,3,5,0,0,2,0 | 298 | 149 | 0.0425328658804 | | 1 | 0.944,1.0,0.952,0. | 0.95,0.928,0.934,0 | 0,028 |
| 33763 | ENSG00000204842 | ATXN2 | chr12 | - | 111552905 | 111552977 | 111552279 | 111552430 | 111554157 | 111554217 | 33763 | 16,6,52,16,18,12,15,16,9,6 | 5,0,2,0,0,2,0,0,1,0,2,0 | 23,24,10,11,4,10,13,16,11, | 4,1,1,0,0,0,1,4,2,3,1,0,0,3,2 | 221 | 149 | 0.0425384823464 | | 1 | 0.683,1.0,0.946,1. | 0.795,0.942,0.871, | -0,016 |
| 31880 | ENSG00000247774 | PCED1B-AS1 | chr12 | - | 47209280 | 47209458 | 47205897 | 47206145 | 47216164 | 47216251 | 31880 | 18,31,51,45,6,5,18,13,32,4 | 2,2,4,22,1,2,3,4,5,4,0,1 | 28,20,21,15,26,23,8,11,3,6 | 2,1,2,0,3,1,1,0,0,0,2,16,5,6 | 298 | 149 | 0.0425438701218 | | 1 | 0.818,0.886,0.864, | 0.875,0.909,0.84,1 | -0,106 |
| 164445 | ENSG00000005700 | IBTK | chr6 | - | 82215911 | 82216250 | 82214226 | 82214829 | 82217959 | 82218137 | 164445 | 7,1,61,49,13,12,9,17,5,7,1, | 0,0,5,0,0,0,0,1,1,0,2,1 | 39,21,16,3,18,15,15,16,17, | 2,5,1,5,1,1,2,0,2,2,0,1,0,0,0 | 298 | 149 | 0.0425606239703 | | 1 | 1.0,1.0,0.859,1.0,1 | 0.907,0.677,0.889, | 0,029 |
| 205268 | ENSG00000106346 | USP42 | chr7 | + | 6116815 | 6116939 | 6115322 | 6115523 | 6135840 | 6135951 | 205268 | 2,2,4,3,1,4,1,5,2,0,1,1 | 6,0,3,1,3,0,0,1,0,4,0,0 | 8,7,3,7,6,1,4,3,1,0,3,4,1,6,2 | 3,5,0,4,0,1,6,4,0,3,2,1,4,4,2 | 273 | 149 | 0.0425909399588 | | 1 | 0.154,1.0,0.421,0. | 0.593,0.433,1.0,0. | 0,206 |
| 410 | ENSG00000186866 | POFUT2 | chr21 | - | 45266021 | 45266227 | 45263927 | 45265635 | 45267589 | 45267713 | 410 | 23,11,11,5,7,10,1,4,3,1,1,1 | 18,4,17,6,5,9,1,2,9,4,3,6 | 0,4,6,2,2,2,5,7,4,5,4,2,8,5,4 | 12,3,8,7,5,2,4,9,3,6,18,9,7, | 298 | 149 | 0.04259231024 | | 1 | 0.39,0.579,0.244,0 | 0.0,0.4,0.273,0.12 | 0,067 |
| 202436 | ENSG00000154222 | CC2D1B | chr1 | - | 52359704 | 52359835 | 52359458 | 52359534 | 52360073 | 52360233 | 202436 | 11,9,28,11,12,16,10,11,13, | 0,2,2,0,2,10,2,6,1,0,0,3 | 18,17,6,10,11,8,11,24,7,10 | 4,4,2,5,4,1,2,5,1,0,6,2,3,1,8 | 280 | 149 | 0.0425990315462 | | 1 | 1.0,0.705,0.882,1. | 0.705,0.693,0.615, | 0,12 |
| 123689 | ENSG00000179532 | DNHD1 | chr11 | + | 6538382 | 6538510 | 6533923 | 6534229 | 6538611 | 6538810 | 123689 | 9,9,18,7,9,16,11,12,9,7,4,1 | 1,1,2,3,0,2,0,1,0,0,4,0 | 12,7,17,26,4,2,3,16,13,20,1 | 0,0,0,1,0,0,0,0,2,0,0,0,0,0,0 | 277 | 149 | 0.0426147127329 | | 1 | 0.829,0.829,0.829, | 1.0,1.0,1.0,0.933,1 | -0,113 |
| 171103 | ENSG00000147403 | RPL10 | chrX | + | 154400557 | 154400626 | 154399882 | 154399941 | 154400701 | 154400811 | 171103 | 491,214,1222,1207,729,52 | 77,31,190,101,52,34,38,52 | 375,314,217,103,317,246,4 | 89,62,38,21,75,21,36,73,20 | 218 | 149 | 0.0426151628617 | | 1 | 0.813,0.825,0.815, | 0.742,0.776,0.796, | 0,025 |
| 145568 | ENSG00000006194 | ZNF263 | chr16 | + | 3285993 | 3286149 | 3283545 | 3284205 | 3288453 | 3288570 | 145568 | 16,2,4,0,5,4,3,2,2,1,1,0 | 3,3,5,5,3,2,0,2,4,4,2,1 | 1,4,3,1,1,1,2,2,2,8,3,2,3,7,4 | 1,1,0,0,1,0,3,3,0,7,0,3,3,2,0 | 298 | 149 | 0.0426307658922 | | 1 | 0.727,0.25,0.286,0 | 0.333,0.667,1.0,1. | -0,25 |
| 56154 | ENSG00000158796 | DEDD | chr1 | - | 161123074 | 161123174 | 161121743 | 161122523 | 161123838 | 161123946 | 56154 | 21,33,23,26,19,22,10,12,25 | 0,7,0,2,3,7,6,10,0,0,0,0 | 20,22,14,19,13,9,18,26,9,2 | 9,4,2,5,3,5,5,4,1,5,0,6,2,0,1 | 249 | 149 | 0.0426711359749 | | 1 | 1.0,0.738,1.0,0.88 | 0.571,0.767,0.807, | 0,083 |
| 46541 | ENSG00000178694 | NSUN3 | chr3 | + | 94063811 | 94063979 | 94063010 | 94063138 | 94064436 | 94064518 | 46541 | 8,0,21,14,9,5,5,12,13,3,7,8 | 36,24,35,24,14,32,15,24,13 | 10,10,2,9,4,4,2,8,2,8,3,6,4, | 25,34,18,11,26,16,17,13,7, | 298 | 149 | 0.0427217399621 | | 1 | 0.1,0.0,0.231,0.22 | 0.167,0.128,0.053, | 0,067 |
| 139668 | ENSG00000124243 | BCAS4 | chr20 | + | 50851941 | 50852120 | 50841765 | 50841900 | 50876485 | 50877177 | 139668 | 6,8,0,2,2,9,3,13,7,4,0,2 | 178,99,227,133,111,94,11 | 3,4,6,6,1,1,0,2,2,7,2,2,2,1,1 | 96,105,96,70,67,92,69,89,4 | 298 | 149 | 0.0427367547969 | | 1 | 0.017,0.039,0.0,0. | 0.015,0.019,0.03,0 | 0,005 |
| 36230 | ENSG00000155229 | MMS19 | chr10 | - | 97468965 | 97469104 | 97467504 | 97467583 | 97469645 | 97469723 | 36230 | 19,14,24,26,20,13,18,21,28 | 0,0,3,2,2,6,3,1,2,1,0,1 | 26,16,7,9,14,5,3,13,11,12,4 | 1,0,3,1,0,0,0,2,0,1,3,2,0,0,5 | 288 | 149 | 0.0427460263143 | | 1 | 1.0,1.0,0.805,0.87 | 0.931,1.0,0.547,0. | 0,032 |
| 221812 | ENSG00000112159 | MDN1 | chr6 | - | 89726674 | 89726715 | 89725198 | 89725396 | 89727832 | 89727955 | 221812 | 2,1,7,3,1,2,0,3,0,2,1,3 | 9,12,14,18,19,5,8,8,16,7,3, | 3,3,0,1,4,1,2,0,3,1,1,2,0,0,1 | 21,11,2,7,7,11,15,24,6,8,7, | 190 | 149 | 0.042752396183 | | 1 | 0.148,0.061,0.282, | 0.101,0.176,0.0,0. | 0,041 |
| 209400 | ENSG00000091490 | SEL1L3 | chr4 | - | 25784498 | 25784562 | 25784227 | 25784290 | 25788223 | 25788364 | 209400 | 8,5,16,5,3,1,4,11,0,0,1,4 | 121,103,247,187,127,95,67 | 3,10,10,14,5,1,2,1,8,8,3,13 | 104,65,92,65,64,64,30,45,6 | 213 | 149 | 0.0427764956294 | | 1 | 0.044,0.033,0.043, | 0.02,0.097,0.071,0 | -0,017 |
| 34436 | ENSG00000166913 | YWHAB | chr20 | + | 44902897 | 44903121 | 44901610 | 44901833 | 44903992 | 44904116 | 34436 | 17,6,33,9,4,11,5,12,7,2,4,1 | 74,56,102,117,130,102,47, | 28,30,10,26,6,8,18,11,6,13 | 111,108,39,47,43,36,43,46 | 298 | 149 | 0.0427935800979 | | 1 | 0.103,0.051,0.139, | 0.112,0.122,0.114, | -0,021 |
| 137339 | ENSG00000196684 | HSH2D | chr19 | + | 16153076 | 16153204 | 16152551 | 16152641 | 16154398 | 16154863 | 137339 | 14,13,49,30,33,30,13,19,17 | 0,0,2,1,0,0,0,3,3,4,0,5 | 23,35,8,13,13,10,5,11,10,1 | 0,1,0,0,0,0,0,1,1,0,0,0,0,0,0 | 277 | 149 | 0.0428178880918 | | 1 | 1.0,1.0,0.929,0.94 | 1.0,0.95,1.0,1.0,1. | -0,078 |
| 194060 | ENSG00000113318 | MSH3 | chr5 | + | 80752159 | 80752256 | 80744505 | 80744615 | 80761545 | 80761678 | 194060 | 1,4,3,2,4,1,0,2,0,0,1,0 | 6,4,19,14,9,9,6,8,3,1,3,5 | 0,3,1,3,0,0,0,4,1,1,0,3,1,0,1 | 8,17,14,6,13,17,6,10,16,9,3 | 246 | 149 | 0.042821758631 | | 1 | 0.092,0.377,0.087, | 0.0,0.097,0.041,0. | 0,045 |
| 23249 | ENSG00000131966 | ACTR10 | chr14 | + | 58213630 | 58213698 | 58208998 | 58209107 | 58215204 | 58215284 | 23249 | 6,1,24,24,10,12,6,3,2,7,3,7 | 1,1,6,11,2,6,2,3,2,0,5,8 | 13,14,7,0,2,3,5,7,3,6,4,10,5 | 10,8,6,4,5,2,0,9,6,2,2,4,8,3 | 217 | 149 | 0.042833802821 | | 1 | 0.805,0.407,0.733, | 0.472,0.546,0.445, | 0,13 |
| 185740 | ENSG00000198160 | MIER1 | chr1 | + | 66972874 | 66972991 | 66971654 | 66971736 | 66976594 | 66976722 | 185740 | 16,13,82,48,16,27,14,26,5, | 3,0,0,2,0,2,3,2,0,0,0,1 | 33,66,19,17,30,13,15,26,19 | 2,1,0,1,0,4,3,6,0,0,5,0,1,3,0 | 266 | 149 | 0.04283995792 | | 1 | 0.749,1.0,1.0,0.93 | 0.902,0.974,1.0,0. | 0,019 |
| 131067 | ENSG00000050426 | LETMD1 | chr12 | + | 51056149 | 51056245 | 51049033 | 51049185 | 51056349 | 51056502 | 131067 | 23,11,44,25,9,13,9,24,4,4,5 | 7,7,6,18,9,9,13,4,15,4,5,14 | 34,21,26,13,13,19,7,18,22, | 6,1,3,4,15,5,3,8,4,15,7,22,1 | 245 | 149 | 0.042860306303 | | 1 | 0.666,0.489,0.817, | 0.775,0.927,0.841, | -0,142 |
| 131063 | ENSG00000050426 | LETMD1 | chr12 | + | 51056143 | 51056245 | 51049033 | 51049185 | 51056349 | 51056428 | 131063 | 25,11,49,26,11,13,9,25,6,4 | 7,7,6,18,9,9,13,4,15,4,5,14 | 36,25,26,13,14,21,7,19,24, | 6,1,3,4,15,5,3,8,4,15,7,22,1 | 251 | 149 | 0.0428876926671 | | 1 | 0.679,0.483,0.829, | 0.781,0.937,0.837, | -0,14 |
| 116068 | ENSG00000165209 | STRBP | chr9 | - | 123173676 | 123173834 | 123169901 | 123170046 | 123179006 | 123179227 | 116068 | 355,150,466,349,204,162,1 | 11,10,19,9,10,12,10,9,3,8,2 | 342,292,293,151,134,110,8 | 16,16,10,26,13,3,2,3,7,1,8, | 298 | 149 | 0.0429324994643 | | 1 | 0.942,0.882,0.925, | 0.914,0.901,0.936, | 0,019 |
| 208608 | ENSG00000285417 | | chr1 | + | 206056178 | 206056463 | 206052844 | 206053265 | 206060400 | 206060449 | 208608 | 9,0,11,24,5,4,8,27,14,10,1, | 11,1,12,7,5,16,7,21,8,18,2, | 0,2,2,8,0,1,2,0,0,4,1,6,8,10 | 2,0,3,6,1,0,0,2,2,3,7,23,10, | 298 | 149 | 0.0429451102824 | | 1 | 0.29,0.0,0.314,0.6 | 0.0,1.0,0.25,0.4,0. | -0,059 |
| 206566 | ENSG00000138029 | HADHB | chr2 | + | 26285406 | 26285489 | 26284882 | 26284957 | 26289917 | 26290465 | 206566 | 36,32,61,45,21,15,10,14,33 | 4,3,15,10,4,12,7,7,6,0,0,1 | 38,29,32,9,28,19,18,24,15, | 15,23,14,4,19,5,9,5,6,2,8,9 | 232 | 149 | 0.0429451617271 | | 1 | 0.853,0.873,0.723, | 0.619,0.447,0.595, | 0,112 |
| 48106 | ENSG00000182866 | LCK | chr1 | + | 32276336 | 32276489 | 32275913 | 32276063 | 32276606 | 32276786 | 48106 | 99,19,93,101,80,29,58,38,9 | 18,1,5,4,4,7,5,4,13,4,3,6 | 69,29,48,5,28,48,23,45,25, | 11,12,6,5,4,5,2,4,2,8,3,4,0, | 298 | 149 | 0.0429629830494 | | 1 | 0.733,0.905,0.903, | 0.758,0.547,0.8,0. | 0,067 |
| 70998 | ENSG00000174606 | ANGEL2 | chr1 | - | 213013092 | 213013418 | 213008209 | 213008466 | 213015612 | 213015867 | 70998 | 46,46,79,44,44,39,35,40,33 | 5,1,6,6,2,2,0,1,1,0,0,0 | 49,52,26,40,27,17,17,40,26 | 0,6,2,2,0,1,2,1,0,0,5,0,1,0,0 | 298 | 149 | 0.0429696424389 | | 1 | 0.821,0.958,0.868, | 1.0,0.812,0.867,0. | -0,006 |
| 73594 | ENSG00000218510 | LINC00339 | chr1 | + | 22027413 | 22027534 | 22025521 | 22025643 | 22030279 | 22030374 | 73594 | 0,0,5,4,1,4,0,1,0,6,0,0 | 5,11,21,19,11,18,2,9,15,4,5 | 2,0,0,0,0,0,1,2,0,2,0,11,0,3 | 5,5,4,8,3,5,5,9,3,13,12,3,4, | 270 | 149 | 0.0429827530999 | | 1 | 0.0,0.0,0.116,0.10 | 0.181,0.0,0.0,0.0,0 | -0,023 |
| 170109 | ENSG00000168936 | TMEM129 | chr4 | - | 1717515 | 1717690 | 1715951 | 1717428 | 1718151 | 1718626 | 170109 | 7,8,20,11,13,5,5,5,8,4,3,4 | 0,1,0,2,3,0,0,0,0,0,0,0 | 4,9,5,3,15,7,3,18,3,11,6,15 | 1,1,0,0,1,1,2,1,0,1,3,4,0,0,3 | 298 | 149 | 0.0430255669372 | | 1 | 1.0,0.8,1.0,0.733,0 | 0.667,0.818,1.0,1. | 0,115 |
| 129875 | ENSG00000145439 | CBR4 | chr4 | - | 169007635 | 169007756 | 169006754 | 169006891 | 169009947 | 169010080 | 129875 | 70,24,79,39,51,34,26,29,43 | 3,2,1,3,0,1,1,3,4,0,0,0 | 48,32,31,18,15,23,21,31,25 | 0,0,0,0,0,0,0,0,0,0,0,0,2,0,0 | 270 | 149 | 0.0430545221692 | | 1 | 0.928,0.869,0.978, | 1.0,1.0,1.0,1.0,1.0, | -0,057 |
| 212069 | ENSG00000079263 | SP140 | chr2 | + | 230253333 | 230253417 | 230248884 | 230248968 | 230255451 | 230255532 | 212069 | 37,22,42,33,23,16,14,13,23 | 95,68,98,67,24,21,28,28,65 | 44,36,21,16,39,29,1,4,16,4 | 32,39,29,33,60,27,10,19,31 | 233 | 149 | 0.0430563312481 | | 1 | 0.199,0.171,0.215, | 0.468,0.371,0.317, | -0,038 |
| 5454 | ENSG00000064607 | SUGP2 | chr19 | - | 19003495 | 19003608 | 19001612 | 19001674 | 19004167 | 19004269 | 5454 | 6,9,6,6,2,7,9,7,0,0,0,4 | 41,37,87,57,38,33,29,33,39 | 1,5,2,2,3,1,2,6,3,2,2,8,4,4,4 | 39,51,11,16,37,15,20,46,38 | 262 | 149 | 0.0430721191576 | | 1 | 0.077,0.122,0.038, | 0.014,0.053,0.094, | 0,017 |
| 251596 | ENSG00000133114 | GPALPP1 | chr13 | + | 45005119 | 45005241 | 45004310 | 45004437 | 45006201 | 45006303 | 251596 | 2,1,2,1,0,0,0,3,0,0,0,0 | 16,14,44,18,20,15,16,14,15 | 3,2,1,2,0,0,3,0,2,1,2,0,2,3,2 | 24,10,10,14,10,5,5,24,11,9 | 271 | 149 | 0.0430761864872 | | 1 | 0.064,0.038,0.024, | 0.064,0.099,0.052, | -0,037 |
| 252870 | ENSG00000179889 | PDXDC1 | chr16 | + | 15001657 | 15001856 | 14998342 | 14998405 | 15004186 | 15004333 | 252870 | 32,17,57,44,34,39,22,52,8, | 7,10,14,8,1,0,12,2,2,0,2,5 | 23,30,11,11,20,16,20,57,24 | 11,8,3,8,5,8,0,4,9,8,4,20,8, | 298 | 149 | 0.0430881431608 | | 1 | 0.696,0.459,0.671, | 0.511,0.652,0.647, | 0,108 |
| 253351 | ENSG00000089022 | MAPKAPK5 | chr12 | + | 111882693 | 111882853 | 111880446 | 111880527 | 111883580 | 111883768 | 253351 | 8,1,9,6,0,3,2,10,2,1,2,8 | 46,37,60,42,29,35,37,32,53 | 3,6,0,10,4,1,11,7,1,2,15,1,4 | 43,23,36,13,22,18,18,32,8, | 298 | 149 | 0.0430915942016 | | 1 | 0.08,0.013,0.07,0. | 0.034,0.115,0.0,0. | -0,033 |
| 247984 | ENSG00000083799 | CYLD | chr16 | + | 50744729 | 50745036 | 50742761 | 50742841 | 50749575 | 50750202 | 247984 | 3,3,6,8,4,4,3,4,0,1,0,4 | 20,15,54,34,16,29,42,30,8, | 7,14,2,5,0,7,7,2,0,2,5,14,9, | 46,52,37,29,24,15,16,34,23 | 298 | 149 | 0.0431716261416 | | 1 | 0.07,0.091,0.053,0 | 0.071,0.119,0.026, | -0,018 |
| 207054 | ENSG00000119866 | BCL11A | chr2 | - | 60487351 | 60487481 | 60468739 | 60468833 | 60545970 | 60546394 | 207054 | 4,2,9,5,1,0,0,1,0,0,2,0 | 52,21,52,46,43,21,20,15,44 | 0,0,1,0,2,1,0,1,0,0,0,1,3,2,0 | 57,32,28,26,18,28,59,42,11 | 279 | 149 | 0.0431754088243 | | 1 | 0.039,0.048,0.085, | 0.0,0.0,0.019,0.0,0 | 0,015 |
| 140448 | ENSG00000089063 | TMEM230 | chr20 | - | 5111495 | 5111605 | 5109331 | 5109445 | 5112960 | 5113076 | 140448 | 0,0,6,2,0,5,0,0,2,6,0,1 | 20,33,33,23,21,26,33,34,29 | 2,3,4,2,0,0,8,6,0,0,6,5,0,2,0 | 17,29,8,4,4,8,17,21,8,20,38 | 259 | 149 | 0.043187037077 | | 1 | 0.0,0.0,0.095,0.04 | 0.063,0.056,0.223, | -0,04 |
| 102515 | ENSG00000086300 | SNX10 | chr7 | + | 26354056 | 26354184 | 26346419 | 26346466 | 26360974 | 26361061 | 102515 | 1,0,11,2,0,7,2,2,0,1,0,0 | 15,4,48,29,15,14,18,25,15, | 7,9,3,4,4,4,1,4,2,3,1,6,2,2,0 | 21,26,12,17,13,9,11,20,17, | 277 | 149 | 0.0432177094408 | | 1 | 0.035,0.0,0.11,0.0 | 0.152,0.157,0.119, | -0,045 |
| 223479 | ENSG00000167380 | ZNF226 | chr19 | + | 44174983 | 44175171 | 44172859 | 44172952 | 44175497 | 44177687 | 223479 | 4,2,34,16,10,3,7,7,1,2,4,5 | 3,2,23,26,5,5,14,21,16,6,3, | 9,6,12,5,9,12,6,4,4,7,4,6,5, | 2,2,7,0,2,4,10,11,7,13,13,3 | 298 | 149 | 0.0432285779642 | | 1 | 0.4,0.333,0.425,0. | 0.692,0.6,0.462,1. | -0,151 |
| 72233 | ENSG00000181722 | ZBTB20 | chr3 | - | 114713975 | 114714015 | 114693527 | 114693575 | 114801100 | 114801174 | 72233 | 4,4,2,2,2,7,2,5,4,0,1,0 | 35,40,17,14,18,22,26,27,26 | 0,1,1,0,2,1,3,0,1,2,1,1,2,2,4 | 17,27,13,13,13,8,11,12,14, | 189 | 149 | 0.0432329742084 | | 1 | 0.083,0.073,0.085, | 0.0,0.028,0.057,0. | -0,006 |
| 19886 | ENSG00000245534 | RORA-AS1 | chr15 | + | 60488523 | 60488591 | 60479151 | 60479398 | 60510407 | 60510533 | 19886 | 8,3,14,7,9,6,3,6,12,2,3,8 | 8,4,5,1,2,3,3,7,5,2,4,1 | 4,8,6,5,0,7,5,4,4,1,10,8,7,4 | 3,0,2,0,3,0,1,0,3,1,3,2,1,4,4 | 217 | 149 | 0.0432766620763 | | 1 | 0.407,0.34,0.658,0 | 0.478,1.0,0.673,1. | -0,116 |
| 39698 | ENSG00000167515 | TRAPPC2L | chr16 | + | 88859344 | 88859750 | 88858618 | 88858789 | 88859892 | 88859972 | 39698 | 164,156,235,164,73,68,79, | 4,0,21,7,0,3,4,10,8,4,0,4 | 144,178,40,8,70,62,68,118 | 8,9,2,2,5,0,5,14,3,2,7,4,7,3 | 298 | 149 | 0.0433418864852 | | 1 | 0.953,1.0,0.848,0. | 0.9,0.908,0.909,0. | 0,054 |
| 86564 | ENSG00000132196 | HSD17B7 | chr1 | + | 162802831 | 162802949 | 162797827 | 162799937 | 162803430 | 162803535 | 86564 | 5,0,1,0,2,0,0,0,0,0,0,0 | 12,8,12,13,13,6,7,12,5,6,8, | 0,0,1,0,0,2,1,2,2,0,2,6,3,4,1 | 9,18,15,2,25,9,7,17,4,23,15 | 267 | 149 | 0.0433770524766 | | 1 | 0.189,0.0,0.044,0. | 0.0,0.0,0.036,0.0,0 | -0,043 |
| 129281 | ENSG00000179889 | PDXDC1 | chr16 | + | 15001775 | 15001856 | 14998339 | 14998405 | 15004186 | 15004333 | 129281 | 65,59,117,97,61,74,64,83,2 | 7,10,14,8,1,0,12,2,2,0,2,5 | 80,72,25,32,50,37,53,121,5 | 11,8,3,8,5,8,0,4,9,8,4,20,8, | 230 | 149 | 0.0434350305445 | | 1 | 0.857,0.793,0.844, | 0.825,0.854,0.844, | 0,052 |
| 185810 | ENSG00000055917 | PUM2 | chr2 | - | 20278391 | 20278819 | 20263192 | 20263460 | 20282946 | 20283231 | 185810 | 9,5,15,5,9,1,5,8,14,7,3,5 | 24,11,44,29,26,44,17,11,20 | 20,20,8,39,2,3,24,17,1,7,7, | 29,44,11,28,16,22,31,16,5, | 298 | 149 | 0.0434511399697 | | 1 | 0.158,0.185,0.146, | 0.256,0.185,0.267, | -0,022 |
| 177986 | ENSG00000182866 | LCK | chr1 | + | 32276336 | 32276479 | 32275913 | 32276063 | 32276606 | 32277173 | 177986 | 62,14,70,79,43,15,37,26,67 | 18,1,5,4,4,7,5,4,13,4,3,6 | 48,20,36,4,18,29,9,26,17,5 | 11,12,6,5,4,5,2,4,2,8,3,4,0, | 292 | 149 | 0.043467754129 | | 1 | 0.637,0.877,0.877, | 0.69,0.46,0.754,0. | 0,081 |
| 13136 | ENSG00000006194 | ZNF263 | chr16 | + | 3285680 | 3285754 | 3283505 | 3284205 | 3286022 | 3286149 | 13136 | 6,3,5,2,2,4,3,1,3,4,1,3 | 2,10,9,5,0,4,4,0,7,4,1,8 | 1,3,0,0,5,2,4,3,2,4,0,2,3,0,1 | 8,7,4,7,4,1,3,2,2,4,10,6,2,6 | 223 | 149 | 0.043508399991 | | 1 | 0.667,0.167,0.271, | 0.077,0.223,0.0,0. | 0,196 |
| 38945 | ENSG00000291091 | SDHAP4 | chr3 | - | 197622186 | 197622330 | 197621703 | 197621868 | 197623219 | 197623382 | 38945 | 22,25,38,24,21,24,13,26,19 | 35,18,20,18,12,15,4,8,7,3,6 | 17,11,21,9,13,18,9,31,16,2 | 21,39,14,16,14,8,9,18,6,11 | 293 | 149 | 0.0435143357308 | | 1 | 0.242,0.414,0.491, | 0.292,0.125,0.433, | 0,093 |
| 40613 | ENSG00000115524 | SF3B1 | chr2 | - | 197409769 | 197410007 | 197408368 | 197408581 | 197416740 | 197416911 | 40613 | 1497,1043,1569,1009,855, | 24,26,9,5,15,8,9,15,11,11,1 | 1017,1154,885,710,698,32 | 22,26,17,7,10,3,11,15,9,11 | 298 | 149 | 0.0435285494536 | | 1 | 0.969,0.953,0.989, | 0.959,0.957,0.963, | 0,004 |
| 44538 | ENSG00000187514 | PTMA | chr2 | + | 231712451 | 231712516 | 231711889 | 231711983 | 231712803 | 231713289 | 44538 | 173,117,262,155,209,183,1 | 19,6,10,8,9,3,12,18,7,7,2,8 | 164,152,138,72,95,56,121, | 9,12,12,15,13,2,7,11,2,7,29 | 214 | 149 | 0.0435468748264 | | 1 | 0.864,0.931,0.948, | 0.927,0.898,0.889, | 0,02 |
| 239010 | ENSG00000167632 | TRAPPC9 | chr8 | - | 140347756 | 140347940 | 140311247 | 140311374 | 140360049 | 140360193 | 239010 | 5,2,3,2,0,11,9,6,3,2,0,0 | 22,17,25,14,17,34,23,31,16 | 2,6,4,15,7,1,6,0,1,3,9,7,0,5 | 21,13,15,22,13,10,15,34,22 | 298 | 149 | 0.0435542187526 | | 1 | 0.102,0.056,0.057, | 0.045,0.188,0.118, | -0,047 |
| 124680 | ENSG00000152556 | PFKM | chr12 | + | 48135290 | 48135383 | 48134942 | 48135038 | 48137720 | 48137815 | 124680 | 29,22,19,29,13,25,18,6,19, | 0,8,0,2,2,2,0,2,0,0,0,1 | 17,18,21,8,16,6,7,21,14,12 | 1,4,1,3,0,0,0,1,1,0,0,7,4,8,3 | 242 | 149 | 0.0435791846316 | | 1 | 1.0,0.629,1.0,0.89 | 0.913,0.735,0.928, | 0,044 |
| 240853 | ENSG00000109689 | STIM2 | chr4 | + | 26966094 | 26966146 | 26957611 | 26957726 | 26995378 | 26995490 | 240853 | 5,4,11,1,4,3,2,3,1,0,0,4 | 123,103,310,138,90,99,88, | 7,3,4,0,4,1,6,4,3,2,8,8,4,1,1 | 129,134,134,90,96,57,72,1 | 201 | 149 | 0.0435812229778 | | 1 | 0.029,0.028,0.026, | 0.039,0.016,0.022, | -0,012 |
| 192823 | ENSG00000096060 | FKBP5 | chr6 | - | 35630676 | 35630749 | 35620131 | 35620274 | 35637013 | 35637158 | 192823 | 2,0,0,2,0,2,0,12,2,0,0,0 | 11,4,23,41,12,67,37,197,59 | 4,3,2,5,2,1,4,6,0,0,9,2,0,1,2 | 27,65,31,37,19,23,45,138,2 | 222 | 149 | 0.0435857065581 | | 1 | 0.109,0.0,0.0,0.03 | 0.09,0.03,0.042,0. | -0,021 |
| 180868 | ENSG00000114841 | DNAH1 | chr3 | + | 52323810 | 52323880 | 52322408 | 52322775 | 52326139 | 52326314 | 180868 | 2,0,8,1,0,3,3,3,0,0,0,2 | 9,10,5,3,6,11,3,7,3,1,1,0 | 5,0,4,1,3,4,0,2,3,3,2,3,5,3,2 | 5,13,3,15,7,1,5,2,9,7,0,0,7, | 219 | 149 | 0.0436262644057 | | 1 | 0.131,0.0,0.521,0. | 0.405,0.0,0.476,0. | -0,143 |
| 233816 | ENSG00000175463 | TBC1D10C | chr11 | + | 67407548 | 67407760 | 67406826 | 67407016 | 67408978 | 67409133 | 233816 | 4,0,7,0,0,0,0,0,0,1,0,0 | 19,19,37,19,25,12,19,22,14 | 0,4,1,2,2,0,0,1,3,1,1,2,1,4,1 | 15,18,10,6,9,5,14,26,13,13 | 298 | 149 | 0.0436330013288 | | 1 | 0.095,0.0,0.086,0. | 0.0,0.1,0.048,0.14 | -0,033 |

| | | | | | | | | | | | | | | | | | | | | | | | |
|---|---|---|---|---|---|---|---|---|---|---|---|---|---|---|---|---|---|---|---|---|---|---|---|
| 207655 | ENSG00000130348 | QRSL1 | chr6 | + | 106654842 | 106654922 | 106652466 | 106652582 | 106655614 | 106655732 | 207655 | 19,16,61,61,13,15,9,18,23, | 4,8,10,11,8,5,2,2,1,2,2,0 | 28,24,16,7,9,32,4,13,13,34 | 5,2,5,2,0,0,0,2,4,11,3,0,0,0 | 229 | 149 | 0.0436348750024 | | 1 | 0.756,0.565,0.799, | 0.785,0.886,0.676, | -0,076 |
| 45514 | ENSG00000155657 | TTN | chr2 | - | 178734327 | 178734606 | 178733613 | 178733892 | 178735510 | 178736074 | 45514 | 111,63,166,75,54,77,79,73 | 0,0,2,1,0,2,2,1,0,0,0,0 | 77,104,77,49,141,81,48,83 | 2,1,6,1,3,0,0,3,0,0,5,0,0,1, | 298 | 149 | 0.0436660802247 | | 1 | 1.0,1.0,0.976,0.97 | 0.951,0.981,0.865, | 0,025 |
| 175733 | ENSG00000155657 | TTN | chr2 | - | 178734327 | 178734815 | 178733613 | 178733892 | 178735510 | 178736074 | 175733 | 111,63,166,75,54,77,79,73 | 0,0,2,1,0,2,2,1,0,0,0,0 | 77,104,77,49,141,81,48,83 | 2,1,6,1,3,0,0,3,0,0,5,0,0,1, | 298 | 149 | 0.0436660802247 | | 1 | 1.0,1.0,0.976,0.97 | 0.951,0.981,0.865, | 0,025 |
| 121603 | ENSG00000166405 | RIC3 | chr11 | - | 8126658 | 8126807 | 8106055 | 8111137 | 8137377 | 8137471 | 121603 | 87,29,128,62,64,73,56,47, | 0,0,7,0,6,2,5,4,0,3,2,2 | 83,81,29,60,23,23,23,46,1 | 2,4,3,3,0,2,2,1,1,3,7,3,5,6, | 298 | 149 | 0.0436698181727 | | 1 | 1.0,1.0,0.901,1.0,0 | 0.954,0.91,0.829,0 | 0,047 |
| 185243 | ENSG00000132185 | FCRLA | chr1 | + | 161712067 | 161712218 | 161707294 | 161707343 | 161713084 | 161713468 | 185243 | 264,212,591,519,396,179, | 10,2,4,2,0,4,2,0,6,12,0,6 | 445,344,121,85,131,131,1 | 3,4,0,2,0,0,2,2,0,0,2,7,2,2, | 298 | 149 | 0.043689490691 | | 1 | 0.93,0.981,0.987,0 | 0.987,0.977,1.0,0. | -0,012 |
| 142901 | ENSG00000154262 | ABCA6 | chr17 | - | 69126385 | 69126634 | 69124887 | 69125035 | 69128618 | 69128804 | 142901 | 10,8,44,27,7,2,20,22,7,3,0, | 89,74,546,380,143,62,243, | 15,10,7,5,13,6,5,10,5,12,6, | 385,356,105,29,128,167,8 | 298 | 149 | 0.0436949416526 | | 1 | 0.053,0.051,0.039, | 0.019,0.014,0.032, | 0,002 |
| 39336 | ENSG00000075826 | SEC31B | chr10 | - | 100501653 | 100501710 | 100500109 | 100500159 | 100502253 | 100502484 | 39336 | 1,0,3,4,1,2,2,2,5,0,2,0 | 5,8,15,14,3,10,12,7,1,2,10, | 0,1,0,0,6,0,1,0,2,4,2,2,1,2, | 4,3,19,7,18,5,7,8,9,15,2,7, | 206 | 149 | 0.0437554635807 | | 1 | 0.126,0.0,0.126,0. | 0.0,0.194,0.0,0.0,0 | -0,01 |
| 79844 | ENSG00000005483 | KMT2E | chr7 | + | 105068474 | 105068545 | 105066726 | 105066807 | 105073618 | 105073677 | 79844 | 9,2,5,7,5,6,6,1,2,1,7,2 | 173,135,196,130,48,71,68, | 12,12,16,11,5,3,5,5,5,3,7,6 | 150,174,110,186,119,54,7 | 220 | 149 | 0.0437624895621 | | 1 | 0.034,0.01,0.017,0 | 0.051,0.045,0.09,0 | -0,001 |
| 216505 | ENSG00000168918 | INPP5D | chr2 | + | 233145267 | 233145515 | 233139841 | 233139929 | 233146161 | 233146242 | 216505 | 10,2,3,2,2,4,4,15,0,4,0,0 | 78,70,108,118,84,81,106,1 | 1,8,5,12,3,6,7,4,1,0,19,4,0, | 82,106,56,41,43,52,38,78, | 298 | 149 | 0.0438029740983 | | 1 | 0.06,0.014,0.014,0 | 0.006,0.036,0.043, | -0,015 |
| 126227 | ENSG00000120539 | MASTL | chr10 | + | 27173120 | 27173259 | 27169943 | 27171083 | 27180952 | 27181066 | 126227 | 21,8,39,19,4,12,16,13,8,2, | 1,0,3,4,2,0,6,6,0,0,3,1 | 11,17,6,24,19,6,15,13,5,11 | 3,2,1,5,4,3,1,5,3,6,1,9,2,2, | 288 | 149 | 0.0438058904894 | | 1 | 0.916,1.0,0.871,0. | 0.655,0.815,0.756, | 0,068 |
| 113279 | ENSG00000064313 | TAF2 | chr8 | - | 119744880 | 119745036 | 119744287 | 119744393 | 119746704 | 119746934 | 113279 | 3,3,12,2,0,2,0,2,2,1,0,0 | 11,10,27,19,7,9,10,10,16,4 | 4,1,2,0,1,2,3,5,1,3,6,0,0,0, | 12,15,16,5,8,6,1,13,6,15,1 | 298 | 149 | 0.0438100728959 | | 1 | 0.12,0.13,0.182,0. | 0.143,0.032,0.059, | -0,044 |
| 202995 | ENSG00000239305 | RNF103 | chr2 | - | 86616716 | 86616791 | 86612158 | 86612274 | 86620329 | 86620469 | 202995 | 0,0,3,2,1,0,0,0,0,0,0,0 | 12,12,42,17,3,8,4,10,14,4, | 3,2,4,3,1,2,2,0,0,0,2,2,1,0, | 19,38,7,8,18,6,20,12,7,6,1 | 224 | 149 | 0.0438273339114 | | 1 | 0.0,0.0,0.045,0.07 | 0.095,0.034,0.275, | -0,06 |
| 4827 | ENSG00000245680 | ZNF585B | chr19 | - | 37199399 | 37199497 | 37190072 | 37190150 | 37207039 | 37207254 | 4827 | 3,2,9,9,5,0,1,8,0,6,1,1 | 0,1,6,5,12,1,1,5,7,3,1,4 | 7,2,1,2,0,1,7,6,0,0,0,2,6,6, | 7,0,0,0,1,0,1,6,2,1,3,1,7,0, | 247 | 149 | 0.0438276913087 | | 1 | 1.0,0.547,0.475,0. | 0.376,1.0,1.0,1.0,0 | -0,183 |
| 111673 | ENSG00000185900 | POMK | chr8 | + | 43102504 | 43102600 | 43097523 | 43097615 | 43103527 | 43103830 | 111673 | 11,0,14,29,26,6,11,1,20,6, | 3,1,7,5,6,5,2,2,2,0,1,2 | 8,2,5,5,4,5,4,9,2,10,5,7,8,1 | 5,3,5,2,1,2,5,3,0,3,0,5,3,10 | 245 | 149 | 0.0438778520661 | | 1 | 0.69,0.0,0.549,0.7 | 0.493,0.288,0.378, | 0,057 |
| 165899 | ENSG00000172869 | DMXL1 | chr5 | + | 119099231 | 119099416 | 119097978 | 119098104 | 119101934 | 119102210 | 165899 | 5,1,11,7,5,2,2,3,1,5,0,4 | 160,127,129,92,50,78,67,7 | 4,3,3,1,1,2,2,0,0,2,6,8,5,0, | 91,115,119,74,73,59,41,65 | 298 | 149 | 0.0439192310184 | | 1 | 0.015,0.004,0.041, | 0.022,0.013,0.012, | 0,009 |
| 201298 | ENSG00000055070 | SZRD1 | chr1 | + | 16391374 | 16391424 | 16380015 | 16380093 | 16393227 | 16393385 | 201298 | 20,13,11,6,3,5,7,11,27,9,4, | 2,0,0,0,0,0,0,0,0,0,0,4 | 11,9,8,2,1,1,7,16,1,3,7,18, | 2,2,1,4,0,0,2,2,1,0,3,3,1,0, | 199 | 149 | 0.0439289705418 | | 1 | 0.882,1.0,1.0,1.0,1 | 0.805,0.771,0.857, | 0,141 |
| 72178 | ENSG00000132680 | KHDC4 | chr1 | - | 155926675 | 155926839 | 155925631 | 155925843 | 155927103 | 155927156 | 72178 | 48,34,78,90,78,47,42,81,6 | 13,2,10,12,2,3,0,2,3,2,1,3 | 41,37,24,16,40,29,19,43,4 | 19,4,4,1,3,2,1,3,7,2,8,5,4,1 | 298 | 149 | 0.0439620813911 | | 1 | 0.649,0.895,0.796, | 0.519,0.822,0.75,0 | 0,064 |
| 200128 | ENSG00000131051 | RBM39 | chr20 | - | 35734945 | 35735081 | 35734185 | 35734257 | 35738967 | 35739017 | 200128 | 11,8,26,10,5,5,1,9,8,0,0,2 | 30,10,27,8,7,7,1,8,10,14,3, | 6,14,5,18,5,5,1,8,8,13,8,14 | 6,2,18,14,13,6,4,3,2,8,3,18 | 285 | 149 | 0.0439623478706 | | 1 | 0.161,0.295,0.335, | 0.343,0.785,0.127, | -0,142 |
| 55465 | ENSG00000132185 | FCRLA | chr1 | + | 161711933 | 161712218 | 161706971 | 161707343 | 161713084 | 161713468 | 55465 | 269,215,602,530,427,178, | 10,2,4,2,0,4,2,0,6,12,0,6 | 464,352,124,90,133,132,2 | 3,4,0,2,0,0,2,2,0,0,2,7,2,2, | 298 | 149 | 0.0439691608159 | | 1 | 0.931,0.982,0.987, | 0.987,0.978,1.0,0. | -0,011 |
| 119401 | ENSG00000100731 | PCNX1 | chr14 | + | 70988566 | 70988699 | 70976941 | 70978648 | 70995740 | 70995843 | 119401 | 6,14,18,4,13,8,10,12,5,7,3, | 3,0,3,4,0,7,0,1,2,2,2,2 | 0,3,7,2,4,6,2,15,5,15,3,6,1 | 4,3,6,0,0,0,3,1,0,0,0,0,0,2, | 282 | 149 | 0.0439845581324 | | 1 | 0.514,1.0,0.76,0.3 | 0.0,0.346,0.381,1. | -0,073 |
| 88301 | ENSG00000205531 | NAP1L4 | chr11 | - | 2954300 | 2954410 | 2951779 | 2951809 | 2954526 | 2954646 | 88301 | 13,3,4,2,4,3,2,4,1,2,0,0 | 82,46,59,40,29,26,23,16,6 | 3,4,3,9,5,1,2,0,2,6,2,4,3,5, | 33,13,29,17,16,17,15,32,6, | 259 | 149 | 0.0440164977594 | | 1 | 0.084,0.036,0.038, | 0.05,0.15,0.056,0. | -0,038 |
| 176055 | ENSG00000116191 | RALGPS2 | chr1 | + | 178902142 | 178902211 | 178897663 | 178897756 | 178906775 | 178906811 | 176055 | 427,311,288,146,105,58,1 | 8,3,5,4,3,5,5,0,0,4,0,2 | 277,246,181,250,124,90,5 | 5,4,1,2,0,3,2,0,1,2,0,0,4,2, | 218 | 149 | 0.0440197578838 | | 1 | 0.973,0.986,0.975, | 0.974,0.977,0.992, | -0,012 |
| 49131 | ENSG00000009307 | CSDE1 | chr1 | - | 114730257 | 114730422 | 114726982 | 114727090 | 114732603 | 114732816 | 49131 | 367,314,795,465,241,235, | 30,36,86,47,12,25,16,41,9, | 424,325,209,187,271,178, | 27,65,39,70,97,22,21,19,3 | 298 | 149 | 0.0440509900795 | | 1 | 0.859,0.813,0.822, | 0.887,0.714,0.728, | 0,052 |
| 143646 | ENSG00000267121 | FMNL1-DT | chr17 | - | 45219933 | 45220075 | 45219349 | 45219632 | 45220267 | 45222222 | 143646 | 11,2,24,23,8,4,7,10,3,7,9,1 | 42,16,85,47,56,42,32,17,2 | 7,15,1,0,3,7,5,11,7,6,2,2,9, | 44,41,22,18,50,21,21,67,3 | 291 | 149 | 0.0440692475211 | | 1 | 0.118,0.06,0.126,0 | 0.075,0.158,0.023, | 0,038 |
| 148544 | ENSG00000122299 | ZC3H7A | chr16 | - | 11769030 | 11769112 | 11768314 | 11768501 | 11770782 | 11770987 | 148544 | 29,28,34,35,9,9,16,28,21,2 | 1,4,6,1,3,2,2,2,2,3,2,1 | 16,45,32,22,15,14,10,10,2 | 0,8,2,8,9,4,6,7,1,3,3,2,0,2, | 231 | 149 | 0.0440799092736 | | 1 | 0.949,0.819,0.785, | 1.0,0.784,0.912,0. | 0,077 |
| 97401 | ENSG00000083168 | KAT6A | chr8 | - | 41942792 | 41943000 | 41940841 | 41941444 | 41943747 | 41943979 | 97401 | 103,78,165,79,76,50,58,73 | 3,0,6,1,6,6,1,0,2,0,5,0 | 129,110,59,137,54,61,74,8 | 2,4,3,9,5,5,0,2,2,3,8,9,4,8, | 298 | 149 | 0.0440844650569 | | 1 | 0.945,1.0,0.932,0. | 0.97,0.932,0.908,0 | 0,029 |
| 61504 | ENSG00000111445 | RFC5 | chr12 | + | 118017689 | 118018053 | 118016721 | 118016892 | 118019071 | 118019136 | 61504 | 11,3,8,7,4,5,6,3,7,1,4,2 | 9,8,15,12,5,6,2,5,11,9,2,2 | 3,6,4,2,3,1,3,8,2,10,9,0,9,6 | 8,10,2,0,0,1,1,9,1,8,7,19,1, | 298 | 149 | 0.044087206624 | | 1 | 0.379,0.158,0.211, | 0.158,0.231,0.5,1. | -0,228 |
| 230681 | ENSG00000225648 | SBDSP1 | chr7 | + | 72833763 | 72833899 | 72831601 | 72831798 | 72836576 | 72836701 | 230681 | 91,30,220,132,45,76,94,10 | 8,0,6,1,0,1,0,9,4,0,2,0 | 123,117,114,123,96,75,29, | 6,0,4,1,3,0,0,1,3,2,4,0,1,0, | 285 | 149 | 0.0441172450675 | | 1 | 0.856,1.0,0.95,0.9 | 0.915,1.0,0.937,0. | -0,023 |
| 108685 | ENSG00000196696 | PDXDC2P-NPIP | chr16 | - | 69976810 | 69978003 | 69976296 | 69976732 | 69978076 | 69978112 | 108685 | 3,2,54,22,4,6,16,13,0,0,3,1 | 22,5,4,0,9,3,5,8,13,5,1,5 | 7,8,7,5,13,2,6,2,15,8,6,2,1 | 6,4,1,3,0,2,4,7,1,3,3,2,0,6, | 298 | 149 | 0.0441309895548 | | 1 | 0.064,0.167,0.871, | 0.368,0.5,0.778,0. | -0,141 |
| 112731 | ENSG00000172269 | DPAGT1 | chr11 | - | 119097854 | 119098043 | 119096522 | 119097551 | 119098402 | 119098487 | 112731 | 9,6,23,26,9,8,18,8,12,4,10, | 0,0,2,0,2,0,2,6,0,0,0,2 | 13,4,8,2,3,6,3,2,11,13,7,21 | 0,0,0,0,0,1,0,1,1,1,2,0,1,1, | 298 | 149 | 0.0441442523662 | | 1 | 1.0,1.0,0.852,1.0,0 | 1.0,1.0,1.0,1.0,1.0, | -0,022 |
| 32116 | ENSG00000119522 | DENND1A | chr9 | - | 123387729 | 123387858 | 123383654 | 123383913 | 123403401 | 123403490 | 32116 | 2,0,1,1,0,0,0,0,0,0,0,0 | 4,5,9,2,1,11,5,7,5,1,2,3 | 0,2,3,1,0,0,0,0,1,2,0,0,4,0, | 5,3,7,9,3,4,1,2,0,2,6,8,3,6, | 278 | 149 | 0.044167150214 | | 1 | 0.211,0.0,0.056,0. | 0.0,0.263,0.187,0. | -0,165 |
| 239705 | ENSG00000095209 | TMEM38B | chr9 | + | 105714021 | 105714505 | 105705674 | 105705753 | 105721536 | 105721721 | 239705 | 0,0,3,2,0,0,3,0,0,0,2,2 | 6,4,15,9,5,5,5,2,4,4,3,4 | 5,3,1,1,1,0,0,1,0,1,4,4,0,2, | 19,5,3,0,4,6,5,9,2,6,7,13,6, | 298 | 149 | 0.0441749046494 | | 1 | 0.0,0.0,0.091,0.1,0 | 0.116,0.231,0.143, | -0,127 |
| 218116 | ENSG00000064933 | PMS1 | chr2 | + | 189817041 | 189817110 | 189805651 | 189805754 | 189818016 | 189818181 | 218116 | 0,0,5,4,2,6,2,3,3,0,1,0 | 17,3,95,60,38,23,18,24,16, | 4,2,0,1,0,1,0,0,0,0,3,1,0,3, | 46,33,17,0,9,10,13,30,16,1 | 218 | 149 | 0.0441798931076 | | 1 | 0.0,0.0,0.035,0.04 | 0.056,0.04,0.0,1.0, | -0,037 |
| 112500 | ENSG00000119397 | CNTRL | chr9 | + | 121141380 | 121141588 | 121140640 | 121140786 | 121142090 | 121142270 | 112500 | 160,73,196,141,102,78,55, | 13,9,24,6,11,16,20,9,4,4,7, | 155,114,103,129,78,68,67, | 10,13,24,18,5,8,7,21,12,23 | 298 | 149 | 0.0442012593974 | | 1 | 0.86,0.802,0.803,0 | 0.886,0.814,0.682, | 0,046 |
| 127550 | ENSG00000107862 | GBF1 | chr10 | + | 102381126 | 102381255 | 102379283 | 102380686 | 102382055 | 102382336 | 127550 | 50,41,55,25,34,19,46,40,6 | 4,3,8,1,0,0,0,1,0,6,3,0 | 23,34,29,23,21,19,29,26,1 | 0,0,2,5,2,0,0,0,0,0,3,0,2,2, | 278 | 149 | 0.0442246962355 | | 1 | 0.87,0.88,0.787,0. | 1.0,1.0,0.886,0.71 | 0 |
| 126791 | ENSG00000066084 | DIP2B | chr12 | + | 50691048 | 50691151 | 50686572 | 50686682 | 50695266 | 50695360 | 126791 | 63,62,49,34,13,22,18,25,4 | 10,5,6,2,1,2,3,10,2,0,0,1 | 36,22,36,30,25,20,14,17,2 | 1,8,5,7,7,3,2,12,7,5,4,0,0,0 | 252 | 149 | 0.0442269157618 | | 1 | 0.788,0.88,0.828,0 | 0.955,0.619,0.81,0 | 0,093 |
| 116814 | ENSG00000108424 | KPNB1 | chr17 | + | 47673489 | 47673561 | 47673017 | 47673165 | 47674637 | 47674755 | 116814 | 185,206,426,316,116,122, | 6,0,0,0,0,2,0,2,0,0,0,0 | 311,365,155,122,185,143, | 2,7,4,11,4,4,0,0,4,0,4,0,3,0 | 221 | 149 | 0.0442324686282 | | 1 | 0.954,1.0,1.0,1.0,1 | 0.991,0.972,0.963, | 0,018 |
| 208323 | ENSG00000049618 | ARID1B | chr6 | + | 157110475 | 157110561 | 157084771 | 157084905 | 157133027 | 157133207 | 208323 | 7,13,32,23,4,10,16,20,4,6, | 0,0,0,0,0,1,1,0,0,0,1,0 | 32,37,12,28,13,15,23,18,2 | 2,0,5,3,0,0,2,0,0,0,4,0,1,1, | 235 | 149 | 0.0442373137207 | | 1 | 1.0,1.0,1.0,1.0,1.0, | 0.91,1.0,0.603,0.8 | 0,055 |
| 255647 | ENSG00000171792 | RHNO1 | chr12 | + | 2885377 | 2885534 | 2877222 | 2877282 | 2887910 | 2887934 | 255647 | 81,61,45,41,34,30,24,31,4 | 3,2,2,7,0,3,3,2,1,0,0,2 | 6,24,32,27,24,12,21,34,9,3 | 7,1,1,0,3,2,1,4,2,0,1,3,4,4, | 298 | 149 | 0.0442950944934 | | 1 | 0.931,0.938,0.918, | 0.3,0.923,0.941,1. | 0,085 |
| 245930 | ENSG00000166405 | RIC3 | chr11 | - | 8137379 | 8137471 | 8135786 | 8135844 | 8138271 | 8138347 | 245930 | 61,26,95,42,48,43,41,42,3 | 1,0,2,0,0,0,1,0,0,0,3,0 | 57,69,60,65,28,18,18,31,1 | 6,1,2,6,0,0,0,3,0,0,4,1,1,0, | 241 | 149 | 0.0443174244172 | | 1 | 0.974,1.0,0.967,1. | 0.855,0.977,0.949, | 0,028 |
| 45054 | ENSG00000111642 | CHD4 | chr12 | - | 6600925 | 6601053 | 6600533 | 6600669 | 6601288 | 6601530 | 45054 | 195,164,278,183,140,167, | 10,2,5,1,3,9,6,10,0,4,0,3 | 170,261,112,61,171,95,85, | 8,11,4,12,4,5,1,14,6,1,11,8 | 277 | 149 | 0.0443425392896 | | 1 | 0.913,0.978,0.968, | 0.92,0.927,0.938,0 | 0,028 |
| 86809 | ENSG00000144134 | RABL2A | chr2 | + | 113640893 | 113641005 | 113635050 | 113635130 | 113641352 | 113641450 | 86809 | 12,4,6,12,4,10,5,9,3,5,1,2 | 5,1,1,1,2,4,4,1,0,0,2,2 | 9,11,6,7,4,4,2,6,4,13,3,3,1 | 0,0,1,2,0,2,0,5,2,5,0,0,0,1, | 261 | 149 | 0.0443478486854 | | 1 | 0.578,0.695,0.774, | 1.0,1.0,0.774,0.66 | -0,14 |
| 178615 | ENSG00000120008 | WDR11 | chr10 | + | 120905833 | 120906021 | 120905318 | 120905416 | 120906775 | 120906855 | 178615 | 185,173,244,156,117,56,6 | 7,3,10,6,5,5,6,4,2,0,1,4 | 146,148,209,125,126,111, | 6,10,1,5,17,2,0,6,6,15,6,4, | 298 | 149 | 0.0443604360924 | | 1 | 0.93,0.966,0.924,0 | 0.924,0.881,0.991, | 0,025 |
| 74298 | ENSG00000099622 | CIRBP | chr19 | + | 1273600 | 1273715 | 1271980 | 1272051 | 1274306 | 1274440 | 74298 | 3,1,20,15,24,7,7,8,11,7,3,3 | 16,7,13,6,8,6,1,0,4,2,1,7 | 3,0,4,1,4,8,5,4,1,7,8,7,17,1 | 10,6,2,1,5,3,3,10,4,9,9,30, | 264 | 149 | 0.0443830251957 | | 1 | 0.096,0.075,0.465, | 0.145,0.0,0.53,0.3 | 0,171 |
| 36684 | ENSG00000182150 | ERCC6L2 | chr9 | + | 95907077 | 95907271 | 95897848 | 95897971 | 95915667 | 95915829 | 36684 | 35,20,103,49,14,21,29,32, | 2,0,4,2,2,2,0,2,0,1,3,1 | 54,42,34,38,24,23,19,34,1 | 4,6,4,5,2,1,1,3,1,1,4,0,1,4, | 298 | 149 | 0.0444111608562 | | 1 | 0.897,1.0,0.928,0. | 0.871,0.778,0.81,0 | 0,004 |
| 77356 | ENSG00000128815 | WDFY4 | chr10 | + | 48817244 | 48817409 | 48813956 | 48814082 | 48820233 | 48820437 | 77356 | 188,140,166,151,221,191, | 0,1,2,2,1,6,0,1,0,0,1,4 | 165,151,50,169,45,59,164, | 2,9,0,5,0,0,1,2,1,0,6,3,0,1, | 298 | 149 | 0.0444232641909 | | 1 | 1.0,0.986,0.976,0. | 0.976,0.893,1.0,0. | 0,008 |
| 88392 | ENSG00000136699 | SMPD4 | chr2 | - | 130161185 | 130161272 | 130157250 | 130157396 | 130164373 | 130164445 | 88392 | 14,7,1,8,4,11,12,12,19,7,1, | 14,2,3,5,4,6,17,8,19,13,4,1 | 2,4,5,13,3,1,3,3,7,5,1,3,1,0 | 7,1,0,9,3,5,0,13,2,16,5,14,1 | 236 | 149 | 0.0444766654293 | | 1 | 0.387,0.688,0.174, | 0.153,0.716,1.0,0. | 0,041 |
| 138232 | ENSG00000125731 | SH2D3A | chr19 | - | 6760637 | 6760984 | 6757329 | 6759670 | 6763679 | 6764481 | 138232 | 33,13,39,30,34,27,11,21,1 | 6,0,1,1,2,2,4,4,0,2,1,0 | 20,19,9,12,17,6,18,9,16,15 | 1,4,0,3,1,0,0,1,0,0,0,2,2,0, | 298 | 149 | 0.0444865088093 | | 1 | 0.733,1.0,0.951,0. | 0.909,0.704,1.0,0. | -0,048 |
| 198166 | ENSG00000126775 | ATG14 | chr14 | - | 55385678 | 55386096 | 55381961 | 55382191 | 55390910 | 55390992 | 198166 | 54,43,57,33,23,42,35,29,1 | 6,0,3,1,0,1,1,2,0,0,0,3 | 46,71,33,47,29,13,13,34,2 | 1,3,0,0,2,0,2,0,1,0,3,0,0,0, | 298 | 149 | 0.0445291101718 | | 1 | 0.818,1.0,0.905,0. | 0.958,0.922,1.0,1. | -0,027 |
| 16770 | ENSG00000122299 | ZC3H7A | chr16 | - | 11769030 | 11769095 | 11768314 | 11768501 | 11770782 | 11770841 | 16770 | 45,41,58,68,18,34,31,60,2 | 1,4,6,1,3,2,2,2,2,3,2,1 | 56,81,53,43,24,30,26,40,4 | 0,8,2,8,9,4,6,7,1,3,3,2,0,2, | 214 | 149 | 0.0445478763509 | | 1 | 0.969,0.877,0.871, | 1.0,0.876,0.949,0. | 0,027 |
| 233322 | ENSG00000215301 | DDX3X | chrX | + | 41346930 | 41347012 | 41346504 | 41346622 | 41347639 | 41347694 | 233322 | 3,1,13,16,9,3,3,7,2,4,1,2 | 3,1,7,4,3,0,2,0,0,1,0,2 | 16,12,1,9,6,5,1,11,6,5,16,1 | 8,5,1,31,0,6,5,8,1,6,10,0,3, | 231 | 149 | 0.0445680887413 | | 1 | 0.392,0.392,0.545, | 0.563,0.608,0.392, | 0,157 |
| 145667 | ENSG00000167978 | SRRM2 | chr16 | + | 2769982 | 2770227 | 2768996 | 2769284 | 2770351 | 2770465 | 145667 | 9,0,7,7,1,2,2,2,7,3,1,0 | 36,23,13,5,6,12,18,13,6,4, | 3,8,2,9,2,7,14,4,1,0,7,3,6,3 | 3,11,4,30,9,1,17,12,10,3,9, | 298 | 149 | 0.0445895482986 | | 1 | 0.111,0.0,0.212,0. | 0.333,0.267,0.2,0. | -0,071 |
| 80654 | ENSG00000109466 | KLHL2 | chr4 | + | 165219643 | 165219722 | 165207822 | 165207902 | 165219933 | 165220059 | 80654 | 0,0,11,6,0,8,1,9,0,0,0,0 | 4,3,10,12,5,3,5,17,1,1,2,10 | 3,8,2,0,3,1,6,7,4,1,6,3,1,3, | 7,2,0,3,2,0,3,7,2,4,4,7,11,5 | 228 | 149 | 0.0446074657034 | | 1 | 0.0,0.0,0.418,0.24 | 0.219,0.723,1.0,0. | -0,258 |
| 135169 | ENSG00000093000 | NUP50 | chr22 | + | 45170987 | 45171116 | 45168165 | 45168246 | 45171599 | 45172206 | 135169 | 2,4,9,2,0,1,3,3,3,1,3,0 | 11,10,13,10,10,8,16,10,25, | 4,3,4,1,4,3,2,3,1,3,2,5,0,5, | 7,10,5,5,6,6,5,5,4,4,7,7,11, | 278 | 149 | 0.0446181544601 | | 1 | 0.089,0.177,0.271, | 0.234,0.139,0.3,0. | -0,066 |
| 8075 | ENSG00000176225 | RTTN | chr18 | - | 70168854 | 70168973 | 70166918 | 70167031 | 70176674 | 70176845 | 8075 | 8,7,28,7,3,4,3,12,2,1,5,8 | 0,0,1,0,0,0,1,0,0,2,0,0 | 11,7,0,8,17,2,12,6,4,5,15,8 | 2,3,2,3,0,0,0,2,0,0,0,0,0,0, | 268 | 149 | 0.0446274728484 | | 1 | 1.0,1.0,0.94,1.0,1. | 0.754,0.565,0.0,0. | 0,096 |
| 102618 | ENSG00000122566 | HNRNPA2B1 | chr7 | - | 26191421 | 26191650 | 26190992 | 26191128 | 26192494 | 26192577 | 102618 | 9,0,11,0,0,4,3,9,0,0,1,6 | 11,5,18,17,12,8,10,28,8,1, | 0,3,4,4,2,3,4,8,0,1,3,0,6,2, | 23,33,13,20,8,14,7,24,21,1 | 298 | 149 | 0.0446541569507 | | 1 | 0.29,0.0,0.234,0.0, | 0.0,0.043,0.133,0. | 0,007 |
| 251049 | ENSG00000139620 | KANSL2 | chr12 | - | 48661022 | 48661243 | 48660467 | 48660619 | 48662559 | 48662592 | 251049 | 7,4,2,8,1,3,3,3,2,3,4,4 | 1,0,2,0,0,2,2,1,0,0,0,0 | 2,7,1,2,2,1,2,1,0,1,0,6,1,0, | 3,1,4,0,0,0,0,1,1,0,1,1,1,3, | 298 | 149 | 0.0447801876807 | | 1 | 0.778,1.0,0.333,1. | 0.25,0.778,0.111,1 | 0,2 |
| 226442 | ENSG00000181090 | EHMT1 | chr9 | + | 137778239 | 137778290 | 137777969 | 137778055 | 137779634 | 137779717 | 226442 | 2,0,9,3,1,3,4,4,2,1,0,4 | 83,60,101,85,67,48,80,65, | 8,9,3,2,2,0,8,2,5,0,0,2,3,4, | 77,62,60,77,27,32,26,50,4 | 200 | 149 | 0.0447808622845 | | 1 | 0.018,0.0,0.062,0. | 0.072,0.098,0.036, | -0,021 |
| 149344 | ENSG00000140157 | NIPA2 | chr15 | + | 22850939 | 22851233 | 22845145 | 22845267 | 22851638 | 22851870 | 149344 | 0,0,13,1,2,2,0,2,1,3,2,0 | 26,11,28,20,11,8,8,7,25,12 | 6,6,0,3,3,3,1,1,2,2,1,3,4,3, | 20,11,22,2,3,6,2,5,11,4,8,8 | 298 | 149 | 0.0448034155951 | | 1 | 0.0,0.0,0.188,0.02 | 0.13,0.214,0.0,0.4 | -0,096 |
| 81144 | ENSG00000166986 | MARS1 | chr12 | + | 57489423 | 57489558 | 57489266 | 57489345 | 57490206 | 57490379 | 81144 | 85,27,99,71,31,46,39,32,1 | 3,0,5,0,2,5,1,2,3,2,0,0 | 58,62,35,23,41,13,25,31,3 | 1,4,0,3,7,0,6,2,0,5,5,5,2,3, | 284 | 149 | 0.0448168863673 | | 1 | 0.937,1.0,0.912,1. | 0.968,0.89,1.0,0.8 | 0,037 |
| 224786 | ENSG00000145725 | PPIP5K2 | chr5 | + | 103133452 | 103133618 | 103129305 | 103129703 | 103136731 | 103136822 | 224786 | 19,7,76,44,34,34,19,22,4,5 | 1,0,4,1,0,3,0,0,1,0,0,0 | 53,29,12,15,24,8,6,24,14,1 | 0,5,4,1,9,0,0,0,1,0,0,0,0,0, | 298 | 149 | 0.0448298570351 | | 1 | 0.905,1.0,0.905,0. | 1.0,0.744,0.6,0.88 | 0,051 |
| 178618 | ENSG00000120008 | WDR11 | chr10 | + | 120905862 | 120906021 | 120905318 | 120905416 | 120906775 | 120906855 | 178618 | 185,173,244,156,117,54,6 | 7,3,10,6,5,5,6,4,2,0,1,4 | 146,148,209,127,126,111, | 6,10,1,5,17,2,0,6,6,15,6,4, | 298 | 149 | 0.0448983707787 | | 1 | 0.93,0.966,0.924,0 | 0.924,0.881,0.991, | 0,025 |
| 139232 | ENSG00000171456 | ASXL1 | chr20 | + | 32371779 | 32371874 | 32369014 | 32369123 | 32428127 | 32428248 | 139232 | 2,4,1,2,3,2,4,4,0,2,3,0 | 38,29,92,58,21,29,29,36,9, | 2,10,1,3,2,0,0,5,2,11,3,5,4, | 36,54,29,43,35,29,10,34,1 | 244 | 149 | 0.0449010561679 | | 1 | 0.031,0.078,0.007, | 0.033,0.102,0.021, | -0,006 |
| 159430 | ENSG00000132780 | NASP | chr1 | + | 45617462 | 45617525 | 45616625 | 45616703 | 45618060 | 45618904 | 159430 | 55,34,48,34,27,14,11,9,28, | 8,2,2,2,0,2,4,4,2,0,0,5 | 35,31,21,12,8,10,11,11,10, | 5,8,2,7,1,0,3,4,0,4,2,5,2,1, | 212 | 149 | 0.0449377074197 | | 1 | 0.829,0.923,0.944, | 0.831,0.731,0.881, | 0,059 |
| 242663 | ENSG00000115459 | ELMOD3 | chr2 | + | 85369465 | 85369830 | 85368685 | 85368754 | 85371085 | 85371104 | 242663 | 13,8,12,16,10,11,7,7,8,5,4, | 1,1,1,0,0,0,1,0,1,0,0,0 | 26,14,6,3,8,2,7,13,8,11,10, | 3,0,2,1,0,0,0,1,1,2,0,0,8,0, | 298 | 149 | 0.0449528717644 | | 1 | 0.867,0.8,0.857,1. | 0.812,1.0,0.6,0.6,1 | 0,082 |
| 96954 | ENSG00000198815 | FOXJ3 | chr1 | - | 42323683 | 42323827 | 42311049 | 42311110 | 42335058 | 42335235 | 96954 | 1,2,4,4,8,1,3,4,0,0,0,7 | 5,3,12,6,9,12,8,12,7,12,7,8 | 0,2,0,3,1,1,0,1,3,0,4,3,0,4, | 4,3,9,10,2,3,10,6,4,2,17,18 | 293 | 149 | 0.0449818744811 | | 1 | 0.092,0.253,0.145, | 0.0,0.253,0.0,0.13 | 0,02 |
| 157512 | ENSG00000078319 | PMS2P1 | chr7 | - | 100335332 | 100335428 | 100332441 | 100332581 | 100335837 | 100335860 | 157512 | 0,0,2,2,0,0,0,1,0,0,0,0 | 33,26,42,52,14,16,14,28,1 | 3,0,1,2,0,0,0,1,0,2,2,13,1,0 | 47,34,25,14,25,6,7,12,18,2 | 245 | 149 | 0.045021490832 | | 1 | 0.0,0.0,0.028,0.02 | 0.037,0.0,0.024,0. | -0,022 |
| 16201 | ENSG00000149923 | PPP4C | chr16 | + | 30082745 | 30082847 | 30082483 | 30082534 | 30083393 | 30083567 | 16201 | 55,42,125,81,49,61,22,58, | 1,1,3,1,0,0,0,1,2,0,0,0 | 66,96,50,28,51,24,23,51,4 | 4,6,3,4,0,0,0,0,0,0,1,1,4,2 | 251 | 149 | 0.0451073238904 | | 1 | 0.97,0.961,0.961,0 | 0.907,0.905,0.908, | 0,023 |
| 144790 | ENSG00000141568 | FOXK2 | chr17 | + | 82584063 | 82584188 | 82571723 | 82571870 | 82585903 | 82586200 | 144790 | 11,10,8,3,11,9,5,6,45,11,5, | 7,11,1,4,6,2,4,3,2,2,1,3 | 6,12,11,2,2,8,4,12,4,10,23, | 1,0,2,1,2,0,0,10,0,1,6,5,2,1 | 274 | 149 | 0.0451084675097 | | 1 | 0.461,0.331,0.813, | 0.765,1.0,0.749,0. | -0,16 |
| 130082 | ENSG00000139350 | NEDD1 | chr12 | + | 96917620 | 96917737 | 96912575 | 96912817 | 96919984 | 96920118 | 130082 | 17,7,33,27,22,9,9,15,18,9, | 1,0,8,1,2,5,1,2,0,0,0,1 | 19,25,27,7,7,0,4,16,3,15,1 | 0,0,1,2,3,1,0,0,3,4,1,3,0,0,1 | 266 | 149 | 0.045124843485 | | 1 | 0.905,1.0,0.698,0. | 1.0,1.0,0.938,0.66 | 0,093 |
| 43067 | ENSG00000163930 | BAP1 | chr3 | - | 52405109 | 52405294 | 52404452 | 52404586 | 52405764 | 52405912 | 43067 | 52,40,43,54,40,28,27,27,3 | 0,6,7,0,2,8,3,5,0,0,0,0 | 49,44,35,25,43,22,22,39,1 | 4,8,3,7,7,0,7,9,4,0,4,5,2,0, | 298 | 149 | 0.0451251926932 | | 1 | 1.0,0.769,0.754,1. | 0.86,0.733,0.854,0 | 0,067 |
| 194741 | ENSG00000276087 | | chr2 | + | 24142642 | 24142746 | 24139369 | 24139450 | 24146747 | 24146835 | 194741 | 3,0,11,8,1,4,3,2,1,0,0,4 | 52,28,68,55,41,41,14,31,1 | 20,5,7,3,0,9,2,2,4,7,4,2,2,4 | 97,65,34,11,22,19,32,68,6, | 253 | 149 | 0.0451256901794 | | 1 | 0.033,0.0,0.087,0. | 0.108,0.043,0.108, | -0,037 |
| 7211 | ENSG00000080839 | RBL1 | chr20 | - | 37003701 | 37003866 | 36996348 | 36998929 | 37007410 | 37007559 | 7211 | 24,12,21,12,7,13,7,7,8,4,1 | 0,2,0,0,0,1,0,0,0,0,0,0 | 16,6,16,5,6,3,5,9,4,14,12,2 | 1,1,2,2,1,0,0,1,3,1,0,4,0,0, | 298 | 149 | 0.0452012752537 | | 1 | 1.0,0.75,1.0,1.0,1. | 0.889,0.75,0.8,0.5 | 0,141 |
| 21806 | ENSG00000100429 | HDAC10 | chr22 | - | 50250426 | 50250523 | 50250062 | 50250160 | 50250770 | 50250905 | 21806 | 13,12,10,14,9,12,8,10,8,6, | 3,4,0,1,1,2,2,2,1,2,2,0 | 15,14,6,5,7,6,7,17,10,27,6, | 2,2,0,0,2,0,1,2,0,5,3,0,1,3, | 246 | 149 | 0.0452457104458 | | 1 | 0.724,0.645,1.0,0. | 0.82,0.809,1.0,1.0, | -0,073 |
| 5481 | ENSG00000167232 | ZNF91 | chr19 | - | 23374637 | 23374764 | 23359402 | 23362725 | 23395324 | 23395446 | 5481 | 61,40,125,98,29,45,36,67, | 2,3,3,7,0,3,2,0,0,0,2,1 | 66,73,44,56,58,60,21,28,4 | 5,0,0,5,3,2,0,0,0,2,0,3,0,0, | 276 | 149 | 0.0453461028377 | | 1 | 0.943,0.878,0.957, | 0.877,1.0,1.0,0.85 | -0,027 |
| 22253 | ENSG00000213463 | SYNJ2BP | chr14 | - | 70388469 | 70388606 | 70375675 | 70375771 | 70416899 | 70417090 | 22253 | 12,17,34,27,21,19,19,12,4, | 1,5,3,1,0,2,0,2,0,4,0,0 | 27,27,15,9,11,19,1,7,12,19 | 2,2,4,0,2,0,0,1,2,0,1,0,0,0, | 286 | 149 | 0.0453518598345 | | 1 | 0.862,0.639,0.855, | 0.876,0.876,0.661, | -0,047 |
| 181925 | ENSG00000163166 | IWS1 | chr2 | - | 127523676 | 127523791 | 127505554 | 127505752 | 127526174 | 127526440 | 181925 | 14,8,26,15,3,10,8,19,10,6, | 5,0,4,0,2,2,0,1,0,1,0,0 | 10,16,2,3,4,5,13,6,2,1,7,16 | 2,0,2,0,0,2,1,0,0,0,2,0,0,0, | 264 | 149 | 0.0453724578757 | | 1 | 0.612,1.0,0.786,1. | 0.738,1.0,0.361,1. | -0,032 |
| 52186 | ENSG00000167110 | GOLGA2 | chr9 | - | 128272784 | 128272865 | 128268419 | 128268524 | 128273849 | 128273972 | 52186 | 9,8,36,14,19,3,5,12,6,2,4,6 | 8,3,28,13,7,18,2,15,7,4,4,6 | 24,21,26,13,19,7,8,26,4,13 | 9,11,1,16,13,6,2,15,4,5,11, | 230 | 149 | 0.0453820350126 | | 1 | 0.422,0.633,0.454, | 0.633,0.553,0.944, | -0,111 |
| 13090 | ENSG00000182628 | SKA2 | chr17 | - | 59119298 | 59119495 | 59112090 | 59112345 | 59131280 | 59131367 | 13090 | 7,2,20,13,5,5,3,2,7,3,2,4 | 6,3,14,9,1,7,1,1,3,2,1,0 | 2,5,3,9,6,3,2,1,3,8,1,10,4,7 | 1,2,4,1,1,1,1,0,0,2,0,6,3,1, | 298 | 149 | 0.0454054261817 | | 1 | 0.368,0.25,0.417,0 | 0.5,0.556,0.273,0. | -0,159 |
| 152004 | ENSG00000171606 | ZNF274 | chr19 | + | 58210221 | 58210430 | 58209960 | 58210073 | 58211559 | 58211686 | 152004 | 16,10,23,2,4,0,6,12,3,5,5,3 | 50,26,59,28,33,32,22,34,8 | 8,8,14,30,5,7,9,5,2,6,16,6, | 36,56,52,25,25,9,15,32,5,6 | 298 | 149 | 0.0454270627873 | | 1 | 0.138,0.161,0.163, | 0.1,0.067,0.119,0. | -0,055 |

| | | | | | | | | | | | | | | | | | | | | | | |
|---|---|---|---|---|---|---|---|---|---|---|---|---|---|---|---|---|---|---|---|---|---|---|
| 126119 | ENSG00000197535 | MYO5A | chr15 | - | 52407291 | 52407399 | 52405107 | 52405393 | 52408058 | 52408136 | 126119 | 76,47,134,110,75,51,39,48 | 3,1,9,3,3,4,5,5,0,0,0,5 | 114,117,47,59,75,49,47,59 | 5,11,5,2,6,2,5,3,4,5,9,15,3, | 257 | 149 | 0.0454296768384 | 1 | 0.936,0.965,0.896, | 0.93,0.86,0.845,0. | 0,037 |
| 164394 | ENSG00000124203 | ZNF831 | chr20 | + | 59199532 | 59199601 | 59195868 | 59196005 | 59206904 | 59207056 | 164394 | 14,1,3,4,1,8,1,17,0,2,0,1 | 15,12,32,29,11,23,8,26,16, | 5,0,1,0,1,0,4,2,1,0,13,0,2,3 | 17,3,9,6,6,10,22,9,1,15,54, | 218 | 149 | 0.0454354778364 | 1 | 0.389,0.054,0.06,0 | 0.167,0.0,0.071,0. | 0,02 |
| 104264 | ENSG00000214021 | TTLL3 | chr3 | + | 9818554 | 9818920 | 9817644 | 9817759 | 9820545 | 9820616 | 104264 | 34,15,20,9,24,22,12,16,15, | 1,1,0,0,1,1,1,0,0,0,0,0 | 23,16,15,34,9,11,11,33,12, | 2,1,1,0,0,1,0,8,0,0,3,0,1,3, | 298 | 149 | 0.0454545439612 | 1 | 0.944,0.882,1.0,1. | 0.852,0.889,0.882, | 0,064 |
| 29211 | ENSG00000011243 | AKAP8L | chr19 | - | 15398629 | 15398703 | 15397713 | 15397855 | 15399301 | 15399410 | 29211 | 7,6,16,6,3,9,5,6,6,5,1,1 | 62,60,67,82,41,25,35,53,97 | 7,3,2,0,4,2,1,1,3,4,4,3,7,5, | 54,75,26,21,35,14,23,43,15 | 223 | 149 | 0.0454611103657 | 1 | 0.07,0.063,0.138,0 | 0.08,0.026,0.049,0 | 0,018 |
| 193246 | ENSG00000070610 | GBA2 | chr9 | - | 35740515 | 35740628 | 35740208 | 35740362 | 35740824 | 35741064 | 193246 | 25,16,35,30,34,29,20,28,2 | 10,0,12,2,1,0,1,3,2,5,0,7 | 20,10,14,10,8,12,8,21,21,2 | 5,6,2,2,4,1,3,3,4,1,6,7,5,4, | 262 | 149 | 0.0455102086048 | 1 | 0.587,1.0,0.624,0. | 0.695,0.487,0.799, | 0,114 |
| 248459 | ENSG00000110713 | NUP98 | chr11 | - | 3703007 | 3703110 | 3702771 | 3702892 | 3705199 | 3705356 | 248459 | 4,0,5,2,1,0,5,10,0,0,1,2 | 52,47,61,35,16,20,17,30,4 | 4,5,3,10,4,2,2,7,0,1,14,1,2, | 40,56,26,17,26,16,39,44,11 | 252 | 149 | 0.0455607508767 | 1 | 0.044,0.0,0.046,0. | 0.056,0.05,0.064,0 | -0,01 |
| 153784 | ENSG00000258461 | | chr15 | + | 42360078 | 42360114 | 42354339 | 42354453 | 42384482 | 42384552 | 153784 | 3,1,3,6,4,5,3,5,7,1,3,7 | 15,25,13,9,23,23,15,18,13, | 3,3,1,1,2,3,4,3,2,3,0,0,3,0, | 13,20,7,2,6,7,15,17,15,19,1 | 185 | 149 | 0.0455633356866 | 1 | 0.139,0.031,0.157, | 0.157,0.108,0.103, | 0,053 |
| 32214 | ENSG00000049192 | ADAMTS6 | chr5 | - | 65297107 | 65297244 | 65291328 | 65291470 | 65299984 | 65300131 | 32214 | 4,1,8,4,10,2,2,4,1,0,4,10 | 1,0,17,15,5,11,6,3,8,3,6,8 | 4,9,0,0,0,2,2,1,0,7,7,0,0,3, | 13,10,4,1,6,8,2,0,8,5,7,7,4, | 286 | 149 | 0.045572070225 | 1 | 0.676,1.0,0.197,0. | 0.138,0.319,0.0,0. | 0,12 |
| 61587 | ENSG00000048740 | CELF2 | chr10 | + | 11328925 | 11329097 | 11325835 | 11325979 | 11330455 | 11330639 | 61587 | 118,65,216,125,73,94,70,1 | 3,0,5,1,0,5,8,6,0,0,0,2 | 158,117,93,57,51,45,70,90 | 5,1,5,8,6,2,0,3,0,2,5,4,1,2, | 298 | 149 | 0.0455853637956 | 1 | 0.952,1.0,0.956,0. | 0.94,0.983,0.903,0 | 0,019 |
| 146640 | ENSG00000103035 | PSMD7 | chr16 | + | 74299571 | 74299745 | 74296828 | 74296988 | 74300114 | 74300206 | 146640 | 4,2,7,0,0,2,0,5,1,2,1,1 | 17,22,70,38,35,27,35,35,2 | 2,3,6,10,2,2,7,1,0,3,0,0,4,4 | 28,56,15,25,32,13,26,46,27 | 298 | 149 | 0.0456441451888 | 1 | 0.105,0.043,0.048, | 0.034,0.026,0.167, | -0,025 |
| 252237 | ENSG00000196151 | WDSUB1 | chr2 | - | 159248371 | 159248503 | 159235797 | 159236190 | 159256195 | 159256375 | 252237 | 6,5,15,6,6,5,7,5,3,2,2,4 | 0,0,2,0,0,0,0,0,1,0,0,1 | 12,5,9,1,6,2,2,15,8,8,12,14 | 1,1,3,3,2,0,1,0,1,5,0,0,0,0, | 281 | 149 | 0.045694603236 | 1 | 1.0,1.0,0.799,1.0,1 | 0.864,0.726,0.614, | 0,127 |
| 182494 | ENSG00000226571 | | chr6 | + | 139442297 | 139442434 | 139436336 | 139436460 | 139451557 | 139451669 | 182494 | 21,6,15,3,0,4,0,0,8,9,0,0 | 40,20,69,28,23,25,16,24,2 | 1,2,0,0,3,5,1,2,0,0,7,2,3,0, | 34,33,6,24,49,10,22,30,33, | 286 | 149 | 0.0457922820388 | 1 | 0.215,0.135,0.102, | 0.015,0.031,0.0,0. | 0,013 |
| 130257 | ENSG00000100982 | PCIF1 | chr20 | + | 45943339 | 45943423 | 45943096 | 45943244 | 45943665 | 45943765 | 130257 | 13,8,24,30,9,10,10,4,8,3,5, | 1,4,4,1,0,1,0,0,0,1,3,1 | 13,15,7,4,9,3,13,8,4,6,6,8, | 2,6,0,3,0,0,4,0,2,0,1,1,0,3, | 233 | 149 | 0.045807533177 | 1 | 0.893,0.561,0.793, | 0.806,0.615,1.0,0. | 0,022 |
| 188362 | ENSG00000100280 | AP1B1 | chr22 | - | 29356425 | 29356535 | 29354649 | 29354871 | 29358725 | 29358971 | 188362 | 39,28,78,81,36,47,44,49,1 | 5,2,0,0,2,0,1,0,2,2,3,2 | 74,64,11,15,35,11,23,28,1 | 1,1,1,0,0,0,2,0,0,0,0,8,1,1, | 259 | 149 | 0.0458253026814 | 1 | 0.818,0.89,1.0,1.0, | 0.977,0.974,0.864, | -0,045 |
| 160287 | ENSG00000166888 | STAT6 | chr12 | - | 57099766 | 57099819 | 57098791 | 57099440 | 57099995 | 57100090 | 160287 | 58,34,139,87,62,59,47,42, | 1,0,17,7,0,2,4,0,2,4,0,1 | 71,42,55,19,29,36,20,47,2 | 2,4,5,1,0,0,2,11,2,1,3,1,2,1 | 202 | 149 | 0.0458259564445 | 1 | 0.977,1.0,0.858,0. | 0.963,0.886,0.89,0 | 0,015 |
| 168123 | ENSG00000155189 | AGPAT5 | chr8 | + | 6709480 | 6709546 | 6708692 | 6708887 | 6724869 | 6724939 | 168123 | 4,0,11,0,2,1,6,4,5,0,0,6 | 11,9,23,19,37,22,12,22,4,6 | 8,2,9,0,4,0,2,3,10,6,5,2,4,2 | 23,33,12,1,13,14,9,14,12,2 | 215 | 149 | 0.0458451845687 | 1 | 0.201,0.0,0.249,0. | 0.194,0.04,0.342,0 | -0,039 |
| 193391 | ENSG00000135596 | MICAL1 | chr6 | - | 109452020 | 109452122 | 109451599 | 109451700 | 109452245 | 109452401 | 193391 | 4,1,6,11,2,2,3,4,1,0,2,1 | 43,39,63,44,36,39,41,43,1 | 2,4,7,2,7,5,3,5,0,7,3,1,2,9, | 85,43,11,16,27,19,13,51,1 | 251 | 149 | 0.0458462584104 | 1 | 0.052,0.015,0.054, | 0.014,0.052,0.274, | -0,037 |
| 191738 | ENSG00000158773 | USF1 | chr1 | - | 161041650 | 161041760 | 161041323 | 161041411 | 161042115 | 161042217 | 191738 | 106,117,95,85,60,51,60,77 | 21,9,23,5,11,13,5,6,17,18, | 80,109,48,71,68,32,26,81, | 26,36,5,18,13,5,7,7,7,22,11 | 259 | 149 | 0.0458522391222 | 1 | 0.744,0.882,0.704, | 0.639,0.635,0.847, | 0,05 |
| 246882 | ENSG00000173933 | RBM4 | chr11 | + | 66657135 | 66657259 | 66643749 | 66644140 | 66665855 | 66668374 | 246882 | 7,5,15,8,8,3,6,0,5,3,2,2 | 2,1,2,0,4,0,2,5,4,1,3,2 | 5,1,4,0,1,4,5,2,3,6,2,0,2,4, | 2,4,1,2,2,3,0,5,0,4,4,8,1,3, | 273 | 149 | 0.045862885862 | 1 | 0.656,0.732,0.804, | 0.577,0.12,0.686,0 | 0,145 |
| 150907 | ENSG00000129003 | VPS13C | chr15 | - | 61905901 | 61905987 | 61890164 | 61890400 | 61907263 | 61907390 | 150907 | 5,5,1,1,5,4,1,1,0,4,2,3 | 61,35,121,65,71,89,55,74, | 4,13,5,8,4,2,5,3,7,0,5,3,4,2 | 64,124,40,101,61,38,72,66 | 235 | 149 | 0.0458659394899 | 1 | 0.049,0.083,0.005, | 0.038,0.062,0.073, | 0,002 |
| 254306 | ENSG00000111596 | CNOT2 | chr12 | + | 70329408 | 70329570 | 70319297 | 70319364 | 70335437 | 70335464 | 254306 | 5,1,2,3,6,0,5,0,0,0,0,2 | 8,5,10,5,5,13,12,5,6,1,6,5 | 0,3,2,7,3,1,0,1,1,1,0,0,0,3, | 13,10,0,5,5,5,7,6,2,2,12,7, | 298 | 149 | 0.0458915538324 | 1 | 0.238,0.091,0.091, | 0.0,0.13,1.0,0.412, | -0,051 |
| 103694 | ENSG00000113013 | HSPA9 | chr5 | - | 138574067 | 138574126 | 138573762 | 138573850 | 138575237 | 138575318 | 103694 | 48,41,94,73,48,60,62,46,4 | 5,0,12,4,2,4,7,4,2,1,3,1 | 57,103,25,31,53,21,48,51, | 5,8,2,17,2,1,2,2,3,1,9,21,4, | 208 | 149 | 0.0458969916625 | 1 | 0.873,1.0,0.849,0. | 0.891,0.902,0.9,0. | 0,037 |
| 96481 | ENSG00000099968 | BCL2L13 | chr22 | + | 17696140 | 17696210 | 17688985 | 17689142 | 17726676 | 17730855 | 96481 | 7,8,53,37,11,11,13,11,11,5 | 3,0,0,6,1,0,2,2,1,2,0,2 | 19,9,11,5,10,6,9,11,4,13,1 | 3,0,0,0,1,0,0,5,0,0,0,0,2,2, | 219 | 149 | 0.0459440535318 | 1 | 0.614,1.0,1.0,0.80 | 0.812,1.0,1.0,1.0,0 | -0,083 |
| 82244 | ENSG00000004897 | CDC27 | chr17 | - | 47171916 | 47172064 | 47158205 | 47158303 | 47181561 | 47181637 | 82244 | 30,17,14,7,9,9,1,17,14,11, | 2,1,3,0,3,0,2,1,0,0,0,3 | 13,7,19,26,10,16,9,8,3,12, | 0,3,3,0,2,1,1,1,0,2,5,7,4,0, | 297 | 149 | 0.0459646460744 | 1 | 0.883,0.895,0.701, | 1.0,0.539,0.761,1. | 0,042 |
| 206553 | ENSG00000138029 | HADHB | chr2 | + | 26260980 | 26261063 | 26254429 | 26254474 | 26263379 | 26263479 | 206553 | 6,4,5,4,0,0,2,4,1,0,1,1 | 52,33,96,63,24,23,20,25,4 | 5,4,2,2,1,0,0,4,5,7,1,1,8,8, | 58,35,48,10,22,22,21,35,1 | 232 | 149 | 0.0459803339436 | 1 | 0.069,0.072,0.032, | 0.052,0.068,0.026, | -0,024 |
| 114179 | ENSG00000153339 | TRAPPC8 | chr18 | - | 31832083 | 31832173 | 31829196 | 31830989 | 31839311 | 31839457 | 114179 | 83,34,134,90,67,42,44,43, | 6,4,6,7,2,6,0,4,1,3,4,2 | 66,42,48,21,27,27,35,52,1 | 2,7,8,10,2,4,2,9,4,1,6,3,2,7 | 239 | 149 | 0.0459866429228 | 1 | 0.896,0.841,0.933, | 0.954,0.789,0.789, | 0,028 |
| 215110 | ENSG00000115355 | CCDC88A | chr2 | - | 55297924 | 55298034 | 55296257 | 55296523 | 55301205 | 55301277 | 215110 | 49,7,102,46,12,4,18,9,6,1, | 5,4,10,12,0,0,3,7,0,0,0,1 | 46,44,12,3,15,11,11,19,9,7 | 13,6,7,0,0,2,0,12,2,1,4,0,9, | 259 | 149 | 0.0459881572948 | 1 | 0.849,0.502,0.854, | 0.671,0.808,0.497, | 0,097 |
| 9098 | ENSG00000108312 | UBTF | chr17 | - | 44208618 | 44209001 | 44207863 | 44207911 | 44209351 | 44209422 | 9098 | 6,2,1,4,4,0,70,41,7,3,2,2 | 79,39,99,76,54,53,29,38,6 | 3,1,1,0,6,5,3,5,1,7,9,5,3,7, | 67,67,45,11,55,41,29,49,1 | 298 | 149 | 0.046011784707 | 1 | 0.037,0.025,0.005, | 0.022,0.007,0.011, | 0,055 |
| 173233 | ENSG00000114316 | USP4 | chr3 | - | 49293510 | 49293667 | 49292509 | 49292598 | 49294406 | 49294598 | 173233 | 1,0,3,2,6,0,0,2,7,4,2,2 | 41,7,31,17,15,15,14,14,29, | 2,4,2,0,2,0,0,2,0,0,1,0,1,4, | 10,36,20,16,14,13,19,8,18, | 298 | 149 | 0.0460160978026 | 1 | 0.012,0.0,0.046,0. | 0.091,0.053,0.048, | 0,011 |
| 161349 | ENSG00000165819 | METTL3 | chr14 | - | 21500494 | 21500567 | 21499763 | 21499802 | 21500912 | 21501129 | 161349 | 78,40,44,37,32,28,22,34,4 | 8,4,15,3,10,5,2,12,2,7,1,4 | 33,28,31,11,27,24,10,47,2 | 6,10,2,1,0,2,2,1,0,0,4,7,2,2 | 222 | 149 | 0.0460369181439 | 1 | 0.867,0.87,0.663,0 | 0.787,0.653,0.912, | -0,045 |
| 199943 | ENSG00000151883 | PARP8 | chr5 | + | 50715630 | 50715667 | 50668070 | 50668125 | 50750150 | 50750188 | 199943 | 8,2,4,2,1,3,2,0,0,2,3,2 | 33,25,57,38,17,27,17,18,2 | 1,2,7,4,0,0,0,1,2,0,2,0,2,0, | 32,20,11,20,14,13,6,26,5,1 | 186 | 149 | 0.0460568260492 | 1 | 0.163,0.06,0.053,0 | 0.024,0.074,0.338, | 0,014 |
| 74028 | ENSG00000089127 | OAS1 | chr12 | + | 112917546 | 112917700 | 112913636 | 112916738 | 112919388 | 112919432 | 74028 | 37,9,36,74,87,19,17,8,13,1 | 0,0,0,0,0,1,0,0,0,0,0,0 | 28,14,35,22,26,63,4,9,18,5 | 0,0,2,2,1,1,0,0,1,4,0,0,2,2, | 298 | 149 | 0.0461405164327 | 1 | 1.0,1.0,1.0,1.0,1.0, | 1.0,1.0,0.897,0.84 | 0,049 |
| 1649 | ENSG00000100138 | SNU13 | chr22 | - | 41681225 | 41681559 | 41680243 | 41680364 | 41682355 | 41682456 | 1649 | 7,1,1,2,2,2,2,3,0,0,0,4 | 14,14,46,41,19,18,14,15,1 | 4,6,7,6,1,3,0,0,0,6,5,5,2,0, | 16,22,17,16,15,10,18,28,1 | 298 | 149 | 0.0461446528649 | 1 | 0.2,0.034,0.011,0. | 0.111,0.12,0.171,0 | -0,028 |
| 235369 | ENSG00000104361 | NIPAL2 | chr8 | - | 98193384 | 98193501 | 98189832 | 98193090 | 98194727 | 98194822 | 235369 | 10,0,13,6,5,5,19,14,0,0,0,0 | 17,14,63,51,13,14,9,5,15,1 | 13,4,12,5,4,6,0,0,7,3,7,19, | 37,21,6,5,3,11,14,25,15,20 | 266 | 149 | 0.0461674967358 | 1 | 0.248,0.0,0.104,0. | 0.164,0.096,0.528, | -0,054 |
| 134800 | ENSG00000242259 | C22orf39 | chr22 | - | 19408273 | 19408407 | 19407183 | 19407274 | 19408482 | 19408593 | 134800 | 6,5,2,8,3,2,1,2,1,0,1,0 | 14,20,11,12,7,9,7,8,1,4,1,4 | 6,11,0,3,4,2,6,7,2,4,1,7,6,2 | 7,18,5,10,19,8,8,10,4,13,7, | 283 | 149 | 0.0461739908335 | 1 | 0.184,0.116,0.087, | 0.311,0.243,0.0,0. | -0,048 |
| 44820 | ENSG00000101596 | SMCHD1 | chr18 | + | 2778168 | 2778239 | 2777805 | 2777915 | 2784449 | 2784621 | 44820 | 541,439,651,360,240,221, | 115,73,107,95,26,44,54,72 | 187,271,347,217,269,217, | 23,76,62,109,168,43,26,42 | 220 | 149 | 0.0461823630097 | 1 | 0.761,0.803,0.805, | 0.846,0.707,0.791, | 0,04 |
| 172309 | ENSG00000066777 | ARFGEF1 | chr8 | - | 67275765 | 67276109 | 67271701 | 67271936 | 67277281 | 67277457 | 172309 | 38,19,92,72,32,35,34,45,3 | 1,0,0,2,1,1,1,7,0,1,1,0 | 58,34,37,19,14,38,17,34,3 | 0,4,1,3,9,0,0,4,3,1,2,0,0,0, | 298 | 149 | 0.0461898690901 | 1 | 0.95,1.0,1.0,0.947, | 1.0,0.81,0.949,0.7 | 0,039 |
| 63261 | ENSG00000119004 | CYP20A1 | chr2 | + | 203246754 | 203246921 | 203245845 | 203245895 | 203251966 | 203252109 | 63261 | 41,46,80,79,42,52,22,52,2 | 7,2,13,3,1,13,5,9,0,1,0,7 | 57,38,22,23,55,32,22,48,1 | 7,13,8,12,4,0,4,5,2,7,7,9,3, | 298 | 149 | 0.0461941419073 | 1 | 0.745,0.92,0.755,0 | 0.803,0.594,0.579, | 0,102 |
| 183520 | ENSG00000163568 | AIM2 | chr1 | - | 159073237 | 159073408 | 159068567 | 159068701 | 159076632 | 159076766 | 183520 | 28,19,88,100,75,68,97,107 | 4,0,5,4,0,1,0,3,1,0,4,2 | 49,31,44,7,38,109,5,42,48, | 3,0,5,0,7,11,0,1,2,4,2,9,0,2 | 298 | 149 | 0.0462179284897 | 1 | 0.778,1.0,0.898,0. | 0.891,1.0,0.815,1. | -0,008 |
| 180353 | ENSG00000151092 | NGLY1 | chr3 | - | 25735969 | 25736149 | 25733871 | 25733982 | 25736308 | 25736400 | 180353 | 6,5,7,3,5,7,4,7,2,1,3,4 | 1,0,3,0,0,3,2,0,1,0,2,1 | 11,6,5,5,13,3,3,9,4,2,5,4,1, | 0,0,1,0,2,0,0,0,0,1,0,0,0,0, | 298 | 149 | 0.046241288848 | 1 | 0.75,1.0,0.538,1.0, | 1.0,1.0,0.714,1.0,0 | -0,146 |
| 39474 | ENSG00000162736 | NCSTN | chr1 | + | 160355654 | 160355958 | 160354117 | 160354290 | 160356259 | 160356347 | 39474 | 74,60,80,64,48,34,42,64,5 | 2,0,2,3,6,1,0,2,3,0,0,5 | 72,78,44,18,51,34,38,54,3 | 1,3,1,10,4,1,1,1,3,1,2,8,2,1 | 298 | 149 | 0.0462608719757 | 1 | 0.949,1.0,0.952,0. | 0.973,0.929,0.957, | 0,058 |
| 23093 | ENSG00000119689 | DLST | chr14 | + | 74894311 | 74894409 | 74893347 | 74893424 | 74898368 | 74898455 | 23093 | 62,33,75,42,36,22,35,23,1 | 1,1,7,0,2,3,5,4,2,0,2,4 | 46,47,18,30,36,16,21,38,2 | 0,3,5,12,5,7,2,3,4,1,5,4,2,0 | 247 | 149 | 0.0462913373992 | 1 | 0.974,0.952,0.866, | 1.0,0.904,0.685,0. | 0,056 |
| 108670 | ENSG00000120860 | WASHC3 | chr12 | - | 102019276 | 102019371 | 102012942 | 102013192 | 102025973 | 102026038 | 108670 | 4,1,2,5,3,0,3,2,4,0,0,1 | 25,16,64,62,32,22,16,17,4 | 4,3,1,4,3,4,0,2,1,4,3,3,5,2, | 31,21,21,12,12,13,2,13,5,2 | 244 | 149 | 0.0462984243376 | 1 | 0.089,0.037,0.019, | 0.073,0.08,0.028,0 | -0,04 |
| 171646 | ENSG00000158411 | MITD1 | chr2 | - | 99171338 | 99171429 | 99170536 | 99170652 | 99171509 | 99171646 | 171646 | 36,9,37,20,24,18,7,13,8,7, | 3,1,2,4,1,1,1,0,3,3,0,0 | 18,17,2,4,2,5,3,10,10,29,9, | 0,0,0,0,3,0,1,0,5,0,0,2,1,1, | 240 | 149 | 0.0463025282289 | 1 | 0.882,0.848,0.92,0 | 1.0,1.0,1.0,1.0,0.2 | -0,023 |
| 237321 | ENSG00000120910 | PPP3CC | chr8 | + | 22539731 | 22539845 | 22532920 | 22533018 | 22540614 | 22541142 | 237321 | 3,1,9,5,4,0,0,1,1,1,0,2 | 32,26,40,20,11,4,11,4,12,6 | 1,0,0,2,1,0,1,2,3,2,3,3,0,2, | 19,19,10,5,16,7,12,11,9,14 | 263 | 149 | 0.0463194330602 | 1 | 0.05,0.021,0.113,0 | 0.029,0.0,0.0,0.18 | -0,009 |
| 253763 | ENSG00000100982 | PCIF1 | chr20 | + | 45945712 | 45945883 | 45945006 | 45945030 | 45946028 | 45946115 | 253763 | 7,10,21,11,13,12,6,10,7,20 | 7,5,6,3,4,5,7,2,0,0,0,2 | 10,26,9,10,7,6,14,7,2,3,15, | 1,10,2,14,5,0,2,14,2,1,17,9 | 298 | 149 | 0.046346407885 | 1 | 0.333,0.5,0.636,0. | 0.833,0.565,0.692, | 0,144 |
| 151016 | ENSG00000170776 | AKAP13 | chr15 | + | 85577762 | 85577870 | 85575130 | 85575329 | 85578929 | 85582107 | 151016 | 24,26,41,19,9,29,11,28,11, | 303,208,227,146,147,150, | 23,40,21,38,56,19,4,11,14, | 194,205,147,158,186,128, | 257 | 149 | 0.0463849962758 | 1 | 0.044,0.068,0.095, | 0.064,0.102,0.076, | -0,011 |
| 220627 | ENSG00000065427 | KARS1 | chr16 | - | 75636453 | 75636543 | 75635911 | 75636098 | 75640183 | 75640349 | 220627 | 15,8,66,44,19,19,3,28,19,4 | 4,0,4,7,0,3,1,5,1,0,1,0 | 50,40,8,6,18,16,12,16,12,2 | 7,1,3,1,5,1,2,3,1,2,3,0,1,1, | 239 | 149 | 0.046391474677 | 1 | 0.7,1.0,0.911,0.79 | 0.817,0.961,0.624, | 0,008 |
| 60270 | ENSG00000138138 | ATAD1 | chr10 | - | 87759700 | 87759813 | 87756788 | 87756922 | 87767672 | 87767723 | 60270 | 2,2,17,9,3,4,2,8,5,0,0,2 | 20,11,26,11,16,15,8,10,8,3 | 3,11,2,0,12,1,0,4,8,6,3,1,0, | 31,40,6,24,14,8,4,12,18,10 | 262 | 149 | 0.0464082961509 | 1 | 0.054,0.094,0.271, | 0.052,0.135,0.159, | 0,039 |
| 44950 | ENSG00000164073 | MFSD8 | chr4 | - | 127939852 | 127939997 | 127938782 | 127938838 | 127942044 | 127942158 | 44950 | 29,21,42,16,26,20,30,25,1 | 0,0,3,4,4,1,2,5,0,2,3,7 | 35,19,14,9,10,17,16,33,8,2 | 5,3,6,1,5,2,1,0,5,6,3,8,6,5, | 294 | 149 | 0.0464220283764 | 1 | 1.0,1.0,0.876,0.67, | 0.78,0.762,0.542,0 | 0,071 |
| 193742 | ENSG00000077254 | USP33 | chr1 | - | 77720261 | 77720586 | 77718595 | 77718641 | 77721171 | 77721738 | 193742 | 8,15,33,21,12,25,11,22,9,5 | 8,8,34,24,25,20,14,15,6,12 | 30,44,7,25,18,15,24,16,14, | 18,20,11,8,8,7,9,19,4,12,2 | 298 | 149 | 0.0464238399499 | 1 | 0.333,0.484,0.327, | 0.455,0.524,0.241, | -0,084 |
| 90050 | ENSG00000102081 | FMR1 | chrX | + | 147924918 | 147925055 | 147921932 | 147921985 | 147925539 | 147925633 | 90050 | 1,1,1,3,0,0,0,1,0,0,0,2 | 13,13,25,8,3,5,4,13,22,3,6, | 0,3,1,2,0,0,1,4,2,2,2,0,3,1, | 10,13,13,9,8,2,7,10,5,26,7, | 286 | 149 | 0.0464591966222 | 1 | 0.039,0.039,0.02,0 | 0.0,0.107,0.039,0. | -0,03 |
| 216281 | ENSG00000163219 | ARHGAP25 | chr2 | + | 68787821 | 68787956 | 68782232 | 68782320 | 68807272 | 68807480 | 216281 | 141,219,135,156,124,79,6 | 35,45,31,31,6,14,10,13,3,1 | 190,346,86,58,98,41,65,76 | 38,77,14,31,29,9,15,7,9,15 | 284 | 149 | 0.0465031878848 | 1 | 0.679,0.719,0.696, | 0.724,0.702,0.763, | 0,066 |
| 49493 | ENSG00000116406 | EDEM3 | chr1 | - | 184723750 | 184723856 | 184718837 | 184721386 | 184726254 | 184726389 | 49493 | 69,39,66,40,15,24,27,17,5 | 2,1,4,1,1,7,0,0,0,0,2,1 | 40,25,31,11,14,24,8,37,14, | 2,2,3,5,3,0,0,4,5,4,2,3,0,0, | 255 | 149 | 0.0465185138295 | 1 | 0.953,0.958,0.906, | 0.921,0.88,0.858,0 | 0,048 |
| 212065 | ENSG00000079263 | SP140 | chr2 | + | 230253316 | 230253417 | 230250980 | 230251061 | 230255451 | 230255532 | 212065 | 39,24,46,38,25,17,13,12,2 | 17,6,10,15,3,2,7,9,0,1,7,5 | 44,39,19,16,39,31,1,3,18,4 | 7,2,1,1,0,1,0,2,4,7,6,13,2,7 | 250 | 149 | 0.04655168007 | 1 | 0.578,0.704,0.733, | 0.789,0.921,0.919, | -0,096 |
| 209177 | ENSG00000170310 | STX8 | chr17 | - | 9437796 | 9437887 | 9378551 | 9378653 | 9491828 | 9491921 | 209177 | 3,2,0,0,1,13,7,3,3,0,0,0 | 36,11,23,28,34,17,21,32,2 | 0,4,2,2,3,0,1,0,1,5,1,3,1,0, | 30,29,18,14,21,12,24,31,7, | 240 | 149 | 0.0465572316709 | 1 | 0.049,0.101,0.0,0. | 0.0,0.079,0.065,0. | 0,019 |
| 43144 | ENSG00000188313 | PLSCR1 | chr3 | - | 146521543 | 146522053 | 146517005 | 146517167 | 146525604 | 146525647 | 43144 | 7,2,4,13,6,5,7,7,9,1,5,9 | 0,7,5,6,1,0,0,4,0,0,0,2 | 9,12,13,3,1,14,4,9,1,7,5,52 | 7,3,3,0,6,2,4,2,1,0,2,15,2,7 | 298 | 149 | 0.0465854499838 | 1 | 1.0,0.125,0.286,0. | 0.391,0.667,0.684, | 0,165 |
| 188798 | ENSG00000082512 | TRAF5 | chr1 | + | 211359911 | 211360146 | 211356366 | 211356468 | 211360701 | 211360779 | 188798 | 43,37,100,61,29,44,39,40, | 2,0,0,2,0,1,0,4,0,0,0,0 | 46,42,33,37,46,37,23,22,4 | 1,0,0,2,2,4,1,2,5,4,0,0,2,0, | 298 | 149 | 0.046593423477 | 1 | 0.915,1.0,1.0,0.93 | 0.958,1.0,1.0,0.90 | 0,032 |
| 72242 | ENSG00000181722 | ZBTB20 | chr3 | - | 114904326 | 114904368 | 114900303 | 114900342 | 114974365 | 114974416 | 72242 | 4,3,2,0,0,1,2,2,2,1,1,0 | 31,12,52,20,23,42,18,58,2 | 1,0,0,0,0,0,1,1,3,0,2,2,1,0, | 44,29,34,39,33,22,14,23,3 | 191 | 149 | 0.0466076712033 | 1 | 0.091,0.163,0.029, | 0.017,0.0,0.0,0.0,0 | 0,032 |
| 95928 | ENSG00000286053 | ASDURF | chr2 | + | 189665421 | 189665451 | 189663900 | 189663954 | 189666040 | 189666401 | 95928 | 25,23,28,23,3,9,9,6,10,11, | 3,0,2,0,1,1,1,6,1,3,0,2 | 15,22,9,3,9,4,7,4,7,12,16,9 | 3,2,1,1,0,0,0,1,0,0,4,0,0,0, | 179 | 149 | 0.0466174556384 | 1 | 0.874,1.0,0.921,1. | 0.806,0.902,0.882, | -0,054 |
| 129360 | ENSG00000188342 | GTF2F2 | chr13 | + | 45220865 | 45220909 | 45207423 | 45207505 | 45252870 | 45252970 | 129360 | 0,0,0,2,0,0,2,0,0,0,0,0 | 67,27,60,57,13,35,14,20,2 | 5,1,3,0,0,0,1,1,0,0,1,3,9,1, | 32,33,28,15,16,14,10,35,1 | 193 | 149 | 0.0466298994879 | 1 | 0.0,0.0,0.0,0.026,0 | 0.108,0.023,0.076, | -0,042 |
| 151569 | ENSG00000077238 | IL4R | chr16 | + | 27356694 | 27356838 | 27355807 | 27355907 | 27358915 | 27358994 | 151569 | 22,2,16,5,0,2,8,11,2,1,8,3 | 149,46,475,272,146,169,1 | 5,8,9,4,2,3,4,1,0,0,4,0,2,3, | 221,111,123,25,34,52,56,4 | 293 | 149 | 0.0466602080069 | 1 | 0.07,0.022,0.017,0 | 0.011,0.035,0.036, | 0,003 |
| 101779 | ENSG00000164896 | FASTK | chr7 | - | 151078922 | 151079021 | 151078561 | 151078701 | 151079499 | 151079922 | 101779 | 52,33,50,42,34,56,20,26,4 | 4,0,3,0,1,2,0,2,0,0,2,0 | 16,7,14,10,18,16,15,32,14, | 2,0,0,1,2,0,2,8,1,3,0,1,0,1, | 248 | 149 | 0.0466644615005 | 1 | 0.886,1.0,0.909,1. | 0.828,1.0,1.0,0.85 | 0,045 |
| 17840 | ENSG00000140374 | ETFA | chr15 | - | 76309931 | 76310062 | 76295590 | 76295737 | 76311349 | 76311418 | 17840 | 0,0,10,0,0,0,0,1,3,5,0,4 | 4,5,14,18,9,13,2,7,8,4,4,15 | 2,1,1,0,2,2,0,0,0,0,5,5,3,0, | 12,4,5,1,1,8,9,11,8,11,11,3 | 280 | 149 | 0.0466652602112 | 1 | 0.0,0.0,0.275,0.0,0 | 0.081,0.117,0.096, | 0,001 |
| 161355 | ENSG00000165819 | METTL3 | chr14 | - | 21500494 | 21500647 | 21499763 | 21499802 | 21500912 | 21501129 | 161355 | 215,91,93,69,68,57,59,70, | 8,4,15,3,10,5,2,12,2,7,1,4 | 70,58,59,26,50,46,25,98,4 | 6,10,2,1,0,2,2,1,0,0,4,7,2,2 | 298 | 149 | 0.0466771639349 | 1 | 0.931,0.919,0.756, | 0.854,0.744,0.937, | -0,034 |
| 16618 | ENSG00000103111 | MON1B | chr16 | + | 77193450 | 77193777 | 77191475 | 77191633 | 77194334 | 77194360 | 16618 | 26,30,21,20,14,11,12,9,17, | 5,0,4,8,0,0,0,0,0,0,1,1 | 5,24,8,9,21,17,8,15,9,3,14, | 2,4,0,1,3,2,0,1,0,1,4,7,0,3, | 298 | 149 | 0.0467178260668 | 1 | 0.722,1.0,0.724,0. | 0.556,0.75,1.0,0.8 | 0,135 |
| 194497 | ENSG00000198624 | CCDC69 | chr5 | - | 151185424 | 151185541 | 151184343 | 151184441 | 151186022 | 151186124 | 194497 | 70,74,173,129,78,52,72,67 | 1,0,3,0,0,0,0,2,0,0,0,2 | 112,125,49,51,58,39,43,75 | 0,8,0,1,2,1,1,0,1,0,3,3,2,2, | 266 | 149 | 0.0467310150248 | 1 | 0.975,1.0,0.97,1.0, | 1.0,0.897,1.0,0.96 | 0,029 |
| 129872 | ENSG00000145439 | CBR4 | chr4 | - | 169005884 | 169005914 | 169001574 | 169002205 | 169006754 | 169006891 | 129872 | 9,2,7,4,3,2,2,8,0,2,0,2 | 27,13,40,27,19,13,18,16,1 | 13,9,8,1,3,8,2,9,0,1,5,7,0,5 | 34,15,11,8,6,12,16,12,8,32 | 179 | 149 | 0.0467607352878 | 1 | 0.217,0.114,0.127, | 0.241,0.333,0.377, | -0,079 |
| 243437 | ENSG00000174106 | LEMD3 | chr12 | + | 65240636 | 65240809 | 65240135 | 65240238 | 65240908 | 65241087 | 243437 | 7,5,22,4,3,5,7,10,2,0,6,7 | 10,16,89,68,23,33,35,63,1 | 5,6,2,7,7,8,4,13,5,3,11,0,2, | 62,63,20,19,33,20,21,29,1 | 298 | 149 | 0.0467617910049 | 1 | 0.259,0.135,0.11,0 | 0.039,0.045,0.048, | 0,025 |
| 61250 | ENSG00000114026 | OGG1 | chr3 | + | 9754703 | 9754885 | 9751769 | 9751949 | 9756470 | 9756621 | 61250 | 9,10,25,39,35,33,16,19,33, | 0,0,2,4,2,0,0,5,2,1,0,0 | 41,29,10,6,16,10,9,15,9,13 | 0,2,0,0,0,2,0,0,0,0,0,0,0,0, | 298 | 149 | 0.0467669036934 | 1 | 1.0,1.0,0.862,0.83, | 1.0,0.879,1.0,1.0,1 | -0,045 |
| 223029 | ENSG00000104133 | SPG11 | chr15 | - | 44662838 | 44662929 | 44660431 | 44660616 | 44663390 | 44663688 | 223029 | 2,0,2,4,1,2,4,7,0,0,3,3 | 11,5,43,12,18,27,16,24,9,7 | 1,2,0,4,4,1,2,0,3,0,3,1,1,0, | 10,22,10,10,9,10,10,15,8,2 | 240 | 149 | 0.0468007676294 | 1 | 0.101,0.0,0.028,0. | 0.058,0.053,0.0,0. | 0,003 |
| 79205 | ENSG00000112078 | KCTD20 | chr6 | + | 36479590 | 36479711 | 36470051 | 36470257 | 36481561 | 36481759 | 79205 | 26,12,58,75,22,31,17,24,2 | 2,3,2,1,0,1,0,0,0,0,2,0 | 74,66,21,23,21,18,7,26,16, | 2,0,0,0,0,0,0,2,0,0,4,12,0,1 | 270 | 149 | 0.0468249236432 | 1 | 0.878,0.688,0.941, | 0.953,1.0,1.0,1.0,1 | -0,015 |
| 64640 | ENSG00000168894 | RNF181 | chr2 | + | 85596856 | 85596931 | 85596518 | 85596649 | 85597103 | 85597178 | 64640 | 44,44,61,59,40,26,31,32,5 | 23,18,24,26,13,14,14,21,5, | 48,43,35,15,23,16,20,27,1 | 32,32,24,10,12,10,8,17,7,9 | 224 | 149 | 0.0468377043457 | 1 | 0.56,0.619,0.628,0 | 0.499,0.472,0.492, | 0,068 |
| 4752 | ENSG00000076662 | ICAM3 | chr19 | - | 10335065 | 10335353 | 10334527 | 10334782 | 10335670 | 10335976 | 4752 | 105,58,110,68,99,70,99,11 | 6,4,8,3,3,2,0,2,3,3,0,1 | 92,95,50,10,22,34,46,62,3 | 5,4,3,3,2,0,0,0,1,0,3,11,6,8 | 298 | 149 | 0.0468416506687 | 1 | 0.897,0.879,0.873, | 0.902,0.922,0.893, | 0,036 |
| 26686 | ENSG00000135506 | OS9 | chr12 | + | 57720098 | 57720304 | 57718992 | 57719182 | 57720405 | 57720518 | 26686 | 52,38,109,108,44,48,39,51 | 49,24,134,85,53,41,41,55, | 59,69,25,17,30,28,22,43,2 | 67,85,51,24,37,28,28,36,3 | 298 | 149 | 0.0468574922027 | 1 | 0.347,0.442,0.289, | 0.306,0.289,0.197, | 0,05 |
| 9752 | ENSG00000108604 | SMARCD2 | chr17 | - | 63833297 | 63833420 | 63832089 | 63833170 | 63833586 | 63833722 | 9752 | 187,180,217,130,97,100,6 | 5,0,3,0,0,3,5,3,2,0,2,0 | 141,164,199,110,122,76,5 | 3,12,3,1,3,5,3,6,0,0,1,0,0,3 | 272 | 149 | 0.0468836755395 | 1 | 0.953,1.0,0.975,1. | 0.963,0.882,0.973, | 0,011 |
| 237331 | ENSG00000120910 | PPP3CC | chr8 | + | 22539811 | 22539936 | 22539468 | 22539498 | 22540614 | 22541142 | 237331 | 28,14,47,47,18,9,8,5,10,27 | 91,45,100,64,53,29,69,74, | 15,22,7,8,7,26,2,19,6,22,9, | 64,54,74,56,43,28,28,67,3 | 274 | 149 | 0.0469234419504 | 1 | 0.143,0.145,0.204, | 0.113,0.181,0.049, | 0,029 |
| 237087 | ENSG00000173064 | HECTD4 | chr12 | - | 112235584 | 112235784 | 112235076 | 112235266 | 112236944 | 112237098 | 237087 | 20,14,32,33,14,24,26,22,4, | 0,2,0,0,0,0,1,2,0,0,0,0 | 11,18,10,8,12,12,11,27,9,1 | 2,4,0,3,0,0,0,2,0,0,4,0,1,3, | 298 | 149 | 0.0469265498954 | 1 | 1.0,0.778,1.0,1.0,1 | 0.733,0.692,1.0,0. | 0,076 |
| 146988 | ENSG00000125124 | BBS2 | chr16 | - | 56501650 | 56502079 | 56501352 | 56501497 | 56502316 | 56502456 | 146988 | 2,1,7,4,1,7,14,9,8,1,1,0 | 15,8,4,16,11,8,12,8,3,11,9, | 10,6,14,8,6,0,5,3,3,7,23,1, | 12,6,13,7,7,1,12,23,4,9,9,4 | 298 | 149 | 0.0469344008615 | 1 | 0.062,0.059,0.467, | 0.294,0.333,0.35,0 | -0,056 |
| 237085 | ENSG00000173064 | HECTD4 | chr12 | - | 112235409 | 112235784 | 112235076 | 112235266 | 112236944 | 112237098 | 237085 | 20,14,32,33,16,24,26,22,4, | 0,2,0,0,0,0,1,2,0,0,0,0 | 11,18,10,8,12,11,11,27,9,1 | 2,4,0,3,0,0,0,2,0,0,4,0,1,3, | 298 | 149 | 0.0469380322603 | 1 | 1.0,0.778,1.0,1.0,1 | 0.733,0.692,1.0,0. | 0,076 |

| | | | | | | | | | | | | | | | | | | | | | | |
|---|---|---|---|---|---|---|---|---|---|---|---|---|---|---|---|---|---|---|---|---|---|---|
| 35757 | ENSG00000162980 | ARL5A | chr2 | - | 151810377 | 151810561 | 151806820 | 151806972 | 151812356 | 151812440 | 35757 | 7,5,2,2,2,3,4,3,6,3,2,1 | 66,57,97,78,50,48,42,69,11 | 2,4,8,6,0,1,0,6,2,7,5,21,1,2 | 41,65,69,14,19,36,41,50,29 | 298 | 149 | 0.0469558479744 | 1 | 0.05,0.042,0.01,0.0 | 0.024,0.03,0.055,0 | -0,009 |
| 234700 | ENSG00000179195 | ZNF664 | chr12 | + | 124004565 | 124004935 | 123988045 | 123988138 | 124011380 | 124011441 | 234700 | 4,4,8,4,12,7,3,6,10,4,1,3 | 28,18,28,18,25,11,16,17,24 | 4,9,1,0,2,3,1,9,4,4,1,3,5,4,3 | 26,8,10,18,7,5,19,22,8,16,1 | 298 | 149 | 0.0469810388446 | 1 | 0.067,0.1,0.125,0.1 | 0.071,0.36,0.048,0 | 0,028 |
| 67120 | ENSG00000143498 | TAF1A | chr1 | - | 222579758 | 222579872 | 222577444 | 222577643 | 222584127 | 222584297 | 67120 | 15,12,23,9,11,7,2,3,10,3,4, | 1,1,6,2,3,4,2,12,0,0,2,4 | 13,9,6,0,1,9,4,0,2,8,13,6,8, | 10,6,1,6,1,4,0,9,2,6,6,4,11, | 263 | 149 | 0.0469829541805 | 1 | 0.895,0.872,0.685, | 0.424,0.459,0.773, | 0,139 |
| 79397 | ENSG00000116809 | ZBTB17 | chr1 | - | 15943807 | 15943895 | 15943598 | 15943715 | 15944299 | 15944358 | 79397 | 4,12,17,10,18,9,10,25,11,7 | 2,0,5,7,3,2,0,0,2,1,0,4 | 12,7,4,6,4,4,7,21,4,3,6,14,9 | 2,1,0,1,4,1,0,0,3,0,2,1,2,2,2 | 237 | 149 | 0.0470355488085 | 1 | 0.557,1.0,0.681,0.4 | 0.79,0.815,1.0,0.79 | -0,003 |
| 244967 | ENSG00000107036 | RIC1 | chr9 | + | 5765413 | 5765480 | 5763139 | 5763868 | 5768969 | 5769674 | 244967 | 17,27,23,29,15,24,22,33,20 | 0,0,2,2,0,2,4,4,0,0,0,0 | 23,24,19,24,30,14,10,17,15 | 6,4,2,1,6,0,1,2,1,2,0,1,0,0,0 | 216 | 149 | 0.0470682713862 | 1 | 1.0,1.0,0.888,0.909 | 0.726,0.805,0.868, | 0,038 |
| 72890 | ENSG00000134186 | PRPF38B | chr1 | + | 108693589 | 108693642 | 108692326 | 108692867 | 108695701 | 108695770 | 72890 | 26,32,37,27,2,14,13,23,7,1 | 61,46,56,22,37,21,26,42,17 | 25,38,15,17,23,14,12,17,16 | 32,39,19,22,13,10,17,25,10 | 202 | 149 | 0.0471099129768 | 1 | 0.239,0.339,0.328, | 0.366,0.418,0.368, | -0,081 |
| 200468 | ENSG00000123064 | DDX54 | chr12 | - | 113179958 | 113180005 | 113179142 | 113179331 | 113180928 | 113181058 | 200468 | 46,32,22,16,4,6,7,8,26,29,3 | 1,5,3,0,0,2,0,2,0,0,0,0 | 13,9,17,13,11,5,8,9,2,10,6, | 1,0,0,2,2,1,3,2,0,4,2,2,0,0,1 | 196 | 149 | 0.0471158402561 | 1 | 0.972,0.83,0.848,1 | 0.908,1.0,1.0,0.83 | 0,07 |
| 26388 | ENSG00000115760 | BIRC6 | chr2 | + | 32477367 | 32477583 | 32476212 | 32476344 | 32478634 | 32478818 | 26388 | 146,112,210,124,109,118,1 | 17,9,20,22,4,21,15,11,10,6 | 199,111,77,62,59,49,81,13 | 19,65,14,41,13,1,14,25,2,6 | 298 | 149 | 0.0471374290232 | 1 | 0.811,0.862,0.84,0 | 0.84,0.461,0.733,0 | 0,061 |
| 118564 | ENSG00000187239 | FNBP1 | chr9 | - | 129923843 | 129924026 | 129908889 | 129908999 | 129924959 | 129925157 | 118564 | 5,1,10,7,7,2,1,4,1,7,5,2 | 128,119,140,104,48,57,38, | 3,2,5,11,4,5,0,0,0,8,0,5,4,6 | 65,64,164,75,85,54,38,61,8 | 298 | 149 | 0.047137801626 | 1 | 0.019,0.004,0.034, | 0.023,0.015,0.015, | 0,016 |
| 33040 | ENSG00000109756 | RAPGEF2 | chr4 | + | 159344035 | 159344059 | 159343280 | 159343404 | 159345105 | 159345329 | 33040 | 7,30,34,21,6,3,3,14,3,19,2, | 19,13,27,4,5,3,2,7,4,5,1,2 | 17,31,12,9,19,7,15,26,4,1,2 | 2,13,1,0,6,3,7,5,3,3,5,6,6,0 | 173 | 149 | 0.0471419949792 | 1 | 0.241,0.665,0.52,0 | 0.88,0.673,0.912,1 | -0,113 |
| 63963 | ENSG00000138674 | SEC31A | chr4 | - | 82880661 | 82880711 | 82878775 | 82878928 | 82880798 | 82880922 | 63963 | 16,7,26,28,14,27,13,16,4,2 | 23,14,125,106,27,49,41,45 | 25,21,12,35,36,6,9,22,13,1 | 81,47,7,21,16,17,30,37,22, | 199 | 149 | 0.047169149597 | 1 | 0.342,0.272,0.135, | 0.188,0.251,0.562, | -0,08 |
| 40391 | ENSG00000131374 | TBC1D5 | chr3 | - | 17562018 | 17562142 | 17508473 | 17508605 | 17623848 | 17623913 | 40391 | 6,2,22,8,4,15,6,3,7,5,0,5 | 29,31,71,38,28,29,31,41,21 | 15,8,4,12,3,5,5,9,5,9,10,14 | 41,41,11,27,26,25,15,12,24 | 273 | 149 | 0.0471939020469 | 1 | 0.101,0.034,0.145, | 0.166,0.096,0.166, | -0,044 |
| 198402 | ENSG00000069248 | NUP133 | chr1 | - | 229503200 | 229503270 | 229501998 | 229502102 | 229506039 | 229506158 | 198402 | 0,1,0,0,0,2,0,3,0,0,0,3 | 13,7,70,27,15,14,17,23,12, | 1,7,1,1,0,0,1,0,0,2,2,6,0,2,0 | 29,47,8,9,22,14,12,13,18,4 | 219 | 149 | 0.0472247163489 | 1 | 0.0,0.089,0.0,0.0,0 | 0.023,0.092,0.078, | -0,015 |
| 119801 | ENSG00000130309 | COLGALT1 | chr19 | + | 17571869 | 17571987 | 17568508 | 17568713 | 17572482 | 17572602 | 119801 | 6,4,6,0,3,7,2,6,1,0,1,0 | 29,14,35,25,16,22,22,21,21 | 6,3,3,7,4,2,4,1,6,4,1,8,7,5,1 | 40,21,14,11,25,13,6,26,14, | 267 | 149 | 0.0472694455881 | 1 | 0.104,0.138,0.087, | 0.077,0.074,0.107, | -0,04 |
| 150721 | ENSG00000114062 | UBE3A | chr15 | - | 25354515 | 25354683 | 25354406 | 25354426 | 25355891 | 25356056 | 150721 | 71,45,186,66,30,43,42,57,3 | 11,7,27,0,1,4,5,8,6,2,7,3 | 37,62,31,16,46,23,23,40,24 | 5,14,14,6,5,4,1,9,0,3,17,5,5 | 298 | 149 | 0.0473068402013 | 1 | 0.763,0.763,0.775, | 0.787,0.689,0.525, | 0,076 |
| 25219 | ENSG00000083457 | ITGAE | chr17 | - | 3723287 | 3723383 | 3720306 | 3720402 | 3723687 | 3723744 | 25219 | 17,8,67,36,8,16,15,29,4,4,9 | 5,4,31,32,11,10,10,8,5,5,1, | 31,18,20,8,7,5,2,18,7,20,20 | 16,22,15,2,15,9,3,6,9,22,5, | 245 | 149 | 0.047334214407 | 1 | 0.674,0.549,0.568, | 0.541,0.332,0.448, | 0,113 |
| 193769 | ENSG00000168894 | RNF181 | chr2 | + | 85596859 | 85596931 | 85596518 | 85596649 | 85597103 | 85597178 | 193769 | 41,45,49,50,34,23,28,30,49 | 23,18,24,26,13,14,14,21,5, | 40,36,29,10,22,15,15,24,15 | 32,32,24,10,12,10,8,17,7,9 | 221 | 149 | 0.0473362738933 | 1 | 0.546,0.628,0.579, | 0.457,0.431,0.449, | 0,07 |
| 59943 | ENSG00000084072 | PPIE | chr1 | + | 39745374 | 39745498 | 39743823 | 39743924 | 39748902 | 39749088 | 59943 | 30,12,62,46,35,31,24,16,26 | 0,0,7,6,1,0,0,0,2,0,0,1 | 35,27,14,3,12,13,19,38,14, | 4,2,0,0,0,0,0,1,1,1,4,0,0,0,5 | 273 | 149 | 0.0474044406594 | 1 | 1.0,1.0,0.829,0.807 | 0.827,0.88,1.0,1.0, | 0,053 |
| 234341 | ENSG00000116754 | SRSF11 | chr1 | + | 70231094 | 70231566 | 70228421 | 70228555 | 70231858 | 70231975 | 234341 | 18,5,8,8,2,7,2,7,3,5,2,3 | 15,7,14,6,1,10,4,7,11,2,2,5 | 3,12,8,54,7,10,15,14,3,3,4, | 8,18,4,24,9,1,10,9,0,4,6,4,5 | 298 | 149 | 0.0474070702499 | 1 | 0.375,0.263,0.222, | 0.158,0.25,0.5,0.52 | -0,165 |
| 226113 | ENSG00000129103 | SUMF2 | chr7 | + | 56074476 | 56074736 | 56074173 | 56074218 | 56076833 | 56076889 | 226113 | 50,33,124,164,78,81,49,83 | 7,4,17,3,5,3,7,9,9,3,3,6 | 61,88,46,25,31,24,42,66,36 | 11,16,8,6,6,5,5,6,4,5,4,12,1 | 298 | 149 | 0.0474308611192 | 1 | 0.781,0.805,0.785, | 0.735,0.733,0.742, | 0,05 |
| 155482 | ENSG00000173575 | CHD2 | chr15 | + | 92888254 | 92888314 | 92886202 | 92886254 | 92892003 | 92892072 | 155482 | 29,6,29,21,11,7,6,23,11,0,4 | 111,55,334,194,108,116,10 | 32,17,14,43,6,25,6,14,12,1 | 224,247,128,109,129,73,40 | 209 | 149 | 0.0474404453371 | 1 | 0.157,0.072,0.058, | 0.092,0.047,0.072, | -0,024 |
| 158731 | ENSG00000198746 | GPATCH3 | chr1 | - | 26897089 | 26897121 | 26894235 | 26894410 | 26897300 | 26897725 | 158731 | 3,0,3,1,0,6,2,3,0,0,1,3 | 16,40,32,5,13,14,20,10,10, | 5,4,2,2,4,3,0,1,1,2,5,4,1,2,0 | 5,15,13,11,19,3,11,12,10,5 | 181 | 149 | 0.0474424564624 | 1 | 0.134,0.0,0.072,0.1 | 0.452,0.18,0.112,0 | -0,049 |
| 220433 | ENSG00000196367 | TRRAP | chr7 | + | 98928792 | 98928916 | 98927166 | 98927366 | 98929988 | 98930206 | 220433 | 4,0,1,2,0,0,4,2,0,0,1,0 | 14,8,13,10,14,21,10,7,7,9,7 | 1,4,1,10,5,1,0,1,0,0,0,1,0,0 | 11,20,1,13,11,21,4,10,13,1 | 273 | 149 | 0.0474561176239 | 1 | 0.135,0.0,0.04,0.09 | 0.047,0.098,0.353, | -0,028 |
| 138029 | ENSG00000198521 | ZNF43 | chr19 | - | 21819094 | 21819407 | 21817887 | 21817986 | 21836035 | 21836187 | 138029 | 24,30,39,36,7,17,11,21,16, | 9,2,6,1,2,4,4,0,12,2,1,4 | 30,24,14,10,8,6,11,25,8,16 | 8,1,0,4,0,0,1,5,1,2,5,0,0,2,3 | 298 | 149 | 0.0475153192698 | 1 | 0.571,0.882,0.765, | 0.652,0.923,1.0,0.5 | -0,133 |
| 199830 | ENSG00000115875 | SRSF7 | chr2 | - | 38747061 | 38747261 | 38746693 | 38746747 | 38748046 | 38748157 | 199830 | 11,1,9,16,13,4,8,8,1,1,3,6 | 112,118,247,101,97,97,94, | 24,6,4,6,2,8,1,21,10,7,20,1 | 164,201,77,38,88,58,80,12 | 298 | 149 | 0.0475356694981 | 1 | 0.047,0.004,0.018, | 0.068,0.015,0.025, | -0,01 |
| 12186 | ENSG00000133030 | MPRIP | chr17 | + | 17136218 | 17136450 | 17131616 | 17131701 | 17137915 | 17138075 | 12186 | 51,46,56,33,15,26,29,31,48 | 3,0,4,1,4,2,0,0,0,1,0,2 | 45,32,29,58,9,15,37,39,19, | 1,0,0,0,0,0,2,0,0,2,0,1,0,0 | 298 | 149 | 0.0475608781735 | 1 | 0.895,1.0,0.875,0.9 | 0.957,1.0,1.0,1.0,1 | -0,064 |
| 116023 | ENSG00000011454 | RABGAP1 | chr9 | + | 122990055 | 122990213 | 122989296 | 122989471 | 122996040 | 122996151 | 116023 | 81,57,120,56,50,50,34,44,4 | 1,0,0,0,0,0,3,1,2,0,0,1 | 75,83,96,85,58,44,39,47,45 | 1,4,2,6,1,0,0,0,2,2,0,0,3,0,2 | 298 | 149 | 0.0475636754341 | 1 | 0.976,1.0,1.0,1.0,1 | 0.974,0.912,0.96,0 | 0,008 |
| 55295 | ENSG00000108654 | DDX5 | chr17 | - | 64501056 | 64501143 | 64500548 | 64500773 | 64502009 | 64502069 | 55295 | 23,3,22,28,10,9,5,10,7,1,1, | 824,625,917,701,474,444,3 | 26,17,33,4,6,16,0,16,7,12,1 | 807,884,754,587,420,483,2 | 236 | 149 | 0.0475879661402 | 1 | 0.017,0.003,0.015, | 0.02,0.012,0.027,0 | -0,004 |
| 213431 | ENSG00000035720 | STAP1 | chr4 | + | 67577206 | 67577259 | 67575384 | 67575498 | 67581304 | 67581471 | 213431 | 77,35,89,67,34,41,37,40,75 | 3,0,4,1,1,0,3,4,1,1,2,2 | 48,37,53,28,38,23,18,32,19 | 2,3,3,11,3,0,2,2,2,0,0,0,3,4 | 202 | 149 | 0.0476055926653 | 1 | 0.95,1.0,0.943,0.98 | 0.947,0.901,0.929, | 0,009 |
| 204432 | ENSG00000257093 | DENND11 | chr7 | - | 141685202 | 141685636 | 141674066 | 141674220 | 141686558 | 141686658 | 204432 | 54,53,53,45,68,63,68,51,50 | 6,7,5,3,6,10,10,4,4,0,8,4 | 54,55,47,41,33,21,49,73,20 | 2,10,2,2,4,1,2,3,4,7,5,1,1,0 | 298 | 149 | 0.0476083570051 | 1 | 0.818,0.791,0.841, | 0.931,0.733,0.922, | -0,041 |
| 204433 | ENSG00000257093 | DENND11 | chr7 | - | 141685482 | 141685636 | 141674066 | 141674220 | 141686558 | 141686658 | 204433 | 54,53,53,45,69,63,68,51,50 | 6,7,5,3,6,10,10,4,4,0,8,4 | 54,55,49,41,33,21,49,73,20 | 2,10,2,2,4,1,2,3,4,7,5,1,1,0 | 298 | 149 | 0.0476280749348 | 1 | 0.818,0.791,0.841, | 0.931,0.733,0.925, | -0,041 |
| 91801 | ENSG00000120802 | TMPO | chr12 | + | 98544950 | 98545061 | 98537474 | 98537572 | 98546358 | 98546447 | 91801 | 78,18,132,66,42,56,39,78,5 | 12,6,43,37,45,29,17,13,26, | 78,74,64,29,21,26,22,72,27 | 30,11,30,6,13,25,6,17,5,19 | 260 | 149 | 0.0476323447181 | 1 | 0.788,0.632,0.638, | 0.598,0.794,0.55,0 | -0,075 |
| 200284 | ENSG00000176444 | CLK2 | chr1 | - | 155263572 | 155263708 | 155262879 | 155263400 | 155263949 | 155264550 | 200284 | 7,10,10,11,6,4,3,7,10,0,4,7 | 47,26,64,37,48,57,34,66,22 | 1,5,4,6,0,0,3,4,3,9,9,2,2,9,1 | 36,60,25,25,21,18,35,39,21 | 285 | 149 | 0.0476448065061 | 1 | 0.072,0.167,0.076, | 0.014,0.042,0.077, | 0,027 |
| 250943 | ENSG00000122376 | SHLD2 | chr10 | + | 87175923 | 87176095 | 87170839 | 87170974 | 87180074 | 87180303 | 250943 | 18,12,18,16,17,12,22,5,13, | 18,9,24,13,8,8,5,10,5,2,1,3 | 16,5,7,5,17,8,3,11,5,21,7,1 | 15,17,10,10,29,2,10,13,5,1 | 298 | 149 | 0.0476534173188 | 1 | 0.333,0.4,0.273,0.3 | 0.348,0.128,0.259, | 0,134 |
| 38834 | ENSG00000047849 | MAP4 | chr3 | - | 47909037 | 47912421 | 47877416 | 47877523 | 47914816 | 47914939 | 38834 | 1,1,2,2,0,0,0,4,1,0,0,0 | 59,39,62,36,60,64,31,42,29 | 5,0,0,5,0,0,4,2,0,1,0,8,2,2,2 | 25,45,34,40,31,21,24,55,15 | 298 | 149 | 0.0476679866199 | 1 | 0.008,0.013,0.016, | 0.091,0.0,0.0,0.059 | -0,018 |
| 161584 | ENSG00000154822 | PLCL2 | chr3 | + | 17042165 | 17042217 | 17014707 | 17014911 | 17042873 | 17042949 | 161584 | 4,3,0,0,0,0,2,6,0,1,0,0 | 115,61,274,138,75,107,51, | 0,0,5,1,0,0,1,4,0,0,16,5,1,1 | 201,143,69,46,82,63,71,72 | 201 | 149 | 0.0476726825282 | 1 | 0.025,0.035,0.0,0.0 | 0.0,0.0,0.051,0.01 | -0,01 |
| 18743 | ENSG00000242028 | HYPK | chr15 | + | 43801131 | 43801187 | 43800494 | 43800784 | 43801517 | 43801569 | 18743 | 152,116,105,89,38,46,29,3 | 15,5,4,12,2,2,1,6,2,2,1,2 | 43,64,48,34,58,29,18,42,31 | 7,4,6,19,0,2,6,3,3,2,8,3,0,2 | 205 | 149 | 0.0476816568396 | 1 | 0.88,0.944,0.95,0.8 | 0.817,0.921,0.853, | 0,049 |
| 134823 | ENSG00000128159 | TUBGCP6 | chr22 | - | 50221509 | 50221874 | 50219956 | 50220015 | 50222027 | 50222102 | 134823 | 52,32,61,27,30,46,38,46,26 | 6,4,5,0,9,3,3,10,7,0,3,2 | 50,30,12,19,20,13,21,45,23 | 3,10,1,4,9,1,1,8,2,4,5,9,4,6 | 298 | 149 | 0.0477041436584 | 1 | 0.812,0.8,0.859,1.0 | 0.893,0.6,0.857,0.7 | 0,079 |
| 119802 | ENSG00000130309 | COLGALT1 | chr19 | + | 17571869 | 17571991 | 17568508 | 17568713 | 17572482 | 17572602 | 119802 | 5,4,10,3,5,7,2,6,0,0,1,0 | 29,14,35,25,16,22,22,21,21 | 4,6,8,3,4,5,5,1,8,8,2,9,12,4 | 40,21,14,11,25,13,6,26,14, | 271 | 149 | 0.0477105228002 | 1 | 0.087,0.136,0.136, | 0.052,0.136,0.239, | -0,055 |
| 117000 | ENSG00000132589 | FLOT2 | chr17 | - | 28885856 | 28885947 | 28884224 | 28884315 | 28888944 | 28889026 | 117000 | 1,0,7,9,2,5,0,5,7,6,2,2 | 56,36,80,83,79,32,67,53,73 | 7,3,3,0,3,1,0,1,0,5,2,1,2,2,5 | 62,42,44,9,31,22,30,41,20, | 240 | 149 | 0.0477166177606 | 1 | 0.011,0.0,0.052,0.0 | 0.066,0.042,0.041, | 0,012 |
| 67248 | ENSG00000168010 | ATG16L2 | chr11 | + | 72823543 | 72823797 | 72822847 | 72822961 | 72824059 | 72824122 | 67248 | 0,1,3,1,2,0,3,8,1,4,2,3 | 49,70,61,47,32,28,23,9,19, | 3,3,2,2,2,1,0,2,1,0,0,0,1,2,1 | 61,52,28,29,18,22,19,30,40 | 298 | 149 | 0.0477309721989 | 1 | 0.0,0.007,0.024,0.0 | 0.024,0.028,0.034, | 0,039 |
| 246595 | ENSG00000054654 | SYNE2 | chr14 | + | 64009940 | 64010116 | 64007042 | 64007222 | 64016472 | 64016631 | 246595 | 87,102,178,121,74,124,118 | 4,1,7,4,6,10,4,10,4,1,6,3 | 76,136,70,99,138,108,14,1 | 4,3,4,13,12,3,0,3,7,3,3,2,0, | 298 | 149 | 0.047753510621 | 1 | 0.916,0.981,0.927, | 0.905,0.958,0.897, | -0,054 |
| 109861 | ENSG00000174197 | MGA | chr15 | + | 41727179 | 41727406 | 41713150 | 41713496 | 41729163 | 41729349 | 109861 | 48,73,85,59,32,45,33,70,54 | 2,0,1,0,0,0,0,0,0,0,1,0 | 67,69,28,24,39,21,26,38,19 | 3,2,5,2,0,0,2,2,2,1,0,0,0,0,4 | 298 | 149 | 0.0477567585861 | 1 | 0.923,1.0,0.977,1.0 | 0.918,0.945,0.737, | 0,055 |
| 229109 | ENSG00000106609 | TMEM248 | chr7 | + | 66953225 | 66953572 | 66950951 | 66951135 | 66955501 | 66958551 | 229109 | 128,101,196,203,58,47,85, | 4,1,11,6,7,2,3,3,0,2,0,2 | 123,164,83,26,67,54,47,88 | 8,6,0,6,0,1,4,4,1,9,10,2,2,3 | 298 | 149 | 0.0478211359079 | 1 | 0.941,0.981,0.899, | 0.885,0.932,1.0,0.6 | 0,041 |
| 198649 | ENSG00000223745 | CCDC18-AS1 | chr1 | - | 93315148 | 93315266 | 93309752 | 93309790 | 93324634 | 93324729 | 198649 | 0,0,0,0,0,0,1,0,0,0,0,2 | 1,4,10,5,3,4,10,11,3,1,7,3 | 0,0,0,0,0,2,0,2,2,5,4,1,0,1,1 | 1,4,2,1,8,1,2,6,3,6,9,0,3,3,0 | 267 | 149 | 0.0478350800612 | 1 | 0.0,0.0,0.0,0.0,0.0, | 0.0,0.0,0.0,0.0,0.0, | -0,2 |
| 8799 | ENSG00000154889 | MPPE1 | chr18 | - | 11886712 | 11886778 | 11886498 | 11886621 | 11886916 | 11887025 | 8799 | 7,8,12,22,5,7,4,10,9,8,4,9 | 2,1,1,7,1,1,0,4,2,0,0,0 | 11,3,3,1,9,13,4,7,6,24,3,11 | 5,4,0,0,1,0,0,1,1,0,0,0,3,0,0 | 215 | 149 | 0.0478788670565 | 1 | 0.708,0.847,0.893, | 0.604,0.342,1.0,1.0 | -0,016 |
| 37287 | ENSG00000251022 | THAP9-AS1 | chr4 | - | 82895691 | 82895774 | 82893008 | 82894847 | 82897988 | 82898062 | 37287 | 14,5,26,23,12,10,1,5,4,1,4, | 2,2,6,7,0,2,2,3,0,0,2,1 | 8,8,12,0,4,1,6,7,3,9,8,9,8,9 | 9,5,0,2,7,1,2,4,1,3,3,0,5,10 | 232 | 149 | 0.047889706036 | 1 | 0.818,0.616,0.736, | 0.363,0.507,1.0,0.0 | 0,15 |
| 213133 | ENSG00000169045 | HNRNPH1 | chr5 | - | 179616898 | 179616922 | 179616125 | 179616218 | 179617020 | 179617110 | 213133 | 0,2,7,4,1,2,1,2,4,1,0,3 | 85,63,38,17,26,27,29,23,68 | 2,5,2,3,1,0,2,2,0,1,4,5,2,1,7 | 26,13,32,30,5,11,6,31,8,33 | 173 | 149 | 0.0479032248394 | 1 | 0.0,0.027,0.137,0.1 | 0.062,0.249,0.051, | -0,043 |
| 107969 | ENSG00000068308 | OTUD5 | chrX | - | 48936190 | 48936358 | 48934953 | 48935018 | 48944189 | 48944283 | 107969 | 36,12,22,17,4,5,13,16,8,3,1 | 169,118,125,76,49,64,33,5 | 21,24,13,83,5,6,8,10,4,4,8, | 57,65,65,31,53,46,35,54,32 | 298 | 149 | 0.0479570925853 | 1 | 0.096,0.048,0.081, | 0.156,0.156,0.091, | -0,04 |
| 77018 | ENSG00000085832 | EPS15 | chr1 | - | 51448045 | 51448135 | 51446959 | 51447105 | 51461090 | 51461150 | 77018 | 125,66,143,123,102,106,73 | 5,5,12,5,3,6,1,6,4,0,0,2 | 132,96,56,33,74,77,30,129 | 9,11,2,7,8,7,5,9,0,0,13,7,5, | 239 | 149 | 0.0479649298107 | 1 | 0.94,0.892,0.881,0 | 0.901,0.845,0.946, | 0,045 |
| 96275 | ENSG00000125779 | PANK2 | chr20 | + | 3912457 | 3912634 | 3910576 | 3910830 | 3916926 | 3917050 | 96275 | 109,54,297,164,103,111,84 | 3,4,7,10,8,1,2,6,6,2,0,1 | 130,143,89,37,102,65,42,6 | 10,1,6,2,4,10,0,4,3,0,15,4,6 | 298 | 149 | 0.04797158314 | 1 | 0.948,0.871,0.955, | 0.867,0.986,0.881, | 0,032 |
| 35274 | ENSG00000104517 | UBR5 | chr8 | - | 102258315 | 102258424 | 102257631 | 102257717 | 102258984 | 102259129 | 35274 | 1,0,12,18,0,3,0,1,2,1,0,2 | 58,58,95,47,82,84,70,69,73 | 2,2,4,2,1,4,1,10,0,5,5,0,0,0 | 73,93,80,88,38,16,57,75,54 | 258 | 149 | 0.048015113865 | 1 | 0.01,0.0,0.068,0.18 | 0.016,0.012,0.028, | 0,006 |
| 131975 | ENSG00000135297 | MTO1 | chr6 | + | 73497735 | 73497896 | 73492233 | 73492352 | 73500573 | 73500923 | 131975 | 86,51,142,63,47,55,47,51,4 | 2,0,2,5,4,0,2,2,2,3,2,2 | 83,73,71,28,48,25,26,40,17 | 3,2,3,4,2,0,0,0,0,2,1,3,0,2,0 | 298 | 149 | 0.0480826412534 | 1 | 0.956,1.0,0.973,0.8 | 0.933,0.948,0.922, | -0,043 |
| 61140 | ENSG00000134698 | AGO4 | chr1 | + | 35832436 | 35832570 | 35832056 | 35832185 | 35833989 | 35834174 | 61140 | 27,16,69,54,28,14,17,16,20 | 0,2,3,0,0,0,0,2,1,0,0,2 | 38,30,27,18,10,17,23,43,16 | 5,4,0,0,2,0,0,4,0,4,1,0,2,0,0 | 283 | 149 | 0.0481974457977 | 1 | 1.0,0.808,0.924,1.0 | 0.8,0.798,1.0,1.0,0 | 0,01 |
| 41949 | ENSG00000137502 | RAB30 | chr11 | - | 82994038 | 82994106 | 82987582 | 82987770 | 82997223 | 82997324 | 41949 | 34,38,39,41,7,9,8,9,29,9,5, | 5,3,6,8,0,1,3,3,2,3,0,4 | 33,74,37,32,35,11,19,28,13 | 3,9,5,13,19,9,14,1,7,1,3,8,2 | 217 | 149 | 0.0482063075409 | 1 | 0.824,0.897,0.817, | 0.883,0.85,0.836,0 | 0,084 |
| 43521 | ENSG00000196455 | PIK3R4 | chr3 | - | 130735868 | 130736002 | 130733547 | 130734130 | 130744485 | 130745264 | 43521 | 10,24,35,52,26,14,14,22,11 | 1,2,3,0,2,0,4,0,0,0,1,0 | 13,42,26,13,17,17,17,14,12 | 0,0,2,3,4,2,4,1,2,0,2,0,2,0,1 | 283 | 149 | 0.0482652393269 | 1 | 0.84,0.863,0.86,1.0 | 1.0,1.0,0.873,0.69 | 0,036 |
| 61687 | ENSG00000270629 | NBPF14 | chr1 | - | 148593597 | 148593700 | 148592551 | 148592766 | 148595542 | 148595717 | 61687 | 49,31,40,26,47,35,30,29,21 | 10,9,8,6,5,14,2,10,3,2,1,7 | 16,21,14,18,9,14,11,12,46, | 7,6,3,10,5,7,2,2,3,4,2,34,3, | 252 | 149 | 0.0482921068562 | 1 | 0.743,0.671,0.747, | 0.575,0.674,0.734, | 0,117 |
| 209488 | ENSG00000110880 | CORO1C | chr12 | - | 108653037 | 108653181 | 108652271 | 108652417 | 108654305 | 108654410 | 209488 | 7,10,60,17,6,9,8,10,12,4,0, | 26,17,48,45,28,20,13,24,15 | 25,18,8,9,7,4,23,20,9,11,24 | 15,25,10,18,27,14,15,10,8, | 293 | 149 | 0.04830756751 | 1 | 0.12,0.23,0.389,0.1 | 0.459,0.268,0.289, | -0,091 |
| 174531 | ENSG00000163755 | HPS3 | chr3 | + | 149157353 | 149157531 | 149155106 | 149155215 | 149158665 | 149158846 | 174531 | 38,22,48,19,39,38,21,37,34 | 5,0,3,0,0,0,3,1,1,4,0,4 | 49,31,32,7,13,30,17,27,26, | 0,0,0,0,0,1,0,2,0,1,2,4,3,0,2 | 298 | 149 | 0.0483080678629 | 1 | 0.792,1.0,0.889,1.0 | 1.0,1.0,1.0,1.0,1.0, | -0,026 |
| 177100 | ENSG00000179912 | R3HDM2 | chr12 | - | 57292571 | 57292690 | 57289008 | 57289040 | 57295398 | 57295594 | 177100 | 11,5,19,11,10,10,6,10,3,7,7 | 3,5,13,4,2,10,2,6,2,5,0,9 | 16,13,7,8,9,4,19,8,8,5,11,1 | 4,0,2,4,3,0,9,0,2,4,2,10,9,3 | 268 | 149 | 0.0483167311736 | 1 | 0.671,0.357,0.448, | 0.69,1.0,0.661,0.52 | -0,134 |
| 210660 | ENSG00000184922 | FMNL1 | chr17 | + | 45245618 | 45245733 | 45245325 | 45245416 | 45245877 | 45245973 | 210660 | 36,26,38,20,33,19,13,21,34 | 9,4,10,5,8,9,8,1,1,3,2,9 | 15,21,14,28,22,8,23,16,21, | 4,12,3,15,10,1,17,21,8,3,7, | 264 | 149 | 0.0483368245373 | 1 | 0.693,0.786,0.682, | 0.679,0.497,0.725, | 0,082 |
| 186602 | ENSG00000112118 | MCM3 | chr6 | - | 52283293 | 52283402 | 52282652 | 52282861 | 52284596 | 52284881 | 186602 | 22,10,49,46,16,22,4,17,20, | 0,4,0,1,0,0,0,3,0,0,0,0 | 31,24,6,0,17,18,8,15,10,27 | 10,1,3,1,0,1,0,0,0,0,1,5,0,1 | 258 | 149 | 0.0483641130972 | 1 | 1.0,0.591,1.0,0.964 | 0.642,0.933,0.536, | 0,125 |
| 51154 | ENSG00000197111 | PCBP2 | chr12 | + | 53465931 | 53465973 | 53464759 | 53464852 | 53467220 | 53467293 | 51154 | 104,78,193,156,86,174,76, | 10,0,7,5,12,8,9,16,0,3,5,5 | 112,212,95,257,114,71,268 | 7,11,12,8,5,0,16,6,2,0,2,4,3 | 191 | 149 | 0.0483838693568 | 1 | 0.89,1.0,0.956,0.9 | 0.926,0.938,0.861, | -0,021 |
| 213598 | ENSG00000272752 | STAG3L5P-PVR | chr7 | + | 100350679 | 100350815 | 100349798 | 100349887 | 100351251 | 100351410 | 213598 | 0,2,9,11,0,0,0,4,4,4,0,0 | 101,44,52,49,67,111,34,59 | 0,0,0,0,7,7,0,0,1,3,0,0,2,0,0 | 36,20,26,33,26,17,18,33,23 | 285 | 149 | 0.0483839129506 | 1 | 0.0,0.023,0.083,0.1 | 0.0,0.0,0.0,0.0,0.1 | 0,003 |
| 139624 | ENSG00000132670 | PTPRA | chr20 | + | 3034900 | 3035133 | 3027706 | 3027841 | 3035584 | 3035710 | 139624 | 7,3,4,3,0,2,3,2,7,0,0,4 | 62,33,70,64,20,27,15,26,37 | 5,0,12,9,3,2,4,2,3,3,3,9,3,0 | 31,38,22,22,33,22,18,34,16 | 298 | 149 | 0.0484522916033 | 1 | 0.053,0.043,0.028, | 0.075,0.0,0.214,0.1 | -0,026 |
| 70182 | ENSG00000242372 | EIF6 | chr20 | - | 35284175 | 35284492 | 35279941 | 35280118 | 35284725 | 35284770 | 70182 | 28,17,69,46,4,4,12,10,13,5 | 6,0,4,1,2,1,0,0,0,1,0,2 | 19,18,21,5,21,7,4,7,3,11,16 | 4,2,0,0,3,3,0,3,3,0,3,3,0,1,0 | 298 | 149 | 0.048471113433 | 1 | 0.7,1.0,0.896,0.958 | 0.704,0.818,1.0,1.0 | 0,057 |
| 178588 | ENSG00000120008 | WDR11 | chr10 | + | 120880362 | 120880562 | 120878352 | 120878459 | 120880825 | 120880901 | 178588 | 5,4,26,10,1,0,3,3,3,0,0,5 | 154,137,314,171,134,71,87 | 9,10,12,11,9,7,1,6,11,10,11 | 173,168,204,140,128,109,6 | 298 | 149 | 0.0484760903052 | 1 | 0.016,0.014,0.04,0 | 0.025,0.029,0.029, | -0,013 |
| 162889 | ENSG00000155329 | ZCCHC10 | chr5 | - | 133022810 | 133022906 | 133006758 | 133006920 | 133026496 | 133026549 | 162889 | 4,3,10,3,0,4,4,6,2,6,3,4 | 0,4,15,8,2,4,4,17,4,2,4,2 | 11,10,6,1,4,0,3,8,1,6,15,6,5 | 0,1,1,6,5,1,4,1,10,4,4,1,10, | 245 | 149 | 0.0485027081536 | 1 | 1.0,0.313,0.288,0.1 | 1.0,0.859,0.785,0.0 | -0,137 |
| 5174 | ENSG00000157540 | DYRK1A | chr21 | + | 37430268 | 37430400 | 37420298 | 37420384 | 37472683 | 37472829 | 5174 | 14,11,25,3,1,10,7,27,4,0,4, | 31,21,32,14,19,28,21,34,41 | 6,46,13,37,16,6,12,19,10,2 | 39,37,32,41,19,17,23,32,9, | 281 | 149 | 0.0485148909616 | 1 | 0.193,0.217,0.293, | 0.075,0.397,0.177, | -0,07 |
| 36845 | ENSG00000166004 | CEP295 | chr11 | + | 93721953 | 93722050 | 93721311 | 93721412 | 93723040 | 93723289 | 36845 | 13,28,25,12,10,1,2,4,7,2,2, | 7,1,8,6,1,4,1,0,4,0,0,3 | 17,18,6,3,2,5,3,7,1,13,3,7,8 | 5,5,1,4,7,2,1,12,2,2,4,3,4,5 | 246 | 149 | 0.048555505548 | 1 | 0.529,0.944,0.654, | 0.673,0.686,0.784, | 0,177 |
| 189223 | ENSG00000118308 | IRAG2 | chr12 | + | 25066364 | 25066516 | 25063719 | 25063816 | 25069349 | 25069431 | 189223 | 517,172,1273,948,458,436 | 12,6,40,45,10,6,8,21,9,10,1 | 592,720,448,183,252,214,1 | 14,12,5,3,6,8,4,6,5,10,6,3,2 | 298 | 149 | 0.0485759866439 | 1 | 0.956,0.935,0.941, | 0.955,0.968,0.978, | -0,013 |
| 44739 | ENSG00000169359 | SLC33A1 | chr3 | - | 155833856 | 155834041 | 155829687 | 155829903 | 155842431 | 155842619 | 44739 | 15,15,15,16,17,12,16,20,22 | 5,4,12,6,1,4,7,4,1,2,0,5 | 14,7,5,1,3,8,2,5,0,18,4,13,1 | 8,15,5,1,3,2,1,2,4,1,5,2,4,2 | 298 | 149 | 0.0486127085587 | 1 | 0.6,0.652,0.385,0.5 | 0.467,0.189,0.333, | 0,177 |
| 135073 | ENSG00000100030 | MAPK1 | chr22 | - | 21785020 | 21785165 | 21772872 | 21772982 | 21788256 | 21788388 | 135073 | 4,4,8,0,3,9,4,4,0,4,2,4 | 102,66,115,96,59,54,59,63 | 7,14,3,6,3,1,2,6,6,5,9,2,5,5 | 97,63,80,51,36,29,38,84,42 | 294 | 149 | 0.0486609793774 | 1 | 0.019,0.03,0.034,0 | 0.035,0.101,0.019, | -0,009 |
| 9425 | ENSG00000118046 | STK11 | chr19 | + | 1216267 | 1216398 | 1206815 | 1207203 | 1218416 | 1218500 | 9425 | 22,14,14,8,1,4,10,17,15,3,3 | 34,29,77,74,33,41,25,27,23 | 4,7,8,5,2,6,2,5,3,10,6,11,7, | 23,25,12,16,11,21,18,50,14 | 280 | 149 | 0.048673878478 | 1 | 0.256,0.204,0.088, | 0.085,0.13,0.262,0 | 0,034 |
| 1579 | ENSG00000128159 | TUBGCP6 | chr22 | - | 50221274 | 50221874 | 50219956 | 50220015 | 50222027 | 50222102 | 1579 | 51,32,61,27,30,46,38,46,26 | 6,4,5,0,9,3,3,10,7,0,3,2 | 50,30,12,19,20,13,21,45,23 | 3,10,1,4,9,1,1,8,2,4,5,9,4,6 | 298 | 149 | 0.0486751395518 | 1 | 0.81,0.8,0.859,1.0, | 0.893,0.6,0.857,0.7 | 0,079 |
| 72847 | ENSG00000162664 | ZNF326 | chr1 | + | 90022249 | 90022345 | 90018684 | 90018784 | 90027353 | 90028226 | 72847 | 42,62,22,15,8,8,9,27,20,12 | 4,2,4,0,1,0,0,0,0,0,0,2 | 18,47,12,21,20,8,3,11,9,23 | 5,1,2,5,2,1,0,0,0,3,0,0,0,1,2 | 245 | 149 | 0.0486816711701 | 1 | 0.865,0.95,0.77,1.0 | 0.686,0.966,0.785, | 0,074 |
| 126544 | ENSG00000136754 | ABI1 | chr10 | - | 26759061 | 26759235 | 26751597 | 26751783 | 26765217 | 26765318 | 126544 | 192,107,151,153,92,95,133 | 34,14,49,62,65,45,22,39,13 | 151,159,174,224,84,57,73, | 42,15,33,3,15,5,20,32,11,3 | 298 | 149 | 0.0487034299812 | 1 | 0.738,0.793,0.606, | 0.643,0.841,0.725, | -0,068 |
| 189224 | ENSG00000118308 | IRAG2 | chr12 | + | 25066364 | 25066745 | 25063719 | 25063816 | 25069349 | 25069431 | 189224 | 517,172,1273,948,458,436 | 12,6,40,45,10,6,8,21,9,10,1 | 592,720,448,183,252,214,1 | 14,12,5,3,6,8,4,6,5,10,6,3,2 | 298 | 149 | 0.0487042223677 | 1 | 0.956,0.935,0.941, | 0.955,0.968,0.978, | -0,013 |
| 54070 | ENSG00000143742 | SRP9 | chr1 | + | 225786861 | 225786985 | 225783299 | 225783368 | 225787970 | 225788266 | 54070 | 3,2,18,22,7,14,1,14,4,1,2,3 | 1,0,14,0,0,0,1,0,1,2,0,3 | 19,17,5,4,4,5,5,2,6,9,7,19,9 | 2,0,1,2,2,0,6,0,0,0,1,1,1,4,3 | 273 | 149 | 0.0487168572333 | 1 | 0.621,1.0,0.412,1.0 | 0.838,1.0,0.732,0.5 | -0,073 |
| 244188 | ENSG00000173120 | KDM2A | chr11 | + | 67149765 | 67149863 | 67121268 | 67121358 | 67180078 | 67180453 | 244188 | 5,2,3,3,0,3,4,2,1,0,0,0 | 67,59,65,39,32,35,38,26,28 | 0,5,1,1,0,1,2,1,0,2,0,0,3,0,1 | 37,38,32,49,20,15,35,41,23 | 247 | 149 | 0.0487457825208 | 1 | 0.043,0.02,0.027,0 | 0.0,0.074,0.019,0.0 | 0,008 |

| | | | | | | | | | | | | | | | | | | | | | | | |
|---|---|---|---|---|---|---|---|---|---|---|---|---|---|---|---|---|---|---|---|---|---|---|---|
| 76071 | ENSG00000111801 | BTN3A3 | chr6 | + | 26443381 | 26443442 | 26440471 | 26440648 | 26443956 | 26444298 | 76071 | 23,25,29,29,7,7,12,10,6,2, | 0,0,0,0,0,0,0,0,0,0,0,0 | 17,9,19,7,6,16,2,9,17,34,1 | 0,0,0,0,0,0,0,0,0,0,0,0,11,9 | 210 | 149 | 0.0487823953607 | | 1 | 1.0,1.0,1.0,1.0,1.0, | 1.0,1.0,1.0,1.0,1.0, | 0,06 |
| 123979 | ENSG00000162241 | SLC25A45 | chr11 | - | 65379375 | 65379561 | 65376817 | 65377076 | 65379866 | 65379938 | 123979 | 30,15,68,32,27,21,33,31,3 | 3,3,5,2,0,2,3,1,0,0,0,2 | 31,26,28,11,16,17,7,21,8,1 | 4,4,0,1,3,1,0,1,1,4,0,8,2,4,1 | 298 | 149 | 0.0488122281945 | | 1 | 0.833,0.714,0.872, | 0.795,0.765,1.0,0. | 0,066 |
| 183712 | ENSG00000105483 | CARD8 | chr19 | - | 48249522 | 48249631 | 48240961 | 48241063 | 48249750 | 48249878 | 183712 | 22,13,34,33,24,12,9,12,21, | 10,10,13,17,6,5,6,7,2,5,5,8 | 14,13,8,7,10,11,9,15,22,48 | 7,13,5,6,12,11,3,15,16,26, | 258 | 149 | 0.0488642764954 | | 1 | 0.56,0.429,0.602,0 | 0.536,0.366,0.48,0 | 0,083 |
| 948 | ENSG00000100083 | GGA1 | chr22 | + | 37624968 | 37625076 | 37623551 | 37623633 | 37629461 | 37629526 | 948 | 18,11,26,20,26,20,11,26,1 | 3,4,4,0,2,4,1,0,6,0,0,3 | 17,17,9,5,16,12,11,15,10,2 | 2,3,1,2,2,1,3,9,2,3,3,10,3,1 | 257 | 149 | 0.0488670969694 | | 1 | 0.777,0.615,0.79,1 | 0.831,0.767,0.839, | 0,086 |
| 133327 | ENSG00000182670 | TTC3 | chr21 | + | 37157988 | 37158203 | 37157160 | 37157214 | 37159698 | 37159745 | 133327 | 2,1,7,9,1,5,0,6,1,0,1,1 | 15,10,28,27,16,27,11,27,7, | 6,8,4,4,5,3,1,3,4,4,2,0,3,4, | 20,26,13,8,21,10,10,24,9,1 | 298 | 149 | 0.0488674640528 | | 1 | 0.062,0.048,0.111, | 0.13,0.133,0.133,0 | -0,04 |
| 256295 | ENSG00000111450 | STX2 | chr12 | - | 130804304 | 130804540 | 130801414 | 130801488 | 130806981 | 130807090 | 256295 | 0,0,0,0,1,5,4,3,0,0,0,0 | 1,1,10,13,11,13,6,8,5,1,4,7 | 0,2,0,0,1,5,1,1,2,0,4,9,4,2, | 3,0,1,14,1,2,2,10,2,5,5,16,1 | 298 | 149 | 0.0488948683403 | | 1 | 0.0,0.0,0.0,0.0,0.0 | 0.0,1.0,0.0,0.0,0.3 | -0,156 |
| 238695 | ENSG00000064313 | TAF2 | chr8 | - | 119767074 | 119767155 | 119762414 | 119762608 | 119778018 | 119778129 | 238695 | 0,0,4,0,1,1,3,3,0,2,0,3 | 20,5,37,13,15,5,10,16,1,2,1 | 1,5,1,0,2,0,0,0,2,1,4,2,0,0, | 10,27,12,9,21,4,7,6,6,23,1 | 230 | 149 | 0.0489014731574 | | 1 | 0.0,0.0,0.065,0.0,0 | 0.061,0.107,0.051, | 0,004 |
| 31980 | ENSG00000118965 | WDR35 | chr2 | - | 19960553 | 19960614 | 19953833 | 19953978 | 19966723 | 19966909 | 31980 | 14,15,49,25,8,31,17,11,19, | 8,0,7,3,0,0,2,1,1,3,0,5 | 17,22,8,4,3,12,7,25,8,10,1 | 5,3,4,9,3,0,1,6,2,2,2,3,6,4, | 210 | 149 | 0.0489069113679 | | 1 | 0.554,1.0,0.832,0. | 0.707,0.839,0.587, | 0,112 |
| 94419 | ENSG00000072121 | ZFYVE26 | chr14 | - | 67805216 | 67805305 | 67804100 | 67804264 | 67805453 | 67805618 | 94419 | 15,15,42,15,21,9,20,23,19, | 2,5,5,2,4,5,7,8,0,2,1,10 | 21,15,18,12,22,5,14,14,15, | 7,16,2,12,8,4,2,10,8,10,6,3 | 238 | 149 | 0.0489765402523 | | 1 | 0.824,0.653,0.84,0 | 0.653,0.37,0.849,0 | 0,092 |
| 185867 | ENSG00000158796 | DEDD | chr1 | - | 161123077 | 161123221 | 161122350 | 161122523 | 161123838 | 161123946 | 185867 | 20,22,30,29,9,18,16,22,23, | 0,7,0,2,3,7,6,10,0,0,0,0 | 24,23,20,17,19,13,14,30,8, | 9,4,2,5,3,5,5,4,1,5,0,6,2,0,1 | 293 | 149 | 0.0489871690995 | | 1 | 1.0,0.615,1.0,0.88 | 0.576,0.745,0.836, | 0,084 |
| 9949 | ENSG00000178852 | EFCAB13 | chr17 | + | 47361377 | 47361521 | 47345015 | 47345098 | 47370436 | 47370508 | 9949 | 38,39,191,105,19,48,71,98 | 6,5,21,15,12,8,1,4,5,5,4,5 | 5,3,0,0,64,49,68,78,75,112 | 14,16,18,27,0,1,6,9,2,2,11, | 293 | 149 | 0.0490325166142 | | 1 | 0.763,0.799,0.822, | 0.154,0.087,0.0,0. | 0,128 |
| 142100 | ENSG00000265257 | | chr18 | + | 9103091 | 9103433 | 9102735 | 9102797 | 9117837 | 9117903 | 142100 | 7,2,17,6,1,2,4,12,1,4,0,3 | 24,26,30,21,10,8,10,13,7,9 | 6,6,5,6,6,0,3,0,4,5,5,10,4,3 | 8,10,15,2,5,5,6,7,4,6,9,20,1 | 298 | 149 | 0.0490518751117 | | 1 | 0.127,0.037,0.221, | 0.273,0.231,0.143, | -0,105 |
| 187878 | ENSG00000118197 | DDX59 | chr1 | - | 200650424 | 200650585 | 200649073 | 200649226 | 200654440 | 200654582 | 187878 | 30,14,40,13,5,15,7,11,10,9 | 2,0,5,1,0,0,0,1,0,0,2,0 | 23,15,8,15,0,11,7,9,5,10,8, | 1,1,0,5,3,0,1,1,0,2,3,2,0,0,1 | 298 | 149 | 0.0490569604241 | | 1 | 0.882,1.0,0.8,0.86 | 0.92,0.882,1.0,0.6, | 0,109 |
| 92684 | ENSG00000136861 | CDK5RAP2 | chr9 | - | 120407011 | 120407248 | 120404035 | 120404113 | 120408346 | 120408468 | 92684 | 23,20,34,32,12,9,6,22,21,5 | 1,0,2,5,3,4,4,2,0,1,2,0 | 19,24,15,36,4,5,26,31,7,23 | 4,7,10,0,4,3,0,5,1,1,0,8,5,1 | 298 | 149 | 0.049067271852 | | 1 | 0.92,1.0,0.895,0.7 | 0.704,0.632,0.429, | 0,082 |
| 189764 | ENSG00000019582 | CD74 | chr5 | - | 150402562 | 150402628 | 150401636 | 150402250 | 150404679 | 150404767 | 189764 | 497,395,571,466,112,62,1 | 17,1,0,5,1,4,2,6,6,0,4,0 | 241,370,218,230,193,125, | 4,2,1,71,1,4,0,1,0,0,0,1,2,2 | 215 | 149 | 0.0490976061121 | | 1 | 0.953,0.996,1.0,0. | 0.977,0.992,0.993, | 0,003 |
| 213086 | ENSG00000197299 | BLM | chr15 | + | 90760150 | 90760279 | 90754810 | 90754938 | 90760593 | 90761255 | 213086 | 3,5,23,18,1,8,9,4,7,1,0,3 | 0,0,2,0,2,0,0,0,0,0,4,2 | 9,19,5,3,4,6,5,1,2,6,15,8,5, | 0,0,0,0,2,0,0,0,2,0,0,0,1,0, | 278 | 149 | 0.0491017678784 | | 1 | 1.0,1.0,0.86,1.0,0. | 1.0,1.0,1.0,1.0,0.5 | -0,106 |
| 109276 | ENSG00000035681 | NSMAF | chr8 | - | 58605926 | 58606035 | 58603209 | 58603386 | 58607768 | 58607840 | 109276 | 72,38,80,51,36,42,35,55,4 | 0,0,1,0,0,2,2,3,4,0,0,2 | 54,58,50,16,29,7,18,36,12, | 2,0,8,0,0,2,0,1,0,0,2,2,4,2, | 258 | 149 | 0.0491390697335 | | 1 | 1.0,1.0,0.979,1.0,1 | 0.94,1.0,0.783,1.0, | 0,033 |
| 91593 | ENSG00000024048 | UBR2 | chr6 | + | 42612170 | 42612291 | 42606588 | 42606651 | 42615070 | 42615178 | 91593 | 104,47,175,101,53,108,67, | 1,0,1,3,0,1,0,2,2,0,0,0 | 133,130,106,69,105,36,45, | 1,1,5,2,1,0,0,0,4,0,3,6,0,2, | 270 | 149 | 0.0491510791931 | | 1 | 0.983,1.0,0.99,0.9 | 0.987,0.986,0.921, | 0,015 |
| 105919 | ENSG00000258529 | | chr11 | - | 111840654 | 111840809 | 111838248 | 111838378 | 111844600 | 111844723 | 105919 | 13,12,22,17,3,6,6,8,12,7,2, | 1,4,7,6,0,0,4,0,5,2,1,2 | 15,19,15,9,11,5,10,9,11,8,1 | 1,4,4,5,0,3,1,0,2,3,0,1,0,0, | 298 | 149 | 0.0491991995881 | | 1 | 0.867,0.6,0.611,0. | 0.882,0.704,0.652, | -0,09 |
| 50061 | ENSG00000168724 | DNAJC21 | chr5 | + | 34949542 | 34949677 | 34945721 | 34945803 | 34950169 | 34950342 | 50061 | 8,2,12,6,4,0,2,0,5,4,0,1 | 34,19,44,48,26,28,18,25,2 | 3,0,3,3,0,5,3,4,1,2,0,9,1,4, | 37,46,24,24,27,7,15,34,13, | 284 | 149 | 0.0492037411348 | | 1 | 0.11,0.052,0.125,0 | 0.041,0.0,0.062,0. | 0,009 |
| 245907 | ENSG00000166405 | RIC3 | chr11 | - | 8126658 | 8126802 | 8106055 | 8111137 | 8137377 | 8137471 | 245907 | 79,25,124,58,59,70,48,45, | 0,0,7,0,6,2,5,4,0,3,2,2 | 74,77,27,57,23,22,18,42,1 | 2,4,3,3,0,2,2,1,1,3,7,3,5,6, | 293 | 149 | 0.0492038294193 | | 1 | 1.0,1.0,0.9,1.0,0.8 | 0.95,0.907,0.821,0 | 0,049 |
| 26529 | ENSG00000107201 | DDX58 | chr9 | - | 32492390 | 32492538 | 32491300 | 32491420 | 32493760 | 32493942 | 26529 | 21,15,53,24,31,13,28,23,4 | 9,4,8,6,6,2,4,1,5,4,1,0 | 33,21,29,28,8,9,8,27,12,65 | 6,1,0,4,3,1,0,1,3,2,6,7,2,7, | 297 | 149 | 0.0492177089771 | | 1 | 0.539,0.653,0.769, | 0.734,0.913,1.0,0. | -0,072 |
| 92623 | ENSG00000184182 | UBE2F | chr2 | + | 238016532 | 238016633 | 237994743 | 237994809 | 238025341 | 238025412 | 92623 | 0,2,7,1,1,1,1,0,1,0,0,1 | 2,1,16,11,5,0,1,3,1,4,1,6 | 2,2,0,1,2,2,0,0,1,4,0,3,3,4, | 9,15,3,3,4,3,2,1,0,4,8,8,2,0 | 250 | 149 | 0.0492967549769 | | 1 | 0.0,0.544,0.207,0. | 0.117,0.074,0.0,0. | -0,027 |
| 232998 | ENSG00000005889 | ZFX | chrX | + | 24173630 | 24173723 | 24172755 | 24172800 | 24179182 | 24179770 | 232998 | 0,1,20,8,0,4,4,2,0,0,1,2 | 16,9,87,41,17,18,8,27,5,4, | 6,3,6,2,1,7,4,2,0,3,5,1,1,1, | 58,71,25,17,32,13,13,20,2 | 242 | 149 | 0.0493048143312 | | 1 | 0.0,0.064,0.124,0. | 0.06,0.025,0.129,0 | 0,001 |
| 80442 | ENSG00000290927 | | chr16 | + | 31702534 | 31702681 | 31700562 | 31700738 | 31703155 | 31703363 | 80442 | 87,60,88,38,29,38,18,24,3 | 15,15,19,12,5,2,7,5,2,8,7,7 | 26,32,29,40,25,10,12,21,1 | 12,8,3,13,9,4,5,7,2,8,5,9,2 | 296 | 149 | 0.0493756463292 | | 1 | 0.745,0.668,0.7,0. | 0.522,0.668,0.83,0 | 0,076 |
| 104101 | ENSG00000291178 | | chr7 | - | 100335295 | 100335428 | 100332441 | 100332576 | 100335837 | 100335932 | 104101 | 0,0,2,4,0,0,0,1,0,0,0,0 | 17,10,15,22,8,13,10,5,16,1 | 3,0,1,2,0,0,0,1,0,0,2,11,1,0 | 11,7,17,9,11,8,14,10,1,14,1 | 282 | 149 | 0.0493800312703 | | 1 | 0.0,0.0,0.066,0.08 | 0.126,0.0,0.03,0.1 | -0,018 |
| 94727 | ENSG00000146247 | PHIP | chr6 | - | 78970048 | 78970173 | 78969834 | 78969917 | 78970780 | 78970888 | 94727 | 42,42,104,42,34,30,27,68, | 3,0,11,8,4,5,2,2,1,2,0,1 | 55,83,43,52,37,24,24,44,2 | 3,10,7,6,5,5,0,6,10,1,7,2,4, | 274 | 149 | 0.0494066627838 | | 1 | 0.884,1.0,0.837,0. | 0.909,0.819,0.77,0 | 0,05 |
| 69812 | ENSG00000223745 | CCDC18-AS1 | chr1 | - | 93318167 | 93318228 | 93264644 | 93264772 | 93324634 | 93324691 | 69812 | 2,0,4,0,3,2,2,1,0,0,0,0 | 7,16,27,4,5,9,3,7,2,2,8,7 | 1,1,3,2,1,0,0,1,2,0,4,3,1,0, | 3,9,4,0,6,8,5,12,14,8,15,7, | 210 | 149 | 0.0494647455614 | | 1 | 0.169,0.0,0.095,0. | 0.191,0.073,0.347, | -0,112 |
| 81895 | ENSG00000168385 | SEPTIN2 | chr2 | + | 241335321 | 241335351 | 241335125 | 241335212 | 241335958 | 241336336 | 81895 | 1,1,1,0,1,1,1,2,0,0,0,1 | 9,1,30,34,7,3,0,3,7,6,2,2 | 2,1,0,0,1,0,1,3,0,1,0,0,6,2, | 1,3,2,1,22,6,0,1,6,5,1,1,3,7 | 179 | 149 | 0.0494692414437 | | 1 | 0.085,0.454,0.027, | 0.625,0.217,0.0,0. | -0,045 |
| 150997 | ENSG00000170776 | AKAP13 | chr15 | + | 85487181 | 85487369 | 85485709 | 85485753 | 85521427 | 85521575 | 150997 | 1,0,28,14,3,0,0,0,0,0,0,2 | 51,40,109,51,53,84,53,71, | 0,4,2,2,9,5,0,2,2,4,4,3,2,2, | 95,65,31,48,62,51,27,43,6 | 298 | 149 | 0.0494757888882 | | 1 | 0.01,0.0,0.114,0.1 | 0.0,0.03,0.031,0.0 | -0,002 |
| 249067 | ENSG00000187147 | RNF220 | chr1 | + | 44635544 | 44635641 | 44632234 | 44632385 | 44636029 | 44636495 | 249067 | 27,36,28,19,18,5,8,9,19,27 | 2,0,4,1,3,0,4,4,0,0,0,3 | 21,18,16,15,25,10,6,11,8,2 | 4,2,0,6,2,0,0,2,7,6,3,5,0,0, | 246 | 149 | 0.0494878642226 | | 1 | 0.891,1.0,0.809,0. | 0.761,0.845,1.0,0. | 0,058 |
| 127435 | ENSG00000012779 | ALOX5 | chr10 | + | 45428617 | 45428764 | 45424959 | 45425132 | 45440429 | 45440633 | 127435 | 872,588,669,605,809,469, | 1,1,6,0,0,1,0,3,3,1,2,3 | 677,630,450,723,421,293, | 12,8,2,7,6,0,5,6,0,1,1,7,0,2 | 296 | 149 | 0.0495359314044 | | 1 | 0.998,0.997,0.982, | 0.966,0.975,0.991, | 0,01 |
| 215017 | ENSG00000003402 | CFLAR | chr2 | + | 201162995 | 201163177 | 201160431 | 201160942 | 201163834 | 201176687 | 215017 | 11,11,51,33,16,20,12,20,1 | 62,40,139,85,64,66,69,106 | 13,23,14,43,15,21,9,35,21, | 68,102,28,26,46,25,41,105 | 298 | 149 | 0.049537360103 | | 1 | 0.081,0.121,0.155, | 0.087,0.101,0.2,0. | -0,046 |
| 160493 | ENSG00000186532 | SMYD4 | chr17 | - | 1783615 | 1784094 | 1783359 | 1783476 | 1784325 | 1784461 | 160493 | 0,1,5,2,2,4,1,5,2,6,0,5 | 8,0,32,9,4,5,3,5,10,0,1,9 | 1,0,0,2,2,0,0,1,0,1,4,1,2,2, | 5,2,5,3,3,1,1,5,6,5,8,13,5,1 | 298 | 149 | 0.0495445957915 | | 1 | 0.0,1.0,0.072,0.1,0 | 0.091,0.0,0.0,0.25, | 0,135 |
| 127427 | ENSG00000012779 | ALOX5 | chr10 | + | 45395854 | 45395936 | 45382482 | 45382681 | 45412190 | 45412313 | 127427 | 606,523,787,566,659,439, | 4,7,19,16,9,3,9,17,2,3,4,0 | 611,660,345,597,459,271,1 | 19,21,7,6,15,7,2,5,17,9,12, | 231 | 149 | 0.0496083686081 | | 1 | 0.99,0.98,0.964,0. | 0.954,0.953,0.97,0 | 0,007 |
| 216279 | ENSG00000163219 | ARHGAP25 | chr2 | + | 68787561 | 68787956 | 68782232 | 68782320 | 68807272 | 68807480 | 216279 | 141,219,135,156,124,79,6 | 35,45,31,31,6,14,10,13,3,1 | 190,346,87,58,98,41,65,76 | 38,77,14,31,29,9,15,7,9,15 | 298 | 149 | 0.0496454124329 | | 1 | 0.668,0.709,0.685, | 0.714,0.692,0.757, | 0,066 |
| 1577 | ENSG00000128159 | TUBGCP6 | chr22 | - | 50220191 | 50221874 | 50219956 | 50220015 | 50222027 | 50222102 | 1577 | 51,36,62,27,30,46,38,46,2 | 6,4,5,0,9,3,3,10,7,0,3,2 | 52,30,12,19,20,14,23,45,2 | 3,10,1,4,9,1,1,8,2,4,5,9,4,6 | 298 | 149 | 0.0496584789817 | | 1 | 0.81,0.818,0.861,1 | 0.897,0.6,0.857,0. | 0,079 |
| 253647 | ENSG00000145781 | COMMD10 | chr5 | + | 116317516 | 116317597 | 116303529 | 116303683 | 116399481 | 116399622 | 253647 | 8,0,9,2,5,3,4,3,1,2,2,1 | 10,2,11,1,11,6,6,2,6,1,0,4 | 7,3,4,3,3,3,6,3,3,5,0,5,4, | 3,6,2,1,5,0,3,5,6,0,2,2,4,5,1 | 230 | 149 | 0.0496652791538 | | 1 | 0.341,0.0,0.346,0. | 0.602,0.245,0.564, | -0,13 |
| 113124 | ENSG00000158669 | GPAT4 | chr8 | + | 41610735 | 41610810 | 41609654 | 41609955 | 41611902 | 41611992 | 113124 | 49,34,43,41,28,29,20,22,4 | 2,0,0,4,2,0,1,2,0,0,3,0 | 45,25,27,13,16,15,5,32,6,3 | 1,3,1,8,1,2,0,0,0,0,0,10,2,0 | 224 | 149 | 0.0496879213769 | | 1 | 0.942,1.0,1.0,0.87 | 0.968,0.847,0.947, | 0,019 |
| 102228 | ENSG00000144815 | NXPE3 | chr3 | + | 101807052 | 101807126 | 101801234 | 101801989 | 101816794 | 101817001 | 102228 | 11,8,35,34,11,10,6,13,12,9 | 0,1,0,0,3,0,4,4,4,0,0,0 | 24,21,7,11,12,14,3,11,10,2 | 0,0,0,0,1,0,0,0,1,0,0,0,2,2, | 223 | 149 | 0.0496962909244 | | 1 | 1.0,0.842,1.0,1.0,0 | 1.0,1.0,1.0,1.0,0.8 | -0,095 |
| 94713 | ENSG00000108021 | TASOR2 | chr10 | + | 5762531 | 5762646 | 5761289 | 5761471 | 5763028 | 5763632 | 94713 | 34,21,62,44,21,32,30,34,1 | 0,0,2,0,0,0,0,0,0,0,0,0 | 33,58,37,32,20,17,23,31,1 | 3,2,4,5,3,0,0,0,2,0,2,0,0,0,1 | 264 | 149 | 0.0497096263723 | | 1 | 1.0,1.0,0.946,1.0,1 | 0.861,0.942,0.839, | 0,063 |
| 83128 | ENSG00000164548 | TRA2A | chr7 | - | 23522353 | 23522432 | 23521706 | 23521840 | 23531788 | 23532023 | 83128 | 72,19,48,13,10,8,22,28,9,1 | 14,7,72,52,14,24,4,4,10,4, | 22,21,22,20,15,14,2,5,2,22 | 36,49,23,11,10,15,5,17,19, | 228 | 149 | 0.0497114286146 | | 1 | 0.771,0.639,0.303, | 0.285,0.219,0.385, | 0,135 |
| 176072 | ENSG00000179262 | RAD23A | chr19 | + | 12949019 | 12949159 | 12948688 | 12948813 | 12949274 | 12949408 | 176072 | 21,9,32,15,2,7,7,6,6,5,3,9 | 1,0,5,1,2,3,2,1,0,0,0,2 | 10,12,6,10,2,4,4,13,4,4,8,2 | 2,3,0,0,0,0,0,4,0,0,4,0,0,0, | 289 | 149 | 0.0497335367149 | | 1 | 0.915,1.0,0.767,0. | 0.721,0.673,1.0,1. | -0,112 |
| 60999 | ENSG00000163781 | TOPBP1 | chr3 | - | 133618212 | 133618433 | 133617159 | 133617326 | 133620154 | 133620347 | 60999 | 73,46,48,33,41,31,27,42,7 | 15,1,7,3,11,14,5,16,20,6,5, | 43,31,31,22,26,15,13,38,8, | 13,2,15,14,11,9,2,10,5,7,8, | 298 | 149 | 0.0497347840638 | | 1 | 0.709,0.958,0.774, | 0.623,0.886,0.508, | 0,094 |
| 168090 | ENSG00000203875 | SNHG5 | chr6 | - | 85677794 | 85677879 | 85677006 | 85677492 | 85677953 | 85678748 | 168090 | 12,4,62,31,3,1,1,3,0,0,5,5 | 1,5,25,43,14,8,6,7,2,3,10,1 | 8,51,14,7,9,9,3,6,15,24,5,4 | 11,9,2,4,15,10,11,15,4,12, | 234 | 149 | 0.0497531965526 | | 1 | 0.884,0.337,0.612, | 0.317,0.783,0.817, | -0,143 |
| 214320 | ENSG00000129472 | RAB2B | chr14 | - | 21466651 | 21466752 | 21463655 | 21463767 | 21468356 | 21468449 | 214320 | 3,2,1,5,0,3,0,1,0,0,1,4 | 42,17,45,43,26,11,8,3,31,1 | 1,0,0,3,1,0,0,0,1,2,3,0,0,2, | 25,16,13,14,26,16,18,25,1 | 250 | 149 | 0.0497553671821 | | 1 | 0.041,0.066,0.013, | 0.023,0.0,0.0,0.11 | 0,028 |
| 116181 | ENSG00000119314 | PTBP3 | chr9 | - | 112250928 | 112251103 | 112234819 | 112234897 | 112252677 | 112252788 | 116181 | 164,103,310,210,149,140, | 1,0,0,1,2,1,1,1,0,0,0,0 | 214,269,206,172,146,98,1 | 5,2,3,2,1,0,0,0,0,0,8,10,2,1 | 298 | 149 | 0.0497711957179 | | 1 | 0.988,1.0,1.0,0.99 | 0.955,0.985,0.972, | 0,016 |
| 167669 | ENSG00000101109 | STK4 | chr20 | + | 44975323 | 44975441 | 44972077 | 44972158 | 44978442 | 44978571 | 167669 | 7,3,31,14,0,5,6,16,2,10,2, | 166,90,299,182,96,74,81,1 | 11,28,6,6,13,4,2,4,2,11,1,5 | 209,231,139,109,89,76,66, | 267 | 149 | 0.0497744968463 | | 1 | 0.023,0.018,0.055, | 0.029,0.063,0.024, | 0,008 |
| 213204 | ENSG00000114978 | MOB1A | chr2 | - | 74159090 | 74159216 | 74152527 | 74156645 | 74165217 | 74165351 | 213204 | 173,149,370,241,172,141, | 0,4,6,2,3,0,4,0,4,5,0,1 | 364,480,135,158,150,102, | 9,11,4,5,3,1,3,3,3,0,5,9,0,3 | 275 | 149 | 0.0497872605837 | | 1 | 1.0,0.953,0.971,0. | 0.956,0.959,0.948, | 0,007 |
| 254746 | ENSG00000228144 | | chr12 | - | 66160735 | 66161172 | 66153339 | 66153448 | 66169854 | 66169985 | 254746 | 2,0,4,0,2,0,0,0,0,0,4,1 | 72,40,132,103,56,65,73,80 | 4,5,4,0,1,0,0,2,0,3,3,4,9,4, | 94,66,65,16,41,35,27,51,1 | 298 | 149 | 0.0498049949922 | | 1 | 0.014,0.0,0.015,0. | 0.021,0.036,0.03,0 | -0,011 |
| 135101 | ENSG00000196236 | XPNPEP3 | chr22 | + | 40883151 | 40883226 | 40881769 | 40882177 | 40886312 | 40886515 | 135101 | 0,0,3,3,0,2,0,3,0,1,2,0 | 6,2,13,14,2,4,5,2,2,0,4,15 | 0,0,2,3,0,0,0,0,1,0,1,0,0,1, | 5,12,6,15,4,3,7,12,9,8,13,1 | 224 | 149 | 0.049816525984 | | 1 | 0.0,0.0,0.133,0.12 | 0.0,0.0,0.181,0.11 | 0,146 |
| 69269 | ENSG00000126775 | ATG14 | chr14 | - | 55385858 | 55386096 | 55381961 | 55382191 | 55390910 | 55390992 | 69269 | 90,69,101,66,40,58,61,54, | 6,0,3,1,0,1,1,2,0,0,0,3 | 77,100,57,78,58,22,32,53, | 1,3,0,0,2,0,2,0,1,0,3,0,0,0, | 298 | 149 | 0.0498233888698 | | 1 | 0.882,1.0,0.944,0. | 0.975,0.943,1.0,1. | -0,014 |
| 95405 | ENSG00000111832 | RWDD1 | chr6 | + | 116574057 | 116574171 | 116571488 | 116571655 | 116580294 | 116580360 | 95405 | 45,16,42,21,17,12,26,15,2 | 34,17,20,29,15,14,18,13,5 | 26,29,31,23,19,13,24,17,1 | 17,18,16,15,15,8,5,27,4,18 | 263 | 149 | 0.049836690675 | | 1 | 0.429,0.348,0.543, | 0.464,0.477,0.523, | -0,091 |
| 224306 | ENSG00000083642 | PDS5B | chr13 | + | 32674609 | 32674745 | 32673215 | 32673356 | 32675843 | 32675959 | 224306 | 5,0,5,2,0,4,0,4,0,1,0,0 | 53,30,31,12,22,20,21,14,9, | 1,9,0,7,1,2,6,1,4,4,0,0,1,3, | 27,37,7,50,17,21,22,22,13, | 285 | 149 | 0.0498446899661 | | 1 | 0.047,0.0,0.078,0. | 0.019,0.113,0.0,0. | -0,023 |
| 29179 | ENSG00000134644 | PUM1 | chr1 | - | 31006993 | 31007102 | 30995053 | 30995220 | 31028795 | 31028864 | 29179 | 89,30,100,50,32,40,26,36, | 7,1,7,5,0,4,0,5,0,3,2,3 | 72,77,40,54,44,41,11,30,2 | 9,2,2,1,7,5,0,3,4,4,7,4,7,6, | 258 | 149 | 0.0498704825972 | | 1 | 0.88,0.945,0.892,0 | 0.822,0.957,0.92,0 | 0,042 |
| 19612 | ENSG00000174485 | DENND4A | chr15 | - | 65700543 | 65700672 | 65697266 | 65697383 | 65701050 | 65701192 | 19612 | 27,13,45,9,6,8,4,9,16,10,9, | 133,159,163,85,38,43,29,5 | 16,17,11,31,13,14,9,16,9,3 | 93,113,49,106,91,37,30,61 | 278 | 149 | 0.0498711757969 | | 1 | 0.098,0.042,0.129, | 0.084,0.075,0.107, | -0,02 |
| 1629 | ENSG00000183765 | CHEK2 | chr22 | - | 28730419 | 28730548 | 28725242 | 28725367 | 28734402 | 28734727 | 1629 | 8,0,5,1,0,2,0,3,0,3,1,0 | 12,8,15,10,7,4,1,2,4,4,0,10 | 6,1,1,2,4,4,0,1,4,6,0,4,6,5, | 11,5,2,0,3,6,3,6,13,18,5,17 | 278 | 149 | 0.0498716459697 | | 1 | 0.263,0.0,0.152,0. | 0.226,0.097,0.211, | -0,014 |
| 247402 | ENSG00000149311 | ATM | chr11 | + | 108244934 | 108245026 | 108243961 | 108244118 | 108246963 | 108247127 | 247402 | 27,9,159,80,13,19,9,21,16, | 4,0,2,0,0,0,0,0,2,0,1,0 | 75,63,7,8,33,34,10,15,36,4 | 9,10,0,0,4,0,0,0,0,1,0,0,0,0 | 241 | 149 | 0.0498783847205 | | 1 | 0.807,1.0,0.98,1.0, | 0.837,0.796,1.0,1. | 0,01 |
| 8967 | ENSG00000182446 | NPLOC4 | chr17 | - | 81563920 | 81563979 | 81559090 | 81559416 | 81565504 | 81565607 | 8967 | 0,0,16,10,6,7,10,7,4,8,5,6 | 13,12,62,57,16,6,53,34,46, | 4,0,0,1,3,0,3,3,2,6,1,0,7,2, | 14,24,14,6,7,12,10,18,8,17 | 208 | 149 | 0.0498835932828 | | 1 | 0.0,0.0,0.156,0.11 | 0.17,0.0,0.0,0.107, | 0,037 |
| 62378 | ENSG00000111877 | MCM9 | chr6 | - | 118826146 | 118826292 | 118813441 | 118816294 | 118826781 | 118826864 | 62378 | 43,21,82,20,17,20,15,22,2 | 22,22,14,3,4,6,5,4,4,15,3,2 | 36,49,18,33,37,7,26,13,24, | 8,8,3,11,5,4,7,3,5,7,4,4,4,3 | 295 | 149 | 0.0499036151605 | | 1 | 0.497,0.325,0.747, | 0.694,0.756,0.752, | -0,084 |
| 143507 | ENSG00000120071 | KANSL1 | chr17 | - | 46162476 | 46162525 | 46094559 | 46094701 | 46170854 | 46172232 | 143507 | 2,0,2,1,1,0,1,2,0,3,0,0 | 72,53,67,46,77,88,35,37,4 | 3,0,3,2,2,0,2,3,3,1,3,2,4,3, | 96,108,70,141,35,31,42,55 | 198 | 149 | 0.0499552597243 | | 1 | 0.02,0.0,0.022,0.0 | 0.023,0.0,0.031,0. | -0,016 |
| 152521 | ENSG00000258643 | BCL2L2-PABPN | chr14 | + | 23323938 | 23324052 | 23323376 | 23323483 | 23324137 | 23324289 | 152521 | 276,158,260,140,89,128,1 | 9,3,3,7,5,2,3,6,16,6,1,7 | 170,139,134,86,115,84,51, | 8,1,4,0,4,1,0,6,4,1,4,6,2,3, | 263 | 149 | 0.0499587833907 | | 1 | 0.946,0.968,0.98,0 | 0.923,0.987,0.95,1 | -0,016 |
| 152000 | ENSG00000171606 | ZNF274 | chr19 | + | 58210166 | 58210430 | 58209960 | 58210073 | 58211559 | 58211686 | 152000 | 16,10,23,2,4,0,6,12,3,5,5,3 | 50,26,59,28,33,32,22,34,8 | 8,8,14,30,5,7,11,5,2,6,16,4 | 36,56,52,25,25,9,15,32,5,6 | 298 | 149 | 0.0499606729778 | | 1 | 0.138,0.161,0.163, | 0.1,0.067,0.119,0. | -0,055 |
| 199250 | ENSG00000076321 | KLHL20 | chr1 | + | 173725703 | 173725858 | 173716002 | 173716066 | 173733712 | 173734513 | 199250 | 11,5,5,5,0,6,1,6,2,2,5,1 | 11,6,28,14,13,16,5,8,1,7,9, | 13,4,3,6,5,0,2,7,2,3,0,0,0,0 | 11,26,2,5,22,5,11,20,1,11,1 | 298 | 149 | 0.0499749794566 | | 1 | 0.333,0.294,0.082, | 0.371,0.071,0.429, | 0,048 |
| 10315 | ENSG00000170836 | PPM1D | chr17 | + | 60656931 | 60657042 | 60656598 | 60656841 | 60662994 | 60664473 | 10315 | 5,2,0,3,0,2,0,2,1,3,2,2 | 5,3,49,10,8,10,13,15,7,6,7, | 7,4,13,8,0,0,3,0,2,6,4,2,4,4 | 11,38,22,16,11,4,22,15,7,2 | 260 | 149 | 0.0499848278385 | | 1 | 0.364,0.276,0.0,0. | 0.267,0.057,0.253, | 0,007 |
